# Mink by mink: stitching together signatures of subspecies adaptation through a pangenome of threatened mustelids

**DOI:** 10.1101/2025.07.25.666817

**Authors:** Aishatu Affini, Hailey Baranowski, Scott Forbes, Elena N. Foust, Kristyn Hatley, Ethan L Ni, Airianna McGuire, Kyle Paist, Mary Rutter, Robin N. Smith, Nataly Vargas, Harshita Akella, Kate Castellano, Nicole Pauloski, Noah Reid, Rachel O’Neill, Teisha King, Elizabeth L. Jockusch, Jill L. Wegrzyn, Paul Hapeman

## Abstract

The American mink (*Neogale vison*), a semi-aquatic Mustelidae carnivoran with broad ecological range across North America, includes several putative subspecies of conservation concern. To investigate the evolutionary history and adaptive signatures of mink subspecies, chromosome-scale genome assemblies were generated for six individuals representing three southern subspecies: *N. vison evergladensis*, *N. vison vulgivaga*, and *N. vison lutensis*. Genomes were assembled using Illumina short reads, scaffolded with Oxford Nanopore long reads, and aligned to the phased *N. vison* reference genome. Assemblies ranged from 75.9% to 97.8% completeness, with five meeting thresholds for pangenome construction. A reference-free pangenome revealed an open architecture, highlighting considerable subspecies diversity. Subspecies-specific gene enrichment reflected adaptation: *N. vison evergladensis* showed enrichment in traits related to reproduction and sensory function; *N. vison vulgivaga* in cytoskeletal remodeling and oxidative stress; and *N. vison lutensis* in neuronal development, synaptic plasticity and cellular stress pathways. Assessment of the mitogenomes resolved *N. vison lutensis* as a distinct lineage, while nuclear data supported broader subspecies divergence but lacked fine scale resolution. *N. vison evergladensis* showed multiple signatures of small population size, including inbreeding coefficients (F_ROH_) above 0.5, and displayed consistent population decline over time via demographic inference. Our findings support *evergladensis* as a distinct subspecies, supported by both the mitogenome phylogeny, and significant functional differentiation. As the first pangenome for Mustelidae, this study demonstrates the power of integrating cross-platform sequencing with natural history specimens to improve the resolution on signatures of adaptation and inform conservation policy and management of threatened populations.

## Introduction

Accurate subspecies delineation is critical for effective conservation management (Zink and Klicka 2022). Differences in habitat, breeding time, and morphology (Burbrink et al. 2022; Walsh et al. 2017) as well as genetic markers (Haig et al. 2006; Martien et al. 2017; Wirgin et al. 2015) have been used to support the delineation of subspecies. However, these methods often prove insufficient to resolve evolutionarily distinct lineages. Inaccurate taxonomic assignment hinders estimation of population size and diversity that aid the development of conservation listing and management.

Advances in sequencing and assembly technologies have made the collection of genomic data more affordable, providing a tool to understand complex population histories and detect fine-scale differentiation for subspecies classifications (du Plessis et al. 2023; Wang et al. 2023). While reference genome assemblies provide a valuable resource for comparative and functional studies, pangenomes offer a more accurate tool for understanding the genetic diversity of putative subspecies by enabling assessments of evolutionary history, adaptation, and population divergence (Kim et al. 2021; Rajput et al. 2023). There is great potential for the use of pangenomes in conservation but its application has largely been limited to comparisons among viruses, bacteria, and domesticated animals (Gong et al. 2023).

The American mink (*Neogale vison*) is a semi-aquatic, mustelid native to wetland and riparian habitats throughout North America except the southwestern United States (Chapman and Feldhamer 1982) (Fig. 1A). Mink have also been introduced to many other parts of the world for fur farming, including Europe and South America (Bonesi and Palazon 2007; Reid et al. 2016). Fifteen subspecies of mink have been reported throughout their native range (Wozencraft 2005), but the subspecies designations have not been thoroughly evaluated. Several subspecies of mink occur in Florida and are of increasing conservation concern due to knowledge gaps in ecological information and threats from climate change and habitat loss (Florida Fish and Wildlife Conservation Commission 2016).

**Fig 1A:**
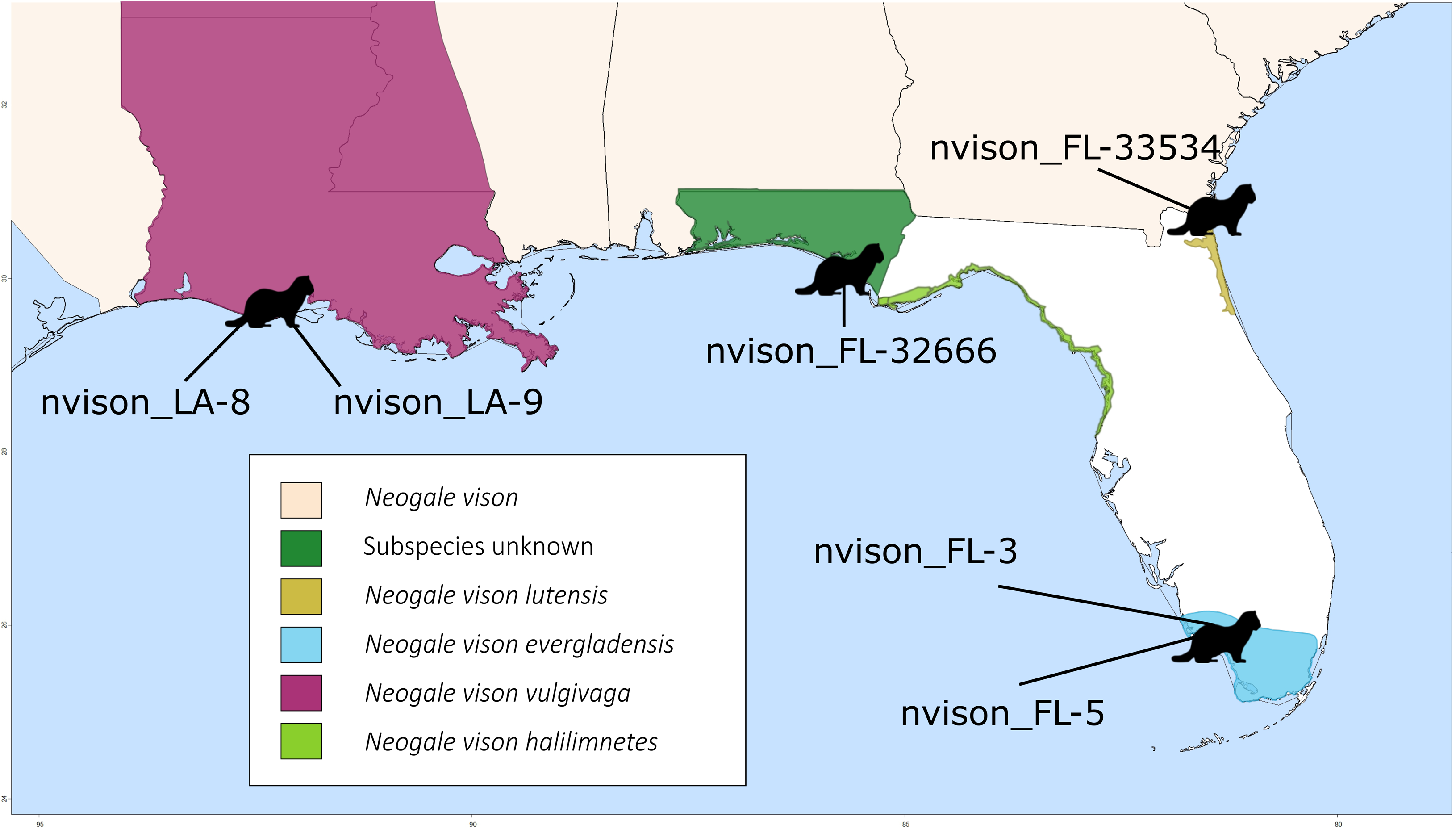
American mink range map with sampling locations and putative subspecies ranges. The map shows the *Neogale vison* species range using data imported from the IUCN (International Union for the Conservation of Nature) Red List database (Reid et al., 2016), as well as subspecies range estimates from Hapeman and Smith (2024). The Florida subspecies estimates are sourced from Hapeman and Smith (2024), and the *vulgivaga* range in Louisiana was adapted from Trani and Chapman (2007). Silhouettes indicate approximate sample locations.

The Everglades mink (*N. v. evergladensis*) of south Florida was originally described as a subspecies based on a single specimen (Hamilton 1948) but a morphometric analysis of mink samples from the southeastern United States concluded that *evergladensis* was a disjunct population of *N. vison* rather than a subspecies (Humphrey and Setzer 1989) (Fig. 1B). A recent study provided support for *evergladensis* as a separate taxonomic unit using mtDNA *Cyt b* and microsatellites (Hapeman and Smith 2024). While *evergladensis* is currently a state-listed threatened subspecies, its taxonomic status requires further investigation.

**Fig 1B:**
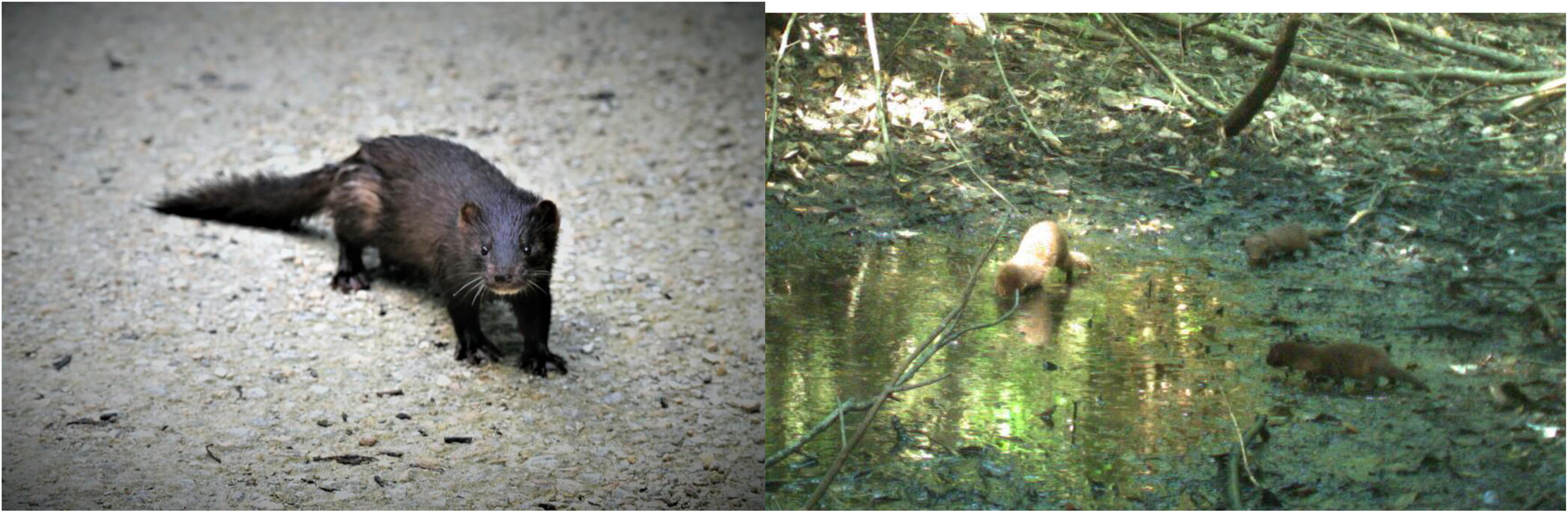
Everglades mink (*Neogale vison evergladensis*). Left: Photograph by user dalempijevic1, taken Aug 14, 2022 (iNaturalist; https://www.inaturalist.org/observations/130841747). Right: Image courtesy of Florida Fish and Wildlife Conservation Commission.

To resolve subspecies of *N. vison*, we assembled whole genomes from samples of American mink collected from populations in Florida and Louisiana and derived a pangenome. With the data, we investigated the evolutionary and phylogenetic relationships among subspecies, characterized demographic history and inbreeding coefficients, and identified genes that may contribute to local adaptation.

## Methods and Materials

### Sample Collection

Tissue samples of American mink (*Neogale vison*) were collected following protocols outlined in Hapeman and Smith (2024). The individuals used in this study include nvison_FL-3 (2004 Collier County, Florida (*evergladensis*; roadkill)); nvison_FL-5, (2005 the same locality and subspecies); nvison_FL-32666, (2000 Bay County, Florida (unknown subspecies), archived at the Florida Museum of Natural History); nvison_FL-33534 (2016 Nassau County, Florida (*lutensis*), archived at the Florida Museum of Natural History; and nvison_LA-8 and nvison_LA-9 (both 2015 Vermillion Parish, Louisiana (*vulgivaga*) using live trapping with assistance from the Louisiana Department of Wildlife and Fisheries). All tissue samples were preserved in ethanol.

### DNA Extraction, Library Preparation, and Sequencing

High molecular weight (HMW) DNA was extracted using a modified phenol-chloroform-isoamyl alcohol protocol (Green et al., 2012) optimized for efficient tissue lysis. DNA was precipitated with ethanol, resuspended in nuclease-free water, purified with Ampure XP beads, and quantified using spectrophotometry (NanoDrop™ One) and fluorimetry (Qubit™ 4). HMW DNA was recovered from each sample at quantities sufficient for library preparation, with the exception of nvison_LA-8 and nvison_FL-32666, which were too low in concentration or degraded, respectively.

Illumina sequencing was conducted at the University of Connecticut’s Center for Genome Innovation. Libraries were prepared using the NEBNext Ultra™ II FS DNA Library Prep Kit (New England Biolabs), with fragmentation times adjusted based on DNA quality to achieve target insert sizes of 300–350 base pairs. For the degraded sample, nvison_FL-32666, a second library was prepared without fragmentation. All libraries were normalized, pooled, and sequenced on the Illumina NovaSeq 6000 S4 platform using v1.5 chemistry with a target depth of approximately 80X.

Oxford Nanopore Technologies (ONT) sequencing was also performed at the Center for Genome Innovation using the PromethION platform with FLO-PRO114M flow cells and the SQK-LSK114 ligation sequencing kit.

Samples nvison_LA-9, nvison_FL-3, nvison_FL-5, and nvison_FL-33534 were sequenced using both Illumina and Oxford Nanopore platforms. Sample nvison_LA-8 and both libraries of nvison_FL-32666 were sequenced using Illumina only. Detailed extraction and library preparation protocols are provided in <u>File S1</u>.

### Genome Assembly, and Scaffolding

We performed a quality assessment of ONT long reads using Nanoplot (v1.33) (De Coster & Rademakers, 2023). Reads were screened for potential contamination using Centrifuge (v1.0.4) (Kim et al., 2016) with a custom database containing fungal, bacterial, archaeal, and viral sequences from NCBI RefSeq. Additional filtering was performed using Seqkit (v2.8.1) (Shen et al., 2024). Genome size estimates were generated using k-mer distributions computed with kmerfreq (v4.0) and GCE (v1.0.2) (Liu et al., 2013).

Illumina short reads were similarly assessed for contamination with Kraken (v2.1.2) (Wood et al., 2019), and filtered using Seqkit. Due to elevated adapter read-through, Fastp (v0.23.2) (Chen et al., 2018) was used to trim reads from nvison_FL-32666. All other samples were used as cleaned but untrimmed reads in accordance with MaSuRCA recommendations.

Genome assemblies were generated using MaSuRCA (v4.1.2) (Zimin et al., 2017) in short-read mode for all samples. The sample nvison_LA-9, which had a more contiguous long-read dataset, was additionally assembled using Flye (v2.9.1) (Kolmogorov et al., 2019). Long-read and short-read assemblies for samples with ONT data (nvison_FL-3, nvison_FL-5, nvison_FL-33534, and nvison_LA-9) were scaffolded to the 15 pseudochromosomes of the phased *Neogale vison* reference genome (haplotype 1, NCBI BioProjectID: PRJEB76007) using RagTag (v2.1.0) (Alonge et al., 2022). Assemblies were further polished and gap-filled using SAMBA (v4.1.1) (Zimin & Salzberg, 2022). For nvison_FL-5, SAMBA was run prior to RagTag due to the high contiguity of the initial assembly.

Assembly quality was assessed using QUAST (v5.2.0) (Gurevich et al., 2013) and Compleasm (v0.2.6) (Li & Huang, 2023) with the carnivora_odb10 database. Genome accuracy was evaluated using Merqury (v1.3) (Rhie et al., 2020) by comparing each chromosome-scale assembly to k-mer profiles derived from the corresponding trimmed Illumina reads.

Final assemblies underwent contamination screening using NCBI’s Foreign Contamination Screen-GX (FCS-GX) and FCS-Adapter (v0.5.4) (Astashyn et al., 2024). Sequences identified as non-host or adapter-derived were hardmasked prior to submission.

### Structural and Functional Genome Annotation

RepeatModeler (v2.0.4) (Smit et al., 2008–2015) was used to identify and classify repeats in each genome and generate repeat libraries, which were used by RepeatMasker (v4.1.5) (Smit et al., 2013–2015) to softmask each assembly prior to gene annotation.

Next, we performed gene annotation using EGAPx (https://github.com/ncbi/egapx), which integrates protein and transcriptomic evidence with hidden Markov models (HMMs) for gene prediction. Up to sixteen publicly available *N. vison* RNA-seq libraries from NCBI were included based on sequencing depth (greater than 20 million paired reads) and tissue diversity. The NCBI taxonomic identifier 452646 enabled automatic selection of relevant protein evidence. We then evaluated the completeness of annotations using Compleasm (v0.2.6) with the carnivora_odb10 database (Li & Huang, 2023), and structural quality using AGAT (Dainat, n.d.). Functional annotations were generated using EnTAP (v2.0.0) (Hart et al., 2020). The bioinformatics workflow is shown in Fig. 1C.

**Fig 1C:**
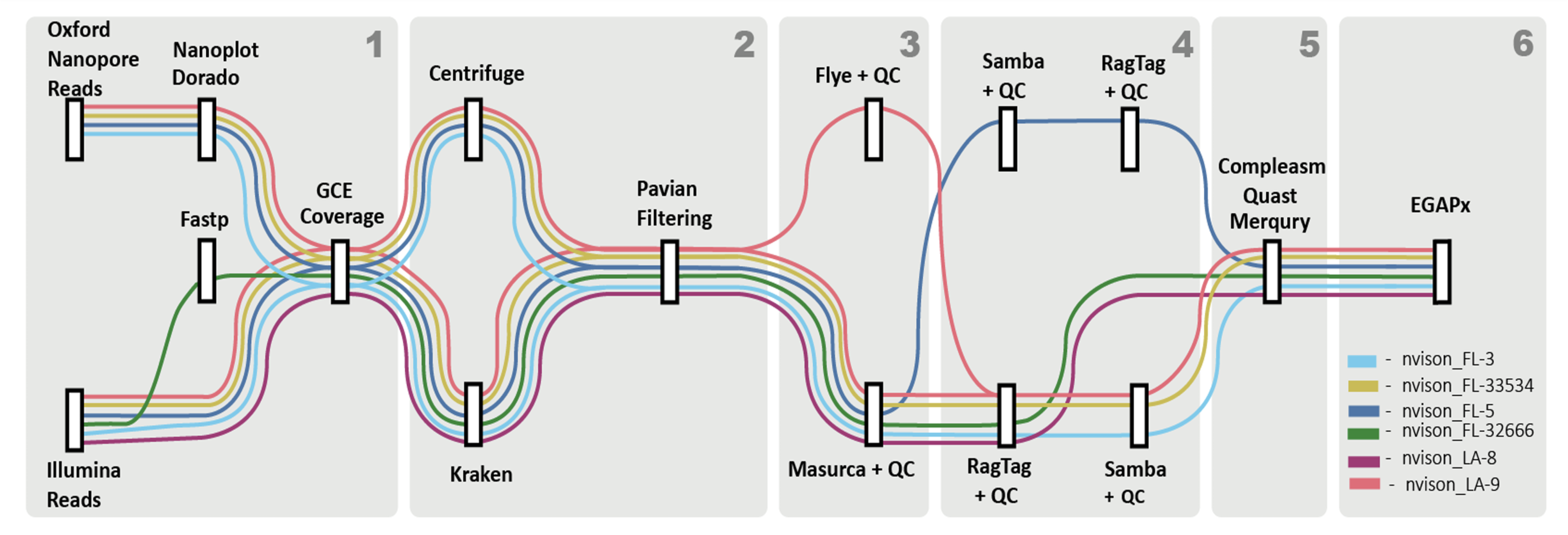
Informatic workflow from sequencing to pangenome construction. The workflow illustrates the bioinformatic steps each sample underwent for pangenome construction. The color coding of the samples reflects their corresponding putative species as follows: *vulgivaga* samples (nvison_LA-8; nvison_LA-9; pink), *evergladensis* samples (nvison_FL-3; nvison_FL-5; blue), unknown sample (nvison_FL-32666; dark green), and the *lutensis* sample (nvison_FL-33534; yellow). The boxes with numerical values in the upper right (1-6) represent the major steps of the workflow: (1) Read quality control and genome size estimation, (2) Contamination identification and filtering, (3) Assembly, (4) Polishing, gap filling and scaffolding, (5) Genome contiguity and completeness, (6) Annotation.

### Genome-wide Synteny

Synteny among assemblies (excluding nvison_FL-33534) was analyzed using MCScanX (v1.0.0) (Wang et al., 2012), and visualized with SynViso (https://github.com/kiranbandi/synvisio). DIAMOND (v2.1.8) (Buchfink et al., 2015) was used to align protein sequences using sample-specific protein databases. For each genome, the longest protein isoforms were used for both alignment and database construction. For subspecies comparisons, chromosomes were anchored to the reference assembly, and gene order in other assemblies was adjusted based on syntenic relationships.

### Gene Family Evolution

Orthologous gene families were identified using OrthoFinder (v2.5.4) (Emms & Kelly, 2019) with all six study genomes, the reference genome nvison_hap1, and 18 additional species of Carnivora across the families Mustelidae, Ursidae, Phocidae, Felidae, Hyaenidae, and Canidae (Table S1). Compleasm (v0.2.6) with the carnivora_odb10 database was used to confirm a minimum of 85% gene space completeness prior to comparative analysis.

OrthoFinder followed default settings using DIAMOND for sequence similarity and MAFFT for multiple sequence alignment. Phylogenetic trees were reconstructed using IQ-TREE (v3.0.0) (Wong *et al*., 2025). Following initial evaluation, OrthoFinder was subsequently re-run with the subspecies tree amended for consistency with the mitogenome.

OrthoFinder output included orthogroups and gene copy number for each species. Functional annotation of gene families was performed using EggNOG (v2.2.12) (Huerta-Cepas et al., 2018). Gene family contraction and expansion were analyzed using CAFE (v5.0) (De Bie et al., 2006), which implements a birth-death model and uses the OrthoFinder-derived phylogeny to estimate evolutionary rates (lambda values) and p-values for each orthogroup. Gene families with unusually high copy numbers were excluded to avoid skewing lambda estimates. CafePlotter (v0.2.0) (Moshiashvili, 2024) was used to visualize gene family evolution across the phylogeny. Functional categories from EggNOG were incorporated into CafePlotter to contextualize gene family evolution.

### Genomic Estimates of Inbreeding and Coalescent Modeling

Coalescent modeling was performed using SMC++ (v1.15.4) (Terhorst et al., 2017) to infer recent demographic history and divergence per individual. This method reconstructs local genealogies and identifies recombination events across the genome (Mather et al., 2019).

Short-read Illumina data from each sample were aligned to the chromosome-scale haplotype 1 reference using BWA-MEM2 (v2.2.1) (M. Vasimuddin, 2013), and read groups were assigned accordingly. BAM files were analyzed with Mosdepth (v0.3.8) (Pedersen & Quinlan, 2017) to assess genome-wide coverage, using 1000 bp windows and a minimum mapping quality of 30. Callable, low quality, and high quality regions were identified using Mosdepth’s “quantize” feature after setting sample dependent thresholds (Table S2). Regions input to the masked bed file used in SMC++ were those deemed as no converge and those where coverage was exceedingly high (Table S2).

Variant sites were called using GATK (v4.3.0.0) (Van der Auwera & O’Connor, 2020), following best practices. Variants were generated using HaplotypeCaller per chromosome and merged into a genome-wide VCF. The X chromosome, derived from the female reference individual, was excluded. Generation time was estimated as two years based on a lifespan of three to four years (Ohio Department of Natural Resources, n.d.) and reproductive parameters reported by Amstislavsky and Ternovskaya (2000). A mutation rate of 1×10⁻⁸ substitutions per site per generation was applied following a previous study of mustelids (Derežanin et al., 2021). SMC++ was run with cubic spline interpolation to reconstruct demographic trajectories. Samples were assigned to a population based on subspecies, for a total of four populations (*vison, evergladensis, vulgivaga,* and *lutensis*). The model for each population was plotted to assess demographic history.

We assessed inbreeding using Runs of Homozygosity (ROH) generated with bcftools (v1.19) (Li et al., 2009) command *roh*, and we retained segments > 500 Kb through filtering. Inbreeding coefficients were generated with PLINK (v1.9.0-b7.7) (Purcell et al., 2007) from the GATK-derived variants from the coalescent analysis. ROH analysis was conducted with bcftools (v1.19).

### Mitogenomes and Phylogenetic Inference

NOVOPlasty (v4.3.5) (Dierckxsens et al., 2017) was used to assemble circular mitochondrial genomes with Illumina short reads, the *N. vison* mitochondrial reference genome (NC_020641), and CO1 seed sequence (OZ067098.1). Assemblies were annotated in Geneious Prime (v2025.1.3) using the Transfer Annotation by Homology function after aligning each sample to NC_020641 and the outgroup *N. frenata* (NC_020640) using MAFFT(v7.490).

A maximum likelihood tree of mitochondrial genomes was inferred in IQ-TREE (v3.0.0) (Wong et al., 2025), with 42 candidate partitions (one for each codon position in each protein coding gene, 16S rRNA, 28S rRNA, and the concatenated tRNAs). Actual partitions and models for each partition were selected using the MFP+MERGE option. Support was calculated using 1000 non-parametric bootstrapping replicates. *Neogale frenata* was included as an outgroup. Mitochondrial diversity metrics and pairwise comparisons were calculated using MEGA11 (Tamura et al., 2021) and R packages ape (Paradis et al., 2004) and pegas (Paradis, 2010).

### Pangenome Construction

Pantools (v4.3.3) (Sheikhizadeh et al., 2016) was used to build a reference-free pangenome using chromosome-scale assemblies and annotated protein-coding genes from all samples, including nvison_hap1. The assembly nvison_FL-33534 was excluded due to low completeness. Subspecies designations were included as phenotypic metadata to track gene presence–absence variation and gene function by group.

Database construction included annotation data and homology group inference. The “optimal-grouping” function was used to determine the best relaxation parameter for clustering homologs based on F1 and precision scores. Relaxation level 3, corresponding to 75% sequence similarity, was selected as the threshold for grouping genes. Compleasm (v0.2.6) with the carnivora_odb10 database was used to classify genes as core (present in all genomes), accessory (present in some genomes), or unique (present in one genome). Gene distributions were also analyzed by subspecies using the “functional_classification” module to identify shared, specific, and exclusive gene content. The structure of the pangenome was examined to determine whether it followed an open or closed model based on the accumulation of novel genes across genomes.

### Gene Ontology (GO) Enrichment

Gene Ontology (GO) enrichment analysis using Pantools identified overrepresented biological processes in each subspecies. Genes were grouped by subspecies, with homology nodes as the foreground set and the remaining pangenome as the background. GO terms with p < 0.01 were retained. Gene ratios were calculated relative to the background, and enrichment status, associated genes, and presence–absence variation were recorded.

## Results

### Sequencing

Illumina (150bp PE) data generated for six samples had initial depths ranging from 96X to 193X coverage. Sample nvison_FL-32666 had two Illumina libraries, sheared and unfragmented, and had the highest Illumina coverage across both libraries (158X and 193X, respectively), while nvison_FL-5 had the lowest coverage (96X) (Table S3). Post-contaminant filtering with Kraken reduced the coverage from 82X to 187X across all samples (Tables S3; Table S4).

We sequenced long reads (ONT) for five of the samples, with initial coverage ranging from 4X to 34X. In all cases, read lengths were low, with N50s 2 Kb or less (435 bp to 2 Kb). Sample nvison_LA-9 and nvison_FL-3 had the highest ONT coverage (34X and 30X, respectively), while nvison_FL-33534 had the lowest (4X). After contaminant filtering with Centrifuge, coverage ranged from 4X to 33X (Table S3).

### Genome Assembly

Initial assemblies using MaSuRCA in short-read mode were highly fragmented, with completeness ranging from 21.6% to 34.7%, N50 values between 1.2 and 6.2 Kb, and contig counts from 593K to over 2.1M. nvison_FL-5 had the best initial assembly (34.7% completeness, N50: 6.2 Kb, 593,378 contigs), while nvison_FL-32666 was the most fragmented (21.6% completeness, N50: 1.2 Kb, 2.16M contigs). A long-read assembly of nvison_LA-9 showed substantial improvement over all short-read assemblies, with 64.7% completeness, a 40 Kb N50, and 109,919 contigs. We scaffolded all assemblies to haplotype 1 of the mink reference, which dramatically improved quality metrics. nvison_LA-8 and nvison_FL-3 showed the greatest gains, with completeness increasing to over 72%, N50s over 220 Mb, and contig counts reduced by over 1 million. nvison_LA-9 reached 74.5% completeness and a 225.7 Mb N50, while nvison_FL-32666 improved to 66.6% completeness and a 247.6 Mb N50. nvison_FL-33534 remained the most fragmented after scaffolding, with 49.1% completeness and a 203.9 Mb N50. Across all samples, 97.7–98.1% of each assembly was placed within the 15 expected pseudo-chromosomes (Table 1; Table S5,). Polishing and gap filling with SAMBA further improved assembly quality. nvison_FL-5, the only sample polished before scaffolding, improved to 63.1% completeness and a 209.2 Mb N50.

**Table 1:**
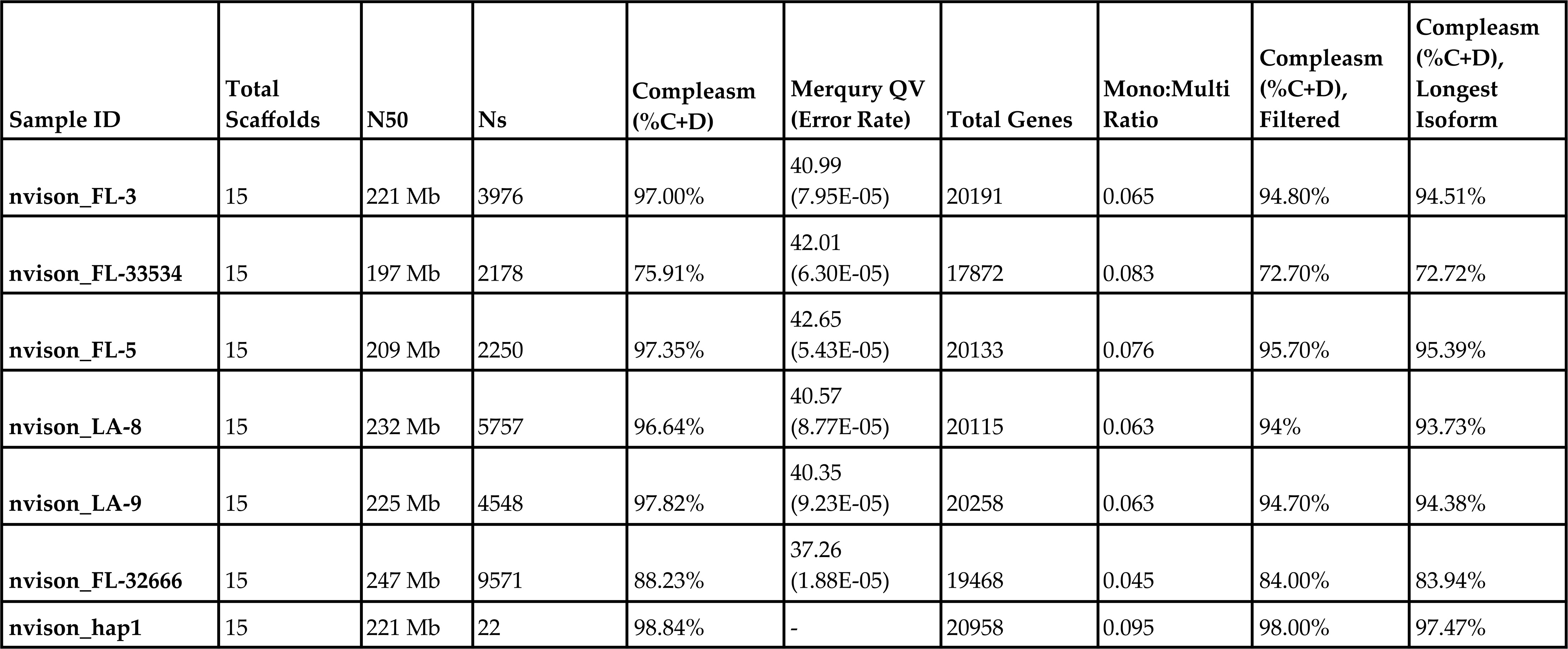
Sequence read, assembly, and structural annotation statistics.

Post-scaffolding polishing increased contiguity and completeness in all other samples, including nvison_FL-3 (+65.1%, 221.5 Mb N50), nvison_FL-33534 (+46.8%, 203.9 Mb N50), and nvison_LA-9 (+24.3%, 217.5 Mb N50).

We evaluated final chromosome-scale assemblies (15 pseudo-chromosomes) for completeness, contiguity, and correctness (Table 1; Table S5). The most complete assembly was from nvison_LA-9 (97.05% completeness, N50: 225 Mb, QV: 40.35, 2.73 Gb total length), though it had relatively large gaps. nvison_FL-3, nvison_FL-5 and nvison_LA-8 all had genome completeness around 96%, N50 values around 200 Mb, and variable gapping. The quality scores of nvison_FL-3 and nvison_LA-8 were similar at 40.99 and 40.56, while nvison_FL-5 was slightly higher at 42.69. Genomes of the museum samples were less complete (76.2–87.6%), with more gaps and lower quality scores (37.52, 42.01). N50 values were similar to the those of the other assemblies (197 Mb, 203 Mb) (Table 1; Table S5). All genome assemblies were highly syntenic with the reference genome (Fig. 1D).

**Fig 1D:**
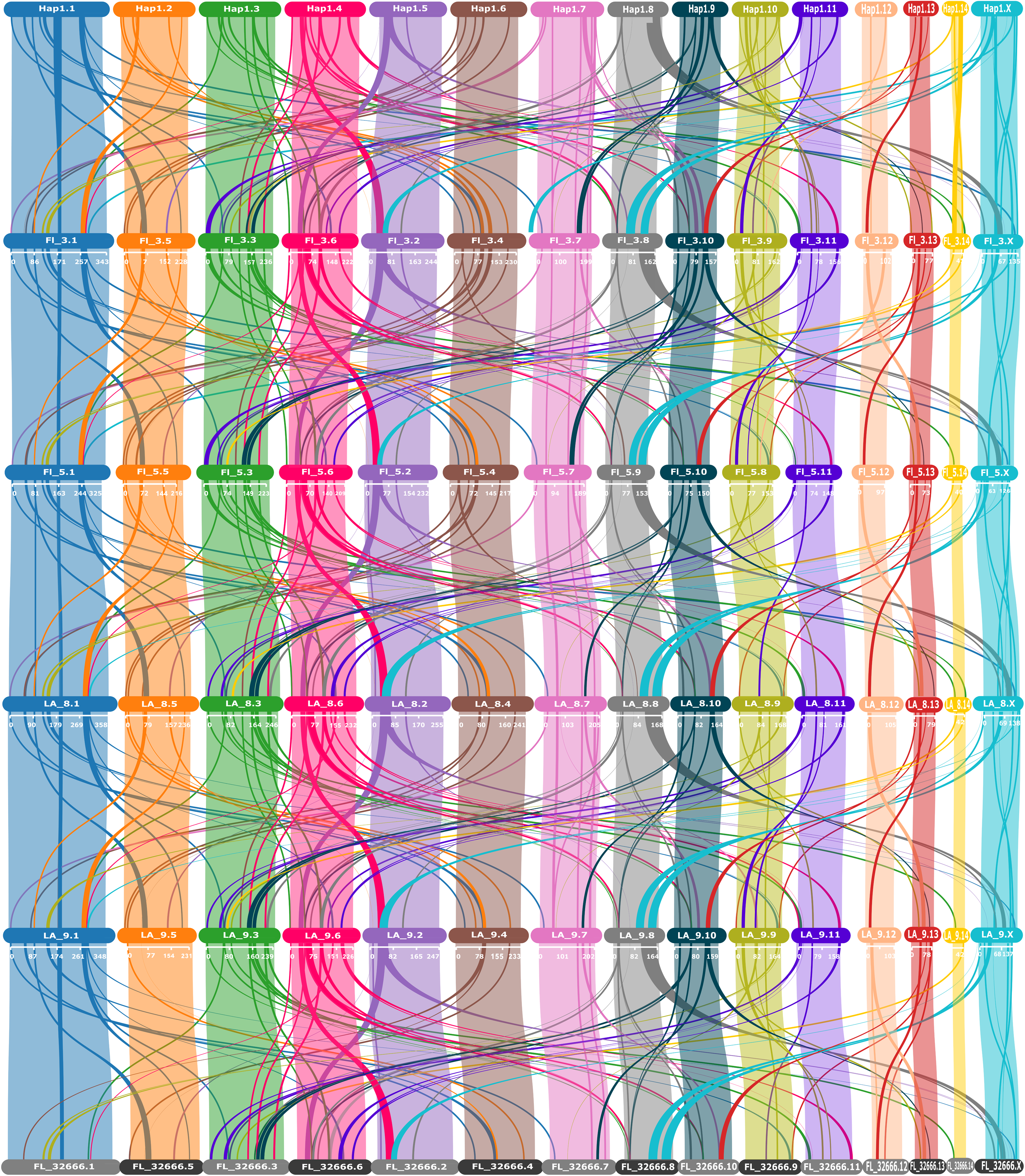
Alignment of 15 pseudo-chromosomes from each sample to the *N. vison* reference genome (*N.* vison_hap1). Each sample’s assembly was anchored to the published *N. vison* reference genome, with 15 pseudo-chromosomes displayed, including the X chromosome. Labels are formatted as sample ID followed by chromosome number (SampleID_1).

The total proportion of hardmasked bases, comprising both adapter and mitochondrial sequences, was minimal across all assemblies: 0.004% (nvison_FL-32666), 0.006% (nvison_FL-33534), and 0.0002% (nvison_LA-8).

### Structural and Functional Annotation

The pseudo-chromosome scale assemblies for all six genomes were softmasked and annotated for repetitive elements, with total masked content ranging from 29.4% to 31.9% (Tables S6 and S7). Retroelements comprised the majority (22.7%–27.7%), followed by DNA transposons (1.26%–1.53%) and unclassified repeats (1.17%–2.92%). We annotated all individuals using EGAPx, with 10 to 16 RNA-Seq libraries aligned per sample and alignment rates between 65.6% and 94.9% (mean: 87.1%) (Table S8). Annotation completeness ranged from 72.7% to 95.4%, with protein-coding gene counts between 17,872 and 20,258 (Table S8). We observed the highest completeness in nvison_FL-5 and the reference genome (nvison_hap1), while lower completeness was associated with museum-preserved samples.

### Coalescence

The demographic plot (Fig. 1E) shows effective population size (N_e_) trajectories inferred from each sample over the past 100,000 years. *vulgivaga* maintained the most stable N_e_, with a gradual decline that plateaued ∼10,000 years ago. *evergladensis* experienced the steepest N_e_ decline, beginning around 6,000 years ago. *lutensis* and *N. vison* showed similar histories, with sharp declines ∼20,000 years ago followed by moderate fluctuations.

**Fig 1E:**
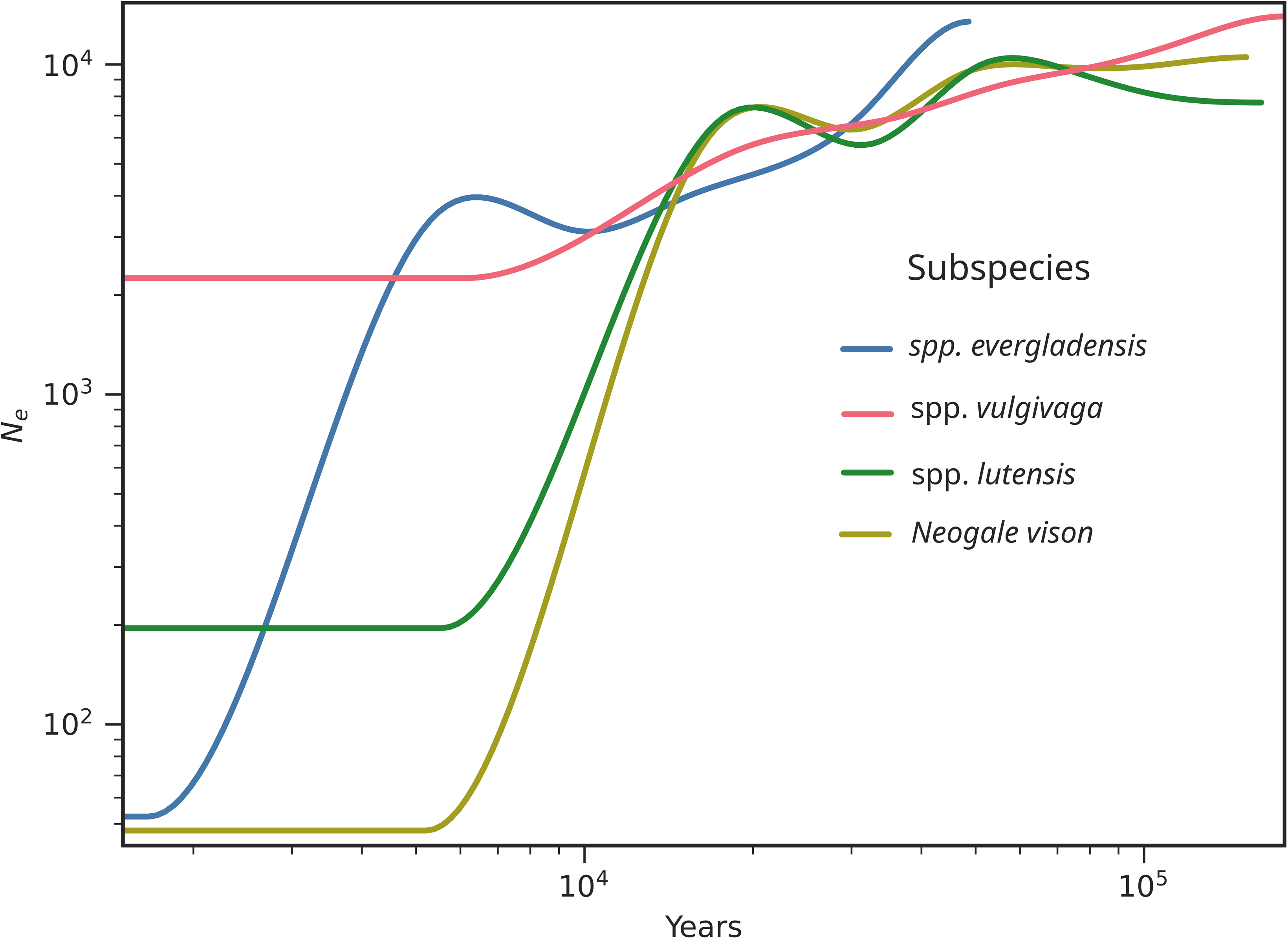
Preliminary demographic history plot. All samples (solid lines) are mapped to a single reference genome, with a generation time of two years and a mutation rate of 1.0e-8. Each curve represents changes in Ne over time for each subspecies.

Inbreeding estimates (Table 2) varied greatly across subspecies, but were relatively consistent within subspecies. Values ranged from low inbreeding (F_ROH_ = 0.0154 in *vulgivaga*, nvison_LA-8) to high inbreeding in *evergladensis* (F_ROH_ = 0.5293 and 0.6504 in nvison_FL-3 and nvison_FL-5). One sample (nvison_FL-32666) showed a negative F_ROH_ value, which could be due to high heterozygosity, but may more likely be sampling error (Purcell et al., 2007) due to the sample’s more gapped genome.

**Table 2:**
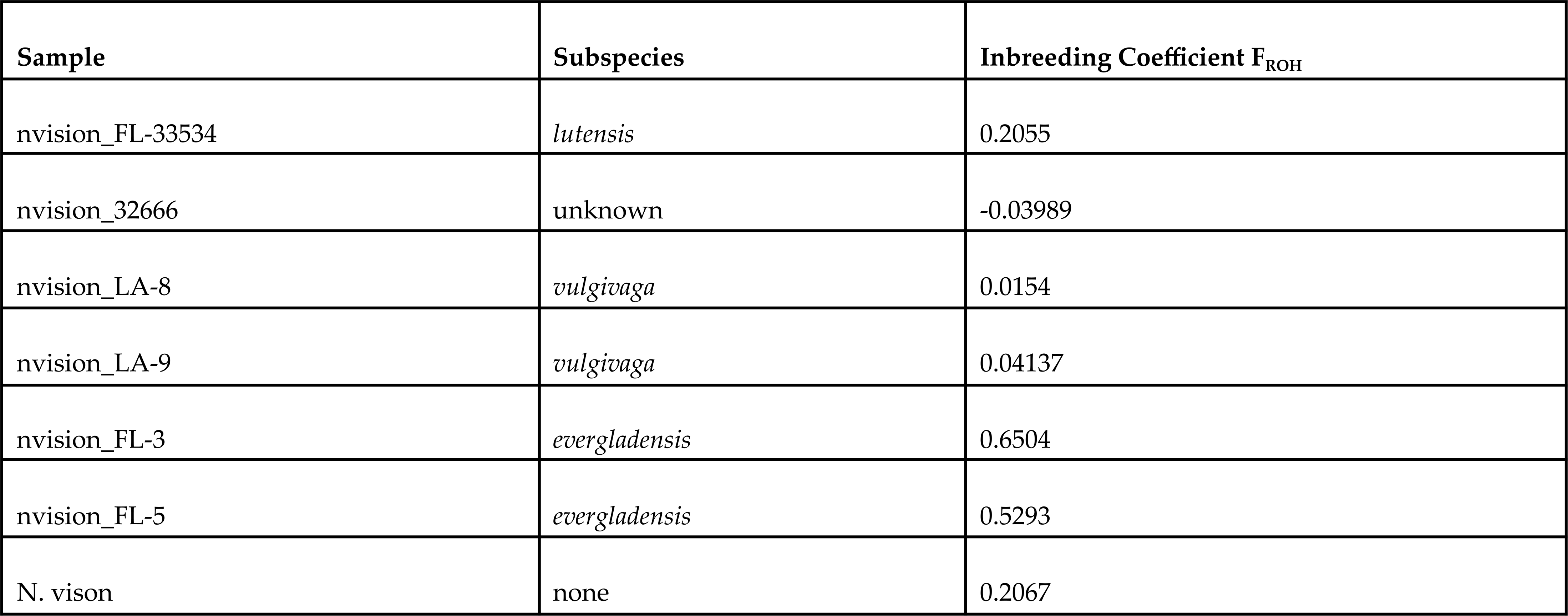
Inbreeding coefficients for subspecies and the American mink reference genome.

Across all genomes, we identified 2,927 ROH > 500 Kb (mean: 859.48 Kb/sample), with the longest at 6.02 Mb (Fig. 1F). ROH were most prevalent in *evergladensis* by far, aligning with its high inbreeding estimate. The *evergladensis* samples also had longer ROH (Fig. 1F). Over 50% of the protein-coding gene space in *evergladensis* overlaps with the ROH (Table S9).

**Fig. 1F:**
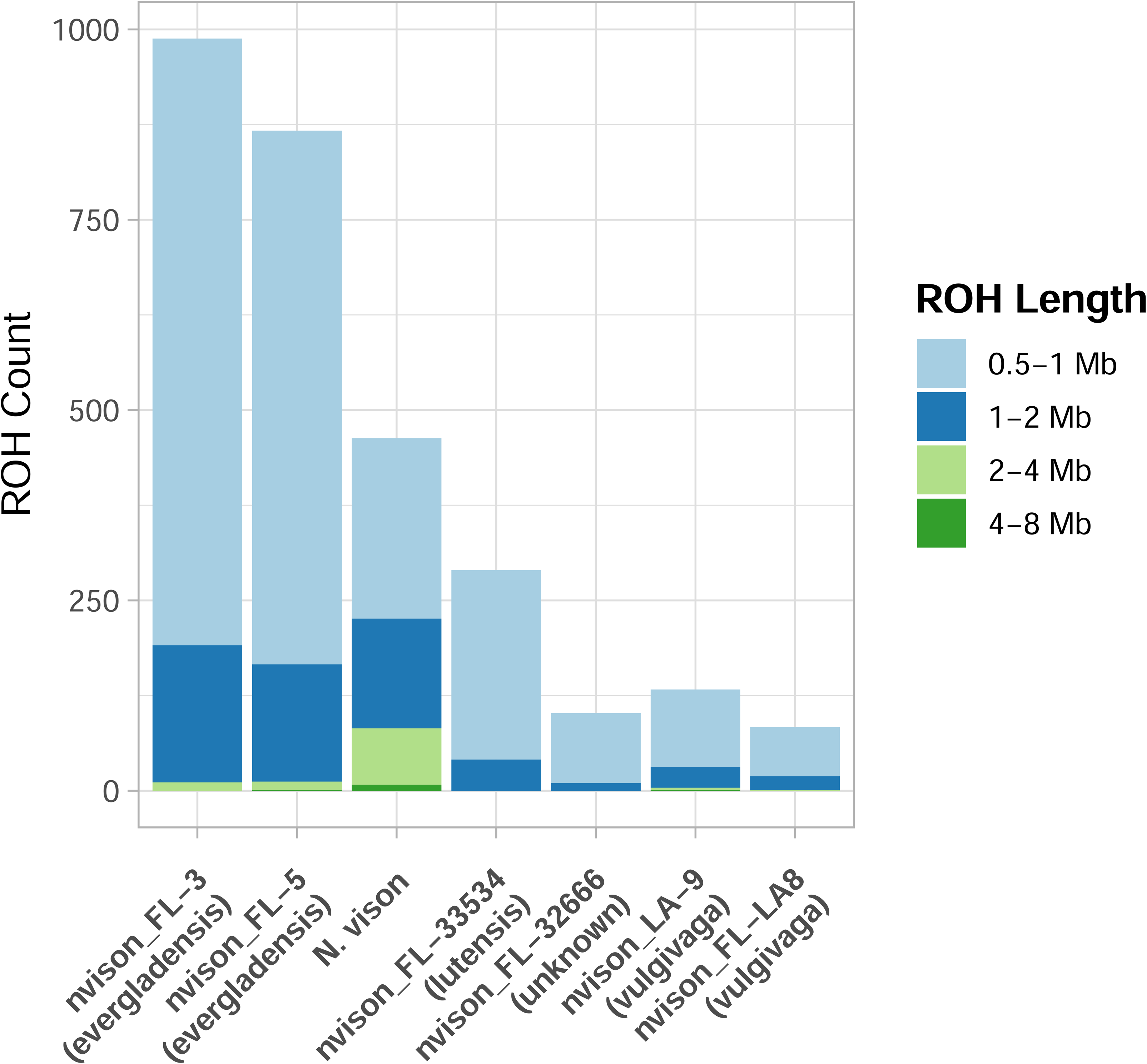
Distribution of length and total counts of runs of homozygosity (ROH) across the subspecies samples. A minimum length of 500 Kb was applied to define the presence of a ROH. Counts of each class of ROH (0.5-1 Mb, 1-2 Mb, 2-4 Mb, and 4-8 Mb) are depicted in the stacked histogram.

### Gene Family Evolution

Gene family evolution between our American mink samples and 18 other species in the order Carnivora (Table S1) was assessed using OrthoFinder. The 448,907 genes were placed into 22,440 gene families. 12,461 gene families were shared among all 24 samples, and 2,209 orthogroups were present in all samples except one of the two mink samples with lower genome completeness. The mink samples also contained unique orthogroups absent from other carnivorans. There were 194 orthogroups conserved among the 5 *N. vison* samples (Fig. 2A) and 228 orthogroups present in 4 of 5 mink samples (likely reflecting, in part, lower genome completeness). An additional 122 orthogroups were present in 2–3 mink samples. Each mink sample also had 1–8 unique multicopy orthogroups and 2–20 unique genes.

**Figure 2A:**
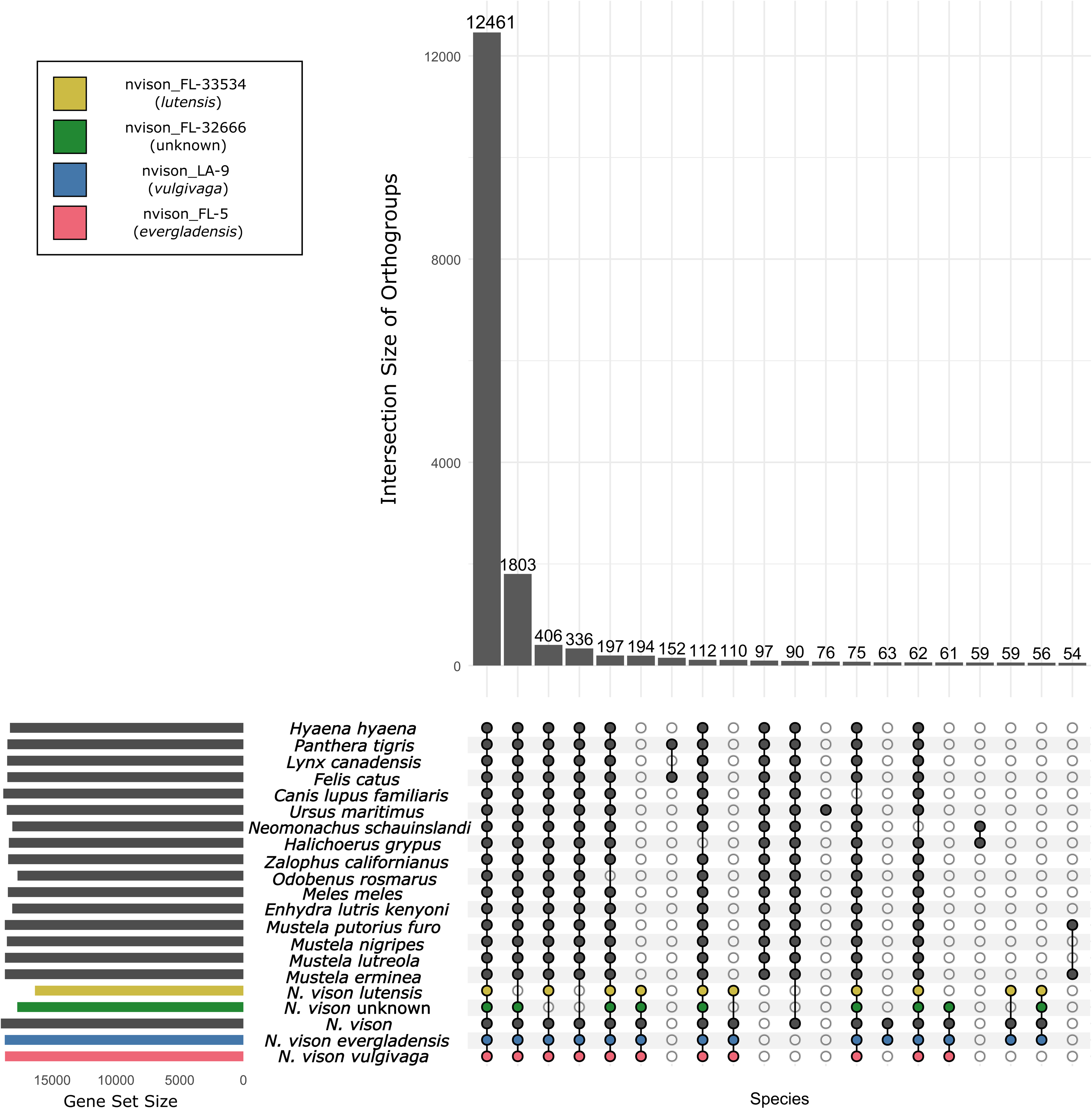
Conserved orthogroups (gene families) among species. This UpsetR plot highlights conserved orthogroups shared among species, as well as species-specific or lineage-specific orthogroups. The horizontal bars (left) represent the total number of orthogroups detected in each species, and the rows representing the mink subspecies are highlighted. The vertical bars indicate the number of shared orthogroups for each combination of species, with filled dots below each bar representing the contributing species.

We utilized CAFE to assess gene family evolution, retaining one sample per mink subspecies. Each *N. vison* lineage showed unique gene family expansions (214–286 families per sample) and contractions (447–2357 families) (Fig. 2B). The unusually high number of contracting gene families observed in nvison_FL-33534 may be an artifact resulting from a substantially lower genome completeness.

**Figure 2B:**
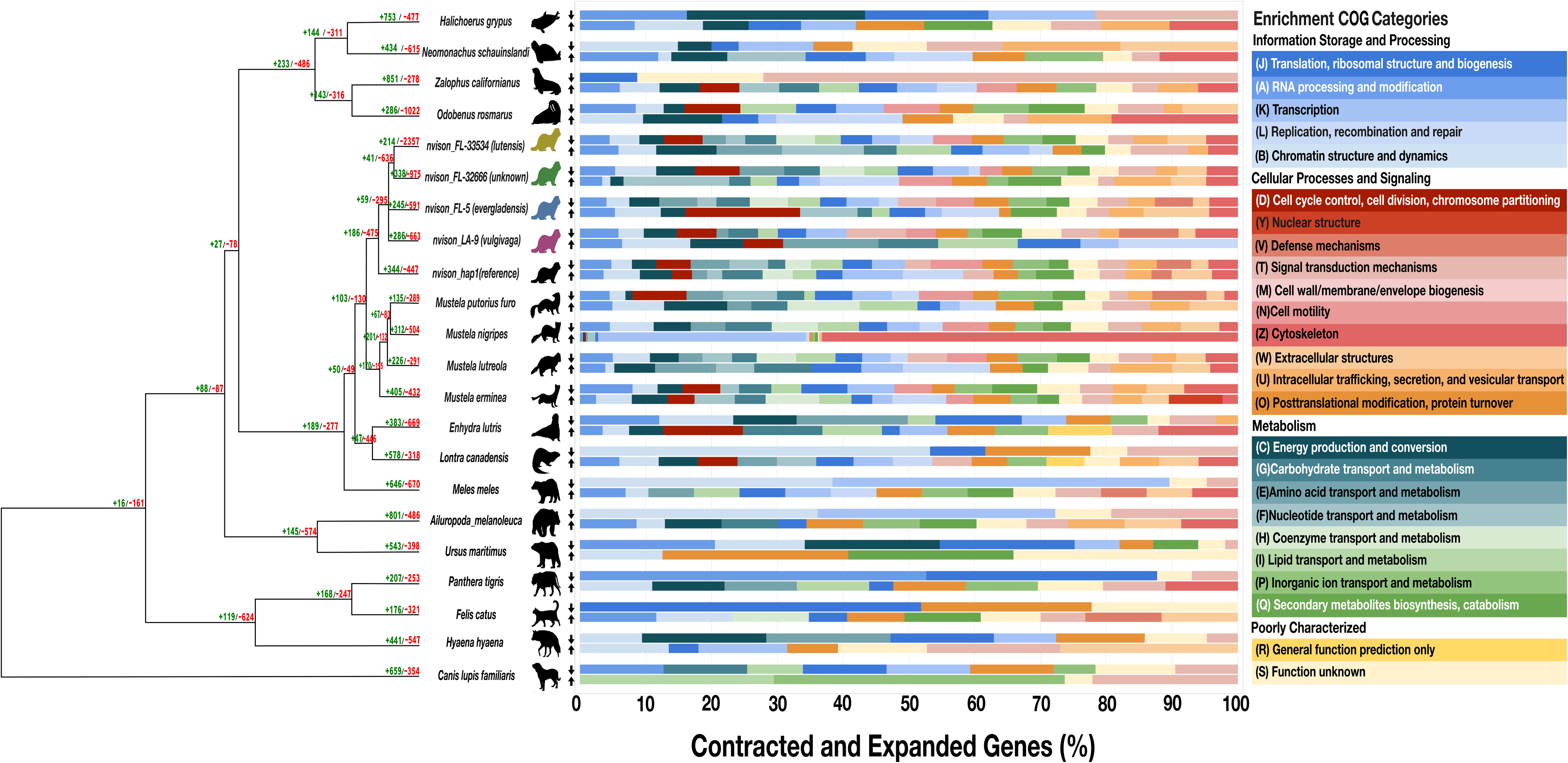
COG (Clusters of Orthologous Groups) category enrichment analysis based on gene family evolution inferred using CAFE (Computational Analysis of gene Family Evolution). The phylogenetic tree on the left shows relationships among mustelids and representative carnivoran taxa. Numbers in green indicate the number of expanded gene families, while numbers in red indicate the number of contracted gene families as identified by CAFE. Animal silhouettes represent sampled species and mink haplotypes, including representatives from putative *N. vison* subspecies (*vulgivaga*, *evergladensis*, *lutensis*, and unknown). The up arrows to the left of the bar graph indicate expanding genes while the down arrows indicate contracting genes in each species and each COG category.

To evaluate how gene family changes influence the biology of mink subspecies, we performed Clusters of Orthologous Groups (COG) category enrichment analysis. In nvison_LA-9 (*vulgivaga*), the category *replication, recombination, and repair* had the highest proportion of expanding genes, while *lipid transport and metabolism*, *translation, ribosomal structure and biogenesis*, *cell wall/membrane/envelope biogenesis*, and *cell motility* showed the greatest proportions of gene contraction. In nvison_FL-5, the top expanding categories were *cell cycle control, cell division, chromosome partitioning*, while *amino acid transport and metabolism*, *coenzyme transport and metabolism*, *inorganic ion transport and metabolism*, and *defense mechanisms* exhibited high levels of gene contraction. For nvison_FL-32666, *nucleotide transport and metabolism* had the highest percentage of gene expansion. In contrast, *cell cycle control, cell division, chromosome partitioning, amino acid transport and metabolism*, and *coenzyme transport and metabolism* had the most contracting genes. Sample nvison_FL-33534 showed no categories with a high proportion of expanding genes. The highest percentages of contraction were observed in *energy production and conversion* and *signal transduction mechanisms*. Additional contraction was noted in *cell cycle control, cell division, chromosome partitioning*, *coenzyme transport and metabolism*, *cell motility*, *inorganic ion transport and metabolism*, and *extracellular structures*. Finally, in nvison_hap1, the categories *transcription* and *replication, recombination, and repair* had the highest percentages of expanding genes, while *cell motility* had the greatest proportion of contracting genes (Fig. 2B).

### Mitogenome Phylogenetic and Diversity Analysis

We used NOVOPlasty to assemble a circularized mitochondrial genome for nvison_LA-8, nvison_FL-3, nvison_FL-5, and nvison_FL-32666. For nvison_LA-9 and nvison_FL-33534, NOVOPlasty generated multiple contigs, manually merged using Geneious Prime (v2025.1.3) (Table S10). Fifteen N’s were inserted into the D-loop repetitive region to represent the assembly gap in samples, nvison_LA-9 and nvison_FL-33534. The genomic length of the mitochondrial reference (NC_020641) is 16,552 bp. The assembled genomes ranged from 16,461 bp (nvison_FL-33534) to 16,573 bp (nvison_LA-8) (Table S10). Differences in length are due to D-loop variation between the genomes. All assembled mitochondrial genomes consisted of 13 protein-coding genes, 22 tRNA genes, 2 rRNA genes, and a repetitive non-coding region (Fig. 3A; Table S11). These results were consistent with the published *N. vision* reference (NC_020641).

**Figure 3A:**
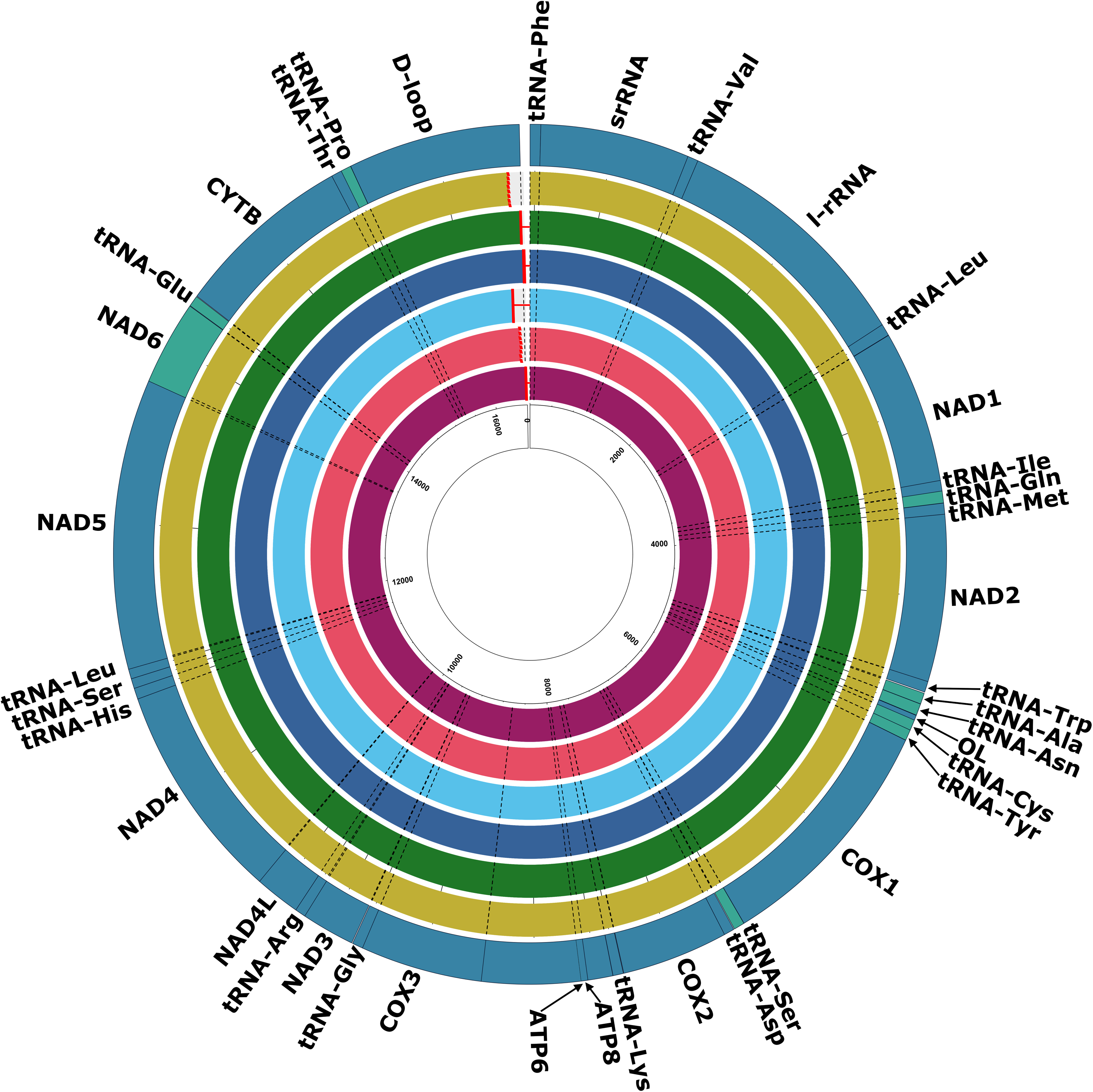
Mitogenome Circos: Circos plot visualising mitochondrial genome features across *N. vison* genomes. Each ring represents one isolate. For the reference genome (NC_020641), genes on the minus (-) strand are shown in a lighter color. The red solid lines connected to the beginning of the genome represent assemblies that successfully assembled the entire repetitive region in the D-loop. The dotted lines indicate species with incomplete D-loop repeat regions, highlighting gene conservation across individuals. Plot generated using R package circlize (v0.4.16).

Within the 13 mitochondrial protein-coding genes, *ATP8* and *NAD3* were the most conserved across the three subspecies while *NAD4* exhibited the greatest nucleotide differences, 11 found in *evergladensis*, 20 in *lutensis*, and 13 in *vulgivaga*. Across all subspecies, *lutensis* exhibited the highest number of nucleotide differences across both coding and non-coding regions. Divergence was lowest between the two *evergladensis* samples; their D-loop region differed by a large indel (Table S11).

The mitogenome tree was rooted using the outgroup *Neogale fermata*. Within *N. vison*, the first diversification event separated the mitochondrial reference genome from all others. This was followed by separation of the clade containing nvison_FL-32666 and nvison_FL-33534 from the clade containing nvison_FL-3, nvison_FL-5, nvison_LA-8 and nvison_LA-9 with strong bootstrap support (95). Clear separation in the mitogenome tree helped identify sample nvison_FL-32666 (unknown) as part of *lutensis*. The two *evergladensis* samples (nvison_FL-3 and nvison_FL-5) were then separated from the two *vulgivaga* samples (nvison_LA-8 and nvison_LA-9) with high confidence (bootstrap = 100) (Fig. 3B).

**Figure 3B:**
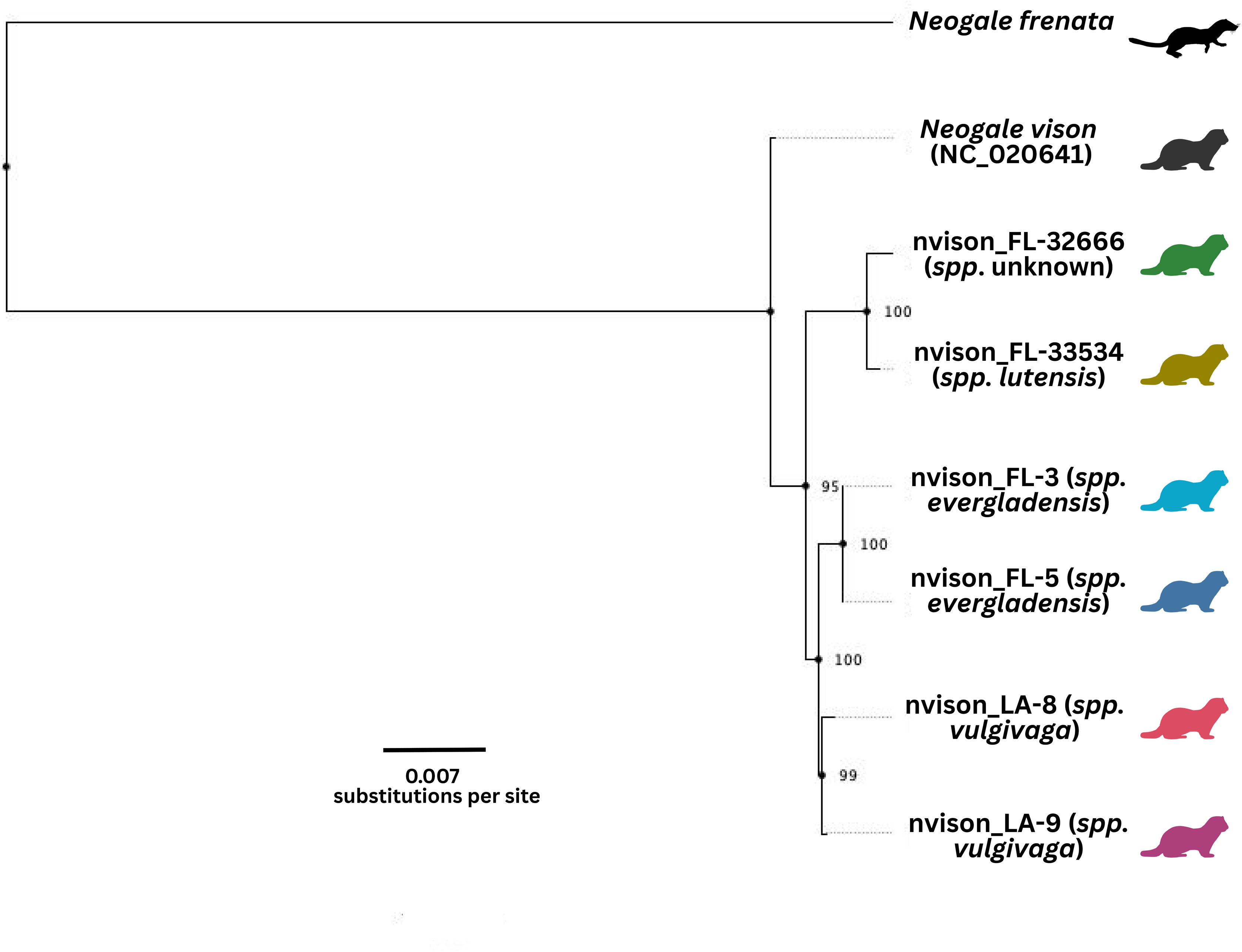
Mitochondrial phylogeny of *Neogale* lineages. A maximum likelihood phylogeny was constructed using mitochondrial genome sequences to assess evolutionary relationships among *Neogale vison* individuals and an outgroup (*Neogale frenata*, long-tailed weasel). The tree includes a reference haplotype (*N.* vison_hap*1*) and representatives of several putative subspecies, including spp. *evergladensis* (*nvison_FL-5; nvison_FL-3*), spp. *lutensis* (*nvison_FL-33534*), spp. *vulgivaga (nvison_LA-8; nvison_LA-9*) and a sample (*nvison_FL-32666*). Branch support values (shown at nodes) are based on 1000 bootstrap replicates and indicate high confidence in most relationships. FigTree (v1.4.4) used to visualize the tree.

### Pangenome

After the initial construction of the pangenome, genes were first categorized into core, accessory, and unique groups to comparatively assess the gene space and genetic variation. Core genes are shared across all genomes, accessory genes are shared by some genomes, and unique genes are specific to individual genomes. Across all sampled genomes, a total of 25,344 homology groups were identified (Fig. 4A). Of these, 14,528 (57.32%) were core genes, 6,090 (24.03%) accessory and 4726 (18.65%) were unique gene groups. Among the core genome, 13,500 (92.9%) were single copy gene families (Fig. 4B). Heap’s Law was calculated to be 0.817, which indicates an open pangenome. Consistent with Heap’s Law, the pangenome accumulation curve generated shows an increase in the number of gene groups with additional genomes added (Fig. 4A). The largest number of unique gene groups was identified in sample nvison_FL-32666 with 1774 (9.44%).

**Figure 4A:**
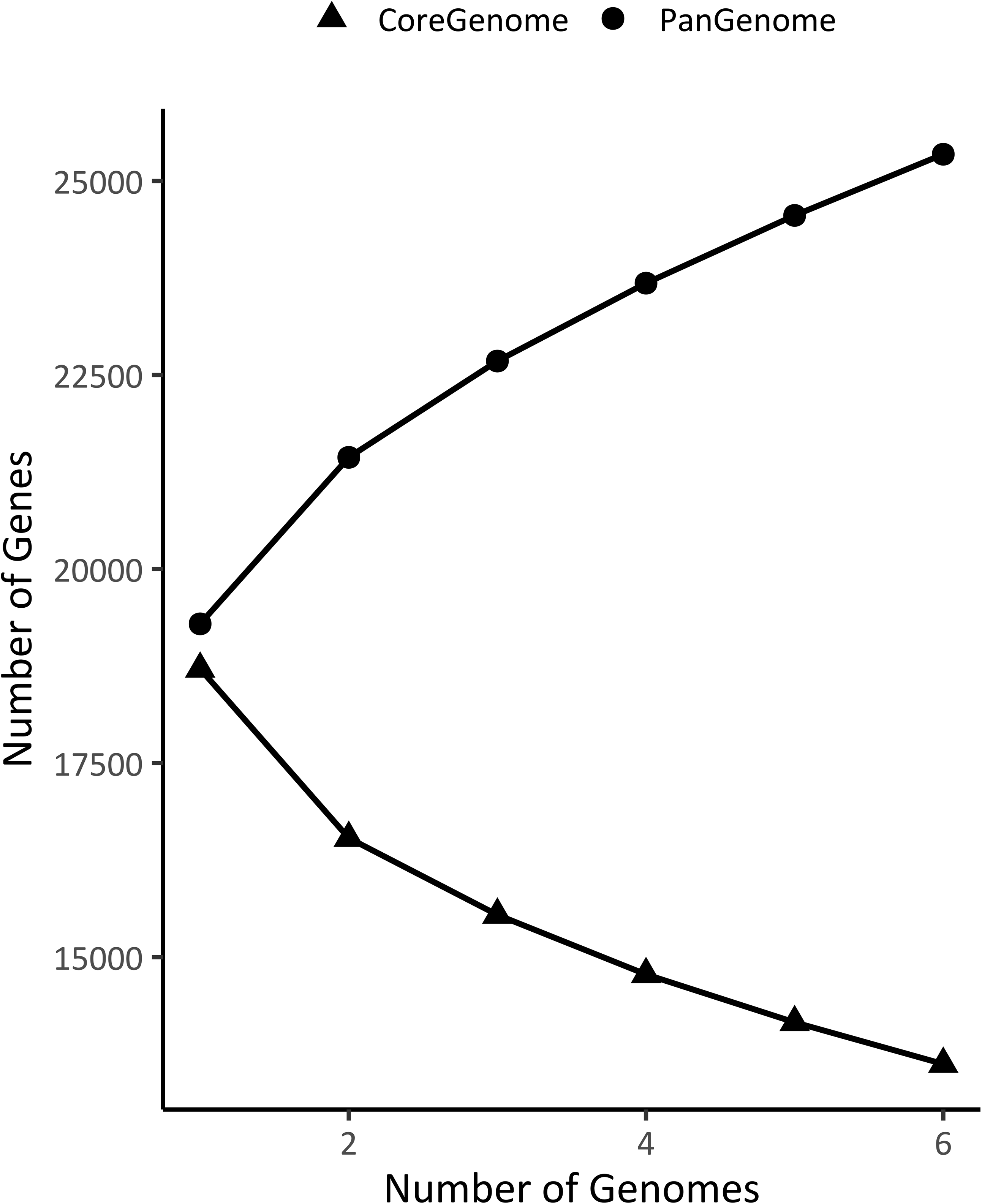
Pangenome completeness assessed across the six assembled and annotated genomes. Each point represents the total number of genes classified as pan-genome (present in at least one genome) and core-genome (present in all genomes to that point). Core genes show a steady decrease, while the pangenome grows with each added genome, consistent with an open pangenome.

**Figure 4B.**
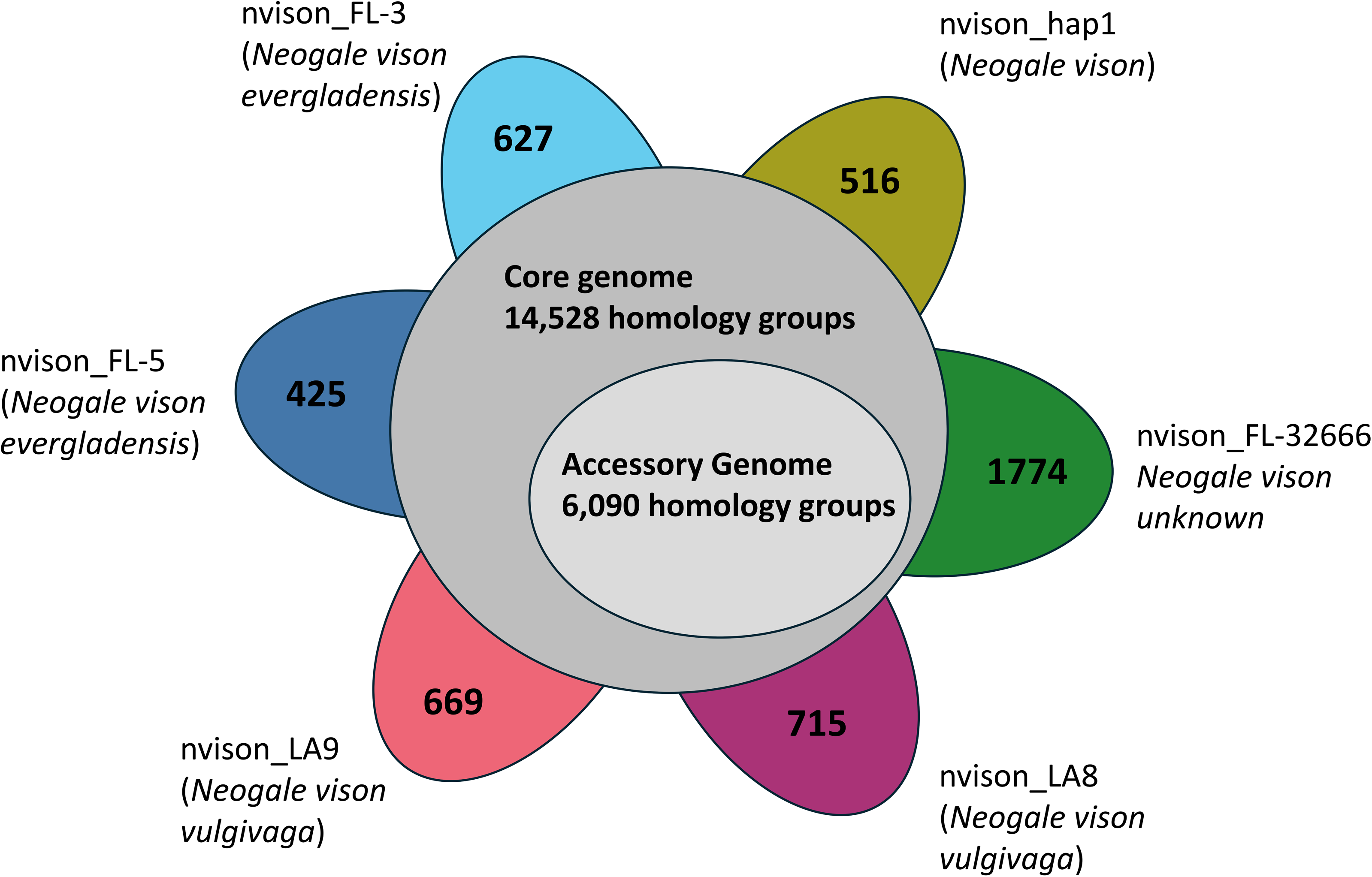
Pangenome homology group statistics. Interior grey circles represent the number of core and accessory homology groups in the pangenome. Exterior bubbles represent the number of unique homology groups in each sample.

### Gene Ontology (GO) Enrichment

For the GO enrichment analysis, a total of 34 significantly enriched GO terms were selected for further analysis across all subspecies based on their potential relevance to environmental adaptation. Overall, 17% of these terms were related to development, 13% to cell differentiation, 13% to behavior and movement, 13% to immune response, and 13% to neurological processes (Table S12). Eight percent were related to cell signaling, 8% to nucleic acid processes, and 8% to enzymatic activity. Five percent were related to cellular stress, and 2% to regulation of biological quality.

In *evergladensis* (nvison_FL-3, nvison_FL-5), enriched GO terms primarily involve cell differentiation, development, and aging. Notably, *lung cell differentiation* (GO:0002065, 4 genes, p = 0.00047) and *aging* (GO:0035315, 6 genes, p = 0.00280) were over-represented. Epithelium-related terms included *epithelial cell development* (4 genes, p = 0.00360) and *epithelial cell differentiation* (GO:0030855, 14 genes, p = 0.00470), both over-represented. *Inner ear receptor cell differentiation* (GO:0060119, 4 genes, p = 0.00880) and *inner ear development* (GO:0042491, 4 genes, p = 0.00540) were also significantly enriched.

In *vulgivaga* (nvison_LA-8, nvison_LA-9), enriched terms spanned *development and movement*. Development-related terms included positive regulation of *developmental processes* (GO:0051094, 2 genes, p = 0.00660; GO:0032502, 34 genes, p = 0.00550) and *anatomical structure development* (GO:0048856, 32 genes, p = 0.00660), all under-represented. *Locomotion* (GO:0040011, 14 genes, p = 0.00650) was also under-represented. Additional under-represented terms included *regulation of cell differentiation* (GO:0045595, 4 genes, p = 0.00450) and *regulation of biological quality* (GO:0065009, 10 genes, p = 0.00660).

In *lutensis* (nvison_FL-32666), enriched terms reflected thermal tolerance, pollutant response, and neural development/signaling. Key terms included *head development* (GO:0060322, 109 genes, p = 0.00750), *o-methyltransferase activity* (GO:0016319, 24 genes, p = 0.00004), *response to pain* (GO:0048265, 16 genes, p = 7.60000e-07), *ATPase activity couple*d to transmembrane transport (GO:0042626, 23 genes, p = 0.00005), *synaptic vesicle uncoating* (GO:0099604, 1 gene, p = 0.00016), *regulation of synaptic plasticity* (GO:0048167, 40 genes, p = 0.00004), *interspecies interactions* (GO:0051703, 17 genes, p = 0.00004), *synapse assembly* (GO:0007416, 32 genes, p = 0.00001), and *neuromuscular synaptic transmission* (GO:0097477, 5 genes, p = 0.00007), all over-represented.

In the primary reference individual (*N. vison*), enrichment patterns focused on immune responses, signal transduction, and stress responses. Under-represented terms included transcription from *RNA polymerase II promoter* (GO:0006366, 6 genes, p = 0.00000), *RNA biosynthetic process* (GO:0032774, 11 genes, p = 0.00005). Over-represented immune-related terms included *B cell activation* (GO:0042113, 18 genes, p = 2.00000e-06), *regulation of antiviral defense re*sponses (GO:0050688, 16 genes, p = 1.00000e-05), and *response to type I interferon* (GO:0034340, 15 genes, p = 1.00000e-05). *C*ellular stress-related terms included over-representation of *cellular response to type I interferon* (GO:0071357, 15 genes, p = 1.00000e-05) and under-representation of regulation of *cellular response to stress* (GO:0080135, 14 genes, p = 0.00740). Additional over-represented terms included *negative regulation of organ growth* (GO:0046621, 10 genes, p = 0.00003), *histone acetyltransferase activity* (GO:0004416, 3 genes, p = 0.00009), *phosphoglucomutase activity* (GO:0008967, 3 genes, p = 0.00009), *modulation of chemokine production* (GO:0035821, 4 genes, p = 4.60000e-06), *p53-mediated signal transduction* (GO:0035499, 3 genes, p = 1.00000e-05), and *interleukin-6 receptor binding* (GO:0005132, 15 genes, p = 1.00000e-05).

## Discussion

Species of conservation concern frequently lack comprehensive genomic tools, particularly at the subspecies level. The American mink (*Neogale vison*), with its broad North American range, economic significance, and phased reference genome (O’Brien et al., 2025), provides a framework to resolve subspecies and population delineation. This study advances subspecies resolution by sequencing six individuals representing three focal subspecies in the southern portion of the native range: *N. vison evergladensis, N. vison lutensis,* and *N. vison vulgivaga*. Notably, all six genomes were generated from natural history collections and field samples, illustrating that such samples can provide valuable genome-scale information with respect to subspecies and population relationships. The reference genome was generated from a wild-caught individual within the introduced range of American mink, which is presumed to be descended from domesticated individuals imported for fur farming (O’Brien et al., 2025). Thus, the new genomes also provide insights into differences between wild and domesticated mink.

### Population decline and loss of genetic diversity in N. v. evergladensis

Based on our mink subspecies genome assemblies, demographic reconstruction revealed a sustained decline in effective population size in *evergladensis*, with reductions beginning approximately 60,000 years ago (Fig. 1E). This pattern, which was not observed in other subspecies, suggests long-term population decline. Complementing these results, the mitogenome phylogeny placed *evergladensis* as more closely related to the *vulgivaga* subspecies from Louisiana than to the geographically proximate *lutensis* subspecies (Fig. 3B), highlighting a complex history of isolation that is not strictly aligned with geography (Hapeman and Smith 2024). Mitochondrial nucleotide diversity statistics support the conclusion that the *evergladensis* population is small.

Signals of inbreeding, supported by high F values, elevated frequency of ROH segments (Table 2; Fig. 1F), and previous support from microsatellite assessments (Hapeman and Smith 2024), further support conservation concern for *evergladensis*. The F_ROH_ value for *evergladensis* exceeds that of many critically endangered mammals, including black and white rhinoceroses (Mellya et al. 2025; Sánchez-Barreiro et al. 2021), Chinese pangolins (Lan et al. 2025) and western lowland gorillas (Alvarez-Estape et al. 2023), as well as species with a well documented population bottleneck (e.g., Hoelzel et al. 2024). Comparable values of F_ROH_ have been documented in several small, isolated populations including southern Indian tigers (Khan et al. 2021) and Appenine brown bears (Clendenin et al. 2025). Comparisons to the mink reference genome also indicate that loss of genetic diversity in *evergladensis* exceeds the loss of genetic diversity associated with domestication followed by introduction.

### Mitogenome phylogeny resolves subspecies relationships

Due to their rapid evolutionary rate, lack of recombination, and maternal inheritance, mitochondrial genomes can improve the resolution of recent divergence and phylogeographic structure exceeding biparentally inherited nuclear markers, particularly in taxa with complex demographic histories (Galtier et al., 2009). A previous study in *N. vison* focused on one mitochondrial gene, cytochrome b (*CYTB*), due to its high variability (Caine et al., 2006). By expanding our analyses to include full mitogenomes, we found well-supported patterns of maternal lineage divergence among the *N. vison* subspecies. Subspecies *evergladensis* and *vulgivaga* are most similar, corroborating the Hapeman and Smith (2024) findings that they share a recent common ancestor. Overall, *lutensis* exhibits higher nucleotide differences than *evergladensis* and *vulgivaga*. In particular, the D-loop region is more conserved for *evergladensis*, consistent with smaller population size. In addition, the resulting mitochondrial phylogeny (Fig. 3B) identified nvison_FL-32666 (unknown) from the Florida panhandle as *lutensis*, a subspecies which also occurs on the Atlantic coast of Florida. This individual was not confidently placed using nuclear genomic data alone.

### Advantages of pangenomes for comparative analysis

Pangenomes offer a powerful alternative to linear reference-based approaches by minimizing reference bias and capturing variation missing from a single genome (Secomandi et al. 2025). In this study, the graph-based tool PanTools enabled subspecies-level comparisons, revealing functionally distinct gene space and adaptive variation. Its homology grouping method has been shown to resolve the same gene families, but with greater precision than OrthoFinder (Sheikhizadeh-Anari et al. 2018), underscoring its utility for fine-scale comparative analysis. Traditional methods for gene family evolution assessment, OrthoFinder and CAFE, were still leveraged for broader assessments of gene evolution across representative species of Carnivora (Table S13; Table S14).

### Resilience through immune defense genes in invasive American mink (Neogale vison)

American mink is an invasive species in Europe and Asia, first introduced through fur farming (Zhang et al. 2021). The *N. vison* phased reference was derived from a wild-caught sample from Arundel, England (O’Brien et al., 2025). Both domestication and introduction into a new environment tend to reduce effective population size relative to the populations of origin, but can also greatly alter patterns of selection. In the pangenome assessment, presence/absence variations (PAV) assessment was significantly enriched for *immune response* functions including regulation of defense response to virus, cellular response to type 1 interferon, interleukin-6 receptor binding and B cell activation (Fig. 4C). Genes associated with these responses include interferon alpha-1 2-like (*IFNA5*), signal transducer and activator of transcription (*STAT5B*), and cardiotrophin-like cytokine factor 1 (*CLCF1*). These genes were present in *N. vison* and absent in other subspecies with *IFNA5* being particularly abundant in *N. vison* (Fig. 4D).

**Figure 4C:**
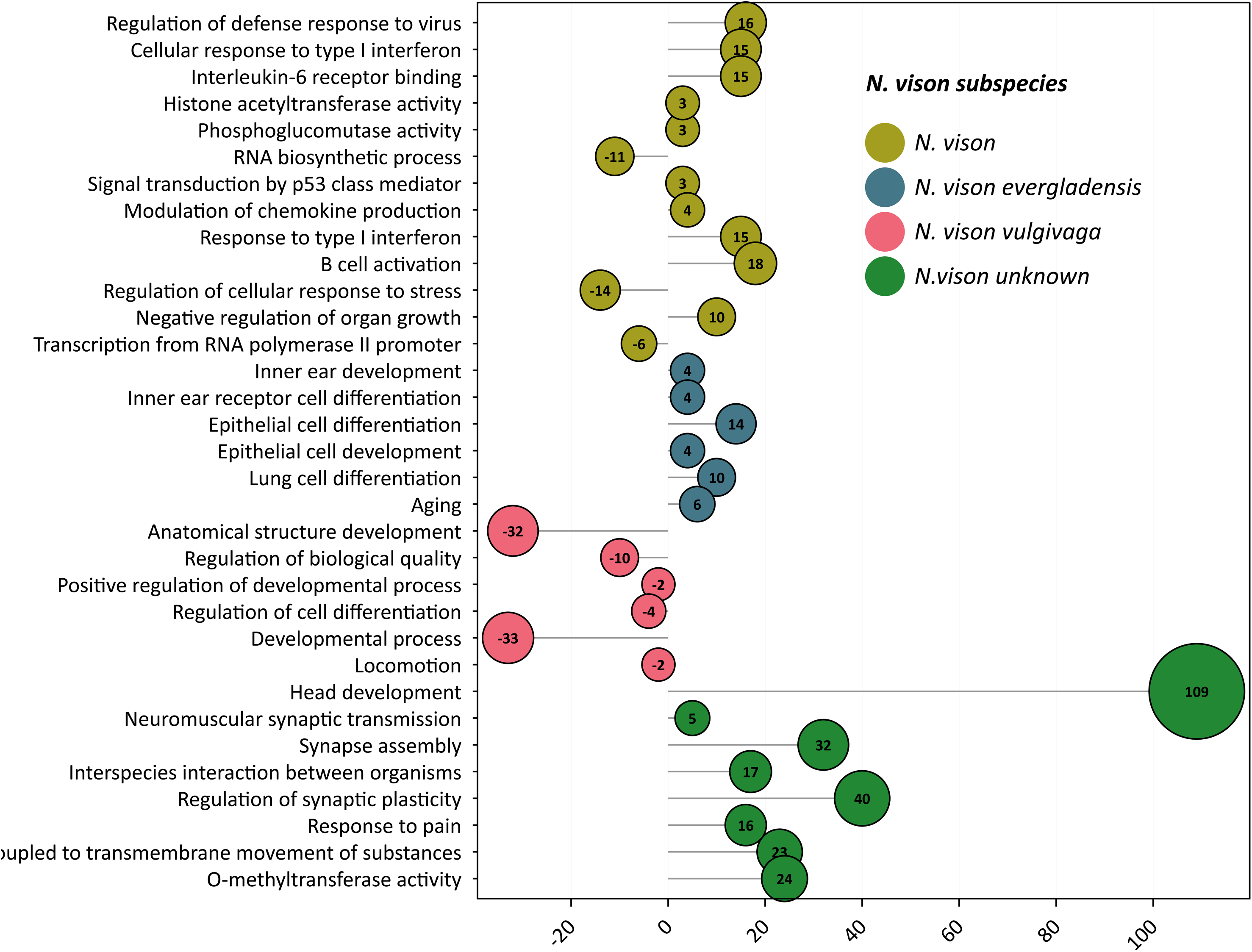
Functional trends of PAV genes with their associated function, for each subspecies. Negative values indicate under-represented genes in a function, while over-represented genes are represented by positive values. The number in each category represents the total number of genes represented.

**Figure 4D.**
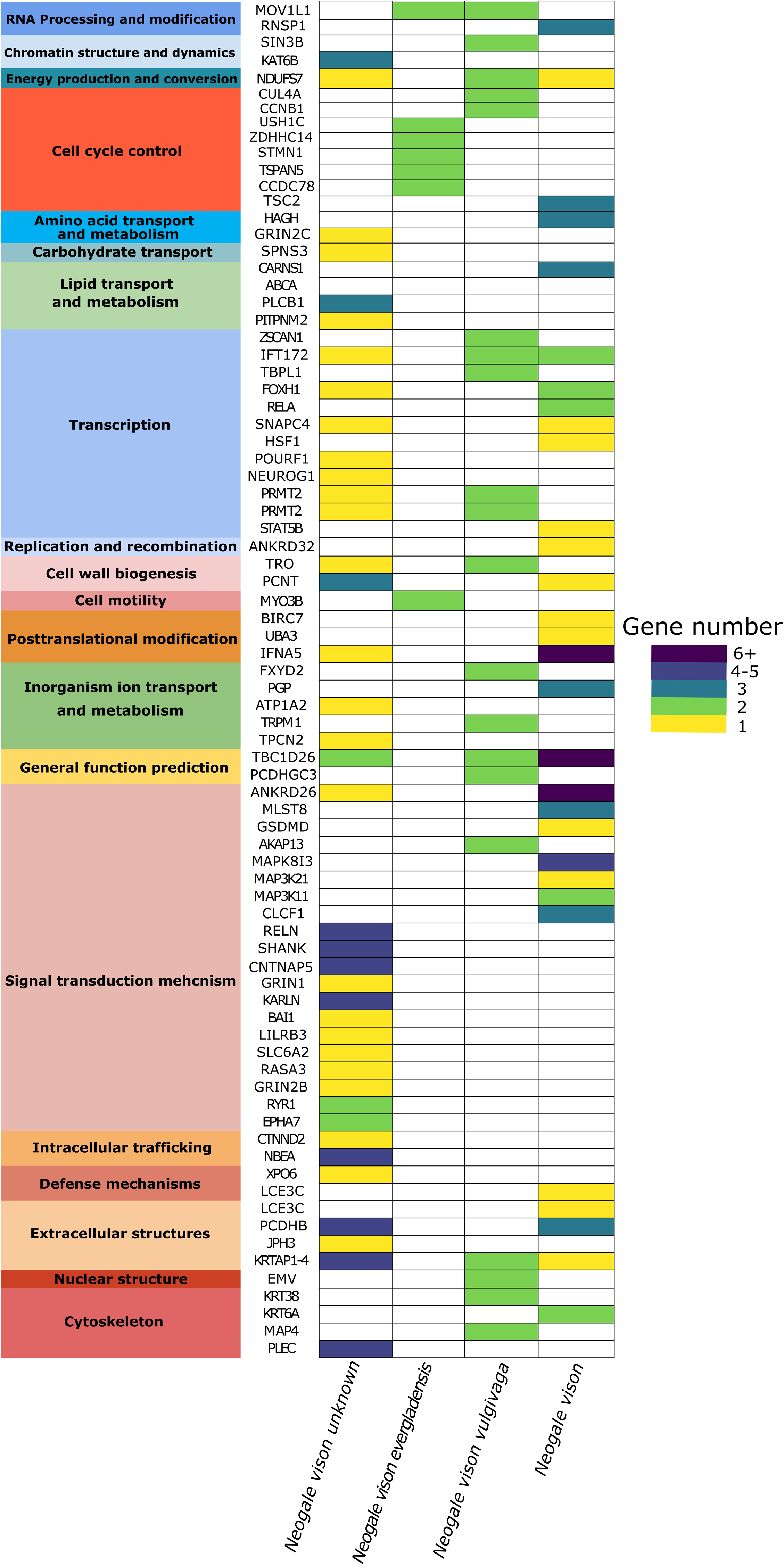
Gene level presence absence variation matrix. Gene names are shown on the y axis and are color coded based on COG function; mink subspecies are shown on the x axis. The color gradient represents the number of genes in each subspecies.

While viral outbreaks are seen in wild populations, they are more frequent in captive populations due to overcrowding and exposure to humans. Aleutian disease and SARS CoV-2 are common diseases that impact mink populations globally and are prevalent in mustelid populations in England (Yamaguchi & Macdonald, 2001; Jahid et al., 2024).

Aleutian disease is particularly known for decreasing reproductive success and increasing offspring mortality which could act as a selection pressure on the immune function related genes observed in *N. vison* compared to the other subspecies.

Selection on immune function is commonly observed during domestication (Hou et al., 2020). Immune adaptation in response to introduction is often species-specific and depends on many ecological variables (Cornet et al., 2016). A previous study investigating immune-associated genes of the American gray squirrel suggested that these genes are energetically costly and may be undergoing negative selection in a new environment where native parasites are no longer abundant (Romeo et al., 2023). Alternatively, there is evidence in house mice that immune-associated genes undergo positive selection in a new environment where exposure to new pathogens can occur (Charbonnel et al., 2020).

Although further research is needed to investigate shifts in immune function within introduced species, the enriched immune genes in *N. vison* have been linked to increased immune functions in other species. For example, B cell activation was downregulated in individuals with severe cases of SARS CoV-2 compared to individuals with mild cases indicating a possible immune advantage of having overrepresentation of this functional category in viral infections (Pence et al., 2022).

### Adaptive capacity as an invasive species in American mink (Neogale vison)

The ankyrin repeat domain-containing protein 26 (ANKRD26) gene, present in eight copies, was overrepresented in pathways associated with the regulation of organ growth.

ANKRD26 is highly conserved across mammals and has been shown to cluster near genomic breakpoints, regions often associated with structural variation and local adaptation (Du et al. 2014). While present across all carnivoran species in our comparative dataset, ANKRD26 is specifically expanded in *N. vison*. Given the ecological breadth of *N. vison* across diverse environments, the expansion of this gene family may reflect selection for developmental plasticity in response to environmental heterogeneity.

### Transcriptional plasticity and neural adaptation in N. v. vulgivaga

The southern estuarine mink (*vulgivaga*) inhabits large diversity of habitats expanding through Louisiana and western Mississippi, mostly concentrated in freshwater marshes, brackish, and coastal marshes. This habitat variability suggests they may have plastic responses that allow them to survive in highly dynamic and fluctuating environments (Vinton et al., 2022). *N. v vulgivaga* displayed enrichment of SIN3 transcription regulator family member B (*SIN3B*), RNA-binding motif protein 44 (*RBM44*), and small nuclear ribonucleoprotein polypeptide C (*SNRPC*) aligning with enhanced transcriptional flexibility, a trait observed in marine mammals exposed to variable environments (Xu et al., 2025).

*N. v. vulgivaga* was also enriched in intraflagellar transport 88 (*IFT88*)which regulates neural, vestibular, and musculoskeletal systems necessary for movement (Wang et al., 2025). kinesin family member 5B (*KIF5B*) was also enriched, which is involved in synaptic plasticity and memory formation(Flex et al., 2023). In aquatic and semi-aquatic mammals, efficient senses are essential for navigating dense vegetation, capturing prey, and avoiding predators in dynamic environments (Mortensen et al., 2021). *IFT88* and *KIF5B* are conserved in carnivora and are only found in *vulgivaga* (Fig. 4D) and are underrepresented (Fig. 4C).

### Enrichment of sensory and reproduction genes may support differences in reproductive behavior in N. v. evergladensis

*N. v. evergladensis* is distinct from other mink species in temperate zones, as some evidence suggests they mate in the autumn, in contrast to northern mink species, which mate in the winter/spring (Hapeman and Smith, 2024). This shift is in response to seasonal flooding during wet seasons in the Everglades (Humphrey and Zinn, 1982), providing a unique hydrological and vegetational habitat for *evergladensis* (Hapeman and Smith, 2024). In our analysis of significant gene enrichment in *evergladensis,* we identified enriched genes involved in sensory perception and reproduction. Sensory genes, such as Usher syndrome 1C (*USH1C*) and Myosin IIIB (*MYO3B*), were over represented and enriched in *evergladensis*. *USH1C* is linked to human vestibular dysfunction disorders, and a study of genes associated with balance in semi-aquatic mammals identified *USH1C* as a potential contributor to adaptation to both land and underwater environments by reducing vertigo (Dong et al., 2025). Enrichment of this gene is significant because *evergladensis* also inhabits varied habitats, exposing it to similar physical balance pressures. Meanwhile, *MYO3B* is present in photoreceptors and cochlear hair cells in mice (Katti et al., 2009). Enrichment of these genes indicate selection on the visual, auditory, and vestibular systems, and collectively might contribute to the adaptation of *evergladensis* to its unique environment.

Likewise, enriched genes involved in reproduction, such as Sperm associated antigen 9 (*SPAG9*), Zinc Finger DHHC-Type Palmitoyltransferase 14 (*ZDHHC14*), and Mov10 Like RNA Helicase (*MOV10L1*), may have a role in the evolution of novel mating behavior.

*MOV10L1* encodes a putative RNA helicase and is expressed in spermatocytes in the development of sperm cells (Vourekas et al., 2015). Similarly, *SPAG9* and *ZDHHC14* are all highly expressed in testis (Jagadish et al., 2005, Yeste-Velasco et al., 2014). *SPAG9* is a gene PAV variant unique to *evergladensis*, and is not represented in other carnivorans in our comparative analysis (Fig. 4D). While potentially implicated in reproduction, *SPAG9* has also been associated with mammalian color and patterning (Hempel et al. 2024). *MOV10L1* is present in only *evergladensis* and *vulgivaga,* and is rapidly expanding in these subspecies (Fig. 2B). This gene is contracting at the phylogenetic node representing the divergence of *lutensis*; this notable trend of gene evolution potentially suggests mating behavior is evolving in mink subspecies (Fig. 2B). Further, *ZDHHC14* is a copy number variant (two copies) only present in *evergladensis,* further supporting its potential role in novel behavior (Fig. 4D).

Varied spermatogenesis activity has been observed with seasonal reproductive changes in mink (Blottner et al., 2006), therefore it is possible that enrichment of these genes contributes to variation in mating behavior in *evergladensis*.

### Strong enrichment of neural development and intracellular stress response may support cognitive plasticity in N. v. lutensis

The *lutensis* subspecies of American mink exhibits several unique genomic signatures associated with neural development, coordination, and craniofacial morphology. Enrichment of genes such as Reelin (*RELN*), Down syndrome cell adhesion molecule 1 (*DSCAM1*), kalirin RhoGEF kinase (*KALRN*), and neurobeachin (*NBEA*) suggests advanced neurodevelopmental capacity in this lineage. *RELN*, a conserved glycoprotein in mammals, is essential for neuronal migration and synaptic plasticity and is linked to behavioral flexibility and brain development (DeSilva et al., 1997). A similar analysis of the southern sea otter found evidence of positive selection for *RELN*, which is believed to facilitate genetic adaptation in learning and memory, influencing prey capture and preference (Beichman et al., 2019). *KALRN* promotes dendritic branching and maturation, with overexpression associated with increased dendritic complexity and potentially brain volume (Ma et al., 2003). *DSCAML1*, a gene highly conserved among vertebrates, is implicated in axon guidance and craniofacial morphology, and has been shown to affect head size in zebrafish (Ma et al., 2019, 2023). *NBEA* regulates protein trafficking in neurons and is essential for synaptic development (Miller et al., 2015). These genes, along with the significant, broad enrichment of the *head development* may explain previously observed differences in skull size and shape in *lutensis*, as described in morphological assessments (Humphrey and Setzer 1989) (Fig. 4C).

The coastal salt marsh ecosystems of northern Florida are known for their environmental instability and sensitivity to climate change. These ecosystems experience wide fluctuations in salinity, temperature, and nutrient availability, and are increasingly threatened by sea-level rise, saltwater intrusion, and anthropogenic disturbances (Jeppesen et al., 2020). In addition, the individual genome of *lutensis* (nvison_FL-32666), originating from Bay County, Florida, may offer further insight. This region has a documented history of industrial pollution (United States Environmental Protection Agency, 1975; Florida Department of Environmental Protection, 2022). Thus, specific neural and cranial features may enhance sensory processing and behavioral flexibility, traits that may be advantageous where rapid environmental change is prevalent.

Several functional groups of genes further support adaptations for environmental resilience, including associations with ATPase activity coupled to transmembrane transport (myosin heavy chain 9 gene), O-methyltransferase activity linked to thermal adaptation, and synaptic vesicle uncoating (Transient Receptor Potential Melastatin 2 gene), associated with response to heat stress (Huson et al., 2011; Liu et al., 2024) (Fig. 4C).

Particularly notable is the enrichment of *ATP1A2*, the alpha-2 subunit of the Na⁺/K⁺-ATPase complex. This gene maintains ionic gradients essential for cellular homeostasis and has been implicated in salinity adaptation in salmonids (Dalziel et al. 2014). While *ATP1A2* is broadly conserved in carnivorans, its specific enrichment in *lutensis* suggests possible adaptation to the variable salinity conditions of its habitat (Fig. 4C).

## Conclusion

This study demonstrates the utility of pangenome approaches in examining adaptive signatures among closely related subspecies, using *N. vison* as a model. By integrating genomic data from natural history collections and leveraging gene presence–absence variation, the research uncovers distinct evolutionary trajectories and adaptive genes influenced by environmental pressures. These findings not only reinforce the value of reference-free pangenomes for conservation genomics but also provide insights into subspecies delineation, informing more targeted and effective management strategies.

## Acknowledgements

Several museum specimens were provided thanks to the University of Florida Board of Trustees – Florida Museum of Natural History Genetic Resources Repository. The authors acknowledge contributions of the HPC resources from the Computational Biology Core and sequencing resources from the Center for Genome Innovation, both within the Institute for Systems Genomics at the University of Connecticut.

## Author contribution statement

AA, HB, SF, ENF, KH, ELN, AM, KP, MR, RNS, NV, HA, NR, and JLW contributed to the genomic analysis. MR, KC, NP, and RJO directed and executed sample preparation, DNA extraction, and sequencing. HA, NR, TK, ELJ, JLW, and PH provided guidance and training on informatic approaches. ELJ, JLW, and PH conceived and designed the study. PH selected study samples and coordinated acquisition. All authors participated in manuscript preparation and editing.

## Data Archiving

All of the Illumina (paired-end) and Oxford Nanopore Technologies (ONT) raw data, and de-novo genome assemblies are available under the BioProject ID PRJNA1249195. The reviewer link for our BioProject is: https://dataview.ncbi.nlm.nih.gov/object/PRJNA1249195?reviewer=spedibf5t7reqojn9kks5cplti. The full workflow and scripts are available here: https://gitlab.com/PlantGenomicsLab/minkconservation_ramp2025

## Supplementary Figures/Tables

File S1 - Sequencing Methods

File S2 - Newick Mitochondria Tree

Table S1 - Comparative Statistics (other genomes)

Table S2 - Coalescence Statistics

Table S3 - Genome Coverage Statistics

Table S4 - Sequence Read Contamination Statistics

Table S5 - Genome Assembly Statistics

Table S6 - Genome Repeat Statistics

Table S7 - Repeat Classification

Table S8 - Genome Annotation Statistics

Table S9 - Protein-coding genes in ROH Regions shared among *evergladensis* individuals

Table S10 - Mitochondrial Genome Statistics

Table S11 - Nucleotide Conservation and Differences Across Mitochondrial Regions in *Neogale vison* Subspecies

Table S12 - Gene Ontology Terms of Interest with Functional Definitions

Table S13 - Hierarchical Orthologous Groups and Associated Genes in Selected Carnivorans

Table S14 - Pangenome Homology Grouping

## References

Alonge, M., Lebeigle, L., Kirsche, M., Jenike, K., Ou, S., Aganezov, S. et al. (2022). Automated assembly scaffolding using RagTag elevates a new tomato system for high-throughput genome editing. Genome Biology, 23(1), 258. 10.1186/s13059-022-02823-7

Amstislavsky, S., & Ternovskaya, Y. (2000). Reproduction in mustelids. Animal Reproduction Science, 60–61, 571–581. 10.1016/S0378-4320(00)00126-3

Astashyn, A., Tvedte, E.S., Sweeney, D., Sapojnikov, V., Bouk, N., Joukov V. et al. (2024). Rapid and sensitive detection of genome contamination at scale with FCS-GX. Genome Biology 25, 60. 10.1186/s13059-024-03198-7

van der Auwera, G., & O’Connor, B. D. (2020). Genomics in the cloud: using Docker, GATK, and WDL in Terra (First edition.). O’Reilly Media.

Beichman, A. C., Koepfli, K.-P., Li, G., Murphy, W., Dobrynin, P., Kliver, S. et al. (2019). Aquatic adaptation and depleted diversity: A deep dive into the genomes of the sea otter and giant otter. Molecular Biology and Evolution, 36(12), 2631–2655. 10.1093/molbev/msz101

Blottner, S., Schön, J., & Jewgenow, K. (2006). Seasonally activated spermatogenesis is correlated with increased testicular production of testosterone and epidermal growth factor in mink (*Mustela vison*). Theriogenology, 66(6-7), 1593–1598. 10.1016/j.theriogenology.2006.01.041

Bonesi L, Palazon S (2007) The American mink in Europe: Status, impacts, and control. Biological Conservation, 134, 470–483. 10.1016/j.biocon.2006.09.006.

Buchfink, B., Xie, C., & Huson, D. H. (2015). Fast and sensitive protein alignment using DIAMOND. Nature Methods, 12(1), 59–60. 10.1038/nmeth.3176

Burbrink FT, Crother BI, Murray CM, Smith BT, Ruane S, Myers EA et al. (2022) Empirical and philosophical problems with the subspecies rank. Ecology and Evolution, 12, e9069.

Cainé, L., Lima, G., Pontes, L., Abrantes, D., Pereira, M., & Pinheiro, M. (2006). Species identification by cytochrome b gene: Casework samples. International Congress Series, 1288, 145–147. 10.1016/j.ics.2005.09.152

Chapman, J. A., & Feldhamer, G. A. (1982). Wild mammals of North America: Biology, Management, and Economics. Johns Hopkins University Press, Baltimore and London.

Charbonnel, N., Galan, M., Tatard, C., Loiseau, A., Diagne, C., Dalecky, A. et al. (2020). Differential immune gene expression associated with contemporary range expansion in two invasive rodents in Senegal. Scientific Reports, 10, 18257. 10.1038/s41598-020-75060-2

Chen, S., Zhou, Y., Chen, Y., & Gu, J. (2018). fastp: an ultra-fast all-in-one FASTQ preprocessor. Bioinformatics, 34(17), i884–i890. 10.1093/bioinformatics/bty560

Clendenin, H. R., Pollard, M. D., & Puckett, E. E. (2025). Linking Measures of Inbreeding and Genetic Load to Demographic Histories Across Three Species of Bears. Evolutionary Applications, 18(7), e70133. 10.1111/eva.70133

Cornet, S., Brouat, C., Diagne, C., & Charbonnel, N. (2016). Eco-immunology and bioinvasion: revisiting the evolution of increased competitive ability hypotheses. Evolutionary Applications, 9(8), 952–962. 10.1111/eva.12406

Dalziel, A. C., Bittman, J., Mandic, M., Ou, M., & Schulte, P. M. (2014). Origins and functional diversification of salinity-responsive Na+, K+ ATPase α1 paralogs in salmonids. Molecular Ecology, 23(14), 3483–3503. 10.1111/mec.12828

Dainat, J. (n.d.). Welcome to AGAT’s documentation! - AGAT. Retrieved April 25, 2025, from https://nbisweden.github.io/AGAT/.

Dong, Y., Wei, Q., Sun, G., Gao, X., Lyu, T., Wang, L., Zhou, S. et al. (2025). Evolutionary analysis of genes associated with the sense of balance in semi-aquatic mammals. BMC Ecology and Evolution, 25, 8. 10.1186/s12862-024-02345-9

DeSilva, U., D’Arcangelo, G., Braden, V. V., Chen, J., Miao, G. G., Curran, T. et al. (1997). The human Reelin gene: Isolation, sequencing, and mapping on chromosome 7. Genome Research, 7(2), 157–164. 10.1101/gr.7.2.157

De Bie, T., Cristianini, N., Demuth, J. P., & Hahn, M. W. (2006). CAFE: A computational tool for the study of gene family evolution. Bioinformatics, 22(10), 1269–1271. 10.1093/bioinformatics/btl097

De Coster, W., & Rademakers, R. (2023). NanoPack2: population-scale evaluation of long-read sequencing data. Bioinformatics, 39(5), btad311. 10.1093/bioinformatics/btad311

Derežanin, L., Blažytė, A., Dobrynin, P., Duchêne, D. A., Horacio Grau, J., Jeon., S. et al. (2022). Multiple types of genomic variation contribute to adaptive traits in the mustelid subfamily Guloninae. Evolutionary Biology, 31(10), 2898–2919. 10.1101/2021.09.27.46165

Dierckxsens, N., Mardulyn, P., & Smits, G. (2016). Novoplasty: de novo assembly of organelle genomes from whole genome data. Nucleic Acids Research, 45(4), e18. 10.1093/nar/gkw955

Du, X., Servin, B., Womack, J. E., & Faraut, T. (2014). An update of the goat genome assembly using dense radiation hybrid maps allows detailed analysis of evolutionary rearrangements in Bovidae. BMC Genomics, 15, 625. 10.1186/1471-2164-15-625

du Plessis SJ, Hong S, Lee B, Koepfli KP, Chadwick EA, Hailer F (2023) Mitochondrial genome-based synthesis and timeline of Eurasian otter (*Lutra lutra*) phylogeography, Animal Cells and Systems, 27, 366–377.

Emms, D.M., Kelly, S. (2019) OrthoFinder: phylogenetic orthology inference for comparative genomics. Genome Biology, 20, 238. 10.1186/s13059-019-1832-y

Florida Fish and Wildlife Conservation Commission. 2016. Florida’s Imperiled Species Management Plan. Tallahassee, Florida.

Flex, E., Albadri, S., Radio, F. C., Cecchetti, S., Lauri, A., Priolo, et al. (2023). Dominantly acting KIF5B variants with pleiotropic cellular consequences cause variable clinical phenotypes. Human Molecular Genetics, 32(3), 473–488. 10.1093/hmg/ddac213

Gong, Y, Li, Y, Liu, X, Ma, Y, Jiang, L. (2023) A review of the pangenome: how it affects our understanding of genomic variation, selection and breeding in domestic animals? Journal of Animal Science and Biotechnology, 14, 73. 10.1186/s40104-023-00860-1

Green, M. R., & Sambrook, J. (2012). Molecular Cloning: a Laboratory Manual (4th ed.). Cold Spring Harbor Laboratory Press.

Gurevich, A., Saveliev, V., Vyahhi, N., & Tesler, G. (2013). QUAST: quality assessment tool for genome assemblies. Bioinformatics, 29(8), 1072–1075. 10.1093/bioinformatics/btt086

Haig, S.M., Beever, E.A., Chambers, S.M., Draheim, H.M, Dugger, B.D., Dunham, S.M., et al. (2006) Taxonomic considerations in listing subspecies under the U.S. Endangered Species Act. Conservation Biology, 20(6), 1584–1594. 10.1111/j.1523-1739.2006.00530.x

Hapeman, P., and Smith, L. M. (2024). Genetics, geography, and subspecies status of American mink in Florida, with an emphasis on *Neogale vison evergladensis*. Systematics and Biodiversity, 22(1), 2330371. 10.1080/14772000.2024.2330371

Hart, A. J., Ginzburg, S., Fisher, C. R., Rahmatpour, N., Mitton, J. B., Paul, R., et al. (2020). EnTAP: Bringing faster and smarter functional annotation to non-model eukaryotic transcriptomes. Molecular Ecology Resources, 20(2), 591–604. 10.1111/1755-0998.13106

Hempel, E., Faith, J. T., Preick, M., De Jager, D., Barish, S., Hartmann, S. et al. (2024). Colonial-driven extinction of the blue antelope despite genomic adaptation to low population size. Current Biology, 34(9), 2020–2029.e6. 10.1016/j.cub.2024.03.051

Hoelzel, A. R., Gkafas, G. A., Kang, H., Sarigol, F., Le Boeuf, B., Costa, D. P. et al. (2024). Genomics of post-bottleneck recovery in the northern elephant seal. Nature Ecology & Evolution, 8(4), 686–694. 10.1038/s41559-024-02337-4

Hou, Y., Qi, F., Bai, X., Ren, T., Shen, X., Chu, Q., et al. (2020). Genome-wide analysis reveals molecular convergence underlying domestication in 7 bird and mammals. BMC Genomics, 21, 204. 10.1186/s12864-020-6613-1

Huang, N., & Li, H. (2023). compleasm: a faster and more accurate reimplementation of BUSCO. Bioinformatics, 39(10), btad595. 10.1093/bioinformatics/btad595

Huerta-Cepas, J., Szklarczyk, D., Heller, D., Hernández-Plaza, A., Forslund, S. K., Cook, H. et al (2018). Eggnog 5.0: A hierarchical, functionally and phylogenetically annotated orthology resource based on 5090 organisms and 2502 viruses. Nucleic Acids Research, 47, D309-D317. 10.1093/nar/gky1085

Humphrey, Steven R., & Setzer, H. W. (1989). Geographic variation and taxonomic revision of Mink (mustela vison) in Florida. Journal of Mammalogy, 70(2), 241–252. 10.2307/1381505

Humphrey, S. R., & Zinn, T. L. (1982). Seasonal habitat use by river otters and Everglades mink in Florida. The Journal of Wildlife Management, 46(2), 375. 10.2307/3808649

Huson, H. J., vonHoldt, B. M., Rimbault, M., Byers, A. M., Runstadler, J. A., Parker, H. G. et al. (2011). Breed-specific ancestry studies and genome-wide association analysis highlight an association between the MYH9 gene and heat tolerance in Alaskan sprint racing sled dogs. Mammalian Genome, 23(1–2), 178–194. 10.1007/s00335-011-9374-y

Jagadish, N., Rana, R., Mishra, D., Kumar, M., & Suri, A. (2005). Sperm associated antigen 9 (SPAG9): A new member of c-Jun NH2-terminal kinase (JNK) interacting protein exclusively expressed in testis. Kobe Journal of Medical Sciences, 54(2), 66–71. 10.2302/kjm.54.66

Jahid, M. J., Bowman, A. S., Nolting, J. M. (2024). SARS-CoV-2 Outbreaks on mink farms-a review of current knowledge on virus infection, spread, spillover, and containment. Viruses, 16(1), 81. doi: 10.3390/v16010081.

Jeppesen, E., Beklioğlu, M., Özkan, K., & Akyürek, Z. (2020). Salinization increase due to climate change will have substantial negative effects on inland waters: A call for multifaceted research at the local and global scale. Innovation 1(2), 100030. 10.1016/j.xinn.2020.100030

Katti, C., Dalal, J. S., Dosé, A. C., Burnside, B., & Battelle, B. (2009). Cloning and distribution of myosin 3B in the mouse retina: Differential distribution in cone outer segments. Experimental Eye Research, 89(2), 224–237. 10.1016/j.exer.2009.03.011

Khan, A., Patel, K., Shukla, H., Viswanathan, A., Borthakur, U., Nigam, P. et al. (2021). Genomic evidence for inbreeding depression and purging of deleterious genetic variation in Indian tigers. Proceedings of the National Academy of Sciences, 118(49), e2023018118. 10.1073/pnas.2023018118

Kim, E., Cho, E.-J., Yang, S.-M., & Kim, H.-Y. (2021). Identification and Monitoring of *Lactobacillus delbrueckii* Subspecies Using Pangenomic-Based Novel Genetic Markers. Journal of Microbiology and Biotechnology, 31(2), 280–289. 10.4014/jmb.2009.09034

Kim, D., Song, L., Breitwieser, F. P., & Salzberg, S. L. (2016). Centrifuge: rapid and sensitive classification of metagenomic sequences. Genome Research, 26(12), 1721–1729. 10.1101/gr.210641.116

Kolmogorov, M., Yuan, J., Lin, Y., & Pevzner, P. A. (2019). Assembly of long, error-prone reads using repeat graphs. Nature Biotechnology, 37(5), 540–546. 10.1038/s41587-019-0072-8

Lan, T., Tian, Y., Shi, M., Liu, B., Lin, Y., Xia, Y. et al. (2025). Enhancing inbreeding estimation and global conservation insights through chromosome-level assemblies of the Chinese and Malayan pangolin. GigaScience, 14, 1–16. 10.1093/gigascience/giaf003

Li, H., Handsaker, B., Wysoker, A., Fennell, T., Ruan, J., Homer, N. et al. (2009). The Sequence Alignment/Map format and SAMtools. Bioinformatics, 25(16), 2078. 10.1093/bioinformatics/btp352

Liu, B., Shi, Y., Yuan, J., Hu, X., Zhang, H., Li, N. et al. (2013). Estimation of genomic characteristics by analyzing k-mer frequency in de novo genome projects. *ArXiv*. https://arxiv.org/abs/1308.2012

Liu, D., Yang, Y., Chen, Z., Fan, Y., Liu, J., Xu, Y. et al. (2024). Temperature adaptation patterns in Chinese cattle revealed by TRPM2 gene mutation analysis. Animal Biotechnology, 35(1), 2299944. 10.1080/10495398.2024.2299944

Ma, M., Huang, J., Wang, Y., Eipper, B. A., & Mains, R. E. (2003). Kalirin, a multifunctional Rho Guanine Nucleotide Exchange Factor, is necessary for maintenance of hippocampal pyramidal neuron dendrites and dendritic spines. The Journal of Neuroscience, 23(33), 10593. 10.1523/JNEUROSCI.23-33-10593.2003

Ma, M., Ramirez, A. D., Wang, T., Roberts, R. L., Harmon, K. E., Schoppik, D. et al. (2019). Zebrafish dscaml1 deficiency impairs retinal patterning and oculomotor function. The Journal of Neuroscience, 40(1), 143–158. 10.1523/jneurosci.1783-19.2019

Ma, M., Brunal, A. A., Clark, K. C., Studtmann, C., Stebbins, K., Higashijima, S., & Pan, Y. A. (2023). Deficiency in the cell-adhesion molecule dscaml1 impairs hypothalamic CRH neuron development and perturbs normal neuroendocrine stress axis function. Frontiers in Cell and Developmental Biology, 11, 1113675. 10.3389/fcell.2023.1113675

Martien, K. K., Leslie, M.S., Taylor, B.L., Morin, P.A., Archer, F.I., Hancock-Hanser, B.L., (2017). Analytical approaches to subspecies delimitation with genetic data. Marine Mammal Science, 33, 27–55. doi: 10.1111/mms.12409.

Mather, N., Traves, S. M., & Ho, S. Y. W. (2019). A practical introduction to sequentially Markovian coalescent methods for estimating demographic history from genomic data. Ecology and Evolution, 10(1), 579–589. 10.1002/ece3.5888

Md, V., Misra, S., Li, H., & Aluru, S. (2019). Efficient architecture-aware acceleration of BWA-MEM for multicore systems. IEEE International Parallel and Distributed Processing Symposium (IPDPS), 2019, 314–324. 10.1109/IPDPS.2019.00041

Mellya, R. V., Hopcraft, J. G., Mwakilema, W., Eblate, E. M., Mduma, S., Mnaya, B. et al. (2025). Natural dispersal is better than translocation for reducing risks of inbreeding depression in eastern black rhinoceros (*Diceros bicornis michaeli*). Proceedings of the National Academy of Sciences, 122(23), e2414412122. 10.1073/pnas.2414412122

Miller, A. C., Voelker, L. H., Shah, A. N., & Moens, C. B. (2015). *Neurobeachin* is required postsynaptically for electrical and chemical synapse formation. Current Biology, 25(1), 16–28. 10.1016/j.cub.2014.10.051

Mortensen, R. M., Reinhardt, S., Hjønnevåg, M. E., Wilson, R. P., & Rosell, F. (2021). Aquatic habitat use in a semi-aquatic mammal: The Eurasian beaver. Animal Biotelemetry, 9(1), 35. 10.1186/s40317-021-00259-7

Moshiashvili, L. (2024). CafePlotter (v0.2.0) [Computer software]. GitHub. https://github.com/moshi4/CafePlotter

O’Brien, M. F., Lopez Colom, R., Natural History Museum Genome Acquisition Lab, Darwin Tree of Life Barcoding collective, Wellcome Sanger Institute Tree of Life Management, Samples and Laboratory team, Wellcome Sanger Institute Scientific Operations: Sequencing Operations, et al. (2025). The genome sequence of the American mink, Neogale vison (Schreber, 1777), version 1. Wellcome Open Research, 10, 103 10.12688/wellcomeopenres.23780.1

Ohio Department of Natural Resources. (n.d.). American mink. Retrieved July 22, 2025, from https://ohiodnr.gov/discover-and-learn/animals/mammals/american-mink

Paradis, E. (2010). Pegas: An R package for population genetics with an integrated–modular approach. Bioinformatics, 26(3), 419–420. 10.1093/bioinformatics/btp696

Paradis, E., Claude, J., & Strimmer, K. (2004). APE: Analyses of phylogenetics and evolution in R language. Bioinformatics, 20(2), 289–290. 10.1093/bioinformatics/btg412

Pedersen, B. S., & Quinlan, A. R. (2017). Mosdepth: Quick coverage calculation for genomes and exomes. Bioinformatics, 34(5), 867–868. 10.1093/bioinformatics/btx699

Pence, S., Caykara, B., Pence, H. H., Tekin, S., Keskin, B. C., Uncu, A. T. et al. (2022). Transcriptomic analysis of asymptomatic and symptomatic severe Turkish patients in SARS-CoV-2 infection. Northern Clinics of Istanbul, 9(2), 122–130. 10.14744/nci.2022.28000

Rajput, A., Chauhan, S. M., Mohite, O. S., Hyun, J. C., Ardalani, O., Jahn, L. J. et al. (2023). Pangenome analysis reveals the genetic basis for taxonomic classification of the Lactobacillaceae family. Food Microbiology, 115, 104334. 10.1016/j.fm.2023.104334

Reid, F., Schiaffini, M. & Schipper, J. 2016. Neovison vison. The IUCN Red List of Threatened Species 2016: e.T41661A45214988. 10.2305/IUCN.UK.2016-1.RLTS.T41661A45214988.en. Accessed on 10 April 2025.

Rhie, A., Walenz, B. P., Koren, S., & Phillippy, A. M. (2020). Merqury: reference-free quality, completeness, and phasing assessment for genome assemblies. Genome Biology, 21(1), 245. 10.1186/s13059-020-02134-9

Romeo, C., Filipe, J., Wauters, L. A., Comazzi, S., Riva, F., & Ferrari, N. (2023). Shifts in immune responses of an invasive alien species: A test of the evolution of increased competitive ability hypothesis using American Eastern gray squirrels in Italy. Science of the Total Environment, 900, 165747. 10.1016/j.scitotenv.2023.165747

Sánchez-Barreiro, F., Gopalakrishnan, S., Ramos-Madrigal, J., Westbury, M. V., Margaryan, A., Ciucani, M. M. et al. (2021). Historical population declines prompted significant genomic erosion in the northern and southern white rhinoceros (Ceratotherium simum). Molecular Ecology, 30(23), 6355–6369. 10.1111/mec.16043

Secomandi, S., Gallo, G. R., Rossi, R., Rodríguez Fernandes, C., Jarvis, E. D., Bonisoli-Alquati, A. et al. (2025). Pangenome graphs and their applications in biodiversity genomics. Nature Genetics, 57(1), 13–26. 10.1038/s41588-024-02029-6

Sheikhizadeh-Anari, S., de Ridder, D., Schranz, M. E., & Smit, S. (2018). Efficient inference of homologs in large eukaryotic pan-proteomes. BMC Bioinformatics, 19, 340. 10.1186/s12859-018-2362-4

Sheikhizadeh, S., Schranz, M. E., Akdel, M., de Ridder, D., & Smit, S. (2016). PanTools: representation, storage and exploration of pan-genomic data. Bioinformatics, 32(17), i487–i493. 10.1093/bioinformatics/btw455

Shen, W., Sipos, B., & Zhao, L. (2024). SeqKit2: A Swiss army knife for sequence and alignment processing. iMeta, 3(3), e191. 10.1002/imt2.191

Smit, A. F. A., Hubley, R., & Green, P. (2013–2015). RepeatMasker Open-4.0. http://www.repeatmasker.org

Smit, A. F. A., & Hubley, R. (2008–2015). RepeatModeler Open-1.0. http://www.repeatmasker.org

Tamura, K., Stecher, G., & Kumar, S. (2021). MEGA11: Molecular Evolutionary Genetics Analysis Version 11. Molecular Biology and Evolution, 38(7), 3022–3027. 10.1093/molbev/msab120

Terhorst, J., Kamm, J. A., & Song, Y. S. (2017). Robust and scalable inference of population history from hundreds of unphased whole genomes. Nature Genetics, 49(2), 303–309. 10.1038/ng.3748

The Florida Department of Environmental Protection. (2022). Panama City and Lynn Haven: How “Enforcement” Turns Into Quiet Acknowledgment of the Inevitable. peer.org. https://peer.org/wp-content/uploads/2022/03/03_3_2022-FDEP-Panama-City-and-Lynn-Haven.pdf

United States Environmental Protection Agency. Water Quality Study: St. Andrew Bay, Florida. EPA, 1975. EPA Report No. 330/2-75-003. EPA National Service Center for Environmental Publications, https://nepis.epa.gov/.

Vinton, A. C., Gascoigne, S. J., Sepil, I., & Salguero-Gómez, R. (2022). Plasticity’s role in adaptive evolution depends on environmental change components. Trends in Ecology & Evolution, 37(12), 1067–1078. 10.1016/j.tree.2022.08.008

Vourekas, A., Zheng, K., Fu, Q., Maragkakis, M., Alexiou, P., Ma, J. et al. (2015). The RNA helicase MOV10L1 binds piRNA precursors to initiate piRNA processing. Genes & Development, 29(6), 617. 10.1101/gad.254631.114

Vuruputoor, V. S., Monyak, D., Fetter, K. C., Webster, C., Bhattarai, A., Shrestha, B. et al. (2023). Welcome to the big leaves: Best practices for improving genome annotation in non-model plant genomes. Applications in Plant Sciences, 11(4), e11533. 10.1002/aps3.11533

Walsh, J., Lovette, I. J., Winder, V., Elphick, C. S., Olsen, B. J., Shriver, G. et al. (2017). Subspecies delineation amid phenotypic, geographic and genetic discordance in a songbird. Molecular Ecology, 26(5), 1242–1255. 10.1111/mec.14010

Wang, Y., Tang, H., DeBarry, J. D., Tan, X., Li, J., Wang, X. et al. (2012). MCScanX: A toolkit for detection and evolutionary analysis of gene synteny and collinearity. Nucleic Acids Research, 40(7), e49. 10.1093/nar/gkr1293

Wang, C., Wu, D., Yuan, H., Yao, C., Han, L., Wu, J. et al. (2023). Population genomic analysis provides evidence of the past success and future potential of South China tiger captive conservation. BMC Biology, 21, 64. 10.1186/s12915-023-01552-y

Wang, X., Yin, G., Yang, Y., & Tian, X. (2025). Ciliary and non-ciliary roles of IFT88 in development and diseases. International Journal of Molecular Sciences, 26(5), 2110. 10.3390/ijms26052110

Wirgin, I., Maceda, L., Waldman, J., Mayack, D. T. (2015). Genetic variation and population structure of American mink Neovison vison from PCB-contaminated and non-contaminated locales in eastern North America. Ecotoxicology, 24, 1961–75. 10.1007/s10646-015-1533-6

Wong, T. K. F., Ly-Trong, N., Ren, H., Baños, H., Roger, A. J., Susko, E., et al. (2025). IQ-TREE 3: Phylogenomic inference software using complex evolutionary models. EcoEvoRxiv. 10.32942/X2P62N

Wood, D. E., Lu, J., & Langmead, B. (2019). Improved metagenomic analysis with Kraken 2. Genome Biology, 20(1), 257. 10.1186/s13059-019-1891-0

Wozencraft, W. C. (2005). Order Carnivora. In: Wilson DE, Reeder DM (eds) Mammal Species of the World: A Taxonomic and Geographic Reference, 3^rd^ edn. Vol I. Johns Hopkins University Press, Baltimore, pp 532–628.

Yamaguchi, N., & Macdonald, D. W. (2001). Detection of Aleutian disease antibodies in feral American mink in southern England. Veterinary Record, 149(16), 485–488. 10.1136/vr.149.16.485

Yeste-Velasco, M., Mao, X., Grose, R., Kudahetti, S. C., Lin, D., Marzec, J. et al. (2014). Identification of ZDHHC14 as a novel human tumour suppressor gene. The Journal of Pathology, 232(5), 566–577. 10.1002/path.4327

Xu, S.; Shan, L.; Tian, R.; Yu, Z.; Sun, D.; Zhang, Z. et al. (2025). Multi-level genomic convergence of secondary aquatic adaptation in marine mammals. Innovation, 6(3), 100798. 10.1016/j.xinn.2025.100798

Zhang, L., Hua, Y., & Wei, S. (2021). High genetic diversity of an invasive alien species: comparison between fur-farmed and feral American Mink (*Neovison vison*) in China. Animals, 11(2), 472. 10.3390/ani11020472

Zimin, A. V., Puiu, D., Luo, M.-C., Zhu, T., Koren, S., Marçais et al. (2017). Hybrid assembly of the large and highly repetitive genome of *Aegilops tauschii*, a progenitor of bread wheat, with the MaSuRCA mega-reads algorithm. Genome Research, 27(5), 787–792. 10.1101/gr.213405.116

Zimin, A. V., & Salzberg, S. L. (2022). The SAMBA tool uses long reads to improve the contiguity of genome assemblies. PLoS Computational Biology, 18(2), e1009860. 10.1371/journal.pcbi.1009860

Zink, R. M., & Klicka, L. B. (2022). The taxonomic basis of subspecies listed as threatened and endangered under the Endangered Species Act. Frontiers in Conservation Science, 3. 10.3389/fcosc.2022.971280

