## Supplementary material for "Mink by mink: stitching together signatures of subspecies adaptation through a pangenome of threatened mustelids": File S1

**DNA Extraction Protocol**

High molecular weight DNA was extracted from mink tissue using a modified Phenol/Chloroform/Isoamyl alcohol protocol from Green *et al*., 2012. All tissues were rinsed twice in PBS and stored at -80 °C before DNA extraction.

Lysis and Digestion:

Tissues were homogenized using a mortal and pestle. Each sample was digested in a modified lysis buffer with the following recipe:

a. 400 mM of 1M Tris-Cl (pH 8.0)
b. 60 mM of 0.5M EDTA (pH 8.0)
c. 150 mM of 5M NaCl
d. 1% SDS from 20% stock

Proteinase K was added to a final concentration of 200 μg/mL (from a 20 mg/mL stock). Samples were incubated at 56 °C for 1 hour with agitation every 15 minutes until fully digested. RNase A was added to a final concentration of 20 μg/mL, followed by a 15-minute incubation at 37 °C.

Extraction and Precipitation:

Equal volumes phenol:chloroform:isoamyl alcohol (25:24:1) was added to each digested sample and vortexed to form an emulsion. Samples were then centrifuged at 5500 x g for 10 minutes at 4 °C, and the supernatant was transferred to a new 1.5mL tube to separate DNA from proteins. DNA was precipitated with:

a. 1/10 volume of 3M Sodium Acetate
b. 2 volumes of -20 °C 100% ethanol

Samples were gently inverted to mix and stored at -20°C overnight. The following day, samples were centrifuged at 10,000 x g for 20 minutes at 4 °C to pellet DNA. The supernatant was then discarded, and pellets were left to air dry for ~30 seconds. Finally, the pellet was resuspended in 100 uL of nuclease-free water (NFW). DNA was stored at 4°C until library preparation.

DNA Purification and QC

Due to low DNA yield in nvison_FL-5, multiple elutes from separate extractions were pooled. RNase A was added again (20ug/mL), followed by a Ampure XP clean-up step. The combined DNA samples were cleaned using Ampure XP beads. The cleanup was done using the following procedure: 2X volume of Ampure XP beads was added to each sample. Samples were then incubated at RT in a Hula mixer at 9 RPM for 10 minutes. Beads were then washed and eluted in 100 uL NFW, followed by a 10-minute incubation at 37 °C.

DNA concentration and purity was measured using Thermo Fisher Scientific's NanoDrop™ One and Qubit™ 4 Fluorometer. DNA was stored at 4 °C.

Illumina Library Preparation and Sequencing

For Library Preparation, the NEBNext Ultra™ II FS DNA Library Prep Kit (New England Biolabs, Ipswich, MA) was used. Fragmentation size was optimized based on sample quality to achieve a target insert length of 300-350 bp. For nvison_FL-32666, which was degraded – two libraries were prepared: one with shearing to 350 bp, and the other without fragmentation. Library fragment sizes and adapter dimer presence were assessed using Agilent TapeStation 4200 D1000 High Sensitivity assay (Agilent Technologies, Santa Clara, CA, USA), and library concentration was quantified using Qubit™ 3.0 (dsDNA HS Assay). Both libraries for nvison_FL-32666 passed QC and were sequenced. The target genome coverage was approximately 80X per sample.

Oxford Nanopore Library Preparation and Sequencing

Adapter ligation, cleanup, priming, and flow cell loading were performed according to the Ligation Sequencing DNA V14 (SQK-LSK114) protocol with input DNA adjusted to a concentration of 400 fmol.

References:

Green, M. R., & Sambrook, J. (2012). Molecular cloning : a laboratory manual (4th ed.). Cold Spring Harbor Laboratory Press.
