## Supplementary material for "Mink by mink: stitching together signatures of subspecies adaptation through a pangenome of threatened mustelids": File S2

('Neogale frenata':0.0669648201,('Neogale vison':3.838107000000007E-4,
((nvison_FL-32666:0.0019760060999999968,nvison_FL-33534:9.631642000000
051E-4):0.004630246500000004,
((nvison_FL-3:1.0017000000028808E-6,nvison_FL-5:1.0017000000028808E-6)
:0.0018858551000000001,
(nvison_LA-8:9.74474000000003E-4,nvison_LA-9:3.452670000000102E-4):2.7
618109999999474E-4):9.542176000000013E-4):0.0026594892999999998):0.0576
99078200000004);
