## Supplementary material for "Mink by mink: stitching together signatures of subspecies adaptation through a pangenome of threatened mustelids": Table S1

| Species | Source | Version | Chromosomes | Diploid/Haploid | Protein-coding genes | Genome Completeness (compleasm) | Protein Completeness (compleasm) | BioProject | Link |
|---|---|---|---|---|---|---|---|---|---|
| ***Protein files from this paper*** | | | | | | | | | |
| Neogale vison (Haplotype-1 American mink assembly) | NCBI | hap1.1 | Yes | Haploid | 19,992 | 98.84% | 97.47% | PRJEB76007 | https://www.ncbi. |
| nvison_FL-32666 | Affini et al., 2025 | v.3.0 | Yes | Haploid | 20,191 | 88.23% | 83.94% | PRJNA1249195 | https://dataview.r |
| nvison_FL-5 | Affini et al., 2025 | v.2.0 | Yes | Haploid | 20,133 | 97.35% | 95.39% | PRJNA1249195 | https://dataview.r |
| nvison_LA-9 | Affini et al., 2025 | v.2.0 | Yes | Haploid | 20,258 | 97.82% | 94.30% | PRJNA1249195 | https://dataview.r |
| ***Mustelidae*** | | | | | | | | | |
| Mustela putorius furo (domestic ferret) | NCBI | 1 | No, contig | Haploid | 20,794 | 99.47% | 98.66% | PRJNA580247 | https://www.ncbi. |
| Mustela nigripes (black-footed ferret) | NCBI | 1 | Yes | Haploid | 20,269 | 99.22% | 98.32% | PRJNA634921 | https://www.ncbi. |
| Mustela lutreola (European mink) | NCBI | 1 | Yes | Haploid | 20,701 | 99.67% | 98.81% | PRJNA984926 | https://www.ncbi. |
| Mustela erminea (ermine) | NCBI | 1 | Yes | Haploid | 20,934 | 99.32 | 98.58% | PRJNA561931 | https://www.ncbi. |
| Mustela nivalis (least weasel) | NCBI | 1 | Yes | Haploid | N/A | 98.86% | N/A | PRJEB83720 | https://www.ncbi. |
| Lontra canadensis (Northern American river otter) | NCBI | 1 | No, scaffold | Haploid | 20,305 | 99.28% | 98.64% | PRJNA578111 | https://www.ncbi. |
| Meles meles (Eurasian badger) | NCBI | 1 | Yes | Haploid | 21,063 | 97.87% | 98.84% | PRJEB49353 | https://www.ncbi. |
| Enhydra lutris kenyoni (Northern Sea Otter) | NCBI | 1 | No, scaffold | Haploid | 19,340 | 98.41% | 99.20% | PRJNA388419 | https://www.ncbi. |
| ***Ursidae*** | | | | | | | | | |
| Ailuropoda melanoleuca (giant panda) | NCBI | 1 | Yes | Haploid | 20,837 | 98.59% | 97.27% | PRJNA588422 | https://www.ncbi. |
| Ursus maritimus (polar bear) | NCBI | 1 | No, scaffold | Haploid | 20,419 | 99.26% | 98.54% | PRJNA669153 | https://www.ncbi. |
| ***Odobenidae*** | | | | | | | | | |
| Odobenus rosmarus divergens (Pacific walrus) | NCBI | 1 | No, scaffold | Haploid | 19,094 | 97.01% | 97.24% | PRJNA167474 | https://www.ncbi. |
| ***Phocidae*** | | | | | | | | | |
| Neomonachus schauinslandi (Hawaiian monk seal) | NCBI | 1 | Yes | Haploid | 19,838 | 97.67% | 98.30% | PRJNA384558 | https://www.ncbi. |
| Leptonychotes weddellii (Weddell seal) | NCBI | 1 | No, scaffold | Haploid | 22,214 | 89.65% | 84.54% | PRJNA68235 | https://www.ncbi. |
| Halichoerus grypus (gray seal) | NCBI | 1 | No, scaffold | Haploid | 21,245 | 96.06% | 94.20% | PRJNA577240 | https://www.ncbi. |
| Zalophus californianus (California sea lion) | NCBI | 1 | Yes | Haploid | 21,397 | 99.47% | 98.45% | PRJNA559673 | https://www.ncbi. |
| ***Felidae*** | | | | | | | | | |
| Felis catus (domestic cat) | NCBI | 1 | Yes | Haploid | 20,453 | 99.39% | 98.95% | PRJNA684600 | https://www.ncbi. |
| Panthera tigris (tiger) | NCBI | 1 | Yes | Haploid | 20,232 | 99.28% | 99% | PRJNA684344 | https://www.ncbi. |
| ***Hyaenidae*** | | | | | | | | | |
| Hyaena hyaena (striped hyena) | NCBI | 1 | Yes | Haploid | 20,024 | 98.49% | 97.63 | PRJNA390068 | https://www.ncbi. |
| ***Canidae*** | | | | | | | | | |
| Canis lupus familiaris (dog) | NCBI | 1 | Yes | Haploid | 21,175 | 97.72% | 98.12% | PRJNA587469 | https://www.ncbi. |
