## Supplementary material for "Mink by mink: stitching together signatures of subspecies adaptation through a pangenome of threatened mustelids": Table S2

**Mapping Rates** | hap1_guided_SMC++/

| Sample | Error rate | Bases mapped | Total # of reads | Alignment rate |
|---|---|---|---|---|
| nvison_FL-32666 | 6.23E-03 | 374806454723 | 387476056437 | 96.73% |
| nvison_FL-33534 | 6.47E-03 | 358886758413 | 371481490451 | 96.61% |
| nvison_LA-8 | 5.40E-03 | 322114493733 | 326073652970 | 98.79% |
| nvison_LA-9 | 4.81E-03 | 327091619248 | 332115592426 | 98.49% |
| nvison_FL-3 | 4.62E-03 | 389913434150 | 394825795140 | 98.76% |
| nvison_FL-5 | 5.58E-03 | 249653330312 | 252722795424 | 98.79% |
| nvison_Haplotype-1 | 5.52E-03 | 175344574869 | 178393439406 | 98.29% |

**Sample Quality Thresholds for Masking**

| Sample | No Coverage Threshold | Low Qualitity Threshold | High Qualitity Threshold | |
|---|---|---|---|---|
| nvison_FL-32666 | 0.00E+00 | 1 - 24 | 25 - 250 | |
| nvison_FL-33534 | 0.00E+00 | 1- 24 | 25 - 150 | |
| nvison_LA-8 | 0.00E+00 | 1 - 59 | 60 - 120 | |
| nvison_LA-9 | 0.00E+00 | 1 - 54 | 55 - 95 | |
| nvison_FL-3 | 0.00E+00 | 1 - 74 | 75 - 105 | |
| nvison_FL-5 | 0.00E+00 | 1 - 44 | 45 - 100 | |
| nvison_Haplotype-1 | 0.00E+00 | 1 - 34 | 35 - 65 | *based on depth coverage |

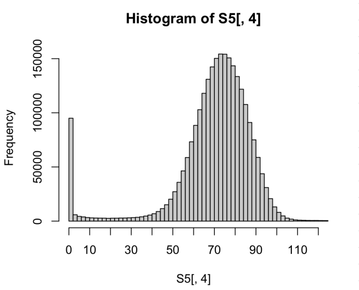


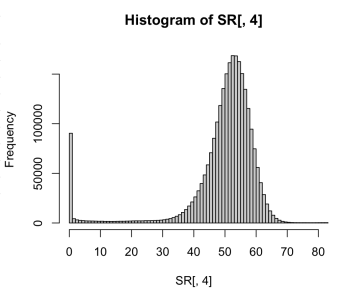


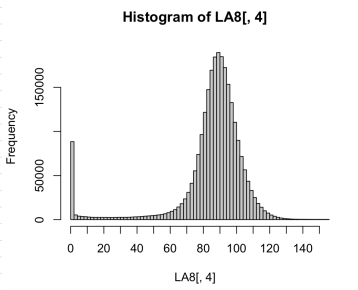


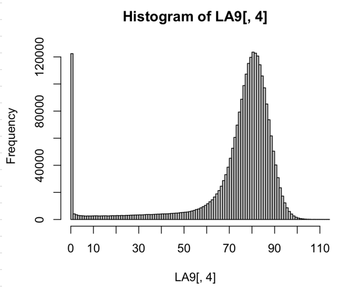


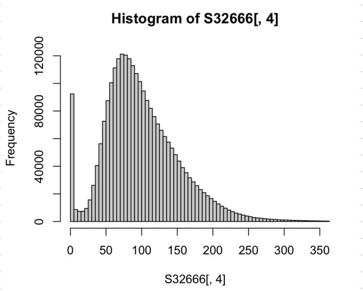


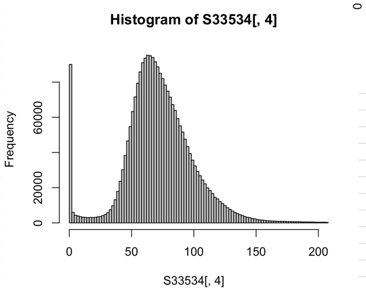
