## Supplementary material for "Mink by mink: stitching together signatures of subspecies adaptation through a pangenome of threatened mustelids": Table S3

| Sample Metadata | | | | | | | | | | |
|---|---|---|---|---|---|---|---|---|---|---|
| **Sample ID** | **ONT/Illumina** | **Coverage** | **Coverage post-contm** | **Read Length N50** | **Basecaller** | **Flowcell** | **NCBI Accession** | **Tissue Type** | **Sex** | **Collection Date** |
| **nvison_FL-3** | Illumina | 158.77X | 124.91X | 150bp PE | | | | Tissue | Male | 2004 |
| | ONT | 29.63X | 27.52X | 832 bp | Dorado (dna_r10.4.1_e8.2_400bps_sup@v5.0.0) | FLO-PRO114M | | | | |
| **nvison_LA-9** | Illumina | 154.67X | 152.07X | 150 bp PE | | | | Tissue | Male | 2015 |
| | ONT | 34.13X | 33.77X | 2020 bp | Dorado (dna_r10.4.1_e8.2_400bps_sup@v5.0.0) | FLO-PRO114M | | | | |
| **nvison_FL-5** | Illumina | 95.6X | 82X | 150bp PE | | | | Tissue | Male | 2004 |
| | ONT | 12.06X | 11.9X | 435 bp | Dorado (dna_r10.4.1_e8.2_400bps_sup@v5.0.0) | FLO-PRO114M | | | | |
| **nvison_LA-8** | Illumina | 129.51X | 105.06X | | | | | Tissue | Male | 2015 |
| **nvison_FL-33534** | ONT | 4.11X | 4.08x | 759 bp | Dorado (dna_r10.4.1_e8.2_400bps_sup@v5.0.0) | FLO-PRO114M | | Tissue | Male | 2016 |
| | Illumina | 151.21X | 132.58x | 150bp PE | | | | | | |
| **nvison_FL-32666 - sheared** | Illumina | 157.83X | 156.26x | 150bp PE | | | | Tissue | Putative Male | 2000 |
| **nvison_FL-32666 - unfragmented** | Illumina | 192.56X | 187.30x | 150bp PE | | | | Tissue | | 2000 |
