## Supplementary material for "Mink by mink: stitching together signatures of subspecies adaptation through a pangenome of threatened mustelids": Table S4

**ContamInation Statistics (Kraken/Centrifuge)**

| **Sample ID:** | **ONT/Illumina** | **Contaminant 1** | **Contaminant 2** | **Contaminant 3** | **Contaminant 4** | **Contaminant 5** |
|---|---|---|---|---|---|---|
| nvison_FL-5 | Illumina | *Xanthomonas euvesicatoria* | *Pseudomonas tolaasii* | *Streptomyces lividans* | *Staphylococcus virus Andhra* | *Fusarium pseudograminearum* |
| nvison_FL-5 | ONT | *Cutibacterium acnes* | *Pseudomonas stutzeri group* | *Pseudomonas putida group* | *Streptococcus suis* | *Streptococcus pneumoniae* |
| nvison_LA-8 | Illumina | *Xanthomonas euvesicatoria* | *Pseudomonas tolaasii* | *Streptomyces lividans* | *Klebsiella pneumoniae* | *Staphylococcus virus Andhra* |
| nvison_FL-32666 - Sheared | Illumina | *Klebsiella pneumoniae* | *Xanthomonas euvesicatoria* | *Pseudomonas tolaasii* | *Streptomyces lividans* | *Staphylococcus virus Andhra* |
| nvison_FL-32666 - Unfragmented | Illumina | *Xanthomonas euvesicatoria* | *Pseudomonas tolaasii* | *Streptomyces lividans* | *Klebsiella pneumoniae* | *Fusarium pseudograminearum* |
| nvison_FL-3 | Illumina | *Streptomyces lividans* | *Xanthomonas euvesicatoria* | *Pseudomonas tolaasii* | *Klebsiella pneumoniae* | *Klebsiella grimontii* |
| nvison_FL-3 | ONT | *Clostridium botulinum* | *Enterococcus faecalis* | *Edwardsiella tarda* | *Clostridium pasteurianum* | *Clostridium perfringens* |
| nvison_FL-33534 | Illumina | *Xanathomonas euvesicatoria* | *Pseudomonas tolaasii* | *Streptomyces lividans* | *Klebsiella pneumoniae* | *Pseudomonas fluorescens* |
| nvison_FL-33534 | ONT | Pseudomonas syringae group | Pseudomonas fluorescens group | *Pseudomonas putida group* | *Pseudomonas chlororaphis group* | *Pseudomonas aeruginosa group* |
| nvison_LA-9 | Illumina | *Xanthomonas euvesicatoria* | *Streptomyces lividans* | *Pseudomonas tolaasii* | *Klebsiella pneumoniae* | *Staphylococcus virus Andhra* |
| nvison_LA-9 | ONT | *Aeromonas hydrophila* | *Cutibacterium acnes* | *Pantoea ananatis* | *Streptococcus suis* | *Aeromonas media* |

| **FCS Statistics - Unplaced Scaffolds** | **Contaminant 1** | **Contaminant 2** | **Contaminant 3** | **# Sequences** | **# Bases** |
|---|---|---|---|---|---|
| nvison_FL-5 | *Homo sapiens* | *Sarcocystis neurona* | *Drosophila melanogaster* | 47 | *62462* |
| nvison_LA-8 | *Sarcocystis neurona* | *Homo sapiens* | *Synthetic* | 267 | 105188 |
| nvison_FL-32666 | *Psychrobacter sp* | *Wohlfahrtiimonas chitiniclastica* | *Metalysinibacillus jejuensis* | 10326 | 7578306 |
| nvison_FL-3 | *Sarcocystis neurona* | *Homo sapiens* | - | 1015 | 721785 |
| nvison_FL-33534 | *Sarcocystis neurona* | *Homo sapiens* | *Rhopapillomavirus* | 267 | 105188 |
| nvison_LA-9 | *synthetic* | - | - | 1 | 6418 |

| will remove - using for git | | | | 1987.166667 | | |
|---|---|---|---|---|---|---|
| **Sample ID:** | **ONT/Illumina** | **Contaminant 1** | **Contaminant 2** | **Contaminant 3** | **Contaminant 4** | **Contaminant 5** |
| nvison_FL-5 | Illumina | *X. euvesicatoria* | *P. tolaasii* | *S. lividans* | *S. virus Andhra* | *F. pseudograminearum* |
| nvison_FL-5 | ONT | *C. acnes* | *P. stutzeri group* | *P. putida group* | *S. suis* | *S. pneumoniae* |
| nvison_LA-8 | Illumina | *X. euvesicatoria* | *P. tolaasii* | *S. lividans* | *K. pneumoniae* | *S. virus Andhra* |
| nvison_FL-32666 - Sheared | Illumina | *K. pneumoniae* | *X. euvesicatoria* | *P. tolaasii* | *S. lividans* | *S.virus Andhra* |
| nvison_FL-32666 - Unfragmented | Illumina | *X. euvesicatoria* | *P. tolaasii* | *S. lividans* | *K. pneumoniae* | *F. pseudograminearum* |
| nvison_FL-3 | Illumina | *S. lividans* | *X. euvesicatoria* | *P. tolaasii* | *K. pneumoniae* | *K. grimontii* |
| nvison_FL-3 | ONT | *C. botulinum* | *E. faecalis* | *E. tarda* | *C. pasteurianum* | *C. perfringens* |
| nvison_FL-33534 | Illumina | *X. euvesicatoria* | *P. tolaasii* | *S. lividans* | *K. pneumoniae* | *P. fluorescens* |
| nvison_FL-33534 | ONT | P. syringae group | P. fluorescens group | *P. putida group* | *P. chlororaphis group* | *P. aeruginosa group* |
| nvison_LA-9 | Illumina | *X. euvesicatoria* | *S. lividans* | *P. tolaasii* | *K. pneumoniae* | *S. virus Andhra* |
| nvison_LA-9 | ONT | *A. hydrophila* | *C. acnes* | *P. ananatis* | *S. suis* | *Aeromonas media* |
