## Supplementary material for "Mink by mink: stitching together signatures of subspecies adaptation through a pangenome of threatened mustelids": Table S5

**Assembly Statistics**

| Sample ID | Assembler/Scaffolder | Total Scaffolds | N50 | Ns | Compleasm (% single copy) | Compleasm (% duplicated) | Merqury QV | Merqury ER |
|---|---|---|---|---|---|---|---|---|
| **nvison_FL-3** | Masurca SR/SAMBA/RagTag | 15 | 221 Mb | 3976 | 96.19% | 0.81% | 40.99 | 7.95E-05 |
| **nvison_FL-33534** | Masurca/SAMBA/RagTag | 15 | 197 Mb | 2178 | 75.15% | 0.82% | 42.01 | 6.30E-05 |
| **nvison_FL-5** | Masurca SR/SAMBA/RagTag | 15 | 209 Mb | 2250 | 96.41% | 0.94% | 42.65 | 5.43E-05 |
| **nvison_LA-8** | Masurca SR/RagTag | 15 | 232 Mb | 5757 | 95.92% | 0.72% | 40.57 | 8.77E-05 |
| **nvison_LA-9** | Masurca SR | 15 | 225 Mb | 4548 | 97.05% | 0.77% | 40.35 | 9.23E-05 |
| **nvison_FL-32666 - sheared and unfragmented** | Masurca SR/RagTag | 15 | 247 Mb | 9571 | 87.64% | 0.59% | 37.26 | 1.88E-05 |
