## Supplementary material for "Mink by mink: stitching together signatures of subspecies adaptation through a pangenome of threatened mustelids": Table S6

| Repeat Statsitics | | | | |
|---|---|---|---|---|
| **Sample ID** | **% Masked** | **Retroelements** | **DNA transposons** | **Unclassified** |
| nv_FL-33534 | 31.90% | 27.70% | 1.53% | 1.26% |
| nv_LA-8 | 29.69% | 25.60% | 1.36% | 1.30% |
| nv_LA-9 | 30.04% | 26.09% | 1.34% | 1.21% |
| nv_FL-3 | 29.39% | 26% | 1.41% | 1.17% |
| nv_FL-5 | 30.38% | 26.48% | 1.47% | 1.03% |
| nv_FL-32666 | 28.07% | 23.59% | 1.34% | 1.84% |
