## Supplementary material for "Mink by mink: stitching together signatures of subspecies adaptation through a pangenome of threatened mustelids": Table S7

| Sample_nvison_FL-33534_ | Number of Elem | Length Occupie | Percentage of Sequence | | Number of Base | Percent Masked |
|---|---|---|---|---|---|---|
| Retroelements | 2751730 | 680398255 | 27.70% | Bases Masked | 783509594 | 31.90% |
| SINEs | 1309652 | 220637929 | 8.98% | | **Base Pairs** | |
| Penelope | 5813 | 805666 | 0.03% | | | |
| LINEs | 1109720 | 379276301 | 15.44% | Total Length | 2456191777 | |
| L1/CIN4 | 992844 | 364231430 | 14.83% | | **Percentage** | |
| Retroviral | 328934 | 79359372 | 3.23% | GC | 40.92% | |
| LTR elements | 332358 | 80484025 | 3.28% | | | |
| DNA transposons | 232610 | 37641030 | 1.53% | | | |
| hobo-Activator | 152741 | 23922472 | 0.97% | | | |
| Tc1-IS630-Pogo | 73476 | 12729778 | 0.52% | | | |
| Unclassified | 151094 | 30842639 | 1.26% | | | |
| Total interspersed repeats | N/A | 749687590 | 30.52% | | | |
| Small RNA: | 1161673 | 200952347 | 8.18% | | | |
| Satellites | 710 | 613670 | 0.02% | | | |
| Simple repeats | 707458 | 26874713 | 1.09% | | | |
| Low complexity | 118007 | 5932853 | 0.24% | | | |

| Sample_nvison_LA-8_Masurc | Number of Elem | Length Occupie | Percentage of Sequence | | Number of Base | Percent Masked |
|---|---|---|---|---|---|---|
| Retroelements | 3089939 | 716348763 | 22.73% | Bases Masked | 830810288 | 29.69 |
| SINEs | 1485807 | 233301609 | 7.05% | | **Base Pairs** | |
| Penelope | 1190 | 443219 | 0.02% | | | |
| LINEs | 1199968 | 397006982 | 12.80% | Total Length | 2798604795 | |
| L1/CIN4 | 1081478 | 378625251 | 12.23% | | **Percentage** | |
| Retroviral | 402981 | 85765818 | 2.88% | GC | 41.45% | |
| LTR elements | 404164 | 86040172 | 2.89% | | | |
| DNA transposons | 248377 | 37973704 | 1.26% | | | |
| hobo-Activator | 158260 | 23659603 | 0.77% | | | |
| Tc1-IS630-Pogo | 89281 | 14006369 | 0.49% | | | |
| Unclassified | 159935 | 36395281 | 2.92% | | | |
| Total interspersed repeats | N/A | 791160967 | 26.91% | | | |
| Small RNA: | 1314924 | 213141251 | 7.62% | | | |
| Satellites | 266 | 58187 | 0.00% | | | |
| Simple repeats | 824110 | 32858084 | 1.17% | | | |
| Low complexity | 129641 | 6471104 | 0.23% | | | |

| Sample_nvison_FL-3_15_I | Number of Elem | Length Occupie | Percentage of Sequence | | Number of Base | Percent Masked |
|---|---|---|---|---|---|---|
| Retroelements | 2953324 | 653119761 | 26.00 % | Bases Masked | 759025152 | 30.21 % |
| SINEs | 1001438 | 143785203 | 5.72 % | | **Base Pairs** | |
| LINEs | 1110735 | 349403971 | 13.91 % | | | |
| L2/CR1/Rex | 103572 | 16286467 | 0.65 % | Total Length | 2512474419 | |
| RTE/Bov-B | 2078 | 320933 | 0.01 % | | **Percentage** | |
| L1/CIN4 | 1005085 | 332796571 | 13.25 % | GC | 41.64 % | |
| LTR elements | 841151 | 159930588 | 6.37 % | | | |
| Ty1/Copia | 151 | 82335 | 0.00 % | | | |
| Retroviral | 343466 | 80639417 | 3.21 % | | | |
| DNA transposons | 215359 | 36004953 | 1.43 % | | | |
| hobo-Activator | 145412 | 22377590 | 0.89 % | | | |
| Tc1-IS630-Pogo | 69777 | 22377590 | 0.54 % | | | |
| Unclassified | 70628 | 29125246 | 1.16 % | | | |
| Total interspersed repeats | N/A | 718249961 | 28.59 % | | | |
| Small RNA: | 852972 | 125518281 | 5.00 % | | | |
| Satellites | 808 | 775143 | 0.03 % | | | |
| Simple repeats | 774410 | 33175574 | 1.32 % | | | |
| Low complexity | 122651 | 6401721 | 0.25 % | | | |

| Sample_nvison_LA-9_flye2 | Number of Elem | Length Occupie | Percentage of Sequence | | Number of Base | Percent Masked |
|---|---|---|---|---|---|---|
| Retroelements | 3008796 | 626345494 | 28.35 | Bases Masked | 710416481 | 32.16 |
| SINEs | 365754 | 42053118 | 1.9 | | **Base Pairs** | |
| Penelope | 1021 | 240314 | 0.01 | | | |
| LINEs | 1779789 | 445743461 | 20.18 | Total Length | 2209197289 | |
| L1/CIN4 | 1679501 | 430705335 | 19.5 | | **Percentage** | |
| Retroviral | 519920 | 97005100 | 4.39 | GC | 41.73% | |
| LTR elements | 863253 | 138548915 | 6.27 | | | |
| DNA transposons | 203976 | 32726777 | 1.48 | | | |
| hobo-Activator | 123037 | 19516932 | 0.88 | | | |
| Tc1-IS630-Pogo | 80216 | 12900617 | 0.58 | | | |
| Unclassified | 34719 | 5908596 | 0.27 | | | |
| Total interspersed repeats | N/A | 665221181 | 30.11 | | | |
| Small RNA: | 211070 | 21486400 | 0.97 | | | |
| Satellites | 606 | 810947 | 0.04 | | | |
| Simple repeats | 662604 | 38602026 | 1.75 | | | |
| Low complexity | 103946 | 5490815 | 0.25 | | | |

| Sample_nvison_FL-5_MAS | Number of Elem | Length Occupie | Percentage of Sequence | | Number of Base | Percent Masked |
|---|---|---|---|---|---|---|
| Retroelements | 2776135 | 675822389 | 26.48% | Bases Masked | 775289861 | 30.38 |
| SINEs | 1356028 | 221234621 | 8.67% | | **Base Pairs** | |
| LINEs | 1037869 | 372395583 | 14.59% | Total Length | 2552017607 | |
| L1/CIN4 | 939586 | 355150280 | 13.92% | | **Percentage** | |
| Retroviral | 378648 | 81104841 | 3.18% | GC | 41.62 | |
| LTR elements | 382238 | 82192185 | 3.22% | | | |
| DNA transposons | 223682 | 37443344 | 1.47% | | | |
| hobo-Activator | 162715 | 25716279 | 1.01% | | | |
| Tc1-IS630-Pogo | 56817 | 11327254 | 0.44% | | | |
| Unclassified | 110765 | 26194130 | 1.03% | | | |
| Total interspersed repeats | N/A | 740478735 | 29.02% | | | |
| Small RNA: | 1197297 | 203028714 | 7.96% | | | |
| Satellites | 268 | 58058 | 0.00% | | | |
| Simple repeats | 741291 | 28275947 | 1.11% | | | |
| Low complexity | 124120 | 6229066 | 0.24% | | | |

| Sample_nvison_FL-32666_Ma | Number of Elem | Length Occupie | Percentage of Sequence | | Number of Base | Percent Masked |
|---|---|---|---|---|---|---|
| Retroelements | 3431487 | 690571203 | 23.59% | Bases Masked | 821570622 | 28.07% |
| SINEs | 1558278 | 220054863 | 7.52% | | **Base Pairs** | |
| Penelope | 6287 | 919300 | 0.03% | | | |
| LINEs | 1381956 | 384825878 | 13.15% | Total Length | 2926869213 | |
| L1/CIN4 | 1248814 | 364606762 | 12.46% | | **Percentage** | |
| Retroviral | 486643 | 84493184 | 2.89% | GC | 41.36 | |
| LTR elements | 491253 | 85690462 | 2.93% | | | |
| DNA transposons | 300832 | 39315306 | 1.34% | | | |
| hobo-Activator | 158288 | 22573823 | 0.77% | | | |
| Tc1-IS630-Pogo | 136977 | 16094145 | 0.55% | | | |
| Unclassified | 446294 | 53878645 | 1.84% | | | |
| Total interspersed repeats | N/A | 784684454 | 26.81% | | | |
| Small RNA: | 1413981 | 204811016 | 7.00% | | | |
| Satellites | 1 | 417 | 0.00% | | | |
| Simple repeats | 771509 | 29579391 | 1.01% | | | |
| Low complexity | 131591 | 6613378 | 0.23% | | | |
