## Supplementary material for "Mink by mink: stitching together signatures of subspecies adaptation through a pangenome of threatened mustelids": Table S8

**Annotation Statistics**

| Sample ID | Total Genes | Seq Similarity Search (%) | Gene Family and/or Seq Simi | Mono/Multi Ratio | Compleasm (Complete %) | Compleasm (Complete %) (Longest Isoform) | Compleasm (Duplicate %) (Longest Isoform) | Mono exonic genes |
|---|---|---|---|---|---|---|---|---|
| Sample 3 | 20191 | 87.99% | 91.02% of total retained | 0.065 | 94.80% | 94.51% | 4.55% | 1235 |
| Sample 5 | 20133 | 88.54% | 92.04% of total retained | 0.076 | 95.70% | 95.39% | 4.68% | 1424 |
| LA8 | 20115 | 88.69% | 91.56% of total retained | 0.063 | 94% | 93.73% | 4.43% | 1195 |
| LA9 | 20258 | 87.97% | 90.92% of total retained | 0.063 | 94.70% | 94.38% | 4.42% | 1203 |
| Sample 32666 | 19468 | 87.99% | 91.72% of total retained | 0.045 | 84.00% | 83.94% | 3.79% | 847 |
| Sample 33534 | 17872 | 84.83% | 89.25% of total retained | 0.083 | 72.70% | 72.72% | 3.63% | 1363 |
| Hap1 | 20958 | 89.32% | 92.49% of total retained | 0.095 | 98.00% | 97.47% | 5.85% | 1820 |

| | |
|---|---|
| **Alignment rates - Sample 5** | |
| ERR216198 | 72.65 |
| SRR18697429 | 88.82 |
| SRR18697436 | 90.44 |
| SRR22999228 | 85.96 |
| SRR22999229 | 85.52 |
| SRR22999230 | 86.29 |
| SRR22999232 | 89.94 |
| SRR22999234 | 91.49 |
| SRR22999235 | 90.57 |
| SRR22999236 | 90.59 |
| **LA8** | |
| SRR18697429 | 90.33 |
| SRR18697436 | 91.36 |
| SRR22999228 | 87.97 |
| SRR22999229 | 87.9 |
| SRR22999230 | 88.32 |
| SRR22999232 | 90.79 |
| SRR22999234 | 91.82 |
| SRR22999235 | 91.25 |
| SRR22999236 | 91.16 |
| **Sample 3** | |
| ERR216198 | 76.01 |
| SRR18697428 | 88.75 |
| SRR18697429 | 88.55 |
| SRR18697430 | 89.3 |
| SRR18697433 | 88.84 |
| SRR18697434 | 88.27 |
| SRR18697436 | 90.04 |
| SRR18697438 | 84.27 |
| SRR18697439 | 87.85 |
| SRR22999228 | 85.84 |
| SRR22999229 | 85.64 |
| SRR22999230 | 86.43 |
| SRR22999232 | 89.36 |
| SRR22999234 | 90.54 |
| SRR22999235 | 89.84 |
| SRR22999236 | 89.7 |
| **Hap1** | |
| ERR216198 | 78.14 |
| SRR18697429 | 92.88 |
| SRR18697436 | 94.46 |
| SRR22999228 | 89.83 |
| SRR22999229 | 89.51 |
| SRR22999230 | 90.38 |
| SRR22999232 | 93.45 |
| SRR22999234 | 94.94 |
| SRR22999235 | 94.16 |
| SRR22999236 | 93.96 |
| **32666** | |
| ERR216198 | 82.2 |
| SRR18697428 | 80.52 |
| SRR18697429 | 81.53 |
| SRR18697430 | 81.32 |
| SRR18697433 | 81.13 |
| SRR18697434 | 79.43 |
| SRR18697436 | 81.4 |
| SRR18697438 | 76.87 |
| SRR18697439 | 79.78 |
| SRR22999228 | 79.85 |
| SRR22999229 | 79.33 |
| SRR22999230 | 79.73 |
| SRR22999232 | 82.13 |
| SRR22999234 | 83.02 |
| SRR22999235 | 82.61 |
| SRR22999236 | 82.54 |
| **33534** | |
| ERR216198 | 73.8 |
| SRR18697428 | 71.32 |
| SRR18697429 | 74.53 |
| SRR18697430 | 72.71 |
| SRR18697433 | 70.91 |
| SRR18697436 | 71.47 |
| SRR22999228 | 73.1 |
| SRR22999229 | 73.27 |
| SRR22999230 | 73.6 |
| SRR22999232 | 74.89 |
| SRR22999234 | 74.01 |
| SRR22999235 | 75.13 |
| SRR22999236 | 74.67 |
| | 73.34 |
| **LA9** | |
| ERR216198 | 75.77 |
| SRR18697428 | 88.37 |
| SRR18697429 | 88.49 |
| SRR18697430 | 88.8 |
| SRR18697433 | 88.18 |
| SRR18697434 | 87.84 |
| SRR18697436 | 89.59 |
| SRR18697438 | 84.75 |
| SRR18697439 | 87.43 |
| SRR22999228 | 85.57 |
| SRR22999229 | 85.27 |
| SRR22999230 | 85.99 |
| SRR22999232 | 89.02 |
| SRR22999234 | 90.31 |
| SRR22999235 | 89.58 |
| SRR22999236 | 89.44 |
| | 87.15 |
