## Supplementary material for "Mink by mink: stitching together signatures of subspecies adaptation through a pangenome of threatened mustelids": Table S9: Table S9 - Protein-coding genes in ROH Regions shared among evergladensis individuals.html

|  | A | B | C | D |
| --- | --- | --- | --- | --- |
| 1 | #CHR | End | Start | Gene |
| 2 | 1 | 590166 | 609603 | mNeoVis.2.1.1.chr1\_hap1.g00013.t01 |
| 3 | 1 | 609749 | 626936 | mNeoVis.2.1.1.chr1\_hap1.g00014.t01 |
| 4 | 1 | 615859 | 618432 | mNeoVis.2.1.1.chr1\_hap1.g00015.t01 |
| 5 | 1 | 651545 | 720527 | mNeoVis.2.1.1.chr1\_hap1.g00016.t01 |
| 6 | 1 | 746056 | 760735 | mNeoVis.2.1.1.chr1\_hap1.g00017.t01 |
| 7 | 1 | 763401 | 778163 | mNeoVis.2.1.1.chr1\_hap1.g00018.t01 |
| 8 | 1 | 780695 | 801246 | mNeoVis.2.1.1.chr1\_hap1.g00019.t01 |
| 9 | 1 | 813559 | 828263 | mNeoVis.2.1.1.chr1\_hap1.g00020.t01 |
| 10 | 1 | 843609 | 877617 | mNeoVis.2.1.1.chr1\_hap1.g00021.t01 |
| 11 | 1 | 893427 | 937083 | mNeoVis.2.1.1.chr1\_hap1.g00022.t01 |
| 12 | 1 | 1813618 | 1818946 | mNeoVis.2.1.1.chr1\_hap1.g00026.t01 |
| 13 | 1 | 2458994 | 2464613 | mNeoVis.2.1.1.chr1\_hap1.g00027.t01 |
| 14 | 1 | 3661499 | 3808438 | mNeoVis.2.1.1.chr1\_hap1.g00028.t01 |
| 15 | 1 | 3881727 | 3938229 | mNeoVis.2.1.1.chr1\_hap1.g00029.t01 |
| 16 | 1 | 3881727 | 3938229 | mNeoVis.2.1.1.chr1\_hap1.g00029.t01 |
| 17 | 1 | 4732966 | 4768773 | mNeoVis.2.1.1.chr1\_hap1.g00031.t01 |
| 18 | 1 | 4822129 | 4850168 | mNeoVis.2.1.1.chr1\_hap1.g00032.t01 |
| 19 | 1 | 4850289 | 4868236 | mNeoVis.2.1.1.chr1\_hap1.g00033.t01 |
| 20 | 1 | 4929560 | 4974795 | mNeoVis.2.1.1.chr1\_hap1.g00034.t01 |
| 21 | 1 | 4929560 | 4974795 | mNeoVis.2.1.1.chr1\_hap1.g00034.t01 |
| 22 | 1 | 5261798 | 5265685 | mNeoVis.2.1.1.chr1\_hap1.g00035.t01 |
| 23 | 1 | 5261798 | 5265685 | mNeoVis.2.1.1.chr1\_hap1.g00035.t01 |
| 24 | 1 | 5301428 | 5306809 | mNeoVis.2.1.1.chr1\_hap1.g00036.t01 |
| 25 | 1 | 5301428 | 5306809 | mNeoVis.2.1.1.chr1\_hap1.g00036.t01 |
| 26 | 1 | 5331202 | 5364639 | mNeoVis.2.1.1.chr1\_hap1.g00037.t01 |
| 27 | 1 | 5331202 | 5364639 | mNeoVis.2.1.1.chr1\_hap1.g00037.t01 |
| 28 | 1 | 5500050 | 5902672 | mNeoVis.2.1.1.chr1\_hap1.g00038.t01 |
| 29 | 1 | 5906281 | 5920959 | mNeoVis.2.1.1.chr1\_hap1.g00039.t01 |
| 30 | 1 | 5945578 | 5955792 | mNeoVis.2.1.1.chr1\_hap1.g00040.t01 |
| 31 | 1 | 5955905 | 5988558 | mNeoVis.2.1.1.chr1\_hap1.g00041.t01 |
| 32 | 1 | 6923108 | 7397133 | mNeoVis.2.1.1.chr1\_hap1.g00042.t01 |
| 33 | 1 | 7470085 | 7470981 | mNeoVis.2.1.1.chr1\_hap1.g00043.t01 |
| 34 | 1 | 7959931 | 7976769 | mNeoVis.2.1.1.chr1\_hap1.g00044.t01 |
| 35 | 1 | 7977135 | 7990587 | mNeoVis.2.1.1.chr1\_hap1.g00045.t01 |
| 36 | 1 | 7994599 | 8017891 | mNeoVis.2.1.1.chr1\_hap1.g00046.t01 |
| 37 | 1 | 8080405 | 8138860 | mNeoVis.2.1.1.chr1\_hap1.g00047.t01 |
| 38 | 1 | 8147155 | 8160471 | mNeoVis.2.1.1.chr1\_hap1.g00048.t01 |
| 39 | 1 | 8222564 | 8298418 | mNeoVis.2.1.1.chr1\_hap1.g00049.t01 |
| 40 | 1 | 8312732 | 8377961 | mNeoVis.2.1.1.chr1\_hap1.g00050.t01 |
| 41 | 1 | 8338438 | 8351574 | mNeoVis.2.1.1.chr1\_hap1.g00051.t01 |
| 42 | 1 | 8575748 | 9481510 | mNeoVis.2.1.1.chr1\_hap1.g00052.t01 |
| 43 | 1 | 10805341 | 10869759 | mNeoVis.2.1.1.chr1\_hap1.g00053.t01 |
| 44 | 1 | 10805341 | 10869759 | mNeoVis.2.1.1.chr1\_hap1.g00053.t01 |
| 45 | 1 | 10908196 | 11126254 | mNeoVis.2.1.1.chr1\_hap1.g00054.t01 |
| 46 | 1 | 11279333 | 11621991 | mNeoVis.2.1.1.chr1\_hap1.g00055.t01 |
| 47 | 1 | 12152901 | 12180071 | mNeoVis.2.1.1.chr1\_hap1.g00059.t01 |
| 48 | 1 | 12370666 | 12742320 | mNeoVis.2.1.1.chr1\_hap1.g00060.t01 |
| 49 | 1 | 12850692 | 12951780 | mNeoVis.2.1.1.chr1\_hap1.g00061.t01 |
| 50 | 1 | 12952005 | 12984242 | mNeoVis.2.1.1.chr1\_hap1.g00062.t01 |
| 51 | 1 | 13058841 | 13162999 | mNeoVis.2.1.1.chr1\_hap1.g00063.t01 |
| 52 | 1 | 13058841 | 13162999 | mNeoVis.2.1.1.chr1\_hap1.g00063.t01 |
| 53 | 1 | 13166412 | 13180703 | mNeoVis.2.1.1.chr1\_hap1.g00064.t01 |
| 54 | 1 | 13166412 | 13180703 | mNeoVis.2.1.1.chr1\_hap1.g00064.t01 |
| 55 | 1 | 13188461 | 13315752 | mNeoVis.2.1.1.chr1\_hap1.g00065.t01 |
| 56 | 1 | 13188461 | 13315752 | mNeoVis.2.1.1.chr1\_hap1.g00065.t01 |
| 57 | 1 | 13340285 | 13553858 | mNeoVis.2.1.1.chr1\_hap1.g00066.t01 |
| 58 | 1 | 13340285 | 13553858 | mNeoVis.2.1.1.chr1\_hap1.g00066.t01 |
| 59 | 1 | 13801377 | 13803112 | mNeoVis.2.1.1.chr1\_hap1.g00067.t01 |
| 60 | 1 | 13801377 | 13803112 | mNeoVis.2.1.1.chr1\_hap1.g00067.t01 |
| 61 | 1 | 14029845 | 14035653 | mNeoVis.2.1.1.chr1\_hap1.g00068.t01 |
| 62 | 1 | 14029845 | 14035653 | mNeoVis.2.1.1.chr1\_hap1.g00068.t01 |
| 63 | 1 | 14035503 | 14037279 | mNeoVis.2.1.1.chr1\_hap1.g00069.t01 |
| 64 | 1 | 14035503 | 14037279 | mNeoVis.2.1.1.chr1\_hap1.g00069.t01 |
| 65 | 1 | 18310932 | 18311754 | mNeoVis.2.1.1.chr1\_hap1.g00072.t01 |
| 66 | 1 | 18310932 | 18311754 | mNeoVis.2.1.1.chr1\_hap1.g00072.t01 |
| 67 | 1 | 19298223 | 19473231 | mNeoVis.2.1.1.chr1\_hap1.g00073.t01 |
| 68 | 1 | 20055315 | 20055630 | mNeoVis.2.1.1.chr1\_hap1.g00074.t01 |
| 69 | 1 | 20339463 | 20392579 | mNeoVis.2.1.1.chr1\_hap1.g00075.t01 |
| 70 | 1 | 21157048 | 21291449 | mNeoVis.2.1.1.chr1\_hap1.g00077.t01 |
| 71 | 1 | 21157048 | 21291449 | mNeoVis.2.1.1.chr1\_hap1.g00077.t01 |
| 72 | 1 | 24138081 | 24646987 | mNeoVis.2.1.1.chr1\_hap1.g00078.t01 |
| 73 | 1 | 24138081 | 24646987 | mNeoVis.2.1.1.chr1\_hap1.g00078.t01 |
| 74 | 1 | 24629695 | 24754581 | mNeoVis.2.1.1.chr1\_hap1.g00079.t01 |
| 75 | 1 | 24629695 | 24754581 | mNeoVis.2.1.1.chr1\_hap1.g00079.t01 |
| 76 | 1 | 24808592 | 24927796 | mNeoVis.2.1.1.chr1\_hap1.g00080.t01 |
| 77 | 1 | 24808592 | 24927796 | mNeoVis.2.1.1.chr1\_hap1.g00080.t01 |
| 78 | 1 | 24927948 | 24949470 | mNeoVis.2.1.1.chr1\_hap1.g00081.t01 |
| 79 | 1 | 25006410 | 25355391 | mNeoVis.2.1.1.chr1\_hap1.g00082.t01 |
| 80 | 1 | 25006410 | 25355391 | mNeoVis.2.1.1.chr1\_hap1.g00082.t01 |
| 81 | 1 | 25372006 | 25434326 | mNeoVis.2.1.1.chr1\_hap1.g00083.t01 |
| 82 | 1 | 25372006 | 25434326 | mNeoVis.2.1.1.chr1\_hap1.g00083.t01 |
| 83 | 1 | 25372006 | 25434326 | mNeoVis.2.1.1.chr1\_hap1.g00083.t01 |
| 84 | 1 | 25372006 | 25434326 | mNeoVis.2.1.1.chr1\_hap1.g00083.t01 |
| 85 | 1 | 25477958 | 25551501 | mNeoVis.2.1.1.chr1\_hap1.g00084.t01 |
| 86 | 1 | 25477958 | 25551501 | mNeoVis.2.1.1.chr1\_hap1.g00084.t01 |
| 87 | 1 | 25477958 | 25551501 | mNeoVis.2.1.1.chr1\_hap1.g00084.t01 |
| 88 | 1 | 25477958 | 25551501 | mNeoVis.2.1.1.chr1\_hap1.g00084.t01 |
| 89 | 1 | 25573233 | 25659703 | mNeoVis.2.1.1.chr1\_hap1.g00085.t01 |
| 90 | 1 | 25573233 | 25659703 | mNeoVis.2.1.1.chr1\_hap1.g00085.t01 |
| 91 | 1 | 25573233 | 25659703 | mNeoVis.2.1.1.chr1\_hap1.g00085.t01 |
| 92 | 1 | 25573233 | 25659703 | mNeoVis.2.1.1.chr1\_hap1.g00085.t01 |
| 93 | 1 | 25750996 | 25768809 | mNeoVis.2.1.1.chr1\_hap1.g00087.t01 |
| 94 | 1 | 25750996 | 25768809 | mNeoVis.2.1.1.chr1\_hap1.g00087.t01 |
| 95 | 1 | 25830856 | 25902513 | mNeoVis.2.1.1.chr1\_hap1.g00088.t01 |
| 96 | 1 | 25830856 | 25902513 | mNeoVis.2.1.1.chr1\_hap1.g00088.t01 |
| 97 | 1 | 26468054 | 26494667 | mNeoVis.2.1.1.chr1\_hap1.g00089.t01 |
| 98 | 1 | 26468054 | 26494667 | mNeoVis.2.1.1.chr1\_hap1.g00089.t01 |
| 99 | 1 | 26525274 | 26814717 | mNeoVis.2.1.1.chr1\_hap1.g00090.t01 |
| 100 | 1 | 26525274 | 26814717 | mNeoVis.2.1.1.chr1\_hap1.g00090.t01 |
| 101 | 1 | 26827239 | 26832819 | mNeoVis.2.1.1.chr1\_hap1.g00091.t01 |
| 102 | 1 | 26827239 | 26832819 | mNeoVis.2.1.1.chr1\_hap1.g00091.t01 |
| 103 | 1 | 26827239 | 26832819 | mNeoVis.2.1.1.chr1\_hap1.g00091.t01 |
| 104 | 1 | 26833212 | 26865146 | mNeoVis.2.1.1.chr1\_hap1.g00092.t01 |
| 105 | 1 | 26833212 | 26865146 | mNeoVis.2.1.1.chr1\_hap1.g00092.t01 |
| 106 | 1 | 26833212 | 26865146 | mNeoVis.2.1.1.chr1\_hap1.g00092.t01 |
| 107 | 1 | 26868687 | 26890553 | mNeoVis.2.1.1.chr1\_hap1.g00093.t01 |
| 108 | 1 | 26868687 | 26890553 | mNeoVis.2.1.1.chr1\_hap1.g00093.t01 |
| 109 | 1 | 26868687 | 26890553 | mNeoVis.2.1.1.chr1\_hap1.g00093.t01 |
| 110 | 1 | 26895070 | 26917033 | mNeoVis.2.1.1.chr1\_hap1.g00094.t01 |
| 111 | 1 | 26895070 | 26917033 | mNeoVis.2.1.1.chr1\_hap1.g00094.t01 |
| 112 | 1 | 26895070 | 26917033 | mNeoVis.2.1.1.chr1\_hap1.g00094.t01 |
| 113 | 1 | 26895070 | 26917033 | mNeoVis.2.1.1.chr1\_hap1.g00094.t01 |
| 114 | 1 | 27172950 | 27228298 | mNeoVis.2.1.1.chr1\_hap1.g00095.t01 |
| 115 | 1 | 27172950 | 27228298 | mNeoVis.2.1.1.chr1\_hap1.g00095.t01 |
| 116 | 1 | 27172950 | 27228298 | mNeoVis.2.1.1.chr1\_hap1.g00095.t01 |
| 117 | 1 | 27172950 | 27228298 | mNeoVis.2.1.1.chr1\_hap1.g00095.t01 |
| 118 | 1 | 27234590 | 27280874 | mNeoVis.2.1.1.chr1\_hap1.g00096.t01 |
| 119 | 1 | 27234590 | 27280874 | mNeoVis.2.1.1.chr1\_hap1.g00096.t01 |
| 120 | 1 | 27234590 | 27280874 | mNeoVis.2.1.1.chr1\_hap1.g00096.t01 |
| 121 | 1 | 27234590 | 27280874 | mNeoVis.2.1.1.chr1\_hap1.g00096.t01 |
| 122 | 1 | 27283894 | 27304821 | mNeoVis.2.1.1.chr1\_hap1.g00097.t01 |
| 123 | 1 | 27283894 | 27304821 | mNeoVis.2.1.1.chr1\_hap1.g00097.t01 |
| 124 | 1 | 27283894 | 27304821 | mNeoVis.2.1.1.chr1\_hap1.g00097.t01 |
| 125 | 1 | 27291993 | 27303701 | mNeoVis.2.1.1.chr1\_hap1.g00098.t01 |
| 126 | 1 | 27291993 | 27303701 | mNeoVis.2.1.1.chr1\_hap1.g00098.t01 |
| 127 | 1 | 27291993 | 27303701 | mNeoVis.2.1.1.chr1\_hap1.g00098.t01 |
| 128 | 1 | 27291993 | 27303701 | mNeoVis.2.1.1.chr1\_hap1.g00098.t01 |
| 129 | 1 | 27305091 | 27325916 | mNeoVis.2.1.1.chr1\_hap1.g00099.t01 |
| 130 | 1 | 27305091 | 27325916 | mNeoVis.2.1.1.chr1\_hap1.g00099.t01 |
| 131 | 1 | 27305091 | 27325916 | mNeoVis.2.1.1.chr1\_hap1.g00099.t01 |
| 132 | 1 | 27349539 | 27385037 | mNeoVis.2.1.1.chr1\_hap1.g00100.t01 |
| 133 | 1 | 27349539 | 27385037 | mNeoVis.2.1.1.chr1\_hap1.g00100.t01 |
| 134 | 1 | 27349539 | 27385037 | mNeoVis.2.1.1.chr1\_hap1.g00100.t01 |
| 135 | 1 | 27349539 | 27385037 | mNeoVis.2.1.1.chr1\_hap1.g00100.t01 |
| 136 | 1 | 27396569 | 27572085 | mNeoVis.2.1.1.chr1\_hap1.g00101.t01 |
| 137 | 1 | 27396569 | 27572085 | mNeoVis.2.1.1.chr1\_hap1.g00101.t01 |
| 138 | 1 | 27832691 | 28013380 | mNeoVis.2.1.1.chr1\_hap1.g00102.t01 |
| 139 | 1 | 27880200 | 27900960 | mNeoVis.2.1.1.chr1\_hap1.g00103.t01 |
| 140 | 1 | 27880200 | 27900960 | mNeoVis.2.1.1.chr1\_hap1.g00103.t01 |
| 141 | 1 | 28045553 | 28073090 | mNeoVis.2.1.1.chr1\_hap1.g00104.t01 |
| 142 | 1 | 28085790 | 28101892 | mNeoVis.2.1.1.chr1\_hap1.g00105.t01 |
| 143 | 1 | 28120876 | 28176690 | mNeoVis.2.1.1.chr1\_hap1.g00106.t01 |
| 144 | 1 | 28204131 | 28254068 | mNeoVis.2.1.1.chr1\_hap1.g00107.t01 |
| 145 | 1 | 28275595 | 28318121 | mNeoVis.2.1.1.chr1\_hap1.g00108.t01 |
| 146 | 1 | 28318288 | 28352332 | mNeoVis.2.1.1.chr1\_hap1.g00109.t01 |
| 147 | 1 | 28368908 | 28412574 | mNeoVis.2.1.1.chr1\_hap1.g00110.t01 |
| 148 | 1 | 28418208 | 28463818 | mNeoVis.2.1.1.chr1\_hap1.g00111.t01 |
| 149 | 1 | 28733303 | 28809807 | mNeoVis.2.1.1.chr1\_hap1.g00112.t01 |
| 150 | 1 | 28733303 | 28809807 | mNeoVis.2.1.1.chr1\_hap1.g00112.t01 |
| 151 | 1 | 28733303 | 28809807 | mNeoVis.2.1.1.chr1\_hap1.g00112.t01 |
| 152 | 1 | 28969521 | 29173979 | mNeoVis.2.1.1.chr1\_hap1.g00113.t01 |
| 153 | 1 | 28969521 | 29173979 | mNeoVis.2.1.1.chr1\_hap1.g00113.t01 |
| 154 | 1 | 28969521 | 29173979 | mNeoVis.2.1.1.chr1\_hap1.g00113.t01 |
| 155 | 1 | 28969521 | 29173979 | mNeoVis.2.1.1.chr1\_hap1.g00113.t01 |
| 156 | 1 | 29352544 | 29418826 | mNeoVis.2.1.1.chr1\_hap1.g00115.t01 |
| 157 | 1 | 29425009 | 29732860 | mNeoVis.2.1.1.chr1\_hap1.g00116.t01 |
| 158 | 1 | 29766933 | 29792327 | mNeoVis.2.1.1.chr1\_hap1.g00117.t01 |
| 159 | 1 | 30189289 | 30368747 | mNeoVis.2.1.1.chr1\_hap1.g00118.t01 |
| 160 | 1 | 30382483 | 30461168 | mNeoVis.2.1.1.chr1\_hap1.g00119.t01 |
| 161 | 1 | 30682014 | 30683217 | mNeoVis.2.1.1.chr1\_hap1.g00120.t01 |
| 162 | 1 | 30702209 | 30755055 | mNeoVis.2.1.1.chr1\_hap1.g00121.t01 |
| 163 | 1 | 30757052 | 30887314 | mNeoVis.2.1.1.chr1\_hap1.g00122.t01 |
| 164 | 1 | 30814512 | 30920407 | mNeoVis.2.1.1.chr1\_hap1.g00123.t01 |
| 165 | 1 | 30907934 | 31059008 | mNeoVis.2.1.1.chr1\_hap1.g00124.t01 |
| 166 | 1 | 30907934 | 31059008 | mNeoVis.2.1.1.chr1\_hap1.g00124.t01 |
| 167 | 1 | 30907934 | 31059008 | mNeoVis.2.1.1.chr1\_hap1.g00124.t01 |
| 168 | 1 | 31081891 | 31131287 | mNeoVis.2.1.1.chr1\_hap1.g00125.t01 |
| 169 | 1 | 31081891 | 31131287 | mNeoVis.2.1.1.chr1\_hap1.g00125.t01 |
| 170 | 1 | 31081891 | 31131287 | mNeoVis.2.1.1.chr1\_hap1.g00125.t01 |
| 171 | 1 | 31142945 | 31305092 | mNeoVis.2.1.1.chr1\_hap1.g00126.t01 |
| 172 | 1 | 31142945 | 31305092 | mNeoVis.2.1.1.chr1\_hap1.g00126.t01 |
| 173 | 1 | 31142945 | 31305092 | mNeoVis.2.1.1.chr1\_hap1.g00126.t01 |
| 174 | 1 | 32155201 | 32168840 | mNeoVis.2.1.1.chr1\_hap1.g00127.t01 |
| 175 | 1 | 32155201 | 32168840 | mNeoVis.2.1.1.chr1\_hap1.g00127.t01 |
| 176 | 1 | 32176319 | 32213300 | mNeoVis.2.1.1.chr1\_hap1.g00128.t01 |
| 177 | 1 | 32176319 | 32213300 | mNeoVis.2.1.1.chr1\_hap1.g00128.t01 |
| 178 | 1 | 32215909 | 32248756 | mNeoVis.2.1.1.chr1\_hap1.g00129.t01 |
| 179 | 1 | 32215909 | 32248756 | mNeoVis.2.1.1.chr1\_hap1.g00129.t01 |
| 180 | 1 | 32270042 | 32272521 | mNeoVis.2.1.1.chr1\_hap1.g00130.t01 |
| 181 | 1 | 32270042 | 32272521 | mNeoVis.2.1.1.chr1\_hap1.g00130.t01 |
| 182 | 1 | 32272827 | 32288814 | mNeoVis.2.1.1.chr1\_hap1.g00131.t01 |
| 183 | 1 | 32272827 | 32288814 | mNeoVis.2.1.1.chr1\_hap1.g00131.t01 |
| 184 | 1 | 32316669 | 32375153 | mNeoVis.2.1.1.chr1\_hap1.g00132.t01 |
| 185 | 1 | 32390133 | 32457635 | mNeoVis.2.1.1.chr1\_hap1.g00133.t01 |
| 186 | 1 | 32390133 | 32457635 | mNeoVis.2.1.1.chr1\_hap1.g00133.t01 |
| 187 | 1 | 32441146 | 32442516 | mNeoVis.2.1.1.chr1\_hap1.g00134.t01 |
| 188 | 1 | 32441146 | 32442516 | mNeoVis.2.1.1.chr1\_hap1.g00134.t01 |
| 189 | 1 | 32512905 | 32568548 | mNeoVis.2.1.1.chr1\_hap1.g00135.t01 |
| 190 | 1 | 32512905 | 32568548 | mNeoVis.2.1.1.chr1\_hap1.g00135.t01 |
| 191 | 1 | 32649646 | 32823510 | mNeoVis.2.1.1.chr1\_hap1.g00136.t01 |
| 192 | 1 | 32649646 | 32823510 | mNeoVis.2.1.1.chr1\_hap1.g00136.t01 |
| 193 | 1 | 33000555 | 33118973 | mNeoVis.2.1.1.chr1\_hap1.g00137.t01 |
| 194 | 1 | 33138034 | 33163514 | mNeoVis.2.1.1.chr1\_hap1.g00138.t01 |
| 195 | 1 | 33163667 | 33176826 | mNeoVis.2.1.1.chr1\_hap1.g00139.t01 |
| 196 | 1 | 33459144 | 33459810 | mNeoVis.2.1.1.chr1\_hap1.g00140.t01 |
| 197 | 1 | 33544844 | 33802354 | mNeoVis.2.1.1.chr1\_hap1.g00141.t01 |
| 198 | 1 | 33827044 | 33858758 | mNeoVis.2.1.1.chr1\_hap1.g00142.t01 |
| 199 | 1 | 33858429 | 33871405 | mNeoVis.2.1.1.chr1\_hap1.g00143.t01 |
| 200 | 1 | 34003007 | 34033239 | mNeoVis.2.1.1.chr1\_hap1.g00144.t01 |
| 201 | 1 | 34003007 | 34033239 | mNeoVis.2.1.1.chr1\_hap1.g00144.t01 |
| 202 | 1 | 34043820 | 34100957 | mNeoVis.2.1.1.chr1\_hap1.g00145.t01 |
| 203 | 1 | 34043820 | 34100957 | mNeoVis.2.1.1.chr1\_hap1.g00145.t01 |
| 204 | 1 | 34104820 | 34124777 | mNeoVis.2.1.1.chr1\_hap1.g00146.t01 |
| 205 | 1 | 34104820 | 34124777 | mNeoVis.2.1.1.chr1\_hap1.g00146.t01 |
| 206 | 1 | 34155054 | 34168245 | mNeoVis.2.1.1.chr1\_hap1.g00147.t01 |
| 207 | 1 | 34155054 | 34168245 | mNeoVis.2.1.1.chr1\_hap1.g00147.t01 |
| 208 | 1 | 34155054 | 34168245 | mNeoVis.2.1.1.chr1\_hap1.g00147.t01 |
| 209 | 1 | 34195411 | 34215273 | mNeoVis.2.1.1.chr1\_hap1.g00148.t01 |
| 210 | 1 | 34195411 | 34215273 | mNeoVis.2.1.1.chr1\_hap1.g00148.t01 |
| 211 | 1 | 34195411 | 34215273 | mNeoVis.2.1.1.chr1\_hap1.g00148.t01 |
| 212 | 1 | 34215624 | 34249397 | mNeoVis.2.1.1.chr1\_hap1.g00149.t01 |
| 213 | 1 | 34215624 | 34249397 | mNeoVis.2.1.1.chr1\_hap1.g00149.t01 |
| 214 | 1 | 34215624 | 34249397 | mNeoVis.2.1.1.chr1\_hap1.g00149.t01 |
| 215 | 1 | 34254936 | 34286118 | mNeoVis.2.1.1.chr1\_hap1.g00150.t01 |
| 216 | 1 | 34254936 | 34286118 | mNeoVis.2.1.1.chr1\_hap1.g00150.t01 |
| 217 | 1 | 34254936 | 34286118 | mNeoVis.2.1.1.chr1\_hap1.g00150.t01 |
| 218 | 1 | 34304380 | 34396969 | mNeoVis.2.1.1.chr1\_hap1.g00151.t01 |
| 219 | 1 | 34304380 | 34396969 | mNeoVis.2.1.1.chr1\_hap1.g00151.t01 |
| 220 | 1 | 34304380 | 34396969 | mNeoVis.2.1.1.chr1\_hap1.g00151.t01 |
| 221 | 1 | 34398456 | 34470496 | mNeoVis.2.1.1.chr1\_hap1.g00152.t01 |
| 222 | 1 | 34398456 | 34470496 | mNeoVis.2.1.1.chr1\_hap1.g00152.t01 |
| 223 | 1 | 34398456 | 34470496 | mNeoVis.2.1.1.chr1\_hap1.g00152.t01 |
| 224 | 1 | 34760646 | 35168987 | mNeoVis.2.1.1.chr1\_hap1.g00153.t01 |
| 225 | 1 | 34760646 | 35168987 | mNeoVis.2.1.1.chr1\_hap1.g00153.t01 |
| 226 | 1 | 34828056 | 34913169 | mNeoVis.2.1.1.chr1\_hap1.g00154.t01 |
| 227 | 1 | 34828056 | 34913169 | mNeoVis.2.1.1.chr1\_hap1.g00154.t01 |
| 228 | 1 | 35218212 | 35233966 | mNeoVis.2.1.1.chr1\_hap1.g00155.t01 |
| 229 | 1 | 35218212 | 35233966 | mNeoVis.2.1.1.chr1\_hap1.g00155.t01 |
| 230 | 1 | 35596331 | 36001850 | mNeoVis.2.1.1.chr1\_hap1.g00156.t01 |
| 231 | 1 | 36329696 | 36376355 | mNeoVis.2.1.1.chr1\_hap1.g00157.t01 |
| 232 | 1 | 36329696 | 36376355 | mNeoVis.2.1.1.chr1\_hap1.g00157.t01 |
| 233 | 1 | 36512278 | 36513779 | mNeoVis.2.1.1.chr1\_hap1.g00158.t01 |
| 234 | 1 | 36512278 | 36513779 | mNeoVis.2.1.1.chr1\_hap1.g00158.t01 |
| 235 | 1 | 36513709 | 36688478 | mNeoVis.2.1.1.chr1\_hap1.g00159.t01 |
| 236 | 1 | 37146639 | 37157593 | mNeoVis.2.1.1.chr1\_hap1.g00160.t01 |
| 237 | 1 | 37146639 | 37157593 | mNeoVis.2.1.1.chr1\_hap1.g00160.t01 |
| 238 | 1 | 38044694 | 38155408 | mNeoVis.2.1.1.chr1\_hap1.g00161.t01 |
| 239 | 1 | 38312141 | 38474114 | mNeoVis.2.1.1.chr1\_hap1.g00162.t01 |
| 240 | 1 | 38312141 | 38474114 | mNeoVis.2.1.1.chr1\_hap1.g00162.t01 |
| 241 | 1 | 38321708 | 38323906 | mNeoVis.2.1.1.chr1\_hap1.g00163.t01 |
| 242 | 1 | 38321708 | 38323906 | mNeoVis.2.1.1.chr1\_hap1.g00163.t01 |
| 243 | 1 | 38498819 | 38608714 | mNeoVis.2.1.1.chr1\_hap1.g00164.t01 |
| 244 | 1 | 38498819 | 38608714 | mNeoVis.2.1.1.chr1\_hap1.g00164.t01 |
| 245 | 1 | 38611178 | 38620040 | mNeoVis.2.1.1.chr1\_hap1.g00165.t01 |
| 246 | 1 | 38611178 | 38620040 | mNeoVis.2.1.1.chr1\_hap1.g00165.t01 |
| 247 | 1 | 38619587 | 38635505 | mNeoVis.2.1.1.chr1\_hap1.g00166.t01 |
| 248 | 1 | 38619587 | 38635505 | mNeoVis.2.1.1.chr1\_hap1.g00166.t01 |
| 249 | 1 | 39779635 | 39850478 | mNeoVis.2.1.1.chr1\_hap1.g00171.t01 |
| 250 | 1 | 39795341 | 39811777 | mNeoVis.2.1.1.chr1\_hap1.g00172.t01 |
| 251 | 1 | 39850613 | 39881209 | mNeoVis.2.1.1.chr1\_hap1.g00173.t01 |
| 252 | 1 | 40070663 | 40133137 | mNeoVis.2.1.1.chr1\_hap1.g00174.t01 |
| 253 | 1 | 40193505 | 40202612 | mNeoVis.2.1.1.chr1\_hap1.g00175.t01 |
| 254 | 1 | 40225857 | 40236065 | mNeoVis.2.1.1.chr1\_hap1.g00176.t01 |
| 255 | 1 | 40275468 | 40353672 | mNeoVis.2.1.1.chr1\_hap1.g00177.t01 |
| 256 | 1 | 40305385 | 40314908 | mNeoVis.2.1.1.chr1\_hap1.g00178.t01 |
| 257 | 1 | 40323369 | 40327102 | mNeoVis.2.1.1.chr1\_hap1.g00179.t01 |
| 258 | 1 | 40335517 | 40341515 | mNeoVis.2.1.1.chr1\_hap1.g00180.t01 |
| 259 | 1 | 40353915 | 40363392 | mNeoVis.2.1.1.chr1\_hap1.g00181.t01 |
| 260 | 1 | 40384660 | 40431049 | mNeoVis.2.1.1.chr1\_hap1.g00182.t01 |
| 261 | 1 | 40473177 | 40622715 | mNeoVis.2.1.1.chr1\_hap1.g00183.t01 |
| 262 | 1 | 40745267 | 40747262 | mNeoVis.2.1.1.chr1\_hap1.g00184.t01 |
| 263 | 1 | 41112871 | 41174032 | mNeoVis.2.1.1.chr1\_hap1.g00187.t01 |
| 264 | 1 | 41176100 | 41185128 | mNeoVis.2.1.1.chr1\_hap1.g00188.t01 |
| 265 | 1 | 41190245 | 41201510 | mNeoVis.2.1.1.chr1\_hap1.g00189.t01 |
| 266 | 1 | 41207376 | 41213064 | mNeoVis.2.1.1.chr1\_hap1.g00190.t01 |
| 267 | 1 | 41309567 | 41351998 | mNeoVis.2.1.1.chr1\_hap1.g00191.t01 |
| 268 | 1 | 41370750 | 41423983 | mNeoVis.2.1.1.chr1\_hap1.g00192.t01 |
| 269 | 1 | 41500057 | 41611610 | mNeoVis.2.1.1.chr1\_hap1.g00193.t01 |
| 270 | 1 | 41500057 | 41611610 | mNeoVis.2.1.1.chr1\_hap1.g00193.t01 |
| 271 | 1 | 41614820 | 41672503 | mNeoVis.2.1.1.chr1\_hap1.g00194.t01 |
| 272 | 1 | 41614820 | 41672503 | mNeoVis.2.1.1.chr1\_hap1.g00194.t01 |
| 273 | 1 | 41685573 | 41703127 | mNeoVis.2.1.1.chr1\_hap1.g00195.t01 |
| 274 | 1 | 41685573 | 41703127 | mNeoVis.2.1.1.chr1\_hap1.g00195.t01 |
| 275 | 1 | 41719152 | 41764450 | mNeoVis.2.1.1.chr1\_hap1.g00196.t01 |
| 276 | 1 | 41719152 | 41764450 | mNeoVis.2.1.1.chr1\_hap1.g00196.t01 |
| 277 | 1 | 41764609 | 41794246 | mNeoVis.2.1.1.chr1\_hap1.g00197.t01 |
| 278 | 1 | 41764609 | 41794246 | mNeoVis.2.1.1.chr1\_hap1.g00197.t01 |
| 279 | 1 | 41813348 | 41849935 | mNeoVis.2.1.1.chr1\_hap1.g00198.t01 |
| 280 | 1 | 41813348 | 41849935 | mNeoVis.2.1.1.chr1\_hap1.g00198.t01 |
| 281 | 1 | 41859812 | 41875878 | mNeoVis.2.1.1.chr1\_hap1.g00199.t01 |
| 282 | 1 | 41859812 | 41875878 | mNeoVis.2.1.1.chr1\_hap1.g00199.t01 |
| 283 | 1 | 41881329 | 41899031 | mNeoVis.2.1.1.chr1\_hap1.g00200.t01 |
| 284 | 1 | 41881329 | 41899031 | mNeoVis.2.1.1.chr1\_hap1.g00200.t01 |
| 285 | 1 | 41905714 | 41966282 | mNeoVis.2.1.1.chr1\_hap1.g00201.t01 |
| 286 | 1 | 41972760 | 41979388 | mNeoVis.2.1.1.chr1\_hap1.g00202.t01 |
| 287 | 1 | 42029085 | 42066489 | mNeoVis.2.1.1.chr1\_hap1.g00203.t01 |
| 288 | 1 | 42067616 | 42076497 | mNeoVis.2.1.1.chr1\_hap1.g00204.t01 |
| 289 | 1 | 42097848 | 42104588 | mNeoVis.2.1.1.chr1\_hap1.g00205.t01 |
| 290 | 1 | 42128162 | 42186464 | mNeoVis.2.1.1.chr1\_hap1.g00206.t01 |
| 291 | 1 | 42217044 | 42269909 | mNeoVis.2.1.1.chr1\_hap1.g00207.t01 |
| 292 | 1 | 42269998 | 42287920 | mNeoVis.2.1.1.chr1\_hap1.g00208.t01 |
| 293 | 1 | 42294367 | 42304711 | mNeoVis.2.1.1.chr1\_hap1.g00209.t01 |
| 294 | 1 | 42311428 | 42342431 | mNeoVis.2.1.1.chr1\_hap1.g00210.t01 |
| 295 | 1 | 42364804 | 42385868 | mNeoVis.2.1.1.chr1\_hap1.g00211.t01 |
| 296 | 1 | 42437985 | 42448677 | mNeoVis.2.1.1.chr1\_hap1.g00212.t01 |
| 297 | 1 | 42465508 | 42477779 | mNeoVis.2.1.1.chr1\_hap1.g00213.t01 |
| 298 | 1 | 42521992 | 42530113 | mNeoVis.2.1.1.chr1\_hap1.g00214.t01 |
| 299 | 1 | 42530617 | 42579601 | mNeoVis.2.1.1.chr1\_hap1.g00215.t01 |
| 300 | 1 | 42590815 | 42637905 | mNeoVis.2.1.1.chr1\_hap1.g00216.t01 |
| 301 | 1 | 43771419 | 43785261 | mNeoVis.2.1.1.chr1\_hap1.g00227.t01 |
| 302 | 1 | 45011627 | 45264433 | mNeoVis.2.1.1.chr1\_hap1.g00230.t01 |
| 303 | 1 | 45279646 | 45312524 | mNeoVis.2.1.1.chr1\_hap1.g00231.t01 |
| 304 | 1 | 45279646 | 45312524 | mNeoVis.2.1.1.chr1\_hap1.g00231.t01 |
| 305 | 1 | 45312462 | 45331705 | mNeoVis.2.1.1.chr1\_hap1.g00232.t01 |
| 306 | 1 | 45312462 | 45331705 | mNeoVis.2.1.1.chr1\_hap1.g00232.t01 |
| 307 | 1 | 45451445 | 45665585 | mNeoVis.2.1.1.chr1\_hap1.g00233.t01 |
| 308 | 1 | 45675571 | 45690809 | mNeoVis.2.1.1.chr1\_hap1.g00234.t01 |
| 309 | 1 | 45677647 | 45680115 | mNeoVis.2.1.1.chr1\_hap1.g00235.t01 |
| 310 | 1 | 45704125 | 45707400 | mNeoVis.2.1.1.chr1\_hap1.g00236.t01 |
| 311 | 1 | 45880739 | 45939906 | mNeoVis.2.1.1.chr1\_hap1.g00237.t01 |
| 312 | 1 | 47865482 | 47898297 | mNeoVis.2.1.1.chr1\_hap1.g00243.t01 |
| 313 | 1 | 47944278 | 47965853 | mNeoVis.2.1.1.chr1\_hap1.g00244.t01 |
| 314 | 1 | 47977201 | 47995684 | mNeoVis.2.1.1.chr1\_hap1.g00245.t01 |
| 315 | 1 | 48009087 | 48016250 | mNeoVis.2.1.1.chr1\_hap1.g00246.t01 |
| 316 | 1 | 48031046 | 48033994 | mNeoVis.2.1.1.chr1\_hap1.g00247.t01 |
| 317 | 1 | 48047102 | 48114347 | mNeoVis.2.1.1.chr1\_hap1.g00248.t01 |
| 318 | 1 | 48194445 | 48200431 | mNeoVis.2.1.1.chr1\_hap1.g00250.t01 |
| 319 | 1 | 48269368 | 48282477 | mNeoVis.2.1.1.chr1\_hap1.g00251.t01 |
| 320 | 1 | 48284273 | 48363892 | mNeoVis.2.1.1.chr1\_hap1.g00252.t01 |
| 321 | 1 | 48408773 | 48444429 | mNeoVis.2.1.1.chr1\_hap1.g00253.t01 |
| 322 | 1 | 48444745 | 48483630 | mNeoVis.2.1.1.chr1\_hap1.g00254.t01 |
| 323 | 1 | 48485353 | 48500651 | mNeoVis.2.1.1.chr1\_hap1.g00255.t01 |
| 324 | 1 | 48527733 | 48587568 | mNeoVis.2.1.1.chr1\_hap1.g00256.t01 |
| 325 | 1 | 49314848 | 49701609 | mNeoVis.2.1.1.chr1\_hap1.g00258.t01 |
| 326 | 1 | 49748099 | 49803592 | mNeoVis.2.1.1.chr1\_hap1.g00259.t01 |
| 327 | 1 | 49818905 | 49846401 | mNeoVis.2.1.1.chr1\_hap1.g00260.t01 |
| 328 | 1 | 49844955 | 49864269 | mNeoVis.2.1.1.chr1\_hap1.g00261.t01 |
| 329 | 1 | 49886676 | 49900879 | mNeoVis.2.1.1.chr1\_hap1.g00262.t01 |
| 330 | 1 | 50364522 | 50412949 | mNeoVis.2.1.1.chr1\_hap1.g00263.t01 |
| 331 | 1 | 50440413 | 50485742 | mNeoVis.2.1.1.chr1\_hap1.g00264.t01 |
| 332 | 1 | 50492927 | 50532846 | mNeoVis.2.1.1.chr1\_hap1.g00265.t01 |
| 333 | 1 | 50604890 | 50621517 | mNeoVis.2.1.1.chr1\_hap1.g00266.t01 |
| 334 | 1 | 50628913 | 50684090 | mNeoVis.2.1.1.chr1\_hap1.g00267.t01 |
| 335 | 1 | 50715869 | 50737997 | mNeoVis.2.1.1.chr1\_hap1.g00268.t01 |
| 336 | 1 | 50761285 | 50776686 | mNeoVis.2.1.1.chr1\_hap1.g00269.t01 |
| 337 | 1 | 50780308 | 50787710 | mNeoVis.2.1.1.chr1\_hap1.g00270.t01 |
| 338 | 1 | 51563885 | 51807199 | mNeoVis.2.1.1.chr1\_hap1.g00272.t01 |
| 339 | 1 | 51937121 | 51952229 | mNeoVis.2.1.1.chr1\_hap1.g00273.t01 |
| 340 | 1 | 51937121 | 51952229 | mNeoVis.2.1.1.chr1\_hap1.g00273.t01 |
| 341 | 1 | 51974610 | 52170920 | mNeoVis.2.1.1.chr1\_hap1.g00274.t01 |
| 342 | 1 | 51974610 | 52170920 | mNeoVis.2.1.1.chr1\_hap1.g00274.t01 |
| 343 | 1 | 52269603 | 52377195 | mNeoVis.2.1.1.chr1\_hap1.g00275.t01 |
| 344 | 1 | 52269603 | 52377195 | mNeoVis.2.1.1.chr1\_hap1.g00275.t01 |
| 345 | 1 | 52269603 | 52377195 | mNeoVis.2.1.1.chr1\_hap1.g00275.t01 |
| 346 | 1 | 53417458 | 53425167 | mNeoVis.2.1.1.chr1\_hap1.g00276.t01 |
| 347 | 1 | 53417458 | 53425167 | mNeoVis.2.1.1.chr1\_hap1.g00276.t01 |
| 348 | 1 | 53417458 | 53425167 | mNeoVis.2.1.1.chr1\_hap1.g00276.t01 |
| 349 | 1 | 53442846 | 53450097 | mNeoVis.2.1.1.chr1\_hap1.g00277.t01 |
| 350 | 1 | 53442846 | 53450097 | mNeoVis.2.1.1.chr1\_hap1.g00277.t01 |
| 351 | 1 | 53442846 | 53450097 | mNeoVis.2.1.1.chr1\_hap1.g00277.t01 |
| 352 | 1 | 53450336 | 53493026 | mNeoVis.2.1.1.chr1\_hap1.g00278.t01 |
| 353 | 1 | 53450336 | 53493026 | mNeoVis.2.1.1.chr1\_hap1.g00278.t01 |
| 354 | 1 | 53450336 | 53493026 | mNeoVis.2.1.1.chr1\_hap1.g00278.t01 |
| 355 | 1 | 53497900 | 53511192 | mNeoVis.2.1.1.chr1\_hap1.g00279.t01 |
| 356 | 1 | 53497900 | 53511192 | mNeoVis.2.1.1.chr1\_hap1.g00279.t01 |
| 357 | 1 | 53497900 | 53511192 | mNeoVis.2.1.1.chr1\_hap1.g00279.t01 |
| 358 | 1 | 53508926 | 53526352 | mNeoVis.2.1.1.chr1\_hap1.g00280.t01 |
| 359 | 1 | 53508926 | 53526352 | mNeoVis.2.1.1.chr1\_hap1.g00280.t01 |
| 360 | 1 | 53508926 | 53526352 | mNeoVis.2.1.1.chr1\_hap1.g00280.t01 |
| 361 | 1 | 54379660 | 54379988 | mNeoVis.2.1.1.chr1\_hap1.g00281.t01 |
| 362 | 1 | 54379660 | 54379988 | mNeoVis.2.1.1.chr1\_hap1.g00281.t01 |
| 363 | 1 | 54379660 | 54379988 | mNeoVis.2.1.1.chr1\_hap1.g00281.t01 |
| 364 | 1 | 56211994 | 56674853 | mNeoVis.2.1.1.chr1\_hap1.g00282.t01 |
| 365 | 1 | 56696175 | 56696613 | mNeoVis.2.1.1.chr1\_hap1.g00283.t01 |
| 366 | 1 | 57973593 | 58125339 | mNeoVis.2.1.1.chr1\_hap1.g00285.t01 |
| 367 | 1 | 58634533 | 58642240 | mNeoVis.2.1.1.chr1\_hap1.g00290.t01 |
| 368 | 1 | 59487228 | 59601595 | mNeoVis.2.1.1.chr1\_hap1.g00293.t01 |
| 369 | 1 | 59705930 | 59709695 | mNeoVis.2.1.1.chr1\_hap1.g00294.t01 |
| 370 | 1 | 59801519 | 59809744 | mNeoVis.2.1.1.chr1\_hap1.g00295.t01 |
| 371 | 1 | 59811601 | 59854081 | mNeoVis.2.1.1.chr1\_hap1.g00296.t01 |
| 372 | 1 | 59920273 | 59929660 | mNeoVis.2.1.1.chr1\_hap1.g00298.t01 |
| 373 | 1 | 60272748 | 60294998 | mNeoVis.2.1.1.chr1\_hap1.g00299.t01 |
| 374 | 1 | 60272748 | 60294998 | mNeoVis.2.1.1.chr1\_hap1.g00299.t01 |
| 375 | 1 | 60335494 | 60654465 | mNeoVis.2.1.1.chr1\_hap1.g00300.t01 |
| 376 | 1 | 60335494 | 60654465 | mNeoVis.2.1.1.chr1\_hap1.g00300.t01 |
| 377 | 1 | 60663950 | 60666122 | mNeoVis.2.1.1.chr1\_hap1.g00301.t01 |
| 378 | 1 | 60663950 | 60666122 | mNeoVis.2.1.1.chr1\_hap1.g00301.t01 |
| 379 | 1 | 60718677 | 60735426 | mNeoVis.2.1.1.chr1\_hap1.g00302.t01 |
| 380 | 1 | 60718677 | 60735426 | mNeoVis.2.1.1.chr1\_hap1.g00302.t01 |
| 381 | 1 | 60743481 | 60776562 | mNeoVis.2.1.1.chr1\_hap1.g00303.t01 |
| 382 | 1 | 60743481 | 60776562 | mNeoVis.2.1.1.chr1\_hap1.g00303.t01 |
| 383 | 1 | 61063575 | 61077326 | mNeoVis.2.1.1.chr1\_hap1.g00304.t01 |
| 384 | 1 | 61063575 | 61077326 | mNeoVis.2.1.1.chr1\_hap1.g00304.t01 |
| 385 | 1 | 61135757 | 61257318 | mNeoVis.2.1.1.chr1\_hap1.g00305.t01 |
| 386 | 1 | 61135757 | 61257318 | mNeoVis.2.1.1.chr1\_hap1.g00305.t01 |
| 387 | 1 | 61308561 | 61429239 | mNeoVis.2.1.1.chr1\_hap1.g00306.t01 |
| 388 | 1 | 61308561 | 61429239 | mNeoVis.2.1.1.chr1\_hap1.g00306.t01 |
| 389 | 1 | 61339870 | 61348057 | mNeoVis.2.1.1.chr1\_hap1.g00307.t01 |
| 390 | 1 | 61339870 | 61348057 | mNeoVis.2.1.1.chr1\_hap1.g00307.t01 |
| 391 | 1 | 61438265 | 61452773 | mNeoVis.2.1.1.chr1\_hap1.g00308.t01 |
| 392 | 1 | 61438265 | 61452773 | mNeoVis.2.1.1.chr1\_hap1.g00308.t01 |
| 393 | 1 | 61447330 | 61511702 | mNeoVis.2.1.1.chr1\_hap1.g00309.t01 |
| 394 | 1 | 61540698 | 61640810 | mNeoVis.2.1.1.chr1\_hap1.g00310.t01 |
| 395 | 1 | 61798673 | 61837090 | mNeoVis.2.1.1.chr1\_hap1.g00311.t01 |
| 396 | 1 | 61798673 | 61837090 | mNeoVis.2.1.1.chr1\_hap1.g00311.t01 |
| 397 | 1 | 61898428 | 61901531 | mNeoVis.2.1.1.chr1\_hap1.g00312.t01 |
| 398 | 1 | 61898428 | 61901531 | mNeoVis.2.1.1.chr1\_hap1.g00312.t01 |
| 399 | 1 | 61952201 | 62059242 | mNeoVis.2.1.1.chr1\_hap1.g00313.t01 |
| 400 | 1 | 61952201 | 62059242 | mNeoVis.2.1.1.chr1\_hap1.g00313.t01 |
| 401 | 1 | 62063878 | 62072100 | mNeoVis.2.1.1.chr1\_hap1.g00314.t01 |
| 402 | 1 | 62063878 | 62072100 | mNeoVis.2.1.1.chr1\_hap1.g00314.t01 |
| 403 | 1 | 62080841 | 62111215 | mNeoVis.2.1.1.chr1\_hap1.g00315.t01 |
| 404 | 1 | 62080841 | 62111215 | mNeoVis.2.1.1.chr1\_hap1.g00315.t01 |
| 405 | 1 | 62123390 | 62179503 | mNeoVis.2.1.1.chr1\_hap1.g00316.t01 |
| 406 | 1 | 62123390 | 62179503 | mNeoVis.2.1.1.chr1\_hap1.g00316.t01 |
| 407 | 1 | 62183952 | 62195610 | mNeoVis.2.1.1.chr1\_hap1.g00317.t01 |
| 408 | 1 | 62183952 | 62195610 | mNeoVis.2.1.1.chr1\_hap1.g00317.t01 |
| 409 | 1 | 62243205 | 62245762 | mNeoVis.2.1.1.chr1\_hap1.g00318.t01 |
| 410 | 1 | 62243205 | 62245762 | mNeoVis.2.1.1.chr1\_hap1.g00318.t01 |
| 411 | 1 | 62321016 | 62336294 | mNeoVis.2.1.1.chr1\_hap1.g00319.t01 |
| 412 | 1 | 62321016 | 62336294 | mNeoVis.2.1.1.chr1\_hap1.g00319.t01 |
| 413 | 1 | 62572626 | 62580096 | mNeoVis.2.1.1.chr1\_hap1.g00320.t01 |
| 414 | 1 | 62572626 | 62580096 | mNeoVis.2.1.1.chr1\_hap1.g00320.t01 |
| 415 | 1 | 62798055 | 62861119 | mNeoVis.2.1.1.chr1\_hap1.g00321.t01 |
| 416 | 1 | 62798055 | 62861119 | mNeoVis.2.1.1.chr1\_hap1.g00321.t01 |
| 417 | 1 | 62798055 | 62861119 | mNeoVis.2.1.1.chr1\_hap1.g00321.t01 |
| 418 | 1 | 62888476 | 62978199 | mNeoVis.2.1.1.chr1\_hap1.g00322.t01 |
| 419 | 1 | 62888476 | 62978199 | mNeoVis.2.1.1.chr1\_hap1.g00322.t01 |
| 420 | 1 | 62888476 | 62978199 | mNeoVis.2.1.1.chr1\_hap1.g00322.t01 |
| 421 | 1 | 63452925 | 63458086 | mNeoVis.2.1.1.chr1\_hap1.g00323.t01 |
| 422 | 1 | 63840905 | 63874929 | mNeoVis.2.1.1.chr1\_hap1.g00324.t01 |
| 423 | 1 | 63897250 | 63935068 | mNeoVis.2.1.1.chr1\_hap1.g00325.t01 |
| 424 | 1 | 64018993 | 64073577 | mNeoVis.2.1.1.chr1\_hap1.g00326.t01 |
| 425 | 1 | 64137430 | 64219220 | mNeoVis.2.1.1.chr1\_hap1.g00327.t01 |
| 426 | 1 | 64271757 | 64292813 | mNeoVis.2.1.1.chr1\_hap1.g00328.t01 |
| 427 | 1 | 64357504 | 64431907 | mNeoVis.2.1.1.chr1\_hap1.g00329.t01 |
| 428 | 1 | 64432019 | 64445792 | mNeoVis.2.1.1.chr1\_hap1.g00330.t01 |
| 429 | 1 | 64455304 | 64464409 | mNeoVis.2.1.1.chr1\_hap1.g00331.t01 |
| 430 | 1 | 64466099 | 64471729 | mNeoVis.2.1.1.chr1\_hap1.g00332.t01 |
| 431 | 1 | 64471103 | 64472042 | mNeoVis.2.1.1.chr1\_hap1.g00333.t01 |
| 432 | 1 | 64474169 | 64494154 | mNeoVis.2.1.1.chr1\_hap1.g00334.t01 |
| 433 | 1 | 64598527 | 64617508 | mNeoVis.2.1.1.chr1\_hap1.g00336.t01 |
| 434 | 1 | 64625108 | 64661906 | mNeoVis.2.1.1.chr1\_hap1.g00337.t01 |
| 435 | 1 | 64662088 | 64672840 | mNeoVis.2.1.1.chr1\_hap1.g00338.t01 |
| 436 | 1 | 64682684 | 64691399 | mNeoVis.2.1.1.chr1\_hap1.g00339.t01 |
| 437 | 1 | 64700831 | 64703824 | mNeoVis.2.1.1.chr1\_hap1.g00340.t01 |
| 438 | 1 | 64705590 | 64712449 | mNeoVis.2.1.1.chr1\_hap1.g00341.t01 |
| 439 | 1 | 64839153 | 64899780 | mNeoVis.2.1.1.chr1\_hap1.g00342.t01 |
| 440 | 1 | 64899751 | 64916842 | mNeoVis.2.1.1.chr1\_hap1.g00343.t01 |
| 441 | 1 | 64923002 | 65054000 | mNeoVis.2.1.1.chr1\_hap1.g00344.t01 |
| 442 | 1 | 65003792 | 65005037 | mNeoVis.2.1.1.chr1\_hap1.g00345.t01 |
| 443 | 1 | 65088032 | 65122527 | mNeoVis.2.1.1.chr1\_hap1.g00346.t01 |
| 444 | 1 | 65135804 | 65146732 | mNeoVis.2.1.1.chr1\_hap1.g00347.t01 |
| 445 | 1 | 65169415 | 65322407 | mNeoVis.2.1.1.chr1\_hap1.g00348.t01 |
| 446 | 1 | 65323630 | 65325912 | mNeoVis.2.1.1.chr1\_hap1.g00349.t01 |
| 447 | 1 | 65325988 | 65329516 | mNeoVis.2.1.1.chr1\_hap1.g00350.t01 |
| 448 | 1 | 65329755 | 65334699 | mNeoVis.2.1.1.chr1\_hap1.g00351.t01 |
| 449 | 1 | 65353259 | 65374065 | mNeoVis.2.1.1.chr1\_hap1.g00352.t01 |
| 450 | 1 | 65377124 | 65393308 | mNeoVis.2.1.1.chr1\_hap1.g00353.t01 |
| 451 | 1 | 65402201 | 65415098 | mNeoVis.2.1.1.chr1\_hap1.g00354.t01 |
| 452 | 1 | 65416166 | 65416954 | mNeoVis.2.1.1.chr1\_hap1.g00355.t01 |
| 453 | 1 | 65417636 | 65425581 | mNeoVis.2.1.1.chr1\_hap1.g00356.t01 |
| 454 | 1 | 65431929 | 65447287 | mNeoVis.2.1.1.chr1\_hap1.g00357.t01 |
| 455 | 1 | 65766543 | 65819482 | mNeoVis.2.1.1.chr1\_hap1.g00362.t01 |
| 456 | 1 | 65844419 | 65981848 | mNeoVis.2.1.1.chr1\_hap1.g00363.t01 |
| 457 | 1 | 65880157 | 65928676 | mNeoVis.2.1.1.chr1\_hap1.g00364.t01 |
| 458 | 1 | 65982007 | 66334076 | mNeoVis.2.1.1.chr1\_hap1.g00365.t01 |
| 459 | 1 | 65982007 | 66334076 | mNeoVis.2.1.1.chr1\_hap1.g00365.t01 |
| 460 | 1 | 66002352 | 66024493 | mNeoVis.2.1.1.chr1\_hap1.g00366.t01 |
| 461 | 1 | 66002352 | 66024493 | mNeoVis.2.1.1.chr1\_hap1.g00366.t01 |
| 462 | 1 | 66387154 | 66429740 | mNeoVis.2.1.1.chr1\_hap1.g00367.t01 |
| 463 | 1 | 66387154 | 66429740 | mNeoVis.2.1.1.chr1\_hap1.g00367.t01 |
| 464 | 1 | 66524024 | 66534699 | mNeoVis.2.1.1.chr1\_hap1.g00368.t01 |
| 465 | 1 | 66921928 | 67025643 | mNeoVis.2.1.1.chr1\_hap1.g00370.t01 |
| 466 | 1 | 66998603 | 67022905 | mNeoVis.2.1.1.chr1\_hap1.g00371.t01 |
| 467 | 1 | 67028549 | 67052260 | mNeoVis.2.1.1.chr1\_hap1.g00372.t01 |
| 468 | 1 | 67083518 | 67330084 | mNeoVis.2.1.1.chr1\_hap1.g00373.t01 |
| 469 | 1 | 68074372 | 68319516 | mNeoVis.2.1.1.chr1\_hap1.g00376.t01 |
| 470 | 1 | 68324404 | 68342137 | mNeoVis.2.1.1.chr1\_hap1.g00377.t01 |
| 471 | 1 | 72257846 | 72273583 | mNeoVis.2.1.1.chr1\_hap1.g00411.t01 |
| 472 | 1 | 72257846 | 72273583 | mNeoVis.2.1.1.chr1\_hap1.g00411.t01 |
| 473 | 1 | 72277925 | 72306672 | mNeoVis.2.1.1.chr1\_hap1.g00412.t01 |
| 474 | 1 | 72277925 | 72306672 | mNeoVis.2.1.1.chr1\_hap1.g00412.t01 |
| 475 | 1 | 72384398 | 72427265 | mNeoVis.2.1.1.chr1\_hap1.g00413.t01 |
| 476 | 1 | 72384398 | 72427265 | mNeoVis.2.1.1.chr1\_hap1.g00413.t01 |
| 477 | 1 | 72427057 | 72441759 | mNeoVis.2.1.1.chr1\_hap1.g00414.t01 |
| 478 | 1 | 72427057 | 72441759 | mNeoVis.2.1.1.chr1\_hap1.g00414.t01 |
| 479 | 1 | 72458955 | 72480371 | mNeoVis.2.1.1.chr1\_hap1.g00415.t01 |
| 480 | 1 | 72458955 | 72480371 | mNeoVis.2.1.1.chr1\_hap1.g00415.t01 |
| 481 | 1 | 72485087 | 72593092 | mNeoVis.2.1.1.chr1\_hap1.g00416.t01 |
| 482 | 1 | 72485087 | 72593092 | mNeoVis.2.1.1.chr1\_hap1.g00416.t01 |
| 483 | 1 | 72596323 | 72611645 | mNeoVis.2.1.1.chr1\_hap1.g00417.t01 |
| 484 | 1 | 72596323 | 72611645 | mNeoVis.2.1.1.chr1\_hap1.g00417.t01 |
| 485 | 1 | 72620131 | 72628131 | mNeoVis.2.1.1.chr1\_hap1.g00418.t01 |
| 486 | 1 | 72620131 | 72628131 | mNeoVis.2.1.1.chr1\_hap1.g00418.t01 |
| 487 | 1 | 72711982 | 72761554 | mNeoVis.2.1.1.chr1\_hap1.g00419.t01 |
| 488 | 1 | 72711982 | 72761554 | mNeoVis.2.1.1.chr1\_hap1.g00419.t01 |
| 489 | 1 | 72783599 | 72811872 | mNeoVis.2.1.1.chr1\_hap1.g00420.t01 |
| 490 | 1 | 72783599 | 72811872 | mNeoVis.2.1.1.chr1\_hap1.g00420.t01 |
| 491 | 1 | 72823365 | 72889578 | mNeoVis.2.1.1.chr1\_hap1.g00421.t01 |
| 492 | 1 | 72867005 | 72867781 | mNeoVis.2.1.1.chr1\_hap1.g00422.t01 |
| 493 | 1 | 72892608 | 72902458 | mNeoVis.2.1.1.chr1\_hap1.g00423.t01 |
| 494 | 1 | 72908180 | 72978940 | mNeoVis.2.1.1.chr1\_hap1.g00424.t01 |
| 495 | 1 | 72985935 | 72996815 | mNeoVis.2.1.1.chr1\_hap1.g00425.t01 |
| 496 | 1 | 73017909 | 73034458 | mNeoVis.2.1.1.chr1\_hap1.g00426.t01 |
| 497 | 1 | 73067095 | 73067942 | mNeoVis.2.1.1.chr1\_hap1.g00428.t01 |
| 498 | 1 | 73094290 | 73123411 | mNeoVis.2.1.1.chr1\_hap1.g00429.t01 |
| 499 | 1 | 73138350 | 73144438 | mNeoVis.2.1.1.chr1\_hap1.g00430.t01 |
| 500 | 1 | 73425603 | 73499841 | mNeoVis.2.1.1.chr1\_hap1.g00431.t01 |
| 501 | 1 | 73541389 | 73546776 | mNeoVis.2.1.1.chr1\_hap1.g00432.t01 |
| 502 | 1 | 73541389 | 73546776 | mNeoVis.2.1.1.chr1\_hap1.g00432.t01 |
| 503 | 1 | 73613987 | 73614730 | mNeoVis.2.1.1.chr1\_hap1.g00433.t01 |
| 504 | 1 | 73613987 | 73614730 | mNeoVis.2.1.1.chr1\_hap1.g00433.t01 |
| 505 | 1 | 73646500 | 73654647 | mNeoVis.2.1.1.chr1\_hap1.g00434.t01 |
| 506 | 1 | 73646500 | 73654647 | mNeoVis.2.1.1.chr1\_hap1.g00434.t01 |
| 507 | 1 | 73786822 | 73792262 | mNeoVis.2.1.1.chr1\_hap1.g00435.t01 |
| 508 | 1 | 73786822 | 73792262 | mNeoVis.2.1.1.chr1\_hap1.g00435.t01 |
| 509 | 1 | 73801204 | 73843681 | mNeoVis.2.1.1.chr1\_hap1.g00436.t01 |
| 510 | 1 | 73801204 | 73843681 | mNeoVis.2.1.1.chr1\_hap1.g00436.t01 |
| 511 | 1 | 73850519 | 73912627 | mNeoVis.2.1.1.chr1\_hap1.g00437.t01 |
| 512 | 1 | 73850519 | 73912627 | mNeoVis.2.1.1.chr1\_hap1.g00437.t01 |
| 513 | 1 | 73854172 | 73857008 | mNeoVis.2.1.1.chr1\_hap1.g00438.t01 |
| 514 | 1 | 73854172 | 73857008 | mNeoVis.2.1.1.chr1\_hap1.g00438.t01 |
| 515 | 1 | 73913018 | 73930111 | mNeoVis.2.1.1.chr1\_hap1.g00439.t01 |
| 516 | 1 | 73913018 | 73930111 | mNeoVis.2.1.1.chr1\_hap1.g00439.t01 |
| 517 | 1 | 73993860 | 74026799 | mNeoVis.2.1.1.chr1\_hap1.g00440.t01 |
| 518 | 1 | 73993860 | 74026799 | mNeoVis.2.1.1.chr1\_hap1.g00440.t01 |
| 519 | 1 | 74081964 | 74136038 | mNeoVis.2.1.1.chr1\_hap1.g00441.t01 |
| 520 | 1 | 74144569 | 74145796 | mNeoVis.2.1.1.chr1\_hap1.g00442.t01 |
| 521 | 1 | 74166039 | 74314056 | mNeoVis.2.1.1.chr1\_hap1.g00443.t01 |
| 522 | 1 | 74848255 | 75345396 | mNeoVis.2.1.1.chr1\_hap1.g00444.t01 |
| 523 | 1 | 75350260 | 75401255 | mNeoVis.2.1.1.chr1\_hap1.g00445.t01 |
| 524 | 1 | 75908982 | 75914423 | mNeoVis.2.1.1.chr1\_hap1.g00452.t01 |
| 525 | 1 | 75937337 | 75959416 | mNeoVis.2.1.1.chr1\_hap1.g00453.t01 |
| 526 | 1 | 75959763 | 75994274 | mNeoVis.2.1.1.chr1\_hap1.g00454.t01 |
| 527 | 1 | 75994397 | 76004704 | mNeoVis.2.1.1.chr1\_hap1.g00455.t01 |
| 528 | 1 | 76005330 | 76024063 | mNeoVis.2.1.1.chr1\_hap1.g00456.t01 |
| 529 | 1 | 76055310 | 76073586 | mNeoVis.2.1.1.chr1\_hap1.g00457.t01 |
| 530 | 1 | 76081618 | 76114648 | mNeoVis.2.1.1.chr1\_hap1.g00458.t01 |
| 531 | 1 | 76114449 | 76119852 | mNeoVis.2.1.1.chr1\_hap1.g00459.t01 |
| 532 | 1 | 76119715 | 76130972 | mNeoVis.2.1.1.chr1\_hap1.g00460.t01 |
| 533 | 1 | 76134237 | 76204071 | mNeoVis.2.1.1.chr1\_hap1.g00461.t01 |
| 534 | 1 | 76207235 | 76211892 | mNeoVis.2.1.1.chr1\_hap1.g00462.t01 |
| 535 | 1 | 76225029 | 76228882 | mNeoVis.2.1.1.chr1\_hap1.g00463.t01 |
| 536 | 1 | 76258620 | 76290142 | mNeoVis.2.1.1.chr1\_hap1.g00464.t01 |
| 537 | 1 | 76303922 | 76322616 | mNeoVis.2.1.1.chr1\_hap1.g00465.t01 |
| 538 | 1 | 76322771 | 76642026 | mNeoVis.2.1.1.chr1\_hap1.g00466.t01 |
| 539 | 1 | 76569971 | 76576506 | mNeoVis.2.1.1.chr1\_hap1.g00467.t01 |
| 540 | 1 | 77631116 | 77642546 | mNeoVis.2.1.1.chr1\_hap1.g00484.t01 |
| 541 | 1 | 77631116 | 77642546 | mNeoVis.2.1.1.chr1\_hap1.g00484.t01 |
| 542 | 1 | 77649021 | 77651863 | mNeoVis.2.1.1.chr1\_hap1.g00485.t01 |
| 543 | 1 | 77649021 | 77651863 | mNeoVis.2.1.1.chr1\_hap1.g00485.t01 |
| 544 | 1 | 77695531 | 77736765 | mNeoVis.2.1.1.chr1\_hap1.g00486.t01 |
| 545 | 1 | 77695531 | 77736765 | mNeoVis.2.1.1.chr1\_hap1.g00486.t01 |
| 546 | 1 | 77757199 | 77840189 | mNeoVis.2.1.1.chr1\_hap1.g00487.t01 |
| 547 | 1 | 77757199 | 77840189 | mNeoVis.2.1.1.chr1\_hap1.g00487.t01 |
| 548 | 1 | 77849048 | 77861415 | mNeoVis.2.1.1.chr1\_hap1.g00488.t01 |
| 549 | 1 | 77849048 | 77861415 | mNeoVis.2.1.1.chr1\_hap1.g00488.t01 |
| 550 | 1 | 77874846 | 77888857 | mNeoVis.2.1.1.chr1\_hap1.g00489.t01 |
| 551 | 1 | 77874846 | 77888857 | mNeoVis.2.1.1.chr1\_hap1.g00489.t01 |
| 552 | 1 | 77912134 | 78038183 | mNeoVis.2.1.1.chr1\_hap1.g00490.t01 |
| 553 | 1 | 77912134 | 78038183 | mNeoVis.2.1.1.chr1\_hap1.g00490.t01 |
| 554 | 1 | 78041038 | 78043855 | mNeoVis.2.1.1.chr1\_hap1.g00491.t01 |
| 555 | 1 | 78041038 | 78043855 | mNeoVis.2.1.1.chr1\_hap1.g00491.t01 |
| 556 | 1 | 78044041 | 78052614 | mNeoVis.2.1.1.chr1\_hap1.g00492.t01 |
| 557 | 1 | 78044041 | 78052614 | mNeoVis.2.1.1.chr1\_hap1.g00492.t01 |
| 558 | 1 | 78051023 | 78056757 | mNeoVis.2.1.1.chr1\_hap1.g00493.t01 |
| 559 | 1 | 78051023 | 78056757 | mNeoVis.2.1.1.chr1\_hap1.g00493.t01 |
| 560 | 1 | 78056832 | 78059306 | mNeoVis.2.1.1.chr1\_hap1.g00494.t01 |
| 561 | 1 | 78056832 | 78059306 | mNeoVis.2.1.1.chr1\_hap1.g00494.t01 |
| 562 | 1 | 78059723 | 78061647 | mNeoVis.2.1.1.chr1\_hap1.g00495.t01 |
| 563 | 1 | 78059723 | 78061647 | mNeoVis.2.1.1.chr1\_hap1.g00495.t01 |
| 564 | 1 | 78061779 | 78071588 | mNeoVis.2.1.1.chr1\_hap1.g00496.t01 |
| 565 | 1 | 78061779 | 78071588 | mNeoVis.2.1.1.chr1\_hap1.g00496.t01 |
| 566 | 1 | 78087015 | 78089454 | mNeoVis.2.1.1.chr1\_hap1.g00497.t01 |
| 567 | 1 | 78087015 | 78089454 | mNeoVis.2.1.1.chr1\_hap1.g00497.t01 |
| 568 | 1 | 78093685 | 78099584 | mNeoVis.2.1.1.chr1\_hap1.g00498.t01 |
| 569 | 1 | 78093685 | 78099584 | mNeoVis.2.1.1.chr1\_hap1.g00498.t01 |
| 570 | 1 | 78099651 | 78101601 | mNeoVis.2.1.1.chr1\_hap1.g00499.t01 |
| 571 | 1 | 78099651 | 78101601 | mNeoVis.2.1.1.chr1\_hap1.g00499.t01 |
| 572 | 1 | 78101762 | 78114987 | mNeoVis.2.1.1.chr1\_hap1.g00500.t01 |
| 573 | 1 | 78101762 | 78114987 | mNeoVis.2.1.1.chr1\_hap1.g00500.t01 |
| 574 | 1 | 78117327 | 78122900 | mNeoVis.2.1.1.chr1\_hap1.g00501.t01 |
| 575 | 1 | 78117327 | 78122900 | mNeoVis.2.1.1.chr1\_hap1.g00501.t01 |
| 576 | 1 | 78123043 | 78125692 | mNeoVis.2.1.1.chr1\_hap1.g00502.t01 |
| 577 | 1 | 78123043 | 78125692 | mNeoVis.2.1.1.chr1\_hap1.g00502.t01 |
| 578 | 1 | 78125840 | 78140076 | mNeoVis.2.1.1.chr1\_hap1.g00503.t01 |
| 579 | 1 | 78125840 | 78140076 | mNeoVis.2.1.1.chr1\_hap1.g00503.t01 |
| 580 | 1 | 78140134 | 78147333 | mNeoVis.2.1.1.chr1\_hap1.g00504.t01 |
| 581 | 1 | 78140134 | 78147333 | mNeoVis.2.1.1.chr1\_hap1.g00504.t01 |
| 582 | 1 | 78147246 | 78154719 | mNeoVis.2.1.1.chr1\_hap1.g00505.t01 |
| 583 | 1 | 78147246 | 78154719 | mNeoVis.2.1.1.chr1\_hap1.g00505.t01 |
| 584 | 1 | 78356029 | 78402826 | mNeoVis.2.1.1.chr1\_hap1.g00506.t01 |
| 585 | 1 | 78440206 | 78442776 | mNeoVis.2.1.1.chr1\_hap1.g00507.t01 |
| 586 | 1 | 78481297 | 78510423 | mNeoVis.2.1.1.chr1\_hap1.g00508.t01 |
| 587 | 1 | 78492262 | 78502946 | mNeoVis.2.1.1.chr1\_hap1.g00509.t01 |
| 588 | 1 | 78511821 | 78521803 | mNeoVis.2.1.1.chr1\_hap1.g00510.t01 |
| 589 | 1 | 78523415 | 78531268 | mNeoVis.2.1.1.chr1\_hap1.g00511.t01 |
| 590 | 1 | 78531784 | 78550025 | mNeoVis.2.1.1.chr1\_hap1.g00512.t01 |
| 591 | 1 | 78534402 | 78540226 | mNeoVis.2.1.1.chr1\_hap1.g00513.t01 |
| 592 | 1 | 78568072 | 78594144 | mNeoVis.2.1.1.chr1\_hap1.g00514.t01 |
| 593 | 1 | 78608222 | 78609802 | mNeoVis.2.1.1.chr1\_hap1.g00515.t01 |
| 594 | 1 | 78617065 | 78619407 | mNeoVis.2.1.1.chr1\_hap1.g00516.t01 |
| 595 | 1 | 78638846 | 78643400 | mNeoVis.2.1.1.chr1\_hap1.g00517.t01 |
| 596 | 1 | 78670898 | 78683585 | mNeoVis.2.1.1.chr1\_hap1.g00518.t01 |
| 597 | 1 | 78683675 | 78819806 | mNeoVis.2.1.1.chr1\_hap1.g00519.t01 |
| 598 | 1 | 78688961 | 78693582 | mNeoVis.2.1.1.chr1\_hap1.g00520.t01 |
| 599 | 1 | 78693863 | 78711250 | mNeoVis.2.1.1.chr1\_hap1.g00521.t01 |
| 600 | 1 | 78700607 | 78706018 | mNeoVis.2.1.1.chr1\_hap1.g00522.t01 |
| 601 | 1 | 78718590 | 78723357 | mNeoVis.2.1.1.chr1\_hap1.g00523.t01 |
| 602 | 1 | 78730519 | 78812071 | mNeoVis.2.1.1.chr1\_hap1.g00524.t01 |
| 603 | 1 | 78733156 | 78736328 | mNeoVis.2.1.1.chr1\_hap1.g00525.t01 |
| 604 | 1 | 78821970 | 78832953 | mNeoVis.2.1.1.chr1\_hap1.g00526.t01 |
| 605 | 1 | 78833093 | 78861258 | mNeoVis.2.1.1.chr1\_hap1.g00527.t01 |
| 606 | 1 | 80438022 | 80439787 | mNeoVis.2.1.1.chr1\_hap1.g00543.t01 |
| 607 | 1 | 81256786 | 81270275 | mNeoVis.2.1.1.chr1\_hap1.g00545.t01 |
| 608 | 1 | 81256786 | 81270275 | mNeoVis.2.1.1.chr1\_hap1.g00545.t01 |
| 609 | 1 | 81270321 | 81569968 | mNeoVis.2.1.1.chr1\_hap1.g00546.t01 |
| 610 | 1 | 82136065 | 82238820 | mNeoVis.2.1.1.chr1\_hap1.g00547.t01 |
| 611 | 1 | 82715819 | 82779495 | mNeoVis.2.1.1.chr1\_hap1.g00548.t01 |
| 612 | 1 | 82715819 | 82779495 | mNeoVis.2.1.1.chr1\_hap1.g00548.t01 |
| 613 | 1 | 82715819 | 82779495 | mNeoVis.2.1.1.chr1\_hap1.g00548.t01 |
| 614 | 1 | 82808441 | 82821435 | mNeoVis.2.1.1.chr1\_hap1.g00549.t01 |
| 615 | 1 | 82808441 | 82821435 | mNeoVis.2.1.1.chr1\_hap1.g00549.t01 |
| 616 | 1 | 82808441 | 82821435 | mNeoVis.2.1.1.chr1\_hap1.g00549.t01 |
| 617 | 1 | 82873044 | 83001379 | mNeoVis.2.1.1.chr1\_hap1.g00550.t01 |
| 618 | 1 | 82873044 | 83001379 | mNeoVis.2.1.1.chr1\_hap1.g00550.t01 |
| 619 | 1 | 82873044 | 83001379 | mNeoVis.2.1.1.chr1\_hap1.g00550.t01 |
| 620 | 1 | 82985314 | 83022261 | mNeoVis.2.1.1.chr1\_hap1.g00551.t01 |
| 621 | 1 | 82985314 | 83022261 | mNeoVis.2.1.1.chr1\_hap1.g00551.t01 |
| 622 | 1 | 82985314 | 83022261 | mNeoVis.2.1.1.chr1\_hap1.g00551.t01 |
| 623 | 1 | 83031053 | 83100896 | mNeoVis.2.1.1.chr1\_hap1.g00552.t01 |
| 624 | 1 | 83031053 | 83100896 | mNeoVis.2.1.1.chr1\_hap1.g00552.t01 |
| 625 | 1 | 83031053 | 83100896 | mNeoVis.2.1.1.chr1\_hap1.g00552.t01 |
| 626 | 1 | 83116214 | 83195933 | mNeoVis.2.1.1.chr1\_hap1.g00553.t01 |
| 627 | 1 | 83116214 | 83195933 | mNeoVis.2.1.1.chr1\_hap1.g00553.t01 |
| 628 | 1 | 83116214 | 83195933 | mNeoVis.2.1.1.chr1\_hap1.g00553.t01 |
| 629 | 1 | 83243903 | 83245235 | mNeoVis.2.1.1.chr1\_hap1.g00554.t01 |
| 630 | 1 | 83243903 | 83245235 | mNeoVis.2.1.1.chr1\_hap1.g00554.t01 |
| 631 | 1 | 83243903 | 83245235 | mNeoVis.2.1.1.chr1\_hap1.g00554.t01 |
| 632 | 1 | 83339697 | 83402236 | mNeoVis.2.1.1.chr1\_hap1.g00555.t01 |
| 633 | 1 | 83339697 | 83402236 | mNeoVis.2.1.1.chr1\_hap1.g00555.t01 |
| 634 | 1 | 83339697 | 83402236 | mNeoVis.2.1.1.chr1\_hap1.g00555.t01 |
| 635 | 1 | 83405181 | 83406440 | mNeoVis.2.1.1.chr1\_hap1.g00556.t01 |
| 636 | 1 | 83405181 | 83406440 | mNeoVis.2.1.1.chr1\_hap1.g00556.t01 |
| 637 | 1 | 83405181 | 83406440 | mNeoVis.2.1.1.chr1\_hap1.g00556.t01 |
| 638 | 1 | 83493388 | 83918030 | mNeoVis.2.1.1.chr1\_hap1.g00557.t01 |
| 639 | 1 | 84048296 | 84183948 | mNeoVis.2.1.1.chr1\_hap1.g00558.t01 |
| 640 | 1 | 84951964 | 84970719 | mNeoVis.2.1.1.chr1\_hap1.g00567.t01 |
| 641 | 1 | 84996935 | 85005250 | mNeoVis.2.1.1.chr1\_hap1.g00568.t01 |
| 642 | 1 | 85006730 | 85053605 | mNeoVis.2.1.1.chr1\_hap1.g00569.t01 |
| 643 | 1 | 85041478 | 85046653 | mNeoVis.2.1.1.chr1\_hap1.g00570.t01 |
| 644 | 1 | 85084238 | 85090719 | mNeoVis.2.1.1.chr1\_hap1.g00571.t01 |
| 645 | 1 | 85095699 | 85098323 | mNeoVis.2.1.1.chr1\_hap1.g00572.t01 |
| 646 | 1 | 85098392 | 85141841 | mNeoVis.2.1.1.chr1\_hap1.g00573.t01 |
| 647 | 1 | 85130892 | 85186811 | mNeoVis.2.1.1.chr1\_hap1.g00574.t01 |
| 648 | 1 | 85130892 | 85186811 | mNeoVis.2.1.1.chr1\_hap1.g00574.t01 |
| 649 | 1 | 85143674 | 85144834 | mNeoVis.2.1.1.chr1\_hap1.g00575.t01 |
| 650 | 1 | 85143674 | 85144834 | mNeoVis.2.1.1.chr1\_hap1.g00575.t01 |
| 651 | 1 | 85186891 | 85294836 | mNeoVis.2.1.1.chr1\_hap1.g00576.t01 |
| 652 | 1 | 85186891 | 85294836 | mNeoVis.2.1.1.chr1\_hap1.g00576.t01 |
| 653 | 1 | 85294793 | 85388331 | mNeoVis.2.1.1.chr1\_hap1.g00577.t01 |
| 654 | 1 | 85294793 | 85388331 | mNeoVis.2.1.1.chr1\_hap1.g00577.t01 |
| 655 | 1 | 85457189 | 85541603 | mNeoVis.2.1.1.chr1\_hap1.g00578.t01 |
| 656 | 1 | 85520204 | 85520641 | mNeoVis.2.1.1.chr1\_hap1.g00579.t01 |
| 657 | 1 | 85520204 | 85520641 | mNeoVis.2.1.1.chr1\_hap1.g00579.t01 |
| 658 | 1 | 85556911 | 85630794 | mNeoVis.2.1.1.chr1\_hap1.g00580.t01 |
| 659 | 1 | 85556911 | 85630794 | mNeoVis.2.1.1.chr1\_hap1.g00580.t01 |
| 660 | 1 | 85634471 | 85726437 | mNeoVis.2.1.1.chr1\_hap1.g00581.t01 |
| 661 | 1 | 85747827 | 85808046 | mNeoVis.2.1.1.chr1\_hap1.g00582.t01 |
| 662 | 1 | 86345833 | 86346788 | mNeoVis.2.1.1.chr1\_hap1.g00584.t01 |
| 663 | 1 | 86489516 | 86570423 | mNeoVis.2.1.1.chr1\_hap1.g00585.t01 |
| 664 | 1 | 86532880 | 86673149 | mNeoVis.2.1.1.chr1\_hap1.g00586.t01 |
| 665 | 1 | 86608469 | 86615748 | mNeoVis.2.1.1.chr1\_hap1.g00587.t01 |
| 666 | 1 | 86810970 | 86888596 | mNeoVis.2.1.1.chr1\_hap1.g00588.t01 |
| 667 | 1 | 87074441 | 87077538 | mNeoVis.2.1.1.chr1\_hap1.g00589.t01 |
| 668 | 1 | 87074441 | 87077538 | mNeoVis.2.1.1.chr1\_hap1.g00589.t01 |
| 669 | 1 | 87915867 | 87959464 | mNeoVis.2.1.1.chr1\_hap1.g00591.t01 |
| 670 | 1 | 87915867 | 87959464 | mNeoVis.2.1.1.chr1\_hap1.g00591.t01 |
| 671 | 1 | 87915867 | 87959464 | mNeoVis.2.1.1.chr1\_hap1.g00591.t01 |
| 672 | 1 | 87919575 | 87925502 | mNeoVis.2.1.1.chr1\_hap1.g00592.t01 |
| 673 | 1 | 87919575 | 87925502 | mNeoVis.2.1.1.chr1\_hap1.g00592.t01 |
| 674 | 1 | 87919575 | 87925502 | mNeoVis.2.1.1.chr1\_hap1.g00592.t01 |
| 675 | 1 | 87925573 | 87927417 | mNeoVis.2.1.1.chr1\_hap1.g00593.t01 |
| 676 | 1 | 87925573 | 87927417 | mNeoVis.2.1.1.chr1\_hap1.g00593.t01 |
| 677 | 1 | 87925573 | 87927417 | mNeoVis.2.1.1.chr1\_hap1.g00593.t01 |
| 678 | 1 | 90436792 | 90445891 | mNeoVis.2.1.1.chr1\_hap1.g00599.t01 |
| 679 | 1 | 90436792 | 90445891 | mNeoVis.2.1.1.chr1\_hap1.g00599.t01 |
| 680 | 1 | 90603201 | 90802825 | mNeoVis.2.1.1.chr1\_hap1.g00600.t01 |
| 681 | 1 | 90603201 | 90802825 | mNeoVis.2.1.1.chr1\_hap1.g00600.t01 |
| 682 | 1 | 90603201 | 90802825 | mNeoVis.2.1.1.chr1\_hap1.g00600.t01 |
| 683 | 1 | 90651475 | 90652231 | mNeoVis.2.1.1.chr1\_hap1.g00601.t01 |
| 684 | 1 | 90651475 | 90652231 | mNeoVis.2.1.1.chr1\_hap1.g00601.t01 |
| 685 | 1 | 90651475 | 90652231 | mNeoVis.2.1.1.chr1\_hap1.g00601.t01 |
| 686 | 1 | 90825152 | 91028328 | mNeoVis.2.1.1.chr1\_hap1.g00602.t01 |
| 687 | 1 | 90825152 | 91028328 | mNeoVis.2.1.1.chr1\_hap1.g00602.t01 |
| 688 | 1 | 92864863 | 92885729 | mNeoVis.2.1.1.chr1\_hap1.g00607.t01 |
| 689 | 1 | 93327666 | 93427215 | mNeoVis.2.1.1.chr1\_hap1.g00609.t01 |
| 690 | 1 | 93327666 | 93427215 | mNeoVis.2.1.1.chr1\_hap1.g00609.t01 |
| 691 | 1 | 93327666 | 93427215 | mNeoVis.2.1.1.chr1\_hap1.g00609.t01 |
| 692 | 1 | 93492619 | 93512487 | mNeoVis.2.1.1.chr1\_hap1.g00610.t01 |
| 693 | 1 | 93492619 | 93512487 | mNeoVis.2.1.1.chr1\_hap1.g00610.t01 |
| 694 | 1 | 93492619 | 93512487 | mNeoVis.2.1.1.chr1\_hap1.g00610.t01 |
| 695 | 1 | 93512632 | 93557247 | mNeoVis.2.1.1.chr1\_hap1.g00611.t01 |
| 696 | 1 | 93512632 | 93557247 | mNeoVis.2.1.1.chr1\_hap1.g00611.t01 |
| 697 | 1 | 93512632 | 93557247 | mNeoVis.2.1.1.chr1\_hap1.g00611.t01 |
| 698 | 1 | 94182705 | 94222675 | mNeoVis.2.1.1.chr1\_hap1.g00614.t01 |
| 699 | 1 | 94222740 | 94243943 | mNeoVis.2.1.1.chr1\_hap1.g00615.t01 |
| 700 | 1 | 94246027 | 94278392 | mNeoVis.2.1.1.chr1\_hap1.g00616.t01 |
| 701 | 1 | 94278978 | 94318304 | mNeoVis.2.1.1.chr1\_hap1.g00617.t01 |
| 702 | 1 | 94278984 | 94283482 | mNeoVis.2.1.1.chr1\_hap1.g00618.t01 |
| 703 | 1 | 94319552 | 94364611 | mNeoVis.2.1.1.chr1\_hap1.g00619.t01 |
| 704 | 1 | 94366038 | 94463518 | mNeoVis.2.1.1.chr1\_hap1.g00620.t01 |
| 705 | 1 | 94550519 | 94551772 | mNeoVis.2.1.1.chr1\_hap1.g00621.t01 |
| 706 | 1 | 94552209 | 94680323 | mNeoVis.2.1.1.chr1\_hap1.g00622.t01 |
| 707 | 1 | 94718945 | 94767061 | mNeoVis.2.1.1.chr1\_hap1.g00623.t01 |
| 708 | 1 | 94718945 | 94767061 | mNeoVis.2.1.1.chr1\_hap1.g00623.t01 |
| 709 | 1 | 94767289 | 94768029 | mNeoVis.2.1.1.chr1\_hap1.g00624.t01 |
| 710 | 1 | 94767289 | 94768029 | mNeoVis.2.1.1.chr1\_hap1.g00624.t01 |
| 711 | 1 | 95080894 | 95634264 | mNeoVis.2.1.1.chr1\_hap1.g00625.t01 |
| 712 | 1 | 95641776 | 95656018 | mNeoVis.2.1.1.chr1\_hap1.g00626.t01 |
| 713 | 1 | 96583521 | 96668988 | mNeoVis.2.1.1.chr1\_hap1.g00628.t01 |
| 714 | 1 | 98103623 | 98105373 | mNeoVis.2.1.1.chr1\_hap1.g00647.t01 |
| 715 | 1 | 98444174 | 98539940 | mNeoVis.2.1.1.chr1\_hap1.g00648.t01 |
| 716 | 1 | 98545839 | 98633286 | mNeoVis.2.1.1.chr1\_hap1.g00649.t01 |
| 717 | 1 | 98633322 | 98669152 | mNeoVis.2.1.1.chr1\_hap1.g00650.t01 |
| 718 | 1 | 98743504 | 98811388 | mNeoVis.2.1.1.chr1\_hap1.g00651.t01 |
| 719 | 1 | 98743504 | 98811388 | mNeoVis.2.1.1.chr1\_hap1.g00651.t01 |
| 720 | 1 | 99067998 | 99162262 | mNeoVis.2.1.1.chr1\_hap1.g00652.t01 |
| 721 | 1 | 99383634 | 99392890 | mNeoVis.2.1.1.chr1\_hap1.g00654.t01 |
| 722 | 1 | 99424695 | 99427134 | mNeoVis.2.1.1.chr1\_hap1.g00655.t01 |
| 723 | 1 | 99520060 | 99661838 | mNeoVis.2.1.1.chr1\_hap1.g00656.t01 |
| 724 | 1 | 99709427 | 99718619 | mNeoVis.2.1.1.chr1\_hap1.g00657.t01 |
| 725 | 1 | 99749040 | 99761755 | mNeoVis.2.1.1.chr1\_hap1.g00658.t01 |
| 726 | 1 | 99761014 | 99778787 | mNeoVis.2.1.1.chr1\_hap1.g00659.t01 |
| 727 | 1 | 99804700 | 100124020 | mNeoVis.2.1.1.chr1\_hap1.g00660.t01 |
| 728 | 1 | 100707330 | 100720040 | mNeoVis.2.1.1.chr1\_hap1.g00661.t01 |
| 729 | 1 | 100761226 | 100786636 | mNeoVis.2.1.1.chr1\_hap1.g00662.t01 |
| 730 | 1 | 101354778 | 101383808 | mNeoVis.2.1.1.chr1\_hap1.g00669.t01 |
| 731 | 1 | 101382811 | 101501776 | mNeoVis.2.1.1.chr1\_hap1.g00670.t01 |
| 732 | 1 | 101501665 | 101569223 | mNeoVis.2.1.1.chr1\_hap1.g00671.t01 |
| 733 | 1 | 101577205 | 101644686 | mNeoVis.2.1.1.chr1\_hap1.g00672.t01 |
| 734 | 1 | 101648140 | 101684665 | mNeoVis.2.1.1.chr1\_hap1.g00673.t01 |
| 735 | 1 | 101648140 | 101684665 | mNeoVis.2.1.1.chr1\_hap1.g00673.t01 |
| 736 | 1 | 101899192 | 102169510 | mNeoVis.2.1.1.chr1\_hap1.g00674.t01 |
| 737 | 1 | 101899192 | 102169510 | mNeoVis.2.1.1.chr1\_hap1.g00674.t01 |
| 738 | 1 | 101899192 | 102169510 | mNeoVis.2.1.1.chr1\_hap1.g00674.t01 |
| 739 | 1 | 102208513 | 102502768 | mNeoVis.2.1.1.chr1\_hap1.g00675.t01 |
| 740 | 1 | 102414306 | 102420878 | mNeoVis.2.1.1.chr1\_hap1.g00676.t01 |
| 741 | 1 | 102414306 | 102420878 | mNeoVis.2.1.1.chr1\_hap1.g00676.t01 |
| 742 | 1 | 102510042 | 102522191 | mNeoVis.2.1.1.chr1\_hap1.g00677.t01 |
| 743 | 1 | 102592007 | 102604397 | mNeoVis.2.1.1.chr1\_hap1.g00678.t01 |
| 744 | 1 | 102612636 | 102629467 | mNeoVis.2.1.1.chr1\_hap1.g00679.t01 |
| 745 | 1 | 102673477 | 102686417 | mNeoVis.2.1.1.chr1\_hap1.g00680.t01 |
| 746 | 1 | 102673477 | 102686417 | mNeoVis.2.1.1.chr1\_hap1.g00680.t01 |
| 747 | 1 | 102725215 | 102846749 | mNeoVis.2.1.1.chr1\_hap1.g00681.t01 |
| 748 | 1 | 102776034 | 102785234 | mNeoVis.2.1.1.chr1\_hap1.g00682.t01 |
| 749 | 1 | 102776034 | 102785234 | mNeoVis.2.1.1.chr1\_hap1.g00682.t01 |
| 750 | 1 | 102892499 | 103131444 | mNeoVis.2.1.1.chr1\_hap1.g00683.t01 |
| 751 | 1 | 102892499 | 103131444 | mNeoVis.2.1.1.chr1\_hap1.g00683.t01 |
| 752 | 1 | 103134050 | 103316766 | mNeoVis.2.1.1.chr1\_hap1.g00684.t01 |
| 753 | 1 | 103328000 | 103336041 | mNeoVis.2.1.1.chr1\_hap1.g00685.t01 |
| 754 | 1 | 103373609 | 103451664 | mNeoVis.2.1.1.chr1\_hap1.g00686.t01 |
| 755 | 1 | 104396028 | 104567486 | mNeoVis.2.1.1.chr1\_hap1.g00691.t01 |
| 756 | 1 | 104549949 | 104551403 | mNeoVis.2.1.1.chr1\_hap1.g00692.t01 |
| 757 | 1 | 104583773 | 104649272 | mNeoVis.2.1.1.chr1\_hap1.g00693.t01 |
| 758 | 1 | 104686405 | 104703277 | mNeoVis.2.1.1.chr1\_hap1.g00694.t01 |
| 759 | 1 | 104829209 | 104829496 | mNeoVis.2.1.1.chr1\_hap1.g00696.t01 |
| 760 | 1 | 104919385 | 105050828 | mNeoVis.2.1.1.chr1\_hap1.g00697.t01 |
| 761 | 1 | 105117421 | 105187425 | mNeoVis.2.1.1.chr1\_hap1.g00698.t01 |
| 762 | 1 | 105190148 | 105280756 | mNeoVis.2.1.1.chr1\_hap1.g00699.t01 |
| 763 | 1 | 105420741 | 105434505 | mNeoVis.2.1.1.chr1\_hap1.g00700.t01 |
| 764 | 1 | 105420741 | 105434505 | mNeoVis.2.1.1.chr1\_hap1.g00700.t01 |
| 765 | 1 | 105478653 | 105522325 | mNeoVis.2.1.1.chr1\_hap1.g00701.t01 |
| 766 | 1 | 105478653 | 105522325 | mNeoVis.2.1.1.chr1\_hap1.g00701.t01 |
| 767 | 1 | 105578859 | 105680386 | mNeoVis.2.1.1.chr1\_hap1.g00702.t01 |
| 768 | 1 | 105717005 | 105718715 | mNeoVis.2.1.1.chr1\_hap1.g00703.t01 |
| 769 | 1 | 105761029 | 105810496 | mNeoVis.2.1.1.chr1\_hap1.g00704.t01 |
| 770 | 1 | 105761029 | 105810496 | mNeoVis.2.1.1.chr1\_hap1.g00704.t01 |
| 771 | 1 | 105824179 | 105857505 | mNeoVis.2.1.1.chr1\_hap1.g00705.t01 |
| 772 | 1 | 105824179 | 105857505 | mNeoVis.2.1.1.chr1\_hap1.g00705.t01 |
| 773 | 1 | 105866009 | 105869160 | mNeoVis.2.1.1.chr1\_hap1.g00706.t01 |
| 774 | 1 | 105866009 | 105869160 | mNeoVis.2.1.1.chr1\_hap1.g00706.t01 |
| 775 | 1 | 105917702 | 105942148 | mNeoVis.2.1.1.chr1\_hap1.g00707.t01 |
| 776 | 1 | 105942303 | 106170427 | mNeoVis.2.1.1.chr1\_hap1.g00708.t01 |
| 777 | 1 | 106428858 | 106460635 | mNeoVis.2.1.1.chr1\_hap1.g00710.t01 |
| 778 | 1 | 106428858 | 106460635 | mNeoVis.2.1.1.chr1\_hap1.g00710.t01 |
| 779 | 1 | 106455328 | 106551105 | mNeoVis.2.1.1.chr1\_hap1.g00711.t01 |
| 780 | 1 | 106455328 | 106551105 | mNeoVis.2.1.1.chr1\_hap1.g00711.t01 |
| 781 | 1 | 106490502 | 106500659 | mNeoVis.2.1.1.chr1\_hap1.g00712.t01 |
| 782 | 1 | 106490502 | 106500659 | mNeoVis.2.1.1.chr1\_hap1.g00712.t01 |
| 783 | 1 | 107961655 | 107980229 | mNeoVis.2.1.1.chr1\_hap1.g00719.t01 |
| 784 | 1 | 107980323 | 108244294 | mNeoVis.2.1.1.chr1\_hap1.g00720.t01 |
| 785 | 1 | 108239024 | 108440207 | mNeoVis.2.1.1.chr1\_hap1.g00721.t01 |
| 786 | 1 | 108493235 | 108571196 | mNeoVis.2.1.1.chr1\_hap1.g00722.t01 |
| 787 | 1 | 108493235 | 108571196 | mNeoVis.2.1.1.chr1\_hap1.g00722.t01 |
| 788 | 1 | 109645408 | 109645707 | mNeoVis.2.1.1.chr1\_hap1.g00724.t01 |
| 789 | 1 | 111082302 | 111084663 | mNeoVis.2.1.1.chr1\_hap1.g00726.t01 |
| 790 | 1 | 111603803 | 111729045 | mNeoVis.2.1.1.chr1\_hap1.g00727.t01 |
| 791 | 1 | 111603803 | 111729045 | mNeoVis.2.1.1.chr1\_hap1.g00727.t01 |
| 792 | 1 | 111726674 | 111752266 | mNeoVis.2.1.1.chr1\_hap1.g00728.t01 |
| 793 | 1 | 111726674 | 111752266 | mNeoVis.2.1.1.chr1\_hap1.g00728.t01 |
| 794 | 1 | 112408339 | 112481925 | mNeoVis.2.1.1.chr1\_hap1.g00729.t01 |
| 795 | 1 | 112408339 | 112481925 | mNeoVis.2.1.1.chr1\_hap1.g00729.t01 |
| 796 | 1 | 112486151 | 112654593 | mNeoVis.2.1.1.chr1\_hap1.g00730.t01 |
| 797 | 1 | 112816138 | 112842358 | mNeoVis.2.1.1.chr1\_hap1.g00731.t01 |
| 798 | 1 | 112816138 | 112842358 | mNeoVis.2.1.1.chr1\_hap1.g00731.t01 |
| 799 | 1 | 112868132 | 113027409 | mNeoVis.2.1.1.chr1\_hap1.g00732.t01 |
| 800 | 1 | 112868132 | 113027409 | mNeoVis.2.1.1.chr1\_hap1.g00732.t01 |
| 801 | 1 | 113032578 | 113363783 | mNeoVis.2.1.1.chr1\_hap1.g00733.t01 |
| 802 | 1 | 114311597 | 114331335 | mNeoVis.2.1.1.chr1\_hap1.g00735.t01 |
| 803 | 1 | 114311597 | 114331335 | mNeoVis.2.1.1.chr1\_hap1.g00735.t01 |
| 804 | 1 | 114311597 | 114331335 | mNeoVis.2.1.1.chr1\_hap1.g00735.t01 |
| 805 | 1 | 114331225 | 114440296 | mNeoVis.2.1.1.chr1\_hap1.g00736.t01 |
| 806 | 1 | 114331225 | 114440296 | mNeoVis.2.1.1.chr1\_hap1.g00736.t01 |
| 807 | 1 | 114379010 | 114394968 | mNeoVis.2.1.1.chr1\_hap1.g00737.t01 |
| 808 | 1 | 114379010 | 114394968 | mNeoVis.2.1.1.chr1\_hap1.g00737.t01 |
| 809 | 1 | 114441144 | 114454241 | mNeoVis.2.1.1.chr1\_hap1.g00738.t01 |
| 810 | 1 | 114441144 | 114454241 | mNeoVis.2.1.1.chr1\_hap1.g00738.t01 |
| 811 | 1 | 116616474 | 116657676 | mNeoVis.2.1.1.chr1\_hap1.g00742.t01 |
| 812 | 1 | 117308910 | 117318530 | mNeoVis.2.1.1.chr1\_hap1.g00743.t01 |
| 813 | 1 | 117342089 | 117780122 | mNeoVis.2.1.1.chr1\_hap1.g00744.t01 |
| 814 | 1 | 117464689 | 117465756 | mNeoVis.2.1.1.chr1\_hap1.g00745.t01 |
| 815 | 1 | 117806674 | 118218248 | mNeoVis.2.1.1.chr1\_hap1.g00746.t01 |
| 816 | 1 | 117806674 | 118218248 | mNeoVis.2.1.1.chr1\_hap1.g00746.t01 |
| 817 | 1 | 118218264 | 118299541 | mNeoVis.2.1.1.chr1\_hap1.g00747.t01 |
| 818 | 1 | 118309560 | 118827524 | mNeoVis.2.1.1.chr1\_hap1.g00748.t01 |
| 819 | 1 | 118309560 | 118827524 | mNeoVis.2.1.1.chr1\_hap1.g00748.t01 |
| 820 | 1 | 119849986 | 119895635 | mNeoVis.2.1.1.chr1\_hap1.g00758.t01 |
| 821 | 1 | 120114095 | 120226699 | mNeoVis.2.1.1.chr1\_hap1.g00759.t01 |
| 822 | 1 | 120214563 | 120225117 | mNeoVis.2.1.1.chr1\_hap1.g00760.t01 |
| 823 | 1 | 120227979 | 120387400 | mNeoVis.2.1.1.chr1\_hap1.g00761.t01 |
| 824 | 1 | 120273469 | 120378303 | mNeoVis.2.1.1.chr1\_hap1.g00762.t01 |
| 825 | 1 | 123415985 | 123440369 | mNeoVis.2.1.1.chr1\_hap1.g00770.t01 |
| 826 | 1 | 123647685 | 123748588 | mNeoVis.2.1.1.chr1\_hap1.g00771.t01 |
| 827 | 1 | 123647685 | 123748588 | mNeoVis.2.1.1.chr1\_hap1.g00771.t01 |
| 828 | 1 | 123825060 | 123827332 | mNeoVis.2.1.1.chr1\_hap1.g00772.t01 |
| 829 | 1 | 123825060 | 123827332 | mNeoVis.2.1.1.chr1\_hap1.g00772.t01 |
| 830 | 1 | 125490930 | 125517175 | mNeoVis.2.1.1.chr1\_hap1.g00777.t01 |
| 831 | 1 | 125522027 | 125596194 | mNeoVis.2.1.1.chr1\_hap1.g00778.t01 |
| 832 | 1 | 125640262 | 125659416 | mNeoVis.2.1.1.chr1\_hap1.g00779.t01 |
| 833 | 1 | 125683583 | 125798587 | mNeoVis.2.1.1.chr1\_hap1.g00780.t01 |
| 834 | 1 | 125798599 | 125902384 | mNeoVis.2.1.1.chr1\_hap1.g00781.t01 |
| 835 | 1 | 125922862 | 125937625 | mNeoVis.2.1.1.chr1\_hap1.g00782.t01 |
| 836 | 1 | 127952629 | 128243545 | mNeoVis.2.1.1.chr1\_hap1.g00783.t01 |
| 837 | 1 | 128551988 | 128785961 | mNeoVis.2.1.1.chr1\_hap1.g00784.t01 |
| 838 | 1 | 128551988 | 128785961 | mNeoVis.2.1.1.chr1\_hap1.g00784.t01 |
| 839 | 1 | 130601389 | 131127650 | mNeoVis.2.1.1.chr1\_hap1.g00787.t01 |
| 840 | 1 | 131237274 | 131310539 | mNeoVis.2.1.1.chr1\_hap1.g00788.t01 |
| 841 | 1 | 131560555 | 131727682 | mNeoVis.2.1.1.chr1\_hap1.g00789.t01 |
| 842 | 1 | 131560555 | 131727682 | mNeoVis.2.1.1.chr1\_hap1.g00789.t01 |
| 843 | 1 | 131560998 | 131567725 | mNeoVis.2.1.1.chr1\_hap1.g00790.t01 |
| 844 | 1 | 131560998 | 131567725 | mNeoVis.2.1.1.chr1\_hap1.g00790.t01 |
| 845 | 1 | 132176496 | 132457070 | mNeoVis.2.1.1.chr1\_hap1.g00791.t01 |
| 846 | 1 | 132465196 | 132485815 | mNeoVis.2.1.1.chr1\_hap1.g00792.t01 |
| 847 | 1 | 132465196 | 132485815 | mNeoVis.2.1.1.chr1\_hap1.g00792.t01 |
| 848 | 1 | 132697519 | 132704435 | mNeoVis.2.1.1.chr1\_hap1.g00793.t01 |
| 849 | 1 | 132697519 | 132704435 | mNeoVis.2.1.1.chr1\_hap1.g00793.t01 |
| 850 | 1 | 132697519 | 132704435 | mNeoVis.2.1.1.chr1\_hap1.g00793.t01 |
| 851 | 1 | 132712340 | 132758202 | mNeoVis.2.1.1.chr1\_hap1.g00794.t01 |
| 852 | 1 | 132712340 | 132758202 | mNeoVis.2.1.1.chr1\_hap1.g00794.t01 |
| 853 | 1 | 132712340 | 132758202 | mNeoVis.2.1.1.chr1\_hap1.g00794.t01 |
| 854 | 1 | 132762033 | 133065831 | mNeoVis.2.1.1.chr1\_hap1.g00795.t01 |
| 855 | 1 | 133064102 | 133093515 | mNeoVis.2.1.1.chr1\_hap1.g00796.t01 |
| 856 | 1 | 133617305 | 133640294 | mNeoVis.2.1.1.chr1\_hap1.g00798.t01 |
| 857 | 1 | 133617305 | 133640294 | mNeoVis.2.1.1.chr1\_hap1.g00798.t01 |
| 858 | 1 | 133643580 | 133898217 | mNeoVis.2.1.1.chr1\_hap1.g00799.t01 |
| 859 | 1 | 133643580 | 133898217 | mNeoVis.2.1.1.chr1\_hap1.g00799.t01 |
| 860 | 1 | 134093001 | 134236686 | mNeoVis.2.1.1.chr1\_hap1.g00800.t01 |
| 861 | 1 | 134246870 | 134254000 | mNeoVis.2.1.1.chr1\_hap1.g00801.t01 |
| 862 | 1 | 134260965 | 134298309 | mNeoVis.2.1.1.chr1\_hap1.g00802.t01 |
| 863 | 1 | 134342273 | 134413854 | mNeoVis.2.1.1.chr1\_hap1.g00803.t01 |
| 864 | 1 | 136484279 | 136808300 | mNeoVis.2.1.1.chr1\_hap1.g00813.t01 |
| 865 | 1 | 136771189 | 136772407 | mNeoVis.2.1.1.chr1\_hap1.g00814.t01 |
| 866 | 1 | 137511592 | 137623933 | mNeoVis.2.1.1.chr1\_hap1.g00815.t01 |
| 867 | 1 | 137716936 | 137796521 | mNeoVis.2.1.1.chr1\_hap1.g00816.t01 |
| 868 | 1 | 138091419 | 138106922 | mNeoVis.2.1.1.chr1\_hap1.g00818.t01 |
| 869 | 1 | 138121123 | 138286288 | mNeoVis.2.1.1.chr1\_hap1.g00819.t01 |
| 870 | 1 | 138305109 | 138363312 | mNeoVis.2.1.1.chr1\_hap1.g00820.t01 |
| 871 | 1 | 139780923 | 139844065 | mNeoVis.2.1.1.chr1\_hap1.g00828.t01 |
| 872 | 1 | 139847218 | 140009254 | mNeoVis.2.1.1.chr1\_hap1.g00829.t01 |
| 873 | 1 | 140146678 | 140148163 | mNeoVis.2.1.1.chr1\_hap1.g00830.t01 |
| 874 | 1 | 142684351 | 142809427 | mNeoVis.2.1.1.chr1\_hap1.g00839.t01 |
| 875 | 1 | 142904114 | 143029856 | mNeoVis.2.1.1.chr1\_hap1.g00840.t01 |
| 876 | 1 | 143483435 | 143483831 | mNeoVis.2.1.1.chr1\_hap1.g00841.t01 |
| 877 | 1 | 143483435 | 143483831 | mNeoVis.2.1.1.chr1\_hap1.g00841.t01 |
| 878 | 1 | 144878106 | 144891743 | mNeoVis.2.1.1.chr1\_hap1.g00842.t01 |
| 879 | 1 | 145241125 | 145367350 | mNeoVis.2.1.1.chr1\_hap1.g00843.t01 |
| 880 | 1 | 146414992 | 146420140 | mNeoVis.2.1.1.chr1\_hap1.g00853.t01 |
| 881 | 1 | 146484368 | 146512406 | mNeoVis.2.1.1.chr1\_hap1.g00854.t01 |
| 882 | 1 | 146515713 | 146572673 | mNeoVis.2.1.1.chr1\_hap1.g00855.t01 |
| 883 | 1 | 146598350 | 146611420 | mNeoVis.2.1.1.chr1\_hap1.g00856.t01 |
| 884 | 1 | 146611561 | 146658867 | mNeoVis.2.1.1.chr1\_hap1.g00857.t01 |
| 885 | 1 | 147144274 | 147530968 | mNeoVis.2.1.1.chr1\_hap1.g00858.t01 |
| 886 | 1 | 148696660 | 148697601 | mNeoVis.2.1.1.chr1\_hap1.g00874.t01 |
| 887 | 1 | 148875787 | 148878388 | mNeoVis.2.1.1.chr1\_hap1.g00875.t01 |
| 888 | 1 | 148875787 | 148878388 | mNeoVis.2.1.1.chr1\_hap1.g00875.t01 |
| 889 | 1 | 148891955 | 148892893 | mNeoVis.2.1.1.chr1\_hap1.g00876.t01 |
| 890 | 1 | 148891955 | 148892893 | mNeoVis.2.1.1.chr1\_hap1.g00876.t01 |
| 891 | 1 | 148910510 | 148925442 | mNeoVis.2.1.1.chr1\_hap1.g00877.t01 |
| 892 | 1 | 148910510 | 148925442 | mNeoVis.2.1.1.chr1\_hap1.g00877.t01 |
| 893 | 1 | 148959661 | 148965221 | mNeoVis.2.1.1.chr1\_hap1.g00878.t01 |
| 894 | 1 | 148959661 | 148965221 | mNeoVis.2.1.1.chr1\_hap1.g00878.t01 |
| 895 | 1 | 148965665 | 148990478 | mNeoVis.2.1.1.chr1\_hap1.g00879.t01 |
| 896 | 1 | 148965665 | 148990478 | mNeoVis.2.1.1.chr1\_hap1.g00879.t01 |
| 897 | 1 | 148996606 | 149009236 | mNeoVis.2.1.1.chr1\_hap1.g00880.t01 |
| 898 | 1 | 148996606 | 149009236 | mNeoVis.2.1.1.chr1\_hap1.g00880.t01 |
| 899 | 1 | 149036162 | 149049651 | mNeoVis.2.1.1.chr1\_hap1.g00881.t01 |
| 900 | 1 | 149036162 | 149049651 | mNeoVis.2.1.1.chr1\_hap1.g00881.t01 |
| 901 | 1 | 149060156 | 149077399 | mNeoVis.2.1.1.chr1\_hap1.g00882.t01 |
| 902 | 1 | 149060156 | 149077399 | mNeoVis.2.1.1.chr1\_hap1.g00882.t01 |
| 903 | 1 | 149091217 | 149310593 | mNeoVis.2.1.1.chr1\_hap1.g00883.t01 |
| 904 | 1 | 149091217 | 149310593 | mNeoVis.2.1.1.chr1\_hap1.g00883.t01 |
| 905 | 1 | 149220863 | 149222275 | mNeoVis.2.1.1.chr1\_hap1.g00884.t01 |
| 906 | 1 | 149220863 | 149222275 | mNeoVis.2.1.1.chr1\_hap1.g00884.t01 |
| 907 | 1 | 154037849 | 154042370 | mNeoVis.2.1.1.chr1\_hap1.g00949.t01 |
| 908 | 1 | 154042580 | 154087953 | mNeoVis.2.1.1.chr1\_hap1.g00950.t01 |
| 909 | 1 | 154126426 | 154137776 | mNeoVis.2.1.1.chr1\_hap1.g00951.t01 |
| 910 | 1 | 154165835 | 154177185 | mNeoVis.2.1.1.chr1\_hap1.g00952.t01 |
| 911 | 1 | 154211585 | 154254021 | mNeoVis.2.1.1.chr1\_hap1.g00953.t01 |
| 912 | 1 | 154269453 | 154283870 | mNeoVis.2.1.1.chr1\_hap1.g00954.t01 |
| 913 | 1 | 154284618 | 154286584 | mNeoVis.2.1.1.chr1\_hap1.g00955.t01 |
| 914 | 1 | 154288115 | 154291315 | mNeoVis.2.1.1.chr1\_hap1.g00956.t01 |
| 915 | 1 | 154291530 | 154299142 | mNeoVis.2.1.1.chr1\_hap1.g00957.t01 |
| 916 | 1 | 154299899 | 154307547 | mNeoVis.2.1.1.chr1\_hap1.g00958.t01 |
| 917 | 1 | 154314203 | 154318998 | mNeoVis.2.1.1.chr1\_hap1.g00959.t01 |
| 918 | 1 | 154329407 | 154465723 | mNeoVis.2.1.1.chr1\_hap1.g00960.t01 |
| 919 | 1 | 154494879 | 154501825 | mNeoVis.2.1.1.chr1\_hap1.g00961.t01 |
| 920 | 1 | 154502695 | 154509225 | mNeoVis.2.1.1.chr1\_hap1.g00962.t01 |
| 921 | 1 | 154516475 | 154518084 | mNeoVis.2.1.1.chr1\_hap1.g00963.t01 |
| 922 | 1 | 154534119 | 154537253 | mNeoVis.2.1.1.chr1\_hap1.g00964.t01 |
| 923 | 1 | 154537457 | 154537980 | mNeoVis.2.1.1.chr1\_hap1.g00965.t01 |
| 924 | 1 | 154542066 | 154546725 | mNeoVis.2.1.1.chr1\_hap1.g00966.t01 |
| 925 | 1 | 154555668 | 154575481 | mNeoVis.2.1.1.chr1\_hap1.g00967.t01 |
| 926 | 1 | 177656213 | 177683587 | mNeoVis.2.1.1.chr1\_hap1.g01093.t01 |
| 927 | 1 | 177748299 | 177832519 | mNeoVis.2.1.1.chr1\_hap1.g01094.t01 |
| 928 | 1 | 177784962 | 177789450 | mNeoVis.2.1.1.chr1\_hap1.g01095.t01 |
| 929 | 1 | 177798433 | 177809771 | mNeoVis.2.1.1.chr1\_hap1.g01096.t01 |
| 930 | 1 | 177843273 | 177850166 | mNeoVis.2.1.1.chr1\_hap1.g01097.t01 |
| 931 | 1 | 177875499 | 177896531 | mNeoVis.2.1.1.chr1\_hap1.g01098.t01 |
| 932 | 1 | 177896542 | 177904534 | mNeoVis.2.1.1.chr1\_hap1.g01099.t01 |
| 933 | 1 | 177898060 | 177929264 | mNeoVis.2.1.1.chr1\_hap1.g01100.t01 |
| 934 | 1 | 177904722 | 177909997 | mNeoVis.2.1.1.chr1\_hap1.g01101.t01 |
| 935 | 1 | 177910157 | 177911236 | mNeoVis.2.1.1.chr1\_hap1.g01102.t01 |
| 936 | 1 | 177949304 | 177950203 | mNeoVis.2.1.1.chr1\_hap1.g01103.t01 |
| 937 | 1 | 177965769 | 177966666 | mNeoVis.2.1.1.chr1\_hap1.g01104.t01 |
| 938 | 1 | 177971784 | 177972988 | mNeoVis.2.1.1.chr1\_hap1.g01105.t01 |
| 939 | 1 | 177984626 | 178040502 | mNeoVis.2.1.1.chr1\_hap1.g01106.t01 |
| 940 | 1 | 178041878 | 178068456 | mNeoVis.2.1.1.chr1\_hap1.g01107.t01 |
| 941 | 1 | 178041878 | 178068456 | mNeoVis.2.1.1.chr1\_hap1.g01107.t01 |
| 942 | 1 | 178083215 | 178086734 | mNeoVis.2.1.1.chr1\_hap1.g01108.t01 |
| 943 | 1 | 178083215 | 178086734 | mNeoVis.2.1.1.chr1\_hap1.g01108.t01 |
| 944 | 1 | 178126385 | 178135699 | mNeoVis.2.1.1.chr1\_hap1.g01109.t01 |
| 945 | 1 | 178126385 | 178135699 | mNeoVis.2.1.1.chr1\_hap1.g01109.t01 |
| 946 | 1 | 178146541 | 178189473 | mNeoVis.2.1.1.chr1\_hap1.g01110.t01 |
| 947 | 1 | 178146541 | 178189473 | mNeoVis.2.1.1.chr1\_hap1.g01110.t01 |
| 948 | 1 | 178160338 | 178176687 | mNeoVis.2.1.1.chr1\_hap1.g01111.t01 |
| 949 | 1 | 178160338 | 178176687 | mNeoVis.2.1.1.chr1\_hap1.g01111.t01 |
| 950 | 1 | 178178976 | 178181399 | mNeoVis.2.1.1.chr1\_hap1.g01112.t01 |
| 951 | 1 | 178178976 | 178181399 | mNeoVis.2.1.1.chr1\_hap1.g01112.t01 |
| 952 | 1 | 178232183 | 178233482 | mNeoVis.2.1.1.chr1\_hap1.g01113.t01 |
| 953 | 1 | 178232183 | 178233482 | mNeoVis.2.1.1.chr1\_hap1.g01113.t01 |
| 954 | 1 | 178233874 | 178238608 | mNeoVis.2.1.1.chr1\_hap1.g01114.t01 |
| 955 | 1 | 178233874 | 178238608 | mNeoVis.2.1.1.chr1\_hap1.g01114.t01 |
| 956 | 1 | 178238981 | 178239373 | mNeoVis.2.1.1.chr1\_hap1.g01115.t01 |
| 957 | 1 | 178238981 | 178239373 | mNeoVis.2.1.1.chr1\_hap1.g01115.t01 |
| 958 | 1 | 178240555 | 178241282 | mNeoVis.2.1.1.chr1\_hap1.g01116.t01 |
| 959 | 1 | 178240555 | 178241282 | mNeoVis.2.1.1.chr1\_hap1.g01116.t01 |
| 960 | 1 | 178245395 | 178249188 | mNeoVis.2.1.1.chr1\_hap1.g01117.t01 |
| 961 | 1 | 178245395 | 178249188 | mNeoVis.2.1.1.chr1\_hap1.g01117.t01 |
| 962 | 1 | 178250105 | 178250515 | mNeoVis.2.1.1.chr1\_hap1.g01118.t01 |
| 963 | 1 | 178250105 | 178250515 | mNeoVis.2.1.1.chr1\_hap1.g01118.t01 |
| 964 | 1 | 178251338 | 178254340 | mNeoVis.2.1.1.chr1\_hap1.g01119.t01 |
| 965 | 1 | 178251338 | 178254340 | mNeoVis.2.1.1.chr1\_hap1.g01119.t01 |
| 966 | 1 | 178261871 | 178262182 | mNeoVis.2.1.1.chr1\_hap1.g01120.t01 |
| 967 | 1 | 178261871 | 178262182 | mNeoVis.2.1.1.chr1\_hap1.g01120.t01 |
| 968 | 1 | 178266192 | 178267694 | mNeoVis.2.1.1.chr1\_hap1.g01121.t01 |
| 969 | 1 | 178266192 | 178267694 | mNeoVis.2.1.1.chr1\_hap1.g01121.t01 |
| 970 | 1 | 178266247 | 178267030 | mNeoVis.2.1.1.chr1\_hap1.g01122.t01 |
| 971 | 1 | 178266247 | 178267030 | mNeoVis.2.1.1.chr1\_hap1.g01122.t01 |
| 972 | 1 | 178274855 | 178276130 | mNeoVis.2.1.1.chr1\_hap1.g01123.t01 |
| 973 | 1 | 178274855 | 178276130 | mNeoVis.2.1.1.chr1\_hap1.g01123.t01 |
| 974 | 1 | 178280645 | 178281789 | mNeoVis.2.1.1.chr1\_hap1.g01124.t01 |
| 975 | 1 | 178280645 | 178281789 | mNeoVis.2.1.1.chr1\_hap1.g01124.t01 |
| 976 | 1 | 178285161 | 178285797 | mNeoVis.2.1.1.chr1\_hap1.g01125.t01 |
| 977 | 1 | 178285161 | 178285797 | mNeoVis.2.1.1.chr1\_hap1.g01125.t01 |
| 978 | 1 | 178286531 | 178287021 | mNeoVis.2.1.1.chr1\_hap1.g01126.t01 |
| 979 | 1 | 178286531 | 178287021 | mNeoVis.2.1.1.chr1\_hap1.g01126.t01 |
| 980 | 1 | 178287316 | 178287823 | mNeoVis.2.1.1.chr1\_hap1.g01127.t01 |
| 981 | 1 | 178287316 | 178287823 | mNeoVis.2.1.1.chr1\_hap1.g01127.t01 |
| 982 | 1 | 178291711 | 178297458 | mNeoVis.2.1.1.chr1\_hap1.g01128.t01 |
| 983 | 1 | 178291711 | 178297458 | mNeoVis.2.1.1.chr1\_hap1.g01128.t01 |
| 984 | 1 | 178292361 | 178294123 | mNeoVis.2.1.1.chr1\_hap1.g01129.t01 |
| 985 | 1 | 178292361 | 178294123 | mNeoVis.2.1.1.chr1\_hap1.g01129.t01 |
| 986 | 1 | 178294326 | 178294828 | mNeoVis.2.1.1.chr1\_hap1.g01130.t01 |
| 987 | 1 | 178294326 | 178294828 | mNeoVis.2.1.1.chr1\_hap1.g01130.t01 |
| 988 | 1 | 178302951 | 178303751 | mNeoVis.2.1.1.chr1\_hap1.g01131.t01 |
| 989 | 1 | 178302951 | 178303751 | mNeoVis.2.1.1.chr1\_hap1.g01131.t01 |
| 990 | 1 | 178306224 | 178309140 | mNeoVis.2.1.1.chr1\_hap1.g01132.t01 |
| 991 | 1 | 178306224 | 178309140 | mNeoVis.2.1.1.chr1\_hap1.g01132.t01 |
| 992 | 1 | 178317944 | 178331299 | mNeoVis.2.1.1.chr1\_hap1.g01133.t01 |
| 993 | 1 | 178317944 | 178331299 | mNeoVis.2.1.1.chr1\_hap1.g01133.t01 |
| 994 | 1 | 178318482 | 178326449 | mNeoVis.2.1.1.chr1\_hap1.g01134.t01 |
| 995 | 1 | 178318482 | 178326449 | mNeoVis.2.1.1.chr1\_hap1.g01134.t01 |
| 996 | 1 | 178327344 | 178329190 | mNeoVis.2.1.1.chr1\_hap1.g01135.t01 |
| 997 | 1 | 178327344 | 178329190 | mNeoVis.2.1.1.chr1\_hap1.g01135.t01 |
| 998 | 1 | 178331346 | 178337890 | mNeoVis.2.1.1.chr1\_hap1.g01136.t01 |
| 999 | 1 | 178331346 | 178337890 | mNeoVis.2.1.1.chr1\_hap1.g01136.t01 |
| 1000 | 1 | 178347970 | 178369565 | mNeoVis.2.1.1.chr1\_hap1.g01137.t01 |
| 1001 | 1 | 178347970 | 178369565 | mNeoVis.2.1.1.chr1\_hap1.g01137.t01 |
| 1002 | 1 | 178348578 | 178355760 | mNeoVis.2.1.1.chr1\_hap1.g01138.t01 |
| 1003 | 1 | 178348578 | 178355760 | mNeoVis.2.1.1.chr1\_hap1.g01138.t01 |
| 1004 | 1 | 178356581 | 178357293 | mNeoVis.2.1.1.chr1\_hap1.g01139.t01 |
| 1005 | 1 | 178356581 | 178357293 | mNeoVis.2.1.1.chr1\_hap1.g01139.t01 |
| 1006 | 1 | 178378839 | 178379326 | mNeoVis.2.1.1.chr1\_hap1.g01140.t01 |
| 1007 | 1 | 178378839 | 178379326 | mNeoVis.2.1.1.chr1\_hap1.g01140.t01 |
| 1008 | 1 | 178379607 | 178394081 | mNeoVis.2.1.1.chr1\_hap1.g01141.t01 |
| 1009 | 1 | 178379607 | 178394081 | mNeoVis.2.1.1.chr1\_hap1.g01141.t01 |
| 1010 | 1 | 178384576 | 178404282 | mNeoVis.2.1.1.chr1\_hap1.g01142.t01 |
| 1011 | 1 | 178384576 | 178404282 | mNeoVis.2.1.1.chr1\_hap1.g01142.t01 |
| 1012 | 1 | 178397586 | 178401948 | mNeoVis.2.1.1.chr1\_hap1.g01143.t01 |
| 1013 | 1 | 178397586 | 178401948 | mNeoVis.2.1.1.chr1\_hap1.g01143.t01 |
| 1014 | 1 | 178410404 | 178419888 | mNeoVis.2.1.1.chr1\_hap1.g01144.t01 |
| 1015 | 1 | 178410404 | 178419888 | mNeoVis.2.1.1.chr1\_hap1.g01144.t01 |
| 1016 | 1 | 178436252 | 178456473 | mNeoVis.2.1.1.chr1\_hap1.g01145.t01 |
| 1017 | 1 | 178436252 | 178456473 | mNeoVis.2.1.1.chr1\_hap1.g01145.t01 |
| 1018 | 1 | 178444545 | 178444970 | mNeoVis.2.1.1.chr1\_hap1.g01146.t01 |
| 1019 | 1 | 178444545 | 178444970 | mNeoVis.2.1.1.chr1\_hap1.g01146.t01 |
| 1020 | 1 | 178445841 | 178446306 | mNeoVis.2.1.1.chr1\_hap1.g01147.t01 |
| 1021 | 1 | 178445841 | 178446306 | mNeoVis.2.1.1.chr1\_hap1.g01147.t01 |
| 1022 | 1 | 178446660 | 178447105 | mNeoVis.2.1.1.chr1\_hap1.g01148.t01 |
| 1023 | 1 | 178446660 | 178447105 | mNeoVis.2.1.1.chr1\_hap1.g01148.t01 |
| 1024 | 1 | 178457416 | 178458112 | mNeoVis.2.1.1.chr1\_hap1.g01149.t01 |
| 1025 | 1 | 178457416 | 178458112 | mNeoVis.2.1.1.chr1\_hap1.g01149.t01 |
| 1026 | 1 | 178460832 | 178462615 | mNeoVis.2.1.1.chr1\_hap1.g01150.t01 |
| 1027 | 1 | 178460832 | 178462615 | mNeoVis.2.1.1.chr1\_hap1.g01150.t01 |
| 1028 | 1 | 178466573 | 178468863 | mNeoVis.2.1.1.chr1\_hap1.g01151.t01 |
| 1029 | 1 | 178466573 | 178468863 | mNeoVis.2.1.1.chr1\_hap1.g01151.t01 |
| 1030 | 1 | 178469196 | 178469652 | mNeoVis.2.1.1.chr1\_hap1.g01152.t01 |
| 1031 | 1 | 178469196 | 178469652 | mNeoVis.2.1.1.chr1\_hap1.g01152.t01 |
| 1032 | 1 | 178502437 | 178521624 | mNeoVis.2.1.1.chr1\_hap1.g01153.t01 |
| 1033 | 1 | 178502437 | 178521624 | mNeoVis.2.1.1.chr1\_hap1.g01153.t01 |
| 1034 | 1 | 178537947 | 178548626 | mNeoVis.2.1.1.chr1\_hap1.g01154.t01 |
| 1035 | 1 | 178537947 | 178548626 | mNeoVis.2.1.1.chr1\_hap1.g01154.t01 |
| 1036 | 1 | 178604135 | 178743326 | mNeoVis.2.1.1.chr1\_hap1.g01156.t01 |
| 1037 | 1 | 178646696 | 178668296 | mNeoVis.2.1.1.chr1\_hap1.g01157.t01 |
| 1038 | 1 | 178646696 | 178668296 | mNeoVis.2.1.1.chr1\_hap1.g01157.t01 |
| 1039 | 1 | 178675321 | 178700158 | mNeoVis.2.1.1.chr1\_hap1.g01158.t01 |
| 1040 | 1 | 178675321 | 178700158 | mNeoVis.2.1.1.chr1\_hap1.g01158.t01 |
| 1041 | 1 | 178675321 | 178700158 | mNeoVis.2.1.1.chr1\_hap1.g01158.t01 |
| 1042 | 1 | 178684590 | 178692021 | mNeoVis.2.1.1.chr1\_hap1.g01159.t01 |
| 1043 | 1 | 178684590 | 178692021 | mNeoVis.2.1.1.chr1\_hap1.g01159.t01 |
| 1044 | 1 | 178684590 | 178692021 | mNeoVis.2.1.1.chr1\_hap1.g01159.t01 |
| 1045 | 1 | 178688352 | 178691101 | mNeoVis.2.1.1.chr1\_hap1.g01160.t01 |
| 1046 | 1 | 178688352 | 178691101 | mNeoVis.2.1.1.chr1\_hap1.g01160.t01 |
| 1047 | 1 | 178688352 | 178691101 | mNeoVis.2.1.1.chr1\_hap1.g01160.t01 |
| 1048 | 1 | 178700407 | 178700970 | mNeoVis.2.1.1.chr1\_hap1.g01161.t01 |
| 1049 | 1 | 178700407 | 178700970 | mNeoVis.2.1.1.chr1\_hap1.g01161.t01 |
| 1050 | 1 | 178700407 | 178700970 | mNeoVis.2.1.1.chr1\_hap1.g01161.t01 |
| 1051 | 1 | 178733450 | 178770615 | mNeoVis.2.1.1.chr1\_hap1.g01162.t01 |
| 1052 | 1 | 178733450 | 178770615 | mNeoVis.2.1.1.chr1\_hap1.g01162.t01 |
| 1053 | 1 | 178733450 | 178770615 | mNeoVis.2.1.1.chr1\_hap1.g01162.t01 |
| 1054 | 1 | 178800143 | 179099215 | mNeoVis.2.1.1.chr1\_hap1.g01163.t01 |
| 1055 | 1 | 178800143 | 179099215 | mNeoVis.2.1.1.chr1\_hap1.g01163.t01 |
| 1056 | 1 | 178800143 | 179099215 | mNeoVis.2.1.1.chr1\_hap1.g01163.t01 |
| 1057 | 1 | 179199964 | 179413543 | mNeoVis.2.1.1.chr1\_hap1.g01164.t01 |
| 1058 | 1 | 179199964 | 179413543 | mNeoVis.2.1.1.chr1\_hap1.g01164.t01 |
| 1059 | 1 | 179415162 | 179417380 | mNeoVis.2.1.1.chr1\_hap1.g01165.t01 |
| 1060 | 1 | 179415162 | 179417380 | mNeoVis.2.1.1.chr1\_hap1.g01165.t01 |
| 1061 | 1 | 179427224 | 179436627 | mNeoVis.2.1.1.chr1\_hap1.g01166.t01 |
| 1062 | 1 | 179427224 | 179436627 | mNeoVis.2.1.1.chr1\_hap1.g01166.t01 |
| 1063 | 1 | 179501057 | 179515538 | mNeoVis.2.1.1.chr1\_hap1.g01167.t01 |
| 1064 | 1 | 179517544 | 179531591 | mNeoVis.2.1.1.chr1\_hap1.g01168.t01 |
| 1065 | 1 | 179517544 | 179531591 | mNeoVis.2.1.1.chr1\_hap1.g01168.t01 |
| 1066 | 1 | 179531673 | 179546676 | mNeoVis.2.1.1.chr1\_hap1.g01169.t01 |
| 1067 | 1 | 179531673 | 179546676 | mNeoVis.2.1.1.chr1\_hap1.g01169.t01 |
| 1068 | 1 | 179590467 | 179642398 | mNeoVis.2.1.1.chr1\_hap1.g01170.t01 |
| 1069 | 1 | 179590467 | 179642398 | mNeoVis.2.1.1.chr1\_hap1.g01170.t01 |
| 1070 | 1 | 179645242 | 179671522 | mNeoVis.2.1.1.chr1\_hap1.g01171.t01 |
| 1071 | 1 | 179645242 | 179671522 | mNeoVis.2.1.1.chr1\_hap1.g01171.t01 |
| 1072 | 1 | 179673953 | 179728328 | mNeoVis.2.1.1.chr1\_hap1.g01172.t01 |
| 1073 | 1 | 179673953 | 179728328 | mNeoVis.2.1.1.chr1\_hap1.g01172.t01 |
| 1074 | 1 | 179728032 | 179768113 | mNeoVis.2.1.1.chr1\_hap1.g01173.t01 |
| 1075 | 1 | 179728032 | 179768113 | mNeoVis.2.1.1.chr1\_hap1.g01173.t01 |
| 1076 | 1 | 179798211 | 179950387 | mNeoVis.2.1.1.chr1\_hap1.g01174.t01 |
| 1077 | 1 | 179972402 | 179992814 | mNeoVis.2.1.1.chr1\_hap1.g01175.t01 |
| 1078 | 1 | 180556484 | 180565732 | mNeoVis.2.1.1.chr1\_hap1.g01176.t01 |
| 1079 | 1 | 180556484 | 180565732 | mNeoVis.2.1.1.chr1\_hap1.g01176.t01 |
| 1080 | 1 | 180556484 | 180565732 | mNeoVis.2.1.1.chr1\_hap1.g01176.t01 |
| 1081 | 1 | 181544328 | 181791720 | mNeoVis.2.1.1.chr1\_hap1.g01178.t01 |
| 1082 | 1 | 181544328 | 181791720 | mNeoVis.2.1.1.chr1\_hap1.g01178.t01 |
| 1083 | 1 | 181586569 | 181596372 | mNeoVis.2.1.1.chr1\_hap1.g01179.t01 |
| 1084 | 1 | 181586569 | 181596372 | mNeoVis.2.1.1.chr1\_hap1.g01179.t01 |
| 1085 | 1 | 181785431 | 181939316 | mNeoVis.2.1.1.chr1\_hap1.g01180.t01 |
| 1086 | 1 | 181785431 | 181939316 | mNeoVis.2.1.1.chr1\_hap1.g01180.t01 |
| 1087 | 1 | 182148358 | 182152737 | mNeoVis.2.1.1.chr1\_hap1.g01181.t01 |
| 1088 | 1 | 182148358 | 182152737 | mNeoVis.2.1.1.chr1\_hap1.g01181.t01 |
| 1089 | 1 | 182195619 | 182210883 | mNeoVis.2.1.1.chr1\_hap1.g01182.t01 |
| 1090 | 1 | 182195619 | 182210883 | mNeoVis.2.1.1.chr1\_hap1.g01182.t01 |
| 1091 | 1 | 183158224 | 183234349 | mNeoVis.2.1.1.chr1\_hap1.g01184.t01 |
| 1092 | 1 | 183405476 | 183515922 | mNeoVis.2.1.1.chr1\_hap1.g01185.t01 |
| 1093 | 1 | 185295993 | 185329059 | mNeoVis.2.1.1.chr1\_hap1.g01190.t01 |
| 1094 | 1 | 185329455 | 185390713 | mNeoVis.2.1.1.chr1\_hap1.g01191.t01 |
| 1095 | 1 | 185390924 | 185415319 | mNeoVis.2.1.1.chr1\_hap1.g01192.t01 |
| 1096 | 1 | 185425190 | 185426621 | mNeoVis.2.1.1.chr1\_hap1.g01193.t01 |
| 1097 | 1 | 185517022 | 185724372 | mNeoVis.2.1.1.chr1\_hap1.g01194.t01 |
| 1098 | 1 | 185734238 | 185734988 | mNeoVis.2.1.1.chr1\_hap1.g01195.t01 |
| 1099 | 1 | 185749210 | 185759062 | mNeoVis.2.1.1.chr1\_hap1.g01196.t01 |
| 1100 | 1 | 185758818 | 185832848 | mNeoVis.2.1.1.chr1\_hap1.g01197.t01 |
| 1101 | 1 | 185835040 | 185844674 | mNeoVis.2.1.1.chr1\_hap1.g01198.t01 |
| 1102 | 1 | 185874287 | 185990831 | mNeoVis.2.1.1.chr1\_hap1.g01199.t01 |
| 1103 | 1 | 185958088 | 185958990 | mNeoVis.2.1.1.chr1\_hap1.g01200.t01 |
| 1104 | 1 | 186085406 | 186097656 | mNeoVis.2.1.1.chr1\_hap1.g01201.t01 |
| 1105 | 1 | 186191256 | 186217165 | mNeoVis.2.1.1.chr1\_hap1.g01202.t01 |
| 1106 | 1 | 186394510 | 186430236 | mNeoVis.2.1.1.chr1\_hap1.g01203.t01 |
| 1107 | 1 | 186486850 | 186882916 | mNeoVis.2.1.1.chr1\_hap1.g01204.t01 |
| 1108 | 1 | 186892185 | 186935480 | mNeoVis.2.1.1.chr1\_hap1.g01205.t01 |
| 1109 | 1 | 186892185 | 186935480 | mNeoVis.2.1.1.chr1\_hap1.g01205.t01 |
| 1110 | 1 | 187043617 | 187064085 | mNeoVis.2.1.1.chr1\_hap1.g01206.t01 |
| 1111 | 1 | 187043617 | 187064085 | mNeoVis.2.1.1.chr1\_hap1.g01206.t01 |
| 1112 | 1 | 187465043 | 187593184 | mNeoVis.2.1.1.chr1\_hap1.g01207.t01 |
| 1113 | 1 | 187593754 | 187854197 | mNeoVis.2.1.1.chr1\_hap1.g01208.t01 |
| 1114 | 1 | 188811654 | 188830338 | mNeoVis.2.1.1.chr1\_hap1.g01209.t01 |
| 1115 | 1 | 188961955 | 189025732 | mNeoVis.2.1.1.chr1\_hap1.g01210.t01 |
| 1116 | 1 | 189393082 | 189499037 | mNeoVis.2.1.1.chr1\_hap1.g01215.t01 |
| 1117 | 1 | 189521201 | 189554446 | mNeoVis.2.1.1.chr1\_hap1.g01216.t01 |
| 1118 | 1 | 189574489 | 190126168 | mNeoVis.2.1.1.chr1\_hap1.g01217.t01 |
| 1119 | 1 | 189574489 | 190126168 | mNeoVis.2.1.1.chr1\_hap1.g01217.t01 |
| 1120 | 1 | 191167310 | 191202666 | mNeoVis.2.1.1.chr1\_hap1.g01222.t01 |
| 1121 | 1 | 191167310 | 191202666 | mNeoVis.2.1.1.chr1\_hap1.g01222.t01 |
| 1122 | 1 | 191399502 | 191515440 | mNeoVis.2.1.1.chr1\_hap1.g01223.t01 |
| 1123 | 1 | 191399502 | 191515440 | mNeoVis.2.1.1.chr1\_hap1.g01223.t01 |
| 1124 | 1 | 191558188 | 191586614 | mNeoVis.2.1.1.chr1\_hap1.g01224.t01 |
| 1125 | 1 | 191558188 | 191586614 | mNeoVis.2.1.1.chr1\_hap1.g01224.t01 |
| 1126 | 1 | 191615412 | 191679209 | mNeoVis.2.1.1.chr1\_hap1.g01225.t01 |
| 1127 | 1 | 194109392 | 194135703 | mNeoVis.2.1.1.chr1\_hap1.g01236.t01 |
| 1128 | 1 | 194415985 | 194436312 | mNeoVis.2.1.1.chr1\_hap1.g01237.t01 |
| 1129 | 1 | 194922229 | 194940935 | mNeoVis.2.1.1.chr1\_hap1.g01241.t01 |
| 1130 | 1 | 194945113 | 194991247 | mNeoVis.2.1.1.chr1\_hap1.g01242.t01 |
| 1131 | 1 | 195080274 | 195113833 | mNeoVis.2.1.1.chr1\_hap1.g01243.t01 |
| 1132 | 1 | 195113959 | 195164242 | mNeoVis.2.1.1.chr1\_hap1.g01244.t01 |
| 1133 | 1 | 195174421 | 195200150 | mNeoVis.2.1.1.chr1\_hap1.g01245.t01 |
| 1134 | 1 | 195226879 | 195404343 | mNeoVis.2.1.1.chr1\_hap1.g01246.t01 |
| 1135 | 1 | 195558026 | 195558635 | mNeoVis.2.1.1.chr1\_hap1.g01247.t01 |
| 1136 | 1 | 195793021 | 195851090 | mNeoVis.2.1.1.chr1\_hap1.g01248.t01 |
| 1137 | 1 | 195793021 | 195851090 | mNeoVis.2.1.1.chr1\_hap1.g01248.t01 |
| 1138 | 1 | 196148245 | 196319556 | mNeoVis.2.1.1.chr1\_hap1.g01249.t01 |
| 1139 | 1 | 196148245 | 196319556 | mNeoVis.2.1.1.chr1\_hap1.g01249.t01 |
| 1140 | 1 | 197461894 | 197471259 | mNeoVis.2.1.1.chr1\_hap1.g01255.t01 |
| 1141 | 1 | 197506655 | 197751368 | mNeoVis.2.1.1.chr1\_hap1.g01256.t01 |
| 1142 | 1 | 197506655 | 197751368 | mNeoVis.2.1.1.chr1\_hap1.g01256.t01 |
| 1143 | 1 | 198294391 | 198315783 | mNeoVis.2.1.1.chr1\_hap1.g01257.t01 |
| 1144 | 1 | 198299652 | 198330105 | mNeoVis.2.1.1.chr1\_hap1.g01258.t01 |
| 1145 | 1 | 198330196 | 198345240 | mNeoVis.2.1.1.chr1\_hap1.g01259.t01 |
| 1146 | 1 | 198348957 | 198387547 | mNeoVis.2.1.1.chr1\_hap1.g01260.t01 |
| 1147 | 1 | 198388687 | 198559134 | mNeoVis.2.1.1.chr1\_hap1.g01261.t01 |
| 1148 | 1 | 198604775 | 198624136 | mNeoVis.2.1.1.chr1\_hap1.g01262.t01 |
| 1149 | 1 | 198703088 | 198730496 | mNeoVis.2.1.1.chr1\_hap1.g01263.t01 |
| 1150 | 1 | 198747607 | 198808319 | mNeoVis.2.1.1.chr1\_hap1.g01264.t01 |
| 1151 | 1 | 198747607 | 198808319 | mNeoVis.2.1.1.chr1\_hap1.g01264.t01 |
| 1152 | 1 | 199002700 | 199174211 | mNeoVis.2.1.1.chr1\_hap1.g01265.t01 |
| 1153 | 1 | 199175869 | 199197563 | mNeoVis.2.1.1.chr1\_hap1.g01266.t01 |
| 1154 | 1 | 199222420 | 199225704 | mNeoVis.2.1.1.chr1\_hap1.g01267.t01 |
| 1155 | 1 | 199273465 | 199277852 | mNeoVis.2.1.1.chr1\_hap1.g01268.t01 |
| 1156 | 1 | 199278355 | 199315560 | mNeoVis.2.1.1.chr1\_hap1.g01269.t01 |
| 1157 | 1 | 199319241 | 199351887 | mNeoVis.2.1.1.chr1\_hap1.g01270.t01 |
| 1158 | 1 | 199403736 | 199421479 | mNeoVis.2.1.1.chr1\_hap1.g01271.t01 |
| 1159 | 1 | 199449706 | 199472349 | mNeoVis.2.1.1.chr1\_hap1.g01272.t01 |
| 1160 | 1 | 199501840 | 199514763 | mNeoVis.2.1.1.chr1\_hap1.g01273.t01 |
| 1161 | 1 | 199560054 | 199569944 | mNeoVis.2.1.1.chr1\_hap1.g01274.t01 |
| 1162 | 1 | 199598836 | 199624036 | mNeoVis.2.1.1.chr1\_hap1.g01275.t01 |
| 1163 | 1 | 199618733 | 199724500 | mNeoVis.2.1.1.chr1\_hap1.g01276.t01 |
| 1164 | 1 | 200117594 | 200747233 | mNeoVis.2.1.1.chr1\_hap1.g01277.t01 |
| 1165 | 1 | 200117594 | 200747233 | mNeoVis.2.1.1.chr1\_hap1.g01277.t01 |
| 1166 | 1 | 200757600 | 200761203 | mNeoVis.2.1.1.chr1\_hap1.g01278.t01 |
| 1167 | 1 | 200757600 | 200761203 | mNeoVis.2.1.1.chr1\_hap1.g01278.t01 |
| 1168 | 1 | 200766021 | 200767228 | mNeoVis.2.1.1.chr1\_hap1.g01279.t01 |
| 1169 | 1 | 200766021 | 200767228 | mNeoVis.2.1.1.chr1\_hap1.g01279.t01 |
| 1170 | 1 | 206440288 | 206457798 | mNeoVis.2.1.1.chr1\_hap1.g01423.t01 |
| 1171 | 1 | 206457816 | 206464564 | mNeoVis.2.1.1.chr1\_hap1.g01424.t01 |
| 1172 | 1 | 206464714 | 206467122 | mNeoVis.2.1.1.chr1\_hap1.g01425.t01 |
| 1173 | 1 | 206467262 | 206475789 | mNeoVis.2.1.1.chr1\_hap1.g01426.t01 |
| 1174 | 1 | 206475975 | 206477279 | mNeoVis.2.1.1.chr1\_hap1.g01427.t01 |
| 1175 | 1 | 206478266 | 206485668 | mNeoVis.2.1.1.chr1\_hap1.g01428.t01 |
| 1176 | 1 | 206486380 | 206496994 | mNeoVis.2.1.1.chr1\_hap1.g01429.t01 |
| 1177 | 1 | 206497135 | 206500191 | mNeoVis.2.1.1.chr1\_hap1.g01430.t01 |
| 1178 | 1 | 206501161 | 206505621 | mNeoVis.2.1.1.chr1\_hap1.g01431.t01 |
| 1179 | 1 | 206506041 | 206512058 | mNeoVis.2.1.1.chr1\_hap1.g01432.t01 |
| 1180 | 1 | 206547340 | 206561562 | mNeoVis.2.1.1.chr1\_hap1.g01433.t01 |
| 1181 | 1 | 206561617 | 206566654 | mNeoVis.2.1.1.chr1\_hap1.g01434.t01 |
| 1182 | 1 | 206566756 | 206568278 | mNeoVis.2.1.1.chr1\_hap1.g01435.t01 |
| 1183 | 1 | 206570455 | 206602110 | mNeoVis.2.1.1.chr1\_hap1.g01436.t01 |
| 1184 | 1 | 206602954 | 206605886 | mNeoVis.2.1.1.chr1\_hap1.g01437.t01 |
| 1185 | 1 | 206698408 | 206711868 | mNeoVis.2.1.1.chr1\_hap1.g01438.t01 |
| 1186 | 1 | 206705967 | 206711176 | mNeoVis.2.1.1.chr1\_hap1.g01439.t01 |
| 1187 | 1 | 206730075 | 206795792 | mNeoVis.2.1.1.chr1\_hap1.g01440.t01 |
| 1188 | 1 | 206796787 | 206810442 | mNeoVis.2.1.1.chr1\_hap1.g01441.t01 |
| 1189 | 1 | 206818120 | 206838969 | mNeoVis.2.1.1.chr1\_hap1.g01442.t01 |
| 1190 | 1 | 206857123 | 206874478 | mNeoVis.2.1.1.chr1\_hap1.g01443.t01 |
| 1191 | 1 | 206881099 | 206886567 | mNeoVis.2.1.1.chr1\_hap1.g01444.t01 |
| 1192 | 1 | 207094688 | 207212374 | mNeoVis.2.1.1.chr1\_hap1.g01445.t01 |
| 1193 | 1 | 207277933 | 207286183 | mNeoVis.2.1.1.chr1\_hap1.g01446.t01 |
| 1194 | 1 | 207281978 | 207290018 | mNeoVis.2.1.1.chr1\_hap1.g01447.t01 |
| 1195 | 1 | 209420149 | 209539259 | mNeoVis.2.1.1.chr1\_hap1.g01479.t01 |
| 1196 | 1 | 209942384 | 209948620 | mNeoVis.2.1.1.chr1\_hap1.g01480.t01 |
| 1197 | 1 | 209958592 | 209964827 | mNeoVis.2.1.1.chr1\_hap1.g01481.t01 |
| 1198 | 1 | 209974800 | 209981066 | mNeoVis.2.1.1.chr1\_hap1.g01482.t01 |
| 1199 | 1 | 209987130 | 209988523 | mNeoVis.2.1.1.chr1\_hap1.g01483.t01 |
| 1200 | 1 | 209989586 | 209995825 | mNeoVis.2.1.1.chr1\_hap1.g01484.t01 |
| 1201 | 1 | 210987784 | 211006032 | mNeoVis.2.1.1.chr1\_hap1.g01491.t01 |
| 1202 | 1 | 211037091 | 211040095 | mNeoVis.2.1.1.chr1\_hap1.g01492.t01 |
| 1203 | 1 | 211058055 | 211094828 | mNeoVis.2.1.1.chr1\_hap1.g01493.t01 |
| 1204 | 1 | 211058076 | 211059668 | mNeoVis.2.1.1.chr1\_hap1.g01494.t01 |
| 1205 | 1 | 211100169 | 211101944 | mNeoVis.2.1.1.chr1\_hap1.g01495.t01 |
| 1206 | 1 | 212068490 | 212069792 | mNeoVis.2.1.1.chr1\_hap1.g01498.t01 |
| 1207 | 1 | 212891197 | 212913457 | mNeoVis.2.1.1.chr1\_hap1.g01499.t01 |
| 1208 | 1 | 212891197 | 212913457 | mNeoVis.2.1.1.chr1\_hap1.g01499.t01 |
| 1209 | 1 | 212891197 | 212913457 | mNeoVis.2.1.1.chr1\_hap1.g01499.t01 |
| 1210 | 1 | 213196495 | 213277586 | mNeoVis.2.1.1.chr1\_hap1.g01500.t01 |
| 1211 | 1 | 213196495 | 213277586 | mNeoVis.2.1.1.chr1\_hap1.g01500.t01 |
| 1212 | 1 | 213277740 | 213442692 | mNeoVis.2.1.1.chr1\_hap1.g01501.t01 |
| 1213 | 1 | 213277740 | 213442692 | mNeoVis.2.1.1.chr1\_hap1.g01501.t01 |
| 1214 | 1 | 213395084 | 213396064 | mNeoVis.2.1.1.chr1\_hap1.g01502.t01 |
| 1215 | 1 | 213395084 | 213396064 | mNeoVis.2.1.1.chr1\_hap1.g01502.t01 |
| 1216 | 1 | 213495122 | 213519662 | mNeoVis.2.1.1.chr1\_hap1.g01503.t01 |
| 1217 | 1 | 213525155 | 213650460 | mNeoVis.2.1.1.chr1\_hap1.g01504.t01 |
| 1218 | 1 | 213777000 | 213841383 | mNeoVis.2.1.1.chr1\_hap1.g01505.t01 |
| 1219 | 1 | 214246769 | 214383304 | mNeoVis.2.1.1.chr1\_hap1.g01507.t01 |
| 1220 | 1 | 214246769 | 214383304 | mNeoVis.2.1.1.chr1\_hap1.g01507.t01 |
| 1221 | 1 | 214809235 | 214896995 | mNeoVis.2.1.1.chr1\_hap1.g01509.t01 |
| 1222 | 1 | 218110220 | 218116278 | mNeoVis.2.1.1.chr1\_hap1.g01510.t01 |
| 1223 | 1 | 219083976 | 219084839 | mNeoVis.2.1.1.chr1\_hap1.g01512.t01 |
| 1224 | 1 | 219083976 | 219084839 | mNeoVis.2.1.1.chr1\_hap1.g01512.t01 |
| 1225 | 1 | 219764918 | 219851215 | mNeoVis.2.1.1.chr1\_hap1.g01513.t01 |
| 1226 | 1 | 219897084 | 219901927 | mNeoVis.2.1.1.chr1\_hap1.g01514.t01 |
| 1227 | 1 | 219902200 | 219905888 | mNeoVis.2.1.1.chr1\_hap1.g01515.t01 |
| 1228 | 1 | 220066920 | 220096960 | mNeoVis.2.1.1.chr1\_hap1.g01516.t01 |
| 1229 | 1 | 220386762 | 220444702 | mNeoVis.2.1.1.chr1\_hap1.g01517.t01 |
| 1230 | 1 | 220451729 | 220698143 | mNeoVis.2.1.1.chr1\_hap1.g01518.t01 |
| 1231 | 1 | 221892199 | 222116478 | mNeoVis.2.1.1.chr1\_hap1.g01520.t01 |
| 1232 | 1 | 221892199 | 222116478 | mNeoVis.2.1.1.chr1\_hap1.g01520.t01 |
| 1233 | 1 | 222080426 | 222104450 | mNeoVis.2.1.1.chr1\_hap1.g01521.t01 |
| 1234 | 1 | 222080426 | 222104450 | mNeoVis.2.1.1.chr1\_hap1.g01521.t01 |
| 1235 | 1 | 222218438 | 222566249 | mNeoVis.2.1.1.chr1\_hap1.g01522.t01 |
| 1236 | 1 | 222605918 | 222652945 | mNeoVis.2.1.1.chr1\_hap1.g01523.t01 |
| 1237 | 1 | 222673378 | 222690936 | mNeoVis.2.1.1.chr1\_hap1.g01524.t01 |
| 1238 | 1 | 222694090 | 222791833 | mNeoVis.2.1.1.chr1\_hap1.g01525.t01 |
| 1239 | 1 | 222800431 | 222830887 | mNeoVis.2.1.1.chr1\_hap1.g01526.t01 |
| 1240 | 1 | 222842036 | 222893394 | mNeoVis.2.1.1.chr1\_hap1.g01527.t01 |
| 1241 | 1 | 222893377 | 223364607 | mNeoVis.2.1.1.chr1\_hap1.g01528.t01 |
| 1242 | 1 | 223928098 | 224043658 | mNeoVis.2.1.1.chr1\_hap1.g01530.t01 |
| 1243 | 1 | 223928098 | 224043658 | mNeoVis.2.1.1.chr1\_hap1.g01530.t01 |
| 1244 | 1 | 224447646 | 224498195 | mNeoVis.2.1.1.chr1\_hap1.g01532.t01 |
| 1245 | 1 | 224527461 | 224633721 | mNeoVis.2.1.1.chr1\_hap1.g01533.t01 |
| 1246 | 1 | 224720479 | 224721086 | mNeoVis.2.1.1.chr1\_hap1.g01534.t01 |
| 1247 | 1 | 224835763 | 224917288 | mNeoVis.2.1.1.chr1\_hap1.g01535.t01 |
| 1248 | 1 | 225408649 | 225558261 | mNeoVis.2.1.1.chr1\_hap1.g01537.t01 |
| 1249 | 1 | 225763395 | 225891182 | mNeoVis.2.1.1.chr1\_hap1.g01538.t01 |
| 1250 | 1 | 225891377 | 226050910 | mNeoVis.2.1.1.chr1\_hap1.g01539.t01 |
| 1251 | 1 | 226143179 | 226191393 | mNeoVis.2.1.1.chr1\_hap1.g01540.t01 |
| 1252 | 1 | 228027134 | 228036271 | mNeoVis.2.1.1.chr1\_hap1.g01559.t01 |
| 1253 | 1 | 229218051 | 229218512 | mNeoVis.2.1.1.chr1\_hap1.g01562.t01 |
| 1254 | 1 | 229218051 | 229218512 | mNeoVis.2.1.1.chr1\_hap1.g01562.t01 |
| 1255 | 1 | 229427472 | 229432283 | mNeoVis.2.1.1.chr1\_hap1.g01563.t01 |
| 1256 | 1 | 229461558 | 229462792 | mNeoVis.2.1.1.chr1\_hap1.g01564.t01 |
| 1257 | 1 | 229554622 | 229579552 | mNeoVis.2.1.1.chr1\_hap1.g01565.t01 |
| 1258 | 1 | 229632544 | 229634192 | mNeoVis.2.1.1.chr1\_hap1.g01566.t01 |
| 1259 | 1 | 229664252 | 229692843 | mNeoVis.2.1.1.chr1\_hap1.g01567.t01 |
| 1260 | 1 | 229716564 | 229738409 | mNeoVis.2.1.1.chr1\_hap1.g01568.t01 |
| 1261 | 1 | 229809873 | 229828883 | mNeoVis.2.1.1.chr1\_hap1.g01569.t01 |
| 1262 | 1 | 230159033 | 230190423 | mNeoVis.2.1.1.chr1\_hap1.g01577.t01 |
| 1263 | 1 | 231434025 | 231614707 | mNeoVis.2.1.1.chr1\_hap1.g01578.t01 |
| 1264 | 1 | 231434025 | 231614707 | mNeoVis.2.1.1.chr1\_hap1.g01578.t01 |
| 1265 | 1 | 231673253 | 231725057 | mNeoVis.2.1.1.chr1\_hap1.g01579.t01 |
| 1266 | 1 | 231673253 | 231725057 | mNeoVis.2.1.1.chr1\_hap1.g01579.t01 |
| 1267 | 1 | 231759769 | 231790601 | mNeoVis.2.1.1.chr1\_hap1.g01580.t01 |
| 1268 | 1 | 231791288 | 231832114 | mNeoVis.2.1.1.chr1\_hap1.g01581.t01 |
| 1269 | 1 | 232262069 | 232374350 | mNeoVis.2.1.1.chr1\_hap1.g01584.t01 |
| 1270 | 1 | 232422620 | 232520650 | mNeoVis.2.1.1.chr1\_hap1.g01585.t01 |
| 1271 | 1 | 232422620 | 232520650 | mNeoVis.2.1.1.chr1\_hap1.g01585.t01 |
| 1272 | 1 | 232531723 | 232568789 | mNeoVis.2.1.1.chr1\_hap1.g01586.t01 |
| 1273 | 1 | 232531723 | 232568789 | mNeoVis.2.1.1.chr1\_hap1.g01586.t01 |
| 1274 | 1 | 232598055 | 232612123 | mNeoVis.2.1.1.chr1\_hap1.g01587.t01 |
| 1275 | 1 | 232598055 | 232612123 | mNeoVis.2.1.1.chr1\_hap1.g01587.t01 |
| 1276 | 1 | 232622842 | 232661435 | mNeoVis.2.1.1.chr1\_hap1.g01588.t01 |
| 1277 | 1 | 232622842 | 232661435 | mNeoVis.2.1.1.chr1\_hap1.g01588.t01 |
| 1278 | 1 | 232664720 | 232677853 | mNeoVis.2.1.1.chr1\_hap1.g01589.t01 |
| 1279 | 1 | 232664720 | 232677853 | mNeoVis.2.1.1.chr1\_hap1.g01589.t01 |
| 1280 | 1 | 232664720 | 232677853 | mNeoVis.2.1.1.chr1\_hap1.g01589.t01 |
| 1281 | 1 | 232687066 | 232714273 | mNeoVis.2.1.1.chr1\_hap1.g01590.t01 |
| 1282 | 1 | 232687066 | 232714273 | mNeoVis.2.1.1.chr1\_hap1.g01590.t01 |
| 1283 | 1 | 232687066 | 232714273 | mNeoVis.2.1.1.chr1\_hap1.g01590.t01 |
| 1284 | 1 | 232714548 | 232806417 | mNeoVis.2.1.1.chr1\_hap1.g01591.t01 |
| 1285 | 1 | 232714548 | 232806417 | mNeoVis.2.1.1.chr1\_hap1.g01591.t01 |
| 1286 | 1 | 232714548 | 232806417 | mNeoVis.2.1.1.chr1\_hap1.g01591.t01 |
| 1287 | 1 | 232854721 | 233124590 | mNeoVis.2.1.1.chr1\_hap1.g01592.t01 |
| 1288 | 1 | 233164788 | 233175432 | mNeoVis.2.1.1.chr1\_hap1.g01593.t01 |
| 1289 | 1 | 233181551 | 233196608 | mNeoVis.2.1.1.chr1\_hap1.g01594.t01 |
| 1290 | 1 | 233296960 | 233396529 | mNeoVis.2.1.1.chr1\_hap1.g01595.t01 |
| 1291 | 1 | 233714619 | 233923628 | mNeoVis.2.1.1.chr1\_hap1.g01596.t01 |
| 1292 | 1 | 234724464 | 234774170 | mNeoVis.2.1.1.chr1\_hap1.g01598.t01 |
| 1293 | 1 | 234780734 | 234805978 | mNeoVis.2.1.1.chr1\_hap1.g01599.t01 |
| 1294 | 1 | 234836587 | 234848992 | mNeoVis.2.1.1.chr1\_hap1.g01600.t01 |
| 1295 | 1 | 234850480 | 234865480 | mNeoVis.2.1.1.chr1\_hap1.g01601.t01 |
| 1296 | 1 | 234867757 | 234873589 | mNeoVis.2.1.1.chr1\_hap1.g01602.t01 |
| 1297 | 1 | 234879345 | 234886881 | mNeoVis.2.1.1.chr1\_hap1.g01603.t01 |
| 1298 | 1 | 234887502 | 234891031 | mNeoVis.2.1.1.chr1\_hap1.g01604.t01 |
| 1299 | 1 | 234891282 | 234897795 | mNeoVis.2.1.1.chr1\_hap1.g01605.t01 |
| 1300 | 1 | 234910421 | 234917591 | mNeoVis.2.1.1.chr1\_hap1.g01606.t01 |
| 1301 | 1 | 234924533 | 234926020 | mNeoVis.2.1.1.chr1\_hap1.g01607.t01 |
| 1302 | 1 | 234954321 | 234955887 | mNeoVis.2.1.1.chr1\_hap1.g01608.t01 |
| 1303 | 1 | 234957508 | 234975648 | mNeoVis.2.1.1.chr1\_hap1.g01609.t01 |
| 1304 | 1 | 234980548 | 235003064 | mNeoVis.2.1.1.chr1\_hap1.g01610.t01 |
| 1305 | 1 | 235013142 | 235016785 | mNeoVis.2.1.1.chr1\_hap1.g01611.t01 |
| 1306 | 1 | 235330293 | 235342392 | mNeoVis.2.1.1.chr1\_hap1.g01612.t01 |
| 1307 | 1 | 235430671 | 235447584 | mNeoVis.2.1.1.chr1\_hap1.g01613.t01 |
| 1308 | 1 | 235447983 | 235461585 | mNeoVis.2.1.1.chr1\_hap1.g01614.t01 |
| 1309 | 1 | 235462436 | 235467907 | mNeoVis.2.1.1.chr1\_hap1.g01615.t01 |
| 1310 | 1 | 235473979 | 235481603 | mNeoVis.2.1.1.chr1\_hap1.g01616.t01 |
| 1311 | 1 | 235483192 | 235518602 | mNeoVis.2.1.1.chr1\_hap1.g01617.t01 |
| 1312 | 1 | 235518007 | 235566102 | mNeoVis.2.1.1.chr1\_hap1.g01618.t01 |
| 1313 | 1 | 235566821 | 235571000 | mNeoVis.2.1.1.chr1\_hap1.g01619.t01 |
| 1314 | 1 | 235571152 | 235575341 | mNeoVis.2.1.1.chr1\_hap1.g01620.t01 |
| 1315 | 1 | 235577047 | 235580790 | mNeoVis.2.1.1.chr1\_hap1.g01621.t01 |
| 1316 | 1 | 235580902 | 235605731 | mNeoVis.2.1.1.chr1\_hap1.g01622.t01 |
| 1317 | 1 | 235619989 | 235624782 | mNeoVis.2.1.1.chr1\_hap1.g01623.t01 |
| 1318 | 1 | 235624934 | 235644419 | mNeoVis.2.1.1.chr1\_hap1.g01624.t01 |
| 1319 | 1 | 235765187 | 235767244 | mNeoVis.2.1.1.chr1\_hap1.g01625.t01 |
| 1320 | 1 | 235823087 | 235851519 | mNeoVis.2.1.1.chr1\_hap1.g01626.t01 |
| 1321 | 1 | 235870827 | 235874290 | mNeoVis.2.1.1.chr1\_hap1.g01627.t01 |
| 1322 | 1 | 235874316 | 235880514 | mNeoVis.2.1.1.chr1\_hap1.g01628.t01 |
| 1323 | 1 | 235888695 | 235929158 | mNeoVis.2.1.1.chr1\_hap1.g01629.t01 |
| 1324 | 1 | 235943670 | 235947262 | mNeoVis.2.1.1.chr1\_hap1.g01630.t01 |
| 1325 | 1 | 235947517 | 235985829 | mNeoVis.2.1.1.chr1\_hap1.g01631.t01 |
| 1326 | 1 | 235986485 | 235996781 | mNeoVis.2.1.1.chr1\_hap1.g01632.t01 |
| 1327 | 1 | 236352217 | 236370146 | mNeoVis.2.1.1.chr1\_hap1.g01648.t01 |
| 1328 | 1 | 236373736 | 236501046 | mNeoVis.2.1.1.chr1\_hap1.g01649.t01 |
| 1329 | 1 | 236613568 | 236816175 | mNeoVis.2.1.1.chr1\_hap1.g01650.t01 |
| 1330 | 1 | 236821550 | 236839969 | mNeoVis.2.1.1.chr1\_hap1.g01651.t01 |
| 1331 | 1 | 236821550 | 236839969 | mNeoVis.2.1.1.chr1\_hap1.g01651.t01 |
| 1332 | 1 | 236847438 | 236856685 | mNeoVis.2.1.1.chr1\_hap1.g01652.t01 |
| 1333 | 1 | 236847438 | 236856685 | mNeoVis.2.1.1.chr1\_hap1.g01652.t01 |
| 1334 | 1 | 236859359 | 236865341 | mNeoVis.2.1.1.chr1\_hap1.g01653.t01 |
| 1335 | 1 | 236859359 | 236865341 | mNeoVis.2.1.1.chr1\_hap1.g01653.t01 |
| 1336 | 1 | 236871354 | 236888573 | mNeoVis.2.1.1.chr1\_hap1.g01654.t01 |
| 1337 | 1 | 236888674 | 236922097 | mNeoVis.2.1.1.chr1\_hap1.g01655.t01 |
| 1338 | 1 | 236943798 | 236965799 | mNeoVis.2.1.1.chr1\_hap1.g01656.t01 |
| 1339 | 1 | 236967054 | 237121567 | mNeoVis.2.1.1.chr1\_hap1.g01657.t01 |
| 1340 | 1 | 237055057 | 237061740 | mNeoVis.2.1.1.chr1\_hap1.g01658.t01 |
| 1341 | 1 | 237062670 | 237073580 | mNeoVis.2.1.1.chr1\_hap1.g01659.t01 |
| 1342 | 1 | 237097359 | 237184036 | mNeoVis.2.1.1.chr1\_hap1.g01660.t01 |
| 1343 | 1 | 237184036 | 237185895 | mNeoVis.2.1.1.chr1\_hap1.g01661.t01 |
| 1344 | 1 | 237185956 | 237189608 | mNeoVis.2.1.1.chr1\_hap1.g01662.t01 |
| 1345 | 1 | 237186399 | 237202142 | mNeoVis.2.1.1.chr1\_hap1.g01663.t01 |
| 1346 | 1 | 237218886 | 237227703 | mNeoVis.2.1.1.chr1\_hap1.g01664.t01 |
| 1347 | 1 | 237228768 | 237281916 | mNeoVis.2.1.1.chr1\_hap1.g01665.t01 |
| 1348 | 1 | 237308952 | 237324886 | mNeoVis.2.1.1.chr1\_hap1.g01666.t01 |
| 1349 | 1 | 237683241 | 237691332 | mNeoVis.2.1.1.chr1\_hap1.g01670.t01 |
| 1350 | 1 | 237694790 | 237703924 | mNeoVis.2.1.1.chr1\_hap1.g01671.t01 |
| 1351 | 1 | 237712657 | 237717218 | mNeoVis.2.1.1.chr1\_hap1.g01672.t01 |
| 1352 | 1 | 237721386 | 237725927 | mNeoVis.2.1.1.chr1\_hap1.g01673.t01 |
| 1353 | 1 | 237735162 | 237776551 | mNeoVis.2.1.1.chr1\_hap1.g01674.t01 |
| 1354 | 1 | 237778140 | 237806832 | mNeoVis.2.1.1.chr1\_hap1.g01675.t01 |
| 1355 | 1 | 237805760 | 237812765 | mNeoVis.2.1.1.chr1\_hap1.g01676.t01 |
| 1356 | 1 | 237814734 | 237826242 | mNeoVis.2.1.1.chr1\_hap1.g01677.t01 |
| 1357 | 1 | 237836216 | 237854501 | mNeoVis.2.1.1.chr1\_hap1.g01678.t01 |
| 1358 | 1 | 239580233 | 239586620 | mNeoVis.2.1.1.chr1\_hap1.g01683.t01 |
| 1359 | 1 | 239588366 | 239602441 | mNeoVis.2.1.1.chr1\_hap1.g01684.t01 |
| 1360 | 1 | 239655468 | 239692993 | mNeoVis.2.1.1.chr1\_hap1.g01685.t01 |
| 1361 | 1 | 239655468 | 239692993 | mNeoVis.2.1.1.chr1\_hap1.g01685.t01 |
| 1362 | 1 | 239754477 | 239769767 | mNeoVis.2.1.1.chr1\_hap1.g01686.t01 |
| 1363 | 1 | 239754477 | 239769767 | mNeoVis.2.1.1.chr1\_hap1.g01686.t01 |
| 1364 | 1 | 239777436 | 239810492 | mNeoVis.2.1.1.chr1\_hap1.g01687.t01 |
| 1365 | 1 | 239777436 | 239810492 | mNeoVis.2.1.1.chr1\_hap1.g01687.t01 |
| 1366 | 1 | 240185668 | 240210451 | mNeoVis.2.1.1.chr1\_hap1.g01689.t01 |
| 1367 | 1 | 240233464 | 240647023 | mNeoVis.2.1.1.chr1\_hap1.g01690.t01 |
| 1368 | 1 | 240664060 | 240996047 | mNeoVis.2.1.1.chr1\_hap1.g01691.t01 |
| 1369 | 1 | 241096449 | 241156988 | mNeoVis.2.1.1.chr1\_hap1.g01692.t01 |
| 1370 | 1 | 241287152 | 241305080 | mNeoVis.2.1.1.chr1\_hap1.g01693.t01 |
| 1371 | 1 | 241302300 | 241355343 | mNeoVis.2.1.1.chr1\_hap1.g01694.t01 |
| 1372 | 1 | 241389758 | 241396853 | mNeoVis.2.1.1.chr1\_hap1.g01695.t01 |
| 1373 | 1 | 241411195 | 241443584 | mNeoVis.2.1.1.chr1\_hap1.g01696.t01 |
| 1374 | 1 | 241458724 | 241533912 | mNeoVis.2.1.1.chr1\_hap1.g01697.t01 |
| 1375 | 1 | 241533994 | 241574622 | mNeoVis.2.1.1.chr1\_hap1.g01698.t01 |
| 1376 | 1 | 241534031 | 241565884 | mNeoVis.2.1.1.chr1\_hap1.g01699.t01 |
| 1377 | 1 | 242566657 | 242568079 | mNeoVis.2.1.1.chr1\_hap1.g01719.t01 |
| 1378 | 1 | 242585323 | 242597051 | mNeoVis.2.1.1.chr1\_hap1.g01720.t01 |
| 1379 | 1 | 242600154 | 242628518 | mNeoVis.2.1.1.chr1\_hap1.g01721.t01 |
| 1380 | 1 | 242630515 | 242676755 | mNeoVis.2.1.1.chr1\_hap1.g01722.t01 |
| 1381 | 1 | 242685553 | 242696959 | mNeoVis.2.1.1.chr1\_hap1.g01723.t01 |
| 1382 | 1 | 242719904 | 242740400 | mNeoVis.2.1.1.chr1\_hap1.g01724.t01 |
| 1383 | 1 | 242769568 | 242832291 | mNeoVis.2.1.1.chr1\_hap1.g01725.t01 |
| 1384 | 1 | 243089071 | 243094929 | mNeoVis.2.1.1.chr1\_hap1.g01726.t01 |
| 1385 | 1 | 243109257 | 243260895 | mNeoVis.2.1.1.chr1\_hap1.g01727.t01 |
| 1386 | 1 | 243261582 | 243327591 | mNeoVis.2.1.1.chr1\_hap1.g01728.t01 |
| 1387 | 1 | 243341526 | 243381718 | mNeoVis.2.1.1.chr1\_hap1.g01729.t01 |
| 1388 | 1 | 243539879 | 243573661 | mNeoVis.2.1.1.chr1\_hap1.g01732.t01 |
| 1389 | 1 | 243581483 | 243971427 | mNeoVis.2.1.1.chr1\_hap1.g01733.t01 |
| 1390 | 1 | 244065402 | 244255045 | mNeoVis.2.1.1.chr1\_hap1.g01734.t01 |
| 1391 | 1 | 244175136 | 244185819 | mNeoVis.2.1.1.chr1\_hap1.g01735.t01 |
| 1392 | 1 | 244390028 | 244482494 | mNeoVis.2.1.1.chr1\_hap1.g01736.t01 |
| 1393 | 1 | 244433825 | 244434010 | mNeoVis.2.1.1.chr1\_hap1.g01737.t01 |
| 1394 | 1 | 244447383 | 244461089 | mNeoVis.2.1.1.chr1\_hap1.g01738.t01 |
| 1395 | 1 | 244730007 | 244896366 | mNeoVis.2.1.1.chr1\_hap1.g01740.t01 |
| 1396 | 1 | 244802030 | 244860344 | mNeoVis.2.1.1.chr1\_hap1.g01741.t01 |
| 1397 | 1 | 246314824 | 246315101 | mNeoVis.2.1.1.chr1\_hap1.g01742.t01 |
| 1398 | 1 | 246341932 | 246384872 | mNeoVis.2.1.1.chr1\_hap1.g01743.t01 |
| 1399 | 1 | 246383458 | 246618658 | mNeoVis.2.1.1.chr1\_hap1.g01744.t01 |
| 1400 | 1 | 246689245 | 246702662 | mNeoVis.2.1.1.chr1\_hap1.g01745.t01 |
| 1401 | 1 | 247580290 | 247613360 | mNeoVis.2.1.1.chr1\_hap1.g01746.t01 |
| 1402 | 1 | 247618056 | 247652429 | mNeoVis.2.1.1.chr1\_hap1.g01747.t01 |
| 1403 | 1 | 247686695 | 247735762 | mNeoVis.2.1.1.chr1\_hap1.g01748.t01 |
| 1404 | 1 | 247748600 | 247828666 | mNeoVis.2.1.1.chr1\_hap1.g01749.t01 |
| 1405 | 1 | 247877883 | 247883670 | mNeoVis.2.1.1.chr1\_hap1.g01750.t01 |
| 1406 | 1 | 247887214 | 247904792 | mNeoVis.2.1.1.chr1\_hap1.g01751.t01 |
| 1407 | 1 | 248170548 | 248610033 | mNeoVis.2.1.1.chr1\_hap1.g01753.t01 |
| 1408 | 1 | 248170548 | 248610033 | mNeoVis.2.1.1.chr1\_hap1.g01753.t01 |
| 1409 | 1 | 249579967 | 249584876 | mNeoVis.2.1.1.chr1\_hap1.g01755.t01 |
| 1410 | 1 | 249579967 | 249584876 | mNeoVis.2.1.1.chr1\_hap1.g01755.t01 |
| 1411 | 1 | 249579967 | 249584876 | mNeoVis.2.1.1.chr1\_hap1.g01755.t01 |
| 1412 | 1 | 250125022 | 250154398 | mNeoVis.2.1.1.chr1\_hap1.g01758.t01 |
| 1413 | 1 | 250547215 | 250559558 | mNeoVis.2.1.1.chr1\_hap1.g01759.t01 |
| 1414 | 1 | 250547215 | 250559558 | mNeoVis.2.1.1.chr1\_hap1.g01759.t01 |
| 1415 | 1 | 250547215 | 250559558 | mNeoVis.2.1.1.chr1\_hap1.g01759.t01 |
| 1416 | 1 | 250573633 | 250734280 | mNeoVis.2.1.1.chr1\_hap1.g01760.t01 |
| 1417 | 1 | 250573633 | 250734280 | mNeoVis.2.1.1.chr1\_hap1.g01760.t01 |
| 1418 | 1 | 250573633 | 250734280 | mNeoVis.2.1.1.chr1\_hap1.g01760.t01 |
| 1419 | 1 | 250749941 | 250769523 | mNeoVis.2.1.1.chr1\_hap1.g01761.t01 |
| 1420 | 1 | 250749941 | 250769523 | mNeoVis.2.1.1.chr1\_hap1.g01761.t01 |
| 1421 | 1 | 250749941 | 250769523 | mNeoVis.2.1.1.chr1\_hap1.g01761.t01 |
| 1422 | 1 | 250777174 | 250842265 | mNeoVis.2.1.1.chr1\_hap1.g01762.t01 |
| 1423 | 1 | 250777174 | 250842265 | mNeoVis.2.1.1.chr1\_hap1.g01762.t01 |
| 1424 | 1 | 250777174 | 250842265 | mNeoVis.2.1.1.chr1\_hap1.g01762.t01 |
| 1425 | 1 | 251055713 | 251080240 | mNeoVis.2.1.1.chr1\_hap1.g01763.t01 |
| 1426 | 1 | 251055713 | 251080240 | mNeoVis.2.1.1.chr1\_hap1.g01763.t01 |
| 1427 | 1 | 251121455 | 251130617 | mNeoVis.2.1.1.chr1\_hap1.g01764.t01 |
| 1428 | 1 | 251121455 | 251130617 | mNeoVis.2.1.1.chr1\_hap1.g01764.t01 |
| 1429 | 1 | 251508448 | 251509252 | mNeoVis.2.1.1.chr1\_hap1.g01765.t01 |
| 1430 | 1 | 251824357 | 251838265 | mNeoVis.2.1.1.chr1\_hap1.g01766.t01 |
| 1431 | 1 | 251838284 | 251927375 | mNeoVis.2.1.1.chr1\_hap1.g01767.t01 |
| 1432 | 1 | 251989368 | 252010801 | mNeoVis.2.1.1.chr1\_hap1.g01768.t01 |
| 1433 | 1 | 252012381 | 252082182 | mNeoVis.2.1.1.chr1\_hap1.g01769.t01 |
| 1434 | 1 | 252163705 | 252197737 | mNeoVis.2.1.1.chr1\_hap1.g01770.t01 |
| 1435 | 1 | 252197782 | 252246132 | mNeoVis.2.1.1.chr1\_hap1.g01771.t01 |
| 1436 | 1 | 252226530 | 252226937 | mNeoVis.2.1.1.chr1\_hap1.g01772.t01 |
| 1437 | 1 | 252315313 | 252333006 | mNeoVis.2.1.1.chr1\_hap1.g01773.t01 |
| 1438 | 1 | 252426953 | 252611832 | mNeoVis.2.1.1.chr1\_hap1.g01774.t01 |
| 1439 | 1 | 253331453 | 253490993 | mNeoVis.2.1.1.chr1\_hap1.g01776.t01 |
| 1440 | 1 | 253331453 | 253490993 | mNeoVis.2.1.1.chr1\_hap1.g01776.t01 |
| 1441 | 1 | 253626095 | 254260742 | mNeoVis.2.1.1.chr1\_hap1.g01777.t01 |
| 1442 | 1 | 254326317 | 254456496 | mNeoVis.2.1.1.chr1\_hap1.g01778.t01 |
| 1443 | 1 | 254572309 | 254597003 | mNeoVis.2.1.1.chr1\_hap1.g01779.t01 |
| 1444 | 1 | 254572309 | 254597003 | mNeoVis.2.1.1.chr1\_hap1.g01779.t01 |
| 1445 | 1 | 254734818 | 254843875 | mNeoVis.2.1.1.chr1\_hap1.g01780.t01 |
| 1446 | 1 | 254734818 | 254843875 | mNeoVis.2.1.1.chr1\_hap1.g01780.t01 |
| 1447 | 1 | 254846932 | 254902743 | mNeoVis.2.1.1.chr1\_hap1.g01781.t01 |
| 1448 | 1 | 254846932 | 254902743 | mNeoVis.2.1.1.chr1\_hap1.g01781.t01 |
| 1449 | 1 | 255001752 | 255074877 | mNeoVis.2.1.1.chr1\_hap1.g01782.t01 |
| 1450 | 1 | 255001752 | 255074877 | mNeoVis.2.1.1.chr1\_hap1.g01782.t01 |
| 1451 | 1 | 255156128 | 255306990 | mNeoVis.2.1.1.chr1\_hap1.g01783.t01 |
| 1452 | 1 | 255156128 | 255306990 | mNeoVis.2.1.1.chr1\_hap1.g01783.t01 |
| 1453 | 1 | 255156128 | 255306990 | mNeoVis.2.1.1.chr1\_hap1.g01783.t01 |
| 1454 | 1 | 255422596 | 255430620 | mNeoVis.2.1.1.chr1\_hap1.g01784.t01 |
| 1455 | 1 | 255422596 | 255430620 | mNeoVis.2.1.1.chr1\_hap1.g01784.t01 |
| 1456 | 1 | 255422596 | 255430620 | mNeoVis.2.1.1.chr1\_hap1.g01784.t01 |
| 1457 | 1 | 255436526 | 255565550 | mNeoVis.2.1.1.chr1\_hap1.g01785.t01 |
| 1458 | 1 | 255436526 | 255565550 | mNeoVis.2.1.1.chr1\_hap1.g01785.t01 |
| 1459 | 1 | 255436526 | 255565550 | mNeoVis.2.1.1.chr1\_hap1.g01785.t01 |
| 1460 | 1 | 255677559 | 255678513 | mNeoVis.2.1.1.chr1\_hap1.g01786.t01 |
| 1461 | 1 | 255677559 | 255678513 | mNeoVis.2.1.1.chr1\_hap1.g01786.t01 |
| 1462 | 1 | 255677559 | 255678513 | mNeoVis.2.1.1.chr1\_hap1.g01786.t01 |
| 1463 | 1 | 255677559 | 255678513 | mNeoVis.2.1.1.chr1\_hap1.g01786.t01 |
| 1464 | 1 | 255688251 | 255841884 | mNeoVis.2.1.1.chr1\_hap1.g01787.t01 |
| 1465 | 1 | 255688251 | 255841884 | mNeoVis.2.1.1.chr1\_hap1.g01787.t01 |
| 1466 | 1 | 255688251 | 255841884 | mNeoVis.2.1.1.chr1\_hap1.g01787.t01 |
| 1467 | 1 | 255688251 | 255841884 | mNeoVis.2.1.1.chr1\_hap1.g01787.t01 |
| 1468 | 1 | 255876697 | 255878000 | mNeoVis.2.1.1.chr1\_hap1.g01788.t01 |
| 1469 | 1 | 255876697 | 255878000 | mNeoVis.2.1.1.chr1\_hap1.g01788.t01 |
| 1470 | 1 | 255876697 | 255878000 | mNeoVis.2.1.1.chr1\_hap1.g01788.t01 |
| 1471 | 1 | 255876697 | 255878000 | mNeoVis.2.1.1.chr1\_hap1.g01788.t01 |
| 1472 | 1 | 255884199 | 255911385 | mNeoVis.2.1.1.chr1\_hap1.g01789.t01 |
| 1473 | 1 | 255884199 | 255911385 | mNeoVis.2.1.1.chr1\_hap1.g01789.t01 |
| 1474 | 1 | 255884199 | 255911385 | mNeoVis.2.1.1.chr1\_hap1.g01789.t01 |
| 1475 | 1 | 256049155 | 256060372 | mNeoVis.2.1.1.chr1\_hap1.g01790.t01 |
| 1476 | 1 | 256049155 | 256060372 | mNeoVis.2.1.1.chr1\_hap1.g01790.t01 |
| 1477 | 1 | 256049155 | 256060372 | mNeoVis.2.1.1.chr1\_hap1.g01790.t01 |
| 1478 | 1 | 256062930 | 256106596 | mNeoVis.2.1.1.chr1\_hap1.g01791.t01 |
| 1479 | 1 | 256062930 | 256106596 | mNeoVis.2.1.1.chr1\_hap1.g01791.t01 |
| 1480 | 1 | 256062930 | 256106596 | mNeoVis.2.1.1.chr1\_hap1.g01791.t01 |
| 1481 | 1 | 256121933 | 256195155 | mNeoVis.2.1.1.chr1\_hap1.g01792.t01 |
| 1482 | 1 | 256121933 | 256195155 | mNeoVis.2.1.1.chr1\_hap1.g01792.t01 |
| 1483 | 1 | 256121933 | 256195155 | mNeoVis.2.1.1.chr1\_hap1.g01792.t01 |
| 1484 | 1 | 256254299 | 256329127 | mNeoVis.2.1.1.chr1\_hap1.g01793.t01 |
| 1485 | 1 | 256254299 | 256329127 | mNeoVis.2.1.1.chr1\_hap1.g01793.t01 |
| 1486 | 1 | 256254299 | 256329127 | mNeoVis.2.1.1.chr1\_hap1.g01793.t01 |
| 1487 | 1 | 256365990 | 256369048 | mNeoVis.2.1.1.chr1\_hap1.g01794.t01 |
| 1488 | 1 | 256365990 | 256369048 | mNeoVis.2.1.1.chr1\_hap1.g01794.t01 |
| 1489 | 1 | 256365990 | 256369048 | mNeoVis.2.1.1.chr1\_hap1.g01794.t01 |
| 1490 | 1 | 256705053 | 256792286 | mNeoVis.2.1.1.chr1\_hap1.g01795.t01 |
| 1491 | 1 | 256705053 | 256792286 | mNeoVis.2.1.1.chr1\_hap1.g01795.t01 |
| 1492 | 1 | 256838792 | 256887553 | mNeoVis.2.1.1.chr1\_hap1.g01796.t01 |
| 1493 | 1 | 256838792 | 256887553 | mNeoVis.2.1.1.chr1\_hap1.g01796.t01 |
| 1494 | 1 | 256915925 | 256916971 | mNeoVis.2.1.1.chr1\_hap1.g01797.t01 |
| 1495 | 1 | 256924899 | 256930837 | mNeoVis.2.1.1.chr1\_hap1.g01798.t01 |
| 1496 | 1 | 256956240 | 256976842 | mNeoVis.2.1.1.chr1\_hap1.g01799.t01 |
| 1497 | 1 | 256985063 | 256986106 | mNeoVis.2.1.1.chr1\_hap1.g01800.t01 |
| 1498 | 1 | 257004421 | 257005464 | mNeoVis.2.1.1.chr1\_hap1.g01802.t01 |
| 1499 | 1 | 257024165 | 257025196 | mNeoVis.2.1.1.chr1\_hap1.g01803.t01 |
| 1500 | 1 | 257043834 | 257091262 | mNeoVis.2.1.1.chr1\_hap1.g01804.t01 |
| 1501 | 1 | 257043834 | 257091262 | mNeoVis.2.1.1.chr1\_hap1.g01804.t01 |
| 1502 | 1 | 257067190 | 257080416 | mNeoVis.2.1.1.chr1\_hap1.g01805.t01 |
| 1503 | 1 | 257067190 | 257080416 | mNeoVis.2.1.1.chr1\_hap1.g01805.t01 |
| 1504 | 1 | 257094293 | 257150538 | mNeoVis.2.1.1.chr1\_hap1.g01806.t01 |
| 1505 | 1 | 257094293 | 257150538 | mNeoVis.2.1.1.chr1\_hap1.g01806.t01 |
| 1506 | 1 | 257189819 | 257215281 | mNeoVis.2.1.1.chr1\_hap1.g01807.t01 |
| 1507 | 1 | 257189819 | 257215281 | mNeoVis.2.1.1.chr1\_hap1.g01807.t01 |
| 1508 | 1 | 257231396 | 257234130 | mNeoVis.2.1.1.chr1\_hap1.g01808.t01 |
| 1509 | 1 | 257231396 | 257234130 | mNeoVis.2.1.1.chr1\_hap1.g01808.t01 |
| 1510 | 1 | 257631343 | 257897947 | mNeoVis.2.1.1.chr1\_hap1.g01809.t01 |
| 1511 | 1 | 257631343 | 257897947 | mNeoVis.2.1.1.chr1\_hap1.g01809.t01 |
| 1512 | 1 | 257631343 | 257897947 | mNeoVis.2.1.1.chr1\_hap1.g01809.t01 |
| 1513 | 1 | 257631343 | 257897947 | mNeoVis.2.1.1.chr1\_hap1.g01809.t01 |
| 1514 | 1 | 258201668 | 258204752 | mNeoVis.2.1.1.chr1\_hap1.g01810.t01 |
| 1515 | 1 | 258201668 | 258204752 | mNeoVis.2.1.1.chr1\_hap1.g01810.t01 |
| 1516 | 1 | 258201668 | 258204752 | mNeoVis.2.1.1.chr1\_hap1.g01810.t01 |
| 1517 | 1 | 258252114 | 258288866 | mNeoVis.2.1.1.chr1\_hap1.g01811.t01 |
| 1518 | 1 | 258252114 | 258288866 | mNeoVis.2.1.1.chr1\_hap1.g01811.t01 |
| 1519 | 1 | 258252114 | 258288866 | mNeoVis.2.1.1.chr1\_hap1.g01811.t01 |
| 1520 | 1 | 258289724 | 258342237 | mNeoVis.2.1.1.chr1\_hap1.g01812.t01 |
| 1521 | 1 | 258289724 | 258342237 | mNeoVis.2.1.1.chr1\_hap1.g01812.t01 |
| 1522 | 1 | 258289724 | 258342237 | mNeoVis.2.1.1.chr1\_hap1.g01812.t01 |
| 1523 | 1 | 258420362 | 258528852 | mNeoVis.2.1.1.chr1\_hap1.g01813.t01 |
| 1524 | 1 | 258420362 | 258528852 | mNeoVis.2.1.1.chr1\_hap1.g01813.t01 |
| 1525 | 1 | 259019425 | 259041172 | mNeoVis.2.1.1.chr1\_hap1.g01814.t01 |
| 1526 | 1 | 259019425 | 259041172 | mNeoVis.2.1.1.chr1\_hap1.g01814.t01 |
| 1527 | 1 | 259019425 | 259041172 | mNeoVis.2.1.1.chr1\_hap1.g01814.t01 |
| 1528 | 1 | 259050222 | 259135118 | mNeoVis.2.1.1.chr1\_hap1.g01815.t01 |
| 1529 | 1 | 259050222 | 259135118 | mNeoVis.2.1.1.chr1\_hap1.g01815.t01 |
| 1530 | 1 | 259050222 | 259135118 | mNeoVis.2.1.1.chr1\_hap1.g01815.t01 |
| 1531 | 1 | 259250234 | 259284812 | mNeoVis.2.1.1.chr1\_hap1.g01816.t01 |
| 1532 | 1 | 259250234 | 259284812 | mNeoVis.2.1.1.chr1\_hap1.g01816.t01 |
| 1533 | 1 | 259340581 | 259555400 | mNeoVis.2.1.1.chr1\_hap1.g01817.t01 |
| 1534 | 1 | 260018119 | 260167998 | mNeoVis.2.1.1.chr1\_hap1.g01818.t01 |
| 1535 | 1 | 260201695 | 260214116 | mNeoVis.2.1.1.chr1\_hap1.g01819.t01 |
| 1536 | 1 | 260225124 | 260251972 | mNeoVis.2.1.1.chr1\_hap1.g01820.t01 |
| 1537 | 1 | 260296638 | 260463544 | mNeoVis.2.1.1.chr1\_hap1.g01821.t01 |
| 1538 | 1 | 260296638 | 260463544 | mNeoVis.2.1.1.chr1\_hap1.g01821.t01 |
| 1539 | 1 | 260441050 | 260446088 | mNeoVis.2.1.1.chr1\_hap1.g01822.t01 |
| 1540 | 1 | 260441050 | 260446088 | mNeoVis.2.1.1.chr1\_hap1.g01822.t01 |
| 1541 | 1 | 260469384 | 260672164 | mNeoVis.2.1.1.chr1\_hap1.g01823.t01 |
| 1542 | 1 | 260469384 | 260672164 | mNeoVis.2.1.1.chr1\_hap1.g01823.t01 |
| 1543 | 1 | 260705795 | 260796721 | mNeoVis.2.1.1.chr1\_hap1.g01824.t01 |
| 1544 | 1 | 260705795 | 260796721 | mNeoVis.2.1.1.chr1\_hap1.g01824.t01 |
| 1545 | 1 | 260803428 | 260807216 | mNeoVis.2.1.1.chr1\_hap1.g01825.t01 |
| 1546 | 1 | 260803428 | 260807216 | mNeoVis.2.1.1.chr1\_hap1.g01825.t01 |
| 1547 | 1 | 261313475 | 261314935 | mNeoVis.2.1.1.chr1\_hap1.g01829.t01 |
| 1548 | 1 | 261313475 | 261314935 | mNeoVis.2.1.1.chr1\_hap1.g01829.t01 |
| 1549 | 1 | 261639403 | 261655034 | mNeoVis.2.1.1.chr1\_hap1.g01830.t01 |
| 1550 | 1 | 261639403 | 261655034 | mNeoVis.2.1.1.chr1\_hap1.g01830.t01 |
| 1551 | 1 | 261813662 | 261826756 | mNeoVis.2.1.1.chr1\_hap1.g01831.t01 |
| 1552 | 1 | 261813662 | 261826756 | mNeoVis.2.1.1.chr1\_hap1.g01831.t01 |
| 1553 | 1 | 261874207 | 262056847 | mNeoVis.2.1.1.chr1\_hap1.g01832.t01 |
| 1554 | 1 | 262101883 | 262106498 | mNeoVis.2.1.1.chr1\_hap1.g01833.t01 |
| 1555 | 1 | 262101883 | 262106498 | mNeoVis.2.1.1.chr1\_hap1.g01833.t01 |
| 1556 | 1 | 262116844 | 262125230 | mNeoVis.2.1.1.chr1\_hap1.g01834.t01 |
| 1557 | 1 | 262116844 | 262125230 | mNeoVis.2.1.1.chr1\_hap1.g01834.t01 |
| 1558 | 1 | 262138649 | 262374173 | mNeoVis.2.1.1.chr1\_hap1.g01835.t01 |
| 1559 | 1 | 262443201 | 262456173 | mNeoVis.2.1.1.chr1\_hap1.g01836.t01 |
| 1560 | 1 | 262458101 | 262532126 | mNeoVis.2.1.1.chr1\_hap1.g01837.t01 |
| 1561 | 1 | 265247995 | 265267047 | mNeoVis.2.1.1.chr1\_hap1.g01844.t01 |
| 1562 | 1 | 265247995 | 265267047 | mNeoVis.2.1.1.chr1\_hap1.g01844.t01 |
| 1563 | 1 | 265247995 | 265267047 | mNeoVis.2.1.1.chr1\_hap1.g01844.t01 |
| 1564 | 1 | 265297767 | 265299570 | mNeoVis.2.1.1.chr1\_hap1.g01845.t01 |
| 1565 | 1 | 265297767 | 265299570 | mNeoVis.2.1.1.chr1\_hap1.g01845.t01 |
| 1566 | 1 | 265297767 | 265299570 | mNeoVis.2.1.1.chr1\_hap1.g01845.t01 |
| 1567 | 1 | 265308638 | 265374398 | mNeoVis.2.1.1.chr1\_hap1.g01846.t01 |
| 1568 | 1 | 265308638 | 265374398 | mNeoVis.2.1.1.chr1\_hap1.g01846.t01 |
| 1569 | 1 | 265308638 | 265374398 | mNeoVis.2.1.1.chr1\_hap1.g01846.t01 |
| 1570 | 1 | 265445758 | 265582392 | mNeoVis.2.1.1.chr1\_hap1.g01847.t01 |
| 1571 | 1 | 265445758 | 265582392 | mNeoVis.2.1.1.chr1\_hap1.g01847.t01 |
| 1572 | 1 | 265847931 | 266047581 | mNeoVis.2.1.1.chr1\_hap1.g01848.t01 |
| 1573 | 1 | 266129578 | 266393517 | mNeoVis.2.1.1.chr1\_hap1.g01849.t01 |
| 1574 | 1 | 266477283 | 266499650 | mNeoVis.2.1.1.chr1\_hap1.g01850.t01 |
| 1575 | 1 | 266477283 | 266499650 | mNeoVis.2.1.1.chr1\_hap1.g01850.t01 |
| 1576 | 1 | 266499718 | 266533555 | mNeoVis.2.1.1.chr1\_hap1.g01851.t01 |
| 1577 | 1 | 266538060 | 266556237 | mNeoVis.2.1.1.chr1\_hap1.g01852.t01 |
| 1578 | 1 | 266592217 | 266853853 | mNeoVis.2.1.1.chr1\_hap1.g01853.t01 |
| 1579 | 1 | 266859873 | 266874413 | mNeoVis.2.1.1.chr1\_hap1.g01854.t01 |
| 1580 | 1 | 266874536 | 266919606 | mNeoVis.2.1.1.chr1\_hap1.g01855.t01 |
| 1581 | 1 | 266921469 | 267029692 | mNeoVis.2.1.1.chr1\_hap1.g01856.t01 |
| 1582 | 1 | 267057582 | 267058323 | mNeoVis.2.1.1.chr1\_hap1.g01857.t01 |
| 1583 | 1 | 267058438 | 267215309 | mNeoVis.2.1.1.chr1\_hap1.g01858.t01 |
| 1584 | 1 | 267836832 | 267837516 | mNeoVis.2.1.1.chr1\_hap1.g01860.t01 |
| 1585 | 1 | 267836832 | 267837516 | mNeoVis.2.1.1.chr1\_hap1.g01860.t01 |
| 1586 | 1 | 267836832 | 267837516 | mNeoVis.2.1.1.chr1\_hap1.g01860.t01 |
| 1587 | 1 | 267836832 | 267837516 | mNeoVis.2.1.1.chr1\_hap1.g01860.t01 |
| 1588 | 1 | 268365994 | 268471629 | mNeoVis.2.1.1.chr1\_hap1.g01861.t01 |
| 1589 | 1 | 268365994 | 268471629 | mNeoVis.2.1.1.chr1\_hap1.g01861.t01 |
| 1590 | 1 | 268365994 | 268471629 | mNeoVis.2.1.1.chr1\_hap1.g01861.t01 |
| 1591 | 1 | 268508846 | 268526540 | mNeoVis.2.1.1.chr1\_hap1.g01862.t01 |
| 1592 | 1 | 268508846 | 268526540 | mNeoVis.2.1.1.chr1\_hap1.g01862.t01 |
| 1593 | 1 | 268575928 | 268665542 | mNeoVis.2.1.1.chr1\_hap1.g01863.t01 |
| 1594 | 1 | 268575928 | 268665542 | mNeoVis.2.1.1.chr1\_hap1.g01863.t01 |
| 1595 | 1 | 268701238 | 269116988 | mNeoVis.2.1.1.chr1\_hap1.g01864.t01 |
| 1596 | 1 | 268701238 | 269116988 | mNeoVis.2.1.1.chr1\_hap1.g01864.t01 |
| 1597 | 1 | 269851774 | 270025501 | mNeoVis.2.1.1.chr1\_hap1.g01867.t01 |
| 1598 | 1 | 270105984 | 270161775 | mNeoVis.2.1.1.chr1\_hap1.g01868.t01 |
| 1599 | 1 | 271223125 | 271517539 | mNeoVis.2.1.1.chr1\_hap1.g01870.t01 |
| 1600 | 1 | 271722439 | 271813674 | mNeoVis.2.1.1.chr1\_hap1.g01871.t01 |
| 1601 | 1 | 271834516 | 271871831 | mNeoVis.2.1.1.chr1\_hap1.g01872.t01 |
| 1602 | 1 | 271888951 | 271923869 | mNeoVis.2.1.1.chr1\_hap1.g01873.t01 |
| 1603 | 1 | 271930530 | 271952462 | mNeoVis.2.1.1.chr1\_hap1.g01874.t01 |
| 1604 | 1 | 271953728 | 271989359 | mNeoVis.2.1.1.chr1\_hap1.g01875.t01 |
| 1605 | 1 | 271995771 | 272025724 | mNeoVis.2.1.1.chr1\_hap1.g01876.t01 |
| 1606 | 1 | 272025600 | 272049047 | mNeoVis.2.1.1.chr1\_hap1.g01877.t01 |
| 1607 | 1 | 272054477 | 272068320 | mNeoVis.2.1.1.chr1\_hap1.g01878.t01 |
| 1608 | 1 | 272071091 | 272116716 | mNeoVis.2.1.1.chr1\_hap1.g01879.t01 |
| 1609 | 1 | 272123237 | 272170179 | mNeoVis.2.1.1.chr1\_hap1.g01880.t01 |
| 1610 | 1 | 272889505 | 272890144 | mNeoVis.2.1.1.chr1\_hap1.g01887.t01 |
| 1611 | 1 | 272946676 | 273096209 | mNeoVis.2.1.1.chr1\_hap1.g01888.t01 |
| 1612 | 1 | 274843181 | 274849222 | mNeoVis.2.1.1.chr1\_hap1.g01891.t01 |
| 1613 | 1 | 275016572 | 275017937 | mNeoVis.2.1.1.chr1\_hap1.g01892.t01 |
| 1614 | 1 | 275401158 | 275419383 | mNeoVis.2.1.1.chr1\_hap1.g01893.t01 |
| 1615 | 1 | 275421241 | 275578146 | mNeoVis.2.1.1.chr1\_hap1.g01894.t01 |
| 1616 | 1 | 275708314 | 275739030 | mNeoVis.2.1.1.chr1\_hap1.g01895.t01 |
| 1617 | 1 | 275967369 | 276063875 | mNeoVis.2.1.1.chr1\_hap1.g01896.t01 |
| 1618 | 1 | 276159930 | 276235565 | mNeoVis.2.1.1.chr1\_hap1.g01897.t01 |
| 1619 | 1 | 276390626 | 276421541 | mNeoVis.2.1.1.chr1\_hap1.g01898.t01 |
| 1620 | 1 | 276421377 | 276500076 | mNeoVis.2.1.1.chr1\_hap1.g01899.t01 |
| 1621 | 1 | 277944006 | 277950748 | mNeoVis.2.1.1.chr1\_hap1.g01903.t01 |
| 1622 | 1 | 277951333 | 278143245 | mNeoVis.2.1.1.chr1\_hap1.g01904.t01 |
| 1623 | 1 | 278575110 | 278580378 | mNeoVis.2.1.1.chr1\_hap1.g01906.t01 |
| 1624 | 1 | 279199093 | 279298991 | mNeoVis.2.1.1.chr1\_hap1.g01908.t01 |
| 1625 | 1 | 279299650 | 279325510 | mNeoVis.2.1.1.chr1\_hap1.g01909.t01 |
| 1626 | 1 | 279318941 | 279327998 | mNeoVis.2.1.1.chr1\_hap1.g01910.t01 |
| 1627 | 1 | 279337363 | 279521910 | mNeoVis.2.1.1.chr1\_hap1.g01911.t01 |
| 1628 | 1 | 279598857 | 279603517 | mNeoVis.2.1.1.chr1\_hap1.g01912.t01 |
| 1629 | 1 | 279598857 | 279603517 | mNeoVis.2.1.1.chr1\_hap1.g01912.t01 |
| 1630 | 1 | 279599878 | 279615180 | mNeoVis.2.1.1.chr1\_hap1.g01913.t01 |
| 1631 | 1 | 279599878 | 279615180 | mNeoVis.2.1.1.chr1\_hap1.g01913.t01 |
| 1632 | 1 | 279626910 | 279774885 | mNeoVis.2.1.1.chr1\_hap1.g01914.t01 |
| 1633 | 1 | 279626910 | 279774885 | mNeoVis.2.1.1.chr1\_hap1.g01914.t01 |
| 1634 | 1 | 279903986 | 279908356 | mNeoVis.2.1.1.chr1\_hap1.g01915.t01 |
| 1635 | 1 | 279910277 | 280002436 | mNeoVis.2.1.1.chr1\_hap1.g01916.t01 |
| 1636 | 1 | 280081358 | 280130473 | mNeoVis.2.1.1.chr1\_hap1.g01917.t01 |
| 1637 | 1 | 280669237 | 280698503 | mNeoVis.2.1.1.chr1\_hap1.g01919.t01 |
| 1638 | 1 | 280669237 | 280698503 | mNeoVis.2.1.1.chr1\_hap1.g01919.t01 |
| 1639 | 1 | 280706966 | 280750354 | mNeoVis.2.1.1.chr1\_hap1.g01920.t01 |
| 1640 | 1 | 280706966 | 280750354 | mNeoVis.2.1.1.chr1\_hap1.g01920.t01 |
| 1641 | 1 | 281135162 | 281219418 | mNeoVis.2.1.1.chr1\_hap1.g01921.t01 |
| 1642 | 1 | 281291868 | 281342083 | mNeoVis.2.1.1.chr1\_hap1.g01922.t01 |
| 1643 | 1 | 281350839 | 281434769 | mNeoVis.2.1.1.chr1\_hap1.g01923.t01 |
| 1644 | 1 | 281458083 | 281483309 | mNeoVis.2.1.1.chr1\_hap1.g01924.t01 |
| 1645 | 1 | 281541064 | 281541703 | mNeoVis.2.1.1.chr1\_hap1.g01926.t01 |
| 1646 | 1 | 282472541 | 282518237 | mNeoVis.2.1.1.chr1\_hap1.g01928.t01 |
| 1647 | 1 | 282542085 | 282631512 | mNeoVis.2.1.1.chr1\_hap1.g01929.t01 |
| 1648 | 1 | 282647629 | 282649663 | mNeoVis.2.1.1.chr1\_hap1.g01930.t01 |
| 1649 | 1 | 282668845 | 282679206 | mNeoVis.2.1.1.chr1\_hap1.g01931.t01 |
| 1650 | 1 | 282683620 | 282701328 | mNeoVis.2.1.1.chr1\_hap1.g01932.t01 |
| 1651 | 1 | 282756130 | 282790103 | mNeoVis.2.1.1.chr1\_hap1.g01933.t01 |
| 1652 | 1 | 282796757 | 282824562 | mNeoVis.2.1.1.chr1\_hap1.g01934.t01 |
| 1653 | 1 | 282825428 | 282884238 | mNeoVis.2.1.1.chr1\_hap1.g01935.t01 |
| 1654 | 1 | 282884281 | 282947553 | mNeoVis.2.1.1.chr1\_hap1.g01936.t01 |
| 1655 | 1 | 282954326 | 282972369 | mNeoVis.2.1.1.chr1\_hap1.g01937.t01 |
| 1656 | 1 | 283311692 | 283332698 | mNeoVis.2.1.1.chr1\_hap1.g01939.t01 |
| 1657 | 1 | 283418628 | 283438661 | mNeoVis.2.1.1.chr1\_hap1.g01940.t01 |
| 1658 | 1 | 283439823 | 283513740 | mNeoVis.2.1.1.chr1\_hap1.g01941.t01 |
| 1659 | 1 | 284278660 | 284294387 | mNeoVis.2.1.1.chr1\_hap1.g01945.t01 |
| 1660 | 1 | 284306206 | 284337739 | mNeoVis.2.1.1.chr1\_hap1.g01946.t01 |
| 1661 | 1 | 284363663 | 284409689 | mNeoVis.2.1.1.chr1\_hap1.g01947.t01 |
| 1662 | 1 | 284414619 | 284439647 | mNeoVis.2.1.1.chr1\_hap1.g01948.t01 |
| 1663 | 1 | 284414619 | 284439647 | mNeoVis.2.1.1.chr1\_hap1.g01948.t01 |
| 1664 | 1 | 284414619 | 284439647 | mNeoVis.2.1.1.chr1\_hap1.g01948.t01 |
| 1665 | 1 | 284414619 | 284439647 | mNeoVis.2.1.1.chr1\_hap1.g01948.t01 |
| 1666 | 1 | 284454395 | 284496192 | mNeoVis.2.1.1.chr1\_hap1.g01949.t01 |
| 1667 | 1 | 284454395 | 284496192 | mNeoVis.2.1.1.chr1\_hap1.g01949.t01 |
| 1668 | 1 | 284454395 | 284496192 | mNeoVis.2.1.1.chr1\_hap1.g01949.t01 |
| 1669 | 1 | 284454395 | 284496192 | mNeoVis.2.1.1.chr1\_hap1.g01949.t01 |
| 1670 | 1 | 284672878 | 284708091 | mNeoVis.2.1.1.chr1\_hap1.g01950.t01 |
| 1671 | 1 | 284672878 | 284708091 | mNeoVis.2.1.1.chr1\_hap1.g01950.t01 |
| 1672 | 1 | 284672878 | 284708091 | mNeoVis.2.1.1.chr1\_hap1.g01950.t01 |
| 1673 | 1 | 284672878 | 284708091 | mNeoVis.2.1.1.chr1\_hap1.g01950.t01 |
| 1674 | 1 | 284708191 | 284715447 | mNeoVis.2.1.1.chr1\_hap1.g01951.t01 |
| 1675 | 1 | 284708191 | 284715447 | mNeoVis.2.1.1.chr1\_hap1.g01951.t01 |
| 1676 | 1 | 284708191 | 284715447 | mNeoVis.2.1.1.chr1\_hap1.g01951.t01 |
| 1677 | 1 | 284708191 | 284715447 | mNeoVis.2.1.1.chr1\_hap1.g01951.t01 |
| 1678 | 1 | 284717444 | 284884761 | mNeoVis.2.1.1.chr1\_hap1.g01952.t01 |
| 1679 | 1 | 284717444 | 284884761 | mNeoVis.2.1.1.chr1\_hap1.g01952.t01 |
| 1680 | 1 | 284717444 | 284884761 | mNeoVis.2.1.1.chr1\_hap1.g01952.t01 |
| 1681 | 1 | 284717444 | 284884761 | mNeoVis.2.1.1.chr1\_hap1.g01952.t01 |
| 1682 | 1 | 284897106 | 284934202 | mNeoVis.2.1.1.chr1\_hap1.g01953.t01 |
| 1683 | 1 | 284897106 | 284934202 | mNeoVis.2.1.1.chr1\_hap1.g01953.t01 |
| 1684 | 1 | 284897106 | 284934202 | mNeoVis.2.1.1.chr1\_hap1.g01953.t01 |
| 1685 | 1 | 284897106 | 284934202 | mNeoVis.2.1.1.chr1\_hap1.g01953.t01 |
| 1686 | 1 | 284929342 | 284951955 | mNeoVis.2.1.1.chr1\_hap1.g01954.t01 |
| 1687 | 1 | 284929342 | 284951955 | mNeoVis.2.1.1.chr1\_hap1.g01954.t01 |
| 1688 | 1 | 284929342 | 284951955 | mNeoVis.2.1.1.chr1\_hap1.g01954.t01 |
| 1689 | 1 | 284929342 | 284951955 | mNeoVis.2.1.1.chr1\_hap1.g01954.t01 |
| 1690 | 1 | 284952345 | 285316203 | mNeoVis.2.1.1.chr1\_hap1.g01955.t01 |
| 1691 | 1 | 285136942 | 285138412 | mNeoVis.2.1.1.chr1\_hap1.g01956.t01 |
| 1692 | 1 | 285136942 | 285138412 | mNeoVis.2.1.1.chr1\_hap1.g01956.t01 |
| 1693 | 1 | 285136942 | 285138412 | mNeoVis.2.1.1.chr1\_hap1.g01956.t01 |
| 1694 | 1 | 285514641 | 285586297 | mNeoVis.2.1.1.chr1\_hap1.g01957.t01 |
| 1695 | 1 | 285514641 | 285586297 | mNeoVis.2.1.1.chr1\_hap1.g01957.t01 |
| 1696 | 1 | 288100297 | 288273492 | mNeoVis.2.1.1.chr1\_hap1.g01958.t01 |
| 1697 | 1 | 290199764 | 290289376 | mNeoVis.2.1.1.chr1\_hap1.g01962.t01 |
| 1698 | 1 | 291723380 | 291853840 | mNeoVis.2.1.1.chr1\_hap1.g01969.t01 |
| 1699 | 1 | 292754268 | 292811412 | mNeoVis.2.1.1.chr1\_hap1.g01971.t01 |
| 1700 | 1 | 292754268 | 292811412 | mNeoVis.2.1.1.chr1\_hap1.g01971.t01 |
| 1701 | 1 | 293367147 | 293371522 | mNeoVis.2.1.1.chr1\_hap1.g01972.t01 |
| 1702 | 1 | 293367147 | 293371522 | mNeoVis.2.1.1.chr1\_hap1.g01972.t01 |
| 1703 | 1 | 293400998 | 293480053 | mNeoVis.2.1.1.chr1\_hap1.g01973.t01 |
| 1704 | 1 | 293400998 | 293480053 | mNeoVis.2.1.1.chr1\_hap1.g01973.t01 |
| 1705 | 1 | 293750740 | 293826961 | mNeoVis.2.1.1.chr1\_hap1.g01974.t01 |
| 1706 | 1 | 293750740 | 293826961 | mNeoVis.2.1.1.chr1\_hap1.g01974.t01 |
| 1707 | 1 | 294755443 | 294827520 | mNeoVis.2.1.1.chr1\_hap1.g01983.t01 |
| 1708 | 1 | 300130492 | 300322346 | mNeoVis.2.1.1.chr1\_hap1.g01995.t01 |
| 1709 | 1 | 300323817 | 300368093 | mNeoVis.2.1.1.chr1\_hap1.g01996.t01 |
| 1710 | 1 | 300368259 | 300425923 | mNeoVis.2.1.1.chr1\_hap1.g01997.t01 |
| 1711 | 1 | 300768280 | 301030297 | mNeoVis.2.1.1.chr1\_hap1.g02001.t01 |
| 1712 | 1 | 301061500 | 301221866 | mNeoVis.2.1.1.chr1\_hap1.g02002.t01 |
| 1713 | 1 | 301259683 | 301356489 | mNeoVis.2.1.1.chr1\_hap1.g02003.t01 |
| 1714 | 1 | 301385367 | 301456382 | mNeoVis.2.1.1.chr1\_hap1.g02004.t01 |
| 1715 | 1 | 301516082 | 301551409 | mNeoVis.2.1.1.chr1\_hap1.g02005.t01 |
| 1716 | 1 | 301641103 | 301661288 | mNeoVis.2.1.1.chr1\_hap1.g02006.t01 |
| 1717 | 1 | 301678979 | 301718960 | mNeoVis.2.1.1.chr1\_hap1.g02007.t01 |
| 1718 | 1 | 301995757 | 302120801 | mNeoVis.2.1.1.chr1\_hap1.g02009.t01 |
| 1719 | 1 | 302126371 | 302153031 | mNeoVis.2.1.1.chr1\_hap1.g02010.t01 |
| 1720 | 1 | 302273803 | 302386461 | mNeoVis.2.1.1.chr1\_hap1.g02011.t01 |
| 1721 | 1 | 304594126 | 304641814 | mNeoVis.2.1.1.chr1\_hap1.g02040.t01 |
| 1722 | 1 | 304672779 | 304882998 | mNeoVis.2.1.1.chr1\_hap1.g02041.t01 |
| 1723 | 1 | 305040765 | 305055550 | mNeoVis.2.1.1.chr1\_hap1.g02042.t01 |
| 1724 | 1 | 305049651 | 305075531 | mNeoVis.2.1.1.chr1\_hap1.g02043.t01 |
| 1725 | 1 | 305075524 | 305087725 | mNeoVis.2.1.1.chr1\_hap1.g02044.t01 |
| 1726 | 1 | 305093472 | 305241228 | mNeoVis.2.1.1.chr1\_hap1.g02045.t01 |
| 1727 | 1 | 305317976 | 305319127 | mNeoVis.2.1.1.chr1\_hap1.g02046.t01 |
| 1728 | 1 | 306614498 | 306619697 | mNeoVis.2.1.1.chr1\_hap1.g02047.t01 |
| 1729 | 1 | 306684863 | 306714566 | mNeoVis.2.1.1.chr1\_hap1.g02048.t01 |
| 1730 | 1 | 306684863 | 306714566 | mNeoVis.2.1.1.chr1\_hap1.g02048.t01 |
| 1731 | 1 | 306773155 | 307037228 | mNeoVis.2.1.1.chr1\_hap1.g02049.t01 |
| 1732 | 1 | 307672190 | 307756846 | mNeoVis.2.1.1.chr1\_hap1.g02050.t01 |
| 1733 | 1 | 310992867 | 311470882 | mNeoVis.2.1.1.chr1\_hap1.g02079.t01 |
| 1734 | 1 | 311761098 | 311770945 | mNeoVis.2.1.1.chr1\_hap1.g02082.t01 |
| 1735 | 1 | 311778802 | 311791842 | mNeoVis.2.1.1.chr1\_hap1.g02083.t01 |
| 1736 | 1 | 311778802 | 311791842 | mNeoVis.2.1.1.chr1\_hap1.g02083.t01 |
| 1737 | 1 | 311798810 | 311833128 | mNeoVis.2.1.1.chr1\_hap1.g02084.t01 |
| 1738 | 1 | 311798810 | 311833128 | mNeoVis.2.1.1.chr1\_hap1.g02084.t01 |
| 1739 | 1 | 312588652 | 312642376 | mNeoVis.2.1.1.chr1\_hap1.g02085.t01 |
| 1740 | 1 | 312687663 | 312874176 | mNeoVis.2.1.1.chr1\_hap1.g02086.t01 |
| 1741 | 1 | 312915601 | 313004094 | mNeoVis.2.1.1.chr1\_hap1.g02087.t01 |
| 1742 | 1 | 312915601 | 313004094 | mNeoVis.2.1.1.chr1\_hap1.g02087.t01 |
| 1743 | 1 | 313003830 | 313006086 | mNeoVis.2.1.1.chr1\_hap1.g02088.t01 |
| 1744 | 1 | 313003830 | 313006086 | mNeoVis.2.1.1.chr1\_hap1.g02088.t01 |
| 1745 | 1 | 313019537 | 313126299 | mNeoVis.2.1.1.chr1\_hap1.g02089.t01 |
| 1746 | 1 | 313184520 | 313278511 | mNeoVis.2.1.1.chr1\_hap1.g02090.t01 |
| 1747 | 1 | 313184520 | 313278511 | mNeoVis.2.1.1.chr1\_hap1.g02090.t01 |
| 1748 | 1 | 313283874 | 313642069 | mNeoVis.2.1.1.chr1\_hap1.g02091.t01 |
| 1749 | 1 | 313283874 | 313642069 | mNeoVis.2.1.1.chr1\_hap1.g02091.t01 |
| 1750 | 1 | 313642020 | 313720766 | mNeoVis.2.1.1.chr1\_hap1.g02092.t01 |
| 1751 | 1 | 313655339 | 313665020 | mNeoVis.2.1.1.chr1\_hap1.g02093.t01 |
| 1752 | 1 | 313655339 | 313665020 | mNeoVis.2.1.1.chr1\_hap1.g02093.t01 |
| 1753 | 1 | 313771999 | 313831557 | mNeoVis.2.1.1.chr1\_hap1.g02094.t01 |
| 1754 | 1 | 313771999 | 313831557 | mNeoVis.2.1.1.chr1\_hap1.g02094.t01 |
| 1755 | 1 | 315008345 | 315453307 | mNeoVis.2.1.1.chr1\_hap1.g02095.t01 |
| 1756 | 1 | 315008345 | 315453307 | mNeoVis.2.1.1.chr1\_hap1.g02095.t01 |
| 1757 | 1 | 315519615 | 315548179 | mNeoVis.2.1.1.chr1\_hap1.g02096.t01 |
| 1758 | 1 | 315519615 | 315548179 | mNeoVis.2.1.1.chr1\_hap1.g02096.t01 |
| 1759 | 1 | 315526819 | 315528547 | mNeoVis.2.1.1.chr1\_hap1.g02097.t01 |
| 1760 | 1 | 315526819 | 315528547 | mNeoVis.2.1.1.chr1\_hap1.g02097.t01 |
| 1761 | 1 | 315593265 | 315849302 | mNeoVis.2.1.1.chr1\_hap1.g02098.t01 |
| 1762 | 1 | 315593265 | 315849302 | mNeoVis.2.1.1.chr1\_hap1.g02098.t01 |
| 1763 | 1 | 316106376 | 316202997 | mNeoVis.2.1.1.chr1\_hap1.g02100.t01 |
| 1764 | 1 | 316225936 | 316281303 | mNeoVis.2.1.1.chr1\_hap1.g02101.t01 |
| 1765 | 1 | 316281369 | 316294139 | mNeoVis.2.1.1.chr1\_hap1.g02102.t01 |
| 1766 | 1 | 316328794 | 316556849 | mNeoVis.2.1.1.chr1\_hap1.g02103.t01 |
| 1767 | 1 | 316586188 | 316657080 | mNeoVis.2.1.1.chr1\_hap1.g02104.t01 |
| 1768 | 1 | 316657791 | 316665450 | mNeoVis.2.1.1.chr1\_hap1.g02105.t01 |
| 1769 | 1 | 316672889 | 316754579 | mNeoVis.2.1.1.chr1\_hap1.g02106.t01 |
| 1770 | 1 | 316914282 | 316935228 | mNeoVis.2.1.1.chr1\_hap1.g02108.t01 |
| 1771 | 1 | 316968851 | 316977833 | mNeoVis.2.1.1.chr1\_hap1.g02109.t01 |
| 1772 | 1 | 317086030 | 317202499 | mNeoVis.2.1.1.chr1\_hap1.g02110.t01 |
| 1773 | 1 | 317283624 | 317320166 | mNeoVis.2.1.1.chr1\_hap1.g02111.t01 |
| 1774 | 1 | 317639043 | 317653141 | mNeoVis.2.1.1.chr1\_hap1.g02115.t01 |
| 1775 | 1 | 317653059 | 317664245 | mNeoVis.2.1.1.chr1\_hap1.g02116.t01 |
| 1776 | 1 | 317664401 | 317670197 | mNeoVis.2.1.1.chr1\_hap1.g02117.t01 |
| 1777 | 1 | 317671717 | 317692810 | mNeoVis.2.1.1.chr1\_hap1.g02118.t01 |
| 1778 | 1 | 317733176 | 317734157 | mNeoVis.2.1.1.chr1\_hap1.g02119.t01 |
| 1779 | 1 | 317803397 | 317899359 | mNeoVis.2.1.1.chr1\_hap1.g02120.t01 |
| 1780 | 1 | 317906569 | 317933120 | mNeoVis.2.1.1.chr1\_hap1.g02121.t01 |
| 1781 | 1 | 317968723 | 318002464 | mNeoVis.2.1.1.chr1\_hap1.g02122.t01 |
| 1782 | 1 | 317968723 | 318002464 | mNeoVis.2.1.1.chr1\_hap1.g02122.t01 |
| 1783 | 1 | 318061732 | 318153538 | mNeoVis.2.1.1.chr1\_hap1.g02123.t01 |
| 1784 | 1 | 318153679 | 318191575 | mNeoVis.2.1.1.chr1\_hap1.g02124.t01 |
| 1785 | 1 | 318153679 | 318191575 | mNeoVis.2.1.1.chr1\_hap1.g02124.t01 |
| 1786 | 1 | 318241665 | 318354828 | mNeoVis.2.1.1.chr1\_hap1.g02125.t01 |
| 1787 | 1 | 318241665 | 318354828 | mNeoVis.2.1.1.chr1\_hap1.g02125.t01 |
| 1788 | 1 | 318360960 | 318480942 | mNeoVis.2.1.1.chr1\_hap1.g02126.t01 |
| 1789 | 1 | 318360960 | 318480942 | mNeoVis.2.1.1.chr1\_hap1.g02126.t01 |
| 1790 | 1 | 319240641 | 319241203 | mNeoVis.2.1.1.chr1\_hap1.g02128.t01 |
| 1791 | 1 | 319240641 | 319241203 | mNeoVis.2.1.1.chr1\_hap1.g02128.t01 |
| 1792 | 1 | 319673828 | 319674064 | mNeoVis.2.1.1.chr1\_hap1.g02129.t01 |
| 1793 | 1 | 319673828 | 319674064 | mNeoVis.2.1.1.chr1\_hap1.g02129.t01 |
| 1794 | 1 | 324751320 | 324776276 | mNeoVis.2.1.1.chr1\_hap1.g02154.t01 |
| 1795 | 1 | 324887762 | 324997085 | mNeoVis.2.1.1.chr1\_hap1.g02155.t01 |
| 1796 | 1 | 324998120 | 325000404 | mNeoVis.2.1.1.chr1\_hap1.g02156.t01 |
| 1797 | 2 | 471784 | 531710 | mNeoVis.2.1.1.chr2\_hap1.g00024.t01 |
| 1798 | 2 | 543100 | 575510 | mNeoVis.2.1.1.chr2\_hap1.g00025.t01 |
| 1799 | 2 | 650936 | 739388 | mNeoVis.2.1.1.chr2\_hap1.g00026.t01 |
| 1800 | 2 | 756244 | 757507 | mNeoVis.2.1.1.chr2\_hap1.g00027.t01 |
| 1801 | 2 | 760241 | 916727 | mNeoVis.2.1.1.chr2\_hap1.g00028.t01 |
| 1802 | 2 | 985378 | 1008078 | mNeoVis.2.1.1.chr2\_hap1.g00029.t01 |
| 1803 | 2 | 1704130 | 1883014 | mNeoVis.2.1.1.chr2\_hap1.g00037.t01 |
| 1804 | 2 | 4337873 | 4364237 | mNeoVis.2.1.1.chr2\_hap1.g00042.t01 |
| 1805 | 2 | 4337873 | 4364237 | mNeoVis.2.1.1.chr2\_hap1.g00042.t01 |
| 1806 | 2 | 4369543 | 4521144 | mNeoVis.2.1.1.chr2\_hap1.g00043.t01 |
| 1807 | 2 | 4369543 | 4521144 | mNeoVis.2.1.1.chr2\_hap1.g00043.t01 |
| 1808 | 2 | 4534759 | 4547620 | mNeoVis.2.1.1.chr2\_hap1.g00044.t01 |
| 1809 | 2 | 4534759 | 4547620 | mNeoVis.2.1.1.chr2\_hap1.g00044.t01 |
| 1810 | 2 | 4630247 | 4631981 | mNeoVis.2.1.1.chr2\_hap1.g00045.t01 |
| 1811 | 2 | 4630247 | 4631981 | mNeoVis.2.1.1.chr2\_hap1.g00045.t01 |
| 1812 | 2 | 4681579 | 4682401 | mNeoVis.2.1.1.chr2\_hap1.g00046.t01 |
| 1813 | 2 | 4767245 | 4770791 | mNeoVis.2.1.1.chr2\_hap1.g00047.t01 |
| 1814 | 2 | 5091314 | 5150378 | mNeoVis.2.1.1.chr2\_hap1.g00049.t01 |
| 1815 | 2 | 6095003 | 6122716 | mNeoVis.2.1.1.chr2\_hap1.g00052.t01 |
| 1816 | 2 | 6189023 | 6245616 | mNeoVis.2.1.1.chr2\_hap1.g00053.t01 |
| 1817 | 2 | 6245573 | 6259881 | mNeoVis.2.1.1.chr2\_hap1.g00054.t01 |
| 1818 | 2 | 6259799 | 6292463 | mNeoVis.2.1.1.chr2\_hap1.g00055.t01 |
| 1819 | 2 | 6295866 | 6302752 | mNeoVis.2.1.1.chr2\_hap1.g00056.t01 |
| 1820 | 2 | 6304478 | 6347258 | mNeoVis.2.1.1.chr2\_hap1.g00057.t01 |
| 1821 | 2 | 6379373 | 6390281 | mNeoVis.2.1.1.chr2\_hap1.g00058.t01 |
| 1822 | 2 | 6379373 | 6390281 | mNeoVis.2.1.1.chr2\_hap1.g00058.t01 |
| 1823 | 2 | 6923701 | 6959661 | mNeoVis.2.1.1.chr2\_hap1.g00059.t01 |
| 1824 | 2 | 6960609 | 6999554 | mNeoVis.2.1.1.chr2\_hap1.g00060.t01 |
| 1825 | 2 | 7077278 | 7103781 | mNeoVis.2.1.1.chr2\_hap1.g00061.t01 |
| 1826 | 2 | 7110404 | 7182199 | mNeoVis.2.1.1.chr2\_hap1.g00062.t01 |
| 1827 | 2 | 7182196 | 7254445 | mNeoVis.2.1.1.chr2\_hap1.g00063.t01 |
| 1828 | 2 | 7258370 | 7378804 | mNeoVis.2.1.1.chr2\_hap1.g00064.t01 |
| 1829 | 2 | 7392865 | 7395437 | mNeoVis.2.1.1.chr2\_hap1.g00065.t01 |
| 1830 | 2 | 7408179 | 7429874 | mNeoVis.2.1.1.chr2\_hap1.g00066.t01 |
| 1831 | 2 | 9024313 | 9073230 | mNeoVis.2.1.1.chr2\_hap1.g00082.t01 |
| 1832 | 2 | 9094190 | 9149757 | mNeoVis.2.1.1.chr2\_hap1.g00083.t01 |
| 1833 | 2 | 9094190 | 9149757 | mNeoVis.2.1.1.chr2\_hap1.g00083.t01 |
| 1834 | 2 | 9181890 | 9223521 | mNeoVis.2.1.1.chr2\_hap1.g00084.t01 |
| 1835 | 2 | 9181890 | 9223521 | mNeoVis.2.1.1.chr2\_hap1.g00084.t01 |
| 1836 | 2 | 9231500 | 9418986 | mNeoVis.2.1.1.chr2\_hap1.g00085.t01 |
| 1837 | 2 | 9231500 | 9418986 | mNeoVis.2.1.1.chr2\_hap1.g00085.t01 |
| 1838 | 2 | 9442910 | 9455433 | mNeoVis.2.1.1.chr2\_hap1.g00086.t01 |
| 1839 | 2 | 9442910 | 9455433 | mNeoVis.2.1.1.chr2\_hap1.g00086.t01 |
| 1840 | 2 | 9458524 | 9464615 | mNeoVis.2.1.1.chr2\_hap1.g00087.t01 |
| 1841 | 2 | 9458524 | 9464615 | mNeoVis.2.1.1.chr2\_hap1.g00087.t01 |
| 1842 | 2 | 9478602 | 9648330 | mNeoVis.2.1.1.chr2\_hap1.g00088.t01 |
| 1843 | 2 | 9478602 | 9648330 | mNeoVis.2.1.1.chr2\_hap1.g00088.t01 |
| 1844 | 2 | 9760322 | 9863438 | mNeoVis.2.1.1.chr2\_hap1.g00089.t01 |
| 1845 | 2 | 9760322 | 9863438 | mNeoVis.2.1.1.chr2\_hap1.g00089.t01 |
| 1846 | 2 | 11760694 | 11769528 | mNeoVis.2.1.1.chr2\_hap1.g00100.t01 |
| 1847 | 2 | 11771563 | 11792768 | mNeoVis.2.1.1.chr2\_hap1.g00101.t01 |
| 1848 | 2 | 11794357 | 11817364 | mNeoVis.2.1.1.chr2\_hap1.g00102.t01 |
| 1849 | 2 | 11835137 | 11873425 | mNeoVis.2.1.1.chr2\_hap1.g00103.t01 |
| 1850 | 2 | 11876025 | 11878703 | mNeoVis.2.1.1.chr2\_hap1.g00104.t01 |
| 1851 | 2 | 12035845 | 12038927 | mNeoVis.2.1.1.chr2\_hap1.g00105.t01 |
| 1852 | 2 | 12066738 | 12142002 | mNeoVis.2.1.1.chr2\_hap1.g00106.t01 |
| 1853 | 2 | 12199762 | 12201247 | mNeoVis.2.1.1.chr2\_hap1.g00107.t01 |
| 1854 | 2 | 12375680 | 12376583 | mNeoVis.2.1.1.chr2\_hap1.g00108.t01 |
| 1855 | 2 | 12437312 | 12469357 | mNeoVis.2.1.1.chr2\_hap1.g00109.t01 |
| 1856 | 2 | 12496533 | 12509768 | mNeoVis.2.1.1.chr2\_hap1.g00110.t01 |
| 1857 | 2 | 12703979 | 12745936 | mNeoVis.2.1.1.chr2\_hap1.g00111.t01 |
| 1858 | 2 | 12703979 | 12745936 | mNeoVis.2.1.1.chr2\_hap1.g00111.t01 |
| 1859 | 2 | 13906157 | 13924777 | mNeoVis.2.1.1.chr2\_hap1.g00121.t01 |
| 1860 | 2 | 13931453 | 13947821 | mNeoVis.2.1.1.chr2\_hap1.g00122.t01 |
| 1861 | 2 | 13957861 | 13976637 | mNeoVis.2.1.1.chr2\_hap1.g00123.t01 |
| 1862 | 2 | 14020584 | 14056804 | mNeoVis.2.1.1.chr2\_hap1.g00124.t01 |
| 1863 | 2 | 14061108 | 14077692 | mNeoVis.2.1.1.chr2\_hap1.g00125.t01 |
| 1864 | 2 | 14101925 | 14157801 | mNeoVis.2.1.1.chr2\_hap1.g00126.t01 |
| 1865 | 2 | 14202441 | 14413517 | mNeoVis.2.1.1.chr2\_hap1.g00127.t01 |
| 1866 | 2 | 15349947 | 15401466 | mNeoVis.2.1.1.chr2\_hap1.g00129.t01 |
| 1867 | 2 | 15423819 | 15467337 | mNeoVis.2.1.1.chr2\_hap1.g00130.t01 |
| 1868 | 2 | 15521883 | 15573684 | mNeoVis.2.1.1.chr2\_hap1.g00131.t01 |
| 1869 | 2 | 15683609 | 15905411 | mNeoVis.2.1.1.chr2\_hap1.g00132.t01 |
| 1870 | 2 | 15929463 | 16027895 | mNeoVis.2.1.1.chr2\_hap1.g00133.t01 |
| 1871 | 2 | 16028362 | 16075887 | mNeoVis.2.1.1.chr2\_hap1.g00134.t01 |
| 1872 | 2 | 16081740 | 16128606 | mNeoVis.2.1.1.chr2\_hap1.g00135.t01 |
| 1873 | 2 | 16378017 | 16440473 | mNeoVis.2.1.1.chr2\_hap1.g00139.t01 |
| 1874 | 2 | 16440737 | 16463479 | mNeoVis.2.1.1.chr2\_hap1.g00140.t01 |
| 1875 | 2 | 16511854 | 16536546 | mNeoVis.2.1.1.chr2\_hap1.g00141.t01 |
| 1876 | 2 | 16511854 | 16536546 | mNeoVis.2.1.1.chr2\_hap1.g00141.t01 |
| 1877 | 2 | 16550853 | 16580032 | mNeoVis.2.1.1.chr2\_hap1.g00142.t01 |
| 1878 | 2 | 16550853 | 16580032 | mNeoVis.2.1.1.chr2\_hap1.g00142.t01 |
| 1879 | 2 | 16585474 | 16655228 | mNeoVis.2.1.1.chr2\_hap1.g00143.t01 |
| 1880 | 2 | 16585474 | 16655228 | mNeoVis.2.1.1.chr2\_hap1.g00143.t01 |
| 1881 | 2 | 16654809 | 16675388 | mNeoVis.2.1.1.chr2\_hap1.g00144.t01 |
| 1882 | 2 | 16654809 | 16675388 | mNeoVis.2.1.1.chr2\_hap1.g00144.t01 |
| 1883 | 2 | 17004790 | 17202661 | mNeoVis.2.1.1.chr2\_hap1.g00145.t01 |
| 1884 | 2 | 17659554 | 17662890 | mNeoVis.2.1.1.chr2\_hap1.g00146.t01 |
| 1885 | 2 | 17659554 | 17662890 | mNeoVis.2.1.1.chr2\_hap1.g00146.t01 |
| 1886 | 2 | 17662753 | 17679226 | mNeoVis.2.1.1.chr2\_hap1.g00147.t01 |
| 1887 | 2 | 17662753 | 17679226 | mNeoVis.2.1.1.chr2\_hap1.g00147.t01 |
| 1888 | 2 | 17680454 | 17707981 | mNeoVis.2.1.1.chr2\_hap1.g00148.t01 |
| 1889 | 2 | 17680454 | 17707981 | mNeoVis.2.1.1.chr2\_hap1.g00148.t01 |
| 1890 | 2 | 17776562 | 17795208 | mNeoVis.2.1.1.chr2\_hap1.g00149.t01 |
| 1891 | 2 | 17776562 | 17795208 | mNeoVis.2.1.1.chr2\_hap1.g00149.t01 |
| 1892 | 2 | 17835314 | 17895864 | mNeoVis.2.1.1.chr2\_hap1.g00150.t01 |
| 1893 | 2 | 17835314 | 17895864 | mNeoVis.2.1.1.chr2\_hap1.g00150.t01 |
| 1894 | 2 | 18457517 | 18478134 | mNeoVis.2.1.1.chr2\_hap1.g00151.t01 |
| 1895 | 2 | 18746687 | 18759944 | mNeoVis.2.1.1.chr2\_hap1.g00152.t01 |
| 1896 | 2 | 18746687 | 18759944 | mNeoVis.2.1.1.chr2\_hap1.g00152.t01 |
| 1897 | 2 | 18797801 | 18800718 | mNeoVis.2.1.1.chr2\_hap1.g00153.t01 |
| 1898 | 2 | 18797801 | 18800718 | mNeoVis.2.1.1.chr2\_hap1.g00153.t01 |
| 1899 | 2 | 18806133 | 18831657 | mNeoVis.2.1.1.chr2\_hap1.g00154.t01 |
| 1900 | 2 | 18806133 | 18831657 | mNeoVis.2.1.1.chr2\_hap1.g00154.t01 |
| 1901 | 2 | 18831866 | 18926736 | mNeoVis.2.1.1.chr2\_hap1.g00155.t01 |
| 1902 | 2 | 18831866 | 18926736 | mNeoVis.2.1.1.chr2\_hap1.g00155.t01 |
| 1903 | 2 | 18926054 | 18940315 | mNeoVis.2.1.1.chr2\_hap1.g00156.t01 |
| 1904 | 2 | 18926054 | 18940315 | mNeoVis.2.1.1.chr2\_hap1.g00156.t01 |
| 1905 | 2 | 18941415 | 18971252 | mNeoVis.2.1.1.chr2\_hap1.g00157.t01 |
| 1906 | 2 | 18941415 | 18971252 | mNeoVis.2.1.1.chr2\_hap1.g00157.t01 |
| 1907 | 2 | 21755397 | 21800624 | mNeoVis.2.1.1.chr2\_hap1.g00167.t01 |
| 1908 | 2 | 21755397 | 21800624 | mNeoVis.2.1.1.chr2\_hap1.g00167.t01 |
| 1909 | 2 | 23343319 | 23474887 | mNeoVis.2.1.1.chr2\_hap1.g00169.t01 |
| 1910 | 2 | 23682133 | 24266297 | mNeoVis.2.1.1.chr2\_hap1.g00170.t01 |
| 1911 | 2 | 24632590 | 24643707 | mNeoVis.2.1.1.chr2\_hap1.g00174.t01 |
| 1912 | 2 | 24657996 | 24758831 | mNeoVis.2.1.1.chr2\_hap1.g00175.t01 |
| 1913 | 2 | 24761590 | 24765675 | mNeoVis.2.1.1.chr2\_hap1.g00176.t01 |
| 1914 | 2 | 24765866 | 24851504 | mNeoVis.2.1.1.chr2\_hap1.g00177.t01 |
| 1915 | 2 | 24853467 | 24917420 | mNeoVis.2.1.1.chr2\_hap1.g00178.t01 |
| 1916 | 2 | 24925932 | 24926475 | mNeoVis.2.1.1.chr2\_hap1.g00179.t01 |
| 1917 | 2 | 24958110 | 25002403 | mNeoVis.2.1.1.chr2\_hap1.g00180.t01 |
| 1918 | 2 | 25269685 | 25352813 | mNeoVis.2.1.1.chr2\_hap1.g00182.t01 |
| 1919 | 2 | 25353381 | 25373769 | mNeoVis.2.1.1.chr2\_hap1.g00183.t01 |
| 1920 | 2 | 25353381 | 25373769 | mNeoVis.2.1.1.chr2\_hap1.g00183.t01 |
| 1921 | 2 | 25385107 | 25388242 | mNeoVis.2.1.1.chr2\_hap1.g00184.t01 |
| 1922 | 2 | 25385107 | 25388242 | mNeoVis.2.1.1.chr2\_hap1.g00184.t01 |
| 1923 | 2 | 25385107 | 25388242 | mNeoVis.2.1.1.chr2\_hap1.g00184.t01 |
| 1924 | 2 | 25385107 | 25388242 | mNeoVis.2.1.1.chr2\_hap1.g00184.t01 |
| 1925 | 2 | 25390812 | 25395540 | mNeoVis.2.1.1.chr2\_hap1.g00185.t01 |
| 1926 | 2 | 25390812 | 25395540 | mNeoVis.2.1.1.chr2\_hap1.g00185.t01 |
| 1927 | 2 | 25390812 | 25395540 | mNeoVis.2.1.1.chr2\_hap1.g00185.t01 |
| 1928 | 2 | 25390812 | 25395540 | mNeoVis.2.1.1.chr2\_hap1.g00185.t01 |
| 1929 | 2 | 25397166 | 25440333 | mNeoVis.2.1.1.chr2\_hap1.g00186.t01 |
| 1930 | 2 | 25397166 | 25440333 | mNeoVis.2.1.1.chr2\_hap1.g00186.t01 |
| 1931 | 2 | 25397166 | 25440333 | mNeoVis.2.1.1.chr2\_hap1.g00186.t01 |
| 1932 | 2 | 25397166 | 25440333 | mNeoVis.2.1.1.chr2\_hap1.g00186.t01 |
| 1933 | 2 | 25440410 | 25446517 | mNeoVis.2.1.1.chr2\_hap1.g00187.t01 |
| 1934 | 2 | 25440410 | 25446517 | mNeoVis.2.1.1.chr2\_hap1.g00187.t01 |
| 1935 | 2 | 25440410 | 25446517 | mNeoVis.2.1.1.chr2\_hap1.g00187.t01 |
| 1936 | 2 | 25440410 | 25446517 | mNeoVis.2.1.1.chr2\_hap1.g00187.t01 |
| 1937 | 2 | 25446665 | 25459778 | mNeoVis.2.1.1.chr2\_hap1.g00188.t01 |
| 1938 | 2 | 25446665 | 25459778 | mNeoVis.2.1.1.chr2\_hap1.g00188.t01 |
| 1939 | 2 | 25446665 | 25459778 | mNeoVis.2.1.1.chr2\_hap1.g00188.t01 |
| 1940 | 2 | 25446665 | 25459778 | mNeoVis.2.1.1.chr2\_hap1.g00188.t01 |
| 1941 | 2 | 25475756 | 25506415 | mNeoVis.2.1.1.chr2\_hap1.g00189.t01 |
| 1942 | 2 | 25475756 | 25506415 | mNeoVis.2.1.1.chr2\_hap1.g00189.t01 |
| 1943 | 2 | 25475756 | 25506415 | mNeoVis.2.1.1.chr2\_hap1.g00189.t01 |
| 1944 | 2 | 25475756 | 25506415 | mNeoVis.2.1.1.chr2\_hap1.g00189.t01 |
| 1945 | 2 | 25514656 | 25527883 | mNeoVis.2.1.1.chr2\_hap1.g00190.t01 |
| 1946 | 2 | 25514656 | 25527883 | mNeoVis.2.1.1.chr2\_hap1.g00190.t01 |
| 1947 | 2 | 25514656 | 25527883 | mNeoVis.2.1.1.chr2\_hap1.g00190.t01 |
| 1948 | 2 | 25514656 | 25527883 | mNeoVis.2.1.1.chr2\_hap1.g00190.t01 |
| 1949 | 2 | 25609731 | 25666076 | mNeoVis.2.1.1.chr2\_hap1.g00191.t01 |
| 1950 | 2 | 25609731 | 25666076 | mNeoVis.2.1.1.chr2\_hap1.g00191.t01 |
| 1951 | 2 | 25609731 | 25666076 | mNeoVis.2.1.1.chr2\_hap1.g00191.t01 |
| 1952 | 2 | 25609731 | 25666076 | mNeoVis.2.1.1.chr2\_hap1.g00191.t01 |
| 1953 | 2 | 25670940 | 25813912 | mNeoVis.2.1.1.chr2\_hap1.g00192.t01 |
| 1954 | 2 | 25670940 | 25813912 | mNeoVis.2.1.1.chr2\_hap1.g00192.t01 |
| 1955 | 2 | 25670940 | 25813912 | mNeoVis.2.1.1.chr2\_hap1.g00192.t01 |
| 1956 | 2 | 25670940 | 25813912 | mNeoVis.2.1.1.chr2\_hap1.g00192.t01 |
| 1957 | 2 | 25828704 | 25845803 | mNeoVis.2.1.1.chr2\_hap1.g00193.t01 |
| 1958 | 2 | 25828704 | 25845803 | mNeoVis.2.1.1.chr2\_hap1.g00193.t01 |
| 1959 | 2 | 25828704 | 25845803 | mNeoVis.2.1.1.chr2\_hap1.g00193.t01 |
| 1960 | 2 | 25828704 | 25845803 | mNeoVis.2.1.1.chr2\_hap1.g00193.t01 |
| 1961 | 2 | 25853446 | 25864116 | mNeoVis.2.1.1.chr2\_hap1.g00194.t01 |
| 1962 | 2 | 25853446 | 25864116 | mNeoVis.2.1.1.chr2\_hap1.g00194.t01 |
| 1963 | 2 | 25853446 | 25864116 | mNeoVis.2.1.1.chr2\_hap1.g00194.t01 |
| 1964 | 2 | 25853446 | 25864116 | mNeoVis.2.1.1.chr2\_hap1.g00194.t01 |
| 1965 | 2 | 25878517 | 25884317 | mNeoVis.2.1.1.chr2\_hap1.g00195.t01 |
| 1966 | 2 | 25878517 | 25884317 | mNeoVis.2.1.1.chr2\_hap1.g00195.t01 |
| 1967 | 2 | 25878517 | 25884317 | mNeoVis.2.1.1.chr2\_hap1.g00195.t01 |
| 1968 | 2 | 25878517 | 25884317 | mNeoVis.2.1.1.chr2\_hap1.g00195.t01 |
| 1969 | 2 | 25891872 | 25952625 | mNeoVis.2.1.1.chr2\_hap1.g00196.t01 |
| 1970 | 2 | 25891872 | 25952625 | mNeoVis.2.1.1.chr2\_hap1.g00196.t01 |
| 1971 | 2 | 25891872 | 25952625 | mNeoVis.2.1.1.chr2\_hap1.g00196.t01 |
| 1972 | 2 | 25891872 | 25952625 | mNeoVis.2.1.1.chr2\_hap1.g00196.t01 |
| 1973 | 2 | 25957776 | 26018422 | mNeoVis.2.1.1.chr2\_hap1.g00197.t01 |
| 1974 | 2 | 25957776 | 26018422 | mNeoVis.2.1.1.chr2\_hap1.g00197.t01 |
| 1975 | 2 | 25957776 | 26018422 | mNeoVis.2.1.1.chr2\_hap1.g00197.t01 |
| 1976 | 2 | 25983351 | 26018728 | mNeoVis.2.1.1.chr2\_hap1.g00198.t01 |
| 1977 | 2 | 25983351 | 26018728 | mNeoVis.2.1.1.chr2\_hap1.g00198.t01 |
| 1978 | 2 | 25983351 | 26018728 | mNeoVis.2.1.1.chr2\_hap1.g00198.t01 |
| 1979 | 2 | 26032842 | 26047129 | mNeoVis.2.1.1.chr2\_hap1.g00199.t01 |
| 1980 | 2 | 26032842 | 26047129 | mNeoVis.2.1.1.chr2\_hap1.g00199.t01 |
| 1981 | 2 | 26058247 | 26169475 | mNeoVis.2.1.1.chr2\_hap1.g00200.t01 |
| 1982 | 2 | 26172045 | 26191393 | mNeoVis.2.1.1.chr2\_hap1.g00201.t01 |
| 1983 | 2 | 26192639 | 26205515 | mNeoVis.2.1.1.chr2\_hap1.g00202.t01 |
| 1984 | 2 | 26207534 | 26212123 | mNeoVis.2.1.1.chr2\_hap1.g00203.t01 |
| 1985 | 2 | 26222599 | 26229549 | mNeoVis.2.1.1.chr2\_hap1.g00204.t01 |
| 1986 | 2 | 26229671 | 26231782 | mNeoVis.2.1.1.chr2\_hap1.g00205.t01 |
| 1987 | 2 | 26234410 | 26249077 | mNeoVis.2.1.1.chr2\_hap1.g00206.t01 |
| 1988 | 2 | 26249164 | 26257533 | mNeoVis.2.1.1.chr2\_hap1.g00207.t01 |
| 1989 | 2 | 27305245 | 27306718 | mNeoVis.2.1.1.chr2\_hap1.g00229.t01 |
| 1990 | 2 | 27342546 | 27347831 | mNeoVis.2.1.1.chr2\_hap1.g00230.t01 |
| 1991 | 2 | 27403967 | 27409385 | mNeoVis.2.1.1.chr2\_hap1.g00231.t01 |
| 1992 | 2 | 27431085 | 27439212 | mNeoVis.2.1.1.chr2\_hap1.g00232.t01 |
| 1993 | 2 | 27440514 | 27459207 | mNeoVis.2.1.1.chr2\_hap1.g00233.t01 |
| 1994 | 2 | 27459685 | 27470012 | mNeoVis.2.1.1.chr2\_hap1.g00234.t01 |
| 1995 | 2 | 27472173 | 27476660 | mNeoVis.2.1.1.chr2\_hap1.g00235.t01 |
| 1996 | 2 | 27477296 | 27478938 | mNeoVis.2.1.1.chr2\_hap1.g00236.t01 |
| 1997 | 2 | 27479564 | 27497345 | mNeoVis.2.1.1.chr2\_hap1.g00237.t01 |
| 1998 | 2 | 27499437 | 27547798 | mNeoVis.2.1.1.chr2\_hap1.g00238.t01 |
| 1999 | 2 | 27628838 | 27708284 | mNeoVis.2.1.1.chr2\_hap1.g00239.t01 |
| 2000 | 2 | 27874891 | 27885490 | mNeoVis.2.1.1.chr2\_hap1.g00240.t01 |
| 2001 | 2 | 27888584 | 27893221 | mNeoVis.2.1.1.chr2\_hap1.g00241.t01 |
| 2002 | 2 | 27897004 | 27927490 | mNeoVis.2.1.1.chr2\_hap1.g00242.t01 |
| 2003 | 2 | 27930603 | 27954787 | mNeoVis.2.1.1.chr2\_hap1.g00243.t01 |
| 2004 | 2 | 28008737 | 28024338 | mNeoVis.2.1.1.chr2\_hap1.g00244.t01 |
| 2005 | 2 | 28060054 | 28090149 | mNeoVis.2.1.1.chr2\_hap1.g00245.t01 |
| 2006 | 2 | 28091416 | 28098261 | mNeoVis.2.1.1.chr2\_hap1.g00246.t01 |
| 2007 | 2 | 28104650 | 28135853 | mNeoVis.2.1.1.chr2\_hap1.g00247.t01 |
| 2008 | 2 | 28138490 | 28181723 | mNeoVis.2.1.1.chr2\_hap1.g00248.t01 |
| 2009 | 2 | 28183906 | 28224400 | mNeoVis.2.1.1.chr2\_hap1.g00249.t01 |
| 2010 | 2 | 28228751 | 28272162 | mNeoVis.2.1.1.chr2\_hap1.g00250.t01 |
| 2011 | 2 | 28287390 | 28391362 | mNeoVis.2.1.1.chr2\_hap1.g00251.t01 |
| 2012 | 2 | 28530114 | 28543699 | mNeoVis.2.1.1.chr2\_hap1.g00254.t01 |
| 2013 | 2 | 28546048 | 28591853 | mNeoVis.2.1.1.chr2\_hap1.g00255.t01 |
| 2014 | 2 | 28610995 | 28621019 | mNeoVis.2.1.1.chr2\_hap1.g00256.t01 |
| 2015 | 2 | 28682815 | 28689118 | mNeoVis.2.1.1.chr2\_hap1.g00257.t01 |
| 2016 | 2 | 28777489 | 28808536 | mNeoVis.2.1.1.chr2\_hap1.g00258.t01 |
| 2017 | 2 | 28810257 | 28870924 | mNeoVis.2.1.1.chr2\_hap1.g00259.t01 |
| 2018 | 2 | 28881453 | 28882845 | mNeoVis.2.1.1.chr2\_hap1.g00260.t01 |
| 2019 | 2 | 29624347 | 29657632 | mNeoVis.2.1.1.chr2\_hap1.g00262.t01 |
| 2020 | 2 | 29649831 | 29678512 | mNeoVis.2.1.1.chr2\_hap1.g00263.t01 |
| 2021 | 2 | 29784960 | 29800598 | mNeoVis.2.1.1.chr2\_hap1.g00264.t01 |
| 2022 | 2 | 29803369 | 29896570 | mNeoVis.2.1.1.chr2\_hap1.g00265.t01 |
| 2023 | 2 | 29989137 | 30125359 | mNeoVis.2.1.1.chr2\_hap1.g00266.t01 |
| 2024 | 2 | 30138381 | 30144682 | mNeoVis.2.1.1.chr2\_hap1.g00267.t01 |
| 2025 | 2 | 30214397 | 30220225 | mNeoVis.2.1.1.chr2\_hap1.g00269.t01 |
| 2026 | 2 | 30867146 | 31000561 | mNeoVis.2.1.1.chr2\_hap1.g00287.t01 |
| 2027 | 2 | 31023738 | 31024148 | mNeoVis.2.1.1.chr2\_hap1.g00288.t01 |
| 2028 | 2 | 31100265 | 31214340 | mNeoVis.2.1.1.chr2\_hap1.g00289.t01 |
| 2029 | 2 | 31218826 | 31287744 | mNeoVis.2.1.1.chr2\_hap1.g00290.t01 |
| 2030 | 2 | 31292089 | 31407156 | mNeoVis.2.1.1.chr2\_hap1.g00291.t01 |
| 2031 | 2 | 31414378 | 31424400 | mNeoVis.2.1.1.chr2\_hap1.g00292.t01 |
| 2032 | 2 | 31414378 | 31424400 | mNeoVis.2.1.1.chr2\_hap1.g00292.t01 |
| 2033 | 2 | 31424553 | 31456312 | mNeoVis.2.1.1.chr2\_hap1.g00293.t01 |
| 2034 | 2 | 31424553 | 31456312 | mNeoVis.2.1.1.chr2\_hap1.g00293.t01 |
| 2035 | 2 | 31485419 | 31559282 | mNeoVis.2.1.1.chr2\_hap1.g00294.t01 |
| 2036 | 2 | 31485419 | 31559282 | mNeoVis.2.1.1.chr2\_hap1.g00294.t01 |
| 2037 | 2 | 31575629 | 31604710 | mNeoVis.2.1.1.chr2\_hap1.g00295.t01 |
| 2038 | 2 | 31575629 | 31604710 | mNeoVis.2.1.1.chr2\_hap1.g00295.t01 |
| 2039 | 2 | 31676939 | 31694371 | mNeoVis.2.1.1.chr2\_hap1.g00296.t01 |
| 2040 | 2 | 31676939 | 31694371 | mNeoVis.2.1.1.chr2\_hap1.g00296.t01 |
| 2041 | 2 | 31702340 | 31809711 | mNeoVis.2.1.1.chr2\_hap1.g00297.t01 |
| 2042 | 2 | 31809680 | 31862771 | mNeoVis.2.1.1.chr2\_hap1.g00298.t01 |
| 2043 | 2 | 31809680 | 31862771 | mNeoVis.2.1.1.chr2\_hap1.g00298.t01 |
| 2044 | 2 | 31866737 | 31921322 | mNeoVis.2.1.1.chr2\_hap1.g00299.t01 |
| 2045 | 2 | 31866737 | 31921322 | mNeoVis.2.1.1.chr2\_hap1.g00299.t01 |
| 2046 | 2 | 31992486 | 32012691 | mNeoVis.2.1.1.chr2\_hap1.g00300.t01 |
| 2047 | 2 | 31992486 | 32012691 | mNeoVis.2.1.1.chr2\_hap1.g00300.t01 |
| 2048 | 2 | 32017806 | 32057772 | mNeoVis.2.1.1.chr2\_hap1.g00301.t01 |
| 2049 | 2 | 32017806 | 32057772 | mNeoVis.2.1.1.chr2\_hap1.g00301.t01 |
| 2050 | 2 | 32090369 | 32318194 | mNeoVis.2.1.1.chr2\_hap1.g00302.t01 |
| 2051 | 2 | 32335555 | 32381718 | mNeoVis.2.1.1.chr2\_hap1.g00303.t01 |
| 2052 | 2 | 32688932 | 32715080 | mNeoVis.2.1.1.chr2\_hap1.g00308.t01 |
| 2053 | 2 | 32719061 | 33048183 | mNeoVis.2.1.1.chr2\_hap1.g00309.t01 |
| 2054 | 2 | 32778393 | 32780221 | mNeoVis.2.1.1.chr2\_hap1.g00310.t01 |
| 2055 | 2 | 33102889 | 33103854 | mNeoVis.2.1.1.chr2\_hap1.g00311.t01 |
| 2056 | 2 | 33102889 | 33103854 | mNeoVis.2.1.1.chr2\_hap1.g00311.t01 |
| 2057 | 2 | 33122660 | 33132662 | mNeoVis.2.1.1.chr2\_hap1.g00312.t01 |
| 2058 | 2 | 33122660 | 33132662 | mNeoVis.2.1.1.chr2\_hap1.g00312.t01 |
| 2059 | 2 | 33209017 | 33246885 | mNeoVis.2.1.1.chr2\_hap1.g00313.t01 |
| 2060 | 2 | 33209017 | 33246885 | mNeoVis.2.1.1.chr2\_hap1.g00313.t01 |
| 2061 | 2 | 33270796 | 33317554 | mNeoVis.2.1.1.chr2\_hap1.g00314.t01 |
| 2062 | 2 | 33270796 | 33317554 | mNeoVis.2.1.1.chr2\_hap1.g00314.t01 |
| 2063 | 2 | 33317693 | 33362582 | mNeoVis.2.1.1.chr2\_hap1.g00315.t01 |
| 2064 | 2 | 33317693 | 33362582 | mNeoVis.2.1.1.chr2\_hap1.g00315.t01 |
| 2065 | 2 | 33377294 | 33385908 | mNeoVis.2.1.1.chr2\_hap1.g00316.t01 |
| 2066 | 2 | 33377294 | 33385908 | mNeoVis.2.1.1.chr2\_hap1.g00316.t01 |
| 2067 | 2 | 33388850 | 33406322 | mNeoVis.2.1.1.chr2\_hap1.g00317.t01 |
| 2068 | 2 | 33388850 | 33406322 | mNeoVis.2.1.1.chr2\_hap1.g00317.t01 |
| 2069 | 2 | 33428174 | 33453095 | mNeoVis.2.1.1.chr2\_hap1.g00318.t01 |
| 2070 | 2 | 33428174 | 33453095 | mNeoVis.2.1.1.chr2\_hap1.g00318.t01 |
| 2071 | 2 | 33463210 | 33623834 | mNeoVis.2.1.1.chr2\_hap1.g00319.t01 |
| 2072 | 2 | 33616290 | 33617528 | mNeoVis.2.1.1.chr2\_hap1.g00320.t01 |
| 2073 | 2 | 34487715 | 34675582 | mNeoVis.2.1.1.chr2\_hap1.g00330.t01 |
| 2074 | 2 | 34700108 | 34797343 | mNeoVis.2.1.1.chr2\_hap1.g00331.t01 |
| 2075 | 2 | 34700108 | 34797343 | mNeoVis.2.1.1.chr2\_hap1.g00331.t01 |
| 2076 | 2 | 34807593 | 34809255 | mNeoVis.2.1.1.chr2\_hap1.g00332.t01 |
| 2077 | 2 | 34807593 | 34809255 | mNeoVis.2.1.1.chr2\_hap1.g00332.t01 |
| 2078 | 2 | 34856494 | 34925014 | mNeoVis.2.1.1.chr2\_hap1.g00333.t01 |
| 2079 | 2 | 34856494 | 34925014 | mNeoVis.2.1.1.chr2\_hap1.g00333.t01 |
| 2080 | 2 | 35043302 | 35048158 | mNeoVis.2.1.1.chr2\_hap1.g00334.t01 |
| 2081 | 2 | 35043302 | 35048158 | mNeoVis.2.1.1.chr2\_hap1.g00334.t01 |
| 2082 | 2 | 35139093 | 35213899 | mNeoVis.2.1.1.chr2\_hap1.g00335.t01 |
| 2083 | 2 | 36926496 | 36973040 | mNeoVis.2.1.1.chr2\_hap1.g00345.t01 |
| 2084 | 2 | 37173701 | 37206530 | mNeoVis.2.1.1.chr2\_hap1.g00346.t01 |
| 2085 | 2 | 37173701 | 37206530 | mNeoVis.2.1.1.chr2\_hap1.g00346.t01 |
| 2086 | 2 | 37215310 | 37269517 | mNeoVis.2.1.1.chr2\_hap1.g00347.t01 |
| 2087 | 2 | 37215310 | 37269517 | mNeoVis.2.1.1.chr2\_hap1.g00347.t01 |
| 2088 | 2 | 37267651 | 37299939 | mNeoVis.2.1.1.chr2\_hap1.g00348.t01 |
| 2089 | 2 | 37267651 | 37299939 | mNeoVis.2.1.1.chr2\_hap1.g00348.t01 |
| 2090 | 2 | 37432339 | 37440378 | mNeoVis.2.1.1.chr2\_hap1.g00349.t01 |
| 2091 | 2 | 37432339 | 37440378 | mNeoVis.2.1.1.chr2\_hap1.g00349.t01 |
| 2092 | 2 | 37452985 | 37473456 | mNeoVis.2.1.1.chr2\_hap1.g00350.t01 |
| 2093 | 2 | 37452985 | 37473456 | mNeoVis.2.1.1.chr2\_hap1.g00350.t01 |
| 2094 | 2 | 37494486 | 37511055 | mNeoVis.2.1.1.chr2\_hap1.g00351.t01 |
| 2095 | 2 | 37494486 | 37511055 | mNeoVis.2.1.1.chr2\_hap1.g00351.t01 |
| 2096 | 2 | 39272355 | 39374682 | mNeoVis.2.1.1.chr2\_hap1.g00376.t01 |
| 2097 | 2 | 39272355 | 39374682 | mNeoVis.2.1.1.chr2\_hap1.g00376.t01 |
| 2098 | 2 | 39340286 | 39340618 | mNeoVis.2.1.1.chr2\_hap1.g00377.t01 |
| 2099 | 2 | 39340286 | 39340618 | mNeoVis.2.1.1.chr2\_hap1.g00377.t01 |
| 2100 | 2 | 39377701 | 39434301 | mNeoVis.2.1.1.chr2\_hap1.g00378.t01 |
| 2101 | 2 | 39377701 | 39434301 | mNeoVis.2.1.1.chr2\_hap1.g00378.t01 |
| 2102 | 2 | 39475650 | 39498348 | mNeoVis.2.1.1.chr2\_hap1.g00379.t01 |
| 2103 | 2 | 39475650 | 39498348 | mNeoVis.2.1.1.chr2\_hap1.g00379.t01 |
| 2104 | 2 | 39499176 | 39529515 | mNeoVis.2.1.1.chr2\_hap1.g00380.t01 |
| 2105 | 2 | 39499176 | 39529515 | mNeoVis.2.1.1.chr2\_hap1.g00380.t01 |
| 2106 | 2 | 39529651 | 39655792 | mNeoVis.2.1.1.chr2\_hap1.g00381.t01 |
| 2107 | 2 | 39529651 | 39655792 | mNeoVis.2.1.1.chr2\_hap1.g00381.t01 |
| 2108 | 2 | 39697322 | 39726165 | mNeoVis.2.1.1.chr2\_hap1.g00382.t01 |
| 2109 | 2 | 39697322 | 39726165 | mNeoVis.2.1.1.chr2\_hap1.g00382.t01 |
| 2110 | 2 | 39744197 | 39755788 | mNeoVis.2.1.1.chr2\_hap1.g00383.t01 |
| 2111 | 2 | 39744197 | 39755788 | mNeoVis.2.1.1.chr2\_hap1.g00383.t01 |
| 2112 | 2 | 39763655 | 39809369 | mNeoVis.2.1.1.chr2\_hap1.g00384.t01 |
| 2113 | 2 | 39763655 | 39809369 | mNeoVis.2.1.1.chr2\_hap1.g00384.t01 |
| 2114 | 2 | 39813242 | 39815152 | mNeoVis.2.1.1.chr2\_hap1.g00385.t01 |
| 2115 | 2 | 39813242 | 39815152 | mNeoVis.2.1.1.chr2\_hap1.g00385.t01 |
| 2116 | 2 | 39852398 | 39926292 | mNeoVis.2.1.1.chr2\_hap1.g00386.t01 |
| 2117 | 2 | 39852398 | 39926292 | mNeoVis.2.1.1.chr2\_hap1.g00386.t01 |
| 2118 | 2 | 39966042 | 39994697 | mNeoVis.2.1.1.chr2\_hap1.g00387.t01 |
| 2119 | 2 | 39966042 | 39994697 | mNeoVis.2.1.1.chr2\_hap1.g00387.t01 |
| 2120 | 2 | 40025185 | 40051299 | mNeoVis.2.1.1.chr2\_hap1.g00388.t01 |
| 2121 | 2 | 40025185 | 40051299 | mNeoVis.2.1.1.chr2\_hap1.g00388.t01 |
| 2122 | 2 | 40152278 | 40187684 | mNeoVis.2.1.1.chr2\_hap1.g00389.t01 |
| 2123 | 2 | 40152278 | 40187684 | mNeoVis.2.1.1.chr2\_hap1.g00389.t01 |
| 2124 | 2 | 40203934 | 40206671 | mNeoVis.2.1.1.chr2\_hap1.g00390.t01 |
| 2125 | 2 | 40203934 | 40206671 | mNeoVis.2.1.1.chr2\_hap1.g00390.t01 |
| 2126 | 2 | 40220322 | 40318208 | mNeoVis.2.1.1.chr2\_hap1.g00391.t01 |
| 2127 | 2 | 40337669 | 40609092 | mNeoVis.2.1.1.chr2\_hap1.g00392.t01 |
| 2128 | 2 | 40337669 | 40609092 | mNeoVis.2.1.1.chr2\_hap1.g00392.t01 |
| 2129 | 2 | 40379647 | 40416875 | mNeoVis.2.1.1.chr2\_hap1.g00393.t01 |
| 2130 | 2 | 40379647 | 40416875 | mNeoVis.2.1.1.chr2\_hap1.g00393.t01 |
| 2131 | 2 | 40636500 | 40804268 | mNeoVis.2.1.1.chr2\_hap1.g00394.t01 |
| 2132 | 2 | 40815269 | 40916265 | mNeoVis.2.1.1.chr2\_hap1.g00395.t01 |
| 2133 | 2 | 40951320 | 41007288 | mNeoVis.2.1.1.chr2\_hap1.g00396.t01 |
| 2134 | 2 | 41007927 | 41031335 | mNeoVis.2.1.1.chr2\_hap1.g00397.t01 |
| 2135 | 2 | 42196983 | 42256272 | mNeoVis.2.1.1.chr2\_hap1.g00417.t01 |
| 2136 | 2 | 42259025 | 42269681 | mNeoVis.2.1.1.chr2\_hap1.g00418.t01 |
| 2137 | 2 | 42344909 | 42430596 | mNeoVis.2.1.1.chr2\_hap1.g00419.t01 |
| 2138 | 2 | 45288910 | 46347967 | mNeoVis.2.1.1.chr2\_hap1.g00427.t01 |
| 2139 | 2 | 46042355 | 46042765 | mNeoVis.2.1.1.chr2\_hap1.g00428.t01 |
| 2140 | 2 | 46042355 | 46042765 | mNeoVis.2.1.1.chr2\_hap1.g00428.t01 |
| 2141 | 2 | 47192421 | 47283309 | mNeoVis.2.1.1.chr2\_hap1.g00429.t01 |
| 2142 | 2 | 47192421 | 47283309 | mNeoVis.2.1.1.chr2\_hap1.g00429.t01 |
| 2143 | 2 | 47303794 | 47357073 | mNeoVis.2.1.1.chr2\_hap1.g00430.t01 |
| 2144 | 2 | 47303794 | 47357073 | mNeoVis.2.1.1.chr2\_hap1.g00430.t01 |
| 2145 | 2 | 47380108 | 47401068 | mNeoVis.2.1.1.chr2\_hap1.g00431.t01 |
| 2146 | 2 | 47380108 | 47401068 | mNeoVis.2.1.1.chr2\_hap1.g00431.t01 |
| 2147 | 2 | 47425790 | 47430566 | mNeoVis.2.1.1.chr2\_hap1.g00432.t01 |
| 2148 | 2 | 47425790 | 47430566 | mNeoVis.2.1.1.chr2\_hap1.g00432.t01 |
| 2149 | 2 | 47434924 | 47449353 | mNeoVis.2.1.1.chr2\_hap1.g00433.t01 |
| 2150 | 2 | 47434924 | 47449353 | mNeoVis.2.1.1.chr2\_hap1.g00433.t01 |
| 2151 | 2 | 47452586 | 47490349 | mNeoVis.2.1.1.chr2\_hap1.g00434.t01 |
| 2152 | 2 | 47452586 | 47490349 | mNeoVis.2.1.1.chr2\_hap1.g00434.t01 |
| 2153 | 2 | 47505413 | 47509923 | mNeoVis.2.1.1.chr2\_hap1.g00435.t01 |
| 2154 | 2 | 47505413 | 47509923 | mNeoVis.2.1.1.chr2\_hap1.g00435.t01 |
| 2155 | 2 | 47514683 | 47519678 | mNeoVis.2.1.1.chr2\_hap1.g00436.t01 |
| 2156 | 2 | 47514683 | 47519678 | mNeoVis.2.1.1.chr2\_hap1.g00436.t01 |
| 2157 | 2 | 47533566 | 47548651 | mNeoVis.2.1.1.chr2\_hap1.g00437.t01 |
| 2158 | 2 | 47533566 | 47548651 | mNeoVis.2.1.1.chr2\_hap1.g00437.t01 |
| 2159 | 2 | 47569169 | 47580148 | mNeoVis.2.1.1.chr2\_hap1.g00438.t01 |
| 2160 | 2 | 47569169 | 47580148 | mNeoVis.2.1.1.chr2\_hap1.g00438.t01 |
| 2161 | 2 | 47591185 | 47598646 | mNeoVis.2.1.1.chr2\_hap1.g00439.t01 |
| 2162 | 2 | 47591185 | 47598646 | mNeoVis.2.1.1.chr2\_hap1.g00439.t01 |
| 2163 | 2 | 47618781 | 47632352 | mNeoVis.2.1.1.chr2\_hap1.g00440.t01 |
| 2164 | 2 | 47618781 | 47632352 | mNeoVis.2.1.1.chr2\_hap1.g00440.t01 |
| 2165 | 2 | 47646176 | 47682907 | mNeoVis.2.1.1.chr2\_hap1.g00441.t01 |
| 2166 | 2 | 47734797 | 47795137 | mNeoVis.2.1.1.chr2\_hap1.g00442.t01 |
| 2167 | 2 | 48997081 | 49001245 | mNeoVis.2.1.1.chr2\_hap1.g00445.t01 |
| 2168 | 2 | 49005033 | 49054013 | mNeoVis.2.1.1.chr2\_hap1.g00446.t01 |
| 2169 | 2 | 49088276 | 49205649 | mNeoVis.2.1.1.chr2\_hap1.g00447.t01 |
| 2170 | 2 | 49088276 | 49205649 | mNeoVis.2.1.1.chr2\_hap1.g00447.t01 |
| 2171 | 2 | 51371695 | 51375784 | mNeoVis.2.1.1.chr2\_hap1.g00450.t01 |
| 2172 | 2 | 51490012 | 51492074 | mNeoVis.2.1.1.chr2\_hap1.g00451.t01 |
| 2173 | 2 | 51494538 | 51506489 | mNeoVis.2.1.1.chr2\_hap1.g00452.t01 |
| 2174 | 2 | 51527343 | 51578219 | mNeoVis.2.1.1.chr2\_hap1.g00453.t01 |
| 2175 | 2 | 51581759 | 51591676 | mNeoVis.2.1.1.chr2\_hap1.g00454.t01 |
| 2176 | 2 | 51610637 | 51645324 | mNeoVis.2.1.1.chr2\_hap1.g00455.t01 |
| 2177 | 2 | 51649975 | 51837480 | mNeoVis.2.1.1.chr2\_hap1.g00456.t01 |
| 2178 | 2 | 52458685 | 52480791 | mNeoVis.2.1.1.chr2\_hap1.g00458.t01 |
| 2179 | 2 | 52482060 | 52599466 | mNeoVis.2.1.1.chr2\_hap1.g00459.t01 |
| 2180 | 2 | 52669514 | 52675395 | mNeoVis.2.1.1.chr2\_hap1.g00460.t01 |
| 2181 | 2 | 52669514 | 52675395 | mNeoVis.2.1.1.chr2\_hap1.g00460.t01 |
| 2182 | 2 | 52704753 | 52755843 | mNeoVis.2.1.1.chr2\_hap1.g00461.t01 |
| 2183 | 2 | 52704753 | 52755843 | mNeoVis.2.1.1.chr2\_hap1.g00461.t01 |
| 2184 | 2 | 52762097 | 52767520 | mNeoVis.2.1.1.chr2\_hap1.g00462.t01 |
| 2185 | 2 | 52762097 | 52767520 | mNeoVis.2.1.1.chr2\_hap1.g00462.t01 |
| 2186 | 2 | 52767644 | 52781869 | mNeoVis.2.1.1.chr2\_hap1.g00463.t01 |
| 2187 | 2 | 52767644 | 52781869 | mNeoVis.2.1.1.chr2\_hap1.g00463.t01 |
| 2188 | 2 | 52783925 | 52785218 | mNeoVis.2.1.1.chr2\_hap1.g00464.t01 |
| 2189 | 2 | 52783925 | 52785218 | mNeoVis.2.1.1.chr2\_hap1.g00464.t01 |
| 2190 | 2 | 52785228 | 52788135 | mNeoVis.2.1.1.chr2\_hap1.g00465.t01 |
| 2191 | 2 | 52785228 | 52788135 | mNeoVis.2.1.1.chr2\_hap1.g00465.t01 |
| 2192 | 2 | 52788951 | 52794753 | mNeoVis.2.1.1.chr2\_hap1.g00466.t01 |
| 2193 | 2 | 52788951 | 52794753 | mNeoVis.2.1.1.chr2\_hap1.g00466.t01 |
| 2194 | 2 | 52794973 | 52818200 | mNeoVis.2.1.1.chr2\_hap1.g00467.t01 |
| 2195 | 2 | 52794973 | 52818200 | mNeoVis.2.1.1.chr2\_hap1.g00467.t01 |
| 2196 | 2 | 52825132 | 52836765 | mNeoVis.2.1.1.chr2\_hap1.g00468.t01 |
| 2197 | 2 | 52825132 | 52836765 | mNeoVis.2.1.1.chr2\_hap1.g00468.t01 |
| 2198 | 2 | 52836444 | 52844068 | mNeoVis.2.1.1.chr2\_hap1.g00469.t01 |
| 2199 | 2 | 52836444 | 52844068 | mNeoVis.2.1.1.chr2\_hap1.g00469.t01 |
| 2200 | 2 | 52851115 | 52969943 | mNeoVis.2.1.1.chr2\_hap1.g00470.t01 |
| 2201 | 2 | 52971730 | 53027069 | mNeoVis.2.1.1.chr2\_hap1.g00471.t01 |
| 2202 | 2 | 52971730 | 53027069 | mNeoVis.2.1.1.chr2\_hap1.g00471.t01 |
| 2203 | 2 | 53033960 | 53075809 | mNeoVis.2.1.1.chr2\_hap1.g00472.t01 |
| 2204 | 2 | 53033960 | 53075809 | mNeoVis.2.1.1.chr2\_hap1.g00472.t01 |
| 2205 | 2 | 53078029 | 53197454 | mNeoVis.2.1.1.chr2\_hap1.g00473.t01 |
| 2206 | 2 | 53198831 | 53203152 | mNeoVis.2.1.1.chr2\_hap1.g00474.t01 |
| 2207 | 2 | 53200578 | 53208065 | mNeoVis.2.1.1.chr2\_hap1.g00475.t01 |
| 2208 | 2 | 53209069 | 53287864 | mNeoVis.2.1.1.chr2\_hap1.g00476.t01 |
| 2209 | 2 | 53288001 | 53319509 | mNeoVis.2.1.1.chr2\_hap1.g00477.t01 |
| 2210 | 2 | 53288001 | 53319509 | mNeoVis.2.1.1.chr2\_hap1.g00477.t01 |
| 2211 | 2 | 53320531 | 53358013 | mNeoVis.2.1.1.chr2\_hap1.g00478.t01 |
| 2212 | 2 | 53320531 | 53358013 | mNeoVis.2.1.1.chr2\_hap1.g00478.t01 |
| 2213 | 2 | 53356882 | 53434707 | mNeoVis.2.1.1.chr2\_hap1.g00479.t01 |
| 2214 | 2 | 53356882 | 53434707 | mNeoVis.2.1.1.chr2\_hap1.g00479.t01 |
| 2215 | 2 | 53485120 | 53508658 | mNeoVis.2.1.1.chr2\_hap1.g00480.t01 |
| 2216 | 2 | 53485120 | 53508658 | mNeoVis.2.1.1.chr2\_hap1.g00480.t01 |
| 2217 | 2 | 53509004 | 53535495 | mNeoVis.2.1.1.chr2\_hap1.g00481.t01 |
| 2218 | 2 | 53509004 | 53535495 | mNeoVis.2.1.1.chr2\_hap1.g00481.t01 |
| 2219 | 2 | 53540490 | 53745925 | mNeoVis.2.1.1.chr2\_hap1.g00482.t01 |
| 2220 | 2 | 53540490 | 53745925 | mNeoVis.2.1.1.chr2\_hap1.g00482.t01 |
| 2221 | 2 | 53645228 | 53646643 | mNeoVis.2.1.1.chr2\_hap1.g00483.t01 |
| 2222 | 2 | 53645228 | 53646643 | mNeoVis.2.1.1.chr2\_hap1.g00483.t01 |
| 2223 | 2 | 54169872 | 54269140 | mNeoVis.2.1.1.chr2\_hap1.g00489.t01 |
| 2224 | 2 | 54233024 | 54234359 | mNeoVis.2.1.1.chr2\_hap1.g00490.t01 |
| 2225 | 2 | 54278847 | 54304829 | mNeoVis.2.1.1.chr2\_hap1.g00491.t01 |
| 2226 | 2 | 54347901 | 54380784 | mNeoVis.2.1.1.chr2\_hap1.g00492.t01 |
| 2227 | 2 | 54437567 | 54471083 | mNeoVis.2.1.1.chr2\_hap1.g00493.t01 |
| 2228 | 2 | 54437567 | 54471083 | mNeoVis.2.1.1.chr2\_hap1.g00493.t01 |
| 2229 | 2 | 54471296 | 54547873 | mNeoVis.2.1.1.chr2\_hap1.g00494.t01 |
| 2230 | 2 | 54471296 | 54547873 | mNeoVis.2.1.1.chr2\_hap1.g00494.t01 |
| 2231 | 2 | 54512929 | 54536459 | mNeoVis.2.1.1.chr2\_hap1.g00495.t01 |
| 2232 | 2 | 54512929 | 54536459 | mNeoVis.2.1.1.chr2\_hap1.g00495.t01 |
| 2233 | 2 | 54545115 | 54629048 | mNeoVis.2.1.1.chr2\_hap1.g00496.t01 |
| 2234 | 2 | 54545115 | 54629048 | mNeoVis.2.1.1.chr2\_hap1.g00496.t01 |
| 2235 | 2 | 54565984 | 54566355 | mNeoVis.2.1.1.chr2\_hap1.g00497.t01 |
| 2236 | 2 | 54565984 | 54566355 | mNeoVis.2.1.1.chr2\_hap1.g00497.t01 |
| 2237 | 2 | 54650125 | 54868757 | mNeoVis.2.1.1.chr2\_hap1.g00498.t01 |
| 2238 | 2 | 54650125 | 54868757 | mNeoVis.2.1.1.chr2\_hap1.g00498.t01 |
| 2239 | 2 | 54895143 | 54933693 | mNeoVis.2.1.1.chr2\_hap1.g00499.t01 |
| 2240 | 2 | 54895143 | 54933693 | mNeoVis.2.1.1.chr2\_hap1.g00499.t01 |
| 2241 | 2 | 54946849 | 55024783 | mNeoVis.2.1.1.chr2\_hap1.g00500.t01 |
| 2242 | 2 | 54946849 | 55024783 | mNeoVis.2.1.1.chr2\_hap1.g00500.t01 |
| 2243 | 2 | 55318055 | 55322277 | mNeoVis.2.1.1.chr2\_hap1.g00501.t01 |
| 2244 | 2 | 55550358 | 55628411 | mNeoVis.2.1.1.chr2\_hap1.g00506.t01 |
| 2245 | 2 | 55650247 | 55657257 | mNeoVis.2.1.1.chr2\_hap1.g00507.t01 |
| 2246 | 2 | 55650247 | 55657257 | mNeoVis.2.1.1.chr2\_hap1.g00507.t01 |
| 2247 | 2 | 55660429 | 55683432 | mNeoVis.2.1.1.chr2\_hap1.g00508.t01 |
| 2248 | 2 | 55660429 | 55683432 | mNeoVis.2.1.1.chr2\_hap1.g00508.t01 |
| 2249 | 2 | 55707185 | 55775966 | mNeoVis.2.1.1.chr2\_hap1.g00509.t01 |
| 2250 | 2 | 55707185 | 55775966 | mNeoVis.2.1.1.chr2\_hap1.g00509.t01 |
| 2251 | 2 | 55806154 | 55832481 | mNeoVis.2.1.1.chr2\_hap1.g00510.t01 |
| 2252 | 2 | 55806154 | 55832481 | mNeoVis.2.1.1.chr2\_hap1.g00510.t01 |
| 2253 | 2 | 55845761 | 55881666 | mNeoVis.2.1.1.chr2\_hap1.g00511.t01 |
| 2254 | 2 | 55845761 | 55881666 | mNeoVis.2.1.1.chr2\_hap1.g00511.t01 |
| 2255 | 2 | 55898639 | 55914735 | mNeoVis.2.1.1.chr2\_hap1.g00512.t01 |
| 2256 | 2 | 55898639 | 55914735 | mNeoVis.2.1.1.chr2\_hap1.g00512.t01 |
| 2257 | 2 | 55917019 | 55923586 | mNeoVis.2.1.1.chr2\_hap1.g00513.t01 |
| 2258 | 2 | 55917019 | 55923586 | mNeoVis.2.1.1.chr2\_hap1.g00513.t01 |
| 2259 | 2 | 55930990 | 55947778 | mNeoVis.2.1.1.chr2\_hap1.g00514.t01 |
| 2260 | 2 | 55930990 | 55947778 | mNeoVis.2.1.1.chr2\_hap1.g00514.t01 |
| 2261 | 2 | 55947806 | 55992320 | mNeoVis.2.1.1.chr2\_hap1.g00515.t01 |
| 2262 | 2 | 55947806 | 55992320 | mNeoVis.2.1.1.chr2\_hap1.g00515.t01 |
| 2263 | 2 | 56038601 | 56117049 | mNeoVis.2.1.1.chr2\_hap1.g00516.t01 |
| 2264 | 2 | 56038601 | 56117049 | mNeoVis.2.1.1.chr2\_hap1.g00516.t01 |
| 2265 | 2 | 56163840 | 56179517 | mNeoVis.2.1.1.chr2\_hap1.g00517.t01 |
| 2266 | 2 | 56163840 | 56179517 | mNeoVis.2.1.1.chr2\_hap1.g00517.t01 |
| 2267 | 2 | 56315617 | 56318842 | mNeoVis.2.1.1.chr2\_hap1.g00518.t01 |
| 2268 | 2 | 56358051 | 56358698 | mNeoVis.2.1.1.chr2\_hap1.g00519.t01 |
| 2269 | 2 | 56402829 | 56451616 | mNeoVis.2.1.1.chr2\_hap1.g00520.t01 |
| 2270 | 2 | 56472948 | 56486235 | mNeoVis.2.1.1.chr2\_hap1.g00521.t01 |
| 2271 | 2 | 56472948 | 56486235 | mNeoVis.2.1.1.chr2\_hap1.g00521.t01 |
| 2272 | 2 | 56487596 | 56603169 | mNeoVis.2.1.1.chr2\_hap1.g00522.t01 |
| 2273 | 2 | 56589762 | 56590696 | mNeoVis.2.1.1.chr2\_hap1.g00523.t01 |
| 2274 | 2 | 56589762 | 56590696 | mNeoVis.2.1.1.chr2\_hap1.g00523.t01 |
| 2275 | 2 | 56604467 | 56647633 | mNeoVis.2.1.1.chr2\_hap1.g00524.t01 |
| 2276 | 2 | 56604467 | 56647633 | mNeoVis.2.1.1.chr2\_hap1.g00524.t01 |
| 2277 | 2 | 56663208 | 56692194 | mNeoVis.2.1.1.chr2\_hap1.g00525.t01 |
| 2278 | 2 | 56663208 | 56692194 | mNeoVis.2.1.1.chr2\_hap1.g00525.t01 |
| 2279 | 2 | 56702843 | 56735252 | mNeoVis.2.1.1.chr2\_hap1.g00526.t01 |
| 2280 | 2 | 56702843 | 56735252 | mNeoVis.2.1.1.chr2\_hap1.g00526.t01 |
| 2281 | 2 | 56753783 | 56765372 | mNeoVis.2.1.1.chr2\_hap1.g00527.t01 |
| 2282 | 2 | 56753783 | 56765372 | mNeoVis.2.1.1.chr2\_hap1.g00527.t01 |
| 2283 | 2 | 56821134 | 56854606 | mNeoVis.2.1.1.chr2\_hap1.g00528.t01 |
| 2284 | 2 | 56821134 | 56854606 | mNeoVis.2.1.1.chr2\_hap1.g00528.t01 |
| 2285 | 2 | 56821134 | 56854606 | mNeoVis.2.1.1.chr2\_hap1.g00528.t01 |
| 2286 | 2 | 56860545 | 56887646 | mNeoVis.2.1.1.chr2\_hap1.g00529.t01 |
| 2287 | 2 | 56860545 | 56887646 | mNeoVis.2.1.1.chr2\_hap1.g00529.t01 |
| 2288 | 2 | 56860545 | 56887646 | mNeoVis.2.1.1.chr2\_hap1.g00529.t01 |
| 2289 | 2 | 56894949 | 56930821 | mNeoVis.2.1.1.chr2\_hap1.g00530.t01 |
| 2290 | 2 | 56894949 | 56930821 | mNeoVis.2.1.1.chr2\_hap1.g00530.t01 |
| 2291 | 2 | 56894949 | 56930821 | mNeoVis.2.1.1.chr2\_hap1.g00530.t01 |
| 2292 | 2 | 56935309 | 56998975 | mNeoVis.2.1.1.chr2\_hap1.g00531.t01 |
| 2293 | 2 | 56935309 | 56998975 | mNeoVis.2.1.1.chr2\_hap1.g00531.t01 |
| 2294 | 2 | 56935309 | 56998975 | mNeoVis.2.1.1.chr2\_hap1.g00531.t01 |
| 2295 | 2 | 57026342 | 57094499 | mNeoVis.2.1.1.chr2\_hap1.g00532.t01 |
| 2296 | 2 | 57026342 | 57094499 | mNeoVis.2.1.1.chr2\_hap1.g00532.t01 |
| 2297 | 2 | 57115648 | 57250766 | mNeoVis.2.1.1.chr2\_hap1.g00533.t01 |
| 2298 | 2 | 57115648 | 57250766 | mNeoVis.2.1.1.chr2\_hap1.g00533.t01 |
| 2299 | 2 | 57288406 | 57334367 | mNeoVis.2.1.1.chr2\_hap1.g00534.t01 |
| 2300 | 2 | 57288406 | 57334367 | mNeoVis.2.1.1.chr2\_hap1.g00534.t01 |
| 2301 | 2 | 57344554 | 57401875 | mNeoVis.2.1.1.chr2\_hap1.g00535.t01 |
| 2302 | 2 | 57344554 | 57401875 | mNeoVis.2.1.1.chr2\_hap1.g00535.t01 |
| 2303 | 2 | 57407018 | 57463113 | mNeoVis.2.1.1.chr2\_hap1.g00536.t01 |
| 2304 | 2 | 57407018 | 57463113 | mNeoVis.2.1.1.chr2\_hap1.g00536.t01 |
| 2305 | 2 | 57463428 | 57475054 | mNeoVis.2.1.1.chr2\_hap1.g00537.t01 |
| 2306 | 2 | 57463428 | 57475054 | mNeoVis.2.1.1.chr2\_hap1.g00537.t01 |
| 2307 | 2 | 57463428 | 57475054 | mNeoVis.2.1.1.chr2\_hap1.g00537.t01 |
| 2308 | 2 | 57494400 | 57516970 | mNeoVis.2.1.1.chr2\_hap1.g00538.t01 |
| 2309 | 2 | 57494400 | 57516970 | mNeoVis.2.1.1.chr2\_hap1.g00538.t01 |
| 2310 | 2 | 57494400 | 57516970 | mNeoVis.2.1.1.chr2\_hap1.g00538.t01 |
| 2311 | 2 | 57550171 | 57551035 | mNeoVis.2.1.1.chr2\_hap1.g00539.t01 |
| 2312 | 2 | 57550171 | 57551035 | mNeoVis.2.1.1.chr2\_hap1.g00539.t01 |
| 2313 | 2 | 57550171 | 57551035 | mNeoVis.2.1.1.chr2\_hap1.g00539.t01 |
| 2314 | 2 | 57552201 | 57690885 | mNeoVis.2.1.1.chr2\_hap1.g00540.t01 |
| 2315 | 2 | 57552201 | 57690885 | mNeoVis.2.1.1.chr2\_hap1.g00540.t01 |
| 2316 | 2 | 57552201 | 57690885 | mNeoVis.2.1.1.chr2\_hap1.g00540.t01 |
| 2317 | 2 | 57690880 | 57806158 | mNeoVis.2.1.1.chr2\_hap1.g00541.t01 |
| 2318 | 2 | 57690880 | 57806158 | mNeoVis.2.1.1.chr2\_hap1.g00541.t01 |
| 2319 | 2 | 57810249 | 57840188 | mNeoVis.2.1.1.chr2\_hap1.g00542.t01 |
| 2320 | 2 | 58682994 | 58854119 | mNeoVis.2.1.1.chr2\_hap1.g00546.t01 |
| 2321 | 2 | 59548636 | 59549515 | mNeoVis.2.1.1.chr2\_hap1.g00547.t01 |
| 2322 | 2 | 59548636 | 59549515 | mNeoVis.2.1.1.chr2\_hap1.g00547.t01 |
| 2323 | 2 | 61924266 | 62023388 | mNeoVis.2.1.1.chr2\_hap1.g00549.t01 |
| 2324 | 2 | 62077300 | 62338989 | mNeoVis.2.1.1.chr2\_hap1.g00550.t01 |
| 2325 | 2 | 62355119 | 62390852 | mNeoVis.2.1.1.chr2\_hap1.g00551.t01 |
| 2326 | 2 | 62646112 | 62652993 | mNeoVis.2.1.1.chr2\_hap1.g00552.t01 |
| 2327 | 2 | 62655726 | 62661206 | mNeoVis.2.1.1.chr2\_hap1.g00553.t01 |
| 2328 | 2 | 63271084 | 63317091 | mNeoVis.2.1.1.chr2\_hap1.g00556.t01 |
| 2329 | 2 | 63385031 | 63448372 | mNeoVis.2.1.1.chr2\_hap1.g00557.t01 |
| 2330 | 2 | 63577881 | 63695388 | mNeoVis.2.1.1.chr2\_hap1.g00558.t01 |
| 2331 | 2 | 63898476 | 63953717 | mNeoVis.2.1.1.chr2\_hap1.g00559.t01 |
| 2332 | 2 | 64333052 | 64454170 | mNeoVis.2.1.1.chr2\_hap1.g00560.t01 |
| 2333 | 2 | 64333052 | 64454170 | mNeoVis.2.1.1.chr2\_hap1.g00560.t01 |
| 2334 | 2 | 64528496 | 64544519 | mNeoVis.2.1.1.chr2\_hap1.g00561.t01 |
| 2335 | 2 | 64946334 | 65279172 | mNeoVis.2.1.1.chr2\_hap1.g00562.t01 |
| 2336 | 2 | 64946334 | 65279172 | mNeoVis.2.1.1.chr2\_hap1.g00562.t01 |
| 2337 | 2 | 65379529 | 65487358 | mNeoVis.2.1.1.chr2\_hap1.g00563.t01 |
| 2338 | 2 | 65379529 | 65487358 | mNeoVis.2.1.1.chr2\_hap1.g00563.t01 |
| 2339 | 2 | 65379529 | 65487358 | mNeoVis.2.1.1.chr2\_hap1.g00563.t01 |
| 2340 | 2 | 65500707 | 65522216 | mNeoVis.2.1.1.chr2\_hap1.g00564.t01 |
| 2341 | 2 | 65500707 | 65522216 | mNeoVis.2.1.1.chr2\_hap1.g00564.t01 |
| 2342 | 2 | 65500707 | 65522216 | mNeoVis.2.1.1.chr2\_hap1.g00564.t01 |
| 2343 | 2 | 65563299 | 65594410 | mNeoVis.2.1.1.chr2\_hap1.g00565.t01 |
| 2344 | 2 | 65563299 | 65594410 | mNeoVis.2.1.1.chr2\_hap1.g00565.t01 |
| 2345 | 2 | 65563299 | 65594410 | mNeoVis.2.1.1.chr2\_hap1.g00565.t01 |
| 2346 | 2 | 65744350 | 65744853 | mNeoVis.2.1.1.chr2\_hap1.g00566.t01 |
| 2347 | 2 | 65758101 | 65758413 | mNeoVis.2.1.1.chr2\_hap1.g00567.t01 |
| 2348 | 2 | 65874558 | 66017723 | mNeoVis.2.1.1.chr2\_hap1.g00568.t01 |
| 2349 | 2 | 66017704 | 66095627 | mNeoVis.2.1.1.chr2\_hap1.g00569.t01 |
| 2350 | 2 | 66541626 | 66812953 | mNeoVis.2.1.1.chr2\_hap1.g00570.t01 |
| 2351 | 2 | 66969679 | 66986451 | mNeoVis.2.1.1.chr2\_hap1.g00571.t01 |
| 2352 | 2 | 67035658 | 67075783 | mNeoVis.2.1.1.chr2\_hap1.g00572.t01 |
| 2353 | 2 | 67142929 | 67158896 | mNeoVis.2.1.1.chr2\_hap1.g00573.t01 |
| 2354 | 2 | 67162415 | 67213523 | mNeoVis.2.1.1.chr2\_hap1.g00574.t01 |
| 2355 | 2 | 71836920 | 71838992 | mNeoVis.2.1.1.chr2\_hap1.g00579.t01 |
| 2356 | 2 | 71836920 | 71838992 | mNeoVis.2.1.1.chr2\_hap1.g00579.t01 |
| 2357 | 2 | 72265895 | 72269124 | mNeoVis.2.1.1.chr2\_hap1.g00580.t01 |
| 2358 | 2 | 72265895 | 72269124 | mNeoVis.2.1.1.chr2\_hap1.g00580.t01 |
| 2359 | 2 | 72867367 | 72871364 | mNeoVis.2.1.1.chr2\_hap1.g00582.t01 |
| 2360 | 2 | 72867367 | 72871364 | mNeoVis.2.1.1.chr2\_hap1.g00582.t01 |
| 2361 | 2 | 72867367 | 72871364 | mNeoVis.2.1.1.chr2\_hap1.g00582.t01 |
| 2362 | 2 | 73791231 | 73901642 | mNeoVis.2.1.1.chr2\_hap1.g00584.t01 |
| 2363 | 2 | 73884525 | 73884967 | mNeoVis.2.1.1.chr2\_hap1.g00585.t01 |
| 2364 | 2 | 73923282 | 74186921 | mNeoVis.2.1.1.chr2\_hap1.g00586.t01 |
| 2365 | 2 | 73923282 | 74186921 | mNeoVis.2.1.1.chr2\_hap1.g00586.t01 |
| 2366 | 2 | 74290725 | 74337852 | mNeoVis.2.1.1.chr2\_hap1.g00587.t01 |
| 2367 | 2 | 74411601 | 74483800 | mNeoVis.2.1.1.chr2\_hap1.g00588.t01 |
| 2368 | 2 | 74411601 | 74483800 | mNeoVis.2.1.1.chr2\_hap1.g00588.t01 |
| 2369 | 2 | 74489572 | 74557112 | mNeoVis.2.1.1.chr2\_hap1.g00589.t01 |
| 2370 | 2 | 74489572 | 74557112 | mNeoVis.2.1.1.chr2\_hap1.g00589.t01 |
| 2371 | 2 | 74571138 | 74573857 | mNeoVis.2.1.1.chr2\_hap1.g00590.t01 |
| 2372 | 2 | 74571138 | 74573857 | mNeoVis.2.1.1.chr2\_hap1.g00590.t01 |
| 2373 | 2 | 74574925 | 74660237 | mNeoVis.2.1.1.chr2\_hap1.g00591.t01 |
| 2374 | 2 | 74574925 | 74660237 | mNeoVis.2.1.1.chr2\_hap1.g00591.t01 |
| 2375 | 2 | 75491765 | 75578749 | mNeoVis.2.1.1.chr2\_hap1.g00598.t01 |
| 2376 | 2 | 75504259 | 75531164 | mNeoVis.2.1.1.chr2\_hap1.g00599.t01 |
| 2377 | 2 | 75583300 | 75622456 | mNeoVis.2.1.1.chr2\_hap1.g00600.t01 |
| 2378 | 2 | 75638111 | 75655127 | mNeoVis.2.1.1.chr2\_hap1.g00601.t01 |
| 2379 | 2 | 75655409 | 75678727 | mNeoVis.2.1.1.chr2\_hap1.g00602.t01 |
| 2380 | 2 | 75703621 | 75752901 | mNeoVis.2.1.1.chr2\_hap1.g00603.t01 |
| 2381 | 2 | 75703621 | 75752901 | mNeoVis.2.1.1.chr2\_hap1.g00603.t01 |
| 2382 | 2 | 75703621 | 75752901 | mNeoVis.2.1.1.chr2\_hap1.g00603.t01 |
| 2383 | 2 | 75760269 | 75777231 | mNeoVis.2.1.1.chr2\_hap1.g00604.t01 |
| 2384 | 2 | 75760269 | 75777231 | mNeoVis.2.1.1.chr2\_hap1.g00604.t01 |
| 2385 | 2 | 75760269 | 75777231 | mNeoVis.2.1.1.chr2\_hap1.g00604.t01 |
| 2386 | 2 | 75812204 | 75836881 | mNeoVis.2.1.1.chr2\_hap1.g00605.t01 |
| 2387 | 2 | 75812204 | 75836881 | mNeoVis.2.1.1.chr2\_hap1.g00605.t01 |
| 2388 | 2 | 75812204 | 75836881 | mNeoVis.2.1.1.chr2\_hap1.g00605.t01 |
| 2389 | 2 | 75991567 | 75995741 | mNeoVis.2.1.1.chr2\_hap1.g00606.t01 |
| 2390 | 2 | 75991567 | 75995741 | mNeoVis.2.1.1.chr2\_hap1.g00606.t01 |
| 2391 | 2 | 75991567 | 75995741 | mNeoVis.2.1.1.chr2\_hap1.g00606.t01 |
| 2392 | 2 | 76000363 | 76031342 | mNeoVis.2.1.1.chr2\_hap1.g00607.t01 |
| 2393 | 2 | 76000363 | 76031342 | mNeoVis.2.1.1.chr2\_hap1.g00607.t01 |
| 2394 | 2 | 76000363 | 76031342 | mNeoVis.2.1.1.chr2\_hap1.g00607.t01 |
| 2395 | 2 | 76061821 | 76082820 | mNeoVis.2.1.1.chr2\_hap1.g00608.t01 |
| 2396 | 2 | 76061821 | 76082820 | mNeoVis.2.1.1.chr2\_hap1.g00608.t01 |
| 2397 | 2 | 76061821 | 76082820 | mNeoVis.2.1.1.chr2\_hap1.g00608.t01 |
| 2398 | 2 | 76095793 | 76140256 | mNeoVis.2.1.1.chr2\_hap1.g00609.t01 |
| 2399 | 2 | 76095793 | 76140256 | mNeoVis.2.1.1.chr2\_hap1.g00609.t01 |
| 2400 | 2 | 76095793 | 76140256 | mNeoVis.2.1.1.chr2\_hap1.g00609.t01 |
| 2401 | 2 | 76171650 | 76192623 | mNeoVis.2.1.1.chr2\_hap1.g00610.t01 |
| 2402 | 2 | 76171650 | 76192623 | mNeoVis.2.1.1.chr2\_hap1.g00610.t01 |
| 2403 | 2 | 76171650 | 76192623 | mNeoVis.2.1.1.chr2\_hap1.g00610.t01 |
| 2404 | 2 | 76204872 | 76226365 | mNeoVis.2.1.1.chr2\_hap1.g00611.t01 |
| 2405 | 2 | 76204872 | 76226365 | mNeoVis.2.1.1.chr2\_hap1.g00611.t01 |
| 2406 | 2 | 76232449 | 76256813 | mNeoVis.2.1.1.chr2\_hap1.g00612.t01 |
| 2407 | 2 | 76325911 | 76356577 | mNeoVis.2.1.1.chr2\_hap1.g00615.t01 |
| 2408 | 2 | 76407684 | 76645397 | mNeoVis.2.1.1.chr2\_hap1.g00616.t01 |
| 2409 | 2 | 77895343 | 78064360 | mNeoVis.2.1.1.chr2\_hap1.g00617.t01 |
| 2410 | 2 | 77895343 | 78064360 | mNeoVis.2.1.1.chr2\_hap1.g00617.t01 |
| 2411 | 2 | 78281724 | 78288621 | mNeoVis.2.1.1.chr2\_hap1.g00618.t01 |
| 2412 | 2 | 78281724 | 78288621 | mNeoVis.2.1.1.chr2\_hap1.g00618.t01 |
| 2413 | 2 | 78319326 | 78319677 | mNeoVis.2.1.1.chr2\_hap1.g00619.t01 |
| 2414 | 2 | 79099793 | 79101358 | mNeoVis.2.1.1.chr2\_hap1.g00621.t01 |
| 2415 | 2 | 79099793 | 79101358 | mNeoVis.2.1.1.chr2\_hap1.g00621.t01 |
| 2416 | 2 | 79186572 | 79295723 | mNeoVis.2.1.1.chr2\_hap1.g00622.t01 |
| 2417 | 2 | 79186572 | 79295723 | mNeoVis.2.1.1.chr2\_hap1.g00622.t01 |
| 2418 | 2 | 79314455 | 79380086 | mNeoVis.2.1.1.chr2\_hap1.g00623.t01 |
| 2419 | 2 | 79314455 | 79380086 | mNeoVis.2.1.1.chr2\_hap1.g00623.t01 |
| 2420 | 2 | 79473004 | 79520578 | mNeoVis.2.1.1.chr2\_hap1.g00624.t01 |
| 2421 | 2 | 79521426 | 79569416 | mNeoVis.2.1.1.chr2\_hap1.g00625.t01 |
| 2422 | 2 | 79603282 | 79665895 | mNeoVis.2.1.1.chr2\_hap1.g00626.t01 |
| 2423 | 2 | 82710904 | 82716154 | mNeoVis.2.1.1.chr2\_hap1.g00642.t01 |
| 2424 | 2 | 82753745 | 82774894 | mNeoVis.2.1.1.chr2\_hap1.g00643.t01 |
| 2425 | 2 | 82777629 | 82798204 | mNeoVis.2.1.1.chr2\_hap1.g00644.t01 |
| 2426 | 2 | 82780205 | 82831676 | mNeoVis.2.1.1.chr2\_hap1.g00645.t01 |
| 2427 | 2 | 82980348 | 83000910 | mNeoVis.2.1.1.chr2\_hap1.g00646.t01 |
| 2428 | 2 | 83062361 | 83064483 | mNeoVis.2.1.1.chr2\_hap1.g00647.t01 |
| 2429 | 2 | 83064050 | 83067683 | mNeoVis.2.1.1.chr2\_hap1.g00648.t01 |
| 2430 | 2 | 84779293 | 84822982 | mNeoVis.2.1.1.chr2\_hap1.g00660.t01 |
| 2431 | 2 | 84827574 | 84843699 | mNeoVis.2.1.1.chr2\_hap1.g00661.t01 |
| 2432 | 2 | 84843819 | 84878834 | mNeoVis.2.1.1.chr2\_hap1.g00662.t01 |
| 2433 | 2 | 84896149 | 84900957 | mNeoVis.2.1.1.chr2\_hap1.g00663.t01 |
| 2434 | 2 | 84901174 | 84914046 | mNeoVis.2.1.1.chr2\_hap1.g00664.t01 |
| 2435 | 2 | 84923512 | 84982084 | mNeoVis.2.1.1.chr2\_hap1.g00665.t01 |
| 2436 | 2 | 85085354 | 85119900 | mNeoVis.2.1.1.chr2\_hap1.g00666.t01 |
| 2437 | 2 | 85123379 | 85125803 | mNeoVis.2.1.1.chr2\_hap1.g00667.t01 |
| 2438 | 2 | 85837472 | 86051738 | mNeoVis.2.1.1.chr2\_hap1.g00669.t01 |
| 2439 | 2 | 86196723 | 86200288 | mNeoVis.2.1.1.chr2\_hap1.g00670.t01 |
| 2440 | 2 | 86403265 | 86711004 | mNeoVis.2.1.1.chr2\_hap1.g00672.t01 |
| 2441 | 2 | 86734477 | 86792039 | mNeoVis.2.1.1.chr2\_hap1.g00673.t01 |
| 2442 | 2 | 86805093 | 86812761 | mNeoVis.2.1.1.chr2\_hap1.g00674.t01 |
| 2443 | 2 | 86984607 | 87027393 | mNeoVis.2.1.1.chr2\_hap1.g00675.t01 |
| 2444 | 2 | 87257633 | 87258902 | mNeoVis.2.1.1.chr2\_hap1.g00677.t01 |
| 2445 | 2 | 87257633 | 87258902 | mNeoVis.2.1.1.chr2\_hap1.g00677.t01 |
| 2446 | 2 | 87375974 | 87377939 | mNeoVis.2.1.1.chr2\_hap1.g00678.t01 |
| 2447 | 2 | 87375974 | 87377939 | mNeoVis.2.1.1.chr2\_hap1.g00678.t01 |
| 2448 | 2 | 87375974 | 87377939 | mNeoVis.2.1.1.chr2\_hap1.g00678.t01 |
| 2449 | 2 | 87375974 | 87377939 | mNeoVis.2.1.1.chr2\_hap1.g00678.t01 |
| 2450 | 2 | 87674521 | 87701180 | mNeoVis.2.1.1.chr2\_hap1.g00679.t01 |
| 2451 | 2 | 87674521 | 87701180 | mNeoVis.2.1.1.chr2\_hap1.g00679.t01 |
| 2452 | 2 | 87674521 | 87701180 | mNeoVis.2.1.1.chr2\_hap1.g00679.t01 |
| 2453 | 2 | 87685724 | 87771645 | mNeoVis.2.1.1.chr2\_hap1.g00680.t01 |
| 2454 | 2 | 87685724 | 87771645 | mNeoVis.2.1.1.chr2\_hap1.g00680.t01 |
| 2455 | 2 | 87685724 | 87771645 | mNeoVis.2.1.1.chr2\_hap1.g00680.t01 |
| 2456 | 2 | 87877987 | 87881965 | mNeoVis.2.1.1.chr2\_hap1.g00681.t01 |
| 2457 | 2 | 87877987 | 87881965 | mNeoVis.2.1.1.chr2\_hap1.g00681.t01 |
| 2458 | 2 | 87877987 | 87881965 | mNeoVis.2.1.1.chr2\_hap1.g00681.t01 |
| 2459 | 2 | 87877987 | 87881965 | mNeoVis.2.1.1.chr2\_hap1.g00681.t01 |
| 2460 | 2 | 88014560 | 88138256 | mNeoVis.2.1.1.chr2\_hap1.g00682.t01 |
| 2461 | 2 | 88014560 | 88138256 | mNeoVis.2.1.1.chr2\_hap1.g00682.t01 |
| 2462 | 2 | 88014560 | 88138256 | mNeoVis.2.1.1.chr2\_hap1.g00682.t01 |
| 2463 | 2 | 88308931 | 88325625 | mNeoVis.2.1.1.chr2\_hap1.g00683.t01 |
| 2464 | 2 | 88308931 | 88325625 | mNeoVis.2.1.1.chr2\_hap1.g00683.t01 |
| 2465 | 2 | 88308931 | 88325625 | mNeoVis.2.1.1.chr2\_hap1.g00683.t01 |
| 2466 | 2 | 88331447 | 88352104 | mNeoVis.2.1.1.chr2\_hap1.g00684.t01 |
| 2467 | 2 | 88331447 | 88352104 | mNeoVis.2.1.1.chr2\_hap1.g00684.t01 |
| 2468 | 2 | 88356056 | 88494926 | mNeoVis.2.1.1.chr2\_hap1.g00685.t01 |
| 2469 | 2 | 88356056 | 88494926 | mNeoVis.2.1.1.chr2\_hap1.g00685.t01 |
| 2470 | 2 | 88495082 | 88506672 | mNeoVis.2.1.1.chr2\_hap1.g00686.t01 |
| 2471 | 2 | 88495082 | 88506672 | mNeoVis.2.1.1.chr2\_hap1.g00686.t01 |
| 2472 | 2 | 88508938 | 88510694 | mNeoVis.2.1.1.chr2\_hap1.g00687.t01 |
| 2473 | 2 | 88508938 | 88510694 | mNeoVis.2.1.1.chr2\_hap1.g00687.t01 |
| 2474 | 2 | 88525633 | 88608081 | mNeoVis.2.1.1.chr2\_hap1.g00688.t01 |
| 2475 | 2 | 88525633 | 88608081 | mNeoVis.2.1.1.chr2\_hap1.g00688.t01 |
| 2476 | 2 | 88609003 | 88666303 | mNeoVis.2.1.1.chr2\_hap1.g00689.t01 |
| 2477 | 2 | 88609003 | 88666303 | mNeoVis.2.1.1.chr2\_hap1.g00689.t01 |
| 2478 | 2 | 88701977 | 88744784 | mNeoVis.2.1.1.chr2\_hap1.g00691.t01 |
| 2479 | 2 | 88774263 | 88963009 | mNeoVis.2.1.1.chr2\_hap1.g00692.t01 |
| 2480 | 2 | 89028706 | 89112279 | mNeoVis.2.1.1.chr2\_hap1.g00693.t01 |
| 2481 | 2 | 89175198 | 89223809 | mNeoVis.2.1.1.chr2\_hap1.g00694.t01 |
| 2482 | 2 | 90813262 | 90910035 | mNeoVis.2.1.1.chr2\_hap1.g00720.t01 |
| 2483 | 2 | 90938544 | 91124991 | mNeoVis.2.1.1.chr2\_hap1.g00721.t01 |
| 2484 | 2 | 91501844 | 91528801 | mNeoVis.2.1.1.chr2\_hap1.g00722.t01 |
| 2485 | 2 | 91529876 | 91560615 | mNeoVis.2.1.1.chr2\_hap1.g00723.t01 |
| 2486 | 2 | 91598511 | 91635301 | mNeoVis.2.1.1.chr2\_hap1.g00724.t01 |
| 2487 | 2 | 91640397 | 91693565 | mNeoVis.2.1.1.chr2\_hap1.g00725.t01 |
| 2488 | 2 | 91700606 | 91703483 | mNeoVis.2.1.1.chr2\_hap1.g00726.t01 |
| 2489 | 2 | 91769736 | 91792245 | mNeoVis.2.1.1.chr2\_hap1.g00727.t01 |
| 2490 | 2 | 91817719 | 91864768 | mNeoVis.2.1.1.chr2\_hap1.g00728.t01 |
| 2491 | 2 | 92298497 | 92308683 | mNeoVis.2.1.1.chr2\_hap1.g00729.t01 |
| 2492 | 2 | 133152745 | 133158243 | mNeoVis.2.1.1.chr2\_hap1.g00757.t01 |
| 2493 | 2 | 133180808 | 133187573 | mNeoVis.2.1.1.chr2\_hap1.g00758.t01 |
| 2494 | 2 | 133225188 | 133233880 | mNeoVis.2.1.1.chr2\_hap1.g00759.t01 |
| 2495 | 2 | 133256768 | 133268050 | mNeoVis.2.1.1.chr2\_hap1.g00760.t01 |
| 2496 | 2 | 133365685 | 133429949 | mNeoVis.2.1.1.chr2\_hap1.g00761.t01 |
| 2497 | 2 | 133365685 | 133429949 | mNeoVis.2.1.1.chr2\_hap1.g00761.t01 |
| 2498 | 2 | 133428024 | 133436131 | mNeoVis.2.1.1.chr2\_hap1.g00762.t01 |
| 2499 | 2 | 133428024 | 133436131 | mNeoVis.2.1.1.chr2\_hap1.g00762.t01 |
| 2500 | 2 | 133443859 | 133446179 | mNeoVis.2.1.1.chr2\_hap1.g00763.t01 |
| 2501 | 2 | 133443859 | 133446179 | mNeoVis.2.1.1.chr2\_hap1.g00763.t01 |
| 2502 | 2 | 133451361 | 133475458 | mNeoVis.2.1.1.chr2\_hap1.g00764.t01 |
| 2503 | 2 | 133451361 | 133475458 | mNeoVis.2.1.1.chr2\_hap1.g00764.t01 |
| 2504 | 2 | 133478241 | 133480703 | mNeoVis.2.1.1.chr2\_hap1.g00765.t01 |
| 2505 | 2 | 133478241 | 133480703 | mNeoVis.2.1.1.chr2\_hap1.g00765.t01 |
| 2506 | 2 | 133488123 | 133505226 | mNeoVis.2.1.1.chr2\_hap1.g00766.t01 |
| 2507 | 2 | 133488123 | 133505226 | mNeoVis.2.1.1.chr2\_hap1.g00766.t01 |
| 2508 | 2 | 133505549 | 133508644 | mNeoVis.2.1.1.chr2\_hap1.g00767.t01 |
| 2509 | 2 | 133505549 | 133508644 | mNeoVis.2.1.1.chr2\_hap1.g00767.t01 |
| 2510 | 2 | 133511722 | 133516076 | mNeoVis.2.1.1.chr2\_hap1.g00768.t01 |
| 2511 | 2 | 133511722 | 133516076 | mNeoVis.2.1.1.chr2\_hap1.g00768.t01 |
| 2512 | 2 | 133534389 | 133553160 | mNeoVis.2.1.1.chr2\_hap1.g00769.t01 |
| 2513 | 2 | 133534389 | 133553160 | mNeoVis.2.1.1.chr2\_hap1.g00769.t01 |
| 2514 | 2 | 133545937 | 133560405 | mNeoVis.2.1.1.chr2\_hap1.g00770.t01 |
| 2515 | 2 | 133545937 | 133560405 | mNeoVis.2.1.1.chr2\_hap1.g00770.t01 |
| 2516 | 2 | 133565581 | 133631613 | mNeoVis.2.1.1.chr2\_hap1.g00771.t01 |
| 2517 | 2 | 133565581 | 133631613 | mNeoVis.2.1.1.chr2\_hap1.g00771.t01 |
| 2518 | 2 | 133655375 | 133704822 | mNeoVis.2.1.1.chr2\_hap1.g00772.t01 |
| 2519 | 2 | 133655375 | 133704822 | mNeoVis.2.1.1.chr2\_hap1.g00772.t01 |
| 2520 | 2 | 133706036 | 133730549 | mNeoVis.2.1.1.chr2\_hap1.g00773.t01 |
| 2521 | 2 | 133706036 | 133730549 | mNeoVis.2.1.1.chr2\_hap1.g00773.t01 |
| 2522 | 2 | 133769353 | 133819500 | mNeoVis.2.1.1.chr2\_hap1.g00774.t01 |
| 2523 | 2 | 133769353 | 133819500 | mNeoVis.2.1.1.chr2\_hap1.g00774.t01 |
| 2524 | 2 | 133820546 | 133822974 | mNeoVis.2.1.1.chr2\_hap1.g00775.t01 |
| 2525 | 2 | 133820546 | 133822974 | mNeoVis.2.1.1.chr2\_hap1.g00775.t01 |
| 2526 | 2 | 133823105 | 133875896 | mNeoVis.2.1.1.chr2\_hap1.g00776.t01 |
| 2527 | 2 | 133823105 | 133875896 | mNeoVis.2.1.1.chr2\_hap1.g00776.t01 |
| 2528 | 2 | 133886350 | 133893197 | mNeoVis.2.1.1.chr2\_hap1.g00777.t01 |
| 2529 | 2 | 133902086 | 133929932 | mNeoVis.2.1.1.chr2\_hap1.g00778.t01 |
| 2530 | 2 | 133929878 | 133939573 | mNeoVis.2.1.1.chr2\_hap1.g00779.t01 |
| 2531 | 2 | 133976245 | 133986545 | mNeoVis.2.1.1.chr2\_hap1.g00780.t01 |
| 2532 | 2 | 133991633 | 134031339 | mNeoVis.2.1.1.chr2\_hap1.g00781.t01 |
| 2533 | 2 | 134035723 | 134047846 | mNeoVis.2.1.1.chr2\_hap1.g00782.t01 |
| 2534 | 2 | 134048553 | 134053409 | mNeoVis.2.1.1.chr2\_hap1.g00783.t01 |
| 2535 | 2 | 134054290 | 134056934 | mNeoVis.2.1.1.chr2\_hap1.g00784.t01 |
| 2536 | 2 | 134062002 | 134066502 | mNeoVis.2.1.1.chr2\_hap1.g00785.t01 |
| 2537 | 2 | 134066503 | 134069305 | mNeoVis.2.1.1.chr2\_hap1.g00786.t01 |
| 2538 | 2 | 134080041 | 134093329 | mNeoVis.2.1.1.chr2\_hap1.g00787.t01 |
| 2539 | 2 | 134134439 | 134145341 | mNeoVis.2.1.1.chr2\_hap1.g00789.t01 |
| 2540 | 2 | 134141309 | 134149343 | mNeoVis.2.1.1.chr2\_hap1.g00790.t01 |
| 2541 | 2 | 134149353 | 134152130 | mNeoVis.2.1.1.chr2\_hap1.g00791.t01 |
| 2542 | 2 | 134151965 | 134160339 | mNeoVis.2.1.1.chr2\_hap1.g00792.t01 |
| 2543 | 2 | 134160876 | 134181912 | mNeoVis.2.1.1.chr2\_hap1.g00793.t01 |
| 2544 | 2 | 134182053 | 134192976 | mNeoVis.2.1.1.chr2\_hap1.g00794.t01 |
| 2545 | 2 | 134193113 | 134202281 | mNeoVis.2.1.1.chr2\_hap1.g00795.t01 |
| 2546 | 2 | 134207970 | 134217260 | mNeoVis.2.1.1.chr2\_hap1.g00796.t01 |
| 2547 | 2 | 134217188 | 134249565 | mNeoVis.2.1.1.chr2\_hap1.g00797.t01 |
| 2548 | 2 | 134256095 | 134289421 | mNeoVis.2.1.1.chr2\_hap1.g00798.t01 |
| 2549 | 2 | 134289843 | 134361244 | mNeoVis.2.1.1.chr2\_hap1.g00799.t01 |
| 2550 | 2 | 134361130 | 134372346 | mNeoVis.2.1.1.chr2\_hap1.g00800.t01 |
| 2551 | 2 | 134386300 | 134407549 | mNeoVis.2.1.1.chr2\_hap1.g00801.t01 |
| 2552 | 2 | 134421058 | 134441653 | mNeoVis.2.1.1.chr2\_hap1.g00802.t01 |
| 2553 | 2 | 134444731 | 134484501 | mNeoVis.2.1.1.chr2\_hap1.g00803.t01 |
| 2554 | 2 | 134497499 | 134504809 | mNeoVis.2.1.1.chr2\_hap1.g00804.t01 |
| 2555 | 2 | 134537510 | 134542684 | mNeoVis.2.1.1.chr2\_hap1.g00805.t01 |
| 2556 | 2 | 134552959 | 134570810 | mNeoVis.2.1.1.chr2\_hap1.g00806.t01 |
| 2557 | 2 | 134601563 | 134607394 | mNeoVis.2.1.1.chr2\_hap1.g00807.t01 |
| 2558 | 2 | 134608040 | 134623841 | mNeoVis.2.1.1.chr2\_hap1.g00808.t01 |
| 2559 | 2 | 134629374 | 134731039 | mNeoVis.2.1.1.chr2\_hap1.g00809.t01 |
| 2560 | 2 | 134740747 | 134763075 | mNeoVis.2.1.1.chr2\_hap1.g00810.t01 |
| 2561 | 2 | 134773424 | 134786808 | mNeoVis.2.1.1.chr2\_hap1.g00811.t01 |
| 2562 | 2 | 134794256 | 134798163 | mNeoVis.2.1.1.chr2\_hap1.g00812.t01 |
| 2563 | 2 | 134799737 | 134805670 | mNeoVis.2.1.1.chr2\_hap1.g00813.t01 |
| 2564 | 2 | 134807534 | 134811205 | mNeoVis.2.1.1.chr2\_hap1.g00814.t01 |
| 2565 | 2 | 134811535 | 134817609 | mNeoVis.2.1.1.chr2\_hap1.g00815.t01 |
| 2566 | 2 | 135394148 | 135416454 | mNeoVis.2.1.1.chr2\_hap1.g00839.t01 |
| 2567 | 2 | 135570529 | 135571902 | mNeoVis.2.1.1.chr2\_hap1.g00840.t01 |
| 2568 | 2 | 135601511 | 135605886 | mNeoVis.2.1.1.chr2\_hap1.g00841.t01 |
| 2569 | 2 | 135618189 | 135622354 | mNeoVis.2.1.1.chr2\_hap1.g00842.t01 |
| 2570 | 2 | 135631882 | 135641663 | mNeoVis.2.1.1.chr2\_hap1.g00843.t01 |
| 2571 | 2 | 135632858 | 135646925 | mNeoVis.2.1.1.chr2\_hap1.g00844.t01 |
| 2572 | 2 | 135648726 | 135670017 | mNeoVis.2.1.1.chr2\_hap1.g00845.t01 |
| 2573 | 2 | 135673825 | 135675574 | mNeoVis.2.1.1.chr2\_hap1.g00846.t01 |
| 2574 | 2 | 135675715 | 135680563 | mNeoVis.2.1.1.chr2\_hap1.g00847.t01 |
| 2575 | 2 | 135680808 | 135686532 | mNeoVis.2.1.1.chr2\_hap1.g00848.t01 |
| 2576 | 2 | 135686668 | 135704614 | mNeoVis.2.1.1.chr2\_hap1.g00849.t01 |
| 2577 | 2 | 135716893 | 135734131 | mNeoVis.2.1.1.chr2\_hap1.g00850.t01 |
| 2578 | 2 | 135739244 | 135747688 | mNeoVis.2.1.1.chr2\_hap1.g00851.t01 |
| 2579 | 2 | 135745253 | 135752212 | mNeoVis.2.1.1.chr2\_hap1.g00852.t01 |
| 2580 | 2 | 135752309 | 135761041 | mNeoVis.2.1.1.chr2\_hap1.g00853.t01 |
| 2581 | 2 | 135761226 | 135833221 | mNeoVis.2.1.1.chr2\_hap1.g00854.t01 |
| 2582 | 2 | 135833324 | 135858266 | mNeoVis.2.1.1.chr2\_hap1.g00855.t01 |
| 2583 | 2 | 135871657 | 135909745 | mNeoVis.2.1.1.chr2\_hap1.g00856.t01 |
| 2584 | 2 | 135910137 | 135972474 | mNeoVis.2.1.1.chr2\_hap1.g00857.t01 |
| 2585 | 2 | 135984980 | 135986302 | mNeoVis.2.1.1.chr2\_hap1.g00858.t01 |
| 2586 | 2 | 135997068 | 136016368 | mNeoVis.2.1.1.chr2\_hap1.g00859.t01 |
| 2587 | 2 | 136102018 | 136119941 | mNeoVis.2.1.1.chr2\_hap1.g00860.t01 |
| 2588 | 2 | 136181315 | 136199506 | mNeoVis.2.1.1.chr2\_hap1.g00861.t01 |
| 2589 | 2 | 136230012 | 136242874 | mNeoVis.2.1.1.chr2\_hap1.g00862.t01 |
| 2590 | 2 | 136242835 | 136312756 | mNeoVis.2.1.1.chr2\_hap1.g00863.t01 |
| 2591 | 2 | 137936404 | 137951074 | mNeoVis.2.1.1.chr2\_hap1.g00877.t01 |
| 2592 | 2 | 137983027 | 138004058 | mNeoVis.2.1.1.chr2\_hap1.g00878.t01 |
| 2593 | 2 | 138007500 | 138039667 | mNeoVis.2.1.1.chr2\_hap1.g00879.t01 |
| 2594 | 2 | 138085062 | 138116094 | mNeoVis.2.1.1.chr2\_hap1.g00880.t01 |
| 2595 | 2 | 138129791 | 138138134 | mNeoVis.2.1.1.chr2\_hap1.g00881.t01 |
| 2596 | 2 | 138166917 | 138198934 | mNeoVis.2.1.1.chr2\_hap1.g00882.t01 |
| 2597 | 2 | 138395082 | 138491578 | mNeoVis.2.1.1.chr2\_hap1.g00883.t01 |
| 2598 | 2 | 138524931 | 138631030 | mNeoVis.2.1.1.chr2\_hap1.g00884.t01 |
| 2599 | 2 | 138524931 | 138631030 | mNeoVis.2.1.1.chr2\_hap1.g00884.t01 |
| 2600 | 2 | 139644514 | 139675920 | mNeoVis.2.1.1.chr2\_hap1.g00887.t01 |
| 2601 | 2 | 139675948 | 139749717 | mNeoVis.2.1.1.chr2\_hap1.g00888.t01 |
| 2602 | 2 | 139952014 | 139973912 | mNeoVis.2.1.1.chr2\_hap1.g00889.t01 |
| 2603 | 2 | 139952014 | 139973912 | mNeoVis.2.1.1.chr2\_hap1.g00889.t01 |
| 2604 | 2 | 140043260 | 140230748 | mNeoVis.2.1.1.chr2\_hap1.g00890.t01 |
| 2605 | 2 | 140043260 | 140230748 | mNeoVis.2.1.1.chr2\_hap1.g00890.t01 |
| 2606 | 2 | 140043260 | 140230748 | mNeoVis.2.1.1.chr2\_hap1.g00890.t01 |
| 2607 | 2 | 140310420 | 140323631 | mNeoVis.2.1.1.chr2\_hap1.g00891.t01 |
| 2608 | 2 | 140310420 | 140323631 | mNeoVis.2.1.1.chr2\_hap1.g00891.t01 |
| 2609 | 2 | 140310420 | 140323631 | mNeoVis.2.1.1.chr2\_hap1.g00891.t01 |
| 2610 | 2 | 140310420 | 140323631 | mNeoVis.2.1.1.chr2\_hap1.g00891.t01 |
| 2611 | 2 | 140310420 | 140323631 | mNeoVis.2.1.1.chr2\_hap1.g00891.t01 |
| 2612 | 2 | 140364963 | 140402426 | mNeoVis.2.1.1.chr2\_hap1.g00892.t01 |
| 2613 | 2 | 140364963 | 140402426 | mNeoVis.2.1.1.chr2\_hap1.g00892.t01 |
| 2614 | 2 | 140364963 | 140402426 | mNeoVis.2.1.1.chr2\_hap1.g00892.t01 |
| 2615 | 2 | 140364963 | 140402426 | mNeoVis.2.1.1.chr2\_hap1.g00892.t01 |
| 2616 | 2 | 140364963 | 140402426 | mNeoVis.2.1.1.chr2\_hap1.g00892.t01 |
| 2617 | 2 | 140366814 | 140425112 | mNeoVis.2.1.1.chr2\_hap1.g00893.t01 |
| 2618 | 2 | 140366814 | 140425112 | mNeoVis.2.1.1.chr2\_hap1.g00893.t01 |
| 2619 | 2 | 140366814 | 140425112 | mNeoVis.2.1.1.chr2\_hap1.g00893.t01 |
| 2620 | 2 | 140366814 | 140425112 | mNeoVis.2.1.1.chr2\_hap1.g00893.t01 |
| 2621 | 2 | 140426386 | 140435308 | mNeoVis.2.1.1.chr2\_hap1.g00894.t01 |
| 2622 | 2 | 140426386 | 140435308 | mNeoVis.2.1.1.chr2\_hap1.g00894.t01 |
| 2623 | 2 | 140426386 | 140435308 | mNeoVis.2.1.1.chr2\_hap1.g00894.t01 |
| 2624 | 2 | 140426386 | 140435308 | mNeoVis.2.1.1.chr2\_hap1.g00894.t01 |
| 2625 | 2 | 140443560 | 140486367 | mNeoVis.2.1.1.chr2\_hap1.g00895.t01 |
| 2626 | 2 | 140443560 | 140486367 | mNeoVis.2.1.1.chr2\_hap1.g00895.t01 |
| 2627 | 2 | 140443560 | 140486367 | mNeoVis.2.1.1.chr2\_hap1.g00895.t01 |
| 2628 | 2 | 140443560 | 140486367 | mNeoVis.2.1.1.chr2\_hap1.g00895.t01 |
| 2629 | 2 | 140505346 | 140542403 | mNeoVis.2.1.1.chr2\_hap1.g00896.t01 |
| 2630 | 2 | 140505346 | 140542403 | mNeoVis.2.1.1.chr2\_hap1.g00896.t01 |
| 2631 | 2 | 140505346 | 140542403 | mNeoVis.2.1.1.chr2\_hap1.g00896.t01 |
| 2632 | 2 | 140552742 | 140636311 | mNeoVis.2.1.1.chr2\_hap1.g00897.t01 |
| 2633 | 2 | 140552742 | 140636311 | mNeoVis.2.1.1.chr2\_hap1.g00897.t01 |
| 2634 | 2 | 140552742 | 140636311 | mNeoVis.2.1.1.chr2\_hap1.g00897.t01 |
| 2635 | 2 | 140552742 | 140636311 | mNeoVis.2.1.1.chr2\_hap1.g00897.t01 |
| 2636 | 2 | 140755855 | 140776912 | mNeoVis.2.1.1.chr2\_hap1.g00898.t01 |
| 2637 | 2 | 140755855 | 140776912 | mNeoVis.2.1.1.chr2\_hap1.g00898.t01 |
| 2638 | 2 | 140755855 | 140776912 | mNeoVis.2.1.1.chr2\_hap1.g00898.t01 |
| 2639 | 2 | 140839132 | 140934459 | mNeoVis.2.1.1.chr2\_hap1.g00899.t01 |
| 2640 | 2 | 140839132 | 140934459 | mNeoVis.2.1.1.chr2\_hap1.g00899.t01 |
| 2641 | 2 | 140839132 | 140934459 | mNeoVis.2.1.1.chr2\_hap1.g00899.t01 |
| 2642 | 2 | 140939174 | 140979426 | mNeoVis.2.1.1.chr2\_hap1.g00900.t01 |
| 2643 | 2 | 140939174 | 140979426 | mNeoVis.2.1.1.chr2\_hap1.g00900.t01 |
| 2644 | 2 | 140939174 | 140979426 | mNeoVis.2.1.1.chr2\_hap1.g00900.t01 |
| 2645 | 2 | 141106010 | 141135044 | mNeoVis.2.1.1.chr2\_hap1.g00901.t01 |
| 2646 | 2 | 141106010 | 141135044 | mNeoVis.2.1.1.chr2\_hap1.g00901.t01 |
| 2647 | 2 | 141106010 | 141135044 | mNeoVis.2.1.1.chr2\_hap1.g00901.t01 |
| 2648 | 2 | 141334024 | 141359196 | mNeoVis.2.1.1.chr2\_hap1.g00902.t01 |
| 2649 | 2 | 141407580 | 141494428 | mNeoVis.2.1.1.chr2\_hap1.g00903.t01 |
| 2650 | 2 | 141597232 | 141597639 | mNeoVis.2.1.1.chr2\_hap1.g00904.t01 |
| 2651 | 2 | 141667415 | 141728932 | mNeoVis.2.1.1.chr2\_hap1.g00905.t01 |
| 2652 | 2 | 141735576 | 141789460 | mNeoVis.2.1.1.chr2\_hap1.g00906.t01 |
| 2653 | 2 | 142007804 | 142059656 | mNeoVis.2.1.1.chr2\_hap1.g00907.t01 |
| 2654 | 2 | 142228205 | 142289628 | mNeoVis.2.1.1.chr2\_hap1.g00908.t01 |
| 2655 | 2 | 142299480 | 142335186 | mNeoVis.2.1.1.chr2\_hap1.g00909.t01 |
| 2656 | 2 | 142340657 | 142478797 | mNeoVis.2.1.1.chr2\_hap1.g00910.t01 |
| 2657 | 2 | 142480922 | 142495674 | mNeoVis.2.1.1.chr2\_hap1.g00911.t01 |
| 2658 | 2 | 142537757 | 142547307 | mNeoVis.2.1.1.chr2\_hap1.g00912.t01 |
| 2659 | 2 | 142555368 | 142595502 | mNeoVis.2.1.1.chr2\_hap1.g00913.t01 |
| 2660 | 2 | 142595639 | 142605622 | mNeoVis.2.1.1.chr2\_hap1.g00914.t01 |
| 2661 | 2 | 142619782 | 142641842 | mNeoVis.2.1.1.chr2\_hap1.g00915.t01 |
| 2662 | 2 | 142798598 | 142912947 | mNeoVis.2.1.1.chr2\_hap1.g00918.t01 |
| 2663 | 2 | 143103307 | 143161237 | mNeoVis.2.1.1.chr2\_hap1.g00919.t01 |
| 2664 | 2 | 143239263 | 143242382 | mNeoVis.2.1.1.chr2\_hap1.g00920.t01 |
| 2665 | 2 | 143243671 | 143302588 | mNeoVis.2.1.1.chr2\_hap1.g00921.t01 |
| 2666 | 2 | 143317250 | 143323477 | mNeoVis.2.1.1.chr2\_hap1.g00922.t01 |
| 2667 | 2 | 143323438 | 143335538 | mNeoVis.2.1.1.chr2\_hap1.g00923.t01 |
| 2668 | 2 | 143336762 | 143344168 | mNeoVis.2.1.1.chr2\_hap1.g00924.t01 |
| 2669 | 2 | 143344251 | 143409871 | mNeoVis.2.1.1.chr2\_hap1.g00925.t01 |
| 2670 | 2 | 143351418 | 143408266 | mNeoVis.2.1.1.chr2\_hap1.g00926.t01 |
| 2671 | 2 | 143409868 | 143458709 | mNeoVis.2.1.1.chr2\_hap1.g00927.t01 |
| 2672 | 2 | 143460017 | 143510923 | mNeoVis.2.1.1.chr2\_hap1.g00928.t01 |
| 2673 | 2 | 143460093 | 143460734 | mNeoVis.2.1.1.chr2\_hap1.g00929.t01 |
| 2674 | 2 | 143537440 | 143783075 | mNeoVis.2.1.1.chr2\_hap1.g00930.t01 |
| 2675 | 2 | 143537440 | 143783075 | mNeoVis.2.1.1.chr2\_hap1.g00930.t01 |
| 2676 | 2 | 144004646 | 144073861 | mNeoVis.2.1.1.chr2\_hap1.g00931.t01 |
| 2677 | 2 | 144004646 | 144073861 | mNeoVis.2.1.1.chr2\_hap1.g00931.t01 |
| 2678 | 2 | 144175432 | 144229824 | mNeoVis.2.1.1.chr2\_hap1.g00932.t01 |
| 2679 | 2 | 144175432 | 144229824 | mNeoVis.2.1.1.chr2\_hap1.g00932.t01 |
| 2680 | 2 | 144434209 | 144457331 | mNeoVis.2.1.1.chr2\_hap1.g00937.t01 |
| 2681 | 2 | 144461679 | 144511500 | mNeoVis.2.1.1.chr2\_hap1.g00938.t01 |
| 2682 | 2 | 144511975 | 144593599 | mNeoVis.2.1.1.chr2\_hap1.g00939.t01 |
| 2683 | 2 | 144511975 | 144593599 | mNeoVis.2.1.1.chr2\_hap1.g00939.t01 |
| 2684 | 2 | 144590630 | 144603192 | mNeoVis.2.1.1.chr2\_hap1.g00940.t01 |
| 2685 | 2 | 144590630 | 144603192 | mNeoVis.2.1.1.chr2\_hap1.g00940.t01 |
| 2686 | 2 | 144590630 | 144603192 | mNeoVis.2.1.1.chr2\_hap1.g00940.t01 |
| 2687 | 2 | 144621766 | 144673121 | mNeoVis.2.1.1.chr2\_hap1.g00941.t01 |
| 2688 | 2 | 144621766 | 144673121 | mNeoVis.2.1.1.chr2\_hap1.g00941.t01 |
| 2689 | 2 | 144621766 | 144673121 | mNeoVis.2.1.1.chr2\_hap1.g00941.t01 |
| 2690 | 2 | 144761881 | 144763493 | mNeoVis.2.1.1.chr2\_hap1.g00942.t01 |
| 2691 | 2 | 144761881 | 144763493 | mNeoVis.2.1.1.chr2\_hap1.g00942.t01 |
| 2692 | 2 | 144761881 | 144763493 | mNeoVis.2.1.1.chr2\_hap1.g00942.t01 |
| 2693 | 2 | 145003382 | 145202738 | mNeoVis.2.1.1.chr2\_hap1.g00943.t01 |
| 2694 | 2 | 145003382 | 145202738 | mNeoVis.2.1.1.chr2\_hap1.g00943.t01 |
| 2695 | 2 | 145211715 | 145222551 | mNeoVis.2.1.1.chr2\_hap1.g00944.t01 |
| 2696 | 2 | 145211715 | 145222551 | mNeoVis.2.1.1.chr2\_hap1.g00944.t01 |
| 2697 | 2 | 145237210 | 145251644 | mNeoVis.2.1.1.chr2\_hap1.g00945.t01 |
| 2698 | 2 | 145237210 | 145251644 | mNeoVis.2.1.1.chr2\_hap1.g00945.t01 |
| 2699 | 2 | 145261594 | 145335664 | mNeoVis.2.1.1.chr2\_hap1.g00946.t01 |
| 2700 | 2 | 145398511 | 145443498 | mNeoVis.2.1.1.chr2\_hap1.g00947.t01 |
| 2701 | 2 | 145398511 | 145454782 | mNeoVis.2.1.1.chr2\_hap1.g00948.t01 |
| 2702 | 2 | 145458967 | 145463759 | mNeoVis.2.1.1.chr2\_hap1.g00949.t01 |
| 2703 | 2 | 145474726 | 145489275 | mNeoVis.2.1.1.chr2\_hap1.g00950.t01 |
| 2704 | 2 | 145489379 | 145497748 | mNeoVis.2.1.1.chr2\_hap1.g00951.t01 |
| 2705 | 2 | 145502580 | 145515301 | mNeoVis.2.1.1.chr2\_hap1.g00952.t01 |
| 2706 | 2 | 145561779 | 145569319 | mNeoVis.2.1.1.chr2\_hap1.g00953.t01 |
| 2707 | 2 | 145574674 | 145588347 | mNeoVis.2.1.1.chr2\_hap1.g00954.t01 |
| 2708 | 2 | 145592098 | 145598364 | mNeoVis.2.1.1.chr2\_hap1.g00955.t01 |
| 2709 | 2 | 146543148 | 146552512 | mNeoVis.2.1.1.chr2\_hap1.g00969.t01 |
| 2710 | 2 | 146641315 | 146662719 | mNeoVis.2.1.1.chr2\_hap1.g00970.t01 |
| 2711 | 2 | 146670778 | 146719499 | mNeoVis.2.1.1.chr2\_hap1.g00971.t01 |
| 2712 | 2 | 146754107 | 146754774 | mNeoVis.2.1.1.chr2\_hap1.g00972.t01 |
| 2713 | 2 | 146796685 | 146807251 | mNeoVis.2.1.1.chr2\_hap1.g00973.t01 |
| 2714 | 2 | 146812066 | 146829539 | mNeoVis.2.1.1.chr2\_hap1.g00974.t01 |
| 2715 | 2 | 146842376 | 146880445 | mNeoVis.2.1.1.chr2\_hap1.g00975.t01 |
| 2716 | 2 | 146924349 | 146943496 | mNeoVis.2.1.1.chr2\_hap1.g00976.t01 |
| 2717 | 2 | 147055214 | 147067987 | mNeoVis.2.1.1.chr2\_hap1.g00977.t01 |
| 2718 | 2 | 147093503 | 147097065 | mNeoVis.2.1.1.chr2\_hap1.g00978.t01 |
| 2719 | 2 | 147093503 | 147097065 | mNeoVis.2.1.1.chr2\_hap1.g00978.t01 |
| 2720 | 2 | 147112741 | 147117818 | mNeoVis.2.1.1.chr2\_hap1.g00979.t01 |
| 2721 | 2 | 147112741 | 147117818 | mNeoVis.2.1.1.chr2\_hap1.g00979.t01 |
| 2722 | 2 | 147121359 | 147128759 | mNeoVis.2.1.1.chr2\_hap1.g00980.t01 |
| 2723 | 2 | 147121359 | 147128759 | mNeoVis.2.1.1.chr2\_hap1.g00980.t01 |
| 2724 | 2 | 147150377 | 147163543 | mNeoVis.2.1.1.chr2\_hap1.g00981.t01 |
| 2725 | 2 | 147150377 | 147163543 | mNeoVis.2.1.1.chr2\_hap1.g00981.t01 |
| 2726 | 2 | 147161830 | 147177564 | mNeoVis.2.1.1.chr2\_hap1.g00982.t01 |
| 2727 | 2 | 147161830 | 147177564 | mNeoVis.2.1.1.chr2\_hap1.g00982.t01 |
| 2728 | 2 | 147185319 | 147222307 | mNeoVis.2.1.1.chr2\_hap1.g00983.t01 |
| 2729 | 2 | 147185319 | 147222307 | mNeoVis.2.1.1.chr2\_hap1.g00983.t01 |
| 2730 | 2 | 147225020 | 147264468 | mNeoVis.2.1.1.chr2\_hap1.g00984.t01 |
| 2731 | 2 | 147225020 | 147264468 | mNeoVis.2.1.1.chr2\_hap1.g00984.t01 |
| 2732 | 2 | 147256471 | 147263809 | mNeoVis.2.1.1.chr2\_hap1.g00985.t01 |
| 2733 | 2 | 147256471 | 147263809 | mNeoVis.2.1.1.chr2\_hap1.g00985.t01 |
| 2734 | 2 | 147267485 | 147273171 | mNeoVis.2.1.1.chr2\_hap1.g00986.t01 |
| 2735 | 2 | 147267485 | 147273171 | mNeoVis.2.1.1.chr2\_hap1.g00986.t01 |
| 2736 | 2 | 147275209 | 147283328 | mNeoVis.2.1.1.chr2\_hap1.g00987.t01 |
| 2737 | 2 | 147275209 | 147283328 | mNeoVis.2.1.1.chr2\_hap1.g00987.t01 |
| 2738 | 2 | 147285037 | 147300391 | mNeoVis.2.1.1.chr2\_hap1.g00988.t01 |
| 2739 | 2 | 147285037 | 147300391 | mNeoVis.2.1.1.chr2\_hap1.g00988.t01 |
| 2740 | 2 | 147348796 | 147374607 | mNeoVis.2.1.1.chr2\_hap1.g00989.t01 |
| 2741 | 2 | 147348796 | 147374607 | mNeoVis.2.1.1.chr2\_hap1.g00989.t01 |
| 2742 | 2 | 147378431 | 147445996 | mNeoVis.2.1.1.chr2\_hap1.g00990.t01 |
| 2743 | 2 | 147378431 | 147445996 | mNeoVis.2.1.1.chr2\_hap1.g00990.t01 |
| 2744 | 2 | 147448649 | 147460490 | mNeoVis.2.1.1.chr2\_hap1.g00991.t01 |
| 2745 | 2 | 147448649 | 147460490 | mNeoVis.2.1.1.chr2\_hap1.g00991.t01 |
| 2746 | 2 | 147465688 | 147472830 | mNeoVis.2.1.1.chr2\_hap1.g00992.t01 |
| 2747 | 2 | 147465688 | 147472830 | mNeoVis.2.1.1.chr2\_hap1.g00992.t01 |
| 2748 | 2 | 147473678 | 147497197 | mNeoVis.2.1.1.chr2\_hap1.g00993.t01 |
| 2749 | 2 | 147473678 | 147497197 | mNeoVis.2.1.1.chr2\_hap1.g00993.t01 |
| 2750 | 2 | 147508004 | 147529010 | mNeoVis.2.1.1.chr2\_hap1.g00994.t01 |
| 2751 | 2 | 147508004 | 147529010 | mNeoVis.2.1.1.chr2\_hap1.g00994.t01 |
| 2752 | 2 | 147537128 | 147606933 | mNeoVis.2.1.1.chr2\_hap1.g00995.t01 |
| 2753 | 2 | 147537128 | 147606933 | mNeoVis.2.1.1.chr2\_hap1.g00995.t01 |
| 2754 | 2 | 147607797 | 147616990 | mNeoVis.2.1.1.chr2\_hap1.g00996.t01 |
| 2755 | 2 | 147607797 | 147616990 | mNeoVis.2.1.1.chr2\_hap1.g00996.t01 |
| 2756 | 2 | 147620276 | 147625782 | mNeoVis.2.1.1.chr2\_hap1.g00997.t01 |
| 2757 | 2 | 147637611 | 147646085 | mNeoVis.2.1.1.chr2\_hap1.g00998.t01 |
| 2758 | 2 | 147654576 | 147717413 | mNeoVis.2.1.1.chr2\_hap1.g00999.t01 |
| 2759 | 2 | 147719100 | 147748531 | mNeoVis.2.1.1.chr2\_hap1.g01000.t01 |
| 2760 | 2 | 147749100 | 147809075 | mNeoVis.2.1.1.chr2\_hap1.g01001.t01 |
| 2761 | 2 | 148900531 | 148962436 | mNeoVis.2.1.1.chr2\_hap1.g01019.t01 |
| 2762 | 2 | 151017832 | 151078851 | mNeoVis.2.1.1.chr2\_hap1.g01026.t01 |
| 2763 | 2 | 152610025 | 152652161 | mNeoVis.2.1.1.chr2\_hap1.g01027.t01 |
| 2764 | 2 | 152651134 | 152676737 | mNeoVis.2.1.1.chr2\_hap1.g01028.t01 |
| 2765 | 2 | 153094602 | 153294064 | mNeoVis.2.1.1.chr2\_hap1.g01029.t01 |
| 2766 | 2 | 154116110 | 154167303 | mNeoVis.2.1.1.chr2\_hap1.g01030.t01 |
| 2767 | 2 | 154538468 | 154539433 | mNeoVis.2.1.1.chr2\_hap1.g01031.t01 |
| 2768 | 2 | 154639065 | 154644284 | mNeoVis.2.1.1.chr2\_hap1.g01032.t01 |
| 2769 | 2 | 154865706 | 154902112 | mNeoVis.2.1.1.chr2\_hap1.g01033.t01 |
| 2770 | 2 | 154865706 | 154902112 | mNeoVis.2.1.1.chr2\_hap1.g01033.t01 |
| 2771 | 2 | 154914001 | 155001338 | mNeoVis.2.1.1.chr2\_hap1.g01034.t01 |
| 2772 | 2 | 154914001 | 155001338 | mNeoVis.2.1.1.chr2\_hap1.g01034.t01 |
| 2773 | 2 | 154914001 | 155001338 | mNeoVis.2.1.1.chr2\_hap1.g01034.t01 |
| 2774 | 2 | 155002240 | 155021961 | mNeoVis.2.1.1.chr2\_hap1.g01035.t01 |
| 2775 | 2 | 155002240 | 155021961 | mNeoVis.2.1.1.chr2\_hap1.g01035.t01 |
| 2776 | 2 | 155002240 | 155021961 | mNeoVis.2.1.1.chr2\_hap1.g01035.t01 |
| 2777 | 2 | 155101043 | 155120021 | mNeoVis.2.1.1.chr2\_hap1.g01036.t01 |
| 2778 | 2 | 155101043 | 155120021 | mNeoVis.2.1.1.chr2\_hap1.g01036.t01 |
| 2779 | 2 | 155101043 | 155120021 | mNeoVis.2.1.1.chr2\_hap1.g01036.t01 |
| 2780 | 2 | 155245949 | 155250055 | mNeoVis.2.1.1.chr2\_hap1.g01037.t01 |
| 2781 | 2 | 155245949 | 155250055 | mNeoVis.2.1.1.chr2\_hap1.g01037.t01 |
| 2782 | 2 | 155290684 | 155442654 | mNeoVis.2.1.1.chr2\_hap1.g01038.t01 |
| 2783 | 2 | 155481375 | 155508025 | mNeoVis.2.1.1.chr2\_hap1.g01039.t01 |
| 2784 | 2 | 155486995 | 155487386 | mNeoVis.2.1.1.chr2\_hap1.g01040.t01 |
| 2785 | 2 | 155733072 | 155771462 | mNeoVis.2.1.1.chr2\_hap1.g01046.t01 |
| 2786 | 2 | 155742058 | 155784037 | mNeoVis.2.1.1.chr2\_hap1.g01047.t01 |
| 2787 | 2 | 155831908 | 155906969 | mNeoVis.2.1.1.chr2\_hap1.g01048.t01 |
| 2788 | 2 | 155907049 | 156015957 | mNeoVis.2.1.1.chr2\_hap1.g01049.t01 |
| 2789 | 2 | 156038169 | 156090206 | mNeoVis.2.1.1.chr2\_hap1.g01050.t01 |
| 2790 | 2 | 156378572 | 156419732 | mNeoVis.2.1.1.chr2\_hap1.g01051.t01 |
| 2791 | 2 | 157289661 | 157445386 | mNeoVis.2.1.1.chr2\_hap1.g01054.t01 |
| 2792 | 2 | 157591509 | 158436051 | mNeoVis.2.1.1.chr2\_hap1.g01055.t01 |
| 2793 | 2 | 158730950 | 158732807 | mNeoVis.2.1.1.chr2\_hap1.g01056.t01 |
| 2794 | 2 | 159133609 | 159322362 | mNeoVis.2.1.1.chr2\_hap1.g01058.t01 |
| 2795 | 2 | 160023534 | 160037397 | mNeoVis.2.1.1.chr2\_hap1.g01059.t01 |
| 2796 | 2 | 160023534 | 160037397 | mNeoVis.2.1.1.chr2\_hap1.g01059.t01 |
| 2797 | 2 | 160070386 | 160171993 | mNeoVis.2.1.1.chr2\_hap1.g01060.t01 |
| 2798 | 2 | 160070386 | 160171993 | mNeoVis.2.1.1.chr2\_hap1.g01060.t01 |
| 2799 | 2 | 160177538 | 160266682 | mNeoVis.2.1.1.chr2\_hap1.g01061.t01 |
| 2800 | 2 | 160177538 | 160266682 | mNeoVis.2.1.1.chr2\_hap1.g01061.t01 |
| 2801 | 2 | 160177538 | 160266682 | mNeoVis.2.1.1.chr2\_hap1.g01061.t01 |
| 2802 | 2 | 160319021 | 160348388 | mNeoVis.2.1.1.chr2\_hap1.g01062.t01 |
| 2803 | 2 | 160319021 | 160348388 | mNeoVis.2.1.1.chr2\_hap1.g01062.t01 |
| 2804 | 2 | 160319021 | 160348388 | mNeoVis.2.1.1.chr2\_hap1.g01062.t01 |
| 2805 | 2 | 160329528 | 160446705 | mNeoVis.2.1.1.chr2\_hap1.g01063.t01 |
| 2806 | 2 | 160329528 | 160446705 | mNeoVis.2.1.1.chr2\_hap1.g01063.t01 |
| 2807 | 2 | 160329528 | 160446705 | mNeoVis.2.1.1.chr2\_hap1.g01063.t01 |
| 2808 | 2 | 160660972 | 160673076 | mNeoVis.2.1.1.chr2\_hap1.g01064.t01 |
| 2809 | 2 | 160660972 | 160673076 | mNeoVis.2.1.1.chr2\_hap1.g01064.t01 |
| 2810 | 2 | 160660972 | 160673076 | mNeoVis.2.1.1.chr2\_hap1.g01064.t01 |
| 2811 | 2 | 160693679 | 160770105 | mNeoVis.2.1.1.chr2\_hap1.g01065.t01 |
| 2812 | 2 | 160693679 | 160770105 | mNeoVis.2.1.1.chr2\_hap1.g01065.t01 |
| 2813 | 2 | 160924155 | 160986998 | mNeoVis.2.1.1.chr2\_hap1.g01066.t01 |
| 2814 | 2 | 161018267 | 161142537 | mNeoVis.2.1.1.chr2\_hap1.g01067.t01 |
| 2815 | 2 | 161177332 | 161199120 | mNeoVis.2.1.1.chr2\_hap1.g01068.t01 |
| 2816 | 2 | 161177332 | 161199120 | mNeoVis.2.1.1.chr2\_hap1.g01068.t01 |
| 2817 | 2 | 161204282 | 161224417 | mNeoVis.2.1.1.chr2\_hap1.g01069.t01 |
| 2818 | 2 | 161204282 | 161224417 | mNeoVis.2.1.1.chr2\_hap1.g01069.t01 |
| 2819 | 2 | 161224569 | 161237994 | mNeoVis.2.1.1.chr2\_hap1.g01070.t01 |
| 2820 | 2 | 161224569 | 161237994 | mNeoVis.2.1.1.chr2\_hap1.g01070.t01 |
| 2821 | 2 | 161370849 | 161480272 | mNeoVis.2.1.1.chr2\_hap1.g01071.t01 |
| 2822 | 2 | 161370849 | 161480272 | mNeoVis.2.1.1.chr2\_hap1.g01071.t01 |
| 2823 | 2 | 161490299 | 161590458 | mNeoVis.2.1.1.chr2\_hap1.g01072.t01 |
| 2824 | 2 | 161603321 | 161603964 | mNeoVis.2.1.1.chr2\_hap1.g01073.t01 |
| 2825 | 2 | 161657702 | 161675016 | mNeoVis.2.1.1.chr2\_hap1.g01074.t01 |
| 2826 | 2 | 161771538 | 161902651 | mNeoVis.2.1.1.chr2\_hap1.g01075.t01 |
| 2827 | 2 | 161903137 | 161919324 | mNeoVis.2.1.1.chr2\_hap1.g01076.t01 |
| 2828 | 2 | 161933178 | 161990335 | mNeoVis.2.1.1.chr2\_hap1.g01077.t01 |
| 2829 | 2 | 162029681 | 162066898 | mNeoVis.2.1.1.chr2\_hap1.g01078.t01 |
| 2830 | 2 | 162064824 | 162193738 | mNeoVis.2.1.1.chr2\_hap1.g01079.t01 |
| 2831 | 2 | 162193974 | 162203694 | mNeoVis.2.1.1.chr2\_hap1.g01080.t01 |
| 2832 | 2 | 162467674 | 162476350 | mNeoVis.2.1.1.chr2\_hap1.g01082.t01 |
| 2833 | 2 | 162506248 | 162677351 | mNeoVis.2.1.1.chr2\_hap1.g01083.t01 |
| 2834 | 2 | 162556334 | 162639727 | mNeoVis.2.1.1.chr2\_hap1.g01084.t01 |
| 2835 | 2 | 162678090 | 162717218 | mNeoVis.2.1.1.chr2\_hap1.g01085.t01 |
| 2836 | 2 | 162753451 | 162867261 | mNeoVis.2.1.1.chr2\_hap1.g01086.t01 |
| 2837 | 2 | 162879564 | 162923558 | mNeoVis.2.1.1.chr2\_hap1.g01087.t01 |
| 2838 | 2 | 163087584 | 163287273 | mNeoVis.2.1.1.chr2\_hap1.g01089.t01 |
| 2839 | 2 | 163407568 | 163434666 | mNeoVis.2.1.1.chr2\_hap1.g01090.t01 |
| 2840 | 2 | 163798563 | 163941119 | mNeoVis.2.1.1.chr2\_hap1.g01093.t01 |
| 2841 | 2 | 163947605 | 164017346 | mNeoVis.2.1.1.chr2\_hap1.g01094.t01 |
| 2842 | 2 | 163947605 | 164017346 | mNeoVis.2.1.1.chr2\_hap1.g01094.t01 |
| 2843 | 2 | 163947605 | 164017346 | mNeoVis.2.1.1.chr2\_hap1.g01094.t01 |
| 2844 | 2 | 164086239 | 164092318 | mNeoVis.2.1.1.chr2\_hap1.g01095.t01 |
| 2845 | 2 | 164086239 | 164092318 | mNeoVis.2.1.1.chr2\_hap1.g01095.t01 |
| 2846 | 2 | 164086239 | 164092318 | mNeoVis.2.1.1.chr2\_hap1.g01095.t01 |
| 2847 | 2 | 164765051 | 164803819 | mNeoVis.2.1.1.chr2\_hap1.g01096.t01 |
| 2848 | 2 | 164844685 | 164845287 | mNeoVis.2.1.1.chr2\_hap1.g01097.t01 |
| 2849 | 2 | 164864254 | 164919764 | mNeoVis.2.1.1.chr2\_hap1.g01098.t01 |
| 2850 | 2 | 165086721 | 165172874 | mNeoVis.2.1.1.chr2\_hap1.g01099.t01 |
| 2851 | 2 | 165227484 | 165291124 | mNeoVis.2.1.1.chr2\_hap1.g01100.t01 |
| 2852 | 2 | 165459188 | 165489209 | mNeoVis.2.1.1.chr2\_hap1.g01101.t01 |
| 2853 | 2 | 165528843 | 165552170 | mNeoVis.2.1.1.chr2\_hap1.g01102.t01 |
| 2854 | 2 | 165647201 | 165659548 | mNeoVis.2.1.1.chr2\_hap1.g01103.t01 |
| 2855 | 2 | 165665055 | 165677062 | mNeoVis.2.1.1.chr2\_hap1.g01104.t01 |
| 2856 | 2 | 165758651 | 165784664 | mNeoVis.2.1.1.chr2\_hap1.g01105.t01 |
| 2857 | 2 | 165963204 | 165996667 | mNeoVis.2.1.1.chr2\_hap1.g01109.t01 |
| 2858 | 2 | 166026177 | 166159835 | mNeoVis.2.1.1.chr2\_hap1.g01110.t01 |
| 2859 | 2 | 166396814 | 166467494 | mNeoVis.2.1.1.chr2\_hap1.g01111.t01 |
| 2860 | 2 | 166396814 | 166467494 | mNeoVis.2.1.1.chr2\_hap1.g01111.t01 |
| 2861 | 2 | 167011563 | 167014446 | mNeoVis.2.1.1.chr2\_hap1.g01112.t01 |
| 2862 | 2 | 167030280 | 167087793 | mNeoVis.2.1.1.chr2\_hap1.g01113.t01 |
| 2863 | 2 | 167672012 | 167850505 | mNeoVis.2.1.1.chr2\_hap1.g01115.t01 |
| 2864 | 2 | 167851019 | 167912054 | mNeoVis.2.1.1.chr2\_hap1.g01116.t01 |
| 2865 | 2 | 167988879 | 168022998 | mNeoVis.2.1.1.chr2\_hap1.g01117.t01 |
| 2866 | 2 | 168888345 | 168981654 | mNeoVis.2.1.1.chr2\_hap1.g01125.t01 |
| 2867 | 2 | 169003701 | 169006637 | mNeoVis.2.1.1.chr2\_hap1.g01126.t01 |
| 2868 | 2 | 169122784 | 169257550 | mNeoVis.2.1.1.chr2\_hap1.g01127.t01 |
| 2869 | 2 | 169122784 | 169257550 | mNeoVis.2.1.1.chr2\_hap1.g01127.t01 |
| 2870 | 2 | 169288678 | 169299532 | mNeoVis.2.1.1.chr2\_hap1.g01128.t01 |
| 2871 | 2 | 169288678 | 169299532 | mNeoVis.2.1.1.chr2\_hap1.g01128.t01 |
| 2872 | 2 | 169305595 | 169311725 | mNeoVis.2.1.1.chr2\_hap1.g01129.t01 |
| 2873 | 2 | 169305595 | 169311725 | mNeoVis.2.1.1.chr2\_hap1.g01129.t01 |
| 2874 | 2 | 169348993 | 169397544 | mNeoVis.2.1.1.chr2\_hap1.g01130.t01 |
| 2875 | 2 | 169423109 | 169490994 | mNeoVis.2.1.1.chr2\_hap1.g01131.t01 |
| 2876 | 2 | 169502433 | 169529049 | mNeoVis.2.1.1.chr2\_hap1.g01132.t01 |
| 2877 | 2 | 169531372 | 169586165 | mNeoVis.2.1.1.chr2\_hap1.g01133.t01 |
| 2878 | 2 | 169808005 | 169852710 | mNeoVis.2.1.1.chr2\_hap1.g01135.t01 |
| 2879 | 2 | 169808005 | 169852710 | mNeoVis.2.1.1.chr2\_hap1.g01135.t01 |
| 2880 | 2 | 169862184 | 169928421 | mNeoVis.2.1.1.chr2\_hap1.g01136.t01 |
| 2881 | 2 | 169862184 | 169928421 | mNeoVis.2.1.1.chr2\_hap1.g01136.t01 |
| 2882 | 2 | 169922165 | 169973555 | mNeoVis.2.1.1.chr2\_hap1.g01137.t01 |
| 2883 | 2 | 169922165 | 169973555 | mNeoVis.2.1.1.chr2\_hap1.g01137.t01 |
| 2884 | 2 | 169973609 | 169981422 | mNeoVis.2.1.1.chr2\_hap1.g01138.t01 |
| 2885 | 2 | 169973609 | 169981422 | mNeoVis.2.1.1.chr2\_hap1.g01138.t01 |
| 2886 | 2 | 169982214 | 170004325 | mNeoVis.2.1.1.chr2\_hap1.g01139.t01 |
| 2887 | 2 | 169982214 | 170004325 | mNeoVis.2.1.1.chr2\_hap1.g01139.t01 |
| 2888 | 2 | 170022906 | 170103013 | mNeoVis.2.1.1.chr2\_hap1.g01140.t01 |
| 2889 | 2 | 170022906 | 170103013 | mNeoVis.2.1.1.chr2\_hap1.g01140.t01 |
| 2890 | 2 | 170057045 | 170072323 | mNeoVis.2.1.1.chr2\_hap1.g01141.t01 |
| 2891 | 2 | 170057045 | 170072323 | mNeoVis.2.1.1.chr2\_hap1.g01141.t01 |
| 2892 | 2 | 170106364 | 170124442 | mNeoVis.2.1.1.chr2\_hap1.g01142.t01 |
| 2893 | 2 | 170106364 | 170124442 | mNeoVis.2.1.1.chr2\_hap1.g01142.t01 |
| 2894 | 2 | 170118422 | 170159938 | mNeoVis.2.1.1.chr2\_hap1.g01143.t01 |
| 2895 | 2 | 170118422 | 170159938 | mNeoVis.2.1.1.chr2\_hap1.g01143.t01 |
| 2896 | 2 | 170206178 | 170281585 | mNeoVis.2.1.1.chr2\_hap1.g01144.t01 |
| 2897 | 2 | 170378411 | 170383096 | mNeoVis.2.1.1.chr2\_hap1.g01145.t01 |
| 2898 | 2 | 170381089 | 170540734 | mNeoVis.2.1.1.chr2\_hap1.g01146.t01 |
| 2899 | 2 | 171120531 | 171129717 | mNeoVis.2.1.1.chr2\_hap1.g01147.t01 |
| 2900 | 2 | 171861359 | 172235971 | mNeoVis.2.1.1.chr2\_hap1.g01148.t01 |
| 2901 | 2 | 172271536 | 172804148 | mNeoVis.2.1.1.chr2\_hap1.g01149.t01 |
| 2902 | 2 | 172461141 | 172463788 | mNeoVis.2.1.1.chr2\_hap1.g01150.t01 |
| 2903 | 2 | 172461141 | 172463788 | mNeoVis.2.1.1.chr2\_hap1.g01150.t01 |
| 2904 | 2 | 172461141 | 172463788 | mNeoVis.2.1.1.chr2\_hap1.g01150.t01 |
| 2905 | 2 | 173179654 | 173375986 | mNeoVis.2.1.1.chr2\_hap1.g01152.t01 |
| 2906 | 2 | 173179654 | 173375986 | mNeoVis.2.1.1.chr2\_hap1.g01152.t01 |
| 2907 | 2 | 173179654 | 173375986 | mNeoVis.2.1.1.chr2\_hap1.g01152.t01 |
| 2908 | 2 | 174861537 | 174971927 | mNeoVis.2.1.1.chr2\_hap1.g01153.t01 |
| 2909 | 2 | 174861537 | 174971927 | mNeoVis.2.1.1.chr2\_hap1.g01153.t01 |
| 2910 | 2 | 175152586 | 175194771 | mNeoVis.2.1.1.chr2\_hap1.g01154.t01 |
| 2911 | 2 | 175152586 | 175194771 | mNeoVis.2.1.1.chr2\_hap1.g01154.t01 |
| 2912 | 2 | 175610085 | 175695315 | mNeoVis.2.1.1.chr2\_hap1.g01156.t01 |
| 2913 | 2 | 175610085 | 175695315 | mNeoVis.2.1.1.chr2\_hap1.g01156.t01 |
| 2914 | 2 | 175718155 | 175764064 | mNeoVis.2.1.1.chr2\_hap1.g01157.t01 |
| 2915 | 2 | 175718155 | 175764064 | mNeoVis.2.1.1.chr2\_hap1.g01157.t01 |
| 2916 | 2 | 175764347 | 175798503 | mNeoVis.2.1.1.chr2\_hap1.g01158.t01 |
| 2917 | 2 | 175764347 | 175798503 | mNeoVis.2.1.1.chr2\_hap1.g01158.t01 |
| 2918 | 2 | 175796199 | 175852457 | mNeoVis.2.1.1.chr2\_hap1.g01159.t01 |
| 2919 | 2 | 175796199 | 175852457 | mNeoVis.2.1.1.chr2\_hap1.g01159.t01 |
| 2920 | 2 | 175858156 | 175948442 | mNeoVis.2.1.1.chr2\_hap1.g01160.t01 |
| 2921 | 2 | 175858156 | 175948442 | mNeoVis.2.1.1.chr2\_hap1.g01160.t01 |
| 2922 | 2 | 175961651 | 176015473 | mNeoVis.2.1.1.chr2\_hap1.g01161.t01 |
| 2923 | 2 | 175961651 | 176015473 | mNeoVis.2.1.1.chr2\_hap1.g01161.t01 |
| 2924 | 2 | 176023415 | 176025069 | mNeoVis.2.1.1.chr2\_hap1.g01162.t01 |
| 2925 | 2 | 176023415 | 176025069 | mNeoVis.2.1.1.chr2\_hap1.g01162.t01 |
| 2926 | 2 | 176024425 | 176149068 | mNeoVis.2.1.1.chr2\_hap1.g01163.t01 |
| 2927 | 2 | 176024425 | 176149068 | mNeoVis.2.1.1.chr2\_hap1.g01163.t01 |
| 2928 | 2 | 176150300 | 176374441 | mNeoVis.2.1.1.chr2\_hap1.g01164.t01 |
| 2929 | 2 | 176150300 | 176374441 | mNeoVis.2.1.1.chr2\_hap1.g01164.t01 |
| 2930 | 2 | 176428696 | 176561362 | mNeoVis.2.1.1.chr2\_hap1.g01165.t01 |
| 2931 | 2 | 176581560 | 176749802 | mNeoVis.2.1.1.chr2\_hap1.g01166.t01 |
| 2932 | 2 | 177712628 | 177731503 | mNeoVis.2.1.1.chr2\_hap1.g01168.t01 |
| 2933 | 2 | 177736369 | 177853276 | mNeoVis.2.1.1.chr2\_hap1.g01169.t01 |
| 2934 | 2 | 177855311 | 177863855 | mNeoVis.2.1.1.chr2\_hap1.g01170.t01 |
| 2935 | 2 | 177882989 | 177907534 | mNeoVis.2.1.1.chr2\_hap1.g01171.t01 |
| 2936 | 2 | 177907829 | 177966377 | mNeoVis.2.1.1.chr2\_hap1.g01172.t01 |
| 2937 | 2 | 179253710 | 179263059 | mNeoVis.2.1.1.chr2\_hap1.g01181.t01 |
| 2938 | 2 | 179253710 | 179263059 | mNeoVis.2.1.1.chr2\_hap1.g01181.t01 |
| 2939 | 2 | 179263150 | 179469831 | mNeoVis.2.1.1.chr2\_hap1.g01182.t01 |
| 2940 | 2 | 179263150 | 179469831 | mNeoVis.2.1.1.chr2\_hap1.g01182.t01 |
| 2941 | 2 | 179280414 | 179280996 | mNeoVis.2.1.1.chr2\_hap1.g01183.t01 |
| 2942 | 2 | 179280414 | 179280996 | mNeoVis.2.1.1.chr2\_hap1.g01183.t01 |
| 2943 | 2 | 180689871 | 180691099 | mNeoVis.2.1.1.chr2\_hap1.g01184.t01 |
| 2944 | 2 | 181194880 | 182054842 | mNeoVis.2.1.1.chr2\_hap1.g01185.t01 |
| 2945 | 2 | 182292256 | 182310415 | mNeoVis.2.1.1.chr2\_hap1.g01186.t01 |
| 2946 | 2 | 182292256 | 182310415 | mNeoVis.2.1.1.chr2\_hap1.g01186.t01 |
| 2947 | 2 | 182325761 | 182524416 | mNeoVis.2.1.1.chr2\_hap1.g01187.t01 |
| 2948 | 2 | 182325761 | 182524416 | mNeoVis.2.1.1.chr2\_hap1.g01187.t01 |
| 2949 | 2 | 182792707 | 182816503 | mNeoVis.2.1.1.chr2\_hap1.g01188.t01 |
| 2950 | 2 | 182792707 | 182816503 | mNeoVis.2.1.1.chr2\_hap1.g01188.t01 |
| 2951 | 2 | 182792707 | 182816503 | mNeoVis.2.1.1.chr2\_hap1.g01188.t01 |
| 2952 | 2 | 182792707 | 182816503 | mNeoVis.2.1.1.chr2\_hap1.g01188.t01 |
| 2953 | 2 | 182862072 | 182960438 | mNeoVis.2.1.1.chr2\_hap1.g01189.t01 |
| 2954 | 2 | 182862072 | 182960438 | mNeoVis.2.1.1.chr2\_hap1.g01189.t01 |
| 2955 | 2 | 182862072 | 182960438 | mNeoVis.2.1.1.chr2\_hap1.g01189.t01 |
| 2956 | 2 | 182862072 | 182960438 | mNeoVis.2.1.1.chr2\_hap1.g01189.t01 |
| 2957 | 2 | 182965142 | 183008983 | mNeoVis.2.1.1.chr2\_hap1.g01190.t01 |
| 2958 | 2 | 182965142 | 183008983 | mNeoVis.2.1.1.chr2\_hap1.g01190.t01 |
| 2959 | 2 | 182965142 | 183008983 | mNeoVis.2.1.1.chr2\_hap1.g01190.t01 |
| 2960 | 2 | 182965142 | 183008983 | mNeoVis.2.1.1.chr2\_hap1.g01190.t01 |
| 2961 | 2 | 183009034 | 183069494 | mNeoVis.2.1.1.chr2\_hap1.g01191.t01 |
| 2962 | 2 | 183009034 | 183069494 | mNeoVis.2.1.1.chr2\_hap1.g01191.t01 |
| 2963 | 2 | 183009034 | 183069494 | mNeoVis.2.1.1.chr2\_hap1.g01191.t01 |
| 2964 | 2 | 183009034 | 183069494 | mNeoVis.2.1.1.chr2\_hap1.g01191.t01 |
| 2965 | 2 | 184575622 | 184594116 | mNeoVis.2.1.1.chr2\_hap1.g01193.t01 |
| 2966 | 2 | 184631065 | 184651620 | mNeoVis.2.1.1.chr2\_hap1.g01194.t01 |
| 2967 | 2 | 184817352 | 184919258 | mNeoVis.2.1.1.chr2\_hap1.g01195.t01 |
| 2968 | 2 | 184957801 | 184976154 | mNeoVis.2.1.1.chr2\_hap1.g01196.t01 |
| 2969 | 2 | 185133220 | 185148540 | mNeoVis.2.1.1.chr2\_hap1.g01198.t01 |
| 2970 | 2 | 185273760 | 185276907 | mNeoVis.2.1.1.chr2\_hap1.g01199.t01 |
| 2971 | 2 | 185484957 | 185500100 | mNeoVis.2.1.1.chr2\_hap1.g01200.t01 |
| 2972 | 2 | 185484957 | 185500100 | mNeoVis.2.1.1.chr2\_hap1.g01200.t01 |
| 2973 | 2 | 185484957 | 185500100 | mNeoVis.2.1.1.chr2\_hap1.g01200.t01 |
| 2974 | 2 | 185505535 | 185586421 | mNeoVis.2.1.1.chr2\_hap1.g01201.t01 |
| 2975 | 2 | 185505535 | 185586421 | mNeoVis.2.1.1.chr2\_hap1.g01201.t01 |
| 2976 | 2 | 185505535 | 185586421 | mNeoVis.2.1.1.chr2\_hap1.g01201.t01 |
| 2977 | 2 | 185629312 | 185690253 | mNeoVis.2.1.1.chr2\_hap1.g01202.t01 |
| 2978 | 2 | 185629312 | 185690253 | mNeoVis.2.1.1.chr2\_hap1.g01202.t01 |
| 2979 | 2 | 185713302 | 185781289 | mNeoVis.2.1.1.chr2\_hap1.g01203.t01 |
| 2980 | 2 | 185713302 | 185781289 | mNeoVis.2.1.1.chr2\_hap1.g01203.t01 |
| 2981 | 2 | 185713302 | 185781289 | mNeoVis.2.1.1.chr2\_hap1.g01203.t01 |
| 2982 | 2 | 185803719 | 185851392 | mNeoVis.2.1.1.chr2\_hap1.g01204.t01 |
| 2983 | 2 | 185803719 | 185851392 | mNeoVis.2.1.1.chr2\_hap1.g01204.t01 |
| 2984 | 2 | 185861723 | 185923126 | mNeoVis.2.1.1.chr2\_hap1.g01205.t01 |
| 2985 | 2 | 185861723 | 185923126 | mNeoVis.2.1.1.chr2\_hap1.g01205.t01 |
| 2986 | 2 | 185923252 | 186004123 | mNeoVis.2.1.1.chr2\_hap1.g01206.t01 |
| 2987 | 2 | 185923252 | 186004123 | mNeoVis.2.1.1.chr2\_hap1.g01206.t01 |
| 2988 | 2 | 186031511 | 186059872 | mNeoVis.2.1.1.chr2\_hap1.g01207.t01 |
| 2989 | 2 | 186031511 | 186059872 | mNeoVis.2.1.1.chr2\_hap1.g01207.t01 |
| 2990 | 2 | 186425508 | 186986439 | mNeoVis.2.1.1.chr2\_hap1.g01209.t01 |
| 2991 | 2 | 187082854 | 187237572 | mNeoVis.2.1.1.chr2\_hap1.g01210.t01 |
| 2992 | 2 | 187246346 | 187262387 | mNeoVis.2.1.1.chr2\_hap1.g01211.t01 |
| 2993 | 2 | 187246346 | 187262387 | mNeoVis.2.1.1.chr2\_hap1.g01211.t01 |
| 2994 | 2 | 187246346 | 187262387 | mNeoVis.2.1.1.chr2\_hap1.g01211.t01 |
| 2995 | 2 | 187268376 | 187407719 | mNeoVis.2.1.1.chr2\_hap1.g01212.t01 |
| 2996 | 2 | 187268376 | 187407719 | mNeoVis.2.1.1.chr2\_hap1.g01212.t01 |
| 2997 | 2 | 187268376 | 187407719 | mNeoVis.2.1.1.chr2\_hap1.g01212.t01 |
| 2998 | 2 | 187455237 | 187492791 | mNeoVis.2.1.1.chr2\_hap1.g01213.t01 |
| 2999 | 2 | 187455237 | 187492791 | mNeoVis.2.1.1.chr2\_hap1.g01213.t01 |
| 3000 | 2 | 187455237 | 187492791 | mNeoVis.2.1.1.chr2\_hap1.g01213.t01 |
| 3001 | 2 | 187645944 | 187770402 | mNeoVis.2.1.1.chr2\_hap1.g01214.t01 |
| 3002 | 2 | 187645944 | 187770402 | mNeoVis.2.1.1.chr2\_hap1.g01214.t01 |
| 3003 | 2 | 187645944 | 187770402 | mNeoVis.2.1.1.chr2\_hap1.g01214.t01 |
| 3004 | 2 | 187771520 | 187862557 | mNeoVis.2.1.1.chr2\_hap1.g01215.t01 |
| 3005 | 2 | 187771520 | 187862557 | mNeoVis.2.1.1.chr2\_hap1.g01215.t01 |
| 3006 | 2 | 187771520 | 187862557 | mNeoVis.2.1.1.chr2\_hap1.g01215.t01 |
| 3007 | 2 | 187911786 | 188119079 | mNeoVis.2.1.1.chr2\_hap1.g01216.t01 |
| 3008 | 2 | 188294515 | 188353318 | mNeoVis.2.1.1.chr2\_hap1.g01217.t01 |
| 3009 | 2 | 188294515 | 188353318 | mNeoVis.2.1.1.chr2\_hap1.g01217.t01 |
| 3010 | 2 | 188353371 | 188376664 | mNeoVis.2.1.1.chr2\_hap1.g01218.t01 |
| 3011 | 2 | 188353371 | 188376664 | mNeoVis.2.1.1.chr2\_hap1.g01218.t01 |
| 3012 | 2 | 188365947 | 188779309 | mNeoVis.2.1.1.chr2\_hap1.g01219.t01 |
| 3013 | 2 | 188365947 | 188779309 | mNeoVis.2.1.1.chr2\_hap1.g01219.t01 |
| 3014 | 2 | 188839191 | 188927774 | mNeoVis.2.1.1.chr2\_hap1.g01220.t01 |
| 3015 | 2 | 188975439 | 189026347 | mNeoVis.2.1.1.chr2\_hap1.g01221.t01 |
| 3016 | 2 | 189026469 | 189092738 | mNeoVis.2.1.1.chr2\_hap1.g01222.t01 |
| 3017 | 2 | 189026469 | 189092738 | mNeoVis.2.1.1.chr2\_hap1.g01222.t01 |
| 3018 | 2 | 189026469 | 189092738 | mNeoVis.2.1.1.chr2\_hap1.g01222.t01 |
| 3019 | 2 | 189092880 | 189160110 | mNeoVis.2.1.1.chr2\_hap1.g01223.t01 |
| 3020 | 2 | 189092880 | 189160110 | mNeoVis.2.1.1.chr2\_hap1.g01223.t01 |
| 3021 | 2 | 189092880 | 189160110 | mNeoVis.2.1.1.chr2\_hap1.g01223.t01 |
| 3022 | 2 | 189193026 | 189195079 | mNeoVis.2.1.1.chr2\_hap1.g01224.t01 |
| 3023 | 2 | 189193026 | 189195079 | mNeoVis.2.1.1.chr2\_hap1.g01224.t01 |
| 3024 | 2 | 189193026 | 189195079 | mNeoVis.2.1.1.chr2\_hap1.g01224.t01 |
| 3025 | 2 | 189643049 | 189724640 | mNeoVis.2.1.1.chr2\_hap1.g01226.t01 |
| 3026 | 2 | 189875219 | 189884907 | mNeoVis.2.1.1.chr2\_hap1.g01227.t01 |
| 3027 | 2 | 190015450 | 190028304 | mNeoVis.2.1.1.chr2\_hap1.g01228.t01 |
| 3028 | 2 | 190131961 | 190200372 | mNeoVis.2.1.1.chr2\_hap1.g01229.t01 |
| 3029 | 2 | 190131961 | 190200372 | mNeoVis.2.1.1.chr2\_hap1.g01229.t01 |
| 3030 | 2 | 190275494 | 190645949 | mNeoVis.2.1.1.chr2\_hap1.g01230.t01 |
| 3031 | 2 | 190275494 | 190645949 | mNeoVis.2.1.1.chr2\_hap1.g01230.t01 |
| 3032 | 2 | 190656483 | 190701869 | mNeoVis.2.1.1.chr2\_hap1.g01231.t01 |
| 3033 | 2 | 190656483 | 190701869 | mNeoVis.2.1.1.chr2\_hap1.g01231.t01 |
| 3034 | 2 | 193415735 | 193418502 | mNeoVis.2.1.1.chr2\_hap1.g01240.t01 |
| 3035 | 2 | 193415735 | 193418502 | mNeoVis.2.1.1.chr2\_hap1.g01240.t01 |
| 3036 | 2 | 193482642 | 193526409 | mNeoVis.2.1.1.chr2\_hap1.g01241.t01 |
| 3037 | 2 | 193482642 | 193526409 | mNeoVis.2.1.1.chr2\_hap1.g01241.t01 |
| 3038 | 2 | 193482642 | 193526409 | mNeoVis.2.1.1.chr2\_hap1.g01241.t01 |
| 3039 | 2 | 193574314 | 193576134 | mNeoVis.2.1.1.chr2\_hap1.g01242.t01 |
| 3040 | 2 | 193574314 | 193576134 | mNeoVis.2.1.1.chr2\_hap1.g01242.t01 |
| 3041 | 2 | 193574314 | 193576134 | mNeoVis.2.1.1.chr2\_hap1.g01242.t01 |
| 3042 | 2 | 193595974 | 193682001 | mNeoVis.2.1.1.chr2\_hap1.g01243.t01 |
| 3043 | 2 | 193595974 | 193682001 | mNeoVis.2.1.1.chr2\_hap1.g01243.t01 |
| 3044 | 2 | 193595974 | 193682001 | mNeoVis.2.1.1.chr2\_hap1.g01243.t01 |
| 3045 | 2 | 195053917 | 195096891 | mNeoVis.2.1.1.chr2\_hap1.g01245.t01 |
| 3046 | 2 | 195053917 | 195096891 | mNeoVis.2.1.1.chr2\_hap1.g01245.t01 |
| 3047 | 2 | 195107621 | 195156552 | mNeoVis.2.1.1.chr2\_hap1.g01246.t01 |
| 3048 | 2 | 195107621 | 195156552 | mNeoVis.2.1.1.chr2\_hap1.g01246.t01 |
| 3049 | 2 | 195211711 | 195347403 | mNeoVis.2.1.1.chr2\_hap1.g01247.t01 |
| 3050 | 2 | 195347582 | 195426470 | mNeoVis.2.1.1.chr2\_hap1.g01248.t01 |
| 3051 | 2 | 196656582 | 196796845 | mNeoVis.2.1.1.chr2\_hap1.g01250.t01 |
| 3052 | 2 | 196656582 | 196796845 | mNeoVis.2.1.1.chr2\_hap1.g01250.t01 |
| 3053 | 2 | 196656582 | 196796845 | mNeoVis.2.1.1.chr2\_hap1.g01250.t01 |
| 3054 | 2 | 196822613 | 196834796 | mNeoVis.2.1.1.chr2\_hap1.g01251.t01 |
| 3055 | 2 | 196822613 | 196834796 | mNeoVis.2.1.1.chr2\_hap1.g01251.t01 |
| 3056 | 2 | 196822613 | 196834796 | mNeoVis.2.1.1.chr2\_hap1.g01251.t01 |
| 3057 | 2 | 196838295 | 196848709 | mNeoVis.2.1.1.chr2\_hap1.g01252.t01 |
| 3058 | 2 | 196838295 | 196848709 | mNeoVis.2.1.1.chr2\_hap1.g01252.t01 |
| 3059 | 2 | 196838295 | 196848709 | mNeoVis.2.1.1.chr2\_hap1.g01252.t01 |
| 3060 | 2 | 196883020 | 196913032 | mNeoVis.2.1.1.chr2\_hap1.g01253.t01 |
| 3061 | 2 | 196883020 | 196913032 | mNeoVis.2.1.1.chr2\_hap1.g01253.t01 |
| 3062 | 2 | 196883020 | 196913032 | mNeoVis.2.1.1.chr2\_hap1.g01253.t01 |
| 3063 | 2 | 196917436 | 196943144 | mNeoVis.2.1.1.chr2\_hap1.g01254.t01 |
| 3064 | 2 | 196917436 | 196943144 | mNeoVis.2.1.1.chr2\_hap1.g01254.t01 |
| 3065 | 2 | 196917436 | 196943144 | mNeoVis.2.1.1.chr2\_hap1.g01254.t01 |
| 3066 | 2 | 196947985 | 196967829 | mNeoVis.2.1.1.chr2\_hap1.g01255.t01 |
| 3067 | 2 | 196947985 | 196967829 | mNeoVis.2.1.1.chr2\_hap1.g01255.t01 |
| 3068 | 2 | 196947985 | 196967829 | mNeoVis.2.1.1.chr2\_hap1.g01255.t01 |
| 3069 | 2 | 196979351 | 196988082 | mNeoVis.2.1.1.chr2\_hap1.g01256.t01 |
| 3070 | 2 | 196979351 | 196988082 | mNeoVis.2.1.1.chr2\_hap1.g01256.t01 |
| 3071 | 2 | 196979351 | 196988082 | mNeoVis.2.1.1.chr2\_hap1.g01256.t01 |
| 3072 | 2 | 196992296 | 197017103 | mNeoVis.2.1.1.chr2\_hap1.g01257.t01 |
| 3073 | 2 | 196992296 | 197017103 | mNeoVis.2.1.1.chr2\_hap1.g01257.t01 |
| 3074 | 2 | 196992296 | 197017103 | mNeoVis.2.1.1.chr2\_hap1.g01257.t01 |
| 3075 | 2 | 197020156 | 197074332 | mNeoVis.2.1.1.chr2\_hap1.g01258.t01 |
| 3076 | 2 | 197020156 | 197074332 | mNeoVis.2.1.1.chr2\_hap1.g01258.t01 |
| 3077 | 2 | 197081818 | 197092641 | mNeoVis.2.1.1.chr2\_hap1.g01259.t01 |
| 3078 | 2 | 197081818 | 197092641 | mNeoVis.2.1.1.chr2\_hap1.g01259.t01 |
| 3079 | 2 | 197092629 | 197135724 | mNeoVis.2.1.1.chr2\_hap1.g01260.t01 |
| 3080 | 2 | 197092629 | 197135724 | mNeoVis.2.1.1.chr2\_hap1.g01260.t01 |
| 3081 | 2 | 197245499 | 197407016 | mNeoVis.2.1.1.chr2\_hap1.g01261.t01 |
| 3082 | 2 | 197245499 | 197407016 | mNeoVis.2.1.1.chr2\_hap1.g01261.t01 |
| 3083 | 2 | 197411040 | 197425964 | mNeoVis.2.1.1.chr2\_hap1.g01262.t01 |
| 3084 | 2 | 197411040 | 197425964 | mNeoVis.2.1.1.chr2\_hap1.g01262.t01 |
| 3085 | 2 | 197426097 | 197447295 | mNeoVis.2.1.1.chr2\_hap1.g01263.t01 |
| 3086 | 2 | 197426097 | 197447295 | mNeoVis.2.1.1.chr2\_hap1.g01263.t01 |
| 3087 | 2 | 197459128 | 197474979 | mNeoVis.2.1.1.chr2\_hap1.g01264.t01 |
| 3088 | 2 | 197459128 | 197474979 | mNeoVis.2.1.1.chr2\_hap1.g01264.t01 |
| 3089 | 2 | 197503956 | 197540277 | mNeoVis.2.1.1.chr2\_hap1.g01265.t01 |
| 3090 | 2 | 197503956 | 197540277 | mNeoVis.2.1.1.chr2\_hap1.g01265.t01 |
| 3091 | 2 | 197540522 | 197567820 | mNeoVis.2.1.1.chr2\_hap1.g01266.t01 |
| 3092 | 2 | 197540522 | 197567820 | mNeoVis.2.1.1.chr2\_hap1.g01266.t01 |
| 3093 | 2 | 197575537 | 197584768 | mNeoVis.2.1.1.chr2\_hap1.g01267.t01 |
| 3094 | 2 | 197575537 | 197584768 | mNeoVis.2.1.1.chr2\_hap1.g01267.t01 |
| 3095 | 2 | 197600566 | 197621558 | mNeoVis.2.1.1.chr2\_hap1.g01268.t01 |
| 3096 | 2 | 197600566 | 197621558 | mNeoVis.2.1.1.chr2\_hap1.g01268.t01 |
| 3097 | 2 | 197621646 | 197655756 | mNeoVis.2.1.1.chr2\_hap1.g01269.t01 |
| 3098 | 2 | 197621646 | 197655756 | mNeoVis.2.1.1.chr2\_hap1.g01269.t01 |
| 3099 | 2 | 197668435 | 197683840 | mNeoVis.2.1.1.chr2\_hap1.g01270.t01 |
| 3100 | 2 | 197668435 | 197683840 | mNeoVis.2.1.1.chr2\_hap1.g01270.t01 |
| 3101 | 2 | 197700963 | 197742666 | mNeoVis.2.1.1.chr2\_hap1.g01271.t01 |
| 3102 | 2 | 197749255 | 197796735 | mNeoVis.2.1.1.chr2\_hap1.g01272.t01 |
| 3103 | 2 | 197749255 | 197796735 | mNeoVis.2.1.1.chr2\_hap1.g01272.t01 |
| 3104 | 2 | 197749255 | 197796735 | mNeoVis.2.1.1.chr2\_hap1.g01272.t01 |
| 3105 | 2 | 197760966 | 197762390 | mNeoVis.2.1.1.chr2\_hap1.g01273.t01 |
| 3106 | 2 | 197760966 | 197762390 | mNeoVis.2.1.1.chr2\_hap1.g01273.t01 |
| 3107 | 2 | 197760966 | 197762390 | mNeoVis.2.1.1.chr2\_hap1.g01273.t01 |
| 3108 | 2 | 197823594 | 198041455 | mNeoVis.2.1.1.chr2\_hap1.g01274.t01 |
| 3109 | 2 | 197823594 | 198041455 | mNeoVis.2.1.1.chr2\_hap1.g01274.t01 |
| 3110 | 2 | 198074281 | 198082292 | mNeoVis.2.1.1.chr2\_hap1.g01275.t01 |
| 3111 | 2 | 198074281 | 198082292 | mNeoVis.2.1.1.chr2\_hap1.g01275.t01 |
| 3112 | 2 | 198094410 | 198097013 | mNeoVis.2.1.1.chr2\_hap1.g01276.t01 |
| 3113 | 2 | 198094410 | 198097013 | mNeoVis.2.1.1.chr2\_hap1.g01276.t01 |
| 3114 | 2 | 198188341 | 198263618 | mNeoVis.2.1.1.chr2\_hap1.g01277.t01 |
| 3115 | 2 | 198188341 | 198263618 | mNeoVis.2.1.1.chr2\_hap1.g01277.t01 |
| 3116 | 2 | 198269836 | 198279939 | mNeoVis.2.1.1.chr2\_hap1.g01278.t01 |
| 3117 | 2 | 198269836 | 198279939 | mNeoVis.2.1.1.chr2\_hap1.g01278.t01 |
| 3118 | 2 | 198282129 | 198289116 | mNeoVis.2.1.1.chr2\_hap1.g01279.t01 |
| 3119 | 2 | 198282129 | 198289116 | mNeoVis.2.1.1.chr2\_hap1.g01279.t01 |
| 3120 | 2 | 198297586 | 198316446 | mNeoVis.2.1.1.chr2\_hap1.g01280.t01 |
| 3121 | 2 | 198297586 | 198316446 | mNeoVis.2.1.1.chr2\_hap1.g01280.t01 |
| 3122 | 2 | 198368362 | 198406834 | mNeoVis.2.1.1.chr2\_hap1.g01281.t01 |
| 3123 | 2 | 198368362 | 198406834 | mNeoVis.2.1.1.chr2\_hap1.g01281.t01 |
| 3124 | 2 | 198408402 | 198431656 | mNeoVis.2.1.1.chr2\_hap1.g01282.t01 |
| 3125 | 2 | 198408402 | 198431656 | mNeoVis.2.1.1.chr2\_hap1.g01282.t01 |
| 3126 | 2 | 199093335 | 199104914 | mNeoVis.2.1.1.chr2\_hap1.g01291.t01 |
| 3127 | 2 | 199105046 | 199135093 | mNeoVis.2.1.1.chr2\_hap1.g01292.t01 |
| 3128 | 2 | 199137581 | 199151695 | mNeoVis.2.1.1.chr2\_hap1.g01293.t01 |
| 3129 | 2 | 199151969 | 199356057 | mNeoVis.2.1.1.chr2\_hap1.g01294.t01 |
| 3130 | 2 | 199400643 | 199422802 | mNeoVis.2.1.1.chr2\_hap1.g01295.t01 |
| 3131 | 2 | 199423830 | 199456726 | mNeoVis.2.1.1.chr2\_hap1.g01296.t01 |
| 3132 | 2 | 199445050 | 199447617 | mNeoVis.2.1.1.chr2\_hap1.g01297.t01 |
| 3133 | 2 | 199586995 | 199702629 | mNeoVis.2.1.1.chr2\_hap1.g01299.t01 |
| 3134 | 2 | 199701778 | 199876295 | mNeoVis.2.1.1.chr2\_hap1.g01300.t01 |
| 3135 | 2 | 199701778 | 199876295 | mNeoVis.2.1.1.chr2\_hap1.g01300.t01 |
| 3136 | 2 | 199857653 | 199859107 | mNeoVis.2.1.1.chr2\_hap1.g01301.t01 |
| 3137 | 2 | 199857653 | 199859107 | mNeoVis.2.1.1.chr2\_hap1.g01301.t01 |
| 3138 | 2 | 199957272 | 200103732 | mNeoVis.2.1.1.chr2\_hap1.g01302.t01 |
| 3139 | 2 | 199957272 | 200103732 | mNeoVis.2.1.1.chr2\_hap1.g01302.t01 |
| 3140 | 2 | 199957272 | 200103732 | mNeoVis.2.1.1.chr2\_hap1.g01302.t01 |
| 3141 | 2 | 199957272 | 200103732 | mNeoVis.2.1.1.chr2\_hap1.g01302.t01 |
| 3142 | 2 | 200032181 | 200032641 | mNeoVis.2.1.1.chr2\_hap1.g01303.t01 |
| 3143 | 2 | 200032181 | 200032641 | mNeoVis.2.1.1.chr2\_hap1.g01303.t01 |
| 3144 | 2 | 200032181 | 200032641 | mNeoVis.2.1.1.chr2\_hap1.g01303.t01 |
| 3145 | 2 | 200032181 | 200032641 | mNeoVis.2.1.1.chr2\_hap1.g01303.t01 |
| 3146 | 2 | 200112774 | 200161183 | mNeoVis.2.1.1.chr2\_hap1.g01304.t01 |
| 3147 | 2 | 200112774 | 200161183 | mNeoVis.2.1.1.chr2\_hap1.g01304.t01 |
| 3148 | 2 | 200112774 | 200161183 | mNeoVis.2.1.1.chr2\_hap1.g01304.t01 |
| 3149 | 2 | 200112774 | 200161183 | mNeoVis.2.1.1.chr2\_hap1.g01304.t01 |
| 3150 | 2 | 200167361 | 200227192 | mNeoVis.2.1.1.chr2\_hap1.g01305.t01 |
| 3151 | 2 | 200167361 | 200227192 | mNeoVis.2.1.1.chr2\_hap1.g01305.t01 |
| 3152 | 2 | 200167361 | 200227192 | mNeoVis.2.1.1.chr2\_hap1.g01305.t01 |
| 3153 | 2 | 200280094 | 200336957 | mNeoVis.2.1.1.chr2\_hap1.g01306.t01 |
| 3154 | 2 | 200280094 | 200336957 | mNeoVis.2.1.1.chr2\_hap1.g01306.t01 |
| 3155 | 2 | 200280094 | 200336957 | mNeoVis.2.1.1.chr2\_hap1.g01306.t01 |
| 3156 | 2 | 200326249 | 200328149 | mNeoVis.2.1.1.chr2\_hap1.g01307.t01 |
| 3157 | 2 | 200326249 | 200328149 | mNeoVis.2.1.1.chr2\_hap1.g01307.t01 |
| 3158 | 2 | 200326249 | 200328149 | mNeoVis.2.1.1.chr2\_hap1.g01307.t01 |
| 3159 | 2 | 200456098 | 200462553 | mNeoVis.2.1.1.chr2\_hap1.g01308.t01 |
| 3160 | 2 | 200959954 | 200965191 | mNeoVis.2.1.1.chr2\_hap1.g01310.t01 |
| 3161 | 2 | 201138679 | 201224191 | mNeoVis.2.1.1.chr2\_hap1.g01311.t01 |
| 3162 | 2 | 202791090 | 202840401 | mNeoVis.2.1.1.chr2\_hap1.g01313.t01 |
| 3163 | 2 | 203062479 | 203212313 | mNeoVis.2.1.1.chr2\_hap1.g01314.t01 |
| 3164 | 2 | 203062479 | 203212313 | mNeoVis.2.1.1.chr2\_hap1.g01314.t01 |
| 3165 | 2 | 203237605 | 203267578 | mNeoVis.2.1.1.chr2\_hap1.g01315.t01 |
| 3166 | 2 | 203237605 | 203267578 | mNeoVis.2.1.1.chr2\_hap1.g01315.t01 |
| 3167 | 2 | 203381265 | 203585294 | mNeoVis.2.1.1.chr2\_hap1.g01316.t01 |
| 3168 | 2 | 203381265 | 203585294 | mNeoVis.2.1.1.chr2\_hap1.g01316.t01 |
| 3169 | 2 | 203381265 | 203585294 | mNeoVis.2.1.1.chr2\_hap1.g01316.t01 |
| 3170 | 2 | 203874005 | 203876566 | mNeoVis.2.1.1.chr2\_hap1.g01317.t01 |
| 3171 | 2 | 203874005 | 203876566 | mNeoVis.2.1.1.chr2\_hap1.g01317.t01 |
| 3172 | 2 | 203874005 | 203876566 | mNeoVis.2.1.1.chr2\_hap1.g01317.t01 |
| 3173 | 2 | 203895688 | 203897115 | mNeoVis.2.1.1.chr2\_hap1.g01318.t01 |
| 3174 | 2 | 203895688 | 203897115 | mNeoVis.2.1.1.chr2\_hap1.g01318.t01 |
| 3175 | 2 | 203895688 | 203897115 | mNeoVis.2.1.1.chr2\_hap1.g01318.t01 |
| 3176 | 2 | 203926520 | 203962533 | mNeoVis.2.1.1.chr2\_hap1.g01319.t01 |
| 3177 | 2 | 203973412 | 204047544 | mNeoVis.2.1.1.chr2\_hap1.g01320.t01 |
| 3178 | 2 | 203973412 | 204047544 | mNeoVis.2.1.1.chr2\_hap1.g01320.t01 |
| 3179 | 2 | 204062382 | 204078139 | mNeoVis.2.1.1.chr2\_hap1.g01321.t01 |
| 3180 | 2 | 204062382 | 204078139 | mNeoVis.2.1.1.chr2\_hap1.g01321.t01 |
| 3181 | 2 | 204097015 | 204102771 | mNeoVis.2.1.1.chr2\_hap1.g01322.t01 |
| 3182 | 2 | 204097015 | 204102771 | mNeoVis.2.1.1.chr2\_hap1.g01322.t01 |
| 3183 | 2 | 204123705 | 204162563 | mNeoVis.2.1.1.chr2\_hap1.g01323.t01 |
| 3184 | 2 | 204123705 | 204162563 | mNeoVis.2.1.1.chr2\_hap1.g01323.t01 |
| 3185 | 2 | 204199515 | 204209644 | mNeoVis.2.1.1.chr2\_hap1.g01324.t01 |
| 3186 | 2 | 204199515 | 204209644 | mNeoVis.2.1.1.chr2\_hap1.g01324.t01 |
| 3187 | 2 | 204229376 | 204241478 | mNeoVis.2.1.1.chr2\_hap1.g01325.t01 |
| 3188 | 2 | 204229376 | 204241478 | mNeoVis.2.1.1.chr2\_hap1.g01325.t01 |
| 3189 | 2 | 204274051 | 204321481 | mNeoVis.2.1.1.chr2\_hap1.g01326.t01 |
| 3190 | 2 | 204274051 | 204321481 | mNeoVis.2.1.1.chr2\_hap1.g01326.t01 |
| 3191 | 2 | 204338962 | 204356425 | mNeoVis.2.1.1.chr2\_hap1.g01327.t01 |
| 3192 | 2 | 204338962 | 204356425 | mNeoVis.2.1.1.chr2\_hap1.g01327.t01 |
| 3193 | 2 | 204404391 | 204440022 | mNeoVis.2.1.1.chr2\_hap1.g01328.t01 |
| 3194 | 2 | 204404391 | 204440022 | mNeoVis.2.1.1.chr2\_hap1.g01328.t01 |
| 3195 | 2 | 204444720 | 204446131 | mNeoVis.2.1.1.chr2\_hap1.g01329.t01 |
| 3196 | 2 | 204444720 | 204446131 | mNeoVis.2.1.1.chr2\_hap1.g01329.t01 |
| 3197 | 2 | 204450271 | 204480646 | mNeoVis.2.1.1.chr2\_hap1.g01330.t01 |
| 3198 | 2 | 204450271 | 204480646 | mNeoVis.2.1.1.chr2\_hap1.g01330.t01 |
| 3199 | 2 | 204496277 | 204504249 | mNeoVis.2.1.1.chr2\_hap1.g01331.t01 |
| 3200 | 2 | 204496277 | 204504249 | mNeoVis.2.1.1.chr2\_hap1.g01331.t01 |
| 3201 | 2 | 204506955 | 204548422 | mNeoVis.2.1.1.chr2\_hap1.g01332.t01 |
| 3202 | 2 | 204506955 | 204548422 | mNeoVis.2.1.1.chr2\_hap1.g01332.t01 |
| 3203 | 2 | 204551736 | 204558519 | mNeoVis.2.1.1.chr2\_hap1.g01333.t01 |
| 3204 | 2 | 204551736 | 204558519 | mNeoVis.2.1.1.chr2\_hap1.g01333.t01 |
| 3205 | 2 | 204568513 | 204569046 | mNeoVis.2.1.1.chr2\_hap1.g01334.t01 |
| 3206 | 2 | 204568513 | 204569046 | mNeoVis.2.1.1.chr2\_hap1.g01334.t01 |
| 3207 | 2 | 204587397 | 204605954 | mNeoVis.2.1.1.chr2\_hap1.g01335.t01 |
| 3208 | 2 | 204587397 | 204605954 | mNeoVis.2.1.1.chr2\_hap1.g01335.t01 |
| 3209 | 2 | 204647970 | 204667570 | mNeoVis.2.1.1.chr2\_hap1.g01336.t01 |
| 3210 | 2 | 204673297 | 204691020 | mNeoVis.2.1.1.chr2\_hap1.g01337.t01 |
| 3211 | 2 | 204705289 | 204722289 | mNeoVis.2.1.1.chr2\_hap1.g01338.t01 |
| 3212 | 2 | 204718891 | 204721614 | mNeoVis.2.1.1.chr2\_hap1.g01339.t01 |
| 3213 | 2 | 204732764 | 204742071 | mNeoVis.2.1.1.chr2\_hap1.g01340.t01 |
| 3214 | 2 | 204742220 | 204759832 | mNeoVis.2.1.1.chr2\_hap1.g01341.t01 |
| 3215 | 2 | 204759754 | 204790166 | mNeoVis.2.1.1.chr2\_hap1.g01342.t01 |
| 3216 | 2 | 204813252 | 204824805 | mNeoVis.2.1.1.chr2\_hap1.g01343.t01 |
| 3217 | 2 | 204829490 | 204838035 | mNeoVis.2.1.1.chr2\_hap1.g01344.t01 |
| 3218 | 2 | 204840273 | 204845998 | mNeoVis.2.1.1.chr2\_hap1.g01345.t01 |
| 3219 | 2 | 205266443 | 205311374 | mNeoVis.2.1.1.chr2\_hap1.g01348.t01 |
| 3220 | 2 | 205312448 | 205320197 | mNeoVis.2.1.1.chr2\_hap1.g01349.t01 |
| 3221 | 2 | 205320463 | 205375931 | mNeoVis.2.1.1.chr2\_hap1.g01350.t01 |
| 3222 | 2 | 205377913 | 205382098 | mNeoVis.2.1.1.chr2\_hap1.g01351.t01 |
| 3223 | 2 | 205382912 | 205427670 | mNeoVis.2.1.1.chr2\_hap1.g01352.t01 |
| 3224 | 2 | 205440380 | 205453025 | mNeoVis.2.1.1.chr2\_hap1.g01353.t01 |
| 3225 | 2 | 205498873 | 205513900 | mNeoVis.2.1.1.chr2\_hap1.g01354.t01 |
| 3226 | 2 | 205515363 | 205519894 | mNeoVis.2.1.1.chr2\_hap1.g01355.t01 |
| 3227 | 2 | 205520021 | 205523694 | mNeoVis.2.1.1.chr2\_hap1.g01356.t01 |
| 3228 | 2 | 205527912 | 205669261 | mNeoVis.2.1.1.chr2\_hap1.g01357.t01 |
| 3229 | 2 | 205577839 | 205578394 | mNeoVis.2.1.1.chr2\_hap1.g01358.t01 |
| 3230 | 2 | 205669168 | 205672760 | mNeoVis.2.1.1.chr2\_hap1.g01359.t01 |
| 3231 | 2 | 205675367 | 205680729 | mNeoVis.2.1.1.chr2\_hap1.g01360.t01 |
| 3232 | 2 | 205680873 | 205683043 | mNeoVis.2.1.1.chr2\_hap1.g01361.t01 |
| 3233 | 2 | 205992118 | 205997798 | mNeoVis.2.1.1.chr2\_hap1.g01363.t01 |
| 3234 | 2 | 205997749 | 206005927 | mNeoVis.2.1.1.chr2\_hap1.g01364.t01 |
| 3235 | 2 | 206005125 | 206140178 | mNeoVis.2.1.1.chr2\_hap1.g01365.t01 |
| 3236 | 2 | 206146507 | 206162449 | mNeoVis.2.1.1.chr2\_hap1.g01366.t01 |
| 3237 | 2 | 206170462 | 206176512 | mNeoVis.2.1.1.chr2\_hap1.g01367.t01 |
| 3238 | 2 | 206176700 | 206178338 | mNeoVis.2.1.1.chr2\_hap1.g01368.t01 |
| 3239 | 2 | 206178350 | 206183341 | mNeoVis.2.1.1.chr2\_hap1.g01369.t01 |
| 3240 | 2 | 206196518 | 206200440 | mNeoVis.2.1.1.chr2\_hap1.g01370.t01 |
| 3241 | 2 | 206196518 | 206200440 | mNeoVis.2.1.1.chr2\_hap1.g01370.t01 |
| 3242 | 2 | 206203471 | 206206298 | mNeoVis.2.1.1.chr2\_hap1.g01371.t01 |
| 3243 | 2 | 206203471 | 206206298 | mNeoVis.2.1.1.chr2\_hap1.g01371.t01 |
| 3244 | 2 | 206216533 | 206237399 | mNeoVis.2.1.1.chr2\_hap1.g01372.t01 |
| 3245 | 2 | 206216533 | 206237399 | mNeoVis.2.1.1.chr2\_hap1.g01372.t01 |
| 3246 | 2 | 206258655 | 206311605 | mNeoVis.2.1.1.chr2\_hap1.g01373.t01 |
| 3247 | 2 | 206258655 | 206311605 | mNeoVis.2.1.1.chr2\_hap1.g01373.t01 |
| 3248 | 2 | 206311680 | 206337013 | mNeoVis.2.1.1.chr2\_hap1.g01374.t01 |
| 3249 | 2 | 206311680 | 206337013 | mNeoVis.2.1.1.chr2\_hap1.g01374.t01 |
| 3250 | 2 | 206336208 | 206561852 | mNeoVis.2.1.1.chr2\_hap1.g01375.t01 |
| 3251 | 2 | 206336208 | 206561852 | mNeoVis.2.1.1.chr2\_hap1.g01375.t01 |
| 3252 | 2 | 206610438 | 206736638 | mNeoVis.2.1.1.chr2\_hap1.g01376.t01 |
| 3253 | 2 | 206610438 | 206736638 | mNeoVis.2.1.1.chr2\_hap1.g01376.t01 |
| 3254 | 2 | 206737022 | 206745704 | mNeoVis.2.1.1.chr2\_hap1.g01377.t01 |
| 3255 | 2 | 206737022 | 206745704 | mNeoVis.2.1.1.chr2\_hap1.g01377.t01 |
| 3256 | 2 | 206784101 | 206790935 | mNeoVis.2.1.1.chr2\_hap1.g01378.t01 |
| 3257 | 2 | 206784101 | 206790935 | mNeoVis.2.1.1.chr2\_hap1.g01378.t01 |
| 3258 | 2 | 206800562 | 206813491 | mNeoVis.2.1.1.chr2\_hap1.g01379.t01 |
| 3259 | 2 | 206800562 | 206813491 | mNeoVis.2.1.1.chr2\_hap1.g01379.t01 |
| 3260 | 2 | 206904814 | 206924589 | mNeoVis.2.1.1.chr2\_hap1.g01380.t01 |
| 3261 | 2 | 206904814 | 206924589 | mNeoVis.2.1.1.chr2\_hap1.g01380.t01 |
| 3262 | 2 | 206924740 | 206928179 | mNeoVis.2.1.1.chr2\_hap1.g01381.t01 |
| 3263 | 2 | 206924740 | 206928179 | mNeoVis.2.1.1.chr2\_hap1.g01381.t01 |
| 3264 | 2 | 206928615 | 206937786 | mNeoVis.2.1.1.chr2\_hap1.g01382.t01 |
| 3265 | 2 | 206928615 | 206937786 | mNeoVis.2.1.1.chr2\_hap1.g01382.t01 |
| 3266 | 2 | 206939403 | 206942689 | mNeoVis.2.1.1.chr2\_hap1.g01383.t01 |
| 3267 | 2 | 206939403 | 206942689 | mNeoVis.2.1.1.chr2\_hap1.g01383.t01 |
| 3268 | 2 | 206944629 | 206948116 | mNeoVis.2.1.1.chr2\_hap1.g01384.t01 |
| 3269 | 2 | 206944629 | 206948116 | mNeoVis.2.1.1.chr2\_hap1.g01384.t01 |
| 3270 | 2 | 206950741 | 206970949 | mNeoVis.2.1.1.chr2\_hap1.g01385.t01 |
| 3271 | 2 | 206950741 | 206970949 | mNeoVis.2.1.1.chr2\_hap1.g01385.t01 |
| 3272 | 2 | 206975537 | 206976637 | mNeoVis.2.1.1.chr2\_hap1.g01386.t01 |
| 3273 | 2 | 206975537 | 206976637 | mNeoVis.2.1.1.chr2\_hap1.g01386.t01 |
| 3274 | 2 | 206981362 | 207175894 | mNeoVis.2.1.1.chr2\_hap1.g01387.t01 |
| 3275 | 2 | 206981362 | 207175894 | mNeoVis.2.1.1.chr2\_hap1.g01387.t01 |
| 3276 | 2 | 207178119 | 207226463 | mNeoVis.2.1.1.chr2\_hap1.g01388.t01 |
| 3277 | 2 | 207178119 | 207226463 | mNeoVis.2.1.1.chr2\_hap1.g01388.t01 |
| 3278 | 2 | 207248033 | 207334361 | mNeoVis.2.1.1.chr2\_hap1.g01389.t01 |
| 3279 | 2 | 207248033 | 207334361 | mNeoVis.2.1.1.chr2\_hap1.g01389.t01 |
| 3280 | 2 | 207388426 | 207391058 | mNeoVis.2.1.1.chr2\_hap1.g01390.t01 |
| 3281 | 2 | 207388426 | 207391058 | mNeoVis.2.1.1.chr2\_hap1.g01390.t01 |
| 3282 | 2 | 207391325 | 207443016 | mNeoVis.2.1.1.chr2\_hap1.g01391.t01 |
| 3283 | 2 | 207391325 | 207443016 | mNeoVis.2.1.1.chr2\_hap1.g01391.t01 |
| 3284 | 2 | 207443118 | 207464131 | mNeoVis.2.1.1.chr2\_hap1.g01392.t01 |
| 3285 | 2 | 207443118 | 207464131 | mNeoVis.2.1.1.chr2\_hap1.g01392.t01 |
| 3286 | 2 | 207464273 | 207469278 | mNeoVis.2.1.1.chr2\_hap1.g01393.t01 |
| 3287 | 2 | 207464273 | 207469278 | mNeoVis.2.1.1.chr2\_hap1.g01393.t01 |
| 3288 | 2 | 207474084 | 207488737 | mNeoVis.2.1.1.chr2\_hap1.g01394.t01 |
| 3289 | 2 | 207474084 | 207488737 | mNeoVis.2.1.1.chr2\_hap1.g01394.t01 |
| 3290 | 2 | 207498908 | 207518939 | mNeoVis.2.1.1.chr2\_hap1.g01395.t01 |
| 3291 | 2 | 207498908 | 207518939 | mNeoVis.2.1.1.chr2\_hap1.g01395.t01 |
| 3292 | 2 | 207538736 | 207542630 | mNeoVis.2.1.1.chr2\_hap1.g01396.t01 |
| 3293 | 2 | 207538736 | 207542630 | mNeoVis.2.1.1.chr2\_hap1.g01396.t01 |
| 3294 | 2 | 207552154 | 207556058 | mNeoVis.2.1.1.chr2\_hap1.g01397.t01 |
| 3295 | 2 | 207552154 | 207556058 | mNeoVis.2.1.1.chr2\_hap1.g01397.t01 |
| 3296 | 2 | 207563234 | 207636036 | mNeoVis.2.1.1.chr2\_hap1.g01398.t01 |
| 3297 | 2 | 207630956 | 207632288 | mNeoVis.2.1.1.chr2\_hap1.g01399.t01 |
| 3298 | 2 | 207630956 | 207632288 | mNeoVis.2.1.1.chr2\_hap1.g01399.t01 |
| 3299 | 2 | 207636105 | 207644255 | mNeoVis.2.1.1.chr2\_hap1.g01400.t01 |
| 3300 | 2 | 207649487 | 207658372 | mNeoVis.2.1.1.chr2\_hap1.g01401.t01 |
| 3301 | 2 | 207658577 | 207659492 | mNeoVis.2.1.1.chr2\_hap1.g01402.t01 |
| 3302 | 2 | 207681513 | 207682460 | mNeoVis.2.1.1.chr2\_hap1.g01404.t01 |
| 3303 | 2 | 207691702 | 207692766 | mNeoVis.2.1.1.chr2\_hap1.g01405.t01 |
| 3304 | 2 | 208039314 | 208045861 | mNeoVis.2.1.1.chr2\_hap1.g01415.t01 |
| 3305 | 2 | 208055445 | 208071098 | mNeoVis.2.1.1.chr2\_hap1.g01416.t01 |
| 3306 | 2 | 208080404 | 208088288 | mNeoVis.2.1.1.chr2\_hap1.g01417.t01 |
| 3307 | 2 | 208088357 | 208107370 | mNeoVis.2.1.1.chr2\_hap1.g01418.t01 |
| 3308 | 2 | 208112310 | 208117272 | mNeoVis.2.1.1.chr2\_hap1.g01419.t01 |
| 3309 | 2 | 208133315 | 208154453 | mNeoVis.2.1.1.chr2\_hap1.g01420.t01 |
| 3310 | 2 | 208133315 | 208154453 | mNeoVis.2.1.1.chr2\_hap1.g01420.t01 |
| 3311 | 2 | 208159264 | 208176305 | mNeoVis.2.1.1.chr2\_hap1.g01421.t01 |
| 3312 | 2 | 208159264 | 208176305 | mNeoVis.2.1.1.chr2\_hap1.g01421.t01 |
| 3313 | 2 | 208178414 | 208243877 | mNeoVis.2.1.1.chr2\_hap1.g01422.t01 |
| 3314 | 2 | 208178414 | 208243877 | mNeoVis.2.1.1.chr2\_hap1.g01422.t01 |
| 3315 | 2 | 208283324 | 208287048 | mNeoVis.2.1.1.chr2\_hap1.g01423.t01 |
| 3316 | 2 | 208283324 | 208287048 | mNeoVis.2.1.1.chr2\_hap1.g01423.t01 |
| 3317 | 2 | 208287166 | 208313439 | mNeoVis.2.1.1.chr2\_hap1.g01424.t01 |
| 3318 | 2 | 208287166 | 208313439 | mNeoVis.2.1.1.chr2\_hap1.g01424.t01 |
| 3319 | 2 | 208323955 | 208354490 | mNeoVis.2.1.1.chr2\_hap1.g01425.t01 |
| 3320 | 2 | 208323955 | 208354490 | mNeoVis.2.1.1.chr2\_hap1.g01425.t01 |
| 3321 | 2 | 208420396 | 208584723 | mNeoVis.2.1.1.chr2\_hap1.g01426.t01 |
| 3322 | 2 | 208420396 | 208584723 | mNeoVis.2.1.1.chr2\_hap1.g01426.t01 |
| 3323 | 2 | 208581731 | 208597314 | mNeoVis.2.1.1.chr2\_hap1.g01427.t01 |
| 3324 | 2 | 208581731 | 208597314 | mNeoVis.2.1.1.chr2\_hap1.g01427.t01 |
| 3325 | 2 | 208581731 | 208597314 | mNeoVis.2.1.1.chr2\_hap1.g01427.t01 |
| 3326 | 2 | 208590989 | 208595804 | mNeoVis.2.1.1.chr2\_hap1.g01428.t01 |
| 3327 | 2 | 208590989 | 208595804 | mNeoVis.2.1.1.chr2\_hap1.g01428.t01 |
| 3328 | 2 | 208590989 | 208595804 | mNeoVis.2.1.1.chr2\_hap1.g01428.t01 |
| 3329 | 2 | 208599172 | 208601344 | mNeoVis.2.1.1.chr2\_hap1.g01429.t01 |
| 3330 | 2 | 208599172 | 208601344 | mNeoVis.2.1.1.chr2\_hap1.g01429.t01 |
| 3331 | 2 | 208599172 | 208601344 | mNeoVis.2.1.1.chr2\_hap1.g01429.t01 |
| 3332 | 2 | 208765212 | 209129714 | mNeoVis.2.1.1.chr2\_hap1.g01430.t01 |
| 3333 | 2 | 209053917 | 209055141 | mNeoVis.2.1.1.chr2\_hap1.g01431.t01 |
| 3334 | 2 | 209150434 | 209156004 | mNeoVis.2.1.1.chr2\_hap1.g01432.t01 |
| 3335 | 2 | 209378863 | 209549890 | mNeoVis.2.1.1.chr2\_hap1.g01434.t01 |
| 3336 | 2 | 209554472 | 209563302 | mNeoVis.2.1.1.chr2\_hap1.g01435.t01 |
| 3337 | 2 | 209562486 | 209593105 | mNeoVis.2.1.1.chr2\_hap1.g01436.t01 |
| 3338 | 2 | 209679134 | 209680480 | mNeoVis.2.1.1.chr2\_hap1.g01437.t01 |
| 3339 | 2 | 209698384 | 209750740 | mNeoVis.2.1.1.chr2\_hap1.g01438.t01 |
| 3340 | 2 | 209762937 | 209804892 | mNeoVis.2.1.1.chr2\_hap1.g01439.t01 |
| 3341 | 2 | 209851395 | 209891646 | mNeoVis.2.1.1.chr2\_hap1.g01440.t01 |
| 3342 | 2 | 209957530 | 209969644 | mNeoVis.2.1.1.chr2\_hap1.g01441.t01 |
| 3343 | 2 | 209975935 | 209982750 | mNeoVis.2.1.1.chr2\_hap1.g01442.t01 |
| 3344 | 2 | 209992665 | 210008525 | mNeoVis.2.1.1.chr2\_hap1.g01443.t01 |
| 3345 | 2 | 210016381 | 210029031 | mNeoVis.2.1.1.chr2\_hap1.g01444.t01 |
| 3346 | 2 | 210049304 | 210098853 | mNeoVis.2.1.1.chr2\_hap1.g01445.t01 |
| 3347 | 2 | 210120801 | 210136332 | mNeoVis.2.1.1.chr2\_hap1.g01446.t01 |
| 3348 | 2 | 210142799 | 210181859 | mNeoVis.2.1.1.chr2\_hap1.g01447.t01 |
| 3349 | 2 | 210189371 | 210192208 | mNeoVis.2.1.1.chr2\_hap1.g01448.t01 |
| 3350 | 2 | 210198644 | 210292336 | mNeoVis.2.1.1.chr2\_hap1.g01449.t01 |
| 3351 | 2 | 210337856 | 210363831 | mNeoVis.2.1.1.chr2\_hap1.g01450.t01 |
| 3352 | 2 | 210364022 | 210391393 | mNeoVis.2.1.1.chr2\_hap1.g01451.t01 |
| 3353 | 2 | 210435262 | 210448854 | mNeoVis.2.1.1.chr2\_hap1.g01452.t01 |
| 3354 | 2 | 211227270 | 211233368 | mNeoVis.2.1.1.chr2\_hap1.g01465.t01 |
| 3355 | 2 | 211244055 | 211256047 | mNeoVis.2.1.1.chr2\_hap1.g01466.t01 |
| 3356 | 2 | 211291583 | 211343634 | mNeoVis.2.1.1.chr2\_hap1.g01467.t01 |
| 3357 | 2 | 211336594 | 211337755 | mNeoVis.2.1.1.chr2\_hap1.g01468.t01 |
| 3358 | 2 | 211348775 | 211351980 | mNeoVis.2.1.1.chr2\_hap1.g01469.t01 |
| 3359 | 2 | 211352264 | 211360197 | mNeoVis.2.1.1.chr2\_hap1.g01470.t01 |
| 3360 | 2 | 211362369 | 211377871 | mNeoVis.2.1.1.chr2\_hap1.g01471.t01 |
| 3361 | 2 | 212040491 | 212043398 | mNeoVis.2.1.1.chr2\_hap1.g01472.t01 |
| 3362 | 2 | 212040491 | 212043398 | mNeoVis.2.1.1.chr2\_hap1.g01472.t01 |
| 3363 | 2 | 212060146 | 212069825 | mNeoVis.2.1.1.chr2\_hap1.g01473.t01 |
| 3364 | 2 | 212060146 | 212069825 | mNeoVis.2.1.1.chr2\_hap1.g01473.t01 |
| 3365 | 2 | 212074898 | 212082473 | mNeoVis.2.1.1.chr2\_hap1.g01474.t01 |
| 3366 | 2 | 212074898 | 212082473 | mNeoVis.2.1.1.chr2\_hap1.g01474.t01 |
| 3367 | 2 | 212086753 | 212109510 | mNeoVis.2.1.1.chr2\_hap1.g01475.t01 |
| 3368 | 2 | 212086753 | 212109510 | mNeoVis.2.1.1.chr2\_hap1.g01475.t01 |
| 3369 | 2 | 212117440 | 212165017 | mNeoVis.2.1.1.chr2\_hap1.g01476.t01 |
| 3370 | 2 | 212117440 | 212165017 | mNeoVis.2.1.1.chr2\_hap1.g01476.t01 |
| 3371 | 2 | 212166514 | 212208134 | mNeoVis.2.1.1.chr2\_hap1.g01477.t01 |
| 3372 | 2 | 212166514 | 212208134 | mNeoVis.2.1.1.chr2\_hap1.g01477.t01 |
| 3373 | 2 | 212208270 | 212209397 | mNeoVis.2.1.1.chr2\_hap1.g01478.t01 |
| 3374 | 2 | 212208270 | 212209397 | mNeoVis.2.1.1.chr2\_hap1.g01478.t01 |
| 3375 | 2 | 212209534 | 212213897 | mNeoVis.2.1.1.chr2\_hap1.g01479.t01 |
| 3376 | 2 | 212209534 | 212213897 | mNeoVis.2.1.1.chr2\_hap1.g01479.t01 |
| 3377 | 2 | 212213881 | 212220143 | mNeoVis.2.1.1.chr2\_hap1.g01480.t01 |
| 3378 | 2 | 212213881 | 212220143 | mNeoVis.2.1.1.chr2\_hap1.g01480.t01 |
| 3379 | 2 | 212243216 | 212284238 | mNeoVis.2.1.1.chr2\_hap1.g01481.t01 |
| 3380 | 2 | 212243216 | 212284238 | mNeoVis.2.1.1.chr2\_hap1.g01481.t01 |
| 3381 | 2 | 212289455 | 212304921 | mNeoVis.2.1.1.chr2\_hap1.g01482.t01 |
| 3382 | 2 | 212289455 | 212304921 | mNeoVis.2.1.1.chr2\_hap1.g01482.t01 |
| 3383 | 2 | 212305093 | 212321028 | mNeoVis.2.1.1.chr2\_hap1.g01483.t01 |
| 3384 | 2 | 212305093 | 212321028 | mNeoVis.2.1.1.chr2\_hap1.g01483.t01 |
| 3385 | 2 | 212353479 | 212372207 | mNeoVis.2.1.1.chr2\_hap1.g01484.t01 |
| 3386 | 2 | 212353479 | 212372207 | mNeoVis.2.1.1.chr2\_hap1.g01484.t01 |
| 3387 | 2 | 212385991 | 212409644 | mNeoVis.2.1.1.chr2\_hap1.g01485.t01 |
| 3388 | 2 | 212385991 | 212409644 | mNeoVis.2.1.1.chr2\_hap1.g01485.t01 |
| 3389 | 2 | 212409609 | 212417492 | mNeoVis.2.1.1.chr2\_hap1.g01486.t01 |
| 3390 | 2 | 212409609 | 212417492 | mNeoVis.2.1.1.chr2\_hap1.g01486.t01 |
| 3391 | 2 | 212415783 | 212436568 | mNeoVis.2.1.1.chr2\_hap1.g01487.t01 |
| 3392 | 2 | 212415783 | 212436568 | mNeoVis.2.1.1.chr2\_hap1.g01487.t01 |
| 3393 | 2 | 212435085 | 212460898 | mNeoVis.2.1.1.chr2\_hap1.g01488.t01 |
| 3394 | 2 | 212435085 | 212460898 | mNeoVis.2.1.1.chr2\_hap1.g01488.t01 |
| 3395 | 2 | 212468769 | 212477526 | mNeoVis.2.1.1.chr2\_hap1.g01489.t01 |
| 3396 | 2 | 212468769 | 212477526 | mNeoVis.2.1.1.chr2\_hap1.g01489.t01 |
| 3397 | 2 | 212833556 | 213059516 | mNeoVis.2.1.1.chr2\_hap1.g01490.t01 |
| 3398 | 2 | 212833556 | 213059516 | mNeoVis.2.1.1.chr2\_hap1.g01490.t01 |
| 3399 | 2 | 213563512 | 213584793 | mNeoVis.2.1.1.chr2\_hap1.g01499.t01 |
| 3400 | 2 | 213590053 | 213601164 | mNeoVis.2.1.1.chr2\_hap1.g01500.t01 |
| 3401 | 2 | 213610190 | 213632696 | mNeoVis.2.1.1.chr2\_hap1.g01501.t01 |
| 3402 | 2 | 213633915 | 213639288 | mNeoVis.2.1.1.chr2\_hap1.g01502.t01 |
| 3403 | 2 | 213640026 | 213643997 | mNeoVis.2.1.1.chr2\_hap1.g01503.t01 |
| 3404 | 2 | 213652729 | 213794740 | mNeoVis.2.1.1.chr2\_hap1.g01504.t01 |
| 3405 | 2 | 213794831 | 213837336 | mNeoVis.2.1.1.chr2\_hap1.g01505.t01 |
| 3406 | 2 | 213794831 | 213837336 | mNeoVis.2.1.1.chr2\_hap1.g01505.t01 |
| 3407 | 2 | 213794831 | 213837336 | mNeoVis.2.1.1.chr2\_hap1.g01505.t01 |
| 3408 | 2 | 213865372 | 213905650 | mNeoVis.2.1.1.chr2\_hap1.g01506.t01 |
| 3409 | 2 | 213865372 | 213905650 | mNeoVis.2.1.1.chr2\_hap1.g01506.t01 |
| 3410 | 2 | 213865372 | 213905650 | mNeoVis.2.1.1.chr2\_hap1.g01506.t01 |
| 3411 | 2 | 213938777 | 213971935 | mNeoVis.2.1.1.chr2\_hap1.g01507.t01 |
| 3412 | 2 | 213938777 | 213971935 | mNeoVis.2.1.1.chr2\_hap1.g01507.t01 |
| 3413 | 2 | 213938777 | 213971935 | mNeoVis.2.1.1.chr2\_hap1.g01507.t01 |
| 3414 | 2 | 213985747 | 213990464 | mNeoVis.2.1.1.chr2\_hap1.g01508.t01 |
| 3415 | 2 | 213985747 | 213990464 | mNeoVis.2.1.1.chr2\_hap1.g01508.t01 |
| 3416 | 2 | 213985747 | 213990464 | mNeoVis.2.1.1.chr2\_hap1.g01508.t01 |
| 3417 | 2 | 214050558 | 214083138 | mNeoVis.2.1.1.chr2\_hap1.g01509.t01 |
| 3418 | 2 | 214050558 | 214083138 | mNeoVis.2.1.1.chr2\_hap1.g01509.t01 |
| 3419 | 2 | 214050558 | 214083138 | mNeoVis.2.1.1.chr2\_hap1.g01509.t01 |
| 3420 | 2 | 214090377 | 214108452 | mNeoVis.2.1.1.chr2\_hap1.g01510.t01 |
| 3421 | 2 | 214090377 | 214108452 | mNeoVis.2.1.1.chr2\_hap1.g01510.t01 |
| 3422 | 2 | 214090377 | 214108452 | mNeoVis.2.1.1.chr2\_hap1.g01510.t01 |
| 3423 | 2 | 214090377 | 214108452 | mNeoVis.2.1.1.chr2\_hap1.g01510.t01 |
| 3424 | 2 | 214114714 | 214123716 | mNeoVis.2.1.1.chr2\_hap1.g01511.t01 |
| 3425 | 2 | 214114714 | 214123716 | mNeoVis.2.1.1.chr2\_hap1.g01511.t01 |
| 3426 | 2 | 214114714 | 214123716 | mNeoVis.2.1.1.chr2\_hap1.g01511.t01 |
| 3427 | 2 | 214114714 | 214123716 | mNeoVis.2.1.1.chr2\_hap1.g01511.t01 |
| 3428 | 2 | 214123628 | 214215093 | mNeoVis.2.1.1.chr2\_hap1.g01512.t01 |
| 3429 | 2 | 214123628 | 214215093 | mNeoVis.2.1.1.chr2\_hap1.g01512.t01 |
| 3430 | 2 | 214123628 | 214215093 | mNeoVis.2.1.1.chr2\_hap1.g01512.t01 |
| 3431 | 2 | 214123628 | 214215093 | mNeoVis.2.1.1.chr2\_hap1.g01512.t01 |
| 3432 | 2 | 214231821 | 214381611 | mNeoVis.2.1.1.chr2\_hap1.g01513.t01 |
| 3433 | 2 | 214231821 | 214381611 | mNeoVis.2.1.1.chr2\_hap1.g01513.t01 |
| 3434 | 2 | 214231821 | 214381611 | mNeoVis.2.1.1.chr2\_hap1.g01513.t01 |
| 3435 | 2 | 214441614 | 214458896 | mNeoVis.2.1.1.chr2\_hap1.g01514.t01 |
| 3436 | 2 | 214441614 | 214458896 | mNeoVis.2.1.1.chr2\_hap1.g01514.t01 |
| 3437 | 2 | 214441614 | 214458896 | mNeoVis.2.1.1.chr2\_hap1.g01514.t01 |
| 3438 | 2 | 214499811 | 214531998 | mNeoVis.2.1.1.chr2\_hap1.g01515.t01 |
| 3439 | 2 | 214499811 | 214531998 | mNeoVis.2.1.1.chr2\_hap1.g01515.t01 |
| 3440 | 2 | 214555570 | 214606664 | mNeoVis.2.1.1.chr2\_hap1.g01516.t01 |
| 3441 | 2 | 214555570 | 214606664 | mNeoVis.2.1.1.chr2\_hap1.g01516.t01 |
| 3442 | 2 | 214608018 | 214618944 | mNeoVis.2.1.1.chr2\_hap1.g01517.t01 |
| 3443 | 2 | 214608018 | 214618944 | mNeoVis.2.1.1.chr2\_hap1.g01517.t01 |
| 3444 | 2 | 214613593 | 214618627 | mNeoVis.2.1.1.chr2\_hap1.g01518.t01 |
| 3445 | 2 | 214613593 | 214618627 | mNeoVis.2.1.1.chr2\_hap1.g01518.t01 |
| 3446 | 2 | 214661124 | 214724665 | mNeoVis.2.1.1.chr2\_hap1.g01519.t01 |
| 3447 | 2 | 214730531 | 214733836 | mNeoVis.2.1.1.chr2\_hap1.g01520.t01 |
| 3448 | 2 | 214785119 | 214787846 | mNeoVis.2.1.1.chr2\_hap1.g01521.t01 |
| 3449 | 2 | 214794316 | 214799602 | mNeoVis.2.1.1.chr2\_hap1.g01522.t01 |
| 3450 | 2 | 214815423 | 214819912 | mNeoVis.2.1.1.chr2\_hap1.g01523.t01 |
| 3451 | 2 | 214820051 | 214823393 | mNeoVis.2.1.1.chr2\_hap1.g01524.t01 |
| 3452 | 2 | 215351811 | 215391495 | mNeoVis.2.1.1.chr2\_hap1.g01525.t01 |
| 3453 | 2 | 215666093 | 215666831 | mNeoVis.2.1.1.chr2\_hap1.g01527.t01 |
| 3454 | 2 | 217978074 | 218113624 | mNeoVis.2.1.1.chr2\_hap1.g01579.t01 |
| 3455 | 2 | 218125724 | 218164021 | mNeoVis.2.1.1.chr2\_hap1.g01580.t01 |
| 3456 | 2 | 218125724 | 218164021 | mNeoVis.2.1.1.chr2\_hap1.g01580.t01 |
| 3457 | 2 | 218125724 | 218164021 | mNeoVis.2.1.1.chr2\_hap1.g01580.t01 |
| 3458 | 2 | 218257987 | 218281864 | mNeoVis.2.1.1.chr2\_hap1.g01581.t01 |
| 3459 | 2 | 218257987 | 218281864 | mNeoVis.2.1.1.chr2\_hap1.g01581.t01 |
| 3460 | 2 | 218257987 | 218281864 | mNeoVis.2.1.1.chr2\_hap1.g01581.t01 |
| 3461 | 2 | 218292005 | 218300944 | mNeoVis.2.1.1.chr2\_hap1.g01582.t01 |
| 3462 | 2 | 218292005 | 218300944 | mNeoVis.2.1.1.chr2\_hap1.g01582.t01 |
| 3463 | 2 | 218292005 | 218300944 | mNeoVis.2.1.1.chr2\_hap1.g01582.t01 |
| 3464 | 2 | 219341173 | 219418299 | mNeoVis.2.1.1.chr2\_hap1.g01583.t01 |
| 3465 | 2 | 219425181 | 219454947 | mNeoVis.2.1.1.chr2\_hap1.g01584.t01 |
| 3466 | 2 | 219461659 | 219489417 | mNeoVis.2.1.1.chr2\_hap1.g01585.t01 |
| 3467 | 2 | 220388871 | 220497477 | mNeoVis.2.1.1.chr2\_hap1.g01602.t01 |
| 3468 | 2 | 220503712 | 220510528 | mNeoVis.2.1.1.chr2\_hap1.g01603.t01 |
| 3469 | 2 | 220510829 | 220535540 | mNeoVis.2.1.1.chr2\_hap1.g01604.t01 |
| 3470 | 2 | 220549833 | 220567922 | mNeoVis.2.1.1.chr2\_hap1.g01605.t01 |
| 3471 | 2 | 220571285 | 220587038 | mNeoVis.2.1.1.chr2\_hap1.g01606.t01 |
| 3472 | 2 | 220606863 | 220626239 | mNeoVis.2.1.1.chr2\_hap1.g01607.t01 |
| 3473 | 2 | 220634396 | 220675771 | mNeoVis.2.1.1.chr2\_hap1.g01608.t01 |
| 3474 | 2 | 220684335 | 220707505 | mNeoVis.2.1.1.chr2\_hap1.g01609.t01 |
| 3475 | 2 | 220727824 | 220731315 | mNeoVis.2.1.1.chr2\_hap1.g01610.t01 |
| 3476 | 2 | 220758280 | 220777884 | mNeoVis.2.1.1.chr2\_hap1.g01611.t01 |
| 3477 | 2 | 220758280 | 220777884 | mNeoVis.2.1.1.chr2\_hap1.g01611.t01 |
| 3478 | 2 | 220785815 | 220850146 | mNeoVis.2.1.1.chr2\_hap1.g01612.t01 |
| 3479 | 2 | 220885480 | 220956363 | mNeoVis.2.1.1.chr2\_hap1.g01613.t01 |
| 3480 | 2 | 220964294 | 220967458 | mNeoVis.2.1.1.chr2\_hap1.g01614.t01 |
| 3481 | 2 | 220973440 | 220978066 | mNeoVis.2.1.1.chr2\_hap1.g01615.t01 |
| 3482 | 2 | 220973440 | 220978066 | mNeoVis.2.1.1.chr2\_hap1.g01615.t01 |
| 3483 | 2 | 220982045 | 220987265 | mNeoVis.2.1.1.chr2\_hap1.g01616.t01 |
| 3484 | 2 | 220982045 | 220987265 | mNeoVis.2.1.1.chr2\_hap1.g01616.t01 |
| 3485 | 2 | 220987513 | 220998655 | mNeoVis.2.1.1.chr2\_hap1.g01617.t01 |
| 3486 | 2 | 220987513 | 220998655 | mNeoVis.2.1.1.chr2\_hap1.g01617.t01 |
| 3487 | 2 | 220998779 | 221008250 | mNeoVis.2.1.1.chr2\_hap1.g01618.t01 |
| 3488 | 2 | 220998779 | 221008250 | mNeoVis.2.1.1.chr2\_hap1.g01618.t01 |
| 3489 | 2 | 221010086 | 221022300 | mNeoVis.2.1.1.chr2\_hap1.g01619.t01 |
| 3490 | 2 | 221010086 | 221022300 | mNeoVis.2.1.1.chr2\_hap1.g01619.t01 |
| 3491 | 2 | 221033186 | 221103351 | mNeoVis.2.1.1.chr2\_hap1.g01620.t01 |
| 3492 | 2 | 221033186 | 221103351 | mNeoVis.2.1.1.chr2\_hap1.g01620.t01 |
| 3493 | 2 | 221155019 | 221205076 | mNeoVis.2.1.1.chr2\_hap1.g01621.t01 |
| 3494 | 2 | 221155019 | 221205076 | mNeoVis.2.1.1.chr2\_hap1.g01621.t01 |
| 3495 | 2 | 221265850 | 221273362 | mNeoVis.2.1.1.chr2\_hap1.g01622.t01 |
| 3496 | 2 | 221265850 | 221273362 | mNeoVis.2.1.1.chr2\_hap1.g01622.t01 |
| 3497 | 2 | 221276290 | 221282522 | mNeoVis.2.1.1.chr2\_hap1.g01623.t01 |
| 3498 | 2 | 221276290 | 221282522 | mNeoVis.2.1.1.chr2\_hap1.g01623.t01 |
| 3499 | 2 | 221307061 | 221315399 | mNeoVis.2.1.1.chr2\_hap1.g01624.t01 |
| 3500 | 2 | 221307061 | 221315399 | mNeoVis.2.1.1.chr2\_hap1.g01624.t01 |
| 3501 | 2 | 221316661 | 221331249 | mNeoVis.2.1.1.chr2\_hap1.g01625.t01 |
| 3502 | 2 | 221316661 | 221331249 | mNeoVis.2.1.1.chr2\_hap1.g01625.t01 |
| 3503 | 2 | 221341651 | 221343177 | mNeoVis.2.1.1.chr2\_hap1.g01626.t01 |
| 3504 | 2 | 221341651 | 221343177 | mNeoVis.2.1.1.chr2\_hap1.g01626.t01 |
| 3505 | 2 | 221350221 | 221357869 | mNeoVis.2.1.1.chr2\_hap1.g01627.t01 |
| 3506 | 2 | 221350221 | 221357869 | mNeoVis.2.1.1.chr2\_hap1.g01627.t01 |
| 3507 | 2 | 221358251 | 221368795 | mNeoVis.2.1.1.chr2\_hap1.g01628.t01 |
| 3508 | 2 | 221358251 | 221368795 | mNeoVis.2.1.1.chr2\_hap1.g01628.t01 |
| 3509 | 2 | 221376895 | 221378217 | mNeoVis.2.1.1.chr2\_hap1.g01629.t01 |
| 3510 | 2 | 221376895 | 221378217 | mNeoVis.2.1.1.chr2\_hap1.g01629.t01 |
| 3511 | 2 | 221382798 | 221409961 | mNeoVis.2.1.1.chr2\_hap1.g01630.t01 |
| 3512 | 2 | 221382798 | 221409961 | mNeoVis.2.1.1.chr2\_hap1.g01630.t01 |
| 3513 | 2 | 221423050 | 221433565 | mNeoVis.2.1.1.chr2\_hap1.g01631.t01 |
| 3514 | 2 | 221423050 | 221433565 | mNeoVis.2.1.1.chr2\_hap1.g01631.t01 |
| 3515 | 2 | 221439641 | 221508966 | mNeoVis.2.1.1.chr2\_hap1.g01632.t01 |
| 3516 | 2 | 221439641 | 221508966 | mNeoVis.2.1.1.chr2\_hap1.g01632.t01 |
| 3517 | 2 | 222658097 | 222663150 | mNeoVis.2.1.1.chr2\_hap1.g01663.t01 |
| 3518 | 2 | 222675033 | 222682196 | mNeoVis.2.1.1.chr2\_hap1.g01664.t01 |
| 3519 | 2 | 222951214 | 222980622 | mNeoVis.2.1.1.chr2\_hap1.g01665.t01 |
| 3520 | 2 | 222951214 | 222980622 | mNeoVis.2.1.1.chr2\_hap1.g01665.t01 |
| 3521 | 2 | 223020397 | 223091802 | mNeoVis.2.1.1.chr2\_hap1.g01666.t01 |
| 3522 | 2 | 223020397 | 223091802 | mNeoVis.2.1.1.chr2\_hap1.g01666.t01 |
| 3523 | 2 | 223137125 | 223168436 | mNeoVis.2.1.1.chr2\_hap1.g01667.t01 |
| 3524 | 2 | 223137125 | 223168436 | mNeoVis.2.1.1.chr2\_hap1.g01667.t01 |
| 3525 | 2 | 223190785 | 223232719 | mNeoVis.2.1.1.chr2\_hap1.g01668.t01 |
| 3526 | 2 | 223190785 | 223232719 | mNeoVis.2.1.1.chr2\_hap1.g01668.t01 |
| 3527 | 2 | 223249551 | 223285957 | mNeoVis.2.1.1.chr2\_hap1.g01669.t01 |
| 3528 | 2 | 223249551 | 223285957 | mNeoVis.2.1.1.chr2\_hap1.g01669.t01 |
| 3529 | 2 | 223301278 | 223353485 | mNeoVis.2.1.1.chr2\_hap1.g01670.t01 |
| 3530 | 2 | 223301278 | 223353485 | mNeoVis.2.1.1.chr2\_hap1.g01670.t01 |
| 3531 | 2 | 223355048 | 223402913 | mNeoVis.2.1.1.chr2\_hap1.g01671.t01 |
| 3532 | 2 | 223355048 | 223402913 | mNeoVis.2.1.1.chr2\_hap1.g01671.t01 |
| 3533 | 2 | 223407758 | 223442420 | mNeoVis.2.1.1.chr2\_hap1.g01672.t01 |
| 3534 | 2 | 223407758 | 223442420 | mNeoVis.2.1.1.chr2\_hap1.g01672.t01 |
| 3535 | 2 | 223541080 | 223570761 | mNeoVis.2.1.1.chr2\_hap1.g01673.t01 |
| 3536 | 2 | 223577792 | 223598245 | mNeoVis.2.1.1.chr2\_hap1.g01674.t01 |
| 3537 | 2 | 223602484 | 223648969 | mNeoVis.2.1.1.chr2\_hap1.g01675.t01 |
| 3538 | 2 | 223710641 | 223720085 | mNeoVis.2.1.1.chr2\_hap1.g01676.t01 |
| 3539 | 2 | 223725034 | 223728742 | mNeoVis.2.1.1.chr2\_hap1.g01677.t01 |
| 3540 | 2 | 223757712 | 223782051 | mNeoVis.2.1.1.chr2\_hap1.g01678.t01 |
| 3541 | 2 | 223757712 | 223782051 | mNeoVis.2.1.1.chr2\_hap1.g01678.t01 |
| 3542 | 2 | 223785189 | 223797459 | mNeoVis.2.1.1.chr2\_hap1.g01679.t01 |
| 3543 | 2 | 223785189 | 223797459 | mNeoVis.2.1.1.chr2\_hap1.g01679.t01 |
| 3544 | 2 | 223801852 | 223816837 | mNeoVis.2.1.1.chr2\_hap1.g01680.t01 |
| 3545 | 2 | 223801852 | 223816837 | mNeoVis.2.1.1.chr2\_hap1.g01680.t01 |
| 3546 | 2 | 223817247 | 223822054 | mNeoVis.2.1.1.chr2\_hap1.g01681.t01 |
| 3547 | 2 | 223817247 | 223822054 | mNeoVis.2.1.1.chr2\_hap1.g01681.t01 |
| 3548 | 2 | 223822104 | 223826663 | mNeoVis.2.1.1.chr2\_hap1.g01682.t01 |
| 3549 | 2 | 223822104 | 223826663 | mNeoVis.2.1.1.chr2\_hap1.g01682.t01 |
| 3550 | 2 | 223828698 | 223835472 | mNeoVis.2.1.1.chr2\_hap1.g01683.t01 |
| 3551 | 2 | 223828698 | 223835472 | mNeoVis.2.1.1.chr2\_hap1.g01683.t01 |
| 3552 | 2 | 223846902 | 223861680 | mNeoVis.2.1.1.chr2\_hap1.g01684.t01 |
| 3553 | 2 | 223846902 | 223861680 | mNeoVis.2.1.1.chr2\_hap1.g01684.t01 |
| 3554 | 2 | 223893784 | 223899357 | mNeoVis.2.1.1.chr2\_hap1.g01685.t01 |
| 3555 | 2 | 223893784 | 223899357 | mNeoVis.2.1.1.chr2\_hap1.g01685.t01 |
| 3556 | 2 | 223987802 | 223989366 | mNeoVis.2.1.1.chr2\_hap1.g01686.t01 |
| 3557 | 2 | 224008027 | 224028953 | mNeoVis.2.1.1.chr2\_hap1.g01687.t01 |
| 3558 | 2 | 224047714 | 224095604 | mNeoVis.2.1.1.chr2\_hap1.g01688.t01 |
| 3559 | 2 | 224098553 | 224134060 | mNeoVis.2.1.1.chr2\_hap1.g01689.t01 |
| 3560 | 2 | 224141057 | 224151982 | mNeoVis.2.1.1.chr2\_hap1.g01690.t01 |
| 3561 | 2 | 224165134 | 224199100 | mNeoVis.2.1.1.chr2\_hap1.g01691.t01 |
| 3562 | 2 | 224241685 | 224249533 | mNeoVis.2.1.1.chr2\_hap1.g01692.t01 |
| 3563 | 2 | 224294780 | 224297586 | mNeoVis.2.1.1.chr2\_hap1.g01693.t01 |
| 3564 | 2 | 224314260 | 224418439 | mNeoVis.2.1.1.chr2\_hap1.g01694.t01 |
| 3565 | 2 | 224419805 | 224480090 | mNeoVis.2.1.1.chr2\_hap1.g01695.t01 |
| 3566 | 2 | 224484650 | 224487382 | mNeoVis.2.1.1.chr2\_hap1.g01696.t01 |
| 3567 | 2 | 224500682 | 224513026 | mNeoVis.2.1.1.chr2\_hap1.g01697.t01 |
| 3568 | 2 | 224500682 | 224513026 | mNeoVis.2.1.1.chr2\_hap1.g01697.t01 |
| 3569 | 2 | 224529815 | 224712414 | mNeoVis.2.1.1.chr2\_hap1.g01698.t01 |
| 3570 | 2 | 224529815 | 224712414 | mNeoVis.2.1.1.chr2\_hap1.g01698.t01 |
| 3571 | 2 | 224557742 | 224559046 | mNeoVis.2.1.1.chr2\_hap1.g01699.t01 |
| 3572 | 2 | 224557742 | 224559046 | mNeoVis.2.1.1.chr2\_hap1.g01699.t01 |
| 3573 | 2 | 224752491 | 224757846 | mNeoVis.2.1.1.chr2\_hap1.g01700.t01 |
| 3574 | 2 | 224752491 | 224757846 | mNeoVis.2.1.1.chr2\_hap1.g01700.t01 |
| 3575 | 2 | 224762067 | 224765178 | mNeoVis.2.1.1.chr2\_hap1.g01701.t01 |
| 3576 | 2 | 224762067 | 224765178 | mNeoVis.2.1.1.chr2\_hap1.g01701.t01 |
| 3577 | 2 | 224768344 | 224771341 | mNeoVis.2.1.1.chr2\_hap1.g01702.t01 |
| 3578 | 2 | 224768344 | 224771341 | mNeoVis.2.1.1.chr2\_hap1.g01702.t01 |
| 3579 | 2 | 224802705 | 224838391 | mNeoVis.2.1.1.chr2\_hap1.g01703.t01 |
| 3580 | 2 | 224802705 | 224838391 | mNeoVis.2.1.1.chr2\_hap1.g01703.t01 |
| 3581 | 2 | 224865706 | 224938599 | mNeoVis.2.1.1.chr2\_hap1.g01704.t01 |
| 3582 | 2 | 224865706 | 224938599 | mNeoVis.2.1.1.chr2\_hap1.g01704.t01 |
| 3583 | 2 | 225191291 | 225218277 | mNeoVis.2.1.1.chr2\_hap1.g01705.t01 |
| 3584 | 2 | 225191291 | 225218277 | mNeoVis.2.1.1.chr2\_hap1.g01705.t01 |
| 3585 | 2 | 225240034 | 225288168 | mNeoVis.2.1.1.chr2\_hap1.g01706.t01 |
| 3586 | 2 | 225312478 | 225318610 | mNeoVis.2.1.1.chr2\_hap1.g01707.t01 |
| 3587 | 2 | 225345479 | 225348090 | mNeoVis.2.1.1.chr2\_hap1.g01708.t01 |
| 3588 | 2 | 225358121 | 225457457 | mNeoVis.2.1.1.chr2\_hap1.g01709.t01 |
| 3589 | 2 | 225457599 | 225465780 | mNeoVis.2.1.1.chr2\_hap1.g01710.t01 |
| 3590 | 2 | 225487837 | 225579927 | mNeoVis.2.1.1.chr2\_hap1.g01711.t01 |
| 3591 | 2 | 225604685 | 225654380 | mNeoVis.2.1.1.chr2\_hap1.g01712.t01 |
| 3592 | 2 | 225667655 | 225730488 | mNeoVis.2.1.1.chr2\_hap1.g01713.t01 |
| 3593 | 2 | 225857417 | 225920098 | mNeoVis.2.1.1.chr2\_hap1.g01714.t01 |
| 3594 | 2 | 225947652 | 225950281 | mNeoVis.2.1.1.chr2\_hap1.g01715.t01 |
| 3595 | 2 | 226244792 | 226245635 | mNeoVis.2.1.1.chr2\_hap1.g01717.t01 |
| 3596 | 2 | 226309544 | 226347264 | mNeoVis.2.1.1.chr2\_hap1.g01718.t01 |
| 3597 | 2 | 226357894 | 226370007 | mNeoVis.2.1.1.chr2\_hap1.g01719.t01 |
| 3598 | 2 | 226371018 | 226411770 | mNeoVis.2.1.1.chr2\_hap1.g01720.t01 |
| 3599 | 2 | 226413543 | 226421667 | mNeoVis.2.1.1.chr2\_hap1.g01721.t01 |
| 3600 | 2 | 226421929 | 226445075 | mNeoVis.2.1.1.chr2\_hap1.g01722.t01 |
| 3601 | 2 | 226454505 | 226477924 | mNeoVis.2.1.1.chr2\_hap1.g01723.t01 |
| 3602 | 2 | 226519318 | 226520806 | mNeoVis.2.1.1.chr2\_hap1.g01724.t01 |
| 3603 | 2 | 226555729 | 226564126 | mNeoVis.2.1.1.chr2\_hap1.g01725.t01 |
| 3604 | 2 | 227068718 | 227069419 | mNeoVis.2.1.1.chr2\_hap1.g01735.t01 |
| 3605 | 2 | 227068718 | 227069419 | mNeoVis.2.1.1.chr2\_hap1.g01735.t01 |
| 3606 | 2 | 227081917 | 227167636 | mNeoVis.2.1.1.chr2\_hap1.g01736.t01 |
| 3607 | 2 | 227081917 | 227167636 | mNeoVis.2.1.1.chr2\_hap1.g01736.t01 |
| 3608 | 2 | 227169143 | 227180611 | mNeoVis.2.1.1.chr2\_hap1.g01737.t01 |
| 3609 | 2 | 227169143 | 227180611 | mNeoVis.2.1.1.chr2\_hap1.g01737.t01 |
| 3610 | 2 | 227186410 | 227197754 | mNeoVis.2.1.1.chr2\_hap1.g01738.t01 |
| 3611 | 2 | 227186410 | 227197754 | mNeoVis.2.1.1.chr2\_hap1.g01738.t01 |
| 3612 | 2 | 227212978 | 227243720 | mNeoVis.2.1.1.chr2\_hap1.g01739.t01 |
| 3613 | 2 | 227212978 | 227243720 | mNeoVis.2.1.1.chr2\_hap1.g01739.t01 |
| 3614 | 2 | 227315899 | 227441378 | mNeoVis.2.1.1.chr2\_hap1.g01740.t01 |
| 3615 | 2 | 227315899 | 227441378 | mNeoVis.2.1.1.chr2\_hap1.g01740.t01 |
| 3616 | 2 | 227443580 | 227464422 | mNeoVis.2.1.1.chr2\_hap1.g01741.t01 |
| 3617 | 2 | 227443580 | 227464422 | mNeoVis.2.1.1.chr2\_hap1.g01741.t01 |
| 3618 | 2 | 227462143 | 227473669 | mNeoVis.2.1.1.chr2\_hap1.g01742.t01 |
| 3619 | 2 | 227462143 | 227473669 | mNeoVis.2.1.1.chr2\_hap1.g01742.t01 |
| 3620 | 2 | 227479747 | 227490980 | mNeoVis.2.1.1.chr2\_hap1.g01743.t01 |
| 3621 | 2 | 227479747 | 227490980 | mNeoVis.2.1.1.chr2\_hap1.g01743.t01 |
| 3622 | 2 | 227491839 | 227498407 | mNeoVis.2.1.1.chr2\_hap1.g01744.t01 |
| 3623 | 2 | 227491839 | 227498407 | mNeoVis.2.1.1.chr2\_hap1.g01744.t01 |
| 3624 | 2 | 227498517 | 227524329 | mNeoVis.2.1.1.chr2\_hap1.g01745.t01 |
| 3625 | 2 | 227498517 | 227524329 | mNeoVis.2.1.1.chr2\_hap1.g01745.t01 |
| 3626 | 2 | 227537376 | 227675143 | mNeoVis.2.1.1.chr2\_hap1.g01746.t01 |
| 3627 | 2 | 227537376 | 227675143 | mNeoVis.2.1.1.chr2\_hap1.g01746.t01 |
| 3628 | 2 | 227771172 | 227818565 | mNeoVis.2.1.1.chr2\_hap1.g01747.t01 |
| 3629 | 2 | 227771172 | 227818565 | mNeoVis.2.1.1.chr2\_hap1.g01747.t01 |
| 3630 | 2 | 227820971 | 227847707 | mNeoVis.2.1.1.chr2\_hap1.g01748.t01 |
| 3631 | 2 | 227820971 | 227847707 | mNeoVis.2.1.1.chr2\_hap1.g01748.t01 |
| 3632 | 2 | 227855298 | 227871805 | mNeoVis.2.1.1.chr2\_hap1.g01749.t01 |
| 3633 | 2 | 227855298 | 227871805 | mNeoVis.2.1.1.chr2\_hap1.g01749.t01 |
| 3634 | 2 | 227948105 | 228045933 | mNeoVis.2.1.1.chr2\_hap1.g01750.t01 |
| 3635 | 2 | 227948105 | 228045933 | mNeoVis.2.1.1.chr2\_hap1.g01750.t01 |
| 3636 | 2 | 228163871 | 228169702 | mNeoVis.2.1.1.chr2\_hap1.g01751.t01 |
| 3637 | 2 | 228163871 | 228169702 | mNeoVis.2.1.1.chr2\_hap1.g01751.t01 |
| 3638 | 2 | 228234483 | 228468133 | mNeoVis.2.1.1.chr2\_hap1.g01752.t01 |
| 3639 | 2 | 228234483 | 228468133 | mNeoVis.2.1.1.chr2\_hap1.g01752.t01 |
| 3640 | 2 | 228698774 | 228716538 | mNeoVis.2.1.1.chr2\_hap1.g01753.t01 |
| 3641 | 2 | 228698774 | 228716538 | mNeoVis.2.1.1.chr2\_hap1.g01753.t01 |
| 3642 | 2 | 228807537 | 228963168 | mNeoVis.2.1.1.chr2\_hap1.g01754.t01 |
| 3643 | 2 | 229006230 | 229031117 | mNeoVis.2.1.1.chr2\_hap1.g01755.t01 |
| 3644 | 2 | 229032309 | 229053259 | mNeoVis.2.1.1.chr2\_hap1.g01756.t01 |
| 3645 | 2 | 229058363 | 229095692 | mNeoVis.2.1.1.chr2\_hap1.g01757.t01 |
| 3646 | 2 | 229118861 | 229147983 | mNeoVis.2.1.1.chr2\_hap1.g01758.t01 |
| 3647 | 2 | 229150316 | 229191005 | mNeoVis.2.1.1.chr2\_hap1.g01759.t01 |
| 3648 | 2 | 229691912 | 229718813 | mNeoVis.2.1.1.chr2\_hap1.g01771.t01 |
| 3649 | 2 | 229761727 | 229768821 | mNeoVis.2.1.1.chr2\_hap1.g01772.t01 |
| 3650 | 2 | 229769269 | 229781269 | mNeoVis.2.1.1.chr2\_hap1.g01773.t01 |
| 3651 | 2 | 229785804 | 229789199 | mNeoVis.2.1.1.chr2\_hap1.g01774.t01 |
| 3652 | 2 | 229796873 | 229799184 | mNeoVis.2.1.1.chr2\_hap1.g01775.t01 |
| 3653 | 2 | 229810399 | 229841216 | mNeoVis.2.1.1.chr2\_hap1.g01776.t01 |
| 3654 | 2 | 229842449 | 229940462 | mNeoVis.2.1.1.chr2\_hap1.g01777.t01 |
| 3655 | 2 | 229994328 | 230015433 | mNeoVis.2.1.1.chr2\_hap1.g01778.t01 |
| 3656 | 2 | 230025801 | 230030377 | mNeoVis.2.1.1.chr2\_hap1.g01779.t01 |
| 3657 | 2 | 230030574 | 230034598 | mNeoVis.2.1.1.chr2\_hap1.g01780.t01 |
| 3658 | 2 | 230036121 | 230074794 | mNeoVis.2.1.1.chr2\_hap1.g01781.t01 |
| 3659 | 2 | 230088457 | 230133094 | mNeoVis.2.1.1.chr2\_hap1.g01782.t01 |
| 3660 | 2 | 230152101 | 230162985 | mNeoVis.2.1.1.chr2\_hap1.g01783.t01 |
| 3661 | 2 | 230163092 | 230199653 | mNeoVis.2.1.1.chr2\_hap1.g01784.t01 |
| 3662 | 2 | 230199802 | 230220068 | mNeoVis.2.1.1.chr2\_hap1.g01785.t01 |
| 3663 | 2 | 230243374 | 230247426 | mNeoVis.2.1.1.chr2\_hap1.g01786.t01 |
| 3664 | 2 | 230253293 | 230269421 | mNeoVis.2.1.1.chr2\_hap1.g01787.t01 |
| 3665 | 2 | 231602024 | 231696962 | mNeoVis.2.1.1.chr2\_hap1.g01792.t01 |
| 3666 | 2 | 231602024 | 231696962 | mNeoVis.2.1.1.chr2\_hap1.g01792.t01 |
| 3667 | 2 | 231748572 | 231776909 | mNeoVis.2.1.1.chr2\_hap1.g01793.t01 |
| 3668 | 2 | 231913410 | 231926533 | mNeoVis.2.1.1.chr2\_hap1.g01795.t01 |
| 3669 | 2 | 231938647 | 231947462 | mNeoVis.2.1.1.chr2\_hap1.g01796.t01 |
| 3670 | 2 | 231972256 | 231986630 | mNeoVis.2.1.1.chr2\_hap1.g01797.t01 |
| 3671 | 2 | 232024219 | 232061737 | mNeoVis.2.1.1.chr2\_hap1.g01798.t01 |
| 3672 | 2 | 232097795 | 232340983 | mNeoVis.2.1.1.chr2\_hap1.g01799.t01 |
| 3673 | 2 | 232355204 | 232388737 | mNeoVis.2.1.1.chr2\_hap1.g01800.t01 |
| 3674 | 2 | 232355204 | 232388737 | mNeoVis.2.1.1.chr2\_hap1.g01800.t01 |
| 3675 | 2 | 232408854 | 232471618 | mNeoVis.2.1.1.chr2\_hap1.g01801.t01 |
| 3676 | 2 | 232408854 | 232471618 | mNeoVis.2.1.1.chr2\_hap1.g01801.t01 |
| 3677 | 2 | 232495738 | 232503157 | mNeoVis.2.1.1.chr2\_hap1.g01802.t01 |
| 3678 | 2 | 232495738 | 232503157 | mNeoVis.2.1.1.chr2\_hap1.g01802.t01 |
| 3679 | 2 | 232510979 | 232538936 | mNeoVis.2.1.1.chr2\_hap1.g01803.t01 |
| 3680 | 2 | 232510979 | 232538936 | mNeoVis.2.1.1.chr2\_hap1.g01803.t01 |
| 3681 | 2 | 232549898 | 232575405 | mNeoVis.2.1.1.chr2\_hap1.g01804.t01 |
| 3682 | 2 | 232580291 | 232586028 | mNeoVis.2.1.1.chr2\_hap1.g01805.t01 |
| 3683 | 2 | 232634767 | 232636155 | mNeoVis.2.1.1.chr2\_hap1.g01806.t01 |
| 3684 | 2 | 232645481 | 232646766 | mNeoVis.2.1.1.chr2\_hap1.g01807.t01 |
| 3685 | 2 | 232646656 | 232680310 | mNeoVis.2.1.1.chr2\_hap1.g01808.t01 |
| 3686 | 2 | 232682669 | 232694533 | mNeoVis.2.1.1.chr2\_hap1.g01809.t01 |
| 3687 | 2 | 232693017 | 232723534 | mNeoVis.2.1.1.chr2\_hap1.g01810.t01 |
| 3688 | 2 | 232731276 | 232737630 | mNeoVis.2.1.1.chr2\_hap1.g01811.t01 |
| 3689 | 2 | 232760957 | 232783915 | mNeoVis.2.1.1.chr2\_hap1.g01812.t01 |
| 3690 | 2 | 234885668 | 234906235 | mNeoVis.2.1.1.chr2\_hap1.g01845.t01 |
| 3691 | 2 | 234937501 | 234962360 | mNeoVis.2.1.1.chr2\_hap1.g01846.t01 |
| 3692 | 2 | 234995212 | 235010279 | mNeoVis.2.1.1.chr2\_hap1.g01847.t01 |
| 3693 | 2 | 235035902 | 235053280 | mNeoVis.2.1.1.chr2\_hap1.g01848.t01 |
| 3694 | 2 | 235053136 | 235480522 | mNeoVis.2.1.1.chr2\_hap1.g01849.t01 |
| 3695 | 2 | 236999707 | 237004077 | mNeoVis.2.1.1.chr2\_hap1.g01866.t01 |
| 3696 | 2 | 237006986 | 237035549 | mNeoVis.2.1.1.chr2\_hap1.g01867.t01 |
| 3697 | 2 | 237040530 | 237043131 | mNeoVis.2.1.1.chr2\_hap1.g01868.t01 |
| 3698 | 2 | 237075830 | 237156040 | mNeoVis.2.1.1.chr2\_hap1.g01869.t01 |
| 3699 | 2 | 237159038 | 237169874 | mNeoVis.2.1.1.chr2\_hap1.g01870.t01 |
| 3700 | 2 | 237171511 | 237180505 | mNeoVis.2.1.1.chr2\_hap1.g01871.t01 |
| 3701 | 2 | 237179805 | 237186481 | mNeoVis.2.1.1.chr2\_hap1.g01872.t01 |
| 3702 | 2 | 237186627 | 237201660 | mNeoVis.2.1.1.chr2\_hap1.g01873.t01 |
| 3703 | 2 | 237205857 | 237214785 | mNeoVis.2.1.1.chr2\_hap1.g01874.t01 |
| 3704 | 2 | 237221548 | 237280323 | mNeoVis.2.1.1.chr2\_hap1.g01875.t01 |
| 3705 | 2 | 237280835 | 237312832 | mNeoVis.2.1.1.chr2\_hap1.g01876.t01 |
| 3706 | 2 | 237361648 | 237467779 | mNeoVis.2.1.1.chr2\_hap1.g01877.t01 |
| 3707 | 3 | 2176345 | 2193686 | mNeoVis.2.1.1.chr3\_hap1.g00013.t01 |
| 3708 | 3 | 3101425 | 3112898 | mNeoVis.2.1.1.chr3\_hap1.g00018.t01 |
| 3709 | 3 | 3623352 | 4246345 | mNeoVis.2.1.1.chr3\_hap1.g00019.t01 |
| 3710 | 3 | 3699675 | 3712526 | mNeoVis.2.1.1.chr3\_hap1.g00020.t01 |
| 3711 | 3 | 3699675 | 3712526 | mNeoVis.2.1.1.chr3\_hap1.g00020.t01 |
| 3712 | 3 | 4253753 | 4286482 | mNeoVis.2.1.1.chr3\_hap1.g00021.t01 |
| 3713 | 3 | 4253753 | 4286482 | mNeoVis.2.1.1.chr3\_hap1.g00021.t01 |
| 3714 | 3 | 4289243 | 4308414 | mNeoVis.2.1.1.chr3\_hap1.g00022.t01 |
| 3715 | 3 | 4329406 | 4347203 | mNeoVis.2.1.1.chr3\_hap1.g00023.t01 |
| 3716 | 3 | 4394480 | 4423081 | mNeoVis.2.1.1.chr3\_hap1.g00024.t01 |
| 3717 | 3 | 4430965 | 4468096 | mNeoVis.2.1.1.chr3\_hap1.g00025.t01 |
| 3718 | 3 | 5768916 | 5777606 | mNeoVis.2.1.1.chr3\_hap1.g00030.t01 |
| 3719 | 3 | 8381745 | 8393194 | mNeoVis.2.1.1.chr3\_hap1.g00035.t01 |
| 3720 | 3 | 8634387 | 8771648 | mNeoVis.2.1.1.chr3\_hap1.g00036.t01 |
| 3721 | 3 | 8634387 | 8771648 | mNeoVis.2.1.1.chr3\_hap1.g00036.t01 |
| 3722 | 3 | 8824227 | 8866759 | mNeoVis.2.1.1.chr3\_hap1.g00037.t01 |
| 3723 | 3 | 8824227 | 8866759 | mNeoVis.2.1.1.chr3\_hap1.g00037.t01 |
| 3724 | 3 | 10356371 | 10585753 | mNeoVis.2.1.1.chr3\_hap1.g00039.t01 |
| 3725 | 3 | 10356371 | 10585753 | mNeoVis.2.1.1.chr3\_hap1.g00039.t01 |
| 3726 | 3 | 12621347 | 12635213 | mNeoVis.2.1.1.chr3\_hap1.g00041.t01 |
| 3727 | 3 | 12641457 | 12664559 | mNeoVis.2.1.1.chr3\_hap1.g00042.t01 |
| 3728 | 3 | 12668099 | 12682915 | mNeoVis.2.1.1.chr3\_hap1.g00043.t01 |
| 3729 | 3 | 12721443 | 12739706 | mNeoVis.2.1.1.chr3\_hap1.g00044.t01 |
| 3730 | 3 | 12745162 | 12811150 | mNeoVis.2.1.1.chr3\_hap1.g00045.t01 |
| 3731 | 3 | 12814548 | 12838441 | mNeoVis.2.1.1.chr3\_hap1.g00046.t01 |
| 3732 | 3 | 12862846 | 12885363 | mNeoVis.2.1.1.chr3\_hap1.g00047.t01 |
| 3733 | 3 | 12906877 | 12917214 | mNeoVis.2.1.1.chr3\_hap1.g00048.t01 |
| 3734 | 3 | 12928516 | 12952933 | mNeoVis.2.1.1.chr3\_hap1.g00049.t01 |
| 3735 | 3 | 12979801 | 13003589 | mNeoVis.2.1.1.chr3\_hap1.g00050.t01 |
| 3736 | 3 | 13043916 | 13074401 | mNeoVis.2.1.1.chr3\_hap1.g00051.t01 |
| 3737 | 3 | 13092512 | 13131546 | mNeoVis.2.1.1.chr3\_hap1.g00052.t01 |
| 3738 | 3 | 13137496 | 13146572 | mNeoVis.2.1.1.chr3\_hap1.g00053.t01 |
| 3739 | 3 | 13421954 | 13631020 | mNeoVis.2.1.1.chr3\_hap1.g00054.t01 |
| 3740 | 3 | 13692209 | 13703134 | mNeoVis.2.1.1.chr3\_hap1.g00055.t01 |
| 3741 | 3 | 13729672 | 13791020 | mNeoVis.2.1.1.chr3\_hap1.g00056.t01 |
| 3742 | 3 | 13898601 | 13948660 | mNeoVis.2.1.1.chr3\_hap1.g00057.t01 |
| 3743 | 3 | 13972231 | 14075236 | mNeoVis.2.1.1.chr3\_hap1.g00058.t01 |
| 3744 | 3 | 14075442 | 14174242 | mNeoVis.2.1.1.chr3\_hap1.g00059.t01 |
| 3745 | 3 | 14188315 | 14362661 | mNeoVis.2.1.1.chr3\_hap1.g00060.t01 |
| 3746 | 3 | 14565830 | 14769059 | mNeoVis.2.1.1.chr3\_hap1.g00061.t01 |
| 3747 | 3 | 15893119 | 15897407 | mNeoVis.2.1.1.chr3\_hap1.g00064.t01 |
| 3748 | 3 | 16125576 | 16144364 | mNeoVis.2.1.1.chr3\_hap1.g00066.t01 |
| 3749 | 3 | 16125576 | 16144364 | mNeoVis.2.1.1.chr3\_hap1.g00066.t01 |
| 3750 | 3 | 16289859 | 16316368 | mNeoVis.2.1.1.chr3\_hap1.g00067.t01 |
| 3751 | 3 | 16319381 | 16341451 | mNeoVis.2.1.1.chr3\_hap1.g00068.t01 |
| 3752 | 3 | 16356516 | 16361632 | mNeoVis.2.1.1.chr3\_hap1.g00069.t01 |
| 3753 | 3 | 16386843 | 16399142 | mNeoVis.2.1.1.chr3\_hap1.g00070.t01 |
| 3754 | 3 | 16429519 | 16445342 | mNeoVis.2.1.1.chr3\_hap1.g00071.t01 |
| 3755 | 3 | 16483520 | 16543927 | mNeoVis.2.1.1.chr3\_hap1.g00072.t01 |
| 3756 | 3 | 16736351 | 16796935 | mNeoVis.2.1.1.chr3\_hap1.g00074.t01 |
| 3757 | 3 | 16872960 | 16944200 | mNeoVis.2.1.1.chr3\_hap1.g00075.t01 |
| 3758 | 3 | 17006712 | 17343600 | mNeoVis.2.1.1.chr3\_hap1.g00076.t01 |
| 3759 | 3 | 19974434 | 20028110 | mNeoVis.2.1.1.chr3\_hap1.g00087.t01 |
| 3760 | 3 | 19974434 | 20028110 | mNeoVis.2.1.1.chr3\_hap1.g00087.t01 |
| 3761 | 3 | 20030295 | 20178562 | mNeoVis.2.1.1.chr3\_hap1.g00088.t01 |
| 3762 | 3 | 20030295 | 20178562 | mNeoVis.2.1.1.chr3\_hap1.g00088.t01 |
| 3763 | 3 | 20250244 | 20273084 | mNeoVis.2.1.1.chr3\_hap1.g00089.t01 |
| 3764 | 3 | 20250244 | 20273084 | mNeoVis.2.1.1.chr3\_hap1.g00089.t01 |
| 3765 | 3 | 20289497 | 20299026 | mNeoVis.2.1.1.chr3\_hap1.g00090.t01 |
| 3766 | 3 | 20289497 | 20299026 | mNeoVis.2.1.1.chr3\_hap1.g00090.t01 |
| 3767 | 3 | 20338121 | 20419849 | mNeoVis.2.1.1.chr3\_hap1.g00091.t01 |
| 3768 | 3 | 20338121 | 20419849 | mNeoVis.2.1.1.chr3\_hap1.g00091.t01 |
| 3769 | 3 | 20439983 | 20455182 | mNeoVis.2.1.1.chr3\_hap1.g00092.t01 |
| 3770 | 3 | 20439983 | 20455182 | mNeoVis.2.1.1.chr3\_hap1.g00092.t01 |
| 3771 | 3 | 20483530 | 20504994 | mNeoVis.2.1.1.chr3\_hap1.g00093.t01 |
| 3772 | 3 | 20483530 | 20504994 | mNeoVis.2.1.1.chr3\_hap1.g00093.t01 |
| 3773 | 3 | 20504892 | 20535618 | mNeoVis.2.1.1.chr3\_hap1.g00094.t01 |
| 3774 | 3 | 20504892 | 20535618 | mNeoVis.2.1.1.chr3\_hap1.g00094.t01 |
| 3775 | 3 | 20538990 | 20562645 | mNeoVis.2.1.1.chr3\_hap1.g00095.t01 |
| 3776 | 3 | 20538990 | 20562645 | mNeoVis.2.1.1.chr3\_hap1.g00095.t01 |
| 3777 | 3 | 20598427 | 20666239 | mNeoVis.2.1.1.chr3\_hap1.g00096.t01 |
| 3778 | 3 | 20598427 | 20666239 | mNeoVis.2.1.1.chr3\_hap1.g00096.t01 |
| 3779 | 3 | 22991230 | 23147990 | mNeoVis.2.1.1.chr3\_hap1.g00098.t01 |
| 3780 | 3 | 23329686 | 23349087 | mNeoVis.2.1.1.chr3\_hap1.g00099.t01 |
| 3781 | 3 | 23382824 | 23385074 | mNeoVis.2.1.1.chr3\_hap1.g00100.t01 |
| 3782 | 3 | 23395833 | 23456055 | mNeoVis.2.1.1.chr3\_hap1.g00101.t01 |
| 3783 | 3 | 23483218 | 23498272 | mNeoVis.2.1.1.chr3\_hap1.g00102.t01 |
| 3784 | 3 | 23510576 | 23572049 | mNeoVis.2.1.1.chr3\_hap1.g00103.t01 |
| 3785 | 3 | 23612243 | 23635673 | mNeoVis.2.1.1.chr3\_hap1.g00104.t01 |
| 3786 | 3 | 23679646 | 23827503 | mNeoVis.2.1.1.chr3\_hap1.g00105.t01 |
| 3787 | 3 | 23916897 | 23944036 | mNeoVis.2.1.1.chr3\_hap1.g00106.t01 |
| 3788 | 3 | 23958452 | 23965409 | mNeoVis.2.1.1.chr3\_hap1.g00107.t01 |
| 3789 | 3 | 23966499 | 23980184 | mNeoVis.2.1.1.chr3\_hap1.g00108.t01 |
| 3790 | 3 | 23980333 | 24016549 | mNeoVis.2.1.1.chr3\_hap1.g00109.t01 |
| 3791 | 3 | 24145614 | 24374946 | mNeoVis.2.1.1.chr3\_hap1.g00110.t01 |
| 3792 | 3 | 24384562 | 24414686 | mNeoVis.2.1.1.chr3\_hap1.g00111.t01 |
| 3793 | 3 | 24560726 | 24582275 | mNeoVis.2.1.1.chr3\_hap1.g00112.t01 |
| 3794 | 3 | 25064005 | 25095055 | mNeoVis.2.1.1.chr3\_hap1.g00116.t01 |
| 3795 | 3 | 25144326 | 25430399 | mNeoVis.2.1.1.chr3\_hap1.g00117.t01 |
| 3796 | 3 | 27355242 | 27377356 | mNeoVis.2.1.1.chr3\_hap1.g00129.t01 |
| 3797 | 3 | 27377496 | 27388759 | mNeoVis.2.1.1.chr3\_hap1.g00130.t01 |
| 3798 | 3 | 27401503 | 27471154 | mNeoVis.2.1.1.chr3\_hap1.g00131.t01 |
| 3799 | 3 | 27482373 | 27586888 | mNeoVis.2.1.1.chr3\_hap1.g00132.t01 |
| 3800 | 3 | 27594395 | 27606476 | mNeoVis.2.1.1.chr3\_hap1.g00133.t01 |
| 3801 | 3 | 27645096 | 27674592 | mNeoVis.2.1.1.chr3\_hap1.g00134.t01 |
| 3802 | 3 | 29178088 | 29413578 | mNeoVis.2.1.1.chr3\_hap1.g00137.t01 |
| 3803 | 3 | 29178088 | 29413578 | mNeoVis.2.1.1.chr3\_hap1.g00137.t01 |
| 3804 | 3 | 29178088 | 29413578 | mNeoVis.2.1.1.chr3\_hap1.g00137.t01 |
| 3805 | 3 | 29813501 | 29823448 | mNeoVis.2.1.1.chr3\_hap1.g00138.t01 |
| 3806 | 3 | 29932154 | 30318757 | mNeoVis.2.1.1.chr3\_hap1.g00139.t01 |
| 3807 | 3 | 30238899 | 30618879 | mNeoVis.2.1.1.chr3\_hap1.g00140.t01 |
| 3808 | 3 | 30921351 | 31064825 | mNeoVis.2.1.1.chr3\_hap1.g00141.t01 |
| 3809 | 3 | 30921351 | 31064825 | mNeoVis.2.1.1.chr3\_hap1.g00141.t01 |
| 3810 | 3 | 30937780 | 30975037 | mNeoVis.2.1.1.chr3\_hap1.g00142.t01 |
| 3811 | 3 | 30937780 | 30975037 | mNeoVis.2.1.1.chr3\_hap1.g00142.t01 |
| 3812 | 3 | 32936077 | 32958893 | mNeoVis.2.1.1.chr3\_hap1.g00143.t01 |
| 3813 | 3 | 33114183 | 33263105 | mNeoVis.2.1.1.chr3\_hap1.g00144.t01 |
| 3814 | 3 | 34444974 | 34566890 | mNeoVis.2.1.1.chr3\_hap1.g00147.t01 |
| 3815 | 3 | 34565130 | 34631655 | mNeoVis.2.1.1.chr3\_hap1.g00148.t01 |
| 3816 | 3 | 34565130 | 34631655 | mNeoVis.2.1.1.chr3\_hap1.g00148.t01 |
| 3817 | 3 | 36220797 | 36277426 | mNeoVis.2.1.1.chr3\_hap1.g00153.t01 |
| 3818 | 3 | 36286988 | 36330732 | mNeoVis.2.1.1.chr3\_hap1.g00154.t01 |
| 3819 | 3 | 36331298 | 36351882 | mNeoVis.2.1.1.chr3\_hap1.g00155.t01 |
| 3820 | 3 | 36361137 | 36362033 | mNeoVis.2.1.1.chr3\_hap1.g00156.t01 |
| 3821 | 3 | 36375068 | 36454187 | mNeoVis.2.1.1.chr3\_hap1.g00157.t01 |
| 3822 | 3 | 36467886 | 36474433 | mNeoVis.2.1.1.chr3\_hap1.g00158.t01 |
| 3823 | 3 | 36890045 | 36906501 | mNeoVis.2.1.1.chr3\_hap1.g00161.t01 |
| 3824 | 3 | 36938958 | 36969519 | mNeoVis.2.1.1.chr3\_hap1.g00162.t01 |
| 3825 | 3 | 36986190 | 36996932 | mNeoVis.2.1.1.chr3\_hap1.g00163.t01 |
| 3826 | 3 | 37007365 | 37027327 | mNeoVis.2.1.1.chr3\_hap1.g00164.t01 |
| 3827 | 3 | 37029235 | 37039154 | mNeoVis.2.1.1.chr3\_hap1.g00165.t01 |
| 3828 | 3 | 37041137 | 37049537 | mNeoVis.2.1.1.chr3\_hap1.g00166.t01 |
| 3829 | 3 | 37142542 | 37240233 | mNeoVis.2.1.1.chr3\_hap1.g00167.t01 |
| 3830 | 3 | 37142542 | 37240233 | mNeoVis.2.1.1.chr3\_hap1.g00167.t01 |
| 3831 | 3 | 37366923 | 37722574 | mNeoVis.2.1.1.chr3\_hap1.g00168.t01 |
| 3832 | 3 | 37918823 | 38323517 | mNeoVis.2.1.1.chr3\_hap1.g00169.t01 |
| 3833 | 3 | 37918823 | 38323517 | mNeoVis.2.1.1.chr3\_hap1.g00169.t01 |
| 3834 | 3 | 38401609 | 38588792 | mNeoVis.2.1.1.chr3\_hap1.g00170.t01 |
| 3835 | 3 | 38401609 | 38588792 | mNeoVis.2.1.1.chr3\_hap1.g00170.t01 |
| 3836 | 3 | 39803275 | 39828051 | mNeoVis.2.1.1.chr3\_hap1.g00174.t01 |
| 3837 | 3 | 39803275 | 39828051 | mNeoVis.2.1.1.chr3\_hap1.g00174.t01 |
| 3838 | 3 | 39865186 | 39902780 | mNeoVis.2.1.1.chr3\_hap1.g00175.t01 |
| 3839 | 3 | 39865186 | 39902780 | mNeoVis.2.1.1.chr3\_hap1.g00175.t01 |
| 3840 | 3 | 39916694 | 39954342 | mNeoVis.2.1.1.chr3\_hap1.g00176.t01 |
| 3841 | 3 | 39916694 | 39954342 | mNeoVis.2.1.1.chr3\_hap1.g00176.t01 |
| 3842 | 3 | 40013776 | 40089871 | mNeoVis.2.1.1.chr3\_hap1.g00177.t01 |
| 3843 | 3 | 40013776 | 40089871 | mNeoVis.2.1.1.chr3\_hap1.g00177.t01 |
| 3844 | 3 | 40225891 | 40293082 | mNeoVis.2.1.1.chr3\_hap1.g00178.t01 |
| 3845 | 3 | 40225891 | 40293082 | mNeoVis.2.1.1.chr3\_hap1.g00178.t01 |
| 3846 | 3 | 40316300 | 40323203 | mNeoVis.2.1.1.chr3\_hap1.g00179.t01 |
| 3847 | 3 | 40316300 | 40323203 | mNeoVis.2.1.1.chr3\_hap1.g00179.t01 |
| 3848 | 3 | 40357711 | 40403508 | mNeoVis.2.1.1.chr3\_hap1.g00180.t01 |
| 3849 | 3 | 40357711 | 40403508 | mNeoVis.2.1.1.chr3\_hap1.g00180.t01 |
| 3850 | 3 | 40418569 | 40452717 | mNeoVis.2.1.1.chr3\_hap1.g00181.t01 |
| 3851 | 3 | 40418569 | 40452717 | mNeoVis.2.1.1.chr3\_hap1.g00181.t01 |
| 3852 | 3 | 40474478 | 40520054 | mNeoVis.2.1.1.chr3\_hap1.g00182.t01 |
| 3853 | 3 | 40474478 | 40520054 | mNeoVis.2.1.1.chr3\_hap1.g00182.t01 |
| 3854 | 3 | 40533196 | 40568411 | mNeoVis.2.1.1.chr3\_hap1.g00183.t01 |
| 3855 | 3 | 40533196 | 40568411 | mNeoVis.2.1.1.chr3\_hap1.g00183.t01 |
| 3856 | 3 | 40702106 | 40734933 | mNeoVis.2.1.1.chr3\_hap1.g00184.t01 |
| 3857 | 3 | 40754172 | 40788613 | mNeoVis.2.1.1.chr3\_hap1.g00185.t01 |
| 3858 | 3 | 43220514 | 43432525 | mNeoVis.2.1.1.chr3\_hap1.g00187.t01 |
| 3859 | 3 | 43220514 | 43432525 | mNeoVis.2.1.1.chr3\_hap1.g00187.t01 |
| 3860 | 3 | 44460402 | 44460632 | mNeoVis.2.1.1.chr3\_hap1.g00190.t01 |
| 3861 | 3 | 44622327 | 44712799 | mNeoVis.2.1.1.chr3\_hap1.g00191.t01 |
| 3862 | 3 | 44778687 | 44907715 | mNeoVis.2.1.1.chr3\_hap1.g00192.t01 |
| 3863 | 3 | 44934045 | 44992495 | mNeoVis.2.1.1.chr3\_hap1.g00193.t01 |
| 3864 | 3 | 44970538 | 44970887 | mNeoVis.2.1.1.chr3\_hap1.g00194.t01 |
| 3865 | 3 | 44993166 | 45083246 | mNeoVis.2.1.1.chr3\_hap1.g00195.t01 |
| 3866 | 3 | 47242097 | 47298542 | mNeoVis.2.1.1.chr3\_hap1.g00205.t01 |
| 3867 | 3 | 47313070 | 47361219 | mNeoVis.2.1.1.chr3\_hap1.g00206.t01 |
| 3868 | 3 | 47361374 | 47379873 | mNeoVis.2.1.1.chr3\_hap1.g00207.t01 |
| 3869 | 3 | 47393101 | 47417263 | mNeoVis.2.1.1.chr3\_hap1.g00208.t01 |
| 3870 | 3 | 47427101 | 47540698 | mNeoVis.2.1.1.chr3\_hap1.g00209.t01 |
| 3871 | 3 | 47566691 | 47678427 | mNeoVis.2.1.1.chr3\_hap1.g00210.t01 |
| 3872 | 3 | 47566691 | 47678427 | mNeoVis.2.1.1.chr3\_hap1.g00210.t01 |
| 3873 | 3 | 47763843 | 47872830 | mNeoVis.2.1.1.chr3\_hap1.g00211.t01 |
| 3874 | 3 | 47937954 | 47943521 | mNeoVis.2.1.1.chr3\_hap1.g00212.t01 |
| 3875 | 3 | 48489680 | 48526365 | mNeoVis.2.1.1.chr3\_hap1.g00213.t01 |
| 3876 | 3 | 49623842 | 49675590 | mNeoVis.2.1.1.chr3\_hap1.g00221.t01 |
| 3877 | 3 | 49675476 | 49728930 | mNeoVis.2.1.1.chr3\_hap1.g00222.t01 |
| 3878 | 3 | 49795393 | 49985338 | mNeoVis.2.1.1.chr3\_hap1.g00223.t01 |
| 3879 | 3 | 51632099 | 51632705 | mNeoVis.2.1.1.chr3\_hap1.g00231.t01 |
| 3880 | 3 | 51632099 | 51632705 | mNeoVis.2.1.1.chr3\_hap1.g00231.t01 |
| 3881 | 3 | 51632099 | 51632705 | mNeoVis.2.1.1.chr3\_hap1.g00231.t01 |
| 3882 | 3 | 51880977 | 51917240 | mNeoVis.2.1.1.chr3\_hap1.g00232.t01 |
| 3883 | 3 | 51880977 | 51917240 | mNeoVis.2.1.1.chr3\_hap1.g00232.t01 |
| 3884 | 3 | 51880977 | 51917240 | mNeoVis.2.1.1.chr3\_hap1.g00232.t01 |
| 3885 | 3 | 51880977 | 51917240 | mNeoVis.2.1.1.chr3\_hap1.g00232.t01 |
| 3886 | 3 | 51917277 | 51967975 | mNeoVis.2.1.1.chr3\_hap1.g00233.t01 |
| 3887 | 3 | 51917277 | 51967975 | mNeoVis.2.1.1.chr3\_hap1.g00233.t01 |
| 3888 | 3 | 51917277 | 51967975 | mNeoVis.2.1.1.chr3\_hap1.g00233.t01 |
| 3889 | 3 | 51917277 | 51967975 | mNeoVis.2.1.1.chr3\_hap1.g00233.t01 |
| 3890 | 3 | 52001658 | 52152001 | mNeoVis.2.1.1.chr3\_hap1.g00234.t01 |
| 3891 | 3 | 52001658 | 52152001 | mNeoVis.2.1.1.chr3\_hap1.g00234.t01 |
| 3892 | 3 | 52001658 | 52152001 | mNeoVis.2.1.1.chr3\_hap1.g00234.t01 |
| 3893 | 3 | 52171207 | 52247609 | mNeoVis.2.1.1.chr3\_hap1.g00235.t01 |
| 3894 | 3 | 52249874 | 52332174 | mNeoVis.2.1.1.chr3\_hap1.g00236.t01 |
| 3895 | 3 | 53270361 | 53340681 | mNeoVis.2.1.1.chr3\_hap1.g00243.t01 |
| 3896 | 3 | 53425940 | 53426366 | mNeoVis.2.1.1.chr3\_hap1.g00244.t01 |
| 3897 | 3 | 54982297 | 54988135 | mNeoVis.2.1.1.chr3\_hap1.g00248.t01 |
| 3898 | 3 | 54988364 | 54994934 | mNeoVis.2.1.1.chr3\_hap1.g00249.t01 |
| 3899 | 3 | 56265676 | 56267146 | mNeoVis.2.1.1.chr3\_hap1.g00253.t01 |
| 3900 | 3 | 56265676 | 56267146 | mNeoVis.2.1.1.chr3\_hap1.g00253.t01 |
| 3901 | 3 | 57470202 | 57495348 | mNeoVis.2.1.1.chr3\_hap1.g00255.t01 |
| 3902 | 3 | 57548958 | 57672831 | mNeoVis.2.1.1.chr3\_hap1.g00256.t01 |
| 3903 | 3 | 59014302 | 59047330 | mNeoVis.2.1.1.chr3\_hap1.g00266.t01 |
| 3904 | 3 | 59082753 | 59106419 | mNeoVis.2.1.1.chr3\_hap1.g00267.t01 |
| 3905 | 3 | 60110595 | 60249811 | mNeoVis.2.1.1.chr3\_hap1.g00269.t01 |
| 3906 | 3 | 60228676 | 60231357 | mNeoVis.2.1.1.chr3\_hap1.g00270.t01 |
| 3907 | 3 | 60249954 | 60270048 | mNeoVis.2.1.1.chr3\_hap1.g00271.t01 |
| 3908 | 3 | 60331106 | 60373811 | mNeoVis.2.1.1.chr3\_hap1.g00272.t01 |
| 3909 | 3 | 62355308 | 62395854 | mNeoVis.2.1.1.chr3\_hap1.g00297.t01 |
| 3910 | 3 | 62400089 | 62452817 | mNeoVis.2.1.1.chr3\_hap1.g00298.t01 |
| 3911 | 3 | 62453658 | 62474804 | mNeoVis.2.1.1.chr3\_hap1.g00299.t01 |
| 3912 | 3 | 62476694 | 62524577 | mNeoVis.2.1.1.chr3\_hap1.g00300.t01 |
| 3913 | 3 | 62546663 | 62551378 | mNeoVis.2.1.1.chr3\_hap1.g00301.t01 |
| 3914 | 3 | 62547436 | 62609178 | mNeoVis.2.1.1.chr3\_hap1.g00302.t01 |
| 3915 | 3 | 62625216 | 62633220 | mNeoVis.2.1.1.chr3\_hap1.g00303.t01 |
| 3916 | 3 | 62625216 | 62633220 | mNeoVis.2.1.1.chr3\_hap1.g00303.t01 |
| 3917 | 3 | 62665600 | 62666775 | mNeoVis.2.1.1.chr3\_hap1.g00304.t01 |
| 3918 | 3 | 62665600 | 62666775 | mNeoVis.2.1.1.chr3\_hap1.g00304.t01 |
| 3919 | 3 | 62666931 | 62674564 | mNeoVis.2.1.1.chr3\_hap1.g00305.t01 |
| 3920 | 3 | 62666931 | 62674564 | mNeoVis.2.1.1.chr3\_hap1.g00305.t01 |
| 3921 | 3 | 62794948 | 62797439 | mNeoVis.2.1.1.chr3\_hap1.g00306.t01 |
| 3922 | 3 | 62794948 | 62797439 | mNeoVis.2.1.1.chr3\_hap1.g00306.t01 |
| 3923 | 3 | 62799887 | 62833323 | mNeoVis.2.1.1.chr3\_hap1.g00307.t01 |
| 3924 | 3 | 62799887 | 62833323 | mNeoVis.2.1.1.chr3\_hap1.g00307.t01 |
| 3925 | 3 | 62833341 | 62859462 | mNeoVis.2.1.1.chr3\_hap1.g00308.t01 |
| 3926 | 3 | 62833341 | 62859462 | mNeoVis.2.1.1.chr3\_hap1.g00308.t01 |
| 3927 | 3 | 62862076 | 62871465 | mNeoVis.2.1.1.chr3\_hap1.g00309.t01 |
| 3928 | 3 | 62862076 | 62871465 | mNeoVis.2.1.1.chr3\_hap1.g00309.t01 |
| 3929 | 3 | 62870202 | 62874218 | mNeoVis.2.1.1.chr3\_hap1.g00310.t01 |
| 3930 | 3 | 62870202 | 62874218 | mNeoVis.2.1.1.chr3\_hap1.g00310.t01 |
| 3931 | 3 | 62875105 | 62962333 | mNeoVis.2.1.1.chr3\_hap1.g00311.t01 |
| 3932 | 3 | 62988878 | 62990311 | mNeoVis.2.1.1.chr3\_hap1.g00312.t01 |
| 3933 | 3 | 62988878 | 62990311 | mNeoVis.2.1.1.chr3\_hap1.g00312.t01 |
| 3934 | 3 | 62992310 | 63089669 | mNeoVis.2.1.1.chr3\_hap1.g00313.t01 |
| 3935 | 3 | 62992310 | 63089669 | mNeoVis.2.1.1.chr3\_hap1.g00313.t01 |
| 3936 | 3 | 63090671 | 63093926 | mNeoVis.2.1.1.chr3\_hap1.g00314.t01 |
| 3937 | 3 | 63090671 | 63093926 | mNeoVis.2.1.1.chr3\_hap1.g00314.t01 |
| 3938 | 3 | 63125811 | 63130041 | mNeoVis.2.1.1.chr3\_hap1.g00315.t01 |
| 3939 | 3 | 63125811 | 63130041 | mNeoVis.2.1.1.chr3\_hap1.g00315.t01 |
| 3940 | 3 | 63130296 | 63139958 | mNeoVis.2.1.1.chr3\_hap1.g00316.t01 |
| 3941 | 3 | 63130296 | 63139958 | mNeoVis.2.1.1.chr3\_hap1.g00316.t01 |
| 3942 | 3 | 63141752 | 63144055 | mNeoVis.2.1.1.chr3\_hap1.g00317.t01 |
| 3943 | 3 | 63141752 | 63144055 | mNeoVis.2.1.1.chr3\_hap1.g00317.t01 |
| 3944 | 3 | 63147482 | 63149031 | mNeoVis.2.1.1.chr3\_hap1.g00318.t01 |
| 3945 | 3 | 63147482 | 63149031 | mNeoVis.2.1.1.chr3\_hap1.g00318.t01 |
| 3946 | 3 | 63151230 | 63242861 | mNeoVis.2.1.1.chr3\_hap1.g00319.t01 |
| 3947 | 3 | 63244182 | 63247402 | mNeoVis.2.1.1.chr3\_hap1.g00320.t01 |
| 3948 | 3 | 63244182 | 63247402 | mNeoVis.2.1.1.chr3\_hap1.g00320.t01 |
| 3949 | 3 | 63247531 | 63249097 | mNeoVis.2.1.1.chr3\_hap1.g00321.t01 |
| 3950 | 3 | 63247531 | 63249097 | mNeoVis.2.1.1.chr3\_hap1.g00321.t01 |
| 3951 | 3 | 63288838 | 63307558 | mNeoVis.2.1.1.chr3\_hap1.g00322.t01 |
| 3952 | 3 | 63288838 | 63307558 | mNeoVis.2.1.1.chr3\_hap1.g00322.t01 |
| 3953 | 3 | 63323313 | 63335597 | mNeoVis.2.1.1.chr3\_hap1.g00323.t01 |
| 3954 | 3 | 63323313 | 63335597 | mNeoVis.2.1.1.chr3\_hap1.g00323.t01 |
| 3955 | 3 | 63339400 | 63375205 | mNeoVis.2.1.1.chr3\_hap1.g00324.t01 |
| 3956 | 3 | 63339400 | 63375205 | mNeoVis.2.1.1.chr3\_hap1.g00324.t01 |
| 3957 | 3 | 63387053 | 63454946 | mNeoVis.2.1.1.chr3\_hap1.g00325.t01 |
| 3958 | 3 | 63387053 | 63454946 | mNeoVis.2.1.1.chr3\_hap1.g00325.t01 |
| 3959 | 3 | 63455578 | 63461127 | mNeoVis.2.1.1.chr3\_hap1.g00326.t01 |
| 3960 | 3 | 63455578 | 63461127 | mNeoVis.2.1.1.chr3\_hap1.g00326.t01 |
| 3961 | 3 | 63480686 | 63503064 | mNeoVis.2.1.1.chr3\_hap1.g00327.t01 |
| 3962 | 3 | 63480686 | 63503064 | mNeoVis.2.1.1.chr3\_hap1.g00327.t01 |
| 3963 | 3 | 63502445 | 63511831 | mNeoVis.2.1.1.chr3\_hap1.g00328.t01 |
| 3964 | 3 | 63502445 | 63511831 | mNeoVis.2.1.1.chr3\_hap1.g00328.t01 |
| 3965 | 3 | 63513247 | 63525115 | mNeoVis.2.1.1.chr3\_hap1.g00329.t01 |
| 3966 | 3 | 63513247 | 63525115 | mNeoVis.2.1.1.chr3\_hap1.g00329.t01 |
| 3967 | 3 | 63539904 | 63545976 | mNeoVis.2.1.1.chr3\_hap1.g00330.t01 |
| 3968 | 3 | 63539904 | 63545976 | mNeoVis.2.1.1.chr3\_hap1.g00330.t01 |
| 3969 | 3 | 63546361 | 63555446 | mNeoVis.2.1.1.chr3\_hap1.g00331.t01 |
| 3970 | 3 | 63546361 | 63555446 | mNeoVis.2.1.1.chr3\_hap1.g00331.t01 |
| 3971 | 3 | 63555711 | 63571794 | mNeoVis.2.1.1.chr3\_hap1.g00332.t01 |
| 3972 | 3 | 63555711 | 63571794 | mNeoVis.2.1.1.chr3\_hap1.g00332.t01 |
| 3973 | 3 | 63576067 | 63585952 | mNeoVis.2.1.1.chr3\_hap1.g00333.t01 |
| 3974 | 3 | 63576067 | 63585952 | mNeoVis.2.1.1.chr3\_hap1.g00333.t01 |
| 3975 | 3 | 63587381 | 63590303 | mNeoVis.2.1.1.chr3\_hap1.g00334.t01 |
| 3976 | 3 | 63587381 | 63590303 | mNeoVis.2.1.1.chr3\_hap1.g00334.t01 |
| 3977 | 3 | 63591020 | 63607375 | mNeoVis.2.1.1.chr3\_hap1.g00335.t01 |
| 3978 | 3 | 63591020 | 63607375 | mNeoVis.2.1.1.chr3\_hap1.g00335.t01 |
| 3979 | 3 | 63608111 | 63620727 | mNeoVis.2.1.1.chr3\_hap1.g00336.t01 |
| 3980 | 3 | 63608111 | 63620727 | mNeoVis.2.1.1.chr3\_hap1.g00336.t01 |
| 3981 | 3 | 63629492 | 63668702 | mNeoVis.2.1.1.chr3\_hap1.g00337.t01 |
| 3982 | 3 | 63629492 | 63668702 | mNeoVis.2.1.1.chr3\_hap1.g00337.t01 |
| 3983 | 3 | 63629492 | 63668702 | mNeoVis.2.1.1.chr3\_hap1.g00337.t01 |
| 3984 | 3 | 63693894 | 63718767 | mNeoVis.2.1.1.chr3\_hap1.g00338.t01 |
| 3985 | 3 | 63693894 | 63718767 | mNeoVis.2.1.1.chr3\_hap1.g00338.t01 |
| 3986 | 3 | 63693894 | 63718767 | mNeoVis.2.1.1.chr3\_hap1.g00338.t01 |
| 3987 | 3 | 63719085 | 63837890 | mNeoVis.2.1.1.chr3\_hap1.g00339.t01 |
| 3988 | 3 | 63719085 | 63837890 | mNeoVis.2.1.1.chr3\_hap1.g00339.t01 |
| 3989 | 3 | 63775717 | 63788652 | mNeoVis.2.1.1.chr3\_hap1.g00340.t01 |
| 3990 | 3 | 63775717 | 63788652 | mNeoVis.2.1.1.chr3\_hap1.g00340.t01 |
| 3991 | 3 | 63775717 | 63788652 | mNeoVis.2.1.1.chr3\_hap1.g00340.t01 |
| 3992 | 3 | 63846520 | 63849852 | mNeoVis.2.1.1.chr3\_hap1.g00341.t01 |
| 3993 | 3 | 63862235 | 63895648 | mNeoVis.2.1.1.chr3\_hap1.g00342.t01 |
| 3994 | 3 | 63913521 | 63919008 | mNeoVis.2.1.1.chr3\_hap1.g00343.t01 |
| 3995 | 3 | 63928556 | 63986045 | mNeoVis.2.1.1.chr3\_hap1.g00344.t01 |
| 3996 | 3 | 63992038 | 63993881 | mNeoVis.2.1.1.chr3\_hap1.g00345.t01 |
| 3997 | 3 | 63996128 | 64001312 | mNeoVis.2.1.1.chr3\_hap1.g00346.t01 |
| 3998 | 3 | 64005771 | 64007897 | mNeoVis.2.1.1.chr3\_hap1.g00347.t01 |
| 3999 | 3 | 64013439 | 64025270 | mNeoVis.2.1.1.chr3\_hap1.g00348.t01 |
| 4000 | 3 | 64027008 | 64053665 | mNeoVis.2.1.1.chr3\_hap1.g00349.t01 |
| 4001 | 3 | 64054920 | 64079109 | mNeoVis.2.1.1.chr3\_hap1.g00350.t01 |
| 4002 | 3 | 64100449 | 64211526 | mNeoVis.2.1.1.chr3\_hap1.g00351.t01 |
| 4003 | 3 | 64247224 | 64275032 | mNeoVis.2.1.1.chr3\_hap1.g00352.t01 |
| 4004 | 3 | 64280192 | 64303880 | mNeoVis.2.1.1.chr3\_hap1.g00353.t01 |
| 4005 | 3 | 67663200 | 67675675 | mNeoVis.2.1.1.chr3\_hap1.g00392.t01 |
| 4006 | 3 | 67706159 | 67840040 | mNeoVis.2.1.1.chr3\_hap1.g00393.t01 |
| 4007 | 3 | 67844759 | 67905181 | mNeoVis.2.1.1.chr3\_hap1.g00394.t01 |
| 4008 | 3 | 67961175 | 67971540 | mNeoVis.2.1.1.chr3\_hap1.g00395.t01 |
| 4009 | 3 | 67961175 | 67971540 | mNeoVis.2.1.1.chr3\_hap1.g00395.t01 |
| 4010 | 3 | 67971558 | 68066341 | mNeoVis.2.1.1.chr3\_hap1.g00396.t01 |
| 4011 | 3 | 67971558 | 68066341 | mNeoVis.2.1.1.chr3\_hap1.g00396.t01 |
| 4012 | 3 | 68077232 | 68121085 | mNeoVis.2.1.1.chr3\_hap1.g00397.t01 |
| 4013 | 3 | 68077232 | 68121085 | mNeoVis.2.1.1.chr3\_hap1.g00397.t01 |
| 4014 | 3 | 68125263 | 68146636 | mNeoVis.2.1.1.chr3\_hap1.g00398.t01 |
| 4015 | 3 | 68125263 | 68146636 | mNeoVis.2.1.1.chr3\_hap1.g00398.t01 |
| 4016 | 3 | 68188106 | 68207314 | mNeoVis.2.1.1.chr3\_hap1.g00399.t01 |
| 4017 | 3 | 68188106 | 68207314 | mNeoVis.2.1.1.chr3\_hap1.g00399.t01 |
| 4018 | 3 | 68230005 | 68244983 | mNeoVis.2.1.1.chr3\_hap1.g00400.t01 |
| 4019 | 3 | 68230005 | 68244983 | mNeoVis.2.1.1.chr3\_hap1.g00400.t01 |
| 4020 | 3 | 68246197 | 68307303 | mNeoVis.2.1.1.chr3\_hap1.g00401.t01 |
| 4021 | 3 | 68246197 | 68307303 | mNeoVis.2.1.1.chr3\_hap1.g00401.t01 |
| 4022 | 3 | 68311593 | 68334480 | mNeoVis.2.1.1.chr3\_hap1.g00402.t01 |
| 4023 | 3 | 68311593 | 68334480 | mNeoVis.2.1.1.chr3\_hap1.g00402.t01 |
| 4024 | 3 | 68362860 | 68371123 | mNeoVis.2.1.1.chr3\_hap1.g00403.t01 |
| 4025 | 3 | 68362860 | 68371123 | mNeoVis.2.1.1.chr3\_hap1.g00403.t01 |
| 4026 | 3 | 68371246 | 68379961 | mNeoVis.2.1.1.chr3\_hap1.g00404.t01 |
| 4027 | 3 | 68371246 | 68379961 | mNeoVis.2.1.1.chr3\_hap1.g00404.t01 |
| 4028 | 3 | 68386997 | 68389686 | mNeoVis.2.1.1.chr3\_hap1.g00405.t01 |
| 4029 | 3 | 68386997 | 68389686 | mNeoVis.2.1.1.chr3\_hap1.g00405.t01 |
| 4030 | 3 | 68389856 | 68412265 | mNeoVis.2.1.1.chr3\_hap1.g00406.t01 |
| 4031 | 3 | 68389856 | 68412265 | mNeoVis.2.1.1.chr3\_hap1.g00406.t01 |
| 4032 | 3 | 68499640 | 68504408 | mNeoVis.2.1.1.chr3\_hap1.g00407.t01 |
| 4033 | 3 | 68499640 | 68504408 | mNeoVis.2.1.1.chr3\_hap1.g00407.t01 |
| 4034 | 3 | 68505301 | 68546944 | mNeoVis.2.1.1.chr3\_hap1.g00408.t01 |
| 4035 | 3 | 68505301 | 68546944 | mNeoVis.2.1.1.chr3\_hap1.g00408.t01 |
| 4036 | 3 | 68561402 | 68720762 | mNeoVis.2.1.1.chr3\_hap1.g00409.t01 |
| 4037 | 3 | 68561402 | 68720762 | mNeoVis.2.1.1.chr3\_hap1.g00409.t01 |
| 4038 | 3 | 68734754 | 68740647 | mNeoVis.2.1.1.chr3\_hap1.g00410.t01 |
| 4039 | 3 | 68734754 | 68740647 | mNeoVis.2.1.1.chr3\_hap1.g00410.t01 |
| 4040 | 3 | 68740770 | 68749700 | mNeoVis.2.1.1.chr3\_hap1.g00411.t01 |
| 4041 | 3 | 68740770 | 68749700 | mNeoVis.2.1.1.chr3\_hap1.g00411.t01 |
| 4042 | 3 | 68761930 | 68768800 | mNeoVis.2.1.1.chr3\_hap1.g00412.t01 |
| 4043 | 3 | 68761930 | 68768800 | mNeoVis.2.1.1.chr3\_hap1.g00412.t01 |
| 4044 | 3 | 68772918 | 68787950 | mNeoVis.2.1.1.chr3\_hap1.g00413.t01 |
| 4045 | 3 | 68772918 | 68787950 | mNeoVis.2.1.1.chr3\_hap1.g00413.t01 |
| 4046 | 3 | 68824857 | 68841833 | mNeoVis.2.1.1.chr3\_hap1.g00414.t01 |
| 4047 | 3 | 68824857 | 68841833 | mNeoVis.2.1.1.chr3\_hap1.g00414.t01 |
| 4048 | 3 | 68850902 | 68861034 | mNeoVis.2.1.1.chr3\_hap1.g00415.t01 |
| 4049 | 3 | 68850902 | 68861034 | mNeoVis.2.1.1.chr3\_hap1.g00415.t01 |
| 4050 | 3 | 68889275 | 68924386 | mNeoVis.2.1.1.chr3\_hap1.g00416.t01 |
| 4051 | 3 | 68934986 | 68938464 | mNeoVis.2.1.1.chr3\_hap1.g00417.t01 |
| 4052 | 3 | 68940154 | 68960022 | mNeoVis.2.1.1.chr3\_hap1.g00418.t01 |
| 4053 | 3 | 68972959 | 68979042 | mNeoVis.2.1.1.chr3\_hap1.g00419.t01 |
| 4054 | 3 | 68981453 | 68993150 | mNeoVis.2.1.1.chr3\_hap1.g00420.t01 |
| 4055 | 3 | 68996222 | 69016334 | mNeoVis.2.1.1.chr3\_hap1.g00421.t01 |
| 4056 | 3 | 69022990 | 69054698 | mNeoVis.2.1.1.chr3\_hap1.g00422.t01 |
| 4057 | 3 | 69057070 | 69061526 | mNeoVis.2.1.1.chr3\_hap1.g00423.t01 |
| 4058 | 3 | 69074554 | 69077403 | mNeoVis.2.1.1.chr3\_hap1.g00424.t01 |
| 4059 | 3 | 69723386 | 69738341 | mNeoVis.2.1.1.chr3\_hap1.g00433.t01 |
| 4060 | 3 | 69739129 | 69774658 | mNeoVis.2.1.1.chr3\_hap1.g00434.t01 |
| 4061 | 3 | 69790095 | 69798772 | mNeoVis.2.1.1.chr3\_hap1.g00435.t01 |
| 4062 | 3 | 69839869 | 69890712 | mNeoVis.2.1.1.chr3\_hap1.g00436.t01 |
| 4063 | 3 | 69901782 | 69904708 | mNeoVis.2.1.1.chr3\_hap1.g00437.t01 |
| 4064 | 3 | 69904854 | 69909366 | mNeoVis.2.1.1.chr3\_hap1.g00438.t01 |
| 4065 | 3 | 69917348 | 69947172 | mNeoVis.2.1.1.chr3\_hap1.g00439.t01 |
| 4066 | 3 | 69947132 | 69952751 | mNeoVis.2.1.1.chr3\_hap1.g00440.t01 |
| 4067 | 3 | 69953488 | 69996913 | mNeoVis.2.1.1.chr3\_hap1.g00441.t01 |
| 4068 | 3 | 70039025 | 70105303 | mNeoVis.2.1.1.chr3\_hap1.g00442.t01 |
| 4069 | 3 | 70112834 | 70115386 | mNeoVis.2.1.1.chr3\_hap1.g00443.t01 |
| 4070 | 3 | 70115526 | 70272692 | mNeoVis.2.1.1.chr3\_hap1.g00444.t01 |
| 4071 | 3 | 70343856 | 70349901 | mNeoVis.2.1.1.chr3\_hap1.g00445.t01 |
| 4072 | 3 | 70353172 | 70366490 | mNeoVis.2.1.1.chr3\_hap1.g00446.t01 |
| 4073 | 3 | 70391212 | 70408956 | mNeoVis.2.1.1.chr3\_hap1.g00448.t01 |
| 4074 | 3 | 70461470 | 71095780 | mNeoVis.2.1.1.chr3\_hap1.g00449.t01 |
| 4075 | 3 | 70846124 | 70846549 | mNeoVis.2.1.1.chr3\_hap1.g00450.t01 |
| 4076 | 3 | 70846124 | 70846549 | mNeoVis.2.1.1.chr3\_hap1.g00450.t01 |
| 4077 | 3 | 71143962 | 71208418 | mNeoVis.2.1.1.chr3\_hap1.g00451.t01 |
| 4078 | 3 | 71143962 | 71208418 | mNeoVis.2.1.1.chr3\_hap1.g00451.t01 |
| 4079 | 3 | 71249285 | 71296464 | mNeoVis.2.1.1.chr3\_hap1.g00452.t01 |
| 4080 | 3 | 72154229 | 72170808 | mNeoVis.2.1.1.chr3\_hap1.g00453.t01 |
| 4081 | 3 | 72188941 | 72195636 | mNeoVis.2.1.1.chr3\_hap1.g00454.t01 |
| 4082 | 3 | 72201133 | 72212669 | mNeoVis.2.1.1.chr3\_hap1.g00455.t01 |
| 4083 | 3 | 72246622 | 72266583 | mNeoVis.2.1.1.chr3\_hap1.g00456.t01 |
| 4084 | 3 | 72273716 | 72288295 | mNeoVis.2.1.1.chr3\_hap1.g00457.t01 |
| 4085 | 3 | 72288268 | 72303280 | mNeoVis.2.1.1.chr3\_hap1.g00458.t01 |
| 4086 | 3 | 72303318 | 72327221 | mNeoVis.2.1.1.chr3\_hap1.g00459.t01 |
| 4087 | 3 | 72334192 | 72346784 | mNeoVis.2.1.1.chr3\_hap1.g00460.t01 |
| 4088 | 3 | 72378550 | 72560192 | mNeoVis.2.1.1.chr3\_hap1.g00461.t01 |
| 4089 | 3 | 72944395 | 73063619 | mNeoVis.2.1.1.chr3\_hap1.g00463.t01 |
| 4090 | 3 | 73131857 | 73140815 | mNeoVis.2.1.1.chr3\_hap1.g00464.t01 |
| 4091 | 3 | 73149706 | 73154020 | mNeoVis.2.1.1.chr3\_hap1.g00465.t01 |
| 4092 | 3 | 73157472 | 73346422 | mNeoVis.2.1.1.chr3\_hap1.g00466.t01 |
| 4093 | 3 | 73358102 | 73363713 | mNeoVis.2.1.1.chr3\_hap1.g00467.t01 |
| 4094 | 3 | 73369297 | 73459182 | mNeoVis.2.1.1.chr3\_hap1.g00468.t01 |
| 4095 | 3 | 73610945 | 73619394 | mNeoVis.2.1.1.chr3\_hap1.g00470.t01 |
| 4096 | 3 | 74306960 | 74312261 | mNeoVis.2.1.1.chr3\_hap1.g00471.t01 |
| 4097 | 3 | 74306960 | 74312261 | mNeoVis.2.1.1.chr3\_hap1.g00471.t01 |
| 4098 | 3 | 74316211 | 74347381 | mNeoVis.2.1.1.chr3\_hap1.g00472.t01 |
| 4099 | 3 | 74316211 | 74347381 | mNeoVis.2.1.1.chr3\_hap1.g00472.t01 |
| 4100 | 3 | 74350278 | 74402893 | mNeoVis.2.1.1.chr3\_hap1.g00473.t01 |
| 4101 | 3 | 74350278 | 74402893 | mNeoVis.2.1.1.chr3\_hap1.g00473.t01 |
| 4102 | 3 | 74413914 | 74429022 | mNeoVis.2.1.1.chr3\_hap1.g00474.t01 |
| 4103 | 3 | 74413914 | 74429022 | mNeoVis.2.1.1.chr3\_hap1.g00474.t01 |
| 4104 | 3 | 74447437 | 74469328 | mNeoVis.2.1.1.chr3\_hap1.g00475.t01 |
| 4105 | 3 | 74447437 | 74469328 | mNeoVis.2.1.1.chr3\_hap1.g00475.t01 |
| 4106 | 3 | 74478802 | 74516012 | mNeoVis.2.1.1.chr3\_hap1.g00476.t01 |
| 4107 | 3 | 74478802 | 74516012 | mNeoVis.2.1.1.chr3\_hap1.g00476.t01 |
| 4108 | 3 | 74520819 | 74575112 | mNeoVis.2.1.1.chr3\_hap1.g00477.t01 |
| 4109 | 3 | 74520819 | 74575112 | mNeoVis.2.1.1.chr3\_hap1.g00477.t01 |
| 4110 | 3 | 74667860 | 74688213 | mNeoVis.2.1.1.chr3\_hap1.g00478.t01 |
| 4111 | 3 | 74667860 | 74688213 | mNeoVis.2.1.1.chr3\_hap1.g00478.t01 |
| 4112 | 3 | 74688558 | 74720876 | mNeoVis.2.1.1.chr3\_hap1.g00479.t01 |
| 4113 | 3 | 74688558 | 74720876 | mNeoVis.2.1.1.chr3\_hap1.g00479.t01 |
| 4114 | 3 | 74716884 | 74767745 | mNeoVis.2.1.1.chr3\_hap1.g00480.t01 |
| 4115 | 3 | 74716884 | 74767745 | mNeoVis.2.1.1.chr3\_hap1.g00480.t01 |
| 4116 | 3 | 74768030 | 74797594 | mNeoVis.2.1.1.chr3\_hap1.g00481.t01 |
| 4117 | 3 | 74768030 | 74797594 | mNeoVis.2.1.1.chr3\_hap1.g00481.t01 |
| 4118 | 3 | 75343204 | 75420841 | mNeoVis.2.1.1.chr3\_hap1.g00491.t01 |
| 4119 | 3 | 75435887 | 75444948 | mNeoVis.2.1.1.chr3\_hap1.g00492.t01 |
| 4120 | 3 | 75453014 | 75467749 | mNeoVis.2.1.1.chr3\_hap1.g00493.t01 |
| 4121 | 3 | 75488168 | 75495342 | mNeoVis.2.1.1.chr3\_hap1.g00494.t01 |
| 4122 | 3 | 75496165 | 75563939 | mNeoVis.2.1.1.chr3\_hap1.g00495.t01 |
| 4123 | 3 | 75534910 | 75546366 | mNeoVis.2.1.1.chr3\_hap1.g00496.t01 |
| 4124 | 3 | 75668487 | 75715551 | mNeoVis.2.1.1.chr3\_hap1.g00497.t01 |
| 4125 | 3 | 75753134 | 75878494 | mNeoVis.2.1.1.chr3\_hap1.g00498.t01 |
| 4126 | 3 | 76578013 | 76588528 | mNeoVis.2.1.1.chr3\_hap1.g00504.t01 |
| 4127 | 3 | 76590279 | 76602098 | mNeoVis.2.1.1.chr3\_hap1.g00505.t01 |
| 4128 | 3 | 76608335 | 76622250 | mNeoVis.2.1.1.chr3\_hap1.g00506.t01 |
| 4129 | 3 | 76634288 | 76656553 | mNeoVis.2.1.1.chr3\_hap1.g00507.t01 |
| 4130 | 3 | 76694953 | 76697688 | mNeoVis.2.1.1.chr3\_hap1.g00508.t01 |
| 4131 | 3 | 76699662 | 76740559 | mNeoVis.2.1.1.chr3\_hap1.g00509.t01 |
| 4132 | 3 | 76745389 | 76764667 | mNeoVis.2.1.1.chr3\_hap1.g00510.t01 |
| 4133 | 3 | 76765807 | 76799895 | mNeoVis.2.1.1.chr3\_hap1.g00511.t01 |
| 4134 | 3 | 76800235 | 76807717 | mNeoVis.2.1.1.chr3\_hap1.g00512.t01 |
| 4135 | 3 | 76808246 | 76818304 | mNeoVis.2.1.1.chr3\_hap1.g00513.t01 |
| 4136 | 3 | 76818293 | 76820399 | mNeoVis.2.1.1.chr3\_hap1.g00514.t01 |
| 4137 | 3 | 76822216 | 76824566 | mNeoVis.2.1.1.chr3\_hap1.g00515.t01 |
| 4138 | 3 | 76886295 | 76909654 | mNeoVis.2.1.1.chr3\_hap1.g00516.t01 |
| 4139 | 3 | 76919634 | 76927082 | mNeoVis.2.1.1.chr3\_hap1.g00517.t01 |
| 4140 | 3 | 76929142 | 76943469 | mNeoVis.2.1.1.chr3\_hap1.g00518.t01 |
| 4141 | 3 | 76966358 | 77015253 | mNeoVis.2.1.1.chr3\_hap1.g00519.t01 |
| 4142 | 3 | 77017196 | 77021341 | mNeoVis.2.1.1.chr3\_hap1.g00520.t01 |
| 4143 | 3 | 77022562 | 77082874 | mNeoVis.2.1.1.chr3\_hap1.g00521.t01 |
| 4144 | 3 | 77093410 | 77110514 | mNeoVis.2.1.1.chr3\_hap1.g00522.t01 |
| 4145 | 3 | 77111129 | 77212564 | mNeoVis.2.1.1.chr3\_hap1.g00523.t01 |
| 4146 | 3 | 77292866 | 77461438 | mNeoVis.2.1.1.chr3\_hap1.g00524.t01 |
| 4147 | 3 | 77471038 | 77481464 | mNeoVis.2.1.1.chr3\_hap1.g00525.t01 |
| 4148 | 3 | 77471038 | 77481464 | mNeoVis.2.1.1.chr3\_hap1.g00525.t01 |
| 4149 | 3 | 77501106 | 77537479 | mNeoVis.2.1.1.chr3\_hap1.g00526.t01 |
| 4150 | 3 | 77501106 | 77537479 | mNeoVis.2.1.1.chr3\_hap1.g00526.t01 |
| 4151 | 3 | 77587389 | 77745852 | mNeoVis.2.1.1.chr3\_hap1.g00527.t01 |
| 4152 | 3 | 77862426 | 77875566 | mNeoVis.2.1.1.chr3\_hap1.g00528.t01 |
| 4153 | 3 | 78592839 | 78593656 | mNeoVis.2.1.1.chr3\_hap1.g00533.t01 |
| 4154 | 3 | 78739868 | 78744734 | mNeoVis.2.1.1.chr3\_hap1.g00534.t01 |
| 4155 | 3 | 78771338 | 78809496 | mNeoVis.2.1.1.chr3\_hap1.g00535.t01 |
| 4156 | 3 | 78817773 | 78830044 | mNeoVis.2.1.1.chr3\_hap1.g00536.t01 |
| 4157 | 3 | 78830531 | 78841509 | mNeoVis.2.1.1.chr3\_hap1.g00537.t01 |
| 4158 | 3 | 78874753 | 79267473 | mNeoVis.2.1.1.chr3\_hap1.g00538.t01 |
| 4159 | 3 | 79360005 | 79466154 | mNeoVis.2.1.1.chr3\_hap1.g00539.t01 |
| 4160 | 3 | 79482909 | 79528200 | mNeoVis.2.1.1.chr3\_hap1.g00540.t01 |
| 4161 | 3 | 79580801 | 79602777 | mNeoVis.2.1.1.chr3\_hap1.g00541.t01 |
| 4162 | 3 | 79607011 | 79731837 | mNeoVis.2.1.1.chr3\_hap1.g00542.t01 |
| 4163 | 3 | 79753618 | 79776672 | mNeoVis.2.1.1.chr3\_hap1.g00543.t01 |
| 4164 | 3 | 80200198 | 80490320 | mNeoVis.2.1.1.chr3\_hap1.g00546.t01 |
| 4165 | 3 | 81470755 | 81474473 | mNeoVis.2.1.1.chr3\_hap1.g00547.t01 |
| 4166 | 3 | 81679600 | 81728627 | mNeoVis.2.1.1.chr3\_hap1.g00549.t01 |
| 4167 | 3 | 82000643 | 82117049 | mNeoVis.2.1.1.chr3\_hap1.g00550.t01 |
| 4168 | 3 | 82563017 | 82586319 | mNeoVis.2.1.1.chr3\_hap1.g00557.t01 |
| 4169 | 3 | 82563017 | 82586319 | mNeoVis.2.1.1.chr3\_hap1.g00557.t01 |
| 4170 | 3 | 82587579 | 82598000 | mNeoVis.2.1.1.chr3\_hap1.g00558.t01 |
| 4171 | 3 | 82587579 | 82598000 | mNeoVis.2.1.1.chr3\_hap1.g00558.t01 |
| 4172 | 3 | 82592678 | 82610837 | mNeoVis.2.1.1.chr3\_hap1.g00559.t01 |
| 4173 | 3 | 82592678 | 82610837 | mNeoVis.2.1.1.chr3\_hap1.g00559.t01 |
| 4174 | 3 | 82610938 | 82617872 | mNeoVis.2.1.1.chr3\_hap1.g00560.t01 |
| 4175 | 3 | 82610938 | 82617872 | mNeoVis.2.1.1.chr3\_hap1.g00560.t01 |
| 4176 | 3 | 82627759 | 82655506 | mNeoVis.2.1.1.chr3\_hap1.g00561.t01 |
| 4177 | 3 | 82627759 | 82655506 | mNeoVis.2.1.1.chr3\_hap1.g00561.t01 |
| 4178 | 3 | 82656739 | 82690979 | mNeoVis.2.1.1.chr3\_hap1.g00562.t01 |
| 4179 | 3 | 82656739 | 82690979 | mNeoVis.2.1.1.chr3\_hap1.g00562.t01 |
| 4180 | 3 | 82708015 | 82726325 | mNeoVis.2.1.1.chr3\_hap1.g00563.t01 |
| 4181 | 3 | 82708015 | 82726325 | mNeoVis.2.1.1.chr3\_hap1.g00563.t01 |
| 4182 | 3 | 82732051 | 82752850 | mNeoVis.2.1.1.chr3\_hap1.g00564.t01 |
| 4183 | 3 | 82732051 | 82752850 | mNeoVis.2.1.1.chr3\_hap1.g00564.t01 |
| 4184 | 3 | 82758361 | 82768389 | mNeoVis.2.1.1.chr3\_hap1.g00565.t01 |
| 4185 | 3 | 82758361 | 82768389 | mNeoVis.2.1.1.chr3\_hap1.g00565.t01 |
| 4186 | 3 | 82773921 | 82872583 | mNeoVis.2.1.1.chr3\_hap1.g00566.t01 |
| 4187 | 3 | 82773921 | 82872583 | mNeoVis.2.1.1.chr3\_hap1.g00566.t01 |
| 4188 | 3 | 83068105 | 83151991 | mNeoVis.2.1.1.chr3\_hap1.g00567.t01 |
| 4189 | 3 | 83068105 | 83151991 | mNeoVis.2.1.1.chr3\_hap1.g00567.t01 |
| 4190 | 3 | 90113031 | 90139753 | mNeoVis.2.1.1.chr3\_hap1.g00664.t01 |
| 4191 | 3 | 90151116 | 90233992 | mNeoVis.2.1.1.chr3\_hap1.g00665.t01 |
| 4192 | 3 | 90281240 | 90805674 | mNeoVis.2.1.1.chr3\_hap1.g00666.t01 |
| 4193 | 3 | 91115263 | 91151674 | mNeoVis.2.1.1.chr3\_hap1.g00668.t01 |
| 4194 | 3 | 91162651 | 91256446 | mNeoVis.2.1.1.chr3\_hap1.g00669.t01 |
| 4195 | 3 | 91414436 | 91446847 | mNeoVis.2.1.1.chr3\_hap1.g00670.t01 |
| 4196 | 3 | 91460272 | 91464519 | mNeoVis.2.1.1.chr3\_hap1.g00671.t01 |
| 4197 | 3 | 138526504 | 138551637 | mNeoVis.2.1.1.chr3\_hap1.g00760.t01 |
| 4198 | 3 | 138618040 | 138710996 | mNeoVis.2.1.1.chr3\_hap1.g00761.t01 |
| 4199 | 3 | 138715930 | 138726800 | mNeoVis.2.1.1.chr3\_hap1.g00762.t01 |
| 4200 | 3 | 138750021 | 138769208 | mNeoVis.2.1.1.chr3\_hap1.g00763.t01 |
| 4201 | 3 | 139917308 | 140267368 | mNeoVis.2.1.1.chr3\_hap1.g00765.t01 |
| 4202 | 3 | 142260660 | 142298209 | mNeoVis.2.1.1.chr3\_hap1.g00768.t01 |
| 4203 | 3 | 142390553 | 142596779 | mNeoVis.2.1.1.chr3\_hap1.g00769.t01 |
| 4204 | 3 | 142596907 | 142639636 | mNeoVis.2.1.1.chr3\_hap1.g00770.t01 |
| 4205 | 3 | 144377685 | 144598547 | mNeoVis.2.1.1.chr3\_hap1.g00773.t01 |
| 4206 | 3 | 144725447 | 144818052 | mNeoVis.2.1.1.chr3\_hap1.g00774.t01 |
| 4207 | 3 | 144828050 | 144869912 | mNeoVis.2.1.1.chr3\_hap1.g00775.t01 |
| 4208 | 3 | 145508239 | 145641783 | mNeoVis.2.1.1.chr3\_hap1.g00779.t01 |
| 4209 | 3 | 145508239 | 145641783 | mNeoVis.2.1.1.chr3\_hap1.g00779.t01 |
| 4210 | 3 | 145656473 | 145713337 | mNeoVis.2.1.1.chr3\_hap1.g00780.t01 |
| 4211 | 3 | 145656473 | 145713337 | mNeoVis.2.1.1.chr3\_hap1.g00780.t01 |
| 4212 | 3 | 145713935 | 145771528 | mNeoVis.2.1.1.chr3\_hap1.g00781.t01 |
| 4213 | 3 | 145785007 | 145820298 | mNeoVis.2.1.1.chr3\_hap1.g00782.t01 |
| 4214 | 3 | 145855594 | 145921664 | mNeoVis.2.1.1.chr3\_hap1.g00783.t01 |
| 4215 | 3 | 148661015 | 148686594 | mNeoVis.2.1.1.chr3\_hap1.g00787.t01 |
| 4216 | 3 | 148756553 | 148871164 | mNeoVis.2.1.1.chr3\_hap1.g00788.t01 |
| 4217 | 3 | 152771056 | 152937549 | mNeoVis.2.1.1.chr3\_hap1.g00791.t01 |
| 4218 | 3 | 152771056 | 152937549 | mNeoVis.2.1.1.chr3\_hap1.g00791.t01 |
| 4219 | 3 | 152986930 | 153594552 | mNeoVis.2.1.1.chr3\_hap1.g00792.t01 |
| 4220 | 3 | 153238215 | 153306369 | mNeoVis.2.1.1.chr3\_hap1.g00793.t01 |
| 4221 | 3 | 153700493 | 154008807 | mNeoVis.2.1.1.chr3\_hap1.g00794.t01 |
| 4222 | 3 | 153857942 | 153860082 | mNeoVis.2.1.1.chr3\_hap1.g00795.t01 |
| 4223 | 3 | 153857942 | 153860082 | mNeoVis.2.1.1.chr3\_hap1.g00795.t01 |
| 4224 | 3 | 158275648 | 158293039 | mNeoVis.2.1.1.chr3\_hap1.g00804.t01 |
| 4225 | 3 | 158275648 | 158293039 | mNeoVis.2.1.1.chr3\_hap1.g00804.t01 |
| 4226 | 3 | 158275648 | 158293039 | mNeoVis.2.1.1.chr3\_hap1.g00804.t01 |
| 4227 | 3 | 158370195 | 158484723 | mNeoVis.2.1.1.chr3\_hap1.g00805.t01 |
| 4228 | 3 | 158370195 | 158484723 | mNeoVis.2.1.1.chr3\_hap1.g00805.t01 |
| 4229 | 3 | 158370195 | 158484723 | mNeoVis.2.1.1.chr3\_hap1.g00805.t01 |
| 4230 | 3 | 158492759 | 158701108 | mNeoVis.2.1.1.chr3\_hap1.g00806.t01 |
| 4231 | 3 | 158492759 | 158701108 | mNeoVis.2.1.1.chr3\_hap1.g00806.t01 |
| 4232 | 3 | 158492759 | 158701108 | mNeoVis.2.1.1.chr3\_hap1.g00806.t01 |
| 4233 | 3 | 158712341 | 158890627 | mNeoVis.2.1.1.chr3\_hap1.g00807.t01 |
| 4234 | 3 | 158712341 | 158890627 | mNeoVis.2.1.1.chr3\_hap1.g00807.t01 |
| 4235 | 3 | 158712341 | 158890627 | mNeoVis.2.1.1.chr3\_hap1.g00807.t01 |
| 4236 | 3 | 158989030 | 159113780 | mNeoVis.2.1.1.chr3\_hap1.g00808.t01 |
| 4237 | 3 | 158989030 | 159113780 | mNeoVis.2.1.1.chr3\_hap1.g00808.t01 |
| 4238 | 3 | 159239143 | 159378434 | mNeoVis.2.1.1.chr3\_hap1.g00809.t01 |
| 4239 | 3 | 159401884 | 159438924 | mNeoVis.2.1.1.chr3\_hap1.g00810.t01 |
| 4240 | 3 | 159401884 | 159438924 | mNeoVis.2.1.1.chr3\_hap1.g00810.t01 |
| 4241 | 3 | 159480298 | 159515056 | mNeoVis.2.1.1.chr3\_hap1.g00811.t01 |
| 4242 | 3 | 159480298 | 159515056 | mNeoVis.2.1.1.chr3\_hap1.g00811.t01 |
| 4243 | 3 | 160793059 | 161080696 | mNeoVis.2.1.1.chr3\_hap1.g00813.t01 |
| 4244 | 3 | 162436609 | 162441916 | mNeoVis.2.1.1.chr3\_hap1.g00815.t01 |
| 4245 | 3 | 162442810 | 162460772 | mNeoVis.2.1.1.chr3\_hap1.g00816.t01 |
| 4246 | 3 | 162484847 | 162587784 | mNeoVis.2.1.1.chr3\_hap1.g00817.t01 |
| 4247 | 3 | 162645865 | 162677105 | mNeoVis.2.1.1.chr3\_hap1.g00818.t01 |
| 4248 | 3 | 162768774 | 162819354 | mNeoVis.2.1.1.chr3\_hap1.g00819.t01 |
| 4249 | 3 | 163014755 | 163359642 | mNeoVis.2.1.1.chr3\_hap1.g00820.t01 |
| 4250 | 3 | 163375129 | 163492135 | mNeoVis.2.1.1.chr3\_hap1.g00821.t01 |
| 4251 | 3 | 163375129 | 163492135 | mNeoVis.2.1.1.chr3\_hap1.g00821.t01 |
| 4252 | 3 | 163375129 | 163492135 | mNeoVis.2.1.1.chr3\_hap1.g00821.t01 |
| 4253 | 3 | 163435423 | 163466219 | mNeoVis.2.1.1.chr3\_hap1.g00822.t01 |
| 4254 | 3 | 163435423 | 163466219 | mNeoVis.2.1.1.chr3\_hap1.g00822.t01 |
| 4255 | 3 | 163435423 | 163466219 | mNeoVis.2.1.1.chr3\_hap1.g00822.t01 |
| 4256 | 3 | 163522063 | 163738433 | mNeoVis.2.1.1.chr3\_hap1.g00823.t01 |
| 4257 | 3 | 163522063 | 163738433 | mNeoVis.2.1.1.chr3\_hap1.g00823.t01 |
| 4258 | 3 | 163522063 | 163738433 | mNeoVis.2.1.1.chr3\_hap1.g00823.t01 |
| 4259 | 3 | 163542715 | 163581787 | mNeoVis.2.1.1.chr3\_hap1.g00824.t01 |
| 4260 | 3 | 163542715 | 163581787 | mNeoVis.2.1.1.chr3\_hap1.g00824.t01 |
| 4261 | 3 | 163542715 | 163581787 | mNeoVis.2.1.1.chr3\_hap1.g00824.t01 |
| 4262 | 3 | 163762172 | 163766219 | mNeoVis.2.1.1.chr3\_hap1.g00825.t01 |
| 4263 | 3 | 163762172 | 163766219 | mNeoVis.2.1.1.chr3\_hap1.g00825.t01 |
| 4264 | 3 | 163762172 | 163766219 | mNeoVis.2.1.1.chr3\_hap1.g00825.t01 |
| 4265 | 3 | 163780492 | 163897983 | mNeoVis.2.1.1.chr3\_hap1.g00826.t01 |
| 4266 | 3 | 163780492 | 163897983 | mNeoVis.2.1.1.chr3\_hap1.g00826.t01 |
| 4267 | 3 | 163894873 | 163975367 | mNeoVis.2.1.1.chr3\_hap1.g00827.t01 |
| 4268 | 3 | 163894873 | 163975367 | mNeoVis.2.1.1.chr3\_hap1.g00827.t01 |
| 4269 | 3 | 165442968 | 165470625 | mNeoVis.2.1.1.chr3\_hap1.g00832.t01 |
| 4270 | 3 | 165461233 | 165863363 | mNeoVis.2.1.1.chr3\_hap1.g00833.t01 |
| 4271 | 3 | 166185399 | 166404025 | mNeoVis.2.1.1.chr3\_hap1.g00835.t01 |
| 4272 | 3 | 166441482 | 166718581 | mNeoVis.2.1.1.chr3\_hap1.g00836.t01 |
| 4273 | 3 | 166722227 | 166766867 | mNeoVis.2.1.1.chr3\_hap1.g00837.t01 |
| 4274 | 3 | 166722227 | 166766867 | mNeoVis.2.1.1.chr3\_hap1.g00837.t01 |
| 4275 | 3 | 166722227 | 166766867 | mNeoVis.2.1.1.chr3\_hap1.g00837.t01 |
| 4276 | 3 | 166768831 | 166779515 | mNeoVis.2.1.1.chr3\_hap1.g00838.t01 |
| 4277 | 3 | 166768831 | 166779515 | mNeoVis.2.1.1.chr3\_hap1.g00838.t01 |
| 4278 | 3 | 166768831 | 166779515 | mNeoVis.2.1.1.chr3\_hap1.g00838.t01 |
| 4279 | 3 | 166782056 | 166788385 | mNeoVis.2.1.1.chr3\_hap1.g00839.t01 |
| 4280 | 3 | 166782056 | 166788385 | mNeoVis.2.1.1.chr3\_hap1.g00839.t01 |
| 4281 | 3 | 166782056 | 166788385 | mNeoVis.2.1.1.chr3\_hap1.g00839.t01 |
| 4282 | 3 | 166790444 | 166810076 | mNeoVis.2.1.1.chr3\_hap1.g00840.t01 |
| 4283 | 3 | 166790444 | 166810076 | mNeoVis.2.1.1.chr3\_hap1.g00840.t01 |
| 4284 | 3 | 166790444 | 166810076 | mNeoVis.2.1.1.chr3\_hap1.g00840.t01 |
| 4285 | 3 | 166837361 | 167044040 | mNeoVis.2.1.1.chr3\_hap1.g00841.t01 |
| 4286 | 3 | 166837361 | 167044040 | mNeoVis.2.1.1.chr3\_hap1.g00841.t01 |
| 4287 | 3 | 166837361 | 167044040 | mNeoVis.2.1.1.chr3\_hap1.g00841.t01 |
| 4288 | 3 | 167096904 | 167115162 | mNeoVis.2.1.1.chr3\_hap1.g00842.t01 |
| 4289 | 3 | 167096904 | 167115162 | mNeoVis.2.1.1.chr3\_hap1.g00842.t01 |
| 4290 | 3 | 167096904 | 167115162 | mNeoVis.2.1.1.chr3\_hap1.g00842.t01 |
| 4291 | 3 | 167119977 | 167141881 | mNeoVis.2.1.1.chr3\_hap1.g00843.t01 |
| 4292 | 3 | 167119977 | 167141881 | mNeoVis.2.1.1.chr3\_hap1.g00843.t01 |
| 4293 | 3 | 167119977 | 167141881 | mNeoVis.2.1.1.chr3\_hap1.g00843.t01 |
| 4294 | 3 | 168803955 | 168872131 | mNeoVis.2.1.1.chr3\_hap1.g00852.t01 |
| 4295 | 3 | 168803955 | 168872131 | mNeoVis.2.1.1.chr3\_hap1.g00852.t01 |
| 4296 | 3 | 168803955 | 168872131 | mNeoVis.2.1.1.chr3\_hap1.g00852.t01 |
| 4297 | 3 | 168943038 | 168944974 | mNeoVis.2.1.1.chr3\_hap1.g00853.t01 |
| 4298 | 3 | 168943038 | 168944974 | mNeoVis.2.1.1.chr3\_hap1.g00853.t01 |
| 4299 | 3 | 168943038 | 168944974 | mNeoVis.2.1.1.chr3\_hap1.g00853.t01 |
| 4300 | 3 | 168957995 | 168974695 | mNeoVis.2.1.1.chr3\_hap1.g00854.t01 |
| 4301 | 3 | 168957995 | 168974695 | mNeoVis.2.1.1.chr3\_hap1.g00854.t01 |
| 4302 | 3 | 168976000 | 168981669 | mNeoVis.2.1.1.chr3\_hap1.g00855.t01 |
| 4303 | 3 | 168976000 | 168981669 | mNeoVis.2.1.1.chr3\_hap1.g00855.t01 |
| 4304 | 3 | 168999710 | 169003414 | mNeoVis.2.1.1.chr3\_hap1.g00856.t01 |
| 4305 | 3 | 168999710 | 169003414 | mNeoVis.2.1.1.chr3\_hap1.g00856.t01 |
| 4306 | 3 | 169007565 | 169009774 | mNeoVis.2.1.1.chr3\_hap1.g00857.t01 |
| 4307 | 3 | 169007565 | 169009774 | mNeoVis.2.1.1.chr3\_hap1.g00857.t01 |
| 4308 | 3 | 169012501 | 169015833 | mNeoVis.2.1.1.chr3\_hap1.g00858.t01 |
| 4309 | 3 | 169012501 | 169015833 | mNeoVis.2.1.1.chr3\_hap1.g00858.t01 |
| 4310 | 3 | 169013342 | 169025215 | mNeoVis.2.1.1.chr3\_hap1.g00859.t01 |
| 4311 | 3 | 169013342 | 169025215 | mNeoVis.2.1.1.chr3\_hap1.g00859.t01 |
| 4312 | 3 | 169031742 | 169032819 | mNeoVis.2.1.1.chr3\_hap1.g00860.t01 |
| 4313 | 3 | 169031742 | 169032819 | mNeoVis.2.1.1.chr3\_hap1.g00860.t01 |
| 4314 | 3 | 169036321 | 169039611 | mNeoVis.2.1.1.chr3\_hap1.g00861.t01 |
| 4315 | 3 | 169048571 | 169052247 | mNeoVis.2.1.1.chr3\_hap1.g00862.t01 |
| 4316 | 3 | 169132891 | 169217404 | mNeoVis.2.1.1.chr3\_hap1.g00863.t01 |
| 4317 | 3 | 169879974 | 169883249 | mNeoVis.2.1.1.chr3\_hap1.g00864.t01 |
| 4318 | 3 | 169893802 | 169978888 | mNeoVis.2.1.1.chr3\_hap1.g00865.t01 |
| 4319 | 3 | 169998112 | 170001495 | mNeoVis.2.1.1.chr3\_hap1.g00866.t01 |
| 4320 | 3 | 170280413 | 170297375 | mNeoVis.2.1.1.chr3\_hap1.g00868.t01 |
| 4321 | 3 | 170280413 | 170297375 | mNeoVis.2.1.1.chr3\_hap1.g00868.t01 |
| 4322 | 3 | 170320861 | 170436984 | mNeoVis.2.1.1.chr3\_hap1.g00869.t01 |
| 4323 | 3 | 170320861 | 170436984 | mNeoVis.2.1.1.chr3\_hap1.g00869.t01 |
| 4324 | 3 | 170488047 | 170533776 | mNeoVis.2.1.1.chr3\_hap1.g00870.t01 |
| 4325 | 3 | 170488047 | 170533776 | mNeoVis.2.1.1.chr3\_hap1.g00870.t01 |
| 4326 | 3 | 170488047 | 170533776 | mNeoVis.2.1.1.chr3\_hap1.g00870.t01 |
| 4327 | 3 | 170533610 | 170565117 | mNeoVis.2.1.1.chr3\_hap1.g00871.t01 |
| 4328 | 3 | 170533610 | 170565117 | mNeoVis.2.1.1.chr3\_hap1.g00871.t01 |
| 4329 | 3 | 170533610 | 170565117 | mNeoVis.2.1.1.chr3\_hap1.g00871.t01 |
| 4330 | 3 | 170565265 | 170607344 | mNeoVis.2.1.1.chr3\_hap1.g00872.t01 |
| 4331 | 3 | 170565265 | 170607344 | mNeoVis.2.1.1.chr3\_hap1.g00872.t01 |
| 4332 | 3 | 170565265 | 170607344 | mNeoVis.2.1.1.chr3\_hap1.g00872.t01 |
| 4333 | 3 | 170617706 | 170620299 | mNeoVis.2.1.1.chr3\_hap1.g00873.t01 |
| 4334 | 3 | 170617706 | 170620299 | mNeoVis.2.1.1.chr3\_hap1.g00873.t01 |
| 4335 | 3 | 170617706 | 170620299 | mNeoVis.2.1.1.chr3\_hap1.g00873.t01 |
| 4336 | 3 | 170682632 | 170860695 | mNeoVis.2.1.1.chr3\_hap1.g00874.t01 |
| 4337 | 3 | 170682632 | 170860695 | mNeoVis.2.1.1.chr3\_hap1.g00874.t01 |
| 4338 | 3 | 170728295 | 170730254 | mNeoVis.2.1.1.chr3\_hap1.g00875.t01 |
| 4339 | 3 | 170728295 | 170730254 | mNeoVis.2.1.1.chr3\_hap1.g00875.t01 |
| 4340 | 3 | 170965132 | 171021564 | mNeoVis.2.1.1.chr3\_hap1.g00876.t01 |
| 4341 | 3 | 170965132 | 171021564 | mNeoVis.2.1.1.chr3\_hap1.g00876.t01 |
| 4342 | 3 | 171039316 | 171043512 | mNeoVis.2.1.1.chr3\_hap1.g00877.t01 |
| 4343 | 3 | 171039316 | 171043512 | mNeoVis.2.1.1.chr3\_hap1.g00877.t01 |
| 4344 | 3 | 171086159 | 171090994 | mNeoVis.2.1.1.chr3\_hap1.g00878.t01 |
| 4345 | 3 | 171086159 | 171090994 | mNeoVis.2.1.1.chr3\_hap1.g00878.t01 |
| 4346 | 3 | 171383176 | 171421520 | mNeoVis.2.1.1.chr3\_hap1.g00879.t01 |
| 4347 | 3 | 171383176 | 171421520 | mNeoVis.2.1.1.chr3\_hap1.g00879.t01 |
| 4348 | 3 | 171413248 | 171417269 | mNeoVis.2.1.1.chr3\_hap1.g00880.t01 |
| 4349 | 3 | 171413248 | 171417269 | mNeoVis.2.1.1.chr3\_hap1.g00880.t01 |
| 4350 | 3 | 171456976 | 171468847 | mNeoVis.2.1.1.chr3\_hap1.g00881.t01 |
| 4351 | 3 | 171456976 | 171468847 | mNeoVis.2.1.1.chr3\_hap1.g00881.t01 |
| 4352 | 3 | 171527054 | 171707030 | mNeoVis.2.1.1.chr3\_hap1.g00882.t01 |
| 4353 | 3 | 171724172 | 172010520 | mNeoVis.2.1.1.chr3\_hap1.g00883.t01 |
| 4354 | 3 | 172548627 | 172551958 | mNeoVis.2.1.1.chr3\_hap1.g00887.t01 |
| 4355 | 3 | 172562108 | 172566087 | mNeoVis.2.1.1.chr3\_hap1.g00888.t01 |
| 4356 | 3 | 172570693 | 172657232 | mNeoVis.2.1.1.chr3\_hap1.g00889.t01 |
| 4357 | 3 | 172673068 | 172731571 | mNeoVis.2.1.1.chr3\_hap1.g00890.t01 |
| 4358 | 3 | 172731849 | 172856421 | mNeoVis.2.1.1.chr3\_hap1.g00891.t01 |
| 4359 | 3 | 172886192 | 172908864 | mNeoVis.2.1.1.chr3\_hap1.g00892.t01 |
| 4360 | 3 | 173014650 | 173114198 | mNeoVis.2.1.1.chr3\_hap1.g00893.t01 |
| 4361 | 3 | 173114350 | 173215428 | mNeoVis.2.1.1.chr3\_hap1.g00894.t01 |
| 4362 | 3 | 173344376 | 173489099 | mNeoVis.2.1.1.chr3\_hap1.g00895.t01 |
| 4363 | 3 | 173514603 | 173552150 | mNeoVis.2.1.1.chr3\_hap1.g00896.t01 |
| 4364 | 3 | 173514603 | 173552150 | mNeoVis.2.1.1.chr3\_hap1.g00896.t01 |
| 4365 | 3 | 173599210 | 173639505 | mNeoVis.2.1.1.chr3\_hap1.g00897.t01 |
| 4366 | 3 | 173599210 | 173639505 | mNeoVis.2.1.1.chr3\_hap1.g00897.t01 |
| 4367 | 3 | 173654905 | 173693775 | mNeoVis.2.1.1.chr3\_hap1.g00898.t01 |
| 4368 | 3 | 173654905 | 173693775 | mNeoVis.2.1.1.chr3\_hap1.g00898.t01 |
| 4369 | 3 | 173734290 | 173736834 | mNeoVis.2.1.1.chr3\_hap1.g00899.t01 |
| 4370 | 3 | 173734290 | 173736834 | mNeoVis.2.1.1.chr3\_hap1.g00899.t01 |
| 4371 | 3 | 174258107 | 174511717 | mNeoVis.2.1.1.chr3\_hap1.g00902.t01 |
| 4372 | 3 | 174534111 | 174541741 | mNeoVis.2.1.1.chr3\_hap1.g00903.t01 |
| 4373 | 3 | 174541442 | 174560996 | mNeoVis.2.1.1.chr3\_hap1.g00904.t01 |
| 4374 | 3 | 175353411 | 175411303 | mNeoVis.2.1.1.chr3\_hap1.g00917.t01 |
| 4375 | 3 | 175429631 | 175749866 | mNeoVis.2.1.1.chr3\_hap1.g00918.t01 |
| 4376 | 3 | 175921210 | 176222904 | mNeoVis.2.1.1.chr3\_hap1.g00919.t01 |
| 4377 | 3 | 176339143 | 176862749 | mNeoVis.2.1.1.chr3\_hap1.g00920.t01 |
| 4378 | 3 | 177725534 | 177894125 | mNeoVis.2.1.1.chr3\_hap1.g00924.t01 |
| 4379 | 3 | 177953172 | 178096399 | mNeoVis.2.1.1.chr3\_hap1.g00925.t01 |
| 4380 | 3 | 178116680 | 178193941 | mNeoVis.2.1.1.chr3\_hap1.g00926.t01 |
| 4381 | 3 | 178205910 | 178264187 | mNeoVis.2.1.1.chr3\_hap1.g00927.t01 |
| 4382 | 3 | 178205910 | 178264187 | mNeoVis.2.1.1.chr3\_hap1.g00927.t01 |
| 4383 | 3 | 178263519 | 178307906 | mNeoVis.2.1.1.chr3\_hap1.g00928.t01 |
| 4384 | 3 | 178263519 | 178307906 | mNeoVis.2.1.1.chr3\_hap1.g00928.t01 |
| 4385 | 3 | 178344944 | 178557715 | mNeoVis.2.1.1.chr3\_hap1.g00929.t01 |
| 4386 | 3 | 178344944 | 178557715 | mNeoVis.2.1.1.chr3\_hap1.g00929.t01 |
| 4387 | 3 | 178344944 | 178557715 | mNeoVis.2.1.1.chr3\_hap1.g00929.t01 |
| 4388 | 3 | 178618097 | 178758364 | mNeoVis.2.1.1.chr3\_hap1.g00930.t01 |
| 4389 | 3 | 178618097 | 178758364 | mNeoVis.2.1.1.chr3\_hap1.g00930.t01 |
| 4390 | 3 | 178618097 | 178758364 | mNeoVis.2.1.1.chr3\_hap1.g00930.t01 |
| 4391 | 3 | 178802291 | 178911880 | mNeoVis.2.1.1.chr3\_hap1.g00931.t01 |
| 4392 | 3 | 178802291 | 178911880 | mNeoVis.2.1.1.chr3\_hap1.g00931.t01 |
| 4393 | 3 | 178802291 | 178911880 | mNeoVis.2.1.1.chr3\_hap1.g00931.t01 |
| 4394 | 3 | 179030705 | 179081342 | mNeoVis.2.1.1.chr3\_hap1.g00932.t01 |
| 4395 | 3 | 179030705 | 179081342 | mNeoVis.2.1.1.chr3\_hap1.g00932.t01 |
| 4396 | 3 | 179030705 | 179081342 | mNeoVis.2.1.1.chr3\_hap1.g00932.t01 |
| 4397 | 3 | 179146579 | 179315656 | mNeoVis.2.1.1.chr3\_hap1.g00933.t01 |
| 4398 | 3 | 179146579 | 179315656 | mNeoVis.2.1.1.chr3\_hap1.g00933.t01 |
| 4399 | 3 | 179374533 | 179500016 | mNeoVis.2.1.1.chr3\_hap1.g00934.t01 |
| 4400 | 3 | 179374533 | 179500016 | mNeoVis.2.1.1.chr3\_hap1.g00934.t01 |
| 4401 | 3 | 180202275 | 180333553 | mNeoVis.2.1.1.chr3\_hap1.g00935.t01 |
| 4402 | 3 | 180202275 | 180333553 | mNeoVis.2.1.1.chr3\_hap1.g00935.t01 |
| 4403 | 3 | 180202275 | 180333553 | mNeoVis.2.1.1.chr3\_hap1.g00935.t01 |
| 4404 | 3 | 181003971 | 181489097 | mNeoVis.2.1.1.chr3\_hap1.g00936.t01 |
| 4405 | 3 | 181499365 | 181522750 | mNeoVis.2.1.1.chr3\_hap1.g00937.t01 |
| 4406 | 3 | 183143441 | 183356414 | mNeoVis.2.1.1.chr3\_hap1.g00948.t01 |
| 4407 | 3 | 183423251 | 183498931 | mNeoVis.2.1.1.chr3\_hap1.g00949.t01 |
| 4408 | 3 | 183520720 | 183631611 | mNeoVis.2.1.1.chr3\_hap1.g00950.t01 |
| 4409 | 3 | 184283743 | 184329306 | mNeoVis.2.1.1.chr3\_hap1.g00955.t01 |
| 4410 | 3 | 184283743 | 184329306 | mNeoVis.2.1.1.chr3\_hap1.g00955.t01 |
| 4411 | 3 | 184331998 | 184566294 | mNeoVis.2.1.1.chr3\_hap1.g00956.t01 |
| 4412 | 3 | 184331998 | 184566294 | mNeoVis.2.1.1.chr3\_hap1.g00956.t01 |
| 4413 | 3 | 184664522 | 184685406 | mNeoVis.2.1.1.chr3\_hap1.g00957.t01 |
| 4414 | 3 | 184664522 | 184685406 | mNeoVis.2.1.1.chr3\_hap1.g00957.t01 |
| 4415 | 3 | 184664522 | 184685406 | mNeoVis.2.1.1.chr3\_hap1.g00957.t01 |
| 4416 | 3 | 184664522 | 184685406 | mNeoVis.2.1.1.chr3\_hap1.g00957.t01 |
| 4417 | 3 | 184785152 | 185033725 | mNeoVis.2.1.1.chr3\_hap1.g00958.t01 |
| 4418 | 3 | 184785152 | 185033725 | mNeoVis.2.1.1.chr3\_hap1.g00958.t01 |
| 4419 | 3 | 184785152 | 185033725 | mNeoVis.2.1.1.chr3\_hap1.g00958.t01 |
| 4420 | 3 | 185300482 | 185334117 | mNeoVis.2.1.1.chr3\_hap1.g00960.t01 |
| 4421 | 3 | 185508730 | 185635361 | mNeoVis.2.1.1.chr3\_hap1.g00961.t01 |
| 4422 | 3 | 185777831 | 185861120 | mNeoVis.2.1.1.chr3\_hap1.g00962.t01 |
| 4423 | 3 | 185936121 | 185967052 | mNeoVis.2.1.1.chr3\_hap1.g00963.t01 |
| 4424 | 3 | 185936121 | 185967052 | mNeoVis.2.1.1.chr3\_hap1.g00963.t01 |
| 4425 | 3 | 186028153 | 186034551 | mNeoVis.2.1.1.chr3\_hap1.g00964.t01 |
| 4426 | 3 | 186028153 | 186034551 | mNeoVis.2.1.1.chr3\_hap1.g00964.t01 |
| 4427 | 3 | 186037716 | 186084773 | mNeoVis.2.1.1.chr3\_hap1.g00965.t01 |
| 4428 | 3 | 186037716 | 186084773 | mNeoVis.2.1.1.chr3\_hap1.g00965.t01 |
| 4429 | 3 | 186648398 | 186745618 | mNeoVis.2.1.1.chr3\_hap1.g00966.t01 |
| 4430 | 3 | 186860511 | 186877748 | mNeoVis.2.1.1.chr3\_hap1.g00967.t01 |
| 4431 | 3 | 188079245 | 188230733 | mNeoVis.2.1.1.chr3\_hap1.g00968.t01 |
| 4432 | 3 | 189257205 | 189282895 | mNeoVis.2.1.1.chr3\_hap1.g00970.t01 |
| 4433 | 3 | 189863412 | 189907959 | mNeoVis.2.1.1.chr3\_hap1.g00971.t01 |
| 4434 | 3 | 189908609 | 189964726 | mNeoVis.2.1.1.chr3\_hap1.g00972.t01 |
| 4435 | 3 | 190441119 | 190494043 | mNeoVis.2.1.1.chr3\_hap1.g00974.t01 |
| 4436 | 3 | 190767067 | 190803764 | mNeoVis.2.1.1.chr3\_hap1.g00976.t01 |
| 4437 | 3 | 190845436 | 191062176 | mNeoVis.2.1.1.chr3\_hap1.g00977.t01 |
| 4438 | 3 | 191144159 | 191144921 | mNeoVis.2.1.1.chr3\_hap1.g00979.t01 |
| 4439 | 3 | 191176011 | 191192008 | mNeoVis.2.1.1.chr3\_hap1.g00980.t01 |
| 4440 | 3 | 191239091 | 191261155 | mNeoVis.2.1.1.chr3\_hap1.g00981.t01 |
| 4441 | 3 | 191270954 | 191278827 | mNeoVis.2.1.1.chr3\_hap1.g00982.t01 |
| 4442 | 3 | 191285022 | 191597196 | mNeoVis.2.1.1.chr3\_hap1.g00983.t01 |
| 4443 | 3 | 191972366 | 191992626 | mNeoVis.2.1.1.chr3\_hap1.g00984.t01 |
| 4444 | 3 | 192055631 | 192055942 | mNeoVis.2.1.1.chr3\_hap1.g00985.t01 |
| 4445 | 3 | 192735028 | 192776553 | mNeoVis.2.1.1.chr3\_hap1.g00987.t01 |
| 4446 | 3 | 192962931 | 192963309 | mNeoVis.2.1.1.chr3\_hap1.g00988.t01 |
| 4447 | 3 | 193039230 | 193178210 | mNeoVis.2.1.1.chr3\_hap1.g00989.t01 |
| 4448 | 3 | 193624025 | 193635616 | mNeoVis.2.1.1.chr3\_hap1.g01000.t01 |
| 4449 | 3 | 193635482 | 193750320 | mNeoVis.2.1.1.chr3\_hap1.g01001.t01 |
| 4450 | 3 | 193753625 | 193786282 | mNeoVis.2.1.1.chr3\_hap1.g01002.t01 |
| 4451 | 3 | 193785873 | 193859147 | mNeoVis.2.1.1.chr3\_hap1.g01003.t01 |
| 4452 | 3 | 193858924 | 193888100 | mNeoVis.2.1.1.chr3\_hap1.g01004.t01 |
| 4453 | 3 | 193890531 | 193919518 | mNeoVis.2.1.1.chr3\_hap1.g01005.t01 |
| 4454 | 3 | 193920352 | 193953648 | mNeoVis.2.1.1.chr3\_hap1.g01006.t01 |
| 4455 | 3 | 193955780 | 193963172 | mNeoVis.2.1.1.chr3\_hap1.g01007.t01 |
| 4456 | 3 | 193963312 | 194047482 | mNeoVis.2.1.1.chr3\_hap1.g01008.t01 |
| 4457 | 3 | 194054303 | 194114173 | mNeoVis.2.1.1.chr3\_hap1.g01009.t01 |
| 4458 | 3 | 194121704 | 194130467 | mNeoVis.2.1.1.chr3\_hap1.g01010.t01 |
| 4459 | 3 | 194137042 | 194146860 | mNeoVis.2.1.1.chr3\_hap1.g01011.t01 |
| 4460 | 3 | 194168548 | 194260859 | mNeoVis.2.1.1.chr3\_hap1.g01012.t01 |
| 4461 | 3 | 194266842 | 194273456 | mNeoVis.2.1.1.chr3\_hap1.g01013.t01 |
| 4462 | 3 | 194289051 | 194294943 | mNeoVis.2.1.1.chr3\_hap1.g01014.t01 |
| 4463 | 3 | 194313401 | 194325366 | mNeoVis.2.1.1.chr3\_hap1.g01015.t01 |
| 4464 | 3 | 194330005 | 194339808 | mNeoVis.2.1.1.chr3\_hap1.g01016.t01 |
| 4465 | 3 | 194340882 | 194344940 | mNeoVis.2.1.1.chr3\_hap1.g01017.t01 |
| 4466 | 3 | 194348172 | 194390598 | mNeoVis.2.1.1.chr3\_hap1.g01018.t01 |
| 4467 | 3 | 194390744 | 194419297 | mNeoVis.2.1.1.chr3\_hap1.g01019.t01 |
| 4468 | 3 | 194611064 | 194612145 | mNeoVis.2.1.1.chr3\_hap1.g01020.t01 |
| 4469 | 3 | 194615614 | 194620235 | mNeoVis.2.1.1.chr3\_hap1.g01021.t01 |
| 4470 | 3 | 194733597 | 194734535 | mNeoVis.2.1.1.chr3\_hap1.g01023.t01 |
| 4471 | 3 | 194741494 | 194742429 | mNeoVis.2.1.1.chr3\_hap1.g01024.t01 |
| 4472 | 3 | 194752046 | 194802838 | mNeoVis.2.1.1.chr3\_hap1.g01025.t01 |
| 4473 | 3 | 195272279 | 195426432 | mNeoVis.2.1.1.chr3\_hap1.g01026.t01 |
| 4474 | 3 | 195272279 | 195426432 | mNeoVis.2.1.1.chr3\_hap1.g01026.t01 |
| 4475 | 3 | 195521167 | 195571538 | mNeoVis.2.1.1.chr3\_hap1.g01027.t01 |
| 4476 | 3 | 195521167 | 195571538 | mNeoVis.2.1.1.chr3\_hap1.g01027.t01 |
| 4477 | 3 | 195864725 | 195885582 | mNeoVis.2.1.1.chr3\_hap1.g01028.t01 |
| 4478 | 3 | 196047013 | 196070708 | mNeoVis.2.1.1.chr3\_hap1.g01032.t01 |
| 4479 | 3 | 196071964 | 196081385 | mNeoVis.2.1.1.chr3\_hap1.g01033.t01 |
| 4480 | 3 | 196086894 | 196097346 | mNeoVis.2.1.1.chr3\_hap1.g01034.t01 |
| 4481 | 3 | 196104377 | 196130179 | mNeoVis.2.1.1.chr3\_hap1.g01035.t01 |
| 4482 | 3 | 196131264 | 196168336 | mNeoVis.2.1.1.chr3\_hap1.g01036.t01 |
| 4483 | 3 | 196179050 | 196233686 | mNeoVis.2.1.1.chr3\_hap1.g01037.t01 |
| 4484 | 3 | 196267237 | 196309443 | mNeoVis.2.1.1.chr3\_hap1.g01038.t01 |
| 4485 | 3 | 196319020 | 196359898 | mNeoVis.2.1.1.chr3\_hap1.g01039.t01 |
| 4486 | 3 | 196374711 | 196501115 | mNeoVis.2.1.1.chr3\_hap1.g01040.t01 |
| 4487 | 3 | 196513296 | 196546343 | mNeoVis.2.1.1.chr3\_hap1.g01041.t01 |
| 4488 | 3 | 196513296 | 196546343 | mNeoVis.2.1.1.chr3\_hap1.g01041.t01 |
| 4489 | 3 | 196547556 | 196548382 | mNeoVis.2.1.1.chr3\_hap1.g01042.t01 |
| 4490 | 3 | 196547556 | 196548382 | mNeoVis.2.1.1.chr3\_hap1.g01042.t01 |
| 4491 | 3 | 196553708 | 196601931 | mNeoVis.2.1.1.chr3\_hap1.g01043.t01 |
| 4492 | 3 | 196553708 | 196601931 | mNeoVis.2.1.1.chr3\_hap1.g01043.t01 |
| 4493 | 3 | 196672264 | 196750191 | mNeoVis.2.1.1.chr3\_hap1.g01044.t01 |
| 4494 | 3 | 196984936 | 196998809 | mNeoVis.2.1.1.chr3\_hap1.g01045.t01 |
| 4495 | 3 | 196984936 | 196998809 | mNeoVis.2.1.1.chr3\_hap1.g01045.t01 |
| 4496 | 3 | 197449604 | 197461046 | mNeoVis.2.1.1.chr3\_hap1.g01046.t01 |
| 4497 | 3 | 197449604 | 197461046 | mNeoVis.2.1.1.chr3\_hap1.g01046.t01 |
| 4498 | 3 | 197507312 | 197658456 | mNeoVis.2.1.1.chr3\_hap1.g01047.t01 |
| 4499 | 3 | 197507312 | 197658456 | mNeoVis.2.1.1.chr3\_hap1.g01047.t01 |
| 4500 | 3 | 197690359 | 197753194 | mNeoVis.2.1.1.chr3\_hap1.g01048.t01 |
| 4501 | 3 | 197690359 | 197753194 | mNeoVis.2.1.1.chr3\_hap1.g01048.t01 |
| 4502 | 3 | 197770588 | 197772703 | mNeoVis.2.1.1.chr3\_hap1.g01049.t01 |
| 4503 | 3 | 197770588 | 197772703 | mNeoVis.2.1.1.chr3\_hap1.g01049.t01 |
| 4504 | 3 | 197803317 | 198176171 | mNeoVis.2.1.1.chr3\_hap1.g01050.t01 |
| 4505 | 3 | 197803317 | 198176171 | mNeoVis.2.1.1.chr3\_hap1.g01050.t01 |
| 4506 | 3 | 197803317 | 198176171 | mNeoVis.2.1.1.chr3\_hap1.g01050.t01 |
| 4507 | 3 | 198629036 | 198712210 | mNeoVis.2.1.1.chr3\_hap1.g01051.t01 |
| 4508 | 3 | 198629036 | 198712210 | mNeoVis.2.1.1.chr3\_hap1.g01051.t01 |
| 4509 | 3 | 199061458 | 199065505 | mNeoVis.2.1.1.chr3\_hap1.g01052.t01 |
| 4510 | 3 | 199061458 | 199065505 | mNeoVis.2.1.1.chr3\_hap1.g01052.t01 |
| 4511 | 3 | 199353046 | 199386071 | mNeoVis.2.1.1.chr3\_hap1.g01053.t01 |
| 4512 | 3 | 199353046 | 199386071 | mNeoVis.2.1.1.chr3\_hap1.g01053.t01 |
| 4513 | 3 | 199394282 | 199470863 | mNeoVis.2.1.1.chr3\_hap1.g01054.t01 |
| 4514 | 3 | 199394282 | 199470863 | mNeoVis.2.1.1.chr3\_hap1.g01054.t01 |
| 4515 | 3 | 199517851 | 199534643 | mNeoVis.2.1.1.chr3\_hap1.g01055.t01 |
| 4516 | 3 | 199517851 | 199534643 | mNeoVis.2.1.1.chr3\_hap1.g01055.t01 |
| 4517 | 3 | 199536743 | 199582242 | mNeoVis.2.1.1.chr3\_hap1.g01056.t01 |
| 4518 | 3 | 199536743 | 199582242 | mNeoVis.2.1.1.chr3\_hap1.g01056.t01 |
| 4519 | 3 | 199586932 | 199615359 | mNeoVis.2.1.1.chr3\_hap1.g01057.t01 |
| 4520 | 3 | 199586932 | 199615359 | mNeoVis.2.1.1.chr3\_hap1.g01057.t01 |
| 4521 | 3 | 199607401 | 199615838 | mNeoVis.2.1.1.chr3\_hap1.g01058.t01 |
| 4522 | 3 | 199607401 | 199615838 | mNeoVis.2.1.1.chr3\_hap1.g01058.t01 |
| 4523 | 3 | 199652965 | 199728923 | mNeoVis.2.1.1.chr3\_hap1.g01059.t01 |
| 4524 | 3 | 199652965 | 199728923 | mNeoVis.2.1.1.chr3\_hap1.g01059.t01 |
| 4525 | 3 | 199734036 | 199847263 | mNeoVis.2.1.1.chr3\_hap1.g01060.t01 |
| 4526 | 3 | 199854131 | 199892608 | mNeoVis.2.1.1.chr3\_hap1.g01061.t01 |
| 4527 | 3 | 199892689 | 199930542 | mNeoVis.2.1.1.chr3\_hap1.g01062.t01 |
| 4528 | 3 | 199892689 | 199930542 | mNeoVis.2.1.1.chr3\_hap1.g01062.t01 |
| 4529 | 3 | 199960151 | 200071457 | mNeoVis.2.1.1.chr3\_hap1.g01063.t01 |
| 4530 | 3 | 199960151 | 200071457 | mNeoVis.2.1.1.chr3\_hap1.g01063.t01 |
| 4531 | 3 | 200084982 | 200168329 | mNeoVis.2.1.1.chr3\_hap1.g01064.t01 |
| 4532 | 3 | 200084982 | 200168329 | mNeoVis.2.1.1.chr3\_hap1.g01064.t01 |
| 4533 | 3 | 200084982 | 200168329 | mNeoVis.2.1.1.chr3\_hap1.g01064.t01 |
| 4534 | 3 | 200105336 | 200205701 | mNeoVis.2.1.1.chr3\_hap1.g01065.t01 |
| 4535 | 3 | 200105336 | 200205701 | mNeoVis.2.1.1.chr3\_hap1.g01065.t01 |
| 4536 | 3 | 200105336 | 200205701 | mNeoVis.2.1.1.chr3\_hap1.g01065.t01 |
| 4537 | 3 | 200207815 | 200214871 | mNeoVis.2.1.1.chr3\_hap1.g01066.t01 |
| 4538 | 3 | 200207815 | 200214871 | mNeoVis.2.1.1.chr3\_hap1.g01066.t01 |
| 4539 | 3 | 200207815 | 200214871 | mNeoVis.2.1.1.chr3\_hap1.g01066.t01 |
| 4540 | 3 | 200227562 | 200376921 | mNeoVis.2.1.1.chr3\_hap1.g01067.t01 |
| 4541 | 3 | 200227562 | 200376921 | mNeoVis.2.1.1.chr3\_hap1.g01067.t01 |
| 4542 | 3 | 200299252 | 200308598 | mNeoVis.2.1.1.chr3\_hap1.g01068.t01 |
| 4543 | 3 | 200299252 | 200308598 | mNeoVis.2.1.1.chr3\_hap1.g01068.t01 |
| 4544 | 3 | 200299252 | 200308598 | mNeoVis.2.1.1.chr3\_hap1.g01068.t01 |
| 4545 | 3 | 200389566 | 200425127 | mNeoVis.2.1.1.chr3\_hap1.g01069.t01 |
| 4546 | 3 | 200483793 | 200500257 | mNeoVis.2.1.1.chr3\_hap1.g01070.t01 |
| 4547 | 3 | 200502328 | 200507902 | mNeoVis.2.1.1.chr3\_hap1.g01071.t01 |
| 4548 | 3 | 200511662 | 200520406 | mNeoVis.2.1.1.chr3\_hap1.g01072.t01 |
| 4549 | 3 | 200521021 | 200526077 | mNeoVis.2.1.1.chr3\_hap1.g01073.t01 |
| 4550 | 3 | 200555264 | 200564232 | mNeoVis.2.1.1.chr3\_hap1.g01074.t01 |
| 4551 | 3 | 200572490 | 200577014 | mNeoVis.2.1.1.chr3\_hap1.g01075.t01 |
| 4552 | 3 | 200585639 | 200952267 | mNeoVis.2.1.1.chr3\_hap1.g01076.t01 |
| 4553 | 3 | 200864408 | 200864923 | mNeoVis.2.1.1.chr3\_hap1.g01077.t01 |
| 4554 | 3 | 200982660 | 200987466 | mNeoVis.2.1.1.chr3\_hap1.g01078.t01 |
| 4555 | 3 | 201036295 | 201061844 | mNeoVis.2.1.1.chr3\_hap1.g01079.t01 |
| 4556 | 3 | 201062035 | 201113839 | mNeoVis.2.1.1.chr3\_hap1.g01080.t01 |
| 4557 | 3 | 201129024 | 201133911 | mNeoVis.2.1.1.chr3\_hap1.g01081.t01 |
| 4558 | 3 | 201229294 | 201230625 | mNeoVis.2.1.1.chr3\_hap1.g01082.t01 |
| 4559 | 3 | 201268376 | 201275228 | mNeoVis.2.1.1.chr3\_hap1.g01083.t01 |
| 4560 | 3 | 201287661 | 201327936 | mNeoVis.2.1.1.chr3\_hap1.g01084.t01 |
| 4561 | 3 | 201327796 | 201338241 | mNeoVis.2.1.1.chr3\_hap1.g01085.t01 |
| 4562 | 3 | 201826411 | 201914896 | mNeoVis.2.1.1.chr3\_hap1.g01095.t01 |
| 4563 | 3 | 201956093 | 201995396 | mNeoVis.2.1.1.chr3\_hap1.g01096.t01 |
| 4564 | 3 | 202001560 | 202103912 | mNeoVis.2.1.1.chr3\_hap1.g01097.t01 |
| 4565 | 3 | 202080576 | 202081557 | mNeoVis.2.1.1.chr3\_hap1.g01098.t01 |
| 4566 | 3 | 202080576 | 202081557 | mNeoVis.2.1.1.chr3\_hap1.g01098.t01 |
| 4567 | 3 | 202114451 | 202187667 | mNeoVis.2.1.1.chr3\_hap1.g01099.t01 |
| 4568 | 3 | 202114451 | 202187667 | mNeoVis.2.1.1.chr3\_hap1.g01099.t01 |
| 4569 | 3 | 202187624 | 202224785 | mNeoVis.2.1.1.chr3\_hap1.g01100.t01 |
| 4570 | 3 | 202187624 | 202224785 | mNeoVis.2.1.1.chr3\_hap1.g01100.t01 |
| 4571 | 3 | 202247792 | 202261827 | mNeoVis.2.1.1.chr3\_hap1.g01101.t01 |
| 4572 | 3 | 202247792 | 202261827 | mNeoVis.2.1.1.chr3\_hap1.g01101.t01 |
| 4573 | 3 | 202299607 | 202329937 | mNeoVis.2.1.1.chr3\_hap1.g01102.t01 |
| 4574 | 3 | 202299607 | 202329937 | mNeoVis.2.1.1.chr3\_hap1.g01102.t01 |
| 4575 | 3 | 202344602 | 202437713 | mNeoVis.2.1.1.chr3\_hap1.g01103.t01 |
| 4576 | 3 | 202437869 | 202583429 | mNeoVis.2.1.1.chr3\_hap1.g01104.t01 |
| 4577 | 3 | 202437869 | 202583429 | mNeoVis.2.1.1.chr3\_hap1.g01104.t01 |
| 4578 | 3 | 202622809 | 202943078 | mNeoVis.2.1.1.chr3\_hap1.g01105.t01 |
| 4579 | 3 | 202622809 | 202943078 | mNeoVis.2.1.1.chr3\_hap1.g01105.t01 |
| 4580 | 3 | 202749598 | 202751581 | mNeoVis.2.1.1.chr3\_hap1.g01106.t01 |
| 4581 | 3 | 202749598 | 202751581 | mNeoVis.2.1.1.chr3\_hap1.g01106.t01 |
| 4582 | 3 | 203005238 | 203236505 | mNeoVis.2.1.1.chr3\_hap1.g01107.t01 |
| 4583 | 3 | 203648513 | 203689015 | mNeoVis.2.1.1.chr3\_hap1.g01108.t01 |
| 4584 | 3 | 203648513 | 203689015 | mNeoVis.2.1.1.chr3\_hap1.g01108.t01 |
| 4585 | 3 | 203724133 | 203725397 | mNeoVis.2.1.1.chr3\_hap1.g01109.t01 |
| 4586 | 3 | 203724133 | 203725397 | mNeoVis.2.1.1.chr3\_hap1.g01109.t01 |
| 4587 | 3 | 203871409 | 204043285 | mNeoVis.2.1.1.chr3\_hap1.g01110.t01 |
| 4588 | 3 | 204086644 | 204129431 | mNeoVis.2.1.1.chr3\_hap1.g01111.t01 |
| 4589 | 3 | 204164726 | 204168091 | mNeoVis.2.1.1.chr3\_hap1.g01112.t01 |
| 4590 | 3 | 204217517 | 204233809 | mNeoVis.2.1.1.chr3\_hap1.g01113.t01 |
| 4591 | 3 | 204236332 | 204267477 | mNeoVis.2.1.1.chr3\_hap1.g01114.t01 |
| 4592 | 3 | 204239113 | 204240077 | mNeoVis.2.1.1.chr3\_hap1.g01115.t01 |
| 4593 | 3 | 204270621 | 204314612 | mNeoVis.2.1.1.chr3\_hap1.g01116.t01 |
| 4594 | 3 | 204315422 | 204322820 | mNeoVis.2.1.1.chr3\_hap1.g01117.t01 |
| 4595 | 3 | 204315422 | 204322820 | mNeoVis.2.1.1.chr3\_hap1.g01117.t01 |
| 4596 | 3 | 204317700 | 204318541 | mNeoVis.2.1.1.chr3\_hap1.g01118.t01 |
| 4597 | 3 | 204317700 | 204318541 | mNeoVis.2.1.1.chr3\_hap1.g01118.t01 |
| 4598 | 3 | 204333644 | 204334505 | mNeoVis.2.1.1.chr3\_hap1.g01119.t01 |
| 4599 | 3 | 204333644 | 204334505 | mNeoVis.2.1.1.chr3\_hap1.g01119.t01 |
| 4600 | 3 | 204367409 | 204443793 | mNeoVis.2.1.1.chr3\_hap1.g01120.t01 |
| 4601 | 3 | 204367409 | 204443793 | mNeoVis.2.1.1.chr3\_hap1.g01120.t01 |
| 4602 | 3 | 204511537 | 204523268 | mNeoVis.2.1.1.chr3\_hap1.g01121.t01 |
| 4603 | 3 | 204511537 | 204523268 | mNeoVis.2.1.1.chr3\_hap1.g01121.t01 |
| 4604 | 3 | 204529614 | 204561359 | mNeoVis.2.1.1.chr3\_hap1.g01122.t01 |
| 4605 | 3 | 204529614 | 204561359 | mNeoVis.2.1.1.chr3\_hap1.g01122.t01 |
| 4606 | 3 | 204568458 | 204694817 | mNeoVis.2.1.1.chr3\_hap1.g01123.t01 |
| 4607 | 3 | 204568458 | 204694817 | mNeoVis.2.1.1.chr3\_hap1.g01123.t01 |
| 4608 | 3 | 204574001 | 204574381 | mNeoVis.2.1.1.chr3\_hap1.g01124.t01 |
| 4609 | 3 | 204574001 | 204574381 | mNeoVis.2.1.1.chr3\_hap1.g01124.t01 |
| 4610 | 3 | 204695197 | 204824766 | mNeoVis.2.1.1.chr3\_hap1.g01125.t01 |
| 4611 | 3 | 204695197 | 204824766 | mNeoVis.2.1.1.chr3\_hap1.g01125.t01 |
| 4612 | 3 | 204845682 | 204948500 | mNeoVis.2.1.1.chr3\_hap1.g01126.t01 |
| 4613 | 3 | 204845682 | 204948500 | mNeoVis.2.1.1.chr3\_hap1.g01126.t01 |
| 4614 | 3 | 204845682 | 204948500 | mNeoVis.2.1.1.chr3\_hap1.g01126.t01 |
| 4615 | 3 | 204983148 | 205042149 | mNeoVis.2.1.1.chr3\_hap1.g01127.t01 |
| 4616 | 3 | 204983148 | 205042149 | mNeoVis.2.1.1.chr3\_hap1.g01127.t01 |
| 4617 | 3 | 204983148 | 205042149 | mNeoVis.2.1.1.chr3\_hap1.g01127.t01 |
| 4618 | 3 | 205920734 | 206177995 | mNeoVis.2.1.1.chr3\_hap1.g01128.t01 |
| 4619 | 3 | 205920734 | 206177995 | mNeoVis.2.1.1.chr3\_hap1.g01128.t01 |
| 4620 | 3 | 206536719 | 206742404 | mNeoVis.2.1.1.chr3\_hap1.g01129.t01 |
| 4621 | 3 | 206536719 | 206742404 | mNeoVis.2.1.1.chr3\_hap1.g01129.t01 |
| 4622 | 3 | 206536719 | 206742404 | mNeoVis.2.1.1.chr3\_hap1.g01129.t01 |
| 4623 | 3 | 206785365 | 206796636 | mNeoVis.2.1.1.chr3\_hap1.g01130.t01 |
| 4624 | 3 | 206785365 | 206796636 | mNeoVis.2.1.1.chr3\_hap1.g01130.t01 |
| 4625 | 3 | 206785365 | 206796636 | mNeoVis.2.1.1.chr3\_hap1.g01130.t01 |
| 4626 | 3 | 206884598 | 206976323 | mNeoVis.2.1.1.chr3\_hap1.g01131.t01 |
| 4627 | 3 | 206884598 | 206976323 | mNeoVis.2.1.1.chr3\_hap1.g01131.t01 |
| 4628 | 3 | 206884598 | 206976323 | mNeoVis.2.1.1.chr3\_hap1.g01131.t01 |
| 4629 | 3 | 207036797 | 207053776 | mNeoVis.2.1.1.chr3\_hap1.g01132.t01 |
| 4630 | 3 | 207036797 | 207053776 | mNeoVis.2.1.1.chr3\_hap1.g01132.t01 |
| 4631 | 3 | 207036797 | 207053776 | mNeoVis.2.1.1.chr3\_hap1.g01132.t01 |
| 4632 | 3 | 207321914 | 207383776 | mNeoVis.2.1.1.chr3\_hap1.g01133.t01 |
| 4633 | 3 | 207321914 | 207383776 | mNeoVis.2.1.1.chr3\_hap1.g01133.t01 |
| 4634 | 3 | 207393767 | 207403454 | mNeoVis.2.1.1.chr3\_hap1.g01134.t01 |
| 4635 | 3 | 207393767 | 207403454 | mNeoVis.2.1.1.chr3\_hap1.g01134.t01 |
| 4636 | 3 | 207408943 | 207452520 | mNeoVis.2.1.1.chr3\_hap1.g01135.t01 |
| 4637 | 3 | 207408943 | 207452520 | mNeoVis.2.1.1.chr3\_hap1.g01135.t01 |
| 4638 | 3 | 207475525 | 207531095 | mNeoVis.2.1.1.chr3\_hap1.g01136.t01 |
| 4639 | 3 | 207475525 | 207531095 | mNeoVis.2.1.1.chr3\_hap1.g01136.t01 |
| 4640 | 3 | 207654711 | 207660070 | mNeoVis.2.1.1.chr3\_hap1.g01137.t01 |
| 4641 | 3 | 207654711 | 207660070 | mNeoVis.2.1.1.chr3\_hap1.g01137.t01 |
| 4642 | 3 | 208103036 | 208106501 | mNeoVis.2.1.1.chr3\_hap1.g01138.t01 |
| 4643 | 3 | 208182603 | 208251330 | mNeoVis.2.1.1.chr3\_hap1.g01139.t01 |
| 4644 | 3 | 208182603 | 208251330 | mNeoVis.2.1.1.chr3\_hap1.g01139.t01 |
| 4645 | 3 | 208365420 | 208389599 | mNeoVis.2.1.1.chr3\_hap1.g01140.t01 |
| 4646 | 3 | 208365420 | 208389599 | mNeoVis.2.1.1.chr3\_hap1.g01140.t01 |
| 4647 | 3 | 208409221 | 208481037 | mNeoVis.2.1.1.chr3\_hap1.g01141.t01 |
| 4648 | 3 | 208409221 | 208481037 | mNeoVis.2.1.1.chr3\_hap1.g01141.t01 |
| 4649 | 3 | 208489056 | 208599985 | mNeoVis.2.1.1.chr3\_hap1.g01142.t01 |
| 4650 | 3 | 208489056 | 208599985 | mNeoVis.2.1.1.chr3\_hap1.g01142.t01 |
| 4651 | 3 | 208677736 | 208680665 | mNeoVis.2.1.1.chr3\_hap1.g01143.t01 |
| 4652 | 3 | 208677736 | 208680665 | mNeoVis.2.1.1.chr3\_hap1.g01143.t01 |
| 4653 | 3 | 208699716 | 208797019 | mNeoVis.2.1.1.chr3\_hap1.g01144.t01 |
| 4654 | 3 | 208699716 | 208797019 | mNeoVis.2.1.1.chr3\_hap1.g01144.t01 |
| 4655 | 3 | 209338052 | 209490196 | mNeoVis.2.1.1.chr3\_hap1.g01145.t01 |
| 4656 | 3 | 209338052 | 209490196 | mNeoVis.2.1.1.chr3\_hap1.g01145.t01 |
| 4657 | 3 | 210997338 | 211013919 | mNeoVis.2.1.1.chr3\_hap1.g01146.t01 |
| 4658 | 3 | 213404474 | 213405050 | mNeoVis.2.1.1.chr3\_hap1.g01196.t01 |
| 4659 | 3 | 213560449 | 213580047 | mNeoVis.2.1.1.chr3\_hap1.g01197.t01 |
| 4660 | 3 | 213586328 | 213610444 | mNeoVis.2.1.1.chr3\_hap1.g01198.t01 |
| 4661 | 3 | 213723101 | 213725912 | mNeoVis.2.1.1.chr3\_hap1.g01199.t01 |
| 4662 | 3 | 213723101 | 213725912 | mNeoVis.2.1.1.chr3\_hap1.g01199.t01 |
| 4663 | 3 | 213734146 | 213795046 | mNeoVis.2.1.1.chr3\_hap1.g01200.t01 |
| 4664 | 3 | 213734146 | 213795046 | mNeoVis.2.1.1.chr3\_hap1.g01200.t01 |
| 4665 | 3 | 213830628 | 213938830 | mNeoVis.2.1.1.chr3\_hap1.g01201.t01 |
| 4666 | 3 | 213830628 | 213938830 | mNeoVis.2.1.1.chr3\_hap1.g01201.t01 |
| 4667 | 3 | 213968729 | 214055749 | mNeoVis.2.1.1.chr3\_hap1.g01202.t01 |
| 4668 | 3 | 213968729 | 214055749 | mNeoVis.2.1.1.chr3\_hap1.g01202.t01 |
| 4669 | 3 | 214060246 | 214074414 | mNeoVis.2.1.1.chr3\_hap1.g01203.t01 |
| 4670 | 3 | 214074509 | 214107255 | mNeoVis.2.1.1.chr3\_hap1.g01204.t01 |
| 4671 | 3 | 214126863 | 214144313 | mNeoVis.2.1.1.chr3\_hap1.g01205.t01 |
| 4672 | 3 | 214198622 | 214249519 | mNeoVis.2.1.1.chr3\_hap1.g01206.t01 |
| 4673 | 3 | 214341079 | 214391987 | mNeoVis.2.1.1.chr3\_hap1.g01209.t01 |
| 4674 | 3 | 214411770 | 214472505 | mNeoVis.2.1.1.chr3\_hap1.g01210.t01 |
| 4675 | 3 | 214898353 | 214965129 | mNeoVis.2.1.1.chr3\_hap1.g01211.t01 |
| 4676 | 3 | 214898353 | 214965129 | mNeoVis.2.1.1.chr3\_hap1.g01211.t01 |
| 4677 | 3 | 214973755 | 215004287 | mNeoVis.2.1.1.chr3\_hap1.g01212.t01 |
| 4678 | 3 | 214973755 | 215004287 | mNeoVis.2.1.1.chr3\_hap1.g01212.t01 |
| 4679 | 3 | 215122872 | 215292787 | mNeoVis.2.1.1.chr3\_hap1.g01213.t01 |
| 4680 | 3 | 215409877 | 215489849 | mNeoVis.2.1.1.chr3\_hap1.g01214.t01 |
| 4681 | 3 | 215409877 | 215489849 | mNeoVis.2.1.1.chr3\_hap1.g01214.t01 |
| 4682 | 3 | 215630734 | 215798452 | mNeoVis.2.1.1.chr3\_hap1.g01215.t01 |
| 4683 | 3 | 216965960 | 217120581 | mNeoVis.2.1.1.chr3\_hap1.g01217.t01 |
| 4684 | 3 | 216965960 | 217120581 | mNeoVis.2.1.1.chr3\_hap1.g01217.t01 |
| 4685 | 3 | 217282264 | 217388073 | mNeoVis.2.1.1.chr3\_hap1.g01218.t01 |
| 4686 | 3 | 217529499 | 218694374 | mNeoVis.2.1.1.chr3\_hap1.g01219.t01 |
| 4687 | 3 | 219329647 | 219513042 | mNeoVis.2.1.1.chr3\_hap1.g01220.t01 |
| 4688 | 3 | 219329647 | 219513042 | mNeoVis.2.1.1.chr3\_hap1.g01220.t01 |
| 4689 | 3 | 219513231 | 219554322 | mNeoVis.2.1.1.chr3\_hap1.g01221.t01 |
| 4690 | 3 | 219513231 | 219554322 | mNeoVis.2.1.1.chr3\_hap1.g01221.t01 |
| 4691 | 3 | 219600654 | 219624456 | mNeoVis.2.1.1.chr3\_hap1.g01222.t01 |
| 4692 | 3 | 219600654 | 219624456 | mNeoVis.2.1.1.chr3\_hap1.g01222.t01 |
| 4693 | 3 | 219674807 | 219714506 | mNeoVis.2.1.1.chr3\_hap1.g01223.t01 |
| 4694 | 3 | 219729574 | 219877151 | mNeoVis.2.1.1.chr3\_hap1.g01224.t01 |
| 4695 | 3 | 219878215 | 219893895 | mNeoVis.2.1.1.chr3\_hap1.g01225.t01 |
| 4696 | 3 | 219895859 | 220119796 | mNeoVis.2.1.1.chr3\_hap1.g01226.t01 |
| 4697 | 3 | 221116590 | 221225028 | mNeoVis.2.1.1.chr3\_hap1.g01229.t01 |
| 4698 | 3 | 221244286 | 221321855 | mNeoVis.2.1.1.chr3\_hap1.g01230.t01 |
| 4699 | 3 | 221328010 | 221344117 | mNeoVis.2.1.1.chr3\_hap1.g01231.t01 |
| 4700 | 3 | 221372988 | 221399300 | mNeoVis.2.1.1.chr3\_hap1.g01232.t01 |
| 4701 | 3 | 221401473 | 221403271 | mNeoVis.2.1.1.chr3\_hap1.g01233.t01 |
| 4702 | 3 | 221413975 | 221415641 | mNeoVis.2.1.1.chr3\_hap1.g01234.t01 |
| 4703 | 3 | 221420023 | 221424586 | mNeoVis.2.1.1.chr3\_hap1.g01235.t01 |
| 4704 | 3 | 221429621 | 221431105 | mNeoVis.2.1.1.chr3\_hap1.g01236.t01 |
| 4705 | 3 | 221438884 | 221441180 | mNeoVis.2.1.1.chr3\_hap1.g01237.t01 |
| 4706 | 3 | 221500798 | 221684686 | mNeoVis.2.1.1.chr3\_hap1.g01238.t01 |
| 4707 | 3 | 221736804 | 221737753 | mNeoVis.2.1.1.chr3\_hap1.g01239.t01 |
| 4708 | 3 | 221737306 | 221744670 | mNeoVis.2.1.1.chr3\_hap1.g01240.t01 |
| 4709 | 3 | 221749493 | 221787659 | mNeoVis.2.1.1.chr3\_hap1.g01241.t01 |
| 4710 | 3 | 221859078 | 221866647 | mNeoVis.2.1.1.chr3\_hap1.g01242.t01 |
| 4711 | 3 | 221873051 | 221936061 | mNeoVis.2.1.1.chr3\_hap1.g01243.t01 |
| 4712 | 3 | 222262029 | 222349128 | mNeoVis.2.1.1.chr3\_hap1.g01244.t01 |
| 4713 | 3 | 222262029 | 222349128 | mNeoVis.2.1.1.chr3\_hap1.g01244.t01 |
| 4714 | 3 | 222382976 | 222431348 | mNeoVis.2.1.1.chr3\_hap1.g01245.t01 |
| 4715 | 3 | 222382976 | 222431348 | mNeoVis.2.1.1.chr3\_hap1.g01245.t01 |
| 4716 | 3 | 222382976 | 222431348 | mNeoVis.2.1.1.chr3\_hap1.g01245.t01 |
| 4717 | 3 | 222425717 | 222477210 | mNeoVis.2.1.1.chr3\_hap1.g01246.t01 |
| 4718 | 3 | 222425717 | 222477210 | mNeoVis.2.1.1.chr3\_hap1.g01246.t01 |
| 4719 | 3 | 222425717 | 222477210 | mNeoVis.2.1.1.chr3\_hap1.g01246.t01 |
| 4720 | 3 | 222572930 | 222591794 | mNeoVis.2.1.1.chr3\_hap1.g01247.t01 |
| 4721 | 3 | 222572930 | 222591794 | mNeoVis.2.1.1.chr3\_hap1.g01247.t01 |
| 4722 | 3 | 222572930 | 222591794 | mNeoVis.2.1.1.chr3\_hap1.g01247.t01 |
| 4723 | 3 | 222591899 | 222618430 | mNeoVis.2.1.1.chr3\_hap1.g01248.t01 |
| 4724 | 3 | 222591899 | 222618430 | mNeoVis.2.1.1.chr3\_hap1.g01248.t01 |
| 4725 | 3 | 222591899 | 222618430 | mNeoVis.2.1.1.chr3\_hap1.g01248.t01 |
| 4726 | 3 | 222617781 | 222677050 | mNeoVis.2.1.1.chr3\_hap1.g01249.t01 |
| 4727 | 3 | 222617781 | 222677050 | mNeoVis.2.1.1.chr3\_hap1.g01249.t01 |
| 4728 | 3 | 222617781 | 222677050 | mNeoVis.2.1.1.chr3\_hap1.g01249.t01 |
| 4729 | 3 | 222680476 | 222731886 | mNeoVis.2.1.1.chr3\_hap1.g01250.t01 |
| 4730 | 3 | 222680476 | 222731886 | mNeoVis.2.1.1.chr3\_hap1.g01250.t01 |
| 4731 | 3 | 222680476 | 222731886 | mNeoVis.2.1.1.chr3\_hap1.g01250.t01 |
| 4732 | 3 | 222735565 | 222759849 | mNeoVis.2.1.1.chr3\_hap1.g01251.t01 |
| 4733 | 3 | 222735565 | 222759849 | mNeoVis.2.1.1.chr3\_hap1.g01251.t01 |
| 4734 | 3 | 222735565 | 222759849 | mNeoVis.2.1.1.chr3\_hap1.g01251.t01 |
| 4735 | 3 | 222977321 | 223008748 | mNeoVis.2.1.1.chr3\_hap1.g01252.t01 |
| 4736 | 3 | 222977321 | 223008748 | mNeoVis.2.1.1.chr3\_hap1.g01252.t01 |
| 4737 | 3 | 223082570 | 223127580 | mNeoVis.2.1.1.chr3\_hap1.g01253.t01 |
| 4738 | 3 | 223136416 | 223139378 | mNeoVis.2.1.1.chr3\_hap1.g01254.t01 |
| 4739 | 3 | 223139493 | 223171965 | mNeoVis.2.1.1.chr3\_hap1.g01255.t01 |
| 4740 | 3 | 223224939 | 223275724 | mNeoVis.2.1.1.chr3\_hap1.g01256.t01 |
| 4741 | 3 | 225181517 | 225233323 | mNeoVis.2.1.1.chr3\_hap1.g01259.t01 |
| 4742 | 3 | 225256912 | 225263339 | mNeoVis.2.1.1.chr3\_hap1.g01260.t01 |
| 4743 | 3 | 225256912 | 225263339 | mNeoVis.2.1.1.chr3\_hap1.g01260.t01 |
| 4744 | 3 | 225375402 | 225408379 | mNeoVis.2.1.1.chr3\_hap1.g01261.t01 |
| 4745 | 3 | 225375402 | 225408379 | mNeoVis.2.1.1.chr3\_hap1.g01261.t01 |
| 4746 | 3 | 225532001 | 225626558 | mNeoVis.2.1.1.chr3\_hap1.g01262.t01 |
| 4747 | 3 | 225532001 | 225626558 | mNeoVis.2.1.1.chr3\_hap1.g01262.t01 |
| 4748 | 3 | 225632024 | 225750125 | mNeoVis.2.1.1.chr3\_hap1.g01263.t01 |
| 4749 | 3 | 225767665 | 225829440 | mNeoVis.2.1.1.chr3\_hap1.g01264.t01 |
| 4750 | 3 | 225856672 | 226052578 | mNeoVis.2.1.1.chr3\_hap1.g01265.t01 |
| 4751 | 3 | 226100458 | 226196651 | mNeoVis.2.1.1.chr3\_hap1.g01266.t01 |
| 4752 | 3 | 226166137 | 226188049 | mNeoVis.2.1.1.chr3\_hap1.g01267.t01 |
| 4753 | 3 | 226196320 | 226277085 | mNeoVis.2.1.1.chr3\_hap1.g01268.t01 |
| 4754 | 3 | 226273199 | 226285397 | mNeoVis.2.1.1.chr3\_hap1.g01269.t01 |
| 4755 | 3 | 226290388 | 226364235 | mNeoVis.2.1.1.chr3\_hap1.g01270.t01 |
| 4756 | 3 | 226410609 | 226622514 | mNeoVis.2.1.1.chr3\_hap1.g01271.t01 |
| 4757 | 3 | 226556069 | 226557881 | mNeoVis.2.1.1.chr3\_hap1.g01272.t01 |
| 4758 | 3 | 226623081 | 226634195 | mNeoVis.2.1.1.chr3\_hap1.g01273.t01 |
| 4759 | 3 | 226673358 | 226707206 | mNeoVis.2.1.1.chr3\_hap1.g01274.t01 |
| 4760 | 3 | 227738098 | 227746714 | mNeoVis.2.1.1.chr3\_hap1.g01289.t01 |
| 4761 | 3 | 227746967 | 227824526 | mNeoVis.2.1.1.chr3\_hap1.g01290.t01 |
| 4762 | 3 | 227831360 | 227876807 | mNeoVis.2.1.1.chr3\_hap1.g01291.t01 |
| 4763 | 3 | 227879598 | 227898781 | mNeoVis.2.1.1.chr3\_hap1.g01292.t01 |
| 4764 | 3 | 227898806 | 227908253 | mNeoVis.2.1.1.chr3\_hap1.g01293.t01 |
| 4765 | 3 | 227914509 | 227924870 | mNeoVis.2.1.1.chr3\_hap1.g01294.t01 |
| 4766 | 3 | 227954228 | 227967176 | mNeoVis.2.1.1.chr3\_hap1.g01295.t01 |
| 4767 | 3 | 227967697 | 227970533 | mNeoVis.2.1.1.chr3\_hap1.g01296.t01 |
| 4768 | 3 | 227979564 | 228058863 | mNeoVis.2.1.1.chr3\_hap1.g01297.t01 |
| 4769 | 3 | 228068137 | 228127014 | mNeoVis.2.1.1.chr3\_hap1.g01298.t01 |
| 4770 | 3 | 228150267 | 228212330 | mNeoVis.2.1.1.chr3\_hap1.g01299.t01 |
| 4771 | 3 | 228150267 | 228212330 | mNeoVis.2.1.1.chr3\_hap1.g01299.t01 |
| 4772 | 3 | 228212451 | 228295508 | mNeoVis.2.1.1.chr3\_hap1.g01300.t01 |
| 4773 | 3 | 228212451 | 228295508 | mNeoVis.2.1.1.chr3\_hap1.g01300.t01 |
| 4774 | 3 | 228309499 | 228351141 | mNeoVis.2.1.1.chr3\_hap1.g01301.t01 |
| 4775 | 3 | 228309499 | 228351141 | mNeoVis.2.1.1.chr3\_hap1.g01301.t01 |
| 4776 | 3 | 228350296 | 228387898 | mNeoVis.2.1.1.chr3\_hap1.g01302.t01 |
| 4777 | 3 | 228350296 | 228387898 | mNeoVis.2.1.1.chr3\_hap1.g01302.t01 |
| 4778 | 3 | 228395366 | 228558428 | mNeoVis.2.1.1.chr3\_hap1.g01303.t01 |
| 4779 | 3 | 228395366 | 228558428 | mNeoVis.2.1.1.chr3\_hap1.g01303.t01 |
| 4780 | 3 | 228834301 | 228843846 | mNeoVis.2.1.1.chr3\_hap1.g01304.t01 |
| 4781 | 3 | 228834301 | 228843846 | mNeoVis.2.1.1.chr3\_hap1.g01304.t01 |
| 4782 | 3 | 228845010 | 228865178 | mNeoVis.2.1.1.chr3\_hap1.g01305.t01 |
| 4783 | 3 | 228845010 | 228865178 | mNeoVis.2.1.1.chr3\_hap1.g01305.t01 |
| 4784 | 3 | 228866996 | 228883035 | mNeoVis.2.1.1.chr3\_hap1.g01306.t01 |
| 4785 | 3 | 228866996 | 228883035 | mNeoVis.2.1.1.chr3\_hap1.g01306.t01 |
| 4786 | 3 | 228933667 | 228961418 | mNeoVis.2.1.1.chr3\_hap1.g01307.t01 |
| 4787 | 3 | 228933667 | 228961418 | mNeoVis.2.1.1.chr3\_hap1.g01307.t01 |
| 4788 | 3 | 229254054 | 229439913 | mNeoVis.2.1.1.chr3\_hap1.g01308.t01 |
| 4789 | 3 | 231793140 | 231877154 | mNeoVis.2.1.1.chr3\_hap1.g01322.t01 |
| 4790 | 3 | 231929037 | 232292282 | mNeoVis.2.1.1.chr3\_hap1.g01323.t01 |
| 4791 | 3 | 232312968 | 232348303 | mNeoVis.2.1.1.chr3\_hap1.g01324.t01 |
| 4792 | 3 | 232397828 | 232649415 | mNeoVis.2.1.1.chr3\_hap1.g01325.t01 |
| 4793 | 3 | 232649420 | 232787127 | mNeoVis.2.1.1.chr3\_hap1.g01326.t01 |
| 4794 | 3 | 232713805 | 232728154 | mNeoVis.2.1.1.chr3\_hap1.g01327.t01 |
| 4795 | 3 | 232912089 | 232913584 | mNeoVis.2.1.1.chr3\_hap1.g01328.t01 |
| 4796 | 3 | 234147360 | 234189329 | mNeoVis.2.1.1.chr3\_hap1.g01330.t01 |
| 4797 | 3 | 235532530 | 235532874 | mNeoVis.2.1.1.chr3\_hap1.g01331.t01 |
| 4798 | 3 | 236059807 | 236072657 | mNeoVis.2.1.1.chr3\_hap1.g01333.t01 |
| 4799 | 3 | 236206120 | 236213178 | mNeoVis.2.1.1.chr3\_hap1.g01334.t01 |
| 4800 | 3 | 236214242 | 236224286 | mNeoVis.2.1.1.chr3\_hap1.g01335.t01 |
| 4801 | 3 | 236442099 | 236621147 | mNeoVis.2.1.1.chr3\_hap1.g01336.t01 |
| 4802 | 3 | 236626662 | 236647246 | mNeoVis.2.1.1.chr3\_hap1.g01337.t01 |
| 4803 | 3 | 236838917 | 236877393 | mNeoVis.2.1.1.chr3\_hap1.g01339.t01 |
| 4804 | 3 | 236881620 | 236964007 | mNeoVis.2.1.1.chr3\_hap1.g01340.t01 |
| 4805 | 3 | 237089847 | 237093191 | mNeoVis.2.1.1.chr3\_hap1.g01341.t01 |
| 4806 | 3 | 237089847 | 237093191 | mNeoVis.2.1.1.chr3\_hap1.g01341.t01 |
| 4807 | 3 | 237150138 | 237194956 | mNeoVis.2.1.1.chr3\_hap1.g01342.t01 |
| 4808 | 3 | 237150138 | 237194956 | mNeoVis.2.1.1.chr3\_hap1.g01342.t01 |
| 4809 | 3 | 237246958 | 237286823 | mNeoVis.2.1.1.chr3\_hap1.g01343.t01 |
| 4810 | 3 | 237246958 | 237286823 | mNeoVis.2.1.1.chr3\_hap1.g01343.t01 |
| 4811 | 3 | 237285305 | 237311032 | mNeoVis.2.1.1.chr3\_hap1.g01344.t01 |
| 4812 | 3 | 237315989 | 237383589 | mNeoVis.2.1.1.chr3\_hap1.g01345.t01 |
| 4813 | 3 | 237413187 | 237426962 | mNeoVis.2.1.1.chr3\_hap1.g01346.t01 |
| 4814 | 3 | 237455778 | 237555146 | mNeoVis.2.1.1.chr3\_hap1.g01347.t01 |
| 4815 | 3 | 237553554 | 237626644 | mNeoVis.2.1.1.chr3\_hap1.g01348.t01 |
| 4816 | 3 | 237625381 | 237626779 | mNeoVis.2.1.1.chr3\_hap1.g01349.t01 |
| 4817 | 3 | 237701661 | 237707250 | mNeoVis.2.1.1.chr3\_hap1.g01350.t01 |
| 4818 | 3 | 237799071 | 237849738 | mNeoVis.2.1.1.chr3\_hap1.g01351.t01 |
| 4819 | 3 | 237850738 | 237937985 | mNeoVis.2.1.1.chr3\_hap1.g01352.t01 |
| 4820 | 3 | 237850738 | 237937985 | mNeoVis.2.1.1.chr3\_hap1.g01352.t01 |
| 4821 | 3 | 237936933 | 237951170 | mNeoVis.2.1.1.chr3\_hap1.g01353.t01 |
| 4822 | 3 | 237936933 | 237951170 | mNeoVis.2.1.1.chr3\_hap1.g01353.t01 |
| 4823 | 3 | 237958652 | 237970156 | mNeoVis.2.1.1.chr3\_hap1.g01354.t01 |
| 4824 | 3 | 237958652 | 237970156 | mNeoVis.2.1.1.chr3\_hap1.g01354.t01 |
| 4825 | 3 | 237958652 | 237970156 | mNeoVis.2.1.1.chr3\_hap1.g01354.t01 |
| 4826 | 3 | 237973080 | 238037793 | mNeoVis.2.1.1.chr3\_hap1.g01355.t01 |
| 4827 | 3 | 237973080 | 238037793 | mNeoVis.2.1.1.chr3\_hap1.g01355.t01 |
| 4828 | 3 | 237973080 | 238037793 | mNeoVis.2.1.1.chr3\_hap1.g01355.t01 |
| 4829 | 3 | 238041598 | 238046522 | mNeoVis.2.1.1.chr3\_hap1.g01356.t01 |
| 4830 | 3 | 238041598 | 238046522 | mNeoVis.2.1.1.chr3\_hap1.g01356.t01 |
| 4831 | 3 | 238041598 | 238046522 | mNeoVis.2.1.1.chr3\_hap1.g01356.t01 |
| 4832 | 3 | 238046476 | 238106661 | mNeoVis.2.1.1.chr3\_hap1.g01357.t01 |
| 4833 | 3 | 238046476 | 238106661 | mNeoVis.2.1.1.chr3\_hap1.g01357.t01 |
| 4834 | 3 | 238046476 | 238106661 | mNeoVis.2.1.1.chr3\_hap1.g01357.t01 |
| 4835 | 3 | 238153540 | 238175090 | mNeoVis.2.1.1.chr3\_hap1.g01358.t01 |
| 4836 | 3 | 238153540 | 238175090 | mNeoVis.2.1.1.chr3\_hap1.g01358.t01 |
| 4837 | 3 | 238153540 | 238175090 | mNeoVis.2.1.1.chr3\_hap1.g01358.t01 |
| 4838 | 3 | 238228884 | 238265107 | mNeoVis.2.1.1.chr3\_hap1.g01359.t01 |
| 4839 | 3 | 238228884 | 238265107 | mNeoVis.2.1.1.chr3\_hap1.g01359.t01 |
| 4840 | 3 | 238228884 | 238265107 | mNeoVis.2.1.1.chr3\_hap1.g01359.t01 |
| 4841 | 3 | 238271319 | 238298233 | mNeoVis.2.1.1.chr3\_hap1.g01360.t01 |
| 4842 | 3 | 238271319 | 238298233 | mNeoVis.2.1.1.chr3\_hap1.g01360.t01 |
| 4843 | 3 | 238271319 | 238298233 | mNeoVis.2.1.1.chr3\_hap1.g01360.t01 |
| 4844 | 3 | 238632294 | 238693588 | mNeoVis.2.1.1.chr3\_hap1.g01361.t01 |
| 4845 | 3 | 238709140 | 238747975 | mNeoVis.2.1.1.chr3\_hap1.g01362.t01 |
| 4846 | 3 | 238789451 | 238803724 | mNeoVis.2.1.1.chr3\_hap1.g01363.t01 |
| 4847 | 3 | 239173276 | 239182541 | mNeoVis.2.1.1.chr3\_hap1.g01365.t01 |
| 4848 | 3 | 239455512 | 239471160 | mNeoVis.2.1.1.chr3\_hap1.g01366.t01 |
| 4849 | 3 | 239740706 | 239779304 | mNeoVis.2.1.1.chr3\_hap1.g01367.t01 |
| 4850 | 3 | 239779190 | 239817064 | mNeoVis.2.1.1.chr3\_hap1.g01368.t01 |
| 4851 | 4 | 166333 | 221494 | mNeoVis.2.1.1.chr4\_hap1.g00001.t01 |
| 4852 | 4 | 166333 | 221494 | mNeoVis.2.1.1.chr4\_hap1.g00001.t01 |
| 4853 | 4 | 295780 | 351658 | mNeoVis.2.1.1.chr4\_hap1.g00002.t01 |
| 4854 | 4 | 295780 | 351658 | mNeoVis.2.1.1.chr4\_hap1.g00002.t01 |
| 4855 | 4 | 359835 | 438679 | mNeoVis.2.1.1.chr4\_hap1.g00003.t01 |
| 4856 | 4 | 359835 | 438679 | mNeoVis.2.1.1.chr4\_hap1.g00003.t01 |
| 4857 | 4 | 453907 | 521201 | mNeoVis.2.1.1.chr4\_hap1.g00004.t01 |
| 4858 | 4 | 453907 | 521201 | mNeoVis.2.1.1.chr4\_hap1.g00004.t01 |
| 4859 | 4 | 600189 | 1230227 | mNeoVis.2.1.1.chr4\_hap1.g00005.t01 |
| 4860 | 4 | 600189 | 1230227 | mNeoVis.2.1.1.chr4\_hap1.g00005.t01 |
| 4861 | 4 | 1286876 | 1346191 | mNeoVis.2.1.1.chr4\_hap1.g00006.t01 |
| 4862 | 4 | 1286876 | 1346191 | mNeoVis.2.1.1.chr4\_hap1.g00006.t01 |
| 4863 | 4 | 1286876 | 1346191 | mNeoVis.2.1.1.chr4\_hap1.g00006.t01 |
| 4864 | 4 | 1396839 | 1501711 | mNeoVis.2.1.1.chr4\_hap1.g00007.t01 |
| 4865 | 4 | 1570040 | 1575182 | mNeoVis.2.1.1.chr4\_hap1.g00008.t01 |
| 4866 | 4 | 1586804 | 1609799 | mNeoVis.2.1.1.chr4\_hap1.g00009.t01 |
| 4867 | 4 | 1816693 | 1856939 | mNeoVis.2.1.1.chr4\_hap1.g00011.t01 |
| 4868 | 4 | 2384024 | 2393454 | mNeoVis.2.1.1.chr4\_hap1.g00012.t01 |
| 4869 | 4 | 2384024 | 2393454 | mNeoVis.2.1.1.chr4\_hap1.g00012.t01 |
| 4870 | 4 | 2393499 | 2543787 | mNeoVis.2.1.1.chr4\_hap1.g00013.t01 |
| 4871 | 4 | 2393499 | 2543787 | mNeoVis.2.1.1.chr4\_hap1.g00013.t01 |
| 4872 | 4 | 2619070 | 2625540 | mNeoVis.2.1.1.chr4\_hap1.g00014.t01 |
| 4873 | 4 | 2619070 | 2625540 | mNeoVis.2.1.1.chr4\_hap1.g00014.t01 |
| 4874 | 4 | 2656459 | 2662629 | mNeoVis.2.1.1.chr4\_hap1.g00015.t01 |
| 4875 | 4 | 2656459 | 2662629 | mNeoVis.2.1.1.chr4\_hap1.g00015.t01 |
| 4876 | 4 | 2742835 | 2753139 | mNeoVis.2.1.1.chr4\_hap1.g00016.t01 |
| 4877 | 4 | 2742835 | 2753139 | mNeoVis.2.1.1.chr4\_hap1.g00016.t01 |
| 4878 | 4 | 2854512 | 2868684 | mNeoVis.2.1.1.chr4\_hap1.g00017.t01 |
| 4879 | 4 | 2854512 | 2868684 | mNeoVis.2.1.1.chr4\_hap1.g00017.t01 |
| 4880 | 4 | 2892250 | 2940036 | mNeoVis.2.1.1.chr4\_hap1.g00018.t01 |
| 4881 | 4 | 2892250 | 2940036 | mNeoVis.2.1.1.chr4\_hap1.g00018.t01 |
| 4882 | 4 | 2962844 | 3621375 | mNeoVis.2.1.1.chr4\_hap1.g00019.t01 |
| 4883 | 4 | 3172840 | 3173352 | mNeoVis.2.1.1.chr4\_hap1.g00020.t01 |
| 4884 | 4 | 3172840 | 3173352 | mNeoVis.2.1.1.chr4\_hap1.g00020.t01 |
| 4885 | 4 | 3172840 | 3173352 | mNeoVis.2.1.1.chr4\_hap1.g00020.t01 |
| 4886 | 4 | 5417413 | 5460985 | mNeoVis.2.1.1.chr4\_hap1.g00029.t01 |
| 4887 | 4 | 5476994 | 5483902 | mNeoVis.2.1.1.chr4\_hap1.g00030.t01 |
| 4888 | 4 | 5497464 | 5519893 | mNeoVis.2.1.1.chr4\_hap1.g00031.t01 |
| 4889 | 4 | 5520031 | 5548277 | mNeoVis.2.1.1.chr4\_hap1.g00032.t01 |
| 4890 | 4 | 5610974 | 5620462 | mNeoVis.2.1.1.chr4\_hap1.g00033.t01 |
| 4891 | 4 | 5620583 | 5626282 | mNeoVis.2.1.1.chr4\_hap1.g00034.t01 |
| 4892 | 4 | 5632712 | 5647449 | mNeoVis.2.1.1.chr4\_hap1.g00035.t01 |
| 4893 | 4 | 5650590 | 5653042 | mNeoVis.2.1.1.chr4\_hap1.g00036.t01 |
| 4894 | 4 | 5654981 | 5668245 | mNeoVis.2.1.1.chr4\_hap1.g00037.t01 |
| 4895 | 4 | 5670270 | 5678597 | mNeoVis.2.1.1.chr4\_hap1.g00038.t01 |
| 4896 | 4 | 5685423 | 5706330 | mNeoVis.2.1.1.chr4\_hap1.g00039.t01 |
| 4897 | 4 | 5706475 | 5734785 | mNeoVis.2.1.1.chr4\_hap1.g00040.t01 |
| 4898 | 4 | 5761782 | 5764315 | mNeoVis.2.1.1.chr4\_hap1.g00041.t01 |
| 4899 | 4 | 5764608 | 5768879 | mNeoVis.2.1.1.chr4\_hap1.g00042.t01 |
| 4900 | 4 | 5768969 | 5784203 | mNeoVis.2.1.1.chr4\_hap1.g00043.t01 |
| 4901 | 4 | 5785909 | 5790128 | mNeoVis.2.1.1.chr4\_hap1.g00044.t01 |
| 4902 | 4 | 5791221 | 5795021 | mNeoVis.2.1.1.chr4\_hap1.g00045.t01 |
| 4903 | 4 | 5797424 | 5813340 | mNeoVis.2.1.1.chr4\_hap1.g00046.t01 |
| 4904 | 4 | 5814188 | 5823759 | mNeoVis.2.1.1.chr4\_hap1.g00047.t01 |
| 4905 | 4 | 5823892 | 5841684 | mNeoVis.2.1.1.chr4\_hap1.g00048.t01 |
| 4906 | 4 | 5855592 | 5887070 | mNeoVis.2.1.1.chr4\_hap1.g00049.t01 |
| 4907 | 4 | 5933590 | 5939407 | mNeoVis.2.1.1.chr4\_hap1.g00050.t01 |
| 4908 | 4 | 5966138 | 5970586 | mNeoVis.2.1.1.chr4\_hap1.g00051.t01 |
| 4909 | 4 | 5970939 | 5980477 | mNeoVis.2.1.1.chr4\_hap1.g00052.t01 |
| 4910 | 4 | 6006622 | 6012953 | mNeoVis.2.1.1.chr4\_hap1.g00053.t01 |
| 4911 | 4 | 6027647 | 6046077 | mNeoVis.2.1.1.chr4\_hap1.g00054.t01 |
| 4912 | 4 | 6073005 | 6086710 | mNeoVis.2.1.1.chr4\_hap1.g00055.t01 |
| 4913 | 4 | 6086854 | 6104938 | mNeoVis.2.1.1.chr4\_hap1.g00056.t01 |
| 4914 | 4 | 6108464 | 6115424 | mNeoVis.2.1.1.chr4\_hap1.g00057.t01 |
| 4915 | 4 | 6130374 | 6133832 | mNeoVis.2.1.1.chr4\_hap1.g00058.t01 |
| 4916 | 4 | 6139216 | 6148277 | mNeoVis.2.1.1.chr4\_hap1.g00059.t01 |
| 4917 | 4 | 6154606 | 6160443 | mNeoVis.2.1.1.chr4\_hap1.g00060.t01 |
| 4918 | 4 | 6160716 | 6165743 | mNeoVis.2.1.1.chr4\_hap1.g00061.t01 |
| 4919 | 4 | 11292668 | 11323986 | mNeoVis.2.1.1.chr4\_hap1.g00104.t01 |
| 4920 | 4 | 11328400 | 11337300 | mNeoVis.2.1.1.chr4\_hap1.g00105.t01 |
| 4921 | 4 | 11338508 | 11341144 | mNeoVis.2.1.1.chr4\_hap1.g00106.t01 |
| 4922 | 4 | 11352096 | 11356536 | mNeoVis.2.1.1.chr4\_hap1.g00107.t01 |
| 4923 | 4 | 11377907 | 11379287 | mNeoVis.2.1.1.chr4\_hap1.g00108.t01 |
| 4924 | 4 | 11401181 | 11402268 | mNeoVis.2.1.1.chr4\_hap1.g00109.t01 |
| 4925 | 4 | 11449138 | 11457534 | mNeoVis.2.1.1.chr4\_hap1.g00110.t01 |
| 4926 | 4 | 11541820 | 11542979 | mNeoVis.2.1.1.chr4\_hap1.g00111.t01 |
| 4927 | 4 | 11561524 | 11562468 | mNeoVis.2.1.1.chr4\_hap1.g00112.t01 |
| 4928 | 4 | 11573643 | 11574587 | mNeoVis.2.1.1.chr4\_hap1.g00113.t01 |
| 4929 | 4 | 11624159 | 11643161 | mNeoVis.2.1.1.chr4\_hap1.g00114.t01 |
| 4930 | 4 | 11643381 | 11644807 | mNeoVis.2.1.1.chr4\_hap1.g00115.t01 |
| 4931 | 4 | 11648912 | 11677429 | mNeoVis.2.1.1.chr4\_hap1.g00116.t01 |
| 4932 | 4 | 11692709 | 11707007 | mNeoVis.2.1.1.chr4\_hap1.g00117.t01 |
| 4933 | 4 | 11707114 | 11720937 | mNeoVis.2.1.1.chr4\_hap1.g00118.t01 |
| 4934 | 4 | 11786739 | 11790402 | mNeoVis.2.1.1.chr4\_hap1.g00119.t01 |
| 4935 | 4 | 11990917 | 11995374 | mNeoVis.2.1.1.chr4\_hap1.g00120.t01 |
| 4936 | 4 | 12010914 | 12014266 | mNeoVis.2.1.1.chr4\_hap1.g00121.t01 |
| 4937 | 4 | 12033466 | 12037938 | mNeoVis.2.1.1.chr4\_hap1.g00122.t01 |
| 4938 | 4 | 12045249 | 12052253 | mNeoVis.2.1.1.chr4\_hap1.g00123.t01 |
| 4939 | 4 | 12059259 | 12066133 | mNeoVis.2.1.1.chr4\_hap1.g00124.t01 |
| 4940 | 4 | 12081108 | 12088468 | mNeoVis.2.1.1.chr4\_hap1.g00125.t01 |
| 4941 | 4 | 12093324 | 12190714 | mNeoVis.2.1.1.chr4\_hap1.g00126.t01 |
| 4942 | 4 | 12198892 | 12285724 | mNeoVis.2.1.1.chr4\_hap1.g00127.t01 |
| 4943 | 4 | 12304411 | 12305415 | mNeoVis.2.1.1.chr4\_hap1.g00128.t01 |
| 4944 | 4 | 12321099 | 12332459 | mNeoVis.2.1.1.chr4\_hap1.g00129.t01 |
| 4945 | 4 | 12338902 | 12339846 | mNeoVis.2.1.1.chr4\_hap1.g00130.t01 |
| 4946 | 4 | 12366144 | 12367088 | mNeoVis.2.1.1.chr4\_hap1.g00131.t01 |
| 4947 | 4 | 12376479 | 12377418 | mNeoVis.2.1.1.chr4\_hap1.g00132.t01 |
| 4948 | 4 | 12393613 | 12394548 | mNeoVis.2.1.1.chr4\_hap1.g00133.t01 |
| 4949 | 4 | 12417955 | 12440570 | mNeoVis.2.1.1.chr4\_hap1.g00134.t01 |
| 4950 | 4 | 12443934 | 12449386 | mNeoVis.2.1.1.chr4\_hap1.g00135.t01 |
| 4951 | 4 | 12461495 | 12462382 | mNeoVis.2.1.1.chr4\_hap1.g00136.t01 |
| 4952 | 4 | 12474971 | 12475873 | mNeoVis.2.1.1.chr4\_hap1.g00137.t01 |
| 4953 | 4 | 12482094 | 12485089 | mNeoVis.2.1.1.chr4\_hap1.g00138.t01 |
| 4954 | 4 | 12484116 | 12510749 | mNeoVis.2.1.1.chr4\_hap1.g00139.t01 |
| 4955 | 4 | 12517085 | 12549842 | mNeoVis.2.1.1.chr4\_hap1.g00140.t01 |
| 4956 | 4 | 12559209 | 12610418 | mNeoVis.2.1.1.chr4\_hap1.g00141.t01 |
| 4957 | 4 | 12605482 | 12678548 | mNeoVis.2.1.1.chr4\_hap1.g00142.t01 |
| 4958 | 4 | 13199642 | 13378685 | mNeoVis.2.1.1.chr4\_hap1.g00145.t01 |
| 4959 | 4 | 13390233 | 13398085 | mNeoVis.2.1.1.chr4\_hap1.g00146.t01 |
| 4960 | 4 | 13402708 | 13418643 | mNeoVis.2.1.1.chr4\_hap1.g00147.t01 |
| 4961 | 4 | 13429657 | 13532972 | mNeoVis.2.1.1.chr4\_hap1.g00148.t01 |
| 4962 | 4 | 13554355 | 13578747 | mNeoVis.2.1.1.chr4\_hap1.g00149.t01 |
| 4963 | 4 | 13586740 | 13597855 | mNeoVis.2.1.1.chr4\_hap1.g00150.t01 |
| 4964 | 4 | 13606635 | 13656345 | mNeoVis.2.1.1.chr4\_hap1.g00151.t01 |
| 4965 | 4 | 13772827 | 13858039 | mNeoVis.2.1.1.chr4\_hap1.g00152.t01 |
| 4966 | 4 | 14301941 | 14315446 | mNeoVis.2.1.1.chr4\_hap1.g00156.t01 |
| 4967 | 4 | 14334266 | 14350051 | mNeoVis.2.1.1.chr4\_hap1.g00157.t01 |
| 4968 | 4 | 14369478 | 14431541 | mNeoVis.2.1.1.chr4\_hap1.g00158.t01 |
| 4969 | 4 | 14442792 | 14447052 | mNeoVis.2.1.1.chr4\_hap1.g00159.t01 |
| 4970 | 4 | 14471112 | 14549065 | mNeoVis.2.1.1.chr4\_hap1.g00160.t01 |
| 4971 | 4 | 14510832 | 14511119 | mNeoVis.2.1.1.chr4\_hap1.g00161.t01 |
| 4972 | 4 | 14567811 | 14613294 | mNeoVis.2.1.1.chr4\_hap1.g00162.t01 |
| 4973 | 4 | 14631112 | 14691130 | mNeoVis.2.1.1.chr4\_hap1.g00163.t01 |
| 4974 | 4 | 14695604 | 14719727 | mNeoVis.2.1.1.chr4\_hap1.g00164.t01 |
| 4975 | 4 | 14724327 | 14859899 | mNeoVis.2.1.1.chr4\_hap1.g00165.t01 |
| 4976 | 4 | 14877079 | 14883776 | mNeoVis.2.1.1.chr4\_hap1.g00166.t01 |
| 4977 | 4 | 14883196 | 14957875 | mNeoVis.2.1.1.chr4\_hap1.g00167.t01 |
| 4978 | 4 | 14959434 | 15035783 | mNeoVis.2.1.1.chr4\_hap1.g00168.t01 |
| 4979 | 4 | 15045990 | 15147124 | mNeoVis.2.1.1.chr4\_hap1.g00169.t01 |
| 4980 | 4 | 15335674 | 15421561 | mNeoVis.2.1.1.chr4\_hap1.g00170.t01 |
| 4981 | 4 | 15421954 | 15528332 | mNeoVis.2.1.1.chr4\_hap1.g00171.t01 |
| 4982 | 4 | 15555434 | 16004334 | mNeoVis.2.1.1.chr4\_hap1.g00172.t01 |
| 4983 | 4 | 16271232 | 16416871 | mNeoVis.2.1.1.chr4\_hap1.g00174.t01 |
| 4984 | 4 | 16271232 | 16416871 | mNeoVis.2.1.1.chr4\_hap1.g00174.t01 |
| 4985 | 4 | 17270602 | 17284303 | mNeoVis.2.1.1.chr4\_hap1.g00176.t01 |
| 4986 | 4 | 17287954 | 17328156 | mNeoVis.2.1.1.chr4\_hap1.g00177.t01 |
| 4987 | 4 | 17331933 | 17350220 | mNeoVis.2.1.1.chr4\_hap1.g00178.t01 |
| 4988 | 4 | 17356956 | 17442444 | mNeoVis.2.1.1.chr4\_hap1.g00179.t01 |
| 4989 | 4 | 17477899 | 17620066 | mNeoVis.2.1.1.chr4\_hap1.g00180.t01 |
| 4990 | 4 | 17645108 | 17712563 | mNeoVis.2.1.1.chr4\_hap1.g00181.t01 |
| 4991 | 4 | 17691748 | 17715099 | mNeoVis.2.1.1.chr4\_hap1.g00182.t01 |
| 4992 | 4 | 17691748 | 17715099 | mNeoVis.2.1.1.chr4\_hap1.g00182.t01 |
| 4993 | 4 | 17724732 | 17754136 | mNeoVis.2.1.1.chr4\_hap1.g00183.t01 |
| 4994 | 4 | 17724732 | 17754136 | mNeoVis.2.1.1.chr4\_hap1.g00183.t01 |
| 4995 | 4 | 17754898 | 17759092 | mNeoVis.2.1.1.chr4\_hap1.g00184.t01 |
| 4996 | 4 | 17754898 | 17759092 | mNeoVis.2.1.1.chr4\_hap1.g00184.t01 |
| 4997 | 4 | 17758574 | 17773472 | mNeoVis.2.1.1.chr4\_hap1.g00185.t01 |
| 4998 | 4 | 17758574 | 17773472 | mNeoVis.2.1.1.chr4\_hap1.g00185.t01 |
| 4999 | 4 | 17785033 | 17841337 | mNeoVis.2.1.1.chr4\_hap1.g00186.t01 |
| 5000 | 4 | 17785033 | 17841337 | mNeoVis.2.1.1.chr4\_hap1.g00186.t01 |
| 5001 | 4 | 17852880 | 17925636 | mNeoVis.2.1.1.chr4\_hap1.g00187.t01 |
| 5002 | 4 | 17852880 | 17925636 | mNeoVis.2.1.1.chr4\_hap1.g00187.t01 |
| 5003 | 4 | 18135140 | 18161665 | mNeoVis.2.1.1.chr4\_hap1.g00188.t01 |
| 5004 | 4 | 18135140 | 18161665 | mNeoVis.2.1.1.chr4\_hap1.g00188.t01 |
| 5005 | 4 | 18180643 | 18189746 | mNeoVis.2.1.1.chr4\_hap1.g00189.t01 |
| 5006 | 4 | 18180643 | 18189746 | mNeoVis.2.1.1.chr4\_hap1.g00189.t01 |
| 5007 | 4 | 18187950 | 18207697 | mNeoVis.2.1.1.chr4\_hap1.g00190.t01 |
| 5008 | 4 | 18187950 | 18207697 | mNeoVis.2.1.1.chr4\_hap1.g00190.t01 |
| 5009 | 4 | 18256832 | 18272377 | mNeoVis.2.1.1.chr4\_hap1.g00191.t01 |
| 5010 | 4 | 18256832 | 18272377 | mNeoVis.2.1.1.chr4\_hap1.g00191.t01 |
| 5011 | 4 | 18379792 | 18380629 | mNeoVis.2.1.1.chr4\_hap1.g00192.t01 |
| 5012 | 4 | 18379792 | 18380629 | mNeoVis.2.1.1.chr4\_hap1.g00192.t01 |
| 5013 | 4 | 18379792 | 18380629 | mNeoVis.2.1.1.chr4\_hap1.g00192.t01 |
| 5014 | 4 | 18385249 | 18426676 | mNeoVis.2.1.1.chr4\_hap1.g00193.t01 |
| 5015 | 4 | 18385249 | 18426676 | mNeoVis.2.1.1.chr4\_hap1.g00193.t01 |
| 5016 | 4 | 18385249 | 18426676 | mNeoVis.2.1.1.chr4\_hap1.g00193.t01 |
| 5017 | 4 | 18497964 | 18569751 | mNeoVis.2.1.1.chr4\_hap1.g00194.t01 |
| 5018 | 4 | 18497964 | 18569751 | mNeoVis.2.1.1.chr4\_hap1.g00194.t01 |
| 5019 | 4 | 18497964 | 18569751 | mNeoVis.2.1.1.chr4\_hap1.g00194.t01 |
| 5020 | 4 | 19476864 | 19755291 | mNeoVis.2.1.1.chr4\_hap1.g00196.t01 |
| 5021 | 4 | 19935456 | 20362370 | mNeoVis.2.1.1.chr4\_hap1.g00197.t01 |
| 5022 | 4 | 20833024 | 20880471 | mNeoVis.2.1.1.chr4\_hap1.g00198.t01 |
| 5023 | 4 | 20833024 | 20880471 | mNeoVis.2.1.1.chr4\_hap1.g00198.t01 |
| 5024 | 4 | 20833024 | 20880471 | mNeoVis.2.1.1.chr4\_hap1.g00198.t01 |
| 5025 | 4 | 20883507 | 21064144 | mNeoVis.2.1.1.chr4\_hap1.g00199.t01 |
| 5026 | 4 | 20883507 | 21064144 | mNeoVis.2.1.1.chr4\_hap1.g00199.t01 |
| 5027 | 4 | 20883507 | 21064144 | mNeoVis.2.1.1.chr4\_hap1.g00199.t01 |
| 5028 | 4 | 21225031 | 21439876 | mNeoVis.2.1.1.chr4\_hap1.g00200.t01 |
| 5029 | 4 | 21225031 | 21439876 | mNeoVis.2.1.1.chr4\_hap1.g00200.t01 |
| 5030 | 4 | 21544706 | 21548028 | mNeoVis.2.1.1.chr4\_hap1.g00201.t01 |
| 5031 | 4 | 21544706 | 21548028 | mNeoVis.2.1.1.chr4\_hap1.g00201.t01 |
| 5032 | 4 | 21544706 | 21548028 | mNeoVis.2.1.1.chr4\_hap1.g00201.t01 |
| 5033 | 4 | 21587723 | 21602986 | mNeoVis.2.1.1.chr4\_hap1.g00202.t01 |
| 5034 | 4 | 21587723 | 21602986 | mNeoVis.2.1.1.chr4\_hap1.g00202.t01 |
| 5035 | 4 | 21587723 | 21602986 | mNeoVis.2.1.1.chr4\_hap1.g00202.t01 |
| 5036 | 4 | 21603136 | 21720192 | mNeoVis.2.1.1.chr4\_hap1.g00203.t01 |
| 5037 | 4 | 21603136 | 21720192 | mNeoVis.2.1.1.chr4\_hap1.g00203.t01 |
| 5038 | 4 | 21603136 | 21720192 | mNeoVis.2.1.1.chr4\_hap1.g00203.t01 |
| 5039 | 4 | 21720140 | 21733458 | mNeoVis.2.1.1.chr4\_hap1.g00204.t01 |
| 5040 | 4 | 21720140 | 21733458 | mNeoVis.2.1.1.chr4\_hap1.g00204.t01 |
| 5041 | 4 | 21720140 | 21733458 | mNeoVis.2.1.1.chr4\_hap1.g00204.t01 |
| 5042 | 4 | 21782044 | 21829429 | mNeoVis.2.1.1.chr4\_hap1.g00205.t01 |
| 5043 | 4 | 21782044 | 21829429 | mNeoVis.2.1.1.chr4\_hap1.g00205.t01 |
| 5044 | 4 | 21782044 | 21829429 | mNeoVis.2.1.1.chr4\_hap1.g00205.t01 |
| 5045 | 4 | 21837742 | 21845614 | mNeoVis.2.1.1.chr4\_hap1.g00206.t01 |
| 5046 | 4 | 21837742 | 21845614 | mNeoVis.2.1.1.chr4\_hap1.g00206.t01 |
| 5047 | 4 | 21837742 | 21845614 | mNeoVis.2.1.1.chr4\_hap1.g00206.t01 |
| 5048 | 4 | 21854279 | 21876036 | mNeoVis.2.1.1.chr4\_hap1.g00207.t01 |
| 5049 | 4 | 21854279 | 21876036 | mNeoVis.2.1.1.chr4\_hap1.g00207.t01 |
| 5050 | 4 | 21854279 | 21876036 | mNeoVis.2.1.1.chr4\_hap1.g00207.t01 |
| 5051 | 4 | 21891059 | 21911708 | mNeoVis.2.1.1.chr4\_hap1.g00208.t01 |
| 5052 | 4 | 21891059 | 21911708 | mNeoVis.2.1.1.chr4\_hap1.g00208.t01 |
| 5053 | 4 | 21891059 | 21911708 | mNeoVis.2.1.1.chr4\_hap1.g00208.t01 |
| 5054 | 4 | 21914615 | 21965952 | mNeoVis.2.1.1.chr4\_hap1.g00209.t01 |
| 5055 | 4 | 21914615 | 21965952 | mNeoVis.2.1.1.chr4\_hap1.g00209.t01 |
| 5056 | 4 | 21914615 | 21965952 | mNeoVis.2.1.1.chr4\_hap1.g00209.t01 |
| 5057 | 4 | 21956208 | 21965204 | mNeoVis.2.1.1.chr4\_hap1.g00210.t01 |
| 5058 | 4 | 21956208 | 21965204 | mNeoVis.2.1.1.chr4\_hap1.g00210.t01 |
| 5059 | 4 | 21956208 | 21965204 | mNeoVis.2.1.1.chr4\_hap1.g00210.t01 |
| 5060 | 4 | 21965986 | 22003485 | mNeoVis.2.1.1.chr4\_hap1.g00211.t01 |
| 5061 | 4 | 21965986 | 22003485 | mNeoVis.2.1.1.chr4\_hap1.g00211.t01 |
| 5062 | 4 | 21965986 | 22003485 | mNeoVis.2.1.1.chr4\_hap1.g00211.t01 |
| 5063 | 4 | 22023670 | 22087302 | mNeoVis.2.1.1.chr4\_hap1.g00212.t01 |
| 5064 | 4 | 22023670 | 22087302 | mNeoVis.2.1.1.chr4\_hap1.g00212.t01 |
| 5065 | 4 | 22023670 | 22087302 | mNeoVis.2.1.1.chr4\_hap1.g00212.t01 |
| 5066 | 4 | 22106050 | 22125148 | mNeoVis.2.1.1.chr4\_hap1.g00213.t01 |
| 5067 | 4 | 22106050 | 22125148 | mNeoVis.2.1.1.chr4\_hap1.g00213.t01 |
| 5068 | 4 | 22191013 | 22294249 | mNeoVis.2.1.1.chr4\_hap1.g00214.t01 |
| 5069 | 4 | 22191013 | 22294249 | mNeoVis.2.1.1.chr4\_hap1.g00214.t01 |
| 5070 | 4 | 22355789 | 22503303 | mNeoVis.2.1.1.chr4\_hap1.g00215.t01 |
| 5071 | 4 | 22355789 | 22503303 | mNeoVis.2.1.1.chr4\_hap1.g00215.t01 |
| 5072 | 4 | 22554279 | 22563039 | mNeoVis.2.1.1.chr4\_hap1.g00216.t01 |
| 5073 | 4 | 22554279 | 22563039 | mNeoVis.2.1.1.chr4\_hap1.g00216.t01 |
| 5074 | 4 | 22554279 | 22563039 | mNeoVis.2.1.1.chr4\_hap1.g00216.t01 |
| 5075 | 4 | 22570770 | 22623000 | mNeoVis.2.1.1.chr4\_hap1.g00217.t01 |
| 5076 | 4 | 22570770 | 22623000 | mNeoVis.2.1.1.chr4\_hap1.g00217.t01 |
| 5077 | 4 | 22627543 | 22794980 | mNeoVis.2.1.1.chr4\_hap1.g00218.t01 |
| 5078 | 4 | 22627543 | 22794980 | mNeoVis.2.1.1.chr4\_hap1.g00218.t01 |
| 5079 | 4 | 22802649 | 22824496 | mNeoVis.2.1.1.chr4\_hap1.g00219.t01 |
| 5080 | 4 | 22802649 | 22824496 | mNeoVis.2.1.1.chr4\_hap1.g00219.t01 |
| 5081 | 4 | 22802649 | 22824496 | mNeoVis.2.1.1.chr4\_hap1.g00219.t01 |
| 5082 | 4 | 22838278 | 22858642 | mNeoVis.2.1.1.chr4\_hap1.g00220.t01 |
| 5083 | 4 | 22838278 | 22858642 | mNeoVis.2.1.1.chr4\_hap1.g00220.t01 |
| 5084 | 4 | 22838278 | 22858642 | mNeoVis.2.1.1.chr4\_hap1.g00220.t01 |
| 5085 | 4 | 22927888 | 23017775 | mNeoVis.2.1.1.chr4\_hap1.g00221.t01 |
| 5086 | 4 | 22927888 | 23017775 | mNeoVis.2.1.1.chr4\_hap1.g00221.t01 |
| 5087 | 4 | 22927888 | 23017775 | mNeoVis.2.1.1.chr4\_hap1.g00221.t01 |
| 5088 | 4 | 23017917 | 23030427 | mNeoVis.2.1.1.chr4\_hap1.g00222.t01 |
| 5089 | 4 | 23017917 | 23030427 | mNeoVis.2.1.1.chr4\_hap1.g00222.t01 |
| 5090 | 4 | 23017917 | 23030427 | mNeoVis.2.1.1.chr4\_hap1.g00222.t01 |
| 5091 | 4 | 23049123 | 23073439 | mNeoVis.2.1.1.chr4\_hap1.g00223.t01 |
| 5092 | 4 | 23049123 | 23073439 | mNeoVis.2.1.1.chr4\_hap1.g00223.t01 |
| 5093 | 4 | 23076685 | 23080151 | mNeoVis.2.1.1.chr4\_hap1.g00224.t01 |
| 5094 | 4 | 23076685 | 23080151 | mNeoVis.2.1.1.chr4\_hap1.g00224.t01 |
| 5095 | 4 | 23081073 | 23083385 | mNeoVis.2.1.1.chr4\_hap1.g00225.t01 |
| 5096 | 4 | 23081073 | 23083385 | mNeoVis.2.1.1.chr4\_hap1.g00225.t01 |
| 5097 | 4 | 23083942 | 23086400 | mNeoVis.2.1.1.chr4\_hap1.g00226.t01 |
| 5098 | 4 | 23083942 | 23086400 | mNeoVis.2.1.1.chr4\_hap1.g00226.t01 |
| 5099 | 4 | 23087809 | 23115240 | mNeoVis.2.1.1.chr4\_hap1.g00227.t01 |
| 5100 | 4 | 23087809 | 23115240 | mNeoVis.2.1.1.chr4\_hap1.g00227.t01 |
| 5101 | 4 | 23120287 | 23146680 | mNeoVis.2.1.1.chr4\_hap1.g00228.t01 |
| 5102 | 4 | 23120287 | 23146680 | mNeoVis.2.1.1.chr4\_hap1.g00228.t01 |
| 5103 | 4 | 23159626 | 23164264 | mNeoVis.2.1.1.chr4\_hap1.g00229.t01 |
| 5104 | 4 | 23159626 | 23164264 | mNeoVis.2.1.1.chr4\_hap1.g00229.t01 |
| 5105 | 4 | 23165012 | 23200914 | mNeoVis.2.1.1.chr4\_hap1.g00230.t01 |
| 5106 | 4 | 23165012 | 23200914 | mNeoVis.2.1.1.chr4\_hap1.g00230.t01 |
| 5107 | 4 | 23202881 | 23217851 | mNeoVis.2.1.1.chr4\_hap1.g00231.t01 |
| 5108 | 4 | 23202881 | 23217851 | mNeoVis.2.1.1.chr4\_hap1.g00231.t01 |
| 5109 | 4 | 23227893 | 23237185 | mNeoVis.2.1.1.chr4\_hap1.g00232.t01 |
| 5110 | 4 | 23227893 | 23237185 | mNeoVis.2.1.1.chr4\_hap1.g00232.t01 |
| 5111 | 4 | 23249851 | 23251815 | mNeoVis.2.1.1.chr4\_hap1.g00233.t01 |
| 5112 | 4 | 23249851 | 23251815 | mNeoVis.2.1.1.chr4\_hap1.g00233.t01 |
| 5113 | 4 | 23287625 | 23304001 | mNeoVis.2.1.1.chr4\_hap1.g00234.t01 |
| 5114 | 4 | 23287625 | 23304001 | mNeoVis.2.1.1.chr4\_hap1.g00234.t01 |
| 5115 | 4 | 23320928 | 23330374 | mNeoVis.2.1.1.chr4\_hap1.g00235.t01 |
| 5116 | 4 | 23320928 | 23330374 | mNeoVis.2.1.1.chr4\_hap1.g00235.t01 |
| 5117 | 4 | 23334451 | 23365400 | mNeoVis.2.1.1.chr4\_hap1.g00236.t01 |
| 5118 | 4 | 23334451 | 23365400 | mNeoVis.2.1.1.chr4\_hap1.g00236.t01 |
| 5119 | 4 | 23410139 | 23425335 | mNeoVis.2.1.1.chr4\_hap1.g00237.t01 |
| 5120 | 4 | 23410139 | 23425335 | mNeoVis.2.1.1.chr4\_hap1.g00237.t01 |
| 5121 | 4 | 23561118 | 23976649 | mNeoVis.2.1.1.chr4\_hap1.g00238.t01 |
| 5122 | 4 | 23561118 | 23976649 | mNeoVis.2.1.1.chr4\_hap1.g00238.t01 |
| 5123 | 4 | 23621141 | 23625424 | mNeoVis.2.1.1.chr4\_hap1.g00239.t01 |
| 5124 | 4 | 23621141 | 23625424 | mNeoVis.2.1.1.chr4\_hap1.g00239.t01 |
| 5125 | 4 | 23976774 | 24013965 | mNeoVis.2.1.1.chr4\_hap1.g00240.t01 |
| 5126 | 4 | 23976774 | 24013965 | mNeoVis.2.1.1.chr4\_hap1.g00240.t01 |
| 5127 | 4 | 24016685 | 24025816 | mNeoVis.2.1.1.chr4\_hap1.g00241.t01 |
| 5128 | 4 | 24016685 | 24025816 | mNeoVis.2.1.1.chr4\_hap1.g00241.t01 |
| 5129 | 4 | 24027225 | 24030356 | mNeoVis.2.1.1.chr4\_hap1.g00242.t01 |
| 5130 | 4 | 24027225 | 24030356 | mNeoVis.2.1.1.chr4\_hap1.g00242.t01 |
| 5131 | 4 | 24033266 | 24038439 | mNeoVis.2.1.1.chr4\_hap1.g00243.t01 |
| 5132 | 4 | 24033266 | 24038439 | mNeoVis.2.1.1.chr4\_hap1.g00243.t01 |
| 5133 | 4 | 24171031 | 24223054 | mNeoVis.2.1.1.chr4\_hap1.g00244.t01 |
| 5134 | 4 | 24171031 | 24223054 | mNeoVis.2.1.1.chr4\_hap1.g00244.t01 |
| 5135 | 4 | 24776585 | 24777377 | mNeoVis.2.1.1.chr4\_hap1.g00246.t01 |
| 5136 | 4 | 26273857 | 26358652 | mNeoVis.2.1.1.chr4\_hap1.g00248.t01 |
| 5137 | 4 | 26273857 | 26358652 | mNeoVis.2.1.1.chr4\_hap1.g00248.t01 |
| 5138 | 4 | 26403332 | 26423060 | mNeoVis.2.1.1.chr4\_hap1.g00249.t01 |
| 5139 | 4 | 26403332 | 26423060 | mNeoVis.2.1.1.chr4\_hap1.g00249.t01 |
| 5140 | 4 | 26843802 | 26848185 | mNeoVis.2.1.1.chr4\_hap1.g00251.t01 |
| 5141 | 4 | 26934133 | 26936512 | mNeoVis.2.1.1.chr4\_hap1.g00252.t01 |
| 5142 | 4 | 26996943 | 27013911 | mNeoVis.2.1.1.chr4\_hap1.g00253.t01 |
| 5143 | 4 | 27028021 | 27063316 | mNeoVis.2.1.1.chr4\_hap1.g00254.t01 |
| 5144 | 4 | 27128701 | 27195705 | mNeoVis.2.1.1.chr4\_hap1.g00255.t01 |
| 5145 | 4 | 27128701 | 27195705 | mNeoVis.2.1.1.chr4\_hap1.g00255.t01 |
| 5146 | 4 | 27207771 | 27216312 | mNeoVis.2.1.1.chr4\_hap1.g00256.t01 |
| 5147 | 4 | 27207771 | 27216312 | mNeoVis.2.1.1.chr4\_hap1.g00256.t01 |
| 5148 | 4 | 27223205 | 27286398 | mNeoVis.2.1.1.chr4\_hap1.g00257.t01 |
| 5149 | 4 | 27223205 | 27286398 | mNeoVis.2.1.1.chr4\_hap1.g00257.t01 |
| 5150 | 4 | 27243232 | 27279698 | mNeoVis.2.1.1.chr4\_hap1.g00258.t01 |
| 5151 | 4 | 27243232 | 27279698 | mNeoVis.2.1.1.chr4\_hap1.g00258.t01 |
| 5152 | 4 | 27281473 | 27300687 | mNeoVis.2.1.1.chr4\_hap1.g00259.t01 |
| 5153 | 4 | 27281473 | 27300687 | mNeoVis.2.1.1.chr4\_hap1.g00259.t01 |
| 5154 | 4 | 27281473 | 27300687 | mNeoVis.2.1.1.chr4\_hap1.g00259.t01 |
| 5155 | 4 | 27309223 | 27359353 | mNeoVis.2.1.1.chr4\_hap1.g00260.t01 |
| 5156 | 4 | 27309223 | 27359353 | mNeoVis.2.1.1.chr4\_hap1.g00260.t01 |
| 5157 | 4 | 27309223 | 27359353 | mNeoVis.2.1.1.chr4\_hap1.g00260.t01 |
| 5158 | 4 | 27505887 | 27591835 | mNeoVis.2.1.1.chr4\_hap1.g00261.t01 |
| 5159 | 4 | 27505887 | 27591835 | mNeoVis.2.1.1.chr4\_hap1.g00261.t01 |
| 5160 | 4 | 27661814 | 27664052 | mNeoVis.2.1.1.chr4\_hap1.g00262.t01 |
| 5161 | 4 | 27661814 | 27664052 | mNeoVis.2.1.1.chr4\_hap1.g00262.t01 |
| 5162 | 4 | 27741379 | 28272106 | mNeoVis.2.1.1.chr4\_hap1.g00263.t01 |
| 5163 | 4 | 27764485 | 27771972 | mNeoVis.2.1.1.chr4\_hap1.g00264.t01 |
| 5164 | 4 | 27764485 | 27771972 | mNeoVis.2.1.1.chr4\_hap1.g00264.t01 |
| 5165 | 4 | 27911452 | 27912762 | mNeoVis.2.1.1.chr4\_hap1.g00265.t01 |
| 5166 | 4 | 27911452 | 27912762 | mNeoVis.2.1.1.chr4\_hap1.g00265.t01 |
| 5167 | 4 | 27914792 | 27916240 | mNeoVis.2.1.1.chr4\_hap1.g00266.t01 |
| 5168 | 4 | 27914792 | 27916240 | mNeoVis.2.1.1.chr4\_hap1.g00266.t01 |
| 5169 | 4 | 28286941 | 28289327 | mNeoVis.2.1.1.chr4\_hap1.g00267.t01 |
| 5170 | 4 | 28414762 | 28467898 | mNeoVis.2.1.1.chr4\_hap1.g00268.t01 |
| 5171 | 4 | 28478615 | 28655123 | mNeoVis.2.1.1.chr4\_hap1.g00269.t01 |
| 5172 | 4 | 28663643 | 28938099 | mNeoVis.2.1.1.chr4\_hap1.g00270.t01 |
| 5173 | 4 | 29001945 | 29052970 | mNeoVis.2.1.1.chr4\_hap1.g00271.t01 |
| 5174 | 4 | 29059234 | 29073032 | mNeoVis.2.1.1.chr4\_hap1.g00272.t01 |
| 5175 | 4 | 29093286 | 29355858 | mNeoVis.2.1.1.chr4\_hap1.g00273.t01 |
| 5176 | 4 | 29364751 | 29391443 | mNeoVis.2.1.1.chr4\_hap1.g00274.t01 |
| 5177 | 4 | 29364751 | 29391443 | mNeoVis.2.1.1.chr4\_hap1.g00274.t01 |
| 5178 | 4 | 29454983 | 29519821 | mNeoVis.2.1.1.chr4\_hap1.g00275.t01 |
| 5179 | 4 | 29454983 | 29519821 | mNeoVis.2.1.1.chr4\_hap1.g00275.t01 |
| 5180 | 4 | 30079304 | 30160759 | mNeoVis.2.1.1.chr4\_hap1.g00277.t01 |
| 5181 | 4 | 31603058 | 31630923 | mNeoVis.2.1.1.chr4\_hap1.g00278.t01 |
| 5182 | 4 | 31631057 | 31637905 | mNeoVis.2.1.1.chr4\_hap1.g00279.t01 |
| 5183 | 4 | 31638154 | 31867292 | mNeoVis.2.1.1.chr4\_hap1.g00280.t01 |
| 5184 | 4 | 31736433 | 31813379 | mNeoVis.2.1.1.chr4\_hap1.g00281.t01 |
| 5185 | 4 | 31914503 | 32066170 | mNeoVis.2.1.1.chr4\_hap1.g00282.t01 |
| 5186 | 4 | 32145033 | 32315725 | mNeoVis.2.1.1.chr4\_hap1.g00283.t01 |
| 5187 | 4 | 32357026 | 32414091 | mNeoVis.2.1.1.chr4\_hap1.g00284.t01 |
| 5188 | 4 | 32440559 | 32487864 | mNeoVis.2.1.1.chr4\_hap1.g00285.t01 |
| 5189 | 4 | 32521159 | 32527477 | mNeoVis.2.1.1.chr4\_hap1.g00286.t01 |
| 5190 | 4 | 32528288 | 32763616 | mNeoVis.2.1.1.chr4\_hap1.g00287.t01 |
| 5191 | 4 | 32792520 | 32854334 | mNeoVis.2.1.1.chr4\_hap1.g00288.t01 |
| 5192 | 4 | 32912707 | 33026770 | mNeoVis.2.1.1.chr4\_hap1.g00289.t01 |
| 5193 | 4 | 33131596 | 33167431 | mNeoVis.2.1.1.chr4\_hap1.g00290.t01 |
| 5194 | 4 | 33183932 | 33191895 | mNeoVis.2.1.1.chr4\_hap1.g00291.t01 |
| 5195 | 4 | 33415671 | 33461867 | mNeoVis.2.1.1.chr4\_hap1.g00292.t01 |
| 5196 | 4 | 33415671 | 33461867 | mNeoVis.2.1.1.chr4\_hap1.g00292.t01 |
| 5197 | 4 | 33415671 | 33461867 | mNeoVis.2.1.1.chr4\_hap1.g00292.t01 |
| 5198 | 4 | 33433413 | 33673487 | mNeoVis.2.1.1.chr4\_hap1.g00293.t01 |
| 5199 | 4 | 33433413 | 33673487 | mNeoVis.2.1.1.chr4\_hap1.g00293.t01 |
| 5200 | 4 | 33433413 | 33673487 | mNeoVis.2.1.1.chr4\_hap1.g00293.t01 |
| 5201 | 4 | 33804136 | 34481509 | mNeoVis.2.1.1.chr4\_hap1.g00294.t01 |
| 5202 | 4 | 33814055 | 33977180 | mNeoVis.2.1.1.chr4\_hap1.g00295.t01 |
| 5203 | 4 | 33814055 | 33977180 | mNeoVis.2.1.1.chr4\_hap1.g00295.t01 |
| 5204 | 4 | 33814055 | 33977180 | mNeoVis.2.1.1.chr4\_hap1.g00295.t01 |
| 5205 | 4 | 33814055 | 33977180 | mNeoVis.2.1.1.chr4\_hap1.g00295.t01 |
| 5206 | 4 | 34232270 | 34251456 | mNeoVis.2.1.1.chr4\_hap1.g00296.t01 |
| 5207 | 4 | 34232270 | 34251456 | mNeoVis.2.1.1.chr4\_hap1.g00296.t01 |
| 5208 | 4 | 34232270 | 34251456 | mNeoVis.2.1.1.chr4\_hap1.g00296.t01 |
| 5209 | 4 | 34232270 | 34251456 | mNeoVis.2.1.1.chr4\_hap1.g00296.t01 |
| 5210 | 4 | 34515727 | 34606850 | mNeoVis.2.1.1.chr4\_hap1.g00297.t01 |
| 5211 | 4 | 34515727 | 34606850 | mNeoVis.2.1.1.chr4\_hap1.g00297.t01 |
| 5212 | 4 | 34515727 | 34606850 | mNeoVis.2.1.1.chr4\_hap1.g00297.t01 |
| 5213 | 4 | 34803638 | 35358279 | mNeoVis.2.1.1.chr4\_hap1.g00298.t01 |
| 5214 | 4 | 35450979 | 35494229 | mNeoVis.2.1.1.chr4\_hap1.g00299.t01 |
| 5215 | 4 | 35450979 | 35494229 | mNeoVis.2.1.1.chr4\_hap1.g00299.t01 |
| 5216 | 4 | 35450979 | 35494229 | mNeoVis.2.1.1.chr4\_hap1.g00299.t01 |
| 5217 | 4 | 36126104 | 36126495 | mNeoVis.2.1.1.chr4\_hap1.g00300.t01 |
| 5218 | 4 | 36168147 | 36170355 | mNeoVis.2.1.1.chr4\_hap1.g00301.t01 |
| 5219 | 4 | 36197410 | 36203130 | mNeoVis.2.1.1.chr4\_hap1.g00302.t01 |
| 5220 | 4 | 36321166 | 36425708 | mNeoVis.2.1.1.chr4\_hap1.g00303.t01 |
| 5221 | 4 | 36321166 | 36425708 | mNeoVis.2.1.1.chr4\_hap1.g00303.t01 |
| 5222 | 4 | 36375899 | 36582035 | mNeoVis.2.1.1.chr4\_hap1.g00304.t01 |
| 5223 | 4 | 36375899 | 36582035 | mNeoVis.2.1.1.chr4\_hap1.g00304.t01 |
| 5224 | 4 | 36444277 | 36477143 | mNeoVis.2.1.1.chr4\_hap1.g00305.t01 |
| 5225 | 4 | 36444277 | 36477143 | mNeoVis.2.1.1.chr4\_hap1.g00305.t01 |
| 5226 | 4 | 36707609 | 36714900 | mNeoVis.2.1.1.chr4\_hap1.g00306.t01 |
| 5227 | 4 | 36707609 | 36714900 | mNeoVis.2.1.1.chr4\_hap1.g00306.t01 |
| 5228 | 4 | 36722934 | 36745011 | mNeoVis.2.1.1.chr4\_hap1.g00307.t01 |
| 5229 | 4 | 36722934 | 36745011 | mNeoVis.2.1.1.chr4\_hap1.g00307.t01 |
| 5230 | 4 | 36744920 | 36746112 | mNeoVis.2.1.1.chr4\_hap1.g00308.t01 |
| 5231 | 4 | 36744920 | 36746112 | mNeoVis.2.1.1.chr4\_hap1.g00308.t01 |
| 5232 | 4 | 36840492 | 36949012 | mNeoVis.2.1.1.chr4\_hap1.g00309.t01 |
| 5233 | 4 | 36949237 | 37380051 | mNeoVis.2.1.1.chr4\_hap1.g00310.t01 |
| 5234 | 4 | 36949237 | 37380051 | mNeoVis.2.1.1.chr4\_hap1.g00310.t01 |
| 5235 | 4 | 37342526 | 37490868 | mNeoVis.2.1.1.chr4\_hap1.g00311.t01 |
| 5236 | 4 | 37490896 | 38359259 | mNeoVis.2.1.1.chr4\_hap1.g00312.t01 |
| 5237 | 4 | 37897001 | 37933088 | mNeoVis.2.1.1.chr4\_hap1.g00313.t01 |
| 5238 | 4 | 37897001 | 37933088 | mNeoVis.2.1.1.chr4\_hap1.g00313.t01 |
| 5239 | 4 | 40099362 | 40106988 | mNeoVis.2.1.1.chr4\_hap1.g00314.t01 |
| 5240 | 4 | 40106567 | 40114846 | mNeoVis.2.1.1.chr4\_hap1.g00315.t01 |
| 5241 | 4 | 40115168 | 40279004 | mNeoVis.2.1.1.chr4\_hap1.g00316.t01 |
| 5242 | 4 | 40138262 | 40139332 | mNeoVis.2.1.1.chr4\_hap1.g00317.t01 |
| 5243 | 4 | 40323775 | 40850549 | mNeoVis.2.1.1.chr4\_hap1.g00318.t01 |
| 5244 | 4 | 40958350 | 41121129 | mNeoVis.2.1.1.chr4\_hap1.g00319.t01 |
| 5245 | 4 | 40958350 | 41121129 | mNeoVis.2.1.1.chr4\_hap1.g00319.t01 |
| 5246 | 4 | 41158481 | 41207427 | mNeoVis.2.1.1.chr4\_hap1.g00320.t01 |
| 5247 | 4 | 41158481 | 41207427 | mNeoVis.2.1.1.chr4\_hap1.g00320.t01 |
| 5248 | 4 | 41207198 | 41284068 | mNeoVis.2.1.1.chr4\_hap1.g00321.t01 |
| 5249 | 4 | 41297185 | 41312241 | mNeoVis.2.1.1.chr4\_hap1.g00322.t01 |
| 5250 | 4 | 41297185 | 41312241 | mNeoVis.2.1.1.chr4\_hap1.g00322.t01 |
| 5251 | 4 | 41297185 | 41312241 | mNeoVis.2.1.1.chr4\_hap1.g00322.t01 |
| 5252 | 4 | 41320388 | 41442608 | mNeoVis.2.1.1.chr4\_hap1.g00323.t01 |
| 5253 | 4 | 41320388 | 41442608 | mNeoVis.2.1.1.chr4\_hap1.g00323.t01 |
| 5254 | 4 | 41320388 | 41442608 | mNeoVis.2.1.1.chr4\_hap1.g00323.t01 |
| 5255 | 4 | 41537232 | 41646659 | mNeoVis.2.1.1.chr4\_hap1.g00324.t01 |
| 5256 | 4 | 41537232 | 41646659 | mNeoVis.2.1.1.chr4\_hap1.g00324.t01 |
| 5257 | 4 | 41537232 | 41646659 | mNeoVis.2.1.1.chr4\_hap1.g00324.t01 |
| 5258 | 4 | 41693441 | 41729920 | mNeoVis.2.1.1.chr4\_hap1.g00325.t01 |
| 5259 | 4 | 41693441 | 41729920 | mNeoVis.2.1.1.chr4\_hap1.g00325.t01 |
| 5260 | 4 | 41693441 | 41729920 | mNeoVis.2.1.1.chr4\_hap1.g00325.t01 |
| 5261 | 4 | 41730164 | 41762735 | mNeoVis.2.1.1.chr4\_hap1.g00326.t01 |
| 5262 | 4 | 41730164 | 41762735 | mNeoVis.2.1.1.chr4\_hap1.g00326.t01 |
| 5263 | 4 | 41730164 | 41762735 | mNeoVis.2.1.1.chr4\_hap1.g00326.t01 |
| 5264 | 4 | 41944604 | 41973034 | mNeoVis.2.1.1.chr4\_hap1.g00327.t01 |
| 5265 | 4 | 41944604 | 41973034 | mNeoVis.2.1.1.chr4\_hap1.g00327.t01 |
| 5266 | 4 | 42069189 | 42130296 | mNeoVis.2.1.1.chr4\_hap1.g00328.t01 |
| 5267 | 4 | 42069189 | 42130296 | mNeoVis.2.1.1.chr4\_hap1.g00328.t01 |
| 5268 | 4 | 42141263 | 42233323 | mNeoVis.2.1.1.chr4\_hap1.g00329.t01 |
| 5269 | 4 | 42264072 | 42358919 | mNeoVis.2.1.1.chr4\_hap1.g00330.t01 |
| 5270 | 4 | 42403162 | 42550911 | mNeoVis.2.1.1.chr4\_hap1.g00331.t01 |
| 5271 | 4 | 42403162 | 42550911 | mNeoVis.2.1.1.chr4\_hap1.g00331.t01 |
| 5272 | 4 | 42766119 | 42846169 | mNeoVis.2.1.1.chr4\_hap1.g00332.t01 |
| 5273 | 4 | 42766119 | 42846169 | mNeoVis.2.1.1.chr4\_hap1.g00332.t01 |
| 5274 | 4 | 42854343 | 43105340 | mNeoVis.2.1.1.chr4\_hap1.g00333.t01 |
| 5275 | 4 | 43308905 | 43309216 | mNeoVis.2.1.1.chr4\_hap1.g00334.t01 |
| 5276 | 4 | 43308905 | 43309216 | mNeoVis.2.1.1.chr4\_hap1.g00334.t01 |
| 5277 | 4 | 43389713 | 43850728 | mNeoVis.2.1.1.chr4\_hap1.g00335.t01 |
| 5278 | 4 | 43850803 | 43876582 | mNeoVis.2.1.1.chr4\_hap1.g00336.t01 |
| 5279 | 4 | 43903451 | 43903921 | mNeoVis.2.1.1.chr4\_hap1.g00337.t01 |
| 5280 | 4 | 43911058 | 43961677 | mNeoVis.2.1.1.chr4\_hap1.g00338.t01 |
| 5281 | 4 | 43966657 | 44007022 | mNeoVis.2.1.1.chr4\_hap1.g00339.t01 |
| 5282 | 4 | 44011117 | 44026605 | mNeoVis.2.1.1.chr4\_hap1.g00340.t01 |
| 5283 | 4 | 44031305 | 44048913 | mNeoVis.2.1.1.chr4\_hap1.g00341.t01 |
| 5284 | 4 | 44073185 | 44082790 | mNeoVis.2.1.1.chr4\_hap1.g00342.t01 |
| 5285 | 4 | 44080560 | 44103535 | mNeoVis.2.1.1.chr4\_hap1.g00343.t01 |
| 5286 | 4 | 44099216 | 44106519 | mNeoVis.2.1.1.chr4\_hap1.g00344.t01 |
| 5287 | 4 | 44103765 | 44165996 | mNeoVis.2.1.1.chr4\_hap1.g00345.t01 |
| 5288 | 4 | 44172434 | 44175508 | mNeoVis.2.1.1.chr4\_hap1.g00346.t01 |
| 5289 | 4 | 44391596 | 44826844 | mNeoVis.2.1.1.chr4\_hap1.g00347.t01 |
| 5290 | 4 | 44834178 | 44852875 | mNeoVis.2.1.1.chr4\_hap1.g00348.t01 |
| 5291 | 4 | 44903435 | 44989076 | mNeoVis.2.1.1.chr4\_hap1.g00349.t01 |
| 5292 | 4 | 45148547 | 45151957 | mNeoVis.2.1.1.chr4\_hap1.g00350.t01 |
| 5293 | 4 | 45310429 | 45367976 | mNeoVis.2.1.1.chr4\_hap1.g00351.t01 |
| 5294 | 4 | 45427929 | 45439424 | mNeoVis.2.1.1.chr4\_hap1.g00352.t01 |
| 5295 | 4 | 45461479 | 45474829 | mNeoVis.2.1.1.chr4\_hap1.g00353.t01 |
| 5296 | 4 | 45495483 | 45595812 | mNeoVis.2.1.1.chr4\_hap1.g00354.t01 |
| 5297 | 4 | 45683927 | 45714411 | mNeoVis.2.1.1.chr4\_hap1.g00355.t01 |
| 5298 | 4 | 45683927 | 45714411 | mNeoVis.2.1.1.chr4\_hap1.g00355.t01 |
| 5299 | 4 | 45683927 | 45714411 | mNeoVis.2.1.1.chr4\_hap1.g00355.t01 |
| 5300 | 4 | 45726005 | 45771081 | mNeoVis.2.1.1.chr4\_hap1.g00356.t01 |
| 5301 | 4 | 45726005 | 45771081 | mNeoVis.2.1.1.chr4\_hap1.g00356.t01 |
| 5302 | 4 | 45726005 | 45771081 | mNeoVis.2.1.1.chr4\_hap1.g00356.t01 |
| 5303 | 4 | 45827352 | 45836288 | mNeoVis.2.1.1.chr4\_hap1.g00357.t01 |
| 5304 | 4 | 45827352 | 45836288 | mNeoVis.2.1.1.chr4\_hap1.g00357.t01 |
| 5305 | 4 | 45827352 | 45836288 | mNeoVis.2.1.1.chr4\_hap1.g00357.t01 |
| 5306 | 4 | 45850422 | 45918578 | mNeoVis.2.1.1.chr4\_hap1.g00358.t01 |
| 5307 | 4 | 45850422 | 45918578 | mNeoVis.2.1.1.chr4\_hap1.g00358.t01 |
| 5308 | 4 | 45850422 | 45918578 | mNeoVis.2.1.1.chr4\_hap1.g00358.t01 |
| 5309 | 4 | 45958020 | 46056513 | mNeoVis.2.1.1.chr4\_hap1.g00359.t01 |
| 5310 | 4 | 45958020 | 46056513 | mNeoVis.2.1.1.chr4\_hap1.g00359.t01 |
| 5311 | 4 | 45958020 | 46056513 | mNeoVis.2.1.1.chr4\_hap1.g00359.t01 |
| 5312 | 4 | 46056542 | 46058352 | mNeoVis.2.1.1.chr4\_hap1.g00360.t01 |
| 5313 | 4 | 46056542 | 46058352 | mNeoVis.2.1.1.chr4\_hap1.g00360.t01 |
| 5314 | 4 | 46056542 | 46058352 | mNeoVis.2.1.1.chr4\_hap1.g00360.t01 |
| 5315 | 4 | 46157667 | 46185824 | mNeoVis.2.1.1.chr4\_hap1.g00361.t01 |
| 5316 | 4 | 46157667 | 46185824 | mNeoVis.2.1.1.chr4\_hap1.g00361.t01 |
| 5317 | 4 | 46221745 | 46278309 | mNeoVis.2.1.1.chr4\_hap1.g00362.t01 |
| 5318 | 4 | 46221745 | 46278309 | mNeoVis.2.1.1.chr4\_hap1.g00362.t01 |
| 5319 | 4 | 46280128 | 46300386 | mNeoVis.2.1.1.chr4\_hap1.g00363.t01 |
| 5320 | 4 | 46280128 | 46300386 | mNeoVis.2.1.1.chr4\_hap1.g00363.t01 |
| 5321 | 4 | 46303581 | 46364823 | mNeoVis.2.1.1.chr4\_hap1.g00364.t01 |
| 5322 | 4 | 46303581 | 46364823 | mNeoVis.2.1.1.chr4\_hap1.g00364.t01 |
| 5323 | 4 | 46367845 | 46445478 | mNeoVis.2.1.1.chr4\_hap1.g00365.t01 |
| 5324 | 4 | 46367845 | 46445478 | mNeoVis.2.1.1.chr4\_hap1.g00365.t01 |
| 5325 | 4 | 46550149 | 46553358 | mNeoVis.2.1.1.chr4\_hap1.g00366.t01 |
| 5326 | 4 | 46550149 | 46553358 | mNeoVis.2.1.1.chr4\_hap1.g00366.t01 |
| 5327 | 4 | 46584912 | 46875049 | mNeoVis.2.1.1.chr4\_hap1.g00367.t01 |
| 5328 | 4 | 46584912 | 46875049 | mNeoVis.2.1.1.chr4\_hap1.g00367.t01 |
| 5329 | 4 | 47104709 | 47109899 | mNeoVis.2.1.1.chr4\_hap1.g00369.t01 |
| 5330 | 4 | 47215213 | 47616920 | mNeoVis.2.1.1.chr4\_hap1.g00370.t01 |
| 5331 | 4 | 47827426 | 48150281 | mNeoVis.2.1.1.chr4\_hap1.g00371.t01 |
| 5332 | 4 | 48150791 | 48233030 | mNeoVis.2.1.1.chr4\_hap1.g00372.t01 |
| 5333 | 4 | 48300306 | 48410642 | mNeoVis.2.1.1.chr4\_hap1.g00373.t01 |
| 5334 | 4 | 48300306 | 48410642 | mNeoVis.2.1.1.chr4\_hap1.g00373.t01 |
| 5335 | 4 | 48300306 | 48410642 | mNeoVis.2.1.1.chr4\_hap1.g00373.t01 |
| 5336 | 4 | 48654013 | 48658883 | mNeoVis.2.1.1.chr4\_hap1.g00374.t01 |
| 5337 | 4 | 48654013 | 48658883 | mNeoVis.2.1.1.chr4\_hap1.g00374.t01 |
| 5338 | 4 | 48654013 | 48658883 | mNeoVis.2.1.1.chr4\_hap1.g00374.t01 |
| 5339 | 4 | 48703411 | 48708767 | mNeoVis.2.1.1.chr4\_hap1.g00375.t01 |
| 5340 | 4 | 48703411 | 48708767 | mNeoVis.2.1.1.chr4\_hap1.g00375.t01 |
| 5341 | 4 | 48703411 | 48708767 | mNeoVis.2.1.1.chr4\_hap1.g00375.t01 |
| 5342 | 4 | 48718309 | 48722051 | mNeoVis.2.1.1.chr4\_hap1.g00376.t01 |
| 5343 | 4 | 48718309 | 48722051 | mNeoVis.2.1.1.chr4\_hap1.g00376.t01 |
| 5344 | 4 | 48718309 | 48722051 | mNeoVis.2.1.1.chr4\_hap1.g00376.t01 |
| 5345 | 4 | 48726447 | 48738984 | mNeoVis.2.1.1.chr4\_hap1.g00377.t01 |
| 5346 | 4 | 48726447 | 48738984 | mNeoVis.2.1.1.chr4\_hap1.g00377.t01 |
| 5347 | 4 | 48726447 | 48738984 | mNeoVis.2.1.1.chr4\_hap1.g00377.t01 |
| 5348 | 4 | 48738093 | 48741504 | mNeoVis.2.1.1.chr4\_hap1.g00378.t01 |
| 5349 | 4 | 48738093 | 48741504 | mNeoVis.2.1.1.chr4\_hap1.g00378.t01 |
| 5350 | 4 | 48738093 | 48741504 | mNeoVis.2.1.1.chr4\_hap1.g00378.t01 |
| 5351 | 4 | 48747246 | 48750315 | mNeoVis.2.1.1.chr4\_hap1.g00379.t01 |
| 5352 | 4 | 48747246 | 48750315 | mNeoVis.2.1.1.chr4\_hap1.g00379.t01 |
| 5353 | 4 | 48747246 | 48750315 | mNeoVis.2.1.1.chr4\_hap1.g00379.t01 |
| 5354 | 4 | 48751715 | 48797459 | mNeoVis.2.1.1.chr4\_hap1.g00380.t01 |
| 5355 | 4 | 48751715 | 48797459 | mNeoVis.2.1.1.chr4\_hap1.g00380.t01 |
| 5356 | 4 | 48753884 | 48758617 | mNeoVis.2.1.1.chr4\_hap1.g00381.t01 |
| 5357 | 4 | 48753884 | 48758617 | mNeoVis.2.1.1.chr4\_hap1.g00381.t01 |
| 5358 | 4 | 48753884 | 48758617 | mNeoVis.2.1.1.chr4\_hap1.g00381.t01 |
| 5359 | 4 | 48760312 | 48763229 | mNeoVis.2.1.1.chr4\_hap1.g00382.t01 |
| 5360 | 4 | 48760312 | 48763229 | mNeoVis.2.1.1.chr4\_hap1.g00382.t01 |
| 5361 | 4 | 48760312 | 48763229 | mNeoVis.2.1.1.chr4\_hap1.g00382.t01 |
| 5362 | 4 | 48773635 | 48775927 | mNeoVis.2.1.1.chr4\_hap1.g00383.t01 |
| 5363 | 4 | 48773635 | 48775927 | mNeoVis.2.1.1.chr4\_hap1.g00383.t01 |
| 5364 | 4 | 48773635 | 48775927 | mNeoVis.2.1.1.chr4\_hap1.g00383.t01 |
| 5365 | 4 | 48801506 | 48804206 | mNeoVis.2.1.1.chr4\_hap1.g00384.t01 |
| 5366 | 4 | 48801506 | 48804206 | mNeoVis.2.1.1.chr4\_hap1.g00384.t01 |
| 5367 | 4 | 48801506 | 48804206 | mNeoVis.2.1.1.chr4\_hap1.g00384.t01 |
| 5368 | 4 | 48806115 | 48811271 | mNeoVis.2.1.1.chr4\_hap1.g00385.t01 |
| 5369 | 4 | 48806115 | 48811271 | mNeoVis.2.1.1.chr4\_hap1.g00385.t01 |
| 5370 | 4 | 48806115 | 48811271 | mNeoVis.2.1.1.chr4\_hap1.g00385.t01 |
| 5371 | 4 | 48815865 | 48982379 | mNeoVis.2.1.1.chr4\_hap1.g00386.t01 |
| 5372 | 4 | 48815865 | 48982379 | mNeoVis.2.1.1.chr4\_hap1.g00386.t01 |
| 5373 | 4 | 48815865 | 48982379 | mNeoVis.2.1.1.chr4\_hap1.g00386.t01 |
| 5374 | 4 | 48983789 | 49154122 | mNeoVis.2.1.1.chr4\_hap1.g00387.t01 |
| 5375 | 4 | 48983789 | 49154122 | mNeoVis.2.1.1.chr4\_hap1.g00387.t01 |
| 5376 | 4 | 48983789 | 49154122 | mNeoVis.2.1.1.chr4\_hap1.g00387.t01 |
| 5377 | 4 | 49269184 | 49274767 | mNeoVis.2.1.1.chr4\_hap1.g00388.t01 |
| 5378 | 4 | 49269184 | 49274767 | mNeoVis.2.1.1.chr4\_hap1.g00388.t01 |
| 5379 | 4 | 50384857 | 50388794 | mNeoVis.2.1.1.chr4\_hap1.g00393.t01 |
| 5380 | 4 | 50384857 | 50388794 | mNeoVis.2.1.1.chr4\_hap1.g00393.t01 |
| 5381 | 4 | 50413648 | 50454411 | mNeoVis.2.1.1.chr4\_hap1.g00394.t01 |
| 5382 | 4 | 50413648 | 50454411 | mNeoVis.2.1.1.chr4\_hap1.g00394.t01 |
| 5383 | 4 | 50463157 | 50465733 | mNeoVis.2.1.1.chr4\_hap1.g00395.t01 |
| 5384 | 4 | 50463157 | 50465733 | mNeoVis.2.1.1.chr4\_hap1.g00395.t01 |
| 5385 | 4 | 50548511 | 50731230 | mNeoVis.2.1.1.chr4\_hap1.g00396.t01 |
| 5386 | 4 | 50548511 | 50731230 | mNeoVis.2.1.1.chr4\_hap1.g00396.t01 |
| 5387 | 4 | 50753401 | 50837906 | mNeoVis.2.1.1.chr4\_hap1.g00397.t01 |
| 5388 | 4 | 50753401 | 50837906 | mNeoVis.2.1.1.chr4\_hap1.g00397.t01 |
| 5389 | 4 | 50847045 | 50934975 | mNeoVis.2.1.1.chr4\_hap1.g00398.t01 |
| 5390 | 4 | 50847045 | 50934975 | mNeoVis.2.1.1.chr4\_hap1.g00398.t01 |
| 5391 | 4 | 51218165 | 51225777 | mNeoVis.2.1.1.chr4\_hap1.g00399.t01 |
| 5392 | 4 | 51624157 | 51712663 | mNeoVis.2.1.1.chr4\_hap1.g00400.t01 |
| 5393 | 4 | 51720661 | 51746700 | mNeoVis.2.1.1.chr4\_hap1.g00401.t01 |
| 5394 | 4 | 51772266 | 51822171 | mNeoVis.2.1.1.chr4\_hap1.g00402.t01 |
| 5395 | 4 | 51879178 | 51903698 | mNeoVis.2.1.1.chr4\_hap1.g00403.t01 |
| 5396 | 4 | 51895407 | 51896275 | mNeoVis.2.1.1.chr4\_hap1.g00404.t01 |
| 5397 | 4 | 51940013 | 52085922 | mNeoVis.2.1.1.chr4\_hap1.g00405.t01 |
| 5398 | 4 | 52086158 | 52096333 | mNeoVis.2.1.1.chr4\_hap1.g00406.t01 |
| 5399 | 4 | 52100003 | 52136029 | mNeoVis.2.1.1.chr4\_hap1.g00407.t01 |
| 5400 | 4 | 52168841 | 52194019 | mNeoVis.2.1.1.chr4\_hap1.g00408.t01 |
| 5401 | 4 | 52200643 | 52251868 | mNeoVis.2.1.1.chr4\_hap1.g00409.t01 |
| 5402 | 4 | 52534798 | 52544504 | mNeoVis.2.1.1.chr4\_hap1.g00411.t01 |
| 5403 | 4 | 52534798 | 52544504 | mNeoVis.2.1.1.chr4\_hap1.g00411.t01 |
| 5404 | 4 | 52534798 | 52544504 | mNeoVis.2.1.1.chr4\_hap1.g00411.t01 |
| 5405 | 4 | 52577016 | 52578323 | mNeoVis.2.1.1.chr4\_hap1.g00412.t01 |
| 5406 | 4 | 52577016 | 52578323 | mNeoVis.2.1.1.chr4\_hap1.g00412.t01 |
| 5407 | 4 | 52577016 | 52578323 | mNeoVis.2.1.1.chr4\_hap1.g00412.t01 |
| 5408 | 4 | 52586833 | 52591261 | mNeoVis.2.1.1.chr4\_hap1.g00413.t01 |
| 5409 | 4 | 52586833 | 52591261 | mNeoVis.2.1.1.chr4\_hap1.g00413.t01 |
| 5410 | 4 | 52586833 | 52591261 | mNeoVis.2.1.1.chr4\_hap1.g00413.t01 |
| 5411 | 4 | 52852185 | 53075478 | mNeoVis.2.1.1.chr4\_hap1.g00414.t01 |
| 5412 | 4 | 52852185 | 53075478 | mNeoVis.2.1.1.chr4\_hap1.g00414.t01 |
| 5413 | 4 | 52852185 | 53075478 | mNeoVis.2.1.1.chr4\_hap1.g00414.t01 |
| 5414 | 4 | 53083914 | 53174756 | mNeoVis.2.1.1.chr4\_hap1.g00415.t01 |
| 5415 | 4 | 53083914 | 53174756 | mNeoVis.2.1.1.chr4\_hap1.g00415.t01 |
| 5416 | 4 | 53220590 | 53222005 | mNeoVis.2.1.1.chr4\_hap1.g00416.t01 |
| 5417 | 4 | 53220590 | 53222005 | mNeoVis.2.1.1.chr4\_hap1.g00416.t01 |
| 5418 | 4 | 53220590 | 53222005 | mNeoVis.2.1.1.chr4\_hap1.g00416.t01 |
| 5419 | 4 | 53221908 | 53266889 | mNeoVis.2.1.1.chr4\_hap1.g00417.t01 |
| 5420 | 4 | 53221908 | 53266889 | mNeoVis.2.1.1.chr4\_hap1.g00417.t01 |
| 5421 | 4 | 53221908 | 53266889 | mNeoVis.2.1.1.chr4\_hap1.g00417.t01 |
| 5422 | 4 | 53267043 | 53591749 | mNeoVis.2.1.1.chr4\_hap1.g00418.t01 |
| 5423 | 4 | 53267043 | 53591749 | mNeoVis.2.1.1.chr4\_hap1.g00418.t01 |
| 5424 | 4 | 53616194 | 53699807 | mNeoVis.2.1.1.chr4\_hap1.g00419.t01 |
| 5425 | 4 | 53616194 | 53699807 | mNeoVis.2.1.1.chr4\_hap1.g00419.t01 |
| 5426 | 4 | 54214093 | 54223191 | mNeoVis.2.1.1.chr4\_hap1.g00420.t01 |
| 5427 | 4 | 54214093 | 54223191 | mNeoVis.2.1.1.chr4\_hap1.g00420.t01 |
| 5428 | 4 | 54255942 | 54258564 | mNeoVis.2.1.1.chr4\_hap1.g00421.t01 |
| 5429 | 4 | 54255942 | 54258564 | mNeoVis.2.1.1.chr4\_hap1.g00421.t01 |
| 5430 | 4 | 54284681 | 54428853 | mNeoVis.2.1.1.chr4\_hap1.g00422.t01 |
| 5431 | 4 | 54586805 | 54667873 | mNeoVis.2.1.1.chr4\_hap1.g00423.t01 |
| 5432 | 4 | 54586805 | 54667873 | mNeoVis.2.1.1.chr4\_hap1.g00423.t01 |
| 5433 | 4 | 54809125 | 54845987 | mNeoVis.2.1.1.chr4\_hap1.g00424.t01 |
| 5434 | 4 | 54809125 | 54845987 | mNeoVis.2.1.1.chr4\_hap1.g00424.t01 |
| 5435 | 4 | 55182014 | 55201482 | mNeoVis.2.1.1.chr4\_hap1.g00426.t01 |
| 5436 | 4 | 55660825 | 55664264 | mNeoVis.2.1.1.chr4\_hap1.g00427.t01 |
| 5437 | 4 | 55696776 | 55698790 | mNeoVis.2.1.1.chr4\_hap1.g00428.t01 |
| 5438 | 4 | 55788366 | 56658635 | mNeoVis.2.1.1.chr4\_hap1.g00429.t01 |
| 5439 | 4 | 56778505 | 56779628 | mNeoVis.2.1.1.chr4\_hap1.g00430.t01 |
| 5440 | 4 | 56843426 | 56979477 | mNeoVis.2.1.1.chr4\_hap1.g00431.t01 |
| 5441 | 4 | 57375194 | 57426407 | mNeoVis.2.1.1.chr4\_hap1.g00433.t01 |
| 5442 | 4 | 57866239 | 57877676 | mNeoVis.2.1.1.chr4\_hap1.g00434.t01 |
| 5443 | 4 | 57884093 | 57917389 | mNeoVis.2.1.1.chr4\_hap1.g00435.t01 |
| 5444 | 4 | 57964048 | 58023678 | mNeoVis.2.1.1.chr4\_hap1.g00436.t01 |
| 5445 | 4 | 58024052 | 58062103 | mNeoVis.2.1.1.chr4\_hap1.g00437.t01 |
| 5446 | 4 | 58065570 | 58107910 | mNeoVis.2.1.1.chr4\_hap1.g00438.t01 |
| 5447 | 4 | 58157552 | 58161749 | mNeoVis.2.1.1.chr4\_hap1.g00439.t01 |
| 5448 | 4 | 58890393 | 58953329 | mNeoVis.2.1.1.chr4\_hap1.g00441.t01 |
| 5449 | 4 | 59021386 | 59394043 | mNeoVis.2.1.1.chr4\_hap1.g00442.t01 |
| 5450 | 4 | 59550733 | 59551597 | mNeoVis.2.1.1.chr4\_hap1.g00443.t01 |
| 5451 | 4 | 59617233 | 60308910 | mNeoVis.2.1.1.chr4\_hap1.g00444.t01 |
| 5452 | 4 | 60460362 | 60553361 | mNeoVis.2.1.1.chr4\_hap1.g00445.t01 |
| 5453 | 4 | 61700115 | 61702439 | mNeoVis.2.1.1.chr4\_hap1.g00446.t01 |
| 5454 | 4 | 61749840 | 61827026 | mNeoVis.2.1.1.chr4\_hap1.g00447.t01 |
| 5455 | 4 | 61749840 | 61827026 | mNeoVis.2.1.1.chr4\_hap1.g00447.t01 |
| 5456 | 4 | 61864366 | 61870222 | mNeoVis.2.1.1.chr4\_hap1.g00448.t01 |
| 5457 | 4 | 61864366 | 61870222 | mNeoVis.2.1.1.chr4\_hap1.g00448.t01 |
| 5458 | 4 | 61881389 | 61882303 | mNeoVis.2.1.1.chr4\_hap1.g00449.t01 |
| 5459 | 4 | 61881389 | 61882303 | mNeoVis.2.1.1.chr4\_hap1.g00449.t01 |
| 5460 | 4 | 61937450 | 62011805 | mNeoVis.2.1.1.chr4\_hap1.g00450.t01 |
| 5461 | 4 | 61937450 | 62011805 | mNeoVis.2.1.1.chr4\_hap1.g00450.t01 |
| 5462 | 4 | 62050549 | 62075976 | mNeoVis.2.1.1.chr4\_hap1.g00451.t01 |
| 5463 | 4 | 62050549 | 62075976 | mNeoVis.2.1.1.chr4\_hap1.g00451.t01 |
| 5464 | 4 | 62050549 | 62075976 | mNeoVis.2.1.1.chr4\_hap1.g00451.t01 |
| 5465 | 4 | 62456793 | 62899853 | mNeoVis.2.1.1.chr4\_hap1.g00452.t01 |
| 5466 | 4 | 62456793 | 62899853 | mNeoVis.2.1.1.chr4\_hap1.g00452.t01 |
| 5467 | 4 | 63119846 | 63336344 | mNeoVis.2.1.1.chr4\_hap1.g00453.t01 |
| 5468 | 4 | 63358110 | 63365551 | mNeoVis.2.1.1.chr4\_hap1.g00454.t01 |
| 5469 | 4 | 65111296 | 65547682 | mNeoVis.2.1.1.chr4\_hap1.g00455.t01 |
| 5470 | 4 | 65546732 | 65691238 | mNeoVis.2.1.1.chr4\_hap1.g00456.t01 |
| 5471 | 4 | 65546732 | 65691238 | mNeoVis.2.1.1.chr4\_hap1.g00456.t01 |
| 5472 | 4 | 65708202 | 65812449 | mNeoVis.2.1.1.chr4\_hap1.g00457.t01 |
| 5473 | 4 | 65913148 | 66135602 | mNeoVis.2.1.1.chr4\_hap1.g00458.t01 |
| 5474 | 4 | 65913148 | 66135602 | mNeoVis.2.1.1.chr4\_hap1.g00458.t01 |
| 5475 | 4 | 66135521 | 66139241 | mNeoVis.2.1.1.chr4\_hap1.g00459.t01 |
| 5476 | 4 | 66135521 | 66139241 | mNeoVis.2.1.1.chr4\_hap1.g00459.t01 |
| 5477 | 4 | 66162796 | 66200593 | mNeoVis.2.1.1.chr4\_hap1.g00460.t01 |
| 5478 | 4 | 66162796 | 66200593 | mNeoVis.2.1.1.chr4\_hap1.g00460.t01 |
| 5479 | 4 | 66216921 | 66370648 | mNeoVis.2.1.1.chr4\_hap1.g00461.t01 |
| 5480 | 4 | 66216921 | 66370648 | mNeoVis.2.1.1.chr4\_hap1.g00461.t01 |
| 5481 | 4 | 66450750 | 66518794 | mNeoVis.2.1.1.chr4\_hap1.g00462.t01 |
| 5482 | 4 | 66450750 | 66518794 | mNeoVis.2.1.1.chr4\_hap1.g00462.t01 |
| 5483 | 4 | 66656347 | 66671195 | mNeoVis.2.1.1.chr4\_hap1.g00463.t01 |
| 5484 | 4 | 66656347 | 66671195 | mNeoVis.2.1.1.chr4\_hap1.g00463.t01 |
| 5485 | 4 | 66767464 | 66775533 | mNeoVis.2.1.1.chr4\_hap1.g00464.t01 |
| 5486 | 4 | 66767464 | 66775533 | mNeoVis.2.1.1.chr4\_hap1.g00464.t01 |
| 5487 | 4 | 66769227 | 67292071 | mNeoVis.2.1.1.chr4\_hap1.g00465.t01 |
| 5488 | 4 | 67207355 | 67271089 | mNeoVis.2.1.1.chr4\_hap1.g00466.t01 |
| 5489 | 4 | 67207355 | 67271089 | mNeoVis.2.1.1.chr4\_hap1.g00466.t01 |
| 5490 | 4 | 67332812 | 67337158 | mNeoVis.2.1.1.chr4\_hap1.g00467.t01 |
| 5491 | 4 | 67332812 | 67337158 | mNeoVis.2.1.1.chr4\_hap1.g00467.t01 |
| 5492 | 4 | 67347007 | 67352072 | mNeoVis.2.1.1.chr4\_hap1.g00468.t01 |
| 5493 | 4 | 67347007 | 67352072 | mNeoVis.2.1.1.chr4\_hap1.g00468.t01 |
| 5494 | 4 | 67608556 | 67634720 | mNeoVis.2.1.1.chr4\_hap1.g00469.t01 |
| 5495 | 4 | 67608556 | 67634720 | mNeoVis.2.1.1.chr4\_hap1.g00469.t01 |
| 5496 | 4 | 67608556 | 67634720 | mNeoVis.2.1.1.chr4\_hap1.g00469.t01 |
| 5497 | 4 | 67919826 | 68101920 | mNeoVis.2.1.1.chr4\_hap1.g00470.t01 |
| 5498 | 4 | 67919826 | 68101920 | mNeoVis.2.1.1.chr4\_hap1.g00470.t01 |
| 5499 | 4 | 67919826 | 68101920 | mNeoVis.2.1.1.chr4\_hap1.g00470.t01 |
| 5500 | 4 | 68118869 | 68404685 | mNeoVis.2.1.1.chr4\_hap1.g00471.t01 |
| 5501 | 4 | 68118869 | 68404685 | mNeoVis.2.1.1.chr4\_hap1.g00471.t01 |
| 5502 | 4 | 68118869 | 68404685 | mNeoVis.2.1.1.chr4\_hap1.g00471.t01 |
| 5503 | 4 | 68553333 | 68568061 | mNeoVis.2.1.1.chr4\_hap1.g00472.t01 |
| 5504 | 4 | 68553333 | 68568061 | mNeoVis.2.1.1.chr4\_hap1.g00472.t01 |
| 5505 | 4 | 68553333 | 68568061 | mNeoVis.2.1.1.chr4\_hap1.g00472.t01 |
| 5506 | 4 | 68598285 | 68662198 | mNeoVis.2.1.1.chr4\_hap1.g00473.t01 |
| 5507 | 4 | 68598285 | 68662198 | mNeoVis.2.1.1.chr4\_hap1.g00473.t01 |
| 5508 | 4 | 68598285 | 68662198 | mNeoVis.2.1.1.chr4\_hap1.g00473.t01 |
| 5509 | 4 | 68710700 | 68778334 | mNeoVis.2.1.1.chr4\_hap1.g00474.t01 |
| 5510 | 4 | 68799099 | 68831578 | mNeoVis.2.1.1.chr4\_hap1.g00475.t01 |
| 5511 | 4 | 68799099 | 68831578 | mNeoVis.2.1.1.chr4\_hap1.g00475.t01 |
| 5512 | 4 | 68833927 | 69138272 | mNeoVis.2.1.1.chr4\_hap1.g00476.t01 |
| 5513 | 4 | 69278543 | 69287475 | mNeoVis.2.1.1.chr4\_hap1.g00477.t01 |
| 5514 | 4 | 69286855 | 69353477 | mNeoVis.2.1.1.chr4\_hap1.g00478.t01 |
| 5515 | 4 | 69379332 | 69437023 | mNeoVis.2.1.1.chr4\_hap1.g00479.t01 |
| 5516 | 4 | 69499565 | 69535229 | mNeoVis.2.1.1.chr4\_hap1.g00480.t01 |
| 5517 | 4 | 69918865 | 69928610 | mNeoVis.2.1.1.chr4\_hap1.g00481.t01 |
| 5518 | 4 | 69930080 | 70295823 | mNeoVis.2.1.1.chr4\_hap1.g00482.t01 |
| 5519 | 4 | 69988698 | 69993381 | mNeoVis.2.1.1.chr4\_hap1.g00483.t01 |
| 5520 | 4 | 70015117 | 70026342 | mNeoVis.2.1.1.chr4\_hap1.g00484.t01 |
| 5521 | 4 | 70333119 | 70483834 | mNeoVis.2.1.1.chr4\_hap1.g00485.t01 |
| 5522 | 4 | 70530774 | 70657636 | mNeoVis.2.1.1.chr4\_hap1.g00486.t01 |
| 5523 | 4 | 70663951 | 70719167 | mNeoVis.2.1.1.chr4\_hap1.g00487.t01 |
| 5524 | 4 | 70743617 | 70764119 | mNeoVis.2.1.1.chr4\_hap1.g00488.t01 |
| 5525 | 4 | 70965134 | 71211439 | mNeoVis.2.1.1.chr4\_hap1.g00489.t01 |
| 5526 | 4 | 71222844 | 71250455 | mNeoVis.2.1.1.chr4\_hap1.g00490.t01 |
| 5527 | 4 | 71398998 | 71423792 | mNeoVis.2.1.1.chr4\_hap1.g00495.t01 |
| 5528 | 4 | 71434048 | 71676799 | mNeoVis.2.1.1.chr4\_hap1.g00496.t01 |
| 5529 | 4 | 71581708 | 71619619 | mNeoVis.2.1.1.chr4\_hap1.g00497.t01 |
| 5530 | 4 | 71625391 | 71660335 | mNeoVis.2.1.1.chr4\_hap1.g00498.t01 |
| 5531 | 4 | 71677239 | 71699414 | mNeoVis.2.1.1.chr4\_hap1.g00499.t01 |
| 5532 | 4 | 71702086 | 71858123 | mNeoVis.2.1.1.chr4\_hap1.g00500.t01 |
| 5533 | 4 | 71897147 | 71903532 | mNeoVis.2.1.1.chr4\_hap1.g00501.t01 |
| 5534 | 4 | 72230815 | 72231602 | mNeoVis.2.1.1.chr4\_hap1.g00502.t01 |
| 5535 | 4 | 73066300 | 73134642 | mNeoVis.2.1.1.chr4\_hap1.g00506.t01 |
| 5536 | 4 | 73166099 | 73176388 | mNeoVis.2.1.1.chr4\_hap1.g00507.t01 |
| 5537 | 4 | 73187535 | 73216774 | mNeoVis.2.1.1.chr4\_hap1.g00508.t01 |
| 5538 | 4 | 73216800 | 73257383 | mNeoVis.2.1.1.chr4\_hap1.g00509.t01 |
| 5539 | 4 | 73264860 | 73316969 | mNeoVis.2.1.1.chr4\_hap1.g00510.t01 |
| 5540 | 4 | 73324529 | 73345221 | mNeoVis.2.1.1.chr4\_hap1.g00511.t01 |
| 5541 | 4 | 73389633 | 73406601 | mNeoVis.2.1.1.chr4\_hap1.g00512.t01 |
| 5542 | 4 | 74254179 | 74775516 | mNeoVis.2.1.1.chr4\_hap1.g00513.t01 |
| 5543 | 4 | 74740555 | 74742395 | mNeoVis.2.1.1.chr4\_hap1.g00514.t01 |
| 5544 | 4 | 74740555 | 74742395 | mNeoVis.2.1.1.chr4\_hap1.g00514.t01 |
| 5545 | 4 | 75207925 | 75234179 | mNeoVis.2.1.1.chr4\_hap1.g00515.t01 |
| 5546 | 4 | 75207925 | 75234179 | mNeoVis.2.1.1.chr4\_hap1.g00515.t01 |
| 5547 | 4 | 75207925 | 75234179 | mNeoVis.2.1.1.chr4\_hap1.g00515.t01 |
| 5548 | 4 | 75274157 | 75289193 | mNeoVis.2.1.1.chr4\_hap1.g00516.t01 |
| 5549 | 4 | 75274157 | 75289193 | mNeoVis.2.1.1.chr4\_hap1.g00516.t01 |
| 5550 | 4 | 75291024 | 75522147 | mNeoVis.2.1.1.chr4\_hap1.g00517.t01 |
| 5551 | 4 | 75291024 | 75522147 | mNeoVis.2.1.1.chr4\_hap1.g00517.t01 |
| 5552 | 4 | 75531198 | 75557897 | mNeoVis.2.1.1.chr4\_hap1.g00518.t01 |
| 5553 | 4 | 75531198 | 75557897 | mNeoVis.2.1.1.chr4\_hap1.g00518.t01 |
| 5554 | 4 | 75558045 | 75611913 | mNeoVis.2.1.1.chr4\_hap1.g00519.t01 |
| 5555 | 4 | 75558045 | 75611913 | mNeoVis.2.1.1.chr4\_hap1.g00519.t01 |
| 5556 | 4 | 75612615 | 75752869 | mNeoVis.2.1.1.chr4\_hap1.g00520.t01 |
| 5557 | 4 | 75612615 | 75752869 | mNeoVis.2.1.1.chr4\_hap1.g00520.t01 |
| 5558 | 4 | 75805187 | 75911360 | mNeoVis.2.1.1.chr4\_hap1.g00521.t01 |
| 5559 | 4 | 75934782 | 76000231 | mNeoVis.2.1.1.chr4\_hap1.g00522.t01 |
| 5560 | 4 | 76002982 | 76053333 | mNeoVis.2.1.1.chr4\_hap1.g00523.t01 |
| 5561 | 4 | 76130904 | 76151452 | mNeoVis.2.1.1.chr4\_hap1.g00524.t01 |
| 5562 | 4 | 76151277 | 76196811 | mNeoVis.2.1.1.chr4\_hap1.g00525.t01 |
| 5563 | 4 | 76304680 | 76479694 | mNeoVis.2.1.1.chr4\_hap1.g00526.t01 |
| 5564 | 4 | 76304680 | 76479694 | mNeoVis.2.1.1.chr4\_hap1.g00526.t01 |
| 5565 | 4 | 76485986 | 76708229 | mNeoVis.2.1.1.chr4\_hap1.g00527.t01 |
| 5566 | 4 | 76485986 | 76708229 | mNeoVis.2.1.1.chr4\_hap1.g00527.t01 |
| 5567 | 4 | 78054396 | 78257476 | mNeoVis.2.1.1.chr4\_hap1.g00528.t01 |
| 5568 | 4 | 78272681 | 78318294 | mNeoVis.2.1.1.chr4\_hap1.g00529.t01 |
| 5569 | 4 | 78272681 | 78318294 | mNeoVis.2.1.1.chr4\_hap1.g00529.t01 |
| 5570 | 4 | 78339014 | 79114581 | mNeoVis.2.1.1.chr4\_hap1.g00530.t01 |
| 5571 | 4 | 78575958 | 78611164 | mNeoVis.2.1.1.chr4\_hap1.g00531.t01 |
| 5572 | 4 | 78575958 | 78611164 | mNeoVis.2.1.1.chr4\_hap1.g00531.t01 |
| 5573 | 4 | 79406166 | 79657875 | mNeoVis.2.1.1.chr4\_hap1.g00532.t01 |
| 5574 | 4 | 79845761 | 80261038 | mNeoVis.2.1.1.chr4\_hap1.g00533.t01 |
| 5575 | 4 | 79845761 | 80261038 | mNeoVis.2.1.1.chr4\_hap1.g00533.t01 |
| 5576 | 4 | 80554427 | 81066714 | mNeoVis.2.1.1.chr4\_hap1.g00534.t01 |
| 5577 | 4 | 81223436 | 81315159 | mNeoVis.2.1.1.chr4\_hap1.g00535.t01 |
| 5578 | 4 | 81223436 | 81315159 | mNeoVis.2.1.1.chr4\_hap1.g00535.t01 |
| 5579 | 4 | 81925463 | 81926855 | mNeoVis.2.1.1.chr4\_hap1.g00536.t01 |
| 5580 | 4 | 81925463 | 81926855 | mNeoVis.2.1.1.chr4\_hap1.g00536.t01 |
| 5581 | 4 | 81925463 | 81926855 | mNeoVis.2.1.1.chr4\_hap1.g00536.t01 |
| 5582 | 4 | 81925463 | 81926855 | mNeoVis.2.1.1.chr4\_hap1.g00536.t01 |
| 5583 | 4 | 82062411 | 82236506 | mNeoVis.2.1.1.chr4\_hap1.g00537.t01 |
| 5584 | 4 | 82062411 | 82236506 | mNeoVis.2.1.1.chr4\_hap1.g00537.t01 |
| 5585 | 4 | 82062411 | 82236506 | mNeoVis.2.1.1.chr4\_hap1.g00537.t01 |
| 5586 | 4 | 82062411 | 82236506 | mNeoVis.2.1.1.chr4\_hap1.g00537.t01 |
| 5587 | 4 | 82317551 | 82398282 | mNeoVis.2.1.1.chr4\_hap1.g00538.t01 |
| 5588 | 4 | 82317551 | 82398282 | mNeoVis.2.1.1.chr4\_hap1.g00538.t01 |
| 5589 | 4 | 82317551 | 82398282 | mNeoVis.2.1.1.chr4\_hap1.g00538.t01 |
| 5590 | 4 | 82317551 | 82398282 | mNeoVis.2.1.1.chr4\_hap1.g00538.t01 |
| 5591 | 4 | 82465362 | 82520754 | mNeoVis.2.1.1.chr4\_hap1.g00539.t01 |
| 5592 | 4 | 82465362 | 82520754 | mNeoVis.2.1.1.chr4\_hap1.g00539.t01 |
| 5593 | 4 | 82465362 | 82520754 | mNeoVis.2.1.1.chr4\_hap1.g00539.t01 |
| 5594 | 4 | 82465362 | 82520754 | mNeoVis.2.1.1.chr4\_hap1.g00539.t01 |
| 5595 | 4 | 82735130 | 82749572 | mNeoVis.2.1.1.chr4\_hap1.g00540.t01 |
| 5596 | 4 | 82770776 | 82853446 | mNeoVis.2.1.1.chr4\_hap1.g00541.t01 |
| 5597 | 4 | 82912307 | 83061706 | mNeoVis.2.1.1.chr4\_hap1.g00542.t01 |
| 5598 | 4 | 83023451 | 83164071 | mNeoVis.2.1.1.chr4\_hap1.g00543.t01 |
| 5599 | 4 | 83464085 | 84828093 | mNeoVis.2.1.1.chr4\_hap1.g00544.t01 |
| 5600 | 4 | 84891552 | 85012240 | mNeoVis.2.1.1.chr4\_hap1.g00545.t01 |
| 5601 | 4 | 85012471 | 85017644 | mNeoVis.2.1.1.chr4\_hap1.g00546.t01 |
| 5602 | 4 | 85025906 | 85095931 | mNeoVis.2.1.1.chr4\_hap1.g00547.t01 |
| 5603 | 4 | 85151005 | 85246164 | mNeoVis.2.1.1.chr4\_hap1.g00548.t01 |
| 5604 | 4 | 85370347 | 85454775 | mNeoVis.2.1.1.chr4\_hap1.g00549.t01 |
| 5605 | 4 | 85524725 | 85530195 | mNeoVis.2.1.1.chr4\_hap1.g00551.t01 |
| 5606 | 4 | 85898240 | 85936365 | mNeoVis.2.1.1.chr4\_hap1.g00555.t01 |
| 5607 | 4 | 85936366 | 85989287 | mNeoVis.2.1.1.chr4\_hap1.g00556.t01 |
| 5608 | 4 | 86030000 | 86044440 | mNeoVis.2.1.1.chr4\_hap1.g00557.t01 |
| 5609 | 4 | 86044523 | 86082584 | mNeoVis.2.1.1.chr4\_hap1.g00558.t01 |
| 5610 | 4 | 86082882 | 86099109 | mNeoVis.2.1.1.chr4\_hap1.g00559.t01 |
| 5611 | 4 | 86099985 | 86160344 | mNeoVis.2.1.1.chr4\_hap1.g00560.t01 |
| 5612 | 4 | 86174178 | 86686334 | mNeoVis.2.1.1.chr4\_hap1.g00561.t01 |
| 5613 | 4 | 86174178 | 86686334 | mNeoVis.2.1.1.chr4\_hap1.g00561.t01 |
| 5614 | 4 | 86807488 | 86903533 | mNeoVis.2.1.1.chr4\_hap1.g00562.t01 |
| 5615 | 4 | 87014761 | 87492208 | mNeoVis.2.1.1.chr4\_hap1.g00563.t01 |
| 5616 | 4 | 87693021 | 87791913 | mNeoVis.2.1.1.chr4\_hap1.g00564.t01 |
| 5617 | 4 | 87794801 | 88035425 | mNeoVis.2.1.1.chr4\_hap1.g00565.t01 |
| 5618 | 4 | 88083056 | 88139939 | mNeoVis.2.1.1.chr4\_hap1.g00566.t01 |
| 5619 | 4 | 88144876 | 88182228 | mNeoVis.2.1.1.chr4\_hap1.g00567.t01 |
| 5620 | 4 | 88181948 | 88192928 | mNeoVis.2.1.1.chr4\_hap1.g00568.t01 |
| 5621 | 4 | 88207365 | 88443414 | mNeoVis.2.1.1.chr4\_hap1.g00569.t01 |
| 5622 | 4 | 88521212 | 88578619 | mNeoVis.2.1.1.chr4\_hap1.g00570.t01 |
| 5623 | 4 | 88635983 | 88646043 | mNeoVis.2.1.1.chr4\_hap1.g00571.t01 |
| 5624 | 4 | 88649155 | 88649708 | mNeoVis.2.1.1.chr4\_hap1.g00572.t01 |
| 5625 | 4 | 88663448 | 88688519 | mNeoVis.2.1.1.chr4\_hap1.g00573.t01 |
| 5626 | 4 | 88818522 | 88856552 | mNeoVis.2.1.1.chr4\_hap1.g00574.t01 |
| 5627 | 4 | 89529388 | 89566307 | mNeoVis.2.1.1.chr4\_hap1.g00577.t01 |
| 5628 | 4 | 89667123 | 89773408 | mNeoVis.2.1.1.chr4\_hap1.g00578.t01 |
| 5629 | 4 | 89781401 | 89810984 | mNeoVis.2.1.1.chr4\_hap1.g00579.t01 |
| 5630 | 4 | 89781401 | 89810984 | mNeoVis.2.1.1.chr4\_hap1.g00579.t01 |
| 5631 | 4 | 89810979 | 90106454 | mNeoVis.2.1.1.chr4\_hap1.g00580.t01 |
| 5632 | 4 | 90024953 | 90030312 | mNeoVis.2.1.1.chr4\_hap1.g00581.t01 |
| 5633 | 4 | 90024953 | 90030312 | mNeoVis.2.1.1.chr4\_hap1.g00581.t01 |
| 5634 | 4 | 90111309 | 90125357 | mNeoVis.2.1.1.chr4\_hap1.g00582.t01 |
| 5635 | 4 | 90111309 | 90125357 | mNeoVis.2.1.1.chr4\_hap1.g00582.t01 |
| 5636 | 4 | 90126066 | 90174609 | mNeoVis.2.1.1.chr4\_hap1.g00583.t01 |
| 5637 | 4 | 90126066 | 90174609 | mNeoVis.2.1.1.chr4\_hap1.g00583.t01 |
| 5638 | 4 | 90194870 | 90248531 | mNeoVis.2.1.1.chr4\_hap1.g00584.t01 |
| 5639 | 4 | 90194870 | 90248531 | mNeoVis.2.1.1.chr4\_hap1.g00584.t01 |
| 5640 | 4 | 90272010 | 90291946 | mNeoVis.2.1.1.chr4\_hap1.g00585.t01 |
| 5641 | 4 | 90272010 | 90291946 | mNeoVis.2.1.1.chr4\_hap1.g00585.t01 |
| 5642 | 4 | 90297862 | 90320344 | mNeoVis.2.1.1.chr4\_hap1.g00586.t01 |
| 5643 | 4 | 90297862 | 90320344 | mNeoVis.2.1.1.chr4\_hap1.g00586.t01 |
| 5644 | 4 | 90400748 | 90427820 | mNeoVis.2.1.1.chr4\_hap1.g00587.t01 |
| 5645 | 4 | 90430388 | 90498577 | mNeoVis.2.1.1.chr4\_hap1.g00588.t01 |
| 5646 | 4 | 90518147 | 90617709 | mNeoVis.2.1.1.chr4\_hap1.g00589.t01 |
| 5647 | 4 | 90655160 | 90922637 | mNeoVis.2.1.1.chr4\_hap1.g00590.t01 |
| 5648 | 4 | 90947990 | 90987059 | mNeoVis.2.1.1.chr4\_hap1.g00591.t01 |
| 5649 | 4 | 91011149 | 91015749 | mNeoVis.2.1.1.chr4\_hap1.g00592.t01 |
| 5650 | 4 | 91025764 | 91102431 | mNeoVis.2.1.1.chr4\_hap1.g00593.t01 |
| 5651 | 4 | 91111218 | 91121355 | mNeoVis.2.1.1.chr4\_hap1.g00594.t01 |
| 5652 | 4 | 91125906 | 91150648 | mNeoVis.2.1.1.chr4\_hap1.g00595.t01 |
| 5653 | 4 | 91346924 | 91486872 | mNeoVis.2.1.1.chr4\_hap1.g00596.t01 |
| 5654 | 4 | 91541042 | 91568429 | mNeoVis.2.1.1.chr4\_hap1.g00597.t01 |
| 5655 | 4 | 93486778 | 93489344 | mNeoVis.2.1.1.chr4\_hap1.g00603.t01 |
| 5656 | 4 | 93495367 | 93506353 | mNeoVis.2.1.1.chr4\_hap1.g00604.t01 |
| 5657 | 4 | 93559182 | 93570801 | mNeoVis.2.1.1.chr4\_hap1.g00605.t01 |
| 5658 | 4 | 93571491 | 93573844 | mNeoVis.2.1.1.chr4\_hap1.g00606.t01 |
| 5659 | 4 | 93583090 | 93590678 | mNeoVis.2.1.1.chr4\_hap1.g00607.t01 |
| 5660 | 4 | 93607613 | 93619755 | mNeoVis.2.1.1.chr4\_hap1.g00608.t01 |
| 5661 | 4 | 93619877 | 93626996 | mNeoVis.2.1.1.chr4\_hap1.g00609.t01 |
| 5662 | 4 | 93636960 | 93638660 | mNeoVis.2.1.1.chr4\_hap1.g00610.t01 |
| 5663 | 4 | 93640784 | 93674867 | mNeoVis.2.1.1.chr4\_hap1.g00611.t01 |
| 5664 | 4 | 93681930 | 93692246 | mNeoVis.2.1.1.chr4\_hap1.g00612.t01 |
| 5665 | 4 | 93695187 | 93774078 | mNeoVis.2.1.1.chr4\_hap1.g00613.t01 |
| 5666 | 4 | 93805206 | 93865409 | mNeoVis.2.1.1.chr4\_hap1.g00614.t01 |
| 5667 | 4 | 93869113 | 93885618 | mNeoVis.2.1.1.chr4\_hap1.g00615.t01 |
| 5668 | 4 | 93896111 | 93911438 | mNeoVis.2.1.1.chr4\_hap1.g00616.t01 |
| 5669 | 4 | 93911588 | 93914061 | mNeoVis.2.1.1.chr4\_hap1.g00617.t01 |
| 5670 | 4 | 93927653 | 93981505 | mNeoVis.2.1.1.chr4\_hap1.g00618.t01 |
| 5671 | 4 | 93988353 | 94002554 | mNeoVis.2.1.1.chr4\_hap1.g00619.t01 |
| 5672 | 4 | 94014250 | 94017269 | mNeoVis.2.1.1.chr4\_hap1.g00620.t01 |
| 5673 | 4 | 94028060 | 94036284 | mNeoVis.2.1.1.chr4\_hap1.g00621.t01 |
| 5674 | 4 | 94054443 | 94056973 | mNeoVis.2.1.1.chr4\_hap1.g00622.t01 |
| 5675 | 4 | 94073093 | 94077117 | mNeoVis.2.1.1.chr4\_hap1.g00623.t01 |
| 5676 | 4 | 94082612 | 94096145 | mNeoVis.2.1.1.chr4\_hap1.g00624.t01 |
| 5677 | 4 | 94097963 | 94100434 | mNeoVis.2.1.1.chr4\_hap1.g00625.t01 |
| 5678 | 4 | 94107001 | 94147805 | mNeoVis.2.1.1.chr4\_hap1.g00626.t01 |
| 5679 | 4 | 94149231 | 94155327 | mNeoVis.2.1.1.chr4\_hap1.g00627.t01 |
| 5680 | 4 | 94169216 | 94177489 | mNeoVis.2.1.1.chr4\_hap1.g00628.t01 |
| 5681 | 4 | 94185069 | 94186809 | mNeoVis.2.1.1.chr4\_hap1.g00629.t01 |
| 5682 | 4 | 94205522 | 94217019 | mNeoVis.2.1.1.chr4\_hap1.g00630.t01 |
| 5683 | 4 | 94346198 | 94439899 | mNeoVis.2.1.1.chr4\_hap1.g00631.t01 |
| 5684 | 4 | 94348533 | 94353813 | mNeoVis.2.1.1.chr4\_hap1.g00632.t01 |
| 5685 | 4 | 94443962 | 94446755 | mNeoVis.2.1.1.chr4\_hap1.g00633.t01 |
| 5686 | 4 | 94462402 | 94466696 | mNeoVis.2.1.1.chr4\_hap1.g00634.t01 |
| 5687 | 4 | 94474087 | 94479504 | mNeoVis.2.1.1.chr4\_hap1.g00635.t01 |
| 5688 | 4 | 95100674 | 95269224 | mNeoVis.2.1.1.chr4\_hap1.g00636.t01 |
| 5689 | 4 | 95404724 | 95484731 | mNeoVis.2.1.1.chr4\_hap1.g00637.t01 |
| 5690 | 4 | 95484740 | 95498283 | mNeoVis.2.1.1.chr4\_hap1.g00638.t01 |
| 5691 | 4 | 95484740 | 95498283 | mNeoVis.2.1.1.chr4\_hap1.g00638.t01 |
| 5692 | 4 | 95502973 | 95514263 | mNeoVis.2.1.1.chr4\_hap1.g00639.t01 |
| 5693 | 4 | 95502973 | 95514263 | mNeoVis.2.1.1.chr4\_hap1.g00639.t01 |
| 5694 | 4 | 95522493 | 95542307 | mNeoVis.2.1.1.chr4\_hap1.g00640.t01 |
| 5695 | 4 | 95522493 | 95542307 | mNeoVis.2.1.1.chr4\_hap1.g00640.t01 |
| 5696 | 4 | 95548306 | 95565214 | mNeoVis.2.1.1.chr4\_hap1.g00641.t01 |
| 5697 | 4 | 95548306 | 95565214 | mNeoVis.2.1.1.chr4\_hap1.g00641.t01 |
| 5698 | 4 | 95568026 | 95592261 | mNeoVis.2.1.1.chr4\_hap1.g00642.t01 |
| 5699 | 4 | 95568026 | 95592261 | mNeoVis.2.1.1.chr4\_hap1.g00642.t01 |
| 5700 | 4 | 95618293 | 95681281 | mNeoVis.2.1.1.chr4\_hap1.g00643.t01 |
| 5701 | 4 | 95618293 | 95681281 | mNeoVis.2.1.1.chr4\_hap1.g00643.t01 |
| 5702 | 4 | 95695928 | 95948715 | mNeoVis.2.1.1.chr4\_hap1.g00644.t01 |
| 5703 | 4 | 95695928 | 95948715 | mNeoVis.2.1.1.chr4\_hap1.g00644.t01 |
| 5704 | 4 | 96302242 | 96375784 | mNeoVis.2.1.1.chr4\_hap1.g00645.t01 |
| 5705 | 4 | 96986041 | 97089891 | mNeoVis.2.1.1.chr4\_hap1.g00646.t01 |
| 5706 | 4 | 97114791 | 97230781 | mNeoVis.2.1.1.chr4\_hap1.g00647.t01 |
| 5707 | 4 | 97233357 | 97296903 | mNeoVis.2.1.1.chr4\_hap1.g00648.t01 |
| 5708 | 4 | 97444469 | 97538192 | mNeoVis.2.1.1.chr4\_hap1.g00649.t01 |
| 5709 | 4 | 97444469 | 97538192 | mNeoVis.2.1.1.chr4\_hap1.g00649.t01 |
| 5710 | 4 | 97567095 | 97572921 | mNeoVis.2.1.1.chr4\_hap1.g00650.t01 |
| 5711 | 4 | 97567095 | 97572921 | mNeoVis.2.1.1.chr4\_hap1.g00650.t01 |
| 5712 | 4 | 97580668 | 97664893 | mNeoVis.2.1.1.chr4\_hap1.g00651.t01 |
| 5713 | 4 | 97676664 | 97854554 | mNeoVis.2.1.1.chr4\_hap1.g00652.t01 |
| 5714 | 4 | 98025972 | 98214992 | mNeoVis.2.1.1.chr4\_hap1.g00653.t01 |
| 5715 | 4 | 98025972 | 98214992 | mNeoVis.2.1.1.chr4\_hap1.g00653.t01 |
| 5716 | 4 | 98303131 | 98303702 | mNeoVis.2.1.1.chr4\_hap1.g00654.t01 |
| 5717 | 4 | 98303131 | 98303702 | mNeoVis.2.1.1.chr4\_hap1.g00654.t01 |
| 5718 | 4 | 98303131 | 98303702 | mNeoVis.2.1.1.chr4\_hap1.g00654.t01 |
| 5719 | 4 | 99874707 | 99876027 | mNeoVis.2.1.1.chr4\_hap1.g00655.t01 |
| 5720 | 4 | 100747359 | 100747926 | mNeoVis.2.1.1.chr4\_hap1.g00656.t01 |
| 5721 | 4 | 100747359 | 100747926 | mNeoVis.2.1.1.chr4\_hap1.g00656.t01 |
| 5722 | 4 | 101388839 | 101417254 | mNeoVis.2.1.1.chr4\_hap1.g00657.t01 |
| 5723 | 4 | 101388839 | 101417254 | mNeoVis.2.1.1.chr4\_hap1.g00657.t01 |
| 5724 | 4 | 101638702 | 101641199 | mNeoVis.2.1.1.chr4\_hap1.g00658.t01 |
| 5725 | 4 | 101638702 | 101641199 | mNeoVis.2.1.1.chr4\_hap1.g00658.t01 |
| 5726 | 4 | 101638702 | 101641199 | mNeoVis.2.1.1.chr4\_hap1.g00658.t01 |
| 5727 | 4 | 101922396 | 102119009 | mNeoVis.2.1.1.chr4\_hap1.g00659.t01 |
| 5728 | 4 | 101922396 | 102119009 | mNeoVis.2.1.1.chr4\_hap1.g00659.t01 |
| 5729 | 4 | 102623566 | 102666583 | mNeoVis.2.1.1.chr4\_hap1.g00664.t01 |
| 5730 | 4 | 103264328 | 103269295 | mNeoVis.2.1.1.chr4\_hap1.g00666.t01 |
| 5731 | 4 | 103269374 | 103278787 | mNeoVis.2.1.1.chr4\_hap1.g00667.t01 |
| 5732 | 4 | 103283827 | 103293734 | mNeoVis.2.1.1.chr4\_hap1.g00668.t01 |
| 5733 | 4 | 103941526 | 104205909 | mNeoVis.2.1.1.chr4\_hap1.g00669.t01 |
| 5734 | 4 | 103941526 | 104205909 | mNeoVis.2.1.1.chr4\_hap1.g00669.t01 |
| 5735 | 4 | 104409529 | 104427908 | mNeoVis.2.1.1.chr4\_hap1.g00670.t01 |
| 5736 | 4 | 104409529 | 104427908 | mNeoVis.2.1.1.chr4\_hap1.g00670.t01 |
| 5737 | 4 | 106115120 | 106119320 | mNeoVis.2.1.1.chr4\_hap1.g00672.t01 |
| 5738 | 4 | 106154391 | 106275816 | mNeoVis.2.1.1.chr4\_hap1.g00673.t01 |
| 5739 | 4 | 106275970 | 106312980 | mNeoVis.2.1.1.chr4\_hap1.g00674.t01 |
| 5740 | 4 | 106378466 | 106450889 | mNeoVis.2.1.1.chr4\_hap1.g00675.t01 |
| 5741 | 4 | 106492394 | 106497755 | mNeoVis.2.1.1.chr4\_hap1.g00676.t01 |
| 5742 | 4 | 106591603 | 107022687 | mNeoVis.2.1.1.chr4\_hap1.g00677.t01 |
| 5743 | 4 | 107154782 | 107336181 | mNeoVis.2.1.1.chr4\_hap1.g00678.t01 |
| 5744 | 4 | 107154782 | 107336181 | mNeoVis.2.1.1.chr4\_hap1.g00678.t01 |
| 5745 | 4 | 107225631 | 107226170 | mNeoVis.2.1.1.chr4\_hap1.g00679.t01 |
| 5746 | 4 | 107225631 | 107226170 | mNeoVis.2.1.1.chr4\_hap1.g00679.t01 |
| 5747 | 4 | 151136377 | 151137651 | mNeoVis.2.1.1.chr4\_hap1.g00688.t01 |
| 5748 | 4 | 151179421 | 151421161 | mNeoVis.2.1.1.chr4\_hap1.g00689.t01 |
| 5749 | 4 | 151421789 | 151441233 | mNeoVis.2.1.1.chr4\_hap1.g00690.t01 |
| 5750 | 4 | 151508230 | 151568140 | mNeoVis.2.1.1.chr4\_hap1.g00691.t01 |
| 5751 | 4 | 152262715 | 152277204 | mNeoVis.2.1.1.chr4\_hap1.g00692.t01 |
| 5752 | 4 | 152262715 | 152277204 | mNeoVis.2.1.1.chr4\_hap1.g00692.t01 |
| 5753 | 4 | 152277305 | 152319548 | mNeoVis.2.1.1.chr4\_hap1.g00693.t01 |
| 5754 | 4 | 152277305 | 152319548 | mNeoVis.2.1.1.chr4\_hap1.g00693.t01 |
| 5755 | 4 | 152488716 | 152492510 | mNeoVis.2.1.1.chr4\_hap1.g00694.t01 |
| 5756 | 4 | 152488716 | 152492510 | mNeoVis.2.1.1.chr4\_hap1.g00694.t01 |
| 5757 | 4 | 152643442 | 152670898 | mNeoVis.2.1.1.chr4\_hap1.g00695.t01 |
| 5758 | 4 | 153346554 | 153347759 | mNeoVis.2.1.1.chr4\_hap1.g00696.t01 |
| 5759 | 4 | 153346554 | 153347759 | mNeoVis.2.1.1.chr4\_hap1.g00696.t01 |
| 5760 | 4 | 155327051 | 155819984 | mNeoVis.2.1.1.chr4\_hap1.g00698.t01 |
| 5761 | 4 | 155826055 | 155890135 | mNeoVis.2.1.1.chr4\_hap1.g00699.t01 |
| 5762 | 4 | 155826055 | 155890135 | mNeoVis.2.1.1.chr4\_hap1.g00699.t01 |
| 5763 | 4 | 155890368 | 156040071 | mNeoVis.2.1.1.chr4\_hap1.g00700.t01 |
| 5764 | 4 | 155890368 | 156040071 | mNeoVis.2.1.1.chr4\_hap1.g00700.t01 |
| 5765 | 4 | 156092860 | 156439553 | mNeoVis.2.1.1.chr4\_hap1.g00701.t01 |
| 5766 | 4 | 156092860 | 156439553 | mNeoVis.2.1.1.chr4\_hap1.g00701.t01 |
| 5767 | 4 | 156092860 | 156439553 | mNeoVis.2.1.1.chr4\_hap1.g00701.t01 |
| 5768 | 4 | 156519915 | 156540250 | mNeoVis.2.1.1.chr4\_hap1.g00702.t01 |
| 5769 | 4 | 156593092 | 156688128 | mNeoVis.2.1.1.chr4\_hap1.g00703.t01 |
| 5770 | 4 | 157979575 | 158056331 | mNeoVis.2.1.1.chr4\_hap1.g00707.t01 |
| 5771 | 4 | 158066890 | 158156044 | mNeoVis.2.1.1.chr4\_hap1.g00708.t01 |
| 5772 | 4 | 158118747 | 158119883 | mNeoVis.2.1.1.chr4\_hap1.g00709.t01 |
| 5773 | 4 | 158173868 | 158180512 | mNeoVis.2.1.1.chr4\_hap1.g00710.t01 |
| 5774 | 4 | 158468167 | 158470933 | mNeoVis.2.1.1.chr4\_hap1.g00711.t01 |
| 5775 | 4 | 158599594 | 158654700 | mNeoVis.2.1.1.chr4\_hap1.g00712.t01 |
| 5776 | 4 | 162022356 | 162048617 | mNeoVis.2.1.1.chr4\_hap1.g00726.t01 |
| 5777 | 4 | 162022356 | 162048617 | mNeoVis.2.1.1.chr4\_hap1.g00726.t01 |
| 5778 | 4 | 162079021 | 162087142 | mNeoVis.2.1.1.chr4\_hap1.g00727.t01 |
| 5779 | 4 | 162079021 | 162087142 | mNeoVis.2.1.1.chr4\_hap1.g00727.t01 |
| 5780 | 4 | 162124810 | 162164021 | mNeoVis.2.1.1.chr4\_hap1.g00728.t01 |
| 5781 | 4 | 162124810 | 162164021 | mNeoVis.2.1.1.chr4\_hap1.g00728.t01 |
| 5782 | 4 | 162163384 | 162223191 | mNeoVis.2.1.1.chr4\_hap1.g00729.t01 |
| 5783 | 4 | 162163384 | 162223191 | mNeoVis.2.1.1.chr4\_hap1.g00729.t01 |
| 5784 | 4 | 166070186 | 166091880 | mNeoVis.2.1.1.chr4\_hap1.g00737.t01 |
| 5785 | 4 | 166070186 | 166091880 | mNeoVis.2.1.1.chr4\_hap1.g00737.t01 |
| 5786 | 4 | 166091773 | 166117665 | mNeoVis.2.1.1.chr4\_hap1.g00738.t01 |
| 5787 | 4 | 166091773 | 166117665 | mNeoVis.2.1.1.chr4\_hap1.g00738.t01 |
| 5788 | 4 | 166107717 | 166126662 | mNeoVis.2.1.1.chr4\_hap1.g00739.t01 |
| 5789 | 4 | 166107717 | 166126662 | mNeoVis.2.1.1.chr4\_hap1.g00739.t01 |
| 5790 | 4 | 166168976 | 166208622 | mNeoVis.2.1.1.chr4\_hap1.g00740.t01 |
| 5791 | 4 | 166168976 | 166208622 | mNeoVis.2.1.1.chr4\_hap1.g00740.t01 |
| 5792 | 4 | 166222976 | 166223254 | mNeoVis.2.1.1.chr4\_hap1.g00741.t01 |
| 5793 | 4 | 166222976 | 166223254 | mNeoVis.2.1.1.chr4\_hap1.g00741.t01 |
| 5794 | 4 | 167202163 | 167213921 | mNeoVis.2.1.1.chr4\_hap1.g00742.t01 |
| 5795 | 4 | 167384002 | 167386902 | mNeoVis.2.1.1.chr4\_hap1.g00743.t01 |
| 5796 | 4 | 167384002 | 167386902 | mNeoVis.2.1.1.chr4\_hap1.g00743.t01 |
| 5797 | 4 | 167394304 | 167568932 | mNeoVis.2.1.1.chr4\_hap1.g00744.t01 |
| 5798 | 4 | 167394304 | 167568932 | mNeoVis.2.1.1.chr4\_hap1.g00744.t01 |
| 5799 | 4 | 175907858 | 175911269 | mNeoVis.2.1.1.chr4\_hap1.g00781.t01 |
| 5800 | 4 | 176086436 | 176151409 | mNeoVis.2.1.1.chr4\_hap1.g00782.t01 |
| 5801 | 4 | 176369546 | 176764620 | mNeoVis.2.1.1.chr4\_hap1.g00783.t01 |
| 5802 | 4 | 176546014 | 176550137 | mNeoVis.2.1.1.chr4\_hap1.g00784.t01 |
| 5803 | 4 | 176845166 | 176879506 | mNeoVis.2.1.1.chr4\_hap1.g00785.t01 |
| 5804 | 4 | 176845166 | 176879506 | mNeoVis.2.1.1.chr4\_hap1.g00785.t01 |
| 5805 | 4 | 176887907 | 176914384 | mNeoVis.2.1.1.chr4\_hap1.g00786.t01 |
| 5806 | 4 | 176887907 | 176914384 | mNeoVis.2.1.1.chr4\_hap1.g00786.t01 |
| 5807 | 4 | 177024234 | 177046504 | mNeoVis.2.1.1.chr4\_hap1.g00787.t01 |
| 5808 | 4 | 177024234 | 177046504 | mNeoVis.2.1.1.chr4\_hap1.g00787.t01 |
| 5809 | 4 | 177051352 | 177055388 | mNeoVis.2.1.1.chr4\_hap1.g00788.t01 |
| 5810 | 4 | 177051352 | 177055388 | mNeoVis.2.1.1.chr4\_hap1.g00788.t01 |
| 5811 | 4 | 177055533 | 177083495 | mNeoVis.2.1.1.chr4\_hap1.g00789.t01 |
| 5812 | 4 | 177055533 | 177083495 | mNeoVis.2.1.1.chr4\_hap1.g00789.t01 |
| 5813 | 4 | 177164526 | 177494330 | mNeoVis.2.1.1.chr4\_hap1.g00790.t01 |
| 5814 | 4 | 177164526 | 177494330 | mNeoVis.2.1.1.chr4\_hap1.g00790.t01 |
| 5815 | 4 | 177534235 | 177614294 | mNeoVis.2.1.1.chr4\_hap1.g00791.t01 |
| 5816 | 4 | 177655021 | 177677120 | mNeoVis.2.1.1.chr4\_hap1.g00792.t01 |
| 5817 | 4 | 177681593 | 177689110 | mNeoVis.2.1.1.chr4\_hap1.g00793.t01 |
| 5818 | 4 | 177695203 | 177724051 | mNeoVis.2.1.1.chr4\_hap1.g00794.t01 |
| 5819 | 4 | 178553383 | 178585704 | mNeoVis.2.1.1.chr4\_hap1.g00797.t01 |
| 5820 | 4 | 178712166 | 178753059 | mNeoVis.2.1.1.chr4\_hap1.g00798.t01 |
| 5821 | 4 | 179143043 | 179169935 | mNeoVis.2.1.1.chr4\_hap1.g00799.t01 |
| 5822 | 4 | 179143043 | 179169935 | mNeoVis.2.1.1.chr4\_hap1.g00799.t01 |
| 5823 | 4 | 180058761 | 180241095 | mNeoVis.2.1.1.chr4\_hap1.g00800.t01 |
| 5824 | 4 | 180058761 | 180241095 | mNeoVis.2.1.1.chr4\_hap1.g00800.t01 |
| 5825 | 4 | 180341621 | 180358816 | mNeoVis.2.1.1.chr4\_hap1.g00801.t01 |
| 5826 | 4 | 180341621 | 180358816 | mNeoVis.2.1.1.chr4\_hap1.g00801.t01 |
| 5827 | 4 | 181565020 | 181565295 | mNeoVis.2.1.1.chr4\_hap1.g00802.t01 |
| 5828 | 4 | 181662634 | 181755649 | mNeoVis.2.1.1.chr4\_hap1.g00803.t01 |
| 5829 | 4 | 181804788 | 181947466 | mNeoVis.2.1.1.chr4\_hap1.g00804.t01 |
| 5830 | 4 | 181870201 | 181916202 | mNeoVis.2.1.1.chr4\_hap1.g00805.t01 |
| 5831 | 4 | 182368133 | 182369106 | mNeoVis.2.1.1.chr4\_hap1.g00806.t01 |
| 5832 | 4 | 182587083 | 182638385 | mNeoVis.2.1.1.chr4\_hap1.g00807.t01 |
| 5833 | 4 | 182716431 | 182720351 | mNeoVis.2.1.1.chr4\_hap1.g00808.t01 |
| 5834 | 4 | 182843855 | 182996538 | mNeoVis.2.1.1.chr4\_hap1.g00809.t01 |
| 5835 | 4 | 182843855 | 182996538 | mNeoVis.2.1.1.chr4\_hap1.g00809.t01 |
| 5836 | 4 | 183128726 | 183294957 | mNeoVis.2.1.1.chr4\_hap1.g00810.t01 |
| 5837 | 4 | 183128726 | 183294957 | mNeoVis.2.1.1.chr4\_hap1.g00810.t01 |
| 5838 | 4 | 183128726 | 183294957 | mNeoVis.2.1.1.chr4\_hap1.g00810.t01 |
| 5839 | 4 | 183296127 | 183329379 | mNeoVis.2.1.1.chr4\_hap1.g00811.t01 |
| 5840 | 4 | 183296127 | 183329379 | mNeoVis.2.1.1.chr4\_hap1.g00811.t01 |
| 5841 | 4 | 183296127 | 183329379 | mNeoVis.2.1.1.chr4\_hap1.g00811.t01 |
| 5842 | 4 | 183397274 | 183400124 | mNeoVis.2.1.1.chr4\_hap1.g00812.t01 |
| 5843 | 4 | 183397274 | 183400124 | mNeoVis.2.1.1.chr4\_hap1.g00812.t01 |
| 5844 | 4 | 183435410 | 183675588 | mNeoVis.2.1.1.chr4\_hap1.g00813.t01 |
| 5845 | 4 | 183758709 | 183897589 | mNeoVis.2.1.1.chr4\_hap1.g00814.t01 |
| 5846 | 4 | 183957447 | 184045719 | mNeoVis.2.1.1.chr4\_hap1.g00815.t01 |
| 5847 | 4 | 183957447 | 184045719 | mNeoVis.2.1.1.chr4\_hap1.g00815.t01 |
| 5848 | 4 | 184045629 | 184049814 | mNeoVis.2.1.1.chr4\_hap1.g00816.t01 |
| 5849 | 4 | 184045629 | 184049814 | mNeoVis.2.1.1.chr4\_hap1.g00816.t01 |
| 5850 | 4 | 184165780 | 184177417 | mNeoVis.2.1.1.chr4\_hap1.g00817.t01 |
| 5851 | 4 | 184165780 | 184177417 | mNeoVis.2.1.1.chr4\_hap1.g00817.t01 |
| 5852 | 4 | 184187217 | 184191307 | mNeoVis.2.1.1.chr4\_hap1.g00818.t01 |
| 5853 | 4 | 184187217 | 184191307 | mNeoVis.2.1.1.chr4\_hap1.g00818.t01 |
| 5854 | 4 | 184201706 | 184205801 | mNeoVis.2.1.1.chr4\_hap1.g00819.t01 |
| 5855 | 4 | 184201706 | 184205801 | mNeoVis.2.1.1.chr4\_hap1.g00819.t01 |
| 5856 | 4 | 184244422 | 184255193 | mNeoVis.2.1.1.chr4\_hap1.g00820.t01 |
| 5857 | 4 | 184244422 | 184255193 | mNeoVis.2.1.1.chr4\_hap1.g00820.t01 |
| 5858 | 4 | 184331447 | 184365939 | mNeoVis.2.1.1.chr4\_hap1.g00821.t01 |
| 5859 | 4 | 184331447 | 184365939 | mNeoVis.2.1.1.chr4\_hap1.g00821.t01 |
| 5860 | 4 | 184371598 | 184375504 | mNeoVis.2.1.1.chr4\_hap1.g00822.t01 |
| 5861 | 4 | 184371598 | 184375504 | mNeoVis.2.1.1.chr4\_hap1.g00822.t01 |
| 5862 | 4 | 184376006 | 184400867 | mNeoVis.2.1.1.chr4\_hap1.g00823.t01 |
| 5863 | 4 | 184376006 | 184400867 | mNeoVis.2.1.1.chr4\_hap1.g00823.t01 |
| 5864 | 4 | 184411712 | 184437422 | mNeoVis.2.1.1.chr4\_hap1.g00824.t01 |
| 5865 | 4 | 184455322 | 184495228 | mNeoVis.2.1.1.chr4\_hap1.g00825.t01 |
| 5866 | 4 | 184494925 | 184649441 | mNeoVis.2.1.1.chr4\_hap1.g00826.t01 |
| 5867 | 4 | 185959731 | 185960660 | mNeoVis.2.1.1.chr4\_hap1.g00827.t01 |
| 5868 | 4 | 187401008 | 187411153 | mNeoVis.2.1.1.chr4\_hap1.g00829.t01 |
| 5869 | 4 | 187434633 | 187471275 | mNeoVis.2.1.1.chr4\_hap1.g00830.t01 |
| 5870 | 4 | 187482795 | 187512209 | mNeoVis.2.1.1.chr4\_hap1.g00831.t01 |
| 5871 | 4 | 187512194 | 187517530 | mNeoVis.2.1.1.chr4\_hap1.g00832.t01 |
| 5872 | 4 | 187535405 | 187565371 | mNeoVis.2.1.1.chr4\_hap1.g00833.t01 |
| 5873 | 4 | 187583643 | 187591634 | mNeoVis.2.1.1.chr4\_hap1.g00834.t01 |
| 5874 | 4 | 187609653 | 187620039 | mNeoVis.2.1.1.chr4\_hap1.g00835.t01 |
| 5875 | 4 | 187693850 | 187702515 | mNeoVis.2.1.1.chr4\_hap1.g00836.t01 |
| 5876 | 4 | 187715171 | 187731342 | mNeoVis.2.1.1.chr4\_hap1.g00837.t01 |
| 5877 | 4 | 188051932 | 188072463 | mNeoVis.2.1.1.chr4\_hap1.g00838.t01 |
| 5878 | 4 | 188051932 | 188072463 | mNeoVis.2.1.1.chr4\_hap1.g00838.t01 |
| 5879 | 4 | 188091735 | 188136854 | mNeoVis.2.1.1.chr4\_hap1.g00839.t01 |
| 5880 | 4 | 188091735 | 188136854 | mNeoVis.2.1.1.chr4\_hap1.g00839.t01 |
| 5881 | 4 | 188171671 | 188183997 | mNeoVis.2.1.1.chr4\_hap1.g00840.t01 |
| 5882 | 4 | 188171671 | 188183997 | mNeoVis.2.1.1.chr4\_hap1.g00840.t01 |
| 5883 | 4 | 188252609 | 188363864 | mNeoVis.2.1.1.chr4\_hap1.g00841.t01 |
| 5884 | 4 | 188252609 | 188363864 | mNeoVis.2.1.1.chr4\_hap1.g00841.t01 |
| 5885 | 4 | 188365180 | 188411101 | mNeoVis.2.1.1.chr4\_hap1.g00842.t01 |
| 5886 | 4 | 188365180 | 188411101 | mNeoVis.2.1.1.chr4\_hap1.g00842.t01 |
| 5887 | 4 | 188416193 | 188465716 | mNeoVis.2.1.1.chr4\_hap1.g00843.t01 |
| 5888 | 4 | 189420675 | 189527947 | mNeoVis.2.1.1.chr4\_hap1.g00846.t01 |
| 5889 | 4 | 189420675 | 189527947 | mNeoVis.2.1.1.chr4\_hap1.g00846.t01 |
| 5890 | 4 | 189420675 | 189527947 | mNeoVis.2.1.1.chr4\_hap1.g00846.t01 |
| 5891 | 4 | 189743242 | 190038372 | mNeoVis.2.1.1.chr4\_hap1.g00847.t01 |
| 5892 | 4 | 191207077 | 191241771 | mNeoVis.2.1.1.chr4\_hap1.g00848.t01 |
| 5893 | 4 | 191317526 | 191342370 | mNeoVis.2.1.1.chr4\_hap1.g00849.t01 |
| 5894 | 4 | 191359202 | 191410440 | mNeoVis.2.1.1.chr4\_hap1.g00850.t01 |
| 5895 | 4 | 191426466 | 191470304 | mNeoVis.2.1.1.chr4\_hap1.g00851.t01 |
| 5896 | 4 | 191475644 | 191497913 | mNeoVis.2.1.1.chr4\_hap1.g00852.t01 |
| 5897 | 4 | 191826514 | 191972981 | mNeoVis.2.1.1.chr4\_hap1.g00853.t01 |
| 5898 | 4 | 192090596 | 192292350 | mNeoVis.2.1.1.chr4\_hap1.g00854.t01 |
| 5899 | 4 | 192293932 | 192332607 | mNeoVis.2.1.1.chr4\_hap1.g00855.t01 |
| 5900 | 4 | 192343776 | 192400632 | mNeoVis.2.1.1.chr4\_hap1.g00856.t01 |
| 5901 | 4 | 192436621 | 192453855 | mNeoVis.2.1.1.chr4\_hap1.g00857.t01 |
| 5902 | 4 | 193197447 | 193333126 | mNeoVis.2.1.1.chr4\_hap1.g00860.t01 |
| 5903 | 4 | 193197447 | 193333126 | mNeoVis.2.1.1.chr4\_hap1.g00860.t01 |
| 5904 | 4 | 193760366 | 193863733 | mNeoVis.2.1.1.chr4\_hap1.g00861.t01 |
| 5905 | 4 | 193760366 | 193863733 | mNeoVis.2.1.1.chr4\_hap1.g00861.t01 |
| 5906 | 4 | 193981820 | 194086472 | mNeoVis.2.1.1.chr4\_hap1.g00862.t01 |
| 5907 | 4 | 193981820 | 194086472 | mNeoVis.2.1.1.chr4\_hap1.g00862.t01 |
| 5908 | 4 | 194411702 | 194532750 | mNeoVis.2.1.1.chr4\_hap1.g00863.t01 |
| 5909 | 4 | 195556370 | 195629185 | mNeoVis.2.1.1.chr4\_hap1.g00875.t01 |
| 5910 | 4 | 195647831 | 195697439 | mNeoVis.2.1.1.chr4\_hap1.g00876.t01 |
| 5911 | 4 | 195697577 | 195712184 | mNeoVis.2.1.1.chr4\_hap1.g00877.t01 |
| 5912 | 4 | 195742726 | 195764563 | mNeoVis.2.1.1.chr4\_hap1.g00878.t01 |
| 5913 | 4 | 195842412 | 195867641 | mNeoVis.2.1.1.chr4\_hap1.g00879.t01 |
| 5914 | 4 | 195909993 | 195984715 | mNeoVis.2.1.1.chr4\_hap1.g00880.t01 |
| 5915 | 4 | 196057526 | 196078615 | mNeoVis.2.1.1.chr4\_hap1.g00881.t01 |
| 5916 | 4 | 196958988 | 196977183 | mNeoVis.2.1.1.chr4\_hap1.g00882.t01 |
| 5917 | 4 | 197034739 | 197039489 | mNeoVis.2.1.1.chr4\_hap1.g00883.t01 |
| 5918 | 4 | 197042891 | 197063138 | mNeoVis.2.1.1.chr4\_hap1.g00884.t01 |
| 5919 | 4 | 197063339 | 197126158 | mNeoVis.2.1.1.chr4\_hap1.g00885.t01 |
| 5920 | 4 | 197258309 | 197361656 | mNeoVis.2.1.1.chr4\_hap1.g00886.t01 |
| 5921 | 4 | 197956412 | 197960905 | mNeoVis.2.1.1.chr4\_hap1.g00888.t01 |
| 5922 | 4 | 198208008 | 198266015 | mNeoVis.2.1.1.chr4\_hap1.g00889.t01 |
| 5923 | 4 | 198401330 | 198543066 | mNeoVis.2.1.1.chr4\_hap1.g00891.t01 |
| 5924 | 4 | 198547742 | 198550735 | mNeoVis.2.1.1.chr4\_hap1.g00892.t01 |
| 5925 | 4 | 198560179 | 198584227 | mNeoVis.2.1.1.chr4\_hap1.g00893.t01 |
| 5926 | 4 | 198560179 | 198584227 | mNeoVis.2.1.1.chr4\_hap1.g00893.t01 |
| 5927 | 4 | 198594513 | 198606468 | mNeoVis.2.1.1.chr4\_hap1.g00894.t01 |
| 5928 | 4 | 198594513 | 198606468 | mNeoVis.2.1.1.chr4\_hap1.g00894.t01 |
| 5929 | 4 | 198606581 | 198646019 | mNeoVis.2.1.1.chr4\_hap1.g00895.t01 |
| 5930 | 4 | 198606581 | 198646019 | mNeoVis.2.1.1.chr4\_hap1.g00895.t01 |
| 5931 | 4 | 198674247 | 198746194 | mNeoVis.2.1.1.chr4\_hap1.g00896.t01 |
| 5932 | 4 | 198674247 | 198746194 | mNeoVis.2.1.1.chr4\_hap1.g00896.t01 |
| 5933 | 4 | 198865044 | 198869364 | mNeoVis.2.1.1.chr4\_hap1.g00897.t01 |
| 5934 | 4 | 198865044 | 198869364 | mNeoVis.2.1.1.chr4\_hap1.g00897.t01 |
| 5935 | 4 | 198890451 | 199173145 | mNeoVis.2.1.1.chr4\_hap1.g00898.t01 |
| 5936 | 4 | 199289474 | 199297168 | mNeoVis.2.1.1.chr4\_hap1.g00899.t01 |
| 5937 | 4 | 199289474 | 199297168 | mNeoVis.2.1.1.chr4\_hap1.g00899.t01 |
| 5938 | 4 | 199363855 | 200147785 | mNeoVis.2.1.1.chr4\_hap1.g00900.t01 |
| 5939 | 4 | 199911630 | 199913326 | mNeoVis.2.1.1.chr4\_hap1.g00901.t01 |
| 5940 | 4 | 199911630 | 199913326 | mNeoVis.2.1.1.chr4\_hap1.g00901.t01 |
| 5941 | 4 | 200128662 | 200137326 | mNeoVis.2.1.1.chr4\_hap1.g00902.t01 |
| 5942 | 4 | 200148460 | 200170120 | mNeoVis.2.1.1.chr4\_hap1.g00903.t01 |
| 5943 | 4 | 200210824 | 200327918 | mNeoVis.2.1.1.chr4\_hap1.g00904.t01 |
| 5944 | 4 | 200343548 | 200357273 | mNeoVis.2.1.1.chr4\_hap1.g00905.t01 |
| 5945 | 4 | 200361023 | 200361934 | mNeoVis.2.1.1.chr4\_hap1.g00906.t01 |
| 5946 | 4 | 200363611 | 200366309 | mNeoVis.2.1.1.chr4\_hap1.g00907.t01 |
| 5947 | 4 | 200369049 | 200439277 | mNeoVis.2.1.1.chr4\_hap1.g00908.t01 |
| 5948 | 4 | 200455167 | 200514721 | mNeoVis.2.1.1.chr4\_hap1.g00909.t01 |
| 5949 | 4 | 200585064 | 200742652 | mNeoVis.2.1.1.chr4\_hap1.g00910.t01 |
| 5950 | 4 | 200747315 | 200812200 | mNeoVis.2.1.1.chr4\_hap1.g00911.t01 |
| 5951 | 4 | 200874104 | 200890265 | mNeoVis.2.1.1.chr4\_hap1.g00912.t01 |
| 5952 | 4 | 200923743 | 200999128 | mNeoVis.2.1.1.chr4\_hap1.g00913.t01 |
| 5953 | 4 | 201021956 | 201053978 | mNeoVis.2.1.1.chr4\_hap1.g00914.t01 |
| 5954 | 4 | 201252010 | 201259561 | mNeoVis.2.1.1.chr4\_hap1.g00915.t01 |
| 5955 | 4 | 201427858 | 201478595 | mNeoVis.2.1.1.chr4\_hap1.g00916.t01 |
| 5956 | 4 | 201469270 | 201470771 | mNeoVis.2.1.1.chr4\_hap1.g00917.t01 |
| 5957 | 4 | 201478370 | 201639108 | mNeoVis.2.1.1.chr4\_hap1.g00918.t01 |
| 5958 | 4 | 202112366 | 202153268 | mNeoVis.2.1.1.chr4\_hap1.g00920.t01 |
| 5959 | 4 | 202112366 | 202153268 | mNeoVis.2.1.1.chr4\_hap1.g00920.t01 |
| 5960 | 4 | 202189876 | 202325174 | mNeoVis.2.1.1.chr4\_hap1.g00921.t01 |
| 5961 | 4 | 202189876 | 202325174 | mNeoVis.2.1.1.chr4\_hap1.g00921.t01 |
| 5962 | 4 | 202433139 | 202441879 | mNeoVis.2.1.1.chr4\_hap1.g00922.t01 |
| 5963 | 4 | 202433139 | 202441879 | mNeoVis.2.1.1.chr4\_hap1.g00922.t01 |
| 5964 | 4 | 202515389 | 202520397 | mNeoVis.2.1.1.chr4\_hap1.g00923.t01 |
| 5965 | 4 | 202515389 | 202520397 | mNeoVis.2.1.1.chr4\_hap1.g00923.t01 |
| 5966 | 4 | 202658309 | 202690248 | mNeoVis.2.1.1.chr4\_hap1.g00924.t01 |
| 5967 | 4 | 202658309 | 202690248 | mNeoVis.2.1.1.chr4\_hap1.g00924.t01 |
| 5968 | 4 | 202658309 | 202690248 | mNeoVis.2.1.1.chr4\_hap1.g00924.t01 |
| 5969 | 4 | 202803952 | 202846598 | mNeoVis.2.1.1.chr4\_hap1.g00925.t01 |
| 5970 | 4 | 202803952 | 202846598 | mNeoVis.2.1.1.chr4\_hap1.g00925.t01 |
| 5971 | 4 | 202803952 | 202846598 | mNeoVis.2.1.1.chr4\_hap1.g00925.t01 |
| 5972 | 4 | 202895969 | 202979105 | mNeoVis.2.1.1.chr4\_hap1.g00926.t01 |
| 5973 | 4 | 202895969 | 202979105 | mNeoVis.2.1.1.chr4\_hap1.g00926.t01 |
| 5974 | 4 | 202895969 | 202979105 | mNeoVis.2.1.1.chr4\_hap1.g00926.t01 |
| 5975 | 4 | 203021566 | 203061860 | mNeoVis.2.1.1.chr4\_hap1.g00927.t01 |
| 5976 | 4 | 203021566 | 203061860 | mNeoVis.2.1.1.chr4\_hap1.g00927.t01 |
| 5977 | 4 | 203089825 | 203101103 | mNeoVis.2.1.1.chr4\_hap1.g00928.t01 |
| 5978 | 4 | 203089825 | 203101103 | mNeoVis.2.1.1.chr4\_hap1.g00928.t01 |
| 5979 | 4 | 203111059 | 203124918 | mNeoVis.2.1.1.chr4\_hap1.g00929.t01 |
| 5980 | 4 | 203111059 | 203124918 | mNeoVis.2.1.1.chr4\_hap1.g00929.t01 |
| 5981 | 4 | 203125163 | 203155644 | mNeoVis.2.1.1.chr4\_hap1.g00930.t01 |
| 5982 | 4 | 203125163 | 203155644 | mNeoVis.2.1.1.chr4\_hap1.g00930.t01 |
| 5983 | 4 | 203224166 | 203815561 | mNeoVis.2.1.1.chr4\_hap1.g00931.t01 |
| 5984 | 4 | 203224166 | 203815561 | mNeoVis.2.1.1.chr4\_hap1.g00931.t01 |
| 5985 | 4 | 203293197 | 203293960 | mNeoVis.2.1.1.chr4\_hap1.g00932.t01 |
| 5986 | 4 | 203293197 | 203293960 | mNeoVis.2.1.1.chr4\_hap1.g00932.t01 |
| 5987 | 4 | 203857669 | 203873982 | mNeoVis.2.1.1.chr4\_hap1.g00933.t01 |
| 5988 | 4 | 203857669 | 203873982 | mNeoVis.2.1.1.chr4\_hap1.g00933.t01 |
| 5989 | 4 | 203857669 | 203873982 | mNeoVis.2.1.1.chr4\_hap1.g00933.t01 |
| 5990 | 4 | 203899358 | 203977346 | mNeoVis.2.1.1.chr4\_hap1.g00934.t01 |
| 5991 | 4 | 203899358 | 203977346 | mNeoVis.2.1.1.chr4\_hap1.g00934.t01 |
| 5992 | 4 | 203899358 | 203977346 | mNeoVis.2.1.1.chr4\_hap1.g00934.t01 |
| 5993 | 4 | 203993391 | 204066695 | mNeoVis.2.1.1.chr4\_hap1.g00935.t01 |
| 5994 | 4 | 203993391 | 204066695 | mNeoVis.2.1.1.chr4\_hap1.g00935.t01 |
| 5995 | 4 | 203993391 | 204066695 | mNeoVis.2.1.1.chr4\_hap1.g00935.t01 |
| 5996 | 4 | 204650415 | 205103128 | mNeoVis.2.1.1.chr4\_hap1.g00936.t01 |
| 5997 | 4 | 204650415 | 205103128 | mNeoVis.2.1.1.chr4\_hap1.g00936.t01 |
| 5998 | 4 | 204650415 | 205103128 | mNeoVis.2.1.1.chr4\_hap1.g00936.t01 |
| 5999 | 4 | 205537318 | 205896481 | mNeoVis.2.1.1.chr4\_hap1.g00937.t01 |
| 6000 | 4 | 205537318 | 205896481 | mNeoVis.2.1.1.chr4\_hap1.g00937.t01 |
| 6001 | 4 | 205904856 | 205946406 | mNeoVis.2.1.1.chr4\_hap1.g00938.t01 |
| 6002 | 4 | 205904856 | 205946406 | mNeoVis.2.1.1.chr4\_hap1.g00938.t01 |
| 6003 | 4 | 206353013 | 206650085 | mNeoVis.2.1.1.chr4\_hap1.g00939.t01 |
| 6004 | 4 | 206353013 | 206650085 | mNeoVis.2.1.1.chr4\_hap1.g00939.t01 |
| 6005 | 4 | 206353013 | 206650085 | mNeoVis.2.1.1.chr4\_hap1.g00939.t01 |
| 6006 | 4 | 206884293 | 207036534 | mNeoVis.2.1.1.chr4\_hap1.g00940.t01 |
| 6007 | 4 | 207117498 | 207162524 | mNeoVis.2.1.1.chr4\_hap1.g00941.t01 |
| 6008 | 4 | 207117498 | 207162524 | mNeoVis.2.1.1.chr4\_hap1.g00941.t01 |
| 6009 | 4 | 207117498 | 207162524 | mNeoVis.2.1.1.chr4\_hap1.g00941.t01 |
| 6010 | 4 | 207356778 | 207399792 | mNeoVis.2.1.1.chr4\_hap1.g00942.t01 |
| 6011 | 4 | 207356778 | 207399792 | mNeoVis.2.1.1.chr4\_hap1.g00942.t01 |
| 6012 | 4 | 207356778 | 207399792 | mNeoVis.2.1.1.chr4\_hap1.g00942.t01 |
| 6013 | 4 | 207356778 | 207399792 | mNeoVis.2.1.1.chr4\_hap1.g00942.t01 |
| 6014 | 4 | 207740651 | 207744291 | mNeoVis.2.1.1.chr4\_hap1.g00943.t01 |
| 6015 | 4 | 207740651 | 207744291 | mNeoVis.2.1.1.chr4\_hap1.g00943.t01 |
| 6016 | 4 | 207740651 | 207744291 | mNeoVis.2.1.1.chr4\_hap1.g00943.t01 |
| 6017 | 4 | 207740651 | 207744291 | mNeoVis.2.1.1.chr4\_hap1.g00943.t01 |
| 6018 | 4 | 207740651 | 207744291 | mNeoVis.2.1.1.chr4\_hap1.g00943.t01 |
| 6019 | 4 | 207985206 | 208021831 | mNeoVis.2.1.1.chr4\_hap1.g00944.t01 |
| 6020 | 4 | 207985206 | 208021831 | mNeoVis.2.1.1.chr4\_hap1.g00944.t01 |
| 6021 | 4 | 207985206 | 208021831 | mNeoVis.2.1.1.chr4\_hap1.g00944.t01 |
| 6022 | 4 | 207985206 | 208021831 | mNeoVis.2.1.1.chr4\_hap1.g00944.t01 |
| 6023 | 4 | 208138268 | 208218939 | mNeoVis.2.1.1.chr4\_hap1.g00945.t01 |
| 6024 | 4 | 208138268 | 208218939 | mNeoVis.2.1.1.chr4\_hap1.g00945.t01 |
| 6025 | 4 | 208138268 | 208218939 | mNeoVis.2.1.1.chr4\_hap1.g00945.t01 |
| 6026 | 4 | 208138268 | 208218939 | mNeoVis.2.1.1.chr4\_hap1.g00945.t01 |
| 6027 | 4 | 208219086 | 208229142 | mNeoVis.2.1.1.chr4\_hap1.g00946.t01 |
| 6028 | 4 | 208219086 | 208229142 | mNeoVis.2.1.1.chr4\_hap1.g00946.t01 |
| 6029 | 4 | 208219086 | 208229142 | mNeoVis.2.1.1.chr4\_hap1.g00946.t01 |
| 6030 | 4 | 208246155 | 208402699 | mNeoVis.2.1.1.chr4\_hap1.g00947.t01 |
| 6031 | 4 | 208246155 | 208402699 | mNeoVis.2.1.1.chr4\_hap1.g00947.t01 |
| 6032 | 4 | 208246155 | 208402699 | mNeoVis.2.1.1.chr4\_hap1.g00947.t01 |
| 6033 | 4 | 208410379 | 208429293 | mNeoVis.2.1.1.chr4\_hap1.g00948.t01 |
| 6034 | 4 | 208410379 | 208429293 | mNeoVis.2.1.1.chr4\_hap1.g00948.t01 |
| 6035 | 4 | 208410379 | 208429293 | mNeoVis.2.1.1.chr4\_hap1.g00948.t01 |
| 6036 | 4 | 208410379 | 208429293 | mNeoVis.2.1.1.chr4\_hap1.g00948.t01 |
| 6037 | 4 | 208438807 | 208513270 | mNeoVis.2.1.1.chr4\_hap1.g00949.t01 |
| 6038 | 4 | 208438807 | 208513270 | mNeoVis.2.1.1.chr4\_hap1.g00949.t01 |
| 6039 | 4 | 208438807 | 208513270 | mNeoVis.2.1.1.chr4\_hap1.g00949.t01 |
| 6040 | 4 | 208438807 | 208513270 | mNeoVis.2.1.1.chr4\_hap1.g00949.t01 |
| 6041 | 4 | 208758379 | 208764809 | mNeoVis.2.1.1.chr4\_hap1.g00950.t01 |
| 6042 | 4 | 208758379 | 208764809 | mNeoVis.2.1.1.chr4\_hap1.g00950.t01 |
| 6043 | 4 | 208846064 | 208847777 | mNeoVis.2.1.1.chr4\_hap1.g00951.t01 |
| 6044 | 4 | 208846064 | 208847777 | mNeoVis.2.1.1.chr4\_hap1.g00951.t01 |
| 6045 | 4 | 209174718 | 209232412 | mNeoVis.2.1.1.chr4\_hap1.g00952.t01 |
| 6046 | 4 | 209174718 | 209232412 | mNeoVis.2.1.1.chr4\_hap1.g00952.t01 |
| 6047 | 4 | 210498067 | 210499489 | mNeoVis.2.1.1.chr4\_hap1.g00953.t01 |
| 6048 | 4 | 210498067 | 210499489 | mNeoVis.2.1.1.chr4\_hap1.g00953.t01 |
| 6049 | 4 | 210498067 | 210499489 | mNeoVis.2.1.1.chr4\_hap1.g00953.t01 |
| 6050 | 4 | 213533887 | 213783887 | mNeoVis.2.1.1.chr4\_hap1.g00956.t01 |
| 6051 | 4 | 213533887 | 213783887 | mNeoVis.2.1.1.chr4\_hap1.g00956.t01 |
| 6052 | 4 | 213761782 | 213763011 | mNeoVis.2.1.1.chr4\_hap1.g00957.t01 |
| 6053 | 4 | 213761782 | 213763011 | mNeoVis.2.1.1.chr4\_hap1.g00957.t01 |
| 6054 | 4 | 214463706 | 214562792 | mNeoVis.2.1.1.chr4\_hap1.g00958.t01 |
| 6055 | 4 | 214463706 | 214562792 | mNeoVis.2.1.1.chr4\_hap1.g00958.t01 |
| 6056 | 4 | 214615549 | 214719307 | mNeoVis.2.1.1.chr4\_hap1.g00959.t01 |
| 6057 | 4 | 214728498 | 214738763 | mNeoVis.2.1.1.chr4\_hap1.g00960.t01 |
| 6058 | 4 | 215447892 | 215469018 | mNeoVis.2.1.1.chr4\_hap1.g00965.t01 |
| 6059 | 4 | 215713337 | 215995542 | mNeoVis.2.1.1.chr4\_hap1.g00966.t01 |
| 6060 | 4 | 216116915 | 216446893 | mNeoVis.2.1.1.chr4\_hap1.g00967.t01 |
| 6061 | 4 | 216713932 | 216742651 | mNeoVis.2.1.1.chr4\_hap1.g00968.t01 |
| 6062 | 4 | 216713932 | 216742651 | mNeoVis.2.1.1.chr4\_hap1.g00968.t01 |
| 6063 | 4 | 216861619 | 216897284 | mNeoVis.2.1.1.chr4\_hap1.g00969.t01 |
| 6064 | 4 | 216861619 | 216897284 | mNeoVis.2.1.1.chr4\_hap1.g00969.t01 |
| 6065 | 4 | 216873464 | 216929867 | mNeoVis.2.1.1.chr4\_hap1.g00970.t01 |
| 6066 | 4 | 216873464 | 216929867 | mNeoVis.2.1.1.chr4\_hap1.g00970.t01 |
| 6067 | 4 | 216974882 | 217000565 | mNeoVis.2.1.1.chr4\_hap1.g00971.t01 |
| 6068 | 4 | 217087120 | 217094569 | mNeoVis.2.1.1.chr4\_hap1.g00972.t01 |
| 6069 | 4 | 217087120 | 217094569 | mNeoVis.2.1.1.chr4\_hap1.g00972.t01 |
| 6070 | 4 | 217165845 | 217273335 | mNeoVis.2.1.1.chr4\_hap1.g00973.t01 |
| 6071 | 4 | 217165845 | 217273335 | mNeoVis.2.1.1.chr4\_hap1.g00973.t01 |
| 6072 | 4 | 217317967 | 217416638 | mNeoVis.2.1.1.chr4\_hap1.g00974.t01 |
| 6073 | 4 | 217417024 | 217432654 | mNeoVis.2.1.1.chr4\_hap1.g00975.t01 |
| 6074 | 4 | 217417024 | 217432654 | mNeoVis.2.1.1.chr4\_hap1.g00975.t01 |
| 6075 | 4 | 217443624 | 217580081 | mNeoVis.2.1.1.chr4\_hap1.g00976.t01 |
| 6076 | 4 | 217443624 | 217580081 | mNeoVis.2.1.1.chr4\_hap1.g00976.t01 |
| 6077 | 4 | 217623531 | 217845013 | mNeoVis.2.1.1.chr4\_hap1.g00977.t01 |
| 6078 | 4 | 217623531 | 217845013 | mNeoVis.2.1.1.chr4\_hap1.g00977.t01 |
| 6079 | 4 | 217645272 | 217683272 | mNeoVis.2.1.1.chr4\_hap1.g00978.t01 |
| 6080 | 4 | 217645272 | 217683272 | mNeoVis.2.1.1.chr4\_hap1.g00978.t01 |
| 6081 | 4 | 217860414 | 217909364 | mNeoVis.2.1.1.chr4\_hap1.g00979.t01 |
| 6082 | 4 | 217860414 | 217909364 | mNeoVis.2.1.1.chr4\_hap1.g00979.t01 |
| 6083 | 4 | 217909568 | 217983947 | mNeoVis.2.1.1.chr4\_hap1.g00980.t01 |
| 6084 | 4 | 218002432 | 218233377 | mNeoVis.2.1.1.chr4\_hap1.g00981.t01 |
| 6085 | 4 | 218937692 | 218970664 | mNeoVis.2.1.1.chr4\_hap1.g00983.t01 |
| 6086 | 4 | 218937692 | 218970664 | mNeoVis.2.1.1.chr4\_hap1.g00983.t01 |
| 6087 | 4 | 219902709 | 220048570 | mNeoVis.2.1.1.chr4\_hap1.g00984.t01 |
| 6088 | 4 | 220083777 | 220112320 | mNeoVis.2.1.1.chr4\_hap1.g00985.t01 |
| 6089 | 4 | 220132385 | 220201460 | mNeoVis.2.1.1.chr4\_hap1.g00986.t01 |
| 6090 | 4 | 220132385 | 220201460 | mNeoVis.2.1.1.chr4\_hap1.g00986.t01 |
| 6091 | 4 | 220227717 | 220247616 | mNeoVis.2.1.1.chr4\_hap1.g00987.t01 |
| 6092 | 4 | 220227717 | 220247616 | mNeoVis.2.1.1.chr4\_hap1.g00987.t01 |
| 6093 | 4 | 220255006 | 220275209 | mNeoVis.2.1.1.chr4\_hap1.g00988.t01 |
| 6094 | 4 | 220255006 | 220275209 | mNeoVis.2.1.1.chr4\_hap1.g00988.t01 |
| 6095 | 4 | 220279920 | 220280479 | mNeoVis.2.1.1.chr4\_hap1.g00989.t01 |
| 6096 | 4 | 220279920 | 220280479 | mNeoVis.2.1.1.chr4\_hap1.g00989.t01 |
| 6097 | 4 | 220285727 | 220309514 | mNeoVis.2.1.1.chr4\_hap1.g00990.t01 |
| 6098 | 4 | 220285727 | 220309514 | mNeoVis.2.1.1.chr4\_hap1.g00990.t01 |
| 6099 | 4 | 220323295 | 220391304 | mNeoVis.2.1.1.chr4\_hap1.g00991.t01 |
| 6100 | 4 | 220323295 | 220391304 | mNeoVis.2.1.1.chr4\_hap1.g00991.t01 |
| 6101 | 4 | 220400858 | 220419588 | mNeoVis.2.1.1.chr4\_hap1.g00992.t01 |
| 6102 | 4 | 220400858 | 220419588 | mNeoVis.2.1.1.chr4\_hap1.g00992.t01 |
| 6103 | 4 | 220470035 | 220499870 | mNeoVis.2.1.1.chr4\_hap1.g00993.t01 |
| 6104 | 4 | 220470035 | 220499870 | mNeoVis.2.1.1.chr4\_hap1.g00993.t01 |
| 6105 | 4 | 220583719 | 220594527 | mNeoVis.2.1.1.chr4\_hap1.g00994.t01 |
| 6106 | 4 | 220583719 | 220594527 | mNeoVis.2.1.1.chr4\_hap1.g00994.t01 |
| 6107 | 4 | 220619030 | 220678119 | mNeoVis.2.1.1.chr4\_hap1.g00995.t01 |
| 6108 | 4 | 220702902 | 220746751 | mNeoVis.2.1.1.chr4\_hap1.g00996.t01 |
| 6109 | 4 | 220824708 | 220948077 | mNeoVis.2.1.1.chr4\_hap1.g00997.t01 |
| 6110 | 4 | 221097159 | 221161558 | mNeoVis.2.1.1.chr4\_hap1.g00998.t01 |
| 6111 | 4 | 221208967 | 221217609 | mNeoVis.2.1.1.chr4\_hap1.g00999.t01 |
| 6112 | 4 | 221227026 | 221239673 | mNeoVis.2.1.1.chr4\_hap1.g01000.t01 |
| 6113 | 4 | 221240188 | 221281062 | mNeoVis.2.1.1.chr4\_hap1.g01001.t01 |
| 6114 | 4 | 221281030 | 221288483 | mNeoVis.2.1.1.chr4\_hap1.g01002.t01 |
| 6115 | 4 | 221667541 | 221727061 | mNeoVis.2.1.1.chr4\_hap1.g01007.t01 |
| 6116 | 4 | 221727100 | 221940486 | mNeoVis.2.1.1.chr4\_hap1.g01008.t01 |
| 6117 | 4 | 221987561 | 221995078 | mNeoVis.2.1.1.chr4\_hap1.g01009.t01 |
| 6118 | 4 | 222520223 | 222520645 | mNeoVis.2.1.1.chr4\_hap1.g01010.t01 |
| 6119 | 4 | 222890606 | 222896469 | mNeoVis.2.1.1.chr4\_hap1.g01011.t01 |
| 6120 | 4 | 222890606 | 222896469 | mNeoVis.2.1.1.chr4\_hap1.g01011.t01 |
| 6121 | 4 | 225337828 | 225539616 | mNeoVis.2.1.1.chr4\_hap1.g01014.t01 |
| 6122 | 4 | 225683624 | 225720283 | mNeoVis.2.1.1.chr4\_hap1.g01015.t01 |
| 6123 | 4 | 225744570 | 225894546 | mNeoVis.2.1.1.chr4\_hap1.g01016.t01 |
| 6124 | 4 | 227372140 | 227489356 | mNeoVis.2.1.1.chr4\_hap1.g01019.t01 |
| 6125 | 4 | 227493937 | 227524074 | mNeoVis.2.1.1.chr4\_hap1.g01020.t01 |
| 6126 | 4 | 227531797 | 227565791 | mNeoVis.2.1.1.chr4\_hap1.g01021.t01 |
| 6127 | 4 | 233380027 | 233464519 | mNeoVis.2.1.1.chr4\_hap1.g01038.t01 |
| 6128 | 4 | 233491714 | 233971847 | mNeoVis.2.1.1.chr4\_hap1.g01039.t01 |
| 6129 | 4 | 233836222 | 233837348 | mNeoVis.2.1.1.chr4\_hap1.g01040.t01 |
| 6130 | 4 | 233836222 | 233837348 | mNeoVis.2.1.1.chr4\_hap1.g01040.t01 |
| 6131 | 4 | 233836222 | 233837348 | mNeoVis.2.1.1.chr4\_hap1.g01040.t01 |
| 6132 | 4 | 233978186 | 233984228 | mNeoVis.2.1.1.chr4\_hap1.g01041.t01 |
| 6133 | 4 | 233978186 | 233984228 | mNeoVis.2.1.1.chr4\_hap1.g01041.t01 |
| 6134 | 4 | 233978186 | 233984228 | mNeoVis.2.1.1.chr4\_hap1.g01041.t01 |
| 6135 | 4 | 234011609 | 234014780 | mNeoVis.2.1.1.chr4\_hap1.g01042.t01 |
| 6136 | 4 | 234011609 | 234014780 | mNeoVis.2.1.1.chr4\_hap1.g01042.t01 |
| 6137 | 4 | 234011609 | 234014780 | mNeoVis.2.1.1.chr4\_hap1.g01042.t01 |
| 6138 | 4 | 234019324 | 234125283 | mNeoVis.2.1.1.chr4\_hap1.g01043.t01 |
| 6139 | 4 | 234019324 | 234125283 | mNeoVis.2.1.1.chr4\_hap1.g01043.t01 |
| 6140 | 4 | 234019324 | 234125283 | mNeoVis.2.1.1.chr4\_hap1.g01043.t01 |
| 6141 | 4 | 234148448 | 234394261 | mNeoVis.2.1.1.chr4\_hap1.g01044.t01 |
| 6142 | 4 | 234470935 | 234521381 | mNeoVis.2.1.1.chr4\_hap1.g01045.t01 |
| 6143 | 4 | 234533727 | 234602623 | mNeoVis.2.1.1.chr4\_hap1.g01046.t01 |
| 6144 | 4 | 234637872 | 234638945 | mNeoVis.2.1.1.chr4\_hap1.g01047.t01 |
| 6145 | 4 | 234665926 | 234697623 | mNeoVis.2.1.1.chr4\_hap1.g01048.t01 |
| 6146 | 4 | 234699684 | 234767479 | mNeoVis.2.1.1.chr4\_hap1.g01049.t01 |
| 6147 | 4 | 234775689 | 234779447 | mNeoVis.2.1.1.chr4\_hap1.g01050.t01 |
| 6148 | 5 | 418477 | 442332 | mNeoVis.2.1.1.chr5\_hap1.g00028.t01 |
| 6149 | 5 | 445900 | 446884 | mNeoVis.2.1.1.chr5\_hap1.g00029.t01 |
| 6150 | 5 | 450880 | 474641 | mNeoVis.2.1.1.chr5\_hap1.g00030.t01 |
| 6151 | 5 | 474571 | 479166 | mNeoVis.2.1.1.chr5\_hap1.g00031.t01 |
| 6152 | 5 | 479742 | 486802 | mNeoVis.2.1.1.chr5\_hap1.g00032.t01 |
| 6153 | 5 | 488389 | 492234 | mNeoVis.2.1.1.chr5\_hap1.g00033.t01 |
| 6154 | 5 | 497054 | 504150 | mNeoVis.2.1.1.chr5\_hap1.g00034.t01 |
| 6155 | 5 | 512305 | 518649 | mNeoVis.2.1.1.chr5\_hap1.g00035.t01 |
| 6156 | 5 | 531290 | 540215 | mNeoVis.2.1.1.chr5\_hap1.g00036.t01 |
| 6157 | 5 | 548578 | 558827 | mNeoVis.2.1.1.chr5\_hap1.g00037.t01 |
| 6158 | 5 | 564206 | 566552 | mNeoVis.2.1.1.chr5\_hap1.g00038.t01 |
| 6159 | 5 | 572449 | 603014 | mNeoVis.2.1.1.chr5\_hap1.g00039.t01 |
| 6160 | 5 | 674957 | 691725 | mNeoVis.2.1.1.chr5\_hap1.g00040.t01 |
| 6161 | 5 | 695840 | 713417 | mNeoVis.2.1.1.chr5\_hap1.g00041.t01 |
| 6162 | 5 | 715990 | 739216 | mNeoVis.2.1.1.chr5\_hap1.g00042.t01 |
| 6163 | 5 | 744259 | 755140 | mNeoVis.2.1.1.chr5\_hap1.g00043.t01 |
| 6164 | 5 | 765648 | 774108 | mNeoVis.2.1.1.chr5\_hap1.g00044.t01 |
| 6165 | 5 | 788669 | 793431 | mNeoVis.2.1.1.chr5\_hap1.g00045.t01 |
| 6166 | 5 | 793229 | 812942 | mNeoVis.2.1.1.chr5\_hap1.g00046.t01 |
| 6167 | 5 | 816773 | 840386 | mNeoVis.2.1.1.chr5\_hap1.g00047.t01 |
| 6168 | 5 | 843202 | 863639 | mNeoVis.2.1.1.chr5\_hap1.g00048.t01 |
| 6169 | 5 | 876495 | 887487 | mNeoVis.2.1.1.chr5\_hap1.g00049.t01 |
| 6170 | 5 | 903336 | 916796 | mNeoVis.2.1.1.chr5\_hap1.g00050.t01 |
| 6171 | 5 | 917054 | 919580 | mNeoVis.2.1.1.chr5\_hap1.g00051.t01 |
| 6172 | 5 | 923349 | 957697 | mNeoVis.2.1.1.chr5\_hap1.g00052.t01 |
| 6173 | 5 | 982404 | 1040634 | mNeoVis.2.1.1.chr5\_hap1.g00053.t01 |
| 6174 | 5 | 1040777 | 1045030 | mNeoVis.2.1.1.chr5\_hap1.g00054.t01 |
| 6175 | 5 | 1045189 | 1055408 | mNeoVis.2.1.1.chr5\_hap1.g00055.t01 |
| 6176 | 5 | 1056000 | 1058948 | mNeoVis.2.1.1.chr5\_hap1.g00056.t01 |
| 6177 | 5 | 1059144 | 1062845 | mNeoVis.2.1.1.chr5\_hap1.g00057.t01 |
| 6178 | 5 | 1070439 | 1072341 | mNeoVis.2.1.1.chr5\_hap1.g00058.t01 |
| 6179 | 5 | 1074517 | 1077822 | mNeoVis.2.1.1.chr5\_hap1.g00059.t01 |
| 6180 | 5 | 1079729 | 1085650 | mNeoVis.2.1.1.chr5\_hap1.g00060.t01 |
| 6181 | 5 | 3472555 | 3474080 | mNeoVis.2.1.1.chr5\_hap1.g00140.t01 |
| 6182 | 5 | 3537155 | 3542505 | mNeoVis.2.1.1.chr5\_hap1.g00141.t01 |
| 6183 | 5 | 3556801 | 3590352 | mNeoVis.2.1.1.chr5\_hap1.g00142.t01 |
| 6184 | 5 | 3589457 | 3626325 | mNeoVis.2.1.1.chr5\_hap1.g00143.t01 |
| 6185 | 5 | 3629829 | 3632077 | mNeoVis.2.1.1.chr5\_hap1.g00144.t01 |
| 6186 | 5 | 3632220 | 3636796 | mNeoVis.2.1.1.chr5\_hap1.g00145.t01 |
| 6187 | 5 | 3639150 | 3640186 | mNeoVis.2.1.1.chr5\_hap1.g00146.t01 |
| 6188 | 5 | 3640275 | 3660373 | mNeoVis.2.1.1.chr5\_hap1.g00147.t01 |
| 6189 | 5 | 3661031 | 3708598 | mNeoVis.2.1.1.chr5\_hap1.g00148.t01 |
| 6190 | 5 | 3710302 | 3712233 | mNeoVis.2.1.1.chr5\_hap1.g00149.t01 |
| 6191 | 5 | 3712576 | 3718114 | mNeoVis.2.1.1.chr5\_hap1.g00150.t01 |
| 6192 | 5 | 3723497 | 3737437 | mNeoVis.2.1.1.chr5\_hap1.g00151.t01 |
| 6193 | 5 | 3746731 | 3750054 | mNeoVis.2.1.1.chr5\_hap1.g00152.t01 |
| 6194 | 5 | 3773496 | 3779158 | mNeoVis.2.1.1.chr5\_hap1.g00153.t01 |
| 6195 | 5 | 3780234 | 3783631 | mNeoVis.2.1.1.chr5\_hap1.g00154.t01 |
| 6196 | 5 | 3785107 | 3786562 | mNeoVis.2.1.1.chr5\_hap1.g00155.t01 |
| 6197 | 5 | 3786643 | 3840226 | mNeoVis.2.1.1.chr5\_hap1.g00156.t01 |
| 6198 | 5 | 3852116 | 3941809 | mNeoVis.2.1.1.chr5\_hap1.g00157.t01 |
| 6199 | 5 | 3959036 | 3991626 | mNeoVis.2.1.1.chr5\_hap1.g00158.t01 |
| 6200 | 5 | 3996422 | 4014820 | mNeoVis.2.1.1.chr5\_hap1.g00159.t01 |
| 6201 | 5 | 4017979 | 4073739 | mNeoVis.2.1.1.chr5\_hap1.g00160.t01 |
| 6202 | 5 | 4091237 | 4108927 | mNeoVis.2.1.1.chr5\_hap1.g00161.t01 |
| 6203 | 5 | 4131855 | 4137289 | mNeoVis.2.1.1.chr5\_hap1.g00162.t01 |
| 6204 | 5 | 4139507 | 4141171 | mNeoVis.2.1.1.chr5\_hap1.g00163.t01 |
| 6205 | 5 | 4141438 | 4142319 | mNeoVis.2.1.1.chr5\_hap1.g00164.t01 |
| 6206 | 5 | 4143663 | 4153055 | mNeoVis.2.1.1.chr5\_hap1.g00165.t01 |
| 6207 | 5 | 4153950 | 4167959 | mNeoVis.2.1.1.chr5\_hap1.g00166.t01 |
| 6208 | 5 | 4167176 | 4178471 | mNeoVis.2.1.1.chr5\_hap1.g00167.t01 |
| 6209 | 5 | 4178697 | 4200242 | mNeoVis.2.1.1.chr5\_hap1.g00168.t01 |
| 6210 | 5 | 4202738 | 4205266 | mNeoVis.2.1.1.chr5\_hap1.g00169.t01 |
| 6211 | 5 | 4205269 | 4213514 | mNeoVis.2.1.1.chr5\_hap1.g00170.t01 |
| 6212 | 5 | 4218037 | 4222977 | mNeoVis.2.1.1.chr5\_hap1.g00171.t01 |
| 6213 | 5 | 4238044 | 4241495 | mNeoVis.2.1.1.chr5\_hap1.g00172.t01 |
| 6214 | 5 | 4278804 | 4281911 | mNeoVis.2.1.1.chr5\_hap1.g00173.t01 |
| 6215 | 5 | 4314495 | 4318138 | mNeoVis.2.1.1.chr5\_hap1.g00174.t01 |
| 6216 | 5 | 4322735 | 4355695 | mNeoVis.2.1.1.chr5\_hap1.g00175.t01 |
| 6217 | 5 | 4361428 | 4367558 | mNeoVis.2.1.1.chr5\_hap1.g00176.t01 |
| 6218 | 5 | 4369658 | 4378760 | mNeoVis.2.1.1.chr5\_hap1.g00177.t01 |
| 6219 | 5 | 4380151 | 4389469 | mNeoVis.2.1.1.chr5\_hap1.g00178.t01 |
| 6220 | 5 | 4391701 | 4440269 | mNeoVis.2.1.1.chr5\_hap1.g00179.t01 |
| 6221 | 5 | 6981107 | 6987722 | mNeoVis.2.1.1.chr5\_hap1.g00266.t01 |
| 6222 | 5 | 6997777 | 7024403 | mNeoVis.2.1.1.chr5\_hap1.g00267.t01 |
| 6223 | 5 | 7049428 | 7053943 | mNeoVis.2.1.1.chr5\_hap1.g00268.t01 |
| 6224 | 5 | 7054088 | 7057853 | mNeoVis.2.1.1.chr5\_hap1.g00269.t01 |
| 6225 | 5 | 7062701 | 7067069 | mNeoVis.2.1.1.chr5\_hap1.g00270.t01 |
| 6226 | 5 | 7067019 | 7069909 | mNeoVis.2.1.1.chr5\_hap1.g00271.t01 |
| 6227 | 5 | 7070012 | 7074285 | mNeoVis.2.1.1.chr5\_hap1.g00272.t01 |
| 6228 | 5 | 7088614 | 7128088 | mNeoVis.2.1.1.chr5\_hap1.g00273.t01 |
| 6229 | 5 | 7152083 | 7160508 | mNeoVis.2.1.1.chr5\_hap1.g00274.t01 |
| 6230 | 5 | 7172359 | 7179377 | mNeoVis.2.1.1.chr5\_hap1.g00275.t01 |
| 6231 | 5 | 7187616 | 7196463 | mNeoVis.2.1.1.chr5\_hap1.g00276.t01 |
| 6232 | 5 | 7196617 | 7197854 | mNeoVis.2.1.1.chr5\_hap1.g00277.t01 |
| 6233 | 5 | 7199386 | 7206086 | mNeoVis.2.1.1.chr5\_hap1.g00278.t01 |
| 6234 | 5 | 7208158 | 7211458 | mNeoVis.2.1.1.chr5\_hap1.g00279.t01 |
| 6235 | 5 | 7211955 | 7216383 | mNeoVis.2.1.1.chr5\_hap1.g00280.t01 |
| 6236 | 5 | 7216429 | 7230805 | mNeoVis.2.1.1.chr5\_hap1.g00281.t01 |
| 6237 | 5 | 7230803 | 7239939 | mNeoVis.2.1.1.chr5\_hap1.g00282.t01 |
| 6238 | 5 | 7240949 | 7259820 | mNeoVis.2.1.1.chr5\_hap1.g00283.t01 |
| 6239 | 5 | 7267063 | 7277670 | mNeoVis.2.1.1.chr5\_hap1.g00284.t01 |
| 6240 | 5 | 7288666 | 7329181 | mNeoVis.2.1.1.chr5\_hap1.g00285.t01 |
| 6241 | 5 | 7342327 | 7350605 | mNeoVis.2.1.1.chr5\_hap1.g00286.t01 |
| 6242 | 5 | 7355724 | 7360145 | mNeoVis.2.1.1.chr5\_hap1.g00287.t01 |
| 6243 | 5 | 7379655 | 7386098 | mNeoVis.2.1.1.chr5\_hap1.g00288.t01 |
| 6244 | 5 | 7385741 | 7394208 | mNeoVis.2.1.1.chr5\_hap1.g00289.t01 |
| 6245 | 5 | 7394633 | 7397218 | mNeoVis.2.1.1.chr5\_hap1.g00290.t01 |
| 6246 | 5 | 7398720 | 7408544 | mNeoVis.2.1.1.chr5\_hap1.g00291.t01 |
| 6247 | 5 | 7415073 | 7446090 | mNeoVis.2.1.1.chr5\_hap1.g00292.t01 |
| 6248 | 5 | 7446600 | 7449771 | mNeoVis.2.1.1.chr5\_hap1.g00293.t01 |
| 6249 | 5 | 7455430 | 7469640 | mNeoVis.2.1.1.chr5\_hap1.g00294.t01 |
| 6250 | 5 | 11937948 | 11940522 | mNeoVis.2.1.1.chr5\_hap1.g00457.t01 |
| 6251 | 5 | 11956027 | 12054054 | mNeoVis.2.1.1.chr5\_hap1.g00458.t01 |
| 6252 | 5 | 12057602 | 12081154 | mNeoVis.2.1.1.chr5\_hap1.g00459.t01 |
| 6253 | 5 | 12081234 | 12098715 | mNeoVis.2.1.1.chr5\_hap1.g00460.t01 |
| 6254 | 5 | 12097641 | 12115975 | mNeoVis.2.1.1.chr5\_hap1.g00461.t01 |
| 6255 | 5 | 12120466 | 12137503 | mNeoVis.2.1.1.chr5\_hap1.g00462.t01 |
| 6256 | 5 | 12139784 | 12191403 | mNeoVis.2.1.1.chr5\_hap1.g00463.t01 |
| 6257 | 5 | 12206216 | 12217195 | mNeoVis.2.1.1.chr5\_hap1.g00464.t01 |
| 6258 | 5 | 12218235 | 12235274 | mNeoVis.2.1.1.chr5\_hap1.g00465.t01 |
| 6259 | 5 | 12253476 | 12316978 | mNeoVis.2.1.1.chr5\_hap1.g00466.t01 |
| 6260 | 5 | 12325049 | 12359203 | mNeoVis.2.1.1.chr5\_hap1.g00467.t01 |
| 6261 | 5 | 12363572 | 12365617 | mNeoVis.2.1.1.chr5\_hap1.g00468.t01 |
| 6262 | 5 | 12368244 | 12438564 | mNeoVis.2.1.1.chr5\_hap1.g00469.t01 |
| 6263 | 5 | 12471544 | 12489704 | mNeoVis.2.1.1.chr5\_hap1.g00470.t01 |
| 6264 | 5 | 12489838 | 12575050 | mNeoVis.2.1.1.chr5\_hap1.g00471.t01 |
| 6265 | 5 | 12576039 | 12579800 | mNeoVis.2.1.1.chr5\_hap1.g00472.t01 |
| 6266 | 5 | 12580095 | 12583959 | mNeoVis.2.1.1.chr5\_hap1.g00473.t01 |
| 6267 | 5 | 12586444 | 12596649 | mNeoVis.2.1.1.chr5\_hap1.g00474.t01 |
| 6268 | 5 | 12600016 | 12608689 | mNeoVis.2.1.1.chr5\_hap1.g00475.t01 |
| 6269 | 5 | 12610397 | 12618332 | mNeoVis.2.1.1.chr5\_hap1.g00476.t01 |
| 6270 | 5 | 12618472 | 12628843 | mNeoVis.2.1.1.chr5\_hap1.g00477.t01 |
| 6271 | 5 | 12625588 | 12631370 | mNeoVis.2.1.1.chr5\_hap1.g00478.t01 |
| 6272 | 5 | 12632326 | 12634996 | mNeoVis.2.1.1.chr5\_hap1.g00479.t01 |
| 6273 | 5 | 12636981 | 12643502 | mNeoVis.2.1.1.chr5\_hap1.g00480.t01 |
| 6274 | 5 | 12645550 | 12655905 | mNeoVis.2.1.1.chr5\_hap1.g00481.t01 |
| 6275 | 5 | 12656477 | 12661007 | mNeoVis.2.1.1.chr5\_hap1.g00482.t01 |
| 6276 | 5 | 12665894 | 12681816 | mNeoVis.2.1.1.chr5\_hap1.g00483.t01 |
| 6277 | 5 | 12691138 | 12693795 | mNeoVis.2.1.1.chr5\_hap1.g00484.t01 |
| 6278 | 5 | 12697333 | 12706707 | mNeoVis.2.1.1.chr5\_hap1.g00485.t01 |
| 6279 | 5 | 12718091 | 12721785 | mNeoVis.2.1.1.chr5\_hap1.g00486.t01 |
| 6280 | 5 | 12729677 | 12736209 | mNeoVis.2.1.1.chr5\_hap1.g00487.t01 |
| 6281 | 5 | 12737933 | 12741398 | mNeoVis.2.1.1.chr5\_hap1.g00488.t01 |
| 6282 | 5 | 12749116 | 12783748 | mNeoVis.2.1.1.chr5\_hap1.g00489.t01 |
| 6283 | 5 | 12790475 | 12798088 | mNeoVis.2.1.1.chr5\_hap1.g00490.t01 |
| 6284 | 5 | 12807090 | 12817980 | mNeoVis.2.1.1.chr5\_hap1.g00491.t01 |
| 6285 | 5 | 13582110 | 13589599 | mNeoVis.2.1.1.chr5\_hap1.g00523.t01 |
| 6286 | 5 | 13590623 | 13594615 | mNeoVis.2.1.1.chr5\_hap1.g00524.t01 |
| 6287 | 5 | 13595808 | 13598640 | mNeoVis.2.1.1.chr5\_hap1.g00525.t01 |
| 6288 | 5 | 13599096 | 13610205 | mNeoVis.2.1.1.chr5\_hap1.g00526.t01 |
| 6289 | 5 | 13616518 | 13621806 | mNeoVis.2.1.1.chr5\_hap1.g00527.t01 |
| 6290 | 5 | 13636997 | 13665410 | mNeoVis.2.1.1.chr5\_hap1.g00528.t01 |
| 6291 | 5 | 13674715 | 13749094 | mNeoVis.2.1.1.chr5\_hap1.g00529.t01 |
| 6292 | 5 | 13754366 | 13762051 | mNeoVis.2.1.1.chr5\_hap1.g00530.t01 |
| 6293 | 5 | 13795796 | 13834006 | mNeoVis.2.1.1.chr5\_hap1.g00531.t01 |
| 6294 | 5 | 13834113 | 13836171 | mNeoVis.2.1.1.chr5\_hap1.g00532.t01 |
| 6295 | 5 | 13836421 | 13853325 | mNeoVis.2.1.1.chr5\_hap1.g00533.t01 |
| 6296 | 5 | 13856509 | 13871234 | mNeoVis.2.1.1.chr5\_hap1.g00534.t01 |
| 6297 | 5 | 13871462 | 13879248 | mNeoVis.2.1.1.chr5\_hap1.g00535.t01 |
| 6298 | 5 | 13879877 | 13887426 | mNeoVis.2.1.1.chr5\_hap1.g00536.t01 |
| 6299 | 5 | 13932733 | 13963015 | mNeoVis.2.1.1.chr5\_hap1.g00537.t01 |
| 6300 | 5 | 13963316 | 13991514 | mNeoVis.2.1.1.chr5\_hap1.g00538.t01 |
| 6301 | 5 | 13995305 | 14012217 | mNeoVis.2.1.1.chr5\_hap1.g00539.t01 |
| 6302 | 5 | 14017341 | 14033994 | mNeoVis.2.1.1.chr5\_hap1.g00540.t01 |
| 6303 | 5 | 14038222 | 14056064 | mNeoVis.2.1.1.chr5\_hap1.g00541.t01 |
| 6304 | 5 | 14063441 | 14072074 | mNeoVis.2.1.1.chr5\_hap1.g00542.t01 |
| 6305 | 5 | 14072204 | 14078123 | mNeoVis.2.1.1.chr5\_hap1.g00543.t01 |
| 6306 | 5 | 14078169 | 14079635 | mNeoVis.2.1.1.chr5\_hap1.g00544.t01 |
| 6307 | 5 | 14084988 | 14108899 | mNeoVis.2.1.1.chr5\_hap1.g00545.t01 |
| 6308 | 5 | 14115292 | 14122086 | mNeoVis.2.1.1.chr5\_hap1.g00546.t01 |
| 6309 | 5 | 14123014 | 14129818 | mNeoVis.2.1.1.chr5\_hap1.g00547.t01 |
| 6310 | 5 | 14132671 | 14164705 | mNeoVis.2.1.1.chr5\_hap1.g00548.t01 |
| 6311 | 5 | 14164902 | 14190798 | mNeoVis.2.1.1.chr5\_hap1.g00549.t01 |
| 6312 | 5 | 14222297 | 14314545 | mNeoVis.2.1.1.chr5\_hap1.g00550.t01 |
| 6313 | 5 | 14319169 | 14320595 | mNeoVis.2.1.1.chr5\_hap1.g00551.t01 |
| 6314 | 5 | 14320734 | 14330309 | mNeoVis.2.1.1.chr5\_hap1.g00552.t01 |
| 6315 | 5 | 14330747 | 14337461 | mNeoVis.2.1.1.chr5\_hap1.g00553.t01 |
| 6316 | 5 | 14339169 | 14348134 | mNeoVis.2.1.1.chr5\_hap1.g00554.t01 |
| 6317 | 5 | 14362638 | 14399056 | mNeoVis.2.1.1.chr5\_hap1.g00555.t01 |
| 6318 | 5 | 14399315 | 14405859 | mNeoVis.2.1.1.chr5\_hap1.g00556.t01 |
| 6319 | 5 | 14406396 | 14416001 | mNeoVis.2.1.1.chr5\_hap1.g00557.t01 |
| 6320 | 5 | 14417560 | 14433980 | mNeoVis.2.1.1.chr5\_hap1.g00558.t01 |
| 6321 | 5 | 14476046 | 14488532 | mNeoVis.2.1.1.chr5\_hap1.g00559.t01 |
| 6322 | 5 | 14502494 | 14508350 | mNeoVis.2.1.1.chr5\_hap1.g00560.t01 |
| 6323 | 5 | 14655203 | 14690303 | mNeoVis.2.1.1.chr5\_hap1.g00564.t01 |
| 6324 | 5 | 14701914 | 14709605 | mNeoVis.2.1.1.chr5\_hap1.g00565.t01 |
| 6325 | 5 | 14737943 | 14796090 | mNeoVis.2.1.1.chr5\_hap1.g00566.t01 |
| 6326 | 5 | 14863135 | 14864806 | mNeoVis.2.1.1.chr5\_hap1.g00567.t01 |
| 6327 | 5 | 14980048 | 15127331 | mNeoVis.2.1.1.chr5\_hap1.g00568.t01 |
| 6328 | 5 | 15150393 | 15212434 | mNeoVis.2.1.1.chr5\_hap1.g00569.t01 |
| 6329 | 5 | 15222533 | 15283633 | mNeoVis.2.1.1.chr5\_hap1.g00570.t01 |
| 6330 | 5 | 16645724 | 16679677 | mNeoVis.2.1.1.chr5\_hap1.g00604.t01 |
| 6331 | 5 | 16685670 | 16856683 | mNeoVis.2.1.1.chr5\_hap1.g00605.t01 |
| 6332 | 5 | 16918642 | 16935517 | mNeoVis.2.1.1.chr5\_hap1.g00606.t01 |
| 6333 | 5 | 16936056 | 16983587 | mNeoVis.2.1.1.chr5\_hap1.g00607.t01 |
| 6334 | 5 | 17172820 | 17200612 | mNeoVis.2.1.1.chr5\_hap1.g00608.t01 |
| 6335 | 5 | 17754905 | 17784881 | mNeoVis.2.1.1.chr5\_hap1.g00629.t01 |
| 6336 | 5 | 17798163 | 17849436 | mNeoVis.2.1.1.chr5\_hap1.g00630.t01 |
| 6337 | 5 | 17858003 | 17872495 | mNeoVis.2.1.1.chr5\_hap1.g00631.t01 |
| 6338 | 5 | 17862369 | 17865715 | mNeoVis.2.1.1.chr5\_hap1.g00632.t01 |
| 6339 | 5 | 17872644 | 17880676 | mNeoVis.2.1.1.chr5\_hap1.g00633.t01 |
| 6340 | 5 | 17880621 | 17884660 | mNeoVis.2.1.1.chr5\_hap1.g00634.t01 |
| 6341 | 5 | 17886125 | 17887620 | mNeoVis.2.1.1.chr5\_hap1.g00635.t01 |
| 6342 | 5 | 17888138 | 17893073 | mNeoVis.2.1.1.chr5\_hap1.g00636.t01 |
| 6343 | 5 | 17893542 | 17948254 | mNeoVis.2.1.1.chr5\_hap1.g00637.t01 |
| 6344 | 5 | 17948393 | 17955624 | mNeoVis.2.1.1.chr5\_hap1.g00638.t01 |
| 6345 | 5 | 17957209 | 17967670 | mNeoVis.2.1.1.chr5\_hap1.g00639.t01 |
| 6346 | 5 | 17967863 | 17979675 | mNeoVis.2.1.1.chr5\_hap1.g00640.t01 |
| 6347 | 5 | 17992488 | 17998770 | mNeoVis.2.1.1.chr5\_hap1.g00641.t01 |
| 6348 | 5 | 18002260 | 18007611 | mNeoVis.2.1.1.chr5\_hap1.g00642.t01 |
| 6349 | 5 | 18008336 | 18021815 | mNeoVis.2.1.1.chr5\_hap1.g00643.t01 |
| 6350 | 5 | 18025503 | 18065137 | mNeoVis.2.1.1.chr5\_hap1.g00644.t01 |
| 6351 | 5 | 18072387 | 18075712 | mNeoVis.2.1.1.chr5\_hap1.g00645.t01 |
| 6352 | 5 | 18076292 | 18128682 | mNeoVis.2.1.1.chr5\_hap1.g00646.t01 |
| 6353 | 5 | 18137682 | 18138842 | mNeoVis.2.1.1.chr5\_hap1.g00647.t01 |
| 6354 | 5 | 18139882 | 18199089 | mNeoVis.2.1.1.chr5\_hap1.g00648.t01 |
| 6355 | 5 | 18199301 | 18202667 | mNeoVis.2.1.1.chr5\_hap1.g00649.t01 |
| 6356 | 5 | 18203681 | 18208311 | mNeoVis.2.1.1.chr5\_hap1.g00650.t01 |
| 6357 | 5 | 18208447 | 18213005 | mNeoVis.2.1.1.chr5\_hap1.g00651.t01 |
| 6358 | 5 | 18248470 | 18318453 | mNeoVis.2.1.1.chr5\_hap1.g00652.t01 |
| 6359 | 5 | 18329745 | 18424802 | mNeoVis.2.1.1.chr5\_hap1.g00653.t01 |
| 6360 | 5 | 18426747 | 18435493 | mNeoVis.2.1.1.chr5\_hap1.g00654.t01 |
| 6361 | 5 | 18435916 | 18440642 | mNeoVis.2.1.1.chr5\_hap1.g00655.t01 |
| 6362 | 5 | 18441419 | 18469579 | mNeoVis.2.1.1.chr5\_hap1.g00656.t01 |
| 6363 | 5 | 18465404 | 18468565 | mNeoVis.2.1.1.chr5\_hap1.g00657.t01 |
| 6364 | 5 | 18469549 | 18471997 | mNeoVis.2.1.1.chr5\_hap1.g00658.t01 |
| 6365 | 5 | 18472296 | 18538673 | mNeoVis.2.1.1.chr5\_hap1.g00659.t01 |
| 6366 | 5 | 18540814 | 18548323 | mNeoVis.2.1.1.chr5\_hap1.g00660.t01 |
| 6367 | 5 | 18552113 | 18553815 | mNeoVis.2.1.1.chr5\_hap1.g00661.t01 |
| 6368 | 5 | 18553846 | 18562612 | mNeoVis.2.1.1.chr5\_hap1.g00662.t01 |
| 6369 | 5 | 18569234 | 18591045 | mNeoVis.2.1.1.chr5\_hap1.g00663.t01 |
| 6370 | 5 | 18592158 | 18604896 | mNeoVis.2.1.1.chr5\_hap1.g00664.t01 |
| 6371 | 5 | 18608267 | 18609812 | mNeoVis.2.1.1.chr5\_hap1.g00665.t01 |
| 6372 | 5 | 18634612 | 18711014 | mNeoVis.2.1.1.chr5\_hap1.g00666.t01 |
| 6373 | 5 | 18718088 | 18743581 | mNeoVis.2.1.1.chr5\_hap1.g00667.t01 |
| 6374 | 5 | 19811817 | 19815503 | mNeoVis.2.1.1.chr5\_hap1.g00691.t01 |
| 6375 | 5 | 19817695 | 19820839 | mNeoVis.2.1.1.chr5\_hap1.g00692.t01 |
| 6376 | 5 | 19824242 | 19908292 | mNeoVis.2.1.1.chr5\_hap1.g00693.t01 |
| 6377 | 5 | 19908432 | 20049865 | mNeoVis.2.1.1.chr5\_hap1.g00694.t01 |
| 6378 | 5 | 19908432 | 20049865 | mNeoVis.2.1.1.chr5\_hap1.g00694.t01 |
| 6379 | 5 | 20049936 | 20080817 | mNeoVis.2.1.1.chr5\_hap1.g00695.t01 |
| 6380 | 5 | 20049936 | 20080817 | mNeoVis.2.1.1.chr5\_hap1.g00695.t01 |
| 6381 | 5 | 20083592 | 20095016 | mNeoVis.2.1.1.chr5\_hap1.g00696.t01 |
| 6382 | 5 | 20083592 | 20095016 | mNeoVis.2.1.1.chr5\_hap1.g00696.t01 |
| 6383 | 5 | 20118315 | 20119002 | mNeoVis.2.1.1.chr5\_hap1.g00697.t01 |
| 6384 | 5 | 20118315 | 20119002 | mNeoVis.2.1.1.chr5\_hap1.g00697.t01 |
| 6385 | 5 | 20142556 | 20144813 | mNeoVis.2.1.1.chr5\_hap1.g00698.t01 |
| 6386 | 5 | 20142556 | 20144813 | mNeoVis.2.1.1.chr5\_hap1.g00698.t01 |
| 6387 | 5 | 20153915 | 20155436 | mNeoVis.2.1.1.chr5\_hap1.g00699.t01 |
| 6388 | 5 | 20153915 | 20155436 | mNeoVis.2.1.1.chr5\_hap1.g00699.t01 |
| 6389 | 5 | 20193686 | 20195497 | mNeoVis.2.1.1.chr5\_hap1.g00700.t01 |
| 6390 | 5 | 20193686 | 20195497 | mNeoVis.2.1.1.chr5\_hap1.g00700.t01 |
| 6391 | 5 | 20211191 | 20218814 | mNeoVis.2.1.1.chr5\_hap1.g00701.t01 |
| 6392 | 5 | 20211191 | 20218814 | mNeoVis.2.1.1.chr5\_hap1.g00701.t01 |
| 6393 | 5 | 20219639 | 20225389 | mNeoVis.2.1.1.chr5\_hap1.g00702.t01 |
| 6394 | 5 | 20219639 | 20225389 | mNeoVis.2.1.1.chr5\_hap1.g00702.t01 |
| 6395 | 5 | 20231096 | 20232208 | mNeoVis.2.1.1.chr5\_hap1.g00703.t01 |
| 6396 | 5 | 20231096 | 20232208 | mNeoVis.2.1.1.chr5\_hap1.g00703.t01 |
| 6397 | 5 | 20232914 | 20242993 | mNeoVis.2.1.1.chr5\_hap1.g00704.t01 |
| 6398 | 5 | 20232914 | 20242993 | mNeoVis.2.1.1.chr5\_hap1.g00704.t01 |
| 6399 | 5 | 20244090 | 20248299 | mNeoVis.2.1.1.chr5\_hap1.g00705.t01 |
| 6400 | 5 | 20244090 | 20248299 | mNeoVis.2.1.1.chr5\_hap1.g00705.t01 |
| 6401 | 5 | 20247501 | 20261001 | mNeoVis.2.1.1.chr5\_hap1.g00706.t01 |
| 6402 | 5 | 20247501 | 20261001 | mNeoVis.2.1.1.chr5\_hap1.g00706.t01 |
| 6403 | 5 | 20263141 | 20265547 | mNeoVis.2.1.1.chr5\_hap1.g00707.t01 |
| 6404 | 5 | 20263141 | 20265547 | mNeoVis.2.1.1.chr5\_hap1.g00707.t01 |
| 6405 | 5 | 20308862 | 20316332 | mNeoVis.2.1.1.chr5\_hap1.g00708.t01 |
| 6406 | 5 | 20308862 | 20316332 | mNeoVis.2.1.1.chr5\_hap1.g00708.t01 |
| 6407 | 5 | 20332546 | 20401196 | mNeoVis.2.1.1.chr5\_hap1.g00709.t01 |
| 6408 | 5 | 20332546 | 20401196 | mNeoVis.2.1.1.chr5\_hap1.g00709.t01 |
| 6409 | 5 | 20426667 | 20440269 | mNeoVis.2.1.1.chr5\_hap1.g00710.t01 |
| 6410 | 5 | 20426667 | 20440269 | mNeoVis.2.1.1.chr5\_hap1.g00710.t01 |
| 6411 | 5 | 20442394 | 20449977 | mNeoVis.2.1.1.chr5\_hap1.g00711.t01 |
| 6412 | 5 | 20442394 | 20449977 | mNeoVis.2.1.1.chr5\_hap1.g00711.t01 |
| 6413 | 5 | 20453027 | 20462592 | mNeoVis.2.1.1.chr5\_hap1.g00712.t01 |
| 6414 | 5 | 20453027 | 20462592 | mNeoVis.2.1.1.chr5\_hap1.g00712.t01 |
| 6415 | 5 | 20471315 | 20476287 | mNeoVis.2.1.1.chr5\_hap1.g00713.t01 |
| 6416 | 5 | 20471315 | 20476287 | mNeoVis.2.1.1.chr5\_hap1.g00713.t01 |
| 6417 | 5 | 20478051 | 20498632 | mNeoVis.2.1.1.chr5\_hap1.g00714.t01 |
| 6418 | 5 | 20478051 | 20498632 | mNeoVis.2.1.1.chr5\_hap1.g00714.t01 |
| 6419 | 5 | 20498776 | 20532571 | mNeoVis.2.1.1.chr5\_hap1.g00715.t01 |
| 6420 | 5 | 20498776 | 20532571 | mNeoVis.2.1.1.chr5\_hap1.g00715.t01 |
| 6421 | 5 | 20552418 | 20627261 | mNeoVis.2.1.1.chr5\_hap1.g00716.t01 |
| 6422 | 5 | 20552418 | 20627261 | mNeoVis.2.1.1.chr5\_hap1.g00716.t01 |
| 6423 | 5 | 20627646 | 20636678 | mNeoVis.2.1.1.chr5\_hap1.g00717.t01 |
| 6424 | 5 | 20627646 | 20636678 | mNeoVis.2.1.1.chr5\_hap1.g00717.t01 |
| 6425 | 5 | 20639466 | 20649492 | mNeoVis.2.1.1.chr5\_hap1.g00718.t01 |
| 6426 | 5 | 20639466 | 20649492 | mNeoVis.2.1.1.chr5\_hap1.g00718.t01 |
| 6427 | 5 | 20657804 | 20669328 | mNeoVis.2.1.1.chr5\_hap1.g00719.t01 |
| 6428 | 5 | 20674236 | 20677361 | mNeoVis.2.1.1.chr5\_hap1.g00720.t01 |
| 6429 | 5 | 20678858 | 20711436 | mNeoVis.2.1.1.chr5\_hap1.g00721.t01 |
| 6430 | 5 | 20713578 | 20741018 | mNeoVis.2.1.1.chr5\_hap1.g00722.t01 |
| 6431 | 5 | 20740877 | 20749524 | mNeoVis.2.1.1.chr5\_hap1.g00723.t01 |
| 6432 | 5 | 20752125 | 20756016 | mNeoVis.2.1.1.chr5\_hap1.g00724.t01 |
| 6433 | 5 | 21375433 | 21435417 | mNeoVis.2.1.1.chr5\_hap1.g00739.t01 |
| 6434 | 5 | 21551595 | 21852763 | mNeoVis.2.1.1.chr5\_hap1.g00740.t01 |
| 6435 | 5 | 21853110 | 21892218 | mNeoVis.2.1.1.chr5\_hap1.g00741.t01 |
| 6436 | 5 | 21892362 | 21907220 | mNeoVis.2.1.1.chr5\_hap1.g00742.t01 |
| 6437 | 5 | 21902140 | 21924820 | mNeoVis.2.1.1.chr5\_hap1.g00743.t01 |
| 6438 | 5 | 21927405 | 21935370 | mNeoVis.2.1.1.chr5\_hap1.g00744.t01 |
| 6439 | 5 | 22842411 | 22852085 | mNeoVis.2.1.1.chr5\_hap1.g00746.t01 |
| 6440 | 5 | 25243437 | 25249456 | mNeoVis.2.1.1.chr5\_hap1.g00758.t01 |
| 6441 | 5 | 25260017 | 25285186 | mNeoVis.2.1.1.chr5\_hap1.g00759.t01 |
| 6442 | 5 | 25308753 | 25371732 | mNeoVis.2.1.1.chr5\_hap1.g00760.t01 |
| 6443 | 5 | 25422177 | 25566135 | mNeoVis.2.1.1.chr5\_hap1.g00761.t01 |
| 6444 | 5 | 25623361 | 25770216 | mNeoVis.2.1.1.chr5\_hap1.g00762.t01 |
| 6445 | 5 | 25776620 | 25778680 | mNeoVis.2.1.1.chr5\_hap1.g00763.t01 |
| 6446 | 5 | 25789188 | 25811330 | mNeoVis.2.1.1.chr5\_hap1.g00764.t01 |
| 6447 | 5 | 25830334 | 25883219 | mNeoVis.2.1.1.chr5\_hap1.g00765.t01 |
| 6448 | 5 | 25893642 | 25904271 | mNeoVis.2.1.1.chr5\_hap1.g00766.t01 |
| 6449 | 5 | 25904311 | 25905780 | mNeoVis.2.1.1.chr5\_hap1.g00767.t01 |
| 6450 | 5 | 25913743 | 25990532 | mNeoVis.2.1.1.chr5\_hap1.g00768.t01 |
| 6451 | 5 | 25991015 | 25997027 | mNeoVis.2.1.1.chr5\_hap1.g00769.t01 |
| 6452 | 5 | 25996375 | 26036152 | mNeoVis.2.1.1.chr5\_hap1.g00770.t01 |
| 6453 | 5 | 26036621 | 26069599 | mNeoVis.2.1.1.chr5\_hap1.g00771.t01 |
| 6454 | 5 | 26076922 | 26129973 | mNeoVis.2.1.1.chr5\_hap1.g00772.t01 |
| 6455 | 5 | 26141073 | 26169400 | mNeoVis.2.1.1.chr5\_hap1.g00773.t01 |
| 6456 | 5 | 26145580 | 26147041 | mNeoVis.2.1.1.chr5\_hap1.g00774.t01 |
| 6457 | 5 | 26206538 | 26493487 | mNeoVis.2.1.1.chr5\_hap1.g00775.t01 |
| 6458 | 5 | 30348795 | 30349564 | mNeoVis.2.1.1.chr5\_hap1.g00782.t01 |
| 6459 | 5 | 30358495 | 30484514 | mNeoVis.2.1.1.chr5\_hap1.g00783.t01 |
| 6460 | 5 | 30818393 | 30838866 | mNeoVis.2.1.1.chr5\_hap1.g00785.t01 |
| 6461 | 5 | 30944304 | 31062149 | mNeoVis.2.1.1.chr5\_hap1.g00786.t01 |
| 6462 | 5 | 30993473 | 31015090 | mNeoVis.2.1.1.chr5\_hap1.g00787.t01 |
| 6463 | 5 | 31062131 | 31152487 | mNeoVis.2.1.1.chr5\_hap1.g00788.t01 |
| 6464 | 5 | 31160511 | 31176059 | mNeoVis.2.1.1.chr5\_hap1.g00789.t01 |
| 6465 | 5 | 31626391 | 31793390 | mNeoVis.2.1.1.chr5\_hap1.g00791.t01 |
| 6466 | 5 | 31673312 | 31696932 | mNeoVis.2.1.1.chr5\_hap1.g00792.t01 |
| 6467 | 5 | 31832649 | 32021820 | mNeoVis.2.1.1.chr5\_hap1.g00793.t01 |
| 6468 | 5 | 31832649 | 32021820 | mNeoVis.2.1.1.chr5\_hap1.g00793.t01 |
| 6469 | 5 | 33256676 | 33384186 | mNeoVis.2.1.1.chr5\_hap1.g00795.t01 |
| 6470 | 5 | 33256676 | 33384186 | mNeoVis.2.1.1.chr5\_hap1.g00795.t01 |
| 6471 | 5 | 33384976 | 33496345 | mNeoVis.2.1.1.chr5\_hap1.g00796.t01 |
| 6472 | 5 | 33384976 | 33496345 | mNeoVis.2.1.1.chr5\_hap1.g00796.t01 |
| 6473 | 5 | 33936072 | 33949150 | mNeoVis.2.1.1.chr5\_hap1.g00797.t01 |
| 6474 | 5 | 33936072 | 33949150 | mNeoVis.2.1.1.chr5\_hap1.g00797.t01 |
| 6475 | 5 | 33966997 | 33980176 | mNeoVis.2.1.1.chr5\_hap1.g00798.t01 |
| 6476 | 5 | 33966997 | 33980176 | mNeoVis.2.1.1.chr5\_hap1.g00798.t01 |
| 6477 | 5 | 35455550 | 35456451 | mNeoVis.2.1.1.chr5\_hap1.g00801.t01 |
| 6478 | 5 | 35455550 | 35456451 | mNeoVis.2.1.1.chr5\_hap1.g00801.t01 |
| 6479 | 5 | 35455550 | 35456451 | mNeoVis.2.1.1.chr5\_hap1.g00801.t01 |
| 6480 | 5 | 35627590 | 35825982 | mNeoVis.2.1.1.chr5\_hap1.g00802.t01 |
| 6481 | 5 | 35867909 | 35906992 | mNeoVis.2.1.1.chr5\_hap1.g00803.t01 |
| 6482 | 5 | 35867909 | 35906992 | mNeoVis.2.1.1.chr5\_hap1.g00803.t01 |
| 6483 | 5 | 35912596 | 35944959 | mNeoVis.2.1.1.chr5\_hap1.g00804.t01 |
| 6484 | 5 | 35912596 | 35944959 | mNeoVis.2.1.1.chr5\_hap1.g00804.t01 |
| 6485 | 5 | 35947331 | 35971834 | mNeoVis.2.1.1.chr5\_hap1.g00805.t01 |
| 6486 | 5 | 35947331 | 35971834 | mNeoVis.2.1.1.chr5\_hap1.g00805.t01 |
| 6487 | 5 | 35975652 | 35997326 | mNeoVis.2.1.1.chr5\_hap1.g00806.t01 |
| 6488 | 5 | 35975652 | 35997326 | mNeoVis.2.1.1.chr5\_hap1.g00806.t01 |
| 6489 | 5 | 36001057 | 36040926 | mNeoVis.2.1.1.chr5\_hap1.g00807.t01 |
| 6490 | 5 | 36001057 | 36040926 | mNeoVis.2.1.1.chr5\_hap1.g00807.t01 |
| 6491 | 5 | 36075022 | 36253017 | mNeoVis.2.1.1.chr5\_hap1.g00808.t01 |
| 6492 | 5 | 36075022 | 36253017 | mNeoVis.2.1.1.chr5\_hap1.g00808.t01 |
| 6493 | 5 | 36582663 | 36801118 | mNeoVis.2.1.1.chr5\_hap1.g00809.t01 |
| 6494 | 5 | 36998743 | 37102765 | mNeoVis.2.1.1.chr5\_hap1.g00810.t01 |
| 6495 | 5 | 36998743 | 37102765 | mNeoVis.2.1.1.chr5\_hap1.g00810.t01 |
| 6496 | 5 | 36998743 | 37102765 | mNeoVis.2.1.1.chr5\_hap1.g00810.t01 |
| 6497 | 5 | 37267629 | 37328116 | mNeoVis.2.1.1.chr5\_hap1.g00811.t01 |
| 6498 | 5 | 37267629 | 37328116 | mNeoVis.2.1.1.chr5\_hap1.g00811.t01 |
| 6499 | 5 | 37267629 | 37328116 | mNeoVis.2.1.1.chr5\_hap1.g00811.t01 |
| 6500 | 5 | 37267629 | 37328116 | mNeoVis.2.1.1.chr5\_hap1.g00811.t01 |
| 6501 | 5 | 37433711 | 37539489 | mNeoVis.2.1.1.chr5\_hap1.g00812.t01 |
| 6502 | 5 | 37433711 | 37539489 | mNeoVis.2.1.1.chr5\_hap1.g00812.t01 |
| 6503 | 5 | 37645294 | 37819723 | mNeoVis.2.1.1.chr5\_hap1.g00813.t01 |
| 6504 | 5 | 37645294 | 37819723 | mNeoVis.2.1.1.chr5\_hap1.g00813.t01 |
| 6505 | 5 | 37645294 | 37819723 | mNeoVis.2.1.1.chr5\_hap1.g00813.t01 |
| 6506 | 5 | 37915908 | 37916519 | mNeoVis.2.1.1.chr5\_hap1.g00814.t01 |
| 6507 | 5 | 37915908 | 37916519 | mNeoVis.2.1.1.chr5\_hap1.g00814.t01 |
| 6508 | 5 | 37915908 | 37916519 | mNeoVis.2.1.1.chr5\_hap1.g00814.t01 |
| 6509 | 5 | 38178183 | 38365382 | mNeoVis.2.1.1.chr5\_hap1.g00815.t01 |
| 6510 | 5 | 38178183 | 38365382 | mNeoVis.2.1.1.chr5\_hap1.g00815.t01 |
| 6511 | 5 | 38394080 | 38396686 | mNeoVis.2.1.1.chr5\_hap1.g00816.t01 |
| 6512 | 5 | 38394080 | 38396686 | mNeoVis.2.1.1.chr5\_hap1.g00816.t01 |
| 6513 | 5 | 38407030 | 38421712 | mNeoVis.2.1.1.chr5\_hap1.g00817.t01 |
| 6514 | 5 | 38407030 | 38421712 | mNeoVis.2.1.1.chr5\_hap1.g00817.t01 |
| 6515 | 5 | 38931795 | 39159885 | mNeoVis.2.1.1.chr5\_hap1.g00818.t01 |
| 6516 | 5 | 39266336 | 39368496 | mNeoVis.2.1.1.chr5\_hap1.g00819.t01 |
| 6517 | 5 | 39395557 | 39485596 | mNeoVis.2.1.1.chr5\_hap1.g00820.t01 |
| 6518 | 5 | 39508070 | 39559534 | mNeoVis.2.1.1.chr5\_hap1.g00821.t01 |
| 6519 | 5 | 39826061 | 40067872 | mNeoVis.2.1.1.chr5\_hap1.g00822.t01 |
| 6520 | 5 | 39826061 | 40067872 | mNeoVis.2.1.1.chr5\_hap1.g00822.t01 |
| 6521 | 5 | 40138855 | 40183975 | mNeoVis.2.1.1.chr5\_hap1.g00823.t01 |
| 6522 | 5 | 40138855 | 40183975 | mNeoVis.2.1.1.chr5\_hap1.g00823.t01 |
| 6523 | 5 | 40138855 | 40183975 | mNeoVis.2.1.1.chr5\_hap1.g00823.t01 |
| 6524 | 5 | 40725841 | 41064233 | mNeoVis.2.1.1.chr5\_hap1.g00824.t01 |
| 6525 | 5 | 41202392 | 41203442 | mNeoVis.2.1.1.chr5\_hap1.g00825.t01 |
| 6526 | 5 | 41792160 | 42010814 | mNeoVis.2.1.1.chr5\_hap1.g00826.t01 |
| 6527 | 5 | 42658664 | 42955390 | mNeoVis.2.1.1.chr5\_hap1.g00827.t01 |
| 6528 | 5 | 42658664 | 42955390 | mNeoVis.2.1.1.chr5\_hap1.g00827.t01 |
| 6529 | 5 | 43544953 | 44493056 | mNeoVis.2.1.1.chr5\_hap1.g00828.t01 |
| 6530 | 5 | 43761672 | 43763102 | mNeoVis.2.1.1.chr5\_hap1.g00829.t01 |
| 6531 | 5 | 43761672 | 43763102 | mNeoVis.2.1.1.chr5\_hap1.g00829.t01 |
| 6532 | 5 | 44498868 | 44569707 | mNeoVis.2.1.1.chr5\_hap1.g00830.t01 |
| 6533 | 5 | 44498868 | 44569707 | mNeoVis.2.1.1.chr5\_hap1.g00830.t01 |
| 6534 | 5 | 44502404 | 44537100 | mNeoVis.2.1.1.chr5\_hap1.g00831.t01 |
| 6535 | 5 | 44502404 | 44537100 | mNeoVis.2.1.1.chr5\_hap1.g00831.t01 |
| 6536 | 5 | 44571381 | 44595959 | mNeoVis.2.1.1.chr5\_hap1.g00832.t01 |
| 6537 | 5 | 44571381 | 44595959 | mNeoVis.2.1.1.chr5\_hap1.g00832.t01 |
| 6538 | 5 | 45200657 | 45239712 | mNeoVis.2.1.1.chr5\_hap1.g00833.t01 |
| 6539 | 5 | 45200657 | 45239712 | mNeoVis.2.1.1.chr5\_hap1.g00833.t01 |
| 6540 | 5 | 45646831 | 45648254 | mNeoVis.2.1.1.chr5\_hap1.g00834.t01 |
| 6541 | 5 | 45646831 | 45648254 | mNeoVis.2.1.1.chr5\_hap1.g00834.t01 |
| 6542 | 5 | 45681271 | 45693133 | mNeoVis.2.1.1.chr5\_hap1.g00835.t01 |
| 6543 | 5 | 45681271 | 45693133 | mNeoVis.2.1.1.chr5\_hap1.g00835.t01 |
| 6544 | 5 | 45718175 | 45825008 | mNeoVis.2.1.1.chr5\_hap1.g00836.t01 |
| 6545 | 5 | 45718175 | 45825008 | mNeoVis.2.1.1.chr5\_hap1.g00836.t01 |
| 6546 | 5 | 45857187 | 46185766 | mNeoVis.2.1.1.chr5\_hap1.g00837.t01 |
| 6547 | 5 | 45857187 | 46185766 | mNeoVis.2.1.1.chr5\_hap1.g00837.t01 |
| 6548 | 5 | 45857187 | 46185766 | mNeoVis.2.1.1.chr5\_hap1.g00837.t01 |
| 6549 | 5 | 46304879 | 46310557 | mNeoVis.2.1.1.chr5\_hap1.g00838.t01 |
| 6550 | 5 | 46304879 | 46310557 | mNeoVis.2.1.1.chr5\_hap1.g00838.t01 |
| 6551 | 5 | 46448431 | 46502679 | mNeoVis.2.1.1.chr5\_hap1.g00839.t01 |
| 6552 | 5 | 46448431 | 46502679 | mNeoVis.2.1.1.chr5\_hap1.g00839.t01 |
| 6553 | 5 | 46510733 | 46538476 | mNeoVis.2.1.1.chr5\_hap1.g00840.t01 |
| 6554 | 5 | 46510733 | 46538476 | mNeoVis.2.1.1.chr5\_hap1.g00840.t01 |
| 6555 | 5 | 46682329 | 47094184 | mNeoVis.2.1.1.chr5\_hap1.g00841.t01 |
| 6556 | 5 | 48503406 | 48505352 | mNeoVis.2.1.1.chr5\_hap1.g00843.t01 |
| 6557 | 5 | 49966657 | 49980013 | mNeoVis.2.1.1.chr5\_hap1.g00846.t01 |
| 6558 | 5 | 49984149 | 50003966 | mNeoVis.2.1.1.chr5\_hap1.g00847.t01 |
| 6559 | 5 | 50125597 | 50225669 | mNeoVis.2.1.1.chr5\_hap1.g00848.t01 |
| 6560 | 5 | 50241991 | 50494826 | mNeoVis.2.1.1.chr5\_hap1.g00849.t01 |
| 6561 | 5 | 50586824 | 50611924 | mNeoVis.2.1.1.chr5\_hap1.g00850.t01 |
| 6562 | 5 | 50586824 | 50611924 | mNeoVis.2.1.1.chr5\_hap1.g00850.t01 |
| 6563 | 5 | 50623377 | 50705978 | mNeoVis.2.1.1.chr5\_hap1.g00851.t01 |
| 6564 | 5 | 50623377 | 50705978 | mNeoVis.2.1.1.chr5\_hap1.g00851.t01 |
| 6565 | 5 | 50722477 | 50750808 | mNeoVis.2.1.1.chr5\_hap1.g00852.t01 |
| 6566 | 5 | 50722477 | 50750808 | mNeoVis.2.1.1.chr5\_hap1.g00852.t01 |
| 6567 | 5 | 50779085 | 50825028 | mNeoVis.2.1.1.chr5\_hap1.g00853.t01 |
| 6568 | 5 | 50779085 | 50825028 | mNeoVis.2.1.1.chr5\_hap1.g00853.t01 |
| 6569 | 5 | 50826301 | 50844660 | mNeoVis.2.1.1.chr5\_hap1.g00854.t01 |
| 6570 | 5 | 50826301 | 50844660 | mNeoVis.2.1.1.chr5\_hap1.g00854.t01 |
| 6571 | 5 | 50847087 | 50862545 | mNeoVis.2.1.1.chr5\_hap1.g00855.t01 |
| 6572 | 5 | 50864116 | 50869405 | mNeoVis.2.1.1.chr5\_hap1.g00856.t01 |
| 6573 | 5 | 50869420 | 50881412 | mNeoVis.2.1.1.chr5\_hap1.g00857.t01 |
| 6574 | 5 | 50895848 | 50907523 | mNeoVis.2.1.1.chr5\_hap1.g00858.t01 |
| 6575 | 5 | 50921569 | 50949517 | mNeoVis.2.1.1.chr5\_hap1.g00859.t01 |
| 6576 | 5 | 50951691 | 50958339 | mNeoVis.2.1.1.chr5\_hap1.g00860.t01 |
| 6577 | 5 | 50962594 | 50972894 | mNeoVis.2.1.1.chr5\_hap1.g00861.t01 |
| 6578 | 5 | 50985118 | 50988369 | mNeoVis.2.1.1.chr5\_hap1.g00862.t01 |
| 6579 | 5 | 50985132 | 51006415 | mNeoVis.2.1.1.chr5\_hap1.g00863.t01 |
| 6580 | 5 | 51006490 | 51019557 | mNeoVis.2.1.1.chr5\_hap1.g00864.t01 |
| 6581 | 5 | 51019592 | 51028363 | mNeoVis.2.1.1.chr5\_hap1.g00865.t01 |
| 6582 | 5 | 51028605 | 51038015 | mNeoVis.2.1.1.chr5\_hap1.g00866.t01 |
| 6583 | 5 | 51028605 | 51038015 | mNeoVis.2.1.1.chr5\_hap1.g00866.t01 |
| 6584 | 5 | 51048236 | 51067987 | mNeoVis.2.1.1.chr5\_hap1.g00867.t01 |
| 6585 | 5 | 51048236 | 51067987 | mNeoVis.2.1.1.chr5\_hap1.g00867.t01 |
| 6586 | 5 | 51068071 | 51129792 | mNeoVis.2.1.1.chr5\_hap1.g00868.t01 |
| 6587 | 5 | 51068071 | 51129792 | mNeoVis.2.1.1.chr5\_hap1.g00868.t01 |
| 6588 | 5 | 51129944 | 51133111 | mNeoVis.2.1.1.chr5\_hap1.g00869.t01 |
| 6589 | 5 | 51129944 | 51133111 | mNeoVis.2.1.1.chr5\_hap1.g00869.t01 |
| 6590 | 5 | 51138256 | 51148018 | mNeoVis.2.1.1.chr5\_hap1.g00870.t01 |
| 6591 | 5 | 51138256 | 51148018 | mNeoVis.2.1.1.chr5\_hap1.g00870.t01 |
| 6592 | 5 | 51161796 | 51170128 | mNeoVis.2.1.1.chr5\_hap1.g00871.t01 |
| 6593 | 5 | 51161796 | 51170128 | mNeoVis.2.1.1.chr5\_hap1.g00871.t01 |
| 6594 | 5 | 51178000 | 51236766 | mNeoVis.2.1.1.chr5\_hap1.g00872.t01 |
| 6595 | 5 | 51178000 | 51236766 | mNeoVis.2.1.1.chr5\_hap1.g00872.t01 |
| 6596 | 5 | 51239296 | 51334502 | mNeoVis.2.1.1.chr5\_hap1.g00873.t01 |
| 6597 | 5 | 51239296 | 51334502 | mNeoVis.2.1.1.chr5\_hap1.g00873.t01 |
| 6598 | 5 | 51271917 | 51290597 | mNeoVis.2.1.1.chr5\_hap1.g00874.t01 |
| 6599 | 5 | 51271917 | 51290597 | mNeoVis.2.1.1.chr5\_hap1.g00874.t01 |
| 6600 | 5 | 51294473 | 51327272 | mNeoVis.2.1.1.chr5\_hap1.g00875.t01 |
| 6601 | 5 | 51294473 | 51327272 | mNeoVis.2.1.1.chr5\_hap1.g00875.t01 |
| 6602 | 5 | 51331424 | 51497037 | mNeoVis.2.1.1.chr5\_hap1.g00876.t01 |
| 6603 | 5 | 51331424 | 51497037 | mNeoVis.2.1.1.chr5\_hap1.g00876.t01 |
| 6604 | 5 | 51331424 | 51497037 | mNeoVis.2.1.1.chr5\_hap1.g00876.t01 |
| 6605 | 5 | 51331424 | 51497037 | mNeoVis.2.1.1.chr5\_hap1.g00876.t01 |
| 6606 | 5 | 51808604 | 51936347 | mNeoVis.2.1.1.chr5\_hap1.g00877.t01 |
| 6607 | 5 | 51808604 | 51936347 | mNeoVis.2.1.1.chr5\_hap1.g00877.t01 |
| 6608 | 5 | 51808604 | 51936347 | mNeoVis.2.1.1.chr5\_hap1.g00877.t01 |
| 6609 | 5 | 52007145 | 52028190 | mNeoVis.2.1.1.chr5\_hap1.g00878.t01 |
| 6610 | 5 | 52007145 | 52028190 | mNeoVis.2.1.1.chr5\_hap1.g00878.t01 |
| 6611 | 5 | 52007145 | 52028190 | mNeoVis.2.1.1.chr5\_hap1.g00878.t01 |
| 6612 | 5 | 52055530 | 52194127 | mNeoVis.2.1.1.chr5\_hap1.g00879.t01 |
| 6613 | 5 | 52055530 | 52194127 | mNeoVis.2.1.1.chr5\_hap1.g00879.t01 |
| 6614 | 5 | 52457165 | 52541895 | mNeoVis.2.1.1.chr5\_hap1.g00882.t01 |
| 6615 | 5 | 52457165 | 52541895 | mNeoVis.2.1.1.chr5\_hap1.g00882.t01 |
| 6616 | 5 | 52659652 | 52711795 | mNeoVis.2.1.1.chr5\_hap1.g00883.t01 |
| 6617 | 5 | 52659652 | 52711795 | mNeoVis.2.1.1.chr5\_hap1.g00883.t01 |
| 6618 | 5 | 52674491 | 52676056 | mNeoVis.2.1.1.chr5\_hap1.g00884.t01 |
| 6619 | 5 | 52674491 | 52676056 | mNeoVis.2.1.1.chr5\_hap1.g00884.t01 |
| 6620 | 5 | 52829878 | 53038537 | mNeoVis.2.1.1.chr5\_hap1.g00885.t01 |
| 6621 | 5 | 52829878 | 53038537 | mNeoVis.2.1.1.chr5\_hap1.g00885.t01 |
| 6622 | 5 | 52998687 | 53004748 | mNeoVis.2.1.1.chr5\_hap1.g00886.t01 |
| 6623 | 5 | 52998687 | 53004748 | mNeoVis.2.1.1.chr5\_hap1.g00886.t01 |
| 6624 | 5 | 53183921 | 53254563 | mNeoVis.2.1.1.chr5\_hap1.g00887.t01 |
| 6625 | 5 | 53183921 | 53254563 | mNeoVis.2.1.1.chr5\_hap1.g00887.t01 |
| 6626 | 5 | 53294402 | 53339666 | mNeoVis.2.1.1.chr5\_hap1.g00888.t01 |
| 6627 | 5 | 53294402 | 53339666 | mNeoVis.2.1.1.chr5\_hap1.g00888.t01 |
| 6628 | 5 | 53346834 | 53352105 | mNeoVis.2.1.1.chr5\_hap1.g00889.t01 |
| 6629 | 5 | 53346834 | 53352105 | mNeoVis.2.1.1.chr5\_hap1.g00889.t01 |
| 6630 | 5 | 53384723 | 53407776 | mNeoVis.2.1.1.chr5\_hap1.g00890.t01 |
| 6631 | 5 | 53384723 | 53407776 | mNeoVis.2.1.1.chr5\_hap1.g00890.t01 |
| 6632 | 5 | 53408862 | 53495873 | mNeoVis.2.1.1.chr5\_hap1.g00891.t01 |
| 6633 | 5 | 53408862 | 53495873 | mNeoVis.2.1.1.chr5\_hap1.g00891.t01 |
| 6634 | 5 | 53537026 | 53590432 | mNeoVis.2.1.1.chr5\_hap1.g00892.t01 |
| 6635 | 5 | 53537026 | 53590432 | mNeoVis.2.1.1.chr5\_hap1.g00892.t01 |
| 6636 | 5 | 53614080 | 53644720 | mNeoVis.2.1.1.chr5\_hap1.g00893.t01 |
| 6637 | 5 | 53614080 | 53644720 | mNeoVis.2.1.1.chr5\_hap1.g00893.t01 |
| 6638 | 5 | 53648640 | 53653833 | mNeoVis.2.1.1.chr5\_hap1.g00894.t01 |
| 6639 | 5 | 53648640 | 53653833 | mNeoVis.2.1.1.chr5\_hap1.g00894.t01 |
| 6640 | 5 | 53655397 | 53675742 | mNeoVis.2.1.1.chr5\_hap1.g00895.t01 |
| 6641 | 5 | 53655397 | 53675742 | mNeoVis.2.1.1.chr5\_hap1.g00895.t01 |
| 6642 | 5 | 53676687 | 53688347 | mNeoVis.2.1.1.chr5\_hap1.g00896.t01 |
| 6643 | 5 | 53676687 | 53688347 | mNeoVis.2.1.1.chr5\_hap1.g00896.t01 |
| 6644 | 5 | 53688503 | 53760438 | mNeoVis.2.1.1.chr5\_hap1.g00897.t01 |
| 6645 | 5 | 53688503 | 53760438 | mNeoVis.2.1.1.chr5\_hap1.g00897.t01 |
| 6646 | 5 | 53767274 | 53772478 | mNeoVis.2.1.1.chr5\_hap1.g00898.t01 |
| 6647 | 5 | 53767274 | 53772478 | mNeoVis.2.1.1.chr5\_hap1.g00898.t01 |
| 6648 | 5 | 53780162 | 53788898 | mNeoVis.2.1.1.chr5\_hap1.g00899.t01 |
| 6649 | 5 | 53780162 | 53788898 | mNeoVis.2.1.1.chr5\_hap1.g00899.t01 |
| 6650 | 5 | 53793119 | 53850366 | mNeoVis.2.1.1.chr5\_hap1.g00900.t01 |
| 6651 | 5 | 53793119 | 53850366 | mNeoVis.2.1.1.chr5\_hap1.g00900.t01 |
| 6652 | 5 | 53885060 | 54140692 | mNeoVis.2.1.1.chr5\_hap1.g00901.t01 |
| 6653 | 5 | 54209636 | 54212137 | mNeoVis.2.1.1.chr5\_hap1.g00902.t01 |
| 6654 | 5 | 54248314 | 54277733 | mNeoVis.2.1.1.chr5\_hap1.g00903.t01 |
| 6655 | 5 | 54283887 | 54297478 | mNeoVis.2.1.1.chr5\_hap1.g00904.t01 |
| 6656 | 5 | 54418376 | 54540944 | mNeoVis.2.1.1.chr5\_hap1.g00906.t01 |
| 6657 | 5 | 54593054 | 54682855 | mNeoVis.2.1.1.chr5\_hap1.g00907.t01 |
| 6658 | 5 | 54682410 | 54711259 | mNeoVis.2.1.1.chr5\_hap1.g00908.t01 |
| 6659 | 5 | 54787173 | 54842141 | mNeoVis.2.1.1.chr5\_hap1.g00909.t01 |
| 6660 | 5 | 55720866 | 55779646 | mNeoVis.2.1.1.chr5\_hap1.g00918.t01 |
| 6661 | 5 | 55720866 | 55779646 | mNeoVis.2.1.1.chr5\_hap1.g00918.t01 |
| 6662 | 5 | 55931646 | 56017978 | mNeoVis.2.1.1.chr5\_hap1.g00919.t01 |
| 6663 | 5 | 55931646 | 56017978 | mNeoVis.2.1.1.chr5\_hap1.g00919.t01 |
| 6664 | 5 | 55931646 | 56017978 | mNeoVis.2.1.1.chr5\_hap1.g00919.t01 |
| 6665 | 5 | 55968329 | 55970245 | mNeoVis.2.1.1.chr5\_hap1.g00920.t01 |
| 6666 | 5 | 55968329 | 55970245 | mNeoVis.2.1.1.chr5\_hap1.g00920.t01 |
| 6667 | 5 | 55968329 | 55970245 | mNeoVis.2.1.1.chr5\_hap1.g00920.t01 |
| 6668 | 5 | 56052672 | 56075314 | mNeoVis.2.1.1.chr5\_hap1.g00921.t01 |
| 6669 | 5 | 56052672 | 56075314 | mNeoVis.2.1.1.chr5\_hap1.g00921.t01 |
| 6670 | 5 | 56052672 | 56075314 | mNeoVis.2.1.1.chr5\_hap1.g00921.t01 |
| 6671 | 5 | 56124534 | 56225967 | mNeoVis.2.1.1.chr5\_hap1.g00922.t01 |
| 6672 | 5 | 56124534 | 56225967 | mNeoVis.2.1.1.chr5\_hap1.g00922.t01 |
| 6673 | 5 | 56124534 | 56225967 | mNeoVis.2.1.1.chr5\_hap1.g00922.t01 |
| 6674 | 5 | 56251820 | 56629068 | mNeoVis.2.1.1.chr5\_hap1.g00923.t01 |
| 6675 | 5 | 56656271 | 56697927 | mNeoVis.2.1.1.chr5\_hap1.g00924.t01 |
| 6676 | 5 | 56656271 | 56697927 | mNeoVis.2.1.1.chr5\_hap1.g00924.t01 |
| 6677 | 5 | 56695933 | 56780953 | mNeoVis.2.1.1.chr5\_hap1.g00925.t01 |
| 6678 | 5 | 56695933 | 56780953 | mNeoVis.2.1.1.chr5\_hap1.g00925.t01 |
| 6679 | 5 | 56780795 | 56822597 | mNeoVis.2.1.1.chr5\_hap1.g00926.t01 |
| 6680 | 5 | 56780795 | 56822597 | mNeoVis.2.1.1.chr5\_hap1.g00926.t01 |
| 6681 | 5 | 56833058 | 56834606 | mNeoVis.2.1.1.chr5\_hap1.g00927.t01 |
| 6682 | 5 | 56833058 | 56834606 | mNeoVis.2.1.1.chr5\_hap1.g00927.t01 |
| 6683 | 5 | 56846372 | 56881531 | mNeoVis.2.1.1.chr5\_hap1.g00928.t01 |
| 6684 | 5 | 56846372 | 56881531 | mNeoVis.2.1.1.chr5\_hap1.g00928.t01 |
| 6685 | 5 | 56887702 | 56916873 | mNeoVis.2.1.1.chr5\_hap1.g00929.t01 |
| 6686 | 5 | 56887702 | 56916873 | mNeoVis.2.1.1.chr5\_hap1.g00929.t01 |
| 6687 | 5 | 56931150 | 56943502 | mNeoVis.2.1.1.chr5\_hap1.g00930.t01 |
| 6688 | 5 | 56931150 | 56943502 | mNeoVis.2.1.1.chr5\_hap1.g00930.t01 |
| 6689 | 5 | 57002271 | 57029413 | mNeoVis.2.1.1.chr5\_hap1.g00931.t01 |
| 6690 | 5 | 57002271 | 57029413 | mNeoVis.2.1.1.chr5\_hap1.g00931.t01 |
| 6691 | 5 | 57030625 | 57053180 | mNeoVis.2.1.1.chr5\_hap1.g00932.t01 |
| 6692 | 5 | 57030625 | 57053180 | mNeoVis.2.1.1.chr5\_hap1.g00932.t01 |
| 6693 | 5 | 57063427 | 57071804 | mNeoVis.2.1.1.chr5\_hap1.g00933.t01 |
| 6694 | 5 | 57063427 | 57071804 | mNeoVis.2.1.1.chr5\_hap1.g00933.t01 |
| 6695 | 5 | 57085156 | 57160282 | mNeoVis.2.1.1.chr5\_hap1.g00934.t01 |
| 6696 | 5 | 57085156 | 57160282 | mNeoVis.2.1.1.chr5\_hap1.g00934.t01 |
| 6697 | 5 | 57218429 | 57244131 | mNeoVis.2.1.1.chr5\_hap1.g00935.t01 |
| 6698 | 5 | 57319753 | 57334275 | mNeoVis.2.1.1.chr5\_hap1.g00936.t01 |
| 6699 | 5 | 57352936 | 57353975 | mNeoVis.2.1.1.chr5\_hap1.g00937.t01 |
| 6700 | 5 | 57404122 | 57654934 | mNeoVis.2.1.1.chr5\_hap1.g00938.t01 |
| 6701 | 5 | 57669598 | 57714611 | mNeoVis.2.1.1.chr5\_hap1.g00939.t01 |
| 6702 | 5 | 57723695 | 57741161 | mNeoVis.2.1.1.chr5\_hap1.g00940.t01 |
| 6703 | 5 | 57779382 | 57862261 | mNeoVis.2.1.1.chr5\_hap1.g00941.t01 |
| 6704 | 5 | 58061726 | 58068966 | mNeoVis.2.1.1.chr5\_hap1.g00943.t01 |
| 6705 | 5 | 58069608 | 58105555 | mNeoVis.2.1.1.chr5\_hap1.g00944.t01 |
| 6706 | 5 | 58118050 | 58139483 | mNeoVis.2.1.1.chr5\_hap1.g00945.t01 |
| 6707 | 5 | 58145190 | 58158976 | mNeoVis.2.1.1.chr5\_hap1.g00946.t01 |
| 6708 | 5 | 58350423 | 58405889 | mNeoVis.2.1.1.chr5\_hap1.g00947.t01 |
| 6709 | 5 | 58632795 | 58682565 | mNeoVis.2.1.1.chr5\_hap1.g00948.t01 |
| 6710 | 5 | 58632795 | 58682565 | mNeoVis.2.1.1.chr5\_hap1.g00948.t01 |
| 6711 | 5 | 58632795 | 58682565 | mNeoVis.2.1.1.chr5\_hap1.g00948.t01 |
| 6712 | 5 | 58632795 | 58682565 | mNeoVis.2.1.1.chr5\_hap1.g00948.t01 |
| 6713 | 5 | 58699756 | 58943655 | mNeoVis.2.1.1.chr5\_hap1.g00949.t01 |
| 6714 | 5 | 58699756 | 58943655 | mNeoVis.2.1.1.chr5\_hap1.g00949.t01 |
| 6715 | 5 | 58699756 | 58943655 | mNeoVis.2.1.1.chr5\_hap1.g00949.t01 |
| 6716 | 5 | 58979867 | 59072903 | mNeoVis.2.1.1.chr5\_hap1.g00950.t01 |
| 6717 | 5 | 59919923 | 59952680 | mNeoVis.2.1.1.chr5\_hap1.g00961.t01 |
| 6718 | 5 | 59952837 | 59994517 | mNeoVis.2.1.1.chr5\_hap1.g00962.t01 |
| 6719 | 5 | 60037766 | 60054499 | mNeoVis.2.1.1.chr5\_hap1.g00963.t01 |
| 6720 | 5 | 60037766 | 60054499 | mNeoVis.2.1.1.chr5\_hap1.g00963.t01 |
| 6721 | 5 | 60078651 | 60090398 | mNeoVis.2.1.1.chr5\_hap1.g00964.t01 |
| 6722 | 5 | 60078651 | 60090398 | mNeoVis.2.1.1.chr5\_hap1.g00964.t01 |
| 6723 | 5 | 60118759 | 60167371 | mNeoVis.2.1.1.chr5\_hap1.g00965.t01 |
| 6724 | 5 | 60118759 | 60167371 | mNeoVis.2.1.1.chr5\_hap1.g00965.t01 |
| 6725 | 5 | 60183889 | 60241914 | mNeoVis.2.1.1.chr5\_hap1.g00966.t01 |
| 6726 | 5 | 60183889 | 60241914 | mNeoVis.2.1.1.chr5\_hap1.g00966.t01 |
| 6727 | 5 | 60243776 | 60362395 | mNeoVis.2.1.1.chr5\_hap1.g00967.t01 |
| 6728 | 5 | 60243776 | 60362395 | mNeoVis.2.1.1.chr5\_hap1.g00967.t01 |
| 6729 | 5 | 60901268 | 60935382 | mNeoVis.2.1.1.chr5\_hap1.g00968.t01 |
| 6730 | 5 | 60901268 | 60935382 | mNeoVis.2.1.1.chr5\_hap1.g00968.t01 |
| 6731 | 5 | 61262611 | 61320163 | mNeoVis.2.1.1.chr5\_hap1.g00970.t01 |
| 6732 | 5 | 61262611 | 61320163 | mNeoVis.2.1.1.chr5\_hap1.g00970.t01 |
| 6733 | 5 | 61611145 | 61791612 | mNeoVis.2.1.1.chr5\_hap1.g00971.t01 |
| 6734 | 5 | 61611145 | 61791612 | mNeoVis.2.1.1.chr5\_hap1.g00971.t01 |
| 6735 | 5 | 61814443 | 61872892 | mNeoVis.2.1.1.chr5\_hap1.g00972.t01 |
| 6736 | 5 | 61814443 | 61872892 | mNeoVis.2.1.1.chr5\_hap1.g00972.t01 |
| 6737 | 5 | 62050662 | 62127794 | mNeoVis.2.1.1.chr5\_hap1.g00974.t01 |
| 6738 | 5 | 62050662 | 62127794 | mNeoVis.2.1.1.chr5\_hap1.g00974.t01 |
| 6739 | 5 | 62133683 | 62158443 | mNeoVis.2.1.1.chr5\_hap1.g00975.t01 |
| 6740 | 5 | 62133683 | 62158443 | mNeoVis.2.1.1.chr5\_hap1.g00975.t01 |
| 6741 | 5 | 62169129 | 62248479 | mNeoVis.2.1.1.chr5\_hap1.g00976.t01 |
| 6742 | 5 | 62169129 | 62248479 | mNeoVis.2.1.1.chr5\_hap1.g00976.t01 |
| 6743 | 5 | 62169761 | 62180819 | mNeoVis.2.1.1.chr5\_hap1.g00977.t01 |
| 6744 | 5 | 62169761 | 62180819 | mNeoVis.2.1.1.chr5\_hap1.g00977.t01 |
| 6745 | 5 | 62275553 | 62279681 | mNeoVis.2.1.1.chr5\_hap1.g00978.t01 |
| 6746 | 5 | 62275553 | 62279681 | mNeoVis.2.1.1.chr5\_hap1.g00978.t01 |
| 6747 | 5 | 62330535 | 62395592 | mNeoVis.2.1.1.chr5\_hap1.g00979.t01 |
| 6748 | 5 | 62330535 | 62395592 | mNeoVis.2.1.1.chr5\_hap1.g00979.t01 |
| 6749 | 5 | 62423648 | 62493374 | mNeoVis.2.1.1.chr5\_hap1.g00980.t01 |
| 6750 | 5 | 62423648 | 62493374 | mNeoVis.2.1.1.chr5\_hap1.g00980.t01 |
| 6751 | 5 | 62553604 | 62629623 | mNeoVis.2.1.1.chr5\_hap1.g00981.t01 |
| 6752 | 5 | 63997149 | 64025368 | mNeoVis.2.1.1.chr5\_hap1.g00982.t01 |
| 6753 | 5 | 63997149 | 64025368 | mNeoVis.2.1.1.chr5\_hap1.g00982.t01 |
| 6754 | 5 | 63997149 | 64025368 | mNeoVis.2.1.1.chr5\_hap1.g00982.t01 |
| 6755 | 5 | 63997149 | 64025368 | mNeoVis.2.1.1.chr5\_hap1.g00982.t01 |
| 6756 | 5 | 64290435 | 64402838 | mNeoVis.2.1.1.chr5\_hap1.g00983.t01 |
| 6757 | 5 | 64290435 | 64402838 | mNeoVis.2.1.1.chr5\_hap1.g00983.t01 |
| 6758 | 5 | 64290435 | 64402838 | mNeoVis.2.1.1.chr5\_hap1.g00983.t01 |
| 6759 | 5 | 64999460 | 65152300 | mNeoVis.2.1.1.chr5\_hap1.g00984.t01 |
| 6760 | 5 | 65270134 | 65305230 | mNeoVis.2.1.1.chr5\_hap1.g00985.t01 |
| 6761 | 5 | 65270134 | 65305230 | mNeoVis.2.1.1.chr5\_hap1.g00985.t01 |
| 6762 | 5 | 65372864 | 65477903 | mNeoVis.2.1.1.chr5\_hap1.g00986.t01 |
| 6763 | 5 | 65372864 | 65477903 | mNeoVis.2.1.1.chr5\_hap1.g00986.t01 |
| 6764 | 5 | 65518900 | 65527084 | mNeoVis.2.1.1.chr5\_hap1.g00987.t01 |
| 6765 | 5 | 65527092 | 65590876 | mNeoVis.2.1.1.chr5\_hap1.g00988.t01 |
| 6766 | 5 | 65586647 | 65703260 | mNeoVis.2.1.1.chr5\_hap1.g00989.t01 |
| 6767 | 5 | 65744476 | 65872936 | mNeoVis.2.1.1.chr5\_hap1.g00990.t01 |
| 6768 | 5 | 65930964 | 66269562 | mNeoVis.2.1.1.chr5\_hap1.g00991.t01 |
| 6769 | 5 | 66680943 | 66729795 | mNeoVis.2.1.1.chr5\_hap1.g00999.t01 |
| 6770 | 5 | 66680943 | 66729795 | mNeoVis.2.1.1.chr5\_hap1.g00999.t01 |
| 6771 | 5 | 66720677 | 66730473 | mNeoVis.2.1.1.chr5\_hap1.g01000.t01 |
| 6772 | 5 | 66720677 | 66730473 | mNeoVis.2.1.1.chr5\_hap1.g01000.t01 |
| 6773 | 5 | 66743847 | 66769484 | mNeoVis.2.1.1.chr5\_hap1.g01001.t01 |
| 6774 | 5 | 66743847 | 66769484 | mNeoVis.2.1.1.chr5\_hap1.g01001.t01 |
| 6775 | 5 | 66792240 | 66858155 | mNeoVis.2.1.1.chr5\_hap1.g01002.t01 |
| 6776 | 5 | 66792240 | 66858155 | mNeoVis.2.1.1.chr5\_hap1.g01002.t01 |
| 6777 | 5 | 66867212 | 66902595 | mNeoVis.2.1.1.chr5\_hap1.g01003.t01 |
| 6778 | 5 | 66867212 | 66902595 | mNeoVis.2.1.1.chr5\_hap1.g01003.t01 |
| 6779 | 5 | 66905835 | 66933256 | mNeoVis.2.1.1.chr5\_hap1.g01004.t01 |
| 6780 | 5 | 66905835 | 66933256 | mNeoVis.2.1.1.chr5\_hap1.g01004.t01 |
| 6781 | 5 | 66905835 | 66933256 | mNeoVis.2.1.1.chr5\_hap1.g01004.t01 |
| 6782 | 5 | 66951333 | 67046149 | mNeoVis.2.1.1.chr5\_hap1.g01005.t01 |
| 6783 | 5 | 66951333 | 67046149 | mNeoVis.2.1.1.chr5\_hap1.g01005.t01 |
| 6784 | 5 | 66951333 | 67046149 | mNeoVis.2.1.1.chr5\_hap1.g01005.t01 |
| 6785 | 5 | 67098206 | 67206757 | mNeoVis.2.1.1.chr5\_hap1.g01006.t01 |
| 6786 | 5 | 67098206 | 67206757 | mNeoVis.2.1.1.chr5\_hap1.g01006.t01 |
| 6787 | 5 | 67253942 | 67423829 | mNeoVis.2.1.1.chr5\_hap1.g01007.t01 |
| 6788 | 5 | 67423798 | 67472311 | mNeoVis.2.1.1.chr5\_hap1.g01008.t01 |
| 6789 | 5 | 67472461 | 67482081 | mNeoVis.2.1.1.chr5\_hap1.g01009.t01 |
| 6790 | 5 | 67482191 | 67515251 | mNeoVis.2.1.1.chr5\_hap1.g01010.t01 |
| 6791 | 5 | 67518069 | 67530252 | mNeoVis.2.1.1.chr5\_hap1.g01011.t01 |
| 6792 | 5 | 67552575 | 67561257 | mNeoVis.2.1.1.chr5\_hap1.g01012.t01 |
| 6793 | 5 | 67618716 | 67619765 | mNeoVis.2.1.1.chr5\_hap1.g01013.t01 |
| 6794 | 5 | 67642565 | 67658127 | mNeoVis.2.1.1.chr5\_hap1.g01014.t01 |
| 6795 | 5 | 72078005 | 72081391 | mNeoVis.2.1.1.chr5\_hap1.g01028.t01 |
| 6796 | 5 | 72145311 | 72178780 | mNeoVis.2.1.1.chr5\_hap1.g01029.t01 |
| 6797 | 5 | 72198520 | 72202455 | mNeoVis.2.1.1.chr5\_hap1.g01030.t01 |
| 6798 | 5 | 72215896 | 72226191 | mNeoVis.2.1.1.chr5\_hap1.g01031.t01 |
| 6799 | 5 | 72228620 | 72260151 | mNeoVis.2.1.1.chr5\_hap1.g01032.t01 |
| 6800 | 5 | 72261702 | 72277830 | mNeoVis.2.1.1.chr5\_hap1.g01033.t01 |
| 6801 | 5 | 72762621 | 72851471 | mNeoVis.2.1.1.chr5\_hap1.g01035.t01 |
| 6802 | 5 | 72762621 | 72851471 | mNeoVis.2.1.1.chr5\_hap1.g01035.t01 |
| 6803 | 5 | 72849472 | 72890123 | mNeoVis.2.1.1.chr5\_hap1.g01036.t01 |
| 6804 | 5 | 72849472 | 72890123 | mNeoVis.2.1.1.chr5\_hap1.g01036.t01 |
| 6805 | 5 | 73663240 | 73777923 | mNeoVis.2.1.1.chr5\_hap1.g01038.t01 |
| 6806 | 5 | 73818086 | 73823737 | mNeoVis.2.1.1.chr5\_hap1.g01039.t01 |
| 6807 | 5 | 73958858 | 74068254 | mNeoVis.2.1.1.chr5\_hap1.g01040.t01 |
| 6808 | 5 | 74073147 | 74105066 | mNeoVis.2.1.1.chr5\_hap1.g01041.t01 |
| 6809 | 5 | 74119959 | 74144366 | mNeoVis.2.1.1.chr5\_hap1.g01042.t01 |
| 6810 | 5 | 74144285 | 74151166 | mNeoVis.2.1.1.chr5\_hap1.g01043.t01 |
| 6811 | 5 | 74164354 | 74225581 | mNeoVis.2.1.1.chr5\_hap1.g01044.t01 |
| 6812 | 5 | 74230753 | 74243628 | mNeoVis.2.1.1.chr5\_hap1.g01045.t01 |
| 6813 | 5 | 74230753 | 74243628 | mNeoVis.2.1.1.chr5\_hap1.g01045.t01 |
| 6814 | 5 | 74257320 | 74264504 | mNeoVis.2.1.1.chr5\_hap1.g01046.t01 |
| 6815 | 5 | 74257320 | 74264504 | mNeoVis.2.1.1.chr5\_hap1.g01046.t01 |
| 6816 | 5 | 74257320 | 74264504 | mNeoVis.2.1.1.chr5\_hap1.g01046.t01 |
| 6817 | 5 | 74264633 | 74277722 | mNeoVis.2.1.1.chr5\_hap1.g01047.t01 |
| 6818 | 5 | 74264633 | 74277722 | mNeoVis.2.1.1.chr5\_hap1.g01047.t01 |
| 6819 | 5 | 74264633 | 74277722 | mNeoVis.2.1.1.chr5\_hap1.g01047.t01 |
| 6820 | 5 | 74277784 | 74331290 | mNeoVis.2.1.1.chr5\_hap1.g01048.t01 |
| 6821 | 5 | 74277784 | 74331290 | mNeoVis.2.1.1.chr5\_hap1.g01048.t01 |
| 6822 | 5 | 74339379 | 74346464 | mNeoVis.2.1.1.chr5\_hap1.g01049.t01 |
| 6823 | 5 | 74339379 | 74346464 | mNeoVis.2.1.1.chr5\_hap1.g01049.t01 |
| 6824 | 5 | 74390149 | 74401244 | mNeoVis.2.1.1.chr5\_hap1.g01050.t01 |
| 6825 | 5 | 74390149 | 74401244 | mNeoVis.2.1.1.chr5\_hap1.g01050.t01 |
| 6826 | 5 | 74407099 | 74408695 | mNeoVis.2.1.1.chr5\_hap1.g01051.t01 |
| 6827 | 5 | 74407099 | 74408695 | mNeoVis.2.1.1.chr5\_hap1.g01051.t01 |
| 6828 | 5 | 74421795 | 74428256 | mNeoVis.2.1.1.chr5\_hap1.g01052.t01 |
| 6829 | 5 | 74421795 | 74428256 | mNeoVis.2.1.1.chr5\_hap1.g01052.t01 |
| 6830 | 5 | 75235204 | 75292347 | mNeoVis.2.1.1.chr5\_hap1.g01058.t01 |
| 6831 | 5 | 75306991 | 75414235 | mNeoVis.2.1.1.chr5\_hap1.g01059.t01 |
| 6832 | 5 | 75474891 | 75537323 | mNeoVis.2.1.1.chr5\_hap1.g01060.t01 |
| 6833 | 5 | 75474891 | 75537323 | mNeoVis.2.1.1.chr5\_hap1.g01060.t01 |
| 6834 | 5 | 75637167 | 75949047 | mNeoVis.2.1.1.chr5\_hap1.g01061.t01 |
| 6835 | 5 | 75966272 | 76062659 | mNeoVis.2.1.1.chr5\_hap1.g01062.t01 |
| 6836 | 5 | 76694561 | 76832166 | mNeoVis.2.1.1.chr5\_hap1.g01063.t01 |
| 6837 | 5 | 76694561 | 76832166 | mNeoVis.2.1.1.chr5\_hap1.g01063.t01 |
| 6838 | 5 | 76861339 | 76951474 | mNeoVis.2.1.1.chr5\_hap1.g01064.t01 |
| 6839 | 5 | 76861339 | 76951474 | mNeoVis.2.1.1.chr5\_hap1.g01064.t01 |
| 6840 | 5 | 77193276 | 77218465 | mNeoVis.2.1.1.chr5\_hap1.g01065.t01 |
| 6841 | 5 | 77193276 | 77218465 | mNeoVis.2.1.1.chr5\_hap1.g01065.t01 |
| 6842 | 5 | 77442896 | 78177743 | mNeoVis.2.1.1.chr5\_hap1.g01066.t01 |
| 6843 | 5 | 78262808 | 78785827 | mNeoVis.2.1.1.chr5\_hap1.g01067.t01 |
| 6844 | 5 | 79769119 | 79877485 | mNeoVis.2.1.1.chr5\_hap1.g01075.t01 |
| 6845 | 5 | 79880285 | 80108331 | mNeoVis.2.1.1.chr5\_hap1.g01076.t01 |
| 6846 | 5 | 80415329 | 80504925 | mNeoVis.2.1.1.chr5\_hap1.g01077.t01 |
| 6847 | 5 | 80843011 | 80919126 | mNeoVis.2.1.1.chr5\_hap1.g01078.t01 |
| 6848 | 5 | 81192295 | 81202450 | mNeoVis.2.1.1.chr5\_hap1.g01079.t01 |
| 6849 | 5 | 81201796 | 81258113 | mNeoVis.2.1.1.chr5\_hap1.g01080.t01 |
| 6850 | 5 | 81309512 | 81369351 | mNeoVis.2.1.1.chr5\_hap1.g01081.t01 |
| 6851 | 5 | 81369402 | 82082858 | mNeoVis.2.1.1.chr5\_hap1.g01082.t01 |
| 6852 | 5 | 82957960 | 82986135 | mNeoVis.2.1.1.chr5\_hap1.g01084.t01 |
| 6853 | 5 | 83038065 | 83042137 | mNeoVis.2.1.1.chr5\_hap1.g01085.t01 |
| 6854 | 5 | 83042225 | 83098069 | mNeoVis.2.1.1.chr5\_hap1.g01086.t01 |
| 6855 | 5 | 83074096 | 83160628 | mNeoVis.2.1.1.chr5\_hap1.g01087.t01 |
| 6856 | 5 | 85080070 | 85375370 | mNeoVis.2.1.1.chr5\_hap1.g01090.t01 |
| 6857 | 5 | 85080070 | 85375370 | mNeoVis.2.1.1.chr5\_hap1.g01090.t01 |
| 6858 | 5 | 85080070 | 85375370 | mNeoVis.2.1.1.chr5\_hap1.g01090.t01 |
| 6859 | 5 | 86408755 | 86409076 | mNeoVis.2.1.1.chr5\_hap1.g01091.t01 |
| 6860 | 5 | 86540220 | 86555403 | mNeoVis.2.1.1.chr5\_hap1.g01092.t01 |
| 6861 | 5 | 86562906 | 86665687 | mNeoVis.2.1.1.chr5\_hap1.g01093.t01 |
| 6862 | 5 | 86684345 | 86705006 | mNeoVis.2.1.1.chr5\_hap1.g01094.t01 |
| 6863 | 5 | 86705162 | 86734689 | mNeoVis.2.1.1.chr5\_hap1.g01095.t01 |
| 6864 | 5 | 86739976 | 86794642 | mNeoVis.2.1.1.chr5\_hap1.g01096.t01 |
| 6865 | 5 | 87125362 | 87510333 | mNeoVis.2.1.1.chr5\_hap1.g01097.t01 |
| 6866 | 5 | 87969515 | 87982311 | mNeoVis.2.1.1.chr5\_hap1.g01098.t01 |
| 6867 | 5 | 87969515 | 87982311 | mNeoVis.2.1.1.chr5\_hap1.g01098.t01 |
| 6868 | 5 | 88135295 | 88235259 | mNeoVis.2.1.1.chr5\_hap1.g01099.t01 |
| 6869 | 5 | 88135295 | 88235259 | mNeoVis.2.1.1.chr5\_hap1.g01099.t01 |
| 6870 | 5 | 88331199 | 88388213 | mNeoVis.2.1.1.chr5\_hap1.g01100.t01 |
| 6871 | 5 | 88331199 | 88388213 | mNeoVis.2.1.1.chr5\_hap1.g01100.t01 |
| 6872 | 5 | 88331199 | 88388213 | mNeoVis.2.1.1.chr5\_hap1.g01100.t01 |
| 6873 | 5 | 88786732 | 89343939 | mNeoVis.2.1.1.chr5\_hap1.g01101.t01 |
| 6874 | 5 | 89403354 | 89593958 | mNeoVis.2.1.1.chr5\_hap1.g01102.t01 |
| 6875 | 5 | 89794815 | 89813344 | mNeoVis.2.1.1.chr5\_hap1.g01103.t01 |
| 6876 | 5 | 89794815 | 89813344 | mNeoVis.2.1.1.chr5\_hap1.g01103.t01 |
| 6877 | 5 | 89794815 | 89813344 | mNeoVis.2.1.1.chr5\_hap1.g01103.t01 |
| 6878 | 5 | 89888482 | 90072768 | mNeoVis.2.1.1.chr5\_hap1.g01104.t01 |
| 6879 | 5 | 89888482 | 90072768 | mNeoVis.2.1.1.chr5\_hap1.g01104.t01 |
| 6880 | 5 | 89888482 | 90072768 | mNeoVis.2.1.1.chr5\_hap1.g01104.t01 |
| 6881 | 5 | 90027790 | 90060118 | mNeoVis.2.1.1.chr5\_hap1.g01105.t01 |
| 6882 | 5 | 90027790 | 90060118 | mNeoVis.2.1.1.chr5\_hap1.g01105.t01 |
| 6883 | 5 | 90027790 | 90060118 | mNeoVis.2.1.1.chr5\_hap1.g01105.t01 |
| 6884 | 5 | 90027790 | 90060118 | mNeoVis.2.1.1.chr5\_hap1.g01105.t01 |
| 6885 | 5 | 90085554 | 90122250 | mNeoVis.2.1.1.chr5\_hap1.g01106.t01 |
| 6886 | 5 | 90085554 | 90122250 | mNeoVis.2.1.1.chr5\_hap1.g01106.t01 |
| 6887 | 5 | 90085554 | 90122250 | mNeoVis.2.1.1.chr5\_hap1.g01106.t01 |
| 6888 | 5 | 90085554 | 90122250 | mNeoVis.2.1.1.chr5\_hap1.g01106.t01 |
| 6889 | 5 | 90122459 | 90129658 | mNeoVis.2.1.1.chr5\_hap1.g01107.t01 |
| 6890 | 5 | 90122459 | 90129658 | mNeoVis.2.1.1.chr5\_hap1.g01107.t01 |
| 6891 | 5 | 90122459 | 90129658 | mNeoVis.2.1.1.chr5\_hap1.g01107.t01 |
| 6892 | 5 | 90122459 | 90129658 | mNeoVis.2.1.1.chr5\_hap1.g01107.t01 |
| 6893 | 5 | 90153310 | 90195788 | mNeoVis.2.1.1.chr5\_hap1.g01108.t01 |
| 6894 | 5 | 90153310 | 90195788 | mNeoVis.2.1.1.chr5\_hap1.g01108.t01 |
| 6895 | 5 | 90153310 | 90195788 | mNeoVis.2.1.1.chr5\_hap1.g01108.t01 |
| 6896 | 5 | 90153310 | 90195788 | mNeoVis.2.1.1.chr5\_hap1.g01108.t01 |
| 6897 | 5 | 90359689 | 90360303 | mNeoVis.2.1.1.chr5\_hap1.g01109.t01 |
| 6898 | 5 | 90359689 | 90360303 | mNeoVis.2.1.1.chr5\_hap1.g01109.t01 |
| 6899 | 5 | 90359689 | 90360303 | mNeoVis.2.1.1.chr5\_hap1.g01109.t01 |
| 6900 | 5 | 90359689 | 90360303 | mNeoVis.2.1.1.chr5\_hap1.g01109.t01 |
| 6901 | 5 | 90491976 | 90627652 | mNeoVis.2.1.1.chr5\_hap1.g01110.t01 |
| 6902 | 5 | 90491976 | 90627652 | mNeoVis.2.1.1.chr5\_hap1.g01110.t01 |
| 6903 | 5 | 90631889 | 90658002 | mNeoVis.2.1.1.chr5\_hap1.g01111.t01 |
| 6904 | 5 | 90631889 | 90658002 | mNeoVis.2.1.1.chr5\_hap1.g01111.t01 |
| 6905 | 5 | 90658094 | 90704653 | mNeoVis.2.1.1.chr5\_hap1.g01112.t01 |
| 6906 | 5 | 90658094 | 90704653 | mNeoVis.2.1.1.chr5\_hap1.g01112.t01 |
| 6907 | 5 | 90740164 | 90795860 | mNeoVis.2.1.1.chr5\_hap1.g01113.t01 |
| 6908 | 5 | 90740164 | 90795860 | mNeoVis.2.1.1.chr5\_hap1.g01113.t01 |
| 6909 | 5 | 90783607 | 90798649 | mNeoVis.2.1.1.chr5\_hap1.g01114.t01 |
| 6910 | 5 | 90783607 | 90798649 | mNeoVis.2.1.1.chr5\_hap1.g01114.t01 |
| 6911 | 5 | 90798075 | 90813287 | mNeoVis.2.1.1.chr5\_hap1.g01115.t01 |
| 6912 | 5 | 90798075 | 90813287 | mNeoVis.2.1.1.chr5\_hap1.g01115.t01 |
| 6913 | 5 | 90854897 | 90927128 | mNeoVis.2.1.1.chr5\_hap1.g01116.t01 |
| 6914 | 5 | 90927902 | 90970502 | mNeoVis.2.1.1.chr5\_hap1.g01117.t01 |
| 6915 | 5 | 91044293 | 91147147 | mNeoVis.2.1.1.chr5\_hap1.g01118.t01 |
| 6916 | 5 | 91228262 | 91379862 | mNeoVis.2.1.1.chr5\_hap1.g01119.t01 |
| 6917 | 5 | 91415156 | 91565903 | mNeoVis.2.1.1.chr5\_hap1.g01120.t01 |
| 6918 | 5 | 91567655 | 91647207 | mNeoVis.2.1.1.chr5\_hap1.g01121.t01 |
| 6919 | 5 | 91753480 | 91930469 | mNeoVis.2.1.1.chr5\_hap1.g01122.t01 |
| 6920 | 5 | 91761514 | 91788666 | mNeoVis.2.1.1.chr5\_hap1.g01123.t01 |
| 6921 | 5 | 91929963 | 91940276 | mNeoVis.2.1.1.chr5\_hap1.g01124.t01 |
| 6922 | 5 | 91940435 | 92196601 | mNeoVis.2.1.1.chr5\_hap1.g01125.t01 |
| 6923 | 5 | 92208551 | 92211273 | mNeoVis.2.1.1.chr5\_hap1.g01126.t01 |
| 6924 | 5 | 92236511 | 92275345 | mNeoVis.2.1.1.chr5\_hap1.g01127.t01 |
| 6925 | 5 | 92281454 | 92329655 | mNeoVis.2.1.1.chr5\_hap1.g01128.t01 |
| 6926 | 5 | 92348229 | 92834052 | mNeoVis.2.1.1.chr5\_hap1.g01129.t01 |
| 6927 | 5 | 92722227 | 92722731 | mNeoVis.2.1.1.chr5\_hap1.g01130.t01 |
| 6928 | 5 | 93064618 | 93111560 | mNeoVis.2.1.1.chr5\_hap1.g01131.t01 |
| 6929 | 5 | 93163315 | 93291470 | mNeoVis.2.1.1.chr5\_hap1.g01132.t01 |
| 6930 | 5 | 93288676 | 93424720 | mNeoVis.2.1.1.chr5\_hap1.g01133.t01 |
| 6931 | 5 | 93368218 | 93374509 | mNeoVis.2.1.1.chr5\_hap1.g01134.t01 |
| 6932 | 5 | 93424939 | 93452867 | mNeoVis.2.1.1.chr5\_hap1.g01135.t01 |
| 6933 | 5 | 93424939 | 93452867 | mNeoVis.2.1.1.chr5\_hap1.g01135.t01 |
| 6934 | 5 | 93441243 | 93490757 | mNeoVis.2.1.1.chr5\_hap1.g01136.t01 |
| 6935 | 5 | 93441243 | 93490757 | mNeoVis.2.1.1.chr5\_hap1.g01136.t01 |
| 6936 | 5 | 93812548 | 94189285 | mNeoVis.2.1.1.chr5\_hap1.g01137.t01 |
| 6937 | 5 | 93812548 | 94189285 | mNeoVis.2.1.1.chr5\_hap1.g01137.t01 |
| 6938 | 5 | 94191553 | 94261568 | mNeoVis.2.1.1.chr5\_hap1.g01138.t01 |
| 6939 | 5 | 94330688 | 94346715 | mNeoVis.2.1.1.chr5\_hap1.g01139.t01 |
| 6940 | 5 | 94379128 | 94447552 | mNeoVis.2.1.1.chr5\_hap1.g01140.t01 |
| 6941 | 5 | 94461075 | 94504438 | mNeoVis.2.1.1.chr5\_hap1.g01141.t01 |
| 6942 | 5 | 94530559 | 94559956 | mNeoVis.2.1.1.chr5\_hap1.g01142.t01 |
| 6943 | 5 | 94563379 | 94593708 | mNeoVis.2.1.1.chr5\_hap1.g01143.t01 |
| 6944 | 5 | 94599102 | 94654700 | mNeoVis.2.1.1.chr5\_hap1.g01144.t01 |
| 6945 | 5 | 94678870 | 94758904 | mNeoVis.2.1.1.chr5\_hap1.g01145.t01 |
| 6946 | 5 | 94802431 | 94804324 | mNeoVis.2.1.1.chr5\_hap1.g01146.t01 |
| 6947 | 5 | 94845974 | 94930623 | mNeoVis.2.1.1.chr5\_hap1.g01147.t01 |
| 6948 | 5 | 95018804 | 95036753 | mNeoVis.2.1.1.chr5\_hap1.g01148.t01 |
| 6949 | 5 | 95115949 | 95147558 | mNeoVis.2.1.1.chr5\_hap1.g01149.t01 |
| 6950 | 5 | 95123306 | 95181405 | mNeoVis.2.1.1.chr5\_hap1.g01150.t01 |
| 6951 | 5 | 95202715 | 95249577 | mNeoVis.2.1.1.chr5\_hap1.g01151.t01 |
| 6952 | 5 | 95371234 | 95792062 | mNeoVis.2.1.1.chr5\_hap1.g01152.t01 |
| 6953 | 5 | 96306872 | 96515831 | mNeoVis.2.1.1.chr5\_hap1.g01153.t01 |
| 6954 | 5 | 96306872 | 96515831 | mNeoVis.2.1.1.chr5\_hap1.g01153.t01 |
| 6955 | 5 | 96516752 | 96557175 | mNeoVis.2.1.1.chr5\_hap1.g01154.t01 |
| 6956 | 5 | 96516752 | 96557175 | mNeoVis.2.1.1.chr5\_hap1.g01154.t01 |
| 6957 | 5 | 96587776 | 96694334 | mNeoVis.2.1.1.chr5\_hap1.g01155.t01 |
| 6958 | 5 | 96587776 | 96694334 | mNeoVis.2.1.1.chr5\_hap1.g01155.t01 |
| 6959 | 5 | 96699594 | 97128603 | mNeoVis.2.1.1.chr5\_hap1.g01156.t01 |
| 6960 | 5 | 96699594 | 97128603 | mNeoVis.2.1.1.chr5\_hap1.g01156.t01 |
| 6961 | 5 | 97188959 | 97230438 | mNeoVis.2.1.1.chr5\_hap1.g01157.t01 |
| 6962 | 5 | 97188959 | 97230438 | mNeoVis.2.1.1.chr5\_hap1.g01157.t01 |
| 6963 | 5 | 97235684 | 97317541 | mNeoVis.2.1.1.chr5\_hap1.g01158.t01 |
| 6964 | 5 | 97235684 | 97317541 | mNeoVis.2.1.1.chr5\_hap1.g01158.t01 |
| 6965 | 5 | 97327516 | 97365351 | mNeoVis.2.1.1.chr5\_hap1.g01159.t01 |
| 6966 | 5 | 97327516 | 97365351 | mNeoVis.2.1.1.chr5\_hap1.g01159.t01 |
| 6967 | 5 | 98074085 | 98075093 | mNeoVis.2.1.1.chr5\_hap1.g01160.t01 |
| 6968 | 5 | 98247147 | 98275735 | mNeoVis.2.1.1.chr5\_hap1.g01161.t01 |
| 6969 | 5 | 98247147 | 98275735 | mNeoVis.2.1.1.chr5\_hap1.g01161.t01 |
| 6970 | 5 | 98247147 | 98275735 | mNeoVis.2.1.1.chr5\_hap1.g01161.t01 |
| 6971 | 5 | 98303780 | 98347536 | mNeoVis.2.1.1.chr5\_hap1.g01162.t01 |
| 6972 | 5 | 98303780 | 98347536 | mNeoVis.2.1.1.chr5\_hap1.g01162.t01 |
| 6973 | 5 | 98303780 | 98347536 | mNeoVis.2.1.1.chr5\_hap1.g01162.t01 |
| 6974 | 5 | 98351764 | 98363682 | mNeoVis.2.1.1.chr5\_hap1.g01163.t01 |
| 6975 | 5 | 98351764 | 98363682 | mNeoVis.2.1.1.chr5\_hap1.g01163.t01 |
| 6976 | 5 | 98351764 | 98363682 | mNeoVis.2.1.1.chr5\_hap1.g01163.t01 |
| 6977 | 5 | 98384081 | 98397968 | mNeoVis.2.1.1.chr5\_hap1.g01164.t01 |
| 6978 | 5 | 98384081 | 98397968 | mNeoVis.2.1.1.chr5\_hap1.g01164.t01 |
| 6979 | 5 | 98384081 | 98397968 | mNeoVis.2.1.1.chr5\_hap1.g01164.t01 |
| 6980 | 5 | 98409849 | 98540121 | mNeoVis.2.1.1.chr5\_hap1.g01165.t01 |
| 6981 | 5 | 98409849 | 98540121 | mNeoVis.2.1.1.chr5\_hap1.g01165.t01 |
| 6982 | 5 | 98409849 | 98540121 | mNeoVis.2.1.1.chr5\_hap1.g01165.t01 |
| 6983 | 5 | 98524495 | 98546246 | mNeoVis.2.1.1.chr5\_hap1.g01166.t01 |
| 6984 | 5 | 98524495 | 98546246 | mNeoVis.2.1.1.chr5\_hap1.g01166.t01 |
| 6985 | 5 | 98524495 | 98546246 | mNeoVis.2.1.1.chr5\_hap1.g01166.t01 |
| 6986 | 5 | 98562724 | 98626598 | mNeoVis.2.1.1.chr5\_hap1.g01167.t01 |
| 6987 | 5 | 98562724 | 98626598 | mNeoVis.2.1.1.chr5\_hap1.g01167.t01 |
| 6988 | 5 | 98562724 | 98626598 | mNeoVis.2.1.1.chr5\_hap1.g01167.t01 |
| 6989 | 5 | 98666123 | 98715530 | mNeoVis.2.1.1.chr5\_hap1.g01168.t01 |
| 6990 | 5 | 98666123 | 98715530 | mNeoVis.2.1.1.chr5\_hap1.g01168.t01 |
| 6991 | 5 | 98666123 | 98715530 | mNeoVis.2.1.1.chr5\_hap1.g01168.t01 |
| 6992 | 5 | 98843735 | 98858170 | mNeoVis.2.1.1.chr5\_hap1.g01170.t01 |
| 6993 | 5 | 98881431 | 99082999 | mNeoVis.2.1.1.chr5\_hap1.g01171.t01 |
| 6994 | 5 | 99000864 | 99002402 | mNeoVis.2.1.1.chr5\_hap1.g01172.t01 |
| 6995 | 5 | 99168741 | 99169874 | mNeoVis.2.1.1.chr5\_hap1.g01173.t01 |
| 6996 | 5 | 99222583 | 99223326 | mNeoVis.2.1.1.chr5\_hap1.g01174.t01 |
| 6997 | 5 | 99229063 | 99230532 | mNeoVis.2.1.1.chr5\_hap1.g01175.t01 |
| 6998 | 5 | 99258777 | 99260764 | mNeoVis.2.1.1.chr5\_hap1.g01176.t01 |
| 6999 | 5 | 99732131 | 99733397 | mNeoVis.2.1.1.chr5\_hap1.g01180.t01 |
| 7000 | 5 | 99740243 | 99764434 | mNeoVis.2.1.1.chr5\_hap1.g01181.t01 |
| 7001 | 5 | 99780869 | 99901179 | mNeoVis.2.1.1.chr5\_hap1.g01182.t01 |
| 7002 | 5 | 100108015 | 100740487 | mNeoVis.2.1.1.chr5\_hap1.g01183.t01 |
| 7003 | 5 | 100812142 | 100834586 | mNeoVis.2.1.1.chr5\_hap1.g01184.t01 |
| 7004 | 5 | 101688616 | 101736352 | mNeoVis.2.1.1.chr5\_hap1.g01192.t01 |
| 7005 | 5 | 101747011 | 101775595 | mNeoVis.2.1.1.chr5\_hap1.g01193.t01 |
| 7006 | 5 | 101794217 | 101861588 | mNeoVis.2.1.1.chr5\_hap1.g01194.t01 |
| 7007 | 5 | 101897999 | 101972690 | mNeoVis.2.1.1.chr5\_hap1.g01195.t01 |
| 7008 | 5 | 102031201 | 102042902 | mNeoVis.2.1.1.chr5\_hap1.g01196.t01 |
| 7009 | 5 | 102055819 | 102079571 | mNeoVis.2.1.1.chr5\_hap1.g01197.t01 |
| 7010 | 5 | 102095355 | 102108738 | mNeoVis.2.1.1.chr5\_hap1.g01198.t01 |
| 7011 | 5 | 102110262 | 102113297 | mNeoVis.2.1.1.chr5\_hap1.g01199.t01 |
| 7012 | 5 | 102117215 | 102188249 | mNeoVis.2.1.1.chr5\_hap1.g01200.t01 |
| 7013 | 5 | 102196549 | 102232689 | mNeoVis.2.1.1.chr5\_hap1.g01201.t01 |
| 7014 | 5 | 102196549 | 102232689 | mNeoVis.2.1.1.chr5\_hap1.g01201.t01 |
| 7015 | 5 | 102232422 | 102242548 | mNeoVis.2.1.1.chr5\_hap1.g01202.t01 |
| 7016 | 5 | 102232422 | 102242548 | mNeoVis.2.1.1.chr5\_hap1.g01202.t01 |
| 7017 | 5 | 102248193 | 102298444 | mNeoVis.2.1.1.chr5\_hap1.g01203.t01 |
| 7018 | 5 | 102248193 | 102298444 | mNeoVis.2.1.1.chr5\_hap1.g01203.t01 |
| 7019 | 5 | 102323221 | 102374606 | mNeoVis.2.1.1.chr5\_hap1.g01204.t01 |
| 7020 | 5 | 102378506 | 102441367 | mNeoVis.2.1.1.chr5\_hap1.g01205.t01 |
| 7021 | 5 | 102442485 | 102511015 | mNeoVis.2.1.1.chr5\_hap1.g01206.t01 |
| 7022 | 5 | 102541536 | 102582082 | mNeoVis.2.1.1.chr5\_hap1.g01207.t01 |
| 7023 | 5 | 102695334 | 102786335 | mNeoVis.2.1.1.chr5\_hap1.g01208.t01 |
| 7024 | 5 | 102695334 | 102786335 | mNeoVis.2.1.1.chr5\_hap1.g01208.t01 |
| 7025 | 5 | 102834090 | 102891264 | mNeoVis.2.1.1.chr5\_hap1.g01209.t01 |
| 7026 | 5 | 102834090 | 102891264 | mNeoVis.2.1.1.chr5\_hap1.g01209.t01 |
| 7027 | 5 | 102896951 | 103044615 | mNeoVis.2.1.1.chr5\_hap1.g01210.t01 |
| 7028 | 5 | 102896951 | 103044615 | mNeoVis.2.1.1.chr5\_hap1.g01210.t01 |
| 7029 | 5 | 103085195 | 103206087 | mNeoVis.2.1.1.chr5\_hap1.g01211.t01 |
| 7030 | 5 | 103232079 | 103409867 | mNeoVis.2.1.1.chr5\_hap1.g01212.t01 |
| 7031 | 5 | 103536268 | 103583089 | mNeoVis.2.1.1.chr5\_hap1.g01213.t01 |
| 7032 | 5 | 103662471 | 104324305 | mNeoVis.2.1.1.chr5\_hap1.g01215.t01 |
| 7033 | 5 | 104330789 | 104361079 | mNeoVis.2.1.1.chr5\_hap1.g01216.t01 |
| 7034 | 5 | 104380778 | 104494892 | mNeoVis.2.1.1.chr5\_hap1.g01217.t01 |
| 7035 | 5 | 104510439 | 104538154 | mNeoVis.2.1.1.chr5\_hap1.g01218.t01 |
| 7036 | 5 | 104563839 | 104651983 | mNeoVis.2.1.1.chr5\_hap1.g01219.t01 |
| 7037 | 5 | 104673488 | 104812076 | mNeoVis.2.1.1.chr5\_hap1.g01220.t01 |
| 7038 | 5 | 104822589 | 104947676 | mNeoVis.2.1.1.chr5\_hap1.g01221.t01 |
| 7039 | 5 | 105011223 | 105081674 | mNeoVis.2.1.1.chr5\_hap1.g01222.t01 |
| 7040 | 5 | 105103699 | 105204751 | mNeoVis.2.1.1.chr5\_hap1.g01223.t01 |
| 7041 | 5 | 105354766 | 105358163 | mNeoVis.2.1.1.chr5\_hap1.g01225.t01 |
| 7042 | 5 | 105358240 | 105407802 | mNeoVis.2.1.1.chr5\_hap1.g01226.t01 |
| 7043 | 5 | 105373963 | 105375640 | mNeoVis.2.1.1.chr5\_hap1.g01227.t01 |
| 7044 | 5 | 105408989 | 105525703 | mNeoVis.2.1.1.chr5\_hap1.g01228.t01 |
| 7045 | 5 | 105530472 | 105563812 | mNeoVis.2.1.1.chr5\_hap1.g01229.t01 |
| 7046 | 5 | 105576245 | 105632715 | mNeoVis.2.1.1.chr5\_hap1.g01230.t01 |
| 7047 | 5 | 105670190 | 105682636 | mNeoVis.2.1.1.chr5\_hap1.g01231.t01 |
| 7048 | 5 | 105688083 | 105744890 | mNeoVis.2.1.1.chr5\_hap1.g01232.t01 |
| 7049 | 5 | 105793467 | 105816577 | mNeoVis.2.1.1.chr5\_hap1.g01233.t01 |
| 7050 | 5 | 105857213 | 105886462 | mNeoVis.2.1.1.chr5\_hap1.g01234.t01 |
| 7051 | 5 | 106292472 | 106308025 | mNeoVis.2.1.1.chr5\_hap1.g01244.t01 |
| 7052 | 5 | 106308271 | 106325063 | mNeoVis.2.1.1.chr5\_hap1.g01245.t01 |
| 7053 | 5 | 106365305 | 106393802 | mNeoVis.2.1.1.chr5\_hap1.g01246.t01 |
| 7054 | 5 | 106432891 | 106438373 | mNeoVis.2.1.1.chr5\_hap1.g01247.t01 |
| 7055 | 5 | 106438289 | 106479853 | mNeoVis.2.1.1.chr5\_hap1.g01248.t01 |
| 7056 | 5 | 106480999 | 106575100 | mNeoVis.2.1.1.chr5\_hap1.g01249.t01 |
| 7057 | 5 | 106598181 | 106664804 | mNeoVis.2.1.1.chr5\_hap1.g01250.t01 |
| 7058 | 5 | 106632430 | 106668359 | mNeoVis.2.1.1.chr5\_hap1.g01251.t01 |
| 7059 | 5 | 106668398 | 106669251 | mNeoVis.2.1.1.chr5\_hap1.g01252.t01 |
| 7060 | 5 | 106672091 | 106696314 | mNeoVis.2.1.1.chr5\_hap1.g01253.t01 |
| 7061 | 5 | 106722747 | 106746496 | mNeoVis.2.1.1.chr5\_hap1.g01254.t01 |
| 7062 | 5 | 106757569 | 107022565 | mNeoVis.2.1.1.chr5\_hap1.g01255.t01 |
| 7063 | 5 | 107178264 | 107210582 | mNeoVis.2.1.1.chr5\_hap1.g01256.t01 |
| 7064 | 5 | 107272437 | 107289790 | mNeoVis.2.1.1.chr5\_hap1.g01257.t01 |
| 7065 | 5 | 107300550 | 107311703 | mNeoVis.2.1.1.chr5\_hap1.g01258.t01 |
| 7066 | 5 | 107311063 | 107340932 | mNeoVis.2.1.1.chr5\_hap1.g01259.t01 |
| 7067 | 5 | 107376794 | 107536551 | mNeoVis.2.1.1.chr5\_hap1.g01260.t01 |
| 7068 | 5 | 107541172 | 107698823 | mNeoVis.2.1.1.chr5\_hap1.g01261.t01 |
| 7069 | 5 | 107999907 | 108005148 | mNeoVis.2.1.1.chr5\_hap1.g01262.t01 |
| 7070 | 5 | 108018180 | 108056311 | mNeoVis.2.1.1.chr5\_hap1.g01263.t01 |
| 7071 | 5 | 108062659 | 108088248 | mNeoVis.2.1.1.chr5\_hap1.g01264.t01 |
| 7072 | 5 | 108165499 | 108257941 | mNeoVis.2.1.1.chr5\_hap1.g01265.t01 |
| 7073 | 5 | 108261612 | 108263518 | mNeoVis.2.1.1.chr5\_hap1.g01266.t01 |
| 7074 | 5 | 108297680 | 108310978 | mNeoVis.2.1.1.chr5\_hap1.g01267.t01 |
| 7075 | 5 | 108319252 | 108321729 | mNeoVis.2.1.1.chr5\_hap1.g01268.t01 |
| 7076 | 5 | 108484658 | 108487212 | mNeoVis.2.1.1.chr5\_hap1.g01269.t01 |
| 7077 | 5 | 108854414 | 108945729 | mNeoVis.2.1.1.chr5\_hap1.g01270.t01 |
| 7078 | 5 | 108854414 | 108945729 | mNeoVis.2.1.1.chr5\_hap1.g01270.t01 |
| 7079 | 5 | 109244120 | 109262387 | mNeoVis.2.1.1.chr5\_hap1.g01273.t01 |
| 7080 | 5 | 109522729 | 109635590 | mNeoVis.2.1.1.chr5\_hap1.g01274.t01 |
| 7081 | 5 | 109522729 | 109635590 | mNeoVis.2.1.1.chr5\_hap1.g01274.t01 |
| 7082 | 5 | 109522729 | 109635590 | mNeoVis.2.1.1.chr5\_hap1.g01274.t01 |
| 7083 | 5 | 109986834 | 110252611 | mNeoVis.2.1.1.chr5\_hap1.g01275.t01 |
| 7084 | 5 | 110946345 | 111017203 | mNeoVis.2.1.1.chr5\_hap1.g01276.t01 |
| 7085 | 5 | 111062658 | 111080773 | mNeoVis.2.1.1.chr5\_hap1.g01277.t01 |
| 7086 | 5 | 111081352 | 111122197 | mNeoVis.2.1.1.chr5\_hap1.g01278.t01 |
| 7087 | 5 | 111400067 | 111410897 | mNeoVis.2.1.1.chr5\_hap1.g01279.t01 |
| 7088 | 5 | 111593121 | 111721944 | mNeoVis.2.1.1.chr5\_hap1.g01280.t01 |
| 7089 | 5 | 111785388 | 111801118 | mNeoVis.2.1.1.chr5\_hap1.g01281.t01 |
| 7090 | 5 | 111813352 | 111889741 | mNeoVis.2.1.1.chr5\_hap1.g01282.t01 |
| 7091 | 5 | 111909322 | 111925616 | mNeoVis.2.1.1.chr5\_hap1.g01283.t01 |
| 7092 | 5 | 112057306 | 112214189 | mNeoVis.2.1.1.chr5\_hap1.g01284.t01 |
| 7093 | 5 | 112057306 | 112214189 | mNeoVis.2.1.1.chr5\_hap1.g01284.t01 |
| 7094 | 5 | 112057306 | 112214189 | mNeoVis.2.1.1.chr5\_hap1.g01284.t01 |
| 7095 | 5 | 112259849 | 112361431 | mNeoVis.2.1.1.chr5\_hap1.g01285.t01 |
| 7096 | 5 | 112259849 | 112361431 | mNeoVis.2.1.1.chr5\_hap1.g01285.t01 |
| 7097 | 5 | 112396984 | 112488204 | mNeoVis.2.1.1.chr5\_hap1.g01286.t01 |
| 7098 | 5 | 112738872 | 112888623 | mNeoVis.2.1.1.chr5\_hap1.g01287.t01 |
| 7099 | 5 | 113019206 | 113019814 | mNeoVis.2.1.1.chr5\_hap1.g01288.t01 |
| 7100 | 5 | 113466072 | 113504785 | mNeoVis.2.1.1.chr5\_hap1.g01290.t01 |
| 7101 | 5 | 113626596 | 113627283 | mNeoVis.2.1.1.chr5\_hap1.g01291.t01 |
| 7102 | 5 | 114183855 | 114206714 | mNeoVis.2.1.1.chr5\_hap1.g01292.t01 |
| 7103 | 5 | 114183855 | 114206714 | mNeoVis.2.1.1.chr5\_hap1.g01292.t01 |
| 7104 | 5 | 114227008 | 114230544 | mNeoVis.2.1.1.chr5\_hap1.g01293.t01 |
| 7105 | 5 | 114227008 | 114230544 | mNeoVis.2.1.1.chr5\_hap1.g01293.t01 |
| 7106 | 5 | 114251300 | 114252753 | mNeoVis.2.1.1.chr5\_hap1.g01294.t01 |
| 7107 | 5 | 114251300 | 114252753 | mNeoVis.2.1.1.chr5\_hap1.g01294.t01 |
| 7108 | 5 | 114485844 | 114507087 | mNeoVis.2.1.1.chr5\_hap1.g01295.t01 |
| 7109 | 5 | 114485844 | 114507087 | mNeoVis.2.1.1.chr5\_hap1.g01295.t01 |
| 7110 | 5 | 114559015 | 114615654 | mNeoVis.2.1.1.chr5\_hap1.g01296.t01 |
| 7111 | 5 | 114559015 | 114615654 | mNeoVis.2.1.1.chr5\_hap1.g01296.t01 |
| 7112 | 5 | 114633318 | 114635827 | mNeoVis.2.1.1.chr5\_hap1.g01297.t01 |
| 7113 | 5 | 114633318 | 114635827 | mNeoVis.2.1.1.chr5\_hap1.g01297.t01 |
| 7114 | 5 | 114914208 | 114924827 | mNeoVis.2.1.1.chr5\_hap1.g01299.t01 |
| 7115 | 5 | 114914208 | 114924827 | mNeoVis.2.1.1.chr5\_hap1.g01299.t01 |
| 7116 | 5 | 114947708 | 114963715 | mNeoVis.2.1.1.chr5\_hap1.g01300.t01 |
| 7117 | 5 | 114947708 | 114963715 | mNeoVis.2.1.1.chr5\_hap1.g01300.t01 |
| 7118 | 5 | 114947708 | 114963715 | mNeoVis.2.1.1.chr5\_hap1.g01300.t01 |
| 7119 | 5 | 114963710 | 114970581 | mNeoVis.2.1.1.chr5\_hap1.g01301.t01 |
| 7120 | 5 | 114963710 | 114970581 | mNeoVis.2.1.1.chr5\_hap1.g01301.t01 |
| 7121 | 5 | 114963710 | 114970581 | mNeoVis.2.1.1.chr5\_hap1.g01301.t01 |
| 7122 | 5 | 115002308 | 115029924 | mNeoVis.2.1.1.chr5\_hap1.g01302.t01 |
| 7123 | 5 | 115002308 | 115029924 | mNeoVis.2.1.1.chr5\_hap1.g01302.t01 |
| 7124 | 5 | 115002308 | 115029924 | mNeoVis.2.1.1.chr5\_hap1.g01302.t01 |
| 7125 | 5 | 115002308 | 115029924 | mNeoVis.2.1.1.chr5\_hap1.g01302.t01 |
| 7126 | 5 | 115031808 | 115041105 | mNeoVis.2.1.1.chr5\_hap1.g01303.t01 |
| 7127 | 5 | 115031808 | 115041105 | mNeoVis.2.1.1.chr5\_hap1.g01303.t01 |
| 7128 | 5 | 115031808 | 115041105 | mNeoVis.2.1.1.chr5\_hap1.g01303.t01 |
| 7129 | 5 | 115031808 | 115041105 | mNeoVis.2.1.1.chr5\_hap1.g01303.t01 |
| 7130 | 5 | 115045021 | 115059087 | mNeoVis.2.1.1.chr5\_hap1.g01304.t01 |
| 7131 | 5 | 115045021 | 115059087 | mNeoVis.2.1.1.chr5\_hap1.g01304.t01 |
| 7132 | 5 | 115045021 | 115059087 | mNeoVis.2.1.1.chr5\_hap1.g01304.t01 |
| 7133 | 5 | 115045021 | 115059087 | mNeoVis.2.1.1.chr5\_hap1.g01304.t01 |
| 7134 | 5 | 115085209 | 115092598 | mNeoVis.2.1.1.chr5\_hap1.g01305.t01 |
| 7135 | 5 | 115085209 | 115092598 | mNeoVis.2.1.1.chr5\_hap1.g01305.t01 |
| 7136 | 5 | 115085209 | 115092598 | mNeoVis.2.1.1.chr5\_hap1.g01305.t01 |
| 7137 | 5 | 115085209 | 115092598 | mNeoVis.2.1.1.chr5\_hap1.g01305.t01 |
| 7138 | 5 | 115109076 | 115128094 | mNeoVis.2.1.1.chr5\_hap1.g01306.t01 |
| 7139 | 5 | 115109076 | 115128094 | mNeoVis.2.1.1.chr5\_hap1.g01306.t01 |
| 7140 | 5 | 115109076 | 115128094 | mNeoVis.2.1.1.chr5\_hap1.g01306.t01 |
| 7141 | 5 | 115109076 | 115128094 | mNeoVis.2.1.1.chr5\_hap1.g01306.t01 |
| 7142 | 5 | 115128283 | 115147624 | mNeoVis.2.1.1.chr5\_hap1.g01307.t01 |
| 7143 | 5 | 115128283 | 115147624 | mNeoVis.2.1.1.chr5\_hap1.g01307.t01 |
| 7144 | 5 | 115128283 | 115147624 | mNeoVis.2.1.1.chr5\_hap1.g01307.t01 |
| 7145 | 5 | 115128283 | 115147624 | mNeoVis.2.1.1.chr5\_hap1.g01307.t01 |
| 7146 | 5 | 115150221 | 115156256 | mNeoVis.2.1.1.chr5\_hap1.g01308.t01 |
| 7147 | 5 | 115150221 | 115156256 | mNeoVis.2.1.1.chr5\_hap1.g01308.t01 |
| 7148 | 5 | 115150221 | 115156256 | mNeoVis.2.1.1.chr5\_hap1.g01308.t01 |
| 7149 | 5 | 115150221 | 115156256 | mNeoVis.2.1.1.chr5\_hap1.g01308.t01 |
| 7150 | 5 | 115294720 | 115504906 | mNeoVis.2.1.1.chr5\_hap1.g01309.t01 |
| 7151 | 5 | 115294720 | 115504906 | mNeoVis.2.1.1.chr5\_hap1.g01309.t01 |
| 7152 | 5 | 115294720 | 115504906 | mNeoVis.2.1.1.chr5\_hap1.g01309.t01 |
| 7153 | 5 | 115294720 | 115504906 | mNeoVis.2.1.1.chr5\_hap1.g01309.t01 |
| 7154 | 5 | 115526977 | 115585209 | mNeoVis.2.1.1.chr5\_hap1.g01310.t01 |
| 7155 | 5 | 115526977 | 115585209 | mNeoVis.2.1.1.chr5\_hap1.g01310.t01 |
| 7156 | 5 | 115526977 | 115585209 | mNeoVis.2.1.1.chr5\_hap1.g01310.t01 |
| 7157 | 5 | 115656785 | 115675849 | mNeoVis.2.1.1.chr5\_hap1.g01311.t01 |
| 7158 | 5 | 115656785 | 115675849 | mNeoVis.2.1.1.chr5\_hap1.g01311.t01 |
| 7159 | 5 | 115952399 | 115986606 | mNeoVis.2.1.1.chr5\_hap1.g01313.t01 |
| 7160 | 5 | 115995778 | 116041256 | mNeoVis.2.1.1.chr5\_hap1.g01314.t01 |
| 7161 | 5 | 116048579 | 116058389 | mNeoVis.2.1.1.chr5\_hap1.g01315.t01 |
| 7162 | 5 | 116058969 | 116230173 | mNeoVis.2.1.1.chr5\_hap1.g01316.t01 |
| 7163 | 5 | 116058969 | 116230173 | mNeoVis.2.1.1.chr5\_hap1.g01316.t01 |
| 7164 | 5 | 116241394 | 116292224 | mNeoVis.2.1.1.chr5\_hap1.g01317.t01 |
| 7165 | 5 | 116241394 | 116292224 | mNeoVis.2.1.1.chr5\_hap1.g01317.t01 |
| 7166 | 5 | 116325046 | 116408148 | mNeoVis.2.1.1.chr5\_hap1.g01318.t01 |
| 7167 | 5 | 116325046 | 116408148 | mNeoVis.2.1.1.chr5\_hap1.g01318.t01 |
| 7168 | 5 | 116763358 | 116765099 | mNeoVis.2.1.1.chr5\_hap1.g01320.t01 |
| 7169 | 5 | 116763358 | 116765099 | mNeoVis.2.1.1.chr5\_hap1.g01320.t01 |
| 7170 | 5 | 116874333 | 116894014 | mNeoVis.2.1.1.chr5\_hap1.g01321.t01 |
| 7171 | 5 | 116874333 | 116894014 | mNeoVis.2.1.1.chr5\_hap1.g01321.t01 |
| 7172 | 5 | 116874333 | 116894014 | mNeoVis.2.1.1.chr5\_hap1.g01321.t01 |
| 7173 | 5 | 116954179 | 116955958 | mNeoVis.2.1.1.chr5\_hap1.g01322.t01 |
| 7174 | 5 | 116954179 | 116955958 | mNeoVis.2.1.1.chr5\_hap1.g01322.t01 |
| 7175 | 5 | 116954179 | 116955958 | mNeoVis.2.1.1.chr5\_hap1.g01322.t01 |
| 7176 | 5 | 117041772 | 117050502 | mNeoVis.2.1.1.chr5\_hap1.g01323.t01 |
| 7177 | 5 | 117041772 | 117050502 | mNeoVis.2.1.1.chr5\_hap1.g01323.t01 |
| 7178 | 5 | 117041772 | 117050502 | mNeoVis.2.1.1.chr5\_hap1.g01323.t01 |
| 7179 | 5 | 117051733 | 117057951 | mNeoVis.2.1.1.chr5\_hap1.g01324.t01 |
| 7180 | 5 | 117051733 | 117057951 | mNeoVis.2.1.1.chr5\_hap1.g01324.t01 |
| 7181 | 5 | 117051733 | 117057951 | mNeoVis.2.1.1.chr5\_hap1.g01324.t01 |
| 7182 | 5 | 117061268 | 117064676 | mNeoVis.2.1.1.chr5\_hap1.g01325.t01 |
| 7183 | 5 | 117061268 | 117064676 | mNeoVis.2.1.1.chr5\_hap1.g01325.t01 |
| 7184 | 5 | 117061268 | 117064676 | mNeoVis.2.1.1.chr5\_hap1.g01325.t01 |
| 7185 | 5 | 117066447 | 117081606 | mNeoVis.2.1.1.chr5\_hap1.g01326.t01 |
| 7186 | 5 | 117066447 | 117081606 | mNeoVis.2.1.1.chr5\_hap1.g01326.t01 |
| 7187 | 5 | 117066447 | 117081606 | mNeoVis.2.1.1.chr5\_hap1.g01326.t01 |
| 7188 | 5 | 117081581 | 117087985 | mNeoVis.2.1.1.chr5\_hap1.g01327.t01 |
| 7189 | 5 | 117081581 | 117087985 | mNeoVis.2.1.1.chr5\_hap1.g01327.t01 |
| 7190 | 5 | 117081581 | 117087985 | mNeoVis.2.1.1.chr5\_hap1.g01327.t01 |
| 7191 | 5 | 117090076 | 117109836 | mNeoVis.2.1.1.chr5\_hap1.g01328.t01 |
| 7192 | 5 | 117090076 | 117109836 | mNeoVis.2.1.1.chr5\_hap1.g01328.t01 |
| 7193 | 5 | 117090076 | 117109836 | mNeoVis.2.1.1.chr5\_hap1.g01328.t01 |
| 7194 | 5 | 117114084 | 117123591 | mNeoVis.2.1.1.chr5\_hap1.g01329.t01 |
| 7195 | 5 | 117114084 | 117123591 | mNeoVis.2.1.1.chr5\_hap1.g01329.t01 |
| 7196 | 5 | 117114084 | 117123591 | mNeoVis.2.1.1.chr5\_hap1.g01329.t01 |
| 7197 | 5 | 117129718 | 117143243 | mNeoVis.2.1.1.chr5\_hap1.g01330.t01 |
| 7198 | 5 | 117129718 | 117143243 | mNeoVis.2.1.1.chr5\_hap1.g01330.t01 |
| 7199 | 5 | 117129718 | 117143243 | mNeoVis.2.1.1.chr5\_hap1.g01330.t01 |
| 7200 | 5 | 117151783 | 117153185 | mNeoVis.2.1.1.chr5\_hap1.g01331.t01 |
| 7201 | 5 | 117151783 | 117153185 | mNeoVis.2.1.1.chr5\_hap1.g01331.t01 |
| 7202 | 5 | 117151783 | 117153185 | mNeoVis.2.1.1.chr5\_hap1.g01331.t01 |
| 7203 | 5 | 117154208 | 117160928 | mNeoVis.2.1.1.chr5\_hap1.g01332.t01 |
| 7204 | 5 | 117154208 | 117160928 | mNeoVis.2.1.1.chr5\_hap1.g01332.t01 |
| 7205 | 5 | 117154208 | 117160928 | mNeoVis.2.1.1.chr5\_hap1.g01332.t01 |
| 7206 | 5 | 117161582 | 117167051 | mNeoVis.2.1.1.chr5\_hap1.g01333.t01 |
| 7207 | 5 | 117161582 | 117167051 | mNeoVis.2.1.1.chr5\_hap1.g01333.t01 |
| 7208 | 5 | 117161582 | 117167051 | mNeoVis.2.1.1.chr5\_hap1.g01333.t01 |
| 7209 | 5 | 117167074 | 117178006 | mNeoVis.2.1.1.chr5\_hap1.g01334.t01 |
| 7210 | 5 | 117167074 | 117178006 | mNeoVis.2.1.1.chr5\_hap1.g01334.t01 |
| 7211 | 5 | 117167074 | 117178006 | mNeoVis.2.1.1.chr5\_hap1.g01334.t01 |
| 7212 | 5 | 117206344 | 117213586 | mNeoVis.2.1.1.chr5\_hap1.g01335.t01 |
| 7213 | 5 | 117206344 | 117213586 | mNeoVis.2.1.1.chr5\_hap1.g01335.t01 |
| 7214 | 5 | 117206344 | 117213586 | mNeoVis.2.1.1.chr5\_hap1.g01335.t01 |
| 7215 | 5 | 117216103 | 117225231 | mNeoVis.2.1.1.chr5\_hap1.g01336.t01 |
| 7216 | 5 | 117216103 | 117225231 | mNeoVis.2.1.1.chr5\_hap1.g01336.t01 |
| 7217 | 5 | 117216103 | 117225231 | mNeoVis.2.1.1.chr5\_hap1.g01336.t01 |
| 7218 | 5 | 117227021 | 117244613 | mNeoVis.2.1.1.chr5\_hap1.g01337.t01 |
| 7219 | 5 | 117227021 | 117244613 | mNeoVis.2.1.1.chr5\_hap1.g01337.t01 |
| 7220 | 5 | 117240506 | 117241399 | mNeoVis.2.1.1.chr5\_hap1.g01338.t01 |
| 7221 | 5 | 117240506 | 117241399 | mNeoVis.2.1.1.chr5\_hap1.g01338.t01 |
| 7222 | 5 | 117251652 | 117260698 | mNeoVis.2.1.1.chr5\_hap1.g01339.t01 |
| 7223 | 5 | 117251652 | 117260698 | mNeoVis.2.1.1.chr5\_hap1.g01339.t01 |
| 7224 | 5 | 117295086 | 117300564 | mNeoVis.2.1.1.chr5\_hap1.g01340.t01 |
| 7225 | 5 | 117295086 | 117300564 | mNeoVis.2.1.1.chr5\_hap1.g01340.t01 |
| 7226 | 5 | 117306805 | 117313460 | mNeoVis.2.1.1.chr5\_hap1.g01341.t01 |
| 7227 | 5 | 117306805 | 117313460 | mNeoVis.2.1.1.chr5\_hap1.g01341.t01 |
| 7228 | 5 | 117320160 | 117325077 | mNeoVis.2.1.1.chr5\_hap1.g01342.t01 |
| 7229 | 5 | 117320160 | 117325077 | mNeoVis.2.1.1.chr5\_hap1.g01342.t01 |
| 7230 | 5 | 117336476 | 117424743 | mNeoVis.2.1.1.chr5\_hap1.g01343.t01 |
| 7231 | 5 | 117446884 | 117499223 | mNeoVis.2.1.1.chr5\_hap1.g01344.t01 |
| 7232 | 5 | 117502401 | 117512546 | mNeoVis.2.1.1.chr5\_hap1.g01345.t01 |
| 7233 | 5 | 117512306 | 117633863 | mNeoVis.2.1.1.chr5\_hap1.g01346.t01 |
| 7234 | 5 | 117658847 | 117708110 | mNeoVis.2.1.1.chr5\_hap1.g01347.t01 |
| 7235 | 5 | 117783730 | 117842529 | mNeoVis.2.1.1.chr5\_hap1.g01348.t01 |
| 7236 | 5 | 118070384 | 118089759 | mNeoVis.2.1.1.chr5\_hap1.g01350.t01 |
| 7237 | 5 | 118177061 | 118212644 | mNeoVis.2.1.1.chr5\_hap1.g01351.t01 |
| 7238 | 5 | 118219094 | 118275915 | mNeoVis.2.1.1.chr5\_hap1.g01352.t01 |
| 7239 | 5 | 118297317 | 118334212 | mNeoVis.2.1.1.chr5\_hap1.g01353.t01 |
| 7240 | 5 | 118376326 | 118493551 | mNeoVis.2.1.1.chr5\_hap1.g01354.t01 |
| 7241 | 5 | 118376326 | 118493551 | mNeoVis.2.1.1.chr5\_hap1.g01354.t01 |
| 7242 | 5 | 119126450 | 119267788 | mNeoVis.2.1.1.chr5\_hap1.g01355.t01 |
| 7243 | 5 | 119412545 | 119412799 | mNeoVis.2.1.1.chr5\_hap1.g01356.t01 |
| 7244 | 5 | 119460687 | 119462281 | mNeoVis.2.1.1.chr5\_hap1.g01357.t01 |
| 7245 | 5 | 120383242 | 120439474 | mNeoVis.2.1.1.chr5\_hap1.g01360.t01 |
| 7246 | 5 | 120383242 | 120439474 | mNeoVis.2.1.1.chr5\_hap1.g01360.t01 |
| 7247 | 5 | 120383242 | 120439474 | mNeoVis.2.1.1.chr5\_hap1.g01360.t01 |
| 7248 | 5 | 120439622 | 120449651 | mNeoVis.2.1.1.chr5\_hap1.g01361.t01 |
| 7249 | 5 | 120439622 | 120449651 | mNeoVis.2.1.1.chr5\_hap1.g01361.t01 |
| 7250 | 5 | 120439622 | 120449651 | mNeoVis.2.1.1.chr5\_hap1.g01361.t01 |
| 7251 | 5 | 121047957 | 121217553 | mNeoVis.2.1.1.chr5\_hap1.g01362.t01 |
| 7252 | 5 | 121047957 | 121217553 | mNeoVis.2.1.1.chr5\_hap1.g01362.t01 |
| 7253 | 5 | 121211326 | 121336200 | mNeoVis.2.1.1.chr5\_hap1.g01363.t01 |
| 7254 | 5 | 121211326 | 121336200 | mNeoVis.2.1.1.chr5\_hap1.g01363.t01 |
| 7255 | 5 | 121356773 | 121371632 | mNeoVis.2.1.1.chr5\_hap1.g01364.t01 |
| 7256 | 5 | 121356773 | 121371632 | mNeoVis.2.1.1.chr5\_hap1.g01364.t01 |
| 7257 | 5 | 121371649 | 121399079 | mNeoVis.2.1.1.chr5\_hap1.g01365.t01 |
| 7258 | 5 | 121371649 | 121399079 | mNeoVis.2.1.1.chr5\_hap1.g01365.t01 |
| 7259 | 5 | 121399214 | 121428467 | mNeoVis.2.1.1.chr5\_hap1.g01366.t01 |
| 7260 | 5 | 121399214 | 121428467 | mNeoVis.2.1.1.chr5\_hap1.g01366.t01 |
| 7261 | 5 | 121503652 | 121561756 | mNeoVis.2.1.1.chr5\_hap1.g01367.t01 |
| 7262 | 5 | 121503652 | 121561756 | mNeoVis.2.1.1.chr5\_hap1.g01367.t01 |
| 7263 | 5 | 121563170 | 121596813 | mNeoVis.2.1.1.chr5\_hap1.g01368.t01 |
| 7264 | 5 | 121563170 | 121596813 | mNeoVis.2.1.1.chr5\_hap1.g01368.t01 |
| 7265 | 5 | 121609432 | 121619304 | mNeoVis.2.1.1.chr5\_hap1.g01369.t01 |
| 7266 | 5 | 121609432 | 121619304 | mNeoVis.2.1.1.chr5\_hap1.g01369.t01 |
| 7267 | 5 | 121685684 | 121789877 | mNeoVis.2.1.1.chr5\_hap1.g01370.t01 |
| 7268 | 5 | 121708239 | 121792269 | mNeoVis.2.1.1.chr5\_hap1.g01371.t01 |
| 7269 | 5 | 121826739 | 121859727 | mNeoVis.2.1.1.chr5\_hap1.g01372.t01 |
| 7270 | 5 | 122016688 | 122248917 | mNeoVis.2.1.1.chr5\_hap1.g01374.t01 |
| 7271 | 5 | 122016688 | 122248917 | mNeoVis.2.1.1.chr5\_hap1.g01374.t01 |
| 7272 | 5 | 122640456 | 122651328 | mNeoVis.2.1.1.chr5\_hap1.g01375.t01 |
| 7273 | 5 | 122640456 | 122651328 | mNeoVis.2.1.1.chr5\_hap1.g01375.t01 |
| 7274 | 5 | 122640456 | 122651328 | mNeoVis.2.1.1.chr5\_hap1.g01375.t01 |
| 7275 | 5 | 123487460 | 123657945 | mNeoVis.2.1.1.chr5\_hap1.g01376.t01 |
| 7276 | 5 | 123487460 | 123657945 | mNeoVis.2.1.1.chr5\_hap1.g01376.t01 |
| 7277 | 5 | 125283902 | 125284577 | mNeoVis.2.1.1.chr5\_hap1.g01378.t01 |
| 7278 | 5 | 125283902 | 125284577 | mNeoVis.2.1.1.chr5\_hap1.g01378.t01 |
| 7279 | 5 | 125531516 | 126113288 | mNeoVis.2.1.1.chr5\_hap1.g01379.t01 |
| 7280 | 5 | 126274634 | 127114654 | mNeoVis.2.1.1.chr5\_hap1.g01380.t01 |
| 7281 | 5 | 127298475 | 127545335 | mNeoVis.2.1.1.chr5\_hap1.g01381.t01 |
| 7282 | 5 | 127609452 | 127672434 | mNeoVis.2.1.1.chr5\_hap1.g01382.t01 |
| 7283 | 5 | 127609452 | 127672434 | mNeoVis.2.1.1.chr5\_hap1.g01382.t01 |
| 7284 | 5 | 127694164 | 127753593 | mNeoVis.2.1.1.chr5\_hap1.g01383.t01 |
| 7285 | 5 | 127694164 | 127753593 | mNeoVis.2.1.1.chr5\_hap1.g01383.t01 |
| 7286 | 5 | 127830954 | 127850147 | mNeoVis.2.1.1.chr5\_hap1.g01384.t01 |
| 7287 | 5 | 127830954 | 127850147 | mNeoVis.2.1.1.chr5\_hap1.g01384.t01 |
| 7288 | 5 | 127879696 | 127884153 | mNeoVis.2.1.1.chr5\_hap1.g01385.t01 |
| 7289 | 5 | 127879696 | 127884153 | mNeoVis.2.1.1.chr5\_hap1.g01385.t01 |
| 7290 | 5 | 127941423 | 128342623 | mNeoVis.2.1.1.chr5\_hap1.g01386.t01 |
| 7291 | 5 | 128414174 | 128435030 | mNeoVis.2.1.1.chr5\_hap1.g01387.t01 |
| 7292 | 5 | 128464289 | 128668685 | mNeoVis.2.1.1.chr5\_hap1.g01388.t01 |
| 7293 | 5 | 128724033 | 128785849 | mNeoVis.2.1.1.chr5\_hap1.g01389.t01 |
| 7294 | 5 | 129200223 | 129232849 | mNeoVis.2.1.1.chr5\_hap1.g01391.t01 |
| 7295 | 5 | 129200223 | 129232849 | mNeoVis.2.1.1.chr5\_hap1.g01391.t01 |
| 7296 | 5 | 129277250 | 129291133 | mNeoVis.2.1.1.chr5\_hap1.g01392.t01 |
| 7297 | 5 | 129277250 | 129291133 | mNeoVis.2.1.1.chr5\_hap1.g01392.t01 |
| 7298 | 5 | 129299348 | 129304925 | mNeoVis.2.1.1.chr5\_hap1.g01393.t01 |
| 7299 | 5 | 129299348 | 129304925 | mNeoVis.2.1.1.chr5\_hap1.g01393.t01 |
| 7300 | 5 | 129429963 | 129433190 | mNeoVis.2.1.1.chr5\_hap1.g01394.t01 |
| 7301 | 5 | 129429963 | 129433190 | mNeoVis.2.1.1.chr5\_hap1.g01394.t01 |
| 7302 | 5 | 129539661 | 129656368 | mNeoVis.2.1.1.chr5\_hap1.g01395.t01 |
| 7303 | 5 | 129539661 | 129656368 | mNeoVis.2.1.1.chr5\_hap1.g01395.t01 |
| 7304 | 5 | 129680215 | 129696332 | mNeoVis.2.1.1.chr5\_hap1.g01396.t01 |
| 7305 | 5 | 129680215 | 129696332 | mNeoVis.2.1.1.chr5\_hap1.g01396.t01 |
| 7306 | 5 | 129712668 | 129743577 | mNeoVis.2.1.1.chr5\_hap1.g01397.t01 |
| 7307 | 5 | 129712668 | 129743577 | mNeoVis.2.1.1.chr5\_hap1.g01397.t01 |
| 7308 | 5 | 129774799 | 129845427 | mNeoVis.2.1.1.chr5\_hap1.g01398.t01 |
| 7309 | 5 | 129872807 | 129950936 | mNeoVis.2.1.1.chr5\_hap1.g01399.t01 |
| 7310 | 5 | 129951659 | 129995141 | mNeoVis.2.1.1.chr5\_hap1.g01400.t01 |
| 7311 | 5 | 130032878 | 130057885 | mNeoVis.2.1.1.chr5\_hap1.g01401.t01 |
| 7312 | 5 | 130058026 | 130184215 | mNeoVis.2.1.1.chr5\_hap1.g01402.t01 |
| 7313 | 5 | 130079501 | 130095940 | mNeoVis.2.1.1.chr5\_hap1.g01403.t01 |
| 7314 | 5 | 130156178 | 130190596 | mNeoVis.2.1.1.chr5\_hap1.g01404.t01 |
| 7315 | 5 | 130195160 | 130219312 | mNeoVis.2.1.1.chr5\_hap1.g01405.t01 |
| 7316 | 5 | 130197803 | 130244838 | mNeoVis.2.1.1.chr5\_hap1.g01406.t01 |
| 7317 | 5 | 130244942 | 130249677 | mNeoVis.2.1.1.chr5\_hap1.g01407.t01 |
| 7318 | 5 | 130316383 | 130863098 | mNeoVis.2.1.1.chr5\_hap1.g01408.t01 |
| 7319 | 5 | 131027762 | 131052792 | mNeoVis.2.1.1.chr5\_hap1.g01409.t01 |
| 7320 | 5 | 131082294 | 131360379 | mNeoVis.2.1.1.chr5\_hap1.g01410.t01 |
| 7321 | 5 | 131631781 | 131641736 | mNeoVis.2.1.1.chr5\_hap1.g01411.t01 |
| 7322 | 5 | 131631781 | 131641736 | mNeoVis.2.1.1.chr5\_hap1.g01411.t01 |
| 7323 | 5 | 131756524 | 131836205 | mNeoVis.2.1.1.chr5\_hap1.g01412.t01 |
| 7324 | 5 | 131756524 | 131836205 | mNeoVis.2.1.1.chr5\_hap1.g01412.t01 |
| 7325 | 5 | 131925729 | 131950690 | mNeoVis.2.1.1.chr5\_hap1.g01413.t01 |
| 7326 | 5 | 132021125 | 132097536 | mNeoVis.2.1.1.chr5\_hap1.g01414.t01 |
| 7327 | 5 | 132098062 | 132141482 | mNeoVis.2.1.1.chr5\_hap1.g01415.t01 |
| 7328 | 5 | 132152830 | 132295768 | mNeoVis.2.1.1.chr5\_hap1.g01416.t01 |
| 7329 | 5 | 132301676 | 132338257 | mNeoVis.2.1.1.chr5\_hap1.g01417.t01 |
| 7330 | 5 | 132391464 | 132515819 | mNeoVis.2.1.1.chr5\_hap1.g01418.t01 |
| 7331 | 5 | 133827907 | 133887655 | mNeoVis.2.1.1.chr5\_hap1.g01419.t01 |
| 7332 | 5 | 134143338 | 134377714 | mNeoVis.2.1.1.chr5\_hap1.g01420.t01 |
| 7333 | 5 | 134391136 | 134400380 | mNeoVis.2.1.1.chr5\_hap1.g01421.t01 |
| 7334 | 5 | 134501164 | 134603518 | mNeoVis.2.1.1.chr5\_hap1.g01422.t01 |
| 7335 | 5 | 137672760 | 137746423 | mNeoVis.2.1.1.chr5\_hap1.g01423.t01 |
| 7336 | 5 | 137773339 | 137808241 | mNeoVis.2.1.1.chr5\_hap1.g01424.t01 |
| 7337 | 5 | 137896197 | 137923364 | mNeoVis.2.1.1.chr5\_hap1.g01425.t01 |
| 7338 | 5 | 138027802 | 138053448 | mNeoVis.2.1.1.chr5\_hap1.g01426.t01 |
| 7339 | 5 | 138065986 | 138367381 | mNeoVis.2.1.1.chr5\_hap1.g01427.t01 |
| 7340 | 5 | 138367335 | 138399749 | mNeoVis.2.1.1.chr5\_hap1.g01428.t01 |
| 7341 | 5 | 138427767 | 138524573 | mNeoVis.2.1.1.chr5\_hap1.g01429.t01 |
| 7342 | 5 | 138444122 | 138444977 | mNeoVis.2.1.1.chr5\_hap1.g01430.t01 |
| 7343 | 5 | 138524733 | 138582884 | mNeoVis.2.1.1.chr5\_hap1.g01431.t01 |
| 7344 | 5 | 138657418 | 138693287 | mNeoVis.2.1.1.chr5\_hap1.g01433.t01 |
| 7345 | 5 | 138693308 | 138807997 | mNeoVis.2.1.1.chr5\_hap1.g01434.t01 |
| 7346 | 5 | 138776530 | 138830057 | mNeoVis.2.1.1.chr5\_hap1.g01435.t01 |
| 7347 | 5 | 138793743 | 138795544 | mNeoVis.2.1.1.chr5\_hap1.g01436.t01 |
| 7348 | 5 | 138973943 | 139008958 | mNeoVis.2.1.1.chr5\_hap1.g01437.t01 |
| 7349 | 5 | 139014220 | 139021380 | mNeoVis.2.1.1.chr5\_hap1.g01438.t01 |
| 7350 | 5 | 140268228 | 140328928 | mNeoVis.2.1.1.chr5\_hap1.g01442.t01 |
| 7351 | 5 | 140306407 | 140306926 | mNeoVis.2.1.1.chr5\_hap1.g01443.t01 |
| 7352 | 5 | 140344413 | 140387889 | mNeoVis.2.1.1.chr5\_hap1.g01444.t01 |
| 7353 | 5 | 140344413 | 140387889 | mNeoVis.2.1.1.chr5\_hap1.g01444.t01 |
| 7354 | 5 | 140357903 | 140365044 | mNeoVis.2.1.1.chr5\_hap1.g01445.t01 |
| 7355 | 5 | 140357903 | 140365044 | mNeoVis.2.1.1.chr5\_hap1.g01445.t01 |
| 7356 | 5 | 140396686 | 140416602 | mNeoVis.2.1.1.chr5\_hap1.g01446.t01 |
| 7357 | 5 | 140396686 | 140416602 | mNeoVis.2.1.1.chr5\_hap1.g01446.t01 |
| 7358 | 5 | 140453165 | 140874334 | mNeoVis.2.1.1.chr5\_hap1.g01447.t01 |
| 7359 | 5 | 140877846 | 140886874 | mNeoVis.2.1.1.chr5\_hap1.g01448.t01 |
| 7360 | 5 | 141475296 | 141725113 | mNeoVis.2.1.1.chr5\_hap1.g01449.t01 |
| 7361 | 5 | 141475296 | 141725113 | mNeoVis.2.1.1.chr5\_hap1.g01449.t01 |
| 7362 | 5 | 141533409 | 141539124 | mNeoVis.2.1.1.chr5\_hap1.g01450.t01 |
| 7363 | 5 | 141533409 | 141539124 | mNeoVis.2.1.1.chr5\_hap1.g01450.t01 |
| 7364 | 5 | 141901387 | 141914544 | mNeoVis.2.1.1.chr5\_hap1.g01451.t01 |
| 7365 | 5 | 141901387 | 141914544 | mNeoVis.2.1.1.chr5\_hap1.g01451.t01 |
| 7366 | 5 | 141901387 | 141914544 | mNeoVis.2.1.1.chr5\_hap1.g01451.t01 |
| 7367 | 5 | 142014058 | 142205894 | mNeoVis.2.1.1.chr5\_hap1.g01452.t01 |
| 7368 | 5 | 142014058 | 142205894 | mNeoVis.2.1.1.chr5\_hap1.g01452.t01 |
| 7369 | 5 | 142326269 | 142358024 | mNeoVis.2.1.1.chr5\_hap1.g01453.t01 |
| 7370 | 5 | 142326269 | 142358024 | mNeoVis.2.1.1.chr5\_hap1.g01453.t01 |
| 7371 | 5 | 142326269 | 142358024 | mNeoVis.2.1.1.chr5\_hap1.g01453.t01 |
| 7372 | 5 | 142439308 | 142460847 | mNeoVis.2.1.1.chr5\_hap1.g01454.t01 |
| 7373 | 5 | 142439308 | 142460847 | mNeoVis.2.1.1.chr5\_hap1.g01454.t01 |
| 7374 | 5 | 142439308 | 142460847 | mNeoVis.2.1.1.chr5\_hap1.g01454.t01 |
| 7375 | 5 | 142439308 | 142460847 | mNeoVis.2.1.1.chr5\_hap1.g01454.t01 |
| 7376 | 5 | 142468865 | 142480921 | mNeoVis.2.1.1.chr5\_hap1.g01455.t01 |
| 7377 | 5 | 142468865 | 142480921 | mNeoVis.2.1.1.chr5\_hap1.g01455.t01 |
| 7378 | 5 | 142468865 | 142480921 | mNeoVis.2.1.1.chr5\_hap1.g01455.t01 |
| 7379 | 5 | 142468865 | 142480921 | mNeoVis.2.1.1.chr5\_hap1.g01455.t01 |
| 7380 | 5 | 142483506 | 142896205 | mNeoVis.2.1.1.chr5\_hap1.g01456.t01 |
| 7381 | 5 | 142483506 | 142896205 | mNeoVis.2.1.1.chr5\_hap1.g01456.t01 |
| 7382 | 5 | 142958496 | 142985485 | mNeoVis.2.1.1.chr5\_hap1.g01457.t01 |
| 7383 | 5 | 142958496 | 142985485 | mNeoVis.2.1.1.chr5\_hap1.g01457.t01 |
| 7384 | 5 | 143161653 | 143188629 | mNeoVis.2.1.1.chr5\_hap1.g01459.t01 |
| 7385 | 5 | 143202861 | 143216885 | mNeoVis.2.1.1.chr5\_hap1.g01460.t01 |
| 7386 | 5 | 143227895 | 143447939 | mNeoVis.2.1.1.chr5\_hap1.g01461.t01 |
| 7387 | 5 | 143589062 | 143657033 | mNeoVis.2.1.1.chr5\_hap1.g01462.t01 |
| 7388 | 5 | 143624780 | 143625359 | mNeoVis.2.1.1.chr5\_hap1.g01463.t01 |
| 7389 | 5 | 143694847 | 143792511 | mNeoVis.2.1.1.chr5\_hap1.g01464.t01 |
| 7390 | 5 | 144148499 | 144149825 | mNeoVis.2.1.1.chr5\_hap1.g01465.t01 |
| 7391 | 5 | 144293261 | 144365176 | mNeoVis.2.1.1.chr5\_hap1.g01466.t01 |
| 7392 | 5 | 144380209 | 144430251 | mNeoVis.2.1.1.chr5\_hap1.g01467.t01 |
| 7393 | 5 | 144437682 | 144564420 | mNeoVis.2.1.1.chr5\_hap1.g01468.t01 |
| 7394 | 5 | 144437682 | 144564420 | mNeoVis.2.1.1.chr5\_hap1.g01468.t01 |
| 7395 | 5 | 145433462 | 145439707 | mNeoVis.2.1.1.chr5\_hap1.g01469.t01 |
| 7396 | 5 | 145439571 | 145444931 | mNeoVis.2.1.1.chr5\_hap1.g01470.t01 |
| 7397 | 5 | 145737294 | 145742840 | mNeoVis.2.1.1.chr5\_hap1.g01471.t01 |
| 7398 | 5 | 145737294 | 145742840 | mNeoVis.2.1.1.chr5\_hap1.g01471.t01 |
| 7399 | 5 | 146064252 | 146263859 | mNeoVis.2.1.1.chr5\_hap1.g01472.t01 |
| 7400 | 5 | 146610397 | 146617519 | mNeoVis.2.1.1.chr5\_hap1.g01473.t01 |
| 7401 | 5 | 146702491 | 146745470 | mNeoVis.2.1.1.chr5\_hap1.g01474.t01 |
| 7402 | 5 | 146702491 | 146745470 | mNeoVis.2.1.1.chr5\_hap1.g01474.t01 |
| 7403 | 5 | 147011081 | 147065406 | mNeoVis.2.1.1.chr5\_hap1.g01475.t01 |
| 7404 | 5 | 147011081 | 147065406 | mNeoVis.2.1.1.chr5\_hap1.g01475.t01 |
| 7405 | 5 | 147069126 | 147383025 | mNeoVis.2.1.1.chr5\_hap1.g01476.t01 |
| 7406 | 5 | 147069126 | 147383025 | mNeoVis.2.1.1.chr5\_hap1.g01476.t01 |
| 7407 | 5 | 147109722 | 147151904 | mNeoVis.2.1.1.chr5\_hap1.g01477.t01 |
| 7408 | 5 | 147109722 | 147151904 | mNeoVis.2.1.1.chr5\_hap1.g01477.t01 |
| 7409 | 5 | 147109722 | 147151904 | mNeoVis.2.1.1.chr5\_hap1.g01477.t01 |
| 7410 | 5 | 147157606 | 147160779 | mNeoVis.2.1.1.chr5\_hap1.g01478.t01 |
| 7411 | 5 | 147157606 | 147160779 | mNeoVis.2.1.1.chr5\_hap1.g01478.t01 |
| 7412 | 5 | 147157606 | 147160779 | mNeoVis.2.1.1.chr5\_hap1.g01478.t01 |
| 7413 | 5 | 147170589 | 147192503 | mNeoVis.2.1.1.chr5\_hap1.g01479.t01 |
| 7414 | 5 | 147170589 | 147192503 | mNeoVis.2.1.1.chr5\_hap1.g01479.t01 |
| 7415 | 5 | 147170589 | 147192503 | mNeoVis.2.1.1.chr5\_hap1.g01479.t01 |
| 7416 | 5 | 147210714 | 147265071 | mNeoVis.2.1.1.chr5\_hap1.g01480.t01 |
| 7417 | 5 | 147210714 | 147265071 | mNeoVis.2.1.1.chr5\_hap1.g01480.t01 |
| 7418 | 5 | 147210714 | 147265071 | mNeoVis.2.1.1.chr5\_hap1.g01480.t01 |
| 7419 | 5 | 147273774 | 147280567 | mNeoVis.2.1.1.chr5\_hap1.g01481.t01 |
| 7420 | 5 | 147273774 | 147280567 | mNeoVis.2.1.1.chr5\_hap1.g01481.t01 |
| 7421 | 5 | 147273774 | 147280567 | mNeoVis.2.1.1.chr5\_hap1.g01481.t01 |
| 7422 | 5 | 147518059 | 147555484 | mNeoVis.2.1.1.chr5\_hap1.g01482.t01 |
| 7423 | 5 | 147518059 | 147555484 | mNeoVis.2.1.1.chr5\_hap1.g01482.t01 |
| 7424 | 5 | 147578721 | 147614532 | mNeoVis.2.1.1.chr5\_hap1.g01483.t01 |
| 7425 | 5 | 147578721 | 147614532 | mNeoVis.2.1.1.chr5\_hap1.g01483.t01 |
| 7426 | 5 | 147683767 | 147701863 | mNeoVis.2.1.1.chr5\_hap1.g01484.t01 |
| 7427 | 5 | 147683767 | 147701863 | mNeoVis.2.1.1.chr5\_hap1.g01484.t01 |
| 7428 | 5 | 147754815 | 147788494 | mNeoVis.2.1.1.chr5\_hap1.g01485.t01 |
| 7429 | 5 | 147754815 | 147788494 | mNeoVis.2.1.1.chr5\_hap1.g01485.t01 |
| 7430 | 5 | 147811681 | 147836716 | mNeoVis.2.1.1.chr5\_hap1.g01486.t01 |
| 7431 | 5 | 147811681 | 147836716 | mNeoVis.2.1.1.chr5\_hap1.g01486.t01 |
| 7432 | 5 | 147853021 | 147890175 | mNeoVis.2.1.1.chr5\_hap1.g01487.t01 |
| 7433 | 5 | 147853021 | 147890175 | mNeoVis.2.1.1.chr5\_hap1.g01487.t01 |
| 7434 | 5 | 147890234 | 147896259 | mNeoVis.2.1.1.chr5\_hap1.g01488.t01 |
| 7435 | 5 | 147890234 | 147896259 | mNeoVis.2.1.1.chr5\_hap1.g01488.t01 |
| 7436 | 5 | 147955820 | 148004678 | mNeoVis.2.1.1.chr5\_hap1.g01489.t01 |
| 7437 | 5 | 147955820 | 148004678 | mNeoVis.2.1.1.chr5\_hap1.g01489.t01 |
| 7438 | 5 | 148054536 | 148084818 | mNeoVis.2.1.1.chr5\_hap1.g01490.t01 |
| 7439 | 5 | 148263466 | 148265033 | mNeoVis.2.1.1.chr5\_hap1.g01491.t01 |
| 7440 | 5 | 148396779 | 148404463 | mNeoVis.2.1.1.chr5\_hap1.g01492.t01 |
| 7441 | 5 | 148407391 | 148562231 | mNeoVis.2.1.1.chr5\_hap1.g01493.t01 |
| 7442 | 5 | 148572588 | 148604435 | mNeoVis.2.1.1.chr5\_hap1.g01494.t01 |
| 7443 | 5 | 148611854 | 148654543 | mNeoVis.2.1.1.chr5\_hap1.g01495.t01 |
| 7444 | 5 | 148676530 | 148818578 | mNeoVis.2.1.1.chr5\_hap1.g01496.t01 |
| 7445 | 5 | 148834542 | 148861176 | mNeoVis.2.1.1.chr5\_hap1.g01497.t01 |
| 7446 | 5 | 148944658 | 148952288 | mNeoVis.2.1.1.chr5\_hap1.g01498.t01 |
| 7447 | 5 | 148997277 | 149025781 | mNeoVis.2.1.1.chr5\_hap1.g01499.t01 |
| 7448 | 5 | 149066732 | 149097680 | mNeoVis.2.1.1.chr5\_hap1.g01500.t01 |
| 7449 | 5 | 149109294 | 149168304 | mNeoVis.2.1.1.chr5\_hap1.g01501.t01 |
| 7450 | 5 | 149158604 | 149198290 | mNeoVis.2.1.1.chr5\_hap1.g01502.t01 |
| 7451 | 5 | 149220337 | 149274353 | mNeoVis.2.1.1.chr5\_hap1.g01503.t01 |
| 7452 | 5 | 149280162 | 149314717 | mNeoVis.2.1.1.chr5\_hap1.g01504.t01 |
| 7453 | 5 | 149406405 | 149437895 | mNeoVis.2.1.1.chr5\_hap1.g01505.t01 |
| 7454 | 5 | 149443473 | 149478283 | mNeoVis.2.1.1.chr5\_hap1.g01506.t01 |
| 7455 | 5 | 149568118 | 149616863 | mNeoVis.2.1.1.chr5\_hap1.g01507.t01 |
| 7456 | 5 | 149568118 | 149616863 | mNeoVis.2.1.1.chr5\_hap1.g01507.t01 |
| 7457 | 5 | 150690392 | 150692552 | mNeoVis.2.1.1.chr5\_hap1.g01508.t01 |
| 7458 | 5 | 150690392 | 150692552 | mNeoVis.2.1.1.chr5\_hap1.g01508.t01 |
| 7459 | 5 | 150690392 | 150692552 | mNeoVis.2.1.1.chr5\_hap1.g01508.t01 |
| 7460 | 5 | 150693436 | 150728153 | mNeoVis.2.1.1.chr5\_hap1.g01509.t01 |
| 7461 | 5 | 150693436 | 150728153 | mNeoVis.2.1.1.chr5\_hap1.g01509.t01 |
| 7462 | 5 | 150693436 | 150728153 | mNeoVis.2.1.1.chr5\_hap1.g01509.t01 |
| 7463 | 5 | 150699493 | 150703539 | mNeoVis.2.1.1.chr5\_hap1.g01510.t01 |
| 7464 | 5 | 150699493 | 150703539 | mNeoVis.2.1.1.chr5\_hap1.g01510.t01 |
| 7465 | 5 | 150699493 | 150703539 | mNeoVis.2.1.1.chr5\_hap1.g01510.t01 |
| 7466 | 5 | 151889733 | 151985186 | mNeoVis.2.1.1.chr5\_hap1.g01515.t01 |
| 7467 | 5 | 151889733 | 151985186 | mNeoVis.2.1.1.chr5\_hap1.g01515.t01 |
| 7468 | 5 | 153451430 | 153601217 | mNeoVis.2.1.1.chr5\_hap1.g01516.t01 |
| 7469 | 5 | 153451430 | 153601217 | mNeoVis.2.1.1.chr5\_hap1.g01516.t01 |
| 7470 | 5 | 153836376 | 153859469 | mNeoVis.2.1.1.chr5\_hap1.g01517.t01 |
| 7471 | 5 | 153836376 | 153859469 | mNeoVis.2.1.1.chr5\_hap1.g01517.t01 |
| 7472 | 5 | 153836376 | 153859469 | mNeoVis.2.1.1.chr5\_hap1.g01517.t01 |
| 7473 | 5 | 154654040 | 154657182 | mNeoVis.2.1.1.chr5\_hap1.g01519.t01 |
| 7474 | 5 | 154654040 | 154657182 | mNeoVis.2.1.1.chr5\_hap1.g01519.t01 |
| 7475 | 5 | 154704957 | 154767412 | mNeoVis.2.1.1.chr5\_hap1.g01520.t01 |
| 7476 | 5 | 154808484 | 154818252 | mNeoVis.2.1.1.chr5\_hap1.g01521.t01 |
| 7477 | 5 | 154808484 | 154818252 | mNeoVis.2.1.1.chr5\_hap1.g01521.t01 |
| 7478 | 5 | 154877418 | 154943341 | mNeoVis.2.1.1.chr5\_hap1.g01522.t01 |
| 7479 | 5 | 154877418 | 154943341 | mNeoVis.2.1.1.chr5\_hap1.g01522.t01 |
| 7480 | 5 | 154952138 | 155025230 | mNeoVis.2.1.1.chr5\_hap1.g01523.t01 |
| 7481 | 5 | 154952138 | 155025230 | mNeoVis.2.1.1.chr5\_hap1.g01523.t01 |
| 7482 | 5 | 155028056 | 155165504 | mNeoVis.2.1.1.chr5\_hap1.g01524.t01 |
| 7483 | 5 | 155028056 | 155165504 | mNeoVis.2.1.1.chr5\_hap1.g01524.t01 |
| 7484 | 5 | 155164497 | 155263145 | mNeoVis.2.1.1.chr5\_hap1.g01525.t01 |
| 7485 | 5 | 155164497 | 155263145 | mNeoVis.2.1.1.chr5\_hap1.g01525.t01 |
| 7486 | 5 | 155263930 | 155371623 | mNeoVis.2.1.1.chr5\_hap1.g01526.t01 |
| 7487 | 5 | 155397468 | 155482619 | mNeoVis.2.1.1.chr5\_hap1.g01527.t01 |
| 7488 | 5 | 155397468 | 155482619 | mNeoVis.2.1.1.chr5\_hap1.g01527.t01 |
| 7489 | 5 | 155489545 | 155672159 | mNeoVis.2.1.1.chr5\_hap1.g01528.t01 |
| 7490 | 5 | 155489545 | 155672159 | mNeoVis.2.1.1.chr5\_hap1.g01528.t01 |
| 7491 | 5 | 155696570 | 155742854 | mNeoVis.2.1.1.chr5\_hap1.g01529.t01 |
| 7492 | 5 | 155696570 | 155742854 | mNeoVis.2.1.1.chr5\_hap1.g01529.t01 |
| 7493 | 5 | 155758425 | 155822732 | mNeoVis.2.1.1.chr5\_hap1.g01530.t01 |
| 7494 | 5 | 155758425 | 155822732 | mNeoVis.2.1.1.chr5\_hap1.g01530.t01 |
| 7495 | 5 | 155852270 | 155856778 | mNeoVis.2.1.1.chr5\_hap1.g01531.t01 |
| 7496 | 5 | 155852270 | 155856778 | mNeoVis.2.1.1.chr5\_hap1.g01531.t01 |
| 7497 | 5 | 155861334 | 155908492 | mNeoVis.2.1.1.chr5\_hap1.g01532.t01 |
| 7498 | 5 | 155861334 | 155908492 | mNeoVis.2.1.1.chr5\_hap1.g01532.t01 |
| 7499 | 5 | 155923466 | 156060332 | mNeoVis.2.1.1.chr5\_hap1.g01533.t01 |
| 7500 | 5 | 155944329 | 155947033 | mNeoVis.2.1.1.chr5\_hap1.g01534.t01 |
| 7501 | 5 | 155944329 | 155947033 | mNeoVis.2.1.1.chr5\_hap1.g01534.t01 |
| 7502 | 5 | 156102127 | 156240191 | mNeoVis.2.1.1.chr5\_hap1.g01535.t01 |
| 7503 | 5 | 156102127 | 156240191 | mNeoVis.2.1.1.chr5\_hap1.g01535.t01 |
| 7504 | 5 | 156250578 | 156271792 | mNeoVis.2.1.1.chr5\_hap1.g01536.t01 |
| 7505 | 5 | 156250578 | 156271792 | mNeoVis.2.1.1.chr5\_hap1.g01536.t01 |
| 7506 | 5 | 156365683 | 156444782 | mNeoVis.2.1.1.chr5\_hap1.g01537.t01 |
| 7507 | 5 | 156365683 | 156444782 | mNeoVis.2.1.1.chr5\_hap1.g01537.t01 |
| 7508 | 5 | 156496388 | 156532479 | mNeoVis.2.1.1.chr5\_hap1.g01538.t01 |
| 7509 | 5 | 156496388 | 156532479 | mNeoVis.2.1.1.chr5\_hap1.g01538.t01 |
| 7510 | 5 | 157400243 | 157430561 | mNeoVis.2.1.1.chr5\_hap1.g01541.t01 |
| 7511 | 5 | 157437572 | 157480024 | mNeoVis.2.1.1.chr5\_hap1.g01542.t01 |
| 7512 | 5 | 157493364 | 157499645 | mNeoVis.2.1.1.chr5\_hap1.g01543.t01 |
| 7513 | 5 | 157604339 | 157657564 | mNeoVis.2.1.1.chr5\_hap1.g01544.t01 |
| 7514 | 5 | 157691587 | 157711671 | mNeoVis.2.1.1.chr5\_hap1.g01545.t01 |
| 7515 | 5 | 157691587 | 157711671 | mNeoVis.2.1.1.chr5\_hap1.g01545.t01 |
| 7516 | 5 | 157741375 | 157835294 | mNeoVis.2.1.1.chr5\_hap1.g01546.t01 |
| 7517 | 5 | 157741375 | 157835294 | mNeoVis.2.1.1.chr5\_hap1.g01546.t01 |
| 7518 | 5 | 157897114 | 158118003 | mNeoVis.2.1.1.chr5\_hap1.g01547.t01 |
| 7519 | 5 | 158120086 | 158129117 | mNeoVis.2.1.1.chr5\_hap1.g01548.t01 |
| 7520 | 5 | 158120086 | 158129117 | mNeoVis.2.1.1.chr5\_hap1.g01548.t01 |
| 7521 | 5 | 158146103 | 158249421 | mNeoVis.2.1.1.chr5\_hap1.g01549.t01 |
| 7522 | 5 | 158261666 | 158269458 | mNeoVis.2.1.1.chr5\_hap1.g01550.t01 |
| 7523 | 5 | 158277329 | 158304666 | mNeoVis.2.1.1.chr5\_hap1.g01551.t01 |
| 7524 | 5 | 158320933 | 158346069 | mNeoVis.2.1.1.chr5\_hap1.g01552.t01 |
| 7525 | 5 | 158355020 | 158362924 | mNeoVis.2.1.1.chr5\_hap1.g01553.t01 |
| 7526 | 5 | 158373802 | 158402516 | mNeoVis.2.1.1.chr5\_hap1.g01554.t01 |
| 7527 | 5 | 158402636 | 158425350 | mNeoVis.2.1.1.chr5\_hap1.g01555.t01 |
| 7528 | 5 | 158428357 | 158459221 | mNeoVis.2.1.1.chr5\_hap1.g01556.t01 |
| 7529 | 5 | 158463311 | 158494807 | mNeoVis.2.1.1.chr5\_hap1.g01557.t01 |
| 7530 | 5 | 158500571 | 158617431 | mNeoVis.2.1.1.chr5\_hap1.g01558.t01 |
| 7531 | 5 | 158649466 | 158679746 | mNeoVis.2.1.1.chr5\_hap1.g01559.t01 |
| 7532 | 5 | 158687017 | 158716857 | mNeoVis.2.1.1.chr5\_hap1.g01560.t01 |
| 7533 | 5 | 158724968 | 158740311 | mNeoVis.2.1.1.chr5\_hap1.g01561.t01 |
| 7534 | 5 | 159436446 | 159436961 | mNeoVis.2.1.1.chr5\_hap1.g01563.t01 |
| 7535 | 5 | 160589028 | 160663741 | mNeoVis.2.1.1.chr5\_hap1.g01564.t01 |
| 7536 | 5 | 160589028 | 160663741 | mNeoVis.2.1.1.chr5\_hap1.g01564.t01 |
| 7537 | 5 | 161284282 | 161933170 | mNeoVis.2.1.1.chr5\_hap1.g01565.t01 |
| 7538 | 5 | 161284282 | 161933170 | mNeoVis.2.1.1.chr5\_hap1.g01565.t01 |
| 7539 | 5 | 161995566 | 162097209 | mNeoVis.2.1.1.chr5\_hap1.g01566.t01 |
| 7540 | 5 | 161995566 | 162097209 | mNeoVis.2.1.1.chr5\_hap1.g01566.t01 |
| 7541 | 5 | 162543611 | 163330626 | mNeoVis.2.1.1.chr5\_hap1.g01567.t01 |
| 7542 | 5 | 162828613 | 162829114 | mNeoVis.2.1.1.chr5\_hap1.g01568.t01 |
| 7543 | 5 | 163080505 | 163192268 | mNeoVis.2.1.1.chr5\_hap1.g01569.t01 |
| 7544 | 5 | 163354497 | 163365731 | mNeoVis.2.1.1.chr5\_hap1.g01570.t01 |
| 7545 | 5 | 163437766 | 163477270 | mNeoVis.2.1.1.chr5\_hap1.g01571.t01 |
| 7546 | 5 | 163510734 | 163535113 | mNeoVis.2.1.1.chr5\_hap1.g01572.t01 |
| 7547 | 5 | 163535324 | 163625852 | mNeoVis.2.1.1.chr5\_hap1.g01573.t01 |
| 7548 | 5 | 163626564 | 163638647 | mNeoVis.2.1.1.chr5\_hap1.g01574.t01 |
| 7549 | 5 | 163637858 | 163640787 | mNeoVis.2.1.1.chr5\_hap1.g01575.t01 |
| 7550 | 5 | 163640922 | 163732288 | mNeoVis.2.1.1.chr5\_hap1.g01576.t01 |
| 7551 | 5 | 163755546 | 163815470 | mNeoVis.2.1.1.chr5\_hap1.g01577.t01 |
| 7552 | 5 | 163816075 | 163852824 | mNeoVis.2.1.1.chr5\_hap1.g01578.t01 |
| 7553 | 5 | 163868995 | 163920019 | mNeoVis.2.1.1.chr5\_hap1.g01579.t01 |
| 7554 | 5 | 164039508 | 164094101 | mNeoVis.2.1.1.chr5\_hap1.g01581.t01 |
| 7555 | 5 | 164097515 | 164269380 | mNeoVis.2.1.1.chr5\_hap1.g01582.t01 |
| 7556 | 5 | 164269533 | 164339492 | mNeoVis.2.1.1.chr5\_hap1.g01583.t01 |
| 7557 | 5 | 164452706 | 164530092 | mNeoVis.2.1.1.chr5\_hap1.g01584.t01 |
| 7558 | 5 | 164501695 | 164513929 | mNeoVis.2.1.1.chr5\_hap1.g01585.t01 |
| 7559 | 5 | 164552081 | 164586429 | mNeoVis.2.1.1.chr5\_hap1.g01586.t01 |
| 7560 | 5 | 164617473 | 164632884 | mNeoVis.2.1.1.chr5\_hap1.g01587.t01 |
| 7561 | 5 | 164661311 | 164677496 | mNeoVis.2.1.1.chr5\_hap1.g01588.t01 |
| 7562 | 5 | 164752710 | 164753384 | mNeoVis.2.1.1.chr5\_hap1.g01589.t01 |
| 7563 | 5 | 164832797 | 164867050 | mNeoVis.2.1.1.chr5\_hap1.g01590.t01 |
| 7564 | 5 | 164878310 | 164900434 | mNeoVis.2.1.1.chr5\_hap1.g01591.t01 |
| 7565 | 5 | 164899868 | 164925519 | mNeoVis.2.1.1.chr5\_hap1.g01592.t01 |
| 7566 | 5 | 164955367 | 164983826 | mNeoVis.2.1.1.chr5\_hap1.g01593.t01 |
| 7567 | 5 | 164997638 | 165034099 | mNeoVis.2.1.1.chr5\_hap1.g01594.t01 |
| 7568 | 5 | 165051740 | 165087701 | mNeoVis.2.1.1.chr5\_hap1.g01595.t01 |
| 7569 | 5 | 165088078 | 165129729 | mNeoVis.2.1.1.chr5\_hap1.g01596.t01 |
| 7570 | 5 | 165112820 | 165113119 | mNeoVis.2.1.1.chr5\_hap1.g01597.t01 |
| 7571 | 5 | 165129860 | 165217708 | mNeoVis.2.1.1.chr5\_hap1.g01598.t01 |
| 7572 | 5 | 165222441 | 165238379 | mNeoVis.2.1.1.chr5\_hap1.g01599.t01 |
| 7573 | 5 | 165242033 | 165269961 | mNeoVis.2.1.1.chr5\_hap1.g01600.t01 |
| 7574 | 5 | 165276359 | 165314838 | mNeoVis.2.1.1.chr5\_hap1.g01601.t01 |
| 7575 | 5 | 165326580 | 165341427 | mNeoVis.2.1.1.chr5\_hap1.g01602.t01 |
| 7576 | 5 | 165348530 | 165365141 | mNeoVis.2.1.1.chr5\_hap1.g01603.t01 |
| 7577 | 5 | 165367697 | 165592472 | mNeoVis.2.1.1.chr5\_hap1.g01604.t01 |
| 7578 | 5 | 165597190 | 165649426 | mNeoVis.2.1.1.chr5\_hap1.g01605.t01 |
| 7579 | 5 | 165665076 | 165758873 | mNeoVis.2.1.1.chr5\_hap1.g01606.t01 |
| 7580 | 5 | 165718773 | 165722483 | mNeoVis.2.1.1.chr5\_hap1.g01607.t01 |
| 7581 | 5 | 166027118 | 166152382 | mNeoVis.2.1.1.chr5\_hap1.g01608.t01 |
| 7582 | 5 | 166027118 | 166152382 | mNeoVis.2.1.1.chr5\_hap1.g01608.t01 |
| 7583 | 5 | 166225738 | 166287258 | mNeoVis.2.1.1.chr5\_hap1.g01609.t01 |
| 7584 | 5 | 166225738 | 166287258 | mNeoVis.2.1.1.chr5\_hap1.g01609.t01 |
| 7585 | 5 | 166225738 | 166287258 | mNeoVis.2.1.1.chr5\_hap1.g01609.t01 |
| 7586 | 5 | 167630829 | 167642111 | mNeoVis.2.1.1.chr5\_hap1.g01611.t01 |
| 7587 | 5 | 167766726 | 167780794 | mNeoVis.2.1.1.chr5\_hap1.g01612.t01 |
| 7588 | 5 | 167800861 | 167977807 | mNeoVis.2.1.1.chr5\_hap1.g01613.t01 |
| 7589 | 5 | 167981718 | 168012791 | mNeoVis.2.1.1.chr5\_hap1.g01614.t01 |
| 7590 | 5 | 168060616 | 168098031 | mNeoVis.2.1.1.chr5\_hap1.g01615.t01 |
| 7591 | 5 | 168147059 | 168149212 | mNeoVis.2.1.1.chr5\_hap1.g01616.t01 |
| 7592 | 5 | 168194866 | 168304224 | mNeoVis.2.1.1.chr5\_hap1.g01617.t01 |
| 7593 | 5 | 168304427 | 168337817 | mNeoVis.2.1.1.chr5\_hap1.g01618.t01 |
| 7594 | 5 | 168376602 | 168538242 | mNeoVis.2.1.1.chr5\_hap1.g01619.t01 |
| 7595 | 5 | 168376602 | 168538242 | mNeoVis.2.1.1.chr5\_hap1.g01619.t01 |
| 7596 | 5 | 168538391 | 168676185 | mNeoVis.2.1.1.chr5\_hap1.g01620.t01 |
| 7597 | 5 | 168538391 | 168676185 | mNeoVis.2.1.1.chr5\_hap1.g01620.t01 |
| 7598 | 5 | 168676401 | 168730645 | mNeoVis.2.1.1.chr5\_hap1.g01621.t01 |
| 7599 | 5 | 168676401 | 168730645 | mNeoVis.2.1.1.chr5\_hap1.g01621.t01 |
| 7600 | 5 | 168804957 | 168850565 | mNeoVis.2.1.1.chr5\_hap1.g01622.t01 |
| 7601 | 5 | 168804957 | 168850565 | mNeoVis.2.1.1.chr5\_hap1.g01622.t01 |
| 7602 | 5 | 168969235 | 168973694 | mNeoVis.2.1.1.chr5\_hap1.g01623.t01 |
| 7603 | 5 | 169089879 | 169176676 | mNeoVis.2.1.1.chr5\_hap1.g01624.t01 |
| 7604 | 5 | 169368322 | 169533710 | mNeoVis.2.1.1.chr5\_hap1.g01625.t01 |
| 7605 | 5 | 169508128 | 169509389 | mNeoVis.2.1.1.chr5\_hap1.g01626.t01 |
| 7606 | 5 | 170786647 | 171003238 | mNeoVis.2.1.1.chr5\_hap1.g01627.t01 |
| 7607 | 5 | 170786647 | 171003238 | mNeoVis.2.1.1.chr5\_hap1.g01627.t01 |
| 7608 | 5 | 171077599 | 171292687 | mNeoVis.2.1.1.chr5\_hap1.g01628.t01 |
| 7609 | 5 | 171077599 | 171292687 | mNeoVis.2.1.1.chr5\_hap1.g01628.t01 |
| 7610 | 5 | 172242102 | 172242644 | mNeoVis.2.1.1.chr5\_hap1.g01629.t01 |
| 7611 | 5 | 173894353 | 174000171 | mNeoVis.2.1.1.chr5\_hap1.g01631.t01 |
| 7612 | 5 | 174510460 | 174521981 | mNeoVis.2.1.1.chr5\_hap1.g01632.t01 |
| 7613 | 5 | 174546580 | 174591108 | mNeoVis.2.1.1.chr5\_hap1.g01633.t01 |
| 7614 | 5 | 174598510 | 174646178 | mNeoVis.2.1.1.chr5\_hap1.g01634.t01 |
| 7615 | 5 | 174635444 | 174649447 | mNeoVis.2.1.1.chr5\_hap1.g01635.t01 |
| 7616 | 5 | 174651574 | 174684365 | mNeoVis.2.1.1.chr5\_hap1.g01636.t01 |
| 7617 | 5 | 174696465 | 174713711 | mNeoVis.2.1.1.chr5\_hap1.g01637.t01 |
| 7618 | 5 | 174717690 | 174722874 | mNeoVis.2.1.1.chr5\_hap1.g01638.t01 |
| 7619 | 5 | 174733388 | 174746807 | mNeoVis.2.1.1.chr5\_hap1.g01639.t01 |
| 7620 | 5 | 174746854 | 174909571 | mNeoVis.2.1.1.chr5\_hap1.g01640.t01 |
| 7621 | 5 | 174912781 | 174991388 | mNeoVis.2.1.1.chr5\_hap1.g01641.t01 |
| 7622 | 5 | 175242969 | 175493751 | mNeoVis.2.1.1.chr5\_hap1.g01642.t01 |
| 7623 | 5 | 175494074 | 175527296 | mNeoVis.2.1.1.chr5\_hap1.g01643.t01 |
| 7624 | 5 | 175540665 | 175620646 | mNeoVis.2.1.1.chr5\_hap1.g01644.t01 |
| 7625 | 5 | 175635923 | 175700176 | mNeoVis.2.1.1.chr5\_hap1.g01645.t01 |
| 7626 | 5 | 175724533 | 175759420 | mNeoVis.2.1.1.chr5\_hap1.g01646.t01 |
| 7627 | 5 | 175759548 | 175784572 | mNeoVis.2.1.1.chr5\_hap1.g01647.t01 |
| 7628 | 5 | 175810602 | 175834991 | mNeoVis.2.1.1.chr5\_hap1.g01648.t01 |
| 7629 | 5 | 175840524 | 175899006 | mNeoVis.2.1.1.chr5\_hap1.g01649.t01 |
| 7630 | 5 | 175933282 | 175958636 | mNeoVis.2.1.1.chr5\_hap1.g01650.t01 |
| 7631 | 5 | 175959818 | 176349753 | mNeoVis.2.1.1.chr5\_hap1.g01651.t01 |
| 7632 | 5 | 176008135 | 176008737 | mNeoVis.2.1.1.chr5\_hap1.g01652.t01 |
| 7633 | 5 | 176037242 | 176343075 | mNeoVis.2.1.1.chr5\_hap1.g01653.t01 |
| 7634 | 5 | 176366427 | 176521361 | mNeoVis.2.1.1.chr5\_hap1.g01654.t01 |
| 7635 | 5 | 177188146 | 177283983 | mNeoVis.2.1.1.chr5\_hap1.g01655.t01 |
| 7636 | 5 | 177276158 | 177344792 | mNeoVis.2.1.1.chr5\_hap1.g01656.t01 |
| 7637 | 5 | 178161042 | 178166835 | mNeoVis.2.1.1.chr5\_hap1.g01675.t01 |
| 7638 | 5 | 178215286 | 178221182 | mNeoVis.2.1.1.chr5\_hap1.g01676.t01 |
| 7639 | 5 | 178226095 | 178227015 | mNeoVis.2.1.1.chr5\_hap1.g01677.t01 |
| 7640 | 5 | 178247335 | 178248255 | mNeoVis.2.1.1.chr5\_hap1.g01678.t01 |
| 7641 | 5 | 178270191 | 178318844 | mNeoVis.2.1.1.chr5\_hap1.g01679.t01 |
| 7642 | 5 | 178331210 | 178364388 | mNeoVis.2.1.1.chr5\_hap1.g01680.t01 |
| 7643 | 5 | 178364317 | 178481677 | mNeoVis.2.1.1.chr5\_hap1.g01681.t01 |
| 7644 | 5 | 178437177 | 178546900 | mNeoVis.2.1.1.chr5\_hap1.g01682.t01 |
| 7645 | 5 | 178501809 | 178546652 | mNeoVis.2.1.1.chr5\_hap1.g01683.t01 |
| 7646 | 5 | 178555004 | 179428355 | mNeoVis.2.1.1.chr5\_hap1.g01684.t01 |
| 7647 | 5 | 178575493 | 178622395 | mNeoVis.2.1.1.chr5\_hap1.g01685.t01 |
| 7648 | 5 | 181670240 | 181726101 | mNeoVis.2.1.1.chr5\_hap1.g01686.t01 |
| 7649 | 5 | 181670240 | 181726101 | mNeoVis.2.1.1.chr5\_hap1.g01686.t01 |
| 7650 | 5 | 181670240 | 181726101 | mNeoVis.2.1.1.chr5\_hap1.g01686.t01 |
| 7651 | 5 | 181731033 | 181806239 | mNeoVis.2.1.1.chr5\_hap1.g01687.t01 |
| 7652 | 5 | 181731033 | 181806239 | mNeoVis.2.1.1.chr5\_hap1.g01687.t01 |
| 7653 | 5 | 181764301 | 181776181 | mNeoVis.2.1.1.chr5\_hap1.g01688.t01 |
| 7654 | 5 | 181764301 | 181776181 | mNeoVis.2.1.1.chr5\_hap1.g01688.t01 |
| 7655 | 5 | 181812264 | 181880071 | mNeoVis.2.1.1.chr5\_hap1.g01689.t01 |
| 7656 | 5 | 181812264 | 181880071 | mNeoVis.2.1.1.chr5\_hap1.g01689.t01 |
| 7657 | 5 | 182502650 | 182862485 | mNeoVis.2.1.1.chr5\_hap1.g01690.t01 |
| 7658 | 5 | 183396446 | 183510859 | mNeoVis.2.1.1.chr5\_hap1.g01691.t01 |
| 7659 | 5 | 183396446 | 183510859 | mNeoVis.2.1.1.chr5\_hap1.g01691.t01 |
| 7660 | 5 | 183676519 | 183689613 | mNeoVis.2.1.1.chr5\_hap1.g01692.t01 |
| 7661 | 5 | 183676519 | 183689613 | mNeoVis.2.1.1.chr5\_hap1.g01692.t01 |
| 7662 | 5 | 183696952 | 183775835 | mNeoVis.2.1.1.chr5\_hap1.g01693.t01 |
| 7663 | 5 | 183696952 | 183775835 | mNeoVis.2.1.1.chr5\_hap1.g01693.t01 |
| 7664 | 5 | 183775992 | 183782869 | mNeoVis.2.1.1.chr5\_hap1.g01694.t01 |
| 7665 | 5 | 183775992 | 183782869 | mNeoVis.2.1.1.chr5\_hap1.g01694.t01 |
| 7666 | 5 | 183839191 | 183986285 | mNeoVis.2.1.1.chr5\_hap1.g01695.t01 |
| 7667 | 5 | 183839191 | 183986285 | mNeoVis.2.1.1.chr5\_hap1.g01695.t01 |
| 7668 | 5 | 184025172 | 184040833 | mNeoVis.2.1.1.chr5\_hap1.g01696.t01 |
| 7669 | 5 | 184025172 | 184040833 | mNeoVis.2.1.1.chr5\_hap1.g01696.t01 |
| 7670 | 5 | 184367145 | 184412451 | mNeoVis.2.1.1.chr5\_hap1.g01697.t01 |
| 7671 | 5 | 184367145 | 184412451 | mNeoVis.2.1.1.chr5\_hap1.g01697.t01 |
| 7672 | 5 | 184367145 | 184412451 | mNeoVis.2.1.1.chr5\_hap1.g01697.t01 |
| 7673 | 5 | 184415029 | 184442344 | mNeoVis.2.1.1.chr5\_hap1.g01698.t01 |
| 7674 | 5 | 184415029 | 184442344 | mNeoVis.2.1.1.chr5\_hap1.g01698.t01 |
| 7675 | 5 | 184557833 | 184792851 | mNeoVis.2.1.1.chr5\_hap1.g01699.t01 |
| 7676 | 5 | 184557833 | 184792851 | mNeoVis.2.1.1.chr5\_hap1.g01699.t01 |
| 7677 | 5 | 185493528 | 186578111 | mNeoVis.2.1.1.chr5\_hap1.g01700.t01 |
| 7678 | 5 | 188794554 | 188806354 | mNeoVis.2.1.1.chr5\_hap1.g01702.t01 |
| 7679 | 5 | 188794554 | 188806354 | mNeoVis.2.1.1.chr5\_hap1.g01702.t01 |
| 7680 | 5 | 189694971 | 189966226 | mNeoVis.2.1.1.chr5\_hap1.g01703.t01 |
| 7681 | 5 | 189936245 | 189965410 | mNeoVis.2.1.1.chr5\_hap1.g01704.t01 |
| 7682 | 5 | 193522843 | 194826187 | mNeoVis.2.1.1.chr5\_hap1.g01706.t01 |
| 7683 | 5 | 195243408 | 195243964 | mNeoVis.2.1.1.chr5\_hap1.g01707.t01 |
| 7684 | 5 | 195243408 | 195243964 | mNeoVis.2.1.1.chr5\_hap1.g01707.t01 |
| 7685 | 5 | 195280319 | 195360883 | mNeoVis.2.1.1.chr5\_hap1.g01708.t01 |
| 7686 | 5 | 195280319 | 195360883 | mNeoVis.2.1.1.chr5\_hap1.g01708.t01 |
| 7687 | 5 | 195360975 | 195374273 | mNeoVis.2.1.1.chr5\_hap1.g01709.t01 |
| 7688 | 5 | 195360975 | 195374273 | mNeoVis.2.1.1.chr5\_hap1.g01709.t01 |
| 7689 | 5 | 195420472 | 195511015 | mNeoVis.2.1.1.chr5\_hap1.g01710.t01 |
| 7690 | 5 | 195520755 | 195532234 | mNeoVis.2.1.1.chr5\_hap1.g01711.t01 |
| 7691 | 5 | 195532454 | 195697637 | mNeoVis.2.1.1.chr5\_hap1.g01712.t01 |
| 7692 | 5 | 195632418 | 195721621 | mNeoVis.2.1.1.chr5\_hap1.g01713.t01 |
| 7693 | 5 | 196020482 | 196121548 | mNeoVis.2.1.1.chr5\_hap1.g01714.t01 |
| 7694 | 5 | 196216378 | 196218980 | mNeoVis.2.1.1.chr5\_hap1.g01715.t01 |
| 7695 | 5 | 196216378 | 196218980 | mNeoVis.2.1.1.chr5\_hap1.g01715.t01 |
| 7696 | 5 | 196733700 | 196868590 | mNeoVis.2.1.1.chr5\_hap1.g01716.t01 |
| 7697 | 5 | 196733700 | 196868590 | mNeoVis.2.1.1.chr5\_hap1.g01716.t01 |
| 7698 | 5 | 196959520 | 197225419 | mNeoVis.2.1.1.chr5\_hap1.g01717.t01 |
| 7699 | 5 | 197147525 | 197397001 | mNeoVis.2.1.1.chr5\_hap1.g01718.t01 |
| 7700 | 5 | 197959338 | 197959960 | mNeoVis.2.1.1.chr5\_hap1.g01720.t01 |
| 7701 | 5 | 197959338 | 197959960 | mNeoVis.2.1.1.chr5\_hap1.g01720.t01 |
| 7702 | 5 | 198170367 | 198171383 | mNeoVis.2.1.1.chr5\_hap1.g01721.t01 |
| 7703 | 5 | 198251220 | 198292106 | mNeoVis.2.1.1.chr5\_hap1.g01722.t01 |
| 7704 | 5 | 198307765 | 198327012 | mNeoVis.2.1.1.chr5\_hap1.g01723.t01 |
| 7705 | 5 | 198449565 | 198482473 | mNeoVis.2.1.1.chr5\_hap1.g01724.t01 |
| 7706 | 5 | 198974676 | 199017803 | mNeoVis.2.1.1.chr5\_hap1.g01726.t01 |
| 7707 | 5 | 199004960 | 199124684 | mNeoVis.2.1.1.chr5\_hap1.g01727.t01 |
| 7708 | 5 | 201489701 | 201511421 | mNeoVis.2.1.1.chr5\_hap1.g01729.t01 |
| 7709 | 5 | 203789687 | 203827788 | mNeoVis.2.1.1.chr5\_hap1.g01730.t01 |
| 7710 | 5 | 206093527 | 206328379 | mNeoVis.2.1.1.chr5\_hap1.g01736.t01 |
| 7711 | 5 | 206632784 | 206736657 | mNeoVis.2.1.1.chr5\_hap1.g01737.t01 |
| 7712 | 5 | 206963003 | 206972368 | mNeoVis.2.1.1.chr5\_hap1.g01738.t01 |
| 7713 | 5 | 207030528 | 207079817 | mNeoVis.2.1.1.chr5\_hap1.g01739.t01 |
| 7714 | 5 | 209900026 | 209902451 | mNeoVis.2.1.1.chr5\_hap1.g01751.t01 |
| 7715 | 5 | 209950739 | 209953145 | mNeoVis.2.1.1.chr5\_hap1.g01752.t01 |
| 7716 | 5 | 210009012 | 210010635 | mNeoVis.2.1.1.chr5\_hap1.g01753.t01 |
| 7717 | 5 | 210017020 | 210039879 | mNeoVis.2.1.1.chr5\_hap1.g01754.t01 |
| 7718 | 5 | 210053677 | 210054540 | mNeoVis.2.1.1.chr5\_hap1.g01755.t01 |
| 7719 | 5 | 210068214 | 210075404 | mNeoVis.2.1.1.chr5\_hap1.g01756.t01 |
| 7720 | 5 | 210101064 | 210101950 | mNeoVis.2.1.1.chr5\_hap1.g01757.t01 |
| 7721 | 5 | 210126198 | 210134508 | mNeoVis.2.1.1.chr5\_hap1.g01758.t01 |
| 7722 | 5 | 210165021 | 210165802 | mNeoVis.2.1.1.chr5\_hap1.g01759.t01 |
| 7723 | 5 | 210171971 | 210172757 | mNeoVis.2.1.1.chr5\_hap1.g01760.t01 |
| 7724 | 5 | 210179484 | 210180372 | mNeoVis.2.1.1.chr5\_hap1.g01761.t01 |
| 7725 | 5 | 210184409 | 210185031 | mNeoVis.2.1.1.chr5\_hap1.g01762.t01 |
| 7726 | 5 | 210244841 | 210245323 | mNeoVis.2.1.1.chr5\_hap1.g01763.t01 |
| 7727 | 5 | 210251504 | 210252025 | mNeoVis.2.1.1.chr5\_hap1.g01764.t01 |
| 7728 | 5 | 210375894 | 210376509 | mNeoVis.2.1.1.chr5\_hap1.g01765.t01 |
| 7729 | 5 | 210409711 | 210410329 | mNeoVis.2.1.1.chr5\_hap1.g01766.t01 |
| 7730 | 5 | 210416119 | 210470678 | mNeoVis.2.1.1.chr5\_hap1.g01767.t01 |
| 7731 | 5 | 210425182 | 210425877 | mNeoVis.2.1.1.chr5\_hap1.g01768.t01 |
| 7732 | 5 | 210449362 | 210450070 | mNeoVis.2.1.1.chr5\_hap1.g01769.t01 |
| 7733 | 5 | 210465078 | 210465559 | mNeoVis.2.1.1.chr5\_hap1.g01770.t01 |
| 7734 | 5 | 210494030 | 210494555 | mNeoVis.2.1.1.chr5\_hap1.g01771.t01 |
| 7735 | 5 | 210584423 | 210594841 | mNeoVis.2.1.1.chr5\_hap1.g01772.t01 |
| 7736 | 5 | 210598750 | 210599424 | mNeoVis.2.1.1.chr5\_hap1.g01773.t01 |
| 7737 | 5 | 210604273 | 210605110 | mNeoVis.2.1.1.chr5\_hap1.g01774.t01 |
| 7738 | 5 | 210611416 | 210616749 | mNeoVis.2.1.1.chr5\_hap1.g01775.t01 |
| 7739 | 5 | 210619882 | 210620554 | mNeoVis.2.1.1.chr5\_hap1.g01776.t01 |
| 7740 | 5 | 211915478 | 211926810 | mNeoVis.2.1.1.chr5\_hap1.g01785.t01 |
| 7741 | 5 | 211943117 | 211959886 | mNeoVis.2.1.1.chr5\_hap1.g01786.t01 |
| 7742 | 5 | 211961167 | 212029289 | mNeoVis.2.1.1.chr5\_hap1.g01787.t01 |
| 7743 | 5 | 212079421 | 212118300 | mNeoVis.2.1.1.chr5\_hap1.g01788.t01 |
| 7744 | 5 | 212153652 | 212164614 | mNeoVis.2.1.1.chr5\_hap1.g01789.t01 |
| 7745 | 5 | 212168644 | 212256501 | mNeoVis.2.1.1.chr5\_hap1.g01790.t01 |
| 7746 | 5 | 212263357 | 212297753 | mNeoVis.2.1.1.chr5\_hap1.g01791.t01 |
| 7747 | 5 | 212299971 | 212306830 | mNeoVis.2.1.1.chr5\_hap1.g01792.t01 |
| 7748 | 5 | 212327289 | 212343627 | mNeoVis.2.1.1.chr5\_hap1.g01793.t01 |
| 7749 | 5 | 212987295 | 213023201 | mNeoVis.2.1.1.chr5\_hap1.g01801.t01 |
| 7750 | 5 | 213022687 | 213055734 | mNeoVis.2.1.1.chr5\_hap1.g01802.t01 |
| 7751 | 5 | 213045958 | 213066334 | mNeoVis.2.1.1.chr5\_hap1.g01803.t01 |
| 7752 | 5 | 213067174 | 213098985 | mNeoVis.2.1.1.chr5\_hap1.g01804.t01 |
| 7753 | 5 | 213135882 | 213281635 | mNeoVis.2.1.1.chr5\_hap1.g01805.t01 |
| 7754 | 5 | 213291727 | 213300140 | mNeoVis.2.1.1.chr5\_hap1.g01806.t01 |
| 7755 | 5 | 213435019 | 213501166 | mNeoVis.2.1.1.chr5\_hap1.g01807.t01 |
| 7756 | 5 | 213456114 | 213468724 | mNeoVis.2.1.1.chr5\_hap1.g01808.t01 |
| 7757 | 5 | 213943645 | 213988314 | mNeoVis.2.1.1.chr5\_hap1.g01815.t01 |
| 7758 | 5 | 214064267 | 214307792 | mNeoVis.2.1.1.chr5\_hap1.g01816.t01 |
| 7759 | 5 | 215248629 | 215277162 | mNeoVis.2.1.1.chr5\_hap1.g01817.t01 |
| 7760 | 5 | 215280489 | 215285172 | mNeoVis.2.1.1.chr5\_hap1.g01818.t01 |
| 7761 | 5 | 215308228 | 215314047 | mNeoVis.2.1.1.chr5\_hap1.g01819.t01 |
| 7762 | 5 | 215323963 | 215333161 | mNeoVis.2.1.1.chr5\_hap1.g01820.t01 |
| 7763 | 5 | 215384917 | 215401645 | mNeoVis.2.1.1.chr5\_hap1.g01821.t01 |
| 7764 | 5 | 217352112 | 217401699 | mNeoVis.2.1.1.chr5\_hap1.g01835.t01 |
| 7765 | 5 | 217461224 | 217704305 | mNeoVis.2.1.1.chr5\_hap1.g01836.t01 |
| 7766 | 5 | 217828795 | 217847061 | mNeoVis.2.1.1.chr5\_hap1.g01837.t01 |
| 7767 | 5 | 220930602 | 220966930 | mNeoVis.2.1.1.chr5\_hap1.g01867.t01 |
| 7768 | 5 | 220981936 | 221001537 | mNeoVis.2.1.1.chr5\_hap1.g01868.t01 |
| 7769 | 5 | 221005376 | 221074587 | mNeoVis.2.1.1.chr5\_hap1.g01869.t01 |
| 7770 | 5 | 221115452 | 221124159 | mNeoVis.2.1.1.chr5\_hap1.g01870.t01 |
| 7771 | 5 | 221128396 | 221209806 | mNeoVis.2.1.1.chr5\_hap1.g01871.t01 |
| 7772 | 5 | 221222933 | 221227095 | mNeoVis.2.1.1.chr5\_hap1.g01872.t01 |
| 7773 | 5 | 221252877 | 221273992 | mNeoVis.2.1.1.chr5\_hap1.g01873.t01 |
| 7774 | 5 | 221286168 | 221297568 | mNeoVis.2.1.1.chr5\_hap1.g01874.t01 |
| 7775 | 5 | 221325316 | 221387698 | mNeoVis.2.1.1.chr5\_hap1.g01875.t01 |
| 7776 | 5 | 221398151 | 221418537 | mNeoVis.2.1.1.chr5\_hap1.g01876.t01 |
| 7777 | 5 | 221427369 | 221440300 | mNeoVis.2.1.1.chr5\_hap1.g01877.t01 |
| 7778 | 6 | 1818780 | 1855660 | mNeoVis.2.1.1.chr6\_hap1.g00020.t01 |
| 7779 | 6 | 2013495 | 2110826 | mNeoVis.2.1.1.chr6\_hap1.g00021.t01 |
| 7780 | 6 | 3739420 | 3754421 | mNeoVis.2.1.1.chr6\_hap1.g00032.t01 |
| 7781 | 6 | 3765867 | 3787695 | mNeoVis.2.1.1.chr6\_hap1.g00033.t01 |
| 7782 | 6 | 4119853 | 4157011 | mNeoVis.2.1.1.chr6\_hap1.g00034.t01 |
| 7783 | 6 | 4144731 | 4266367 | mNeoVis.2.1.1.chr6\_hap1.g00035.t01 |
| 7784 | 6 | 6111972 | 6423774 | mNeoVis.2.1.1.chr6\_hap1.g00049.t01 |
| 7785 | 6 | 6415804 | 6507703 | mNeoVis.2.1.1.chr6\_hap1.g00050.t01 |
| 7786 | 6 | 6510059 | 6514298 | mNeoVis.2.1.1.chr6\_hap1.g00051.t01 |
| 7787 | 6 | 6525362 | 6557314 | mNeoVis.2.1.1.chr6\_hap1.g00052.t01 |
| 7788 | 6 | 6579014 | 6649849 | mNeoVis.2.1.1.chr6\_hap1.g00053.t01 |
| 7789 | 6 | 6653064 | 6688659 | mNeoVis.2.1.1.chr6\_hap1.g00054.t01 |
| 7790 | 6 | 7163887 | 7251768 | mNeoVis.2.1.1.chr6\_hap1.g00064.t01 |
| 7791 | 6 | 7257937 | 7312907 | mNeoVis.2.1.1.chr6\_hap1.g00065.t01 |
| 7792 | 6 | 7326036 | 7327951 | mNeoVis.2.1.1.chr6\_hap1.g00066.t01 |
| 7793 | 6 | 7353706 | 7369180 | mNeoVis.2.1.1.chr6\_hap1.g00067.t01 |
| 7794 | 6 | 7369204 | 7387605 | mNeoVis.2.1.1.chr6\_hap1.g00068.t01 |
| 7795 | 6 | 7388128 | 7499784 | mNeoVis.2.1.1.chr6\_hap1.g00069.t01 |
| 7796 | 6 | 7598427 | 7602798 | mNeoVis.2.1.1.chr6\_hap1.g00070.t01 |
| 7797 | 6 | 7932098 | 7979265 | mNeoVis.2.1.1.chr6\_hap1.g00072.t01 |
| 7798 | 6 | 8214358 | 8227021 | mNeoVis.2.1.1.chr6\_hap1.g00073.t01 |
| 7799 | 6 | 8277188 | 8284502 | mNeoVis.2.1.1.chr6\_hap1.g00074.t01 |
| 7800 | 6 | 8332429 | 8340199 | mNeoVis.2.1.1.chr6\_hap1.g00075.t01 |
| 7801 | 6 | 8344932 | 8477892 | mNeoVis.2.1.1.chr6\_hap1.g00076.t01 |
| 7802 | 6 | 8504324 | 8506229 | mNeoVis.2.1.1.chr6\_hap1.g00077.t01 |
| 7803 | 6 | 8514043 | 8516063 | mNeoVis.2.1.1.chr6\_hap1.g00078.t01 |
| 7804 | 6 | 8609161 | 8611337 | mNeoVis.2.1.1.chr6\_hap1.g00079.t01 |
| 7805 | 6 | 8626544 | 8627434 | mNeoVis.2.1.1.chr6\_hap1.g00080.t01 |
| 7806 | 6 | 8632594 | 8633441 | mNeoVis.2.1.1.chr6\_hap1.g00081.t01 |
| 7807 | 6 | 8644621 | 8646724 | mNeoVis.2.1.1.chr6\_hap1.g00082.t01 |
| 7808 | 6 | 8736400 | 8739878 | mNeoVis.2.1.1.chr6\_hap1.g00083.t01 |
| 7809 | 6 | 8845423 | 8889826 | mNeoVis.2.1.1.chr6\_hap1.g00084.t01 |
| 7810 | 6 | 8897659 | 8944542 | mNeoVis.2.1.1.chr6\_hap1.g00085.t01 |
| 7811 | 6 | 8952773 | 8974236 | mNeoVis.2.1.1.chr6\_hap1.g00086.t01 |
| 7812 | 6 | 8982933 | 9003466 | mNeoVis.2.1.1.chr6\_hap1.g00087.t01 |
| 7813 | 6 | 9017635 | 9034737 | mNeoVis.2.1.1.chr6\_hap1.g00088.t01 |
| 7814 | 6 | 9146544 | 9310760 | mNeoVis.2.1.1.chr6\_hap1.g00089.t01 |
| 7815 | 6 | 9315783 | 9328220 | mNeoVis.2.1.1.chr6\_hap1.g00090.t01 |
| 7816 | 6 | 9686856 | 9944231 | mNeoVis.2.1.1.chr6\_hap1.g00091.t01 |
| 7817 | 6 | 10062507 | 10075000 | mNeoVis.2.1.1.chr6\_hap1.g00092.t01 |
| 7818 | 6 | 10062507 | 10075000 | mNeoVis.2.1.1.chr6\_hap1.g00092.t01 |
| 7819 | 6 | 10355610 | 10390527 | mNeoVis.2.1.1.chr6\_hap1.g00093.t01 |
| 7820 | 6 | 10355610 | 10390527 | mNeoVis.2.1.1.chr6\_hap1.g00093.t01 |
| 7821 | 6 | 10519223 | 10870020 | mNeoVis.2.1.1.chr6\_hap1.g00094.t01 |
| 7822 | 6 | 10519223 | 10870020 | mNeoVis.2.1.1.chr6\_hap1.g00094.t01 |
| 7823 | 6 | 10964760 | 10994539 | mNeoVis.2.1.1.chr6\_hap1.g00095.t01 |
| 7824 | 6 | 10964760 | 10994539 | mNeoVis.2.1.1.chr6\_hap1.g00095.t01 |
| 7825 | 6 | 10967471 | 10982783 | mNeoVis.2.1.1.chr6\_hap1.g00096.t01 |
| 7826 | 6 | 10967471 | 10982783 | mNeoVis.2.1.1.chr6\_hap1.g00096.t01 |
| 7827 | 6 | 10997541 | 11019847 | mNeoVis.2.1.1.chr6\_hap1.g00097.t01 |
| 7828 | 6 | 10997541 | 11019847 | mNeoVis.2.1.1.chr6\_hap1.g00097.t01 |
| 7829 | 6 | 11100549 | 11114436 | mNeoVis.2.1.1.chr6\_hap1.g00098.t01 |
| 7830 | 6 | 11100549 | 11114436 | mNeoVis.2.1.1.chr6\_hap1.g00098.t01 |
| 7831 | 6 | 11125439 | 11209727 | mNeoVis.2.1.1.chr6\_hap1.g00099.t01 |
| 7832 | 6 | 11221296 | 11225844 | mNeoVis.2.1.1.chr6\_hap1.g00100.t01 |
| 7833 | 6 | 11250726 | 11259804 | mNeoVis.2.1.1.chr6\_hap1.g00101.t01 |
| 7834 | 6 | 11277688 | 11291354 | mNeoVis.2.1.1.chr6\_hap1.g00102.t01 |
| 7835 | 6 | 11310551 | 11323637 | mNeoVis.2.1.1.chr6\_hap1.g00103.t01 |
| 7836 | 6 | 11361460 | 11375367 | mNeoVis.2.1.1.chr6\_hap1.g00104.t01 |
| 7837 | 6 | 11399555 | 11416199 | mNeoVis.2.1.1.chr6\_hap1.g00105.t01 |
| 7838 | 6 | 11544147 | 11562183 | mNeoVis.2.1.1.chr6\_hap1.g00107.t01 |
| 7839 | 6 | 11576555 | 11578717 | mNeoVis.2.1.1.chr6\_hap1.g00108.t01 |
| 7840 | 6 | 11576555 | 11578717 | mNeoVis.2.1.1.chr6\_hap1.g00108.t01 |
| 7841 | 6 | 11646588 | 11665698 | mNeoVis.2.1.1.chr6\_hap1.g00109.t01 |
| 7842 | 6 | 11646588 | 11665698 | mNeoVis.2.1.1.chr6\_hap1.g00109.t01 |
| 7843 | 6 | 11689763 | 11699040 | mNeoVis.2.1.1.chr6\_hap1.g00110.t01 |
| 7844 | 6 | 11689763 | 11699040 | mNeoVis.2.1.1.chr6\_hap1.g00110.t01 |
| 7845 | 6 | 11750705 | 11765114 | mNeoVis.2.1.1.chr6\_hap1.g00111.t01 |
| 7846 | 6 | 11750705 | 11765114 | mNeoVis.2.1.1.chr6\_hap1.g00111.t01 |
| 7847 | 6 | 11862138 | 11862416 | mNeoVis.2.1.1.chr6\_hap1.g00112.t01 |
| 7848 | 6 | 11862138 | 11862416 | mNeoVis.2.1.1.chr6\_hap1.g00112.t01 |
| 7849 | 6 | 11909851 | 11917770 | mNeoVis.2.1.1.chr6\_hap1.g00113.t01 |
| 7850 | 6 | 11909851 | 11917770 | mNeoVis.2.1.1.chr6\_hap1.g00113.t01 |
| 7851 | 6 | 11927393 | 11937281 | mNeoVis.2.1.1.chr6\_hap1.g00114.t01 |
| 7852 | 6 | 11927393 | 11937281 | mNeoVis.2.1.1.chr6\_hap1.g00114.t01 |
| 7853 | 6 | 11997291 | 12020902 | mNeoVis.2.1.1.chr6\_hap1.g00115.t01 |
| 7854 | 6 | 11997291 | 12020902 | mNeoVis.2.1.1.chr6\_hap1.g00115.t01 |
| 7855 | 6 | 12031909 | 12066689 | mNeoVis.2.1.1.chr6\_hap1.g00116.t01 |
| 7856 | 6 | 12031909 | 12066689 | mNeoVis.2.1.1.chr6\_hap1.g00116.t01 |
| 7857 | 6 | 12076662 | 12115790 | mNeoVis.2.1.1.chr6\_hap1.g00117.t01 |
| 7858 | 6 | 12076662 | 12115790 | mNeoVis.2.1.1.chr6\_hap1.g00117.t01 |
| 7859 | 6 | 12171594 | 12236083 | mNeoVis.2.1.1.chr6\_hap1.g00118.t01 |
| 7860 | 6 | 12481115 | 12517444 | mNeoVis.2.1.1.chr6\_hap1.g00121.t01 |
| 7861 | 6 | 12580432 | 12668168 | mNeoVis.2.1.1.chr6\_hap1.g00122.t01 |
| 7862 | 6 | 12669622 | 12698600 | mNeoVis.2.1.1.chr6\_hap1.g00123.t01 |
| 7863 | 6 | 12710823 | 12803962 | mNeoVis.2.1.1.chr6\_hap1.g00124.t01 |
| 7864 | 6 | 12822612 | 12855623 | mNeoVis.2.1.1.chr6\_hap1.g00125.t01 |
| 7865 | 6 | 12822612 | 12855623 | mNeoVis.2.1.1.chr6\_hap1.g00125.t01 |
| 7866 | 6 | 12866451 | 12918083 | mNeoVis.2.1.1.chr6\_hap1.g00126.t01 |
| 7867 | 6 | 12866451 | 12918083 | mNeoVis.2.1.1.chr6\_hap1.g00126.t01 |
| 7868 | 6 | 12938846 | 12987534 | mNeoVis.2.1.1.chr6\_hap1.g00127.t01 |
| 7869 | 6 | 12938846 | 12987534 | mNeoVis.2.1.1.chr6\_hap1.g00127.t01 |
| 7870 | 6 | 13069812 | 13086454 | mNeoVis.2.1.1.chr6\_hap1.g00128.t01 |
| 7871 | 6 | 13069812 | 13086454 | mNeoVis.2.1.1.chr6\_hap1.g00128.t01 |
| 7872 | 6 | 13102342 | 13189054 | mNeoVis.2.1.1.chr6\_hap1.g00129.t01 |
| 7873 | 6 | 13102342 | 13189054 | mNeoVis.2.1.1.chr6\_hap1.g00129.t01 |
| 7874 | 6 | 13201846 | 13249228 | mNeoVis.2.1.1.chr6\_hap1.g00130.t01 |
| 7875 | 6 | 13201846 | 13249228 | mNeoVis.2.1.1.chr6\_hap1.g00130.t01 |
| 7876 | 6 | 13249104 | 13345517 | mNeoVis.2.1.1.chr6\_hap1.g00131.t01 |
| 7877 | 6 | 15052621 | 15402703 | mNeoVis.2.1.1.chr6\_hap1.g00132.t01 |
| 7878 | 6 | 15052621 | 15402703 | mNeoVis.2.1.1.chr6\_hap1.g00132.t01 |
| 7879 | 6 | 17647380 | 17661546 | mNeoVis.2.1.1.chr6\_hap1.g00134.t01 |
| 7880 | 6 | 17647380 | 17661546 | mNeoVis.2.1.1.chr6\_hap1.g00134.t01 |
| 7881 | 6 | 17725490 | 17769110 | mNeoVis.2.1.1.chr6\_hap1.g00135.t01 |
| 7882 | 6 | 17725490 | 17769110 | mNeoVis.2.1.1.chr6\_hap1.g00135.t01 |
| 7883 | 6 | 17982748 | 18781332 | mNeoVis.2.1.1.chr6\_hap1.g00136.t01 |
| 7884 | 6 | 18870629 | 18884614 | mNeoVis.2.1.1.chr6\_hap1.g00137.t01 |
| 7885 | 6 | 19711032 | 19711553 | mNeoVis.2.1.1.chr6\_hap1.g00138.t01 |
| 7886 | 6 | 21915697 | 22374473 | mNeoVis.2.1.1.chr6\_hap1.g00139.t01 |
| 7887 | 6 | 21915697 | 22374473 | mNeoVis.2.1.1.chr6\_hap1.g00139.t01 |
| 7888 | 6 | 22475029 | 22552736 | mNeoVis.2.1.1.chr6\_hap1.g00140.t01 |
| 7889 | 6 | 22552648 | 22598366 | mNeoVis.2.1.1.chr6\_hap1.g00141.t01 |
| 7890 | 6 | 22598468 | 22612186 | mNeoVis.2.1.1.chr6\_hap1.g00142.t01 |
| 7891 | 6 | 22847249 | 22877082 | mNeoVis.2.1.1.chr6\_hap1.g00146.t01 |
| 7892 | 6 | 22847249 | 22877082 | mNeoVis.2.1.1.chr6\_hap1.g00146.t01 |
| 7893 | 6 | 22908137 | 22957190 | mNeoVis.2.1.1.chr6\_hap1.g00147.t01 |
| 7894 | 6 | 22908137 | 22957190 | mNeoVis.2.1.1.chr6\_hap1.g00147.t01 |
| 7895 | 6 | 22965064 | 22981356 | mNeoVis.2.1.1.chr6\_hap1.g00148.t01 |
| 7896 | 6 | 22965064 | 22981356 | mNeoVis.2.1.1.chr6\_hap1.g00148.t01 |
| 7897 | 6 | 22982793 | 23014645 | mNeoVis.2.1.1.chr6\_hap1.g00149.t01 |
| 7898 | 6 | 22982793 | 23014645 | mNeoVis.2.1.1.chr6\_hap1.g00149.t01 |
| 7899 | 6 | 23019641 | 23069639 | mNeoVis.2.1.1.chr6\_hap1.g00150.t01 |
| 7900 | 6 | 23019641 | 23069639 | mNeoVis.2.1.1.chr6\_hap1.g00150.t01 |
| 7901 | 6 | 23045652 | 23081430 | mNeoVis.2.1.1.chr6\_hap1.g00151.t01 |
| 7902 | 6 | 23045652 | 23081430 | mNeoVis.2.1.1.chr6\_hap1.g00151.t01 |
| 7903 | 6 | 23091877 | 23130077 | mNeoVis.2.1.1.chr6\_hap1.g00152.t01 |
| 7904 | 6 | 23091877 | 23130077 | mNeoVis.2.1.1.chr6\_hap1.g00152.t01 |
| 7905 | 6 | 23136011 | 23273785 | mNeoVis.2.1.1.chr6\_hap1.g00153.t01 |
| 7906 | 6 | 23136011 | 23273785 | mNeoVis.2.1.1.chr6\_hap1.g00153.t01 |
| 7907 | 6 | 23345664 | 23355404 | mNeoVis.2.1.1.chr6\_hap1.g00154.t01 |
| 7908 | 6 | 23430881 | 23510540 | mNeoVis.2.1.1.chr6\_hap1.g00155.t01 |
| 7909 | 6 | 23538423 | 23600569 | mNeoVis.2.1.1.chr6\_hap1.g00156.t01 |
| 7910 | 6 | 23621159 | 23637909 | mNeoVis.2.1.1.chr6\_hap1.g00157.t01 |
| 7911 | 6 | 24098535 | 24113187 | mNeoVis.2.1.1.chr6\_hap1.g00162.t01 |
| 7912 | 6 | 24270348 | 24313978 | mNeoVis.2.1.1.chr6\_hap1.g00163.t01 |
| 7913 | 6 | 25190870 | 25251713 | mNeoVis.2.1.1.chr6\_hap1.g00168.t01 |
| 7914 | 6 | 25568378 | 25749870 | mNeoVis.2.1.1.chr6\_hap1.g00169.t01 |
| 7915 | 6 | 25750652 | 25827911 | mNeoVis.2.1.1.chr6\_hap1.g00170.t01 |
| 7916 | 6 | 26181610 | 26215868 | mNeoVis.2.1.1.chr6\_hap1.g00172.t01 |
| 7917 | 6 | 26290183 | 26294829 | mNeoVis.2.1.1.chr6\_hap1.g00173.t01 |
| 7918 | 6 | 26440309 | 26496658 | mNeoVis.2.1.1.chr6\_hap1.g00174.t01 |
| 7919 | 6 | 27083933 | 27087289 | mNeoVis.2.1.1.chr6\_hap1.g00179.t01 |
| 7920 | 6 | 27114007 | 27168423 | mNeoVis.2.1.1.chr6\_hap1.g00180.t01 |
| 7921 | 6 | 27230637 | 27246344 | mNeoVis.2.1.1.chr6\_hap1.g00181.t01 |
| 7922 | 6 | 27250584 | 27277572 | mNeoVis.2.1.1.chr6\_hap1.g00182.t01 |
| 7923 | 6 | 27413745 | 27581026 | mNeoVis.2.1.1.chr6\_hap1.g00183.t01 |
| 7924 | 6 | 27581157 | 27588790 | mNeoVis.2.1.1.chr6\_hap1.g00184.t01 |
| 7925 | 6 | 27589535 | 27596218 | mNeoVis.2.1.1.chr6\_hap1.g00185.t01 |
| 7926 | 6 | 27638649 | 27729815 | mNeoVis.2.1.1.chr6\_hap1.g00186.t01 |
| 7927 | 6 | 27775688 | 27818075 | mNeoVis.2.1.1.chr6\_hap1.g00187.t01 |
| 7928 | 6 | 27848607 | 27940577 | mNeoVis.2.1.1.chr6\_hap1.g00188.t01 |
| 7929 | 6 | 27942828 | 28002810 | mNeoVis.2.1.1.chr6\_hap1.g00189.t01 |
| 7930 | 6 | 28021457 | 28048516 | mNeoVis.2.1.1.chr6\_hap1.g00190.t01 |
| 7931 | 6 | 28021457 | 28048516 | mNeoVis.2.1.1.chr6\_hap1.g00190.t01 |
| 7932 | 6 | 28066342 | 28095853 | mNeoVis.2.1.1.chr6\_hap1.g00191.t01 |
| 7933 | 6 | 28066342 | 28095853 | mNeoVis.2.1.1.chr6\_hap1.g00191.t01 |
| 7934 | 6 | 28114724 | 28191335 | mNeoVis.2.1.1.chr6\_hap1.g00192.t01 |
| 7935 | 6 | 28114724 | 28191335 | mNeoVis.2.1.1.chr6\_hap1.g00192.t01 |
| 7936 | 6 | 28198966 | 28469559 | mNeoVis.2.1.1.chr6\_hap1.g00193.t01 |
| 7937 | 6 | 28470199 | 28573064 | mNeoVis.2.1.1.chr6\_hap1.g00194.t01 |
| 7938 | 6 | 28592702 | 28607689 | mNeoVis.2.1.1.chr6\_hap1.g00195.t01 |
| 7939 | 6 | 29567534 | 29690282 | mNeoVis.2.1.1.chr6\_hap1.g00200.t01 |
| 7940 | 6 | 29786971 | 29856959 | mNeoVis.2.1.1.chr6\_hap1.g00201.t01 |
| 7941 | 6 | 30924278 | 30948430 | mNeoVis.2.1.1.chr6\_hap1.g00202.t01 |
| 7942 | 6 | 30924278 | 30948430 | mNeoVis.2.1.1.chr6\_hap1.g00202.t01 |
| 7943 | 6 | 30951139 | 30974915 | mNeoVis.2.1.1.chr6\_hap1.g00203.t01 |
| 7944 | 6 | 30951139 | 30974915 | mNeoVis.2.1.1.chr6\_hap1.g00203.t01 |
| 7945 | 6 | 30999745 | 31028433 | mNeoVis.2.1.1.chr6\_hap1.g00204.t01 |
| 7946 | 6 | 30999745 | 31028433 | mNeoVis.2.1.1.chr6\_hap1.g00204.t01 |
| 7947 | 6 | 31184351 | 31442776 | mNeoVis.2.1.1.chr6\_hap1.g00205.t01 |
| 7948 | 6 | 32363660 | 32492575 | mNeoVis.2.1.1.chr6\_hap1.g00206.t01 |
| 7949 | 6 | 32683206 | 32914792 | mNeoVis.2.1.1.chr6\_hap1.g00207.t01 |
| 7950 | 6 | 32920968 | 32925877 | mNeoVis.2.1.1.chr6\_hap1.g00208.t01 |
| 7951 | 6 | 33117920 | 33158613 | mNeoVis.2.1.1.chr6\_hap1.g00209.t01 |
| 7952 | 6 | 33180228 | 33270974 | mNeoVis.2.1.1.chr6\_hap1.g00210.t01 |
| 7953 | 6 | 33294890 | 33307476 | mNeoVis.2.1.1.chr6\_hap1.g00211.t01 |
| 7954 | 6 | 33905050 | 33919147 | mNeoVis.2.1.1.chr6\_hap1.g00214.t01 |
| 7955 | 6 | 33919873 | 33956662 | mNeoVis.2.1.1.chr6\_hap1.g00215.t01 |
| 7956 | 6 | 33962716 | 34337284 | mNeoVis.2.1.1.chr6\_hap1.g00216.t01 |
| 7957 | 6 | 34496614 | 34654117 | mNeoVis.2.1.1.chr6\_hap1.g00218.t01 |
| 7958 | 6 | 34616385 | 34641288 | mNeoVis.2.1.1.chr6\_hap1.g00219.t01 |
| 7959 | 6 | 34705226 | 34716288 | mNeoVis.2.1.1.chr6\_hap1.g00220.t01 |
| 7960 | 6 | 34785734 | 34832260 | mNeoVis.2.1.1.chr6\_hap1.g00221.t01 |
| 7961 | 6 | 36059376 | 36064547 | mNeoVis.2.1.1.chr6\_hap1.g00236.t01 |
| 7962 | 6 | 36067138 | 36092212 | mNeoVis.2.1.1.chr6\_hap1.g00237.t01 |
| 7963 | 6 | 36146874 | 36183737 | mNeoVis.2.1.1.chr6\_hap1.g00238.t01 |
| 7964 | 6 | 36594868 | 36814258 | mNeoVis.2.1.1.chr6\_hap1.g00239.t01 |
| 7965 | 6 | 36848686 | 36884225 | mNeoVis.2.1.1.chr6\_hap1.g00240.t01 |
| 7966 | 6 | 36893364 | 36940492 | mNeoVis.2.1.1.chr6\_hap1.g00241.t01 |
| 7967 | 6 | 37061337 | 37091211 | mNeoVis.2.1.1.chr6\_hap1.g00242.t01 |
| 7968 | 6 | 37094273 | 37121519 | mNeoVis.2.1.1.chr6\_hap1.g00243.t01 |
| 7969 | 6 | 37151398 | 37333792 | mNeoVis.2.1.1.chr6\_hap1.g00244.t01 |
| 7970 | 6 | 38139967 | 38207921 | mNeoVis.2.1.1.chr6\_hap1.g00245.t01 |
| 7971 | 6 | 38139967 | 38207921 | mNeoVis.2.1.1.chr6\_hap1.g00245.t01 |
| 7972 | 6 | 38246766 | 38434425 | mNeoVis.2.1.1.chr6\_hap1.g00246.t01 |
| 7973 | 6 | 38246766 | 38434425 | mNeoVis.2.1.1.chr6\_hap1.g00246.t01 |
| 7974 | 6 | 39513091 | 39519824 | mNeoVis.2.1.1.chr6\_hap1.g00247.t01 |
| 7975 | 6 | 40494370 | 40495557 | mNeoVis.2.1.1.chr6\_hap1.g00249.t01 |
| 7976 | 6 | 48249574 | 48285316 | mNeoVis.2.1.1.chr6\_hap1.g00264.t01 |
| 7977 | 6 | 48359648 | 48387789 | mNeoVis.2.1.1.chr6\_hap1.g00265.t01 |
| 7978 | 6 | 48451332 | 48560377 | mNeoVis.2.1.1.chr6\_hap1.g00266.t01 |
| 7979 | 6 | 48563875 | 48568977 | mNeoVis.2.1.1.chr6\_hap1.g00267.t01 |
| 7980 | 6 | 48742159 | 48800042 | mNeoVis.2.1.1.chr6\_hap1.g00268.t01 |
| 7981 | 6 | 48742159 | 48800042 | mNeoVis.2.1.1.chr6\_hap1.g00268.t01 |
| 7982 | 6 | 48742159 | 48800042 | mNeoVis.2.1.1.chr6\_hap1.g00268.t01 |
| 7983 | 6 | 49187467 | 49489158 | mNeoVis.2.1.1.chr6\_hap1.g00269.t01 |
| 7984 | 6 | 49187467 | 49489158 | mNeoVis.2.1.1.chr6\_hap1.g00269.t01 |
| 7985 | 6 | 49768499 | 49884703 | mNeoVis.2.1.1.chr6\_hap1.g00270.t01 |
| 7986 | 6 | 49838403 | 49848465 | mNeoVis.2.1.1.chr6\_hap1.g00271.t01 |
| 7987 | 6 | 50300448 | 50416539 | mNeoVis.2.1.1.chr6\_hap1.g00272.t01 |
| 7988 | 6 | 50300448 | 50416539 | mNeoVis.2.1.1.chr6\_hap1.g00272.t01 |
| 7989 | 6 | 50539434 | 50667898 | mNeoVis.2.1.1.chr6\_hap1.g00273.t01 |
| 7990 | 6 | 50539434 | 50667898 | mNeoVis.2.1.1.chr6\_hap1.g00273.t01 |
| 7991 | 6 | 52176778 | 52196216 | mNeoVis.2.1.1.chr6\_hap1.g00275.t01 |
| 7992 | 6 | 52252276 | 52610059 | mNeoVis.2.1.1.chr6\_hap1.g00276.t01 |
| 7993 | 6 | 57407634 | 57479709 | mNeoVis.2.1.1.chr6\_hap1.g00297.t01 |
| 7994 | 6 | 58743294 | 58798339 | mNeoVis.2.1.1.chr6\_hap1.g00300.t01 |
| 7995 | 6 | 58818418 | 58821034 | mNeoVis.2.1.1.chr6\_hap1.g00301.t01 |
| 7996 | 6 | 58876738 | 58989789 | mNeoVis.2.1.1.chr6\_hap1.g00302.t01 |
| 7997 | 6 | 59011591 | 59084724 | mNeoVis.2.1.1.chr6\_hap1.g00303.t01 |
| 7998 | 6 | 59524326 | 59595373 | mNeoVis.2.1.1.chr6\_hap1.g00304.t01 |
| 7999 | 6 | 60644024 | 60710761 | mNeoVis.2.1.1.chr6\_hap1.g00305.t01 |
| 8000 | 6 | 61417221 | 61552412 | mNeoVis.2.1.1.chr6\_hap1.g00306.t01 |
| 8001 | 6 | 61571116 | 61595125 | mNeoVis.2.1.1.chr6\_hap1.g00307.t01 |
| 8002 | 6 | 62364739 | 62477476 | mNeoVis.2.1.1.chr6\_hap1.g00316.t01 |
| 8003 | 6 | 62733837 | 62737986 | mNeoVis.2.1.1.chr6\_hap1.g00317.t01 |
| 8004 | 6 | 62862279 | 62866300 | mNeoVis.2.1.1.chr6\_hap1.g00318.t01 |
| 8005 | 6 | 63489215 | 63551176 | mNeoVis.2.1.1.chr6\_hap1.g00322.t01 |
| 8006 | 6 | 63489215 | 63551176 | mNeoVis.2.1.1.chr6\_hap1.g00322.t01 |
| 8007 | 6 | 63556131 | 63633165 | mNeoVis.2.1.1.chr6\_hap1.g00323.t01 |
| 8008 | 6 | 63556131 | 63633165 | mNeoVis.2.1.1.chr6\_hap1.g00323.t01 |
| 8009 | 6 | 63637885 | 63747361 | mNeoVis.2.1.1.chr6\_hap1.g00324.t01 |
| 8010 | 6 | 63637885 | 63747361 | mNeoVis.2.1.1.chr6\_hap1.g00324.t01 |
| 8011 | 6 | 63747511 | 63875730 | mNeoVis.2.1.1.chr6\_hap1.g00325.t01 |
| 8012 | 6 | 63747511 | 63875730 | mNeoVis.2.1.1.chr6\_hap1.g00325.t01 |
| 8013 | 6 | 63874911 | 63920193 | mNeoVis.2.1.1.chr6\_hap1.g00326.t01 |
| 8014 | 6 | 64324635 | 64326731 | mNeoVis.2.1.1.chr6\_hap1.g00331.t01 |
| 8015 | 6 | 64346673 | 64352124 | mNeoVis.2.1.1.chr6\_hap1.g00332.t01 |
| 8016 | 6 | 64366495 | 64369165 | mNeoVis.2.1.1.chr6\_hap1.g00333.t01 |
| 8017 | 6 | 64372340 | 64374570 | mNeoVis.2.1.1.chr6\_hap1.g00334.t01 |
| 8018 | 6 | 64380145 | 64381624 | mNeoVis.2.1.1.chr6\_hap1.g00335.t01 |
| 8019 | 6 | 64435626 | 64517687 | mNeoVis.2.1.1.chr6\_hap1.g00336.t01 |
| 8020 | 6 | 64536556 | 64634760 | mNeoVis.2.1.1.chr6\_hap1.g00337.t01 |
| 8021 | 6 | 64640005 | 64642669 | mNeoVis.2.1.1.chr6\_hap1.g00338.t01 |
| 8022 | 6 | 64640005 | 64642669 | mNeoVis.2.1.1.chr6\_hap1.g00338.t01 |
| 8023 | 6 | 64643092 | 64657910 | mNeoVis.2.1.1.chr6\_hap1.g00339.t01 |
| 8024 | 6 | 64643092 | 64657910 | mNeoVis.2.1.1.chr6\_hap1.g00339.t01 |
| 8025 | 6 | 64659482 | 64667654 | mNeoVis.2.1.1.chr6\_hap1.g00340.t01 |
| 8026 | 6 | 64659482 | 64667654 | mNeoVis.2.1.1.chr6\_hap1.g00340.t01 |
| 8027 | 6 | 64746434 | 65170827 | mNeoVis.2.1.1.chr6\_hap1.g00341.t01 |
| 8028 | 6 | 64746434 | 65170827 | mNeoVis.2.1.1.chr6\_hap1.g00341.t01 |
| 8029 | 6 | 65175677 | 65310600 | mNeoVis.2.1.1.chr6\_hap1.g00342.t01 |
| 8030 | 6 | 65175677 | 65310600 | mNeoVis.2.1.1.chr6\_hap1.g00342.t01 |
| 8031 | 6 | 65320530 | 65425331 | mNeoVis.2.1.1.chr6\_hap1.g00343.t01 |
| 8032 | 6 | 65320530 | 65425331 | mNeoVis.2.1.1.chr6\_hap1.g00343.t01 |
| 8033 | 6 | 65476926 | 65525059 | mNeoVis.2.1.1.chr6\_hap1.g00344.t01 |
| 8034 | 6 | 65476926 | 65525059 | mNeoVis.2.1.1.chr6\_hap1.g00344.t01 |
| 8035 | 6 | 65549842 | 65579504 | mNeoVis.2.1.1.chr6\_hap1.g00345.t01 |
| 8036 | 6 | 65549842 | 65579504 | mNeoVis.2.1.1.chr6\_hap1.g00345.t01 |
| 8037 | 6 | 65620508 | 65666011 | mNeoVis.2.1.1.chr6\_hap1.g00346.t01 |
| 8038 | 6 | 65620508 | 65666011 | mNeoVis.2.1.1.chr6\_hap1.g00346.t01 |
| 8039 | 6 | 65679192 | 65702323 | mNeoVis.2.1.1.chr6\_hap1.g00347.t01 |
| 8040 | 6 | 65679192 | 65702323 | mNeoVis.2.1.1.chr6\_hap1.g00347.t01 |
| 8041 | 6 | 65702004 | 65717863 | mNeoVis.2.1.1.chr6\_hap1.g00348.t01 |
| 8042 | 6 | 65756794 | 65778505 | mNeoVis.2.1.1.chr6\_hap1.g00349.t01 |
| 8043 | 6 | 65958570 | 65963137 | mNeoVis.2.1.1.chr6\_hap1.g00350.t01 |
| 8044 | 6 | 66085052 | 66087177 | mNeoVis.2.1.1.chr6\_hap1.g00351.t01 |
| 8045 | 6 | 66244967 | 66257207 | mNeoVis.2.1.1.chr6\_hap1.g00352.t01 |
| 8046 | 6 | 66266541 | 66308548 | mNeoVis.2.1.1.chr6\_hap1.g00353.t01 |
| 8047 | 6 | 66310174 | 66415342 | mNeoVis.2.1.1.chr6\_hap1.g00354.t01 |
| 8048 | 6 | 66436051 | 66457312 | mNeoVis.2.1.1.chr6\_hap1.g00355.t01 |
| 8049 | 6 | 66452562 | 66591346 | mNeoVis.2.1.1.chr6\_hap1.g00356.t01 |
| 8050 | 6 | 66602639 | 66700543 | mNeoVis.2.1.1.chr6\_hap1.g00357.t01 |
| 8051 | 6 | 66700364 | 66719562 | mNeoVis.2.1.1.chr6\_hap1.g00358.t01 |
| 8052 | 6 | 66720089 | 66724472 | mNeoVis.2.1.1.chr6\_hap1.g00359.t01 |
| 8053 | 6 | 66725143 | 66809687 | mNeoVis.2.1.1.chr6\_hap1.g00360.t01 |
| 8054 | 6 | 66822507 | 66840800 | mNeoVis.2.1.1.chr6\_hap1.g00361.t01 |
| 8055 | 6 | 66880590 | 66922489 | mNeoVis.2.1.1.chr6\_hap1.g00362.t01 |
| 8056 | 6 | 66931766 | 67097892 | mNeoVis.2.1.1.chr6\_hap1.g00363.t01 |
| 8057 | 6 | 67155403 | 67193361 | mNeoVis.2.1.1.chr6\_hap1.g00365.t01 |
| 8058 | 6 | 67239835 | 67302149 | mNeoVis.2.1.1.chr6\_hap1.g00366.t01 |
| 8059 | 6 | 67310535 | 67324539 | mNeoVis.2.1.1.chr6\_hap1.g00367.t01 |
| 8060 | 6 | 67329902 | 67346558 | mNeoVis.2.1.1.chr6\_hap1.g00368.t01 |
| 8061 | 6 | 67346456 | 67488374 | mNeoVis.2.1.1.chr6\_hap1.g00369.t01 |
| 8062 | 6 | 67492882 | 67499524 | mNeoVis.2.1.1.chr6\_hap1.g00370.t01 |
| 8063 | 6 | 67505778 | 67508103 | mNeoVis.2.1.1.chr6\_hap1.g00371.t01 |
| 8064 | 6 | 67542773 | 67562447 | mNeoVis.2.1.1.chr6\_hap1.g00372.t01 |
| 8065 | 6 | 67562895 | 67626830 | mNeoVis.2.1.1.chr6\_hap1.g00373.t01 |
| 8066 | 6 | 67675711 | 67715573 | mNeoVis.2.1.1.chr6\_hap1.g00374.t01 |
| 8067 | 6 | 67719554 | 67735509 | mNeoVis.2.1.1.chr6\_hap1.g00375.t01 |
| 8068 | 6 | 67779268 | 67792118 | mNeoVis.2.1.1.chr6\_hap1.g00376.t01 |
| 8069 | 6 | 67792260 | 67797775 | mNeoVis.2.1.1.chr6\_hap1.g00377.t01 |
| 8070 | 6 | 67798593 | 67829523 | mNeoVis.2.1.1.chr6\_hap1.g00378.t01 |
| 8071 | 6 | 67831236 | 67861853 | mNeoVis.2.1.1.chr6\_hap1.g00379.t01 |
| 8072 | 6 | 68038329 | 68041809 | mNeoVis.2.1.1.chr6\_hap1.g00380.t01 |
| 8073 | 6 | 68046790 | 68082304 | mNeoVis.2.1.1.chr6\_hap1.g00381.t01 |
| 8074 | 6 | 68091582 | 68123847 | mNeoVis.2.1.1.chr6\_hap1.g00382.t01 |
| 8075 | 6 | 68169349 | 68194744 | mNeoVis.2.1.1.chr6\_hap1.g00383.t01 |
| 8076 | 6 | 68220117 | 68224160 | mNeoVis.2.1.1.chr6\_hap1.g00384.t01 |
| 8077 | 6 | 68234242 | 68253869 | mNeoVis.2.1.1.chr6\_hap1.g00385.t01 |
| 8078 | 6 | 68255845 | 68290812 | mNeoVis.2.1.1.chr6\_hap1.g00386.t01 |
| 8079 | 6 | 68311142 | 68323874 | mNeoVis.2.1.1.chr6\_hap1.g00387.t01 |
| 8080 | 6 | 68338934 | 68353656 | mNeoVis.2.1.1.chr6\_hap1.g00388.t01 |
| 8081 | 6 | 68348940 | 68358353 | mNeoVis.2.1.1.chr6\_hap1.g00389.t01 |
| 8082 | 6 | 68362554 | 68375170 | mNeoVis.2.1.1.chr6\_hap1.g00390.t01 |
| 8083 | 6 | 68374627 | 68400680 | mNeoVis.2.1.1.chr6\_hap1.g00391.t01 |
| 8084 | 6 | 68400792 | 68456680 | mNeoVis.2.1.1.chr6\_hap1.g00392.t01 |
| 8085 | 6 | 68468763 | 68503009 | mNeoVis.2.1.1.chr6\_hap1.g00393.t01 |
| 8086 | 6 | 68521096 | 68573754 | mNeoVis.2.1.1.chr6\_hap1.g00394.t01 |
| 8087 | 6 | 68573948 | 68592596 | mNeoVis.2.1.1.chr6\_hap1.g00395.t01 |
| 8088 | 6 | 68592736 | 68596085 | mNeoVis.2.1.1.chr6\_hap1.g00396.t01 |
| 8089 | 6 | 68599769 | 68601516 | mNeoVis.2.1.1.chr6\_hap1.g00397.t01 |
| 8090 | 6 | 68603383 | 68636173 | mNeoVis.2.1.1.chr6\_hap1.g00398.t01 |
| 8091 | 6 | 68656440 | 68661939 | mNeoVis.2.1.1.chr6\_hap1.g00399.t01 |
| 8092 | 6 | 68674654 | 68721281 | mNeoVis.2.1.1.chr6\_hap1.g00400.t01 |
| 8093 | 6 | 70103882 | 70186492 | mNeoVis.2.1.1.chr6\_hap1.g00402.t01 |
| 8094 | 6 | 70583330 | 70617783 | mNeoVis.2.1.1.chr6\_hap1.g00403.t01 |
| 8095 | 6 | 70674880 | 70941601 | mNeoVis.2.1.1.chr6\_hap1.g00404.t01 |
| 8096 | 6 | 71038731 | 71160689 | mNeoVis.2.1.1.chr6\_hap1.g00405.t01 |
| 8097 | 6 | 71183544 | 71194488 | mNeoVis.2.1.1.chr6\_hap1.g00406.t01 |
| 8098 | 6 | 71197533 | 71228763 | mNeoVis.2.1.1.chr6\_hap1.g00407.t01 |
| 8099 | 6 | 71298974 | 71359950 | mNeoVis.2.1.1.chr6\_hap1.g00409.t01 |
| 8100 | 6 | 71364808 | 71401643 | mNeoVis.2.1.1.chr6\_hap1.g00410.t01 |
| 8101 | 6 | 71560744 | 71965293 | mNeoVis.2.1.1.chr6\_hap1.g00411.t01 |
| 8102 | 6 | 71969424 | 71992957 | mNeoVis.2.1.1.chr6\_hap1.g00412.t01 |
| 8103 | 6 | 72083191 | 72129921 | mNeoVis.2.1.1.chr6\_hap1.g00413.t01 |
| 8104 | 6 | 77104349 | 77111584 | mNeoVis.2.1.1.chr6\_hap1.g00424.t01 |
| 8105 | 6 | 78019806 | 78190156 | mNeoVis.2.1.1.chr6\_hap1.g00426.t01 |
| 8106 | 6 | 78224561 | 78225114 | mNeoVis.2.1.1.chr6\_hap1.g00427.t01 |
| 8107 | 6 | 81655321 | 81678076 | mNeoVis.2.1.1.chr6\_hap1.g00428.t01 |
| 8108 | 6 | 82819826 | 82868938 | mNeoVis.2.1.1.chr6\_hap1.g00437.t01 |
| 8109 | 6 | 82870983 | 82871549 | mNeoVis.2.1.1.chr6\_hap1.g00438.t01 |
| 8110 | 6 | 82877661 | 82902788 | mNeoVis.2.1.1.chr6\_hap1.g00439.t01 |
| 8111 | 6 | 82908080 | 82987397 | mNeoVis.2.1.1.chr6\_hap1.g00440.t01 |
| 8112 | 6 | 85028974 | 85207751 | mNeoVis.2.1.1.chr6\_hap1.g00441.t01 |
| 8113 | 6 | 86795569 | 86797991 | mNeoVis.2.1.1.chr6\_hap1.g00443.t01 |
| 8114 | 6 | 86884576 | 86889912 | mNeoVis.2.1.1.chr6\_hap1.g00444.t01 |
| 8115 | 6 | 88976777 | 89076074 | mNeoVis.2.1.1.chr6\_hap1.g00463.t01 |
| 8116 | 6 | 89029734 | 89068762 | mNeoVis.2.1.1.chr6\_hap1.g00464.t01 |
| 8117 | 6 | 89117869 | 89159529 | mNeoVis.2.1.1.chr6\_hap1.g00465.t01 |
| 8118 | 6 | 89234421 | 89453904 | mNeoVis.2.1.1.chr6\_hap1.g00466.t01 |
| 8119 | 6 | 94367446 | 94642086 | mNeoVis.2.1.1.chr6\_hap1.g00484.t01 |
| 8120 | 6 | 94761229 | 94852111 | mNeoVis.2.1.1.chr6\_hap1.g00485.t01 |
| 8121 | 6 | 94974780 | 94976436 | mNeoVis.2.1.1.chr6\_hap1.g00486.t01 |
| 8122 | 6 | 95546641 | 95891669 | mNeoVis.2.1.1.chr6\_hap1.g00488.t01 |
| 8123 | 6 | 95994886 | 96221974 | mNeoVis.2.1.1.chr6\_hap1.g00489.t01 |
| 8124 | 6 | 96302615 | 96315276 | mNeoVis.2.1.1.chr6\_hap1.g00490.t01 |
| 8125 | 6 | 99418923 | 99512748 | mNeoVis.2.1.1.chr6\_hap1.g00514.t01 |
| 8126 | 6 | 100554446 | 100597730 | mNeoVis.2.1.1.chr6\_hap1.g00519.t01 |
| 8127 | 6 | 100554446 | 100597730 | mNeoVis.2.1.1.chr6\_hap1.g00519.t01 |
| 8128 | 6 | 100554446 | 100597730 | mNeoVis.2.1.1.chr6\_hap1.g00519.t01 |
| 8129 | 6 | 100599033 | 100720644 | mNeoVis.2.1.1.chr6\_hap1.g00520.t01 |
| 8130 | 6 | 100599033 | 100720644 | mNeoVis.2.1.1.chr6\_hap1.g00520.t01 |
| 8131 | 6 | 100599033 | 100720644 | mNeoVis.2.1.1.chr6\_hap1.g00520.t01 |
| 8132 | 6 | 100700218 | 100780578 | mNeoVis.2.1.1.chr6\_hap1.g00521.t01 |
| 8133 | 6 | 100700218 | 100780578 | mNeoVis.2.1.1.chr6\_hap1.g00521.t01 |
| 8134 | 6 | 100700218 | 100780578 | mNeoVis.2.1.1.chr6\_hap1.g00521.t01 |
| 8135 | 6 | 100780673 | 100799749 | mNeoVis.2.1.1.chr6\_hap1.g00522.t01 |
| 8136 | 6 | 100780673 | 100799749 | mNeoVis.2.1.1.chr6\_hap1.g00522.t01 |
| 8137 | 6 | 100780673 | 100799749 | mNeoVis.2.1.1.chr6\_hap1.g00522.t01 |
| 8138 | 6 | 100794839 | 100816961 | mNeoVis.2.1.1.chr6\_hap1.g00523.t01 |
| 8139 | 6 | 100794839 | 100816961 | mNeoVis.2.1.1.chr6\_hap1.g00523.t01 |
| 8140 | 6 | 100794839 | 100816961 | mNeoVis.2.1.1.chr6\_hap1.g00523.t01 |
| 8141 | 6 | 100835873 | 100921354 | mNeoVis.2.1.1.chr6\_hap1.g00524.t01 |
| 8142 | 6 | 100835873 | 100921354 | mNeoVis.2.1.1.chr6\_hap1.g00524.t01 |
| 8143 | 6 | 100849656 | 100850447 | mNeoVis.2.1.1.chr6\_hap1.g00525.t01 |
| 8144 | 6 | 100849656 | 100850447 | mNeoVis.2.1.1.chr6\_hap1.g00525.t01 |
| 8145 | 6 | 100849656 | 100850447 | mNeoVis.2.1.1.chr6\_hap1.g00525.t01 |
| 8146 | 6 | 100909639 | 100920398 | mNeoVis.2.1.1.chr6\_hap1.g00526.t01 |
| 8147 | 6 | 100909639 | 100920398 | mNeoVis.2.1.1.chr6\_hap1.g00526.t01 |
| 8148 | 6 | 101660515 | 101763449 | mNeoVis.2.1.1.chr6\_hap1.g00528.t01 |
| 8149 | 6 | 102574323 | 102656536 | mNeoVis.2.1.1.chr6\_hap1.g00532.t01 |
| 8150 | 6 | 102697341 | 102822867 | mNeoVis.2.1.1.chr6\_hap1.g00533.t01 |
| 8151 | 6 | 102824950 | 102845230 | mNeoVis.2.1.1.chr6\_hap1.g00534.t01 |
| 8152 | 6 | 102851148 | 102984176 | mNeoVis.2.1.1.chr6\_hap1.g00535.t01 |
| 8153 | 6 | 103056817 | 103141199 | mNeoVis.2.1.1.chr6\_hap1.g00536.t01 |
| 8154 | 6 | 103221286 | 103359745 | mNeoVis.2.1.1.chr6\_hap1.g00537.t01 |
| 8155 | 6 | 103914102 | 103940573 | mNeoVis.2.1.1.chr6\_hap1.g00538.t01 |
| 8156 | 6 | 103914102 | 103940573 | mNeoVis.2.1.1.chr6\_hap1.g00538.t01 |
| 8157 | 6 | 104562790 | 104640048 | mNeoVis.2.1.1.chr6\_hap1.g00539.t01 |
| 8158 | 6 | 105007614 | 105029939 | mNeoVis.2.1.1.chr6\_hap1.g00541.t01 |
| 8159 | 6 | 105021642 | 105032016 | mNeoVis.2.1.1.chr6\_hap1.g00542.t01 |
| 8160 | 6 | 105031934 | 105081930 | mNeoVis.2.1.1.chr6\_hap1.g00543.t01 |
| 8161 | 6 | 105087599 | 105150950 | mNeoVis.2.1.1.chr6\_hap1.g00544.t01 |
| 8162 | 6 | 105170306 | 105186597 | mNeoVis.2.1.1.chr6\_hap1.g00545.t01 |
| 8163 | 6 | 105232364 | 105257983 | mNeoVis.2.1.1.chr6\_hap1.g00546.t01 |
| 8164 | 6 | 105285059 | 105409166 | mNeoVis.2.1.1.chr6\_hap1.g00547.t01 |
| 8165 | 6 | 105423238 | 105542974 | mNeoVis.2.1.1.chr6\_hap1.g00548.t01 |
| 8166 | 6 | 105423238 | 105542974 | mNeoVis.2.1.1.chr6\_hap1.g00548.t01 |
| 8167 | 6 | 105659537 | 105742911 | mNeoVis.2.1.1.chr6\_hap1.g00549.t01 |
| 8168 | 6 | 105881178 | 106182403 | mNeoVis.2.1.1.chr6\_hap1.g00550.t01 |
| 8169 | 6 | 107258004 | 107366770 | mNeoVis.2.1.1.chr6\_hap1.g00552.t01 |
| 8170 | 6 | 107258004 | 107366770 | mNeoVis.2.1.1.chr6\_hap1.g00552.t01 |
| 8171 | 6 | 107531312 | 107536321 | mNeoVis.2.1.1.chr6\_hap1.g00553.t01 |
| 8172 | 6 | 107531312 | 107536321 | mNeoVis.2.1.1.chr6\_hap1.g00553.t01 |
| 8173 | 6 | 107653375 | 107667972 | mNeoVis.2.1.1.chr6\_hap1.g00554.t01 |
| 8174 | 6 | 107718268 | 107720497 | mNeoVis.2.1.1.chr6\_hap1.g00555.t01 |
| 8175 | 6 | 107721940 | 107779002 | mNeoVis.2.1.1.chr6\_hap1.g00556.t01 |
| 8176 | 6 | 107779076 | 107793512 | mNeoVis.2.1.1.chr6\_hap1.g00557.t01 |
| 8177 | 6 | 107795049 | 107857560 | mNeoVis.2.1.1.chr6\_hap1.g00558.t01 |
| 8178 | 6 | 107953367 | 107972836 | mNeoVis.2.1.1.chr6\_hap1.g00559.t01 |
| 8179 | 6 | 107985716 | 108043040 | mNeoVis.2.1.1.chr6\_hap1.g00560.t01 |
| 8180 | 6 | 108043118 | 108060581 | mNeoVis.2.1.1.chr6\_hap1.g00561.t01 |
| 8181 | 6 | 108068361 | 108101087 | mNeoVis.2.1.1.chr6\_hap1.g00562.t01 |
| 8182 | 6 | 108218279 | 108234098 | mNeoVis.2.1.1.chr6\_hap1.g00563.t01 |
| 8183 | 6 | 108275560 | 108288072 | mNeoVis.2.1.1.chr6\_hap1.g00564.t01 |
| 8184 | 6 | 108338491 | 108354562 | mNeoVis.2.1.1.chr6\_hap1.g00565.t01 |
| 8185 | 6 | 108370421 | 108410462 | mNeoVis.2.1.1.chr6\_hap1.g00566.t01 |
| 8186 | 6 | 108428567 | 108443971 | mNeoVis.2.1.1.chr6\_hap1.g00567.t01 |
| 8187 | 6 | 108834193 | 109010287 | mNeoVis.2.1.1.chr6\_hap1.g00570.t01 |
| 8188 | 6 | 109024374 | 109183971 | mNeoVis.2.1.1.chr6\_hap1.g00571.t01 |
| 8189 | 6 | 109024374 | 109183971 | mNeoVis.2.1.1.chr6\_hap1.g00571.t01 |
| 8190 | 6 | 111322071 | 111323474 | mNeoVis.2.1.1.chr6\_hap1.g00573.t01 |
| 8191 | 6 | 111322071 | 111323474 | mNeoVis.2.1.1.chr6\_hap1.g00573.t01 |
| 8192 | 6 | 111322071 | 111323474 | mNeoVis.2.1.1.chr6\_hap1.g00573.t01 |
| 8193 | 6 | 111548558 | 111908204 | mNeoVis.2.1.1.chr6\_hap1.g00574.t01 |
| 8194 | 6 | 111915785 | 112019932 | mNeoVis.2.1.1.chr6\_hap1.g00575.t01 |
| 8195 | 6 | 112700504 | 112729567 | mNeoVis.2.1.1.chr6\_hap1.g00578.t01 |
| 8196 | 6 | 112733757 | 112809298 | mNeoVis.2.1.1.chr6\_hap1.g00579.t01 |
| 8197 | 6 | 113097495 | 113098559 | mNeoVis.2.1.1.chr6\_hap1.g00580.t01 |
| 8198 | 6 | 115835345 | 115853425 | mNeoVis.2.1.1.chr6\_hap1.g00583.t01 |
| 8199 | 6 | 115835345 | 115853425 | mNeoVis.2.1.1.chr6\_hap1.g00583.t01 |
| 8200 | 6 | 115835345 | 115853425 | mNeoVis.2.1.1.chr6\_hap1.g00583.t01 |
| 8201 | 6 | 117540455 | 117609553 | mNeoVis.2.1.1.chr6\_hap1.g00587.t01 |
| 8202 | 6 | 117723432 | 117729368 | mNeoVis.2.1.1.chr6\_hap1.g00588.t01 |
| 8203 | 6 | 118508422 | 118535166 | mNeoVis.2.1.1.chr6\_hap1.g00596.t01 |
| 8204 | 6 | 118526041 | 118527367 | mNeoVis.2.1.1.chr6\_hap1.g00597.t01 |
| 8205 | 6 | 118558622 | 118719887 | mNeoVis.2.1.1.chr6\_hap1.g00598.t01 |
| 8206 | 6 | 118731042 | 118793691 | mNeoVis.2.1.1.chr6\_hap1.g00599.t01 |
| 8207 | 6 | 118815014 | 118823515 | mNeoVis.2.1.1.chr6\_hap1.g00600.t01 |
| 8208 | 6 | 118829934 | 118905565 | mNeoVis.2.1.1.chr6\_hap1.g00601.t01 |
| 8209 | 6 | 118965958 | 118975653 | mNeoVis.2.1.1.chr6\_hap1.g00602.t01 |
| 8210 | 6 | 118987675 | 119001706 | mNeoVis.2.1.1.chr6\_hap1.g00603.t01 |
| 8211 | 6 | 119335142 | 119364415 | mNeoVis.2.1.1.chr6\_hap1.g00607.t01 |
| 8212 | 6 | 119372535 | 119412909 | mNeoVis.2.1.1.chr6\_hap1.g00608.t01 |
| 8213 | 6 | 119421234 | 119439152 | mNeoVis.2.1.1.chr6\_hap1.g00609.t01 |
| 8214 | 6 | 119505813 | 119562940 | mNeoVis.2.1.1.chr6\_hap1.g00610.t01 |
| 8215 | 6 | 119596693 | 119704110 | mNeoVis.2.1.1.chr6\_hap1.g00611.t01 |
| 8216 | 6 | 124182004 | 124182844 | mNeoVis.2.1.1.chr6\_hap1.g00630.t01 |
| 8217 | 6 | 124182004 | 124182844 | mNeoVis.2.1.1.chr6\_hap1.g00630.t01 |
| 8218 | 6 | 124190868 | 124448903 | mNeoVis.2.1.1.chr6\_hap1.g00631.t01 |
| 8219 | 6 | 124190868 | 124448903 | mNeoVis.2.1.1.chr6\_hap1.g00631.t01 |
| 8220 | 6 | 124749869 | 124788774 | mNeoVis.2.1.1.chr6\_hap1.g00638.t01 |
| 8221 | 6 | 124976053 | 124977274 | mNeoVis.2.1.1.chr6\_hap1.g00639.t01 |
| 8222 | 6 | 125126809 | 125144054 | mNeoVis.2.1.1.chr6\_hap1.g00640.t01 |
| 8223 | 6 | 125141919 | 125150585 | mNeoVis.2.1.1.chr6\_hap1.g00641.t01 |
| 8224 | 6 | 125246146 | 125281302 | mNeoVis.2.1.1.chr6\_hap1.g00642.t01 |
| 8225 | 6 | 125672758 | 125721070 | mNeoVis.2.1.1.chr6\_hap1.g00644.t01 |
| 8226 | 6 | 125756803 | 125858465 | mNeoVis.2.1.1.chr6\_hap1.g00645.t01 |
| 8227 | 6 | 125813977 | 125831645 | mNeoVis.2.1.1.chr6\_hap1.g00646.t01 |
| 8228 | 6 | 126625458 | 126824933 | mNeoVis.2.1.1.chr6\_hap1.g00647.t01 |
| 8229 | 6 | 126625458 | 126824933 | mNeoVis.2.1.1.chr6\_hap1.g00647.t01 |
| 8230 | 6 | 126910024 | 127010698 | mNeoVis.2.1.1.chr6\_hap1.g00648.t01 |
| 8231 | 6 | 126910024 | 127010698 | mNeoVis.2.1.1.chr6\_hap1.g00648.t01 |
| 8232 | 6 | 127051595 | 127220266 | mNeoVis.2.1.1.chr6\_hap1.g00649.t01 |
| 8233 | 6 | 127952527 | 128029730 | mNeoVis.2.1.1.chr6\_hap1.g00650.t01 |
| 8234 | 6 | 128025902 | 128101518 | mNeoVis.2.1.1.chr6\_hap1.g00651.t01 |
| 8235 | 6 | 128276345 | 128380427 | mNeoVis.2.1.1.chr6\_hap1.g00652.t01 |
| 8236 | 6 | 128276345 | 128380427 | mNeoVis.2.1.1.chr6\_hap1.g00652.t01 |
| 8237 | 6 | 128382427 | 128387925 | mNeoVis.2.1.1.chr6\_hap1.g00653.t01 |
| 8238 | 6 | 128388103 | 128417430 | mNeoVis.2.1.1.chr6\_hap1.g00654.t01 |
| 8239 | 6 | 128417954 | 128430472 | mNeoVis.2.1.1.chr6\_hap1.g00655.t01 |
| 8240 | 6 | 128467542 | 128603649 | mNeoVis.2.1.1.chr6\_hap1.g00656.t01 |
| 8241 | 6 | 128611090 | 128693777 | mNeoVis.2.1.1.chr6\_hap1.g00657.t01 |
| 8242 | 6 | 128752803 | 128994140 | mNeoVis.2.1.1.chr6\_hap1.g00658.t01 |
| 8243 | 6 | 132803122 | 132824723 | mNeoVis.2.1.1.chr6\_hap1.g00662.t01 |
| 8244 | 6 | 132824371 | 132829513 | mNeoVis.2.1.1.chr6\_hap1.g00663.t01 |
| 8245 | 6 | 132952079 | 132954710 | mNeoVis.2.1.1.chr6\_hap1.g00664.t01 |
| 8246 | 6 | 132969298 | 133005304 | mNeoVis.2.1.1.chr6\_hap1.g00665.t01 |
| 8247 | 6 | 133836972 | 133950029 | mNeoVis.2.1.1.chr6\_hap1.g00667.t01 |
| 8248 | 6 | 134221652 | 134357502 | mNeoVis.2.1.1.chr6\_hap1.g00668.t01 |
| 8249 | 6 | 138085081 | 138504315 | mNeoVis.2.1.1.chr6\_hap1.g00688.t01 |
| 8250 | 6 | 138544403 | 138566351 | mNeoVis.2.1.1.chr6\_hap1.g00689.t01 |
| 8251 | 6 | 139668502 | 139782134 | mNeoVis.2.1.1.chr6\_hap1.g00695.t01 |
| 8252 | 6 | 139982930 | 140023852 | mNeoVis.2.1.1.chr6\_hap1.g00696.t01 |
| 8253 | 6 | 140022294 | 140041508 | mNeoVis.2.1.1.chr6\_hap1.g00697.t01 |
| 8254 | 6 | 140041765 | 140146534 | mNeoVis.2.1.1.chr6\_hap1.g00698.t01 |
| 8255 | 6 | 140146704 | 140156197 | mNeoVis.2.1.1.chr6\_hap1.g00699.t01 |
| 8256 | 6 | 140146704 | 140156197 | mNeoVis.2.1.1.chr6\_hap1.g00699.t01 |
| 8257 | 6 | 140229401 | 140269646 | mNeoVis.2.1.1.chr6\_hap1.g00700.t01 |
| 8258 | 6 | 140229401 | 140269646 | mNeoVis.2.1.1.chr6\_hap1.g00700.t01 |
| 8259 | 6 | 140348703 | 140578705 | mNeoVis.2.1.1.chr6\_hap1.g00701.t01 |
| 8260 | 6 | 140348703 | 140578705 | mNeoVis.2.1.1.chr6\_hap1.g00701.t01 |
| 8261 | 6 | 140832781 | 140847739 | mNeoVis.2.1.1.chr6\_hap1.g00702.t01 |
| 8262 | 6 | 141070260 | 141162070 | mNeoVis.2.1.1.chr6\_hap1.g00703.t01 |
| 8263 | 6 | 141716101 | 141720414 | mNeoVis.2.1.1.chr6\_hap1.g00709.t01 |
| 8264 | 6 | 142250876 | 142251955 | mNeoVis.2.1.1.chr6\_hap1.g00711.t01 |
| 8265 | 6 | 146062225 | 146063903 | mNeoVis.2.1.1.chr6\_hap1.g00721.t01 |
| 8266 | 6 | 146143847 | 146401102 | mNeoVis.2.1.1.chr6\_hap1.g00722.t01 |
| 8267 | 6 | 146422432 | 146433370 | mNeoVis.2.1.1.chr6\_hap1.g00723.t01 |
| 8268 | 6 | 146453914 | 146467851 | mNeoVis.2.1.1.chr6\_hap1.g00724.t01 |
| 8269 | 6 | 147642658 | 147737204 | mNeoVis.2.1.1.chr6\_hap1.g00733.t01 |
| 8270 | 6 | 148126358 | 148166608 | mNeoVis.2.1.1.chr6\_hap1.g00734.t01 |
| 8271 | 6 | 148309161 | 148390929 | mNeoVis.2.1.1.chr6\_hap1.g00735.t01 |
| 8272 | 6 | 148420046 | 148460879 | mNeoVis.2.1.1.chr6\_hap1.g00736.t01 |
| 8273 | 6 | 148512049 | 148636132 | mNeoVis.2.1.1.chr6\_hap1.g00737.t01 |
| 8274 | 6 | 148635833 | 148680245 | mNeoVis.2.1.1.chr6\_hap1.g00738.t01 |
| 8275 | 6 | 149777504 | 149795817 | mNeoVis.2.1.1.chr6\_hap1.g00746.t01 |
| 8276 | 6 | 149825082 | 149851393 | mNeoVis.2.1.1.chr6\_hap1.g00747.t01 |
| 8277 | 6 | 149842505 | 149843796 | mNeoVis.2.1.1.chr6\_hap1.g00748.t01 |
| 8278 | 6 | 149863462 | 149977159 | mNeoVis.2.1.1.chr6\_hap1.g00749.t01 |
| 8279 | 6 | 149979179 | 149989158 | mNeoVis.2.1.1.chr6\_hap1.g00750.t01 |
| 8280 | 6 | 150005893 | 150068495 | mNeoVis.2.1.1.chr6\_hap1.g00751.t01 |
| 8281 | 6 | 150074766 | 150105238 | mNeoVis.2.1.1.chr6\_hap1.g00752.t01 |
| 8282 | 6 | 150113067 | 150129653 | mNeoVis.2.1.1.chr6\_hap1.g00753.t01 |
| 8283 | 6 | 151220320 | 151402562 | mNeoVis.2.1.1.chr6\_hap1.g00767.t01 |
| 8284 | 6 | 151422790 | 151618375 | mNeoVis.2.1.1.chr6\_hap1.g00768.t01 |
| 8285 | 6 | 152329723 | 152343154 | mNeoVis.2.1.1.chr6\_hap1.g00770.t01 |
| 8286 | 6 | 152360776 | 152392636 | mNeoVis.2.1.1.chr6\_hap1.g00771.t01 |
| 8287 | 6 | 152407203 | 152426856 | mNeoVis.2.1.1.chr6\_hap1.g00772.t01 |
| 8288 | 6 | 152496545 | 152591084 | mNeoVis.2.1.1.chr6\_hap1.g00773.t01 |
| 8289 | 6 | 152590052 | 152626075 | mNeoVis.2.1.1.chr6\_hap1.g00774.t01 |
| 8290 | 6 | 152649514 | 152656157 | mNeoVis.2.1.1.chr6\_hap1.g00775.t01 |
| 8291 | 6 | 152691689 | 152786632 | mNeoVis.2.1.1.chr6\_hap1.g00776.t01 |
| 8292 | 6 | 152847319 | 152930477 | mNeoVis.2.1.1.chr6\_hap1.g00777.t01 |
| 8293 | 6 | 153236774 | 153294110 | mNeoVis.2.1.1.chr6\_hap1.g00779.t01 |
| 8294 | 6 | 153306674 | 153323924 | mNeoVis.2.1.1.chr6\_hap1.g00780.t01 |
| 8295 | 6 | 153309907 | 153310140 | mNeoVis.2.1.1.chr6\_hap1.g00781.t01 |
| 8296 | 6 | 153380277 | 153422502 | mNeoVis.2.1.1.chr6\_hap1.g00782.t01 |
| 8297 | 6 | 153454708 | 153527973 | mNeoVis.2.1.1.chr6\_hap1.g00783.t01 |
| 8298 | 6 | 153771171 | 153792180 | mNeoVis.2.1.1.chr6\_hap1.g00786.t01 |
| 8299 | 6 | 153796109 | 153815113 | mNeoVis.2.1.1.chr6\_hap1.g00787.t01 |
| 8300 | 6 | 153955138 | 153958090 | mNeoVis.2.1.1.chr6\_hap1.g00788.t01 |
| 8301 | 6 | 154306040 | 154311191 | mNeoVis.2.1.1.chr6\_hap1.g00789.t01 |
| 8302 | 6 | 154918932 | 154948150 | mNeoVis.2.1.1.chr6\_hap1.g00793.t01 |
| 8303 | 6 | 155099824 | 155124043 | mNeoVis.2.1.1.chr6\_hap1.g00794.t01 |
| 8304 | 6 | 155144754 | 155191788 | mNeoVis.2.1.1.chr6\_hap1.g00795.t01 |
| 8305 | 6 | 155879281 | 155881942 | mNeoVis.2.1.1.chr6\_hap1.g00796.t01 |
| 8306 | 6 | 156022004 | 156046494 | mNeoVis.2.1.1.chr6\_hap1.g00797.t01 |
| 8307 | 6 | 156022004 | 156046494 | mNeoVis.2.1.1.chr6\_hap1.g00797.t01 |
| 8308 | 6 | 156055099 | 156060651 | mNeoVis.2.1.1.chr6\_hap1.g00798.t01 |
| 8309 | 6 | 156055099 | 156060651 | mNeoVis.2.1.1.chr6\_hap1.g00798.t01 |
| 8310 | 6 | 156077134 | 156090266 | mNeoVis.2.1.1.chr6\_hap1.g00799.t01 |
| 8311 | 6 | 156077134 | 156090266 | mNeoVis.2.1.1.chr6\_hap1.g00799.t01 |
| 8312 | 6 | 156090581 | 156093281 | mNeoVis.2.1.1.chr6\_hap1.g00800.t01 |
| 8313 | 6 | 156090581 | 156093281 | mNeoVis.2.1.1.chr6\_hap1.g00800.t01 |
| 8314 | 6 | 156101241 | 156244823 | mNeoVis.2.1.1.chr6\_hap1.g00801.t01 |
| 8315 | 6 | 156101241 | 156244823 | mNeoVis.2.1.1.chr6\_hap1.g00801.t01 |
| 8316 | 6 | 156139320 | 156139676 | mNeoVis.2.1.1.chr6\_hap1.g00802.t01 |
| 8317 | 6 | 156139320 | 156139676 | mNeoVis.2.1.1.chr6\_hap1.g00802.t01 |
| 8318 | 6 | 159303777 | 159305488 | mNeoVis.2.1.1.chr6\_hap1.g00804.t01 |
| 8319 | 6 | 159367419 | 159475738 | mNeoVis.2.1.1.chr6\_hap1.g00805.t01 |
| 8320 | 6 | 159545129 | 159571604 | mNeoVis.2.1.1.chr6\_hap1.g00806.t01 |
| 8321 | 6 | 159580832 | 159581404 | mNeoVis.2.1.1.chr6\_hap1.g00807.t01 |
| 8322 | 6 | 159775266 | 159865838 | mNeoVis.2.1.1.chr6\_hap1.g00808.t01 |
| 8323 | 6 | 159775266 | 159865838 | mNeoVis.2.1.1.chr6\_hap1.g00808.t01 |
| 8324 | 6 | 160064633 | 160256866 | mNeoVis.2.1.1.chr6\_hap1.g00809.t01 |
| 8325 | 6 | 160064633 | 160256866 | mNeoVis.2.1.1.chr6\_hap1.g00809.t01 |
| 8326 | 6 | 160546005 | 160594030 | mNeoVis.2.1.1.chr6\_hap1.g00810.t01 |
| 8327 | 6 | 160594810 | 160638720 | mNeoVis.2.1.1.chr6\_hap1.g00811.t01 |
| 8328 | 6 | 160642734 | 160647298 | mNeoVis.2.1.1.chr6\_hap1.g00812.t01 |
| 8329 | 6 | 160650329 | 160857525 | mNeoVis.2.1.1.chr6\_hap1.g00813.t01 |
| 8330 | 6 | 161599306 | 161692601 | mNeoVis.2.1.1.chr6\_hap1.g00817.t01 |
| 8331 | 6 | 161718522 | 161735676 | mNeoVis.2.1.1.chr6\_hap1.g00818.t01 |
| 8332 | 6 | 161772816 | 161845848 | mNeoVis.2.1.1.chr6\_hap1.g00819.t01 |
| 8333 | 6 | 162386846 | 162399260 | mNeoVis.2.1.1.chr6\_hap1.g00820.t01 |
| 8334 | 6 | 162386846 | 162399260 | mNeoVis.2.1.1.chr6\_hap1.g00820.t01 |
| 8335 | 6 | 162543728 | 162547558 | mNeoVis.2.1.1.chr6\_hap1.g00821.t01 |
| 8336 | 6 | 162543728 | 162547558 | mNeoVis.2.1.1.chr6\_hap1.g00821.t01 |
| 8337 | 6 | 162543728 | 162547558 | mNeoVis.2.1.1.chr6\_hap1.g00821.t01 |
| 8338 | 6 | 162576014 | 162580370 | mNeoVis.2.1.1.chr6\_hap1.g00822.t01 |
| 8339 | 6 | 162576014 | 162580370 | mNeoVis.2.1.1.chr6\_hap1.g00822.t01 |
| 8340 | 6 | 162576014 | 162580370 | mNeoVis.2.1.1.chr6\_hap1.g00822.t01 |
| 8341 | 6 | 162683695 | 162720443 | mNeoVis.2.1.1.chr6\_hap1.g00823.t01 |
| 8342 | 6 | 162683695 | 162720443 | mNeoVis.2.1.1.chr6\_hap1.g00823.t01 |
| 8343 | 6 | 162683695 | 162720443 | mNeoVis.2.1.1.chr6\_hap1.g00823.t01 |
| 8344 | 6 | 162786504 | 162822727 | mNeoVis.2.1.1.chr6\_hap1.g00824.t01 |
| 8345 | 6 | 162786504 | 162822727 | mNeoVis.2.1.1.chr6\_hap1.g00824.t01 |
| 8346 | 6 | 162786504 | 162822727 | mNeoVis.2.1.1.chr6\_hap1.g00824.t01 |
| 8347 | 6 | 162846328 | 162934447 | mNeoVis.2.1.1.chr6\_hap1.g00825.t01 |
| 8348 | 6 | 162846328 | 162934447 | mNeoVis.2.1.1.chr6\_hap1.g00825.t01 |
| 8349 | 6 | 162846328 | 162934447 | mNeoVis.2.1.1.chr6\_hap1.g00825.t01 |
| 8350 | 6 | 162935339 | 162954717 | mNeoVis.2.1.1.chr6\_hap1.g00826.t01 |
| 8351 | 6 | 162935339 | 162954717 | mNeoVis.2.1.1.chr6\_hap1.g00826.t01 |
| 8352 | 6 | 162935339 | 162954717 | mNeoVis.2.1.1.chr6\_hap1.g00826.t01 |
| 8353 | 6 | 163002760 | 163016204 | mNeoVis.2.1.1.chr6\_hap1.g00827.t01 |
| 8354 | 6 | 163002760 | 163016204 | mNeoVis.2.1.1.chr6\_hap1.g00827.t01 |
| 8355 | 6 | 163002760 | 163016204 | mNeoVis.2.1.1.chr6\_hap1.g00827.t01 |
| 8356 | 6 | 163074759 | 163092419 | mNeoVis.2.1.1.chr6\_hap1.g00828.t01 |
| 8357 | 6 | 163074759 | 163092419 | mNeoVis.2.1.1.chr6\_hap1.g00828.t01 |
| 8358 | 6 | 164847443 | 164848825 | mNeoVis.2.1.1.chr6\_hap1.g00855.t01 |
| 8359 | 6 | 165624888 | 165675183 | mNeoVis.2.1.1.chr6\_hap1.g00856.t01 |
| 8360 | 6 | 165693901 | 165719594 | mNeoVis.2.1.1.chr6\_hap1.g00857.t01 |
| 8361 | 6 | 165693901 | 165719594 | mNeoVis.2.1.1.chr6\_hap1.g00857.t01 |
| 8362 | 6 | 165739025 | 165772421 | mNeoVis.2.1.1.chr6\_hap1.g00858.t01 |
| 8363 | 6 | 165739025 | 165772421 | mNeoVis.2.1.1.chr6\_hap1.g00858.t01 |
| 8364 | 6 | 166075168 | 166095652 | mNeoVis.2.1.1.chr6\_hap1.g00859.t01 |
| 8365 | 6 | 172099728 | 172117884 | mNeoVis.2.1.1.chr6\_hap1.g00901.t01 |
| 8366 | 6 | 172184517 | 172189013 | mNeoVis.2.1.1.chr6\_hap1.g00902.t01 |
| 8367 | 6 | 172251930 | 172290779 | mNeoVis.2.1.1.chr6\_hap1.g00903.t01 |
| 8368 | 6 | 172290976 | 172312229 | mNeoVis.2.1.1.chr6\_hap1.g00904.t01 |
| 8369 | 6 | 172477092 | 172479579 | mNeoVis.2.1.1.chr6\_hap1.g00905.t01 |
| 8370 | 6 | 172525644 | 172532367 | mNeoVis.2.1.1.chr6\_hap1.g00906.t01 |
| 8371 | 6 | 172532457 | 172559768 | mNeoVis.2.1.1.chr6\_hap1.g00907.t01 |
| 8372 | 6 | 172714382 | 172732539 | mNeoVis.2.1.1.chr6\_hap1.g00908.t01 |
| 8373 | 6 | 172788062 | 172792553 | mNeoVis.2.1.1.chr6\_hap1.g00909.t01 |
| 8374 | 6 | 172788062 | 172792553 | mNeoVis.2.1.1.chr6\_hap1.g00909.t01 |
| 8375 | 6 | 172798115 | 172848361 | mNeoVis.2.1.1.chr6\_hap1.g00910.t01 |
| 8376 | 6 | 172798115 | 172848361 | mNeoVis.2.1.1.chr6\_hap1.g00910.t01 |
| 8377 | 6 | 173070409 | 173072898 | mNeoVis.2.1.1.chr6\_hap1.g00911.t01 |
| 8378 | 6 | 173070409 | 173072898 | mNeoVis.2.1.1.chr6\_hap1.g00911.t01 |
| 8379 | 6 | 177685660 | 177687571 | mNeoVis.2.1.1.chr6\_hap1.g00913.t01 |
| 8380 | 6 | 177719829 | 177794921 | mNeoVis.2.1.1.chr6\_hap1.g00914.t01 |
| 8381 | 6 | 178823075 | 178889249 | mNeoVis.2.1.1.chr6\_hap1.g00916.t01 |
| 8382 | 6 | 179691115 | 179702588 | mNeoVis.2.1.1.chr6\_hap1.g00925.t01 |
| 8383 | 6 | 179749977 | 179767768 | mNeoVis.2.1.1.chr6\_hap1.g00926.t01 |
| 8384 | 6 | 179781153 | 179801062 | mNeoVis.2.1.1.chr6\_hap1.g00927.t01 |
| 8385 | 6 | 179925691 | 179937734 | mNeoVis.2.1.1.chr6\_hap1.g00928.t01 |
| 8386 | 6 | 179925691 | 179937734 | mNeoVis.2.1.1.chr6\_hap1.g00928.t01 |
| 8387 | 6 | 179937971 | 180044392 | mNeoVis.2.1.1.chr6\_hap1.g00929.t01 |
| 8388 | 6 | 179937971 | 180044392 | mNeoVis.2.1.1.chr6\_hap1.g00929.t01 |
| 8389 | 6 | 180053147 | 180066542 | mNeoVis.2.1.1.chr6\_hap1.g00930.t01 |
| 8390 | 6 | 180053147 | 180066542 | mNeoVis.2.1.1.chr6\_hap1.g00930.t01 |
| 8391 | 6 | 180080063 | 180083224 | mNeoVis.2.1.1.chr6\_hap1.g00931.t01 |
| 8392 | 6 | 180080063 | 180083224 | mNeoVis.2.1.1.chr6\_hap1.g00931.t01 |
| 8393 | 6 | 180084882 | 180112395 | mNeoVis.2.1.1.chr6\_hap1.g00932.t01 |
| 8394 | 6 | 180084882 | 180112395 | mNeoVis.2.1.1.chr6\_hap1.g00932.t01 |
| 8395 | 6 | 180128419 | 180186791 | mNeoVis.2.1.1.chr6\_hap1.g00933.t01 |
| 8396 | 6 | 180241952 | 180265997 | mNeoVis.2.1.1.chr6\_hap1.g00934.t01 |
| 8397 | 6 | 180268756 | 180285936 | mNeoVis.2.1.1.chr6\_hap1.g00935.t01 |
| 8398 | 6 | 180378953 | 180490673 | mNeoVis.2.1.1.chr6\_hap1.g00936.t01 |
| 8399 | 6 | 180378953 | 180490673 | mNeoVis.2.1.1.chr6\_hap1.g00936.t01 |
| 8400 | 6 | 180525064 | 180736035 | mNeoVis.2.1.1.chr6\_hap1.g00937.t01 |
| 8401 | 6 | 180525064 | 180736035 | mNeoVis.2.1.1.chr6\_hap1.g00937.t01 |
| 8402 | 6 | 180738688 | 180742769 | mNeoVis.2.1.1.chr6\_hap1.g00938.t01 |
| 8403 | 6 | 180738688 | 180742769 | mNeoVis.2.1.1.chr6\_hap1.g00938.t01 |
| 8404 | 6 | 180760237 | 180795089 | mNeoVis.2.1.1.chr6\_hap1.g00939.t01 |
| 8405 | 6 | 180760237 | 180795089 | mNeoVis.2.1.1.chr6\_hap1.g00939.t01 |
| 8406 | 6 | 180824613 | 180837460 | mNeoVis.2.1.1.chr6\_hap1.g00940.t01 |
| 8407 | 6 | 180824613 | 180837460 | mNeoVis.2.1.1.chr6\_hap1.g00940.t01 |
| 8408 | 6 | 180849302 | 180866385 | mNeoVis.2.1.1.chr6\_hap1.g00941.t01 |
| 8409 | 6 | 180849302 | 180866385 | mNeoVis.2.1.1.chr6\_hap1.g00941.t01 |
| 8410 | 6 | 181035136 | 181080743 | mNeoVis.2.1.1.chr6\_hap1.g00942.t01 |
| 8411 | 6 | 181035136 | 181080743 | mNeoVis.2.1.1.chr6\_hap1.g00942.t01 |
| 8412 | 6 | 181246786 | 181274863 | mNeoVis.2.1.1.chr6\_hap1.g00943.t01 |
| 8413 | 6 | 181246786 | 181274863 | mNeoVis.2.1.1.chr6\_hap1.g00943.t01 |
| 8414 | 6 | 181384772 | 181636054 | mNeoVis.2.1.1.chr6\_hap1.g00944.t01 |
| 8415 | 6 | 183789759 | 183837761 | mNeoVis.2.1.1.chr6\_hap1.g00959.t01 |
| 8416 | 6 | 183837684 | 183859903 | mNeoVis.2.1.1.chr6\_hap1.g00960.t01 |
| 8417 | 6 | 183859530 | 183984911 | mNeoVis.2.1.1.chr6\_hap1.g00961.t01 |
| 8418 | 6 | 183984905 | 184021596 | mNeoVis.2.1.1.chr6\_hap1.g00962.t01 |
| 8419 | 6 | 183989801 | 183997479 | mNeoVis.2.1.1.chr6\_hap1.g00963.t01 |
| 8420 | 6 | 184035270 | 184060232 | mNeoVis.2.1.1.chr6\_hap1.g00964.t01 |
| 8421 | 6 | 184060484 | 184071041 | mNeoVis.2.1.1.chr6\_hap1.g00965.t01 |
| 8422 | 6 | 184071178 | 184081534 | mNeoVis.2.1.1.chr6\_hap1.g00966.t01 |
| 8423 | 6 | 184081699 | 184088345 | mNeoVis.2.1.1.chr6\_hap1.g00967.t01 |
| 8424 | 6 | 184091056 | 184119816 | mNeoVis.2.1.1.chr6\_hap1.g00968.t01 |
| 8425 | 6 | 184135872 | 184155653 | mNeoVis.2.1.1.chr6\_hap1.g00969.t01 |
| 8426 | 6 | 184186969 | 184194382 | mNeoVis.2.1.1.chr6\_hap1.g00970.t01 |
| 8427 | 6 | 184205489 | 184210027 | mNeoVis.2.1.1.chr6\_hap1.g00971.t01 |
| 8428 | 6 | 184245124 | 184277156 | mNeoVis.2.1.1.chr6\_hap1.g00972.t01 |
| 8429 | 6 | 184283853 | 184288207 | mNeoVis.2.1.1.chr6\_hap1.g00973.t01 |
| 8430 | 6 | 184288411 | 184323564 | mNeoVis.2.1.1.chr6\_hap1.g00974.t01 |
| 8431 | 6 | 184339272 | 184355260 | mNeoVis.2.1.1.chr6\_hap1.g00975.t01 |
| 8432 | 6 | 184360492 | 184377949 | mNeoVis.2.1.1.chr6\_hap1.g00976.t01 |
| 8433 | 6 | 184407418 | 184411730 | mNeoVis.2.1.1.chr6\_hap1.g00977.t01 |
| 8434 | 6 | 184407418 | 184411730 | mNeoVis.2.1.1.chr6\_hap1.g00977.t01 |
| 8435 | 6 | 185087871 | 185247272 | mNeoVis.2.1.1.chr6\_hap1.g00978.t01 |
| 8436 | 6 | 185087871 | 185247272 | mNeoVis.2.1.1.chr6\_hap1.g00978.t01 |
| 8437 | 6 | 185087871 | 185247272 | mNeoVis.2.1.1.chr6\_hap1.g00978.t01 |
| 8438 | 6 | 185264959 | 185373480 | mNeoVis.2.1.1.chr6\_hap1.g00979.t01 |
| 8439 | 6 | 185264959 | 185373480 | mNeoVis.2.1.1.chr6\_hap1.g00979.t01 |
| 8440 | 6 | 185264959 | 185373480 | mNeoVis.2.1.1.chr6\_hap1.g00979.t01 |
| 8441 | 6 | 186023655 | 186556591 | mNeoVis.2.1.1.chr6\_hap1.g00981.t01 |
| 8442 | 6 | 186023655 | 186556591 | mNeoVis.2.1.1.chr6\_hap1.g00981.t01 |
| 8443 | 6 | 188133199 | 188138597 | mNeoVis.2.1.1.chr6\_hap1.g00983.t01 |
| 8444 | 6 | 188144016 | 188160672 | mNeoVis.2.1.1.chr6\_hap1.g00984.t01 |
| 8445 | 6 | 188178342 | 188189549 | mNeoVis.2.1.1.chr6\_hap1.g00985.t01 |
| 8446 | 6 | 188178342 | 188189549 | mNeoVis.2.1.1.chr6\_hap1.g00985.t01 |
| 8447 | 6 | 188187364 | 188201671 | mNeoVis.2.1.1.chr6\_hap1.g00986.t01 |
| 8448 | 6 | 188187364 | 188201671 | mNeoVis.2.1.1.chr6\_hap1.g00986.t01 |
| 8449 | 6 | 188216568 | 188290234 | mNeoVis.2.1.1.chr6\_hap1.g00987.t01 |
| 8450 | 6 | 188216568 | 188290234 | mNeoVis.2.1.1.chr6\_hap1.g00987.t01 |
| 8451 | 6 | 188767904 | 188893091 | mNeoVis.2.1.1.chr6\_hap1.g00988.t01 |
| 8452 | 6 | 188767904 | 188893091 | mNeoVis.2.1.1.chr6\_hap1.g00988.t01 |
| 8453 | 6 | 188767904 | 188893091 | mNeoVis.2.1.1.chr6\_hap1.g00988.t01 |
| 8454 | 6 | 188893095 | 188893558 | mNeoVis.2.1.1.chr6\_hap1.g00989.t01 |
| 8455 | 6 | 188893095 | 188893558 | mNeoVis.2.1.1.chr6\_hap1.g00989.t01 |
| 8456 | 6 | 188893095 | 188893558 | mNeoVis.2.1.1.chr6\_hap1.g00989.t01 |
| 8457 | 6 | 189012726 | 189036778 | mNeoVis.2.1.1.chr6\_hap1.g00990.t01 |
| 8458 | 6 | 189012726 | 189036778 | mNeoVis.2.1.1.chr6\_hap1.g00990.t01 |
| 8459 | 6 | 190291958 | 190442048 | mNeoVis.2.1.1.chr6\_hap1.g00991.t01 |
| 8460 | 6 | 190442198 | 190448498 | mNeoVis.2.1.1.chr6\_hap1.g00992.t01 |
| 8461 | 6 | 190459959 | 190503889 | mNeoVis.2.1.1.chr6\_hap1.g00993.t01 |
| 8462 | 6 | 190491084 | 190504286 | mNeoVis.2.1.1.chr6\_hap1.g00994.t01 |
| 8463 | 6 | 190504203 | 190510457 | mNeoVis.2.1.1.chr6\_hap1.g00995.t01 |
| 8464 | 6 | 190648277 | 190708026 | mNeoVis.2.1.1.chr6\_hap1.g00996.t01 |
| 8465 | 6 | 190723910 | 190758000 | mNeoVis.2.1.1.chr6\_hap1.g00997.t01 |
| 8466 | 6 | 190775795 | 190800704 | mNeoVis.2.1.1.chr6\_hap1.g00998.t01 |
| 8467 | 6 | 190810759 | 190859787 | mNeoVis.2.1.1.chr6\_hap1.g00999.t01 |
| 8468 | 6 | 190888226 | 190969415 | mNeoVis.2.1.1.chr6\_hap1.g01000.t01 |
| 8469 | 6 | 190974644 | 190995843 | mNeoVis.2.1.1.chr6\_hap1.g01001.t01 |
| 8470 | 6 | 192436117 | 192437492 | mNeoVis.2.1.1.chr6\_hap1.g01005.t01 |
| 8471 | 6 | 192436117 | 192437492 | mNeoVis.2.1.1.chr6\_hap1.g01005.t01 |
| 8472 | 6 | 192448437 | 192449351 | mNeoVis.2.1.1.chr6\_hap1.g01006.t01 |
| 8473 | 6 | 192448437 | 192449351 | mNeoVis.2.1.1.chr6\_hap1.g01006.t01 |
| 8474 | 6 | 192541964 | 192768801 | mNeoVis.2.1.1.chr6\_hap1.g01007.t01 |
| 8475 | 6 | 192578707 | 192585370 | mNeoVis.2.1.1.chr6\_hap1.g01008.t01 |
| 8476 | 6 | 192578707 | 192585370 | mNeoVis.2.1.1.chr6\_hap1.g01008.t01 |
| 8477 | 6 | 193930430 | 194047605 | mNeoVis.2.1.1.chr6\_hap1.g01009.t01 |
| 8478 | 6 | 194351164 | 194370849 | mNeoVis.2.1.1.chr6\_hap1.g01010.t01 |
| 8479 | 6 | 194351164 | 194370849 | mNeoVis.2.1.1.chr6\_hap1.g01010.t01 |
| 8480 | 6 | 194351164 | 194370849 | mNeoVis.2.1.1.chr6\_hap1.g01010.t01 |
| 8481 | 6 | 194385849 | 194435096 | mNeoVis.2.1.1.chr6\_hap1.g01011.t01 |
| 8482 | 6 | 194385849 | 194435096 | mNeoVis.2.1.1.chr6\_hap1.g01011.t01 |
| 8483 | 6 | 194385849 | 194435096 | mNeoVis.2.1.1.chr6\_hap1.g01011.t01 |
| 8484 | 6 | 194437486 | 194522855 | mNeoVis.2.1.1.chr6\_hap1.g01012.t01 |
| 8485 | 6 | 194437486 | 194522855 | mNeoVis.2.1.1.chr6\_hap1.g01012.t01 |
| 8486 | 6 | 194437486 | 194522855 | mNeoVis.2.1.1.chr6\_hap1.g01012.t01 |
| 8487 | 6 | 194522300 | 194591301 | mNeoVis.2.1.1.chr6\_hap1.g01013.t01 |
| 8488 | 6 | 194522300 | 194591301 | mNeoVis.2.1.1.chr6\_hap1.g01013.t01 |
| 8489 | 6 | 194522300 | 194591301 | mNeoVis.2.1.1.chr6\_hap1.g01013.t01 |
| 8490 | 6 | 194641720 | 194808972 | mNeoVis.2.1.1.chr6\_hap1.g01014.t01 |
| 8491 | 6 | 194641720 | 194808972 | mNeoVis.2.1.1.chr6\_hap1.g01014.t01 |
| 8492 | 6 | 194704505 | 194705152 | mNeoVis.2.1.1.chr6\_hap1.g01015.t01 |
| 8493 | 6 | 194704505 | 194705152 | mNeoVis.2.1.1.chr6\_hap1.g01015.t01 |
| 8494 | 6 | 194704505 | 194705152 | mNeoVis.2.1.1.chr6\_hap1.g01015.t01 |
| 8495 | 6 | 194809341 | 194894186 | mNeoVis.2.1.1.chr6\_hap1.g01016.t01 |
| 8496 | 6 | 194809341 | 194894186 | mNeoVis.2.1.1.chr6\_hap1.g01016.t01 |
| 8497 | 6 | 194903837 | 194974711 | mNeoVis.2.1.1.chr6\_hap1.g01017.t01 |
| 8498 | 6 | 195048197 | 195155953 | mNeoVis.2.1.1.chr6\_hap1.g01018.t01 |
| 8499 | 6 | 195172757 | 195228155 | mNeoVis.2.1.1.chr6\_hap1.g01019.t01 |
| 8500 | 6 | 195228274 | 195246604 | mNeoVis.2.1.1.chr6\_hap1.g01020.t01 |
| 8501 | 6 | 195232624 | 195321459 | mNeoVis.2.1.1.chr6\_hap1.g01021.t01 |
| 8502 | 6 | 195349345 | 195453463 | mNeoVis.2.1.1.chr6\_hap1.g01022.t01 |
| 8503 | 6 | 195475843 | 195485288 | mNeoVis.2.1.1.chr6\_hap1.g01023.t01 |
| 8504 | 6 | 196329828 | 196413847 | mNeoVis.2.1.1.chr6\_hap1.g01025.t01 |
| 8505 | 6 | 196762791 | 197039198 | mNeoVis.2.1.1.chr6\_hap1.g01026.t01 |
| 8506 | 6 | 197626703 | 197627097 | mNeoVis.2.1.1.chr6\_hap1.g01027.t01 |
| 8507 | 6 | 197659086 | 198468897 | mNeoVis.2.1.1.chr6\_hap1.g01028.t01 |
| 8508 | 6 | 197660153 | 197685750 | mNeoVis.2.1.1.chr6\_hap1.g01029.t01 |
| 8509 | 6 | 198823689 | 198825022 | mNeoVis.2.1.1.chr6\_hap1.g01030.t01 |
| 8510 | 6 | 198941349 | 198943678 | mNeoVis.2.1.1.chr6\_hap1.g01031.t01 |
| 8511 | 6 | 199471457 | 199500161 | mNeoVis.2.1.1.chr6\_hap1.g01033.t01 |
| 8512 | 6 | 199471457 | 199500161 | mNeoVis.2.1.1.chr6\_hap1.g01033.t01 |
| 8513 | 6 | 199683734 | 199722463 | mNeoVis.2.1.1.chr6\_hap1.g01034.t01 |
| 8514 | 6 | 199683734 | 199722463 | mNeoVis.2.1.1.chr6\_hap1.g01034.t01 |
| 8515 | 6 | 199709261 | 199834013 | mNeoVis.2.1.1.chr6\_hap1.g01035.t01 |
| 8516 | 6 | 199852127 | 199899425 | mNeoVis.2.1.1.chr6\_hap1.g01036.t01 |
| 8517 | 6 | 200161447 | 200163004 | mNeoVis.2.1.1.chr6\_hap1.g01037.t01 |
| 8518 | 6 | 200169073 | 200210090 | mNeoVis.2.1.1.chr6\_hap1.g01038.t01 |
| 8519 | 6 | 200252936 | 200357249 | mNeoVis.2.1.1.chr6\_hap1.g01039.t01 |
| 8520 | 6 | 200434788 | 200476403 | mNeoVis.2.1.1.chr6\_hap1.g01040.t01 |
| 8521 | 6 | 200558460 | 200570106 | mNeoVis.2.1.1.chr6\_hap1.g01041.t01 |
| 8522 | 6 | 200558460 | 200570106 | mNeoVis.2.1.1.chr6\_hap1.g01041.t01 |
| 8523 | 6 | 200898572 | 201113395 | mNeoVis.2.1.1.chr6\_hap1.g01042.t01 |
| 8524 | 6 | 201105284 | 201113411 | mNeoVis.2.1.1.chr6\_hap1.g01043.t01 |
| 8525 | 6 | 202470390 | 202841616 | mNeoVis.2.1.1.chr6\_hap1.g01058.t01 |
| 8526 | 6 | 202821350 | 202824424 | mNeoVis.2.1.1.chr6\_hap1.g01059.t01 |
| 8527 | 6 | 202886538 | 202916601 | mNeoVis.2.1.1.chr6\_hap1.g01060.t01 |
| 8528 | 6 | 202930388 | 202948953 | mNeoVis.2.1.1.chr6\_hap1.g01061.t01 |
| 8529 | 6 | 203006295 | 203008488 | mNeoVis.2.1.1.chr6\_hap1.g01064.t01 |
| 8530 | 6 | 203008316 | 203056348 | mNeoVis.2.1.1.chr6\_hap1.g01065.t01 |
| 8531 | 6 | 203051511 | 203174723 | mNeoVis.2.1.1.chr6\_hap1.g01066.t01 |
| 8532 | 6 | 203182029 | 203217684 | mNeoVis.2.1.1.chr6\_hap1.g01067.t01 |
| 8533 | 6 | 203228372 | 203232467 | mNeoVis.2.1.1.chr6\_hap1.g01068.t01 |
| 8534 | 6 | 203333872 | 203345630 | mNeoVis.2.1.1.chr6\_hap1.g01069.t01 |
| 8535 | 6 | 203473582 | 203572354 | mNeoVis.2.1.1.chr6\_hap1.g01070.t01 |
| 8536 | 6 | 203584503 | 203620241 | mNeoVis.2.1.1.chr6\_hap1.g01071.t01 |
| 8537 | 6 | 203620957 | 203751273 | mNeoVis.2.1.1.chr6\_hap1.g01072.t01 |
| 8538 | 6 | 203686746 | 203709891 | mNeoVis.2.1.1.chr6\_hap1.g01073.t01 |
| 8539 | 6 | 203770096 | 203771894 | mNeoVis.2.1.1.chr6\_hap1.g01074.t01 |
| 8540 | 6 | 203819878 | 203901465 | mNeoVis.2.1.1.chr6\_hap1.g01075.t01 |
| 8541 | 6 | 204013344 | 204120930 | mNeoVis.2.1.1.chr6\_hap1.g01076.t01 |
| 8542 | 6 | 204013344 | 204120930 | mNeoVis.2.1.1.chr6\_hap1.g01076.t01 |
| 8543 | 6 | 204180250 | 204389331 | mNeoVis.2.1.1.chr6\_hap1.g01077.t01 |
| 8544 | 6 | 204180250 | 204389331 | mNeoVis.2.1.1.chr6\_hap1.g01077.t01 |
| 8545 | 6 | 204435507 | 204480846 | mNeoVis.2.1.1.chr6\_hap1.g01078.t01 |
| 8546 | 6 | 204435507 | 204480846 | mNeoVis.2.1.1.chr6\_hap1.g01078.t01 |
| 8547 | 6 | 204481079 | 204496992 | mNeoVis.2.1.1.chr6\_hap1.g01079.t01 |
| 8548 | 6 | 204481079 | 204496992 | mNeoVis.2.1.1.chr6\_hap1.g01079.t01 |
| 8549 | 6 | 204481079 | 204496992 | mNeoVis.2.1.1.chr6\_hap1.g01079.t01 |
| 8550 | 6 | 204588905 | 204595641 | mNeoVis.2.1.1.chr6\_hap1.g01080.t01 |
| 8551 | 6 | 204588905 | 204595641 | mNeoVis.2.1.1.chr6\_hap1.g01080.t01 |
| 8552 | 6 | 204588905 | 204595641 | mNeoVis.2.1.1.chr6\_hap1.g01080.t01 |
| 8553 | 6 | 204634960 | 204638740 | mNeoVis.2.1.1.chr6\_hap1.g01081.t01 |
| 8554 | 6 | 204634960 | 204638740 | mNeoVis.2.1.1.chr6\_hap1.g01081.t01 |
| 8555 | 6 | 204634960 | 204638740 | mNeoVis.2.1.1.chr6\_hap1.g01081.t01 |
| 8556 | 6 | 204732773 | 204733565 | mNeoVis.2.1.1.chr6\_hap1.g01082.t01 |
| 8557 | 6 | 204732773 | 204733565 | mNeoVis.2.1.1.chr6\_hap1.g01082.t01 |
| 8558 | 6 | 204735093 | 204736016 | mNeoVis.2.1.1.chr6\_hap1.g01083.t01 |
| 8559 | 6 | 204735093 | 204736016 | mNeoVis.2.1.1.chr6\_hap1.g01083.t01 |
| 8560 | 6 | 204738230 | 204941013 | mNeoVis.2.1.1.chr6\_hap1.g01084.t01 |
| 8561 | 6 | 204738230 | 204941013 | mNeoVis.2.1.1.chr6\_hap1.g01084.t01 |
| 8562 | 6 | 205026083 | 205068309 | mNeoVis.2.1.1.chr6\_hap1.g01085.t01 |
| 8563 | 6 | 205026083 | 205068309 | mNeoVis.2.1.1.chr6\_hap1.g01085.t01 |
| 8564 | 6 | 205026083 | 205068309 | mNeoVis.2.1.1.chr6\_hap1.g01085.t01 |
| 8565 | 6 | 205137258 | 205971838 | mNeoVis.2.1.1.chr6\_hap1.g01086.t01 |
| 8566 | 6 | 207198857 | 207629090 | mNeoVis.2.1.1.chr6\_hap1.g01087.t01 |
| 8567 | 6 | 208102656 | 208250509 | mNeoVis.2.1.1.chr6\_hap1.g01088.t01 |
| 8568 | 6 | 210104375 | 210114858 | mNeoVis.2.1.1.chr6\_hap1.g01090.t01 |
| 8569 | 6 | 210180378 | 210227720 | mNeoVis.2.1.1.chr6\_hap1.g01091.t01 |
| 8570 | 6 | 211087449 | 211096113 | mNeoVis.2.1.1.chr6\_hap1.g01094.t01 |
| 8571 | 6 | 211096145 | 211161550 | mNeoVis.2.1.1.chr6\_hap1.g01095.t01 |
| 8572 | 6 | 211131361 | 211178269 | mNeoVis.2.1.1.chr6\_hap1.g01096.t01 |
| 8573 | 6 | 211184800 | 211197511 | mNeoVis.2.1.1.chr6\_hap1.g01097.t01 |
| 8574 | 6 | 211199861 | 211307241 | mNeoVis.2.1.1.chr6\_hap1.g01098.t01 |
| 8575 | 6 | 211324873 | 211325770 | mNeoVis.2.1.1.chr6\_hap1.g01099.t01 |
| 8576 | 6 | 218451310 | 218453755 | mNeoVis.2.1.1.chr6\_hap1.g01118.t01 |
| 8577 | 6 | 218451310 | 218453755 | mNeoVis.2.1.1.chr6\_hap1.g01118.t01 |
| 8578 | 7 | 645470 | 649842 | mNeoVis.2.1.1.chr7\_hap1.g00025.t01 |
| 8579 | 7 | 650086 | 669957 | mNeoVis.2.1.1.chr7\_hap1.g00026.t01 |
| 8580 | 7 | 683167 | 723253 | mNeoVis.2.1.1.chr7\_hap1.g00027.t01 |
| 8581 | 7 | 797012 | 870560 | mNeoVis.2.1.1.chr7\_hap1.g00028.t01 |
| 8582 | 7 | 884407 | 888345 | mNeoVis.2.1.1.chr7\_hap1.g00029.t01 |
| 8583 | 7 | 888355 | 912898 | mNeoVis.2.1.1.chr7\_hap1.g00030.t01 |
| 8584 | 7 | 912858 | 915496 | mNeoVis.2.1.1.chr7\_hap1.g00031.t01 |
| 8585 | 7 | 915933 | 920301 | mNeoVis.2.1.1.chr7\_hap1.g00032.t01 |
| 8586 | 7 | 933173 | 979620 | mNeoVis.2.1.1.chr7\_hap1.g00033.t01 |
| 8587 | 7 | 979809 | 987238 | mNeoVis.2.1.1.chr7\_hap1.g00034.t01 |
| 8588 | 7 | 987373 | 994531 | mNeoVis.2.1.1.chr7\_hap1.g00035.t01 |
| 8589 | 7 | 1000104 | 1006264 | mNeoVis.2.1.1.chr7\_hap1.g00036.t01 |
| 8590 | 7 | 1023667 | 1031846 | mNeoVis.2.1.1.chr7\_hap1.g00037.t01 |
| 8591 | 7 | 1033067 | 1040533 | mNeoVis.2.1.1.chr7\_hap1.g00038.t01 |
| 8592 | 7 | 1042335 | 1046581 | mNeoVis.2.1.1.chr7\_hap1.g00039.t01 |
| 8593 | 7 | 1047976 | 1108823 | mNeoVis.2.1.1.chr7\_hap1.g00040.t01 |
| 8594 | 7 | 1143567 | 1207896 | mNeoVis.2.1.1.chr7\_hap1.g00041.t01 |
| 8595 | 7 | 1215669 | 1253564 | mNeoVis.2.1.1.chr7\_hap1.g00042.t01 |
| 8596 | 7 | 1450510 | 1548463 | mNeoVis.2.1.1.chr7\_hap1.g00043.t01 |
| 8597 | 7 | 1561173 | 1580702 | mNeoVis.2.1.1.chr7\_hap1.g00044.t01 |
| 8598 | 7 | 1595107 | 1626118 | mNeoVis.2.1.1.chr7\_hap1.g00045.t01 |
| 8599 | 7 | 1659661 | 1706014 | mNeoVis.2.1.1.chr7\_hap1.g00046.t01 |
| 8600 | 7 | 1713043 | 1781492 | mNeoVis.2.1.1.chr7\_hap1.g00047.t01 |
| 8601 | 7 | 1726323 | 1730109 | mNeoVis.2.1.1.chr7\_hap1.g00048.t01 |
| 8602 | 7 | 1850883 | 1901715 | mNeoVis.2.1.1.chr7\_hap1.g00049.t01 |
| 8603 | 7 | 1906685 | 1919594 | mNeoVis.2.1.1.chr7\_hap1.g00050.t01 |
| 8604 | 7 | 2465996 | 2469222 | mNeoVis.2.1.1.chr7\_hap1.g00054.t01 |
| 8605 | 7 | 2465996 | 2469222 | mNeoVis.2.1.1.chr7\_hap1.g00054.t01 |
| 8606 | 7 | 2477567 | 2480164 | mNeoVis.2.1.1.chr7\_hap1.g00055.t01 |
| 8607 | 7 | 2477567 | 2480164 | mNeoVis.2.1.1.chr7\_hap1.g00055.t01 |
| 8608 | 7 | 2477567 | 2480164 | mNeoVis.2.1.1.chr7\_hap1.g00055.t01 |
| 8609 | 7 | 2495535 | 2518167 | mNeoVis.2.1.1.chr7\_hap1.g00056.t01 |
| 8610 | 7 | 2495535 | 2518167 | mNeoVis.2.1.1.chr7\_hap1.g00056.t01 |
| 8611 | 7 | 2495535 | 2518167 | mNeoVis.2.1.1.chr7\_hap1.g00056.t01 |
| 8612 | 7 | 2528772 | 2532642 | mNeoVis.2.1.1.chr7\_hap1.g00057.t01 |
| 8613 | 7 | 2528772 | 2532642 | mNeoVis.2.1.1.chr7\_hap1.g00057.t01 |
| 8614 | 7 | 2528772 | 2532642 | mNeoVis.2.1.1.chr7\_hap1.g00057.t01 |
| 8615 | 7 | 2534280 | 2927157 | mNeoVis.2.1.1.chr7\_hap1.g00058.t01 |
| 8616 | 7 | 2534280 | 2927157 | mNeoVis.2.1.1.chr7\_hap1.g00058.t01 |
| 8617 | 7 | 2534280 | 2927157 | mNeoVis.2.1.1.chr7\_hap1.g00058.t01 |
| 8618 | 7 | 2950379 | 2971039 | mNeoVis.2.1.1.chr7\_hap1.g00059.t01 |
| 8619 | 7 | 2950379 | 2971039 | mNeoVis.2.1.1.chr7\_hap1.g00059.t01 |
| 8620 | 7 | 2950379 | 2971039 | mNeoVis.2.1.1.chr7\_hap1.g00059.t01 |
| 8621 | 7 | 3033922 | 3041845 | mNeoVis.2.1.1.chr7\_hap1.g00060.t01 |
| 8622 | 7 | 3033922 | 3041845 | mNeoVis.2.1.1.chr7\_hap1.g00060.t01 |
| 8623 | 7 | 3033922 | 3041845 | mNeoVis.2.1.1.chr7\_hap1.g00060.t01 |
| 8624 | 7 | 3041972 | 3059970 | mNeoVis.2.1.1.chr7\_hap1.g00061.t01 |
| 8625 | 7 | 3041972 | 3059970 | mNeoVis.2.1.1.chr7\_hap1.g00061.t01 |
| 8626 | 7 | 3041972 | 3059970 | mNeoVis.2.1.1.chr7\_hap1.g00061.t01 |
| 8627 | 7 | 3083883 | 3113082 | mNeoVis.2.1.1.chr7\_hap1.g00062.t01 |
| 8628 | 7 | 3083883 | 3113082 | mNeoVis.2.1.1.chr7\_hap1.g00062.t01 |
| 8629 | 7 | 3125179 | 3138496 | mNeoVis.2.1.1.chr7\_hap1.g00063.t01 |
| 8630 | 7 | 6278860 | 6304320 | mNeoVis.2.1.1.chr7\_hap1.g00093.t01 |
| 8631 | 7 | 6311069 | 6533978 | mNeoVis.2.1.1.chr7\_hap1.g00094.t01 |
| 8632 | 7 | 6569557 | 6629358 | mNeoVis.2.1.1.chr7\_hap1.g00095.t01 |
| 8633 | 7 | 6650448 | 6757982 | mNeoVis.2.1.1.chr7\_hap1.g00096.t01 |
| 8634 | 7 | 6650448 | 6757982 | mNeoVis.2.1.1.chr7\_hap1.g00096.t01 |
| 8635 | 7 | 6704009 | 6794411 | mNeoVis.2.1.1.chr7\_hap1.g00097.t01 |
| 8636 | 7 | 6704009 | 6794411 | mNeoVis.2.1.1.chr7\_hap1.g00097.t01 |
| 8637 | 7 | 6798587 | 6812224 | mNeoVis.2.1.1.chr7\_hap1.g00098.t01 |
| 8638 | 7 | 6798587 | 6812224 | mNeoVis.2.1.1.chr7\_hap1.g00098.t01 |
| 8639 | 7 | 6818916 | 6831592 | mNeoVis.2.1.1.chr7\_hap1.g00099.t01 |
| 8640 | 7 | 6818916 | 6831592 | mNeoVis.2.1.1.chr7\_hap1.g00099.t01 |
| 8641 | 7 | 6838432 | 6853504 | mNeoVis.2.1.1.chr7\_hap1.g00100.t01 |
| 8642 | 7 | 6838432 | 6853504 | mNeoVis.2.1.1.chr7\_hap1.g00100.t01 |
| 8643 | 7 | 6856413 | 6878873 | mNeoVis.2.1.1.chr7\_hap1.g00101.t01 |
| 8644 | 7 | 6856413 | 6878873 | mNeoVis.2.1.1.chr7\_hap1.g00101.t01 |
| 8645 | 7 | 6879696 | 6895910 | mNeoVis.2.1.1.chr7\_hap1.g00102.t01 |
| 8646 | 7 | 6879696 | 6895910 | mNeoVis.2.1.1.chr7\_hap1.g00102.t01 |
| 8647 | 7 | 7232344 | 7241646 | mNeoVis.2.1.1.chr7\_hap1.g00104.t01 |
| 8648 | 7 | 8058568 | 8065041 | mNeoVis.2.1.1.chr7\_hap1.g00105.t01 |
| 8649 | 7 | 9370062 | 9377165 | mNeoVis.2.1.1.chr7\_hap1.g00107.t01 |
| 8650 | 7 | 9410182 | 9547597 | mNeoVis.2.1.1.chr7\_hap1.g00108.t01 |
| 8651 | 7 | 9586168 | 9596229 | mNeoVis.2.1.1.chr7\_hap1.g00109.t01 |
| 8652 | 7 | 10594730 | 11115852 | mNeoVis.2.1.1.chr7\_hap1.g00114.t01 |
| 8653 | 7 | 10594730 | 11115852 | mNeoVis.2.1.1.chr7\_hap1.g00114.t01 |
| 8654 | 7 | 12425634 | 12446792 | mNeoVis.2.1.1.chr7\_hap1.g00141.t01 |
| 8655 | 7 | 12459076 | 12480202 | mNeoVis.2.1.1.chr7\_hap1.g00142.t01 |
| 8656 | 7 | 12489816 | 12540589 | mNeoVis.2.1.1.chr7\_hap1.g00143.t01 |
| 8657 | 7 | 12540936 | 12572815 | mNeoVis.2.1.1.chr7\_hap1.g00144.t01 |
| 8658 | 7 | 12573122 | 12598981 | mNeoVis.2.1.1.chr7\_hap1.g00145.t01 |
| 8659 | 7 | 12599227 | 12642122 | mNeoVis.2.1.1.chr7\_hap1.g00146.t01 |
| 8660 | 7 | 12698599 | 12707457 | mNeoVis.2.1.1.chr7\_hap1.g00147.t01 |
| 8661 | 7 | 12707947 | 12729780 | mNeoVis.2.1.1.chr7\_hap1.g00148.t01 |
| 8662 | 7 | 12731073 | 12829936 | mNeoVis.2.1.1.chr7\_hap1.g00149.t01 |
| 8663 | 7 | 13223114 | 13235954 | mNeoVis.2.1.1.chr7\_hap1.g00151.t01 |
| 8664 | 7 | 13281675 | 13310532 | mNeoVis.2.1.1.chr7\_hap1.g00152.t01 |
| 8665 | 7 | 13336403 | 13346160 | mNeoVis.2.1.1.chr7\_hap1.g00153.t01 |
| 8666 | 7 | 13348332 | 13363940 | mNeoVis.2.1.1.chr7\_hap1.g00154.t01 |
| 8667 | 7 | 13365214 | 13396823 | mNeoVis.2.1.1.chr7\_hap1.g00155.t01 |
| 8668 | 7 | 13428504 | 13437565 | mNeoVis.2.1.1.chr7\_hap1.g00156.t01 |
| 8669 | 7 | 13470629 | 13518558 | mNeoVis.2.1.1.chr7\_hap1.g00157.t01 |
| 8670 | 7 | 13518830 | 13532193 | mNeoVis.2.1.1.chr7\_hap1.g00158.t01 |
| 8671 | 7 | 13908106 | 13912521 | mNeoVis.2.1.1.chr7\_hap1.g00166.t01 |
| 8672 | 7 | 13914506 | 13922767 | mNeoVis.2.1.1.chr7\_hap1.g00167.t01 |
| 8673 | 7 | 13922919 | 13940942 | mNeoVis.2.1.1.chr7\_hap1.g00168.t01 |
| 8674 | 7 | 13944600 | 13987792 | mNeoVis.2.1.1.chr7\_hap1.g00169.t01 |
| 8675 | 7 | 15744706 | 15749755 | mNeoVis.2.1.1.chr7\_hap1.g00171.t01 |
| 8676 | 7 | 15744706 | 15749755 | mNeoVis.2.1.1.chr7\_hap1.g00171.t01 |
| 8677 | 7 | 15938865 | 15947144 | mNeoVis.2.1.1.chr7\_hap1.g00172.t01 |
| 8678 | 7 | 15960539 | 16108726 | mNeoVis.2.1.1.chr7\_hap1.g00173.t01 |
| 8679 | 7 | 16115425 | 16123335 | mNeoVis.2.1.1.chr7\_hap1.g00174.t01 |
| 8680 | 7 | 16141354 | 16153951 | mNeoVis.2.1.1.chr7\_hap1.g00175.t01 |
| 8681 | 7 | 16158585 | 16273789 | mNeoVis.2.1.1.chr7\_hap1.g00176.t01 |
| 8682 | 7 | 17013212 | 17057983 | mNeoVis.2.1.1.chr7\_hap1.g00190.t01 |
| 8683 | 7 | 17128849 | 17154567 | mNeoVis.2.1.1.chr7\_hap1.g00191.t01 |
| 8684 | 7 | 17228990 | 17309821 | mNeoVis.2.1.1.chr7\_hap1.g00192.t01 |
| 8685 | 7 | 17309737 | 17357800 | mNeoVis.2.1.1.chr7\_hap1.g00193.t01 |
| 8686 | 7 | 17357840 | 17369136 | mNeoVis.2.1.1.chr7\_hap1.g00194.t01 |
| 8687 | 7 | 17369399 | 17402041 | mNeoVis.2.1.1.chr7\_hap1.g00195.t01 |
| 8688 | 7 | 17404393 | 17415217 | mNeoVis.2.1.1.chr7\_hap1.g00196.t01 |
| 8689 | 7 | 17417948 | 17432003 | mNeoVis.2.1.1.chr7\_hap1.g00197.t01 |
| 8690 | 7 | 17429333 | 17561609 | mNeoVis.2.1.1.chr7\_hap1.g00198.t01 |
| 8691 | 7 | 20097379 | 20247902 | mNeoVis.2.1.1.chr7\_hap1.g00264.t01 |
| 8692 | 7 | 20097379 | 20247902 | mNeoVis.2.1.1.chr7\_hap1.g00264.t01 |
| 8693 | 7 | 20138960 | 20143805 | mNeoVis.2.1.1.chr7\_hap1.g00265.t01 |
| 8694 | 7 | 20138960 | 20143805 | mNeoVis.2.1.1.chr7\_hap1.g00265.t01 |
| 8695 | 7 | 26238313 | 26258870 | mNeoVis.2.1.1.chr7\_hap1.g00285.t01 |
| 8696 | 7 | 26262026 | 26279842 | mNeoVis.2.1.1.chr7\_hap1.g00286.t01 |
| 8697 | 7 | 26282682 | 26304074 | mNeoVis.2.1.1.chr7\_hap1.g00287.t01 |
| 8698 | 7 | 26306098 | 26344032 | mNeoVis.2.1.1.chr7\_hap1.g00288.t01 |
| 8699 | 7 | 26369291 | 26390906 | mNeoVis.2.1.1.chr7\_hap1.g00289.t01 |
| 8700 | 7 | 26390683 | 26421480 | mNeoVis.2.1.1.chr7\_hap1.g00290.t01 |
| 8701 | 7 | 26431940 | 26444844 | mNeoVis.2.1.1.chr7\_hap1.g00291.t01 |
| 8702 | 7 | 26444796 | 26457879 | mNeoVis.2.1.1.chr7\_hap1.g00292.t01 |
| 8703 | 7 | 26459528 | 26473216 | mNeoVis.2.1.1.chr7\_hap1.g00293.t01 |
| 8704 | 7 | 26473418 | 26488750 | mNeoVis.2.1.1.chr7\_hap1.g00294.t01 |
| 8705 | 7 | 26497562 | 26499900 | mNeoVis.2.1.1.chr7\_hap1.g00295.t01 |
| 8706 | 7 | 26517398 | 26528779 | mNeoVis.2.1.1.chr7\_hap1.g00296.t01 |
| 8707 | 7 | 26536073 | 26541979 | mNeoVis.2.1.1.chr7\_hap1.g00297.t01 |
| 8708 | 7 | 26591905 | 26610420 | mNeoVis.2.1.1.chr7\_hap1.g00298.t01 |
| 8709 | 7 | 26612551 | 26618902 | mNeoVis.2.1.1.chr7\_hap1.g00299.t01 |
| 8710 | 7 | 26623433 | 26667895 | mNeoVis.2.1.1.chr7\_hap1.g00300.t01 |
| 8711 | 7 | 26674729 | 26704454 | mNeoVis.2.1.1.chr7\_hap1.g00301.t01 |
| 8712 | 7 | 26710054 | 26754960 | mNeoVis.2.1.1.chr7\_hap1.g00302.t01 |
| 8713 | 7 | 26762442 | 26847069 | mNeoVis.2.1.1.chr7\_hap1.g00303.t01 |
| 8714 | 7 | 26872288 | 26884789 | mNeoVis.2.1.1.chr7\_hap1.g00304.t01 |
| 8715 | 7 | 27249744 | 27290319 | mNeoVis.2.1.1.chr7\_hap1.g00315.t01 |
| 8716 | 7 | 27292686 | 27458978 | mNeoVis.2.1.1.chr7\_hap1.g00316.t01 |
| 8717 | 7 | 27719279 | 27747555 | mNeoVis.2.1.1.chr7\_hap1.g00317.t01 |
| 8718 | 7 | 27719279 | 27747555 | mNeoVis.2.1.1.chr7\_hap1.g00317.t01 |
| 8719 | 7 | 27759360 | 27790506 | mNeoVis.2.1.1.chr7\_hap1.g00318.t01 |
| 8720 | 7 | 27759360 | 27790506 | mNeoVis.2.1.1.chr7\_hap1.g00318.t01 |
| 8721 | 7 | 29099337 | 29471691 | mNeoVis.2.1.1.chr7\_hap1.g00326.t01 |
| 8722 | 7 | 29471610 | 29684542 | mNeoVis.2.1.1.chr7\_hap1.g00327.t01 |
| 8723 | 7 | 29471610 | 29684542 | mNeoVis.2.1.1.chr7\_hap1.g00327.t01 |
| 8724 | 7 | 29632580 | 29707105 | mNeoVis.2.1.1.chr7\_hap1.g00328.t01 |
| 8725 | 7 | 29632580 | 29707105 | mNeoVis.2.1.1.chr7\_hap1.g00328.t01 |
| 8726 | 7 | 29641611 | 29651212 | mNeoVis.2.1.1.chr7\_hap1.g00329.t01 |
| 8727 | 7 | 29641611 | 29651212 | mNeoVis.2.1.1.chr7\_hap1.g00329.t01 |
| 8728 | 7 | 32357724 | 32395252 | mNeoVis.2.1.1.chr7\_hap1.g00337.t01 |
| 8729 | 7 | 32395554 | 32436048 | mNeoVis.2.1.1.chr7\_hap1.g00338.t01 |
| 8730 | 7 | 32462552 | 32532054 | mNeoVis.2.1.1.chr7\_hap1.g00339.t01 |
| 8731 | 7 | 32540854 | 32612225 | mNeoVis.2.1.1.chr7\_hap1.g00340.t01 |
| 8732 | 7 | 32629978 | 32649164 | mNeoVis.2.1.1.chr7\_hap1.g00341.t01 |
| 8733 | 7 | 32647118 | 32741317 | mNeoVis.2.1.1.chr7\_hap1.g00342.t01 |
| 8734 | 7 | 35281216 | 35352092 | mNeoVis.2.1.1.chr7\_hap1.g00354.t01 |
| 8735 | 7 | 35411013 | 35433895 | mNeoVis.2.1.1.chr7\_hap1.g00355.t01 |
| 8736 | 7 | 35456080 | 35489690 | mNeoVis.2.1.1.chr7\_hap1.g00356.t01 |
| 8737 | 7 | 35456080 | 35489690 | mNeoVis.2.1.1.chr7\_hap1.g00356.t01 |
| 8738 | 7 | 35569410 | 35579533 | mNeoVis.2.1.1.chr7\_hap1.g00357.t01 |
| 8739 | 7 | 35569410 | 35579533 | mNeoVis.2.1.1.chr7\_hap1.g00357.t01 |
| 8740 | 7 | 35579676 | 35612842 | mNeoVis.2.1.1.chr7\_hap1.g00358.t01 |
| 8741 | 7 | 35579676 | 35612842 | mNeoVis.2.1.1.chr7\_hap1.g00358.t01 |
| 8742 | 7 | 35647465 | 35693679 | mNeoVis.2.1.1.chr7\_hap1.g00359.t01 |
| 8743 | 7 | 35647465 | 35693679 | mNeoVis.2.1.1.chr7\_hap1.g00359.t01 |
| 8744 | 7 | 35696328 | 35703638 | mNeoVis.2.1.1.chr7\_hap1.g00360.t01 |
| 8745 | 7 | 35696328 | 35703638 | mNeoVis.2.1.1.chr7\_hap1.g00360.t01 |
| 8746 | 7 | 35703764 | 35732908 | mNeoVis.2.1.1.chr7\_hap1.g00361.t01 |
| 8747 | 7 | 35703764 | 35732908 | mNeoVis.2.1.1.chr7\_hap1.g00361.t01 |
| 8748 | 7 | 35775332 | 35805705 | mNeoVis.2.1.1.chr7\_hap1.g00362.t01 |
| 8749 | 7 | 38951078 | 39020814 | mNeoVis.2.1.1.chr7\_hap1.g00375.t01 |
| 8750 | 7 | 39713752 | 39832728 | mNeoVis.2.1.1.chr7\_hap1.g00376.t01 |
| 8751 | 7 | 39834409 | 39872522 | mNeoVis.2.1.1.chr7\_hap1.g00377.t01 |
| 8752 | 7 | 39888072 | 39957369 | mNeoVis.2.1.1.chr7\_hap1.g00378.t01 |
| 8753 | 7 | 40120009 | 40185238 | mNeoVis.2.1.1.chr7\_hap1.g00383.t01 |
| 8754 | 7 | 40185378 | 40207023 | mNeoVis.2.1.1.chr7\_hap1.g00384.t01 |
| 8755 | 7 | 40208905 | 40279280 | mNeoVis.2.1.1.chr7\_hap1.g00385.t01 |
| 8756 | 7 | 40255201 | 40255740 | mNeoVis.2.1.1.chr7\_hap1.g00386.t01 |
| 8757 | 7 | 40279523 | 40283158 | mNeoVis.2.1.1.chr7\_hap1.g00387.t01 |
| 8758 | 7 | 40284228 | 40342273 | mNeoVis.2.1.1.chr7\_hap1.g00388.t01 |
| 8759 | 7 | 40351253 | 40394886 | mNeoVis.2.1.1.chr7\_hap1.g00389.t01 |
| 8760 | 7 | 40396052 | 40436869 | mNeoVis.2.1.1.chr7\_hap1.g00390.t01 |
| 8761 | 7 | 40457674 | 40469411 | mNeoVis.2.1.1.chr7\_hap1.g00391.t01 |
| 8762 | 7 | 40469406 | 40485802 | mNeoVis.2.1.1.chr7\_hap1.g00392.t01 |
| 8763 | 7 | 40545135 | 40547736 | mNeoVis.2.1.1.chr7\_hap1.g00393.t01 |
| 8764 | 7 | 41275633 | 41339393 | mNeoVis.2.1.1.chr7\_hap1.g00398.t01 |
| 8765 | 7 | 41358969 | 41441073 | mNeoVis.2.1.1.chr7\_hap1.g00399.t01 |
| 8766 | 7 | 41448011 | 41473978 | mNeoVis.2.1.1.chr7\_hap1.g00400.t01 |
| 8767 | 7 | 41479589 | 41504355 | mNeoVis.2.1.1.chr7\_hap1.g00401.t01 |
| 8768 | 7 | 41508638 | 41548071 | mNeoVis.2.1.1.chr7\_hap1.g00402.t01 |
| 8769 | 7 | 41555861 | 41582267 | mNeoVis.2.1.1.chr7\_hap1.g00403.t01 |
| 8770 | 7 | 41594683 | 41652208 | mNeoVis.2.1.1.chr7\_hap1.g00404.t01 |
| 8771 | 7 | 41601329 | 41602879 | mNeoVis.2.1.1.chr7\_hap1.g00405.t01 |
| 8772 | 7 | 41606807 | 41609063 | mNeoVis.2.1.1.chr7\_hap1.g00406.t01 |
| 8773 | 7 | 41758721 | 41792500 | mNeoVis.2.1.1.chr7\_hap1.g00409.t01 |
| 8774 | 7 | 41810027 | 41819339 | mNeoVis.2.1.1.chr7\_hap1.g00410.t01 |
| 8775 | 7 | 41819978 | 41874378 | mNeoVis.2.1.1.chr7\_hap1.g00411.t01 |
| 8776 | 7 | 41877145 | 41884994 | mNeoVis.2.1.1.chr7\_hap1.g00412.t01 |
| 8777 | 7 | 41886379 | 41902994 | mNeoVis.2.1.1.chr7\_hap1.g00413.t01 |
| 8778 | 7 | 41920013 | 41925933 | mNeoVis.2.1.1.chr7\_hap1.g00414.t01 |
| 8779 | 7 | 41926818 | 41934729 | mNeoVis.2.1.1.chr7\_hap1.g00415.t01 |
| 8780 | 7 | 41937690 | 41941986 | mNeoVis.2.1.1.chr7\_hap1.g00416.t01 |
| 8781 | 7 | 41942174 | 41951618 | mNeoVis.2.1.1.chr7\_hap1.g00417.t01 |
| 8782 | 7 | 41951914 | 41963057 | mNeoVis.2.1.1.chr7\_hap1.g00418.t01 |
| 8783 | 7 | 42005985 | 42027316 | mNeoVis.2.1.1.chr7\_hap1.g00419.t01 |
| 8784 | 7 | 42038014 | 42052077 | mNeoVis.2.1.1.chr7\_hap1.g00420.t01 |
| 8785 | 7 | 42052731 | 42063545 | mNeoVis.2.1.1.chr7\_hap1.g00421.t01 |
| 8786 | 7 | 42065500 | 42067044 | mNeoVis.2.1.1.chr7\_hap1.g00422.t01 |
| 8787 | 7 | 42070286 | 42083889 | mNeoVis.2.1.1.chr7\_hap1.g00423.t01 |
| 8788 | 7 | 42096144 | 42107723 | mNeoVis.2.1.1.chr7\_hap1.g00424.t01 |
| 8789 | 7 | 42111645 | 42112544 | mNeoVis.2.1.1.chr7\_hap1.g00425.t01 |
| 8790 | 7 | 42116116 | 42117958 | mNeoVis.2.1.1.chr7\_hap1.g00426.t01 |
| 8791 | 7 | 42138777 | 42139766 | mNeoVis.2.1.1.chr7\_hap1.g00427.t01 |
| 8792 | 7 | 42170282 | 42173115 | mNeoVis.2.1.1.chr7\_hap1.g00428.t01 |
| 8793 | 7 | 42203534 | 42206615 | mNeoVis.2.1.1.chr7\_hap1.g00429.t01 |
| 8794 | 7 | 42215256 | 42229704 | mNeoVis.2.1.1.chr7\_hap1.g00430.t01 |
| 8795 | 7 | 42235496 | 42240232 | mNeoVis.2.1.1.chr7\_hap1.g00431.t01 |
| 8796 | 7 | 42244178 | 42253838 | mNeoVis.2.1.1.chr7\_hap1.g00432.t01 |
| 8797 | 7 | 42254028 | 42255796 | mNeoVis.2.1.1.chr7\_hap1.g00433.t01 |
| 8798 | 7 | 42257882 | 42334685 | mNeoVis.2.1.1.chr7\_hap1.g00434.t01 |
| 8799 | 7 | 42301939 | 42302996 | mNeoVis.2.1.1.chr7\_hap1.g00435.t01 |
| 8800 | 7 | 42334698 | 42342472 | mNeoVis.2.1.1.chr7\_hap1.g00436.t01 |
| 8801 | 7 | 42341492 | 42351155 | mNeoVis.2.1.1.chr7\_hap1.g00437.t01 |
| 8802 | 7 | 42352404 | 42372319 | mNeoVis.2.1.1.chr7\_hap1.g00438.t01 |
| 8803 | 7 | 42372776 | 42375411 | mNeoVis.2.1.1.chr7\_hap1.g00439.t01 |
| 8804 | 7 | 42375560 | 42377850 | mNeoVis.2.1.1.chr7\_hap1.g00440.t01 |
| 8805 | 7 | 42377928 | 42379327 | mNeoVis.2.1.1.chr7\_hap1.g00441.t01 |
| 8806 | 7 | 42380291 | 42385565 | mNeoVis.2.1.1.chr7\_hap1.g00442.t01 |
| 8807 | 7 | 42385622 | 42388237 | mNeoVis.2.1.1.chr7\_hap1.g00443.t01 |
| 8808 | 7 | 42390586 | 42397630 | mNeoVis.2.1.1.chr7\_hap1.g00444.t01 |
| 8809 | 7 | 42401277 | 42413618 | mNeoVis.2.1.1.chr7\_hap1.g00445.t01 |
| 8810 | 7 | 42413664 | 42421450 | mNeoVis.2.1.1.chr7\_hap1.g00446.t01 |
| 8811 | 7 | 42441358 | 42469444 | mNeoVis.2.1.1.chr7\_hap1.g00447.t01 |
| 8812 | 7 | 42472932 | 42481036 | mNeoVis.2.1.1.chr7\_hap1.g00448.t01 |
| 8813 | 7 | 42482232 | 42491123 | mNeoVis.2.1.1.chr7\_hap1.g00449.t01 |
| 8814 | 7 | 44096222 | 44119907 | mNeoVis.2.1.1.chr7\_hap1.g00494.t01 |
| 8815 | 7 | 44144430 | 44164401 | mNeoVis.2.1.1.chr7\_hap1.g00495.t01 |
| 8816 | 7 | 44164665 | 44192581 | mNeoVis.2.1.1.chr7\_hap1.g00496.t01 |
| 8817 | 7 | 44181675 | 44197789 | mNeoVis.2.1.1.chr7\_hap1.g00497.t01 |
| 8818 | 7 | 44197945 | 44435546 | mNeoVis.2.1.1.chr7\_hap1.g00498.t01 |
| 8819 | 7 | 44437814 | 44446569 | mNeoVis.2.1.1.chr7\_hap1.g00499.t01 |
| 8820 | 7 | 44465275 | 44471796 | mNeoVis.2.1.1.chr7\_hap1.g00500.t01 |
| 8821 | 7 | 44479551 | 44511360 | mNeoVis.2.1.1.chr7\_hap1.g00501.t01 |
| 8822 | 7 | 44519090 | 44521761 | mNeoVis.2.1.1.chr7\_hap1.g00502.t01 |
| 8823 | 7 | 44521906 | 44529026 | mNeoVis.2.1.1.chr7\_hap1.g00503.t01 |
| 8824 | 7 | 44532380 | 44543870 | mNeoVis.2.1.1.chr7\_hap1.g00504.t01 |
| 8825 | 7 | 44532380 | 44543870 | mNeoVis.2.1.1.chr7\_hap1.g00504.t01 |
| 8826 | 7 | 44545181 | 44571303 | mNeoVis.2.1.1.chr7\_hap1.g00505.t01 |
| 8827 | 7 | 44545181 | 44571303 | mNeoVis.2.1.1.chr7\_hap1.g00505.t01 |
| 8828 | 7 | 44577724 | 44584502 | mNeoVis.2.1.1.chr7\_hap1.g00506.t01 |
| 8829 | 7 | 44577724 | 44584502 | mNeoVis.2.1.1.chr7\_hap1.g00506.t01 |
| 8830 | 7 | 44585360 | 44589394 | mNeoVis.2.1.1.chr7\_hap1.g00507.t01 |
| 8831 | 7 | 44585360 | 44589394 | mNeoVis.2.1.1.chr7\_hap1.g00507.t01 |
| 8832 | 7 | 44591496 | 44600092 | mNeoVis.2.1.1.chr7\_hap1.g00508.t01 |
| 8833 | 7 | 44591496 | 44600092 | mNeoVis.2.1.1.chr7\_hap1.g00508.t01 |
| 8834 | 7 | 44600935 | 44604729 | mNeoVis.2.1.1.chr7\_hap1.g00509.t01 |
| 8835 | 7 | 44600935 | 44604729 | mNeoVis.2.1.1.chr7\_hap1.g00509.t01 |
| 8836 | 7 | 44605772 | 44618413 | mNeoVis.2.1.1.chr7\_hap1.g00510.t01 |
| 8837 | 7 | 44605772 | 44618413 | mNeoVis.2.1.1.chr7\_hap1.g00510.t01 |
| 8838 | 7 | 44624031 | 44734812 | mNeoVis.2.1.1.chr7\_hap1.g00511.t01 |
| 8839 | 7 | 44734911 | 44750752 | mNeoVis.2.1.1.chr7\_hap1.g00512.t01 |
| 8840 | 7 | 44751670 | 44764950 | mNeoVis.2.1.1.chr7\_hap1.g00513.t01 |
| 8841 | 7 | 44769879 | 44838653 | mNeoVis.2.1.1.chr7\_hap1.g00514.t01 |
| 8842 | 7 | 44838316 | 44849581 | mNeoVis.2.1.1.chr7\_hap1.g00515.t01 |
| 8843 | 7 | 44867354 | 44869363 | mNeoVis.2.1.1.chr7\_hap1.g00516.t01 |
| 8844 | 7 | 44878868 | 44880953 | mNeoVis.2.1.1.chr7\_hap1.g00517.t01 |
| 8845 | 7 | 44886106 | 44886578 | mNeoVis.2.1.1.chr7\_hap1.g00518.t01 |
| 8846 | 7 | 44887714 | 44896714 | mNeoVis.2.1.1.chr7\_hap1.g00519.t01 |
| 8847 | 7 | 44898000 | 44908619 | mNeoVis.2.1.1.chr7\_hap1.g00520.t01 |
| 8848 | 7 | 44911855 | 44924150 | mNeoVis.2.1.1.chr7\_hap1.g00521.t01 |
| 8849 | 7 | 44944571 | 44951803 | mNeoVis.2.1.1.chr7\_hap1.g00522.t01 |
| 8850 | 7 | 44951899 | 44968866 | mNeoVis.2.1.1.chr7\_hap1.g00523.t01 |
| 8851 | 7 | 44968914 | 44978444 | mNeoVis.2.1.1.chr7\_hap1.g00524.t01 |
| 8852 | 7 | 44978612 | 44982516 | mNeoVis.2.1.1.chr7\_hap1.g00525.t01 |
| 8853 | 7 | 44982671 | 44992984 | mNeoVis.2.1.1.chr7\_hap1.g00526.t01 |
| 8854 | 7 | 44992105 | 44995897 | mNeoVis.2.1.1.chr7\_hap1.g00527.t01 |
| 8855 | 7 | 44996037 | 45025313 | mNeoVis.2.1.1.chr7\_hap1.g00528.t01 |
| 8856 | 7 | 45055203 | 45062294 | mNeoVis.2.1.1.chr7\_hap1.g00529.t01 |
| 8857 | 7 | 45094796 | 45111500 | mNeoVis.2.1.1.chr7\_hap1.g00530.t01 |
| 8858 | 7 | 45121560 | 45163843 | mNeoVis.2.1.1.chr7\_hap1.g00531.t01 |
| 8859 | 7 | 45176516 | 45179615 | mNeoVis.2.1.1.chr7\_hap1.g00532.t01 |
| 8860 | 7 | 45180743 | 45183809 | mNeoVis.2.1.1.chr7\_hap1.g00533.t01 |
| 8861 | 7 | 45203556 | 45207637 | mNeoVis.2.1.1.chr7\_hap1.g00534.t01 |
| 8862 | 7 | 45207351 | 45209322 | mNeoVis.2.1.1.chr7\_hap1.g00535.t01 |
| 8863 | 7 | 45220116 | 45222537 | mNeoVis.2.1.1.chr7\_hap1.g00536.t01 |
| 8864 | 7 | 45256712 | 45258652 | mNeoVis.2.1.1.chr7\_hap1.g00537.t01 |
| 8865 | 7 | 45264281 | 45290449 | mNeoVis.2.1.1.chr7\_hap1.g00538.t01 |
| 8866 | 7 | 45282356 | 45290883 | mNeoVis.2.1.1.chr7\_hap1.g00539.t01 |
| 8867 | 7 | 45305487 | 45330366 | mNeoVis.2.1.1.chr7\_hap1.g00540.t01 |
| 8868 | 7 | 45330541 | 45335084 | mNeoVis.2.1.1.chr7\_hap1.g00541.t01 |
| 8869 | 7 | 45335580 | 45339423 | mNeoVis.2.1.1.chr7\_hap1.g00542.t01 |
| 8870 | 7 | 45344801 | 45347439 | mNeoVis.2.1.1.chr7\_hap1.g00543.t01 |
| 8871 | 7 | 47852052 | 47866694 | mNeoVis.2.1.1.chr7\_hap1.g00642.t01 |
| 8872 | 7 | 47871112 | 47878235 | mNeoVis.2.1.1.chr7\_hap1.g00643.t01 |
| 8873 | 7 | 47888828 | 47907611 | mNeoVis.2.1.1.chr7\_hap1.g00644.t01 |
| 8874 | 7 | 47921675 | 47931885 | mNeoVis.2.1.1.chr7\_hap1.g00645.t01 |
| 8875 | 7 | 47943868 | 47969538 | mNeoVis.2.1.1.chr7\_hap1.g00646.t01 |
| 8876 | 7 | 47998624 | 48015315 | mNeoVis.2.1.1.chr7\_hap1.g00647.t01 |
| 8877 | 7 | 47998624 | 48015315 | mNeoVis.2.1.1.chr7\_hap1.g00647.t01 |
| 8878 | 7 | 48029001 | 48047707 | mNeoVis.2.1.1.chr7\_hap1.g00648.t01 |
| 8879 | 7 | 48029001 | 48047707 | mNeoVis.2.1.1.chr7\_hap1.g00648.t01 |
| 8880 | 7 | 48050999 | 48054757 | mNeoVis.2.1.1.chr7\_hap1.g00649.t01 |
| 8881 | 7 | 48050999 | 48054757 | mNeoVis.2.1.1.chr7\_hap1.g00649.t01 |
| 8882 | 7 | 48055333 | 48084676 | mNeoVis.2.1.1.chr7\_hap1.g00650.t01 |
| 8883 | 7 | 48055333 | 48084676 | mNeoVis.2.1.1.chr7\_hap1.g00650.t01 |
| 8884 | 7 | 48083401 | 48107431 | mNeoVis.2.1.1.chr7\_hap1.g00651.t01 |
| 8885 | 7 | 48083401 | 48107431 | mNeoVis.2.1.1.chr7\_hap1.g00651.t01 |
| 8886 | 7 | 48275697 | 48322011 | mNeoVis.2.1.1.chr7\_hap1.g00652.t01 |
| 8887 | 7 | 48275697 | 48322011 | mNeoVis.2.1.1.chr7\_hap1.g00652.t01 |
| 8888 | 7 | 48338424 | 48354372 | mNeoVis.2.1.1.chr7\_hap1.g00653.t01 |
| 8889 | 7 | 48338424 | 48354372 | mNeoVis.2.1.1.chr7\_hap1.g00653.t01 |
| 8890 | 7 | 48368300 | 48387391 | mNeoVis.2.1.1.chr7\_hap1.g00654.t01 |
| 8891 | 7 | 48368300 | 48387391 | mNeoVis.2.1.1.chr7\_hap1.g00654.t01 |
| 8892 | 7 | 48406268 | 48407194 | mNeoVis.2.1.1.chr7\_hap1.g00655.t01 |
| 8893 | 7 | 48406268 | 48407194 | mNeoVis.2.1.1.chr7\_hap1.g00655.t01 |
| 8894 | 7 | 48441159 | 48442148 | mNeoVis.2.1.1.chr7\_hap1.g00656.t01 |
| 8895 | 7 | 48441159 | 48442148 | mNeoVis.2.1.1.chr7\_hap1.g00656.t01 |
| 8896 | 7 | 48450604 | 48451569 | mNeoVis.2.1.1.chr7\_hap1.g00657.t01 |
| 8897 | 7 | 48450604 | 48451569 | mNeoVis.2.1.1.chr7\_hap1.g00657.t01 |
| 8898 | 7 | 48461146 | 48462123 | mNeoVis.2.1.1.chr7\_hap1.g00658.t01 |
| 8899 | 7 | 48461146 | 48462123 | mNeoVis.2.1.1.chr7\_hap1.g00658.t01 |
| 8900 | 7 | 48472011 | 48472956 | mNeoVis.2.1.1.chr7\_hap1.g00659.t01 |
| 8901 | 7 | 48601886 | 48604022 | mNeoVis.2.1.1.chr7\_hap1.g00662.t01 |
| 8902 | 7 | 48601886 | 48604022 | mNeoVis.2.1.1.chr7\_hap1.g00662.t01 |
| 8903 | 7 | 48610167 | 48616045 | mNeoVis.2.1.1.chr7\_hap1.g00663.t01 |
| 8904 | 7 | 48610167 | 48616045 | mNeoVis.2.1.1.chr7\_hap1.g00663.t01 |
| 8905 | 7 | 48619437 | 48620536 | mNeoVis.2.1.1.chr7\_hap1.g00664.t01 |
| 8906 | 7 | 48619437 | 48620536 | mNeoVis.2.1.1.chr7\_hap1.g00664.t01 |
| 8907 | 7 | 48648004 | 48666214 | mNeoVis.2.1.1.chr7\_hap1.g00665.t01 |
| 8908 | 7 | 48648004 | 48666214 | mNeoVis.2.1.1.chr7\_hap1.g00665.t01 |
| 8909 | 7 | 48672057 | 48687723 | mNeoVis.2.1.1.chr7\_hap1.g00666.t01 |
| 8910 | 7 | 48672057 | 48687723 | mNeoVis.2.1.1.chr7\_hap1.g00666.t01 |
| 8911 | 7 | 48692004 | 48713549 | mNeoVis.2.1.1.chr7\_hap1.g00667.t01 |
| 8912 | 7 | 48692004 | 48713549 | mNeoVis.2.1.1.chr7\_hap1.g00667.t01 |
| 8913 | 7 | 48719275 | 48776247 | mNeoVis.2.1.1.chr7\_hap1.g00668.t01 |
| 8914 | 7 | 48719275 | 48776247 | mNeoVis.2.1.1.chr7\_hap1.g00668.t01 |
| 8915 | 7 | 48787705 | 48801552 | mNeoVis.2.1.1.chr7\_hap1.g00669.t01 |
| 8916 | 7 | 48787705 | 48801552 | mNeoVis.2.1.1.chr7\_hap1.g00669.t01 |
| 8917 | 7 | 48845479 | 48872188 | mNeoVis.2.1.1.chr7\_hap1.g00670.t01 |
| 8918 | 7 | 48845479 | 48872188 | mNeoVis.2.1.1.chr7\_hap1.g00670.t01 |
| 8919 | 7 | 48897320 | 48905709 | mNeoVis.2.1.1.chr7\_hap1.g00671.t01 |
| 8920 | 7 | 48897320 | 48905709 | mNeoVis.2.1.1.chr7\_hap1.g00671.t01 |
| 8921 | 7 | 48926992 | 48931543 | mNeoVis.2.1.1.chr7\_hap1.g00672.t01 |
| 8922 | 7 | 48926992 | 48931543 | mNeoVis.2.1.1.chr7\_hap1.g00672.t01 |
| 8923 | 7 | 48942089 | 48956134 | mNeoVis.2.1.1.chr7\_hap1.g00673.t01 |
| 8924 | 7 | 48942089 | 48956134 | mNeoVis.2.1.1.chr7\_hap1.g00673.t01 |
| 8925 | 7 | 48958500 | 48974694 | mNeoVis.2.1.1.chr7\_hap1.g00674.t01 |
| 8926 | 7 | 48958500 | 48974694 | mNeoVis.2.1.1.chr7\_hap1.g00674.t01 |
| 8927 | 7 | 48987270 | 49009266 | mNeoVis.2.1.1.chr7\_hap1.g00675.t01 |
| 8928 | 7 | 48987270 | 49009266 | mNeoVis.2.1.1.chr7\_hap1.g00675.t01 |
| 8929 | 7 | 49020812 | 49049035 | mNeoVis.2.1.1.chr7\_hap1.g00676.t01 |
| 8930 | 7 | 49020812 | 49049035 | mNeoVis.2.1.1.chr7\_hap1.g00676.t01 |
| 8931 | 7 | 49057598 | 49085128 | mNeoVis.2.1.1.chr7\_hap1.g00677.t01 |
| 8932 | 7 | 49057598 | 49085128 | mNeoVis.2.1.1.chr7\_hap1.g00677.t01 |
| 8933 | 7 | 49132929 | 49144918 | mNeoVis.2.1.1.chr7\_hap1.g00678.t01 |
| 8934 | 7 | 49132929 | 49144918 | mNeoVis.2.1.1.chr7\_hap1.g00678.t01 |
| 8935 | 7 | 54810367 | 54824950 | mNeoVis.2.1.1.chr7\_hap1.g00850.t01 |
| 8936 | 7 | 54829401 | 54889958 | mNeoVis.2.1.1.chr7\_hap1.g00851.t01 |
| 8937 | 7 | 54866047 | 54910291 | mNeoVis.2.1.1.chr7\_hap1.g00852.t01 |
| 8938 | 7 | 54891657 | 54893894 | mNeoVis.2.1.1.chr7\_hap1.g00853.t01 |
| 8939 | 7 | 54894071 | 54898126 | mNeoVis.2.1.1.chr7\_hap1.g00854.t01 |
| 8940 | 7 | 54915636 | 54922460 | mNeoVis.2.1.1.chr7\_hap1.g00855.t01 |
| 8941 | 7 | 54922659 | 54927710 | mNeoVis.2.1.1.chr7\_hap1.g00856.t01 |
| 8942 | 7 | 54928182 | 54952662 | mNeoVis.2.1.1.chr7\_hap1.g00857.t01 |
| 8943 | 7 | 54954339 | 54960441 | mNeoVis.2.1.1.chr7\_hap1.g00858.t01 |
| 8944 | 7 | 54962187 | 54985002 | mNeoVis.2.1.1.chr7\_hap1.g00859.t01 |
| 8945 | 7 | 54986322 | 54992877 | mNeoVis.2.1.1.chr7\_hap1.g00860.t01 |
| 8946 | 7 | 55004279 | 55014045 | mNeoVis.2.1.1.chr7\_hap1.g00861.t01 |
| 8947 | 7 | 55017997 | 55034341 | mNeoVis.2.1.1.chr7\_hap1.g00862.t01 |
| 8948 | 7 | 55034485 | 55115398 | mNeoVis.2.1.1.chr7\_hap1.g00863.t01 |
| 8949 | 7 | 55151494 | 55162141 | mNeoVis.2.1.1.chr7\_hap1.g00864.t01 |
| 8950 | 7 | 55168780 | 55181449 | mNeoVis.2.1.1.chr7\_hap1.g00865.t01 |
| 8951 | 7 | 55194298 | 55208368 | mNeoVis.2.1.1.chr7\_hap1.g00866.t01 |
| 8952 | 7 | 55207301 | 55238946 | mNeoVis.2.1.1.chr7\_hap1.g00867.t01 |
| 8953 | 7 | 55245204 | 55252610 | mNeoVis.2.1.1.chr7\_hap1.g00868.t01 |
| 8954 | 7 | 55264355 | 55268660 | mNeoVis.2.1.1.chr7\_hap1.g00869.t01 |
| 8955 | 7 | 55268461 | 55292491 | mNeoVis.2.1.1.chr7\_hap1.g00870.t01 |
| 8956 | 7 | 55268463 | 55306007 | mNeoVis.2.1.1.chr7\_hap1.g00871.t01 |
| 8957 | 7 | 55306274 | 55317398 | mNeoVis.2.1.1.chr7\_hap1.g00872.t01 |
| 8958 | 7 | 55318289 | 55326132 | mNeoVis.2.1.1.chr7\_hap1.g00873.t01 |
| 8959 | 7 | 55326592 | 55332856 | mNeoVis.2.1.1.chr7\_hap1.g00874.t01 |
| 8960 | 7 | 57531291 | 57541368 | mNeoVis.2.1.1.chr7\_hap1.g00982.t01 |
| 8961 | 7 | 57583109 | 57594932 | mNeoVis.2.1.1.chr7\_hap1.g00983.t01 |
| 8962 | 7 | 57598864 | 57608935 | mNeoVis.2.1.1.chr7\_hap1.g00984.t01 |
| 8963 | 7 | 57608567 | 57633808 | mNeoVis.2.1.1.chr7\_hap1.g00985.t01 |
| 8964 | 7 | 57636477 | 57668941 | mNeoVis.2.1.1.chr7\_hap1.g00986.t01 |
| 8965 | 7 | 57675638 | 57685050 | mNeoVis.2.1.1.chr7\_hap1.g00987.t01 |
| 8966 | 7 | 57693331 | 57694953 | mNeoVis.2.1.1.chr7\_hap1.g00988.t01 |
| 8967 | 7 | 57702135 | 57705617 | mNeoVis.2.1.1.chr7\_hap1.g00989.t01 |
| 8968 | 7 | 57710062 | 57719475 | mNeoVis.2.1.1.chr7\_hap1.g00990.t01 |
| 8969 | 7 | 57758766 | 57762714 | mNeoVis.2.1.1.chr7\_hap1.g00991.t01 |
| 8970 | 7 | 57765371 | 57770679 | mNeoVis.2.1.1.chr7\_hap1.g00992.t01 |
| 8971 | 7 | 57781486 | 57785750 | mNeoVis.2.1.1.chr7\_hap1.g00993.t01 |
| 8972 | 7 | 57807525 | 57812025 | mNeoVis.2.1.1.chr7\_hap1.g00994.t01 |
| 8973 | 7 | 57847506 | 57868265 | mNeoVis.2.1.1.chr7\_hap1.g00995.t01 |
| 8974 | 7 | 57869417 | 57893588 | mNeoVis.2.1.1.chr7\_hap1.g00996.t01 |
| 8975 | 7 | 57944320 | 57949400 | mNeoVis.2.1.1.chr7\_hap1.g00997.t01 |
| 8976 | 7 | 57955664 | 58005169 | mNeoVis.2.1.1.chr7\_hap1.g00998.t01 |
| 8977 | 7 | 57972653 | 57983420 | mNeoVis.2.1.1.chr7\_hap1.g00999.t01 |
| 8978 | 7 | 58020129 | 58043983 | mNeoVis.2.1.1.chr7\_hap1.g01000.t01 |
| 8979 | 7 | 58073961 | 58075462 | mNeoVis.2.1.1.chr7\_hap1.g01001.t01 |
| 8980 | 7 | 58264477 | 58271983 | mNeoVis.2.1.1.chr7\_hap1.g01014.t01 |
| 8981 | 7 | 58280113 | 58304957 | mNeoVis.2.1.1.chr7\_hap1.g01015.t01 |
| 8982 | 7 | 58321002 | 58331744 | mNeoVis.2.1.1.chr7\_hap1.g01016.t01 |
| 8983 | 7 | 58333486 | 58379746 | mNeoVis.2.1.1.chr7\_hap1.g01017.t01 |
| 8984 | 7 | 58406023 | 58418917 | mNeoVis.2.1.1.chr7\_hap1.g01018.t01 |
| 8985 | 7 | 58423983 | 58425626 | mNeoVis.2.1.1.chr7\_hap1.g01019.t01 |
| 8986 | 7 | 58426727 | 58436554 | mNeoVis.2.1.1.chr7\_hap1.g01020.t01 |
| 8987 | 7 | 58442561 | 58449517 | mNeoVis.2.1.1.chr7\_hap1.g01021.t01 |
| 8988 | 7 | 58486597 | 58500833 | mNeoVis.2.1.1.chr7\_hap1.g01022.t01 |
| 8989 | 7 | 58500921 | 58503294 | mNeoVis.2.1.1.chr7\_hap1.g01023.t01 |
| 8990 | 7 | 58503595 | 58520849 | mNeoVis.2.1.1.chr7\_hap1.g01024.t01 |
| 8991 | 7 | 58503595 | 58520849 | mNeoVis.2.1.1.chr7\_hap1.g01024.t01 |
| 8992 | 7 | 58527956 | 58543035 | mNeoVis.2.1.1.chr7\_hap1.g01025.t01 |
| 8993 | 7 | 58527956 | 58543035 | mNeoVis.2.1.1.chr7\_hap1.g01025.t01 |
| 8994 | 7 | 58542519 | 58550763 | mNeoVis.2.1.1.chr7\_hap1.g01026.t01 |
| 8995 | 7 | 58542519 | 58550763 | mNeoVis.2.1.1.chr7\_hap1.g01026.t01 |
| 8996 | 7 | 58559744 | 58569170 | mNeoVis.2.1.1.chr7\_hap1.g01027.t01 |
| 8997 | 7 | 58559744 | 58569170 | mNeoVis.2.1.1.chr7\_hap1.g01027.t01 |
| 8998 | 7 | 58572040 | 58596258 | mNeoVis.2.1.1.chr7\_hap1.g01028.t01 |
| 8999 | 7 | 58572040 | 58596258 | mNeoVis.2.1.1.chr7\_hap1.g01028.t01 |
| 9000 | 7 | 58605466 | 58626603 | mNeoVis.2.1.1.chr7\_hap1.g01029.t01 |
| 9001 | 7 | 58605466 | 58626603 | mNeoVis.2.1.1.chr7\_hap1.g01029.t01 |
| 9002 | 7 | 58673030 | 58676060 | mNeoVis.2.1.1.chr7\_hap1.g01030.t01 |
| 9003 | 7 | 58673030 | 58676060 | mNeoVis.2.1.1.chr7\_hap1.g01030.t01 |
| 9004 | 7 | 58676177 | 58686470 | mNeoVis.2.1.1.chr7\_hap1.g01031.t01 |
| 9005 | 7 | 58676177 | 58686470 | mNeoVis.2.1.1.chr7\_hap1.g01031.t01 |
| 9006 | 7 | 58686541 | 58696366 | mNeoVis.2.1.1.chr7\_hap1.g01032.t01 |
| 9007 | 7 | 58686541 | 58696366 | mNeoVis.2.1.1.chr7\_hap1.g01032.t01 |
| 9008 | 7 | 58697956 | 58737735 | mNeoVis.2.1.1.chr7\_hap1.g01033.t01 |
| 9009 | 7 | 58697956 | 58737735 | mNeoVis.2.1.1.chr7\_hap1.g01033.t01 |
| 9010 | 7 | 58738045 | 58741877 | mNeoVis.2.1.1.chr7\_hap1.g01034.t01 |
| 9011 | 7 | 58738045 | 58741877 | mNeoVis.2.1.1.chr7\_hap1.g01034.t01 |
| 9012 | 7 | 58752584 | 58756390 | mNeoVis.2.1.1.chr7\_hap1.g01035.t01 |
| 9013 | 7 | 58752584 | 58756390 | mNeoVis.2.1.1.chr7\_hap1.g01035.t01 |
| 9014 | 7 | 58756856 | 58781586 | mNeoVis.2.1.1.chr7\_hap1.g01036.t01 |
| 9015 | 7 | 58756856 | 58781586 | mNeoVis.2.1.1.chr7\_hap1.g01036.t01 |
| 9016 | 7 | 58781764 | 58809101 | mNeoVis.2.1.1.chr7\_hap1.g01037.t01 |
| 9017 | 7 | 58781764 | 58809101 | mNeoVis.2.1.1.chr7\_hap1.g01037.t01 |
| 9018 | 7 | 58813249 | 58841084 | mNeoVis.2.1.1.chr7\_hap1.g01038.t01 |
| 9019 | 7 | 58813249 | 58841084 | mNeoVis.2.1.1.chr7\_hap1.g01038.t01 |
| 9020 | 7 | 58845778 | 58847310 | mNeoVis.2.1.1.chr7\_hap1.g01039.t01 |
| 9021 | 7 | 58845778 | 58847310 | mNeoVis.2.1.1.chr7\_hap1.g01039.t01 |
| 9022 | 7 | 58847904 | 58849794 | mNeoVis.2.1.1.chr7\_hap1.g01040.t01 |
| 9023 | 7 | 58847904 | 58849794 | mNeoVis.2.1.1.chr7\_hap1.g01040.t01 |
| 9024 | 7 | 58856329 | 58858818 | mNeoVis.2.1.1.chr7\_hap1.g01041.t01 |
| 9025 | 7 | 58856329 | 58858818 | mNeoVis.2.1.1.chr7\_hap1.g01041.t01 |
| 9026 | 7 | 58861569 | 58864414 | mNeoVis.2.1.1.chr7\_hap1.g01042.t01 |
| 9027 | 7 | 58861569 | 58864414 | mNeoVis.2.1.1.chr7\_hap1.g01042.t01 |
| 9028 | 7 | 58866236 | 58876515 | mNeoVis.2.1.1.chr7\_hap1.g01043.t01 |
| 9029 | 7 | 58866236 | 58876515 | mNeoVis.2.1.1.chr7\_hap1.g01043.t01 |
| 9030 | 7 | 58877912 | 58904051 | mNeoVis.2.1.1.chr7\_hap1.g01044.t01 |
| 9031 | 7 | 58877912 | 58904051 | mNeoVis.2.1.1.chr7\_hap1.g01044.t01 |
| 9032 | 7 | 58909996 | 58919421 | mNeoVis.2.1.1.chr7\_hap1.g01045.t01 |
| 9033 | 7 | 58909996 | 58919421 | mNeoVis.2.1.1.chr7\_hap1.g01045.t01 |
| 9034 | 7 | 58922157 | 58940555 | mNeoVis.2.1.1.chr7\_hap1.g01046.t01 |
| 9035 | 7 | 58922157 | 58940555 | mNeoVis.2.1.1.chr7\_hap1.g01046.t01 |
| 9036 | 7 | 58951300 | 58961026 | mNeoVis.2.1.1.chr7\_hap1.g01047.t01 |
| 9037 | 7 | 58951300 | 58961026 | mNeoVis.2.1.1.chr7\_hap1.g01047.t01 |
| 9038 | 7 | 58984104 | 58991457 | mNeoVis.2.1.1.chr7\_hap1.g01048.t01 |
| 9039 | 7 | 58984104 | 58991457 | mNeoVis.2.1.1.chr7\_hap1.g01048.t01 |
| 9040 | 7 | 59030085 | 59045106 | mNeoVis.2.1.1.chr7\_hap1.g01049.t01 |
| 9041 | 7 | 59030085 | 59045106 | mNeoVis.2.1.1.chr7\_hap1.g01049.t01 |
| 9042 | 7 | 59102216 | 59155563 | mNeoVis.2.1.1.chr7\_hap1.g01050.t01 |
| 9043 | 7 | 59157883 | 59172399 | mNeoVis.2.1.1.chr7\_hap1.g01051.t01 |
| 9044 | 7 | 59197038 | 59215923 | mNeoVis.2.1.1.chr7\_hap1.g01052.t01 |
| 9045 | 7 | 59220271 | 59242179 | mNeoVis.2.1.1.chr7\_hap1.g01053.t01 |
| 9046 | 7 | 59707952 | 59723728 | mNeoVis.2.1.1.chr7\_hap1.g01066.t01 |
| 9047 | 7 | 59733771 | 59753120 | mNeoVis.2.1.1.chr7\_hap1.g01067.t01 |
| 9048 | 7 | 59753126 | 59759317 | mNeoVis.2.1.1.chr7\_hap1.g01068.t01 |
| 9049 | 7 | 59759704 | 59762821 | mNeoVis.2.1.1.chr7\_hap1.g01069.t01 |
| 9050 | 7 | 59768905 | 59789435 | mNeoVis.2.1.1.chr7\_hap1.g01070.t01 |
| 9051 | 7 | 59796559 | 59809029 | mNeoVis.2.1.1.chr7\_hap1.g01071.t01 |
| 9052 | 7 | 59816574 | 59830398 | mNeoVis.2.1.1.chr7\_hap1.g01072.t01 |
| 9053 | 7 | 59835622 | 59838317 | mNeoVis.2.1.1.chr7\_hap1.g01073.t01 |
| 9054 | 7 | 59835743 | 59841661 | mNeoVis.2.1.1.chr7\_hap1.g01074.t01 |
| 9055 | 7 | 59838557 | 59858896 | mNeoVis.2.1.1.chr7\_hap1.g01075.t01 |
| 9056 | 7 | 59847263 | 59851277 | mNeoVis.2.1.1.chr7\_hap1.g01076.t01 |
| 9057 | 7 | 59861259 | 59867184 | mNeoVis.2.1.1.chr7\_hap1.g01077.t01 |
| 9058 | 7 | 59866603 | 59917340 | mNeoVis.2.1.1.chr7\_hap1.g01078.t01 |
| 9059 | 7 | 59891348 | 59896650 | mNeoVis.2.1.1.chr7\_hap1.g01079.t01 |
| 9060 | 7 | 59918120 | 59938695 | mNeoVis.2.1.1.chr7\_hap1.g01080.t01 |
| 9061 | 7 | 59943572 | 59947727 | mNeoVis.2.1.1.chr7\_hap1.g01081.t01 |
| 9062 | 7 | 59955545 | 60012642 | mNeoVis.2.1.1.chr7\_hap1.g01082.t01 |
| 9063 | 7 | 59978498 | 59981813 | mNeoVis.2.1.1.chr7\_hap1.g01083.t01 |
| 9064 | 7 | 60032988 | 60044267 | mNeoVis.2.1.1.chr7\_hap1.g01084.t01 |
| 9065 | 7 | 60049881 | 60056095 | mNeoVis.2.1.1.chr7\_hap1.g01085.t01 |
| 9066 | 7 | 60098830 | 60106484 | mNeoVis.2.1.1.chr7\_hap1.g01086.t01 |
| 9067 | 7 | 60113290 | 60119227 | mNeoVis.2.1.1.chr7\_hap1.g01087.t01 |
| 9068 | 7 | 60120335 | 60124774 | mNeoVis.2.1.1.chr7\_hap1.g01088.t01 |
| 9069 | 7 | 60129101 | 60138246 | mNeoVis.2.1.1.chr7\_hap1.g01089.t01 |
| 9070 | 7 | 60141702 | 60145216 | mNeoVis.2.1.1.chr7\_hap1.g01090.t01 |
| 9071 | 7 | 60146328 | 60164361 | mNeoVis.2.1.1.chr7\_hap1.g01091.t01 |
| 9072 | 7 | 60164639 | 60182350 | mNeoVis.2.1.1.chr7\_hap1.g01092.t01 |
| 9073 | 7 | 60203475 | 60206464 | mNeoVis.2.1.1.chr7\_hap1.g01093.t01 |
| 9074 | 7 | 60209643 | 60230807 | mNeoVis.2.1.1.chr7\_hap1.g01094.t01 |
| 9075 | 7 | 60235098 | 60290652 | mNeoVis.2.1.1.chr7\_hap1.g01095.t01 |
| 9076 | 7 | 60290652 | 60296387 | mNeoVis.2.1.1.chr7\_hap1.g01096.t01 |
| 9077 | 7 | 60305463 | 60337833 | mNeoVis.2.1.1.chr7\_hap1.g01097.t01 |
| 9078 | 7 | 97798727 | 97849299 | mNeoVis.2.1.1.chr7\_hap1.g01130.t01 |
| 9079 | 7 | 97853268 | 97866242 | mNeoVis.2.1.1.chr7\_hap1.g01131.t01 |
| 9080 | 7 | 97862591 | 97886613 | mNeoVis.2.1.1.chr7\_hap1.g01132.t01 |
| 9081 | 7 | 97894066 | 97982671 | mNeoVis.2.1.1.chr7\_hap1.g01133.t01 |
| 9082 | 7 | 98005638 | 98034264 | mNeoVis.2.1.1.chr7\_hap1.g01134.t01 |
| 9083 | 7 | 98076422 | 98125699 | mNeoVis.2.1.1.chr7\_hap1.g01135.t01 |
| 9084 | 7 | 98165982 | 98170581 | mNeoVis.2.1.1.chr7\_hap1.g01136.t01 |
| 9085 | 7 | 98180231 | 98180677 | mNeoVis.2.1.1.chr7\_hap1.g01137.t01 |
| 9086 | 7 | 99629864 | 99651863 | mNeoVis.2.1.1.chr7\_hap1.g01143.t01 |
| 9087 | 7 | 99651973 | 99690066 | mNeoVis.2.1.1.chr7\_hap1.g01144.t01 |
| 9088 | 7 | 99651973 | 99690066 | mNeoVis.2.1.1.chr7\_hap1.g01144.t01 |
| 9089 | 7 | 99724113 | 99730455 | mNeoVis.2.1.1.chr7\_hap1.g01145.t01 |
| 9090 | 7 | 99724113 | 99730455 | mNeoVis.2.1.1.chr7\_hap1.g01145.t01 |
| 9091 | 7 | 100103872 | 100173189 | mNeoVis.2.1.1.chr7\_hap1.g01147.t01 |
| 9092 | 7 | 105110877 | 105418846 | mNeoVis.2.1.1.chr7\_hap1.g01192.t01 |
| 9093 | 7 | 105168474 | 105170484 | mNeoVis.2.1.1.chr7\_hap1.g01193.t01 |
| 9094 | 7 | 105168474 | 105170484 | mNeoVis.2.1.1.chr7\_hap1.g01193.t01 |
| 9095 | 7 | 105446578 | 105488905 | mNeoVis.2.1.1.chr7\_hap1.g01194.t01 |
| 9096 | 7 | 105446578 | 105488905 | mNeoVis.2.1.1.chr7\_hap1.g01194.t01 |
| 9097 | 7 | 105500587 | 105515373 | mNeoVis.2.1.1.chr7\_hap1.g01195.t01 |
| 9098 | 7 | 105500587 | 105515373 | mNeoVis.2.1.1.chr7\_hap1.g01195.t01 |
| 9099 | 7 | 105517551 | 105577455 | mNeoVis.2.1.1.chr7\_hap1.g01196.t01 |
| 9100 | 7 | 105517551 | 105577455 | mNeoVis.2.1.1.chr7\_hap1.g01196.t01 |
| 9101 | 7 | 105722385 | 105737686 | mNeoVis.2.1.1.chr7\_hap1.g01197.t01 |
| 9102 | 7 | 105722385 | 105737686 | mNeoVis.2.1.1.chr7\_hap1.g01197.t01 |
| 9103 | 7 | 105755646 | 105788674 | mNeoVis.2.1.1.chr7\_hap1.g01198.t01 |
| 9104 | 7 | 105755646 | 105788674 | mNeoVis.2.1.1.chr7\_hap1.g01198.t01 |
| 9105 | 7 | 105784049 | 105794665 | mNeoVis.2.1.1.chr7\_hap1.g01199.t01 |
| 9106 | 7 | 105784049 | 105794665 | mNeoVis.2.1.1.chr7\_hap1.g01199.t01 |
| 9107 | 7 | 106311746 | 106316803 | mNeoVis.2.1.1.chr7\_hap1.g01206.t01 |
| 9108 | 7 | 106316934 | 106345700 | mNeoVis.2.1.1.chr7\_hap1.g01207.t01 |
| 9109 | 7 | 106345625 | 106348273 | mNeoVis.2.1.1.chr7\_hap1.g01208.t01 |
| 9110 | 7 | 106356348 | 106366239 | mNeoVis.2.1.1.chr7\_hap1.g01209.t01 |
| 9111 | 7 | 106541347 | 106558371 | mNeoVis.2.1.1.chr7\_hap1.g01210.t01 |
| 9112 | 7 | 107066503 | 107391850 | mNeoVis.2.1.1.chr7\_hap1.g01211.t01 |
| 9113 | 7 | 108403828 | 108418539 | mNeoVis.2.1.1.chr7\_hap1.g01213.t01 |
| 9114 | 7 | 108424743 | 108452341 | mNeoVis.2.1.1.chr7\_hap1.g01214.t01 |
| 9115 | 7 | 108454133 | 108462276 | mNeoVis.2.1.1.chr7\_hap1.g01215.t01 |
| 9116 | 7 | 108463523 | 108467960 | mNeoVis.2.1.1.chr7\_hap1.g01216.t01 |
| 9117 | 7 | 108483284 | 108485728 | mNeoVis.2.1.1.chr7\_hap1.g01217.t01 |
| 9118 | 7 | 108490788 | 108493282 | mNeoVis.2.1.1.chr7\_hap1.g01218.t01 |
| 9119 | 7 | 108495540 | 108497639 | mNeoVis.2.1.1.chr7\_hap1.g01219.t01 |
| 9120 | 7 | 108496984 | 108751032 | mNeoVis.2.1.1.chr7\_hap1.g01220.t01 |
| 9121 | 7 | 108795715 | 108821663 | mNeoVis.2.1.1.chr7\_hap1.g01221.t01 |
| 9122 | 7 | 108829706 | 108845391 | mNeoVis.2.1.1.chr7\_hap1.g01222.t01 |
| 9123 | 7 | 108847239 | 108853101 | mNeoVis.2.1.1.chr7\_hap1.g01223.t01 |
| 9124 | 7 | 108853352 | 108878833 | mNeoVis.2.1.1.chr7\_hap1.g01224.t01 |
| 9125 | 7 | 108879049 | 108923732 | mNeoVis.2.1.1.chr7\_hap1.g01225.t01 |
| 9126 | 7 | 108925315 | 108947860 | mNeoVis.2.1.1.chr7\_hap1.g01226.t01 |
| 9127 | 7 | 108955596 | 109017540 | mNeoVis.2.1.1.chr7\_hap1.g01227.t01 |
| 9128 | 7 | 109424273 | 109453992 | mNeoVis.2.1.1.chr7\_hap1.g01229.t01 |
| 9129 | 7 | 109480110 | 109508796 | mNeoVis.2.1.1.chr7\_hap1.g01230.t01 |
| 9130 | 7 | 109545534 | 109557870 | mNeoVis.2.1.1.chr7\_hap1.g01231.t01 |
| 9131 | 7 | 109557922 | 109670804 | mNeoVis.2.1.1.chr7\_hap1.g01232.t01 |
| 9132 | 7 | 109622068 | 109659933 | mNeoVis.2.1.1.chr7\_hap1.g01233.t01 |
| 9133 | 7 | 109680719 | 109700464 | mNeoVis.2.1.1.chr7\_hap1.g01234.t01 |
| 9134 | 7 | 109709929 | 109721903 | mNeoVis.2.1.1.chr7\_hap1.g01235.t01 |
| 9135 | 7 | 109730851 | 109783042 | mNeoVis.2.1.1.chr7\_hap1.g01236.t01 |
| 9136 | 7 | 109783559 | 109796831 | mNeoVis.2.1.1.chr7\_hap1.g01237.t01 |
| 9137 | 7 | 109826765 | 109837896 | mNeoVis.2.1.1.chr7\_hap1.g01238.t01 |
| 9138 | 7 | 109842128 | 109848348 | mNeoVis.2.1.1.chr7\_hap1.g01239.t01 |
| 9139 | 7 | 109849682 | 109860143 | mNeoVis.2.1.1.chr7\_hap1.g01240.t01 |
| 9140 | 7 | 109864075 | 109905914 | mNeoVis.2.1.1.chr7\_hap1.g01241.t01 |
| 9141 | 7 | 109908089 | 109915856 | mNeoVis.2.1.1.chr7\_hap1.g01242.t01 |
| 9142 | 7 | 109943848 | 110022720 | mNeoVis.2.1.1.chr7\_hap1.g01243.t01 |
| 9143 | 7 | 110023255 | 110026725 | mNeoVis.2.1.1.chr7\_hap1.g01244.t01 |
| 9144 | 7 | 110027032 | 110031491 | mNeoVis.2.1.1.chr7\_hap1.g01245.t01 |
| 9145 | 7 | 110040650 | 110055597 | mNeoVis.2.1.1.chr7\_hap1.g01246.t01 |
| 9146 | 7 | 110061519 | 110089004 | mNeoVis.2.1.1.chr7\_hap1.g01247.t01 |
| 9147 | 7 | 110093985 | 110139154 | mNeoVis.2.1.1.chr7\_hap1.g01248.t01 |
| 9148 | 7 | 110139367 | 110155228 | mNeoVis.2.1.1.chr7\_hap1.g01249.t01 |
| 9149 | 7 | 110155688 | 110169095 | mNeoVis.2.1.1.chr7\_hap1.g01250.t01 |
| 9150 | 7 | 110458514 | 110468325 | mNeoVis.2.1.1.chr7\_hap1.g01265.t01 |
| 9151 | 7 | 110471570 | 110484187 | mNeoVis.2.1.1.chr7\_hap1.g01266.t01 |
| 9152 | 7 | 110487226 | 110506552 | mNeoVis.2.1.1.chr7\_hap1.g01267.t01 |
| 9153 | 7 | 110513767 | 110525492 | mNeoVis.2.1.1.chr7\_hap1.g01268.t01 |
| 9154 | 7 | 110526151 | 110530706 | mNeoVis.2.1.1.chr7\_hap1.g01269.t01 |
| 9155 | 7 | 110531492 | 110540778 | mNeoVis.2.1.1.chr7\_hap1.g01270.t01 |
| 9156 | 7 | 110540999 | 110622845 | mNeoVis.2.1.1.chr7\_hap1.g01271.t01 |
| 9157 | 7 | 110623158 | 110630718 | mNeoVis.2.1.1.chr7\_hap1.g01272.t01 |
| 9158 | 7 | 110623158 | 110630718 | mNeoVis.2.1.1.chr7\_hap1.g01272.t01 |
| 9159 | 7 | 110654938 | 110657735 | mNeoVis.2.1.1.chr7\_hap1.g01273.t01 |
| 9160 | 7 | 110654938 | 110657735 | mNeoVis.2.1.1.chr7\_hap1.g01273.t01 |
| 9161 | 7 | 110659405 | 110661881 | mNeoVis.2.1.1.chr7\_hap1.g01274.t01 |
| 9162 | 7 | 110659405 | 110661881 | mNeoVis.2.1.1.chr7\_hap1.g01274.t01 |
| 9163 | 7 | 110661894 | 110666789 | mNeoVis.2.1.1.chr7\_hap1.g01275.t01 |
| 9164 | 7 | 110661894 | 110666789 | mNeoVis.2.1.1.chr7\_hap1.g01275.t01 |
| 9165 | 7 | 110671656 | 110697230 | mNeoVis.2.1.1.chr7\_hap1.g01276.t01 |
| 9166 | 7 | 110671656 | 110697230 | mNeoVis.2.1.1.chr7\_hap1.g01276.t01 |
| 9167 | 7 | 110712181 | 110718205 | mNeoVis.2.1.1.chr7\_hap1.g01277.t01 |
| 9168 | 7 | 110712181 | 110718205 | mNeoVis.2.1.1.chr7\_hap1.g01277.t01 |
| 9169 | 7 | 110927219 | 110989822 | mNeoVis.2.1.1.chr7\_hap1.g01278.t01 |
| 9170 | 7 | 110927219 | 110989822 | mNeoVis.2.1.1.chr7\_hap1.g01278.t01 |
| 9171 | 7 | 111328508 | 111353489 | mNeoVis.2.1.1.chr7\_hap1.g01279.t01 |
| 9172 | 7 | 111328508 | 111353489 | mNeoVis.2.1.1.chr7\_hap1.g01279.t01 |
| 9173 | 7 | 111418312 | 111436042 | mNeoVis.2.1.1.chr7\_hap1.g01280.t01 |
| 9174 | 7 | 111445386 | 111522466 | mNeoVis.2.1.1.chr7\_hap1.g01281.t01 |
| 9175 | 7 | 111445386 | 111522466 | mNeoVis.2.1.1.chr7\_hap1.g01281.t01 |
| 9176 | 7 | 111528860 | 111536753 | mNeoVis.2.1.1.chr7\_hap1.g01282.t01 |
| 9177 | 7 | 111528860 | 111536753 | mNeoVis.2.1.1.chr7\_hap1.g01282.t01 |
| 9178 | 7 | 111539568 | 111682824 | mNeoVis.2.1.1.chr7\_hap1.g01283.t01 |
| 9179 | 7 | 111539568 | 111682824 | mNeoVis.2.1.1.chr7\_hap1.g01283.t01 |
| 9180 | 7 | 112412887 | 112425042 | mNeoVis.2.1.1.chr7\_hap1.g01287.t01 |
| 9181 | 7 | 112412887 | 112425042 | mNeoVis.2.1.1.chr7\_hap1.g01287.t01 |
| 9182 | 7 | 112536733 | 112687288 | mNeoVis.2.1.1.chr7\_hap1.g01288.t01 |
| 9183 | 7 | 112536733 | 112687288 | mNeoVis.2.1.1.chr7\_hap1.g01288.t01 |
| 9184 | 7 | 113467686 | 113477534 | mNeoVis.2.1.1.chr7\_hap1.g01289.t01 |
| 9185 | 7 | 113467686 | 113477534 | mNeoVis.2.1.1.chr7\_hap1.g01289.t01 |
| 9186 | 7 | 113552479 | 113689350 | mNeoVis.2.1.1.chr7\_hap1.g01290.t01 |
| 9187 | 7 | 113552479 | 113689350 | mNeoVis.2.1.1.chr7\_hap1.g01290.t01 |
| 9188 | 7 | 113696160 | 113725545 | mNeoVis.2.1.1.chr7\_hap1.g01291.t01 |
| 9189 | 7 | 113696160 | 113725545 | mNeoVis.2.1.1.chr7\_hap1.g01291.t01 |
| 9190 | 7 | 113739604 | 113794456 | mNeoVis.2.1.1.chr7\_hap1.g01292.t01 |
| 9191 | 7 | 113739604 | 113794456 | mNeoVis.2.1.1.chr7\_hap1.g01292.t01 |
| 9192 | 7 | 113739604 | 113794456 | mNeoVis.2.1.1.chr7\_hap1.g01292.t01 |
| 9193 | 7 | 113807986 | 113841795 | mNeoVis.2.1.1.chr7\_hap1.g01293.t01 |
| 9194 | 7 | 113807986 | 113841795 | mNeoVis.2.1.1.chr7\_hap1.g01293.t01 |
| 9195 | 7 | 113877489 | 113882096 | mNeoVis.2.1.1.chr7\_hap1.g01294.t01 |
| 9196 | 7 | 113877489 | 113882096 | mNeoVis.2.1.1.chr7\_hap1.g01294.t01 |
| 9197 | 7 | 113877489 | 113882096 | mNeoVis.2.1.1.chr7\_hap1.g01294.t01 |
| 9198 | 7 | 113884844 | 113985864 | mNeoVis.2.1.1.chr7\_hap1.g01295.t01 |
| 9199 | 7 | 113884844 | 113985864 | mNeoVis.2.1.1.chr7\_hap1.g01295.t01 |
| 9200 | 7 | 113884844 | 113985864 | mNeoVis.2.1.1.chr7\_hap1.g01295.t01 |
| 9201 | 7 | 114211374 | 114381766 | mNeoVis.2.1.1.chr7\_hap1.g01296.t01 |
| 9202 | 7 | 114211374 | 114381766 | mNeoVis.2.1.1.chr7\_hap1.g01296.t01 |
| 9203 | 7 | 114385779 | 114407574 | mNeoVis.2.1.1.chr7\_hap1.g01297.t01 |
| 9204 | 7 | 114385779 | 114407574 | mNeoVis.2.1.1.chr7\_hap1.g01297.t01 |
| 9205 | 7 | 114477402 | 114494064 | mNeoVis.2.1.1.chr7\_hap1.g01298.t01 |
| 9206 | 7 | 114502578 | 114503513 | mNeoVis.2.1.1.chr7\_hap1.g01299.t01 |
| 9207 | 7 | 114577354 | 114589483 | mNeoVis.2.1.1.chr7\_hap1.g01300.t01 |
| 9208 | 7 | 114589027 | 114590001 | mNeoVis.2.1.1.chr7\_hap1.g01301.t01 |
| 9209 | 7 | 115341132 | 115342151 | mNeoVis.2.1.1.chr7\_hap1.g01338.t01 |
| 9210 | 7 | 115359702 | 115360640 | mNeoVis.2.1.1.chr7\_hap1.g01339.t01 |
| 9211 | 7 | 115390208 | 115391143 | mNeoVis.2.1.1.chr7\_hap1.g01340.t01 |
| 9212 | 7 | 115506696 | 115513469 | mNeoVis.2.1.1.chr7\_hap1.g01341.t01 |
| 9213 | 7 | 115516646 | 115527127 | mNeoVis.2.1.1.chr7\_hap1.g01342.t01 |
| 9214 | 7 | 115529903 | 115570920 | mNeoVis.2.1.1.chr7\_hap1.g01343.t01 |
| 9215 | 7 | 115570945 | 115584379 | mNeoVis.2.1.1.chr7\_hap1.g01344.t01 |
| 9216 | 7 | 115625656 | 115632603 | mNeoVis.2.1.1.chr7\_hap1.g01345.t01 |
| 9217 | 7 | 115632867 | 115637917 | mNeoVis.2.1.1.chr7\_hap1.g01346.t01 |
| 9218 | 7 | 115638934 | 115647463 | mNeoVis.2.1.1.chr7\_hap1.g01347.t01 |
| 9219 | 7 | 115651635 | 115685581 | mNeoVis.2.1.1.chr7\_hap1.g01348.t01 |
| 9220 | 7 | 115730838 | 115742982 | mNeoVis.2.1.1.chr7\_hap1.g01349.t01 |
| 9221 | 7 | 115745286 | 115759267 | mNeoVis.2.1.1.chr7\_hap1.g01350.t01 |
| 9222 | 7 | 115770103 | 115785800 | mNeoVis.2.1.1.chr7\_hap1.g01351.t01 |
| 9223 | 7 | 115799825 | 115879828 | mNeoVis.2.1.1.chr7\_hap1.g01352.t01 |
| 9224 | 7 | 115900380 | 115923408 | mNeoVis.2.1.1.chr7\_hap1.g01353.t01 |
| 9225 | 7 | 115929820 | 115946200 | mNeoVis.2.1.1.chr7\_hap1.g01354.t01 |
| 9226 | 7 | 116941623 | 116952869 | mNeoVis.2.1.1.chr7\_hap1.g01373.t01 |
| 9227 | 7 | 116952877 | 117002815 | mNeoVis.2.1.1.chr7\_hap1.g01374.t01 |
| 9228 | 7 | 117003063 | 117009007 | mNeoVis.2.1.1.chr7\_hap1.g01375.t01 |
| 9229 | 7 | 117009232 | 117018425 | mNeoVis.2.1.1.chr7\_hap1.g01376.t01 |
| 9230 | 7 | 117022021 | 117036174 | mNeoVis.2.1.1.chr7\_hap1.g01377.t01 |
| 9231 | 7 | 117041713 | 117080831 | mNeoVis.2.1.1.chr7\_hap1.g01378.t01 |
| 9232 | 7 | 117088843 | 117123026 | mNeoVis.2.1.1.chr7\_hap1.g01379.t01 |
| 9233 | 7 | 117125930 | 117661680 | mNeoVis.2.1.1.chr7\_hap1.g01380.t01 |
| 9234 | 7 | 119209226 | 119339158 | mNeoVis.2.1.1.chr7\_hap1.g01382.t01 |
| 9235 | 7 | 119434497 | 119557054 | mNeoVis.2.1.1.chr7\_hap1.g01383.t01 |
| 9236 | 7 | 119587619 | 119617015 | mNeoVis.2.1.1.chr7\_hap1.g01384.t01 |
| 9237 | 7 | 119659058 | 119663568 | mNeoVis.2.1.1.chr7\_hap1.g01385.t01 |
| 9238 | 7 | 119711688 | 119916548 | mNeoVis.2.1.1.chr7\_hap1.g01386.t01 |
| 9239 | 7 | 120091725 | 120166270 | mNeoVis.2.1.1.chr7\_hap1.g01387.t01 |
| 9240 | 7 | 120424139 | 120461736 | mNeoVis.2.1.1.chr7\_hap1.g01388.t01 |
| 9241 | 7 | 124190927 | 124192335 | mNeoVis.2.1.1.chr7\_hap1.g01401.t01 |
| 9242 | 7 | 124190927 | 124192335 | mNeoVis.2.1.1.chr7\_hap1.g01401.t01 |
| 9243 | 7 | 124311349 | 124317215 | mNeoVis.2.1.1.chr7\_hap1.g01402.t01 |
| 9244 | 7 | 124311349 | 124317215 | mNeoVis.2.1.1.chr7\_hap1.g01402.t01 |
| 9245 | 7 | 124358231 | 124405300 | mNeoVis.2.1.1.chr7\_hap1.g01403.t01 |
| 9246 | 7 | 124358231 | 124405300 | mNeoVis.2.1.1.chr7\_hap1.g01403.t01 |
| 9247 | 7 | 131260998 | 131385829 | mNeoVis.2.1.1.chr7\_hap1.g01420.t01 |
| 9248 | 7 | 131390534 | 131431205 | mNeoVis.2.1.1.chr7\_hap1.g01421.t01 |
| 9249 | 7 | 131431743 | 131461710 | mNeoVis.2.1.1.chr7\_hap1.g01422.t01 |
| 9250 | 7 | 131604274 | 131817953 | mNeoVis.2.1.1.chr7\_hap1.g01423.t01 |
| 9251 | 7 | 131935584 | 132009299 | mNeoVis.2.1.1.chr7\_hap1.g01424.t01 |
| 9252 | 7 | 132045483 | 132073006 | mNeoVis.2.1.1.chr7\_hap1.g01425.t01 |
| 9253 | 7 | 132100409 | 132102412 | mNeoVis.2.1.1.chr7\_hap1.g01426.t01 |
| 9254 | 7 | 132112590 | 132143853 | mNeoVis.2.1.1.chr7\_hap1.g01427.t01 |
| 9255 | 7 | 132143725 | 132154293 | mNeoVis.2.1.1.chr7\_hap1.g01428.t01 |
| 9256 | 7 | 132210539 | 132303778 | mNeoVis.2.1.1.chr7\_hap1.g01429.t01 |
| 9257 | 7 | 132430510 | 132475611 | mNeoVis.2.1.1.chr7\_hap1.g01430.t01 |
| 9258 | 7 | 132510680 | 132514618 | mNeoVis.2.1.1.chr7\_hap1.g01431.t01 |
| 9259 | 7 | 132530218 | 132546009 | mNeoVis.2.1.1.chr7\_hap1.g01432.t01 |
| 9260 | 7 | 132566098 | 132572801 | mNeoVis.2.1.1.chr7\_hap1.g01433.t01 |
| 9261 | 7 | 132819486 | 132908595 | mNeoVis.2.1.1.chr7\_hap1.g01438.t01 |
| 9262 | 7 | 133035495 | 133078849 | mNeoVis.2.1.1.chr7\_hap1.g01439.t01 |
| 9263 | 7 | 133080245 | 133101452 | mNeoVis.2.1.1.chr7\_hap1.g01440.t01 |
| 9264 | 7 | 133106417 | 133130055 | mNeoVis.2.1.1.chr7\_hap1.g01441.t01 |
| 9265 | 7 | 133130106 | 133140348 | mNeoVis.2.1.1.chr7\_hap1.g01442.t01 |
| 9266 | 7 | 133140309 | 133195910 | mNeoVis.2.1.1.chr7\_hap1.g01443.t01 |
| 9267 | 7 | 133333740 | 133345279 | mNeoVis.2.1.1.chr7\_hap1.g01444.t01 |
| 9268 | 7 | 133369469 | 133453019 | mNeoVis.2.1.1.chr7\_hap1.g01445.t01 |
| 9269 | 7 | 133693473 | 133705561 | mNeoVis.2.1.1.chr7\_hap1.g01447.t01 |
| 9270 | 7 | 136344621 | 136368949 | mNeoVis.2.1.1.chr7\_hap1.g01449.t01 |
| 9271 | 7 | 136376296 | 136445035 | mNeoVis.2.1.1.chr7\_hap1.g01450.t01 |
| 9272 | 7 | 136476558 | 136480361 | mNeoVis.2.1.1.chr7\_hap1.g01451.t01 |
| 9273 | 7 | 136511160 | 136574660 | mNeoVis.2.1.1.chr7\_hap1.g01452.t01 |
| 9274 | 7 | 136601464 | 136681655 | mNeoVis.2.1.1.chr7\_hap1.g01453.t01 |
| 9275 | 7 | 136731564 | 136983634 | mNeoVis.2.1.1.chr7\_hap1.g01454.t01 |
| 9276 | 7 | 136916801 | 137030966 | mNeoVis.2.1.1.chr7\_hap1.g01455.t01 |
| 9277 | 7 | 138908975 | 138915744 | mNeoVis.2.1.1.chr7\_hap1.g01462.t01 |
| 9278 | 7 | 138908975 | 138915744 | mNeoVis.2.1.1.chr7\_hap1.g01462.t01 |
| 9279 | 7 | 138950200 | 138957780 | mNeoVis.2.1.1.chr7\_hap1.g01463.t01 |
| 9280 | 7 | 138950200 | 138957780 | mNeoVis.2.1.1.chr7\_hap1.g01463.t01 |
| 9281 | 7 | 138957462 | 138966626 | mNeoVis.2.1.1.chr7\_hap1.g01464.t01 |
| 9282 | 7 | 138957462 | 138966626 | mNeoVis.2.1.1.chr7\_hap1.g01464.t01 |
| 9283 | 7 | 139048593 | 139069931 | mNeoVis.2.1.1.chr7\_hap1.g01465.t01 |
| 9284 | 7 | 139048593 | 139069931 | mNeoVis.2.1.1.chr7\_hap1.g01465.t01 |
| 9285 | 7 | 139153679 | 139160876 | mNeoVis.2.1.1.chr7\_hap1.g01466.t01 |
| 9286 | 7 | 139153679 | 139160876 | mNeoVis.2.1.1.chr7\_hap1.g01466.t01 |
| 9287 | 7 | 146541485 | 146684151 | mNeoVis.2.1.1.chr7\_hap1.g01486.t01 |
| 9288 | 7 | 146541485 | 146684151 | mNeoVis.2.1.1.chr7\_hap1.g01486.t01 |
| 9289 | 7 | 146685080 | 146876661 | mNeoVis.2.1.1.chr7\_hap1.g01487.t01 |
| 9290 | 7 | 146685080 | 146876661 | mNeoVis.2.1.1.chr7\_hap1.g01487.t01 |
| 9291 | 7 | 146875723 | 146929789 | mNeoVis.2.1.1.chr7\_hap1.g01488.t01 |
| 9292 | 7 | 146929981 | 146945163 | mNeoVis.2.1.1.chr7\_hap1.g01489.t01 |
| 9293 | 7 | 146993005 | 147018992 | mNeoVis.2.1.1.chr7\_hap1.g01490.t01 |
| 9294 | 7 | 147035059 | 147045621 | mNeoVis.2.1.1.chr7\_hap1.g01491.t01 |
| 9295 | 7 | 147046505 | 147049125 | mNeoVis.2.1.1.chr7\_hap1.g01492.t01 |
| 9296 | 7 | 147065256 | 147080976 | mNeoVis.2.1.1.chr7\_hap1.g01493.t01 |
| 9297 | 7 | 147092324 | 147206151 | mNeoVis.2.1.1.chr7\_hap1.g01494.t01 |
| 9298 | 7 | 147212477 | 147253973 | mNeoVis.2.1.1.chr7\_hap1.g01495.t01 |
| 9299 | 7 | 147431750 | 147456050 | mNeoVis.2.1.1.chr7\_hap1.g01497.t01 |
| 9300 | 7 | 147461162 | 147477845 | mNeoVis.2.1.1.chr7\_hap1.g01498.t01 |
| 9301 | 7 | 147572720 | 147722408 | mNeoVis.2.1.1.chr7\_hap1.g01499.t01 |
| 9302 | 7 | 150198995 | 150210054 | mNeoVis.2.1.1.chr7\_hap1.g01532.t01 |
| 9303 | 7 | 150251191 | 150306901 | mNeoVis.2.1.1.chr7\_hap1.g01533.t01 |
| 9304 | 7 | 150323591 | 150362273 | mNeoVis.2.1.1.chr7\_hap1.g01534.t01 |
| 9305 | 7 | 150371307 | 150435869 | mNeoVis.2.1.1.chr7\_hap1.g01535.t01 |
| 9306 | 7 | 150435948 | 150567149 | mNeoVis.2.1.1.chr7\_hap1.g01536.t01 |
| 9307 | 7 | 150586551 | 150598452 | mNeoVis.2.1.1.chr7\_hap1.g01537.t01 |
| 9308 | 7 | 150608435 | 150615861 | mNeoVis.2.1.1.chr7\_hap1.g01538.t01 |
| 9309 | 7 | 150624840 | 150638475 | mNeoVis.2.1.1.chr7\_hap1.g01539.t01 |
| 9310 | 7 | 150631926 | 150648528 | mNeoVis.2.1.1.chr7\_hap1.g01540.t01 |
| 9311 | 7 | 150648683 | 150692574 | mNeoVis.2.1.1.chr7\_hap1.g01541.t01 |
| 9312 | 7 | 150692630 | 150695490 | mNeoVis.2.1.1.chr7\_hap1.g01542.t01 |
| 9313 | 7 | 151971660 | 152114210 | mNeoVis.2.1.1.chr7\_hap1.g01556.t01 |
| 9314 | 7 | 152144477 | 152149004 | mNeoVis.2.1.1.chr7\_hap1.g01557.t01 |
| 9315 | 7 | 152149131 | 152163739 | mNeoVis.2.1.1.chr7\_hap1.g01558.t01 |
| 9316 | 7 | 152165576 | 152170817 | mNeoVis.2.1.1.chr7\_hap1.g01559.t01 |
| 9317 | 7 | 152192675 | 152200375 | mNeoVis.2.1.1.chr7\_hap1.g01560.t01 |
| 9318 | 7 | 152203264 | 152206977 | mNeoVis.2.1.1.chr7\_hap1.g01561.t01 |
| 9319 | 7 | 152227984 | 152231115 | mNeoVis.2.1.1.chr7\_hap1.g01562.t01 |
| 9320 | 7 | 152231715 | 152234137 | mNeoVis.2.1.1.chr7\_hap1.g01563.t01 |
| 9321 | 7 | 152236693 | 152242113 | mNeoVis.2.1.1.chr7\_hap1.g01564.t01 |
| 9322 | 7 | 152242834 | 152256234 | mNeoVis.2.1.1.chr7\_hap1.g01565.t01 |
| 9323 | 7 | 152256324 | 152319092 | mNeoVis.2.1.1.chr7\_hap1.g01566.t01 |
| 9324 | 7 | 152319202 | 152323370 | mNeoVis.2.1.1.chr7\_hap1.g01567.t01 |
| 9325 | 7 | 152324505 | 152411505 | mNeoVis.2.1.1.chr7\_hap1.g01568.t01 |
| 9326 | 7 | 152411624 | 152430941 | mNeoVis.2.1.1.chr7\_hap1.g01569.t01 |
| 9327 | 7 | 152433190 | 152451100 | mNeoVis.2.1.1.chr7\_hap1.g01570.t01 |
| 9328 | 7 | 152451833 | 152454665 | mNeoVis.2.1.1.chr7\_hap1.g01571.t01 |
| 9329 | 7 | 152458058 | 152467485 | mNeoVis.2.1.1.chr7\_hap1.g01572.t01 |
| 9330 | 7 | 152467600 | 152473030 | mNeoVis.2.1.1.chr7\_hap1.g01573.t01 |
| 9331 | 7 | 154555041 | 154555976 | mNeoVis.2.1.1.chr7\_hap1.g01671.t01 |
| 9332 | 7 | 154559871 | 154560809 | mNeoVis.2.1.1.chr7\_hap1.g01672.t01 |
| 9333 | 7 | 154569943 | 154570875 | mNeoVis.2.1.1.chr7\_hap1.g01673.t01 |
| 9334 | 7 | 154578935 | 154579873 | mNeoVis.2.1.1.chr7\_hap1.g01674.t01 |
| 9335 | 7 | 154593230 | 154594168 | mNeoVis.2.1.1.chr7\_hap1.g01675.t01 |
| 9336 | 7 | 154646070 | 154652679 | mNeoVis.2.1.1.chr7\_hap1.g01676.t01 |
| 9337 | 7 | 154674780 | 154675733 | mNeoVis.2.1.1.chr7\_hap1.g01677.t01 |
| 9338 | 7 | 154686122 | 154721650 | mNeoVis.2.1.1.chr7\_hap1.g01678.t01 |
| 9339 | 7 | 154702885 | 154704160 | mNeoVis.2.1.1.chr7\_hap1.g01679.t01 |
| 9340 | 7 | 154713026 | 154718792 | mNeoVis.2.1.1.chr7\_hap1.g01680.t01 |
| 9341 | 7 | 154732774 | 154733756 | mNeoVis.2.1.1.chr7\_hap1.g01681.t01 |
| 9342 | 7 | 154742942 | 154744470 | mNeoVis.2.1.1.chr7\_hap1.g01682.t01 |
| 9343 | 7 | 154752329 | 154753273 | mNeoVis.2.1.1.chr7\_hap1.g01683.t01 |
| 9344 | 7 | 154769173 | 154770117 | mNeoVis.2.1.1.chr7\_hap1.g01684.t01 |
| 9345 | 7 | 154788738 | 154790184 | mNeoVis.2.1.1.chr7\_hap1.g01685.t01 |
| 9346 | 7 | 154805408 | 154806364 | mNeoVis.2.1.1.chr7\_hap1.g01686.t01 |
| 9347 | 7 | 154822868 | 154823928 | mNeoVis.2.1.1.chr7\_hap1.g01687.t01 |
| 9348 | 7 | 154827644 | 154829587 | mNeoVis.2.1.1.chr7\_hap1.g01688.t01 |
| 9349 | 7 | 154838523 | 154841238 | mNeoVis.2.1.1.chr7\_hap1.g01689.t01 |
| 9350 | 7 | 154844984 | 154847280 | mNeoVis.2.1.1.chr7\_hap1.g01690.t01 |
| 9351 | 7 | 154849530 | 154855330 | mNeoVis.2.1.1.chr7\_hap1.g01691.t01 |
| 9352 | 7 | 154857291 | 154858310 | mNeoVis.2.1.1.chr7\_hap1.g01692.t01 |
| 9353 | 7 | 154867881 | 154868828 | mNeoVis.2.1.1.chr7\_hap1.g01693.t01 |
| 9354 | 7 | 154887969 | 154888910 | mNeoVis.2.1.1.chr7\_hap1.g01694.t01 |
| 9355 | 7 | 154903128 | 154904078 | mNeoVis.2.1.1.chr7\_hap1.g01695.t01 |
| 9356 | 7 | 154918411 | 154920472 | mNeoVis.2.1.1.chr7\_hap1.g01696.t01 |
| 9357 | 7 | 154944307 | 154990581 | mNeoVis.2.1.1.chr7\_hap1.g01697.t01 |
| 9358 | 7 | 155019255 | 155032296 | mNeoVis.2.1.1.chr7\_hap1.g01698.t01 |
| 9359 | 7 | 155066969 | 155067910 | mNeoVis.2.1.1.chr7\_hap1.g01699.t01 |
| 9360 | 7 | 155079080 | 155080030 | mNeoVis.2.1.1.chr7\_hap1.g01700.t01 |
| 9361 | 7 | 155092797 | 155106188 | mNeoVis.2.1.1.chr7\_hap1.g01701.t01 |
| 9362 | 7 | 155094031 | 155094981 | mNeoVis.2.1.1.chr7\_hap1.g01702.t01 |
| 9363 | 7 | 155134901 | 155135890 | mNeoVis.2.1.1.chr7\_hap1.g01703.t01 |
| 9364 | 7 | 155160255 | 155161220 | mNeoVis.2.1.1.chr7\_hap1.g01704.t01 |
| 9365 | 7 | 155169838 | 155170821 | mNeoVis.2.1.1.chr7\_hap1.g01705.t01 |
| 9366 | 7 | 155184490 | 155185449 | mNeoVis.2.1.1.chr7\_hap1.g01706.t01 |
| 9367 | 7 | 155198801 | 155203332 | mNeoVis.2.1.1.chr7\_hap1.g01707.t01 |
| 9368 | 7 | 155226830 | 155227801 | mNeoVis.2.1.1.chr7\_hap1.g01708.t01 |
| 9369 | 7 | 157034676 | 157189387 | mNeoVis.2.1.1.chr7\_hap1.g01776.t01 |
| 9370 | 7 | 157255191 | 157280148 | mNeoVis.2.1.1.chr7\_hap1.g01777.t01 |
| 9371 | 7 | 157282854 | 157427986 | mNeoVis.2.1.1.chr7\_hap1.g01778.t01 |
| 9372 | 7 | 157430595 | 157439194 | mNeoVis.2.1.1.chr7\_hap1.g01779.t01 |
| 9373 | 7 | 157452005 | 157469854 | mNeoVis.2.1.1.chr7\_hap1.g01780.t01 |
| 9374 | 7 | 157491430 | 157493737 | mNeoVis.2.1.1.chr7\_hap1.g01781.t01 |
| 9375 | 7 | 158783104 | 158810505 | mNeoVis.2.1.1.chr7\_hap1.g01805.t01 |
| 9376 | 7 | 158868489 | 158924301 | mNeoVis.2.1.1.chr7\_hap1.g01806.t01 |
| 9377 | 7 | 158936023 | 158997991 | mNeoVis.2.1.1.chr7\_hap1.g01807.t01 |
| 9378 | 7 | 159020632 | 159038278 | mNeoVis.2.1.1.chr7\_hap1.g01808.t01 |
| 9379 | 7 | 159084314 | 159152513 | mNeoVis.2.1.1.chr7\_hap1.g01809.t01 |
| 9380 | 7 | 161085441 | 161197246 | mNeoVis.2.1.1.chr7\_hap1.g01822.t01 |
| 9381 | 7 | 161085441 | 161197246 | mNeoVis.2.1.1.chr7\_hap1.g01822.t01 |
| 9382 | 7 | 161201428 | 161247080 | mNeoVis.2.1.1.chr7\_hap1.g01823.t01 |
| 9383 | 7 | 161201428 | 161247080 | mNeoVis.2.1.1.chr7\_hap1.g01823.t01 |
| 9384 | 7 | 162166841 | 162168686 | mNeoVis.2.1.1.chr7\_hap1.g01828.t01 |
| 9385 | 7 | 162399843 | 162497962 | mNeoVis.2.1.1.chr7\_hap1.g01829.t01 |
| 9386 | 7 | 162498107 | 162547900 | mNeoVis.2.1.1.chr7\_hap1.g01830.t01 |
| 9387 | 7 | 162590267 | 162643841 | mNeoVis.2.1.1.chr7\_hap1.g01831.t01 |
| 9388 | 7 | 162695115 | 162695453 | mNeoVis.2.1.1.chr7\_hap1.g01832.t01 |
| 9389 | 7 | 162710731 | 162778211 | mNeoVis.2.1.1.chr7\_hap1.g01833.t01 |
| 9390 | 7 | 162788840 | 162823613 | mNeoVis.2.1.1.chr7\_hap1.g01834.t01 |
| 9391 | 7 | 163232302 | 163303679 | mNeoVis.2.1.1.chr7\_hap1.g01836.t01 |
| 9392 | 7 | 163232302 | 163303679 | mNeoVis.2.1.1.chr7\_hap1.g01836.t01 |
| 9393 | 7 | 163232302 | 163303679 | mNeoVis.2.1.1.chr7\_hap1.g01836.t01 |
| 9394 | 7 | 163377068 | 163418473 | mNeoVis.2.1.1.chr7\_hap1.g01837.t01 |
| 9395 | 7 | 163377068 | 163418473 | mNeoVis.2.1.1.chr7\_hap1.g01837.t01 |
| 9396 | 7 | 163377068 | 163418473 | mNeoVis.2.1.1.chr7\_hap1.g01837.t01 |
| 9397 | 7 | 163424352 | 163436278 | mNeoVis.2.1.1.chr7\_hap1.g01838.t01 |
| 9398 | 7 | 163424352 | 163436278 | mNeoVis.2.1.1.chr7\_hap1.g01838.t01 |
| 9399 | 7 | 163424352 | 163436278 | mNeoVis.2.1.1.chr7\_hap1.g01838.t01 |
| 9400 | 7 | 163577893 | 163739524 | mNeoVis.2.1.1.chr7\_hap1.g01839.t01 |
| 9401 | 7 | 163577893 | 163739524 | mNeoVis.2.1.1.chr7\_hap1.g01839.t01 |
| 9402 | 7 | 163577893 | 163739524 | mNeoVis.2.1.1.chr7\_hap1.g01839.t01 |
| 9403 | 7 | 163769885 | 163793148 | mNeoVis.2.1.1.chr7\_hap1.g01840.t01 |
| 9404 | 7 | 165780219 | 165783706 | mNeoVis.2.1.1.chr7\_hap1.g01847.t01 |
| 9405 | 7 | 165791739 | 165896901 | mNeoVis.2.1.1.chr7\_hap1.g01848.t01 |
| 9406 | 7 | 165895552 | 165935867 | mNeoVis.2.1.1.chr7\_hap1.g01849.t01 |
| 9407 | 7 | 165956094 | 166002451 | mNeoVis.2.1.1.chr7\_hap1.g01850.t01 |
| 9408 | 7 | 166012894 | 166033695 | mNeoVis.2.1.1.chr7\_hap1.g01851.t01 |
| 9409 | 7 | 166038042 | 166041723 | mNeoVis.2.1.1.chr7\_hap1.g01852.t01 |
| 9410 | 7 | 166045069 | 166118601 | mNeoVis.2.1.1.chr7\_hap1.g01853.t01 |
| 9411 | 7 | 166138889 | 166181944 | mNeoVis.2.1.1.chr7\_hap1.g01854.t01 |
| 9412 | 7 | 166184669 | 166262887 | mNeoVis.2.1.1.chr7\_hap1.g01855.t01 |
| 9413 | 7 | 166671880 | 166700677 | mNeoVis.2.1.1.chr7\_hap1.g01860.t01 |
| 9414 | 7 | 166704263 | 166706425 | mNeoVis.2.1.1.chr7\_hap1.g01861.t01 |
| 9415 | 7 | 166734368 | 166739283 | mNeoVis.2.1.1.chr7\_hap1.g01862.t01 |
| 9416 | 7 | 166751379 | 166799393 | mNeoVis.2.1.1.chr7\_hap1.g01863.t01 |
| 9417 | 7 | 166799457 | 166839266 | mNeoVis.2.1.1.chr7\_hap1.g01864.t01 |
| 9418 | 7 | 166866904 | 166878600 | mNeoVis.2.1.1.chr7\_hap1.g01865.t01 |
| 9419 | 7 | 166885372 | 166930364 | mNeoVis.2.1.1.chr7\_hap1.g01866.t01 |
| 9420 | 7 | 166938633 | 166981574 | mNeoVis.2.1.1.chr7\_hap1.g01867.t01 |
| 9421 | 7 | 166983159 | 167040838 | mNeoVis.2.1.1.chr7\_hap1.g01868.t01 |
| 9422 | 7 | 167049420 | 167051848 | mNeoVis.2.1.1.chr7\_hap1.g01869.t01 |
| 9423 | 7 | 167049420 | 167051848 | mNeoVis.2.1.1.chr7\_hap1.g01869.t01 |
| 9424 | 7 | 167058488 | 167078972 | mNeoVis.2.1.1.chr7\_hap1.g01870.t01 |
| 9425 | 7 | 167058488 | 167078972 | mNeoVis.2.1.1.chr7\_hap1.g01870.t01 |
| 9426 | 7 | 167100836 | 167110010 | mNeoVis.2.1.1.chr7\_hap1.g01871.t01 |
| 9427 | 7 | 167100836 | 167110010 | mNeoVis.2.1.1.chr7\_hap1.g01871.t01 |
| 9428 | 7 | 167111016 | 167130160 | mNeoVis.2.1.1.chr7\_hap1.g01872.t01 |
| 9429 | 7 | 167111016 | 167130160 | mNeoVis.2.1.1.chr7\_hap1.g01872.t01 |
| 9430 | 7 | 167131447 | 167175848 | mNeoVis.2.1.1.chr7\_hap1.g01873.t01 |
| 9431 | 7 | 167131447 | 167175848 | mNeoVis.2.1.1.chr7\_hap1.g01873.t01 |
| 9432 | 7 | 167230372 | 167232072 | mNeoVis.2.1.1.chr7\_hap1.g01874.t01 |
| 9433 | 7 | 167230372 | 167232072 | mNeoVis.2.1.1.chr7\_hap1.g01874.t01 |
| 9434 | 7 | 167279703 | 167328917 | mNeoVis.2.1.1.chr7\_hap1.g01875.t01 |
| 9435 | 7 | 167279703 | 167328917 | mNeoVis.2.1.1.chr7\_hap1.g01875.t01 |
| 9436 | 7 | 167335338 | 167354364 | mNeoVis.2.1.1.chr7\_hap1.g01876.t01 |
| 9437 | 7 | 167335338 | 167354364 | mNeoVis.2.1.1.chr7\_hap1.g01876.t01 |
| 9438 | 7 | 167377293 | 167394994 | mNeoVis.2.1.1.chr7\_hap1.g01877.t01 |
| 9439 | 7 | 167377293 | 167394994 | mNeoVis.2.1.1.chr7\_hap1.g01877.t01 |
| 9440 | 7 | 167810566 | 168207438 | mNeoVis.2.1.1.chr7\_hap1.g01878.t01 |
| 9441 | 7 | 168231648 | 168235713 | mNeoVis.2.1.1.chr7\_hap1.g01879.t01 |
| 9442 | 7 | 168238917 | 168355447 | mNeoVis.2.1.1.chr7\_hap1.g01880.t01 |
| 9443 | 7 | 168416191 | 168431749 | mNeoVis.2.1.1.chr7\_hap1.g01881.t01 |
| 9444 | 7 | 168416191 | 168431749 | mNeoVis.2.1.1.chr7\_hap1.g01881.t01 |
| 9445 | 7 | 168434723 | 168563453 | mNeoVis.2.1.1.chr7\_hap1.g01882.t01 |
| 9446 | 7 | 168434723 | 168563453 | mNeoVis.2.1.1.chr7\_hap1.g01882.t01 |
| 9447 | 7 | 168621299 | 168674031 | mNeoVis.2.1.1.chr7\_hap1.g01883.t01 |
| 9448 | 7 | 168621299 | 168674031 | mNeoVis.2.1.1.chr7\_hap1.g01883.t01 |
| 9449 | 7 | 168849793 | 168850347 | mNeoVis.2.1.1.chr7\_hap1.g01885.t01 |
| 9450 | 7 | 168849793 | 168850347 | mNeoVis.2.1.1.chr7\_hap1.g01885.t01 |
| 9451 | 7 | 169236036 | 169237689 | mNeoVis.2.1.1.chr7\_hap1.g01886.t01 |
| 9452 | 7 | 169236036 | 169237689 | mNeoVis.2.1.1.chr7\_hap1.g01886.t01 |
| 9453 | 7 | 169968901 | 170008524 | mNeoVis.2.1.1.chr7\_hap1.g01887.t01 |
| 9454 | 7 | 169968901 | 170008524 | mNeoVis.2.1.1.chr7\_hap1.g01887.t01 |
| 9455 | 7 | 170007893 | 170109845 | mNeoVis.2.1.1.chr7\_hap1.g01888.t01 |
| 9456 | 7 | 170007893 | 170109845 | mNeoVis.2.1.1.chr7\_hap1.g01888.t01 |
| 9457 | 7 | 170133831 | 170174054 | mNeoVis.2.1.1.chr7\_hap1.g01889.t01 |
| 9458 | 7 | 170133831 | 170174054 | mNeoVis.2.1.1.chr7\_hap1.g01889.t01 |
| 9459 | 7 | 170133831 | 170174054 | mNeoVis.2.1.1.chr7\_hap1.g01889.t01 |
| 9460 | 7 | 170384601 | 170386104 | mNeoVis.2.1.1.chr7\_hap1.g01890.t01 |
| 9461 | 7 | 170384601 | 170386104 | mNeoVis.2.1.1.chr7\_hap1.g01890.t01 |
| 9462 | 7 | 170409153 | 170535670 | mNeoVis.2.1.1.chr7\_hap1.g01891.t01 |
| 9463 | 7 | 170409153 | 170535670 | mNeoVis.2.1.1.chr7\_hap1.g01891.t01 |
| 9464 | 7 | 170538045 | 170547493 | mNeoVis.2.1.1.chr7\_hap1.g01892.t01 |
| 9465 | 7 | 170538045 | 170547493 | mNeoVis.2.1.1.chr7\_hap1.g01892.t01 |
| 9466 | 7 | 170556979 | 170567223 | mNeoVis.2.1.1.chr7\_hap1.g01893.t01 |
| 9467 | 7 | 170556979 | 170567223 | mNeoVis.2.1.1.chr7\_hap1.g01893.t01 |
| 9468 | 7 | 171252793 | 171305974 | mNeoVis.2.1.1.chr7\_hap1.g01894.t01 |
| 9469 | 7 | 171252793 | 171305974 | mNeoVis.2.1.1.chr7\_hap1.g01894.t01 |
| 9470 | 7 | 173281455 | 173754736 | mNeoVis.2.1.1.chr7\_hap1.g01896.t01 |
| 9471 | 7 | 173681898 | 173694649 | mNeoVis.2.1.1.chr7\_hap1.g01897.t01 |
| 9472 | 7 | 173681898 | 173694649 | mNeoVis.2.1.1.chr7\_hap1.g01897.t01 |
| 9473 | 7 | 173762976 | 173810026 | mNeoVis.2.1.1.chr7\_hap1.g01898.t01 |
| 9474 | 7 | 173762976 | 173810026 | mNeoVis.2.1.1.chr7\_hap1.g01898.t01 |
| 9475 | 7 | 173762976 | 173810026 | mNeoVis.2.1.1.chr7\_hap1.g01898.t01 |
| 9476 | 7 | 174022229 | 174024330 | mNeoVis.2.1.1.chr7\_hap1.g01899.t01 |
| 9477 | 7 | 174022229 | 174024330 | mNeoVis.2.1.1.chr7\_hap1.g01899.t01 |
| 9478 | 7 | 174022229 | 174024330 | mNeoVis.2.1.1.chr7\_hap1.g01899.t01 |
| 9479 | 7 | 174065428 | 174125397 | mNeoVis.2.1.1.chr7\_hap1.g01900.t01 |
| 9480 | 7 | 174065428 | 174125397 | mNeoVis.2.1.1.chr7\_hap1.g01900.t01 |
| 9481 | 7 | 174210479 | 174338267 | mNeoVis.2.1.1.chr7\_hap1.g01901.t01 |
| 9482 | 7 | 174210479 | 174338267 | mNeoVis.2.1.1.chr7\_hap1.g01901.t01 |
| 9483 | 7 | 174343127 | 174447305 | mNeoVis.2.1.1.chr7\_hap1.g01902.t01 |
| 9484 | 7 | 174462087 | 174473549 | mNeoVis.2.1.1.chr7\_hap1.g01903.t01 |
| 9485 | 7 | 174703557 | 174705633 | mNeoVis.2.1.1.chr7\_hap1.g01905.t01 |
| 9486 | 7 | 174703557 | 174705633 | mNeoVis.2.1.1.chr7\_hap1.g01905.t01 |
| 9487 | 7 | 174776364 | 174893796 | mNeoVis.2.1.1.chr7\_hap1.g01906.t01 |
| 9488 | 7 | 174776364 | 174893796 | mNeoVis.2.1.1.chr7\_hap1.g01906.t01 |
| 9489 | 7 | 174914017 | 174993582 | mNeoVis.2.1.1.chr7\_hap1.g01907.t01 |
| 9490 | 7 | 174914017 | 174993582 | mNeoVis.2.1.1.chr7\_hap1.g01907.t01 |
| 9491 | 7 | 174993599 | 175185670 | mNeoVis.2.1.1.chr7\_hap1.g01908.t01 |
| 9492 | 7 | 174993599 | 175185670 | mNeoVis.2.1.1.chr7\_hap1.g01908.t01 |
| 9493 | 7 | 175207478 | 175207696 | mNeoVis.2.1.1.chr7\_hap1.g01909.t01 |
| 9494 | 7 | 175207478 | 175207696 | mNeoVis.2.1.1.chr7\_hap1.g01909.t01 |
| 9495 | 7 | 176748504 | 176756062 | mNeoVis.2.1.1.chr7\_hap1.g01910.t01 |
| 9496 | 7 | 176939732 | 176955966 | mNeoVis.2.1.1.chr7\_hap1.g01911.t01 |
| 9497 | 7 | 176939732 | 176955966 | mNeoVis.2.1.1.chr7\_hap1.g01911.t01 |
| 9498 | 7 | 177035715 | 177055132 | mNeoVis.2.1.1.chr7\_hap1.g01912.t01 |
| 9499 | 7 | 177035715 | 177055132 | mNeoVis.2.1.1.chr7\_hap1.g01912.t01 |
| 9500 | 7 | 177127375 | 177302016 | mNeoVis.2.1.1.chr7\_hap1.g01913.t01 |
| 9501 | 7 | 177127375 | 177302016 | mNeoVis.2.1.1.chr7\_hap1.g01913.t01 |
| 9502 | 7 | 177535271 | 177982878 | mNeoVis.2.1.1.chr7\_hap1.g01914.t01 |
| 9503 | 7 | 177535271 | 177982878 | mNeoVis.2.1.1.chr7\_hap1.g01914.t01 |
| 9504 | 7 | 177535271 | 177982878 | mNeoVis.2.1.1.chr7\_hap1.g01914.t01 |
| 9505 | 7 | 177982820 | 178050637 | mNeoVis.2.1.1.chr7\_hap1.g01915.t01 |
| 9506 | 7 | 177982820 | 178050637 | mNeoVis.2.1.1.chr7\_hap1.g01915.t01 |
| 9507 | 7 | 178049128 | 178131408 | mNeoVis.2.1.1.chr7\_hap1.g01916.t01 |
| 9508 | 7 | 178049128 | 178131408 | mNeoVis.2.1.1.chr7\_hap1.g01916.t01 |
| 9509 | 7 | 178131466 | 178374238 | mNeoVis.2.1.1.chr7\_hap1.g01917.t01 |
| 9510 | 7 | 178131466 | 178374238 | mNeoVis.2.1.1.chr7\_hap1.g01917.t01 |
| 9511 | 7 | 178379899 | 178397767 | mNeoVis.2.1.1.chr7\_hap1.g01918.t01 |
| 9512 | 7 | 178379899 | 178397767 | mNeoVis.2.1.1.chr7\_hap1.g01918.t01 |
| 9513 | 7 | 178446415 | 178446723 | mNeoVis.2.1.1.chr7\_hap1.g01919.t01 |
| 9514 | 7 | 178661705 | 178674338 | mNeoVis.2.1.1.chr7\_hap1.g01920.t01 |
| 9515 | 7 | 178661705 | 178674338 | mNeoVis.2.1.1.chr7\_hap1.g01920.t01 |
| 9516 | 7 | 178679569 | 178857204 | mNeoVis.2.1.1.chr7\_hap1.g01921.t01 |
| 9517 | 7 | 178679569 | 178857204 | mNeoVis.2.1.1.chr7\_hap1.g01921.t01 |
| 9518 | 7 | 178894489 | 178943418 | mNeoVis.2.1.1.chr7\_hap1.g01922.t01 |
| 9519 | 7 | 178894489 | 178943418 | mNeoVis.2.1.1.chr7\_hap1.g01922.t01 |
| 9520 | 7 | 178894489 | 178943418 | mNeoVis.2.1.1.chr7\_hap1.g01922.t01 |
| 9521 | 7 | 178894489 | 178943418 | mNeoVis.2.1.1.chr7\_hap1.g01922.t01 |
| 9522 | 7 | 179049774 | 179101868 | mNeoVis.2.1.1.chr7\_hap1.g01923.t01 |
| 9523 | 7 | 179049774 | 179101868 | mNeoVis.2.1.1.chr7\_hap1.g01923.t01 |
| 9524 | 7 | 179049774 | 179101868 | mNeoVis.2.1.1.chr7\_hap1.g01923.t01 |
| 9525 | 7 | 179049774 | 179101868 | mNeoVis.2.1.1.chr7\_hap1.g01923.t01 |
| 9526 | 7 | 179102024 | 179249420 | mNeoVis.2.1.1.chr7\_hap1.g01924.t01 |
| 9527 | 7 | 179279032 | 179294346 | mNeoVis.2.1.1.chr7\_hap1.g01925.t01 |
| 9528 | 7 | 179317865 | 179392632 | mNeoVis.2.1.1.chr7\_hap1.g01926.t01 |
| 9529 | 7 | 179417567 | 179435540 | mNeoVis.2.1.1.chr7\_hap1.g01927.t01 |
| 9530 | 7 | 179440497 | 179476999 | mNeoVis.2.1.1.chr7\_hap1.g01928.t01 |
| 9531 | 7 | 179482965 | 179551364 | mNeoVis.2.1.1.chr7\_hap1.g01929.t01 |
| 9532 | 7 | 179634970 | 179731301 | mNeoVis.2.1.1.chr7\_hap1.g01930.t01 |
| 9533 | 7 | 179892549 | 180004931 | mNeoVis.2.1.1.chr7\_hap1.g01931.t01 |
| 9534 | 7 | 180020615 | 180022778 | mNeoVis.2.1.1.chr7\_hap1.g01932.t01 |
| 9535 | 7 | 180033920 | 180054675 | mNeoVis.2.1.1.chr7\_hap1.g01933.t01 |
| 9536 | 7 | 180057449 | 180088879 | mNeoVis.2.1.1.chr7\_hap1.g01934.t01 |
| 9537 | 7 | 180624637 | 180665885 | mNeoVis.2.1.1.chr7\_hap1.g01939.t01 |
| 9538 | 7 | 180659036 | 180665156 | mNeoVis.2.1.1.chr7\_hap1.g01940.t01 |
| 9539 | 7 | 180671607 | 180714331 | mNeoVis.2.1.1.chr7\_hap1.g01941.t01 |
| 9540 | 7 | 180833595 | 180862536 | mNeoVis.2.1.1.chr7\_hap1.g01942.t01 |
| 9541 | 7 | 181027614 | 181047608 | mNeoVis.2.1.1.chr7\_hap1.g01943.t01 |
| 9542 | 7 | 181047774 | 181120708 | mNeoVis.2.1.1.chr7\_hap1.g01944.t01 |
| 9543 | 7 | 181156725 | 181214646 | mNeoVis.2.1.1.chr7\_hap1.g01945.t01 |
| 9544 | 7 | 181260387 | 181348157 | mNeoVis.2.1.1.chr7\_hap1.g01946.t01 |
| 9545 | 7 | 181260387 | 181348157 | mNeoVis.2.1.1.chr7\_hap1.g01946.t01 |
| 9546 | 7 | 181366230 | 181424825 | mNeoVis.2.1.1.chr7\_hap1.g01947.t01 |
| 9547 | 7 | 181366230 | 181424825 | mNeoVis.2.1.1.chr7\_hap1.g01947.t01 |
| 9548 | 7 | 181524492 | 181596029 | mNeoVis.2.1.1.chr7\_hap1.g01948.t01 |
| 9549 | 7 | 181524492 | 181596029 | mNeoVis.2.1.1.chr7\_hap1.g01948.t01 |
| 9550 | 7 | 181674957 | 181677649 | mNeoVis.2.1.1.chr7\_hap1.g01949.t01 |
| 9551 | 7 | 181674957 | 181677649 | mNeoVis.2.1.1.chr7\_hap1.g01949.t01 |
| 9552 | 7 | 181718445 | 181829458 | mNeoVis.2.1.1.chr7\_hap1.g01950.t01 |
| 9553 | 7 | 183874149 | 184276433 | mNeoVis.2.1.1.chr7\_hap1.g01957.t01 |
| 9554 | 7 | 183948635 | 183999982 | mNeoVis.2.1.1.chr7\_hap1.g01958.t01 |
| 9555 | 7 | 184185952 | 184187100 | mNeoVis.2.1.1.chr7\_hap1.g01959.t01 |
| 9556 | 7 | 186812671 | 186817686 | mNeoVis.2.1.1.chr7\_hap1.g01961.t01 |
| 9557 | 7 | 186812671 | 186817686 | mNeoVis.2.1.1.chr7\_hap1.g01961.t01 |
| 9558 | 7 | 187992115 | 188021284 | mNeoVis.2.1.1.chr7\_hap1.g01962.t01 |
| 9559 | 7 | 187992115 | 188021284 | mNeoVis.2.1.1.chr7\_hap1.g01962.t01 |
| 9560 | 7 | 188031303 | 188150142 | mNeoVis.2.1.1.chr7\_hap1.g01963.t01 |
| 9561 | 7 | 188031303 | 188150142 | mNeoVis.2.1.1.chr7\_hap1.g01963.t01 |
| 9562 | 7 | 188304794 | 188465190 | mNeoVis.2.1.1.chr7\_hap1.g01964.t01 |
| 9563 | 7 | 188304794 | 188465190 | mNeoVis.2.1.1.chr7\_hap1.g01964.t01 |
| 9564 | 7 | 188478851 | 188516270 | mNeoVis.2.1.1.chr7\_hap1.g01965.t01 |
| 9565 | 7 | 188478851 | 188516270 | mNeoVis.2.1.1.chr7\_hap1.g01965.t01 |
| 9566 | 7 | 188535460 | 188536759 | mNeoVis.2.1.1.chr7\_hap1.g01966.t01 |
| 9567 | 7 | 188535460 | 188536759 | mNeoVis.2.1.1.chr7\_hap1.g01966.t01 |
| 9568 | 7 | 188615949 | 188633802 | mNeoVis.2.1.1.chr7\_hap1.g01967.t01 |
| 9569 | 7 | 188615949 | 188633802 | mNeoVis.2.1.1.chr7\_hap1.g01967.t01 |
| 9570 | 7 | 188637564 | 188659422 | mNeoVis.2.1.1.chr7\_hap1.g01968.t01 |
| 9571 | 7 | 188637564 | 188659422 | mNeoVis.2.1.1.chr7\_hap1.g01968.t01 |
| 9572 | 7 | 188666081 | 188806425 | mNeoVis.2.1.1.chr7\_hap1.g01969.t01 |
| 9573 | 7 | 188666081 | 188806425 | mNeoVis.2.1.1.chr7\_hap1.g01969.t01 |
| 9574 | 7 | 188823978 | 188867973 | mNeoVis.2.1.1.chr7\_hap1.g01970.t01 |
| 9575 | 7 | 188823978 | 188867973 | mNeoVis.2.1.1.chr7\_hap1.g01970.t01 |
| 9576 | 7 | 188951281 | 188951508 | mNeoVis.2.1.1.chr7\_hap1.g01971.t01 |
| 9577 | 7 | 189081230 | 189122237 | mNeoVis.2.1.1.chr7\_hap1.g01972.t01 |
| 9578 | 7 | 189205729 | 189383925 | mNeoVis.2.1.1.chr7\_hap1.g01973.t01 |
| 9579 | 7 | 189384059 | 189400519 | mNeoVis.2.1.1.chr7\_hap1.g01974.t01 |
| 9580 | 7 | 189561832 | 189640201 | mNeoVis.2.1.1.chr7\_hap1.g01975.t01 |
| 9581 | 7 | 189652339 | 189695444 | mNeoVis.2.1.1.chr7\_hap1.g01976.t01 |
| 9582 | 7 | 189980668 | 190010695 | mNeoVis.2.1.1.chr7\_hap1.g01977.t01 |
| 9583 | 7 | 190133932 | 190140660 | mNeoVis.2.1.1.chr7\_hap1.g01978.t01 |
| 9584 | 7 | 190168317 | 190204495 | mNeoVis.2.1.1.chr7\_hap1.g01979.t01 |
| 9585 | 7 | 190207109 | 190225944 | mNeoVis.2.1.1.chr7\_hap1.g01980.t01 |
| 9586 | 7 | 190225990 | 190226795 | mNeoVis.2.1.1.chr7\_hap1.g01981.t01 |
| 9587 | 7 | 190229541 | 190238719 | mNeoVis.2.1.1.chr7\_hap1.g01982.t01 |
| 9588 | 7 | 190239163 | 190244811 | mNeoVis.2.1.1.chr7\_hap1.g01983.t01 |
| 9589 | 7 | 190244979 | 190437205 | mNeoVis.2.1.1.chr7\_hap1.g01984.t01 |
| 9590 | 7 | 190567615 | 190602624 | mNeoVis.2.1.1.chr7\_hap1.g01985.t01 |
| 9591 | 7 | 190621254 | 190651906 | mNeoVis.2.1.1.chr7\_hap1.g01986.t01 |
| 9592 | 7 | 190652149 | 190655249 | mNeoVis.2.1.1.chr7\_hap1.g01987.t01 |
| 9593 | 7 | 190655178 | 190657965 | mNeoVis.2.1.1.chr7\_hap1.g01988.t01 |
| 9594 | 7 | 190667415 | 190848166 | mNeoVis.2.1.1.chr7\_hap1.g01989.t01 |
| 9595 | 7 | 190854296 | 190864684 | mNeoVis.2.1.1.chr7\_hap1.g01990.t01 |
| 9596 | 7 | 190864632 | 190921717 | mNeoVis.2.1.1.chr7\_hap1.g01991.t01 |
| 9597 | 7 | 190924036 | 190942379 | mNeoVis.2.1.1.chr7\_hap1.g01992.t01 |
| 9598 | 7 | 190942377 | 190947249 | mNeoVis.2.1.1.chr7\_hap1.g01993.t01 |
| 9599 | 7 | 190949285 | 190967689 | mNeoVis.2.1.1.chr7\_hap1.g01994.t01 |
| 9600 | 7 | 191386552 | 191394286 | mNeoVis.2.1.1.chr7\_hap1.g02001.t01 |
| 9601 | 7 | 191394378 | 191406845 | mNeoVis.2.1.1.chr7\_hap1.g02002.t01 |
| 9602 | 7 | 191407997 | 191447969 | mNeoVis.2.1.1.chr7\_hap1.g02003.t01 |
| 9603 | 7 | 191449274 | 191464682 | mNeoVis.2.1.1.chr7\_hap1.g02004.t01 |
| 9604 | 7 | 191466615 | 191481790 | mNeoVis.2.1.1.chr7\_hap1.g02005.t01 |
| 9605 | 7 | 191504808 | 191513701 | mNeoVis.2.1.1.chr7\_hap1.g02006.t01 |
| 9606 | 7 | 191515885 | 191522072 | mNeoVis.2.1.1.chr7\_hap1.g02007.t01 |
| 9607 | 7 | 191523669 | 191533394 | mNeoVis.2.1.1.chr7\_hap1.g02008.t01 |
| 9608 | 7 | 191545868 | 191602796 | mNeoVis.2.1.1.chr7\_hap1.g02009.t01 |
| 9609 | 7 | 191638537 | 191657060 | mNeoVis.2.1.1.chr7\_hap1.g02010.t01 |
| 9610 | 7 | 191644989 | 191651874 | mNeoVis.2.1.1.chr7\_hap1.g02011.t01 |
| 9611 | 7 | 191657207 | 191660456 | mNeoVis.2.1.1.chr7\_hap1.g02012.t01 |
| 9612 | 7 | 191661123 | 191662330 | mNeoVis.2.1.1.chr7\_hap1.g02013.t01 |
| 9613 | 7 | 191687182 | 191706298 | mNeoVis.2.1.1.chr7\_hap1.g02014.t01 |
| 9614 | 7 | 191727073 | 191772202 | mNeoVis.2.1.1.chr7\_hap1.g02015.t01 |
| 9615 | 7 | 191773732 | 191816728 | mNeoVis.2.1.1.chr7\_hap1.g02016.t01 |
| 9616 | 7 | 191827155 | 191881202 | mNeoVis.2.1.1.chr7\_hap1.g02017.t01 |
| 9617 | 7 | 191969853 | 192133523 | mNeoVis.2.1.1.chr7\_hap1.g02018.t01 |
| 9618 | 7 | 192155711 | 192156690 | mNeoVis.2.1.1.chr7\_hap1.g02019.t01 |
| 9619 | 7 | 192178348 | 192179265 | mNeoVis.2.1.1.chr7\_hap1.g02020.t01 |
| 9620 | 7 | 192876664 | 192877581 | mNeoVis.2.1.1.chr7\_hap1.g02052.t01 |
| 9621 | 7 | 192897996 | 192899433 | mNeoVis.2.1.1.chr7\_hap1.g02053.t01 |
| 9622 | 7 | 192932073 | 192932975 | mNeoVis.2.1.1.chr7\_hap1.g02054.t01 |
| 9623 | 7 | 192955543 | 192956573 | mNeoVis.2.1.1.chr7\_hap1.g02055.t01 |
| 9624 | 7 | 192964715 | 192965629 | mNeoVis.2.1.1.chr7\_hap1.g02056.t01 |
| 9625 | 7 | 192984603 | 192985505 | mNeoVis.2.1.1.chr7\_hap1.g02057.t01 |
| 9626 | 7 | 193012120 | 193013052 | mNeoVis.2.1.1.chr7\_hap1.g02058.t01 |
| 9627 | 7 | 193030740 | 193031666 | mNeoVis.2.1.1.chr7\_hap1.g02059.t01 |
| 9628 | 7 | 193060294 | 193061223 | mNeoVis.2.1.1.chr7\_hap1.g02060.t01 |
| 9629 | 7 | 193076684 | 193077764 | mNeoVis.2.1.1.chr7\_hap1.g02061.t01 |
| 9630 | 7 | 193097745 | 193098659 | mNeoVis.2.1.1.chr7\_hap1.g02062.t01 |
| 9631 | 7 | 193106990 | 193107916 | mNeoVis.2.1.1.chr7\_hap1.g02063.t01 |
| 9632 | 7 | 193136400 | 193137332 | mNeoVis.2.1.1.chr7\_hap1.g02064.t01 |
| 9633 | 7 | 193158785 | 193159711 | mNeoVis.2.1.1.chr7\_hap1.g02065.t01 |
| 9634 | 7 | 193182231 | 193183163 | mNeoVis.2.1.1.chr7\_hap1.g02066.t01 |
| 9635 | 7 | 193192392 | 193193378 | mNeoVis.2.1.1.chr7\_hap1.g02067.t01 |
| 9636 | 7 | 193207336 | 193208268 | mNeoVis.2.1.1.chr7\_hap1.g02068.t01 |
| 9637 | 7 | 193231528 | 193232460 | mNeoVis.2.1.1.chr7\_hap1.g02069.t01 |
| 9638 | 7 | 193250645 | 193251577 | mNeoVis.2.1.1.chr7\_hap1.g02070.t01 |
| 9639 | 7 | 193269718 | 193286366 | mNeoVis.2.1.1.chr7\_hap1.g02071.t01 |
| 9640 | 7 | 193296060 | 193296992 | mNeoVis.2.1.1.chr7\_hap1.g02072.t01 |
| 9641 | 7 | 193339041 | 193340173 | mNeoVis.2.1.1.chr7\_hap1.g02073.t01 |
| 9642 | 7 | 193376427 | 193409690 | mNeoVis.2.1.1.chr7\_hap1.g02074.t01 |
| 9643 | 7 | 193397194 | 193420401 | mNeoVis.2.1.1.chr7\_hap1.g02075.t01 |
| 9644 | 7 | 193442640 | 193443572 | mNeoVis.2.1.1.chr7\_hap1.g02076.t01 |
| 9645 | 7 | 193455673 | 193456605 | mNeoVis.2.1.1.chr7\_hap1.g02077.t01 |
| 9646 | 7 | 196057573 | 196094810 | mNeoVis.2.1.1.chr7\_hap1.g02198.t01 |
| 9647 | 7 | 196067407 | 196068360 | mNeoVis.2.1.1.chr7\_hap1.g02199.t01 |
| 9648 | 7 | 196108019 | 196108963 | mNeoVis.2.1.1.chr7\_hap1.g02200.t01 |
| 9649 | 7 | 196117119 | 196118057 | mNeoVis.2.1.1.chr7\_hap1.g02201.t01 |
| 9650 | 7 | 196152773 | 196153717 | mNeoVis.2.1.1.chr7\_hap1.g02202.t01 |
| 9651 | 7 | 196164566 | 196165504 | mNeoVis.2.1.1.chr7\_hap1.g02203.t01 |
| 9652 | 7 | 196195103 | 196196041 | mNeoVis.2.1.1.chr7\_hap1.g02204.t01 |
| 9653 | 7 | 196250790 | 196251762 | mNeoVis.2.1.1.chr7\_hap1.g02205.t01 |
| 9654 | 7 | 196270683 | 196309293 | mNeoVis.2.1.1.chr7\_hap1.g02206.t01 |
| 9655 | 7 | 196310179 | 196360323 | mNeoVis.2.1.1.chr7\_hap1.g02207.t01 |
| 9656 | 7 | 196362302 | 196367938 | mNeoVis.2.1.1.chr7\_hap1.g02208.t01 |
| 9657 | 7 | 196412702 | 196441501 | mNeoVis.2.1.1.chr7\_hap1.g02209.t01 |
| 9658 | 7 | 196455814 | 196484065 | mNeoVis.2.1.1.chr7\_hap1.g02210.t01 |
| 9659 | 7 | 196490440 | 196497426 | mNeoVis.2.1.1.chr7\_hap1.g02211.t01 |
| 9660 | 7 | 196530061 | 196540655 | mNeoVis.2.1.1.chr7\_hap1.g02212.t01 |
| 9661 | 7 | 196553404 | 196562633 | mNeoVis.2.1.1.chr7\_hap1.g02213.t01 |
| 9662 | 7 | 196994893 | 196995828 | mNeoVis.2.1.1.chr7\_hap1.g02230.t01 |
| 9663 | 7 | 197006903 | 197007841 | mNeoVis.2.1.1.chr7\_hap1.g02231.t01 |
| 9664 | 7 | 197016898 | 197017833 | mNeoVis.2.1.1.chr7\_hap1.g02232.t01 |
| 9665 | 7 | 197027536 | 197028486 | mNeoVis.2.1.1.chr7\_hap1.g02233.t01 |
| 9666 | 7 | 197054150 | 197087726 | mNeoVis.2.1.1.chr7\_hap1.g02234.t01 |
| 9667 | 7 | 197106673 | 197138021 | mNeoVis.2.1.1.chr7\_hap1.g02235.t01 |
| 9668 | 7 | 197143221 | 197146116 | mNeoVis.2.1.1.chr7\_hap1.g02236.t01 |
| 9669 | 7 | 197182649 | 197183578 | mNeoVis.2.1.1.chr7\_hap1.g02237.t01 |
| 9670 | 7 | 197214023 | 197215153 | mNeoVis.2.1.1.chr7\_hap1.g02238.t01 |
| 9671 | 7 | 197226107 | 197261246 | mNeoVis.2.1.1.chr7\_hap1.g02239.t01 |
| 9672 | 7 | 197268668 | 197274401 | mNeoVis.2.1.1.chr7\_hap1.g02240.t01 |
| 9673 | 7 | 197274404 | 197362258 | mNeoVis.2.1.1.chr7\_hap1.g02241.t01 |
| 9674 | 7 | 197279801 | 197301431 | mNeoVis.2.1.1.chr7\_hap1.g02242.t01 |
| 9675 | 7 | 197306758 | 197320734 | mNeoVis.2.1.1.chr7\_hap1.g02243.t01 |
| 9676 | 7 | 197372694 | 197377916 | mNeoVis.2.1.1.chr7\_hap1.g02244.t01 |
| 9677 | 7 | 197390876 | 197398710 | mNeoVis.2.1.1.chr7\_hap1.g02245.t01 |
| 9678 | 7 | 197414865 | 197421004 | mNeoVis.2.1.1.chr7\_hap1.g02246.t01 |
| 9679 | 7 | 197452089 | 197464991 | mNeoVis.2.1.1.chr7\_hap1.g02247.t01 |
| 9680 | 7 | 197482224 | 197486145 | mNeoVis.2.1.1.chr7\_hap1.g02248.t01 |
| 9681 | 7 | 197497213 | 197506952 | mNeoVis.2.1.1.chr7\_hap1.g02249.t01 |
| 9682 | 7 | 197515009 | 197527017 | mNeoVis.2.1.1.chr7\_hap1.g02250.t01 |
| 9683 | 7 | 197550736 | 197752908 | mNeoVis.2.1.1.chr7\_hap1.g02251.t01 |
| 9684 | 7 | 197596684 | 197618592 | mNeoVis.2.1.1.chr7\_hap1.g02252.t01 |
| 9685 | 7 | 197623514 | 197648312 | mNeoVis.2.1.1.chr7\_hap1.g02253.t01 |
| 9686 | 7 | 197656788 | 197678665 | mNeoVis.2.1.1.chr7\_hap1.g02254.t01 |
| 9687 | 7 | 197683407 | 197695587 | mNeoVis.2.1.1.chr7\_hap1.g02255.t01 |
| 9688 | 7 | 197734189 | 197746183 | mNeoVis.2.1.1.chr7\_hap1.g02256.t01 |
| 9689 | 7 | 197752960 | 197768291 | mNeoVis.2.1.1.chr7\_hap1.g02257.t01 |
| 9690 | 7 | 200976344 | 200987899 | mNeoVis.2.1.1.chr7\_hap1.g02373.t01 |
| 9691 | 7 | 200993357 | 201006416 | mNeoVis.2.1.1.chr7\_hap1.g02374.t01 |
| 9692 | 7 | 201014759 | 201032052 | mNeoVis.2.1.1.chr7\_hap1.g02375.t01 |
| 9693 | 7 | 201032402 | 201038703 | mNeoVis.2.1.1.chr7\_hap1.g02376.t01 |
| 9694 | 7 | 201047833 | 201066913 | mNeoVis.2.1.1.chr7\_hap1.g02377.t01 |
| 9695 | 7 | 201074691 | 201096348 | mNeoVis.2.1.1.chr7\_hap1.g02378.t01 |
| 9696 | 7 | 201109916 | 201129400 | mNeoVis.2.1.1.chr7\_hap1.g02379.t01 |
| 9697 | 7 | 201134603 | 201142429 | mNeoVis.2.1.1.chr7\_hap1.g02380.t01 |
| 9698 | 7 | 201142580 | 201145530 | mNeoVis.2.1.1.chr7\_hap1.g02381.t01 |
| 9699 | 7 | 201145656 | 201173585 | mNeoVis.2.1.1.chr7\_hap1.g02382.t01 |
| 9700 | 7 | 201189279 | 201206477 | mNeoVis.2.1.1.chr7\_hap1.g02383.t01 |
| 9701 | 7 | 201207325 | 201212395 | mNeoVis.2.1.1.chr7\_hap1.g02384.t01 |
| 9702 | 7 | 201214896 | 201227516 | mNeoVis.2.1.1.chr7\_hap1.g02385.t01 |
| 9703 | 7 | 201227904 | 201230373 | mNeoVis.2.1.1.chr7\_hap1.g02386.t01 |
| 9704 | 7 | 201230520 | 201242669 | mNeoVis.2.1.1.chr7\_hap1.g02387.t01 |
| 9705 | 7 | 201251910 | 201264712 | mNeoVis.2.1.1.chr7\_hap1.g02388.t01 |
| 9706 | 7 | 201264798 | 201268871 | mNeoVis.2.1.1.chr7\_hap1.g02389.t01 |
| 9707 | 7 | 201268842 | 201272575 | mNeoVis.2.1.1.chr7\_hap1.g02390.t01 |
| 9708 | 7 | 201273501 | 201290196 | mNeoVis.2.1.1.chr7\_hap1.g02391.t01 |
| 9709 | 7 | 201290258 | 201294879 | mNeoVis.2.1.1.chr7\_hap1.g02392.t01 |
| 9710 | 7 | 201295064 | 201299067 | mNeoVis.2.1.1.chr7\_hap1.g02393.t01 |
| 9711 | 7 | 201299384 | 201300959 | mNeoVis.2.1.1.chr7\_hap1.g02394.t01 |
| 9712 | 7 | 201300981 | 201305127 | mNeoVis.2.1.1.chr7\_hap1.g02395.t01 |
| 9713 | 7 | 201306032 | 201312536 | mNeoVis.2.1.1.chr7\_hap1.g02396.t01 |
| 9714 | 7 | 201312353 | 201343943 | mNeoVis.2.1.1.chr7\_hap1.g02397.t01 |
| 9715 | 7 | 201352025 | 201374097 | mNeoVis.2.1.1.chr7\_hap1.g02398.t01 |
| 9716 | 7 | 201375439 | 201400213 | mNeoVis.2.1.1.chr7\_hap1.g02399.t01 |
| 9717 | 7 | 201418686 | 201432045 | mNeoVis.2.1.1.chr7\_hap1.g02400.t01 |
| 9718 | 7 | 201456594 | 201484000 | mNeoVis.2.1.1.chr7\_hap1.g02401.t01 |
| 9719 | 7 | 201494711 | 201500802 | mNeoVis.2.1.1.chr7\_hap1.g02402.t01 |
| 9720 | 7 | 201510263 | 201524221 | mNeoVis.2.1.1.chr7\_hap1.g02403.t01 |
| 9721 | 7 | 201524174 | 201530074 | mNeoVis.2.1.1.chr7\_hap1.g02404.t01 |
| 9722 | 7 | 201536790 | 201542643 | mNeoVis.2.1.1.chr7\_hap1.g02405.t01 |
| 9723 | 7 | 201536790 | 201542643 | mNeoVis.2.1.1.chr7\_hap1.g02405.t01 |
| 9724 | 7 | 201552502 | 201553750 | mNeoVis.2.1.1.chr7\_hap1.g02406.t01 |
| 9725 | 7 | 201553890 | 201556179 | mNeoVis.2.1.1.chr7\_hap1.g02407.t01 |
| 9726 | 7 | 201558209 | 201574342 | mNeoVis.2.1.1.chr7\_hap1.g02408.t01 |
| 9727 | 7 | 201574764 | 201577877 | mNeoVis.2.1.1.chr7\_hap1.g02409.t01 |
| 9728 | 7 | 201578980 | 201593808 | mNeoVis.2.1.1.chr7\_hap1.g02410.t01 |
| 9729 | 7 | 201594735 | 201614221 | mNeoVis.2.1.1.chr7\_hap1.g02411.t01 |
| 9730 | 7 | 201615004 | 201626451 | mNeoVis.2.1.1.chr7\_hap1.g02412.t01 |
| 9731 | 7 | 201628268 | 201637004 | mNeoVis.2.1.1.chr7\_hap1.g02413.t01 |
| 9732 | 7 | 201670521 | 201677768 | mNeoVis.2.1.1.chr7\_hap1.g02414.t01 |
| 9733 | 7 | 201675335 | 201679372 | mNeoVis.2.1.1.chr7\_hap1.g02415.t01 |
| 9734 | 7 | 201700965 | 201706866 | mNeoVis.2.1.1.chr7\_hap1.g02416.t01 |
| 9735 | 7 | 201711661 | 201723393 | mNeoVis.2.1.1.chr7\_hap1.g02417.t01 |
| 9736 | 7 | 201713770 | 201752421 | mNeoVis.2.1.1.chr7\_hap1.g02418.t01 |
| 9737 | 7 | 201752855 | 201765027 | mNeoVis.2.1.1.chr7\_hap1.g02419.t01 |
| 9738 | 7 | 201766045 | 201769823 | mNeoVis.2.1.1.chr7\_hap1.g02420.t01 |
| 9739 | 7 | 201769825 | 201777558 | mNeoVis.2.1.1.chr7\_hap1.g02421.t01 |
| 9740 | 7 | 201778225 | 201784200 | mNeoVis.2.1.1.chr7\_hap1.g02422.t01 |
| 9741 | 7 | 201786963 | 201791392 | mNeoVis.2.1.1.chr7\_hap1.g02423.t01 |
| 9742 | 7 | 201791521 | 201795825 | mNeoVis.2.1.1.chr7\_hap1.g02424.t01 |
| 9743 | 7 | 201797389 | 201799540 | mNeoVis.2.1.1.chr7\_hap1.g02425.t01 |
| 9744 | 7 | 201799678 | 201804638 | mNeoVis.2.1.1.chr7\_hap1.g02426.t01 |
| 9745 | 7 | 201818780 | 201821069 | mNeoVis.2.1.1.chr7\_hap1.g02427.t01 |
| 9746 | 7 | 201821205 | 201823528 | mNeoVis.2.1.1.chr7\_hap1.g02428.t01 |
| 9747 | 7 | 201848380 | 201856942 | mNeoVis.2.1.1.chr7\_hap1.g02429.t01 |
| 9748 | 7 | 201857883 | 201873800 | mNeoVis.2.1.1.chr7\_hap1.g02430.t01 |
| 9749 | 7 | 201896529 | 201899557 | mNeoVis.2.1.1.chr7\_hap1.g02431.t01 |
| 9750 | 7 | 201899816 | 201901473 | mNeoVis.2.1.1.chr7\_hap1.g02432.t01 |
| 9751 | 7 | 201907535 | 201909002 | mNeoVis.2.1.1.chr7\_hap1.g02433.t01 |
| 9752 | 7 | 201911015 | 201920602 | mNeoVis.2.1.1.chr7\_hap1.g02434.t01 |
| 9753 | 7 | 201936326 | 201937574 | mNeoVis.2.1.1.chr7\_hap1.g02435.t01 |
| 9754 | 7 | 201938340 | 201940004 | mNeoVis.2.1.1.chr7\_hap1.g02436.t01 |
| 9755 | 7 | 201942036 | 201958256 | mNeoVis.2.1.1.chr7\_hap1.g02437.t01 |
| 9756 | 7 | 201958550 | 201961649 | mNeoVis.2.1.1.chr7\_hap1.g02438.t01 |
| 9757 | 7 | 201958550 | 201961649 | mNeoVis.2.1.1.chr7\_hap1.g02438.t01 |
| 9758 | 7 | 201962763 | 201976932 | mNeoVis.2.1.1.chr7\_hap1.g02439.t01 |
| 9759 | 7 | 201962763 | 201976932 | mNeoVis.2.1.1.chr7\_hap1.g02439.t01 |
| 9760 | 7 | 201978528 | 201998055 | mNeoVis.2.1.1.chr7\_hap1.g02440.t01 |
| 9761 | 7 | 201978528 | 201998055 | mNeoVis.2.1.1.chr7\_hap1.g02440.t01 |
| 9762 | 7 | 202000915 | 202010398 | mNeoVis.2.1.1.chr7\_hap1.g02441.t01 |
| 9763 | 7 | 202000915 | 202010398 | mNeoVis.2.1.1.chr7\_hap1.g02441.t01 |
| 9764 | 7 | 202012104 | 202020841 | mNeoVis.2.1.1.chr7\_hap1.g02442.t01 |
| 9765 | 7 | 202012104 | 202020841 | mNeoVis.2.1.1.chr7\_hap1.g02442.t01 |
| 9766 | 7 | 202054329 | 202061575 | mNeoVis.2.1.1.chr7\_hap1.g02443.t01 |
| 9767 | 7 | 202054329 | 202061575 | mNeoVis.2.1.1.chr7\_hap1.g02443.t01 |
| 9768 | 7 | 202061771 | 202063180 | mNeoVis.2.1.1.chr7\_hap1.g02444.t01 |
| 9769 | 7 | 202061771 | 202063180 | mNeoVis.2.1.1.chr7\_hap1.g02444.t01 |
| 9770 | 7 | 202082362 | 202088271 | mNeoVis.2.1.1.chr7\_hap1.g02445.t01 |
| 9771 | 7 | 202082362 | 202088271 | mNeoVis.2.1.1.chr7\_hap1.g02445.t01 |
| 9772 | 7 | 202093068 | 202104786 | mNeoVis.2.1.1.chr7\_hap1.g02446.t01 |
| 9773 | 7 | 202093068 | 202104786 | mNeoVis.2.1.1.chr7\_hap1.g02446.t01 |
| 9774 | 7 | 202095153 | 202133829 | mNeoVis.2.1.1.chr7\_hap1.g02447.t01 |
| 9775 | 7 | 202095153 | 202133829 | mNeoVis.2.1.1.chr7\_hap1.g02447.t01 |
| 9776 | 7 | 202134265 | 202146434 | mNeoVis.2.1.1.chr7\_hap1.g02448.t01 |
| 9777 | 7 | 202134265 | 202146434 | mNeoVis.2.1.1.chr7\_hap1.g02448.t01 |
| 9778 | 7 | 202147448 | 202151208 | mNeoVis.2.1.1.chr7\_hap1.g02449.t01 |
| 9779 | 7 | 202147448 | 202151208 | mNeoVis.2.1.1.chr7\_hap1.g02449.t01 |
| 9780 | 7 | 202151051 | 202158977 | mNeoVis.2.1.1.chr7\_hap1.g02450.t01 |
| 9781 | 7 | 202151051 | 202158977 | mNeoVis.2.1.1.chr7\_hap1.g02450.t01 |
| 9782 | 7 | 202159644 | 202165615 | mNeoVis.2.1.1.chr7\_hap1.g02451.t01 |
| 9783 | 7 | 202159644 | 202165615 | mNeoVis.2.1.1.chr7\_hap1.g02451.t01 |
| 9784 | 7 | 202167385 | 202172812 | mNeoVis.2.1.1.chr7\_hap1.g02452.t01 |
| 9785 | 7 | 202167385 | 202172812 | mNeoVis.2.1.1.chr7\_hap1.g02452.t01 |
| 9786 | 7 | 202172992 | 202177240 | mNeoVis.2.1.1.chr7\_hap1.g02453.t01 |
| 9787 | 7 | 202178746 | 202180959 | mNeoVis.2.1.1.chr7\_hap1.g02454.t01 |
| 9788 | 7 | 202181097 | 202186058 | mNeoVis.2.1.1.chr7\_hap1.g02455.t01 |
| 9789 | 7 | 202200193 | 202202482 | mNeoVis.2.1.1.chr7\_hap1.g02456.t01 |
| 9790 | 7 | 202202618 | 202204941 | mNeoVis.2.1.1.chr7\_hap1.g02457.t01 |
| 9791 | 7 | 202229799 | 202238343 | mNeoVis.2.1.1.chr7\_hap1.g02458.t01 |
| 9792 | 7 | 202239280 | 202254781 | mNeoVis.2.1.1.chr7\_hap1.g02459.t01 |
| 9793 | 7 | 202276003 | 202280742 | mNeoVis.2.1.1.chr7\_hap1.g02460.t01 |
| 9794 | 7 | 202281009 | 202282654 | mNeoVis.2.1.1.chr7\_hap1.g02461.t01 |
| 9795 | 7 | 202288726 | 202290190 | mNeoVis.2.1.1.chr7\_hap1.g02462.t01 |
| 9796 | 7 | 202292190 | 202301778 | mNeoVis.2.1.1.chr7\_hap1.g02463.t01 |
| 9797 | 7 | 202324240 | 202332879 | mNeoVis.2.1.1.chr7\_hap1.g02464.t01 |
| 9798 | 7 | 202335795 | 202350187 | mNeoVis.2.1.1.chr7\_hap1.g02465.t01 |
| 9799 | 7 | 202351748 | 202493925 | mNeoVis.2.1.1.chr7\_hap1.g02466.t01 |
| 9800 | 7 | 202505433 | 202515356 | mNeoVis.2.1.1.chr7\_hap1.g02467.t01 |
| 9801 | 7 | 202516846 | 202523675 | mNeoVis.2.1.1.chr7\_hap1.g02468.t01 |
| 9802 | 7 | 202524536 | 202530415 | mNeoVis.2.1.1.chr7\_hap1.g02469.t01 |
| 9803 | 7 | 202530779 | 202535161 | mNeoVis.2.1.1.chr7\_hap1.g02470.t01 |
| 9804 | 7 | 202537515 | 202542276 | mNeoVis.2.1.1.chr7\_hap1.g02471.t01 |
| 9805 | 7 | 202554426 | 202557037 | mNeoVis.2.1.1.chr7\_hap1.g02472.t01 |
| 9806 | 7 | 202566344 | 202573139 | mNeoVis.2.1.1.chr7\_hap1.g02473.t01 |
| 9807 | 7 | 202573549 | 202580752 | mNeoVis.2.1.1.chr7\_hap1.g02474.t01 |
| 9808 | 7 | 202581005 | 202584423 | mNeoVis.2.1.1.chr7\_hap1.g02475.t01 |
| 9809 | 7 | 203366067 | 203385153 | mNeoVis.2.1.1.chr7\_hap1.g02498.t01 |
| 9810 | 7 | 203387857 | 203397957 | mNeoVis.2.1.1.chr7\_hap1.g02499.t01 |
| 9811 | 7 | 203398985 | 203406348 | mNeoVis.2.1.1.chr7\_hap1.g02500.t01 |
| 9812 | 7 | 203443817 | 203503201 | mNeoVis.2.1.1.chr7\_hap1.g02501.t01 |
| 9813 | 7 | 203505407 | 203516195 | mNeoVis.2.1.1.chr7\_hap1.g02502.t01 |
| 9814 | 7 | 203518511 | 203522644 | mNeoVis.2.1.1.chr7\_hap1.g02503.t01 |
| 9815 | 7 | 203523886 | 203540510 | mNeoVis.2.1.1.chr7\_hap1.g02504.t01 |
| 9816 | 7 | 203536842 | 203558743 | mNeoVis.2.1.1.chr7\_hap1.g02505.t01 |
| 9817 | 7 | 203576870 | 203581057 | mNeoVis.2.1.1.chr7\_hap1.g02506.t01 |
| 9818 | 7 | 203587091 | 203596021 | mNeoVis.2.1.1.chr7\_hap1.g02507.t01 |
| 9819 | 7 | 203597296 | 203599325 | mNeoVis.2.1.1.chr7\_hap1.g02508.t01 |
| 9820 | 7 | 203599475 | 203605014 | mNeoVis.2.1.1.chr7\_hap1.g02509.t01 |
| 9821 | 7 | 203605406 | 203610693 | mNeoVis.2.1.1.chr7\_hap1.g02510.t01 |
| 9822 | 7 | 203624536 | 203627481 | mNeoVis.2.1.1.chr7\_hap1.g02511.t01 |
| 9823 | 7 | 203656070 | 203658210 | mNeoVis.2.1.1.chr7\_hap1.g02512.t01 |
| 9824 | 7 | 203666287 | 203670119 | mNeoVis.2.1.1.chr7\_hap1.g02513.t01 |
| 9825 | 7 | 203675801 | 203677555 | mNeoVis.2.1.1.chr7\_hap1.g02514.t01 |
| 9826 | 7 | 203678715 | 203692209 | mNeoVis.2.1.1.chr7\_hap1.g02515.t01 |
| 9827 | 7 | 203692807 | 203698962 | mNeoVis.2.1.1.chr7\_hap1.g02516.t01 |
| 9828 | 7 | 203707173 | 203712078 | mNeoVis.2.1.1.chr7\_hap1.g02517.t01 |
| 9829 | 7 | 203715394 | 203719665 | mNeoVis.2.1.1.chr7\_hap1.g02518.t01 |
| 9830 | 7 | 203720485 | 203723361 | mNeoVis.2.1.1.chr7\_hap1.g02519.t01 |
| 9831 | 7 | 203727651 | 203732840 | mNeoVis.2.1.1.chr7\_hap1.g02520.t01 |
| 9832 | 7 | 203732985 | 203735339 | mNeoVis.2.1.1.chr7\_hap1.g02521.t01 |
| 9833 | 7 | 203735450 | 203741063 | mNeoVis.2.1.1.chr7\_hap1.g02522.t01 |
| 9834 | 7 | 203742207 | 203750144 | mNeoVis.2.1.1.chr7\_hap1.g02523.t01 |
| 9835 | 7 | 203756580 | 203762302 | mNeoVis.2.1.1.chr7\_hap1.g02524.t01 |
| 9836 | 7 | 203763924 | 203767146 | mNeoVis.2.1.1.chr7\_hap1.g02525.t01 |
| 9837 | 7 | 203766925 | 203772690 | mNeoVis.2.1.1.chr7\_hap1.g02526.t01 |
| 9838 | 7 | 203784495 | 203794023 | mNeoVis.2.1.1.chr7\_hap1.g02527.t01 |
| 9839 | 7 | 203801029 | 203802821 | mNeoVis.2.1.1.chr7\_hap1.g02528.t01 |
| 9840 | 7 | 204005813 | 204007773 | mNeoVis.2.1.1.chr7\_hap1.g02548.t01 |
| 9841 | 7 | 204008956 | 204017441 | mNeoVis.2.1.1.chr7\_hap1.g02549.t01 |
| 9842 | 7 | 204020446 | 204028259 | mNeoVis.2.1.1.chr7\_hap1.g02550.t01 |
| 9843 | 7 | 204046378 | 204068677 | mNeoVis.2.1.1.chr7\_hap1.g02551.t01 |
| 9844 | 7 | 204064921 | 204081643 | mNeoVis.2.1.1.chr7\_hap1.g02552.t01 |
| 9845 | 7 | 204082890 | 204087020 | mNeoVis.2.1.1.chr7\_hap1.g02553.t01 |
| 9846 | 7 | 204089355 | 204100123 | mNeoVis.2.1.1.chr7\_hap1.g02554.t01 |
| 9847 | 7 | 204102327 | 204161668 | mNeoVis.2.1.1.chr7\_hap1.g02555.t01 |
| 9848 | 7 | 204172328 | 204174120 | mNeoVis.2.1.1.chr7\_hap1.g02556.t01 |
| 9849 | 7 | 204181126 | 204190654 | mNeoVis.2.1.1.chr7\_hap1.g02557.t01 |
| 9850 | 7 | 204202459 | 204208224 | mNeoVis.2.1.1.chr7\_hap1.g02558.t01 |
| 9851 | 7 | 204208003 | 204211225 | mNeoVis.2.1.1.chr7\_hap1.g02559.t01 |
| 9852 | 7 | 204212802 | 204218569 | mNeoVis.2.1.1.chr7\_hap1.g02560.t01 |
| 9853 | 7 | 204223346 | 204232942 | mNeoVis.2.1.1.chr7\_hap1.g02561.t01 |
| 9854 | 7 | 204234086 | 204239699 | mNeoVis.2.1.1.chr7\_hap1.g02562.t01 |
| 9855 | 7 | 204239810 | 204242171 | mNeoVis.2.1.1.chr7\_hap1.g02563.t01 |
| 9856 | 7 | 204242309 | 204247498 | mNeoVis.2.1.1.chr7\_hap1.g02564.t01 |
| 9857 | 7 | 204251788 | 204254664 | mNeoVis.2.1.1.chr7\_hap1.g02565.t01 |
| 9858 | 7 | 204255484 | 204259755 | mNeoVis.2.1.1.chr7\_hap1.g02566.t01 |
| 9859 | 7 | 204263071 | 204267981 | mNeoVis.2.1.1.chr7\_hap1.g02567.t01 |
| 9860 | 7 | 204276264 | 204282342 | mNeoVis.2.1.1.chr7\_hap1.g02568.t01 |
| 9861 | 7 | 204282940 | 204296432 | mNeoVis.2.1.1.chr7\_hap1.g02569.t01 |
| 9862 | 7 | 204297594 | 204299355 | mNeoVis.2.1.1.chr7\_hap1.g02570.t01 |
| 9863 | 7 | 204305030 | 204308862 | mNeoVis.2.1.1.chr7\_hap1.g02571.t01 |
| 9864 | 7 | 204316939 | 204319079 | mNeoVis.2.1.1.chr7\_hap1.g02572.t01 |
| 9865 | 7 | 204347665 | 204350613 | mNeoVis.2.1.1.chr7\_hap1.g02573.t01 |
| 9866 | 7 | 204364453 | 204369743 | mNeoVis.2.1.1.chr7\_hap1.g02574.t01 |
| 9867 | 7 | 204370135 | 204375674 | mNeoVis.2.1.1.chr7\_hap1.g02575.t01 |
| 9868 | 7 | 204375824 | 204377853 | mNeoVis.2.1.1.chr7\_hap1.g02576.t01 |
| 9869 | 7 | 204379128 | 204388058 | mNeoVis.2.1.1.chr7\_hap1.g02577.t01 |
| 9870 | 7 | 204394092 | 204398279 | mNeoVis.2.1.1.chr7\_hap1.g02578.t01 |
| 9871 | 7 | 204416406 | 204438309 | mNeoVis.2.1.1.chr7\_hap1.g02579.t01 |
| 9872 | 7 | 204434549 | 204451263 | mNeoVis.2.1.1.chr7\_hap1.g02580.t01 |
| 9873 | 7 | 204452526 | 204456638 | mNeoVis.2.1.1.chr7\_hap1.g02581.t01 |
| 9874 | 7 | 204458954 | 204469742 | mNeoVis.2.1.1.chr7\_hap1.g02582.t01 |
| 9875 | 7 | 204471948 | 204531332 | mNeoVis.2.1.1.chr7\_hap1.g02583.t01 |
| 9876 | 7 | 204601482 | 204612143 | mNeoVis.2.1.1.chr7\_hap1.g02584.t01 |
| 9877 | 7 | 205059408 | 205071483 | mNeoVis.2.1.1.chr7\_hap1.g02590.t01 |
| 9878 | 7 | 205071438 | 205097020 | mNeoVis.2.1.1.chr7\_hap1.g02591.t01 |
| 9879 | 7 | 205113886 | 205123118 | mNeoVis.2.1.1.chr7\_hap1.g02592.t01 |
| 9880 | 7 | 205128503 | 205137319 | mNeoVis.2.1.1.chr7\_hap1.g02593.t01 |
| 9881 | 7 | 205155697 | 205184310 | mNeoVis.2.1.1.chr7\_hap1.g02594.t01 |
| 9882 | 7 | 205231237 | 205235106 | mNeoVis.2.1.1.chr7\_hap1.g02595.t01 |
| 9883 | 7 | 205591890 | 205604456 | mNeoVis.2.1.1.chr7\_hap1.g02596.t01 |
| 9884 | 7 | 205609457 | 205617397 | mNeoVis.2.1.1.chr7\_hap1.g02597.t01 |
| 9885 | 7 | 205628117 | 205633081 | mNeoVis.2.1.1.chr7\_hap1.g02598.t01 |
| 9886 | 7 | 205679681 | 205681444 | mNeoVis.2.1.1.chr7\_hap1.g02599.t01 |
| 9887 | 7 | 205706883 | 205714051 | mNeoVis.2.1.1.chr7\_hap1.g02600.t01 |
| 9888 | 7 | 205835039 | 205970065 | mNeoVis.2.1.1.chr7\_hap1.g02601.t01 |
| 9889 | 7 | 205982099 | 205988074 | mNeoVis.2.1.1.chr7\_hap1.g02602.t01 |
| 9890 | 7 | 206013852 | 206096904 | mNeoVis.2.1.1.chr7\_hap1.g02603.t01 |
| 9891 | 7 | 206106630 | 206136530 | mNeoVis.2.1.1.chr7\_hap1.g02604.t01 |
| 9892 | 8 | 171473 | 185522 | mNeoVis.2.1.1.chr8\_hap1.g00002.t01 |
| 9893 | 8 | 206301 | 279028 | mNeoVis.2.1.1.chr8\_hap1.g00003.t01 |
| 9894 | 8 | 250533 | 256194 | mNeoVis.2.1.1.chr8\_hap1.g00004.t01 |
| 9895 | 8 | 279018 | 436941 | mNeoVis.2.1.1.chr8\_hap1.g00005.t01 |
| 9896 | 8 | 385502 | 388669 | mNeoVis.2.1.1.chr8\_hap1.g00006.t01 |
| 9897 | 8 | 438239 | 445460 | mNeoVis.2.1.1.chr8\_hap1.g00007.t01 |
| 9898 | 8 | 449765 | 457968 | mNeoVis.2.1.1.chr8\_hap1.g00008.t01 |
| 9899 | 8 | 457915 | 466865 | mNeoVis.2.1.1.chr8\_hap1.g00009.t01 |
| 9900 | 8 | 469833 | 496021 | mNeoVis.2.1.1.chr8\_hap1.g00010.t01 |
| 9901 | 8 | 500152 | 542545 | mNeoVis.2.1.1.chr8\_hap1.g00011.t01 |
| 9902 | 8 | 542880 | 609660 | mNeoVis.2.1.1.chr8\_hap1.g00012.t01 |
| 9903 | 8 | 619104 | 648362 | mNeoVis.2.1.1.chr8\_hap1.g00013.t01 |
| 9904 | 8 | 654496 | 660425 | mNeoVis.2.1.1.chr8\_hap1.g00014.t01 |
| 9905 | 8 | 660567 | 676641 | mNeoVis.2.1.1.chr8\_hap1.g00015.t01 |
| 9906 | 8 | 676872 | 692286 | mNeoVis.2.1.1.chr8\_hap1.g00016.t01 |
| 9907 | 8 | 697700 | 700627 | mNeoVis.2.1.1.chr8\_hap1.g00017.t01 |
| 9908 | 8 | 708335 | 710051 | mNeoVis.2.1.1.chr8\_hap1.g00018.t01 |
| 9909 | 8 | 856458 | 869496 | mNeoVis.2.1.1.chr8\_hap1.g00020.t01 |
| 9910 | 8 | 871224 | 874761 | mNeoVis.2.1.1.chr8\_hap1.g00021.t01 |
| 9911 | 8 | 902660 | 943247 | mNeoVis.2.1.1.chr8\_hap1.g00022.t01 |
| 9912 | 8 | 934808 | 941115 | mNeoVis.2.1.1.chr8\_hap1.g00023.t01 |
| 9913 | 8 | 952590 | 1042748 | mNeoVis.2.1.1.chr8\_hap1.g00024.t01 |
| 9914 | 8 | 1044469 | 1063388 | mNeoVis.2.1.1.chr8\_hap1.g00025.t01 |
| 9915 | 8 | 1070669 | 1076478 | mNeoVis.2.1.1.chr8\_hap1.g00026.t01 |
| 9916 | 8 | 1076833 | 1081772 | mNeoVis.2.1.1.chr8\_hap1.g00027.t01 |
| 9917 | 8 | 1082123 | 1086075 | mNeoVis.2.1.1.chr8\_hap1.g00028.t01 |
| 9918 | 8 | 1103278 | 1105335 | mNeoVis.2.1.1.chr8\_hap1.g00029.t01 |
| 9919 | 8 | 1107975 | 1109946 | mNeoVis.2.1.1.chr8\_hap1.g00030.t01 |
| 9920 | 8 | 1111813 | 1114234 | mNeoVis.2.1.1.chr8\_hap1.g00031.t01 |
| 9921 | 8 | 1114768 | 1122599 | mNeoVis.2.1.1.chr8\_hap1.g00032.t01 |
| 9922 | 8 | 1122762 | 1126143 | mNeoVis.2.1.1.chr8\_hap1.g00033.t01 |
| 9923 | 8 | 1127086 | 1150911 | mNeoVis.2.1.1.chr8\_hap1.g00034.t01 |
| 9924 | 8 | 1162882 | 1170588 | mNeoVis.2.1.1.chr8\_hap1.g00035.t01 |
| 9925 | 8 | 1175625 | 1177651 | mNeoVis.2.1.1.chr8\_hap1.g00036.t01 |
| 9926 | 8 | 1179961 | 1184518 | mNeoVis.2.1.1.chr8\_hap1.g00037.t01 |
| 9927 | 8 | 1188848 | 1197849 | mNeoVis.2.1.1.chr8\_hap1.g00038.t01 |
| 9928 | 8 | 1197943 | 1204064 | mNeoVis.2.1.1.chr8\_hap1.g00039.t01 |
| 9929 | 8 | 1204413 | 1211099 | mNeoVis.2.1.1.chr8\_hap1.g00040.t01 |
| 9930 | 8 | 1213010 | 1213573 | mNeoVis.2.1.1.chr8\_hap1.g00041.t01 |
| 9931 | 8 | 1216044 | 1223377 | mNeoVis.2.1.1.chr8\_hap1.g00042.t01 |
| 9932 | 8 | 1222334 | 1226083 | mNeoVis.2.1.1.chr8\_hap1.g00043.t01 |
| 9933 | 8 | 1237007 | 1241590 | mNeoVis.2.1.1.chr8\_hap1.g00044.t01 |
| 9934 | 8 | 1246431 | 1255989 | mNeoVis.2.1.1.chr8\_hap1.g00045.t01 |
| 9935 | 8 | 1259781 | 1261568 | mNeoVis.2.1.1.chr8\_hap1.g00046.t01 |
| 9936 | 8 | 1262071 | 1271150 | mNeoVis.2.1.1.chr8\_hap1.g00047.t01 |
| 9937 | 8 | 1274092 | 1281867 | mNeoVis.2.1.1.chr8\_hap1.g00048.t01 |
| 9938 | 8 | 1302736 | 1309315 | mNeoVis.2.1.1.chr8\_hap1.g00049.t01 |
| 9939 | 8 | 1312193 | 1316247 | mNeoVis.2.1.1.chr8\_hap1.g00050.t01 |
| 9940 | 8 | 1317248 | 1331236 | mNeoVis.2.1.1.chr8\_hap1.g00051.t01 |
| 9941 | 8 | 1331692 | 1333845 | mNeoVis.2.1.1.chr8\_hap1.g00052.t01 |
| 9942 | 8 | 1334818 | 1340896 | mNeoVis.2.1.1.chr8\_hap1.g00053.t01 |
| 9943 | 8 | 1340445 | 1347580 | mNeoVis.2.1.1.chr8\_hap1.g00054.t01 |
| 9944 | 8 | 1341088 | 1342503 | mNeoVis.2.1.1.chr8\_hap1.g00055.t01 |
| 9945 | 8 | 1348208 | 1354724 | mNeoVis.2.1.1.chr8\_hap1.g00056.t01 |
| 9946 | 8 | 1359065 | 1412760 | mNeoVis.2.1.1.chr8\_hap1.g00057.t01 |
| 9947 | 8 | 1412913 | 1434284 | mNeoVis.2.1.1.chr8\_hap1.g00058.t01 |
| 9948 | 8 | 1416737 | 1426105 | mNeoVis.2.1.1.chr8\_hap1.g00059.t01 |
| 9949 | 8 | 1446895 | 1449695 | mNeoVis.2.1.1.chr8\_hap1.g00060.t01 |
| 9950 | 8 | 1484663 | 1545136 | mNeoVis.2.1.1.chr8\_hap1.g00061.t01 |
| 9951 | 8 | 1624597 | 1664190 | mNeoVis.2.1.1.chr8\_hap1.g00062.t01 |
| 9952 | 8 | 1666337 | 1668797 | mNeoVis.2.1.1.chr8\_hap1.g00063.t01 |
| 9953 | 8 | 1668921 | 1679815 | mNeoVis.2.1.1.chr8\_hap1.g00064.t01 |
| 9954 | 8 | 1682071 | 1699194 | mNeoVis.2.1.1.chr8\_hap1.g00065.t01 |
| 9955 | 8 | 1703789 | 1706795 | mNeoVis.2.1.1.chr8\_hap1.g00066.t01 |
| 9956 | 8 | 2613159 | 2619519 | mNeoVis.2.1.1.chr8\_hap1.g00081.t01 |
| 9957 | 8 | 2653009 | 2655642 | mNeoVis.2.1.1.chr8\_hap1.g00082.t01 |
| 9958 | 8 | 2661298 | 2670732 | mNeoVis.2.1.1.chr8\_hap1.g00083.t01 |
| 9959 | 8 | 2682031 | 2682422 | mNeoVis.2.1.1.chr8\_hap1.g00084.t01 |
| 9960 | 8 | 2698899 | 2705964 | mNeoVis.2.1.1.chr8\_hap1.g00085.t01 |
| 9961 | 8 | 2717965 | 2743045 | mNeoVis.2.1.1.chr8\_hap1.g00086.t01 |
| 9962 | 8 | 2758218 | 3181721 | mNeoVis.2.1.1.chr8\_hap1.g00087.t01 |
| 9963 | 8 | 3181870 | 3188488 | mNeoVis.2.1.1.chr8\_hap1.g00088.t01 |
| 9964 | 8 | 3204626 | 3223095 | mNeoVis.2.1.1.chr8\_hap1.g00089.t01 |
| 9965 | 8 | 3229889 | 3258922 | mNeoVis.2.1.1.chr8\_hap1.g00090.t01 |
| 9966 | 8 | 3285196 | 3330177 | mNeoVis.2.1.1.chr8\_hap1.g00091.t01 |
| 9967 | 8 | 3296709 | 3306552 | mNeoVis.2.1.1.chr8\_hap1.g00092.t01 |
| 9968 | 8 | 3337568 | 3375535 | mNeoVis.2.1.1.chr8\_hap1.g00093.t01 |
| 9969 | 8 | 3421860 | 3462806 | mNeoVis.2.1.1.chr8\_hap1.g00094.t01 |
| 9970 | 8 | 3492133 | 3583180 | mNeoVis.2.1.1.chr8\_hap1.g00095.t01 |
| 9971 | 8 | 3582229 | 3617754 | mNeoVis.2.1.1.chr8\_hap1.g00096.t01 |
| 9972 | 8 | 3629798 | 3633099 | mNeoVis.2.1.1.chr8\_hap1.g00097.t01 |
| 9973 | 8 | 3706789 | 3712566 | mNeoVis.2.1.1.chr8\_hap1.g00098.t01 |
| 9974 | 8 | 3713568 | 3718399 | mNeoVis.2.1.1.chr8\_hap1.g00099.t01 |
| 9975 | 8 | 3724342 | 3732459 | mNeoVis.2.1.1.chr8\_hap1.g00100.t01 |
| 9976 | 8 | 3736440 | 3752700 | mNeoVis.2.1.1.chr8\_hap1.g00101.t01 |
| 9977 | 8 | 3752911 | 3763659 | mNeoVis.2.1.1.chr8\_hap1.g00102.t01 |
| 9978 | 8 | 3764970 | 3768875 | mNeoVis.2.1.1.chr8\_hap1.g00103.t01 |
| 9979 | 8 | 3769458 | 3780678 | mNeoVis.2.1.1.chr8\_hap1.g00104.t01 |
| 9980 | 8 | 3782577 | 3792164 | mNeoVis.2.1.1.chr8\_hap1.g00105.t01 |
| 9981 | 8 | 3792655 | 3809849 | mNeoVis.2.1.1.chr8\_hap1.g00106.t01 |
| 9982 | 8 | 3810174 | 3957179 | mNeoVis.2.1.1.chr8\_hap1.g00107.t01 |
| 9983 | 8 | 4265004 | 4348666 | mNeoVis.2.1.1.chr8\_hap1.g00108.t01 |
| 9984 | 8 | 6977758 | 7223454 | mNeoVis.2.1.1.chr8\_hap1.g00197.t01 |
| 9985 | 8 | 7243428 | 7263726 | mNeoVis.2.1.1.chr8\_hap1.g00198.t01 |
| 9986 | 8 | 7275748 | 7282537 | mNeoVis.2.1.1.chr8\_hap1.g00199.t01 |
| 9987 | 8 | 7287452 | 7298682 | mNeoVis.2.1.1.chr8\_hap1.g00200.t01 |
| 9988 | 8 | 7298593 | 7315259 | mNeoVis.2.1.1.chr8\_hap1.g00201.t01 |
| 9989 | 8 | 7315871 | 7328791 | mNeoVis.2.1.1.chr8\_hap1.g00202.t01 |
| 9990 | 8 | 7348927 | 7354942 | mNeoVis.2.1.1.chr8\_hap1.g00203.t01 |
| 9991 | 8 | 7422000 | 7724372 | mNeoVis.2.1.1.chr8\_hap1.g00204.t01 |
| 9992 | 8 | 9872751 | 9883332 | mNeoVis.2.1.1.chr8\_hap1.g00210.t01 |
| 9993 | 8 | 9911490 | 9966145 | mNeoVis.2.1.1.chr8\_hap1.g00211.t01 |
| 9994 | 8 | 10329815 | 10361687 | mNeoVis.2.1.1.chr8\_hap1.g00212.t01 |
| 9995 | 8 | 10372709 | 10492298 | mNeoVis.2.1.1.chr8\_hap1.g00213.t01 |
| 9996 | 8 | 10546858 | 10617571 | mNeoVis.2.1.1.chr8\_hap1.g00214.t01 |
| 9997 | 8 | 10632880 | 10650930 | mNeoVis.2.1.1.chr8\_hap1.g00215.t01 |
| 9998 | 8 | 10707397 | 10766800 | mNeoVis.2.1.1.chr8\_hap1.g00216.t01 |
| 9999 | 8 | 10782028 | 10847578 | mNeoVis.2.1.1.chr8\_hap1.g00217.t01 |
| 10000 | 8 | 10866389 | 10934382 | mNeoVis.2.1.1.chr8\_hap1.g00218.t01 |
| 10001 | 8 | 11018858 | 11038355 | mNeoVis.2.1.1.chr8\_hap1.g00219.t01 |
| 10002 | 8 | 11100966 | 11148289 | mNeoVis.2.1.1.chr8\_hap1.g00220.t01 |
| 10003 | 8 | 11152348 | 11173270 | mNeoVis.2.1.1.chr8\_hap1.g00221.t01 |
| 10004 | 8 | 11211910 | 11244280 | mNeoVis.2.1.1.chr8\_hap1.g00222.t01 |
| 10005 | 8 | 11244376 | 11349089 | mNeoVis.2.1.1.chr8\_hap1.g00223.t01 |
| 10006 | 8 | 11324815 | 11341742 | mNeoVis.2.1.1.chr8\_hap1.g00224.t01 |
| 10007 | 8 | 11348292 | 11357535 | mNeoVis.2.1.1.chr8\_hap1.g00225.t01 |
| 10008 | 8 | 11382632 | 11423639 | mNeoVis.2.1.1.chr8\_hap1.g00226.t01 |
| 10009 | 8 | 11476340 | 11545652 | mNeoVis.2.1.1.chr8\_hap1.g00227.t01 |
| 10010 | 8 | 11764603 | 11796105 | mNeoVis.2.1.1.chr8\_hap1.g00228.t01 |
| 10011 | 8 | 12281759 | 12314517 | mNeoVis.2.1.1.chr8\_hap1.g00230.t01 |
| 10012 | 8 | 13193932 | 13222890 | mNeoVis.2.1.1.chr8\_hap1.g00236.t01 |
| 10013 | 8 | 13234366 | 13489550 | mNeoVis.2.1.1.chr8\_hap1.g00237.t01 |
| 10014 | 8 | 13507869 | 13527140 | mNeoVis.2.1.1.chr8\_hap1.g00238.t01 |
| 10015 | 8 | 13507869 | 13527140 | mNeoVis.2.1.1.chr8\_hap1.g00238.t01 |
| 10016 | 8 | 13586252 | 13745157 | mNeoVis.2.1.1.chr8\_hap1.g00239.t01 |
| 10017 | 8 | 13586252 | 13745157 | mNeoVis.2.1.1.chr8\_hap1.g00239.t01 |
| 10018 | 8 | 13748093 | 13755506 | mNeoVis.2.1.1.chr8\_hap1.g00240.t01 |
| 10019 | 8 | 13748093 | 13755506 | mNeoVis.2.1.1.chr8\_hap1.g00240.t01 |
| 10020 | 8 | 13771224 | 13784771 | mNeoVis.2.1.1.chr8\_hap1.g00241.t01 |
| 10021 | 8 | 13771224 | 13784771 | mNeoVis.2.1.1.chr8\_hap1.g00241.t01 |
| 10022 | 8 | 13851017 | 13858288 | mNeoVis.2.1.1.chr8\_hap1.g00242.t01 |
| 10023 | 8 | 13851017 | 13858288 | mNeoVis.2.1.1.chr8\_hap1.g00242.t01 |
| 10024 | 8 | 13858146 | 13976928 | mNeoVis.2.1.1.chr8\_hap1.g00243.t01 |
| 10025 | 8 | 13858146 | 13976928 | mNeoVis.2.1.1.chr8\_hap1.g00243.t01 |
| 10026 | 8 | 13985604 | 14026562 | mNeoVis.2.1.1.chr8\_hap1.g00244.t01 |
| 10027 | 8 | 13985604 | 14026562 | mNeoVis.2.1.1.chr8\_hap1.g00244.t01 |
| 10028 | 8 | 14026607 | 14033945 | mNeoVis.2.1.1.chr8\_hap1.g00245.t01 |
| 10029 | 8 | 14026607 | 14033945 | mNeoVis.2.1.1.chr8\_hap1.g00245.t01 |
| 10030 | 8 | 14036067 | 14054771 | mNeoVis.2.1.1.chr8\_hap1.g00246.t01 |
| 10031 | 8 | 14036067 | 14054771 | mNeoVis.2.1.1.chr8\_hap1.g00246.t01 |
| 10032 | 8 | 14042227 | 14074870 | mNeoVis.2.1.1.chr8\_hap1.g00247.t01 |
| 10033 | 8 | 14042227 | 14074870 | mNeoVis.2.1.1.chr8\_hap1.g00247.t01 |
| 10034 | 8 | 14112542 | 14162235 | mNeoVis.2.1.1.chr8\_hap1.g00248.t01 |
| 10035 | 8 | 14112542 | 14162235 | mNeoVis.2.1.1.chr8\_hap1.g00248.t01 |
| 10036 | 8 | 14824097 | 14843362 | mNeoVis.2.1.1.chr8\_hap1.g00269.t01 |
| 10037 | 8 | 14867170 | 14878233 | mNeoVis.2.1.1.chr8\_hap1.g00270.t01 |
| 10038 | 8 | 14880007 | 15250120 | mNeoVis.2.1.1.chr8\_hap1.g00271.t01 |
| 10039 | 8 | 15370973 | 15455229 | mNeoVis.2.1.1.chr8\_hap1.g00272.t01 |
| 10040 | 8 | 15370973 | 15455229 | mNeoVis.2.1.1.chr8\_hap1.g00272.t01 |
| 10041 | 8 | 15462033 | 15519194 | mNeoVis.2.1.1.chr8\_hap1.g00273.t01 |
| 10042 | 8 | 15462033 | 15519194 | mNeoVis.2.1.1.chr8\_hap1.g00273.t01 |
| 10043 | 8 | 15529983 | 15647580 | mNeoVis.2.1.1.chr8\_hap1.g00274.t01 |
| 10044 | 8 | 15677295 | 15690077 | mNeoVis.2.1.1.chr8\_hap1.g00275.t01 |
| 10045 | 8 | 16211545 | 16262765 | mNeoVis.2.1.1.chr8\_hap1.g00277.t01 |
| 10046 | 8 | 16211545 | 16262765 | mNeoVis.2.1.1.chr8\_hap1.g00277.t01 |
| 10047 | 8 | 16281612 | 16310069 | mNeoVis.2.1.1.chr8\_hap1.g00278.t01 |
| 10048 | 8 | 16281612 | 16310069 | mNeoVis.2.1.1.chr8\_hap1.g00278.t01 |
| 10049 | 8 | 16310214 | 16383362 | mNeoVis.2.1.1.chr8\_hap1.g00279.t01 |
| 10050 | 8 | 16310214 | 16383362 | mNeoVis.2.1.1.chr8\_hap1.g00279.t01 |
| 10051 | 8 | 16402199 | 16581170 | mNeoVis.2.1.1.chr8\_hap1.g00280.t01 |
| 10052 | 8 | 16402199 | 16581170 | mNeoVis.2.1.1.chr8\_hap1.g00280.t01 |
| 10053 | 8 | 16582344 | 16653168 | mNeoVis.2.1.1.chr8\_hap1.g00281.t01 |
| 10054 | 8 | 16582344 | 16653168 | mNeoVis.2.1.1.chr8\_hap1.g00281.t01 |
| 10055 | 8 | 16582344 | 16653168 | mNeoVis.2.1.1.chr8\_hap1.g00281.t01 |
| 10056 | 8 | 16709503 | 16711749 | mNeoVis.2.1.1.chr8\_hap1.g00282.t01 |
| 10057 | 8 | 16709503 | 16711749 | mNeoVis.2.1.1.chr8\_hap1.g00282.t01 |
| 10058 | 8 | 16709503 | 16711749 | mNeoVis.2.1.1.chr8\_hap1.g00282.t01 |
| 10059 | 8 | 16721170 | 16766608 | mNeoVis.2.1.1.chr8\_hap1.g00283.t01 |
| 10060 | 8 | 16721170 | 16766608 | mNeoVis.2.1.1.chr8\_hap1.g00283.t01 |
| 10061 | 8 | 16721170 | 16766608 | mNeoVis.2.1.1.chr8\_hap1.g00283.t01 |
| 10062 | 8 | 16893902 | 16913317 | mNeoVis.2.1.1.chr8\_hap1.g00284.t01 |
| 10063 | 8 | 16893902 | 16913317 | mNeoVis.2.1.1.chr8\_hap1.g00284.t01 |
| 10064 | 8 | 16893902 | 16913317 | mNeoVis.2.1.1.chr8\_hap1.g00284.t01 |
| 10065 | 8 | 16921632 | 17070019 | mNeoVis.2.1.1.chr8\_hap1.g00285.t01 |
| 10066 | 8 | 16921632 | 17070019 | mNeoVis.2.1.1.chr8\_hap1.g00285.t01 |
| 10067 | 8 | 16921632 | 17070019 | mNeoVis.2.1.1.chr8\_hap1.g00285.t01 |
| 10068 | 8 | 17075384 | 17121197 | mNeoVis.2.1.1.chr8\_hap1.g00286.t01 |
| 10069 | 8 | 17075384 | 17121197 | mNeoVis.2.1.1.chr8\_hap1.g00286.t01 |
| 10070 | 8 | 17075384 | 17121197 | mNeoVis.2.1.1.chr8\_hap1.g00286.t01 |
| 10071 | 8 | 17144423 | 17164977 | mNeoVis.2.1.1.chr8\_hap1.g00287.t01 |
| 10072 | 8 | 17144423 | 17164977 | mNeoVis.2.1.1.chr8\_hap1.g00287.t01 |
| 10073 | 8 | 17144423 | 17164977 | mNeoVis.2.1.1.chr8\_hap1.g00287.t01 |
| 10074 | 8 | 17198514 | 17216973 | mNeoVis.2.1.1.chr8\_hap1.g00288.t01 |
| 10075 | 8 | 17198514 | 17216973 | mNeoVis.2.1.1.chr8\_hap1.g00288.t01 |
| 10076 | 8 | 17198514 | 17216973 | mNeoVis.2.1.1.chr8\_hap1.g00288.t01 |
| 10077 | 8 | 17242069 | 17242880 | mNeoVis.2.1.1.chr8\_hap1.g00289.t01 |
| 10078 | 8 | 17242069 | 17242880 | mNeoVis.2.1.1.chr8\_hap1.g00289.t01 |
| 10079 | 8 | 17242069 | 17242880 | mNeoVis.2.1.1.chr8\_hap1.g00289.t01 |
| 10080 | 8 | 17265542 | 17273516 | mNeoVis.2.1.1.chr8\_hap1.g00290.t01 |
| 10081 | 8 | 17265542 | 17273516 | mNeoVis.2.1.1.chr8\_hap1.g00290.t01 |
| 10082 | 8 | 17275371 | 17285456 | mNeoVis.2.1.1.chr8\_hap1.g00291.t01 |
| 10083 | 8 | 17275371 | 17285456 | mNeoVis.2.1.1.chr8\_hap1.g00291.t01 |
| 10084 | 8 | 17287724 | 17316298 | mNeoVis.2.1.1.chr8\_hap1.g00292.t01 |
| 10085 | 8 | 17287724 | 17316298 | mNeoVis.2.1.1.chr8\_hap1.g00292.t01 |
| 10086 | 8 | 17346273 | 17392329 | mNeoVis.2.1.1.chr8\_hap1.g00293.t01 |
| 10087 | 8 | 17393645 | 17424869 | mNeoVis.2.1.1.chr8\_hap1.g00294.t01 |
| 10088 | 8 | 17482757 | 17526204 | mNeoVis.2.1.1.chr8\_hap1.g00295.t01 |
| 10089 | 8 | 17527086 | 17541321 | mNeoVis.2.1.1.chr8\_hap1.g00296.t01 |
| 10090 | 8 | 17543206 | 17568541 | mNeoVis.2.1.1.chr8\_hap1.g00297.t01 |
| 10091 | 8 | 17601172 | 17605033 | mNeoVis.2.1.1.chr8\_hap1.g00298.t01 |
| 10092 | 8 | 17615715 | 17617889 | mNeoVis.2.1.1.chr8\_hap1.g00299.t01 |
| 10093 | 8 | 17621909 | 17628305 | mNeoVis.2.1.1.chr8\_hap1.g00300.t01 |
| 10094 | 8 | 17629913 | 17648180 | mNeoVis.2.1.1.chr8\_hap1.g00301.t01 |
| 10095 | 8 | 17653334 | 17683777 | mNeoVis.2.1.1.chr8\_hap1.g00302.t01 |
| 10096 | 8 | 17683143 | 17711716 | mNeoVis.2.1.1.chr8\_hap1.g00303.t01 |
| 10097 | 8 | 17753541 | 17760437 | mNeoVis.2.1.1.chr8\_hap1.g00304.t01 |
| 10098 | 8 | 17768596 | 17788121 | mNeoVis.2.1.1.chr8\_hap1.g00305.t01 |
| 10099 | 8 | 17795926 | 17805533 | mNeoVis.2.1.1.chr8\_hap1.g00306.t01 |
| 10100 | 8 | 17805674 | 17807471 | mNeoVis.2.1.1.chr8\_hap1.g00307.t01 |
| 10101 | 8 | 17807608 | 17811894 | mNeoVis.2.1.1.chr8\_hap1.g00308.t01 |
| 10102 | 8 | 17812045 | 17854926 | mNeoVis.2.1.1.chr8\_hap1.g00309.t01 |
| 10103 | 8 | 17854997 | 17861343 | mNeoVis.2.1.1.chr8\_hap1.g00310.t01 |
| 10104 | 8 | 17866346 | 17908769 | mNeoVis.2.1.1.chr8\_hap1.g00311.t01 |
| 10105 | 8 | 17939320 | 17946266 | mNeoVis.2.1.1.chr8\_hap1.g00312.t01 |
| 10106 | 8 | 17961169 | 18009598 | mNeoVis.2.1.1.chr8\_hap1.g00313.t01 |
| 10107 | 8 | 18009947 | 18021372 | mNeoVis.2.1.1.chr8\_hap1.g00314.t01 |
| 10108 | 8 | 18009947 | 18021372 | mNeoVis.2.1.1.chr8\_hap1.g00314.t01 |
| 10109 | 8 | 18023460 | 18038043 | mNeoVis.2.1.1.chr8\_hap1.g00315.t01 |
| 10110 | 8 | 18023460 | 18038043 | mNeoVis.2.1.1.chr8\_hap1.g00315.t01 |
| 10111 | 8 | 18096829 | 18112991 | mNeoVis.2.1.1.chr8\_hap1.g00316.t01 |
| 10112 | 8 | 18096829 | 18112991 | mNeoVis.2.1.1.chr8\_hap1.g00316.t01 |
| 10113 | 8 | 18154325 | 18268540 | mNeoVis.2.1.1.chr8\_hap1.g00317.t01 |
| 10114 | 8 | 18154325 | 18268540 | mNeoVis.2.1.1.chr8\_hap1.g00317.t01 |
| 10115 | 8 | 18272126 | 18285941 | mNeoVis.2.1.1.chr8\_hap1.g00318.t01 |
| 10116 | 8 | 18272126 | 18285941 | mNeoVis.2.1.1.chr8\_hap1.g00318.t01 |
| 10117 | 8 | 18289047 | 18299546 | mNeoVis.2.1.1.chr8\_hap1.g00319.t01 |
| 10118 | 8 | 18289047 | 18299546 | mNeoVis.2.1.1.chr8\_hap1.g00319.t01 |
| 10119 | 8 | 18299627 | 18306355 | mNeoVis.2.1.1.chr8\_hap1.g00320.t01 |
| 10120 | 8 | 18299627 | 18306355 | mNeoVis.2.1.1.chr8\_hap1.g00320.t01 |
| 10121 | 8 | 18323134 | 18328138 | mNeoVis.2.1.1.chr8\_hap1.g00321.t01 |
| 10122 | 8 | 18323134 | 18328138 | mNeoVis.2.1.1.chr8\_hap1.g00321.t01 |
| 10123 | 8 | 18328767 | 18337856 | mNeoVis.2.1.1.chr8\_hap1.g00322.t01 |
| 10124 | 8 | 18328767 | 18337856 | mNeoVis.2.1.1.chr8\_hap1.g00322.t01 |
| 10125 | 8 | 18368070 | 18379896 | mNeoVis.2.1.1.chr8\_hap1.g00323.t01 |
| 10126 | 8 | 18368070 | 18379896 | mNeoVis.2.1.1.chr8\_hap1.g00323.t01 |
| 10127 | 8 | 18395589 | 18409609 | mNeoVis.2.1.1.chr8\_hap1.g00324.t01 |
| 10128 | 8 | 18395589 | 18409609 | mNeoVis.2.1.1.chr8\_hap1.g00324.t01 |
| 10129 | 8 | 18417083 | 18424712 | mNeoVis.2.1.1.chr8\_hap1.g00325.t01 |
| 10130 | 8 | 18417083 | 18424712 | mNeoVis.2.1.1.chr8\_hap1.g00325.t01 |
| 10131 | 8 | 18424795 | 18428939 | mNeoVis.2.1.1.chr8\_hap1.g00326.t01 |
| 10132 | 8 | 18424795 | 18428939 | mNeoVis.2.1.1.chr8\_hap1.g00326.t01 |
| 10133 | 8 | 18434485 | 18445150 | mNeoVis.2.1.1.chr8\_hap1.g00327.t01 |
| 10134 | 8 | 18434485 | 18445150 | mNeoVis.2.1.1.chr8\_hap1.g00327.t01 |
| 10135 | 8 | 18447074 | 18462039 | mNeoVis.2.1.1.chr8\_hap1.g00328.t01 |
| 10136 | 8 | 18447074 | 18462039 | mNeoVis.2.1.1.chr8\_hap1.g00328.t01 |
| 10137 | 8 | 18493246 | 18517485 | mNeoVis.2.1.1.chr8\_hap1.g00329.t01 |
| 10138 | 8 | 18493246 | 18517485 | mNeoVis.2.1.1.chr8\_hap1.g00329.t01 |
| 10139 | 8 | 18519439 | 18524181 | mNeoVis.2.1.1.chr8\_hap1.g00330.t01 |
| 10140 | 8 | 18519439 | 18524181 | mNeoVis.2.1.1.chr8\_hap1.g00330.t01 |
| 10141 | 8 | 18526342 | 18548546 | mNeoVis.2.1.1.chr8\_hap1.g00331.t01 |
| 10142 | 8 | 18526342 | 18548546 | mNeoVis.2.1.1.chr8\_hap1.g00331.t01 |
| 10143 | 8 | 18560941 | 18564408 | mNeoVis.2.1.1.chr8\_hap1.g00332.t01 |
| 10144 | 8 | 18560941 | 18564408 | mNeoVis.2.1.1.chr8\_hap1.g00332.t01 |
| 10145 | 8 | 18565346 | 18569457 | mNeoVis.2.1.1.chr8\_hap1.g00333.t01 |
| 10146 | 8 | 18565346 | 18569457 | mNeoVis.2.1.1.chr8\_hap1.g00333.t01 |
| 10147 | 8 | 18602359 | 18607834 | mNeoVis.2.1.1.chr8\_hap1.g00334.t01 |
| 10148 | 8 | 18602359 | 18607834 | mNeoVis.2.1.1.chr8\_hap1.g00334.t01 |
| 10149 | 8 | 18611363 | 18616374 | mNeoVis.2.1.1.chr8\_hap1.g00335.t01 |
| 10150 | 8 | 18629481 | 18656265 | mNeoVis.2.1.1.chr8\_hap1.g00336.t01 |
| 10151 | 8 | 18665192 | 18678226 | mNeoVis.2.1.1.chr8\_hap1.g00337.t01 |
| 10152 | 8 | 18693610 | 18718223 | mNeoVis.2.1.1.chr8\_hap1.g00338.t01 |
| 10153 | 8 | 18725038 | 18728164 | mNeoVis.2.1.1.chr8\_hap1.g00339.t01 |
| 10154 | 8 | 19821536 | 19842299 | mNeoVis.2.1.1.chr8\_hap1.g00383.t01 |
| 10155 | 8 | 19897000 | 19921417 | mNeoVis.2.1.1.chr8\_hap1.g00384.t01 |
| 10156 | 8 | 19925925 | 19929881 | mNeoVis.2.1.1.chr8\_hap1.g00385.t01 |
| 10157 | 8 | 19931814 | 19932967 | mNeoVis.2.1.1.chr8\_hap1.g00386.t01 |
| 10158 | 8 | 19935448 | 19952830 | mNeoVis.2.1.1.chr8\_hap1.g00387.t01 |
| 10159 | 8 | 19954462 | 19979573 | mNeoVis.2.1.1.chr8\_hap1.g00388.t01 |
| 10160 | 8 | 19980338 | 19980884 | mNeoVis.2.1.1.chr8\_hap1.g00389.t01 |
| 10161 | 8 | 19982113 | 19989789 | mNeoVis.2.1.1.chr8\_hap1.g00390.t01 |
| 10162 | 8 | 19990481 | 19998608 | mNeoVis.2.1.1.chr8\_hap1.g00391.t01 |
| 10163 | 8 | 20022361 | 20051644 | mNeoVis.2.1.1.chr8\_hap1.g00392.t01 |
| 10164 | 8 | 20030051 | 20042163 | mNeoVis.2.1.1.chr8\_hap1.g00393.t01 |
| 10165 | 8 | 20051784 | 20058129 | mNeoVis.2.1.1.chr8\_hap1.g00394.t01 |
| 10166 | 8 | 20058455 | 20087131 | mNeoVis.2.1.1.chr8\_hap1.g00395.t01 |
| 10167 | 8 | 20088569 | 20095306 | mNeoVis.2.1.1.chr8\_hap1.g00396.t01 |
| 10168 | 8 | 20096643 | 20122332 | mNeoVis.2.1.1.chr8\_hap1.g00397.t01 |
| 10169 | 8 | 20131926 | 20176122 | mNeoVis.2.1.1.chr8\_hap1.g00398.t01 |
| 10170 | 8 | 20176425 | 20188046 | mNeoVis.2.1.1.chr8\_hap1.g00399.t01 |
| 10171 | 8 | 20188217 | 20194457 | mNeoVis.2.1.1.chr8\_hap1.g00400.t01 |
| 10172 | 8 | 20194552 | 20204002 | mNeoVis.2.1.1.chr8\_hap1.g00401.t01 |
| 10173 | 8 | 20205699 | 20213920 | mNeoVis.2.1.1.chr8\_hap1.g00402.t01 |
| 10174 | 8 | 20214991 | 20221914 | mNeoVis.2.1.1.chr8\_hap1.g00403.t01 |
| 10175 | 8 | 20232242 | 20255306 | mNeoVis.2.1.1.chr8\_hap1.g00404.t01 |
| 10176 | 8 | 20266088 | 20278697 | mNeoVis.2.1.1.chr8\_hap1.g00405.t01 |
| 10177 | 8 | 20279708 | 20281379 | mNeoVis.2.1.1.chr8\_hap1.g00406.t01 |
| 10178 | 8 | 20297478 | 20299595 | mNeoVis.2.1.1.chr8\_hap1.g00407.t01 |
| 10179 | 8 | 20338680 | 20348233 | mNeoVis.2.1.1.chr8\_hap1.g00408.t01 |
| 10180 | 8 | 20350118 | 20355257 | mNeoVis.2.1.1.chr8\_hap1.g00409.t01 |
| 10181 | 8 | 20361192 | 20389546 | mNeoVis.2.1.1.chr8\_hap1.g00410.t01 |
| 10182 | 8 | 20399476 | 20403645 | mNeoVis.2.1.1.chr8\_hap1.g00411.t01 |
| 10183 | 8 | 20436702 | 20441051 | mNeoVis.2.1.1.chr8\_hap1.g00412.t01 |
| 10184 | 8 | 20445726 | 20450060 | mNeoVis.2.1.1.chr8\_hap1.g00413.t01 |
| 10185 | 8 | 20457375 | 20461351 | mNeoVis.2.1.1.chr8\_hap1.g00414.t01 |
| 10186 | 8 | 20917821 | 20919317 | mNeoVis.2.1.1.chr8\_hap1.g00437.t01 |
| 10187 | 8 | 20922353 | 20923095 | mNeoVis.2.1.1.chr8\_hap1.g00438.t01 |
| 10188 | 8 | 20927623 | 20928407 | mNeoVis.2.1.1.chr8\_hap1.g00439.t01 |
| 10189 | 8 | 20935529 | 20936234 | mNeoVis.2.1.1.chr8\_hap1.g00440.t01 |
| 10190 | 8 | 20939843 | 20946920 | mNeoVis.2.1.1.chr8\_hap1.g00441.t01 |
| 10191 | 8 | 20957563 | 20961445 | mNeoVis.2.1.1.chr8\_hap1.g00442.t01 |
| 10192 | 8 | 20975751 | 20976358 | mNeoVis.2.1.1.chr8\_hap1.g00443.t01 |
| 10193 | 8 | 20988803 | 20989514 | mNeoVis.2.1.1.chr8\_hap1.g00444.t01 |
| 10194 | 8 | 20988803 | 20989514 | mNeoVis.2.1.1.chr8\_hap1.g00444.t01 |
| 10195 | 8 | 20995614 | 20996353 | mNeoVis.2.1.1.chr8\_hap1.g00445.t01 |
| 10196 | 8 | 20995614 | 20996353 | mNeoVis.2.1.1.chr8\_hap1.g00445.t01 |
| 10197 | 8 | 21008482 | 21015038 | mNeoVis.2.1.1.chr8\_hap1.g00446.t01 |
| 10198 | 8 | 21008482 | 21015038 | mNeoVis.2.1.1.chr8\_hap1.g00446.t01 |
| 10199 | 8 | 21023032 | 21034138 | mNeoVis.2.1.1.chr8\_hap1.g00447.t01 |
| 10200 | 8 | 21023032 | 21034138 | mNeoVis.2.1.1.chr8\_hap1.g00447.t01 |
| 10201 | 8 | 21051138 | 21070605 | mNeoVis.2.1.1.chr8\_hap1.g00448.t01 |
| 10202 | 8 | 21051138 | 21070605 | mNeoVis.2.1.1.chr8\_hap1.g00448.t01 |
| 10203 | 8 | 21107866 | 21115695 | mNeoVis.2.1.1.chr8\_hap1.g00449.t01 |
| 10204 | 8 | 21107866 | 21115695 | mNeoVis.2.1.1.chr8\_hap1.g00449.t01 |
| 10205 | 8 | 21123403 | 21129805 | mNeoVis.2.1.1.chr8\_hap1.g00450.t01 |
| 10206 | 8 | 21123403 | 21129805 | mNeoVis.2.1.1.chr8\_hap1.g00450.t01 |
| 10207 | 8 | 21163008 | 21167660 | mNeoVis.2.1.1.chr8\_hap1.g00451.t01 |
| 10208 | 8 | 21163008 | 21167660 | mNeoVis.2.1.1.chr8\_hap1.g00451.t01 |
| 10209 | 8 | 21175284 | 21183170 | mNeoVis.2.1.1.chr8\_hap1.g00452.t01 |
| 10210 | 8 | 21175284 | 21183170 | mNeoVis.2.1.1.chr8\_hap1.g00452.t01 |
| 10211 | 8 | 21199260 | 21203943 | mNeoVis.2.1.1.chr8\_hap1.g00453.t01 |
| 10212 | 8 | 21199260 | 21203943 | mNeoVis.2.1.1.chr8\_hap1.g00453.t01 |
| 10213 | 8 | 21209482 | 21215671 | mNeoVis.2.1.1.chr8\_hap1.g00454.t01 |
| 10214 | 8 | 21209482 | 21215671 | mNeoVis.2.1.1.chr8\_hap1.g00454.t01 |
| 10215 | 8 | 21220277 | 21228034 | mNeoVis.2.1.1.chr8\_hap1.g00455.t01 |
| 10216 | 8 | 21220277 | 21228034 | mNeoVis.2.1.1.chr8\_hap1.g00455.t01 |
| 10217 | 8 | 21246720 | 21251130 | mNeoVis.2.1.1.chr8\_hap1.g00456.t01 |
| 10218 | 8 | 21246720 | 21251130 | mNeoVis.2.1.1.chr8\_hap1.g00456.t01 |
| 10219 | 8 | 21280146 | 21292209 | mNeoVis.2.1.1.chr8\_hap1.g00457.t01 |
| 10220 | 8 | 21280146 | 21292209 | mNeoVis.2.1.1.chr8\_hap1.g00457.t01 |
| 10221 | 8 | 21294798 | 21315309 | mNeoVis.2.1.1.chr8\_hap1.g00458.t01 |
| 10222 | 8 | 21294798 | 21315309 | mNeoVis.2.1.1.chr8\_hap1.g00458.t01 |
| 10223 | 8 | 21356675 | 21370468 | mNeoVis.2.1.1.chr8\_hap1.g00459.t01 |
| 10224 | 8 | 21356675 | 21370468 | mNeoVis.2.1.1.chr8\_hap1.g00459.t01 |
| 10225 | 8 | 21414040 | 21435950 | mNeoVis.2.1.1.chr8\_hap1.g00460.t01 |
| 10226 | 8 | 21414040 | 21435950 | mNeoVis.2.1.1.chr8\_hap1.g00460.t01 |
| 10227 | 8 | 21445418 | 21458616 | mNeoVis.2.1.1.chr8\_hap1.g00461.t01 |
| 10228 | 8 | 21445418 | 21458616 | mNeoVis.2.1.1.chr8\_hap1.g00461.t01 |
| 10229 | 8 | 21479528 | 21509918 | mNeoVis.2.1.1.chr8\_hap1.g00462.t01 |
| 10230 | 8 | 21517812 | 21521923 | mNeoVis.2.1.1.chr8\_hap1.g00463.t01 |
| 10231 | 8 | 21525661 | 21540176 | mNeoVis.2.1.1.chr8\_hap1.g00464.t01 |
| 10232 | 8 | 21574984 | 21596771 | mNeoVis.2.1.1.chr8\_hap1.g00465.t01 |
| 10233 | 8 | 21596657 | 21642009 | mNeoVis.2.1.1.chr8\_hap1.g00466.t01 |
| 10234 | 8 | 21655859 | 21669673 | mNeoVis.2.1.1.chr8\_hap1.g00467.t01 |
| 10235 | 8 | 21672852 | 21693775 | mNeoVis.2.1.1.chr8\_hap1.g00468.t01 |
| 10236 | 8 | 21695874 | 21708326 | mNeoVis.2.1.1.chr8\_hap1.g00469.t01 |
| 10237 | 8 | 21729350 | 21735463 | mNeoVis.2.1.1.chr8\_hap1.g00470.t01 |
| 10238 | 8 | 21735645 | 21758801 | mNeoVis.2.1.1.chr8\_hap1.g00471.t01 |
| 10239 | 8 | 21766202 | 21791754 | mNeoVis.2.1.1.chr8\_hap1.g00472.t01 |
| 10240 | 8 | 21793345 | 21797021 | mNeoVis.2.1.1.chr8\_hap1.g00473.t01 |
| 10241 | 8 | 21801035 | 21814975 | mNeoVis.2.1.1.chr8\_hap1.g00474.t01 |
| 10242 | 8 | 21817430 | 21830937 | mNeoVis.2.1.1.chr8\_hap1.g00475.t01 |
| 10243 | 8 | 21839791 | 21841455 | mNeoVis.2.1.1.chr8\_hap1.g00476.t01 |
| 10244 | 8 | 21851897 | 21858296 | mNeoVis.2.1.1.chr8\_hap1.g00477.t01 |
| 10245 | 8 | 21861396 | 21869575 | mNeoVis.2.1.1.chr8\_hap1.g00478.t01 |
| 10246 | 8 | 21892782 | 21901037 | mNeoVis.2.1.1.chr8\_hap1.g00479.t01 |
| 10247 | 8 | 21904704 | 21998015 | mNeoVis.2.1.1.chr8\_hap1.g00480.t01 |
| 10248 | 8 | 22007446 | 22016229 | mNeoVis.2.1.1.chr8\_hap1.g00481.t01 |
| 10249 | 8 | 22021533 | 22022919 | mNeoVis.2.1.1.chr8\_hap1.g00482.t01 |
| 10250 | 8 | 22101875 | 22110589 | mNeoVis.2.1.1.chr8\_hap1.g00488.t01 |
| 10251 | 8 | 22128846 | 22132452 | mNeoVis.2.1.1.chr8\_hap1.g00489.t01 |
| 10252 | 8 | 22193108 | 22248828 | mNeoVis.2.1.1.chr8\_hap1.g00490.t01 |
| 10253 | 8 | 22255541 | 22285751 | mNeoVis.2.1.1.chr8\_hap1.g00491.t01 |
| 10254 | 8 | 22287382 | 22396297 | mNeoVis.2.1.1.chr8\_hap1.g00492.t01 |
| 10255 | 8 | 22419504 | 22431997 | mNeoVis.2.1.1.chr8\_hap1.g00493.t01 |
| 10256 | 8 | 22437040 | 22442006 | mNeoVis.2.1.1.chr8\_hap1.g00494.t01 |
| 10257 | 8 | 22444595 | 22464588 | mNeoVis.2.1.1.chr8\_hap1.g00495.t01 |
| 10258 | 8 | 22467866 | 22475245 | mNeoVis.2.1.1.chr8\_hap1.g00496.t01 |
| 10259 | 8 | 22478070 | 22535115 | mNeoVis.2.1.1.chr8\_hap1.g00497.t01 |
| 10260 | 8 | 22575235 | 22602562 | mNeoVis.2.1.1.chr8\_hap1.g00498.t01 |
| 10261 | 8 | 22609648 | 22646757 | mNeoVis.2.1.1.chr8\_hap1.g00499.t01 |
| 10262 | 8 | 22656768 | 22661898 | mNeoVis.2.1.1.chr8\_hap1.g00500.t01 |
| 10263 | 8 | 22665699 | 22669833 | mNeoVis.2.1.1.chr8\_hap1.g00501.t01 |
| 10264 | 8 | 22684323 | 22708952 | mNeoVis.2.1.1.chr8\_hap1.g00502.t01 |
| 10265 | 8 | 22709955 | 22737304 | mNeoVis.2.1.1.chr8\_hap1.g00503.t01 |
| 10266 | 8 | 22738355 | 22747015 | mNeoVis.2.1.1.chr8\_hap1.g00504.t01 |
| 10267 | 8 | 22750654 | 22761953 | mNeoVis.2.1.1.chr8\_hap1.g00505.t01 |
| 10268 | 8 | 22763297 | 22766521 | mNeoVis.2.1.1.chr8\_hap1.g00506.t01 |
| 10269 | 8 | 22770637 | 22789383 | mNeoVis.2.1.1.chr8\_hap1.g00507.t01 |
| 10270 | 8 | 22811779 | 22815002 | mNeoVis.2.1.1.chr8\_hap1.g00508.t01 |
| 10271 | 8 | 22858219 | 22920837 | mNeoVis.2.1.1.chr8\_hap1.g00509.t01 |
| 10272 | 8 | 22933490 | 23004218 | mNeoVis.2.1.1.chr8\_hap1.g00510.t01 |
| 10273 | 8 | 22996504 | 23006106 | mNeoVis.2.1.1.chr8\_hap1.g00511.t01 |
| 10274 | 8 | 23021388 | 23053611 | mNeoVis.2.1.1.chr8\_hap1.g00512.t01 |
| 10275 | 8 | 23060051 | 23076658 | mNeoVis.2.1.1.chr8\_hap1.g00513.t01 |
| 10276 | 8 | 23080824 | 23090527 | mNeoVis.2.1.1.chr8\_hap1.g00514.t01 |
| 10277 | 8 | 23120067 | 23206491 | mNeoVis.2.1.1.chr8\_hap1.g00515.t01 |
| 10278 | 8 | 23239762 | 23264587 | mNeoVis.2.1.1.chr8\_hap1.g00516.t01 |
| 10279 | 8 | 23274522 | 23292085 | mNeoVis.2.1.1.chr8\_hap1.g00517.t01 |
| 10280 | 8 | 23312614 | 23324043 | mNeoVis.2.1.1.chr8\_hap1.g00518.t01 |
| 10281 | 8 | 23372641 | 23387511 | mNeoVis.2.1.1.chr8\_hap1.g00519.t01 |
| 10282 | 8 | 23389671 | 23397796 | mNeoVis.2.1.1.chr8\_hap1.g00520.t01 |
| 10283 | 8 | 23395910 | 23400850 | mNeoVis.2.1.1.chr8\_hap1.g00521.t01 |
| 10284 | 8 | 23401351 | 23404761 | mNeoVis.2.1.1.chr8\_hap1.g00522.t01 |
| 10285 | 8 | 23408840 | 23414929 | mNeoVis.2.1.1.chr8\_hap1.g00523.t01 |
| 10286 | 8 | 23452934 | 23477753 | mNeoVis.2.1.1.chr8\_hap1.g00524.t01 |
| 10287 | 8 | 23484777 | 23491920 | mNeoVis.2.1.1.chr8\_hap1.g00525.t01 |
| 10288 | 8 | 23493976 | 23499405 | mNeoVis.2.1.1.chr8\_hap1.g00526.t01 |
| 10289 | 8 | 23506482 | 23515254 | mNeoVis.2.1.1.chr8\_hap1.g00527.t01 |
| 10290 | 8 | 24016141 | 24017552 | mNeoVis.2.1.1.chr8\_hap1.g00533.t01 |
| 10291 | 8 | 24028045 | 24030882 | mNeoVis.2.1.1.chr8\_hap1.g00534.t01 |
| 10292 | 8 | 24035849 | 24038013 | mNeoVis.2.1.1.chr8\_hap1.g00535.t01 |
| 10293 | 8 | 24062486 | 24064287 | mNeoVis.2.1.1.chr8\_hap1.g00536.t01 |
| 10294 | 8 | 24077103 | 24080171 | mNeoVis.2.1.1.chr8\_hap1.g00537.t01 |
| 10295 | 8 | 24081660 | 24090875 | mNeoVis.2.1.1.chr8\_hap1.g00538.t01 |
| 10296 | 8 | 24093151 | 24096656 | mNeoVis.2.1.1.chr8\_hap1.g00539.t01 |
| 10297 | 8 | 24097954 | 24111816 | mNeoVis.2.1.1.chr8\_hap1.g00540.t01 |
| 10298 | 8 | 24107090 | 24112259 | mNeoVis.2.1.1.chr8\_hap1.g00541.t01 |
| 10299 | 8 | 24177754 | 24180393 | mNeoVis.2.1.1.chr8\_hap1.g00542.t01 |
| 10300 | 8 | 24182816 | 24186390 | mNeoVis.2.1.1.chr8\_hap1.g00543.t01 |
| 10301 | 8 | 24214321 | 24252861 | mNeoVis.2.1.1.chr8\_hap1.g00544.t01 |
| 10302 | 8 | 28732090 | 28829959 | mNeoVis.2.1.1.chr8\_hap1.g00599.t01 |
| 10303 | 8 | 29370720 | 29372179 | mNeoVis.2.1.1.chr8\_hap1.g00600.t01 |
| 10304 | 8 | 29410551 | 29477101 | mNeoVis.2.1.1.chr8\_hap1.g00601.t01 |
| 10305 | 8 | 29476979 | 29484408 | mNeoVis.2.1.1.chr8\_hap1.g00602.t01 |
| 10306 | 8 | 29484394 | 29658208 | mNeoVis.2.1.1.chr8\_hap1.g00603.t01 |
| 10307 | 8 | 29757180 | 29809695 | mNeoVis.2.1.1.chr8\_hap1.g00604.t01 |
| 10308 | 8 | 29856437 | 29883959 | mNeoVis.2.1.1.chr8\_hap1.g00605.t01 |
| 10309 | 8 | 29941548 | 29960700 | mNeoVis.2.1.1.chr8\_hap1.g00606.t01 |
| 10310 | 8 | 30148174 | 30151721 | mNeoVis.2.1.1.chr8\_hap1.g00607.t01 |
| 10311 | 8 | 30179811 | 30205175 | mNeoVis.2.1.1.chr8\_hap1.g00608.t01 |
| 10312 | 8 | 30190363 | 30210570 | mNeoVis.2.1.1.chr8\_hap1.g00609.t01 |
| 10313 | 8 | 30210462 | 30832281 | mNeoVis.2.1.1.chr8\_hap1.g00610.t01 |
| 10314 | 8 | 30900940 | 30902762 | mNeoVis.2.1.1.chr8\_hap1.g00611.t01 |
| 10315 | 8 | 31100525 | 31118861 | mNeoVis.2.1.1.chr8\_hap1.g00612.t01 |
| 10316 | 8 | 31130654 | 31155989 | mNeoVis.2.1.1.chr8\_hap1.g00613.t01 |
| 10317 | 8 | 31170615 | 31191594 | mNeoVis.2.1.1.chr8\_hap1.g00614.t01 |
| 10318 | 8 | 31221068 | 31238239 | mNeoVis.2.1.1.chr8\_hap1.g00615.t01 |
| 10319 | 8 | 31242313 | 31274732 | mNeoVis.2.1.1.chr8\_hap1.g00616.t01 |
| 10320 | 8 | 31354400 | 31372747 | mNeoVis.2.1.1.chr8\_hap1.g00617.t01 |
| 10321 | 8 | 31915339 | 31916790 | mNeoVis.2.1.1.chr8\_hap1.g00619.t01 |
| 10322 | 8 | 31988496 | 31993843 | mNeoVis.2.1.1.chr8\_hap1.g00620.t01 |
| 10323 | 8 | 32003372 | 32086228 | mNeoVis.2.1.1.chr8\_hap1.g00621.t01 |
| 10324 | 8 | 32101707 | 32118017 | mNeoVis.2.1.1.chr8\_hap1.g00622.t01 |
| 10325 | 8 | 32133742 | 32137765 | mNeoVis.2.1.1.chr8\_hap1.g00623.t01 |
| 10326 | 8 | 32137207 | 32212034 | mNeoVis.2.1.1.chr8\_hap1.g00624.t01 |
| 10327 | 8 | 32137210 | 32151713 | mNeoVis.2.1.1.chr8\_hap1.g00625.t01 |
| 10328 | 8 | 32267963 | 32268898 | mNeoVis.2.1.1.chr8\_hap1.g00626.t01 |
| 10329 | 8 | 32277151 | 32278086 | mNeoVis.2.1.1.chr8\_hap1.g00627.t01 |
| 10330 | 8 | 32288767 | 32289839 | mNeoVis.2.1.1.chr8\_hap1.g00628.t01 |
| 10331 | 8 | 32306297 | 32307326 | mNeoVis.2.1.1.chr8\_hap1.g00629.t01 |
| 10332 | 8 | 32320843 | 32330058 | mNeoVis.2.1.1.chr8\_hap1.g00630.t01 |
| 10333 | 8 | 32330620 | 32342318 | mNeoVis.2.1.1.chr8\_hap1.g00631.t01 |
| 10334 | 8 | 32360574 | 32380427 | mNeoVis.2.1.1.chr8\_hap1.g00632.t01 |
| 10335 | 8 | 32381419 | 32391136 | mNeoVis.2.1.1.chr8\_hap1.g00633.t01 |
| 10336 | 8 | 32404645 | 32430551 | mNeoVis.2.1.1.chr8\_hap1.g00634.t01 |
| 10337 | 8 | 32445634 | 32451585 | mNeoVis.2.1.1.chr8\_hap1.g00635.t01 |
| 10338 | 8 | 32453041 | 32516534 | mNeoVis.2.1.1.chr8\_hap1.g00636.t01 |
| 10339 | 8 | 32518979 | 32565703 | mNeoVis.2.1.1.chr8\_hap1.g00637.t01 |
| 10340 | 8 | 32566677 | 32591663 | mNeoVis.2.1.1.chr8\_hap1.g00638.t01 |
| 10341 | 8 | 32593192 | 32616600 | mNeoVis.2.1.1.chr8\_hap1.g00639.t01 |
| 10342 | 8 | 32616694 | 32683308 | mNeoVis.2.1.1.chr8\_hap1.g00640.t01 |
| 10343 | 8 | 32814298 | 33017606 | mNeoVis.2.1.1.chr8\_hap1.g00643.t01 |
| 10344 | 8 | 33030458 | 33171802 | mNeoVis.2.1.1.chr8\_hap1.g00644.t01 |
| 10345 | 8 | 33175610 | 33218020 | mNeoVis.2.1.1.chr8\_hap1.g00645.t01 |
| 10346 | 8 | 33218212 | 33240364 | mNeoVis.2.1.1.chr8\_hap1.g00646.t01 |
| 10347 | 8 | 33244447 | 33249509 | mNeoVis.2.1.1.chr8\_hap1.g00647.t01 |
| 10348 | 8 | 33256892 | 33313320 | mNeoVis.2.1.1.chr8\_hap1.g00648.t01 |
| 10349 | 8 | 33355105 | 33421709 | mNeoVis.2.1.1.chr8\_hap1.g00649.t01 |
| 10350 | 8 | 35403199 | 35445017 | mNeoVis.2.1.1.chr8\_hap1.g00669.t01 |
| 10351 | 8 | 35516047 | 35577256 | mNeoVis.2.1.1.chr8\_hap1.g00670.t01 |
| 10352 | 8 | 35516047 | 35577256 | mNeoVis.2.1.1.chr8\_hap1.g00670.t01 |
| 10353 | 8 | 35555909 | 35557222 | mNeoVis.2.1.1.chr8\_hap1.g00671.t01 |
| 10354 | 8 | 35555909 | 35557222 | mNeoVis.2.1.1.chr8\_hap1.g00671.t01 |
| 10355 | 8 | 35591989 | 35601287 | mNeoVis.2.1.1.chr8\_hap1.g00672.t01 |
| 10356 | 8 | 35591989 | 35601287 | mNeoVis.2.1.1.chr8\_hap1.g00672.t01 |
| 10357 | 8 | 35601368 | 35839054 | mNeoVis.2.1.1.chr8\_hap1.g00673.t01 |
| 10358 | 8 | 35601368 | 35839054 | mNeoVis.2.1.1.chr8\_hap1.g00673.t01 |
| 10359 | 8 | 35840107 | 35842191 | mNeoVis.2.1.1.chr8\_hap1.g00674.t01 |
| 10360 | 8 | 35840107 | 35842191 | mNeoVis.2.1.1.chr8\_hap1.g00674.t01 |
| 10361 | 8 | 35859512 | 35865278 | mNeoVis.2.1.1.chr8\_hap1.g00675.t01 |
| 10362 | 8 | 35859512 | 35865278 | mNeoVis.2.1.1.chr8\_hap1.g00675.t01 |
| 10363 | 8 | 35868036 | 35885168 | mNeoVis.2.1.1.chr8\_hap1.g00676.t01 |
| 10364 | 8 | 35868036 | 35885168 | mNeoVis.2.1.1.chr8\_hap1.g00676.t01 |
| 10365 | 8 | 35899988 | 35907879 | mNeoVis.2.1.1.chr8\_hap1.g00677.t01 |
| 10366 | 8 | 35907757 | 35940928 | mNeoVis.2.1.1.chr8\_hap1.g00678.t01 |
| 10367 | 8 | 35941034 | 35945073 | mNeoVis.2.1.1.chr8\_hap1.g00679.t01 |
| 10368 | 8 | 35951847 | 35984545 | mNeoVis.2.1.1.chr8\_hap1.g00680.t01 |
| 10369 | 8 | 35986384 | 35994514 | mNeoVis.2.1.1.chr8\_hap1.g00681.t01 |
| 10370 | 8 | 35995337 | 35999400 | mNeoVis.2.1.1.chr8\_hap1.g00682.t01 |
| 10371 | 8 | 36274511 | 36282415 | mNeoVis.2.1.1.chr8\_hap1.g00683.t01 |
| 10372 | 8 | 36274511 | 36282415 | mNeoVis.2.1.1.chr8\_hap1.g00683.t01 |
| 10373 | 8 | 36286657 | 36387148 | mNeoVis.2.1.1.chr8\_hap1.g00684.t01 |
| 10374 | 8 | 36410177 | 36698216 | mNeoVis.2.1.1.chr8\_hap1.g00685.t01 |
| 10375 | 8 | 36410177 | 36698216 | mNeoVis.2.1.1.chr8\_hap1.g00685.t01 |
| 10376 | 8 | 36663657 | 36677233 | mNeoVis.2.1.1.chr8\_hap1.g00686.t01 |
| 10377 | 8 | 36663657 | 36677233 | mNeoVis.2.1.1.chr8\_hap1.g00686.t01 |
| 10378 | 8 | 36698568 | 36751295 | mNeoVis.2.1.1.chr8\_hap1.g00687.t01 |
| 10379 | 8 | 36698568 | 36751295 | mNeoVis.2.1.1.chr8\_hap1.g00687.t01 |
| 10380 | 8 | 36762605 | 36789834 | mNeoVis.2.1.1.chr8\_hap1.g00688.t01 |
| 10381 | 8 | 36762605 | 36789834 | mNeoVis.2.1.1.chr8\_hap1.g00688.t01 |
| 10382 | 8 | 36884949 | 36906947 | mNeoVis.2.1.1.chr8\_hap1.g00690.t01 |
| 10383 | 8 | 36962709 | 37008535 | mNeoVis.2.1.1.chr8\_hap1.g00691.t01 |
| 10384 | 8 | 37094244 | 37126544 | mNeoVis.2.1.1.chr8\_hap1.g00692.t01 |
| 10385 | 8 | 37178849 | 37180750 | mNeoVis.2.1.1.chr8\_hap1.g00693.t01 |
| 10386 | 8 | 37190372 | 37192241 | mNeoVis.2.1.1.chr8\_hap1.g00694.t01 |
| 10387 | 8 | 37206084 | 37207367 | mNeoVis.2.1.1.chr8\_hap1.g00695.t01 |
| 10388 | 8 | 37238866 | 37244670 | mNeoVis.2.1.1.chr8\_hap1.g00696.t01 |
| 10389 | 8 | 37274110 | 37277753 | mNeoVis.2.1.1.chr8\_hap1.g00697.t01 |
| 10390 | 8 | 37287211 | 37290224 | mNeoVis.2.1.1.chr8\_hap1.g00698.t01 |
| 10391 | 8 | 37292277 | 37295806 | mNeoVis.2.1.1.chr8\_hap1.g00699.t01 |
| 10392 | 8 | 37317819 | 37323915 | mNeoVis.2.1.1.chr8\_hap1.g00700.t01 |
| 10393 | 8 | 37325229 | 37337721 | mNeoVis.2.1.1.chr8\_hap1.g00701.t01 |
| 10394 | 8 | 37345277 | 37373939 | mNeoVis.2.1.1.chr8\_hap1.g00702.t01 |
| 10395 | 8 | 37384179 | 37409573 | mNeoVis.2.1.1.chr8\_hap1.g00703.t01 |
| 10396 | 8 | 38377504 | 38386269 | mNeoVis.2.1.1.chr8\_hap1.g00719.t01 |
| 10397 | 8 | 38387873 | 38389758 | mNeoVis.2.1.1.chr8\_hap1.g00720.t01 |
| 10398 | 8 | 38397873 | 38399585 | mNeoVis.2.1.1.chr8\_hap1.g00721.t01 |
| 10399 | 8 | 38427122 | 38428948 | mNeoVis.2.1.1.chr8\_hap1.g00722.t01 |
| 10400 | 8 | 38436699 | 38438677 | mNeoVis.2.1.1.chr8\_hap1.g00723.t01 |
| 10401 | 8 | 39231493 | 39491777 | mNeoVis.2.1.1.chr8\_hap1.g00724.t01 |
| 10402 | 8 | 39503326 | 39510968 | mNeoVis.2.1.1.chr8\_hap1.g00725.t01 |
| 10403 | 8 | 39503326 | 39510968 | mNeoVis.2.1.1.chr8\_hap1.g00725.t01 |
| 10404 | 8 | 39545317 | 39557529 | mNeoVis.2.1.1.chr8\_hap1.g00726.t01 |
| 10405 | 8 | 39545317 | 39557529 | mNeoVis.2.1.1.chr8\_hap1.g00726.t01 |
| 10406 | 8 | 39545317 | 39557529 | mNeoVis.2.1.1.chr8\_hap1.g00726.t01 |
| 10407 | 8 | 39567328 | 39582412 | mNeoVis.2.1.1.chr8\_hap1.g00727.t01 |
| 10408 | 8 | 39567328 | 39582412 | mNeoVis.2.1.1.chr8\_hap1.g00727.t01 |
| 10409 | 8 | 39567328 | 39582412 | mNeoVis.2.1.1.chr8\_hap1.g00727.t01 |
| 10410 | 8 | 39582151 | 39927249 | mNeoVis.2.1.1.chr8\_hap1.g00728.t01 |
| 10411 | 8 | 39582151 | 39927249 | mNeoVis.2.1.1.chr8\_hap1.g00728.t01 |
| 10412 | 8 | 39928534 | 39932878 | mNeoVis.2.1.1.chr8\_hap1.g00729.t01 |
| 10413 | 8 | 39928534 | 39932878 | mNeoVis.2.1.1.chr8\_hap1.g00729.t01 |
| 10414 | 8 | 39938143 | 39965738 | mNeoVis.2.1.1.chr8\_hap1.g00730.t01 |
| 10415 | 8 | 39938143 | 39965738 | mNeoVis.2.1.1.chr8\_hap1.g00730.t01 |
| 10416 | 8 | 40024671 | 40051957 | mNeoVis.2.1.1.chr8\_hap1.g00731.t01 |
| 10417 | 8 | 40056184 | 40067521 | mNeoVis.2.1.1.chr8\_hap1.g00732.t01 |
| 10418 | 8 | 40072784 | 40121906 | mNeoVis.2.1.1.chr8\_hap1.g00733.t01 |
| 10419 | 8 | 40163234 | 40256979 | mNeoVis.2.1.1.chr8\_hap1.g00734.t01 |
| 10420 | 8 | 40163234 | 40256979 | mNeoVis.2.1.1.chr8\_hap1.g00734.t01 |
| 10421 | 8 | 40264624 | 40293021 | mNeoVis.2.1.1.chr8\_hap1.g00735.t01 |
| 10422 | 8 | 40264624 | 40293021 | mNeoVis.2.1.1.chr8\_hap1.g00735.t01 |
| 10423 | 8 | 40306365 | 40315312 | mNeoVis.2.1.1.chr8\_hap1.g00736.t01 |
| 10424 | 8 | 40306365 | 40315312 | mNeoVis.2.1.1.chr8\_hap1.g00736.t01 |
| 10425 | 8 | 40317645 | 40367810 | mNeoVis.2.1.1.chr8\_hap1.g00737.t01 |
| 10426 | 8 | 40317645 | 40367810 | mNeoVis.2.1.1.chr8\_hap1.g00737.t01 |
| 10427 | 8 | 40376494 | 40414760 | mNeoVis.2.1.1.chr8\_hap1.g00738.t01 |
| 10428 | 8 | 40376494 | 40414760 | mNeoVis.2.1.1.chr8\_hap1.g00738.t01 |
| 10429 | 8 | 40426556 | 40469688 | mNeoVis.2.1.1.chr8\_hap1.g00739.t01 |
| 10430 | 8 | 40426556 | 40469688 | mNeoVis.2.1.1.chr8\_hap1.g00739.t01 |
| 10431 | 8 | 40497298 | 40522731 | mNeoVis.2.1.1.chr8\_hap1.g00740.t01 |
| 10432 | 8 | 40497298 | 40522731 | mNeoVis.2.1.1.chr8\_hap1.g00740.t01 |
| 10433 | 8 | 40525248 | 40531478 | mNeoVis.2.1.1.chr8\_hap1.g00741.t01 |
| 10434 | 8 | 40525248 | 40531478 | mNeoVis.2.1.1.chr8\_hap1.g00741.t01 |
| 10435 | 8 | 40734128 | 40740609 | mNeoVis.2.1.1.chr8\_hap1.g00742.t01 |
| 10436 | 8 | 40895813 | 41101468 | mNeoVis.2.1.1.chr8\_hap1.g00744.t01 |
| 10437 | 8 | 40941699 | 40946179 | mNeoVis.2.1.1.chr8\_hap1.g00745.t01 |
| 10438 | 8 | 40948823 | 40958331 | mNeoVis.2.1.1.chr8\_hap1.g00746.t01 |
| 10439 | 8 | 40966424 | 40969408 | mNeoVis.2.1.1.chr8\_hap1.g00747.t01 |
| 10440 | 8 | 40966424 | 40969408 | mNeoVis.2.1.1.chr8\_hap1.g00747.t01 |
| 10441 | 8 | 41148897 | 41152472 | mNeoVis.2.1.1.chr8\_hap1.g00748.t01 |
| 10442 | 8 | 41148897 | 41152472 | mNeoVis.2.1.1.chr8\_hap1.g00748.t01 |
| 10443 | 8 | 41148897 | 41152472 | mNeoVis.2.1.1.chr8\_hap1.g00748.t01 |
| 10444 | 8 | 41264875 | 41283002 | mNeoVis.2.1.1.chr8\_hap1.g00749.t01 |
| 10445 | 8 | 41264875 | 41283002 | mNeoVis.2.1.1.chr8\_hap1.g00749.t01 |
| 10446 | 8 | 41264875 | 41283002 | mNeoVis.2.1.1.chr8\_hap1.g00749.t01 |
| 10447 | 8 | 41287991 | 41299182 | mNeoVis.2.1.1.chr8\_hap1.g00750.t01 |
| 10448 | 8 | 41287991 | 41299182 | mNeoVis.2.1.1.chr8\_hap1.g00750.t01 |
| 10449 | 8 | 41287991 | 41299182 | mNeoVis.2.1.1.chr8\_hap1.g00750.t01 |
| 10450 | 8 | 41365310 | 41515212 | mNeoVis.2.1.1.chr8\_hap1.g00751.t01 |
| 10451 | 8 | 42087665 | 42116102 | mNeoVis.2.1.1.chr8\_hap1.g00765.t01 |
| 10452 | 8 | 42126307 | 42147619 | mNeoVis.2.1.1.chr8\_hap1.g00766.t01 |
| 10453 | 8 | 42151774 | 42157563 | mNeoVis.2.1.1.chr8\_hap1.g00767.t01 |
| 10454 | 8 | 42158404 | 42171124 | mNeoVis.2.1.1.chr8\_hap1.g00768.t01 |
| 10455 | 8 | 42173714 | 42177572 | mNeoVis.2.1.1.chr8\_hap1.g00769.t01 |
| 10456 | 8 | 42178179 | 42200699 | mNeoVis.2.1.1.chr8\_hap1.g00770.t01 |
| 10457 | 8 | 42204228 | 42212857 | mNeoVis.2.1.1.chr8\_hap1.g00771.t01 |
| 10458 | 8 | 42213007 | 42245638 | mNeoVis.2.1.1.chr8\_hap1.g00772.t01 |
| 10459 | 8 | 42246419 | 42257964 | mNeoVis.2.1.1.chr8\_hap1.g00773.t01 |
| 10460 | 8 | 42258128 | 42291954 | mNeoVis.2.1.1.chr8\_hap1.g00774.t01 |
| 10461 | 8 | 42292998 | 42300325 | mNeoVis.2.1.1.chr8\_hap1.g00775.t01 |
| 10462 | 8 | 42304539 | 42309337 | mNeoVis.2.1.1.chr8\_hap1.g00776.t01 |
| 10463 | 8 | 42310515 | 42314082 | mNeoVis.2.1.1.chr8\_hap1.g00777.t01 |
| 10464 | 8 | 42314188 | 42316326 | mNeoVis.2.1.1.chr8\_hap1.g00778.t01 |
| 10465 | 8 | 42318679 | 42328475 | mNeoVis.2.1.1.chr8\_hap1.g00779.t01 |
| 10466 | 8 | 42329644 | 42337724 | mNeoVis.2.1.1.chr8\_hap1.g00780.t01 |
| 10467 | 8 | 42337869 | 42422157 | mNeoVis.2.1.1.chr8\_hap1.g00781.t01 |
| 10468 | 8 | 42430522 | 42435072 | mNeoVis.2.1.1.chr8\_hap1.g00782.t01 |
| 10469 | 8 | 42452878 | 42468483 | mNeoVis.2.1.1.chr8\_hap1.g00783.t01 |
| 10470 | 8 | 42468691 | 42473433 | mNeoVis.2.1.1.chr8\_hap1.g00784.t01 |
| 10471 | 8 | 42475785 | 42518202 | mNeoVis.2.1.1.chr8\_hap1.g00785.t01 |
| 10472 | 8 | 42523624 | 42567134 | mNeoVis.2.1.1.chr8\_hap1.g00786.t01 |
| 10473 | 8 | 42574857 | 42617742 | mNeoVis.2.1.1.chr8\_hap1.g00787.t01 |
| 10474 | 8 | 42633467 | 42729018 | mNeoVis.2.1.1.chr8\_hap1.g00788.t01 |
| 10475 | 8 | 43977213 | 43992231 | mNeoVis.2.1.1.chr8\_hap1.g00806.t01 |
| 10476 | 8 | 43999223 | 43999885 | mNeoVis.2.1.1.chr8\_hap1.g00807.t01 |
| 10477 | 8 | 44053891 | 44231534 | mNeoVis.2.1.1.chr8\_hap1.g00808.t01 |
| 10478 | 8 | 44223742 | 44238595 | mNeoVis.2.1.1.chr8\_hap1.g00809.t01 |
| 10479 | 8 | 44248235 | 44402765 | mNeoVis.2.1.1.chr8\_hap1.g00810.t01 |
| 10480 | 8 | 44405901 | 44414162 | mNeoVis.2.1.1.chr8\_hap1.g00811.t01 |
| 10481 | 8 | 44405901 | 44414162 | mNeoVis.2.1.1.chr8\_hap1.g00811.t01 |
| 10482 | 8 | 44414195 | 44433680 | mNeoVis.2.1.1.chr8\_hap1.g00812.t01 |
| 10483 | 8 | 44414195 | 44433680 | mNeoVis.2.1.1.chr8\_hap1.g00812.t01 |
| 10484 | 8 | 44437472 | 44438681 | mNeoVis.2.1.1.chr8\_hap1.g00813.t01 |
| 10485 | 8 | 44437472 | 44438681 | mNeoVis.2.1.1.chr8\_hap1.g00813.t01 |
| 10486 | 8 | 44438722 | 44445424 | mNeoVis.2.1.1.chr8\_hap1.g00814.t01 |
| 10487 | 8 | 44438722 | 44445424 | mNeoVis.2.1.1.chr8\_hap1.g00814.t01 |
| 10488 | 8 | 44446473 | 44454447 | mNeoVis.2.1.1.chr8\_hap1.g00815.t01 |
| 10489 | 8 | 44446473 | 44454447 | mNeoVis.2.1.1.chr8\_hap1.g00815.t01 |
| 10490 | 8 | 44471697 | 44608138 | mNeoVis.2.1.1.chr8\_hap1.g00816.t01 |
| 10491 | 8 | 44622894 | 44628692 | mNeoVis.2.1.1.chr8\_hap1.g00817.t01 |
| 10492 | 8 | 44635567 | 44638562 | mNeoVis.2.1.1.chr8\_hap1.g00818.t01 |
| 10493 | 8 | 44661274 | 44803211 | mNeoVis.2.1.1.chr8\_hap1.g00819.t01 |
| 10494 | 8 | 44826708 | 44856754 | mNeoVis.2.1.1.chr8\_hap1.g00820.t01 |
| 10495 | 8 | 44874861 | 44932950 | mNeoVis.2.1.1.chr8\_hap1.g00821.t01 |
| 10496 | 8 | 47910507 | 47914077 | mNeoVis.2.1.1.chr8\_hap1.g00865.t01 |
| 10497 | 8 | 47934799 | 47958077 | mNeoVis.2.1.1.chr8\_hap1.g00866.t01 |
| 10498 | 8 | 47981835 | 47987176 | mNeoVis.2.1.1.chr8\_hap1.g00867.t01 |
| 10499 | 8 | 48005137 | 48024208 | mNeoVis.2.1.1.chr8\_hap1.g00868.t01 |
| 10500 | 8 | 48024550 | 48045789 | mNeoVis.2.1.1.chr8\_hap1.g00869.t01 |
| 10501 | 8 | 48073635 | 48074370 | mNeoVis.2.1.1.chr8\_hap1.g00870.t01 |
| 10502 | 8 | 48076562 | 48144288 | mNeoVis.2.1.1.chr8\_hap1.g00871.t01 |
| 10503 | 8 | 48145985 | 48182804 | mNeoVis.2.1.1.chr8\_hap1.g00872.t01 |
| 10504 | 8 | 48292322 | 48298405 | mNeoVis.2.1.1.chr8\_hap1.g00873.t01 |
| 10505 | 8 | 48326038 | 48329213 | mNeoVis.2.1.1.chr8\_hap1.g00874.t01 |
| 10506 | 8 | 48329299 | 48334410 | mNeoVis.2.1.1.chr8\_hap1.g00875.t01 |
| 10507 | 8 | 48363304 | 48376074 | mNeoVis.2.1.1.chr8\_hap1.g00876.t01 |
| 10508 | 8 | 48380577 | 48462566 | mNeoVis.2.1.1.chr8\_hap1.g00877.t01 |
| 10509 | 8 | 48520789 | 48544110 | mNeoVis.2.1.1.chr8\_hap1.g00878.t01 |
| 10510 | 8 | 48545245 | 48604642 | mNeoVis.2.1.1.chr8\_hap1.g00879.t01 |
| 10511 | 8 | 48561233 | 48589634 | mNeoVis.2.1.1.chr8\_hap1.g00880.t01 |
| 10512 | 8 | 48706515 | 48730982 | mNeoVis.2.1.1.chr8\_hap1.g00882.t01 |
| 10513 | 8 | 48732694 | 48744109 | mNeoVis.2.1.1.chr8\_hap1.g00883.t01 |
| 10514 | 8 | 48744404 | 48770386 | mNeoVis.2.1.1.chr8\_hap1.g00884.t01 |
| 10515 | 8 | 48776588 | 48781632 | mNeoVis.2.1.1.chr8\_hap1.g00885.t01 |
| 10516 | 8 | 48782844 | 48795946 | mNeoVis.2.1.1.chr8\_hap1.g00886.t01 |
| 10517 | 8 | 48795915 | 48812739 | mNeoVis.2.1.1.chr8\_hap1.g00887.t01 |
| 10518 | 8 | 48820015 | 48844839 | mNeoVis.2.1.1.chr8\_hap1.g00888.t01 |
| 10519 | 8 | 48847221 | 48899766 | mNeoVis.2.1.1.chr8\_hap1.g00889.t01 |
| 10520 | 8 | 48908337 | 48927355 | mNeoVis.2.1.1.chr8\_hap1.g00890.t01 |
| 10521 | 8 | 48937366 | 48962172 | mNeoVis.2.1.1.chr8\_hap1.g00891.t01 |
| 10522 | 8 | 48962517 | 48981825 | mNeoVis.2.1.1.chr8\_hap1.g00892.t01 |
| 10523 | 8 | 48982915 | 49014696 | mNeoVis.2.1.1.chr8\_hap1.g00893.t01 |
| 10524 | 8 | 49014913 | 49140199 | mNeoVis.2.1.1.chr8\_hap1.g00894.t01 |
| 10525 | 8 | 49148027 | 49170683 | mNeoVis.2.1.1.chr8\_hap1.g00895.t01 |
| 10526 | 8 | 49550172 | 49586686 | mNeoVis.2.1.1.chr8\_hap1.g00900.t01 |
| 10527 | 8 | 49550172 | 49586686 | mNeoVis.2.1.1.chr8\_hap1.g00900.t01 |
| 10528 | 8 | 49550172 | 49586686 | mNeoVis.2.1.1.chr8\_hap1.g00900.t01 |
| 10529 | 8 | 49610184 | 49635519 | mNeoVis.2.1.1.chr8\_hap1.g00901.t01 |
| 10530 | 8 | 49610184 | 49635519 | mNeoVis.2.1.1.chr8\_hap1.g00901.t01 |
| 10531 | 8 | 49610184 | 49635519 | mNeoVis.2.1.1.chr8\_hap1.g00901.t01 |
| 10532 | 8 | 49640068 | 49658833 | mNeoVis.2.1.1.chr8\_hap1.g00902.t01 |
| 10533 | 8 | 49640068 | 49658833 | mNeoVis.2.1.1.chr8\_hap1.g00902.t01 |
| 10534 | 8 | 49640068 | 49658833 | mNeoVis.2.1.1.chr8\_hap1.g00902.t01 |
| 10535 | 8 | 49679289 | 49684017 | mNeoVis.2.1.1.chr8\_hap1.g00903.t01 |
| 10536 | 8 | 49679289 | 49684017 | mNeoVis.2.1.1.chr8\_hap1.g00903.t01 |
| 10537 | 8 | 49679289 | 49684017 | mNeoVis.2.1.1.chr8\_hap1.g00903.t01 |
| 10538 | 8 | 49689607 | 49835489 | mNeoVis.2.1.1.chr8\_hap1.g00904.t01 |
| 10539 | 8 | 49689607 | 49835489 | mNeoVis.2.1.1.chr8\_hap1.g00904.t01 |
| 10540 | 8 | 49689607 | 49835489 | mNeoVis.2.1.1.chr8\_hap1.g00904.t01 |
| 10541 | 8 | 49904519 | 49947111 | mNeoVis.2.1.1.chr8\_hap1.g00905.t01 |
| 10542 | 8 | 49904519 | 49947111 | mNeoVis.2.1.1.chr8\_hap1.g00905.t01 |
| 10543 | 8 | 49904519 | 49947111 | mNeoVis.2.1.1.chr8\_hap1.g00905.t01 |
| 10544 | 8 | 49959332 | 50008222 | mNeoVis.2.1.1.chr8\_hap1.g00906.t01 |
| 10545 | 8 | 49959332 | 50008222 | mNeoVis.2.1.1.chr8\_hap1.g00906.t01 |
| 10546 | 8 | 53436293 | 53458242 | mNeoVis.2.1.1.chr8\_hap1.g00949.t01 |
| 10547 | 8 | 53452110 | 53473956 | mNeoVis.2.1.1.chr8\_hap1.g00950.t01 |
| 10548 | 8 | 53474076 | 53505654 | mNeoVis.2.1.1.chr8\_hap1.g00951.t01 |
| 10549 | 8 | 53506869 | 53662114 | mNeoVis.2.1.1.chr8\_hap1.g00952.t01 |
| 10550 | 8 | 53662646 | 53732789 | mNeoVis.2.1.1.chr8\_hap1.g00953.t01 |
| 10551 | 8 | 53748193 | 53760906 | mNeoVis.2.1.1.chr8\_hap1.g00954.t01 |
| 10552 | 8 | 53795438 | 53797587 | mNeoVis.2.1.1.chr8\_hap1.g00955.t01 |
| 10553 | 8 | 53820173 | 53838090 | mNeoVis.2.1.1.chr8\_hap1.g00956.t01 |
| 10554 | 8 | 53842183 | 53876498 | mNeoVis.2.1.1.chr8\_hap1.g00957.t01 |
| 10555 | 8 | 53921480 | 53940339 | mNeoVis.2.1.1.chr8\_hap1.g00958.t01 |
| 10556 | 8 | 53967232 | 54008920 | mNeoVis.2.1.1.chr8\_hap1.g00959.t01 |
| 10557 | 8 | 54023763 | 54045253 | mNeoVis.2.1.1.chr8\_hap1.g00960.t01 |
| 10558 | 8 | 54256699 | 54281339 | mNeoVis.2.1.1.chr8\_hap1.g00965.t01 |
| 10559 | 8 | 54347741 | 54378959 | mNeoVis.2.1.1.chr8\_hap1.g00966.t01 |
| 10560 | 8 | 54448583 | 54486878 | mNeoVis.2.1.1.chr8\_hap1.g00967.t01 |
| 10561 | 8 | 54527870 | 54558374 | mNeoVis.2.1.1.chr8\_hap1.g00968.t01 |
| 10562 | 8 | 55353563 | 55394245 | mNeoVis.2.1.1.chr8\_hap1.g00969.t01 |
| 10563 | 8 | 55353563 | 55394245 | mNeoVis.2.1.1.chr8\_hap1.g00969.t01 |
| 10564 | 8 | 55353563 | 55394245 | mNeoVis.2.1.1.chr8\_hap1.g00969.t01 |
| 10565 | 8 | 55429240 | 55560740 | mNeoVis.2.1.1.chr8\_hap1.g00970.t01 |
| 10566 | 8 | 55429240 | 55560740 | mNeoVis.2.1.1.chr8\_hap1.g00970.t01 |
| 10567 | 8 | 55429240 | 55560740 | mNeoVis.2.1.1.chr8\_hap1.g00970.t01 |
| 10568 | 8 | 55942578 | 56035383 | mNeoVis.2.1.1.chr8\_hap1.g00971.t01 |
| 10569 | 8 | 56464556 | 56484962 | mNeoVis.2.1.1.chr8\_hap1.g00972.t01 |
| 10570 | 8 | 56484931 | 56652712 | mNeoVis.2.1.1.chr8\_hap1.g00973.t01 |
| 10571 | 8 | 56663867 | 56762179 | mNeoVis.2.1.1.chr8\_hap1.g00974.t01 |
| 10572 | 8 | 57202695 | 57335609 | mNeoVis.2.1.1.chr8\_hap1.g00975.t01 |
| 10573 | 8 | 57202695 | 57335609 | mNeoVis.2.1.1.chr8\_hap1.g00975.t01 |
| 10574 | 8 | 57359390 | 57388814 | mNeoVis.2.1.1.chr8\_hap1.g00976.t01 |
| 10575 | 8 | 57749535 | 58026152 | mNeoVis.2.1.1.chr8\_hap1.g00978.t01 |
| 10576 | 8 | 59157753 | 59181142 | mNeoVis.2.1.1.chr8\_hap1.g00994.t01 |
| 10577 | 8 | 59185309 | 59221297 | mNeoVis.2.1.1.chr8\_hap1.g00995.t01 |
| 10578 | 8 | 59224393 | 59274940 | mNeoVis.2.1.1.chr8\_hap1.g00996.t01 |
| 10579 | 8 | 59276857 | 59313693 | mNeoVis.2.1.1.chr8\_hap1.g00997.t01 |
| 10580 | 8 | 59314296 | 59570777 | mNeoVis.2.1.1.chr8\_hap1.g00998.t01 |
| 10581 | 8 | 59778487 | 59838779 | mNeoVis.2.1.1.chr8\_hap1.g01000.t01 |
| 10582 | 8 | 59848578 | 59897843 | mNeoVis.2.1.1.chr8\_hap1.g01001.t01 |
| 10583 | 8 | 59898973 | 59901105 | mNeoVis.2.1.1.chr8\_hap1.g01002.t01 |
| 10584 | 8 | 59919048 | 59936892 | mNeoVis.2.1.1.chr8\_hap1.g01003.t01 |
| 10585 | 8 | 60009031 | 60121596 | mNeoVis.2.1.1.chr8\_hap1.g01004.t01 |
| 10586 | 8 | 60126822 | 60157028 | mNeoVis.2.1.1.chr8\_hap1.g01005.t01 |
| 10587 | 8 | 60190472 | 60198488 | mNeoVis.2.1.1.chr8\_hap1.g01006.t01 |
| 10588 | 8 | 60202109 | 60207476 | mNeoVis.2.1.1.chr8\_hap1.g01007.t01 |
| 10589 | 8 | 60207566 | 60216374 | mNeoVis.2.1.1.chr8\_hap1.g01008.t01 |
| 10590 | 8 | 60231569 | 60235080 | mNeoVis.2.1.1.chr8\_hap1.g01009.t01 |
| 10591 | 8 | 60243166 | 60253795 | mNeoVis.2.1.1.chr8\_hap1.g01010.t01 |
| 10592 | 8 | 60368873 | 60373187 | mNeoVis.2.1.1.chr8\_hap1.g01019.t01 |
| 10593 | 8 | 60382149 | 60385980 | mNeoVis.2.1.1.chr8\_hap1.g01020.t01 |
| 10594 | 8 | 60387137 | 60402696 | mNeoVis.2.1.1.chr8\_hap1.g01021.t01 |
| 10595 | 8 | 60411687 | 60415728 | mNeoVis.2.1.1.chr8\_hap1.g01022.t01 |
| 10596 | 8 | 60418933 | 60440289 | mNeoVis.2.1.1.chr8\_hap1.g01023.t01 |
| 10597 | 8 | 60446756 | 60460110 | mNeoVis.2.1.1.chr8\_hap1.g01024.t01 |
| 10598 | 8 | 60469874 | 60478737 | mNeoVis.2.1.1.chr8\_hap1.g01025.t01 |
| 10599 | 8 | 60498978 | 60511394 | mNeoVis.2.1.1.chr8\_hap1.g01026.t01 |
| 10600 | 8 | 60545411 | 60565872 | mNeoVis.2.1.1.chr8\_hap1.g01027.t01 |
| 10601 | 8 | 60565757 | 60567527 | mNeoVis.2.1.1.chr8\_hap1.g01028.t01 |
| 10602 | 8 | 60568525 | 60574541 | mNeoVis.2.1.1.chr8\_hap1.g01029.t01 |
| 10603 | 8 | 60578801 | 60582128 | mNeoVis.2.1.1.chr8\_hap1.g01030.t01 |
| 10604 | 8 | 60582936 | 60607424 | mNeoVis.2.1.1.chr8\_hap1.g01031.t01 |
| 10605 | 8 | 60606942 | 60631733 | mNeoVis.2.1.1.chr8\_hap1.g01032.t01 |
| 10606 | 8 | 60628946 | 60630479 | mNeoVis.2.1.1.chr8\_hap1.g01033.t01 |
| 10607 | 8 | 60632190 | 60633197 | mNeoVis.2.1.1.chr8\_hap1.g01034.t01 |
| 10608 | 8 | 60633427 | 60655137 | mNeoVis.2.1.1.chr8\_hap1.g01035.t01 |
| 10609 | 8 | 60655582 | 60737398 | mNeoVis.2.1.1.chr8\_hap1.g01036.t01 |
| 10610 | 8 | 60743389 | 60749349 | mNeoVis.2.1.1.chr8\_hap1.g01037.t01 |
| 10611 | 8 | 60751062 | 60761305 | mNeoVis.2.1.1.chr8\_hap1.g01038.t01 |
| 10612 | 8 | 60762266 | 60775902 | mNeoVis.2.1.1.chr8\_hap1.g01039.t01 |
| 10613 | 8 | 60780997 | 60787212 | mNeoVis.2.1.1.chr8\_hap1.g01040.t01 |
| 10614 | 8 | 60795007 | 60797925 | mNeoVis.2.1.1.chr8\_hap1.g01041.t01 |
| 10615 | 8 | 60799174 | 60802308 | mNeoVis.2.1.1.chr8\_hap1.g01042.t01 |
| 10616 | 8 | 60808731 | 60823456 | mNeoVis.2.1.1.chr8\_hap1.g01043.t01 |
| 10617 | 8 | 60824731 | 60826356 | mNeoVis.2.1.1.chr8\_hap1.g01044.t01 |
| 10618 | 8 | 60826825 | 60829907 | mNeoVis.2.1.1.chr8\_hap1.g01045.t01 |
| 10619 | 8 | 60831480 | 60834028 | mNeoVis.2.1.1.chr8\_hap1.g01046.t01 |
| 10620 | 8 | 60834183 | 60840161 | mNeoVis.2.1.1.chr8\_hap1.g01047.t01 |
| 10621 | 8 | 60840833 | 60848670 | mNeoVis.2.1.1.chr8\_hap1.g01048.t01 |
| 10622 | 8 | 60849580 | 60852490 | mNeoVis.2.1.1.chr8\_hap1.g01049.t01 |
| 10623 | 8 | 60852797 | 60859722 | mNeoVis.2.1.1.chr8\_hap1.g01050.t01 |
| 10624 | 8 | 60967754 | 60969754 | mNeoVis.2.1.1.chr8\_hap1.g01058.t01 |
| 10625 | 8 | 60971591 | 60972956 | mNeoVis.2.1.1.chr8\_hap1.g01059.t01 |
| 10626 | 8 | 60973404 | 60988317 | mNeoVis.2.1.1.chr8\_hap1.g01060.t01 |
| 10627 | 8 | 60989040 | 60990165 | mNeoVis.2.1.1.chr8\_hap1.g01061.t01 |
| 10628 | 8 | 60989040 | 60990165 | mNeoVis.2.1.1.chr8\_hap1.g01061.t01 |
| 10629 | 8 | 60995878 | 61000308 | mNeoVis.2.1.1.chr8\_hap1.g01062.t01 |
| 10630 | 8 | 60995878 | 61000308 | mNeoVis.2.1.1.chr8\_hap1.g01062.t01 |
| 10631 | 8 | 61000765 | 61007175 | mNeoVis.2.1.1.chr8\_hap1.g01063.t01 |
| 10632 | 8 | 61000765 | 61007175 | mNeoVis.2.1.1.chr8\_hap1.g01063.t01 |
| 10633 | 8 | 61015679 | 61017722 | mNeoVis.2.1.1.chr8\_hap1.g01064.t01 |
| 10634 | 8 | 61015679 | 61017722 | mNeoVis.2.1.1.chr8\_hap1.g01064.t01 |
| 10635 | 8 | 61017872 | 61021162 | mNeoVis.2.1.1.chr8\_hap1.g01065.t01 |
| 10636 | 8 | 61017872 | 61021162 | mNeoVis.2.1.1.chr8\_hap1.g01065.t01 |
| 10637 | 8 | 61021280 | 61033002 | mNeoVis.2.1.1.chr8\_hap1.g01066.t01 |
| 10638 | 8 | 61021280 | 61033002 | mNeoVis.2.1.1.chr8\_hap1.g01066.t01 |
| 10639 | 8 | 61033121 | 61036700 | mNeoVis.2.1.1.chr8\_hap1.g01067.t01 |
| 10640 | 8 | 61033121 | 61036700 | mNeoVis.2.1.1.chr8\_hap1.g01067.t01 |
| 10641 | 8 | 61037631 | 61048589 | mNeoVis.2.1.1.chr8\_hap1.g01068.t01 |
| 10642 | 8 | 61037631 | 61048589 | mNeoVis.2.1.1.chr8\_hap1.g01068.t01 |
| 10643 | 8 | 61048736 | 61052013 | mNeoVis.2.1.1.chr8\_hap1.g01069.t01 |
| 10644 | 8 | 61048736 | 61052013 | mNeoVis.2.1.1.chr8\_hap1.g01069.t01 |
| 10645 | 8 | 61051640 | 61055635 | mNeoVis.2.1.1.chr8\_hap1.g01070.t01 |
| 10646 | 8 | 61051640 | 61055635 | mNeoVis.2.1.1.chr8\_hap1.g01070.t01 |
| 10647 | 8 | 61071878 | 61077539 | mNeoVis.2.1.1.chr8\_hap1.g01071.t01 |
| 10648 | 8 | 61071878 | 61077539 | mNeoVis.2.1.1.chr8\_hap1.g01071.t01 |
| 10649 | 8 | 61078117 | 61083733 | mNeoVis.2.1.1.chr8\_hap1.g01072.t01 |
| 10650 | 8 | 61078117 | 61083733 | mNeoVis.2.1.1.chr8\_hap1.g01072.t01 |
| 10651 | 8 | 61091878 | 61094552 | mNeoVis.2.1.1.chr8\_hap1.g01073.t01 |
| 10652 | 8 | 61091878 | 61094552 | mNeoVis.2.1.1.chr8\_hap1.g01073.t01 |
| 10653 | 8 | 61094825 | 61101077 | mNeoVis.2.1.1.chr8\_hap1.g01074.t01 |
| 10654 | 8 | 61094825 | 61101077 | mNeoVis.2.1.1.chr8\_hap1.g01074.t01 |
| 10655 | 8 | 61101402 | 61107285 | mNeoVis.2.1.1.chr8\_hap1.g01075.t01 |
| 10656 | 8 | 61101402 | 61107285 | mNeoVis.2.1.1.chr8\_hap1.g01075.t01 |
| 10657 | 8 | 61108622 | 61110689 | mNeoVis.2.1.1.chr8\_hap1.g01076.t01 |
| 10658 | 8 | 61108622 | 61110689 | mNeoVis.2.1.1.chr8\_hap1.g01076.t01 |
| 10659 | 8 | 61111608 | 61115899 | mNeoVis.2.1.1.chr8\_hap1.g01077.t01 |
| 10660 | 8 | 61111608 | 61115899 | mNeoVis.2.1.1.chr8\_hap1.g01077.t01 |
| 10661 | 8 | 61116436 | 61123833 | mNeoVis.2.1.1.chr8\_hap1.g01078.t01 |
| 10662 | 8 | 61116436 | 61123833 | mNeoVis.2.1.1.chr8\_hap1.g01078.t01 |
| 10663 | 8 | 61123904 | 61129244 | mNeoVis.2.1.1.chr8\_hap1.g01079.t01 |
| 10664 | 8 | 61123904 | 61129244 | mNeoVis.2.1.1.chr8\_hap1.g01079.t01 |
| 10665 | 8 | 61131624 | 61152560 | mNeoVis.2.1.1.chr8\_hap1.g01080.t01 |
| 10666 | 8 | 61131624 | 61152560 | mNeoVis.2.1.1.chr8\_hap1.g01080.t01 |
| 10667 | 8 | 61159869 | 61163617 | mNeoVis.2.1.1.chr8\_hap1.g01081.t01 |
| 10668 | 8 | 61159869 | 61163617 | mNeoVis.2.1.1.chr8\_hap1.g01081.t01 |
| 10669 | 8 | 61188267 | 61219973 | mNeoVis.2.1.1.chr8\_hap1.g01082.t01 |
| 10670 | 8 | 61188267 | 61219973 | mNeoVis.2.1.1.chr8\_hap1.g01082.t01 |
| 10671 | 8 | 61238828 | 61241407 | mNeoVis.2.1.1.chr8\_hap1.g01083.t01 |
| 10672 | 8 | 61238828 | 61241407 | mNeoVis.2.1.1.chr8\_hap1.g01083.t01 |
| 10673 | 8 | 61241800 | 61246370 | mNeoVis.2.1.1.chr8\_hap1.g01084.t01 |
| 10674 | 8 | 61241800 | 61246370 | mNeoVis.2.1.1.chr8\_hap1.g01084.t01 |
| 10675 | 8 | 61249387 | 61253514 | mNeoVis.2.1.1.chr8\_hap1.g01085.t01 |
| 10676 | 8 | 61249387 | 61253514 | mNeoVis.2.1.1.chr8\_hap1.g01085.t01 |
| 10677 | 8 | 61259180 | 61261926 | mNeoVis.2.1.1.chr8\_hap1.g01086.t01 |
| 10678 | 8 | 61259180 | 61261926 | mNeoVis.2.1.1.chr8\_hap1.g01086.t01 |
| 10679 | 8 | 61262132 | 61263756 | mNeoVis.2.1.1.chr8\_hap1.g01087.t01 |
| 10680 | 8 | 61262132 | 61263756 | mNeoVis.2.1.1.chr8\_hap1.g01087.t01 |
| 10681 | 8 | 61264835 | 61276788 | mNeoVis.2.1.1.chr8\_hap1.g01088.t01 |
| 10682 | 8 | 61264835 | 61276788 | mNeoVis.2.1.1.chr8\_hap1.g01088.t01 |
| 10683 | 8 | 61355510 | 61361327 | mNeoVis.2.1.1.chr8\_hap1.g01089.t01 |
| 10684 | 8 | 61355510 | 61361327 | mNeoVis.2.1.1.chr8\_hap1.g01089.t01 |
| 10685 | 8 | 61405739 | 61413862 | mNeoVis.2.1.1.chr8\_hap1.g01090.t01 |
| 10686 | 8 | 61405739 | 61413862 | mNeoVis.2.1.1.chr8\_hap1.g01090.t01 |
| 10687 | 8 | 61425669 | 61426063 | mNeoVis.2.1.1.chr8\_hap1.g01091.t01 |
| 10688 | 8 | 61425669 | 61426063 | mNeoVis.2.1.1.chr8\_hap1.g01091.t01 |
| 10689 | 8 | 61430893 | 61431167 | mNeoVis.2.1.1.chr8\_hap1.g01092.t01 |
| 10690 | 8 | 61430893 | 61431167 | mNeoVis.2.1.1.chr8\_hap1.g01092.t01 |
| 10691 | 8 | 61444761 | 61463820 | mNeoVis.2.1.1.chr8\_hap1.g01093.t01 |
| 10692 | 8 | 61444761 | 61463820 | mNeoVis.2.1.1.chr8\_hap1.g01093.t01 |
| 10693 | 8 | 61470043 | 61477996 | mNeoVis.2.1.1.chr8\_hap1.g01094.t01 |
| 10694 | 8 | 61470043 | 61477996 | mNeoVis.2.1.1.chr8\_hap1.g01094.t01 |
| 10695 | 8 | 61483181 | 61485455 | mNeoVis.2.1.1.chr8\_hap1.g01095.t01 |
| 10696 | 8 | 61483181 | 61485455 | mNeoVis.2.1.1.chr8\_hap1.g01095.t01 |
| 10697 | 8 | 61485237 | 61489233 | mNeoVis.2.1.1.chr8\_hap1.g01096.t01 |
| 10698 | 8 | 61485237 | 61489233 | mNeoVis.2.1.1.chr8\_hap1.g01096.t01 |
| 10699 | 8 | 61489337 | 61494617 | mNeoVis.2.1.1.chr8\_hap1.g01097.t01 |
| 10700 | 8 | 61489337 | 61494617 | mNeoVis.2.1.1.chr8\_hap1.g01097.t01 |
| 10701 | 8 | 61514700 | 61520869 | mNeoVis.2.1.1.chr8\_hap1.g01098.t01 |
| 10702 | 8 | 61521728 | 61527996 | mNeoVis.2.1.1.chr8\_hap1.g01099.t01 |
| 10703 | 8 | 61524790 | 61526376 | mNeoVis.2.1.1.chr8\_hap1.g01100.t01 |
| 10704 | 8 | 61531705 | 61533669 | mNeoVis.2.1.1.chr8\_hap1.g01101.t01 |
| 10705 | 8 | 61536228 | 61536867 | mNeoVis.2.1.1.chr8\_hap1.g01102.t01 |
| 10706 | 8 | 61542835 | 61554847 | mNeoVis.2.1.1.chr8\_hap1.g01103.t01 |
| 10707 | 8 | 61560036 | 61602607 | mNeoVis.2.1.1.chr8\_hap1.g01104.t01 |
| 10708 | 8 | 61603956 | 61625540 | mNeoVis.2.1.1.chr8\_hap1.g01105.t01 |
| 10709 | 8 | 61627285 | 61630024 | mNeoVis.2.1.1.chr8\_hap1.g01106.t01 |
| 10710 | 8 | 61630173 | 61634675 | mNeoVis.2.1.1.chr8\_hap1.g01107.t01 |
| 10711 | 8 | 61635263 | 61638705 | mNeoVis.2.1.1.chr8\_hap1.g01108.t01 |
| 10712 | 8 | 61638738 | 61644872 | mNeoVis.2.1.1.chr8\_hap1.g01109.t01 |
| 10713 | 8 | 61645439 | 61649167 | mNeoVis.2.1.1.chr8\_hap1.g01110.t01 |
| 10714 | 8 | 61649712 | 61663620 | mNeoVis.2.1.1.chr8\_hap1.g01111.t01 |
| 10715 | 8 | 61664421 | 61669712 | mNeoVis.2.1.1.chr8\_hap1.g01112.t01 |
| 10716 | 8 | 61670109 | 61690488 | mNeoVis.2.1.1.chr8\_hap1.g01113.t01 |
| 10717 | 8 | 61709545 | 61721222 | mNeoVis.2.1.1.chr8\_hap1.g01114.t01 |
| 10718 | 8 | 61750051 | 61767034 | mNeoVis.2.1.1.chr8\_hap1.g01115.t01 |
| 10719 | 8 | 61802036 | 61892351 | mNeoVis.2.1.1.chr8\_hap1.g01116.t01 |
| 10720 | 8 | 61893974 | 61914186 | mNeoVis.2.1.1.chr8\_hap1.g01117.t01 |
| 10721 | 8 | 61914312 | 61921290 | mNeoVis.2.1.1.chr8\_hap1.g01118.t01 |
| 10722 | 8 | 61921198 | 61925960 | mNeoVis.2.1.1.chr8\_hap1.g01119.t01 |
| 10723 | 8 | 61928247 | 61941877 | mNeoVis.2.1.1.chr8\_hap1.g01120.t01 |
| 10724 | 8 | 61947133 | 61957770 | mNeoVis.2.1.1.chr8\_hap1.g01121.t01 |
| 10725 | 8 | 61957907 | 61960396 | mNeoVis.2.1.1.chr8\_hap1.g01122.t01 |
| 10726 | 8 | 61966394 | 61968317 | mNeoVis.2.1.1.chr8\_hap1.g01123.t01 |
| 10727 | 8 | 62254597 | 62287664 | mNeoVis.2.1.1.chr8\_hap1.g01125.t01 |
| 10728 | 8 | 62422178 | 62432374 | mNeoVis.2.1.1.chr8\_hap1.g01126.t01 |
| 10729 | 8 | 62432613 | 62438025 | mNeoVis.2.1.1.chr8\_hap1.g01127.t01 |
| 10730 | 8 | 62438185 | 62495749 | mNeoVis.2.1.1.chr8\_hap1.g01128.t01 |
| 10731 | 8 | 62504994 | 62542201 | mNeoVis.2.1.1.chr8\_hap1.g01129.t01 |
| 10732 | 8 | 62542893 | 62546074 | mNeoVis.2.1.1.chr8\_hap1.g01130.t01 |
| 10733 | 8 | 62546671 | 62548684 | mNeoVis.2.1.1.chr8\_hap1.g01131.t01 |
| 10734 | 8 | 62549854 | 62556209 | mNeoVis.2.1.1.chr8\_hap1.g01132.t01 |
| 10735 | 8 | 62573520 | 62597372 | mNeoVis.2.1.1.chr8\_hap1.g01133.t01 |
| 10736 | 8 | 62619992 | 62625287 | mNeoVis.2.1.1.chr8\_hap1.g01134.t01 |
| 10737 | 8 | 62632772 | 62641735 | mNeoVis.2.1.1.chr8\_hap1.g01135.t01 |
| 10738 | 8 | 62657878 | 62674214 | mNeoVis.2.1.1.chr8\_hap1.g01136.t01 |
| 10739 | 8 | 62665873 | 62672469 | mNeoVis.2.1.1.chr8\_hap1.g01137.t01 |
| 10740 | 8 | 62693146 | 62696921 | mNeoVis.2.1.1.chr8\_hap1.g01138.t01 |
| 10741 | 8 | 62698083 | 62710660 | mNeoVis.2.1.1.chr8\_hap1.g01139.t01 |
| 10742 | 8 | 62710526 | 62740272 | mNeoVis.2.1.1.chr8\_hap1.g01140.t01 |
| 10743 | 8 | 62746453 | 62779783 | mNeoVis.2.1.1.chr8\_hap1.g01141.t01 |
| 10744 | 8 | 62814824 | 62923598 | mNeoVis.2.1.1.chr8\_hap1.g01142.t01 |
| 10745 | 8 | 62928146 | 63000785 | mNeoVis.2.1.1.chr8\_hap1.g01143.t01 |
| 10746 | 8 | 63002739 | 63014381 | mNeoVis.2.1.1.chr8\_hap1.g01144.t01 |
| 10747 | 8 | 63014808 | 63113505 | mNeoVis.2.1.1.chr8\_hap1.g01145.t01 |
| 10748 | 8 | 68154535 | 68231089 | mNeoVis.2.1.1.chr8\_hap1.g01146.t01 |
| 10749 | 8 | 68236609 | 68253934 | mNeoVis.2.1.1.chr8\_hap1.g01147.t01 |
| 10750 | 8 | 80741698 | 80796760 | mNeoVis.2.1.1.chr8\_hap1.g01160.t01 |
| 10751 | 8 | 80829662 | 80907404 | mNeoVis.2.1.1.chr8\_hap1.g01161.t01 |
| 10752 | 8 | 80935286 | 80985145 | mNeoVis.2.1.1.chr8\_hap1.g01162.t01 |
| 10753 | 8 | 81056431 | 81174430 | mNeoVis.2.1.1.chr8\_hap1.g01163.t01 |
| 10754 | 8 | 81207663 | 81211050 | mNeoVis.2.1.1.chr8\_hap1.g01164.t01 |
| 10755 | 8 | 81219469 | 81238941 | mNeoVis.2.1.1.chr8\_hap1.g01165.t01 |
| 10756 | 8 | 81271636 | 81277502 | mNeoVis.2.1.1.chr8\_hap1.g01166.t01 |
| 10757 | 8 | 81547604 | 81704115 | mNeoVis.2.1.1.chr8\_hap1.g01168.t01 |
| 10758 | 8 | 81736519 | 81891698 | mNeoVis.2.1.1.chr8\_hap1.g01169.t01 |
| 10759 | 8 | 81902565 | 81922922 | mNeoVis.2.1.1.chr8\_hap1.g01170.t01 |
| 10760 | 8 | 81926628 | 81953245 | mNeoVis.2.1.1.chr8\_hap1.g01171.t01 |
| 10761 | 8 | 82081183 | 82085167 | mNeoVis.2.1.1.chr8\_hap1.g01173.t01 |
| 10762 | 8 | 82140188 | 82213767 | mNeoVis.2.1.1.chr8\_hap1.g01174.t01 |
| 10763 | 8 | 82287465 | 82295243 | mNeoVis.2.1.1.chr8\_hap1.g01175.t01 |
| 10764 | 8 | 82302225 | 82321716 | mNeoVis.2.1.1.chr8\_hap1.g01176.t01 |
| 10765 | 8 | 82321784 | 82322599 | mNeoVis.2.1.1.chr8\_hap1.g01177.t01 |
| 10766 | 8 | 82352946 | 82420639 | mNeoVis.2.1.1.chr8\_hap1.g01178.t01 |
| 10767 | 8 | 82695518 | 82729889 | mNeoVis.2.1.1.chr8\_hap1.g01180.t01 |
| 10768 | 8 | 84168808 | 84326317 | mNeoVis.2.1.1.chr8\_hap1.g01181.t01 |
| 10769 | 8 | 84336611 | 84436129 | mNeoVis.2.1.1.chr8\_hap1.g01182.t01 |
| 10770 | 8 | 84336611 | 84436129 | mNeoVis.2.1.1.chr8\_hap1.g01182.t01 |
| 10771 | 8 | 84336611 | 84436129 | mNeoVis.2.1.1.chr8\_hap1.g01182.t01 |
| 10772 | 8 | 84576826 | 84667700 | mNeoVis.2.1.1.chr8\_hap1.g01183.t01 |
| 10773 | 8 | 84576826 | 84667700 | mNeoVis.2.1.1.chr8\_hap1.g01183.t01 |
| 10774 | 8 | 84576826 | 84667700 | mNeoVis.2.1.1.chr8\_hap1.g01183.t01 |
| 10775 | 8 | 84722214 | 84893233 | mNeoVis.2.1.1.chr8\_hap1.g01184.t01 |
| 10776 | 8 | 84722214 | 84893233 | mNeoVis.2.1.1.chr8\_hap1.g01184.t01 |
| 10777 | 8 | 84899628 | 84906522 | mNeoVis.2.1.1.chr8\_hap1.g01185.t01 |
| 10778 | 8 | 84899628 | 84906522 | mNeoVis.2.1.1.chr8\_hap1.g01185.t01 |
| 10779 | 8 | 84899628 | 84906522 | mNeoVis.2.1.1.chr8\_hap1.g01185.t01 |
| 10780 | 8 | 84913996 | 85253593 | mNeoVis.2.1.1.chr8\_hap1.g01186.t01 |
| 10781 | 8 | 84913996 | 85253593 | mNeoVis.2.1.1.chr8\_hap1.g01186.t01 |
| 10782 | 8 | 85176104 | 85177817 | mNeoVis.2.1.1.chr8\_hap1.g01187.t01 |
| 10783 | 8 | 85176104 | 85177817 | mNeoVis.2.1.1.chr8\_hap1.g01187.t01 |
| 10784 | 8 | 85350407 | 85352974 | mNeoVis.2.1.1.chr8\_hap1.g01188.t01 |
| 10785 | 8 | 85350407 | 85352974 | mNeoVis.2.1.1.chr8\_hap1.g01188.t01 |
| 10786 | 8 | 85421520 | 85463438 | mNeoVis.2.1.1.chr8\_hap1.g01189.t01 |
| 10787 | 8 | 87143526 | 87150520 | mNeoVis.2.1.1.chr8\_hap1.g01214.t01 |
| 10788 | 8 | 87231245 | 87507938 | mNeoVis.2.1.1.chr8\_hap1.g01215.t01 |
| 10789 | 8 | 87548431 | 87557976 | mNeoVis.2.1.1.chr8\_hap1.g01216.t01 |
| 10790 | 8 | 87576641 | 87689997 | mNeoVis.2.1.1.chr8\_hap1.g01217.t01 |
| 10791 | 8 | 87802360 | 87941011 | mNeoVis.2.1.1.chr8\_hap1.g01218.t01 |
| 10792 | 8 | 87986691 | 87989869 | mNeoVis.2.1.1.chr8\_hap1.g01219.t01 |
| 10793 | 8 | 89305581 | 89307231 | mNeoVis.2.1.1.chr8\_hap1.g01239.t01 |
| 10794 | 8 | 89305581 | 89348943 | mNeoVis.2.1.1.chr8\_hap1.g01240.t01 |
| 10795 | 8 | 89444390 | 89445626 | mNeoVis.2.1.1.chr8\_hap1.g01241.t01 |
| 10796 | 8 | 89444390 | 89445626 | mNeoVis.2.1.1.chr8\_hap1.g01241.t01 |
| 10797 | 8 | 89562794 | 89569521 | mNeoVis.2.1.1.chr8\_hap1.g01242.t01 |
| 10798 | 8 | 89562794 | 89569521 | mNeoVis.2.1.1.chr8\_hap1.g01242.t01 |
| 10799 | 8 | 89600442 | 89608037 | mNeoVis.2.1.1.chr8\_hap1.g01243.t01 |
| 10800 | 8 | 89600442 | 89608037 | mNeoVis.2.1.1.chr8\_hap1.g01243.t01 |
| 10801 | 8 | 89616535 | 89626582 | mNeoVis.2.1.1.chr8\_hap1.g01244.t01 |
| 10802 | 8 | 89616535 | 89626582 | mNeoVis.2.1.1.chr8\_hap1.g01244.t01 |
| 10803 | 8 | 89645301 | 89720518 | mNeoVis.2.1.1.chr8\_hap1.g01245.t01 |
| 10804 | 8 | 89645301 | 89720518 | mNeoVis.2.1.1.chr8\_hap1.g01245.t01 |
| 10805 | 8 | 89763949 | 89895587 | mNeoVis.2.1.1.chr8\_hap1.g01246.t01 |
| 10806 | 8 | 89901094 | 90078455 | mNeoVis.2.1.1.chr8\_hap1.g01247.t01 |
| 10807 | 8 | 90231413 | 90249435 | mNeoVis.2.1.1.chr8\_hap1.g01248.t01 |
| 10808 | 8 | 90258096 | 90276137 | mNeoVis.2.1.1.chr8\_hap1.g01249.t01 |
| 10809 | 8 | 90273474 | 90305612 | mNeoVis.2.1.1.chr8\_hap1.g01250.t01 |
| 10810 | 8 | 93337419 | 93686467 | mNeoVis.2.1.1.chr8\_hap1.g01266.t01 |
| 10811 | 8 | 93738483 | 93746408 | mNeoVis.2.1.1.chr8\_hap1.g01267.t01 |
| 10812 | 8 | 93738483 | 93746408 | mNeoVis.2.1.1.chr8\_hap1.g01267.t01 |
| 10813 | 8 | 93747591 | 93806003 | mNeoVis.2.1.1.chr8\_hap1.g01268.t01 |
| 10814 | 8 | 93747591 | 93806003 | mNeoVis.2.1.1.chr8\_hap1.g01268.t01 |
| 10815 | 8 | 93808032 | 93844972 | mNeoVis.2.1.1.chr8\_hap1.g01269.t01 |
| 10816 | 8 | 93808032 | 93844972 | mNeoVis.2.1.1.chr8\_hap1.g01269.t01 |
| 10817 | 8 | 93852440 | 93925408 | mNeoVis.2.1.1.chr8\_hap1.g01270.t01 |
| 10818 | 8 | 93852440 | 93925408 | mNeoVis.2.1.1.chr8\_hap1.g01270.t01 |
| 10819 | 8 | 93956633 | 94154107 | mNeoVis.2.1.1.chr8\_hap1.g01271.t01 |
| 10820 | 8 | 93956633 | 94154107 | mNeoVis.2.1.1.chr8\_hap1.g01271.t01 |
| 10821 | 8 | 94352948 | 94393952 | mNeoVis.2.1.1.chr8\_hap1.g01272.t01 |
| 10822 | 8 | 94424556 | 94607109 | mNeoVis.2.1.1.chr8\_hap1.g01273.t01 |
| 10823 | 8 | 94816784 | 94905290 | mNeoVis.2.1.1.chr8\_hap1.g01274.t01 |
| 10824 | 8 | 95069965 | 95091636 | mNeoVis.2.1.1.chr8\_hap1.g01275.t01 |
| 10825 | 8 | 96037364 | 96704748 | mNeoVis.2.1.1.chr8\_hap1.g01276.t01 |
| 10826 | 8 | 96516754 | 96519092 | mNeoVis.2.1.1.chr8\_hap1.g01277.t01 |
| 10827 | 8 | 96516754 | 96519092 | mNeoVis.2.1.1.chr8\_hap1.g01277.t01 |
| 10828 | 8 | 96796086 | 97126535 | mNeoVis.2.1.1.chr8\_hap1.g01278.t01 |
| 10829 | 8 | 98479203 | 98513307 | mNeoVis.2.1.1.chr8\_hap1.g01289.t01 |
| 10830 | 8 | 98556100 | 98761191 | mNeoVis.2.1.1.chr8\_hap1.g01290.t01 |
| 10831 | 8 | 99113485 | 99125697 | mNeoVis.2.1.1.chr8\_hap1.g01291.t01 |
| 10832 | 8 | 99387068 | 99547114 | mNeoVis.2.1.1.chr8\_hap1.g01292.t01 |
| 10833 | 8 | 99552657 | 99573715 | mNeoVis.2.1.1.chr8\_hap1.g01293.t01 |
| 10834 | 8 | 99583679 | 99611498 | mNeoVis.2.1.1.chr8\_hap1.g01294.t01 |
| 10835 | 8 | 99611929 | 99624544 | mNeoVis.2.1.1.chr8\_hap1.g01295.t01 |
| 10836 | 8 | 99772492 | 99773625 | mNeoVis.2.1.1.chr8\_hap1.g01296.t01 |
| 10837 | 8 | 99872630 | 100110503 | mNeoVis.2.1.1.chr8\_hap1.g01297.t01 |
| 10838 | 8 | 100147893 | 100240391 | mNeoVis.2.1.1.chr8\_hap1.g01298.t01 |
| 10839 | 8 | 100332953 | 100381558 | mNeoVis.2.1.1.chr8\_hap1.g01299.t01 |
| 10840 | 8 | 100336910 | 100337824 | mNeoVis.2.1.1.chr8\_hap1.g01300.t01 |
| 10841 | 8 | 100947662 | 101046731 | mNeoVis.2.1.1.chr8\_hap1.g01301.t01 |
| 10842 | 8 | 101086691 | 101121880 | mNeoVis.2.1.1.chr8\_hap1.g01302.t01 |
| 10843 | 8 | 101704080 | 101726766 | mNeoVis.2.1.1.chr8\_hap1.g01309.t01 |
| 10844 | 8 | 101730146 | 101795978 | mNeoVis.2.1.1.chr8\_hap1.g01310.t01 |
| 10845 | 8 | 101796114 | 101811910 | mNeoVis.2.1.1.chr8\_hap1.g01311.t01 |
| 10846 | 8 | 101812162 | 101884415 | mNeoVis.2.1.1.chr8\_hap1.g01312.t01 |
| 10847 | 8 | 101943526 | 101950526 | mNeoVis.2.1.1.chr8\_hap1.g01313.t01 |
| 10848 | 8 | 101943526 | 101950526 | mNeoVis.2.1.1.chr8\_hap1.g01313.t01 |
| 10849 | 8 | 101981390 | 102004773 | mNeoVis.2.1.1.chr8\_hap1.g01314.t01 |
| 10850 | 8 | 101981390 | 102004773 | mNeoVis.2.1.1.chr8\_hap1.g01314.t01 |
| 10851 | 8 | 102008029 | 102017704 | mNeoVis.2.1.1.chr8\_hap1.g01315.t01 |
| 10852 | 8 | 102008029 | 102017704 | mNeoVis.2.1.1.chr8\_hap1.g01315.t01 |
| 10853 | 8 | 102040850 | 102223145 | mNeoVis.2.1.1.chr8\_hap1.g01316.t01 |
| 10854 | 8 | 102040850 | 102223145 | mNeoVis.2.1.1.chr8\_hap1.g01316.t01 |
| 10855 | 8 | 102270750 | 102314448 | mNeoVis.2.1.1.chr8\_hap1.g01317.t01 |
| 10856 | 8 | 102270750 | 102314448 | mNeoVis.2.1.1.chr8\_hap1.g01317.t01 |
| 10857 | 8 | 102315215 | 102428715 | mNeoVis.2.1.1.chr8\_hap1.g01318.t01 |
| 10858 | 8 | 102409178 | 102441334 | mNeoVis.2.1.1.chr8\_hap1.g01319.t01 |
| 10859 | 8 | 102529122 | 102634368 | mNeoVis.2.1.1.chr8\_hap1.g01320.t01 |
| 10860 | 8 | 102703421 | 102725550 | mNeoVis.2.1.1.chr8\_hap1.g01321.t01 |
| 10861 | 8 | 102768186 | 102799204 | mNeoVis.2.1.1.chr8\_hap1.g01322.t01 |
| 10862 | 8 | 103824940 | 103830816 | mNeoVis.2.1.1.chr8\_hap1.g01325.t01 |
| 10863 | 8 | 103831741 | 103848932 | mNeoVis.2.1.1.chr8\_hap1.g01326.t01 |
| 10864 | 8 | 103861932 | 103863660 | mNeoVis.2.1.1.chr8\_hap1.g01327.t01 |
| 10865 | 8 | 103892148 | 103909232 | mNeoVis.2.1.1.chr8\_hap1.g01328.t01 |
| 10866 | 8 | 103990939 | 104094154 | mNeoVis.2.1.1.chr8\_hap1.g01329.t01 |
| 10867 | 8 | 104099713 | 104131776 | mNeoVis.2.1.1.chr8\_hap1.g01330.t01 |
| 10868 | 8 | 104099713 | 104131776 | mNeoVis.2.1.1.chr8\_hap1.g01330.t01 |
| 10869 | 8 | 104156423 | 104159123 | mNeoVis.2.1.1.chr8\_hap1.g01331.t01 |
| 10870 | 8 | 104156423 | 104159123 | mNeoVis.2.1.1.chr8\_hap1.g01331.t01 |
| 10871 | 8 | 104183611 | 104235728 | mNeoVis.2.1.1.chr8\_hap1.g01332.t01 |
| 10872 | 8 | 104183611 | 104235728 | mNeoVis.2.1.1.chr8\_hap1.g01332.t01 |
| 10873 | 8 | 104244462 | 104338470 | mNeoVis.2.1.1.chr8\_hap1.g01333.t01 |
| 10874 | 8 | 104244462 | 104338470 | mNeoVis.2.1.1.chr8\_hap1.g01333.t01 |
| 10875 | 8 | 104339003 | 104460547 | mNeoVis.2.1.1.chr8\_hap1.g01334.t01 |
| 10876 | 8 | 104481894 | 104526767 | mNeoVis.2.1.1.chr8\_hap1.g01335.t01 |
| 10877 | 8 | 104544965 | 104547670 | mNeoVis.2.1.1.chr8\_hap1.g01336.t01 |
| 10878 | 8 | 104561200 | 104745747 | mNeoVis.2.1.1.chr8\_hap1.g01337.t01 |
| 10879 | 8 | 104797360 | 104812008 | mNeoVis.2.1.1.chr8\_hap1.g01338.t01 |
| 10880 | 8 | 104954120 | 105051991 | mNeoVis.2.1.1.chr8\_hap1.g01339.t01 |
| 10881 | 8 | 105059079 | 105105436 | mNeoVis.2.1.1.chr8\_hap1.g01340.t01 |
| 10882 | 8 | 105123146 | 105168292 | mNeoVis.2.1.1.chr8\_hap1.g01341.t01 |
| 10883 | 8 | 105123146 | 105168292 | mNeoVis.2.1.1.chr8\_hap1.g01341.t01 |
| 10884 | 8 | 105167082 | 105338681 | mNeoVis.2.1.1.chr8\_hap1.g01342.t01 |
| 10885 | 8 | 105167082 | 105338681 | mNeoVis.2.1.1.chr8\_hap1.g01342.t01 |
| 10886 | 8 | 105234905 | 105237005 | mNeoVis.2.1.1.chr8\_hap1.g01343.t01 |
| 10887 | 8 | 105234905 | 105237005 | mNeoVis.2.1.1.chr8\_hap1.g01343.t01 |
| 10888 | 8 | 105390585 | 105417896 | mNeoVis.2.1.1.chr8\_hap1.g01344.t01 |
| 10889 | 8 | 105390585 | 105417896 | mNeoVis.2.1.1.chr8\_hap1.g01344.t01 |
| 10890 | 8 | 105533370 | 105554514 | mNeoVis.2.1.1.chr8\_hap1.g01345.t01 |
| 10891 | 8 | 105533370 | 105554514 | mNeoVis.2.1.1.chr8\_hap1.g01345.t01 |
| 10892 | 8 | 105563451 | 105573171 | mNeoVis.2.1.1.chr8\_hap1.g01346.t01 |
| 10893 | 8 | 105563451 | 105573171 | mNeoVis.2.1.1.chr8\_hap1.g01346.t01 |
| 10894 | 8 | 105575525 | 105629481 | mNeoVis.2.1.1.chr8\_hap1.g01347.t01 |
| 10895 | 8 | 105575525 | 105629481 | mNeoVis.2.1.1.chr8\_hap1.g01347.t01 |
| 10896 | 8 | 106660951 | 106722577 | mNeoVis.2.1.1.chr8\_hap1.g01348.t01 |
| 10897 | 8 | 106698815 | 106751728 | mNeoVis.2.1.1.chr8\_hap1.g01349.t01 |
| 10898 | 8 | 106755189 | 106781008 | mNeoVis.2.1.1.chr8\_hap1.g01350.t01 |
| 10899 | 8 | 106809146 | 106996538 | mNeoVis.2.1.1.chr8\_hap1.g01351.t01 |
| 10900 | 8 | 107086266 | 107128158 | mNeoVis.2.1.1.chr8\_hap1.g01352.t01 |
| 10901 | 8 | 107156896 | 107198585 | mNeoVis.2.1.1.chr8\_hap1.g01353.t01 |
| 10902 | 8 | 107227473 | 107283196 | mNeoVis.2.1.1.chr8\_hap1.g01354.t01 |
| 10903 | 8 | 107295658 | 107359080 | mNeoVis.2.1.1.chr8\_hap1.g01355.t01 |
| 10904 | 8 | 107359722 | 107474676 | mNeoVis.2.1.1.chr8\_hap1.g01356.t01 |
| 10905 | 8 | 107586186 | 107657467 | mNeoVis.2.1.1.chr8\_hap1.g01357.t01 |
| 10906 | 8 | 107702395 | 107715067 | mNeoVis.2.1.1.chr8\_hap1.g01358.t01 |
| 10907 | 8 | 109015594 | 109034115 | mNeoVis.2.1.1.chr8\_hap1.g01371.t01 |
| 10908 | 8 | 109270085 | 109282106 | mNeoVis.2.1.1.chr8\_hap1.g01372.t01 |
| 10909 | 8 | 109282129 | 109325693 | mNeoVis.2.1.1.chr8\_hap1.g01373.t01 |
| 10910 | 8 | 109374921 | 109951811 | mNeoVis.2.1.1.chr8\_hap1.g01374.t01 |
| 10911 | 8 | 109903054 | 110174366 | mNeoVis.2.1.1.chr8\_hap1.g01375.t01 |
| 10912 | 8 | 110056032 | 110142456 | mNeoVis.2.1.1.chr8\_hap1.g01376.t01 |
| 10913 | 8 | 110170475 | 110257486 | mNeoVis.2.1.1.chr8\_hap1.g01377.t01 |
| 10914 | 8 | 110379694 | 110388072 | mNeoVis.2.1.1.chr8\_hap1.g01378.t01 |
| 10915 | 8 | 110509516 | 110509887 | mNeoVis.2.1.1.chr8\_hap1.g01379.t01 |
| 10916 | 8 | 110510002 | 110511106 | mNeoVis.2.1.1.chr8\_hap1.g01380.t01 |
| 10917 | 8 | 110553019 | 110584861 | mNeoVis.2.1.1.chr8\_hap1.g01381.t01 |
| 10918 | 8 | 110794778 | 110839587 | mNeoVis.2.1.1.chr8\_hap1.g01382.t01 |
| 10919 | 8 | 110794778 | 110839587 | mNeoVis.2.1.1.chr8\_hap1.g01382.t01 |
| 10920 | 8 | 110868802 | 111065967 | mNeoVis.2.1.1.chr8\_hap1.g01383.t01 |
| 10921 | 8 | 111146017 | 111475312 | mNeoVis.2.1.1.chr8\_hap1.g01384.t01 |
| 10922 | 8 | 111456678 | 111508337 | mNeoVis.2.1.1.chr8\_hap1.g01385.t01 |
| 10923 | 8 | 111522619 | 111546510 | mNeoVis.2.1.1.chr8\_hap1.g01386.t01 |
| 10924 | 8 | 111568023 | 111636431 | mNeoVis.2.1.1.chr8\_hap1.g01387.t01 |
| 10925 | 8 | 111655515 | 111712673 | mNeoVis.2.1.1.chr8\_hap1.g01388.t01 |
| 10926 | 8 | 111655515 | 111712673 | mNeoVis.2.1.1.chr8\_hap1.g01388.t01 |
| 10927 | 8 | 111716846 | 111721068 | mNeoVis.2.1.1.chr8\_hap1.g01389.t01 |
| 10928 | 8 | 111716846 | 111721068 | mNeoVis.2.1.1.chr8\_hap1.g01389.t01 |
| 10929 | 8 | 111793785 | 111798565 | mNeoVis.2.1.1.chr8\_hap1.g01390.t01 |
| 10930 | 8 | 111793785 | 111798565 | mNeoVis.2.1.1.chr8\_hap1.g01390.t01 |
| 10931 | 8 | 111830050 | 111898014 | mNeoVis.2.1.1.chr8\_hap1.g01391.t01 |
| 10932 | 8 | 111830050 | 111898014 | mNeoVis.2.1.1.chr8\_hap1.g01391.t01 |
| 10933 | 8 | 111898540 | 112014363 | mNeoVis.2.1.1.chr8\_hap1.g01392.t01 |
| 10934 | 8 | 111898540 | 112014363 | mNeoVis.2.1.1.chr8\_hap1.g01392.t01 |
| 10935 | 8 | 112018290 | 112059863 | mNeoVis.2.1.1.chr8\_hap1.g01393.t01 |
| 10936 | 8 | 112018290 | 112059863 | mNeoVis.2.1.1.chr8\_hap1.g01393.t01 |
| 10937 | 8 | 112086705 | 112111649 | mNeoVis.2.1.1.chr8\_hap1.g01394.t01 |
| 10938 | 8 | 112086705 | 112111649 | mNeoVis.2.1.1.chr8\_hap1.g01394.t01 |
| 10939 | 8 | 112208085 | 112212836 | mNeoVis.2.1.1.chr8\_hap1.g01395.t01 |
| 10940 | 8 | 112382660 | 112429445 | mNeoVis.2.1.1.chr8\_hap1.g01396.t01 |
| 10941 | 8 | 128829308 | 129220061 | mNeoVis.2.1.1.chr8\_hap1.g01409.t01 |
| 10942 | 8 | 129798152 | 129883985 | mNeoVis.2.1.1.chr8\_hap1.g01410.t01 |
| 10943 | 8 | 129798152 | 129883985 | mNeoVis.2.1.1.chr8\_hap1.g01410.t01 |
| 10944 | 8 | 129798152 | 129883985 | mNeoVis.2.1.1.chr8\_hap1.g01410.t01 |
| 10945 | 8 | 129798152 | 129883985 | mNeoVis.2.1.1.chr8\_hap1.g01410.t01 |
| 10946 | 8 | 129967684 | 129988004 | mNeoVis.2.1.1.chr8\_hap1.g01411.t01 |
| 10947 | 8 | 129967684 | 129988004 | mNeoVis.2.1.1.chr8\_hap1.g01411.t01 |
| 10948 | 8 | 129967684 | 129988004 | mNeoVis.2.1.1.chr8\_hap1.g01411.t01 |
| 10949 | 8 | 129967684 | 129988004 | mNeoVis.2.1.1.chr8\_hap1.g01411.t01 |
| 10950 | 8 | 129988119 | 130012565 | mNeoVis.2.1.1.chr8\_hap1.g01412.t01 |
| 10951 | 8 | 129988119 | 130012565 | mNeoVis.2.1.1.chr8\_hap1.g01412.t01 |
| 10952 | 8 | 129988119 | 130012565 | mNeoVis.2.1.1.chr8\_hap1.g01412.t01 |
| 10953 | 8 | 129988119 | 130012565 | mNeoVis.2.1.1.chr8\_hap1.g01412.t01 |
| 10954 | 8 | 130012827 | 130042641 | mNeoVis.2.1.1.chr8\_hap1.g01413.t01 |
| 10955 | 8 | 130012827 | 130042641 | mNeoVis.2.1.1.chr8\_hap1.g01413.t01 |
| 10956 | 8 | 130012827 | 130042641 | mNeoVis.2.1.1.chr8\_hap1.g01413.t01 |
| 10957 | 8 | 130012827 | 130042641 | mNeoVis.2.1.1.chr8\_hap1.g01413.t01 |
| 10958 | 8 | 130042906 | 130240918 | mNeoVis.2.1.1.chr8\_hap1.g01414.t01 |
| 10959 | 8 | 130042906 | 130240918 | mNeoVis.2.1.1.chr8\_hap1.g01414.t01 |
| 10960 | 8 | 130042906 | 130240918 | mNeoVis.2.1.1.chr8\_hap1.g01414.t01 |
| 10961 | 8 | 130275900 | 130293666 | mNeoVis.2.1.1.chr8\_hap1.g01415.t01 |
| 10962 | 8 | 130275900 | 130293666 | mNeoVis.2.1.1.chr8\_hap1.g01415.t01 |
| 10963 | 8 | 130275900 | 130293666 | mNeoVis.2.1.1.chr8\_hap1.g01415.t01 |
| 10964 | 8 | 132845125 | 133119224 | mNeoVis.2.1.1.chr8\_hap1.g01422.t01 |
| 10965 | 8 | 133431437 | 133435769 | mNeoVis.2.1.1.chr8\_hap1.g01423.t01 |
| 10966 | 8 | 133666213 | 133674543 | mNeoVis.2.1.1.chr8\_hap1.g01424.t01 |
| 10967 | 8 | 133677088 | 133739057 | mNeoVis.2.1.1.chr8\_hap1.g01425.t01 |
| 10968 | 8 | 133712541 | 133722825 | mNeoVis.2.1.1.chr8\_hap1.g01426.t01 |
| 10969 | 8 | 133739192 | 133748302 | mNeoVis.2.1.1.chr8\_hap1.g01427.t01 |
| 10970 | 8 | 133751085 | 133772026 | mNeoVis.2.1.1.chr8\_hap1.g01428.t01 |
| 10971 | 8 | 133787390 | 134070152 | mNeoVis.2.1.1.chr8\_hap1.g01429.t01 |
| 10972 | 8 | 134293931 | 134400718 | mNeoVis.2.1.1.chr8\_hap1.g01430.t01 |
| 10973 | 8 | 134429639 | 134487746 | mNeoVis.2.1.1.chr8\_hap1.g01431.t01 |
| 10974 | 8 | 134668110 | 134692291 | mNeoVis.2.1.1.chr8\_hap1.g01432.t01 |
| 10975 | 8 | 134668110 | 134692291 | mNeoVis.2.1.1.chr8\_hap1.g01432.t01 |
| 10976 | 8 | 135233650 | 135257180 | mNeoVis.2.1.1.chr8\_hap1.g01433.t01 |
| 10977 | 8 | 135259414 | 135263863 | mNeoVis.2.1.1.chr8\_hap1.g01434.t01 |
| 10978 | 8 | 135273417 | 135317725 | mNeoVis.2.1.1.chr8\_hap1.g01435.t01 |
| 10979 | 8 | 136669648 | 136674448 | mNeoVis.2.1.1.chr8\_hap1.g01439.t01 |
| 10980 | 8 | 139598815 | 139603140 | mNeoVis.2.1.1.chr8\_hap1.g01440.t01 |
| 10981 | 8 | 140475757 | 140639540 | mNeoVis.2.1.1.chr8\_hap1.g01441.t01 |
| 10982 | 8 | 140475757 | 140639540 | mNeoVis.2.1.1.chr8\_hap1.g01441.t01 |
| 10983 | 8 | 140605614 | 140788762 | mNeoVis.2.1.1.chr8\_hap1.g01442.t01 |
| 10984 | 8 | 140605614 | 140788762 | mNeoVis.2.1.1.chr8\_hap1.g01442.t01 |
| 10985 | 8 | 141196493 | 141203528 | mNeoVis.2.1.1.chr8\_hap1.g01443.t01 |
| 10986 | 8 | 141196493 | 141203528 | mNeoVis.2.1.1.chr8\_hap1.g01443.t01 |
| 10987 | 8 | 142665884 | 142673571 | mNeoVis.2.1.1.chr8\_hap1.g01444.t01 |
| 10988 | 8 | 144651878 | 144877434 | mNeoVis.2.1.1.chr8\_hap1.g01448.t01 |
| 10989 | 8 | 144906014 | 145051960 | mNeoVis.2.1.1.chr8\_hap1.g01449.t01 |
| 10990 | 8 | 144906014 | 145051960 | mNeoVis.2.1.1.chr8\_hap1.g01449.t01 |
| 10991 | 8 | 145023915 | 145098869 | mNeoVis.2.1.1.chr8\_hap1.g01450.t01 |
| 10992 | 8 | 145023915 | 145098869 | mNeoVis.2.1.1.chr8\_hap1.g01450.t01 |
| 10993 | 8 | 148746209 | 148780377 | mNeoVis.2.1.1.chr8\_hap1.g01454.t01 |
| 10994 | 8 | 148746209 | 148780377 | mNeoVis.2.1.1.chr8\_hap1.g01454.t01 |
| 10995 | 8 | 148848709 | 148865571 | mNeoVis.2.1.1.chr8\_hap1.g01455.t01 |
| 10996 | 8 | 148848709 | 148865571 | mNeoVis.2.1.1.chr8\_hap1.g01455.t01 |
| 10997 | 8 | 148867676 | 148896275 | mNeoVis.2.1.1.chr8\_hap1.g01456.t01 |
| 10998 | 8 | 148867676 | 148896275 | mNeoVis.2.1.1.chr8\_hap1.g01456.t01 |
| 10999 | 8 | 148971117 | 148974574 | mNeoVis.2.1.1.chr8\_hap1.g01457.t01 |
| 11000 | 8 | 148971117 | 148974574 | mNeoVis.2.1.1.chr8\_hap1.g01457.t01 |
| 11001 | 8 | 149219770 | 149463960 | mNeoVis.2.1.1.chr8\_hap1.g01458.t01 |
| 11002 | 8 | 149219770 | 149463960 | mNeoVis.2.1.1.chr8\_hap1.g01458.t01 |
| 11003 | 8 | 149627607 | 149652789 | mNeoVis.2.1.1.chr8\_hap1.g01459.t01 |
| 11004 | 8 | 149654782 | 149706805 | mNeoVis.2.1.1.chr8\_hap1.g01460.t01 |
| 11005 | 8 | 149729823 | 149790809 | mNeoVis.2.1.1.chr8\_hap1.g01461.t01 |
| 11006 | 8 | 149769828 | 149787094 | mNeoVis.2.1.1.chr8\_hap1.g01462.t01 |
| 11007 | 8 | 149794990 | 149992757 | mNeoVis.2.1.1.chr8\_hap1.g01463.t01 |
| 11008 | 8 | 150032949 | 150679976 | mNeoVis.2.1.1.chr8\_hap1.g01464.t01 |
| 11009 | 8 | 150084055 | 150084787 | mNeoVis.2.1.1.chr8\_hap1.g01465.t01 |
| 11010 | 8 | 150785985 | 150793338 | mNeoVis.2.1.1.chr8\_hap1.g01466.t01 |
| 11011 | 8 | 151982125 | 152047834 | mNeoVis.2.1.1.chr8\_hap1.g01471.t01 |
| 11012 | 8 | 152169784 | 152221070 | mNeoVis.2.1.1.chr8\_hap1.g01472.t01 |
| 11013 | 8 | 152230379 | 152488617 | mNeoVis.2.1.1.chr8\_hap1.g01473.t01 |
| 11014 | 8 | 153787948 | 153843698 | mNeoVis.2.1.1.chr8\_hap1.g01482.t01 |
| 11015 | 8 | 153856010 | 153919677 | mNeoVis.2.1.1.chr8\_hap1.g01483.t01 |
| 11016 | 8 | 154181374 | 154497318 | mNeoVis.2.1.1.chr8\_hap1.g01484.t01 |
| 11017 | 8 | 154727957 | 155342446 | mNeoVis.2.1.1.chr8\_hap1.g01486.t01 |
| 11018 | 8 | 155344333 | 155365163 | mNeoVis.2.1.1.chr8\_hap1.g01487.t01 |
| 11019 | 8 | 155374107 | 155374618 | mNeoVis.2.1.1.chr8\_hap1.g01488.t01 |
| 11020 | 8 | 155454374 | 155524372 | mNeoVis.2.1.1.chr8\_hap1.g01489.t01 |
| 11021 | 8 | 155530395 | 155540933 | mNeoVis.2.1.1.chr8\_hap1.g01490.t01 |
| 11022 | 8 | 155574695 | 155604921 | mNeoVis.2.1.1.chr8\_hap1.g01491.t01 |
| 11023 | 8 | 155609810 | 155616406 | mNeoVis.2.1.1.chr8\_hap1.g01492.t01 |
| 11024 | 8 | 155873267 | 155893402 | mNeoVis.2.1.1.chr8\_hap1.g01497.t01 |
| 11025 | 8 | 158878016 | 158922114 | mNeoVis.2.1.1.chr8\_hap1.g01498.t01 |
| 11026 | 8 | 158929227 | 158954992 | mNeoVis.2.1.1.chr8\_hap1.g01499.t01 |
| 11027 | 8 | 159219299 | 159249409 | mNeoVis.2.1.1.chr8\_hap1.g01500.t01 |
| 11028 | 8 | 160344163 | 160380145 | mNeoVis.2.1.1.chr8\_hap1.g01505.t01 |
| 11029 | 8 | 162330882 | 162450818 | mNeoVis.2.1.1.chr8\_hap1.g01516.t01 |
| 11030 | 8 | 162330882 | 162450818 | mNeoVis.2.1.1.chr8\_hap1.g01516.t01 |
| 11031 | 8 | 162456095 | 162472418 | mNeoVis.2.1.1.chr8\_hap1.g01517.t01 |
| 11032 | 8 | 162456095 | 162472418 | mNeoVis.2.1.1.chr8\_hap1.g01517.t01 |
| 11033 | 8 | 162456095 | 162472418 | mNeoVis.2.1.1.chr8\_hap1.g01517.t01 |
| 11034 | 8 | 162819368 | 162862842 | mNeoVis.2.1.1.chr8\_hap1.g01518.t01 |
| 11035 | 8 | 162819368 | 162862842 | mNeoVis.2.1.1.chr8\_hap1.g01518.t01 |
| 11036 | 8 | 162819368 | 162862842 | mNeoVis.2.1.1.chr8\_hap1.g01518.t01 |
| 11037 | 8 | 163006990 | 163008174 | mNeoVis.2.1.1.chr8\_hap1.g01519.t01 |
| 11038 | 8 | 163032750 | 163034013 | mNeoVis.2.1.1.chr8\_hap1.g01520.t01 |
| 11039 | 8 | 163034119 | 163066567 | mNeoVis.2.1.1.chr8\_hap1.g01521.t01 |
| 11040 | 8 | 163777095 | 163781684 | mNeoVis.2.1.1.chr8\_hap1.g01532.t01 |
| 11041 | 8 | 163788431 | 163801980 | mNeoVis.2.1.1.chr8\_hap1.g01533.t01 |
| 11042 | 8 | 163803495 | 163829949 | mNeoVis.2.1.1.chr8\_hap1.g01534.t01 |
| 11043 | 8 | 163830197 | 163868118 | mNeoVis.2.1.1.chr8\_hap1.g01535.t01 |
| 11044 | 8 | 163891613 | 163934317 | mNeoVis.2.1.1.chr8\_hap1.g01536.t01 |
| 11045 | 8 | 163935310 | 163941507 | mNeoVis.2.1.1.chr8\_hap1.g01537.t01 |
| 11046 | 8 | 163947558 | 163961236 | mNeoVis.2.1.1.chr8\_hap1.g01538.t01 |
| 11047 | 8 | 163971431 | 164008297 | mNeoVis.2.1.1.chr8\_hap1.g01539.t01 |
| 11048 | 8 | 164013724 | 164044794 | mNeoVis.2.1.1.chr8\_hap1.g01540.t01 |
| 11049 | 8 | 164055366 | 164063304 | mNeoVis.2.1.1.chr8\_hap1.g01541.t01 |
| 11050 | 8 | 164110337 | 164119321 | mNeoVis.2.1.1.chr8\_hap1.g01542.t01 |
| 11051 | 8 | 164121881 | 164234791 | mNeoVis.2.1.1.chr8\_hap1.g01543.t01 |
| 11052 | 8 | 164315284 | 164344412 | mNeoVis.2.1.1.chr8\_hap1.g01544.t01 |
| 11053 | 9 | 2679542 | 2767905 | mNeoVis.2.1.1.chr9\_hap1.g00003.t01 |
| 11054 | 9 | 2679542 | 2767905 | mNeoVis.2.1.1.chr9\_hap1.g00003.t01 |
| 11055 | 9 | 2730472 | 2732671 | mNeoVis.2.1.1.chr9\_hap1.g00004.t01 |
| 11056 | 9 | 2730472 | 2732671 | mNeoVis.2.1.1.chr9\_hap1.g00004.t01 |
| 11057 | 9 | 2829258 | 2848990 | mNeoVis.2.1.1.chr9\_hap1.g00005.t01 |
| 11058 | 9 | 2858763 | 2861141 | mNeoVis.2.1.1.chr9\_hap1.g00006.t01 |
| 11059 | 9 | 2863277 | 2911486 | mNeoVis.2.1.1.chr9\_hap1.g00007.t01 |
| 11060 | 9 | 2950744 | 3299177 | mNeoVis.2.1.1.chr9\_hap1.g00008.t01 |
| 11061 | 9 | 3631798 | 3632396 | mNeoVis.2.1.1.chr9\_hap1.g00009.t01 |
| 11062 | 9 | 3631798 | 3632396 | mNeoVis.2.1.1.chr9\_hap1.g00009.t01 |
| 11063 | 9 | 3996567 | 4008075 | mNeoVis.2.1.1.chr9\_hap1.g00010.t01 |
| 11064 | 9 | 4291306 | 4348422 | mNeoVis.2.1.1.chr9\_hap1.g00011.t01 |
| 11065 | 9 | 4291306 | 4348422 | mNeoVis.2.1.1.chr9\_hap1.g00011.t01 |
| 11066 | 9 | 4572178 | 4623437 | mNeoVis.2.1.1.chr9\_hap1.g00012.t01 |
| 11067 | 9 | 4572178 | 4623437 | mNeoVis.2.1.1.chr9\_hap1.g00012.t01 |
| 11068 | 9 | 4624172 | 4661894 | mNeoVis.2.1.1.chr9\_hap1.g00013.t01 |
| 11069 | 9 | 4624172 | 4661894 | mNeoVis.2.1.1.chr9\_hap1.g00013.t01 |
| 11070 | 9 | 6116114 | 6118078 | mNeoVis.2.1.1.chr9\_hap1.g00015.t01 |
| 11071 | 9 | 6131643 | 6143587 | mNeoVis.2.1.1.chr9\_hap1.g00016.t01 |
| 11072 | 9 | 7260823 | 7430040 | mNeoVis.2.1.1.chr9\_hap1.g00033.t01 |
| 11073 | 9 | 13327239 | 13333651 | mNeoVis.2.1.1.chr9\_hap1.g00050.t01 |
| 11074 | 9 | 13372968 | 13452105 | mNeoVis.2.1.1.chr9\_hap1.g00051.t01 |
| 11075 | 9 | 13628776 | 13950132 | mNeoVis.2.1.1.chr9\_hap1.g00052.t01 |
| 11076 | 9 | 13753955 | 13755088 | mNeoVis.2.1.1.chr9\_hap1.g00053.t01 |
| 11077 | 9 | 14034417 | 14218668 | mNeoVis.2.1.1.chr9\_hap1.g00054.t01 |
| 11078 | 9 | 14284204 | 14351329 | mNeoVis.2.1.1.chr9\_hap1.g00055.t01 |
| 11079 | 9 | 14471197 | 14507644 | mNeoVis.2.1.1.chr9\_hap1.g00056.t01 |
| 11080 | 9 | 14485436 | 14486163 | mNeoVis.2.1.1.chr9\_hap1.g00057.t01 |
| 11081 | 9 | 16373501 | 16447983 | mNeoVis.2.1.1.chr9\_hap1.g00076.t01 |
| 11082 | 9 | 16451689 | 16686994 | mNeoVis.2.1.1.chr9\_hap1.g00077.t01 |
| 11083 | 9 | 17735277 | 17745422 | mNeoVis.2.1.1.chr9\_hap1.g00080.t01 |
| 11084 | 9 | 17745726 | 17768910 | mNeoVis.2.1.1.chr9\_hap1.g00081.t01 |
| 11085 | 9 | 17797260 | 17890282 | mNeoVis.2.1.1.chr9\_hap1.g00082.t01 |
| 11086 | 9 | 19937736 | 20000388 | mNeoVis.2.1.1.chr9\_hap1.g00092.t01 |
| 11087 | 9 | 20014757 | 20024381 | mNeoVis.2.1.1.chr9\_hap1.g00093.t01 |
| 11088 | 9 | 20025727 | 20029664 | mNeoVis.2.1.1.chr9\_hap1.g00094.t01 |
| 11089 | 9 | 20308998 | 20531326 | mNeoVis.2.1.1.chr9\_hap1.g00095.t01 |
| 11090 | 9 | 20423181 | 20423939 | mNeoVis.2.1.1.chr9\_hap1.g00096.t01 |
| 11091 | 9 | 20461334 | 20542512 | mNeoVis.2.1.1.chr9\_hap1.g00097.t01 |
| 11092 | 9 | 21321438 | 21435245 | mNeoVis.2.1.1.chr9\_hap1.g00102.t01 |
| 11093 | 9 | 21808713 | 22287650 | mNeoVis.2.1.1.chr9\_hap1.g00103.t01 |
| 11094 | 9 | 21817242 | 21822207 | mNeoVis.2.1.1.chr9\_hap1.g00104.t01 |
| 11095 | 9 | 22384513 | 23165585 | mNeoVis.2.1.1.chr9\_hap1.g00105.t01 |
| 11096 | 9 | 22670285 | 22672053 | mNeoVis.2.1.1.chr9\_hap1.g00106.t01 |
| 11097 | 9 | 23548911 | 23579949 | mNeoVis.2.1.1.chr9\_hap1.g00107.t01 |
| 11098 | 9 | 23584938 | 23662191 | mNeoVis.2.1.1.chr9\_hap1.g00108.t01 |
| 11099 | 9 | 23594813 | 23597672 | mNeoVis.2.1.1.chr9\_hap1.g00109.t01 |
| 11100 | 9 | 23660388 | 24047506 | mNeoVis.2.1.1.chr9\_hap1.g00110.t01 |
| 11101 | 9 | 24131954 | 24221511 | mNeoVis.2.1.1.chr9\_hap1.g00111.t01 |
| 11102 | 9 | 24243859 | 24345957 | mNeoVis.2.1.1.chr9\_hap1.g00112.t01 |
| 11103 | 9 | 24367801 | 24531040 | mNeoVis.2.1.1.chr9\_hap1.g00113.t01 |
| 11104 | 9 | 24549379 | 24566678 | mNeoVis.2.1.1.chr9\_hap1.g00114.t01 |
| 11105 | 9 | 24869406 | 24877261 | mNeoVis.2.1.1.chr9\_hap1.g00116.t01 |
| 11106 | 9 | 24903441 | 24962474 | mNeoVis.2.1.1.chr9\_hap1.g00117.t01 |
| 11107 | 9 | 24977856 | 25237912 | mNeoVis.2.1.1.chr9\_hap1.g00118.t01 |
| 11108 | 9 | 25242706 | 25428994 | mNeoVis.2.1.1.chr9\_hap1.g00119.t01 |
| 11109 | 9 | 25484192 | 25492210 | mNeoVis.2.1.1.chr9\_hap1.g00120.t01 |
| 11110 | 9 | 25493079 | 25562222 | mNeoVis.2.1.1.chr9\_hap1.g00121.t01 |
| 11111 | 9 | 26175776 | 26268703 | mNeoVis.2.1.1.chr9\_hap1.g00122.t01 |
| 11112 | 9 | 26274316 | 26275154 | mNeoVis.2.1.1.chr9\_hap1.g00123.t01 |
| 11113 | 9 | 26336961 | 26511751 | mNeoVis.2.1.1.chr9\_hap1.g00124.t01 |
| 11114 | 9 | 27924193 | 28527690 | mNeoVis.2.1.1.chr9\_hap1.g00142.t01 |
| 11115 | 9 | 28939312 | 28957564 | mNeoVis.2.1.1.chr9\_hap1.g00148.t01 |
| 11116 | 9 | 28987672 | 28997564 | mNeoVis.2.1.1.chr9\_hap1.g00149.t01 |
| 11117 | 9 | 28988921 | 29048829 | mNeoVis.2.1.1.chr9\_hap1.g00150.t01 |
| 11118 | 9 | 29064849 | 29110889 | mNeoVis.2.1.1.chr9\_hap1.g00151.t01 |
| 11119 | 9 | 29185807 | 29314292 | mNeoVis.2.1.1.chr9\_hap1.g00152.t01 |
| 11120 | 9 | 29404027 | 29834816 | mNeoVis.2.1.1.chr9\_hap1.g00153.t01 |
| 11121 | 9 | 29404027 | 29834816 | mNeoVis.2.1.1.chr9\_hap1.g00153.t01 |
| 11122 | 9 | 29910782 | 29966201 | mNeoVis.2.1.1.chr9\_hap1.g00154.t01 |
| 11123 | 9 | 29910782 | 29966201 | mNeoVis.2.1.1.chr9\_hap1.g00154.t01 |
| 11124 | 9 | 29966475 | 29991357 | mNeoVis.2.1.1.chr9\_hap1.g00155.t01 |
| 11125 | 9 | 29966475 | 29991357 | mNeoVis.2.1.1.chr9\_hap1.g00155.t01 |
| 11126 | 9 | 29993858 | 30021446 | mNeoVis.2.1.1.chr9\_hap1.g00156.t01 |
| 11127 | 9 | 29993858 | 30021446 | mNeoVis.2.1.1.chr9\_hap1.g00156.t01 |
| 11128 | 9 | 30024686 | 30081031 | mNeoVis.2.1.1.chr9\_hap1.g00157.t01 |
| 11129 | 9 | 30024686 | 30081031 | mNeoVis.2.1.1.chr9\_hap1.g00157.t01 |
| 11130 | 9 | 30789659 | 30890223 | mNeoVis.2.1.1.chr9\_hap1.g00161.t01 |
| 11131 | 9 | 31013468 | 31197138 | mNeoVis.2.1.1.chr9\_hap1.g00162.t01 |
| 11132 | 9 | 31013468 | 31197138 | mNeoVis.2.1.1.chr9\_hap1.g00162.t01 |
| 11133 | 9 | 31439871 | 31630075 | mNeoVis.2.1.1.chr9\_hap1.g00163.t01 |
| 11134 | 9 | 31648425 | 31650164 | mNeoVis.2.1.1.chr9\_hap1.g00164.t01 |
| 11135 | 9 | 31648425 | 31650164 | mNeoVis.2.1.1.chr9\_hap1.g00164.t01 |
| 11136 | 9 | 31845279 | 31955303 | mNeoVis.2.1.1.chr9\_hap1.g00165.t01 |
| 11137 | 9 | 32043960 | 32089979 | mNeoVis.2.1.1.chr9\_hap1.g00166.t01 |
| 11138 | 9 | 32243273 | 32309019 | mNeoVis.2.1.1.chr9\_hap1.g00167.t01 |
| 11139 | 9 | 32901366 | 32947509 | mNeoVis.2.1.1.chr9\_hap1.g00169.t01 |
| 11140 | 9 | 33118690 | 33255191 | mNeoVis.2.1.1.chr9\_hap1.g00170.t01 |
| 11141 | 9 | 33118690 | 33255191 | mNeoVis.2.1.1.chr9\_hap1.g00170.t01 |
| 11142 | 9 | 33118690 | 33255191 | mNeoVis.2.1.1.chr9\_hap1.g00170.t01 |
| 11143 | 9 | 33925989 | 34584805 | mNeoVis.2.1.1.chr9\_hap1.g00171.t01 |
| 11144 | 9 | 34734371 | 34902421 | mNeoVis.2.1.1.chr9\_hap1.g00172.t01 |
| 11145 | 9 | 34902798 | 34972868 | mNeoVis.2.1.1.chr9\_hap1.g00173.t01 |
| 11146 | 9 | 35091945 | 35315540 | mNeoVis.2.1.1.chr9\_hap1.g00174.t01 |
| 11147 | 9 | 35091945 | 35315540 | mNeoVis.2.1.1.chr9\_hap1.g00174.t01 |
| 11148 | 9 | 35103403 | 35103982 | mNeoVis.2.1.1.chr9\_hap1.g00175.t01 |
| 11149 | 9 | 35103403 | 35103982 | mNeoVis.2.1.1.chr9\_hap1.g00175.t01 |
| 11150 | 9 | 35344910 | 35349240 | mNeoVis.2.1.1.chr9\_hap1.g00176.t01 |
| 11151 | 9 | 35344910 | 35349240 | mNeoVis.2.1.1.chr9\_hap1.g00176.t01 |
| 11152 | 9 | 35377233 | 35380862 | mNeoVis.2.1.1.chr9\_hap1.g00177.t01 |
| 11153 | 9 | 35377233 | 35380862 | mNeoVis.2.1.1.chr9\_hap1.g00177.t01 |
| 11154 | 9 | 37666641 | 37667621 | mNeoVis.2.1.1.chr9\_hap1.g00179.t01 |
| 11155 | 9 | 41240819 | 41304002 | mNeoVis.2.1.1.chr9\_hap1.g00180.t01 |
| 11156 | 9 | 41240819 | 41304002 | mNeoVis.2.1.1.chr9\_hap1.g00180.t01 |
| 11157 | 9 | 41320797 | 41339687 | mNeoVis.2.1.1.chr9\_hap1.g00181.t01 |
| 11158 | 9 | 41320797 | 41339687 | mNeoVis.2.1.1.chr9\_hap1.g00181.t01 |
| 11159 | 9 | 41355015 | 41365235 | mNeoVis.2.1.1.chr9\_hap1.g00182.t01 |
| 11160 | 9 | 41355015 | 41365235 | mNeoVis.2.1.1.chr9\_hap1.g00182.t01 |
| 11161 | 9 | 41437697 | 41635703 | mNeoVis.2.1.1.chr9\_hap1.g00183.t01 |
| 11162 | 9 | 41437697 | 41635703 | mNeoVis.2.1.1.chr9\_hap1.g00183.t01 |
| 11163 | 9 | 41657476 | 41704648 | mNeoVis.2.1.1.chr9\_hap1.g00184.t01 |
| 11164 | 9 | 41657476 | 41704648 | mNeoVis.2.1.1.chr9\_hap1.g00184.t01 |
| 11165 | 9 | 41706183 | 41791715 | mNeoVis.2.1.1.chr9\_hap1.g00185.t01 |
| 11166 | 9 | 41706183 | 41791715 | mNeoVis.2.1.1.chr9\_hap1.g00185.t01 |
| 11167 | 9 | 41706183 | 41791715 | mNeoVis.2.1.1.chr9\_hap1.g00185.t01 |
| 11168 | 9 | 41800803 | 41816301 | mNeoVis.2.1.1.chr9\_hap1.g00186.t01 |
| 11169 | 9 | 41800803 | 41816301 | mNeoVis.2.1.1.chr9\_hap1.g00186.t01 |
| 11170 | 9 | 41800803 | 41816301 | mNeoVis.2.1.1.chr9\_hap1.g00186.t01 |
| 11171 | 9 | 41849591 | 42024949 | mNeoVis.2.1.1.chr9\_hap1.g00187.t01 |
| 11172 | 9 | 41849591 | 42024949 | mNeoVis.2.1.1.chr9\_hap1.g00187.t01 |
| 11173 | 9 | 41849591 | 42024949 | mNeoVis.2.1.1.chr9\_hap1.g00187.t01 |
| 11174 | 9 | 42069216 | 42087143 | mNeoVis.2.1.1.chr9\_hap1.g00188.t01 |
| 11175 | 9 | 42069216 | 42087143 | mNeoVis.2.1.1.chr9\_hap1.g00188.t01 |
| 11176 | 9 | 42069216 | 42087143 | mNeoVis.2.1.1.chr9\_hap1.g00188.t01 |
| 11177 | 9 | 42140474 | 42175093 | mNeoVis.2.1.1.chr9\_hap1.g00189.t01 |
| 11178 | 9 | 42140474 | 42175093 | mNeoVis.2.1.1.chr9\_hap1.g00189.t01 |
| 11179 | 9 | 42140474 | 42175093 | mNeoVis.2.1.1.chr9\_hap1.g00189.t01 |
| 11180 | 9 | 42189858 | 42209621 | mNeoVis.2.1.1.chr9\_hap1.g00190.t01 |
| 11181 | 9 | 42189858 | 42209621 | mNeoVis.2.1.1.chr9\_hap1.g00190.t01 |
| 11182 | 9 | 42189858 | 42209621 | mNeoVis.2.1.1.chr9\_hap1.g00190.t01 |
| 11183 | 9 | 42234792 | 42239383 | mNeoVis.2.1.1.chr9\_hap1.g00191.t01 |
| 11184 | 9 | 42234792 | 42239383 | mNeoVis.2.1.1.chr9\_hap1.g00191.t01 |
| 11185 | 9 | 42234792 | 42239383 | mNeoVis.2.1.1.chr9\_hap1.g00191.t01 |
| 11186 | 9 | 42243075 | 42267965 | mNeoVis.2.1.1.chr9\_hap1.g00192.t01 |
| 11187 | 9 | 42243075 | 42267965 | mNeoVis.2.1.1.chr9\_hap1.g00192.t01 |
| 11188 | 9 | 42243075 | 42267965 | mNeoVis.2.1.1.chr9\_hap1.g00192.t01 |
| 11189 | 9 | 42304789 | 42931160 | mNeoVis.2.1.1.chr9\_hap1.g00193.t01 |
| 11190 | 9 | 42304789 | 42931160 | mNeoVis.2.1.1.chr9\_hap1.g00193.t01 |
| 11191 | 9 | 42985670 | 43039732 | mNeoVis.2.1.1.chr9\_hap1.g00194.t01 |
| 11192 | 9 | 42985670 | 43039732 | mNeoVis.2.1.1.chr9\_hap1.g00194.t01 |
| 11193 | 9 | 42985670 | 43039732 | mNeoVis.2.1.1.chr9\_hap1.g00194.t01 |
| 11194 | 9 | 42985670 | 43039732 | mNeoVis.2.1.1.chr9\_hap1.g00194.t01 |
| 11195 | 9 | 43070057 | 43183306 | mNeoVis.2.1.1.chr9\_hap1.g00195.t01 |
| 11196 | 9 | 43070057 | 43183306 | mNeoVis.2.1.1.chr9\_hap1.g00195.t01 |
| 11197 | 9 | 43070057 | 43183306 | mNeoVis.2.1.1.chr9\_hap1.g00195.t01 |
| 11198 | 9 | 43070057 | 43183306 | mNeoVis.2.1.1.chr9\_hap1.g00195.t01 |
| 11199 | 9 | 43144900 | 43161411 | mNeoVis.2.1.1.chr9\_hap1.g00196.t01 |
| 11200 | 9 | 43144900 | 43161411 | mNeoVis.2.1.1.chr9\_hap1.g00196.t01 |
| 11201 | 9 | 43144900 | 43161411 | mNeoVis.2.1.1.chr9\_hap1.g00196.t01 |
| 11202 | 9 | 43144900 | 43161411 | mNeoVis.2.1.1.chr9\_hap1.g00196.t01 |
| 11203 | 9 | 43185621 | 43280276 | mNeoVis.2.1.1.chr9\_hap1.g00197.t01 |
| 11204 | 9 | 43185621 | 43280276 | mNeoVis.2.1.1.chr9\_hap1.g00197.t01 |
| 11205 | 9 | 43185621 | 43280276 | mNeoVis.2.1.1.chr9\_hap1.g00197.t01 |
| 11206 | 9 | 43337908 | 43350902 | mNeoVis.2.1.1.chr9\_hap1.g00198.t01 |
| 11207 | 9 | 43337908 | 43350902 | mNeoVis.2.1.1.chr9\_hap1.g00198.t01 |
| 11208 | 9 | 43337908 | 43350902 | mNeoVis.2.1.1.chr9\_hap1.g00198.t01 |
| 11209 | 9 | 43384378 | 43386318 | mNeoVis.2.1.1.chr9\_hap1.g00199.t01 |
| 11210 | 9 | 43384378 | 43386318 | mNeoVis.2.1.1.chr9\_hap1.g00199.t01 |
| 11211 | 9 | 43384378 | 43386318 | mNeoVis.2.1.1.chr9\_hap1.g00199.t01 |
| 11212 | 9 | 43489347 | 43489759 | mNeoVis.2.1.1.chr9\_hap1.g00201.t01 |
| 11213 | 9 | 43489347 | 43489759 | mNeoVis.2.1.1.chr9\_hap1.g00201.t01 |
| 11214 | 9 | 43489347 | 43489759 | mNeoVis.2.1.1.chr9\_hap1.g00201.t01 |
| 11215 | 9 | 44088532 | 44126624 | mNeoVis.2.1.1.chr9\_hap1.g00202.t01 |
| 11216 | 9 | 44088532 | 44126624 | mNeoVis.2.1.1.chr9\_hap1.g00202.t01 |
| 11217 | 9 | 44117292 | 44117857 | mNeoVis.2.1.1.chr9\_hap1.g00203.t01 |
| 11218 | 9 | 44117292 | 44117857 | mNeoVis.2.1.1.chr9\_hap1.g00203.t01 |
| 11219 | 9 | 44126284 | 44276053 | mNeoVis.2.1.1.chr9\_hap1.g00204.t01 |
| 11220 | 9 | 44126284 | 44276053 | mNeoVis.2.1.1.chr9\_hap1.g00204.t01 |
| 11221 | 9 | 44353819 | 44432350 | mNeoVis.2.1.1.chr9\_hap1.g00205.t01 |
| 11222 | 9 | 44353819 | 44432350 | mNeoVis.2.1.1.chr9\_hap1.g00205.t01 |
| 11223 | 9 | 44433109 | 44524656 | mNeoVis.2.1.1.chr9\_hap1.g00206.t01 |
| 11224 | 9 | 44433109 | 44524656 | mNeoVis.2.1.1.chr9\_hap1.g00206.t01 |
| 11225 | 9 | 44536252 | 44561794 | mNeoVis.2.1.1.chr9\_hap1.g00207.t01 |
| 11226 | 9 | 44536252 | 44561794 | mNeoVis.2.1.1.chr9\_hap1.g00207.t01 |
| 11227 | 9 | 44574370 | 44673440 | mNeoVis.2.1.1.chr9\_hap1.g00208.t01 |
| 11228 | 9 | 44677550 | 44733374 | mNeoVis.2.1.1.chr9\_hap1.g00209.t01 |
| 11229 | 9 | 44739546 | 44764401 | mNeoVis.2.1.1.chr9\_hap1.g00210.t01 |
| 11230 | 9 | 44773405 | 44875938 | mNeoVis.2.1.1.chr9\_hap1.g00211.t01 |
| 11231 | 9 | 44876319 | 44927025 | mNeoVis.2.1.1.chr9\_hap1.g00212.t01 |
| 11232 | 9 | 44927138 | 44963171 | mNeoVis.2.1.1.chr9\_hap1.g00213.t01 |
| 11233 | 9 | 44975851 | 45056275 | mNeoVis.2.1.1.chr9\_hap1.g00214.t01 |
| 11234 | 9 | 45079015 | 45087734 | mNeoVis.2.1.1.chr9\_hap1.g00215.t01 |
| 11235 | 9 | 45093647 | 45105612 | mNeoVis.2.1.1.chr9\_hap1.g00216.t01 |
| 11236 | 9 | 45123746 | 45132577 | mNeoVis.2.1.1.chr9\_hap1.g00217.t01 |
| 11237 | 9 | 45132712 | 45145766 | mNeoVis.2.1.1.chr9\_hap1.g00218.t01 |
| 11238 | 9 | 45132712 | 45145766 | mNeoVis.2.1.1.chr9\_hap1.g00218.t01 |
| 11239 | 9 | 45145780 | 45153084 | mNeoVis.2.1.1.chr9\_hap1.g00219.t01 |
| 11240 | 9 | 45145780 | 45153084 | mNeoVis.2.1.1.chr9\_hap1.g00219.t01 |
| 11241 | 9 | 45154877 | 45220511 | mNeoVis.2.1.1.chr9\_hap1.g00220.t01 |
| 11242 | 9 | 45154877 | 45220511 | mNeoVis.2.1.1.chr9\_hap1.g00220.t01 |
| 11243 | 9 | 45257961 | 45297585 | mNeoVis.2.1.1.chr9\_hap1.g00221.t01 |
| 11244 | 9 | 45257961 | 45297585 | mNeoVis.2.1.1.chr9\_hap1.g00221.t01 |
| 11245 | 9 | 45425952 | 45509832 | mNeoVis.2.1.1.chr9\_hap1.g00222.t01 |
| 11246 | 9 | 45425952 | 45509832 | mNeoVis.2.1.1.chr9\_hap1.g00222.t01 |
| 11247 | 9 | 45513175 | 45579079 | mNeoVis.2.1.1.chr9\_hap1.g00223.t01 |
| 11248 | 9 | 45513175 | 45579079 | mNeoVis.2.1.1.chr9\_hap1.g00223.t01 |
| 11249 | 9 | 45693444 | 45799307 | mNeoVis.2.1.1.chr9\_hap1.g00224.t01 |
| 11250 | 9 | 45693444 | 45799307 | mNeoVis.2.1.1.chr9\_hap1.g00224.t01 |
| 11251 | 9 | 45813313 | 45923135 | mNeoVis.2.1.1.chr9\_hap1.g00225.t01 |
| 11252 | 9 | 45813313 | 45923135 | mNeoVis.2.1.1.chr9\_hap1.g00225.t01 |
| 11253 | 9 | 46005584 | 46050762 | mNeoVis.2.1.1.chr9\_hap1.g00226.t01 |
| 11254 | 9 | 46005584 | 46050762 | mNeoVis.2.1.1.chr9\_hap1.g00226.t01 |
| 11255 | 9 | 46433014 | 46462639 | mNeoVis.2.1.1.chr9\_hap1.g00227.t01 |
| 11256 | 9 | 46433014 | 46462639 | mNeoVis.2.1.1.chr9\_hap1.g00227.t01 |
| 11257 | 9 | 46477523 | 46507805 | mNeoVis.2.1.1.chr9\_hap1.g00228.t01 |
| 11258 | 9 | 46477523 | 46507805 | mNeoVis.2.1.1.chr9\_hap1.g00228.t01 |
| 11259 | 9 | 46477523 | 46507805 | mNeoVis.2.1.1.chr9\_hap1.g00228.t01 |
| 11260 | 9 | 46502934 | 46504416 | mNeoVis.2.1.1.chr9\_hap1.g00229.t01 |
| 11261 | 9 | 46502934 | 46504416 | mNeoVis.2.1.1.chr9\_hap1.g00229.t01 |
| 11262 | 9 | 46502934 | 46504416 | mNeoVis.2.1.1.chr9\_hap1.g00229.t01 |
| 11263 | 9 | 46516760 | 46528002 | mNeoVis.2.1.1.chr9\_hap1.g00230.t01 |
| 11264 | 9 | 46516760 | 46528002 | mNeoVis.2.1.1.chr9\_hap1.g00230.t01 |
| 11265 | 9 | 46516760 | 46528002 | mNeoVis.2.1.1.chr9\_hap1.g00230.t01 |
| 11266 | 9 | 46581801 | 46597714 | mNeoVis.2.1.1.chr9\_hap1.g00231.t01 |
| 11267 | 9 | 46581801 | 46597714 | mNeoVis.2.1.1.chr9\_hap1.g00231.t01 |
| 11268 | 9 | 46638346 | 46679319 | mNeoVis.2.1.1.chr9\_hap1.g00232.t01 |
| 11269 | 9 | 46638346 | 46679319 | mNeoVis.2.1.1.chr9\_hap1.g00232.t01 |
| 11270 | 9 | 46679486 | 46710784 | mNeoVis.2.1.1.chr9\_hap1.g00233.t01 |
| 11271 | 9 | 46712894 | 46761280 | mNeoVis.2.1.1.chr9\_hap1.g00234.t01 |
| 11272 | 9 | 46762804 | 46792981 | mNeoVis.2.1.1.chr9\_hap1.g00235.t01 |
| 11273 | 9 | 46795796 | 46831580 | mNeoVis.2.1.1.chr9\_hap1.g00236.t01 |
| 11274 | 9 | 46868992 | 46926007 | mNeoVis.2.1.1.chr9\_hap1.g00237.t01 |
| 11275 | 9 | 47640816 | 47708072 | mNeoVis.2.1.1.chr9\_hap1.g00240.t01 |
| 11276 | 9 | 47640816 | 47708072 | mNeoVis.2.1.1.chr9\_hap1.g00240.t01 |
| 11277 | 9 | 47773811 | 48114596 | mNeoVis.2.1.1.chr9\_hap1.g00241.t01 |
| 11278 | 9 | 47773811 | 48114596 | mNeoVis.2.1.1.chr9\_hap1.g00241.t01 |
| 11279 | 9 | 48118195 | 48254434 | mNeoVis.2.1.1.chr9\_hap1.g00242.t01 |
| 11280 | 9 | 48118195 | 48254434 | mNeoVis.2.1.1.chr9\_hap1.g00242.t01 |
| 11281 | 9 | 48239486 | 48241234 | mNeoVis.2.1.1.chr9\_hap1.g00243.t01 |
| 11282 | 9 | 48239486 | 48241234 | mNeoVis.2.1.1.chr9\_hap1.g00243.t01 |
| 11283 | 9 | 48311458 | 48344380 | mNeoVis.2.1.1.chr9\_hap1.g00244.t01 |
| 11284 | 9 | 48311458 | 48344380 | mNeoVis.2.1.1.chr9\_hap1.g00244.t01 |
| 11285 | 9 | 48407227 | 48424027 | mNeoVis.2.1.1.chr9\_hap1.g00245.t01 |
| 11286 | 9 | 48407227 | 48424027 | mNeoVis.2.1.1.chr9\_hap1.g00245.t01 |
| 11287 | 9 | 48424217 | 48437302 | mNeoVis.2.1.1.chr9\_hap1.g00246.t01 |
| 11288 | 9 | 48424217 | 48437302 | mNeoVis.2.1.1.chr9\_hap1.g00246.t01 |
| 11289 | 9 | 48440947 | 48467451 | mNeoVis.2.1.1.chr9\_hap1.g00247.t01 |
| 11290 | 9 | 48474328 | 48480974 | mNeoVis.2.1.1.chr9\_hap1.g00248.t01 |
| 11291 | 9 | 48491416 | 48566281 | mNeoVis.2.1.1.chr9\_hap1.g00249.t01 |
| 11292 | 9 | 48549964 | 48557935 | mNeoVis.2.1.1.chr9\_hap1.g00250.t01 |
| 11293 | 9 | 48580919 | 48583049 | mNeoVis.2.1.1.chr9\_hap1.g00251.t01 |
| 11294 | 9 | 48583122 | 48613744 | mNeoVis.2.1.1.chr9\_hap1.g00252.t01 |
| 11295 | 9 | 48616411 | 48620696 | mNeoVis.2.1.1.chr9\_hap1.g00253.t01 |
| 11296 | 9 | 48621561 | 48633806 | mNeoVis.2.1.1.chr9\_hap1.g00254.t01 |
| 11297 | 9 | 48634295 | 48647621 | mNeoVis.2.1.1.chr9\_hap1.g00255.t01 |
| 11298 | 9 | 48648459 | 48679589 | mNeoVis.2.1.1.chr9\_hap1.g00256.t01 |
| 11299 | 9 | 48687496 | 48699988 | mNeoVis.2.1.1.chr9\_hap1.g00257.t01 |
| 11300 | 9 | 48701422 | 48710455 | mNeoVis.2.1.1.chr9\_hap1.g00258.t01 |
| 11301 | 9 | 48711019 | 48718068 | mNeoVis.2.1.1.chr9\_hap1.g00259.t01 |
| 11302 | 9 | 48720185 | 48740480 | mNeoVis.2.1.1.chr9\_hap1.g00260.t01 |
| 11303 | 9 | 48744451 | 48750996 | mNeoVis.2.1.1.chr9\_hap1.g00261.t01 |
| 11304 | 9 | 48773998 | 48799508 | mNeoVis.2.1.1.chr9\_hap1.g00262.t01 |
| 11305 | 9 | 48799625 | 48804306 | mNeoVis.2.1.1.chr9\_hap1.g00263.t01 |
| 11306 | 9 | 48815061 | 48819930 | mNeoVis.2.1.1.chr9\_hap1.g00264.t01 |
| 11307 | 9 | 48821383 | 48827770 | mNeoVis.2.1.1.chr9\_hap1.g00265.t01 |
| 11308 | 9 | 48837153 | 48878268 | mNeoVis.2.1.1.chr9\_hap1.g00266.t01 |
| 11309 | 9 | 48878327 | 48884050 | mNeoVis.2.1.1.chr9\_hap1.g00267.t01 |
| 11310 | 9 | 48884920 | 48895756 | mNeoVis.2.1.1.chr9\_hap1.g00268.t01 |
| 11311 | 9 | 48895855 | 48900871 | mNeoVis.2.1.1.chr9\_hap1.g00269.t01 |
| 11312 | 9 | 48903764 | 48905929 | mNeoVis.2.1.1.chr9\_hap1.g00270.t01 |
| 11313 | 9 | 48906488 | 48919784 | mNeoVis.2.1.1.chr9\_hap1.g00271.t01 |
| 11314 | 9 | 48920547 | 48924085 | mNeoVis.2.1.1.chr9\_hap1.g00272.t01 |
| 11315 | 9 | 48924193 | 48926648 | mNeoVis.2.1.1.chr9\_hap1.g00273.t01 |
| 11316 | 9 | 48942191 | 48948393 | mNeoVis.2.1.1.chr9\_hap1.g00274.t01 |
| 11317 | 9 | 48948696 | 48957032 | mNeoVis.2.1.1.chr9\_hap1.g00275.t01 |
| 11318 | 9 | 50412436 | 50432440 | mNeoVis.2.1.1.chr9\_hap1.g00304.t01 |
| 11319 | 9 | 50412436 | 50432440 | mNeoVis.2.1.1.chr9\_hap1.g00304.t01 |
| 11320 | 9 | 50463562 | 50625664 | mNeoVis.2.1.1.chr9\_hap1.g00305.t01 |
| 11321 | 9 | 50463562 | 50625664 | mNeoVis.2.1.1.chr9\_hap1.g00305.t01 |
| 11322 | 9 | 50766408 | 50784390 | mNeoVis.2.1.1.chr9\_hap1.g00306.t01 |
| 11323 | 9 | 50766408 | 50784390 | mNeoVis.2.1.1.chr9\_hap1.g00306.t01 |
| 11324 | 9 | 51147503 | 51155308 | mNeoVis.2.1.1.chr9\_hap1.g00307.t01 |
| 11325 | 9 | 51645308 | 51902425 | mNeoVis.2.1.1.chr9\_hap1.g00311.t01 |
| 11326 | 9 | 51955868 | 52066927 | mNeoVis.2.1.1.chr9\_hap1.g00312.t01 |
| 11327 | 9 | 51955868 | 52066927 | mNeoVis.2.1.1.chr9\_hap1.g00312.t01 |
| 11328 | 9 | 51993967 | 51995026 | mNeoVis.2.1.1.chr9\_hap1.g00313.t01 |
| 11329 | 9 | 51993967 | 51995026 | mNeoVis.2.1.1.chr9\_hap1.g00313.t01 |
| 11330 | 9 | 52110286 | 52222897 | mNeoVis.2.1.1.chr9\_hap1.g00314.t01 |
| 11331 | 9 | 52110286 | 52222897 | mNeoVis.2.1.1.chr9\_hap1.g00314.t01 |
| 11332 | 9 | 52273200 | 52765538 | mNeoVis.2.1.1.chr9\_hap1.g00315.t01 |
| 11333 | 9 | 54399403 | 54582572 | mNeoVis.2.1.1.chr9\_hap1.g00316.t01 |
| 11334 | 9 | 54399403 | 54582572 | mNeoVis.2.1.1.chr9\_hap1.g00316.t01 |
| 11335 | 9 | 54569760 | 54570844 | mNeoVis.2.1.1.chr9\_hap1.g00317.t01 |
| 11336 | 9 | 54569760 | 54570844 | mNeoVis.2.1.1.chr9\_hap1.g00317.t01 |
| 11337 | 9 | 54569760 | 54570844 | mNeoVis.2.1.1.chr9\_hap1.g00317.t01 |
| 11338 | 9 | 56305865 | 56336642 | mNeoVis.2.1.1.chr9\_hap1.g00336.t01 |
| 11339 | 9 | 56305865 | 56336642 | mNeoVis.2.1.1.chr9\_hap1.g00336.t01 |
| 11340 | 9 | 56340215 | 56354200 | mNeoVis.2.1.1.chr9\_hap1.g00337.t01 |
| 11341 | 9 | 56340215 | 56354200 | mNeoVis.2.1.1.chr9\_hap1.g00337.t01 |
| 11342 | 9 | 56354357 | 56363567 | mNeoVis.2.1.1.chr9\_hap1.g00338.t01 |
| 11343 | 9 | 56354357 | 56363567 | mNeoVis.2.1.1.chr9\_hap1.g00338.t01 |
| 11344 | 9 | 56362956 | 56367903 | mNeoVis.2.1.1.chr9\_hap1.g00339.t01 |
| 11345 | 9 | 56362956 | 56367903 | mNeoVis.2.1.1.chr9\_hap1.g00339.t01 |
| 11346 | 9 | 56367842 | 56385766 | mNeoVis.2.1.1.chr9\_hap1.g00340.t01 |
| 11347 | 9 | 56367842 | 56385766 | mNeoVis.2.1.1.chr9\_hap1.g00340.t01 |
| 11348 | 9 | 56389938 | 56398155 | mNeoVis.2.1.1.chr9\_hap1.g00341.t01 |
| 11349 | 9 | 56389938 | 56398155 | mNeoVis.2.1.1.chr9\_hap1.g00341.t01 |
| 11350 | 9 | 56398306 | 56409173 | mNeoVis.2.1.1.chr9\_hap1.g00342.t01 |
| 11351 | 9 | 56398306 | 56409173 | mNeoVis.2.1.1.chr9\_hap1.g00342.t01 |
| 11352 | 9 | 56411362 | 56413213 | mNeoVis.2.1.1.chr9\_hap1.g00343.t01 |
| 11353 | 9 | 56411362 | 56413213 | mNeoVis.2.1.1.chr9\_hap1.g00343.t01 |
| 11354 | 9 | 56415917 | 56431995 | mNeoVis.2.1.1.chr9\_hap1.g00344.t01 |
| 11355 | 9 | 56415917 | 56431995 | mNeoVis.2.1.1.chr9\_hap1.g00344.t01 |
| 11356 | 9 | 56431293 | 56595746 | mNeoVis.2.1.1.chr9\_hap1.g00345.t01 |
| 11357 | 9 | 56431293 | 56595746 | mNeoVis.2.1.1.chr9\_hap1.g00345.t01 |
| 11358 | 9 | 56595623 | 56604062 | mNeoVis.2.1.1.chr9\_hap1.g00346.t01 |
| 11359 | 9 | 56595623 | 56604062 | mNeoVis.2.1.1.chr9\_hap1.g00346.t01 |
| 11360 | 9 | 56609611 | 56612201 | mNeoVis.2.1.1.chr9\_hap1.g00347.t01 |
| 11361 | 9 | 56609611 | 56612201 | mNeoVis.2.1.1.chr9\_hap1.g00347.t01 |
| 11362 | 9 | 56612378 | 56617379 | mNeoVis.2.1.1.chr9\_hap1.g00348.t01 |
| 11363 | 9 | 56612378 | 56617379 | mNeoVis.2.1.1.chr9\_hap1.g00348.t01 |
| 11364 | 9 | 56619938 | 56629339 | mNeoVis.2.1.1.chr9\_hap1.g00349.t01 |
| 11365 | 9 | 56619938 | 56629339 | mNeoVis.2.1.1.chr9\_hap1.g00349.t01 |
| 11366 | 9 | 56632924 | 56712664 | mNeoVis.2.1.1.chr9\_hap1.g00350.t01 |
| 11367 | 9 | 56632924 | 56712664 | mNeoVis.2.1.1.chr9\_hap1.g00350.t01 |
| 11368 | 9 | 56728260 | 56741875 | mNeoVis.2.1.1.chr9\_hap1.g00351.t01 |
| 11369 | 9 | 56728260 | 56741875 | mNeoVis.2.1.1.chr9\_hap1.g00351.t01 |
| 11370 | 9 | 56741815 | 56747980 | mNeoVis.2.1.1.chr9\_hap1.g00352.t01 |
| 11371 | 9 | 56741815 | 56747980 | mNeoVis.2.1.1.chr9\_hap1.g00352.t01 |
| 11372 | 9 | 56756577 | 56782229 | mNeoVis.2.1.1.chr9\_hap1.g00353.t01 |
| 11373 | 9 | 56756577 | 56782229 | mNeoVis.2.1.1.chr9\_hap1.g00353.t01 |
| 11374 | 9 | 56786298 | 56806890 | mNeoVis.2.1.1.chr9\_hap1.g00354.t01 |
| 11375 | 9 | 56786298 | 56806890 | mNeoVis.2.1.1.chr9\_hap1.g00354.t01 |
| 11376 | 9 | 56810850 | 56820236 | mNeoVis.2.1.1.chr9\_hap1.g00355.t01 |
| 11377 | 9 | 56810850 | 56820236 | mNeoVis.2.1.1.chr9\_hap1.g00355.t01 |
| 11378 | 9 | 56825972 | 56834759 | mNeoVis.2.1.1.chr9\_hap1.g00356.t01 |
| 11379 | 9 | 56825972 | 56834759 | mNeoVis.2.1.1.chr9\_hap1.g00356.t01 |
| 11380 | 9 | 56834867 | 56838625 | mNeoVis.2.1.1.chr9\_hap1.g00357.t01 |
| 11381 | 9 | 56834867 | 56838625 | mNeoVis.2.1.1.chr9\_hap1.g00357.t01 |
| 11382 | 9 | 56847244 | 56856054 | mNeoVis.2.1.1.chr9\_hap1.g00358.t01 |
| 11383 | 9 | 56847244 | 56856054 | mNeoVis.2.1.1.chr9\_hap1.g00358.t01 |
| 11384 | 9 | 56859149 | 56869420 | mNeoVis.2.1.1.chr9\_hap1.g00359.t01 |
| 11385 | 9 | 56859149 | 56869420 | mNeoVis.2.1.1.chr9\_hap1.g00359.t01 |
| 11386 | 9 | 56946021 | 57011903 | mNeoVis.2.1.1.chr9\_hap1.g00360.t01 |
| 11387 | 9 | 56946021 | 57011903 | mNeoVis.2.1.1.chr9\_hap1.g00360.t01 |
| 11388 | 9 | 56982555 | 57006237 | mNeoVis.2.1.1.chr9\_hap1.g00361.t01 |
| 11389 | 9 | 56982555 | 57006237 | mNeoVis.2.1.1.chr9\_hap1.g00361.t01 |
| 11390 | 9 | 57011847 | 57026450 | mNeoVis.2.1.1.chr9\_hap1.g00362.t01 |
| 11391 | 9 | 57011847 | 57026450 | mNeoVis.2.1.1.chr9\_hap1.g00362.t01 |
| 11392 | 9 | 57055045 | 57076151 | mNeoVis.2.1.1.chr9\_hap1.g00363.t01 |
| 11393 | 9 | 57055045 | 57076151 | mNeoVis.2.1.1.chr9\_hap1.g00363.t01 |
| 11394 | 9 | 57085128 | 57091394 | mNeoVis.2.1.1.chr9\_hap1.g00364.t01 |
| 11395 | 9 | 57085128 | 57091394 | mNeoVis.2.1.1.chr9\_hap1.g00364.t01 |
| 11396 | 9 | 57093085 | 57128232 | mNeoVis.2.1.1.chr9\_hap1.g00365.t01 |
| 11397 | 9 | 57093085 | 57128232 | mNeoVis.2.1.1.chr9\_hap1.g00365.t01 |
| 11398 | 9 | 57133394 | 57225079 | mNeoVis.2.1.1.chr9\_hap1.g00366.t01 |
| 11399 | 9 | 57133394 | 57225079 | mNeoVis.2.1.1.chr9\_hap1.g00366.t01 |
| 11400 | 9 | 57225354 | 57241257 | mNeoVis.2.1.1.chr9\_hap1.g00367.t01 |
| 11401 | 9 | 57225354 | 57241257 | mNeoVis.2.1.1.chr9\_hap1.g00367.t01 |
| 11402 | 9 | 57241399 | 57242537 | mNeoVis.2.1.1.chr9\_hap1.g00368.t01 |
| 11403 | 9 | 57241399 | 57242537 | mNeoVis.2.1.1.chr9\_hap1.g00368.t01 |
| 11404 | 9 | 57255521 | 57260267 | mNeoVis.2.1.1.chr9\_hap1.g00369.t01 |
| 11405 | 9 | 57255521 | 57260267 | mNeoVis.2.1.1.chr9\_hap1.g00369.t01 |
| 11406 | 9 | 57260543 | 57288008 | mNeoVis.2.1.1.chr9\_hap1.g00370.t01 |
| 11407 | 9 | 57260543 | 57288008 | mNeoVis.2.1.1.chr9\_hap1.g00370.t01 |
| 11408 | 9 | 57295599 | 57296911 | mNeoVis.2.1.1.chr9\_hap1.g00371.t01 |
| 11409 | 9 | 57297502 | 57316803 | mNeoVis.2.1.1.chr9\_hap1.g00372.t01 |
| 11410 | 9 | 57318081 | 57347042 | mNeoVis.2.1.1.chr9\_hap1.g00373.t01 |
| 11411 | 9 | 57323389 | 57338520 | mNeoVis.2.1.1.chr9\_hap1.g00374.t01 |
| 11412 | 9 | 57338937 | 57342104 | mNeoVis.2.1.1.chr9\_hap1.g00375.t01 |
| 11413 | 9 | 57347559 | 57375546 | mNeoVis.2.1.1.chr9\_hap1.g00376.t01 |
| 11414 | 9 | 57376284 | 57379783 | mNeoVis.2.1.1.chr9\_hap1.g00377.t01 |
| 11415 | 9 | 57385259 | 57403468 | mNeoVis.2.1.1.chr9\_hap1.g00378.t01 |
| 11416 | 9 | 57403617 | 57409881 | mNeoVis.2.1.1.chr9\_hap1.g00379.t01 |
| 11417 | 9 | 57410058 | 57415667 | mNeoVis.2.1.1.chr9\_hap1.g00380.t01 |
| 11418 | 9 | 57414718 | 57447663 | mNeoVis.2.1.1.chr9\_hap1.g00381.t01 |
| 11419 | 9 | 57458129 | 57479265 | mNeoVis.2.1.1.chr9\_hap1.g00382.t01 |
| 11420 | 9 | 57480621 | 57483630 | mNeoVis.2.1.1.chr9\_hap1.g00383.t01 |
| 11421 | 9 | 57484093 | 57489134 | mNeoVis.2.1.1.chr9\_hap1.g00384.t01 |
| 11422 | 9 | 57502646 | 57516892 | mNeoVis.2.1.1.chr9\_hap1.g00385.t01 |
| 11423 | 9 | 57521089 | 57526438 | mNeoVis.2.1.1.chr9\_hap1.g00386.t01 |
| 11424 | 9 | 60171163 | 60179060 | mNeoVis.2.1.1.chr9\_hap1.g00455.t01 |
| 11425 | 9 | 60180033 | 60207394 | mNeoVis.2.1.1.chr9\_hap1.g00456.t01 |
| 11426 | 9 | 60213742 | 60252935 | mNeoVis.2.1.1.chr9\_hap1.g00457.t01 |
| 11427 | 9 | 60256538 | 60262083 | mNeoVis.2.1.1.chr9\_hap1.g00458.t01 |
| 11428 | 9 | 60271577 | 60287329 | mNeoVis.2.1.1.chr9\_hap1.g00459.t01 |
| 11429 | 9 | 60318999 | 60340072 | mNeoVis.2.1.1.chr9\_hap1.g00460.t01 |
| 11430 | 9 | 60350276 | 60364268 | mNeoVis.2.1.1.chr9\_hap1.g00461.t01 |
| 11431 | 9 | 60367840 | 60369859 | mNeoVis.2.1.1.chr9\_hap1.g00462.t01 |
| 11432 | 9 | 60380472 | 60399831 | mNeoVis.2.1.1.chr9\_hap1.g00463.t01 |
| 11433 | 9 | 60418428 | 60425538 | mNeoVis.2.1.1.chr9\_hap1.g00464.t01 |
| 11434 | 9 | 60428130 | 60434640 | mNeoVis.2.1.1.chr9\_hap1.g00465.t01 |
| 11435 | 9 | 60435005 | 60472554 | mNeoVis.2.1.1.chr9\_hap1.g00466.t01 |
| 11436 | 9 | 60508066 | 60514656 | mNeoVis.2.1.1.chr9\_hap1.g00467.t01 |
| 11437 | 9 | 60631074 | 60669899 | mNeoVis.2.1.1.chr9\_hap1.g00468.t01 |
| 11438 | 9 | 60651031 | 60653866 | mNeoVis.2.1.1.chr9\_hap1.g00469.t01 |
| 11439 | 9 | 60656019 | 60667426 | mNeoVis.2.1.1.chr9\_hap1.g00470.t01 |
| 11440 | 9 | 60673766 | 60677333 | mNeoVis.2.1.1.chr9\_hap1.g00471.t01 |
| 11441 | 9 | 60683883 | 60686498 | mNeoVis.2.1.1.chr9\_hap1.g00472.t01 |
| 11442 | 9 | 60696481 | 60701655 | mNeoVis.2.1.1.chr9\_hap1.g00473.t01 |
| 11443 | 9 | 60711479 | 60713635 | mNeoVis.2.1.1.chr9\_hap1.g00474.t01 |
| 11444 | 9 | 60730207 | 60731995 | mNeoVis.2.1.1.chr9\_hap1.g00475.t01 |
| 11445 | 9 | 60740158 | 60748150 | mNeoVis.2.1.1.chr9\_hap1.g00476.t01 |
| 11446 | 9 | 60958268 | 60975116 | mNeoVis.2.1.1.chr9\_hap1.g00478.t01 |
| 11447 | 9 | 61001771 | 61010889 | mNeoVis.2.1.1.chr9\_hap1.g00479.t01 |
| 11448 | 9 | 61057819 | 61146716 | mNeoVis.2.1.1.chr9\_hap1.g00480.t01 |
| 11449 | 9 | 61147706 | 61152044 | mNeoVis.2.1.1.chr9\_hap1.g00481.t01 |
| 11450 | 9 | 61152236 | 61157529 | mNeoVis.2.1.1.chr9\_hap1.g00482.t01 |
| 11451 | 9 | 61164754 | 61174354 | mNeoVis.2.1.1.chr9\_hap1.g00483.t01 |
| 11452 | 9 | 61178898 | 61197933 | mNeoVis.2.1.1.chr9\_hap1.g00484.t01 |
| 11453 | 9 | 61208996 | 61212277 | mNeoVis.2.1.1.chr9\_hap1.g00485.t01 |
| 11454 | 9 | 61220659 | 61226426 | mNeoVis.2.1.1.chr9\_hap1.g00486.t01 |
| 11455 | 9 | 61232020 | 61278134 | mNeoVis.2.1.1.chr9\_hap1.g00487.t01 |
| 11456 | 9 | 61318122 | 61325543 | mNeoVis.2.1.1.chr9\_hap1.g00488.t01 |
| 11457 | 9 | 61329749 | 61344894 | mNeoVis.2.1.1.chr9\_hap1.g00489.t01 |
| 11458 | 9 | 61345176 | 61352042 | mNeoVis.2.1.1.chr9\_hap1.g00490.t01 |
| 11459 | 9 | 61352268 | 61355885 | mNeoVis.2.1.1.chr9\_hap1.g00491.t01 |
| 11460 | 9 | 61356847 | 61381168 | mNeoVis.2.1.1.chr9\_hap1.g00492.t01 |
| 11461 | 9 | 61387113 | 61389468 | mNeoVis.2.1.1.chr9\_hap1.g00493.t01 |
| 11462 | 9 | 61395997 | 61396655 | mNeoVis.2.1.1.chr9\_hap1.g00494.t01 |
| 11463 | 9 | 61405578 | 61426629 | mNeoVis.2.1.1.chr9\_hap1.g00495.t01 |
| 11464 | 9 | 61428763 | 61444747 | mNeoVis.2.1.1.chr9\_hap1.g00496.t01 |
| 11465 | 9 | 61445201 | 61454798 | mNeoVis.2.1.1.chr9\_hap1.g00497.t01 |
| 11466 | 9 | 61453826 | 61481708 | mNeoVis.2.1.1.chr9\_hap1.g00498.t01 |
| 11467 | 9 | 61499201 | 61508993 | mNeoVis.2.1.1.chr9\_hap1.g00499.t01 |
| 11468 | 9 | 61509932 | 61513947 | mNeoVis.2.1.1.chr9\_hap1.g00500.t01 |
| 11469 | 9 | 62083078 | 62088676 | mNeoVis.2.1.1.chr9\_hap1.g00517.t01 |
| 11470 | 9 | 62109525 | 62120012 | mNeoVis.2.1.1.chr9\_hap1.g00518.t01 |
| 11471 | 9 | 62121682 | 62134371 | mNeoVis.2.1.1.chr9\_hap1.g00519.t01 |
| 11472 | 9 | 62137150 | 62148600 | mNeoVis.2.1.1.chr9\_hap1.g00520.t01 |
| 11473 | 9 | 62156998 | 62165991 | mNeoVis.2.1.1.chr9\_hap1.g00521.t01 |
| 11474 | 9 | 62180416 | 62187934 | mNeoVis.2.1.1.chr9\_hap1.g00522.t01 |
| 11475 | 9 | 62192770 | 62207072 | mNeoVis.2.1.1.chr9\_hap1.g00523.t01 |
| 11476 | 9 | 62221512 | 62228390 | mNeoVis.2.1.1.chr9\_hap1.g00524.t01 |
| 11477 | 9 | 62236832 | 62243888 | mNeoVis.2.1.1.chr9\_hap1.g00525.t01 |
| 11478 | 9 | 62253840 | 62259382 | mNeoVis.2.1.1.chr9\_hap1.g00526.t01 |
| 11479 | 9 | 62264208 | 62269355 | mNeoVis.2.1.1.chr9\_hap1.g00527.t01 |
| 11480 | 9 | 62277099 | 62280039 | mNeoVis.2.1.1.chr9\_hap1.g00528.t01 |
| 11481 | 9 | 62282599 | 62289766 | mNeoVis.2.1.1.chr9\_hap1.g00529.t01 |
| 11482 | 9 | 62291161 | 62305331 | mNeoVis.2.1.1.chr9\_hap1.g00530.t01 |
| 11483 | 9 | 62310907 | 62313000 | mNeoVis.2.1.1.chr9\_hap1.g00531.t01 |
| 11484 | 9 | 62343760 | 62366653 | mNeoVis.2.1.1.chr9\_hap1.g00532.t01 |
| 11485 | 9 | 62405956 | 62409438 | mNeoVis.2.1.1.chr9\_hap1.g00533.t01 |
| 11486 | 9 | 62411233 | 62417555 | mNeoVis.2.1.1.chr9\_hap1.g00534.t01 |
| 11487 | 9 | 62426490 | 62436320 | mNeoVis.2.1.1.chr9\_hap1.g00535.t01 |
| 11488 | 9 | 62468780 | 62476405 | mNeoVis.2.1.1.chr9\_hap1.g00536.t01 |
| 11489 | 9 | 62484235 | 62520119 | mNeoVis.2.1.1.chr9\_hap1.g00537.t01 |
| 11490 | 9 | 62549834 | 62564557 | mNeoVis.2.1.1.chr9\_hap1.g00538.t01 |
| 11491 | 9 | 62576209 | 62579026 | mNeoVis.2.1.1.chr9\_hap1.g00539.t01 |
| 11492 | 9 | 62634952 | 62641901 | mNeoVis.2.1.1.chr9\_hap1.g00540.t01 |
| 11493 | 9 | 62642978 | 62645715 | mNeoVis.2.1.1.chr9\_hap1.g00541.t01 |
| 11494 | 9 | 63218809 | 63234550 | mNeoVis.2.1.1.chr9\_hap1.g00551.t01 |
| 11495 | 9 | 63235006 | 63246919 | mNeoVis.2.1.1.chr9\_hap1.g00552.t01 |
| 11496 | 9 | 63243430 | 63300140 | mNeoVis.2.1.1.chr9\_hap1.g00553.t01 |
| 11497 | 9 | 63303120 | 63343091 | mNeoVis.2.1.1.chr9\_hap1.g00554.t01 |
| 11498 | 9 | 63357662 | 63386500 | mNeoVis.2.1.1.chr9\_hap1.g00555.t01 |
| 11499 | 9 | 63386930 | 63432046 | mNeoVis.2.1.1.chr9\_hap1.g00556.t01 |
| 11500 | 9 | 63405038 | 63495928 | mNeoVis.2.1.1.chr9\_hap1.g00557.t01 |
| 11501 | 9 | 63505514 | 63507467 | mNeoVis.2.1.1.chr9\_hap1.g00558.t01 |
| 11502 | 9 | 63505514 | 63507467 | mNeoVis.2.1.1.chr9\_hap1.g00558.t01 |
| 11503 | 9 | 63507983 | 63735343 | mNeoVis.2.1.1.chr9\_hap1.g00559.t01 |
| 11504 | 9 | 63507983 | 63735343 | mNeoVis.2.1.1.chr9\_hap1.g00559.t01 |
| 11505 | 9 | 63564146 | 63565011 | mNeoVis.2.1.1.chr9\_hap1.g00560.t01 |
| 11506 | 9 | 63564146 | 63565011 | mNeoVis.2.1.1.chr9\_hap1.g00560.t01 |
| 11507 | 9 | 63750757 | 63832986 | mNeoVis.2.1.1.chr9\_hap1.g00561.t01 |
| 11508 | 9 | 63836163 | 63927104 | mNeoVis.2.1.1.chr9\_hap1.g00562.t01 |
| 11509 | 9 | 63965162 | 64058379 | mNeoVis.2.1.1.chr9\_hap1.g00563.t01 |
| 11510 | 9 | 64061306 | 64091670 | mNeoVis.2.1.1.chr9\_hap1.g00564.t01 |
| 11511 | 9 | 64103088 | 64108037 | mNeoVis.2.1.1.chr9\_hap1.g00565.t01 |
| 11512 | 9 | 64108554 | 64116354 | mNeoVis.2.1.1.chr9\_hap1.g00566.t01 |
| 11513 | 9 | 64118772 | 64131444 | mNeoVis.2.1.1.chr9\_hap1.g00567.t01 |
| 11514 | 9 | 64133148 | 64141630 | mNeoVis.2.1.1.chr9\_hap1.g00568.t01 |
| 11515 | 9 | 64180948 | 64207293 | mNeoVis.2.1.1.chr9\_hap1.g00569.t01 |
| 11516 | 9 | 64217828 | 64222731 | mNeoVis.2.1.1.chr9\_hap1.g00570.t01 |
| 11517 | 9 | 64229480 | 64233272 | mNeoVis.2.1.1.chr9\_hap1.g00571.t01 |
| 11518 | 9 | 64240696 | 64245851 | mNeoVis.2.1.1.chr9\_hap1.g00572.t01 |
| 11519 | 9 | 64293823 | 64327291 | mNeoVis.2.1.1.chr9\_hap1.g00573.t01 |
| 11520 | 9 | 64344478 | 64350340 | mNeoVis.2.1.1.chr9\_hap1.g00574.t01 |
| 11521 | 9 | 66005776 | 66006717 | mNeoVis.2.1.1.chr9\_hap1.g00627.t01 |
| 11522 | 9 | 66027291 | 66031474 | mNeoVis.2.1.1.chr9\_hap1.g00628.t01 |
| 11523 | 9 | 66044133 | 66050256 | mNeoVis.2.1.1.chr9\_hap1.g00629.t01 |
| 11524 | 9 | 66051012 | 66083973 | mNeoVis.2.1.1.chr9\_hap1.g00630.t01 |
| 11525 | 9 | 66085016 | 66127623 | mNeoVis.2.1.1.chr9\_hap1.g00631.t01 |
| 11526 | 9 | 66129073 | 66178490 | mNeoVis.2.1.1.chr9\_hap1.g00632.t01 |
| 11527 | 9 | 66217889 | 66248874 | mNeoVis.2.1.1.chr9\_hap1.g00633.t01 |
| 11528 | 9 | 66268826 | 66273694 | mNeoVis.2.1.1.chr9\_hap1.g00634.t01 |
| 11529 | 9 | 66344044 | 66376716 | mNeoVis.2.1.1.chr9\_hap1.g00635.t01 |
| 11530 | 9 | 66397228 | 66432726 | mNeoVis.2.1.1.chr9\_hap1.g00636.t01 |
| 11531 | 9 | 66433744 | 66441382 | mNeoVis.2.1.1.chr9\_hap1.g00637.t01 |
| 11532 | 9 | 66454893 | 66478196 | mNeoVis.2.1.1.chr9\_hap1.g00638.t01 |
| 11533 | 9 | 66488373 | 66503165 | mNeoVis.2.1.1.chr9\_hap1.g00639.t01 |
| 11534 | 9 | 66956729 | 66960012 | mNeoVis.2.1.1.chr9\_hap1.g00643.t01 |
| 11535 | 9 | 67152794 | 67215841 | mNeoVis.2.1.1.chr9\_hap1.g00644.t01 |
| 11536 | 9 | 67152794 | 67215841 | mNeoVis.2.1.1.chr9\_hap1.g00644.t01 |
| 11537 | 9 | 67296176 | 67312752 | mNeoVis.2.1.1.chr9\_hap1.g00645.t01 |
| 11538 | 9 | 67296176 | 67312752 | mNeoVis.2.1.1.chr9\_hap1.g00645.t01 |
| 11539 | 9 | 67530414 | 67544426 | mNeoVis.2.1.1.chr9\_hap1.g00646.t01 |
| 11540 | 9 | 67530414 | 67544426 | mNeoVis.2.1.1.chr9\_hap1.g00646.t01 |
| 11541 | 9 | 67587123 | 67655565 | mNeoVis.2.1.1.chr9\_hap1.g00647.t01 |
| 11542 | 9 | 67869918 | 68048453 | mNeoVis.2.1.1.chr9\_hap1.g00649.t01 |
| 11543 | 9 | 68367268 | 68551184 | mNeoVis.2.1.1.chr9\_hap1.g00650.t01 |
| 11544 | 9 | 68701905 | 68734687 | mNeoVis.2.1.1.chr9\_hap1.g00651.t01 |
| 11545 | 9 | 68701905 | 68734687 | mNeoVis.2.1.1.chr9\_hap1.g00651.t01 |
| 11546 | 9 | 71126499 | 71299470 | mNeoVis.2.1.1.chr9\_hap1.g00660.t01 |
| 11547 | 9 | 71266863 | 71282581 | mNeoVis.2.1.1.chr9\_hap1.g00661.t01 |
| 11548 | 9 | 71305724 | 71308913 | mNeoVis.2.1.1.chr9\_hap1.g00662.t01 |
| 11549 | 9 | 71330574 | 71408991 | mNeoVis.2.1.1.chr9\_hap1.g00663.t01 |
| 11550 | 9 | 71417539 | 71475553 | mNeoVis.2.1.1.chr9\_hap1.g00664.t01 |
| 11551 | 9 | 71782127 | 71782563 | mNeoVis.2.1.1.chr9\_hap1.g00665.t01 |
| 11552 | 9 | 71901780 | 72257637 | mNeoVis.2.1.1.chr9\_hap1.g00666.t01 |
| 11553 | 9 | 72360783 | 72717369 | mNeoVis.2.1.1.chr9\_hap1.g00667.t01 |
| 11554 | 9 | 72360783 | 72717369 | mNeoVis.2.1.1.chr9\_hap1.g00667.t01 |
| 11555 | 9 | 72360783 | 72717369 | mNeoVis.2.1.1.chr9\_hap1.g00667.t01 |
| 11556 | 9 | 72815542 | 72839377 | mNeoVis.2.1.1.chr9\_hap1.g00668.t01 |
| 11557 | 9 | 72815542 | 72839377 | mNeoVis.2.1.1.chr9\_hap1.g00668.t01 |
| 11558 | 9 | 72815542 | 72839377 | mNeoVis.2.1.1.chr9\_hap1.g00668.t01 |
| 11559 | 9 | 73058857 | 73065204 | mNeoVis.2.1.1.chr9\_hap1.g00669.t01 |
| 11560 | 9 | 73142772 | 73280084 | mNeoVis.2.1.1.chr9\_hap1.g00670.t01 |
| 11561 | 9 | 73326214 | 73383407 | mNeoVis.2.1.1.chr9\_hap1.g00671.t01 |
| 11562 | 9 | 73326214 | 73383407 | mNeoVis.2.1.1.chr9\_hap1.g00671.t01 |
| 11563 | 9 | 73402437 | 73777308 | mNeoVis.2.1.1.chr9\_hap1.g00672.t01 |
| 11564 | 9 | 73402437 | 73777308 | mNeoVis.2.1.1.chr9\_hap1.g00672.t01 |
| 11565 | 9 | 73662962 | 73663181 | mNeoVis.2.1.1.chr9\_hap1.g00673.t01 |
| 11566 | 9 | 73662962 | 73663181 | mNeoVis.2.1.1.chr9\_hap1.g00673.t01 |
| 11567 | 9 | 73811332 | 73911781 | mNeoVis.2.1.1.chr9\_hap1.g00674.t01 |
| 11568 | 9 | 73911835 | 73979821 | mNeoVis.2.1.1.chr9\_hap1.g00675.t01 |
| 11569 | 9 | 73911835 | 73979821 | mNeoVis.2.1.1.chr9\_hap1.g00675.t01 |
| 11570 | 9 | 74077254 | 74230368 | mNeoVis.2.1.1.chr9\_hap1.g00676.t01 |
| 11571 | 9 | 74077254 | 74230368 | mNeoVis.2.1.1.chr9\_hap1.g00676.t01 |
| 11572 | 9 | 74575095 | 74793058 | mNeoVis.2.1.1.chr9\_hap1.g00677.t01 |
| 11573 | 9 | 75087795 | 75098593 | mNeoVis.2.1.1.chr9\_hap1.g00678.t01 |
| 11574 | 9 | 75087795 | 75098593 | mNeoVis.2.1.1.chr9\_hap1.g00678.t01 |
| 11575 | 9 | 75466696 | 75468132 | mNeoVis.2.1.1.chr9\_hap1.g00679.t01 |
| 11576 | 9 | 75466696 | 75468132 | mNeoVis.2.1.1.chr9\_hap1.g00679.t01 |
| 11577 | 9 | 75466696 | 75468132 | mNeoVis.2.1.1.chr9\_hap1.g00679.t01 |
| 11578 | 9 | 75556269 | 75622996 | mNeoVis.2.1.1.chr9\_hap1.g00680.t01 |
| 11579 | 9 | 75556269 | 75622996 | mNeoVis.2.1.1.chr9\_hap1.g00680.t01 |
| 11580 | 9 | 75556269 | 75622996 | mNeoVis.2.1.1.chr9\_hap1.g00680.t01 |
| 11581 | 9 | 75833452 | 76075946 | mNeoVis.2.1.1.chr9\_hap1.g00681.t01 |
| 11582 | 9 | 75833452 | 76075946 | mNeoVis.2.1.1.chr9\_hap1.g00681.t01 |
| 11583 | 9 | 75833452 | 76075946 | mNeoVis.2.1.1.chr9\_hap1.g00681.t01 |
| 11584 | 9 | 76109646 | 76134378 | mNeoVis.2.1.1.chr9\_hap1.g00682.t01 |
| 11585 | 9 | 76109646 | 76134378 | mNeoVis.2.1.1.chr9\_hap1.g00682.t01 |
| 11586 | 9 | 76109646 | 76134378 | mNeoVis.2.1.1.chr9\_hap1.g00682.t01 |
| 11587 | 9 | 76131216 | 76178749 | mNeoVis.2.1.1.chr9\_hap1.g00683.t01 |
| 11588 | 9 | 76131216 | 76178749 | mNeoVis.2.1.1.chr9\_hap1.g00683.t01 |
| 11589 | 9 | 76131216 | 76178749 | mNeoVis.2.1.1.chr9\_hap1.g00683.t01 |
| 11590 | 9 | 76207840 | 76307944 | mNeoVis.2.1.1.chr9\_hap1.g00684.t01 |
| 11591 | 9 | 76207840 | 76307944 | mNeoVis.2.1.1.chr9\_hap1.g00684.t01 |
| 11592 | 9 | 76207840 | 76307944 | mNeoVis.2.1.1.chr9\_hap1.g00684.t01 |
| 11593 | 9 | 76207840 | 76307944 | mNeoVis.2.1.1.chr9\_hap1.g00684.t01 |
| 11594 | 9 | 76429803 | 76651939 | mNeoVis.2.1.1.chr9\_hap1.g00685.t01 |
| 11595 | 9 | 76429803 | 76651939 | mNeoVis.2.1.1.chr9\_hap1.g00685.t01 |
| 11596 | 9 | 76429803 | 76651939 | mNeoVis.2.1.1.chr9\_hap1.g00685.t01 |
| 11597 | 9 | 76474202 | 76517604 | mNeoVis.2.1.1.chr9\_hap1.g00686.t01 |
| 11598 | 9 | 76474202 | 76517604 | mNeoVis.2.1.1.chr9\_hap1.g00686.t01 |
| 11599 | 9 | 76474202 | 76517604 | mNeoVis.2.1.1.chr9\_hap1.g00686.t01 |
| 11600 | 9 | 76765557 | 76793943 | mNeoVis.2.1.1.chr9\_hap1.g00687.t01 |
| 11601 | 9 | 76765557 | 76793943 | mNeoVis.2.1.1.chr9\_hap1.g00687.t01 |
| 11602 | 9 | 77558051 | 77618318 | mNeoVis.2.1.1.chr9\_hap1.g00696.t01 |
| 11603 | 9 | 77618464 | 77632782 | mNeoVis.2.1.1.chr9\_hap1.g00697.t01 |
| 11604 | 9 | 77637783 | 77649956 | mNeoVis.2.1.1.chr9\_hap1.g00698.t01 |
| 11605 | 9 | 77637783 | 77649956 | mNeoVis.2.1.1.chr9\_hap1.g00698.t01 |
| 11606 | 9 | 77649842 | 77701442 | mNeoVis.2.1.1.chr9\_hap1.g00699.t01 |
| 11607 | 9 | 77649842 | 77701442 | mNeoVis.2.1.1.chr9\_hap1.g00699.t01 |
| 11608 | 9 | 77729676 | 77751916 | mNeoVis.2.1.1.chr9\_hap1.g00700.t01 |
| 11609 | 9 | 77729676 | 77751916 | mNeoVis.2.1.1.chr9\_hap1.g00700.t01 |
| 11610 | 9 | 77846266 | 77853664 | mNeoVis.2.1.1.chr9\_hap1.g00701.t01 |
| 11611 | 9 | 77846266 | 77853664 | mNeoVis.2.1.1.chr9\_hap1.g00701.t01 |
| 11612 | 9 | 77867743 | 78006809 | mNeoVis.2.1.1.chr9\_hap1.g00702.t01 |
| 11613 | 9 | 77867743 | 78006809 | mNeoVis.2.1.1.chr9\_hap1.g00702.t01 |
| 11614 | 9 | 78077981 | 78098351 | mNeoVis.2.1.1.chr9\_hap1.g00703.t01 |
| 11615 | 9 | 78077981 | 78098351 | mNeoVis.2.1.1.chr9\_hap1.g00703.t01 |
| 11616 | 9 | 78077981 | 78098351 | mNeoVis.2.1.1.chr9\_hap1.g00703.t01 |
| 11617 | 9 | 78112193 | 78305305 | mNeoVis.2.1.1.chr9\_hap1.g00704.t01 |
| 11618 | 9 | 78112193 | 78305305 | mNeoVis.2.1.1.chr9\_hap1.g00704.t01 |
| 11619 | 9 | 78112193 | 78305305 | mNeoVis.2.1.1.chr9\_hap1.g00704.t01 |
| 11620 | 9 | 78383223 | 78482096 | mNeoVis.2.1.1.chr9\_hap1.g00705.t01 |
| 11621 | 9 | 78383223 | 78482096 | mNeoVis.2.1.1.chr9\_hap1.g00705.t01 |
| 11622 | 9 | 78383223 | 78482096 | mNeoVis.2.1.1.chr9\_hap1.g00705.t01 |
| 11623 | 9 | 78559358 | 78630056 | mNeoVis.2.1.1.chr9\_hap1.g00706.t01 |
| 11624 | 9 | 78559358 | 78630056 | mNeoVis.2.1.1.chr9\_hap1.g00706.t01 |
| 11625 | 9 | 78647039 | 78845218 | mNeoVis.2.1.1.chr9\_hap1.g00707.t01 |
| 11626 | 9 | 78647039 | 78845218 | mNeoVis.2.1.1.chr9\_hap1.g00707.t01 |
| 11627 | 9 | 80542946 | 80619829 | mNeoVis.2.1.1.chr9\_hap1.g00709.t01 |
| 11628 | 9 | 80542946 | 80619829 | mNeoVis.2.1.1.chr9\_hap1.g00709.t01 |
| 11629 | 9 | 80718461 | 80765186 | mNeoVis.2.1.1.chr9\_hap1.g00710.t01 |
| 11630 | 9 | 80718461 | 80765186 | mNeoVis.2.1.1.chr9\_hap1.g00710.t01 |
| 11631 | 9 | 80795907 | 80829591 | mNeoVis.2.1.1.chr9\_hap1.g00711.t01 |
| 11632 | 9 | 80795907 | 80829591 | mNeoVis.2.1.1.chr9\_hap1.g00711.t01 |
| 11633 | 9 | 80829729 | 80892086 | mNeoVis.2.1.1.chr9\_hap1.g00712.t01 |
| 11634 | 9 | 80829729 | 80892086 | mNeoVis.2.1.1.chr9\_hap1.g00712.t01 |
| 11635 | 9 | 80892158 | 80898849 | mNeoVis.2.1.1.chr9\_hap1.g00713.t01 |
| 11636 | 9 | 80892158 | 80898849 | mNeoVis.2.1.1.chr9\_hap1.g00713.t01 |
| 11637 | 9 | 80904093 | 81044453 | mNeoVis.2.1.1.chr9\_hap1.g00714.t01 |
| 11638 | 9 | 80904093 | 81044453 | mNeoVis.2.1.1.chr9\_hap1.g00714.t01 |
| 11639 | 9 | 81605828 | 81742638 | mNeoVis.2.1.1.chr9\_hap1.g00716.t01 |
| 11640 | 9 | 81780108 | 81784802 | mNeoVis.2.1.1.chr9\_hap1.g00717.t01 |
| 11641 | 9 | 81780108 | 81784802 | mNeoVis.2.1.1.chr9\_hap1.g00717.t01 |
| 11642 | 9 | 81860009 | 81896940 | mNeoVis.2.1.1.chr9\_hap1.g00718.t01 |
| 11643 | 9 | 81860009 | 81896940 | mNeoVis.2.1.1.chr9\_hap1.g00718.t01 |
| 11644 | 9 | 81972950 | 82472746 | mNeoVis.2.1.1.chr9\_hap1.g00719.t01 |
| 11645 | 9 | 82549938 | 82580179 | mNeoVis.2.1.1.chr9\_hap1.g00720.t01 |
| 11646 | 9 | 82549938 | 82580179 | mNeoVis.2.1.1.chr9\_hap1.g00720.t01 |
| 11647 | 9 | 82580448 | 82611303 | mNeoVis.2.1.1.chr9\_hap1.g00721.t01 |
| 11648 | 9 | 82580448 | 82611303 | mNeoVis.2.1.1.chr9\_hap1.g00721.t01 |
| 11649 | 9 | 82618861 | 82653755 | mNeoVis.2.1.1.chr9\_hap1.g00722.t01 |
| 11650 | 9 | 82618861 | 82653755 | mNeoVis.2.1.1.chr9\_hap1.g00722.t01 |
| 11651 | 9 | 82792208 | 82875179 | mNeoVis.2.1.1.chr9\_hap1.g00724.t01 |
| 11652 | 9 | 82884422 | 82982301 | mNeoVis.2.1.1.chr9\_hap1.g00725.t01 |
| 11653 | 9 | 83037103 | 83066581 | mNeoVis.2.1.1.chr9\_hap1.g00726.t01 |
| 11654 | 9 | 83037103 | 83066581 | mNeoVis.2.1.1.chr9\_hap1.g00726.t01 |
| 11655 | 9 | 83088849 | 83262949 | mNeoVis.2.1.1.chr9\_hap1.g00727.t01 |
| 11656 | 9 | 83088849 | 83262949 | mNeoVis.2.1.1.chr9\_hap1.g00727.t01 |
| 11657 | 9 | 83088849 | 83262949 | mNeoVis.2.1.1.chr9\_hap1.g00727.t01 |
| 11658 | 9 | 83168148 | 83170929 | mNeoVis.2.1.1.chr9\_hap1.g00728.t01 |
| 11659 | 9 | 83168148 | 83170929 | mNeoVis.2.1.1.chr9\_hap1.g00728.t01 |
| 11660 | 9 | 83168148 | 83170929 | mNeoVis.2.1.1.chr9\_hap1.g00728.t01 |
| 11661 | 9 | 83263090 | 83264382 | mNeoVis.2.1.1.chr9\_hap1.g00729.t01 |
| 11662 | 9 | 83263090 | 83264382 | mNeoVis.2.1.1.chr9\_hap1.g00729.t01 |
| 11663 | 9 | 83263090 | 83264382 | mNeoVis.2.1.1.chr9\_hap1.g00729.t01 |
| 11664 | 9 | 83270768 | 83313296 | mNeoVis.2.1.1.chr9\_hap1.g00730.t01 |
| 11665 | 9 | 83270768 | 83313296 | mNeoVis.2.1.1.chr9\_hap1.g00730.t01 |
| 11666 | 9 | 83270768 | 83313296 | mNeoVis.2.1.1.chr9\_hap1.g00730.t01 |
| 11667 | 9 | 83322116 | 83331753 | mNeoVis.2.1.1.chr9\_hap1.g00731.t01 |
| 11668 | 9 | 83322116 | 83331753 | mNeoVis.2.1.1.chr9\_hap1.g00731.t01 |
| 11669 | 9 | 83322116 | 83331753 | mNeoVis.2.1.1.chr9\_hap1.g00731.t01 |
| 11670 | 9 | 83336646 | 83356669 | mNeoVis.2.1.1.chr9\_hap1.g00732.t01 |
| 11671 | 9 | 83336646 | 83356669 | mNeoVis.2.1.1.chr9\_hap1.g00732.t01 |
| 11672 | 9 | 83336646 | 83356669 | mNeoVis.2.1.1.chr9\_hap1.g00732.t01 |
| 11673 | 9 | 83383820 | 83562465 | mNeoVis.2.1.1.chr9\_hap1.g00733.t01 |
| 11674 | 9 | 83383820 | 83562465 | mNeoVis.2.1.1.chr9\_hap1.g00733.t01 |
| 11675 | 9 | 83383820 | 83562465 | mNeoVis.2.1.1.chr9\_hap1.g00733.t01 |
| 11676 | 9 | 83499537 | 83500434 | mNeoVis.2.1.1.chr9\_hap1.g00734.t01 |
| 11677 | 9 | 83499537 | 83500434 | mNeoVis.2.1.1.chr9\_hap1.g00734.t01 |
| 11678 | 9 | 83499537 | 83500434 | mNeoVis.2.1.1.chr9\_hap1.g00734.t01 |
| 11679 | 9 | 83521092 | 83586790 | mNeoVis.2.1.1.chr9\_hap1.g00735.t01 |
| 11680 | 9 | 83521092 | 83586790 | mNeoVis.2.1.1.chr9\_hap1.g00735.t01 |
| 11681 | 9 | 83725092 | 83745681 | mNeoVis.2.1.1.chr9\_hap1.g00736.t01 |
| 11682 | 9 | 83725092 | 83745681 | mNeoVis.2.1.1.chr9\_hap1.g00736.t01 |
| 11683 | 9 | 83745024 | 84092370 | mNeoVis.2.1.1.chr9\_hap1.g00737.t01 |
| 11684 | 9 | 84557534 | 84699703 | mNeoVis.2.1.1.chr9\_hap1.g00738.t01 |
| 11685 | 9 | 84557534 | 84699703 | mNeoVis.2.1.1.chr9\_hap1.g00738.t01 |
| 11686 | 9 | 84611007 | 84611402 | mNeoVis.2.1.1.chr9\_hap1.g00739.t01 |
| 11687 | 9 | 84611007 | 84611402 | mNeoVis.2.1.1.chr9\_hap1.g00739.t01 |
| 11688 | 9 | 84702375 | 84741474 | mNeoVis.2.1.1.chr9\_hap1.g00740.t01 |
| 11689 | 9 | 84702375 | 84741474 | mNeoVis.2.1.1.chr9\_hap1.g00740.t01 |
| 11690 | 9 | 84753640 | 84809396 | mNeoVis.2.1.1.chr9\_hap1.g00741.t01 |
| 11691 | 9 | 84753640 | 84809396 | mNeoVis.2.1.1.chr9\_hap1.g00741.t01 |
| 11692 | 9 | 84826135 | 84833270 | mNeoVis.2.1.1.chr9\_hap1.g00742.t01 |
| 11693 | 9 | 84826135 | 84833270 | mNeoVis.2.1.1.chr9\_hap1.g00742.t01 |
| 11694 | 9 | 84835146 | 85096544 | mNeoVis.2.1.1.chr9\_hap1.g00743.t01 |
| 11695 | 9 | 85838241 | 85887440 | mNeoVis.2.1.1.chr9\_hap1.g00747.t01 |
| 11696 | 9 | 85950632 | 86017431 | mNeoVis.2.1.1.chr9\_hap1.g00748.t01 |
| 11697 | 9 | 86045068 | 86247173 | mNeoVis.2.1.1.chr9\_hap1.g00749.t01 |
| 11698 | 9 | 86658606 | 86687174 | mNeoVis.2.1.1.chr9\_hap1.g00754.t01 |
| 11699 | 9 | 86658606 | 86687174 | mNeoVis.2.1.1.chr9\_hap1.g00754.t01 |
| 11700 | 9 | 86711211 | 86830258 | mNeoVis.2.1.1.chr9\_hap1.g00755.t01 |
| 11701 | 9 | 86711211 | 86830258 | mNeoVis.2.1.1.chr9\_hap1.g00755.t01 |
| 11702 | 9 | 86725492 | 86771215 | mNeoVis.2.1.1.chr9\_hap1.g00756.t01 |
| 11703 | 9 | 86725492 | 86771215 | mNeoVis.2.1.1.chr9\_hap1.g00756.t01 |
| 11704 | 9 | 86846139 | 86892170 | mNeoVis.2.1.1.chr9\_hap1.g00757.t01 |
| 11705 | 9 | 86846139 | 86892170 | mNeoVis.2.1.1.chr9\_hap1.g00757.t01 |
| 11706 | 9 | 86925483 | 86985492 | mNeoVis.2.1.1.chr9\_hap1.g00758.t01 |
| 11707 | 9 | 86925483 | 86985492 | mNeoVis.2.1.1.chr9\_hap1.g00758.t01 |
| 11708 | 9 | 86995871 | 87317760 | mNeoVis.2.1.1.chr9\_hap1.g00759.t01 |
| 11709 | 9 | 86995871 | 87317760 | mNeoVis.2.1.1.chr9\_hap1.g00759.t01 |
| 11710 | 9 | 87479891 | 87532657 | mNeoVis.2.1.1.chr9\_hap1.g00760.t01 |
| 11711 | 9 | 88118224 | 88155157 | mNeoVis.2.1.1.chr9\_hap1.g00761.t01 |
| 11712 | 9 | 88118224 | 88155157 | mNeoVis.2.1.1.chr9\_hap1.g00761.t01 |
| 11713 | 9 | 88183552 | 88421478 | mNeoVis.2.1.1.chr9\_hap1.g00762.t01 |
| 11714 | 9 | 88183552 | 88421478 | mNeoVis.2.1.1.chr9\_hap1.g00762.t01 |
| 11715 | 9 | 88476492 | 88476770 | mNeoVis.2.1.1.chr9\_hap1.g00763.t01 |
| 11716 | 9 | 88476492 | 88476770 | mNeoVis.2.1.1.chr9\_hap1.g00763.t01 |
| 11717 | 9 | 88592345 | 88592519 | mNeoVis.2.1.1.chr9\_hap1.g00764.t01 |
| 11718 | 9 | 88752152 | 88811920 | mNeoVis.2.1.1.chr9\_hap1.g00765.t01 |
| 11719 | 9 | 88762320 | 88763385 | mNeoVis.2.1.1.chr9\_hap1.g00766.t01 |
| 11720 | 9 | 88838196 | 89203577 | mNeoVis.2.1.1.chr9\_hap1.g00767.t01 |
| 11721 | 9 | 89214570 | 89346387 | mNeoVis.2.1.1.chr9\_hap1.g00768.t01 |
| 11722 | 9 | 89214570 | 89346387 | mNeoVis.2.1.1.chr9\_hap1.g00768.t01 |
| 11723 | 9 | 89214570 | 89346387 | mNeoVis.2.1.1.chr9\_hap1.g00768.t01 |
| 11724 | 9 | 89398998 | 89400991 | mNeoVis.2.1.1.chr9\_hap1.g00769.t01 |
| 11725 | 9 | 89398998 | 89400991 | mNeoVis.2.1.1.chr9\_hap1.g00769.t01 |
| 11726 | 9 | 89398998 | 89400991 | mNeoVis.2.1.1.chr9\_hap1.g00769.t01 |
| 11727 | 9 | 89711042 | 89736122 | mNeoVis.2.1.1.chr9\_hap1.g00770.t01 |
| 11728 | 9 | 90277677 | 90383728 | mNeoVis.2.1.1.chr9\_hap1.g00771.t01 |
| 11729 | 9 | 90277677 | 90383728 | mNeoVis.2.1.1.chr9\_hap1.g00771.t01 |
| 11730 | 9 | 90277677 | 90383728 | mNeoVis.2.1.1.chr9\_hap1.g00771.t01 |
| 11731 | 9 | 90547406 | 90601493 | mNeoVis.2.1.1.chr9\_hap1.g00772.t01 |
| 11732 | 9 | 90547406 | 90601493 | mNeoVis.2.1.1.chr9\_hap1.g00772.t01 |
| 11733 | 9 | 90547406 | 90601493 | mNeoVis.2.1.1.chr9\_hap1.g00772.t01 |
| 11734 | 9 | 90766905 | 90788668 | mNeoVis.2.1.1.chr9\_hap1.g00773.t01 |
| 11735 | 9 | 90766905 | 90788668 | mNeoVis.2.1.1.chr9\_hap1.g00773.t01 |
| 11736 | 9 | 90766905 | 90788668 | mNeoVis.2.1.1.chr9\_hap1.g00773.t01 |
| 11737 | 9 | 90849376 | 90927054 | mNeoVis.2.1.1.chr9\_hap1.g00774.t01 |
| 11738 | 9 | 90849376 | 90927054 | mNeoVis.2.1.1.chr9\_hap1.g00774.t01 |
| 11739 | 9 | 90849376 | 90927054 | mNeoVis.2.1.1.chr9\_hap1.g00774.t01 |
| 11740 | 9 | 90952746 | 90953442 | mNeoVis.2.1.1.chr9\_hap1.g00775.t01 |
| 11741 | 9 | 90952746 | 90953442 | mNeoVis.2.1.1.chr9\_hap1.g00775.t01 |
| 11742 | 9 | 90952746 | 90953442 | mNeoVis.2.1.1.chr9\_hap1.g00775.t01 |
| 11743 | 9 | 91075861 | 91191877 | mNeoVis.2.1.1.chr9\_hap1.g00776.t01 |
| 11744 | 9 | 91075861 | 91191877 | mNeoVis.2.1.1.chr9\_hap1.g00776.t01 |
| 11745 | 9 | 91195074 | 91212447 | mNeoVis.2.1.1.chr9\_hap1.g00777.t01 |
| 11746 | 9 | 91195074 | 91212447 | mNeoVis.2.1.1.chr9\_hap1.g00777.t01 |
| 11747 | 9 | 91291392 | 91474643 | mNeoVis.2.1.1.chr9\_hap1.g00778.t01 |
| 11748 | 9 | 91291392 | 91474643 | mNeoVis.2.1.1.chr9\_hap1.g00778.t01 |
| 11749 | 9 | 91291392 | 91474643 | mNeoVis.2.1.1.chr9\_hap1.g00778.t01 |
| 11750 | 9 | 91487857 | 91599968 | mNeoVis.2.1.1.chr9\_hap1.g00779.t01 |
| 11751 | 9 | 91487857 | 91599968 | mNeoVis.2.1.1.chr9\_hap1.g00779.t01 |
| 11752 | 9 | 91487857 | 91599968 | mNeoVis.2.1.1.chr9\_hap1.g00779.t01 |
| 11753 | 9 | 91818133 | 91936122 | mNeoVis.2.1.1.chr9\_hap1.g00780.t01 |
| 11754 | 9 | 91818133 | 91936122 | mNeoVis.2.1.1.chr9\_hap1.g00780.t01 |
| 11755 | 9 | 91818133 | 91936122 | mNeoVis.2.1.1.chr9\_hap1.g00780.t01 |
| 11756 | 9 | 92052878 | 92070017 | mNeoVis.2.1.1.chr9\_hap1.g00781.t01 |
| 11757 | 9 | 92075849 | 92096504 | mNeoVis.2.1.1.chr9\_hap1.g00782.t01 |
| 11758 | 9 | 92096559 | 92115752 | mNeoVis.2.1.1.chr9\_hap1.g00783.t01 |
| 11759 | 9 | 92150772 | 92154136 | mNeoVis.2.1.1.chr9\_hap1.g00784.t01 |
| 11760 | 9 | 92201060 | 92214559 | mNeoVis.2.1.1.chr9\_hap1.g00785.t01 |
| 11761 | 9 | 92229906 | 92244200 | mNeoVis.2.1.1.chr9\_hap1.g00786.t01 |
| 11762 | 9 | 92233253 | 92242298 | mNeoVis.2.1.1.chr9\_hap1.g00787.t01 |
| 11763 | 9 | 92244890 | 92260698 | mNeoVis.2.1.1.chr9\_hap1.g00788.t01 |
| 11764 | 9 | 92272711 | 92275069 | mNeoVis.2.1.1.chr9\_hap1.g00789.t01 |
| 11765 | 9 | 92278499 | 92279221 | mNeoVis.2.1.1.chr9\_hap1.g00791.t01 |
| 11766 | 9 | 92454843 | 92509462 | mNeoVis.2.1.1.chr9\_hap1.g00792.t01 |
| 11767 | 9 | 92454843 | 92509462 | mNeoVis.2.1.1.chr9\_hap1.g00792.t01 |
| 11768 | 9 | 92477147 | 92477682 | mNeoVis.2.1.1.chr9\_hap1.g00793.t01 |
| 11769 | 9 | 92477147 | 92477682 | mNeoVis.2.1.1.chr9\_hap1.g00793.t01 |
| 11770 | 9 | 92515016 | 92604663 | mNeoVis.2.1.1.chr9\_hap1.g00794.t01 |
| 11771 | 9 | 92515016 | 92604663 | mNeoVis.2.1.1.chr9\_hap1.g00794.t01 |
| 11772 | 9 | 92901458 | 93308629 | mNeoVis.2.1.1.chr9\_hap1.g00795.t01 |
| 11773 | 9 | 92901458 | 93308629 | mNeoVis.2.1.1.chr9\_hap1.g00795.t01 |
| 11774 | 9 | 95017422 | 95036709 | mNeoVis.2.1.1.chr9\_hap1.g00814.t01 |
| 11775 | 9 | 95017422 | 95036709 | mNeoVis.2.1.1.chr9\_hap1.g00814.t01 |
| 11776 | 9 | 95140672 | 95158195 | mNeoVis.2.1.1.chr9\_hap1.g00815.t01 |
| 11777 | 9 | 95140672 | 95158195 | mNeoVis.2.1.1.chr9\_hap1.g00815.t01 |
| 11778 | 9 | 95181629 | 95182128 | mNeoVis.2.1.1.chr9\_hap1.g00816.t01 |
| 11779 | 9 | 95181629 | 95182128 | mNeoVis.2.1.1.chr9\_hap1.g00816.t01 |
| 11780 | 9 | 95187970 | 95188941 | mNeoVis.2.1.1.chr9\_hap1.g00817.t01 |
| 11781 | 9 | 95187970 | 95188941 | mNeoVis.2.1.1.chr9\_hap1.g00817.t01 |
| 11782 | 9 | 95196061 | 95196996 | mNeoVis.2.1.1.chr9\_hap1.g00818.t01 |
| 11783 | 9 | 95196061 | 95196996 | mNeoVis.2.1.1.chr9\_hap1.g00818.t01 |
| 11784 | 9 | 95206192 | 95208359 | mNeoVis.2.1.1.chr9\_hap1.g00819.t01 |
| 11785 | 9 | 95206192 | 95208359 | mNeoVis.2.1.1.chr9\_hap1.g00819.t01 |
| 11786 | 9 | 95235662 | 95236656 | mNeoVis.2.1.1.chr9\_hap1.g00820.t01 |
| 11787 | 9 | 95235662 | 95236656 | mNeoVis.2.1.1.chr9\_hap1.g00820.t01 |
| 11788 | 9 | 95256655 | 95257578 | mNeoVis.2.1.1.chr9\_hap1.g00821.t01 |
| 11789 | 9 | 95256655 | 95257578 | mNeoVis.2.1.1.chr9\_hap1.g00821.t01 |
| 11790 | 9 | 95274401 | 95275357 | mNeoVis.2.1.1.chr9\_hap1.g00822.t01 |
| 11791 | 9 | 95274401 | 95275357 | mNeoVis.2.1.1.chr9\_hap1.g00822.t01 |
| 11792 | 9 | 95295083 | 95296054 | mNeoVis.2.1.1.chr9\_hap1.g00823.t01 |
| 11793 | 9 | 95295083 | 95296054 | mNeoVis.2.1.1.chr9\_hap1.g00823.t01 |
| 11794 | 9 | 95306767 | 95307762 | mNeoVis.2.1.1.chr9\_hap1.g00824.t01 |
| 11795 | 9 | 95306767 | 95307762 | mNeoVis.2.1.1.chr9\_hap1.g00824.t01 |
| 11796 | 9 | 95309576 | 95310498 | mNeoVis.2.1.1.chr9\_hap1.g00825.t01 |
| 11797 | 9 | 95309576 | 95310498 | mNeoVis.2.1.1.chr9\_hap1.g00825.t01 |
| 11798 | 9 | 95314680 | 95315618 | mNeoVis.2.1.1.chr9\_hap1.g00826.t01 |
| 11799 | 9 | 95314680 | 95315618 | mNeoVis.2.1.1.chr9\_hap1.g00826.t01 |
| 11800 | 9 | 96237285 | 96246385 | mNeoVis.2.1.1.chr9\_hap1.g00842.t01 |
| 11801 | 9 | 96237285 | 96246385 | mNeoVis.2.1.1.chr9\_hap1.g00842.t01 |
| 11802 | 9 | 96265396 | 96275589 | mNeoVis.2.1.1.chr9\_hap1.g00843.t01 |
| 11803 | 9 | 96265396 | 96275589 | mNeoVis.2.1.1.chr9\_hap1.g00843.t01 |
| 11804 | 9 | 96281029 | 96290498 | mNeoVis.2.1.1.chr9\_hap1.g00844.t01 |
| 11805 | 9 | 96281029 | 96290498 | mNeoVis.2.1.1.chr9\_hap1.g00844.t01 |
| 11806 | 9 | 96310847 | 96331128 | mNeoVis.2.1.1.chr9\_hap1.g00845.t01 |
| 11807 | 9 | 96310847 | 96331128 | mNeoVis.2.1.1.chr9\_hap1.g00845.t01 |
| 11808 | 9 | 96335333 | 96359690 | mNeoVis.2.1.1.chr9\_hap1.g00846.t01 |
| 11809 | 9 | 96335333 | 96359690 | mNeoVis.2.1.1.chr9\_hap1.g00846.t01 |
| 11810 | 9 | 96362074 | 96376999 | mNeoVis.2.1.1.chr9\_hap1.g00847.t01 |
| 11811 | 9 | 96362074 | 96376999 | mNeoVis.2.1.1.chr9\_hap1.g00847.t01 |
| 11812 | 9 | 96378689 | 96388219 | mNeoVis.2.1.1.chr9\_hap1.g00848.t01 |
| 11813 | 9 | 96378689 | 96388219 | mNeoVis.2.1.1.chr9\_hap1.g00848.t01 |
| 11814 | 9 | 96396753 | 96405042 | mNeoVis.2.1.1.chr9\_hap1.g00849.t01 |
| 11815 | 9 | 96396753 | 96405042 | mNeoVis.2.1.1.chr9\_hap1.g00849.t01 |
| 11816 | 9 | 96430460 | 96454693 | mNeoVis.2.1.1.chr9\_hap1.g00850.t01 |
| 11817 | 9 | 96430460 | 96454693 | mNeoVis.2.1.1.chr9\_hap1.g00850.t01 |
| 11818 | 9 | 96475412 | 96484898 | mNeoVis.2.1.1.chr9\_hap1.g00851.t01 |
| 11819 | 9 | 96475412 | 96484898 | mNeoVis.2.1.1.chr9\_hap1.g00851.t01 |
| 11820 | 9 | 96494899 | 96503531 | mNeoVis.2.1.1.chr9\_hap1.g00852.t01 |
| 11821 | 9 | 96494899 | 96503531 | mNeoVis.2.1.1.chr9\_hap1.g00852.t01 |
| 11822 | 9 | 96505398 | 96523404 | mNeoVis.2.1.1.chr9\_hap1.g00853.t01 |
| 11823 | 9 | 96505398 | 96523404 | mNeoVis.2.1.1.chr9\_hap1.g00853.t01 |
| 11824 | 9 | 96506894 | 96507365 | mNeoVis.2.1.1.chr9\_hap1.g00854.t01 |
| 11825 | 9 | 96506894 | 96507365 | mNeoVis.2.1.1.chr9\_hap1.g00854.t01 |
| 11826 | 9 | 96523650 | 96537116 | mNeoVis.2.1.1.chr9\_hap1.g00855.t01 |
| 11827 | 9 | 96523650 | 96537116 | mNeoVis.2.1.1.chr9\_hap1.g00855.t01 |
| 11828 | 9 | 96545073 | 96577128 | mNeoVis.2.1.1.chr9\_hap1.g00856.t01 |
| 11829 | 9 | 96545073 | 96577128 | mNeoVis.2.1.1.chr9\_hap1.g00856.t01 |
| 11830 | 9 | 96577272 | 96590297 | mNeoVis.2.1.1.chr9\_hap1.g00857.t01 |
| 11831 | 9 | 96577272 | 96590297 | mNeoVis.2.1.1.chr9\_hap1.g00857.t01 |
| 11832 | 9 | 96622621 | 96632073 | mNeoVis.2.1.1.chr9\_hap1.g00858.t01 |
| 11833 | 9 | 96622621 | 96632073 | mNeoVis.2.1.1.chr9\_hap1.g00858.t01 |
| 11834 | 9 | 96660327 | 96680960 | mNeoVis.2.1.1.chr9\_hap1.g00859.t01 |
| 11835 | 9 | 96660327 | 96680960 | mNeoVis.2.1.1.chr9\_hap1.g00859.t01 |
| 11836 | 9 | 96714732 | 96727007 | mNeoVis.2.1.1.chr9\_hap1.g00860.t01 |
| 11837 | 9 | 96726909 | 96763472 | mNeoVis.2.1.1.chr9\_hap1.g00861.t01 |
| 11838 | 9 | 96789214 | 96799051 | mNeoVis.2.1.1.chr9\_hap1.g00862.t01 |
| 11839 | 9 | 96831956 | 96877038 | mNeoVis.2.1.1.chr9\_hap1.g00863.t01 |
| 11840 | 9 | 96914572 | 96921791 | mNeoVis.2.1.1.chr9\_hap1.g00864.t01 |
| 11841 | 9 | 97286018 | 97326505 | mNeoVis.2.1.1.chr9\_hap1.g00870.t01 |
| 11842 | 9 | 97326139 | 97396334 | mNeoVis.2.1.1.chr9\_hap1.g00871.t01 |
| 11843 | 9 | 97402674 | 97412638 | mNeoVis.2.1.1.chr9\_hap1.g00872.t01 |
| 11844 | 9 | 97413359 | 97433041 | mNeoVis.2.1.1.chr9\_hap1.g00873.t01 |
| 11845 | 9 | 97454691 | 97496470 | mNeoVis.2.1.1.chr9\_hap1.g00874.t01 |
| 11846 | 9 | 97543408 | 97699169 | mNeoVis.2.1.1.chr9\_hap1.g00875.t01 |
| 11847 | 9 | 97633029 | 97647365 | mNeoVis.2.1.1.chr9\_hap1.g00876.t01 |
| 11848 | 9 | 97668619 | 97679036 | mNeoVis.2.1.1.chr9\_hap1.g00877.t01 |
| 11849 | 9 | 97709834 | 97718336 | mNeoVis.2.1.1.chr9\_hap1.g00878.t01 |
| 11850 | 9 | 97735256 | 97772624 | mNeoVis.2.1.1.chr9\_hap1.g00879.t01 |
| 11851 | 9 | 97772674 | 97777935 | mNeoVis.2.1.1.chr9\_hap1.g00880.t01 |
| 11852 | 9 | 97879741 | 97880262 | mNeoVis.2.1.1.chr9\_hap1.g00881.t01 |
| 11853 | 9 | 98245611 | 98303974 | mNeoVis.2.1.1.chr9\_hap1.g00886.t01 |
| 11854 | 9 | 98323634 | 98407321 | mNeoVis.2.1.1.chr9\_hap1.g00887.t01 |
| 11855 | 9 | 98323634 | 98407321 | mNeoVis.2.1.1.chr9\_hap1.g00887.t01 |
| 11856 | 9 | 98554606 | 98622500 | mNeoVis.2.1.1.chr9\_hap1.g00888.t01 |
| 11857 | 9 | 98554606 | 98622500 | mNeoVis.2.1.1.chr9\_hap1.g00888.t01 |
| 11858 | 9 | 98554606 | 98622500 | mNeoVis.2.1.1.chr9\_hap1.g00888.t01 |
| 11859 | 9 | 98684509 | 98762989 | mNeoVis.2.1.1.chr9\_hap1.g00889.t01 |
| 11860 | 9 | 98684509 | 98762989 | mNeoVis.2.1.1.chr9\_hap1.g00889.t01 |
| 11861 | 9 | 98684509 | 98762989 | mNeoVis.2.1.1.chr9\_hap1.g00889.t01 |
| 11862 | 9 | 98684509 | 98762989 | mNeoVis.2.1.1.chr9\_hap1.g00889.t01 |
| 11863 | 9 | 98774069 | 98937021 | mNeoVis.2.1.1.chr9\_hap1.g00890.t01 |
| 11864 | 9 | 98774069 | 98937021 | mNeoVis.2.1.1.chr9\_hap1.g00890.t01 |
| 11865 | 9 | 98774069 | 98937021 | mNeoVis.2.1.1.chr9\_hap1.g00890.t01 |
| 11866 | 9 | 98774069 | 98937021 | mNeoVis.2.1.1.chr9\_hap1.g00890.t01 |
| 11867 | 9 | 99242353 | 99331602 | mNeoVis.2.1.1.chr9\_hap1.g00891.t01 |
| 11868 | 9 | 99242353 | 99331602 | mNeoVis.2.1.1.chr9\_hap1.g00891.t01 |
| 11869 | 9 | 99242353 | 99331602 | mNeoVis.2.1.1.chr9\_hap1.g00891.t01 |
| 11870 | 9 | 99527051 | 99532854 | mNeoVis.2.1.1.chr9\_hap1.g00892.t01 |
| 11871 | 9 | 99527051 | 99532854 | mNeoVis.2.1.1.chr9\_hap1.g00892.t01 |
| 11872 | 9 | 99527051 | 99532854 | mNeoVis.2.1.1.chr9\_hap1.g00892.t01 |
| 11873 | 9 | 99527051 | 99532854 | mNeoVis.2.1.1.chr9\_hap1.g00892.t01 |
| 11874 | 9 | 99538052 | 99565719 | mNeoVis.2.1.1.chr9\_hap1.g00893.t01 |
| 11875 | 9 | 99538052 | 99565719 | mNeoVis.2.1.1.chr9\_hap1.g00893.t01 |
| 11876 | 9 | 99538052 | 99565719 | mNeoVis.2.1.1.chr9\_hap1.g00893.t01 |
| 11877 | 9 | 99538052 | 99565719 | mNeoVis.2.1.1.chr9\_hap1.g00893.t01 |
| 11878 | 9 | 99578612 | 99625748 | mNeoVis.2.1.1.chr9\_hap1.g00894.t01 |
| 11879 | 9 | 99578612 | 99625748 | mNeoVis.2.1.1.chr9\_hap1.g00894.t01 |
| 11880 | 9 | 99578612 | 99625748 | mNeoVis.2.1.1.chr9\_hap1.g00894.t01 |
| 11881 | 9 | 99578612 | 99625748 | mNeoVis.2.1.1.chr9\_hap1.g00894.t01 |
| 11882 | 9 | 99662869 | 99693261 | mNeoVis.2.1.1.chr9\_hap1.g00895.t01 |
| 11883 | 9 | 99662869 | 99693261 | mNeoVis.2.1.1.chr9\_hap1.g00895.t01 |
| 11884 | 9 | 99662869 | 99693261 | mNeoVis.2.1.1.chr9\_hap1.g00895.t01 |
| 11885 | 9 | 99662869 | 99693261 | mNeoVis.2.1.1.chr9\_hap1.g00895.t01 |
| 11886 | 9 | 99693721 | 99939218 | mNeoVis.2.1.1.chr9\_hap1.g00896.t01 |
| 11887 | 9 | 99693721 | 99939218 | mNeoVis.2.1.1.chr9\_hap1.g00896.t01 |
| 11888 | 9 | 100578416 | 100620402 | mNeoVis.2.1.1.chr9\_hap1.g00897.t01 |
| 11889 | 9 | 100578416 | 100620402 | mNeoVis.2.1.1.chr9\_hap1.g00897.t01 |
| 11890 | 9 | 100620936 | 100783055 | mNeoVis.2.1.1.chr9\_hap1.g00898.t01 |
| 11891 | 9 | 100646665 | 100685076 | mNeoVis.2.1.1.chr9\_hap1.g00899.t01 |
| 11892 | 9 | 100646665 | 100685076 | mNeoVis.2.1.1.chr9\_hap1.g00899.t01 |
| 11893 | 9 | 100699559 | 100701785 | mNeoVis.2.1.1.chr9\_hap1.g00900.t01 |
| 11894 | 9 | 100699559 | 100701785 | mNeoVis.2.1.1.chr9\_hap1.g00900.t01 |
| 11895 | 9 | 100919345 | 100933123 | mNeoVis.2.1.1.chr9\_hap1.g00901.t01 |
| 11896 | 9 | 101363856 | 101368805 | mNeoVis.2.1.1.chr9\_hap1.g00902.t01 |
| 11897 | 9 | 101452467 | 101456926 | mNeoVis.2.1.1.chr9\_hap1.g00904.t01 |
| 11898 | 9 | 101489017 | 101522596 | mNeoVis.2.1.1.chr9\_hap1.g00905.t01 |
| 11899 | 9 | 101749816 | 101793951 | mNeoVis.2.1.1.chr9\_hap1.g00906.t01 |
| 11900 | 9 | 101813213 | 101858010 | mNeoVis.2.1.1.chr9\_hap1.g00907.t01 |
| 11901 | 9 | 101865478 | 101881243 | mNeoVis.2.1.1.chr9\_hap1.g00908.t01 |
| 11902 | 9 | 101914019 | 101943516 | mNeoVis.2.1.1.chr9\_hap1.g00909.t01 |
| 11903 | 9 | 101914019 | 101943516 | mNeoVis.2.1.1.chr9\_hap1.g00909.t01 |
| 11904 | 9 | 101946763 | 102022991 | mNeoVis.2.1.1.chr9\_hap1.g00910.t01 |
| 11905 | 9 | 101946763 | 102022991 | mNeoVis.2.1.1.chr9\_hap1.g00910.t01 |
| 11906 | 9 | 102242983 | 102277435 | mNeoVis.2.1.1.chr9\_hap1.g00911.t01 |
| 11907 | 9 | 102242983 | 102277435 | mNeoVis.2.1.1.chr9\_hap1.g00911.t01 |
| 11908 | 9 | 102242983 | 102277435 | mNeoVis.2.1.1.chr9\_hap1.g00911.t01 |
| 11909 | 9 | 102344578 | 102349752 | mNeoVis.2.1.1.chr9\_hap1.g00912.t01 |
| 11910 | 9 | 102344578 | 102349752 | mNeoVis.2.1.1.chr9\_hap1.g00912.t01 |
| 11911 | 9 | 102344578 | 102349752 | mNeoVis.2.1.1.chr9\_hap1.g00912.t01 |
| 11912 | 9 | 102362958 | 102383309 | mNeoVis.2.1.1.chr9\_hap1.g00913.t01 |
| 11913 | 9 | 102362958 | 102383309 | mNeoVis.2.1.1.chr9\_hap1.g00913.t01 |
| 11914 | 9 | 102362958 | 102383309 | mNeoVis.2.1.1.chr9\_hap1.g00913.t01 |
| 11915 | 9 | 102445368 | 102451512 | mNeoVis.2.1.1.chr9\_hap1.g00914.t01 |
| 11916 | 9 | 102445368 | 102451512 | mNeoVis.2.1.1.chr9\_hap1.g00914.t01 |
| 11917 | 9 | 102445368 | 102451512 | mNeoVis.2.1.1.chr9\_hap1.g00914.t01 |
| 11918 | 9 | 102543964 | 102572950 | mNeoVis.2.1.1.chr9\_hap1.g00915.t01 |
| 11919 | 9 | 102543964 | 102572950 | mNeoVis.2.1.1.chr9\_hap1.g00915.t01 |
| 11920 | 9 | 102543964 | 102572950 | mNeoVis.2.1.1.chr9\_hap1.g00915.t01 |
| 11921 | 9 | 102576624 | 102600120 | mNeoVis.2.1.1.chr9\_hap1.g00916.t01 |
| 11922 | 9 | 102576624 | 102600120 | mNeoVis.2.1.1.chr9\_hap1.g00916.t01 |
| 11923 | 9 | 102576624 | 102600120 | mNeoVis.2.1.1.chr9\_hap1.g00916.t01 |
| 11924 | 9 | 102598707 | 102599660 | mNeoVis.2.1.1.chr9\_hap1.g00917.t01 |
| 11925 | 9 | 102598707 | 102599660 | mNeoVis.2.1.1.chr9\_hap1.g00917.t01 |
| 11926 | 9 | 102598707 | 102599660 | mNeoVis.2.1.1.chr9\_hap1.g00917.t01 |
| 11927 | 9 | 102617876 | 102696772 | mNeoVis.2.1.1.chr9\_hap1.g00918.t01 |
| 11928 | 9 | 102617876 | 102696772 | mNeoVis.2.1.1.chr9\_hap1.g00918.t01 |
| 11929 | 9 | 102617876 | 102696772 | mNeoVis.2.1.1.chr9\_hap1.g00918.t01 |
| 11930 | 9 | 102708622 | 102744714 | mNeoVis.2.1.1.chr9\_hap1.g00919.t01 |
| 11931 | 9 | 102708622 | 102744714 | mNeoVis.2.1.1.chr9\_hap1.g00919.t01 |
| 11932 | 9 | 102708622 | 102744714 | mNeoVis.2.1.1.chr9\_hap1.g00919.t01 |
| 11933 | 9 | 102755517 | 102880244 | mNeoVis.2.1.1.chr9\_hap1.g00920.t01 |
| 11934 | 9 | 102755517 | 102880244 | mNeoVis.2.1.1.chr9\_hap1.g00920.t01 |
| 11935 | 9 | 102755517 | 102880244 | mNeoVis.2.1.1.chr9\_hap1.g00920.t01 |
| 11936 | 9 | 102889510 | 103091041 | mNeoVis.2.1.1.chr9\_hap1.g00921.t01 |
| 11937 | 9 | 102889510 | 103091041 | mNeoVis.2.1.1.chr9\_hap1.g00921.t01 |
| 11938 | 9 | 102889510 | 103091041 | mNeoVis.2.1.1.chr9\_hap1.g00921.t01 |
| 11939 | 9 | 103091029 | 103210110 | mNeoVis.2.1.1.chr9\_hap1.g00922.t01 |
| 11940 | 9 | 103091029 | 103210110 | mNeoVis.2.1.1.chr9\_hap1.g00922.t01 |
| 11941 | 9 | 103223115 | 103282007 | mNeoVis.2.1.1.chr9\_hap1.g00923.t01 |
| 11942 | 9 | 103223115 | 103282007 | mNeoVis.2.1.1.chr9\_hap1.g00923.t01 |
| 11943 | 9 | 103305587 | 103422116 | mNeoVis.2.1.1.chr9\_hap1.g00924.t01 |
| 11944 | 9 | 103305587 | 103422116 | mNeoVis.2.1.1.chr9\_hap1.g00924.t01 |
| 11945 | 9 | 103422732 | 103657878 | mNeoVis.2.1.1.chr9\_hap1.g00925.t01 |
| 11946 | 9 | 103422732 | 103657878 | mNeoVis.2.1.1.chr9\_hap1.g00925.t01 |
| 11947 | 9 | 103830745 | 103917385 | mNeoVis.2.1.1.chr9\_hap1.g00926.t01 |
| 11948 | 9 | 103830745 | 103917385 | mNeoVis.2.1.1.chr9\_hap1.g00926.t01 |
| 11949 | 9 | 103830745 | 103917385 | mNeoVis.2.1.1.chr9\_hap1.g00926.t01 |
| 11950 | 9 | 104062719 | 104188381 | mNeoVis.2.1.1.chr9\_hap1.g00927.t01 |
| 11951 | 9 | 104062719 | 104188381 | mNeoVis.2.1.1.chr9\_hap1.g00927.t01 |
| 11952 | 9 | 104209896 | 104261714 | mNeoVis.2.1.1.chr9\_hap1.g00928.t01 |
| 11953 | 9 | 104209896 | 104261714 | mNeoVis.2.1.1.chr9\_hap1.g00928.t01 |
| 11954 | 9 | 104261856 | 104272044 | mNeoVis.2.1.1.chr9\_hap1.g00929.t01 |
| 11955 | 9 | 104261856 | 104272044 | mNeoVis.2.1.1.chr9\_hap1.g00929.t01 |
| 11956 | 9 | 104278234 | 104297516 | mNeoVis.2.1.1.chr9\_hap1.g00930.t01 |
| 11957 | 9 | 104278234 | 104297516 | mNeoVis.2.1.1.chr9\_hap1.g00930.t01 |
| 11958 | 9 | 104310943 | 104312013 | mNeoVis.2.1.1.chr9\_hap1.g00931.t01 |
| 11959 | 9 | 104310943 | 104312013 | mNeoVis.2.1.1.chr9\_hap1.g00931.t01 |
| 11960 | 9 | 104332583 | 104367178 | mNeoVis.2.1.1.chr9\_hap1.g00932.t01 |
| 11961 | 9 | 104332583 | 104367178 | mNeoVis.2.1.1.chr9\_hap1.g00932.t01 |
| 11962 | 9 | 104409027 | 104482990 | mNeoVis.2.1.1.chr9\_hap1.g00933.t01 |
| 11963 | 9 | 104409027 | 104482990 | mNeoVis.2.1.1.chr9\_hap1.g00933.t01 |
| 11964 | 9 | 104483961 | 104604518 | mNeoVis.2.1.1.chr9\_hap1.g00934.t01 |
| 11965 | 9 | 104483961 | 104604518 | mNeoVis.2.1.1.chr9\_hap1.g00934.t01 |
| 11966 | 9 | 104823789 | 105200888 | mNeoVis.2.1.1.chr9\_hap1.g00935.t01 |
| 11967 | 9 | 105216337 | 105315734 | mNeoVis.2.1.1.chr9\_hap1.g00936.t01 |
| 11968 | 9 | 105216337 | 105315734 | mNeoVis.2.1.1.chr9\_hap1.g00936.t01 |
| 11969 | 9 | 106615491 | 106619185 | mNeoVis.2.1.1.chr9\_hap1.g00937.t01 |
| 11970 | 9 | 106615491 | 106619185 | mNeoVis.2.1.1.chr9\_hap1.g00937.t01 |
| 11971 | 9 | 106615491 | 106619185 | mNeoVis.2.1.1.chr9\_hap1.g00937.t01 |
| 11972 | 9 | 106829118 | 106851780 | mNeoVis.2.1.1.chr9\_hap1.g00938.t01 |
| 11973 | 9 | 106829118 | 106851780 | mNeoVis.2.1.1.chr9\_hap1.g00938.t01 |
| 11974 | 9 | 106829118 | 106851780 | mNeoVis.2.1.1.chr9\_hap1.g00938.t01 |
| 11975 | 9 | 106949127 | 107136204 | mNeoVis.2.1.1.chr9\_hap1.g00939.t01 |
| 11976 | 9 | 106949127 | 107136204 | mNeoVis.2.1.1.chr9\_hap1.g00939.t01 |
| 11977 | 9 | 107183226 | 107232640 | mNeoVis.2.1.1.chr9\_hap1.g00940.t01 |
| 11978 | 9 | 107183226 | 107232640 | mNeoVis.2.1.1.chr9\_hap1.g00940.t01 |
| 11979 | 9 | 107239710 | 107267154 | mNeoVis.2.1.1.chr9\_hap1.g00941.t01 |
| 11980 | 9 | 107239710 | 107267154 | mNeoVis.2.1.1.chr9\_hap1.g00941.t01 |
| 11981 | 9 | 107273838 | 107305250 | mNeoVis.2.1.1.chr9\_hap1.g00942.t01 |
| 11982 | 9 | 107317092 | 107362088 | mNeoVis.2.1.1.chr9\_hap1.g00943.t01 |
| 11983 | 9 | 107358723 | 107376344 | mNeoVis.2.1.1.chr9\_hap1.g00944.t01 |
| 11984 | 9 | 107358723 | 107376344 | mNeoVis.2.1.1.chr9\_hap1.g00944.t01 |
| 11985 | 9 | 107778407 | 107830094 | mNeoVis.2.1.1.chr9\_hap1.g00945.t01 |
| 11986 | 9 | 107778407 | 107830094 | mNeoVis.2.1.1.chr9\_hap1.g00945.t01 |
| 11987 | 9 | 107837634 | 107871968 | mNeoVis.2.1.1.chr9\_hap1.g00946.t01 |
| 11988 | 9 | 107837634 | 107871968 | mNeoVis.2.1.1.chr9\_hap1.g00946.t01 |
| 11989 | 9 | 108051467 | 108056056 | mNeoVis.2.1.1.chr9\_hap1.g00947.t01 |
| 11990 | 9 | 108057023 | 108214374 | mNeoVis.2.1.1.chr9\_hap1.g00948.t01 |
| 11991 | 9 | 108365960 | 108484114 | mNeoVis.2.1.1.chr9\_hap1.g00949.t01 |
| 11992 | 9 | 108458685 | 108477197 | mNeoVis.2.1.1.chr9\_hap1.g00950.t01 |
| 11993 | 9 | 108606184 | 108645943 | mNeoVis.2.1.1.chr9\_hap1.g00951.t01 |
| 11994 | 9 | 108647603 | 108988435 | mNeoVis.2.1.1.chr9\_hap1.g00952.t01 |
| 11995 | 9 | 109389108 | 109774319 | mNeoVis.2.1.1.chr9\_hap1.g00953.t01 |
| 11996 | 9 | 110405442 | 110948381 | mNeoVis.2.1.1.chr9\_hap1.g00954.t01 |
| 11997 | 9 | 110405442 | 110948381 | mNeoVis.2.1.1.chr9\_hap1.g00954.t01 |
| 11998 | 9 | 110405442 | 110948381 | mNeoVis.2.1.1.chr9\_hap1.g00954.t01 |
| 11999 | 9 | 111019753 | 111134801 | mNeoVis.2.1.1.chr9\_hap1.g00955.t01 |
| 12000 | 9 | 111019753 | 111134801 | mNeoVis.2.1.1.chr9\_hap1.g00955.t01 |
| 12001 | 9 | 111019753 | 111134801 | mNeoVis.2.1.1.chr9\_hap1.g00955.t01 |
| 12002 | 9 | 111019753 | 111134801 | mNeoVis.2.1.1.chr9\_hap1.g00955.t01 |
| 12003 | 9 | 111191329 | 111340915 | mNeoVis.2.1.1.chr9\_hap1.g00956.t01 |
| 12004 | 9 | 111191329 | 111340915 | mNeoVis.2.1.1.chr9\_hap1.g00956.t01 |
| 12005 | 9 | 111191329 | 111340915 | mNeoVis.2.1.1.chr9\_hap1.g00956.t01 |
| 12006 | 9 | 111592227 | 111722401 | mNeoVis.2.1.1.chr9\_hap1.g00957.t01 |
| 12007 | 9 | 111592227 | 111722401 | mNeoVis.2.1.1.chr9\_hap1.g00957.t01 |
| 12008 | 9 | 111749820 | 111986029 | mNeoVis.2.1.1.chr9\_hap1.g00958.t01 |
| 12009 | 9 | 111749820 | 111986029 | mNeoVis.2.1.1.chr9\_hap1.g00958.t01 |
| 12010 | 9 | 112009842 | 112011711 | mNeoVis.2.1.1.chr9\_hap1.g00959.t01 |
| 12011 | 9 | 112009842 | 112011711 | mNeoVis.2.1.1.chr9\_hap1.g00959.t01 |
| 12012 | 9 | 112009842 | 112011711 | mNeoVis.2.1.1.chr9\_hap1.g00959.t01 |
| 12013 | 9 | 112019277 | 112021856 | mNeoVis.2.1.1.chr9\_hap1.g00960.t01 |
| 12014 | 9 | 112019277 | 112021856 | mNeoVis.2.1.1.chr9\_hap1.g00960.t01 |
| 12015 | 9 | 112019277 | 112021856 | mNeoVis.2.1.1.chr9\_hap1.g00960.t01 |
| 12016 | 9 | 112072099 | 112202374 | mNeoVis.2.1.1.chr9\_hap1.g00961.t01 |
| 12017 | 9 | 112072099 | 112202374 | mNeoVis.2.1.1.chr9\_hap1.g00961.t01 |
| 12018 | 9 | 112202024 | 112213542 | mNeoVis.2.1.1.chr9\_hap1.g00962.t01 |
| 12019 | 9 | 112202024 | 112213542 | mNeoVis.2.1.1.chr9\_hap1.g00962.t01 |
| 12020 | 9 | 112202024 | 112213542 | mNeoVis.2.1.1.chr9\_hap1.g00962.t01 |
| 12021 | 9 | 112319157 | 112477890 | mNeoVis.2.1.1.chr9\_hap1.g00963.t01 |
| 12022 | 9 | 112319157 | 112477890 | mNeoVis.2.1.1.chr9\_hap1.g00963.t01 |
| 12023 | 9 | 112319157 | 112477890 | mNeoVis.2.1.1.chr9\_hap1.g00963.t01 |
| 12024 | 9 | 112478588 | 112964359 | mNeoVis.2.1.1.chr9\_hap1.g00964.t01 |
| 12025 | 9 | 113389783 | 113396577 | mNeoVis.2.1.1.chr9\_hap1.g00965.t01 |
| 12026 | 9 | 113396617 | 113519653 | mNeoVis.2.1.1.chr9\_hap1.g00966.t01 |
| 12027 | 9 | 113681616 | 113684324 | mNeoVis.2.1.1.chr9\_hap1.g00967.t01 |
| 12028 | 9 | 113681616 | 113684324 | mNeoVis.2.1.1.chr9\_hap1.g00967.t01 |
| 12029 | 9 | 113684502 | 114132030 | mNeoVis.2.1.1.chr9\_hap1.g00968.t01 |
| 12030 | 9 | 115319615 | 115741392 | mNeoVis.2.1.1.chr9\_hap1.g00969.t01 |
| 12031 | 9 | 115319615 | 115741392 | mNeoVis.2.1.1.chr9\_hap1.g00969.t01 |
| 12032 | 9 | 115529309 | 115529656 | mNeoVis.2.1.1.chr9\_hap1.g00970.t01 |
| 12033 | 9 | 115529309 | 115529656 | mNeoVis.2.1.1.chr9\_hap1.g00970.t01 |
| 12034 | 9 | 116078889 | 116152629 | mNeoVis.2.1.1.chr9\_hap1.g00972.t01 |
| 12035 | 9 | 116152656 | 116407955 | mNeoVis.2.1.1.chr9\_hap1.g00973.t01 |
| 12036 | 9 | 116857297 | 116885510 | mNeoVis.2.1.1.chr9\_hap1.g00974.t01 |
| 12037 | 9 | 116857297 | 116885510 | mNeoVis.2.1.1.chr9\_hap1.g00974.t01 |
| 12038 | 9 | 116857297 | 116885510 | mNeoVis.2.1.1.chr9\_hap1.g00974.t01 |
| 12039 | 9 | 116913021 | 116925556 | mNeoVis.2.1.1.chr9\_hap1.g00975.t01 |
| 12040 | 9 | 116913021 | 116925556 | mNeoVis.2.1.1.chr9\_hap1.g00975.t01 |
| 12041 | 9 | 116913021 | 116925556 | mNeoVis.2.1.1.chr9\_hap1.g00975.t01 |
| 12042 | 9 | 117005990 | 117227293 | mNeoVis.2.1.1.chr9\_hap1.g00976.t01 |
| 12043 | 9 | 117005990 | 117227293 | mNeoVis.2.1.1.chr9\_hap1.g00976.t01 |
| 12044 | 9 | 118074602 | 118075149 | mNeoVis.2.1.1.chr9\_hap1.g00977.t01 |
| 12045 | 9 | 118074602 | 118075149 | mNeoVis.2.1.1.chr9\_hap1.g00977.t01 |
| 12046 | 9 | 118074602 | 118075149 | mNeoVis.2.1.1.chr9\_hap1.g00977.t01 |
| 12047 | 9 | 118074602 | 118075149 | mNeoVis.2.1.1.chr9\_hap1.g00977.t01 |
| 12048 | 9 | 118695113 | 118730101 | mNeoVis.2.1.1.chr9\_hap1.g00978.t01 |
| 12049 | 9 | 118695113 | 118730101 | mNeoVis.2.1.1.chr9\_hap1.g00978.t01 |
| 12050 | 9 | 118695113 | 118730101 | mNeoVis.2.1.1.chr9\_hap1.g00978.t01 |
| 12051 | 9 | 118733662 | 118745380 | mNeoVis.2.1.1.chr9\_hap1.g00979.t01 |
| 12052 | 9 | 118733662 | 118745380 | mNeoVis.2.1.1.chr9\_hap1.g00979.t01 |
| 12053 | 9 | 118733662 | 118745380 | mNeoVis.2.1.1.chr9\_hap1.g00979.t01 |
| 12054 | 9 | 118745899 | 118843452 | mNeoVis.2.1.1.chr9\_hap1.g00980.t01 |
| 12055 | 9 | 118745899 | 118843452 | mNeoVis.2.1.1.chr9\_hap1.g00980.t01 |
| 12056 | 9 | 118745899 | 118843452 | mNeoVis.2.1.1.chr9\_hap1.g00980.t01 |
| 12057 | 9 | 118843654 | 118891156 | mNeoVis.2.1.1.chr9\_hap1.g00981.t01 |
| 12058 | 9 | 118843654 | 118891156 | mNeoVis.2.1.1.chr9\_hap1.g00981.t01 |
| 12059 | 9 | 118843654 | 118891156 | mNeoVis.2.1.1.chr9\_hap1.g00981.t01 |
| 12060 | 9 | 118903613 | 119091225 | mNeoVis.2.1.1.chr9\_hap1.g00982.t01 |
| 12061 | 9 | 119130322 | 119220168 | mNeoVis.2.1.1.chr9\_hap1.g00983.t01 |
| 12062 | 9 | 119130322 | 119220168 | mNeoVis.2.1.1.chr9\_hap1.g00983.t01 |
| 12063 | 9 | 119130322 | 119220168 | mNeoVis.2.1.1.chr9\_hap1.g00983.t01 |
| 12064 | 9 | 119234760 | 119394889 | mNeoVis.2.1.1.chr9\_hap1.g00984.t01 |
| 12065 | 9 | 119234760 | 119394889 | mNeoVis.2.1.1.chr9\_hap1.g00984.t01 |
| 12066 | 9 | 119234760 | 119394889 | mNeoVis.2.1.1.chr9\_hap1.g00984.t01 |
| 12067 | 9 | 119856178 | 119860893 | mNeoVis.2.1.1.chr9\_hap1.g00985.t01 |
| 12068 | 9 | 119856178 | 119860893 | mNeoVis.2.1.1.chr9\_hap1.g00985.t01 |
| 12069 | 9 | 119917394 | 120017120 | mNeoVis.2.1.1.chr9\_hap1.g00986.t01 |
| 12070 | 9 | 120007665 | 120015695 | mNeoVis.2.1.1.chr9\_hap1.g00987.t01 |
| 12071 | 9 | 120007665 | 120015695 | mNeoVis.2.1.1.chr9\_hap1.g00987.t01 |
| 12072 | 9 | 120079890 | 120203970 | mNeoVis.2.1.1.chr9\_hap1.g00988.t01 |
| 12073 | 9 | 120079890 | 120203970 | mNeoVis.2.1.1.chr9\_hap1.g00988.t01 |
| 12074 | 9 | 120079890 | 120203970 | mNeoVis.2.1.1.chr9\_hap1.g00988.t01 |
| 12075 | 9 | 120291250 | 120321829 | mNeoVis.2.1.1.chr9\_hap1.g00989.t01 |
| 12076 | 9 | 120291250 | 120321829 | mNeoVis.2.1.1.chr9\_hap1.g00989.t01 |
| 12077 | 9 | 120291250 | 120321829 | mNeoVis.2.1.1.chr9\_hap1.g00989.t01 |
| 12078 | 9 | 120498830 | 120629913 | mNeoVis.2.1.1.chr9\_hap1.g00990.t01 |
| 12079 | 9 | 120498830 | 120629913 | mNeoVis.2.1.1.chr9\_hap1.g00990.t01 |
| 12080 | 9 | 121207406 | 121210111 | mNeoVis.2.1.1.chr9\_hap1.g00991.t01 |
| 12081 | 9 | 121215726 | 121260320 | mNeoVis.2.1.1.chr9\_hap1.g00992.t01 |
| 12082 | 9 | 121298118 | 121305980 | mNeoVis.2.1.1.chr9\_hap1.g00993.t01 |
| 12083 | 9 | 121348178 | 121355594 | mNeoVis.2.1.1.chr9\_hap1.g00994.t01 |
| 12084 | 9 | 121390021 | 121431341 | mNeoVis.2.1.1.chr9\_hap1.g00995.t01 |
| 12085 | 9 | 122229172 | 122231895 | mNeoVis.2.1.1.chr9\_hap1.g00996.t01 |
| 12086 | 9 | 122229172 | 122231895 | mNeoVis.2.1.1.chr9\_hap1.g00996.t01 |
| 12087 | 9 | 122229172 | 122231895 | mNeoVis.2.1.1.chr9\_hap1.g00996.t01 |
| 12088 | 9 | 122683988 | 122709950 | mNeoVis.2.1.1.chr9\_hap1.g00997.t01 |
| 12089 | 9 | 122683988 | 122709950 | mNeoVis.2.1.1.chr9\_hap1.g00997.t01 |
| 12090 | 9 | 122710971 | 122712824 | mNeoVis.2.1.1.chr9\_hap1.g00998.t01 |
| 12091 | 9 | 122710971 | 122712824 | mNeoVis.2.1.1.chr9\_hap1.g00998.t01 |
| 12092 | 9 | 122736101 | 122855536 | mNeoVis.2.1.1.chr9\_hap1.g00999.t01 |
| 12093 | 9 | 122736101 | 122855536 | mNeoVis.2.1.1.chr9\_hap1.g00999.t01 |
| 12094 | 9 | 122891541 | 123223371 | mNeoVis.2.1.1.chr9\_hap1.g01000.t01 |
| 12095 | 9 | 122891541 | 123223371 | mNeoVis.2.1.1.chr9\_hap1.g01000.t01 |
| 12096 | 9 | 123122676 | 123166139 | mNeoVis.2.1.1.chr9\_hap1.g01001.t01 |
| 12097 | 9 | 123122676 | 123166139 | mNeoVis.2.1.1.chr9\_hap1.g01001.t01 |
| 12098 | 9 | 123223892 | 123240064 | mNeoVis.2.1.1.chr9\_hap1.g01002.t01 |
| 12099 | 9 | 123223892 | 123240064 | mNeoVis.2.1.1.chr9\_hap1.g01002.t01 |
| 12100 | 9 | 123243285 | 123264756 | mNeoVis.2.1.1.chr9\_hap1.g01003.t01 |
| 12101 | 9 | 123300074 | 123333502 | mNeoVis.2.1.1.chr9\_hap1.g01004.t01 |
| 12102 | 9 | 123384097 | 123388369 | mNeoVis.2.1.1.chr9\_hap1.g01005.t01 |
| 12103 | 9 | 123437556 | 123700476 | mNeoVis.2.1.1.chr9\_hap1.g01006.t01 |
| 12104 | 9 | 123879161 | 124023369 | mNeoVis.2.1.1.chr9\_hap1.g01007.t01 |
| 12105 | 9 | 124019351 | 124161869 | mNeoVis.2.1.1.chr9\_hap1.g01008.t01 |
| 12106 | 9 | 124248672 | 124300520 | mNeoVis.2.1.1.chr9\_hap1.g01009.t01 |
| 12107 | 9 | 124604789 | 124629250 | mNeoVis.2.1.1.chr9\_hap1.g01010.t01 |
| 12108 | 9 | 124604789 | 124629250 | mNeoVis.2.1.1.chr9\_hap1.g01010.t01 |
| 12109 | 9 | 124642919 | 124687745 | mNeoVis.2.1.1.chr9\_hap1.g01011.t01 |
| 12110 | 9 | 124642919 | 124687745 | mNeoVis.2.1.1.chr9\_hap1.g01011.t01 |
| 12111 | 9 | 124692794 | 124826590 | mNeoVis.2.1.1.chr9\_hap1.g01012.t01 |
| 12112 | 9 | 124692794 | 124826590 | mNeoVis.2.1.1.chr9\_hap1.g01012.t01 |
| 12113 | 9 | 124826691 | 124889891 | mNeoVis.2.1.1.chr9\_hap1.g01013.t01 |
| 12114 | 9 | 124826691 | 124889891 | mNeoVis.2.1.1.chr9\_hap1.g01013.t01 |
| 12115 | 9 | 124899679 | 125028514 | mNeoVis.2.1.1.chr9\_hap1.g01014.t01 |
| 12116 | 9 | 125028568 | 125060950 | mNeoVis.2.1.1.chr9\_hap1.g01015.t01 |
| 12117 | 9 | 125176058 | 125291042 | mNeoVis.2.1.1.chr9\_hap1.g01017.t01 |
| 12118 | 9 | 125335334 | 125335993 | mNeoVis.2.1.1.chr9\_hap1.g01018.t01 |
| 12119 | 9 | 125340478 | 125347460 | mNeoVis.2.1.1.chr9\_hap1.g01019.t01 |
| 12120 | 9 | 125348643 | 125397162 | mNeoVis.2.1.1.chr9\_hap1.g01020.t01 |
| 12121 | 9 | 125397612 | 125413110 | mNeoVis.2.1.1.chr9\_hap1.g01021.t01 |
| 12122 | 9 | 125416820 | 125444086 | mNeoVis.2.1.1.chr9\_hap1.g01022.t01 |
| 12123 | 9 | 125448593 | 125481955 | mNeoVis.2.1.1.chr9\_hap1.g01023.t01 |
| 12124 | 9 | 125606886 | 125676469 | mNeoVis.2.1.1.chr9\_hap1.g01024.t01 |
| 12125 | 9 | 125651654 | 125795838 | mNeoVis.2.1.1.chr9\_hap1.g01025.t01 |
| 12126 | 9 | 125862738 | 126185254 | mNeoVis.2.1.1.chr9\_hap1.g01026.t01 |
| 12127 | 9 | 126222992 | 126275233 | mNeoVis.2.1.1.chr9\_hap1.g01027.t01 |
| 12128 | 9 | 126222992 | 126275233 | mNeoVis.2.1.1.chr9\_hap1.g01027.t01 |
| 12129 | 9 | 126359782 | 126360374 | mNeoVis.2.1.1.chr9\_hap1.g01028.t01 |
| 12130 | 9 | 126359782 | 126360374 | mNeoVis.2.1.1.chr9\_hap1.g01028.t01 |
| 12131 | 9 | 127171811 | 127614041 | mNeoVis.2.1.1.chr9\_hap1.g01029.t01 |
| 12132 | 9 | 128691305 | 128694011 | mNeoVis.2.1.1.chr9\_hap1.g01035.t01 |
| 12133 | 9 | 128691305 | 128694011 | mNeoVis.2.1.1.chr9\_hap1.g01035.t01 |
| 12134 | 9 | 129020693 | 129126164 | mNeoVis.2.1.1.chr9\_hap1.g01036.t01 |
| 12135 | 9 | 129020693 | 129126164 | mNeoVis.2.1.1.chr9\_hap1.g01036.t01 |
| 12136 | 9 | 129133451 | 129155989 | mNeoVis.2.1.1.chr9\_hap1.g01037.t01 |
| 12137 | 9 | 129133451 | 129155989 | mNeoVis.2.1.1.chr9\_hap1.g01037.t01 |
| 12138 | 9 | 129156826 | 129158412 | mNeoVis.2.1.1.chr9\_hap1.g01038.t01 |
| 12139 | 9 | 129156826 | 129158412 | mNeoVis.2.1.1.chr9\_hap1.g01038.t01 |
| 12140 | 9 | 129158526 | 129180882 | mNeoVis.2.1.1.chr9\_hap1.g01039.t01 |
| 12141 | 9 | 129158526 | 129180882 | mNeoVis.2.1.1.chr9\_hap1.g01039.t01 |
| 12142 | 9 | 129180340 | 129242099 | mNeoVis.2.1.1.chr9\_hap1.g01040.t01 |
| 12143 | 9 | 129254457 | 129302338 | mNeoVis.2.1.1.chr9\_hap1.g01041.t01 |
| 12144 | 9 | 129388068 | 129432173 | mNeoVis.2.1.1.chr9\_hap1.g01042.t01 |
| 12145 | 9 | 129388068 | 129432173 | mNeoVis.2.1.1.chr9\_hap1.g01042.t01 |
| 12146 | 9 | 129438341 | 129479189 | mNeoVis.2.1.1.chr9\_hap1.g01043.t01 |
| 12147 | 9 | 129438341 | 129479189 | mNeoVis.2.1.1.chr9\_hap1.g01043.t01 |
| 12148 | 9 | 129498318 | 129498619 | mNeoVis.2.1.1.chr9\_hap1.g01044.t01 |
| 12149 | 9 | 129498318 | 129498619 | mNeoVis.2.1.1.chr9\_hap1.g01044.t01 |
| 12150 | 9 | 129637293 | 130018155 | mNeoVis.2.1.1.chr9\_hap1.g01045.t01 |
| 12151 | 9 | 130032880 | 130100802 | mNeoVis.2.1.1.chr9\_hap1.g01046.t01 |
| 12152 | 9 | 130032880 | 130100802 | mNeoVis.2.1.1.chr9\_hap1.g01046.t01 |
| 12153 | 9 | 130170898 | 130265496 | mNeoVis.2.1.1.chr9\_hap1.g01047.t01 |
| 12154 | 9 | 130170898 | 130265496 | mNeoVis.2.1.1.chr9\_hap1.g01047.t01 |
| 12155 | 9 | 130248396 | 130279106 | mNeoVis.2.1.1.chr9\_hap1.g01048.t01 |
| 12156 | 9 | 130248396 | 130279106 | mNeoVis.2.1.1.chr9\_hap1.g01048.t01 |
| 12157 | 9 | 130278534 | 130334428 | mNeoVis.2.1.1.chr9\_hap1.g01049.t01 |
| 12158 | 9 | 130278534 | 130334428 | mNeoVis.2.1.1.chr9\_hap1.g01049.t01 |
| 12159 | 9 | 130393241 | 130481926 | mNeoVis.2.1.1.chr9\_hap1.g01050.t01 |
| 12160 | 9 | 130497327 | 130530566 | mNeoVis.2.1.1.chr9\_hap1.g01051.t01 |
| 12161 | 9 | 130530153 | 130541020 | mNeoVis.2.1.1.chr9\_hap1.g01052.t01 |
| 12162 | 9 | 130557031 | 130635847 | mNeoVis.2.1.1.chr9\_hap1.g01053.t01 |
| 12163 | 9 | 130665899 | 130705861 | mNeoVis.2.1.1.chr9\_hap1.g01054.t01 |
| 12164 | 9 | 130665899 | 130705861 | mNeoVis.2.1.1.chr9\_hap1.g01054.t01 |
| 12165 | 9 | 130789488 | 130844352 | mNeoVis.2.1.1.chr9\_hap1.g01055.t01 |
| 12166 | 9 | 130789488 | 130844352 | mNeoVis.2.1.1.chr9\_hap1.g01055.t01 |
| 12167 | 9 | 130856485 | 130901104 | mNeoVis.2.1.1.chr9\_hap1.g01056.t01 |
| 12168 | 9 | 130856485 | 130901104 | mNeoVis.2.1.1.chr9\_hap1.g01056.t01 |
| 12169 | 9 | 130900987 | 131150720 | mNeoVis.2.1.1.chr9\_hap1.g01057.t01 |
| 12170 | 9 | 130900987 | 131150720 | mNeoVis.2.1.1.chr9\_hap1.g01057.t01 |
| 12171 | 9 | 130970871 | 130975242 | mNeoVis.2.1.1.chr9\_hap1.g01058.t01 |
| 12172 | 9 | 130970871 | 130975242 | mNeoVis.2.1.1.chr9\_hap1.g01058.t01 |
| 12173 | 9 | 131161673 | 131217523 | mNeoVis.2.1.1.chr9\_hap1.g01059.t01 |
| 12174 | 9 | 131161673 | 131217523 | mNeoVis.2.1.1.chr9\_hap1.g01059.t01 |
| 12175 | 9 | 131220825 | 131391322 | mNeoVis.2.1.1.chr9\_hap1.g01060.t01 |
| 12176 | 9 | 131533762 | 131609117 | mNeoVis.2.1.1.chr9\_hap1.g01061.t01 |
| 12177 | 9 | 131641009 | 131643328 | mNeoVis.2.1.1.chr9\_hap1.g01062.t01 |
| 12178 | 9 | 132445849 | 132464543 | mNeoVis.2.1.1.chr9\_hap1.g01068.t01 |
| 12179 | 9 | 132468797 | 132477936 | mNeoVis.2.1.1.chr9\_hap1.g01069.t01 |
| 12180 | 9 | 132481695 | 132507372 | mNeoVis.2.1.1.chr9\_hap1.g01070.t01 |
| 12181 | 9 | 132507492 | 132540315 | mNeoVis.2.1.1.chr9\_hap1.g01071.t01 |
| 12182 | 9 | 132553148 | 132601769 | mNeoVis.2.1.1.chr9\_hap1.g01072.t01 |
| 12183 | 9 | 132601920 | 132618734 | mNeoVis.2.1.1.chr9\_hap1.g01073.t01 |
| 12184 | 9 | 132625440 | 132625856 | mNeoVis.2.1.1.chr9\_hap1.g01074.t01 |
| 12185 | 9 | 132669588 | 132796650 | mNeoVis.2.1.1.chr9\_hap1.g01075.t01 |
| 12186 | 9 | 132738301 | 132739815 | mNeoVis.2.1.1.chr9\_hap1.g01076.t01 |
| 12187 | 9 | 133002661 | 133003378 | mNeoVis.2.1.1.chr9\_hap1.g01078.t01 |
| 12188 | 9 | 135177494 | 135178575 | mNeoVis.2.1.1.chr9\_hap1.g01095.t01 |
| 12189 | 9 | 135267232 | 135556850 | mNeoVis.2.1.1.chr9\_hap1.g01096.t01 |
| 12190 | 9 | 135576404 | 135601099 | mNeoVis.2.1.1.chr9\_hap1.g01097.t01 |
| 12191 | 9 | 135602195 | 135604048 | mNeoVis.2.1.1.chr9\_hap1.g01098.t01 |
| 12192 | 9 | 135610123 | 135610514 | mNeoVis.2.1.1.chr9\_hap1.g01099.t01 |
| 12193 | 9 | 135703904 | 135725063 | mNeoVis.2.1.1.chr9\_hap1.g01102.t01 |
| 12194 | 9 | 135726608 | 135777783 | mNeoVis.2.1.1.chr9\_hap1.g01103.t01 |
| 12195 | 9 | 135777816 | 135801952 | mNeoVis.2.1.1.chr9\_hap1.g01104.t01 |
| 12196 | 9 | 135801960 | 136262002 | mNeoVis.2.1.1.chr9\_hap1.g01105.t01 |
| 12197 | 9 | 136043219 | 136043406 | mNeoVis.2.1.1.chr9\_hap1.g01106.t01 |
| 12198 | 9 | 136050757 | 136107676 | mNeoVis.2.1.1.chr9\_hap1.g01107.t01 |
| 12199 | 9 | 136163664 | 136165593 | mNeoVis.2.1.1.chr9\_hap1.g01108.t01 |
| 12200 | 9 | 137801119 | 137819051 | mNeoVis.2.1.1.chr9\_hap1.g01112.t01 |
| 12201 | 9 | 137837132 | 137838494 | mNeoVis.2.1.1.chr9\_hap1.g01113.t01 |
| 12202 | 9 | 137958616 | 137990229 | mNeoVis.2.1.1.chr9\_hap1.g01114.t01 |
| 12203 | 9 | 137994042 | 138036844 | mNeoVis.2.1.1.chr9\_hap1.g01115.t01 |
| 12204 | 9 | 138073030 | 138079350 | mNeoVis.2.1.1.chr9\_hap1.g01116.t01 |
| 12205 | 9 | 138081648 | 138101074 | mNeoVis.2.1.1.chr9\_hap1.g01117.t01 |
| 12206 | 9 | 138178040 | 138189778 | mNeoVis.2.1.1.chr9\_hap1.g01118.t01 |
| 12207 | 9 | 138270413 | 138283954 | mNeoVis.2.1.1.chr9\_hap1.g01120.t01 |
| 12208 | 9 | 138297068 | 138313373 | mNeoVis.2.1.1.chr9\_hap1.g01121.t01 |
| 12209 | 9 | 138327127 | 138594925 | mNeoVis.2.1.1.chr9\_hap1.g01122.t01 |
| 12210 | 9 | 138651497 | 138665522 | mNeoVis.2.1.1.chr9\_hap1.g01123.t01 |
| 12211 | 9 | 138651497 | 138665522 | mNeoVis.2.1.1.chr9\_hap1.g01123.t01 |
| 12212 | 9 | 138668526 | 138678450 | mNeoVis.2.1.1.chr9\_hap1.g01124.t01 |
| 12213 | 9 | 138668526 | 138678450 | mNeoVis.2.1.1.chr9\_hap1.g01124.t01 |
| 12214 | 9 | 138690754 | 138696032 | mNeoVis.2.1.1.chr9\_hap1.g01125.t01 |
| 12215 | 9 | 138690754 | 138696032 | mNeoVis.2.1.1.chr9\_hap1.g01125.t01 |
| 12216 | 9 | 138704374 | 138794078 | mNeoVis.2.1.1.chr9\_hap1.g01126.t01 |
| 12217 | 9 | 138704374 | 138794078 | mNeoVis.2.1.1.chr9\_hap1.g01126.t01 |
| 12218 | 9 | 138865802 | 138878037 | mNeoVis.2.1.1.chr9\_hap1.g01127.t01 |
| 12219 | 9 | 138865802 | 138878037 | mNeoVis.2.1.1.chr9\_hap1.g01127.t01 |
| 12220 | 9 | 138902927 | 138917315 | mNeoVis.2.1.1.chr9\_hap1.g01128.t01 |
| 12221 | 9 | 138902927 | 138917315 | mNeoVis.2.1.1.chr9\_hap1.g01128.t01 |
| 12222 | 9 | 138919887 | 138933604 | mNeoVis.2.1.1.chr9\_hap1.g01129.t01 |
| 12223 | 9 | 138919887 | 138933604 | mNeoVis.2.1.1.chr9\_hap1.g01129.t01 |
| 12224 | 9 | 138960824 | 139070462 | mNeoVis.2.1.1.chr9\_hap1.g01130.t01 |
| 12225 | 9 | 139091949 | 139101645 | mNeoVis.2.1.1.chr9\_hap1.g01131.t01 |
| 12226 | 9 | 139113786 | 139132021 | mNeoVis.2.1.1.chr9\_hap1.g01132.t01 |
| 12227 | 9 | 139148778 | 139165101 | mNeoVis.2.1.1.chr9\_hap1.g01133.t01 |
| 12228 | 9 | 139207821 | 139226954 | mNeoVis.2.1.1.chr9\_hap1.g01134.t01 |
| 12229 | 9 | 139207821 | 139226954 | mNeoVis.2.1.1.chr9\_hap1.g01134.t01 |
| 12230 | 9 | 139260399 | 139285788 | mNeoVis.2.1.1.chr9\_hap1.g01135.t01 |
| 12231 | 9 | 139260399 | 139285788 | mNeoVis.2.1.1.chr9\_hap1.g01135.t01 |
| 12232 | 9 | 139315923 | 139344084 | mNeoVis.2.1.1.chr9\_hap1.g01136.t01 |
| 12233 | 9 | 139315923 | 139344084 | mNeoVis.2.1.1.chr9\_hap1.g01136.t01 |
| 12234 | 9 | 139346608 | 139358984 | mNeoVis.2.1.1.chr9\_hap1.g01137.t01 |
| 12235 | 9 | 139346608 | 139358984 | mNeoVis.2.1.1.chr9\_hap1.g01137.t01 |
| 12236 | 9 | 139359267 | 139366393 | mNeoVis.2.1.1.chr9\_hap1.g01138.t01 |
| 12237 | 9 | 139359267 | 139366393 | mNeoVis.2.1.1.chr9\_hap1.g01138.t01 |
| 12238 | 9 | 139377749 | 139387330 | mNeoVis.2.1.1.chr9\_hap1.g01139.t01 |
| 12239 | 9 | 139377749 | 139387330 | mNeoVis.2.1.1.chr9\_hap1.g01139.t01 |
| 12240 | 9 | 139389109 | 139419225 | mNeoVis.2.1.1.chr9\_hap1.g01140.t01 |
| 12241 | 9 | 139389109 | 139419225 | mNeoVis.2.1.1.chr9\_hap1.g01140.t01 |
| 12242 | 9 | 139435129 | 139450525 | mNeoVis.2.1.1.chr9\_hap1.g01141.t01 |
| 12243 | 9 | 139435129 | 139450525 | mNeoVis.2.1.1.chr9\_hap1.g01141.t01 |
| 12244 | 9 | 139471761 | 139480041 | mNeoVis.2.1.1.chr9\_hap1.g01142.t01 |
| 12245 | 9 | 139471761 | 139480041 | mNeoVis.2.1.1.chr9\_hap1.g01142.t01 |
| 12246 | 9 | 139487831 | 139496825 | mNeoVis.2.1.1.chr9\_hap1.g01143.t01 |
| 12247 | 9 | 139487831 | 139496825 | mNeoVis.2.1.1.chr9\_hap1.g01143.t01 |
| 12248 | 9 | 139508322 | 139524150 | mNeoVis.2.1.1.chr9\_hap1.g01144.t01 |
| 12249 | 9 | 139508322 | 139524150 | mNeoVis.2.1.1.chr9\_hap1.g01144.t01 |
| 12250 | 9 | 139547332 | 139576992 | mNeoVis.2.1.1.chr9\_hap1.g01145.t01 |
| 12251 | 9 | 139547332 | 139576992 | mNeoVis.2.1.1.chr9\_hap1.g01145.t01 |
| 12252 | 9 | 139621621 | 139678408 | mNeoVis.2.1.1.chr9\_hap1.g01146.t01 |
| 12253 | 9 | 139621621 | 139678408 | mNeoVis.2.1.1.chr9\_hap1.g01146.t01 |
| 12254 | 9 | 139714855 | 139730440 | mNeoVis.2.1.1.chr9\_hap1.g01147.t01 |
| 12255 | 9 | 139733211 | 139758323 | mNeoVis.2.1.1.chr9\_hap1.g01148.t01 |
| 12256 | 9 | 139787597 | 139795519 | mNeoVis.2.1.1.chr9\_hap1.g01149.t01 |
| 12257 | 9 | 139795811 | 139799246 | mNeoVis.2.1.1.chr9\_hap1.g01150.t01 |
| 12258 | 9 | 139813213 | 139831600 | mNeoVis.2.1.1.chr9\_hap1.g01151.t01 |
| 12259 | 9 | 139843497 | 139869228 | mNeoVis.2.1.1.chr9\_hap1.g01152.t01 |
| 12260 | 9 | 139870454 | 139878286 | mNeoVis.2.1.1.chr9\_hap1.g01153.t01 |
| 12261 | 9 | 139878331 | 139889528 | mNeoVis.2.1.1.chr9\_hap1.g01154.t01 |
| 12262 | 9 | 139893152 | 139903198 | mNeoVis.2.1.1.chr9\_hap1.g01155.t01 |
| 12263 | 9 | 139966983 | 139986709 | mNeoVis.2.1.1.chr9\_hap1.g01156.t01 |
| 12264 | 9 | 139988240 | 139993807 | mNeoVis.2.1.1.chr9\_hap1.g01157.t01 |
| 12265 | 9 | 139995062 | 140020006 | mNeoVis.2.1.1.chr9\_hap1.g01158.t01 |
| 12266 | 9 | 140020433 | 140081863 | mNeoVis.2.1.1.chr9\_hap1.g01159.t01 |
| 12267 | 9 | 140092880 | 140102967 | mNeoVis.2.1.1.chr9\_hap1.g01160.t01 |
| 12268 | 9 | 140104582 | 140148974 | mNeoVis.2.1.1.chr9\_hap1.g01161.t01 |
| 12269 | 9 | 140159725 | 140193231 | mNeoVis.2.1.1.chr9\_hap1.g01162.t01 |
| 12270 | 9 | 140221029 | 140243682 | mNeoVis.2.1.1.chr9\_hap1.g01163.t01 |
| 12271 | 9 | 140254555 | 140266547 | mNeoVis.2.1.1.chr9\_hap1.g01164.t01 |
| 12272 | 9 | 140266545 | 140286991 | mNeoVis.2.1.1.chr9\_hap1.g01165.t01 |
| 12273 | 9 | 140297086 | 140346669 | mNeoVis.2.1.1.chr9\_hap1.g01166.t01 |
| 12274 | 9 | 140350544 | 140430349 | mNeoVis.2.1.1.chr9\_hap1.g01167.t01 |
| 12275 | 9 | 140434661 | 140445854 | mNeoVis.2.1.1.chr9\_hap1.g01168.t01 |
| 12276 | 9 | 140452378 | 140454344 | mNeoVis.2.1.1.chr9\_hap1.g01169.t01 |
| 12277 | 9 | 140455296 | 140474148 | mNeoVis.2.1.1.chr9\_hap1.g01170.t01 |
| 12278 | 9 | 140474265 | 140499592 | mNeoVis.2.1.1.chr9\_hap1.g01171.t01 |
| 12279 | 9 | 140501735 | 140520850 | mNeoVis.2.1.1.chr9\_hap1.g01172.t01 |
| 12280 | 9 | 140540111 | 140555573 | mNeoVis.2.1.1.chr9\_hap1.g01173.t01 |
| 12281 | 9 | 140564411 | 140587082 | mNeoVis.2.1.1.chr9\_hap1.g01174.t01 |
| 12282 | 9 | 140602076 | 140613044 | mNeoVis.2.1.1.chr9\_hap1.g01175.t01 |
| 12283 | 9 | 140964918 | 140999993 | mNeoVis.2.1.1.chr9\_hap1.g01184.t01 |
| 12284 | 9 | 141024111 | 141058081 | mNeoVis.2.1.1.chr9\_hap1.g01185.t01 |
| 12285 | 9 | 141073263 | 141075608 | mNeoVis.2.1.1.chr9\_hap1.g01186.t01 |
| 12286 | 9 | 141081053 | 141087194 | mNeoVis.2.1.1.chr9\_hap1.g01187.t01 |
| 12287 | 9 | 141090656 | 141092772 | mNeoVis.2.1.1.chr9\_hap1.g01188.t01 |
| 12288 | 9 | 141097930 | 141101313 | mNeoVis.2.1.1.chr9\_hap1.g01189.t01 |
| 12289 | 9 | 141124708 | 141236629 | mNeoVis.2.1.1.chr9\_hap1.g01190.t01 |
| 12290 | 9 | 141275174 | 141377567 | mNeoVis.2.1.1.chr9\_hap1.g01191.t01 |
| 12291 | 9 | 141382354 | 141450756 | mNeoVis.2.1.1.chr9\_hap1.g01192.t01 |
| 12292 | 9 | 141466651 | 141500635 | mNeoVis.2.1.1.chr9\_hap1.g01193.t01 |
| 12293 | 9 | 141512158 | 141764588 | mNeoVis.2.1.1.chr9\_hap1.g01194.t01 |
| 12294 | 9 | 141785493 | 141804360 | mNeoVis.2.1.1.chr9\_hap1.g01195.t01 |
| 12295 | 9 | 141821586 | 141869703 | mNeoVis.2.1.1.chr9\_hap1.g01196.t01 |
| 12296 | 9 | 141869837 | 141929728 | mNeoVis.2.1.1.chr9\_hap1.g01197.t01 |
| 12297 | 9 | 141951193 | 142143573 | mNeoVis.2.1.1.chr9\_hap1.g01198.t01 |
| 12298 | 9 | 142103877 | 142107122 | mNeoVis.2.1.1.chr9\_hap1.g01199.t01 |
| 12299 | 9 | 142170506 | 142212611 | mNeoVis.2.1.1.chr9\_hap1.g01200.t01 |
| 12300 | 9 | 142218135 | 142252113 | mNeoVis.2.1.1.chr9\_hap1.g01201.t01 |
| 12301 | 9 | 142252462 | 142333390 | mNeoVis.2.1.1.chr9\_hap1.g01202.t01 |
| 12302 | 9 | 142258417 | 142259591 | mNeoVis.2.1.1.chr9\_hap1.g01203.t01 |
| 12303 | 9 | 142355736 | 142371214 | mNeoVis.2.1.1.chr9\_hap1.g01204.t01 |
| 12304 | 9 | 142477288 | 142558801 | mNeoVis.2.1.1.chr9\_hap1.g01205.t01 |
| 12305 | 9 | 142614043 | 142632172 | mNeoVis.2.1.1.chr9\_hap1.g01206.t01 |
| 12306 | 9 | 142632325 | 142638970 | mNeoVis.2.1.1.chr9\_hap1.g01207.t01 |
| 12307 | 9 | 142647914 | 142675384 | mNeoVis.2.1.1.chr9\_hap1.g01208.t01 |
| 12308 | 9 | 142686075 | 142742758 | mNeoVis.2.1.1.chr9\_hap1.g01209.t01 |
| 12309 | 9 | 142759097 | 142774548 | mNeoVis.2.1.1.chr9\_hap1.g01210.t01 |
| 12310 | 9 | 142795480 | 142808920 | mNeoVis.2.1.1.chr9\_hap1.g01211.t01 |
| 12311 | 9 | 142816675 | 142826646 | mNeoVis.2.1.1.chr9\_hap1.g01212.t01 |
| 12312 | 9 | 142826677 | 142879381 | mNeoVis.2.1.1.chr9\_hap1.g01213.t01 |
| 12313 | 9 | 142880479 | 142892773 | mNeoVis.2.1.1.chr9\_hap1.g01214.t01 |
| 12314 | 9 | 142903909 | 142916033 | mNeoVis.2.1.1.chr9\_hap1.g01215.t01 |
| 12315 | 9 | 142916290 | 142926438 | mNeoVis.2.1.1.chr9\_hap1.g01216.t01 |
| 12316 | 9 | 142935065 | 142955074 | mNeoVis.2.1.1.chr9\_hap1.g01217.t01 |
| 12317 | 9 | 142955179 | 142982119 | mNeoVis.2.1.1.chr9\_hap1.g01218.t01 |
| 12318 | 9 | 142987599 | 142998980 | mNeoVis.2.1.1.chr9\_hap1.g01219.t01 |
| 12319 | 9 | 143001307 | 143007683 | mNeoVis.2.1.1.chr9\_hap1.g01220.t01 |
| 12320 | 9 | 143011964 | 143091161 | mNeoVis.2.1.1.chr9\_hap1.g01221.t01 |
| 12321 | 9 | 143092181 | 143120330 | mNeoVis.2.1.1.chr9\_hap1.g01222.t01 |
| 12322 | 9 | 143111158 | 143168823 | mNeoVis.2.1.1.chr9\_hap1.g01223.t01 |
| 12323 | 9 | 143171948 | 143176823 | mNeoVis.2.1.1.chr9\_hap1.g01224.t01 |
| 12324 | 9 | 143184763 | 143187743 | mNeoVis.2.1.1.chr9\_hap1.g01225.t01 |
| 12325 | 9 | 143195660 | 143206527 | mNeoVis.2.1.1.chr9\_hap1.g01226.t01 |
| 12326 | 9 | 143211840 | 143215839 | mNeoVis.2.1.1.chr9\_hap1.g01227.t01 |
| 12327 | 9 | 143222815 | 143249112 | mNeoVis.2.1.1.chr9\_hap1.g01228.t01 |
| 12328 | 9 | 143250687 | 143287116 | mNeoVis.2.1.1.chr9\_hap1.g01229.t01 |
| 12329 | 9 | 143312615 | 143327819 | mNeoVis.2.1.1.chr9\_hap1.g01230.t01 |
| 12330 | 9 | 143316287 | 143327446 | mNeoVis.2.1.1.chr9\_hap1.g01231.t01 |
| 12331 | 9 | 143327857 | 143332171 | mNeoVis.2.1.1.chr9\_hap1.g01232.t01 |
| 12332 | 9 | 143333387 | 143340995 | mNeoVis.2.1.1.chr9\_hap1.g01233.t01 |
| 12333 | 9 | 143351009 | 143359566 | mNeoVis.2.1.1.chr9\_hap1.g01234.t01 |
| 12334 | 9 | 143361005 | 143365588 | mNeoVis.2.1.1.chr9\_hap1.g01235.t01 |
| 12335 | 9 | 143370978 | 143478317 | mNeoVis.2.1.1.chr9\_hap1.g01236.t01 |
| 12336 | 9 | 143576798 | 143610507 | mNeoVis.2.1.1.chr9\_hap1.g01237.t01 |
| 12337 | 9 | 143698328 | 143727524 | mNeoVis.2.1.1.chr9\_hap1.g01238.t01 |
| 12338 | 9 | 143727663 | 143735905 | mNeoVis.2.1.1.chr9\_hap1.g01239.t01 |
| 12339 | 9 | 143742613 | 143765683 | mNeoVis.2.1.1.chr9\_hap1.g01240.t01 |
| 12340 | 9 | 143770264 | 143799401 | mNeoVis.2.1.1.chr9\_hap1.g01241.t01 |
| 12341 | 9 | 143838604 | 143863025 | mNeoVis.2.1.1.chr9\_hap1.g01242.t01 |
| 12342 | 9 | 143904032 | 143950440 | mNeoVis.2.1.1.chr9\_hap1.g01243.t01 |
| 12343 | 9 | 143961212 | 143989889 | mNeoVis.2.1.1.chr9\_hap1.g01244.t01 |
| 12344 | 9 | 143961212 | 143989889 | mNeoVis.2.1.1.chr9\_hap1.g01244.t01 |
| 12345 | 9 | 144113528 | 144144215 | mNeoVis.2.1.1.chr9\_hap1.g01245.t01 |
| 12346 | 9 | 144113528 | 144144215 | mNeoVis.2.1.1.chr9\_hap1.g01245.t01 |
| 12347 | 9 | 144169003 | 144187230 | mNeoVis.2.1.1.chr9\_hap1.g01246.t01 |
| 12348 | 9 | 144169003 | 144187230 | mNeoVis.2.1.1.chr9\_hap1.g01246.t01 |
| 12349 | 9 | 144188963 | 144198045 | mNeoVis.2.1.1.chr9\_hap1.g01247.t01 |
| 12350 | 9 | 144188963 | 144198045 | mNeoVis.2.1.1.chr9\_hap1.g01247.t01 |
| 12351 | 9 | 144203155 | 144213521 | mNeoVis.2.1.1.chr9\_hap1.g01248.t01 |
| 12352 | 9 | 144203155 | 144213521 | mNeoVis.2.1.1.chr9\_hap1.g01248.t01 |
| 12353 | 9 | 144215619 | 144232398 | mNeoVis.2.1.1.chr9\_hap1.g01249.t01 |
| 12354 | 9 | 144215619 | 144232398 | mNeoVis.2.1.1.chr9\_hap1.g01249.t01 |
| 12355 | 9 | 144256434 | 144390832 | mNeoVis.2.1.1.chr9\_hap1.g01250.t01 |
| 12356 | 9 | 144256434 | 144390832 | mNeoVis.2.1.1.chr9\_hap1.g01250.t01 |
| 12357 | 9 | 144460190 | 144521550 | mNeoVis.2.1.1.chr9\_hap1.g01251.t01 |
| 12358 | 9 | 144460190 | 144521550 | mNeoVis.2.1.1.chr9\_hap1.g01251.t01 |
| 12359 | 9 | 144533530 | 144751309 | mNeoVis.2.1.1.chr9\_hap1.g01252.t01 |
| 12360 | 9 | 144761823 | 144794214 | mNeoVis.2.1.1.chr9\_hap1.g01253.t01 |
| 12361 | 9 | 144761823 | 144794214 | mNeoVis.2.1.1.chr9\_hap1.g01253.t01 |
| 12362 | 9 | 144761823 | 144794214 | mNeoVis.2.1.1.chr9\_hap1.g01253.t01 |
| 12363 | 9 | 144803383 | 144814695 | mNeoVis.2.1.1.chr9\_hap1.g01254.t01 |
| 12364 | 9 | 144803383 | 144814695 | mNeoVis.2.1.1.chr9\_hap1.g01254.t01 |
| 12365 | 9 | 144803383 | 144814695 | mNeoVis.2.1.1.chr9\_hap1.g01254.t01 |
| 12366 | 9 | 144851331 | 144867586 | mNeoVis.2.1.1.chr9\_hap1.g01255.t01 |
| 12367 | 9 | 144851331 | 144867586 | mNeoVis.2.1.1.chr9\_hap1.g01255.t01 |
| 12368 | 9 | 144851331 | 144867586 | mNeoVis.2.1.1.chr9\_hap1.g01255.t01 |
| 12369 | 9 | 144870275 | 144904803 | mNeoVis.2.1.1.chr9\_hap1.g01256.t01 |
| 12370 | 9 | 144870275 | 144904803 | mNeoVis.2.1.1.chr9\_hap1.g01256.t01 |
| 12371 | 9 | 144870275 | 144904803 | mNeoVis.2.1.1.chr9\_hap1.g01256.t01 |
| 12372 | 9 | 144921509 | 145014830 | mNeoVis.2.1.1.chr9\_hap1.g01257.t01 |
| 12373 | 9 | 144921509 | 145014830 | mNeoVis.2.1.1.chr9\_hap1.g01257.t01 |
| 12374 | 9 | 145017058 | 145039462 | mNeoVis.2.1.1.chr9\_hap1.g01258.t01 |
| 12375 | 9 | 145017058 | 145039462 | mNeoVis.2.1.1.chr9\_hap1.g01258.t01 |
| 12376 | 9 | 145093098 | 145133269 | mNeoVis.2.1.1.chr9\_hap1.g01259.t01 |
| 12377 | 9 | 145093098 | 145133269 | mNeoVis.2.1.1.chr9\_hap1.g01259.t01 |
| 12378 | 9 | 145248673 | 145254483 | mNeoVis.2.1.1.chr9\_hap1.g01260.t01 |
| 12379 | 9 | 145248673 | 145254483 | mNeoVis.2.1.1.chr9\_hap1.g01260.t01 |
| 12380 | 9 | 145330457 | 145346128 | mNeoVis.2.1.1.chr9\_hap1.g01261.t01 |
| 12381 | 9 | 145330457 | 145346128 | mNeoVis.2.1.1.chr9\_hap1.g01261.t01 |
| 12382 | 9 | 145389796 | 145416310 | mNeoVis.2.1.1.chr9\_hap1.g01262.t01 |
| 12383 | 9 | 145429122 | 145498076 | mNeoVis.2.1.1.chr9\_hap1.g01263.t01 |
| 12384 | 9 | 145503649 | 145586792 | mNeoVis.2.1.1.chr9\_hap1.g01264.t01 |
| 12385 | 9 | 145700975 | 145718271 | mNeoVis.2.1.1.chr9\_hap1.g01265.t01 |
| 12386 | 9 | 145727079 | 145759402 | mNeoVis.2.1.1.chr9\_hap1.g01266.t01 |
| 12387 | 9 | 146322245 | 146370103 | mNeoVis.2.1.1.chr9\_hap1.g01273.t01 |
| 12388 | 9 | 146471993 | 146495564 | mNeoVis.2.1.1.chr9\_hap1.g01274.t01 |
| 12389 | 9 | 146521435 | 146587195 | mNeoVis.2.1.1.chr9\_hap1.g01275.t01 |
| 12390 | 9 | 146587440 | 146592047 | mNeoVis.2.1.1.chr9\_hap1.g01276.t01 |
| 12391 | 9 | 146594677 | 146600678 | mNeoVis.2.1.1.chr9\_hap1.g01277.t01 |
| 12392 | 9 | 146605589 | 146613382 | mNeoVis.2.1.1.chr9\_hap1.g01278.t01 |
| 12393 | 9 | 146623635 | 146631127 | mNeoVis.2.1.1.chr9\_hap1.g01279.t01 |
| 12394 | 9 | 146645290 | 146650219 | mNeoVis.2.1.1.chr9\_hap1.g01281.t01 |
| 12395 | 9 | 146656206 | 146664008 | mNeoVis.2.1.1.chr9\_hap1.g01282.t01 |
| 12396 | 9 | 146667088 | 146688753 | mNeoVis.2.1.1.chr9\_hap1.g01283.t01 |
| 12397 | 9 | 146691034 | 146711349 | mNeoVis.2.1.1.chr9\_hap1.g01284.t01 |
| 12398 | 9 | 146715454 | 146734174 | mNeoVis.2.1.1.chr9\_hap1.g01285.t01 |
| 12399 | 9 | 146737260 | 146867662 | mNeoVis.2.1.1.chr9\_hap1.g01286.t01 |
| 12400 | 9 | 146900347 | 146972687 | mNeoVis.2.1.1.chr9\_hap1.g01287.t01 |
| 12401 | 9 | 146976073 | 147015010 | mNeoVis.2.1.1.chr9\_hap1.g01288.t01 |
| 12402 | 9 | 147056604 | 147333279 | mNeoVis.2.1.1.chr9\_hap1.g01289.t01 |
| 12403 | 9 | 148236574 | 148402802 | mNeoVis.2.1.1.chr9\_hap1.g01290.t01 |
| 12404 | 9 | 149072623 | 149113238 | mNeoVis.2.1.1.chr9\_hap1.g01291.t01 |
| 12405 | 9 | 149219406 | 149222518 | mNeoVis.2.1.1.chr9\_hap1.g01295.t01 |
| 12406 | 9 | 149247278 | 149268497 | mNeoVis.2.1.1.chr9\_hap1.g01296.t01 |
| 12407 | 9 | 149270859 | 149313248 | mNeoVis.2.1.1.chr9\_hap1.g01297.t01 |
| 12408 | 9 | 149314483 | 149337213 | mNeoVis.2.1.1.chr9\_hap1.g01298.t01 |
| 12409 | 9 | 149336929 | 149401942 | mNeoVis.2.1.1.chr9\_hap1.g01299.t01 |
| 12410 | 9 | 149407844 | 149415902 | mNeoVis.2.1.1.chr9\_hap1.g01300.t01 |
| 12411 | 9 | 149419707 | 149427604 | mNeoVis.2.1.1.chr9\_hap1.g01301.t01 |
| 12412 | 9 | 149428580 | 149431372 | mNeoVis.2.1.1.chr9\_hap1.g01302.t01 |
| 12413 | 9 | 149433240 | 149445006 | mNeoVis.2.1.1.chr9\_hap1.g01303.t01 |
| 12414 | 9 | 149444107 | 149465019 | mNeoVis.2.1.1.chr9\_hap1.g01304.t01 |
| 12415 | 9 | 149470469 | 149476157 | mNeoVis.2.1.1.chr9\_hap1.g01305.t01 |
| 12416 | 9 | 149477356 | 149486560 | mNeoVis.2.1.1.chr9\_hap1.g01306.t01 |
| 12417 | 9 | 149488510 | 149495093 | mNeoVis.2.1.1.chr9\_hap1.g01307.t01 |
| 12418 | 9 | 149499422 | 149528752 | mNeoVis.2.1.1.chr9\_hap1.g01308.t01 |
| 12419 | 9 | 149531999 | 149542500 | mNeoVis.2.1.1.chr9\_hap1.g01309.t01 |
| 12420 | 9 | 149553144 | 149637436 | mNeoVis.2.1.1.chr9\_hap1.g01310.t01 |
| 12421 | 9 | 149637578 | 149659050 | mNeoVis.2.1.1.chr9\_hap1.g01311.t01 |
| 12422 | 9 | 149668507 | 149671035 | mNeoVis.2.1.1.chr9\_hap1.g01312.t01 |
| 12423 | 9 | 149673637 | 149676143 | mNeoVis.2.1.1.chr9\_hap1.g01313.t01 |
| 12424 | 9 | 149685044 | 149689598 | mNeoVis.2.1.1.chr9\_hap1.g01314.t01 |
| 12425 | 9 | 149689754 | 149703265 | mNeoVis.2.1.1.chr9\_hap1.g01315.t01 |
| 12426 | 9 | 149703399 | 149705390 | mNeoVis.2.1.1.chr9\_hap1.g01316.t01 |
| 12427 | 9 | 149705880 | 149708948 | mNeoVis.2.1.1.chr9\_hap1.g01317.t01 |
| 12428 | 9 | 149708966 | 149714342 | mNeoVis.2.1.1.chr9\_hap1.g01318.t01 |
| 12429 | 9 | 149723534 | 149726218 | mNeoVis.2.1.1.chr9\_hap1.g01319.t01 |
| 12430 | 9 | 149726284 | 149729019 | mNeoVis.2.1.1.chr9\_hap1.g01320.t01 |
| 12431 | 9 | 149729311 | 149758321 | mNeoVis.2.1.1.chr9\_hap1.g01321.t01 |
| 12432 | 9 | 149761243 | 149769162 | mNeoVis.2.1.1.chr9\_hap1.g01322.t01 |
| 12433 | 9 | 149769715 | 149773075 | mNeoVis.2.1.1.chr9\_hap1.g01323.t01 |
| 12434 | 9 | 149789006 | 149799631 | mNeoVis.2.1.1.chr9\_hap1.g01324.t01 |
| 12435 | 9 | 149809154 | 149812369 | mNeoVis.2.1.1.chr9\_hap1.g01325.t01 |
| 12436 | 9 | 149832841 | 149884311 | mNeoVis.2.1.1.chr9\_hap1.g01326.t01 |
| 12437 | 9 | 149889064 | 149894620 | mNeoVis.2.1.1.chr9\_hap1.g01327.t01 |
| 12438 | 10 | 2109144 | 2214782 | mNeoVis.2.1.1.chr10\_hap1.g00036.t01 |
| 12439 | 10 | 2203624 | 2204557 | mNeoVis.2.1.1.chr10\_hap1.g00037.t01 |
| 12440 | 10 | 2215772 | 2223167 | mNeoVis.2.1.1.chr10\_hap1.g00038.t01 |
| 12441 | 10 | 2270189 | 2360603 | mNeoVis.2.1.1.chr10\_hap1.g00039.t01 |
| 12442 | 10 | 2360848 | 2372425 | mNeoVis.2.1.1.chr10\_hap1.g00040.t01 |
| 12443 | 10 | 2372893 | 2384554 | mNeoVis.2.1.1.chr10\_hap1.g00041.t01 |
| 12444 | 10 | 2384978 | 2440131 | mNeoVis.2.1.1.chr10\_hap1.g00042.t01 |
| 12445 | 10 | 2461622 | 2486326 | mNeoVis.2.1.1.chr10\_hap1.g00043.t01 |
| 12446 | 10 | 2461622 | 2486326 | mNeoVis.2.1.1.chr10\_hap1.g00043.t01 |
| 12447 | 10 | 2486448 | 2489796 | mNeoVis.2.1.1.chr10\_hap1.g00044.t01 |
| 12448 | 10 | 2486448 | 2489796 | mNeoVis.2.1.1.chr10\_hap1.g00044.t01 |
| 12449 | 10 | 2509416 | 2519693 | mNeoVis.2.1.1.chr10\_hap1.g00045.t01 |
| 12450 | 10 | 2509416 | 2519693 | mNeoVis.2.1.1.chr10\_hap1.g00045.t01 |
| 12451 | 10 | 2522593 | 2525772 | mNeoVis.2.1.1.chr10\_hap1.g00046.t01 |
| 12452 | 10 | 2522593 | 2525772 | mNeoVis.2.1.1.chr10\_hap1.g00046.t01 |
| 12453 | 10 | 2539257 | 2661185 | mNeoVis.2.1.1.chr10\_hap1.g00047.t01 |
| 12454 | 10 | 2539257 | 2661185 | mNeoVis.2.1.1.chr10\_hap1.g00047.t01 |
| 12455 | 10 | 2599991 | 2601714 | mNeoVis.2.1.1.chr10\_hap1.g00048.t01 |
| 12456 | 10 | 2599991 | 2601714 | mNeoVis.2.1.1.chr10\_hap1.g00048.t01 |
| 12457 | 10 | 2694909 | 2704351 | mNeoVis.2.1.1.chr10\_hap1.g00049.t01 |
| 12458 | 10 | 2694909 | 2704351 | mNeoVis.2.1.1.chr10\_hap1.g00049.t01 |
| 12459 | 10 | 2704297 | 2743005 | mNeoVis.2.1.1.chr10\_hap1.g00050.t01 |
| 12460 | 10 | 2704297 | 2743005 | mNeoVis.2.1.1.chr10\_hap1.g00050.t01 |
| 12461 | 10 | 2758775 | 2760007 | mNeoVis.2.1.1.chr10\_hap1.g00051.t01 |
| 12462 | 10 | 2758775 | 2760007 | mNeoVis.2.1.1.chr10\_hap1.g00051.t01 |
| 12463 | 10 | 2811927 | 2823508 | mNeoVis.2.1.1.chr10\_hap1.g00052.t01 |
| 12464 | 10 | 2832141 | 2843765 | mNeoVis.2.1.1.chr10\_hap1.g00053.t01 |
| 12465 | 10 | 2847861 | 2852730 | mNeoVis.2.1.1.chr10\_hap1.g00054.t01 |
| 12466 | 10 | 2868341 | 2969255 | mNeoVis.2.1.1.chr10\_hap1.g00055.t01 |
| 12467 | 10 | 2987366 | 2997403 | mNeoVis.2.1.1.chr10\_hap1.g00056.t01 |
| 12468 | 10 | 5112590 | 5118561 | mNeoVis.2.1.1.chr10\_hap1.g00079.t01 |
| 12469 | 10 | 5128708 | 5141952 | mNeoVis.2.1.1.chr10\_hap1.g00080.t01 |
| 12470 | 10 | 5151987 | 5182544 | mNeoVis.2.1.1.chr10\_hap1.g00081.t01 |
| 12471 | 10 | 5235090 | 5257216 | mNeoVis.2.1.1.chr10\_hap1.g00082.t01 |
| 12472 | 10 | 5256992 | 5361321 | mNeoVis.2.1.1.chr10\_hap1.g00083.t01 |
| 12473 | 10 | 5415334 | 5541758 | mNeoVis.2.1.1.chr10\_hap1.g00084.t01 |
| 12474 | 10 | 5600630 | 5634085 | mNeoVis.2.1.1.chr10\_hap1.g00085.t01 |
| 12475 | 10 | 5639122 | 5663505 | mNeoVis.2.1.1.chr10\_hap1.g00086.t01 |
| 12476 | 10 | 5693578 | 5767795 | mNeoVis.2.1.1.chr10\_hap1.g00087.t01 |
| 12477 | 10 | 5768215 | 5796153 | mNeoVis.2.1.1.chr10\_hap1.g00088.t01 |
| 12478 | 10 | 5768215 | 5796153 | mNeoVis.2.1.1.chr10\_hap1.g00088.t01 |
| 12479 | 10 | 5796330 | 5876375 | mNeoVis.2.1.1.chr10\_hap1.g00089.t01 |
| 12480 | 10 | 5796330 | 5876375 | mNeoVis.2.1.1.chr10\_hap1.g00089.t01 |
| 12481 | 10 | 5962549 | 6072307 | mNeoVis.2.1.1.chr10\_hap1.g00090.t01 |
| 12482 | 10 | 5962549 | 6072307 | mNeoVis.2.1.1.chr10\_hap1.g00090.t01 |
| 12483 | 10 | 6157065 | 6221296 | mNeoVis.2.1.1.chr10\_hap1.g00091.t01 |
| 12484 | 10 | 11703142 | 11734987 | mNeoVis.2.1.1.chr10\_hap1.g00142.t01 |
| 12485 | 10 | 11777534 | 11851211 | mNeoVis.2.1.1.chr10\_hap1.g00143.t01 |
| 12486 | 10 | 11904496 | 12064411 | mNeoVis.2.1.1.chr10\_hap1.g00144.t01 |
| 12487 | 10 | 12106471 | 12113722 | mNeoVis.2.1.1.chr10\_hap1.g00145.t01 |
| 12488 | 10 | 12761926 | 13320069 | mNeoVis.2.1.1.chr10\_hap1.g00147.t01 |
| 12489 | 10 | 14546736 | 14577562 | mNeoVis.2.1.1.chr10\_hap1.g00163.t01 |
| 12490 | 10 | 14578861 | 14621620 | mNeoVis.2.1.1.chr10\_hap1.g00164.t01 |
| 12491 | 10 | 14622232 | 14636506 | mNeoVis.2.1.1.chr10\_hap1.g00165.t01 |
| 12492 | 10 | 14636486 | 14649521 | mNeoVis.2.1.1.chr10\_hap1.g00166.t01 |
| 12493 | 10 | 14652935 | 14702763 | mNeoVis.2.1.1.chr10\_hap1.g00167.t01 |
| 12494 | 10 | 14702842 | 14758387 | mNeoVis.2.1.1.chr10\_hap1.g00168.t01 |
| 12495 | 10 | 14737767 | 14760295 | mNeoVis.2.1.1.chr10\_hap1.g00169.t01 |
| 12496 | 10 | 14760529 | 14782026 | mNeoVis.2.1.1.chr10\_hap1.g00170.t01 |
| 12497 | 10 | 14916327 | 14932080 | mNeoVis.2.1.1.chr10\_hap1.g00171.t01 |
| 12498 | 10 | 14945565 | 14961801 | mNeoVis.2.1.1.chr10\_hap1.g00172.t01 |
| 12499 | 10 | 14996903 | 14999023 | mNeoVis.2.1.1.chr10\_hap1.g00173.t01 |
| 12500 | 10 | 16786288 | 16856574 | mNeoVis.2.1.1.chr10\_hap1.g00200.t01 |
| 12501 | 10 | 17051807 | 17071887 | mNeoVis.2.1.1.chr10\_hap1.g00201.t01 |
| 12502 | 10 | 17051807 | 17071887 | mNeoVis.2.1.1.chr10\_hap1.g00201.t01 |
| 12503 | 10 | 17074414 | 17102189 | mNeoVis.2.1.1.chr10\_hap1.g00202.t01 |
| 12504 | 10 | 17074414 | 17102189 | mNeoVis.2.1.1.chr10\_hap1.g00202.t01 |
| 12505 | 10 | 17115921 | 17149720 | mNeoVis.2.1.1.chr10\_hap1.g00203.t01 |
| 12506 | 10 | 17115921 | 17149720 | mNeoVis.2.1.1.chr10\_hap1.g00203.t01 |
| 12507 | 10 | 17227450 | 17227842 | mNeoVis.2.1.1.chr10\_hap1.g00204.t01 |
| 12508 | 10 | 17227450 | 17227842 | mNeoVis.2.1.1.chr10\_hap1.g00204.t01 |
| 12509 | 10 | 17227450 | 17227842 | mNeoVis.2.1.1.chr10\_hap1.g00204.t01 |
| 12510 | 10 | 17269710 | 17274631 | mNeoVis.2.1.1.chr10\_hap1.g00205.t01 |
| 12511 | 10 | 17269710 | 17274631 | mNeoVis.2.1.1.chr10\_hap1.g00205.t01 |
| 12512 | 10 | 17269710 | 17274631 | mNeoVis.2.1.1.chr10\_hap1.g00205.t01 |
| 12513 | 10 | 17329134 | 17349265 | mNeoVis.2.1.1.chr10\_hap1.g00206.t01 |
| 12514 | 10 | 17329134 | 17349265 | mNeoVis.2.1.1.chr10\_hap1.g00206.t01 |
| 12515 | 10 | 17329134 | 17349265 | mNeoVis.2.1.1.chr10\_hap1.g00206.t01 |
| 12516 | 10 | 17363470 | 17374292 | mNeoVis.2.1.1.chr10\_hap1.g00207.t01 |
| 12517 | 10 | 17363470 | 17374292 | mNeoVis.2.1.1.chr10\_hap1.g00207.t01 |
| 12518 | 10 | 17363470 | 17374292 | mNeoVis.2.1.1.chr10\_hap1.g00207.t01 |
| 12519 | 10 | 17453236 | 17453628 | mNeoVis.2.1.1.chr10\_hap1.g00208.t01 |
| 12520 | 10 | 17453236 | 17453628 | mNeoVis.2.1.1.chr10\_hap1.g00208.t01 |
| 12521 | 10 | 17453236 | 17453628 | mNeoVis.2.1.1.chr10\_hap1.g00208.t01 |
| 12522 | 10 | 17495415 | 17500114 | mNeoVis.2.1.1.chr10\_hap1.g00209.t01 |
| 12523 | 10 | 17495415 | 17500114 | mNeoVis.2.1.1.chr10\_hap1.g00209.t01 |
| 12524 | 10 | 17495415 | 17500114 | mNeoVis.2.1.1.chr10\_hap1.g00209.t01 |
| 12525 | 10 | 17554827 | 17574973 | mNeoVis.2.1.1.chr10\_hap1.g00210.t01 |
| 12526 | 10 | 17554827 | 17574973 | mNeoVis.2.1.1.chr10\_hap1.g00210.t01 |
| 12527 | 10 | 17554827 | 17574973 | mNeoVis.2.1.1.chr10\_hap1.g00210.t01 |
| 12528 | 10 | 17589146 | 17599971 | mNeoVis.2.1.1.chr10\_hap1.g00211.t01 |
| 12529 | 10 | 17589146 | 17599971 | mNeoVis.2.1.1.chr10\_hap1.g00211.t01 |
| 12530 | 10 | 17589146 | 17599971 | mNeoVis.2.1.1.chr10\_hap1.g00211.t01 |
| 12531 | 10 | 20401656 | 20415652 | mNeoVis.2.1.1.chr10\_hap1.g00228.t01 |
| 12532 | 10 | 20668933 | 21073217 | mNeoVis.2.1.1.chr10\_hap1.g00229.t01 |
| 12533 | 10 | 20668933 | 21073217 | mNeoVis.2.1.1.chr10\_hap1.g00229.t01 |
| 12534 | 10 | 21085615 | 21249234 | mNeoVis.2.1.1.chr10\_hap1.g00230.t01 |
| 12535 | 10 | 21085615 | 21249234 | mNeoVis.2.1.1.chr10\_hap1.g00230.t01 |
| 12536 | 10 | 21286215 | 21332422 | mNeoVis.2.1.1.chr10\_hap1.g00231.t01 |
| 12537 | 10 | 21286215 | 21332422 | mNeoVis.2.1.1.chr10\_hap1.g00231.t01 |
| 12538 | 10 | 21360981 | 21504275 | mNeoVis.2.1.1.chr10\_hap1.g00232.t01 |
| 12539 | 10 | 21360981 | 21504275 | mNeoVis.2.1.1.chr10\_hap1.g00232.t01 |
| 12540 | 10 | 21530527 | 21584891 | mNeoVis.2.1.1.chr10\_hap1.g00233.t01 |
| 12541 | 10 | 21530527 | 21584891 | mNeoVis.2.1.1.chr10\_hap1.g00233.t01 |
| 12542 | 10 | 22745470 | 22749785 | mNeoVis.2.1.1.chr10\_hap1.g00234.t01 |
| 12543 | 10 | 27027334 | 27086832 | mNeoVis.2.1.1.chr10\_hap1.g00237.t01 |
| 12544 | 10 | 27086800 | 27113915 | mNeoVis.2.1.1.chr10\_hap1.g00238.t01 |
| 12545 | 10 | 27245527 | 27371150 | mNeoVis.2.1.1.chr10\_hap1.g00239.t01 |
| 12546 | 10 | 28085085 | 28484050 | mNeoVis.2.1.1.chr10\_hap1.g00245.t01 |
| 12547 | 10 | 28149674 | 28169245 | mNeoVis.2.1.1.chr10\_hap1.g00246.t01 |
| 12548 | 10 | 28665112 | 28772390 | mNeoVis.2.1.1.chr10\_hap1.g00247.t01 |
| 12549 | 10 | 28774115 | 28864989 | mNeoVis.2.1.1.chr10\_hap1.g00248.t01 |
| 12550 | 10 | 29565311 | 29738843 | mNeoVis.2.1.1.chr10\_hap1.g00255.t01 |
| 12551 | 10 | 29643573 | 29644391 | mNeoVis.2.1.1.chr10\_hap1.g00256.t01 |
| 12552 | 10 | 29779898 | 29855986 | mNeoVis.2.1.1.chr10\_hap1.g00257.t01 |
| 12553 | 10 | 29857107 | 29886637 | mNeoVis.2.1.1.chr10\_hap1.g00258.t01 |
| 12554 | 10 | 30063115 | 30099361 | mNeoVis.2.1.1.chr10\_hap1.g00260.t01 |
| 12555 | 10 | 30063115 | 30099361 | mNeoVis.2.1.1.chr10\_hap1.g00260.t01 |
| 12556 | 10 | 30137987 | 30251365 | mNeoVis.2.1.1.chr10\_hap1.g00261.t01 |
| 12557 | 10 | 30137987 | 30251365 | mNeoVis.2.1.1.chr10\_hap1.g00261.t01 |
| 12558 | 10 | 30290685 | 30331720 | mNeoVis.2.1.1.chr10\_hap1.g00262.t01 |
| 12559 | 10 | 30290685 | 30331720 | mNeoVis.2.1.1.chr10\_hap1.g00262.t01 |
| 12560 | 10 | 30363414 | 30436821 | mNeoVis.2.1.1.chr10\_hap1.g00263.t01 |
| 12561 | 10 | 30363414 | 30436821 | mNeoVis.2.1.1.chr10\_hap1.g00263.t01 |
| 12562 | 10 | 30450109 | 30517221 | mNeoVis.2.1.1.chr10\_hap1.g00264.t01 |
| 12563 | 10 | 30450109 | 30517221 | mNeoVis.2.1.1.chr10\_hap1.g00264.t01 |
| 12564 | 10 | 30526378 | 30563317 | mNeoVis.2.1.1.chr10\_hap1.g00265.t01 |
| 12565 | 10 | 30526378 | 30563317 | mNeoVis.2.1.1.chr10\_hap1.g00265.t01 |
| 12566 | 10 | 30567446 | 30573366 | mNeoVis.2.1.1.chr10\_hap1.g00266.t01 |
| 12567 | 10 | 30567446 | 30573366 | mNeoVis.2.1.1.chr10\_hap1.g00266.t01 |
| 12568 | 10 | 30573549 | 30606379 | mNeoVis.2.1.1.chr10\_hap1.g00267.t01 |
| 12569 | 10 | 30573549 | 30606379 | mNeoVis.2.1.1.chr10\_hap1.g00267.t01 |
| 12570 | 10 | 30734385 | 30911955 | mNeoVis.2.1.1.chr10\_hap1.g00268.t01 |
| 12571 | 10 | 30930590 | 31043292 | mNeoVis.2.1.1.chr10\_hap1.g00269.t01 |
| 12572 | 10 | 31053912 | 31105186 | mNeoVis.2.1.1.chr10\_hap1.g00270.t01 |
| 12573 | 10 | 31157688 | 31197698 | mNeoVis.2.1.1.chr10\_hap1.g00271.t01 |
| 12574 | 10 | 31199180 | 31226611 | mNeoVis.2.1.1.chr10\_hap1.g00272.t01 |
| 12575 | 10 | 31298537 | 31437244 | mNeoVis.2.1.1.chr10\_hap1.g00273.t01 |
| 12576 | 10 | 31452506 | 31603144 | mNeoVis.2.1.1.chr10\_hap1.g00274.t01 |
| 12577 | 10 | 31452506 | 31603144 | mNeoVis.2.1.1.chr10\_hap1.g00274.t01 |
| 12578 | 10 | 31659451 | 31680443 | mNeoVis.2.1.1.chr10\_hap1.g00275.t01 |
| 12579 | 10 | 31659451 | 31680443 | mNeoVis.2.1.1.chr10\_hap1.g00275.t01 |
| 12580 | 10 | 31683125 | 31688215 | mNeoVis.2.1.1.chr10\_hap1.g00276.t01 |
| 12581 | 10 | 31683125 | 31688215 | mNeoVis.2.1.1.chr10\_hap1.g00276.t01 |
| 12582 | 10 | 31730031 | 31825371 | mNeoVis.2.1.1.chr10\_hap1.g00277.t01 |
| 12583 | 10 | 31730031 | 31825371 | mNeoVis.2.1.1.chr10\_hap1.g00277.t01 |
| 12584 | 10 | 31861182 | 31864812 | mNeoVis.2.1.1.chr10\_hap1.g00278.t01 |
| 12585 | 10 | 31861182 | 31864812 | mNeoVis.2.1.1.chr10\_hap1.g00278.t01 |
| 12586 | 10 | 31895321 | 31925436 | mNeoVis.2.1.1.chr10\_hap1.g00279.t01 |
| 12587 | 10 | 31895321 | 31925436 | mNeoVis.2.1.1.chr10\_hap1.g00279.t01 |
| 12588 | 10 | 31935689 | 32000710 | mNeoVis.2.1.1.chr10\_hap1.g00280.t01 |
| 12589 | 10 | 32000993 | 32019083 | mNeoVis.2.1.1.chr10\_hap1.g00281.t01 |
| 12590 | 10 | 32042377 | 32136869 | mNeoVis.2.1.1.chr10\_hap1.g00282.t01 |
| 12591 | 10 | 32144540 | 32210597 | mNeoVis.2.1.1.chr10\_hap1.g00283.t01 |
| 12592 | 10 | 32253404 | 32351630 | mNeoVis.2.1.1.chr10\_hap1.g00284.t01 |
| 12593 | 10 | 32418601 | 32515920 | mNeoVis.2.1.1.chr10\_hap1.g00285.t01 |
| 12594 | 10 | 32493489 | 32494432 | mNeoVis.2.1.1.chr10\_hap1.g00286.t01 |
| 12595 | 10 | 32560970 | 32566701 | mNeoVis.2.1.1.chr10\_hap1.g00287.t01 |
| 12596 | 10 | 33044393 | 33068787 | mNeoVis.2.1.1.chr10\_hap1.g00289.t01 |
| 12597 | 10 | 33437715 | 33503972 | mNeoVis.2.1.1.chr10\_hap1.g00290.t01 |
| 12598 | 10 | 33437715 | 33503972 | mNeoVis.2.1.1.chr10\_hap1.g00290.t01 |
| 12599 | 10 | 33522217 | 33532492 | mNeoVis.2.1.1.chr10\_hap1.g00291.t01 |
| 12600 | 10 | 33522217 | 33532492 | mNeoVis.2.1.1.chr10\_hap1.g00291.t01 |
| 12601 | 10 | 33559272 | 33574782 | mNeoVis.2.1.1.chr10\_hap1.g00292.t01 |
| 12602 | 10 | 33577974 | 33608305 | mNeoVis.2.1.1.chr10\_hap1.g00293.t01 |
| 12603 | 10 | 33657548 | 33664749 | mNeoVis.2.1.1.chr10\_hap1.g00295.t01 |
| 12604 | 10 | 33665714 | 33715633 | mNeoVis.2.1.1.chr10\_hap1.g00296.t01 |
| 12605 | 10 | 33736869 | 33777351 | mNeoVis.2.1.1.chr10\_hap1.g00297.t01 |
| 12606 | 10 | 33828146 | 33850048 | mNeoVis.2.1.1.chr10\_hap1.g00298.t01 |
| 12607 | 10 | 33850195 | 33866981 | mNeoVis.2.1.1.chr10\_hap1.g00299.t01 |
| 12608 | 10 | 33867159 | 33898588 | mNeoVis.2.1.1.chr10\_hap1.g00300.t01 |
| 12609 | 10 | 33898680 | 33976478 | mNeoVis.2.1.1.chr10\_hap1.g00301.t01 |
| 12610 | 10 | 33986773 | 34017680 | mNeoVis.2.1.1.chr10\_hap1.g00302.t01 |
| 12611 | 10 | 35234463 | 35235057 | mNeoVis.2.1.1.chr10\_hap1.g00305.t01 |
| 12612 | 10 | 35295022 | 35613117 | mNeoVis.2.1.1.chr10\_hap1.g00306.t01 |
| 12613 | 10 | 35821585 | 35846110 | mNeoVis.2.1.1.chr10\_hap1.g00307.t01 |
| 12614 | 10 | 35857555 | 35924787 | mNeoVis.2.1.1.chr10\_hap1.g00308.t01 |
| 12615 | 10 | 35934780 | 35956810 | mNeoVis.2.1.1.chr10\_hap1.g00309.t01 |
| 12616 | 10 | 35959556 | 35962871 | mNeoVis.2.1.1.chr10\_hap1.g00310.t01 |
| 12617 | 10 | 35969198 | 35983637 | mNeoVis.2.1.1.chr10\_hap1.g00311.t01 |
| 12618 | 10 | 37036992 | 37074778 | mNeoVis.2.1.1.chr10\_hap1.g00325.t01 |
| 12619 | 10 | 37055843 | 37114114 | mNeoVis.2.1.1.chr10\_hap1.g00326.t01 |
| 12620 | 10 | 37112087 | 37153245 | mNeoVis.2.1.1.chr10\_hap1.g00327.t01 |
| 12621 | 10 | 37112087 | 37153245 | mNeoVis.2.1.1.chr10\_hap1.g00327.t01 |
| 12622 | 10 | 37181503 | 37327587 | mNeoVis.2.1.1.chr10\_hap1.g00328.t01 |
| 12623 | 10 | 37181503 | 37327587 | mNeoVis.2.1.1.chr10\_hap1.g00328.t01 |
| 12624 | 10 | 37389176 | 37393972 | mNeoVis.2.1.1.chr10\_hap1.g00329.t01 |
| 12625 | 10 | 37405747 | 37484940 | mNeoVis.2.1.1.chr10\_hap1.g00330.t01 |
| 12626 | 10 | 37692963 | 37851615 | mNeoVis.2.1.1.chr10\_hap1.g00332.t01 |
| 12627 | 10 | 38426570 | 38462483 | mNeoVis.2.1.1.chr10\_hap1.g00333.t01 |
| 12628 | 10 | 38437970 | 38571582 | mNeoVis.2.1.1.chr10\_hap1.g00334.t01 |
| 12629 | 10 | 38707112 | 38743535 | mNeoVis.2.1.1.chr10\_hap1.g00335.t01 |
| 12630 | 10 | 38755693 | 38796616 | mNeoVis.2.1.1.chr10\_hap1.g00336.t01 |
| 12631 | 10 | 39874025 | 39967628 | mNeoVis.2.1.1.chr10\_hap1.g00344.t01 |
| 12632 | 10 | 39878576 | 39880409 | mNeoVis.2.1.1.chr10\_hap1.g00345.t01 |
| 12633 | 10 | 40057452 | 40101099 | mNeoVis.2.1.1.chr10\_hap1.g00346.t01 |
| 12634 | 10 | 40168918 | 40192058 | mNeoVis.2.1.1.chr10\_hap1.g00347.t01 |
| 12635 | 10 | 40168918 | 40192058 | mNeoVis.2.1.1.chr10\_hap1.g00347.t01 |
| 12636 | 10 | 40192207 | 40223418 | mNeoVis.2.1.1.chr10\_hap1.g00348.t01 |
| 12637 | 10 | 40192207 | 40223418 | mNeoVis.2.1.1.chr10\_hap1.g00348.t01 |
| 12638 | 10 | 40208839 | 40209624 | mNeoVis.2.1.1.chr10\_hap1.g00349.t01 |
| 12639 | 10 | 40208839 | 40209624 | mNeoVis.2.1.1.chr10\_hap1.g00349.t01 |
| 12640 | 10 | 40223329 | 40376219 | mNeoVis.2.1.1.chr10\_hap1.g00350.t01 |
| 12641 | 10 | 40223329 | 40376219 | mNeoVis.2.1.1.chr10\_hap1.g00350.t01 |
| 12642 | 10 | 40404735 | 40629901 | mNeoVis.2.1.1.chr10\_hap1.g00351.t01 |
| 12643 | 10 | 40647387 | 40731332 | mNeoVis.2.1.1.chr10\_hap1.g00352.t01 |
| 12644 | 10 | 40716474 | 40743973 | mNeoVis.2.1.1.chr10\_hap1.g00353.t01 |
| 12645 | 10 | 40716474 | 40743973 | mNeoVis.2.1.1.chr10\_hap1.g00353.t01 |
| 12646 | 10 | 40744006 | 40757235 | mNeoVis.2.1.1.chr10\_hap1.g00354.t01 |
| 12647 | 10 | 40763203 | 40767788 | mNeoVis.2.1.1.chr10\_hap1.g00355.t01 |
| 12648 | 10 | 40809448 | 40974515 | mNeoVis.2.1.1.chr10\_hap1.g00356.t01 |
| 12649 | 10 | 41499531 | 41510040 | mNeoVis.2.1.1.chr10\_hap1.g00357.t01 |
| 12650 | 10 | 41499531 | 41510040 | mNeoVis.2.1.1.chr10\_hap1.g00357.t01 |
| 12651 | 10 | 41577621 | 41709360 | mNeoVis.2.1.1.chr10\_hap1.g00358.t01 |
| 12652 | 10 | 41577621 | 41709360 | mNeoVis.2.1.1.chr10\_hap1.g00358.t01 |
| 12653 | 10 | 41709905 | 41727583 | mNeoVis.2.1.1.chr10\_hap1.g00359.t01 |
| 12654 | 10 | 41709905 | 41727583 | mNeoVis.2.1.1.chr10\_hap1.g00359.t01 |
| 12655 | 10 | 41727528 | 41735170 | mNeoVis.2.1.1.chr10\_hap1.g00360.t01 |
| 12656 | 10 | 41727528 | 41735170 | mNeoVis.2.1.1.chr10\_hap1.g00360.t01 |
| 12657 | 10 | 41745071 | 41811575 | mNeoVis.2.1.1.chr10\_hap1.g00361.t01 |
| 12658 | 10 | 41745071 | 41811575 | mNeoVis.2.1.1.chr10\_hap1.g00361.t01 |
| 12659 | 10 | 41745071 | 41811575 | mNeoVis.2.1.1.chr10\_hap1.g00361.t01 |
| 12660 | 10 | 41834368 | 41861664 | mNeoVis.2.1.1.chr10\_hap1.g00362.t01 |
| 12661 | 10 | 41834368 | 41861664 | mNeoVis.2.1.1.chr10\_hap1.g00362.t01 |
| 12662 | 10 | 41834368 | 41861664 | mNeoVis.2.1.1.chr10\_hap1.g00362.t01 |
| 12663 | 10 | 41870342 | 41900431 | mNeoVis.2.1.1.chr10\_hap1.g00363.t01 |
| 12664 | 10 | 41870342 | 41900431 | mNeoVis.2.1.1.chr10\_hap1.g00363.t01 |
| 12665 | 10 | 41870342 | 41900431 | mNeoVis.2.1.1.chr10\_hap1.g00363.t01 |
| 12666 | 10 | 41938412 | 42071011 | mNeoVis.2.1.1.chr10\_hap1.g00364.t01 |
| 12667 | 10 | 42206870 | 42411898 | mNeoVis.2.1.1.chr10\_hap1.g00365.t01 |
| 12668 | 10 | 42515369 | 42537280 | mNeoVis.2.1.1.chr10\_hap1.g00367.t01 |
| 12669 | 10 | 42537253 | 42564287 | mNeoVis.2.1.1.chr10\_hap1.g00368.t01 |
| 12670 | 10 | 42606990 | 42632049 | mNeoVis.2.1.1.chr10\_hap1.g00369.t01 |
| 12671 | 10 | 42632121 | 42695149 | mNeoVis.2.1.1.chr10\_hap1.g00370.t01 |
| 12672 | 10 | 43017044 | 43040642 | mNeoVis.2.1.1.chr10\_hap1.g00371.t01 |
| 12673 | 10 | 43033836 | 43037711 | mNeoVis.2.1.1.chr10\_hap1.g00372.t01 |
| 12674 | 10 | 43033836 | 43037711 | mNeoVis.2.1.1.chr10\_hap1.g00372.t01 |
| 12675 | 10 | 43172391 | 43240937 | mNeoVis.2.1.1.chr10\_hap1.g00373.t01 |
| 12676 | 10 | 43172391 | 43240937 | mNeoVis.2.1.1.chr10\_hap1.g00373.t01 |
| 12677 | 10 | 44035939 | 44143944 | mNeoVis.2.1.1.chr10\_hap1.g00375.t01 |
| 12678 | 10 | 44244984 | 44275640 | mNeoVis.2.1.1.chr10\_hap1.g00376.t01 |
| 12679 | 10 | 44277309 | 44314922 | mNeoVis.2.1.1.chr10\_hap1.g00377.t01 |
| 12680 | 10 | 44322424 | 44406621 | mNeoVis.2.1.1.chr10\_hap1.g00378.t01 |
| 12681 | 10 | 44322424 | 44406621 | mNeoVis.2.1.1.chr10\_hap1.g00378.t01 |
| 12682 | 10 | 44472440 | 44511034 | mNeoVis.2.1.1.chr10\_hap1.g00379.t01 |
| 12683 | 10 | 44472440 | 44511034 | mNeoVis.2.1.1.chr10\_hap1.g00379.t01 |
| 12684 | 10 | 44514304 | 44531819 | mNeoVis.2.1.1.chr10\_hap1.g00380.t01 |
| 12685 | 10 | 44583510 | 44597250 | mNeoVis.2.1.1.chr10\_hap1.g00381.t01 |
| 12686 | 10 | 44601642 | 44624444 | mNeoVis.2.1.1.chr10\_hap1.g00382.t01 |
| 12687 | 10 | 44624500 | 44690865 | mNeoVis.2.1.1.chr10\_hap1.g00383.t01 |
| 12688 | 10 | 44726944 | 44772063 | mNeoVis.2.1.1.chr10\_hap1.g00384.t01 |
| 12689 | 10 | 44726944 | 44772063 | mNeoVis.2.1.1.chr10\_hap1.g00384.t01 |
| 12690 | 10 | 44748138 | 44749247 | mNeoVis.2.1.1.chr10\_hap1.g00385.t01 |
| 12691 | 10 | 44748138 | 44749247 | mNeoVis.2.1.1.chr10\_hap1.g00385.t01 |
| 12692 | 10 | 44814809 | 45021045 | mNeoVis.2.1.1.chr10\_hap1.g00386.t01 |
| 12693 | 10 | 44814809 | 45021045 | mNeoVis.2.1.1.chr10\_hap1.g00386.t01 |
| 12694 | 10 | 45044108 | 45074526 | mNeoVis.2.1.1.chr10\_hap1.g00387.t01 |
| 12695 | 10 | 45044108 | 45074526 | mNeoVis.2.1.1.chr10\_hap1.g00387.t01 |
| 12696 | 10 | 45044108 | 45074526 | mNeoVis.2.1.1.chr10\_hap1.g00387.t01 |
| 12697 | 10 | 45074585 | 45098310 | mNeoVis.2.1.1.chr10\_hap1.g00388.t01 |
| 12698 | 10 | 45074585 | 45098310 | mNeoVis.2.1.1.chr10\_hap1.g00388.t01 |
| 12699 | 10 | 45074585 | 45098310 | mNeoVis.2.1.1.chr10\_hap1.g00388.t01 |
| 12700 | 10 | 45107692 | 45125766 | mNeoVis.2.1.1.chr10\_hap1.g00389.t01 |
| 12701 | 10 | 45107692 | 45125766 | mNeoVis.2.1.1.chr10\_hap1.g00389.t01 |
| 12702 | 10 | 45107692 | 45125766 | mNeoVis.2.1.1.chr10\_hap1.g00389.t01 |
| 12703 | 10 | 45127411 | 45142621 | mNeoVis.2.1.1.chr10\_hap1.g00390.t01 |
| 12704 | 10 | 45127411 | 45142621 | mNeoVis.2.1.1.chr10\_hap1.g00390.t01 |
| 12705 | 10 | 45127411 | 45142621 | mNeoVis.2.1.1.chr10\_hap1.g00390.t01 |
| 12706 | 10 | 45489525 | 45496594 | mNeoVis.2.1.1.chr10\_hap1.g00391.t01 |
| 12707 | 10 | 46002956 | 46003964 | mNeoVis.2.1.1.chr10\_hap1.g00392.t01 |
| 12708 | 10 | 46002956 | 46003964 | mNeoVis.2.1.1.chr10\_hap1.g00392.t01 |
| 12709 | 10 | 46002956 | 46003964 | mNeoVis.2.1.1.chr10\_hap1.g00392.t01 |
| 12710 | 10 | 46184397 | 46267996 | mNeoVis.2.1.1.chr10\_hap1.g00393.t01 |
| 12711 | 10 | 46184397 | 46267996 | mNeoVis.2.1.1.chr10\_hap1.g00393.t01 |
| 12712 | 10 | 46184397 | 46267996 | mNeoVis.2.1.1.chr10\_hap1.g00393.t01 |
| 12713 | 10 | 46325161 | 46416065 | mNeoVis.2.1.1.chr10\_hap1.g00394.t01 |
| 12714 | 10 | 46325161 | 46416065 | mNeoVis.2.1.1.chr10\_hap1.g00394.t01 |
| 12715 | 10 | 46444742 | 46445476 | mNeoVis.2.1.1.chr10\_hap1.g00395.t01 |
| 12716 | 10 | 46444742 | 46445476 | mNeoVis.2.1.1.chr10\_hap1.g00395.t01 |
| 12717 | 10 | 46565387 | 46587170 | mNeoVis.2.1.1.chr10\_hap1.g00398.t01 |
| 12718 | 10 | 46597209 | 46643672 | mNeoVis.2.1.1.chr10\_hap1.g00399.t01 |
| 12719 | 10 | 46645734 | 46709236 | mNeoVis.2.1.1.chr10\_hap1.g00400.t01 |
| 12720 | 10 | 46709779 | 46816894 | mNeoVis.2.1.1.chr10\_hap1.g00401.t01 |
| 12721 | 10 | 46937674 | 46951146 | mNeoVis.2.1.1.chr10\_hap1.g00402.t01 |
| 12722 | 10 | 46937674 | 46951146 | mNeoVis.2.1.1.chr10\_hap1.g00402.t01 |
| 12723 | 10 | 46989869 | 46999254 | mNeoVis.2.1.1.chr10\_hap1.g00403.t01 |
| 12724 | 10 | 46989869 | 46999254 | mNeoVis.2.1.1.chr10\_hap1.g00403.t01 |
| 12725 | 10 | 47194464 | 47257128 | mNeoVis.2.1.1.chr10\_hap1.g00404.t01 |
| 12726 | 10 | 47194464 | 47257128 | mNeoVis.2.1.1.chr10\_hap1.g00404.t01 |
| 12727 | 10 | 47194464 | 47257128 | mNeoVis.2.1.1.chr10\_hap1.g00404.t01 |
| 12728 | 10 | 47256846 | 47272463 | mNeoVis.2.1.1.chr10\_hap1.g00405.t01 |
| 12729 | 10 | 47256846 | 47272463 | mNeoVis.2.1.1.chr10\_hap1.g00405.t01 |
| 12730 | 10 | 47256846 | 47272463 | mNeoVis.2.1.1.chr10\_hap1.g00405.t01 |
| 12731 | 10 | 47299847 | 47414160 | mNeoVis.2.1.1.chr10\_hap1.g00406.t01 |
| 12732 | 10 | 47299847 | 47414160 | mNeoVis.2.1.1.chr10\_hap1.g00406.t01 |
| 12733 | 10 | 47513827 | 47535880 | mNeoVis.2.1.1.chr10\_hap1.g00407.t01 |
| 12734 | 10 | 47513827 | 47535880 | mNeoVis.2.1.1.chr10\_hap1.g00407.t01 |
| 12735 | 10 | 47513827 | 47535880 | mNeoVis.2.1.1.chr10\_hap1.g00407.t01 |
| 12736 | 10 | 47537623 | 47746253 | mNeoVis.2.1.1.chr10\_hap1.g00408.t01 |
| 12737 | 10 | 47537623 | 47746253 | mNeoVis.2.1.1.chr10\_hap1.g00408.t01 |
| 12738 | 10 | 47947198 | 47960861 | mNeoVis.2.1.1.chr10\_hap1.g00409.t01 |
| 12739 | 10 | 47947198 | 47960861 | mNeoVis.2.1.1.chr10\_hap1.g00409.t01 |
| 12740 | 10 | 48068044 | 48175433 | mNeoVis.2.1.1.chr10\_hap1.g00410.t01 |
| 12741 | 10 | 48068044 | 48175433 | mNeoVis.2.1.1.chr10\_hap1.g00410.t01 |
| 12742 | 10 | 48207075 | 48249473 | mNeoVis.2.1.1.chr10\_hap1.g00411.t01 |
| 12743 | 10 | 48249303 | 48278769 | mNeoVis.2.1.1.chr10\_hap1.g00412.t01 |
| 12744 | 10 | 48561773 | 48670033 | mNeoVis.2.1.1.chr10\_hap1.g00416.t01 |
| 12745 | 10 | 48675380 | 48684571 | mNeoVis.2.1.1.chr10\_hap1.g00417.t01 |
| 12746 | 10 | 48678637 | 48737649 | mNeoVis.2.1.1.chr10\_hap1.g00418.t01 |
| 12747 | 10 | 48738384 | 48750300 | mNeoVis.2.1.1.chr10\_hap1.g00419.t01 |
| 12748 | 10 | 48750382 | 48790564 | mNeoVis.2.1.1.chr10\_hap1.g00420.t01 |
| 12749 | 10 | 48794008 | 48877056 | mNeoVis.2.1.1.chr10\_hap1.g00421.t01 |
| 12750 | 10 | 48877817 | 48902990 | mNeoVis.2.1.1.chr10\_hap1.g00422.t01 |
| 12751 | 10 | 49060911 | 49064371 | mNeoVis.2.1.1.chr10\_hap1.g00423.t01 |
| 12752 | 10 | 49097254 | 49148633 | mNeoVis.2.1.1.chr10\_hap1.g00424.t01 |
| 12753 | 10 | 49149461 | 49160681 | mNeoVis.2.1.1.chr10\_hap1.g00425.t01 |
| 12754 | 10 | 49160831 | 49162103 | mNeoVis.2.1.1.chr10\_hap1.g00426.t01 |
| 12755 | 10 | 49166398 | 49218551 | mNeoVis.2.1.1.chr10\_hap1.g00427.t01 |
| 12756 | 10 | 49225827 | 49254484 | mNeoVis.2.1.1.chr10\_hap1.g00428.t01 |
| 12757 | 10 | 49254676 | 49267023 | mNeoVis.2.1.1.chr10\_hap1.g00429.t01 |
| 12758 | 10 | 49267160 | 49271094 | mNeoVis.2.1.1.chr10\_hap1.g00430.t01 |
| 12759 | 10 | 49271212 | 49272661 | mNeoVis.2.1.1.chr10\_hap1.g00431.t01 |
| 12760 | 10 | 49274564 | 49285162 | mNeoVis.2.1.1.chr10\_hap1.g00432.t01 |
| 12761 | 10 | 49291562 | 49293402 | mNeoVis.2.1.1.chr10\_hap1.g00433.t01 |
| 12762 | 10 | 52072410 | 52073216 | mNeoVis.2.1.1.chr10\_hap1.g00435.t01 |
| 12763 | 10 | 52072410 | 52073216 | mNeoVis.2.1.1.chr10\_hap1.g00435.t01 |
| 12764 | 10 | 52072410 | 52073216 | mNeoVis.2.1.1.chr10\_hap1.g00435.t01 |
| 12765 | 10 | 53132236 | 53133003 | mNeoVis.2.1.1.chr10\_hap1.g00437.t01 |
| 12766 | 10 | 53163776 | 53215300 | mNeoVis.2.1.1.chr10\_hap1.g00438.t01 |
| 12767 | 10 | 56567923 | 56798582 | mNeoVis.2.1.1.chr10\_hap1.g00447.t01 |
| 12768 | 10 | 56605450 | 56606615 | mNeoVis.2.1.1.chr10\_hap1.g00448.t01 |
| 12769 | 10 | 58703566 | 58741185 | mNeoVis.2.1.1.chr10\_hap1.g00450.t01 |
| 12770 | 10 | 58754112 | 58839941 | mNeoVis.2.1.1.chr10\_hap1.g00451.t01 |
| 12771 | 10 | 58862641 | 58869915 | mNeoVis.2.1.1.chr10\_hap1.g00452.t01 |
| 12772 | 10 | 58874177 | 58891467 | mNeoVis.2.1.1.chr10\_hap1.g00453.t01 |
| 12773 | 10 | 58895672 | 58920747 | mNeoVis.2.1.1.chr10\_hap1.g00454.t01 |
| 12774 | 10 | 58921390 | 58923519 | mNeoVis.2.1.1.chr10\_hap1.g00455.t01 |
| 12775 | 10 | 58932005 | 58998818 | mNeoVis.2.1.1.chr10\_hap1.g00456.t01 |
| 12776 | 10 | 59929885 | 59975437 | mNeoVis.2.1.1.chr10\_hap1.g00460.t01 |
| 12777 | 10 | 60026539 | 60252840 | mNeoVis.2.1.1.chr10\_hap1.g00461.t01 |
| 12778 | 10 | 60267722 | 60271781 | mNeoVis.2.1.1.chr10\_hap1.g00462.t01 |
| 12779 | 10 | 60274555 | 60282158 | mNeoVis.2.1.1.chr10\_hap1.g00463.t01 |
| 12780 | 10 | 60645956 | 61143922 | mNeoVis.2.1.1.chr10\_hap1.g00465.t01 |
| 12781 | 10 | 61142017 | 61160630 | mNeoVis.2.1.1.chr10\_hap1.g00466.t01 |
| 12782 | 10 | 61228788 | 61230179 | mNeoVis.2.1.1.chr10\_hap1.g00467.t01 |
| 12783 | 10 | 61240677 | 61241291 | mNeoVis.2.1.1.chr10\_hap1.g00468.t01 |
| 12784 | 10 | 61295872 | 61299998 | mNeoVis.2.1.1.chr10\_hap1.g00469.t01 |
| 12785 | 10 | 61462135 | 61482885 | mNeoVis.2.1.1.chr10\_hap1.g00470.t01 |
| 12786 | 10 | 61857441 | 61891487 | mNeoVis.2.1.1.chr10\_hap1.g00471.t01 |
| 12787 | 10 | 61906808 | 62173404 | mNeoVis.2.1.1.chr10\_hap1.g00472.t01 |
| 12788 | 10 | 62175029 | 62177311 | mNeoVis.2.1.1.chr10\_hap1.g00473.t01 |
| 12789 | 10 | 62177197 | 62231829 | mNeoVis.2.1.1.chr10\_hap1.g00474.t01 |
| 12790 | 10 | 62265268 | 62267023 | mNeoVis.2.1.1.chr10\_hap1.g00475.t01 |
| 12791 | 10 | 62267224 | 62291179 | mNeoVis.2.1.1.chr10\_hap1.g00476.t01 |
| 12792 | 10 | 62292985 | 62312574 | mNeoVis.2.1.1.chr10\_hap1.g00477.t01 |
| 12793 | 10 | 62297487 | 62300219 | mNeoVis.2.1.1.chr10\_hap1.g00478.t01 |
| 12794 | 10 | 62320056 | 62361888 | mNeoVis.2.1.1.chr10\_hap1.g00479.t01 |
| 12795 | 10 | 62361987 | 62398132 | mNeoVis.2.1.1.chr10\_hap1.g00480.t01 |
| 12796 | 10 | 62447404 | 62566742 | mNeoVis.2.1.1.chr10\_hap1.g00481.t01 |
| 12797 | 10 | 62580706 | 62582017 | mNeoVis.2.1.1.chr10\_hap1.g00482.t01 |
| 12798 | 10 | 62586777 | 62592417 | mNeoVis.2.1.1.chr10\_hap1.g00483.t01 |
| 12799 | 10 | 62652096 | 62717314 | mNeoVis.2.1.1.chr10\_hap1.g00484.t01 |
| 12800 | 10 | 62757193 | 62780980 | mNeoVis.2.1.1.chr10\_hap1.g00485.t01 |
| 12801 | 10 | 62837920 | 62861956 | mNeoVis.2.1.1.chr10\_hap1.g00486.t01 |
| 12802 | 10 | 63292149 | 63320174 | mNeoVis.2.1.1.chr10\_hap1.g00488.t01 |
| 12803 | 10 | 63394858 | 64242409 | mNeoVis.2.1.1.chr10\_hap1.g00489.t01 |
| 12804 | 10 | 64320756 | 64743556 | mNeoVis.2.1.1.chr10\_hap1.g00490.t01 |
| 12805 | 10 | 64861135 | 64958853 | mNeoVis.2.1.1.chr10\_hap1.g00491.t01 |
| 12806 | 10 | 66844900 | 67175237 | mNeoVis.2.1.1.chr10\_hap1.g00504.t01 |
| 12807 | 10 | 67555052 | 67894357 | mNeoVis.2.1.1.chr10\_hap1.g00505.t01 |
| 12808 | 10 | 67880160 | 67888425 | mNeoVis.2.1.1.chr10\_hap1.g00506.t01 |
| 12809 | 10 | 67902131 | 67904613 | mNeoVis.2.1.1.chr10\_hap1.g00507.t01 |
| 12810 | 10 | 68336721 | 68337860 | mNeoVis.2.1.1.chr10\_hap1.g00508.t01 |
| 12811 | 10 | 71812672 | 71815889 | mNeoVis.2.1.1.chr10\_hap1.g00514.t01 |
| 12812 | 10 | 71831988 | 71838101 | mNeoVis.2.1.1.chr10\_hap1.g00515.t01 |
| 12813 | 10 | 71846858 | 71895816 | mNeoVis.2.1.1.chr10\_hap1.g00516.t01 |
| 12814 | 10 | 71920975 | 71924617 | mNeoVis.2.1.1.chr10\_hap1.g00517.t01 |
| 12815 | 10 | 71934908 | 71938514 | mNeoVis.2.1.1.chr10\_hap1.g00518.t01 |
| 12816 | 10 | 71947962 | 71970218 | mNeoVis.2.1.1.chr10\_hap1.g00519.t01 |
| 12817 | 10 | 71990111 | 72000118 | mNeoVis.2.1.1.chr10\_hap1.g00520.t01 |
| 12818 | 10 | 72025897 | 72029346 | mNeoVis.2.1.1.chr10\_hap1.g00521.t01 |
| 12819 | 10 | 72030513 | 72045754 | mNeoVis.2.1.1.chr10\_hap1.g00522.t01 |
| 12820 | 10 | 72047520 | 72049244 | mNeoVis.2.1.1.chr10\_hap1.g00523.t01 |
| 12821 | 10 | 72049678 | 72054855 | mNeoVis.2.1.1.chr10\_hap1.g00524.t01 |
| 12822 | 10 | 72053162 | 72057784 | mNeoVis.2.1.1.chr10\_hap1.g00525.t01 |
| 12823 | 10 | 72057767 | 72063122 | mNeoVis.2.1.1.chr10\_hap1.g00526.t01 |
| 12824 | 10 | 72063193 | 72069899 | mNeoVis.2.1.1.chr10\_hap1.g00527.t01 |
| 12825 | 10 | 72093288 | 72099458 | mNeoVis.2.1.1.chr10\_hap1.g00528.t01 |
| 12826 | 10 | 72099655 | 72116615 | mNeoVis.2.1.1.chr10\_hap1.g00529.t01 |
| 12827 | 10 | 72125187 | 72128818 | mNeoVis.2.1.1.chr10\_hap1.g00530.t01 |
| 12828 | 10 | 72128708 | 72135826 | mNeoVis.2.1.1.chr10\_hap1.g00531.t01 |
| 12829 | 10 | 72135828 | 72146988 | mNeoVis.2.1.1.chr10\_hap1.g00532.t01 |
| 12830 | 10 | 72147703 | 72149478 | mNeoVis.2.1.1.chr10\_hap1.g00533.t01 |
| 12831 | 10 | 72149854 | 72154110 | mNeoVis.2.1.1.chr10\_hap1.g00534.t01 |
| 12832 | 10 | 72155629 | 72158131 | mNeoVis.2.1.1.chr10\_hap1.g00535.t01 |
| 12833 | 10 | 72164132 | 72168951 | mNeoVis.2.1.1.chr10\_hap1.g00536.t01 |
| 12834 | 10 | 72169224 | 72178173 | mNeoVis.2.1.1.chr10\_hap1.g00537.t01 |
| 12835 | 10 | 72178214 | 72185725 | mNeoVis.2.1.1.chr10\_hap1.g00538.t01 |
| 12836 | 10 | 72193016 | 72198369 | mNeoVis.2.1.1.chr10\_hap1.g00539.t01 |
| 12837 | 10 | 72540731 | 72550170 | mNeoVis.2.1.1.chr10\_hap1.g00553.t01 |
| 12838 | 10 | 72550315 | 72558861 | mNeoVis.2.1.1.chr10\_hap1.g00554.t01 |
| 12839 | 10 | 72559093 | 72562278 | mNeoVis.2.1.1.chr10\_hap1.g00555.t01 |
| 12840 | 10 | 72564629 | 72595280 | mNeoVis.2.1.1.chr10\_hap1.g00556.t01 |
| 12841 | 10 | 72631017 | 72639012 | mNeoVis.2.1.1.chr10\_hap1.g00557.t01 |
| 12842 | 10 | 72657184 | 72679074 | mNeoVis.2.1.1.chr10\_hap1.g00558.t01 |
| 12843 | 10 | 72684246 | 72708455 | mNeoVis.2.1.1.chr10\_hap1.g00559.t01 |
| 12844 | 10 | 72710585 | 72713219 | mNeoVis.2.1.1.chr10\_hap1.g00560.t01 |
| 12845 | 10 | 72713640 | 72739349 | mNeoVis.2.1.1.chr10\_hap1.g00561.t01 |
| 12846 | 10 | 72725218 | 72733121 | mNeoVis.2.1.1.chr10\_hap1.g00562.t01 |
| 12847 | 10 | 72736057 | 72742411 | mNeoVis.2.1.1.chr10\_hap1.g00563.t01 |
| 12848 | 10 | 72749795 | 72766264 | mNeoVis.2.1.1.chr10\_hap1.g00564.t01 |
| 12849 | 10 | 72770181 | 72776213 | mNeoVis.2.1.1.chr10\_hap1.g00565.t01 |
| 12850 | 10 | 72777043 | 72799312 | mNeoVis.2.1.1.chr10\_hap1.g00566.t01 |
| 12851 | 10 | 72791151 | 72817817 | mNeoVis.2.1.1.chr10\_hap1.g00567.t01 |
| 12852 | 10 | 72841446 | 72891335 | mNeoVis.2.1.1.chr10\_hap1.g00568.t01 |
| 12853 | 10 | 72896440 | 72898060 | mNeoVis.2.1.1.chr10\_hap1.g00569.t01 |
| 12854 | 10 | 72906686 | 72911814 | mNeoVis.2.1.1.chr10\_hap1.g00570.t01 |
| 12855 | 10 | 72911938 | 72914323 | mNeoVis.2.1.1.chr10\_hap1.g00571.t01 |
| 12856 | 10 | 72915107 | 72949750 | mNeoVis.2.1.1.chr10\_hap1.g00572.t01 |
| 12857 | 10 | 72950740 | 72960679 | mNeoVis.2.1.1.chr10\_hap1.g00573.t01 |
| 12858 | 10 | 72964349 | 72965209 | mNeoVis.2.1.1.chr10\_hap1.g00574.t01 |
| 12859 | 10 | 72971281 | 72978772 | mNeoVis.2.1.1.chr10\_hap1.g00575.t01 |
| 12860 | 10 | 72991606 | 73015691 | mNeoVis.2.1.1.chr10\_hap1.g00576.t01 |
| 12861 | 10 | 73017658 | 73028395 | mNeoVis.2.1.1.chr10\_hap1.g00577.t01 |
| 12862 | 10 | 73696205 | 73698777 | mNeoVis.2.1.1.chr10\_hap1.g00591.t01 |
| 12863 | 10 | 74422621 | 74641646 | mNeoVis.2.1.1.chr10\_hap1.g00592.t01 |
| 12864 | 10 | 75343291 | 75387213 | mNeoVis.2.1.1.chr10\_hap1.g00594.t01 |
| 12865 | 10 | 75343291 | 75387213 | mNeoVis.2.1.1.chr10\_hap1.g00594.t01 |
| 12866 | 10 | 75474666 | 75544256 | mNeoVis.2.1.1.chr10\_hap1.g00595.t01 |
| 12867 | 10 | 75474666 | 75544256 | mNeoVis.2.1.1.chr10\_hap1.g00595.t01 |
| 12868 | 10 | 75509934 | 75510515 | mNeoVis.2.1.1.chr10\_hap1.g00596.t01 |
| 12869 | 10 | 75509934 | 75510515 | mNeoVis.2.1.1.chr10\_hap1.g00596.t01 |
| 12870 | 10 | 75551540 | 75572799 | mNeoVis.2.1.1.chr10\_hap1.g00597.t01 |
| 12871 | 10 | 75551540 | 75572799 | mNeoVis.2.1.1.chr10\_hap1.g00597.t01 |
| 12872 | 10 | 75589976 | 75787361 | mNeoVis.2.1.1.chr10\_hap1.g00598.t01 |
| 12873 | 10 | 75589976 | 75787361 | mNeoVis.2.1.1.chr10\_hap1.g00598.t01 |
| 12874 | 10 | 75797109 | 75891311 | mNeoVis.2.1.1.chr10\_hap1.g00599.t01 |
| 12875 | 10 | 75797109 | 75891311 | mNeoVis.2.1.1.chr10\_hap1.g00599.t01 |
| 12876 | 10 | 75898637 | 75944037 | mNeoVis.2.1.1.chr10\_hap1.g00600.t01 |
| 12877 | 10 | 75898637 | 75944037 | mNeoVis.2.1.1.chr10\_hap1.g00600.t01 |
| 12878 | 10 | 75950180 | 76006082 | mNeoVis.2.1.1.chr10\_hap1.g00601.t01 |
| 12879 | 10 | 75950180 | 76006082 | mNeoVis.2.1.1.chr10\_hap1.g00601.t01 |
| 12880 | 10 | 76031552 | 76071825 | mNeoVis.2.1.1.chr10\_hap1.g00602.t01 |
| 12881 | 10 | 76031552 | 76071825 | mNeoVis.2.1.1.chr10\_hap1.g00602.t01 |
| 12882 | 10 | 76268666 | 76338708 | mNeoVis.2.1.1.chr10\_hap1.g00606.t01 |
| 12883 | 10 | 76316960 | 76338294 | mNeoVis.2.1.1.chr10\_hap1.g00607.t01 |
| 12884 | 10 | 76338770 | 76355619 | mNeoVis.2.1.1.chr10\_hap1.g00608.t01 |
| 12885 | 10 | 76358436 | 76395263 | mNeoVis.2.1.1.chr10\_hap1.g00609.t01 |
| 12886 | 10 | 76463599 | 76622211 | mNeoVis.2.1.1.chr10\_hap1.g00610.t01 |
| 12887 | 10 | 76649494 | 76711972 | mNeoVis.2.1.1.chr10\_hap1.g00611.t01 |
| 12888 | 10 | 76649494 | 76711972 | mNeoVis.2.1.1.chr10\_hap1.g00611.t01 |
| 12889 | 10 | 76738661 | 76852101 | mNeoVis.2.1.1.chr10\_hap1.g00612.t01 |
| 12890 | 10 | 76955842 | 76992492 | mNeoVis.2.1.1.chr10\_hap1.g00613.t01 |
| 12891 | 10 | 77013117 | 77073294 | mNeoVis.2.1.1.chr10\_hap1.g00614.t01 |
| 12892 | 10 | 77073537 | 77166581 | mNeoVis.2.1.1.chr10\_hap1.g00615.t01 |
| 12893 | 10 | 77181400 | 77234179 | mNeoVis.2.1.1.chr10\_hap1.g00616.t01 |
| 12894 | 10 | 78250044 | 78303044 | mNeoVis.2.1.1.chr10\_hap1.g00627.t01 |
| 12895 | 10 | 78250044 | 78303044 | mNeoVis.2.1.1.chr10\_hap1.g00627.t01 |
| 12896 | 10 | 78375940 | 78480226 | mNeoVis.2.1.1.chr10\_hap1.g00628.t01 |
| 12897 | 10 | 78375940 | 78480226 | mNeoVis.2.1.1.chr10\_hap1.g00628.t01 |
| 12898 | 10 | 78375940 | 78480226 | mNeoVis.2.1.1.chr10\_hap1.g00628.t01 |
| 12899 | 10 | 78487664 | 78516627 | mNeoVis.2.1.1.chr10\_hap1.g00629.t01 |
| 12900 | 10 | 78487664 | 78516627 | mNeoVis.2.1.1.chr10\_hap1.g00629.t01 |
| 12901 | 10 | 78487664 | 78516627 | mNeoVis.2.1.1.chr10\_hap1.g00629.t01 |
| 12902 | 10 | 78579050 | 78641255 | mNeoVis.2.1.1.chr10\_hap1.g00630.t01 |
| 12903 | 10 | 78579050 | 78641255 | mNeoVis.2.1.1.chr10\_hap1.g00630.t01 |
| 12904 | 10 | 78579050 | 78641255 | mNeoVis.2.1.1.chr10\_hap1.g00630.t01 |
| 12905 | 10 | 78668637 | 78806249 | mNeoVis.2.1.1.chr10\_hap1.g00631.t01 |
| 12906 | 10 | 78668637 | 78806249 | mNeoVis.2.1.1.chr10\_hap1.g00631.t01 |
| 12907 | 10 | 78751188 | 78753161 | mNeoVis.2.1.1.chr10\_hap1.g00632.t01 |
| 12908 | 10 | 78751188 | 78753161 | mNeoVis.2.1.1.chr10\_hap1.g00632.t01 |
| 12909 | 10 | 78817837 | 78912600 | mNeoVis.2.1.1.chr10\_hap1.g00633.t01 |
| 12910 | 10 | 79227452 | 79299983 | mNeoVis.2.1.1.chr10\_hap1.g00635.t01 |
| 12911 | 10 | 79227452 | 79299983 | mNeoVis.2.1.1.chr10\_hap1.g00635.t01 |
| 12912 | 10 | 79307537 | 79320450 | mNeoVis.2.1.1.chr10\_hap1.g00636.t01 |
| 12913 | 10 | 79307537 | 79320450 | mNeoVis.2.1.1.chr10\_hap1.g00636.t01 |
| 12914 | 10 | 79343715 | 79433670 | mNeoVis.2.1.1.chr10\_hap1.g00637.t01 |
| 12915 | 10 | 79343715 | 79433670 | mNeoVis.2.1.1.chr10\_hap1.g00637.t01 |
| 12916 | 10 | 79437487 | 79439370 | mNeoVis.2.1.1.chr10\_hap1.g00638.t01 |
| 12917 | 10 | 79437487 | 79439370 | mNeoVis.2.1.1.chr10\_hap1.g00638.t01 |
| 12918 | 10 | 79462179 | 79495188 | mNeoVis.2.1.1.chr10\_hap1.g00639.t01 |
| 12919 | 10 | 79462179 | 79495188 | mNeoVis.2.1.1.chr10\_hap1.g00639.t01 |
| 12920 | 10 | 79495741 | 79524335 | mNeoVis.2.1.1.chr10\_hap1.g00640.t01 |
| 12921 | 10 | 79495741 | 79524335 | mNeoVis.2.1.1.chr10\_hap1.g00640.t01 |
| 12922 | 10 | 82448516 | 82481275 | mNeoVis.2.1.1.chr10\_hap1.g00651.t01 |
| 12923 | 10 | 82481279 | 82490666 | mNeoVis.2.1.1.chr10\_hap1.g00652.t01 |
| 12924 | 10 | 82490968 | 82497505 | mNeoVis.2.1.1.chr10\_hap1.g00653.t01 |
| 12925 | 10 | 82495592 | 82514293 | mNeoVis.2.1.1.chr10\_hap1.g00654.t01 |
| 12926 | 10 | 82539208 | 82571911 | mNeoVis.2.1.1.chr10\_hap1.g00655.t01 |
| 12927 | 10 | 82610219 | 82629815 | mNeoVis.2.1.1.chr10\_hap1.g00656.t01 |
| 12928 | 10 | 82610219 | 82629815 | mNeoVis.2.1.1.chr10\_hap1.g00656.t01 |
| 12929 | 10 | 82635779 | 82671585 | mNeoVis.2.1.1.chr10\_hap1.g00657.t01 |
| 12930 | 10 | 82635779 | 82671585 | mNeoVis.2.1.1.chr10\_hap1.g00657.t01 |
| 12931 | 10 | 82702098 | 82707929 | mNeoVis.2.1.1.chr10\_hap1.g00658.t01 |
| 12932 | 10 | 82702098 | 82707929 | mNeoVis.2.1.1.chr10\_hap1.g00658.t01 |
| 12933 | 10 | 82729629 | 82734579 | mNeoVis.2.1.1.chr10\_hap1.g00659.t01 |
| 12934 | 10 | 82729629 | 82734579 | mNeoVis.2.1.1.chr10\_hap1.g00659.t01 |
| 12935 | 10 | 82736600 | 82832267 | mNeoVis.2.1.1.chr10\_hap1.g00660.t01 |
| 12936 | 10 | 82736600 | 82832267 | mNeoVis.2.1.1.chr10\_hap1.g00660.t01 |
| 12937 | 10 | 82800491 | 82803118 | mNeoVis.2.1.1.chr10\_hap1.g00661.t01 |
| 12938 | 10 | 82800491 | 82803118 | mNeoVis.2.1.1.chr10\_hap1.g00661.t01 |
| 12939 | 10 | 82832847 | 82857438 | mNeoVis.2.1.1.chr10\_hap1.g00662.t01 |
| 12940 | 10 | 82832847 | 82857438 | mNeoVis.2.1.1.chr10\_hap1.g00662.t01 |
| 12941 | 10 | 82832847 | 82857438 | mNeoVis.2.1.1.chr10\_hap1.g00662.t01 |
| 12942 | 10 | 82857347 | 82858167 | mNeoVis.2.1.1.chr10\_hap1.g00663.t01 |
| 12943 | 10 | 82857347 | 82858167 | mNeoVis.2.1.1.chr10\_hap1.g00663.t01 |
| 12944 | 10 | 82857347 | 82858167 | mNeoVis.2.1.1.chr10\_hap1.g00663.t01 |
| 12945 | 10 | 82868234 | 82952427 | mNeoVis.2.1.1.chr10\_hap1.g00664.t01 |
| 12946 | 10 | 82868234 | 82952427 | mNeoVis.2.1.1.chr10\_hap1.g00664.t01 |
| 12947 | 10 | 82868234 | 82952427 | mNeoVis.2.1.1.chr10\_hap1.g00664.t01 |
| 12948 | 10 | 82960591 | 83067240 | mNeoVis.2.1.1.chr10\_hap1.g00665.t01 |
| 12949 | 10 | 82960591 | 83067240 | mNeoVis.2.1.1.chr10\_hap1.g00665.t01 |
| 12950 | 10 | 83159346 | 83468468 | mNeoVis.2.1.1.chr10\_hap1.g00666.t01 |
| 12951 | 10 | 83179821 | 83180665 | mNeoVis.2.1.1.chr10\_hap1.g00667.t01 |
| 12952 | 10 | 83179821 | 83180665 | mNeoVis.2.1.1.chr10\_hap1.g00667.t01 |
| 12953 | 10 | 83451486 | 83536136 | mNeoVis.2.1.1.chr10\_hap1.g00668.t01 |
| 12954 | 10 | 83536123 | 83551314 | mNeoVis.2.1.1.chr10\_hap1.g00669.t01 |
| 12955 | 10 | 83552590 | 83554472 | mNeoVis.2.1.1.chr10\_hap1.g00670.t01 |
| 12956 | 10 | 83554487 | 83559765 | mNeoVis.2.1.1.chr10\_hap1.g00671.t01 |
| 12957 | 10 | 83559908 | 83561740 | mNeoVis.2.1.1.chr10\_hap1.g00672.t01 |
| 12958 | 10 | 83561893 | 83582238 | mNeoVis.2.1.1.chr10\_hap1.g00673.t01 |
| 12959 | 10 | 83582282 | 83586737 | mNeoVis.2.1.1.chr10\_hap1.g00674.t01 |
| 12960 | 10 | 83586873 | 83610914 | mNeoVis.2.1.1.chr10\_hap1.g00675.t01 |
| 12961 | 10 | 84102682 | 84248806 | mNeoVis.2.1.1.chr10\_hap1.g00692.t01 |
| 12962 | 10 | 84272872 | 84435777 | mNeoVis.2.1.1.chr10\_hap1.g00693.t01 |
| 12963 | 10 | 84437707 | 84450825 | mNeoVis.2.1.1.chr10\_hap1.g00694.t01 |
| 12964 | 10 | 84452375 | 84575954 | mNeoVis.2.1.1.chr10\_hap1.g00695.t01 |
| 12965 | 10 | 84587991 | 84603292 | mNeoVis.2.1.1.chr10\_hap1.g00696.t01 |
| 12966 | 10 | 84603187 | 84609703 | mNeoVis.2.1.1.chr10\_hap1.g00697.t01 |
| 12967 | 10 | 84613273 | 84657668 | mNeoVis.2.1.1.chr10\_hap1.g00698.t01 |
| 12968 | 10 | 84639929 | 84641116 | mNeoVis.2.1.1.chr10\_hap1.g00699.t01 |
| 12969 | 10 | 84658740 | 84730491 | mNeoVis.2.1.1.chr10\_hap1.g00700.t01 |
| 12970 | 10 | 84736270 | 84786626 | mNeoVis.2.1.1.chr10\_hap1.g00701.t01 |
| 12971 | 10 | 84794325 | 84866728 | mNeoVis.2.1.1.chr10\_hap1.g00702.t01 |
| 12972 | 10 | 85678393 | 85694239 | mNeoVis.2.1.1.chr10\_hap1.g00718.t01 |
| 12973 | 10 | 85695828 | 85729739 | mNeoVis.2.1.1.chr10\_hap1.g00719.t01 |
| 12974 | 10 | 85729848 | 85747903 | mNeoVis.2.1.1.chr10\_hap1.g00720.t01 |
| 12975 | 10 | 85779028 | 85826813 | mNeoVis.2.1.1.chr10\_hap1.g00721.t01 |
| 12976 | 10 | 85827136 | 85862351 | mNeoVis.2.1.1.chr10\_hap1.g00722.t01 |
| 12977 | 10 | 85906539 | 86044120 | mNeoVis.2.1.1.chr10\_hap1.g00723.t01 |
| 12978 | 10 | 86065093 | 86067467 | mNeoVis.2.1.1.chr10\_hap1.g00724.t01 |
| 12979 | 10 | 86081075 | 86090081 | mNeoVis.2.1.1.chr10\_hap1.g00725.t01 |
| 12980 | 10 | 86113122 | 86122140 | mNeoVis.2.1.1.chr10\_hap1.g00726.t01 |
| 12981 | 10 | 86138898 | 86141241 | mNeoVis.2.1.1.chr10\_hap1.g00727.t01 |
| 12982 | 10 | 86153823 | 86167351 | mNeoVis.2.1.1.chr10\_hap1.g00728.t01 |
| 12983 | 10 | 86167512 | 86192583 | mNeoVis.2.1.1.chr10\_hap1.g00729.t01 |
| 12984 | 10 | 86193391 | 86200457 | mNeoVis.2.1.1.chr10\_hap1.g00730.t01 |
| 12985 | 10 | 86200532 | 86235108 | mNeoVis.2.1.1.chr10\_hap1.g00731.t01 |
| 12986 | 10 | 86230611 | 86234426 | mNeoVis.2.1.1.chr10\_hap1.g00732.t01 |
| 12987 | 10 | 86235323 | 86239045 | mNeoVis.2.1.1.chr10\_hap1.g00733.t01 |
| 12988 | 10 | 86245232 | 86262604 | mNeoVis.2.1.1.chr10\_hap1.g00734.t01 |
| 12989 | 10 | 86262742 | 86292683 | mNeoVis.2.1.1.chr10\_hap1.g00735.t01 |
| 12990 | 10 | 89831588 | 89843668 | mNeoVis.2.1.1.chr10\_hap1.g00766.t01 |
| 12991 | 10 | 89904656 | 90113052 | mNeoVis.2.1.1.chr10\_hap1.g00767.t01 |
| 12992 | 10 | 89904656 | 90113052 | mNeoVis.2.1.1.chr10\_hap1.g00767.t01 |
| 12993 | 10 | 90585028 | 90774003 | mNeoVis.2.1.1.chr10\_hap1.g00768.t01 |
| 12994 | 10 | 90585028 | 90774003 | mNeoVis.2.1.1.chr10\_hap1.g00768.t01 |
| 12995 | 10 | 90838763 | 91121853 | mNeoVis.2.1.1.chr10\_hap1.g00769.t01 |
| 12996 | 10 | 90838763 | 91121853 | mNeoVis.2.1.1.chr10\_hap1.g00769.t01 |
| 12997 | 10 | 90838763 | 91121853 | mNeoVis.2.1.1.chr10\_hap1.g00769.t01 |
| 12998 | 10 | 91121881 | 91297939 | mNeoVis.2.1.1.chr10\_hap1.g00770.t01 |
| 12999 | 10 | 91612342 | 91622470 | mNeoVis.2.1.1.chr10\_hap1.g00771.t01 |
| 13000 | 10 | 91624102 | 91719272 | mNeoVis.2.1.1.chr10\_hap1.g00772.t01 |
| 13001 | 10 | 91765720 | 91771012 | mNeoVis.2.1.1.chr10\_hap1.g00773.t01 |
| 13002 | 10 | 91765720 | 91771012 | mNeoVis.2.1.1.chr10\_hap1.g00773.t01 |
| 13003 | 10 | 91801552 | 91804938 | mNeoVis.2.1.1.chr10\_hap1.g00774.t01 |
| 13004 | 10 | 91801552 | 91804938 | mNeoVis.2.1.1.chr10\_hap1.g00774.t01 |
| 13005 | 10 | 91935572 | 91956279 | mNeoVis.2.1.1.chr10\_hap1.g00775.t01 |
| 13006 | 10 | 91935572 | 91956279 | mNeoVis.2.1.1.chr10\_hap1.g00775.t01 |
| 13007 | 10 | 91935572 | 91956279 | mNeoVis.2.1.1.chr10\_hap1.g00775.t01 |
| 13008 | 10 | 91959167 | 92012691 | mNeoVis.2.1.1.chr10\_hap1.g00776.t01 |
| 13009 | 10 | 91959167 | 92012691 | mNeoVis.2.1.1.chr10\_hap1.g00776.t01 |
| 13010 | 10 | 91959167 | 92012691 | mNeoVis.2.1.1.chr10\_hap1.g00776.t01 |
| 13011 | 10 | 92036210 | 92048339 | mNeoVis.2.1.1.chr10\_hap1.g00777.t01 |
| 13012 | 10 | 92036210 | 92048339 | mNeoVis.2.1.1.chr10\_hap1.g00777.t01 |
| 13013 | 10 | 92036210 | 92048339 | mNeoVis.2.1.1.chr10\_hap1.g00777.t01 |
| 13014 | 10 | 92079195 | 92493444 | mNeoVis.2.1.1.chr10\_hap1.g00778.t01 |
| 13015 | 10 | 92079195 | 92493444 | mNeoVis.2.1.1.chr10\_hap1.g00778.t01 |
| 13016 | 10 | 92527837 | 92536083 | mNeoVis.2.1.1.chr10\_hap1.g00779.t01 |
| 13017 | 10 | 92527837 | 92536083 | mNeoVis.2.1.1.chr10\_hap1.g00779.t01 |
| 13018 | 10 | 92527837 | 92536083 | mNeoVis.2.1.1.chr10\_hap1.g00779.t01 |
| 13019 | 10 | 92588739 | 93116688 | mNeoVis.2.1.1.chr10\_hap1.g00780.t01 |
| 13020 | 10 | 93116798 | 93303950 | mNeoVis.2.1.1.chr10\_hap1.g00781.t01 |
| 13021 | 10 | 93116798 | 93303950 | mNeoVis.2.1.1.chr10\_hap1.g00781.t01 |
| 13022 | 10 | 93116798 | 93303950 | mNeoVis.2.1.1.chr10\_hap1.g00781.t01 |
| 13023 | 10 | 93318494 | 93320092 | mNeoVis.2.1.1.chr10\_hap1.g00782.t01 |
| 13024 | 10 | 93318494 | 93320092 | mNeoVis.2.1.1.chr10\_hap1.g00782.t01 |
| 13025 | 10 | 93318494 | 93320092 | mNeoVis.2.1.1.chr10\_hap1.g00782.t01 |
| 13026 | 10 | 93338669 | 93351477 | mNeoVis.2.1.1.chr10\_hap1.g00783.t01 |
| 13027 | 10 | 93338669 | 93351477 | mNeoVis.2.1.1.chr10\_hap1.g00783.t01 |
| 13028 | 10 | 93338669 | 93351477 | mNeoVis.2.1.1.chr10\_hap1.g00783.t01 |
| 13029 | 10 | 93386190 | 93423459 | mNeoVis.2.1.1.chr10\_hap1.g00784.t01 |
| 13030 | 10 | 93386190 | 93423459 | mNeoVis.2.1.1.chr10\_hap1.g00784.t01 |
| 13031 | 10 | 93386190 | 93423459 | mNeoVis.2.1.1.chr10\_hap1.g00784.t01 |
| 13032 | 10 | 93437309 | 93442862 | mNeoVis.2.1.1.chr10\_hap1.g00785.t01 |
| 13033 | 10 | 93437309 | 93442862 | mNeoVis.2.1.1.chr10\_hap1.g00785.t01 |
| 13034 | 10 | 93437309 | 93442862 | mNeoVis.2.1.1.chr10\_hap1.g00785.t01 |
| 13035 | 10 | 93442708 | 93532418 | mNeoVis.2.1.1.chr10\_hap1.g00786.t01 |
| 13036 | 10 | 93442708 | 93532418 | mNeoVis.2.1.1.chr10\_hap1.g00786.t01 |
| 13037 | 10 | 95666567 | 95667547 | mNeoVis.2.1.1.chr10\_hap1.g00856.t01 |
| 13038 | 10 | 95696804 | 95716372 | mNeoVis.2.1.1.chr10\_hap1.g00857.t01 |
| 13039 | 10 | 95719661 | 95728695 | mNeoVis.2.1.1.chr10\_hap1.g00858.t01 |
| 13040 | 10 | 95747658 | 95756867 | mNeoVis.2.1.1.chr10\_hap1.g00859.t01 |
| 13041 | 10 | 95761212 | 95800051 | mNeoVis.2.1.1.chr10\_hap1.g00860.t01 |
| 13042 | 10 | 95813060 | 95825188 | mNeoVis.2.1.1.chr10\_hap1.g00861.t01 |
| 13043 | 10 | 95834205 | 95840388 | mNeoVis.2.1.1.chr10\_hap1.g00862.t01 |
| 13044 | 10 | 95840681 | 95843755 | mNeoVis.2.1.1.chr10\_hap1.g00863.t01 |
| 13045 | 10 | 95843837 | 95847529 | mNeoVis.2.1.1.chr10\_hap1.g00864.t01 |
| 13046 | 10 | 95860756 | 95869480 | mNeoVis.2.1.1.chr10\_hap1.g00865.t01 |
| 13047 | 10 | 95881340 | 95887225 | mNeoVis.2.1.1.chr10\_hap1.g00866.t01 |
| 13048 | 10 | 95911048 | 95916978 | mNeoVis.2.1.1.chr10\_hap1.g00867.t01 |
| 13049 | 10 | 95935157 | 95935783 | mNeoVis.2.1.1.chr10\_hap1.g00868.t01 |
| 13050 | 10 | 95940353 | 95940861 | mNeoVis.2.1.1.chr10\_hap1.g00869.t01 |
| 13051 | 10 | 95981512 | 95982725 | mNeoVis.2.1.1.chr10\_hap1.g00870.t01 |
| 13052 | 10 | 96044583 | 96079736 | mNeoVis.2.1.1.chr10\_hap1.g00871.t01 |
| 13053 | 10 | 96044590 | 96063138 | mNeoVis.2.1.1.chr10\_hap1.g00872.t01 |
| 13054 | 10 | 96056529 | 96057080 | mNeoVis.2.1.1.chr10\_hap1.g00873.t01 |
| 13055 | 10 | 96143186 | 96143745 | mNeoVis.2.1.1.chr10\_hap1.g00874.t01 |
| 13056 | 10 | 96208008 | 96216448 | mNeoVis.2.1.1.chr10\_hap1.g00875.t01 |
| 13057 | 10 | 96296833 | 96298738 | mNeoVis.2.1.1.chr10\_hap1.g00876.t01 |
| 13058 | 10 | 96332104 | 96333512 | mNeoVis.2.1.1.chr10\_hap1.g00877.t01 |
| 13059 | 10 | 97347741 | 97349753 | mNeoVis.2.1.1.chr10\_hap1.g00907.t01 |
| 13060 | 10 | 97398120 | 97459450 | mNeoVis.2.1.1.chr10\_hap1.g00908.t01 |
| 13061 | 10 | 97502661 | 97621738 | mNeoVis.2.1.1.chr10\_hap1.g00909.t01 |
| 13062 | 10 | 97728702 | 97759591 | mNeoVis.2.1.1.chr10\_hap1.g00910.t01 |
| 13063 | 10 | 97728702 | 97759591 | mNeoVis.2.1.1.chr10\_hap1.g00910.t01 |
| 13064 | 10 | 97803764 | 97829760 | mNeoVis.2.1.1.chr10\_hap1.g00911.t01 |
| 13065 | 10 | 97803764 | 97829760 | mNeoVis.2.1.1.chr10\_hap1.g00911.t01 |
| 13066 | 10 | 97821582 | 97822273 | mNeoVis.2.1.1.chr10\_hap1.g00912.t01 |
| 13067 | 10 | 97821582 | 97822273 | mNeoVis.2.1.1.chr10\_hap1.g00912.t01 |
| 13068 | 10 | 97935805 | 97969756 | mNeoVis.2.1.1.chr10\_hap1.g00913.t01 |
| 13069 | 10 | 98000867 | 98127732 | mNeoVis.2.1.1.chr10\_hap1.g00914.t01 |
| 13070 | 10 | 98000867 | 98127732 | mNeoVis.2.1.1.chr10\_hap1.g00914.t01 |
| 13071 | 10 | 98250406 | 98361661 | mNeoVis.2.1.1.chr10\_hap1.g00915.t01 |
| 13072 | 10 | 98368803 | 98384319 | mNeoVis.2.1.1.chr10\_hap1.g00916.t01 |
| 13073 | 10 | 98410934 | 98432684 | mNeoVis.2.1.1.chr10\_hap1.g00917.t01 |
| 13074 | 10 | 98434604 | 98443964 | mNeoVis.2.1.1.chr10\_hap1.g00918.t01 |
| 13075 | 10 | 98730937 | 98739643 | mNeoVis.2.1.1.chr10\_hap1.g00920.t01 |
| 13076 | 10 | 98757712 | 99003091 | mNeoVis.2.1.1.chr10\_hap1.g00921.t01 |
| 13077 | 10 | 99017170 | 99022534 | mNeoVis.2.1.1.chr10\_hap1.g00922.t01 |
| 13078 | 10 | 99017170 | 99022534 | mNeoVis.2.1.1.chr10\_hap1.g00922.t01 |
| 13079 | 10 | 99042176 | 99234867 | mNeoVis.2.1.1.chr10\_hap1.g00923.t01 |
| 13080 | 10 | 99042176 | 99234867 | mNeoVis.2.1.1.chr10\_hap1.g00923.t01 |
| 13081 | 10 | 99234958 | 99286079 | mNeoVis.2.1.1.chr10\_hap1.g00924.t01 |
| 13082 | 10 | 99234958 | 99286079 | mNeoVis.2.1.1.chr10\_hap1.g00924.t01 |
| 13083 | 10 | 99292290 | 99408942 | mNeoVis.2.1.1.chr10\_hap1.g00925.t01 |
| 13084 | 10 | 99641659 | 99717370 | mNeoVis.2.1.1.chr10\_hap1.g00926.t01 |
| 13085 | 10 | 99833516 | 99846841 | mNeoVis.2.1.1.chr10\_hap1.g00927.t01 |
| 13086 | 10 | 99846837 | 99883781 | mNeoVis.2.1.1.chr10\_hap1.g00928.t01 |
| 13087 | 10 | 99883988 | 99888720 | mNeoVis.2.1.1.chr10\_hap1.g00929.t01 |
| 13088 | 10 | 99890449 | 99893996 | mNeoVis.2.1.1.chr10\_hap1.g00930.t01 |
| 13089 | 10 | 99895651 | 99973879 | mNeoVis.2.1.1.chr10\_hap1.g00931.t01 |
| 13090 | 10 | 100000726 | 100017900 | mNeoVis.2.1.1.chr10\_hap1.g00932.t01 |
| 13091 | 10 | 100019419 | 100052743 | mNeoVis.2.1.1.chr10\_hap1.g00933.t01 |
| 13092 | 10 | 100052880 | 100094344 | mNeoVis.2.1.1.chr10\_hap1.g00934.t01 |
| 13093 | 10 | 100443800 | 100464007 | mNeoVis.2.1.1.chr10\_hap1.g00936.t01 |
| 13094 | 10 | 100443800 | 100464007 | mNeoVis.2.1.1.chr10\_hap1.g00936.t01 |
| 13095 | 10 | 100536479 | 100667847 | mNeoVis.2.1.1.chr10\_hap1.g00937.t01 |
| 13096 | 10 | 100536479 | 100667847 | mNeoVis.2.1.1.chr10\_hap1.g00937.t01 |
| 13097 | 10 | 100667827 | 100696750 | mNeoVis.2.1.1.chr10\_hap1.g00938.t01 |
| 13098 | 10 | 100667827 | 100696750 | mNeoVis.2.1.1.chr10\_hap1.g00938.t01 |
| 13099 | 10 | 100708314 | 100739409 | mNeoVis.2.1.1.chr10\_hap1.g00939.t01 |
| 13100 | 10 | 100708314 | 100739409 | mNeoVis.2.1.1.chr10\_hap1.g00939.t01 |
| 13101 | 10 | 100739850 | 100780532 | mNeoVis.2.1.1.chr10\_hap1.g00940.t01 |
| 13102 | 10 | 100739850 | 100780532 | mNeoVis.2.1.1.chr10\_hap1.g00940.t01 |
| 13103 | 10 | 100792180 | 100860155 | mNeoVis.2.1.1.chr10\_hap1.g00941.t01 |
| 13104 | 10 | 100792180 | 100860155 | mNeoVis.2.1.1.chr10\_hap1.g00941.t01 |
| 13105 | 10 | 100877646 | 100917328 | mNeoVis.2.1.1.chr10\_hap1.g00942.t01 |
| 13106 | 10 | 101221012 | 101224887 | mNeoVis.2.1.1.chr10\_hap1.g00948.t01 |
| 13107 | 10 | 101230482 | 101243804 | mNeoVis.2.1.1.chr10\_hap1.g00949.t01 |
| 13108 | 10 | 101243865 | 101247748 | mNeoVis.2.1.1.chr10\_hap1.g00950.t01 |
| 13109 | 10 | 101262373 | 101285946 | mNeoVis.2.1.1.chr10\_hap1.g00951.t01 |
| 13110 | 10 | 101295166 | 101313441 | mNeoVis.2.1.1.chr10\_hap1.g00952.t01 |
| 13111 | 10 | 101313588 | 101359081 | mNeoVis.2.1.1.chr10\_hap1.g00953.t01 |
| 13112 | 10 | 101371896 | 101445580 | mNeoVis.2.1.1.chr10\_hap1.g00954.t01 |
| 13113 | 10 | 101403563 | 101426057 | mNeoVis.2.1.1.chr10\_hap1.g00955.t01 |
| 13114 | 10 | 101475950 | 101502333 | mNeoVis.2.1.1.chr10\_hap1.g00956.t01 |
| 13115 | 10 | 101527625 | 101545754 | mNeoVis.2.1.1.chr10\_hap1.g00957.t01 |
| 13116 | 10 | 101547233 | 101778492 | mNeoVis.2.1.1.chr10\_hap1.g00958.t01 |
| 13117 | 10 | 101783698 | 101852088 | mNeoVis.2.1.1.chr10\_hap1.g00959.t01 |
| 13118 | 10 | 101855727 | 101867573 | mNeoVis.2.1.1.chr10\_hap1.g00960.t01 |
| 13119 | 10 | 101873737 | 102072109 | mNeoVis.2.1.1.chr10\_hap1.g00961.t01 |
| 13120 | 10 | 102074945 | 102080784 | mNeoVis.2.1.1.chr10\_hap1.g00962.t01 |
| 13121 | 10 | 102080924 | 102084433 | mNeoVis.2.1.1.chr10\_hap1.g00963.t01 |
| 13122 | 10 | 102097471 | 102131983 | mNeoVis.2.1.1.chr10\_hap1.g00964.t01 |
| 13123 | 10 | 102133800 | 102151459 | mNeoVis.2.1.1.chr10\_hap1.g00965.t01 |
| 13124 | 10 | 102352364 | 102537337 | mNeoVis.2.1.1.chr10\_hap1.g00967.t01 |
| 13125 | 10 | 102371485 | 102372246 | mNeoVis.2.1.1.chr10\_hap1.g00968.t01 |
| 13126 | 10 | 102542035 | 102545936 | mNeoVis.2.1.1.chr10\_hap1.g00969.t01 |
| 13127 | 10 | 102548640 | 102656682 | mNeoVis.2.1.1.chr10\_hap1.g00970.t01 |
| 13128 | 10 | 102782165 | 102880928 | mNeoVis.2.1.1.chr10\_hap1.g00972.t01 |
| 13129 | 10 | 102886368 | 102945837 | mNeoVis.2.1.1.chr10\_hap1.g00973.t01 |
| 13130 | 10 | 102974363 | 102977985 | mNeoVis.2.1.1.chr10\_hap1.g00974.t01 |
| 13131 | 10 | 102985961 | 103003950 | mNeoVis.2.1.1.chr10\_hap1.g00975.t01 |
| 13132 | 10 | 103048610 | 103070799 | mNeoVis.2.1.1.chr10\_hap1.g00976.t01 |
| 13133 | 10 | 103056638 | 103084590 | mNeoVis.2.1.1.chr10\_hap1.g00977.t01 |
| 13134 | 10 | 103940969 | 103943354 | mNeoVis.2.1.1.chr10\_hap1.g00981.t01 |
| 13135 | 10 | 103948824 | 104129754 | mNeoVis.2.1.1.chr10\_hap1.g00982.t01 |
| 13136 | 10 | 104347435 | 104742754 | mNeoVis.2.1.1.chr10\_hap1.g00983.t01 |
| 13137 | 10 | 105349784 | 105452013 | mNeoVis.2.1.1.chr10\_hap1.g00984.t01 |
| 13138 | 10 | 105458696 | 105522021 | mNeoVis.2.1.1.chr10\_hap1.g00985.t01 |
| 13139 | 10 | 105458696 | 105522021 | mNeoVis.2.1.1.chr10\_hap1.g00985.t01 |
| 13140 | 10 | 105543030 | 105586783 | mNeoVis.2.1.1.chr10\_hap1.g00986.t01 |
| 13141 | 10 | 105543030 | 105586783 | mNeoVis.2.1.1.chr10\_hap1.g00986.t01 |
| 13142 | 10 | 105792628 | 105794362 | mNeoVis.2.1.1.chr10\_hap1.g00987.t01 |
| 13143 | 10 | 105926117 | 105927116 | mNeoVis.2.1.1.chr10\_hap1.g00988.t01 |
| 13144 | 10 | 106217904 | 106246415 | mNeoVis.2.1.1.chr10\_hap1.g00989.t01 |
| 13145 | 10 | 106246283 | 106266903 | mNeoVis.2.1.1.chr10\_hap1.g00990.t01 |
| 13146 | 10 | 106283006 | 106312092 | mNeoVis.2.1.1.chr10\_hap1.g00991.t01 |
| 13147 | 10 | 106409831 | 106468468 | mNeoVis.2.1.1.chr10\_hap1.g00992.t01 |
| 13148 | 10 | 107946893 | 107955240 | mNeoVis.2.1.1.chr10\_hap1.g01002.t01 |
| 13149 | 10 | 107970946 | 108059899 | mNeoVis.2.1.1.chr10\_hap1.g01003.t01 |
| 13150 | 10 | 108097536 | 108257110 | mNeoVis.2.1.1.chr10\_hap1.g01004.t01 |
| 13151 | 10 | 109443787 | 109580338 | mNeoVis.2.1.1.chr10\_hap1.g01010.t01 |
| 13152 | 10 | 109683259 | 109761675 | mNeoVis.2.1.1.chr10\_hap1.g01011.t01 |
| 13153 | 10 | 109830523 | 109958756 | mNeoVis.2.1.1.chr10\_hap1.g01012.t01 |
| 13154 | 10 | 109973053 | 109995823 | mNeoVis.2.1.1.chr10\_hap1.g01013.t01 |
| 13155 | 10 | 110026252 | 110081780 | mNeoVis.2.1.1.chr10\_hap1.g01014.t01 |
| 13156 | 10 | 110026252 | 110081780 | mNeoVis.2.1.1.chr10\_hap1.g01014.t01 |
| 13157 | 10 | 110087541 | 110096536 | mNeoVis.2.1.1.chr10\_hap1.g01015.t01 |
| 13158 | 10 | 110087541 | 110096536 | mNeoVis.2.1.1.chr10\_hap1.g01015.t01 |
| 13159 | 10 | 110096738 | 110135685 | mNeoVis.2.1.1.chr10\_hap1.g01016.t01 |
| 13160 | 10 | 110096738 | 110135685 | mNeoVis.2.1.1.chr10\_hap1.g01016.t01 |
| 13161 | 10 | 110135676 | 110176982 | mNeoVis.2.1.1.chr10\_hap1.g01017.t01 |
| 13162 | 10 | 110135676 | 110176982 | mNeoVis.2.1.1.chr10\_hap1.g01017.t01 |
| 13163 | 10 | 110177057 | 110179020 | mNeoVis.2.1.1.chr10\_hap1.g01018.t01 |
| 13164 | 10 | 110177057 | 110179020 | mNeoVis.2.1.1.chr10\_hap1.g01018.t01 |
| 13165 | 10 | 110186751 | 110265304 | mNeoVis.2.1.1.chr10\_hap1.g01019.t01 |
| 13166 | 10 | 110186751 | 110265304 | mNeoVis.2.1.1.chr10\_hap1.g01019.t01 |
| 13167 | 10 | 110271676 | 110288499 | mNeoVis.2.1.1.chr10\_hap1.g01020.t01 |
| 13168 | 10 | 110271676 | 110288499 | mNeoVis.2.1.1.chr10\_hap1.g01020.t01 |
| 13169 | 10 | 110444326 | 110445068 | mNeoVis.2.1.1.chr10\_hap1.g01021.t01 |
| 13170 | 10 | 110444326 | 110445068 | mNeoVis.2.1.1.chr10\_hap1.g01021.t01 |
| 13171 | 10 | 110630004 | 110631267 | mNeoVis.2.1.1.chr10\_hap1.g01022.t01 |
| 13172 | 10 | 110969292 | 111566543 | mNeoVis.2.1.1.chr10\_hap1.g01023.t01 |
| 13173 | 10 | 112475945 | 112637878 | mNeoVis.2.1.1.chr10\_hap1.g01025.t01 |
| 13174 | 10 | 112475945 | 112637878 | mNeoVis.2.1.1.chr10\_hap1.g01025.t01 |
| 13175 | 10 | 112638203 | 112650887 | mNeoVis.2.1.1.chr10\_hap1.g01026.t01 |
| 13176 | 10 | 112638203 | 112650887 | mNeoVis.2.1.1.chr10\_hap1.g01026.t01 |
| 13177 | 10 | 112678846 | 112699248 | mNeoVis.2.1.1.chr10\_hap1.g01027.t01 |
| 13178 | 10 | 112678846 | 112699248 | mNeoVis.2.1.1.chr10\_hap1.g01027.t01 |
| 13179 | 10 | 112699356 | 112719228 | mNeoVis.2.1.1.chr10\_hap1.g01028.t01 |
| 13180 | 10 | 112699356 | 112719228 | mNeoVis.2.1.1.chr10\_hap1.g01028.t01 |
| 13181 | 10 | 112750731 | 112838443 | mNeoVis.2.1.1.chr10\_hap1.g01029.t01 |
| 13182 | 10 | 113288830 | 113335830 | mNeoVis.2.1.1.chr10\_hap1.g01030.t01 |
| 13183 | 10 | 113901200 | 113906529 | mNeoVis.2.1.1.chr10\_hap1.g01032.t01 |
| 13184 | 10 | 113910862 | 114202914 | mNeoVis.2.1.1.chr10\_hap1.g01033.t01 |
| 13185 | 10 | 113965450 | 113968309 | mNeoVis.2.1.1.chr10\_hap1.g01034.t01 |
| 13186 | 10 | 114201977 | 114220151 | mNeoVis.2.1.1.chr10\_hap1.g01035.t01 |
| 13187 | 10 | 114236452 | 114271555 | mNeoVis.2.1.1.chr10\_hap1.g01036.t01 |
| 13188 | 10 | 114272705 | 114297241 | mNeoVis.2.1.1.chr10\_hap1.g01037.t01 |
| 13189 | 10 | 114306296 | 114335066 | mNeoVis.2.1.1.chr10\_hap1.g01038.t01 |
| 13190 | 10 | 114345157 | 114375019 | mNeoVis.2.1.1.chr10\_hap1.g01039.t01 |
| 13191 | 10 | 114398176 | 114425134 | mNeoVis.2.1.1.chr10\_hap1.g01040.t01 |
| 13192 | 10 | 114444862 | 114454905 | mNeoVis.2.1.1.chr10\_hap1.g01041.t01 |
| 13193 | 10 | 114476201 | 114481493 | mNeoVis.2.1.1.chr10\_hap1.g01042.t01 |
| 13194 | 10 | 116343869 | 116357661 | mNeoVis.2.1.1.chr10\_hap1.g01068.t01 |
| 13195 | 10 | 116357875 | 116435946 | mNeoVis.2.1.1.chr10\_hap1.g01069.t01 |
| 13196 | 10 | 116438718 | 116445204 | mNeoVis.2.1.1.chr10\_hap1.g01070.t01 |
| 13197 | 10 | 116463709 | 116468218 | mNeoVis.2.1.1.chr10\_hap1.g01071.t01 |
| 13198 | 10 | 116497976 | 116516841 | mNeoVis.2.1.1.chr10\_hap1.g01072.t01 |
| 13199 | 10 | 116525222 | 116534251 | mNeoVis.2.1.1.chr10\_hap1.g01073.t01 |
| 13200 | 10 | 116535116 | 116540398 | mNeoVis.2.1.1.chr10\_hap1.g01074.t01 |
| 13201 | 10 | 116545286 | 116552330 | mNeoVis.2.1.1.chr10\_hap1.g01075.t01 |
| 13202 | 10 | 116552890 | 116565390 | mNeoVis.2.1.1.chr10\_hap1.g01076.t01 |
| 13203 | 10 | 116577883 | 116586271 | mNeoVis.2.1.1.chr10\_hap1.g01077.t01 |
| 13204 | 10 | 116587326 | 116593882 | mNeoVis.2.1.1.chr10\_hap1.g01078.t01 |
| 13205 | 10 | 116609470 | 116624932 | mNeoVis.2.1.1.chr10\_hap1.g01079.t01 |
| 13206 | 10 | 116631853 | 116638083 | mNeoVis.2.1.1.chr10\_hap1.g01080.t01 |
| 13207 | 10 | 116640174 | 116678076 | mNeoVis.2.1.1.chr10\_hap1.g01081.t01 |
| 13208 | 10 | 116678956 | 116703575 | mNeoVis.2.1.1.chr10\_hap1.g01082.t01 |
| 13209 | 10 | 116816660 | 116831211 | mNeoVis.2.1.1.chr10\_hap1.g01083.t01 |
| 13210 | 10 | 116867258 | 116873572 | mNeoVis.2.1.1.chr10\_hap1.g01084.t01 |
| 13211 | 10 | 116875856 | 116880932 | mNeoVis.2.1.1.chr10\_hap1.g01085.t01 |
| 13212 | 10 | 123099345 | 123820897 | mNeoVis.2.1.1.chr10\_hap1.g01118.t01 |
| 13213 | 10 | 123512815 | 123513484 | mNeoVis.2.1.1.chr10\_hap1.g01119.t01 |
| 13214 | 10 | 123848305 | 123933029 | mNeoVis.2.1.1.chr10\_hap1.g01120.t01 |
| 13215 | 10 | 128442696 | 128493171 | mNeoVis.2.1.1.chr10\_hap1.g01131.t01 |
| 13216 | 10 | 128695382 | 128769260 | mNeoVis.2.1.1.chr10\_hap1.g01132.t01 |
| 13217 | 10 | 128778731 | 128830962 | mNeoVis.2.1.1.chr10\_hap1.g01133.t01 |
| 13218 | 10 | 128828159 | 128865069 | mNeoVis.2.1.1.chr10\_hap1.g01134.t01 |
| 13219 | 10 | 128873065 | 128892709 | mNeoVis.2.1.1.chr10\_hap1.g01135.t01 |
| 13220 | 10 | 128893034 | 128923835 | mNeoVis.2.1.1.chr10\_hap1.g01136.t01 |
| 13221 | 10 | 128986667 | 129237370 | mNeoVis.2.1.1.chr10\_hap1.g01137.t01 |
| 13222 | 10 | 129379673 | 129381002 | mNeoVis.2.1.1.chr10\_hap1.g01139.t01 |
| 13223 | 10 | 130200810 | 130261311 | mNeoVis.2.1.1.chr10\_hap1.g01141.t01 |
| 13224 | 10 | 130263018 | 130273642 | mNeoVis.2.1.1.chr10\_hap1.g01142.t01 |
| 13225 | 10 | 130422582 | 130581341 | mNeoVis.2.1.1.chr10\_hap1.g01143.t01 |
| 13226 | 10 | 130422582 | 130581341 | mNeoVis.2.1.1.chr10\_hap1.g01143.t01 |
| 13227 | 10 | 130588675 | 130603162 | mNeoVis.2.1.1.chr10\_hap1.g01144.t01 |
| 13228 | 10 | 130588675 | 130603162 | mNeoVis.2.1.1.chr10\_hap1.g01144.t01 |
| 13229 | 10 | 130608929 | 130617469 | mNeoVis.2.1.1.chr10\_hap1.g01145.t01 |
| 13230 | 10 | 130608929 | 130617469 | mNeoVis.2.1.1.chr10\_hap1.g01145.t01 |
| 13231 | 10 | 130656279 | 130727627 | mNeoVis.2.1.1.chr10\_hap1.g01146.t01 |
| 13232 | 10 | 130656279 | 130727627 | mNeoVis.2.1.1.chr10\_hap1.g01146.t01 |
| 13233 | 10 | 130808019 | 130931324 | mNeoVis.2.1.1.chr10\_hap1.g01147.t01 |
| 13234 | 10 | 130934181 | 130970895 | mNeoVis.2.1.1.chr10\_hap1.g01148.t01 |
| 13235 | 10 | 131171107 | 131220561 | mNeoVis.2.1.1.chr10\_hap1.g01149.t01 |
| 13236 | 10 | 131220825 | 131243163 | mNeoVis.2.1.1.chr10\_hap1.g01150.t01 |
| 13237 | 10 | 131253899 | 131369726 | mNeoVis.2.1.1.chr10\_hap1.g01151.t01 |
| 13238 | 10 | 131788915 | 131823682 | mNeoVis.2.1.1.chr10\_hap1.g01153.t01 |
| 13239 | 10 | 131788915 | 131823682 | mNeoVis.2.1.1.chr10\_hap1.g01153.t01 |
| 13240 | 10 | 131839510 | 132001600 | mNeoVis.2.1.1.chr10\_hap1.g01154.t01 |
| 13241 | 10 | 131839510 | 132001600 | mNeoVis.2.1.1.chr10\_hap1.g01154.t01 |
| 13242 | 10 | 131844721 | 131851517 | mNeoVis.2.1.1.chr10\_hap1.g01155.t01 |
| 13243 | 10 | 131844721 | 131851517 | mNeoVis.2.1.1.chr10\_hap1.g01155.t01 |
| 13244 | 10 | 132162150 | 132324385 | mNeoVis.2.1.1.chr10\_hap1.g01156.t01 |
| 13245 | 10 | 132162150 | 132324385 | mNeoVis.2.1.1.chr10\_hap1.g01156.t01 |
| 13246 | 10 | 132162150 | 132324385 | mNeoVis.2.1.1.chr10\_hap1.g01156.t01 |
| 13247 | 10 | 132162150 | 132324385 | mNeoVis.2.1.1.chr10\_hap1.g01156.t01 |
| 13248 | 10 | 132333025 | 132425636 | mNeoVis.2.1.1.chr10\_hap1.g01157.t01 |
| 13249 | 10 | 132333025 | 132425636 | mNeoVis.2.1.1.chr10\_hap1.g01157.t01 |
| 13250 | 10 | 132333025 | 132425636 | mNeoVis.2.1.1.chr10\_hap1.g01157.t01 |
| 13251 | 10 | 132333025 | 132425636 | mNeoVis.2.1.1.chr10\_hap1.g01157.t01 |
| 13252 | 10 | 132427722 | 132455151 | mNeoVis.2.1.1.chr10\_hap1.g01158.t01 |
| 13253 | 10 | 132427722 | 132455151 | mNeoVis.2.1.1.chr10\_hap1.g01158.t01 |
| 13254 | 10 | 132427722 | 132455151 | mNeoVis.2.1.1.chr10\_hap1.g01158.t01 |
| 13255 | 10 | 132427722 | 132455151 | mNeoVis.2.1.1.chr10\_hap1.g01158.t01 |
| 13256 | 10 | 132621705 | 132922656 | mNeoVis.2.1.1.chr10\_hap1.g01159.t01 |
| 13257 | 10 | 132926039 | 132927685 | mNeoVis.2.1.1.chr10\_hap1.g01160.t01 |
| 13258 | 10 | 132926039 | 132927685 | mNeoVis.2.1.1.chr10\_hap1.g01160.t01 |
| 13259 | 10 | 132970548 | 133058781 | mNeoVis.2.1.1.chr10\_hap1.g01161.t01 |
| 13260 | 10 | 132970548 | 133058781 | mNeoVis.2.1.1.chr10\_hap1.g01161.t01 |
| 13261 | 10 | 133366528 | 133407424 | mNeoVis.2.1.1.chr10\_hap1.g01162.t01 |
| 13262 | 10 | 133465153 | 133478990 | mNeoVis.2.1.1.chr10\_hap1.g01163.t01 |
| 13263 | 10 | 133516747 | 133522752 | mNeoVis.2.1.1.chr10\_hap1.g01164.t01 |
| 13264 | 10 | 134978170 | 134985631 | mNeoVis.2.1.1.chr10\_hap1.g01165.t01 |
| 13265 | 10 | 134978170 | 134985631 | mNeoVis.2.1.1.chr10\_hap1.g01165.t01 |
| 13266 | 10 | 134986650 | 135163844 | mNeoVis.2.1.1.chr10\_hap1.g01166.t01 |
| 13267 | 10 | 134986650 | 135163844 | mNeoVis.2.1.1.chr10\_hap1.g01166.t01 |
| 13268 | 10 | 136721877 | 136967149 | mNeoVis.2.1.1.chr10\_hap1.g01167.t01 |
| 13269 | 10 | 136721877 | 136967149 | mNeoVis.2.1.1.chr10\_hap1.g01167.t01 |
| 13270 | 10 | 136971824 | 136973720 | mNeoVis.2.1.1.chr10\_hap1.g01168.t01 |
| 13271 | 10 | 136971824 | 136973720 | mNeoVis.2.1.1.chr10\_hap1.g01168.t01 |
| 13272 | 10 | 137075214 | 137088716 | mNeoVis.2.1.1.chr10\_hap1.g01169.t01 |
| 13273 | 10 | 137641587 | 137881513 | mNeoVis.2.1.1.chr10\_hap1.g01170.t01 |
| 13274 | 10 | 137918461 | 137963225 | mNeoVis.2.1.1.chr10\_hap1.g01171.t01 |
| 13275 | 10 | 137974601 | 138085568 | mNeoVis.2.1.1.chr10\_hap1.g01172.t01 |
| 13276 | 10 | 138092052 | 138105516 | mNeoVis.2.1.1.chr10\_hap1.g01173.t01 |
| 13277 | 10 | 138260570 | 138287929 | mNeoVis.2.1.1.chr10\_hap1.g01174.t01 |
| 13278 | 10 | 138415357 | 138478735 | mNeoVis.2.1.1.chr10\_hap1.g01175.t01 |
| 13279 | 10 | 138744552 | 139045793 | mNeoVis.2.1.1.chr10\_hap1.g01176.t01 |
| 13280 | 10 | 139536868 | 139725644 | mNeoVis.2.1.1.chr10\_hap1.g01177.t01 |
| 13281 | 10 | 139777163 | 139818306 | mNeoVis.2.1.1.chr10\_hap1.g01178.t01 |
| 13282 | 10 | 139853463 | 140175698 | mNeoVis.2.1.1.chr10\_hap1.g01179.t01 |
| 13283 | 10 | 140128721 | 140138625 | mNeoVis.2.1.1.chr10\_hap1.g01180.t01 |
| 13284 | 10 | 140239922 | 140334796 | mNeoVis.2.1.1.chr10\_hap1.g01181.t01 |
| 13285 | 10 | 140532244 | 141246973 | mNeoVis.2.1.1.chr10\_hap1.g01182.t01 |
| 13286 | 10 | 140532244 | 141246973 | mNeoVis.2.1.1.chr10\_hap1.g01182.t01 |
| 13287 | 10 | 140567375 | 140577001 | mNeoVis.2.1.1.chr10\_hap1.g01183.t01 |
| 13288 | 10 | 140567375 | 140577001 | mNeoVis.2.1.1.chr10\_hap1.g01183.t01 |
| 13289 | 10 | 141282718 | 141345234 | mNeoVis.2.1.1.chr10\_hap1.g01184.t01 |
| 13290 | 10 | 141282718 | 141345234 | mNeoVis.2.1.1.chr10\_hap1.g01184.t01 |
| 13291 | 10 | 141593903 | 141995711 | mNeoVis.2.1.1.chr10\_hap1.g01185.t01 |
| 13292 | 10 | 142036537 | 142073079 | mNeoVis.2.1.1.chr10\_hap1.g01186.t01 |
| 13293 | 10 | 142073077 | 142437232 | mNeoVis.2.1.1.chr10\_hap1.g01187.t01 |
| 13294 | 10 | 142073077 | 142437232 | mNeoVis.2.1.1.chr10\_hap1.g01187.t01 |
| 13295 | 10 | 142073077 | 142437232 | mNeoVis.2.1.1.chr10\_hap1.g01187.t01 |
| 13296 | 10 | 142436685 | 142627750 | mNeoVis.2.1.1.chr10\_hap1.g01188.t01 |
| 13297 | 10 | 142436685 | 142627750 | mNeoVis.2.1.1.chr10\_hap1.g01188.t01 |
| 13298 | 10 | 142436685 | 142627750 | mNeoVis.2.1.1.chr10\_hap1.g01188.t01 |
| 13299 | 10 | 142627832 | 142638229 | mNeoVis.2.1.1.chr10\_hap1.g01189.t01 |
| 13300 | 10 | 142627832 | 142638229 | mNeoVis.2.1.1.chr10\_hap1.g01189.t01 |
| 13301 | 10 | 142627832 | 142638229 | mNeoVis.2.1.1.chr10\_hap1.g01189.t01 |
| 13302 | 10 | 142669450 | 142691131 | mNeoVis.2.1.1.chr10\_hap1.g01190.t01 |
| 13303 | 10 | 142669450 | 142691131 | mNeoVis.2.1.1.chr10\_hap1.g01190.t01 |
| 13304 | 10 | 142669450 | 142691131 | mNeoVis.2.1.1.chr10\_hap1.g01190.t01 |
| 13305 | 10 | 142735114 | 142743860 | mNeoVis.2.1.1.chr10\_hap1.g01191.t01 |
| 13306 | 10 | 142746856 | 142757803 | mNeoVis.2.1.1.chr10\_hap1.g01192.t01 |
| 13307 | 10 | 142911094 | 142947338 | mNeoVis.2.1.1.chr10\_hap1.g01193.t01 |
| 13308 | 10 | 142948839 | 142953110 | mNeoVis.2.1.1.chr10\_hap1.g01194.t01 |
| 13309 | 10 | 142966582 | 142992524 | mNeoVis.2.1.1.chr10\_hap1.g01195.t01 |
| 13310 | 10 | 143063264 | 143161168 | mNeoVis.2.1.1.chr10\_hap1.g01196.t01 |
| 13311 | 10 | 143167029 | 143176282 | mNeoVis.2.1.1.chr10\_hap1.g01197.t01 |
| 13312 | 10 | 143186392 | 143269534 | mNeoVis.2.1.1.chr10\_hap1.g01198.t01 |
| 13313 | 10 | 143269684 | 143286878 | mNeoVis.2.1.1.chr10\_hap1.g01199.t01 |
| 13314 | 10 | 143481917 | 143524776 | mNeoVis.2.1.1.chr10\_hap1.g01202.t01 |
| 13315 | 10 | 143526172 | 143552723 | mNeoVis.2.1.1.chr10\_hap1.g01203.t01 |
| 13316 | 10 | 143558456 | 143574094 | mNeoVis.2.1.1.chr10\_hap1.g01204.t01 |
| 13317 | 10 | 143576746 | 143586545 | mNeoVis.2.1.1.chr10\_hap1.g01205.t01 |
| 13318 | 10 | 143586622 | 143631113 | mNeoVis.2.1.1.chr10\_hap1.g01206.t01 |
| 13319 | 10 | 143636491 | 143654920 | mNeoVis.2.1.1.chr10\_hap1.g01207.t01 |
| 13320 | 10 | 143654790 | 143657139 | mNeoVis.2.1.1.chr10\_hap1.g01208.t01 |
| 13321 | 10 | 143662703 | 143687753 | mNeoVis.2.1.1.chr10\_hap1.g01209.t01 |
| 13322 | 10 | 143708555 | 143761867 | mNeoVis.2.1.1.chr10\_hap1.g01210.t01 |
| 13323 | 10 | 143764157 | 143776627 | mNeoVis.2.1.1.chr10\_hap1.g01211.t01 |
| 13324 | 10 | 143887238 | 143958007 | mNeoVis.2.1.1.chr10\_hap1.g01212.t01 |
| 13325 | 10 | 143959282 | 143977084 | mNeoVis.2.1.1.chr10\_hap1.g01213.t01 |
| 13326 | 10 | 144030988 | 144079689 | mNeoVis.2.1.1.chr10\_hap1.g01214.t01 |
| 13327 | 10 | 144070824 | 144074411 | mNeoVis.2.1.1.chr10\_hap1.g01215.t01 |
| 13328 | 10 | 144082894 | 144095585 | mNeoVis.2.1.1.chr10\_hap1.g01216.t01 |
| 13329 | 10 | 144095681 | 144101486 | mNeoVis.2.1.1.chr10\_hap1.g01217.t01 |
| 13330 | 10 | 144106121 | 144128044 | mNeoVis.2.1.1.chr10\_hap1.g01218.t01 |
| 13331 | 10 | 144130661 | 144156471 | mNeoVis.2.1.1.chr10\_hap1.g01219.t01 |
| 13332 | 10 | 144169074 | 144181071 | mNeoVis.2.1.1.chr10\_hap1.g01220.t01 |
| 13333 | 10 | 144181385 | 144200136 | mNeoVis.2.1.1.chr10\_hap1.g01221.t01 |
| 13334 | 10 | 144198023 | 144199679 | mNeoVis.2.1.1.chr10\_hap1.g01222.t01 |
| 13335 | 10 | 144206189 | 144224667 | mNeoVis.2.1.1.chr10\_hap1.g01223.t01 |
| 13336 | 10 | 144916757 | 144961357 | mNeoVis.2.1.1.chr10\_hap1.g01236.t01 |
| 13337 | 10 | 144973604 | 145018163 | mNeoVis.2.1.1.chr10\_hap1.g01237.t01 |
| 13338 | 10 | 145048597 | 145196191 | mNeoVis.2.1.1.chr10\_hap1.g01238.t01 |
| 13339 | 10 | 145048597 | 145196191 | mNeoVis.2.1.1.chr10\_hap1.g01238.t01 |
| 13340 | 10 | 145236212 | 145238861 | mNeoVis.2.1.1.chr10\_hap1.g01239.t01 |
| 13341 | 10 | 145236212 | 145238861 | mNeoVis.2.1.1.chr10\_hap1.g01239.t01 |
| 13342 | 10 | 145240609 | 145336877 | mNeoVis.2.1.1.chr10\_hap1.g01240.t01 |
| 13343 | 10 | 145240609 | 145336877 | mNeoVis.2.1.1.chr10\_hap1.g01240.t01 |
| 13344 | 10 | 145347983 | 145377116 | mNeoVis.2.1.1.chr10\_hap1.g01241.t01 |
| 13345 | 10 | 145347983 | 145377116 | mNeoVis.2.1.1.chr10\_hap1.g01241.t01 |
| 13346 | 10 | 145377902 | 145381785 | mNeoVis.2.1.1.chr10\_hap1.g01242.t01 |
| 13347 | 10 | 145377902 | 145381785 | mNeoVis.2.1.1.chr10\_hap1.g01242.t01 |
| 13348 | 10 | 145407068 | 145452363 | mNeoVis.2.1.1.chr10\_hap1.g01243.t01 |
| 13349 | 10 | 145407068 | 145452363 | mNeoVis.2.1.1.chr10\_hap1.g01243.t01 |
| 13350 | 10 | 145451701 | 145468336 | mNeoVis.2.1.1.chr10\_hap1.g01244.t01 |
| 13351 | 10 | 145451701 | 145468336 | mNeoVis.2.1.1.chr10\_hap1.g01244.t01 |
| 13352 | 10 | 145482334 | 145560473 | mNeoVis.2.1.1.chr10\_hap1.g01245.t01 |
| 13353 | 10 | 145482334 | 145560473 | mNeoVis.2.1.1.chr10\_hap1.g01245.t01 |
| 13354 | 10 | 145591164 | 145595607 | mNeoVis.2.1.1.chr10\_hap1.g01246.t01 |
| 13355 | 10 | 145591164 | 145595607 | mNeoVis.2.1.1.chr10\_hap1.g01246.t01 |
| 13356 | 10 | 145603370 | 145609559 | mNeoVis.2.1.1.chr10\_hap1.g01247.t01 |
| 13357 | 10 | 145603370 | 145609559 | mNeoVis.2.1.1.chr10\_hap1.g01247.t01 |
| 13358 | 10 | 145609699 | 145648330 | mNeoVis.2.1.1.chr10\_hap1.g01248.t01 |
| 13359 | 10 | 145609699 | 145648330 | mNeoVis.2.1.1.chr10\_hap1.g01248.t01 |
| 13360 | 10 | 145649107 | 145710031 | mNeoVis.2.1.1.chr10\_hap1.g01249.t01 |
| 13361 | 10 | 145649107 | 145710031 | mNeoVis.2.1.1.chr10\_hap1.g01249.t01 |
| 13362 | 10 | 145683231 | 145710236 | mNeoVis.2.1.1.chr10\_hap1.g01250.t01 |
| 13363 | 10 | 145683231 | 145710236 | mNeoVis.2.1.1.chr10\_hap1.g01250.t01 |
| 13364 | 10 | 145711607 | 145786128 | mNeoVis.2.1.1.chr10\_hap1.g01251.t01 |
| 13365 | 10 | 145711607 | 145786128 | mNeoVis.2.1.1.chr10\_hap1.g01251.t01 |
| 13366 | 10 | 146504853 | 146522777 | mNeoVis.2.1.1.chr10\_hap1.g01254.t01 |
| 13367 | 10 | 146504853 | 146522777 | mNeoVis.2.1.1.chr10\_hap1.g01254.t01 |
| 13368 | 10 | 146522849 | 146639067 | mNeoVis.2.1.1.chr10\_hap1.g01255.t01 |
| 13369 | 10 | 146522849 | 146639067 | mNeoVis.2.1.1.chr10\_hap1.g01255.t01 |
| 13370 | 10 | 146696482 | 146701303 | mNeoVis.2.1.1.chr10\_hap1.g01256.t01 |
| 13371 | 10 | 146696482 | 146701303 | mNeoVis.2.1.1.chr10\_hap1.g01256.t01 |
| 13372 | 10 | 147173140 | 147211064 | mNeoVis.2.1.1.chr10\_hap1.g01257.t01 |
| 13373 | 10 | 147173140 | 147211064 | mNeoVis.2.1.1.chr10\_hap1.g01257.t01 |
| 13374 | 10 | 147216508 | 147224402 | mNeoVis.2.1.1.chr10\_hap1.g01258.t01 |
| 13375 | 10 | 147216508 | 147224402 | mNeoVis.2.1.1.chr10\_hap1.g01258.t01 |
| 13376 | 10 | 147227421 | 147321499 | mNeoVis.2.1.1.chr10\_hap1.g01259.t01 |
| 13377 | 10 | 147227421 | 147321499 | mNeoVis.2.1.1.chr10\_hap1.g01259.t01 |
| 13378 | 10 | 147344483 | 147353736 | mNeoVis.2.1.1.chr10\_hap1.g01260.t01 |
| 13379 | 10 | 147424175 | 147426074 | mNeoVis.2.1.1.chr10\_hap1.g01261.t01 |
| 13380 | 10 | 147436220 | 147459608 | mNeoVis.2.1.1.chr10\_hap1.g01262.t01 |
| 13381 | 10 | 147459438 | 147585776 | mNeoVis.2.1.1.chr10\_hap1.g01263.t01 |
| 13382 | 10 | 147601578 | 147644537 | mNeoVis.2.1.1.chr10\_hap1.g01264.t01 |
| 13383 | 10 | 147708599 | 147850306 | mNeoVis.2.1.1.chr10\_hap1.g01265.t01 |
| 13384 | 10 | 147955589 | 147956404 | mNeoVis.2.1.1.chr10\_hap1.g01266.t01 |
| 13385 | 10 | 147955589 | 147956404 | mNeoVis.2.1.1.chr10\_hap1.g01266.t01 |
| 13386 | 10 | 147966812 | 147990191 | mNeoVis.2.1.1.chr10\_hap1.g01267.t01 |
| 13387 | 10 | 147966812 | 147990191 | mNeoVis.2.1.1.chr10\_hap1.g01267.t01 |
| 13388 | 10 | 148002026 | 148067233 | mNeoVis.2.1.1.chr10\_hap1.g01268.t01 |
| 13389 | 10 | 148002026 | 148067233 | mNeoVis.2.1.1.chr10\_hap1.g01268.t01 |
| 13390 | 10 | 148066482 | 148115255 | mNeoVis.2.1.1.chr10\_hap1.g01269.t01 |
| 13391 | 10 | 148066482 | 148115255 | mNeoVis.2.1.1.chr10\_hap1.g01269.t01 |
| 13392 | 10 | 148128686 | 148163270 | mNeoVis.2.1.1.chr10\_hap1.g01270.t01 |
| 13393 | 10 | 148128686 | 148163270 | mNeoVis.2.1.1.chr10\_hap1.g01270.t01 |
| 13394 | 10 | 148350318 | 148361270 | mNeoVis.2.1.1.chr10\_hap1.g01271.t01 |
| 13395 | 10 | 148350318 | 148361270 | mNeoVis.2.1.1.chr10\_hap1.g01271.t01 |
| 13396 | 10 | 148413594 | 148417504 | mNeoVis.2.1.1.chr10\_hap1.g01272.t01 |
| 13397 | 10 | 148413594 | 148417504 | mNeoVis.2.1.1.chr10\_hap1.g01272.t01 |
| 13398 | 10 | 148425190 | 148426802 | mNeoVis.2.1.1.chr10\_hap1.g01273.t01 |
| 13399 | 10 | 148425190 | 148426802 | mNeoVis.2.1.1.chr10\_hap1.g01273.t01 |
| 13400 | 10 | 148550862 | 148637837 | mNeoVis.2.1.1.chr10\_hap1.g01274.t01 |
| 13401 | 10 | 148653647 | 148666546 | mNeoVis.2.1.1.chr10\_hap1.g01275.t01 |
| 13402 | 10 | 148683516 | 148701920 | mNeoVis.2.1.1.chr10\_hap1.g01276.t01 |
| 13403 | 10 | 148734082 | 148773670 | mNeoVis.2.1.1.chr10\_hap1.g01277.t01 |
| 13404 | 10 | 148777680 | 148834863 | mNeoVis.2.1.1.chr10\_hap1.g01278.t01 |
| 13405 | 10 | 148843385 | 148861209 | mNeoVis.2.1.1.chr10\_hap1.g01279.t01 |
| 13406 | 10 | 148852583 | 148865817 | mNeoVis.2.1.1.chr10\_hap1.g01280.t01 |
| 13407 | 10 | 148875820 | 148889197 | mNeoVis.2.1.1.chr10\_hap1.g01281.t01 |
| 13408 | 10 | 148889979 | 148949678 | mNeoVis.2.1.1.chr10\_hap1.g01282.t01 |
| 13409 | 10 | 148949794 | 148963585 | mNeoVis.2.1.1.chr10\_hap1.g01283.t01 |
| 13410 | 10 | 148964262 | 148980989 | mNeoVis.2.1.1.chr10\_hap1.g01284.t01 |
| 13411 | 10 | 149003328 | 149009709 | mNeoVis.2.1.1.chr10\_hap1.g01285.t01 |
| 13412 | 10 | 149057647 | 149100304 | mNeoVis.2.1.1.chr10\_hap1.g01286.t01 |
| 13413 | 10 | 149101442 | 149122686 | mNeoVis.2.1.1.chr10\_hap1.g01287.t01 |
| 13414 | 10 | 149124279 | 149131723 | mNeoVis.2.1.1.chr10\_hap1.g01288.t01 |
| 13415 | 10 | 149145417 | 149212828 | mNeoVis.2.1.1.chr10\_hap1.g01289.t01 |
| 13416 | 10 | 149213002 | 149230211 | mNeoVis.2.1.1.chr10\_hap1.g01290.t01 |
| 13417 | 10 | 149232181 | 149243624 | mNeoVis.2.1.1.chr10\_hap1.g01291.t01 |
| 13418 | 11 | 96276 | 119267 | mNeoVis.2.1.1.chr11\_hap1.g00002.t01 |
| 13419 | 11 | 130511 | 166518 | mNeoVis.2.1.1.chr11\_hap1.g00003.t01 |
| 13420 | 11 | 182294 | 231335 | mNeoVis.2.1.1.chr11\_hap1.g00004.t01 |
| 13421 | 11 | 233832 | 253669 | mNeoVis.2.1.1.chr11\_hap1.g00005.t01 |
| 13422 | 11 | 247678 | 249724 | mNeoVis.2.1.1.chr11\_hap1.g00006.t01 |
| 13423 | 11 | 253110 | 259586 | mNeoVis.2.1.1.chr11\_hap1.g00007.t01 |
| 13424 | 11 | 260339 | 268242 | mNeoVis.2.1.1.chr11\_hap1.g00008.t01 |
| 13425 | 11 | 278852 | 280633 | mNeoVis.2.1.1.chr11\_hap1.g00009.t01 |
| 13426 | 11 | 289561 | 291524 | mNeoVis.2.1.1.chr11\_hap1.g00010.t01 |
| 13427 | 11 | 294050 | 333641 | mNeoVis.2.1.1.chr11\_hap1.g00011.t01 |
| 13428 | 11 | 334546 | 339641 | mNeoVis.2.1.1.chr11\_hap1.g00012.t01 |
| 13429 | 11 | 341913 | 349762 | mNeoVis.2.1.1.chr11\_hap1.g00013.t01 |
| 13430 | 11 | 353112 | 364580 | mNeoVis.2.1.1.chr11\_hap1.g00014.t01 |
| 13431 | 11 | 369566 | 410626 | mNeoVis.2.1.1.chr11\_hap1.g00015.t01 |
| 13432 | 11 | 409148 | 415240 | mNeoVis.2.1.1.chr11\_hap1.g00016.t01 |
| 13433 | 11 | 415358 | 432535 | mNeoVis.2.1.1.chr11\_hap1.g00017.t01 |
| 13434 | 11 | 434054 | 435879 | mNeoVis.2.1.1.chr11\_hap1.g00018.t01 |
| 13435 | 11 | 459115 | 517488 | mNeoVis.2.1.1.chr11\_hap1.g00019.t01 |
| 13436 | 11 | 518533 | 523213 | mNeoVis.2.1.1.chr11\_hap1.g00020.t01 |
| 13437 | 11 | 523924 | 526814 | mNeoVis.2.1.1.chr11\_hap1.g00021.t01 |
| 13438 | 11 | 526939 | 537108 | mNeoVis.2.1.1.chr11\_hap1.g00022.t01 |
| 13439 | 11 | 539569 | 544421 | mNeoVis.2.1.1.chr11\_hap1.g00023.t01 |
| 13440 | 11 | 544658 | 574681 | mNeoVis.2.1.1.chr11\_hap1.g00024.t01 |
| 13441 | 11 | 574805 | 586464 | mNeoVis.2.1.1.chr11\_hap1.g00025.t01 |
| 13442 | 11 | 593233 | 618821 | mNeoVis.2.1.1.chr11\_hap1.g00026.t01 |
| 13443 | 11 | 629599 | 644260 | mNeoVis.2.1.1.chr11\_hap1.g00027.t01 |
| 13444 | 11 | 646785 | 649249 | mNeoVis.2.1.1.chr11\_hap1.g00028.t01 |
| 13445 | 11 | 654878 | 664219 | mNeoVis.2.1.1.chr11\_hap1.g00029.t01 |
| 13446 | 11 | 661961 | 671947 | mNeoVis.2.1.1.chr11\_hap1.g00030.t01 |
| 13447 | 11 | 672099 | 673583 | mNeoVis.2.1.1.chr11\_hap1.g00031.t01 |
| 13448 | 11 | 683794 | 699705 | mNeoVis.2.1.1.chr11\_hap1.g00032.t01 |
| 13449 | 11 | 686196 | 690886 | mNeoVis.2.1.1.chr11\_hap1.g00033.t01 |
| 13450 | 11 | 711461 | 763657 | mNeoVis.2.1.1.chr11\_hap1.g00034.t01 |
| 13451 | 11 | 788557 | 801744 | mNeoVis.2.1.1.chr11\_hap1.g00035.t01 |
| 13452 | 11 | 808261 | 821627 | mNeoVis.2.1.1.chr11\_hap1.g00036.t01 |
| 13453 | 11 | 827788 | 844332 | mNeoVis.2.1.1.chr11\_hap1.g00037.t01 |
| 13454 | 11 | 857646 | 868107 | mNeoVis.2.1.1.chr11\_hap1.g00038.t01 |
| 13455 | 11 | 868276 | 874075 | mNeoVis.2.1.1.chr11\_hap1.g00039.t01 |
| 13456 | 11 | 883661 | 894800 | mNeoVis.2.1.1.chr11\_hap1.g00040.t01 |
| 13457 | 11 | 1023916 | 1026202 | mNeoVis.2.1.1.chr11\_hap1.g00041.t01 |
| 13458 | 11 | 1052390 | 1066061 | mNeoVis.2.1.1.chr11\_hap1.g00042.t01 |
| 13459 | 11 | 1067074 | 1077481 | mNeoVis.2.1.1.chr11\_hap1.g00043.t01 |
| 13460 | 11 | 1077675 | 1126537 | mNeoVis.2.1.1.chr11\_hap1.g00044.t01 |
| 13461 | 11 | 1132974 | 1141943 | mNeoVis.2.1.1.chr11\_hap1.g00045.t01 |
| 13462 | 11 | 1147530 | 1163778 | mNeoVis.2.1.1.chr11\_hap1.g00046.t01 |
| 13463 | 11 | 1167677 | 1175691 | mNeoVis.2.1.1.chr11\_hap1.g00047.t01 |
| 13464 | 11 | 1179593 | 1184576 | mNeoVis.2.1.1.chr11\_hap1.g00048.t01 |
| 13465 | 11 | 1206341 | 1245290 | mNeoVis.2.1.1.chr11\_hap1.g00049.t01 |
| 13466 | 11 | 1263860 | 1285928 | mNeoVis.2.1.1.chr11\_hap1.g00050.t01 |
| 13467 | 11 | 1353715 | 1359926 | mNeoVis.2.1.1.chr11\_hap1.g00051.t01 |
| 13468 | 11 | 1397532 | 1408211 | mNeoVis.2.1.1.chr11\_hap1.g00052.t01 |
| 13469 | 11 | 1428777 | 1446297 | mNeoVis.2.1.1.chr11\_hap1.g00053.t01 |
| 13470 | 11 | 1448132 | 1461185 | mNeoVis.2.1.1.chr11\_hap1.g00054.t01 |
| 13471 | 11 | 1462393 | 1463514 | mNeoVis.2.1.1.chr11\_hap1.g00055.t01 |
| 13472 | 11 | 1764957 | 2261780 | mNeoVis.2.1.1.chr11\_hap1.g00065.t01 |
| 13473 | 11 | 3159159 | 3200009 | mNeoVis.2.1.1.chr11\_hap1.g00066.t01 |
| 13474 | 11 | 3207171 | 3214995 | mNeoVis.2.1.1.chr11\_hap1.g00067.t01 |
| 13475 | 11 | 3215375 | 3216504 | mNeoVis.2.1.1.chr11\_hap1.g00068.t01 |
| 13476 | 11 | 3220143 | 3292316 | mNeoVis.2.1.1.chr11\_hap1.g00069.t01 |
| 13477 | 11 | 3300377 | 3499889 | mNeoVis.2.1.1.chr11\_hap1.g00070.t01 |
| 13478 | 11 | 3685657 | 3708291 | mNeoVis.2.1.1.chr11\_hap1.g00071.t01 |
| 13479 | 11 | 3685657 | 3708291 | mNeoVis.2.1.1.chr11\_hap1.g00071.t01 |
| 13480 | 11 | 3713894 | 3851712 | mNeoVis.2.1.1.chr11\_hap1.g00072.t01 |
| 13481 | 11 | 3713894 | 3851712 | mNeoVis.2.1.1.chr11\_hap1.g00072.t01 |
| 13482 | 11 | 3943428 | 3952253 | mNeoVis.2.1.1.chr11\_hap1.g00073.t01 |
| 13483 | 11 | 3943428 | 3952253 | mNeoVis.2.1.1.chr11\_hap1.g00073.t01 |
| 13484 | 11 | 3953621 | 3957513 | mNeoVis.2.1.1.chr11\_hap1.g00074.t01 |
| 13485 | 11 | 3953621 | 3957513 | mNeoVis.2.1.1.chr11\_hap1.g00074.t01 |
| 13486 | 11 | 3961332 | 3966766 | mNeoVis.2.1.1.chr11\_hap1.g00075.t01 |
| 13487 | 11 | 3961332 | 3966766 | mNeoVis.2.1.1.chr11\_hap1.g00075.t01 |
| 13488 | 11 | 3978787 | 3988439 | mNeoVis.2.1.1.chr11\_hap1.g00076.t01 |
| 13489 | 11 | 3978787 | 3988439 | mNeoVis.2.1.1.chr11\_hap1.g00076.t01 |
| 13490 | 11 | 3988493 | 4000900 | mNeoVis.2.1.1.chr11\_hap1.g00077.t01 |
| 13491 | 11 | 3988493 | 4000900 | mNeoVis.2.1.1.chr11\_hap1.g00077.t01 |
| 13492 | 11 | 4063502 | 4070317 | mNeoVis.2.1.1.chr11\_hap1.g00078.t01 |
| 13493 | 11 | 4073234 | 4082137 | mNeoVis.2.1.1.chr11\_hap1.g00079.t01 |
| 13494 | 11 | 4073234 | 4082137 | mNeoVis.2.1.1.chr11\_hap1.g00079.t01 |
| 13495 | 11 | 4074664 | 4130400 | mNeoVis.2.1.1.chr11\_hap1.g00080.t01 |
| 13496 | 11 | 4074664 | 4130400 | mNeoVis.2.1.1.chr11\_hap1.g00080.t01 |
| 13497 | 11 | 4238089 | 4259445 | mNeoVis.2.1.1.chr11\_hap1.g00081.t01 |
| 13498 | 11 | 4238089 | 4259445 | mNeoVis.2.1.1.chr11\_hap1.g00081.t01 |
| 13499 | 11 | 4269848 | 4295423 | mNeoVis.2.1.1.chr11\_hap1.g00082.t01 |
| 13500 | 11 | 4269848 | 4295423 | mNeoVis.2.1.1.chr11\_hap1.g00082.t01 |
| 13501 | 11 | 4423328 | 4473312 | mNeoVis.2.1.1.chr11\_hap1.g00083.t01 |
| 13502 | 11 | 4489622 | 4540605 | mNeoVis.2.1.1.chr11\_hap1.g00084.t01 |
| 13503 | 11 | 4558445 | 4614943 | mNeoVis.2.1.1.chr11\_hap1.g00085.t01 |
| 13504 | 11 | 4614845 | 4695589 | mNeoVis.2.1.1.chr11\_hap1.g00086.t01 |
| 13505 | 11 | 4695928 | 4708971 | mNeoVis.2.1.1.chr11\_hap1.g00087.t01 |
| 13506 | 11 | 4753030 | 4761052 | mNeoVis.2.1.1.chr11\_hap1.g00088.t01 |
| 13507 | 11 | 6159951 | 6160922 | mNeoVis.2.1.1.chr11\_hap1.g00106.t01 |
| 13508 | 11 | 6345292 | 6350944 | mNeoVis.2.1.1.chr11\_hap1.g00107.t01 |
| 13509 | 11 | 6628259 | 6922718 | mNeoVis.2.1.1.chr11\_hap1.g00108.t01 |
| 13510 | 11 | 7061939 | 7355220 | mNeoVis.2.1.1.chr11\_hap1.g00109.t01 |
| 13511 | 11 | 7410363 | 7564954 | mNeoVis.2.1.1.chr11\_hap1.g00110.t01 |
| 13512 | 11 | 8229779 | 8263477 | mNeoVis.2.1.1.chr11\_hap1.g00114.t01 |
| 13513 | 11 | 8815105 | 9148143 | mNeoVis.2.1.1.chr11\_hap1.g00116.t01 |
| 13514 | 11 | 8815105 | 9148143 | mNeoVis.2.1.1.chr11\_hap1.g00116.t01 |
| 13515 | 11 | 9395949 | 9426480 | mNeoVis.2.1.1.chr11\_hap1.g00117.t01 |
| 13516 | 11 | 9395949 | 9426480 | mNeoVis.2.1.1.chr11\_hap1.g00117.t01 |
| 13517 | 11 | 9724673 | 9744674 | mNeoVis.2.1.1.chr11\_hap1.g00118.t01 |
| 13518 | 11 | 9929524 | 10069463 | mNeoVis.2.1.1.chr11\_hap1.g00120.t01 |
| 13519 | 11 | 10327150 | 10346681 | mNeoVis.2.1.1.chr11\_hap1.g00121.t01 |
| 13520 | 11 | 10382380 | 10384583 | mNeoVis.2.1.1.chr11\_hap1.g00122.t01 |
| 13521 | 11 | 10384860 | 10404098 | mNeoVis.2.1.1.chr11\_hap1.g00123.t01 |
| 13522 | 11 | 10402223 | 10438727 | mNeoVis.2.1.1.chr11\_hap1.g00124.t01 |
| 13523 | 11 | 10644938 | 10997808 | mNeoVis.2.1.1.chr11\_hap1.g00128.t01 |
| 13524 | 11 | 10976227 | 10978217 | mNeoVis.2.1.1.chr11\_hap1.g00129.t01 |
| 13525 | 11 | 10998349 | 11021618 | mNeoVis.2.1.1.chr11\_hap1.g00130.t01 |
| 13526 | 11 | 11030367 | 11060675 | mNeoVis.2.1.1.chr11\_hap1.g00131.t01 |
| 13527 | 11 | 11066213 | 11081754 | mNeoVis.2.1.1.chr11\_hap1.g00132.t01 |
| 13528 | 11 | 11125014 | 11131038 | mNeoVis.2.1.1.chr11\_hap1.g00133.t01 |
| 13529 | 11 | 11148176 | 11165510 | mNeoVis.2.1.1.chr11\_hap1.g00134.t01 |
| 13530 | 11 | 11178018 | 11189376 | mNeoVis.2.1.1.chr11\_hap1.g00135.t01 |
| 13531 | 11 | 11192463 | 11193035 | mNeoVis.2.1.1.chr11\_hap1.g00136.t01 |
| 13532 | 11 | 11199394 | 11270141 | mNeoVis.2.1.1.chr11\_hap1.g00137.t01 |
| 13533 | 11 | 11366986 | 11413767 | mNeoVis.2.1.1.chr11\_hap1.g00138.t01 |
| 13534 | 11 | 18205514 | 18210904 | mNeoVis.2.1.1.chr11\_hap1.g00226.t01 |
| 13535 | 11 | 18211061 | 18253307 | mNeoVis.2.1.1.chr11\_hap1.g00227.t01 |
| 13536 | 11 | 18252237 | 18362851 | mNeoVis.2.1.1.chr11\_hap1.g00228.t01 |
| 13537 | 11 | 18368194 | 18403651 | mNeoVis.2.1.1.chr11\_hap1.g00229.t01 |
| 13538 | 11 | 18412908 | 18513343 | mNeoVis.2.1.1.chr11\_hap1.g00230.t01 |
| 13539 | 11 | 20112398 | 20256743 | mNeoVis.2.1.1.chr11\_hap1.g00232.t01 |
| 13540 | 11 | 20263152 | 20282937 | mNeoVis.2.1.1.chr11\_hap1.g00233.t01 |
| 13541 | 11 | 20285181 | 20377287 | mNeoVis.2.1.1.chr11\_hap1.g00234.t01 |
| 13542 | 11 | 20406384 | 20433112 | mNeoVis.2.1.1.chr11\_hap1.g00235.t01 |
| 13543 | 11 | 20406384 | 20433112 | mNeoVis.2.1.1.chr11\_hap1.g00235.t01 |
| 13544 | 11 | 20441546 | 20446265 | mNeoVis.2.1.1.chr11\_hap1.g00236.t01 |
| 13545 | 11 | 20441546 | 20446265 | mNeoVis.2.1.1.chr11\_hap1.g00236.t01 |
| 13546 | 11 | 20506093 | 20529975 | mNeoVis.2.1.1.chr11\_hap1.g00237.t01 |
| 13547 | 11 | 20506093 | 20529975 | mNeoVis.2.1.1.chr11\_hap1.g00237.t01 |
| 13548 | 11 | 20535772 | 20542988 | mNeoVis.2.1.1.chr11\_hap1.g00238.t01 |
| 13549 | 11 | 20535772 | 20542988 | mNeoVis.2.1.1.chr11\_hap1.g00238.t01 |
| 13550 | 11 | 20535772 | 20542988 | mNeoVis.2.1.1.chr11\_hap1.g00238.t01 |
| 13551 | 11 | 20545381 | 20626382 | mNeoVis.2.1.1.chr11\_hap1.g00239.t01 |
| 13552 | 11 | 20545381 | 20626382 | mNeoVis.2.1.1.chr11\_hap1.g00239.t01 |
| 13553 | 11 | 20545381 | 20626382 | mNeoVis.2.1.1.chr11\_hap1.g00239.t01 |
| 13554 | 11 | 20663043 | 20685461 | mNeoVis.2.1.1.chr11\_hap1.g00240.t01 |
| 13555 | 11 | 20663043 | 20685461 | mNeoVis.2.1.1.chr11\_hap1.g00240.t01 |
| 13556 | 11 | 20663043 | 20685461 | mNeoVis.2.1.1.chr11\_hap1.g00240.t01 |
| 13557 | 11 | 20735611 | 20758883 | mNeoVis.2.1.1.chr11\_hap1.g00241.t01 |
| 13558 | 11 | 20735611 | 20758883 | mNeoVis.2.1.1.chr11\_hap1.g00241.t01 |
| 13559 | 11 | 20735611 | 20758883 | mNeoVis.2.1.1.chr11\_hap1.g00241.t01 |
| 13560 | 11 | 20768606 | 20795888 | mNeoVis.2.1.1.chr11\_hap1.g00242.t01 |
| 13561 | 11 | 20768606 | 20795888 | mNeoVis.2.1.1.chr11\_hap1.g00242.t01 |
| 13562 | 11 | 20821181 | 20861310 | mNeoVis.2.1.1.chr11\_hap1.g00243.t01 |
| 13563 | 11 | 20821181 | 20861310 | mNeoVis.2.1.1.chr11\_hap1.g00243.t01 |
| 13564 | 11 | 20899871 | 20933364 | mNeoVis.2.1.1.chr11\_hap1.g00244.t01 |
| 13565 | 11 | 20899871 | 20933364 | mNeoVis.2.1.1.chr11\_hap1.g00244.t01 |
| 13566 | 11 | 20958902 | 21013005 | mNeoVis.2.1.1.chr11\_hap1.g00245.t01 |
| 13567 | 11 | 21531167 | 21581801 | mNeoVis.2.1.1.chr11\_hap1.g00251.t01 |
| 13568 | 11 | 21602623 | 21625966 | mNeoVis.2.1.1.chr11\_hap1.g00252.t01 |
| 13569 | 11 | 21635489 | 21654553 | mNeoVis.2.1.1.chr11\_hap1.g00253.t01 |
| 13570 | 11 | 21659277 | 21715152 | mNeoVis.2.1.1.chr11\_hap1.g00254.t01 |
| 13571 | 11 | 21714795 | 21739726 | mNeoVis.2.1.1.chr11\_hap1.g00255.t01 |
| 13572 | 11 | 21742953 | 21769358 | mNeoVis.2.1.1.chr11\_hap1.g00256.t01 |
| 13573 | 11 | 21782333 | 21850911 | mNeoVis.2.1.1.chr11\_hap1.g00257.t01 |
| 13574 | 11 | 21867600 | 21927060 | mNeoVis.2.1.1.chr11\_hap1.g00258.t01 |
| 13575 | 11 | 21927255 | 21940513 | mNeoVis.2.1.1.chr11\_hap1.g00259.t01 |
| 13576 | 11 | 21950833 | 22020686 | mNeoVis.2.1.1.chr11\_hap1.g00260.t01 |
| 13577 | 11 | 22023420 | 22040085 | mNeoVis.2.1.1.chr11\_hap1.g00261.t01 |
| 13578 | 11 | 22042878 | 22122524 | mNeoVis.2.1.1.chr11\_hap1.g00262.t01 |
| 13579 | 11 | 22125725 | 22154086 | mNeoVis.2.1.1.chr11\_hap1.g00263.t01 |
| 13580 | 11 | 22154228 | 22163200 | mNeoVis.2.1.1.chr11\_hap1.g00264.t01 |
| 13581 | 11 | 23767683 | 23776950 | mNeoVis.2.1.1.chr11\_hap1.g00298.t01 |
| 13582 | 11 | 23799566 | 23893518 | mNeoVis.2.1.1.chr11\_hap1.g00299.t01 |
| 13583 | 11 | 23838247 | 23875267 | mNeoVis.2.1.1.chr11\_hap1.g00300.t01 |
| 13584 | 11 | 23893731 | 23895359 | mNeoVis.2.1.1.chr11\_hap1.g00301.t01 |
| 13585 | 11 | 23916622 | 23922910 | mNeoVis.2.1.1.chr11\_hap1.g00302.t01 |
| 13586 | 11 | 23938610 | 24071920 | mNeoVis.2.1.1.chr11\_hap1.g00303.t01 |
| 13587 | 11 | 24131339 | 24144371 | mNeoVis.2.1.1.chr11\_hap1.g00304.t01 |
| 13588 | 11 | 24144488 | 24300165 | mNeoVis.2.1.1.chr11\_hap1.g00305.t01 |
| 13589 | 11 | 24300243 | 24320448 | mNeoVis.2.1.1.chr11\_hap1.g00306.t01 |
| 13590 | 11 | 24331073 | 24413159 | mNeoVis.2.1.1.chr11\_hap1.g00307.t01 |
| 13591 | 11 | 24526847 | 24578381 | mNeoVis.2.1.1.chr11\_hap1.g00308.t01 |
| 13592 | 11 | 24590347 | 24635856 | mNeoVis.2.1.1.chr11\_hap1.g00309.t01 |
| 13593 | 11 | 24642432 | 24658539 | mNeoVis.2.1.1.chr11\_hap1.g00310.t01 |
| 13594 | 11 | 24670392 | 24681010 | mNeoVis.2.1.1.chr11\_hap1.g00311.t01 |
| 13595 | 11 | 24682388 | 24698863 | mNeoVis.2.1.1.chr11\_hap1.g00312.t01 |
| 13596 | 11 | 24688020 | 24689693 | mNeoVis.2.1.1.chr11\_hap1.g00313.t01 |
| 13597 | 11 | 24692363 | 24693711 | mNeoVis.2.1.1.chr11\_hap1.g00314.t01 |
| 13598 | 11 | 24922019 | 24951282 | mNeoVis.2.1.1.chr11\_hap1.g00316.t01 |
| 13599 | 11 | 24959410 | 24997189 | mNeoVis.2.1.1.chr11\_hap1.g00317.t01 |
| 13600 | 11 | 25008002 | 25019068 | mNeoVis.2.1.1.chr11\_hap1.g00318.t01 |
| 13601 | 11 | 25043480 | 25053883 | mNeoVis.2.1.1.chr11\_hap1.g00319.t01 |
| 13602 | 11 | 25058149 | 25078172 | mNeoVis.2.1.1.chr11\_hap1.g00320.t01 |
| 13603 | 11 | 25058149 | 25078172 | mNeoVis.2.1.1.chr11\_hap1.g00320.t01 |
| 13604 | 11 | 25099527 | 25106076 | mNeoVis.2.1.1.chr11\_hap1.g00321.t01 |
| 13605 | 11 | 25099527 | 25106076 | mNeoVis.2.1.1.chr11\_hap1.g00321.t01 |
| 13606 | 11 | 25113445 | 25123616 | mNeoVis.2.1.1.chr11\_hap1.g00322.t01 |
| 13607 | 11 | 25113445 | 25123616 | mNeoVis.2.1.1.chr11\_hap1.g00322.t01 |
| 13608 | 11 | 25166431 | 25170764 | mNeoVis.2.1.1.chr11\_hap1.g00323.t01 |
| 13609 | 11 | 25166431 | 25170764 | mNeoVis.2.1.1.chr11\_hap1.g00323.t01 |
| 13610 | 11 | 25181086 | 25189422 | mNeoVis.2.1.1.chr11\_hap1.g00324.t01 |
| 13611 | 11 | 25181086 | 25189422 | mNeoVis.2.1.1.chr11\_hap1.g00324.t01 |
| 13612 | 11 | 25216614 | 25245307 | mNeoVis.2.1.1.chr11\_hap1.g00325.t01 |
| 13613 | 11 | 25216614 | 25245307 | mNeoVis.2.1.1.chr11\_hap1.g00325.t01 |
| 13614 | 11 | 25250524 | 25266334 | mNeoVis.2.1.1.chr11\_hap1.g00326.t01 |
| 13615 | 11 | 25250524 | 25266334 | mNeoVis.2.1.1.chr11\_hap1.g00326.t01 |
| 13616 | 11 | 25273246 | 25285334 | mNeoVis.2.1.1.chr11\_hap1.g00327.t01 |
| 13617 | 11 | 25273246 | 25285334 | mNeoVis.2.1.1.chr11\_hap1.g00327.t01 |
| 13618 | 11 | 25290994 | 25309144 | mNeoVis.2.1.1.chr11\_hap1.g00328.t01 |
| 13619 | 11 | 25290994 | 25309144 | mNeoVis.2.1.1.chr11\_hap1.g00328.t01 |
| 13620 | 11 | 25312214 | 25329841 | mNeoVis.2.1.1.chr11\_hap1.g00329.t01 |
| 13621 | 11 | 25312214 | 25329841 | mNeoVis.2.1.1.chr11\_hap1.g00329.t01 |
| 13622 | 11 | 25330474 | 25330872 | mNeoVis.2.1.1.chr11\_hap1.g00330.t01 |
| 13623 | 11 | 25330474 | 25330872 | mNeoVis.2.1.1.chr11\_hap1.g00330.t01 |
| 13624 | 11 | 25348102 | 25409801 | mNeoVis.2.1.1.chr11\_hap1.g00331.t01 |
| 13625 | 11 | 25348102 | 25409801 | mNeoVis.2.1.1.chr11\_hap1.g00331.t01 |
| 13626 | 11 | 25415750 | 25444451 | mNeoVis.2.1.1.chr11\_hap1.g00332.t01 |
| 13627 | 11 | 25415750 | 25444451 | mNeoVis.2.1.1.chr11\_hap1.g00332.t01 |
| 13628 | 11 | 25451687 | 25475955 | mNeoVis.2.1.1.chr11\_hap1.g00333.t01 |
| 13629 | 11 | 25451687 | 25475955 | mNeoVis.2.1.1.chr11\_hap1.g00333.t01 |
| 13630 | 11 | 25513491 | 25533921 | mNeoVis.2.1.1.chr11\_hap1.g00334.t01 |
| 13631 | 11 | 25513491 | 25533921 | mNeoVis.2.1.1.chr11\_hap1.g00334.t01 |
| 13632 | 11 | 25763000 | 25841203 | mNeoVis.2.1.1.chr11\_hap1.g00336.t01 |
| 13633 | 11 | 25868689 | 25914132 | mNeoVis.2.1.1.chr11\_hap1.g00337.t01 |
| 13634 | 11 | 25938754 | 25965140 | mNeoVis.2.1.1.chr11\_hap1.g00338.t01 |
| 13635 | 11 | 25965228 | 25977075 | mNeoVis.2.1.1.chr11\_hap1.g00339.t01 |
| 13636 | 11 | 25979114 | 25984384 | mNeoVis.2.1.1.chr11\_hap1.g00340.t01 |
| 13637 | 11 | 25997492 | 26042610 | mNeoVis.2.1.1.chr11\_hap1.g00341.t01 |
| 13638 | 11 | 26048402 | 26090782 | mNeoVis.2.1.1.chr11\_hap1.g00342.t01 |
| 13639 | 11 | 26107652 | 26123780 | mNeoVis.2.1.1.chr11\_hap1.g00343.t01 |
| 13640 | 11 | 26136385 | 26159784 | mNeoVis.2.1.1.chr11\_hap1.g00344.t01 |
| 13641 | 11 | 26174748 | 26180643 | mNeoVis.2.1.1.chr11\_hap1.g00345.t01 |
| 13642 | 11 | 26182257 | 26223373 | mNeoVis.2.1.1.chr11\_hap1.g00346.t01 |
| 13643 | 11 | 26231058 | 26240965 | mNeoVis.2.1.1.chr11\_hap1.g00347.t01 |
| 13644 | 11 | 26242776 | 26252091 | mNeoVis.2.1.1.chr11\_hap1.g00348.t01 |
| 13645 | 11 | 26252261 | 26254938 | mNeoVis.2.1.1.chr11\_hap1.g00349.t01 |
| 13646 | 11 | 26264183 | 26277394 | mNeoVis.2.1.1.chr11\_hap1.g00350.t01 |
| 13647 | 11 | 26290826 | 26351232 | mNeoVis.2.1.1.chr11\_hap1.g00351.t01 |
| 13648 | 11 | 26351325 | 26416004 | mNeoVis.2.1.1.chr11\_hap1.g00352.t01 |
| 13649 | 11 | 26433310 | 26439633 | mNeoVis.2.1.1.chr11\_hap1.g00353.t01 |
| 13650 | 11 | 26463053 | 26464260 | mNeoVis.2.1.1.chr11\_hap1.g00354.t01 |
| 13651 | 11 | 26492343 | 26533537 | mNeoVis.2.1.1.chr11\_hap1.g00355.t01 |
| 13652 | 11 | 26558352 | 26567899 | mNeoVis.2.1.1.chr11\_hap1.g00356.t01 |
| 13653 | 11 | 26568086 | 26573009 | mNeoVis.2.1.1.chr11\_hap1.g00357.t01 |
| 13654 | 11 | 26581478 | 26589106 | mNeoVis.2.1.1.chr11\_hap1.g00358.t01 |
| 13655 | 11 | 29963830 | 29987402 | mNeoVis.2.1.1.chr11\_hap1.g00427.t01 |
| 13656 | 11 | 29987795 | 30000931 | mNeoVis.2.1.1.chr11\_hap1.g00428.t01 |
| 13657 | 11 | 30007024 | 30014474 | mNeoVis.2.1.1.chr11\_hap1.g00429.t01 |
| 13658 | 11 | 30014578 | 30017476 | mNeoVis.2.1.1.chr11\_hap1.g00430.t01 |
| 13659 | 11 | 30025160 | 30034902 | mNeoVis.2.1.1.chr11\_hap1.g00431.t01 |
| 13660 | 11 | 30036681 | 30046646 | mNeoVis.2.1.1.chr11\_hap1.g00432.t01 |
| 13661 | 11 | 30065378 | 30090097 | mNeoVis.2.1.1.chr11\_hap1.g00433.t01 |
| 13662 | 11 | 30101251 | 30137272 | mNeoVis.2.1.1.chr11\_hap1.g00434.t01 |
| 13663 | 11 | 30145443 | 30248618 | mNeoVis.2.1.1.chr11\_hap1.g00435.t01 |
| 13664 | 11 | 30301247 | 30394458 | mNeoVis.2.1.1.chr11\_hap1.g00436.t01 |
| 13665 | 11 | 30397319 | 30417167 | mNeoVis.2.1.1.chr11\_hap1.g00437.t01 |
| 13666 | 11 | 32277679 | 32477026 | mNeoVis.2.1.1.chr11\_hap1.g00455.t01 |
| 13667 | 11 | 32437530 | 32450649 | mNeoVis.2.1.1.chr11\_hap1.g00456.t01 |
| 13668 | 11 | 33425948 | 33464941 | mNeoVis.2.1.1.chr11\_hap1.g00457.t01 |
| 13669 | 11 | 33425948 | 33464941 | mNeoVis.2.1.1.chr11\_hap1.g00457.t01 |
| 13670 | 11 | 33425948 | 33464941 | mNeoVis.2.1.1.chr11\_hap1.g00457.t01 |
| 13671 | 11 | 33483923 | 33533879 | mNeoVis.2.1.1.chr11\_hap1.g00458.t01 |
| 13672 | 11 | 33483923 | 33533879 | mNeoVis.2.1.1.chr11\_hap1.g00458.t01 |
| 13673 | 11 | 33483923 | 33533879 | mNeoVis.2.1.1.chr11\_hap1.g00458.t01 |
| 13674 | 11 | 33560530 | 33603636 | mNeoVis.2.1.1.chr11\_hap1.g00459.t01 |
| 13675 | 11 | 33560530 | 33603636 | mNeoVis.2.1.1.chr11\_hap1.g00459.t01 |
| 13676 | 11 | 33560530 | 33603636 | mNeoVis.2.1.1.chr11\_hap1.g00459.t01 |
| 13677 | 11 | 33709141 | 34393495 | mNeoVis.2.1.1.chr11\_hap1.g00460.t01 |
| 13678 | 11 | 35537026 | 35567644 | mNeoVis.2.1.1.chr11\_hap1.g00464.t01 |
| 13679 | 11 | 35537026 | 35567644 | mNeoVis.2.1.1.chr11\_hap1.g00464.t01 |
| 13680 | 11 | 35720561 | 35805835 | mNeoVis.2.1.1.chr11\_hap1.g00465.t01 |
| 13681 | 11 | 35720561 | 35805835 | mNeoVis.2.1.1.chr11\_hap1.g00465.t01 |
| 13682 | 11 | 35763960 | 35764687 | mNeoVis.2.1.1.chr11\_hap1.g00466.t01 |
| 13683 | 11 | 35763960 | 35764687 | mNeoVis.2.1.1.chr11\_hap1.g00466.t01 |
| 13684 | 11 | 35888670 | 35897821 | mNeoVis.2.1.1.chr11\_hap1.g00467.t01 |
| 13685 | 11 | 35888670 | 35897821 | mNeoVis.2.1.1.chr11\_hap1.g00467.t01 |
| 13686 | 11 | 35897967 | 35898145 | mNeoVis.2.1.1.chr11\_hap1.g00468.t01 |
| 13687 | 11 | 35897967 | 35898145 | mNeoVis.2.1.1.chr11\_hap1.g00468.t01 |
| 13688 | 11 | 35904887 | 35918793 | mNeoVis.2.1.1.chr11\_hap1.g00469.t01 |
| 13689 | 11 | 35904887 | 35918793 | mNeoVis.2.1.1.chr11\_hap1.g00469.t01 |
| 13690 | 11 | 36875092 | 36875608 | mNeoVis.2.1.1.chr11\_hap1.g00472.t01 |
| 13691 | 11 | 37354617 | 37390162 | mNeoVis.2.1.1.chr11\_hap1.g00473.t01 |
| 13692 | 11 | 38314502 | 38458969 | mNeoVis.2.1.1.chr11\_hap1.g00474.t01 |
| 13693 | 11 | 38508899 | 38583309 | mNeoVis.2.1.1.chr11\_hap1.g00475.t01 |
| 13694 | 11 | 38648090 | 38924855 | mNeoVis.2.1.1.chr11\_hap1.g00476.t01 |
| 13695 | 11 | 38982385 | 39044928 | mNeoVis.2.1.1.chr11\_hap1.g00477.t01 |
| 13696 | 11 | 38982385 | 39044928 | mNeoVis.2.1.1.chr11\_hap1.g00477.t01 |
| 13697 | 11 | 39044956 | 39073370 | mNeoVis.2.1.1.chr11\_hap1.g00478.t01 |
| 13698 | 11 | 39044956 | 39073370 | mNeoVis.2.1.1.chr11\_hap1.g00478.t01 |
| 13699 | 11 | 39081105 | 39217193 | mNeoVis.2.1.1.chr11\_hap1.g00479.t01 |
| 13700 | 11 | 39081105 | 39217193 | mNeoVis.2.1.1.chr11\_hap1.g00479.t01 |
| 13701 | 11 | 39280165 | 39396304 | mNeoVis.2.1.1.chr11\_hap1.g00481.t01 |
| 13702 | 11 | 39531966 | 39544216 | mNeoVis.2.1.1.chr11\_hap1.g00482.t01 |
| 13703 | 11 | 41475455 | 41686062 | mNeoVis.2.1.1.chr11\_hap1.g00483.t01 |
| 13704 | 11 | 41475455 | 41686062 | mNeoVis.2.1.1.chr11\_hap1.g00483.t01 |
| 13705 | 11 | 41830926 | 41842416 | mNeoVis.2.1.1.chr11\_hap1.g00484.t01 |
| 13706 | 11 | 41830926 | 41842416 | mNeoVis.2.1.1.chr11\_hap1.g00484.t01 |
| 13707 | 11 | 42500053 | 42548504 | mNeoVis.2.1.1.chr11\_hap1.g00489.t01 |
| 13708 | 11 | 42500053 | 42548504 | mNeoVis.2.1.1.chr11\_hap1.g00489.t01 |
| 13709 | 11 | 42582094 | 42621485 | mNeoVis.2.1.1.chr11\_hap1.g00490.t01 |
| 13710 | 11 | 42582094 | 42621485 | mNeoVis.2.1.1.chr11\_hap1.g00490.t01 |
| 13711 | 11 | 42596173 | 42596384 | mNeoVis.2.1.1.chr11\_hap1.g00491.t01 |
| 13712 | 11 | 42596173 | 42596384 | mNeoVis.2.1.1.chr11\_hap1.g00491.t01 |
| 13713 | 11 | 42626501 | 42690894 | mNeoVis.2.1.1.chr11\_hap1.g00492.t01 |
| 13714 | 11 | 42626501 | 42690894 | mNeoVis.2.1.1.chr11\_hap1.g00492.t01 |
| 13715 | 11 | 42706269 | 42741581 | mNeoVis.2.1.1.chr11\_hap1.g00493.t01 |
| 13716 | 11 | 42706269 | 42741581 | mNeoVis.2.1.1.chr11\_hap1.g00493.t01 |
| 13717 | 11 | 42907068 | 42936658 | mNeoVis.2.1.1.chr11\_hap1.g00494.t01 |
| 13718 | 11 | 43229124 | 43292491 | mNeoVis.2.1.1.chr11\_hap1.g00500.t01 |
| 13719 | 11 | 43292740 | 43308406 | mNeoVis.2.1.1.chr11\_hap1.g00501.t01 |
| 13720 | 11 | 43311367 | 43326760 | mNeoVis.2.1.1.chr11\_hap1.g00502.t01 |
| 13721 | 11 | 43327002 | 43373770 | mNeoVis.2.1.1.chr11\_hap1.g00503.t01 |
| 13722 | 11 | 43381569 | 43382879 | mNeoVis.2.1.1.chr11\_hap1.g00504.t01 |
| 13723 | 11 | 43393969 | 43568999 | mNeoVis.2.1.1.chr11\_hap1.g00505.t01 |
| 13724 | 11 | 43615124 | 43615833 | mNeoVis.2.1.1.chr11\_hap1.g00506.t01 |
| 13725 | 11 | 43829854 | 43831123 | mNeoVis.2.1.1.chr11\_hap1.g00507.t01 |
| 13726 | 11 | 45121674 | 45123212 | mNeoVis.2.1.1.chr11\_hap1.g00514.t01 |
| 13727 | 11 | 45161762 | 45467369 | mNeoVis.2.1.1.chr11\_hap1.g00515.t01 |
| 13728 | 11 | 46432747 | 46560627 | mNeoVis.2.1.1.chr11\_hap1.g00518.t01 |
| 13729 | 11 | 46432747 | 46560627 | mNeoVis.2.1.1.chr11\_hap1.g00518.t01 |
| 13730 | 11 | 46607949 | 46699563 | mNeoVis.2.1.1.chr11\_hap1.g00519.t01 |
| 13731 | 11 | 46607949 | 46699563 | mNeoVis.2.1.1.chr11\_hap1.g00519.t01 |
| 13732 | 11 | 46705850 | 46707367 | mNeoVis.2.1.1.chr11\_hap1.g00520.t01 |
| 13733 | 11 | 46705850 | 46707367 | mNeoVis.2.1.1.chr11\_hap1.g00520.t01 |
| 13734 | 11 | 46809217 | 46811891 | mNeoVis.2.1.1.chr11\_hap1.g00521.t01 |
| 13735 | 11 | 46809217 | 46811891 | mNeoVis.2.1.1.chr11\_hap1.g00521.t01 |
| 13736 | 11 | 46995609 | 47006149 | mNeoVis.2.1.1.chr11\_hap1.g00522.t01 |
| 13737 | 11 | 46995609 | 47006149 | mNeoVis.2.1.1.chr11\_hap1.g00522.t01 |
| 13738 | 11 | 48263625 | 48266633 | mNeoVis.2.1.1.chr11\_hap1.g00524.t01 |
| 13739 | 11 | 48276577 | 48280430 | mNeoVis.2.1.1.chr11\_hap1.g00525.t01 |
| 13740 | 11 | 48305086 | 48323375 | mNeoVis.2.1.1.chr11\_hap1.g00526.t01 |
| 13741 | 11 | 49662494 | 49856686 | mNeoVis.2.1.1.chr11\_hap1.g00537.t01 |
| 13742 | 11 | 56375661 | 56404262 | mNeoVis.2.1.1.chr11\_hap1.g00546.t01 |
| 13743 | 11 | 56490983 | 56503297 | mNeoVis.2.1.1.chr11\_hap1.g00547.t01 |
| 13744 | 11 | 56558896 | 56728778 | mNeoVis.2.1.1.chr11\_hap1.g00548.t01 |
| 13745 | 11 | 56739624 | 56759752 | mNeoVis.2.1.1.chr11\_hap1.g00549.t01 |
| 13746 | 11 | 56763959 | 56855710 | mNeoVis.2.1.1.chr11\_hap1.g00550.t01 |
| 13747 | 11 | 56858628 | 56916781 | mNeoVis.2.1.1.chr11\_hap1.g00551.t01 |
| 13748 | 11 | 56936236 | 56979833 | mNeoVis.2.1.1.chr11\_hap1.g00552.t01 |
| 13749 | 11 | 59016495 | 59044153 | mNeoVis.2.1.1.chr11\_hap1.g00564.t01 |
| 13750 | 11 | 59073911 | 59119017 | mNeoVis.2.1.1.chr11\_hap1.g00565.t01 |
| 13751 | 11 | 59358072 | 59382130 | mNeoVis.2.1.1.chr11\_hap1.g00566.t01 |
| 13752 | 11 | 59838453 | 59840422 | mNeoVis.2.1.1.chr11\_hap1.g00567.t01 |
| 13753 | 11 | 59838453 | 59840422 | mNeoVis.2.1.1.chr11\_hap1.g00567.t01 |
| 13754 | 11 | 60790012 | 60792441 | mNeoVis.2.1.1.chr11\_hap1.g00569.t01 |
| 13755 | 11 | 60816253 | 60827073 | mNeoVis.2.1.1.chr11\_hap1.g00570.t01 |
| 13756 | 11 | 60867719 | 60961159 | mNeoVis.2.1.1.chr11\_hap1.g00571.t01 |
| 13757 | 11 | 60972561 | 60981823 | mNeoVis.2.1.1.chr11\_hap1.g00572.t01 |
| 13758 | 11 | 61012831 | 61015153 | mNeoVis.2.1.1.chr11\_hap1.g00573.t01 |
| 13759 | 11 | 61136209 | 61285665 | mNeoVis.2.1.1.chr11\_hap1.g00574.t01 |
| 13760 | 11 | 61552901 | 61621879 | mNeoVis.2.1.1.chr11\_hap1.g00575.t01 |
| 13761 | 11 | 61611318 | 61642194 | mNeoVis.2.1.1.chr11\_hap1.g00576.t01 |
| 13762 | 11 | 61649327 | 61723612 | mNeoVis.2.1.1.chr11\_hap1.g00577.t01 |
| 13763 | 11 | 61736842 | 61739025 | mNeoVis.2.1.1.chr11\_hap1.g00578.t01 |
| 13764 | 11 | 61873468 | 61941489 | mNeoVis.2.1.1.chr11\_hap1.g00579.t01 |
| 13765 | 11 | 61873468 | 61941489 | mNeoVis.2.1.1.chr11\_hap1.g00579.t01 |
| 13766 | 11 | 61873468 | 61941489 | mNeoVis.2.1.1.chr11\_hap1.g00579.t01 |
| 13767 | 11 | 62088590 | 62090089 | mNeoVis.2.1.1.chr11\_hap1.g00580.t01 |
| 13768 | 11 | 62088590 | 62090089 | mNeoVis.2.1.1.chr11\_hap1.g00580.t01 |
| 13769 | 11 | 62088590 | 62090089 | mNeoVis.2.1.1.chr11\_hap1.g00580.t01 |
| 13770 | 11 | 63892475 | 63950519 | mNeoVis.2.1.1.chr11\_hap1.g00581.t01 |
| 13771 | 11 | 63892475 | 63950519 | mNeoVis.2.1.1.chr11\_hap1.g00581.t01 |
| 13772 | 11 | 63973100 | 63989869 | mNeoVis.2.1.1.chr11\_hap1.g00582.t01 |
| 13773 | 11 | 63973100 | 63989869 | mNeoVis.2.1.1.chr11\_hap1.g00582.t01 |
| 13774 | 11 | 63991339 | 64094313 | mNeoVis.2.1.1.chr11\_hap1.g00583.t01 |
| 13775 | 11 | 63991339 | 64094313 | mNeoVis.2.1.1.chr11\_hap1.g00583.t01 |
| 13776 | 11 | 64170465 | 64234669 | mNeoVis.2.1.1.chr11\_hap1.g00584.t01 |
| 13777 | 11 | 64170465 | 64234669 | mNeoVis.2.1.1.chr11\_hap1.g00584.t01 |
| 13778 | 11 | 64267557 | 64291567 | mNeoVis.2.1.1.chr11\_hap1.g00585.t01 |
| 13779 | 11 | 64307962 | 64346496 | mNeoVis.2.1.1.chr11\_hap1.g00586.t01 |
| 13780 | 11 | 64350611 | 64369015 | mNeoVis.2.1.1.chr11\_hap1.g00587.t01 |
| 13781 | 11 | 64484267 | 64519858 | mNeoVis.2.1.1.chr11\_hap1.g00588.t01 |
| 13782 | 11 | 64484267 | 64519858 | mNeoVis.2.1.1.chr11\_hap1.g00588.t01 |
| 13783 | 11 | 64531971 | 64605232 | mNeoVis.2.1.1.chr11\_hap1.g00589.t01 |
| 13784 | 11 | 64531971 | 64605232 | mNeoVis.2.1.1.chr11\_hap1.g00589.t01 |
| 13785 | 11 | 64686329 | 64719606 | mNeoVis.2.1.1.chr11\_hap1.g00590.t01 |
| 13786 | 11 | 64686329 | 64719606 | mNeoVis.2.1.1.chr11\_hap1.g00590.t01 |
| 13787 | 11 | 64804139 | 64992275 | mNeoVis.2.1.1.chr11\_hap1.g00591.t01 |
| 13788 | 11 | 64804139 | 64992275 | mNeoVis.2.1.1.chr11\_hap1.g00591.t01 |
| 13789 | 11 | 65178291 | 65242580 | mNeoVis.2.1.1.chr11\_hap1.g00592.t01 |
| 13790 | 11 | 65178291 | 65242580 | mNeoVis.2.1.1.chr11\_hap1.g00592.t01 |
| 13791 | 11 | 65178291 | 65242580 | mNeoVis.2.1.1.chr11\_hap1.g00592.t01 |
| 13792 | 11 | 65258360 | 65310557 | mNeoVis.2.1.1.chr11\_hap1.g00593.t01 |
| 13793 | 11 | 65258360 | 65310557 | mNeoVis.2.1.1.chr11\_hap1.g00593.t01 |
| 13794 | 11 | 65258360 | 65310557 | mNeoVis.2.1.1.chr11\_hap1.g00593.t01 |
| 13795 | 11 | 65343081 | 65371715 | mNeoVis.2.1.1.chr11\_hap1.g00594.t01 |
| 13796 | 11 | 65343081 | 65371715 | mNeoVis.2.1.1.chr11\_hap1.g00594.t01 |
| 13797 | 11 | 65376300 | 65395724 | mNeoVis.2.1.1.chr11\_hap1.g00595.t01 |
| 13798 | 11 | 65376300 | 65395724 | mNeoVis.2.1.1.chr11\_hap1.g00595.t01 |
| 13799 | 11 | 65450329 | 65557064 | mNeoVis.2.1.1.chr11\_hap1.g00596.t01 |
| 13800 | 11 | 65450329 | 65557064 | mNeoVis.2.1.1.chr11\_hap1.g00596.t01 |
| 13801 | 11 | 65558022 | 65566516 | mNeoVis.2.1.1.chr11\_hap1.g00597.t01 |
| 13802 | 11 | 65558022 | 65566516 | mNeoVis.2.1.1.chr11\_hap1.g00597.t01 |
| 13803 | 11 | 65571921 | 65728505 | mNeoVis.2.1.1.chr11\_hap1.g00598.t01 |
| 13804 | 11 | 65571921 | 65728505 | mNeoVis.2.1.1.chr11\_hap1.g00598.t01 |
| 13805 | 11 | 65942306 | 65951493 | mNeoVis.2.1.1.chr11\_hap1.g00599.t01 |
| 13806 | 11 | 65990300 | 66020204 | mNeoVis.2.1.1.chr11\_hap1.g00600.t01 |
| 13807 | 11 | 66053738 | 66073654 | mNeoVis.2.1.1.chr11\_hap1.g00601.t01 |
| 13808 | 11 | 66104542 | 66154798 | mNeoVis.2.1.1.chr11\_hap1.g00602.t01 |
| 13809 | 11 | 66344125 | 66885870 | mNeoVis.2.1.1.chr11\_hap1.g00603.t01 |
| 13810 | 11 | 66344125 | 66885870 | mNeoVis.2.1.1.chr11\_hap1.g00603.t01 |
| 13811 | 11 | 66949676 | 67009955 | mNeoVis.2.1.1.chr11\_hap1.g00604.t01 |
| 13812 | 11 | 66949676 | 67009955 | mNeoVis.2.1.1.chr11\_hap1.g00604.t01 |
| 13813 | 11 | 66949676 | 67009955 | mNeoVis.2.1.1.chr11\_hap1.g00604.t01 |
| 13814 | 11 | 66949676 | 67009955 | mNeoVis.2.1.1.chr11\_hap1.g00604.t01 |
| 13815 | 11 | 67010093 | 67090727 | mNeoVis.2.1.1.chr11\_hap1.g00605.t01 |
| 13816 | 11 | 67010093 | 67090727 | mNeoVis.2.1.1.chr11\_hap1.g00605.t01 |
| 13817 | 11 | 67090901 | 67104703 | mNeoVis.2.1.1.chr11\_hap1.g00606.t01 |
| 13818 | 11 | 67090901 | 67104703 | mNeoVis.2.1.1.chr11\_hap1.g00606.t01 |
| 13819 | 11 | 67576756 | 67593527 | mNeoVis.2.1.1.chr11\_hap1.g00607.t01 |
| 13820 | 11 | 67576756 | 67593527 | mNeoVis.2.1.1.chr11\_hap1.g00607.t01 |
| 13821 | 11 | 67597852 | 67638281 | mNeoVis.2.1.1.chr11\_hap1.g00608.t01 |
| 13822 | 11 | 67597852 | 67638281 | mNeoVis.2.1.1.chr11\_hap1.g00608.t01 |
| 13823 | 11 | 67658906 | 67767326 | mNeoVis.2.1.1.chr11\_hap1.g00609.t01 |
| 13824 | 11 | 67658906 | 67767326 | mNeoVis.2.1.1.chr11\_hap1.g00609.t01 |
| 13825 | 11 | 67793755 | 67803888 | mNeoVis.2.1.1.chr11\_hap1.g00610.t01 |
| 13826 | 11 | 67793755 | 67803888 | mNeoVis.2.1.1.chr11\_hap1.g00610.t01 |
| 13827 | 11 | 67918539 | 67981027 | mNeoVis.2.1.1.chr11\_hap1.g00611.t01 |
| 13828 | 11 | 67918539 | 67981027 | mNeoVis.2.1.1.chr11\_hap1.g00611.t01 |
| 13829 | 11 | 68047189 | 68057627 | mNeoVis.2.1.1.chr11\_hap1.g00612.t01 |
| 13830 | 11 | 68047189 | 68057627 | mNeoVis.2.1.1.chr11\_hap1.g00612.t01 |
| 13831 | 11 | 68057889 | 68093409 | mNeoVis.2.1.1.chr11\_hap1.g00613.t01 |
| 13832 | 11 | 68057889 | 68093409 | mNeoVis.2.1.1.chr11\_hap1.g00613.t01 |
| 13833 | 11 | 68093189 | 68128142 | mNeoVis.2.1.1.chr11\_hap1.g00614.t01 |
| 13834 | 11 | 68093189 | 68128142 | mNeoVis.2.1.1.chr11\_hap1.g00614.t01 |
| 13835 | 11 | 68142551 | 68150525 | mNeoVis.2.1.1.chr11\_hap1.g00615.t01 |
| 13836 | 11 | 68142551 | 68150525 | mNeoVis.2.1.1.chr11\_hap1.g00615.t01 |
| 13837 | 11 | 68176320 | 68178304 | mNeoVis.2.1.1.chr11\_hap1.g00616.t01 |
| 13838 | 11 | 68176320 | 68178304 | mNeoVis.2.1.1.chr11\_hap1.g00616.t01 |
| 13839 | 11 | 68245004 | 68328114 | mNeoVis.2.1.1.chr11\_hap1.g00617.t01 |
| 13840 | 11 | 68245004 | 68328114 | mNeoVis.2.1.1.chr11\_hap1.g00617.t01 |
| 13841 | 11 | 68411783 | 68892822 | mNeoVis.2.1.1.chr11\_hap1.g00618.t01 |
| 13842 | 11 | 68932689 | 69131044 | mNeoVis.2.1.1.chr11\_hap1.g00619.t01 |
| 13843 | 11 | 68932689 | 69131044 | mNeoVis.2.1.1.chr11\_hap1.g00619.t01 |
| 13844 | 11 | 68932689 | 69131044 | mNeoVis.2.1.1.chr11\_hap1.g00619.t01 |
| 13845 | 11 | 69455653 | 69459483 | mNeoVis.2.1.1.chr11\_hap1.g00620.t01 |
| 13846 | 11 | 69509608 | 69513486 | mNeoVis.2.1.1.chr11\_hap1.g00621.t01 |
| 13847 | 11 | 69666202 | 70038223 | mNeoVis.2.1.1.chr11\_hap1.g00622.t01 |
| 13848 | 11 | 70038389 | 70084127 | mNeoVis.2.1.1.chr11\_hap1.g00623.t01 |
| 13849 | 11 | 70083969 | 70115257 | mNeoVis.2.1.1.chr11\_hap1.g00624.t01 |
| 13850 | 11 | 70149206 | 70229430 | mNeoVis.2.1.1.chr11\_hap1.g00625.t01 |
| 13851 | 11 | 70149206 | 70229430 | mNeoVis.2.1.1.chr11\_hap1.g00625.t01 |
| 13852 | 11 | 70204254 | 70205544 | mNeoVis.2.1.1.chr11\_hap1.g00626.t01 |
| 13853 | 11 | 70204254 | 70205544 | mNeoVis.2.1.1.chr11\_hap1.g00626.t01 |
| 13854 | 11 | 70383734 | 70487002 | mNeoVis.2.1.1.chr11\_hap1.g00627.t01 |
| 13855 | 11 | 70383734 | 70487002 | mNeoVis.2.1.1.chr11\_hap1.g00627.t01 |
| 13856 | 11 | 70495108 | 70513330 | mNeoVis.2.1.1.chr11\_hap1.g00628.t01 |
| 13857 | 11 | 70513573 | 70541441 | mNeoVis.2.1.1.chr11\_hap1.g00629.t01 |
| 13858 | 11 | 70521421 | 70588659 | mNeoVis.2.1.1.chr11\_hap1.g00630.t01 |
| 13859 | 11 | 70593810 | 70713426 | mNeoVis.2.1.1.chr11\_hap1.g00631.t01 |
| 13860 | 11 | 70679227 | 70680160 | mNeoVis.2.1.1.chr11\_hap1.g00632.t01 |
| 13861 | 11 | 70685929 | 70688246 | mNeoVis.2.1.1.chr11\_hap1.g00633.t01 |
| 13862 | 11 | 70736508 | 71063658 | mNeoVis.2.1.1.chr11\_hap1.g00634.t01 |
| 13863 | 11 | 70736508 | 71063658 | mNeoVis.2.1.1.chr11\_hap1.g00634.t01 |
| 13864 | 11 | 71051413 | 71051964 | mNeoVis.2.1.1.chr11\_hap1.g00635.t01 |
| 13865 | 11 | 71051413 | 71051964 | mNeoVis.2.1.1.chr11\_hap1.g00635.t01 |
| 13866 | 11 | 71067216 | 71071521 | mNeoVis.2.1.1.chr11\_hap1.g00636.t01 |
| 13867 | 11 | 71067216 | 71071521 | mNeoVis.2.1.1.chr11\_hap1.g00636.t01 |
| 13868 | 11 | 71402711 | 71406805 | mNeoVis.2.1.1.chr11\_hap1.g00637.t01 |
| 13869 | 11 | 71419341 | 71431396 | mNeoVis.2.1.1.chr11\_hap1.g00638.t01 |
| 13870 | 11 | 71433099 | 71435290 | mNeoVis.2.1.1.chr11\_hap1.g00639.t01 |
| 13871 | 11 | 71439879 | 71484254 | mNeoVis.2.1.1.chr11\_hap1.g00640.t01 |
| 13872 | 11 | 71494399 | 71550282 | mNeoVis.2.1.1.chr11\_hap1.g00641.t01 |
| 13873 | 11 | 71551257 | 71578393 | mNeoVis.2.1.1.chr11\_hap1.g00642.t01 |
| 13874 | 11 | 71596847 | 71635308 | mNeoVis.2.1.1.chr11\_hap1.g00643.t01 |
| 13875 | 11 | 73589763 | 73664190 | mNeoVis.2.1.1.chr11\_hap1.g00669.t01 |
| 13876 | 11 | 73686036 | 73737661 | mNeoVis.2.1.1.chr11\_hap1.g00670.t01 |
| 13877 | 11 | 73879693 | 73903827 | mNeoVis.2.1.1.chr11\_hap1.g00671.t01 |
| 13878 | 11 | 73903794 | 73994221 | mNeoVis.2.1.1.chr11\_hap1.g00672.t01 |
| 13879 | 11 | 73994347 | 74005097 | mNeoVis.2.1.1.chr11\_hap1.g00673.t01 |
| 13880 | 11 | 74310829 | 74378340 | mNeoVis.2.1.1.chr11\_hap1.g00674.t01 |
| 13881 | 11 | 74443687 | 74513509 | mNeoVis.2.1.1.chr11\_hap1.g00675.t01 |
| 13882 | 11 | 74536739 | 74564747 | mNeoVis.2.1.1.chr11\_hap1.g00676.t01 |
| 13883 | 11 | 74536739 | 74564747 | mNeoVis.2.1.1.chr11\_hap1.g00676.t01 |
| 13884 | 11 | 74580885 | 74637490 | mNeoVis.2.1.1.chr11\_hap1.g00677.t01 |
| 13885 | 11 | 74580885 | 74637490 | mNeoVis.2.1.1.chr11\_hap1.g00677.t01 |
| 13886 | 11 | 74641324 | 74697716 | mNeoVis.2.1.1.chr11\_hap1.g00678.t01 |
| 13887 | 11 | 74641324 | 74697716 | mNeoVis.2.1.1.chr11\_hap1.g00678.t01 |
| 13888 | 11 | 74852732 | 75268031 | mNeoVis.2.1.1.chr11\_hap1.g00679.t01 |
| 13889 | 11 | 74862221 | 75027719 | mNeoVis.2.1.1.chr11\_hap1.g00680.t01 |
| 13890 | 11 | 75697256 | 75907718 | mNeoVis.2.1.1.chr11\_hap1.g00681.t01 |
| 13891 | 11 | 75697256 | 75907718 | mNeoVis.2.1.1.chr11\_hap1.g00681.t01 |
| 13892 | 11 | 76247090 | 76316138 | mNeoVis.2.1.1.chr11\_hap1.g00682.t01 |
| 13893 | 11 | 76247090 | 76316138 | mNeoVis.2.1.1.chr11\_hap1.g00682.t01 |
| 13894 | 11 | 76247090 | 76316138 | mNeoVis.2.1.1.chr11\_hap1.g00682.t01 |
| 13895 | 11 | 76743187 | 76850217 | mNeoVis.2.1.1.chr11\_hap1.g00683.t01 |
| 13896 | 11 | 78687063 | 78930671 | mNeoVis.2.1.1.chr11\_hap1.g00684.t01 |
| 13897 | 11 | 79076259 | 79186438 | mNeoVis.2.1.1.chr11\_hap1.g00685.t01 |
| 13898 | 11 | 79161316 | 79171343 | mNeoVis.2.1.1.chr11\_hap1.g00686.t01 |
| 13899 | 11 | 79186499 | 79214550 | mNeoVis.2.1.1.chr11\_hap1.g00687.t01 |
| 13900 | 11 | 79226002 | 79234294 | mNeoVis.2.1.1.chr11\_hap1.g00688.t01 |
| 13901 | 11 | 79662779 | 79700687 | mNeoVis.2.1.1.chr11\_hap1.g00692.t01 |
| 13902 | 11 | 79803379 | 80000937 | mNeoVis.2.1.1.chr11\_hap1.g00693.t01 |
| 13903 | 11 | 80008082 | 80321622 | mNeoVis.2.1.1.chr11\_hap1.g00694.t01 |
| 13904 | 11 | 80320813 | 80394193 | mNeoVis.2.1.1.chr11\_hap1.g00695.t01 |
| 13905 | 11 | 80526747 | 80567991 | mNeoVis.2.1.1.chr11\_hap1.g00696.t01 |
| 13906 | 11 | 80567306 | 80569744 | mNeoVis.2.1.1.chr11\_hap1.g00697.t01 |
| 13907 | 11 | 80570648 | 80593902 | mNeoVis.2.1.1.chr11\_hap1.g00698.t01 |
| 13908 | 11 | 80818084 | 80849300 | mNeoVis.2.1.1.chr11\_hap1.g00701.t01 |
| 13909 | 11 | 80851555 | 80921348 | mNeoVis.2.1.1.chr11\_hap1.g00702.t01 |
| 13910 | 11 | 80933218 | 80979638 | mNeoVis.2.1.1.chr11\_hap1.g00703.t01 |
| 13911 | 11 | 81115948 | 81174005 | mNeoVis.2.1.1.chr11\_hap1.g00704.t01 |
| 13912 | 11 | 81376696 | 81405901 | mNeoVis.2.1.1.chr11\_hap1.g00705.t01 |
| 13913 | 11 | 81376696 | 81405901 | mNeoVis.2.1.1.chr11\_hap1.g00705.t01 |
| 13914 | 11 | 81419793 | 81620593 | mNeoVis.2.1.1.chr11\_hap1.g00706.t01 |
| 13915 | 11 | 81419793 | 81620593 | mNeoVis.2.1.1.chr11\_hap1.g00706.t01 |
| 13916 | 11 | 83019678 | 83135414 | mNeoVis.2.1.1.chr11\_hap1.g00707.t01 |
| 13917 | 11 | 84642210 | 84843873 | mNeoVis.2.1.1.chr11\_hap1.g00709.t01 |
| 13918 | 11 | 85050468 | 85091452 | mNeoVis.2.1.1.chr11\_hap1.g00710.t01 |
| 13919 | 11 | 85250498 | 85347474 | mNeoVis.2.1.1.chr11\_hap1.g00711.t01 |
| 13920 | 11 | 85521417 | 85551333 | mNeoVis.2.1.1.chr11\_hap1.g00712.t01 |
| 13921 | 11 | 85631985 | 85662093 | mNeoVis.2.1.1.chr11\_hap1.g00713.t01 |
| 13922 | 11 | 85662231 | 85739013 | mNeoVis.2.1.1.chr11\_hap1.g00714.t01 |
| 13923 | 11 | 85739586 | 85765214 | mNeoVis.2.1.1.chr11\_hap1.g00715.t01 |
| 13924 | 11 | 85770745 | 85775307 | mNeoVis.2.1.1.chr11\_hap1.g00716.t01 |
| 13925 | 11 | 85775359 | 85830914 | mNeoVis.2.1.1.chr11\_hap1.g00717.t01 |
| 13926 | 11 | 85861748 | 85864369 | mNeoVis.2.1.1.chr11\_hap1.g00718.t01 |
| 13927 | 11 | 87404233 | 87404550 | mNeoVis.2.1.1.chr11\_hap1.g00730.t01 |
| 13928 | 11 | 87562972 | 87569860 | mNeoVis.2.1.1.chr11\_hap1.g00731.t01 |
| 13929 | 11 | 88211488 | 88212465 | mNeoVis.2.1.1.chr11\_hap1.g00734.t01 |
| 13930 | 11 | 88404246 | 88472497 | mNeoVis.2.1.1.chr11\_hap1.g00735.t01 |
| 13931 | 11 | 88404246 | 88472497 | mNeoVis.2.1.1.chr11\_hap1.g00735.t01 |
| 13932 | 11 | 88472806 | 88571947 | mNeoVis.2.1.1.chr11\_hap1.g00736.t01 |
| 13933 | 11 | 88472806 | 88571947 | mNeoVis.2.1.1.chr11\_hap1.g00736.t01 |
| 13934 | 11 | 88627879 | 88685436 | mNeoVis.2.1.1.chr11\_hap1.g00737.t01 |
| 13935 | 11 | 88627879 | 88685436 | mNeoVis.2.1.1.chr11\_hap1.g00737.t01 |
| 13936 | 11 | 88956657 | 88964335 | mNeoVis.2.1.1.chr11\_hap1.g00738.t01 |
| 13937 | 11 | 89449948 | 89476656 | mNeoVis.2.1.1.chr11\_hap1.g00742.t01 |
| 13938 | 11 | 89476793 | 89507671 | mNeoVis.2.1.1.chr11\_hap1.g00743.t01 |
| 13939 | 11 | 89603897 | 89606630 | mNeoVis.2.1.1.chr11\_hap1.g00744.t01 |
| 13940 | 11 | 89616661 | 89655147 | mNeoVis.2.1.1.chr11\_hap1.g00745.t01 |
| 13941 | 11 | 89689667 | 89806125 | mNeoVis.2.1.1.chr11\_hap1.g00746.t01 |
| 13942 | 11 | 89861154 | 89874885 | mNeoVis.2.1.1.chr11\_hap1.g00747.t01 |
| 13943 | 11 | 91676696 | 91691179 | mNeoVis.2.1.1.chr11\_hap1.g00750.t01 |
| 13944 | 11 | 91676696 | 91691179 | mNeoVis.2.1.1.chr11\_hap1.g00750.t01 |
| 13945 | 11 | 92234375 | 92252264 | mNeoVis.2.1.1.chr11\_hap1.g00751.t01 |
| 13946 | 11 | 92293135 | 92315904 | mNeoVis.2.1.1.chr11\_hap1.g00752.t01 |
| 13947 | 11 | 92316216 | 92325527 | mNeoVis.2.1.1.chr11\_hap1.g00753.t01 |
| 13948 | 11 | 92439215 | 92455097 | mNeoVis.2.1.1.chr11\_hap1.g00755.t01 |
| 13949 | 11 | 92499159 | 92525415 | mNeoVis.2.1.1.chr11\_hap1.g00756.t01 |
| 13950 | 11 | 92567075 | 92570196 | mNeoVis.2.1.1.chr11\_hap1.g00757.t01 |
| 13951 | 11 | 92570321 | 92648999 | mNeoVis.2.1.1.chr11\_hap1.g00758.t01 |
| 13952 | 11 | 92676996 | 92690991 | mNeoVis.2.1.1.chr11\_hap1.g00759.t01 |
| 13953 | 11 | 92777166 | 92819319 | mNeoVis.2.1.1.chr11\_hap1.g00760.t01 |
| 13954 | 11 | 92852709 | 92858253 | mNeoVis.2.1.1.chr11\_hap1.g00761.t01 |
| 13955 | 11 | 92905093 | 92950477 | mNeoVis.2.1.1.chr11\_hap1.g00762.t01 |
| 13956 | 11 | 92931789 | 92937811 | mNeoVis.2.1.1.chr11\_hap1.g00763.t01 |
| 13957 | 11 | 92970544 | 93192420 | mNeoVis.2.1.1.chr11\_hap1.g00764.t01 |
| 13958 | 11 | 93245014 | 93304034 | mNeoVis.2.1.1.chr11\_hap1.g00765.t01 |
| 13959 | 11 | 93245014 | 93304034 | mNeoVis.2.1.1.chr11\_hap1.g00765.t01 |
| 13960 | 11 | 93314101 | 93337562 | mNeoVis.2.1.1.chr11\_hap1.g00766.t01 |
| 13961 | 11 | 93314101 | 93337562 | mNeoVis.2.1.1.chr11\_hap1.g00766.t01 |
| 13962 | 11 | 93351402 | 93525461 | mNeoVis.2.1.1.chr11\_hap1.g00767.t01 |
| 13963 | 11 | 93615722 | 93692493 | mNeoVis.2.1.1.chr11\_hap1.g00768.t01 |
| 13964 | 11 | 93695159 | 93734192 | mNeoVis.2.1.1.chr11\_hap1.g00769.t01 |
| 13965 | 11 | 93741136 | 93752994 | mNeoVis.2.1.1.chr11\_hap1.g00770.t01 |
| 13966 | 11 | 93759726 | 93787228 | mNeoVis.2.1.1.chr11\_hap1.g00771.t01 |
| 13967 | 11 | 93805942 | 93807565 | mNeoVis.2.1.1.chr11\_hap1.g00772.t01 |
| 13968 | 11 | 93982469 | 94017307 | mNeoVis.2.1.1.chr11\_hap1.g00774.t01 |
| 13969 | 11 | 94028816 | 94063131 | mNeoVis.2.1.1.chr11\_hap1.g00775.t01 |
| 13970 | 11 | 94071073 | 94091014 | mNeoVis.2.1.1.chr11\_hap1.g00776.t01 |
| 13971 | 11 | 94092424 | 94103317 | mNeoVis.2.1.1.chr11\_hap1.g00777.t01 |
| 13972 | 11 | 94103935 | 94108797 | mNeoVis.2.1.1.chr11\_hap1.g00778.t01 |
| 13973 | 11 | 94206493 | 94313410 | mNeoVis.2.1.1.chr11\_hap1.g00779.t01 |
| 13974 | 11 | 95493300 | 95512321 | mNeoVis.2.1.1.chr11\_hap1.g00794.t01 |
| 13975 | 11 | 95540422 | 96162493 | mNeoVis.2.1.1.chr11\_hap1.g00795.t01 |
| 13976 | 11 | 95540422 | 96162493 | mNeoVis.2.1.1.chr11\_hap1.g00795.t01 |
| 13977 | 11 | 96195539 | 96202028 | mNeoVis.2.1.1.chr11\_hap1.g00796.t01 |
| 13978 | 11 | 96195539 | 96202028 | mNeoVis.2.1.1.chr11\_hap1.g00796.t01 |
| 13979 | 11 | 96230239 | 96249464 | mNeoVis.2.1.1.chr11\_hap1.g00797.t01 |
| 13980 | 11 | 96257977 | 96368534 | mNeoVis.2.1.1.chr11\_hap1.g00798.t01 |
| 13981 | 11 | 96370346 | 96406597 | mNeoVis.2.1.1.chr11\_hap1.g00799.t01 |
| 13982 | 11 | 96858852 | 96860492 | mNeoVis.2.1.1.chr11\_hap1.g00807.t01 |
| 13983 | 11 | 96867155 | 96873870 | mNeoVis.2.1.1.chr11\_hap1.g00808.t01 |
| 13984 | 11 | 96883221 | 96908790 | mNeoVis.2.1.1.chr11\_hap1.g00809.t01 |
| 13985 | 11 | 96904960 | 96913238 | mNeoVis.2.1.1.chr11\_hap1.g00810.t01 |
| 13986 | 11 | 96926084 | 96952777 | mNeoVis.2.1.1.chr11\_hap1.g00811.t01 |
| 13987 | 11 | 96981908 | 97003628 | mNeoVis.2.1.1.chr11\_hap1.g00812.t01 |
| 13988 | 11 | 97007519 | 97036696 | mNeoVis.2.1.1.chr11\_hap1.g00813.t01 |
| 13989 | 11 | 97061108 | 97154327 | mNeoVis.2.1.1.chr11\_hap1.g00814.t01 |
| 13990 | 11 | 97182079 | 97192226 | mNeoVis.2.1.1.chr11\_hap1.g00815.t01 |
| 13991 | 11 | 97199486 | 97217558 | mNeoVis.2.1.1.chr11\_hap1.g00816.t01 |
| 13992 | 11 | 97217706 | 97351703 | mNeoVis.2.1.1.chr11\_hap1.g00817.t01 |
| 13993 | 11 | 97231634 | 97243592 | mNeoVis.2.1.1.chr11\_hap1.g00818.t01 |
| 13994 | 11 | 97366892 | 97385584 | mNeoVis.2.1.1.chr11\_hap1.g00819.t01 |
| 13995 | 11 | 97390032 | 97419699 | mNeoVis.2.1.1.chr11\_hap1.g00820.t01 |
| 13996 | 11 | 97403359 | 97409811 | mNeoVis.2.1.1.chr11\_hap1.g00821.t01 |
| 13997 | 11 | 97409802 | 97415895 | mNeoVis.2.1.1.chr11\_hap1.g00822.t01 |
| 13998 | 11 | 97420032 | 97435215 | mNeoVis.2.1.1.chr11\_hap1.g00823.t01 |
| 13999 | 11 | 97443888 | 97447887 | mNeoVis.2.1.1.chr11\_hap1.g00824.t01 |
| 14000 | 11 | 97448028 | 97450826 | mNeoVis.2.1.1.chr11\_hap1.g00825.t01 |
| 14001 | 11 | 97450963 | 97455309 | mNeoVis.2.1.1.chr11\_hap1.g00826.t01 |
| 14002 | 11 | 97455835 | 97458504 | mNeoVis.2.1.1.chr11\_hap1.g00827.t01 |
| 14003 | 11 | 97459514 | 97466516 | mNeoVis.2.1.1.chr11\_hap1.g00828.t01 |
| 14004 | 11 | 97466541 | 97473540 | mNeoVis.2.1.1.chr11\_hap1.g00829.t01 |
| 14005 | 11 | 97477430 | 97485441 | mNeoVis.2.1.1.chr11\_hap1.g00830.t01 |
| 14006 | 11 | 97485551 | 97489364 | mNeoVis.2.1.1.chr11\_hap1.g00831.t01 |
| 14007 | 11 | 97493587 | 97495277 | mNeoVis.2.1.1.chr11\_hap1.g00832.t01 |
| 14008 | 11 | 97497245 | 97503310 | mNeoVis.2.1.1.chr11\_hap1.g00833.t01 |
| 14009 | 11 | 97504205 | 97507288 | mNeoVis.2.1.1.chr11\_hap1.g00834.t01 |
| 14010 | 11 | 97507467 | 97510693 | mNeoVis.2.1.1.chr11\_hap1.g00835.t01 |
| 14011 | 11 | 97510641 | 97528291 | mNeoVis.2.1.1.chr11\_hap1.g00836.t01 |
| 14012 | 11 | 97528735 | 97532032 | mNeoVis.2.1.1.chr11\_hap1.g00837.t01 |
| 14013 | 11 | 97532066 | 97605015 | mNeoVis.2.1.1.chr11\_hap1.g00838.t01 |
| 14014 | 11 | 97606453 | 97633900 | mNeoVis.2.1.1.chr11\_hap1.g00839.t01 |
| 14015 | 11 | 97758244 | 97818198 | mNeoVis.2.1.1.chr11\_hap1.g00840.t01 |
| 14016 | 11 | 97849934 | 97869851 | mNeoVis.2.1.1.chr11\_hap1.g00841.t01 |
| 14017 | 11 | 97924993 | 98075743 | mNeoVis.2.1.1.chr11\_hap1.g00842.t01 |
| 14018 | 11 | 98279902 | 98351597 | mNeoVis.2.1.1.chr11\_hap1.g00843.t01 |
| 14019 | 11 | 98367438 | 98371065 | mNeoVis.2.1.1.chr11\_hap1.g00844.t01 |
| 14020 | 11 | 98389439 | 98401130 | mNeoVis.2.1.1.chr11\_hap1.g00845.t01 |
| 14021 | 11 | 98430729 | 98476501 | mNeoVis.2.1.1.chr11\_hap1.g00846.t01 |
| 14022 | 11 | 98657313 | 98786348 | mNeoVis.2.1.1.chr11\_hap1.g00847.t01 |
| 14023 | 11 | 99329419 | 100131154 | mNeoVis.2.1.1.chr11\_hap1.g00848.t01 |
| 14024 | 11 | 101077257 | 101336188 | mNeoVis.2.1.1.chr11\_hap1.g00849.t01 |
| 14025 | 11 | 101077257 | 101336188 | mNeoVis.2.1.1.chr11\_hap1.g00849.t01 |
| 14026 | 11 | 101380339 | 101382570 | mNeoVis.2.1.1.chr11\_hap1.g00850.t01 |
| 14027 | 11 | 101380339 | 101382570 | mNeoVis.2.1.1.chr11\_hap1.g00850.t01 |
| 14028 | 11 | 101808311 | 102814040 | mNeoVis.2.1.1.chr11\_hap1.g00851.t01 |
| 14029 | 11 | 102424485 | 102427160 | mNeoVis.2.1.1.chr11\_hap1.g00852.t01 |
| 14030 | 11 | 102424485 | 102427160 | mNeoVis.2.1.1.chr11\_hap1.g00852.t01 |
| 14031 | 11 | 103373224 | 103777095 | mNeoVis.2.1.1.chr11\_hap1.g00853.t01 |
| 14032 | 11 | 103373224 | 103777095 | mNeoVis.2.1.1.chr11\_hap1.g00853.t01 |
| 14033 | 11 | 103777690 | 104058966 | mNeoVis.2.1.1.chr11\_hap1.g00854.t01 |
| 14034 | 11 | 105956944 | 105989368 | mNeoVis.2.1.1.chr11\_hap1.g00855.t01 |
| 14035 | 11 | 105956944 | 105989368 | mNeoVis.2.1.1.chr11\_hap1.g00855.t01 |
| 14036 | 11 | 105956944 | 105989368 | mNeoVis.2.1.1.chr11\_hap1.g00855.t01 |
| 14037 | 11 | 105956944 | 105989368 | mNeoVis.2.1.1.chr11\_hap1.g00855.t01 |
| 14038 | 11 | 106022529 | 106242902 | mNeoVis.2.1.1.chr11\_hap1.g00856.t01 |
| 14039 | 11 | 106022529 | 106242902 | mNeoVis.2.1.1.chr11\_hap1.g00856.t01 |
| 14040 | 11 | 106022529 | 106242902 | mNeoVis.2.1.1.chr11\_hap1.g00856.t01 |
| 14041 | 11 | 106169123 | 106298438 | mNeoVis.2.1.1.chr11\_hap1.g00857.t01 |
| 14042 | 11 | 106169123 | 106298438 | mNeoVis.2.1.1.chr11\_hap1.g00857.t01 |
| 14043 | 11 | 106169123 | 106298438 | mNeoVis.2.1.1.chr11\_hap1.g00857.t01 |
| 14044 | 11 | 106326123 | 106327213 | mNeoVis.2.1.1.chr11\_hap1.g00858.t01 |
| 14045 | 11 | 106326123 | 106327213 | mNeoVis.2.1.1.chr11\_hap1.g00858.t01 |
| 14046 | 11 | 106326123 | 106327213 | mNeoVis.2.1.1.chr11\_hap1.g00858.t01 |
| 14047 | 11 | 106379352 | 106454468 | mNeoVis.2.1.1.chr11\_hap1.g00859.t01 |
| 14048 | 11 | 106379352 | 106454468 | mNeoVis.2.1.1.chr11\_hap1.g00859.t01 |
| 14049 | 11 | 106379352 | 106454468 | mNeoVis.2.1.1.chr11\_hap1.g00859.t01 |
| 14050 | 11 | 106512431 | 106666819 | mNeoVis.2.1.1.chr11\_hap1.g00860.t01 |
| 14051 | 11 | 106512431 | 106666819 | mNeoVis.2.1.1.chr11\_hap1.g00860.t01 |
| 14052 | 11 | 106512431 | 106666819 | mNeoVis.2.1.1.chr11\_hap1.g00860.t01 |
| 14053 | 11 | 106673691 | 106681601 | mNeoVis.2.1.1.chr11\_hap1.g00861.t01 |
| 14054 | 11 | 106673691 | 106681601 | mNeoVis.2.1.1.chr11\_hap1.g00861.t01 |
| 14055 | 11 | 106673691 | 106681601 | mNeoVis.2.1.1.chr11\_hap1.g00861.t01 |
| 14056 | 11 | 106681593 | 106694333 | mNeoVis.2.1.1.chr11\_hap1.g00862.t01 |
| 14057 | 11 | 106681593 | 106694333 | mNeoVis.2.1.1.chr11\_hap1.g00862.t01 |
| 14058 | 11 | 106681593 | 106694333 | mNeoVis.2.1.1.chr11\_hap1.g00862.t01 |
| 14059 | 11 | 106695440 | 106721400 | mNeoVis.2.1.1.chr11\_hap1.g00863.t01 |
| 14060 | 11 | 106695440 | 106721400 | mNeoVis.2.1.1.chr11\_hap1.g00863.t01 |
| 14061 | 11 | 106695440 | 106721400 | mNeoVis.2.1.1.chr11\_hap1.g00863.t01 |
| 14062 | 11 | 106723455 | 106735313 | mNeoVis.2.1.1.chr11\_hap1.g00864.t01 |
| 14063 | 11 | 106723455 | 106735313 | mNeoVis.2.1.1.chr11\_hap1.g00864.t01 |
| 14064 | 11 | 106723455 | 106735313 | mNeoVis.2.1.1.chr11\_hap1.g00864.t01 |
| 14065 | 11 | 106753117 | 106760341 | mNeoVis.2.1.1.chr11\_hap1.g00865.t01 |
| 14066 | 11 | 106753117 | 106760341 | mNeoVis.2.1.1.chr11\_hap1.g00865.t01 |
| 14067 | 11 | 106753117 | 106760341 | mNeoVis.2.1.1.chr11\_hap1.g00865.t01 |
| 14068 | 11 | 106826861 | 106833960 | mNeoVis.2.1.1.chr11\_hap1.g00866.t01 |
| 14069 | 11 | 106826861 | 106833960 | mNeoVis.2.1.1.chr11\_hap1.g00866.t01 |
| 14070 | 11 | 106826861 | 106833960 | mNeoVis.2.1.1.chr11\_hap1.g00866.t01 |
| 14071 | 11 | 106837357 | 106847270 | mNeoVis.2.1.1.chr11\_hap1.g00867.t01 |
| 14072 | 11 | 106837357 | 106847270 | mNeoVis.2.1.1.chr11\_hap1.g00867.t01 |
| 14073 | 11 | 106837357 | 106847270 | mNeoVis.2.1.1.chr11\_hap1.g00867.t01 |
| 14074 | 11 | 106856856 | 106860240 | mNeoVis.2.1.1.chr11\_hap1.g00868.t01 |
| 14075 | 11 | 106856856 | 106860240 | mNeoVis.2.1.1.chr11\_hap1.g00868.t01 |
| 14076 | 11 | 106856856 | 106860240 | mNeoVis.2.1.1.chr11\_hap1.g00868.t01 |
| 14077 | 11 | 106861923 | 106869763 | mNeoVis.2.1.1.chr11\_hap1.g00869.t01 |
| 14078 | 11 | 106861923 | 106869763 | mNeoVis.2.1.1.chr11\_hap1.g00869.t01 |
| 14079 | 11 | 106861923 | 106869763 | mNeoVis.2.1.1.chr11\_hap1.g00869.t01 |
| 14080 | 11 | 106872143 | 106874023 | mNeoVis.2.1.1.chr11\_hap1.g00870.t01 |
| 14081 | 11 | 106872143 | 106874023 | mNeoVis.2.1.1.chr11\_hap1.g00870.t01 |
| 14082 | 11 | 106872143 | 106874023 | mNeoVis.2.1.1.chr11\_hap1.g00870.t01 |
| 14083 | 11 | 106874688 | 106880260 | mNeoVis.2.1.1.chr11\_hap1.g00871.t01 |
| 14084 | 11 | 106874688 | 106880260 | mNeoVis.2.1.1.chr11\_hap1.g00871.t01 |
| 14085 | 11 | 106874688 | 106880260 | mNeoVis.2.1.1.chr11\_hap1.g00871.t01 |
| 14086 | 11 | 106881009 | 106886461 | mNeoVis.2.1.1.chr11\_hap1.g00872.t01 |
| 14087 | 11 | 106881009 | 106886461 | mNeoVis.2.1.1.chr11\_hap1.g00872.t01 |
| 14088 | 11 | 106881009 | 106886461 | mNeoVis.2.1.1.chr11\_hap1.g00872.t01 |
| 14089 | 11 | 106888324 | 106915230 | mNeoVis.2.1.1.chr11\_hap1.g00873.t01 |
| 14090 | 11 | 106888324 | 106915230 | mNeoVis.2.1.1.chr11\_hap1.g00873.t01 |
| 14091 | 11 | 106888324 | 106915230 | mNeoVis.2.1.1.chr11\_hap1.g00873.t01 |
| 14092 | 11 | 106923376 | 106931320 | mNeoVis.2.1.1.chr11\_hap1.g00874.t01 |
| 14093 | 11 | 106923376 | 106931320 | mNeoVis.2.1.1.chr11\_hap1.g00874.t01 |
| 14094 | 11 | 106923376 | 106931320 | mNeoVis.2.1.1.chr11\_hap1.g00874.t01 |
| 14095 | 11 | 106952615 | 106956369 | mNeoVis.2.1.1.chr11\_hap1.g00875.t01 |
| 14096 | 11 | 106952615 | 106956369 | mNeoVis.2.1.1.chr11\_hap1.g00875.t01 |
| 14097 | 11 | 107004357 | 107034978 | mNeoVis.2.1.1.chr11\_hap1.g00876.t01 |
| 14098 | 11 | 107004357 | 107034978 | mNeoVis.2.1.1.chr11\_hap1.g00876.t01 |
| 14099 | 11 | 107075390 | 107130104 | mNeoVis.2.1.1.chr11\_hap1.g00877.t01 |
| 14100 | 11 | 107075390 | 107130104 | mNeoVis.2.1.1.chr11\_hap1.g00877.t01 |
| 14101 | 11 | 107075390 | 107130104 | mNeoVis.2.1.1.chr11\_hap1.g00877.t01 |
| 14102 | 11 | 107214414 | 107285176 | mNeoVis.2.1.1.chr11\_hap1.g00878.t01 |
| 14103 | 11 | 107214414 | 107285176 | mNeoVis.2.1.1.chr11\_hap1.g00878.t01 |
| 14104 | 11 | 107214414 | 107285176 | mNeoVis.2.1.1.chr11\_hap1.g00878.t01 |
| 14105 | 11 | 107285439 | 107319391 | mNeoVis.2.1.1.chr11\_hap1.g00879.t01 |
| 14106 | 11 | 107285439 | 107319391 | mNeoVis.2.1.1.chr11\_hap1.g00879.t01 |
| 14107 | 11 | 107285439 | 107319391 | mNeoVis.2.1.1.chr11\_hap1.g00879.t01 |
| 14108 | 11 | 107319540 | 107366440 | mNeoVis.2.1.1.chr11\_hap1.g00880.t01 |
| 14109 | 11 | 107319540 | 107366440 | mNeoVis.2.1.1.chr11\_hap1.g00880.t01 |
| 14110 | 11 | 107319540 | 107366440 | mNeoVis.2.1.1.chr11\_hap1.g00880.t01 |
| 14111 | 11 | 107370469 | 107382044 | mNeoVis.2.1.1.chr11\_hap1.g00881.t01 |
| 14112 | 11 | 107370469 | 107382044 | mNeoVis.2.1.1.chr11\_hap1.g00881.t01 |
| 14113 | 11 | 107370469 | 107382044 | mNeoVis.2.1.1.chr11\_hap1.g00881.t01 |
| 14114 | 11 | 107377723 | 107492036 | mNeoVis.2.1.1.chr11\_hap1.g00882.t01 |
| 14115 | 11 | 107377723 | 107492036 | mNeoVis.2.1.1.chr11\_hap1.g00882.t01 |
| 14116 | 11 | 107377723 | 107492036 | mNeoVis.2.1.1.chr11\_hap1.g00882.t01 |
| 14117 | 11 | 107557607 | 107610534 | mNeoVis.2.1.1.chr11\_hap1.g00883.t01 |
| 14118 | 11 | 107557607 | 107610534 | mNeoVis.2.1.1.chr11\_hap1.g00883.t01 |
| 14119 | 11 | 107716729 | 107737749 | mNeoVis.2.1.1.chr11\_hap1.g00884.t01 |
| 14120 | 11 | 107818613 | 107882158 | mNeoVis.2.1.1.chr11\_hap1.g00885.t01 |
| 14121 | 11 | 107818613 | 107882158 | mNeoVis.2.1.1.chr11\_hap1.g00885.t01 |
| 14122 | 11 | 107890111 | 107918553 | mNeoVis.2.1.1.chr11\_hap1.g00886.t01 |
| 14123 | 11 | 107890111 | 107918553 | mNeoVis.2.1.1.chr11\_hap1.g00886.t01 |
| 14124 | 11 | 107890111 | 107918553 | mNeoVis.2.1.1.chr11\_hap1.g00886.t01 |
| 14125 | 11 | 107933951 | 107949528 | mNeoVis.2.1.1.chr11\_hap1.g00887.t01 |
| 14126 | 11 | 107933951 | 107949528 | mNeoVis.2.1.1.chr11\_hap1.g00887.t01 |
| 14127 | 11 | 107933951 | 107949528 | mNeoVis.2.1.1.chr11\_hap1.g00887.t01 |
| 14128 | 11 | 107967229 | 107984513 | mNeoVis.2.1.1.chr11\_hap1.g00888.t01 |
| 14129 | 11 | 107967229 | 107984513 | mNeoVis.2.1.1.chr11\_hap1.g00888.t01 |
| 14130 | 11 | 107967229 | 107984513 | mNeoVis.2.1.1.chr11\_hap1.g00888.t01 |
| 14131 | 11 | 107992744 | 108032473 | mNeoVis.2.1.1.chr11\_hap1.g00889.t01 |
| 14132 | 11 | 107992744 | 108032473 | mNeoVis.2.1.1.chr11\_hap1.g00889.t01 |
| 14133 | 11 | 108043425 | 108048538 | mNeoVis.2.1.1.chr11\_hap1.g00890.t01 |
| 14134 | 11 | 108043425 | 108048538 | mNeoVis.2.1.1.chr11\_hap1.g00890.t01 |
| 14135 | 11 | 108075899 | 108090986 | mNeoVis.2.1.1.chr11\_hap1.g00891.t01 |
| 14136 | 11 | 108075899 | 108090986 | mNeoVis.2.1.1.chr11\_hap1.g00891.t01 |
| 14137 | 11 | 108247774 | 108253779 | mNeoVis.2.1.1.chr11\_hap1.g00892.t01 |
| 14138 | 11 | 108247774 | 108253779 | mNeoVis.2.1.1.chr11\_hap1.g00892.t01 |
| 14139 | 11 | 108313134 | 108317367 | mNeoVis.2.1.1.chr11\_hap1.g00893.t01 |
| 14140 | 11 | 108313134 | 108317367 | mNeoVis.2.1.1.chr11\_hap1.g00893.t01 |
| 14141 | 11 | 108338091 | 108402568 | mNeoVis.2.1.1.chr11\_hap1.g00894.t01 |
| 14142 | 11 | 108444290 | 108514588 | mNeoVis.2.1.1.chr11\_hap1.g00895.t01 |
| 14143 | 11 | 109319824 | 109376180 | mNeoVis.2.1.1.chr11\_hap1.g00900.t01 |
| 14144 | 11 | 109386943 | 109422426 | mNeoVis.2.1.1.chr11\_hap1.g00901.t01 |
| 14145 | 11 | 109434554 | 109439743 | mNeoVis.2.1.1.chr11\_hap1.g00902.t01 |
| 14146 | 11 | 109482067 | 109489001 | mNeoVis.2.1.1.chr11\_hap1.g00903.t01 |
| 14147 | 11 | 109492045 | 109497576 | mNeoVis.2.1.1.chr11\_hap1.g00904.t01 |
| 14148 | 11 | 109521118 | 109527155 | mNeoVis.2.1.1.chr11\_hap1.g00905.t01 |
| 14149 | 11 | 109537111 | 109703939 | mNeoVis.2.1.1.chr11\_hap1.g00906.t01 |
| 14150 | 11 | 109641792 | 109648447 | mNeoVis.2.1.1.chr11\_hap1.g00907.t01 |
| 14151 | 11 | 109687462 | 109696985 | mNeoVis.2.1.1.chr11\_hap1.g00908.t01 |
| 14152 | 11 | 109687462 | 109696985 | mNeoVis.2.1.1.chr11\_hap1.g00908.t01 |
| 14153 | 11 | 109704093 | 109735360 | mNeoVis.2.1.1.chr11\_hap1.g00909.t01 |
| 14154 | 11 | 109704093 | 109735360 | mNeoVis.2.1.1.chr11\_hap1.g00909.t01 |
| 14155 | 11 | 109786135 | 109801879 | mNeoVis.2.1.1.chr11\_hap1.g00910.t01 |
| 14156 | 11 | 109850641 | 109855200 | mNeoVis.2.1.1.chr11\_hap1.g00911.t01 |
| 14157 | 11 | 109859841 | 109893002 | mNeoVis.2.1.1.chr11\_hap1.g00912.t01 |
| 14158 | 11 | 109897429 | 109931864 | mNeoVis.2.1.1.chr11\_hap1.g00913.t01 |
| 14159 | 11 | 109941843 | 109969409 | mNeoVis.2.1.1.chr11\_hap1.g00914.t01 |
| 14160 | 11 | 109941843 | 109969409 | mNeoVis.2.1.1.chr11\_hap1.g00914.t01 |
| 14161 | 11 | 109974046 | 110038794 | mNeoVis.2.1.1.chr11\_hap1.g00915.t01 |
| 14162 | 11 | 109974046 | 110038794 | mNeoVis.2.1.1.chr11\_hap1.g00915.t01 |
| 14163 | 11 | 110045224 | 110077776 | mNeoVis.2.1.1.chr11\_hap1.g00916.t01 |
| 14164 | 11 | 110045224 | 110077776 | mNeoVis.2.1.1.chr11\_hap1.g00916.t01 |
| 14165 | 11 | 110087933 | 110137770 | mNeoVis.2.1.1.chr11\_hap1.g00917.t01 |
| 14166 | 11 | 110087933 | 110137770 | mNeoVis.2.1.1.chr11\_hap1.g00917.t01 |
| 14167 | 11 | 110087933 | 110137770 | mNeoVis.2.1.1.chr11\_hap1.g00917.t01 |
| 14168 | 11 | 110146689 | 110198991 | mNeoVis.2.1.1.chr11\_hap1.g00918.t01 |
| 14169 | 11 | 110146689 | 110198991 | mNeoVis.2.1.1.chr11\_hap1.g00918.t01 |
| 14170 | 11 | 110146689 | 110198991 | mNeoVis.2.1.1.chr11\_hap1.g00918.t01 |
| 14171 | 11 | 110213856 | 110327412 | mNeoVis.2.1.1.chr11\_hap1.g00919.t01 |
| 14172 | 11 | 110213856 | 110327412 | mNeoVis.2.1.1.chr11\_hap1.g00919.t01 |
| 14173 | 11 | 110213856 | 110327412 | mNeoVis.2.1.1.chr11\_hap1.g00919.t01 |
| 14174 | 11 | 110350043 | 110350557 | mNeoVis.2.1.1.chr11\_hap1.g00920.t01 |
| 14175 | 11 | 110350043 | 110350557 | mNeoVis.2.1.1.chr11\_hap1.g00920.t01 |
| 14176 | 11 | 110350043 | 110350557 | mNeoVis.2.1.1.chr11\_hap1.g00920.t01 |
| 14177 | 11 | 110352898 | 110454652 | mNeoVis.2.1.1.chr11\_hap1.g00921.t01 |
| 14178 | 11 | 110352898 | 110454652 | mNeoVis.2.1.1.chr11\_hap1.g00921.t01 |
| 14179 | 11 | 110352898 | 110454652 | mNeoVis.2.1.1.chr11\_hap1.g00921.t01 |
| 14180 | 11 | 110446824 | 110449518 | mNeoVis.2.1.1.chr11\_hap1.g00922.t01 |
| 14181 | 11 | 110446824 | 110449518 | mNeoVis.2.1.1.chr11\_hap1.g00922.t01 |
| 14182 | 11 | 110446824 | 110449518 | mNeoVis.2.1.1.chr11\_hap1.g00922.t01 |
| 14183 | 11 | 110446824 | 110449518 | mNeoVis.2.1.1.chr11\_hap1.g00922.t01 |
| 14184 | 11 | 110460142 | 110466961 | mNeoVis.2.1.1.chr11\_hap1.g00923.t01 |
| 14185 | 11 | 110460142 | 110466961 | mNeoVis.2.1.1.chr11\_hap1.g00923.t01 |
| 14186 | 11 | 110460142 | 110466961 | mNeoVis.2.1.1.chr11\_hap1.g00923.t01 |
| 14187 | 11 | 110460142 | 110466961 | mNeoVis.2.1.1.chr11\_hap1.g00923.t01 |
| 14188 | 11 | 110966313 | 111009719 | mNeoVis.2.1.1.chr11\_hap1.g00924.t01 |
| 14189 | 11 | 110966313 | 111009719 | mNeoVis.2.1.1.chr11\_hap1.g00924.t01 |
| 14190 | 11 | 110966313 | 111009719 | mNeoVis.2.1.1.chr11\_hap1.g00924.t01 |
| 14191 | 11 | 111009303 | 111012037 | mNeoVis.2.1.1.chr11\_hap1.g00925.t01 |
| 14192 | 11 | 111009303 | 111012037 | mNeoVis.2.1.1.chr11\_hap1.g00925.t01 |
| 14193 | 11 | 111009303 | 111012037 | mNeoVis.2.1.1.chr11\_hap1.g00925.t01 |
| 14194 | 11 | 111012153 | 111344491 | mNeoVis.2.1.1.chr11\_hap1.g00926.t01 |
| 14195 | 11 | 111012153 | 111344491 | mNeoVis.2.1.1.chr11\_hap1.g00926.t01 |
| 14196 | 11 | 111360783 | 111404886 | mNeoVis.2.1.1.chr11\_hap1.g00927.t01 |
| 14197 | 11 | 111360783 | 111404886 | mNeoVis.2.1.1.chr11\_hap1.g00927.t01 |
| 14198 | 11 | 111360783 | 111404886 | mNeoVis.2.1.1.chr11\_hap1.g00927.t01 |
| 14199 | 11 | 111409841 | 111444457 | mNeoVis.2.1.1.chr11\_hap1.g00928.t01 |
| 14200 | 11 | 111409841 | 111444457 | mNeoVis.2.1.1.chr11\_hap1.g00928.t01 |
| 14201 | 11 | 111409841 | 111444457 | mNeoVis.2.1.1.chr11\_hap1.g00928.t01 |
| 14202 | 11 | 111466148 | 111467960 | mNeoVis.2.1.1.chr11\_hap1.g00929.t01 |
| 14203 | 11 | 111466148 | 111467960 | mNeoVis.2.1.1.chr11\_hap1.g00929.t01 |
| 14204 | 11 | 111466148 | 111467960 | mNeoVis.2.1.1.chr11\_hap1.g00929.t01 |
| 14205 | 11 | 111473791 | 111532060 | mNeoVis.2.1.1.chr11\_hap1.g00930.t01 |
| 14206 | 11 | 111544288 | 111572752 | mNeoVis.2.1.1.chr11\_hap1.g00931.t01 |
| 14207 | 11 | 111544288 | 111572752 | mNeoVis.2.1.1.chr11\_hap1.g00931.t01 |
| 14208 | 11 | 111600909 | 111625487 | mNeoVis.2.1.1.chr11\_hap1.g00932.t01 |
| 14209 | 11 | 111600909 | 111625487 | mNeoVis.2.1.1.chr11\_hap1.g00932.t01 |
| 14210 | 11 | 111650438 | 111671574 | mNeoVis.2.1.1.chr11\_hap1.g00933.t01 |
| 14211 | 11 | 111650438 | 111671574 | mNeoVis.2.1.1.chr11\_hap1.g00933.t01 |
| 14212 | 11 | 111679346 | 111691505 | mNeoVis.2.1.1.chr11\_hap1.g00934.t01 |
| 14213 | 11 | 111679346 | 111691505 | mNeoVis.2.1.1.chr11\_hap1.g00934.t01 |
| 14214 | 11 | 111699621 | 111713651 | mNeoVis.2.1.1.chr11\_hap1.g00935.t01 |
| 14215 | 11 | 111699621 | 111713651 | mNeoVis.2.1.1.chr11\_hap1.g00935.t01 |
| 14216 | 11 | 111734689 | 111744864 | mNeoVis.2.1.1.chr11\_hap1.g00936.t01 |
| 14217 | 11 | 111769861 | 111783481 | mNeoVis.2.1.1.chr11\_hap1.g00937.t01 |
| 14218 | 11 | 111789285 | 111869186 | mNeoVis.2.1.1.chr11\_hap1.g00938.t01 |
| 14219 | 11 | 111879108 | 111901603 | mNeoVis.2.1.1.chr11\_hap1.g00939.t01 |
| 14220 | 11 | 111901722 | 111914386 | mNeoVis.2.1.1.chr11\_hap1.g00940.t01 |
| 14221 | 11 | 111979245 | 111982872 | mNeoVis.2.1.1.chr11\_hap1.g00941.t01 |
| 14222 | 11 | 111979245 | 111982872 | mNeoVis.2.1.1.chr11\_hap1.g00941.t01 |
| 14223 | 11 | 111982526 | 112003418 | mNeoVis.2.1.1.chr11\_hap1.g00942.t01 |
| 14224 | 11 | 111982526 | 112003418 | mNeoVis.2.1.1.chr11\_hap1.g00942.t01 |
| 14225 | 11 | 112008213 | 112010535 | mNeoVis.2.1.1.chr11\_hap1.g00943.t01 |
| 14226 | 11 | 112008213 | 112010535 | mNeoVis.2.1.1.chr11\_hap1.g00943.t01 |
| 14227 | 11 | 112038548 | 112059825 | mNeoVis.2.1.1.chr11\_hap1.g00944.t01 |
| 14228 | 11 | 112038548 | 112059825 | mNeoVis.2.1.1.chr11\_hap1.g00944.t01 |
| 14229 | 11 | 112068580 | 112081069 | mNeoVis.2.1.1.chr11\_hap1.g00945.t01 |
| 14230 | 11 | 112068580 | 112081069 | mNeoVis.2.1.1.chr11\_hap1.g00945.t01 |
| 14231 | 11 | 112081210 | 112087138 | mNeoVis.2.1.1.chr11\_hap1.g00946.t01 |
| 14232 | 11 | 112081210 | 112087138 | mNeoVis.2.1.1.chr11\_hap1.g00946.t01 |
| 14233 | 11 | 112088556 | 112118431 | mNeoVis.2.1.1.chr11\_hap1.g00947.t01 |
| 14234 | 11 | 112088556 | 112118431 | mNeoVis.2.1.1.chr11\_hap1.g00947.t01 |
| 14235 | 11 | 112127967 | 112163058 | mNeoVis.2.1.1.chr11\_hap1.g00948.t01 |
| 14236 | 11 | 112127967 | 112163058 | mNeoVis.2.1.1.chr11\_hap1.g00948.t01 |
| 14237 | 11 | 112166962 | 112170641 | mNeoVis.2.1.1.chr11\_hap1.g00949.t01 |
| 14238 | 11 | 112166962 | 112170641 | mNeoVis.2.1.1.chr11\_hap1.g00949.t01 |
| 14239 | 11 | 112223254 | 112236531 | mNeoVis.2.1.1.chr11\_hap1.g00950.t01 |
| 14240 | 11 | 112261466 | 112274087 | mNeoVis.2.1.1.chr11\_hap1.g00951.t01 |
| 14241 | 11 | 112293057 | 112454493 | mNeoVis.2.1.1.chr11\_hap1.g00952.t01 |
| 14242 | 11 | 112897142 | 112907820 | mNeoVis.2.1.1.chr11\_hap1.g00956.t01 |
| 14243 | 11 | 114199418 | 114585777 | mNeoVis.2.1.1.chr11\_hap1.g00958.t01 |
| 14244 | 11 | 114858208 | 114898846 | mNeoVis.2.1.1.chr11\_hap1.g00959.t01 |
| 14245 | 11 | 114915961 | 114965220 | mNeoVis.2.1.1.chr11\_hap1.g00960.t01 |
| 14246 | 11 | 115126538 | 115162890 | mNeoVis.2.1.1.chr11\_hap1.g00961.t01 |
| 14247 | 11 | 115126538 | 115162890 | mNeoVis.2.1.1.chr11\_hap1.g00961.t01 |
| 14248 | 11 | 115159125 | 115161236 | mNeoVis.2.1.1.chr11\_hap1.g00962.t01 |
| 14249 | 11 | 115159125 | 115161236 | mNeoVis.2.1.1.chr11\_hap1.g00962.t01 |
| 14250 | 11 | 115161377 | 115352171 | mNeoVis.2.1.1.chr11\_hap1.g00963.t01 |
| 14251 | 11 | 115161377 | 115352171 | mNeoVis.2.1.1.chr11\_hap1.g00963.t01 |
| 14252 | 11 | 115354329 | 115399314 | mNeoVis.2.1.1.chr11\_hap1.g00964.t01 |
| 14253 | 11 | 115354329 | 115399314 | mNeoVis.2.1.1.chr11\_hap1.g00964.t01 |
| 14254 | 11 | 115455079 | 115600402 | mNeoVis.2.1.1.chr11\_hap1.g00965.t01 |
| 14255 | 11 | 115455079 | 115600402 | mNeoVis.2.1.1.chr11\_hap1.g00965.t01 |
| 14256 | 11 | 115604099 | 115678586 | mNeoVis.2.1.1.chr11\_hap1.g00966.t01 |
| 14257 | 11 | 115604099 | 115678586 | mNeoVis.2.1.1.chr11\_hap1.g00966.t01 |
| 14258 | 11 | 115620759 | 115657787 | mNeoVis.2.1.1.chr11\_hap1.g00967.t01 |
| 14259 | 11 | 115620759 | 115657787 | mNeoVis.2.1.1.chr11\_hap1.g00967.t01 |
| 14260 | 11 | 115657704 | 115684297 | mNeoVis.2.1.1.chr11\_hap1.g00968.t01 |
| 14261 | 11 | 115657704 | 115684297 | mNeoVis.2.1.1.chr11\_hap1.g00968.t01 |
| 14262 | 11 | 115686199 | 115691361 | mNeoVis.2.1.1.chr11\_hap1.g00969.t01 |
| 14263 | 11 | 115686199 | 115691361 | mNeoVis.2.1.1.chr11\_hap1.g00969.t01 |
| 14264 | 11 | 115760745 | 115765086 | mNeoVis.2.1.1.chr11\_hap1.g00971.t01 |
| 14265 | 11 | 115773416 | 115789866 | mNeoVis.2.1.1.chr11\_hap1.g00972.t01 |
| 14266 | 11 | 115790176 | 115807612 | mNeoVis.2.1.1.chr11\_hap1.g00973.t01 |
| 14267 | 11 | 115809508 | 115861059 | mNeoVis.2.1.1.chr11\_hap1.g00974.t01 |
| 14268 | 11 | 115918687 | 115957124 | mNeoVis.2.1.1.chr11\_hap1.g00975.t01 |
| 14269 | 11 | 116107988 | 116162972 | mNeoVis.2.1.1.chr11\_hap1.g00976.t01 |
| 14270 | 11 | 117115375 | 117128168 | mNeoVis.2.1.1.chr11\_hap1.g00987.t01 |
| 14271 | 11 | 117115375 | 117128168 | mNeoVis.2.1.1.chr11\_hap1.g00987.t01 |
| 14272 | 11 | 117118302 | 117122031 | mNeoVis.2.1.1.chr11\_hap1.g00988.t01 |
| 14273 | 11 | 117118302 | 117122031 | mNeoVis.2.1.1.chr11\_hap1.g00988.t01 |
| 14274 | 11 | 117131951 | 117136890 | mNeoVis.2.1.1.chr11\_hap1.g00989.t01 |
| 14275 | 11 | 117131951 | 117136890 | mNeoVis.2.1.1.chr11\_hap1.g00989.t01 |
| 14276 | 11 | 117152083 | 117323498 | mNeoVis.2.1.1.chr11\_hap1.g00990.t01 |
| 14277 | 11 | 117152083 | 117323498 | mNeoVis.2.1.1.chr11\_hap1.g00990.t01 |
| 14278 | 11 | 117866926 | 117973653 | mNeoVis.2.1.1.chr11\_hap1.g00992.t01 |
| 14279 | 11 | 117991840 | 118009222 | mNeoVis.2.1.1.chr11\_hap1.g00993.t01 |
| 14280 | 11 | 119606708 | 119652631 | mNeoVis.2.1.1.chr11\_hap1.g00995.t01 |
| 14281 | 11 | 120143172 | 120236438 | mNeoVis.2.1.1.chr11\_hap1.g00996.t01 |
| 14282 | 11 | 120143172 | 120236438 | mNeoVis.2.1.1.chr11\_hap1.g00996.t01 |
| 14283 | 11 | 120280274 | 120470086 | mNeoVis.2.1.1.chr11\_hap1.g00997.t01 |
| 14284 | 11 | 120471284 | 120514276 | mNeoVis.2.1.1.chr11\_hap1.g00998.t01 |
| 14285 | 11 | 124693028 | 124716752 | mNeoVis.2.1.1.chr11\_hap1.g01036.t01 |
| 14286 | 11 | 124766841 | 125235275 | mNeoVis.2.1.1.chr11\_hap1.g01037.t01 |
| 14287 | 11 | 125173240 | 125256317 | mNeoVis.2.1.1.chr11\_hap1.g01038.t01 |
| 14288 | 11 | 125258120 | 125268010 | mNeoVis.2.1.1.chr11\_hap1.g01039.t01 |
| 14289 | 11 | 125514389 | 125549539 | mNeoVis.2.1.1.chr11\_hap1.g01051.t01 |
| 14290 | 11 | 125549614 | 125551234 | mNeoVis.2.1.1.chr11\_hap1.g01052.t01 |
| 14291 | 11 | 125551360 | 125562620 | mNeoVis.2.1.1.chr11\_hap1.g01053.t01 |
| 14292 | 11 | 125580262 | 125598364 | mNeoVis.2.1.1.chr11\_hap1.g01054.t01 |
| 14293 | 11 | 125598956 | 125602444 | mNeoVis.2.1.1.chr11\_hap1.g01055.t01 |
| 14294 | 11 | 125602544 | 125608687 | mNeoVis.2.1.1.chr11\_hap1.g01056.t01 |
| 14295 | 11 | 125609115 | 125613591 | mNeoVis.2.1.1.chr11\_hap1.g01057.t01 |
| 14296 | 11 | 125617613 | 125641332 | mNeoVis.2.1.1.chr11\_hap1.g01058.t01 |
| 14297 | 11 | 125643970 | 125654834 | mNeoVis.2.1.1.chr11\_hap1.g01059.t01 |
| 14298 | 11 | 125654975 | 125656524 | mNeoVis.2.1.1.chr11\_hap1.g01060.t01 |
| 14299 | 11 | 125656572 | 125678278 | mNeoVis.2.1.1.chr11\_hap1.g01061.t01 |
| 14300 | 11 | 125679328 | 125683392 | mNeoVis.2.1.1.chr11\_hap1.g01062.t01 |
| 14301 | 11 | 125693963 | 125702160 | mNeoVis.2.1.1.chr11\_hap1.g01063.t01 |
| 14302 | 11 | 125710654 | 125734012 | mNeoVis.2.1.1.chr11\_hap1.g01064.t01 |
| 14303 | 11 | 125734265 | 125739785 | mNeoVis.2.1.1.chr11\_hap1.g01065.t01 |
| 14304 | 11 | 125739302 | 125751689 | mNeoVis.2.1.1.chr11\_hap1.g01066.t01 |
| 14305 | 11 | 125788974 | 125793698 | mNeoVis.2.1.1.chr11\_hap1.g01067.t01 |
| 14306 | 11 | 125805695 | 125808315 | mNeoVis.2.1.1.chr11\_hap1.g01068.t01 |
| 14307 | 11 | 125811706 | 125815783 | mNeoVis.2.1.1.chr11\_hap1.g01069.t01 |
| 14308 | 11 | 125815807 | 125821183 | mNeoVis.2.1.1.chr11\_hap1.g01070.t01 |
| 14309 | 11 | 125837349 | 125839594 | mNeoVis.2.1.1.chr11\_hap1.g01071.t01 |
| 14310 | 11 | 125840676 | 125852247 | mNeoVis.2.1.1.chr11\_hap1.g01072.t01 |
| 14311 | 11 | 125852598 | 125860384 | mNeoVis.2.1.1.chr11\_hap1.g01073.t01 |
| 14312 | 11 | 125866154 | 125867193 | mNeoVis.2.1.1.chr11\_hap1.g01074.t01 |
| 14313 | 11 | 125866215 | 125884628 | mNeoVis.2.1.1.chr11\_hap1.g01075.t01 |
| 14314 | 11 | 125890926 | 125899303 | mNeoVis.2.1.1.chr11\_hap1.g01076.t01 |
| 14315 | 11 | 125903857 | 125915590 | mNeoVis.2.1.1.chr11\_hap1.g01077.t01 |
| 14316 | 11 | 125973371 | 126065633 | mNeoVis.2.1.1.chr11\_hap1.g01078.t01 |
| 14317 | 11 | 126082358 | 126084440 | mNeoVis.2.1.1.chr11\_hap1.g01079.t01 |
| 14318 | 11 | 126223566 | 126229028 | mNeoVis.2.1.1.chr11\_hap1.g01084.t01 |
| 14319 | 11 | 126223566 | 126229028 | mNeoVis.2.1.1.chr11\_hap1.g01084.t01 |
| 14320 | 11 | 126232371 | 126246387 | mNeoVis.2.1.1.chr11\_hap1.g01085.t01 |
| 14321 | 11 | 126232371 | 126246387 | mNeoVis.2.1.1.chr11\_hap1.g01085.t01 |
| 14322 | 11 | 126265991 | 126304611 | mNeoVis.2.1.1.chr11\_hap1.g01086.t01 |
| 14323 | 11 | 126265991 | 126304611 | mNeoVis.2.1.1.chr11\_hap1.g01086.t01 |
| 14324 | 11 | 126347646 | 126396108 | mNeoVis.2.1.1.chr11\_hap1.g01087.t01 |
| 14325 | 11 | 126347646 | 126396108 | mNeoVis.2.1.1.chr11\_hap1.g01087.t01 |
| 14326 | 11 | 126397608 | 126412072 | mNeoVis.2.1.1.chr11\_hap1.g01088.t01 |
| 14327 | 11 | 126397608 | 126412072 | mNeoVis.2.1.1.chr11\_hap1.g01088.t01 |
| 14328 | 11 | 126415583 | 126507121 | mNeoVis.2.1.1.chr11\_hap1.g01089.t01 |
| 14329 | 11 | 126415583 | 126507121 | mNeoVis.2.1.1.chr11\_hap1.g01089.t01 |
| 14330 | 11 | 126508444 | 126550602 | mNeoVis.2.1.1.chr11\_hap1.g01090.t01 |
| 14331 | 11 | 126508444 | 126550602 | mNeoVis.2.1.1.chr11\_hap1.g01090.t01 |
| 14332 | 11 | 126555099 | 126596135 | mNeoVis.2.1.1.chr11\_hap1.g01091.t01 |
| 14333 | 11 | 126555099 | 126596135 | mNeoVis.2.1.1.chr11\_hap1.g01091.t01 |
| 14334 | 11 | 126623939 | 126633914 | mNeoVis.2.1.1.chr11\_hap1.g01092.t01 |
| 14335 | 11 | 126623939 | 126633914 | mNeoVis.2.1.1.chr11\_hap1.g01092.t01 |
| 14336 | 11 | 126645777 | 126679348 | mNeoVis.2.1.1.chr11\_hap1.g01093.t01 |
| 14337 | 11 | 126645777 | 126679348 | mNeoVis.2.1.1.chr11\_hap1.g01093.t01 |
| 14338 | 11 | 126679493 | 126725740 | mNeoVis.2.1.1.chr11\_hap1.g01094.t01 |
| 14339 | 11 | 126679493 | 126725740 | mNeoVis.2.1.1.chr11\_hap1.g01094.t01 |
| 14340 | 11 | 126726952 | 126741670 | mNeoVis.2.1.1.chr11\_hap1.g01095.t01 |
| 14341 | 11 | 126726952 | 126741670 | mNeoVis.2.1.1.chr11\_hap1.g01095.t01 |
| 14342 | 11 | 126769394 | 126858490 | mNeoVis.2.1.1.chr11\_hap1.g01096.t01 |
| 14343 | 11 | 126912039 | 126948564 | mNeoVis.2.1.1.chr11\_hap1.g01097.t01 |
| 14344 | 11 | 126987088 | 127122122 | mNeoVis.2.1.1.chr11\_hap1.g01098.t01 |
| 14345 | 11 | 127052510 | 127052979 | mNeoVis.2.1.1.chr11\_hap1.g01099.t01 |
| 14346 | 11 | 127169339 | 127404784 | mNeoVis.2.1.1.chr11\_hap1.g01100.t01 |
| 14347 | 11 | 127436135 | 127532077 | mNeoVis.2.1.1.chr11\_hap1.g01101.t01 |
| 14348 | 11 | 127436135 | 127532077 | mNeoVis.2.1.1.chr11\_hap1.g01101.t01 |
| 14349 | 11 | 127567370 | 127582505 | mNeoVis.2.1.1.chr11\_hap1.g01102.t01 |
| 14350 | 11 | 127567370 | 127582505 | mNeoVis.2.1.1.chr11\_hap1.g01102.t01 |
| 14351 | 11 | 127585053 | 127636512 | mNeoVis.2.1.1.chr11\_hap1.g01103.t01 |
| 14352 | 11 | 127585053 | 127636512 | mNeoVis.2.1.1.chr11\_hap1.g01103.t01 |
| 14353 | 11 | 127717074 | 127751057 | mNeoVis.2.1.1.chr11\_hap1.g01104.t01 |
| 14354 | 11 | 127717074 | 127751057 | mNeoVis.2.1.1.chr11\_hap1.g01104.t01 |
| 14355 | 11 | 127770249 | 127849681 | mNeoVis.2.1.1.chr11\_hap1.g01105.t01 |
| 14356 | 11 | 127770249 | 127849681 | mNeoVis.2.1.1.chr11\_hap1.g01105.t01 |
| 14357 | 11 | 127849312 | 127861195 | mNeoVis.2.1.1.chr11\_hap1.g01106.t01 |
| 14358 | 11 | 127849312 | 127861195 | mNeoVis.2.1.1.chr11\_hap1.g01106.t01 |
| 14359 | 11 | 127861158 | 127865211 | mNeoVis.2.1.1.chr11\_hap1.g01107.t01 |
| 14360 | 11 | 127861158 | 127865211 | mNeoVis.2.1.1.chr11\_hap1.g01107.t01 |
| 14361 | 11 | 127878864 | 128118299 | mNeoVis.2.1.1.chr11\_hap1.g01108.t01 |
| 14362 | 11 | 127878864 | 128118299 | mNeoVis.2.1.1.chr11\_hap1.g01108.t01 |
| 14363 | 11 | 128248433 | 128386984 | mNeoVis.2.1.1.chr11\_hap1.g01109.t01 |
| 14364 | 11 | 128248433 | 128386984 | mNeoVis.2.1.1.chr11\_hap1.g01109.t01 |
| 14365 | 11 | 128248433 | 128386984 | mNeoVis.2.1.1.chr11\_hap1.g01109.t01 |
| 14366 | 11 | 128390840 | 128420705 | mNeoVis.2.1.1.chr11\_hap1.g01110.t01 |
| 14367 | 11 | 128390840 | 128420705 | mNeoVis.2.1.1.chr11\_hap1.g01110.t01 |
| 14368 | 11 | 128390840 | 128420705 | mNeoVis.2.1.1.chr11\_hap1.g01110.t01 |
| 14369 | 11 | 128421100 | 128427057 | mNeoVis.2.1.1.chr11\_hap1.g01111.t01 |
| 14370 | 11 | 128421100 | 128427057 | mNeoVis.2.1.1.chr11\_hap1.g01111.t01 |
| 14371 | 11 | 128421100 | 128427057 | mNeoVis.2.1.1.chr11\_hap1.g01111.t01 |
| 14372 | 11 | 128427260 | 128437210 | mNeoVis.2.1.1.chr11\_hap1.g01112.t01 |
| 14373 | 11 | 128427260 | 128437210 | mNeoVis.2.1.1.chr11\_hap1.g01112.t01 |
| 14374 | 11 | 128427260 | 128437210 | mNeoVis.2.1.1.chr11\_hap1.g01112.t01 |
| 14375 | 11 | 128437647 | 128452642 | mNeoVis.2.1.1.chr11\_hap1.g01113.t01 |
| 14376 | 11 | 128437647 | 128452642 | mNeoVis.2.1.1.chr11\_hap1.g01113.t01 |
| 14377 | 11 | 128437647 | 128452642 | mNeoVis.2.1.1.chr11\_hap1.g01113.t01 |
| 14378 | 11 | 128455716 | 128467028 | mNeoVis.2.1.1.chr11\_hap1.g01114.t01 |
| 14379 | 11 | 128455716 | 128467028 | mNeoVis.2.1.1.chr11\_hap1.g01114.t01 |
| 14380 | 11 | 128455716 | 128467028 | mNeoVis.2.1.1.chr11\_hap1.g01114.t01 |
| 14381 | 11 | 128468705 | 128494345 | mNeoVis.2.1.1.chr11\_hap1.g01115.t01 |
| 14382 | 11 | 128468705 | 128494345 | mNeoVis.2.1.1.chr11\_hap1.g01115.t01 |
| 14383 | 11 | 128468705 | 128494345 | mNeoVis.2.1.1.chr11\_hap1.g01115.t01 |
| 14384 | 11 | 128496452 | 128509369 | mNeoVis.2.1.1.chr11\_hap1.g01116.t01 |
| 14385 | 11 | 128496452 | 128509369 | mNeoVis.2.1.1.chr11\_hap1.g01116.t01 |
| 14386 | 11 | 128496452 | 128509369 | mNeoVis.2.1.1.chr11\_hap1.g01116.t01 |
| 14387 | 11 | 128512106 | 128562598 | mNeoVis.2.1.1.chr11\_hap1.g01117.t01 |
| 14388 | 11 | 128755172 | 129051455 | mNeoVis.2.1.1.chr11\_hap1.g01119.t01 |
| 14389 | 11 | 130629543 | 130630165 | mNeoVis.2.1.1.chr11\_hap1.g01120.t01 |
| 14390 | 11 | 131062566 | 131098738 | mNeoVis.2.1.1.chr11\_hap1.g01121.t01 |
| 14391 | 11 | 131165708 | 131203079 | mNeoVis.2.1.1.chr11\_hap1.g01122.t01 |
| 14392 | 11 | 131328597 | 131437984 | mNeoVis.2.1.1.chr11\_hap1.g01123.t01 |
| 14393 | 11 | 131447775 | 131455618 | mNeoVis.2.1.1.chr11\_hap1.g01124.t01 |
| 14394 | 11 | 131469148 | 131498762 | mNeoVis.2.1.1.chr11\_hap1.g01125.t01 |
| 14395 | 11 | 131631529 | 131633951 | mNeoVis.2.1.1.chr11\_hap1.g01126.t01 |
| 14396 | 11 | 131639378 | 131639596 | mNeoVis.2.1.1.chr11\_hap1.g01127.t01 |
| 14397 | 11 | 131663766 | 131673626 | mNeoVis.2.1.1.chr11\_hap1.g01128.t01 |
| 14398 | 11 | 131841274 | 131865422 | mNeoVis.2.1.1.chr11\_hap1.g01130.t01 |
| 14399 | 11 | 131996843 | 132014486 | mNeoVis.2.1.1.chr11\_hap1.g01131.t01 |
| 14400 | 11 | 132024848 | 132046263 | mNeoVis.2.1.1.chr11\_hap1.g01132.t01 |
| 14401 | 11 | 132048333 | 132113043 | mNeoVis.2.1.1.chr11\_hap1.g01133.t01 |
| 14402 | 11 | 132113574 | 132124689 | mNeoVis.2.1.1.chr11\_hap1.g01134.t01 |
| 14403 | 11 | 132562192 | 132573845 | mNeoVis.2.1.1.chr11\_hap1.g01135.t01 |
| 14404 | 11 | 132874541 | 132876143 | mNeoVis.2.1.1.chr11\_hap1.g01136.t01 |
| 14405 | 11 | 133241625 | 133263778 | mNeoVis.2.1.1.chr11\_hap1.g01137.t01 |
| 14406 | 11 | 133266609 | 133271481 | mNeoVis.2.1.1.chr11\_hap1.g01138.t01 |
| 14407 | 11 | 133611363 | 133656071 | mNeoVis.2.1.1.chr11\_hap1.g01139.t01 |
| 14408 | 11 | 133611363 | 133656071 | mNeoVis.2.1.1.chr11\_hap1.g01139.t01 |
| 14409 | 11 | 133756587 | 133796841 | mNeoVis.2.1.1.chr11\_hap1.g01140.t01 |
| 14410 | 11 | 133756587 | 133796841 | mNeoVis.2.1.1.chr11\_hap1.g01140.t01 |
| 14411 | 11 | 133779031 | 133787453 | mNeoVis.2.1.1.chr11\_hap1.g01141.t01 |
| 14412 | 11 | 133779031 | 133787453 | mNeoVis.2.1.1.chr11\_hap1.g01141.t01 |
| 14413 | 11 | 133824930 | 133825511 | mNeoVis.2.1.1.chr11\_hap1.g01142.t01 |
| 14414 | 11 | 133824930 | 133825511 | mNeoVis.2.1.1.chr11\_hap1.g01142.t01 |
| 14415 | 11 | 133852578 | 133885522 | mNeoVis.2.1.1.chr11\_hap1.g01143.t01 |
| 14416 | 11 | 133852578 | 133885522 | mNeoVis.2.1.1.chr11\_hap1.g01143.t01 |
| 14417 | 11 | 133885660 | 133957383 | mNeoVis.2.1.1.chr11\_hap1.g01144.t01 |
| 14418 | 11 | 133885660 | 133957383 | mNeoVis.2.1.1.chr11\_hap1.g01144.t01 |
| 14419 | 11 | 133964457 | 134031074 | mNeoVis.2.1.1.chr11\_hap1.g01145.t01 |
| 14420 | 11 | 133964457 | 134031074 | mNeoVis.2.1.1.chr11\_hap1.g01145.t01 |
| 14421 | 11 | 134089273 | 134096957 | mNeoVis.2.1.1.chr11\_hap1.g01146.t01 |
| 14422 | 11 | 134160594 | 134182212 | mNeoVis.2.1.1.chr11\_hap1.g01147.t01 |
| 14423 | 11 | 134280225 | 134297095 | mNeoVis.2.1.1.chr11\_hap1.g01148.t01 |
| 14424 | 11 | 134765660 | 134870169 | mNeoVis.2.1.1.chr11\_hap1.g01149.t01 |
| 14425 | 11 | 134765660 | 134870169 | mNeoVis.2.1.1.chr11\_hap1.g01149.t01 |
| 14426 | 11 | 135572098 | 135604065 | mNeoVis.2.1.1.chr11\_hap1.g01151.t01 |
| 14427 | 11 | 135627837 | 135951589 | mNeoVis.2.1.1.chr11\_hap1.g01152.t01 |
| 14428 | 11 | 136382206 | 136389216 | mNeoVis.2.1.1.chr11\_hap1.g01153.t01 |
| 14429 | 11 | 137827615 | 137852966 | mNeoVis.2.1.1.chr11\_hap1.g01154.t01 |
| 14430 | 11 | 140062519 | 140133417 | mNeoVis.2.1.1.chr11\_hap1.g01172.t01 |
| 14431 | 11 | 140283181 | 140320036 | mNeoVis.2.1.1.chr11\_hap1.g01173.t01 |
| 14432 | 11 | 140343126 | 140395636 | mNeoVis.2.1.1.chr11\_hap1.g01174.t01 |
| 14433 | 11 | 140395766 | 140405678 | mNeoVis.2.1.1.chr11\_hap1.g01175.t01 |
| 14434 | 11 | 140408215 | 140451723 | mNeoVis.2.1.1.chr11\_hap1.g01176.t01 |
| 14435 | 11 | 140451811 | 140464191 | mNeoVis.2.1.1.chr11\_hap1.g01177.t01 |
| 14436 | 11 | 140769075 | 140890288 | mNeoVis.2.1.1.chr11\_hap1.g01179.t01 |
| 14437 | 11 | 140907242 | 141000570 | mNeoVis.2.1.1.chr11\_hap1.g01180.t01 |
| 14438 | 11 | 141026813 | 141029234 | mNeoVis.2.1.1.chr11\_hap1.g01181.t01 |
| 14439 | 11 | 142895408 | 142912710 | mNeoVis.2.1.1.chr11\_hap1.g01187.t01 |
| 14440 | 11 | 144766891 | 144797615 | mNeoVis.2.1.1.chr11\_hap1.g01190.t01 |
| 14441 | 11 | 144803173 | 144838142 | mNeoVis.2.1.1.chr11\_hap1.g01191.t01 |
| 14442 | 11 | 144847234 | 145010826 | mNeoVis.2.1.1.chr11\_hap1.g01192.t01 |
| 14443 | 11 | 144861792 | 144876744 | mNeoVis.2.1.1.chr11\_hap1.g01193.t01 |
| 14444 | 11 | 144900592 | 144907367 | mNeoVis.2.1.1.chr11\_hap1.g01194.t01 |
| 14445 | 11 | 144913902 | 144989282 | mNeoVis.2.1.1.chr11\_hap1.g01195.t01 |
| 14446 | 11 | 144992459 | 145117718 | mNeoVis.2.1.1.chr11\_hap1.g01196.t01 |
| 14447 | 11 | 145629781 | 146064374 | mNeoVis.2.1.1.chr11\_hap1.g01197.t01 |
| 14448 | 11 | 146160142 | 146192864 | mNeoVis.2.1.1.chr11\_hap1.g01199.t01 |
| 14449 | 11 | 146208526 | 146219261 | mNeoVis.2.1.1.chr11\_hap1.g01200.t01 |
| 14450 | 11 | 146236360 | 146242373 | mNeoVis.2.1.1.chr11\_hap1.g01201.t01 |
| 14451 | 11 | 146479596 | 146487534 | mNeoVis.2.1.1.chr11\_hap1.g01202.t01 |
| 14452 | 11 | 146488826 | 146502762 | mNeoVis.2.1.1.chr11\_hap1.g01203.t01 |
| 14453 | 11 | 146504354 | 146539013 | mNeoVis.2.1.1.chr11\_hap1.g01204.t01 |
| 14454 | 11 | 146630919 | 146643875 | mNeoVis.2.1.1.chr11\_hap1.g01205.t01 |
| 14455 | 13 | 4613952 | 4635192 | mNeoVis.2.1.1.chr13\_hap1.g00142.t01 |
| 14456 | 13 | 4638450 | 4693802 | mNeoVis.2.1.1.chr13\_hap1.g00143.t01 |
| 14457 | 13 | 4731431 | 4764563 | mNeoVis.2.1.1.chr13\_hap1.g00144.t01 |
| 14458 | 13 | 4776589 | 4913318 | mNeoVis.2.1.1.chr13\_hap1.g00145.t01 |
| 14459 | 13 | 4913390 | 4914278 | mNeoVis.2.1.1.chr13\_hap1.g00146.t01 |
| 14460 | 13 | 4917276 | 4927389 | mNeoVis.2.1.1.chr13\_hap1.g00147.t01 |
| 14461 | 13 | 4934541 | 4947872 | mNeoVis.2.1.1.chr13\_hap1.g00148.t01 |
| 14462 | 13 | 4964843 | 5014878 | mNeoVis.2.1.1.chr13\_hap1.g00149.t01 |
| 14463 | 13 | 5067575 | 5119588 | mNeoVis.2.1.1.chr13\_hap1.g00150.t01 |
| 14464 | 13 | 5772252 | 5778455 | mNeoVis.2.1.1.chr13\_hap1.g00161.t01 |
| 14465 | 13 | 5778594 | 5789716 | mNeoVis.2.1.1.chr13\_hap1.g00162.t01 |
| 14466 | 13 | 5789795 | 5797260 | mNeoVis.2.1.1.chr13\_hap1.g00163.t01 |
| 14467 | 13 | 6107770 | 6110428 | mNeoVis.2.1.1.chr13\_hap1.g00164.t01 |
| 14468 | 13 | 6130606 | 6159507 | mNeoVis.2.1.1.chr13\_hap1.g00165.t01 |
| 14469 | 13 | 6159941 | 6173755 | mNeoVis.2.1.1.chr13\_hap1.g00166.t01 |
| 14470 | 13 | 6176007 | 6185541 | mNeoVis.2.1.1.chr13\_hap1.g00167.t01 |
| 14471 | 13 | 6193572 | 6220084 | mNeoVis.2.1.1.chr13\_hap1.g00168.t01 |
| 14472 | 13 | 6223415 | 6238680 | mNeoVis.2.1.1.chr13\_hap1.g00169.t01 |
| 14473 | 13 | 6239779 | 6287729 | mNeoVis.2.1.1.chr13\_hap1.g00170.t01 |
| 14474 | 13 | 6287795 | 6289859 | mNeoVis.2.1.1.chr13\_hap1.g00171.t01 |
| 14475 | 13 | 6292400 | 6303272 | mNeoVis.2.1.1.chr13\_hap1.g00172.t01 |
| 14476 | 13 | 6306110 | 6334266 | mNeoVis.2.1.1.chr13\_hap1.g00173.t01 |
| 14477 | 13 | 6336893 | 6367034 | mNeoVis.2.1.1.chr13\_hap1.g00174.t01 |
| 14478 | 13 | 6371470 | 6378514 | mNeoVis.2.1.1.chr13\_hap1.g00175.t01 |
| 14479 | 13 | 6378538 | 6382780 | mNeoVis.2.1.1.chr13\_hap1.g00176.t01 |
| 14480 | 13 | 6382931 | 6399876 | mNeoVis.2.1.1.chr13\_hap1.g00177.t01 |
| 14481 | 13 | 6409410 | 6439996 | mNeoVis.2.1.1.chr13\_hap1.g00178.t01 |
| 14482 | 13 | 6443338 | 6446740 | mNeoVis.2.1.1.chr13\_hap1.g00179.t01 |
| 14483 | 13 | 6446895 | 6457549 | mNeoVis.2.1.1.chr13\_hap1.g00180.t01 |
| 14484 | 13 | 6462494 | 6468720 | mNeoVis.2.1.1.chr13\_hap1.g00181.t01 |
| 14485 | 13 | 6491184 | 6509580 | mNeoVis.2.1.1.chr13\_hap1.g00182.t01 |
| 14486 | 13 | 6509579 | 6571617 | mNeoVis.2.1.1.chr13\_hap1.g00183.t01 |
| 14487 | 13 | 6577522 | 6602243 | mNeoVis.2.1.1.chr13\_hap1.g00184.t01 |
| 14488 | 13 | 6604312 | 6638953 | mNeoVis.2.1.1.chr13\_hap1.g00185.t01 |
| 14489 | 13 | 6647043 | 6657461 | mNeoVis.2.1.1.chr13\_hap1.g00186.t01 |
| 14490 | 13 | 6678669 | 6693814 | mNeoVis.2.1.1.chr13\_hap1.g00187.t01 |
| 14491 | 13 | 6701243 | 6714831 | mNeoVis.2.1.1.chr13\_hap1.g00188.t01 |
| 14492 | 13 | 6720718 | 6729257 | mNeoVis.2.1.1.chr13\_hap1.g00189.t01 |
| 14493 | 13 | 6729331 | 6739143 | mNeoVis.2.1.1.chr13\_hap1.g00190.t01 |
| 14494 | 13 | 6762445 | 6768895 | mNeoVis.2.1.1.chr13\_hap1.g00191.t01 |
| 14495 | 13 | 6762445 | 6768895 | mNeoVis.2.1.1.chr13\_hap1.g00191.t01 |
| 14496 | 13 | 6769036 | 6787147 | mNeoVis.2.1.1.chr13\_hap1.g00192.t01 |
| 14497 | 13 | 6769036 | 6787147 | mNeoVis.2.1.1.chr13\_hap1.g00192.t01 |
| 14498 | 13 | 6789739 | 6834069 | mNeoVis.2.1.1.chr13\_hap1.g00193.t01 |
| 14499 | 13 | 6789739 | 6834069 | mNeoVis.2.1.1.chr13\_hap1.g00193.t01 |
| 14500 | 13 | 6838465 | 6854992 | mNeoVis.2.1.1.chr13\_hap1.g00194.t01 |
| 14501 | 13 | 6838465 | 6854992 | mNeoVis.2.1.1.chr13\_hap1.g00194.t01 |
| 14502 | 13 | 6855922 | 6859517 | mNeoVis.2.1.1.chr13\_hap1.g00195.t01 |
| 14503 | 13 | 6855922 | 6859517 | mNeoVis.2.1.1.chr13\_hap1.g00195.t01 |
| 14504 | 13 | 6863959 | 6867525 | mNeoVis.2.1.1.chr13\_hap1.g00196.t01 |
| 14505 | 13 | 6863959 | 6867525 | mNeoVis.2.1.1.chr13\_hap1.g00196.t01 |
| 14506 | 13 | 6878469 | 6883462 | mNeoVis.2.1.1.chr13\_hap1.g00197.t01 |
| 14507 | 13 | 6878469 | 6883462 | mNeoVis.2.1.1.chr13\_hap1.g00197.t01 |
| 14508 | 13 | 6891240 | 6925801 | mNeoVis.2.1.1.chr13\_hap1.g00198.t01 |
| 14509 | 13 | 6891240 | 6925801 | mNeoVis.2.1.1.chr13\_hap1.g00198.t01 |
| 14510 | 13 | 6926792 | 6931459 | mNeoVis.2.1.1.chr13\_hap1.g00199.t01 |
| 14511 | 13 | 6926792 | 6931459 | mNeoVis.2.1.1.chr13\_hap1.g00199.t01 |
| 14512 | 13 | 6974704 | 7009251 | mNeoVis.2.1.1.chr13\_hap1.g00200.t01 |
| 14513 | 13 | 6974704 | 7009251 | mNeoVis.2.1.1.chr13\_hap1.g00200.t01 |
| 14514 | 13 | 7011095 | 7013957 | mNeoVis.2.1.1.chr13\_hap1.g00201.t01 |
| 14515 | 13 | 7011095 | 7013957 | mNeoVis.2.1.1.chr13\_hap1.g00201.t01 |
| 14516 | 13 | 7016128 | 7024419 | mNeoVis.2.1.1.chr13\_hap1.g00202.t01 |
| 14517 | 13 | 7016128 | 7024419 | mNeoVis.2.1.1.chr13\_hap1.g00202.t01 |
| 14518 | 13 | 7024776 | 7038348 | mNeoVis.2.1.1.chr13\_hap1.g00203.t01 |
| 14519 | 13 | 7024776 | 7038348 | mNeoVis.2.1.1.chr13\_hap1.g00203.t01 |
| 14520 | 13 | 7045494 | 7056948 | mNeoVis.2.1.1.chr13\_hap1.g00204.t01 |
| 14521 | 13 | 7045494 | 7056948 | mNeoVis.2.1.1.chr13\_hap1.g00204.t01 |
| 14522 | 13 | 7066718 | 7072271 | mNeoVis.2.1.1.chr13\_hap1.g00205.t01 |
| 14523 | 13 | 7066718 | 7072271 | mNeoVis.2.1.1.chr13\_hap1.g00205.t01 |
| 14524 | 13 | 7080275 | 7112362 | mNeoVis.2.1.1.chr13\_hap1.g00206.t01 |
| 14525 | 13 | 7080275 | 7112362 | mNeoVis.2.1.1.chr13\_hap1.g00206.t01 |
| 14526 | 13 | 7112503 | 7122198 | mNeoVis.2.1.1.chr13\_hap1.g00207.t01 |
| 14527 | 13 | 7112503 | 7122198 | mNeoVis.2.1.1.chr13\_hap1.g00207.t01 |
| 14528 | 13 | 7137297 | 7141397 | mNeoVis.2.1.1.chr13\_hap1.g00208.t01 |
| 14529 | 13 | 7137297 | 7141397 | mNeoVis.2.1.1.chr13\_hap1.g00208.t01 |
| 14530 | 13 | 7148244 | 7177402 | mNeoVis.2.1.1.chr13\_hap1.g00209.t01 |
| 14531 | 13 | 7148244 | 7177402 | mNeoVis.2.1.1.chr13\_hap1.g00209.t01 |
| 14532 | 13 | 7178642 | 7182505 | mNeoVis.2.1.1.chr13\_hap1.g00210.t01 |
| 14533 | 13 | 7178642 | 7182505 | mNeoVis.2.1.1.chr13\_hap1.g00210.t01 |
| 14534 | 13 | 7182443 | 7194482 | mNeoVis.2.1.1.chr13\_hap1.g00211.t01 |
| 14535 | 13 | 7182443 | 7194482 | mNeoVis.2.1.1.chr13\_hap1.g00211.t01 |
| 14536 | 13 | 7194745 | 7196576 | mNeoVis.2.1.1.chr13\_hap1.g00212.t01 |
| 14537 | 13 | 7194745 | 7196576 | mNeoVis.2.1.1.chr13\_hap1.g00212.t01 |
| 14538 | 13 | 7196532 | 7204829 | mNeoVis.2.1.1.chr13\_hap1.g00213.t01 |
| 14539 | 13 | 7196532 | 7204829 | mNeoVis.2.1.1.chr13\_hap1.g00213.t01 |
| 14540 | 13 | 7214772 | 7283746 | mNeoVis.2.1.1.chr13\_hap1.g00214.t01 |
| 14541 | 13 | 7214772 | 7283746 | mNeoVis.2.1.1.chr13\_hap1.g00214.t01 |
| 14542 | 13 | 7318800 | 7370391 | mNeoVis.2.1.1.chr13\_hap1.g00215.t01 |
| 14543 | 13 | 7318800 | 7370391 | mNeoVis.2.1.1.chr13\_hap1.g00215.t01 |
| 14544 | 13 | 7370540 | 7411529 | mNeoVis.2.1.1.chr13\_hap1.g00216.t01 |
| 14545 | 13 | 7370540 | 7411529 | mNeoVis.2.1.1.chr13\_hap1.g00216.t01 |
| 14546 | 13 | 7411567 | 7415140 | mNeoVis.2.1.1.chr13\_hap1.g00217.t01 |
| 14547 | 13 | 7411567 | 7415140 | mNeoVis.2.1.1.chr13\_hap1.g00217.t01 |
| 14548 | 13 | 7416876 | 7436606 | mNeoVis.2.1.1.chr13\_hap1.g00218.t01 |
| 14549 | 13 | 7416876 | 7436606 | mNeoVis.2.1.1.chr13\_hap1.g00218.t01 |
| 14550 | 13 | 7436744 | 7445363 | mNeoVis.2.1.1.chr13\_hap1.g00219.t01 |
| 14551 | 13 | 7436744 | 7445363 | mNeoVis.2.1.1.chr13\_hap1.g00219.t01 |
| 14552 | 13 | 7448390 | 7593460 | mNeoVis.2.1.1.chr13\_hap1.g00220.t01 |
| 14553 | 13 | 7600649 | 7820488 | mNeoVis.2.1.1.chr13\_hap1.g00221.t01 |
| 14554 | 13 | 7694413 | 7729415 | mNeoVis.2.1.1.chr13\_hap1.g00222.t01 |
| 14555 | 13 | 7891429 | 7915693 | mNeoVis.2.1.1.chr13\_hap1.g00223.t01 |
| 14556 | 13 | 7935659 | 7994224 | mNeoVis.2.1.1.chr13\_hap1.g00224.t01 |
| 14557 | 13 | 8041599 | 8121170 | mNeoVis.2.1.1.chr13\_hap1.g00225.t01 |
| 14558 | 13 | 8208080 | 8396172 | mNeoVis.2.1.1.chr13\_hap1.g00226.t01 |
| 14559 | 13 | 8208080 | 8396172 | mNeoVis.2.1.1.chr13\_hap1.g00226.t01 |
| 14560 | 13 | 8680719 | 8892303 | mNeoVis.2.1.1.chr13\_hap1.g00227.t01 |
| 14561 | 13 | 8922199 | 9167864 | mNeoVis.2.1.1.chr13\_hap1.g00228.t01 |
| 14562 | 13 | 8922199 | 9167864 | mNeoVis.2.1.1.chr13\_hap1.g00228.t01 |
| 14563 | 13 | 9216324 | 9290884 | mNeoVis.2.1.1.chr13\_hap1.g00229.t01 |
| 14564 | 13 | 9216324 | 9290884 | mNeoVis.2.1.1.chr13\_hap1.g00229.t01 |
| 14565 | 13 | 9292985 | 9302257 | mNeoVis.2.1.1.chr13\_hap1.g00230.t01 |
| 14566 | 13 | 9292985 | 9302257 | mNeoVis.2.1.1.chr13\_hap1.g00230.t01 |
| 14567 | 13 | 9302417 | 9307129 | mNeoVis.2.1.1.chr13\_hap1.g00231.t01 |
| 14568 | 13 | 9302417 | 9307129 | mNeoVis.2.1.1.chr13\_hap1.g00231.t01 |
| 14569 | 13 | 9309087 | 9331058 | mNeoVis.2.1.1.chr13\_hap1.g00232.t01 |
| 14570 | 13 | 9309087 | 9331058 | mNeoVis.2.1.1.chr13\_hap1.g00232.t01 |
| 14571 | 13 | 9337190 | 9374127 | mNeoVis.2.1.1.chr13\_hap1.g00233.t01 |
| 14572 | 13 | 9337190 | 9374127 | mNeoVis.2.1.1.chr13\_hap1.g00233.t01 |
| 14573 | 13 | 9377709 | 9510387 | mNeoVis.2.1.1.chr13\_hap1.g00234.t01 |
| 14574 | 13 | 9377709 | 9510387 | mNeoVis.2.1.1.chr13\_hap1.g00234.t01 |
| 14575 | 13 | 9520437 | 9568192 | mNeoVis.2.1.1.chr13\_hap1.g00235.t01 |
| 14576 | 13 | 9520437 | 9568192 | mNeoVis.2.1.1.chr13\_hap1.g00235.t01 |
| 14577 | 13 | 9569208 | 9576269 | mNeoVis.2.1.1.chr13\_hap1.g00236.t01 |
| 14578 | 13 | 9569208 | 9576269 | mNeoVis.2.1.1.chr13\_hap1.g00236.t01 |
| 14579 | 13 | 9581013 | 9585320 | mNeoVis.2.1.1.chr13\_hap1.g00237.t01 |
| 14580 | 13 | 9581013 | 9585320 | mNeoVis.2.1.1.chr13\_hap1.g00237.t01 |
| 14581 | 13 | 9585327 | 9590112 | mNeoVis.2.1.1.chr13\_hap1.g00238.t01 |
| 14582 | 13 | 9585327 | 9590112 | mNeoVis.2.1.1.chr13\_hap1.g00238.t01 |
| 14583 | 13 | 9622270 | 9650656 | mNeoVis.2.1.1.chr13\_hap1.g00239.t01 |
| 14584 | 13 | 9622270 | 9650656 | mNeoVis.2.1.1.chr13\_hap1.g00239.t01 |
| 14585 | 13 | 9656349 | 9884850 | mNeoVis.2.1.1.chr13\_hap1.g00240.t01 |
| 14586 | 13 | 9898904 | 9924350 | mNeoVis.2.1.1.chr13\_hap1.g00241.t01 |
| 14587 | 13 | 9926585 | 9945543 | mNeoVis.2.1.1.chr13\_hap1.g00242.t01 |
| 14588 | 13 | 9974828 | 10039063 | mNeoVis.2.1.1.chr13\_hap1.g00243.t01 |
| 14589 | 13 | 10040084 | 10121444 | mNeoVis.2.1.1.chr13\_hap1.g00244.t01 |
| 14590 | 13 | 10272401 | 10292799 | mNeoVis.2.1.1.chr13\_hap1.g00245.t01 |
| 14591 | 13 | 10272401 | 10292799 | mNeoVis.2.1.1.chr13\_hap1.g00245.t01 |
| 14592 | 13 | 10358101 | 10852077 | mNeoVis.2.1.1.chr13\_hap1.g00246.t01 |
| 14593 | 13 | 10852224 | 10868995 | mNeoVis.2.1.1.chr13\_hap1.g00247.t01 |
| 14594 | 13 | 11030204 | 11102456 | mNeoVis.2.1.1.chr13\_hap1.g00248.t01 |
| 14595 | 13 | 11030204 | 11102456 | mNeoVis.2.1.1.chr13\_hap1.g00248.t01 |
| 14596 | 13 | 11093551 | 11098319 | mNeoVis.2.1.1.chr13\_hap1.g00249.t01 |
| 14597 | 13 | 11093551 | 11098319 | mNeoVis.2.1.1.chr13\_hap1.g00249.t01 |
| 14598 | 13 | 11105424 | 11262861 | mNeoVis.2.1.1.chr13\_hap1.g00250.t01 |
| 14599 | 13 | 11105424 | 11262861 | mNeoVis.2.1.1.chr13\_hap1.g00250.t01 |
| 14600 | 13 | 11166852 | 11172966 | mNeoVis.2.1.1.chr13\_hap1.g00251.t01 |
| 14601 | 13 | 11166852 | 11172966 | mNeoVis.2.1.1.chr13\_hap1.g00251.t01 |
| 14602 | 13 | 11277890 | 11283962 | mNeoVis.2.1.1.chr13\_hap1.g00252.t01 |
| 14603 | 13 | 11277890 | 11283962 | mNeoVis.2.1.1.chr13\_hap1.g00252.t01 |
| 14604 | 13 | 11284033 | 11289790 | mNeoVis.2.1.1.chr13\_hap1.g00253.t01 |
| 14605 | 13 | 11284033 | 11289790 | mNeoVis.2.1.1.chr13\_hap1.g00253.t01 |
| 14606 | 13 | 11290964 | 11341737 | mNeoVis.2.1.1.chr13\_hap1.g00254.t01 |
| 14607 | 13 | 11290964 | 11341737 | mNeoVis.2.1.1.chr13\_hap1.g00254.t01 |
| 14608 | 13 | 11354492 | 11363215 | mNeoVis.2.1.1.chr13\_hap1.g00255.t01 |
| 14609 | 13 | 11354492 | 11363215 | mNeoVis.2.1.1.chr13\_hap1.g00255.t01 |
| 14610 | 13 | 11369214 | 11382226 | mNeoVis.2.1.1.chr13\_hap1.g00256.t01 |
| 14611 | 13 | 11373609 | 11374559 | mNeoVis.2.1.1.chr13\_hap1.g00257.t01 |
| 14612 | 13 | 11404524 | 11407480 | mNeoVis.2.1.1.chr13\_hap1.g00258.t01 |
| 14613 | 13 | 11421400 | 11422931 | mNeoVis.2.1.1.chr13\_hap1.g00259.t01 |
| 14614 | 13 | 11428446 | 11430022 | mNeoVis.2.1.1.chr13\_hap1.g00260.t01 |
| 14615 | 13 | 11435547 | 11437059 | mNeoVis.2.1.1.chr13\_hap1.g00261.t01 |
| 14616 | 13 | 11447308 | 11448249 | mNeoVis.2.1.1.chr13\_hap1.g00262.t01 |
| 14617 | 13 | 11456249 | 11457178 | mNeoVis.2.1.1.chr13\_hap1.g00263.t01 |
| 14618 | 13 | 12527955 | 12581879 | mNeoVis.2.1.1.chr13\_hap1.g00292.t01 |
| 14619 | 13 | 12591741 | 12607126 | mNeoVis.2.1.1.chr13\_hap1.g00293.t01 |
| 14620 | 13 | 12659817 | 12687154 | mNeoVis.2.1.1.chr13\_hap1.g00294.t01 |
| 14621 | 13 | 12690060 | 12716750 | mNeoVis.2.1.1.chr13\_hap1.g00295.t01 |
| 14622 | 13 | 12795429 | 12820237 | mNeoVis.2.1.1.chr13\_hap1.g00296.t01 |
| 14623 | 13 | 12821911 | 12912321 | mNeoVis.2.1.1.chr13\_hap1.g00297.t01 |
| 14624 | 13 | 13337239 | 13506217 | mNeoVis.2.1.1.chr13\_hap1.g00306.t01 |
| 14625 | 13 | 14347166 | 14518489 | mNeoVis.2.1.1.chr13\_hap1.g00307.t01 |
| 14626 | 13 | 14347166 | 14518489 | mNeoVis.2.1.1.chr13\_hap1.g00307.t01 |
| 14627 | 13 | 14607422 | 14656498 | mNeoVis.2.1.1.chr13\_hap1.g00308.t01 |
| 14628 | 13 | 14607422 | 14656498 | mNeoVis.2.1.1.chr13\_hap1.g00308.t01 |
| 14629 | 13 | 14984321 | 14992814 | mNeoVis.2.1.1.chr13\_hap1.g00309.t01 |
| 14630 | 13 | 14984321 | 14992814 | mNeoVis.2.1.1.chr13\_hap1.g00309.t01 |
| 14631 | 13 | 14984321 | 14992814 | mNeoVis.2.1.1.chr13\_hap1.g00309.t01 |
| 14632 | 13 | 14984321 | 14992814 | mNeoVis.2.1.1.chr13\_hap1.g00309.t01 |
| 14633 | 13 | 15853754 | 15867364 | mNeoVis.2.1.1.chr13\_hap1.g00310.t01 |
| 14634 | 13 | 15853754 | 15867364 | mNeoVis.2.1.1.chr13\_hap1.g00310.t01 |
| 14635 | 13 | 15853754 | 15867364 | mNeoVis.2.1.1.chr13\_hap1.g00310.t01 |
| 14636 | 13 | 16124894 | 16982882 | mNeoVis.2.1.1.chr13\_hap1.g00311.t01 |
| 14637 | 13 | 16768048 | 16780427 | mNeoVis.2.1.1.chr13\_hap1.g00312.t01 |
| 14638 | 13 | 17012593 | 17248392 | mNeoVis.2.1.1.chr13\_hap1.g00313.t01 |
| 14639 | 13 | 18096523 | 18192533 | mNeoVis.2.1.1.chr13\_hap1.g00314.t01 |
| 14640 | 13 | 18356629 | 18372446 | mNeoVis.2.1.1.chr13\_hap1.g00316.t01 |
| 14641 | 13 | 18457114 | 18466038 | mNeoVis.2.1.1.chr13\_hap1.g00317.t01 |
| 14642 | 13 | 18478831 | 18506017 | mNeoVis.2.1.1.chr13\_hap1.g00318.t01 |
| 14643 | 13 | 18513105 | 18520539 | mNeoVis.2.1.1.chr13\_hap1.g00319.t01 |
| 14644 | 13 | 18568439 | 18659129 | mNeoVis.2.1.1.chr13\_hap1.g00320.t01 |
| 14645 | 13 | 18671282 | 18712659 | mNeoVis.2.1.1.chr13\_hap1.g00321.t01 |
| 14646 | 13 | 18714766 | 18722347 | mNeoVis.2.1.1.chr13\_hap1.g00322.t01 |
| 14647 | 13 | 18726831 | 18733665 | mNeoVis.2.1.1.chr13\_hap1.g00323.t01 |
| 14648 | 13 | 18738796 | 18742006 | mNeoVis.2.1.1.chr13\_hap1.g00324.t01 |
| 14649 | 13 | 18750420 | 18886740 | mNeoVis.2.1.1.chr13\_hap1.g00325.t01 |
| 14650 | 13 | 18929541 | 18959714 | mNeoVis.2.1.1.chr13\_hap1.g00326.t01 |
| 14651 | 13 | 18962598 | 19148504 | mNeoVis.2.1.1.chr13\_hap1.g00327.t01 |
| 14652 | 13 | 19381938 | 19513765 | mNeoVis.2.1.1.chr13\_hap1.g00328.t01 |
| 14653 | 13 | 19381938 | 19513765 | mNeoVis.2.1.1.chr13\_hap1.g00328.t01 |
| 14654 | 13 | 19516525 | 19535355 | mNeoVis.2.1.1.chr13\_hap1.g00329.t01 |
| 14655 | 13 | 19516525 | 19535355 | mNeoVis.2.1.1.chr13\_hap1.g00329.t01 |
| 14656 | 13 | 19535495 | 19537928 | mNeoVis.2.1.1.chr13\_hap1.g00330.t01 |
| 14657 | 13 | 19535495 | 19537928 | mNeoVis.2.1.1.chr13\_hap1.g00330.t01 |
| 14658 | 13 | 19547526 | 19558326 | mNeoVis.2.1.1.chr13\_hap1.g00331.t01 |
| 14659 | 13 | 19547526 | 19558326 | mNeoVis.2.1.1.chr13\_hap1.g00331.t01 |
| 14660 | 13 | 19563142 | 19565779 | mNeoVis.2.1.1.chr13\_hap1.g00332.t01 |
| 14661 | 13 | 19563142 | 19565779 | mNeoVis.2.1.1.chr13\_hap1.g00332.t01 |
| 14662 | 13 | 19567811 | 19587043 | mNeoVis.2.1.1.chr13\_hap1.g00333.t01 |
| 14663 | 13 | 19623586 | 19632813 | mNeoVis.2.1.1.chr13\_hap1.g00335.t01 |
| 14664 | 13 | 19634133 | 19652069 | mNeoVis.2.1.1.chr13\_hap1.g00336.t01 |
| 14665 | 13 | 19652124 | 19661119 | mNeoVis.2.1.1.chr13\_hap1.g00337.t01 |
| 14666 | 13 | 19663652 | 19671390 | mNeoVis.2.1.1.chr13\_hap1.g00338.t01 |
| 14667 | 13 | 19698744 | 19754045 | mNeoVis.2.1.1.chr13\_hap1.g00339.t01 |
| 14668 | 13 | 19755958 | 19764890 | mNeoVis.2.1.1.chr13\_hap1.g00340.t01 |
| 14669 | 13 | 19810847 | 19824620 | mNeoVis.2.1.1.chr13\_hap1.g00341.t01 |
| 14670 | 13 | 19810847 | 19824620 | mNeoVis.2.1.1.chr13\_hap1.g00341.t01 |
| 14671 | 13 | 19842023 | 19843267 | mNeoVis.2.1.1.chr13\_hap1.g00342.t01 |
| 14672 | 13 | 19842023 | 19843267 | mNeoVis.2.1.1.chr13\_hap1.g00342.t01 |
| 14673 | 13 | 19844178 | 19845701 | mNeoVis.2.1.1.chr13\_hap1.g00343.t01 |
| 14674 | 13 | 19844178 | 19845701 | mNeoVis.2.1.1.chr13\_hap1.g00343.t01 |
| 14675 | 13 | 19856846 | 19864563 | mNeoVis.2.1.1.chr13\_hap1.g00344.t01 |
| 14676 | 13 | 19856846 | 19864563 | mNeoVis.2.1.1.chr13\_hap1.g00344.t01 |
| 14677 | 13 | 19928576 | 19938091 | mNeoVis.2.1.1.chr13\_hap1.g00345.t01 |
| 14678 | 13 | 19928576 | 19938091 | mNeoVis.2.1.1.chr13\_hap1.g00345.t01 |
| 14679 | 13 | 19928576 | 19938091 | mNeoVis.2.1.1.chr13\_hap1.g00345.t01 |
| 14680 | 13 | 19941892 | 20052904 | mNeoVis.2.1.1.chr13\_hap1.g00346.t01 |
| 14681 | 13 | 19941892 | 20052904 | mNeoVis.2.1.1.chr13\_hap1.g00346.t01 |
| 14682 | 13 | 19941892 | 20052904 | mNeoVis.2.1.1.chr13\_hap1.g00346.t01 |
| 14683 | 13 | 20082028 | 20100154 | mNeoVis.2.1.1.chr13\_hap1.g00347.t01 |
| 14684 | 13 | 20082028 | 20100154 | mNeoVis.2.1.1.chr13\_hap1.g00347.t01 |
| 14685 | 13 | 20082028 | 20100154 | mNeoVis.2.1.1.chr13\_hap1.g00347.t01 |
| 14686 | 13 | 20120605 | 20266449 | mNeoVis.2.1.1.chr13\_hap1.g00348.t01 |
| 14687 | 13 | 20120605 | 20266449 | mNeoVis.2.1.1.chr13\_hap1.g00348.t01 |
| 14688 | 13 | 20287036 | 20347567 | mNeoVis.2.1.1.chr13\_hap1.g00349.t01 |
| 14689 | 13 | 20287036 | 20347567 | mNeoVis.2.1.1.chr13\_hap1.g00349.t01 |
| 14690 | 13 | 20372707 | 20495385 | mNeoVis.2.1.1.chr13\_hap1.g00350.t01 |
| 14691 | 13 | 20372707 | 20495385 | mNeoVis.2.1.1.chr13\_hap1.g00350.t01 |
| 14692 | 13 | 20522801 | 20653341 | mNeoVis.2.1.1.chr13\_hap1.g00351.t01 |
| 14693 | 13 | 20751346 | 20751878 | mNeoVis.2.1.1.chr13\_hap1.g00352.t01 |
| 14694 | 13 | 20755989 | 20794830 | mNeoVis.2.1.1.chr13\_hap1.g00353.t01 |
| 14695 | 13 | 20866116 | 20947060 | mNeoVis.2.1.1.chr13\_hap1.g00354.t01 |
| 14696 | 13 | 20970053 | 20997417 | mNeoVis.2.1.1.chr13\_hap1.g00356.t01 |
| 14697 | 13 | 20997498 | 21038195 | mNeoVis.2.1.1.chr13\_hap1.g00357.t01 |
| 14698 | 13 | 21052665 | 21067004 | mNeoVis.2.1.1.chr13\_hap1.g00358.t01 |
| 14699 | 13 | 21076933 | 21172901 | mNeoVis.2.1.1.chr13\_hap1.g00359.t01 |
| 14700 | 13 | 21190777 | 21191727 | mNeoVis.2.1.1.chr13\_hap1.g00360.t01 |
| 14701 | 13 | 21307622 | 21528278 | mNeoVis.2.1.1.chr13\_hap1.g00361.t01 |
| 14702 | 13 | 21576293 | 21666602 | mNeoVis.2.1.1.chr13\_hap1.g00362.t01 |
| 14703 | 13 | 21576293 | 21666602 | mNeoVis.2.1.1.chr13\_hap1.g00362.t01 |
| 14704 | 13 | 21728976 | 21908772 | mNeoVis.2.1.1.chr13\_hap1.g00363.t01 |
| 14705 | 13 | 21728976 | 21908772 | mNeoVis.2.1.1.chr13\_hap1.g00363.t01 |
| 14706 | 13 | 21728976 | 21908772 | mNeoVis.2.1.1.chr13\_hap1.g00363.t01 |
| 14707 | 13 | 21934460 | 21958609 | mNeoVis.2.1.1.chr13\_hap1.g00364.t01 |
| 14708 | 13 | 21934460 | 21958609 | mNeoVis.2.1.1.chr13\_hap1.g00364.t01 |
| 14709 | 13 | 21934460 | 21958609 | mNeoVis.2.1.1.chr13\_hap1.g00364.t01 |
| 14710 | 13 | 21982681 | 21996273 | mNeoVis.2.1.1.chr13\_hap1.g00365.t01 |
| 14711 | 13 | 21982681 | 21996273 | mNeoVis.2.1.1.chr13\_hap1.g00365.t01 |
| 14712 | 13 | 21982681 | 21996273 | mNeoVis.2.1.1.chr13\_hap1.g00365.t01 |
| 14713 | 13 | 22028597 | 22034859 | mNeoVis.2.1.1.chr13\_hap1.g00366.t01 |
| 14714 | 13 | 22028597 | 22034859 | mNeoVis.2.1.1.chr13\_hap1.g00366.t01 |
| 14715 | 13 | 22576208 | 22721956 | mNeoVis.2.1.1.chr13\_hap1.g00368.t01 |
| 14716 | 13 | 22576208 | 22721956 | mNeoVis.2.1.1.chr13\_hap1.g00368.t01 |
| 14717 | 13 | 22576208 | 22721956 | mNeoVis.2.1.1.chr13\_hap1.g00368.t01 |
| 14718 | 13 | 22761065 | 22888740 | mNeoVis.2.1.1.chr13\_hap1.g00369.t01 |
| 14719 | 13 | 22761065 | 22888740 | mNeoVis.2.1.1.chr13\_hap1.g00369.t01 |
| 14720 | 13 | 22761065 | 22888740 | mNeoVis.2.1.1.chr13\_hap1.g00369.t01 |
| 14721 | 13 | 22892217 | 22913978 | mNeoVis.2.1.1.chr13\_hap1.g00370.t01 |
| 14722 | 13 | 22892217 | 22913978 | mNeoVis.2.1.1.chr13\_hap1.g00370.t01 |
| 14723 | 13 | 22892217 | 22913978 | mNeoVis.2.1.1.chr13\_hap1.g00370.t01 |
| 14724 | 13 | 22925951 | 23024587 | mNeoVis.2.1.1.chr13\_hap1.g00371.t01 |
| 14725 | 13 | 22925951 | 23024587 | mNeoVis.2.1.1.chr13\_hap1.g00371.t01 |
| 14726 | 13 | 22925951 | 23024587 | mNeoVis.2.1.1.chr13\_hap1.g00371.t01 |
| 14727 | 13 | 23026423 | 23088517 | mNeoVis.2.1.1.chr13\_hap1.g00372.t01 |
| 14728 | 13 | 23026423 | 23088517 | mNeoVis.2.1.1.chr13\_hap1.g00372.t01 |
| 14729 | 13 | 23026423 | 23088517 | mNeoVis.2.1.1.chr13\_hap1.g00372.t01 |
| 14730 | 13 | 23089687 | 23097106 | mNeoVis.2.1.1.chr13\_hap1.g00373.t01 |
| 14731 | 13 | 23089687 | 23097106 | mNeoVis.2.1.1.chr13\_hap1.g00373.t01 |
| 14732 | 13 | 23089687 | 23097106 | mNeoVis.2.1.1.chr13\_hap1.g00373.t01 |
| 14733 | 13 | 23097415 | 23158690 | mNeoVis.2.1.1.chr13\_hap1.g00374.t01 |
| 14734 | 13 | 23162544 | 23164045 | mNeoVis.2.1.1.chr13\_hap1.g00375.t01 |
| 14735 | 13 | 23168578 | 23170008 | mNeoVis.2.1.1.chr13\_hap1.g00376.t01 |
| 14736 | 13 | 23417073 | 23417480 | mNeoVis.2.1.1.chr13\_hap1.g00377.t01 |
| 14737 | 13 | 23436837 | 23442255 | mNeoVis.2.1.1.chr13\_hap1.g00378.t01 |
| 14738 | 13 | 24445258 | 24488662 | mNeoVis.2.1.1.chr13\_hap1.g00380.t01 |
| 14739 | 13 | 24445258 | 24488662 | mNeoVis.2.1.1.chr13\_hap1.g00380.t01 |
| 14740 | 13 | 24459811 | 24487580 | mNeoVis.2.1.1.chr13\_hap1.g00381.t01 |
| 14741 | 13 | 24459811 | 24487580 | mNeoVis.2.1.1.chr13\_hap1.g00381.t01 |
| 14742 | 13 | 24692746 | 24831439 | mNeoVis.2.1.1.chr13\_hap1.g00382.t01 |
| 14743 | 13 | 24692746 | 24831439 | mNeoVis.2.1.1.chr13\_hap1.g00382.t01 |
| 14744 | 13 | 25376942 | 25576527 | mNeoVis.2.1.1.chr13\_hap1.g00383.t01 |
| 14745 | 13 | 25713975 | 25769582 | mNeoVis.2.1.1.chr13\_hap1.g00384.t01 |
| 14746 | 13 | 25802765 | 25819115 | mNeoVis.2.1.1.chr13\_hap1.g00385.t01 |
| 14747 | 13 | 27094954 | 27098778 | mNeoVis.2.1.1.chr13\_hap1.g00406.t01 |
| 14748 | 13 | 27127741 | 27129218 | mNeoVis.2.1.1.chr13\_hap1.g00407.t01 |
| 14749 | 13 | 27235973 | 27300361 | mNeoVis.2.1.1.chr13\_hap1.g00408.t01 |
| 14750 | 13 | 28200172 | 28235250 | mNeoVis.2.1.1.chr13\_hap1.g00409.t01 |
| 14751 | 13 | 29040932 | 29192491 | mNeoVis.2.1.1.chr13\_hap1.g00410.t01 |
| 14752 | 13 | 29040932 | 29192491 | mNeoVis.2.1.1.chr13\_hap1.g00410.t01 |
| 14753 | 13 | 29168479 | 29169393 | mNeoVis.2.1.1.chr13\_hap1.g00411.t01 |
| 14754 | 13 | 29168479 | 29169393 | mNeoVis.2.1.1.chr13\_hap1.g00411.t01 |
| 14755 | 13 | 29198378 | 29224462 | mNeoVis.2.1.1.chr13\_hap1.g00412.t01 |
| 14756 | 13 | 29198378 | 29224462 | mNeoVis.2.1.1.chr13\_hap1.g00412.t01 |
| 14757 | 13 | 29246851 | 29273556 | mNeoVis.2.1.1.chr13\_hap1.g00413.t01 |
| 14758 | 13 | 29297276 | 29313806 | mNeoVis.2.1.1.chr13\_hap1.g00414.t01 |
| 14759 | 13 | 29321562 | 29335839 | mNeoVis.2.1.1.chr13\_hap1.g00415.t01 |
| 14760 | 13 | 29336049 | 29339807 | mNeoVis.2.1.1.chr13\_hap1.g00416.t01 |
| 14761 | 13 | 29363938 | 29379599 | mNeoVis.2.1.1.chr13\_hap1.g00417.t01 |
| 14762 | 13 | 29363938 | 29379599 | mNeoVis.2.1.1.chr13\_hap1.g00417.t01 |
| 14763 | 13 | 29396427 | 29964147 | mNeoVis.2.1.1.chr13\_hap1.g00418.t01 |
| 14764 | 13 | 29797285 | 29798897 | mNeoVis.2.1.1.chr13\_hap1.g00419.t01 |
| 14765 | 13 | 29797285 | 29798897 | mNeoVis.2.1.1.chr13\_hap1.g00419.t01 |
| 14766 | 13 | 29981687 | 29991455 | mNeoVis.2.1.1.chr13\_hap1.g00420.t01 |
| 14767 | 13 | 29992931 | 30078198 | mNeoVis.2.1.1.chr13\_hap1.g00421.t01 |
| 14768 | 13 | 30095341 | 30125537 | mNeoVis.2.1.1.chr13\_hap1.g00422.t01 |
| 14769 | 13 | 30195439 | 30261713 | mNeoVis.2.1.1.chr13\_hap1.g00423.t01 |
| 14770 | 13 | 30262395 | 30432512 | mNeoVis.2.1.1.chr13\_hap1.g00424.t01 |
| 14771 | 13 | 30432670 | 30528342 | mNeoVis.2.1.1.chr13\_hap1.g00425.t01 |
| 14772 | 13 | 30799048 | 31091872 | mNeoVis.2.1.1.chr13\_hap1.g00428.t01 |
| 14773 | 13 | 31195911 | 31202741 | mNeoVis.2.1.1.chr13\_hap1.g00429.t01 |
| 14774 | 13 | 31202868 | 31208069 | mNeoVis.2.1.1.chr13\_hap1.g00430.t01 |
| 14775 | 13 | 31251541 | 31309018 | mNeoVis.2.1.1.chr13\_hap1.g00431.t01 |
| 14776 | 13 | 31338765 | 31441411 | mNeoVis.2.1.1.chr13\_hap1.g00432.t01 |
| 14777 | 13 | 31571940 | 31627629 | mNeoVis.2.1.1.chr13\_hap1.g00434.t01 |
| 14778 | 13 | 31643132 | 31982697 | mNeoVis.2.1.1.chr13\_hap1.g00435.t01 |
| 14779 | 13 | 31994578 | 32046465 | mNeoVis.2.1.1.chr13\_hap1.g00436.t01 |
| 14780 | 13 | 32057457 | 32104703 | mNeoVis.2.1.1.chr13\_hap1.g00437.t01 |
| 14781 | 13 | 32106260 | 32134126 | mNeoVis.2.1.1.chr13\_hap1.g00438.t01 |
| 14782 | 13 | 32106260 | 32134126 | mNeoVis.2.1.1.chr13\_hap1.g00438.t01 |
| 14783 | 13 | 32136870 | 32160522 | mNeoVis.2.1.1.chr13\_hap1.g00439.t01 |
| 14784 | 13 | 32136870 | 32160522 | mNeoVis.2.1.1.chr13\_hap1.g00439.t01 |
| 14785 | 13 | 32174233 | 32200408 | mNeoVis.2.1.1.chr13\_hap1.g00440.t01 |
| 14786 | 13 | 32174233 | 32200408 | mNeoVis.2.1.1.chr13\_hap1.g00440.t01 |
| 14787 | 13 | 32223669 | 32248736 | mNeoVis.2.1.1.chr13\_hap1.g00441.t01 |
| 14788 | 13 | 32223669 | 32248736 | mNeoVis.2.1.1.chr13\_hap1.g00441.t01 |
| 14789 | 13 | 32254278 | 32270989 | mNeoVis.2.1.1.chr13\_hap1.g00442.t01 |
| 14790 | 13 | 32254278 | 32270989 | mNeoVis.2.1.1.chr13\_hap1.g00442.t01 |
| 14791 | 13 | 32310123 | 32314208 | mNeoVis.2.1.1.chr13\_hap1.g00443.t01 |
| 14792 | 13 | 32310123 | 32314208 | mNeoVis.2.1.1.chr13\_hap1.g00443.t01 |
| 14793 | 13 | 32342024 | 32342894 | mNeoVis.2.1.1.chr13\_hap1.g00444.t01 |
| 14794 | 13 | 32342024 | 32342894 | mNeoVis.2.1.1.chr13\_hap1.g00444.t01 |
| 14795 | 13 | 32461960 | 32484087 | mNeoVis.2.1.1.chr13\_hap1.g00445.t01 |
| 14796 | 13 | 32461960 | 32484087 | mNeoVis.2.1.1.chr13\_hap1.g00445.t01 |
| 14797 | 13 | 32487372 | 32526637 | mNeoVis.2.1.1.chr13\_hap1.g00446.t01 |
| 14798 | 13 | 32487372 | 32526637 | mNeoVis.2.1.1.chr13\_hap1.g00446.t01 |
| 14799 | 13 | 32526934 | 32558703 | mNeoVis.2.1.1.chr13\_hap1.g00447.t01 |
| 14800 | 13 | 32526934 | 32558703 | mNeoVis.2.1.1.chr13\_hap1.g00447.t01 |
| 14801 | 13 | 32550519 | 32634847 | mNeoVis.2.1.1.chr13\_hap1.g00448.t01 |
| 14802 | 13 | 32550519 | 32634847 | mNeoVis.2.1.1.chr13\_hap1.g00448.t01 |
| 14803 | 13 | 32639077 | 32712503 | mNeoVis.2.1.1.chr13\_hap1.g00449.t01 |
| 14804 | 13 | 32639077 | 32712503 | mNeoVis.2.1.1.chr13\_hap1.g00449.t01 |
| 14805 | 13 | 32727874 | 32790660 | mNeoVis.2.1.1.chr13\_hap1.g00450.t01 |
| 14806 | 13 | 32727874 | 32790660 | mNeoVis.2.1.1.chr13\_hap1.g00450.t01 |
| 14807 | 13 | 32795193 | 32847835 | mNeoVis.2.1.1.chr13\_hap1.g00451.t01 |
| 14808 | 13 | 32795193 | 32847835 | mNeoVis.2.1.1.chr13\_hap1.g00451.t01 |
| 14809 | 13 | 32857674 | 32873524 | mNeoVis.2.1.1.chr13\_hap1.g00452.t01 |
| 14810 | 13 | 32857674 | 32873524 | mNeoVis.2.1.1.chr13\_hap1.g00452.t01 |
| 14811 | 13 | 32879952 | 32884724 | mNeoVis.2.1.1.chr13\_hap1.g00453.t01 |
| 14812 | 13 | 32879952 | 32884724 | mNeoVis.2.1.1.chr13\_hap1.g00453.t01 |
| 14813 | 13 | 33123286 | 33258421 | mNeoVis.2.1.1.chr13\_hap1.g00454.t01 |
| 14814 | 13 | 33123286 | 33258421 | mNeoVis.2.1.1.chr13\_hap1.g00454.t01 |
| 14815 | 13 | 33274947 | 33294507 | mNeoVis.2.1.1.chr13\_hap1.g00455.t01 |
| 14816 | 13 | 33274947 | 33294507 | mNeoVis.2.1.1.chr13\_hap1.g00455.t01 |
| 14817 | 13 | 33300183 | 33346081 | mNeoVis.2.1.1.chr13\_hap1.g00456.t01 |
| 14818 | 13 | 33300183 | 33346081 | mNeoVis.2.1.1.chr13\_hap1.g00456.t01 |
| 14819 | 13 | 33364319 | 33370289 | mNeoVis.2.1.1.chr13\_hap1.g00457.t01 |
| 14820 | 13 | 33364319 | 33370289 | mNeoVis.2.1.1.chr13\_hap1.g00457.t01 |
| 14821 | 13 | 33371629 | 33387723 | mNeoVis.2.1.1.chr13\_hap1.g00458.t01 |
| 14822 | 13 | 33371629 | 33387723 | mNeoVis.2.1.1.chr13\_hap1.g00458.t01 |
| 14823 | 13 | 33391764 | 33534959 | mNeoVis.2.1.1.chr13\_hap1.g00459.t01 |
| 14824 | 13 | 33391764 | 33534959 | mNeoVis.2.1.1.chr13\_hap1.g00459.t01 |
| 14825 | 13 | 33545559 | 33555803 | mNeoVis.2.1.1.chr13\_hap1.g00460.t01 |
| 14826 | 13 | 33545559 | 33555803 | mNeoVis.2.1.1.chr13\_hap1.g00460.t01 |
| 14827 | 13 | 33549925 | 33619547 | mNeoVis.2.1.1.chr13\_hap1.g00461.t01 |
| 14828 | 13 | 33549925 | 33619547 | mNeoVis.2.1.1.chr13\_hap1.g00461.t01 |
| 14829 | 13 | 33636326 | 33652534 | mNeoVis.2.1.1.chr13\_hap1.g00462.t01 |
| 14830 | 13 | 33636326 | 33652534 | mNeoVis.2.1.1.chr13\_hap1.g00462.t01 |
| 14831 | 13 | 33663164 | 33676855 | mNeoVis.2.1.1.chr13\_hap1.g00463.t01 |
| 14832 | 13 | 33663164 | 33676855 | mNeoVis.2.1.1.chr13\_hap1.g00463.t01 |
| 14833 | 13 | 33677948 | 33687522 | mNeoVis.2.1.1.chr13\_hap1.g00464.t01 |
| 14834 | 13 | 33677948 | 33687522 | mNeoVis.2.1.1.chr13\_hap1.g00464.t01 |
| 14835 | 13 | 33748233 | 33997224 | mNeoVis.2.1.1.chr13\_hap1.g00465.t01 |
| 14836 | 13 | 34036003 | 34226497 | mNeoVis.2.1.1.chr13\_hap1.g00466.t01 |
| 14837 | 13 | 34363564 | 34462588 | mNeoVis.2.1.1.chr13\_hap1.g00467.t01 |
| 14838 | 13 | 34532471 | 34673236 | mNeoVis.2.1.1.chr13\_hap1.g00468.t01 |
| 14839 | 13 | 34687696 | 34735733 | mNeoVis.2.1.1.chr13\_hap1.g00469.t01 |
| 14840 | 13 | 34709341 | 34757533 | mNeoVis.2.1.1.chr13\_hap1.g00470.t01 |
| 14841 | 13 | 34719554 | 34720930 | mNeoVis.2.1.1.chr13\_hap1.g00471.t01 |
| 14842 | 13 | 34764603 | 34784842 | mNeoVis.2.1.1.chr13\_hap1.g00472.t01 |
| 14843 | 13 | 34806442 | 34808404 | mNeoVis.2.1.1.chr13\_hap1.g00473.t01 |
| 14844 | 13 | 34812273 | 34831583 | mNeoVis.2.1.1.chr13\_hap1.g00474.t01 |
| 14845 | 13 | 34846131 | 34922106 | mNeoVis.2.1.1.chr13\_hap1.g00475.t01 |
| 14846 | 13 | 34938044 | 34949678 | mNeoVis.2.1.1.chr13\_hap1.g00476.t01 |
| 14847 | 13 | 34948309 | 34949265 | mNeoVis.2.1.1.chr13\_hap1.g00477.t01 |
| 14848 | 13 | 34954582 | 34955538 | mNeoVis.2.1.1.chr13\_hap1.g00478.t01 |
| 14849 | 13 | 34968865 | 34969824 | mNeoVis.2.1.1.chr13\_hap1.g00479.t01 |
| 14850 | 13 | 34978216 | 34979175 | mNeoVis.2.1.1.chr13\_hap1.g00480.t01 |
| 14851 | 13 | 34992002 | 34992961 | mNeoVis.2.1.1.chr13\_hap1.g00481.t01 |
| 14852 | 13 | 35023943 | 35024902 | mNeoVis.2.1.1.chr13\_hap1.g00482.t01 |
| 14853 | 13 | 35045273 | 35046232 | mNeoVis.2.1.1.chr13\_hap1.g00483.t01 |
| 14854 | 13 | 35045273 | 35046232 | mNeoVis.2.1.1.chr13\_hap1.g00483.t01 |
| 14855 | 13 | 35066969 | 35067928 | mNeoVis.2.1.1.chr13\_hap1.g00484.t01 |
| 14856 | 13 | 35066969 | 35067928 | mNeoVis.2.1.1.chr13\_hap1.g00484.t01 |
| 14857 | 13 | 35084911 | 35107396 | mNeoVis.2.1.1.chr13\_hap1.g00485.t01 |
| 14858 | 13 | 35084911 | 35107396 | mNeoVis.2.1.1.chr13\_hap1.g00485.t01 |
| 14859 | 13 | 35084921 | 35088314 | mNeoVis.2.1.1.chr13\_hap1.g00486.t01 |
| 14860 | 13 | 35084921 | 35088314 | mNeoVis.2.1.1.chr13\_hap1.g00486.t01 |
| 14861 | 13 | 35127469 | 35129544 | mNeoVis.2.1.1.chr13\_hap1.g00487.t01 |
| 14862 | 13 | 35127469 | 35129544 | mNeoVis.2.1.1.chr13\_hap1.g00487.t01 |
| 14863 | 13 | 35156809 | 35158906 | mNeoVis.2.1.1.chr13\_hap1.g00488.t01 |
| 14864 | 13 | 35156809 | 35158906 | mNeoVis.2.1.1.chr13\_hap1.g00488.t01 |
| 14865 | 13 | 35162829 | 35170352 | mNeoVis.2.1.1.chr13\_hap1.g00489.t01 |
| 14866 | 13 | 35162829 | 35170352 | mNeoVis.2.1.1.chr13\_hap1.g00489.t01 |
| 14867 | 13 | 35170853 | 35197952 | mNeoVis.2.1.1.chr13\_hap1.g00490.t01 |
| 14868 | 13 | 35170853 | 35197952 | mNeoVis.2.1.1.chr13\_hap1.g00490.t01 |
| 14869 | 13 | 35198625 | 35208490 | mNeoVis.2.1.1.chr13\_hap1.g00491.t01 |
| 14870 | 13 | 35198625 | 35208490 | mNeoVis.2.1.1.chr13\_hap1.g00491.t01 |
| 14871 | 13 | 35212016 | 35214138 | mNeoVis.2.1.1.chr13\_hap1.g00492.t01 |
| 14872 | 13 | 35212016 | 35214138 | mNeoVis.2.1.1.chr13\_hap1.g00492.t01 |
| 14873 | 13 | 35214885 | 35217130 | mNeoVis.2.1.1.chr13\_hap1.g00493.t01 |
| 14874 | 13 | 35214885 | 35217130 | mNeoVis.2.1.1.chr13\_hap1.g00493.t01 |
| 14875 | 13 | 35217222 | 35235397 | mNeoVis.2.1.1.chr13\_hap1.g00494.t01 |
| 14876 | 13 | 35217222 | 35235397 | mNeoVis.2.1.1.chr13\_hap1.g00494.t01 |
| 14877 | 13 | 35270070 | 35271105 | mNeoVis.2.1.1.chr13\_hap1.g00495.t01 |
| 14878 | 13 | 35270070 | 35271105 | mNeoVis.2.1.1.chr13\_hap1.g00495.t01 |
| 14879 | 13 | 35271155 | 35274471 | mNeoVis.2.1.1.chr13\_hap1.g00496.t01 |
| 14880 | 13 | 35271155 | 35274471 | mNeoVis.2.1.1.chr13\_hap1.g00496.t01 |
| 14881 | 13 | 35274346 | 35286988 | mNeoVis.2.1.1.chr13\_hap1.g00497.t01 |
| 14882 | 13 | 35274346 | 35286988 | mNeoVis.2.1.1.chr13\_hap1.g00497.t01 |
| 14883 | 13 | 35286855 | 35292813 | mNeoVis.2.1.1.chr13\_hap1.g00498.t01 |
| 14884 | 13 | 35286855 | 35292813 | mNeoVis.2.1.1.chr13\_hap1.g00498.t01 |
| 14885 | 13 | 35292879 | 35325371 | mNeoVis.2.1.1.chr13\_hap1.g00499.t01 |
| 14886 | 13 | 35292879 | 35325371 | mNeoVis.2.1.1.chr13\_hap1.g00499.t01 |
| 14887 | 13 | 35292879 | 35325371 | mNeoVis.2.1.1.chr13\_hap1.g00499.t01 |
| 14888 | 13 | 35330396 | 35337763 | mNeoVis.2.1.1.chr13\_hap1.g00500.t01 |
| 14889 | 13 | 35330396 | 35337763 | mNeoVis.2.1.1.chr13\_hap1.g00500.t01 |
| 14890 | 13 | 35330396 | 35337763 | mNeoVis.2.1.1.chr13\_hap1.g00500.t01 |
| 14891 | 13 | 35338714 | 35344797 | mNeoVis.2.1.1.chr13\_hap1.g00501.t01 |
| 14892 | 13 | 35338714 | 35344797 | mNeoVis.2.1.1.chr13\_hap1.g00501.t01 |
| 14893 | 13 | 35338714 | 35344797 | mNeoVis.2.1.1.chr13\_hap1.g00501.t01 |
| 14894 | 13 | 35350897 | 35357193 | mNeoVis.2.1.1.chr13\_hap1.g00502.t01 |
| 14895 | 13 | 35350897 | 35357193 | mNeoVis.2.1.1.chr13\_hap1.g00502.t01 |
| 14896 | 13 | 35350897 | 35357193 | mNeoVis.2.1.1.chr13\_hap1.g00502.t01 |
| 14897 | 13 | 35355076 | 35357804 | mNeoVis.2.1.1.chr13\_hap1.g00503.t01 |
| 14898 | 13 | 35355076 | 35357804 | mNeoVis.2.1.1.chr13\_hap1.g00503.t01 |
| 14899 | 13 | 35355076 | 35357804 | mNeoVis.2.1.1.chr13\_hap1.g00503.t01 |
| 14900 | 13 | 35359504 | 35366829 | mNeoVis.2.1.1.chr13\_hap1.g00504.t01 |
| 14901 | 13 | 35359504 | 35366829 | mNeoVis.2.1.1.chr13\_hap1.g00504.t01 |
| 14902 | 13 | 35359504 | 35366829 | mNeoVis.2.1.1.chr13\_hap1.g00504.t01 |
| 14903 | 13 | 35374080 | 35381309 | mNeoVis.2.1.1.chr13\_hap1.g00505.t01 |
| 14904 | 13 | 35374080 | 35381309 | mNeoVis.2.1.1.chr13\_hap1.g00505.t01 |
| 14905 | 13 | 35374080 | 35381309 | mNeoVis.2.1.1.chr13\_hap1.g00505.t01 |
| 14906 | 13 | 35381448 | 35385882 | mNeoVis.2.1.1.chr13\_hap1.g00506.t01 |
| 14907 | 13 | 35381448 | 35385882 | mNeoVis.2.1.1.chr13\_hap1.g00506.t01 |
| 14908 | 13 | 35381448 | 35385882 | mNeoVis.2.1.1.chr13\_hap1.g00506.t01 |
| 14909 | 13 | 35387355 | 35404205 | mNeoVis.2.1.1.chr13\_hap1.g00507.t01 |
| 14910 | 13 | 35387355 | 35404205 | mNeoVis.2.1.1.chr13\_hap1.g00507.t01 |
| 14911 | 13 | 35387355 | 35404205 | mNeoVis.2.1.1.chr13\_hap1.g00507.t01 |
| 14912 | 13 | 35404266 | 35476242 | mNeoVis.2.1.1.chr13\_hap1.g00508.t01 |
| 14913 | 13 | 35404266 | 35476242 | mNeoVis.2.1.1.chr13\_hap1.g00508.t01 |
| 14914 | 13 | 35404266 | 35476242 | mNeoVis.2.1.1.chr13\_hap1.g00508.t01 |
| 14915 | 13 | 35482050 | 35610398 | mNeoVis.2.1.1.chr13\_hap1.g00509.t01 |
| 14916 | 13 | 35482050 | 35610398 | mNeoVis.2.1.1.chr13\_hap1.g00509.t01 |
| 14917 | 13 | 35611802 | 35674383 | mNeoVis.2.1.1.chr13\_hap1.g00510.t01 |
| 14918 | 13 | 35611802 | 35674383 | mNeoVis.2.1.1.chr13\_hap1.g00510.t01 |
| 14919 | 13 | 35762533 | 35762996 | mNeoVis.2.1.1.chr13\_hap1.g00511.t01 |
| 14920 | 13 | 35762533 | 35762996 | mNeoVis.2.1.1.chr13\_hap1.g00511.t01 |
| 14921 | 13 | 35867735 | 35880090 | mNeoVis.2.1.1.chr13\_hap1.g00512.t01 |
| 14922 | 13 | 35881010 | 35884350 | mNeoVis.2.1.1.chr13\_hap1.g00513.t01 |
| 14923 | 13 | 35889322 | 35896951 | mNeoVis.2.1.1.chr13\_hap1.g00514.t01 |
| 14924 | 13 | 35901143 | 35908323 | mNeoVis.2.1.1.chr13\_hap1.g00515.t01 |
| 14925 | 13 | 35908781 | 35924399 | mNeoVis.2.1.1.chr13\_hap1.g00516.t01 |
| 14926 | 13 | 35929589 | 35933779 | mNeoVis.2.1.1.chr13\_hap1.g00517.t01 |
| 14927 | 13 | 35950157 | 35959433 | mNeoVis.2.1.1.chr13\_hap1.g00518.t01 |
| 14928 | 13 | 35966025 | 35996733 | mNeoVis.2.1.1.chr13\_hap1.g00519.t01 |
| 14929 | 13 | 35995562 | 35998785 | mNeoVis.2.1.1.chr13\_hap1.g00520.t01 |
| 14930 | 13 | 36085336 | 36091935 | mNeoVis.2.1.1.chr13\_hap1.g00521.t01 |
| 14931 | 13 | 36117212 | 36123826 | mNeoVis.2.1.1.chr13\_hap1.g00522.t01 |
| 14932 | 13 | 36191563 | 36192743 | mNeoVis.2.1.1.chr13\_hap1.g00523.t01 |
| 14933 | 13 | 36204025 | 36205818 | mNeoVis.2.1.1.chr13\_hap1.g00524.t01 |
| 14934 | 13 | 36221787 | 36238020 | mNeoVis.2.1.1.chr13\_hap1.g00525.t01 |
| 14935 | 13 | 36224410 | 36225404 | mNeoVis.2.1.1.chr13\_hap1.g00526.t01 |
| 14936 | 13 | 36224434 | 36228898 | mNeoVis.2.1.1.chr13\_hap1.g00527.t01 |
| 14937 | 13 | 36239723 | 36243037 | mNeoVis.2.1.1.chr13\_hap1.g00528.t01 |
| 14938 | 13 | 36250230 | 36252279 | mNeoVis.2.1.1.chr13\_hap1.g00529.t01 |
| 14939 | 13 | 36261241 | 36268945 | mNeoVis.2.1.1.chr13\_hap1.g00530.t01 |
| 14940 | 13 | 36269684 | 36275843 | mNeoVis.2.1.1.chr13\_hap1.g00531.t01 |
| 14941 | 13 | 36277314 | 36279663 | mNeoVis.2.1.1.chr13\_hap1.g00532.t01 |
| 14942 | 13 | 36298881 | 36336674 | mNeoVis.2.1.1.chr13\_hap1.g00533.t01 |
| 14943 | 13 | 36356544 | 36358513 | mNeoVis.2.1.1.chr13\_hap1.g00534.t01 |
| 14944 | 13 | 36358572 | 36416791 | mNeoVis.2.1.1.chr13\_hap1.g00535.t01 |
| 14945 | 13 | 36416893 | 36472959 | mNeoVis.2.1.1.chr13\_hap1.g00536.t01 |
| 14946 | 13 | 36473332 | 36497637 | mNeoVis.2.1.1.chr13\_hap1.g00537.t01 |
| 14947 | 13 | 36502893 | 36514882 | mNeoVis.2.1.1.chr13\_hap1.g00538.t01 |
| 14948 | 13 | 36515031 | 36598081 | mNeoVis.2.1.1.chr13\_hap1.g00539.t01 |
| 14949 | 13 | 36599223 | 36700147 | mNeoVis.2.1.1.chr13\_hap1.g00540.t01 |
| 14950 | 13 | 36700289 | 36731763 | mNeoVis.2.1.1.chr13\_hap1.g00541.t01 |
| 14951 | 13 | 36784511 | 36921897 | mNeoVis.2.1.1.chr13\_hap1.g00542.t01 |
| 14952 | 13 | 36925857 | 37045126 | mNeoVis.2.1.1.chr13\_hap1.g00543.t01 |
| 14953 | 13 | 37054654 | 37067670 | mNeoVis.2.1.1.chr13\_hap1.g00544.t01 |
| 14954 | 13 | 37077143 | 37083628 | mNeoVis.2.1.1.chr13\_hap1.g00545.t01 |
| 14955 | 13 | 37118945 | 37130559 | mNeoVis.2.1.1.chr13\_hap1.g00546.t01 |
| 14956 | 13 | 37141024 | 37221808 | mNeoVis.2.1.1.chr13\_hap1.g00547.t01 |
| 14957 | 13 | 37231363 | 37249363 | mNeoVis.2.1.1.chr13\_hap1.g00548.t01 |
| 14958 | 13 | 37249678 | 37261413 | mNeoVis.2.1.1.chr13\_hap1.g00549.t01 |
| 14959 | 13 | 37265269 | 37287884 | mNeoVis.2.1.1.chr13\_hap1.g00550.t01 |
| 14960 | 13 | 37342077 | 37393507 | mNeoVis.2.1.1.chr13\_hap1.g00551.t01 |
| 14961 | 13 | 37449599 | 37459855 | mNeoVis.2.1.1.chr13\_hap1.g00553.t01 |
| 14962 | 13 | 37482102 | 37556573 | mNeoVis.2.1.1.chr13\_hap1.g00554.t01 |
| 14963 | 13 | 37636753 | 37642871 | mNeoVis.2.1.1.chr13\_hap1.g00555.t01 |
| 14964 | 13 | 37786328 | 37797238 | mNeoVis.2.1.1.chr13\_hap1.g00556.t01 |
| 14965 | 13 | 37786328 | 37797238 | mNeoVis.2.1.1.chr13\_hap1.g00556.t01 |
| 14966 | 13 | 37797730 | 37798520 | mNeoVis.2.1.1.chr13\_hap1.g00557.t01 |
| 14967 | 13 | 37797730 | 37798520 | mNeoVis.2.1.1.chr13\_hap1.g00557.t01 |
| 14968 | 13 | 37798978 | 37809003 | mNeoVis.2.1.1.chr13\_hap1.g00558.t01 |
| 14969 | 13 | 37798978 | 37809003 | mNeoVis.2.1.1.chr13\_hap1.g00558.t01 |
| 14970 | 13 | 37818956 | 37857484 | mNeoVis.2.1.1.chr13\_hap1.g00559.t01 |
| 14971 | 13 | 37818956 | 37857484 | mNeoVis.2.1.1.chr13\_hap1.g00559.t01 |
| 14972 | 13 | 37861646 | 37918077 | mNeoVis.2.1.1.chr13\_hap1.g00560.t01 |
| 14973 | 13 | 37861646 | 37918077 | mNeoVis.2.1.1.chr13\_hap1.g00560.t01 |
| 14974 | 13 | 38794552 | 38795829 | mNeoVis.2.1.1.chr13\_hap1.g00561.t01 |
| 14975 | 13 | 40127979 | 40465694 | mNeoVis.2.1.1.chr13\_hap1.g00562.t01 |
| 14976 | 13 | 40545070 | 41738783 | mNeoVis.2.1.1.chr13\_hap1.g00563.t01 |
| 14977 | 13 | 41897419 | 42068011 | mNeoVis.2.1.1.chr13\_hap1.g00564.t01 |
| 14978 | 13 | 42068052 | 42092671 | mNeoVis.2.1.1.chr13\_hap1.g00565.t01 |
| 14979 | 13 | 42068052 | 42092671 | mNeoVis.2.1.1.chr13\_hap1.g00565.t01 |
| 14980 | 13 | 42068052 | 42092671 | mNeoVis.2.1.1.chr13\_hap1.g00565.t01 |
| 14981 | 13 | 42112347 | 42113985 | mNeoVis.2.1.1.chr13\_hap1.g00566.t01 |
| 14982 | 13 | 42112347 | 42113985 | mNeoVis.2.1.1.chr13\_hap1.g00566.t01 |
| 14983 | 13 | 42112347 | 42113985 | mNeoVis.2.1.1.chr13\_hap1.g00566.t01 |
| 14984 | 13 | 42113561 | 42316362 | mNeoVis.2.1.1.chr13\_hap1.g00567.t01 |
| 14985 | 13 | 42113561 | 42316362 | mNeoVis.2.1.1.chr13\_hap1.g00567.t01 |
| 14986 | 13 | 42113561 | 42316362 | mNeoVis.2.1.1.chr13\_hap1.g00567.t01 |
| 14987 | 13 | 42118482 | 42119278 | mNeoVis.2.1.1.chr13\_hap1.g00568.t01 |
| 14988 | 13 | 42118482 | 42119278 | mNeoVis.2.1.1.chr13\_hap1.g00568.t01 |
| 14989 | 13 | 42118482 | 42119278 | mNeoVis.2.1.1.chr13\_hap1.g00568.t01 |
| 14990 | 13 | 42127838 | 42129045 | mNeoVis.2.1.1.chr13\_hap1.g00569.t01 |
| 14991 | 13 | 42127838 | 42129045 | mNeoVis.2.1.1.chr13\_hap1.g00569.t01 |
| 14992 | 13 | 42127838 | 42129045 | mNeoVis.2.1.1.chr13\_hap1.g00569.t01 |
| 14993 | 13 | 42329478 | 42342088 | mNeoVis.2.1.1.chr13\_hap1.g00570.t01 |
| 14994 | 13 | 42329478 | 42342088 | mNeoVis.2.1.1.chr13\_hap1.g00570.t01 |
| 14995 | 13 | 42329478 | 42342088 | mNeoVis.2.1.1.chr13\_hap1.g00570.t01 |
| 14996 | 13 | 42357853 | 42392909 | mNeoVis.2.1.1.chr13\_hap1.g00571.t01 |
| 14997 | 13 | 42357853 | 42392909 | mNeoVis.2.1.1.chr13\_hap1.g00571.t01 |
| 14998 | 13 | 42357853 | 42392909 | mNeoVis.2.1.1.chr13\_hap1.g00571.t01 |
| 14999 | 13 | 42388376 | 42488117 | mNeoVis.2.1.1.chr13\_hap1.g00572.t01 |
| 15000 | 13 | 42388376 | 42488117 | mNeoVis.2.1.1.chr13\_hap1.g00572.t01 |
| 15001 | 13 | 42388376 | 42488117 | mNeoVis.2.1.1.chr13\_hap1.g00572.t01 |
| 15002 | 13 | 42523928 | 42606260 | mNeoVis.2.1.1.chr13\_hap1.g00573.t01 |
| 15003 | 13 | 42523928 | 42606260 | mNeoVis.2.1.1.chr13\_hap1.g00573.t01 |
| 15004 | 13 | 42523928 | 42606260 | mNeoVis.2.1.1.chr13\_hap1.g00573.t01 |
| 15005 | 13 | 42577494 | 42584150 | mNeoVis.2.1.1.chr13\_hap1.g00574.t01 |
| 15006 | 13 | 42577494 | 42584150 | mNeoVis.2.1.1.chr13\_hap1.g00574.t01 |
| 15007 | 13 | 42577494 | 42584150 | mNeoVis.2.1.1.chr13\_hap1.g00574.t01 |
| 15008 | 13 | 42608553 | 42633125 | mNeoVis.2.1.1.chr13\_hap1.g00575.t01 |
| 15009 | 13 | 42608553 | 42633125 | mNeoVis.2.1.1.chr13\_hap1.g00575.t01 |
| 15010 | 13 | 42608553 | 42633125 | mNeoVis.2.1.1.chr13\_hap1.g00575.t01 |
| 15011 | 13 | 42643771 | 42703910 | mNeoVis.2.1.1.chr13\_hap1.g00576.t01 |
| 15012 | 13 | 42643771 | 42703910 | mNeoVis.2.1.1.chr13\_hap1.g00576.t01 |
| 15013 | 13 | 42643771 | 42703910 | mNeoVis.2.1.1.chr13\_hap1.g00576.t01 |
| 15014 | 13 | 42860577 | 42873967 | mNeoVis.2.1.1.chr13\_hap1.g00577.t01 |
| 15015 | 13 | 42860577 | 42873967 | mNeoVis.2.1.1.chr13\_hap1.g00577.t01 |
| 15016 | 13 | 43663303 | 43664707 | mNeoVis.2.1.1.chr13\_hap1.g00578.t01 |
| 15017 | 13 | 43663303 | 43664707 | mNeoVis.2.1.1.chr13\_hap1.g00578.t01 |
| 15018 | 13 | 44610161 | 44612830 | mNeoVis.2.1.1.chr13\_hap1.g00579.t01 |
| 15019 | 13 | 45187838 | 45333636 | mNeoVis.2.1.1.chr13\_hap1.g00580.t01 |
| 15020 | 13 | 46319741 | 46324779 | mNeoVis.2.1.1.chr13\_hap1.g00581.t01 |
| 15021 | 13 | 46319741 | 46324779 | mNeoVis.2.1.1.chr13\_hap1.g00581.t01 |
| 15022 | 13 | 46319741 | 46324779 | mNeoVis.2.1.1.chr13\_hap1.g00581.t01 |
| 15023 | 13 | 46319741 | 46324779 | mNeoVis.2.1.1.chr13\_hap1.g00581.t01 |
| 15024 | 13 | 46539299 | 46646856 | mNeoVis.2.1.1.chr13\_hap1.g00582.t01 |
| 15025 | 13 | 46539299 | 46646856 | mNeoVis.2.1.1.chr13\_hap1.g00582.t01 |
| 15026 | 13 | 46539299 | 46646856 | mNeoVis.2.1.1.chr13\_hap1.g00582.t01 |
| 15027 | 13 | 46647407 | 46655938 | mNeoVis.2.1.1.chr13\_hap1.g00583.t01 |
| 15028 | 13 | 46647407 | 46655938 | mNeoVis.2.1.1.chr13\_hap1.g00583.t01 |
| 15029 | 13 | 46647407 | 46655938 | mNeoVis.2.1.1.chr13\_hap1.g00583.t01 |
| 15030 | 13 | 46683297 | 46689603 | mNeoVis.2.1.1.chr13\_hap1.g00584.t01 |
| 15031 | 13 | 46683297 | 46689603 | mNeoVis.2.1.1.chr13\_hap1.g00584.t01 |
| 15032 | 13 | 46683297 | 46689603 | mNeoVis.2.1.1.chr13\_hap1.g00584.t01 |
| 15033 | 13 | 46711318 | 46755971 | mNeoVis.2.1.1.chr13\_hap1.g00585.t01 |
| 15034 | 13 | 46711318 | 46755971 | mNeoVis.2.1.1.chr13\_hap1.g00585.t01 |
| 15035 | 13 | 46711318 | 46755971 | mNeoVis.2.1.1.chr13\_hap1.g00585.t01 |
| 15036 | 13 | 47036792 | 47037631 | mNeoVis.2.1.1.chr13\_hap1.g00587.t01 |
| 15037 | 13 | 47036792 | 47037631 | mNeoVis.2.1.1.chr13\_hap1.g00587.t01 |
| 15038 | 13 | 47050166 | 47050753 | mNeoVis.2.1.1.chr13\_hap1.g00588.t01 |
| 15039 | 13 | 47050166 | 47050753 | mNeoVis.2.1.1.chr13\_hap1.g00588.t01 |
| 15040 | 13 | 47063753 | 47064388 | mNeoVis.2.1.1.chr13\_hap1.g00589.t01 |
| 15041 | 13 | 47069693 | 47070280 | mNeoVis.2.1.1.chr13\_hap1.g00590.t01 |
| 15042 | 13 | 47075587 | 47082553 | mNeoVis.2.1.1.chr13\_hap1.g00591.t01 |
| 15043 | 13 | 47093892 | 47094455 | mNeoVis.2.1.1.chr13\_hap1.g00592.t01 |
| 15044 | 13 | 47099788 | 47100351 | mNeoVis.2.1.1.chr13\_hap1.g00593.t01 |
| 15045 | 13 | 47114593 | 47115308 | mNeoVis.2.1.1.chr13\_hap1.g00594.t01 |
| 15046 | 13 | 47129872 | 47130549 | mNeoVis.2.1.1.chr13\_hap1.g00595.t01 |
| 15047 | 13 | 47148850 | 47149485 | mNeoVis.2.1.1.chr13\_hap1.g00596.t01 |
| 15048 | 13 | 47182475 | 47186557 | mNeoVis.2.1.1.chr13\_hap1.g00597.t01 |
| 15049 | 13 | 47206268 | 47206936 | mNeoVis.2.1.1.chr13\_hap1.g00598.t01 |
| 15050 | 13 | 47222823 | 47223386 | mNeoVis.2.1.1.chr13\_hap1.g00599.t01 |
| 15051 | 13 | 47228924 | 47229559 | mNeoVis.2.1.1.chr13\_hap1.g00600.t01 |
| 15052 | 13 | 47243692 | 47244369 | mNeoVis.2.1.1.chr13\_hap1.g00601.t01 |
| 15053 | 13 | 47249804 | 47250367 | mNeoVis.2.1.1.chr13\_hap1.g00602.t01 |
| 15054 | 13 | 47254654 | 47255313 | mNeoVis.2.1.1.chr13\_hap1.g00603.t01 |
| 15055 | 13 | 47261047 | 47261633 | mNeoVis.2.1.1.chr13\_hap1.g00604.t01 |
| 15056 | 13 | 47286623 | 47287180 | mNeoVis.2.1.1.chr13\_hap1.g00605.t01 |
| 15057 | 13 | 47286623 | 47287180 | mNeoVis.2.1.1.chr13\_hap1.g00605.t01 |
| 15058 | 13 | 47310093 | 47310653 | mNeoVis.2.1.1.chr13\_hap1.g00606.t01 |
| 15059 | 13 | 47310093 | 47310653 | mNeoVis.2.1.1.chr13\_hap1.g00606.t01 |
| 15060 | 13 | 47337201 | 47391079 | mNeoVis.2.1.1.chr13\_hap1.g00607.t01 |
| 15061 | 13 | 47337201 | 47391079 | mNeoVis.2.1.1.chr13\_hap1.g00607.t01 |
| 15062 | 13 | 47394730 | 47701002 | mNeoVis.2.1.1.chr13\_hap1.g00608.t01 |
| 15063 | 13 | 47764459 | 48052355 | mNeoVis.2.1.1.chr13\_hap1.g00609.t01 |
| 15064 | 13 | 47764459 | 48052355 | mNeoVis.2.1.1.chr13\_hap1.g00609.t01 |
| 15065 | 13 | 48201319 | 48202595 | mNeoVis.2.1.1.chr13\_hap1.g00610.t01 |
| 15066 | 13 | 48201319 | 48202595 | mNeoVis.2.1.1.chr13\_hap1.g00610.t01 |
| 15067 | 13 | 48201319 | 48202595 | mNeoVis.2.1.1.chr13\_hap1.g00610.t01 |
| 15068 | 13 | 48451291 | 48698390 | mNeoVis.2.1.1.chr13\_hap1.g00611.t01 |
| 15069 | 13 | 48740538 | 48773672 | mNeoVis.2.1.1.chr13\_hap1.g00612.t01 |
| 15070 | 13 | 48740538 | 48773672 | mNeoVis.2.1.1.chr13\_hap1.g00612.t01 |
| 15071 | 13 | 48740538 | 48773672 | mNeoVis.2.1.1.chr13\_hap1.g00612.t01 |
| 15072 | 13 | 48798370 | 48802675 | mNeoVis.2.1.1.chr13\_hap1.g00613.t01 |
| 15073 | 13 | 48798370 | 48802675 | mNeoVis.2.1.1.chr13\_hap1.g00613.t01 |
| 15074 | 13 | 48798370 | 48802675 | mNeoVis.2.1.1.chr13\_hap1.g00613.t01 |
| 15075 | 13 | 48802966 | 48941191 | mNeoVis.2.1.1.chr13\_hap1.g00614.t01 |
| 15076 | 13 | 48802966 | 48941191 | mNeoVis.2.1.1.chr13\_hap1.g00614.t01 |
| 15077 | 13 | 48802966 | 48941191 | mNeoVis.2.1.1.chr13\_hap1.g00614.t01 |
| 15078 | 13 | 49027948 | 49046195 | mNeoVis.2.1.1.chr13\_hap1.g00615.t01 |
| 15079 | 13 | 49027948 | 49046195 | mNeoVis.2.1.1.chr13\_hap1.g00615.t01 |
| 15080 | 13 | 49027948 | 49046195 | mNeoVis.2.1.1.chr13\_hap1.g00615.t01 |
| 15081 | 13 | 49035570 | 49095367 | mNeoVis.2.1.1.chr13\_hap1.g00616.t01 |
| 15082 | 13 | 49035570 | 49095367 | mNeoVis.2.1.1.chr13\_hap1.g00616.t01 |
| 15083 | 13 | 49099826 | 49101437 | mNeoVis.2.1.1.chr13\_hap1.g00617.t01 |
| 15084 | 13 | 49099826 | 49101437 | mNeoVis.2.1.1.chr13\_hap1.g00617.t01 |
| 15085 | 13 | 49099826 | 49101437 | mNeoVis.2.1.1.chr13\_hap1.g00617.t01 |
| 15086 | 13 | 49110586 | 49217546 | mNeoVis.2.1.1.chr13\_hap1.g00618.t01 |
| 15087 | 13 | 49110586 | 49217546 | mNeoVis.2.1.1.chr13\_hap1.g00618.t01 |
| 15088 | 13 | 49110586 | 49217546 | mNeoVis.2.1.1.chr13\_hap1.g00618.t01 |
| 15089 | 13 | 49175696 | 49176641 | mNeoVis.2.1.1.chr13\_hap1.g00619.t01 |
| 15090 | 13 | 49175696 | 49176641 | mNeoVis.2.1.1.chr13\_hap1.g00619.t01 |
| 15091 | 13 | 49175696 | 49176641 | mNeoVis.2.1.1.chr13\_hap1.g00619.t01 |
| 15092 | 13 | 49234533 | 49625019 | mNeoVis.2.1.1.chr13\_hap1.g00620.t01 |
| 15093 | 13 | 49234533 | 49625019 | mNeoVis.2.1.1.chr13\_hap1.g00620.t01 |
| 15094 | 13 | 50238267 | 50268559 | mNeoVis.2.1.1.chr13\_hap1.g00621.t01 |
| 15095 | 13 | 50238267 | 50268559 | mNeoVis.2.1.1.chr13\_hap1.g00621.t01 |
| 15096 | 13 | 50278138 | 50375212 | mNeoVis.2.1.1.chr13\_hap1.g00622.t01 |
| 15097 | 13 | 50278138 | 50375212 | mNeoVis.2.1.1.chr13\_hap1.g00622.t01 |
| 15098 | 13 | 50563194 | 50854411 | mNeoVis.2.1.1.chr13\_hap1.g00623.t01 |
| 15099 | 13 | 50563194 | 50854411 | mNeoVis.2.1.1.chr13\_hap1.g00623.t01 |
| 15100 | 13 | 51151976 | 51504393 | mNeoVis.2.1.1.chr13\_hap1.g00624.t01 |
| 15101 | 13 | 51151976 | 51504393 | mNeoVis.2.1.1.chr13\_hap1.g00624.t01 |
| 15102 | 13 | 51151976 | 51504393 | mNeoVis.2.1.1.chr13\_hap1.g00624.t01 |
| 15103 | 13 | 51151976 | 51504393 | mNeoVis.2.1.1.chr13\_hap1.g00624.t01 |
| 15104 | 13 | 51864195 | 52162252 | mNeoVis.2.1.1.chr13\_hap1.g00625.t01 |
| 15105 | 13 | 52200024 | 52246507 | mNeoVis.2.1.1.chr13\_hap1.g00626.t01 |
| 15106 | 13 | 52317219 | 52317756 | mNeoVis.2.1.1.chr13\_hap1.g00628.t01 |
| 15107 | 13 | 52420519 | 52524112 | mNeoVis.2.1.1.chr13\_hap1.g00629.t01 |
| 15108 | 13 | 52421240 | 52422484 | mNeoVis.2.1.1.chr13\_hap1.g00630.t01 |
| 15109 | 13 | 52654314 | 52843089 | mNeoVis.2.1.1.chr13\_hap1.g00631.t01 |
| 15110 | 13 | 52857039 | 52859600 | mNeoVis.2.1.1.chr13\_hap1.g00632.t01 |
| 15111 | 13 | 52857039 | 52859600 | mNeoVis.2.1.1.chr13\_hap1.g00632.t01 |
| 15112 | 13 | 52876342 | 52943264 | mNeoVis.2.1.1.chr13\_hap1.g00633.t01 |
| 15113 | 13 | 52876342 | 52943264 | mNeoVis.2.1.1.chr13\_hap1.g00633.t01 |
| 15114 | 13 | 53210998 | 53444626 | mNeoVis.2.1.1.chr13\_hap1.g00634.t01 |
| 15115 | 13 | 53839689 | 53913139 | mNeoVis.2.1.1.chr13\_hap1.g00635.t01 |
| 15116 | 13 | 53839689 | 53913139 | mNeoVis.2.1.1.chr13\_hap1.g00635.t01 |
| 15117 | 13 | 53839689 | 53913139 | mNeoVis.2.1.1.chr13\_hap1.g00635.t01 |
| 15118 | 13 | 54202039 | 54357195 | mNeoVis.2.1.1.chr13\_hap1.g00636.t01 |
| 15119 | 13 | 54202039 | 54357195 | mNeoVis.2.1.1.chr13\_hap1.g00636.t01 |
| 15120 | 13 | 54202039 | 54357195 | mNeoVis.2.1.1.chr13\_hap1.g00636.t01 |
| 15121 | 13 | 54202039 | 54357195 | mNeoVis.2.1.1.chr13\_hap1.g00636.t01 |
| 15122 | 13 | 54632629 | 54683804 | mNeoVis.2.1.1.chr13\_hap1.g00637.t01 |
| 15123 | 13 | 54632629 | 54683804 | mNeoVis.2.1.1.chr13\_hap1.g00637.t01 |
| 15124 | 13 | 54632629 | 54683804 | mNeoVis.2.1.1.chr13\_hap1.g00637.t01 |
| 15125 | 13 | 54632629 | 54683804 | mNeoVis.2.1.1.chr13\_hap1.g00637.t01 |
| 15126 | 13 | 54745356 | 54763806 | mNeoVis.2.1.1.chr13\_hap1.g00638.t01 |
| 15127 | 13 | 54745356 | 54763806 | mNeoVis.2.1.1.chr13\_hap1.g00638.t01 |
| 15128 | 13 | 54745356 | 54763806 | mNeoVis.2.1.1.chr13\_hap1.g00638.t01 |
| 15129 | 13 | 61150916 | 61193496 | mNeoVis.2.1.1.chr13\_hap1.g00642.t01 |
| 15130 | 13 | 61212736 | 61308419 | mNeoVis.2.1.1.chr13\_hap1.g00643.t01 |
| 15131 | 13 | 61327378 | 61412111 | mNeoVis.2.1.1.chr13\_hap1.g00644.t01 |
| 15132 | 13 | 61534863 | 61593241 | mNeoVis.2.1.1.chr13\_hap1.g00645.t01 |
| 15133 | 13 | 61665570 | 61670168 | mNeoVis.2.1.1.chr13\_hap1.g00646.t01 |
| 15134 | 13 | 68541171 | 68568362 | mNeoVis.2.1.1.chr13\_hap1.g00656.t01 |
| 15135 | 13 | 68633070 | 68643316 | mNeoVis.2.1.1.chr13\_hap1.g00657.t01 |
| 15136 | 13 | 68633070 | 68643316 | mNeoVis.2.1.1.chr13\_hap1.g00657.t01 |
| 15137 | 13 | 68666143 | 68760552 | mNeoVis.2.1.1.chr13\_hap1.g00658.t01 |
| 15138 | 13 | 68666143 | 68760552 | mNeoVis.2.1.1.chr13\_hap1.g00658.t01 |
| 15139 | 13 | 68772882 | 68812288 | mNeoVis.2.1.1.chr13\_hap1.g00659.t01 |
| 15140 | 13 | 68772882 | 68812288 | mNeoVis.2.1.1.chr13\_hap1.g00659.t01 |
| 15141 | 13 | 68869136 | 68892215 | mNeoVis.2.1.1.chr13\_hap1.g00660.t01 |
| 15142 | 13 | 68869136 | 68892215 | mNeoVis.2.1.1.chr13\_hap1.g00660.t01 |
| 15143 | 13 | 68894982 | 68944795 | mNeoVis.2.1.1.chr13\_hap1.g00661.t01 |
| 15144 | 13 | 68894982 | 68944795 | mNeoVis.2.1.1.chr13\_hap1.g00661.t01 |
| 15145 | 13 | 68964869 | 69005426 | mNeoVis.2.1.1.chr13\_hap1.g00662.t01 |
| 15146 | 13 | 68964869 | 69005426 | mNeoVis.2.1.1.chr13\_hap1.g00662.t01 |
| 15147 | 13 | 68964869 | 69005426 | mNeoVis.2.1.1.chr13\_hap1.g00662.t01 |
| 15148 | 13 | 69219553 | 69355026 | mNeoVis.2.1.1.chr13\_hap1.g00663.t01 |
| 15149 | 13 | 69219553 | 69355026 | mNeoVis.2.1.1.chr13\_hap1.g00663.t01 |
| 15150 | 13 | 69689312 | 69760343 | mNeoVis.2.1.1.chr13\_hap1.g00664.t01 |
| 15151 | 13 | 69689312 | 69760343 | mNeoVis.2.1.1.chr13\_hap1.g00664.t01 |
| 15152 | 13 | 69833501 | 70090718 | mNeoVis.2.1.1.chr13\_hap1.g00665.t01 |
| 15153 | 13 | 69833501 | 70090718 | mNeoVis.2.1.1.chr13\_hap1.g00665.t01 |
| 15154 | 13 | 69833501 | 70090718 | mNeoVis.2.1.1.chr13\_hap1.g00665.t01 |
| 15155 | 13 | 70112033 | 70448076 | mNeoVis.2.1.1.chr13\_hap1.g00666.t01 |
| 15156 | 13 | 70530878 | 70556318 | mNeoVis.2.1.1.chr13\_hap1.g00667.t01 |
| 15157 | 13 | 70568774 | 70601321 | mNeoVis.2.1.1.chr13\_hap1.g00668.t01 |
| 15158 | 13 | 70635043 | 70638379 | mNeoVis.2.1.1.chr13\_hap1.g00669.t01 |
| 15159 | 13 | 70675947 | 70680992 | mNeoVis.2.1.1.chr13\_hap1.g00670.t01 |
| 15160 | 13 | 70735812 | 70739160 | mNeoVis.2.1.1.chr13\_hap1.g00671.t01 |
| 15161 | 13 | 70782834 | 70783940 | mNeoVis.2.1.1.chr13\_hap1.g00672.t01 |
| 15162 | 13 | 71439970 | 71441532 | mNeoVis.2.1.1.chr13\_hap1.g00674.t01 |
| 15163 | 13 | 71439970 | 71441532 | mNeoVis.2.1.1.chr13\_hap1.g00674.t01 |
| 15164 | 13 | 71520725 | 71571898 | mNeoVis.2.1.1.chr13\_hap1.g00675.t01 |
| 15165 | 13 | 71520725 | 71571898 | mNeoVis.2.1.1.chr13\_hap1.g00675.t01 |
| 15166 | 13 | 71520725 | 71571898 | mNeoVis.2.1.1.chr13\_hap1.g00675.t01 |
| 15167 | 13 | 71520725 | 71571898 | mNeoVis.2.1.1.chr13\_hap1.g00675.t01 |
| 15168 | 13 | 71936770 | 71997174 | mNeoVis.2.1.1.chr13\_hap1.g00676.t01 |
| 15169 | 13 | 71936770 | 71997174 | mNeoVis.2.1.1.chr13\_hap1.g00676.t01 |
| 15170 | 13 | 71936770 | 71997174 | mNeoVis.2.1.1.chr13\_hap1.g00676.t01 |
| 15171 | 13 | 72001005 | 72012185 | mNeoVis.2.1.1.chr13\_hap1.g00677.t01 |
| 15172 | 13 | 72001005 | 72012185 | mNeoVis.2.1.1.chr13\_hap1.g00677.t01 |
| 15173 | 13 | 72001005 | 72012185 | mNeoVis.2.1.1.chr13\_hap1.g00677.t01 |
| 15174 | 13 | 72012336 | 72043291 | mNeoVis.2.1.1.chr13\_hap1.g00678.t01 |
| 15175 | 13 | 72012336 | 72043291 | mNeoVis.2.1.1.chr13\_hap1.g00678.t01 |
| 15176 | 13 | 72012336 | 72043291 | mNeoVis.2.1.1.chr13\_hap1.g00678.t01 |
| 15177 | 13 | 72058831 | 72196964 | mNeoVis.2.1.1.chr13\_hap1.g00679.t01 |
| 15178 | 13 | 72058831 | 72196964 | mNeoVis.2.1.1.chr13\_hap1.g00679.t01 |
| 15179 | 13 | 72197639 | 72301041 | mNeoVis.2.1.1.chr13\_hap1.g00680.t01 |
| 15180 | 13 | 72384985 | 72544901 | mNeoVis.2.1.1.chr13\_hap1.g00681.t01 |
| 15181 | 13 | 72384985 | 72544901 | mNeoVis.2.1.1.chr13\_hap1.g00681.t01 |
| 15182 | 13 | 72384985 | 72544901 | mNeoVis.2.1.1.chr13\_hap1.g00681.t01 |
| 15183 | 13 | 72793098 | 72826739 | mNeoVis.2.1.1.chr13\_hap1.g00682.t01 |
| 15184 | 13 | 72793098 | 72826739 | mNeoVis.2.1.1.chr13\_hap1.g00682.t01 |
| 15185 | 13 | 72793098 | 72826739 | mNeoVis.2.1.1.chr13\_hap1.g00682.t01 |
| 15186 | 13 | 72793098 | 72826739 | mNeoVis.2.1.1.chr13\_hap1.g00682.t01 |
| 15187 | 13 | 72932927 | 73257376 | mNeoVis.2.1.1.chr13\_hap1.g00683.t01 |
| 15188 | 13 | 72932927 | 73257376 | mNeoVis.2.1.1.chr13\_hap1.g00683.t01 |
| 15189 | 13 | 72932927 | 73257376 | mNeoVis.2.1.1.chr13\_hap1.g00683.t01 |
| 15190 | 13 | 72932927 | 73257376 | mNeoVis.2.1.1.chr13\_hap1.g00683.t01 |
| 15191 | 13 | 73434234 | 73703355 | mNeoVis.2.1.1.chr13\_hap1.g00684.t01 |
| 15192 | 13 | 73537354 | 73596843 | mNeoVis.2.1.1.chr13\_hap1.g00685.t01 |
| 15193 | 13 | 73537354 | 73596843 | mNeoVis.2.1.1.chr13\_hap1.g00685.t01 |
| 15194 | 13 | 73537354 | 73596843 | mNeoVis.2.1.1.chr13\_hap1.g00685.t01 |
| 15195 | 13 | 73836651 | 73856838 | mNeoVis.2.1.1.chr13\_hap1.g00686.t01 |
| 15196 | 13 | 73856144 | 73867702 | mNeoVis.2.1.1.chr13\_hap1.g00687.t01 |
| 15197 | 13 | 73884173 | 73901094 | mNeoVis.2.1.1.chr13\_hap1.g00688.t01 |
| 15198 | 13 | 73956685 | 74020769 | mNeoVis.2.1.1.chr13\_hap1.g00690.t01 |
| 15199 | 13 | 73956685 | 74020769 | mNeoVis.2.1.1.chr13\_hap1.g00690.t01 |
| 15200 | 13 | 74020924 | 74102782 | mNeoVis.2.1.1.chr13\_hap1.g00691.t01 |
| 15201 | 13 | 74020924 | 74102782 | mNeoVis.2.1.1.chr13\_hap1.g00691.t01 |
| 15202 | 13 | 74131681 | 74184884 | mNeoVis.2.1.1.chr13\_hap1.g00692.t01 |
| 15203 | 13 | 74131681 | 74184884 | mNeoVis.2.1.1.chr13\_hap1.g00692.t01 |
| 15204 | 13 | 74131681 | 74184884 | mNeoVis.2.1.1.chr13\_hap1.g00692.t01 |
| 15205 | 13 | 74196691 | 74208390 | mNeoVis.2.1.1.chr13\_hap1.g00693.t01 |
| 15206 | 13 | 74196691 | 74208390 | mNeoVis.2.1.1.chr13\_hap1.g00693.t01 |
| 15207 | 13 | 74196691 | 74208390 | mNeoVis.2.1.1.chr13\_hap1.g00693.t01 |
| 15208 | 13 | 74259373 | 74581698 | mNeoVis.2.1.1.chr13\_hap1.g00694.t01 |
| 15209 | 13 | 74259373 | 74581698 | mNeoVis.2.1.1.chr13\_hap1.g00694.t01 |
| 15210 | 13 | 74259373 | 74581698 | mNeoVis.2.1.1.chr13\_hap1.g00694.t01 |
| 15211 | 13 | 75923510 | 75999853 | mNeoVis.2.1.1.chr13\_hap1.g00705.t01 |
| 15212 | 13 | 76489535 | 76492527 | mNeoVis.2.1.1.chr13\_hap1.g00706.t01 |
| 15213 | 13 | 77269082 | 77355647 | mNeoVis.2.1.1.chr13\_hap1.g00709.t01 |
| 15214 | 13 | 77269082 | 77355647 | mNeoVis.2.1.1.chr13\_hap1.g00709.t01 |
| 15215 | 13 | 77269082 | 77355647 | mNeoVis.2.1.1.chr13\_hap1.g00709.t01 |
| 15216 | 13 | 77269082 | 77355647 | mNeoVis.2.1.1.chr13\_hap1.g00709.t01 |
| 15217 | 13 | 78107416 | 78820094 | mNeoVis.2.1.1.chr13\_hap1.g00710.t01 |
| 15218 | 13 | 78422183 | 78424777 | mNeoVis.2.1.1.chr13\_hap1.g00711.t01 |
| 15219 | 13 | 78437085 | 78439264 | mNeoVis.2.1.1.chr13\_hap1.g00712.t01 |
| 15220 | 13 | 78439356 | 78443828 | mNeoVis.2.1.1.chr13\_hap1.g00713.t01 |
| 15221 | 13 | 78897727 | 79074353 | mNeoVis.2.1.1.chr13\_hap1.g00714.t01 |
| 15222 | 13 | 78897727 | 79074353 | mNeoVis.2.1.1.chr13\_hap1.g00714.t01 |
| 15223 | 13 | 79806986 | 79807495 | mNeoVis.2.1.1.chr13\_hap1.g00715.t01 |
| 15224 | 13 | 79806986 | 79807495 | mNeoVis.2.1.1.chr13\_hap1.g00715.t01 |
| 15225 | 13 | 80158254 | 80187818 | mNeoVis.2.1.1.chr13\_hap1.g00716.t01 |
| 15226 | 13 | 80158254 | 80187818 | mNeoVis.2.1.1.chr13\_hap1.g00716.t01 |
| 15227 | 13 | 80220263 | 80255274 | mNeoVis.2.1.1.chr13\_hap1.g00717.t01 |
| 15228 | 13 | 80220263 | 80255274 | mNeoVis.2.1.1.chr13\_hap1.g00717.t01 |
| 15229 | 13 | 84601704 | 84618610 | mNeoVis.2.1.1.chr13\_hap1.g00733.t01 |
| 15230 | 13 | 84601704 | 84618610 | mNeoVis.2.1.1.chr13\_hap1.g00733.t01 |
| 15231 | 13 | 84601704 | 84618610 | mNeoVis.2.1.1.chr13\_hap1.g00733.t01 |
| 15232 | 13 | 84705181 | 84738120 | mNeoVis.2.1.1.chr13\_hap1.g00734.t01 |
| 15233 | 13 | 84705181 | 84738120 | mNeoVis.2.1.1.chr13\_hap1.g00734.t01 |
| 15234 | 13 | 84705181 | 84738120 | mNeoVis.2.1.1.chr13\_hap1.g00734.t01 |
| 15235 | 13 | 84722255 | 84841815 | mNeoVis.2.1.1.chr13\_hap1.g00735.t01 |
| 15236 | 13 | 84722255 | 84841815 | mNeoVis.2.1.1.chr13\_hap1.g00735.t01 |
| 15237 | 13 | 84722255 | 84841815 | mNeoVis.2.1.1.chr13\_hap1.g00735.t01 |
| 15238 | 13 | 84738214 | 84791058 | mNeoVis.2.1.1.chr13\_hap1.g00736.t01 |
| 15239 | 13 | 84738214 | 84791058 | mNeoVis.2.1.1.chr13\_hap1.g00736.t01 |
| 15240 | 13 | 84738214 | 84791058 | mNeoVis.2.1.1.chr13\_hap1.g00736.t01 |
| 15241 | 13 | 85813995 | 85900860 | mNeoVis.2.1.1.chr13\_hap1.g00744.t01 |
| 15242 | 13 | 85888212 | 85889363 | mNeoVis.2.1.1.chr13\_hap1.g00745.t01 |
| 15243 | 13 | 86003678 | 86019228 | mNeoVis.2.1.1.chr13\_hap1.g00746.t01 |
| 15244 | 13 | 86977476 | 86999103 | mNeoVis.2.1.1.chr13\_hap1.g00748.t01 |
| 15245 | 13 | 87178850 | 87321724 | mNeoVis.2.1.1.chr13\_hap1.g00750.t01 |
| 15246 | 13 | 87440764 | 87516547 | mNeoVis.2.1.1.chr13\_hap1.g00751.t01 |
| 15247 | 13 | 87558690 | 87608375 | mNeoVis.2.1.1.chr13\_hap1.g00752.t01 |
| 15248 | 13 | 87600525 | 87631445 | mNeoVis.2.1.1.chr13\_hap1.g00753.t01 |
| 15249 | 13 | 88078805 | 88079229 | mNeoVis.2.1.1.chr13\_hap1.g00757.t01 |
| 15250 | 13 | 88122527 | 88141846 | mNeoVis.2.1.1.chr13\_hap1.g00758.t01 |
| 15251 | 13 | 88227501 | 88369270 | mNeoVis.2.1.1.chr13\_hap1.g00759.t01 |
| 15252 | 13 | 88421725 | 88425379 | mNeoVis.2.1.1.chr13\_hap1.g00760.t01 |
| 15253 | 13 | 88641857 | 88648298 | mNeoVis.2.1.1.chr13\_hap1.g00761.t01 |
| 15254 | 13 | 88652613 | 88852769 | mNeoVis.2.1.1.chr13\_hap1.g00762.t01 |
| 15255 | 13 | 88891669 | 88893369 | mNeoVis.2.1.1.chr13\_hap1.g00763.t01 |
| 15256 | 13 | 90552805 | 90733858 | mNeoVis.2.1.1.chr13\_hap1.g00770.t01 |
| 15257 | 13 | 91073843 | 91103788 | mNeoVis.2.1.1.chr13\_hap1.g00771.t01 |
| 15258 | 13 | 91171031 | 91183549 | mNeoVis.2.1.1.chr13\_hap1.g00772.t01 |
| 15259 | 13 | 91224528 | 91281318 | mNeoVis.2.1.1.chr13\_hap1.g00773.t01 |
| 15260 | 13 | 91318425 | 91595050 | mNeoVis.2.1.1.chr13\_hap1.g00774.t01 |
| 15261 | 13 | 91628880 | 91865458 | mNeoVis.2.1.1.chr13\_hap1.g00775.t01 |
| 15262 | 13 | 91628880 | 91865458 | mNeoVis.2.1.1.chr13\_hap1.g00775.t01 |
| 15263 | 13 | 92138554 | 92579250 | mNeoVis.2.1.1.chr13\_hap1.g00776.t01 |
| 15264 | 13 | 92138554 | 92579250 | mNeoVis.2.1.1.chr13\_hap1.g00776.t01 |
| 15265 | 13 | 92760888 | 92837015 | mNeoVis.2.1.1.chr13\_hap1.g00777.t01 |
| 15266 | 13 | 92760888 | 92837015 | mNeoVis.2.1.1.chr13\_hap1.g00777.t01 |
| 15267 | 13 | 92800729 | 92905385 | mNeoVis.2.1.1.chr13\_hap1.g00778.t01 |
| 15268 | 13 | 92900854 | 92903778 | mNeoVis.2.1.1.chr13\_hap1.g00779.t01 |
| 15269 | 13 | 92918150 | 92944348 | mNeoVis.2.1.1.chr13\_hap1.g00780.t01 |
| 15270 | 13 | 92925377 | 92965385 | mNeoVis.2.1.1.chr13\_hap1.g00781.t01 |
| 15271 | 13 | 93341527 | 93357764 | mNeoVis.2.1.1.chr13\_hap1.g00784.t01 |
| 15272 | 13 | 93357827 | 93534104 | mNeoVis.2.1.1.chr13\_hap1.g00785.t01 |
| 15273 | 13 | 93385902 | 93391295 | mNeoVis.2.1.1.chr13\_hap1.g00786.t01 |
| 15274 | 13 | 93412818 | 93498636 | mNeoVis.2.1.1.chr13\_hap1.g00787.t01 |
| 15275 | 13 | 93563950 | 93609474 | mNeoVis.2.1.1.chr13\_hap1.g00788.t01 |
| 15276 | 13 | 93668359 | 93817158 | mNeoVis.2.1.1.chr13\_hap1.g00789.t01 |
| 15277 | 13 | 93725246 | 93726696 | mNeoVis.2.1.1.chr13\_hap1.g00790.t01 |
| 15278 | 13 | 93815920 | 93859218 | mNeoVis.2.1.1.chr13\_hap1.g00791.t01 |
| 15279 | 13 | 94282263 | 94305988 | mNeoVis.2.1.1.chr13\_hap1.g00798.t01 |
| 15280 | 13 | 94313038 | 94323439 | mNeoVis.2.1.1.chr13\_hap1.g00799.t01 |
| 15281 | 13 | 94340721 | 94365775 | mNeoVis.2.1.1.chr13\_hap1.g00800.t01 |
| 15282 | 13 | 94384224 | 94417306 | mNeoVis.2.1.1.chr13\_hap1.g00801.t01 |
| 15283 | 13 | 94464034 | 94509846 | mNeoVis.2.1.1.chr13\_hap1.g00802.t01 |
| 15284 | 13 | 94541005 | 94584106 | mNeoVis.2.1.1.chr13\_hap1.g00803.t01 |
| 15285 | 13 | 94639668 | 94672312 | mNeoVis.2.1.1.chr13\_hap1.g00804.t01 |
| 15286 | 13 | 94685339 | 94698428 | mNeoVis.2.1.1.chr13\_hap1.g00805.t01 |
| 15287 | 13 | 94702706 | 94717848 | mNeoVis.2.1.1.chr13\_hap1.g00806.t01 |
| 15288 | 13 | 94725530 | 94734358 | mNeoVis.2.1.1.chr13\_hap1.g00807.t01 |
| 15289 | 13 | 94992311 | 95090959 | mNeoVis.2.1.1.chr13\_hap1.g00809.t01 |
| 15290 | 13 | 95099723 | 95167761 | mNeoVis.2.1.1.chr13\_hap1.g00810.t01 |
| 15291 | 13 | 95309188 | 95312333 | mNeoVis.2.1.1.chr13\_hap1.g00811.t01 |
| 15292 | 13 | 95347201 | 95408754 | mNeoVis.2.1.1.chr13\_hap1.g00812.t01 |
| 15293 | 13 | 95821354 | 95868852 | mNeoVis.2.1.1.chr13\_hap1.g00819.t01 |
| 15294 | 13 | 96079297 | 96157263 | mNeoVis.2.1.1.chr13\_hap1.g00820.t01 |
| 15295 | 13 | 96079297 | 96157263 | mNeoVis.2.1.1.chr13\_hap1.g00820.t01 |
| 15296 | 13 | 96169085 | 96567412 | mNeoVis.2.1.1.chr13\_hap1.g00821.t01 |
| 15297 | 13 | 96185589 | 96193185 | mNeoVis.2.1.1.chr13\_hap1.g00822.t01 |
| 15298 | 13 | 96185589 | 96193185 | mNeoVis.2.1.1.chr13\_hap1.g00822.t01 |
| 15299 | 13 | 96414953 | 96427033 | mNeoVis.2.1.1.chr13\_hap1.g00823.t01 |
| 15300 | 13 | 96414953 | 96427033 | mNeoVis.2.1.1.chr13\_hap1.g00823.t01 |
| 15301 | 13 | 96427105 | 96522456 | mNeoVis.2.1.1.chr13\_hap1.g00824.t01 |
| 15302 | 13 | 96427105 | 96522456 | mNeoVis.2.1.1.chr13\_hap1.g00824.t01 |
| 15303 | 13 | 96568101 | 96573130 | mNeoVis.2.1.1.chr13\_hap1.g00825.t01 |
| 15304 | 13 | 96568101 | 96573130 | mNeoVis.2.1.1.chr13\_hap1.g00825.t01 |
| 15305 | 13 | 96574329 | 96634120 | mNeoVis.2.1.1.chr13\_hap1.g00826.t01 |
| 15306 | 13 | 96574329 | 96634120 | mNeoVis.2.1.1.chr13\_hap1.g00826.t01 |
| 15307 | 13 | 96631949 | 96705646 | mNeoVis.2.1.1.chr13\_hap1.g00827.t01 |
| 15308 | 13 | 96631949 | 96705646 | mNeoVis.2.1.1.chr13\_hap1.g00827.t01 |
| 15309 | 13 | 96756386 | 96757903 | mNeoVis.2.1.1.chr13\_hap1.g00828.t01 |
| 15310 | 13 | 96756386 | 96757903 | mNeoVis.2.1.1.chr13\_hap1.g00828.t01 |
| 15311 | 13 | 96756386 | 96757903 | mNeoVis.2.1.1.chr13\_hap1.g00828.t01 |
| 15312 | 13 | 97791066 | 97803969 | mNeoVis.2.1.1.chr13\_hap1.g00833.t01 |
| 15313 | 13 | 97965519 | 98118714 | mNeoVis.2.1.1.chr13\_hap1.g00834.t01 |
| 15314 | 13 | 98158567 | 98203344 | mNeoVis.2.1.1.chr13\_hap1.g00835.t01 |
| 15315 | 14 | 257811 | 327233 | mNeoVis.2.1.1.chr14\_hap1.g00004.t01 |
| 15316 | 14 | 333923 | 360276 | mNeoVis.2.1.1.chr14\_hap1.g00005.t01 |
| 15317 | 14 | 360042 | 495353 | mNeoVis.2.1.1.chr14\_hap1.g00006.t01 |
| 15318 | 14 | 360914 | 647520 | mNeoVis.2.1.1.chr14\_hap1.g00007.t01 |
| 15319 | 14 | 499555 | 544526 | mNeoVis.2.1.1.chr14\_hap1.g00008.t01 |
| 15320 | 14 | 583562 | 611322 | mNeoVis.2.1.1.chr14\_hap1.g00009.t01 |
| 15321 | 14 | 672322 | 697514 | mNeoVis.2.1.1.chr14\_hap1.g00010.t01 |
| 15322 | 14 | 793877 | 869179 | mNeoVis.2.1.1.chr14\_hap1.g00011.t01 |
| 15323 | 14 | 1642947 | 1664621 | mNeoVis.2.1.1.chr14\_hap1.g00015.t01 |
| 15324 | 14 | 1642947 | 1664621 | mNeoVis.2.1.1.chr14\_hap1.g00015.t01 |
| 15325 | 14 | 1669781 | 1703941 | mNeoVis.2.1.1.chr14\_hap1.g00016.t01 |
| 15326 | 14 | 1669781 | 1703941 | mNeoVis.2.1.1.chr14\_hap1.g00016.t01 |
| 15327 | 14 | 1723624 | 1742596 | mNeoVis.2.1.1.chr14\_hap1.g00017.t01 |
| 15328 | 14 | 1723624 | 1742596 | mNeoVis.2.1.1.chr14\_hap1.g00017.t01 |
| 15329 | 14 | 1769831 | 1835681 | mNeoVis.2.1.1.chr14\_hap1.g00018.t01 |
| 15330 | 14 | 1769831 | 1835681 | mNeoVis.2.1.1.chr14\_hap1.g00018.t01 |
| 15331 | 14 | 1842800 | 1881071 | mNeoVis.2.1.1.chr14\_hap1.g00019.t01 |
| 15332 | 14 | 1842800 | 1881071 | mNeoVis.2.1.1.chr14\_hap1.g00019.t01 |
| 15333 | 14 | 1910806 | 1931759 | mNeoVis.2.1.1.chr14\_hap1.g00020.t01 |
| 15334 | 14 | 1910806 | 1931759 | mNeoVis.2.1.1.chr14\_hap1.g00020.t01 |
| 15335 | 14 | 1933111 | 1965536 | mNeoVis.2.1.1.chr14\_hap1.g00021.t01 |
| 15336 | 14 | 1933111 | 1965536 | mNeoVis.2.1.1.chr14\_hap1.g00021.t01 |
| 15337 | 14 | 1938820 | 1949313 | mNeoVis.2.1.1.chr14\_hap1.g00022.t01 |
| 15338 | 14 | 1938820 | 1949313 | mNeoVis.2.1.1.chr14\_hap1.g00022.t01 |
| 15339 | 14 | 1988079 | 1990612 | mNeoVis.2.1.1.chr14\_hap1.g00023.t01 |
| 15340 | 14 | 1988079 | 1990612 | mNeoVis.2.1.1.chr14\_hap1.g00023.t01 |
| 15341 | 14 | 1990518 | 2039824 | mNeoVis.2.1.1.chr14\_hap1.g00024.t01 |
| 15342 | 14 | 1990518 | 2039824 | mNeoVis.2.1.1.chr14\_hap1.g00024.t01 |
| 15343 | 14 | 2038910 | 2081793 | mNeoVis.2.1.1.chr14\_hap1.g00025.t01 |
| 15344 | 14 | 2038910 | 2081793 | mNeoVis.2.1.1.chr14\_hap1.g00025.t01 |
| 15345 | 14 | 2317113 | 2358771 | mNeoVis.2.1.1.chr14\_hap1.g00027.t01 |
| 15346 | 14 | 2356217 | 2374823 | mNeoVis.2.1.1.chr14\_hap1.g00028.t01 |
| 15347 | 14 | 2381506 | 2394825 | mNeoVis.2.1.1.chr14\_hap1.g00029.t01 |
| 15348 | 14 | 2634421 | 2653474 | mNeoVis.2.1.1.chr14\_hap1.g00030.t01 |
| 15349 | 14 | 2738207 | 2812546 | mNeoVis.2.1.1.chr14\_hap1.g00031.t01 |
| 15350 | 14 | 2742326 | 2756249 | mNeoVis.2.1.1.chr14\_hap1.g00032.t01 |
| 15351 | 14 | 2959301 | 3040782 | mNeoVis.2.1.1.chr14\_hap1.g00033.t01 |
| 15352 | 14 | 3081029 | 3103645 | mNeoVis.2.1.1.chr14\_hap1.g00034.t01 |
| 15353 | 14 | 3122814 | 3142584 | mNeoVis.2.1.1.chr14\_hap1.g00035.t01 |
| 15354 | 14 | 3154749 | 3181980 | mNeoVis.2.1.1.chr14\_hap1.g00036.t01 |
| 15355 | 14 | 3187916 | 3248508 | mNeoVis.2.1.1.chr14\_hap1.g00037.t01 |
| 15356 | 14 | 3332095 | 3427562 | mNeoVis.2.1.1.chr14\_hap1.g00038.t01 |
| 15357 | 14 | 3446524 | 3457461 | mNeoVis.2.1.1.chr14\_hap1.g00039.t01 |
| 15358 | 14 | 3493010 | 3517535 | mNeoVis.2.1.1.chr14\_hap1.g00040.t01 |
| 15359 | 14 | 3522464 | 3536008 | mNeoVis.2.1.1.chr14\_hap1.g00041.t01 |
| 15360 | 14 | 3620227 | 3623124 | mNeoVis.2.1.1.chr14\_hap1.g00043.t01 |
| 15361 | 14 | 4108373 | 4122899 | mNeoVis.2.1.1.chr14\_hap1.g00044.t01 |
| 15362 | 14 | 4116791 | 4159436 | mNeoVis.2.1.1.chr14\_hap1.g00045.t01 |
| 15363 | 14 | 4123041 | 4125501 | mNeoVis.2.1.1.chr14\_hap1.g00046.t01 |
| 15364 | 14 | 4207568 | 4302659 | mNeoVis.2.1.1.chr14\_hap1.g00047.t01 |
| 15365 | 14 | 4316530 | 4332639 | mNeoVis.2.1.1.chr14\_hap1.g00048.t01 |
| 15366 | 14 | 4345135 | 4353107 | mNeoVis.2.1.1.chr14\_hap1.g00049.t01 |
| 15367 | 14 | 4719913 | 4730317 | mNeoVis.2.1.1.chr14\_hap1.g00050.t01 |
| 15368 | 14 | 4794477 | 4816940 | mNeoVis.2.1.1.chr14\_hap1.g00051.t01 |
| 15369 | 14 | 5021696 | 5090170 | mNeoVis.2.1.1.chr14\_hap1.g00052.t01 |
| 15370 | 14 | 5140526 | 5316133 | mNeoVis.2.1.1.chr14\_hap1.g00053.t01 |
| 15371 | 14 | 5266823 | 5267183 | mNeoVis.2.1.1.chr14\_hap1.g00054.t01 |
| 15372 | 14 | 5398463 | 5466055 | mNeoVis.2.1.1.chr14\_hap1.g00055.t01 |
| 15373 | 14 | 5433768 | 5481758 | mNeoVis.2.1.1.chr14\_hap1.g00056.t01 |
| 15374 | 14 | 5515032 | 5552424 | mNeoVis.2.1.1.chr14\_hap1.g00057.t01 |
| 15375 | 14 | 5515032 | 5552424 | mNeoVis.2.1.1.chr14\_hap1.g00057.t01 |
| 15376 | 14 | 5567513 | 5628105 | mNeoVis.2.1.1.chr14\_hap1.g00058.t01 |
| 15377 | 14 | 5567513 | 5628105 | mNeoVis.2.1.1.chr14\_hap1.g00058.t01 |
| 15378 | 14 | 5661868 | 5679051 | mNeoVis.2.1.1.chr14\_hap1.g00059.t01 |
| 15379 | 14 | 5661868 | 5679051 | mNeoVis.2.1.1.chr14\_hap1.g00059.t01 |
| 15380 | 14 | 5680332 | 5694621 | mNeoVis.2.1.1.chr14\_hap1.g00060.t01 |
| 15381 | 14 | 5680332 | 5694621 | mNeoVis.2.1.1.chr14\_hap1.g00060.t01 |
| 15382 | 14 | 5752264 | 6118026 | mNeoVis.2.1.1.chr14\_hap1.g00061.t01 |
| 15383 | 14 | 5752264 | 6118026 | mNeoVis.2.1.1.chr14\_hap1.g00061.t01 |
| 15384 | 14 | 6107453 | 6158350 | mNeoVis.2.1.1.chr14\_hap1.g00062.t01 |
| 15385 | 14 | 6107453 | 6158350 | mNeoVis.2.1.1.chr14\_hap1.g00062.t01 |
| 15386 | 14 | 6165281 | 6277770 | mNeoVis.2.1.1.chr14\_hap1.g00063.t01 |
| 15387 | 14 | 6165281 | 6277770 | mNeoVis.2.1.1.chr14\_hap1.g00063.t01 |
| 15388 | 14 | 6248120 | 6249573 | mNeoVis.2.1.1.chr14\_hap1.g00064.t01 |
| 15389 | 14 | 6248120 | 6249573 | mNeoVis.2.1.1.chr14\_hap1.g00064.t01 |
| 15390 | 14 | 6427041 | 6921088 | mNeoVis.2.1.1.chr14\_hap1.g00065.t01 |
| 15391 | 14 | 7024357 | 7051578 | mNeoVis.2.1.1.chr14\_hap1.g00066.t01 |
| 15392 | 14 | 7024357 | 7051578 | mNeoVis.2.1.1.chr14\_hap1.g00066.t01 |
| 15393 | 14 | 7024357 | 7051578 | mNeoVis.2.1.1.chr14\_hap1.g00066.t01 |
| 15394 | 14 | 7051779 | 7154643 | mNeoVis.2.1.1.chr14\_hap1.g00067.t01 |
| 15395 | 14 | 7051779 | 7154643 | mNeoVis.2.1.1.chr14\_hap1.g00067.t01 |
| 15396 | 14 | 7051779 | 7154643 | mNeoVis.2.1.1.chr14\_hap1.g00067.t01 |
| 15397 | 14 | 7121853 | 7160752 | mNeoVis.2.1.1.chr14\_hap1.g00068.t01 |
| 15398 | 14 | 7121853 | 7160752 | mNeoVis.2.1.1.chr14\_hap1.g00068.t01 |
| 15399 | 14 | 7121853 | 7160752 | mNeoVis.2.1.1.chr14\_hap1.g00068.t01 |
| 15400 | 14 | 7169046 | 7302975 | mNeoVis.2.1.1.chr14\_hap1.g00069.t01 |
| 15401 | 14 | 7169046 | 7302975 | mNeoVis.2.1.1.chr14\_hap1.g00069.t01 |
| 15402 | 14 | 7169046 | 7302975 | mNeoVis.2.1.1.chr14\_hap1.g00069.t01 |
| 15403 | 14 | 7340086 | 7376001 | mNeoVis.2.1.1.chr14\_hap1.g00070.t01 |
| 15404 | 14 | 7340086 | 7376001 | mNeoVis.2.1.1.chr14\_hap1.g00070.t01 |
| 15405 | 14 | 7340086 | 7376001 | mNeoVis.2.1.1.chr14\_hap1.g00070.t01 |
| 15406 | 14 | 7415710 | 7417269 | mNeoVis.2.1.1.chr14\_hap1.g00071.t01 |
| 15407 | 14 | 7415710 | 7417269 | mNeoVis.2.1.1.chr14\_hap1.g00071.t01 |
| 15408 | 14 | 7415710 | 7417269 | mNeoVis.2.1.1.chr14\_hap1.g00071.t01 |
| 15409 | 14 | 7440915 | 7449978 | mNeoVis.2.1.1.chr14\_hap1.g00072.t01 |
| 15410 | 14 | 7440915 | 7449978 | mNeoVis.2.1.1.chr14\_hap1.g00072.t01 |
| 15411 | 14 | 7440915 | 7449978 | mNeoVis.2.1.1.chr14\_hap1.g00072.t01 |
| 15412 | 14 | 7506237 | 7893488 | mNeoVis.2.1.1.chr14\_hap1.g00073.t01 |
| 15413 | 14 | 8342481 | 8466645 | mNeoVis.2.1.1.chr14\_hap1.g00074.t01 |
| 15414 | 14 | 8342481 | 8466645 | mNeoVis.2.1.1.chr14\_hap1.g00074.t01 |
| 15415 | 14 | 8467315 | 8696195 | mNeoVis.2.1.1.chr14\_hap1.g00075.t01 |
| 15416 | 14 | 8467315 | 8696195 | mNeoVis.2.1.1.chr14\_hap1.g00075.t01 |
| 15417 | 14 | 8706927 | 8999381 | mNeoVis.2.1.1.chr14\_hap1.g00076.t01 |
| 15418 | 14 | 8706927 | 8999381 | mNeoVis.2.1.1.chr14\_hap1.g00076.t01 |
| 15419 | 14 | 8706927 | 8999381 | mNeoVis.2.1.1.chr14\_hap1.g00076.t01 |
| 15420 | 14 | 9039321 | 9132561 | mNeoVis.2.1.1.chr14\_hap1.g00077.t01 |
| 15421 | 14 | 9039321 | 9132561 | mNeoVis.2.1.1.chr14\_hap1.g00077.t01 |
| 15422 | 14 | 9039321 | 9132561 | mNeoVis.2.1.1.chr14\_hap1.g00077.t01 |
| 15423 | 14 | 9173697 | 9182072 | mNeoVis.2.1.1.chr14\_hap1.g00078.t01 |
| 15424 | 14 | 9173697 | 9182072 | mNeoVis.2.1.1.chr14\_hap1.g00078.t01 |
| 15425 | 14 | 9173697 | 9182072 | mNeoVis.2.1.1.chr14\_hap1.g00078.t01 |
| 15426 | 14 | 9173697 | 9182072 | mNeoVis.2.1.1.chr14\_hap1.g00078.t01 |
| 15427 | 14 | 9439735 | 9471047 | mNeoVis.2.1.1.chr14\_hap1.g00079.t01 |
| 15428 | 14 | 9439735 | 9471047 | mNeoVis.2.1.1.chr14\_hap1.g00079.t01 |
| 15429 | 14 | 9439735 | 9471047 | mNeoVis.2.1.1.chr14\_hap1.g00079.t01 |
| 15430 | 14 | 9473697 | 9509504 | mNeoVis.2.1.1.chr14\_hap1.g00080.t01 |
| 15431 | 14 | 9473697 | 9509504 | mNeoVis.2.1.1.chr14\_hap1.g00080.t01 |
| 15432 | 14 | 9473697 | 9509504 | mNeoVis.2.1.1.chr14\_hap1.g00080.t01 |
| 15433 | 14 | 9511115 | 9638706 | mNeoVis.2.1.1.chr14\_hap1.g00081.t01 |
| 15434 | 14 | 9511115 | 9638706 | mNeoVis.2.1.1.chr14\_hap1.g00081.t01 |
| 15435 | 14 | 9511115 | 9638706 | mNeoVis.2.1.1.chr14\_hap1.g00081.t01 |
| 15436 | 14 | 9643874 | 9649210 | mNeoVis.2.1.1.chr14\_hap1.g00082.t01 |
| 15437 | 14 | 9643874 | 9649210 | mNeoVis.2.1.1.chr14\_hap1.g00082.t01 |
| 15438 | 14 | 9643874 | 9649210 | mNeoVis.2.1.1.chr14\_hap1.g00082.t01 |
| 15439 | 14 | 9647889 | 9692144 | mNeoVis.2.1.1.chr14\_hap1.g00083.t01 |
| 15440 | 14 | 9647889 | 9692144 | mNeoVis.2.1.1.chr14\_hap1.g00083.t01 |
| 15441 | 14 | 9647889 | 9692144 | mNeoVis.2.1.1.chr14\_hap1.g00083.t01 |
| 15442 | 14 | 9708231 | 9709036 | mNeoVis.2.1.1.chr14\_hap1.g00084.t01 |
| 15443 | 14 | 9708231 | 9709036 | mNeoVis.2.1.1.chr14\_hap1.g00084.t01 |
| 15444 | 14 | 9708231 | 9709036 | mNeoVis.2.1.1.chr14\_hap1.g00084.t01 |
| 15445 | 14 | 9714244 | 9716807 | mNeoVis.2.1.1.chr14\_hap1.g00085.t01 |
| 15446 | 14 | 9714244 | 9716807 | mNeoVis.2.1.1.chr14\_hap1.g00085.t01 |
| 15447 | 14 | 9714244 | 9716807 | mNeoVis.2.1.1.chr14\_hap1.g00085.t01 |
| 15448 | 14 | 9752206 | 9760260 | mNeoVis.2.1.1.chr14\_hap1.g00086.t01 |
| 15449 | 14 | 9752206 | 9760260 | mNeoVis.2.1.1.chr14\_hap1.g00086.t01 |
| 15450 | 14 | 9752206 | 9760260 | mNeoVis.2.1.1.chr14\_hap1.g00086.t01 |
| 15451 | 14 | 9769050 | 9779407 | mNeoVis.2.1.1.chr14\_hap1.g00087.t01 |
| 15452 | 14 | 9866041 | 9977923 | mNeoVis.2.1.1.chr14\_hap1.g00088.t01 |
| 15453 | 14 | 10020597 | 10436725 | mNeoVis.2.1.1.chr14\_hap1.g00089.t01 |
| 15454 | 14 | 10471098 | 11161585 | mNeoVis.2.1.1.chr14\_hap1.g00090.t01 |
| 15455 | 14 | 10471098 | 11161585 | mNeoVis.2.1.1.chr14\_hap1.g00090.t01 |
| 15456 | 14 | 11177167 | 11195916 | mNeoVis.2.1.1.chr14\_hap1.g00091.t01 |
| 15457 | 14 | 11177167 | 11195916 | mNeoVis.2.1.1.chr14\_hap1.g00091.t01 |
| 15458 | 14 | 11196062 | 11306027 | mNeoVis.2.1.1.chr14\_hap1.g00092.t01 |
| 15459 | 14 | 11196062 | 11306027 | mNeoVis.2.1.1.chr14\_hap1.g00092.t01 |
| 15460 | 14 | 11340977 | 11375294 | mNeoVis.2.1.1.chr14\_hap1.g00093.t01 |
| 15461 | 14 | 11340977 | 11375294 | mNeoVis.2.1.1.chr14\_hap1.g00093.t01 |
| 15462 | 14 | 11340977 | 11375294 | mNeoVis.2.1.1.chr14\_hap1.g00093.t01 |
| 15463 | 14 | 11377653 | 11448489 | mNeoVis.2.1.1.chr14\_hap1.g00094.t01 |
| 15464 | 14 | 11377653 | 11448489 | mNeoVis.2.1.1.chr14\_hap1.g00094.t01 |
| 15465 | 14 | 11448289 | 11530273 | mNeoVis.2.1.1.chr14\_hap1.g00095.t01 |
| 15466 | 14 | 11448289 | 11530273 | mNeoVis.2.1.1.chr14\_hap1.g00095.t01 |
| 15467 | 14 | 11666439 | 11976164 | mNeoVis.2.1.1.chr14\_hap1.g00096.t01 |
| 15468 | 14 | 11666439 | 11976164 | mNeoVis.2.1.1.chr14\_hap1.g00096.t01 |
| 15469 | 14 | 11977512 | 12013505 | mNeoVis.2.1.1.chr14\_hap1.g00097.t01 |
| 15470 | 14 | 11977512 | 12013505 | mNeoVis.2.1.1.chr14\_hap1.g00097.t01 |
| 15471 | 14 | 12036298 | 12057246 | mNeoVis.2.1.1.chr14\_hap1.g00098.t01 |
| 15472 | 14 | 12036298 | 12057246 | mNeoVis.2.1.1.chr14\_hap1.g00098.t01 |
| 15473 | 14 | 12090726 | 12134587 | mNeoVis.2.1.1.chr14\_hap1.g00099.t01 |
| 15474 | 14 | 12090726 | 12134587 | mNeoVis.2.1.1.chr14\_hap1.g00099.t01 |
| 15475 | 14 | 12163578 | 12181377 | mNeoVis.2.1.1.chr14\_hap1.g00100.t01 |
| 15476 | 14 | 12163578 | 12181377 | mNeoVis.2.1.1.chr14\_hap1.g00100.t01 |
| 15477 | 14 | 12239090 | 12247009 | mNeoVis.2.1.1.chr14\_hap1.g00101.t01 |
| 15478 | 14 | 12239090 | 12247009 | mNeoVis.2.1.1.chr14\_hap1.g00101.t01 |
| 15479 | 14 | 12319557 | 12331948 | mNeoVis.2.1.1.chr14\_hap1.g00102.t01 |
| 15480 | 14 | 12368251 | 12452420 | mNeoVis.2.1.1.chr14\_hap1.g00103.t01 |
| 15481 | 14 | 12527302 | 12618898 | mNeoVis.2.1.1.chr14\_hap1.g00104.t01 |
| 15482 | 14 | 12619095 | 12680097 | mNeoVis.2.1.1.chr14\_hap1.g00105.t01 |
| 15483 | 14 | 12790436 | 12855573 | mNeoVis.2.1.1.chr14\_hap1.g00106.t01 |
| 15484 | 14 | 12857808 | 12894943 | mNeoVis.2.1.1.chr14\_hap1.g00107.t01 |
| 15485 | 14 | 12901230 | 12911159 | mNeoVis.2.1.1.chr14\_hap1.g00108.t01 |
| 15486 | 14 | 18247980 | 18269246 | mNeoVis.2.1.1.chr14\_hap1.g00147.t01 |
| 15487 | 14 | 18276091 | 18281931 | mNeoVis.2.1.1.chr14\_hap1.g00148.t01 |
| 15488 | 14 | 18291079 | 18345630 | mNeoVis.2.1.1.chr14\_hap1.g00149.t01 |
| 15489 | 14 | 18452270 | 18487055 | mNeoVis.2.1.1.chr14\_hap1.g00150.t01 |
| 15490 | 14 | 18503986 | 18554830 | mNeoVis.2.1.1.chr14\_hap1.g00151.t01 |
| 15491 | 14 | 18562055 | 18596453 | mNeoVis.2.1.1.chr14\_hap1.g00152.t01 |
| 15492 | 14 | 18652890 | 18779151 | mNeoVis.2.1.1.chr14\_hap1.g00153.t01 |
| 15493 | 14 | 18810171 | 18869432 | mNeoVis.2.1.1.chr14\_hap1.g00154.t01 |
| 15494 | 14 | 19063767 | 19156426 | mNeoVis.2.1.1.chr14\_hap1.g00156.t01 |
| 15495 | 14 | 19157575 | 19188993 | mNeoVis.2.1.1.chr14\_hap1.g00157.t01 |
| 15496 | 14 | 19221578 | 19230295 | mNeoVis.2.1.1.chr14\_hap1.g00158.t01 |
| 15497 | 14 | 19230502 | 19481128 | mNeoVis.2.1.1.chr14\_hap1.g00159.t01 |
| 15498 | 14 | 19241345 | 19243800 | mNeoVis.2.1.1.chr14\_hap1.g00160.t01 |
| 15499 | 14 | 19881485 | 20115498 | mNeoVis.2.1.1.chr14\_hap1.g00164.t01 |
| 15500 | 14 | 20152641 | 20155930 | mNeoVis.2.1.1.chr14\_hap1.g00165.t01 |
| 15501 | 14 | 20173017 | 20245184 | mNeoVis.2.1.1.chr14\_hap1.g00166.t01 |
| 15502 | 14 | 20454806 | 20455974 | mNeoVis.2.1.1.chr14\_hap1.g00168.t01 |
| 15503 | 14 | 20474021 | 20475564 | mNeoVis.2.1.1.chr14\_hap1.g00169.t01 |
| 15504 | 14 | 20474021 | 20475564 | mNeoVis.2.1.1.chr14\_hap1.g00169.t01 |
| 15505 | 14 | 20485385 | 20530507 | mNeoVis.2.1.1.chr14\_hap1.g00170.t01 |
| 15506 | 14 | 20485385 | 20530507 | mNeoVis.2.1.1.chr14\_hap1.g00170.t01 |
| 15507 | 14 | 20541107 | 20586435 | mNeoVis.2.1.1.chr14\_hap1.g00171.t01 |
| 15508 | 14 | 20541107 | 20586435 | mNeoVis.2.1.1.chr14\_hap1.g00171.t01 |
| 15509 | 14 | 20586524 | 20691256 | mNeoVis.2.1.1.chr14\_hap1.g00172.t01 |
| 15510 | 14 | 20586524 | 20691256 | mNeoVis.2.1.1.chr14\_hap1.g00172.t01 |
| 15511 | 14 | 20697162 | 20716761 | mNeoVis.2.1.1.chr14\_hap1.g00173.t01 |
| 15512 | 14 | 20697162 | 20716761 | mNeoVis.2.1.1.chr14\_hap1.g00173.t01 |
| 15513 | 14 | 21638752 | 21655817 | mNeoVis.2.1.1.chr14\_hap1.g00175.t01 |
| 15514 | 14 | 21655818 | 21721794 | mNeoVis.2.1.1.chr14\_hap1.g00176.t01 |
| 15515 | 14 | 21721832 | 21761590 | mNeoVis.2.1.1.chr14\_hap1.g00177.t01 |
| 15516 | 14 | 21788093 | 21802892 | mNeoVis.2.1.1.chr14\_hap1.g00179.t01 |
| 15517 | 14 | 21989175 | 21996786 | mNeoVis.2.1.1.chr14\_hap1.g00180.t01 |
| 15518 | 14 | 22974410 | 23091405 | mNeoVis.2.1.1.chr14\_hap1.g00187.t01 |
| 15519 | 14 | 23093970 | 23111090 | mNeoVis.2.1.1.chr14\_hap1.g00188.t01 |
| 15520 | 14 | 23113646 | 23156556 | mNeoVis.2.1.1.chr14\_hap1.g00189.t01 |
| 15521 | 14 | 23239661 | 23399560 | mNeoVis.2.1.1.chr14\_hap1.g00190.t01 |
| 15522 | 14 | 23282433 | 23306286 | mNeoVis.2.1.1.chr14\_hap1.g00191.t01 |
| 15523 | 14 | 23566211 | 23575682 | mNeoVis.2.1.1.chr14\_hap1.g00192.t01 |
| 15524 | 14 | 23590558 | 23606853 | mNeoVis.2.1.1.chr14\_hap1.g00193.t01 |
| 15525 | 14 | 25812816 | 25825856 | mNeoVis.2.1.1.chr14\_hap1.g00202.t01 |
| 15526 | 14 | 25825778 | 26070473 | mNeoVis.2.1.1.chr14\_hap1.g00203.t01 |
| 15527 | 14 | 26478300 | 26620281 | mNeoVis.2.1.1.chr14\_hap1.g00204.t01 |
| 15528 | 14 | 27765852 | 27851450 | mNeoVis.2.1.1.chr14\_hap1.g00212.t01 |
| 15529 | 14 | 27867993 | 27878266 | mNeoVis.2.1.1.chr14\_hap1.g00213.t01 |
| 15530 | 14 | 27878683 | 27905323 | mNeoVis.2.1.1.chr14\_hap1.g00214.t01 |
| 15531 | 14 | 27915264 | 27942801 | mNeoVis.2.1.1.chr14\_hap1.g00215.t01 |
| 15532 | 14 | 27943932 | 27956870 | mNeoVis.2.1.1.chr14\_hap1.g00216.t01 |
| 15533 | 14 | 27962766 | 28025591 | mNeoVis.2.1.1.chr14\_hap1.g00217.t01 |
| 15534 | 14 | 28041252 | 28093610 | mNeoVis.2.1.1.chr14\_hap1.g00218.t01 |
| 15535 | 14 | 28070808 | 28073137 | mNeoVis.2.1.1.chr14\_hap1.g00219.t01 |
| 15536 | 14 | 28096013 | 28200311 | mNeoVis.2.1.1.chr14\_hap1.g00220.t01 |
| 15537 | 14 | 28201634 | 28214430 | mNeoVis.2.1.1.chr14\_hap1.g00221.t01 |
| 15538 | 14 | 30514232 | 30535075 | mNeoVis.2.1.1.chr14\_hap1.g00249.t01 |
| 15539 | 14 | 30603827 | 30605809 | mNeoVis.2.1.1.chr14\_hap1.g00250.t01 |
| 15540 | 14 | 30618973 | 30703949 | mNeoVis.2.1.1.chr14\_hap1.g00251.t01 |
| 15541 | 14 | 30746500 | 30752774 | mNeoVis.2.1.1.chr14\_hap1.g00252.t01 |
| 15542 | 14 | 30787306 | 30962246 | mNeoVis.2.1.1.chr14\_hap1.g00253.t01 |
| 15543 | 14 | 32109518 | 32123377 | mNeoVis.2.1.1.chr14\_hap1.g00258.t01 |
| 15544 | 14 | 32200310 | 32394065 | mNeoVis.2.1.1.chr14\_hap1.g00259.t01 |
| 15545 | 14 | 32466713 | 32491361 | mNeoVis.2.1.1.chr14\_hap1.g00260.t01 |
| 15546 | 14 | 32513266 | 32532978 | mNeoVis.2.1.1.chr14\_hap1.g00261.t01 |
| 15547 | 14 | 32533988 | 32535719 | mNeoVis.2.1.1.chr14\_hap1.g00262.t01 |
| 15548 | 14 | 32537139 | 32559428 | mNeoVis.2.1.1.chr14\_hap1.g00263.t01 |
| 15549 | 14 | 32569613 | 32633287 | mNeoVis.2.1.1.chr14\_hap1.g00264.t01 |
| 15550 | 14 | 32627578 | 32705013 | mNeoVis.2.1.1.chr14\_hap1.g00265.t01 |
| 15551 | 14 | 32640002 | 32641019 | mNeoVis.2.1.1.chr14\_hap1.g00266.t01 |
| 15552 | 14 | 32706182 | 32734611 | mNeoVis.2.1.1.chr14\_hap1.g00267.t01 |
| 15553 | 14 | 32706182 | 32734611 | mNeoVis.2.1.1.chr14\_hap1.g00267.t01 |
| 15554 | 14 | 34531914 | 34567490 | mNeoVis.2.1.1.chr14\_hap1.g00274.t01 |
| 15555 | 14 | 34622270 | 34650011 | mNeoVis.2.1.1.chr14\_hap1.g00275.t01 |
| 15556 | 14 | 34729506 | 34738100 | mNeoVis.2.1.1.chr14\_hap1.g00276.t01 |
| 15557 | 14 | 34729506 | 34738100 | mNeoVis.2.1.1.chr14\_hap1.g00276.t01 |
| 15558 | 14 | 34751055 | 34758825 | mNeoVis.2.1.1.chr14\_hap1.g00277.t01 |
| 15559 | 14 | 34751055 | 34758825 | mNeoVis.2.1.1.chr14\_hap1.g00277.t01 |
| 15560 | 14 | 34769506 | 34792352 | mNeoVis.2.1.1.chr14\_hap1.g00278.t01 |
| 15561 | 14 | 34769506 | 34792352 | mNeoVis.2.1.1.chr14\_hap1.g00278.t01 |
| 15562 | 14 | 34805385 | 34814906 | mNeoVis.2.1.1.chr14\_hap1.g00279.t01 |
| 15563 | 14 | 34805385 | 34814906 | mNeoVis.2.1.1.chr14\_hap1.g00279.t01 |
| 15564 | 14 | 34813532 | 34844660 | mNeoVis.2.1.1.chr14\_hap1.g00280.t01 |
| 15565 | 14 | 34813532 | 34844660 | mNeoVis.2.1.1.chr14\_hap1.g00280.t01 |
| 15566 | 14 | 34846179 | 34853702 | mNeoVis.2.1.1.chr14\_hap1.g00281.t01 |
| 15567 | 14 | 34846179 | 34853702 | mNeoVis.2.1.1.chr14\_hap1.g00281.t01 |
| 15568 | 14 | 34846542 | 34878162 | mNeoVis.2.1.1.chr14\_hap1.g00282.t01 |
| 15569 | 14 | 34846542 | 34878162 | mNeoVis.2.1.1.chr14\_hap1.g00282.t01 |
| 15570 | 14 | 34927157 | 34939207 | mNeoVis.2.1.1.chr14\_hap1.g00283.t01 |
| 15571 | 14 | 34927157 | 34939207 | mNeoVis.2.1.1.chr14\_hap1.g00283.t01 |
| 15572 | 14 | 34947962 | 34965996 | mNeoVis.2.1.1.chr14\_hap1.g00284.t01 |
| 15573 | 14 | 34947962 | 34965996 | mNeoVis.2.1.1.chr14\_hap1.g00284.t01 |
| 15574 | 14 | 34971881 | 34989188 | mNeoVis.2.1.1.chr14\_hap1.g00285.t01 |
| 15575 | 14 | 34971881 | 34989188 | mNeoVis.2.1.1.chr14\_hap1.g00285.t01 |
| 15576 | 14 | 34989322 | 34992840 | mNeoVis.2.1.1.chr14\_hap1.g00286.t01 |
| 15577 | 14 | 34989322 | 34992840 | mNeoVis.2.1.1.chr14\_hap1.g00286.t01 |
| 15578 | 14 | 35022940 | 35025501 | mNeoVis.2.1.1.chr14\_hap1.g00287.t01 |
| 15579 | 14 | 35022940 | 35025501 | mNeoVis.2.1.1.chr14\_hap1.g00287.t01 |
| 15580 | 14 | 35039762 | 35044645 | mNeoVis.2.1.1.chr14\_hap1.g00288.t01 |
| 15581 | 14 | 35039762 | 35044645 | mNeoVis.2.1.1.chr14\_hap1.g00288.t01 |
| 15582 | 14 | 35105980 | 35111386 | mNeoVis.2.1.1.chr14\_hap1.g00289.t01 |
| 15583 | 14 | 35105980 | 35111386 | mNeoVis.2.1.1.chr14\_hap1.g00289.t01 |
| 15584 | 14 | 35497251 | 35570980 | mNeoVis.2.1.1.chr14\_hap1.g00295.t01 |
| 15585 | 14 | 35548615 | 35817088 | mNeoVis.2.1.1.chr14\_hap1.g00296.t01 |
| 15586 | 14 | 35828995 | 36091846 | mNeoVis.2.1.1.chr14\_hap1.g00297.t01 |
| 15587 | 14 | 35828995 | 36091846 | mNeoVis.2.1.1.chr14\_hap1.g00297.t01 |
| 15588 | 14 | 36193573 | 36202081 | mNeoVis.2.1.1.chr14\_hap1.g00298.t01 |
| 15589 | 14 | 36193573 | 36202081 | mNeoVis.2.1.1.chr14\_hap1.g00298.t01 |
| 15590 | 14 | 36193573 | 36202081 | mNeoVis.2.1.1.chr14\_hap1.g00298.t01 |
| 15591 | 14 | 36211991 | 36223932 | mNeoVis.2.1.1.chr14\_hap1.g00299.t01 |
| 15592 | 14 | 36211991 | 36223932 | mNeoVis.2.1.1.chr14\_hap1.g00299.t01 |
| 15593 | 14 | 36211991 | 36223932 | mNeoVis.2.1.1.chr14\_hap1.g00299.t01 |
| 15594 | 14 | 36251526 | 36392782 | mNeoVis.2.1.1.chr14\_hap1.g00300.t01 |
| 15595 | 14 | 36251526 | 36392782 | mNeoVis.2.1.1.chr14\_hap1.g00300.t01 |
| 15596 | 14 | 36273537 | 36297749 | mNeoVis.2.1.1.chr14\_hap1.g00301.t01 |
| 15597 | 14 | 36273537 | 36297749 | mNeoVis.2.1.1.chr14\_hap1.g00301.t01 |
| 15598 | 14 | 36273537 | 36297749 | mNeoVis.2.1.1.chr14\_hap1.g00301.t01 |
| 15599 | 14 | 36406196 | 36558999 | mNeoVis.2.1.1.chr14\_hap1.g00302.t01 |
| 15600 | 14 | 36406196 | 36558999 | mNeoVis.2.1.1.chr14\_hap1.g00302.t01 |
| 15601 | 14 | 36730559 | 36750467 | mNeoVis.2.1.1.chr14\_hap1.g00303.t01 |
| 15602 | 14 | 36835609 | 36961446 | mNeoVis.2.1.1.chr14\_hap1.g00304.t01 |
| 15603 | 14 | 36975260 | 36982928 | mNeoVis.2.1.1.chr14\_hap1.g00305.t01 |
| 15604 | 14 | 38110596 | 38250380 | mNeoVis.2.1.1.chr14\_hap1.g00306.t01 |
| 15605 | 14 | 38110596 | 38250380 | mNeoVis.2.1.1.chr14\_hap1.g00306.t01 |
| 15606 | 14 | 38442943 | 38448098 | mNeoVis.2.1.1.chr14\_hap1.g00307.t01 |
| 15607 | 14 | 38570102 | 38624289 | mNeoVis.2.1.1.chr14\_hap1.g00308.t01 |
| 15608 | 14 | 38650974 | 38672731 | mNeoVis.2.1.1.chr14\_hap1.g00309.t01 |
| 15609 | 14 | 38878263 | 38880272 | mNeoVis.2.1.1.chr14\_hap1.g00311.t01 |
| 15610 | 14 | 38898382 | 38919162 | mNeoVis.2.1.1.chr14\_hap1.g00312.t01 |
| 15611 | 14 | 38919774 | 38951615 | mNeoVis.2.1.1.chr14\_hap1.g00313.t01 |
| 15612 | 14 | 38955442 | 39003576 | mNeoVis.2.1.1.chr14\_hap1.g00314.t01 |
| 15613 | 14 | 39015466 | 39084117 | mNeoVis.2.1.1.chr14\_hap1.g00315.t01 |
| 15614 | 14 | 39015466 | 39084117 | mNeoVis.2.1.1.chr14\_hap1.g00315.t01 |
| 15615 | 14 | 39085754 | 39087479 | mNeoVis.2.1.1.chr14\_hap1.g00316.t01 |
| 15616 | 14 | 39085754 | 39087479 | mNeoVis.2.1.1.chr14\_hap1.g00316.t01 |
| 15617 | 14 | 39085754 | 39087479 | mNeoVis.2.1.1.chr14\_hap1.g00316.t01 |
| 15618 | 14 | 39103576 | 39123039 | mNeoVis.2.1.1.chr14\_hap1.g00317.t01 |
| 15619 | 14 | 39103576 | 39123039 | mNeoVis.2.1.1.chr14\_hap1.g00317.t01 |
| 15620 | 14 | 39103576 | 39123039 | mNeoVis.2.1.1.chr14\_hap1.g00317.t01 |
| 15621 | 14 | 39154215 | 39184858 | mNeoVis.2.1.1.chr14\_hap1.g00318.t01 |
| 15622 | 14 | 39154215 | 39184858 | mNeoVis.2.1.1.chr14\_hap1.g00318.t01 |
| 15623 | 14 | 39154215 | 39184858 | mNeoVis.2.1.1.chr14\_hap1.g00318.t01 |
| 15624 | 14 | 39227232 | 39277219 | mNeoVis.2.1.1.chr14\_hap1.g00319.t01 |
| 15625 | 14 | 39227232 | 39277219 | mNeoVis.2.1.1.chr14\_hap1.g00319.t01 |
| 15626 | 14 | 39227232 | 39277219 | mNeoVis.2.1.1.chr14\_hap1.g00319.t01 |
| 15627 | 14 | 39297941 | 39312064 | mNeoVis.2.1.1.chr14\_hap1.g00320.t01 |
| 15628 | 14 | 39297941 | 39312064 | mNeoVis.2.1.1.chr14\_hap1.g00320.t01 |
| 15629 | 14 | 39297941 | 39312064 | mNeoVis.2.1.1.chr14\_hap1.g00320.t01 |
| 15630 | 14 | 39321308 | 39338184 | mNeoVis.2.1.1.chr14\_hap1.g00321.t01 |
| 15631 | 14 | 39321308 | 39338184 | mNeoVis.2.1.1.chr14\_hap1.g00321.t01 |
| 15632 | 14 | 39321308 | 39338184 | mNeoVis.2.1.1.chr14\_hap1.g00321.t01 |
| 15633 | 14 | 39346757 | 39359011 | mNeoVis.2.1.1.chr14\_hap1.g00322.t01 |
| 15634 | 14 | 39346757 | 39359011 | mNeoVis.2.1.1.chr14\_hap1.g00322.t01 |
| 15635 | 14 | 39346757 | 39359011 | mNeoVis.2.1.1.chr14\_hap1.g00322.t01 |
| 15636 | 14 | 39390545 | 39393474 | mNeoVis.2.1.1.chr14\_hap1.g00323.t01 |
| 15637 | 14 | 39390545 | 39393474 | mNeoVis.2.1.1.chr14\_hap1.g00323.t01 |
| 15638 | 14 | 39390545 | 39393474 | mNeoVis.2.1.1.chr14\_hap1.g00323.t01 |
| 15639 | 14 | 39409794 | 39433112 | mNeoVis.2.1.1.chr14\_hap1.g00324.t01 |
| 15640 | 14 | 39409794 | 39433112 | mNeoVis.2.1.1.chr14\_hap1.g00324.t01 |
| 15641 | 14 | 39409794 | 39433112 | mNeoVis.2.1.1.chr14\_hap1.g00324.t01 |
| 15642 | 14 | 39456265 | 39702543 | mNeoVis.2.1.1.chr14\_hap1.g00325.t01 |
| 15643 | 14 | 39456265 | 39702543 | mNeoVis.2.1.1.chr14\_hap1.g00325.t01 |
| 15644 | 14 | 39456265 | 39702543 | mNeoVis.2.1.1.chr14\_hap1.g00325.t01 |
| 15645 | 14 | 39710260 | 39766708 | mNeoVis.2.1.1.chr14\_hap1.g00326.t01 |
| 15646 | 14 | 39710260 | 39766708 | mNeoVis.2.1.1.chr14\_hap1.g00326.t01 |
| 15647 | 14 | 39710260 | 39766708 | mNeoVis.2.1.1.chr14\_hap1.g00326.t01 |
| 15648 | 14 | 39776087 | 39780094 | mNeoVis.2.1.1.chr14\_hap1.g00327.t01 |
| 15649 | 14 | 39776087 | 39780094 | mNeoVis.2.1.1.chr14\_hap1.g00327.t01 |
| 15650 | 14 | 39776087 | 39780094 | mNeoVis.2.1.1.chr14\_hap1.g00327.t01 |
| 15651 | 14 | 39785373 | 39826102 | mNeoVis.2.1.1.chr14\_hap1.g00328.t01 |
| 15652 | 14 | 39785373 | 39826102 | mNeoVis.2.1.1.chr14\_hap1.g00328.t01 |
| 15653 | 14 | 39833320 | 39847443 | mNeoVis.2.1.1.chr14\_hap1.g00329.t01 |
| 15654 | 14 | 39833320 | 39847443 | mNeoVis.2.1.1.chr14\_hap1.g00329.t01 |
| 15655 | 14 | 39866585 | 39883824 | mNeoVis.2.1.1.chr14\_hap1.g00330.t01 |
| 15656 | 14 | 39866585 | 39883824 | mNeoVis.2.1.1.chr14\_hap1.g00330.t01 |
| 15657 | 14 | 39884468 | 39892113 | mNeoVis.2.1.1.chr14\_hap1.g00331.t01 |
| 15658 | 14 | 39884468 | 39892113 | mNeoVis.2.1.1.chr14\_hap1.g00331.t01 |
| 15659 | 14 | 40394799 | 40435857 | mNeoVis.2.1.1.chr14\_hap1.g00339.t01 |
| 15660 | 14 | 40518938 | 40614699 | mNeoVis.2.1.1.chr14\_hap1.g00340.t01 |
| 15661 | 14 | 40621160 | 40631015 | mNeoVis.2.1.1.chr14\_hap1.g00341.t01 |
| 15662 | 14 | 40623321 | 40623956 | mNeoVis.2.1.1.chr14\_hap1.g00342.t01 |
| 15663 | 14 | 40653278 | 40661298 | mNeoVis.2.1.1.chr14\_hap1.g00343.t01 |
| 15664 | 14 | 40667162 | 40678797 | mNeoVis.2.1.1.chr14\_hap1.g00344.t01 |
| 15665 | 14 | 40681247 | 40712128 | mNeoVis.2.1.1.chr14\_hap1.g00345.t01 |
| 15666 | 14 | 40719069 | 40734762 | mNeoVis.2.1.1.chr14\_hap1.g00346.t01 |
| 15667 | 14 | 40738649 | 40743818 | mNeoVis.2.1.1.chr14\_hap1.g00347.t01 |
| 15668 | 14 | 40778492 | 40788632 | mNeoVis.2.1.1.chr14\_hap1.g00348.t01 |
| 15669 | 14 | 40799346 | 40815098 | mNeoVis.2.1.1.chr14\_hap1.g00349.t01 |
| 15670 | 14 | 40816300 | 40865625 | mNeoVis.2.1.1.chr14\_hap1.g00350.t01 |
| 15671 | 14 | 40869190 | 40872044 | mNeoVis.2.1.1.chr14\_hap1.g00351.t01 |
| 15672 | 14 | 40911674 | 40943597 | mNeoVis.2.1.1.chr14\_hap1.g00352.t01 |
| 15673 | 14 | 40944755 | 40952342 | mNeoVis.2.1.1.chr14\_hap1.g00353.t01 |
| 15674 | 14 | 40955531 | 40962732 | mNeoVis.2.1.1.chr14\_hap1.g00354.t01 |
| 15675 | 14 | 40990166 | 41004303 | mNeoVis.2.1.1.chr14\_hap1.g00355.t01 |
| 15676 | 14 | 41054589 | 41058786 | mNeoVis.2.1.1.chr14\_hap1.g00356.t01 |
| 15677 | 14 | 41080983 | 41091197 | mNeoVis.2.1.1.chr14\_hap1.g00357.t01 |
| 15678 | 14 | 41193378 | 41199596 | mNeoVis.2.1.1.chr14\_hap1.g00358.t01 |
| 15679 | 14 | 41203920 | 41210071 | mNeoVis.2.1.1.chr14\_hap1.g00359.t01 |
| 15680 | 14 | 41307882 | 41403805 | mNeoVis.2.1.1.chr14\_hap1.g00360.t01 |
| 15681 | 14 | 41408896 | 41421607 | mNeoVis.2.1.1.chr14\_hap1.g00361.t01 |
| 15682 | 14 | 41447013 | 41447111 | mNeoVis.2.1.1.chr14\_hap1.g00362.t01 |
| 15683 | 14 | 41456545 | 41483503 | mNeoVis.2.1.1.chr14\_hap1.g00363.t01 |
| 15684 | 14 | 41492607 | 41498282 | mNeoVis.2.1.1.chr14\_hap1.g00364.t01 |
| 15685 | 14 | 41655536 | 41700993 | mNeoVis.2.1.1.chr14\_hap1.g00365.t01 |
| 15686 | 14 | 41701135 | 41721292 | mNeoVis.2.1.1.chr14\_hap1.g00366.t01 |
| 15687 | 14 | 41723331 | 41733952 | mNeoVis.2.1.1.chr14\_hap1.g00367.t01 |
| 15688 | 14 | 41749014 | 41754504 | mNeoVis.2.1.1.chr14\_hap1.g00368.t01 |
| 15689 | 14 | 41755098 | 41769381 | mNeoVis.2.1.1.chr14\_hap1.g00369.t01 |
| 15690 | 14 | 41774141 | 41915199 | mNeoVis.2.1.1.chr14\_hap1.g00370.t01 |
| 15691 | 14 | 41937694 | 41947770 | mNeoVis.2.1.1.chr14\_hap1.g00371.t01 |
| 15692 | 14 | 41954357 | 42019044 | mNeoVis.2.1.1.chr14\_hap1.g00372.t01 |
| 15693 | 14 | 42036456 | 42082094 | mNeoVis.2.1.1.chr14\_hap1.g00373.t01 |
| 15694 | 14 | 42131591 | 42193717 | mNeoVis.2.1.1.chr14\_hap1.g00374.t01 |
| 15695 | 14 | 42431580 | 42504608 | mNeoVis.2.1.1.chr14\_hap1.g00375.t01 |
| 15696 | 14 | 42431580 | 42504608 | mNeoVis.2.1.1.chr14\_hap1.g00375.t01 |
| 15697 | 14 | 42522980 | 42555523 | mNeoVis.2.1.1.chr14\_hap1.g00376.t01 |
| 15698 | 14 | 42522980 | 42555523 | mNeoVis.2.1.1.chr14\_hap1.g00376.t01 |
| 15699 | 14 | 42558991 | 42561390 | mNeoVis.2.1.1.chr14\_hap1.g00377.t01 |
| 15700 | 14 | 42558991 | 42561390 | mNeoVis.2.1.1.chr14\_hap1.g00377.t01 |
| 15701 | 14 | 42561981 | 42600259 | mNeoVis.2.1.1.chr14\_hap1.g00378.t01 |
| 15702 | 14 | 42561981 | 42600259 | mNeoVis.2.1.1.chr14\_hap1.g00378.t01 |
| 15703 | 14 | 42620544 | 42669560 | mNeoVis.2.1.1.chr14\_hap1.g00379.t01 |
| 15704 | 14 | 42620544 | 42669560 | mNeoVis.2.1.1.chr14\_hap1.g00379.t01 |
| 15705 | 14 | 42688366 | 42726283 | mNeoVis.2.1.1.chr14\_hap1.g00380.t01 |
| 15706 | 14 | 42688366 | 42726283 | mNeoVis.2.1.1.chr14\_hap1.g00380.t01 |
| 15707 | 14 | 42748606 | 42766324 | mNeoVis.2.1.1.chr14\_hap1.g00381.t01 |
| 15708 | 14 | 42748606 | 42766324 | mNeoVis.2.1.1.chr14\_hap1.g00381.t01 |
| 15709 | 14 | 42777934 | 42785275 | mNeoVis.2.1.1.chr14\_hap1.g00382.t01 |
| 15710 | 14 | 42777934 | 42785275 | mNeoVis.2.1.1.chr14\_hap1.g00382.t01 |
| 15711 | 14 | 42817719 | 42844294 | mNeoVis.2.1.1.chr14\_hap1.g00383.t01 |
| 15712 | 14 | 42817719 | 42844294 | mNeoVis.2.1.1.chr14\_hap1.g00383.t01 |
| 15713 | 14 | 42863893 | 42863994 | mNeoVis.2.1.1.chr14\_hap1.g00384.t01 |
| 15714 | 14 | 42863893 | 42863994 | mNeoVis.2.1.1.chr14\_hap1.g00384.t01 |
| 15715 | 14 | 42935384 | 42944630 | mNeoVis.2.1.1.chr14\_hap1.g00385.t01 |
| 15716 | 14 | 42935384 | 42944630 | mNeoVis.2.1.1.chr14\_hap1.g00385.t01 |
| 15717 | 14 | 42972906 | 43000893 | mNeoVis.2.1.1.chr14\_hap1.g00386.t01 |
| 15718 | 14 | 43007026 | 43025931 | mNeoVis.2.1.1.chr14\_hap1.g00387.t01 |
| 15719 | 14 | 43043283 | 43069512 | mNeoVis.2.1.1.chr14\_hap1.g00388.t01 |
| 15720 | 14 | 43093697 | 43126386 | mNeoVis.2.1.1.chr14\_hap1.g00389.t01 |
| 15721 | 14 | 43133085 | 43141087 | mNeoVis.2.1.1.chr14\_hap1.g00390.t01 |
| 15722 | 14 | 43156301 | 43178494 | mNeoVis.2.1.1.chr14\_hap1.g00391.t01 |
| 15723 | 14 | 43192465 | 43213072 | mNeoVis.2.1.1.chr14\_hap1.g00392.t01 |
| 15724 | 14 | 43221721 | 43250129 | mNeoVis.2.1.1.chr14\_hap1.g00393.t01 |
| 15725 | 14 | 43282158 | 43306446 | mNeoVis.2.1.1.chr14\_hap1.g00394.t01 |
| 15726 | 14 | 43315858 | 43328109 | mNeoVis.2.1.1.chr14\_hap1.g00395.t01 |
| 15727 | 14 | 49986907 | 50080805 | mNeoVis.2.1.1.chr14\_hap1.g00407.t01 |
| 15728 | 14 | 52202710 | 52228338 | mNeoVis.2.1.1.chr14\_hap1.g00417.t01 |
| 15729 | 14 | 52202710 | 52228338 | mNeoVis.2.1.1.chr14\_hap1.g00417.t01 |
| 15730 | 14 | 52287190 | 52291670 | mNeoVis.2.1.1.chr14\_hap1.g00418.t01 |
| 15731 | 14 | 52287190 | 52291670 | mNeoVis.2.1.1.chr14\_hap1.g00418.t01 |
| 15732 | 14 | 52483044 | 52502037 | mNeoVis.2.1.1.chr14\_hap1.g00419.t01 |
| 15733 | 14 | 52483044 | 52502037 | mNeoVis.2.1.1.chr14\_hap1.g00419.t01 |
| 15734 | 14 | 52642877 | 52668081 | mNeoVis.2.1.1.chr14\_hap1.g00420.t01 |
| 15735 | 14 | 52642877 | 52668081 | mNeoVis.2.1.1.chr14\_hap1.g00420.t01 |
| 15736 | 14 | 53660015 | 53660886 | mNeoVis.2.1.1.chr14\_hap1.g00421.t01 |
| 15737 | 14 | 53660015 | 53660886 | mNeoVis.2.1.1.chr14\_hap1.g00421.t01 |
| 15738 | 14 | 54110728 | 54510805 | mNeoVis.2.1.1.chr14\_hap1.g00422.t01 |
| 15739 | 14 | 56951076 | 57002541 | mNeoVis.2.1.1.chr14\_hap1.g00424.t01 |
| 15740 | 14 | 57007105 | 57700421 | mNeoVis.2.1.1.chr14\_hap1.g00425.t01 |
| 15741 | 14 | 57781190 | 58012073 | mNeoVis.2.1.1.chr14\_hap1.g00426.t01 |
| 15742 | 14 | 57781190 | 58012073 | mNeoVis.2.1.1.chr14\_hap1.g00426.t01 |
| 15743 | 14 | 58495272 | 58647129 | mNeoVis.2.1.1.chr14\_hap1.g00427.t01 |
| 15744 | 14 | 58650318 | 58819818 | mNeoVis.2.1.1.chr14\_hap1.g00428.t01 |
| 15745 | 14 | 58844626 | 58995784 | mNeoVis.2.1.1.chr14\_hap1.g00429.t01 |
| 15746 | 14 | 59344479 | 59388178 | mNeoVis.2.1.1.chr14\_hap1.g00430.t01 |
| 15747 | 14 | 60044598 | 60080034 | mNeoVis.2.1.1.chr14\_hap1.g00432.t01 |
| 15748 | 14 | 60639020 | 60656730 | mNeoVis.2.1.1.chr14\_hap1.g00433.t01 |
| 15749 | 14 | 60688566 | 60754482 | mNeoVis.2.1.1.chr14\_hap1.g00434.t01 |
| 15750 | 14 | 60790670 | 60846191 | mNeoVis.2.1.1.chr14\_hap1.g00436.t01 |
| 15751 | 14 | 60846657 | 60944622 | mNeoVis.2.1.1.chr14\_hap1.g00437.t01 |
| 15752 | 14 | 61144694 | 61259275 | mNeoVis.2.1.1.chr14\_hap1.g00438.t01 |
| 15753 | 14 | 61988473 | 62069343 | mNeoVis.2.1.1.chr14\_hap1.g00443.t01 |
| 15754 | 14 | 62081910 | 62121583 | mNeoVis.2.1.1.chr14\_hap1.g00444.t01 |
| 15755 | 14 | 62659223 | 62683950 | mNeoVis.2.1.1.chr14\_hap1.g00445.t01 |
| 15756 | 14 | 62692630 | 62722163 | mNeoVis.2.1.1.chr14\_hap1.g00446.t01 |
| 15757 | 14 | 62747990 | 62780666 | mNeoVis.2.1.1.chr14\_hap1.g00447.t01 |
| 15758 | 14 | 62782568 | 62805385 | mNeoVis.2.1.1.chr14\_hap1.g00448.t01 |
| 15759 | 14 | 62807156 | 62846508 | mNeoVis.2.1.1.chr14\_hap1.g00449.t01 |
| 15760 | 14 | 62847474 | 62867754 | mNeoVis.2.1.1.chr14\_hap1.g00450.t01 |
| 15761 | 14 | 62847474 | 62867754 | mNeoVis.2.1.1.chr14\_hap1.g00450.t01 |
| 15762 | 14 | 62847474 | 62867754 | mNeoVis.2.1.1.chr14\_hap1.g00450.t01 |
| 15763 | 14 | 62876105 | 62880898 | mNeoVis.2.1.1.chr14\_hap1.g00451.t01 |
| 15764 | 14 | 62876105 | 62880898 | mNeoVis.2.1.1.chr14\_hap1.g00451.t01 |
| 15765 | 14 | 62876105 | 62880898 | mNeoVis.2.1.1.chr14\_hap1.g00451.t01 |
| 15766 | 14 | 62912500 | 62939029 | mNeoVis.2.1.1.chr14\_hap1.g00452.t01 |
| 15767 | 14 | 62912500 | 62939029 | mNeoVis.2.1.1.chr14\_hap1.g00452.t01 |
| 15768 | 14 | 62912500 | 62939029 | mNeoVis.2.1.1.chr14\_hap1.g00452.t01 |
| 15769 | 14 | 63173289 | 63261823 | mNeoVis.2.1.1.chr14\_hap1.g00453.t01 |
| 15770 | 14 | 63173289 | 63261823 | mNeoVis.2.1.1.chr14\_hap1.g00453.t01 |
| 15771 | 14 | 63362048 | 63436606 | mNeoVis.2.1.1.chr14\_hap1.g00454.t01 |
| 15772 | 14 | 63362048 | 63436606 | mNeoVis.2.1.1.chr14\_hap1.g00454.t01 |
| 15773 | 14 | 63436690 | 63446436 | mNeoVis.2.1.1.chr14\_hap1.g00455.t01 |
| 15774 | 14 | 63436690 | 63446436 | mNeoVis.2.1.1.chr14\_hap1.g00455.t01 |
| 15775 | 14 | 63465333 | 63508518 | mNeoVis.2.1.1.chr14\_hap1.g00456.t01 |
| 15776 | 14 | 63465333 | 63508518 | mNeoVis.2.1.1.chr14\_hap1.g00456.t01 |
| 15777 | 14 | 63522696 | 63568557 | mNeoVis.2.1.1.chr14\_hap1.g00457.t01 |
| 15778 | 14 | 63522696 | 63568557 | mNeoVis.2.1.1.chr14\_hap1.g00457.t01 |
| 15779 | 14 | 63548519 | 63554389 | mNeoVis.2.1.1.chr14\_hap1.g00458.t01 |
| 15780 | 14 | 63548519 | 63554389 | mNeoVis.2.1.1.chr14\_hap1.g00458.t01 |
| 15781 | 14 | 63634704 | 63654288 | mNeoVis.2.1.1.chr14\_hap1.g00459.t01 |
| 15782 | 14 | 63634704 | 63654288 | mNeoVis.2.1.1.chr14\_hap1.g00459.t01 |
| 15783 | 14 | 63740494 | 63781687 | mNeoVis.2.1.1.chr14\_hap1.g00460.t01 |
| 15784 | 14 | 63784307 | 63789791 | mNeoVis.2.1.1.chr14\_hap1.g00461.t01 |
| 15785 | 14 | 63820207 | 63976817 | mNeoVis.2.1.1.chr14\_hap1.g00462.t01 |
| 15786 | 14 | 64259310 | 64540164 | mNeoVis.2.1.1.chr14\_hap1.g00463.t01 |
| 15787 | 14 | 65404458 | 65433055 | mNeoVis.2.1.1.chr14\_hap1.g00464.t01 |
| 15788 | 14 | 65404458 | 65433055 | mNeoVis.2.1.1.chr14\_hap1.g00464.t01 |
| 15789 | 14 | 65432558 | 65438020 | mNeoVis.2.1.1.chr14\_hap1.g00465.t01 |
| 15790 | 14 | 65432558 | 65438020 | mNeoVis.2.1.1.chr14\_hap1.g00465.t01 |
| 15791 | 14 | 65510144 | 65560063 | mNeoVis.2.1.1.chr14\_hap1.g00466.t01 |
| 15792 | 14 | 65510144 | 65560063 | mNeoVis.2.1.1.chr14\_hap1.g00466.t01 |
| 15793 | 14 | 65601601 | 65608776 | mNeoVis.2.1.1.chr14\_hap1.g00467.t01 |
| 15794 | 14 | 65601601 | 65608776 | mNeoVis.2.1.1.chr14\_hap1.g00467.t01 |
| 15795 | 14 | 65762037 | 65769706 | mNeoVis.2.1.1.chr14\_hap1.g00468.t01 |
| 15796 | 14 | 65762037 | 65769706 | mNeoVis.2.1.1.chr14\_hap1.g00468.t01 |
| 15797 | 14 | 65798595 | 65821534 | mNeoVis.2.1.1.chr14\_hap1.g00469.t01 |
| 15798 | 14 | 65798595 | 65821534 | mNeoVis.2.1.1.chr14\_hap1.g00469.t01 |
| 15799 | 14 | 65825312 | 65896335 | mNeoVis.2.1.1.chr14\_hap1.g00470.t01 |
| 15800 | 14 | 65825312 | 65896335 | mNeoVis.2.1.1.chr14\_hap1.g00470.t01 |
| 15801 | 14 | 66016297 | 66052677 | mNeoVis.2.1.1.chr14\_hap1.g00471.t01 |
| 15802 | 14 | 66058265 | 66074531 | mNeoVis.2.1.1.chr14\_hap1.g00472.t01 |
| 15803 | 14 | 66081800 | 66104875 | mNeoVis.2.1.1.chr14\_hap1.g00473.t01 |
| 15804 | 14 | 66127466 | 66151261 | mNeoVis.2.1.1.chr14\_hap1.g00474.t01 |
| 15805 | 14 | 66166852 | 66169475 | mNeoVis.2.1.1.chr14\_hap1.g00476.t01 |
| 15806 | 14 | 66472864 | 66505275 | mNeoVis.2.1.1.chr14\_hap1.g00477.t01 |
| 15807 | 14 | 66525326 | 66739567 | mNeoVis.2.1.1.chr14\_hap1.g00478.t01 |
| 15808 | 14 | 66607548 | 66609009 | mNeoVis.2.1.1.chr14\_hap1.g00479.t01 |
| 15809 | 14 | 66747385 | 66755681 | mNeoVis.2.1.1.chr14\_hap1.g00480.t01 |
| 15810 | 14 | 66774487 | 66778597 | mNeoVis.2.1.1.chr14\_hap1.g00481.t01 |
| 15811 | 14 | 66786102 | 66792926 | mNeoVis.2.1.1.chr14\_hap1.g00482.t01 |
| 15812 | 14 | 66814380 | 66835172 | mNeoVis.2.1.1.chr14\_hap1.g00483.t01 |
| 15813 | 14 | 66872706 | 66879267 | mNeoVis.2.1.1.chr14\_hap1.g00484.t01 |
| 15814 | 14 | 67182592 | 67185617 | mNeoVis.2.1.1.chr14\_hap1.g00485.t01 |
| 15815 | 14 | 67187187 | 67220078 | mNeoVis.2.1.1.chr14\_hap1.g00486.t01 |
| 15816 | 14 | 67224326 | 67229330 | mNeoVis.2.1.1.chr14\_hap1.g00487.t01 |
| 15817 | 14 | 67243581 | 67265712 | mNeoVis.2.1.1.chr14\_hap1.g00488.t01 |
| 15818 | 14 | 67276364 | 67280345 | mNeoVis.2.1.1.chr14\_hap1.g00489.t01 |
| 15819 | 14 | 67280362 | 67284161 | mNeoVis.2.1.1.chr14\_hap1.g00490.t01 |
| 15820 | 14 | 67286434 | 67288053 | mNeoVis.2.1.1.chr14\_hap1.g00491.t01 |
| 15821 | 14 | 67289975 | 67293200 | mNeoVis.2.1.1.chr14\_hap1.g00492.t01 |
| 15822 | 14 | 67293729 | 67303882 | mNeoVis.2.1.1.chr14\_hap1.g00493.t01 |
| 15823 | 14 | 67527544 | 67555899 | mNeoVis.2.1.1.chr14\_hap1.g00508.t01 |
| 15824 | 14 | 67557568 | 67579830 | mNeoVis.2.1.1.chr14\_hap1.g00509.t01 |
| 15825 | 14 | 67586561 | 67596919 | mNeoVis.2.1.1.chr14\_hap1.g00510.t01 |
| 15826 | 14 | 67603399 | 67656086 | mNeoVis.2.1.1.chr14\_hap1.g00511.t01 |
| 15827 | 14 | 67630525 | 67656425 | mNeoVis.2.1.1.chr14\_hap1.g00512.t01 |
| 15828 | 14 | 67679219 | 67679377 | mNeoVis.2.1.1.chr14\_hap1.g00513.t01 |
| 15829 | 14 | 67682098 | 67714012 | mNeoVis.2.1.1.chr14\_hap1.g00514.t01 |
| 15830 | 14 | 67723126 | 67748605 | mNeoVis.2.1.1.chr14\_hap1.g00515.t01 |
| 15831 | 14 | 67756034 | 67786783 | mNeoVis.2.1.1.chr14\_hap1.g00516.t01 |
| 15832 | 14 | 67834277 | 67859054 | mNeoVis.2.1.1.chr14\_hap1.g00517.t01 |
| 15833 | 14 | 67849238 | 67855284 | mNeoVis.2.1.1.chr14\_hap1.g00518.t01 |
| 15834 | 14 | 67882169 | 67882570 | mNeoVis.2.1.1.chr14\_hap1.g00519.t01 |
| 15835 | 14 | 67892838 | 67907498 | mNeoVis.2.1.1.chr14\_hap1.g00520.t01 |
| 15836 | 14 | 67907553 | 67953320 | mNeoVis.2.1.1.chr14\_hap1.g00521.t01 |
| 15837 | 14 | 67957188 | 67963197 | mNeoVis.2.1.1.chr14\_hap1.g00522.t01 |
| 15838 | 14 | 67974806 | 68015142 | mNeoVis.2.1.1.chr14\_hap1.g00523.t01 |
| 15839 | 14 | 68016346 | 68025540 | mNeoVis.2.1.1.chr14\_hap1.g00524.t01 |
| 15840 | 14 | 68028197 | 68036526 | mNeoVis.2.1.1.chr14\_hap1.g00525.t01 |
| 15841 | 14 | 68040566 | 68069307 | mNeoVis.2.1.1.chr14\_hap1.g00526.t01 |
| 15842 | 14 | 68075293 | 68098986 | mNeoVis.2.1.1.chr14\_hap1.g00527.t01 |
| 15843 | 14 | 68111453 | 68118358 | mNeoVis.2.1.1.chr14\_hap1.g00528.t01 |
| 15844 | 14 | 68119727 | 68127207 | mNeoVis.2.1.1.chr14\_hap1.g00529.t01 |
| 15845 | 14 | 68134392 | 68163241 | mNeoVis.2.1.1.chr14\_hap1.g00530.t01 |
| 15846 | 14 | 68169566 | 68171696 | mNeoVis.2.1.1.chr14\_hap1.g00531.t01 |
| 15847 | 14 | 68207087 | 68218935 | mNeoVis.2.1.1.chr14\_hap1.g00532.t01 |
| 15848 | 14 | 68230111 | 68241355 | mNeoVis.2.1.1.chr14\_hap1.g00533.t01 |
| 15849 | 14 | 68242810 | 68249931 | mNeoVis.2.1.1.chr14\_hap1.g00534.t01 |
| 15850 | 14 | 68264181 | 68286969 | mNeoVis.2.1.1.chr14\_hap1.g00535.t01 |
| 15851 | 14 | 68295785 | 68303012 | mNeoVis.2.1.1.chr14\_hap1.g00536.t01 |
| 15852 | 14 | 68314537 | 68325592 | mNeoVis.2.1.1.chr14\_hap1.g00537.t01 |
| 15853 | 14 | 68332857 | 68341039 | mNeoVis.2.1.1.chr14\_hap1.g00538.t01 |
| 15854 | 14 | 68367185 | 68368518 | mNeoVis.2.1.1.chr14\_hap1.g00539.t01 |
| 15855 | 14 | 68508820 | 68756954 | mNeoVis.2.1.1.chr14\_hap1.g00540.t01 |
| 15856 | 14 | 68518444 | 68519248 | mNeoVis.2.1.1.chr14\_hap1.g00541.t01 |
| 15857 | 14 | 68581184 | 68582644 | mNeoVis.2.1.1.chr14\_hap1.g00542.t01 |
| 15858 | 14 | 68636846 | 68637795 | mNeoVis.2.1.1.chr14\_hap1.g00543.t01 |
| 15859 | 14 | 68645419 | 68646408 | mNeoVis.2.1.1.chr14\_hap1.g00544.t01 |
| 15860 | 14 | 68706349 | 68707290 | mNeoVis.2.1.1.chr14\_hap1.g00545.t01 |
| 15861 | 14 | 68728072 | 68729663 | mNeoVis.2.1.1.chr14\_hap1.g00546.t01 |
| 15862 | 14 | 68734686 | 68740226 | mNeoVis.2.1.1.chr14\_hap1.g00547.t01 |
| 15863 | 14 | 68760950 | 68762105 | mNeoVis.2.1.1.chr14\_hap1.g00548.t01 |
| 15864 | 14 | 68819311 | 68822056 | mNeoVis.2.1.1.chr14\_hap1.g00549.t01 |
| 15865 | 14 | 69438918 | 69442051 | mNeoVis.2.1.1.chr14\_hap1.g00576.t01 |
| 15866 | 14 | 69465121 | 69467888 | mNeoVis.2.1.1.chr14\_hap1.g00577.t01 |
| 15867 | 14 | 69482220 | 69485702 | mNeoVis.2.1.1.chr14\_hap1.g00578.t01 |
| 15868 | 14 | 69497674 | 69501144 | mNeoVis.2.1.1.chr14\_hap1.g00579.t01 |
| 15869 | 14 | 69512139 | 69531404 | mNeoVis.2.1.1.chr14\_hap1.g00580.t01 |
| 15870 | 14 | 69558496 | 69564185 | mNeoVis.2.1.1.chr14\_hap1.g00581.t01 |
| 15871 | 14 | 69585408 | 69586445 | mNeoVis.2.1.1.chr14\_hap1.g00582.t01 |
| 15872 | 14 | 69604611 | 69692455 | mNeoVis.2.1.1.chr14\_hap1.g00583.t01 |
| 15873 | 14 | 69615014 | 69619840 | mNeoVis.2.1.1.chr14\_hap1.g00584.t01 |
| 15874 | 14 | 69722568 | 69744603 | mNeoVis.2.1.1.chr14\_hap1.g00585.t01 |
| 15875 | 14 | 69777704 | 69786518 | mNeoVis.2.1.1.chr14\_hap1.g00586.t01 |
| 15876 | 14 | 69798752 | 69808317 | mNeoVis.2.1.1.chr14\_hap1.g00587.t01 |
| 15877 | 14 | 69816807 | 69875857 | mNeoVis.2.1.1.chr14\_hap1.g00588.t01 |
| 15878 | 14 | 69904384 | 69926758 | mNeoVis.2.1.1.chr14\_hap1.g00589.t01 |
| 15879 | 14 | 69940147 | 69973221 | mNeoVis.2.1.1.chr14\_hap1.g00590.t01 |
| 15880 | 14 | 70207718 | 70221466 | mNeoVis.2.1.1.chr14\_hap1.g00596.t01 |
| 15881 | 14 | 70239792 | 70244281 | mNeoVis.2.1.1.chr14\_hap1.g00597.t01 |
| 15882 | 14 | 70262202 | 70339655 | mNeoVis.2.1.1.chr14\_hap1.g00598.t01 |
| 15883 | 14 | 70340959 | 70350996 | mNeoVis.2.1.1.chr14\_hap1.g00599.t01 |
| 15884 | 14 | 70351936 | 70370901 | mNeoVis.2.1.1.chr14\_hap1.g00600.t01 |
| 15885 | 14 | 70382183 | 70399441 | mNeoVis.2.1.1.chr14\_hap1.g00601.t01 |
| 15886 | 14 | 70401616 | 70416203 | mNeoVis.2.1.1.chr14\_hap1.g00602.t01 |
| 15887 | 14 | 70425940 | 70433860 | mNeoVis.2.1.1.chr14\_hap1.g00603.t01 |
| 15888 | 14 | 70438179 | 70461008 | mNeoVis.2.1.1.chr14\_hap1.g00604.t01 |
| 15889 | 14 | 70473433 | 70482247 | mNeoVis.2.1.1.chr14\_hap1.g00605.t01 |
| 15890 | 14 | 70483094 | 70487308 | mNeoVis.2.1.1.chr14\_hap1.g00606.t01 |
| 15891 | 14 | 70487601 | 70493705 | mNeoVis.2.1.1.chr14\_hap1.g00607.t01 |
| 15892 | 14 | 70493999 | 70498448 | mNeoVis.2.1.1.chr14\_hap1.g00608.t01 |
| 15893 | 14 | 70511240 | 70515679 | mNeoVis.2.1.1.chr14\_hap1.g00609.t01 |
| 15894 | 14 | 70528746 | 70537580 | mNeoVis.2.1.1.chr14\_hap1.g00610.t01 |
| 15895 | 14 | 70544906 | 70557533 | mNeoVis.2.1.1.chr14\_hap1.g00611.t01 |
| 15896 | 14 | 70572737 | 70576959 | mNeoVis.2.1.1.chr14\_hap1.g00612.t01 |
| 15897 | 14 | 70585187 | 70591430 | mNeoVis.2.1.1.chr14\_hap1.g00613.t01 |
| 15898 | 14 | 70591758 | 70593963 | mNeoVis.2.1.1.chr14\_hap1.g00614.t01 |
| 15899 | 14 | 70597302 | 70602991 | mNeoVis.2.1.1.chr14\_hap1.g00615.t01 |
| 15900 | 14 | 70607575 | 70642332 | mNeoVis.2.1.1.chr14\_hap1.g00616.t01 |
| 15901 | 14 | 70663419 | 70693717 | mNeoVis.2.1.1.chr14\_hap1.g00617.t01 |
| 15902 | 14 | 70756334 | 70766102 | mNeoVis.2.1.1.chr14\_hap1.g00618.t01 |
| 15903 | 14 | 70775846 | 70781611 | mNeoVis.2.1.1.chr14\_hap1.g00619.t01 |
| 15904 | 14 | 70781911 | 70799634 | mNeoVis.2.1.1.chr14\_hap1.g00620.t01 |
| 15905 | 14 | 70802883 | 70805501 | mNeoVis.2.1.1.chr14\_hap1.g00621.t01 |
| 15906 | 14 | 70805720 | 70811968 | mNeoVis.2.1.1.chr14\_hap1.g00622.t01 |
| 15907 | 14 | 70814859 | 70837107 | mNeoVis.2.1.1.chr14\_hap1.g00623.t01 |
| 15908 | 14 | 70838568 | 70842277 | mNeoVis.2.1.1.chr14\_hap1.g00624.t01 |
| 15909 | 14 | 70842370 | 70843181 | mNeoVis.2.1.1.chr14\_hap1.g00625.t01 |
| 15910 | 14 | 70847845 | 70871921 | mNeoVis.2.1.1.chr14\_hap1.g00626.t01 |
| 15911 | 14 | 70877943 | 70895020 | mNeoVis.2.1.1.chr14\_hap1.g00627.t01 |
| 15912 | 14 | 70904636 | 70921046 | mNeoVis.2.1.1.chr14\_hap1.g00628.t01 |
| 15913 | 14 | 70942475 | 70952779 | mNeoVis.2.1.1.chr14\_hap1.g00629.t01 |
| 15914 | 14 | 70953555 | 70959310 | mNeoVis.2.1.1.chr14\_hap1.g00630.t01 |
| 15915 | 14 | 70960366 | 70963408 | mNeoVis.2.1.1.chr14\_hap1.g00631.t01 |
| 15916 | 14 | 70964103 | 70978627 | mNeoVis.2.1.1.chr14\_hap1.g00632.t01 |
| 15917 | 14 | 70986882 | 70992529 | mNeoVis.2.1.1.chr14\_hap1.g00633.t01 |
| 15918 | 14 | 71001893 | 71037657 | mNeoVis.2.1.1.chr14\_hap1.g00634.t01 |
| 15919 | 14 | 71054142 | 71072298 | mNeoVis.2.1.1.chr14\_hap1.g00635.t01 |
| 15920 | 14 | 71073075 | 71078586 | mNeoVis.2.1.1.chr14\_hap1.g00636.t01 |
| 15921 | 14 | 71088219 | 71105541 | mNeoVis.2.1.1.chr14\_hap1.g00637.t01 |
| 15922 | 14 | 71106005 | 71172590 | mNeoVis.2.1.1.chr14\_hap1.g00638.t01 |
| 15923 | 14 | 71172732 | 71176747 | mNeoVis.2.1.1.chr14\_hap1.g00639.t01 |
| 15924 | 14 | 71184698 | 71210983 | mNeoVis.2.1.1.chr14\_hap1.g00640.t01 |
| 15925 | 14 | 71211362 | 71390787 | mNeoVis.2.1.1.chr14\_hap1.g00641.t01 |
| 15926 | 14 | 71967779 | 71970870 | mNeoVis.2.1.1.chr14\_hap1.g00678.t01 |
| 15927 | 14 | 71975028 | 71979748 | mNeoVis.2.1.1.chr14\_hap1.g00679.t01 |
| 15928 | 14 | 71981526 | 71982796 | mNeoVis.2.1.1.chr14\_hap1.g00680.t01 |
| 15929 | 14 | 71982979 | 72068314 | mNeoVis.2.1.1.chr14\_hap1.g00681.t01 |
| 15930 | 14 | 72069129 | 72107959 | mNeoVis.2.1.1.chr14\_hap1.g00682.t01 |
| 15931 | 14 | 72113691 | 72115403 | mNeoVis.2.1.1.chr14\_hap1.g00683.t01 |
| 15932 | 14 | 72115505 | 72158252 | mNeoVis.2.1.1.chr14\_hap1.g00684.t01 |
| 15933 | 14 | 72159449 | 72162189 | mNeoVis.2.1.1.chr14\_hap1.g00685.t01 |
| 15934 | 14 | 72167540 | 72171724 | mNeoVis.2.1.1.chr14\_hap1.g00686.t01 |
| 15935 | 14 | 72175606 | 72191670 | mNeoVis.2.1.1.chr14\_hap1.g00687.t01 |
| 15936 | 14 | 72214945 | 72250154 | mNeoVis.2.1.1.chr14\_hap1.g00688.t01 |
| 15937 | 14 | 72259183 | 72274482 | mNeoVis.2.1.1.chr14\_hap1.g00689.t01 |
| 15938 | 14 | 72306877 | 72313963 | mNeoVis.2.1.1.chr14\_hap1.g00690.t01 |
| 15939 | 14 | 72321687 | 72334061 | mNeoVis.2.1.1.chr14\_hap1.g00691.t01 |
| 15940 | 14 | 72333914 | 72355337 | mNeoVis.2.1.1.chr14\_hap1.g00692.t01 |
| 15941 | 14 | 72405092 | 72522781 | mNeoVis.2.1.1.chr14\_hap1.g00693.t01 |
| 15942 | 14 | 72549750 | 72557788 | mNeoVis.2.1.1.chr14\_hap1.g00694.t01 |
| 15943 | 14 | 72563967 | 72573609 | mNeoVis.2.1.1.chr14\_hap1.g00695.t01 |
| 15944 | 14 | 72574369 | 72578118 | mNeoVis.2.1.1.chr14\_hap1.g00696.t01 |
| 15945 | 14 | 72578482 | 72586183 | mNeoVis.2.1.1.chr14\_hap1.g00697.t01 |
| 15946 | 14 | 72588739 | 72591949 | mNeoVis.2.1.1.chr14\_hap1.g00698.t01 |
| 15947 | 14 | 72594656 | 72600120 | mNeoVis.2.1.1.chr14\_hap1.g00699.t01 |
| 15948 | 14 | 72600506 | 72601709 | mNeoVis.2.1.1.chr14\_hap1.g00700.t01 |
| 15949 | 14 | 72608334 | 72623399 | mNeoVis.2.1.1.chr14\_hap1.g00701.t01 |
| 15950 | 14 | 72623413 | 72632286 | mNeoVis.2.1.1.chr14\_hap1.g00702.t01 |
| 15951 | 14 | 72632356 | 72645181 | mNeoVis.2.1.1.chr14\_hap1.g00703.t01 |
| 15952 | 14 | 72645468 | 72655238 | mNeoVis.2.1.1.chr14\_hap1.g00704.t01 |
| 15953 | 15 | 8703961 | 8704686 | mNeoVis.2.1.1.chr15\_hap1.g00006.t01 |
| 15954 | 15 | 8706906 | 8741937 | mNeoVis.2.1.1.chr15\_hap1.g00007.t01 |
| 15955 | 15 | 8716743 | 8717244 | mNeoVis.2.1.1.chr15\_hap1.g00008.t01 |
| 15956 | 15 | 8752545 | 8790977 | mNeoVis.2.1.1.chr15\_hap1.g00009.t01 |
| 15957 | 15 | 8816271 | 8860244 | mNeoVis.2.1.1.chr15\_hap1.g00010.t01 |
| 15958 | 15 | 8910027 | 8917631 | mNeoVis.2.1.1.chr15\_hap1.g00011.t01 |
| 15959 | 15 | 8926274 | 8927620 | mNeoVis.2.1.1.chr15\_hap1.g00012.t01 |
| 15960 | 15 | 8937372 | 8947423 | mNeoVis.2.1.1.chr15\_hap1.g00013.t01 |
| 15961 | 15 | 8970075 | 8980907 | mNeoVis.2.1.1.chr15\_hap1.g00014.t01 |
| 15962 | 15 | 8981675 | 8987272 | mNeoVis.2.1.1.chr15\_hap1.g00015.t01 |
| 15963 | 15 | 8987235 | 8998635 | mNeoVis.2.1.1.chr15\_hap1.g00016.t01 |
| 15964 | 15 | 9001755 | 9006029 | mNeoVis.2.1.1.chr15\_hap1.g00017.t01 |
| 15965 | 15 | 9015263 | 9017367 | mNeoVis.2.1.1.chr15\_hap1.g00018.t01 |
| 15966 | 15 | 9018632 | 9023765 | mNeoVis.2.1.1.chr15\_hap1.g00019.t01 |
| 15967 | 15 | 9024005 | 9032280 | mNeoVis.2.1.1.chr15\_hap1.g00020.t01 |
| 15968 | 15 | 9033043 | 9042167 | mNeoVis.2.1.1.chr15\_hap1.g00021.t01 |
| 15969 | 15 | 9060587 | 9066577 | mNeoVis.2.1.1.chr15\_hap1.g00022.t01 |
| 15970 | 15 | 9078608 | 9093258 | mNeoVis.2.1.1.chr15\_hap1.g00023.t01 |
| 15971 | 15 | 9095716 | 9099131 | mNeoVis.2.1.1.chr15\_hap1.g00024.t01 |
| 15972 | 15 | 9100270 | 9123243 | mNeoVis.2.1.1.chr15\_hap1.g00025.t01 |
| 15973 | 15 | 9123379 | 9128421 | mNeoVis.2.1.1.chr15\_hap1.g00026.t01 |
| 15974 | 15 | 9128960 | 9147473 | mNeoVis.2.1.1.chr15\_hap1.g00027.t01 |
| 15975 | 15 | 9170061 | 9172874 | mNeoVis.2.1.1.chr15\_hap1.g00028.t01 |
| 15976 | 15 | 9173306 | 9177016 | mNeoVis.2.1.1.chr15\_hap1.g00029.t01 |
| 15977 | 15 | 9179391 | 9183330 | mNeoVis.2.1.1.chr15\_hap1.g00030.t01 |
| 15978 | 15 | 9243524 | 9268803 | mNeoVis.2.1.1.chr15\_hap1.g00031.t01 |
| 15979 | 15 | 9271891 | 9283860 | mNeoVis.2.1.1.chr15\_hap1.g00032.t01 |
| 15980 | 15 | 9283917 | 9287350 | mNeoVis.2.1.1.chr15\_hap1.g00033.t01 |
| 15981 | 15 | 9287360 | 9304656 | mNeoVis.2.1.1.chr15\_hap1.g00034.t01 |
| 15982 | 15 | 9305088 | 9337683 | mNeoVis.2.1.1.chr15\_hap1.g00035.t01 |
| 15983 | 15 | 9365860 | 9378862 | mNeoVis.2.1.1.chr15\_hap1.g00036.t01 |
| 15984 | 15 | 9380058 | 9385236 | mNeoVis.2.1.1.chr15\_hap1.g00037.t01 |
| 15985 | 15 | 10105076 | 10116315 | mNeoVis.2.1.1.chr15\_hap1.g00088.t01 |
| 15986 | 15 | 10127917 | 10145963 | mNeoVis.2.1.1.chr15\_hap1.g00089.t01 |
| 15987 | 15 | 10178156 | 10191485 | mNeoVis.2.1.1.chr15\_hap1.g00090.t01 |
| 15988 | 15 | 10193228 | 10196660 | mNeoVis.2.1.1.chr15\_hap1.g00091.t01 |
| 15989 | 15 | 10209987 | 10216856 | mNeoVis.2.1.1.chr15\_hap1.g00092.t01 |
| 15990 | 15 | 10220867 | 10235842 | mNeoVis.2.1.1.chr15\_hap1.g00093.t01 |
| 15991 | 15 | 10235995 | 10248715 | mNeoVis.2.1.1.chr15\_hap1.g00094.t01 |
| 15992 | 15 | 10250866 | 10257005 | mNeoVis.2.1.1.chr15\_hap1.g00095.t01 |
| 15993 | 15 | 10263750 | 10273847 | mNeoVis.2.1.1.chr15\_hap1.g00096.t01 |
| 15994 | 15 | 10279106 | 10286970 | mNeoVis.2.1.1.chr15\_hap1.g00097.t01 |
| 15995 | 15 | 10287577 | 10291573 | mNeoVis.2.1.1.chr15\_hap1.g00098.t01 |
| 15996 | 15 | 10319863 | 10367963 | mNeoVis.2.1.1.chr15\_hap1.g00099.t01 |
| 15997 | 15 | 10319863 | 10367963 | mNeoVis.2.1.1.chr15\_hap1.g00099.t01 |
| 15998 | 15 | 10394392 | 10493586 | mNeoVis.2.1.1.chr15\_hap1.g00100.t01 |
| 15999 | 15 | 10394392 | 10493586 | mNeoVis.2.1.1.chr15\_hap1.g00100.t01 |
| 16000 | 15 | 10394392 | 10493586 | mNeoVis.2.1.1.chr15\_hap1.g00100.t01 |
| 16001 | 15 | 10517853 | 10714239 | mNeoVis.2.1.1.chr15\_hap1.g00101.t01 |
| 16002 | 15 | 10517853 | 10714239 | mNeoVis.2.1.1.chr15\_hap1.g00101.t01 |
| 16003 | 15 | 10718586 | 10892073 | mNeoVis.2.1.1.chr15\_hap1.g00102.t01 |
| 16004 | 15 | 10718586 | 10892073 | mNeoVis.2.1.1.chr15\_hap1.g00102.t01 |
| 16005 | 15 | 10892206 | 10960854 | mNeoVis.2.1.1.chr15\_hap1.g00103.t01 |
| 16006 | 15 | 10968008 | 11001000 | mNeoVis.2.1.1.chr15\_hap1.g00104.t01 |
| 16007 | 15 | 11032048 | 11067557 | mNeoVis.2.1.1.chr15\_hap1.g00105.t01 |
| 16008 | 15 | 11032048 | 11067557 | mNeoVis.2.1.1.chr15\_hap1.g00105.t01 |
| 16009 | 15 | 11110813 | 11147227 | mNeoVis.2.1.1.chr15\_hap1.g00106.t01 |
| 16010 | 15 | 11110813 | 11147227 | mNeoVis.2.1.1.chr15\_hap1.g00106.t01 |
| 16011 | 15 | 11147745 | 11162465 | mNeoVis.2.1.1.chr15\_hap1.g00107.t01 |
| 16012 | 15 | 11147745 | 11162465 | mNeoVis.2.1.1.chr15\_hap1.g00107.t01 |
| 16013 | 15 | 11358770 | 11436480 | mNeoVis.2.1.1.chr15\_hap1.g00108.t01 |
| 16014 | 15 | 11358770 | 11436480 | mNeoVis.2.1.1.chr15\_hap1.g00108.t01 |
| 16015 | 15 | 12726184 | 12742746 | mNeoVis.2.1.1.chr15\_hap1.g00110.t01 |
| 16016 | 15 | 12747499 | 12754491 | mNeoVis.2.1.1.chr15\_hap1.g00111.t01 |
| 16017 | 15 | 12777495 | 12835127 | mNeoVis.2.1.1.chr15\_hap1.g00112.t01 |
| 16018 | 15 | 12875078 | 12890532 | mNeoVis.2.1.1.chr15\_hap1.g00113.t01 |
| 16019 | 15 | 12888796 | 12934223 | mNeoVis.2.1.1.chr15\_hap1.g00114.t01 |
| 16020 | 15 | 12903472 | 12917533 | mNeoVis.2.1.1.chr15\_hap1.g00115.t01 |
| 16021 | 15 | 12931334 | 12945937 | mNeoVis.2.1.1.chr15\_hap1.g00116.t01 |
| 16022 | 15 | 12931334 | 12973269 | mNeoVis.2.1.1.chr15\_hap1.g00117.t01 |
| 16023 | 15 | 12987074 | 13001132 | mNeoVis.2.1.1.chr15\_hap1.g00118.t01 |
| 16024 | 15 | 13014936 | 13056813 | mNeoVis.2.1.1.chr15\_hap1.g00119.t01 |
| 16025 | 15 | 13015630 | 13029829 | mNeoVis.2.1.1.chr15\_hap1.g00120.t01 |
| 16026 | 15 | 13613302 | 13652678 | mNeoVis.2.1.1.chr15\_hap1.g00134.t01 |
| 16027 | 15 | 13666380 | 13693112 | mNeoVis.2.1.1.chr15\_hap1.g00135.t01 |
| 16028 | 15 | 13755077 | 13803052 | mNeoVis.2.1.1.chr15\_hap1.g00136.t01 |
| 16029 | 15 | 13788911 | 13802338 | mNeoVis.2.1.1.chr15\_hap1.g00137.t01 |
| 16030 | 15 | 13815068 | 13886166 | mNeoVis.2.1.1.chr15\_hap1.g00138.t01 |
| 16031 | 15 | 14077553 | 14163758 | mNeoVis.2.1.1.chr15\_hap1.g00139.t01 |
| 16032 | 15 | 14077553 | 14163758 | mNeoVis.2.1.1.chr15\_hap1.g00139.t01 |
| 16033 | 15 | 14138120 | 14163807 | mNeoVis.2.1.1.chr15\_hap1.g00140.t01 |
| 16034 | 15 | 14138120 | 14163807 | mNeoVis.2.1.1.chr15\_hap1.g00140.t01 |
| 16035 | 15 | 14282595 | 14348059 | mNeoVis.2.1.1.chr15\_hap1.g00141.t01 |
| 16036 | 15 | 14282595 | 14348059 | mNeoVis.2.1.1.chr15\_hap1.g00141.t01 |
| 16037 | 15 | 14374001 | 14399720 | mNeoVis.2.1.1.chr15\_hap1.g00142.t01 |
| 16038 | 15 | 14374001 | 14399720 | mNeoVis.2.1.1.chr15\_hap1.g00142.t01 |
| 16039 | 15 | 14460810 | 14518421 | mNeoVis.2.1.1.chr15\_hap1.g00143.t01 |
| 16040 | 15 | 14521332 | 14602847 | mNeoVis.2.1.1.chr15\_hap1.g00144.t01 |
| 16041 | 15 | 14547219 | 14549416 | mNeoVis.2.1.1.chr15\_hap1.g00145.t01 |
| 16042 | 15 | 14606485 | 14647026 | mNeoVis.2.1.1.chr15\_hap1.g00146.t01 |
| 16043 | 15 | 14651596 | 14678362 | mNeoVis.2.1.1.chr15\_hap1.g00147.t01 |
| 16044 | 15 | 14678482 | 14685394 | mNeoVis.2.1.1.chr15\_hap1.g00148.t01 |
| 16045 | 15 | 14678482 | 14685394 | mNeoVis.2.1.1.chr15\_hap1.g00148.t01 |
| 16046 | 15 | 14701415 | 14710932 | mNeoVis.2.1.1.chr15\_hap1.g00149.t01 |
| 16047 | 15 | 14701415 | 14710932 | mNeoVis.2.1.1.chr15\_hap1.g00149.t01 |
| 16048 | 15 | 14716631 | 14742657 | mNeoVis.2.1.1.chr15\_hap1.g00150.t01 |
| 16049 | 15 | 14716631 | 14742657 | mNeoVis.2.1.1.chr15\_hap1.g00150.t01 |
| 16050 | 15 | 14742782 | 14762929 | mNeoVis.2.1.1.chr15\_hap1.g00151.t01 |
| 16051 | 15 | 14742782 | 14762929 | mNeoVis.2.1.1.chr15\_hap1.g00151.t01 |
| 16052 | 15 | 14776250 | 14786134 | mNeoVis.2.1.1.chr15\_hap1.g00152.t01 |
| 16053 | 15 | 14776250 | 14786134 | mNeoVis.2.1.1.chr15\_hap1.g00152.t01 |
| 16054 | 15 | 14786258 | 14801459 | mNeoVis.2.1.1.chr15\_hap1.g00153.t01 |
| 16055 | 15 | 14786258 | 14801459 | mNeoVis.2.1.1.chr15\_hap1.g00153.t01 |
| 16056 | 15 | 14821158 | 14827875 | mNeoVis.2.1.1.chr15\_hap1.g00154.t01 |
| 16057 | 15 | 14821158 | 14827875 | mNeoVis.2.1.1.chr15\_hap1.g00154.t01 |
| 16058 | 15 | 14897810 | 15219785 | mNeoVis.2.1.1.chr15\_hap1.g00155.t01 |
| 16059 | 15 | 18775141 | 18827893 | mNeoVis.2.1.1.chr15\_hap1.g00200.t01 |
| 16060 | 15 | 18834621 | 18849710 | mNeoVis.2.1.1.chr15\_hap1.g00201.t01 |
| 16061 | 15 | 18871007 | 18877591 | mNeoVis.2.1.1.chr15\_hap1.g00202.t01 |
| 16062 | 15 | 18923996 | 19058211 | mNeoVis.2.1.1.chr15\_hap1.g00203.t01 |
| 16063 | 15 | 19137241 | 19246041 | mNeoVis.2.1.1.chr15\_hap1.g00204.t01 |
| 16064 | 15 | 19248388 | 19256977 | mNeoVis.2.1.1.chr15\_hap1.g00205.t01 |
| 16065 | 15 | 19263814 | 19271494 | mNeoVis.2.1.1.chr15\_hap1.g00206.t01 |
| 16066 | 15 | 19681900 | 19976367 | mNeoVis.2.1.1.chr15\_hap1.g00208.t01 |
| 16067 | 15 | 20674879 | 20730012 | mNeoVis.2.1.1.chr15\_hap1.g00209.t01 |
| 16068 | 15 | 20674879 | 20730012 | mNeoVis.2.1.1.chr15\_hap1.g00209.t01 |
| 16069 | 15 | 20733065 | 20783728 | mNeoVis.2.1.1.chr15\_hap1.g00210.t01 |
| 16070 | 15 | 20733065 | 20783728 | mNeoVis.2.1.1.chr15\_hap1.g00210.t01 |
| 16071 | 15 | 20783870 | 20912947 | mNeoVis.2.1.1.chr15\_hap1.g00211.t01 |
| 16072 | 15 | 20946770 | 20965054 | mNeoVis.2.1.1.chr15\_hap1.g00212.t01 |
| 16073 | 15 | 20970741 | 21083695 | mNeoVis.2.1.1.chr15\_hap1.g00213.t01 |
| 16074 | 15 | 21084700 | 21117701 | mNeoVis.2.1.1.chr15\_hap1.g00214.t01 |
| 16075 | 15 | 21122042 | 21161598 | mNeoVis.2.1.1.chr15\_hap1.g00215.t01 |
| 16076 | 15 | 21163348 | 21276795 | mNeoVis.2.1.1.chr15\_hap1.g00216.t01 |
| 16077 | 15 | 21305505 | 21317076 | mNeoVis.2.1.1.chr15\_hap1.g00217.t01 |
| 16078 | 15 | 21334677 | 21387669 | mNeoVis.2.1.1.chr15\_hap1.g00218.t01 |
| 16079 | 15 | 21386855 | 21399405 | mNeoVis.2.1.1.chr15\_hap1.g00219.t01 |
| 16080 | 15 | 21400979 | 21431237 | mNeoVis.2.1.1.chr15\_hap1.g00220.t01 |
| 16081 | 15 | 21440894 | 21448047 | mNeoVis.2.1.1.chr15\_hap1.g00221.t01 |
| 16082 | 15 | 21858277 | 22116982 | mNeoVis.2.1.1.chr15\_hap1.g00227.t01 |
| 16083 | 15 | 21858277 | 22116982 | mNeoVis.2.1.1.chr15\_hap1.g00227.t01 |
| 16084 | 15 | 22117004 | 22156088 | mNeoVis.2.1.1.chr15\_hap1.g00228.t01 |
| 16085 | 15 | 22117004 | 22156088 | mNeoVis.2.1.1.chr15\_hap1.g00228.t01 |
| 16086 | 15 | 22739854 | 23016818 | mNeoVis.2.1.1.chr15\_hap1.g00229.t01 |
| 16087 | 15 | 23098329 | 23200589 | mNeoVis.2.1.1.chr15\_hap1.g00230.t01 |
| 16088 | 15 | 30981613 | 30985351 | mNeoVis.2.1.1.chr15\_hap1.g00316.t01 |
| 16089 | 15 | 30988537 | 30997681 | mNeoVis.2.1.1.chr15\_hap1.g00317.t01 |
| 16090 | 15 | 31012686 | 31019918 | mNeoVis.2.1.1.chr15\_hap1.g00318.t01 |
| 16091 | 15 | 31019976 | 31022948 | mNeoVis.2.1.1.chr15\_hap1.g00319.t01 |
| 16092 | 15 | 31023214 | 31024551 | mNeoVis.2.1.1.chr15\_hap1.g00320.t01 |
| 16093 | 15 | 31024796 | 31026869 | mNeoVis.2.1.1.chr15\_hap1.g00321.t01 |
| 16094 | 15 | 31027593 | 31030209 | mNeoVis.2.1.1.chr15\_hap1.g00322.t01 |
| 16095 | 15 | 31030473 | 31032333 | mNeoVis.2.1.1.chr15\_hap1.g00323.t01 |
| 16096 | 15 | 31036542 | 31038497 | mNeoVis.2.1.1.chr15\_hap1.g00324.t01 |
| 16097 | 15 | 31040733 | 31052567 | mNeoVis.2.1.1.chr15\_hap1.g00325.t01 |
| 16098 | 15 | 31057528 | 31068046 | mNeoVis.2.1.1.chr15\_hap1.g00326.t01 |
| 16099 | 15 | 31068191 | 31071561 | mNeoVis.2.1.1.chr15\_hap1.g00327.t01 |
| 16100 | 15 | 31072519 | 31076603 | mNeoVis.2.1.1.chr15\_hap1.g00328.t01 |
| 16101 | 15 | 31085199 | 31107470 | mNeoVis.2.1.1.chr15\_hap1.g00329.t01 |
| 16102 | 15 | 31114464 | 31123254 | mNeoVis.2.1.1.chr15\_hap1.g00330.t01 |
| 16103 | 15 | 31148658 | 31166864 | mNeoVis.2.1.1.chr15\_hap1.g00331.t01 |
| 16104 | 15 | 31167350 | 31175774 | mNeoVis.2.1.1.chr15\_hap1.g00332.t01 |
| 16105 | 15 | 31176939 | 31188835 | mNeoVis.2.1.1.chr15\_hap1.g00333.t01 |
| 16106 | 15 | 31186107 | 31199108 | mNeoVis.2.1.1.chr15\_hap1.g00334.t01 |
| 16107 | 15 | 31195747 | 31226402 | mNeoVis.2.1.1.chr15\_hap1.g00335.t01 |
| 16108 | 15 | 31227323 | 31232780 | mNeoVis.2.1.1.chr15\_hap1.g00336.t01 |
| 16109 | 15 | 31253052 | 31259205 | mNeoVis.2.1.1.chr15\_hap1.g00337.t01 |
| 16110 | 15 | 31262844 | 31286521 | mNeoVis.2.1.1.chr15\_hap1.g00338.t01 |
| 16111 | 15 | 31286809 | 31368900 | mNeoVis.2.1.1.chr15\_hap1.g00339.t01 |
| 16112 | 15 | 31374857 | 31384419 | mNeoVis.2.1.1.chr15\_hap1.g00340.t01 |
| 16113 | 15 | 31384560 | 31389954 | mNeoVis.2.1.1.chr15\_hap1.g00341.t01 |
| 16114 | 15 | 31400573 | 31407706 | mNeoVis.2.1.1.chr15\_hap1.g00342.t01 |
| 16115 | 15 | 31400573 | 31407706 | mNeoVis.2.1.1.chr15\_hap1.g00342.t01 |
| 16116 | 15 | 31423898 | 31428204 | mNeoVis.2.1.1.chr15\_hap1.g00343.t01 |
| 16117 | 15 | 31423898 | 31428204 | mNeoVis.2.1.1.chr15\_hap1.g00343.t01 |
| 16118 | 15 | 31433768 | 31438200 | mNeoVis.2.1.1.chr15\_hap1.g00344.t01 |
| 16119 | 15 | 31433768 | 31438200 | mNeoVis.2.1.1.chr15\_hap1.g00344.t01 |
| 16120 | 15 | 31439269 | 31458163 | mNeoVis.2.1.1.chr15\_hap1.g00345.t01 |
| 16121 | 15 | 31439269 | 31458163 | mNeoVis.2.1.1.chr15\_hap1.g00345.t01 |
| 16122 | 15 | 31460560 | 31526932 | mNeoVis.2.1.1.chr15\_hap1.g00346.t01 |
| 16123 | 15 | 31460560 | 31526932 | mNeoVis.2.1.1.chr15\_hap1.g00346.t01 |
| 16124 | 15 | 31526842 | 31566155 | mNeoVis.2.1.1.chr15\_hap1.g00347.t01 |
| 16125 | 15 | 31526842 | 31566155 | mNeoVis.2.1.1.chr15\_hap1.g00347.t01 |
| 16126 | 15 | 31526842 | 31566155 | mNeoVis.2.1.1.chr15\_hap1.g00347.t01 |
| 16127 | 15 | 31526842 | 31566155 | mNeoVis.2.1.1.chr15\_hap1.g00347.t01 |
| 16128 | 15 | 31526842 | 31566155 | mNeoVis.2.1.1.chr15\_hap1.g00347.t01 |
| 16129 | 15 | 31572259 | 31581491 | mNeoVis.2.1.1.chr15\_hap1.g00348.t01 |
| 16130 | 15 | 31572259 | 31581491 | mNeoVis.2.1.1.chr15\_hap1.g00348.t01 |
| 16131 | 15 | 31572259 | 31581491 | mNeoVis.2.1.1.chr15\_hap1.g00348.t01 |
| 16132 | 15 | 31572259 | 31581491 | mNeoVis.2.1.1.chr15\_hap1.g00348.t01 |
| 16133 | 15 | 31572259 | 31581491 | mNeoVis.2.1.1.chr15\_hap1.g00348.t01 |
| 16134 | 15 | 31581678 | 31590512 | mNeoVis.2.1.1.chr15\_hap1.g00349.t01 |
| 16135 | 15 | 31581678 | 31590512 | mNeoVis.2.1.1.chr15\_hap1.g00349.t01 |
| 16136 | 15 | 31581678 | 31590512 | mNeoVis.2.1.1.chr15\_hap1.g00349.t01 |
| 16137 | 15 | 31581678 | 31590512 | mNeoVis.2.1.1.chr15\_hap1.g00349.t01 |
| 16138 | 15 | 31581678 | 31590512 | mNeoVis.2.1.1.chr15\_hap1.g00349.t01 |
| 16139 | 15 | 31591711 | 31594000 | mNeoVis.2.1.1.chr15\_hap1.g00350.t01 |
| 16140 | 15 | 31591711 | 31594000 | mNeoVis.2.1.1.chr15\_hap1.g00350.t01 |
| 16141 | 15 | 31591711 | 31594000 | mNeoVis.2.1.1.chr15\_hap1.g00350.t01 |
| 16142 | 15 | 31591711 | 31594000 | mNeoVis.2.1.1.chr15\_hap1.g00350.t01 |
| 16143 | 15 | 31591711 | 31594000 | mNeoVis.2.1.1.chr15\_hap1.g00350.t01 |
| 16144 | 15 | 31594757 | 31604114 | mNeoVis.2.1.1.chr15\_hap1.g00351.t01 |
| 16145 | 15 | 31594757 | 31604114 | mNeoVis.2.1.1.chr15\_hap1.g00351.t01 |
| 16146 | 15 | 31594757 | 31604114 | mNeoVis.2.1.1.chr15\_hap1.g00351.t01 |
| 16147 | 15 | 31594757 | 31604114 | mNeoVis.2.1.1.chr15\_hap1.g00351.t01 |
| 16148 | 15 | 31594757 | 31604114 | mNeoVis.2.1.1.chr15\_hap1.g00351.t01 |
| 16149 | 15 | 31608722 | 31609865 | mNeoVis.2.1.1.chr15\_hap1.g00352.t01 |
| 16150 | 15 | 31608722 | 31609865 | mNeoVis.2.1.1.chr15\_hap1.g00352.t01 |
| 16151 | 15 | 31608722 | 31609865 | mNeoVis.2.1.1.chr15\_hap1.g00352.t01 |
| 16152 | 15 | 31608722 | 31609865 | mNeoVis.2.1.1.chr15\_hap1.g00352.t01 |
| 16153 | 15 | 31608722 | 31609865 | mNeoVis.2.1.1.chr15\_hap1.g00352.t01 |
| 16154 | 15 | 31610558 | 31614081 | mNeoVis.2.1.1.chr15\_hap1.g00353.t01 |
| 16155 | 15 | 31610558 | 31614081 | mNeoVis.2.1.1.chr15\_hap1.g00353.t01 |
| 16156 | 15 | 31610558 | 31614081 | mNeoVis.2.1.1.chr15\_hap1.g00353.t01 |
| 16157 | 15 | 31610558 | 31614081 | mNeoVis.2.1.1.chr15\_hap1.g00353.t01 |
| 16158 | 15 | 31610558 | 31614081 | mNeoVis.2.1.1.chr15\_hap1.g00353.t01 |
| 16159 | 15 | 31614088 | 31617831 | mNeoVis.2.1.1.chr15\_hap1.g00354.t01 |
| 16160 | 15 | 31614088 | 31617831 | mNeoVis.2.1.1.chr15\_hap1.g00354.t01 |
| 16161 | 15 | 31614088 | 31617831 | mNeoVis.2.1.1.chr15\_hap1.g00354.t01 |
| 16162 | 15 | 31614088 | 31617831 | mNeoVis.2.1.1.chr15\_hap1.g00354.t01 |
| 16163 | 15 | 31614088 | 31617831 | mNeoVis.2.1.1.chr15\_hap1.g00354.t01 |
| 16164 | 15 | 31624670 | 31641764 | mNeoVis.2.1.1.chr15\_hap1.g00355.t01 |
| 16165 | 15 | 31624670 | 31641764 | mNeoVis.2.1.1.chr15\_hap1.g00355.t01 |
| 16166 | 15 | 31624670 | 31641764 | mNeoVis.2.1.1.chr15\_hap1.g00355.t01 |
| 16167 | 15 | 31624670 | 31641764 | mNeoVis.2.1.1.chr15\_hap1.g00355.t01 |
| 16168 | 15 | 31624670 | 31641764 | mNeoVis.2.1.1.chr15\_hap1.g00355.t01 |
| 16169 | 15 | 31641953 | 31658906 | mNeoVis.2.1.1.chr15\_hap1.g00356.t01 |
| 16170 | 15 | 31641953 | 31658906 | mNeoVis.2.1.1.chr15\_hap1.g00356.t01 |
| 16171 | 15 | 31641953 | 31658906 | mNeoVis.2.1.1.chr15\_hap1.g00356.t01 |
| 16172 | 15 | 31641953 | 31658906 | mNeoVis.2.1.1.chr15\_hap1.g00356.t01 |
| 16173 | 15 | 31641953 | 31658906 | mNeoVis.2.1.1.chr15\_hap1.g00356.t01 |
| 16174 | 15 | 31663819 | 31669583 | mNeoVis.2.1.1.chr15\_hap1.g00357.t01 |
| 16175 | 15 | 31663819 | 31669583 | mNeoVis.2.1.1.chr15\_hap1.g00357.t01 |
| 16176 | 15 | 31663819 | 31669583 | mNeoVis.2.1.1.chr15\_hap1.g00357.t01 |
| 16177 | 15 | 31663819 | 31669583 | mNeoVis.2.1.1.chr15\_hap1.g00357.t01 |
| 16178 | 15 | 31663819 | 31669583 | mNeoVis.2.1.1.chr15\_hap1.g00357.t01 |
| 16179 | 15 | 31680110 | 31721783 | mNeoVis.2.1.1.chr15\_hap1.g00358.t01 |
| 16180 | 15 | 31680110 | 31721783 | mNeoVis.2.1.1.chr15\_hap1.g00358.t01 |
| 16181 | 15 | 31680110 | 31721783 | mNeoVis.2.1.1.chr15\_hap1.g00358.t01 |
| 16182 | 15 | 31680110 | 31721783 | mNeoVis.2.1.1.chr15\_hap1.g00358.t01 |
| 16183 | 15 | 31680110 | 31721783 | mNeoVis.2.1.1.chr15\_hap1.g00358.t01 |
| 16184 | 15 | 31721785 | 31753747 | mNeoVis.2.1.1.chr15\_hap1.g00359.t01 |
| 16185 | 15 | 31721785 | 31753747 | mNeoVis.2.1.1.chr15\_hap1.g00359.t01 |
| 16186 | 15 | 31721785 | 31753747 | mNeoVis.2.1.1.chr15\_hap1.g00359.t01 |
| 16187 | 15 | 31721785 | 31753747 | mNeoVis.2.1.1.chr15\_hap1.g00359.t01 |
| 16188 | 15 | 31721785 | 31753747 | mNeoVis.2.1.1.chr15\_hap1.g00359.t01 |
| 16189 | 15 | 31753850 | 31759161 | mNeoVis.2.1.1.chr15\_hap1.g00360.t01 |
| 16190 | 15 | 31753850 | 31759161 | mNeoVis.2.1.1.chr15\_hap1.g00360.t01 |
| 16191 | 15 | 31753850 | 31759161 | mNeoVis.2.1.1.chr15\_hap1.g00360.t01 |
| 16192 | 15 | 31753850 | 31759161 | mNeoVis.2.1.1.chr15\_hap1.g00360.t01 |
| 16193 | 15 | 31753850 | 31759161 | mNeoVis.2.1.1.chr15\_hap1.g00360.t01 |
| 16194 | 15 | 31759915 | 31771697 | mNeoVis.2.1.1.chr15\_hap1.g00361.t01 |
| 16195 | 15 | 31759915 | 31771697 | mNeoVis.2.1.1.chr15\_hap1.g00361.t01 |
| 16196 | 15 | 31759915 | 31771697 | mNeoVis.2.1.1.chr15\_hap1.g00361.t01 |
| 16197 | 15 | 31759915 | 31771697 | mNeoVis.2.1.1.chr15\_hap1.g00361.t01 |
| 16198 | 15 | 31759915 | 31771697 | mNeoVis.2.1.1.chr15\_hap1.g00361.t01 |
| 16199 | 15 | 31776610 | 31784546 | mNeoVis.2.1.1.chr15\_hap1.g00362.t01 |
| 16200 | 15 | 31776610 | 31784546 | mNeoVis.2.1.1.chr15\_hap1.g00362.t01 |
| 16201 | 15 | 31776610 | 31784546 | mNeoVis.2.1.1.chr15\_hap1.g00362.t01 |
| 16202 | 15 | 31776610 | 31784546 | mNeoVis.2.1.1.chr15\_hap1.g00362.t01 |
| 16203 | 15 | 31776610 | 31784546 | mNeoVis.2.1.1.chr15\_hap1.g00362.t01 |
| 16204 | 15 | 31784811 | 31795501 | mNeoVis.2.1.1.chr15\_hap1.g00363.t01 |
| 16205 | 15 | 31784811 | 31795501 | mNeoVis.2.1.1.chr15\_hap1.g00363.t01 |
| 16206 | 15 | 31784811 | 31795501 | mNeoVis.2.1.1.chr15\_hap1.g00363.t01 |
| 16207 | 15 | 31784811 | 31795501 | mNeoVis.2.1.1.chr15\_hap1.g00363.t01 |
| 16208 | 15 | 31784811 | 31795501 | mNeoVis.2.1.1.chr15\_hap1.g00363.t01 |
| 16209 | 15 | 31798309 | 31803184 | mNeoVis.2.1.1.chr15\_hap1.g00364.t01 |
| 16210 | 15 | 31798309 | 31803184 | mNeoVis.2.1.1.chr15\_hap1.g00364.t01 |
| 16211 | 15 | 31798309 | 31803184 | mNeoVis.2.1.1.chr15\_hap1.g00364.t01 |
| 16212 | 15 | 31798309 | 31803184 | mNeoVis.2.1.1.chr15\_hap1.g00364.t01 |
| 16213 | 15 | 31798309 | 31803184 | mNeoVis.2.1.1.chr15\_hap1.g00364.t01 |
| 16214 | 15 | 31806187 | 31808199 | mNeoVis.2.1.1.chr15\_hap1.g00365.t01 |
| 16215 | 15 | 31806187 | 31808199 | mNeoVis.2.1.1.chr15\_hap1.g00365.t01 |
| 16216 | 15 | 31806187 | 31808199 | mNeoVis.2.1.1.chr15\_hap1.g00365.t01 |
| 16217 | 15 | 31806187 | 31808199 | mNeoVis.2.1.1.chr15\_hap1.g00365.t01 |
| 16218 | 15 | 31806187 | 31808199 | mNeoVis.2.1.1.chr15\_hap1.g00365.t01 |
| 16219 | 15 | 31810192 | 31813091 | mNeoVis.2.1.1.chr15\_hap1.g00366.t01 |
| 16220 | 15 | 31810192 | 31813091 | mNeoVis.2.1.1.chr15\_hap1.g00366.t01 |
| 16221 | 15 | 31810192 | 31813091 | mNeoVis.2.1.1.chr15\_hap1.g00366.t01 |
| 16222 | 15 | 31810192 | 31813091 | mNeoVis.2.1.1.chr15\_hap1.g00366.t01 |
| 16223 | 15 | 31810192 | 31813091 | mNeoVis.2.1.1.chr15\_hap1.g00366.t01 |
| 16224 | 15 | 31813157 | 31819597 | mNeoVis.2.1.1.chr15\_hap1.g00367.t01 |
| 16225 | 15 | 31813157 | 31819597 | mNeoVis.2.1.1.chr15\_hap1.g00367.t01 |
| 16226 | 15 | 31813157 | 31819597 | mNeoVis.2.1.1.chr15\_hap1.g00367.t01 |
| 16227 | 15 | 31813157 | 31819597 | mNeoVis.2.1.1.chr15\_hap1.g00367.t01 |
| 16228 | 15 | 31813157 | 31819597 | mNeoVis.2.1.1.chr15\_hap1.g00367.t01 |
| 16229 | 15 | 31820678 | 31824813 | mNeoVis.2.1.1.chr15\_hap1.g00368.t01 |
| 16230 | 15 | 31820678 | 31824813 | mNeoVis.2.1.1.chr15\_hap1.g00368.t01 |
| 16231 | 15 | 31820678 | 31824813 | mNeoVis.2.1.1.chr15\_hap1.g00368.t01 |
| 16232 | 15 | 31820678 | 31824813 | mNeoVis.2.1.1.chr15\_hap1.g00368.t01 |
| 16233 | 15 | 31820678 | 31824813 | mNeoVis.2.1.1.chr15\_hap1.g00368.t01 |
| 16234 | 15 | 31824810 | 31826689 | mNeoVis.2.1.1.chr15\_hap1.g00369.t01 |
| 16235 | 15 | 31824810 | 31826689 | mNeoVis.2.1.1.chr15\_hap1.g00369.t01 |
| 16236 | 15 | 31824810 | 31826689 | mNeoVis.2.1.1.chr15\_hap1.g00369.t01 |
| 16237 | 15 | 31824810 | 31826689 | mNeoVis.2.1.1.chr15\_hap1.g00369.t01 |
| 16238 | 15 | 31824810 | 31826689 | mNeoVis.2.1.1.chr15\_hap1.g00369.t01 |
| 16239 | 15 | 31826683 | 31829538 | mNeoVis.2.1.1.chr15\_hap1.g00370.t01 |
| 16240 | 15 | 31826683 | 31829538 | mNeoVis.2.1.1.chr15\_hap1.g00370.t01 |
| 16241 | 15 | 31826683 | 31829538 | mNeoVis.2.1.1.chr15\_hap1.g00370.t01 |
| 16242 | 15 | 31826683 | 31829538 | mNeoVis.2.1.1.chr15\_hap1.g00370.t01 |
| 16243 | 15 | 31826683 | 31829538 | mNeoVis.2.1.1.chr15\_hap1.g00370.t01 |
| 16244 | 15 | 31828786 | 31841401 | mNeoVis.2.1.1.chr15\_hap1.g00371.t01 |
| 16245 | 15 | 31828786 | 31841401 | mNeoVis.2.1.1.chr15\_hap1.g00371.t01 |
| 16246 | 15 | 31828786 | 31841401 | mNeoVis.2.1.1.chr15\_hap1.g00371.t01 |
| 16247 | 15 | 31828786 | 31841401 | mNeoVis.2.1.1.chr15\_hap1.g00371.t01 |
| 16248 | 15 | 31828786 | 31841401 | mNeoVis.2.1.1.chr15\_hap1.g00371.t01 |
| 16249 | 15 | 31853758 | 31859810 | mNeoVis.2.1.1.chr15\_hap1.g00372.t01 |
| 16250 | 15 | 31853758 | 31859810 | mNeoVis.2.1.1.chr15\_hap1.g00372.t01 |
| 16251 | 15 | 31853758 | 31859810 | mNeoVis.2.1.1.chr15\_hap1.g00372.t01 |
| 16252 | 15 | 31853758 | 31859810 | mNeoVis.2.1.1.chr15\_hap1.g00372.t01 |
| 16253 | 15 | 31853758 | 31859810 | mNeoVis.2.1.1.chr15\_hap1.g00372.t01 |
| 16254 | 15 | 31891943 | 31924910 | mNeoVis.2.1.1.chr15\_hap1.g00373.t01 |
| 16255 | 15 | 31891943 | 31924910 | mNeoVis.2.1.1.chr15\_hap1.g00373.t01 |
| 16256 | 15 | 31891943 | 31924910 | mNeoVis.2.1.1.chr15\_hap1.g00373.t01 |
| 16257 | 15 | 31891943 | 31924910 | mNeoVis.2.1.1.chr15\_hap1.g00373.t01 |
| 16258 | 15 | 31891943 | 31924910 | mNeoVis.2.1.1.chr15\_hap1.g00373.t01 |
| 16259 | 15 | 31931971 | 31934091 | mNeoVis.2.1.1.chr15\_hap1.g00374.t01 |
| 16260 | 15 | 31931971 | 31934091 | mNeoVis.2.1.1.chr15\_hap1.g00374.t01 |
| 16261 | 15 | 31931971 | 31934091 | mNeoVis.2.1.1.chr15\_hap1.g00374.t01 |
| 16262 | 15 | 31931971 | 31934091 | mNeoVis.2.1.1.chr15\_hap1.g00374.t01 |
| 16263 | 15 | 31931971 | 31934091 | mNeoVis.2.1.1.chr15\_hap1.g00374.t01 |
| 16264 | 15 | 31934328 | 31953298 | mNeoVis.2.1.1.chr15\_hap1.g00375.t01 |
| 16265 | 15 | 31934328 | 31953298 | mNeoVis.2.1.1.chr15\_hap1.g00375.t01 |
| 16266 | 15 | 31934328 | 31953298 | mNeoVis.2.1.1.chr15\_hap1.g00375.t01 |
| 16267 | 15 | 31934328 | 31953298 | mNeoVis.2.1.1.chr15\_hap1.g00375.t01 |
| 16268 | 15 | 31934328 | 31953298 | mNeoVis.2.1.1.chr15\_hap1.g00375.t01 |
| 16269 | 15 | 31953439 | 31963610 | mNeoVis.2.1.1.chr15\_hap1.g00376.t01 |
| 16270 | 15 | 31953439 | 31963610 | mNeoVis.2.1.1.chr15\_hap1.g00376.t01 |
| 16271 | 15 | 31953439 | 31963610 | mNeoVis.2.1.1.chr15\_hap1.g00376.t01 |
| 16272 | 15 | 31953439 | 31963610 | mNeoVis.2.1.1.chr15\_hap1.g00376.t01 |
| 16273 | 15 | 31953439 | 31963610 | mNeoVis.2.1.1.chr15\_hap1.g00376.t01 |
| 16274 | 15 | 31963605 | 31965722 | mNeoVis.2.1.1.chr15\_hap1.g00377.t01 |
| 16275 | 15 | 31963605 | 31965722 | mNeoVis.2.1.1.chr15\_hap1.g00377.t01 |
| 16276 | 15 | 31963605 | 31965722 | mNeoVis.2.1.1.chr15\_hap1.g00377.t01 |
| 16277 | 15 | 31963605 | 31965722 | mNeoVis.2.1.1.chr15\_hap1.g00377.t01 |
| 16278 | 15 | 31963605 | 31965722 | mNeoVis.2.1.1.chr15\_hap1.g00377.t01 |
| 16279 | 15 | 31966712 | 31979361 | mNeoVis.2.1.1.chr15\_hap1.g00378.t01 |
| 16280 | 15 | 31966712 | 31979361 | mNeoVis.2.1.1.chr15\_hap1.g00378.t01 |
| 16281 | 15 | 31966712 | 31979361 | mNeoVis.2.1.1.chr15\_hap1.g00378.t01 |
| 16282 | 15 | 31966712 | 31979361 | mNeoVis.2.1.1.chr15\_hap1.g00378.t01 |
| 16283 | 15 | 31966712 | 31979361 | mNeoVis.2.1.1.chr15\_hap1.g00378.t01 |
| 16284 | 15 | 31970414 | 31975951 | mNeoVis.2.1.1.chr15\_hap1.g00379.t01 |
| 16285 | 15 | 31970414 | 31975951 | mNeoVis.2.1.1.chr15\_hap1.g00379.t01 |
| 16286 | 15 | 31970414 | 31975951 | mNeoVis.2.1.1.chr15\_hap1.g00379.t01 |
| 16287 | 15 | 31970414 | 31975951 | mNeoVis.2.1.1.chr15\_hap1.g00379.t01 |
| 16288 | 15 | 31970414 | 31975951 | mNeoVis.2.1.1.chr15\_hap1.g00379.t01 |
| 16289 | 15 | 31979330 | 31980849 | mNeoVis.2.1.1.chr15\_hap1.g00380.t01 |
| 16290 | 15 | 31979330 | 31980849 | mNeoVis.2.1.1.chr15\_hap1.g00380.t01 |
| 16291 | 15 | 31979330 | 31980849 | mNeoVis.2.1.1.chr15\_hap1.g00380.t01 |
| 16292 | 15 | 31979330 | 31980849 | mNeoVis.2.1.1.chr15\_hap1.g00380.t01 |
| 16293 | 15 | 31979330 | 31980849 | mNeoVis.2.1.1.chr15\_hap1.g00380.t01 |
| 16294 | 15 | 31980492 | 31981542 | mNeoVis.2.1.1.chr15\_hap1.g00381.t01 |
| 16295 | 15 | 31980492 | 31981542 | mNeoVis.2.1.1.chr15\_hap1.g00381.t01 |
| 16296 | 15 | 31980492 | 31981542 | mNeoVis.2.1.1.chr15\_hap1.g00381.t01 |
| 16297 | 15 | 31980492 | 31981542 | mNeoVis.2.1.1.chr15\_hap1.g00381.t01 |
| 16298 | 15 | 31980492 | 31981542 | mNeoVis.2.1.1.chr15\_hap1.g00381.t01 |
| 16299 | 15 | 31981591 | 32027445 | mNeoVis.2.1.1.chr15\_hap1.g00382.t01 |
| 16300 | 15 | 31981591 | 32027445 | mNeoVis.2.1.1.chr15\_hap1.g00382.t01 |
| 16301 | 15 | 31981591 | 32027445 | mNeoVis.2.1.1.chr15\_hap1.g00382.t01 |
| 16302 | 15 | 31981591 | 32027445 | mNeoVis.2.1.1.chr15\_hap1.g00382.t01 |
| 16303 | 15 | 31981591 | 32027445 | mNeoVis.2.1.1.chr15\_hap1.g00382.t01 |
| 16304 | 15 | 32033367 | 32043596 | mNeoVis.2.1.1.chr15\_hap1.g00383.t01 |
| 16305 | 15 | 32033367 | 32043596 | mNeoVis.2.1.1.chr15\_hap1.g00383.t01 |
| 16306 | 15 | 32033367 | 32043596 | mNeoVis.2.1.1.chr15\_hap1.g00383.t01 |
| 16307 | 15 | 32033367 | 32043596 | mNeoVis.2.1.1.chr15\_hap1.g00383.t01 |
| 16308 | 15 | 32033367 | 32043596 | mNeoVis.2.1.1.chr15\_hap1.g00383.t01 |
| 16309 | 15 | 32033367 | 32043596 | mNeoVis.2.1.1.chr15\_hap1.g00383.t01 |
| 16310 | 15 | 32043272 | 32082496 | mNeoVis.2.1.1.chr15\_hap1.g00384.t01 |
| 16311 | 15 | 32043272 | 32082496 | mNeoVis.2.1.1.chr15\_hap1.g00384.t01 |
| 16312 | 15 | 32043272 | 32082496 | mNeoVis.2.1.1.chr15\_hap1.g00384.t01 |
| 16313 | 15 | 32043272 | 32082496 | mNeoVis.2.1.1.chr15\_hap1.g00384.t01 |
| 16314 | 15 | 32043272 | 32082496 | mNeoVis.2.1.1.chr15\_hap1.g00384.t01 |
| 16315 | 15 | 32043272 | 32082496 | mNeoVis.2.1.1.chr15\_hap1.g00384.t01 |
| 16316 | 15 | 32088601 | 32097832 | mNeoVis.2.1.1.chr15\_hap1.g00385.t01 |
| 16317 | 15 | 32088601 | 32097832 | mNeoVis.2.1.1.chr15\_hap1.g00385.t01 |
| 16318 | 15 | 32088601 | 32097832 | mNeoVis.2.1.1.chr15\_hap1.g00385.t01 |
| 16319 | 15 | 32088601 | 32097832 | mNeoVis.2.1.1.chr15\_hap1.g00385.t01 |
| 16320 | 15 | 32088601 | 32097832 | mNeoVis.2.1.1.chr15\_hap1.g00385.t01 |
| 16321 | 15 | 32088601 | 32097832 | mNeoVis.2.1.1.chr15\_hap1.g00385.t01 |
| 16322 | 15 | 32098019 | 32106853 | mNeoVis.2.1.1.chr15\_hap1.g00386.t01 |
| 16323 | 15 | 32098019 | 32106853 | mNeoVis.2.1.1.chr15\_hap1.g00386.t01 |
| 16324 | 15 | 32098019 | 32106853 | mNeoVis.2.1.1.chr15\_hap1.g00386.t01 |
| 16325 | 15 | 32098019 | 32106853 | mNeoVis.2.1.1.chr15\_hap1.g00386.t01 |
| 16326 | 15 | 32098019 | 32106853 | mNeoVis.2.1.1.chr15\_hap1.g00386.t01 |
| 16327 | 15 | 32098019 | 32106853 | mNeoVis.2.1.1.chr15\_hap1.g00386.t01 |
| 16328 | 15 | 32108052 | 32110341 | mNeoVis.2.1.1.chr15\_hap1.g00387.t01 |
| 16329 | 15 | 32108052 | 32110341 | mNeoVis.2.1.1.chr15\_hap1.g00387.t01 |
| 16330 | 15 | 32108052 | 32110341 | mNeoVis.2.1.1.chr15\_hap1.g00387.t01 |
| 16331 | 15 | 32108052 | 32110341 | mNeoVis.2.1.1.chr15\_hap1.g00387.t01 |
| 16332 | 15 | 32108052 | 32110341 | mNeoVis.2.1.1.chr15\_hap1.g00387.t01 |
| 16333 | 15 | 32108052 | 32110341 | mNeoVis.2.1.1.chr15\_hap1.g00387.t01 |
| 16334 | 15 | 32111098 | 32120455 | mNeoVis.2.1.1.chr15\_hap1.g00388.t01 |
| 16335 | 15 | 32111098 | 32120455 | mNeoVis.2.1.1.chr15\_hap1.g00388.t01 |
| 16336 | 15 | 32111098 | 32120455 | mNeoVis.2.1.1.chr15\_hap1.g00388.t01 |
| 16337 | 15 | 32111098 | 32120455 | mNeoVis.2.1.1.chr15\_hap1.g00388.t01 |
| 16338 | 15 | 32111098 | 32120455 | mNeoVis.2.1.1.chr15\_hap1.g00388.t01 |
| 16339 | 15 | 32111098 | 32120455 | mNeoVis.2.1.1.chr15\_hap1.g00388.t01 |
| 16340 | 15 | 32124982 | 32126206 | mNeoVis.2.1.1.chr15\_hap1.g00389.t01 |
| 16341 | 15 | 32124982 | 32126206 | mNeoVis.2.1.1.chr15\_hap1.g00389.t01 |
| 16342 | 15 | 32124982 | 32126206 | mNeoVis.2.1.1.chr15\_hap1.g00389.t01 |
| 16343 | 15 | 32124982 | 32126206 | mNeoVis.2.1.1.chr15\_hap1.g00389.t01 |
| 16344 | 15 | 32124982 | 32126206 | mNeoVis.2.1.1.chr15\_hap1.g00389.t01 |
| 16345 | 15 | 32124982 | 32126206 | mNeoVis.2.1.1.chr15\_hap1.g00389.t01 |
| 16346 | 15 | 32126899 | 32130422 | mNeoVis.2.1.1.chr15\_hap1.g00390.t01 |
| 16347 | 15 | 32126899 | 32130422 | mNeoVis.2.1.1.chr15\_hap1.g00390.t01 |
| 16348 | 15 | 32126899 | 32130422 | mNeoVis.2.1.1.chr15\_hap1.g00390.t01 |
| 16349 | 15 | 32126899 | 32130422 | mNeoVis.2.1.1.chr15\_hap1.g00390.t01 |
| 16350 | 15 | 32126899 | 32130422 | mNeoVis.2.1.1.chr15\_hap1.g00390.t01 |
| 16351 | 15 | 32126899 | 32130422 | mNeoVis.2.1.1.chr15\_hap1.g00390.t01 |
| 16352 | 15 | 32130429 | 32134172 | mNeoVis.2.1.1.chr15\_hap1.g00391.t01 |
| 16353 | 15 | 32130429 | 32134172 | mNeoVis.2.1.1.chr15\_hap1.g00391.t01 |
| 16354 | 15 | 32130429 | 32134172 | mNeoVis.2.1.1.chr15\_hap1.g00391.t01 |
| 16355 | 15 | 32130429 | 32134172 | mNeoVis.2.1.1.chr15\_hap1.g00391.t01 |
| 16356 | 15 | 32130429 | 32134172 | mNeoVis.2.1.1.chr15\_hap1.g00391.t01 |
| 16357 | 15 | 32130429 | 32134172 | mNeoVis.2.1.1.chr15\_hap1.g00391.t01 |
| 16358 | 15 | 32141011 | 32158105 | mNeoVis.2.1.1.chr15\_hap1.g00392.t01 |
| 16359 | 15 | 32141011 | 32158105 | mNeoVis.2.1.1.chr15\_hap1.g00392.t01 |
| 16360 | 15 | 32141011 | 32158105 | mNeoVis.2.1.1.chr15\_hap1.g00392.t01 |
| 16361 | 15 | 32141011 | 32158105 | mNeoVis.2.1.1.chr15\_hap1.g00392.t01 |
| 16362 | 15 | 32141011 | 32158105 | mNeoVis.2.1.1.chr15\_hap1.g00392.t01 |
| 16363 | 15 | 32141011 | 32158105 | mNeoVis.2.1.1.chr15\_hap1.g00392.t01 |
| 16364 | 15 | 32158294 | 32175247 | mNeoVis.2.1.1.chr15\_hap1.g00393.t01 |
| 16365 | 15 | 32158294 | 32175247 | mNeoVis.2.1.1.chr15\_hap1.g00393.t01 |
| 16366 | 15 | 32158294 | 32175247 | mNeoVis.2.1.1.chr15\_hap1.g00393.t01 |
| 16367 | 15 | 32158294 | 32175247 | mNeoVis.2.1.1.chr15\_hap1.g00393.t01 |
| 16368 | 15 | 32158294 | 32175247 | mNeoVis.2.1.1.chr15\_hap1.g00393.t01 |
| 16369 | 15 | 32158294 | 32175247 | mNeoVis.2.1.1.chr15\_hap1.g00393.t01 |
| 16370 | 15 | 32180160 | 32185924 | mNeoVis.2.1.1.chr15\_hap1.g00394.t01 |
| 16371 | 15 | 32180160 | 32185924 | mNeoVis.2.1.1.chr15\_hap1.g00394.t01 |
| 16372 | 15 | 32180160 | 32185924 | mNeoVis.2.1.1.chr15\_hap1.g00394.t01 |
| 16373 | 15 | 32180160 | 32185924 | mNeoVis.2.1.1.chr15\_hap1.g00394.t01 |
| 16374 | 15 | 32180160 | 32185924 | mNeoVis.2.1.1.chr15\_hap1.g00394.t01 |
| 16375 | 15 | 32180160 | 32185924 | mNeoVis.2.1.1.chr15\_hap1.g00394.t01 |
| 16376 | 15 | 32196451 | 32238124 | mNeoVis.2.1.1.chr15\_hap1.g00395.t01 |
| 16377 | 15 | 32196451 | 32238124 | mNeoVis.2.1.1.chr15\_hap1.g00395.t01 |
| 16378 | 15 | 32196451 | 32238124 | mNeoVis.2.1.1.chr15\_hap1.g00395.t01 |
| 16379 | 15 | 32196451 | 32238124 | mNeoVis.2.1.1.chr15\_hap1.g00395.t01 |
| 16380 | 15 | 32196451 | 32238124 | mNeoVis.2.1.1.chr15\_hap1.g00395.t01 |
| 16381 | 15 | 32196451 | 32238124 | mNeoVis.2.1.1.chr15\_hap1.g00395.t01 |
| 16382 | 15 | 32238126 | 32270088 | mNeoVis.2.1.1.chr15\_hap1.g00396.t01 |
| 16383 | 15 | 32238126 | 32270088 | mNeoVis.2.1.1.chr15\_hap1.g00396.t01 |
| 16384 | 15 | 32238126 | 32270088 | mNeoVis.2.1.1.chr15\_hap1.g00396.t01 |
| 16385 | 15 | 32238126 | 32270088 | mNeoVis.2.1.1.chr15\_hap1.g00396.t01 |
| 16386 | 15 | 32238126 | 32270088 | mNeoVis.2.1.1.chr15\_hap1.g00396.t01 |
| 16387 | 15 | 32238126 | 32270088 | mNeoVis.2.1.1.chr15\_hap1.g00396.t01 |
| 16388 | 15 | 32270191 | 32275502 | mNeoVis.2.1.1.chr15\_hap1.g00397.t01 |
| 16389 | 15 | 32270191 | 32275502 | mNeoVis.2.1.1.chr15\_hap1.g00397.t01 |
| 16390 | 15 | 32270191 | 32275502 | mNeoVis.2.1.1.chr15\_hap1.g00397.t01 |
| 16391 | 15 | 32270191 | 32275502 | mNeoVis.2.1.1.chr15\_hap1.g00397.t01 |
| 16392 | 15 | 32270191 | 32275502 | mNeoVis.2.1.1.chr15\_hap1.g00397.t01 |
| 16393 | 15 | 32270191 | 32275502 | mNeoVis.2.1.1.chr15\_hap1.g00397.t01 |
| 16394 | 15 | 32276256 | 32288038 | mNeoVis.2.1.1.chr15\_hap1.g00398.t01 |
| 16395 | 15 | 32276256 | 32288038 | mNeoVis.2.1.1.chr15\_hap1.g00398.t01 |
| 16396 | 15 | 32276256 | 32288038 | mNeoVis.2.1.1.chr15\_hap1.g00398.t01 |
| 16397 | 15 | 32276256 | 32288038 | mNeoVis.2.1.1.chr15\_hap1.g00398.t01 |
| 16398 | 15 | 32276256 | 32288038 | mNeoVis.2.1.1.chr15\_hap1.g00398.t01 |
| 16399 | 15 | 32276256 | 32288038 | mNeoVis.2.1.1.chr15\_hap1.g00398.t01 |
| 16400 | 15 | 32292951 | 32300887 | mNeoVis.2.1.1.chr15\_hap1.g00399.t01 |
| 16401 | 15 | 32292951 | 32300887 | mNeoVis.2.1.1.chr15\_hap1.g00399.t01 |
| 16402 | 15 | 32292951 | 32300887 | mNeoVis.2.1.1.chr15\_hap1.g00399.t01 |
| 16403 | 15 | 32292951 | 32300887 | mNeoVis.2.1.1.chr15\_hap1.g00399.t01 |
| 16404 | 15 | 32292951 | 32300887 | mNeoVis.2.1.1.chr15\_hap1.g00399.t01 |
| 16405 | 15 | 32292951 | 32300887 | mNeoVis.2.1.1.chr15\_hap1.g00399.t01 |
| 16406 | 15 | 32301152 | 32311842 | mNeoVis.2.1.1.chr15\_hap1.g00400.t01 |
| 16407 | 15 | 32301152 | 32311842 | mNeoVis.2.1.1.chr15\_hap1.g00400.t01 |
| 16408 | 15 | 32301152 | 32311842 | mNeoVis.2.1.1.chr15\_hap1.g00400.t01 |
| 16409 | 15 | 32301152 | 32311842 | mNeoVis.2.1.1.chr15\_hap1.g00400.t01 |
| 16410 | 15 | 32301152 | 32311842 | mNeoVis.2.1.1.chr15\_hap1.g00400.t01 |
| 16411 | 15 | 32301152 | 32311842 | mNeoVis.2.1.1.chr15\_hap1.g00400.t01 |
| 16412 | 15 | 32314650 | 32319525 | mNeoVis.2.1.1.chr15\_hap1.g00401.t01 |
| 16413 | 15 | 32314650 | 32319525 | mNeoVis.2.1.1.chr15\_hap1.g00401.t01 |
| 16414 | 15 | 32314650 | 32319525 | mNeoVis.2.1.1.chr15\_hap1.g00401.t01 |
| 16415 | 15 | 32314650 | 32319525 | mNeoVis.2.1.1.chr15\_hap1.g00401.t01 |
| 16416 | 15 | 32314650 | 32319525 | mNeoVis.2.1.1.chr15\_hap1.g00401.t01 |
| 16417 | 15 | 32314650 | 32319525 | mNeoVis.2.1.1.chr15\_hap1.g00401.t01 |
| 16418 | 15 | 32322528 | 32324944 | mNeoVis.2.1.1.chr15\_hap1.g00402.t01 |
| 16419 | 15 | 32322528 | 32324944 | mNeoVis.2.1.1.chr15\_hap1.g00402.t01 |
| 16420 | 15 | 32322528 | 32324944 | mNeoVis.2.1.1.chr15\_hap1.g00402.t01 |
| 16421 | 15 | 32322528 | 32324944 | mNeoVis.2.1.1.chr15\_hap1.g00402.t01 |
| 16422 | 15 | 32322528 | 32324944 | mNeoVis.2.1.1.chr15\_hap1.g00402.t01 |
| 16423 | 15 | 32322528 | 32324944 | mNeoVis.2.1.1.chr15\_hap1.g00402.t01 |
| 16424 | 15 | 32326533 | 32329432 | mNeoVis.2.1.1.chr15\_hap1.g00403.t01 |
| 16425 | 15 | 32326533 | 32329432 | mNeoVis.2.1.1.chr15\_hap1.g00403.t01 |
| 16426 | 15 | 32326533 | 32329432 | mNeoVis.2.1.1.chr15\_hap1.g00403.t01 |
| 16427 | 15 | 32326533 | 32329432 | mNeoVis.2.1.1.chr15\_hap1.g00403.t01 |
| 16428 | 15 | 32326533 | 32329432 | mNeoVis.2.1.1.chr15\_hap1.g00403.t01 |
| 16429 | 15 | 32326533 | 32329432 | mNeoVis.2.1.1.chr15\_hap1.g00403.t01 |
| 16430 | 15 | 32329498 | 32335938 | mNeoVis.2.1.1.chr15\_hap1.g00404.t01 |
| 16431 | 15 | 32329498 | 32335938 | mNeoVis.2.1.1.chr15\_hap1.g00404.t01 |
| 16432 | 15 | 32329498 | 32335938 | mNeoVis.2.1.1.chr15\_hap1.g00404.t01 |
| 16433 | 15 | 32329498 | 32335938 | mNeoVis.2.1.1.chr15\_hap1.g00404.t01 |
| 16434 | 15 | 32329498 | 32335938 | mNeoVis.2.1.1.chr15\_hap1.g00404.t01 |
| 16435 | 15 | 32329498 | 32335938 | mNeoVis.2.1.1.chr15\_hap1.g00404.t01 |
| 16436 | 15 | 32337019 | 32341154 | mNeoVis.2.1.1.chr15\_hap1.g00405.t01 |
| 16437 | 15 | 32337019 | 32341154 | mNeoVis.2.1.1.chr15\_hap1.g00405.t01 |
| 16438 | 15 | 32337019 | 32341154 | mNeoVis.2.1.1.chr15\_hap1.g00405.t01 |
| 16439 | 15 | 32337019 | 32341154 | mNeoVis.2.1.1.chr15\_hap1.g00405.t01 |
| 16440 | 15 | 32337019 | 32341154 | mNeoVis.2.1.1.chr15\_hap1.g00405.t01 |
| 16441 | 15 | 32337019 | 32341154 | mNeoVis.2.1.1.chr15\_hap1.g00405.t01 |
| 16442 | 15 | 32340012 | 32343030 | mNeoVis.2.1.1.chr15\_hap1.g00406.t01 |
| 16443 | 15 | 32340012 | 32343030 | mNeoVis.2.1.1.chr15\_hap1.g00406.t01 |
| 16444 | 15 | 32340012 | 32343030 | mNeoVis.2.1.1.chr15\_hap1.g00406.t01 |
| 16445 | 15 | 32340012 | 32343030 | mNeoVis.2.1.1.chr15\_hap1.g00406.t01 |
| 16446 | 15 | 32340012 | 32343030 | mNeoVis.2.1.1.chr15\_hap1.g00406.t01 |
| 16447 | 15 | 32340012 | 32343030 | mNeoVis.2.1.1.chr15\_hap1.g00406.t01 |
| 16448 | 15 | 32343024 | 32345879 | mNeoVis.2.1.1.chr15\_hap1.g00407.t01 |
| 16449 | 15 | 32343024 | 32345879 | mNeoVis.2.1.1.chr15\_hap1.g00407.t01 |
| 16450 | 15 | 32343024 | 32345879 | mNeoVis.2.1.1.chr15\_hap1.g00407.t01 |
| 16451 | 15 | 32343024 | 32345879 | mNeoVis.2.1.1.chr15\_hap1.g00407.t01 |
| 16452 | 15 | 32343024 | 32345879 | mNeoVis.2.1.1.chr15\_hap1.g00407.t01 |
| 16453 | 15 | 32343024 | 32345879 | mNeoVis.2.1.1.chr15\_hap1.g00407.t01 |
| 16454 | 15 | 32345127 | 32357742 | mNeoVis.2.1.1.chr15\_hap1.g00408.t01 |
| 16455 | 15 | 32345127 | 32357742 | mNeoVis.2.1.1.chr15\_hap1.g00408.t01 |
| 16456 | 15 | 32345127 | 32357742 | mNeoVis.2.1.1.chr15\_hap1.g00408.t01 |
| 16457 | 15 | 32345127 | 32357742 | mNeoVis.2.1.1.chr15\_hap1.g00408.t01 |
| 16458 | 15 | 32345127 | 32357742 | mNeoVis.2.1.1.chr15\_hap1.g00408.t01 |
| 16459 | 15 | 32345127 | 32357742 | mNeoVis.2.1.1.chr15\_hap1.g00408.t01 |
| 16460 | 15 | 32370099 | 32376151 | mNeoVis.2.1.1.chr15\_hap1.g00409.t01 |
| 16461 | 15 | 32370099 | 32376151 | mNeoVis.2.1.1.chr15\_hap1.g00409.t01 |
| 16462 | 15 | 32370099 | 32376151 | mNeoVis.2.1.1.chr15\_hap1.g00409.t01 |
| 16463 | 15 | 32370099 | 32376151 | mNeoVis.2.1.1.chr15\_hap1.g00409.t01 |
| 16464 | 15 | 32370099 | 32376151 | mNeoVis.2.1.1.chr15\_hap1.g00409.t01 |
| 16465 | 15 | 32370099 | 32376151 | mNeoVis.2.1.1.chr15\_hap1.g00409.t01 |
| 16466 | 15 | 32408284 | 32441251 | mNeoVis.2.1.1.chr15\_hap1.g00410.t01 |
| 16467 | 15 | 32408284 | 32441251 | mNeoVis.2.1.1.chr15\_hap1.g00410.t01 |
| 16468 | 15 | 32408284 | 32441251 | mNeoVis.2.1.1.chr15\_hap1.g00410.t01 |
| 16469 | 15 | 32408284 | 32441251 | mNeoVis.2.1.1.chr15\_hap1.g00410.t01 |
| 16470 | 15 | 32408284 | 32441251 | mNeoVis.2.1.1.chr15\_hap1.g00410.t01 |
| 16471 | 15 | 32408284 | 32441251 | mNeoVis.2.1.1.chr15\_hap1.g00410.t01 |
| 16472 | 15 | 32448312 | 32450432 | mNeoVis.2.1.1.chr15\_hap1.g00411.t01 |
| 16473 | 15 | 32448312 | 32450432 | mNeoVis.2.1.1.chr15\_hap1.g00411.t01 |
| 16474 | 15 | 32448312 | 32450432 | mNeoVis.2.1.1.chr15\_hap1.g00411.t01 |
| 16475 | 15 | 32448312 | 32450432 | mNeoVis.2.1.1.chr15\_hap1.g00411.t01 |
| 16476 | 15 | 32448312 | 32450432 | mNeoVis.2.1.1.chr15\_hap1.g00411.t01 |
| 16477 | 15 | 32448312 | 32450432 | mNeoVis.2.1.1.chr15\_hap1.g00411.t01 |
| 16478 | 15 | 32450669 | 32469639 | mNeoVis.2.1.1.chr15\_hap1.g00412.t01 |
| 16479 | 15 | 32450669 | 32469639 | mNeoVis.2.1.1.chr15\_hap1.g00412.t01 |
| 16480 | 15 | 32450669 | 32469639 | mNeoVis.2.1.1.chr15\_hap1.g00412.t01 |
| 16481 | 15 | 32450669 | 32469639 | mNeoVis.2.1.1.chr15\_hap1.g00412.t01 |
| 16482 | 15 | 32450669 | 32469639 | mNeoVis.2.1.1.chr15\_hap1.g00412.t01 |
| 16483 | 15 | 32450669 | 32469639 | mNeoVis.2.1.1.chr15\_hap1.g00412.t01 |
| 16484 | 15 | 32469780 | 32479951 | mNeoVis.2.1.1.chr15\_hap1.g00413.t01 |
| 16485 | 15 | 32469780 | 32479951 | mNeoVis.2.1.1.chr15\_hap1.g00413.t01 |
| 16486 | 15 | 32469780 | 32479951 | mNeoVis.2.1.1.chr15\_hap1.g00413.t01 |
| 16487 | 15 | 32469780 | 32479951 | mNeoVis.2.1.1.chr15\_hap1.g00413.t01 |
| 16488 | 15 | 32469780 | 32479951 | mNeoVis.2.1.1.chr15\_hap1.g00413.t01 |
| 16489 | 15 | 32469780 | 32479951 | mNeoVis.2.1.1.chr15\_hap1.g00413.t01 |
| 16490 | 15 | 32479946 | 32482063 | mNeoVis.2.1.1.chr15\_hap1.g00414.t01 |
| 16491 | 15 | 32479946 | 32482063 | mNeoVis.2.1.1.chr15\_hap1.g00414.t01 |
| 16492 | 15 | 32479946 | 32482063 | mNeoVis.2.1.1.chr15\_hap1.g00414.t01 |
| 16493 | 15 | 32479946 | 32482063 | mNeoVis.2.1.1.chr15\_hap1.g00414.t01 |
| 16494 | 15 | 32479946 | 32482063 | mNeoVis.2.1.1.chr15\_hap1.g00414.t01 |
| 16495 | 15 | 32479946 | 32482063 | mNeoVis.2.1.1.chr15\_hap1.g00414.t01 |
| 16496 | 15 | 32483053 | 32495702 | mNeoVis.2.1.1.chr15\_hap1.g00415.t01 |
| 16497 | 15 | 32483053 | 32495702 | mNeoVis.2.1.1.chr15\_hap1.g00415.t01 |
| 16498 | 15 | 32483053 | 32495702 | mNeoVis.2.1.1.chr15\_hap1.g00415.t01 |
| 16499 | 15 | 32483053 | 32495702 | mNeoVis.2.1.1.chr15\_hap1.g00415.t01 |
| 16500 | 15 | 32483053 | 32495702 | mNeoVis.2.1.1.chr15\_hap1.g00415.t01 |
| 16501 | 15 | 32483053 | 32495702 | mNeoVis.2.1.1.chr15\_hap1.g00415.t01 |
| 16502 | 15 | 32486747 | 32492292 | mNeoVis.2.1.1.chr15\_hap1.g00416.t01 |
| 16503 | 15 | 32486747 | 32492292 | mNeoVis.2.1.1.chr15\_hap1.g00416.t01 |
| 16504 | 15 | 32486747 | 32492292 | mNeoVis.2.1.1.chr15\_hap1.g00416.t01 |
| 16505 | 15 | 32486747 | 32492292 | mNeoVis.2.1.1.chr15\_hap1.g00416.t01 |
| 16506 | 15 | 32486747 | 32492292 | mNeoVis.2.1.1.chr15\_hap1.g00416.t01 |
| 16507 | 15 | 32486747 | 32492292 | mNeoVis.2.1.1.chr15\_hap1.g00416.t01 |
| 16508 | 15 | 32495671 | 32496752 | mNeoVis.2.1.1.chr15\_hap1.g00417.t01 |
| 16509 | 15 | 32495671 | 32496752 | mNeoVis.2.1.1.chr15\_hap1.g00417.t01 |
| 16510 | 15 | 32495671 | 32496752 | mNeoVis.2.1.1.chr15\_hap1.g00417.t01 |
| 16511 | 15 | 32495671 | 32496752 | mNeoVis.2.1.1.chr15\_hap1.g00417.t01 |
| 16512 | 15 | 32495671 | 32496752 | mNeoVis.2.1.1.chr15\_hap1.g00417.t01 |
| 16513 | 15 | 32495671 | 32496752 | mNeoVis.2.1.1.chr15\_hap1.g00417.t01 |
| 16514 | 15 | 32496833 | 32497979 | mNeoVis.2.1.1.chr15\_hap1.g00418.t01 |
| 16515 | 15 | 32496833 | 32497979 | mNeoVis.2.1.1.chr15\_hap1.g00418.t01 |
| 16516 | 15 | 32496833 | 32497979 | mNeoVis.2.1.1.chr15\_hap1.g00418.t01 |
| 16517 | 15 | 32496833 | 32497979 | mNeoVis.2.1.1.chr15\_hap1.g00418.t01 |
| 16518 | 15 | 32496833 | 32497979 | mNeoVis.2.1.1.chr15\_hap1.g00418.t01 |
| 16519 | 15 | 32496833 | 32497979 | mNeoVis.2.1.1.chr15\_hap1.g00418.t01 |
| 16520 | 15 | 32497932 | 32543786 | mNeoVis.2.1.1.chr15\_hap1.g00419.t01 |
| 16521 | 15 | 32497932 | 32543786 | mNeoVis.2.1.1.chr15\_hap1.g00419.t01 |
| 16522 | 15 | 32497932 | 32543786 | mNeoVis.2.1.1.chr15\_hap1.g00419.t01 |
| 16523 | 15 | 32497932 | 32543786 | mNeoVis.2.1.1.chr15\_hap1.g00419.t01 |
| 16524 | 15 | 32497932 | 32543786 | mNeoVis.2.1.1.chr15\_hap1.g00419.t01 |
| 16525 | 15 | 32497932 | 32543786 | mNeoVis.2.1.1.chr15\_hap1.g00419.t01 |
| 16526 | 15 | 32543851 | 32554589 | mNeoVis.2.1.1.chr15\_hap1.g00420.t01 |
| 16527 | 15 | 32543851 | 32554589 | mNeoVis.2.1.1.chr15\_hap1.g00420.t01 |
| 16528 | 15 | 32543851 | 32554589 | mNeoVis.2.1.1.chr15\_hap1.g00420.t01 |
| 16529 | 15 | 32543851 | 32554589 | mNeoVis.2.1.1.chr15\_hap1.g00420.t01 |
| 16530 | 15 | 32555871 | 32598494 | mNeoVis.2.1.1.chr15\_hap1.g00421.t01 |
| 16531 | 15 | 32555871 | 32598494 | mNeoVis.2.1.1.chr15\_hap1.g00421.t01 |
| 16532 | 15 | 32555871 | 32598494 | mNeoVis.2.1.1.chr15\_hap1.g00421.t01 |
| 16533 | 15 | 32555871 | 32598494 | mNeoVis.2.1.1.chr15\_hap1.g00421.t01 |
| 16534 | 15 | 32599705 | 32608948 | mNeoVis.2.1.1.chr15\_hap1.g00422.t01 |
| 16535 | 15 | 32599705 | 32608948 | mNeoVis.2.1.1.chr15\_hap1.g00422.t01 |
| 16536 | 15 | 32599705 | 32608948 | mNeoVis.2.1.1.chr15\_hap1.g00422.t01 |
| 16537 | 15 | 32609036 | 32617971 | mNeoVis.2.1.1.chr15\_hap1.g00423.t01 |
| 16538 | 15 | 32609036 | 32617971 | mNeoVis.2.1.1.chr15\_hap1.g00423.t01 |
| 16539 | 15 | 32609036 | 32617971 | mNeoVis.2.1.1.chr15\_hap1.g00423.t01 |
| 16540 | 15 | 32619170 | 32621460 | mNeoVis.2.1.1.chr15\_hap1.g00424.t01 |
| 16541 | 15 | 32619170 | 32621460 | mNeoVis.2.1.1.chr15\_hap1.g00424.t01 |
| 16542 | 15 | 32619170 | 32621460 | mNeoVis.2.1.1.chr15\_hap1.g00424.t01 |
| 16543 | 15 | 32622217 | 32631584 | mNeoVis.2.1.1.chr15\_hap1.g00425.t01 |
| 16544 | 15 | 32622217 | 32631584 | mNeoVis.2.1.1.chr15\_hap1.g00425.t01 |
| 16545 | 15 | 32622217 | 32631584 | mNeoVis.2.1.1.chr15\_hap1.g00425.t01 |
| 16546 | 15 | 32636109 | 32637333 | mNeoVis.2.1.1.chr15\_hap1.g00426.t01 |
| 16547 | 15 | 32636109 | 32637333 | mNeoVis.2.1.1.chr15\_hap1.g00426.t01 |
| 16548 | 15 | 32636109 | 32637333 | mNeoVis.2.1.1.chr15\_hap1.g00426.t01 |
| 16549 | 15 | 32639242 | 32641550 | mNeoVis.2.1.1.chr15\_hap1.g00427.t01 |
| 16550 | 15 | 32639242 | 32641550 | mNeoVis.2.1.1.chr15\_hap1.g00427.t01 |
| 16551 | 15 | 32639242 | 32641550 | mNeoVis.2.1.1.chr15\_hap1.g00427.t01 |
| 16552 | 15 | 32641557 | 32645300 | mNeoVis.2.1.1.chr15\_hap1.g00428.t01 |
| 16553 | 15 | 32641557 | 32645300 | mNeoVis.2.1.1.chr15\_hap1.g00428.t01 |
| 16554 | 15 | 32641557 | 32645300 | mNeoVis.2.1.1.chr15\_hap1.g00428.t01 |
| 16555 | 15 | 32652133 | 32669199 | mNeoVis.2.1.1.chr15\_hap1.g00429.t01 |
| 16556 | 15 | 32652133 | 32669199 | mNeoVis.2.1.1.chr15\_hap1.g00429.t01 |
| 16557 | 15 | 32652133 | 32669199 | mNeoVis.2.1.1.chr15\_hap1.g00429.t01 |
| 16558 | 15 | 32669388 | 32686320 | mNeoVis.2.1.1.chr15\_hap1.g00430.t01 |
| 16559 | 15 | 32669388 | 32686320 | mNeoVis.2.1.1.chr15\_hap1.g00430.t01 |
| 16560 | 15 | 32691652 | 32697020 | mNeoVis.2.1.1.chr15\_hap1.g00431.t01 |
| 16561 | 15 | 32707546 | 32749228 | mNeoVis.2.1.1.chr15\_hap1.g00432.t01 |
| 16562 | 15 | 32749225 | 32781135 | mNeoVis.2.1.1.chr15\_hap1.g00433.t01 |
| 16563 | 15 | 32781238 | 32786550 | mNeoVis.2.1.1.chr15\_hap1.g00434.t01 |
| 16564 | 15 | 32787304 | 32799097 | mNeoVis.2.1.1.chr15\_hap1.g00435.t01 |
| 16565 | 15 | 32803998 | 32811934 | mNeoVis.2.1.1.chr15\_hap1.g00436.t01 |
| 16566 | 15 | 32812199 | 32822889 | mNeoVis.2.1.1.chr15\_hap1.g00437.t01 |
| 16567 | 15 | 32825696 | 32830571 | mNeoVis.2.1.1.chr15\_hap1.g00438.t01 |
| 16568 | 15 | 32833574 | 32835990 | mNeoVis.2.1.1.chr15\_hap1.g00439.t01 |
| 16569 | 15 | 32836049 | 32840499 | mNeoVis.2.1.1.chr15\_hap1.g00440.t01 |
| 16570 | 15 | 32840572 | 32847080 | mNeoVis.2.1.1.chr15\_hap1.g00441.t01 |
| 16571 | 15 | 32848083 | 32852218 | mNeoVis.2.1.1.chr15\_hap1.g00442.t01 |
| 16572 | 15 | 32851043 | 32854094 | mNeoVis.2.1.1.chr15\_hap1.g00443.t01 |
| 16573 | 15 | 32854088 | 32856944 | mNeoVis.2.1.1.chr15\_hap1.g00444.t01 |
| 16574 | 15 | 32859806 | 32868805 | mNeoVis.2.1.1.chr15\_hap1.g00445.t01 |
| 16575 | 15 | 32881145 | 32887197 | mNeoVis.2.1.1.chr15\_hap1.g00446.t01 |
| 16576 | 15 | 32919307 | 32952272 | mNeoVis.2.1.1.chr15\_hap1.g00447.t01 |
| 16577 | 15 | 32959328 | 32961193 | mNeoVis.2.1.1.chr15\_hap1.g00448.t01 |
| 16578 | 15 | 32961678 | 32979949 | mNeoVis.2.1.1.chr15\_hap1.g00449.t01 |
| 16579 | 15 | 32980789 | 32990960 | mNeoVis.2.1.1.chr15\_hap1.g00450.t01 |
| 16580 | 15 | 32990955 | 32993922 | mNeoVis.2.1.1.chr15\_hap1.g00451.t01 |
| 16581 | 15 | 32994064 | 33006706 | mNeoVis.2.1.1.chr15\_hap1.g00452.t01 |
| 16582 | 15 | 32997777 | 33003322 | mNeoVis.2.1.1.chr15\_hap1.g00453.t01 |
| 16583 | 15 | 33006675 | 33007756 | mNeoVis.2.1.1.chr15\_hap1.g00454.t01 |
| 16584 | 15 | 33007837 | 33008879 | mNeoVis.2.1.1.chr15\_hap1.g00455.t01 |
| 16585 | 15 | 33008928 | 33054778 | mNeoVis.2.1.1.chr15\_hap1.g00456.t01 |
| 16586 | 15 | 33058140 | 33067488 | mNeoVis.2.1.1.chr15\_hap1.g00457.t01 |
| 16587 | 15 | 33071261 | 33084998 | mNeoVis.2.1.1.chr15\_hap1.g00458.t01 |
| 16588 | 15 | 33085298 | 33140643 | mNeoVis.2.1.1.chr15\_hap1.g00459.t01 |
| 16589 | 15 | 33143649 | 33214444 | mNeoVis.2.1.1.chr15\_hap1.g00460.t01 |
| 16590 | 15 | 33183149 | 33199387 | mNeoVis.2.1.1.chr15\_hap1.g00461.t01 |
| 16591 | 15 | 33214577 | 33224990 | mNeoVis.2.1.1.chr15\_hap1.g00462.t01 |
| 16592 | 15 | 33228803 | 33234459 | mNeoVis.2.1.1.chr15\_hap1.g00463.t01 |
| 16593 | 15 | 33237624 | 33259724 | mNeoVis.2.1.1.chr15\_hap1.g00464.t01 |
| 16594 | 15 | 33261101 | 33268092 | mNeoVis.2.1.1.chr15\_hap1.g00465.t01 |
| 16595 | 15 | 33268640 | 33269997 | mNeoVis.2.1.1.chr15\_hap1.g00466.t01 |
| 16596 | 15 | 33277018 | 33278627 | mNeoVis.2.1.1.chr15\_hap1.g00467.t01 |
| 16597 | 15 | 33280638 | 33322502 | mNeoVis.2.1.1.chr15\_hap1.g00468.t01 |
| 16598 | 15 | 33308619 | 33344217 | mNeoVis.2.1.1.chr15\_hap1.g00469.t01 |
| 16599 | 15 | 33343953 | 33346480 | mNeoVis.2.1.1.chr15\_hap1.g00470.t01 |
| 16600 | 15 | 33347330 | 33361872 | mNeoVis.2.1.1.chr15\_hap1.g00471.t01 |
| 16601 | 15 | 33364739 | 33378780 | mNeoVis.2.1.1.chr15\_hap1.g00472.t01 |
| 16602 | 15 | 33519134 | 33525201 | mNeoVis.2.1.1.chr15\_hap1.g00477.t01 |
| 16603 | 15 | 33526156 | 33531400 | mNeoVis.2.1.1.chr15\_hap1.g00478.t01 |
| 16604 | 15 | 33538178 | 33539275 | mNeoVis.2.1.1.chr15\_hap1.g00479.t01 |
| 16605 | 15 | 33602805 | 33607828 | mNeoVis.2.1.1.chr15\_hap1.g00480.t01 |
| 16606 | 15 | 33617881 | 33702822 | mNeoVis.2.1.1.chr15\_hap1.g00481.t01 |
| 16607 | 15 | 33687646 | 33712754 | mNeoVis.2.1.1.chr15\_hap1.g00482.t01 |
| 16608 | 15 | 33746415 | 33749209 | mNeoVis.2.1.1.chr15\_hap1.g00483.t01 |
| 16609 | 15 | 33749182 | 33757575 | mNeoVis.2.1.1.chr15\_hap1.g00484.t01 |
| 16610 | 15 | 33757760 | 33760713 | mNeoVis.2.1.1.chr15\_hap1.g00485.t01 |
| 16611 | 15 | 33761447 | 33769805 | mNeoVis.2.1.1.chr15\_hap1.g00486.t01 |
| 16612 | 15 | 33770305 | 33773426 | mNeoVis.2.1.1.chr15\_hap1.g00487.t01 |
| 16613 | 15 | 33786282 | 33793790 | mNeoVis.2.1.1.chr15\_hap1.g00488.t01 |
| 16614 | 15 | 33793825 | 33796652 | mNeoVis.2.1.1.chr15\_hap1.g00489.t01 |
| 16615 | 15 | 33796749 | 33801244 | mNeoVis.2.1.1.chr15\_hap1.g00490.t01 |
| 16616 | 15 | 33801241 | 33802543 | mNeoVis.2.1.1.chr15\_hap1.g00491.t01 |
| 16617 | 15 | 33814430 | 33825839 | mNeoVis.2.1.1.chr15\_hap1.g00492.t01 |
| 16618 | 15 | 33826925 | 33832048 | mNeoVis.2.1.1.chr15\_hap1.g00493.t01 |
| 16619 | 15 | 33832260 | 33834572 | mNeoVis.2.1.1.chr15\_hap1.g00494.t01 |
| 16620 | 15 | 33833889 | 33836335 | mNeoVis.2.1.1.chr15\_hap1.g00495.t01 |
| 16621 | 15 | 33837768 | 33840396 | mNeoVis.2.1.1.chr15\_hap1.g00496.t01 |
| 16622 | 15 | 33842109 | 33846982 | mNeoVis.2.1.1.chr15\_hap1.g00497.t01 |
| 16623 | 15 | 33847222 | 33863671 | mNeoVis.2.1.1.chr15\_hap1.g00498.t01 |
| 16624 | 15 | 33864657 | 33869840 | mNeoVis.2.1.1.chr15\_hap1.g00499.t01 |
| 16625 | 15 | 33875696 | 33877796 | mNeoVis.2.1.1.chr15\_hap1.g00500.t01 |
| 16626 | 15 | 33878149 | 33881712 | mNeoVis.2.1.1.chr15\_hap1.g00501.t01 |
| 16627 | 15 | 33882494 | 33909756 | mNeoVis.2.1.1.chr15\_hap1.g00502.t01 |
| 16628 | 15 | 33915829 | 33928731 | mNeoVis.2.1.1.chr15\_hap1.g00503.t01 |
| 16629 | 15 | 33933430 | 33935615 | mNeoVis.2.1.1.chr15\_hap1.g00504.t01 |
| 16630 | 15 | 33944571 | 33968152 | mNeoVis.2.1.1.chr15\_hap1.g00505.t01 |
| 16631 | 15 | 34600405 | 34608348 | mNeoVis.2.1.1.chr15\_hap1.g00532.t01 |
| 16632 | 15 | 34600405 | 34608348 | mNeoVis.2.1.1.chr15\_hap1.g00532.t01 |
| 16633 | 15 | 34632196 | 34669830 | mNeoVis.2.1.1.chr15\_hap1.g00533.t01 |
| 16634 | 15 | 34632196 | 34669830 | mNeoVis.2.1.1.chr15\_hap1.g00533.t01 |
| 16635 | 15 | 34678408 | 34679745 | mNeoVis.2.1.1.chr15\_hap1.g00534.t01 |
| 16636 | 15 | 34678408 | 34679745 | mNeoVis.2.1.1.chr15\_hap1.g00534.t01 |
| 16637 | 15 | 34786218 | 34806240 | mNeoVis.2.1.1.chr15\_hap1.g00535.t01 |
| 16638 | 15 | 34786218 | 34806240 | mNeoVis.2.1.1.chr15\_hap1.g00535.t01 |
| 16639 | 15 | 34828543 | 34895791 | mNeoVis.2.1.1.chr15\_hap1.g00536.t01 |
| 16640 | 15 | 34828543 | 34895791 | mNeoVis.2.1.1.chr15\_hap1.g00536.t01 |
| 16641 | 15 | 34908617 | 34950263 | mNeoVis.2.1.1.chr15\_hap1.g00537.t01 |
| 16642 | 15 | 34908617 | 34950263 | mNeoVis.2.1.1.chr15\_hap1.g00537.t01 |
| 16643 | 15 | 34955358 | 34959387 | mNeoVis.2.1.1.chr15\_hap1.g00538.t01 |
| 16644 | 15 | 34955358 | 34959387 | mNeoVis.2.1.1.chr15\_hap1.g00538.t01 |
| 16645 | 15 | 35027187 | 35038039 | mNeoVis.2.1.1.chr15\_hap1.g00539.t01 |
| 16646 | 15 | 35027187 | 35038039 | mNeoVis.2.1.1.chr15\_hap1.g00539.t01 |
| 16647 | 15 | 35044691 | 35166563 | mNeoVis.2.1.1.chr15\_hap1.g00540.t01 |
| 16648 | 15 | 35044691 | 35166563 | mNeoVis.2.1.1.chr15\_hap1.g00540.t01 |
| 16649 | 15 | 35086946 | 35089065 | mNeoVis.2.1.1.chr15\_hap1.g00541.t01 |
| 16650 | 15 | 35086946 | 35089065 | mNeoVis.2.1.1.chr15\_hap1.g00541.t01 |
| 16651 | 15 | 35098933 | 35107693 | mNeoVis.2.1.1.chr15\_hap1.g00542.t01 |
| 16652 | 15 | 35098933 | 35107693 | mNeoVis.2.1.1.chr15\_hap1.g00542.t01 |
| 16653 | 15 | 35116464 | 35131898 | mNeoVis.2.1.1.chr15\_hap1.g00543.t01 |
| 16654 | 15 | 35116464 | 35131898 | mNeoVis.2.1.1.chr15\_hap1.g00543.t01 |
| 16655 | 15 | 35171502 | 35176904 | mNeoVis.2.1.1.chr15\_hap1.g00544.t01 |
| 16656 | 15 | 35171502 | 35176904 | mNeoVis.2.1.1.chr15\_hap1.g00544.t01 |
| 16657 | 15 | 35184296 | 35192554 | mNeoVis.2.1.1.chr15\_hap1.g00545.t01 |
| 16658 | 15 | 35184296 | 35192554 | mNeoVis.2.1.1.chr15\_hap1.g00545.t01 |
| 16659 | 15 | 35207124 | 35258013 | mNeoVis.2.1.1.chr15\_hap1.g00546.t01 |
| 16660 | 15 | 35258802 | 35277682 | mNeoVis.2.1.1.chr15\_hap1.g00547.t01 |
| 16661 | 15 | 35258802 | 35277682 | mNeoVis.2.1.1.chr15\_hap1.g00547.t01 |
| 16662 | 15 | 35279067 | 35334633 | mNeoVis.2.1.1.chr15\_hap1.g00548.t01 |
| 16663 | 15 | 35279067 | 35334633 | mNeoVis.2.1.1.chr15\_hap1.g00548.t01 |
| 16664 | 15 | 35359579 | 35381879 | mNeoVis.2.1.1.chr15\_hap1.g00549.t01 |
| 16665 | 15 | 35382497 | 35406329 | mNeoVis.2.1.1.chr15\_hap1.g00550.t01 |
| 16666 | 15 | 35432788 | 35488249 | mNeoVis.2.1.1.chr15\_hap1.g00551.t01 |
| 16667 | 15 | 35489628 | 35508503 | mNeoVis.2.1.1.chr15\_hap1.g00552.t01 |
| 16668 | 15 | 35509417 | 35560229 | mNeoVis.2.1.1.chr15\_hap1.g00553.t01 |
| 16669 | 15 | 35574833 | 35582998 | mNeoVis.2.1.1.chr15\_hap1.g00554.t01 |
| 16670 | 15 | 35590482 | 35595840 | mNeoVis.2.1.1.chr15\_hap1.g00555.t01 |
| 16671 | 15 | 35600822 | 35722459 | mNeoVis.2.1.1.chr15\_hap1.g00556.t01 |
| 16672 | 15 | 35635480 | 35653009 | mNeoVis.2.1.1.chr15\_hap1.g00557.t01 |
| 16673 | 15 | 35659676 | 35668503 | mNeoVis.2.1.1.chr15\_hap1.g00558.t01 |
| 16674 | 15 | 35675670 | 35680432 | mNeoVis.2.1.1.chr15\_hap1.g00559.t01 |
| 16675 | 15 | 35729368 | 35740209 | mNeoVis.2.1.1.chr15\_hap1.g00560.t01 |
| 16676 | 15 | 35807767 | 35811796 | mNeoVis.2.1.1.chr15\_hap1.g00561.t01 |
| 16677 | 15 | 36225021 | 36540251 | mNeoVis.2.1.1.chr15\_hap1.g00568.t01 |
| 16678 | 15 | 36541164 | 36546644 | mNeoVis.2.1.1.chr15\_hap1.g00569.t01 |
| 16679 | 15 | 36546675 | 36557155 | mNeoVis.2.1.1.chr15\_hap1.g00570.t01 |
| 16680 | 15 | 36557461 | 36596746 | mNeoVis.2.1.1.chr15\_hap1.g00571.t01 |
| 16681 | 15 | 36620466 | 36642473 | mNeoVis.2.1.1.chr15\_hap1.g00572.t01 |
| 16682 | 15 | 36643650 | 36674532 | mNeoVis.2.1.1.chr15\_hap1.g00573.t01 |
| 16683 | 15 | 36686999 | 36706546 | mNeoVis.2.1.1.chr15\_hap1.g00574.t01 |
| 16684 | 15 | 36738365 | 36739187 | mNeoVis.2.1.1.chr15\_hap1.g00575.t01 |
| 16685 | 15 | 38495055 | 38503685 | mNeoVis.2.1.1.chr15\_hap1.g00594.t01 |
| 16686 | 15 | 38495055 | 38503685 | mNeoVis.2.1.1.chr15\_hap1.g00594.t01 |
| 16687 | 15 | 38857496 | 38914136 | mNeoVis.2.1.1.chr15\_hap1.g00595.t01 |
| 16688 | 15 | 38924308 | 38934827 | mNeoVis.2.1.1.chr15\_hap1.g00596.t01 |
| 16689 | 15 | 38984405 | 38987137 | mNeoVis.2.1.1.chr15\_hap1.g00598.t01 |
| 16690 | 15 | 40383683 | 40467053 | mNeoVis.2.1.1.chr15\_hap1.g00629.t01 |
| 16691 | 15 | 40469024 | 40471987 | mNeoVis.2.1.1.chr15\_hap1.g00630.t01 |
| 16692 | 15 | 40472120 | 40496048 | mNeoVis.2.1.1.chr15\_hap1.g00631.t01 |
| 16693 | 15 | 40510547 | 40526067 | mNeoVis.2.1.1.chr15\_hap1.g00632.t01 |
| 16694 | 15 | 40526357 | 40619060 | mNeoVis.2.1.1.chr15\_hap1.g00633.t01 |
| 16695 | 15 | 40766939 | 40778426 | mNeoVis.2.1.1.chr15\_hap1.g00634.t01 |
| 16696 | 15 | 42503733 | 42507164 | mNeoVis.2.1.1.chr15\_hap1.g00691.t01 |
| 16697 | 15 | 42528159 | 42535377 | mNeoVis.2.1.1.chr15\_hap1.g00692.t01 |
| 16698 | 15 | 42544807 | 42550931 | mNeoVis.2.1.1.chr15\_hap1.g00693.t01 |
| 16699 | 15 | 42551742 | 42556739 | mNeoVis.2.1.1.chr15\_hap1.g00694.t01 |
| 16700 | 15 | 42560035 | 42564951 | mNeoVis.2.1.1.chr15\_hap1.g00695.t01 |
| 16701 | 15 | 42578210 | 42582936 | mNeoVis.2.1.1.chr15\_hap1.g00696.t01 |
| 16702 | 15 | 42586545 | 42594415 | mNeoVis.2.1.1.chr15\_hap1.g00697.t01 |
| 16703 | 15 | 42594526 | 42607295 | mNeoVis.2.1.1.chr15\_hap1.g00698.t01 |
| 16704 | 15 | 42611068 | 42615674 | mNeoVis.2.1.1.chr15\_hap1.g00699.t01 |
| 16705 | 15 | 42617632 | 42621753 | mNeoVis.2.1.1.chr15\_hap1.g00700.t01 |
| 16706 | 15 | 42671009 | 42678826 | mNeoVis.2.1.1.chr15\_hap1.g00701.t01 |
| 16707 | 15 | 42702792 | 42824445 | mNeoVis.2.1.1.chr15\_hap1.g00702.t01 |
| 16708 | 15 | 42862582 | 42903527 | mNeoVis.2.1.1.chr15\_hap1.g00703.t01 |
| 16709 | 15 | 42862582 | 42903527 | mNeoVis.2.1.1.chr15\_hap1.g00703.t01 |
| 16710 | 15 | 42998580 | 43332210 | mNeoVis.2.1.1.chr15\_hap1.g00704.t01 |
| 16711 | 15 | 43349885 | 43373230 | mNeoVis.2.1.1.chr15\_hap1.g00705.t01 |
| 16712 | 15 | 43386226 | 43410582 | mNeoVis.2.1.1.chr15\_hap1.g00706.t01 |
| 16713 | 15 | 43415227 | 43429016 | mNeoVis.2.1.1.chr15\_hap1.g00707.t01 |
| 16714 | 15 | 43431536 | 43437588 | mNeoVis.2.1.1.chr15\_hap1.g00708.t01 |
| 16715 | 15 | 43437638 | 43444019 | mNeoVis.2.1.1.chr15\_hap1.g00709.t01 |
| 16716 | 15 | 43444452 | 43448288 | mNeoVis.2.1.1.chr15\_hap1.g00710.t01 |
| 16717 | 15 | 43449893 | 43471014 | mNeoVis.2.1.1.chr15\_hap1.g00711.t01 |
| 16718 | 15 | 43484859 | 43488719 | mNeoVis.2.1.1.chr15\_hap1.g00712.t01 |
| 16719 | 15 | 43499371 | 43503744 | mNeoVis.2.1.1.chr15\_hap1.g00713.t01 |
| 16720 | 15 | 43505042 | 43550090 | mNeoVis.2.1.1.chr15\_hap1.g00714.t01 |
| 16721 | 15 | 43550152 | 43556661 | mNeoVis.2.1.1.chr15\_hap1.g00715.t01 |
| 16722 | 15 | 43550367 | 43575251 | mNeoVis.2.1.1.chr15\_hap1.g00716.t01 |
| 16723 | 15 | 43559133 | 43559563 | mNeoVis.2.1.1.chr15\_hap1.g00717.t01 |
| 16724 | 15 | 43593918 | 43626624 | mNeoVis.2.1.1.chr15\_hap1.g00718.t01 |
| 16725 | 15 | 43639894 | 43641367 | mNeoVis.2.1.1.chr15\_hap1.g00719.t01 |
| 16726 | 15 | 43647685 | 43702159 | mNeoVis.2.1.1.chr15\_hap1.g00720.t01 |
| 16727 | 15 | 43722715 | 43738018 | mNeoVis.2.1.1.chr15\_hap1.g00721.t01 |
| 16728 | 15 | 43737950 | 43799283 | mNeoVis.2.1.1.chr15\_hap1.g00722.t01 |
| 16729 | 15 | 43799552 | 43805579 | mNeoVis.2.1.1.chr15\_hap1.g00723.t01 |
| 16730 | 15 | 43805561 | 43864406 | mNeoVis.2.1.1.chr15\_hap1.g00724.t01 |
| 16731 | 15 | 43885550 | 43897154 | mNeoVis.2.1.1.chr15\_hap1.g00725.t01 |
| 16732 | 15 | 43931753 | 43948592 | mNeoVis.2.1.1.chr15\_hap1.g00726.t01 |
| 16733 | 15 | 43960488 | 43964591 | mNeoVis.2.1.1.chr15\_hap1.g00727.t01 |
| 16734 | 15 | 43988257 | 44120749 | mNeoVis.2.1.1.chr15\_hap1.g00728.t01 |
| 16735 | 15 | 44154628 | 44182805 | mNeoVis.2.1.1.chr15\_hap1.g00729.t01 |
| 16736 | 15 | 44182797 | 44193021 | mNeoVis.2.1.1.chr15\_hap1.g00730.t01 |
| 16737 | 15 | 44197387 | 44209441 | mNeoVis.2.1.1.chr15\_hap1.g00731.t01 |
| 16738 | 15 | 44212973 | 44235685 | mNeoVis.2.1.1.chr15\_hap1.g00732.t01 |
| 16739 | 15 | 44247481 | 44266713 | mNeoVis.2.1.1.chr15\_hap1.g00733.t01 |
| 16740 | 15 | 44277636 | 44280007 | mNeoVis.2.1.1.chr15\_hap1.g00734.t01 |
| 16741 | 15 | 44283504 | 44349811 | mNeoVis.2.1.1.chr15\_hap1.g00735.t01 |
| 16742 | 15 | 44357552 | 44377074 | mNeoVis.2.1.1.chr15\_hap1.g00736.t01 |
| 16743 | 15 | 44389911 | 44397702 | mNeoVis.2.1.1.chr15\_hap1.g00737.t01 |
| 16744 | 15 | 44402479 | 44420862 | mNeoVis.2.1.1.chr15\_hap1.g00738.t01 |
| 16745 | 15 | 44442799 | 44446678 | mNeoVis.2.1.1.chr15\_hap1.g00739.t01 |
| 16746 | 15 | 44451057 | 44453807 | mNeoVis.2.1.1.chr15\_hap1.g00740.t01 |
| 16747 | 15 | 44453804 | 44465247 | mNeoVis.2.1.1.chr15\_hap1.g00741.t01 |
| 16748 | 15 | 44465893 | 44482483 | mNeoVis.2.1.1.chr15\_hap1.g00742.t01 |
| 16749 | 15 | 44490365 | 44493421 | mNeoVis.2.1.1.chr15\_hap1.g00743.t01 |
| 16750 | 15 | 44493363 | 44495675 | mNeoVis.2.1.1.chr15\_hap1.g00744.t01 |
| 16751 | 15 | 44512073 | 44513443 | mNeoVis.2.1.1.chr15\_hap1.g00745.t01 |
| 16752 | 15 | 44515957 | 44516249 | mNeoVis.2.1.1.chr15\_hap1.g00746.t01 |
| 16753 | 15 | 44546617 | 44548285 | mNeoVis.2.1.1.chr15\_hap1.g00747.t01 |
| 16754 | 15 | 44549042 | 44552706 | mNeoVis.2.1.1.chr15\_hap1.g00748.t01 |
| 16755 | 15 | 44561606 | 44566627 | mNeoVis.2.1.1.chr15\_hap1.g00749.t01 |
| 16756 | 15 | 44609611 | 44647872 | mNeoVis.2.1.1.chr15\_hap1.g00750.t01 |
| 16757 | 15 | 44691858 | 44722605 | mNeoVis.2.1.1.chr15\_hap1.g00751.t01 |
| 16758 | 15 | 44730484 | 44755716 | mNeoVis.2.1.1.chr15\_hap1.g00752.t01 |
| 16759 | 15 | 44782714 | 44807294 | mNeoVis.2.1.1.chr15\_hap1.g00753.t01 |
| 16760 | 15 | 44813857 | 44826929 | mNeoVis.2.1.1.chr15\_hap1.g00754.t01 |
| 16761 | 15 | 44827978 | 44842729 | mNeoVis.2.1.1.chr15\_hap1.g00755.t01 |
| 16762 | 15 | 44875024 | 44940009 | mNeoVis.2.1.1.chr15\_hap1.g00756.t01 |
| 16763 | 15 | 44966373 | 45060705 | mNeoVis.2.1.1.chr15\_hap1.g00757.t01 |
| 16764 | 15 | 45096992 | 45196929 | mNeoVis.2.1.1.chr15\_hap1.g00758.t01 |
| 16765 | 15 | 45204412 | 45217276 | mNeoVis.2.1.1.chr15\_hap1.g00759.t01 |
| 16766 | 15 | 45219463 | 45251894 | mNeoVis.2.1.1.chr15\_hap1.g00760.t01 |
| 16767 | 15 | 45260599 | 45281019 | mNeoVis.2.1.1.chr15\_hap1.g00761.t01 |
| 16768 | 15 | 45286029 | 45332807 | mNeoVis.2.1.1.chr15\_hap1.g00762.t01 |
| 16769 | 15 | 49148601 | 49349893 | mNeoVis.2.1.1.chr15\_hap1.g00767.t01 |
| 16770 | 15 | 49349900 | 49356249 | mNeoVis.2.1.1.chr15\_hap1.g00768.t01 |
| 16771 | 15 | 49365469 | 49383618 | mNeoVis.2.1.1.chr15\_hap1.g00769.t01 |
| 16772 | 15 | 49395165 | 49449169 | mNeoVis.2.1.1.chr15\_hap1.g00770.t01 |
