## Supplementary material for "Mink by mink: stitching together signatures of subspecies adaptation through a pangenome of threatened mustelids": Table S10

| Mitochondrial Basic Genome Statistics | | | | | | |
|---|---|---|---|---|---|---|
| | nvison_FL-32666 | nvison_FL-33534 | nvison_FL-3 | nvison_FL-5 | nvison_LA-8 | nvison_LA-9 |
| Genome Size | 16,538 | 16,461 | 16,431 | 16,573 | 16,550 | 16,493 |
| kmer Size | 41 | 41 | 41 | 41 | 41 | 31 |
| N's | 911 | 1,426 | 1,534 | 713 | 839 | 1,258 |
| Protein Coding | 13 | 13 | 13 | 13 | 13 | 13 |
| on-protein Codin | 25 | 25 | 25 | 25 | 25 | 25 |
