## Supplementary material for "Mink by mink: stitching together signatures of subspecies adaptation through a pangenome of threatened mustelids": Table S11

Preliminary Nucleotide Conservation and Differences Across Mitochondrial Regions in *Neogale vison* subspecies.

| | ***N. vison ssp. evergladensis*** | | | | | ***n. vison ssp. lutensis*** | | | | | ***n. vison ssp. vulgivaga*** | | | | |
|---|---|---|---|---|---|---|---|---|---|---|---|---|---|---|---|
| **Coding Region** | Nn | Nc | Nd | h | π | Nn | Nc | Nd | h | π | Nn | Nc | Nd | h | π |
| ATP6 | 681 | 677 | 4 | 0.6666667 | 0.00391581 | 681 | 674 | 7 | 1 | 0.006852668 | 681 | 675 | 6 | 1 | 0.005873715 |
| ATP8 | 204 | 203 | 1 | 0.6666667 | 0.003267974 | 204 | 201 | 3 | 1 | 0.009803922 | 204 | 203 | 1 | 0.6666667 | 0.003267974 |
| COX1 | 1545 | 1541 | 4 | 0.6666667 | 0.001725998 | 1545 | 1531 | 14 | 1 | 0.006040992 | 1545 | 1540 | 5 | 1 | 0.002157497 |
| COX2 | 684 | 680 | 4 | 0.6666667 | 0.003898635 | 684 | 676 | 8 | 1 | 0.007797271 | 684 | 682 | 2 | 0.6666667 | 0.001949318 |
| COX3 | 784 | 779 | 5 | 0.6666667 | 0.004251701 | 784 | 777 | 7 | 1 | 0.005952381 | 784 | 776 | 8 | 1 | 0.006802721 |
| CYTB | 1140 | 1136 | 4 | 0.6666667 | 0.002339181 | 1140 | 1131 | 9 | 1 | 0.005263158 | 1140 | 1133 | 7 | 1 | 0.004093567 |
| NAD1 | 955 | 948 | 7 | 0.6666667 | 0.004886562 | 955 | 944 | 11 | 1 | 0.007678883 | 955 | 948 | 7 | 1 | 0.004886562 |
| NAD2 | 1042 | 1038 | 4 | 0.6666667 | 0.002559181 | 1042 | 1028 | 14 | 1 | 0.008957134 | 1042 | 1037 | 5 | 1 | 0.003198976 |
| NAD3 | 347 | 346 | 1 | 0.6666667 | 0.00192123 | 347 | 346 | 1 | 0.6666667 | 0.00192123 | 347 | 344 | 3 | 1 | 0.005763689 |
| NAD4 | 1378 | 1367 | 11 | 0.6666667 | 0.005321722 | 1378 | 1358 | 20 | 1 | 0.009675859 | 1378 | 1365 | 13 | 1 | 0.006289308 |
| NAD4L | 297 | 293 | 4 | 0.6666667 | 0.008978676 | 297 | 292 | 5 | 0.6666667 | 0.01122334 | 297 | 292 | 5 | 1 | 0.01122334 |
| NAD5 | 1830 | 1821 | 9 | 0.6666667 | 0.003278689 | 1830 | 1816 | 14 | 1 | 0.005100182 | 1830 | 1822 | 8 | 1 | 0.00291439 |
| NAD6 | 534 | 528 | 7 | 0.6666667 | 0.008739076 | 534 | 528 | 6 | 1 | 0.007490637 | 534 | 527 | 7 | 1 | 0.008739076 |
| **Non-coding** | | | | | | | | | | | | | | | |
| D-loop | 1134 | 1034 | 77 | [redacted] | | 1134 | 989 | 106 | [redacted] | | 1134 | 1024 | 66 | [redacted] | |
| **rRNA** | | | | | | | | | | | | | | | |
| srRNA | 963 | 958 | 2 | 0.6666667 | 0.001388889 | 963 | 956 | 5 | 1 | 0.003472222 | 963 | 958 | 2 | 1 | 0.002083333 |
| lrRNA | 1571 | 1562 | 7 | 0.6666667 | 0.002974294 | 1571 | 1561 | 8 | 1 | 0.003399193 | 1571 | 1567 | 2 | 1 | 0.0008497982 |

Total number of nucleotides (Nn), conserved nucleotides (Nc), and nucleotide differences (Nd), h (haplotype diversity), and π ( nucleotide diversity)
