## Supplementary material for "Mink by mink: stitching together signatures of subspecies adaptation through a pangenome of threatened mustelids": Table S12

| Subspecies | GO Term | Associated Gene Count | P-value | Over/Under (ratic | Name | Definition |
|---|---|---|---|---|---|---|
| lutensis | GO:0060322 | 109 | 0.007535907 | Over | Head development | The biological process whose specific outcome is the progression of a head from an initial condition to its mature state. The head is the anterior-most division of the body. |
| lutensis | GO:0097477 | 5 | 0.000072027 | Over | Neuromuscular synaptic transmission | The orderly movement of a lateral motor column neuron from one site to another. A lateral motor column neuron is a motor neuron that is generated only on limb levels and send axons into the limb mesenchym |
| lutensis | GO:0007416 | 32 | 0.000013979 | Over | Synapse assembly | The aggregation, arrangement and bonding together of a set of components to form a synapse. This process ends when the synapse is mature (functional). |
| lutensis | GO:0051703 | 17 | 0.000042981 | Over | Interspecies interaction between organisms | Biological process involved in intraspecies interaction between organisms |
| lutensis | GO:0048167 | 40 | 0.000036774 | Over | Regulation of synaptic plasticity | A process that modulates synaptic plasticity, the ability of synapses to change as circumstances require. They may alter function, such as increasing or decreasing their sensitivity, or they may increase or decrease in actual numbers |
| lutensis | GO:0048265 | 16 | 0.000000758 | Over | Response to pain | Any process that results in a change in state or activity of a cell or an organism (in terms of movement, secretion, enzyme production, gene expression, etc.) as a result of a pain stimulus. Pain stimuli cause activation of nociceptors, peripheral receptors for pain, include receptors which are sensitive to painful mechanical stimuli, extreme heat or cold, a |
| lutensis | GO:0042626 | 23 | 0.000049907 | Over | ATPase activity coupled to transmembrane movement of s | ATPase-coupled transmembrane transporter activity |
| lutensis | GO:0016319 | 24 | 0.000036794 | Over | O-methyltransferase activity | mushroom body development |
| lutensis | GO:0099604 | 1 | 0.000156394 | Over | Synaptic vesicle uncoating | ligand-gated calcium channel activity |
| evergladensis | GO:0035315 | 6 | 0.002777295* | Over | Aging | A developmental process that is associated with progression towards the end of the lifespan of an organism. |
| evergladensis | GO:0002065 | 10 | 0.000474455* | Over | Lung cell differentiation | The process in which a relatively unspecialized cell acquires the specialized structural and/or functional features of a cell in the lung. |
| evergladensis | GO:0060122 | 4 | 0.003556875* | Over | Epithelial cell development | The process whose specific outcome is the progression of an epithelial cell over time, from its formation to the mature state. |
| evergladensis | GO:0030855 | 14 | 0.004683567* | Over | Epithelial cell differentiation | The process in which a relatively unspecialized cell acquires the specialized structural and/or functional features of an epithelial cell. |
| evergladensis | GO:0060119 | 4 | 0.008800984* | Over | Inner ear receptor cell differentiation | The process in which a relatively unspecialized cell acquires the specialized structural and/or functional features of an inner ear receptor cell. |
| evergladensis | GO:0042491 | 4 | 0.005366545* | Over | Inner ear development | The process whose specific outcome is the progression of the inner ear over time, from its formation to the mature structure. |
| vulgivaga | GO:0040011 | 14 | 0.0064911565* | Under | Locomotion | Self-propelled movement of a cell or organism from one location to another. |
| vulgivaga | GO:0032502 | 34 | 0.0054504865* | Under | Developmental process | The biological process whose specific outcome is the progression of an organism over time from an initial condition to a later condition. |
| vulgivaga | GO:0045595 | 4 | 0.0045248095* | Under | Regulation of cell differentiation | Any process that modulates the frequency, rate or extent of cell differentiation. |
| vulgivaga | GO:0051094 | 2 | 0.0065999985* | Under | Positive regulation of developmental process | Any process that activates or increases the frequency, rate or extent of development. |
| vulgivaga | GO:0065009 | 10 | 0.0065735965* | Under | Regulation of biological quality | Any process that modulates a qualitative or quantitative trait of a biological quality, such as size, mass, shape, color, etc. |
| vulgivaga | GO:0048856 | 32 | 0.0065999985* | Under | Anatomical structure development | The biological process whose specific outcome is the progression of an anatomical structure from an initial condition to its mature state. |
| hap1 | GO:0006366 | 6 | 0.000004869 | Under | Transcription from RNA polymerase II promoter | The synthesis of RNA from a DNA template by RNA polymerase II, originating at an RNA polymerase II promoter. |
| hap1 | GO:0046621 | 10 | 0.000025229 | Over | Negative regulation of organ growth | Any process that stops, prevents, or reduces the frequency, rate or extent of organ growth. |
| hap1 | GO:0080135 | 14 | 0.007386786 | Under | Regulation of cellular response to stress | Any process that modulates the frequency, rate or extent of a cell's response to stress. |
| hap1 | GO:0004416 | 3 | 0.00009196 | Over | Histone acetyltransferase activity | Catalysis of the transfer of an acetyl group to a histone. |
| hap1 | GO:0008967 | 3 | 0.00009196 | Over | Phosphoglucomutase activity | Catalysis of the interconversion of glucose-1-phosphate and glucose-6-phosphate. |
| hap1 | GO:0042113 | 18 | 0.000002019 | Over | B cell activation | The change in morphology and behavior of a B cell resulting from exposure to a stimulus such as an antigen or cytokine. |
| hap1 | GO:0034340 | 15 | 0 | Over | Response to type I interferon | Any process that results in a change in state or activity of a cell or an organism as a result of a type I interferon stimulus. |
| hap1 | GO:0035821 | 4 | 0.000004591 | Over | Modulation of chemokine production | Any process that regulates the frequency, rate, or extent of chemokine production. |
| hap1 | GO:0035499 | 3 | 0.000023667 | Over | Signal transduction by p53 class mediator | A series of molecular signals mediated by the p53 class of proteins, leading to a change in cell state or activity. |
| hap1 | GO:0032774 | 11 | 0.000052751 | Under | RNA biosynthetic process | The chemical reactions and pathways resulting in the formation of RNA, including transcription from an RNA polymerase. |
| hap1 | GO:0005132 | 15 | 0 | Over | Interleukin-6 receptor binding | Interacting selectively and non-covalently with the interleukin-6 receptor. |
| hap1 | GO:0071357 | 15 | 0 | Over | Cellular response to type I interferon | Any process that results in a change in state or activity of a cell as a result of a type I interferon stimulus. |
| hap1 | GO:0050688 | 16 | 0.000000108 | Over | Regulation of defense response to virus | Any process that modulates the frequency, rate, or extent of the defense response to a virus. |

* indicate a p-value, averaged among the individuals in a particular subspecies
