## Supplementary material for "Mink by mink: stitching together signatures of subspecies adaptation through a pangenome of threatened mustelids": Table S13: Table S13 - Hierarchical Orthologous Groups and Associated Genes in Selected Carnivorans.html

  
        A  B  C  D  E  F  G  H  I  J  K  L  M  N  O  P  Q  R  S  T  U  V  W  X  Y       1      OG   Gene Tree Parent Clade    Ailuropoda_melanoleuca    Canis_lupis_familiaris   Enhydra_lutris  Felis_catus   Halichoerus_grypus   Hyaena_hyaena  Lynx_canadensis  Meles_meles   Mustela_erminea   Mustela_lutreola   Mustela_nigripes    Mustela_putorius_furo    Neogale_vison_Haplotype_1    Neomonachus_schauinslandi    Odobenus_rosmarus   Panthera_tigris  Ursus_maritimus   Zalophus_californianus   nvison_FL-32666  nvison_FL-33534  nvison_FL-5  nvison_LA-9     2    N0.HOG0000000   OG0000000  n2   rna-XM_011235175.3        rna-XM_019818494.3      rna-XM_039224732.1          rna-XM_059151232.1          rna-XM_044911348.1      rna-XM_042970218.1      rna-XM_027618329.2              3    N0.HOG0000001   OG0000000  n10   rna-XM_034650423.1    rna-XM_038527997.1                    rna-XM_059382684.1, rna-XM_059382685.1    rna-XM_045077458.1    mNeoVis.2.1.1.chr7_hap1.g00785.t01              mNeoVis.3.1.1.chr7.g00726.t01, mNeoVis.3.1.1.chr7.g00732.t01    rna-gnl|WGS_ZZZZ|egapxtmp_013318-R1, rna-gnl|WGS_ZZZZ|egapxtmp_013319-R1    mNeoVis.evergladensis.2.1.chr7.g00772.t01    mNeoVis.vulgivaga.2.1.chr7.g00776.t01, mNeoVis.vulgivaga.2.1.chr7.g00777.t01      4    N0.HOG0000002   OG0000000  n22   rna-XM_034650420.1    rna-XM_038527966.1        rna-XM_036076329.1, rna-XM_036076336.1      rna-XM_032840500.1, rna-XM_032850969.1, rna-XM_032850970.1    rna-XM_045988254.1              rna-XM_044911347.1    rna-XM_004409806.1                    5    N0.HOG0000003   OG0000000  n31   rna-XM_034639133.1    rna-XM_038527968.1    rna-XM_022524121.1      rna-XM_036111863.1      rna-XM_032839414.1    rna-XM_045988276.1    rna-XM_032324142.1    rna-XM_059150670.1    rna-XM_059382641.1    rna-XM_013049652.2    mNeoVis.2.1.1.chr7_hap1.g00784.t01          rna-XM_008684783.2        rna-gnl|WGS_ZZZZ|egapxtmp_013310-R3    mNeoVis.evergladensis.2.1.chr7.g00771.t01        6    N0.HOG0000004   OG0000000  n45   rna-XM_019808296.2, rna-XM_034639069.1    rna-XM_038527960.1      rna-XM_019818583.3, rna-XM_023244968.2    rna-XM_036076332.1, rna-XM_036076342.1    rna-XM_039220146.1, rna-XM_039220163.1    rna-XM_032839411.1    rna-XM_045986780.1, rna-XM_045988251.1    rna-XM_032324133.1    rna-XM_059150667.1    rna-XM_059382657.1    rna-XM_013049656.2, rna-XM_045077277.1    mNeoVis.2.1.1.chr7_hap1.g00792.t01    rna-XM_044911349.1, rna-XM_044912283.1      rna-XM_015542559.2, rna-XM_042970237.1    rna-XM_008684774.2, rna-XM_008684780.2      mNeoVis.3.1.1.chr7.g00734.t01    rna-gnl|WGS_ZZZZ|egapxtmp_012307-R1    mNeoVis.evergladensis.2.1.chr7.g00779.t01    mNeoVis.vulgivaga.2.1.chr7.g00785.t01      7    N0.HOG0000005   OG0000000  n73   rna-XM_019808289.2, rna-XM_034639078.1    rna-XM_038527967.1    rna-XM_022524126.1    rna-XM_019818524.3, rna-XM_023249705.2, rna-XM_045046190.1, rna-XM_045046205.1, rna-XM_045046211.1, rna-XM_045046221.1, rna-XM_045046222.1, rna-XM_045046231.1, rna-XM_045046234.1    rna-XM_036076330.1, rna-XM_036076333.1    rna-XM_039220162.1, rna-XM_039224305.1, rna-XM_039224308.1, rna-XM_039224313.1    rna-XM_032839404.1    rna-XM_045988226.1    rna-XM_032324124.1    rna-XM_059150657.1      rna-XM_045077386.1    mNeoVis.2.1.1.chr7_hap1.g00791.t01    rna-XM_044911350.1, rna-XM_044912407.1, rna-XM_044921996.1    rna-XM_004409810.1    rna-XM_042970023.1, rna-XM_042970026.1, rna-XM_042970219.1, rna-XM_042970220.1, rna-XM_042970221.1, rna-XM_042970222.1, rna-XM_042970223.1, rna-XM_042970228.1, rna-XM_042970230.1    rna-XM_040632919.1, rna-XM_040632921.1    rna-XM_027618284.2, rna-XM_027618301.1, rna-XM_035725229.1      rna-gnl|WGS_ZZZZ|egapxtmp_013309-R1    mNeoVis.evergladensis.2.1.chr7.g00778.t01    mNeoVis.vulgivaga.2.1.chr7.g00784.t01      8    N0.HOG0000006   OG0000000  n119             rna-XM_039224307.1    rna-XM_032839391.1                                    9    N0.HOG0000007   OG0000000  n121   rna-XM_019792542.2, rna-XM_019809349.2    rna-XM_038584456.1    rna-XM_022490629.1, rna-XM_022525123.1    rna-XM_023246073.2, rna-XM_023254666.2, rna-XM_045047143.1    rna-XM_036077281.1, rna-XM_036077292.1, rna-XM_036115502.1, rna-XM_036120500.1    rna-XM_039222030.1, rna-XM_039244152.1, rna-XM_039257561.1    rna-XM_032847836.1    rna-XM_045986824.1, rna-XM_045987592.1, rna-XM_046004744.1    rna-XM_032324737.1    rna-XM_059176162.1    rna-XM_059399631.1    rna-XM_045073960.1    mNeoVis.2.1.1.chr1_hap1.g01041.t01, mNeoVis.2.1.1.chr7_hap1.g00665.t01      rna-XM_004405089.1, rna-XM_004411306.1    rna-XM_042970683.1, rna-XM_042985390.1    rna-XM_040634374.1, rna-XM_040644630.1, rna-XM_040644881.1    rna-XM_027601067.2, rna-XM_027617502.2, rna-XM_027619313.2, rna-XM_035724933.1    mNeoVis.3.1.1.chr1.g01035.t01, mNeoVis.3.1.1.chr7.g00628.t01    rna-gnl|WGS_ZZZZ|egapxtmp_014134-R1, rna-gnl|WGS_ZZZZ|egapxtmp_021101-R1    mNeoVis.evergladensis.2.1.chr1.g01033.t01, mNeoVis.evergladensis.2.1.chr7.g00660.t01    mNeoVis.vulgivaga.2.1.chr1.g01043.t01, mNeoVis.vulgivaga.2.1.chr7.g00663.t01, mNeoVis.vulgivaga.2.1.chr7.g00783.t01      10    N0.HOG0000008   OG0000000  n168   rna-XM_019792535.2, rna-XM_034639493.1    rna-XM_038527612.1    rna-XM_022491841.1    rna-XM_023246060.2, rna-XM_045046736.1    rna-XM_036096589.1, rna-XM_036096611.1    rna-XM_039244112.1, rna-XM_039244115.1    rna-XM_032848560.1    rna-XM_045987580.1, rna-XM_046010714.1    rna-XM_032324631.1, rna-XM_032324820.1    rna-XM_059152283.1, rna-XM_059152357.1, rna-XM_059152437.1    rna-XM_059383158.1    rna-XM_004776939.3, rna-XM_013048037.2    mNeoVis.2.1.1.chr7_hap1.g00667.t01, mNeoVis.2.1.1.chr7_hap1.g00668.t01    rna-XM_044911290.1, rna-XM_044911332.1    rna-XM_004405062.1    rna-XM_042970644.1, rna-XM_042970653.1    rna-XM_040621957.1, rna-XM_040633727.1    rna-XM_027617487.2    mNeoVis.3.1.1.chr7.g00630.t01, mNeoVis.3.1.1.chr7.g00631.t01    rna-gnl|WGS_ZZZZ|egapxtmp_012266-R1, rna-gnl|WGS_ZZZZ|egapxtmp_014128-R1, rna-gnl|WGS_ZZZZ|egapxtmp_014132-R1    mNeoVis.evergladensis.2.1.chr7.g00662.t01, mNeoVis.evergladensis.2.1.chr7.g00663.t01, mNeoVis.evergladensis.2.1.chr7.g00665.t01    mNeoVis.vulgivaga.2.1.chr7.g00664.t01, mNeoVis.vulgivaga.2.1.chr7.g00665.t01      11    N0.HOG0000009   OG0000000     rna-XM_011237978.3, rna-XM_034638428.1    rna-XM_038527629.1      rna-XM_019819072.3    rna-XM_036096602.1, rna-XM_036120128.1    rna-XM_039227408.1, rna-XM_039244126.1    rna-XM_032848561.1    rna-XM_045987594.1    rna-XM_032326142.1    rna-XM_059152442.1    rna-XM_059383203.1    rna-XM_004776936.3    mNeoVis.2.1.1.chr7_hap1.g00669.t01    rna-XM_021700967.2, rna-XM_044911333.1    rna-XM_004405064.1    rna-XM_042968892.1, rna-XM_042970654.1    rna-XM_040633774.1    rna-XM_027617523.2, rna-XM_035725298.1    mNeoVis.3.1.1.chr7.g00632.t01    rna-gnl|WGS_ZZZZ|egapxtmp_014141-R1    mNeoVis.evergladensis.2.1.chr7.g00666.t01        12    N0.HOG0000010   OG0000000  n240   rna-XM_034638440.1    rna-XM_038529187.1    rna-XM_022525373.1      rna-XM_036120106.1    rna-XM_039217743.1    rna-XM_032841893.1    rna-XM_045989370.1      rna-XM_059153713.1      rna-XM_013047688.2    mNeoVis.2.1.1.chr7_hap1.g00492.t01    rna-XM_021700820.1    rna-XM_004396431.1    rna-XM_042968939.1    rna-XM_040633907.1    rna-XM_027617709.1    mNeoVis.3.1.1.chr7.g00474.t01    rna-gnl|WGS_ZZZZ|egapxtmp_012232-R1    mNeoVis.evergladensis.2.1.chr7.g00495.t01        13    N0.HOG0000011   OG0000000  n257     rna-XM_038529038.1      rna-XM_011289854.4                    mNeoVis.2.1.1.chr7_hap1.g00494.t01    rna-XM_021700966.2    rna-XM_004396500.1    rna-XM_007096876.3      rna-XM_035725290.1      rna-gnl|WGS_ZZZZ|egapxtmp_012087-R1      mNeoVis.vulgivaga.2.1.chr7.g00499.t01      14    N0.HOG0000012   OG0000000  n267   rna-XM_011217247.3    rna-NM_001003331.2    rna-XM_022491860.1    rna-XM_011289606.4    rna-XM_036065450.1    rna-XM_039244073.1    rna-XM_032848643.1    rna-XM_045987956.1    rna-XM_032324535.1    rna-XM_059151423.1    rna-XM_059383329.1    rna-XM_045076986.1    mNeoVis.2.1.1.chr7_hap1.g00769.t01    rna-XM_044922001.1    rna-XM_004409538.1    rna-XM_007087127.3    rna-XM_040632954.1    rna-XM_027618369.2      rna-gnl|WGS_ZZZZ|egapxtmp_009902-R9    mNeoVis.evergladensis.2.1.chr7.g00756.t01    mNeoVis.vulgivaga.2.1.chr7.g00759.t01      15    N0.HOG0000013   OG0000000  n286   rna-XM_011237748.3    rna-XM_038528663.1    rna-XM_022491823.1    rna-XM_045045706.1    rna-XM_036093648.1    rna-XM_039248057.1    rna-XM_032838931.1    rna-XM_045987271.1    rna-XM_032326059.1    rna-XM_059151250.1    rna-XM_059383401.1    rna-XM_045077288.1    mNeoVis.2.1.1.chr7_hap1.g01063.t01    rna-XM_021681317.2    rna-XM_004401017.1    rna-XM_042968363.1    rna-XM_008684428.2    rna-XM_027618699.2      rna-gnl|WGS_ZZZZ|egapxtmp_012380-R1    mNeoVis.evergladensis.2.1.chr7.g01055.t01    mNeoVis.vulgivaga.2.1.chr7.g01064.t01      16    N0.HOG0000014   OG0000000  n306   rna-XM_019804311.2    rna-XM_038528889.1    rna-XM_022525171.1    rna-XM_019819525.3    rna-XM_036079702.1, rna-XM_036095035.1      rna-XM_032841845.1    rna-XM_045988615.1    rna-XM_032325096.1    rna-XM_059152071.1    rna-XM_059381870.1    rna-XM_013047729.2    mNeoVis.2.1.1.chr7_hap1.g00556.t01    rna-XM_044911373.1    rna-XM_004396488.1    rna-XM_042969427.1    rna-XM_040633975.1    rna-XM_027618963.2    mNeoVis.3.1.1.chr7.g00539.t01    rna-gnl|WGS_ZZZZ|egapxtmp_012249-R1    mNeoVis.evergladensis.2.1.chr7.g00557.t01    mNeoVis.vulgivaga.2.1.chr7.g00559.t01      17    N0.HOG0000015   OG0000000  n326   rna-XM_034638303.1, rna-XM_034638365.1    rna-XM_038525560.1      rna-XM_045045445.1    rna-XM_036114369.1    rna-XM_039249628.1    rna-XM_032842099.1    rna-XM_045988715.1    rna-XM_032325611.1, rna-XM_032325658.1    rna-XM_059151487.1, rna-XM_059151527.1    rna-XM_059382968.1    rna-XM_004764978.3, rna-XM_045076545.1    mNeoVis.2.1.1.chr7_hap1.g00480.t01, mNeoVis.2.1.1.chr7_hap1.g00481.t01      rna-XM_004396503.1    rna-XM_007096859.3    rna-XM_040632815.1    rna-XM_027617636.2      rna-gnl|WGS_ZZZZ|egapxtmp_012067-R6    mNeoVis.evergladensis.2.1.chr7.g00485.t01    mNeoVis.vulgivaga.2.1.chr7.g00488.t01      18    N0.HOG0000016   OG0000000  n349   rna-XM_011217262.3    rna-XM_038529141.1      rna-XM_045047010.1    rna-XM_036068053.1    rna-XM_039249711.1    rna-XM_032841838.1    rna-XM_045986752.1, rna-XM_045988761.1    rna-XM_032326160.1    rna-XM_059151517.1    rna-XM_059382975.1    rna-XM_013062795.2    mNeoVis.2.1.1.chr7_hap1.g00475.t01, mNeoVis.2.1.1.chr7_hap1.g00476.t01      rna-XM_004396438.1    rna-XM_042970114.1    rna-XM_040632804.1    rna-XM_027617804.2    mNeoVis.3.1.1.chr7.g00461.t01    rna-gnl|WGS_ZZZZ|egapxtmp_012064-R2, rna-gnl|WGS_ZZZZ|egapxtmp_012069-R1    mNeoVis.evergladensis.2.1.chr7.g00481.t01, mNeoVis.evergladensis.2.1.chr7.g00482.t01    mNeoVis.vulgivaga.2.1.chr7.g00482.t01, mNeoVis.vulgivaga.2.1.chr7.g00483.t01      19    N0.HOG0000017   OG0000000  n370                   rna-XM_032325587.1        rna-XM_045076529.1                          20    N0.HOG0000018   OG0000000  n372   rna-XM_034638281.1    rna-XM_038529047.1    rna-XM_022525120.1                rna-XM_059151461.1    rna-XM_059382870.1      mNeoVis.2.1.1.chr7_hap1.g00485.t01    rna-XM_021700836.2      rna-XM_042968933.1          rna-gnl|WGS_ZZZZ|egapxtmp_012056-R2      mNeoVis.vulgivaga.2.1.chr7.g00492.t01      21    N0.HOG0000019   OG0000000  n380   rna-XM_034638310.1    rna-XM_038529216.1      rna-XM_045045446.1    rna-XM_036120117.1      rna-XM_032842124.1        rna-XM_059151466.1    rna-XM_059382898.1      mNeoVis.2.1.1.chr7_hap1.g00484.t01    rna-XM_044911379.1    rna-XM_004396432.1    rna-XM_042968958.1    rna-XM_040632771.1    rna-XM_035725285.1    mNeoVis.3.1.1.chr7.g00466.t01    rna-gnl|WGS_ZZZZ|egapxtmp_012059-R4          22    N0.HOG0000020   OG0000000  n395   rna-XM_011227147.3    rna-XM_038529132.1    rna-XM_022525111.1    rna-XM_006941444.5    rna-XM_036068027.1    rna-XM_039249716.1    rna-XM_032842102.1    rna-XM_045988756.1    rna-XM_032325637.1    rna-XM_059151508.1    rna-XM_059382925.1    rna-XM_004765008.3    mNeoVis.2.1.1.chr7_hap1.g00470.t01    rna-XM_021700828.1    rna-XM_004396447.1    rna-XM_007096826.3    rna-XM_040632808.1    rna-XM_027617836.2    mNeoVis.3.1.1.chr7.g00456.t01    rna-gnl|WGS_ZZZZ|egapxtmp_012065-R1    mNeoVis.evergladensis.2.1.chr7.g00478.t01        23    N0.HOG0000021   OG0000000  n415   rna-XM_034639017.1    rna-XM_038525524.1        rna-XM_036068035.1, rna-XM_036104162.1    rna-XM_039249737.1, rna-XM_039249738.1      rna-XM_045986777.1    rna-XM_032325568.1    rna-XM_059151448.1    rna-XM_059418532.1    rna-XM_013062735.2    mNeoVis.2.1.1.chr7_hap1.g00477.t01      rna-XM_004405070.1      rna-XM_040632800.1    rna-XM_035725255.1    mNeoVis.3.1.1.chr7.g00462.t01, mNeoVis.3.1.1.chr7.g00467.t01, mNeoVis.3.1.1.chr7.g00746.t01, mNeoVis.3.1.1.chr7.g00993.t01    rna-gnl|WGS_ZZZZ|egapxtmp_010029-R1, rna-gnl|WGS_ZZZZ|egapxtmp_012084-R7    mNeoVis.evergladensis.2.1.chr7.g00483.t01    mNeoVis.vulgivaga.2.1.chr7.g00477.t01, mNeoVis.vulgivaga.2.1.chr7.g00484.t01, mNeoVis.vulgivaga.2.1.chr7.g01063.t01      24    N0.HOG0000022   OG0000000  n445     rna-XM_038529189.1      rna-XM_045045514.1    rna-XM_036068071.1      rna-XM_032842132.1      rna-XM_032325675.1      rna-XM_059382970.1    rna-XM_045076553.1    mNeoVis.2.1.1.chr7_hap1.g00474.t01        rna-XM_042969057.1    rna-XM_008711145.2, rna-XM_008712212.2        rna-gnl|WGS_ZZZZ|egapxtmp_012073-R1          25    N0.HOG0000023   OG0000000  n456   rna-XM_034639018.1    rna-XM_038529161.1        rna-XM_036068060.1            rna-XM_059151552.1                rna-XM_040632817.1    rna-XM_035724895.1              26    N0.HOG0000024   OG0000000  n462   rna-XM_034638391.1    rna-XM_038529195.1      rna-XM_019818795.3    rna-XM_036068076.1    rna-XM_039249627.1    rna-XM_032842134.1    rna-XM_045988797.1    rna-XM_032325681.1    rna-XM_059151563.1    rna-XM_059382958.1    rna-XM_045076556.1    mNeoVis.2.1.1.chr7_hap1.g00478.t01        rna-XM_042969040.1    rna-XM_008711155.2    rna-XM_027617868.2    mNeoVis.3.1.1.chr7.g00463.t01    rna-gnl|WGS_ZZZZ|egapxtmp_012075-R1      mNeoVis.vulgivaga.2.1.chr7.g00485.t01      27    N0.HOG0000025   OG0000000  n479   rna-XM_019793387.2    rna-XM_038529185.1    rna-XM_022525113.1    rna-XM_045045524.1    rna-XM_036120131.1    rna-XM_039217746.1    rna-XM_032842127.1    rna-XM_045988784.1    rna-XM_032325687.1    rna-XM_059153783.1    rna-XM_059382888.1    rna-XM_013047707.2    mNeoVis.2.1.1.chr7_hap1.g00490.t01        rna-XM_042969033.1    rna-XM_008688657.2      mNeoVis.3.1.1.chr7.g00472.t01    rna-gnl|WGS_ZZZZ|egapxtmp_012081-R3    mNeoVis.evergladensis.2.1.chr7.g00493.t01    mNeoVis.vulgivaga.2.1.chr7.g00497.t01      28    N0.HOG0000026   OG0000000  n496   rna-XM_034650262.1, rna-XM_034650371.1                                                29    N0.HOG0000027   OG0000000  n502         rna-XM_019818864.3, rna-XM_019819176.3, rna-XM_023244884.2, rna-XM_045046283.1, rna-XM_045047126.1, rna-XM_045047128.1, rna-XM_045047131.1, rna-XM_045047141.1, rna-XM_045047160.1, rna-XM_045047165.1    rna-XM_036096619.1    rna-XM_039219739.1, rna-XM_039219740.1, rna-XM_039219748.1, rna-XM_039219749.1, rna-XM_039244144.1        rna-XM_032324153.1, rna-XM_032324833.1                rna-XM_015536422.1, rna-XM_042967971.1, rna-XM_042967974.1, rna-XM_042967975.1, rna-XM_042967978.1, rna-XM_042967980.1, rna-XM_042969146.1, rna-XM_042969973.1, rna-XM_042969997.1, rna-XM_042969999.1, rna-XM_042970000.1, rna-XM_042970001.1, rna-XM_042970002.1, rna-XM_042970003.1, rna-XM_042970004.1, rna-XM_042970007.1, rna-XM_042970019.1              mNeoVis.vulgivaga.2.1.chr7.g00766.t01      30    N0.HOG0000028   OG0000000  n538             rna-XM_039224246.1, rna-XM_039225719.1      rna-XM_045986790.1, rna-XM_045986793.1, rna-XM_045986796.1                  rna-XM_015541924.2                  31    N0.HOG0000029   OG0000000  n544         rna-XM_023244975.2      rna-XM_039219735.1, rna-XM_039219737.1, rna-XM_039219738.1, rna-XM_039219741.1, rna-XM_039219746.1, rna-XM_039219747.1, rna-XM_039224247.1, rna-XM_039225499.1                      rna-XM_042969143.1, rna-XM_042970016.1                  32    N0.HOG0000030   OG0000000  n555     rna-XM_038525532.1    rna-XM_022524115.1      rna-XM_036111862.1        rna-XM_045988228.1      rna-XM_059150660.1    rna-XM_059382642.1    rna-XM_045077388.1    mNeoVis.2.1.1.chr7_hap1.g00781.t01    rna-XM_044922006.1        rna-XM_008701596.2    rna-XM_035725235.1      rna-gnl|WGS_ZZZZ|egapxtmp_013307-R2      mNeoVis.vulgivaga.2.1.chr7.g00770.t01      33    N0.HOG0000031   OG0000000  n568   rna-XM_011218170.2, rna-XM_034638121.1    rna-XM_038527913.1    rna-XM_022492624.1      rna-XM_036083760.1      rna-XM_032839055.1    rna-XM_045988239.1    rna-XM_032324127.1    rna-XM_059150669.1    rna-XM_059382669.1    rna-XM_045077406.1      rna-XM_044911345.1, rna-XM_044912549.1    rna-XM_004409804.1      rna-XM_008701601.2    rna-XM_027618283.2        mNeoVis.evergladensis.2.1.chr7.g00767.t01    mNeoVis.vulgivaga.2.1.chr7.g00769.t01      34    N0.HOG0000032   OG0000000  n585   rna-XM_034639104.1    rna-XM_038527914.1    rna-XM_022492549.1, rna-XM_022492643.1, rna-XM_022524128.1      rna-XM_036070078.1, rna-XM_036094458.1      rna-XM_032839428.1    rna-XM_045988234.1    rna-XM_032324126.1, rna-XM_032324162.1    rna-XM_059150676.1    rna-XM_059382653.1    rna-XM_045077436.1      rna-XM_021682264.2, rna-XM_021682399.2, rna-XM_044920740.1        rna-XM_040632930.1        rna-gnl|WGS_ZZZZ|egapxtmp_013308-R2    mNeoVis.evergladensis.2.1.chr7.g00766.t01        35    N0.HOG0000033   OG0000000  n606     rna-XM_038525533.1, rna-XM_038525534.1, rna-XM_038527901.1, rna-XM_038527902.1, rna-XM_038527904.1, rna-XM_038527920.1, rna-XM_038527927.1    rna-XM_022524112.1      rna-XM_036111861.1      rna-XM_032839401.1    rna-XM_045988210.1      rna-XM_059150653.1    rna-XM_059382640.1      mNeoVis.2.1.1.chr7_hap1.g00782.t01    rna-XM_044922005.1    rna-XM_004409805.1      rna-XM_040633066.1    rna-XM_035725257.1    mNeoVis.3.1.1.chr7.g00723.t01      mNeoVis.evergladensis.2.1.chr7.g00769.t01    mNeoVis.vulgivaga.2.1.chr7.g00773.t01      36    N0.HOG0000034   OG0000000  n626   rna-XM_034638112.1, rna-XM_034650395.1    rna-XM_038525530.1    rna-XM_022491896.1, rna-XM_022492511.1, rna-XM_022492512.1, rna-XM_022492513.1, rna-XM_022492517.1, rna-XM_022492562.1, rna-XM_022492573.1, rna-XM_022492594.1, rna-XM_022492611.1, rna-XM_022492754.1, rna-XM_022524114.1      rna-XM_036065444.1, rna-XM_036083753.1, rna-XM_036083754.1      rna-XM_032848717.1, rna-XM_032848718.1, rna-XM_032848743.1, rna-XM_032848744.1, rna-XM_032850617.1, rna-XM_032850618.1, rna-XM_032850737.1, rna-XM_032850738.1, rna-XM_032850880.1, rna-XM_032850881.1, rna-XM_032851131.1, rna-XM_032851176.1    rna-XM_045986787.1, rna-XM_045986791.1, rna-XM_045986792.1, rna-XM_045986794.1, rna-XM_045986795.1, rna-XM_045986798.1, rna-XM_045986800.1, rna-XM_045986801.1, rna-XM_045986802.1, rna-XM_045986804.1, rna-XM_045986806.1, rna-XM_045986808.1, rna-XM_045986810.1, rna-XM_045988204.1, rna-XM_045988248.1, rna-XM_045988372.1, rna-XM_045988375.1, rna-XM_045988376.1, rna-XM_045988377.1, rna-XM_045988378.1, rna-XM_045988379.1    rna-XM_032302414.1, rna-XM_032324104.1, rna-XM_032324105.1, rna-XM_032324112.1, rna-XM_032324131.1, rna-XM_032324132.1, rna-XM_032324134.1, rna-XM_032324143.1, rna-XM_032324152.1, rna-XM_032324156.1, rna-XM_032324161.1, rna-XM_032324163.1, rna-XM_032324164.1, rna-XM_032324167.1, rna-XM_032326038.1, rna-XM_032326052.1, rna-XM_032326096.1, rna-XM_032326099.1, rna-XM_032326101.1, rna-XM_032326103.1, rna-XM_032326104.1    rna-XM_059150634.1, rna-XM_059150654.1, rna-XM_059150662.1, rna-XM_059150686.1, rna-XM_059151210.1, rna-XM_059151211.1, rna-XM_059151218.1, rna-XM_059151223.1, rna-XM_059151224.1, rna-XM_059151227.1, rna-XM_059151228.1, rna-XM_059151818.1, rna-XM_059152623.1, rna-XM_059152752.1, rna-XM_059152758.1    rna-XM_059382026.1, rna-XM_059382637.1, rna-XM_059382638.1, rna-XM_059382788.1, rna-XM_059383339.1, rna-XM_059383340.1, rna-XM_059383341.1, rna-XM_059383342.1, rna-XM_059383343.1, rna-XM_059383345.1, rna-XM_059383348.1, rna-XM_059383349.1, rna-XM_059383353.1, rna-XM_059383354.1, rna-XM_059383356.1, rna-XM_059383358.1, rna-XM_059387222.1, rna-XM_059387278.1    rna-XM_004779824.3, rna-XM_013049565.2, rna-XM_013050028.2, rna-XM_013050032.2, rna-XM_013050043.2, rna-XM_045076990.1, rna-XM_045076992.1, rna-XM_045076993.1, rna-XM_045076995.1, rna-XM_045076997.1, rna-XM_045077266.1, rna-XM_045077267.1, rna-XM_045077268.1, rna-XM_045077269.1, rna-XM_045077271.1, rna-XM_045077272.1, rna-XM_045077274.1, rna-XM_045077275.1, rna-XM_045077373.1, rna-XM_045077405.1, rna-XM_045077442.1, rna-XM_045077454.1, rna-XM_045077457.1, rna-XM_045077464.1, rna-XM_045077465.1    mNeoVis.2.1.1.chr7_hap1.g00770.t01, mNeoVis.2.1.1.chr7_hap1.g00772.t01, mNeoVis.2.1.1.chr7_hap1.g00776.t01, mNeoVis.2.1.1.chr7_hap1.g00778.t01, mNeoVis.2.1.1.chr7_hap1.g00779.t01, mNeoVis.2.1.1.chr7_hap1.g00780.t01    rna-XM_044911317.1, rna-XM_044911318.1, rna-XM_044911342.1, rna-XM_044911343.1, rna-XM_044911344.1, rna-XM_044922022.1    rna-XM_004409555.1      rna-XM_008701622.2, rna-XM_040634401.1    rna-XM_035724913.1, rna-XM_035724914.1, rna-XM_035724915.1    mNeoVis.3.1.1.chr7.g00722.t01    rna-gnl|WGS_ZZZZ|egapxtmp_012297-R1, rna-gnl|WGS_ZZZZ|egapxtmp_012300-R1, rna-gnl|WGS_ZZZZ|egapxtmp_012301-R1, rna-gnl|WGS_ZZZZ|egapxtmp_013305-R1, rna-gnl|WGS_ZZZZ|egapxtmp_014209-R4    mNeoVis.evergladensis.2.1.chr7.g00757.t01, mNeoVis.evergladensis.2.1.chr7.g00758.t01, mNeoVis.evergladensis.2.1.chr7.g00760.t01, mNeoVis.evergladensis.2.1.chr7.g00762.t01, mNeoVis.evergladensis.2.1.chr7.g00763.t01, mNeoVis.evergladensis.2.1.chr7.g00765.t01    mNeoVis.vulgivaga.2.1.chr7.g00761.t01, mNeoVis.vulgivaga.2.1.chr7.g00767.t01      37    N0.HOG0000035   OG0000000  n787   rna-XM_019793917.2, rna-XM_034638125.1, rna-XM_034639057.1, rna-XM_034639067.1, rna-XM_034639126.1    rna-XM_038527912.1, rna-XM_038527924.1, rna-XM_038527925.1    rna-XM_022492732.1, rna-XM_022523770.1      rna-XM_036111857.1      rna-XM_032839054.1, rna-XM_032839403.1, rna-XM_032839424.1, rna-XM_032839429.1, rna-XM_032850961.1    rna-XM_045986789.1, rna-XM_045988222.1, rna-XM_045988304.1, rna-XM_045995399.1    rna-XM_032324114.1, rna-XM_032324125.1, rna-XM_032326039.1, rna-XM_032326095.1    rna-XM_059150656.1, rna-XM_059151230.1, rna-XM_059151231.1    rna-XM_059382639.1, rna-XM_059382670.1, rna-XM_059383320.1    rna-XM_045077276.1, rna-XM_045077449.1      rna-XM_021682266.1, rna-XM_021682401.1, rna-XM_044912170.1, rna-XM_044922020.1    rna-XM_004409802.1      rna-XM_008684786.2, rna-XM_040632916.1, rna-XM_040632918.1, rna-XM_040632936.1, rna-XM_040633067.1, rna-XM_040633068.1, rna-XM_040633069.1, rna-XM_040633070.1    rna-XM_035724912.1, rna-XM_035725256.1    mNeoVis.3.1.1.chr7.g00725.t01    rna-gnl|WGS_ZZZZ|egapxtmp_013306-R1    mNeoVis.evergladensis.2.1.chr7.g00768.t01    mNeoVis.vulgivaga.2.1.chr7.g00774.t01, mNeoVis.vulgivaga.2.1.chr7.g00775.t01      38    N0.HOG0000036   OG0000000  n838   rna-XM_034650316.1                                  rna-XM_040632913.1                39    N0.HOG0000037   OG0000000  n840         rna-XM_045046183.1, rna-XM_045047127.1                          rna-XM_015536390.2, rna-XM_042970017.1, rna-XM_042970020.1, rna-XM_042970021.1, rna-XM_042970022.1                  40    N0.HOG0000038   OG0000000  n847   rna-XM_034639071.1, rna-XM_034639072.1, rna-XM_034639191.1    rna-XM_038525531.1    rna-XM_022491917.1, rna-XM_022524136.1          rna-XM_032850866.1    rna-XM_045986797.1    rna-XM_032326093.1    rna-XM_059151220.1    rna-XM_059383346.1, rna-XM_059383355.1              rna-XM_008701603.2, rna-XM_040632997.1, rna-XM_040634402.1    rna-XM_027619365.1, rna-XM_027619370.2    mNeoVis.3.1.1.chr7.g00724.t01        mNeoVis.vulgivaga.2.1.chr7.g00772.t01      41    N0.HOG0000039   OG0000000  n865   rna-XM_034639175.1    rna-XM_038527922.1    rna-XM_022523768.1      rna-XM_036083759.1            rna-XM_059151229.1          rna-XM_021682263.2        rna-XM_040634403.1                42    N0.HOG0000040   OG0000001  n2         rna-XM_003981988.2, rna-XM_003981995.4, rna-XM_045042151.1, rna-XM_045042186.1      rna-XM_039223674.1, rna-XM_039223679.1                      rna-XM_007088318.2, rna-XM_007088595.3, rna-XM_007088603.3, rna-XM_007094196.2                  43    N0.HOG0000041   OG0000001  n12       rna-XM_022497945.1              rna-XM_032303493.1    rna-XM_059159085.1, rna-XM_059160273.1    rna-XM_059388249.1, rna-XM_059389247.1    rna-XM_004748604.1, rna-XM_004781407.2    mNeoVis.2.1.1.chr5_hap1.g00382.t01, mNeoVis.2.1.1.chr5_hap1.g00386.t01              mNeoVis.3.1.1.chr2.g00315.t01, mNeoVis.3.1.1.chr2.g00320.t01    rna-gnl|WGS_ZZZZ|egapxtmp_034878-R1, rna-gnl|WGS_ZZZZ|egapxtmp_034884-R1    mNeoVis.evergladensis.2.1.chr2.g00383.t01, mNeoVis.evergladensis.2.1.chr2.g00399.t01    mNeoVis.vulgivaga.2.1.chr2.g00378.t01, mNeoVis.vulgivaga.2.1.chr2.g00383.t01, mNeoVis.vulgivaga.2.1.chr2.g00392.t01      44    N0.HOG0000042   OG0000001  n31   rna-XM_002929855.2, rna-XM_002931198.2, rna-XM_019793710.2, rna-XM_034650233.1, rna-XM_034653242.1, rna-XM_034653345.1, rna-XM_034653358.1, rna-XM_034657178.1    rna-NM_001388890.1, rna-XM_038565372.1        rna-XM_036076195.1, rna-XM_036089113.1, rna-XM_036089117.1      rna-XM_032883489.1    rna-XM_045989931.1, rna-XM_045991054.1, rna-XM_045991206.1, rna-XM_045991420.1, rna-XM_045991624.1    rna-XM_032303579.1, rna-XM_032303843.1    rna-XM_059160269.1, rna-XM_059165144.1    rna-XM_059387737.1, rna-XM_059387753.1, rna-XM_059388256.1, rna-XM_059388257.1    rna-XM_004761192.2, rna-XM_004781401.1, rna-XM_004781410.1    mNeoVis.2.1.1.chr5_hap1.g00396.t01, mNeoVis.2.1.1.chr5_hap1.g00397.t01, mNeoVis.2.1.1.chr5_hap1.g00410.t01    rna-XM_021683215.1, rna-XM_021683216.1    rna-XM_004408388.1      rna-XM_008706449.2, rna-XM_008711160.2, rna-XM_008711161.2, rna-XM_008711163.1, rna-XM_008711182.2, rna-XM_008711185.1, rna-XM_008711752.2, rna-XM_040628179.1, rna-XM_040628180.1    rna-XM_027581952.2, rna-XM_035726601.1    mNeoVis.3.1.1.chr2.g00328.t01, mNeoVis.3.1.1.chr2.g00341.t01    rna-gnl|WGS_ZZZZ|egapxtmp_034897-R1, rna-gnl|WGS_ZZZZ|egapxtmp_038058-R1, rna-gnl|WGS_ZZZZ|egapxtmp_038070-R1    mNeoVis.evergladensis.2.1.chr2.g00398.t01    mNeoVis.vulgivaga.2.1.chr2.g00386.t01, mNeoVis.vulgivaga.2.1.chr2.g00393.t01, mNeoVis.vulgivaga.2.1.chr2.g00394.t01, mNeoVis.vulgivaga.2.1.chr2.g00411.t01      45    N0.HOG0000043   OG0000001  n86                               rna-XM_004414856.1        rna-XM_027581955.2              46    N0.HOG0000044   OG0000001  n88   rna-XM_019792673.2, rna-XM_034653380.1, rna-XM_034653390.1          rna-XM_036087756.1          rna-XM_032303504.1    rna-XM_059159086.1    rna-XM_059388250.1    rna-XM_004748603.3                          47    N0.HOG0000045   OG0000001  n97   rna-XM_002929860.4, rna-XM_034657184.1    rna-XM_038565364.1, rna-XM_038565365.1            rna-XM_032868474.1        rna-XM_059159101.1      rna-XM_004761199.2                          48    N0.HOG0000046   OG0000001  n103   rna-XM_034653355.1, rna-XM_034657232.1    rna-XM_038565374.1        rna-XM_036089066.1      rna-XM_032868469.1, rna-XM_032868473.1    rna-XM_045989936.1, rna-XM_045989939.1, rna-XM_045989942.1, rna-XM_045989944.1    rna-XM_032303534.1, rna-XM_032303653.1, rna-XM_032303686.1, rna-XM_032303725.1, rna-XM_032303736.1, rna-XM_032303744.1    rna-XM_059159088.1, rna-XM_059159093.1, rna-XM_059159095.1, rna-XM_059159096.1, rna-XM_059159097.1, rna-XM_059165130.1    rna-XM_059388251.1, rna-XM_059388255.1, rna-XM_059388258.1, rna-XM_059388259.1, rna-XM_059388260.1, rna-XM_059388261.1    rna-XM_004781399.1, rna-XM_004781409.1, rna-XM_004782125.1, rna-XM_004782127.1, rna-XM_004782384.1, rna-XM_004782394.3    mNeoVis.2.1.1.chr5_hap1.g00387.t01, mNeoVis.2.1.1.chr5_hap1.g00398.t01, mNeoVis.2.1.1.chr5_hap1.g00404.t01          rna-XM_008706427.2, rna-XM_008711166.1, rna-XM_008711168.2, rna-XM_008711180.2, rna-XM_008711183.2      mNeoVis.3.1.1.chr2.g00321.t01, mNeoVis.3.1.1.chr2.g00327.t01, mNeoVis.3.1.1.chr2.g00329.t01, mNeoVis.3.1.1.chr2.g00333.t01    rna-gnl|WGS_ZZZZ|egapxtmp_034876-R1, rna-gnl|WGS_ZZZZ|egapxtmp_034911-R1, rna-gnl|WGS_ZZZZ|egapxtmp_038057-R1, rna-gnl|WGS_ZZZZ|egapxtmp_038059-R1    mNeoVis.evergladensis.2.1.chr2.g00388.t01, mNeoVis.evergladensis.2.1.chr2.g00403.t01    mNeoVis.vulgivaga.2.1.chr2.g00382.t01, mNeoVis.vulgivaga.2.1.chr2.g00395.t01, mNeoVis.vulgivaga.2.1.chr2.g00400.t01, mNeoVis.vulgivaga.2.1.chr2.g00403.t01, mNeoVis.vulgivaga.2.1.chr2.g00404.t01      49    N0.HOG0000047   OG0000001  n158         rna-XM_004001239.4, rna-XM_011290174.2, rna-XM_019824930.3, rna-XM_019826092.3, rna-XM_045047285.1, rna-XM_045047290.1, rna-XM_045047311.1      rna-XM_039223673.1, rna-XM_039223676.1, rna-XM_039223677.1, rna-XM_039223680.1                      rna-XM_007088980.2, rna-XM_007088981.2, rna-XM_015544715.2, rna-XM_042976863.1, rna-XM_042976886.1, rna-XM_042976918.1, rna-XM_042976958.1, rna-XM_042977310.1, rna-XM_042977311.1, rna-XM_042977312.1, rna-XM_042977313.1, rna-XM_042977314.1, rna-XM_042978420.1                  50    N0.HOG0000048   OG0000001  n184         rna-XM_004001250.2, rna-XM_045047282.1, rna-XM_045047307.1, rna-XM_045047331.1, rna-XM_045047334.1      rna-XM_039223678.1                      rna-XM_007088596.2, rna-XM_007088983.2, rna-XM_007094194.1, rna-XM_042976901.1, rna-XM_042977308.1, rna-XM_042977309.1                  51    N0.HOG0000049   OG0000001  n196   rna-XM_034653383.1    rna-XM_038565368.1      rna-XM_045042170.1    rna-XM_036091756.1    rna-XM_039223666.1    rna-XM_032868466.1    rna-XM_045989935.1    rna-XM_032313760.1    rna-XM_059159092.1    rna-XM_059392587.1    rna-XM_004781404.1          rna-XM_007088599.2        mNeoVis.3.1.1.chr2.g00324.t01    rna-gnl|WGS_ZZZZ|egapxtmp_035361-R1    mNeoVis.evergladensis.2.1.chr2.g00392.t01        52    N0.HOG0000050   OG0000001  n213   rna-XM_002930436.4                                  rna-XM_008684223.2                53    N0.HOG0000051   OG0000001  n216   rna-XM_034650295.1, rna-XM_034653382.1, rna-XM_034657228.1                rna-XM_045991868.1                    rna-XM_008684230.2, rna-XM_008712015.2                54    N0.HOG0000052   OG0000001  n222   rna-XM_002930437.4    rna-NM_001388664.1                            rna-XM_004408384.1      rna-XM_008684222.2                55    N0.HOG0000053   OG0000001  n226                             rna-XM_021683212.2    rna-XM_004408389.1        rna-XM_027585078.1, rna-XM_035728622.1              56    N0.HOG0000054   OG0000001  n231     rna-XM_038565311.1              rna-XM_045991925.1    rna-XM_032303599.1    rna-XM_059160270.1    rna-XM_059387739.1    rna-XM_004781402.1    mNeoVis.2.1.1.chr5_hap1.g00390.t01          rna-XM_008711159.2, rna-XM_008711494.2, rna-XM_008712023.1      mNeoVis.3.1.1.chr2.g00323.t01    rna-gnl|WGS_ZZZZ|egapxtmp_035359-R1    mNeoVis.evergladensis.2.1.chr2.g00391.t01    mNeoVis.vulgivaga.2.1.chr2.g00387.t01      57    N0.HOG0000055   OG0000001  n244   rna-XM_034657189.1    rna-NM_001388660.1              rna-XM_045991613.1    rna-XM_032303819.1    rna-XM_059165139.1    rna-XM_059387747.1    rna-XM_004761195.3    mNeoVis.2.1.1.chr5_hap1.g00408.t01          rna-XM_008684224.2      mNeoVis.3.1.1.chr2.g00340.t01      mNeoVis.evergladensis.2.1.chr2.g00409.t01    mNeoVis.vulgivaga.2.1.chr2.g00408.t01      58    N0.HOG0000056   OG0000001  n255   rna-XM_034653334.1                            rna-XM_044917731.1          rna-XM_035726545.1              59    N0.HOG0000057   OG0000001  n258   rna-XM_034657183.1        rna-XM_045047280.1, rna-XM_045047335.1      rna-XM_039223675.1                      rna-XM_007088598.2    rna-XM_008684219.1                60    N0.HOG0000058   OG0000001  n267   rna-XM_034657190.1    rna-NM_001389028.1        rna-XM_036089092.1          rna-XM_032303841.1    rna-XM_059165141.1    rna-XM_059387750.1    rna-XM_004761194.1    mNeoVis.2.1.1.chr5_hap1.g00401.t01    rna-XM_021683211.1        rna-XM_008706426.2    rna-XM_027585037.2, rna-XM_035728891.1      rna-gnl|WGS_ZZZZ|egapxtmp_034913-R1    mNeoVis.evergladensis.2.1.chr2.g00410.t01    mNeoVis.vulgivaga.2.1.chr2.g00409.t01      61    N0.HOG0000059   OG0000001  n279   rna-XM_034653336.1, rna-XM_034653337.1, rna-XM_034653346.1, rna-XM_034657169.1, rna-XM_034657171.1, rna-XM_034657174.1, rna-XM_034657177.1    rna-NM_001388671.1        rna-XM_036076196.1, rna-XM_036089068.1      rna-XM_032868470.1, rna-XM_032868471.1    rna-XM_045989661.1, rna-XM_045991205.1, rna-XM_045991232.1, rna-XM_045991240.1    rna-XM_032303690.1, rna-XM_032303704.1    rna-XM_059160274.1, rna-XM_059165125.1, rna-XM_059165131.1    rna-XM_059389248.1, rna-XM_059389249.1    rna-XM_004781408.1, rna-XM_004782128.2, rna-XM_045088841.1    mNeoVis.2.1.1.chr5_hap1.g00395.t01, mNeoVis.2.1.1.chr5_hap1.g00399.t01, mNeoVis.2.1.1.chr5_hap1.g00400.t01      rna-XM_004408386.1, rna-XM_004408387.1, rna-XM_004414854.1, rna-XM_004414855.1      rna-XM_008711162.1, rna-XM_008711186.1, rna-XM_008711756.2    rna-XM_027575108.2, rna-XM_027575112.2, rna-XM_027584653.2, rna-XM_027584764.2, rna-XM_027617015.2, rna-XM_027620377.1, rna-XM_035724367.1, rna-XM_035724395.1, rna-XM_035724567.1, rna-XM_035728263.1    mNeoVis.3.1.1.chr2.g00330.t01, mNeoVis.3.1.1.chr2.g00332.t01    rna-gnl|WGS_ZZZZ|egapxtmp_038061-R1    mNeoVis.evergladensis.2.1.chr2.g00396.t01, mNeoVis.evergladensis.2.1.chr2.g00400.t01    mNeoVis.vulgivaga.2.1.chr2.g00396.t01, mNeoVis.vulgivaga.2.1.chr2.g00397.t01      62    N0.HOG0000060   OG0000001  n329   rna-XM_034653330.1, rna-XM_034653341.1, rna-XM_034653384.1, rna-XM_034657170.1, rna-XM_034657172.1, rna-XM_034657173.1      rna-XM_022526818.1              rna-XM_032303448.1, rna-XM_032303519.1    rna-XM_059159087.1    rna-XM_059387738.1    rna-XM_004781398.1            rna-XM_008684225.2        rna-gnl|WGS_ZZZZ|egapxtmp_034910-R1    mNeoVis.evergladensis.2.1.chr2.g00389.t01    mNeoVis.vulgivaga.2.1.chr2.g00384.t01      63    N0.HOG0000061   OG0000001  n344   rna-XM_034657175.1, rna-XM_034657176.1                rna-XM_045991278.1    rna-XM_032303730.1    rna-XM_059165132.1    rna-XM_059387743.1    rna-XM_004782126.1    mNeoVis.2.1.1.chr5_hap1.g00402.t01    rna-XM_021683218.1        rna-XM_008711188.1      mNeoVis.3.1.1.chr2.g00334.t01    rna-gnl|WGS_ZZZZ|egapxtmp_034914-R1    mNeoVis.evergladensis.2.1.chr2.g00401.t01    mNeoVis.vulgivaga.2.1.chr2.g00401.t01      64    N0.HOG0000062   OG0000001  n359   rna-XM_034653361.1    rna-NM_001388657.1    rna-XM_022498198.1    rna-XM_003981997.2      rna-XM_039243415.1    rna-XM_032851082.1    rna-XM_045991882.1, rna-XM_045991885.1    rna-XM_032303844.1    rna-XM_059165146.1      rna-XM_013060460.1    mNeoVis.2.1.1.chr5_hap1.g00412.t01      rna-XM_004408391.1    rna-XM_015540307.1      rna-XM_027585421.1, rna-XM_027585486.2    mNeoVis.3.1.1.chr2.g00343.t01    rna-gnl|WGS_ZZZZ|egapxtmp_034890-R1    mNeoVis.evergladensis.2.1.chr2.g00413.t01    mNeoVis.vulgivaga.2.1.chr2.g00412.t01      65    N0.HOG0000063   OG0000001  n378   rna-XM_002928175.4    rna-NM_001388656.1      rna-XM_003981998.4    rna-XM_036089071.1    rna-XM_039243384.1    rna-XM_032867473.1    rna-XM_045992307.1    rna-XM_032313829.1    rna-XM_059160279.1    rna-XM_059389255.1    rna-XM_004761149.3    mNeoVis.2.1.1.chr5_hap1.g00414.t01    rna-XM_021683209.1      rna-XM_007088605.2    rna-XM_008706448.2      mNeoVis.3.1.1.chr2.g00345.t01    rna-gnl|WGS_ZZZZ|egapxtmp_038071-R1    mNeoVis.evergladensis.2.1.chr2.g00412.t01    mNeoVis.vulgivaga.2.1.chr2.g00414.t01      66    N0.HOG0000064   OG0000001  n397   rna-XM_002929861.4, rna-XM_034650208.1    rna-XM_038565370.1      rna-XM_003981990.3      rna-XM_039223681.1      rna-XM_045991923.1    rna-XM_032360514.1    rna-XM_059165137.1    rna-XM_059387421.1    rna-XM_004761197.1          rna-XM_007088597.2    rna-XM_008711169.2      mNeoVis.3.1.1.chr2.g00339.t01    rna-gnl|WGS_ZZZZ|egapxtmp_038063-R1    mNeoVis.evergladensis.2.1.chr2.g00407.t01    mNeoVis.vulgivaga.2.1.chr2.g00399.t01      67    N0.HOG0000065   OG0000001  n412   rna-XM_002929857.4    rna-XM_038565371.1        rna-XM_036089116.1        rna-XM_045991842.1      rna-XM_059165135.1    rna-XM_059387745.1    rna-XM_004761200.1      rna-XM_021683214.1        rna-XM_008711173.2                68    N0.HOG0000066   OG0000001  n422   rna-XM_002921891.3    rna-XM_038567390.1    rna-XM_022526826.1    rna-XM_003981816.3    rna-XM_036120223.1    rna-XM_039224717.1, rna-XM_039224719.1    rna-XM_032883910.1    rna-XM_045990546.1    rna-XM_032310232.1, rna-XM_032310242.1    rna-XM_059159869.1, rna-XM_059159870.1    rna-XM_059387647.1, rna-XM_059388929.1    rna-XM_004771741.2, rna-XM_045084714.1    mNeoVis.2.1.1.chr5_hap1.g00245.t01    rna-XM_021705186.2      rna-XM_007079387.2    rna-XM_008705518.1    rna-XM_027583020.2    mNeoVis.3.1.1.chr2.g00189.t01    rna-gnl|WGS_ZZZZ|egapxtmp_038121-R1    mNeoVis.evergladensis.2.1.chr2.g00243.t01    mNeoVis.vulgivaga.2.1.chr2.g00241.t01      69    N0.HOG0000067   OG0000001  n447   rna-XM_002921410.3    rna-NM_001388988.1      rna-XM_003981818.4    rna-XM_036119660.1    rna-XM_039251019.1    rna-XM_032883478.1    rna-XM_045991811.1              rna-XM_021705370.2      rna-XM_007079381.1    rna-XM_008705516.2      mNeoVis.3.1.1.chr2.g00182.t01, mNeoVis.3.1.1.chr2.g00186.t01            70    N0.HOG0000068   OG0000001  n458   rna-XM_002921916.4, rna-XM_034650236.1, rna-XM_034650237.1    rna-XM_038565383.1    rna-XM_022527044.1, rna-XM_022527047.1    rna-XM_003981839.6, rna-XM_011289531.3    rna-XM_036095271.1, rna-XM_036120285.1    rna-XM_039224720.1, rna-XM_039224721.1    rna-XM_032883474.1, rna-XM_032883911.1    rna-XM_045990534.1, rna-XM_045990549.1    rna-XM_032310253.1, rna-XM_032310328.1, rna-XM_032358127.1, rna-XM_032361629.1    rna-XM_059159069.1, rna-XM_059159880.1, rna-XM_059159881.1    rna-XM_059387646.1, rna-XM_059387648.1, rna-XM_059388937.1    rna-XM_004771769.1, rna-XM_045084715.1, rna-XM_045084757.1    mNeoVis.2.1.1.chr5_hap1.g00244.t01, mNeoVis.2.1.1.chr5_hap1.g00246.t01, mNeoVis.2.1.1.chr5_hap1.g00249.t01    rna-XM_021705187.1    rna-XM_004398837.1    rna-XM_007079386.2, rna-XM_007079388.3, rna-XM_007095834.2, rna-XM_042969207.1    rna-XM_008705490.2, rna-XM_008705492.2    rna-XM_027583036.2, rna-XM_027583037.2    mNeoVis.3.1.1.chr2.g00187.t01, mNeoVis.3.1.1.chr2.g00191.t01    rna-gnl|WGS_ZZZZ|egapxtmp_038123-R1, rna-gnl|WGS_ZZZZ|egapxtmp_038125-R1, rna-gnl|WGS_ZZZZ|egapxtmp_038131-R1    mNeoVis.evergladensis.2.1.chr2.g00242.t01, mNeoVis.evergladensis.2.1.chr2.g00244.t01, mNeoVis.evergladensis.2.1.chr2.g00247.t01    mNeoVis.vulgivaga.2.1.chr2.g00240.t01      71    N0.HOG0000069   OG0000001  n504   rna-XM_019793347.2, rna-XM_034653331.1, rna-XM_034653332.1, rna-XM_034657168.1    rna-NM_001388904.1    rna-XM_022497944.1, rna-XM_022526819.1    rna-XM_011280078.3, rna-XM_045041506.1, rna-XM_045041518.1, rna-XM_045041534.1, rna-XM_045047244.1, rna-XM_045047249.1, rna-XM_045047250.1, rna-XM_045047259.1    rna-XM_036091758.1, rna-XM_036091760.1    rna-XM_039223670.1, rna-XM_039252229.1    rna-XM_032883487.1    rna-XM_045989926.1, rna-XM_045989928.1    rna-XM_032303435.1, rna-XM_032303459.1, rna-XM_032303469.1, rna-XM_032303484.1    rna-XM_059159081.1, rna-XM_059159082.1, rna-XM_059159083.1    rna-XM_059388245.1, rna-XM_059388246.1, rna-XM_059388247.1, rna-XM_059388248.1    rna-XM_004748605.3, rna-XM_004748606.1, rna-XM_045077623.1    mNeoVis.2.1.1.chr5_hap1.g00384.t01, mNeoVis.2.1.1.chr5_hap1.g00385.t01, mNeoVis.2.1.1.chr5_hap1.g00388.t01    rna-XM_021683224.1      rna-XM_007098165.3, rna-XM_042977306.1, rna-XM_042977307.1    rna-XM_008711687.2, rna-XM_008711760.2, rna-XM_008711761.1, rna-XM_008711762.2, rna-XM_008711763.2, rna-XM_008711765.1, rna-XM_040629154.1    rna-XM_027585085.2    mNeoVis.3.1.1.chr2.g00318.t01, mNeoVis.3.1.1.chr2.g00319.t01    rna-gnl|WGS_ZZZZ|egapxtmp_035356-R1, rna-gnl|WGS_ZZZZ|egapxtmp_035357-R1    mNeoVis.evergladensis.2.1.chr2.g00386.t01, mNeoVis.evergladensis.2.1.chr2.g00387.t01    mNeoVis.vulgivaga.2.1.chr2.g00381.t01      72    N0.HOG0000070   OG0000001  n561         rna-XM_006928537.3, rna-XM_045047287.1, rna-XM_045047317.1, rna-XM_045047329.1      rna-XM_039223672.1                      rna-XM_007088601.2, rna-XM_007088984.2, rna-XM_007093065.2, rna-XM_015540304.2, rna-XM_042976909.1, rna-XM_042976949.1                  73    N0.HOG0000071   OG0000001  n573   rna-XM_002930922.4    rna-NM_001389186.1            rna-XM_032883488.1    rna-XM_045989930.1    rna-XM_032303566.1    rna-XM_059160271.1    rna-XM_059387736.1      mNeoVis.2.1.1.chr5_hap1.g00389.t01          rna-XM_008711757.2      mNeoVis.3.1.1.chr2.g00322.t01    rna-gnl|WGS_ZZZZ|egapxtmp_035358-R1    mNeoVis.evergladensis.2.1.chr2.g00390.t01    mNeoVis.vulgivaga.2.1.chr2.g00385.t01      74    N0.HOG0000072   OG0000001  n583   rna-XM_019793229.2, rna-XM_034653353.1    rna-NM_001388672.1, rna-NM_001388901.1, rna-XM_038565369.1        rna-XM_036091764.1      rna-XM_032850974.1, rna-XM_032850975.1, rna-XM_032868467.1    rna-XM_045991169.1    rna-XM_032303556.1, rna-XM_032303622.1    rna-XM_059159089.1    rna-XM_059387740.1, rna-XM_059388252.1    rna-XM_004781400.3, rna-XM_004781405.1    mNeoVis.2.1.1.chr5_hap1.g00392.t01    rna-XM_021683222.1    rna-XM_004414858.1      rna-XM_008684228.2    rna-XM_027585089.2      rna-gnl|WGS_ZZZZ|egapxtmp_035363-R1    mNeoVis.evergladensis.2.1.chr2.g00393.t01    mNeoVis.vulgivaga.2.1.chr2.g00389.t01      75    N0.HOG0000073   OG0000001  n610   rna-XM_002930953.2, rna-XM_034657235.1    rna-NM_001388888.1    rna-XM_022498088.1            rna-XM_045991044.1      rna-XM_059165138.1, rna-XM_059165143.1    rna-XM_059387746.1, rna-XM_059387752.1    rna-XM_004761193.1, rna-XM_004761196.1    mNeoVis.2.1.1.chr5_hap1.g00409.t01    rna-XM_021683210.2    rna-XM_004408390.1, rna-XM_004414860.1        rna-XM_027585386.1, rna-XM_035728496.1    mNeoVis.3.1.1.chr2.g00331.t01    rna-gnl|WGS_ZZZZ|egapxtmp_034898-R1, rna-gnl|WGS_ZZZZ|egapxtmp_038068-R1    mNeoVis.evergladensis.2.1.chr2.g00408.t01, mNeoVis.evergladensis.2.1.chr2.g00411.t01    mNeoVis.vulgivaga.2.1.chr2.g00407.t01, mNeoVis.vulgivaga.2.1.chr2.g00410.t01      76    N0.HOG0000074   OG0000001  n633   rna-XM_011217950.3, rna-XM_034653362.1, rna-XM_034653386.1, rna-XM_034653387.1    rna-XM_038567028.1            rna-XM_032868502.1    rna-XM_045989677.1      rna-XM_059160272.1    rna-XM_059389246.1    rna-XM_004781406.1    mNeoVis.2.1.1.chr5_hap1.g00394.t01    rna-XM_021683220.1, rna-XM_044912961.1, rna-XM_044917732.1    rna-XM_004408385.1      rna-XM_008684215.2, rna-XM_008684221.1    rna-XM_027607262.2, rna-XM_035724573.1    mNeoVis.3.1.1.chr2.g00326.t01    rna-gnl|WGS_ZZZZ|egapxtmp_034899-R2    mNeoVis.evergladensis.2.1.chr2.g00395.t01    mNeoVis.vulgivaga.2.1.chr2.g00391.t01      77    N0.HOG0000075   OG0000001  n656   rna-XM_034650587.1    rna-NM_001388665.1              rna-XM_045989929.1                rna-XM_004414859.1      rna-XM_008684227.1                78    N0.HOG0000076   OG0000001  n660     rna-NM_001388667.1                                rna-XM_008684232.2                79    N0.HOG0000077   OG0000001  n661   rna-XM_034653354.1, rna-XM_034653356.1, rna-XM_034657180.1              rna-XM_032868476.1    rna-XM_045991073.1, rna-XM_045991123.1, rna-XM_045991135.1, rna-XM_045991168.1, rna-XM_045991173.1    rna-XM_032303663.1, rna-XM_032303756.1    rna-XM_059165127.1, rna-XM_059165133.1, rna-XM_059165136.1    rna-XM_059387742.1, rna-XM_059387744.1    rna-XM_004761198.1, rna-XM_004761202.1, rna-XM_004781411.1    mNeoVis.2.1.1.chr5_hap1.g00406.t01          rna-XM_008711178.2, rna-XM_008711179.1      mNeoVis.3.1.1.chr2.g00338.t01      mNeoVis.evergladensis.2.1.chr2.g00397.t01, mNeoVis.evergladensis.2.1.chr2.g00406.t01        80    N0.HOG0000078   OG0000001  n686         rna-XM_011290109.3                          rna-XM_007088602.3                  81    N0.HOG0000079   OG0000001  n688   rna-XM_019793227.2, rna-XM_019793817.2, rna-XM_034650357.1, rna-XM_034650483.1, rna-XM_034653217.1, rna-XM_034653359.1, rna-XM_034653389.1, rna-XM_034657181.1, rna-XM_034657185.1      rna-XM_022498159.1    rna-XM_045042163.1, rna-XM_045042175.1    rna-XM_036091762.1, rna-XM_036091763.1    rna-XM_039223682.1    rna-XM_032868465.1, rna-XM_032868475.1    rna-XM_045989934.1, rna-XM_045989946.1, rna-XM_045989949.1      rna-XM_059159100.1    rna-XM_059388254.1, rna-XM_059388264.1    rna-XM_013060462.2    mNeoVis.2.1.1.chr5_hap1.g00391.t01, mNeoVis.2.1.1.chr5_hap1.g00407.t01    rna-XM_021683189.1    rna-XM_004414857.1    rna-XM_007088600.1    rna-XM_008684214.2, rna-XM_008684229.1, rna-XM_008711172.2, rna-XM_008711175.1, rna-XM_008711176.2, rna-XM_008712020.1, rna-XM_008712024.2      mNeoVis.3.1.1.chr2.g00337.t01    rna-gnl|WGS_ZZZZ|egapxtmp_035360-R1, rna-gnl|WGS_ZZZZ|egapxtmp_038065-R1    mNeoVis.evergladensis.2.1.chr2.g00405.t01    mNeoVis.vulgivaga.2.1.chr2.g00388.t01, mNeoVis.vulgivaga.2.1.chr2.g00406.t01      82    N0.HOG0000080   OG0000001  n731         rna-XM_003981996.2, rna-XM_011290016.4      rna-XM_039223671.1                      rna-XM_007088604.2, rna-XM_015542963.2                  83    N0.HOG0000081   OG0000001  n738   rna-XM_002931105.4, rna-XM_034653335.1    rna-XM_038567029.1    rna-XM_022498148.1            rna-XM_045991136.1                    rna-XM_008712018.2                84    N0.HOG0000082   OG0000001  n743   rna-XM_034651639.1, rna-XM_034657234.1    rna-NM_001388658.1    rna-XM_022498189.1          rna-XM_032868504.1    rna-XM_045991065.1    rna-XM_032360527.1    rna-XM_059165145.1    rna-XM_059387754.1    rna-XM_004761154.2    mNeoVis.2.1.1.chr5_hap1.g00411.t01              mNeoVis.3.1.1.chr2.g00342.t01            85    N0.HOG0000083   OG0000001  n755   rna-XM_034657188.1, rna-XM_034657927.1                rna-XM_045989938.1, rna-XM_045989940.1, rna-XM_045989943.1, rna-XM_045989945.1      rna-XM_059159098.1    rna-XM_059388262.1    rna-XM_004782385.2    mNeoVis.2.1.1.chr5_hap1.g00405.t01          rna-XM_008711165.2, rna-XM_008711167.2      mNeoVis.3.1.1.chr2.g00336.t01    rna-gnl|WGS_ZZZZ|egapxtmp_038064-R1    mNeoVis.evergladensis.2.1.chr2.g00404.t01    mNeoVis.vulgivaga.2.1.chr2.g00405.t01      86    N0.HOG0000084   OG0000001  n771   rna-XM_034650419.1, rna-XM_034657229.1              rna-XM_032868472.1      rna-XM_032303720.1            rna-XM_021683217.2        rna-XM_008684218.2, rna-XM_008684234.2            mNeoVis.vulgivaga.2.1.chr2.g00398.t01      87    N0.HOG0000085   OG0000001  n779               rna-XM_032868501.1      rna-XM_032360490.1    rna-XM_059165124.1    rna-XM_059387741.1    rna-XM_004781395.2    mNeoVis.2.1.1.chr5_hap1.g00393.t01              mNeoVis.3.1.1.chr2.g00325.t01    rna-gnl|WGS_ZZZZ|egapxtmp_034874-R1    mNeoVis.evergladensis.2.1.chr2.g00394.t01    mNeoVis.vulgivaga.2.1.chr2.g00390.t01      88    N0.HOG0000086   OG0000001  n786   rna-XM_034657227.1    rna-NM_001013422.1                                              89    N0.HOG0000087   OG0000001  n787   rna-XM_002930438.2    rna-XM_038565366.1                                rna-XM_008684217.1                90    N0.HOG0000088   OG0000002  n2   rna-XM_002930681.2    rna-NM_001389118.1, rna-NM_001389170.1, rna-XM_038549518.1    rna-XM_022492486.1    rna-XM_023257761.2      rna-XM_039224457.1, rna-XM_039225522.1    rna-XM_032850725.1    rna-XM_046011122.1, rna-XM_046012940.1    rna-XM_032347862.1, rna-XM_032347867.1    rna-XM_059183521.1, rna-XM_059183546.1    rna-XM_059404893.1, rna-XM_059404905.1    rna-XM_013050505.2, rna-XM_013050826.1, rna-XM_045073256.1    mNeoVis.2.1.1.chr9_hap1.g00428.t01          rna-XM_008711586.2, rna-XM_008711600.1    rna-XM_027594279.2    mNeoVis.3.1.1.chr10.g00412.t01    rna-gnl|WGS_ZZZZ|egapxtmp_022954-R1    mNeoVis.evergladensis.2.1.chr10.g00422.t01, mNeoVis.evergladensis.2.1.chr10.g00433.t01        91    N0.HOG0000089   OG0000002  n28   rna-XM_019793986.2, rna-XM_034643050.1, rna-XM_034646201.1    rna-NM_001389043.1, rna-NM_001389046.1    rna-XM_022492491.1, rna-XM_022492703.1, rna-XM_022519627.1, rna-XM_022519652.1    rna-XM_011284062.3    rna-XM_036082214.1, rna-XM_036114219.1      rna-XM_032850661.1, rna-XM_032850848.1, rna-XM_032850849.1, rna-XM_032851128.1    rna-XM_046013223.1, rna-XM_046013246.1, rna-XM_046013302.1    rna-XM_032345649.1, rna-XM_032345650.1, rna-XM_032347864.1    rna-XM_059183531.1, rna-XM_059183535.1, rna-XM_059183536.1, rna-XM_059183537.1, rna-XM_059183545.1    rna-XM_059404894.1, rna-XM_059404897.1, rna-XM_059404898.1, rna-XM_059404906.1, rna-XM_059404910.1    rna-XM_004781551.3, rna-XM_004782210.2, rna-XM_004782211.3, rna-XM_013050508.2, rna-XM_045072836.1, rna-XM_045073247.1, rna-XM_045073248.1, rna-XM_045073351.1, rna-XM_045073355.1    mNeoVis.2.1.1.chr9_hap1.g00426.t01, mNeoVis.2.1.1.chr9_hap1.g00436.t01, mNeoVis.2.1.1.chr9_hap1.g00437.t01, mNeoVis.2.1.1.chr9_hap1.g00438.t01    rna-XM_021686804.2    rna-XM_004416007.1, rna-XM_004416011.1    rna-XM_015537235.2    rna-XM_008710059.1, rna-XM_008711585.2, rna-XM_008711599.2    rna-XM_027593763.1, rna-XM_027593848.2    mNeoVis.3.1.1.chr10.g00410.t01, mNeoVis.3.1.1.chr10.g00411.t01, mNeoVis.3.1.1.chr10.g00416.t01, mNeoVis.3.1.1.chr10.g00417.t01    rna-gnl|WGS_ZZZZ|egapxtmp_022962-R1, rna-gnl|WGS_ZZZZ|egapxtmp_022963-R1, rna-gnl|WGS_ZZZZ|egapxtmp_023225-R1    mNeoVis.evergladensis.2.1.chr10.g00421.t01, mNeoVis.evergladensis.2.1.chr10.g00430.t01, mNeoVis.evergladensis.2.1.chr10.g00431.t01, mNeoVis.evergladensis.2.1.chr10.g00432.t01    mNeoVis.vulgivaga.2.1.chr10.g00414.t01, mNeoVis.vulgivaga.2.1.chr10.g00423.t01, mNeoVis.vulgivaga.2.1.chr10.g00424.t01      92    N0.HOG0000090   OG0000002  n89   rna-XM_002930991.2      rna-XM_022523060.1, rna-XM_022523091.1      rna-XM_036086543.1    rna-XM_039224443.1, rna-XM_039225156.1, rna-XM_039225616.1    rna-XM_032874350.1    rna-XM_046011123.1, rna-XM_046012361.1    rna-XM_032345668.1, rna-XM_032345671.1    rna-XM_059183512.1    rna-XM_059404915.1, rna-XM_059404916.1    rna-XM_004781539.2, rna-XM_004781548.1    mNeoVis.2.1.1.chr9_hap1.g00419.t01, mNeoVis.2.1.1.chr9_hap1.g00422.t01    rna-XM_021687218.2      rna-XM_015537238.2    rna-XM_008683987.2, rna-XM_008711942.2, rna-XM_040623740.1        rna-gnl|WGS_ZZZZ|egapxtmp_022967-R1    mNeoVis.evergladensis.2.1.chr10.g00413.t01, mNeoVis.evergladensis.2.1.chr10.g00417.t01    mNeoVis.vulgivaga.2.1.chr10.g00408.t01      93    N0.HOG0000091   OG0000002  n117   rna-XM_002930906.1      rna-XM_022523063.1    rna-XM_011284054.2      rna-XM_039225485.1      rna-XM_046012586.1                  rna-XM_007081450.2    rna-XM_008708335.2, rna-XM_008709394.2                94    N0.HOG0000092   OG0000002  n128   rna-XM_002930317.1, rna-XM_034646433.1, rna-XM_034646434.1    rna-NM_001388752.1    rna-XM_022492489.1    rna-XM_011284052.2    rna-XM_036082220.1    rna-XM_039216833.1, rna-XM_039217020.1, rna-XM_039217024.1, rna-XM_039225823.1    rna-XM_032876523.1    rna-XM_046012324.1, rna-XM_046013751.1, rna-XM_046013844.1    rna-XM_032345646.1, rna-XM_032345654.1, rna-XM_032347876.1    rna-XM_059183525.1, rna-XM_059183549.1, rna-XM_059187473.1    rna-XM_059403339.1, rna-XM_059404537.1, rna-XM_059404891.1    rna-XM_004773288.1, rna-XM_004782209.3, rna-XM_004782402.2    mNeoVis.2.1.1.chr9_hap1.g00430.t01, mNeoVis.2.1.1.chr9_hap1.g00434.t01    rna-XM_021686803.2      rna-XM_007081443.2, rna-XM_007081444.2, rna-XM_007081583.1    rna-XM_008684339.1, rna-XM_008711601.2      mNeoVis.3.1.1.chr10.g00400.t01    rna-gnl|WGS_ZZZZ|egapxtmp_022949-R1    mNeoVis.evergladensis.2.1.chr10.g00406.t01, mNeoVis.evergladensis.2.1.chr10.g00437.t01    mNeoVis.vulgivaga.2.1.chr10.g00427.t01      95    N0.HOG0000093   OG0000002  n168     rna-NM_001388819.1    rna-XM_022492628.1, rna-XM_022522966.1    rna-XM_003988848.2, rna-XM_003988904.4    rna-XM_036075356.1    rna-XM_039216834.1, rna-XM_039216888.1    rna-XM_032874325.1    rna-XM_046011841.1, rna-XM_046013579.1    rna-XM_032345680.1, rna-XM_032347875.1    rna-XM_059183502.1, rna-XM_059187474.1    rna-XM_059404535.1    rna-XM_004773031.1, rna-XM_004773287.1    mNeoVis.2.1.1.chr9_hap1.g00409.t01, mNeoVis.2.1.1.chr9_hap1.g00410.t01      rna-XM_004401764.1    rna-XM_007081452.2, rna-XM_007081582.2    rna-XM_008684338.1, rna-XM_008708339.2, rna-XM_040621919.1    rna-XM_027594286.1    mNeoVis.3.1.1.chr10.g00402.t01    rna-gnl|WGS_ZZZZ|egapxtmp_022977-R1    mNeoVis.evergladensis.2.1.chr10.g00407.t01, mNeoVis.evergladensis.2.1.chr10.g00408.t01        96    N0.HOG0000094   OG0000002  n198   rna-XM_002930821.3            rna-XM_039216890.1                      rna-XM_015537242.1    rna-XM_008708338.1    rna-XM_027593891.1              97    N0.HOG0000095   OG0000002  n204         rna-XM_003988849.2, rna-XM_011284046.2      rna-XM_039216893.1, rna-XM_039225435.1                      rna-XM_007081453.2, rna-XM_015537241.2                  98    N0.HOG0000096   OG0000002  n210   rna-XM_002930314.1, rna-XM_034644306.1    rna-NM_001388521.1    rna-XM_022523112.1          rna-XM_032874360.1    rna-XM_046013589.1    rna-XM_032345681.1    rna-XM_059183501.1    rna-XM_059404536.1    rna-XM_004773032.1            rna-XM_008711950.1      mNeoVis.3.1.1.chr10.g00401.t01        mNeoVis.vulgivaga.2.1.chr10.g00402.t01      99    N0.HOG0000097   OG0000002  n221     rna-NM_001313886.1    rna-XM_022523090.1          rna-XM_032874357.1    rna-XM_046013696.1    rna-XM_032345679.1    rna-XM_059183504.1    rna-XM_059404920.1    rna-XM_004773030.1    mNeoVis.2.1.1.chr9_hap1.g00413.t01                rna-gnl|WGS_ZZZZ|egapxtmp_022971-R1    mNeoVis.evergladensis.2.1.chr10.g00411.t01    mNeoVis.vulgivaga.2.1.chr10.g00403.t01      100    N0.HOG0000098   OG0000002  n232           rna-XM_036075353.1, rna-XM_036086540.1, rna-XM_036122305.1                    rna-XM_021687231.2    rna-XM_004401622.1, rna-XM_004401623.1        rna-XM_027593840.1              101    N0.HOG0000099   OG0000002  n243   rna-XM_034646508.1, rna-XM_034650555.1    rna-NM_001389152.1, rna-XM_038588977.1    rna-XM_022492659.1    rna-XM_003988900.1      rna-XM_039225159.1      rna-XM_046012319.1    rna-XM_032345670.1          mNeoVis.2.1.1.chr9_hap1.g00420.t01        rna-XM_007081578.1    rna-XM_008683986.2, rna-XM_008711937.1, rna-XM_008711938.2        rna-gnl|WGS_ZZZZ|egapxtmp_022964-R1    mNeoVis.evergladensis.2.1.chr10.g00414.t01    mNeoVis.vulgivaga.2.1.chr10.g00409.t01      102    N0.HOG0000100   OG0000002  n258   rna-XM_002930685.3, rna-XM_034653232.1    rna-XM_038533680.1, rna-XM_038551554.1, rna-XM_038588938.1, rna-XM_038588976.1, rna-XM_038589084.1    rna-XM_022522967.1      rna-XM_036086538.1      rna-XM_032874324.1    rna-XM_046011834.1, rna-XM_046011837.1, rna-XM_046012358.1    rna-XM_032347871.1    rna-XM_059187475.1    rna-XM_059404914.1    rna-XM_004781547.1    mNeoVis.2.1.1.chr9_hap1.g00421.t01    rna-XM_021686816.1        rna-XM_008683988.2, rna-XM_008690848.2      mNeoVis.3.1.1.chr10.g00408.t01    rna-gnl|WGS_ZZZZ|egapxtmp_022950-R1    mNeoVis.evergladensis.2.1.chr10.g00415.t01    mNeoVis.vulgivaga.2.1.chr10.g00410.t01      103    N0.HOG0000101   OG0000002  n282   rna-XM_002930823.3    rna-NM_001389175.1    rna-XM_022523049.1    rna-XM_003988846.1      rna-XM_039225431.1    rna-XM_032874356.1    rna-XM_046012903.1    rna-XM_032345677.1    rna-XM_059183505.1              rna-XM_007081451.2    rna-XM_008708336.1, rna-XM_008709391.2                104    N0.HOG0000102   OG0000002  n294     rna-NM_001388522.1, rna-NM_001388947.1          rna-XM_039225433.1                        rna-XM_008709389.1                105    N0.HOG0000103   OG0000002  n297   rna-XM_011218265.2    rna-NM_001388949.1, rna-NM_001389076.1, rna-XM_038589061.1          rna-XM_039225434.1                                      106    N0.HOG0000104   OG0000002  n303   rna-XM_019793758.2    rna-NM_001388942.1, rna-NM_001389149.1, rna-XM_038588975.1    rna-XM_022492492.1, rna-XM_022523094.1    rna-XM_011284056.2      rna-XM_039224456.1, rna-XM_039225162.1, rna-XM_039225615.1    rna-XM_032874355.1    rna-XM_046013034.1, rna-XM_046013051.1, rna-XM_046013057.1    rna-XM_032345672.1, rna-XM_032345676.1, rna-XM_032347865.1    rna-XM_059183506.1, rna-XM_059183511.1, rna-XM_059187481.1    rna-XM_059404526.1, rna-XM_059404918.1, rna-XM_059404919.1    rna-XM_004781546.2, rna-XM_004782372.1, rna-XM_045073220.1    mNeoVis.2.1.1.chr9_hap1.g00414.t01, mNeoVis.2.1.1.chr9_hap1.g00432.t01        rna-XM_007081448.1    rna-XM_008683979.2, rna-XM_008710513.2, rna-XM_008711939.2, rna-XM_008711940.2      mNeoVis.3.1.1.chr10.g00405.t01    rna-gnl|WGS_ZZZZ|egapxtmp_022947-R1    mNeoVis.evergladensis.2.1.chr10.g00412.t01, mNeoVis.evergladensis.2.1.chr10.g00426.t01    mNeoVis.vulgivaga.2.1.chr10.g00406.t01, mNeoVis.vulgivaga.2.1.chr10.g00407.t01      107    N0.HOG0000105   OG0000002  n339   rna-XM_002930822.3, rna-XM_002931311.4, rna-XM_011217814.2, rna-XM_034643067.1, rna-XM_034644642.1, rna-XM_034652813.1    rna-NM_001388940.1, rna-NM_001388983.1    rna-XM_022523113.1, rna-XM_022523114.1    rna-XM_003988902.3, rna-XM_006933821.1, rna-XM_011284047.1, rna-XM_011284055.1    rna-XM_036075352.1, rna-XM_036086542.1, rna-XM_036086545.1, rna-XM_036122304.1    rna-XM_039216835.1, rna-XM_039225155.1, rna-XM_039225157.1, rna-XM_039225158.1, rna-XM_039225160.1, rna-XM_039225161.1, rna-XM_039225487.1, rna-XM_039225488.1    rna-XM_032874351.1, rna-XM_032874353.1, rna-XM_032874354.1, rna-XM_032874358.1, rna-XM_032874359.1    rna-XM_046011131.1, rna-XM_046011839.1, rna-XM_046011840.1, rna-XM_046012320.1, rna-XM_046012321.1, rna-XM_046013020.1, rna-XM_046013021.1, rna-XM_046013054.1, rna-XM_046013056.1, rna-XM_046013059.1, rna-XM_046013060.1, rna-XM_046013187.1, rna-XM_046013337.1    rna-XM_032345674.1, rna-XM_032345675.1, rna-XM_032347872.1, rna-XM_032347873.1    rna-XM_059183503.1, rna-XM_059183508.1, rna-XM_059183509.1, rna-XM_059183510.1, rna-XM_059183542.1, rna-XM_059183544.1    rna-XM_059387550.1, rna-XM_059404532.1, rna-XM_059404533.1, rna-XM_059404667.1, rna-XM_059404895.1, rna-XM_059404896.1    rna-XM_004748591.2, rna-XM_004773285.1, rna-XM_004773286.1, rna-XM_004782485.2, rna-XM_013050864.1, rna-XM_045071792.1, rna-XM_045071793.1, rna-XM_045073268.1    mNeoVis.2.1.1.chr9_hap1.g00411.t01, mNeoVis.2.1.1.chr9_hap1.g00415.t01, mNeoVis.2.1.1.chr9_hap1.g00416.t01, mNeoVis.2.1.1.chr9_hap1.g00431.t01    rna-XM_021687219.1, rna-XM_021687230.1    rna-XM_004401621.1, rna-XM_004401763.1    rna-XM_007081449.1, rna-XM_007081579.3, rna-XM_007081581.2, rna-XM_015537240.2    rna-XM_008683976.2, rna-XM_008683977.2, rna-XM_008683978.2, rna-XM_008683985.2, rna-XM_008708337.2, rna-XM_008708340.2, rna-XM_008709961.2, rna-XM_008709999.2, rna-XM_040626577.1, rna-XM_040626578.1    rna-XM_027593858.1, rna-XM_027594029.1    mNeoVis.3.1.1.chr10.g00403.t01, mNeoVis.3.1.1.chr10.g00414.t01    rna-gnl|WGS_ZZZZ|egapxtmp_024655-R1, rna-gnl|WGS_ZZZZ|egapxtmp_024657-R1    mNeoVis.evergladensis.2.1.chr10.g00409.t01, mNeoVis.evergladensis.2.1.chr10.g00424.t01, mNeoVis.evergladensis.2.1.chr10.g00425.t01    mNeoVis.vulgivaga.2.1.chr10.g00404.t01, mNeoVis.vulgivaga.2.1.chr10.g00418.t01, mNeoVis.vulgivaga.2.1.chr10.g00419.t01      108    N0.HOG0000106   OG0000002  n440                           mNeoVis.2.1.1.chr9_hap1.g00439.t01                rna-gnl|WGS_ZZZZ|egapxtmp_024658-R1          109    N0.HOG0000107   OG0000002  n442   rna-XM_011218059.3    rna-NM_001256457.1    rna-XM_022492494.1, rna-XM_022519577.1    rna-XM_003988841.3      rna-XM_039225519.1, rna-XM_039225523.1    rna-XM_032850660.1, rna-XM_032850723.1    rna-XM_046011127.1, rna-XM_046013668.1, rna-XM_046013803.1    rna-XM_032345648.1, rna-XM_032345657.1    rna-XM_059183522.1, rna-XM_059183534.1    rna-XM_059404892.1, rna-XM_059404904.1    rna-XM_004781543.1, rna-XM_004782208.1          rna-XM_007081445.1, rna-XM_042990848.1    rna-XM_008711587.1    rna-XM_035729236.1, rna-XM_035729256.1    mNeoVis.3.1.1.chr10.g00418.t01    rna-gnl|WGS_ZZZZ|egapxtmp_022960-R1    mNeoVis.evergladensis.2.1.chr10.g00423.t01, mNeoVis.evergladensis.2.1.chr10.g00434.t01    mNeoVis.vulgivaga.2.1.chr10.g00416.t01, mNeoVis.vulgivaga.2.1.chr10.g00425.t01      110    N0.HOG0000108   OG0000002  n471     rna-XM_038540995.1    rna-XM_022492488.1, rna-XM_022519639.1            rna-XM_046012325.1, rna-XM_046012858.1, rna-XM_046012987.1    rna-XM_032345656.1    rna-XM_059183523.1, rna-XM_059183547.1    rna-XM_059404903.1    rna-XM_004781544.1, rna-XM_004782427.1    mNeoVis.2.1.1.chr9_hap1.g00429.t01                    mNeoVis.vulgivaga.2.1.chr10.g00417.t01      111    N0.HOG0000109   OG0000002  n484   rna-XM_034646480.1    rna-NM_001256498.1, rna-XM_038588953.1, rna-XM_038588955.1    rna-XM_022519673.1        rna-XM_039225520.1    rna-XM_032876504.1    rna-XM_046013006.1    rna-XM_032345647.1    rna-XM_059183532.1, rna-XM_059183548.1    rna-XM_059403337.1    rna-XM_045073194.1, rna-XM_045073221.1    mNeoVis.2.1.1.chr9_hap1.g00441.t01          rna-XM_008711590.1      mNeoVis.3.1.1.chr10.g00420.t01      mNeoVis.evergladensis.2.1.chr10.g00436.t01    mNeoVis.vulgivaga.2.1.chr10.g00426.t01      112    N0.HOG0000110   OG0000002  n501   rna-XM_002931108.3    rna-NM_001313836.1    rna-XM_022492487.1    rna-XM_011284064.3      rna-XM_039225518.1    rna-XM_032850722.1    rna-XM_046011130.1    rna-XM_032345655.1    rna-XM_059183524.1      rna-XM_004781545.1    mNeoVis.2.1.1.chr9_hap1.g00427.t01, mNeoVis.2.1.1.chr9_hap1.g00440.t01        rna-XM_007081569.2    rna-XM_008711589.1      mNeoVis.3.1.1.chr10.g00419.t01    rna-gnl|WGS_ZZZZ|egapxtmp_022974-R1    mNeoVis.evergladensis.2.1.chr10.g00435.t01        113    N0.HOG0000111   OG0000002  n519   rna-XM_011217111.2    rna-NM_001388754.1, rna-NM_001389060.1, rna-NM_001389064.1      rna-XM_003988829.2      rna-XM_039216999.1    rna-XM_032876529.1      rna-XM_032345645.1    rna-XM_059183554.1    rna-XM_059404886.1      mNeoVis.2.1.1.chr9_hap1.g00447.t01        rna-XM_007081566.2    rna-XM_008711594.2        rna-gnl|WGS_ZZZZ|egapxtmp_022965-R1    mNeoVis.evergladensis.2.1.chr10.g00443.t01    mNeoVis.vulgivaga.2.1.chr10.g00432.t01      114    N0.HOG0000112   OG0000002  n533     rna-XM_038549521.1    rna-XM_022519599.1    rna-XM_011284048.1    rna-XM_036099946.1    rna-XM_039224446.1    rna-XM_032874349.1    rna-XM_046011124.1, rna-XM_046012588.1    rna-XM_032345667.1    rna-XM_059183514.1    rna-XM_059404913.1    rna-XM_004782497.1      rna-XM_021686813.1    rna-XM_004416002.1    rna-XM_015537237.1    rna-XM_008690850.2    rna-XM_027593776.1        mNeoVis.evergladensis.2.1.chr10.g00418.t01    mNeoVis.vulgivaga.2.1.chr13.g00631.t01      115    N0.HOG0000113   OG0000002  n553   rna-XM_002930683.2, rna-XM_034653006.1    rna-XM_038576194.1    rna-XM_022492677.1, rna-XM_022519393.1    rna-XM_023257123.2    rna-XM_036095969.1, rna-XM_036095971.1, rna-XM_036099953.1    rna-XM_039224454.1    rna-XM_032850662.1    rna-XM_046013237.1, rna-XM_046013590.1    rna-XM_032345652.1, rna-XM_032345658.1    rna-XM_059183528.1, rna-XM_059183540.1    rna-XM_059404901.1, rna-XM_059404907.1    rna-XM_004782568.1      rna-XM_021686805.2, rna-XM_021686807.2    rna-XM_004416004.1, rna-XM_004416006.1    rna-XM_015537236.2    rna-XM_008683984.2, rna-XM_008710058.2, rna-XM_008710120.2    rna-XM_035729251.1      rna-gnl|WGS_ZZZZ|egapxtmp_022975-R1    mNeoVis.evergladensis.2.1.chr10.g00427.t01    mNeoVis.vulgivaga.2.1.chr10.g00421.t01      116    N0.HOG0000114   OG0000002  n586   rna-XM_002930680.3    rna-XM_038549519.1    rna-XM_022492553.1    rna-XM_011284057.3, rna-XM_011284059.3, rna-XM_023257122.2    rna-XM_036099947.1, rna-XM_036099950.1    rna-XM_039224447.1, rna-XM_039224449.1    rna-XM_032849960.1, rna-XM_032850680.1, rna-XM_032874345.1, rna-XM_032874348.1    rna-XM_046011125.1    rna-XM_032345662.1, rna-XM_032345666.1    rna-XM_059183515.1, rna-XM_059183517.1    rna-XM_059404909.1, rna-XM_059404912.1    rna-XM_004781541.2, rna-XM_004781550.2    mNeoVis.2.1.1.chr9_hap1.g00425.t01    rna-XM_021686812.1    rna-XM_004416003.1    rna-XM_007081573.2, rna-XM_007081575.3, rna-XM_042990851.1    rna-XM_008683982.2, rna-XM_040627240.1    rna-XM_035729266.1, rna-XM_035729268.1      rna-gnl|WGS_ZZZZ|egapxtmp_022957-R1          117    N0.HOG0000115   OG0000002  n618   rna-XM_002930678.2    rna-NM_001388939.1, rna-XM_038550062.1, rna-XM_038589059.1    rna-XM_022519580.1    rna-XM_006933820.2, rna-XM_011284049.1    rna-XM_036099949.1, rna-XM_036099952.1    rna-XM_039224439.1, rna-XM_039224441.1    rna-XM_032850679.1, rna-XM_032874346.1    rna-XM_046012680.1, rna-XM_046013170.1    rna-XM_032345664.1, rna-XM_032348507.1    rna-XM_059183516.1    rna-XM_059403340.1, rna-XM_059404911.1    rna-XM_004781540.2, rna-XM_045073266.1    mNeoVis.2.1.1.chr9_hap1.g00433.t01    rna-XM_021686809.1      rna-XM_007081447.2, rna-XM_007081571.2, rna-XM_042993697.1      rna-XM_027593827.1    mNeoVis.3.1.1.chr10.g00406.t01        mNeoVis.vulgivaga.2.1.chr10.g00412.t01      118    N0.HOG0000116   OG0000002  n646   rna-XM_011217535.3, rna-XM_034646220.1    rna-NM_001256450.1, rna-NM_001389143.1, rna-XM_038588939.1, rna-XM_038589060.1, rna-XM_038589577.1, rna-XM_038589695.1    rna-XM_022492566.1, rna-XM_022492620.1, rna-XM_022519563.1, rna-XM_022519578.1    rna-XM_011284058.2, rna-XM_023257121.2, rna-XM_023257125.2, rna-XM_023257802.1    rna-XM_036095970.1, rna-XM_036099948.1, rna-XM_036099951.1    rna-XM_039224440.1, rna-XM_039224450.1, rna-XM_039224452.1, rna-XM_039224453.1    rna-XM_032849961.1, rna-XM_032874343.1, rna-XM_032874347.1    rna-XM_046011126.1, rna-XM_046012881.1, rna-XM_046012887.1, rna-XM_046012900.1, rna-XM_046013131.1    rna-XM_032345651.1, rna-XM_032345659.1, rna-XM_032345660.1, rna-XM_032345661.1, rna-XM_032345665.1    rna-XM_059183518.1, rna-XM_059183519.1, rna-XM_059183527.1, rna-XM_059183529.1, rna-XM_059183538.1, rna-XM_059183541.1, rna-XM_059185068.1, rna-XM_059187478.1    rna-XM_059387591.1, rna-XM_059404528.1, rna-XM_059404900.1, rna-XM_059404902.1    rna-XM_045071797.1, rna-XM_045073156.1, rna-XM_045073163.1    mNeoVis.2.1.1.chr9_hap1.g00423.t01, mNeoVis.2.1.1.chr9_hap1.g00424.t01, mNeoVis.2.1.1.chr9_hap1.g00435.t01    rna-XM_021686806.2, rna-XM_021686811.2    rna-XM_004416005.1    rna-XM_007081446.1, rna-XM_007081570.2, rna-XM_007081572.3, rna-XM_007081574.1, rna-XM_042990852.1    rna-XM_008683983.1, rna-XM_040622772.1, rna-XM_040625417.1, rna-XM_040641937.1    rna-XM_027593719.1, rna-XM_035729311.1, rna-XM_035729312.1    mNeoVis.3.1.1.chr10.g00409.t01, mNeoVis.3.1.1.chr10.g00415.t01    rna-gnl|WGS_ZZZZ|egapxtmp_022956-R1, rna-gnl|WGS_ZZZZ|egapxtmp_022958-R1, rna-gnl|WGS_ZZZZ|egapxtmp_022961-R2, rna-gnl|WGS_ZZZZ|egapxtmp_023224-R1, rna-gnl|WGS_ZZZZ|egapxtmp_023229-R1    mNeoVis.evergladensis.2.1.chr10.g00419.t01, mNeoVis.evergladensis.2.1.chr10.g00428.t01, mNeoVis.evergladensis.2.1.chr10.g00429.t01    mNeoVis.vulgivaga.2.1.chr10.g00413.t01, mNeoVis.vulgivaga.2.1.chr10.g00415.t01, mNeoVis.vulgivaga.2.1.chr10.g00420.t01, mNeoVis.vulgivaga.2.1.chr10.g00422.t01      119    N0.HOG0000117   OG0000002  n724   rna-XM_002930684.4    rna-XM_038588892.1, rna-XM_038588973.1, rna-XM_038589062.1, rna-XM_038589083.1    rna-XM_022519636.1, rna-XM_022523092.1    rna-XM_003988844.1    rna-XM_036086544.1, rna-XM_036099945.1    rna-XM_039224444.1, rna-XM_039224445.1, rna-XM_039224451.1    rna-XM_032849963.1, rna-XM_032849964.1    rna-XM_046011129.1, rna-XM_046012318.1, rna-XM_046012359.1, rna-XM_046012618.1, rna-XM_046012779.1, rna-XM_046012918.1    rna-XM_032345716.1, rna-XM_032347869.1, rna-XM_032347870.1    rna-XM_059187477.1    rna-XM_059404530.1    rna-XM_004781549.1    mNeoVis.2.1.1.chr9_hap1.g00417.t01, mNeoVis.2.1.1.chr9_hap1.g00418.t01    rna-XM_021686814.1, rna-XM_021686815.1      rna-XM_007081577.2    rna-XM_008683980.1, rna-XM_008690806.2, rna-XM_008690849.2, rna-XM_008710510.2, rna-XM_040623741.1      mNeoVis.3.1.1.chr10.g00407.t01    rna-gnl|WGS_ZZZZ|egapxtmp_022953-R1    mNeoVis.evergladensis.2.1.chr10.g00416.t01    mNeoVis.vulgivaga.2.1.chr10.g00411.t01      120    N0.HOG0000118   OG0000003  n1   rna-XM_019793867.2    rna-XM_038527358.1    rna-XM_022512790.1    rna-XM_045043859.1    rna-XM_036113703.1    rna-XM_039221445.1    rna-XM_032866714.1    rna-XM_046023306.1    rna-XM_032307327.1    rna-XM_059141176.1    rna-XM_059412064.1    rna-XM_013063353.2    mNeoVis.2.1.1.chr13_hap1.g00813.t01    rna-XM_021694350.1    rna-XM_004393589.1    rna-XM_042963348.1    rna-XM_008690880.2    rna-XM_027615011.2    mNeoVis.3.1.1.chr13.g00736.t01, mNeoVis.3.1.1.chr13.g00737.t01    rna-gnl|WGS_ZZZZ|egapxtmp_030022-R1    mNeoVis.evergladensis.2.1.chr13.g00778.t01    mNeoVis.vulgivaga.2.1.chr13.g00776.t01      121    N0.HOG0000119   OG0000003  n24   rna-XM_034638435.1    rna-XM_038529045.1      rna-XM_045045523.1    rna-XM_036120108.1    rna-XM_039217742.1    rna-XM_032842083.1    rna-XM_045988727.1    rna-XM_032325599.1    rna-XM_059153775.1    rna-XM_059382895.1    rna-XM_004776263.3    mNeoVis.2.1.1.chr7_hap1.g00489.t01      rna-XM_004396501.1    rna-XM_015544101.2    rna-XM_040634410.1    rna-XM_035724896.1    mNeoVis.3.1.1.chr7.g00471.t01    rna-gnl|WGS_ZZZZ|egapxtmp_012057-R1    mNeoVis.evergladensis.2.1.chr7.g00492.t01    mNeoVis.vulgivaga.2.1.chr7.g00496.t01      122    N0.HOG0000120   OG0000003  n43   rna-XM_034639552.1              rna-XM_032850910.1, rna-XM_032850914.1                                    123    N0.HOG0000121   OG0000003  n48   rna-XM_034642829.1    rna-XM_038575929.1        rna-XM_036104273.1    rna-XM_039242936.1                  rna-XM_021689484.1    rna-XM_004415473.1      rna-XM_040619569.1                124    N0.HOG0000122   OG0000003  n54   rna-XM_034662879.1    rna-XM_038578108.1    rna-XM_022499636.1    rna-XM_045040364.1    rna-XM_036087057.1    rna-XM_039257745.1    rna-XM_032877554.1    rna-XM_046027370.1    rna-XM_032312203.1    rna-XM_059169570.1    rna-XM_059397698.1        rna-XM_021700240.2    rna-XM_004410239.1    rna-XM_042959939.1    rna-XM_008696204.2    rna-XM_027589577.2      rna-gnl|WGS_ZZZZ|egapxtmp_008186-R1      mNeoVis.vulgivaga.2.1.chr5.g00355.t01      125    N0.HOG0000123   OG0000003  n74   rna-XM_011237512.3    rna-XM_038573925.1    rna-XM_022498963.1    rna-XM_045055144.1    rna-XM_036117491.1    rna-XM_039225127.1    rna-XM_032869798.1    rna-XM_045997917.1    rna-XM_032329362.1    rna-XM_059155922.1    rna-XM_059384444.1    rna-XM_045086214.1    mNeoVis.2.1.1.chr6_hap1.g01129.t01    rna-XM_044912794.1    rna-XM_004416619.1    rna-XM_015543529.2    rna-XM_008708856.2    rna-XM_027597693.2        mNeoVis.evergladensis.2.1.chr4.g01061.t01    mNeoVis.vulgivaga.2.1.chr4.g01096.t01      126    N0.HOG0000124   OG0000003  n93   rna-XM_034647542.1    rna-XM_038560520.1    rna-XM_022499007.1    rna-XM_006930542.4    rna-XM_036099816.1    rna-XM_039217208.1    rna-XM_032869823.1    rna-XM_045997042.1    rna-XM_032328963.1    rna-XM_059155939.1    rna-XM_059384137.1    rna-XM_004775553.3    mNeoVis.2.1.1.chr6_hap1.g01102.t01    rna-XM_021694143.2      rna-XM_042982885.1    rna-XM_008708866.2    rna-XM_027598588.2, rna-XM_035726369.1        mNeoVis.evergladensis.2.1.chr4.g01037.t01    mNeoVis.vulgivaga.2.1.chr4.g01068.t01, mNeoVis.vulgivaga.2.1.chr4.g01069.t01      127    N0.HOG0000125   OG0000003  n112   rna-XM_019810160.2    rna-XM_038573935.1    rna-XM_022492804.1, rna-XM_022498818.1    rna-XM_019828173.3, rna-XM_023252259.2, rna-XM_045055135.1    rna-XM_036117485.1    rna-XM_039225129.1, rna-XM_039225132.1    rna-XM_032869797.1    rna-XM_045988719.1    rna-XM_032329354.1    rna-XM_059155914.1    rna-XM_059384019.1    rna-XM_045086207.1    mNeoVis.2.1.1.chr6_hap1.g01128.t01      rna-XM_004416618.1    rna-XM_042982784.1, rna-XM_042982798.1, rna-XM_042982805.1    rna-XM_040643346.1    rna-XM_035726131.1      rna-gnl|WGS_ZZZZ|egapxtmp_043180-R2    mNeoVis.evergladensis.2.1.chr4.g01060.t01    mNeoVis.vulgivaga.2.1.chr4.g01094.t01      128    N0.HOG0000126   OG0000003  n135     rna-XM_038560523.1      rna-XM_045042095.1, rna-XM_045055183.1    rna-XM_036081947.1    rna-XM_039217211.1    rna-XM_032869825.1        rna-XM_059155943.1      rna-XM_004775549.3    mNeoVis.2.1.1.chr6_hap1.g01103.t01      rna-XM_004405728.1    rna-XM_042982879.1    rna-XM_040643181.1    rna-XM_027598590.2    mNeoVis.3.1.1.chr4.g01053.t01    rna-gnl|WGS_ZZZZ|egapxtmp_043179-R2    mNeoVis.evergladensis.2.1.chr4.g01039.t01    mNeoVis.vulgivaga.2.1.chr4.g01071.t01      129    N0.HOG0000127   OG0000003  n155   rna-XM_034669774.1    rna-XM_038539499.1    rna-XM_022501647.1    rna-XM_011290426.4    rna-XM_036103019.1    rna-XM_039226053.1    rna-XM_032872735.1    rna-XM_045992864.1    rna-XM_032326977.1    rna-XM_059155643.1    rna-XM_059414362.1    rna-XM_045075142.1    mNeoVis.2.1.1.chr15_hap1.g00608.t01    rna-XM_044915734.1    rna-XM_004394209.1    rna-XM_042972509.1    rna-XM_040634744.1    rna-XM_027612952.2    mNeoVis.3.1.1.chr15.g00392.t01    rna-gnl|WGS_ZZZZ|egapxtmp_033299-R1    mNeoVis.evergladensis.2.1.chr15.g00430.t01    mNeoVis.vulgivaga.2.1.chr15.g00438.t01      130    N0.HOG0000128   OG0000003  n175   rna-XM_034669769.1    rna-XM_038539540.1, rna-XM_038589314.1, rna-XM_038589621.1    rna-XM_022501645.1    rna-XM_006941901.5    rna-XM_036103016.1    rna-XM_039226054.1    rna-XM_032872734.1    rna-XM_045992859.1    rna-XM_032326975.1    rna-XM_059155642.1    rna-XM_059414364.1    rna-XM_045075141.1    mNeoVis.2.1.1.chr15_hap1.g00606.t01    rna-XM_044915287.1      rna-XM_007081115.3    rna-XM_008698320.2    rna-XM_027612959.2    mNeoVis.3.1.1.chr15.g00390.t01    rna-gnl|WGS_ZZZZ|egapxtmp_033298-R1, rna-gnl|WGS_ZZZZ|egapxtmp_033656-R1    mNeoVis.evergladensis.2.1.chr15.g00428.t01    mNeoVis.vulgivaga.2.1.chr15.g00436.t01      131    N0.HOG0000129   OG0000003  n198               rna-XM_032849243.1                                mNeoVis.vulgivaga.2.1.chr3.g00708.t01      132    N0.HOG0000130   OG0000003  n202   rna-XM_011218637.3    rna-XM_038533783.1    rna-XM_022499128.1    rna-XM_023240338.2    rna-XM_036079377.1    rna-XM_039222022.1    rna-XM_032883117.1      rna-XM_032313634.1, rna-XM_032313640.1, rna-XM_032313968.1, rna-XM_032313976.1    rna-XM_059172464.1, rna-XM_059172521.1      rna-XM_045073829.1, rna-XM_045087810.1, rna-XM_045087820.1, rna-XM_045087935.1, rna-XM_045087938.1, rna-XM_045087939.1    mNeoVis.2.1.1.chr2_hap1.g00611.t01, mNeoVis.2.1.1.chr2_hap1.g00710.t01, mNeoVis.2.1.1.chr2_hap1.g00715.t01    rna-XM_044916004.1      rna-XM_042959813.1    rna-XM_040624748.1    rna-XM_027594092.2    mNeoVis.3.1.1.chr5.g00609.t01        mNeoVis.vulgivaga.2.1.chr5.g00610.t01      133    N0.HOG0000131   OG0000003  n229     rna-XM_038533811.1    rna-XM_022499537.1      rna-XM_036066830.1    rna-XM_039219767.1    rna-XM_032883070.1    rna-XM_046027476.1    rna-XM_032313498.1    rna-XM_059133538.1, rna-XM_059172518.1    rna-XM_059387529.1    rna-XM_045087367.1    mNeoVis.2.1.1.chr2_hap1.g00615.t01    rna-XM_044916457.1    rna-XM_004412999.1      rna-XM_008689632.2                134    N0.HOG0000132   OG0000003  n243   rna-XM_034663565.1    rna-XM_038533788.1    rna-XM_022499536.1    rna-XM_045040950.1, rna-XM_045040955.1, rna-XM_045041015.1    rna-XM_036066812.1    rna-XM_039219768.1, rna-XM_039222024.1    rna-XM_032883065.1    rna-XM_046027145.1    rna-XM_032313321.1, rna-XM_032313762.1    rna-XM_059170502.1    rna-XM_059387327.1    rna-XM_045087364.1    mNeoVis.2.1.1.chr2_hap1.g00614.t01    rna-XM_044916456.1      rna-XM_042959780.1, rna-XM_042959783.1, rna-XM_042959801.1    rna-XM_008689631.2    rna-XM_035724294.1    mNeoVis.3.1.1.chr5.g00612.t01    rna-gnl|WGS_ZZZZ|egapxtmp_006903-R1    mNeoVis.evergladensis.2.1.chr5.g00617.t01, mNeoVis.evergladensis.2.1.chr5.g00618.t01    mNeoVis.vulgivaga.2.1.chr5.g00613.t01      135    N0.HOG0000133   OG0000003  n270   rna-XM_011236575.3    rna-XM_038579214.1    rna-XM_022499827.1    rna-XM_045040916.1, rna-XM_045040924.1    rna-XM_036110350.1    rna-XM_039225515.1, rna-XM_039240815.1    rna-XM_032882185.1    rna-XM_046026619.1    rna-XM_032312652.1    rna-XM_059172713.1, rna-XM_059172715.1    rna-XM_059397826.1, rna-XM_059398921.1    rna-XM_013050902.2, rna-XM_045068587.1    mNeoVis.2.1.1.chr2_hap1.g00003.t01    rna-XM_021697280.2    rna-XM_004394982.1    rna-XM_042960769.1    rna-XM_040624620.1    rna-XM_035724352.1, rna-XM_035724353.1    mNeoVis.3.1.1.chr5.g00002.t01    rna-gnl|WGS_ZZZZ|egapxtmp_008140-R1          136    N0.HOG0000134   OG0000003  n295   rna-XM_019794011.2, rna-XM_034647039.1          rna-XM_036084129.1    rna-XM_039230857.1                    rna-XM_004408004.1        rna-XM_035723483.1              137    N0.HOG0000135   OG0000003  n301   rna-XM_019798119.2    rna-XM_038587286.1    rna-XM_022507798.1    rna-XM_045050944.1    rna-XM_036089384.1    rna-XM_039216787.1    rna-XM_032842980.1    rna-XM_045995709.1    rna-XM_032330463.1    rna-XM_059158067.1    rna-XM_059384784.1    rna-XM_013057582.2      rna-XM_044911616.1    rna-XM_004396758.1    rna-XM_007081604.3    rna-XM_040632381.1    rna-XM_027607375.1    mNeoVis.3.1.1.chr12.g00583.t01    rna-gnl|WGS_ZZZZ|egapxtmp_028970-R1, rna-gnl|WGS_ZZZZ|egapxtmp_029082-R1      mNeoVis.vulgivaga.2.1.chr12.g00593.t01      138    N0.HOG0000136   OG0000003  n320   rna-XM_034669525.1    rna-XM_038539498.1    rna-XM_022501644.1    rna-XM_045048242.1    rna-XM_036103015.1    rna-XM_039226162.1    rna-XM_032872315.1    rna-XM_045992853.1      rna-XM_059155640.1      rna-XM_013047083.2    mNeoVis.2.1.1.chr15_hap1.g00609.t01    rna-XM_044915286.1    rna-XM_004394210.1    rna-XM_042972498.1    rna-XM_008698324.2    rna-XM_027612949.2    mNeoVis.3.1.1.chr15.g00393.t01    rna-gnl|WGS_ZZZZ|egapxtmp_033403-R1    mNeoVis.evergladensis.2.1.chr15.g00431.t01        139    N0.HOG0000137   OG0000003  n338   rna-XM_034641916.1                            rna-XM_044919047.1        rna-XM_040638419.1                140    N0.HOG0000138   OG0000003  n342   rna-XM_019793323.2    rna-XM_038529201.1        rna-XM_036068010.1    rna-XM_039249729.1    rna-XM_032842146.1    rna-XM_045988808.1    rna-XM_032325684.1          mNeoVis.2.1.1.chr7_hap1.g00479.t01    rna-XM_021700802.1      rna-XM_042969053.1    rna-XM_008711153.2    rna-XM_027617879.2    mNeoVis.3.1.1.chr7.g00464.t01    rna-gnl|WGS_ZZZZ|egapxtmp_012230-R1    mNeoVis.evergladensis.2.1.chr7.g00484.t01    mNeoVis.vulgivaga.2.1.chr7.g00486.t01      141    N0.HOG0000139   OG0000003  n355   rna-XM_034669779.1    rna-XM_038539532.1    rna-XM_022501651.1    rna-XM_023246240.2    rna-XM_036103020.1    rna-XM_039226117.1    rna-XM_032872738.1    rna-XM_045992870.1    rna-XM_032326979.1    rna-XM_059155645.1    rna-XM_059414365.1    rna-XM_013047077.2    mNeoVis.2.1.1.chr15_hap1.g00605.t01    rna-XM_021680055.2      rna-XM_042972511.1    rna-XM_040634751.1    rna-XM_027612969.1    mNeoVis.3.1.1.chr15.g00389.t01    rna-gnl|WGS_ZZZZ|egapxtmp_033657-R1    mNeoVis.evergladensis.2.1.chr15.g00427.t01    mNeoVis.vulgivaga.2.1.chr15.g00435.t01      142    N0.HOG0000140   OG0000003  n380   rna-XM_034639090.1    rna-XM_038527957.1    rna-XM_022523765.1    rna-XM_023244971.2    rna-XM_036104098.1    rna-XM_039220161.1      rna-XM_045988242.1        rna-XM_059383369.1    rna-XM_045077280.1    mNeoVis.2.1.1.chr7_hap1.g00803.t01        rna-XM_042970229.1    rna-XM_008684755.2    rna-XM_027618328.2    mNeoVis.3.1.1.chr7.g00747.t01    rna-gnl|WGS_ZZZZ|egapxtmp_010922-R6      mNeoVis.vulgivaga.2.1.chr7.g00796.t01      143    N0.HOG0000141   OG0000003  n394   rna-XM_011235654.3    rna-XM_038527969.1    rna-XM_022524125.1    rna-XM_045046245.1        rna-XM_032839413.1    rna-XM_045988257.1    rna-XM_032324024.1    rna-XM_059150668.1    rna-XM_059382663.1    rna-XM_045077420.1    mNeoVis.2.1.1.chr7_hap1.g00804.t01        rna-XM_042970236.1    rna-XM_008684760.2      mNeoVis.3.1.1.chr7.g00748.t01    rna-gnl|WGS_ZZZZ|egapxtmp_010919-R1    mNeoVis.evergladensis.2.1.chr7.g00792.t01, mNeoVis.evergladensis.2.1.chr7.g00793.t01    mNeoVis.vulgivaga.2.1.chr7.g00798.t01      144    N0.HOG0000142   OG0000003  n412   rna-XM_034639111.1        rna-XM_045046194.1    rna-XM_036104041.1        rna-XM_045986776.1    rna-XM_032326074.1        rna-XM_045077281.1          rna-XM_042970226.1    rna-XM_008684757.2                145    N0.HOG0000143   OG0000003  n420   rna-XM_011235184.3    rna-XM_038525537.1      rna-XM_019818578.3, rna-XM_045046210.1    rna-XM_036104058.1, rna-XM_036104110.1    rna-XM_039220156.1, rna-XM_039220168.1    rna-XM_032839051.1    rna-XM_045988215.1    rna-XM_032324135.1    rna-XM_059151237.1    rna-XM_059383368.1      mNeoVis.2.1.1.chr7_hap1.g00798.t01, mNeoVis.2.1.1.chr7_hap1.g00799.t01    rna-XM_021681460.2, rna-XM_044921995.1    rna-XM_004414226.1, rna-XM_004415487.1, rna-XM_004415489.1    rna-XM_015539248.2, rna-XM_042970235.1    rna-XM_040632943.1    rna-XM_027603124.2, rna-XM_035725238.1, rna-XM_035725245.1      rna-gnl|WGS_ZZZZ|egapxtmp_012310-R1    mNeoVis.evergladensis.2.1.chr7.g00786.t01, mNeoVis.evergladensis.2.1.chr7.g00787.t01    mNeoVis.vulgivaga.2.1.chr7.g00789.t01, mNeoVis.vulgivaga.2.1.chr7.g00790.t01, mNeoVis.vulgivaga.2.1.chr7.g00791.t01      146    N0.HOG0000144   OG0000003  n452   rna-XM_019808666.2    rna-XM_038527945.1      rna-XM_045046217.1    rna-XM_036104076.1      rna-XM_032839408.1    rna-XM_045988232.1    rna-XM_032324015.1    rna-XM_059150664.1      rna-XM_045077393.1    mNeoVis.2.1.1.chr7_hap1.g00802.t01    rna-XM_044911351.1    rna-XM_004415494.1    rna-XM_042970224.1    rna-XM_040632929.1    rna-XM_027618306.2    mNeoVis.3.1.1.chr7.g00745.t01    rna-gnl|WGS_ZZZZ|egapxtmp_012312-R1    mNeoVis.evergladensis.2.1.chr7.g00790.t01    mNeoVis.vulgivaga.2.1.chr7.g00795.t01      147    N0.HOG0000145   OG0000003  n470   rna-XM_019808677.2, rna-XM_034639122.1    rna-XM_038527956.1      rna-XM_045046220.1    rna-XM_036104086.1, rna-XM_036104087.1    rna-XM_039220157.1    rna-XM_032839050.1    rna-XM_045988380.1    rna-XM_032324123.1    rna-XM_059150687.1    rna-XM_059382652.1    rna-XM_004781901.3    mNeoVis.2.1.1.chr7_hap1.g00800.t01, mNeoVis.2.1.1.chr7_hap1.g00801.t01    rna-XM_044922003.1, rna-XM_044922008.1    rna-XM_004415495.1    rna-XM_042970268.1    rna-XM_008684766.2    rna-XM_035725258.1    mNeoVis.3.1.1.chr7.g00744.t01    rna-gnl|WGS_ZZZZ|egapxtmp_012311-R1, rna-gnl|WGS_ZZZZ|egapxtmp_014220-R4, rna-gnl|WGS_ZZZZ|egapxtmp_014222-R2    mNeoVis.evergladensis.2.1.chr7.g00788.t01, mNeoVis.evergladensis.2.1.chr7.g00789.t01    mNeoVis.vulgivaga.2.1.chr7.g00792.t01, mNeoVis.vulgivaga.2.1.chr7.g00793.t01, mNeoVis.vulgivaga.2.1.chr7.g00794.t01      148    N0.HOG0000146   OG0000003  n499   rna-XM_034639083.1    rna-XM_038527937.1    rna-XM_022524113.1    rna-XM_045046214.1    rna-XM_036104067.1      rna-XM_032839405.1    rna-XM_045988231.1    rna-XM_032324014.1    rna-XM_059151040.1    rna-XM_059382644.1    rna-XM_013050698.2    mNeoVis.2.1.1.chr7_hap1.g00805.t01        rna-XM_042970344.1    rna-XM_008684748.2    rna-XM_027618303.2    mNeoVis.3.1.1.chr7.g00749.t01    rna-gnl|WGS_ZZZZ|egapxtmp_009697-R2    mNeoVis.evergladensis.2.1.chr7.g00794.t01    mNeoVis.vulgivaga.2.1.chr7.g00799.t01      149    N0.HOG0000147   OG0000003  n517   rna-XM_019803737.2    rna-XM_038574593.1      rna-XM_011287675.4    rna-XM_036081298.1    rna-XM_039222793.1    rna-XM_032849229.1    rna-XM_046024616.1    rna-XM_032309901.1    rna-XM_059139011.1    rna-XM_059408107.1    rna-XM_045087696.1    mNeoVis.2.1.1.chr3_hap1.g00691.t01    rna-XM_021692275.1    rna-XM_004414949.1    rna-XM_042961634.1    rna-XM_040620087.1    rna-XM_027577212.2    mNeoVis.3.1.1.chr3.g00681.t01    rna-gnl|WGS_ZZZZ|egapxtmp_041157-R1    mNeoVis.evergladensis.2.1.chr3.g00699.t01    mNeoVis.vulgivaga.2.1.chr3.g00704.t01      150    N0.HOG0000148   OG0000003  n537   rna-XM_034672341.1      rna-XM_022492145.1    rna-XM_045042327.1    rna-XM_036081464.1    rna-XM_039222772.1    rna-XM_032849237.1    rna-XM_046024422.1    rna-XM_032309910.1    rna-XM_059139965.1      rna-XM_004779588.3    mNeoVis.2.1.1.chr3_hap1.g00692.t01    rna-XM_044921047.1    rna-XM_004414961.1    rna-XM_042961642.1    rna-XM_040620078.1    rna-XM_027577227.1    mNeoVis.3.1.1.chr3.g00683.t01    rna-gnl|WGS_ZZZZ|egapxtmp_042168-R1    mNeoVis.evergladensis.2.1.chr3.g00700.t01    mNeoVis.vulgivaga.2.1.chr3.g00705.t01      151    N0.HOG0000149   OG0000003  n556                             rna-XM_044922197.1    rna-XM_004396437.1                mNeoVis.vulgivaga.2.1.chr7.g00498.t01      152    N0.HOG0000150   OG0000003  n559   rna-XM_034664311.1    rna-XM_038553128.1        rna-XM_036121993.1      rna-XM_032866345.1                rna-XM_044920424.1        rna-XM_040631742.1    rna-XM_035723470.1              153    N0.HOG0000151   OG0000003  n566   rna-XM_034656863.1      rna-XM_022492955.1    rna-XM_019810527.3    rna-XM_036090651.1    rna-XM_039239494.1                    rna-XM_004402850.1    rna-XM_015542581.2    rna-XM_008692496.2    rna-XM_027605750.1              154    N0.HOG0000152   OG0000003  n575   rna-XM_019798463.2    rna-XM_038535004.1      rna-XM_045036759.1    rna-XM_036090646.1    rna-XM_039239464.1    rna-XM_032851578.1    rna-XM_045999784.1    rna-XM_032337282.1    rna-XM_059174064.1    rna-XM_059417182.1    rna-XM_004737801.3    mNeoVis.2.1.1.chr1_hap1.g00212.t01    rna-XM_021700949.2    rna-XM_004402829.1    rna-XM_042994141.1    rna-XM_040630395.1    rna-XM_027605477.1    mNeoVis.3.1.1.chr1.g00213.t01    rna-gnl|WGS_ZZZZ|egapxtmp_027160-R1, rna-gnl|WGS_ZZZZ|egapxtmp_039745-R1    mNeoVis.evergladensis.2.1.chr1.g00207.t01, mNeoVis.evergladensis.2.1.chr1.g00208.t01    mNeoVis.vulgivaga.2.1.chr1.g00210.t01, mNeoVis.vulgivaga.2.1.chr1.g00211.t01      155    N0.HOG0000153   OG0000003  n597   rna-XM_034656855.1    rna-XM_038534996.1, rna-XM_038556376.1    rna-XM_022493332.1    rna-XM_045036715.1    rna-XM_036090640.1    rna-XM_039239455.1    rna-XM_032851581.1    rna-XM_045999790.1    rna-XM_032337287.1    rna-XM_059174067.1    rna-XM_059416530.1    rna-XM_013050318.2    mNeoVis.2.1.1.chr1_hap1.g00213.t01    rna-XM_021700936.1      rna-XM_042994074.1    rna-XM_040630397.1    rna-XM_027605469.1    mNeoVis.3.1.1.chr1.g00214.t01    rna-gnl|WGS_ZZZZ|egapxtmp_039137-R1    mNeoVis.evergladensis.2.1.chr1.g00209.t01    mNeoVis.vulgivaga.2.1.chr1.g00212.t01      156    N0.HOG0000154   OG0000004  n2   rna-XM_011232070.3    rna-XM_038575837.1, rna-XM_038589660.1    rna-XM_022495964.1    rna-XM_003994989.4    rna-XM_036076482.1    rna-XM_039242856.1    rna-XM_032875659.1    rna-XM_046025140.1    rna-XM_032309695.1    rna-XM_059140094.1    rna-XM_059408994.1    rna-XM_004742511.3    mNeoVis.2.1.1.chr3_hap1.g00355.t01    rna-XM_021687899.1    rna-XM_004400560.1    rna-XM_007091094.3      rna-XM_027577415.2    mNeoVis.3.1.1.chr3.g00348.t01    rna-gnl|WGS_ZZZZ|egapxtmp_039464-R1    mNeoVis.evergladensis.2.1.chr3.g00364.t01    mNeoVis.vulgivaga.2.1.chr3.g00358.t01      157    N0.HOG0000155   OG0000004  n23   id-LOC117797918   id-LOC612168   id-LOC111160013    id-LOC111559116, id-LOC123381270        id-LOC116882713    id-LOC123953864    id-LOC116571510, id-LOC116571514, id-LOC116571516    id-LOC131810980, id-LOC131810984    id-LOC132023362, id-LOC132023767    id-LOC101685424, rna-XM_004781370.2          id-LOC102949523, id-LOC102949826, id-LOC102951354, id-LOC102966738, id-LOC107180566, id-LOC122232371, id-LOC122232563, id-LOC122232575, id-LOC122232695, id-LOC122232698, id-LOC122232722, id-LOC122232808, id-LOC122232814, id-LOC122236241, id-LOC122236255, id-LOC122236258    id-LOC121103215                158    N0.HOG0000156   OG0000004  n56   rna-XM_002931370.4   id-LOC612106       id-LOC118550639    id-LOC120227879    id-LOC116882706      id-LOC116571513    id-LOC131810988, id-LOC131810989    id-LOC132023763, id-LOC132023764, id-LOC132023768        id-LOC123326629        id-LOC103677149                159    N0.HOG0000157   OG0000004  n68   id-LOC109488500, id-LOC117797794, id-LOC117797797, id-LOC117797853, rna-XM_002931191.3    id-LOC119869141    id-LOC111139919, id-LOC111140504    id-LOC111557360    id-LOC118524392, id-LOC118542734, id-LOC118550640, rna-XM_036085195.1    id-LOC120227877, id-LOC120227882    id-LOC116882710, id-LOC116882712, id-LOC116882716, id-LOC116882740      id-LOC116572280, rna-XM_032311106.1    id-LOC131811248, id-LOC131811391    id-LOC132023769, id-LOC132023854    id-LOC101681270, rna-XM_013050433.2      id-LOC110583295    rna-XM_004400101.1, rna-XM_004400102.1, rna-XM_004400567.1, rna-XM_004415472.1, rna-XM_004416340.1, rna-XM_004416897.1, rna-XM_004417292.1, rna-XM_004417460.1, rna-XM_004417461.1, rna-XM_004417474.1, rna-XM_004417486.1, rna-XM_004417509.1, rna-XM_004417599.1, rna-XM_004417617.1    id-LOC102964194, id-LOC102964770, id-LOC107181091, id-LOC122232696, id-LOC122232697, id-LOC122232730, id-LOC122236238, id-LOC122236242, id-LOC122236254, id-LOC122236260    id-LOC121101162, id-LOC121103217, rna-XM_040623744.1    id-LOC113912339, id-LOC113912343, id-LOC113912344, id-LOC113912346, id-LOC113912575, id-LOC113912978, id-LOC113933927, id-LOC113935278, id-LOC118355958, id-LOC118356240, id-LOC118356276, rna-XM_035723965.1              160    N0.HOG0000158   OG0000004  n135   id-LOC117795912, rna-XM_019805753.2    rna-XM_038575836.1    rna-XM_022496042.1    id-LOC101088367, id-LOC109496970, id-LOC111557359, id-LOC123381273, id-LOC123381368, id-LOC123381370    rna-XM_036076481.1    id-LOC120226906, id-LOC120226909, rna-XM_039225080.1, rna-XM_039225081.1, rna-XM_039242900.1      rna-XM_046024340.1    rna-XM_032309694.1    rna-XM_059140095.1    rna-XM_059408993.1    rna-XM_004742510.3      rna-XM_044921031.1    rna-XM_004400519.1    id-LOC102962176, id-LOC102968121    rna-XM_040619396.1    id-LOC113913255, rna-XM_027575939.2              161    N0.HOG0000159   OG0000004  n163     id-LOC119866080, id-LOC119877604, id-LOC607582, id-LOC608238, id-LOC608248        id-LOC118526458, id-LOC118526467, id-LOC118526519      id-LOC116863727    id-LOC123954343, id-LOC123954344, id-LOC123954347, id-LOC123954348, id-LOC123954349, id-LOC123954351, id-LOC123954352    id-LOC116572248, id-LOC116572249, id-LOC116572250    id-LOC131811746      id-LOC101689439, id-LOC123387657                  rna-gnl|WGS_ZZZZ|egapxtmp_041074-R1          162    N0.HOG0000160   OG0000004  n185   id-LOC105235181, id-LOC105235281, id-LOC109488360, id-LOC117796044, id-LOC117797806, id-LOC117797934    id-LOC119877621    id-LOC111139903, id-LOC111139904, id-LOC111139905, id-LOC111142666      id-LOC118538191, id-LOC118543526, id-LOC118543527, id-LOC118543538, rna-XM_036077763.1    id-LOC120227888    id-LOC116863063, id-LOC116863067, id-LOC116882727, id-LOC116882731, rna-XM_032876049.1    rna-XM_046024735.1    id-LOC116571651, id-LOC116572251, id-LOC116572252    rna-XM_059140505.1    id-LOC132023367, id-LOC132023722, id-LOC132023725, id-LOC132023726    id-LOC101675839, id-LOC101682133, id-LOC101682403, id-LOC106004691      id-LOC110578359, id-LOC110578361, id-LOC110578363    id-LOC101364392, id-LOC101364687, rna-XM_004400550.1    id-LOC122232574    id-LOC103675572          mNeoVis.evergladensis.2.1.chr3.g00392.t01        163    N0.HOG0000161   OG0000004  n228   id-LOC105235224    id-LOC119877602    id-LOC111139907, id-LOC111139908, id-LOC111139911, id-LOC111140366, id-LOC111142460      id-LOC118538195, id-LOC118543537    id-LOC120227887, id-LOC120227890    id-LOC116862418, id-LOC116862421, id-LOC116882693, id-LOC116882728, id-LOC116882732      id-LOC116571649    id-LOC131811717, id-LOC131811747    id-LOC132023361, id-LOC132023366, id-LOC132023721, id-LOC132023723    id-LOC123387653    mNeoVis.2.1.1.chr3_hap1.g00385.t01    id-LOC110578362    id-LOC101363520, id-LOC101364098    id-LOC122232565, id-LOC122232569, id-LOC122232578    id-LOC103675571, id-LOC103675684                164    N0.HOG0000162   OG0000004  n260   id-LOC109488718, rna-XM_034650256.1    id-LOC119866063                            id-LOC101363807      id-LOC121103019                165    N0.HOG0000163   OG0000004  n264   rna-XM_034642024.1    rna-XM_038575763.1, rna-XM_038575771.1    rna-XM_022495628.1    rna-XM_045041898.1, rna-XM_045041905.1          rna-XM_046024733.1    rna-XM_032309767.1, rna-XM_032309779.1    rna-XM_059140502.1      rna-XM_045065742.1                          166    N0.HOG0000164   OG0000004  n275                 rna-XM_046024732.1          rna-XM_045065721.1                          167    N0.HOG0000165   OG0000004  n278                               id-LOC101364993, id-LOC101365604, id-LOC101365906, id-LOC101379126                    168    N0.HOG0000166   OG0000004  n282         rna-XM_045041880.1, rna-XM_045041886.1                          rna-XM_042962209.1, rna-XM_042962213.1, rna-XM_042962215.1                  169    N0.HOG0000167   OG0000004  n290   id-LOC105234852, id-LOC105234853, id-LOC105235062, id-LOC105235063, id-LOC105235206, id-LOC117797864, id-LOC117797993, id-LOC117797997, id-LOC117798029    id-LOC100685445, id-LOC102155737, id-LOC119866111, id-LOC119868992, id-LOC119869118, id-LOC119869145, id-LOC119878461, id-LOC119879337, rna-XM_038574507.1, rna-XM_038575772.1    id-LOC111139912, id-LOC111139914, id-LOC111160019    id-LOC111557377    id-LOC118531235, id-LOC118543539    id-LOC120227479    id-LOC116882692, id-LOC116882724, id-LOC116882725    id-LOC123953859    id-LOC116571650, id-LOC116572253, id-LOC116572255    id-LOC131810976          rna-XM_044921033.1    id-LOC101365294, id-LOC101386660    id-LOC102963331, id-LOC102966159, id-LOC107180567, id-LOC122232370, id-LOC122232566, id-LOC122232570, id-LOC122232610, id-LOC122232729, id-LOC122236237    id-LOC103675687, id-LOC103675688, id-LOC103675689, id-LOC121105381                170    N0.HOG0000168   OG0000004  n340               id-LOC116882721        rna-XM_059138983.1                              171    N0.HOG0000169   OG0000004  n342    id-LOC486373       id-LOC118531231                    id-LOC110591627          id-LOC118356274              172    N0.HOG0000170   OG0000004  n345   id-LOC105235026, id-LOC117798365, id-LOC117798366    id-LOC119869140        id-LOC118531223                          id-LOC121103017                173    N0.HOG0000171   OG0000004  n350     id-LOC119866055, id-LOC119868854, id-LOC119869112, id-LOC612177    id-LOC111160015              id-LOC116571515    id-LOC131810985      id-LOC101686302              id-LOC118356272              174    N0.HOG0000172   OG0000004  n358     id-LOC119877625, id-LOC606893      id-LOC109493262, id-LOC111557373              id-LOC131811503      id-LOC101679978          id-LOC102966456, id-LOC122232726, rna-XM_042962848.1                  175    N0.HOG0000173   OG0000004  n366   id-LOC105235235, id-LOC109488775, id-LOC117797786, id-LOC117797798, id-LOC117797799, id-LOC117797804, id-LOC117797812, id-LOC117797917, id-LOC117797938, id-LOC117798015, id-LOC117798016, id-LOC117798018, id-LOC117798022, id-LOC117800354    id-LOC119863953, id-LOC119863955, id-LOC119866054, id-LOC119866222, id-LOC119869100, id-LOC119869101, id-LOC119869121, id-LOC119869125, id-LOC119877559, id-LOC119877603, id-LOC119877620, id-LOC119878349, id-LOC119878465, id-LOC119878467, id-LOC119878923, id-LOC486377, id-LOC612173, rna-XM_038574505.1    id-LOC111160016, id-LOC111160018    id-LOC123381274, id-LOC123381352, id-LOC123381354    id-LOC118531225, id-LOC118531226, id-LOC118532663, id-LOC118532665, id-LOC118533552, id-LOC118533554, id-LOC118533555, id-LOC118542698, id-LOC118542699, id-LOC118542731, id-LOC118542733, id-LOC118546749, id-LOC118550635, id-LOC118550641, id-LOC118554150, id-LOC118554158, id-LOC118554348, id-LOC118555066    id-LOC120227489, id-LOC120227880    id-LOC116882702, id-LOC116882703, id-LOC116882709, id-LOC116882718, id-LOC116882719, rna-XM_032883243.1    id-LOC123953860, id-LOC123953861      id-LOC131810983, id-LOC131810987    id-LOC132023766, id-LOC132023770    id-LOC101670366, rna-XM_013050898.2      id-LOC110578043, id-LOC110578192, id-LOC110578224, id-LOC110578926, id-LOC110580005, id-LOC110584173    id-LOC101363121, id-LOC101367944, id-LOC101368241, id-LOC101375066, id-LOC101376715, id-LOC101379416, id-LOC101380000, id-LOC101380763, id-LOC101380823, id-LOC101381360, id-LOC101381406, id-LOC101382568, rna-XM_004411257.1, rna-XM_004417185.1    id-LOC102951050, id-LOC102967593, id-LOC107180790, id-LOC122232571, id-LOC122232572, id-LOC122232573, id-LOC122232720, id-LOC122232723, id-LOC122232724, id-LOC122232725, id-LOC122232806, id-LOC122232807, id-LOC122236239, id-LOC122236240, id-LOC122236243, id-LOC122236244, id-LOC122236245, id-LOC122236246, id-LOC122236248, id-LOC122236251, id-LOC122236256, id-LOC122236257, id-LOC122236259    id-LOC103656290, id-LOC103656291, id-LOC103656293, id-LOC121099736, id-LOC121101163    id-LOC118356257              176    N0.HOG0000174   OG0000004  n484     id-LOC119866215      id-LOC101099449          id-LOC123954073, id-LOC123954074, id-LOC123954078                  id-LOC102964473      id-LOC113915726              177    N0.HOG0000175   OG0000004  n490   id-LOC117798030, rna-XM_002931227.4    id-LOC119877601    id-LOC111139920, id-LOC111142461          id-LOC116878498      id-LOC116571639, id-LOC116571646, rna-XM_032309737.1    id-LOC131811713    id-LOC132023856, id-LOC132023857    id-LOC101681541, rna-XM_045065711.1            rna-XM_040619436.1        id-egapxtmp_042432          178    N0.HOG0000176   OG0000005  n2   rna-XM_002929999.4    rna-NM_001388785.1    rna-XM_022513090.1    rna-XM_045038472.1    rna-XM_036123258.1    rna-XM_039250199.1    rna-XM_032869048.1    rna-XM_046016507.1    rna-XM_032358120.1    rna-XM_059173045.1    rna-XM_059403338.1    rna-XM_013054798.1    mNeoVis.2.1.1.chr7_hap1.g01338.t01    rna-XM_021696225.1      rna-XM_042958686.1    rna-XM_008688231.1    rna-XM_027580992.1    mNeoVis.3.1.1.chr7.g01248.t01    rna-gnl|WGS_ZZZZ|egapxtmp_010616-R1          179    N0.HOG0000177   OG0000005  n20   rna-XM_002930001.3, rna-XM_034667713.1    rna-XM_038535820.1    rna-XM_022513031.1, rna-XM_022513067.1    rna-XM_045038467.1, rna-XM_045038471.1    rna-XM_036123257.1, rna-XM_036123262.1    rna-XM_039250172.1, rna-XM_039250198.1    rna-XM_032869045.1, rna-XM_032869046.1    rna-XM_046016848.1, rna-XM_046017018.1, rna-XM_046017153.1    rna-XM_032360239.1    rna-XM_059156308.1    rna-XM_059392310.1    rna-XM_004750146.3    mNeoVis.2.1.1.chr7_hap1.g01337.t01    rna-XM_021696223.1, rna-XM_021696366.1    rna-XM_004413692.1, rna-XM_004413708.1    rna-XM_007074046.2, rna-XM_007074047.1    rna-XM_008688228.2, rna-XM_008688230.2    rna-XM_027580994.1      rna-gnl|WGS_ZZZZ|egapxtmp_013359-R1    mNeoVis.evergladensis.2.1.chr7.g01321.t01    mNeoVis.vulgivaga.2.1.chr7.g01334.t01      180    N0.HOG0000178   OG0000005  n52   rna-XM_002930000.4    rna-NM_001388799.1, rna-NM_001388878.1    rna-XM_022513124.1    rna-XM_006936748.4, rna-XM_011286461.3    rna-XM_036081750.1    rna-XM_039250194.1, rna-XM_039250227.1    rna-XM_032869024.1      rna-XM_032358777.1        rna-XM_013054803.2    mNeoVis.2.1.1.chr7_hap1.g01309.t01    rna-XM_021696419.1      rna-XM_007074033.2, rna-XM_007074081.2    rna-XM_008688229.1, rna-XM_008689931.2      mNeoVis.3.1.1.chr7.g01231.t01    rna-gnl|WGS_ZZZZ|egapxtmp_013361-R1    mNeoVis.evergladensis.2.1.chr7.g01297.t01    mNeoVis.vulgivaga.2.1.chr7.g01308.t01      181    N0.HOG0000179   OG0000005  n74   rna-XM_002928187.4    rna-NM_001388789.1    rna-XM_022513051.1    rna-XM_003992514.4      rna-XM_039250175.1    rna-XM_032869036.1    rna-XM_046016490.1    rna-XM_032359088.1    rna-XM_059156228.1    rna-XM_059392396.1    rna-XM_004750258.1    mNeoVis.2.1.1.chr7_hap1.g01326.t01    rna-XM_021696639.1      rna-XM_007074041.2    rna-XM_008683618.1      mNeoVis.3.1.1.chr7.g01245.t01    rna-gnl|WGS_ZZZZ|egapxtmp_012435-R1      mNeoVis.vulgivaga.2.1.chr7.g01326.t01      182    N0.HOG0000180   OG0000005  n93     rna-NM_001389090.1    rna-XM_022513346.1    rna-XM_006936754.5    rna-XM_036075357.1    rna-XM_039250211.1      rna-XM_046015963.1, rna-XM_046015964.1    rna-XM_032358111.1, rna-XM_032358112.1    rna-XM_059172085.1, rna-XM_059172211.1    rna-XM_059404135.1    rna-XM_004750255.1    mNeoVis.2.1.1.chr7_hap1.g01321.t01    rna-XM_021687299.2    rna-XM_004413705.1    rna-XM_007074078.2    rna-XM_008683634.2, rna-XM_008683635.2    rna-XM_027580032.2    mNeoVis.3.1.1.chr7.g01239.t01    rna-gnl|WGS_ZZZZ|egapxtmp_012431-R1, rna-gnl|WGS_ZZZZ|egapxtmp_012433-R1    mNeoVis.evergladensis.2.1.chr7.g01306.t01        183    N0.HOG0000181   OG0000005  n117   rna-XM_002928199.3    rna-XM_038535495.1    rna-XM_022513081.1    rna-XM_003992512.5    rna-XM_036075358.1, rna-XM_036113951.1    rna-XM_039250201.1    rna-XM_032869033.1    rna-XM_046016725.1    rna-XM_032359100.1    rna-XM_059156193.1    rna-XM_059392449.1    rna-XM_004750141.3        rna-XM_004413689.1    rna-XM_015538936.1    rna-XM_008683623.2    rna-XM_027580003.2      rna-gnl|WGS_ZZZZ|egapxtmp_013371-R1    mNeoVis.evergladensis.2.1.chr7.g01308.t01    mNeoVis.vulgivaga.2.1.chr7.g01321.t01      184    N0.HOG0000182   OG0000005  n136   rna-XM_002928201.3    rna-NM_001389157.1    rna-XM_022513347.1    rna-XM_023239695.2      rna-XM_039250231.1    rna-XM_032869032.1    rna-XM_046015965.1    rna-XM_032358110.1    rna-XM_059171968.1      rna-XM_004750254.3    mNeoVis.2.1.1.chr7_hap1.g01320.t01    rna-XM_021696636.1      rna-XM_007074077.3    rna-XM_008683636.2      mNeoVis.3.1.1.chr7.g01240.t01    rna-gnl|WGS_ZZZZ|egapxtmp_013350-R1    mNeoVis.evergladensis.2.1.chr7.g01307.t01    mNeoVis.vulgivaga.2.1.chr7.g01322.t01      185    N0.HOG0000183   OG0000005  n155   rna-XM_002928217.4    rna-NM_001389109.1    rna-XM_022513194.1    rna-XM_023238983.2    rna-XM_036081748.1    rna-XM_039250236.1    rna-XM_032868993.1    rna-XM_046017597.1    rna-XM_032359055.1    rna-XM_059156068.1    rna-XM_059392577.1    rna-XM_004750116.2    mNeoVis.2.1.1.chr7_hap1.g01304.t01    rna-XM_021696518.2      rna-XM_042959186.1    rna-XM_008690168.1    rna-XM_027577780.1    mNeoVis.3.1.1.chr7.g01227.t01    rna-gnl|WGS_ZZZZ|egapxtmp_013352-R1    mNeoVis.evergladensis.2.1.chr7.g01292.t01    mNeoVis.vulgivaga.2.1.chr7.g01304.t01      186    N0.HOG0000184   OG0000005  n175   rna-XM_019804619.1    rna-NM_001388781.1    rna-XM_022513126.1    rna-XM_003992522.4, rna-XM_006936772.5, rna-XM_023239244.2    rna-XM_036123255.1    rna-XM_039250208.1, rna-XM_039250209.1, rna-XM_039250218.1    rna-XM_032869052.1    rna-XM_046017118.1    rna-XM_032360020.1    rna-XM_059156351.1    rna-XM_059392250.1    rna-XM_004750150.3      rna-XM_021696473.2    rna-XM_004413695.1    rna-XM_007074085.3, rna-XM_007074087.2, rna-XM_015538968.2    rna-XM_008688237.2    rna-XM_035722934.1    mNeoVis.3.1.1.chr7.g01250.t01    rna-gnl|WGS_ZZZZ|egapxtmp_013362-R1          187    N0.HOG0000185   OG0000005  n201   rna-XM_002928188.2    rna-XM_038535492.1    rna-XM_022513109.1    rna-XM_003992487.4    rna-XM_036113949.1    rna-XM_039250197.1    rna-XM_032869035.1    rna-XM_046016755.1    rna-XM_032359596.1    rna-XM_059156214.1    rna-XM_059392411.1    rna-XM_013054805.2    mNeoVis.2.1.1.chr7_hap1.g01325.t01    rna-XM_021696638.2      rna-XM_007074079.2        mNeoVis.3.1.1.chr7.g01244.t01    rna-gnl|WGS_ZZZZ|egapxtmp_013363-R1    mNeoVis.evergladensis.2.1.chr7.g01312.t01    mNeoVis.vulgivaga.2.1.chr7.g01325.t01      188    N0.HOG0000186   OG0000005  n219   rna-XM_002928189.2    rna-XM_038535493.1    rna-XM_022513089.1    rna-XM_011279812.3    rna-XM_036113948.1    rna-XM_039250195.1      rna-XM_046016794.1    rna-XM_032358882.1    rna-XM_059156206.1    rna-XM_059392439.1    rna-XM_013054804.1    mNeoVis.2.1.1.chr7_hap1.g01324.t01    rna-XM_044919401.1      rna-XM_007074040.2        mNeoVis.3.1.1.chr7.g01243.t01    rna-gnl|WGS_ZZZZ|egapxtmp_013364-R1    mNeoVis.evergladensis.2.1.chr7.g01311.t01    mNeoVis.vulgivaga.2.1.chr7.g01324.t01      189    N0.HOG0000187   OG0000005  n236   rna-XM_002929998.2    rna-NM_001388784.1    rna-XM_022513023.1    rna-XM_019811473.3    rna-XM_036123252.1    rna-XM_039250177.1    rna-XM_032869049.1    rna-XM_046016282.1    rna-XM_032360240.1    rna-XM_059156319.1    rna-XM_059392296.1    rna-XM_004750147.2    mNeoVis.2.1.1.chr7_hap1.g01339.t01    rna-XM_021696370.1    rna-XM_004413693.1    rna-XM_007074048.2    rna-XM_008688234.2    rna-XM_027580993.1        mNeoVis.evergladensis.2.1.chr7.g01322.t01    mNeoVis.vulgivaga.2.1.chr7.g01335.t01      190    N0.HOG0000188   OG0000005  n256                                       mNeoVis.3.1.1.chr7.g01249.t01    rna-gnl|WGS_ZZZZ|egapxtmp_010617-R1      mNeoVis.vulgivaga.2.1.chr7.g01336.t01      191    N0.HOG0000189   OG0000005  n259   rna-XM_002923997.4    rna-NM_001388782.1    rna-XM_022513340.1    rna-XM_006936758.5    rna-XM_036123251.1    rna-XM_039250212.1, rna-XM_039250215.1    rna-XM_032850988.1, rna-XM_032850991.1    rna-XM_046015956.1    rna-XM_032359279.1    rna-XM_059173169.1    rna-XM_059392261.1    rna-XM_004750149.2        rna-XM_004413709.1    rna-XM_007074086.2    rna-XM_008688236.1                192    N0.HOG0000190   OG0000005  n276     rna-NM_001389062.1    rna-XM_022513125.1    rna-XM_003992490.2      rna-XM_039250216.1    rna-XM_032869050.1    rna-XM_046016850.1    rna-XM_032359257.1    rna-XM_059156327.1    rna-XM_059392283.1    rna-XM_045080530.1    mNeoVis.2.1.1.chr7_hap1.g01340.t01      rna-XM_004413694.1    rna-XM_007074084.2    rna-XM_008688235.1    rna-XM_027579982.2        mNeoVis.evergladensis.2.1.chr7.g01323.t01        193    N0.HOG0000191   OG0000005  n291     rna-XM_038535490.1    rna-XM_022513111.1    rna-XM_003992491.2      rna-XM_039250213.1    rna-XM_032869051.1    rna-XM_046015957.1    rna-XM_032359267.1    rna-XM_059156339.1    rna-XM_059392271.1    rna-XM_045080528.1          rna-XM_007074049.1                  194    N0.HOG0000192   OG0000005  n301   rna-XM_034650339.1, rna-XM_034652185.1    rna-XM_038535566.1    rna-XM_022513341.1, rna-XM_022513342.1    rna-XM_045038469.1, rna-XM_045038470.1    rna-XM_036123250.1    rna-XM_039250169.1, rna-XM_039250173.1    rna-XM_032869001.1, rna-XM_032869002.1    rna-XM_046015958.1, rna-XM_046015960.1, rna-XM_046016230.1    rna-XM_032358116.1, rna-XM_032358117.1    rna-XM_059172677.1, rna-XM_059172911.1    rna-XM_059387224.1, rna-XM_059387225.1, rna-XM_059403476.1    rna-XM_045080312.1, rna-XM_045080313.1    mNeoVis.2.1.1.chr7_hap1.g01335.t01, mNeoVis.2.1.1.chr7_hap1.g01336.t01    rna-XM_021696222.1, rna-XM_021696405.1      rna-XM_007074082.2, rna-XM_042958200.1    rna-XM_008688394.2, rna-XM_040630647.1    rna-XM_027580955.1      rna-gnl|WGS_ZZZZ|egapxtmp_010615-R1    mNeoVis.evergladensis.2.1.chr7.g01320.t01    mNeoVis.vulgivaga.2.1.chr7.g01333.t01      195    N0.HOG0000193   OG0000005  n337         rna-XM_045038466.1      rna-XM_039250168.1                      rna-XM_007074045.1                  196    N0.HOG0000194   OG0000005  n341         rna-XM_003992517.2      rna-XM_039250167.1, rna-XM_039250171.1                      rna-XM_007074044.1                  197    N0.HOG0000195   OG0000005  n345   rna-XM_011216697.3, rna-XM_034650101.1, rna-XM_034650220.1, rna-XM_034651776.1          rna-XM_036123253.1, rna-XM_036123342.1                    rna-XM_021696347.1, rna-XM_021696395.2, rna-XM_044919303.1    rna-XM_004413691.1      rna-XM_008683617.2, rna-XM_008683631.2, rna-XM_008688227.2    rna-XM_027577938.1              198    N0.HOG0000196   OG0000005  n360   rna-XM_002931102.4, rna-XM_019792818.2    rna-NM_001389039.1, rna-XM_038535491.1    rna-XM_022513007.1, rna-XM_022513150.1, rna-XM_022513344.1          rna-XM_032851197.1    rna-XM_046016200.1    rna-XM_032358632.1, rna-XM_032359111.1    rna-XM_059156283.1, rna-XM_059156294.1      rna-XM_004750260.1, rna-XM_004750261.2, rna-XM_045080531.1    mNeoVis.2.1.1.chr7_hap1.g01334.t01          rna-XM_008688226.2, rna-XM_008688392.2      mNeoVis.3.1.1.chr7.g01241.t01    rna-gnl|WGS_ZZZZ|egapxtmp_013356-R1    mNeoVis.evergladensis.2.1.chr7.g01319.t01    mNeoVis.vulgivaga.2.1.chr7.g01332.t01      199    N0.HOG0000197   OG0000005  n384     rna-NM_001389084.1    rna-XM_022513066.1, rna-XM_022513154.1          rna-XM_032850925.1, rna-XM_032869042.1, rna-XM_032869043.1, rna-XM_032869044.1    rna-XM_046016161.1, rna-XM_046016166.1, rna-XM_046016280.1, rna-XM_046016286.1, rna-XM_046016294.1, rna-XM_046016397.1    rna-XM_032359087.1, rna-XM_032359091.1, rna-XM_032359092.1    rna-XM_059156252.1, rna-XM_059156260.1    rna-XM_059392333.1, rna-XM_059392347.1    rna-XM_004750145.2, rna-XM_004750259.1    mNeoVis.2.1.1.chr7_hap1.g01332.t01, mNeoVis.2.1.1.chr7_hap1.g01333.t01              mNeoVis.3.1.1.chr7.g01246.t01, mNeoVis.3.1.1.chr7.g01247.t01    rna-gnl|WGS_ZZZZ|egapxtmp_010612-R1, rna-gnl|WGS_ZZZZ|egapxtmp_013355-R1, rna-gnl|WGS_ZZZZ|egapxtmp_013357-R1    mNeoVis.evergladensis.2.1.chr7.g01317.t01, mNeoVis.evergladensis.2.1.chr7.g01318.t01    mNeoVis.vulgivaga.2.1.chr7.g01329.t01, mNeoVis.vulgivaga.2.1.chr7.g01330.t01, mNeoVis.vulgivaga.2.1.chr7.g01331.t01      200    N0.HOG0000198   OG0000005  n417   rna-XM_002928186.4, rna-XM_011218037.3, rna-XM_011236046.3, rna-XM_019793844.2, rna-XM_034650084.1, rna-XM_034650085.1, rna-XM_034650086.1, rna-XM_034667711.1, rna-XM_034667712.1    rna-NM_001388787.1, rna-NM_001389050.1, rna-NM_001389079.1    rna-XM_022513058.1      rna-XM_036123254.1      rna-XM_032869041.1    rna-XM_046016607.1            mNeoVis.2.1.1.chr7_hap1.g01330.t01    rna-XM_021696221.2    rna-XM_004413707.1      rna-XM_008683611.2, rna-XM_008683612.2, rna-XM_008683613.2, rna-XM_008683614.1, rna-XM_008683615.2, rna-XM_008683629.2, rna-XM_008683630.2, rna-XM_008688225.2, rna-XM_040630695.1, rna-XM_040630696.1    rna-XM_027579880.1        mNeoVis.evergladensis.2.1.chr7.g01316.t01        201    N0.HOG0000199   OG0000005  n447       rna-XM_022513064.1          rna-XM_032869040.1    rna-XM_046015959.1, rna-XM_046016529.1            mNeoVis.2.1.1.chr7_hap1.g01329.t01                rna-gnl|WGS_ZZZZ|egapxtmp_012436-R1    mNeoVis.evergladensis.2.1.chr7.g01315.t01    mNeoVis.vulgivaga.2.1.chr7.g01328.t01      202    N0.HOG0000200   OG0000005  n454     rna-NM_001388788.1    rna-XM_022513061.1, rna-XM_022513062.1          rna-XM_032869037.1, rna-XM_032869039.1    rna-XM_046015961.1, rna-XM_046016244.1, rna-XM_046016284.1, rna-XM_046016285.1, rna-XM_046016322.1, rna-XM_046016354.1, rna-XM_046016385.1, rna-XM_046016463.1    rna-XM_032358850.1, rna-XM_032359082.1, rna-XM_032359086.1    rna-XM_059156234.1, rna-XM_059156238.1, rna-XM_059156244.1    rna-XM_059392359.1, rna-XM_059392370.1, rna-XM_059392384.1    rna-XM_004750142.1, rna-XM_004750143.1, rna-XM_004750144.2    mNeoVis.2.1.1.chr7_hap1.g01327.t01, mNeoVis.2.1.1.chr7_hap1.g01328.t01, mNeoVis.2.1.1.chr7_hap1.g01331.t01          rna-XM_008683616.2        rna-gnl|WGS_ZZZZ|egapxtmp_013354-R1, rna-gnl|WGS_ZZZZ|egapxtmp_013360-R1    mNeoVis.evergladensis.2.1.chr7.g01313.t01, mNeoVis.evergladensis.2.1.chr7.g01314.t01    mNeoVis.vulgivaga.2.1.chr7.g01327.t01      203    N0.HOG0000201   OG0000005  n487         rna-XM_006936755.4, rna-XM_006936756.2, rna-XM_019811480.3      rna-XM_039250176.1, rna-XM_039250228.1                      rna-XM_007074042.2, rna-XM_007074043.2, rna-XM_015538957.2                  204    N0.HOG0000202   OG0000006  n1   rna-XM_019796054.1    rna-XM_038530378.1    rna-XM_022493746.1    rna-XM_003980856.4    rna-XM_036123201.1    rna-XM_039236138.1    rna-XM_032850005.1    rna-XM_045998651.1    rna-XM_032336449.1    rna-XM_059173472.1    rna-XM_059416612.1    rna-XM_013052319.2    mNeoVis.2.1.1.chr1_hap1.g00507.t01    rna-XM_021702040.1    rna-XM_004397762.1    rna-XM_007077907.2    rna-XM_040621930.1    rna-XM_027606366.2    mNeoVis.3.1.1.chr1.g00511.t01    rna-gnl|WGS_ZZZZ|egapxtmp_031723-R1    mNeoVis.evergladensis.2.1.chr1.g00512.t01    mNeoVis.vulgivaga.2.1.chr1.g00522.t01      205    N0.HOG0000203   OG0000006  n24       rna-XM_022518067.1, rna-XM_022518071.1          rna-XM_032849949.1, rna-XM_032849954.1      rna-XM_032336377.1    rna-XM_059174843.1    rna-XM_059417769.1    rna-XM_045075368.1      rna-XM_021701844.1    rna-XM_004397749.1, rna-XM_004417769.1    rna-XM_042994849.1    rna-XM_040622331.1      mNeoVis.3.1.1.chr1.g00541.t01    rna-gnl|WGS_ZZZZ|egapxtmp_032516-R1    mNeoVis.evergladensis.2.1.chr1.g00532.t01    mNeoVis.vulgivaga.2.1.chr1.g00541.t01      206    N0.HOG0000204   OG0000006  n41   rna-XM_034652742.1, rna-XM_034655700.1, rna-XM_034655701.1, rna-XM_034656174.1      rna-XM_022492630.1, rna-XM_022492661.1, rna-XM_022492708.1, rna-XM_022492725.1, rna-XM_022492736.1, rna-XM_022493370.1, rna-XM_022493372.1      rna-XM_036123256.1, rna-XM_036123503.1    rna-XM_039225806.1, rna-XM_039225814.1, rna-XM_039236151.1, rna-XM_039236173.1, rna-XM_039236188.1      rna-XM_045999641.1, rna-XM_046000760.1, rna-XM_046000761.1        rna-XM_059387392.1, rna-XM_059387399.1, rna-XM_059387423.1, rna-XM_059387595.1, rna-XM_059387598.1, rna-XM_059416755.1    rna-XM_013052399.2, rna-XM_045075369.1, rna-XM_045075370.1, rna-XM_045075372.1    mNeoVis.2.1.1.chr1_hap1.g00518.t01, mNeoVis.2.1.1.chr1_hap1.g00520.t01, mNeoVis.2.1.1.chr1_hap1.g00522.t01, mNeoVis.2.1.1.chr1_hap1.g00523.t01, mNeoVis.2.1.1.chr1_hap1.g00524.t01    rna-XM_021687650.1, rna-XM_021688716.2, rna-XM_021693640.1, rna-XM_044912344.1, rna-XM_044912451.1, rna-XM_044912468.1, rna-XM_044912472.1, rna-XM_044912545.1    rna-XM_004397751.1, rna-XM_004397855.1, rna-XM_004397857.1, rna-XM_004397859.1, rna-XM_004397862.1, rna-XM_004397864.1, rna-XM_004397865.1    rna-XM_042985611.1, rna-XM_042985656.1, rna-XM_042994471.1    rna-XM_040621955.1    rna-XM_027606365.2    mNeoVis.3.1.1.chr1.g00527.t01, mNeoVis.3.1.1.chr1.g00531.t01, mNeoVis.3.1.1.chr1.g00532.t01, mNeoVis.3.1.1.chr1.g00534.t01, mNeoVis.3.1.1.chr1.g00535.t01, mNeoVis.3.1.1.chr1.g00536.t01, mNeoVis.3.1.1.chr1.g00538.t01, mNeoVis.3.1.1.chr1.g00539.t01    rna-gnl|WGS_ZZZZ|egapxtmp_006414-R1, rna-gnl|WGS_ZZZZ|egapxtmp_006427-R1, rna-gnl|WGS_ZZZZ|egapxtmp_006482-R1, rna-gnl|WGS_ZZZZ|egapxtmp_028371-R1, rna-gnl|WGS_ZZZZ|egapxtmp_028380-R1, rna-gnl|WGS_ZZZZ|egapxtmp_028385-R1, rna-gnl|WGS_ZZZZ|egapxtmp_028392-R1, rna-gnl|WGS_ZZZZ|egapxtmp_032437-R1    mNeoVis.evergladensis.2.1.chr1.g00521.t01, mNeoVis.evergladensis.2.1.chr1.g00523.t01, mNeoVis.evergladensis.2.1.chr1.g00526.t01, mNeoVis.evergladensis.2.1.chr1.g00527.t01, mNeoVis.evergladensis.2.1.chr1.g00529.t01, mNeoVis.evergladensis.2.1.chr1.g00530.t01    mNeoVis.vulgivaga.2.1.chr1.g00535.t01, mNeoVis.vulgivaga.2.1.chr1.g00536.t01, mNeoVis.vulgivaga.2.1.chr1.g00538.t01, mNeoVis.vulgivaga.2.1.chr1.g00539.t01      207    N0.HOG0000205   OG0000006  n125   rna-XM_034651155.1, rna-XM_034656173.1    rna-XM_038530436.1    rna-XM_022492707.1, rna-XM_022493195.1, rna-XM_022493375.1    rna-XM_019833431.3      rna-XM_039236172.1, rna-XM_039236174.1, rna-XM_039236187.1, rna-XM_039236189.1    rna-XM_032850853.1, rna-XM_032850854.1      rna-XM_032336431.1      rna-XM_059387404.1, rna-XM_059387447.1, rna-XM_059387477.1, rna-XM_059416362.1, rna-XM_059416363.1    rna-XM_013052386.2, rna-XM_045075654.1    mNeoVis.2.1.1.chr1_hap1.g00517.t01    rna-XM_021693187.2, rna-XM_021702071.2, rna-XM_021702158.2, rna-XM_044912387.1, rna-XM_044912405.1    rna-XM_004397753.1, rna-XM_004397754.1, rna-XM_004397860.1, rna-XM_004397861.1, rna-XM_004397863.1, rna-XM_004397866.1    rna-XM_007077909.3, rna-XM_042985607.1, rna-XM_042985623.1    rna-XM_040622332.1    rna-XM_035727490.1    mNeoVis.3.1.1.chr1.g00524.t01, mNeoVis.3.1.1.chr1.g00525.t01, mNeoVis.3.1.1.chr1.g00526.t01, mNeoVis.3.1.1.chr1.g00529.t01, mNeoVis.3.1.1.chr1.g00530.t01    rna-gnl|WGS_ZZZZ|egapxtmp_028077-R1, rna-gnl|WGS_ZZZZ|egapxtmp_028339-R1, rna-gnl|WGS_ZZZZ|egapxtmp_028353-R4, rna-gnl|WGS_ZZZZ|egapxtmp_032142-R1    mNeoVis.evergladensis.2.1.chr1.g00519.t01, mNeoVis.evergladensis.2.1.chr1.g00520.t01, mNeoVis.evergladensis.2.1.chr1.g00524.t01, mNeoVis.evergladensis.2.1.chr1.g00525.t01    mNeoVis.vulgivaga.2.1.chr1.g00532.t01, mNeoVis.vulgivaga.2.1.chr1.g00533.t01      208    N0.HOG0000206   OG0000006  n177   rna-XM_034652743.1, rna-XM_034656145.1, rna-XM_034656170.1    rna-XM_038530401.1    rna-XM_022492660.1, rna-XM_022493371.1        rna-XM_039236152.1      rna-XM_046000742.1        rna-XM_059387589.1      mNeoVis.2.1.1.chr1_hap1.g00521.t01    rna-XM_021701905.2, rna-XM_044912469.1    rna-XM_004397856.1, rna-XM_004397858.1, rna-XM_004417768.1        rna-XM_035727458.1    mNeoVis.3.1.1.chr1.g00533.t01, mNeoVis.3.1.1.chr1.g00537.t01    rna-gnl|WGS_ZZZZ|egapxtmp_006440-R1          209    N0.HOG0000207   OG0000006  n195   rna-XM_034656171.1      rna-XM_022493374.1        rna-XM_039236186.1            rna-XM_059387420.1        rna-XM_021694353.1    rna-XM_004397752.1    rna-XM_042959081.1        mNeoVis.3.1.1.chr1.g00528.t01    rna-gnl|WGS_ZZZZ|egapxtmp_028511-R1    mNeoVis.evergladensis.2.1.chr1.g00522.t01    mNeoVis.vulgivaga.2.1.chr1.g00534.t01      210    N0.HOG0000208   OG0000006  n205   rna-XM_034656172.1      rna-XM_022492629.1, rna-XM_022492745.1            rna-XM_046000762.1        rna-XM_059387446.1    rna-XM_045075371.1    mNeoVis.2.1.1.chr1_hap1.g00519.t01, mNeoVis.2.1.1.chr1_hap1.g00525.t01    rna-XM_021693265.1, rna-XM_021694253.2    rna-XM_004397854.1, rna-XM_004417686.1, rna-XM_004417694.1          mNeoVis.3.1.1.chr1.g00540.t01    rna-gnl|WGS_ZZZZ|egapxtmp_006496-R1    mNeoVis.evergladensis.2.1.chr1.g00531.t01    mNeoVis.vulgivaga.2.1.chr1.g00540.t01      211    N0.HOG0000209   OG0000006  n223   rna-XM_002912550.4, rna-XM_002912551.4, rna-XM_019795555.2, rna-XM_034656140.1, rna-XM_034656144.1    rna-XM_038530384.1    rna-XM_022492721.1    rna-XM_019833383.3, rna-XM_023256016.2, rna-XM_023256043.2    rna-XM_036123467.1, rna-XM_036123566.1    rna-XM_039236144.1, rna-XM_039236194.1    rna-XM_032851057.1, rna-XM_032851132.1    rna-XM_045998865.1, rna-XM_045998866.1, rna-XM_045998875.1    rna-XM_032336444.1, rna-XM_032336448.1, rna-XM_032336455.1    rna-XM_059174827.1, rna-XM_059174834.1, rna-XM_059174836.1    rna-XM_059387419.1, rna-XM_059416359.1    rna-XM_045075377.1, rna-XM_045075379.1, rna-XM_045075386.1    mNeoVis.2.1.1.chr1_hap1.g00511.t01, mNeoVis.2.1.1.chr1_hap1.g00512.t01      rna-XM_004397871.1, rna-XM_004417627.1    rna-XM_042985523.1, rna-XM_042985594.1    rna-XM_008691481.2, rna-XM_040621929.1, rna-XM_040622335.1    rna-XM_027604453.2, rna-XM_035727768.1    mNeoVis.3.1.1.chr1.g00513.t01, mNeoVis.3.1.1.chr1.g00517.t01, mNeoVis.3.1.1.chr1.g00518.t01, mNeoVis.3.1.1.chr1.g00519.t01, mNeoVis.3.1.1.chr1.g00520.t01    rna-gnl|WGS_ZZZZ|egapxtmp_031890-R1, rna-gnl|WGS_ZZZZ|egapxtmp_031987-R1    mNeoVis.evergladensis.2.1.chr1.g00515.t01    mNeoVis.vulgivaga.2.1.chr1.g00526.t01      212    N0.HOG0000210   OG0000006  n274   rna-XM_034651165.1    rna-XM_038530397.1    rna-XM_022492712.1    rna-XM_023256049.2    rna-XM_036123564.1      rna-XM_032851113.1    rna-XM_046000667.1    rna-XM_032336437.1    rna-XM_059174841.1    rna-XM_059416361.1    rna-XM_013050971.2        rna-XM_004397867.1    rna-XM_042985600.1    rna-XM_008691651.2    rna-XM_035727500.1              213    N0.HOG0000211   OG0000006  n288   rna-XM_034656143.1    rna-XM_038530388.1      rna-XM_023256044.2    rna-XM_036123513.1    rna-XM_039225768.1    rna-XM_032851130.1    rna-XM_045998652.1    rna-XM_032336435.1    rna-XM_059174840.1    rna-XM_059387426.1    rna-XM_013052402.2    mNeoVis.2.1.1.chr1_hap1.g00514.t01        rna-XM_042985534.1    rna-XM_008691650.2    rna-XM_035727492.1      rna-gnl|WGS_ZZZZ|egapxtmp_006401-R1      mNeoVis.vulgivaga.2.1.chr1.g00529.t01      214    N0.HOG0000212   OG0000006  n305   rna-XM_002912555.4    rna-XM_038530380.1    rna-XM_022493422.1    rna-XM_023255991.2, rna-XM_045039938.1    rna-XM_036123552.1    rna-XM_039236140.1    rna-XM_032849996.1    rna-XM_046001208.1    rna-XM_032336452.1    rna-XM_059174828.1    rna-XM_059416358.1    rna-XM_045075384.1      rna-XM_021702107.1    rna-XM_004397760.1    rna-XM_015535615.2, rna-XM_042985499.1    rna-XM_008691704.2      mNeoVis.3.1.1.chr1.g00514.t01    rna-gnl|WGS_ZZZZ|egapxtmp_031790-R1      mNeoVis.vulgivaga.2.1.chr1.g00523.t01      215    N0.HOG0000213   OG0000006  n326   rna-XM_034656138.1    rna-XM_038530382.1    rna-XM_022493380.1    rna-XM_045060255.1      rna-XM_039236143.1    rna-XM_032850003.1    rna-XM_045998667.1    rna-XM_032336454.1    rna-XM_059174829.1      rna-XM_045075383.1    mNeoVis.2.1.1.chr1_hap1.g00509.t01    rna-XM_021702108.1    rna-XM_004397759.1    rna-XM_042985505.1    rna-XM_008691477.2    rna-XM_035727499.1        mNeoVis.evergladensis.2.1.chr1.g00513.t01    mNeoVis.vulgivaga.2.1.chr1.g00524.t01      216    N0.HOG0000214   OG0000006  n343   rna-XM_002912552.4, rna-XM_002912553.4    rna-XM_038530375.1    rna-XM_022492737.1    rna-XM_045060315.1    rna-XM_036123481.1    rna-XM_039236131.1, rna-XM_039236137.1    rna-XM_032849999.1    rna-XM_045998864.1, rna-XM_046001185.1    rna-XM_032336442.1, rna-XM_032336443.1, rna-XM_032336450.1    rna-XM_059174831.1    rna-XM_059387437.1    rna-XM_045075380.1, rna-XM_045075382.1    mNeoVis.2.1.1.chr1_hap1.g00508.t01      rna-XM_004397870.1    rna-XM_042985487.1    rna-XM_040622334.1    rna-XM_035727491.1, rna-XM_035727494.1    mNeoVis.3.1.1.chr1.g00515.t01, mNeoVis.3.1.1.chr1.g00516.t01      mNeoVis.evergladensis.2.1.chr1.g00514.t01    mNeoVis.vulgivaga.2.1.chr1.g00525.t01      217    N0.HOG0000215   OG0000006  n371   rna-XM_002912549.4        rna-XM_023256035.2    rna-XM_036123247.1    rna-XM_039236132.1    rna-XM_032850994.1    rna-XM_046001085.1    rna-XM_032336440.1        rna-XM_045075376.1        rna-XM_004397758.1    rna-XM_042985565.1    rna-XM_008691489.2    rna-XM_035727496.1              218    N0.HOG0000216   OG0000006  n383   rna-XM_002912603.4, rna-XM_034656139.1    rna-XM_038530386.1    rna-XM_022493379.1    rna-XM_023256012.2, rna-XM_023256023.2    rna-XM_036123527.1, rna-XM_036123563.1    rna-XM_039236141.1, rna-XM_039236191.1    rna-XM_032851099.1, rna-XM_032851150.1    rna-XM_045998861.1    rna-XM_032336441.1, rna-XM_032336446.1, rna-XM_032336447.1    rna-XM_059174833.1, rna-XM_059174835.1    rna-XM_059387398.1, rna-XM_059387458.1    rna-XM_045075381.1    mNeoVis.2.1.1.chr1_hap1.g00510.t01, mNeoVis.2.1.1.chr1_hap1.g00513.t01    rna-XM_044917055.1    rna-XM_004397869.1, rna-XM_004417721.1    rna-XM_042985513.1, rna-XM_042985572.1    rna-XM_008691480.2, rna-XM_008691482.2    rna-XM_035727498.1      rna-gnl|WGS_ZZZZ|egapxtmp_032044-R1    mNeoVis.evergladensis.2.1.chr1.g00516.t01    mNeoVis.vulgivaga.2.1.chr1.g00527.t01      219    N0.HOG0000217   OG0000006  n417   rna-XM_019795563.2    rna-XM_038530377.1    rna-XM_022493421.1    rna-XM_011282592.4    rna-XM_036123532.1    rna-XM_039236139.1    rna-XM_032849998.1    rna-XM_045998766.1    rna-XM_032336451.1    rna-XM_059174826.1    rna-XM_059416357.1    rna-XM_045075385.1        rna-XM_004397761.1    rna-XM_042985493.1    rna-XM_008691496.2    rna-XM_035727767.1    mNeoVis.3.1.1.chr1.g00512.t01            220    N0.HOG0000218   OG0000006  n434   rna-XM_034656142.1    rna-XM_038530396.1    rna-XM_022493376.1    rna-XM_023256060.2    rna-XM_036123544.1    rna-XM_039236193.1    rna-XM_032850633.1    rna-XM_046000666.1    rna-XM_032336438.1    rna-XM_059174838.1    rna-XM_059416360.1    rna-XM_013052301.2      rna-XM_021702110.2    rna-XM_004397868.1      rna-XM_008691485.2    rna-XM_035727497.1              221    N0.HOG0000219   OG0000006  n449   rna-XM_034656141.1    rna-XM_038530387.1    rna-XM_022493378.1    rna-XM_019833357.3      rna-XM_039236142.1    rna-XM_032850987.1      rna-XM_032336439.1    rna-XM_059174837.1      rna-XM_045075375.1        rna-XM_004397757.1    rna-XM_042985560.1    rna-XM_008691484.2    rna-XM_035727495.1          mNeoVis.vulgivaga.2.1.chr1.g00528.t01      222    N0.HOG0000220   OG0000007  n1   rna-XM_002930631.2    rna-NM_001388636.1, rna-XM_038562486.1    rna-XM_022507201.1    rna-XM_045039292.1    rna-XM_036073548.1    rna-XM_039226790.1    rna-XM_032844617.1    rna-XM_046015841.1    rna-XM_032359821.1    rna-XM_059132841.1    rna-XM_059387138.1, rna-XM_059387145.1    rna-XM_004782060.3    mNeoVis.2.1.1.chr7_hap1.g02035.t01    rna-XM_021685189.1    rna-XM_004399504.1    rna-XM_007082203.3    rna-XM_008700939.1    rna-XM_035722905.1    mNeoVis.3.1.1.chr7.g01899.t01    rna-gnl|WGS_ZZZZ|egapxtmp_013546-R1    mNeoVis.evergladensis.2.1.chr7.g01994.t01    mNeoVis.vulgivaga.2.1.chr7.g02016.t01      223    N0.HOG0000221   OG0000007  n24   rna-XM_002930630.3    rna-NM_001388640.1, rna-XM_038562483.1    rna-XM_022506759.1    rna-XM_045039291.1      rna-XM_039226788.1    rna-XM_032844557.1    rna-XM_046016016.1    rna-XM_032358257.1    rna-XM_059158710.1    rna-XM_059386874.1, rna-XM_059386884.1    rna-XM_004782056.1    mNeoVis.2.1.1.chr7_hap1.g02033.t01        rna-XM_007082178.3    rna-XM_008700941.2      mNeoVis.3.1.1.chr7.g01897.t01    rna-gnl|WGS_ZZZZ|egapxtmp_013542-R1      mNeoVis.vulgivaga.2.1.chr7.g02014.t01      224   N0.HOG0000222  OG0000007  n43     rna-NM_001388910.1    rna-XM_022507199.1    rna-XM_011286841.3      rna-XM_039226760.1    rna-XM_032844593.1    rna-XM_046017056.1    rna-XM_032359309.1    rna-XM_059133426.1      rna-XM_013050573.1          rna-XM_007095407.2    rna-XM_008701072.2      mNeoVis.3.1.1.chr7.g01908.t01      mNeoVis.evergladensis.2.1.chr7.g02003.t01    mNeoVis.vulgivaga.2.1.chr7.g02025.t01      225   N0.HOG0000223  OG0000007  n55   rna-XM_034646213.1    rna-NM_001388909.1    rna-XM_022506902.1    rna-XM_019812963.3    rna-XM_036073542.1    rna-XM_039226801.1    rna-XM_032844581.1    rna-XM_046016697.1    rna-XM_032359209.1, rna-XM_032359631.1    rna-XM_059158771.1, rna-XM_059158778.1    rna-XM_059387344.1, rna-XM_059387561.1    rna-XM_004781648.2, rna-XM_004782460.2    mNeoVis.2.1.1.chr7_hap1.g02041.t01        rna-XM_007095401.1    rna-XM_008700931.1      mNeoVis.3.1.1.chr7.g01905.t01      mNeoVis.evergladensis.2.1.chr7.g02000.t01    mNeoVis.vulgivaga.2.1.chr7.g02023.t01      226    N0.HOG0000224   OG0000007  n77   rna-XM_002930388.2, rna-XM_011217181.3, rna-XM_011217186.2, rna-XM_034644807.1, rna-XM_034646283.1, rna-XM_034646377.1, rna-XM_034651354.1, rna-XM_034651808.1    rna-XM_038562485.1, rna-XM_038562487.1    rna-XM_022507203.1, rna-XM_022507204.1    rna-XM_006944301.2, rna-XM_023249654.2      rna-XM_039226800.1    rna-XM_032844532.1, rna-XM_032844615.1, rna-XM_032844616.1    rna-XM_046015843.1, rna-XM_046015844.1, rna-XM_046015846.1, rna-XM_046015848.1, rna-XM_046015849.1, rna-XM_046017270.1    rna-XM_032358330.1, rna-XM_032359813.1, rna-XM_032359814.1, rna-XM_032359815.1, rna-XM_032359816.1, rna-XM_032359817.1, rna-XM_032359820.1, rna-XM_032359823.1    rna-XM_059187540.1, rna-XM_059187629.1, rna-XM_059187645.1, rna-XM_059187742.1, rna-XM_059188229.1, rna-XM_059188358.1, rna-XM_059188478.1, rna-XM_059188611.1    rna-XM_059387133.1, rna-XM_059387134.1, rna-XM_059387136.1, rna-XM_059387137.1, rna-XM_059387139.1, rna-XM_059387140.1, rna-XM_059387142.1, rna-XM_059387143.1, rna-XM_059388816.1    rna-XM_004763222.2, rna-XM_004763223.1, rna-XM_004763224.2, rna-XM_004782061.2, rna-XM_045067383.1, rna-XM_045067387.1, rna-XM_045067395.1, rna-XM_045067826.1    mNeoVis.2.1.1.chr7_hap1.g02028.t01, mNeoVis.2.1.1.chr7_hap1.g02029.t01, mNeoVis.2.1.1.chr7_hap1.g02031.t01, mNeoVis.2.1.1.chr7_hap1.g02032.t01        rna-XM_007082201.2    rna-XM_008700943.2, rna-XM_008700944.2, rna-XM_008700947.2, rna-XM_008700948.2, rna-XM_008700949.1, rna-XM_008700951.2, rna-XM_008700952.1, rna-XM_008701078.2, rna-XM_008701079.2, rna-XM_040638708.1    rna-XM_027579746.1    mNeoVis.3.1.1.chr7.g01894.t01, mNeoVis.3.1.1.chr7.g01895.t01, mNeoVis.3.1.1.chr7.g01896.t01    rna-gnl|WGS_ZZZZ|egapxtmp_010295-R1, rna-gnl|WGS_ZZZZ|egapxtmp_012608-R1, rna-gnl|WGS_ZZZZ|egapxtmp_012609-R1, rna-gnl|WGS_ZZZZ|egapxtmp_012611-R1, rna-gnl|WGS_ZZZZ|egapxtmp_012612-R1    mNeoVis.evergladensis.2.1.chr7.g01991.t01, mNeoVis.evergladensis.2.1.chr7.g01992.t01    mNeoVis.vulgivaga.2.1.chr7.g02011.t01, mNeoVis.vulgivaga.2.1.chr7.g02012.t01, mNeoVis.vulgivaga.2.1.chr7.g02013.t01      227    N0.HOG0000225   OG0000007  n158   rna-XM_019793192.2    rna-XM_038562484.1    rna-XM_022507666.1    rna-XM_003993292.5      rna-XM_039226779.1      rna-XM_046015845.1    rna-XM_032359819.1, rna-XM_032359822.1    rna-XM_059187845.1    rna-XM_059387135.1, rna-XM_059387141.1, rna-XM_059387144.1    rna-XM_013050961.2    mNeoVis.2.1.1.chr7_hap1.g02030.t01        rna-XM_007082202.2    rna-XM_008700945.2, rna-XM_008700950.2, rna-XM_008701080.2, rna-XM_040638709.1    rna-XM_027580559.1      rna-gnl|WGS_ZZZZ|egapxtmp_012610-R1    mNeoVis.evergladensis.2.1.chr7.g01990.t01    mNeoVis.vulgivaga.2.1.chr7.g02002.t01      228    N0.HOG0000226   OG0000007  n179   rna-XM_002930241.4, rna-XM_002931231.2, rna-XM_034644809.1    rna-NM_001388633.1, rna-XM_038563261.1    rna-XM_022506912.1, rna-XM_022507197.1, rna-XM_022507242.1    rna-XM_003993304.4, rna-XM_023239013.2      rna-XM_039226830.1    rna-XM_032844622.1, rna-XM_032844626.1    rna-XM_046015838.1, rna-XM_046017445.1, rna-XM_046017751.1    rna-XM_032358741.1, rna-XM_032359826.1, rna-XM_032359832.1, rna-XM_032359833.1    rna-XM_059133497.1, rna-XM_059133979.1, rna-XM_059158772.1, rna-XM_059158854.1    rna-XM_059386847.1, rna-XM_059387045.1, rna-XM_059387216.1, rna-XM_059388689.1    rna-XM_004781643.3, rna-XM_013050576.1, rna-XM_013050700.1    mNeoVis.2.1.1.chr7_hap1.g02044.t01, mNeoVis.2.1.1.chr7_hap1.g02053.t01, mNeoVis.2.1.1.chr7_hap1.g02063.t01        rna-XM_007095402.1, rna-XM_015543476.2    rna-XM_008700916.1, rna-XM_008701070.2, rna-XM_008701071.2    rna-XM_035722972.1    mNeoVis.3.1.1.chr7.g01909.t01, mNeoVis.3.1.1.chr7.g01916.t01, mNeoVis.3.1.1.chr7.g01920.t01    rna-gnl|WGS_ZZZZ|egapxtmp_012617-R1    mNeoVis.evergladensis.2.1.chr7.g02004.t01, mNeoVis.evergladensis.2.1.chr7.g02019.t01    mNeoVis.vulgivaga.2.1.chr7.g02026.t01, mNeoVis.vulgivaga.2.1.chr7.g02031.t01, mNeoVis.vulgivaga.2.1.chr7.g02038.t01      229   N0.HOG0000227  OG0000007  n227   rna-XM_002930383.4    rna-NM_001388822.1    rna-XM_022506980.1    rna-XM_045039293.1      rna-XM_039226793.1    rna-XM_032844562.1    rna-XM_046016648.1    rna-XM_032359166.1    rna-XM_059158668.1    rna-XM_059386654.1, rna-XM_059386927.1    rna-XM_004763028.1    mNeoVis.2.1.1.chr7_hap1.g02026.t01        rna-XM_007082177.1    rna-XM_008700954.2      mNeoVis.3.1.1.chr7.g01892.t01    rna-gnl|WGS_ZZZZ|egapxtmp_010294-R1    mNeoVis.evergladensis.2.1.chr7.g01988.t01    mNeoVis.vulgivaga.2.1.chr7.g02009.t01      230    N0.HOG0000228   OG0000007  n241   rna-XM_011217182.2    rna-NM_001389204.1    rna-XM_022506973.1    rna-XM_045039294.1      rna-XM_039226789.1    rna-XM_032844559.1    rna-XM_046016968.1    rna-XM_032359188.1    rna-XM_059158683.1      rna-XM_004763027.3    mNeoVis.2.1.1.chr7_hap1.g02027.t01        rna-XM_007082200.3    rna-XM_008700953.2      mNeoVis.3.1.1.chr7.g01893.t01    rna-gnl|WGS_ZZZZ|egapxtmp_010300-R1    mNeoVis.evergladensis.2.1.chr7.g01989.t01    mNeoVis.vulgivaga.2.1.chr7.g02010.t01      231    N0.HOG0000229   OG0000007  n255   rna-XM_034645999.1    rna-XM_038562480.1    rna-XM_022506742.1    rna-XM_019813046.3      rna-XM_039226693.1    rna-XM_032844556.1    rna-XM_046016626.1    rna-XM_032359101.1    rna-XM_059158842.1    rna-XM_059386837.1, rna-XM_059386924.1    rna-XM_013050572.2    mNeoVis.2.1.1.chr7_hap1.g02055.t01        rna-XM_015543496.2    rna-XM_008700923.2      mNeoVis.3.1.1.chr7.g01917.t01    rna-gnl|WGS_ZZZZ|egapxtmp_010310-R1    mNeoVis.evergladensis.2.1.chr7.g02013.t01    mNeoVis.vulgivaga.2.1.chr7.g02033.t01      232    N0.HOG0000230   OG0000007  n272   rna-XM_002930629.1    rna-NM_001388637.1, rna-NM_001388639.1    rna-XM_022506775.1    rna-XM_045039295.1      rna-XM_039226787.1      rna-XM_046016017.1    rna-XM_032358256.1    rna-XM_059158716.1    rna-XM_059386878.1, rna-XM_059386888.1    rna-XM_004782055.1    mNeoVis.2.1.1.chr7_hap1.g02034.t01      rna-XM_004399505.1    rna-XM_007082179.2    rna-XM_008700940.2      mNeoVis.3.1.1.chr7.g01898.t01    rna-gnl|WGS_ZZZZ|egapxtmp_013544-R1    mNeoVis.evergladensis.2.1.chr7.g01993.t01    mNeoVis.vulgivaga.2.1.chr7.g02015.t01      233    N0.HOG0000231   OG0000007  n289   rna-XM_002929984.2, rna-XM_034646190.1    rna-XM_038562473.1, rna-XM_038562474.1, rna-XM_038588901.1, rna-XM_038589276.1, rna-XM_038589635.1, rna-XM_038589674.1    rna-XM_022506969.1    rna-XM_003993261.2      rna-XM_039225297.1, rna-XM_039225298.1    rna-XM_032844579.1, rna-XM_032844583.1    rna-XM_046016363.1, rna-XM_046016705.1, rna-XM_046016707.1    rna-XM_032359700.1, rna-XM_032360075.1, rna-XM_032360077.1    rna-XM_059158859.1, rna-XM_059158871.1, rna-XM_059158873.1    rna-XM_059388272.1, rna-XM_059388340.1    rna-XM_004781915.1, rna-XM_004782614.2, rna-XM_045068083.1    mNeoVis.2.1.1.chr7_hap1.g02067.t01, mNeoVis.2.1.1.chr7_hap1.g02078.t01    rna-XM_021684824.1    rna-XM_004411930.1, rna-XM_004411981.1    rna-XM_007095371.2    rna-XM_008700911.2, rna-XM_040639811.1    rna-XM_027580066.2    mNeoVis.3.1.1.chr7.g01922.t01, mNeoVis.3.1.1.chr7.g01927.t01, mNeoVis.3.1.1.chr7.g01928.t01    rna-gnl|WGS_ZZZZ|egapxtmp_010289-R1, rna-gnl|WGS_ZZZZ|egapxtmp_013545-R1    mNeoVis.evergladensis.2.1.chr7.g02023.t01, mNeoVis.evergladensis.2.1.chr7.g02033.t01    mNeoVis.vulgivaga.2.1.chr7.g02041.t01, mNeoVis.vulgivaga.2.1.chr7.g02050.t01      234   N0.HOG0000232  OG0000007  n330   rna-XM_002931341.2, rna-XM_034646223.1, rna-XM_034650532.1, rna-XM_034651907.1, rna-XM_034653265.1    rna-NM_001388624.1, rna-NM_001388625.1    rna-XM_022506982.1, rna-XM_022507186.1    rna-XM_003993268.2    rna-XM_036073535.1    rna-XM_039225295.1, rna-XM_039225296.1    rna-XM_032846465.1, rna-XM_032846473.1, rna-XM_032846566.1, rna-XM_032846567.1    rna-XM_046015674.1, rna-XM_046015675.1, rna-XM_046015677.1, rna-XM_046015678.1, rna-XM_046015816.1, rna-XM_046016413.1, rna-XM_046016485.1, rna-XM_046016500.1, rna-XM_046016570.1, rna-XM_046016578.1, rna-XM_046016651.1, rna-XM_046016750.1, rna-XM_046016784.1, rna-XM_046017154.1    rna-XM_032358789.1, rna-XM_032358878.1, rna-XM_032359629.1, rna-XM_032359683.1, rna-XM_032359982.1, rna-XM_032360367.1    rna-XM_059158878.1, rna-XM_059158879.1, rna-XM_059158882.1, rna-XM_059158883.1, rna-XM_059158884.1, rna-XM_059158886.1    rna-XM_059388180.1, rna-XM_059388184.1, rna-XM_059388190.1, rna-XM_059388201.1, rna-XM_059388221.1, rna-XM_059388229.1    rna-XM_004782168.1, rna-XM_004782169.1, rna-XM_004782170.1, rna-XM_004782171.2, rna-XM_004782175.1, rna-XM_004782177.2    mNeoVis.2.1.1.chr7_hap1.g02088.t01, mNeoVis.2.1.1.chr7_hap1.g02090.t01, mNeoVis.2.1.1.chr7_hap1.g02092.t01, mNeoVis.2.1.1.chr7_hap1.g02093.t01, mNeoVis.2.1.1.chr7_hap1.g02095.t01    rna-XM_021684827.1    rna-XM_004411979.1, rna-XM_004411980.1, rna-XM_004412004.1    rna-XM_015540516.2, rna-XM_042959251.1    rna-XM_008694060.2, rna-XM_008694063.2, rna-XM_008694066.2, rna-XM_008694067.2, rna-XM_008694068.1, rna-XM_008694207.2    rna-XM_027579677.1, rna-XM_027580031.2, rna-XM_027580060.2, rna-XM_027580061.1, rna-XM_027580062.2, rna-XM_027580075.2    mNeoVis.3.1.1.chr7.g01938.t01    rna-gnl|WGS_ZZZZ|egapxtmp_013541-R1, rna-gnl|WGS_ZZZZ|egapxtmp_013543-R1, rna-gnl|WGS_ZZZZ|egapxtmp_013549-R1, rna-gnl|WGS_ZZZZ|egapxtmp_013553-R1, rna-gnl|WGS_ZZZZ|egapxtmp_013554-R1    mNeoVis.evergladensis.2.1.chr7.g02042.t01, mNeoVis.evergladensis.2.1.chr7.g02043.t01, mNeoVis.evergladensis.2.1.chr7.g02045.t01    mNeoVis.vulgivaga.2.1.chr7.g02060.t01, mNeoVis.vulgivaga.2.1.chr7.g02062.t01, mNeoVis.vulgivaga.2.1.chr7.g02063.t01, mNeoVis.vulgivaga.2.1.chr7.g02064.t01, mNeoVis.vulgivaga.2.1.chr7.g02065.t01, mNeoVis.vulgivaga.2.1.chr7.g02066.t01, mNeoVis.vulgivaga.2.1.chr7.g02067.t01      235   N0.HOG0000233  OG0000008  n1   rna-XM_034643892.1, rna-XM_034645559.1    rna-XM_038577729.1    rna-XM_022519356.1    rna-XM_019834974.3    rna-XM_036096407.1    rna-XM_039241396.1    rna-XM_032876730.1    rna-XM_046010565.1    rna-XM_032348274.1    rna-XM_059185280.1    rna-XM_059403181.1    rna-XM_004774693.3    mNeoVis.2.1.1.chr9_hap1.g00530.t01    rna-XM_021704935.1    rna-XM_004400796.1    rna-XM_042992354.1    rna-XM_008709422.1, rna-XM_040641953.1    rna-XM_027592388.2    mNeoVis.3.1.1.chr10.g00501.t01    rna-gnl|WGS_ZZZZ|egapxtmp_022629-R1    mNeoVis.evergladensis.2.1.chr10.g00522.t01    mNeoVis.vulgivaga.2.1.chr10.g00509.t01      236    N0.HOG0000234   OG0000008  n23   rna-XM_034645321.1    rna-XM_038577738.1    rna-XM_022519472.1    rna-XM_006933719.3    rna-XM_036096395.1    rna-XM_039224947.1    rna-XM_032876718.1    rna-XM_046012316.1    rna-XM_032348296.1    rna-XM_059185272.1    rna-XM_059403178.1    rna-XM_004774686.3    mNeoVis.2.1.1.chr9_hap1.g00519.t01    rna-XM_021686665.1      rna-XM_007073056.2    rna-XM_040642505.1    rna-XM_027592386.2    mNeoVis.3.1.1.chr10.g00492.t01    rna-gnl|WGS_ZZZZ|egapxtmp_022622-R1    mNeoVis.evergladensis.2.1.chr10.g00511.t01        237    N0.HOG0000235   OG0000008  n45   rna-XM_034652631.1, rna-XM_034652685.1                                                238    N0.HOG0000236   OG0000008  n47   rna-XM_002923246.4    rna-NM_001346042.1    rna-XM_022519515.1    rna-XM_003988743.5    rna-XM_036090530.1    rna-XM_039255091.1    rna-XM_032876699.1    rna-XM_046012470.1    rna-XM_032348314.1    rna-XM_059183568.1    rna-XM_059404866.1    rna-XM_004774674.3    mNeoVis.2.1.1.chr9_hap1.g00507.t01    rna-XM_021686674.2    rna-XM_004406247.1    rna-XM_007081360.3    rna-XM_040642511.1    rna-XM_027592396.1    mNeoVis.3.1.1.chr10.g00479.t01    rna-gnl|WGS_ZZZZ|egapxtmp_024671-R1      mNeoVis.vulgivaga.2.1.chr10.g00488.t01      239   N0.HOG0000237  OG0000008  n66   rna-XM_011229904.3    rna-NM_001346015.1    rna-XM_022519495.1    rna-XM_003988742.5    rna-XM_036090691.1    rna-XM_039255082.1    rna-XM_032876717.1    rna-XM_046012427.1    rna-XM_032348315.1    rna-XM_059185257.1    rna-XM_059404865.1    rna-XM_004774675.3      rna-XM_021686673.1    rna-XM_004406248.1    rna-XM_007081545.3    rna-XM_008704618.2    rna-XM_027592398.2    mNeoVis.3.1.1.chr10.g00480.t01    rna-gnl|WGS_ZZZZ|egapxtmp_022619-R1    mNeoVis.evergladensis.2.1.chr10.g00499.t01    mNeoVis.vulgivaga.2.1.chr10.g00489.t01      240    N0.HOG0000238   OG0000008  n87   rna-XM_011229956.3    rna-XM_038577739.1    rna-XM_022519451.1    rna-XM_011283953.4    rna-XM_036096421.1    rna-XM_039255080.1    rna-XM_032876715.1    rna-XM_046012271.1    rna-XM_032348298.1    rna-XM_059185261.1    rna-XM_059403188.1    rna-XM_004774680.1      rna-XM_021686752.1    rna-XM_004406254.1    rna-XM_042992359.1    rna-XM_040642507.1    rna-XM_027592379.2    mNeoVis.3.1.1.chr10.g00487.t01      mNeoVis.evergladensis.2.1.chr10.g00505.t01    mNeoVis.vulgivaga.2.1.chr10.g00497.t01      241    N0.HOG0000239   OG0000008  n107   rna-XM_011229902.3    rna-XM_038577740.1    rna-XM_022519454.1    rna-NM_001278857.1    rna-XM_036096422.1    rna-XM_039255077.1    rna-XM_032876720.1    rna-XM_046012340.1    rna-XM_032348297.1    rna-XM_059183574.1    rna-XM_059404856.1    rna-XM_004774746.3      rna-XM_021686751.1    rna-XM_004406255.1    rna-XM_007081354.2    rna-XM_008704615.1    rna-XM_027592378.2      rna-gnl|WGS_ZZZZ|egapxtmp_022621-R1    mNeoVis.evergladensis.2.1.chr10.g00506.t01    mNeoVis.vulgivaga.2.1.chr10.g00498.t01      242    N0.HOG0000240   OG0000008  n128   rna-XM_002923244.4    rna-NM_001197029.1    rna-XM_022519474.1    rna-NM_001195239.1    rna-XM_036096368.1    rna-XM_039255078.1    rna-XM_032876721.1    rna-XM_046012357.1    rna-XM_032348304.1    rna-XM_059183576.1    rna-XM_059404854.1    rna-XM_045073244.1    mNeoVis.2.1.1.chr9_hap1.g00513.t01    rna-XM_044914834.1    rna-XM_004406256.1    rna-XM_015537244.2    rna-XM_040642506.1    rna-XM_027592380.1        mNeoVis.evergladensis.2.1.chr10.g00507.t01    mNeoVis.vulgivaga.2.1.chr10.g00499.t01      243    N0.HOG0000241   OG0000008  n146   rna-XM_011229901.3    rna-NM_001346046.1    rna-XM_022519455.1    rna-NM_001290235.1    rna-XM_036096419.1    rna-XM_039255076.1    rna-XM_032876719.1    rna-XM_046012339.1    rna-XM_032348294.1    rna-XM_059185263.1    rna-XM_059404857.1    rna-XM_004774679.2      rna-XM_021686755.1    rna-XM_004406266.1    rna-XM_007081352.3    rna-XM_008704617.2    rna-XM_035729201.1    mNeoVis.3.1.1.chr10.g00486.t01      mNeoVis.evergladensis.2.1.chr10.g00504.t01    mNeoVis.vulgivaga.2.1.chr10.g00496.t01      244    N0.HOG0000242   OG0000008  n167   rna-XM_034644789.1    rna-NM_001346012.1    rna-XM_022519417.1    rna-XM_023257101.2    rna-XM_036090349.1    rna-XM_039255072.1    rna-XM_032876703.1    rna-XM_046012246.1    rna-XM_032348313.1    rna-XM_059183569.1    rna-XM_059404862.1    rna-XM_045073070.1    mNeoVis.2.1.1.chr9_hap1.g00508.t01    rna-XM_021686758.1    rna-XM_004406249.1    rna-XM_042990832.1    rna-XM_040641928.1    rna-XM_027592377.1    mNeoVis.3.1.1.chr10.g00481.t01        mNeoVis.vulgivaga.2.1.chr10.g00490.t01      245    N0.HOG0000243   OG0000008  n186   rna-XM_011229898.3    rna-NM_001346020.1    rna-XM_022519460.1    rna-XM_045061845.1    rna-XM_036090229.1    rna-XM_039255073.1    rna-XM_032876709.1    rna-XM_046012257.1    rna-XM_032348308.1    rna-XM_059183572.1    rna-XM_059404861.1    rna-XM_045073072.1    mNeoVis.2.1.1.chr9_hap1.g00509.t01    rna-XM_021686757.1    rna-XM_004417733.1    rna-XM_042990831.1    rna-XM_040641926.1    rna-XM_027592375.1    mNeoVis.3.1.1.chr10.g00482.t01    rna-gnl|WGS_ZZZZ|egapxtmp_022616-R1    mNeoVis.evergladensis.2.1.chr10.g00500.t01    mNeoVis.vulgivaga.2.1.chr10.g00491.t01      246    N0.HOG0000244   OG0000008  n208     rna-XM_038577929.1    rna-XM_022519390.1      rna-XM_036087042.1, rna-XM_036096371.1    rna-XM_039255100.1    rna-XM_032876506.1              mNeoVis.2.1.1.chr9_hap1.g00510.t01    rna-XM_021686737.1        rna-XM_040641957.1    rna-XM_027594273.1    mNeoVis.3.1.1.chr10.g00483.t01    rna-gnl|WGS_ZZZZ|egapxtmp_024672-R1    mNeoVis.evergladensis.2.1.chr10.g00501.t01        247    N0.HOG0000245   OG0000008  n220   rna-XM_034644788.1    rna-XM_038577928.1    rna-XM_022519389.1    rna-XM_045062668.1    rna-XM_036096369.1    rna-XM_039255102.1    rna-XM_032876507.1    rna-XM_046011827.1    rna-XM_032347854.1    rna-XM_059187488.1    rna-XM_059404521.1    rna-XM_013046867.1      rna-XM_021686736.1      rna-XM_042993692.1    rna-XM_040641956.1    rna-XM_035729316.1      rna-gnl|WGS_ZZZZ|egapxtmp_022615-R1    mNeoVis.evergladensis.2.1.chr10.g00502.t01    mNeoVis.vulgivaga.2.1.chr10.g00492.t01      248    N0.HOG0000246   OG0000008  n240   rna-XM_002923245.4    rna-NM_001003386.2    rna-XM_022519456.1    rna-XM_003988739.4    rna-XM_036096417.1    rna-XM_039255069.1    rna-XM_032876725.1    rna-XM_046012353.1    rna-XM_032348305.1    rna-XM_059183573.1    rna-XM_059404858.1    rna-XM_004774678.1      rna-XM_021686671.2    rna-XM_004406252.1    rna-XM_042990830.1    rna-XM_040642508.1    rna-XM_027592367.2    mNeoVis.3.1.1.chr10.g00485.t01    rna-gnl|WGS_ZZZZ|egapxtmp_024673-R1      mNeoVis.vulgivaga.2.1.chr10.g00495.t01      249    N0.HOG0000247   OG0000008  n260   rna-XM_034644901.1    rna-XM_038577741.1    rna-XM_022519471.1    rna-XM_003988741.5    rna-XM_036096423.1    rna-XM_039255075.1    rna-XM_032876708.1    rna-XM_046012331.1    rna-XM_032348312.1    rna-XM_059185258.1    rna-XM_059403189.1    rna-XM_045073071.1    mNeoVis.2.1.1.chr9_hap1.g00511.t01    rna-XM_021686756.1    rna-XM_004406250.1    rna-XM_007081357.3    rna-XM_008704773.2    rna-XM_027592372.1    mNeoVis.3.1.1.chr10.g00484.t01    rna-gnl|WGS_ZZZZ|egapxtmp_022618-R1      mNeoVis.vulgivaga.2.1.chr10.g00493.t01      250    N0.HOG0000248   OG0000008  n280   rna-XM_019803803.2    rna-NM_001003392.1    rna-XM_022519450.1    rna-XM_003988740.5    rna-XM_036096414.1    rna-XM_039255070.1    rna-XM_032876702.1    rna-XM_046012333.1    rna-XM_032348310.1    rna-XM_059185260.1    rna-XM_059404859.1    rna-XM_004774677.3    mNeoVis.2.1.1.chr9_hap1.g00512.t01    rna-XM_021686672.2    rna-XM_004406251.1    rna-XM_007081356.3    rna-XM_040642510.1    rna-XM_027592369.2      rna-gnl|WGS_ZZZZ|egapxtmp_022614-R1      mNeoVis.vulgivaga.2.1.chr10.g00494.t01      251    N0.HOG0000249   OG0000008  n301   rna-XM_034645317.1    rna-XM_038576171.1    rna-XM_022519473.1    rna-XM_003988736.5    rna-XM_036096410.1    rna-XM_039224946.1    rna-XM_032876714.1    rna-XM_046011750.1    rna-XM_032348292.1    rna-XM_059185268.1    rna-XM_059403187.1    rna-XM_004774684.1    mNeoVis.2.1.1.chr9_hap1.g00518.t01    rna-XM_021686663.1    rna-XM_004417506.1    rna-XM_042992357.1    rna-XM_008704767.2    rna-XM_027592382.2    mNeoVis.3.1.1.chr10.g00491.t01    rna-gnl|WGS_ZZZZ|egapxtmp_022620-R1    mNeoVis.evergladensis.2.1.chr10.g00510.t01    mNeoVis.vulgivaga.2.1.chr10.g00503.t01      252    N0.HOG0000250   OG0000008  n321   rna-XM_011229903.3    rna-NM_001346035.1    rna-XM_022519414.1    rna-XM_004001288.6    rna-XM_036096412.1    rna-XM_039255074.1    rna-XM_032876710.1    rna-XM_046012256.1    rna-XM_032348290.1    rna-XM_059185265.1    rna-XM_059403185.1    rna-XM_004774683.2    mNeoVis.2.1.1.chr9_hap1.g00514.t01    rna-XM_044915404.1    rna-XM_004406257.1    rna-XM_042990829.1    rna-XM_008704770.2    rna-XM_027592373.2    mNeoVis.3.1.1.chr10.g00488.t01    rna-gnl|WGS_ZZZZ|egapxtmp_024676-R1    mNeoVis.evergladensis.2.1.chr10.g00508.t01    mNeoVis.vulgivaga.2.1.chr10.g00500.t01      253    N0.HOG0000251   OG0000008  n340   rna-XM_002923243.4, rna-XM_034651777.1, rna-XM_034652339.1    rna-XM_038576173.1, rna-XM_038576175.1    rna-XM_022492730.1    rna-XM_011279741.4, rna-XM_023257100.2, rna-XM_045061842.1    rna-XM_036096411.1, rna-XM_036096413.1    rna-XM_039224950.1    rna-XM_032876711.1, rna-XM_032876712.1    rna-XM_046011748.1, rna-XM_046012289.1    rna-XM_032348291.1, rna-XM_032348303.1    rna-XM_059183577.1, rna-XM_059185266.1    rna-XM_059387425.1    rna-XM_004774685.3, rna-XM_045073161.1    mNeoVis.2.1.1.chr9_hap1.g00515.t01, mNeoVis.2.1.1.chr9_hap1.g00517.t01    rna-XM_021686668.2, rna-XM_021686669.2    rna-XM_004417687.1, rna-XM_004417699.1    rna-XM_007073101.3, rna-XM_042990827.1, rna-XM_042990828.1    rna-XM_008704611.1, rna-XM_040641955.1    rna-XM_035729203.1, rna-XM_035729204.1              254    N0.HOG0000252   OG0000009  n3   rna-XM_034650240.1    rna-NM_001389101.1, rna-NM_001389133.1, rna-XM_038579869.1    rna-XM_022525634.1    rna-XM_003987369.3      rna-XM_039221961.1    rna-XM_032857149.1    rna-XM_046009287.1    rna-XM_032341680.1, rna-XM_032343160.1    rna-XM_059179621.1    rna-XM_059371122.1    rna-XM_013050800.2, rna-XM_045067484.1    mNeoVis.2.1.1.chr10_hap1.g00824.t01, mNeoVis.2.1.1.chr10_hap1.g00830.t01      rna-XM_004413507.1    rna-XM_015541672.2    rna-XM_008710648.2, rna-XM_008710655.2    rna-XM_027571485.2      rna-gnl|WGS_ZZZZ|egapxtmp_017319-R1    mNeoVis.evergladensis.2.1.chr8.g00802.t01    mNeoVis.vulgivaga.2.1.chr9.g00799.t01      255    N0.HOG0000253   OG0000009  n27   rna-XM_034653432.1, rna-XM_034660155.1    rna-XM_038581099.1, rna-XM_038588951.1, rna-XM_038589476.1, rna-XM_038589681.1    rna-XM_022525762.1    rna-XM_003987367.3    rna-XM_036122919.1    rna-XM_039221962.1    rna-XM_032857143.1, rna-XM_032857147.1    rna-XM_046009228.1, rna-XM_046009271.1    rna-XM_032341676.1, rna-XM_032341684.1    rna-XM_059179623.1    rna-XM_059371124.1, rna-XM_059371130.1    rna-XM_004781923.3, rna-XM_004782138.3      rna-XM_021695404.1      rna-XM_007091392.1    rna-XM_008684017.2, rna-XM_008710646.2, rna-XM_008710653.2    rna-XM_027571561.2      rna-gnl|WGS_ZZZZ|egapxtmp_017360-R1, rna-gnl|WGS_ZZZZ|egapxtmp_017361-R1    mNeoVis.evergladensis.2.1.chr8.g00797.t01, mNeoVis.evergladensis.2.1.chr8.g00804.t01    mNeoVis.vulgivaga.2.1.chr9.g00795.t01, mNeoVis.vulgivaga.2.1.chr9.g00798.t01      256    N0.HOG0000254   OG0000009  n57   rna-XM_002931279.2, rna-XM_034661719.1    rna-NM_001389078.1, rna-NM_001389087.1    rna-XM_022525397.1, rna-XM_022526052.1    rna-XM_003987368.4, rna-XM_045059611.1      rna-XM_039221964.1, rna-XM_039221966.1    rna-XM_032857086.1, rna-XM_032857148.1    rna-XM_046009264.1, rna-XM_046009278.1, rna-XM_046009349.1    rna-XM_032341681.1, rna-XM_032341764.1, rna-XM_032343161.1    rna-XM_059179622.1, rna-XM_059183388.1    rna-XM_059371123.1    rna-XM_004781740.2    mNeoVis.2.1.1.chr10_hap1.g00823.t01, mNeoVis.2.1.1.chr10_hap1.g00829.t01      rna-XM_004413508.1, rna-XM_004413513.1    rna-XM_007091391.2, rna-XM_042987726.1    rna-XM_008710647.1, rna-XM_008710654.2    rna-XM_027571372.2, rna-XM_027571494.2    mNeoVis.3.1.1.chr9.g00771.t01    rna-gnl|WGS_ZZZZ|egapxtmp_017342-R1    mNeoVis.evergladensis.2.1.chr8.g00801.t01        257    N0.HOG0000255   OG0000009  n94     rna-NM_001388523.1, rna-XM_038550063.1    rna-XM_022524757.1    rna-XM_003987318.5    rna-XM_036076877.1    rna-XM_039225472.1    rna-XM_032857078.1, rna-XM_032857080.1    rna-XM_046007841.1    rna-XM_032341771.1    rna-XM_059183394.1    rna-XM_059372100.1    rna-XM_004781929.2    mNeoVis.2.1.1.chr10_hap1.g00816.t01      rna-XM_004413509.1    rna-XM_007087482.3      rna-XM_027571329.2, rna-XM_027571330.2    mNeoVis.3.1.1.chr9.g00767.t01    rna-gnl|WGS_ZZZZ|egapxtmp_017364-R1    mNeoVis.evergladensis.2.1.chr8.g00794.t01    mNeoVis.vulgivaga.2.1.chr9.g00792.t01      258    N0.HOG0000256   OG0000009  n114   rna-XM_034653403.1, rna-XM_034653433.1      rna-XM_022525481.1    rna-XM_003987370.3    rna-XM_036076880.1    rna-XM_039221954.1    rna-XM_032857089.1    rna-XM_046009293.1    rna-XM_032341675.1    rna-XM_059179616.1    rna-XM_059371118.1    rna-XM_004781739.1    mNeoVis.2.1.1.chr10_hap1.g00832.t01    rna-XM_044918163.1      rna-XM_007091390.2    rna-XM_008684211.2, rna-XM_008710649.2    rna-XM_027570335.2    mNeoVis.3.1.1.chr9.g00777.t01    rna-gnl|WGS_ZZZZ|egapxtmp_017324-R1    mNeoVis.evergladensis.2.1.chr8.g00806.t01    mNeoVis.vulgivaga.2.1.chr9.g00803.t01      259    N0.HOG0000257   OG0000009  n134     rna-NM_001388941.1    rna-XM_022521673.1    rna-XM_023256232.2    rna-XM_036076879.1    rna-XM_039221974.1    rna-XM_032857088.1    rna-XM_046009505.1    rna-XM_032341762.1    rna-XM_059179619.1    rna-XM_059371119.1    rna-XM_004781751.1    mNeoVis.2.1.1.chr10_hap1.g00831.t01    rna-XM_021695552.1    rna-XM_004415201.1    rna-XM_007091418.1    rna-XM_008684213.1, rna-XM_008710652.2    rna-XM_027571862.2    mNeoVis.3.1.1.chr9.g00772.t01      mNeoVis.evergladensis.2.1.chr8.g00805.t01    mNeoVis.vulgivaga.2.1.chr9.g00802.t01      260    N0.HOG0000258   OG0000009  n152   rna-XM_002931305.4    rna-XM_038581098.1    rna-XM_022521667.1    rna-XM_023256233.2    rna-XM_036076885.1    rna-XM_039221953.1    rna-XM_032857090.1    rna-XM_046007835.1    rna-XM_032341761.1    rna-XM_059183386.1    rna-XM_059372091.1    rna-XM_004781750.1    mNeoVis.2.1.1.chr10_hap1.g00833.t01    rna-XM_021695713.2      rna-XM_042989298.1    rna-XM_008684212.2, rna-XM_008709356.2, rna-XM_008710442.2    rna-XM_027570336.1    mNeoVis.3.1.1.chr9.g00778.t01    rna-gnl|WGS_ZZZZ|egapxtmp_016934-R1    mNeoVis.evergladensis.2.1.chr8.g00807.t01    mNeoVis.vulgivaga.2.1.chr9.g00804.t01      261    N0.HOG0000259   OG0000009  n172   rna-XM_011218372.3, rna-XM_034648307.1      rna-XM_022521680.1, rna-XM_022526536.1    rna-XM_003987404.5, rna-XM_006932705.4      rna-XM_039221946.1, rna-XM_039221947.1    rna-XM_032857087.1, rna-XM_032857150.1    rna-XM_046007836.1, rna-XM_046009353.1    rna-XM_032341679.1, rna-XM_032341763.1, rna-XM_032343162.1    rna-XM_059179620.1, rna-XM_059183387.1    rna-XM_059371121.1, rna-XM_059372093.1, rna-XM_059372095.1    rna-XM_004781752.3, rna-XM_004782142.1, rna-XM_013050610.1    mNeoVis.2.1.1.chr10_hap1.g00819.t01, mNeoVis.2.1.1.chr10_hap1.g00825.t01        rna-XM_007091419.2, rna-XM_015541674.2    rna-XM_008710650.2, rna-XM_008710657.2, rna-XM_008710658.2      mNeoVis.3.1.1.chr9.g00773.t01, mNeoVis.3.1.1.chr9.g00776.t01    rna-gnl|WGS_ZZZZ|egapxtmp_015404-R1, rna-gnl|WGS_ZZZZ|egapxtmp_016933-R1    mNeoVis.evergladensis.2.1.chr8.g00792.t01        262    N0.HOG0000260   OG0000009  n205       rna-XM_022525748.1    rna-XM_023256231.2    rna-XM_036082592.1    rna-XM_039218337.1    rna-XM_032857141.1    rna-XM_046009237.1    rna-XM_032341686.1    rna-XM_059179629.1    rna-XM_059371133.1    rna-XM_004781928.1    mNeoVis.2.1.1.chr10_hap1.g00815.t01    rna-XM_021695718.1      rna-XM_042987725.1    rna-XM_008684030.2    rna-XM_027571297.1    mNeoVis.3.1.1.chr9.g00766.t01    rna-gnl|WGS_ZZZZ|egapxtmp_017337-R1    mNeoVis.evergladensis.2.1.chr8.g00793.t01        263    N0.HOG0000261   OG0000009  n223   rna-XM_011217839.1, rna-XM_034652810.1, rna-XM_034652811.1, rna-XM_034653427.1, rna-XM_034653430.1    rna-NM_001388922.1    rna-XM_022490742.1, rna-XM_022521705.1, rna-XM_022521743.1, rna-XM_022521752.1, rna-XM_022526006.1, rna-XM_022527065.1    rna-XM_003987360.3, rna-XM_003987361.3, rna-XM_003987365.4, rna-XM_006932682.4, rna-XM_006932683.5, rna-XM_023256174.2    rna-XM_036076883.1, rna-XM_036082593.1, rna-XM_036082605.1, rna-XM_036082608.1    rna-XM_039218331.1, rna-XM_039218333.1, rna-XM_039218334.1, rna-XM_039221980.1, rna-XM_039225473.1, rna-XM_039225474.1    rna-XM_032857075.1, rna-XM_032857076.1, rna-XM_032857081.1, rna-XM_032857083.1, rna-XM_032857145.1    rna-XM_046007838.1, rna-XM_046007842.1, rna-XM_046009227.1, rna-XM_046009272.1, rna-XM_046009435.1, rna-XM_046009442.1    rna-XM_032341677.1, rna-XM_032341682.1, rna-XM_032341687.1, rna-XM_032341690.1, rna-XM_032341769.1, rna-XM_032341773.1    rna-XM_059179625.1, rna-XM_059179630.1, rna-XM_059179633.1, rna-XM_059183391.1, rna-XM_059183395.1    rna-XM_059371125.1, rna-XM_059371127.1, rna-XM_059371134.1, rna-XM_059371136.1, rna-XM_059372098.1, rna-XM_059372101.1    rna-XM_004781921.2, rna-XM_004781925.2, rna-XM_004782245.2, rna-XM_004782246.1, rna-XM_004782389.2, rna-XM_045072834.1, rna-XM_045072835.1    mNeoVis.2.1.1.chr10_hap1.g00811.t01, mNeoVis.2.1.1.chr10_hap1.g00812.t01, mNeoVis.2.1.1.chr10_hap1.g00814.t01, mNeoVis.2.1.1.chr10_hap1.g00821.t01, mNeoVis.2.1.1.chr10_hap1.g00822.t01, mNeoVis.2.1.1.chr10_hap1.g00826.t01    rna-XM_021695714.1, rna-XM_021695719.1, rna-XM_021695720.1    rna-XM_004413516.1    rna-XM_007087481.3, rna-XM_007087485.2, rna-XM_007087486.2, rna-XM_015539943.2, rna-XM_015539944.2, rna-XM_015539946.2    rna-XM_008684019.2, rna-XM_008684024.2, rna-XM_008684025.1, rna-XM_008684031.2, rna-XM_008709993.1, rna-XM_040620959.1, rna-XM_040625511.1, rna-XM_040637050.1    rna-XM_027570328.2, rna-XM_027570331.2, rna-XM_027570332.1, rna-XM_027570947.1, rna-XM_027571632.2    mNeoVis.3.1.1.chr9.g00764.t01, mNeoVis.3.1.1.chr9.g00774.t01    rna-gnl|WGS_ZZZZ|egapxtmp_015403-R1, rna-gnl|WGS_ZZZZ|egapxtmp_016927-R1, rna-gnl|WGS_ZZZZ|egapxtmp_016932-R1, rna-gnl|WGS_ZZZZ|egapxtmp_017334-R1, rna-gnl|WGS_ZZZZ|egapxtmp_017336-R1, rna-gnl|WGS_ZZZZ|egapxtmp_017338-R1    mNeoVis.evergladensis.2.1.chr8.g00789.t01, mNeoVis.evergladensis.2.1.chr8.g00790.t01, mNeoVis.evergladensis.2.1.chr8.g00799.t01, mNeoVis.evergladensis.2.1.chr8.g00800.t01    mNeoVis.vulgivaga.2.1.chr9.g00789.t01, mNeoVis.vulgivaga.2.1.chr9.g00791.t01, mNeoVis.vulgivaga.2.1.chr9.g00797.t01      264    N0.HOG0000262   OG0000009  n329     rna-XM_038529586.1    rna-XM_022521696.1, rna-XM_022521734.1    rna-XM_003987366.3, rna-XM_045060731.1    rna-XM_036082553.1    rna-XM_039218335.1    rna-XM_032857084.1    rna-XM_046007837.1    rna-XM_032341766.1    rna-XM_059183390.1    rna-XM_059387312.1    rna-XM_004782140.2    mNeoVis.2.1.1.chr10_hap1.g00827.t01    rna-XM_021695716.1    rna-XM_004413515.1, rna-XM_004413517.1    rna-XM_007087479.3, rna-XM_015539945.2    rna-XM_008684028.2, rna-XM_008710334.2    rna-XM_027570333.2, rna-XM_027571536.2      rna-gnl|WGS_ZZZZ|egapxtmp_017335-R1    mNeoVis.evergladensis.2.1.chr8.g00803.t01    mNeoVis.vulgivaga.2.1.chr9.g00800.t01      265    N0.HOG0000263   OG0000010  n3   rna-XM_002930269.2, rna-XM_011230996.3    rna-NM_001388804.1    rna-XM_022500612.1    rna-XM_003985673.4, rna-XM_003998507.3    rna-XM_036111044.1    rna-XM_039257549.1    rna-XM_032847832.1, rna-XM_032872627.1    rna-XM_045992653.1, rna-XM_045992654.1, rna-XM_045993704.1, rna-XM_046004078.1    rna-XM_032328305.1, rna-XM_032338288.1    rna-XM_059153864.1, rna-XM_059176290.1    rna-XM_059399689.1, rna-XM_059416026.1    rna-XM_004780002.3, rna-XM_013047116.2    mNeoVis.2.1.1.chr1_hap1.g01058.t01    rna-XM_021680039.2, rna-XM_021697188.2    rna-XM_004394221.1, rna-XM_004416712.1    rna-XM_007081120.2, rna-XM_007092695.1    rna-XM_008698382.2, rna-XM_008711578.2    rna-XM_027602255.1    mNeoVis.3.1.1.chr1.g01052.t01, mNeoVis.3.1.1.chr15.g00434.t01    rna-gnl|WGS_ZZZZ|egapxtmp_021171-R1, rna-gnl|WGS_ZZZZ|egapxtmp_033311-R1      mNeoVis.vulgivaga.2.1.chr1.g01066.t01, mNeoVis.vulgivaga.2.1.chr15.g00483.t01      266    N0.HOG0000264   OG0000010  n37   rna-XM_019805587.2, rna-XM_034657971.1, rna-XM_034669578.1, rna-XM_034670519.1    rna-NM_001388519.1, rna-NM_001388647.1    rna-XM_022492700.1, rna-XM_022517045.1    rna-XM_011290658.4, rna-XM_023246286.2, rna-XM_023246287.2    rna-XM_036089118.1, rna-XM_036090600.1, rna-XM_036090609.1, rna-XM_036095395.1    rna-XM_039225743.1, rna-XM_039239316.1, rna-XM_039243408.1    rna-XM_032850056.1, rna-XM_032850973.1, rna-XM_032851046.1    rna-XM_045991810.1, rna-XM_045991845.1, rna-XM_045992964.1, rna-XM_045993717.1, rna-XM_045993730.1    rna-XM_032303848.1, rna-XM_032303849.1, rna-XM_032327978.1, rna-XM_032328630.1, rna-XM_032328631.1    rna-XM_059154191.1, rna-XM_059154816.1, rna-XM_059155521.1, rna-XM_059160334.1, rna-XM_059165162.1    rna-XM_059387773.1, rna-XM_059415895.1, rna-XM_059415987.1    rna-XM_004750978.3, rna-XM_004750979.2, rna-XM_004750980.2, rna-XM_004761182.2, rna-XM_004761183.2    mNeoVis.2.1.1.chr15_hap1.g00311.t01, mNeoVis.2.1.1.chr15_hap1.g00312.t01, mNeoVis.2.1.1.chr15_hap1.g00313.t01, mNeoVis.2.1.1.chr5_hap1.g00449.t01, mNeoVis.2.1.1.chr5_hap1.g00450.t01    rna-XM_021683206.2, rna-XM_021698003.1, rna-XM_021698004.1, rna-XM_044917625.1    rna-XM_004404704.1    rna-XM_015535714.2, rna-XM_042971438.1, rna-XM_042972244.1    rna-XM_008696784.2, rna-XM_008696787.2, rna-XM_008706399.2, rna-XM_008709417.2    rna-XM_035722298.1, rna-XM_035722299.1, rna-XM_035726753.1, rna-XM_035726765.1    mNeoVis.3.1.1.chr15.g00282.t01, mNeoVis.3.1.1.chr15.g00283.t01, mNeoVis.3.1.1.chr15.g00284.t01    rna-gnl|WGS_ZZZZ|egapxtmp_033988-R1, rna-gnl|WGS_ZZZZ|egapxtmp_034151-R8, rna-gnl|WGS_ZZZZ|egapxtmp_034886-R1    mNeoVis.evergladensis.2.1.chr15.g00287.t01, mNeoVis.evergladensis.2.1.chr15.g00288.t01, mNeoVis.evergladensis.2.1.chr15.g00289.t01, mNeoVis.evergladensis.2.1.chr2.g00444.t01    mNeoVis.vulgivaga.2.1.chr15.g00292.t01, mNeoVis.vulgivaga.2.1.chr2.g00446.t01      267    N0.HOG0000265   OG0000010  n113   rna-XM_002928314.2, rna-XM_002930671.2, rna-XM_002931230.3, rna-XM_002931256.2, rna-XM_002931284.3, rna-XM_002931317.4, rna-XM_011217520.3, rna-XM_011218352.3, rna-XM_011236465.3, rna-XM_019793497.2, rna-XM_034640613.1, rna-XM_034663715.1, rna-XM_034664456.1    rna-NM_001014287.1    rna-XM_022501902.1, rna-XM_022501905.1, rna-XM_022501909.1, rna-XM_022509790.1, rna-XM_022521084.1    rna-XM_003995829.2, rna-XM_003996400.5, rna-XM_011288460.3, rna-XM_019816493.3, rna-XM_023242312.1, rna-XM_023243352.2, rna-XM_023243769.2, rna-XM_023243770.2, rna-XM_045043135.1, rna-XM_045043958.1, rna-XM_045043960.1    rna-XM_036092849.1, rna-XM_036092870.1, rna-XM_036107741.1, rna-XM_036107842.1, rna-XM_036107847.1, rna-XM_036107907.1, rna-XM_036121833.1    rna-XM_039223031.1, rna-XM_039224353.1, rna-XM_039224360.1, rna-XM_039224361.1, rna-XM_039224363.1, rna-XM_039224364.1, rna-XM_039224365.1, rna-XM_039253410.1, rna-XM_039253488.1, rna-XM_039253605.1    rna-XM_032846724.1, rna-XM_032849012.1, rna-XM_032849013.1, rna-XM_032849014.1, rna-XM_032865788.1, rna-XM_032865789.1, rna-XM_032865790.1, rna-XM_032866367.1    rna-XM_045986102.1, rna-XM_045986236.1, rna-XM_045986238.1, rna-XM_045996800.1, rna-XM_045996801.1, rna-XM_045996802.1, rna-XM_045996993.1, rna-XM_046021814.1, rna-XM_046021816.1, rna-XM_046021817.1, rna-XM_046021818.1, rna-XM_046023009.1, rna-XM_046023117.1, rna-XM_046023963.1, rna-XM_046023964.1    rna-XM_032307566.1, rna-XM_032307567.1, rna-XM_032307955.1, rna-XM_032308523.1, rna-XM_032308524.1, rna-XM_032308526.1, rna-XM_032308629.1, rna-XM_032308630.1, rna-XM_032308631.1, rna-XM_032320381.1, rna-XM_032320651.1, rna-XM_032321707.1    rna-XM_059141777.1, rna-XM_059142127.1, rna-XM_059142582.1, rna-XM_059142583.1, rna-XM_059142585.1, rna-XM_059142586.1, rna-XM_059143092.1, rna-XM_059143093.1, rna-XM_059147424.1, rna-XM_059147568.1, rna-XM_059148588.1, rna-XM_059148779.1    rna-XM_059378453.1, rna-XM_059378575.1, rna-XM_059378584.1, rna-XM_059412156.1, rna-XM_059412158.1, rna-XM_059412159.1, rna-XM_059412164.1, rna-XM_059412165.1, rna-XM_059412206.1, rna-XM_059412208.1, rna-XM_059412209.1, rna-XM_059412210.1, rna-XM_059412212.1    rna-XM_004782106.1, rna-XM_004782109.2, rna-XM_004782334.1, rna-XM_013050785.2, rna-XM_013050786.2, rna-XM_013050788.2, rna-XM_013050789.2, rna-XM_013050790.2, rna-XM_013050792.2, rna-XM_013053472.2, rna-XM_045073824.1, rna-XM_045082900.1, rna-XM_045082901.1, rna-XM_045085476.1, rna-XM_045085477.1    mNeoVis.2.1.1.chr13_hap1.g00273.t01, mNeoVis.2.1.1.chr13_hap1.g00274.t01, mNeoVis.2.1.1.chr13_hap1.g00275.t01, mNeoVis.2.1.1.chr13_hap1.g00276.t01, mNeoVis.2.1.1.chr13_hap1.g00277.t01, mNeoVis.2.1.1.chr13_hap1.g00278.t01, mNeoVis.2.1.1.chr13_hap1.g00279.t01, mNeoVis.2.1.1.chr13_hap1.g00280.t01, mNeoVis.2.1.1.chr13_hap1.g00281.t01, mNeoVis.2.1.1.chr8_hap1.g00853.t01, mNeoVis.2.1.1.chr8_hap1.g00914.t01, mNeoVis.2.1.1.chr8_hap1.g00915.t01, mNeoVis.2.1.1.chr8_hap1.g00923.t01, mNeoVis.2.1.1.chr8_hap1.g00925.t01    rna-XM_021691589.1, rna-XM_021691590.2, rna-XM_021691591.1, rna-XM_021691592.1, rna-XM_021691700.1, rna-XM_021704358.1, rna-XM_021704432.1    rna-XM_004392624.1, rna-XM_004392625.1, rna-XM_004392644.1, rna-XM_004392646.1, rna-XM_004392647.1, rna-XM_004417000.1    rna-XM_007094370.2, rna-XM_007094371.2, rna-XM_007094514.2, rna-XM_007094515.2, rna-XM_007094519.2, rna-XM_007094520.2, rna-XM_015534845.1, rna-XM_015534884.2, rna-XM_015543014.2, rna-XM_015543015.2, rna-XM_042964737.1, rna-XM_042964738.1, rna-XM_042966510.1, rna-XM_042967238.1    rna-XM_008695527.2, rna-XM_008698802.2, rna-XM_008698804.2, rna-XM_008698929.2, rna-XM_008698931.2, rna-XM_008698934.2, rna-XM_008698935.2, rna-XM_008698936.2, rna-XM_008698938.2, rna-XM_008698940.2, rna-XM_040639913.1    rna-XM_027614686.1, rna-XM_027614688.1, rna-XM_027615135.1, rna-XM_027624688.1, rna-XM_027626527.1, rna-XM_035723466.1, rna-XM_035723467.1, rna-XM_035723794.1    mNeoVis.3.1.1.chr13.g00216.t01, mNeoVis.3.1.1.chr13.g00217.t01, mNeoVis.3.1.1.chr13.g00218.t01, mNeoVis.3.1.1.chr13.g00219.t01, mNeoVis.3.1.1.chr13.g00220.t01, mNeoVis.3.1.1.chr13.g00221.t01, mNeoVis.3.1.1.chr13.g00222.t01, mNeoVis.3.1.1.chr13.g00223.t01, mNeoVis.3.1.1.chr8.g00801.t01    rna-gnl|WGS_ZZZZ|egapxtmp_018660-R1, rna-gnl|WGS_ZZZZ|egapxtmp_019113-R1, rna-gnl|WGS_ZZZZ|egapxtmp_030148-R1, rna-gnl|WGS_ZZZZ|egapxtmp_030152-R1, rna-gnl|WGS_ZZZZ|egapxtmp_030155-R1, rna-gnl|WGS_ZZZZ|egapxtmp_030156-R1, rna-gnl|WGS_ZZZZ|egapxtmp_030157-R1, rna-gnl|WGS_ZZZZ|egapxtmp_030159-R1, rna-gnl|WGS_ZZZZ|egapxtmp_030163-R1, rna-gnl|WGS_ZZZZ|egapxtmp_030165-R1, rna-gnl|WGS_ZZZZ|egapxtmp_030995-R1    mNeoVis.evergladensis.2.1.chr13.g00264.t01, mNeoVis.evergladensis.2.1.chr13.g00265.t01, mNeoVis.evergladensis.2.1.chr13.g00266.t01, mNeoVis.evergladensis.2.1.chr13.g00267.t01, mNeoVis.evergladensis.2.1.chr13.g00268.t01, mNeoVis.evergladensis.2.1.chr13.g00269.t01, mNeoVis.evergladensis.2.1.chr13.g00270.t01, mNeoVis.evergladensis.2.1.chr13.g00271.t01, mNeoVis.evergladensis.2.1.chr9.g00843.t01, mNeoVis.evergladensis.2.1.chr9.g00900.t01    mNeoVis.vulgivaga.2.1.chr13.g00261.t01, mNeoVis.vulgivaga.2.1.chr13.g00262.t01, mNeoVis.vulgivaga.2.1.chr13.g00263.t01, mNeoVis.vulgivaga.2.1.chr13.g00264.t01, mNeoVis.vulgivaga.2.1.chr13.g00265.t01, mNeoVis.vulgivaga.2.1.chr13.g00266.t01, mNeoVis.vulgivaga.2.1.chr13.g00267.t01, mNeoVis.vulgivaga.2.1.chr13.g00268.t01, mNeoVis.vulgivaga.2.1.chr8.g00845.t01      268    N0.HOG0000266   OG0000010  n322   rna-XM_002931218.4, rna-XM_002931229.4    rna-XM_038546156.1    rna-XM_022509920.1    rna-XM_003995842.3, rna-XM_003996403.4    rna-XM_036076645.1    rna-XM_039224362.1, rna-XM_039253501.1    rna-XM_032846777.1, rna-XM_032866353.1    rna-XM_045984141.1, rna-XM_045986656.1, rna-XM_046023597.1    rna-XM_032322095.1    rna-XM_059149075.1    rna-XM_059380166.1    rna-XM_004781931.3    mNeoVis.2.1.1.chr8_hap1.g00860.t01    rna-XM_021704284.1    rna-XM_004392645.1    rna-XM_015534883.2    rna-XM_008695263.2, rna-XM_008698930.1    rna-XM_027568188.1    mNeoVis.3.1.1.chr8.g00806.t01    rna-gnl|WGS_ZZZZ|egapxtmp_018663-R1    mNeoVis.evergladensis.2.1.chr9.g00848.t01    mNeoVis.vulgivaga.2.1.chr8.g00850.t01      269    N0.HOG0000267   OG0000011  n1   rna-XM_011218315.3, rna-XM_034648664.1, rna-XM_034648665.1    rna-XM_038557641.1    rna-XM_022521624.1    rna-XM_023256178.2    rna-XM_036118042.1    rna-XM_039221949.1    rna-XM_032857093.1    rna-XM_046007830.1    rna-XM_032341755.1    rna-XM_059183381.1    rna-XM_059372088.1    rna-XM_004781743.2    mNeoVis.2.1.1.chr10_hap1.g00843.t01    rna-XM_021695708.1    rna-XM_004415202.1    rna-XM_007091413.2    rna-XM_040621911.1    rna-XM_027571400.1    mNeoVis.3.1.1.chr9.g00783.t01    rna-gnl|WGS_ZZZZ|egapxtmp_017316-R1    mNeoVis.evergladensis.2.1.chr8.g00817.t01    mNeoVis.vulgivaga.2.1.chr9.g00812.t01      270    N0.HOG0000268   OG0000011  n24     rna-XM_038551556.1      rna-XM_003987402.3, rna-XM_045060657.1      rna-XM_039221945.1    rna-XM_032857079.1, rna-XM_032857085.1    rna-XM_046007839.1    rna-XM_032341765.1, rna-XM_032341767.1, rna-XM_032341770.1    rna-XM_059183389.1, rna-XM_059183393.1    rna-XM_059372094.1, rna-XM_059372096.1, rna-XM_059372099.1    rna-XM_004782141.1, rna-XM_004782247.3, rna-XM_013050703.1    mNeoVis.2.1.1.chr10_hap1.g00818.t01, mNeoVis.2.1.1.chr10_hap1.g00828.t01    rna-XM_021695715.1    rna-XM_004413514.1    rna-XM_007091421.2    rna-XM_008684027.2    rna-XM_027570334.1    mNeoVis.3.1.1.chr9.g00769.t01, mNeoVis.3.1.1.chr9.g00775.t01    rna-gnl|WGS_ZZZZ|egapxtmp_015405-R1, rna-gnl|WGS_ZZZZ|egapxtmp_016930-R1    mNeoVis.evergladensis.2.1.chr8.g00796.t01    mNeoVis.vulgivaga.2.1.chr9.g00794.t01, mNeoVis.vulgivaga.2.1.chr9.g00801.t01      271    N0.HOG0000269   OG0000011  n59   rna-XM_011217966.2, rna-XM_011218158.2    rna-XM_038557643.1    rna-XM_022526871.1    rna-XM_003987413.3    rna-XM_036118053.1    rna-XM_039221952.1    rna-XM_032857158.1    rna-XM_046009453.1    rna-XM_032341668.1    rna-XM_059181062.1    rna-XM_059371107.1    rna-XM_004782194.2    mNeoVis.2.1.1.chr10_hap1.g00849.t01    rna-XM_021695507.1    rna-XM_004413040.1    rna-XM_007091408.1    rna-XM_008711102.2    rna-XM_027571398.1    mNeoVis.3.1.1.chr9.g00787.t01    rna-gnl|WGS_ZZZZ|egapxtmp_017327-R1    mNeoVis.evergladensis.2.1.chr8.g00823.t01    mNeoVis.vulgivaga.2.1.chr9.g00817.t01      272    N0.HOG0000270   OG0000011  n80   rna-XM_011217964.3, rna-XM_011218157.3, rna-XM_019793979.2, rna-XM_034653428.1    rna-NM_001313835.1    rna-XM_022521603.1, rna-XM_022527075.1    rna-XM_003987412.4    rna-XM_036118040.1, rna-XM_036118054.1    rna-XM_039221960.1, rna-XM_039221972.1    rna-XM_032857157.1    rna-XM_046007827.1, rna-XM_046009498.1, rna-XM_046009594.1    rna-XM_032341669.1, rna-XM_032341670.1, rna-XM_032341678.1, rna-XM_032341752.1    rna-XM_059181058.1, rna-XM_059181063.1, rna-XM_059183377.1    rna-XM_059371108.1, rna-XM_059371110.1, rna-XM_059372086.1    rna-XM_004782192.2, rna-XM_004782193.1, rna-XM_045067488.1    mNeoVis.2.1.1.chr10_hap1.g00846.t01, mNeoVis.2.1.1.chr10_hap1.g00847.t01, mNeoVis.2.1.1.chr10_hap1.g00848.t01    rna-XM_021695706.1    rna-XM_004413041.1, rna-XM_004415204.1    rna-XM_007091409.2, rna-XM_007091410.2    rna-XM_008684007.2, rna-XM_008684008.1, rna-XM_008712297.2, rna-XM_008712298.2    rna-XM_027571481.1    mNeoVis.3.1.1.chr9.g00785.t01, mNeoVis.3.1.1.chr9.g00786.t01    rna-gnl|WGS_ZZZZ|egapxtmp_017320-R1, rna-gnl|WGS_ZZZZ|egapxtmp_017321-R1, rna-gnl|WGS_ZZZZ|egapxtmp_017340-R1    mNeoVis.evergladensis.2.1.chr8.g00819.t01, mNeoVis.evergladensis.2.1.chr8.g00820.t01, mNeoVis.evergladensis.2.1.chr8.g00821.t01    mNeoVis.vulgivaga.2.1.chr9.g00814.t01, mNeoVis.vulgivaga.2.1.chr9.g00815.t01, mNeoVis.vulgivaga.2.1.chr9.g00816.t01      273    N0.HOG0000271   OG0000011  n129   rna-XM_019794037.2, rna-XM_034652771.1, rna-XM_034652817.1, rna-XM_034653396.1    rna-XM_038557642.1    rna-XM_022492761.1, rna-XM_022521612.1    rna-XM_023256179.2, rna-XM_045059614.1    rna-XM_036118041.1    rna-XM_039221950.1, rna-XM_039221951.1    rna-XM_032857094.1, rna-XM_032857095.1    rna-XM_046007828.1, rna-XM_046007829.1    rna-XM_032341753.1, rna-XM_032341754.1    rna-XM_059183379.1, rna-XM_059183380.1    rna-XM_059372087.1    rna-XM_004781741.2, rna-XM_004781742.1    mNeoVis.2.1.1.chr10_hap1.g00844.t01, mNeoVis.2.1.1.chr10_hap1.g00845.t01    rna-XM_044918282.1    rna-XM_004415203.1    rna-XM_007091411.2, rna-XM_007091412.2    rna-XM_008710430.2, rna-XM_008711101.2, rna-XM_040620672.1, rna-XM_040621909.1    rna-XM_027571633.1    mNeoVis.3.1.1.chr9.g00784.t01    rna-gnl|WGS_ZZZZ|egapxtmp_017322-R1    mNeoVis.evergladensis.2.1.chr8.g00818.t01    mNeoVis.vulgivaga.2.1.chr9.g00813.t01      274   N0.HOG0000272  OG0000011  n165   rna-XM_011218254.3, rna-XM_011218333.3, rna-XM_034648659.1    rna-XM_038557595.1    rna-XM_022521637.1, rna-XM_022521646.1    rna-XM_023256177.2, rna-XM_023256236.2    rna-XM_036118044.1    rna-XM_039221948.1, rna-XM_039221968.1    rna-XM_032857091.1, rna-XM_032857092.1    rna-XM_046007831.1, rna-XM_046007832.1    rna-XM_032341756.1, rna-XM_032341758.1    rna-XM_059183382.1, rna-XM_059183383.1    rna-XM_059372089.1, rna-XM_059387320.1    rna-XM_004781744.2, rna-XM_004781746.2, rna-XM_013050801.1    mNeoVis.2.1.1.chr10_hap1.g00839.t01, mNeoVis.2.1.1.chr10_hap1.g00841.t01    rna-XM_021695709.1      rna-XM_007091387.3, rna-XM_007091414.1    rna-XM_008683974.2, rna-XM_008710432.2, rna-XM_008710437.2    rna-XM_027570340.1    mNeoVis.3.1.1.chr9.g00781.t01    rna-gnl|WGS_ZZZZ|egapxtmp_016938-R1, rna-gnl|WGS_ZZZZ|egapxtmp_016941-R1    mNeoVis.evergladensis.2.1.chr8.g00813.t01, mNeoVis.evergladensis.2.1.chr8.g00815.t01    mNeoVis.vulgivaga.2.1.chr9.g00808.t01, mNeoVis.vulgivaga.2.1.chr9.g00810.t01      275   N0.HOG0000273  OG0000011  n203   rna-XM_011218341.2, rna-XM_034648547.1    rna-XM_038557596.1    rna-XM_022526945.1    rna-XM_011283361.3      rna-XM_039224139.1    rna-XM_032850785.1    rna-XM_046007826.1    rna-XM_032341662.1    rna-XM_059183376.1    rna-XM_059372085.1    rna-XM_013050818.2    mNeoVis.2.1.1.chr10_hap1.g00854.t01      rna-XM_004413030.1    rna-XM_015541670.2    rna-XM_008711103.2, rna-XM_040625517.1    rna-XM_027571483.1      rna-gnl|WGS_ZZZZ|egapxtmp_017352-R1    mNeoVis.evergladensis.2.1.chr8.g00828.t01    mNeoVis.vulgivaga.2.1.chr9.g00819.t01      276    N0.HOG0000274   OG0000011  n222         rna-XM_011283360.2      rna-XM_039224149.1                      rna-XM_015541671.2                  277    N0.HOG0000275   OG0000011  n225     rna-XM_038557598.1    rna-XM_022490781.1    rna-XM_011283099.2    rna-XM_036118039.1    rna-XM_039224147.1    rna-XM_032857165.1    rna-XM_046009592.1    rna-XM_032341659.1    rna-XM_059181056.1    rna-XM_059371100.1    rna-XM_004781471.1    mNeoVis.2.1.1.chr10_hap1.g00856.t01    rna-XM_021695705.1      rna-XM_015541675.2    rna-XM_008683653.1      mNeoVis.3.1.1.chr9.g00791.t01    rna-gnl|WGS_ZZZZ|egapxtmp_017323-R1    mNeoVis.evergladensis.2.1.chr8.g00830.t01    mNeoVis.vulgivaga.2.1.chr9.g00820.t01      278    N0.HOG0000276   OG0000011  n241     rna-XM_038557597.1    rna-XM_022526978.1    rna-XM_023255546.2    rna-XM_036118049.1    rna-XM_039224141.1    rna-XM_032857163.1    rna-XM_046009502.1    rna-XM_032341661.1      rna-XM_059371101.1    rna-XM_004782197.3    mNeoVis.2.1.1.chr10_hap1.g00855.t01    rna-XM_021695524.1    rna-XM_004413038.1    rna-XM_015541673.2    rna-XM_008709742.2    rna-XM_027571535.1    mNeoVis.3.1.1.chr9.g00790.t01    rna-gnl|WGS_ZZZZ|egapxtmp_016946-R1    mNeoVis.evergladensis.2.1.chr8.g00829.t01        279   N0.HOG0000277  OG0000011  n259   rna-XM_019793784.2, rna-XM_034650323.1, rna-XM_034653294.1    rna-NM_001256439.1, rna-NM_001256458.1    rna-XM_022526524.1    rna-XM_045059616.1, rna-XM_045060658.1    rna-XM_036118052.1    rna-XM_039221971.1, rna-XM_039224140.1    rna-XM_032857160.1, rna-XM_032857164.1    rna-XM_046009452.1, rna-XM_046009480.1    rna-XM_032341667.1, rna-XM_032341750.1    rna-XM_059181060.1    rna-XM_059371105.1    rna-XM_004782189.2, rna-XM_045067485.1, rna-XM_045083049.1    mNeoVis.2.1.1.chr10_hap1.g00850.t01, mNeoVis.2.1.1.chr10_hap1.g00851.t01      rna-XM_004413032.1    rna-XM_007091384.2    rna-XM_008709960.2, rna-XM_008711098.2, rna-XM_040625516.1    rna-XM_027570341.1, rna-XM_027571373.1    mNeoVis.3.1.1.chr9.g00788.t01    rna-gnl|WGS_ZZZZ|egapxtmp_016944-R1, rna-gnl|WGS_ZZZZ|egapxtmp_017343-R1    mNeoVis.evergladensis.2.1.chr8.g00824.t01, mNeoVis.evergladensis.2.1.chr8.g00825.t01        280    N0.HOG0000278   OG0000011  n295     rna-NM_001389054.1    rna-XM_022526934.1    rna-XM_011283097.4      rna-XM_039221975.1, rna-XM_039221976.1    rna-XM_032857162.1    rna-XM_046009478.1    rna-XM_032341665.1    rna-XM_059181065.1    rna-XM_059371103.1    rna-XM_045067487.1    mNeoVis.2.1.1.chr10_hap1.g00852.t01      rna-XM_004413039.1    rna-XM_007091407.2    rna-XM_008709739.2, rna-XM_008709740.2, rna-XM_008711100.2    rna-XM_027571458.1    mNeoVis.3.1.1.chr9.g00789.t01    rna-gnl|WGS_ZZZZ|egapxtmp_017325-R1    mNeoVis.evergladensis.2.1.chr8.g00826.t01        281    N0.HOG0000279   OG0000011  n314     rna-NM_001313814.1    rna-XM_022526843.1    rna-XM_006932684.4    rna-XM_036118051.1    rna-XM_039221970.1    rna-XM_032857161.1      rna-XM_032341666.1      rna-XM_059371104.1    rna-XM_013050819.2        rna-XM_004413031.1    rna-XM_007091386.2      rna-XM_027571387.1      rna-gnl|WGS_ZZZZ|egapxtmp_016943-R1          282    N0.HOG0000280   OG0000011  n326   rna-XM_002931351.2    rna-NM_001388556.1    rna-XM_022526921.1    rna-XM_006932685.5    rna-XM_036118050.1    rna-XM_039224146.1    rna-XM_032850784.1    rna-XM_046009479.1    rna-XM_032341663.1    rna-XM_059181064.1    rna-XM_059371102.1    rna-XM_004782190.3    mNeoVis.2.1.1.chr10_hap1.g00853.t01    rna-XM_021695514.2      rna-XM_007091385.2    rna-XM_008711099.2    rna-XM_027571419.1        mNeoVis.evergladensis.2.1.chr8.g00827.t01    mNeoVis.vulgivaga.2.1.chr9.g00818.t01      283    N0.HOG0000281   OG0000012  n1   rna-XM_011220444.3    rna-XM_038584309.1    rna-XM_022490571.1, rna-XM_022490621.1    rna-XM_003985738.6, rna-XM_003985771.5    rna-XM_036112333.1    rna-XM_039257313.1    rna-XM_032861716.1, rna-XM_032861793.1    rna-XM_046004098.1    rna-XM_032338384.1, rna-XM_032341278.1    rna-XM_059176411.1, rna-XM_059179505.1    rna-XM_059398950.1, rna-XM_059398953.1    rna-XM_004778781.1, rna-XM_045082222.1    mNeoVis.2.1.1.chr1_hap1.g01159.t01, mNeoVis.2.1.1.chr1_hap1.g01161.t01    rna-XM_044917650.1, rna-XM_044917671.1    rna-XM_004406496.1    rna-XM_007074486.3, rna-XM_042985570.1    rna-XM_008693079.2    rna-XM_035728679.1    mNeoVis.3.1.1.chr1.g01118.t01    rna-gnl|WGS_ZZZZ|egapxtmp_018211-R1, rna-gnl|WGS_ZZZZ|egapxtmp_018452-R1    mNeoVis.evergladensis.2.1.chr1.g01132.t01, mNeoVis.evergladensis.2.1.chr1.g01134.t01    mNeoVis.vulgivaga.2.1.chr1.g01130.t01, mNeoVis.vulgivaga.2.1.chr1.g01132.t01      284    N0.HOG0000282   OG0000012  n34       rna-XM_022490581.1          rna-XM_032861725.1    rna-XM_046006285.1      rna-XM_059176269.1    rna-XM_059398962.1    rna-XM_045082274.1    mNeoVis.2.1.1.chr1_hap1.g01098.t01                  mNeoVis.evergladensis.2.1.chr1.g01096.t01        285    N0.HOG0000283   OG0000012  n39     rna-XM_038562072.1    rna-XM_022492772.1    rna-XM_003990554.6    rna-XM_036098392.1, rna-XM_036098394.1      rna-XM_032851114.1    rna-XM_046012497.1, rna-XM_046012508.1    rna-XM_032303389.1, rna-XM_032303392.1    rna-XM_059137256.1, rna-XM_059137257.1      rna-XM_004782573.3    mNeoVis.2.1.1.chr2_hap1.g00829.t01, mNeoVis.2.1.1.chr2_hap1.g00832.t01      rna-XM_004404171.1, rna-XM_004404174.1      rna-XM_008695745.2, rna-XM_008695751.2    rna-XM_027571471.2, rna-XM_027622608.2              286    N0.HOG0000284   OG0000012  n41   rna-XM_002930223.4    rna-XM_038562074.1    rna-XM_022526073.1    rna-XM_003990556.6      rna-XM_039246244.1    rna-XM_032843521.1    rna-XM_046013796.1      rna-XM_059136607.1      rna-XM_004776832.3    mNeoVis.2.1.1.chr2_hap1.g00826.t01    rna-XM_044914136.1    rna-XM_004404177.1    rna-XM_042993990.1      rna-XM_027607137.1          mNeoVis.vulgivaga.2.1.chr5.g00776.t01      287    N0.HOG0000285   OG0000012  n45   rna-XM_002928374.4    rna-XM_038576872.1    rna-XM_022495087.1    rna-XM_003988442.6    rna-XM_036093770.1    rna-XM_039239908.1    rna-XM_032855052.1    rna-XM_046013405.1    rna-XM_032347412.1    rna-XM_059185994.1    rna-XM_059402650.1    rna-XM_013057874.2      rna-XM_021690519.1    rna-XM_004406990.1    rna-XM_007082765.3    rna-XM_008707027.2    rna-XM_027594188.1          mNeoVis.vulgivaga.2.1.chr10.g00767.t01      288    N0.HOG0000286   OG0000012  n50   rna-XM_034643187.1    rna-XM_038562073.1    rna-XM_022526071.1    rna-XM_003990557.6    rna-XM_036098395.1    rna-XM_039246243.1    rna-XM_032843520.1    rna-XM_046013784.1    rna-XM_032303393.1    rna-XM_059136606.1      rna-XM_004776830.3    mNeoVis.2.1.1.chr2_hap1.g00825.t01    rna-XM_044914135.1    rna-XM_004404178.1    rna-XM_007088035.3    rna-XM_008695744.2    rna-XM_027607123.1    mNeoVis.3.1.1.chr5.g00772.t01      mNeoVis.evergladensis.2.1.chr5.g00782.t01    mNeoVis.vulgivaga.2.1.chr5.g00775.t01      289    N0.HOG0000287   OG0000012  n54   rna-XM_034660187.1    rna-NM_001374525.1, rna-XM_038584346.1                rna-XM_032335062.1          mNeoVis.2.1.1.chr1_hap1.g00964.t01          rna-XM_008708270.2        rna-gnl|WGS_ZZZZ|egapxtmp_021220-R1    mNeoVis.evergladensis.2.1.chr1.g01119.t01, mNeoVis.evergladensis.2.1.chr5.g00783.t01    mNeoVis.vulgivaga.2.1.chr1.g01095.t01      290    N0.HOG0000288   OG0000012  n65                               rna-XM_004413644.1        rna-XM_027604115.2              291    N0.HOG0000289   OG0000012  n70     rna-XM_038584347.1    rna-XM_022490552.1    rna-XM_003985719.5    rna-XM_036112334.1, rna-XM_036112340.1    rna-XM_039257282.1    rna-XM_032861718.1    rna-XM_046006272.1    rna-XM_032338261.1    rna-XM_059176265.1      rna-XM_045082267.1    mNeoVis.2.1.1.chr1_hap1.g01140.t01      rna-XM_004406520.1      rna-XM_040645174.1    rna-XM_035728425.1        mNeoVis.evergladensis.2.1.chr1.g01110.t01        292    N0.HOG0000290   OG0000012  n74                                     rna-XM_027581663.2, rna-XM_035728426.1              293    N0.HOG0000291   OG0000012  n76     rna-XM_038556408.1, rna-XM_038584411.1      rna-XM_003985699.6, rna-XM_011282084.4            rna-XM_032338257.1    rna-XM_059176257.1      rna-XM_045082265.1          rna-XM_015540722.2, rna-XM_042985546.1                  294    N0.HOG0000292   OG0000012  n82   rna-XM_019809261.2, rna-XM_034660121.1, rna-XM_034660122.1, rna-XM_034660123.1, rna-XM_034660124.1, rna-XM_034660125.1    rna-XM_038584348.1, rna-XM_038584433.1, rna-XM_038584434.1      rna-XM_003985691.5, rna-XM_003985693.3, rna-XM_011282063.3, rna-XM_011282066.4, rna-XM_011282078.3, rna-XM_019831689.3    rna-XM_036112327.1, rna-XM_036112335.1, rna-XM_036112338.1, rna-XM_036112339.1, rna-XM_036121559.1, rna-XM_036121561.1, rna-XM_036121562.1, rna-XM_036121564.1    rna-XM_039223538.1, rna-XM_039223540.1, rna-XM_039223544.1, rna-XM_039223546.1, rna-XM_039223554.1, rna-XM_039224913.1, rna-XM_039224917.1, rna-XM_039225804.1, rna-XM_039257280.1, rna-XM_039257283.1    rna-XM_032861723.1        rna-XM_059176263.1, rna-XM_059176264.1      rna-XM_045082235.1, rna-XM_045082269.1, rna-XM_045082270.1    mNeoVis.2.1.1.chr1_hap1.g01115.t01    rna-XM_021687357.2, rna-XM_021697186.1, rna-XM_044917304.1, rna-XM_044917585.1, rna-XM_044917648.1    rna-XM_004413627.1, rna-XM_004413631.1, rna-XM_004413635.1, rna-XM_004413643.1    rna-XM_007074452.3, rna-XM_007074463.3, rna-XM_007074475.3, rna-XM_042985501.1, rna-XM_042985502.1, rna-XM_042985544.1, rna-XM_042985547.1    rna-XM_008692971.2, rna-XM_008692998.2    rna-XM_027575118.2, rna-XM_035728413.1, rna-XM_035728418.1, rna-XM_035728428.1    mNeoVis.3.1.1.chr1.g01088.t01      mNeoVis.evergladensis.2.1.chr1.g01091.t01        295    N0.HOG0000293   OG0000012  n94   rna-XM_002927085.3    rna-XM_038584412.1                rna-XM_032338260.1                  rna-XM_008692978.2, rna-XM_008692984.2, rna-XM_008693004.2, rna-XM_008693021.2                296    N0.HOG0000294   OG0000012  n95   rna-XM_002927087.4, rna-XM_002927089.3, rna-XM_002927091.4    rna-XM_038584408.1                                rna-XM_008693099.2, rna-XM_040645161.1                297    N0.HOG0000295   OG0000012  n103   rna-XM_034655740.1    rna-XM_038584349.1    rna-XM_022490550.1, rna-XM_022490553.1, rna-XM_022490555.1, rna-XM_022490557.1, rna-XM_022490559.1, rna-XM_022491656.1    rna-XM_003985705.5, rna-XM_006927460.4, rna-XM_045058513.1    rna-XM_036112336.1, rna-XM_036112337.1, rna-XM_036119163.1    rna-XM_039220271.1, rna-XM_039257278.1, rna-XM_039257279.1    rna-XM_032848523.1, rna-XM_032861717.1, rna-XM_032861719.1, rna-XM_032861720.1, rna-XM_032861722.1, rna-XM_032861724.1    rna-XM_046000660.1, rna-XM_046006273.1, rna-XM_046006275.1, rna-XM_046006276.1, rna-XM_046006277.1, rna-XM_046006278.1, rna-XM_046006279.1, rna-XM_046006280.1, rna-XM_046006281.1, rna-XM_046006283.1, rna-XM_046006284.1    rna-XM_032338249.1, rna-XM_032338258.1, rna-XM_032338259.1, rna-XM_032338263.1, rna-XM_032338264.1, rna-XM_032338265.1, rna-XM_032338266.1, rna-XM_032338267.1, rna-XM_032338268.1    rna-XM_059176078.1, rna-XM_059176248.1, rna-XM_059176256.1, rna-XM_059176259.1, rna-XM_059176260.1, rna-XM_059176261.1, rna-XM_059176262.1, rna-XM_059176267.1, rna-XM_059176268.1    rna-XM_059399595.1, rna-XM_059399686.1, rna-XM_059416306.1    rna-XM_013050073.2, rna-XM_013050500.2, rna-XM_045082264.1, rna-XM_045082266.1, rna-XM_045082271.1, rna-XM_045082272.1    mNeoVis.2.1.1.chr1_hap1.g01076.t01, mNeoVis.2.1.1.chr1_hap1.g01084.t01, mNeoVis.2.1.1.chr1_hap1.g01087.t01, mNeoVis.2.1.1.chr1_hap1.g01101.t01, mNeoVis.2.1.1.chr1_hap1.g01126.t01, mNeoVis.2.1.1.chr1_hap1.g01128.t01, mNeoVis.2.1.1.chr1_hap1.g01145.t01, mNeoVis.2.1.1.chr1_hap1.g01147.t01    rna-XM_021696845.2    rna-XM_004406515.1, rna-XM_004407068.1, rna-XM_004413658.1    rna-XM_007074492.3, rna-XM_007093442.3    rna-XM_008692953.2, rna-XM_040645173.1    rna-XM_027602240.2, rna-XM_027624356.2    mNeoVis.3.1.1.chr1.g01106.t01, mNeoVis.3.1.1.chr5.g00773.t01    rna-gnl|WGS_ZZZZ|egapxtmp_010164-R1, rna-gnl|WGS_ZZZZ|egapxtmp_018225-R1    mNeoVis.evergladensis.2.1.chr1.g01061.t01, mNeoVis.evergladensis.2.1.chr1.g01067.t01, mNeoVis.evergladensis.2.1.chr1.g01068.t01, mNeoVis.evergladensis.2.1.chr1.g01081.t01, mNeoVis.evergladensis.2.1.chr1.g01099.t01, mNeoVis.evergladensis.2.1.chr1.g01116.t01, mNeoVis.evergladensis.2.1.chr1.g01118.t01    mNeoVis.vulgivaga.2.1.chr1.g00971.t01, mNeoVis.vulgivaga.2.1.chr1.g01081.t01, mNeoVis.vulgivaga.2.1.chr1.g01084.t01      298    N0.HOG0000296   OG0000012  n127   rna-XM_002928522.4    rna-XM_038584413.1          rna-XM_039257337.1    rna-XM_032851224.1    rna-XM_046006286.1    rna-XM_032338269.1    rna-XM_059176270.1      rna-XM_013050499.2    mNeoVis.2.1.1.chr1_hap1.g01080.t01, mNeoVis.2.1.1.chr1_hap1.g01089.t01        rna-XM_042985402.1, rna-XM_042985543.1      rna-XM_035728424.1      rna-gnl|WGS_ZZZZ|egapxtmp_010058-R1      mNeoVis.vulgivaga.2.1.chr1.g01117.t01      299    N0.HOG0000297   OG0000012  n138                               rna-XM_004406516.1, rna-XM_004413655.1, rna-XM_004413657.1      rna-XM_008692949.2    rna-XM_035728427.1              300    N0.HOG0000298   OG0000012  n141             rna-XM_039257284.1                        rna-XM_008692970.2                301    N0.HOG0000299   OG0000013  n2   rna-XM_002919480.3    rna-XM_038586474.1    rna-XM_022507998.1    rna-XM_006943623.5    rna-XM_036114899.1    rna-XM_039225337.1, rna-XM_039225739.1    rna-XM_032842307.1    rna-XM_045996088.1    rna-XM_032331241.1    rna-XM_059157849.1    rna-XM_059385290.1    rna-XM_013064892.2    mNeoVis.2.1.1.chr12_hap1.g00721.t01    rna-XM_021705730.1    rna-XM_004417057.1    rna-XM_007084790.2    rna-XM_008696494.2    rna-XM_027609609.1    mNeoVis.3.1.1.chr12.g00659.t01      mNeoVis.evergladensis.2.1.chr12.g00653.t01    mNeoVis.vulgivaga.2.1.chr12.g00676.t01, mNeoVis.vulgivaga.2.1.chr12.g00678.t01      302    N0.HOG0000300   OG0000013  n24   rna-XM_034649913.1    rna-XM_038586473.1    rna-XM_022508027.1    rna-XM_011291657.2    rna-XM_036099555.1      rna-XM_032842309.1    rna-XM_045996130.1    rna-XM_032331244.1    rna-XM_059157850.1    rna-XM_059385291.1    rna-XM_013064894.2    mNeoVis.2.1.1.chr12_hap1.g00722.t01    rna-XM_021677680.1    rna-XM_004417058.1    rna-XM_042974596.1    rna-XM_040626257.1    rna-XM_027609611.1    mNeoVis.3.1.1.chr12.g00658.t01    rna-gnl|WGS_ZZZZ|egapxtmp_028387-R1    mNeoVis.evergladensis.2.1.chr12.g00654.t01    mNeoVis.vulgivaga.2.1.chr12.g00677.t01      303    N0.HOG0000301   OG0000013  n44   rna-XM_019797282.1    rna-XM_038586481.1    rna-XM_022507812.1    rna-XM_023249163.2    rna-XM_036095618.1    rna-XM_039255357.1    rna-XM_032844345.1    rna-XM_045995554.1    rna-XM_032331004.1    rna-XM_059157656.1    rna-XM_059385282.1    rna-XM_045089213.1    mNeoVis.2.1.1.chr12_hap1.g00711.t01    rna-XM_021677783.1    rna-XM_004417161.1    rna-XM_042975008.1    rna-XM_008696500.1    rna-XM_027608547.1      rna-gnl|WGS_ZZZZ|egapxtmp_029380-R1    mNeoVis.evergladensis.2.1.chr12.g00641.t01    mNeoVis.vulgivaga.2.1.chr12.g00664.t01      304    N0.HOG0000302   OG0000013  n65   rna-XM_019807465.2    rna-XM_038586479.1    rna-XM_022508026.1    rna-XM_006943637.4    rna-XM_036086952.1    rna-XM_039225088.1    rna-XM_032844357.1      rna-XM_032331589.1    rna-XM_059156782.1    rna-XM_059385287.1    rna-XM_045089230.1    mNeoVis.2.1.1.chr12_hap1.g00716.t01    rna-XM_044911849.1    rna-XM_004416693.1    rna-XM_042974372.1    rna-XM_008696449.2    rna-XM_027608682.1    mNeoVis.3.1.1.chr12.g00654.t01    rna-gnl|WGS_ZZZZ|egapxtmp_028251-R1    mNeoVis.evergladensis.2.1.chr12.g00648.t01    mNeoVis.vulgivaga.2.1.chr12.g00670.t01      305    N0.HOG0000303   OG0000013  n84   rna-XM_011234171.2, rna-XM_019807464.2    rna-NM_001003116.1    rna-XM_022508004.1, rna-XM_022508007.1    rna-XM_004000376.5, rna-XM_023249039.2    rna-XM_036086949.1, rna-XM_036086950.1    rna-XM_039225086.1, rna-XM_039225094.1, rna-XM_039225095.1    rna-XM_032844347.1, rna-XM_032844353.1    rna-XM_045994735.1, rna-XM_045996090.1    rna-XM_032331253.1, rna-XM_032331588.1    rna-XM_059156781.1, rna-XM_059157961.1    rna-XM_059385288.1, rna-XM_059385592.1    rna-XM_004761866.2, rna-XM_013061110.2    mNeoVis.2.1.1.chr12_hap1.g00717.t01    rna-XM_021705755.1, rna-XM_044912339.1    rna-XM_004416695.1    rna-XM_015538733.2, rna-XM_015538740.2    rna-XM_008696495.1, rna-XM_008696496.2    rna-XM_027607307.2, rna-XM_035725455.1    mNeoVis.3.1.1.chr12.g00655.t01    rna-gnl|WGS_ZZZZ|egapxtmp_028889-R1, rna-gnl|WGS_ZZZZ|egapxtmp_028890-R1    mNeoVis.evergladensis.2.1.chr12.g00649.t01    mNeoVis.vulgivaga.2.1.chr12.g00671.t01, mNeoVis.vulgivaga.2.1.chr12.g00672.t01      306    N0.HOG0000304   OG0000013  n123   rna-XM_034649752.1    rna-XM_038587117.1    rna-XM_022508005.1    rna-XM_045050420.1    rna-XM_036099558.1    rna-XM_039223228.1    rna-XM_032842303.1    rna-XM_045996089.1    rna-XM_032331135.1    rna-XM_059157691.1    rna-XM_059385570.1    rna-XM_004769146.3    mNeoVis.2.1.1.chr12_hap1.g00725.t01    rna-XM_021705718.1    rna-XM_004416460.1    rna-XM_042974173.1    rna-XM_040633311.1    rna-XM_027609702.2    mNeoVis.3.1.1.chr12.g00660.t01    rna-gnl|WGS_ZZZZ|egapxtmp_028388-R1    mNeoVis.evergladensis.2.1.chr12.g00656.t01, mNeoVis.evergladensis.2.1.chr12.g00657.t01    mNeoVis.vulgivaga.2.1.chr12.g00680.t01      307    N0.HOG0000305   OG0000013  n146   rna-XM_002920360.3    rna-XM_038586621.1    rna-XM_022521317.1    rna-XM_004000964.5    rna-XM_036071227.1    rna-XM_039220956.1    rna-XM_032841404.1    rna-XM_045996127.1    rna-XM_032331116.1    rna-XM_059157651.1    rna-XM_059385971.1    rna-XM_004773647.3    mNeoVis.2.1.1.chr12_hap1.g00138.t01    rna-XM_021684624.1    rna-XM_004405918.1    rna-XM_042974346.1    rna-XM_008701559.2    rna-XM_027608451.2    mNeoVis.3.1.1.chr12.g00108.t01    rna-gnl|WGS_ZZZZ|egapxtmp_029290-R1    mNeoVis.evergladensis.2.1.chr12.g00114.t01    mNeoVis.vulgivaga.2.1.chr12.g00115.t01      308    N0.HOG0000306   OG0000013  n166   rna-XM_034649648.1    rna-XM_038588457.1, rna-XM_038588844.1    rna-XM_022521310.1, rna-XM_022521316.1    rna-XM_045050576.1    rna-XM_036106982.1, rna-XM_036118617.1    rna-XM_039218514.1    rna-XM_032841399.1    rna-XM_045995679.1    rna-XM_032329732.1, rna-XM_032331747.1    rna-XM_059157601.1    rna-XM_059385960.1, rna-XM_059385967.1    rna-XM_004781568.3    mNeoVis.2.1.1.chr12_hap1.g00125.t01, mNeoVis.2.1.1.chr12_hap1.g00129.t01    rna-XM_021684928.1, rna-XM_044911914.1    rna-XM_004417180.1    rna-XM_042975178.1    rna-XM_008701563.1, rna-XM_008701565.2    rna-XM_027607638.1, rna-XM_027608296.1    mNeoVis.3.1.1.chr12.g00097.t01, mNeoVis.3.1.1.chr12.g00100.t01    rna-gnl|WGS_ZZZZ|egapxtmp_027979-R5, rna-gnl|WGS_ZZZZ|egapxtmp_029191-R1    mNeoVis.evergladensis.2.1.chr12.g00099.t01, mNeoVis.evergladensis.2.1.chr12.g00104.t01    mNeoVis.vulgivaga.2.1.chr12.g00101.t01, mNeoVis.vulgivaga.2.1.chr12.g00105.t01      309    N0.HOG0000307   OG0000013  n201   rna-XM_011229366.1    rna-XM_038586486.1    rna-XM_022508021.1    rna-XM_006943647.4    rna-XM_036072047.1, rna-XM_036072048.1    rna-XM_039255907.1, rna-XM_039255913.1    rna-XM_032883158.1    rna-XM_045994743.1, rna-XM_045994744.1    rna-XM_032331269.1, rna-XM_032331306.1    rna-XM_059156786.1, rna-XM_059157848.1    rna-XM_059385275.1    rna-XM_004779856.3, rna-XM_004779858.2    mNeoVis.2.1.1.chr12_hap1.g00695.t01, mNeoVis.2.1.1.chr12_hap1.g00696.t01    rna-XM_021681254.1    rna-XM_004404970.1    rna-XM_007090628.3    rna-XM_008696510.1    rna-XM_027608483.1      rna-gnl|WGS_ZZZZ|egapxtmp_028475-R1    mNeoVis.evergladensis.2.1.chr12.g00627.t01        310    N0.HOG0000308   OG0000013  n226   rna-XM_002922808.4    rna-XM_038586485.1    rna-XM_022507814.1      rna-XM_036092632.1    rna-XM_039256302.1    rna-XM_032883173.1    rna-XM_045994741.1    rna-XM_032329686.1        rna-XM_045089219.1    mNeoVis.2.1.1.chr12_hap1.g00697.t01    rna-XM_021681231.1    rna-XM_004404969.1      rna-XM_008696508.2    rna-XM_027608707.1    mNeoVis.3.1.1.chr12.g00640.t01    rna-gnl|WGS_ZZZZ|egapxtmp_029448-R1, rna-gnl|WGS_ZZZZ|egapxtmp_029451-R1    mNeoVis.evergladensis.2.1.chr12.g00628.t01    mNeoVis.vulgivaga.2.1.chr12.g00650.t01      311    N0.HOG0000309   OG0000013  n243   rna-XM_019800609.2      rna-XM_022518446.1    rna-XM_006943597.3    rna-XM_036083132.1    rna-XM_039233965.1    rna-XM_032840802.1    rna-XM_045995971.1    rna-XM_032331285.1    rna-XM_059158510.1    rna-XM_059385315.1    rna-XM_013059579.1    mNeoVis.2.1.1.chr12_hap1.g00776.t01    rna-XM_021681060.1    rna-XM_004415696.1    rna-XM_007092264.3    rna-XM_040634027.1    rna-XM_027608921.1    mNeoVis.3.1.1.chr12.g00708.t01    rna-gnl|WGS_ZZZZ|egapxtmp_028053-R1    mNeoVis.evergladensis.2.1.chr12.g00703.t01    mNeoVis.vulgivaga.2.1.chr12.g00731.t01, mNeoVis.vulgivaga.2.1.chr12.g00732.t01      312    N0.HOG0000310   OG0000013  n265   rna-XM_034649912.1    rna-XM_038588773.1, rna-XM_038588775.1, rna-XM_038588776.1    rna-XM_022507807.1, rna-XM_022507808.1    rna-XM_011291656.4, rna-XM_023248955.2    rna-XM_036099551.1, rna-XM_036099554.1, rna-XM_036099563.1      rna-XM_032842316.1, rna-XM_032842317.1    rna-XM_045994732.1, rna-XM_045994733.1, rna-XM_045994734.1    rna-XM_032331240.1, rna-XM_032331245.1, rna-XM_032331246.1    rna-XM_059157851.1, rna-XM_059157852.1, rna-XM_059157853.1    rna-XM_059385292.1    rna-XM_013064896.2, rna-XM_045089229.1    mNeoVis.2.1.1.chr12_hap1.g00723.t01, mNeoVis.2.1.1.chr12_hap1.g00724.t01    rna-XM_021677658.1, rna-XM_021677669.1, rna-XM_044911621.1    rna-XM_004417059.1    rna-XM_042973935.1, rna-XM_042973936.1    rna-XM_040636761.1    rna-XM_035725378.1, rna-XM_035725652.1, rna-XM_035725825.1      rna-gnl|WGS_ZZZZ|egapxtmp_028389-R1    mNeoVis.evergladensis.2.1.chr12.g00655.t01    mNeoVis.vulgivaga.2.1.chr12.g00679.t01      313    N0.HOG0000311   OG0000013  n304   rna-XM_034649918.1    rna-XM_038586482.1    rna-XM_022507999.1    rna-XM_004000391.6    rna-XM_036095616.1, rna-XM_036095617.1    rna-XM_039255355.1    rna-XM_032844344.1    rna-XM_045995553.1    rna-XM_032331039.1    rna-XM_059157672.1    rna-XM_059385281.1    rna-XM_004761859.2    mNeoVis.2.1.1.chr12_hap1.g00710.t01      rna-XM_004417160.1    rna-XM_042974260.1    rna-XM_008696501.2    rna-XM_027608438.1      rna-gnl|WGS_ZZZZ|egapxtmp_029382-R1    mNeoVis.evergladensis.2.1.chr12.g00640.t01    mNeoVis.vulgivaga.2.1.chr12.g00663.t01      314    N0.HOG0000312   OG0000013  n323   rna-XM_019797283.2    rna-XM_038586480.1    rna-XM_022507997.1    rna-XM_023249165.2      rna-XM_039255354.1    rna-XM_032844341.1    rna-XM_045995775.1    rna-XM_032330887.1    rna-XM_059157570.1    rna-XM_059385283.1    rna-XM_045089212.1    mNeoVis.2.1.1.chr12_hap1.g00712.t01    rna-XM_021677770.2    rna-XM_004417162.1    rna-XM_042975011.1    rna-XM_008696499.2    rna-XM_027608435.1, rna-XM_027609549.1    mNeoVis.3.1.1.chr12.g00651.t01    rna-gnl|WGS_ZZZZ|egapxtmp_029381-R1    mNeoVis.evergladensis.2.1.chr12.g00642.t01    mNeoVis.vulgivaga.2.1.chr12.g00665.t01      315    N0.HOG0000313   OG0000014  n1   rna-XM_019799301.2    rna-NM_001388872.1, rna-NM_001388917.1, rna-XM_038551634.1    rna-XM_022492614.1, rna-XM_022501390.1    rna-XM_011288557.3, rna-XM_045043550.1    rna-XM_036081205.1    rna-XM_039218628.1, rna-XM_039218629.1, rna-XM_039218630.1, rna-XM_039225700.1    rna-XM_032860856.1, rna-XM_032860857.1, rna-XM_032860858.1    rna-XM_046023126.1, rna-XM_046023199.1, rna-XM_046023210.1, rna-XM_046023231.1    rna-XM_032308272.1, rna-XM_032308273.1, rna-XM_032308277.1, rna-XM_032308278.1    rna-XM_059141690.1, rna-XM_059141694.1, rna-XM_059141696.1, rna-XM_059142169.1    rna-XM_059387505.1, rna-XM_059387507.1, rna-XM_059387537.1, rna-XM_059412260.1    rna-XM_004747575.3, rna-XM_004747576.3, rna-XM_004747879.3, rna-XM_004747881.3    mNeoVis.2.1.1.chr13_hap1.g00481.t01, mNeoVis.2.1.1.chr13_hap1.g00482.t01, mNeoVis.2.1.1.chr13_hap1.g00483.t01, mNeoVis.2.1.1.chr13_hap1.g00484.t01    rna-XM_021691401.1    rna-XM_004392270.1    rna-XM_042964174.1    rna-XM_040641226.1, rna-XM_040641227.1    rna-XM_027614265.1, rna-XM_035723655.1, rna-XM_035723656.1    mNeoVis.3.1.1.chr13.g00417.t01, mNeoVis.3.1.1.chr13.g00418.t01, mNeoVis.3.1.1.chr13.g00419.t01    rna-gnl|WGS_ZZZZ|egapxtmp_030863-R1, rna-gnl|WGS_ZZZZ|egapxtmp_030867-R1, rna-gnl|WGS_ZZZZ|egapxtmp_030868-R1, rna-gnl|WGS_ZZZZ|egapxtmp_030870-R1    mNeoVis.evergladensis.2.1.chr13.g00462.t01, mNeoVis.evergladensis.2.1.chr13.g00463.t01, mNeoVis.evergladensis.2.1.chr13.g00464.t01, mNeoVis.evergladensis.2.1.chr13.g00465.t01    mNeoVis.vulgivaga.2.1.chr13.g00464.t01, mNeoVis.vulgivaga.2.1.chr13.g00465.t01, mNeoVis.vulgivaga.2.1.chr13.g00466.t01, mNeoVis.vulgivaga.2.1.chr13.g00467.t01      316    N0.HOG0000314   OG0000014  n60   rna-XM_019799304.2    rna-NM_001388918.1    rna-XM_022501198.1    rna-XM_045043453.1    rna-XM_036081203.1    rna-XM_039234829.1    rna-XM_032860863.1    rna-XM_046023124.1    rna-XM_032308220.1    rna-XM_059142292.1    rna-XM_059412258.1    rna-XM_013053700.2    mNeoVis.2.1.1.chr13_hap1.g00476.t01    rna-XM_021691402.1    rna-XM_004392273.1    rna-XM_015535049.2    rna-XM_040631398.1    rna-XM_027616088.1    mNeoVis.3.1.1.chr13.g00413.t01    rna-gnl|WGS_ZZZZ|egapxtmp_031216-R1    mNeoVis.evergladensis.2.1.chr13.g00457.t01    mNeoVis.vulgivaga.2.1.chr13.g00460.t01      317    N0.HOG0000315   OG0000014  n79     rna-NM_001388527.1      rna-XM_011288555.1      rna-XM_039234831.1    rna-XM_032860859.1    rna-XM_046023120.1    rna-XM_032308274.1    rna-XM_059142262.1    rna-XM_059387451.1, rna-XM_059387490.1    rna-XM_004747577.2    mNeoVis.2.1.1.chr13_hap1.g00480.t01      rna-XM_004392272.1    rna-XM_007076481.1    rna-XM_008707813.1    rna-XM_027616084.1      rna-gnl|WGS_ZZZZ|egapxtmp_030864-R1    mNeoVis.evergladensis.2.1.chr13.g00461.t01    mNeoVis.vulgivaga.2.1.chr13.g00463.t01      318    N0.HOG0000316   OG0000014  n96   rna-XM_002918886.2    rna-XM_038551635.1    rna-XM_022501195.1    rna-XM_003995617.1    rna-XM_036081208.1    rna-XM_039234832.1    rna-XM_032860861.1    rna-XM_046023116.1    rna-XM_032308276.1    rna-XM_059142249.1    rna-XM_059412259.1    rna-XM_004747578.2    mNeoVis.2.1.1.chr13_hap1.g00479.t01    rna-XM_021691406.1      rna-XM_007076480.2    rna-XM_008708003.1    rna-XM_027616078.1    mNeoVis.3.1.1.chr13.g00416.t01    rna-gnl|WGS_ZZZZ|egapxtmp_030862-R1    mNeoVis.evergladensis.2.1.chr13.g00460.t01    mNeoVis.vulgivaga.2.1.chr13.g00462.t01      319    N0.HOG0000317   OG0000014  n115   rna-XM_034663708.1    rna-NM_001389006.1    rna-XM_022501336.1    rna-XM_003995711.4    rna-XM_036111429.1    rna-XM_039220418.1    rna-XM_032860873.1    rna-XM_046023230.1, rna-XM_046023262.1    rna-XM_032308530.1    rna-XM_059142879.1    rna-XM_059387475.1    rna-XM_004747894.1    mNeoVis.2.1.1.chr13_hap1.g00401.t01    rna-XM_044920562.1    rna-XM_004416753.1    rna-XM_007094472.1    rna-XM_008707903.2    rna-XM_027616225.1    mNeoVis.3.1.1.chr13.g00341.t01    rna-gnl|WGS_ZZZZ|egapxtmp_030090-R1    mNeoVis.evergladensis.2.1.chr13.g00381.t01    mNeoVis.vulgivaga.2.1.chr13.g00388.t01      320    N0.HOG0000318   OG0000014  n136   rna-XM_011237621.3    rna-NM_001256444.1      rna-XM_045042508.1      rna-XM_039240693.1    rna-XM_032860875.1    rna-XM_046023277.1      rna-XM_059142579.1    rna-XM_059412242.1    rna-XM_004747897.2    mNeoVis.2.1.1.chr13_hap1.g00396.t01    rna-XM_044920605.1      rna-XM_007094475.2    rna-XM_008708030.2      mNeoVis.3.1.1.chr13.g00337.t01    rna-gnl|WGS_ZZZZ|egapxtmp_030089-R1    mNeoVis.evergladensis.2.1.chr13.g00383.t01    mNeoVis.vulgivaga.2.1.chr13.g00383.t01      321    N0.HOG0000319   OG0000014  n152   rna-XM_002929570.3    rna-NM_001389020.1        rna-XM_036111458.1                          rna-XM_008708031.2                322    N0.HOG0000320   OG0000014  n156   rna-XM_034663711.1, rna-XM_034665566.1, rna-XM_034665568.1    rna-NM_001389024.1      rna-XM_003995715.5    rna-XM_036111427.1    rna-XM_039240646.1, rna-XM_039241071.1    rna-XM_032860877.1    rna-XM_046023326.1    rna-XM_032306477.1    rna-XM_059142608.1    rna-XM_059412241.1    rna-XM_004747898.2    mNeoVis.2.1.1.chr13_hap1.g00394.t01      rna-XM_004416762.1    rna-XM_007094260.3, rna-XM_007094476.2    rna-XM_008708032.2            mNeoVis.vulgivaga.2.1.chr13.g00381.t01      323    N0.HOG0000321   OG0000014  n175   rna-XM_034663712.1    rna-NM_001388895.1    rna-XM_022501426.1, rna-XM_022501464.1      rna-XM_036111450.1      rna-XM_032860876.1, rna-XM_032860878.1    rna-XM_046023315.1, rna-XM_046023316.1    rna-XM_032306478.1, rna-XM_032307883.1    rna-XM_059142590.1    rna-XM_059412240.1    rna-XM_004747899.1    mNeoVis.2.1.1.chr13_hap1.g00393.t01, mNeoVis.2.1.1.chr13_hap1.g00395.t01    rna-XM_021691539.2, rna-XM_044920603.1    rna-XM_004416755.1      rna-XM_008707900.2    rna-XM_027615137.2    mNeoVis.3.1.1.chr13.g00336.t01    rna-gnl|WGS_ZZZZ|egapxtmp_029898-R1    mNeoVis.evergladensis.2.1.chr13.g00382.t01    mNeoVis.vulgivaga.2.1.chr13.g00382.t01      324   N0.HOG0000322  OG0000014  n201   rna-XM_034663707.1    rna-XM_038551640.1    rna-XM_022501610.1    rna-XM_045043305.1    rna-XM_036111447.1    rna-XM_039220419.1    rna-XM_032860896.1    rna-XM_046021829.1    rna-XM_032306483.1    rna-XM_059143003.1    rna-XM_059410115.1    rna-XM_013053693.2    mNeoVis.2.1.1.chr13_hap1.g00404.t01    rna-XM_021691549.1      rna-XM_042965356.1    rna-XM_008708027.2    rna-XM_035723678.1    mNeoVis.3.1.1.chr13.g00342.t01    rna-gnl|WGS_ZZZZ|egapxtmp_030999-R1    mNeoVis.evergladensis.2.1.chr13.g00388.t01    mNeoVis.vulgivaga.2.1.chr13.g00390.t01      325   N0.HOG0000323  OG0000014  n219   rna-XM_002929574.4    rna-NM_001388529.1    rna-XM_022501309.1    rna-XM_023242524.2    rna-XM_036111448.1    rna-XM_039220415.1    rna-XM_032860897.1    rna-XM_046023181.1    rna-XM_032306482.1    rna-XM_059142880.1    rna-XM_059412245.1    rna-XM_004747893.3    mNeoVis.2.1.1.chr13_hap1.g00403.t01    rna-XM_021691618.2      rna-XM_007094471.2    rna-XM_008707904.2    rna-XM_035723676.1      rna-gnl|WGS_ZZZZ|egapxtmp_030997-R1    mNeoVis.evergladensis.2.1.chr13.g00387.t01    mNeoVis.vulgivaga.2.1.chr13.g00389.t01      326    N0.HOG0000324   OG0000014  n239   rna-XM_002929568.2, rna-XM_034663710.1    rna-NM_001388531.1      rna-XM_045042503.1      rna-XM_039220417.1    rna-XM_032860900.1    rna-XM_046023180.1, rna-XM_046023198.1, rna-XM_046023208.1    rna-XM_032308459.1    rna-XM_059142412.1      rna-XM_004747729.1    mNeoVis.2.1.1.chr13_hap1.g00397.t01      rna-XM_004416754.1    rna-XM_007094474.2    rna-XM_008707901.2      mNeoVis.3.1.1.chr13.g00338.t01    rna-gnl|WGS_ZZZZ|egapxtmp_030092-R1      mNeoVis.vulgivaga.2.1.chr13.g00384.t01      327    N0.HOG0000325   OG0000014  n257   rna-XM_002929569.2    rna-NM_001388530.1    rna-XM_022501328.1    rna-XM_045042511.1, rna-XM_045042512.1    rna-XM_036111449.1    rna-XM_039220413.1, rna-XM_039220416.1    rna-XM_032860874.1    rna-XM_046023201.1    rna-XM_032308457.1    rna-XM_059142384.1    rna-XM_059412243.1    rna-XM_004747728.1    mNeoVis.2.1.1.chr13_hap1.g00398.t01    rna-XM_044920565.1      rna-XM_007094256.2, rna-XM_007094258.2    rna-XM_008707902.2    rna-XM_027616222.1    mNeoVis.3.1.1.chr13.g00339.t01    rna-gnl|WGS_ZZZZ|egapxtmp_030091-R1      mNeoVis.vulgivaga.2.1.chr13.g00385.t01      328    N0.HOG0000326   OG0000014  n281           rna-XM_036081210.1                    rna-XM_021691403.1    rna-XM_004392356.1        rna-XM_027616082.1              329   N0.HOG0000327  OG0000014  n285     rna-NM_001388528.1    rna-XM_022501200.1    rna-XM_003995618.3      rna-XM_039234827.1      rna-XM_046023121.1    rna-XM_032308279.1    rna-XM_059142277.1      rna-XM_045081786.1    mNeoVis.2.1.1.chr13_hap1.g00478.t01        rna-XM_007076602.2    rna-XM_008708004.2      mNeoVis.3.1.1.chr13.g00415.t01    rna-gnl|WGS_ZZZZ|egapxtmp_031003-R1    mNeoVis.evergladensis.2.1.chr13.g00459.t01    mNeoVis.vulgivaga.2.1.chr13.g00461.t01      330    N0.HOG0000328   OG0000014  n296     rna-NM_001388934.1    rna-XM_022501223.1    rna-XM_003995619.4          rna-XM_046023125.1    rna-XM_032308211.1    rna-XM_059142290.1      rna-XM_004747579.3    mNeoVis.2.1.1.chr13_hap1.g00477.t01        rna-XM_007076479.3    rna-XM_008707814.1      mNeoVis.3.1.1.chr13.g00414.t01    rna-gnl|WGS_ZZZZ|egapxtmp_030869-R1    mNeoVis.evergladensis.2.1.chr13.g00458.t01        331    N0.HOG0000329   OG0000014  n308   rna-XM_034663705.1      rna-XM_022492613.1, rna-XM_022501199.1          rna-XM_032851175.1, rna-XM_032860855.1    rna-XM_046023206.1, rna-XM_046023229.1    rna-XM_032308270.1, rna-XM_032308271.1, rna-XM_032308280.1    rna-XM_059141691.1, rna-XM_059141692.1, rna-XM_059141693.1, rna-XM_059141695.1    rna-XM_059387445.1, rna-XM_059387494.1    rna-XM_004747574.3, rna-XM_004747878.3, rna-XM_004747880.2    mNeoVis.2.1.1.chr13_hap1.g00485.t01, mNeoVis.2.1.1.chr13_hap1.g00486.t01          rna-XM_008695229.2      mNeoVis.3.1.1.chr13.g00420.t01    rna-gnl|WGS_ZZZZ|egapxtmp_030866-R1    mNeoVis.evergladensis.2.1.chr13.g00466.t01    mNeoVis.vulgivaga.2.1.chr13.g00468.t01      332    N0.HOG0000330   OG0000015  n2   rna-XM_002930869.1    rna-NM_001388772.1    rna-XM_022490598.1    rna-XM_011279699.3, rna-XM_045057030.1, rna-XM_045058486.1    rna-XM_036111279.1    rna-XM_039225728.1, rna-XM_039225821.1    rna-XM_032861778.1    rna-XM_045997475.1, rna-XM_046005956.1    rna-XM_032328922.1, rna-XM_032341394.1    rna-XM_059155976.1, rna-XM_059179377.1    rna-XM_059401508.1    rna-XM_004766797.1    mNeoVis.2.1.1.chr1_hap1.g01070.t01      rna-XM_004408227.1, rna-XM_004411298.1    rna-XM_042985415.1, rna-XM_042985480.1, rna-XM_042985481.1, rna-XM_042985482.1, rna-XM_042985483.1    rna-XM_008693101.2, rna-XM_008708890.1, rna-XM_008709369.2, rna-XM_008711559.2, rna-XM_008711580.1, rna-XM_008712044.2    rna-XM_027598371.2, rna-XM_027604119.2          mNeoVis.vulgivaga.2.1.chr1.g01074.t01      333    N0.HOG0000331   OG0000015  n36   rna-XM_002931047.1, rna-XM_002931096.4, rna-XM_002931337.4, rna-XM_019807832.2, rna-XM_034650212.1, rna-XM_034653392.1, rna-XM_034653405.1, rna-XM_034653406.1, rna-XM_034653407.1, rna-XM_034653438.1, rna-XM_034653441.1, rna-XM_034660180.1    rna-XM_038583980.1    rna-XM_022490593.1, rna-XM_022492243.1, rna-XM_022492247.1, rna-XM_022492701.1, rna-XM_022492714.1, rna-XM_022492731.1, rna-XM_022527110.1, rna-XM_022527111.1, rna-XM_022527113.1, rna-XM_022527115.1, rna-XM_022527117.1, rna-XM_022527118.1, rna-XM_022527119.1    rna-XM_003980600.2, rna-XM_003985620.5, rna-XM_003985635.4, rna-XM_003985681.2, rna-XM_011282023.2, rna-XM_023249594.2, rna-XM_045058459.1, rna-XM_045058460.1, rna-XM_045058461.1, rna-XM_045058462.1, rna-XM_045058463.1, rna-XM_045058464.1, rna-XM_045058465.1    rna-XM_036066375.1, rna-XM_036070837.1, rna-XM_036074501.1, rna-XM_036074503.1, rna-XM_036074504.1, rna-XM_036074506.1, rna-XM_036077518.1, rna-XM_036111011.1, rna-XM_036111039.1, rna-XM_036111041.1, rna-XM_036111042.1, rna-XM_036121605.1    rna-XM_039218790.1, rna-XM_039223548.1, rna-XM_039223556.1, rna-XM_039224941.1, rna-XM_039225543.1, rna-XM_039225544.1, rna-XM_039225606.1, rna-XM_039225612.1, rna-XM_039225613.1, rna-XM_039225714.1, rna-XM_039257539.1, rna-XM_039257589.1    rna-XM_032847584.1, rna-XM_032847807.1, rna-XM_032847809.1, rna-XM_032847830.1, rna-XM_032850664.1, rna-XM_032850665.1, rna-XM_032861794.1, rna-XM_032861796.1, rna-XM_032861798.1, rna-XM_032861799.1, rna-XM_032869977.1    rna-XM_046000314.1, rna-XM_046004195.1, rna-XM_046004231.1, rna-XM_046005843.1, rna-XM_046005941.1, rna-XM_046005946.1, rna-XM_046005948.1, rna-XM_046005957.1, rna-XM_046005966.1, rna-XM_046006014.1, rna-XM_046006015.1, rna-XM_046006119.1, rna-XM_046006120.1, rna-XM_046006121.1, rna-XM_046006142.1, rna-XM_046006170.1, rna-XM_046006206.1, rna-XM_046006207.1, rna-XM_046006371.1, rna-XM_046006372.1, rna-XM_046006443.1, rna-XM_046006453.1, rna-XM_046006624.1    rna-XM_032328808.1, rna-XM_032328809.1, rna-XM_032335033.1, rna-XM_032338216.1, rna-XM_032338217.1, rna-XM_032338218.1, rna-XM_032338219.1, rna-XM_032338220.1, rna-XM_032338222.1, rna-XM_032338223.1, rna-XM_032338224.1, rna-XM_032338225.1, rna-XM_032338226.1, rna-XM_032338227.1, rna-XM_032338228.1, rna-XM_032338390.1, rna-XM_032338391.1    rna-XM_059173405.1, rna-XM_059176219.1, rna-XM_059176220.1, rna-XM_059176221.1, rna-XM_059176223.1, rna-XM_059176224.1, rna-XM_059176225.1, rna-XM_059176226.1, rna-XM_059176227.1, rna-XM_059176228.1, rna-XM_059176425.1    rna-XM_059384383.1, rna-XM_059387518.1, rna-XM_059387592.1, rna-XM_059398971.1, rna-XM_059399668.1, rna-XM_059399669.1, rna-XM_059399670.1, rna-XM_059399672.1, rna-XM_059399673.1, rna-XM_059416764.1    rna-XM_004766798.2, rna-XM_004778435.2, rna-XM_004779986.2, rna-XM_004779987.3, rna-XM_004780003.3, rna-XM_004780004.2, rna-XM_004781517.2, rna-XM_004781518.1, rna-XM_004781520.2, rna-XM_004781528.2, rna-XM_004781529.1, rna-XM_004781530.2, rna-XM_004782417.1, rna-XM_045073944.1, rna-XM_045073981.1    mNeoVis.2.1.1.chr1_hap1.g01002.t01, mNeoVis.2.1.1.chr1_hap1.g01059.t01, mNeoVis.2.1.1.chr1_hap1.g01060.t01, mNeoVis.2.1.1.chr1_hap1.g01062.t01, mNeoVis.2.1.1.chr1_hap1.g01063.t01, mNeoVis.2.1.1.chr1_hap1.g01064.t01, mNeoVis.2.1.1.chr1_hap1.g01065.t01, mNeoVis.2.1.1.chr1_hap1.g01066.t01, mNeoVis.2.1.1.chr1_hap1.g01067.t01, mNeoVis.2.1.1.chr1_hap1.g01068.t01, mNeoVis.2.1.1.chr1_hap1.g01069.t01, mNeoVis.2.1.1.chr1_hap1.g01073.t01, mNeoVis.2.1.1.chr6_hap1.g01133.t01    rna-XM_021689899.1, rna-XM_021697180.1, rna-XM_021697183.1    rna-XM_004411290.1, rna-XM_004416699.1, rna-XM_004416711.1, rna-XM_004417385.1, rna-XM_004417558.1    rna-XM_007074502.2, rna-XM_007084993.2, rna-XM_007085915.1, rna-XM_007085918.3, rna-XM_007085939.2, rna-XM_007092692.2, rna-XM_007092693.2, rna-XM_007092694.2, rna-XM_007092724.2, rna-XM_007093190.2, rna-XM_007093192.2, rna-XM_007093999.2, rna-XM_015540809.2, rna-XM_042985394.1, rna-XM_042985485.1, rna-XM_042985486.1, rna-XM_042985488.1, rna-XM_042985489.1, rna-XM_042985490.1, rna-XM_042985491.1, rna-XM_042985492.1    rna-XM_008684010.2, rna-XM_008684011.2, rna-XM_008693024.2, rna-XM_008693025.2, rna-XM_008708863.1, rna-XM_008710378.2, rna-XM_008711504.1, rna-XM_008711546.2, rna-XM_008711547.2, rna-XM_008711579.2, rna-XM_008712043.1, rna-XM_040644631.1, rna-XM_040645158.1    rna-XM_027624393.1, rna-XM_035728408.1, rna-XM_035728409.1, rna-XM_035728410.1    mNeoVis.3.1.1.chr1.g01053.t01, mNeoVis.3.1.1.chr1.g01055.t01, mNeoVis.3.1.1.chr1.g01056.t01, mNeoVis.3.1.1.chr1.g01057.t01, mNeoVis.3.1.1.chr1.g01058.t01, mNeoVis.3.1.1.chr1.g01059.t01, mNeoVis.3.1.1.chr1.g01062.t01, mNeoVis.3.1.1.chr1.g01064.t01, mNeoVis.3.1.1.chr4.g01079.t01    rna-gnl|WGS_ZZZZ|egapxtmp_003803-R1, rna-gnl|WGS_ZZZZ|egapxtmp_010048-R1, rna-gnl|WGS_ZZZZ|egapxtmp_010054-R1, rna-gnl|WGS_ZZZZ|egapxtmp_014440-R3, rna-gnl|WGS_ZZZZ|egapxtmp_014470-R1, rna-gnl|WGS_ZZZZ|egapxtmp_014473-R1, rna-gnl|WGS_ZZZZ|egapxtmp_014482-R1, rna-gnl|WGS_ZZZZ|egapxtmp_021181-R1, rna-gnl|WGS_ZZZZ|egapxtmp_021194-R1, rna-gnl|WGS_ZZZZ|egapxtmp_021201-R1, rna-gnl|WGS_ZZZZ|egapxtmp_044008-R1    mNeoVis.evergladensis.2.1.chr1.g00994.t01, mNeoVis.evergladensis.2.1.chr1.g01050.t01, mNeoVis.evergladensis.2.1.chr1.g01051.t01, mNeoVis.evergladensis.2.1.chr1.g01053.t01, mNeoVis.evergladensis.2.1.chr1.g01054.t01, mNeoVis.evergladensis.2.1.chr1.g01055.t01, mNeoVis.evergladensis.2.1.chr1.g01056.t01, mNeoVis.evergladensis.2.1.chr1.g01058.t01, mNeoVis.evergladensis.2.1.chr1.g01060.t01, mNeoVis.evergladensis.2.1.chr4.g01065.t01, mNeoVis.evergladensis.2.1.chr4.g01066.t01    mNeoVis.vulgivaga.2.1.chr1.g01008.t01, mNeoVis.vulgivaga.2.1.chr1.g01067.t01, mNeoVis.vulgivaga.2.1.chr1.g01068.t01, mNeoVis.vulgivaga.2.1.chr1.g01069.t01, mNeoVis.vulgivaga.2.1.chr1.g01070.t01, mNeoVis.vulgivaga.2.1.chr1.g01071.t01, mNeoVis.vulgivaga.2.1.chr1.g01072.t01, mNeoVis.vulgivaga.2.1.chr1.g01073.t01, mNeoVis.vulgivaga.2.1.chr1.g01077.t01, mNeoVis.vulgivaga.2.1.chr1.g01079.t01, mNeoVis.vulgivaga.2.1.chr4.g01100.t01      334   N0.HOG0000332  OG0000015  n274   rna-XM_019793777.2    rna-NM_001388862.1, rna-XM_038556366.1, rna-XM_038589397.1    rna-XM_022491989.1, rna-XM_022492242.1, rna-XM_022492244.1    rna-XM_006927435.3, rna-XM_011281246.2, rna-XM_011283251.3    rna-XM_036078722.1, rna-XM_036078723.1, rna-XM_036092320.1    rna-XM_039218787.1, rna-XM_039218788.1, rna-XM_039220853.1    rna-XM_032847585.1, rna-XM_032848094.1    rna-XM_046000317.1, rna-XM_046001116.1, rna-XM_046001125.1    rna-XM_032335034.1, rna-XM_032335043.1, rna-XM_032335165.1    rna-XM_059173388.1, rna-XM_059173404.1, rna-XM_059173506.1    rna-XM_059416412.1, rna-XM_059416728.1, rna-XM_059416761.1, rna-XM_059416763.1    rna-XM_004778436.2, rna-XM_004781507.1, rna-XM_004781565.3    mNeoVis.2.1.1.chr1_hap1.g00938.t01, mNeoVis.2.1.1.chr1_hap1.g01003.t01, mNeoVis.2.1.1.chr1_hap1.g01010.t01    rna-XM_021689894.1, rna-XM_021690948.1, rna-XM_044916825.1    rna-XM_004417166.1    rna-XM_007084991.2, rna-XM_007084992.1, rna-XM_007092957.1    rna-XM_008683745.1, rna-XM_008712041.1, rna-XM_008712042.1    rna-XM_027606796.2, rna-XM_027624383.1, rna-XM_027624385.2    mNeoVis.3.1.1.chr1.g00951.t01, mNeoVis.3.1.1.chr1.g01003.t01, mNeoVis.3.1.1.chr1.g01011.t01    rna-gnl|WGS_ZZZZ|egapxtmp_003850-R1, rna-gnl|WGS_ZZZZ|egapxtmp_020931-R1, rna-gnl|WGS_ZZZZ|egapxtmp_039723-R1    mNeoVis.evergladensis.2.1.chr1.g00934.t01, mNeoVis.evergladensis.2.1.chr1.g00995.t01, mNeoVis.evergladensis.2.1.chr1.g01003.t01    mNeoVis.vulgivaga.2.1.chr1.g00945.t01, mNeoVis.vulgivaga.2.1.chr1.g01009.t01      335   N0.HOG0000333  OG0000016  n1   rna-XM_034643203.1    rna-XM_038562075.1    rna-XM_022490570.1, rna-XM_022526254.1    rna-XM_006935069.5    rna-XM_036098398.1    rna-XM_039246209.1, rna-XM_039257290.1    rna-XM_032843714.1    rna-XM_046017958.1    rna-XM_032303385.1    rna-XM_059137253.1    rna-XM_059376007.1    rna-XM_004776835.3    mNeoVis.2.1.1.chr2_hap1.g00836.t01    rna-XM_021688070.1    rna-XM_004404170.1, rna-XM_004413662.1    rna-XM_007088031.3    rna-XM_008692968.2    rna-XM_035728442.1    mNeoVis.3.1.1.chr5.g00775.t01    rna-gnl|WGS_ZZZZ|egapxtmp_006208-R1, rna-gnl|WGS_ZZZZ|egapxtmp_021208-R1, rna-gnl|WGS_ZZZZ|egapxtmp_021244-R1    mNeoVis.evergladensis.2.1.chr5.g00790.t01    mNeoVis.vulgivaga.2.1.chr1.g01088.t01      336    N0.HOG0000334   OG0000016  n15   rna-XM_011216961.3, rna-XM_034655739.1    rna-XM_038556407.1    rna-XM_022491657.1, rna-XM_022491663.1    rna-XM_019830161.3    rna-XM_036119164.1    rna-XM_039220268.1, rna-XM_039257287.1, rna-XM_039257291.1, rna-XM_039257294.1, rna-XM_039257295.1    rna-XM_032848497.1, rna-XM_032848524.1      rna-XM_032335020.1, rna-XM_032335063.1    rna-XM_059173430.1, rna-XM_059176079.1    rna-XM_059416307.1, rna-XM_059416430.1    rna-XM_004780621.3, rna-XM_013050060.2    mNeoVis.2.1.1.chr1_hap1.g00965.t01      rna-XM_004407069.1, rna-XM_004407070.1    rna-XM_007093441.3, rna-XM_042985533.1    rna-XM_008708267.2, rna-XM_008708269.2, rna-XM_040645182.1    rna-XM_027624357.1, rna-XM_027624359.2    mNeoVis.3.1.1.chr1.g00974.t01, mNeoVis.3.1.1.chr1.g01079.t01, mNeoVis.3.1.1.chr1.g01102.t01      mNeoVis.evergladensis.2.1.chr1.g00960.t01    mNeoVis.vulgivaga.2.1.chr1.g00972.t01      337    N0.HOG0000335   OG0000016  n37   rna-XM_034643200.1    rna-XM_038562076.1, rna-XM_038562077.1, rna-XM_038562078.1    rna-XM_022492773.1, rna-XM_022526074.1    rna-XM_003990555.6    rna-XM_036098396.1, rna-XM_036098399.1, rna-XM_036098400.1    rna-XM_039246246.1    rna-XM_032843522.1, rna-XM_032851115.1    rna-XM_046012516.1, rna-XM_046012529.1, rna-XM_046013807.1    rna-XM_032303394.1    rna-XM_059136608.1      rna-XM_045083145.1    mNeoVis.2.1.1.chr2_hap1.g00827.t01, mNeoVis.2.1.1.chr2_hap1.g00830.t01, mNeoVis.2.1.1.chr2_hap1.g00831.t01    rna-XM_044912331.1, rna-XM_044914613.1    rna-XM_004404172.1, rna-XM_004404173.1, rna-XM_004404176.1    rna-XM_007088033.3, rna-XM_042993989.1, rna-XM_042993991.1    rna-XM_008695750.2, rna-XM_040626815.1, rna-XM_040626816.1    rna-XM_027571484.2, rna-XM_027607148.1, rna-XM_027622620.2        mNeoVis.evergladensis.2.1.chr5.g00784.t01    mNeoVis.vulgivaga.2.1.chr5.g00777.t01      338    N0.HOG0000336   OG0000016  n38   rna-XM_034660129.1    rna-XM_038584377.1    rna-XM_022490565.1    rna-XM_003985725.6    rna-XM_036112362.1    rna-XM_039257293.1    rna-XM_032861746.1    rna-XM_046006304.1    rna-XM_032338284.1    rna-XM_059176287.1    rna-XM_059399596.1    rna-XM_045082286.1    mNeoVis.2.1.1.chr1_hap1.g01148.t01    rna-XM_044917293.1      rna-XM_007074439.3    rna-XM_040645168.1    rna-XM_035728441.1    mNeoVis.3.1.1.chr1.g01107.t01    rna-gnl|WGS_ZZZZ|egapxtmp_018284-R1    mNeoVis.evergladensis.2.1.chr1.g01120.t01    mNeoVis.vulgivaga.2.1.chr1.g01116.t01      339   N0.HOG0000337  OG0000016  n43   rna-XM_002927101.4, rna-XM_002928525.4, rna-XM_002928532.4, rna-XM_002928539.4, rna-XM_002928552.3, rna-XM_002928555.4, rna-XM_034660114.1, rna-XM_034660117.1, rna-XM_034660127.1, rna-XM_034660128.1    rna-XM_038584350.1, rna-XM_038584351.1, rna-XM_038584366.1, rna-XM_038584367.1, rna-XM_038584368.1, rna-XM_038584369.1, rna-XM_038584405.1, rna-XM_038584425.1, rna-XM_038584426.1, rna-XM_038584437.1    rna-XM_022490561.1, rna-XM_022490562.1, rna-XM_022490563.1, rna-XM_022490564.1    rna-XM_003985682.6, rna-XM_003985690.6, rna-XM_003985694.3, rna-XM_003985706.6, rna-XM_003985716.6, rna-XM_003985720.5, rna-XM_045058515.1, rna-XM_045058520.1, rna-XM_045058521.1    rna-XM_036112345.1, rna-XM_036112354.1, rna-XM_036112355.1, rna-XM_036112357.1, rna-XM_036112360.1, rna-XM_036112361.1, rna-XM_036121570.1, rna-XM_036121573.1, rna-XM_036121579.1    rna-XM_039223539.1, rna-XM_039223555.1, rna-XM_039257285.1, rna-XM_039257286.1, rna-XM_039257289.1    rna-XM_032851090.1, rna-XM_032861732.1, rna-XM_032861736.1, rna-XM_032861743.1, rna-XM_032861744.1    rna-XM_046005614.1, rna-XM_046006290.1, rna-XM_046006294.1, rna-XM_046006296.1, rna-XM_046006298.1, rna-XM_046006299.1, rna-XM_046006300.1, rna-XM_046006301.1, rna-XM_046006303.1    rna-XM_032338271.1, rna-XM_032338272.1, rna-XM_032338274.1, rna-XM_032338276.1, rna-XM_032338278.1, rna-XM_032338282.1, rna-XM_032338283.1, rna-XM_032338286.1, rna-XM_032338287.1    rna-XM_059176271.1, rna-XM_059176272.1, rna-XM_059176274.1, rna-XM_059176275.1, rna-XM_059176281.1, rna-XM_059176283.1, rna-XM_059176285.1, rna-XM_059176286.1, rna-XM_059176288.1      rna-XM_045082236.1, rna-XM_045082275.1, rna-XM_045082279.1, rna-XM_045082281.1, rna-XM_045082285.1, rna-XM_045082287.1    mNeoVis.2.1.1.chr1_hap1.g01075.t01, mNeoVis.2.1.1.chr1_hap1.g01083.t01, mNeoVis.2.1.1.chr1_hap1.g01090.t01, mNeoVis.2.1.1.chr1_hap1.g01114.t01, mNeoVis.2.1.1.chr1_hap1.g01117.t01, mNeoVis.2.1.1.chr1_hap1.g01127.t01, mNeoVis.2.1.1.chr1_hap1.g01132.t01, mNeoVis.2.1.1.chr1_hap1.g01134.t01, mNeoVis.2.1.1.chr1_hap1.g01136.t01, mNeoVis.2.1.1.chr1_hap1.g01141.t01    rna-XM_021696848.2, rna-XM_044917586.1, rna-XM_044917587.1    rna-XM_004413647.1, rna-XM_004413652.1, rna-XM_004413659.1    rna-XM_007074443.3, rna-XM_007074448.3, rna-XM_007074450.3, rna-XM_007074471.3, rna-XM_007074501.3, rna-XM_007096007.3, rna-XM_015540538.2, rna-XM_015540585.2, rna-XM_015540673.2    rna-XM_008692958.2, rna-XM_008692985.2, rna-XM_008693017.2, rna-XM_040645185.1    rna-XM_027602247.2, rna-XM_027602283.2, rna-XM_035728431.1, rna-XM_035728443.1, rna-XM_035728444.1, rna-XM_035728445.1, rna-XM_035728446.1, rna-XM_035728447.1      rna-gnl|WGS_ZZZZ|egapxtmp_018274-R1, rna-gnl|WGS_ZZZZ|egapxtmp_018443-R1    mNeoVis.evergladensis.2.1.chr1.g01102.t01, mNeoVis.evergladensis.2.1.chr1.g01105.t01, mNeoVis.evergladensis.2.1.chr1.g01108.t01, mNeoVis.evergladensis.2.1.chr1.g01111.t01    mNeoVis.vulgivaga.2.1.chr1.g01080.t01, mNeoVis.vulgivaga.2.1.chr1.g01109.t01      340    N0.HOG0000338   OG0000016  n45             rna-XM_039223533.1                        rna-XM_008693022.2                341    N0.HOG0000339   OG0000016  n48   rna-XM_011236404.3    rna-XM_038584360.1, rna-XM_038584376.1      rna-XM_045058518.1, rna-XM_045058519.1    rna-XM_036112349.1, rna-XM_036112352.1    rna-XM_039257288.1    rna-XM_032861739.1    rna-XM_046006288.1    rna-XM_032338277.1, rna-XM_032338280.1    rna-XM_059176280.1      rna-XM_045082282.1, rna-XM_045082283.1    mNeoVis.2.1.1.chr1_hap1.g01138.t01      rna-XM_004406522.1    rna-XM_007074444.3, rna-XM_007096009.3    rna-XM_008692965.2    rna-XM_035728438.1    mNeoVis.3.1.1.chr1.g01100.t01      mNeoVis.evergladensis.2.1.chr1.g01103.t01    mNeoVis.vulgivaga.2.1.chr1.g01113.t01      342    N0.HOG0000340   OG0000016  n50   rna-XM_019807833.2, rna-XM_019807839.2    rna-XM_038584406.1, rna-XM_038584436.1    rna-XM_022490566.1, rna-XM_022490569.1, rna-XM_022490599.1    rna-XM_023254699.2, rna-XM_045058511.1, rna-XM_045058516.1    rna-XM_036112356.1, rna-XM_036112358.1, rna-XM_036121565.1, rna-XM_036121572.1    rna-XM_039224919.1    rna-XM_032861734.1, rna-XM_032861740.1, rna-XM_032861742.1    rna-XM_046006293.1, rna-XM_046006295.1, rna-XM_046006302.1    rna-XM_032338275.1, rna-XM_032338279.1, rna-XM_032338285.1, rna-XM_032341392.1    rna-XM_059176276.1, rna-XM_059176284.1    rna-XM_059399611.1, rna-XM_059399688.1    rna-XM_045082277.1, rna-XM_045082278.1, rna-XM_045082284.1    mNeoVis.2.1.1.chr1_hap1.g01082.t01, mNeoVis.2.1.1.chr1_hap1.g01099.t01, mNeoVis.2.1.1.chr1_hap1.g01102.t01    rna-XM_021696903.2, rna-XM_021697187.1    rna-XM_004406524.1, rna-XM_004413632.1, rna-XM_004413633.1, rna-XM_004413645.1, rna-XM_004413660.1    rna-XM_007074464.3, rna-XM_007074474.3    rna-XM_008692979.2, rna-XM_008692980.2    rna-XM_027602276.2, rna-XM_035728432.1, rna-XM_035728435.1, rna-XM_035728439.1, rna-XM_035728440.1    mNeoVis.3.1.1.chr1.g01068.t01    rna-gnl|WGS_ZZZZ|egapxtmp_021225-R1    mNeoVis.evergladensis.2.1.chr1.g01066.t01, mNeoVis.evergladensis.2.1.chr1.g01082.t01    mNeoVis.vulgivaga.2.1.chr1.g01096.t01      343    N0.HOG0000341   OG0000016  n61   rna-XM_034660130.1    rna-XM_038584424.1      rna-XM_023249598.2      rna-XM_039224920.1          rna-XM_059176278.1      rna-XM_045082306.1          rna-XM_042985549.1    rna-XM_008693005.2                344    N0.HOG0000342   OG0000017  n1   rna-XM_034639338.1    rna-XM_038527520.1    rna-XM_022492078.1    rna-XM_019818993.3    rna-XM_036091931.1    rna-XM_039224605.1    rna-XM_032847696.1    rna-XM_045989575.1    rna-XM_032324789.1    rna-XM_059152423.1    rna-XM_059381965.1    rna-XM_045077342.1    mNeoVis.2.1.1.chr7_hap1.g00632.t01    rna-XM_021682330.2    rna-XM_004410915.1    rna-XM_042969925.1    rna-XM_008706496.2    rna-XM_035725115.1    mNeoVis.3.1.1.chr7.g00606.t01    rna-gnl|WGS_ZZZZ|egapxtmp_010585-R1    mNeoVis.evergladensis.2.1.chr7.g00631.t01    mNeoVis.vulgivaga.2.1.chr7.g00633.t01      345    N0.HOG0000343   OG0000017  n21   rna-XM_034638037.1        rna-XM_045046896.1, rna-XM_045047320.1    rna-XM_036091954.1, rna-XM_036112096.1            rna-XM_059152470.1    rna-XM_059381993.1            rna-XM_042969514.1      rna-XM_027619725.2              346    N0.HOG0000344   OG0000017  n33   rna-XM_019807812.2, rna-XM_034650411.1, rna-XM_034672840.1    rna-XM_038527466.1    rna-XM_022492278.1, rna-XM_022492306.1    rna-XM_045045754.1    rna-XM_036075104.1, rna-XM_036075233.1    rna-XM_039225535.1    rna-XM_032847642.1, rna-XM_032847716.1    rna-XM_045987396.1, rna-XM_045987433.1    rna-XM_032324831.1, rna-XM_032324841.1    rna-XM_059152356.1, rna-XM_059152455.1    rna-XM_059381934.1, rna-XM_059381988.1      mNeoVis.2.1.1.chr7_hap1.g00610.t01    rna-XM_021691860.1, rna-XM_021691861.1      rna-XM_042969836.1    rna-XM_008706458.2, rna-XM_008706461.2    rna-XM_027619661.2, rna-XM_027619699.2      rna-gnl|WGS_ZZZZ|egapxtmp_013819-R1    mNeoVis.evergladensis.2.1.chr7.g00612.t01        347    N0.HOG0000345   OG0000017  n63   rna-XM_034639509.1    rna-XM_038527588.1    rna-XM_022492103.1    rna-XM_045046743.1    rna-XM_036091919.1    rna-XM_039219034.1    rna-XM_032847596.1    rna-XM_045987932.1    rna-XM_032324790.1    rna-XM_059152446.1    rna-XM_059381971.1    rna-XM_013048118.2    mNeoVis.2.1.1.chr7_hap1.g00644.t01    rna-XM_021682275.2      rna-XM_007089519.3    rna-XM_040633745.1    rna-XM_027618147.2    mNeoVis.3.1.1.chr7.g00613.t01      mNeoVis.evergladensis.2.1.chr7.g00641.t01        348    N0.HOG0000346   OG0000017  n81   rna-XM_011231683.3    rna-XM_038527587.1    rna-XM_022492080.1    rna-XM_006940784.5    rna-XM_036091912.1, rna-XM_036091964.1    rna-XM_039219036.1    rna-XM_032847697.1    rna-XM_045987939.1    rna-XM_032324765.1    rna-XM_059152397.1    rna-XM_059381966.1    rna-XM_013048095.2    mNeoVis.2.1.1.chr7_hap1.g00642.t01    rna-XM_021682345.1    rna-XM_004410914.1    rna-XM_042970687.1    rna-XM_008706531.2    rna-XM_027618148.2    mNeoVis.3.1.1.chr7.g00611.t01    rna-gnl|WGS_ZZZZ|egapxtmp_013914-R1    mNeoVis.evergladensis.2.1.chr7.g00639.t01    mNeoVis.vulgivaga.2.1.chr7.g00642.t01      349    N0.HOG0000347   OG0000017  n103   rna-XM_034639337.1    rna-XM_038525522.1      rna-XM_019819014.3    rna-XM_036091932.1    rna-XM_039219007.1, rna-XM_039219012.1      rna-XM_045989596.1        rna-XM_059383445.1        rna-XM_021682331.2, rna-XM_044911328.1    rna-XM_004410918.1    rna-XM_042969509.1, rna-XM_042969520.1    rna-XM_008706519.2    rna-XM_027620310.2, rna-XM_027620319.2              350    N0.HOG0000348   OG0000017  n118   rna-XM_034638051.1    rna-XM_038527576.1    rna-XM_022492074.1    rna-XM_019819030.3    rna-XM_036091950.1    rna-XM_039219005.1                  rna-XM_021682334.2    rna-XM_004410920.1      rna-XM_040634379.1    rna-XM_035724927.1              351    N0.HOG0000349   OG0000017  n130   rna-XM_034639345.1    rna-XM_038527580.1, rna-XM_038527581.1      rna-XM_023244875.2, rna-XM_045047315.1    rna-XM_036091927.1, rna-XM_036091928.1    rna-XM_039219061.1    rna-XM_032847670.1    rna-XM_045989563.1    rna-XM_032324719.1, rna-XM_032347855.1    rna-XM_059152378.1    rna-XM_059381925.1    rna-XM_045077315.1      rna-XM_021682329.2      rna-XM_042969510.1, rna-XM_042969922.1    rna-XM_040633775.1, rna-XM_040634380.1    rna-XM_035725109.1, rna-XM_035725114.1          mNeoVis.vulgivaga.2.1.chr7.g00635.t01      352    N0.HOG0000350   OG0000017  n152   rna-XM_034639364.1    rna-XM_038525685.1          rna-XM_039224497.1                  rna-XM_021691866.1                      353    N0.HOG0000351   OG0000017  n157   rna-XM_034638041.1        rna-XM_045046895.1    rna-XM_036091966.1                        rna-XM_042969533.1        mNeoVis.3.1.1.chr7.g00603.t01    rna-gnl|WGS_ZZZZ|egapxtmp_012125-R3    mNeoVis.evergladensis.2.1.chr7.g00630.t01        354    N0.HOG0000352   OG0000017  n163     rna-XM_038527517.1      rna-XM_003997358.6    rna-XM_036091968.1      rna-XM_032847671.1    rna-XM_045989557.1    rna-XM_032324734.1    rna-XM_059152379.1    rna-XM_059381924.1    rna-XM_004776882.3    mNeoVis.2.1.1.chr7_hap1.g00633.t01      rna-XM_004410916.1    rna-XM_042969904.1    rna-XM_040633784.1    rna-XM_027620311.1    mNeoVis.3.1.1.chr7.g00605.t01    rna-gnl|WGS_ZZZZ|egapxtmp_012118-R1    mNeoVis.evergladensis.2.1.chr7.g00632.t01    mNeoVis.vulgivaga.2.1.chr7.g00632.t01      355    N0.HOG0000353   OG0000017  n180   rna-XM_034650433.1    rna-XM_038527567.1      rna-XM_019818994.3    rna-XM_036091893.1      rna-XM_032847608.1    rna-XM_045989556.1    rna-XM_032324805.1    rna-XM_059152445.1    rna-XM_059383448.1    rna-XM_045077257.1    mNeoVis.2.1.1.chr7_hap1.g00634.t01        rna-XM_042969878.1    rna-XM_008706494.2    rna-XM_035725108.1    mNeoVis.3.1.1.chr7.g00607.t01    rna-gnl|WGS_ZZZZ|egapxtmp_012120-R1    mNeoVis.evergladensis.2.1.chr7.g00633.t01    mNeoVis.vulgivaga.2.1.chr7.g00634.t01      356    N0.HOG0000354   OG0000017  n196   rna-XM_011231654.3        rna-XM_023244873.2    rna-XM_036091914.1    rna-XM_039219058.1    rna-XM_032847659.1    rna-XM_045989551.1    rna-XM_032324684.1    rna-XM_059152304.1    rna-XM_059381923.1    rna-XM_004776879.3    mNeoVis.2.1.1.chr7_hap1.g00636.t01    rna-XM_044911295.1    rna-XM_004410909.1    rna-XM_042969507.1    rna-XM_040633778.1    rna-XM_035725189.1      rna-gnl|WGS_ZZZZ|egapxtmp_014115-R1    mNeoVis.evergladensis.2.1.chr7.g00635.t01    mNeoVis.vulgivaga.2.1.chr7.g00636.t01      357    N0.HOG0000355   OG0000017  n214   rna-XM_034639332.1    rna-XM_038527462.1    rna-XM_022492093.1    rna-XM_045045546.1    rna-XM_036091882.1    rna-XM_039224603.1    rna-XM_032847617.1    rna-XM_045989523.1    rna-XM_032324598.1    rna-XM_059152260.1    rna-XM_059381876.1    rna-XM_004776864.3    mNeoVis.2.1.1.chr7_hap1.g00630.t01      rna-XM_004405691.1    rna-XM_042969823.1    rna-XM_040633787.1    rna-XM_027620304.2    mNeoVis.3.1.1.chr7.g00602.t01    rna-gnl|WGS_ZZZZ|egapxtmp_012260-R1    mNeoVis.evergladensis.2.1.chr7.g00629.t01    mNeoVis.vulgivaga.2.1.chr7.g00631.t01      358    N0.HOG0000356   OG0000017  n234   rna-XM_034639330.1    rna-XM_038527460.1    rna-XM_022492096.1    rna-XM_045045543.1    rna-XM_036091871.1    rna-XM_039224601.1      rna-XM_045989517.1    rna-XM_032324586.1    rna-XM_059152249.1    rna-XM_059381871.1    rna-XM_045077296.1    mNeoVis.2.1.1.chr7_hap1.g00631.t01      rna-XM_004405675.1    rna-XM_042969818.1    rna-XM_040633786.1    rna-XM_027620295.2              359    N0.HOG0000357   OG0000017  n250         rna-XM_011289510.4, rna-XM_023245667.2, rna-XM_045046678.1, rna-XM_045046679.1      rna-XM_039224469.1, rna-XM_039224597.1, rna-XM_039224599.1, rna-XM_039225751.1                      rna-XM_042969817.1, rna-XM_042969819.1, rna-XM_042969821.1, rna-XM_042969833.1                  360    N0.HOG0000358   OG0000017  n262   rna-XM_034639361.1    rna-XM_038527463.1            rna-XM_032847628.1    rna-XM_045989535.1, rna-XM_045989600.1    rna-XM_032324619.1    rna-XM_059152280.1, rna-XM_059152467.1    rna-XM_059381895.1    rna-XM_004780506.3    mNeoVis.2.1.1.chr7_hap1.g00625.t01, mNeoVis.2.1.1.chr7_hap1.g00627.t01    rna-XM_044922242.1    rna-XM_004405697.1      rna-XM_008706511.2    rna-XM_035725160.1      rna-gnl|WGS_ZZZZ|egapxtmp_012115-R1      mNeoVis.vulgivaga.2.1.chr7.g00630.t01      361    N0.HOG0000359   OG0000017  n279   rna-XM_034638657.1    rna-XM_038525519.1    rna-XM_022492305.1, rna-XM_022492702.1      rna-XM_036087163.1, rna-XM_036091888.1, rna-XM_036114216.1      rna-XM_032847609.1    rna-XM_045989519.1, rna-XM_045989520.1, rna-XM_045989521.1    rna-XM_032324594.1, rna-XM_032324596.1, rna-XM_032324599.1    rna-XM_059152255.1, rna-XM_059152256.1, rna-XM_059152259.1    rna-XM_059387322.1    rna-XM_013050915.2, rna-XM_045077256.1, rna-XM_045077297.1, rna-XM_045077301.1, rna-XM_045077307.1    mNeoVis.2.1.1.chr7_hap1.g00628.t01, mNeoVis.2.1.1.chr7_hap1.g00629.t01      rna-XM_004405692.1, rna-XM_004405693.1      rna-XM_040633788.1, rna-XM_040633792.1    rna-XM_035725104.1, rna-XM_035725157.1      rna-gnl|WGS_ZZZZ|egapxtmp_012121-R1    mNeoVis.evergladensis.2.1.chr7.g00627.t01, mNeoVis.evergladensis.2.1.chr7.g00628.t01        362    N0.HOG0000360   OG0000017  n312     rna-XM_038527457.1        rna-XM_036088368.1, rna-XM_036114210.1, rna-XM_036114214.1      rna-XM_032847615.1    rna-XM_045986839.1, rna-XM_045989514.1    rna-XM_032324588.1    rna-XM_059152240.1      rna-XM_013049984.2    mNeoVis.2.1.1.chr7_hap1.g00626.t01    rna-XM_044911327.1, rna-XM_044912258.1    rna-XM_004405694.1      rna-XM_040633789.1, rna-XM_040633791.1    rna-XM_035725137.1, rna-XM_035725148.1, rna-XM_035725149.1              363    N0.HOG0000361   OG0000018  n2       rna-XM_022522018.1          rna-XM_032853609.1    rna-XM_046016525.1    rna-XM_032358166.1    rna-XM_059156845.1    rna-XM_059390603.1    rna-XM_004768209.2    mNeoVis.2.1.1.chr7_hap1.g01630.t01                rna-gnl|WGS_ZZZZ|egapxtmp_012841-R1    mNeoVis.evergladensis.2.1.chr7.g01601.t01        364    N0.HOG0000362   OG0000018  n13   rna-XM_002930702.3    rna-NM_001256448.1    rna-XM_022522026.1          rna-XM_032853613.1    rna-XM_046016600.1                    rna-XM_008710718.1                365    N0.HOG0000363   OG0000018  n18     rna-NM_001389141.1    rna-XM_022522044.1, rna-XM_022522055.1    rna-XM_003992840.3    rna-XM_036114418.1    rna-XM_039223177.1    rna-XM_032853608.1    rna-XM_046016398.1, rna-XM_046016669.1    rna-XM_032359739.1, rna-XM_032359744.1    rna-XM_059156836.1, rna-XM_059156883.1    rna-XM_059390526.1, rna-XM_059390633.1    rna-XM_004768242.1, rna-XM_045071490.1    mNeoVis.2.1.1.chr7_hap1.g01627.t01    rna-XM_021685667.2      rna-XM_007089399.2    rna-XM_008710705.2, rna-XM_008710713.2      mNeoVis.3.1.1.chr7.g01528.t01    rna-gnl|WGS_ZZZZ|egapxtmp_010634-R1, rna-gnl|WGS_ZZZZ|egapxtmp_012859-R1    mNeoVis.evergladensis.2.1.chr7.g01606.t01    mNeoVis.vulgivaga.2.1.chr7.g01618.t01      366    N0.HOG0000364   OG0000018  n45   rna-XM_019793524.2    rna-NM_001389042.1          rna-XM_039223179.1    rna-XM_032852681.1    rna-XM_046016399.1    rna-XM_032359745.1    rna-XM_059156929.1    rna-XM_059390488.1    rna-XM_004768249.1            rna-XM_008710716.1                367    N0.HOG0000365   OG0000018  n61     rna-NM_001389121.1          rna-XM_039223172.1                                      368    N0.HOG0000366   OG0000018  n62     rna-NM_001389031.1    rna-XM_022522063.1    rna-XM_045038818.1      rna-XM_039223171.1    rna-XM_032853610.1    rna-XM_046016276.1, rna-XM_046016407.1    rna-XM_032359081.1, rna-XM_032359454.1    rna-XM_059156855.1, rna-XM_059179341.1    rna-XM_059390566.1    rna-XM_004768244.1, rna-XM_013064293.1    mNeoVis.2.1.1.chr7_hap1.g01632.t01        rna-XM_015540751.2      rna-XM_027579462.1      rna-gnl|WGS_ZZZZ|egapxtmp_012888-R1    mNeoVis.evergladensis.2.1.chr7.g01604.t01    mNeoVis.vulgivaga.2.1.chr7.g01623.t01      369    N0.HOG0000367   OG0000018  n81       rna-XM_022522032.1    rna-XM_011286581.3    rna-XM_036114419.1        rna-XM_046016136.1    rna-XM_032358993.1      rna-XM_059390539.1    rna-XM_004768247.3    mNeoVis.2.1.1.chr7_hap1.g01635.t01    rna-XM_021685668.1      rna-XM_015540750.2    rna-XM_008710712.2        rna-gnl|WGS_ZZZZ|egapxtmp_012856-R1          370    N0.HOG0000368   OG0000018  n93   rna-XM_034653121.1    rna-NM_001388843.1    rna-XM_022522015.1, rna-XM_022522041.1    rna-XM_003992838.3    rna-XM_036081414.1    rna-XM_039223178.1    rna-XM_032852675.1, rna-XM_032852677.1, rna-XM_032852680.1    rna-XM_046015914.1, rna-XM_046015917.1, rna-XM_046016301.1    rna-XM_032359347.1, rna-XM_032359742.1, rna-XM_032360022.1    rna-XM_059156841.1, rna-XM_059156871.1    rna-XM_059390553.1, rna-XM_059390618.1    rna-XM_004768245.2, rna-XM_045071487.1    mNeoVis.2.1.1.chr7_hap1.g01628.t01, mNeoVis.2.1.1.chr7_hap1.g01634.t01      rna-XM_004407570.1, rna-XM_004407571.1    rna-XM_007089400.1    rna-XM_008710706.2, rna-XM_008710711.2, rna-XM_008710714.2    rna-XM_027577911.1    mNeoVis.3.1.1.chr7.g01532.t01    rna-gnl|WGS_ZZZZ|egapxtmp_012513-R1, rna-gnl|WGS_ZZZZ|egapxtmp_012831-R1    mNeoVis.evergladensis.2.1.chr7.g01599.t01, mNeoVis.evergladensis.2.1.chr7.g01605.t01    mNeoVis.vulgivaga.2.1.chr7.g01619.t01      371    N0.HOG0000369   OG0000018  n129     rna-XM_038568845.1    rna-XM_022521972.1        rna-XM_039223170.1    rna-XM_032853612.1    rna-XM_046016201.1    rna-XM_032359027.1    rna-XM_059156916.1    rna-XM_059390501.1    rna-XM_004768212.2    mNeoVis.2.1.1.chr7_hap1.g01636.t01          rna-XM_008710715.1      mNeoVis.3.1.1.chr7.g01538.t01    rna-gnl|WGS_ZZZZ|egapxtmp_012840-R1    mNeoVis.evergladensis.2.1.chr7.g01607.t01    mNeoVis.vulgivaga.2.1.chr7.g01625.t01      372    N0.HOG0000370   OG0000018  n143   rna-XM_002930584.4, rna-XM_002930586.4    rna-NM_001388718.1, rna-NM_001388720.1    rna-XM_022521984.1, rna-XM_022521985.1    rna-XM_003992911.6, rna-XM_003992912.6      rna-XM_039218841.1, rna-XM_039218873.1    rna-XM_032853663.1, rna-XM_032853665.1    rna-XM_046016295.1, rna-XM_046016299.1    rna-XM_032359217.1, rna-XM_032359416.1, rna-XM_032360030.1    rna-XM_059157708.1, rna-XM_059157892.1    rna-XM_059387439.1    rna-XM_004782499.2, rna-XM_045078265.1    mNeoVis.2.1.1.chr7_hap1.g01720.t01, mNeoVis.2.1.1.chr7_hap1.g01722.t01        rna-XM_015537935.2, rna-XM_042958485.1, rna-XM_042958486.1    rna-XM_008709560.2, rna-XM_008709561.2      mNeoVis.3.1.1.chr7.g01604.t01    rna-gnl|WGS_ZZZZ|egapxtmp_012858-R1    mNeoVis.evergladensis.2.1.chr7.g01689.t01        373    N0.HOG0000371   OG0000018  n175   rna-XM_002930583.1, rna-XM_002930588.1    rna-XM_038568859.1    rna-XM_022522006.1, rna-XM_022522016.1    rna-XM_003992878.2, rna-XM_003992879.2      rna-XM_039218845.1, rna-XM_039224510.1    rna-XM_032853661.1, rna-XM_032853662.1    rna-XM_046016577.1, rna-XM_046016580.1    rna-XM_032359590.1, rna-XM_032359606.1    rna-XM_059157832.1, rna-XM_059157844.1    rna-XM_059389640.1, rna-XM_059389653.1    rna-XM_004778986.1, rna-XM_004778987.1    mNeoVis.2.1.1.chr7_hap1.g01717.t01        rna-XM_007083265.1    rna-XM_008709562.1, rna-XM_008709623.1    rna-XM_027580116.1    mNeoVis.3.1.1.chr7.g01599.t01, mNeoVis.3.1.1.chr7.g01600.t01    rna-gnl|WGS_ZZZZ|egapxtmp_012532-R1, rna-gnl|WGS_ZZZZ|egapxtmp_012857-R1    mNeoVis.evergladensis.2.1.chr7.g01686.t01    mNeoVis.vulgivaga.2.1.chr7.g01701.t01      374   N0.HOG0000372  OG0000018  n207   rna-XM_002930585.3    rna-NM_001388719.1    rna-XM_022522005.1    rna-XM_045038132.1      rna-XM_039218843.1    rna-XM_032853664.1    rna-XM_046016612.1    rna-XM_032359312.1    rna-XM_059157882.1    rna-XM_059389588.1    rna-XM_013049241.2, rna-XM_045088009.1    mNeoVis.2.1.1.chr7_hap1.g01721.t01        rna-XM_015537936.2, rna-XM_042959612.1    rna-XM_008709619.2      mNeoVis.3.1.1.chr7.g01603.t01    rna-gnl|WGS_ZZZZ|egapxtmp_012855-R1    mNeoVis.evergladensis.2.1.chr7.g01688.t01    mNeoVis.vulgivaga.2.1.chr7.g01703.t01      375   N0.HOG0000373  OG0000018  n227   rna-XM_011217418.3    rna-XM_038568860.1    rna-XM_022521698.1    rna-XM_003992910.6, rna-XM_004001181.4      rna-XM_039218842.1    rna-XM_032852717.1    rna-XM_046016575.1    rna-XM_032359785.1    rna-XM_059157854.1    rna-XM_059389627.1    rna-XM_004778985.2, rna-XM_045088011.1    mNeoVis.2.1.1.chr7_hap1.g01718.t01        rna-XM_007083266.1, rna-XM_007083267.3    rna-XM_008709622.2    rna-XM_035723022.1    mNeoVis.3.1.1.chr7.g01601.t01    rna-gnl|WGS_ZZZZ|egapxtmp_012871-R1          376    N0.HOG0000374   OG0000018  n246     rna-NM_001388717.1, rna-NM_001389001.1    rna-XM_022522019.1            rna-XM_046016497.1    rna-XM_032359780.1    rna-XM_059157722.1, rna-XM_059157866.1    rna-XM_059389615.1, rna-XM_059389759.1    rna-XM_004779045.2, rna-XM_045088010.1    mNeoVis.2.1.1.chr7_hap1.g01719.t01          rna-XM_008709620.2, rna-XM_008709621.2    rna-XM_027580626.2    mNeoVis.3.1.1.chr7.g01602.t01      mNeoVis.evergladensis.2.1.chr7.g01687.t01    mNeoVis.vulgivaga.2.1.chr7.g01702.t01      377    N0.HOG0000375   OG0000018  n263   rna-XM_002930589.3, rna-XM_002931186.2    rna-NM_001388721.1    rna-XM_022522022.1    rna-XM_003992913.1      rna-XM_039218848.1    rna-XM_032853667.1    rna-XM_046015880.1, rna-XM_046016609.1    rna-XM_032359049.1    rna-XM_059157903.1    rna-XM_059389574.1    rna-XM_004779039.1, rna-XM_045088008.1    mNeoVis.2.1.1.chr7_hap1.g01723.t01        rna-XM_042958794.1    rna-XM_008709558.1, rna-XM_008709618.1      mNeoVis.3.1.1.chr7.g01605.t01, mNeoVis.3.1.1.chr7.g01606.t01    rna-gnl|WGS_ZZZZ|egapxtmp_012833-R1    mNeoVis.evergladensis.2.1.chr7.g01690.t01    mNeoVis.vulgivaga.2.1.chr7.g01704.t01      378    N0.HOG0000376   OG0000018  n285   rna-XM_002931131.4, rna-XM_002931132.4                                  rna-XM_008710708.2, rna-XM_040631303.1                379   N0.HOG0000377  OG0000018  n289     rna-XM_038568156.1    rna-XM_022522020.1, rna-XM_022522070.1    rna-XM_003992804.2      rna-XM_039223174.1    rna-XM_032851135.1, rna-XM_032852676.1, rna-XM_032853604.1    rna-XM_046016527.1, rna-XM_046016528.1    rna-XM_032359375.1, rna-XM_032359461.1    rna-XM_059156814.1, rna-XM_059156852.1, rna-XM_059156862.1    rna-XM_059387413.1, rna-XM_059390580.1, rna-XM_059390706.1    rna-XM_004768207.1, rna-XM_004768211.1    mNeoVis.2.1.1.chr7_hap1.g01624.t01, mNeoVis.2.1.1.chr7_hap1.g01631.t01, mNeoVis.2.1.1.chr7_hap1.g01633.t01    rna-XM_021685672.1      rna-XM_007089344.2        mNeoVis.3.1.1.chr7.g01534.t01, mNeoVis.3.1.1.chr7.g01536.t01    rna-gnl|WGS_ZZZZ|egapxtmp_012515-R1, rna-gnl|WGS_ZZZZ|egapxtmp_012854-R1    mNeoVis.evergladensis.2.1.chr7.g01595.t01, mNeoVis.evergladensis.2.1.chr7.g01602.t01, mNeoVis.evergladensis.2.1.chr7.g01603.t01    mNeoVis.vulgivaga.2.1.chr7.g01614.t01, mNeoVis.vulgivaga.2.1.chr7.g01621.t01, mNeoVis.vulgivaga.2.1.chr7.g01622.t01      380    N0.HOG0000378   OG0000018  n321     rna-NM_001388844.1      rna-XM_003992803.3    rna-XM_036081402.1    rna-XM_039223176.1                  rna-XM_021685671.1    rna-XM_004407554.1, rna-XM_004407555.1    rna-XM_007089345.2      rna-XM_027580237.1, rna-XM_027580333.1              381    N0.HOG0000379   OG0000019  n1   rna-XM_034638974.1    rna-XM_038528105.1    rna-XM_022523880.1    rna-XM_045046318.1    rna-XM_036068129.1    rna-XM_039247143.1    rna-XM_032838892.1    rna-XM_045988200.1    rna-XM_032325766.1    rna-XM_059151009.1    rna-XM_059382073.1    rna-XM_045077361.1    mNeoVis.2.1.1.chr7_hap1.g00863.t01    rna-XM_044911268.1    rna-XM_004409916.1    rna-XM_042970328.1    rna-XM_040634342.1    rna-XM_035724865.1    mNeoVis.3.1.1.chr7.g00805.t01    rna-gnl|WGS_ZZZZ|egapxtmp_012330-R1    mNeoVis.evergladensis.2.1.chr7.g00856.t01    mNeoVis.vulgivaga.2.1.chr7.g00861.t01      382    N0.HOG0000380   OG0000019  n24   rna-XM_019801732.2    rna-XM_038536839.1    rna-XM_022524562.1    rna-XM_023244684.2    rna-XM_036094490.1    rna-XM_039231495.1    rna-XM_032840002.1    rna-XM_045984913.1    rna-XM_032320752.1    rna-XM_059150553.1    rna-XM_059380380.1    rna-XM_045083358.1    mNeoVis.2.1.1.chr8_hap1.g01004.t01    rna-XM_021694815.2    rna-XM_004398449.1    rna-XM_042966028.1    rna-XM_040630018.1    rna-XM_027625680.2    mNeoVis.3.1.1.chr8.g00937.t01    rna-gnl|WGS_ZZZZ|egapxtmp_017557-R1    mNeoVis.evergladensis.2.1.chr9.g00978.t01    mNeoVis.vulgivaga.2.1.chr8.g00979.t01      383    N0.HOG0000381   OG0000019  n45   rna-XM_034670322.1    rna-XM_038540265.1      rna-XM_023246707.2    rna-XM_036115497.1    rna-XM_039233549.1    rna-XM_032836229.1      rna-XM_032328679.1    rna-XM_059155623.1    rna-XM_059415254.1    rna-XM_045072131.1    mNeoVis.2.1.1.chr15_hap1.g00213.t01    rna-XM_044915630.1    rna-XM_004408934.1    rna-XM_042972375.1    rna-XM_040635288.1    rna-XM_027614097.2    mNeoVis.3.1.1.chr15.g00187.t01    rna-gnl|WGS_ZZZZ|egapxtmp_033223-R1    mNeoVis.evergladensis.2.1.chr15.g00192.t01    mNeoVis.vulgivaga.2.1.chr15.g00193.t01      384    N0.HOG0000382   OG0000019  n64   rna-XM_011218801.3    rna-NM_001110767.1    rna-XM_022516497.1, rna-XM_022519126.1    rna-XM_023257448.2    rna-XM_036108739.1    rna-XM_039246115.1    rna-XM_032836576.1    rna-XM_045993786.1, rna-XM_046011458.1    rna-XM_032346410.1    rna-XM_059186854.1    rna-XM_059402106.1    rna-XM_045083927.1    mNeoVis.2.1.1.chr9_hap1.g01126.t01    rna-XM_044915335.1    rna-XM_004417338.1    rna-XM_042993168.1    rna-XM_040637468.1    rna-XM_027592045.2    mNeoVis.3.1.1.chr10.g01080.t01    rna-gnl|WGS_ZZZZ|egapxtmp_033957-R1    mNeoVis.evergladensis.2.1.chr10.g01114.t01    mNeoVis.vulgivaga.2.1.chr10.g01105.t01      385    N0.HOG0000383   OG0000019  n90   rna-XM_002921186.4    rna-NM_001077235.1    rna-XM_022524686.1    rna-XM_011288842.3    rna-XM_036100104.1    rna-XM_039231617.1    rna-XM_032839930.1    rna-XM_045984550.1    rna-XM_032320456.1    rna-XM_059147308.1      rna-XM_045083396.1    mNeoVis.2.1.1.chr8_hap1.g00988.t01    rna-XM_021694820.1    rna-XM_004398427.1    rna-XM_007085947.3    rna-XM_040630039.1    rna-XM_027625156.2    mNeoVis.3.1.1.chr8.g00921.t01    rna-gnl|WGS_ZZZZ|egapxtmp_019265-R1    mNeoVis.evergladensis.2.1.chr9.g00962.t01    mNeoVis.vulgivaga.2.1.chr8.g00962.t01      386    N0.HOG0000384   OG0000019  n111   rna-XM_034647010.1    rna-NM_001076794.1    rna-XM_022524683.1    rna-XM_045045331.1      rna-XM_039231624.1    rna-XM_032839809.1    rna-XM_045984554.1    rna-XM_032321684.1    rna-XM_059147306.1    rna-XM_059379461.1    rna-XM_045083397.1    mNeoVis.2.1.1.chr8_hap1.g00987.t01    rna-XM_044921615.1    rna-XM_004398426.1    rna-XM_007085948.3    rna-XM_008688017.2    rna-XM_027625154.2    mNeoVis.3.1.1.chr8.g00920.t01    rna-gnl|WGS_ZZZZ|egapxtmp_019262-R3    mNeoVis.evergladensis.2.1.chr9.g00961.t01    mNeoVis.vulgivaga.2.1.chr8.g00961.t01      387    N0.HOG0000385   OG0000019  n132   rna-XM_034647204.1, rna-XM_034652165.1, rna-XM_034652848.1, rna-XM_034652903.1    rna-NM_001113712.1    rna-XM_022524690.1    rna-XM_011288844.4      rna-XM_039231628.1      rna-XM_045984549.1    rna-XM_032320454.1    rna-XM_059147309.1    rna-XM_059380099.1    rna-XM_004760163.3    mNeoVis.2.1.1.chr8_hap1.g00984.t01    rna-XM_044921863.1    rna-XM_004398424.1    rna-XM_007085945.3    rna-XM_040630041.1    rna-XM_035724598.1    mNeoVis.3.1.1.chr8.g00917.t01    rna-gnl|WGS_ZZZZ|egapxtmp_019263-R1    mNeoVis.evergladensis.2.1.chr9.g00958.t01    mNeoVis.vulgivaga.2.1.chr8.g00958.t01      388    N0.HOG0000386   OG0000019  n152   rna-XM_034647280.1, rna-XM_034653012.1    rna-XM_038536881.1    rna-XM_022524574.1    rna-XM_045045308.1    rna-XM_036100106.1    rna-XM_039231631.1    rna-XM_032839808.1    rna-XM_045984910.1    rna-XM_032320457.1    rna-XM_059147305.1    rna-XM_059380391.1    rna-XM_045083393.1    mNeoVis.2.1.1.chr8_hap1.g00989.t01    rna-XM_021694529.1    rna-XM_004398618.1    rna-XM_042965988.1    rna-XM_008688010.2    rna-XM_027625162.2    mNeoVis.3.1.1.chr8.g00922.t01    rna-gnl|WGS_ZZZZ|egapxtmp_017548-R1, rna-gnl|WGS_ZZZZ|egapxtmp_019266-R1    mNeoVis.evergladensis.2.1.chr9.g00963.t01    mNeoVis.vulgivaga.2.1.chr8.g00963.t01, mNeoVis.vulgivaga.2.1.chr8.g00964.t01      389    N0.HOG0000387   OG0000019  n179   rna-XM_034657888.1    rna-XM_038582731.1    rna-XM_022502027.1    rna-XM_011285970.4    rna-XM_036068334.1    rna-XM_039244987.1      rna-XM_046001581.1    rna-XM_032344388.1    rna-XM_059165068.1    rna-XM_059388137.1      mNeoVis.2.1.1.chr5_hap1.g01619.t01    rna-XM_044919496.1    rna-XM_004391698.1    rna-XM_007098585.3    rna-XM_008706610.2    rna-XM_027587172.2    mNeoVis.3.1.1.chr2.g01534.t01    rna-gnl|WGS_ZZZZ|egapxtmp_035655-R1    mNeoVis.evergladensis.2.1.chr2.g01606.t01    mNeoVis.vulgivaga.2.1.chr2.g01620.t01      390    N0.HOG0000388   OG0000019  n198   rna-XM_034639917.1    rna-XM_038572080.1    rna-XM_022494369.1    rna-XM_003983604.6    rna-XM_036092264.1    rna-XM_039248695.1    rna-XM_032858813.1    rna-XM_045980379.1    rna-XM_032350873.1    rna-XM_059133434.1    rna-XM_059406855.1    rna-XM_013053365.2    mNeoVis.2.1.1.chr11_hap1.g00293.t01    rna-XM_044918603.1    rna-XM_004393093.1    rna-XM_042980545.1    rna-XM_040638228.1    rna-XM_027620790.2    mNeoVis.3.1.1.chr11.g00264.t01      mNeoVis.evergladensis.2.1.chr11.g00290.t01    mNeoVis.vulgivaga.2.1.chr11.g00292.t01      391    N0.HOG0000389   OG0000019  n218   rna-XM_011231026.3    rna-XM_038539442.1    rna-XM_022500962.1    rna-NM_001009221.1    rna-XM_036064887.1    rna-XM_039226092.1    rna-XM_032872658.1    rna-XM_045993531.1    rna-XM_032328465.1    rna-XM_059155043.1    rna-XM_059415920.1    rna-XM_004774950.2    mNeoVis.2.1.1.chr15_hap1.g00639.t01    rna-XM_021680105.1    rna-XM_004394183.1    rna-XM_042970958.1    rna-XM_040634785.1    rna-XM_027612024.2    mNeoVis.3.1.1.chr15.g00420.t01    rna-gnl|WGS_ZZZZ|egapxtmp_033906-R1    mNeoVis.evergladensis.2.1.chr15.g00461.t01    mNeoVis.vulgivaga.2.1.chr15.g00468.t01      392    N0.HOG0000390   OG0000019  n239   rna-XM_034648669.1, rna-XM_034651190.1    rna-NM_001113711.1    rna-XM_022492780.1, rna-XM_022521116.1    rna-XM_006932746.5    rna-XM_036114770.1    rna-XM_039225829.1, rna-XM_039254680.1    rna-XM_032856354.1    rna-XM_046008445.1    rna-XM_032343855.1    rna-XM_059181679.1    rna-XM_059372910.1    rna-XM_045067767.1    mNeoVis.2.1.1.chr10_hap1.g00558.t01    rna-XM_044912263.1, rna-XM_044912269.1    rna-XM_004402157.1    rna-XM_042989416.1    rna-XM_040620902.1    rna-XM_035728000.1    mNeoVis.3.1.1.chr9.g00516.t01    rna-gnl|WGS_ZZZZ|egapxtmp_016854-R1, rna-gnl|WGS_ZZZZ|egapxtmp_016855-R1    mNeoVis.evergladensis.2.1.chr8.g00540.t01    mNeoVis.vulgivaga.2.1.chr9.g00542.t01      393    N0.HOG0000391   OG0000019  n265   rna-XM_034648522.1    rna-XM_038544221.1    rna-XM_022491983.1, rna-XM_022492751.1    rna-XM_019832816.3      rna-XM_039245149.1, rna-XM_039254701.1    rna-XM_032856356.1    rna-XM_046008446.1    rna-XM_032343854.1    rna-XM_059181677.1    rna-XM_059372137.1    rna-XM_045067765.1    mNeoVis.2.1.1.chr10_hap1.g00559.t01    rna-XM_044912264.1    rna-XM_004402158.1    rna-XM_042989418.1    rna-XM_040620754.1      mNeoVis.3.1.1.chr9.g00517.t01    rna-gnl|WGS_ZZZZ|egapxtmp_030019-R1    mNeoVis.evergladensis.2.1.chr8.g00541.t01    mNeoVis.vulgivaga.2.1.chr9.g00543.t01      394    N0.HOG0000392   OG0000019  n286   rna-XM_034647013.1    rna-NM_001113717.1    rna-XM_022492779.1    rna-XM_023243774.2          rna-XM_045984548.1    rna-XM_032320455.1    rna-XM_059147303.1    rna-XM_059387431.1    rna-XM_045083395.1    mNeoVis.2.1.1.chr8_hap1.g00986.t01    rna-XM_044921798.1    rna-XM_004398617.1    rna-XM_007085949.2    rna-XM_008688018.2    rna-XM_035724543.1    mNeoVis.3.1.1.chr8.g00918.t01    rna-gnl|WGS_ZZZZ|egapxtmp_019115-R1          395    N0.HOG0000393   OG0000019  n302       rna-XM_022492581.1        rna-XM_039231622.1                                      396    N0.HOG0000394   OG0000019  n303   rna-XM_034647012.1    rna-NM_001076795.1    rna-XM_022524575.1, rna-XM_022524576.1    rna-XM_006939791.5    rna-XM_036100100.1    rna-XM_039231618.1    rna-XM_032839802.1    rna-XM_045984546.1    rna-XM_032320452.1    rna-XM_059147302.1    rna-XM_059378589.1, rna-XM_059387456.1    rna-XM_013060371.2    mNeoVis.2.1.1.chr8_hap1.g00985.t01    rna-XM_044921799.1    rna-XM_004398425.1    rna-XM_042965991.1    rna-XM_040628231.1    rna-XM_035724572.1    mNeoVis.3.1.1.chr8.g00919.t01    rna-gnl|WGS_ZZZZ|egapxtmp_019114-R1    mNeoVis.evergladensis.2.1.chr9.g00960.t01    mNeoVis.vulgivaga.2.1.chr8.g00959.t01      397    N0.HOG0000395   OG0000020  n0   id-LOC100482805, id-LOC105235222, id-LOC105235261, id-LOC109488358, id-LOC109488764, id-LOC117798506, id-LOC117798537, id-LOC117798617, id-LOC117798737, id-LOC117798740, id-LOC117798759, id-LOC117798776, id-LOC117798806, id-LOC117798827, id-LOC117798844, id-LOC117800014    id-LOC106559121, id-LOC111097181, id-LOC119869010, id-LOC119869011, id-LOC119869014, id-LOC119869015, id-LOC119869016, id-LOC119869025, id-LOC119869032, id-LOC119869037, id-LOC119869049, id-LOC119869050, id-LOC119869051, id-LOC119869066, id-LOC119872942, id-LOC119872961, id-LOC119875768, id-LOC119875780, id-LOC119876514, id-LOC119876515, id-LOC119876517, id-LOC119876518, id-LOC119876522, id-LOC119876533, id-LOC119876534, id-LOC119876536, id-LOC119876540, id-LOC119876541, id-LOC119876542, id-LOC119876543, id-LOC119876544, id-LOC119876548, id-LOC119876551, id-LOC119876552, id-LOC119876553, id-LOC119876555, id-LOC119876556, id-LOC119876557, id-LOC119876558, id-LOC119876561, id-LOC119876563, id-LOC119876564, id-LOC119876565, id-LOC119876566, id-LOC119876567, id-LOC119876570, id-LOC119876573, id-LOC119876576, id-LOC119876577, id-LOC119876578, id-LOC607467, rna-XM_038545965.1    id-LOC111139941, id-LOC111139942, id-LOC111139944, id-LOC111139945, id-LOC111139947, id-LOC111139950, id-LOC111139953, id-LOC111139954, id-LOC111139955, id-LOC111140130, id-LOC111140131, id-LOC111140132, id-LOC111140135, id-LOC111140261, id-LOC111140362, id-LOC111140438, id-LOC111140471, id-LOC111140537, id-LOC111140544, id-LOC111140567    id-LOC101093027, id-LOC101093530, id-LOC101093781, id-LOC105260306, id-LOC109492277, id-LOC109497166, id-LOC111559009, id-LOC111559020, id-LOC123386228, id-LOC123386257, id-LOC123386258, id-LOC123386262, id-LOC123386263, id-LOC123386264, id-LOC123386266, id-LOC123386270, id-LOC123386272, id-LOC123386273, id-LOC123386275, id-LOC123386279, id-LOC123386299, id-LOC123386300, id-LOC123386301, id-LOC123386302, id-LOC123386303, id-LOC123386305, id-LOC123386306, id-LOC123386307, id-LOC123386308, id-LOC123386310    id-LOC118518698, id-LOC118527461, id-LOC118527462, id-LOC118535942, id-LOC118543833, id-LOC118543836, id-LOC118543838, id-LOC118544684, id-LOC118550599, id-LOC118550614, id-LOC118550616, id-LOC118550618, id-LOC118555003, id-LOC118555004, rna-XM_036115834.1    id-LOC120225928, id-LOC120225929, id-LOC120225930, id-LOC120225931, id-LOC120225932, id-LOC120225934, id-LOC120225936, id-LOC120225937, id-LOC120225978, id-LOC120225980, id-LOC120225981, id-LOC120225982, id-LOC120225983, id-LOC120225985, id-LOC120225986, id-LOC120225987, id-LOC120227645    id-LOC116862288, id-LOC116862289, id-LOC116862296, id-LOC116862300, id-LOC116862302, id-LOC116862304, id-LOC116862305, id-LOC116863562, id-LOC116863635, id-LOC116863732, id-LOC116863817, id-LOC116863853, id-LOC116863892    id-LOC123943490, id-LOC123943491, id-LOC123943493, id-LOC123943494, id-LOC123943701, id-LOC123943702, id-LOC123943703, id-LOC123943705, id-LOC123943706, id-LOC123943709, id-LOC123943711, id-LOC123943713, id-LOC123943792, id-LOC123943793, id-LOC123943794, id-LOC123943795, id-LOC123943797, id-LOC123943798, id-LOC123943799, id-LOC123943800, id-LOC123943801, id-LOC123943802, id-LOC123943803, id-LOC123943805, id-LOC123943806, id-LOC123943809, id-LOC123943810    id-LOC116589874, id-LOC116589880, id-LOC116589881, id-LOC116589882, id-LOC116590248, id-LOC116590249, id-LOC116590250, id-LOC116590251, id-LOC116590252, id-LOC116590259, id-LOC116591961, id-LOC116591964, id-LOC116591966, id-LOC116591969, rna-XM_032341836.1    id-LOC131835176, id-LOC131835177, id-LOC131835183, id-LOC131835184, id-LOC131835185, id-LOC131837084, id-LOC131837085, id-LOC131837275, id-LOC131837276, id-LOC131837277, id-LOC131837278, id-LOC131837279, id-LOC131837280, id-LOC131837282, id-LOC131837284, id-LOC131837286, id-LOC131837287, id-LOC131837288, id-LOC131837289, id-LOC131837291, id-LOC131837292    id-LOC131999473, id-LOC131999474, id-LOC131999476, id-LOC131999477, id-LOC131999478, id-LOC131999482, id-LOC131999483, id-LOC131999485, id-LOC131999486, id-LOC131999487, id-LOC131999489, id-LOC132000341, id-LOC132008642, id-LOC132008819, id-LOC132008910, id-LOC132009152, id-LOC132009220, id-LOC132009225, id-LOC132009241, id-LOC132009249, id-LOC132009301, rna-XM_059372175.1    id-LOC101678893, id-LOC101680397, id-LOC101680710, id-LOC101684716, id-LOC101685012, id-LOC101685311, id-LOC101685623, id-LOC101685910, id-LOC101686202, id-LOC101692463, id-LOC101692737, id-LOC101693682, id-LOC101693963, id-LOC106004701, id-LOC123386916, id-LOC123390516, id-LOC123390518, id-LOC123392246, id-LOC123392247, id-LOC123394068, id-LOC123394188, id-LOC123394233, rna-XM_045080620.1      id-LOC110580926, id-LOC110580936, id-LOC110580954, id-LOC110580972    id-LOC101367247, id-LOC101368101, id-LOC101369209, id-LOC101369953, id-LOC101370087, id-LOC101370174, id-LOC101370252, id-LOC101370382, id-LOC101370550, id-LOC101370666, id-LOC101370696, id-LOC101371730, id-LOC101372028, id-LOC101372145, id-LOC101372322, id-LOC101372452, id-LOC101375713, id-LOC101376579, id-LOC101377458, id-LOC101377736, id-LOC101380190, id-LOC101385190, id-LOC101385490, rna-XM_004413010.1, rna-XM_004413011.1, rna-XM_004413280.1    id-LOC122239533, id-LOC122239536, id-LOC122239538, id-LOC122239542, id-LOC122239735, id-LOC122239736, id-LOC122239739, id-LOC122239743, id-LOC122239746, id-LOC122239747, rna-XM_042989290.1    id-LOC103656545, id-LOC103656546, id-LOC103656549, id-LOC121100627, id-LOC121100629, id-LOC121103216    id-LOC113909372, id-LOC113909941      id-egapxtmp_014751, id-egapxtmp_015248, id-egapxtmp_015436, id-egapxtmp_015438, rna-gnl|WGS_ZZZZ|egapxtmp_015437-R1    mNeoVis.evergladensis.2.1.chr8.g00005.t01        398    N0.HOG0000396   OG0000021  n2   rna-XM_002929980.2, rna-XM_002931159.2, rna-XM_034651896.1    rna-NM_001389049.1, rna-NM_001389055.1, rna-NM_001389124.1    rna-XM_022507039.1, rna-XM_022507100.1    rna-XM_003993265.2    rna-XM_036073551.1    rna-XM_039225637.1, rna-XM_039225692.1    rna-XM_032844591.1, rna-XM_032844597.1, rna-XM_032844609.1    rna-XM_046016627.1, rna-XM_046016765.1, rna-XM_046016860.1, rna-XM_046017087.1    rna-XM_032359684.1, rna-XM_032359715.1, rna-XM_032360043.1    rna-XM_059158845.1, rna-XM_059158849.1, rna-XM_059158851.1, rna-XM_059158855.1    rna-XM_059386466.1, rna-XM_059386467.1, rna-XM_059388353.1    rna-XM_004781912.1, rna-XM_045067955.1, rna-XM_045067971.1, rna-XM_045067974.1    mNeoVis.2.1.1.chr7_hap1.g02058.t01, mNeoVis.2.1.1.chr7_hap1.g02060.t01, mNeoVis.2.1.1.chr7_hap1.g02064.t01    rna-XM_021684821.1    rna-XM_004411931.1    rna-XM_007095368.3    rna-XM_008700915.2, rna-XM_008700920.1        rna-gnl|WGS_ZZZZ|egapxtmp_013534-R1, rna-gnl|WGS_ZZZZ|egapxtmp_013548-R1    mNeoVis.evergladensis.2.1.chr7.g02015.t01, mNeoVis.evergladensis.2.1.chr7.g02016.t01, mNeoVis.evergladensis.2.1.chr7.g02020.t01    mNeoVis.vulgivaga.2.1.chr7.g02036.t01      399    N0.HOG0000397   OG0000021  n47   rna-XM_002930239.2, rna-XM_002931168.2    rna-NM_001388629.1, rna-NM_001389053.1, rna-XM_038562475.1, rna-XM_038562476.1, rna-XM_038562477.1, rna-XM_038562478.1, rna-XM_038563262.1, rna-XM_038563263.1    rna-XM_022506916.1, rna-XM_022507006.1, rna-XM_022507064.1    rna-XM_003993305.2, rna-XM_006937306.3, rna-XM_045038145.1    rna-XM_036073541.1    rna-XM_039225603.1, rna-XM_039225604.1, rna-XM_039225605.1, rna-XM_039225776.1    rna-XM_032844561.1, rna-XM_032844563.1, rna-XM_032844625.1    rna-XM_046016396.1, rna-XM_046016449.1, rna-XM_046016496.1    rna-XM_032359311.1, rna-XM_032359417.1, rna-XM_032359624.1    rna-XM_059158844.1, rna-XM_059158847.1, rna-XM_059158853.1    rna-XM_059386834.1, rna-XM_059386881.1, rna-XM_059387396.1, rna-XM_059387600.1    rna-XM_004781646.2, rna-XM_004781911.2, rna-XM_045067952.1, rna-XM_045067991.1    mNeoVis.2.1.1.chr7_hap1.g02054.t01, mNeoVis.2.1.1.chr7_hap1.g02057.t01, mNeoVis.2.1.1.chr7_hap1.g02062.t01      rna-XM_004417277.1    rna-XM_007095366.2, rna-XM_007095367.2, rna-XM_007095374.2, rna-XM_007095408.3, rna-XM_007095409.3, rna-XM_015543475.2, rna-XM_042958525.1, rna-XM_042958681.1, rna-XM_042959625.1    rna-XM_008700917.2, rna-XM_008700921.2, rna-XM_008700926.1      mNeoVis.3.1.1.chr7.g01919.t01    rna-gnl|WGS_ZZZZ|egapxtmp_010287-R1, rna-gnl|WGS_ZZZZ|egapxtmp_010299-R1    mNeoVis.evergladensis.2.1.chr7.g02012.t01, mNeoVis.evergladensis.2.1.chr7.g02014.t01, mNeoVis.evergladensis.2.1.chr7.g02018.t01    mNeoVis.vulgivaga.2.1.chr7.g02032.t01, mNeoVis.vulgivaga.2.1.chr7.g02035.t01, mNeoVis.vulgivaga.2.1.chr7.g02037.t01      400    N0.HOG0000398   OG0000021  n112   rna-XM_002930237.4, rna-XM_002930238.2, rna-XM_002930242.4, rna-XM_002930243.4    rna-XM_038562479.1    rna-XM_022507099.1, rna-XM_022507142.1    rna-XM_019813047.2    rna-XM_036073553.1    rna-XM_039226910.1    rna-XM_032844582.1, rna-XM_032844595.1, rna-XM_032844611.1    rna-XM_046017149.1, rna-XM_046017203.1, rna-XM_046017260.1, rna-XM_046017317.1, rna-XM_046017323.1, rna-XM_046017579.1    rna-XM_032358742.1, rna-XM_032359099.1, rna-XM_032359212.1, rna-XM_032359379.1, rna-XM_032359422.1    rna-XM_059158774.1, rna-XM_059158785.1, rna-XM_059158841.1, rna-XM_059158843.1    rna-XM_059386840.1, rna-XM_059386842.1, rna-XM_059386843.1, rna-XM_059386930.1, rna-XM_059386943.1, rna-XM_059386944.1    rna-XM_004781637.1, rna-XM_004781642.2, rna-XM_004781644.1, rna-XM_013050574.1    mNeoVis.2.1.1.chr7_hap1.g02045.t01, mNeoVis.2.1.1.chr7_hap1.g02047.t01, mNeoVis.2.1.1.chr7_hap1.g02056.t01      rna-XM_004417278.1    rna-XM_007095403.1    rna-XM_008700922.2, rna-XM_008700924.2, rna-XM_008700925.1, rna-XM_008700927.2, rna-XM_008700930.2    rna-XM_027577873.2    mNeoVis.3.1.1.chr7.g01907.t01, mNeoVis.3.1.1.chr7.g01910.t01    rna-gnl|WGS_ZZZZ|egapxtmp_010283-R1, rna-gnl|WGS_ZZZZ|egapxtmp_010284-R1, rna-gnl|WGS_ZZZZ|egapxtmp_010285-R1    mNeoVis.evergladensis.2.1.chr7.g02006.t01, mNeoVis.evergladensis.2.1.chr7.g02011.t01    mNeoVis.vulgivaga.2.1.chr7.g02034.t01      401    N0.HOG0000399   OG0000021  n167     rna-NM_001389030.1, rna-XM_038562470.1    rna-XM_022507008.1    rna-XM_011286840.2      rna-XM_039225300.1    rna-XM_032844586.1, rna-XM_032844596.1    rna-XM_046015828.1, rna-XM_046015837.1, rna-XM_046016572.1, rna-XM_046016585.1, rna-XM_046016617.1, rna-XM_046016658.1, rna-XM_046016835.1    rna-XM_032358560.1, rna-XM_032359601.1, rna-XM_032360040.1, rna-XM_032360183.1, rna-XM_032360188.1    rna-XM_059158860.1, rna-XM_059158861.1, rna-XM_059158863.1, rna-XM_059158866.1    rna-XM_059388293.1, rna-XM_059388323.1, rna-XM_059388334.1    rna-XM_004781916.2, rna-XM_004782270.1, rna-XM_013050701.2, rna-XM_045068037.1    mNeoVis.2.1.1.chr7_hap1.g02068.t01, mNeoVis.2.1.1.chr7_hap1.g02069.t01, mNeoVis.2.1.1.chr7_hap1.g02070.t01, mNeoVis.2.1.1.chr7_hap1.g02075.t01        rna-XM_007095370.2        mNeoVis.3.1.1.chr7.g01923.t01, mNeoVis.3.1.1.chr7.g01926.t01    rna-gnl|WGS_ZZZZ|egapxtmp_010288-R1, rna-gnl|WGS_ZZZZ|egapxtmp_013524-R1, rna-gnl|WGS_ZZZZ|egapxtmp_013527-R1, rna-gnl|WGS_ZZZZ|egapxtmp_013529-R1    mNeoVis.evergladensis.2.1.chr7.g02024.t01, mNeoVis.evergladensis.2.1.chr7.g02025.t01, mNeoVis.evergladensis.2.1.chr7.g02030.t01    mNeoVis.vulgivaga.2.1.chr7.g02042.t01, mNeoVis.vulgivaga.2.1.chr7.g02046.t01      402    N0.HOG0000400   OG0000021  n214   rna-XM_002929979.2, rna-XM_034646313.1                                  rna-XM_008694056.1, rna-XM_008694198.2                403    N0.HOG0000401   OG0000021  n218   rna-XM_002929976.4, rna-XM_002929982.1, rna-XM_002929983.4, rna-XM_002929985.4, rna-XM_002931192.2, rna-XM_034644683.1, rna-XM_034650426.1    rna-NM_001389041.1    rna-XM_022507221.1          rna-XM_032844587.1, rna-XM_032844629.1    rna-XM_046015834.1, rna-XM_046016523.1, rna-XM_046016613.1, rna-XM_046016625.1, rna-XM_046016760.1, rna-XM_046016761.1, rna-XM_046016767.1, rna-XM_046016980.1, rna-XM_046016992.1, rna-XM_046017201.1    rna-XM_032359835.1, rna-XM_032359983.1, rna-XM_032360087.1, rna-XM_032360181.1, rna-XM_032360182.1, rna-XM_032360184.1    rna-XM_059158858.1, rna-XM_059158865.1, rna-XM_059158867.1, rna-XM_059158870.1    rna-XM_059388281.1, rna-XM_059388289.1, rna-XM_059388300.1, rna-XM_059388309.1, rna-XM_059388312.1, rna-XM_059388342.1    rna-XM_004781913.1, rna-XM_004781919.1, rna-XM_004781920.2, rna-XM_004782269.2, rna-XM_004782271.1, rna-XM_004782272.1    mNeoVis.2.1.1.chr7_hap1.g02066.t01, mNeoVis.2.1.1.chr7_hap1.g02072.t01, mNeoVis.2.1.1.chr7_hap1.g02073.t01, mNeoVis.2.1.1.chr7_hap1.g02074.t01, mNeoVis.2.1.1.chr7_hap1.g02076.t01      rna-XM_004417245.1, rna-XM_004417248.1      rna-XM_008694058.1, rna-XM_008694196.1, rna-XM_008694199.1, rna-XM_008694200.1, rna-XM_008700910.1, rna-XM_008700912.1, rna-XM_008700913.1, rna-XM_008700918.2, rna-XM_040638710.1, rna-XM_040639810.1    rna-XM_027580101.1    mNeoVis.3.1.1.chr7.g01925.t01    rna-gnl|WGS_ZZZZ|egapxtmp_013525-R1, rna-gnl|WGS_ZZZZ|egapxtmp_013526-R1, rna-gnl|WGS_ZZZZ|egapxtmp_013528-R1, rna-gnl|WGS_ZZZZ|egapxtmp_013530-R1    mNeoVis.evergladensis.2.1.chr7.g02022.t01, mNeoVis.evergladensis.2.1.chr7.g02027.t01, mNeoVis.evergladensis.2.1.chr7.g02028.t01, mNeoVis.evergladensis.2.1.chr7.g02029.t01, mNeoVis.evergladensis.2.1.chr7.g02031.t01    mNeoVis.vulgivaga.2.1.chr7.g02040.t01, mNeoVis.vulgivaga.2.1.chr7.g02044.t01, mNeoVis.vulgivaga.2.1.chr7.g02047.t01, mNeoVis.vulgivaga.2.1.chr7.g02048.t01      404    N0.HOG0000402   OG0000021  n289   rna-XM_002929977.2    rna-NM_001388919.1, rna-NM_001389038.1, rna-NM_001389116.1, rna-NM_001389137.1, rna-XM_038562471.1, rna-XM_038562472.1, rna-XM_038589277.1, rna-XM_038589278.1, rna-XM_038589547.1, rna-XM_038589634.1        rna-XM_036073538.1      rna-XM_032844628.1    rna-XM_046016601.1    rna-XM_032359933.1    rna-XM_059158864.1    rna-XM_059388317.1    rna-XM_004781918.1    mNeoVis.2.1.1.chr7_hap1.g02071.t01    rna-XM_021684823.1, rna-XM_021684826.2    rna-XM_004411932.1, rna-XM_004417246.1      rna-XM_008710506.2    rna-XM_035722952.1    mNeoVis.3.1.1.chr7.g01918.t01, mNeoVis.3.1.1.chr7.g01924.t01    rna-gnl|WGS_ZZZZ|egapxtmp_013522-R1    mNeoVis.evergladensis.2.1.chr7.g02026.t01    mNeoVis.vulgivaga.2.1.chr7.g02043.t01      405    N0.HOG0000403   OG0000022  n3     rna-NM_001388676.1    rna-XM_022526827.1    rna-XM_006928381.5      rna-XM_039251603.1    rna-XM_032883473.1    rna-XM_045990548.1    rna-XM_032310207.1    rna-XM_059159867.1    rna-XM_059388926.1    rna-XM_004771768.1          rna-XM_007079389.3            mNeoVis.evergladensis.2.1.chr2.g00240.t01        406    N0.HOG0000404   OG0000022  n16   rna-XM_002921914.4    rna-NM_001388673.1    rna-XM_022492585.1, rna-XM_022492758.1, rna-XM_022526825.1    rna-XM_004001087.4    rna-XM_036095273.1    rna-XM_039224716.1    rna-XM_032883922.1    rna-XM_045990536.1, rna-XM_045990538.1, rna-XM_045990539.1, rna-XM_045990540.1, rna-XM_045990541.1, rna-XM_045990542.1, rna-XM_045990543.1, rna-XM_045990547.1, rna-XM_045990550.1    rna-XM_032310277.1, rna-XM_032310301.1, rna-XM_032310314.1, rna-XM_032310371.1, rna-XM_032361545.1, rna-XM_032361679.1    rna-XM_059159070.1, rna-XM_059159072.1, rna-XM_059159073.1, rna-XM_059159074.1, rna-XM_059159075.1, rna-XM_059159873.1, rna-XM_059159874.1    rna-XM_059387291.1, rna-XM_059387381.1, rna-XM_059387649.1, rna-XM_059387650.1, rna-XM_059387651.1, rna-XM_059387653.1, rna-XM_059388931.1    rna-XM_004771770.3, rna-XM_004771773.3, rna-XM_045077727.1, rna-XM_045077739.1, rna-XM_045077741.1, rna-XM_045077762.1, rna-XM_045084738.1    mNeoVis.2.1.1.chr5_hap1.g00247.t01, mNeoVis.2.1.1.chr5_hap1.g00248.t01, mNeoVis.2.1.1.chr5_hap1.g00250.t01, mNeoVis.2.1.1.chr5_hap1.g00251.t01, mNeoVis.2.1.1.chr5_hap1.g00252.t01    rna-XM_021705190.1      rna-XM_007079385.2, rna-XM_007095833.2, rna-XM_042970188.1    rna-XM_008705494.2    rna-XM_027583028.1    mNeoVis.3.1.1.chr2.g00190.t01, mNeoVis.3.1.1.chr2.g00192.t01, mNeoVis.3.1.1.chr2.g00193.t01    rna-gnl|WGS_ZZZZ|egapxtmp_038052-R1, rna-gnl|WGS_ZZZZ|egapxtmp_038053-R1, rna-gnl|WGS_ZZZZ|egapxtmp_038055-R1, rna-gnl|WGS_ZZZZ|egapxtmp_038112-R1, rna-gnl|WGS_ZZZZ|egapxtmp_038124-R1    mNeoVis.evergladensis.2.1.chr2.g00245.t01, mNeoVis.evergladensis.2.1.chr2.g00246.t01, mNeoVis.evergladensis.2.1.chr2.g00248.t01    mNeoVis.vulgivaga.2.1.chr2.g00242.t01, mNeoVis.vulgivaga.2.1.chr2.g00243.t01      407    N0.HOG0000405   OG0000022  n84   rna-XM_034660032.1    rna-XM_038565382.1    rna-XM_022527048.1    rna-XM_023240976.2    rna-XM_036095268.1    rna-XM_039224715.1    rna-XM_032883924.1    rna-XM_045990551.1    rna-XM_032310341.1    rna-XM_059159883.1, rna-XM_059159884.1    rna-XM_059388939.1, rna-XM_059388940.1    rna-XM_013045312.1    mNeoVis.2.1.1.chr5_hap1.g00253.t01    rna-XM_021705466.1      rna-XM_042978105.1, rna-XM_042978106.1, rna-XM_042978107.1    rna-XM_008705519.2        rna-gnl|WGS_ZZZZ|egapxtmp_038134-R1    mNeoVis.evergladensis.2.1.chr2.g00249.t01    mNeoVis.vulgivaga.2.1.chr2.g00244.t01      408    N0.HOG0000406   OG0000022  n104   rna-XM_002931205.4    rna-XM_038565381.1    rna-XM_022526824.1    rna-XM_023240979.2    rna-XM_036095269.1    rna-XM_039224713.1    rna-XM_032883918.1    rna-XM_045990374.1      rna-XM_059159076.1    rna-XM_059387654.1    rna-XM_013050721.2    mNeoVis.2.1.1.chr5_hap1.g00254.t01    rna-XM_021705304.1      rna-XM_015536310.2    rna-XM_008705495.2    rna-XM_027583027.2    mNeoVis.3.1.1.chr2.g00194.t01    rna-gnl|WGS_ZZZZ|egapxtmp_038128-R1    mNeoVis.evergladensis.2.1.chr2.g00250.t01    mNeoVis.vulgivaga.2.1.chr2.g00245.t01      409    N0.HOG0000407   OG0000022  n123   rna-XM_034657725.1    rna-XM_038567389.1    rna-XM_022527046.1    rna-XM_003981817.2      rna-XM_039224712.1    rna-XM_032883919.1    rna-XM_045990533.1    rna-XM_032361776.1    rna-XM_059159863.1    rna-XM_059388922.1    rna-XM_004781977.2          rna-XM_007079371.2    rna-XM_008705499.2      mNeoVis.3.1.1.chr2.g00195.t01      mNeoVis.evergladensis.2.1.chr2.g00251.t01    mNeoVis.vulgivaga.2.1.chr2.g00246.t01      410    N0.HOG0000408   OG0000022  n140   rna-XM_002921923.4    rna-NM_001389188.1      rna-XM_003981831.6    rna-XM_036120262.1    rna-XM_039251615.1    rna-XM_032883926.1    rna-XM_045991589.1    rna-XM_032360414.1      rna-XM_059388941.1    rna-XM_004771738.3        rna-XM_004398834.1    rna-XM_007090674.2    rna-XM_008705470.1    rna-XM_027583038.2          mNeoVis.vulgivaga.2.1.chr2.g00233.t01      411    N0.HOG0000409   OG0000022  n154   rna-XM_011228336.2    rna-XM_038565328.1    rna-XM_022526902.1    rna-XM_023240954.2    rna-XM_036120242.1    rna-XM_039251604.1    rna-XM_032883913.1    rna-XM_045991640.1    rna-XM_032360433.1    rna-XM_059159879.1    rna-XM_059388936.1    rna-XM_013045274.2          rna-XM_042978095.1    rna-XM_008705477.2          mNeoVis.evergladensis.2.1.chr2.g00239.t01    mNeoVis.vulgivaga.2.1.chr2.g00237.t01      412    N0.HOG0000410   OG0000022  n171   rna-XM_002921925.4      rna-XM_022526830.1    rna-XM_011288856.3      rna-XM_039251636.1    rna-XM_032883925.1    rna-XM_045991474.1    rna-XM_032360379.1        rna-XM_013045276.2          rna-XM_007090713.2    rna-XM_008705469.2          mNeoVis.evergladensis.2.1.chr2.g00233.t01        413    N0.HOG0000411   OG0000022  n182   rna-XM_002921926.4    rna-NM_001388680.1    rna-XM_022526941.1    rna-XM_011288858.2      rna-XM_039251617.1      rna-XM_045991593.1      rna-XM_059159877.1    rna-XM_059388934.1    rna-XM_004771762.3    mNeoVis.2.1.1.chr5_hap1.g00235.t01        rna-XM_015541336.2    rna-XM_008705487.2    rna-XM_027583030.1    mNeoVis.3.1.1.chr2.g00181.t01    rna-gnl|WGS_ZZZZ|egapxtmp_038130-R1    mNeoVis.evergladensis.2.1.chr2.g00232.t01    mNeoVis.vulgivaga.2.1.chr2.g00231.t01      414    N0.HOG0000412   OG0000022  n198   rna-XM_002921922.4    rna-NM_001388679.1    rna-XM_022526946.1    rna-XM_003981832.5      rna-XM_039251623.1    rna-XM_032883906.1, rna-XM_032883907.1    rna-XM_045991476.1, rna-XM_045991681.1    rna-XM_032360391.1    rna-XM_059159876.1, rna-XM_059159878.1    rna-XM_059388932.1, rna-XM_059388935.1    rna-XM_004771763.3, rna-XM_004771764.3    mNeoVis.2.1.1.chr5_hap1.g00236.t01, mNeoVis.2.1.1.chr5_hap1.g00237.t01    rna-XM_044918136.1    rna-XM_004398993.1    rna-XM_007090712.2    rna-XM_008705471.1, rna-XM_008705488.2      mNeoVis.3.1.1.chr2.g00183.t01    rna-gnl|WGS_ZZZZ|egapxtmp_038122-R1, rna-gnl|WGS_ZZZZ|egapxtmp_038129-R1    mNeoVis.evergladensis.2.1.chr2.g00234.t01, mNeoVis.evergladensis.2.1.chr2.g00235.t01    mNeoVis.vulgivaga.2.1.chr2.g00232.t01, mNeoVis.vulgivaga.2.1.chr2.g00234.t01      415    N0.HOG0000413   OG0000022  n228         rna-XM_045053125.1                          rna-XM_007090709.2                  416    N0.HOG0000414   OG0000022  n232   rna-XM_034657224.1    rna-XM_038565329.1    rna-XM_022526947.1    rna-XM_019817043.3      rna-XM_039225831.1, rna-XM_039251583.1    rna-XM_032883923.1    rna-XM_045991859.1    rna-XM_032303354.1    rna-XM_059159067.1    rna-XM_059387645.1    rna-XM_004771767.3    mNeoVis.2.1.1.chr5_hap1.g00241.t01        rna-XM_015541335.2    rna-XM_008705476.1    rna-XM_035726254.1        mNeoVis.evergladensis.2.1.chr2.g00238.t01        417    N0.HOG0000415   OG0000022  n248   rna-XM_002921919.4    rna-XM_038565330.1    rna-XM_022526953.1          rna-XM_032850976.1, rna-XM_032850977.1    rna-XM_045991787.1    rna-XM_032360423.1    rna-XM_059159868.1    rna-XM_059388928.1    rna-XM_004771739.2        rna-XM_004398835.1      rna-XM_008705474.2        rna-gnl|WGS_ZZZZ|egapxtmp_038117-R2    mNeoVis.evergladensis.2.1.chr2.g00237.t01        418    N0.HOG0000416   OG0000022  n260           rna-XM_036120243.1                    rna-XM_021705188.1                      419    N0.HOG0000417   OG0000022  n261   rna-XM_002921920.4, rna-XM_002921921.4    rna-NM_001388678.1, rna-NM_001389145.1    rna-XM_022526829.1    rna-XM_003981833.5, rna-XM_011289502.3    rna-XM_036120254.1    rna-XM_039251602.1, rna-XM_039251621.1    rna-XM_032883470.1, rna-XM_032883471.1    rna-XM_045991477.1, rna-XM_045991644.1    rna-XM_032303332.1, rna-XM_032303345.1    rna-XM_059159066.1, rna-XM_059159875.1    rna-XM_059387385.1, rna-XM_059387643.1    rna-XM_004771765.3, rna-XM_004771766.3    mNeoVis.2.1.1.chr5_hap1.g00238.t01, mNeoVis.2.1.1.chr5_hap1.g00239.t01, mNeoVis.2.1.1.chr5_hap1.g00240.t01    rna-XM_021705492.2, rna-XM_044913096.1    rna-XM_004398994.1    rna-XM_007090710.2, rna-XM_007090711.2    rna-XM_008705472.2, rna-XM_008705473.2    rna-XM_035726271.1, rna-XM_035726275.1      rna-gnl|WGS_ZZZZ|egapxtmp_034644-R1, rna-gnl|WGS_ZZZZ|egapxtmp_034645-R1, rna-gnl|WGS_ZZZZ|egapxtmp_038119-R1    mNeoVis.evergladensis.2.1.chr2.g00236.t01    mNeoVis.vulgivaga.2.1.chr2.g00235.t01      420    N0.HOG0000418   OG0000023  n1   rna-XM_034645215.1, rna-XM_034645221.1    rna-XM_038557849.1    rna-XM_022497294.1    rna-XM_023258494.2    rna-XM_036119055.1    rna-XM_039244490.1    rna-XM_032877302.1    rna-XM_045997228.1    rna-XM_032302584.1    rna-XM_059136966.1    rna-XM_059377315.1    rna-XM_045064215.1    mNeoVis.2.1.1.chr2_hap1.g01481.t01    rna-XM_021683728.1    rna-XM_004402641.1    rna-XM_042996623.1    rna-XM_040619288.1    rna-XM_027613379.2    mNeoVis.3.1.1.chr5.g01402.t01    rna-gnl|WGS_ZZZZ|egapxtmp_006398-R1    mNeoVis.evergladensis.2.1.chr5.g01418.t01    mNeoVis.vulgivaga.2.1.chr5.g01419.t01      421    N0.HOG0000419   OG0000023  n23   rna-XM_002920297.4    rna-XM_038554742.1    rna-XM_022521054.1    rna-XM_011282457.4    rna-XM_036114105.1    rna-XM_039223793.1    rna-XM_032858143.1    rna-XM_046006222.1    rna-XM_032339032.1    rna-XM_059177055.1    rna-XM_059401074.1    rna-XM_004766887.3    mNeoVis.2.1.1.chr1_hap1.g01958.t01    rna-XM_021693107.1    rna-XM_004415003.1    rna-XM_007084905.3    rna-XM_008705038.1    rna-XM_027603117.2    mNeoVis.3.1.1.chr1.g01908.t01, mNeoVis.3.1.1.chr1.g01909.t01, mNeoVis.3.1.1.chr1.g01910.t01    rna-gnl|WGS_ZZZZ|egapxtmp_033937-R4    mNeoVis.evergladensis.2.1.chr1.g01923.t01    mNeoVis.vulgivaga.2.1.chr1.g01921.t01      422    N0.HOG0000420   OG0000023  n45   rna-XM_034643534.1    rna-XM_038531663.1    rna-XM_022514745.1    rna-XM_003989639.6    rna-XM_036093084.1    rna-XM_039232127.1    rna-XM_032864065.1    rna-XM_046013660.1    rna-XM_032303308.1    rna-XM_059137593.1    rna-XM_059376584.1    rna-XM_004741580.2    mNeoVis.2.1.1.chr2_hap1.g01703.t01    rna-XM_021684391.2    rna-XM_004411731.1    rna-XM_042997041.1    rna-XM_040619372.1    rna-XM_027583061.2    mNeoVis.3.1.1.chr5.g01625.t01    rna-gnl|WGS_ZZZZ|egapxtmp_007142-R1    mNeoVis.evergladensis.2.1.chr5.g01642.t01    mNeoVis.vulgivaga.2.1.chr5.g01644.t01      423    N0.HOG0000421   OG0000023  n68   rna-XM_034657115.1    rna-XM_038582514.1    rna-XM_022504843.1    rna-XM_019839578.3    rna-XM_036070278.1    rna-XM_039221986.1, rna-XM_039223137.1    rna-XM_032882241.1    rna-XM_046001266.1    rna-XM_032351607.1    rna-XM_059164402.1    rna-XM_059392124.1    rna-XM_004773009.2    mNeoVis.2.1.1.chr5_hap1.g01684.t01    rna-XM_021687948.1    rna-XM_004391681.1    rna-XM_042998967.1    rna-XM_040621133.1    rna-XM_027586192.2    mNeoVis.3.1.1.chr2.g01595.t01    rna-gnl|WGS_ZZZZ|egapxtmp_036049-R3    mNeoVis.evergladensis.2.1.chr2.g01667.t01    mNeoVis.vulgivaga.2.1.chr2.g01684.t01      424    N0.HOG0000422   OG0000023  n91   rna-XM_034661964.1    rna-XM_038570948.1    rna-XM_022514925.1    rna-XM_023260542.2    rna-XM_036077060.1, rna-XM_036110255.1    rna-XM_039243855.1, rna-XM_039256473.1    rna-XM_032869646.1    rna-XM_046003565.1    rna-XM_032333747.1    rna-XM_059162673.1    rna-XM_059390963.1    rna-XM_004762462.3    mNeoVis.2.1.1.chr5_hap1.g01152.t01    rna-XM_044915538.1    rna-XM_004417390.1    rna-XM_042999022.1    rna-XM_040636595.1, rna-XM_040636596.1    rna-XM_027586952.2    mNeoVis.3.1.1.chr2.g01063.t01    rna-gnl|WGS_ZZZZ|egapxtmp_036306-R1    mNeoVis.evergladensis.2.1.chr2.g01142.t01    mNeoVis.vulgivaga.2.1.chr2.g01148.t01, mNeoVis.vulgivaga.2.1.chr2.g01149.t01      425    N0.HOG0000423   OG0000023  n112   rna-XM_002919539.4    rna-XM_038531657.1    rna-XM_022514489.1    rna-XM_045035271.1    rna-XM_036093078.1    rna-XM_039233725.1    rna-XM_032863701.1    rna-XM_046023935.1    rna-XM_032302009.1    rna-XM_059136587.1    rna-XM_059376596.1    rna-XM_045063934.1    mNeoVis.2.1.1.chr2_hap1.g01698.t01    rna-XM_021683535.1    rna-XM_004394780.1    rna-XM_007083711.3    rna-XM_008694872.2    rna-XM_027581421.2    mNeoVis.3.1.1.chr5.g01619.t01    rna-gnl|WGS_ZZZZ|egapxtmp_007140-R1    mNeoVis.evergladensis.2.1.chr5.g01638.t01        426    N0.HOG0000424   OG0000023  n131   rna-XM_034661745.1    rna-XM_038583600.1    rna-XM_022503938.1    rna-XM_045037186.1    rna-XM_036122529.1    rna-XM_039241727.1    rna-XM_032875544.1    rna-XM_046003498.1    rna-XM_032338629.1    rna-XM_059163257.1    rna-XM_059391375.1    rna-XM_004745167.3    mNeoVis.2.1.1.chr5_hap1.g01321.t01    rna-XM_044914630.1    rna-XM_004406750.1    rna-XM_042956653.1    rna-XM_040621493.1    rna-XM_027587743.1    mNeoVis.3.1.1.chr2.g01241.t01      mNeoVis.evergladensis.2.1.chr2.g01308.t01    mNeoVis.vulgivaga.2.1.chr2.g01315.t01      427    N0.HOG0000425   OG0000023  n150   rna-XM_019808018.2    rna-XM_038539236.1    rna-XM_022500645.1    rna-XM_003998497.5    rna-XM_036073371.1    rna-XM_039221857.1    rna-XM_032872533.1    rna-XM_045993837.1    rna-XM_032327209.1    rna-XM_059154710.1    rna-XM_059414579.1    rna-XM_004761554.3    mNeoVis.2.1.1.chr15_hap1.g00683.t01    rna-XM_021680177.1    rna-XM_004399042.1    rna-XM_042971820.1    rna-XM_040634871.1    rna-XM_027613340.2    mNeoVis.3.1.1.chr15.g00464.t01      mNeoVis.evergladensis.2.1.chr15.g00505.t01    mNeoVis.vulgivaga.2.1.chr15.g00514.t01      428    N0.HOG0000426   OG0000023  n170   rna-XM_011230979.3    rna-XM_038559291.1    rna-XM_022510370.1    rna-XM_023250669.2    rna-XM_036095660.1    rna-XM_039243079.1    rna-XM_032866982.1    rna-XM_046021217.1    rna-XM_032304912.1    rna-XM_059172118.1    rna-XM_059396224.1    rna-XM_004772072.2    mNeoVis.2.1.1.chr4_hap1.g00101.t01    rna-XM_021692889.2    rna-XM_004414585.1    rna-XM_042974153.1    rna-XM_040643880.1    rna-XM_027573839.1    mNeoVis.3.1.1.chr6.g00101.t01    rna-gnl|WGS_ZZZZ|egapxtmp_003143-R2    mNeoVis.evergladensis.2.1.chr6.g00103.t01    mNeoVis.vulgivaga.2.1.chr6.g00100.t01      429    N0.HOG0000427   OG0000023  n191   rna-XM_034671255.1    rna-XM_038531880.1    rna-XM_022514463.1    rna-XM_045035142.1    rna-XM_036067030.1    rna-XM_039229095.1    rna-XM_032863517.1    rna-XM_046016360.1    rna-XM_032301956.1    rna-XM_059136344.1    rna-XM_059376314.1    rna-XM_004741396.3    mNeoVis.2.1.1.chr2_hap1.g01771.t01    rna-XM_021684054.2    rna-XM_004397093.1    rna-XM_042996573.1    rna-XM_040645830.1    rna-XM_027621389.1    mNeoVis.3.1.1.chr5.g01696.t01    rna-gnl|WGS_ZZZZ|egapxtmp_007170-R1    mNeoVis.evergladensis.2.1.chr5.g01711.t01    mNeoVis.vulgivaga.2.1.chr5.g01715.t01      430    N0.HOG0000428   OG0000023  n211   rna-XM_002928416.4    rna-XM_038582501.1, rna-XM_038583245.1    rna-XM_022504614.1    rna-XM_011285988.4    rna-XM_036097323.1, rna-XM_036103817.1    rna-XM_039222850.1    rna-XM_032882231.1    rna-XM_046003410.1    rna-XM_032351763.1    rna-XM_059164416.1    rna-XM_059392132.1    rna-XM_004777945.3    mNeoVis.2.1.1.chr5_hap1.g01690.t01    rna-XM_021689408.2    rna-XM_004391688.1    rna-XM_042956715.1    rna-XM_008704448.2    rna-XM_027587500.2    mNeoVis.3.1.1.chr2.g01600.t01    rna-gnl|WGS_ZZZZ|egapxtmp_034357-R1    mNeoVis.evergladensis.2.1.chr2.g01673.t01    mNeoVis.vulgivaga.2.1.chr2.g01689.t01      431    N0.HOG0000429   OG0000023  n231   rna-XM_011223454.3    rna-XM_038556193.1    rna-XM_022505840.1    rna-XM_023253073.2    rna-XM_036098842.1, rna-XM_036103434.1    rna-XM_039222255.1    rna-XM_032848637.1    rna-XM_045997834.1    rna-XM_032334442.1    rna-XM_059160224.1    rna-XM_059384693.1    rna-XM_004764298.3    mNeoVis.2.1.1.chr6_hap1.g00132.t01    rna-XM_021702245.1    rna-XM_004392867.1    rna-XM_042984445.1    rna-XM_040625400.1    rna-XM_027599098.2    mNeoVis.3.1.1.chr4.g00127.t01    rna-gnl|WGS_ZZZZ|egapxtmp_044311-R1    mNeoVis.evergladensis.2.1.chr4.g00134.t01, mNeoVis.evergladensis.2.1.chr4.g00135.t01    mNeoVis.vulgivaga.2.1.chr4.g00128.t01      432    N0.HOG0000430   OG0000023  n253   rna-XM_002925177.4    rna-XM_038585866.1    rna-XM_022508374.1    rna-XM_019838727.2    rna-XM_036098676.1    rna-XM_039256908.1    rna-XM_032857318.1    rna-XM_046019653.1    rna-XM_032354942.1    rna-XM_059167923.1    rna-XM_059393574.1    rna-XM_013061619.2    mNeoVis.2.1.1.chr3_hap1.g01145.t01    rna-XM_044913762.1    rna-XM_004407479.1    rna-XM_042995412.1    rna-XM_008690419.1    rna-XM_027589823.1    mNeoVis.3.1.1.chr3.g01133.t01    rna-gnl|WGS_ZZZZ|egapxtmp_039929-R1    mNeoVis.evergladensis.2.1.chr3.g01144.t01    mNeoVis.vulgivaga.2.1.chr3.g01159.t01      433    N0.HOG0000431   OG0000024  n6     rna-XM_038582131.1                                  rna-XM_035727343.1              434    N0.HOG0000432   OG0000024  n7   rna-XM_034654784.1    rna-XM_038578239.1, rna-XM_038584948.1      rna-XM_011285395.3    rna-XM_036099180.1      rna-XM_032852428.1, rna-XM_032855969.1    rna-XM_046018938.1    rna-XM_032356027.1    rna-XM_059166954.1    rna-XM_059394225.1    rna-XM_045066688.1    mNeoVis.2.1.1.chr3_hap1.g00850.t01    rna-XM_021702842.2    rna-XM_004403724.1    rna-XM_042994758.1    rna-XM_040643524.1    rna-XM_027588643.2    mNeoVis.3.1.1.chr3.g00833.t01    rna-gnl|WGS_ZZZZ|egapxtmp_042689-R1    mNeoVis.evergladensis.2.1.chr3.g00847.t01        435    N0.HOG0000433   OG0000024  n13             rna-XM_039233984.1          rna-XM_059142840.1    rna-XM_059412217.1    rna-XM_004833879.3                          436    N0.HOG0000434   OG0000024  n18                           mNeoVis.2.1.1.chr5_hap1.g01227.t01              mNeoVis.3.1.1.chr2.g01145.t01      mNeoVis.evergladensis.2.1.chr2.g01215.t01    mNeoVis.vulgivaga.2.1.chr2.g01221.t01      437    N0.HOG0000435   OG0000024  n24   rna-XM_034650813.1    rna-XM_038577914.1                                              438    N0.HOG0000436   OG0000024  n25   rna-XM_034654838.1    rna-XM_038540532.1    rna-XM_022522727.1    rna-XM_045056166.1, rna-XM_045056942.1    rna-XM_036109407.1, rna-XM_036119672.1      rna-XM_032841390.1, rna-XM_032882982.1      rna-XM_032316516.1          mNeoVis.2.1.1.chr12_hap1.g00337.t01, mNeoVis.2.1.1.chr12_hap1.g00806.t01, mNeoVis.2.1.1.chr2_hap1.g00840.t01, mNeoVis.2.1.1.chr2_hap1.g01301.t01, mNeoVis.2.1.1.chr4_hap1.g01040.t01, mNeoVis.2.1.1.chr8_hap1.g01296.t01    rna-XM_044913304.1, rna-XM_044918119.1      rna-XM_042985105.1, rna-XM_042987635.1        mNeoVis.3.1.1.chr5.g01226.t01, mNeoVis.3.1.1.chr8.g01193.t01    rna-gnl|WGS_ZZZZ|egapxtmp_006676-R1, rna-gnl|WGS_ZZZZ|egapxtmp_018783-R1    mNeoVis.evergladensis.2.1.chr5.g00794.t01, mNeoVis.evergladensis.2.1.chr6.g01046.t01, mNeoVis.evergladensis.2.1.chr9.g01244.t01    mNeoVis.vulgivaga.2.1.chr6.g01038.t01      439    N0.HOG0000437   OG0000024  n54               rna-XM_032853138.1              mNeoVis.2.1.1.chr6_hap1.g00626.t01                rna-gnl|WGS_ZZZZ|egapxtmp_003004-R1, rna-gnl|WGS_ZZZZ|egapxtmp_043912-R1          440    N0.HOG0000438   OG0000024  n60                 rna-XM_046024506.1      rna-XM_059164063.1    rna-XM_059405119.1                rna-XM_035726233.1, rna-XM_035727964.1              441    N0.HOG0000439   OG0000024  n66                       rna-XM_059376758.1    rna-XM_045064768.1                          442    N0.HOG0000440   OG0000024  n68                     rna-XM_059171049.1    rna-XM_059395758.1                            443    N0.HOG0000441   OG0000024  n72     rna-XM_038553361.1                          rna-XM_021692816.1          rna-XM_027589443.2, rna-XM_035724339.1, rna-XM_035726917.1    mNeoVis.3.1.1.chr11.g00874.t01    rna-gnl|WGS_ZZZZ|egapxtmp_016827-R1, rna-gnl|WGS_ZZZZ|egapxtmp_036879-R1    mNeoVis.evergladensis.2.1.chr2.g01702.t01    mNeoVis.vulgivaga.2.1.chr7.g00480.t01      444    N0.HOG0000442   OG0000024  n84   rna-XM_034645564.1      rna-XM_022496583.1, rna-XM_022498814.1    rna-XM_019835032.2      rna-XM_039223483.1, rna-XM_039225010.1, rna-XM_039227374.1, rna-XM_039230561.1, rna-XM_039231688.1, rna-XM_039239649.1      rna-XM_045997987.1, rna-XM_046009794.1, rna-XM_046016562.1, rna-XM_046016564.1    rna-XM_032301509.1, rna-XM_032302313.1, rna-XM_032325404.1, rna-XM_032330683.1, rna-XM_032331560.1    rna-XM_059137474.1, rna-XM_059158521.1, rna-XM_059166677.1, rna-XM_059173489.1, rna-XM_059173705.1, rna-XM_059182727.1, rna-XM_059188690.1    rna-XM_059391786.1, rna-XM_059392617.1, rna-XM_059409980.1    rna-XM_045080523.1, rna-XM_045080534.1, rna-XM_045088641.1, rna-XM_045089204.1    mNeoVis.2.1.1.chr10_hap1.g00987.t01, mNeoVis.2.1.1.chr10_hap1.g01022.t01, mNeoVis.2.1.1.chr13_hap1.g00630.t01, mNeoVis.2.1.1.chr1_hap1.g01502.t01, mNeoVis.2.1.1.chr3_hap1.g00384.t01, mNeoVis.2.1.1.chr5_hap1.g01180.t01, mNeoVis.2.1.1.chr6_hap1.g00748.t01, mNeoVis.2.1.1.chr6_hap1.g01074.t01    rna-XM_021703181.1    rna-XM_004392888.1, rna-XM_004393316.1    rna-XM_007081419.3      rna-XM_035724862.1        mNeoVis.evergladensis.2.1.chr1.g01466.t01        445    N0.HOG0000443   OG0000024  n131                 rna-XM_045989609.1            mNeoVis.2.1.1.chr10_hap1.g00510.t01          rna-XM_008684270.2    rna-XM_035727365.1              446    N0.HOG0000444   OG0000024  n137   rna-XM_034645486.1    rna-XM_038577848.1    rna-XM_022519625.1      rna-XM_036124227.1        rna-XM_046013395.1    rna-XM_032348451.1    rna-XM_059185106.1    rna-XM_059403312.1    rna-XM_045073068.1    mNeoVis.2.1.1.chr9_hap1.g00465.t01    rna-XM_021685323.2, rna-XM_021686726.1        rna-XM_008704683.2                447    N0.HOG0000445   OG0000024  n139   rna-XM_019809931.2, rna-XM_034661816.1    rna-XM_038534821.1, rna-XM_038565830.1, rna-XM_038577412.1      rna-XM_011281059.4, rna-XM_011282500.3, rna-XM_019832933.3, rna-XM_023250258.2, rna-XM_045038965.1    rna-XM_036073062.1, rna-XM_036078931.1, rna-XM_036089255.1, rna-XM_036091367.1, rna-XM_036093058.1, rna-XM_036094279.1, rna-XM_036102225.1, rna-XM_036106767.1    rna-XM_039239393.1, rna-XM_039246571.1, rna-XM_039253365.1    rna-XM_032850971.1, rna-XM_032872168.1    rna-XM_046009973.1, rna-XM_046017252.1, rna-XM_046020250.1    rna-XM_032304629.1, rna-XM_032304635.1, rna-XM_032329263.1, rna-XM_032334970.1, rna-XM_032348854.1, rna-XM_032352203.1, rna-XM_032354761.1    rna-XM_059140209.1, rna-XM_059163800.1, rna-XM_059165748.1, rna-XM_059173341.1, rna-XM_059177631.1, rna-XM_059184135.1    rna-XM_059376940.1, rna-XM_059378174.1, rna-XM_059388088.1, rna-XM_059392946.1, rna-XM_059410049.1, rna-XM_059416509.1    rna-XM_013047006.2, rna-XM_013050255.2, rna-XM_013053449.2, rna-XM_045064027.1, rna-XM_045066464.1, rna-XM_045069113.1, rna-XM_045088507.1    mNeoVis.2.1.1.chr12_hap1.g00221.t01, mNeoVis.2.1.1.chr12_hap1.g00494.t01, mNeoVis.2.1.1.chr1_hap1.g00575.t01, mNeoVis.2.1.1.chr6_hap1.g01030.t01    rna-XM_021678540.2, rna-XM_021679331.1, rna-XM_021692073.1, rna-XM_021704735.1    rna-XM_004394537.1    rna-XM_015544613.2      rna-XM_027589148.1      rna-gnl|WGS_ZZZZ|egapxtmp_001170-R1    mNeoVis.evergladensis.2.1.chr10.g00459.t01, mNeoVis.evergladensis.2.1.chr2.g01683.t01    mNeoVis.vulgivaga.2.1.chr2.g01699.t01      448    N0.HOG0000446   OG0000024  n209     rna-XM_038571209.1    rna-XM_022502222.1, rna-XM_022514026.1                rna-XM_059135549.1, rna-XM_059163151.1, rna-XM_059163527.1    rna-XM_059380833.1      mNeoVis.2.1.1.chr10_hap1.g00482.t01, mNeoVis.2.1.1.chr7_hap1.g01401.t01    rna-XM_021689193.2, rna-XM_021701908.2, rna-XM_021704032.2                  mNeoVis.vulgivaga.2.1.chr10.g00448.t01      449    N0.HOG0000447   OG0000024  n214             rna-XM_039230835.1        rna-XM_032304041.1, rna-XM_032312504.1            rna-XM_021684039.2    rna-XM_004406184.1                    450    N0.HOG0000448   OG0000024  n218       rna-XM_022515571.1              rna-XM_032358961.1    rna-XM_059178949.1    rna-XM_059384090.1    rna-XM_045065160.1    mNeoVis.2.1.1.chr12_hap1.g00516.t01                        451    N0.HOG0000449   OG0000024  n223               rna-XM_032857047.1                        rna-XM_027569863.2              452    N0.HOG0000450   OG0000024  n224                       rna-XM_059387967.1, rna-XM_059396374.1, rna-XM_059399139.1    rna-XM_013055590.2, rna-XM_045070204.1, rna-XM_045088178.1    mNeoVis.2.1.1.chr11_hap1.g00734.t01                    mNeoVis.vulgivaga.2.1.chr11.g00736.t01      453    N0.HOG0000451   OG0000024  n231                       rna-XM_059392824.1, rna-XM_059399253.1    rna-XM_013061396.2, rna-XM_045084245.1                mNeoVis.3.1.1.chr4.g00692.t01    rna-gnl|WGS_ZZZZ|egapxtmp_004383-R1          454    N0.HOG0000452   OG0000024  n236   rna-XM_034652112.1    rna-XM_038532079.1        rna-XM_036101548.1          rna-XM_032346810.1    rna-XM_059156417.1, rna-XM_059158372.1      rna-XM_045085593.1    mNeoVis.2.1.1.chr1_hap1.g01092.t01              mNeoVis.3.1.1.chr4.g00048.t01    rna-gnl|WGS_ZZZZ|egapxtmp_041072-R1    mNeoVis.evergladensis.2.1.chr13.g00605.t01, mNeoVis.evergladensis.2.1.chr2.g01168.t01    mNeoVis.vulgivaga.2.1.chr13.g00602.t01, mNeoVis.vulgivaga.2.1.chr3.g00388.t01      455    N0.HOG0000453   OG0000024  n249                 rna-XM_046020663.1                        mNeoVis.3.1.1.chr5.g01023.t01            456    N0.HOG0000454   OG0000025  n1   id-LOC117797283    id-LOC106559087    id-LOC111139586, id-LOC111139598    id-LOC101080387, id-LOC101082664, id-LOC101082918, id-LOC109500742, id-LOC123386008    id-LOC118550023, id-LOC118550122    id-LOC120240432, id-LOC120240454, id-LOC120240465, id-LOC120240470, id-LOC120240471, id-LOC120240472    id-LOC116867075    id-LOC123943627, id-LOC123943628    id-LOC116590189, id-LOC116591095    id-LOC131836321    id-LOC131999778, id-LOC131999781    id-LOC123388327      id-LOC123325717, rna-XM_021695697.1    id-LOC101375374, rna-XM_004402165.1    id-LOC102964273, id-LOC102964843, id-LOC102966868, id-LOC102968300, id-LOC102968892, id-LOC102969501, rna-XM_042989229.1    id-LOC103680421, id-LOC103680431, id-LOC103681025    id-LOC113908991, id-LOC113909263      id-egapxtmp_015282, id-egapxtmp_015724, id-egapxtmp_016170          457    N0.HOG0000455   OG0000025  n45   id-LOC105234897, id-LOC105234920, id-LOC105235009, id-LOC109488739, id-LOC109490671, id-LOC117797298, id-LOC117797300, id-LOC117797710, id-LOC117797716, rna-XM_034648712.1    id-LOC102156003, id-LOC119872979, id-LOC119875827, id-LOC119875829, id-LOC119875830, id-LOC119875833, id-LOC119875839, id-LOC119875840, id-LOC119876476, id-LOC119876483, id-LOC119879437, rna-XM_038544072.1, rna-XM_038544140.1, rna-XM_038589713.1    id-LOC111138978, id-LOC111139290, id-LOC111139382, id-LOC111139435, id-LOC111139445, id-LOC111139447, id-LOC111139476, id-LOC111139492, id-LOC111139515, id-LOC111139536, id-LOC111139557, id-LOC111139566, id-LOC111139573, id-LOC111140231, id-LOC111140233    id-LOC101081900, id-LOC101082160, id-LOC101082412, id-LOC109501106, id-LOC111560853, id-LOC123386007, rna-XM_045059665.1    id-LOC118522991, id-LOC118522993, id-LOC118522995, id-LOC118550022, id-LOC118550108, id-LOC118550123    id-LOC120240428, id-LOC120240429, id-LOC120240458, id-LOC120240466, id-LOC120240467, id-LOC120240468, id-LOC120240469, id-LOC120240475    id-LOC116867076, id-LOC116867084, id-LOC116867086, id-LOC116867362, id-LOC116867365, id-LOC116867368, id-LOC116867370, id-LOC116867371, id-LOC116867377, id-LOC116867380, id-LOC116867381, id-LOC116867388    id-LOC123943601, id-LOC123943603, id-LOC123943605, id-LOC123943607, id-LOC123943609, id-LOC123943613, id-LOC123943614, id-LOC123943615, id-LOC123943617, id-LOC123943620, id-LOC123943622, id-LOC123943623, id-LOC123943624, id-LOC123943625, id-LOC123943626, id-LOC123943630    id-LOC116590167, id-LOC116590168, id-LOC116590172, id-LOC116590173, id-LOC116590178, id-LOC116590181, id-LOC116590183, id-LOC116590184, id-LOC116590185, id-LOC116590187, id-LOC116590188, id-LOC116591106, id-LOC116591110, id-LOC116591112    id-LOC131836312, id-LOC131836316, id-LOC131836318, id-LOC131836331, id-LOC131836336, id-LOC131837183, id-LOC131837184, id-LOC131837188, id-LOC131837189, id-LOC131837192, id-LOC131837193, id-LOC131837194, id-LOC131837197, id-LOC131837200, id-LOC131837203, rna-XM_059183414.1    id-LOC131998744, id-LOC131999372, id-LOC131999373, id-LOC131999376, id-LOC131999377, id-LOC131999378, id-LOC131999381, id-LOC131999384, id-LOC131999388, id-LOC131999395, id-LOC131999396, id-LOC131999398, id-LOC131999400, id-LOC131999402, id-LOC131999426, id-LOC131999776, id-LOC131999777, id-LOC131999779, id-LOC131999784, id-LOC131999785, id-LOC132008787, id-LOC132015211    id-LOC101671122, id-LOC101671423, id-LOC101679175, id-LOC101680371, id-LOC101682112, id-LOC101682693, id-LOC101684438, id-LOC101685296, id-LOC101685891, id-LOC101686748, id-LOC101687901, id-LOC101688192, id-LOC101688475, id-LOC101689313, id-LOC101690182, id-LOC101691049, id-LOC106003341, id-LOC106004720, id-LOC123388328, id-LOC123388333, id-LOC123388334, id-LOC123388363, id-LOC123391338, id-LOC123391341, id-LOC123391345    mNeoVis.2.1.1.chr10_hap1.g00591.t01    id-LOC123323681, id-LOC123323682, id-LOC123323683, id-LOC123325713, id-LOC123325715, rna-XM_044918279.1, rna-XM_044918383.1    id-LOC101365714, id-LOC101366874, id-LOC101367157, id-LOC101367452, id-LOC101368342, id-LOC101369220, id-LOC101370998, id-LOC101371585, id-LOC101372188, id-LOC101372488, id-LOC101372780, id-LOC101373646, id-LOC101375655    id-LOC102965428, id-LOC102966000, id-LOC102967158, id-LOC102967721, id-LOC102968602, id-LOC102969974, id-LOC107181158, id-LOC107181171, id-LOC122239504    id-LOC103679632, id-LOC103680211, id-LOC103680320, id-LOC103680321, id-LOC103680424, id-LOC103680425, id-LOC103680427, id-LOC103680430, id-LOC103680446, id-LOC103680944, id-LOC103680950, id-LOC103681907, id-LOC103681908, id-LOC103682250, id-LOC103682251, id-LOC103682253, id-LOC103682256, id-LOC103682260, id-LOC103682261, id-LOC121100618, id-LOC121100619, id-LOC121100626, id-LOC121102722, id-LOC121103625    id-LOC113908999, id-LOC113909245, id-LOC113909250, id-LOC113909251, id-LOC113909252, id-LOC113909253, id-LOC113909254, id-LOC113909255, id-LOC113909259, id-LOC113909266, id-LOC118357070    mNeoVis.3.1.1.chr9.g00547.t01    id-egapxtmp_016048, rna-gnl|WGS_ZZZZ|egapxtmp_015367-R1    mNeoVis.evergladensis.2.1.chr8.g00574.t01        458    N0.HOG0000456   OG0000026  n4         rna-XM_003993315.2                          rna-XM_007088794.2                  459    N0.HOG0000457   OG0000026  n6   rna-XM_011216740.1    rna-NM_001389089.1    rna-XM_022507180.1      rna-XM_036073549.1    rna-XM_039224973.1    rna-XM_032846564.1    rna-XM_046017265.1    rna-XM_032359853.1    rna-XM_059158890.1    rna-XM_059388151.1    rna-XM_004782258.2    mNeoVis.2.1.1.chr7_hap1.g02082.t01    rna-XM_021685196.1    rna-XM_004412002.1      rna-XM_008694219.1    rna-XM_027580071.1          mNeoVis.vulgivaga.2.1.chr7.g02055.t01      460    N0.HOG0000458   OG0000026  n22   rna-XM_011216737.2    rna-XM_038564482.1    rna-XM_022507181.1        rna-XM_039224964.1      rna-XM_046015810.1    rna-XM_032359852.1    rna-XM_059136365.1    rna-XM_059387606.1              rna-XM_008694074.2        rna-gnl|WGS_ZZZZ|egapxtmp_010751-R1          461    N0.HOG0000459   OG0000026  n33   rna-XM_002930040.4    rna-XM_038564483.1    rna-XM_022507184.1, rna-XM_022507192.1    rna-XM_003993313.2    rna-XM_036073532.1    rna-XM_039224968.1    rna-XM_032846565.1    rna-XM_046015812.1, rna-XM_046015824.1, rna-XM_046015832.1    rna-XM_032358887.1, rna-XM_032359843.1, rna-XM_032359850.1    rna-XM_059135466.1, rna-XM_059135596.1, rna-XM_059136253.1    rna-XM_059387221.1, rna-XM_059387614.1, rna-XM_059388113.1    rna-XM_004782261.2, rna-XM_045067506.1, rna-XM_045067512.1, rna-XM_045074285.1    mNeoVis.2.1.1.chr7_hap1.g02083.t01, mNeoVis.2.1.1.chr7_hap1.g02100.t01        rna-XM_007088792.2    rna-XM_008694073.2        rna-gnl|WGS_ZZZZ|egapxtmp_010293-R1, rna-gnl|WGS_ZZZZ|egapxtmp_010743-R1, rna-gnl|WGS_ZZZZ|egapxtmp_013552-R1    mNeoVis.evergladensis.2.1.chr7.g02038.t01, mNeoVis.evergladensis.2.1.chr7.g02049.t01    mNeoVis.vulgivaga.2.1.chr7.g02056.t01, mNeoVis.vulgivaga.2.1.chr7.g02072.t01      462    N0.HOG0000460   OG0000026  n65   rna-XM_002930039.4    rna-XM_038562469.1    rna-XM_022507185.1, rna-XM_022507191.1    rna-XM_003993311.2    rna-XM_036073533.1    rna-XM_039224967.1      rna-XM_046015813.1, rna-XM_046015833.1    rna-XM_032359849.1    rna-XM_059136118.1    rna-XM_059387705.1    rna-XM_045067528.1    mNeoVis.2.1.1.chr7_hap1.g02098.t01        rna-XM_007088790.1    rna-XM_008694070.1      mNeoVis.3.1.1.chr7.g01934.t01    rna-gnl|WGS_ZZZZ|egapxtmp_013555-R1    mNeoVis.evergladensis.2.1.chr7.g02048.t01    mNeoVis.vulgivaga.2.1.chr7.g02070.t01      463    N0.HOG0000461   OG0000026  n85   rna-XM_002930041.2    rna-NM_001389094.1    rna-XM_022507182.1    rna-XM_003993314.3      rna-XM_039224965.1    rna-XM_032846481.1    rna-XM_046015811.1    rna-XM_032359851.1    rna-XM_059158889.1      rna-XM_004782260.1      rna-XM_021685195.1      rna-XM_007088793.1    rna-XM_008694218.1      mNeoVis.3.1.1.chr7.g01941.t01      mNeoVis.evergladensis.2.1.chr7.g02050.t01    mNeoVis.vulgivaga.2.1.chr7.g02073.t01      464    N0.HOG0000462   OG0000026  n104     rna-NM_001389110.1, rna-NM_001389136.1    rna-XM_022507195.1    rna-XM_011286884.2      rna-XM_039225293.1      rna-XM_046015825.1    rna-XM_032359842.1    rna-XM_059135212.1    rna-XM_059387219.1, rna-XM_059388187.1    rna-XM_004782360.1    mNeoVis.2.1.1.chr7_hap1.g02081.t01          rna-XM_008694061.2      mNeoVis.3.1.1.chr7.g01929.t01      mNeoVis.evergladensis.2.1.chr7.g02037.t01    mNeoVis.vulgivaga.2.1.chr7.g02054.t01      465    N0.HOG0000463   OG0000026  n119   rna-XM_002930038.2, rna-XM_011217780.2, rna-XM_034646479.1, rna-XM_034651596.1, rna-XM_034651789.1, rna-XM_034651905.1    rna-NM_001388623.1    rna-XM_022507189.1, rna-XM_022507194.1, rna-XM_022507377.1, rna-XM_022507448.1, rna-XM_022507505.1, rna-XM_022507549.1    rna-XM_003993312.2, rna-XM_023239015.2      rna-XM_039224966.1, rna-XM_039224970.1    rna-XM_032846482.1, rna-XM_032846509.1, rna-XM_032846518.1, rna-XM_032850676.1    rna-XM_046015670.1, rna-XM_046015671.1, rna-XM_046015673.1, rna-XM_046015822.1, rna-XM_046015823.1, rna-XM_046016721.1, rna-XM_046016764.1, rna-XM_046016816.1, rna-XM_046016900.1, rna-XM_046016962.1, rna-XM_046017028.1, rna-XM_046017049.1, rna-XM_046017222.1, rna-XM_046017510.1    rna-XM_032358886.1, rna-XM_032359498.1, rna-XM_032359840.1, rna-XM_032359847.1, rna-XM_032359981.1, rna-XM_032360459.1, rna-XM_032360463.1    rna-XM_059158875.1, rna-XM_059158877.1, rna-XM_059158887.1, rna-XM_059158888.1    rna-XM_059388170.1, rna-XM_059388176.1, rna-XM_059388235.1, rna-XM_059388265.1    rna-XM_004782172.2, rna-XM_004782179.2, rna-XM_004782181.2, rna-XM_004782262.2, rna-XM_045068114.1, rna-XM_045074286.1    mNeoVis.2.1.1.chr7_hap1.g02080.t01, mNeoVis.2.1.1.chr7_hap1.g02084.t01, mNeoVis.2.1.1.chr7_hap1.g02086.t01, mNeoVis.2.1.1.chr7_hap1.g02097.t01, mNeoVis.2.1.1.chr7_hap1.g02099.t01        rna-XM_007088789.1, rna-XM_015540526.2    rna-XM_008694069.2, rna-XM_008694071.1, rna-XM_008694203.1, rna-XM_008694206.2, rna-XM_008694209.1, rna-XM_008694212.2, rna-XM_008694214.2    rna-XM_035722994.1    mNeoVis.3.1.1.chr7.g01931.t01, mNeoVis.3.1.1.chr7.g01933.t01, mNeoVis.3.1.1.chr7.g01935.t01, mNeoVis.3.1.1.chr7.g01940.t01    rna-gnl|WGS_ZZZZ|egapxtmp_010280-R1, rna-gnl|WGS_ZZZZ|egapxtmp_010719-R1, rna-gnl|WGS_ZZZZ|egapxtmp_013510-R2, rna-gnl|WGS_ZZZZ|egapxtmp_013514-R1, rna-gnl|WGS_ZZZZ|egapxtmp_013519-R1    mNeoVis.evergladensis.2.1.chr7.g02036.t01, mNeoVis.evergladensis.2.1.chr7.g02039.t01, mNeoVis.evergladensis.2.1.chr7.g02041.t01, mNeoVis.evergladensis.2.1.chr7.g02047.t01    mNeoVis.vulgivaga.2.1.chr7.g02053.t01, mNeoVis.vulgivaga.2.1.chr7.g02057.t01, mNeoVis.vulgivaga.2.1.chr7.g02058.t01, mNeoVis.vulgivaga.2.1.chr7.g02069.t01, mNeoVis.vulgivaga.2.1.chr7.g02071.t01      466    N0.HOG0000464   OG0000026  n204   rna-XM_011217781.2, rna-XM_011217981.3, rna-XM_011218257.2, rna-XM_034644810.1, rna-XM_034651801.1, rna-XM_034651934.1    rna-NM_001388626.1    rna-XM_022507579.1    rna-XM_003993308.2    rna-XM_036073534.1, rna-XM_036073537.1    rna-XM_039225301.1    rna-XM_032846502.1, rna-XM_032846506.1, rna-XM_032850675.1    rna-XM_046015664.1, rna-XM_046015665.1, rna-XM_046015667.1, rna-XM_046015672.1, rna-XM_046015817.1, rna-XM_046016483.1, rna-XM_046016865.1, rna-XM_046016875.1, rna-XM_046017031.1, rna-XM_046017050.1, rna-XM_046017097.1, rna-XM_046017760.1    rna-XM_032359839.1, rna-XM_032359845.1, rna-XM_032359846.1    rna-XM_059158874.1, rna-XM_059158880.1, rna-XM_059158885.1    rna-XM_059388253.1    rna-XM_004782174.2, rna-XM_045068151.1, rna-XM_045074287.1    mNeoVis.2.1.1.chr7_hap1.g02089.t01, mNeoVis.2.1.1.chr7_hap1.g02091.t01    rna-XM_021685192.2, rna-XM_021685193.2    rna-XM_004412003.1, rna-XM_004412005.1, rna-XM_004412006.1, rna-XM_004412007.1    rna-XM_042958279.1, rna-XM_042959238.1    rna-XM_008694062.2, rna-XM_008694204.1, rna-XM_008694205.2, rna-XM_008694208.1, rna-XM_008694210.1, rna-XM_008694211.2, rna-XM_008694215.1, rna-XM_008694216.2    rna-XM_027577872.2, rna-XM_027580375.1, rna-XM_027580380.2, rna-XM_027580410.2, rna-XM_027580537.1    mNeoVis.3.1.1.chr7.g01937.t01    rna-gnl|WGS_ZZZZ|egapxtmp_010710-R1, rna-gnl|WGS_ZZZZ|egapxtmp_010745-R1    mNeoVis.evergladensis.2.1.chr7.g02044.t01    mNeoVis.vulgivaga.2.1.chr7.g02051.t01, mNeoVis.vulgivaga.2.1.chr7.g02061.t01      467    N0.HOG0000465   OG0000027  n1   rna-XM_002919851.4, rna-XM_002926695.4, rna-XM_034667222.1    rna-XM_038542821.1, rna-XM_038546578.1    rna-XM_022508154.1, rna-XM_022523255.1    rna-XM_003997160.5, rna-XM_023247314.2    rna-XM_036088675.1, rna-XM_036107113.1, rna-XM_036118073.1    rna-XM_039226319.1, rna-XM_039230190.1, rna-XM_039235801.1, rna-XM_039257207.1    rna-XM_032838730.1, rna-XM_032870606.1    rna-XM_045982578.1, rna-XM_045985685.1    rna-XM_032318232.1, rna-XM_032322627.1, rna-XM_032339497.1    rna-XM_059145945.1, rna-XM_059150348.1    rna-XM_059379999.1, rna-XM_059412898.1    rna-XM_004748891.2, rna-XM_045069566.1    mNeoVis.2.1.1.chr14_hap1.g00228.t01, mNeoVis.2.1.1.chr8_hap1.g00146.t01    rna-XM_021679507.2, rna-XM_021691828.2    rna-XM_004394327.1    rna-XM_007091267.3, rna-XM_007095092.3, rna-XM_042975894.1    rna-XM_008693776.2    rna-XM_027610973.2, rna-XM_027625006.2, rna-XM_035726399.1    mNeoVis.3.1.1.chr14.g00230.t01, mNeoVis.3.1.1.chr8.g00115.t01    rna-gnl|WGS_ZZZZ|egapxtmp_042344-R1    mNeoVis.evergladensis.2.1.chr14.g00230.t01, mNeoVis.evergladensis.2.1.chr9.g00143.t01    mNeoVis.vulgivaga.2.1.chr14.g00231.t01      468    N0.HOG0000466   OG0000027  n9                   rna-XM_032331456.1    rna-XM_059158308.1    rna-XM_059385340.1    rna-XM_013061254.2    mNeoVis.2.1.1.chr12_hap1.g00229.t01    rna-XM_021695554.1, rna-XM_021698992.1          rna-XM_027595180.2    mNeoVis.3.1.1.chr12.g00198.t01    rna-gnl|WGS_ZZZZ|egapxtmp_028655-R1    mNeoVis.evergladensis.2.1.chr12.g00199.t01    mNeoVis.vulgivaga.2.1.chr12.g00204.t01      469    N0.HOG0000467   OG0000027  n23           rna-XM_036098407.1    rna-XM_039246230.1      rna-XM_046017959.1    rna-XM_032302301.1              rna-XM_004393561.1, rna-XM_004404287.1      rna-XM_040626715.1    rna-XM_035727061.1        mNeoVis.evergladensis.2.1.chr1.g01100.t01, mNeoVis.evergladensis.2.1.chr5.g00787.t01    mNeoVis.vulgivaga.2.1.chr1.g01106.t01      470    N0.HOG0000468   OG0000027  n35     rna-NM_001374526.1      rna-XM_003985765.5, rna-XM_023259102.2            rna-XM_032341387.1                rna-XM_042985403.1          rna-gnl|WGS_ZZZZ|egapxtmp_010065-R1, rna-gnl|WGS_ZZZZ|egapxtmp_010178-R1    mNeoVis.evergladensis.2.1.chr1.g01064.t01, mNeoVis.evergladensis.2.1.chr1.g01121.t01        471    N0.HOG0000469   OG0000027  n44                                           mNeoVis.evergladensis.2.1.chr1.g01062.t01, mNeoVis.evergladensis.2.1.chr1.g01092.t01, mNeoVis.evergladensis.2.1.chr1.g01098.t01, mNeoVis.evergladensis.2.1.chr1.g01117.t01    mNeoVis.vulgivaga.2.1.chr1.g01087.t01, mNeoVis.vulgivaga.2.1.chr1.g01122.t01      472    N0.HOG0000470   OG0000027  n46   rna-XM_002928529.4, rna-XM_019807840.2, rna-XM_019809264.2, rna-XM_034660113.1, rna-XM_034660119.1, rna-XM_034660120.1, rna-XM_034660139.1    rna-NM_001374527.1, rna-NM_001374585.1, rna-NM_001374590.1, rna-XM_038562067.1, rna-XM_038584343.1, rna-XM_038584344.1, rna-XM_038584409.1, rna-XM_038584410.1, rna-XM_038584432.1    rna-XM_022490546.1, rna-XM_022490547.1, rna-XM_022490548.1, rna-XM_022490549.1    rna-NM_001374571.1, rna-NM_001374572.1, rna-NM_001374573.1, rna-NM_001374576.1, rna-XM_003985692.5, rna-XM_003985724.4, rna-XM_006931365.4, rna-XM_006931390.4, rna-XM_019831695.2    rna-XM_036112329.1, rna-XM_036112330.1, rna-XM_036112332.1, rna-XM_036121560.1    rna-XM_039223543.1, rna-XM_039257276.1, rna-XM_039257277.1, rna-XM_039257303.1    rna-XM_032861713.1, rna-XM_032861715.1    rna-XM_046006263.1, rna-XM_046006264.1, rna-XM_046006265.1, rna-XM_046006267.1, rna-XM_046006268.1, rna-XM_046006269.1, rna-XM_046006270.1    rna-XM_032338251.1, rna-XM_032338252.1, rna-XM_032338253.1, rna-XM_032338254.1, rna-XM_032338256.1    rna-XM_059176251.1, rna-XM_059176252.1, rna-XM_059176253.1, rna-XM_059176254.1, rna-XM_059176255.1, rna-XM_059179530.1      rna-XM_045082257.1, rna-XM_045082258.1, rna-XM_045082259.1, rna-XM_045082261.1, rna-XM_045082262.1, rna-XM_045082263.1    mNeoVis.2.1.1.chr1_hap1.g01077.t01, mNeoVis.2.1.1.chr1_hap1.g01079.t01, mNeoVis.2.1.1.chr1_hap1.g01088.t01, mNeoVis.2.1.1.chr1_hap1.g01116.t01, mNeoVis.2.1.1.chr1_hap1.g01118.t01, mNeoVis.2.1.1.chr1_hap1.g01123.t01, mNeoVis.2.1.1.chr1_hap1.g01125.t01, mNeoVis.2.1.1.chr1_hap1.g01129.t01, mNeoVis.2.1.1.chr1_hap1.g01146.t01, mNeoVis.2.1.1.chr1_hap1.g01149.t01, mNeoVis.2.1.1.chr1_hap1.g01152.t01    rna-XM_021696844.2, rna-XM_044917303.1, rna-XM_044917582.1, rna-XM_044917583.1    rna-XM_004406514.1, rna-XM_004406532.1, rna-XM_004413628.1, rna-XM_004413651.1    rna-XM_007074479.3, rna-XM_007074488.2, rna-XM_042985500.1, rna-XM_042985536.1, rna-XM_042985537.1, rna-XM_042985538.1, rna-XM_042985539.1, rna-XM_042985541.1, rna-XM_042985542.1    rna-XM_008692974.2, rna-XM_008693020.2, rna-XM_008693098.2, rna-XM_040645175.1    rna-XM_027607102.1, rna-XM_035728420.1, rna-XM_035728421.1, rna-XM_035728422.1, rna-XM_035728453.1      rna-gnl|WGS_ZZZZ|egapxtmp_020930-R1    mNeoVis.evergladensis.2.1.chr1.g01071.t01, mNeoVis.evergladensis.2.1.chr1.g01124.t01    mNeoVis.vulgivaga.2.1.chr1.g01082.t01      473    N0.HOG0000471   OG0000027  n55         rna-XM_011281294.4                          rna-XM_015542587.2                  474    N0.HOG0000472   OG0000027  n57   rna-XM_034660182.1, rna-XM_034660184.1    rna-NM_001376928.1    rna-XM_022490618.1    rna-XM_011282071.4      rna-XM_039257338.1      rna-XM_046006116.1    rna-XM_032341391.1      rna-XM_059387521.1, rna-XM_059387568.1    rna-XM_045082305.1      rna-XM_044917238.1      rna-XM_042985535.1    rna-XM_040644633.1    rna-XM_027604113.2, rna-XM_027624400.2      rna-gnl|WGS_ZZZZ|egapxtmp_010150-R1, rna-gnl|WGS_ZZZZ|egapxtmp_021307-R1, rna-gnl|WGS_ZZZZ|egapxtmp_021337-R1      mNeoVis.vulgivaga.2.1.chr5.g00779.t01      475    N0.HOG0000473   OG0000027  n76   rna-XM_019793752.2    rna-XM_038562066.1, rna-XM_038562071.1    rna-XM_022526273.1      rna-XM_036098401.1    rna-XM_039246205.1    rna-XM_032843668.1, rna-XM_032851116.1    rna-XM_046012482.1    rna-XM_032303388.1    rna-XM_059137255.1    rna-XM_059376006.1    rna-XM_045083144.1    mNeoVis.2.1.1.chr2_hap1.g00828.t01, mNeoVis.2.1.1.chr2_hap1.g00833.t01, mNeoVis.2.1.1.chr2_hap1.g00835.t01    rna-XM_021688069.1    rna-XM_004404175.1    rna-XM_042993986.1, rna-XM_042993987.1, rna-XM_042993988.1    rna-XM_008695746.2        rna-gnl|WGS_ZZZZ|egapxtmp_005758-R1    mNeoVis.evergladensis.2.1.chr5.g00789.t01        476    N0.HOG0000474   OG0000027  n78   rna-XM_002930205.2      rna-XM_022491651.1      rna-XM_036119162.1    rna-XM_039220267.1    rna-XM_032848522.1      rna-XM_032335064.1    rna-XM_059176080.1    rna-XM_059416740.1    rna-XM_004780619.2    mNeoVis.2.1.1.chr1_hap1.g00963.t01    rna-XM_021689909.2        rna-XM_008708271.2      mNeoVis.3.1.1.chr1.g00972.t01    rna-gnl|WGS_ZZZZ|egapxtmp_035642-R1    mNeoVis.evergladensis.2.1.chr1.g00958.t01    mNeoVis.vulgivaga.2.1.chr1.g00970.t01      477    N0.HOG0000475   OG0000027  n82   rna-XM_034660183.1          rna-XM_036112167.1, rna-XM_036121603.1    rna-XM_039257339.1          rna-XM_059138551.1      rna-XM_045083410.1        rna-XM_004413638.1      rna-XM_008692954.2    rna-XM_035728357.1    mNeoVis.3.1.1.chr1.g01094.t01, mNeoVis.3.1.1.chr1.g01108.t01    rna-gnl|WGS_ZZZZ|egapxtmp_010190-R1      mNeoVis.vulgivaga.2.1.chr1.g01118.t01      478    N0.HOG0000476   OG0000027  n94     rna-XM_038525936.1                            rna-XM_004408220.1                    479    N0.HOG0000477   OG0000027  n95               rna-XM_032857643.1                  rna-XM_004415134.1        rna-XM_027611280.2          mNeoVis.vulgivaga.2.1.chr8.g00146.t01      480    N0.HOG0000478   OG0000028  n2       rna-XM_022495838.1          rna-XM_032873888.1      rna-XM_032315095.1    rna-XM_059143988.1, rna-XM_059143989.1    rna-XM_059377779.1    rna-XM_045076703.1, rna-XM_045076704.1, rna-XM_045080317.1                          481    N0.HOG0000479   OG0000028  n11         rna-XM_045039066.1, rna-XM_045042572.1, rna-XM_045043356.1, rna-XM_045045853.1, rna-XM_045046349.1, rna-XM_045050396.1, rna-XM_045050723.1, rna-XM_045050968.1, rna-XM_045052565.1, rna-XM_045052621.1, rna-XM_045052631.1, rna-XM_045056137.1, rna-XM_045056941.1, rna-XM_045057239.1, rna-XM_045057241.1, rna-XM_045057698.1, rna-XM_045057699.1, rna-XM_045057860.1, rna-XM_045058249.1, rna-XM_045060801.1, rna-XM_045060805.1, rna-XM_045061375.1, rna-XM_045061376.1, rna-XM_045061951.1, rna-XM_045062665.1          rna-XM_045979031.1, rna-XM_045986838.1, rna-XM_045994771.1, rna-XM_045994786.1, rna-XM_045996839.1, rna-XM_045996840.1, rna-XM_045996846.1, rna-XM_045996851.1, rna-XM_045996868.1, rna-XM_045996870.1, rna-XM_045996916.1, rna-XM_045997028.1, rna-XM_045997386.1, rna-XM_045997444.1, rna-XM_045999757.1, rna-XM_046001161.1, rna-XM_046011859.1, rna-XM_046024370.1, rna-XM_046024371.1, rna-XM_046026576.1, rna-XM_046027262.1    rna-XM_032304417.1, rna-XM_032309462.1, rna-XM_032321784.1, rna-XM_032321785.1, rna-XM_032321791.1, rna-XM_032323445.1, rna-XM_032326056.1, rna-XM_032331958.1, rna-XM_032331969.1, rna-XM_032335026.1, rna-XM_032337926.1, rna-XM_032341371.1, rna-XM_032341373.1, rna-XM_032341437.1, rna-XM_032344823.1, rna-XM_032347828.1, rna-XM_032354339.1    rna-XM_059140040.1, rna-XM_059151251.1, rna-XM_059156834.1, rna-XM_059158478.1, rna-XM_059172542.1, rna-XM_059172543.1, rna-XM_059173396.1, rna-XM_059173408.1, rna-XM_059175980.1    rna-XM_059376002.1, rna-XM_059377787.1, rna-XM_059388318.1, rna-XM_059397495.1, rna-XM_059397527.1, rna-XM_059416426.1    rna-XM_013047985.2, rna-XM_045062994.1, rna-XM_045067219.1, rna-XM_045071937.1, rna-XM_045083147.1, rna-XM_045086936.1, rna-XM_045087974.1, rna-XM_045087975.1, rna-XM_045087977.1, rna-XM_045088144.1, rna-XM_045089730.1    mNeoVis.2.1.1.chr7_hap1.g02168.t01, mNeoVis.2.1.1.chr9_hap1.g00882.t01          rna-XM_040622882.1, rna-XM_040626731.1, rna-XM_040628262.1      mNeoVis.3.1.1.chr5.g00939.t01    rna-gnl|WGS_ZZZZ|egapxtmp_015407-R1, rna-gnl|WGS_ZZZZ|egapxtmp_033523-R1    mNeoVis.evergladensis.2.1.chr10.g00866.t01, mNeoVis.evergladensis.2.1.chr8.g00876.t01    mNeoVis.vulgivaga.2.1.chr8.g01289.t01      482    N0.HOG0000480   OG0000028  n111     rna-XM_038567190.1              rna-XM_046014700.1                  rna-XM_042987054.1      rna-XM_027575883.2              483    N0.HOG0000481   OG0000028  n114     rna-XM_038532571.1, rna-XM_038541764.1            rna-XM_032847371.1              mNeoVis.2.1.1.chr5_hap1.g01792.t01                  mNeoVis.evergladensis.2.1.chr2.g01776.t01        484    N0.HOG0000482   OG0000028  n120         rna-XM_045035194.1, rna-XM_045042021.1, rna-XM_045043170.1, rna-XM_045047659.1, rna-XM_045048764.1, rna-XM_045050382.1, rna-XM_045054178.1, rna-XM_045056098.1, rna-XM_045057760.1, rna-XM_045058643.1, rna-XM_045062639.1                          rna-XM_042967007.1, rna-XM_042968077.1, rna-XM_042968082.1, rna-XM_042974696.1, rna-XM_042974785.1, rna-XM_042984737.1, rna-XM_042985302.1, rna-XM_042987674.1, rna-XM_042993673.1, rna-XM_042998366.1                  485    N0.HOG0000483   OG0000028  n142       rna-XM_022515689.1, rna-XM_022526543.1    rna-XM_045041791.1, rna-XM_045058458.1      rna-XM_039224778.1, rna-XM_039225779.1, rna-XM_039241279.1    rna-XM_032843116.1, rna-XM_032843117.1, rna-XM_032851985.1, rna-XM_032861956.1, rna-XM_032861957.1, rna-XM_032866623.1, rna-XM_032870291.1, rna-XM_032881267.1    rna-XM_046011774.1    rna-XM_032304419.1, rna-XM_032328625.1, rna-XM_032329707.1, rna-XM_032329708.1, rna-XM_032337946.1, rna-XM_032337947.1, rna-XM_032337948.1, rna-XM_032341397.1, rna-XM_032341409.1    rna-XM_059165027.1, rna-XM_059165029.1, rna-XM_059172538.1, rna-XM_059173526.1, rna-XM_059176190.1    rna-XM_059372071.1, rna-XM_059388314.1, rna-XM_059404531.1, rna-XM_059416772.1    rna-XM_013059435.2, rna-XM_045088148.1, rna-XM_045088149.1, rna-XM_045089504.1, rna-XM_045089725.1          rna-XM_042956886.1, rna-XM_042959623.1, rna-XM_042962823.1, rna-XM_042966405.1, rna-XM_042968078.1, rna-XM_042968079.1, rna-XM_042970018.1, rna-XM_042971198.1, rna-XM_042972888.1, rna-XM_042977111.1, rna-XM_042977192.1, rna-XM_042977206.1, rna-XM_042977211.1, rna-XM_042989204.1, rna-XM_042989205.1, rna-XM_042991860.1, rna-XM_042992075.1, rna-XM_042993667.1, rna-XM_042993685.1, rna-XM_042993906.1, rna-XM_042998819.1              mNeoVis.vulgivaga.2.1.chr3.g00146.t01      486    N0.HOG0000484   OG0000028  n203         rna-XM_023252086.2, rna-XM_045050372.1, rna-XM_045051006.1, rna-XM_045055376.1, rna-XM_045055377.1, rna-XM_045059122.1                          rna-XM_042956409.1, rna-XM_042957130.1, rna-XM_042962226.1, rna-XM_042972707.1, rna-XM_042972893.1, rna-XM_042973950.1, rna-XM_042973968.1, rna-XM_042987612.1, rna-XM_042997572.1    rna-XM_040624738.1                487    N0.HOG0000485   OG0000028  n219     rna-XM_038551600.1              rna-XM_045997434.1    rna-XM_032329693.1    rna-XM_059168864.1, rna-XM_059168865.1                              488    N0.HOG0000486   OG0000028  n223     rna-XM_038529555.1, rna-XM_038538826.1, rna-XM_038544376.1, rna-XM_038550868.1, rna-XM_038551619.1, rna-XM_038559167.1, rna-XM_038565380.1, rna-XM_038572976.1, rna-XM_038584403.1, rna-XM_038588788.1, rna-XM_038588791.1, rna-XM_038588828.1      rna-XM_045033692.1, rna-XM_045062675.1                          rna-XM_042973939.1        mNeoVis.3.1.1.chr7.g00095.t01            489    N0.HOG0000487   OG0000028  n238             rna-XM_039217806.1, rna-XM_039217807.1, rna-XM_039219306.1, rna-XM_039221313.1, rna-XM_039225366.1, rna-XM_039243279.1, rna-XM_039243342.1, rna-XM_039248941.1, rna-XM_039249330.1, rna-XM_039251291.1                      rna-XM_042969094.1                  490    N0.HOG0000488   OG0000028  n248   rna-XM_034643869.1    rna-XM_038551964.1, rna-XM_038562446.1      rna-XM_045041692.1, rna-XM_045052147.1, rna-XM_045052177.1, rna-XM_045053127.1, rna-XM_045053933.1        rna-XM_032879949.1      rna-XM_032307683.1, rna-XM_032353935.1          mNeoVis.2.1.1.chr1_hap1.g00772.t01        rna-XM_042961729.1, rna-XM_042970240.1, rna-XM_042986555.1, rna-XM_042988308.1, rna-XM_042993878.1    rna-XM_040639366.1      mNeoVis.3.1.1.chr3.g00174.t01            491    N0.HOG0000489   OG0000029  n2   rna-XM_011216996.2    rna-NM_001388634.1    rna-XM_022506838.1    rna-XM_003993298.5    rna-XM_036073554.1    rna-XM_039226813.1    rna-XM_032844621.1    rna-XM_046015839.1    rna-XM_032359825.1, rna-XM_032359829.1    rna-XM_059133080.1, rna-XM_059133299.1    rna-XM_059386853.1, rna-XM_059386854.1, rna-XM_059387147.1, rna-XM_059387148.1    rna-XM_004781650.3, rna-XM_004782057.3    mNeoVis.2.1.1.chr7_hap1.g02043.t01    rna-XM_021685190.1    rna-XM_004417544.1    rna-XM_007095406.3    rna-XM_008701073.2    rna-XM_027577876.1        mNeoVis.evergladensis.2.1.chr7.g02002.t01    mNeoVis.vulgivaga.2.1.chr7.g02024.t01      492    N0.HOG0000490   OG0000029  n27   rna-XM_019793062.1    rna-NM_001388950.1, rna-NM_001389174.1    rna-XM_022506849.1    rna-XM_006937314.4      rna-XM_039226797.1    rna-XM_032844619.1    rna-XM_046015840.1    rna-XM_032359216.1, rna-XM_032359828.1    rna-XM_059158770.1    rna-XM_059386838.1, rna-XM_059386923.1    rna-XM_004782051.3    mNeoVis.2.1.1.chr7_hap1.g02042.t01        rna-XM_007095405.3    rna-XM_040639113.1      mNeoVis.3.1.1.chr7.g01906.t01    rna-gnl|WGS_ZZZZ|egapxtmp_010303-R2    mNeoVis.evergladensis.2.1.chr7.g02001.t01    mNeoVis.vulgivaga.2.1.chr7.g02022.t01      493    N0.HOG0000491   OG0000029  n45   rna-XM_034646064.1, rna-XM_034646065.1    rna-NM_001388635.1, rna-XM_038562481.1    rna-XM_022506660.1, rna-XM_022506672.1, rna-XM_022506773.1, rna-XM_022506839.1, rna-XM_022507200.1    rna-XM_011286888.2, rna-XM_023249662.2    rna-XM_036073546.1    rna-XM_039226798.1, rna-XM_039226803.1, rna-XM_039226806.1    rna-XM_032844537.1, rna-XM_032844538.1, rna-XM_032844545.1, rna-XM_032844546.1, rna-XM_032844555.1    rna-XM_046016394.1, rna-XM_046016423.1, rna-XM_046016445.1, rna-XM_046016495.1, rna-XM_046016534.1, rna-XM_046016536.1    rna-XM_032358706.1, rna-XM_032358752.1, rna-XM_032358793.1, rna-XM_032359079.1, rna-XM_032359213.1, rna-XM_032359215.1, rna-XM_032359831.1    rna-XM_059158767.1, rna-XM_059158769.1, rna-XM_059158777.1, rna-XM_059158825.1, rna-XM_059158838.1, rna-XM_059158840.1    rna-XM_059386831.1, rna-XM_059386832.1, rna-XM_059386833.1, rna-XM_059386839.1, rna-XM_059386870.1, rna-XM_059386883.1, rna-XM_059386886.1, rna-XM_059386921.1    rna-XM_004781647.1, rna-XM_004781651.2, rna-XM_004781652.3, rna-XM_004782058.3, rna-XM_004782059.2, rna-XM_013050566.2, rna-XM_045067932.1, rna-XM_045067935.1    mNeoVis.2.1.1.chr7_hap1.g02039.t01, mNeoVis.2.1.1.chr7_hap1.g02040.t01, mNeoVis.2.1.1.chr7_hap1.g02046.t01, mNeoVis.2.1.1.chr7_hap1.g02050.t01, mNeoVis.2.1.1.chr7_hap1.g02052.t01    rna-XM_021684819.1, rna-XM_021684820.1    rna-XM_004417543.1, rna-XM_004417668.1    rna-XM_007082206.3, rna-XM_007082207.3, rna-XM_007095404.3    rna-XM_008701075.2, rna-XM_008701076.2    rna-XM_027579659.1    mNeoVis.3.1.1.chr7.g01904.t01, mNeoVis.3.1.1.chr7.g01913.t01, mNeoVis.3.1.1.chr7.g01915.t01    rna-gnl|WGS_ZZZZ|egapxtmp_010277-R1, rna-gnl|WGS_ZZZZ|egapxtmp_010282-R2, rna-gnl|WGS_ZZZZ|egapxtmp_010290-R1, rna-gnl|WGS_ZZZZ|egapxtmp_012615-R1    mNeoVis.evergladensis.2.1.chr7.g01998.t01, mNeoVis.evergladensis.2.1.chr7.g01999.t01, mNeoVis.evergladensis.2.1.chr7.g02005.t01, mNeoVis.evergladensis.2.1.chr7.g02010.t01    mNeoVis.vulgivaga.2.1.chr7.g02021.t01, mNeoVis.vulgivaga.2.1.chr7.g02028.t01      494    N0.HOG0000492   OG0000029  n128                       rna-XM_059386852.1, rna-XM_059387149.1                        mNeoVis.vulgivaga.2.1.chr7.g02027.t01      495    N0.HOG0000493   OG0000029  n131   rna-XM_019793643.2, rna-XM_034646613.1, rna-XM_034653136.1    rna-NM_001388955.1, rna-XM_038562482.1    rna-XM_022506872.1, rna-XM_022506994.1    rna-XM_003993263.3, rna-XM_045038140.1, rna-XM_045038142.1, rna-XM_045038144.1    rna-XM_036073547.1    rna-XM_039226695.1, rna-XM_039226802.1, rna-XM_039226821.1, rna-XM_039226907.1    rna-XM_032844565.1, rna-XM_032844608.1    rna-XM_046016715.1, rna-XM_046016826.1, rna-XM_046016828.1    rna-XM_032359076.1, rna-XM_032359114.1, rna-XM_032359377.1    rna-XM_059158729.1, rna-XM_059158764.1, rna-XM_059158834.1    rna-XM_059386841.1, rna-XM_059386846.1, rna-XM_059386889.1, rna-XM_059386925.1, rna-XM_059386965.1, rna-XM_059386969.1    rna-XM_004781639.2, rna-XM_004782052.2, rna-XM_004782054.3    mNeoVis.2.1.1.chr7_hap1.g02036.t01, mNeoVis.2.1.1.chr7_hap1.g02038.t01, mNeoVis.2.1.1.chr7_hap1.g02051.t01    rna-XM_021684818.1      rna-XM_007082181.2, rna-XM_007082204.2, rna-XM_007082205.2, rna-XM_007095400.1, rna-XM_042959624.1    rna-XM_008700934.2, rna-XM_008700935.2, rna-XM_008701077.2    rna-XM_027579953.1    mNeoVis.3.1.1.chr7.g01900.t01, mNeoVis.3.1.1.chr7.g01903.t01    rna-gnl|WGS_ZZZZ|egapxtmp_013551-R1, rna-gnl|WGS_ZZZZ|egapxtmp_013560-R1, rna-gnl|WGS_ZZZZ|egapxtmp_013565-R1    mNeoVis.evergladensis.2.1.chr7.g01995.t01, mNeoVis.evergladensis.2.1.chr7.g01997.t01, mNeoVis.evergladensis.2.1.chr7.g02009.t01    mNeoVis.vulgivaga.2.1.chr7.g02017.t01, mNeoVis.vulgivaga.2.1.chr7.g02020.t01, mNeoVis.vulgivaga.2.1.chr7.g02030.t01      496    N0.HOG0000494   OG0000029  n178         rna-XM_045038141.1      rna-XM_039226815.1                      rna-XM_007082180.2                  497    N0.HOG0000495   OG0000029  n182   rna-XM_034650305.1    rna-NM_001388952.1              rna-XM_046016712.1, rna-XM_046016757.1      rna-XM_059158775.1    rna-XM_059386836.1, rna-XM_059386849.1, rna-XM_059386885.1, rna-XM_059387100.1    rna-XM_045067866.1                mNeoVis.3.1.1.chr7.g01902.t01    rna-gnl|WGS_ZZZZ|egapxtmp_012614-R1      mNeoVis.vulgivaga.2.1.chr7.g02019.t01      498    N0.HOG0000496   OG0000029  n194   rna-XM_034646538.1    rna-NM_001388959.1    rna-XM_022506992.1, rna-XM_022506999.1          rna-XM_032844610.1    rna-XM_046016734.1    rna-XM_032359208.1, rna-XM_032359628.1    rna-XM_059158753.1, rna-XM_059158804.1    rna-XM_059386850.1, rna-XM_059387064.1    rna-XM_004781640.1, rna-XM_004782053.2    mNeoVis.2.1.1.chr7_hap1.g02049.t01          rna-XM_040639107.1      mNeoVis.3.1.1.chr7.g01911.t01, mNeoVis.3.1.1.chr7.g01912.t01    rna-gnl|WGS_ZZZZ|egapxtmp_010318-R1    mNeoVis.evergladensis.2.1.chr7.g02008.t01        499    N0.HOG0000497   OG0000029  n213   rna-XM_034644808.1, rna-XM_034645874.1, rna-XM_034650404.1      rna-XM_022506806.1, rna-XM_022506914.1          rna-XM_032844567.1, rna-XM_032844589.1    rna-XM_046016659.1, rna-XM_046016706.1, rna-XM_046016771.1    rna-XM_032359169.1, rna-XM_032359681.1    rna-XM_059158744.1, rna-XM_059158794.1    rna-XM_059386845.1, rna-XM_059386897.1, rna-XM_059386932.1, rna-XM_059386959.1    rna-XM_004781641.1    mNeoVis.2.1.1.chr7_hap1.g02037.t01, mNeoVis.2.1.1.chr7_hap1.g02048.t01      rna-XM_004399598.1      rna-XM_008700936.1, rna-XM_008700938.2      mNeoVis.3.1.1.chr7.g01901.t01    rna-gnl|WGS_ZZZZ|egapxtmp_010320-R1, rna-gnl|WGS_ZZZZ|egapxtmp_010322-R1    mNeoVis.evergladensis.2.1.chr7.g01996.t01, mNeoVis.evergladensis.2.1.chr7.g02007.t01    mNeoVis.vulgivaga.2.1.chr7.g02018.t01, mNeoVis.vulgivaga.2.1.chr7.g02029.t01      500    N0.HOG0000498   OG0000030  n1   rna-XM_034649576.1    rna-XM_038587648.1    rna-XM_022524480.1    rna-XM_004000597.5    rna-XM_036075367.1    rna-XM_039253359.1    rna-XM_032847763.1    rna-XM_045995462.1    rna-XM_032331549.1    rna-XM_059156762.1    rna-XM_059384563.1    rna-XM_004777205.3    mNeoVis.2.1.1.chr12_hap1.g00491.t01, mNeoVis.2.1.1.chr4_hap1.g00072.t01    rna-XM_021680402.1    rna-XM_004396611.1    rna-XM_007097573.3    rna-XM_008712262.2    rna-XM_027608425.1    mNeoVis.3.1.1.chr12.g00448.t01, mNeoVis.3.1.1.chr6.g00076.t01    rna-gnl|WGS_ZZZZ|egapxtmp_002908-R1, rna-gnl|WGS_ZZZZ|egapxtmp_029692-R1    mNeoVis.evergladensis.2.1.chr6.g00077.t01    mNeoVis.vulgivaga.2.1.chr12.g00453.t01, mNeoVis.vulgivaga.2.1.chr6.g00076.t01      501    N0.HOG0000499   OG0000030  n26   rna-XM_002930455.4    rna-XM_038527905.1    rna-XM_022491920.1      rna-XM_036065386.1    rna-XM_039244062.1    rna-XM_032848742.1    rna-XM_045986811.1    rna-XM_032326106.1    rna-XM_059151208.1    rna-XM_059383338.1    rna-XM_013049566.2    mNeoVis.2.1.1.chr7_hap1.g00768.t01    rna-XM_044911340.1    rna-XM_004409556.1      rna-XM_008701619.1    rna-XM_027619355.1    mNeoVis.3.1.1.chr7.g00718.t01    rna-gnl|WGS_ZZZZ|egapxtmp_012294-R1    mNeoVis.evergladensis.2.1.chr7.g00755.t01    mNeoVis.vulgivaga.2.1.chr7.g00758.t01      502    N0.HOG0000500   OG0000030  n46     rna-XM_038576188.1      rna-XM_045061878.1    rna-XM_036124181.1    rna-XM_039255099.1    rna-XM_032876541.1    rna-XM_046013536.1    rna-XM_032345633.1    rna-XM_059185136.1    rna-XM_059403290.1    rna-XM_045073197.1    mNeoVis.2.1.1.chr9_hap1.g00477.t01    rna-XM_044915196.1      rna-XM_042993693.1      rna-XM_035729315.1    mNeoVis.3.1.1.chr10.g00451.t01    rna-gnl|WGS_ZZZZ|egapxtmp_024301-R1    mNeoVis.evergladensis.2.1.chr10.g00471.t01    mNeoVis.vulgivaga.2.1.chr10.g00460.t01      503    N0.HOG0000501   OG0000030  n61     rna-XM_038577003.1      rna-XM_045061502.1      rna-XM_039225154.1                      rna-XM_042991866.1    rna-XM_040642211.1                504    N0.HOG0000502   OG0000030  n66             rna-XM_039238599.1                        rna-XM_040642219.1                505    N0.HOG0000503   OG0000030  n74   rna-XM_034646511.1    rna-XM_038576148.1, rna-XM_038576988.1                                rna-XM_040642208.1                506    N0.HOG0000504   OG0000030  n77     rna-XM_038577909.1, rna-XM_038577911.1, rna-XM_038589380.1                                rna-XM_040642215.1                507    N0.HOG0000505   OG0000030  n80   rna-XM_002930638.4    rna-XM_038576149.1, rna-XM_038576999.1, rna-XM_038577002.1        rna-XM_036109258.1                    rna-XM_021690733.1        rna-XM_040641910.1, rna-XM_040642214.1                508    N0.HOG0000506   OG0000030  n87             rna-XM_039225153.1                      rna-XM_042991867.1                  509    N0.HOG0000507   OG0000030  n88     rna-XM_038576987.1      rna-XM_045061520.1    rna-XM_036109257.1    rna-XM_039223588.1    rna-XM_032854646.1, rna-XM_032854649.1    rna-XM_046013019.1    rna-XM_032347509.1    rna-XM_059185894.1    rna-XM_059402698.1    rna-XM_045073231.1    mNeoVis.2.1.1.chr9_hap1.g00750.t01    rna-XM_044912508.1        rna-XM_040641897.1      mNeoVis.3.1.1.chr10.g00718.t01      mNeoVis.evergladensis.2.1.chr10.g00741.t01    mNeoVis.vulgivaga.2.1.chr10.g00732.t01      510    N0.HOG0000508   OG0000030  n104     rna-XM_038576984.1    rna-XM_022492126.1    rna-XM_045061512.1      rna-XM_039224815.1    rna-XM_032854644.1, rna-XM_032855432.1    rna-XM_046013602.1, rna-XM_046013667.1    rna-XM_032347502.1    rna-XM_059185901.1    rna-XM_059404781.1    rna-XM_045073154.1, rna-XM_045073245.1          rna-XM_042991879.1                  511    N0.HOG0000509   OG0000030  n118     rna-XM_038576993.1      rna-XM_045061523.1    rna-XM_036109256.1    rna-XM_039224816.1                      rna-XM_042991901.1                  512    N0.HOG0000510   OG0000030  n123   rna-XM_034645967.1, rna-XM_034646519.1, rna-XM_034646520.1, rna-XM_034646521.1, rna-XM_034650996.1, rna-XM_034651056.1, rna-XM_034651941.1, rna-XM_034653211.1, rna-XM_034653270.1    rna-XM_038576147.1, rna-XM_038576151.1, rna-XM_038576153.1, rna-XM_038577005.1      rna-XM_045061515.1    rna-XM_036079380.1, rna-XM_036109260.1    rna-XM_039223587.1, rna-XM_039238589.1    rna-XM_032854647.1, rna-XM_032854648.1, rna-XM_032854656.1, rna-XM_032854657.1, rna-XM_032854658.1, rna-XM_032855431.1    rna-XM_046011808.1, rna-XM_046011809.1, rna-XM_046011810.1, rna-XM_046012192.1, rna-XM_046012206.1, rna-XM_046012211.1, rna-XM_046012214.1, rna-XM_046012218.1, rna-XM_046012219.1, rna-XM_046012220.1, rna-XM_046012221.1, rna-XM_046013390.1, rna-XM_046013443.1, rna-XM_046013529.1, rna-XM_046013564.1, rna-XM_046013798.1    rna-XM_032347507.1, rna-XM_032347713.1, rna-XM_032347775.1, rna-XM_032347822.1, rna-XM_032347823.1    rna-XM_059185711.1, rna-XM_059185712.1, rna-XM_059185895.1, rna-XM_059185896.1, rna-XM_059185899.1, rna-XM_059185900.1, rna-XM_059187514.1    rna-XM_059387280.1, rna-XM_059387281.1, rna-XM_059404485.1, rna-XM_059404495.1, rna-XM_059404795.1    rna-XM_045073118.1, rna-XM_045073193.1, rna-XM_045073255.1, rna-XM_045073338.1, rna-XM_045073462.1    mNeoVis.2.1.1.chr9_hap1.g00690.t01, mNeoVis.2.1.1.chr9_hap1.g00691.t01, mNeoVis.2.1.1.chr9_hap1.g00692.t01, mNeoVis.2.1.1.chr9_hap1.g00695.t01, mNeoVis.2.1.1.chr9_hap1.g00751.t01, mNeoVis.2.1.1.chr9_hap1.g00752.t01    rna-XM_044915216.1    rna-XM_004405506.1    rna-XM_042991893.1    rna-XM_040619393.1, rna-XM_040621915.1, rna-XM_040621917.1, rna-XM_040641911.1, rna-XM_040642213.1, rna-XM_040642218.1, rna-XM_040642231.1    rna-XM_035729304.1    mNeoVis.3.1.1.chr10.g00657.t01, mNeoVis.3.1.1.chr10.g00658.t01, mNeoVis.3.1.1.chr10.g00659.t01, mNeoVis.3.1.1.chr10.g00720.t01, mNeoVis.3.1.1.chr10.g00722.t01    rna-gnl|WGS_ZZZZ|egapxtmp_023742-R1, rna-gnl|WGS_ZZZZ|egapxtmp_023911-R1, rna-gnl|WGS_ZZZZ|egapxtmp_023912-R1, rna-gnl|WGS_ZZZZ|egapxtmp_023913-R1, rna-gnl|WGS_ZZZZ|egapxtmp_024738-R1    mNeoVis.evergladensis.2.1.chr10.g00685.t01, mNeoVis.evergladensis.2.1.chr10.g00686.t01, mNeoVis.evergladensis.2.1.chr10.g00688.t01, mNeoVis.evergladensis.2.1.chr10.g00742.t01, mNeoVis.evergladensis.2.1.chr10.g00743.t01    mNeoVis.vulgivaga.2.1.chr10.g00672.t01, mNeoVis.vulgivaga.2.1.chr10.g00673.t01, mNeoVis.vulgivaga.2.1.chr10.g00674.t01, mNeoVis.vulgivaga.2.1.chr10.g00675.t01, mNeoVis.vulgivaga.2.1.chr10.g00733.t01      513    N0.HOG0000511   OG0000030  n220   rna-XM_034646509.1, rna-XM_034646525.1, rna-XM_034651916.1    rna-XM_038577910.1, rna-XM_038577912.1      rna-XM_045061501.1, rna-XM_045061508.1, rna-XM_045061513.1, rna-XM_045061516.1, rna-XM_045061521.1, rna-XM_045061524.1, rna-XM_045061525.1, rna-XM_045062661.1, rna-XM_045062662.1      rna-XM_039223589.1, rna-XM_039224813.1, rna-XM_039224814.1, rna-XM_039224819.1, rna-XM_039225150.1, rna-XM_039225152.1, rna-XM_039238561.1, rna-XM_039238564.1, rna-XM_039238565.1, rna-XM_039238566.1, rna-XM_039238567.1, rna-XM_039238580.1                      rna-XM_042991877.1, rna-XM_042991888.1, rna-XM_042991892.1, rna-XM_042991894.1, rna-XM_042991895.1, rna-XM_042991896.1, rna-XM_042993682.1    rna-XM_040642206.1, rna-XM_040642210.1        rna-gnl|WGS_ZZZZ|egapxtmp_024736-R1          514    N0.HOG0000512   OG0000031  n1   rna-XM_034643579.1    rna-XM_038529612.1    rna-XM_022520800.1    rna-XM_045060820.1    rna-XM_036086168.1    rna-XM_039240353.1    rna-XM_032881664.1    rna-XM_046010517.1    rna-XM_032349852.1    rna-XM_059183712.1    rna-XM_059403994.1    rna-XM_004757921.3    mNeoVis.2.1.1.chr9_hap1.g00019.t01    rna-XM_044915551.1    rna-XM_004411830.1    rna-XM_015537136.2    rna-XM_040645465.1    rna-XM_027592821.2    mNeoVis.3.1.1.chr10.g00018.t01    rna-gnl|WGS_ZZZZ|egapxtmp_023665-R1    mNeoVis.evergladensis.2.1.chr10.g00021.t01    mNeoVis.vulgivaga.2.1.chr10.g00020.t01      515    N0.HOG0000513   OG0000031  n25   rna-XM_002926724.4    rna-XM_038576202.1    rna-XM_022522289.1    rna-XM_004001072.4    rna-XM_036102458.1    rna-XM_039250012.1    rna-XM_032844407.1    rna-XM_046012554.1    rna-XM_032349279.1    rna-XM_059184296.1    rna-XM_059403962.1    rna-XM_045071679.1    mNeoVis.2.1.1.chr9_hap1.g00181.t01    rna-XM_021690472.2    rna-XM_004406904.1    rna-XM_007092789.3    rna-XM_008704809.2    rna-XM_027592581.1    mNeoVis.3.1.1.chr10.g00173.t01    rna-gnl|WGS_ZZZZ|egapxtmp_024328-R1    mNeoVis.evergladensis.2.1.chr10.g00176.t01    mNeoVis.vulgivaga.2.1.chr10.g00173.t01      516    N0.HOG0000514   OG0000031  n40   rna-XM_002925427.4    rna-XM_038541844.1    rna-XM_022508814.1    rna-XM_006942867.5    rna-XM_036079609.1    rna-XM_039227588.1    rna-XM_032878292.1    rna-XM_045982118.1    rna-XM_032319351.1    rna-XM_059146003.1    rna-XM_059413124.1    rna-XM_004756320.3    mNeoVis.2.1.1.chr14_hap1.g00343.t01    rna-XM_021682705.1    rna-XM_004392774.1    rna-XM_007086470.3    rna-XM_008685131.2    rna-XM_035722565.1, rna-XM_035729052.1, rna-XM_035729054.1    mNeoVis.3.1.1.chr14.g00341.t01    rna-gnl|WGS_ZZZZ|egapxtmp_032421-R1    mNeoVis.evergladensis.2.1.chr14.g00343.t01    mNeoVis.vulgivaga.2.1.chr14.g00344.t01      517    N0.HOG0000515   OG0000031  n57     rna-XM_038574482.1    rna-XM_022500444.1, rna-XM_022518032.1          rna-XM_032860882.1    rna-XM_045979041.1, rna-XM_046001163.1, rna-XM_046002249.1, rna-XM_046007796.1, rna-XM_046021826.1    rna-XM_032306961.1    rna-XM_059156375.1    rna-XM_059380170.1, rna-XM_059392597.1, rna-XM_059408052.1    rna-XM_045081789.1    mNeoVis.2.1.1.chr6_hap1.g00271.t01        rna-XM_042961652.1, rna-XM_042962224.1, rna-XM_042962251.1          rna-gnl|WGS_ZZZZ|egapxtmp_043817-R1    mNeoVis.evergladensis.2.1.chr4.g00277.t01    mNeoVis.vulgivaga.2.1.chr4.g00277.t01      518    N0.HOG0000516   OG0000031  n83                               rna-XM_004400859.1        rna-XM_035729162.1              519    N0.HOG0000517   OG0000031  n88   rna-XM_034642067.1, rna-XM_034642679.1    rna-XM_038575954.1, rna-XM_038575966.1    rna-XM_022495015.1, rna-XM_022495399.1, rna-XM_022495405.1, rna-XM_022521181.1    rna-XM_011287925.4    rna-XM_036104318.1, rna-XM_036104326.1    rna-XM_039242808.1, rna-XM_039242858.1    rna-XM_032857992.1, rna-XM_032864012.1, rna-XM_032875946.1, rna-XM_032875958.1    rna-XM_046024379.1, rna-XM_046025397.1    rna-XM_032309601.1, rna-XM_032310951.1    rna-XM_059138649.1, rna-XM_059139545.1, rna-XM_059162282.1    rna-XM_059390711.1, rna-XM_059409027.1    rna-XM_004833880.3, rna-XM_013051315.2, rna-XM_045065283.1    mNeoVis.2.1.1.chr3_hap1.g00329.t01, mNeoVis.2.1.1.chr3_hap1.g00333.t01    rna-XM_021687882.2, rna-XM_021687883.2    rna-XM_004400493.1, rna-XM_004400495.1    rna-XM_042961982.1, rna-XM_042977129.1, rna-XM_042980374.1    rna-XM_008697557.2, rna-XM_008697561.2    rna-XM_035723849.1, rna-XM_035723850.1, rna-XM_035723956.1, rna-XM_035729295.1    mNeoVis.3.1.1.chr3.g00327.t01    rna-gnl|WGS_ZZZZ|egapxtmp_041908-R1    mNeoVis.evergladensis.2.1.chr10.g00582.t01, mNeoVis.evergladensis.2.1.chr3.g00341.t01    mNeoVis.vulgivaga.2.1.chr3.g00332.t01, mNeoVis.vulgivaga.2.1.chr3.g00336.t01      520    N0.HOG0000518   OG0000031  n115     rna-NM_001314090.1    rna-XM_022519351.1    rna-XM_045061749.1    rna-XM_036121122.1      rna-XM_032846471.1, rna-XM_032871280.1, rna-XM_032876873.1    rna-XM_046011466.1    rna-XM_032348094.1    rna-XM_059185462.1    rna-XM_059403026.1    rna-XM_045073177.1    mNeoVis.2.1.1.chr9_hap1.g00588.t01    rna-XM_021704822.2, rna-XM_021704888.1      rna-XM_007072976.3      rna-XM_035723395.1, rna-XM_035729161.1              521    N0.HOG0000519   OG0000031  n116   rna-XM_034644636.1, rna-XM_034651162.1    rna-XM_038564501.1          rna-XM_039221068.1, rna-XM_039241347.1    rna-XM_032862221.1                  rna-XM_004400860.1, rna-XM_004400861.1      rna-XM_008704544.2      mNeoVis.3.1.1.chr5.g01011.t01    rna-gnl|WGS_ZZZZ|egapxtmp_009074-R1    mNeoVis.evergladensis.2.1.chr5.g01032.t01        522    N0.HOG0000520   OG0000031  n128   rna-XM_034655224.1    rna-XM_038585815.1    rna-XM_022496645.1    rna-XM_045034511.1    rna-XM_036083009.1    rna-XM_039225622.1    rna-XM_032857269.1    rna-XM_046018377.1    rna-XM_032354871.1    rna-XM_059167997.1      rna-XM_045078760.1        rna-XM_004402745.1    rna-XM_042995371.1    rna-XM_008690431.2    rna-XM_035726515.1    mNeoVis.3.1.1.chr3.g01150.t01      mNeoVis.evergladensis.2.1.chr3.g01165.t01    mNeoVis.vulgivaga.2.1.chr3.g01179.t01      523    N0.HOG0000521   OG0000031  n138   rna-XM_034652041.1    rna-XM_038585813.1          rna-XM_039225629.1            rna-XM_059393525.1, rna-XM_059395476.1                  mNeoVis.3.1.1.chr8.g01303.t01    rna-gnl|WGS_ZZZZ|egapxtmp_017654-R1      mNeoVis.vulgivaga.2.1.chr4.g00743.t01      524    N0.HOG0000522   OG0000031  n145     rna-XM_038577535.1    rna-XM_022519350.1    rna-XM_003988644.6    rna-XM_036121126.1      rna-XM_032876875.1    rna-XM_046011939.1    rna-XM_032348093.1    rna-XM_059185459.1    rna-XM_059403025.1    rna-XM_045072973.1    mNeoVis.2.1.1.chr9_hap1.g00589.t01    rna-XM_021704821.2      rna-XM_007072974.3    rna-XM_008704542.2    rna-XM_035729104.1        mNeoVis.evergladensis.2.1.chr10.g00583.t01        525    N0.HOG0000523   OG0000031  n146   rna-XM_011231789.2    rna-XM_038577536.1    rna-XM_022499624.1, rna-XM_022519430.1    rna-XM_011283901.4    rna-XM_036085023.1, rna-XM_036106010.1, rna-XM_036109682.1, rna-XM_036121124.1    rna-XM_039225345.1, rna-XM_039241349.1    rna-XM_032845603.1, rna-XM_032857270.1, rna-XM_032867263.1, rna-XM_032876874.1    rna-XM_046001145.1, rna-XM_046011467.1    rna-XM_032348095.1    rna-XM_059185463.1    rna-XM_059403027.1    rna-XM_013049159.2    mNeoVis.2.1.1.chr9_hap1.g00587.t01, mNeoVis.2.1.1.chr9_hap1.g00775.t01    rna-XM_021686649.1, rna-XM_021689552.2, rna-XM_021703064.2, rna-XM_021704799.2, rna-XM_044913624.1      rna-XM_042992207.1    rna-XM_008704543.2      mNeoVis.3.1.1.chr10.g00557.t01    rna-gnl|WGS_ZZZZ|egapxtmp_023005-R1    mNeoVis.evergladensis.2.1.chr10.g00580.t01, mNeoVis.evergladensis.2.1.chr10.g00765.t01, mNeoVis.evergladensis.2.1.chr2.g01498.t01    mNeoVis.vulgivaga.2.1.chr10.g00568.t01, mNeoVis.vulgivaga.2.1.chr10.g00569.t01      526    N0.HOG0000524   OG0000032  n1   rna-XM_002930301.3, rna-XM_002930302.1    rna-XM_038589477.1    rna-XM_022526039.1, rna-XM_022526767.1    rna-XM_023256230.2    rna-XM_036082566.1, rna-XM_036082570.1    rna-XM_039218336.1, rna-XM_039218347.1, rna-XM_039218353.1    rna-XM_032857125.1, rna-XM_032857126.1, rna-XM_032857132.1    rna-XM_046009416.1, rna-XM_046009483.1, rna-XM_046009515.1    rna-XM_032341694.1, rna-XM_032341701.1, rna-XM_032341702.1    rna-XM_059179642.1, rna-XM_059179651.1, rna-XM_059179652.1    rna-XM_059371145.1, rna-XM_059371155.1, rna-XM_059371156.1    rna-XM_004751362.1, rna-XM_004751363.3, rna-XM_004782266.2    mNeoVis.2.1.1.chr10_hap1.g00793.t01, mNeoVis.2.1.1.chr10_hap1.g00794.t01, mNeoVis.2.1.1.chr10_hap1.g00800.t01    rna-XM_021695515.1, rna-XM_021695518.1    rna-XM_004414376.1    rna-XM_007087469.3    rna-XM_008702302.1, rna-XM_008702303.1    rna-XM_027570950.1, rna-XM_027571397.2    mNeoVis.3.1.1.chr9.g00754.t01    rna-gnl|WGS_ZZZZ|egapxtmp_017331-R1, rna-gnl|WGS_ZZZZ|egapxtmp_017366-R1    mNeoVis.evergladensis.2.1.chr8.g00775.t01, mNeoVis.evergladensis.2.1.chr8.g00776.t01, mNeoVis.evergladensis.2.1.chr8.g00781.t01    mNeoVis.vulgivaga.2.1.chr9.g00774.t01, mNeoVis.vulgivaga.2.1.chr9.g00775.t01, mNeoVis.vulgivaga.2.1.chr9.g00776.t01      527    N0.HOG0000525   OG0000032  n47   rna-XM_002930300.3      rna-XM_022526378.1    rna-XM_023256229.2    rna-XM_036082561.1    rna-XM_039218346.1    rna-XM_032857124.1    rna-XM_046009477.1    rna-XM_032341703.1    rna-XM_059179654.1    rna-XM_059371158.1    rna-XM_004751011.1    mNeoVis.2.1.1.chr10_hap1.g00792.t01    rna-XM_021695517.1    rna-XM_004414377.1    rna-XM_007087470.3    rna-XM_008702301.1    rna-XM_027570949.1    mNeoVis.3.1.1.chr9.g00753.t01    rna-gnl|WGS_ZZZZ|egapxtmp_017341-R1      mNeoVis.vulgivaga.2.1.chr9.g00773.t01      528    N0.HOG0000526   OG0000032  n65   rna-XM_002931329.4, rna-XM_034652762.1, rna-XM_034653426.1    rna-NM_001388969.1    rna-XM_022524812.1, rna-XM_022527031.1    rna-XM_003987359.3, rna-XM_023255544.2    rna-XM_036082555.1, rna-XM_036082556.1    rna-XM_039218343.1    rna-XM_032857074.1, rna-XM_032857138.1    rna-XM_046009223.1, rna-XM_046009417.1, rna-XM_046009563.1    rna-XM_032341691.1, rna-XM_032341692.1    rna-XM_059179634.1, rna-XM_059179635.1    rna-XM_059371137.1, rna-XM_059371139.1    rna-XM_004782096.3, rna-XM_045067478.1, rna-XM_045072833.1    mNeoVis.2.1.1.chr10_hap1.g00809.t01    rna-XM_021695560.1, rna-XM_021695576.2    rna-XM_004413511.1, rna-XM_004413518.1    rna-XM_007087487.1, rna-XM_007087488.2    rna-XM_008684033.2, rna-XM_008684034.2    rna-XM_027571299.2    mNeoVis.3.1.1.chr9.g00757.t01, mNeoVis.3.1.1.chr9.g00762.t01    rna-gnl|WGS_ZZZZ|egapxtmp_017326-R1, rna-gnl|WGS_ZZZZ|egapxtmp_017330-R1    mNeoVis.evergladensis.2.1.chr8.g00782.t01, mNeoVis.evergladensis.2.1.chr8.g00787.t01    mNeoVis.vulgivaga.2.1.chr9.g00786.t01, mNeoVis.vulgivaga.2.1.chr9.g00788.t01      529    N0.HOG0000527   OG0000032  n108   rna-XM_011217079.3    rna-NM_001388977.1    rna-XM_022525324.1    rna-XM_011283341.4    rna-XM_036082563.1    rna-XM_039218344.1    rna-XM_032857064.1    rna-XM_046009261.1    rna-XM_032341793.1    rna-XM_059179656.1    rna-XM_059371159.1    rna-XM_004751364.3    mNeoVis.2.1.1.chr10_hap1.g00791.t01    rna-XM_021695726.1    rna-XM_004414395.1    rna-XM_007087496.2    rna-XM_008702300.1    rna-XM_027570322.1    mNeoVis.3.1.1.chr9.g00752.t01    rna-gnl|WGS_ZZZZ|egapxtmp_017318-R1    mNeoVis.evergladensis.2.1.chr8.g00773.t01    mNeoVis.vulgivaga.2.1.chr9.g00772.t01      530    N0.HOG0000528   OG0000032  n126   rna-XM_002930299.2    rna-NM_001389032.1    rna-XM_022525384.1    rna-XM_003987310.4    rna-XM_036082564.1    rna-XM_039218341.1    rna-XM_032857123.1    rna-XM_046009300.1    rna-XM_032341705.1    rna-XM_059179657.1    rna-XM_059371160.1    rna-XM_004751365.1      rna-XM_021695520.1    rna-XM_004414396.1    rna-XM_007087497.1    rna-XM_008702343.2    rna-XM_027570321.1        mNeoVis.evergladensis.2.1.chr8.g00772.t01        531    N0.HOG0000529   OG0000032  n144   rna-XM_002930303.2, rna-XM_011217953.2, rna-XM_011218319.2    rna-NM_001388766.1    rna-XM_022526432.1, rna-XM_022526907.1    rna-XM_003987312.3, rna-XM_003987313.3, rna-XM_011283091.3, rna-XM_011283092.3, rna-XM_023249605.2    rna-XM_036082568.1, rna-XM_036082609.1    rna-XM_039218327.1, rna-XM_039218339.1, rna-XM_039218345.1, rna-XM_039218355.1    rna-XM_032857127.1, rna-XM_032857129.1, rna-XM_032857133.1    rna-XM_046009390.1, rna-XM_046009434.1, rna-XM_046009440.1, rna-XM_046009441.1, rna-XM_046009482.1    rna-XM_032341698.1, rna-XM_032341699.1, rna-XM_032341700.1    rna-XM_059179648.1, rna-XM_059179649.1, rna-XM_059179650.1    rna-XM_059371152.1, rna-XM_059371153.1, rna-XM_059371154.1    rna-XM_004751010.1, rna-XM_004751361.2, rna-XM_013055356.2    mNeoVis.2.1.1.chr10_hap1.g00795.t01    rna-XM_021695519.1, rna-XM_021695523.1    rna-XM_004414373.1, rna-XM_004414393.1, rna-XM_004414394.1    rna-XM_007087465.2, rna-XM_007087466.2, rna-XM_007087467.2, rna-XM_007087468.1, rna-XM_042989285.1, rna-XM_042989286.1    rna-XM_008702304.2, rna-XM_008702305.2, rna-XM_008709972.2    rna-XM_027572940.2, rna-XM_035728163.1      rna-gnl|WGS_ZZZZ|egapxtmp_017351-R1    mNeoVis.evergladensis.2.1.chr8.g00774.t01    mNeoVis.vulgivaga.2.1.chr9.g00779.t01      532    N0.HOG0000530   OG0000032  n200   rna-XM_002931255.4, rna-XM_034651690.1, rna-XM_034660159.1    rna-XM_038579866.1    rna-XM_022490587.1, rna-XM_022490755.1    rna-XM_003987355.3, rna-XM_011283093.3, rna-XM_023253390.1    rna-XM_036082558.1, rna-XM_036082560.1    rna-XM_039218349.1, rna-XM_039218354.1    rna-XM_032857130.1, rna-XM_032857131.1    rna-XM_046009437.1, rna-XM_046009454.1    rna-XM_032341693.1, rna-XM_032341697.1, rna-XM_032341792.1    rna-XM_059179641.1, rna-XM_059179646.1, rna-XM_059179647.1    rna-XM_059371144.1, rna-XM_059371151.1    rna-XM_004751009.2, rna-XM_004782265.1, rna-XM_045067931.1    mNeoVis.2.1.1.chr10_hap1.g00801.t01    rna-XM_021695537.1    rna-XM_004414371.1, rna-XM_004414372.1    rna-XM_007087463.2, rna-XM_007087464.2    rna-XM_008702306.2, rna-XM_008710152.2    rna-XM_027571368.1    mNeoVis.3.1.1.chr9.g00751.t01    rna-gnl|WGS_ZZZZ|egapxtmp_017315-R1    mNeoVis.evergladensis.2.1.chr8.g00777.t01    mNeoVis.vulgivaga.2.1.chr9.g00780.t01      533    N0.HOG0000531   OG0000033  n2         rna-XM_045061054.1      rna-XM_039254766.1                      rna-XM_042991290.1                  534    N0.HOG0000532   OG0000033  n5     rna-XM_038529956.1, rna-XM_038532067.1, rna-XM_038589007.1, rna-XM_038589071.1, rna-XM_038589198.1, rna-XM_038589505.1                          rna-XM_044912452.1        rna-XM_040645301.1                535    N0.HOG0000533   OG0000033  n11   rna-XM_034643993.1    rna-XM_038529955.1, rna-XM_038589031.1    rna-XM_022491573.1      rna-XM_036088825.1      rna-XM_032847139.1    rna-XM_046012769.1    rna-XM_032349730.1    rna-XM_059183830.1    rna-XM_059404994.1    rna-XM_045087408.1    mNeoVis.2.1.1.chr9_hap1.g00054.t01    rna-XM_021686525.1    rna-XM_004417346.1      rna-XM_008683879.2    rna-XM_027593224.2    mNeoVis.3.1.1.chr10.g00058.t01    rna-gnl|WGS_ZZZZ|egapxtmp_023871-R1    mNeoVis.evergladensis.2.1.chr10.g00060.t01    mNeoVis.vulgivaga.2.1.chr10.g00057.t01      536    N0.HOG0000534   OG0000033  n29   rna-XM_034650620.1, rna-XM_034651593.1    rna-XM_038532069.1    rna-XM_022491551.1, rna-XM_022492515.1, rna-XM_022492589.1, rna-XM_022492644.1      rna-XM_036088282.1, rna-XM_036088283.1, rna-XM_036088820.1, rna-XM_036091850.1, rna-XM_036103024.1, rna-XM_036108791.1, rna-XM_036108922.1, rna-XM_036108926.1, rna-XM_036116092.1, rna-XM_036116480.1, rna-XM_036119344.1, rna-XM_036119392.1, rna-XM_036123683.1      rna-XM_032850462.1, rna-XM_032850695.1, rna-XM_032850696.1, rna-XM_032850879.1, rna-XM_032851059.1, rna-XM_032851183.1, rna-XM_032865157.1, rna-XM_032874332.1    rna-XM_046011087.1, rna-XM_046011843.1, rna-XM_046011871.1, rna-XM_046011876.1, rna-XM_046011878.1, rna-XM_046011880.1, rna-XM_046011883.1    rna-XM_032331970.1, rna-XM_032345631.1, rna-XM_032347900.1, rna-XM_032347901.1, rna-XM_032347915.1, rna-XM_032349496.1, rna-XM_032349687.1, rna-XM_032349715.1, rna-XM_032349716.1, rna-XM_032349722.1    rna-XM_059137569.1, rna-XM_059183370.1, rna-XM_059183812.1, rna-XM_059183837.1, rna-XM_059183849.1, rna-XM_059183853.1, rna-XM_059184052.1, rna-XM_059184283.1, rna-XM_059184287.1, rna-XM_059184301.1, rna-XM_059187451.1, rna-XM_059187463.1, rna-XM_059187612.1    rna-XM_059372076.1, rna-XM_059375972.1, rna-XM_059387418.1, rna-XM_059404557.1, rna-XM_059404558.1    rna-XM_045069189.1, rna-XM_045084934.1, rna-XM_045084936.1, rna-XM_045084945.1, rna-XM_045085035.1, rna-XM_045087403.1, rna-XM_045087405.1, rna-XM_045087424.1, rna-XM_045087983.1, rna-XM_045087984.1    mNeoVis.2.1.1.chr9_hap1.g00048.t01, mNeoVis.2.1.1.chr9_hap1.g00060.t01, mNeoVis.2.1.1.chr9_hap1.g00062.t01, mNeoVis.2.1.1.chr9_hap1.g00063.t01, mNeoVis.2.1.1.chr9_hap1.g00064.t01, mNeoVis.2.1.1.chr9_hap1.g00065.t01, mNeoVis.2.1.1.chr9_hap1.g00066.t01, mNeoVis.2.1.1.chr9_hap1.g00067.t01, mNeoVis.2.1.1.chr9_hap1.g00078.t01, mNeoVis.2.1.1.chr9_hap1.g00129.t01, mNeoVis.2.1.1.chr9_hap1.g00132.t01, mNeoVis.2.1.1.chr9_hap1.g00133.t01    rna-XM_044912107.1, rna-XM_044912137.1, rna-XM_044912138.1, rna-XM_044912173.1, rna-XM_044912176.1, rna-XM_044912232.1, rna-XM_044912248.1, rna-XM_044912272.1, rna-XM_044912291.1, rna-XM_044912318.1, rna-XM_044912340.1, rna-XM_044912349.1, rna-XM_044912364.1, rna-XM_044912385.1, rna-XM_044912414.1, rna-XM_044912447.1, rna-XM_044912453.1, rna-XM_044912455.1, rna-XM_044912480.1, rna-XM_044915232.1, rna-XM_044915233.1, rna-XM_044915234.1, rna-XM_044915235.1, rna-XM_044915236.1, rna-XM_044915237.1, rna-XM_044915238.1        rna-XM_008709982.2, rna-XM_040641901.1    rna-XM_035721905.1, rna-XM_035721910.1, rna-XM_035721914.1, rna-XM_035729321.1, rna-XM_035729337.1, rna-XM_035729339.1    mNeoVis.3.1.1.chr10.g00063.t01, mNeoVis.3.1.1.chr10.g00064.t01, mNeoVis.3.1.1.chr10.g00074.t01    rna-gnl|WGS_ZZZZ|egapxtmp_021883-R1, rna-gnl|WGS_ZZZZ|egapxtmp_024168-R1, rna-gnl|WGS_ZZZZ|egapxtmp_024575-R1    mNeoVis.evergladensis.2.1.chr10.g00052.t01, mNeoVis.evergladensis.2.1.chr10.g00065.t01, mNeoVis.evergladensis.2.1.chr10.g00077.t01    mNeoVis.vulgivaga.2.1.chr10.g00062.t01, mNeoVis.vulgivaga.2.1.chr10.g00072.t01      537    N0.HOG0000535   OG0000033  n160   rna-XM_034644369.1, rna-XM_034651501.1, rna-XM_034651598.1, rna-XM_034651604.1                                  rna-XM_040645236.1, rna-XM_040645302.1                538    N0.HOG0000536   OG0000033  n167     rna-XM_038529926.1, rna-XM_038529959.1, rna-XM_038573338.1, rna-XM_038589649.1                                              539    N0.HOG0000537   OG0000033  n172     rna-XM_038589006.1, rna-XM_038589279.1, rna-XM_038589280.1, rna-XM_038589281.1, rna-XM_038589315.1, rna-XM_038589409.1, rna-XM_038589411.1, rna-XM_038589599.1, rna-XM_038589628.1, rna-XM_038589636.1                                              540    N0.HOG0000538   OG0000033  n182     rna-XM_038529728.1, rna-XM_038529729.1, rna-XM_038532070.1, rna-XM_038588967.1, rna-XM_038589009.1, rna-XM_038589154.1, rna-XM_038589250.1, rna-XM_038589320.1, rna-XM_038589367.1, rna-XM_038589427.1, rna-XM_038589544.1, rna-XM_038589591.1, rna-XM_038589710.1        rna-XM_036099593.1                                        541    N0.HOG0000539   OG0000033  n195     rna-XM_038529935.1, rna-XM_038532055.1, rna-XM_038532056.1, rna-XM_038588960.1, rna-XM_038588969.1, rna-XM_038589008.1, rna-XM_038589151.1, rna-XM_038589153.1, rna-XM_038589176.1, rna-XM_038589249.1, rna-XM_038589310.1, rna-XM_038589379.1, rna-XM_038589412.1, rna-XM_038589449.1, rna-XM_038589507.1, rna-XM_038589555.1, rna-XM_038589753.1      rna-XM_045049253.1, rna-XM_045051636.1                                          542    N0.HOG0000540   OG0000033  n213         rna-XM_045041236.1, rna-XM_045041250.1, rna-XM_045041252.1, rna-XM_045041637.1, rna-XM_045041649.1, rna-XM_045042080.1, rna-XM_045051557.1, rna-XM_045051580.1, rna-XM_045051604.1, rna-XM_045061053.1      rna-XM_039220490.1                      rna-XM_042975465.1, rna-XM_042976432.1, rna-XM_042976435.1, rna-XM_042976439.1, rna-XM_042976451.1, rna-XM_042976452.1, rna-XM_042977069.1, rna-XM_042977115.1, rna-XM_042977116.1, rna-XM_042977121.1, rna-XM_042977122.1, rna-XM_042977124.1, rna-XM_042977126.1, rna-XM_042977160.1, rna-XM_042977171.1, rna-XM_042977187.1, rna-XM_042977188.1, rna-XM_042977189.1, rna-XM_042977190.1, rna-XM_042977191.1, rna-XM_042980881.1, rna-XM_042983310.1, rna-XM_042991276.1                  543    N0.HOG0000541   OG0000034  n2   id-LOC117802009   id-LOC483072   id-LOC111140027, id-LOC111140034, id-LOC111140256, id-LOC111140273, id-LOC111140351, id-LOC111140542, id-LOC111140552, id-LOC111147944, id-LOC111147971, id-LOC111147974, id-LOC111158401, rna-XM_022492409.1, rna-XM_022492411.1, rna-XM_022504288.1    id-LOC109498252    id-LOC118535279    id-LOC120226972, id-LOC120226973, id-LOC120226974, id-LOC120226975, id-LOC120226976, id-LOC120226977, id-LOC120226978, id-LOC120226979, id-LOC120227141, id-LOC120227142, id-LOC120227143, id-LOC120227144, id-LOC120227146, id-LOC120227613, id-LOC120227614, id-LOC120227629, id-LOC120245533    id-LOC116858917, id-LOC116858952, id-LOC116858956, id-LOC116863120, id-LOC116863493, id-LOC116863525, rna-XM_032843916.1    id-LOC123925660, id-LOC123925805    id-LOC116584029, id-LOC116596443, id-LOC116596444, id-LOC116596445, id-LOC116596447, id-LOC116596448, id-LOC116596449, id-LOC116596455    id-LOC131807770, id-LOC131807775, id-LOC131807777, id-LOC131807778, id-LOC131807779, id-LOC131807780, id-LOC131807791, id-LOC131807792, id-LOC131808419, id-LOC131808420, id-LOC131808421    id-LOC132022184, id-LOC132022185, id-LOC132022187, id-LOC132022188, id-LOC132022325, id-LOC132022326    id-LOC106004692, id-LOC106004802, id-LOC106004804, id-LOC106004805, id-LOC106004806, id-LOC106004811, rna-XM_045079327.1      id-LOC110587504        id-LOC103671487    id-LOC113937262      id-egapxtmp_026751, id-egapxtmp_026899, id-egapxtmp_026901          544    N0.HOG0000542   OG0000034  n83   id-LOC100481011, id-LOC105240710, id-LOC105240715, id-LOC117802011   id-LOC608379   id-LOC111140015, id-LOC111140018, id-LOC111140021, id-LOC111140030, id-LOC111140032, id-LOC111140252, id-LOC111140297, id-LOC111147972, id-LOC111158400, id-LOC111158402, id-LOC111158453    id-LOC101100802, id-LOC111559716    id-LOC118535280    id-LOC120226971, id-LOC120231520, id-LOC120231546, id-LOC120245532    id-LOC116858921, id-LOC116858922, id-LOC116858965, id-LOC116863644, id-LOC116881082    id-LOC123925659, id-LOC123925803    id-LOC116596446, id-LOC116596450, id-LOC116596452    id-LOC131807771, id-LOC131807784, id-LOC131807787, id-LOC131807795    id-LOC132021371, id-LOC132022186    id-LOC101690499, id-LOC101690781, id-LOC101691089      id-LOC123326013, rna-XM_044918533.1    id-LOC101368783, id-LOC101369686    id-LOC102965596, id-LOC102972940, id-LOC122237377    id-LOC103671486, id-LOC103671488, id-LOC103679829            mNeoVis.vulgivaga.2.1.chr11.g00902.t01      545    N0.HOG0000543   OG0000034  n139             id-LOC120226969, id-LOC120226970, id-LOC120231547, id-LOC120231551, id-LOC120231552, id-LOC120231553, id-LOC120231555, id-LOC120231556, id-LOC120231557, id-LOC120231558                                      546    N0.HOG0000544   OG0000034  n150                     id-LOC131807785      id-LOC123391790                          547    N0.HOG0000545   OG0000034  n152   id-LOC117800782, id-LOC117800798      id-LOC111158396          id-LOC116858950        id-LOC131807772, id-LOC131807773, id-LOC131807776      rna-XM_045079328.1      id-LOC110587503    id-LOC101369084      id-LOC103671482                548    N0.HOG0000546   OG0000034  n163   id-LOC117800810      id-LOC111158394, id-LOC111158397, id-LOC111158398, id-LOC111158498          id-LOC116858918, id-LOC116858919, id-LOC116858920, id-LOC116858948, id-LOC116858951      id-LOC116596441    id-LOC131807781, id-LOC131807782, id-LOC131807789      id-LOC106004809        id-LOC101369380        id-LOC118357291, id-LOC118357292              549    N0.HOG0000547   OG0000034  n181    id-LOC483070               id-LOC116596442    id-LOC131808425      id-LOC101673031                          550    N0.HOG0000548   OG0000034  n184   id-LOC105235173    id-LOC119877168, id-LOC119877169, id-LOC608379-2, id-LOC608379-3, id-LOC608379-4    id-LOC111140013, id-LOC111140020, id-LOC111140022, id-LOC111140023, id-LOC111140028, id-LOC111147973, id-LOC111148004, id-LOC111158404, id-LOC111158405, rna-XM_022520398.1      rna-XM_036091612.1      id-LOC116858954, id-LOC116858957    id-LOC123925796, id-LOC123925806    id-LOC116584030, id-LOC116596398, id-LOC116596438, id-LOC116596457, id-LOC116596458    id-LOC131807768, id-LOC131807769, id-LOC131807788, id-LOC131808415, id-LOC131808416, id-LOC131808417, id-LOC131808418, id-LOC131808422, id-LOC131808423    id-LOC132022178, id-LOC132022182, id-LOC132022190, id-LOC132022324, id-LOC132022327, id-LOC132022328, id-LOC132022329, id-LOC132022331    id-LOC101672758, id-LOC106004753, id-LOC106004754, id-LOC106004755, id-LOC106004808, id-LOC106004812, id-LOC123391781, id-LOC123391789, rna-XM_045079326.1        id-LOC101366157      id-LOC103671857    rna-XM_027621286.1      id-egapxtmp_026897          551    N0.HOG0000549   OG0000034  n238           id-LOC118535278    id-LOC120231530, id-LOC120231562, id-LOC120231563, id-LOC120231564, id-LOC120231565                                      552    N0.HOG0000550   OG0000035  n1   rna-XM_034657064.1    rna-XM_038535417.1    rna-XM_022512452.1    rna-XM_019812450.3    rna-XM_036081906.1    rna-XM_039225006.1    rna-XM_032861908.1    rna-XM_045998702.1    rna-XM_032337714.1    rna-XM_059173699.1    rna-XM_059416471.1    rna-XM_004778670.3    mNeoVis.2.1.1.chr1_hap1.g00071.t01    rna-XM_021695843.1    rna-XM_004406050.1    rna-XM_007075935.3    rna-XM_040630527.1    rna-XM_027605549.2    mNeoVis.3.1.1.chr1.g00075.t01    rna-gnl|WGS_ZZZZ|egapxtmp_021105-R1    mNeoVis.evergladensis.2.1.chr1.g00069.t01    mNeoVis.vulgivaga.2.1.chr1.g00068.t01      553    N0.HOG0000551   OG0000035  n25   rna-XM_034642847.1    rna-NM_001286988.1    rna-XM_022513747.1    rna-XM_023242055.2    rna-XM_036089875.1    rna-XM_039230486.1    rna-XM_032872301.1    rna-XM_046025152.1    rna-XM_032310722.1    rna-XM_059140126.1    rna-XM_059409764.1    rna-XM_004745586.3    mNeoVis.2.1.1.chr3_hap1.g00061.t01    rna-XM_021686396.1    rna-XM_004398055.1    rna-XM_007080575.3    rna-XM_008705403.2    rna-XM_027577077.1    mNeoVis.3.1.1.chr3.g00062.t01    rna-gnl|WGS_ZZZZ|egapxtmp_042106-R1    mNeoVis.evergladensis.2.1.chr3.g00064.t01    mNeoVis.vulgivaga.2.1.chr3.g00058.t01      554    N0.HOG0000552   OG0000035  n47   rna-XM_002922054.4    rna-XM_038525831.1    rna-XM_022513753.1    rna-XM_045041116.1    rna-XM_036100307.1    rna-XM_039238749.1    rna-XM_032872261.1    rna-XM_046026096.1    rna-XM_032311017.1    rna-XM_059140272.1    rna-XM_059410092.1    rna-XM_004745613.3    mNeoVis.2.1.1.chr3_hap1.g00039.t01    rna-XM_021686407.1    rna-XM_004416278.1    rna-XM_007096080.3    rna-XM_008704804.2    rna-XM_027576791.2    mNeoVis.3.1.1.chr3.g00042.t01    rna-gnl|WGS_ZZZZ|egapxtmp_039428-R2    mNeoVis.evergladensis.2.1.chr3.g00045.t01    mNeoVis.vulgivaga.2.1.chr3.g00039.t01      555    N0.HOG0000553   OG0000035  n68   rna-XM_002924875.4    rna-XM_038525835.1    rna-XM_022513744.1    rna-XM_019815403.3    rna-XM_036120071.1    rna-XM_039238744.1    rna-XM_032872262.1    rna-XM_046024025.1    rna-XM_032311015.1    rna-XM_059140328.1    rna-XM_059410091.1    rna-XM_004745610.3    mNeoVis.2.1.1.chr3_hap1.g00040.t01    rna-XM_021686406.1    rna-XM_004398073.1    rna-XM_042962757.1    rna-XM_008704831.1    rna-XM_027576285.2    mNeoVis.3.1.1.chr3.g00043.t01    rna-gnl|WGS_ZZZZ|egapxtmp_040703-R1    mNeoVis.evergladensis.2.1.chr3.g00046.t01    mNeoVis.vulgivaga.2.1.chr3.g00040.t01      556    N0.HOG0000554   OG0000035  n87   rna-XM_019802760.2    rna-XM_038535420.1    rna-XM_022512443.1    rna-XM_045038559.1    rna-XM_036064734.1    rna-XM_039219788.1    rna-XM_032861903.1    rna-XM_046001013.1    rna-XM_032337720.1    rna-XM_059173691.1    rna-XM_059416934.1    rna-XM_045082036.1    mNeoVis.2.1.1.chr1_hap1.g00070.t01    rna-XM_021702503.1    rna-XM_004408332.1    rna-XM_042997273.1    rna-XM_008689887.2    rna-XM_027607141.2    mNeoVis.3.1.1.chr1.g00074.t01    rna-gnl|WGS_ZZZZ|egapxtmp_021344-R3    mNeoVis.evergladensis.2.1.chr1.g00067.t01    mNeoVis.vulgivaga.2.1.chr1.g00067.t01      557    N0.HOG0000555   OG0000035  n107   rna-XM_034638672.1    rna-XM_038538793.1    rna-XM_022518712.1    rna-XM_019819837.3    rna-XM_036091706.1    rna-XM_039218320.1    rna-XM_032854312.1    rna-XM_045989291.1    rna-XM_032325835.1    rna-XM_059152871.1    rna-XM_059381567.1    rna-XM_004744298.2    mNeoVis.2.1.1.chr7_hap1.g00266.t01    rna-XM_021704193.1    rna-XM_004408585.1    rna-XM_042970108.1    rna-XM_040633089.1    rna-XM_035725208.1    mNeoVis.3.1.1.chr7.g00249.t01    rna-gnl|WGS_ZZZZ|egapxtmp_011547-R1    mNeoVis.evergladensis.2.1.chr7.g00266.t01    mNeoVis.vulgivaga.2.1.chr7.g00268.t01      558    N0.HOG0000556   OG0000035  n126   rna-XM_034672331.1    rna-XM_038538790.1    rna-XM_022511679.1    rna-XM_023245550.2    rna-XM_036095796.1    rna-XM_039247652.1    rna-XM_032854311.1    rna-XM_045989064.1    rna-XM_032323554.1    rna-XM_059151437.1    rna-XM_059383692.1    rna-XM_004744300.3    mNeoVis.2.1.1.chr7_hap1.g00264.t01    rna-XM_021704156.1    rna-XM_004404766.1    rna-XM_042968688.1    rna-XM_040633392.1    rna-XM_027619395.2    mNeoVis.3.1.1.chr7.g00247.t01    rna-gnl|WGS_ZZZZ|egapxtmp_010009-R1    mNeoVis.evergladensis.2.1.chr7.g00265.t01    mNeoVis.vulgivaga.2.1.chr7.g00267.t01      559    N0.HOG0000557   OG0000035  n140   rna-XM_034648561.1    rna-XM_038544189.1    rna-XM_022492854.1    rna-XM_011283135.4    rna-XM_036114779.1    rna-XM_039245250.1    rna-XM_032856385.1    rna-XM_046007091.1    rna-XM_032343825.1    rna-XM_059181645.1    rna-XM_059372902.1    rna-XM_013057592.2    mNeoVis.2.1.1.chr10_hap1.g00573.t01    rna-XM_021695423.1    rna-XM_004402160.1    rna-XM_042989385.1    rna-XM_008710573.2    rna-XM_027568702.2    mNeoVis.3.1.1.chr9.g00528.t01    rna-gnl|WGS_ZZZZ|egapxtmp_014387-R2    mNeoVis.evergladensis.2.1.chr8.g00557.t01    mNeoVis.vulgivaga.2.1.chr9.g00558.t01      560    N0.HOG0000558   OG0000035  n162   rna-XM_002917048.2    rna-XM_038533757.1    rna-XM_022512446.1    rna-XM_019812404.3    rna-XM_036106048.1    rna-XM_039220139.1    rna-XM_032861913.1    rna-XM_045999950.1    rna-XM_032337710.1    rna-XM_059173704.1    rna-XM_059416476.1    rna-XM_004738086.3    mNeoVis.2.1.1.chr1_hap1.g00077.t01    rna-XM_021695884.1    rna-XM_004408666.1    rna-XM_042997256.1    rna-XM_040630212.1    rna-XM_027605879.1    mNeoVis.3.1.1.chr1.g00078.t01    rna-gnl|WGS_ZZZZ|egapxtmp_005076-R1    mNeoVis.evergladensis.2.1.chr1.g00076.t01    mNeoVis.vulgivaga.2.1.chr1.g00072.t01      561    N0.HOG0000559   OG0000035  n182   rna-XM_002919906.4    rna-XM_038535409.1    rna-XM_022493225.1    rna-XM_019812349.3    rna-XM_036088432.1    rna-XM_039256414.1    rna-XM_032861916.1    rna-XM_046000245.1    rna-XM_032337708.1    rna-XM_059173706.1    rna-XM_059416941.1    rna-XM_004738085.3    mNeoVis.2.1.1.chr1_hap1.g00079.t01    rna-XM_021695920.1    rna-XM_004411529.1    rna-XM_042997219.1    rna-XM_008692676.1    rna-XM_027606750.2    mNeoVis.3.1.1.chr1.g00080.t01    rna-gnl|WGS_ZZZZ|egapxtmp_020744-R1    mNeoVis.evergladensis.2.1.chr1.g00078.t01    mNeoVis.vulgivaga.2.1.chr1.g00073.t01      562    N0.HOG0000560   OG0000035  n201   rna-XM_034657062.1    rna-XM_038535412.1    rna-XM_022512451.1    rna-XM_045038516.1    rna-XM_036106760.1    rna-XM_039222856.1    rna-XM_032861911.1    rna-XM_045998781.1    rna-XM_032337712.1    rna-XM_059173703.1    rna-XM_059416473.1    rna-XM_004738087.3    mNeoVis.2.1.1.chr1_hap1.g00073.t01    rna-XM_021695870.1    rna-XM_004416461.1    rna-XM_042960623.1    rna-XM_040630525.1    rna-XM_027607154.2    mNeoVis.3.1.1.chr1.g00077.t01    rna-gnl|WGS_ZZZZ|egapxtmp_026693-R1    mNeoVis.evergladensis.2.1.chr1.g00071.t01    mNeoVis.vulgivaga.2.1.chr1.g00070.t01      563    N0.HOG0000561   OG0000036  n1   rna-NM_001304856.1, rna-XM_034651602.1    rna-XM_038571597.1    rna-XM_022509748.1    rna-XM_003996479.4, rna-XM_045033238.1    rna-XM_036070143.1, rna-XM_036077812.1, rna-XM_036122060.1    rna-XM_039251880.1    rna-XM_032838029.1, rna-XM_032838468.1, rna-XM_032852784.1, rna-XM_032858614.1, rna-XM_032861864.1, rna-XM_032863537.1, rna-XM_032865617.1, rna-XM_032867782.1, rna-XM_032873268.1, rna-XM_032876336.1, rna-XM_032883165.1    rna-XM_045986589.1, rna-XM_045995589.1, rna-XM_045997840.1, rna-XM_046000936.1, rna-XM_046006168.1, rna-XM_046009301.1, rna-XM_046010327.1, rna-XM_046016727.1, rna-XM_046018574.1, rna-XM_046025559.1    rna-XM_032310706.1, rna-XM_032312497.1, rna-XM_032314939.1, rna-XM_032318597.1, rna-XM_032329295.1, rna-XM_032330502.1, rna-XM_032331672.1, rna-XM_032338315.1, rna-XM_032338380.1, rna-XM_032345409.1, rna-XM_032346467.1, rna-XM_032350523.1, rna-XM_032358254.1    rna-XM_059133503.1, rna-XM_059137796.1, rna-XM_059138361.1, rna-XM_059141803.1, rna-XM_059142869.1, rna-XM_059145660.1, rna-XM_059145831.1, rna-XM_059149422.1, rna-XM_059155195.1, rna-XM_059157788.1, rna-XM_059158746.1, rna-XM_059169817.1, rna-XM_059170930.1, rna-XM_059172661.1, rna-XM_059176380.1, rna-XM_059183610.1    rna-XM_059380496.1, rna-XM_059384823.1, rna-XM_059385132.1, rna-XM_059394067.1, rna-XM_059395847.1, rna-XM_059396357.1, rna-XM_059399044.1, rna-XM_059399127.1, rna-XM_059407024.1, rna-XM_059407965.1, rna-XM_059412178.1, rna-XM_059414105.1    rna-XM_013053546.2, rna-XM_045068342.1, rna-XM_045079339.1, rna-XM_045079701.1, rna-XM_045081199.1, rna-XM_045081200.1, rna-XM_045081205.1, rna-XM_045082906.1, rna-XM_045083586.1, rna-XM_045084748.1, rna-XM_045089225.1    mNeoVis.2.1.1.chr10_hap1.g00048.t01, mNeoVis.2.1.1.chr10_hap1.g00596.t01, mNeoVis.2.1.1.chr11_hap1.g00920.t01, mNeoVis.2.1.1.chr12_hap1.g00121.t01, mNeoVis.2.1.1.chr13_hap1.g00628.t01, mNeoVis.2.1.1.chr1_hap1.g00811.t01, mNeoVis.2.1.1.chr2_hap1.g01699.t01, mNeoVis.2.1.1.chr4_hap1.g00247.t01, mNeoVis.2.1.1.chr4_hap1.g00337.t01, mNeoVis.2.1.1.chr5_hap1.g01720.t01, mNeoVis.2.1.1.chr6_hap1.g00138.t01, mNeoVis.2.1.1.chr6_hap1.g00438.t01, mNeoVis.2.1.1.chr7_hap1.g00113.t01, mNeoVis.2.1.1.chr7_hap1.g00386.t01, mNeoVis.2.1.1.chr7_hap1.g00777.t01, mNeoVis.2.1.1.chr8_hap1.g00777.t01, mNeoVis.2.1.1.chr9_hap1.g00175.t01, mNeoVis.2.1.1.chr9_hap1.g00203.t01, mNeoVis.2.1.1.chr9_hap1.g00560.t01    rna-XM_021695772.2, rna-XM_044912456.1, rna-XM_044912606.1, rna-XM_044914220.1, rna-XM_044917495.1, rna-XM_044919206.1, rna-XM_044921787.1    rna-XM_004396093.1, rna-XM_004399667.1, rna-XM_004405620.1    rna-NM_001301052.1, rna-XM_042971814.1    rna-XM_008695334.2    rna-XM_027625199.1, rna-XM_035723737.1, rna-XM_035724276.1, rna-XM_035728221.1, rna-XM_035728889.1    mNeoVis.3.1.1.chr1.g00837.t01, mNeoVis.3.1.1.chr1.g01512.t01, mNeoVis.3.1.1.chr12.g00095.t01, mNeoVis.3.1.1.chr2.g00258.t01, mNeoVis.3.1.1.chr5.g00684.t01, mNeoVis.3.1.1.chr5.g01620.t01, mNeoVis.3.1.1.chr6.g00246.t01, mNeoVis.3.1.1.chr7.g00721.t01, mNeoVis.3.1.1.chr8.g00727.t01    rna-gnl|WGS_ZZZZ|egapxtmp_001807-R1, rna-gnl|WGS_ZZZZ|egapxtmp_002544-R1, rna-gnl|WGS_ZZZZ|egapxtmp_002917-R1, rna-gnl|WGS_ZZZZ|egapxtmp_008313-R1, rna-gnl|WGS_ZZZZ|egapxtmp_010566-R1, rna-gnl|WGS_ZZZZ|egapxtmp_010576-R1, rna-gnl|WGS_ZZZZ|egapxtmp_011759-R1, rna-gnl|WGS_ZZZZ|egapxtmp_023002-R1, rna-gnl|WGS_ZZZZ|egapxtmp_029400-R1, rna-gnl|WGS_ZZZZ|egapxtmp_043622-R1    mNeoVis.evergladensis.2.1.chr1.g00818.t01, mNeoVis.evergladensis.2.1.chr10.g00170.t01, mNeoVis.evergladensis.2.1.chr10.g00199.t01, mNeoVis.evergladensis.2.1.chr10.g00553.t01, mNeoVis.evergladensis.2.1.chr11.g00899.t01, mNeoVis.evergladensis.2.1.chr12.g00096.t01, mNeoVis.evergladensis.2.1.chr13.g00603.t01, mNeoVis.evergladensis.2.1.chr2.g00319.t01, mNeoVis.evergladensis.2.1.chr4.g00143.t01, mNeoVis.evergladensis.2.1.chr6.g00246.t01, mNeoVis.evergladensis.2.1.chr6.g00341.t01, mNeoVis.evergladensis.2.1.chr7.g00761.t01, mNeoVis.evergladensis.2.1.chr8.g00047.t01, mNeoVis.evergladensis.2.1.chr9.g00764.t01    mNeoVis.vulgivaga.2.1.chr1.g00833.t01, mNeoVis.vulgivaga.2.1.chr10.g00540.t01, mNeoVis.vulgivaga.2.1.chr11.g00924.t01, mNeoVis.vulgivaga.2.1.chr13.g00605.t01, mNeoVis.vulgivaga.2.1.chr2.g00316.t01, mNeoVis.vulgivaga.2.1.chr4.g00135.t01, mNeoVis.vulgivaga.2.1.chr6.g00247.t01, mNeoVis.vulgivaga.2.1.chr6.g00335.t01, mNeoVis.vulgivaga.2.1.chr7.g00390.t01, mNeoVis.vulgivaga.2.1.chr7.g00765.t01, mNeoVis.vulgivaga.2.1.chr8.g00768.t01      564    N0.HOG0000562   OG0000036  n125   rna-XM_034640272.1    rna-XM_038535266.1      rna-XM_045048870.1, rna-XM_045054619.1, rna-XM_045059126.1, rna-XM_045060971.1    rna-XM_036070793.1, rna-XM_036090130.1, rna-XM_036106388.1    rna-XM_039223091.1, rna-XM_039241100.1, rna-XM_039254259.1    rna-XM_032841173.1, rna-XM_032852655.1, rna-XM_032853715.1, rna-XM_032866129.1, rna-XM_032870300.1, rna-XM_032882629.1    rna-XM_045977881.1, rna-XM_045997684.1, rna-XM_046008204.1, rna-XM_046009664.1, rna-XM_046015510.1    rna-XM_032323817.1, rna-XM_032326623.1, rna-XM_032328859.1, rna-XM_032330634.1, rna-XM_032338319.1, rna-XM_032345587.1    rna-XM_059134278.1, rna-XM_059151170.1, rna-XM_059154949.1, rna-XM_059163841.1, rna-XM_059165244.1, rna-XM_059183644.1, rna-XM_059187505.1    rna-XM_059390967.1, rna-XM_059394560.1, rna-XM_059404496.1, rna-XM_059404652.1, rna-XM_059412296.1    rna-XM_045071381.1, rna-XM_045073243.1, rna-XM_045076515.1, rna-XM_045079424.1, rna-XM_045084853.1    mNeoVis.2.1.1.chr11_hap1.g00268.t01, mNeoVis.2.1.1.chr11_hap1.g00635.t01, mNeoVis.2.1.1.chr1_hap1.g00936.t01, mNeoVis.2.1.1.chr4_hap1.g00989.t01, mNeoVis.2.1.1.chr5_hap1.g00320.t01    rna-XM_044911619.1, rna-XM_044920728.1, rna-XM_044921188.1    rna-XM_004397828.1, rna-XM_004413141.1    rna-XM_042973314.1, rna-XM_042976841.1, rna-XM_042982053.1, rna-XM_042987745.1, rna-XM_042988986.1      rna-XM_027620749.2, rna-XM_035723815.1, rna-XM_035726365.1, rna-XM_035728069.1    mNeoVis.3.1.1.chr1.g01916.t01, mNeoVis.3.1.1.chr6.g00511.t01, mNeoVis.3.1.1.chr7.g00248.t01, mNeoVis.3.1.1.chr7.g00251.t01, mNeoVis.3.1.1.chr7.g01378.t01    rna-gnl|WGS_ZZZZ|egapxtmp_023212-R1, rna-gnl|WGS_ZZZZ|egapxtmp_023500-R1, rna-gnl|WGS_ZZZZ|egapxtmp_035807-R1    mNeoVis.evergladensis.2.1.chr1.g00931.t01    mNeoVis.vulgivaga.2.1.chr7.g00270.t01      565    N0.HOG0000563   OG0000037  n1     rna-XM_038588954.1        rna-XM_036066134.1                                        566    N0.HOG0000564   OG0000037  n2   rna-XM_034651543.1, rna-XM_034652573.1, rna-XM_034653478.1, rna-XM_034653479.1, rna-XM_034653486.1    rna-XM_038530872.1, rna-XM_038549515.1, rna-XM_038555094.1, rna-XM_038556389.1, rna-XM_038557593.1, rna-XM_038559196.1, rna-XM_038559201.1, rna-XM_038559207.1, rna-XM_038559209.1, rna-XM_038559214.1, rna-XM_038559225.1, rna-XM_038559231.1, rna-XM_038559237.1, rna-XM_038559238.1, rna-XM_038559251.1, rna-XM_038559253.1, rna-XM_038559607.1, rna-XM_038560235.1, rna-XM_038560240.1, rna-XM_038560241.1, rna-XM_038560311.1, rna-XM_038560469.1, rna-XM_038560471.1, rna-XM_038560472.1, rna-XM_038560473.1, rna-XM_038560508.1, rna-XM_038560532.1, rna-XM_038560559.1, rna-XM_038560572.1, rna-XM_038562390.1, rna-XM_038576126.1, rna-XM_038583286.1, rna-XM_038583793.1, rna-XM_038588876.1, rna-XM_038588877.1, rna-XM_038588880.1, rna-XM_038588896.1, rna-XM_038588897.1, rna-XM_038588900.1, rna-XM_038588913.1, rna-XM_038588916.1, rna-XM_038588918.1, rna-XM_038588920.1, rna-XM_038588952.1, rna-XM_038588958.1, rna-XM_038588962.1, rna-XM_038589012.1, rna-XM_038589087.1, rna-XM_038589088.1, rna-XM_038589090.1, rna-XM_038589092.1, rna-XM_038589129.1, rna-XM_038589142.1, rna-XM_038589145.1, rna-XM_038589161.1, rna-XM_038589162.1, rna-XM_038589174.1, rna-XM_038589188.1, rna-XM_038589195.1, rna-XM_038589244.1, rna-XM_038589245.1, rna-XM_038589247.1, rna-XM_038589248.1, rna-XM_038589298.1, rna-XM_038589426.1, rna-XM_038589444.1, rna-XM_038589454.1, rna-XM_038589464.1, rna-XM_038589468.1, rna-XM_038589469.1, rna-XM_038589470.1, rna-XM_038589471.1, rna-XM_038589474.1, rna-XM_038589495.1, rna-XM_038589496.1, rna-XM_038589497.1, rna-XM_038589498.1, rna-XM_038589506.1, rna-XM_038589524.1, rna-XM_038589525.1, rna-XM_038589536.1, rna-XM_038589539.1, rna-XM_038589598.1, rna-XM_038589602.1, rna-XM_038589616.1, rna-XM_038589644.1, rna-XM_038589646.1, rna-XM_038589652.1, rna-XM_038589678.1, rna-XM_038589683.1, rna-XM_038589685.1, rna-XM_038589686.1, rna-XM_038589705.1, rna-XM_038589714.1, rna-XM_038589747.1      rna-XM_019824799.3, rna-XM_023242320.2, rna-XM_045051430.1, rna-XM_045051434.1, rna-XM_045051435.1, rna-XM_045051448.1, rna-XM_045051451.1, rna-XM_045051452.1, rna-XM_045051456.1, rna-XM_045051457.1, rna-XM_045051458.1, rna-XM_045051463.1, rna-XM_045051466.1, rna-XM_045051470.1, rna-XM_045051482.1, rna-XM_045051486.1, rna-XM_045051581.1, rna-XM_045052151.1, rna-XM_045052155.1, rna-XM_045052159.1, rna-XM_045052169.1, rna-XM_045056897.1    rna-XM_036096139.1, rna-XM_036097120.1        rna-XM_045978102.1, rna-XM_045978530.1, rna-XM_045978531.1    rna-XM_032308847.1, rna-XM_032314621.1, rna-XM_032314623.1, rna-XM_032314624.1, rna-XM_032315281.1, rna-XM_032315282.1, rna-XM_032315288.1, rna-XM_032315290.1, rna-XM_032315291.1, rna-XM_032315313.1, rna-XM_032315318.1, rna-XM_032315330.1    rna-XM_059143730.1, rna-XM_059143740.1, rna-XM_059143754.1, rna-XM_059143760.1, rna-XM_059143766.1, rna-XM_059143800.1, rna-XM_059143803.1, rna-XM_059144076.1, rna-XM_059144079.1, rna-XM_059144081.1, rna-XM_059144082.1, rna-XM_059144083.1, rna-XM_059144207.1, rna-XM_059144283.1, rna-XM_059144310.1, rna-XM_059144365.1, rna-XM_059144366.1, rna-XM_059144369.1, rna-XM_059144375.1, rna-XM_059144778.1, rna-XM_059144780.1, rna-XM_059144786.1, rna-XM_059144799.1, rna-XM_059144803.1    rna-XM_059377813.1, rna-XM_059387201.1, rna-XM_059387566.1    rna-XM_045069673.1, rna-XM_045069687.1, rna-XM_045069701.1, rna-XM_045069706.1, rna-XM_045069707.1, rna-XM_045069718.1, rna-XM_045072407.1, rna-XM_045072434.1, rna-XM_045072435.1, rna-XM_045072436.1, rna-XM_045072438.1, rna-XM_045072441.1, rna-XM_045072442.1, rna-XM_045072444.1, rna-XM_045072447.1, rna-XM_045072448.1, rna-XM_045087800.1, rna-XM_045087801.1, rna-XM_045087804.1, rna-XM_045087805.1, rna-XM_045087813.1, rna-XM_045087816.1, rna-XM_045087818.1    mNeoVis.2.1.1.chr13_hap1.g00116.t01, mNeoVis.2.1.1.chr2_hap1.g00735.t01, mNeoVis.2.1.1.chr3_hap1.g00697.t01, mNeoVis.2.1.1.chr3_hap1.g00699.t01, mNeoVis.2.1.1.chr4_hap1.g00685.t01, mNeoVis.2.1.1.chr8_hap1.g01191.t01, mNeoVis.2.1.1.chr8_hap1.g01194.t01, mNeoVis.2.1.1.chr8_hap1.g01195.t01, mNeoVis.2.1.1.chr8_hap1.g01201.t01, mNeoVis.2.1.1.chr8_hap1.g01202.t01, mNeoVis.2.1.1.chr8_hap1.g01206.t01, mNeoVis.2.1.1.chr8_hap1.g01209.t01, mNeoVis.2.1.1.chr8_hap1.g01210.t01, mNeoVis.2.1.1.chr8_hap1.g01220.t01, mNeoVis.2.1.1.chr8_hap1.g01221.t01, mNeoVis.2.1.1.chr8_hap1.g01223.t01, mNeoVis.2.1.1.chr8_hap1.g01229.t01, mNeoVis.2.1.1.chr8_hap1.g01230.t01, mNeoVis.2.1.1.chr8_hap1.g01231.t01, mNeoVis.2.1.1.chr8_hap1.g01234.t01, mNeoVis.2.1.1.chr8_hap1.g01237.t01        rna-XM_042981409.1, rna-XM_042981422.1, rna-XM_042981485.1, rna-XM_042999259.1    rna-XM_008709386.2, rna-XM_040620404.1    rna-XM_027570344.1    mNeoVis.3.1.1.chr8.g01127.t01    rna-gnl|WGS_ZZZZ|egapxtmp_013785-R3, rna-gnl|WGS_ZZZZ|egapxtmp_017595-R1, rna-gnl|WGS_ZZZZ|egapxtmp_018012-R1, rna-gnl|WGS_ZZZZ|egapxtmp_018013-R1, rna-gnl|WGS_ZZZZ|egapxtmp_018159-R1    mNeoVis.evergladensis.2.1.chr6.g00688.t01, mNeoVis.evergladensis.2.1.chr9.g01163.t01, mNeoVis.evergladensis.2.1.chr9.g01166.t01, mNeoVis.evergladensis.2.1.chr9.g01167.t01, mNeoVis.evergladensis.2.1.chr9.g01185.t01    mNeoVis.vulgivaga.2.1.chr13.g00110.t01, mNeoVis.vulgivaga.2.1.chr8.g01163.t01, mNeoVis.vulgivaga.2.1.chr8.g01164.t01, mNeoVis.vulgivaga.2.1.chr8.g01165.t01, mNeoVis.vulgivaga.2.1.chr8.g01178.t01, mNeoVis.vulgivaga.2.1.chr8.g01179.t01, mNeoVis.vulgivaga.2.1.chr8.g01187.t01      567    N0.HOG0000565   OG0000038  n2   rna-XM_002926850.4, rna-XM_002926867.4, rna-XM_034644897.1    rna-XM_038562467.1    rna-XM_022507671.1, rna-XM_022507690.1    rna-XM_045038154.1, rna-XM_045038155.1    rna-XM_036092782.1, rna-XM_036092785.1    rna-XM_039224008.1, rna-XM_039224021.1    rna-XM_032846575.1    rna-XM_046014425.1, rna-XM_046017481.1    rna-XM_032356258.1, rna-XM_032360538.1, rna-XM_032360540.1    rna-XM_059137527.1    rna-XM_059388028.1, rna-XM_059388039.1    rna-XM_004781688.2, rna-XM_004781689.2, rna-XM_045068279.1    mNeoVis.2.1.1.chr7_hap1.g02118.t01, mNeoVis.2.1.1.chr7_hap1.g02119.t01    rna-XM_021684833.1    rna-XM_004411994.1    rna-XM_007088806.2    rna-XM_008694093.2, rna-XM_008694227.1, rna-XM_040638717.1    rna-XM_027577866.2      rna-gnl|WGS_ZZZZ|egapxtmp_012625-R1    mNeoVis.evergladensis.2.1.chr7.g02065.t01        568    N0.HOG0000566   OG0000038  n36     rna-NM_001388749.1    rna-XM_022521688.1    rna-XM_003993081.4    rna-XM_036101512.1    rna-XM_039244589.1    rna-XM_032852726.1    rna-XM_046014269.1    rna-XM_032359802.1    rna-XM_059158302.1    rna-XM_059389070.1    rna-XM_004780181.2    mNeoVis.2.1.1.chr7_hap1.g01784.t01    rna-XM_021685664.2      rna-XM_007083287.2    rna-XM_040630679.1    rna-XM_027579019.2    mNeoVis.3.1.1.chr7.g01655.t01    rna-gnl|WGS_ZZZZ|egapxtmp_014009-R1      mNeoVis.vulgivaga.2.1.chr7.g01764.t01      569    N0.HOG0000567   OG0000038  n54   rna-XM_011234342.2    rna-NM_001388954.1      rna-XM_003993346.2    rna-XM_036107505.1    rna-XM_039223509.1      rna-XM_046016576.1    rna-XM_032359874.1    rna-XM_059158934.1    rna-XM_059387959.1    rna-XM_004782242.2    mNeoVis.2.1.1.chr7_hap1.g02146.t01        rna-XM_007088782.1    rna-XM_008694109.1    rna-XM_027581216.2    mNeoVis.3.1.1.chr7.g01972.t01    rna-gnl|WGS_ZZZZ|egapxtmp_013481-R1    mNeoVis.evergladensis.2.1.chr7.g02093.t01    mNeoVis.vulgivaga.2.1.chr7.g02114.t01      570    N0.HOG0000568   OG0000038  n70   rna-XM_002926887.2    rna-NM_001388980.1    rna-XM_022507098.1    rna-XM_003993278.2      rna-XM_039223495.1    rna-XM_032846462.1    rna-XM_046016362.1    rna-XM_032358814.1    rna-XM_059158932.1    rna-XM_059387965.1    rna-XM_004782233.1          rna-XM_007088781.2    rna-XM_008694107.1    rna-XM_027580111.2    mNeoVis.3.1.1.chr7.g01971.t01    rna-gnl|WGS_ZZZZ|egapxtmp_013477-R1    mNeoVis.evergladensis.2.1.chr7.g02092.t01    mNeoVis.vulgivaga.2.1.chr7.g02113.t01      571    N0.HOG0000569   OG0000038  n86   rna-XM_011234351.2                                  rna-XM_008694247.1                572    N0.HOG0000570   OG0000038  n90   rna-XM_002926883.2    rna-XM_038562465.1    rna-XM_022506648.1    rna-XM_003993277.2    rna-XM_036092776.1    rna-XM_039223501.1      rna-XM_046016357.1    rna-XM_032359872.1    rna-XM_059138742.1      rna-XM_004782240.2    mNeoVis.2.1.1.chr7_hap1.g02142.t01        rna-XM_007088824.2    rna-XM_008694105.2        rna-gnl|WGS_ZZZZ|egapxtmp_012630-R1    mNeoVis.evergladensis.2.1.chr7.g02089.t01        573    N0.HOG0000571   OG0000038  n104     rna-XM_038564476.1    rna-XM_022506649.1        rna-XM_039223494.1      rna-XM_046016466.1    rna-XM_032359871.1    rna-XM_059138615.1      rna-XM_004782239.1    mNeoVis.2.1.1.chr7_hap1.g02141.t01              mNeoVis.3.1.1.chr7.g01968.t01    rna-gnl|WGS_ZZZZ|egapxtmp_013492-R1    mNeoVis.evergladensis.2.1.chr7.g02088.t01    mNeoVis.vulgivaga.2.1.chr7.g02110.t01      574    N0.HOG0000572   OG0000038  n115       rna-XM_022507173.1          rna-XM_032846550.1    rna-XM_046015797.1                                  575    N0.HOG0000573   OG0000038  n118                     rna-XM_059158914.1    rna-XM_059405372.1                            576    N0.HOG0000574   OG0000038  n121                   rna-XM_032359867.1          mNeoVis.2.1.1.chr7_hap1.g02129.t01                rna-gnl|WGS_ZZZZ|egapxtmp_012628-R1      mNeoVis.vulgivaga.2.1.chr7.g02098.t01      577    N0.HOG0000575   OG0000038  n125       rna-XM_022507171.1          rna-XM_032846466.1, rna-XM_032846472.1    rna-XM_046016563.1, rna-XM_046016586.1    rna-XM_032359413.1, rna-XM_032360028.1    rna-XM_059158915.1, rna-XM_059158917.1    rna-XM_059405354.1, rna-XM_059405383.1    rna-XM_004781665.2    mNeoVis.2.1.1.chr7_hap1.g02131.t01, mNeoVis.2.1.1.chr7_hap1.g02133.t01      rna-XM_004411974.1        rna-XM_027581067.1, rna-XM_027581068.2    mNeoVis.3.1.1.chr7.g01961.t01    rna-gnl|WGS_ZZZZ|egapxtmp_013482-R1, rna-gnl|WGS_ZZZZ|egapxtmp_013489-R1    mNeoVis.evergladensis.2.1.chr7.g02075.t01, mNeoVis.evergladensis.2.1.chr7.g02078.t01    mNeoVis.vulgivaga.2.1.chr7.g02101.t01      578    N0.HOG0000576   OG0000038  n149   rna-XM_002926858.4    rna-NM_001388986.1    rna-XM_022507170.1    rna-XM_003993276.2, rna-XM_023239020.1      rna-XM_039223498.1, rna-XM_039223504.1, rna-XM_039223510.1    rna-XM_032846547.1    rna-XM_046016279.1    rna-XM_032359870.1    rna-XM_059158929.1    rna-XM_059416749.1    rna-XM_004782238.1    mNeoVis.2.1.1.chr7_hap1.g02140.t01        rna-XM_007088780.2, rna-XM_007088815.2, rna-XM_007088817.2, rna-XM_015540511.2    rna-XM_008694104.2      mNeoVis.3.1.1.chr7.g01969.t01    rna-gnl|WGS_ZZZZ|egapxtmp_012629-R1    mNeoVis.evergladensis.2.1.chr7.g02087.t01    mNeoVis.vulgivaga.2.1.chr7.g02109.t01      579    N0.HOG0000577   OG0000038  n170   rna-XM_002926854.2, rna-XM_002926855.2    rna-NM_001388515.1, rna-NM_001388518.1, rna-NM_001388889.1    rna-XM_022507096.1          rna-XM_032846467.1    rna-XM_046016482.1    rna-XM_032360027.1    rna-XM_059158920.1    rna-XM_059405364.1    rna-XM_004781678.1    mNeoVis.2.1.1.chr7_hap1.g02134.t01          rna-XM_008694099.2, rna-XM_008694100.2        rna-gnl|WGS_ZZZZ|egapxtmp_013488-R1    mNeoVis.evergladensis.2.1.chr7.g02080.t01    mNeoVis.vulgivaga.2.1.chr7.g02104.t01      580    N0.HOG0000578   OG0000038  n186   rna-XM_019807642.2                                  rna-XM_008694098.2, rna-XM_040638758.1, rna-XM_040639115.1                581    N0.HOG0000579   OG0000038  n190           rna-XM_036092779.1                            rna-XM_027581045.2              582    N0.HOG0000580   OG0000038  n193       rna-XM_022507482.1          rna-XM_032846520.1    rna-XM_046017133.1    rna-XM_032358815.1    rna-XM_059158931.1    rna-XM_059387971.1    rna-XM_004782241.1    mNeoVis.2.1.1.chr7_hap1.g02144.t01              mNeoVis.3.1.1.chr7.g01970.t01    rna-gnl|WGS_ZZZZ|egapxtmp_013535-R1    mNeoVis.evergladensis.2.1.chr7.g02091.t01    mNeoVis.vulgivaga.2.1.chr7.g02112.t01      583    N0.HOG0000581   OG0000038  n205   rna-XM_002926886.2    rna-XM_038562464.1          rna-XM_039223514.1                        rna-XM_008694108.1                584    N0.HOG0000582   OG0000038  n208   rna-XM_011234352.1    rna-NM_001388964.1      rna-XM_023239739.2      rna-XM_039223485.1                      rna-XM_007088825.1    rna-XM_008694248.1                585    N0.HOG0000583   OG0000038  n213   rna-XM_002926884.3      rna-XM_022506647.1          rna-XM_032846458.1    rna-XM_046016872.1    rna-XM_032359934.1    rna-XM_059158930.1    rna-XM_059387977.1    rna-XM_004782232.2    mNeoVis.2.1.1.chr7_hap1.g02143.t01          rna-XM_008694106.2          mNeoVis.evergladensis.2.1.chr7.g02090.t01    mNeoVis.vulgivaga.2.1.chr7.g02111.t01      586    N0.HOG0000584   OG0000039  n0   rna-XM_002928528.4, rna-XM_002928543.4, rna-XM_011217917.3, rna-XM_011234659.3, rna-XM_019807841.2, rna-XM_019809268.2, rna-XM_034660131.1, rna-XM_034660132.1, rna-XM_034660133.1, rna-XM_034660134.1    rna-NM_001374580.1, rna-NM_001374588.1, rna-NM_001374589.1, rna-NM_001374591.1, rna-XM_038576140.1, rna-XM_038584378.1, rna-XM_038584427.1    rna-XM_022490542.1, rna-XM_022490572.1, rna-XM_022490574.1, rna-XM_022490575.1, rna-XM_022495204.1, rna-XM_022526238.1, rna-XM_022527121.1    rna-NM_001374574.1, rna-NM_001374578.1, rna-XM_003985683.3, rna-XM_003985688.5, rna-XM_006931391.3, rna-XM_006931394.5, rna-XM_006933534.4, rna-XM_006935068.5, rna-XM_011282072.4, rna-XM_011282077.4, rna-XM_023253583.2, rna-XM_023254700.2, rna-XM_045058522.1, rna-XM_045058523.1    rna-XM_036093771.1, rna-XM_036112170.1, rna-XM_036112363.1, rna-XM_036112364.1, rna-XM_036112365.1, rna-XM_036112366.1, rna-XM_036112373.1, rna-XM_036121576.1, rna-XM_036121577.1    rna-XM_039223547.1, rna-XM_039223561.1, rna-XM_039224921.1, rna-XM_039246211.1, rna-XM_039257296.1, rna-XM_039257298.1, rna-XM_039257299.1, rna-XM_039257302.1, rna-XM_039257304.1, rna-XM_039257305.1    rna-XM_032843566.1, rna-XM_032850995.1, rna-XM_032851031.1, rna-XM_032855419.1, rna-XM_032861704.1, rna-XM_032861747.1, rna-XM_032861748.1, rna-XM_032861760.1, rna-XM_032861761.1, rna-XM_032861762.1    rna-XM_046006114.1, rna-XM_046006307.1, rna-XM_046006309.1, rna-XM_046006310.1, rna-XM_046006311.1, rna-XM_046006316.1, rna-XM_046006317.1, rna-XM_046006321.1, rna-XM_046006322.1, rna-XM_046012557.1, rna-XM_046013946.1    rna-XM_032338291.1, rna-XM_032338292.1, rna-XM_032338293.1, rna-XM_032338294.1, rna-XM_032338295.1, rna-XM_032338296.1, rna-XM_032338297.1, rna-XM_032338298.1, rna-XM_032338299.1, rna-XM_032338300.1, rna-XM_032341385.1, rna-XM_032341386.1, rna-XM_032347413.1    rna-XM_059176292.1, rna-XM_059176293.1, rna-XM_059176294.1, rna-XM_059176295.1, rna-XM_059176296.1, rna-XM_059176302.1, rna-XM_059176304.1, rna-XM_059176305.1, rna-XM_059176306.1, rna-XM_059185996.1    rna-XM_059377120.1, rna-XM_059399604.1, rna-XM_059399613.1, rna-XM_059402651.1    rna-XM_004755789.3, rna-XM_004778774.3, rna-XM_004781521.3, rna-XM_045082288.1, rna-XM_045082289.1, rna-XM_045082291.1, rna-XM_045082292.1, rna-XM_045082293.1, rna-XM_045082299.1, rna-XM_045082301.1, rna-XM_045082302.1    mNeoVis.2.1.1.chr1_hap1.g01078.t01, mNeoVis.2.1.1.chr1_hap1.g01085.t01, mNeoVis.2.1.1.chr1_hap1.g01086.t01, mNeoVis.2.1.1.chr1_hap1.g01100.t01, mNeoVis.2.1.1.chr1_hap1.g01113.t01, mNeoVis.2.1.1.chr1_hap1.g01119.t01, mNeoVis.2.1.1.chr1_hap1.g01120.t01, mNeoVis.2.1.1.chr1_hap1.g01124.t01, mNeoVis.2.1.1.chr1_hap1.g01131.t01, mNeoVis.2.1.1.chr1_hap1.g01133.t01, mNeoVis.2.1.1.chr1_hap1.g01142.t01, mNeoVis.2.1.1.chr1_hap1.g01150.t01, mNeoVis.2.1.1.chr1_hap1.g01151.t01, mNeoVis.2.1.1.chr2_hap1.g00834.t01, mNeoVis.2.1.1.chr9_hap1.g00791.t01    rna-XM_021688068.1, rna-XM_021690607.1, rna-XM_021696850.1, rna-XM_044912454.1, rna-XM_044912461.1, rna-XM_044917302.1, rna-XM_044917580.1, rna-XM_044917590.1, rna-XM_044917591.1, rna-XM_044917592.1    rna-XM_004406517.1, rna-XM_004406525.1, rna-XM_004406991.1, rna-XM_004413629.1, rna-XM_004413634.1, rna-XM_004413646.1, rna-XM_004413650.1, rna-XM_004413653.1, rna-XM_004413661.1, rna-XM_004413663.1    rna-XM_007074436.3, rna-XM_007088032.3, rna-XM_015540513.2, rna-XM_015540675.2, rna-XM_015540747.2, rna-XM_042985396.1, rna-XM_042985555.1, rna-XM_042985557.1, rna-XM_042985558.1, rna-XM_042985559.1, rna-XM_042985561.1, rna-XM_042985562.1, rna-XM_042991806.1    rna-XM_008692955.2, rna-XM_008692967.2, rna-XM_008693003.2, rna-XM_008693091.2, rna-XM_008695756.2, rna-XM_008707026.2, rna-XM_040644635.1, rna-XM_040645171.1, rna-XM_040645187.1, rna-XM_040645188.1, rna-XM_040645189.1, rna-XM_040645190.1, rna-XM_040645191.1, rna-XM_040645197.1    rna-XM_027594572.2, rna-XM_027604114.2, rna-XM_027608948.2, rna-XM_035728352.1, rna-XM_035728448.1, rna-XM_035728449.1, rna-XM_035728450.1, rna-XM_035728451.1, rna-XM_035728452.1, rna-XM_035728455.1    mNeoVis.3.1.1.chr1.g01065.t01, mNeoVis.3.1.1.chr1.g01069.t01, mNeoVis.3.1.1.chr1.g01070.t01, mNeoVis.3.1.1.chr1.g01089.t01, mNeoVis.3.1.1.chr1.g01090.t01, mNeoVis.3.1.1.chr1.g01096.t01, mNeoVis.3.1.1.chr1.g01098.t01, mNeoVis.3.1.1.chr1.g01109.t01, mNeoVis.3.1.1.chr1.g01111.t01, mNeoVis.3.1.1.chr10.g00755.t01, mNeoVis.3.1.1.chr5.g00774.t01    rna-gnl|WGS_ZZZZ|egapxtmp_006456-R1, rna-gnl|WGS_ZZZZ|egapxtmp_010108-R1, rna-gnl|WGS_ZZZZ|egapxtmp_014067-R1, rna-gnl|WGS_ZZZZ|egapxtmp_018327-R1, rna-gnl|WGS_ZZZZ|egapxtmp_018338-R1, rna-gnl|WGS_ZZZZ|egapxtmp_018465-R1, rna-gnl|WGS_ZZZZ|egapxtmp_021260-R1, rna-gnl|WGS_ZZZZ|egapxtmp_021312-R1, rna-gnl|WGS_ZZZZ|egapxtmp_021324-R1, rna-gnl|WGS_ZZZZ|egapxtmp_021328-R1    mNeoVis.evergladensis.2.1.chr1.g01063.t01, mNeoVis.evergladensis.2.1.chr1.g01069.t01, mNeoVis.evergladensis.2.1.chr1.g01070.t01, mNeoVis.evergladensis.2.1.chr1.g01080.t01, mNeoVis.evergladensis.2.1.chr1.g01090.t01, mNeoVis.evergladensis.2.1.chr1.g01093.t01, mNeoVis.evergladensis.2.1.chr1.g01094.t01, mNeoVis.evergladensis.2.1.chr1.g01097.t01, mNeoVis.evergladensis.2.1.chr1.g01101.t01, mNeoVis.evergladensis.2.1.chr1.g01107.t01, mNeoVis.evergladensis.2.1.chr1.g01112.t01, mNeoVis.evergladensis.2.1.chr1.g01122.t01, mNeoVis.evergladensis.2.1.chr1.g01123.t01, mNeoVis.evergladensis.2.1.chr10.g00781.t01, mNeoVis.evergladensis.2.1.chr5.g00788.t01    mNeoVis.vulgivaga.2.1.chr1.g01085.t01, mNeoVis.vulgivaga.2.1.chr1.g01086.t01, mNeoVis.vulgivaga.2.1.chr1.g01104.t01, mNeoVis.vulgivaga.2.1.chr1.g01105.t01, mNeoVis.vulgivaga.2.1.chr1.g01108.t01, mNeoVis.vulgivaga.2.1.chr1.g01110.t01, mNeoVis.vulgivaga.2.1.chr1.g01119.t01, mNeoVis.vulgivaga.2.1.chr1.g01121.t01, mNeoVis.vulgivaga.2.1.chr10.g00768.t01, mNeoVis.vulgivaga.2.1.chr5.g00780.t01      587    N0.HOG0000585   OG0000040  n1   rna-XM_002929675.2, rna-XM_002929677.2, rna-XM_002930259.2, rna-XM_002930260.4, rna-XM_002930261.2, rna-XM_002931323.4, rna-XM_011218094.3, rna-XM_011237720.3, rna-XM_019793089.2, rna-XM_034653250.1, rna-XM_034653296.1, rna-XM_034653297.1    rna-NM_001388542.1, rna-XM_038556363.1    rna-XM_022491969.1, rna-XM_022491978.1, rna-XM_022492027.1    rna-XM_003980544.2, rna-XM_011283070.3, rna-XM_045062378.1    rna-XM_036100216.1, rna-XM_036100218.1, rna-XM_036100232.1    rna-XM_039225468.1, rna-XM_039225645.1, rna-XM_039225760.1, rna-XM_039225787.1    rna-XM_032848084.1, rna-XM_032848086.1, rna-XM_032848099.1    rna-XM_045996830.1, rna-XM_045996998.1, rna-XM_045997016.1, rna-XM_046000832.1, rna-XM_046000868.1, rna-XM_046001135.1    rna-XM_032337890.1    rna-XM_059173393.1    rna-XM_059416415.1    rna-XM_004781850.1    mNeoVis.2.1.1.chr1_hap1.g00945.t01    rna-XM_021690933.1, rna-XM_021690946.1    rna-XM_004413497.1    rna-XM_007092933.1, rna-XM_015542384.2    rna-XM_008683721.2, rna-XM_008683722.2, rna-XM_008683723.2, rna-XM_008683740.2, rna-XM_008683758.2, rna-XM_008683759.2, rna-XM_008709964.2, rna-XM_008709965.2, rna-XM_008710275.1, rna-XM_008710276.2, rna-XM_008710280.2, rna-XM_008710281.1, rna-XM_008710286.2    rna-XM_027604220.2, rna-XM_027606709.1, rna-XM_027606833.1    mNeoVis.3.1.1.chr1.g00957.t01    rna-gnl|WGS_ZZZZ|egapxtmp_040262-R1    mNeoVis.evergladensis.2.1.chr1.g00940.t01    mNeoVis.vulgivaga.2.1.chr1.g00951.t01      588    N0.HOG0000586   OG0000040  n68   rna-XM_002931315.2    rna-NM_001388958.1    rna-XM_022491979.1    rna-XM_003980678.2    rna-XM_036096463.1    rna-XM_039224560.1    rna-XM_032848100.1    rna-XM_046000288.1, rna-XM_046000361.1    rna-XM_032337898.1, rna-XM_032337899.1    rna-XM_059173498.1    rna-XM_059416719.1    rna-XM_004780236.2    mNeoVis.2.1.1.chr1_hap1.g00924.t01    rna-XM_021690952.1, rna-XM_021690953.1      rna-XM_015537894.1    rna-XM_008683760.2    rna-XM_027606693.2    mNeoVis.3.1.1.chr1.g00939.t01    rna-gnl|WGS_ZZZZ|egapxtmp_039035-R1    mNeoVis.evergladensis.2.1.chr1.g00920.t01    mNeoVis.vulgivaga.2.1.chr1.g00932.t01      589    N0.HOG0000587   OG0000040  n90   rna-XM_011234385.2, rna-XM_011237132.2, rna-XM_034653321.1    rna-XM_038557527.1    rna-XM_022491682.1    rna-XM_045053656.1      rna-XM_039225411.1    rna-XM_032848499.1    rna-XM_045999692.1, rna-XM_045999694.1, rna-XM_045999696.1, rna-XM_045999698.1    rna-XM_032335013.1, rna-XM_032335015.1    rna-XM_059173425.1, rna-XM_059176071.1    rna-XM_059416435.1    rna-XM_004780633.1, rna-XM_004782328.1    mNeoVis.2.1.1.chr1_hap1.g00971.t01    rna-XM_044912396.1, rna-XM_044912410.1, rna-XM_044912443.1, rna-XM_044912486.1, rna-XM_044912503.1, rna-XM_044912514.1, rna-XM_044912534.1, rna-XM_044912544.1, rna-XM_044912548.1, rna-XM_044917024.1      rna-XM_007092952.2, rna-XM_007092976.2    rna-XM_008709881.2, rna-XM_008709908.1, rna-XM_008709909.2, rna-XM_008709913.2, rna-XM_008709916.2      mNeoVis.3.1.1.chr1.g00979.t01    rna-gnl|WGS_ZZZZ|egapxtmp_018718-R1    mNeoVis.evergladensis.2.1.chr1.g00966.t01    mNeoVis.vulgivaga.2.1.chr1.g00979.t01      590    N0.HOG0000588   OG0000040  n130   rna-XM_011237719.2    rna-NM_001389015.1          rna-XM_039225799.1                        rna-XM_008710277.1                591    N0.HOG0000589   OG0000040  n133   rna-XM_002929136.4, rna-XM_002929137.4, rna-XM_019809817.2, rna-XM_034653247.1    rna-NM_001388539.1, rna-XM_038556360.1, rna-XM_038556361.1, rna-XM_038556362.1    rna-XM_022491628.1, rna-XM_022492028.1    rna-XM_003980583.3, rna-XM_045056451.1    rna-XM_036100233.1    rna-XM_039218804.1, rna-XM_039222548.1    rna-XM_032844366.1, rna-XM_032849853.1, rna-XM_032849956.1    rna-XM_045980068.1, rna-XM_045996806.1, rna-XM_045997024.1, rna-XM_045998726.1, rna-XM_046001097.1, rna-XM_046001133.1    rna-XM_032334920.1    rna-XM_059173407.1, rna-XM_059173504.1    rna-XM_059416727.1    rna-XM_004781964.1    mNeoVis.2.1.1.chr1_hap1.g00946.t01, mNeoVis.2.1.1.chr1_hap1.g00997.t01, mNeoVis.2.1.1.chr1_hap1.g01008.t01    rna-XM_021690932.1, rna-XM_021690939.1    rna-XM_004403492.1, rna-XM_004403493.1, rna-XM_004406934.1    rna-XM_007092965.2, rna-XM_007092966.2    rna-XM_008709880.2, rna-XM_008709886.2, rna-XM_008710279.1, rna-XM_008710284.1, rna-XM_008710288.2, rna-XM_008710289.2, rna-XM_040622682.1    rna-XM_027606536.1, rna-XM_027606541.1, rna-XM_027606565.1    mNeoVis.3.1.1.chr1.g00958.t01, mNeoVis.3.1.1.chr1.g01008.t01    rna-gnl|WGS_ZZZZ|egapxtmp_010044-R1, rna-gnl|WGS_ZZZZ|egapxtmp_037290-R1, rna-gnl|WGS_ZZZZ|egapxtmp_040326-R1    mNeoVis.evergladensis.2.1.chr1.g00941.t01, mNeoVis.evergladensis.2.1.chr1.g01000.t01    mNeoVis.vulgivaga.2.1.chr1.g00952.t01, mNeoVis.vulgivaga.2.1.chr1.g01001.t01, mNeoVis.vulgivaga.2.1.chr1.g01014.t01      592    N0.HOG0000590   OG0000040  n189   rna-XM_002931248.4, rna-XM_002931252.1    rna-NM_001388851.1    rna-XM_022492011.1, rna-XM_022492013.1    rna-XM_003980636.1, rna-XM_011281402.3      rna-XM_039224544.1, rna-XM_039224561.1    rna-XM_032848082.1, rna-XM_032848083.1    rna-XM_046000644.1, rna-XM_046000729.1    rna-XM_032334927.1, rna-XM_032334929.1    rna-XM_059173379.1, rna-XM_059173496.1    rna-XM_059416404.1, rna-XM_059416718.1    rna-XM_004780213.1, rna-XM_013049799.2    mNeoVis.2.1.1.chr1_hap1.g00921.t01, mNeoVis.2.1.1.chr1_hap1.g00923.t01    rna-XM_021689945.1      rna-XM_007083054.1    rna-XM_008683724.2, rna-XM_008683726.2, rna-XM_040630634.1    rna-XM_027604219.1    mNeoVis.3.1.1.chr1.g00937.t01, mNeoVis.3.1.1.chr1.g00938.t01    rna-gnl|WGS_ZZZZ|egapxtmp_035595-R1    mNeoVis.evergladensis.2.1.chr1.g00918.t01, mNeoVis.evergladensis.2.1.chr1.g00919.t01    mNeoVis.vulgivaga.2.1.chr1.g00930.t01, mNeoVis.vulgivaga.2.1.chr1.g00931.t01      593    N0.HOG0000591   OG0000041  n1   rna-XM_002923800.4    rna-XM_038584717.1    rna-XM_022495337.1    rna-XM_019838209.3    rna-XM_036090408.1    rna-XM_039250145.1    rna-XM_032880362.1    rna-XM_046019407.1    rna-XM_032355837.1    rna-XM_059167122.1    rna-XM_059395488.1    rna-XM_004744002.3    mNeoVis.2.1.1.chr3_hap1.g00922.t01    rna-XM_021702900.1    rna-XM_004396921.1    rna-XM_042994623.1    rna-XM_008687059.2    rna-XM_027591106.1    mNeoVis.3.1.1.chr3.g00908.t01    rna-gnl|WGS_ZZZZ|egapxtmp_038936-R5    mNeoVis.evergladensis.2.1.chr3.g00918.t01    mNeoVis.vulgivaga.2.1.chr3.g00935.t01      594    N0.HOG0000592   OG0000041  n23   rna-XM_034654677.1    rna-XM_038584713.1    rna-XM_022514231.1    rna-XM_019838208.3    rna-XM_036090406.1    rna-XM_039250141.1    rna-XM_032880361.1    rna-XM_046018497.1    rna-XM_032355819.1    rna-XM_059167130.1    rna-XM_059394112.1    rna-XM_045066692.1    mNeoVis.2.1.1.chr3_hap1.g00924.t01    rna-XM_021703139.1    rna-XM_004396924.1    rna-XM_042994620.1    rna-XM_008687055.2    rna-XM_035726822.1    mNeoVis.3.1.1.chr3.g00910.t01    rna-gnl|WGS_ZZZZ|egapxtmp_038940-R1    mNeoVis.evergladensis.2.1.chr3.g00919.t01    mNeoVis.vulgivaga.2.1.chr3.g00937.t01      595    N0.HOG0000593   OG0000041  n47   rna-XM_019805183.2    rna-XM_038546882.1    rna-XM_022519876.1    rna-XM_006940491.4    rna-XM_036116166.1    rna-XM_039235960.1    rna-XM_032870846.1    rna-XM_045985260.1    rna-XM_032320278.1    rna-XM_059147479.1    rna-XM_059381148.1    rna-XM_004749158.3    mNeoVis.2.1.1.chr8_hap1.g00253.t01    rna-XM_021678411.1    rna-XM_004405862.1    rna-XM_042966949.1    rna-XM_040640403.1    rna-XM_027626061.1    mNeoVis.3.1.1.chr8.g00210.t01    rna-gnl|WGS_ZZZZ|egapxtmp_019362-R1    mNeoVis.evergladensis.2.1.chr9.g00242.t01    mNeoVis.vulgivaga.2.1.chr8.g00250.t01      596    N0.HOG0000594   OG0000041  n69   rna-XM_034662225.1    rna-NM_001002994.1    rna-XM_022521391.1    rna-XM_045037851.1    rna-XM_036094215.1    rna-XM_039237226.1    rna-XM_032861173.1    rna-XM_046003336.1    rna-XM_032350129.1    rna-XM_059162056.1    rna-XM_059390533.1    rna-XM_004759336.3    mNeoVis.2.1.1.chr5_hap1.g01005.t01    rna-XM_021705867.1    rna-XM_004412302.1    rna-XM_042999168.1    rna-XM_040636947.1    rna-XM_027582855.2    mNeoVis.3.1.1.chr2.g00924.t01    rna-gnl|WGS_ZZZZ|egapxtmp_035514-R1    mNeoVis.evergladensis.2.1.chr2.g01001.t01    mNeoVis.vulgivaga.2.1.chr2.g01006.t01      597    N0.HOG0000595   OG0000041  n91   rna-XM_002930005.4    rna-XM_038570472.1    rna-XM_022521410.1    rna-XM_045037845.1    rna-XM_036106753.1    rna-XM_039237174.1    rna-XM_032861164.1    rna-XM_046003934.1    rna-XM_032350823.1    rna-XM_059162050.1    rna-XM_059390530.1    rna-XM_045089156.1    mNeoVis.2.1.1.chr5_hap1.g01007.t01    rna-XM_021705863.1    rna-XM_004412115.1, rna-XM_004412331.1    rna-XM_042999271.1    rna-XM_008710137.2    rna-XM_027584593.2    mNeoVis.3.1.1.chr2.g00926.t01    rna-gnl|WGS_ZZZZ|egapxtmp_038456-R1    mNeoVis.evergladensis.2.1.chr2.g01003.t01    mNeoVis.vulgivaga.2.1.chr2.g01008.t01      598    N0.HOG0000596   OG0000041  n113   rna-XM_034661768.1    rna-NM_001003203.1    rna-XM_022521407.1    rna-XM_045037849.1    rna-XM_036094248.1    rna-XM_039237171.1    rna-XM_032861166.1    rna-XM_046003843.1    rna-XM_032351968.1    rna-XM_059162051.1    rna-XM_059387943.1    rna-XM_045089152.1    mNeoVis.2.1.1.chr5_hap1.g01006.t01    rna-XM_021705864.1    rna-XM_004412301.1    rna-XM_007081019.2    rna-XM_008710144.2    rna-XM_027584950.1    mNeoVis.3.1.1.chr2.g00925.t01    rna-gnl|WGS_ZZZZ|egapxtmp_038454-R1    mNeoVis.evergladensis.2.1.chr2.g01002.t01    mNeoVis.vulgivaga.2.1.chr2.g01007.t01      599    N0.HOG0000597   OG0000041  n134   rna-XM_034645208.1    rna-XM_038577714.1    rna-XM_022519536.1    rna-XM_023257081.2    rna-XM_036098857.1    rna-XM_039241351.1    rna-XM_032876747.1    rna-XM_046012736.1    rna-XM_032348254.1    rna-XM_059185302.1    rna-XM_059403160.1    rna-XM_045072918.1    mNeoVis.2.1.1.chr9_hap1.g00542.t01    rna-XM_044914839.1    rna-XM_004400808.1    rna-XM_042992327.1    rna-XM_040642471.1    rna-XM_027593501.2    mNeoVis.3.1.1.chr10.g00511.t01    rna-gnl|WGS_ZZZZ|egapxtmp_022296-R1    mNeoVis.evergladensis.2.1.chr10.g00535.t01    mNeoVis.vulgivaga.2.1.chr10.g00521.t01      600    N0.HOG0000598   OG0000041  n154   rna-XM_034654674.1    rna-XM_038584708.1    rna-XM_022507393.1    rna-XM_019838205.3    rna-XM_036090394.1    rna-XM_039250140.1    rna-XM_032880353.1    rna-XM_046018491.1    rna-XM_032355795.1    rna-XM_059167125.1    rna-XM_059394109.1    rna-XM_004743995.3    mNeoVis.2.1.1.chr3_hap1.g00925.t01    rna-XM_021703128.1    rna-XM_004396923.1    rna-XM_007074624.3    rna-XM_008687048.1    rna-XM_035726815.1    mNeoVis.3.1.1.chr3.g00911.t01, mNeoVis.3.1.1.chr3.g00912.t01, mNeoVis.3.1.1.chr3.g00913.t01    rna-gnl|WGS_ZZZZ|egapxtmp_041213-R1    mNeoVis.evergladensis.2.1.chr3.g00920.t01    mNeoVis.vulgivaga.2.1.chr3.g00938.t01      601    N0.HOG0000599   OG0000041  n176   rna-XM_034654667.1    rna-XM_038584698.1    rna-XM_022509291.1    rna-XM_023259411.2    rna-XM_036090431.1    rna-XM_039241998.1    rna-XM_032880338.1    rna-XM_046019434.1    rna-XM_032355807.1    rna-XM_059167139.1    rna-XM_059394099.1    rna-XM_004743983.3    mNeoVis.2.1.1.chr3_hap1.g00930.t01    rna-XM_044913665.1    rna-XM_004396930.1    rna-XM_042997260.1    rna-XM_008687038.1    rna-XM_035726810.1    mNeoVis.3.1.1.chr3.g00919.t01    rna-gnl|WGS_ZZZZ|egapxtmp_042505-R2    mNeoVis.evergladensis.2.1.chr3.g00925.t01    mNeoVis.vulgivaga.2.1.chr3.g00943.t01      602    N0.HOG0000600   OG0000041  n195   rna-XM_002926919.4    rna-XM_038584693.1    rna-XM_022498669.1    rna-XM_019838191.3    rna-XM_036090433.1    rna-XM_039242011.1    rna-XM_032880329.1    rna-XM_046019657.1    rna-XM_032355810.1    rna-XM_059167145.1    rna-XM_059394095.1    rna-XM_004743978.3    mNeoVis.2.1.1.chr3_hap1.g00931.t01    rna-XM_021703129.1    rna-XM_004396933.1    rna-XM_042994594.1    rna-XM_040644404.1    rna-XM_035726804.1    mNeoVis.3.1.1.chr3.g00920.t01    rna-gnl|WGS_ZZZZ|egapxtmp_041215-R1    mNeoVis.evergladensis.2.1.chr3.g00926.t01    mNeoVis.vulgivaga.2.1.chr3.g00944.t01      603    N0.HOG0000601   OG0000042  n0   rna-XM_002918532.4, rna-XM_002918533.4, rna-XM_002918535.4, rna-XM_034663699.1    rna-NM_001006647.1, rna-NM_001006654.1, rna-XM_038551620.1, rna-XM_038551622.1, rna-XM_038552515.1, rna-XM_038552516.1, rna-XM_038552517.1, rna-XM_038552518.1, rna-XM_038552519.1, rna-XM_038552520.1, rna-XM_038552521.1, rna-XM_038552522.1, rna-XM_038552523.1, rna-XM_038552524.1, rna-XM_038552525.1, rna-XM_038553281.1, rna-XM_038553282.1    rna-XM_022492612.1, rna-XM_022492617.1, rna-XM_022492618.1, rna-XM_022492728.1, rna-XM_022492798.1, rna-XM_022517578.1, rna-XM_022517579.1, rna-XM_022517594.1, rna-XM_022517596.1, rna-XM_022517601.1, rna-XM_022522506.1, rna-XM_022522509.1, rna-XM_022522510.1, rna-XM_022522511.1, rna-XM_022522512.1    rna-NM_001009193.1, rna-NM_001009197.2, rna-XM_004001268.3, rna-XM_023243268.2, rna-XM_023243269.2    rna-XM_036082643.1, rna-XM_036082644.1, rna-XM_036082647.1, rna-XM_036082648.1, rna-XM_036082650.1, rna-XM_036082651.1, rna-XM_036082653.1, rna-XM_036082654.1, rna-XM_036082655.1, rna-XM_036082688.1, rna-XM_036082689.1, rna-XM_036082691.1, rna-XM_036082692.1    rna-XM_039221261.1, rna-XM_039221262.1, rna-XM_039244417.1    rna-XM_032851158.1, rna-XM_032880490.1, rna-XM_032880491.1, rna-XM_032880492.1, rna-XM_032880493.1    rna-XM_046022417.1, rna-XM_046022418.1, rna-XM_046022419.1, rna-XM_046022420.1, rna-XM_046022422.1, rna-XM_046022424.1, rna-XM_046022425.1, rna-XM_046022426.1, rna-XM_046022427.1, rna-XM_046022428.1, rna-XM_046022429.1, rna-XM_046022430.1, rna-XM_046022431.1, rna-XM_046023135.1, rna-XM_046023166.1, rna-XM_046023173.1    rna-XM_032306491.1, rna-XM_032306987.1, rna-XM_032306989.1, rna-XM_032306990.1, rna-XM_032306992.1, rna-XM_032306994.1, rna-XM_032306995.1, rna-XM_032306996.1, rna-XM_032306997.1, rna-XM_032306999.1, rna-XM_032307000.1, rna-XM_032307001.1, rna-XM_032307002.1, rna-XM_032307004.1, rna-XM_032307005.1, rna-XM_032307006.1    rna-XM_059141232.1, rna-XM_059142053.1, rna-XM_059142170.1, rna-XM_059142201.1, rna-XM_059142202.1, rna-XM_059142203.1, rna-XM_059142204.1, rna-XM_059142251.1, rna-XM_059142289.1, rna-XM_059142296.1, rna-XM_059142319.1, rna-XM_059142378.1, rna-XM_059142382.1, rna-XM_059142383.1, rna-XM_059142464.1, rna-XM_059142465.1, rna-XM_059142786.1    rna-XM_059387422.1, rna-XM_059387484.1, rna-XM_059387509.1, rna-XM_059387528.1, rna-XM_059387572.1, rna-XM_059410130.1, rna-XM_059410132.1, rna-XM_059410134.1, rna-XM_059412284.1, rna-XM_059412285.1, rna-XM_059412286.1, rna-XM_059412287.1, rna-XM_059412288.1, rna-XM_059412290.1    rna-NM_001310172.2, rna-NM_001310195.1, rna-XM_004761809.3, rna-XM_004761811.3, rna-XM_004761812.3, rna-XM_004761815.2, rna-XM_004761817.3, rna-XM_004761833.3, rna-XM_004761834.3, rna-XM_004761835.2, rna-XM_004761836.3, rna-XM_004761838.3, rna-XM_004761840.3, rna-XM_045079672.1, rna-XM_045079694.1, rna-XM_045079695.1, rna-XM_045079697.1, rna-XM_045079698.1, rna-XM_045079699.1, rna-XM_045079700.1    mNeoVis.2.1.1.chr13_hap1.g00588.t01, mNeoVis.2.1.1.chr13_hap1.g00589.t01, mNeoVis.2.1.1.chr13_hap1.g00590.t01, mNeoVis.2.1.1.chr13_hap1.g00591.t01, mNeoVis.2.1.1.chr13_hap1.g00592.t01, mNeoVis.2.1.1.chr13_hap1.g00593.t01, mNeoVis.2.1.1.chr13_hap1.g00594.t01, mNeoVis.2.1.1.chr13_hap1.g00595.t01, mNeoVis.2.1.1.chr13_hap1.g00596.t01, mNeoVis.2.1.1.chr13_hap1.g00598.t01, mNeoVis.2.1.1.chr13_hap1.g00599.t01, mNeoVis.2.1.1.chr13_hap1.g00600.t01, mNeoVis.2.1.1.chr13_hap1.g00601.t01, mNeoVis.2.1.1.chr13_hap1.g00602.t01, mNeoVis.2.1.1.chr13_hap1.g00603.t01, mNeoVis.2.1.1.chr13_hap1.g00604.t01    rna-XM_021697285.1, rna-XM_044920310.1, rna-XM_044920504.1    rna-XM_004405654.1, rna-XM_004412911.1, rna-XM_004412913.1    rna-XM_042963643.1, rna-XM_042963645.1, rna-XM_042964304.1, rna-XM_042964317.1, rna-XM_042964318.1, rna-XM_042964319.1, rna-XM_042964349.1    rna-XM_008689794.2, rna-XM_008689795.2, rna-XM_008695066.2, rna-XM_008709917.1, rna-XM_008709918.2, rna-XM_040631390.1    rna-XM_027616076.2, rna-XM_027616123.2, rna-XM_035723646.1, rna-XM_035723652.1, rna-XM_035723657.1    mNeoVis.3.1.1.chr13.g00526.t01, mNeoVis.3.1.1.chr13.g00528.t01    rna-gnl|WGS_ZZZZ|egapxtmp_029939-R1, rna-gnl|WGS_ZZZZ|egapxtmp_029940-R1, rna-gnl|WGS_ZZZZ|egapxtmp_029941-R1, rna-gnl|WGS_ZZZZ|egapxtmp_029943-R1, rna-gnl|WGS_ZZZZ|egapxtmp_029944-R1, rna-gnl|WGS_ZZZZ|egapxtmp_029945-R1, rna-gnl|WGS_ZZZZ|egapxtmp_029946-R1, rna-gnl|WGS_ZZZZ|egapxtmp_029947-R1, rna-gnl|WGS_ZZZZ|egapxtmp_029948-R1, rna-gnl|WGS_ZZZZ|egapxtmp_029949-R1    mNeoVis.evergladensis.2.1.chr13.g00570.t01, mNeoVis.evergladensis.2.1.chr13.g00571.t01, mNeoVis.evergladensis.2.1.chr13.g00572.t01, mNeoVis.evergladensis.2.1.chr13.g00573.t01, mNeoVis.evergladensis.2.1.chr13.g00575.t01, mNeoVis.evergladensis.2.1.chr13.g00576.t01, mNeoVis.evergladensis.2.1.chr13.g00577.t01, mNeoVis.evergladensis.2.1.chr13.g00578.t01, mNeoVis.evergladensis.2.1.chr13.g00579.t01, mNeoVis.evergladensis.2.1.chr13.g00580.t01, mNeoVis.evergladensis.2.1.chr13.g00581.t01    mNeoVis.vulgivaga.2.1.chr13.g00572.t01, mNeoVis.vulgivaga.2.1.chr13.g00573.t01, mNeoVis.vulgivaga.2.1.chr13.g00575.t01, mNeoVis.vulgivaga.2.1.chr13.g00576.t01, mNeoVis.vulgivaga.2.1.chr13.g00577.t01, mNeoVis.vulgivaga.2.1.chr13.g00578.t01, mNeoVis.vulgivaga.2.1.chr13.g00579.t01, mNeoVis.vulgivaga.2.1.chr13.g00580.t01      604    N0.HOG0000602   OG0000043  n1   rna-XM_002929995.3    rna-NM_001388763.1    rna-XM_022519378.1    rna-XM_023257755.2    rna-XM_036103753.1    rna-XM_039221093.1    rna-XM_032876516.1    rna-XM_046013265.1, rna-XM_046013301.1    rna-XM_032347838.1    rna-XM_059183605.1    rna-XM_059404816.1    rna-XM_004781959.1    mNeoVis.2.1.1.chr9_hap1.g00623.t01    rna-XM_021704855.1    rna-XM_004396014.1    rna-XM_007073070.3    rna-XM_008704725.1    rna-XM_027594261.1    mNeoVis.3.1.1.chr10.g00588.t01    rna-gnl|WGS_ZZZZ|egapxtmp_022660-R1    mNeoVis.evergladensis.2.1.chr10.g00616.t01    mNeoVis.vulgivaga.2.1.chr10.g00600.t01      605    N0.HOG0000603   OG0000043  n20         rna-XM_011284012.3      rna-XM_039221064.1                      rna-XM_015544328.2                  606    N0.HOG0000604   OG0000043  n23   rna-XM_011216694.3    rna-NM_001388762.1    rna-XM_022519764.1    rna-XM_045061688.1    rna-XM_036103758.1    rna-XM_039221050.1    rna-XM_032876554.1    rna-XM_046013250.1    rna-XM_032345621.1    rna-XM_059183604.1    rna-XM_059404817.1    rna-XM_004781958.1    mNeoVis.2.1.1.chr9_hap1.g00622.t01    rna-XM_021704901.1    rna-XM_004396013.1    rna-XM_007073071.1    rna-XM_008704511.1    rna-XM_027594363.1      rna-gnl|WGS_ZZZZ|egapxtmp_022661-R1    mNeoVis.evergladensis.2.1.chr10.g00615.t01    mNeoVis.vulgivaga.2.1.chr10.g00599.t01      607    N0.HOG0000605   OG0000043  n42   rna-XM_002927220.4, rna-XM_034652225.1    rna-NM_001389052.1, rna-XM_038576160.1    rna-XM_022519749.1    rna-XM_003988654.2    rna-XM_036103756.1    rna-XM_039221091.1      rna-XM_046013137.1    rna-XM_032345624.1    rna-XM_059183598.1    rna-XM_059404819.1    rna-XM_004778936.1    mNeoVis.2.1.1.chr9_hap1.g00617.t01        rna-XM_007072951.1    rna-XM_008704514.1      mNeoVis.3.1.1.chr10.g00583.t01    rna-gnl|WGS_ZZZZ|egapxtmp_022659-R1    mNeoVis.evergladensis.2.1.chr10.g00610.t01    mNeoVis.vulgivaga.2.1.chr10.g00593.t01      608    N0.HOG0000606   OG0000043  n58   rna-XM_002927218.2    rna-NM_001388760.1      rna-XM_003988656.2    rna-XM_036103760.1    rna-XM_039221048.1      rna-XM_046013138.1    rna-XM_032347843.1    rna-XM_059183596.1    rna-XM_059404820.1    rna-XM_004778934.1    mNeoVis.2.1.1.chr9_hap1.g00615.t01        rna-XM_007073074.2    rna-XM_008710122.2        rna-gnl|WGS_ZZZZ|egapxtmp_024694-R1    mNeoVis.evergladensis.2.1.chr10.g00608.t01    mNeoVis.vulgivaga.2.1.chr10.g00592.t01      609    N0.HOG0000607   OG0000043  n74   rna-XM_002927217.1, rna-XM_011234763.2, rna-XM_034651793.1    rna-NM_001388759.1    rna-XM_022519381.1    rna-XM_003988657.2    rna-XM_036103755.1    rna-XM_039221087.1    rna-XM_032876514.1    rna-XM_046013134.1, rna-XM_046013141.1    rna-XM_032347844.1    rna-XM_059183595.1    rna-XM_059404821.1    rna-XM_004778933.1    mNeoVis.2.1.1.chr9_hap1.g00614.t01    rna-XM_021704854.1    rna-XM_004400906.1    rna-XM_007073075.2    rna-XM_040641914.1, rna-XM_040641915.1      mNeoVis.3.1.1.chr10.g00582.t01    rna-gnl|WGS_ZZZZ|egapxtmp_022654-R1    mNeoVis.evergladensis.2.1.chr10.g00607.t01        610    N0.HOG0000608   OG0000043  n99     rna-NM_001388758.1      rna-XM_003988658.4      rna-XM_039221065.1                      rna-XM_007073076.2                  611    N0.HOG0000609   OG0000043  n102   rna-XM_002927215.3, rna-XM_002927219.2    rna-NM_001389047.1, rna-XM_038576161.1    rna-XM_022519380.1    rna-XM_003988655.2      rna-XM_039221086.1    rna-XM_032876515.1      rna-XM_032347842.1, rna-XM_032347845.1    rna-XM_059187495.1    rna-XM_059404508.1    rna-XM_004778935.1    mNeoVis.2.1.1.chr9_hap1.g00616.t01    rna-XM_044914866.1, rna-XM_044915206.1    rna-XM_004400905.1    rna-XM_007072952.1    rna-XM_008704727.2, rna-XM_008704730.2        rna-gnl|WGS_ZZZZ|egapxtmp_024695-R1    mNeoVis.evergladensis.2.1.chr10.g00609.t01        612    N0.HOG0000610   OG0000043  n123   rna-XM_002930847.3    rna-NM_001388761.1    rna-XM_022519750.1    rna-XM_011284015.2      rna-XM_039221088.1        rna-XM_032347840.1    rna-XM_059183599.1    rna-XM_059404818.1    rna-XM_004778937.1    mNeoVis.2.1.1.chr9_hap1.g00618.t01        rna-XM_007073073.2    rna-XM_008704513.2      mNeoVis.3.1.1.chr10.g00584.t01    rna-gnl|WGS_ZZZZ|egapxtmp_022657-R1    mNeoVis.evergladensis.2.1.chr10.g00611.t01    mNeoVis.vulgivaga.2.1.chr10.g00594.t01      613    N0.HOG0000611   OG0000043  n138   rna-XM_002930846.2, rna-XM_034644782.1      rna-XM_022519379.1    rna-XM_003988653.2    rna-XM_036103757.1    rna-XM_039221090.1      rna-XM_046011820.1    rna-XM_032345622.1, rna-XM_032347839.1    rna-XM_059183600.1    rna-XM_059387345.1    rna-XM_045073341.1, rna-XM_045073342.1    mNeoVis.2.1.1.chr9_hap1.g00619.t01    rna-XM_044914867.1      rna-XM_007073072.2    rna-XM_008704726.1      mNeoVis.3.1.1.chr10.g00585.t01    rna-gnl|WGS_ZZZZ|egapxtmp_024696-R1    mNeoVis.evergladensis.2.1.chr10.g00612.t01    mNeoVis.vulgivaga.2.1.chr10.g00595.t01      614    N0.HOG0000612   OG0000043  n158   rna-XM_002927208.2    rna-NM_001388870.1    rna-XM_022519756.1    rna-XM_045061728.1      rna-XM_039221066.1    rna-XM_032876549.1    rna-XM_046013152.1, rna-XM_046013154.1    rna-XM_032345625.1    rna-XM_059183594.1    rna-XM_059404823.1    rna-XM_013049146.2    mNeoVis.2.1.1.chr9_hap1.g00613.t01          rna-XM_008704515.1      mNeoVis.3.1.1.chr10.g00581.t01        mNeoVis.vulgivaga.2.1.chr10.g00591.t01      615    N0.HOG0000613   OG0000043  n173   rna-XM_002927214.4, rna-XM_002927216.2    rna-XM_038577887.1, rna-XM_038577889.1, rna-XM_038577924.1    rna-XM_022492667.1    rna-XM_023257796.2    rna-XM_036119702.1    rna-XM_039221089.1    rna-XM_032876548.1    rna-XM_046013135.1    rna-XM_032345626.1    rna-XM_059183593.1    rna-XM_059404824.1    rna-XM_004778931.2    mNeoVis.2.1.1.chr9_hap1.g00612.t01    rna-XM_021704853.1      rna-XM_007072953.2, rna-XM_007073077.2    rna-XM_008704729.1, rna-XM_008704732.1      mNeoVis.3.1.1.chr10.g00580.t01    rna-gnl|WGS_ZZZZ|egapxtmp_022658-R1    mNeoVis.evergladensis.2.1.chr10.g00606.t01        616    N0.HOG0000614   OG0000044  n2   rna-XM_002931117.3, rna-XM_034661799.1    rna-XM_038579225.1    rna-XM_022492590.1    rna-XM_023240683.2      rna-XM_039233666.1    rna-XM_032877707.1    rna-XM_045996976.1, rna-XM_046027819.1, rna-XM_046027834.1    rna-XM_032312915.1    rna-XM_059172611.1    rna-XM_059395887.1    rna-XM_004765892.1    mNeoVis.2.1.1.chr2_hap1.g00372.t01        rna-XM_007080085.2    rna-XM_008696187.1    rna-XM_035724323.1    mNeoVis.3.1.1.chr5.g00355.t01    rna-gnl|WGS_ZZZZ|egapxtmp_006842-R1    mNeoVis.evergladensis.2.1.chr5.g00371.t01    mNeoVis.vulgivaga.2.1.chr5.g00371.t01      617    N0.HOG0000615   OG0000044  n23   rna-XM_019793857.2    rna-NM_001256445.1, rna-XM_038589354.1    rna-XM_022500251.1, rna-XM_022500285.1    rna-XM_045040335.1    rna-XM_036113932.1    rna-XM_039233665.1    rna-XM_032842961.1, rna-XM_032877690.1, rna-XM_032877703.1    rna-XM_045996948.1, rna-XM_045996975.1, rna-XM_046027814.1, rna-XM_046027823.1, rna-XM_046027950.1, rna-XM_046027966.1    rna-XM_032311833.1, rna-XM_032312914.1, rna-XM_032313157.1    rna-XM_059172601.1, rna-XM_059172605.1, rna-XM_059172607.1, rna-XM_059172609.1    rna-XM_059387299.1, rna-XM_059387541.1, rna-XM_059387548.1, rna-XM_059395890.1, rna-XM_059397695.1    rna-XM_004749764.3, rna-XM_004765893.2, rna-XM_045068442.1    mNeoVis.2.1.1.chr2_hap1.g00365.t01, mNeoVis.2.1.1.chr2_hap1.g00369.t01, mNeoVis.2.1.1.chr2_hap1.g00371.t01    rna-XM_021699970.1    rna-XM_004408621.1    rna-XM_015536598.2    rna-XM_008696338.1    rna-XM_027590671.1      rna-gnl|WGS_ZZZZ|egapxtmp_006840-R1, rna-gnl|WGS_ZZZZ|egapxtmp_006841-R1, rna-gnl|WGS_ZZZZ|egapxtmp_009440-R1    mNeoVis.evergladensis.2.1.chr5.g00366.t01, mNeoVis.evergladensis.2.1.chr5.g00369.t01, mNeoVis.evergladensis.2.1.chr5.g00370.t01    mNeoVis.vulgivaga.2.1.chr5.g00366.t01, mNeoVis.vulgivaga.2.1.chr5.g00369.t01      618    N0.HOG0000616   OG0000044  n70       rna-XM_022500261.1          rna-XM_032877604.1    rna-XM_046027963.1    rna-XM_032312067.1    rna-XM_059172598.1    rna-XM_059395892.1    rna-XM_013054564.1    mNeoVis.2.1.1.chr2_hap1.g00361.t01                rna-gnl|WGS_ZZZZ|egapxtmp_006839-R1    mNeoVis.evergladensis.2.1.chr5.g00362.t01        619    N0.HOG0000617   OG0000044  n81         rna-XM_019814073.3, rna-XM_019824705.3, rna-XM_023240311.2      rna-XM_039233656.1, rna-XM_039233664.1                      rna-XM_007080050.2, rna-XM_007080051.2, rna-XM_007080087.2                  620    N0.HOG0000618   OG0000044  n89   rna-XM_002930897.3, rna-XM_002930898.3, rna-XM_002930899.3, rna-XM_011218067.2    rna-XM_038589353.1    rna-XM_022499633.1, rna-XM_022499634.1, rna-XM_022500259.1, rna-XM_022500272.1, rna-XM_022500278.1, rna-XM_022500281.1, rna-XM_022500282.1, rna-XM_022500286.1      rna-XM_036113930.1      rna-XM_032842936.1, rna-XM_032877590.1, rna-XM_032877616.1, rna-XM_032877629.1, rna-XM_032877640.1, rna-XM_032877652.1, rna-XM_032877666.1, rna-XM_032877677.1    rna-XM_045996946.1, rna-XM_045996947.1, rna-XM_046026595.1, rna-XM_046027889.1, rna-XM_046027946.1, rna-XM_046027947.1, rna-XM_046027948.1, rna-XM_046027949.1, rna-XM_046027965.1    rna-XM_032312032.1, rna-XM_032312038.1, rna-XM_032312041.1, rna-XM_032312050.1, rna-XM_032312066.1    rna-XM_059172597.1, rna-XM_059172599.1, rna-XM_059172603.1, rna-XM_059172604.1    rna-XM_059395888.1, rna-XM_059395889.1, rna-XM_059395891.1, rna-XM_059395893.1    rna-XM_004749737.1, rna-XM_004749738.1, rna-XM_004749740.1    mNeoVis.2.1.1.chr2_hap1.g00360.t01, mNeoVis.2.1.1.chr2_hap1.g00363.t01, mNeoVis.2.1.1.chr2_hap1.g00364.t01, mNeoVis.2.1.1.chr2_hap1.g00366.t01, mNeoVis.2.1.1.chr2_hap1.g00368.t01    rna-XM_021700175.2, rna-XM_044912246.1, rna-XM_044912352.1    rna-XM_004408622.1, rna-XM_004417006.1, rna-XM_004417007.1      rna-XM_008696188.2, rna-XM_008696339.1, rna-XM_008696341.2, rna-XM_008696342.2, rna-XM_040623836.1, rna-XM_040624818.1    rna-XM_027596875.1    mNeoVis.3.1.1.chr5.g00350.t01, mNeoVis.3.1.1.chr5.g00354.t01, mNeoVis.3.1.1.chr5.g00356.t01, mNeoVis.3.1.1.chr5.g00357.t01    rna-gnl|WGS_ZZZZ|egapxtmp_007681-R1, rna-gnl|WGS_ZZZZ|egapxtmp_007682-R1, rna-gnl|WGS_ZZZZ|egapxtmp_007683-R1, rna-gnl|WGS_ZZZZ|egapxtmp_007684-R1, rna-gnl|WGS_ZZZZ|egapxtmp_007685-R1, rna-gnl|WGS_ZZZZ|egapxtmp_009441-R1    mNeoVis.evergladensis.2.1.chr5.g00360.t01, mNeoVis.evergladensis.2.1.chr5.g00361.t01, mNeoVis.evergladensis.2.1.chr5.g00364.t01, mNeoVis.evergladensis.2.1.chr5.g00365.t01, mNeoVis.evergladensis.2.1.chr5.g00367.t01, mNeoVis.evergladensis.2.1.chr5.g00368.t01    mNeoVis.vulgivaga.2.1.chr5.g00362.t01, mNeoVis.vulgivaga.2.1.chr5.g00364.t01, mNeoVis.vulgivaga.2.1.chr5.g00365.t01, mNeoVis.vulgivaga.2.1.chr5.g00367.t01, mNeoVis.vulgivaga.2.1.chr5.g00368.t01      621    N0.HOG0000619   OG0000044  n169   rna-XM_034650402.1, rna-XM_034651274.1, rna-XM_034663551.1    rna-XM_038579226.1    rna-XM_022500262.1          rna-XM_032877577.1      rna-XM_032312051.1, rna-XM_032312201.1    rna-XM_059172490.1, rna-XM_059172595.1    rna-XM_059395894.1, rna-XM_059397697.1    rna-XM_004749763.3, rna-XM_013054565.2    mNeoVis.2.1.1.chr2_hap1.g00359.t01, mNeoVis.2.1.1.chr2_hap1.g00362.t01          rna-XM_008696189.2, rna-XM_008696192.2, rna-XM_008696343.2      mNeoVis.3.1.1.chr5.g00353.t01    rna-gnl|WGS_ZZZZ|egapxtmp_007238-R1    mNeoVis.evergladensis.2.1.chr5.g00359.t01, mNeoVis.evergladensis.2.1.chr5.g00363.t01    mNeoVis.vulgivaga.2.1.chr5.g00361.t01      622    N0.HOG0000620   OG0000044  n191       rna-XM_022500277.1      rna-XM_036113931.1          rna-XM_032311918.1    rna-XM_059172600.1    rna-XM_059397696.1    rna-XM_004749739.3        rna-XM_004417580.1        rna-XM_027596823.1    mNeoVis.3.1.1.chr5.g00351.t01        mNeoVis.vulgivaga.2.1.chr5.g00363.t01      623    N0.HOG0000621   OG0000045  n2   rna-XM_002922559.4    rna-XM_038531373.1, rna-XM_038535516.1, rna-XM_038549540.1, rna-XM_038576109.1, rna-XM_038581115.1, rna-XM_038588190.1, rna-XM_038589394.1, rna-XM_038589708.1    rna-XM_022518612.1    rna-XM_006943968.4    rna-XM_036066381.1, rna-XM_036098700.1    rna-XM_039219830.1, rna-XM_039238817.1, rna-XM_039247874.1    rna-XM_032845572.1, rna-XM_032877771.1, rna-XM_032882284.1    rna-XM_045995065.1, rna-XM_046003059.1, rna-XM_046016158.1    rna-XM_032330439.1, rna-XM_032334942.1, rna-XM_032334948.1    rna-XM_059157893.1, rna-XM_059165272.1, rna-XM_059171603.1, rna-XM_059186687.1    rna-XM_059385322.1, rna-XM_059385458.1, rna-XM_059385508.1, rna-XM_059404640.1, rna-XM_059404995.1    rna-XM_004762077.3, rna-XM_004833876.3, rna-XM_013047540.2, rna-XM_013051527.2, rna-XM_045089220.1    mNeoVis.2.1.1.chr12_hap1.g00243.t01, mNeoVis.2.1.1.chr15_hap1.g00746.t01, mNeoVis.2.1.1.chr1_hap1.g00990.t01, mNeoVis.2.1.1.chr8_hap1.g01092.t01    rna-XM_021686196.1, rna-XM_021697428.2, rna-XM_021699399.2      rna-XM_042974151.1    rna-XM_040628576.1    rna-XM_027572731.2, rna-XM_027599595.1, rna-XM_027608226.2, rna-XM_035723960.1, rna-XM_035725275.1, rna-XM_035726831.1    mNeoVis.3.1.1.chr12.g00210.t01    rna-gnl|WGS_ZZZZ|egapxtmp_017917-R1, rna-gnl|WGS_ZZZZ|egapxtmp_028657-R1, rna-gnl|WGS_ZZZZ|egapxtmp_041451-R1    mNeoVis.evergladensis.2.1.chr1.g00984.t01, mNeoVis.evergladensis.2.1.chr12.g00212.t01, mNeoVis.evergladensis.2.1.chr15.g00568.t01, mNeoVis.evergladensis.2.1.chr9.g01068.t01    mNeoVis.vulgivaga.2.1.chr12.g00217.t01, mNeoVis.vulgivaga.2.1.chr8.g01068.t01      624    N0.HOG0000622   OG0000045  n5                       rna-XM_059399109.1    rna-XM_045071130.1      rna-XM_044916392.1                      625    N0.HOG0000623   OG0000045  n8   rna-XM_002930768.1    rna-XM_038550869.1, rna-XM_038572339.1        rna-XM_036079267.1    rna-XM_039225693.1        rna-XM_032331685.1    rna-XM_059172640.1    rna-XM_059395782.1    rna-XM_045089497.1      rna-XM_021691086.1    rna-XM_004405237.1      rna-XM_008703896.1    rna-XM_035723769.1              626    N0.HOG0000624   OG0000045  n20     rna-XM_038572212.1        rna-XM_036102263.1      rna-XM_032865581.1        rna-XM_059157797.1    rna-XM_059385471.1    rna-XM_045088022.1      rna-XM_044915801.1          rna-XM_035726879.1              627    N0.HOG0000625   OG0000045  n27                           mNeoVis.2.1.1.chr1_hap1.g01701.t01              mNeoVis.3.1.1.chr1.g01656.t01    rna-gnl|WGS_ZZZZ|egapxtmp_022353-R1    mNeoVis.evergladensis.2.1.chr1.g01668.t01    mNeoVis.vulgivaga.2.1.chr1.g01664.t01      628    N0.HOG0000626   OG0000045  n32                             rna-XM_044911831.1          rna-XM_035728191.1              629    N0.HOG0000627   OG0000045  n35                 rna-XM_046000325.1                      rna-XM_035724050.1              630    N0.HOG0000628   OG0000045  n37                       rna-XM_059404989.1    rna-XM_045087399.1                          631    N0.HOG0000629   OG0000045  n39                   rna-XM_032338352.1            rna-XM_044913431.1                      632    N0.HOG0000630   OG0000045  n41                   rna-XM_032307439.1                    rna-XM_035722586.1              633    N0.HOG0000631   OG0000045  n43     rna-XM_038575426.1    rna-XM_022495348.1    rna-XM_045057155.1    rna-XM_036084679.1, rna-XM_036096922.1      rna-XM_032836352.1    rna-XM_046006570.1    rna-XM_032338576.1, rna-XM_032345639.1    rna-XM_059183611.1    rna-XM_059404777.1    rna-XM_013057933.1    mNeoVis.2.1.1.chr10_hap1.g00493.t01, mNeoVis.2.1.1.chr1_hap1.g00283.t01, mNeoVis.2.1.1.chr1_hap1.g01742.t01, mNeoVis.2.1.1.chr1_hap1.g02112.t01, mNeoVis.2.1.1.chr9_hap1.g00764.t01, mNeoVis.2.1.1.chr9_hap1.g01106.t01        rna-XM_042985262.1    rna-XM_040641913.1    rna-XM_035722690.1, rna-XM_035725226.1, rna-XM_035728741.1, rna-XM_035728948.1    mNeoVis.3.1.1.chr1.g00287.t01, mNeoVis.3.1.1.chr1.g01696.t01, mNeoVis.3.1.1.chr1.g02067.t01, mNeoVis.3.1.1.chr10.g00731.t01, mNeoVis.3.1.1.chr10.g01063.t01    rna-gnl|WGS_ZZZZ|egapxtmp_016044-R1, rna-gnl|WGS_ZZZZ|egapxtmp_020983-R1, rna-gnl|WGS_ZZZZ|egapxtmp_023331-R1, rna-gnl|WGS_ZZZZ|egapxtmp_042051-R1    mNeoVis.evergladensis.2.1.chr1.g00282.t01, mNeoVis.evergladensis.2.1.chr1.g01710.t01, mNeoVis.evergladensis.2.1.chr1.g02082.t01, mNeoVis.evergladensis.2.1.chr10.g00755.t01, mNeoVis.evergladensis.2.1.chr10.g01096.t01, mNeoVis.evergladensis.2.1.chr8.g00473.t01    mNeoVis.vulgivaga.2.1.chr1.g00286.t01, mNeoVis.vulgivaga.2.1.chr1.g01706.t01, mNeoVis.vulgivaga.2.1.chr10.g00743.t01, mNeoVis.vulgivaga.2.1.chr9.g00476.t01      634    N0.HOG0000632   OG0000045  n71     rna-XM_038585974.1        rna-XM_036111502.1, rna-XM_036112101.1                  mNeoVis.2.1.1.chr4_hap1.g00656.t01    rna-XM_044918149.1, rna-XM_044918225.1    rna-XM_004399816.1        rna-XM_035727860.1, rna-XM_035728758.1    mNeoVis.3.1.1.chr6.g00667.t01    rna-gnl|WGS_ZZZZ|egapxtmp_003412-R1    mNeoVis.evergladensis.2.1.chr6.g00663.t01    mNeoVis.vulgivaga.2.1.chr6.g00660.t01      635    N0.HOG0000633   OG0000045  n80                 rna-XM_046003564.1      rna-XM_059168919.1    rna-XM_059392888.1    rna-XM_045063129.1                          636    N0.HOG0000634   OG0000045  n87     rna-XM_038559014.1      rna-XM_045036231.1    rna-XM_036070722.1, rna-XM_036104969.1      rna-XM_032843442.1, rna-XM_032856977.1                rna-XM_021699244.1    rna-XM_004410978.1    rna-XM_015536188.2        mNeoVis.3.1.1.chr12.g00633.t01            637    N0.HOG0000635   OG0000045  n96   rna-XM_002928913.4, rna-XM_034661790.1    rna-XM_038586469.1        rna-XM_036089276.1, rna-XM_036117508.1      rna-XM_032853363.1      rna-XM_032358478.1    rna-XM_059134938.1    rna-XM_059389156.1    rna-XM_045065873.1    mNeoVis.2.1.1.chr7_hap1.g01774.t01, mNeoVis.2.1.1.chr9_hap1.g00763.t01    rna-XM_044919573.1        rna-XM_040630739.1    rna-XM_035722959.1      rna-gnl|WGS_ZZZZ|egapxtmp_013872-R7    mNeoVis.evergladensis.2.1.chr10.g00754.t01, mNeoVis.evergladensis.2.1.chr7.g01735.t01    mNeoVis.vulgivaga.2.1.chr7.g01751.t01      638    N0.HOG0000636   OG0000045  n98     rna-XM_038538844.1                        mNeoVis.2.1.1.chr13_hap1.g00569.t01            rna-XM_035724744.1, rna-XM_035727791.1      rna-gnl|WGS_ZZZZ|egapxtmp_031609-R1    mNeoVis.evergladensis.2.1.chr13.g00553.t01    mNeoVis.vulgivaga.2.1.chr13.g00555.t01      639    N0.HOG0000637   OG0000046  n1   rna-XM_002922114.4    rna-NM_001253741.1    rna-XM_022509195.1    rna-XM_003996860.6    rna-XM_036069120.1    rna-XM_039256744.1    rna-XM_032871613.1    rna-XM_045985375.1    rna-XM_032320598.1    rna-XM_059149434.1    rna-XM_059380604.1    rna-XM_004772745.3    mNeoVis.2.1.1.chr8_hap1.g00410.t01    rna-XM_021681821.2    rna-XM_004410500.1    rna-XM_042966065.1    rna-XM_040640637.1    rna-XM_027568180.1    mNeoVis.3.1.1.chr8.g00374.t01    rna-gnl|WGS_ZZZZ|egapxtmp_019444-R1    mNeoVis.evergladensis.2.1.chr9.g00408.t01    mNeoVis.vulgivaga.2.1.chr8.g00415.t01      640    N0.HOG0000638   OG0000046  n22   rna-XM_002922161.4    rna-XM_038547352.1    rna-XM_022509196.1    rna-XM_003996861.6    rna-XM_036069121.1    rna-XM_039256762.1    rna-XM_032871622.1    rna-XM_045985411.1    rna-XM_032320593.1    rna-XM_059149441.1    rna-XM_059380700.1    rna-XM_045080686.1      rna-XM_021681718.1    rna-XM_004410604.1    rna-XM_042966115.1    rna-XM_040640638.1    rna-XM_027568179.1    mNeoVis.3.1.1.chr6.g00775.t01, mNeoVis.3.1.1.chr8.g00372.t01      mNeoVis.evergladensis.2.1.chr9.g00406.t01    mNeoVis.vulgivaga.2.1.chr8.g00414.t01      641    N0.HOG0000639   OG0000046  n43   rna-XM_034640778.1    rna-XM_038547353.1    rna-XM_022509197.1    rna-XM_003996862.5    rna-XM_036069122.1    rna-XM_039256763.1    rna-XM_032871617.1    rna-XM_045985369.1      rna-XM_059149433.1    rna-XM_059380488.1      mNeoVis.2.1.1.chr8_hap1.g00409.t01    rna-XM_021681820.2    rna-XM_004410501.1    rna-XM_042966116.1    rna-XM_040639878.1    rna-XM_027626146.2    mNeoVis.3.1.1.chr8.g00373.t01    rna-gnl|WGS_ZZZZ|egapxtmp_019794-R1    mNeoVis.evergladensis.2.1.chr9.g00407.t01    mNeoVis.vulgivaga.2.1.chr8.g00413.t01      642    N0.HOG0000640   OG0000046  n61   rna-XM_034640779.1    rna-XM_038547350.1    rna-XM_022509317.1    rna-XM_003996938.4    rna-XM_036069123.1    rna-XM_039256761.1    rna-XM_032871615.1    rna-XM_045985388.1    rna-XM_032320604.1    rna-XM_059149437.1    rna-XM_059380222.1    rna-XM_004772742.3    mNeoVis.2.1.1.chr8_hap1.g00408.t01    rna-XM_044921719.1    rna-XM_004410605.1    rna-XM_015540021.2    rna-XM_040639877.1    rna-XM_027626143.1          mNeoVis.vulgivaga.2.1.chr8.g00412.t01      643    N0.HOG0000641   OG0000046  n81   rna-XM_011228648.3    rna-NM_001253742.1, rna-XM_038556174.1      rna-XM_003996859.5    rna-XM_036069117.1    rna-XM_039256801.1    rna-XM_032871618.1, rna-XM_032877162.1    rna-XM_045985997.1    rna-XM_032320603.1    rna-XM_059149440.1    rna-XM_059378881.1    rna-XM_004772748.3    mNeoVis.2.1.1.chr8_hap1.g00412.t01    rna-XM_021681823.1    rna-XM_004410498.1    rna-XM_042967234.1    rna-XM_040639881.1    rna-XM_027567966.2    mNeoVis.3.1.1.chr8.g00376.t01    rna-gnl|WGS_ZZZZ|egapxtmp_019806-R1    mNeoVis.evergladensis.2.1.chr9.g00410.t01    mNeoVis.vulgivaga.2.1.chr8.g00417.t01      644    N0.HOG0000642   OG0000046  n102   rna-XM_034639721.1, rna-XM_034652617.1, rna-XM_034652696.1    rna-XM_038547368.1    rna-XM_022509128.1      rna-XM_036081479.1    rna-XM_039223296.1    rna-XM_032871586.1    rna-XM_045983810.1    rna-XM_032321082.1    rna-XM_059147938.1    rna-XM_059379542.1    rna-XM_045070153.1    mNeoVis.2.1.1.chr8_hap1.g00451.t01    rna-XM_021703908.2    rna-XM_004395002.1    rna-XM_042965594.1    rna-XM_040640646.1    rna-XM_027624838.2    mNeoVis.3.1.1.chr8.g00407.t01    rna-gnl|WGS_ZZZZ|egapxtmp_019792-R1    mNeoVis.evergladensis.2.1.chr9.g00442.t01    mNeoVis.vulgivaga.2.1.chr8.g00447.t01      645    N0.HOG0000643   OG0000046  n122   rna-XM_002922160.4    rna-XM_038546100.1    rna-XM_022509501.1    rna-XM_003996937.6    rna-XM_036069196.1    rna-XM_039256800.1    rna-XM_032871815.1    rna-XM_045986265.1    rna-XM_032319487.1    rna-XM_059149655.1    rna-XM_059380223.1    rna-XM_045080646.1    mNeoVis.2.1.1.chr8_hap1.g00411.t01      rna-XM_004410603.1    rna-XM_042966389.1    rna-XM_040639879.1    rna-XM_027624663.2    mNeoVis.3.1.1.chr8.g00375.t01    rna-gnl|WGS_ZZZZ|egapxtmp_019445-R1    mNeoVis.evergladensis.2.1.chr9.g00409.t01    mNeoVis.vulgivaga.2.1.chr8.g00416.t01      646    N0.HOG0000644   OG0000046  n143   rna-XM_019802703.2    rna-NM_001346369.1    rna-XM_022509222.1    rna-XM_019817881.3    rna-XM_036069113.1    rna-XM_039256767.1    rna-XM_032871607.1    rna-XM_045985413.1    rna-XM_032320602.1    rna-XM_059149436.1    rna-XM_059380820.1    rna-XM_045080684.1    mNeoVis.2.1.1.chr8_hap1.g00414.t01    rna-XM_021681824.1    rna-XM_004410497.1    rna-XM_007087706.3    rna-XM_040641222.1    rna-XM_027626227.2    mNeoVis.3.1.1.chr8.g00378.t01    rna-gnl|WGS_ZZZZ|egapxtmp_019818-R1    mNeoVis.evergladensis.2.1.chr9.g00412.t01    mNeoVis.vulgivaga.2.1.chr8.g00419.t01      647    N0.HOG0000645   OG0000046  n164   rna-XM_034641051.1    rna-NM_001253744.1    rna-XM_022509219.1    rna-XM_006940345.4    rna-XM_036069119.1    rna-XM_039256764.1    rna-XM_032871614.1    rna-XM_045985996.1    rna-XM_032320601.1    rna-XM_059140345.1, rna-XM_059149435.1    rna-XM_059378880.1    rna-XM_013063491.2, rna-XM_045070176.1, rna-XM_045080687.1    mNeoVis.2.1.1.chr8_hap1.g00413.t01, mNeoVis.2.1.1.chr9_hap1.g00811.t01    rna-XM_021681822.2    rna-XM_004410499.1    rna-XM_042966950.1    rna-XM_040639880.1    rna-XM_027567965.2    mNeoVis.3.1.1.chr10.g00776.t01, mNeoVis.3.1.1.chr3.g00392.t01, mNeoVis.3.1.1.chr8.g00377.t01    rna-gnl|WGS_ZZZZ|egapxtmp_019800-R1, rna-gnl|WGS_ZZZZ|egapxtmp_023763-R1    mNeoVis.evergladensis.2.1.chr10.g00801.t01, mNeoVis.evergladensis.2.1.chr9.g00411.t01    mNeoVis.vulgivaga.2.1.chr10.g00789.t01, mNeoVis.vulgivaga.2.1.chr8.g00418.t01      648    N0.HOG0000646   OG0000047  n3                           mNeoVis.2.1.1.chr4_hap1.g00412.t01              mNeoVis.3.1.1.chr10.g00664.t01      mNeoVis.evergladensis.2.1.chr6.g00415.t01    mNeoVis.vulgivaga.2.1.chr13.g00396.t01      649    N0.HOG0000647   OG0000047  n8   rna-XM_034652152.1            rna-XM_039224104.1            rna-XM_059409134.1    rna-XM_045065875.1                          650    N0.HOG0000648   OG0000047  n12                   rna-XM_032347709.1    rna-XM_059185715.1    rna-XM_059402842.1    rna-NM_001310188.1                          651    N0.HOG0000649   OG0000047  n16       rna-XM_022524914.1            rna-XM_046013448.1                                  652    N0.HOG0000650   OG0000047  n21   rna-XM_034652654.1    rna-XM_038526101.1        rna-XM_036113549.1, rna-XM_036117138.1, rna-XM_036119206.1                    rna-XM_021690600.1, rna-XM_044918728.1, rna-XM_044920186.1    rna-XM_004405498.1, rna-XM_004407105.1, rna-XM_004412399.1      rna-XM_040645364.1    rna-XM_027594944.2, rna-XM_027612532.2, rna-XM_027618471.2, rna-XM_035723303.1, rna-XM_035723310.1, rna-XM_035728562.1, rna-XM_035728720.1              653    N0.HOG0000651   OG0000047  n39   rna-XM_002928987.4    rna-NM_001252164.1                                rna-XM_008693114.2    rna-XM_027605785.2              654    N0.HOG0000652   OG0000047  n42         rna-XM_003988487.4                          rna-XM_007084046.2                  655    N0.HOG0000653   OG0000047  n43             rna-XM_039221223.1, rna-XM_039238576.1    rna-XM_032855308.1, rna-XM_032869859.1                                    656    N0.HOG0000654   OG0000047  n46               rna-XM_032859386.1              mNeoVis.2.1.1.chr9_hap1.g00074.t01, mNeoVis.2.1.1.chr9_hap1.g00700.t01        rna-XM_015535400.2        mNeoVis.3.1.1.chr11.g00399.t01    rna-gnl|WGS_ZZZZ|egapxtmp_003222-R1, rna-gnl|WGS_ZZZZ|egapxtmp_021794-R1    mNeoVis.evergladensis.2.1.chr10.g00072.t01, mNeoVis.evergladensis.2.1.chr10.g00693.t01    mNeoVis.vulgivaga.2.1.chr10.g00681.t01      657    N0.HOG0000655   OG0000047  n53                     rna-XM_059158328.1          rna-XM_021698298.1        rna-XM_008705354.2                658    N0.HOG0000656   OG0000047  n59   rna-XM_011233500.3    rna-NM_001271792.1    rna-XM_022506696.1    rna-XM_011286740.4    rna-XM_036077581.1    rna-XM_039236814.1    rna-XM_032881170.1    rna-XM_046016100.1    rna-XM_032358578.1    rna-XM_059136575.1, rna-XM_059137815.1    rna-XM_059377352.1, rna-XM_059388710.1    rna-XM_004775165.2, rna-XM_045066149.1    mNeoVis.2.1.1.chr7_hap1.g01866.t01    rna-XM_021685325.2    rna-XM_004415661.1    rna-XM_007098438.3    rna-XM_008686923.1    rna-XM_027579359.1    mNeoVis.3.1.1.chr7.g01734.t01    rna-gnl|WGS_ZZZZ|egapxtmp_009804-R2    mNeoVis.evergladensis.2.1.chr7.g01828.t01        659    N0.HOG0000657   OG0000047  n81   rna-XM_002915119.4, rna-XM_011233514.3    rna-XM_038526422.1    rna-XM_022507213.1, rna-XM_022520894.1    rna-XM_045039046.1, rna-XM_045049904.1, rna-XM_045058288.1    rna-XM_036084195.1    rna-XM_039234114.1    rna-XM_032836432.1, rna-XM_032869091.1    rna-XM_046004934.1    rna-XM_032316201.1, rna-XM_032338630.1    rna-XM_059176671.1    rna-XM_059399291.1    rna-XM_013044968.2, rna-XM_045066148.1    mNeoVis.2.1.1.chr1_hap1.g02097.t01    rna-XM_044917287.1    rna-XM_004403889.1, rna-XM_004415660.1    rna-XM_042987389.1    rna-XM_008688522.2, rna-XM_040625787.1, rna-XM_040638883.1    rna-XM_027579574.1, rna-XM_027601969.2, rna-XM_027622357.2, rna-XM_035727424.1    mNeoVis.3.1.1.chr1.g02052.t01, mNeoVis.3.1.1.chr6.g00851.t01    rna-gnl|WGS_ZZZZ|egapxtmp_002969-R1, rna-gnl|WGS_ZZZZ|egapxtmp_035515-R1    mNeoVis.evergladensis.2.1.chr1.g02067.t01, mNeoVis.evergladensis.2.1.chr6.g00846.t01    mNeoVis.vulgivaga.2.1.chr1.g02060.t01, mNeoVis.vulgivaga.2.1.chr6.g00841.t01      660    N0.HOG0000658   OG0000047  n121               rna-XM_032855364.1, rna-XM_032860314.1, rna-XM_032866630.1                                    661    N0.HOG0000659   OG0000047  n124                           mNeoVis.2.1.1.chr5_hap1.g00884.t01              mNeoVis.3.1.1.chr9.g00937.t01            662    N0.HOG0000660   OG0000047  n126       rna-XM_022518541.1            rna-XM_045978638.1                        mNeoVis.3.1.1.chr4.g00248.t01    rna-gnl|WGS_ZZZZ|egapxtmp_043953-R1      mNeoVis.vulgivaga.2.1.chr4.g00255.t01      663    N0.HOG0000661   OG0000047  n132                 rna-XM_045984895.1    rna-XM_032324526.1                                664    N0.HOG0000662   OG0000047  n134                   rna-XM_032315023.1, rna-XM_032358574.1    rna-XM_059137308.1, rna-XM_059137837.1, rna-XM_059158195.1    rna-XM_059385995.1, rna-XM_059407453.1    rna-XM_013050082.2, rna-XM_045065780.1                          665    N0.HOG0000663   OG0000047  n146   rna-XM_034651363.1, rna-XM_034654505.1    rna-XM_038544034.1, rna-XM_038569365.1, rna-XM_038572995.1, rna-XM_038573059.1, rna-XM_038579855.1      rna-XM_019812152.3    rna-XM_036092554.1, rna-XM_036108852.1, rna-XM_036121659.1    rna-XM_039236747.1                  rna-XM_021701226.1, rna-XM_044914532.1, rna-XM_044919055.1, rna-XM_044919488.1, rna-XM_044920270.1    rna-XM_004413550.1    rna-XM_042958262.1, rna-XM_042962568.1      rna-XM_027574892.1, rna-XM_027590404.2, rna-XM_027590694.1, rna-XM_027599084.2              666    N0.HOG0000664   OG0000047  n169   rna-XM_034645235.1    rna-XM_038569203.1                            rna-XM_004400253.1, rna-XM_004409845.1, rna-XM_004415553.1, rna-XM_004415664.1, rna-XM_004415952.1      rna-XM_008686924.2    rna-XM_027576408.2, rna-XM_027579937.2, rna-XM_027607476.2, rna-XM_035721833.1, rna-XM_035722329.1, rna-XM_035723884.1, rna-XM_035727205.1, rna-XM_035729360.1              667    N0.HOG0000665   OG0000047  n184                 rna-XM_046003325.1, rna-XM_046016181.1, rna-XM_046021507.1                                  668    N0.HOG0000666   OG0000047  n186               rna-XM_032881169.1    rna-XM_045988308.1, rna-XM_045988310.1            mNeoVis.2.1.1.chr2_hap1.g00942.t01                    mNeoVis.vulgivaga.2.1.chr2.g00886.t01      669    N0.HOG0000667   OG0000047  n190   rna-XM_034648283.1                          mNeoVis.2.1.1.chr7_hap1.g01865.t01                rna-gnl|WGS_ZZZZ|egapxtmp_009803-R1      mNeoVis.vulgivaga.2.1.chr7.g01844.t01      670    N0.HOG0000668   OG0000047  n192   rna-XM_034657317.1                rna-XM_045997200.1                      rna-XM_035724890.1              671    N0.HOG0000669   OG0000048  n3   rna-XM_002930716.3    rna-NM_001388648.1    rna-XM_022492652.1, rna-XM_022492723.1      rna-XM_036089095.1, rna-XM_036089110.1      rna-XM_032867518.1, rna-XM_032867524.1, rna-XM_032867526.1      rna-XM_032314398.1    rna-XM_059160328.1, rna-XM_059165159.1, rna-XM_059165160.1, rna-XM_059165161.1    rna-XM_059387770.1, rna-XM_059387771.1, rna-XM_059387772.1    rna-XM_004761185.1, rna-XM_004761186.1, rna-XM_045082350.1    mNeoVis.2.1.1.chr5_hap1.g00444.t01, mNeoVis.2.1.1.chr5_hap1.g00445.t01, mNeoVis.2.1.1.chr5_hap1.g00446.t01    rna-XM_021683166.1    rna-XM_004414878.1      rna-XM_040628171.1    rna-XM_027575122.2, rna-XM_035722618.1    mNeoVis.3.1.1.chr2.g00370.t01, mNeoVis.3.1.1.chr2.g00371.t01    rna-gnl|WGS_ZZZZ|egapxtmp_034879-R1, rna-gnl|WGS_ZZZZ|egapxtmp_034882-R1, rna-gnl|WGS_ZZZZ|egapxtmp_034888-R1    mNeoVis.evergladensis.2.1.chr2.g00439.t01, mNeoVis.evergladensis.2.1.chr2.g00440.t01, mNeoVis.evergladensis.2.1.chr2.g00441.t01    mNeoVis.vulgivaga.2.1.chr2.g00441.t01, mNeoVis.vulgivaga.2.1.chr2.g00442.t01, mNeoVis.vulgivaga.2.1.chr2.g00443.t01      672    N0.HOG0000670   OG0000048  n37   rna-XM_002930605.4    rna-NM_001388652.1    rna-XM_022498426.1      rna-XM_036089111.1      rna-XM_032851063.1    rna-XM_045991814.1    rna-XM_032314314.1    rna-XM_059165152.1    rna-XM_059387767.1    rna-XM_004761107.3    mNeoVis.2.1.1.chr5_hap1.g00437.t01    rna-XM_021683167.1    rna-XM_004408383.1      rna-XM_008706408.1    rna-XM_035722667.1    mNeoVis.3.1.1.chr2.g00366.t01    rna-gnl|WGS_ZZZZ|egapxtmp_034880-R1    mNeoVis.evergladensis.2.1.chr2.g00434.t01    mNeoVis.vulgivaga.2.1.chr2.g00436.t01      673    N0.HOG0000671   OG0000048  n54         rna-XM_023241386.2, rna-XM_045041614.1      rna-XM_039243419.1, rna-XM_039243423.1, rna-XM_039243426.1                      rna-XM_007088328.2, rna-XM_007088623.2                  674    N0.HOG0000672   OG0000048  n61   rna-XM_002930606.4    rna-XM_038565308.1    rna-XM_022498425.1    rna-XM_045041609.1    rna-XM_036089112.1    rna-XM_039243425.1    rna-XM_032868483.1    rna-XM_045991813.1    rna-XM_032314326.1    rna-XM_059165153.1    rna-XM_059387765.1    rna-XM_004761108.1    mNeoVis.2.1.1.chr5_hap1.g00436.t01    rna-XM_021683168.1    rna-XM_004408382.1    rna-XM_042977323.1    rna-XM_040629128.1    rna-XM_035722609.1    mNeoVis.3.1.1.chr2.g00365.t01    rna-gnl|WGS_ZZZZ|egapxtmp_034885-R1      mNeoVis.vulgivaga.2.1.chr2.g00435.t01      675    N0.HOG0000673   OG0000048  n82       rna-XM_022492671.1, rna-XM_022497961.1      rna-XM_036089063.1                    rna-XM_044917634.1        rna-XM_008706433.2    rna-XM_027586287.1              676    N0.HOG0000674   OG0000048  n88       rna-XM_022497962.1            rna-XM_045991748.1    rna-XM_032314379.1    rna-XM_059165155.1      rna-XM_013060478.2    mNeoVis.2.1.1.chr5_hap1.g00440.t01                rna-gnl|WGS_ZZZZ|egapxtmp_034895-R1    mNeoVis.evergladensis.2.1.chr2.g00435.t01    mNeoVis.vulgivaga.2.1.chr2.g00437.t01      677    N0.HOG0000675   OG0000048  n97   rna-XM_002930608.4, rna-XM_002930713.4                                  rna-XM_008706407.1, rna-XM_008709976.2                678    N0.HOG0000676   OG0000048  n101   rna-XM_034657195.1      rna-XM_022498434.1      rna-XM_036089059.1      rna-XM_032868509.1    rna-XM_045991746.1, rna-XM_045991747.1            mNeoVis.2.1.1.chr5_hap1.g00442.t01      rna-XM_004408397.1      rna-XM_008709975.2    rna-XM_035722683.1    mNeoVis.3.1.1.chr2.g00368.t01    rna-gnl|WGS_ZZZZ|egapxtmp_034894-R1    mNeoVis.evergladensis.2.1.chr2.g00437.t01    mNeoVis.vulgivaga.2.1.chr2.g00439.t01      679    N0.HOG0000677   OG0000048  n114           rna-XM_036089087.1                      rna-XM_004408395.1        rna-XM_027586214.2              680    N0.HOG0000678   OG0000048  n117   rna-XM_034657197.1    rna-NM_001388650.1, rna-NM_001388927.1    rna-XM_022492696.1    rna-XM_003982014.2, rna-XM_003982015.3, rna-XM_023241202.2, rna-XM_045041619.1, rna-XM_045041622.1, rna-XM_045041644.1    rna-XM_036089057.1    rna-XM_039243398.1, rna-XM_039243424.1, rna-XM_039243427.1, rna-XM_039243429.1, rna-XM_039243430.1    rna-XM_032868511.1    rna-XM_045991750.1, rna-XM_045991752.1, rna-XM_045991846.1    rna-XM_032314365.1    rna-XM_059165158.1    rna-XM_059387769.1    rna-XM_013060464.2    mNeoVis.2.1.1.chr5_hap1.g00443.t01      rna-XM_004414877.1    rna-XM_007088329.1, rna-XM_007088617.3, rna-XM_007088619.3, rna-XM_007088622.2, rna-XM_015540316.1, rna-XM_015540380.2    rna-XM_008706404.2    rna-XM_027586190.2    mNeoVis.3.1.1.chr2.g00369.t01    rna-gnl|WGS_ZZZZ|egapxtmp_034893-R1    mNeoVis.evergladensis.2.1.chr2.g00438.t01    mNeoVis.vulgivaga.2.1.chr2.g00440.t01      681    N0.HOG0000679   OG0000048  n153   rna-XM_002930607.4, rna-XM_019793538.2, rna-XM_034653338.1, rna-XM_034657196.1    rna-NM_001388928.1, rna-NM_001389200.1, rna-XM_038565306.1, rna-XM_038565307.1    rna-XM_022492763.1    rna-XM_045041631.1          rna-XM_045991744.1    rna-XM_032314339.1, rna-XM_032314353.1    rna-XM_059165154.1    rna-XM_059387397.1    rna-XM_004782558.3    mNeoVis.2.1.1.chr5_hap1.g00438.t01, mNeoVis.2.1.1.chr5_hap1.g00439.t01      rna-XM_004408398.1    rna-XM_015540315.1    rna-XM_008706405.2, rna-XM_008706406.2, rna-XM_008709973.2    rna-XM_035722715.1              682    N0.HOG0000680   OG0000048  n176       rna-XM_022498433.1      rna-XM_036089058.1, rna-XM_036089060.1      rna-XM_032868510.1    rna-XM_045991749.1, rna-XM_045991751.1    rna-XM_032314388.1    rna-XM_059165157.1    rna-XM_059387768.1    rna-XM_013060463.2    mNeoVis.2.1.1.chr5_hap1.g00441.t01      rna-XM_004408396.1        rna-XM_027586232.2    mNeoVis.3.1.1.chr2.g00367.t01    rna-gnl|WGS_ZZZZ|egapxtmp_034892-R1    mNeoVis.evergladensis.2.1.chr2.g00436.t01    mNeoVis.vulgivaga.2.1.chr2.g00438.t01      683    N0.HOG0000681   OG0000049  n2   rna-XM_034647954.1, rna-XM_034660420.1, rna-XM_034670040.1    rna-NM_001197116.2    rna-XM_022510968.1    rna-XM_006931840.5    rna-XM_036067704.1    rna-XM_039234028.1, rna-XM_039248918.1    rna-XM_032864360.1, rna-XM_032865368.1, rna-XM_032869159.1, rna-XM_032875112.1, rna-XM_032880940.1    rna-XM_046005737.1, rna-XM_046009229.1, rna-XM_046013349.1, rna-XM_046017025.1    rna-XM_032340475.1    rna-XM_059178463.1    rna-XM_059400150.1    rna-XM_045074789.1    mNeoVis.2.1.1.chr1_hap1.g01488.t01    rna-XM_021693084.2      rna-XM_007090112.3    rna-XM_040643670.1    rna-XM_035726674.1, rna-XM_035727426.1              684    N0.HOG0000682   OG0000049  n3   rna-XM_034650077.1, rna-XM_034651468.1    rna-XM_038538178.1    rna-XM_022514918.1      rna-XM_036115221.1    rna-XM_039225110.1, rna-XM_039254272.1    rna-XM_032851145.1, rna-XM_032853943.1    rna-XM_046025550.1, rna-XM_046025867.1    rna-XM_032321573.1, rna-XM_032331763.1, rna-XM_032334453.1, rna-XM_032338954.1, rna-XM_032350919.1, rna-XM_032359048.1    rna-XM_059158473.1, rna-XM_059168954.1, rna-XM_059174378.1, rna-XM_059178137.1    rna-XM_059376167.1, rna-XM_059408919.1    rna-XM_045077962.1, rna-XM_045079926.1    mNeoVis.2.1.1.chr8_hap1.g00600.t01    rna-XM_044915348.1, rna-XM_044916576.1, rna-XM_044917139.1    rna-XM_004414434.1    rna-XM_042970846.1      rna-XM_035723924.1, rna-XM_035726883.1, rna-XM_035726923.1, rna-XM_035728419.1, rna-XM_035728565.1, rna-XM_035728571.1, rna-XM_035728697.1    mNeoVis.3.1.1.chr4.g00246.t01    rna-gnl|WGS_ZZZZ|egapxtmp_013694-R1, rna-gnl|WGS_ZZZZ|egapxtmp_017448-R1, rna-gnl|WGS_ZZZZ|egapxtmp_021837-R1    mNeoVis.evergladensis.2.1.chr10.g00146.t01    mNeoVis.vulgivaga.2.1.chr1.g01451.t01, mNeoVis.vulgivaga.2.1.chr4.g00253.t01, mNeoVis.vulgivaga.2.1.chr5.g00828.t01, mNeoVis.vulgivaga.2.1.chr6.g00684.t01, mNeoVis.vulgivaga.2.1.chr7.g01403.t01      685    N0.HOG0000683   OG0000049  n52                     rna-XM_059175419.1                  rna-XM_035728583.1              686    N0.HOG0000684   OG0000049  n55               rna-XM_032851055.1, rna-XM_032851056.1, rna-XM_032851149.1                                    687    N0.HOG0000685   OG0000049  n59   rna-XM_034641559.1, rna-XM_034652147.1, rna-XM_034652439.1, rna-XM_034652879.1, rna-XM_034662592.1    rna-XM_038557376.1      rna-XM_045037852.1, rna-XM_045056066.1    rna-XM_036100643.1    rna-XM_039222051.1, rna-XM_039238560.1    rna-XM_032839820.1, rna-XM_032846536.1, rna-XM_032851233.1, rna-XM_032868279.1, rna-XM_032868981.1    rna-XM_045990081.1, rna-XM_045994767.1, rna-XM_045997452.1, rna-XM_046001169.1, rna-XM_046004013.1    rna-XM_032307679.1    rna-XM_059179144.1    rna-XM_059401608.1    rna-XM_045067018.1    mNeoVis.2.1.1.chr4_hap1.g00681.t01, mNeoVis.2.1.1.chr4_hap1.g00682.t01, mNeoVis.2.1.1.chr5_hap1.g01172.t01, mNeoVis.2.1.1.chr8_hap1.g01148.t01    rna-XM_044913486.1, rna-XM_044915207.1, rna-XM_044915526.1, rna-XM_044918999.1    rna-XM_004396883.1, rna-XM_004401272.1, rna-XM_004405648.1, rna-XM_004407125.1, rna-XM_004409573.1, rna-XM_004417657.1, rna-XM_004417664.1      rna-XM_008689818.2, rna-XM_008697945.2, rna-XM_040620862.1, rna-XM_040625233.1, rna-XM_040625449.1, rna-XM_040633733.1    rna-XM_035727699.1    mNeoVis.3.1.1.chr1.g01453.t01, mNeoVis.3.1.1.chr10.g00296.t01, mNeoVis.3.1.1.chr6.g00689.t01      mNeoVis.evergladensis.2.1.chr2.g01162.t01    mNeoVis.vulgivaga.2.1.chr1.g00108.t01, mNeoVis.vulgivaga.2.1.chr10.g00299.t01, mNeoVis.vulgivaga.2.1.chr11.g00106.t01, mNeoVis.vulgivaga.2.1.chr13.g00669.t01, mNeoVis.vulgivaga.2.1.chr2.g01169.t01      688    N0.HOG0000686   OG0000049  n114     rna-XM_038535535.1          rna-XM_039223892.1                          rna-XM_035728576.1              689    N0.HOG0000687   OG0000049  n119   rna-XM_034650499.1    rna-XM_038572871.1    rna-XM_022493858.1    rna-XM_023250968.2    rna-XM_036085031.1    rna-XM_039236923.1    rna-XM_032878782.1    rna-XM_045981247.1    rna-XM_032353556.1    rna-XM_059133028.1    rna-XM_059407420.1    rna-XM_045070689.1    mNeoVis.2.1.1.chr11_hap1.g00032.t01    rna-XM_044918868.1    rna-XM_004410119.1    rna-XM_042979939.1    rna-XM_040637861.1    rna-XM_027623482.1        mNeoVis.evergladensis.2.1.chr11.g00034.t01    mNeoVis.vulgivaga.2.1.chr11.g00032.t01      690    N0.HOG0000688   OG0000049  n128   rna-XM_002913271.4, rna-XM_034652222.1, rna-XM_034652899.1    rna-XM_038531629.1      rna-XM_045035282.1, rna-XM_045038084.1    rna-XM_036089021.1    rna-XM_039222361.1, rna-XM_039231933.1            rna-XM_059387369.1        rna-XM_021683543.1, rna-XM_044915351.1    rna-XM_004394765.1, rna-XM_004403874.1, rna-XM_004413092.1    rna-XM_007083696.3, rna-XM_042958077.1    rna-XM_008694616.1    rna-XM_027575522.2, rna-XM_035729027.1    mNeoVis.3.1.1.chr1.g00115.t01      mNeoVis.evergladensis.2.1.chr1.g01848.t01    mNeoVis.vulgivaga.2.1.chr3.g00416.t01, mNeoVis.vulgivaga.2.1.chr8.g00269.t01      691    N0.HOG0000689   OG0000049  n151               rna-XM_032874117.1    rna-XM_046028123.1          rna-XM_045083048.1    mNeoVis.2.1.1.chr8_hap1.g01327.t01    rna-XM_044911311.1        rna-XM_040643209.1    rna-XM_035728614.1    mNeoVis.3.1.1.chr1.g01431.t01            692    N0.HOG0000690   OG0000050  n1   rna-XM_034645094.1    rna-NM_001388582.1    rna-XM_022507436.1    rna-XM_023239023.2    rna-XM_036076602.1    rna-XM_039245089.1    rna-XM_032849702.1    rna-XM_046016818.1    rna-XM_032358738.1    rna-XM_059158982.1    rna-XM_059387611.1    rna-XM_004777671.3    mNeoVis.2.1.1.chr7_hap1.g02205.t01    rna-XM_021685223.2    rna-XM_004409318.1    rna-XM_015542260.2    rna-XM_008694165.2    rna-XM_027577849.1    mNeoVis.3.1.1.chr7.g02028.t01    rna-gnl|WGS_ZZZZ|egapxtmp_013474-R1    mNeoVis.evergladensis.2.1.chr7.g02154.t01    mNeoVis.vulgivaga.2.1.chr7.g02176.t01      693    N0.HOG0000691   OG0000050  n22   rna-XM_002928225.4    rna-NM_001388583.1    rna-XM_022507219.1    rna-XM_003993400.3    rna-XM_036076603.1    rna-XM_039245069.1    rna-XM_032849697.1    rna-XM_046017043.1    rna-XM_032359314.1    rna-XM_059158981.1    rna-XM_059387612.1    rna-XM_004777613.2      rna-XM_021684865.1    rna-XM_004409317.1    rna-XM_007092640.2    rna-XM_008694282.2    rna-XM_027580272.1    mNeoVis.3.1.1.chr7.g02027.t01    rna-gnl|WGS_ZZZZ|egapxtmp_013550-R1          694    N0.HOG0000692   OG0000050  n42   rna-XM_002928232.4    rna-NM_001388584.1    rna-XM_022506625.1    rna-XM_003993461.3        rna-XM_032849718.1    rna-XM_046016526.1    rna-XM_032359897.1    rna-XM_059158980.1    rna-XM_059387613.1    rna-XM_004777670.2    mNeoVis.2.1.1.chr7_hap1.g02204.t01        rna-XM_007092673.2    rna-XM_008694281.2    rna-XM_027577850.1    mNeoVis.3.1.1.chr7.g02026.t01    rna-gnl|WGS_ZZZZ|egapxtmp_012650-R1    mNeoVis.evergladensis.2.1.chr7.g02153.t01    mNeoVis.vulgivaga.2.1.chr7.g02173.t01      695    N0.HOG0000693   OG0000050  n58   rna-XM_011229262.2    rna-NM_001388588.1    rna-XM_022507155.1    rna-XM_003993456.2      rna-XM_039245066.1    rna-XM_032849698.1    rna-XM_046016524.1    rna-XM_032359380.1    rna-XM_059158971.1    rna-XM_059387679.1    rna-XM_004777663.2    mNeoVis.2.1.1.chr7_hap1.g02199.t01    rna-XM_021684862.1      rna-XM_007092668.1    rna-XM_008694276.1      mNeoVis.3.1.1.chr7.g02022.t01    rna-gnl|WGS_ZZZZ|egapxtmp_013483-R1    mNeoVis.evergladensis.2.1.chr7.g02148.t01    mNeoVis.vulgivaga.2.1.chr7.g02171.t01      696    N0.HOG0000694   OG0000050  n77   rna-XM_002930995.2            rna-XM_039245048.1      rna-XM_046016553.1                    rna-XM_008694279.2      mNeoVis.3.1.1.chr7.g02023.t01            697    N0.HOG0000695   OG0000050  n82   rna-XM_002930993.3    rna-NM_001389168.1    rna-XM_022507161.1    rna-XM_045039333.1      rna-XM_039245047.1    rna-XM_032849699.1    rna-XM_046016561.1    rna-XM_032359475.1    rna-XM_059158977.1              rna-XM_007092650.2    rna-XM_008694162.2    rna-XM_027580148.1              698    N0.HOG0000696   OG0000050  n95   rna-XM_002930997.4    rna-NM_001389167.1, rna-NM_001389185.1      rna-XM_003993458.3, rna-XM_003993460.3      rna-XM_039245052.1, rna-XM_039245071.1      rna-XM_046016554.1, rna-XM_046016555.1, rna-XM_046016556.1    rna-XM_032359423.1, rna-XM_032359482.1    rna-XM_059158974.1, rna-XM_059158979.1    rna-XM_059387616.1, rna-XM_059387652.1    rna-XM_004777666.1, rna-XM_004777669.1    mNeoVis.2.1.1.chr7_hap1.g02201.t01, mNeoVis.2.1.1.chr7_hap1.g02203.t01      rna-XM_004409316.1    rna-XM_007092671.2, rna-XM_007092672.2    rna-XM_008694161.2, rna-XM_008694163.2    rna-XM_027580132.1    mNeoVis.3.1.1.chr7.g02025.t01    rna-gnl|WGS_ZZZZ|egapxtmp_013506-R1, rna-gnl|WGS_ZZZZ|egapxtmp_013569-R1    mNeoVis.evergladensis.2.1.chr7.g02150.t01, mNeoVis.evergladensis.2.1.chr7.g02152.t01    mNeoVis.vulgivaga.2.1.chr7.g02172.t01, mNeoVis.vulgivaga.2.1.chr7.g02175.t01      699    N0.HOG0000697   OG0000050  n127   rna-XM_002922702.2    rna-NM_001388586.1, rna-NM_001388587.1    rna-XM_022507157.1    rna-XM_045038160.1      rna-XM_039245072.1      rna-XM_046016602.1              rna-XM_021684864.1    rna-XM_004409314.1, rna-XM_004409315.1    rna-XM_007092669.2    rna-XM_008694278.2    rna-XM_027580135.2              700    N0.HOG0000698   OG0000050  n139   rna-XM_002930994.4, rna-XM_002930996.1    rna-NM_001388585.1    rna-XM_022507160.1, rna-XM_022507223.1    rna-XM_011286947.1      rna-XM_039245046.1    rna-XM_032849700.1    rna-XM_046016544.1    rna-XM_032359896.1    rna-XM_059158975.1, rna-XM_059158978.1    rna-XM_059387627.1, rna-XM_059387644.1    rna-XM_004777667.1, rna-XM_004777668.1    mNeoVis.2.1.1.chr7_hap1.g02202.t01        rna-XM_007092651.1    rna-XM_008694280.1    rna-XM_027579781.1    mNeoVis.3.1.1.chr7.g02024.t01    rna-gnl|WGS_ZZZZ|egapxtmp_013494-R1    mNeoVis.evergladensis.2.1.chr7.g02151.t01    mNeoVis.vulgivaga.2.1.chr7.g02174.t01      701    N0.HOG0000699   OG0000050  n159   rna-XM_002928233.2, rna-XM_034651828.1    rna-NM_001389181.1    rna-XM_022506626.1, rna-XM_022506627.1    rna-XM_003993399.5, rna-XM_003993457.5      rna-XM_039245034.1    rna-XM_032849719.1, rna-XM_032849720.1    rna-XM_046015779.1, rna-XM_046016770.1    rna-XM_032359425.1    rna-XM_059158972.1, rna-XM_059158973.1    rna-XM_059387660.1, rna-XM_059387670.1    rna-XM_004777664.2, rna-XM_004777665.1    mNeoVis.2.1.1.chr7_hap1.g02198.t01, mNeoVis.2.1.1.chr7_hap1.g02200.t01        rna-XM_007092670.2, rna-XM_015542258.2    rna-XM_008694160.2, rna-XM_040638727.1        rna-gnl|WGS_ZZZZ|egapxtmp_012648-R1    mNeoVis.evergladensis.2.1.chr7.g02149.t01    mNeoVis.vulgivaga.2.1.chr7.g02170.t01      702    N0.HOG0000700   OG0000051  n1   rna-XM_034639619.1    rna-XM_038528155.1    rna-XM_022524188.1    rna-XM_011289698.4    rna-XM_036068148.1    rna-XM_039245068.1    rna-XM_032839039.1    rna-XM_045988619.1    rna-XM_032326040.1    rna-XM_059151055.1    rna-XM_059382232.1    rna-XM_004767237.3    mNeoVis.2.1.1.chr7_hap1.g00868.t01    rna-XM_044911356.1    rna-XM_004409964.1    rna-XM_042970159.1    rna-XM_008684564.2    rna-XM_027617294.2    mNeoVis.3.1.1.chr7.g00808.t01    rna-gnl|WGS_ZZZZ|egapxtmp_013651-R1    mNeoVis.evergladensis.2.1.chr7.g00861.t01    mNeoVis.vulgivaga.2.1.chr7.g00866.t01      703    N0.HOG0000701   OG0000051  n23   rna-XM_002927927.4, rna-XM_034639142.1    rna-XM_038527963.1, rna-XM_038527994.1    rna-XM_022523767.1    rna-XM_023245219.2, rna-XM_023245222.2    rna-XM_036104115.1, rna-XM_036104119.1    rna-XM_039220160.1, rna-XM_039220176.1, rna-XM_039249136.1    rna-XM_032839430.1, rna-XM_032850989.1, rna-XM_032850990.1    rna-XM_045988240.1, rna-XM_045988288.1    rna-XM_032324021.1, rna-XM_032324051.1    rna-XM_059151053.1, rna-XM_059151084.1    rna-XM_059382658.1    rna-XM_045077394.1, rna-XM_045077408.1, rna-XM_045077441.1    mNeoVis.2.1.1.chr7_hap1.g00809.t01    rna-XM_021681401.2, rna-XM_044911385.1    rna-XM_004415486.1, rna-XM_004415490.1    rna-XM_042970351.1, rna-XM_042970362.1    rna-XM_008684805.2, rna-XM_040632961.1    rna-XM_035725246.1, rna-XM_035725248.1    mNeoVis.3.1.1.chr7.g00754.t01, mNeoVis.3.1.1.chr7.g00755.t01      mNeoVis.evergladensis.2.1.chr7.g00798.t01    mNeoVis.vulgivaga.2.1.chr7.g00803.t01, mNeoVis.vulgivaga.2.1.chr7.g00804.t01      704    N0.HOG0000702   OG0000051  n64   rna-XM_034639075.1, rna-XM_034639130.1    rna-XM_038527998.1    rna-XM_022524010.1, rna-XM_022524122.1    rna-XM_011289724.4, rna-XM_045046233.1    rna-XM_036104052.1    rna-XM_039249149.1, rna-XM_039250683.1    rna-XM_032839402.1, rna-XM_032839412.1    rna-XM_045988227.1, rna-XM_045988265.1    rna-XM_032323999.1, rna-XM_032324030.1    rna-XM_059151031.1, rna-XM_059151058.1    rna-XM_059382327.1, rna-XM_059383947.1    rna-XM_013064054.2, rna-XM_013064115.2    mNeoVis.2.1.1.chr7_hap1.g00813.t01, mNeoVis.2.1.1.chr7_hap1.g00814.t01    rna-XM_021681106.1, rna-XM_021681448.2    rna-XM_004409961.1, rna-XM_004409976.1    rna-XM_042970334.1, rna-XM_042970352.1    rna-XM_040632926.1, rna-XM_040632939.1    rna-XM_035724850.1, rna-XM_035725098.1    mNeoVis.3.1.1.chr6.g01057.t01    rna-gnl|WGS_ZZZZ|egapxtmp_013152-R2    mNeoVis.evergladensis.2.1.chr7.g00802.t01, mNeoVis.evergladensis.2.1.chr7.g00803.t01    mNeoVis.vulgivaga.2.1.chr7.g00807.t01, mNeoVis.vulgivaga.2.1.chr7.g00809.t01      705    N0.HOG0000703   OG0000051  n105   rna-XM_011237874.3    rna-XM_038525539.1    rna-XM_022523862.1    rna-XM_023245546.2, rna-XM_045046309.1    rna-XM_036104117.1    rna-XM_039246944.1    rna-XM_032839005.1    rna-XM_045988381.1    rna-XM_032324990.1    rna-XM_059151060.1    rna-XM_059383477.1    rna-XM_013064129.2    mNeoVis.2.1.1.chr7_hap1.g00827.t01    rna-XM_021681399.1    rna-XM_004409972.1    rna-XM_042970356.1, rna-XM_042970378.1    rna-XM_008684720.2    rna-XM_027619934.1    mNeoVis.3.1.1.chr7.g00771.t01      mNeoVis.evergladensis.2.1.chr7.g00816.t01    mNeoVis.vulgivaga.2.1.chr7.g00822.t01      706    N0.HOG0000704   OG0000051  n128   rna-XM_034639127.1, rna-XM_034639185.1    rna-XM_038528006.1    rna-XM_022523861.1, rna-XM_022524011.1    rna-XM_006940936.5    rna-XM_036104101.1    rna-XM_039249855.1, rna-XM_039250691.1    rna-XM_032838976.1, rna-XM_032839422.1    rna-XM_045988252.1, rna-XM_045988305.1    rna-XM_032324989.1    rna-XM_059151056.1, rna-XM_059151087.1    rna-XM_059383371.1, rna-XM_059383476.1    rna-XM_004767148.3, rna-XM_045077424.1        rna-XM_004409973.1    rna-XM_007074205.3, rna-XM_007074217.3    rna-XM_008684718.2    rna-XM_027619933.2, rna-XM_035725186.1    mNeoVis.3.1.1.chr3.g01196.t01, mNeoVis.3.1.1.chr7.g00768.t01    rna-gnl|WGS_ZZZZ|egapxtmp_010596-R1    mNeoVis.evergladensis.2.1.chr7.g00813.t01    mNeoVis.vulgivaga.2.1.chr7.g00808.t01      707    N0.HOG0000705   OG0000051  n158   rna-XM_002927928.4, rna-XM_034639146.1    rna-XM_038528027.1    rna-XM_022523863.1      rna-XM_036104127.1, rna-XM_036104131.1    rna-XM_039246920.1, rna-XM_039249864.1    rna-XM_032838978.1, rna-XM_032838981.1    rna-XM_045988298.1, rna-XM_045988325.1    rna-XM_032324997.1, rna-XM_032324999.1    rna-XM_059151093.1, rna-XM_059151112.1    rna-XM_059383372.1, rna-XM_059387523.1, rna-XM_059387562.1    rna-XM_004767147.3, rna-XM_045077432.1    mNeoVis.2.1.1.chr7_hap1.g00824.t01, mNeoVis.2.1.1.chr7_hap1.g00825.t01    rna-XM_044911443.1      rna-XM_042970373.1    rna-XM_040632944.1, rna-XM_040632968.1    rna-XM_035725127.1, rna-XM_035725132.1    mNeoVis.3.1.1.chr7.g00769.t01    rna-gnl|WGS_ZZZZ|egapxtmp_012317-R1, rna-gnl|WGS_ZZZZ|egapxtmp_012318-R1    mNeoVis.evergladensis.2.1.chr7.g00814.t01    mNeoVis.vulgivaga.2.1.chr7.g00819.t01, mNeoVis.vulgivaga.2.1.chr7.g00820.t01      708    N0.HOG0000706   OG0000052  n1   rna-XM_011235187.1    rna-XM_038525535.1    rna-XM_022523764.1    rna-NM_001167489.1    rna-XM_036065456.1, rna-XM_036076341.1    rna-XM_039244080.1    rna-XM_032839019.1    rna-XM_045986781.1    rna-XM_032326085.1    rna-XM_059151216.1, rna-XM_059151236.1    rna-XM_059383323.1    rna-XM_004779913.2    mNeoVis.2.1.1.chr7_hap1.g00790.t01    rna-XM_021682324.1, rna-XM_044911341.1    rna-XM_004409809.1    rna-XM_007087123.2, rna-XM_015539764.2    rna-XM_008684810.1    rna-XM_027619274.1    mNeoVis.3.1.1.chr7.g00731.t01    rna-gnl|WGS_ZZZZ|egapxtmp_013312-R1    mNeoVis.evergladensis.2.1.chr7.g00777.t01    mNeoVis.vulgivaga.2.1.chr7.g00782.t01      709    N0.HOG0000707   OG0000052  n24   rna-XM_034638043.1, rna-XM_034638056.1    rna-NM_001360006.1    rna-XM_022492089.1, rna-XM_022492105.1    rna-NM_001167473.1, rna-NM_001167476.1, rna-NM_001167482.1, rna-NM_001167483.1, rna-XM_023245895.2    rna-XM_036091967.1    rna-XM_039219006.1, rna-XM_039219017.1, rna-XM_039251082.1    rna-XM_032847598.1, rna-XM_032847602.1    rna-XM_045986833.1    rna-XM_032324823.1    rna-XM_059151188.1, rna-XM_059152739.1    rna-XM_059381977.1, rna-XM_059383443.1    rna-XM_013048029.1    mNeoVis.2.1.1.chr7_hap1.g00640.t01, mNeoVis.2.1.1.chr7_hap1.g01064.t01        rna-XM_007086617.1, rna-XM_007089504.2, rna-XM_007089510.2, rna-XM_007089534.2    rna-XM_008688766.2, rna-XM_008706571.2    rna-XM_027618055.1    mNeoVis.3.1.1.chr7.g00610.t01    rna-gnl|WGS_ZZZZ|egapxtmp_012261-R1, rna-gnl|WGS_ZZZZ|egapxtmp_012381-R1    mNeoVis.evergladensis.2.1.chr7.g00638.t01, mNeoVis.evergladensis.2.1.chr7.g01056.t01    mNeoVis.vulgivaga.2.1.chr7.g00637.t01, mNeoVis.vulgivaga.2.1.chr7.g00640.t01, mNeoVis.vulgivaga.2.1.chr7.g01066.t01      710    N0.HOG0000708   OG0000052  n62   rna-XM_002927586.2, rna-XM_011235182.3, rna-XM_011235191.3, rna-XM_019808282.1, rna-XM_034638111.1, rna-XM_034638971.1    rna-NM_001013852.1    rna-XM_022523762.1, rna-XM_022524131.1, rna-XM_022524132.1    rna-NM_001167478.1, rna-NM_001167480.1, rna-NM_001167484.1, rna-NM_001167485.1, rna-XM_003997553.5, rna-XM_023244982.2, rna-XM_045045404.1, rna-XM_045045410.1, rna-XM_045046284.1    rna-XM_036076327.1    rna-XM_039220147.1, rna-XM_039220149.1, rna-XM_039220150.1, rna-XM_039220151.1, rna-XM_039220152.1, rna-XM_039220153.1, rna-XM_039220169.1, rna-XM_039220170.1, rna-XM_039224314.1, rna-XM_039224737.1, rna-XM_039244079.1    rna-XM_032839020.1, rna-XM_032839021.1, rna-XM_032839052.1, rna-XM_032839421.1, rna-XM_032839425.1, rna-XM_032839427.1    rna-XM_045986779.1, rna-XM_045986782.1, rna-XM_045986784.1, rna-XM_045986785.1, rna-XM_045988324.1    rna-XM_032324146.1, rna-XM_032324147.1, rna-XM_032324148.1, rna-XM_032324159.1, rna-XM_032324160.1, rna-XM_032326087.1, rna-XM_032326091.1, rna-XM_032326092.1    rna-XM_059150672.1, rna-XM_059150674.1, rna-XM_059150675.1, rna-XM_059150685.1, rna-XM_059151233.1    rna-XM_059382675.1, rna-XM_059382677.1, rna-XM_059382678.1, rna-XM_059382679.1, rna-XM_059383322.1, rna-XM_059383360.1    rna-XM_004779902.3, rna-XM_004779905.2, rna-XM_004779906.2, rna-XM_004779911.2, rna-XM_004781910.1, rna-XM_045077279.1    mNeoVis.2.1.1.chr7_hap1.g00786.t01, mNeoVis.2.1.1.chr7_hap1.g00788.t01, mNeoVis.2.1.1.chr7_hap1.g00793.t01, mNeoVis.2.1.1.chr7_hap1.g00795.t01, mNeoVis.2.1.1.chr7_hap1.g00796.t01, mNeoVis.2.1.1.chr7_hap1.g00797.t01    rna-XM_021681457.1, rna-XM_021681458.1    rna-XM_004415496.1    rna-XM_007074324.2, rna-XM_007074325.1, rna-XM_007091061.3, rna-XM_007091062.2, rna-XM_007093341.2, rna-XM_015534648.2, rna-XM_015541551.2, rna-XM_042970044.1, rna-XM_042970238.1, rna-XM_042970239.1, rna-XM_042970242.1, rna-XM_042970243.1, rna-XM_042970244.1, rna-XM_042970251.1, rna-XM_042970260.1, rna-XM_042970261.1, rna-XM_042970262.1, rna-XM_042970263.1, rna-XM_042970267.1    rna-XM_008684776.2, rna-XM_008684811.1, rna-XM_040632979.1, rna-XM_040634398.1, rna-XM_040634399.1, rna-XM_040634405.1    rna-XM_027618450.2, rna-XM_027619276.1, rna-XM_035724916.1    mNeoVis.3.1.1.chr7.g00729.t01, mNeoVis.3.1.1.chr7.g00736.t01, mNeoVis.3.1.1.chr7.g00739.t01, mNeoVis.3.1.1.chr7.g00740.t01, mNeoVis.3.1.1.chr7.g00741.t01    rna-gnl|WGS_ZZZZ|egapxtmp_010592-R1, rna-gnl|WGS_ZZZZ|egapxtmp_010593-R1, rna-gnl|WGS_ZZZZ|egapxtmp_012304-R1, rna-gnl|WGS_ZZZZ|egapxtmp_012306-R1, rna-gnl|WGS_ZZZZ|egapxtmp_012309-R1, rna-gnl|WGS_ZZZZ|egapxtmp_013314-R1, rna-gnl|WGS_ZZZZ|egapxtmp_013315-R1    mNeoVis.evergladensis.2.1.chr7.g00773.t01, mNeoVis.evergladensis.2.1.chr7.g00775.t01, mNeoVis.evergladensis.2.1.chr7.g00780.t01, mNeoVis.evergladensis.2.1.chr7.g00781.t01, mNeoVis.evergladensis.2.1.chr7.g00783.t01, mNeoVis.evergladensis.2.1.chr7.g00784.t01, mNeoVis.evergladensis.2.1.chr7.g00785.t01    mNeoVis.vulgivaga.2.1.chr7.g00780.t01, mNeoVis.vulgivaga.2.1.chr7.g00788.t01      711    N0.HOG0000709   OG0000053  n2   rna-XM_002924988.4, rna-XM_002925007.4    rna-NM_001388723.1    rna-XM_022522048.1    rna-XM_045038856.1, rna-XM_045038857.1    rna-XM_036075987.1    rna-XM_039218854.1    rna-XM_032853669.1    rna-XM_046016847.1    rna-XM_032359788.1    rna-XM_059157936.1    rna-XM_059389538.1    rna-XM_045081432.1    mNeoVis.2.1.1.chr7_hap1.g01726.t01    rna-XM_021685688.2    rna-XM_004416465.1    rna-XM_007083134.1, rna-XM_042958786.1, rna-XM_042958830.1    rna-XM_008709978.2, rna-XM_008711010.2    rna-XM_027580539.1    mNeoVis.3.1.1.chr7.g01608.t01      mNeoVis.evergladensis.2.1.chr7.g01694.t01    mNeoVis.vulgivaga.2.1.chr7.g01707.t01      712    N0.HOG0000710   OG0000053  n26   rna-XM_002924987.4        rna-XM_003992916.2        rna-XM_032852719.1    rna-XM_046015877.1    rna-XM_032359790.1    rna-XM_059183412.1      rna-XM_004779034.2    mNeoVis.2.1.1.chr7_hap1.g01728.t01    rna-XM_021685641.1        rna-XM_008710989.1      mNeoVis.3.1.1.chr7.g01609.t01    rna-gnl|WGS_ZZZZ|egapxtmp_012533-R1    mNeoVis.evergladensis.2.1.chr7.g01696.t01    mNeoVis.vulgivaga.2.1.chr7.g01709.t01      713    N0.HOG0000711   OG0000053  n40     rna-NM_001256500.1, rna-NM_001388725.1                                              714    N0.HOG0000712   OG0000053  n44   rna-XM_002925006.4    rna-XM_038568177.1, rna-XM_038568178.1    rna-XM_022521694.1            rna-XM_046015878.1    rna-XM_032359789.1    rna-XM_059183354.1    rna-XM_059392833.1    rna-XM_045081430.1    mNeoVis.2.1.1.chr7_hap1.g01727.t01          rna-XM_008710990.2        rna-gnl|WGS_ZZZZ|egapxtmp_010656-R1    mNeoVis.evergladensis.2.1.chr7.g01695.t01    mNeoVis.vulgivaga.2.1.chr7.g01708.t01      715    N0.HOG0000713   OG0000053  n56   rna-XM_034666478.1    rna-NM_001388722.1, rna-XM_038568176.1    rna-XM_022521695.1          rna-XM_032852718.1    rna-XM_046015879.1    rna-XM_032359786.1    rna-XM_059183226.1    rna-XM_059392900.1    rna-XM_004779037.2    mNeoVis.2.1.1.chr7_hap1.g01725.t01          rna-XM_008709556.1      mNeoVis.3.1.1.chr7.g01607.t01    rna-gnl|WGS_ZZZZ|egapxtmp_010654-R1    mNeoVis.evergladensis.2.1.chr7.g01693.t01        716    N0.HOG0000714   OG0000053  n69         rna-XM_003992914.2, rna-XM_003992915.5, rna-XM_045038851.1      rna-XM_039218850.1, rna-XM_039218851.1                      rna-XM_007083269.2                  717    N0.HOG0000715   OG0000053  n74   rna-XM_034666477.1    rna-NM_001388724.1    rna-XM_022522061.1    rna-XM_045038852.1, rna-XM_045038853.1, rna-XM_045038854.1, rna-XM_045038855.1      rna-XM_039218853.1, rna-XM_039218856.1, rna-XM_039218872.1    rna-XM_032853668.1    rna-XM_046016838.1    rna-XM_032360088.1    rna-XM_059157926.1    rna-XM_059389550.1    rna-XM_004779038.1          rna-XM_007083135.2, rna-XM_007083268.1, rna-XM_007083270.2, rna-XM_042958774.1    rna-XM_008711009.1          mNeoVis.evergladensis.2.1.chr7.g01692.t01    mNeoVis.vulgivaga.2.1.chr7.g01706.t01      718    N0.HOG0000716   OG0000053  n98   rna-XM_002925005.4        rna-XM_003992947.3      rna-XM_039218846.1    rna-XM_032853670.1    rna-XM_046016758.1    rna-XM_032359936.1    rna-XM_059157946.1      rna-XM_004779033.1    mNeoVis.2.1.1.chr7_hap1.g01729.t01    rna-XM_021685690.1        rna-XM_008710988.2    rna-XM_027580004.1    mNeoVis.3.1.1.chr7.g01610.t01    rna-gnl|WGS_ZZZZ|egapxtmp_012534-R1    mNeoVis.evergladensis.2.1.chr7.g01697.t01    mNeoVis.vulgivaga.2.1.chr7.g01710.t01      719    N0.HOG0000717   OG0000053  n114                               rna-XM_004414461.1, rna-XM_004416466.1, rna-XM_004416467.1        rna-XM_035722997.1, rna-XM_035722999.1              720    N0.HOG0000718   OG0000053  n120     rna-NM_001388728.1    rna-XM_022521693.1            rna-XM_046015876.1                                  721    N0.HOG0000719   OG0000053  n122   rna-XM_002925003.4    rna-NM_001388907.1    rna-XM_022522059.1    rna-XM_045038858.1    rna-XM_036076013.1    rna-XM_039218857.1    rna-XM_032853674.1    rna-XM_046016836.1    rna-XM_032360304.1    rna-XM_059157980.1    rna-XM_059389499.1, rna-XM_059391910.1    rna-XM_004778981.2    mNeoVis.2.1.1.chr7_hap1.g01732.t01    rna-XM_021685691.1        rna-XM_008711006.2, rna-XM_008711007.2        rna-gnl|WGS_ZZZZ|egapxtmp_012837-R1    mNeoVis.evergladensis.2.1.chr7.g01700.t01        722    N0.HOG0000720   OG0000053  n142           rna-XM_036075986.1                    rna-XM_044919492.1                      723    N0.HOG0000721   OG0000053  n145     rna-XM_038568179.1    rna-XM_022522053.1    rna-XM_006937040.5      rna-XM_039218855.1, rna-XM_039218858.1    rna-XM_032853672.1    rna-XM_046016841.1    rna-XM_032360260.1    rna-XM_059157966.1    rna-XM_059389511.1    rna-XM_045078267.1    mNeoVis.2.1.1.chr7_hap1.g01731.t01        rna-XM_042959034.1    rna-XM_040630675.1        rna-gnl|WGS_ZZZZ|egapxtmp_012899-R1    mNeoVis.evergladensis.2.1.chr7.g01699.t01    mNeoVis.vulgivaga.2.1.chr7.g01712.t01      724   N0.HOG0000722  OG0000053  n160     rna-NM_001389018.1      rna-XM_003992880.2                          rna-XM_042958775.1    rna-XM_008709557.2                725   N0.HOG0000723  OG0000053  n163       rna-XM_022521697.1            rna-XM_046016869.1    rna-XM_032360360.1    rna-XM_059157915.1    rna-XM_059389561.1    rna-XM_004778984.1    mNeoVis.2.1.1.chr7_hap1.g01724.t01                  mNeoVis.evergladensis.2.1.chr7.g01691.t01    mNeoVis.vulgivaga.2.1.chr7.g01705.t01      726    N0.HOG0000724   OG0000053  n171   rna-XM_002924986.4    rna-NM_001388726.1    rna-XM_022522060.1          rna-XM_032853671.1    rna-XM_046016837.1    rna-XM_032360190.1    rna-XM_059157957.1    rna-XM_059389524.1    rna-XM_004778983.2    mNeoVis.2.1.1.chr7_hap1.g01730.t01          rna-XM_008711008.2      mNeoVis.3.1.1.chr7.g01611.t01    rna-gnl|WGS_ZZZZ|egapxtmp_012836-R1    mNeoVis.evergladensis.2.1.chr7.g01698.t01    mNeoVis.vulgivaga.2.1.chr7.g01711.t01      727    N0.HOG0000725   OG0000054  n2   rna-XM_019793558.2    rna-NM_001128836.1    rna-XM_022491113.1    rna-XM_045048993.1    rna-XM_036069800.1    rna-XM_039218489.1      rna-XM_045983111.1    rna-XM_032317138.1    rna-XM_059146464.1    rna-XM_059414169.1    rna-XM_045086624.1    mNeoVis.2.1.1.chr14_hap1.g00574.t01    rna-XM_021681850.1    rna-XM_004415673.1    rna-XM_007084269.2    rna-XM_008700744.2    rna-XM_027613259.1    mNeoVis.3.1.1.chr14.g00561.t01    rna-gnl|WGS_ZZZZ|egapxtmp_031956-R1    mNeoVis.evergladensis.2.1.chr14.g00573.t01    mNeoVis.vulgivaga.2.1.chr14.g00569.t01      728    N0.HOG0000726   OG0000054  n23   rna-XM_034667345.1    rna-XM_038586406.1    rna-XM_022491035.1    rna-XM_045048851.1    rna-XM_036069845.1    rna-XM_039218497.1    rna-XM_032845689.1    rna-XM_045983100.1    rna-XM_032317114.1    rna-XM_059145445.1    rna-XM_059413564.1    rna-XM_045086639.1    mNeoVis.2.1.1.chr14_hap1.g00581.t01    rna-XM_021681851.2    rna-XM_004415524.1    rna-XM_042975728.1    rna-XM_040635873.1    rna-XM_027611960.2    mNeoVis.3.1.1.chr14.g00569.t01      mNeoVis.evergladensis.2.1.chr14.g00579.t01    mNeoVis.vulgivaga.2.1.chr14.g00576.t01      729   N0.HOG0000727  OG0000054  n43   rna-XM_011217598.3    rna-XM_038585936.1    rna-XM_022490730.1    rna-XM_006943058.5    rna-XM_036069789.1    rna-XM_039218468.1    rna-XM_032845652.1    rna-XM_045983102.1    rna-XM_032317113.1    rna-XM_059146604.1    rna-XM_059414166.1    rna-XM_004775804.3    mNeoVis.2.1.1.chr14_hap1.g00573.t01    rna-XM_021681944.1    rna-XM_004415682.1    rna-XM_015538472.2    rna-XM_008700806.2    rna-XM_027613791.2    mNeoVis.3.1.1.chr14.g00560.t01      mNeoVis.evergladensis.2.1.chr14.g00571.t01    mNeoVis.vulgivaga.2.1.chr14.g00570.t01      730    N0.HOG0000728   OG0000054  n63   rna-XM_019793557.2    rna-NM_001130830.1    rna-XM_022490964.1    rna-XM_003999680.6    rna-XM_036069790.1    rna-XM_039218491.1    rna-XM_032845651.1    rna-XM_045983316.1    rna-XM_032317130.1    rna-XM_059145661.1    rna-XM_059414168.1    rna-XM_045086621.1      rna-XM_044916018.1    rna-XM_004415674.1    rna-XM_007084268.3    rna-XM_040635885.1    rna-XM_027611959.2    mNeoVis.3.1.1.chr14.g00562.t01      mNeoVis.evergladensis.2.1.chr14.g00572.t01        731    N0.HOG0000729   OG0000054  n82   rna-XM_034667339.1    rna-NM_001128837.1    rna-XM_022490999.1    rna-XM_011291099.4, rna-XM_019822350.3, rna-XM_045048759.1, rna-XM_045048763.1, rna-XM_045048765.1      rna-XM_039218487.1, rna-XM_039218495.1, rna-XM_039218496.1, rna-XM_039218498.1        rna-XM_032317136.1, rna-XM_032317137.1, rna-XM_032317142.1    rna-XM_059145883.1, rna-XM_059146625.1    rna-XM_059414170.1, rna-XM_059414176.1    rna-XM_004775796.3, rna-XM_045086547.1    mNeoVis.2.1.1.chr14_hap1.g00577.t01, mNeoVis.2.1.1.chr14_hap1.g00579.t01        rna-XM_007084270.3, rna-XM_007084271.3, rna-XM_042975879.1, rna-XM_042976108.1, rna-XM_042976109.1, rna-XM_042976502.1, rna-XM_042976710.1    rna-XM_040634609.1      mNeoVis.3.1.1.chr14.g00565.t01, mNeoVis.3.1.1.chr14.g00566.t01    rna-gnl|WGS_ZZZZ|egapxtmp_033041-R1    mNeoVis.evergladensis.2.1.chr14.g00576.t01    mNeoVis.vulgivaga.2.1.chr14.g00574.t01      732    N0.HOG0000730   OG0000054  n118   rna-XM_034667335.1      rna-XM_022490990.1      rna-XM_036069796.1        rna-XM_045983103.1    rna-XM_032317133.1    rna-XM_059146546.1    rna-XM_059414171.1    rna-XM_045086625.1    mNeoVis.2.1.1.chr14_hap1.g00578.t01    rna-XM_021681946.1    rna-XM_004415522.1      rna-XM_040635878.1    rna-XM_027611577.2      rna-gnl|WGS_ZZZZ|egapxtmp_031958-R1    mNeoVis.evergladensis.2.1.chr14.g00577.t01    mNeoVis.vulgivaga.2.1.chr14.g00573.t01      733    N0.HOG0000731   OG0000054  n134           rna-XM_036069792.1, rna-XM_036069801.1                    rna-XM_021681945.1    rna-XM_004415681.1        rna-XM_027613257.2              734   N0.HOG0000732  OG0000054  n139   rna-XM_034650069.1, rna-XM_034667340.1    rna-XM_038586402.1            rna-XM_032845683.1      rna-XM_032317125.1, rna-XM_032317129.1    rna-XM_059146568.1    rna-XM_059413968.1, rna-XM_059414179.1    rna-XM_013047489.2, rna-XM_045086548.1            rna-XM_008700755.2, rna-XM_040635876.1, rna-XM_040635877.1      mNeoVis.3.1.1.chr14.g00568.t01      mNeoVis.evergladensis.2.1.chr14.g00578.t01        735   N0.HOG0000733  OG0000054  n155     rna-XM_038585940.1            rna-XM_032845684.1, rna-XM_032845686.1      rna-XM_032317584.1      rna-XM_059413967.1      mNeoVis.2.1.1.chr14_hap1.g00576.t01              mNeoVis.3.1.1.chr14.g00564.t01    rna-gnl|WGS_ZZZZ|egapxtmp_033057-R1    mNeoVis.evergladensis.2.1.chr14.g00575.t01    mNeoVis.vulgivaga.2.1.chr14.g00572.t01      736    N0.HOG0000734   OG0000054  n164   rna-XM_034667338.1    rna-NM_001128838.1    rna-XM_022490919.1, rna-XM_022490988.1, rna-XM_022490991.1          rna-XM_032845679.1, rna-XM_032845687.1    rna-XM_045983109.1, rna-XM_045983112.1    rna-XM_032317126.1, rna-XM_032317132.1, rna-XM_032317140.1    rna-XM_059146327.1, rna-XM_059146535.1    rna-XM_059413560.1, rna-XM_059414178.1    rna-XM_004775791.3, rna-XM_004775799.3    mNeoVis.2.1.1.chr14_hap1.g00575.t01, mNeoVis.2.1.1.chr14_hap1.g00580.t01          rna-XM_040635881.1      mNeoVis.3.1.1.chr14.g00563.t01    rna-gnl|WGS_ZZZZ|egapxtmp_031957-R1, rna-gnl|WGS_ZZZZ|egapxtmp_033040-R1    mNeoVis.evergladensis.2.1.chr14.g00574.t01    mNeoVis.vulgivaga.2.1.chr14.g00571.t01, mNeoVis.vulgivaga.2.1.chr14.g00575.t01      737    N0.HOG0000735   OG0000055  n1   rna-XM_034643153.1    rna-XM_038543910.1    rna-XM_022495747.1    rna-XM_003995004.5    rna-XM_036121712.1    rna-XM_039217235.1    rna-XM_032875788.1    rna-XM_046026094.1    rna-XM_032309209.1    rna-XM_059140797.1    rna-XM_059409161.1    rna-XM_004742685.3    mNeoVis.2.1.1.chr3_hap1.g00274.t01    rna-XM_021683004.2    rna-XM_004408475.1    rna-XM_042961911.1    rna-XM_040619630.1    rna-XM_027577094.1    mNeoVis.3.1.1.chr3.g00277.t01    rna-gnl|WGS_ZZZZ|egapxtmp_039454-R1    mNeoVis.evergladensis.2.1.chr3.g00284.t01    mNeoVis.vulgivaga.2.1.chr3.g00277.t01      738    N0.HOG0000736   OG0000055  n10   rna-XM_034639053.1    rna-XM_038537984.1    rna-XM_022511442.1      rna-XM_036119740.1    rna-XM_039227820.1    rna-XM_032854594.1    rna-XM_045988019.1    rna-XM_032323491.1    rna-XM_059152902.1    rna-XM_059381860.1    rna-XM_045076586.1    mNeoVis.2.1.1.chr7_hap1.g00009.t01    rna-XM_021705043.1    rna-XM_004392169.1    rna-XM_042968096.1    rna-XM_040633074.1    rna-XM_027617422.2    mNeoVis.3.1.1.chr7.g00010.t01      mNeoVis.evergladensis.2.1.chr7.g00010.t01        739   N0.HOG0000737  OG0000055  n25   rna-XM_002915516.4    rna-XM_038572739.1    rna-XM_022494456.1    rna-NM_001134989.1    rna-XM_036094390.1    rna-XM_039236877.1    rna-XM_032878883.1    rna-XM_045981215.1    rna-XM_032350191.1    rna-XM_059133242.1    rna-XM_059408032.1    rna-XM_004759946.3    mNeoVis.2.1.1.chr11_hap1.g00075.t01    rna-XM_021677795.2    rna-XM_004413990.1    rna-XM_042980035.1    rna-XM_008699496.2    rna-XM_027622346.1    mNeoVis.3.1.1.chr11.g00049.t01    rna-gnl|WGS_ZZZZ|egapxtmp_026783-R1    mNeoVis.evergladensis.2.1.chr11.g00074.t01    mNeoVis.vulgivaga.2.1.chr11.g00074.t01      740    N0.HOG0000738   OG0000055  n46   rna-XM_034660444.1    rna-XM_038553445.1    rna-XM_022510987.1    rna-XM_003985886.5    rna-XM_036120419.1    rna-XM_039220569.1    rna-XM_032847250.1, rna-XM_032860342.1    rna-XM_045996955.1, rna-XM_046004403.1, rna-XM_046022953.1, rna-XM_046023133.1, rna-XM_046023384.1, rna-XM_046023453.1    rna-XM_032338098.1, rna-XM_032339228.1    rna-XM_059179033.1, rna-XM_059180265.1    rna-XM_059399762.1    rna-XM_004781589.3    mNeoVis.2.1.1.chr1_hap1.g01303.t01    rna-XM_021696941.1    rna-XM_004395260.1, rna-XM_004409052.1    rna-XM_007090589.3    rna-XM_008692913.2    rna-XM_035728456.1    mNeoVis.3.1.1.chr1.g01256.t01    rna-gnl|WGS_ZZZZ|egapxtmp_006854-R1    mNeoVis.evergladensis.2.1.chr1.g01274.t01, mNeoVis.evergladensis.2.1.chr5.g00444.t01    mNeoVis.vulgivaga.2.1.chr1.g01273.t01      741    N0.HOG0000739   OG0000055  n63   rna-XM_034657626.1    rna-XM_038567541.1    rna-XM_022525663.1    rna-XM_023244250.2    rna-XM_036120658.1    rna-XM_039221346.1    rna-XM_032883765.1    rna-XM_045992372.1    rna-XM_032308027.1    rna-XM_059159674.1      rna-XM_004771595.3    mNeoVis.2.1.1.chr5_hap1.g00171.t01    rna-XM_021705298.2    rna-XM_004395279.1    rna-XM_042977953.1    rna-XM_040629479.1    rna-XM_027583276.2    mNeoVis.3.1.1.chr2.g00120.t01    rna-gnl|WGS_ZZZZ|egapxtmp_037730-R1    mNeoVis.evergladensis.2.1.chr2.g00167.t01    mNeoVis.vulgivaga.2.1.chr2.g00166.t01      742    N0.HOG0000740   OG0000055  n67       rna-XM_022501961.1                        rna-XM_044920476.1                      743    N0.HOG0000741   OG0000055  n69                       rna-XM_059391801.1                        mNeoVis.vulgivaga.2.1.chr7.g00009.t01      744    N0.HOG0000742   OG0000055  n73           rna-XM_036099603.1      rna-XM_032837885.1                                    745    N0.HOG0000743   OG0000055  n76   rna-XM_034643166.1    rna-XM_038525475.1      rna-XM_045049236.1          rna-XM_045982286.1, rna-XM_046020415.1    rna-XM_032317562.1                rna-XM_015535335.1    rna-XM_040644564.1                746    N0.HOG0000744   OG0000055  n83     rna-XM_038585040.1              rna-XM_046026586.1      rna-XM_059156415.1                  rna-XM_027623853.1    mNeoVis.3.1.1.chr3.g00717.t01      mNeoVis.evergladensis.2.1.chr3.g00736.t01    mNeoVis.vulgivaga.2.1.chr3.g00743.t01      747    N0.HOG0000745   OG0000055  n89   rna-XM_002926120.4        rna-XM_011288741.4    rna-XM_036064930.1    rna-XM_039220719.1    rna-XM_032866320.1    rna-XM_046023793.1    rna-XM_032309009.1    rna-XM_059141652.1    rna-XM_059410213.1    rna-XM_045082798.1    mNeoVis.2.1.1.chr13_hap1.g00015.t01      rna-XM_004416661.1    rna-XM_042963615.1    rna-XM_040632268.1    rna-XM_027615963.2        mNeoVis.evergladensis.2.1.chr13.g00017.t01    mNeoVis.vulgivaga.2.1.chr13.g00016.t01      748    N0.HOG0000746   OG0000055  n92     rna-XM_038584033.1    rna-XM_022517691.1    rna-XM_023253720.2    rna-XM_036085819.1    rna-XM_039246527.1    rna-XM_032861475.1    rna-XM_046004089.1      rna-XM_059179134.1    rna-XM_059399349.1    rna-XM_045067036.1    mNeoVis.2.1.1.chr1_hap1.g01268.t01    rna-XM_044917526.1    rna-XM_004408403.1    rna-XM_042985849.1    rna-XM_040645004.1    rna-XM_035728659.1          mNeoVis.vulgivaga.2.1.chr1.g01240.t01      749    N0.HOG0000747   OG0000055  n98     rna-XM_038584041.1    rna-XM_022517801.1    rna-XM_003985847.5    rna-XM_036085821.1      rna-XM_032861473.1    rna-XM_046004933.1    rna-XM_032340975.1    rna-XM_059179135.1    rna-XM_059399353.1    rna-XM_045067034.1    mNeoVis.2.1.1.chr1_hap1.g01267.t01    rna-XM_044917601.1    rna-XM_004408402.1    rna-XM_042985848.1      rna-XM_035728502.1      rna-gnl|WGS_ZZZZ|egapxtmp_039598-R1    mNeoVis.evergladensis.2.1.chr1.g01239.t01        750    N0.HOG0000748   OG0000056  n2   rna-XM_002929841.1    rna-XM_038557534.1    rna-XM_022491970.1    rna-XM_003980567.2    rna-XM_036074842.1    rna-XM_039220847.1    rna-XM_032848088.1    rna-XM_046001127.1    rna-XM_032337891.1    rna-XM_059173391.1    rna-XM_059416413.1    rna-XM_004781566.1    mNeoVis.2.1.1.chr1_hap1.g00942.t01    rna-XM_021690947.1        rna-XM_008683741.1    rna-XM_027604479.1    mNeoVis.3.1.1.chr1.g00955.t01    rna-gnl|WGS_ZZZZ|egapxtmp_040127-R1    mNeoVis.evergladensis.2.1.chr1.g00938.t01    mNeoVis.vulgivaga.2.1.chr1.g00949.t01      751    N0.HOG0000749   OG0000056  n19   rna-XM_034655680.1    rna-NM_001388544.1    rna-XM_022491988.1    rna-XM_003980566.4      rna-XM_039220845.1    rna-XM_032848089.1    rna-XM_045981637.1, rna-XM_046000731.1    rna-XM_032334922.1    rna-XM_059173392.1    rna-XM_059416414.1    rna-XM_013050519.2    mNeoVis.2.1.1.chr1_hap1.g00943.t01    rna-XM_021690929.1      rna-XM_042980926.1    rna-XM_008683700.2      mNeoVis.3.1.1.chr1.g00956.t01    rna-gnl|WGS_ZZZZ|egapxtmp_021057-R1    mNeoVis.evergladensis.2.1.chr1.g00939.t01    mNeoVis.vulgivaga.2.1.chr1.g00950.t01      752    N0.HOG0000750   OG0000056  n36   rna-XM_002929842.4    rna-NM_001388545.1    rna-XM_022491987.1    rna-XM_003980546.3      rna-XM_039220846.1    rna-XM_032848053.1    rna-XM_045981636.1, rna-XM_045996999.1, rna-XM_045997014.1, rna-XM_046000627.1    rna-XM_032334924.1    rna-XM_059173499.1    rna-XM_059416722.1    rna-XM_004781564.2    mNeoVis.2.1.1.chr1_hap1.g00941.t01        rna-XM_007092934.2    rna-XM_008683701.2    rna-XM_027606258.1    mNeoVis.3.1.1.chr1.g00954.t01    rna-gnl|WGS_ZZZZ|egapxtmp_021059-R1    mNeoVis.evergladensis.2.1.chr1.g00937.t01    mNeoVis.vulgivaga.2.1.chr1.g00948.t01      753    N0.HOG0000751   OG0000056  n57   rna-XM_002929843.1    rna-XM_038557533.1      rna-XM_003980565.2      rna-XM_039220844.1                      rna-XM_007092955.1    rna-XM_008683742.1                754    N0.HOG0000752   OG0000056  n62   rna-XM_002929839.1    rna-NM_001256452.1    rna-XM_022491986.1    rna-XM_011281170.2    rna-XM_036074844.1    rna-XM_039220851.1    rna-XM_032848092.1    rna-XM_045996997.1, rna-XM_045997000.1, rna-XM_045997013.1, rna-XM_045997015.1, rna-XM_046000636.1              rna-XM_021690927.2      rna-XM_015542392.1    rna-XM_008683703.1                755    N0.HOG0000753   OG0000056  n78     rna-NM_001389081.1, rna-XM_038589021.1                                              756    N0.HOG0000754   OG0000056  n80   rna-XM_002929840.2    rna-XM_038556365.1, rna-XM_038589253.1, rna-XM_038589254.1    rna-XM_022491994.1    rna-XM_003980568.1    rna-XM_036112159.1    rna-XM_039220848.1      rna-XM_045980091.1, rna-XM_045980092.1, rna-XM_045996805.1, rna-XM_045996828.1, rna-XM_045996986.1, rna-XM_045996987.1, rna-XM_045997004.1, rna-XM_045997005.1, rna-XM_045997006.1, rna-XM_045997022.1, rna-XM_045997023.1, rna-XM_046000505.1    rna-XM_032334921.1    rna-XM_059173500.1, rna-XM_059173501.1    rna-XM_059416723.1    rna-XM_004782615.2, rna-XM_045069327.1, rna-XM_045069331.1    mNeoVis.2.1.1.chr1_hap1.g00944.t01        rna-XM_007092936.1    rna-XM_040621942.1                757    N0.HOG0000755   OG0000056  n101     rna-NM_001388543.1, rna-XM_038556368.1, rna-XM_038589017.1    rna-XM_022491992.1      rna-XM_036100215.1    rna-XM_039220852.1    rna-XM_032848054.1    rna-XM_045980356.1, rna-XM_045996826.1, rna-XM_045997020.1, rna-XM_046000906.1, rna-XM_046001132.1        rna-XM_059416724.1              rna-XM_008683698.1                758    N0.HOG0000756   OG0000056  n113   rna-XM_002930914.2, rna-XM_002931217.2, rna-XM_034653262.1, rna-XM_034653263.1, rna-XM_034653274.1    rna-NM_001388547.1, rna-XM_038556364.1      rna-XM_003980547.3, rna-XM_003980548.2, rna-XM_006927434.3    rna-XM_036074845.1, rna-XM_036096908.1, rna-XM_036115877.1    rna-XM_039220843.1, rna-XM_039220849.1    rna-XM_032848093.1    rna-XM_045981565.1, rna-XM_045996815.1, rna-XM_045996841.1, rna-XM_045996844.1, rna-XM_045996865.1, rna-XM_045996878.1, rna-XM_045996903.1, rna-XM_045996905.1, rna-XM_045996906.1, rna-XM_045996907.1, rna-XM_045996908.1, rna-XM_045996967.1, rna-XM_045997002.1, rna-XM_045997008.1, rna-XM_045997036.1, rna-XM_046000393.1, rna-XM_046000427.1, rna-XM_046000504.1, rna-XM_046000540.1, rna-XM_046000556.1, rna-XM_046000557.1, rna-XM_046000558.1, rna-XM_046000568.1    rna-XM_032337894.1      rna-XM_059416721.1    rna-XM_045069314.1    mNeoVis.2.1.1.chr1_hap1.g00940.t01    rna-XM_021690926.2, rna-XM_044912123.1    rna-XM_004403494.1    rna-XM_007092937.1, rna-XM_007092954.3, rna-XM_042958368.1    rna-XM_008683704.1, rna-XM_008683705.1, rna-XM_008683706.2, rna-XM_008683707.2, rna-XM_008683708.2, rna-XM_008683743.2, rna-XM_008683744.2    rna-XM_027606469.1, rna-XM_035727739.1    mNeoVis.3.1.1.chr1.g00953.t01    rna-gnl|WGS_ZZZZ|egapxtmp_039906-R1    mNeoVis.evergladensis.2.1.chr1.g00936.t01    mNeoVis.vulgivaga.2.1.chr1.g00947.t01      759    N0.HOG0000757   OG0000057  n2   rna-XM_002926067.1    rna-XM_038530640.1    rna-XM_022518131.1    rna-XM_003980976.4    rna-XM_036072284.1    rna-XM_039225107.1    rna-XM_032850216.1    rna-XM_046000969.1    rna-XM_032334949.1    rna-XM_059175033.1    rna-XM_059417905.1    rna-XM_004744623.1    mNeoVis.2.1.1.chr1_hap1.g00589.t01    rna-XM_021702180.1    rna-XM_004408752.1    rna-XM_007079922.2    rna-XM_008691397.1    rna-XM_027606573.1    mNeoVis.3.1.1.chr1.g00604.t01    rna-gnl|WGS_ZZZZ|egapxtmp_009226-R1    mNeoVis.evergladensis.2.1.chr1.g00594.t01    mNeoVis.vulgivaga.2.1.chr1.g00603.t01      760    N0.HOG0000758   OG0000057  n17   rna-XM_034652566.1, rna-XM_034652615.1, rna-XM_034652843.1, rna-XM_034662529.1    rna-XM_038533952.1    rna-XM_022523621.1    rna-XM_023240445.2    rna-XM_036123368.1    rna-XM_039241274.1    rna-XM_032841109.1    rna-XM_046027083.1    rna-XM_032311861.1    rna-XM_059170518.1    rna-XM_059397268.1    rna-XM_045087090.1    mNeoVis.2.1.1.chr11_hap1.g00812.t01, mNeoVis.2.1.1.chr2_hap1.g00726.t01    rna-XM_021692290.1    rna-XM_004404526.1, rna-XM_004404847.1, rna-XM_004417005.1    rna-XM_042960251.1    rna-XM_040623787.1, rna-XM_040624031.1, rna-XM_040625514.1    rna-XM_035724221.1    mNeoVis.3.1.1.chr5.g00676.t01    rna-gnl|WGS_ZZZZ|egapxtmp_016913-R1    mNeoVis.evergladensis.2.1.chr11.g00795.t01, mNeoVis.evergladensis.2.1.chr5.g00685.t01    mNeoVis.vulgivaga.2.1.chr11.g00814.t01, mNeoVis.vulgivaga.2.1.chr5.g00676.t01      761    N0.HOG0000759   OG0000057  n30   rna-XM_034661201.1    rna-XM_038579968.1    rna-XM_022520656.1    rna-XM_023255528.2    rna-XM_036109144.1    rna-XM_039247272.1    rna-XM_032856906.1    rna-XM_046008702.1    rna-XM_032343216.1    rna-XM_059181101.1    rna-XM_059372462.1    rna-XM_045067785.1    mNeoVis.2.1.1.chr10_hap1.g00773.t01    rna-XM_021695334.1    rna-XM_004409028.1    rna-XM_042989234.1    rna-XM_040620666.1    rna-XM_027572738.2    mNeoVis.3.1.1.chr9.g00732.t01      mNeoVis.evergladensis.2.1.chr8.g00754.t01    mNeoVis.vulgivaga.2.1.chr9.g00753.t01      762    N0.HOG0000760   OG0000057  n34   rna-XM_002921486.4    rna-XM_038561741.1    rna-XM_022504323.1    rna-XM_003984129.6    rna-XM_036084442.1, rna-XM_036087655.1    rna-XM_039240132.1    rna-XM_032850289.1, rna-XM_032850293.1    rna-XM_045979366.1    rna-XM_032352484.1    rna-XM_059188078.1    rna-XM_059405823.1    rna-XM_045079029.1      rna-XM_021699106.1    rna-XM_004398703.1    rna-XM_007083445.3    rna-XM_008700004.2    rna-XM_027623026.1    mNeoVis.3.1.1.chr11.g00775.t01            763    N0.HOG0000761   OG0000057  n38   rna-XM_011218625.3      rna-XM_022499351.1    rna-XM_003993871.6    rna-XM_036079350.1    rna-XM_039222029.1    rna-XM_032883057.1    rna-XM_046027760.1    rna-XM_032312746.1    rna-XM_059172472.1    rna-XM_059397397.1    rna-XM_004780765.3    mNeoVis.2.1.1.chr2_hap1.g00604.t01    rna-XM_021696133.2    rna-XM_004409684.1    rna-XM_007079268.3    rna-XM_008689605.2    rna-XM_027596306.1    mNeoVis.3.1.1.chr5.g00598.t01    rna-gnl|WGS_ZZZZ|egapxtmp_009186-R1    mNeoVis.evergladensis.2.1.chr5.g00606.t01    mNeoVis.vulgivaga.2.1.chr5.g00601.t01      764    N0.HOG0000762   OG0000057  n40   rna-XM_034644851.1    rna-XM_038563623.1    rna-XM_022506403.1      rna-XM_036067815.1    rna-XM_039235346.1    rna-XM_032879084.1    rna-XM_046014581.1    rna-XM_032358099.1    rna-XM_059142357.1      rna-XM_004759628.2    mNeoVis.2.1.1.chr7_hap1.g02606.t01    rna-XM_044919541.1    rna-XM_004394040.1    rna-XM_042959416.1        mNeoVis.3.1.1.chr7.g02270.t01    rna-gnl|WGS_ZZZZ|egapxtmp_010527-R2    mNeoVis.evergladensis.2.1.chr7.g02413.t01    mNeoVis.vulgivaga.2.1.chr7.g02435.t01      765    N0.HOG0000763   OG0000057  n57   rna-XM_034651314.1    rna-XM_038576075.1                                      rna-gnl|WGS_ZZZZ|egapxtmp_025302-R1          766    N0.HOG0000764   OG0000057  n59   rna-XM_034652013.1, rna-XM_034652895.1                              rna-XM_004417672.1                    767    N0.HOG0000765   OG0000057  n61   rna-XM_011218081.2, rna-XM_034652713.1    rna-NM_001003349.2    rna-XM_022523173.1    rna-XM_023244304.2    rna-XM_036094117.1    rna-XM_039226620.1    rna-XM_032870402.1    rna-XM_045984447.1    rna-XM_032304853.1, rna-XM_032335143.1, rna-XM_032344260.1    rna-XM_059148578.1    rna-XM_059379467.1    rna-XM_045085705.1    mNeoVis.2.1.1.chr8_hap1.g00060.t01    rna-XM_021680914.2      rna-XM_042965902.1    rna-XM_040640029.1    rna-XM_027626408.1    mNeoVis.3.1.1.chr8.g00035.t01      mNeoVis.evergladensis.2.1.chr9.g00057.t01    mNeoVis.vulgivaga.2.1.chr8.g00057.t01      768    N0.HOG0000766   OG0000057  n67   rna-XM_034652944.1    rna-NM_001195845.2    rna-XM_022500897.1    rna-XM_006941899.4    rna-XM_036075969.1    rna-XM_039226147.1    rna-XM_032872751.1    rna-XM_045993018.1    rna-XM_032327689.1    rna-XM_059155638.1    rna-XM_059414358.1    rna-XM_045075150.1    mNeoVis.2.1.1.chr15_hap1.g00603.t01    rna-XM_021680249.2    rna-XM_004394150.1    rna-XM_042971108.1    rna-XM_040634733.1    rna-XM_027612302.2    mNeoVis.3.1.1.chr3.g00902.t01    rna-gnl|WGS_ZZZZ|egapxtmp_041389-R1    mNeoVis.evergladensis.2.1.chr3.g00913.t01    mNeoVis.vulgivaga.2.1.chr3.g00929.t01      769    N0.HOG0000767   OG0000058  n1   rna-XM_011221121.3, rna-XM_034651231.1, rna-XM_034651770.1, rna-XM_034652325.1    rna-XM_038549365.1    rna-XM_022501367.1    rna-XM_006939458.5    rna-XM_036112916.1    rna-XM_039239035.1    rna-XM_032865850.1    rna-XM_046008845.1, rna-XM_046009154.1, rna-XM_046023457.1    rna-XM_032308725.1    rna-XM_059141628.1    rna-XM_059410813.1    rna-XM_004757328.3    mNeoVis.2.1.1.chr13_hap1.g00231.t01    rna-XM_021691729.1    rna-XM_004392593.1    rna-XM_007094395.3    rna-XM_008698773.2    rna-XM_027615070.2    mNeoVis.3.1.1.chr13.g00182.t01    rna-gnl|WGS_ZZZZ|egapxtmp_031106-R1    mNeoVis.evergladensis.2.1.chr13.g00229.t01    mNeoVis.vulgivaga.2.1.chr13.g00226.t01      770    N0.HOG0000768   OG0000058  n13   rna-XM_019793821.2    rna-XM_038553570.1    rna-XM_022510697.1    rna-XM_011282307.4    rna-XM_036108228.1    rna-XM_039219548.1    rna-XM_032847288.1    rna-XM_046006596.1    rna-XM_032340834.1    rna-XM_059178865.1    rna-XM_059399889.1    rna-XM_004780935.3    mNeoVis.2.1.1.chr1_hap1.g01358.t01    rna-XM_021697168.1    rna-XM_004415251.1    rna-XM_042986021.1    rna-XM_008711880.2    rna-XM_027603821.2    mNeoVis.3.1.1.chr1.g01307.t01    rna-gnl|WGS_ZZZZ|egapxtmp_012055-R1    mNeoVis.evergladensis.2.1.chr1.g01325.t01    mNeoVis.vulgivaga.2.1.chr1.g01326.t01      771    N0.HOG0000769   OG0000058  n32   rna-XM_002931069.4, rna-XM_034652616.1    rna-NM_001003067.2    rna-XM_022492646.1, rna-XM_022510817.1    rna-XM_019830386.3    rna-XM_036108237.1, rna-XM_036108238.1    rna-XM_039219549.1, rna-XM_039225803.1    rna-XM_032851192.1    rna-XM_046006598.1, rna-XM_046006599.1    rna-XM_032340831.1, rna-XM_032340835.1, rna-XM_032340837.1    rna-XM_059178866.1, rna-XM_059178867.1    rna-XM_059399891.1, rna-XM_059399892.1    rna-XM_004780931.3, rna-XM_004780932.3, rna-XM_045065138.1    mNeoVis.2.1.1.chr10_hap1.g00455.t01, mNeoVis.2.1.1.chr1_hap1.g01359.t01, mNeoVis.2.1.1.chr1_hap1.g01360.t01    rna-XM_021697153.2, rna-XM_044912519.1    rna-XM_004415252.1    rna-XM_042986020.1, rna-XM_042986022.1    rna-XM_008711877.2, rna-XM_008711879.2, rna-XM_040644808.1    rna-XM_035728497.1, rna-XM_035728573.1    mNeoVis.3.1.1.chr1.g01308.t01            772    N0.HOG0000770   OG0000058  n62   rna-XM_034643019.1    rna-XM_038544921.1    rna-XM_022523511.1    rna-XM_006932911.5    rna-XM_036120230.1    rna-XM_039242121.1    rna-XM_032875214.1    rna-XM_046010093.1    rna-XM_032344481.1    rna-XM_059182316.1    rna-XM_059373387.1    rna-XM_004738927.3    mNeoVis.2.1.1.chr10_hap1.g00315.t01    rna-XM_021701309.1    rna-XM_004409366.1    rna-XM_042989879.1    rna-XM_008700919.2    rna-XM_027568738.2    mNeoVis.3.1.1.chr9.g00288.t01      mNeoVis.evergladensis.2.1.chr8.g00299.t01    mNeoVis.vulgivaga.2.1.chr9.g00300.t01      773    N0.HOG0000771   OG0000058  n78       rna-XM_022508233.1                            rna-XM_042975793.1        mNeoVis.3.1.1.chr1.g00073.t01    rna-gnl|WGS_ZZZZ|egapxtmp_010482-R1    mNeoVis.evergladensis.2.1.chr3.g01145.t01        774   N0.HOG0000772  OG0000058  n85   rna-XM_011216931.3, rna-XM_034650288.1, rna-XM_034667260.1, rna-XM_034672111.1    rna-XM_038535851.1    rna-XM_022512849.1    rna-XM_003992478.4    rna-XM_036081702.1, rna-XM_036091286.1, rna-XM_036097328.1, rna-XM_036102981.1, rna-XM_036103546.1    rna-XM_039217173.1, rna-XM_039217699.1, rna-XM_039219640.1, rna-XM_039236405.1, rna-XM_039244667.1, rna-XM_039245959.1, rna-XM_039250756.1, rna-XM_039257512.1    rna-XM_032850097.1, rna-XM_032864169.1, rna-XM_032868771.1    rna-XM_046004928.1, rna-XM_046016489.1    rna-XM_032304872.1, rna-XM_032314963.1, rna-XM_032346479.1, rna-XM_032356457.1    rna-XM_059163843.1, rna-XM_059177758.1, rna-XM_059183482.1    rna-XM_059391783.1, rna-XM_059414982.1    rna-XM_004750096.3, rna-XM_045080307.1, rna-XM_045088486.1    mNeoVis.2.1.1.chr7_hap1.g01294.t01    rna-XM_044916024.1, rna-XM_044917425.1    rna-XM_004403473.1, rna-XM_004413537.1, rna-XM_004414319.1    rna-XM_042958335.1, rna-XM_042959602.1    rna-XM_008689947.2, rna-XM_040634484.1    rna-XM_035722677.1, rna-XM_035725514.1, rna-XM_035726278.1, rna-XM_035726951.1, rna-XM_035727524.1          mNeoVis.vulgivaga.2.1.chr5.g01139.t01      775   N0.HOG0000773  OG0000058  n121     rna-XM_038551914.1    rna-XM_022522255.1    rna-XM_045036760.1, rna-XM_045056007.1                          rna-XM_042984030.1      rna-XM_035728189.1    mNeoVis.3.1.1.chr14.g00422.t01    rna-gnl|WGS_ZZZZ|egapxtmp_031923-R1          776    N0.HOG0000774   OG0000058  n128               rna-XM_032874231.1    rna-XM_046023915.1        rna-XM_059394373.1        rna-XM_044920158.1                      777    N0.HOG0000775   OG0000058  n131       rna-XM_022517794.1              rna-XM_032336828.1    rna-XM_059174483.1    rna-XM_059416587.1    rna-XM_045075550.1      rna-XM_044919593.1                  mNeoVis.vulgivaga.2.1.chr7.g01295.t01      778    N0.HOG0000776   OG0000058  n137   rna-XM_034653055.1              rna-XM_032866493.1                              mNeoVis.evergladensis.2.1.chr7.g01283.t01    mNeoVis.vulgivaga.2.1.chr12.g00654.t01      779   N0.HOG0000777  OG0000059  n1   rna-XM_002931090.2, rna-XM_002931189.2, rna-XM_002931190.2, rna-XM_011218101.2, rna-XM_034650081.1    rna-NM_001388885.1    rna-XM_022521713.1, rna-XM_022522094.1, rna-XM_022522100.1    rna-XM_003992856.3, rna-XM_011286657.2, rna-XM_023239715.2    rna-XM_036081408.1    rna-XM_039225650.1, rna-XM_039225737.1, rna-XM_039225778.1    rna-XM_032852699.1, rna-XM_032853621.1, rna-XM_032853622.1    rna-XM_046015906.1, rna-XM_046015907.1, rna-XM_046015908.1, rna-XM_046016823.1, rna-XM_046016824.1    rna-XM_032359767.1, rna-XM_032359769.1, rna-XM_032360298.1, rna-XM_032360299.1    rna-XM_059157203.1, rna-XM_059157217.1, rna-XM_059181052.1, rna-XM_059181161.1    rna-XM_059390239.1, rna-XM_059390254.1    rna-XM_004779031.2, rna-XM_004779067.2, rna-XM_045071500.1, rna-XM_045071507.1    mNeoVis.2.1.1.chr7_hap1.g01655.t01, mNeoVis.2.1.1.chr7_hap1.g01656.t01, mNeoVis.2.1.1.chr7_hap1.g01657.t01    rna-XM_044919768.1    rna-XM_004407579.1    rna-XM_007089380.2, rna-XM_007089381.2, rna-XM_007089382.2    rna-XM_008710268.2, rna-XM_008710269.2, rna-XM_008710270.2    rna-XM_027577906.1, rna-XM_027580542.2    mNeoVis.3.1.1.chr7.g01554.t01    rna-gnl|WGS_ZZZZ|egapxtmp_012814-R1    mNeoVis.evergladensis.2.1.chr7.g01630.t01, mNeoVis.evergladensis.2.1.chr7.g01631.t01    mNeoVis.vulgivaga.2.1.chr7.g01646.t01, mNeoVis.vulgivaga.2.1.chr7.g01647.t01      780    N0.HOG0000778   OG0000059  n58   rna-XM_002931242.4, rna-XM_034650891.1    rna-NM_001388693.1, rna-NM_001388830.1, rna-NM_001388831.1, rna-NM_001388832.1, rna-XM_038568160.1, rna-XM_038568162.1    rna-XM_022522083.1, rna-XM_022522141.1    rna-XM_006937030.3, rna-XM_011286661.3          rna-XM_046017075.1    rna-XM_032356277.1, rna-XM_032356278.1, rna-XM_032356279.1    rna-XM_059157114.1, rna-XM_059157124.1    rna-XM_059390296.1    rna-XM_004768269.2, rna-XM_045071513.1, rna-XM_045071517.1    mNeoVis.2.1.1.chr7_hap1.g01651.t01    rna-XM_021685677.1      rna-XM_007089385.2, rna-XM_007089386.2    rna-XM_008710770.2    rna-XM_035723000.1    mNeoVis.3.1.1.chr7.g01537.t01    rna-gnl|WGS_ZZZZ|egapxtmp_012868-R1, rna-gnl|WGS_ZZZZ|egapxtmp_012887-R1, rna-gnl|WGS_ZZZZ|egapxtmp_012889-R1    mNeoVis.evergladensis.2.1.chr7.g01626.t01, mNeoVis.evergladensis.2.1.chr7.g01627.t01    mNeoVis.vulgivaga.2.1.chr7.g01643.t01, mNeoVis.vulgivaga.2.1.chr7.g01644.t01      781    N0.HOG0000779   OG0000059  n93   rna-XM_002926085.4, rna-XM_002931045.2, rna-XM_011217984.2, rna-XM_034650186.1, rna-XM_034650187.1    rna-NM_001388695.1    rna-XM_022522091.1, rna-XM_022522095.1    rna-XM_006944313.4, rna-XM_011286578.2, rna-XM_023238993.2, rna-XM_045038121.1, rna-XM_045038122.1    rna-XM_036081409.1, rna-XM_036081422.1    rna-XM_039225675.1, rna-XM_039225676.1    rna-XM_032853623.1, rna-XM_032853624.1, rna-XM_032853625.1, rna-XM_032853627.1, rna-XM_032853628.1    rna-XM_046016756.1, rna-XM_046016820.1, rna-XM_046016867.1, rna-XM_046017094.1, rna-XM_046017096.1, rna-XM_046017115.1    rna-XM_032359502.1, rna-XM_032360300.1, rna-XM_032360314.1, rna-XM_032360316.1, rna-XM_032360497.1    rna-XM_059157241.1, rna-XM_059157254.1, rna-XM_059157280.1, rna-XM_059157307.1, rna-XM_059157326.1    rna-XM_059387565.1, rna-XM_059390202.1, rna-XM_059390210.1, rna-XM_059390223.1, rna-XM_059395260.1    rna-XM_004779027.2, rna-XM_004779028.2, rna-XM_004779029.2, rna-XM_004779030.2, rna-XM_004782528.2, rna-XM_045071510.1    mNeoVis.2.1.1.chr7_hap1.g01658.t01, mNeoVis.2.1.1.chr7_hap1.g01659.t01, mNeoVis.2.1.1.chr7_hap1.g01660.t01, mNeoVis.2.1.1.chr7_hap1.g01663.t01, mNeoVis.2.1.1.chr7_hap1.g01665.t01, mNeoVis.2.1.1.chr7_hap1.g01666.t01    rna-XM_021685626.1, rna-XM_021685678.1    rna-XM_004407557.1    rna-XM_007092231.3, rna-XM_015540752.2, rna-XM_042958927.1, rna-XM_042959030.1, rna-XM_042959057.1, rna-XM_042959070.1, rna-XM_042959089.1    rna-XM_008709614.2, rna-XM_008709617.2, rna-XM_008710266.2, rna-XM_008710267.1, rna-XM_008711584.1    rna-XM_027577905.1, rna-XM_027580554.2    mNeoVis.3.1.1.chr7.g01555.t01, mNeoVis.3.1.1.chr7.g01556.t01, mNeoVis.3.1.1.chr7.g01557.t01, mNeoVis.3.1.1.chr7.g01558.t01, mNeoVis.3.1.1.chr7.g01561.t01    rna-gnl|WGS_ZZZZ|egapxtmp_010641-R1, rna-gnl|WGS_ZZZZ|egapxtmp_010642-R1, rna-gnl|WGS_ZZZZ|egapxtmp_012842-R1, rna-gnl|WGS_ZZZZ|egapxtmp_012847-R1, rna-gnl|WGS_ZZZZ|egapxtmp_012850-R1, rna-gnl|WGS_ZZZZ|egapxtmp_012863-R1, rna-gnl|WGS_ZZZZ|egapxtmp_012882-R1    mNeoVis.evergladensis.2.1.chr7.g01632.t01, mNeoVis.evergladensis.2.1.chr7.g01633.t01, mNeoVis.evergladensis.2.1.chr7.g01635.t01, mNeoVis.evergladensis.2.1.chr7.g01636.t01, mNeoVis.evergladensis.2.1.chr7.g01637.t01    mNeoVis.vulgivaga.2.1.chr7.g01648.t01, mNeoVis.vulgivaga.2.1.chr7.g01649.t01, mNeoVis.vulgivaga.2.1.chr7.g01651.t01, mNeoVis.vulgivaga.2.1.chr7.g01653.t01      782    N0.HOG0000780   OG0000060  n2           rna-XM_036109348.1                              mNeoVis.3.1.1.chr11.g00425.t01        mNeoVis.vulgivaga.2.1.chr11.g00461.t01      783    N0.HOG0000781   OG0000060  n6   rna-XM_019793823.2    rna-NM_001313846.1    rna-XM_022491984.1      rna-XM_036078065.1    rna-XM_039224545.1    rna-XM_032848032.1    rna-XM_045999985.1    rna-XM_032335234.1    rna-XM_059175991.1    rna-XM_059416214.1    rna-XM_013049805.2    mNeoVis.2.1.1.chr1_hap1.g00915.t01    rna-XM_021689937.1    rna-XM_004413495.1      rna-XM_008683735.2    rna-XM_027606231.1      rna-gnl|WGS_ZZZZ|egapxtmp_038957-R1    mNeoVis.evergladensis.2.1.chr1.g00912.t01    mNeoVis.vulgivaga.2.1.chr1.g00924.t01      784    N0.HOG0000782   OG0000060  n26     rna-XM_038551690.1    rna-XM_022521177.1          rna-XM_032850958.1    rna-XM_046000183.1, rna-XM_046000199.1, rna-XM_046000294.1, rna-XM_046000304.1    rna-XM_032334930.1, rna-XM_032334931.1      rna-XM_059416717.1    rna-XM_013059668.1                          785    N0.HOG0000783   OG0000060  n36   rna-XM_034655712.1, rna-XM_034655810.1    rna-XM_038551700.1    rna-XM_022521258.1, rna-XM_022521276.1          rna-XM_032848038.1, rna-XM_032851105.1    rna-XM_045999435.1, rna-XM_045999464.1, rna-XM_046000158.1    rna-XM_032335245.1    rna-XM_059175971.1    rna-XM_059416716.1    rna-XM_045069326.1    mNeoVis.2.1.1.chr1_hap1.g00906.t01, mNeoVis.2.1.1.chr1_hap1.g00910.t01      rna-XM_004416223.1      rna-XM_008710139.2, rna-XM_040622644.1    rna-XM_027606961.1              786    N0.HOG0000784   OG0000060  n57   rna-XM_019804576.2    rna-NM_001286980.1    rna-XM_022491820.1    rna-XM_006941177.4    rna-XM_036107360.1    rna-XM_039245216.1    rna-XM_032839382.1    rna-XM_045989320.1    rna-XM_032326314.1    rna-XM_059150977.1    rna-XM_059381460.1    rna-XM_004767693.1    mNeoVis.2.1.1.chr7_hap1.g01069.t01    rna-XM_021681129.1    rna-XM_004408184.1    rna-XM_042969722.1    rna-XM_008684584.2    rna-XM_027619078.1      rna-gnl|WGS_ZZZZ|egapxtmp_010033-R2    mNeoVis.evergladensis.2.1.chr7.g01061.t01    mNeoVis.vulgivaga.2.1.chr7.g01071.t01      787    N0.HOG0000785   OG0000060  n75   rna-XM_034639010.1    rna-XM_038528687.1    rna-XM_022491804.1    rna-XM_003997740.5    rna-XM_036107363.1    rna-XM_039249451.1    rna-XM_032839386.1    rna-XM_045987518.1    rna-XM_032323899.1    rna-XM_059150979.1    rna-XM_059383402.1    rna-XM_004767683.3    mNeoVis.2.1.1.chr7_hap1.g01068.t01    rna-XM_021681053.1      rna-XM_042969726.1    rna-XM_040634420.1        rna-gnl|WGS_ZZZZ|egapxtmp_010035-R2    mNeoVis.evergladensis.2.1.chr7.g01060.t01    mNeoVis.vulgivaga.2.1.chr7.g01070.t01      788    N0.HOG0000786   OG0000060  n93     rna-NM_001313803.1    rna-XM_022521164.1        rna-XM_039225809.1    rna-XM_032848102.1, rna-XM_032850833.1    rna-XM_046000784.1, rna-XM_046000809.1, rna-XM_046000812.1, rna-XM_046000814.1            mNeoVis.2.1.1.chr1_hap1.g00908.t01, mNeoVis.2.1.1.chr1_hap1.g00913.t01      rna-XM_004413500.1, rna-XM_004413502.1      rna-XM_008683739.2, rna-XM_008710132.2    rna-XM_027606226.2, rna-XM_027606227.1    mNeoVis.3.1.1.chr1.g00930.t01        mNeoVis.vulgivaga.2.1.chr1.g00922.t01, mNeoVis.vulgivaga.2.1.chr15.g00596.t01      789    N0.HOG0000787   OG0000060  n110   rna-XM_011217040.3, rna-XM_019793103.2, rna-XM_034653328.1    rna-XM_038551675.1    rna-XM_022521162.1, rna-XM_022521189.1        rna-XM_039224316.1, rna-XM_039224320.1, rna-XM_039225480.1, rna-XM_039225482.1, rna-XM_039225717.1    rna-XM_032846844.1, rna-XM_032848036.1, rna-XM_032851053.1    rna-XM_045998770.1, rna-XM_045998821.1, rna-XM_045999979.1, rna-XM_045999981.1, rna-XM_046000182.1, rna-XM_046000198.1, rna-XM_046000293.1, rna-XM_046000303.1    rna-XM_032334932.1, rna-XM_032334973.1, rna-XM_032335232.1, rna-XM_032335241.1    rna-XM_059173376.1, rna-XM_059173377.1    rna-XM_059416402.1, rna-XM_059416403.1    rna-XM_013049804.2, rna-XM_013059667.2    mNeoVis.2.1.1.chr1_hap1.g00907.t01, mNeoVis.2.1.1.chr1_hap1.g00914.t01      rna-XM_004413501.1    rna-XM_015537908.2    rna-XM_008710138.2    rna-XM_027606223.1, rna-XM_027607138.1    mNeoVis.3.1.1.chr1.g00927.t01, mNeoVis.3.1.1.chr1.g00929.t01, mNeoVis.3.1.1.chr1.g00931.t01    rna-gnl|WGS_ZZZZ|egapxtmp_006741-R1, rna-gnl|WGS_ZZZZ|egapxtmp_011858-R1, rna-gnl|WGS_ZZZZ|egapxtmp_011859-R1, rna-gnl|WGS_ZZZZ|egapxtmp_038890-R1    mNeoVis.evergladensis.2.1.chr1.g00908.t01, mNeoVis.evergladensis.2.1.chr1.g00910.t01, mNeoVis.evergladensis.2.1.chr1.g00911.t01    mNeoVis.vulgivaga.2.1.chr1.g00918.t01, mNeoVis.vulgivaga.2.1.chr1.g00920.t01, mNeoVis.vulgivaga.2.1.chr1.g00921.t01, mNeoVis.vulgivaga.2.1.chr1.g00923.t01      790    N0.HOG0000788   OG0000060  n161   rna-XM_002931290.4    rna-NM_001313802.1    rna-XM_022521163.1, rna-XM_022521225.1        rna-XM_039224315.1, rna-XM_039225479.1    rna-XM_032851137.1    rna-XM_046000950.1, rna-XM_046000966.1, rna-XM_046001087.1    rna-XM_032334979.1, rna-XM_032335243.1          mNeoVis.2.1.1.chr1_hap1.g00909.t01, mNeoVis.2.1.1.chr1_hap1.g00912.t01      rna-XM_004416222.1      rna-XM_008683738.2, rna-XM_008710129.2    rna-XM_027607230.1              791    N0.HOG0000789   OG0000061  n1   rna-XM_002930262.2, rna-XM_002930270.2, rna-XM_002930271.3, rna-XM_011217031.2, rna-XM_011218296.2, rna-XM_034650605.1, rna-XM_034650622.1, rna-XM_034650625.1, rna-XM_034650627.1, rna-XM_034650777.1, rna-XM_034653324.1, rna-XM_034653325.1    rna-XM_038556367.1    rna-XM_022491968.1, rna-XM_022492385.1, rna-XM_022492388.1    rna-XM_003980597.2, rna-XM_003980603.2, rna-XM_003980604.2, rna-XM_003980605.1, rna-XM_011283236.2, rna-XM_019831422.2, rna-XM_019832928.3    rna-XM_036065929.1, rna-XM_036109027.1, rna-XM_036109028.1, rna-XM_036114482.1, rna-XM_036114484.1    rna-XM_039222992.1, rna-XM_039222997.1, rna-XM_039222998.1, rna-XM_039224229.1    rna-XM_032844361.1, rna-XM_032844362.1, rna-XM_032844363.1, rna-XM_032844371.1, rna-XM_032844372.1, rna-XM_032848087.1, rna-XM_032850957.1    rna-XM_045981642.1, rna-XM_045996827.1, rna-XM_045999187.1    rna-XM_032334917.1, rna-XM_032334994.1, rna-XM_032334996.1, rna-XM_032335022.1, rna-XM_032335028.1, rna-XM_032335044.1, rna-XM_032335046.1    rna-XM_059158835.1, rna-XM_059158836.1, rna-XM_059173406.1, rna-XM_059173433.1    rna-XM_059416423.1, rna-XM_059416449.1, rna-XM_059416451.1    rna-XM_013050659.2, rna-XM_045069237.1, rna-XM_045069239.1, rna-XM_045069241.1    mNeoVis.2.1.1.chr11_hap1.g00542.t01, mNeoVis.2.1.1.chr11_hap1.g00544.t01, mNeoVis.2.1.1.chr1_hap1.g00947.t01, mNeoVis.2.1.1.chr1_hap1.g00991.t01, mNeoVis.2.1.1.chr1_hap1.g00995.t01, mNeoVis.2.1.1.chr1_hap1.g00998.t01, mNeoVis.2.1.1.chr1_hap1.g00999.t01, mNeoVis.2.1.1.chr1_hap1.g01000.t01, mNeoVis.2.1.1.chr9_hap1.g00178.t01    rna-XM_021686917.2, rna-XM_021689890.2, rna-XM_021689900.1, rna-XM_021692322.2, rna-XM_021692379.1, rna-XM_044917037.1, rna-XM_044917038.1    rna-XM_004416787.1, rna-XM_004417083.1, rna-XM_004417084.1    rna-XM_007084983.2, rna-XM_007084996.1, rna-XM_007084997.2, rna-XM_007084998.2, rna-XM_007084999.2, rna-XM_015542383.1, rna-XM_015542388.2, rna-XM_015542389.2    rna-XM_008709370.2, rna-XM_008709433.2, rna-XM_008710075.2, rna-XM_008710079.2, rna-XM_008710080.2, rna-XM_008710100.1, rna-XM_008710101.1, rna-XM_008710102.2, rna-XM_008710104.2, rna-XM_008710107.2, rna-XM_008710109.2, rna-XM_008710110.2, rna-XM_008710111.1, rna-XM_008710113.2, rna-XM_008710114.1, rna-XM_008710116.2, rna-XM_008710117.2, rna-XM_040622722.1, rna-XM_040622730.1, rna-XM_040622731.1, rna-XM_040622732.1    rna-XM_027604486.1, rna-XM_027604487.1, rna-XM_027604489.2, rna-XM_027604592.1, rna-XM_027604641.1    mNeoVis.3.1.1.chr1.g00959.t01, mNeoVis.3.1.1.chr1.g00995.t01, mNeoVis.3.1.1.chr1.g00996.t01, mNeoVis.3.1.1.chr1.g00999.t01, mNeoVis.3.1.1.chr1.g01000.t01, mNeoVis.3.1.1.chr1.g01001.t01, mNeoVis.3.1.1.chr11.g00504.t01, mNeoVis.3.1.1.chr2.g00939.t01    rna-gnl|WGS_ZZZZ|egapxtmp_003817-R1, rna-gnl|WGS_ZZZZ|egapxtmp_006876-R1, rna-gnl|WGS_ZZZZ|egapxtmp_006887-R1, rna-gnl|WGS_ZZZZ|egapxtmp_009772-R1, rna-gnl|WGS_ZZZZ|egapxtmp_010043-R1, rna-gnl|WGS_ZZZZ|egapxtmp_020598-R1, rna-gnl|WGS_ZZZZ|egapxtmp_025212-R1, rna-gnl|WGS_ZZZZ|egapxtmp_025214-R1    mNeoVis.evergladensis.2.1.chr1.g00942.t01, mNeoVis.evergladensis.2.1.chr1.g00985.t01, mNeoVis.evergladensis.2.1.chr1.g00988.t01, mNeoVis.evergladensis.2.1.chr1.g00989.t01, mNeoVis.evergladensis.2.1.chr1.g00990.t01, mNeoVis.evergladensis.2.1.chr1.g00992.t01, mNeoVis.evergladensis.2.1.chr11.g00531.t01    mNeoVis.vulgivaga.2.1.chr1.g00953.t01, mNeoVis.vulgivaga.2.1.chr1.g00998.t01, mNeoVis.vulgivaga.2.1.chr1.g01002.t01, mNeoVis.vulgivaga.2.1.chr1.g01003.t01, mNeoVis.vulgivaga.2.1.chr1.g01004.t01, mNeoVis.vulgivaga.2.1.chr1.g01006.t01, mNeoVis.vulgivaga.2.1.chr11.g00539.t01      792    N0.HOG0000790   OG0000061  n135   rna-XM_002931048.2    rna-XM_038583984.1    rna-XM_022492391.1    rna-XM_003985633.4, rna-XM_045061427.1      rna-XM_039222994.1, rna-XM_039224772.1    rna-XM_032869979.1    rna-XM_045997474.1    rna-XM_032329007.1, rna-XM_032334992.1, rna-XM_032334998.1, rna-XM_032335000.1    rna-XM_059155908.1, rna-XM_059158837.1, rna-XM_059173414.1    rna-XM_059384534.1, rna-XM_059416447.1, rna-XM_059416448.1    rna-XM_004766787.2, rna-XM_013050914.1    mNeoVis.2.1.1.chr11_hap1.g00541.t01, mNeoVis.2.1.1.chr1_hap1.g00986.t01, mNeoVis.2.1.1.chr6_hap1.g01132.t01      rna-XM_004416835.1, rna-XM_004417455.1    rna-XM_007092978.2, rna-XM_015539176.2    rna-XM_008708888.2, rna-XM_008710097.1      mNeoVis.3.1.1.chr1.g00991.t01, mNeoVis.3.1.1.chr11.g00502.t01, mNeoVis.3.1.1.chr4.g01078.t01    rna-gnl|WGS_ZZZZ|egapxtmp_000928-R1, rna-gnl|WGS_ZZZZ|egapxtmp_020174-R1, rna-gnl|WGS_ZZZZ|egapxtmp_025209-R1    mNeoVis.evergladensis.2.1.chr1.g00980.t01, mNeoVis.evergladensis.2.1.chr11.g00529.t01, mNeoVis.evergladensis.2.1.chr4.g01064.t01    mNeoVis.vulgivaga.2.1.chr1.g00993.t01, mNeoVis.vulgivaga.2.1.chr11.g00538.t01, mNeoVis.vulgivaga.2.1.chr4.g01099.t01      793    N0.HOG0000791   OG0000062  n3   rna-XM_002915725.4    rna-XM_038558030.1    rna-XM_022496567.1    rna-XM_019837005.3, rna-XM_045051065.1      rna-XM_039237872.1    rna-XM_032844100.1    rna-XM_046005934.1, rna-XM_046022216.1    rna-XM_032361538.1    rna-XM_059135057.1    rna-XM_059374735.1    rna-XM_004774469.3    mNeoVis.2.1.1.chr12_hap1.g00388.t01, mNeoVis.2.1.1.chr2_hap1.g01295.t01    rna-XM_021684310.1    rna-XM_004393584.1, rna-XM_004400314.1    rna-XM_007077208.3    rna-XM_040627311.1    rna-XM_027598544.2, rna-XM_035726899.1    mNeoVis.3.1.1.chr12.g00350.t01, mNeoVis.3.1.1.chr5.g01220.t01    rna-gnl|WGS_ZZZZ|egapxtmp_005540-R2, rna-gnl|WGS_ZZZZ|egapxtmp_029300-R1    mNeoVis.evergladensis.2.1.chr12.g00346.t01, mNeoVis.evergladensis.2.1.chr5.g01238.t01    mNeoVis.vulgivaga.2.1.chr5.g01232.t01      794    N0.HOG0000792   OG0000062  n11     rna-XM_038530753.1                            rna-XM_004394656.1    rna-XM_042968068.1                  795    N0.HOG0000793   OG0000062  n15                   rna-XM_032328239.1              rna-XM_004415158.1                    796    N0.HOG0000794   OG0000062  n18                 rna-XM_046027295.1                      rna-XM_035723690.1    mNeoVis.3.1.1.chr5.g01587.t01    rna-gnl|WGS_ZZZZ|egapxtmp_007527-R1    mNeoVis.evergladensis.2.1.chr5.g01605.t01        797    N0.HOG0000795   OG0000062  n23                                     rna-XM_027585962.2, rna-XM_035724311.1              798    N0.HOG0000796   OG0000062  n25                           mNeoVis.2.1.1.chr12_hap1.g00405.t01                    mNeoVis.vulgivaga.2.1.chr12.g00369.t01      799    N0.HOG0000797   OG0000062  n31               rna-XM_032874092.1        rna-XM_059175224.1    rna-XM_059418050.1                  mNeoVis.3.1.1.chr1.g00674.t01      mNeoVis.evergladensis.2.1.chr1.g00662.t01, mNeoVis.evergladensis.2.1.chr1.g01475.t01        800    N0.HOG0000798   OG0000062  n37   rna-XM_002929523.4        rna-XM_023257367.2      rna-XM_039242097.1    rna-XM_032848616.1    rna-XM_046001054.1    rna-XM_032335986.1        rna-XM_004766873.3    mNeoVis.2.1.1.chr1_hap1.g00657.t01      rna-XM_004403440.1    rna-XM_042988264.1              mNeoVis.vulgivaga.2.1.chr1.g00674.t01      801    N0.HOG0000799   OG0000062  n44           rna-XM_036116744.1      rna-XM_032856046.1                rna-XM_044916911.1, rna-XM_044917642.1    rna-XM_004410257.1        rna-XM_035724329.1, rna-XM_035728591.1, rna-XM_035728663.1              802    N0.HOG0000800   OG0000062  n52               rna-XM_032880066.1          rna-XM_059412315.1                            803    N0.HOG0000801   OG0000062  n54                             rna-XM_044913642.1    rna-XM_004399697.1                    804    N0.HOG0000802   OG0000062  n56                 rna-XM_046027417.1    rna-XM_032343611.1    rna-XM_059181454.1    rna-XM_059371196.1    rna-XM_045067864.1        rna-XM_004406017.1                    805    N0.HOG0000803   OG0000062  n66   rna-XM_034653429.1    rna-XM_038574956.1        rna-XM_036070057.1, rna-XM_036078803.1, rna-XM_036092407.1, rna-XM_036108851.1, rna-XM_036118637.1, rna-XM_036118810.1, rna-XM_036119145.1      rna-XM_032855395.1    rna-XM_045979320.1        rna-XM_059392768.1        rna-XM_021685807.1, rna-XM_044911760.1, rna-XM_044912573.1, rna-XM_044913732.1, rna-XM_044914765.1, rna-XM_044918512.1      rna-XM_042989237.1      rna-XM_035728888.1, rna-XM_035729297.1    mNeoVis.3.1.1.chr2.g00756.t01            806    N0.HOG0000804   OG0000062  n87     rna-XM_038532441.1      rna-XM_045033263.1, rna-XM_045033334.1    rna-XM_036076343.1, rna-XM_036091252.1    rna-XM_039221196.1      rna-XM_046019205.1    rna-XM_032314686.1            rna-XM_044921589.1          rna-XM_035722090.1, rna-XM_035725731.1    mNeoVis.3.1.1.chr2.g01003.t01            807    N0.HOG0000805   OG0000062  n98                           mNeoVis.2.1.1.chr9_hap1.g00734.t01                rna-gnl|WGS_ZZZZ|egapxtmp_021971-R1      mNeoVis.vulgivaga.2.1.chr10.g00717.t01      808    N0.HOG0000806   OG0000062  n100               rna-XM_032858558.1      rna-XM_032336827.1    rna-XM_059162469.1    rna-XM_059387961.1    rna-XM_045088978.1    mNeoVis.2.1.1.chr2_hap1.g01040.t01        rna-XM_042991354.1        mNeoVis.3.1.1.chr5.g00966.t01        mNeoVis.vulgivaga.2.1.chr5.g00976.t01      809    N0.HOG0000807   OG0000062  n108           rna-XM_036113214.1                      rna-XM_004411705.1        rna-XM_035726544.1              810    N0.HOG0000808   OG0000062  n110                                     rna-XM_035726237.1, rna-XM_035728645.1              811    N0.HOG0000809   OG0000062  n111   rna-XM_034652189.1    rna-NM_001252133.1            rna-XM_032880124.1        rna-XM_059168495.1            rna-XM_004411859.1        rna-XM_035723414.1              812    N0.HOG0000810   OG0000062  n116   rna-XM_034652218.1    rna-XM_038544017.1    rna-XM_022520412.1      rna-XM_036080100.1, rna-XM_036092993.1    rna-XM_039218036.1    rna-XM_032838798.1, rna-XM_032841415.1, rna-XM_032866806.1      rna-XM_032325861.1, rna-XM_032340305.1    rna-XM_059140322.1      rna-XM_045074434.1      rna-XM_021702175.1, rna-XM_044911983.1          rna-XM_035724038.1      rna-gnl|WGS_ZZZZ|egapxtmp_021354-R1          813    N0.HOG0000811   OG0000062  n117       rna-XM_022504886.1      rna-XM_036084456.1, rna-XM_036101323.1    rna-XM_039240510.1    rna-XM_032852105.1        rna-XM_059176322.1, rna-XM_059185579.1    rna-XM_059374547.1, rna-XM_059399257.1    rna-XM_045083069.1    mNeoVis.2.1.1.chr1_hap1.g01512.t01    rna-XM_044920782.1    rna-XM_004410214.1        rna-XM_027587214.1, rna-XM_035723503.1, rna-XM_035726864.1, rna-XM_035728962.1      rna-gnl|WGS_ZZZZ|egapxtmp_037223-R1          814    N0.HOG0000812   OG0000062  n134               rna-XM_032852423.1                rna-XM_044918744.1                      815    N0.HOG0000813   OG0000062  n135                   rna-XM_032332897.1            rna-XM_044919435.1            mNeoVis.3.1.1.chr8.g00593.t01            816    N0.HOG0000814   OG0000063  n1   rna-XM_011218070.3, rna-XM_034650290.1, rna-XM_034650337.1    rna-XM_038527815.1    rna-XM_022491865.1    rna-XM_045045881.1, rna-XM_045045885.1    rna-XM_036082359.1    rna-XM_039244179.1, rna-XM_039244184.1    rna-XM_032848624.1, rna-XM_032848629.1    rna-XM_045987718.1, rna-XM_045988118.1, rna-XM_045988195.1    rna-XM_032324456.1, rna-XM_032324495.1    rna-XM_059153624.1, rna-XM_059153628.1, rna-XM_059153641.1, rna-XM_059153653.1    rna-XM_059381784.1, rna-XM_059381792.1, rna-XM_059383336.1, rna-XM_059387331.1    rna-XM_004779775.3, rna-XM_004779817.3, rna-XM_013049569.2, rna-XM_013049578.2    mNeoVis.2.1.1.chr7_hap1.g00735.t01, mNeoVis.2.1.1.chr7_hap1.g00742.t01, mNeoVis.2.1.1.chr7_hap1.g00744.t01, mNeoVis.2.1.1.chr7_hap1.g00747.t01    rna-XM_021682313.2, rna-XM_044912068.1    rna-XM_004409567.1    rna-XM_007087138.3, rna-XM_042969996.1    rna-XM_008701611.2, rna-XM_008709825.2, rna-XM_040633845.1    rna-XM_035724919.1, rna-XM_035725045.1, rna-XM_035725046.1    mNeoVis.3.1.1.chr7.g00688.t01, mNeoVis.3.1.1.chr7.g00694.t01, mNeoVis.3.1.1.chr7.g00695.t01, mNeoVis.3.1.1.chr7.g00698.t01    rna-gnl|WGS_ZZZZ|egapxtmp_012283-R1, rna-gnl|WGS_ZZZZ|egapxtmp_012285-R1, rna-gnl|WGS_ZZZZ|egapxtmp_014162-R1    mNeoVis.evergladensis.2.1.chr7.g00725.t01, mNeoVis.evergladensis.2.1.chr7.g00732.t01, mNeoVis.evergladensis.2.1.chr7.g00735.t01    mNeoVis.vulgivaga.2.1.chr7.g00727.t01, mNeoVis.vulgivaga.2.1.chr7.g00735.t01, mNeoVis.vulgivaga.2.1.chr7.g00738.t01      817    N0.HOG0000815   OG0000063  n56   rna-XM_034638105.1, rna-XM_034650363.1, rna-XM_034672561.1, rna-XM_034672571.1, rna-XM_034672601.1, rna-XM_034672612.1    rna-XM_038525529.1, rna-XM_038527805.1    rna-XM_022491863.1, rna-XM_022492416.1, rna-XM_022492426.1, rna-XM_022492442.1, rna-XM_022492697.1, rna-XM_022492735.1, rna-XM_022492794.1, rna-XM_022492795.1    rna-XM_006940890.5, rna-XM_045045866.1, rna-XM_045045891.1, rna-XM_045045970.1, rna-XM_045046033.1, rna-XM_045046915.1    rna-XM_036082336.1, rna-XM_036082337.1, rna-XM_036082356.1, rna-XM_036092044.1, rna-XM_036092049.1, rna-XM_036092063.1, rna-XM_036092079.1, rna-XM_036092087.1    rna-XM_039244084.1, rna-XM_039244174.1, rna-XM_039244180.1, rna-XM_039244192.1, rna-XM_039244218.1    rna-XM_032848642.1, rna-XM_032848651.1, rna-XM_032848710.1, rna-XM_032848722.1, rna-XM_032848723.1, rna-XM_032848738.1, rna-XM_032848739.1, rna-XM_032851200.1, rna-XM_032851201.1    rna-XM_045986813.1, rna-XM_045987290.1, rna-XM_045987291.1, rna-XM_045987729.1, rna-XM_045987730.1    rna-XM_032324437.1, rna-XM_032324451.1, rna-XM_032324464.1, rna-XM_032324487.1, rna-XM_032324500.1, rna-XM_032324501.1, rna-XM_032324529.1, rna-XM_032324533.1, rna-XM_032326053.1, rna-XM_032326054.1    rna-XM_059151206.1, rna-XM_059153614.1, rna-XM_059153615.1, rna-XM_059153620.1, rna-XM_059153627.1, rna-XM_059153630.1, rna-XM_059153650.1, rna-XM_059153654.1    rna-XM_059381511.1, rna-XM_059381789.1, rna-XM_059381798.1, rna-XM_059383332.1, rna-XM_059383333.1, rna-XM_059383335.1    rna-XM_004779804.3, rna-XM_013050769.2, rna-XM_045076940.1, rna-XM_045076944.1, rna-XM_045076945.1, rna-XM_045076952.1, rna-XM_045076962.1, rna-XM_045076976.1, rna-XM_045077264.1    mNeoVis.2.1.1.chr7_hap1.g00734.t01, mNeoVis.2.1.1.chr7_hap1.g00741.t01, mNeoVis.2.1.1.chr7_hap1.g00743.t01, mNeoVis.2.1.1.chr7_hap1.g00745.t01    rna-XM_021682394.2, rna-XM_044911335.1, rna-XM_044912551.1    rna-XM_004405023.1, rna-XM_004405024.1, rna-XM_004409568.1, rna-XM_004415863.1, rna-XM_004415867.1, rna-XM_004415870.1    rna-XM_007087142.3, rna-XM_007087167.3, rna-XM_015539794.2, rna-XM_042968214.1    rna-XM_008709816.2, rna-XM_040633843.1, rna-XM_040633850.1, rna-XM_040633863.1, rna-XM_040633868.1, rna-XM_040633884.1    rna-XM_035724918.1, rna-XM_035725040.1, rna-XM_035725041.1, rna-XM_035725044.1, rna-XM_035725055.1, rna-XM_035725056.1    mNeoVis.3.1.1.chr7.g00687.t01, mNeoVis.3.1.1.chr7.g00693.t01, mNeoVis.3.1.1.chr7.g00696.t01    rna-gnl|WGS_ZZZZ|egapxtmp_012284-R1, rna-gnl|WGS_ZZZZ|egapxtmp_014158-R1    mNeoVis.evergladensis.2.1.chr1.g00973.t01, mNeoVis.evergladensis.2.1.chr7.g00724.t01, mNeoVis.evergladensis.2.1.chr7.g00731.t01, mNeoVis.evergladensis.2.1.chr7.g00733.t01    mNeoVis.vulgivaga.2.1.chr7.g00726.t01, mNeoVis.vulgivaga.2.1.chr7.g00733.t01, mNeoVis.vulgivaga.2.1.chr7.g00734.t01, mNeoVis.vulgivaga.2.1.chr7.g00736.t01      818    N0.HOG0000816   OG0000064  n2   rna-XM_034645293.1    rna-NM_001003080.1, rna-NM_001193656.1    rna-XM_022506606.1    rna-NM_001048151.1    rna-XM_036086657.1, rna-XM_036096209.1    rna-XM_039253525.1    rna-XM_032879219.1    rna-XM_046010376.1, rna-XM_046016272.1    rna-XM_032358507.1    rna-XM_059144331.1, rna-XM_059180691.1    rna-XM_059397526.1, rna-XM_059404164.1, rna-XM_059412233.1    rna-XM_004770459.3, rna-XM_045081782.1    mNeoVis.2.1.1.chr6_hap1.g00740.t01, mNeoVis.2.1.1.chr7_hap1.g02296.t01, mNeoVis.2.1.1.chr8_hap1.g01445.t01    rna-XM_021684915.2, rna-XM_044913153.1    rna-XM_004398753.1, rna-XM_004410046.1, rna-XM_004411229.1, rna-XM_004416568.1, rna-XM_004416735.1    rna-XM_042958542.1    rna-XM_040621101.1, rna-XM_040638768.1    rna-XM_035722804.1, rna-XM_035723144.1, rna-XM_035723476.1, rna-XM_035726382.1, rna-XM_035726786.1, rna-XM_035727225.1      rna-gnl|WGS_ZZZZ|egapxtmp_010109-R1    mNeoVis.evergladensis.2.1.chr7.g02243.t01, mNeoVis.evergladensis.2.1.chr9.g01392.t01    mNeoVis.vulgivaga.2.1.chr7.g02261.t01      819    N0.HOG0000817   OG0000064  n32     rna-XM_038570267.1, rna-XM_038582460.1                            rna-XM_004399763.1        rna-XM_035727538.1, rna-XM_035728552.1    mNeoVis.3.1.1.chr7.g02114.t01            820    N0.HOG0000818   OG0000064  n37   rna-XM_034649713.1, rna-XM_034649919.1, rna-XM_034650297.1, rna-XM_034651822.1, rna-XM_034652324.1    rna-XM_038581722.1      rna-XM_011291686.4, rna-XM_045050451.1, rna-XM_045050510.1, rna-XM_045050616.1, rna-XM_045050619.1, rna-XM_045050697.1, rna-XM_045051102.1    rna-XM_036077816.1, rna-XM_036086155.1, rna-XM_036089445.1, rna-XM_036119353.1, rna-XM_036121378.1, rna-XM_036122258.1    rna-XM_039223095.1, rna-XM_039225052.1, rna-XM_039248784.1, rna-XM_039256074.1, rna-XM_039256082.1, rna-XM_039256241.1, rna-XM_039256403.1, rna-XM_039256409.1    rna-XM_032883137.1, rna-XM_032883153.1, rna-XM_032883157.1, rna-XM_032883159.1, rna-XM_032883160.1, rna-XM_032883161.1, rna-XM_032883162.1    rna-XM_045994737.1, rna-XM_045994749.1, rna-XM_045994751.1, rna-XM_045994752.1, rna-XM_045995516.1, rna-XM_045995780.1    rna-XM_032304625.1, rna-XM_032329687.1, rna-XM_032329860.1, rna-XM_032330454.1, rna-XM_032330504.1, rna-XM_032330621.1, rna-XM_032330635.1, rna-XM_032330779.1, rna-XM_032341788.1    rna-XM_059156746.1, rna-XM_059157233.1, rna-XM_059157261.1, rna-XM_059157308.1, rna-XM_059157315.1, rna-XM_059157439.1, rna-XM_059157494.1    rna-XM_059371150.1, rna-XM_059384594.1, rna-XM_059384961.1, rna-XM_059385272.1, rna-XM_059385279.1    rna-XM_045089216.1, rna-XM_045089217.1, rna-XM_045089221.1, rna-XM_045089222.1, rna-XM_045089224.1, rna-XM_045089227.1, rna-XM_045089445.1    mNeoVis.2.1.1.chr10_hap1.g00395.t01, mNeoVis.2.1.1.chr12_hap1.g00685.t01, mNeoVis.2.1.1.chr12_hap1.g00686.t01, mNeoVis.2.1.1.chr12_hap1.g00688.t01, mNeoVis.2.1.1.chr12_hap1.g00690.t01, mNeoVis.2.1.1.chr12_hap1.g00691.t01, mNeoVis.2.1.1.chr12_hap1.g00704.t01    rna-XM_021681562.1, rna-XM_044911644.1, rna-XM_044911645.1, rna-XM_044911646.1, rna-XM_044911647.1, rna-XM_044911657.1, rna-XM_044912662.1, rna-XM_044918633.1    rna-XM_004402012.1, rna-XM_004404975.1, rna-XM_004407438.1, rna-XM_004409592.1    rna-XM_015541301.2, rna-XM_042974292.1, rna-XM_042974296.1, rna-XM_042974324.1, rna-XM_042974398.1, rna-XM_042974468.1, rna-XM_042974819.1    rna-XM_040625336.1, rna-XM_040625405.1, rna-XM_040625558.1, rna-XM_040625885.1, rna-XM_040626137.1, rna-XM_040633087.1, rna-XM_040633101.1    rna-XM_027608306.2, rna-XM_035723385.1, rna-XM_035725488.1, rna-XM_035725489.1, rna-XM_035725490.1, rna-XM_035725620.1, rna-XM_035725624.1, rna-XM_035725625.1, rna-XM_035725673.1, rna-XM_035729239.1    mNeoVis.3.1.1.chr1.g01490.t01, mNeoVis.3.1.1.chr12.g00636.t01, mNeoVis.3.1.1.chr12.g00637.t01, mNeoVis.3.1.1.chr12.g00644.t01    rna-gnl|WGS_ZZZZ|egapxtmp_015336-R1, rna-gnl|WGS_ZZZZ|egapxtmp_028232-R1, rna-gnl|WGS_ZZZZ|egapxtmp_028233-R1, rna-gnl|WGS_ZZZZ|egapxtmp_028819-R1, rna-gnl|WGS_ZZZZ|egapxtmp_029083-R1, rna-gnl|WGS_ZZZZ|egapxtmp_029575-R1    mNeoVis.evergladensis.2.1.chr12.g00622.t01, mNeoVis.evergladensis.2.1.chr12.g00625.t01, mNeoVis.evergladensis.2.1.chr12.g00634.t01, mNeoVis.evergladensis.2.1.chr8.g00377.t01    mNeoVis.vulgivaga.2.1.chr12.g00642.t01, mNeoVis.vulgivaga.2.1.chr12.g00644.t01, mNeoVis.vulgivaga.2.1.chr12.g00646.t01, mNeoVis.vulgivaga.2.1.chr12.g00647.t01, mNeoVis.vulgivaga.2.1.chr12.g00652.t01, mNeoVis.vulgivaga.2.1.chr12.g00657.t01      821    N0.HOG0000819   OG0000065  n1   rna-XM_019801456.2    rna-XM_038529331.1    rna-XM_022525230.1    rna-XM_023245481.2    rna-XM_036067881.1    rna-XM_039249633.1    rna-XM_032841912.1    rna-XM_045988814.1    rna-XM_032323700.1    rna-XM_059152910.1    rna-XM_059381574.1    rna-XM_045075941.1    mNeoVis.2.1.1.chr7_hap1.g00432.t01    rna-XM_044922209.1    rna-XM_004417091.1    rna-XM_042969063.1    rna-XM_008712221.2    rna-XM_027617876.2    mNeoVis.3.1.1.chr7.g00419.t01    rna-gnl|WGS_ZZZZ|egapxtmp_012218-R1    mNeoVis.evergladensis.2.1.chr7.g00436.t01    mNeoVis.vulgivaga.2.1.chr7.g00434.t01      822    N0.HOG0000820   OG0000065  n27   rna-XM_002924114.4                                  rna-XM_040626565.1    rna-XM_035722862.1, rna-XM_035727258.1          mNeoVis.vulgivaga.2.1.chr10.g00968.t01      823    N0.HOG0000821   OG0000065  n32       rna-XM_022525866.1                      mNeoVis.2.1.1.chr10_hap1.g00478.t01                        824    N0.HOG0000822   OG0000065  n36     rna-XM_038542135.1, rna-XM_038549480.1, rna-XM_038550986.1, rna-XM_038557297.1, rna-XM_038586521.1                                              825    N0.HOG0000823   OG0000065  n45           rna-XM_036111776.1          rna-XM_032323229.1                    rna-XM_027570303.2, rna-XM_027609556.2              826    N0.HOG0000824   OG0000065  n49     rna-XM_038569028.1                          rna-XM_044911413.1                      827    N0.HOG0000825   OG0000065  n52           rna-XM_036084337.1                    rna-XM_021679801.2                      828    N0.HOG0000826   OG0000065  n55                     rna-XM_059176821.1    rna-XM_059399276.1    rna-XM_013056646.2                          829    N0.HOG0000827   OG0000065  n58     rna-XM_038579969.1    rna-XM_022508696.1, rna-XM_022524039.1    rna-XM_011287646.4          rna-XM_045995588.1            mNeoVis.2.1.1.chr12_hap1.g00475.t01              mNeoVis.3.1.1.chr12.g00433.t01    rna-gnl|WGS_ZZZZ|egapxtmp_013173-R1      mNeoVis.vulgivaga.2.1.chr12.g00436.t01, mNeoVis.vulgivaga.2.1.chr7.g00858.t01      830    N0.HOG0000828   OG0000065  n67     rna-XM_038569099.1                rna-XM_032347714.1              rna-XM_004397109.1        rna-XM_027585087.2              831    N0.HOG0000829   OG0000065  n71     rna-XM_038532098.1, rna-XM_038552130.1                                              832    N0.HOG0000830   OG0000065  n73     rna-XM_038543859.1, rna-XM_038586433.1                          rna-XM_021684379.2      rna-XM_015537902.2      rna-XM_035724409.1              833    N0.HOG0000831   OG0000065  n77     rna-XM_038574522.1      rna-XM_011287405.2    rna-XM_036108820.1                                        834    N0.HOG0000832   OG0000065  n79   rna-XM_034638851.1    rna-XM_038581887.1    rna-XM_022492777.1, rna-XM_022522270.1              rna-XM_032335758.1                    rna-XM_027591095.2, rna-XM_035723184.1              835    N0.HOG0000833   OG0000065  n85   rna-NM_001304846.1                    rna-XM_059184619.1    rna-XM_059403732.1    rna-NM_001310173.1, rna-XM_004766704.3        rna-XM_004413892.1                    836    N0.HOG0000834   OG0000065  n93     rna-XM_038532102.1        rna-XM_036096160.1            rna-XM_059148305.1    rna-XM_059380795.1    rna-XM_045069975.1    mNeoVis.2.1.1.chr6_hap1.g00636.t01          rna-XM_040632464.1    rna-XM_027595349.1, rna-XM_027605538.2, rna-XM_027608159.2    mNeoVis.3.1.1.chr4.g00641.t01    rna-gnl|WGS_ZZZZ|egapxtmp_008896-R1, rna-gnl|WGS_ZZZZ|egapxtmp_042897-R1    mNeoVis.evergladensis.2.1.chr4.g00644.t01    mNeoVis.vulgivaga.2.1.chr4.g00652.t01      837    N0.HOG0000835   OG0000065  n112     rna-XM_038551655.1, rna-XM_038553380.1, rna-XM_038586276.1                        mNeoVis.2.1.1.chr2_hap1.g01461.t01    rna-XM_021686026.2        rna-XM_008684315.2    rna-XM_035724129.1        mNeoVis.evergladensis.2.1.chr5.g01399.t01        838    N0.HOG0000836   OG0000065  n123     rna-XM_038545974.1    rna-XM_022492412.1, rna-XM_022512167.1            rna-XM_046003650.1    rna-XM_032361168.1    rna-XM_059165038.1, rna-XM_059179658.1    rna-XM_059372483.1, rna-XM_059388325.1, rna-XM_059401483.1    rna-XM_013055396.2, rna-XM_045065585.1, rna-XM_045088137.1    mNeoVis.2.1.1.chr4_hap1.g00745.t01, mNeoVis.2.1.1.chr4_hap1.g00765.t01            rna-XM_035722871.1, rna-XM_035727537.1    mNeoVis.3.1.1.chr10.g00246.t01    rna-gnl|WGS_ZZZZ|egapxtmp_004519-R1, rna-gnl|WGS_ZZZZ|egapxtmp_035842-R1    mNeoVis.evergladensis.2.1.chr6.g00519.t01    mNeoVis.vulgivaga.2.1.chr5.g01399.t01      839    N0.HOG0000837   OG0000065  n144     rna-XM_038565882.1    rna-XM_022492776.1, rna-XM_022512724.1      rna-XM_036115787.1                  mNeoVis.2.1.1.chr8_hap1.g01193.t01    rna-XM_021701476.2          rna-XM_027589721.2, rna-XM_027591086.2      rna-gnl|WGS_ZZZZ|egapxtmp_019117-R1      mNeoVis.vulgivaga.2.1.chr9.g00744.t01      840    N0.HOG0000838   OG0000065  n153                                     rna-XM_035727806.1, rna-XM_035727820.1              841    N0.HOG0000839   OG0000065  n154     rna-XM_038564463.1                rna-XM_032316020.1        rna-XM_045074271.1              rna-XM_035723088.1              842    N0.HOG0000840   OG0000065  n157           rna-XM_036117685.1      rna-XM_032874781.1    rna-XM_046011043.1    rna-XM_032346517.1, rna-XM_032349015.1          mNeoVis.2.1.1.chr9_hap1.g00253.t01    rna-XM_021677889.2, rna-XM_044914949.1        rna-XM_040642080.1    rna-XM_027594890.1        mNeoVis.evergladensis.2.1.chr5.g00462.t01    mNeoVis.vulgivaga.2.1.chr5.g00461.t01      843    N0.HOG0000841   OG0000065  n160                 rna-XM_045984381.1, rna-XM_046006628.1                rna-XM_004403791.1        rna-XM_027590889.2        mNeoVis.evergladensis.2.1.chr10.g00251.t01        844    N0.HOG0000842   OG0000065  n164     rna-NM_001003142.2, rna-XM_038572377.1                              rna-NM_001290607.1                  845    N0.HOG0000843   OG0000065  n166         rna-XM_006933438.4                          rna-XM_042994205.1                  846    N0.HOG0000844   OG0000066  n1   rna-XM_002924109.4    rna-XM_038579338.1    rna-XM_022515514.1    rna-XM_019822692.3    rna-XM_036106088.1    rna-XM_039220288.1    rna-XM_032882578.1    rna-XM_046004116.1    rna-XM_032316655.1    rna-XM_059166422.1    rna-XM_059394598.1    rna-XM_045075770.1    mNeoVis.2.1.1.chr4_hap1.g00716.t01    rna-XM_021688587.1    rna-XM_004414441.1    rna-XM_042972813.1    rna-XM_040625726.1    rna-XM_027577290.2    mNeoVis.3.1.1.chr6.g00722.t01    rna-gnl|WGS_ZZZZ|egapxtmp_003961-R1    mNeoVis.evergladensis.2.1.chr6.g00722.t01    mNeoVis.vulgivaga.2.1.chr6.g00717.t01      847    N0.HOG0000845   OG0000066  n17   rna-XM_011222248.3    rna-XM_038554980.1    rna-XM_022520200.1    rna-XM_023254266.2    rna-XM_036105329.1    rna-XM_039252207.1    rna-XM_032858340.1    rna-XM_046004796.1    rna-XM_032337998.1    rna-XM_059176823.1    rna-XM_059401223.1    rna-XM_004753526.3    mNeoVis.2.1.1.chr1_hap1.g02041.t01    rna-XM_021693674.2    rna-XM_004406721.1    rna-XM_042987006.1    rna-XM_040627650.1    rna-XM_027603145.2    mNeoVis.3.1.1.chr1.g01995.t01    rna-gnl|WGS_ZZZZ|egapxtmp_020693-R1    mNeoVis.evergladensis.2.1.chr1.g02008.t01    mNeoVis.vulgivaga.2.1.chr1.g02001.t01      848    N0.HOG0000846   OG0000066  n30   rna-XM_034639960.1    rna-XM_038572270.1    rna-XM_022494196.1    rna-XM_045053647.1    rna-XM_036101556.1    rna-XM_039227147.1    rna-XM_032845736.1    rna-XM_045980801.1    rna-XM_032350711.1    rna-XM_059133268.1    rna-XM_059406976.1    rna-XM_045084186.1    mNeoVis.2.1.1.chr11_hap1.g00251.t01    rna-XM_021689261.1    rna-XM_004393166.1    rna-XM_007077699.3    rna-XM_040638295.1    rna-XM_027622491.2    mNeoVis.3.1.1.chr11.g00220.t01      mNeoVis.evergladensis.2.1.chr11.g00250.t01    mNeoVis.vulgivaga.2.1.chr11.g00251.t01      849    N0.HOG0000847   OG0000066  n40   rna-XM_034642471.1    rna-NM_001003239.2    rna-XM_022495848.1    rna-XM_045041729.1    rna-XM_036070699.1    rna-XM_039218242.1    rna-XM_032859808.1    rna-XM_046017404.1, rna-XM_046025671.1    rna-XM_032309926.1    rna-XM_059139249.1    rna-XM_059409298.1    rna-XM_045065418.1    mNeoVis.2.1.1.chr3_hap1.g00228.t01    rna-XM_021686044.2    rna-XM_004411198.1    rna-XM_007075938.3    rna-XM_040632682.1    rna-XM_027576216.2    mNeoVis.3.1.1.chr3.g00230.t01    rna-gnl|WGS_ZZZZ|egapxtmp_042415-R1    mNeoVis.evergladensis.2.1.chr3.g00239.t01    mNeoVis.vulgivaga.2.1.chr3.g00231.t01      850    N0.HOG0000848   OG0000066  n56   rna-XM_002913321.4    rna-XM_038531401.1    rna-XM_022496207.1    rna-XM_011284511.4    rna-XM_036114292.1    rna-XM_039233263.1    rna-XM_032863894.1    rna-XM_046026634.1    rna-XM_032301871.1    rna-XM_059136421.1    rna-XM_059376833.1    rna-XM_013047110.2    mNeoVis.2.1.1.chr2_hap1.g01611.t01    rna-XM_021683605.2    rna-XM_004394677.1    rna-XM_042995803.1    rna-XM_040619094.1    rna-XM_027573761.1    mNeoVis.3.1.1.chr5.g01532.t01    rna-gnl|WGS_ZZZZ|egapxtmp_005718-R2    mNeoVis.evergladensis.2.1.chr5.g01550.t01    mNeoVis.vulgivaga.2.1.chr5.g01553.t01      851    N0.HOG0000849   OG0000066  n74   rna-XM_019802460.2    rna-XM_038531235.1    rna-XM_022496368.1    rna-XM_019836589.3    rna-XM_036065179.1    rna-XM_039233102.1    rna-XM_032877068.1    rna-XM_045978520.1    rna-XM_032360910.1    rna-XM_059136867.1    rna-XM_059377027.1    rna-XM_004740918.3    mNeoVis.2.1.1.chr2_hap1.g01553.t01    rna-XM_021683654.1    rna-XM_004406417.1    rna-XM_007092415.3    rna-XM_040619183.1    rna-XM_027611089.1    mNeoVis.3.1.1.chr5.g01474.t01    rna-gnl|WGS_ZZZZ|egapxtmp_007113-R1    mNeoVis.evergladensis.2.1.chr5.g01491.t01    mNeoVis.vulgivaga.2.1.chr5.g01495.t01      852    N0.HOG0000850   OG0000066  n89   rna-XM_034661363.1    rna-XM_038573650.1    rna-XM_022514427.1, rna-XM_022514449.1    rna-XM_003984842.5    rna-XM_036092353.1    rna-XM_039219730.1    rna-XM_032855745.1    rna-XM_045985315.1    rna-XM_032332259.1    rna-XM_059177314.1    rna-XM_059372154.1    rna-XM_004766072.3    mNeoVis.2.1.1.chr6_hap1.g00783.t01    rna-XM_021698849.1    rna-XM_004395256.1    rna-XM_042983580.1    rna-XM_008705072.2    rna-XM_027600249.2    mNeoVis.3.1.1.chr4.g00792.t01    rna-gnl|WGS_ZZZZ|egapxtmp_000846-R1    mNeoVis.evergladensis.2.1.chr4.g00786.t01    mNeoVis.vulgivaga.2.1.chr4.g00805.t01      853    N0.HOG0000851   OG0000066  n109   rna-XM_011223725.3    rna-XM_038571947.1    rna-XM_022494114.1    rna-XM_006929858.5    rna-XM_036093235.1    rna-XM_039237744.1    rna-XM_032858923.1    rna-XM_045980361.1    rna-XM_032351027.1    rna-XM_059133595.1    rna-XM_059406762.1    rna-XM_004746663.3    mNeoVis.2.1.1.chr11_hap1.g00342.t01    rna-XM_021688939.1    rna-XM_004393036.1    rna-XM_042980666.1    rna-XM_040638137.1    rna-XM_027623208.2    mNeoVis.3.1.1.chr11.g00312.t01    rna-gnl|WGS_ZZZZ|egapxtmp_027419-R1    mNeoVis.evergladensis.2.1.chr11.g00339.t01    mNeoVis.vulgivaga.2.1.chr11.g00343.t01      854    N0.HOG0000852   OG0000067  n1   rna-XM_011226623.3    rna-NM_001003364.1    rna-XM_022497654.1    rna-XM_011291675.4    rna-XM_036092014.1    rna-XM_039228192.1    rna-XM_032868108.1    rna-XM_045990688.1    rna-XM_032320460.1    rna-XM_059160990.1    rna-XM_059389778.1    rna-XM_004760587.3    mNeoVis.2.1.1.chr5_hap1.g00671.t01    rna-XM_021686868.2    rna-XM_004399271.1    rna-XM_007088557.3    rna-XM_008703762.2    rna-XM_027585219.2    mNeoVis.3.1.1.chr2.g00588.t01    rna-gnl|WGS_ZZZZ|egapxtmp_035910-R1    mNeoVis.evergladensis.2.1.chr2.g00668.t01    mNeoVis.vulgivaga.2.1.chr2.g00671.t01      855    N0.HOG0000853   OG0000067  n9   rna-XM_011223692.3    rna-XM_038562833.1    rna-XM_022515455.1    rna-XM_023250305.2    rna-XM_036106167.1    rna-XM_039252468.1    rna-XM_032873258.1    rna-XM_046020959.1    rna-XM_032304762.1    rna-XM_059170952.1    rna-XM_059396957.1    rna-XM_004739610.3    mNeoVis.2.1.1.chr4_hap1.g00541.t01    rna-XM_044919933.1    rna-XM_004393626.1    rna-XM_042965276.1    rna-XM_040644319.1    rna-XM_027573991.2    mNeoVis.3.1.1.chr6.g00550.t01    rna-gnl|WGS_ZZZZ|egapxtmp_001981-R1    mNeoVis.evergladensis.2.1.chr6.g00547.t01    mNeoVis.vulgivaga.2.1.chr6.g00550.t01      856    N0.HOG0000854   OG0000067  n16   rna-XM_034654035.1    rna-XM_038541067.1    rna-XM_022497021.1    rna-XM_003990409.6    rna-XM_036109866.1    rna-XM_039251085.1    rna-XM_032881528.1    rna-XM_045996289.1, rna-XM_046007492.1    rna-XM_032303610.1    rna-XM_059137686.1    rna-XM_059375662.1    rna-XM_004769707.3    mNeoVis.2.1.1.chr2_hap1.g00983.t01    rna-XM_021690316.2    rna-XM_004407180.1    rna-XM_007076419.3    rna-XM_008704020.2    rna-XM_027607919.1    mNeoVis.3.1.1.chr5.g00916.t01    rna-gnl|WGS_ZZZZ|egapxtmp_008355-R1    mNeoVis.evergladensis.2.1.chr5.g00936.t01    mNeoVis.vulgivaga.2.1.chr5.g00923.t01      857    N0.HOG0000855   OG0000067  n25   rna-XM_011231771.3    rna-XM_038531014.1    rna-XM_022518717.1    rna-XM_011290154.4    rna-XM_036068939.1    rna-XM_039232634.1    rna-XM_032854439.1    rna-XM_045987871.1    rna-XM_032325123.1    rna-XM_059152961.1    rna-XM_059383069.1    rna-XM_004744208.2    mNeoVis.2.1.1.chr7_hap1.g00316.t01    rna-XM_021705817.1    rna-XM_004395813.1    rna-XM_007090181.3    rna-XM_040633319.1    rna-XM_027618512.2    mNeoVis.3.1.1.chr7.g00302.t01    rna-gnl|WGS_ZZZZ|egapxtmp_011297-R1    mNeoVis.evergladensis.2.1.chr7.g00316.t01    mNeoVis.vulgivaga.2.1.chr7.g00320.t01      858    N0.HOG0000856   OG0000067  n32   rna-XM_002929538.4    rna-XM_038575774.1    rna-XM_022495689.1, rna-XM_022495690.1    rna-XM_023241548.2    rna-XM_036077821.1    rna-XM_039222166.1    rna-XM_032876042.1    rna-XM_046024925.1    rna-XM_032310211.1    rna-XM_059139235.1    rna-XM_059408924.1    rna-XM_045065235.1    mNeoVis.2.1.1.chr3_hap1.g00383.t01    rna-XM_021687889.1    rna-XM_004400548.1    rna-XM_042961952.1    rna-XM_040619481.1    rna-XM_027577503.2    mNeoVis.3.1.1.chr3.g00376.t01    rna-gnl|WGS_ZZZZ|egapxtmp_040040-R1    mNeoVis.evergladensis.2.1.chr3.g00391.t01    mNeoVis.vulgivaga.2.1.chr3.g00387.t01      859    N0.HOG0000857   OG0000067  n37   rna-XM_002920527.4    rna-NM_001003068.3    rna-XM_022498085.1    rna-XM_003982253.5    rna-XM_036091973.1    rna-XM_039228374.1    rna-XM_032868105.1    rna-XM_045991322.1    rna-XM_032320356.1    rna-XM_059160963.1    rna-XM_059389776.1    rna-XM_004760594.3    mNeoVis.2.1.1.chr5_hap1.g00669.t01    rna-XM_021687017.1    rna-XM_004399268.1    rna-XM_042977346.1    rna-XM_040628150.1    rna-XM_027582211.2    mNeoVis.3.1.1.chr2.g00586.t01      mNeoVis.evergladensis.2.1.chr2.g00666.t01    mNeoVis.vulgivaga.2.1.chr2.g00669.t01      860    N0.HOG0000858   OG0000067  n45   rna-XM_002919208.4    rna-XM_038541069.1    rna-XM_022496403.1    rna-XM_045033264.1    rna-XM_036109869.1    rna-XM_039251108.1    rna-XM_032881525.1    rna-XM_046002281.1    rna-XM_032303611.1    rna-XM_059137687.1    rna-XM_059374235.1    rna-XM_004769709.3    mNeoVis.2.1.1.chr2_hap1.g00982.t01    rna-XM_021690021.2    rna-XM_004407178.1    rna-XM_007076418.3    rna-XM_008704022.2    rna-XM_027607933.2    mNeoVis.3.1.1.chr5.g00915.t01    rna-gnl|WGS_ZZZZ|egapxtmp_008353-R1    mNeoVis.evergladensis.2.1.chr5.g00935.t01    mNeoVis.vulgivaga.2.1.chr5.g00922.t01      861    N0.HOG0000859   OG0000067  n60   rna-XM_002918261.4    rna-XM_038562445.1    rna-XM_022515445.1    rna-XM_003982705.5    rna-XM_036087129.1    rna-XM_039252481.1    rna-XM_032873257.1    rna-XM_046021041.1    rna-XM_032304641.1    rna-XM_059170959.1    rna-XM_059395765.1    rna-XM_004739613.3    mNeoVis.2.1.1.chr4_hap1.g00539.t01    rna-XM_021689523.1    rna-XM_004393625.1    rna-XM_007076783.3    rna-XM_008686177.1    rna-XM_027574314.1    mNeoVis.3.1.1.chr6.g00548.t01    rna-gnl|WGS_ZZZZ|egapxtmp_003693-R1    mNeoVis.evergladensis.2.1.chr6.g00545.t01    mNeoVis.vulgivaga.2.1.chr6.g00548.t01      862    N0.HOG0000860   OG0000068  n2   rna-XM_002926891.3    rna-NM_001388607.1    rna-XM_022507294.1    rna-XM_003993349.3      rna-XM_039224046.1      rna-XM_046016615.1    rna-XM_032359701.1    rna-XM_059158941.1    rna-XM_059387898.1    rna-XM_013050834.1    mNeoVis.2.1.1.chr7_hap1.g02151.t01        rna-XM_007088784.1    rna-XM_008694117.2      mNeoVis.3.1.1.chr7.g01979.t01    rna-gnl|WGS_ZZZZ|egapxtmp_012632-R1    mNeoVis.evergladensis.2.1.chr7.g02098.t01    mNeoVis.vulgivaga.2.1.chr7.g02119.t01      863    N0.HOG0000861   OG0000068  n18   rna-XM_002926861.1, rna-XM_002926892.3, rna-XM_011234356.2    rna-XM_038563254.1, rna-XM_038564474.1    rna-XM_022507045.1, rna-XM_022507427.1    rna-XM_004001251.5, rna-XM_011286829.3, rna-XM_045039298.1, rna-XM_045039300.1    rna-XM_036107504.1    rna-XM_039224041.1, rna-XM_039224045.1, rna-XM_039224047.1    rna-XM_032846487.1, rna-XM_032846494.1, rna-XM_032846545.1    rna-XM_046015791.1, rna-XM_046015793.1, rna-XM_046016402.1, rna-XM_046017041.1    rna-XM_032359876.1, rna-XM_032360320.1, rna-XM_032360376.1    rna-XM_059138870.1, rna-XM_059158944.1, rna-XM_059158945.1    rna-XM_059387876.1, rna-XM_059415962.1    rna-XM_004777638.2, rna-XM_004777639.2, rna-XM_045067626.1    mNeoVis.2.1.1.chr7_hap1.g02145.t01, mNeoVis.2.1.1.chr7_hap1.g02152.t01    rna-XM_021685209.1    rna-XM_004411973.1    rna-XM_007088830.2, rna-XM_007088831.2, rna-XM_007088832.2, rna-XM_015540512.2    rna-XM_008694120.2, rna-XM_008694252.1    rna-XM_027577862.1, rna-XM_027580430.2    mNeoVis.3.1.1.chr7.g01978.t01    rna-gnl|WGS_ZZZZ|egapxtmp_010296-R1          864    N0.HOG0000862   OG0000068  n67   rna-XM_011234355.3    rna-NM_001388886.1      rna-XM_003993279.3      rna-XM_039223513.1, rna-XM_039224043.1      rna-XM_046015794.1    rna-XM_032359875.1    rna-XM_059158937.1, rna-XM_059158939.1    rna-XM_059387922.1, rna-XM_059387928.1    rna-XM_004782244.2    mNeoVis.2.1.1.chr7_hap1.g02150.t01        rna-XM_007088827.3    rna-XM_008694250.2      mNeoVis.3.1.1.chr7.g01977.t01    rna-gnl|WGS_ZZZZ|egapxtmp_012631-R1    mNeoVis.evergladensis.2.1.chr7.g02097.t01    mNeoVis.vulgivaga.2.1.chr7.g02118.t01      865    N0.HOG0000863   OG0000068  n87     rna-NM_001388608.1            rna-XM_032846514.1    rna-XM_046016963.1                    rna-XM_008694113.2, rna-XM_008694115.2      mNeoVis.3.1.1.chr7.g01975.t01            866    N0.HOG0000864   OG0000068  n92   rna-XM_002926888.4    rna-NM_001388610.1    rna-XM_022507502.1    rna-XM_003993347.3      rna-XM_039223496.1    rna-XM_032846511.1    rna-XM_046017017.1    rna-XM_032360311.1    rna-XM_059158935.1    rna-XM_059387951.1    rna-XM_004782243.1    mNeoVis.2.1.1.chr7_hap1.g02147.t01        rna-XM_007088826.2    rna-XM_008694110.1      mNeoVis.3.1.1.chr7.g01973.t01    rna-gnl|WGS_ZZZZ|egapxtmp_013537-R1    mNeoVis.evergladensis.2.1.chr7.g02094.t01    mNeoVis.vulgivaga.2.1.chr7.g02115.t01      867    N0.HOG0000865   OG0000068  n106   rna-XM_002926859.4    rna-NM_001388609.1    rna-XM_022507048.1, rna-XM_022507426.1      rna-XM_036107497.1    rna-XM_039223515.1    rna-XM_032846459.1, rna-XM_032846495.1    rna-XM_046015795.1, rna-XM_046017024.1    rna-XM_032360294.1, rna-XM_032360310.1    rna-XM_059158936.1, rna-XM_059158938.1    rna-XM_059387910.1, rna-XM_059387946.1    rna-XM_004782234.2, rna-XM_004782235.1    mNeoVis.2.1.1.chr7_hap1.g02148.t01, mNeoVis.2.1.1.chr7_hap1.g02149.t01        rna-XM_007088783.2    rna-XM_008694112.1, rna-XM_008694114.2      mNeoVis.3.1.1.chr7.g01974.t01, mNeoVis.3.1.1.chr7.g01976.t01    rna-gnl|WGS_ZZZZ|egapxtmp_013536-R1, rna-gnl|WGS_ZZZZ|egapxtmp_013538-R1    mNeoVis.evergladensis.2.1.chr7.g02095.t01, mNeoVis.evergladensis.2.1.chr7.g02096.t01    mNeoVis.vulgivaga.2.1.chr7.g02116.t01, mNeoVis.vulgivaga.2.1.chr7.g02117.t01      868    N0.HOG0000866   OG0000068  n135   rna-XM_034644815.1    rna-NM_001389207.1    rna-XM_022506645.1      rna-XM_036107502.1        rna-XM_046016400.1        rna-XM_059387880.1    rna-XM_013048478.2      rna-XM_021685210.2        rna-XM_008694251.2    rna-XM_035723138.1    mNeoVis.3.1.1.chr7.g01980.t01    rna-gnl|WGS_ZZZZ|egapxtmp_012634-R1      mNeoVis.vulgivaga.2.1.chr7.g02122.t01      869    N0.HOG0000867   OG0000068  n147   rna-XM_002926860.4    rna-NM_001388938.1, rna-NM_001389209.1    rna-XM_022506646.1    rna-XM_011279811.4      rna-XM_039224042.1, rna-XM_039224044.1    rna-XM_032846456.1    rna-XM_046016781.1, rna-XM_046016792.1    rna-XM_032359050.1, rna-XM_032359944.1    rna-XM_059158942.1, rna-XM_059158943.1    rna-XM_059387888.1    rna-XM_004777636.3, rna-XM_004777637.2    mNeoVis.2.1.1.chr7_hap1.g02153.t01        rna-XM_007088828.2, rna-XM_007088829.2    rna-XM_008694119.2        rna-gnl|WGS_ZZZZ|egapxtmp_012633-R1    mNeoVis.evergladensis.2.1.chr7.g02099.t01, mNeoVis.evergladensis.2.1.chr7.g02100.t01    mNeoVis.vulgivaga.2.1.chr7.g02120.t01      870    N0.HOG0000868   OG0000069  n0   rna-XM_034649019.1, rna-XM_034649716.1, rna-XM_034650335.1, rna-XM_034650574.1, rna-XM_034650767.1, rna-XM_034653924.1, rna-XM_034654611.1, rna-XM_034654632.1, rna-XM_034654644.1    rna-XM_038581153.1, rna-XM_038581623.1, rna-XM_038581624.1, rna-XM_038581629.1, rna-XM_038581630.1, rna-XM_038581631.1, rna-XM_038581632.1, rna-XM_038581719.1, rna-XM_038581720.1, rna-XM_038581721.1    rna-XM_022515947.1, rna-XM_022515948.1, rna-XM_022516145.1, rna-XM_022516146.1, rna-XM_022516149.1, rna-XM_022516150.1, rna-XM_022516196.1    rna-XM_019811142.3, rna-XM_019839299.3, rna-XM_019839300.2, rna-XM_023238643.2, rna-XM_023238644.2, rna-XM_023238645.2, rna-XM_023238646.2, rna-XM_045036509.1, rna-XM_045036510.1, rna-XM_045036511.1, rna-XM_045036512.1    rna-XM_036097255.1, rna-XM_036097260.1, rna-XM_036097261.1, rna-XM_036097262.1, rna-XM_036097263.1, rna-XM_036097278.1, rna-XM_036107997.1, rna-XM_036112722.1    rna-XM_039245426.1, rna-XM_039245482.1, rna-XM_039245519.1, rna-XM_039257096.1    rna-XM_032865236.1, rna-XM_032865237.1, rna-XM_032865242.1, rna-XM_032865511.1, rna-XM_032865512.1, rna-XM_032865543.1, rna-XM_032865580.1    rna-XM_046001429.1, rna-XM_046001430.1, rna-XM_046001431.1, rna-XM_046002682.1, rna-XM_046002779.1, rna-XM_046002780.1, rna-XM_046003149.1, rna-XM_046003192.1, rna-XM_046003717.1, rna-XM_046003722.1    rna-XM_032302938.1, rna-XM_032356927.1, rna-XM_032356949.1, rna-XM_032356990.1, rna-XM_032356999.1, rna-XM_032357013.1, rna-XM_032357029.1, rna-XM_032357040.1, rna-XM_032357052.1    rna-XM_059164909.1, rna-XM_059164911.1, rna-XM_059164912.1, rna-XM_059164914.1, rna-XM_059164915.1, rna-XM_059164916.1, rna-XM_059164917.1, rna-XM_059164918.1, rna-XM_059164919.1, rna-XM_059164921.1, rna-XM_059165299.1, rna-XM_059165315.1, rna-XM_059165316.1    rna-XM_059387527.1, rna-XM_059392493.1, rna-XM_059392494.1, rna-XM_059392495.1, rna-XM_059392496.1, rna-XM_059392497.1, rna-XM_059392601.1    rna-XM_045084344.1, rna-XM_045084347.1, rna-XM_045084349.1, rna-XM_045084351.1, rna-XM_045084352.1, rna-XM_045084353.1, rna-XM_045084354.1, rna-XM_045084355.1, rna-XM_045084356.1, rna-XM_045084544.1, rna-XM_045084545.1, rna-XM_045084557.1, rna-XM_045087821.1    mNeoVis.2.1.1.chr5_hap1.g01853.t01, mNeoVis.2.1.1.chr5_hap1.g01897.t01, mNeoVis.2.1.1.chr5_hap1.g01898.t01, mNeoVis.2.1.1.chr5_hap1.g01899.t01, mNeoVis.2.1.1.chr5_hap1.g01900.t01, mNeoVis.2.1.1.chr5_hap1.g01901.t01, mNeoVis.2.1.1.chr5_hap1.g01902.t01, mNeoVis.2.1.1.chr5_hap1.g01906.t01    rna-XM_044913282.1, rna-XM_044918966.1    rna-XM_004410651.1, rna-XM_004410652.1, rna-XM_004410653.1, rna-XM_004410654.1    rna-XM_042956385.1, rna-XM_042956586.1, rna-XM_042956587.1, rna-XM_042998848.1, rna-XM_042998853.1, rna-XM_042998854.1, rna-XM_042998855.1, rna-XM_042998856.1, rna-XM_042998858.1, rna-XM_042998865.1    rna-XM_008710226.2, rna-XM_008710256.2, rna-XM_040621669.1, rna-XM_040621671.1, rna-XM_040621672.1, rna-XM_040621675.1, rna-XM_040621757.1, rna-XM_040621821.1    rna-XM_027585072.1, rna-XM_027585075.1, rna-XM_027586123.1, rna-XM_035728723.1, rna-XM_035728724.1, rna-XM_035728725.1, rna-XM_035728726.1, rna-XM_035728729.1, rna-XM_035728730.1    mNeoVis.3.1.1.chr2.g01754.t01, mNeoVis.3.1.1.chr2.g01790.t01, mNeoVis.3.1.1.chr2.g01791.t01, mNeoVis.3.1.1.chr2.g01792.t01, mNeoVis.3.1.1.chr2.g01793.t01, mNeoVis.3.1.1.chr2.g01794.t01    rna-gnl|WGS_ZZZZ|egapxtmp_034618-R1, rna-gnl|WGS_ZZZZ|egapxtmp_034620-R1, rna-gnl|WGS_ZZZZ|egapxtmp_034621-R1, rna-gnl|WGS_ZZZZ|egapxtmp_035736-R1, rna-gnl|WGS_ZZZZ|egapxtmp_035737-R1, rna-gnl|WGS_ZZZZ|egapxtmp_038694-R1    mNeoVis.evergladensis.2.1.chr2.g01877.t01, mNeoVis.evergladensis.2.1.chr2.g01878.t01, mNeoVis.evergladensis.2.1.chr2.g01879.t01, mNeoVis.evergladensis.2.1.chr2.g01880.t01, mNeoVis.evergladensis.2.1.chr2.g01881.t01, mNeoVis.evergladensis.2.1.chr2.g01882.t01, mNeoVis.evergladensis.2.1.chr2.g01884.t01, mNeoVis.evergladensis.2.1.chr2.g01885.t01    mNeoVis.vulgivaga.2.1.chr2.g01851.t01, mNeoVis.vulgivaga.2.1.chr2.g01892.t01, mNeoVis.vulgivaga.2.1.chr2.g01893.t01, mNeoVis.vulgivaga.2.1.chr2.g01894.t01, mNeoVis.vulgivaga.2.1.chr2.g01895.t01, mNeoVis.vulgivaga.2.1.chr2.g01896.t01, mNeoVis.vulgivaga.2.1.chr2.g01898.t01      871    N0.HOG0000869   OG0000070  n2   rna-XM_034643629.1    rna-XM_038549512.1      rna-XM_045050695.1    rna-XM_036083232.1, rna-XM_036084954.1, rna-XM_036086000.1, rna-XM_036090550.1, rna-XM_036092101.1, rna-XM_036095180.1, rna-XM_036123565.1    rna-XM_039230147.1, rna-XM_039255681.1    rna-XM_032854898.1    rna-XM_045998996.1      rna-XM_059135516.1, rna-XM_059157374.1, rna-XM_059170934.1    rna-XM_059384946.1, rna-XM_059388986.1, rna-XM_059416521.1    rna-XM_045065853.1, rna-XM_045068267.1, rna-XM_045079560.1    mNeoVis.2.1.1.chr5_hap1.g01443.t01    rna-XM_044912599.1, rna-XM_044913083.1, rna-XM_044913233.1, rna-XM_044913263.1, rna-XM_044916779.1, rna-XM_044918756.1, rna-XM_044919432.1, rna-XM_044922068.1    rna-XM_004397371.1, rna-XM_004399721.1, rna-XM_004401061.1, rna-XM_004415694.1    rna-XM_015537344.2, rna-XM_042966461.1      rna-XM_035725714.1      rna-gnl|WGS_ZZZZ|egapxtmp_004454-R1, rna-gnl|WGS_ZZZZ|egapxtmp_036115-R1    mNeoVis.evergladensis.2.1.chr13.g00678.t01, mNeoVis.evergladensis.2.1.chr2.g01431.t01    mNeoVis.vulgivaga.2.1.chr2.g01443.t01      872    N0.HOG0000870   OG0000070  n48               rna-XM_032844340.1                        rna-XM_035723894.1              873    N0.HOG0000871   OG0000070  n50     rna-XM_038535410.1                rna-XM_032311659.1    rna-XM_059139749.1    rna-XM_059409968.1    rna-XM_045081544.1    mNeoVis.2.1.1.chr3_hap1.g00450.t01    rna-XM_044916780.1              rna-gnl|WGS_ZZZZ|egapxtmp_040832-R1    mNeoVis.evergladensis.2.1.chr3.g00458.t01    mNeoVis.vulgivaga.2.1.chr3.g00456.t01      874    N0.HOG0000872   OG0000070  n59     rna-XM_038559015.1                          rna-XM_044916389.1                      875    N0.HOG0000873   OG0000070  n61                           mNeoVis.2.1.1.chr9_hap1.g00881.t01            rna-XM_035727766.1              876    N0.HOG0000874   OG0000070  n63           rna-XM_036106310.1        rna-XM_045989962.1                                  877    N0.HOG0000875   OG0000070  n65           rna-XM_036064581.1                    rna-XM_044918261.1    rna-XM_004392613.1                    878    N0.HOG0000876   OG0000070  n71                 rna-XM_046025188.1                              mNeoVis.vulgivaga.2.1.chr4.g00152.t01      879    N0.HOG0000877   OG0000070  n73     rna-NM_001252321.1    rna-XM_022499730.1                        rna-XM_044916501.1        rna-XM_008696092.2                880    N0.HOG0000878   OG0000070  n74     rna-XM_038543501.1, rna-XM_038586556.1      rna-XM_003994089.6, rna-XM_045056723.1, rna-XM_045057494.1    rna-XM_036080425.1, rna-XM_036087466.1        rna-XM_046002770.1, rna-XM_046027628.1    rna-XM_032305547.1, rna-XM_032311835.1, rna-XM_032329223.1, rna-XM_032359653.1    rna-XM_059145620.1, rna-XM_059172638.1    rna-XM_059377491.1, rna-XM_059381851.1, rna-XM_059395785.1    rna-XM_013062925.2, rna-XM_045076304.1    mNeoVis.2.1.1.chr4_hap1.g00572.t01, mNeoVis.2.1.1.chr4_hap1.g00679.t01    rna-XM_044911719.1      rna-XM_042984734.1      rna-XM_035722372.1, rna-XM_035726137.1, rna-XM_035726754.1      rna-gnl|WGS_ZZZZ|egapxtmp_003074-R1    mNeoVis.evergladensis.2.1.chr6.g00686.t01        881    N0.HOG0000879   OG0000070  n99         rna-XM_045053329.1        rna-XM_032853418.1                                    882    N0.HOG0000880   OG0000070  n105           rna-XM_036072389.1, rna-XM_036076375.1    rna-XM_039245646.1      rna-XM_045991565.1                  rna-XM_042992099.1      rna-XM_035726287.1, rna-XM_035728279.1              883    N0.HOG0000881   OG0000070  n114   rna-XM_011223819.3          rna-XM_036089769.1, rna-XM_036091018.1, rna-XM_036099278.1    rna-XM_039236041.1, rna-XM_039240538.1                      rna-XM_042960087.1                  884    N0.HOG0000882   OG0000070  n116     rna-XM_038575651.1        rna-XM_036118964.1, rna-XM_036121384.1      rna-XM_032838982.1, rna-XM_032864850.1    rna-XM_046014687.1    rna-XM_032331252.1    rna-XM_059140104.1, rna-XM_059155726.1, rna-XM_059169787.1    rna-XM_059387409.1, rna-XM_059388375.1    rna-XM_045066011.1, rna-XM_045075213.1    mNeoVis.2.1.1.chr2_hap1.g00179.t01, mNeoVis.2.1.1.chr2_hap1.g00445.t01    rna-XM_044911668.1, rna-XM_044922205.1    rna-XM_004394818.1, rna-XM_004395262.1, rna-XM_004401059.1, rna-XM_004405001.1        rna-XM_035726285.1, rna-XM_035726424.1    mNeoVis.3.1.1.chr5.g00436.t01    rna-gnl|WGS_ZZZZ|egapxtmp_006530-R1, rna-gnl|WGS_ZZZZ|egapxtmp_007860-R1, rna-gnl|WGS_ZZZZ|egapxtmp_008906-R1, rna-gnl|WGS_ZZZZ|egapxtmp_031031-R1    mNeoVis.evergladensis.2.1.chr5.g00183.t01, mNeoVis.evergladensis.2.1.chr5.g00445.t01    mNeoVis.vulgivaga.2.1.chr5.g00179.t01, mNeoVis.vulgivaga.2.1.chr5.g00444.t01      885    N0.HOG0000883   OG0000070  n146   rna-XM_034651263.1    rna-XM_038545966.1            rna-XM_032855555.1, rna-XM_032864827.1        rna-XM_059153228.1    rna-XM_059375165.1, rna-XM_059381349.1    rna-XM_045077174.1, rna-XM_045079412.1, rna-XM_045089712.1    mNeoVis.2.1.1.chr9_hap1.g00793.t01    rna-XM_044913806.1, rna-XM_044918514.1, rna-XM_044918746.1, rna-XM_044919399.1    rna-XM_004406230.1        rna-XM_035723034.1              886    N0.HOG0000884   OG0000071  n2     rna-XM_038528665.1    rna-XM_022491773.1      rna-XM_036093641.1    rna-XM_039249897.1    rna-XM_032839075.1    rna-XM_045987285.1    rna-XM_032326601.1    rna-XM_059153469.1    rna-XM_059382377.1    rna-XM_004767669.3    mNeoVis.2.1.1.chr7_hap1.g01060.t01    rna-XM_021681056.1          rna-XM_027618696.2      rna-gnl|WGS_ZZZZ|egapxtmp_010473-R1    mNeoVis.evergladensis.2.1.chr7.g01052.t01    mNeoVis.vulgivaga.2.1.chr7.g01061.t01      887    N0.HOG0000885   OG0000071  n15   rna-XM_034639242.1    rna-XM_038528660.1    rna-XM_022491767.1    rna-XM_045045705.1    rna-XM_036093635.1    rna-XM_039249298.1    rna-XM_032839070.1            rna-XM_004767671.3    mNeoVis.2.1.1.chr7_hap1.g01059.t01    rna-XM_044911421.1    rna-XM_004401041.1    rna-XM_042970084.1    rna-XM_008684431.2    rna-XM_027618698.2    mNeoVis.3.1.1.chr7.g00990.t01    rna-gnl|WGS_ZZZZ|egapxtmp_010474-R1    mNeoVis.evergladensis.2.1.chr7.g01051.t01        888    N0.HOG0000886   OG0000071  n34   rna-XM_034639254.1, rna-XM_034672888.1    rna-XM_038528654.1    rna-XM_022491772.1, rna-XM_022524075.1    rna-XM_045046936.1    rna-XM_036081878.1, rna-XM_036093620.1, rna-XM_036093640.1    rna-XM_039247998.1, rna-XM_039249888.1    rna-XM_032839024.1, rna-XM_032839068.1    rna-XM_045988136.1    rna-XM_032325322.1, rna-XM_032326581.1    rna-XM_059151249.1, rna-XM_059153448.1    rna-XM_059382366.1, rna-XM_059383400.1    rna-XM_013063879.2, rna-XM_045077287.1    mNeoVis.2.1.1.chr7_hap1.g01056.t01, mNeoVis.2.1.1.chr7_hap1.g01061.t01    rna-XM_021681057.2, rna-XM_044922171.1    rna-XM_004401042.1    rna-XM_015539500.2    rna-XM_040634112.1, rna-XM_040634123.1    rna-XM_027618688.2, rna-XM_027619410.1    mNeoVis.3.1.1.chr7.g00988.t01, mNeoVis.3.1.1.chr7.g00991.t01    rna-gnl|WGS_ZZZZ|egapxtmp_012379-R1, rna-gnl|WGS_ZZZZ|egapxtmp_014236-R6    mNeoVis.evergladensis.2.1.chr7.g01048.t01, mNeoVis.evergladensis.2.1.chr7.g01053.t01    mNeoVis.vulgivaga.2.1.chr7.g01058.t01, mNeoVis.vulgivaga.2.1.chr7.g01062.t01      889    N0.HOG0000887   OG0000071  n71   rna-XM_019804881.2    rna-XM_038528640.1    rna-XM_022523855.1    rna-XM_023245161.2    rna-XM_036093631.1      rna-XM_032839260.1    rna-XM_045988423.1    rna-XM_032325354.1    rna-XM_059152174.1    rna-XM_059383391.1    rna-XM_013064039.2    mNeoVis.2.1.1.chr7_hap1.g01053.t01    rna-XM_044911369.1        rna-XM_008684440.2    rna-XM_027618047.2      rna-gnl|WGS_ZZZZ|egapxtmp_010229-R1    mNeoVis.evergladensis.2.1.chr7.g01046.t01    mNeoVis.vulgivaga.2.1.chr7.g01055.t01      890    N0.HOG0000888   OG0000071  n89   rna-XM_034637773.1    rna-XM_038528627.1    rna-XM_022523789.1      rna-XM_036093628.1    rna-XM_039246236.1    rna-XM_032839096.1    rna-XM_045987777.1    rna-XM_032325338.1    rna-XM_059152014.1    rna-XM_059382047.1    rna-XM_013064037.2    mNeoVis.2.1.1.chr7_hap1.g01051.t01    rna-XM_044911368.1    rna-XM_004401011.1    rna-XM_042970718.1    rna-XM_040634125.1    rna-XM_035724905.1      rna-gnl|WGS_ZZZZ|egapxtmp_013211-R1    mNeoVis.evergladensis.2.1.chr7.g01044.t01    mNeoVis.vulgivaga.2.1.chr7.g01052.t01      891    N0.HOG0000889   OG0000071  n108   rna-XM_019804880.2    rna-XM_038528635.1      rna-XM_045046575.1    rna-XM_036093591.1    rna-XM_039248500.1    rna-XM_032839265.1    rna-XM_045987774.1    rna-XM_032325352.1    rna-XM_059152172.1    rna-XM_059382046.1    rna-XM_013064035.2    mNeoVis.2.1.1.chr7_hap1.g01052.t01      rna-XM_004401039.1    rna-XM_042970707.1    rna-XM_040634124.1      mNeoVis.3.1.1.chr7.g00986.t01    rna-gnl|WGS_ZZZZ|egapxtmp_011337-R1, rna-gnl|WGS_ZZZZ|egapxtmp_012378-R1    mNeoVis.evergladensis.2.1.chr7.g01045.t01    mNeoVis.vulgivaga.2.1.chr7.g01053.t01      892    N0.HOG0000890   OG0000071  n127   rna-XM_011231088.3    rna-XM_038528641.1      rna-XM_019819424.3    rna-XM_036093584.1    rna-XM_039250051.1    rna-XM_032838863.1    rna-XM_045988585.1    rna-XM_032325286.1    rna-XM_059152032.1    rna-XM_059383582.1    rna-XM_045077188.1    mNeoVis.2.1.1.chr7_hap1.g01054.t01    rna-XM_021681137.2    rna-XM_004401040.1    rna-XM_007086632.3    rna-XM_040634121.1    rna-XM_027618674.2    mNeoVis.3.1.1.chr7.g00987.t01    rna-gnl|WGS_ZZZZ|egapxtmp_010964-R1    mNeoVis.evergladensis.2.1.chr7.g01047.t01    mNeoVis.vulgivaga.2.1.chr7.g01056.t01      893    N0.HOG0000891   OG0000071  n147   rna-XM_002929693.4    rna-XM_038528651.1      rna-XM_045045700.1    rna-XM_036093610.1    rna-XM_039249287.1    rna-XM_032839066.1    rna-XM_045987282.1    rna-XM_032326579.1    rna-XM_059151402.1    rna-XM_059382347.1    rna-XM_004767676.3    mNeoVis.2.1.1.chr7_hap1.g01058.t01    rna-XM_021681316.1    rna-XM_004401014.1    rna-XM_015539499.2      rna-XM_027618681.2    mNeoVis.3.1.1.chr7.g00989.t01      mNeoVis.evergladensis.2.1.chr7.g01050.t01    mNeoVis.vulgivaga.2.1.chr7.g01060.t01      894    N0.HOG0000892   OG0000072  n3   rna-XM_002926882.3    rna-NM_001388967.1    rna-XM_022507116.1    rna-XM_023239839.2      rna-XM_039223503.1    rna-XM_032846449.1    rna-XM_046016498.1    rna-XM_032359869.1    rna-XM_059158928.1    rna-XM_059387995.1    rna-XM_004782237.1    mNeoVis.2.1.1.chr7_hap1.g02139.t01      rna-XM_004411984.1    rna-XM_007088823.1    rna-XM_008694102.1      mNeoVis.3.1.1.chr7.g01967.t01    rna-gnl|WGS_ZZZZ|egapxtmp_013503-R3    mNeoVis.evergladensis.2.1.chr7.g02086.t01    mNeoVis.vulgivaga.2.1.chr7.g02108.t01      895    N0.HOG0000893   OG0000072  n21   rna-XM_002926877.4, rna-XM_011234349.3, rna-XM_011234350.2, rna-XM_034646497.1    rna-NM_001389163.1, rna-XM_038563257.1    rna-XM_022507172.1, rna-XM_022507264.1    rna-XM_003993340.1      rna-XM_039223497.1, rna-XM_039223506.1    rna-XM_032846470.1, rna-XM_032846474.1, rna-XM_032846497.1, rna-XM_032846549.1    rna-XM_046016539.1, rna-XM_046016584.1, rna-XM_046016608.1, rna-XM_046016645.1    rna-XM_032358849.1, rna-XM_032359472.1, rna-XM_032360023.1, rna-XM_032360029.1    rna-XM_059138464.1, rna-XM_059158916.1, rna-XM_059158919.1, rna-XM_059158921.1    rna-XM_059372434.1, rna-XM_059405436.1, rna-XM_059405451.1, rna-XM_059405463.1    rna-XM_004781679.2, rna-XM_004781680.2, rna-XM_004781681.3, rna-XM_045068328.1    mNeoVis.2.1.1.chr7_hap1.g02130.t01, mNeoVis.2.1.1.chr7_hap1.g02132.t01    rna-XM_021684835.2    rna-XM_004411987.1, rna-XM_004411988.1    rna-XM_007088814.2, rna-XM_007088818.2    rna-XM_008694236.1, rna-XM_008694238.1, rna-XM_008694239.1, rna-XM_008694240.1    rna-XM_035723076.1, rna-XM_035723136.1    mNeoVis.3.1.1.chr7.g01962.t01, mNeoVis.3.1.1.chr7.g01963.t01, mNeoVis.3.1.1.chr7.g01965.t01    rna-gnl|WGS_ZZZZ|egapxtmp_010315-R1, rna-gnl|WGS_ZZZZ|egapxtmp_010755-R1    mNeoVis.evergladensis.2.1.chr7.g02076.t01, mNeoVis.evergladensis.2.1.chr7.g02077.t01, mNeoVis.evergladensis.2.1.chr7.g02079.t01    mNeoVis.vulgivaga.2.1.chr7.g02099.t01, mNeoVis.vulgivaga.2.1.chr7.g02100.t01, mNeoVis.vulgivaga.2.1.chr7.g02102.t01      896    N0.HOG0000894   OG0000072  n80     rna-NM_001389153.1    rna-XM_022507166.1    rna-XM_011286831.2      rna-XM_039223505.1, rna-XM_039223508.1, rna-XM_039223511.1, rna-XM_039223512.1    rna-XM_032846475.1    rna-XM_046016537.1, rna-XM_046016573.1    rna-XM_032359455.1    rna-XM_059158922.1    rna-XM_059405423.1    rna-XM_004781664.2, rna-XM_045068330.1    mNeoVis.2.1.1.chr7_hap1.g02137.t01        rna-XM_007088779.2, rna-XM_015540519.1    rna-XM_008694242.1          mNeoVis.evergladensis.2.1.chr7.g02081.t01    mNeoVis.vulgivaga.2.1.chr7.g02106.t01      897    N0.HOG0000895   OG0000072  n101   rna-XM_002926856.4    rna-NM_001389105.1, rna-NM_001389150.1    rna-XM_022507097.1    rna-XM_045039296.1      rna-XM_039223500.1    rna-XM_032846463.1    rna-XM_046016530.1    rna-XM_032358502.1    rna-XM_059158923.1    rna-XM_059405410.1    rna-XM_004782453.1          rna-XM_015540517.2    rna-XM_008694243.1            mNeoVis.vulgivaga.2.1.chr7.g02103.t01      898    N0.HOG0000896   OG0000072  n115   rna-XM_002926857.2    rna-XM_038562466.1    rna-XM_022492784.1, rna-XM_022507036.1    rna-XM_023239738.2      rna-XM_039223492.1      rna-XM_046016416.1, rna-XM_046016472.1    rna-XM_032358500.1, rna-XM_032358501.1    rna-XM_059158924.1, rna-XM_059158926.1    rna-XM_059405397.1    rna-XM_004782452.1, rna-XM_045068348.1    mNeoVis.2.1.1.chr7_hap1.g02135.t01      rna-XM_004411986.1    rna-XM_007088820.1    rna-XM_008694101.2    rna-XM_027580115.1    mNeoVis.3.1.1.chr7.g01964.t01    rna-gnl|WGS_ZZZZ|egapxtmp_013566-R1    mNeoVis.evergladensis.2.1.chr7.g02084.t01        899    N0.HOG0000897   OG0000072  n137   rna-XM_002926881.3    rna-NM_001388970.1    rna-XM_022507041.1    rna-XM_023239841.2      rna-XM_039223499.1    rna-XM_032846450.1    rna-XM_046016395.1    rna-XM_032358877.1    rna-XM_059158927.1    rna-XM_059388006.1    rna-XM_004782236.3        rna-XM_004411985.1    rna-XM_007088822.2    rna-XM_008694246.1      mNeoVis.3.1.1.chr7.g01966.t01      mNeoVis.evergladensis.2.1.chr7.g02083.t01        900    N0.HOG0000898   OG0000072  n152   rna-XM_002926878.3        rna-XM_003993341.1      rna-XM_039223491.1    rna-XM_032846548.1    rna-XM_046015796.1    rna-XM_032359868.1    rna-XM_059138535.1        mNeoVis.2.1.1.chr7_hap1.g02136.t01    rna-XM_021685208.1      rna-XM_007088816.2, rna-XM_042959626.1    rna-XM_040638759.1        rna-gnl|WGS_ZZZZ|egapxtmp_010756-R1    mNeoVis.evergladensis.2.1.chr7.g02082.t01    mNeoVis.vulgivaga.2.1.chr7.g02105.t01      901    N0.HOG0000899   OG0000073  n2   rna-XM_002920040.2    rna-XM_038582471.1    rna-XM_022502232.1    rna-XM_011286019.3    rna-XM_036070315.1    rna-XM_039221985.1    rna-XM_032882256.1      rna-XM_032344708.1    rna-XM_059165079.1    rna-XM_059388372.1    rna-XM_004773029.1    mNeoVis.2.1.1.chr5_hap1.g01675.t01    rna-XM_021687960.1    rna-XM_004391708.1    rna-XM_007095985.2    rna-XM_008689690.1    rna-XM_027586276.1    mNeoVis.3.1.1.chr2.g01588.t01    rna-gnl|WGS_ZZZZ|egapxtmp_038473-R1    mNeoVis.evergladensis.2.1.chr2.g01658.t01    mNeoVis.vulgivaga.2.1.chr2.g01677.t01      902    N0.HOG0000900   OG0000073  n21   rna-XM_002920036.1, rna-XM_034664072.1    rna-NM_001388974.1    rna-XM_022508700.1, rna-XM_022508712.1    rna-XM_011285985.2, rna-XM_011286021.3    rna-XM_036070320.1, rna-XM_036070321.1      rna-XM_032882261.1, rna-XM_032882262.1    rna-XM_046003407.1, rna-XM_046003408.1    rna-XM_032302035.1, rna-XM_032351500.1    rna-XM_059165267.1, rna-XM_059165268.1    rna-XM_059388164.1, rna-XM_059388165.1    rna-XM_004772997.2, rna-XM_004772998.2    mNeoVis.2.1.1.chr5_hap1.g01677.t01, mNeoVis.2.1.1.chr5_hap1.g01678.t01        rna-XM_007095937.1, rna-XM_007095984.2    rna-XM_008689679.2, rna-XM_008689680.2      mNeoVis.3.1.1.chr2.g01590.t01    rna-gnl|WGS_ZZZZ|egapxtmp_037315-R1    mNeoVis.evergladensis.2.1.chr2.g01660.t01, mNeoVis.evergladensis.2.1.chr2.g01661.t01    mNeoVis.vulgivaga.2.1.chr2.g01678.t01      903    N0.HOG0000901   OG0000073  n52   rna-XM_011225963.3    rna-NM_001388769.1    rna-XM_022508725.1    rna-XM_003991557.2      rna-XM_039221992.1    rna-XM_032882263.1    rna-XM_046003413.1    rna-XM_032302026.1    rna-XM_059165266.1    rna-XM_059388163.1    rna-XM_013046112.2    mNeoVis.2.1.1.chr5_hap1.g01676.t01    rna-XM_021687951.1      rna-XM_015543715.2    rna-XM_008689681.1    rna-XM_027585874.1    mNeoVis.3.1.1.chr2.g01589.t01    rna-gnl|WGS_ZZZZ|egapxtmp_037306-R1    mNeoVis.evergladensis.2.1.chr2.g01659.t01    mNeoVis.vulgivaga.2.1.chr2.g01674.t01      904    N0.HOG0000902   OG0000073  n72         rna-XM_011286020.3                          rna-XM_015543713.2                  905    N0.HOG0000903   OG0000073  n74   rna-XM_019793614.2, rna-XM_034650851.1    rna-XM_038582475.1      rna-XM_006935979.5      rna-XM_039225331.1    rna-XM_032852640.1    rna-XM_046003257.1    rna-XM_032301948.1    rna-XM_059165259.1      rna-XM_004772993.1    mNeoVis.2.1.1.chr5_hap1.g01669.t01        rna-XM_007095939.2    rna-XM_008689684.2, rna-XM_008689686.2      mNeoVis.3.1.1.chr2.g01584.t01    rna-gnl|WGS_ZZZZ|egapxtmp_037314-R1    mNeoVis.evergladensis.2.1.chr2.g01653.t01        906    N0.HOG0000904   OG0000073  n91   rna-XM_034651954.1, rna-XM_034664280.1    rna-NM_001388982.1      rna-XM_003991567.2    rna-XM_036070306.1    rna-XM_039221990.1      rna-XM_046003153.1    rna-XM_032302010.1    rna-XM_059165265.1    rna-XM_059388371.1    rna-XM_013046115.1    mNeoVis.2.1.1.chr5_hap1.g01674.t01        rna-XM_007095986.1    rna-XM_008689691.2    rna-XM_027586281.2    mNeoVis.3.1.1.chr2.g01587.t01    rna-gnl|WGS_ZZZZ|egapxtmp_037319-R1    mNeoVis.evergladensis.2.1.chr2.g01657.t01        907    N0.HOG0000905   OG0000073  n109       rna-XM_022507707.1          rna-XM_032882264.1    rna-XM_046003155.1, rna-XM_046003258.1    rna-XM_032301990.1, rna-XM_032301999.1    rna-XM_059165263.1, rna-XM_059165264.1    rna-XM_059388162.1    rna-XM_013046113.1, rna-XM_013046114.1    mNeoVis.2.1.1.chr5_hap1.g01673.t01              mNeoVis.3.1.1.chr2.g01586.t01    rna-gnl|WGS_ZZZZ|egapxtmp_037318-R1    mNeoVis.evergladensis.2.1.chr2.g01655.t01, mNeoVis.evergladensis.2.1.chr2.g01656.t01        908    N0.HOG0000906   OG0000073  n125   rna-XM_011218321.2, rna-XM_034650312.1, rna-XM_034650336.1                                  rna-XM_008689682.2, rna-XM_008689692.2, rna-XM_040621714.1                909    N0.HOG0000907   OG0000073  n131   rna-XM_011217727.2, rna-XM_034650367.1, rna-XM_034668467.1, rna-XM_034668519.1    rna-NM_001388863.1, rna-XM_038582474.1      rna-XM_006935982.4, rna-XM_011286018.3      rna-XM_039221991.1, rna-XM_039225330.1    rna-XM_032882257.1    rna-XM_046003687.1      rna-XM_059165076.1            rna-XM_004391705.1    rna-XM_007095938.1, rna-XM_007095987.2    rna-XM_008689694.2, rna-XM_008689695.2, rna-XM_008689698.2                910    N0.HOG0000908   OG0000073  n149   rna-XM_034668363.1    rna-XM_038582473.1    rna-XM_022502173.1      rna-XM_036070317.1      rna-XM_032882266.1    rna-XM_046003256.1    rna-XM_032301980.1    rna-XM_059165262.1    rna-XM_059388161.1    rna-XM_004772996.2    mNeoVis.2.1.1.chr5_hap1.g01672.t01    rna-XM_021687961.1    rna-XM_004391707.1      rna-XM_008689693.2    rna-XM_027581794.2      rna-gnl|WGS_ZZZZ|egapxtmp_037309-R1    mNeoVis.evergladensis.2.1.chr2.g01654.t01        911    N0.HOG0000909   OG0000074  n1   rna-XM_002923687.4    rna-NM_001006652.1    rna-XM_022525180.1    rna-XM_023245329.2, rna-XM_023249708.2    rna-XM_036094948.1    rna-XM_039249562.1    rna-XM_032841808.1    rna-XM_045987391.1    rna-XM_032323223.1    rna-XM_059151283.1    rna-XM_059383294.1    rna-XM_013047668.2    mNeoVis.2.1.1.chr7_hap1.g00584.t01    rna-XM_021701021.1    rna-XM_004396333.1    rna-XM_042969658.1, rna-XM_042969960.1    rna-XM_040633045.1    rna-XM_027619306.2    mNeoVis.3.1.1.chr7.g00567.t01    rna-gnl|WGS_ZZZZ|egapxtmp_012255-R1    mNeoVis.evergladensis.2.1.chr7.g00586.t01    mNeoVis.vulgivaga.2.1.chr7.g00589.t01      912    N0.HOG0000910   OG0000074  n25   rna-XM_019804276.2      rna-XM_022525264.1      rna-XM_036094853.1      rna-XM_032841811.1, rna-XM_032841963.1    rna-XM_045987393.1    rna-XM_032323225.1    rna-XM_059151285.1    rna-XM_059383297.1    rna-XM_045076839.1    mNeoVis.2.1.1.chr7_hap1.g00583.t01          rna-XM_008706651.1    rna-XM_027619420.1    mNeoVis.3.1.1.chr7.g00566.t01      mNeoVis.evergladensis.2.1.chr7.g00585.t01    mNeoVis.vulgivaga.2.1.chr7.g00588.t01      913    N0.HOG0000911   OG0000074  n42   rna-XM_034672321.1    rna-XM_038528822.1    rna-XM_022525327.1    rna-XM_011289895.4    rna-XM_036095027.1    rna-XM_039244896.1    rna-XM_032841961.1    rna-XM_045989507.1    rna-XM_032323220.1    rna-XM_059150898.1    rna-XM_059382470.1    rna-XM_004776079.3    mNeoVis.2.1.1.chr7_hap1.g00585.t01    rna-XM_021700945.1      rna-XM_042968772.1    rna-XM_040634011.1    rna-XM_027619419.2    mNeoVis.3.1.1.chr7.g00568.t01    rna-gnl|WGS_ZZZZ|egapxtmp_012256-R1    mNeoVis.evergladensis.2.1.chr7.g00587.t01    mNeoVis.vulgivaga.2.1.chr7.g00590.t01      914    N0.HOG0000912   OG0000074  n63   rna-XM_019809407.2, rna-XM_034652999.1    rna-XM_038579212.1        rna-XM_036110371.1                      rna-XM_004394967.1      rna-XM_040624282.1    rna-XM_027596311.1              915    N0.HOG0000913   OG0000074  n70     rna-XM_038528824.1    rna-XM_022525064.1    rna-XM_045046996.1    rna-XM_036095005.1    rna-XM_039248367.1    rna-XM_032841846.1    rna-XM_045986763.1    rna-XM_032324073.1    rna-XM_059151909.1    rna-XM_059383968.1    rna-XM_004776265.3    mNeoVis.2.1.1.chr7_hap1.g00579.t01    rna-XM_021700946.1    rna-XM_004396485.1    rna-XM_015539850.2    rna-XM_040634006.1    rna-XM_027618720.2    mNeoVis.3.1.1.chr7.g00562.t01      mNeoVis.evergladensis.2.1.chr7.g00581.t01    mNeoVis.vulgivaga.2.1.chr7.g00584.t01      916    N0.HOG0000914   OG0000074  n87   rna-XM_002923690.4    rna-XM_038528826.1    rna-XM_022525281.1    rna-XM_006941235.5    rna-XM_036095025.1    rna-XM_039248312.1    rna-XM_032841902.1    rna-XM_045989610.1    rna-XM_032324076.1    rna-XM_059151572.1    rna-XM_059381703.1    rna-XM_004776086.3    mNeoVis.2.1.1.chr7_hap1.g00580.t01    rna-XM_021701109.1    rna-XM_004396335.1    rna-XM_042969807.1    rna-XM_008688728.2    rna-XM_027618839.1    mNeoVis.3.1.1.chr7.g00563.t01      mNeoVis.evergladensis.2.1.chr7.g00582.t01    mNeoVis.vulgivaga.2.1.chr7.g00585.t01      917    N0.HOG0000915   OG0000074  n106   rna-XM_002923688.1    rna-XM_038525669.1    rna-XM_022525142.1      rna-XM_036094846.1    rna-XM_039248011.1    rna-XM_032841800.1    rna-XM_045988055.1    rna-XM_032326149.1    rna-XM_059151920.1    rna-XM_059383751.1    rna-XM_004776085.3      rna-XM_021700796.1    rna-XM_004396334.1    rna-XM_007087241.2    rna-XM_040626189.1    rna-XM_027619102.1    mNeoVis.3.1.1.chr7.g00565.t01      mNeoVis.evergladensis.2.1.chr7.g00584.t01        918    N0.HOG0000916   OG0000074  n122   rna-XM_019804318.2, rna-XM_034638787.1    rna-NM_001037345.1    rna-XM_022525072.1, rna-XM_022525186.1    rna-XM_003997781.6    rna-XM_036094998.1, rna-XM_036095004.1    rna-XM_039244245.1    rna-XM_032850934.1    rna-XM_045989382.1, rna-XM_045989450.1, rna-XM_045989471.1    rna-XM_032324075.1, rna-XM_032325860.1    rna-XM_059152729.1, rna-XM_059152909.1    rna-XM_059381537.1, rna-XM_059383945.1    rna-XM_045076836.1, rna-XM_045076837.1    mNeoVis.2.1.1.chr7_hap1.g00581.t01, mNeoVis.2.1.1.chr7_hap1.g00582.t01    rna-XM_044911389.1, rna-XM_044912327.1      rna-XM_015539851.2    rna-XM_040630632.1, rna-XM_040634009.1, rna-XM_040634418.1, rna-XM_040641894.1    rna-XM_035725101.1, rna-XM_035725342.1    mNeoVis.3.1.1.chr7.g00564.t01      mNeoVis.evergladensis.2.1.chr7.g00583.t01    mNeoVis.vulgivaga.2.1.chr7.g00586.t01, mNeoVis.vulgivaga.2.1.chr7.g00587.t01      919    N0.HOG0000917   OG0000074  n157     rna-NM_001048027.1      rna-XM_011289955.4      rna-XM_039247257.1                      rna-XM_007087253.3                  920    N0.HOG0000918   OG0000075  n2   rna-XM_002928579.2, rna-XM_034650075.1    rna-NM_001388960.1    rna-XM_022521730.1, rna-XM_022522138.1    rna-XM_011286584.4, rna-XM_011286670.2    rna-XM_036114409.1, rna-XM_036114424.1      rna-XM_032850793.1, rna-XM_032853565.1    rna-XM_046015923.1, rna-XM_046016964.1, rna-XM_046016966.1    rna-XM_032359703.1, rna-XM_032359731.1, rna-XM_032360301.1    rna-XM_059156771.1, rna-XM_059156785.1, rna-XM_059178208.1    rna-XM_059390830.1, rna-XM_059397788.1    rna-XM_004768236.1, rna-XM_004782218.2, rna-XM_004782581.1    mNeoVis.2.1.1.chr7_hap1.g01606.t01, mNeoVis.2.1.1.chr7_hap1.g01611.t01, mNeoVis.2.1.1.chr7_hap1.g01613.t01    rna-XM_021685618.2, rna-XM_021685660.1      rna-XM_007089351.2, rna-XM_007089411.2    rna-XM_008709377.1, rna-XM_008710693.1, rna-XM_008710695.2, rna-XM_008710736.2    rna-XM_027580947.2, rna-XM_027580948.2    mNeoVis.3.1.1.chr7.g01516.t01, mNeoVis.3.1.1.chr7.g01519.t01    rna-gnl|WGS_ZZZZ|egapxtmp_012797-R1      mNeoVis.vulgivaga.2.1.chr7.g01600.t01, mNeoVis.vulgivaga.2.1.chr7.g01602.t01      921    N0.HOG0000919   OG0000075  n44   rna-XM_034650182.1, rna-XM_034666457.1    rna-NM_001313854.1    rna-XM_022521677.1    rna-XM_045038815.1    rna-XM_036114423.1    rna-XM_039223167.1    rna-XM_032853599.1    rna-XM_046014227.1    rna-XM_032357027.1    rna-XM_059156803.1    rna-XM_059390748.1    rna-XM_004782215.3    mNeoVis.2.1.1.chr7_hap1.g01619.t01    rna-XM_021685665.1    rna-XM_004407552.1    rna-XM_015540754.2    rna-XM_008710697.1      mNeoVis.3.1.1.chr7.g01523.t01    rna-gnl|WGS_ZZZZ|egapxtmp_012785-R1    mNeoVis.evergladensis.2.1.chr7.g01590.t01    mNeoVis.vulgivaga.2.1.chr7.g01609.t01      922    N0.HOG0000920   OG0000075  n63   rna-XM_034650076.1, rna-XM_034650189.1, rna-XM_034667720.1    rna-NM_001388834.1    rna-XM_022521780.1    rna-XM_003992806.1      rna-XM_039225379.1    rna-XM_032853619.1    rna-XM_046014406.1    rna-XM_032359761.1    rna-XM_059180160.1    rna-XM_059395647.1    rna-XM_004768264.1          rna-XM_015540755.2    rna-XM_008710729.2, rna-XM_008710766.1      mNeoVis.3.1.1.chr7.g01551.t01    rna-gnl|WGS_ZZZZ|egapxtmp_012520-R1      mNeoVis.vulgivaga.2.1.chr7.g01640.t01      923    N0.HOG0000921   OG0000075  n82   rna-XM_002928577.2    rna-XM_038568147.1    rna-XM_022522103.1    rna-XM_011286671.4      rna-XM_039225764.1    rna-XM_032853590.1    rna-XM_046016633.1    rna-XM_032359245.1    rna-XM_059156721.1    rna-XM_059390884.1    rna-XM_045071461.1    mNeoVis.2.1.1.chr7_hap1.g01602.t01    rna-XM_021685615.1      rna-XM_007089413.2    rna-XM_008710734.1      mNeoVis.3.1.1.chr7.g01512.t01    rna-gnl|WGS_ZZZZ|egapxtmp_012902-R1    mNeoVis.evergladensis.2.1.chr7.g01580.t01    mNeoVis.vulgivaga.2.1.chr7.g01596.t01      924    N0.HOG0000922   OG0000075  n100   rna-XM_002928591.2, rna-XM_002928592.2, rna-XM_034665604.1    rna-NM_001388692.1    rna-XM_022521731.1, rna-XM_022522010.1, rna-XM_022522137.1, rna-XM_022522150.1    rna-XM_011286672.3, rna-XM_019811924.3    rna-XM_036071355.1, rna-XM_036103043.1, rna-XM_036103044.1    rna-XM_039225765.1, rna-XM_039225840.1    rna-XM_032853271.1, rna-XM_032853563.1, rna-XM_032853564.1, rna-XM_032853591.1    rna-XM_046015925.1, rna-XM_046015926.1, rna-XM_046016427.1, rna-XM_046017117.1    rna-XM_032359244.1, rna-XM_032359729.1, rna-XM_032359730.1, rna-XM_032360179.1    rna-XM_059134314.1, rna-XM_059177826.1, rna-XM_059177950.1, rna-XM_059178088.1    rna-XM_059390871.1, rna-XM_059390906.1, rna-XM_059397920.1, rna-XM_059398057.1    rna-XM_004768233.1, rna-XM_004768234.1, rna-XM_004768235.2, rna-XM_013064394.2    mNeoVis.2.1.1.chr7_hap1.g01601.t01, mNeoVis.2.1.1.chr7_hap1.g01603.t01, mNeoVis.2.1.1.chr7_hap1.g01604.t01, mNeoVis.2.1.1.chr7_hap1.g01605.t01    rna-XM_021685614.2, rna-XM_021685617.1      rna-XM_007089412.2, rna-XM_042958674.1    rna-XM_008710691.1, rna-XM_008710733.2, rna-XM_008710735.2, rna-XM_040631300.1    rna-XM_035723126.1    mNeoVis.3.1.1.chr7.g01511.t01, mNeoVis.3.1.1.chr7.g01513.t01, mNeoVis.3.1.1.chr7.g01514.t01, mNeoVis.3.1.1.chr7.g01515.t01    rna-gnl|WGS_ZZZZ|egapxtmp_012821-R1, rna-gnl|WGS_ZZZZ|egapxtmp_012822-R1    mNeoVis.evergladensis.2.1.chr7.g01581.t01    mNeoVis.vulgivaga.2.1.chr7.g01595.t01, mNeoVis.vulgivaga.2.1.chr7.g01597.t01, mNeoVis.vulgivaga.2.1.chr7.g01598.t01, mNeoVis.vulgivaga.2.1.chr7.g01599.t01      925    N0.HOG0000923   OG0000076  n1   rna-XM_002924962.4    rna-NM_001346099.1    rna-XM_022509154.1    rna-XM_003996843.6    rna-XM_036068543.1    rna-XM_039223290.1    rna-XM_032871599.1    rna-XM_045985263.1    rna-XM_032321090.1    rna-XM_059147940.1    rna-XM_059378553.1    rna-XM_045080653.1    mNeoVis.2.1.1.chr8_hap1.g00446.t01    rna-XM_021704262.1    rna-XM_004394998.1    rna-XM_007085821.3    rna-XM_040640649.1    rna-XM_027626413.2    mNeoVis.3.1.1.chr8.g00402.t01    rna-gnl|WGS_ZZZZ|egapxtmp_019803-R1    mNeoVis.evergladensis.2.1.chr9.g00437.t01    mNeoVis.vulgivaga.2.1.chr8.g00442.t01      926    N0.HOG0000924   OG0000076  n21   rna-XM_002922111.4    rna-XM_038546022.1    rna-XM_022509397.1    rna-XM_003996856.6    rna-XM_036069199.1    rna-XM_039256774.1    rna-XM_032871794.1    rna-XM_045985541.1    rna-XM_032321719.1    rna-XM_059148313.1    rna-XM_059380696.1    rna-XM_004772751.3    mNeoVis.2.1.1.chr8_hap1.g00417.t01    rna-XM_021681747.1    rna-XM_004410495.1    rna-XM_042966682.1    rna-XM_008701262.2    rna-XM_027626244.1    mNeoVis.3.1.1.chr8.g00381.t01    rna-gnl|WGS_ZZZZ|egapxtmp_019802-R1    mNeoVis.evergladensis.2.1.chr9.g00414.t01    mNeoVis.vulgivaga.2.1.chr8.g00422.t01      927    N0.HOG0000925   OG0000076  n42   rna-XM_002922112.4    rna-XM_038546102.1    rna-XM_022509340.1    rna-XM_003996857.6    rna-XM_036069116.1    rna-XM_039256829.1    rna-XM_032871795.1    rna-XM_045985294.1    rna-XM_032322480.1    rna-XM_059148300.1    rna-XM_059380822.1    rna-XM_004772750.3    mNeoVis.2.1.1.chr8_hap1.g00416.t01    rna-XM_021681720.1    rna-XM_004410496.1    rna-XM_042966680.1    rna-XM_008701265.2    rna-XM_027626164.1    mNeoVis.3.1.1.chr8.g00380.t01        mNeoVis.vulgivaga.2.1.chr8.g00421.t01      928    N0.HOG0000926   OG0000076  n61   rna-XM_002922113.4    rna-XM_038546101.1    rna-XM_022509327.1    rna-XM_045044396.1    rna-XM_036069115.1    rna-XM_039256766.1    rna-XM_032871610.1    rna-XM_045985425.1    rna-XM_032321079.1    rna-XM_059148228.1    rna-XM_059380792.1    rna-XM_004772749.3    mNeoVis.2.1.1.chr8_hap1.g00415.t01    rna-XM_021681719.1    rna-XM_004410602.1    rna-XM_007087705.3    rna-XM_008701264.2    rna-XM_027626149.2    mNeoVis.3.1.1.chr8.g00379.t01      mNeoVis.evergladensis.2.1.chr9.g00413.t01    mNeoVis.vulgivaga.2.1.chr8.g00420.t01      929    N0.HOG0000927   OG0000076  n82   rna-XM_034648398.1    rna-XM_038546104.1    rna-XM_022509172.1    rna-XM_006940342.4    rna-XM_036069108.1    rna-XM_039256723.1    rna-XM_032871793.1    rna-XM_045985333.1    rna-XM_032321715.1    rna-XM_059148135.1    rna-XM_059387552.1    rna-XM_045080976.1    mNeoVis.2.1.1.chr8_hap1.g00421.t01      rna-XM_004410493.1    rna-XM_007087637.2    rna-XM_008701254.2    rna-XM_027626046.1    mNeoVis.3.1.1.chr8.g00385.t01    rna-gnl|WGS_ZZZZ|egapxtmp_019804-R1    mNeoVis.evergladensis.2.1.chr9.g00418.t01    mNeoVis.vulgivaga.2.1.chr8.g00426.t01      930    N0.HOG0000928   OG0000076  n99   rna-XM_034648682.1    rna-XM_038546105.1    rna-XM_022509171.1    rna-XM_006940343.5    rna-XM_036069109.1    rna-XM_039256722.1    rna-XM_032871792.1    rna-XM_045985356.1    rna-XM_032321717.1    rna-XM_059148141.1    rna-XM_059387533.1    rna-XM_004772754.3    mNeoVis.2.1.1.chr8_hap1.g00422.t01      rna-XM_004410600.1    rna-XM_042966012.1    rna-XM_008701256.2    rna-XM_035724616.1    mNeoVis.3.1.1.chr8.g00386.t01, mNeoVis.3.1.1.chr8.g00387.t01    rna-gnl|WGS_ZZZZ|egapxtmp_017414-R1      mNeoVis.vulgivaga.2.1.chr8.g00427.t01      931    N0.HOG0000929   OG0000076  n117   rna-XM_034641239.1, rna-XM_034641240.1    rna-NM_001346094.1, rna-NM_001346095.1    rna-XM_022509156.1    rna-XM_003996855.5, rna-XM_045044400.1    rna-XM_036069110.1, rna-XM_036069197.1    rna-XM_039256724.1, rna-XM_039256743.1    rna-XM_032871790.1, rna-XM_032871791.1    rna-XM_045985332.1, rna-XM_045985387.1    rna-XM_032321716.1, rna-XM_032321718.1    rna-XM_059148142.1, rna-XM_059148180.1    rna-XM_059380227.1, rna-XM_059387459.1, rna-XM_059387478.1    rna-XM_004772755.3, rna-XM_004772756.2    mNeoVis.2.1.1.chr8_hap1.g00424.t01    rna-XM_044912482.1, rna-XM_044912483.1    rna-XM_004410491.1, rna-XM_004410492.1    rna-XM_015540020.2, rna-XM_042966093.1    rna-XM_008701253.2, rna-XM_008701257.2    rna-XM_027625808.2, rna-XM_035724615.1      rna-gnl|WGS_ZZZZ|egapxtmp_019805-R1    mNeoVis.evergladensis.2.1.chr9.g00419.t01, mNeoVis.evergladensis.2.1.chr9.g00420.t01    mNeoVis.vulgivaga.2.1.chr8.g00429.t01      932    N0.HOG0000930   OG0000077  n2   rna-XM_002931288.2, rna-XM_034648662.1    rna-NM_001388554.1, rna-XM_038557594.1    rna-XM_022521644.1, rna-XM_022521658.1    rna-XM_003987373.2    rna-XM_036118046.1    rna-XM_039221978.1    rna-XM_032857155.1    rna-XM_046009991.1, rna-XM_046010291.1    rna-XM_032341757.1, rna-XM_032341759.1    rna-XM_059179610.1, rna-XM_059183384.1    rna-XM_059371111.1, rna-XM_059372090.1    rna-XM_004781745.2, rna-XM_004781748.2    mNeoVis.2.1.1.chr10_hap1.g00837.t01, mNeoVis.2.1.1.chr10_hap1.g00840.t01    rna-XM_021695712.2      rna-XM_007091416.3    rna-XM_008710295.2, rna-XM_008710440.2, rna-XM_008710441.2    rna-XM_027570339.2    mNeoVis.3.1.1.chr9.g00780.t01    rna-gnl|WGS_ZZZZ|egapxtmp_016935-R1, rna-gnl|WGS_ZZZZ|egapxtmp_016939-R1    mNeoVis.evergladensis.2.1.chr8.g00811.t01, mNeoVis.evergladensis.2.1.chr8.g00814.t01    mNeoVis.vulgivaga.2.1.chr9.g00809.t01      933    N0.HOG0000931   OG0000077  n34   rna-XM_034646236.1    rna-NM_001388644.1    rna-XM_022506939.1    rna-XM_045039290.1      rna-XM_039226791.1    rna-XM_032844585.1    rna-XM_046016724.1    rna-XM_032359440.1    rna-XM_059158634.1    rna-XM_059386639.1, rna-XM_059386920.1    rna-XM_004763030.2    mNeoVis.2.1.1.chr7_hap1.g02022.t01      rna-XM_004399506.1    rna-XM_007082174.1    rna-XM_008701081.2      mNeoVis.3.1.1.chr7.g01888.t01    rna-gnl|WGS_ZZZZ|egapxtmp_010317-R1    mNeoVis.evergladensis.2.1.chr7.g01985.t01    mNeoVis.vulgivaga.2.1.chr7.g02005.t01      934    N0.HOG0000932   OG0000077  n53   rna-XM_002929024.4, rna-XM_002929025.4, rna-XM_034651671.1    rna-NM_001017519.1    rna-XM_022507042.1    rna-XM_011286891.4      rna-XM_039226796.1    rna-XM_032844584.1    rna-XM_046016675.1    rna-XM_032359625.1          mNeoVis.2.1.1.chr7_hap1.g02019.t01        rna-XM_007082198.2    rna-XM_008700961.2, rna-XM_008700962.2    rna-XM_027579844.2    mNeoVis.3.1.1.chr7.g01885.t01    rna-gnl|WGS_ZZZZ|egapxtmp_012604-R1          935    N0.HOG0000933   OG0000077  n71   rna-XM_002929981.3    rna-NM_001388923.1    rna-XM_022507105.1    rna-XM_003993266.3    rna-XM_036073539.1    rna-XM_039225294.1    rna-XM_032844630.1    rna-XM_046015829.1    rna-XM_032359836.1    rna-XM_059134551.1    rna-XM_059388374.1    rna-XM_004782273.1    mNeoVis.2.1.1.chr7_hap1.g02077.t01    rna-XM_021684825.1    rna-XM_004417247.1    rna-XM_007095369.2    rna-XM_008694059.2        rna-gnl|WGS_ZZZZ|egapxtmp_012618-R1    mNeoVis.evergladensis.2.1.chr7.g02032.t01    mNeoVis.vulgivaga.2.1.chr7.g02049.t01      936    N0.HOG0000934   OG0000077  n89   rna-XM_002930385.4    rna-NM_001388641.1, rna-NM_001388643.1    rna-XM_022507607.1    rna-XM_045038139.1      rna-XM_039226909.1    rna-XM_032844744.1    rna-XM_046015850.1    rna-XM_032359812.1          mNeoVis.2.1.1.chr7_hap1.g02024.t01        rna-XM_007082176.1    rna-XM_008700957.1      mNeoVis.3.1.1.chr7.g01890.t01    rna-gnl|WGS_ZZZZ|egapxtmp_012606-R1      mNeoVis.vulgivaga.2.1.chr7.g02007.t01      937    N0.HOG0000935   OG0000077  n104   rna-XM_002929028.4    rna-NM_001388646.1    rna-XM_022507085.1    rna-XM_045039286.1      rna-XM_039226794.1    rna-XM_032844607.1    rna-XM_046016726.1    rna-XM_032359658.1    rna-XM_059158607.1    rna-XM_059386642.1, rna-XM_059386931.1    rna-XM_004763226.1    mNeoVis.2.1.1.chr7_hap1.g02020.t01        rna-XM_007082172.2    rna-XM_008700960.2    rna-XM_027580056.1    mNeoVis.3.1.1.chr7.g01886.t01    rna-gnl|WGS_ZZZZ|egapxtmp_010278-R1    mNeoVis.evergladensis.2.1.chr7.g01983.t01    mNeoVis.vulgivaga.2.1.chr7.g02003.t01      938    N0.HOG0000936   OG0000077  n120     rna-NM_001388642.1    rna-XM_022507162.1    rna-XM_003993290.3      rna-XM_039226795.1    rna-XM_032844592.1    rna-XM_046016698.1    rna-XM_032359570.1    rna-XM_059158656.1    rna-XM_059386653.1, rna-XM_059386928.1    rna-XM_004763029.1    mNeoVis.2.1.1.chr7_hap1.g02025.t01        rna-XM_042958303.1    rna-XM_008700955.1      mNeoVis.3.1.1.chr7.g01891.t01    rna-gnl|WGS_ZZZZ|egapxtmp_013547-R1    mNeoVis.evergladensis.2.1.chr7.g01987.t01    mNeoVis.vulgivaga.2.1.chr7.g02008.t01      939    N0.HOG0000937   OG0000077  n134   rna-XM_034646256.1, rna-XM_034650264.1    rna-NM_001388645.1      rna-XM_045039288.1      rna-XM_039226819.1    rna-XM_032844590.1, rna-XM_032851081.1    rna-XM_046016710.1    rna-XM_032359591.1    rna-XM_059158622.1    rna-XM_059386612.1, rna-XM_059386887.1    rna-XM_004763031.1    mNeoVis.2.1.1.chr7_hap1.g02021.t01      rna-XM_004399507.1    rna-XM_042959653.1    rna-XM_008700959.2      mNeoVis.3.1.1.chr7.g01887.t01    rna-gnl|WGS_ZZZZ|egapxtmp_010302-R2    mNeoVis.evergladensis.2.1.chr7.g01984.t01    mNeoVis.vulgivaga.2.1.chr7.g02004.t01      940    N0.HOG0000938   OG0000078  n1   rna-XM_011229861.3    rna-XM_038559246.1, rna-XM_038559509.1, rna-XM_038588921.1    rna-XM_022518267.1    rna-XM_006929062.5    rna-XM_036066746.1    rna-XM_039242968.1    rna-XM_032875026.1    rna-XM_046020936.1    rna-XM_032304450.1    rna-XM_059170663.1    rna-XM_059397153.1    rna-XM_013059650.2    mNeoVis.2.1.1.chr4_hap1.g00631.t01    rna-XM_044920111.1    rna-XM_004409439.1    rna-XM_042963061.1    rna-XM_040644490.1    rna-XM_027574144.2    mNeoVis.3.1.1.chr6.g00642.t01    rna-gnl|WGS_ZZZZ|egapxtmp_004474-R1    mNeoVis.evergladensis.2.1.chr6.g00640.t01    mNeoVis.vulgivaga.2.1.chr6.g00637.t01      941    N0.HOG0000939   OG0000078  n26   rna-XM_002917618.3    rna-XM_038555532.1    rna-XM_022502313.1    rna-XM_004000120.5    rna-XM_036067356.1    rna-XM_039219469.1    rna-XM_032873519.1    rna-XM_045978186.1    rna-XM_032316631.1    rna-XM_059165609.1    rna-XM_059395174.1    rna-XM_004743566.3    mNeoVis.2.1.1.chr4_hap1.g01018.t01    rna-XM_021688464.1    rna-XM_004409835.1    rna-XM_042973080.1    rna-XM_040626047.1    rna-XM_027615207.2    mNeoVis.3.1.1.chr6.g01037.t01    rna-gnl|WGS_ZZZZ|egapxtmp_003793-R1    mNeoVis.evergladensis.2.1.chr6.g01023.t01    mNeoVis.vulgivaga.2.1.chr6.g01015.t01      942    N0.HOG0000940   OG0000078  n46   rna-XM_034637813.1    rna-XM_038531076.1    rna-XM_022518650.1    rna-XM_045047182.1    rna-XM_036119933.1    rna-XM_039241937.1    rna-XM_032854516.1    rna-XM_045987785.1    rna-XM_032326451.1    rna-XM_059153380.1    rna-XM_059382303.1    rna-XM_045076037.1    mNeoVis.2.1.1.chr7_hap1.g00338.t01    rna-XM_021704786.2    rna-XM_004408523.1    rna-XM_042968160.1    rna-XM_040633283.1    rna-XM_035725303.1    mNeoVis.3.1.1.chr7.g00325.t01    rna-gnl|WGS_ZZZZ|egapxtmp_012193-R1    mNeoVis.evergladensis.2.1.chr7.g00339.t01    mNeoVis.vulgivaga.2.1.chr7.g00342.t01      943    N0.HOG0000941   OG0000078  n68   rna-XM_011218389.3, rna-XM_034648697.1    rna-XM_038544325.1    rna-XM_022526806.1    rna-XM_045059721.1    rna-XM_036114708.1    rna-XM_039254690.1    rna-XM_032856257.1    rna-XM_046009346.1    rna-XM_032343949.1    rna-XM_059181768.1    rna-XM_059373002.1    rna-XM_004755198.3    mNeoVis.2.1.1.chr10_hap1.g00522.t01    rna-XM_021695691.1    rna-XM_004402030.1    rna-XM_042989463.1    rna-XM_040620785.1    rna-XM_027569809.1    mNeoVis.3.1.1.chr9.g00482.t01    rna-gnl|WGS_ZZZZ|egapxtmp_014624-R1    mNeoVis.evergladensis.2.1.chr8.g00504.t01    mNeoVis.vulgivaga.2.1.chr9.g00504.t01      944    N0.HOG0000942   OG0000078  n87   rna-XM_034657099.1    rna-XM_038583952.1, rna-XM_038589447.1    rna-XM_022512437.1    rna-XM_023239649.2    rna-XM_036104287.1    rna-XM_039223317.1, rna-XM_039224564.1    rna-XM_032861854.1    rna-XM_046000685.1    rna-XM_032337793.1    rna-XM_059173610.1    rna-XM_059416859.1    rna-XM_045081990.1    mNeoVis.2.1.1.chr1_hap1.g00038.t01    rna-XM_021702488.1    rna-XM_004415371.1    rna-XM_042997717.1    rna-XM_040630559.1    rna-XM_027607059.2    mNeoVis.3.1.1.chr1.g00036.t01, mNeoVis.3.1.1.chr1.g00037.t01, mNeoVis.3.1.1.chr1.g00038.t01    rna-gnl|WGS_ZZZZ|egapxtmp_022347-R1    mNeoVis.evergladensis.2.1.chr1.g00036.t01    mNeoVis.vulgivaga.2.1.chr1.g00037.t01      945    N0.HOG0000943   OG0000078  n114   rna-XM_019807931.2    rna-NM_001195147.1    rna-XM_022519493.1    rna-XM_019834866.3    rna-XM_036121158.1    rna-XM_039221095.1    rna-XM_032876903.1    rna-XM_046011448.1    rna-XM_032348050.1    rna-XM_059185504.1    rna-XM_059403007.1    rna-XM_045073232.1    mNeoVis.2.1.1.chr9_hap1.g00604.t01    rna-XM_021704926.2    rna-XM_004400882.1    rna-XM_042992192.1    rna-XM_040642393.1    rna-XM_027594193.2    mNeoVis.3.1.1.chr10.g00572.t01    rna-gnl|WGS_ZZZZ|egapxtmp_024691-R1    mNeoVis.evergladensis.2.1.chr10.g00598.t01    mNeoVis.vulgivaga.2.1.chr10.g00584.t01      946    N0.HOG0000944   OG0000078  n134   rna-XM_034670712.1    rna-NM_001168460.1, rna-XM_038583046.1    rna-XM_022515072.1    rna-XM_019839852.3    rna-XM_036111499.1, rna-XM_036119886.1    rna-XM_039240624.1    rna-XM_032869501.1    rna-XM_046002127.1    rna-XM_032335217.1    rna-XM_059162861.1    rna-XM_059391080.1    rna-XM_004745509.3    mNeoVis.2.1.1.chr5_hap1.g01210.t01    rna-XM_044920387.1    rna-XM_004391934.1    rna-XM_015544794.2    rna-XM_008708247.2    rna-XM_027585035.2    mNeoVis.3.1.1.chr2.g01122.t01    rna-gnl|WGS_ZZZZ|egapxtmp_035563-R1    mNeoVis.evergladensis.2.1.chr2.g01198.t01    mNeoVis.vulgivaga.2.1.chr2.g01204.t01      947    N0.HOG0000945   OG0000079  n1   rna-XM_002929133.2    rna-NM_001388867.1    rna-XM_022510283.1    rna-XM_045052644.1    rna-XM_036087239.1    rna-XM_039243026.1    rna-XM_032866870.1    rna-XM_046020977.1    rna-XM_032304792.1    rna-XM_059172142.1    rna-XM_059395567.1    rna-XM_004772106.1    mNeoVis.2.1.1.chr4_hap1.g00087.t01    rna-XM_021692936.1    rna-XM_004409608.1    rna-XM_007090901.1    rna-XM_008697399.2    rna-XM_027574244.1    mNeoVis.3.1.1.chr6.g00088.t01    rna-gnl|WGS_ZZZZ|egapxtmp_003139-R1    mNeoVis.evergladensis.2.1.chr6.g00089.t01    mNeoVis.vulgivaga.2.1.chr6.g00087.t01      948    N0.HOG0000946   OG0000079  n23   rna-XM_002921816.2    rna-NM_001388563.1    rna-XM_022510213.1    rna-XM_003983176.3    rna-XM_036087235.1    rna-XM_039243057.1    rna-XM_032865087.1    rna-XM_046020990.1    rna-XM_032305816.1    rna-XM_059172560.1    rna-XM_059395977.1    rna-XM_004772102.1    mNeoVis.2.1.1.chr4_hap1.g00089.t01    rna-XM_021692867.1    rna-XM_004409602.1    rna-XM_007090855.1    rna-XM_008697282.2    rna-XM_027573985.1    mNeoVis.3.1.1.chr6.g00090.t01    rna-gnl|WGS_ZZZZ|egapxtmp_003137-R1    mNeoVis.evergladensis.2.1.chr6.g00092.t01    mNeoVis.vulgivaga.2.1.chr6.g00089.t01      949    N0.HOG0000947   OG0000079  n46   rna-XM_002921817.2    rna-NM_001388513.1    rna-XM_022510333.1    rna-XM_003983175.3    rna-XM_036087242.1    rna-XM_039243058.1    rna-XM_032865099.1    rna-XM_046021124.1    rna-XM_032305554.1    rna-XM_059172707.1    rna-XM_059395571.1    rna-XM_004772079.1      rna-XM_021692946.1    rna-XM_004409603.1    rna-XM_007090854.2    rna-XM_008697283.2    rna-XM_027574273.1    mNeoVis.3.1.1.chr6.g00091.t01            950    N0.HOG0000948   OG0000079  n64   rna-XM_002921829.4    rna-NM_001388968.1    rna-XM_022510294.1    rna-XM_003983185.2    rna-XM_036087248.1      rna-XM_032865064.1    rna-XM_046020988.1    rna-XM_032305725.1    rna-XM_059172708.1    rna-XM_059395570.1    rna-XM_004772080.1    mNeoVis.2.1.1.chr4_hap1.g00088.t01        rna-XM_007090856.1    rna-XM_008697281.2    rna-XM_035723417.1    mNeoVis.3.1.1.chr6.g00089.t01        mNeoVis.vulgivaga.2.1.chr6.g00088.t01      951    N0.HOG0000949   OG0000079  n79   rna-XM_034653915.1    rna-NM_001388856.1    rna-XM_022510293.1    rna-XM_003983186.2    rna-XM_036087245.1    rna-XM_039243055.1    rna-XM_032865051.1    rna-XM_046020992.1    rna-XM_032305724.1    rna-XM_059172709.1    rna-XM_059395569.1    rna-XM_045089519.1        rna-XM_004409609.1    rna-XM_007090900.1    rna-XM_008697402.2    rna-XM_027574990.2      rna-gnl|WGS_ZZZZ|egapxtmp_003138-R1    mNeoVis.evergladensis.2.1.chr6.g00091.t01        952    N0.HOG0000950   OG0000079  n98     rna-NM_001388561.1          rna-XM_039243027.1                      rna-XM_042977223.1                  953    N0.HOG0000951   OG0000079  n100   rna-XM_002921827.4    rna-NM_001388562.1    rna-XM_022510104.1    rna-XM_003983177.2    rna-XM_036087243.1    rna-XM_039243028.1    rna-XM_032866241.1    rna-XM_046020989.1    rna-XM_032304793.1    rna-XM_059172712.1    rna-XM_059395568.1    rna-XM_045089521.1          rna-XM_007090857.1    rna-XM_008697400.2        rna-gnl|WGS_ZZZZ|egapxtmp_003087-R1    mNeoVis.evergladensis.2.1.chr6.g00090.t01        954    N0.HOG0000952   OG0000079  n115     rna-NM_001388821.1                rna-XM_032304791.1    rna-XM_059172711.1    rna-XM_059387412.1    rna-XM_045089520.1    mNeoVis.2.1.1.chr4_hap1.g00085.t01          rna-XM_008697401.2      mNeoVis.3.1.1.chr6.g00086.t01    rna-gnl|WGS_ZZZZ|egapxtmp_003141-R1          955    N0.HOG0000953   OG0000079  n123   rna-XM_034648599.1, rna-XM_034648651.1    rna-NM_001388517.1, rna-NM_001388560.1    rna-XM_022510249.1, rna-XM_022510272.1    rna-XM_003983178.3, rna-XM_011279609.2    rna-XM_036087238.1, rna-XM_036087250.1    rna-XM_039243056.1, rna-XM_039243059.1    rna-XM_032866214.1, rna-XM_032866856.1    rna-XM_046020963.1, rna-XM_046020964.1    rna-XM_032304788.1, rna-XM_032304789.1    rna-XM_059172140.1, rna-XM_059172141.1    rna-XM_059395565.1, rna-XM_059395566.1    rna-XM_004772107.2, rna-XM_004772108.3    mNeoVis.2.1.1.chr4_hap1.g00084.t01, mNeoVis.2.1.1.chr4_hap1.g00086.t01    rna-XM_021692819.2    rna-XM_004409601.1    rna-XM_007090858.1, rna-XM_007090902.1    rna-XM_008697396.2, rna-XM_008697397.2    rna-XM_027573986.1    mNeoVis.3.1.1.chr6.g00085.t01, mNeoVis.3.1.1.chr6.g00087.t01    rna-gnl|WGS_ZZZZ|egapxtmp_003132-R1    mNeoVis.evergladensis.2.1.chr6.g00088.t01    mNeoVis.vulgivaga.2.1.chr6.g00086.t01      956    N0.HOG0000954   OG0000080  n3     id-LOC119877116    id-LOC111151887, id-LOC111151890    rna-XM_023250642.2    id-LOC118532192    id-LOC120232974    id-LOC116872846    id-LOC123951619    rna-XM_032305828.1            id-LOC110582932    id-LOC101363293    id-LOC102964941, id-LOC102966099    id-LOC103669100                957    N0.HOG0000955   OG0000080  n17     id-LOC102156776          id-LOC120238965    id-LOC116872800                            id-egapxtmp_002333          958    N0.HOG0000956   OG0000080  n20     id-LOC111090221        id-LOC118532191    id-LOC120238964                  id-LOC123323180                      959    N0.HOG0000957   OG0000080  n23   id-LOC105239732      id-LOC111151970      id-LOC118532190    id-LOC120232975    id-LOC116872865      id-LOC116569535    id-LOC131829872    id-LOC132015504    id-LOC106004551            id-LOC103669099                960    N0.HOG0000958   OG0000080  n34                 id-LOC123951618              id-LOC123326385        id-LOC103669091                961    N0.HOG0000959   OG0000080  n38   id-LOC105239743    id-LOC119869605            id-LOC116873447                  id-LOC101364470                    962    N0.HOG0000960   OG0000080  n41   id-LOC109490375    id-LOC119874694            id-LOC116872807                  id-LOC101364764                    963    N0.HOG0000961   OG0000080  n45   id-LOC109490377                              id-LOC101364171                    964    N0.HOG0000962   OG0000080  n47     id-LOC119869607      id-LOC111559551      id-LOC120238966    id-LOC116873431        id-LOC131829870      id-LOC106004547                          965    N0.HOG0000963   OG0000080  n56   id-LOC117795149    id-LOC119874695, id-LOC119874696      id-LOC123384060                      id-LOC123326383      id-LOC122235317    rna-XM_040643905.1                966    N0.HOG0000964   OG0000080  n62   id-LOC117797080    id-LOC119869603    id-LOC111151889    id-LOC109497660, id-LOC123384059      id-LOC120232976, id-LOC120233003    id-LOC116872826, id-LOC116872837, id-LOC116872855, id-LOC116872876      id-LOC116569534        id-LOC123387055      id-LOC110582812, id-LOC123326382      id-LOC102965232, id-LOC102965532    id-LOC103669097, id-LOC103669098    id-LOC113910957              967    N0.HOG0000965   OG0000080  n82    id-LOC609004   id-LOC111151888    id-LOC109497661    id-LOC118532193    id-LOC120232977    id-LOC116873458    id-LOC123951616    id-LOC116569692    id-LOC131829874    id-LOC132015627    id-LOC101682998          id-LOC122235318                  968    N0.HOG0000966   OG0000080  n93     id-LOC111090225    id-LOC111151891    id-LOC111557462    id-LOC118532194    id-LOC120232973    id-LOC116873494      id-LOC116569537                id-LOC122236348    id-LOC121105885    id-LOC113910958              969    N0.HOG0000967   OG0000080  n103   id-LOC100478092, id-LOC109490378    id-LOC480788, id-LOC609053      rna-XM_045052894.1    id-LOC118532169    id-LOC120238963, id-LOC120238968    id-LOC116863595, id-LOC116863597, id-LOC116873419    id-LOC123951612, id-LOC123951625    id-LOC116569683, id-LOC116569684    id-LOC131829868    id-LOC132015625, id-LOC132015626    id-LOC101684255, rna-XM_013050448.1        rna-XM_004414591.1    id-LOC102966676, id-LOC122235315    id-LOC103669001, rna-XM_040643899.1    id-LOC113910955, id-LOC113911840              970    N0.HOG0000968   OG0000080  n129     id-LOC119874697    id-LOC111151886      id-LOC118532187      id-LOC116872815    id-LOC123951617    id-LOC116569533    id-LOC131829867            rna-XM_004414590.1    id-LOC102966391                  971    N0.HOG0000969   OG0000080  n138         id-LOC101095329, id-LOC105260394      id-LOC120233013, id-LOC120233015                      id-LOC102964653                  972    N0.HOG0000970   OG0000080  n143     id-LOC102157186, id-LOC119874700        id-LOC118532189, id-LOC118532196      id-LOC116872884, id-LOC116873470        id-LOC131829871, id-LOC131830221    id-LOC132015505, id-LOC132015506    id-LOC123387054        rna-XM_004414589.1                    973    N0.HOG0000971   OG0000080  n154     id-LOC111090224, id-LOC119874702                  id-LOC131829875      id-LOC101686578        id-LOC101362405      id-LOC103669101, id-LOC121105886, rna-XM_040644588.1                974    N0.HOG0000972   OG0000081  n1   rna-XM_002930043.1, rna-XM_034652995.1    rna-NM_001388620.1    rna-XM_022507449.1    rna-XM_003993318.2    rna-XM_036073531.1, rna-XM_036104532.1    rna-XM_039224972.1    rna-XM_032846496.1    rna-XM_046017039.1    rna-XM_032360295.1    rna-XM_059158894.1    rna-XM_059388131.1    rna-XM_004782256.2    mNeoVis.2.1.1.chr7_hap1.g02102.t01    rna-XM_021684828.1    rna-XM_004412001.1    rna-XM_015540528.2    rna-XM_008694075.1    rna-XM_027580583.1    mNeoVis.3.1.1.chr7.g01944.t01    rna-gnl|WGS_ZZZZ|egapxtmp_013486-R1    mNeoVis.evergladensis.2.1.chr7.g02051.t01    mNeoVis.vulgivaga.2.1.chr7.g02075.t01      975    N0.HOG0000973   OG0000081  n26   rna-XM_034644811.1    rna-NM_001388621.1      rna-XM_045038150.1      rna-XM_039224971.1    rna-XM_032846562.1    rna-XM_046016752.1    rna-XM_032360034.1    rna-XM_059158891.1    rna-XM_059388139.1    rna-XM_045068209.1    mNeoVis.2.1.1.chr7_hap1.g02101.t01        rna-XM_007088795.2, rna-XM_042958819.1    rna-XM_008694220.2      mNeoVis.3.1.1.chr7.g01943.t01    rna-gnl|WGS_ZZZZ|egapxtmp_012619-R1      mNeoVis.vulgivaga.2.1.chr7.g02074.t01      976    N0.HOG0000974   OG0000081  n42   rna-XM_002930044.2    rna-NM_001388905.1      rna-XM_003993319.2    rna-XM_036104531.1    rna-XM_039224962.1, rna-XM_039224963.1    rna-XM_032846489.1    rna-XM_046015809.1    rna-XM_032359854.1    rna-XM_059136490.1    rna-XM_059386988.1    rna-XM_004782255.2          rna-XM_007088797.1, rna-XM_015540510.2    rna-XM_008694076.1                977    N0.HOG0000975   OG0000081  n57   rna-XM_002930046.2, rna-XM_034644813.1    rna-NM_001388618.1    rna-XM_022507050.1    rna-XM_003993269.2, rna-XM_006937317.3    rna-XM_036104521.1    rna-XM_039224010.1    rna-XM_032846480.1    rna-XM_046016579.1    rna-XM_032359280.1    rna-XM_059158895.1    rna-XM_059388114.1    rna-XM_004782252.2    mNeoVis.2.1.1.chr7_hap1.g02104.t01    rna-XM_021684829.1      rna-XM_007088769.1, rna-XM_007088800.2    rna-XM_008694078.1    rna-XM_027580024.2      rna-gnl|WGS_ZZZZ|egapxtmp_013501-R1    mNeoVis.evergladensis.2.1.chr7.g02052.t01    mNeoVis.vulgivaga.2.1.chr7.g02077.t01      978    N0.HOG0000976   OG0000081  n80   rna-XM_002930045.3    rna-XM_038564481.1    rna-XM_022507179.1      rna-XM_036104522.1    rna-XM_039224961.1    rna-XM_032846484.1    rna-XM_046015808.1    rna-XM_032359855.1    rna-XM_059136611.1    rna-XM_059385939.1    rna-XM_045067540.1    mNeoVis.2.1.1.chr7_hap1.g02103.t01    rna-XM_021685197.1        rna-XM_008694077.2    rna-XM_035723134.1      rna-gnl|WGS_ZZZZ|egapxtmp_010301-R1      mNeoVis.vulgivaga.2.1.chr7.g02076.t01      979    N0.HOG0000977   OG0000081  n96   rna-XM_002930047.4, rna-XM_002930049.2, rna-XM_011216738.3, rna-XM_034646528.1, rna-XM_034646537.1    rna-NM_001256459.1, rna-NM_001388617.1, rna-XM_038564480.1    rna-XM_022507051.1, rna-XM_022507178.1    rna-XM_003993270.2, rna-XM_011286881.4, rna-XM_019812967.2, rna-XM_045038151.1    rna-XM_036104529.1, rna-XM_036104530.1    rna-XM_039224004.1, rna-XM_039224005.1    rna-XM_032846503.1, rna-XM_032846561.1    rna-XM_046015807.1, rna-XM_046016839.1, rna-XM_046016876.1, rna-XM_046016877.1    rna-XM_032358798.1, rna-XM_032358799.1, rna-XM_032359673.1    rna-XM_059136728.1, rna-XM_059136844.1, rna-XM_059136964.1, rna-XM_059158896.1    rna-XM_059383927.1, rna-XM_059385001.1, rna-XM_059388102.1    rna-XM_004782251.2, rna-XM_004782253.3, rna-XM_004782254.3    mNeoVis.2.1.1.chr7_hap1.g02105.t01, mNeoVis.2.1.1.chr7_hap1.g02106.t01, mNeoVis.2.1.1.chr7_hap1.g02107.t01, mNeoVis.2.1.1.chr7_hap1.g02108.t01    rna-XM_021685198.2    rna-XM_004411978.1, rna-XM_004412000.1    rna-XM_007088798.2, rna-XM_007088799.2, rna-XM_015540529.1    rna-XM_008694081.2, rna-XM_008694082.2, rna-XM_040638714.1, rna-XM_040638715.1    rna-XM_027577869.1, rna-XM_027577870.2    mNeoVis.3.1.1.chr7.g01945.t01, mNeoVis.3.1.1.chr7.g01946.t01    rna-gnl|WGS_ZZZZ|egapxtmp_013496-R1, rna-gnl|WGS_ZZZZ|egapxtmp_013512-R1, rna-gnl|WGS_ZZZZ|egapxtmp_013517-R1    mNeoVis.evergladensis.2.1.chr7.g02053.t01, mNeoVis.evergladensis.2.1.chr7.g02054.t01, mNeoVis.evergladensis.2.1.chr7.g02055.t01, mNeoVis.evergladensis.2.1.chr7.g02056.t01    mNeoVis.vulgivaga.2.1.chr7.g02078.t01, mNeoVis.vulgivaga.2.1.chr7.g02079.t01, mNeoVis.vulgivaga.2.1.chr7.g02080.t01      980    N0.HOG0000978   OG0000082  n1   rna-XM_002918601.1, rna-XM_002926848.2, rna-XM_034643659.1    rna-NM_001389035.1    rna-XM_022507424.1, rna-XM_022522945.1, rna-XM_022522965.1, rna-XM_022526338.1    rna-XM_003988850.2, rna-XM_003988906.1, rna-XM_003993326.5, rna-XM_003993327.5, rna-XM_004000921.2    rna-XM_036086572.1, rna-XM_036086573.1, rna-XM_036092784.1    rna-XM_039216808.1, rna-XM_039216809.1, rna-XM_039224014.1, rna-XM_039224016.1    rna-XM_032846501.1, rna-XM_032874327.1, rna-XM_032874328.1    rna-XM_045994981.1, rna-XM_046010611.1, rna-XM_046010670.1, rna-XM_046017113.1, rna-XM_046017132.1    rna-XM_032329722.1, rna-XM_032347877.1, rna-XM_032360331.1    rna-XM_059156843.1, rna-XM_059158904.1, rna-XM_059187471.1    rna-XM_059385170.1, rna-XM_059388059.1, rna-XM_059404538.1    rna-XM_004773289.1, rna-XM_004778732.2, rna-XM_004781672.3    mNeoVis.2.1.1.chr12_hap1.g00186.t01, mNeoVis.2.1.1.chr9_hap1.g00408.t01    rna-XM_021684832.1, rna-XM_021685084.1, rna-XM_021687279.1    rna-XM_004411996.1    rna-XM_007081584.1, rna-XM_007081585.1, rna-XM_007088773.2, rna-XM_007088774.2, rna-XM_042973971.1    rna-XM_008684341.2, rna-XM_008694090.1, rna-XM_008701675.2, rna-XM_008711959.2    rna-XM_027580573.1, rna-XM_027591828.2    mNeoVis.3.1.1.chr12.g00155.t01, mNeoVis.3.1.1.chr7.g01952.t01    rna-gnl|WGS_ZZZZ|egapxtmp_012623-R1, rna-gnl|WGS_ZZZZ|egapxtmp_022966-R1, rna-gnl|WGS_ZZZZ|egapxtmp_028094-R1    mNeoVis.evergladensis.2.1.chr10.g00405.t01, mNeoVis.evergladensis.2.1.chr12.g00159.t01, mNeoVis.evergladensis.2.1.chr7.g02063.t01    mNeoVis.vulgivaga.2.1.chr10.g00401.t01, mNeoVis.vulgivaga.2.1.chr12.g00162.t01, mNeoVis.vulgivaga.2.1.chr12.g00163.t01, mNeoVis.vulgivaga.2.1.chr7.g02087.t01      981    N0.HOG0000979   OG0000082  n67   rna-XM_002926866.1    rna-NM_001389025.1    rna-XM_022507419.1    rna-XM_003993330.6      rna-XM_039224019.1    rna-XM_032846499.1    rna-XM_046017120.1, rna-XM_046017816.1    rna-XM_032359861.1    rna-XM_059158906.1    rna-XM_059388045.1    rna-XM_004781690.2        rna-XM_004411995.1    rna-XM_007088776.3    rna-XM_008694092.1    rna-XM_035723012.1    mNeoVis.3.1.1.chr7.g01954.t01        mNeoVis.vulgivaga.2.1.chr7.g02090.t01      982    N0.HOG0000980   OG0000082  n86   rna-XM_011234338.2    rna-NM_001389033.1    rna-XM_022507176.1    rna-XM_003993328.6    rna-XM_036092783.1    rna-XM_039224015.1    rna-XM_032846557.1    rna-XM_046017119.1    rna-XM_032360327.1    rna-XM_059158905.1    rna-XM_059388052.1    rna-XM_013050582.2          rna-XM_007088775.3    rna-XM_008694091.2    rna-XM_027577867.2      rna-gnl|WGS_ZZZZ|egapxtmp_012624-R1    mNeoVis.evergladensis.2.1.chr7.g02064.t01    mNeoVis.vulgivaga.2.1.chr7.g02088.t01      983    N0.HOG0000981   OG0000082  n103   rna-XM_002926847.2    rna-NM_001256446.1    rna-XM_022507421.1    rna-XM_023239019.2      rna-XM_039224007.1    rna-XM_032846500.1    rna-XM_046017134.1, rna-XM_046017250.1    rna-XM_032360328.1    rna-XM_059158903.1    rna-XM_059388063.1    rna-XM_013050585.2    mNeoVis.2.1.1.chr7_hap1.g02117.t01    rna-XM_044919567.1      rna-XM_015540523.1    rna-XM_040638716.1    rna-XM_027580030.1    mNeoVis.3.1.1.chr7.g01951.t01    rna-gnl|WGS_ZZZZ|egapxtmp_013567-R1      mNeoVis.vulgivaga.2.1.chr7.g02086.t01      984    N0.HOG0000982   OG0000082  n124   rna-XM_002926864.2    rna-NM_001389126.1    rna-XM_022507384.1          rna-XM_032846558.1    rna-XM_046016902.1, rna-XM_046017038.1    rna-XM_032359859.1    rna-XM_059137298.1      rna-XM_004781693.2    mNeoVis.2.1.1.chr7_hap1.g02116.t01          rna-XM_008694225.2    rna-XM_035722935.1    mNeoVis.3.1.1.chr7.g01950.t01    rna-gnl|WGS_ZZZZ|egapxtmp_010311-R1      mNeoVis.vulgivaga.2.1.chr7.g02085.t01      985    N0.HOG0000983   OG0000082  n138     rna-XM_038563259.1                rna-XM_032359858.1                                986    N0.HOG0000984   OG0000082  n139   rna-XM_011234336.3    rna-NM_001389127.1    rna-XM_022507423.1    rna-XM_003993324.6    rna-XM_036104524.1    rna-XM_039224017.1    rna-XM_032846498.1    rna-XM_046017224.1, rna-XM_046017271.1      rna-XM_059158902.1      rna-XM_004781673.2    mNeoVis.2.1.1.chr7_hap1.g02115.t01    rna-XM_021684831.1    rna-XM_004411997.1    rna-XM_007088772.2    rna-XM_008694088.2    rna-XM_027580580.1      rna-gnl|WGS_ZZZZ|egapxtmp_013495-R1    mNeoVis.evergladensis.2.1.chr7.g02062.t01        987    N0.HOG0000985   OG0000083  n1   rna-XM_002914900.4    rna-NM_001252331.1    rna-XM_022504660.1    rna-XM_003984098.4, rna-XM_045056455.1    rna-XM_036067029.1, rna-XM_036088068.1, rna-XM_036094673.1, rna-XM_036097025.1, rna-XM_036099454.1, rna-XM_036118561.1    rna-XM_039216428.1, rna-XM_039239557.1    rna-XM_032845397.1, rna-XM_032848422.1, rna-XM_032849823.1, rna-XM_032849851.1, rna-XM_032873693.1    rna-XM_045979252.1, rna-XM_045979253.1, rna-XM_045986240.1, rna-XM_046001250.1, rna-XM_046007087.1    rna-XM_032309426.1, rna-XM_032326606.1, rna-XM_032332211.1, rna-XM_032335967.1, rna-XM_032336224.1, rna-XM_032343477.1, rna-XM_032344424.1, rna-XM_032352384.1    rna-XM_059165278.1, rna-XM_059168933.1, rna-XM_059175256.1, rna-XM_059182264.1, rna-XM_059187986.1    rna-XM_059388189.1, rna-XM_059392836.1, rna-XM_059405902.1, rna-XM_059416625.1, rna-XM_059416644.1    rna-XM_045063272.1, rna-XM_045066540.1, rna-XM_045074431.1, rna-XM_045075322.1, rna-XM_045085201.1    mNeoVis.2.1.1.chr1_hap1.g00407.t01, mNeoVis.2.1.1.chr1_hap1.g00579.t01, mNeoVis.2.1.1.chr1_hap1.g00666.t01, mNeoVis.2.1.1.chr3_hap1.g00988.t01, mNeoVis.2.1.1.chr5_hap1.g01091.t01    rna-XM_021694004.1, rna-XM_021699209.1, rna-XM_044911827.1, rna-XM_044914998.1, rna-XM_044921757.1    rna-XM_004398649.1, rna-XM_004404542.1, rna-XM_004417569.1    rna-XM_007083414.3    rna-XM_008699967.2    rna-XM_027622042.1, rna-XM_035722509.1, rna-XM_035725357.1, rna-XM_035727144.1, rna-XM_035729232.1    mNeoVis.3.1.1.chr1.g00683.t01    rna-gnl|WGS_ZZZZ|egapxtmp_020950-R1, rna-gnl|WGS_ZZZZ|egapxtmp_030248-R1, rna-gnl|WGS_ZZZZ|egapxtmp_036599-R1, rna-gnl|WGS_ZZZZ|egapxtmp_038598-R1    mNeoVis.evergladensis.2.1.chr1.g00584.t01, mNeoVis.evergladensis.2.1.chr1.g00672.t01, mNeoVis.evergladensis.2.1.chr2.g01081.t01    mNeoVis.vulgivaga.2.1.chr1.g00593.t01, mNeoVis.vulgivaga.2.1.chr1.g00685.t01, mNeoVis.vulgivaga.2.1.chr11.g00269.t01, mNeoVis.vulgivaga.2.1.chr2.g01088.t01, mNeoVis.vulgivaga.2.1.chr3.g01004.t01      988    N0.HOG0000986   OG0000083  n49   rna-XM_034661798.1    rna-XM_038553915.1      rna-XM_045033271.1    rna-XM_036066654.1, rna-XM_036075637.1, rna-XM_036079077.1, rna-XM_036082457.1, rna-XM_036100277.1, rna-XM_036100826.1, rna-XM_036106651.1    rna-XM_039222726.1, rna-XM_039232466.1, rna-XM_039249856.1    rna-XM_032859833.1, rna-XM_032873821.1    rna-XM_045978406.1, rna-XM_045979144.1, rna-XM_045981578.1, rna-XM_045986786.1, rna-XM_046021418.1    rna-XM_032315219.1    rna-XM_059144170.1, rna-XM_059161605.1    rna-XM_059376183.1, rna-XM_059377811.1    rna-XM_045064095.1, rna-XM_045070883.1, rna-XM_045078992.1    mNeoVis.2.1.1.chr11_hap1.g00271.t01, mNeoVis.2.1.1.chr11_hap1.g00730.t01, mNeoVis.2.1.1.chr11_hap1.g00777.t01, mNeoVis.2.1.1.chr13_hap1.g00757.t01, mNeoVis.2.1.1.chr1_hap1.g00345.t01, mNeoVis.2.1.1.chr8_hap1.g00084.t01    rna-XM_044911826.1, rna-XM_044912668.1, rna-XM_044914037.1, rna-XM_044914244.1, rna-XM_044914554.1, rna-XM_044914828.1, rna-XM_044915767.1, rna-XM_044916108.1, rna-XM_044919710.1    rna-XM_004392406.1    rna-XM_042984700.1, rna-XM_042998160.1    rna-XM_040636591.1    rna-XM_035721937.1, rna-XM_035722941.1, rna-XM_035723772.1, rna-XM_035724202.1, rna-XM_035724310.1, rna-XM_035724795.1, rna-XM_035725322.1, rna-XM_035725766.1, rna-XM_035725771.1, rna-XM_035726165.1, rna-XM_035726910.1, rna-XM_035726935.1, rna-XM_035727307.1, rna-XM_035728288.1, rna-XM_035728610.1    mNeoVis.3.1.1.chr11.g00692.t01, mNeoVis.3.1.1.chr11.g00693.t01, mNeoVis.3.1.1.chr11.g00741.t01, mNeoVis.3.1.1.chr5.g01498.t01    rna-gnl|WGS_ZZZZ|egapxtmp_025911-R1, rna-gnl|WGS_ZZZZ|egapxtmp_026436-R1, rna-gnl|WGS_ZZZZ|egapxtmp_027916-R1, rna-gnl|WGS_ZZZZ|egapxtmp_030234-R1, rna-gnl|WGS_ZZZZ|egapxtmp_030813-R1    mNeoVis.evergladensis.2.1.chr1.g00343.t01, mNeoVis.evergladensis.2.1.chr11.g00268.t01, mNeoVis.evergladensis.2.1.chr11.g00713.t01, mNeoVis.evergladensis.2.1.chr11.g00759.t01, mNeoVis.evergladensis.2.1.chr13.g00721.t01, mNeoVis.evergladensis.2.1.chr9.g00081.t01    mNeoVis.vulgivaga.2.1.chr1.g00349.t01, mNeoVis.vulgivaga.2.1.chr11.g00730.t01, mNeoVis.vulgivaga.2.1.chr11.g00779.t01, mNeoVis.vulgivaga.2.1.chr8.g00084.t01      989    N0.HOG0000987   OG0000084  n1   rna-XM_034667619.1    rna-NM_001389107.1    rna-XM_022522153.1    rna-XM_003992961.3    rna-XM_036095961.1    rna-XM_039224088.1    rna-XM_032852722.1    rna-XM_046016735.1    rna-XM_032359798.1    rna-XM_059158167.1    rna-XM_059389252.1    rna-XM_004782508.2    mNeoVis.2.1.1.chr7_hap1.g01764.t01    rna-XM_021685646.1    rna-XM_004394618.1    rna-XM_007083279.1    rna-XM_008710993.2    rna-XM_027580630.1    mNeoVis.3.1.1.chr7.g01639.t01    rna-gnl|WGS_ZZZZ|egapxtmp_010081-R1    mNeoVis.evergladensis.2.1.chr7.g01726.t01    mNeoVis.vulgivaga.2.1.chr7.g01741.t01      990    N0.HOG0000988   OG0000084  n22   rna-XM_002924989.2    rna-NM_001313821.1      rna-XM_003992960.1    rna-XM_036095924.1    rna-XM_039224089.1    rna-XM_032853685.1    rna-XM_046016701.1, rna-XM_046017280.1    rna-XM_032356406.1    rna-XM_059158156.1    rna-XM_059389263.1    rna-XM_004782455.2    mNeoVis.2.1.1.chr7_hap1.g01763.t01    rna-XM_021685720.1    rna-XM_004394617.1    rna-XM_007083278.1    rna-XM_008710994.2    rna-XM_027580709.2    mNeoVis.3.1.1.chr7.g01638.t01    rna-gnl|WGS_ZZZZ|egapxtmp_010077-R1    mNeoVis.evergladensis.2.1.chr7.g01725.t01    mNeoVis.vulgivaga.2.1.chr7.g01740.t01      991    N0.HOG0000989   OG0000084  n41   rna-XM_034665591.1    rna-NM_001313856.1    rna-XM_022490734.1    rna-XM_003999609.3    rna-XM_036069853.1    rna-XM_039218476.1    rna-XM_032845701.1    rna-XM_045983456.1    rna-XM_032317170.1    rna-XM_059146870.1    rna-XM_059414159.1    rna-XM_004775807.2    mNeoVis.2.1.1.chr14_hap1.g00563.t01    rna-XM_021681939.1    rna-XM_004415678.1    rna-XM_007084274.1    rna-XM_008700737.2    rna-XM_027613346.2    mNeoVis.3.1.1.chr14.g00554.t01    rna-gnl|WGS_ZZZZ|egapxtmp_033045-R1    mNeoVis.evergladensis.2.1.chr14.g00563.t01    mNeoVis.vulgivaga.2.1.chr14.g00561.t01      992    N0.HOG0000990   OG0000084  n60   rna-XM_002931221.2    rna-NM_001388738.1    rna-XM_022522116.1    rna-XM_011286699.4    rna-XM_036095962.1    rna-XM_039224084.1    rna-XM_032853686.1    rna-XM_046016670.1, rna-XM_046016695.1    rna-XM_032359664.1    rna-XM_059158177.1    rna-XM_059389241.1    rna-XM_004782593.2    mNeoVis.2.1.1.chr7_hap1.g01765.t01    rna-XM_021685721.1      rna-XM_007083165.1    rna-XM_008710961.2    rna-XM_027577896.2    mNeoVis.3.1.1.chr7.g01640.t01    rna-gnl|WGS_ZZZZ|egapxtmp_010075-R1    mNeoVis.evergladensis.2.1.chr7.g01727.t01    mNeoVis.vulgivaga.2.1.chr7.g01742.t01      993    N0.HOG0000991   OG0000084  n81   rna-XM_002928699.2, rna-XM_034663653.1    rna-XM_038577992.1    rna-XM_022499662.1    rna-XM_003994551.6    rna-XM_036103313.1    rna-XM_039252438.1    rna-XM_032882189.1    rna-XM_046026616.1    rna-XM_032312211.1        rna-XM_013050895.2    mNeoVis.2.1.1.chr2_hap1.g00007.t01        rna-XM_007079748.2    rna-XM_040623968.1, rna-XM_040624285.1      mNeoVis.3.1.1.chr5.g00007.t01      mNeoVis.evergladensis.2.1.chr5.g00006.t01    mNeoVis.vulgivaga.2.1.chr5.g00005.t01      994    N0.HOG0000992   OG0000084  n97   rna-XM_002921818.3    rna-NM_001388564.1    rna-XM_022510416.1    rna-XM_006929462.2      rna-XM_039243018.1    rna-XM_032865113.1    rna-XM_046021389.1    rna-XM_032305766.1    rna-XM_059172706.1    rna-XM_059395572.1    rna-XM_013045639.1    mNeoVis.2.1.1.chr4_hap1.g00090.t01      rna-XM_004409604.1    rna-XM_007090899.1    rna-XM_008697284.2    rna-XM_027574272.2      rna-gnl|WGS_ZZZZ|egapxtmp_003092-R1    mNeoVis.evergladensis.2.1.chr6.g00093.t01    mNeoVis.vulgivaga.2.1.chr6.g00090.t01      995    N0.HOG0000993   OG0000084  n115   rna-XM_002922468.4, rna-XM_002922473.4    rna-NM_001388864.1, rna-NM_001389146.1    rna-XM_022500263.1, rna-XM_022500397.1    rna-XM_045034705.1    rna-XM_036117628.1    rna-XM_039228573.1    rna-XM_032857980.1    rna-XM_046019703.1, rna-XM_046019704.1    rna-XM_032354824.1, rna-XM_032354826.1    rna-XM_059168966.1, rna-XM_059168967.1    rna-XM_059392734.1, rna-XM_059392735.1    rna-XM_004743743.1, rna-XM_045066926.1    mNeoVis.2.1.1.chr3_hap1.g01023.t01, mNeoVis.2.1.1.chr3_hap1.g01024.t01    rna-XM_021678982.1    rna-XM_004417445.1    rna-XM_007085053.1    rna-XM_008690276.2, rna-XM_008690277.1    rna-XM_027590940.2    mNeoVis.3.1.1.chr3.g01011.t01    rna-gnl|WGS_ZZZZ|egapxtmp_041236-R1    mNeoVis.evergladensis.2.1.chr3.g01021.t01, mNeoVis.evergladensis.2.1.chr3.g01022.t01    mNeoVis.vulgivaga.2.1.chr3.g01039.t01      996    N0.HOG0000994   OG0000085  n2   rna-XM_002921314.3, rna-XM_034670526.1, rna-XM_034671377.1    rna-XM_038555093.1        rna-XM_036071074.1, rna-XM_036079698.1, rna-XM_036097683.1, rna-XM_036113049.1, rna-XM_036120685.1    rna-XM_039235236.1, rna-XM_039236743.1    rna-XM_032836051.1              mNeoVis.2.1.1.chr2_hap1.g01175.t01    rna-XM_044918764.1        rna-XM_040626250.1, rna-XM_040638974.1    rna-XM_035726252.1, rna-XM_035728923.1, rna-XM_035728926.1    mNeoVis.3.1.1.chr5.g01091.t01    rna-gnl|WGS_ZZZZ|egapxtmp_008936-R3    mNeoVis.evergladensis.2.1.chr5.g01118.t01    mNeoVis.vulgivaga.2.1.chr4.g00558.t01, mNeoVis.vulgivaga.2.1.chr5.g01104.t01, mNeoVis.vulgivaga.2.1.chr9.g01102.t01      997    N0.HOG0000995   OG0000085  n25     rna-XM_038555507.1, rna-XM_038561889.1      rna-XM_045034320.1    rna-XM_036074496.1, rna-XM_036081052.1, rna-XM_036098831.1, rna-XM_036101032.1, rna-XM_036119994.1    rna-XM_039225686.1, rna-XM_039229947.1    rna-XM_032882582.1      rna-XM_032328309.1, rna-XM_032334545.1    rna-XM_059153874.1    rna-XM_059415923.1    rna-XM_045075080.1    mNeoVis.2.1.1.chr15_hap1.g00660.t01      rna-XM_004408932.1    rna-XM_015540921.2      rna-XM_035721939.1, rna-XM_035722756.1, rna-XM_035723665.1, rna-XM_035727837.1, rna-XM_035727928.1, rna-XM_035728976.1    mNeoVis.3.1.1.chr15.g00444.t01    rna-gnl|WGS_ZZZZ|egapxtmp_033972-R1    mNeoVis.evergladensis.2.1.chr15.g00482.t01, mNeoVis.evergladensis.2.1.chr8.g00470.t01    mNeoVis.vulgivaga.2.1.chr15.g00492.t01      998    N0.HOG0000996   OG0000085  n52             rna-XM_039254765.1        rna-XM_032315536.1                                999    N0.HOG0000997   OG0000085  n55     rna-XM_038571020.1                              rna-XM_042968006.1      rna-XM_035727261.1              1000    N0.HOG0000998   OG0000085  n59     rna-XM_038569038.1                          rna-XM_044914211.1    rna-XM_004402578.1        rna-XM_035726961.1              1001    N0.HOG0000999   OG0000085  n62     rna-XM_038551469.1                          rna-XM_044920150.1          rna-XM_035722403.1              1002    N0.HOG0001000   OG0000085  n67   rna-XM_019808522.2    rna-XM_038543982.1      rna-NM_001128842.1    rna-XM_036069620.1, rna-XM_036078203.1    rna-XM_039228844.1    rna-XM_032847976.1, rna-XM_032853131.1, rna-XM_032864257.1, rna-XM_032882542.1    rna-XM_046001469.1, rna-XM_046025521.1    rna-XM_032310428.1, rna-XM_032359564.1    rna-XM_059139714.1    rna-XM_059409628.1    rna-XM_013055904.2, rna-XM_045065822.1    mNeoVis.2.1.1.chr3_hap1.g00113.t01, mNeoVis.2.1.1.chr3_hap1.g00751.t01    rna-XM_021686113.1, rna-XM_021701118.1    rna-XM_004410468.1    rna-XM_007080538.2, rna-XM_015544710.2    rna-XM_040632522.1    rna-XM_027576815.1    mNeoVis.3.1.1.chr3.g00117.t01    rna-gnl|WGS_ZZZZ|egapxtmp_042358-R1    mNeoVis.evergladensis.2.1.chr3.g00117.t01    mNeoVis.vulgivaga.2.1.chr3.g00117.t01      1003    N0.HOG0001001   OG0000085  n69                           mNeoVis.2.1.1.chr8_hap1.g01401.t01                rna-gnl|WGS_ZZZZ|egapxtmp_039603-R1          1004    N0.HOG0001002   OG0000085  n71         rna-XM_045047908.1                    mNeoVis.2.1.1.chr5_hap1.g01652.t01                  mNeoVis.evergladensis.2.1.chr2.g01639.t01        1005    N0.HOG0001003   OG0000085  n74   rna-XM_034651476.1    rna-XM_038550501.1    rna-XM_022495727.1      rna-XM_036071318.1, rna-XM_036081299.1, rna-XM_036098355.1, rna-XM_036100236.1, rna-XM_036119311.1    rna-XM_039240794.1, rna-XM_039247055.1    rna-XM_032845273.1, rna-XM_032875405.1    rna-XM_045978265.1, rna-XM_046003300.1    rna-XM_032301678.1, rna-XM_032316959.1    rna-XM_059163255.1, rna-XM_059171288.1    rna-XM_059388037.1, rna-XM_059395711.1    rna-XM_045062745.1, rna-XM_045080990.1, rna-XM_045088708.1    mNeoVis.2.1.1.chr1_hap1.g01306.t01, mNeoVis.2.1.1.chr4_hap1.g00654.t01    rna-XM_021697834.1, rna-XM_044914146.1, rna-XM_044916430.1    rna-XM_004392692.1    rna-XM_042974717.1, rna-XM_042987218.1      rna-XM_035723647.1, rna-XM_035726030.1, rna-XM_035726175.1, rna-XM_035726431.1, rna-XM_035728527.1, rna-XM_035728892.1      rna-gnl|WGS_ZZZZ|egapxtmp_002219-R1, rna-gnl|WGS_ZZZZ|egapxtmp_025571-R1    mNeoVis.evergladensis.2.1.chr1.g01277.t01, mNeoVis.evergladensis.2.1.chr6.g00661.t01    mNeoVis.vulgivaga.2.1.chr1.g01276.t01, mNeoVis.vulgivaga.2.1.chr4.g00020.t01, mNeoVis.vulgivaga.2.1.chr6.g00658.t01      1006    N0.HOG0001004   OG0000085  n115     rna-XM_038559065.1                  rna-XM_059137667.1              rna-XM_042960730.1                  1007    N0.HOG0001005   OG0000085  n117                   rna-XM_032316756.1      rna-XM_059386483.1, rna-XM_059407925.1        rna-XM_044911909.1      rna-XM_015543274.2      rna-XM_035722368.1              1008    N0.HOG0001006   OG0000086  n3   rna-XM_002926914.4, rna-XM_011218338.2, rna-XM_034650586.1, rna-XM_034653320.1    rna-XM_038557525.1    rna-XM_022491661.1      rna-XM_036081043.1, rna-XM_036081044.1    rna-XM_039225180.1    rna-XM_032848500.1, rna-XM_032849958.1    rna-XM_045996819.1, rna-XM_045996834.1, rna-XM_045996862.1, rna-XM_045996932.1, rna-XM_045999693.1, rna-XM_045999697.1, rna-XM_045999763.1    rna-XM_032335012.1, rna-XM_032335014.1, rna-XM_032335024.1    rna-XM_059173424.1, rna-XM_059176072.1    rna-XM_059416436.1, rna-XM_059416437.1    rna-XM_004780634.1, rna-XM_004782327.1    mNeoVis.2.1.1.chr1_hap1.g00970.t01    rna-XM_021688915.2, rna-XM_021690940.1, rna-XM_021692403.1        rna-XM_008709878.1, rna-XM_008709912.2, rna-XM_008709914.2, rna-XM_008709915.2, rna-XM_008710071.1, rna-XM_008710092.2, rna-XM_008710093.2    rna-XM_035727707.1    mNeoVis.3.1.1.chr1.g00978.t01    rna-gnl|WGS_ZZZZ|egapxtmp_019751-R1    mNeoVis.evergladensis.2.1.chr1.g00965.t01    mNeoVis.vulgivaga.2.1.chr1.g00978.t01      1009    N0.HOG0001007   OG0000086  n41   rna-XM_002930731.4    rna-XM_038557532.1    rna-XM_022492026.1        rna-XM_039225497.1      rna-XM_045980311.1, rna-XM_045996825.1, rna-XM_045996996.1, rna-XM_045997010.1, rna-XM_045998725.1, rna-XM_045998981.1, rna-XM_046001129.1                    rna-XM_008710287.1    rna-XM_027606546.1              1010    N0.HOG0001008   OG0000086  n53   rna-XM_002930729.2, rna-XM_002931112.4, rna-XM_034653283.1    rna-NM_001388948.1, rna-NM_001389023.1      rna-XM_003980575.3      rna-XM_039225643.1, rna-XM_039225688.1, rna-XM_039225689.1    rna-XM_032844365.1    rna-XM_045980087.1, rna-XM_046000783.1    rna-XM_032334910.1    rna-XM_059158830.1, rna-XM_059173412.1    rna-XM_059387368.1      mNeoVis.2.1.1.chr1_hap1.g00989.t01        rna-XM_007092948.2, rna-XM_007092949.2    rna-XM_008710065.1, rna-XM_008710069.1, rna-XM_008710090.2, rna-XM_008710282.1    rna-XM_027605755.1    mNeoVis.3.1.1.chr1.g00993.t01    rna-gnl|WGS_ZZZZ|egapxtmp_027617-R1    mNeoVis.evergladensis.2.1.chr1.g00983.t01    mNeoVis.vulgivaga.2.1.chr1.g00997.t01      1011    N0.HOG0001009   OG0000086  n80     rna-NM_001388537.1      rna-XM_011281225.2      rna-XM_039224230.1    rna-XM_032847581.1    rna-XM_045996838.1, rna-XM_045996856.1, rna-XM_045996880.1, rna-XM_045996910.1, rna-XM_045997037.1, rna-XM_046000770.1, rna-XM_046000846.1, rna-XM_046001215.1                rna-XM_004406927.1    rna-XM_007092950.1, rna-XM_007092951.2    rna-XM_008709867.1    rna-XM_027606539.1, rna-XM_035727455.1              1012    N0.HOG0001010   OG0000086  n93         rna-XM_003980595.2      rna-XM_039225412.1, rna-XM_039225710.1                      rna-XM_007092947.2                  1013    N0.HOG0001011   OG0000086  n98   rna-XM_011217583.2    rna-NM_001388538.1                                rna-XM_008710283.1                1014    N0.HOG0001012   OG0000086  n100     rna-NM_001388534.1    rna-XM_022491681.1          rna-XM_032848496.1    rna-XM_045996836.1, rna-XM_045996854.1, rna-XM_045996867.1, rna-XM_045996935.1, rna-XM_045997033.1, rna-XM_045999725.1, rna-XM_045999773.1, rna-XM_045999794.1, rna-XM_046000825.1, rna-XM_046001103.1                    rna-XM_008684235.1, rna-XM_008710066.2, rna-XM_008710067.1, rna-XM_008710068.1, rna-XM_008710086.1, rna-XM_008710087.1, rna-XM_008710088.2, rna-XM_008710089.1, rna-XM_008712033.1                1015    N0.HOG0001013   OG0000086  n118   rna-XM_002929134.3                rna-XM_045996835.1, rna-XM_045996853.1, rna-XM_045996881.1, rna-XM_045996909.1, rna-XM_045996943.1, rna-XM_045997032.1, rna-XM_045999724.1, rna-XM_045999795.1, rna-XM_046001102.1    rna-XM_032334909.1, rna-XM_032334912.1    rna-XM_059158832.1, rna-XM_059173509.1, rna-XM_059173512.1    rna-XM_059416746.1, rna-XM_059416750.1    rna-XM_004781733.1    mNeoVis.2.1.1.chr1_hap1.g00984.t01          rna-XM_008709866.2      mNeoVis.3.1.1.chr1.g00989.t01    rna-gnl|WGS_ZZZZ|egapxtmp_021561-R1    mNeoVis.evergladensis.2.1.chr1.g00977.t01, mNeoVis.evergladensis.2.1.chr1.g00978.t01    mNeoVis.vulgivaga.2.1.chr1.g00989.t01, mNeoVis.vulgivaga.2.1.chr1.g00990.t01      1016    N0.HOG0001014   OG0000087  n2   rna-XM_034656137.1    rna-XM_038530373.1, rna-XM_038589553.1      rna-XM_045060225.1        rna-XM_032851075.1    rna-XM_046001158.1        rna-XM_059416355.1    rna-XM_045075392.1      rna-XM_044912441.1    rna-XM_004397873.1    rna-XM_042985418.1    rna-XM_008691498.2    rna-XM_035727682.1      rna-gnl|WGS_ZZZZ|egapxtmp_031560-R1    mNeoVis.evergladensis.2.1.chr1.g00507.t01        1017    N0.HOG0001015   OG0000087  n17     rna-XM_038530372.1      rna-XM_045060177.1    rna-XM_036123723.1    rna-XM_039252756.1    rna-XM_032851049.1    rna-XM_045999329.1    rna-XM_032336465.1    rna-XM_059174821.1      rna-XM_045075397.1      rna-XM_044912419.1      rna-XM_042956384.1      rna-XM_035727644.1    mNeoVis.3.1.1.chr1.g00504.t01    rna-gnl|WGS_ZZZZ|egapxtmp_031344-R1      mNeoVis.vulgivaga.2.1.chr1.g00515.t01      1018    N0.HOG0001016   OG0000087  n32     rna-XM_038530376.1    rna-XM_022493383.1    rna-XM_023255929.2    rna-XM_036123737.1    rna-XM_039252755.1, rna-XM_039252758.1    rna-XM_032847544.1, rna-XM_032851080.1    rna-XM_045999326.1      rna-XM_059174822.1    rna-XM_059416351.1, rna-XM_059416352.1    rna-XM_045075395.1, rna-XM_045075398.1      rna-XM_044917053.1    rna-XM_004397878.1    rna-XM_042956398.1, rna-XM_042985412.1    rna-XM_040621932.1, rna-XM_040622339.1      mNeoVis.3.1.1.chr1.g00503.t01, mNeoVis.3.1.1.chr1.g00506.t01    rna-gnl|WGS_ZZZZ|egapxtmp_031259-R1    mNeoVis.evergladensis.2.1.chr1.g00502.t01, mNeoVis.evergladensis.2.1.chr1.g00503.t01    mNeoVis.vulgivaga.2.1.chr1.g00514.t01      1019    N0.HOG0001017   OG0000087  n58   rna-XM_034653092.1, rna-XM_034656135.1    rna-XM_038589551.1, rna-XM_038589552.1    rna-XM_022492655.1    rna-XM_045060179.1, rna-XM_045060231.1    rna-XM_036123611.1    rna-XM_039252759.1, rna-XM_039252762.1    rna-XM_032851023.1, rna-XM_032851230.1, rna-XM_032851232.1    rna-XM_045999330.1, rna-XM_046001157.1    rna-XM_032336466.1    rna-XM_059174823.1      rna-XM_013052307.2, rna-XM_045075390.1      rna-XM_021702039.2, rna-XM_044916785.1    rna-XM_004397876.1    rna-XM_042956408.1    rna-XM_040622699.1    rna-XM_035727645.1, rna-XM_035727679.1      rna-gnl|WGS_ZZZZ|egapxtmp_031426-R1    mNeoVis.evergladensis.2.1.chr1.g00504.t01    mNeoVis.vulgivaga.2.1.chr1.g00516.t01, mNeoVis.vulgivaga.2.1.chr1.g00518.t01      1020    N0.HOG0001018   OG0000087  n88   rna-XM_019795580.2      rna-XM_022492681.1        rna-XM_039225286.1    rna-XM_032850932.1          rna-XM_059416356.1    rna-XM_045075387.1      rna-XM_044917054.1        rna-XM_040622337.1    rna-XM_035727646.1    mNeoVis.3.1.1.chr1.g00508.t01      mNeoVis.evergladensis.2.1.chr1.g00509.t01    mNeoVis.vulgivaga.2.1.chr1.g00520.t01      1021    N0.HOG0001019   OG0000087  n100     rna-XM_038530383.1, rna-XM_038589550.1    rna-XM_022492720.1    rna-XM_045060214.1      rna-XM_039225844.1    rna-XM_032850933.1    rna-XM_045999328.1        rna-XM_059387411.1    rna-XM_045075389.1      rna-XM_044912471.1    rna-XM_004397872.1      rna-XM_040621931.1    rna-XM_035727647.1    mNeoVis.3.1.1.chr1.g00507.t01      mNeoVis.evergladensis.2.1.chr1.g00508.t01    mNeoVis.vulgivaga.2.1.chr1.g00519.t01      1022    N0.HOG0001020   OG0000087  n115   rna-XM_034655628.1    rna-XM_038530371.1, rna-XM_038589554.1          rna-XM_039252761.1    rna-XM_032851078.1    rna-XM_046001159.1        rna-XM_059387388.1    rna-XM_045075393.1      rna-XM_044912429.1        rna-XM_008691711.2    rna-XM_035727648.1        mNeoVis.evergladensis.2.1.chr1.g00506.t01        1023    N0.HOG0001021   OG0000087  n126   rna-XM_034653093.1    rna-XM_038529574.1    rna-XM_022493382.1    rna-XM_045033454.1, rna-XM_045033497.1, rna-XM_045060168.1    rna-XM_036123189.1    rna-XM_039252757.1, rna-XM_039252760.1    rna-XM_032851001.1    rna-XM_045999327.1    rna-XM_032336467.1    rna-XM_059174824.1    rna-XM_059416354.1    rna-XM_013052308.2, rna-XM_045075394.1      rna-XM_044912365.1, rna-XM_044912440.1    rna-XM_004397874.1, rna-XM_004397877.1    rna-XM_042956392.1, rna-XM_042956418.1    rna-XM_040622338.1, rna-XM_040622700.1    rna-XM_035727678.1, rna-XM_035727680.1    mNeoVis.3.1.1.chr1.g00505.t01    rna-gnl|WGS_ZZZZ|egapxtmp_031494-R1    mNeoVis.evergladensis.2.1.chr1.g00505.t01    mNeoVis.vulgivaga.2.1.chr1.g00517.t01      1024    N0.HOG0001022   OG0000088  n1   rna-XM_034638501.1    rna-XM_038538261.1    rna-XM_022511606.1    rna-XM_023245046.2    rna-XM_036074807.1    rna-XM_039227582.1    rna-XM_032853930.1    rna-XM_045989410.1    rna-XM_032325430.1    rna-XM_059152231.1    rna-XM_059381303.1    rna-XM_004753962.3    mNeoVis.2.1.1.chr7_hap1.g00114.t01    rna-XM_044922064.1, rna-XM_044922066.1    rna-XM_004416563.1    rna-XM_042969646.1    rna-XM_008689726.2    rna-XM_027620159.2    mNeoVis.3.1.1.chr7.g00098.t01    rna-gnl|WGS_ZZZZ|egapxtmp_010880-R7    mNeoVis.evergladensis.2.1.chr7.g00114.t01    mNeoVis.vulgivaga.2.1.chr7.g00113.t01      1025    N0.HOG0001023   OG0000088  n25   rna-XM_034654380.1    rna-NM_001048108.1    rna-XM_022524243.1    rna-XM_011285199.4    rna-XM_036091701.1, rna-XM_036096912.1    rna-XM_039218225.1    rna-XM_032843937.1    rna-XM_046019554.1    rna-XM_032353926.1    rna-XM_059166598.1    rna-XM_059394484.1    rna-XM_045066473.1    mNeoVis.2.1.1.chr3_hap1.g00736.t01    rna-XM_044913255.1    rna-XM_004397277.1    rna-XM_042994206.1    rna-XM_040619104.1    rna-XM_027590999.2    mNeoVis.3.1.1.chr3.g00718.t01, mNeoVis.3.1.1.chr3.g00719.t01, mNeoVis.3.1.1.chr3.g00720.t01    rna-gnl|WGS_ZZZZ|egapxtmp_041767-R1    mNeoVis.evergladensis.2.1.chr3.g00737.t01    mNeoVis.vulgivaga.2.1.chr3.g00744.t01      1026    N0.HOG0001024   OG0000088  n50   rna-XM_034639817.1    rna-XM_038547250.1    rna-XM_022509973.1    rna-XM_003996899.6    rna-XM_036095435.1    rna-XM_039256819.1    rna-XM_032871688.1    rna-XM_045984092.1    rna-XM_032322802.1    rna-XM_059149544.1    rna-XM_059380773.1    rna-XM_004772664.3    mNeoVis.2.1.1.chr8_hap1.g00369.t01    rna-XM_021681791.1    rna-XM_004410558.1    rna-XM_015540025.2    rna-XM_040641220.1    rna-XM_027625454.2    mNeoVis.3.1.1.chr8.g00335.t01      mNeoVis.evergladensis.2.1.chr9.g00367.t01    mNeoVis.vulgivaga.2.1.chr8.g00374.t01      1027    N0.HOG0001025   OG0000088  n70   rna-XM_034648877.1    rna-XM_038559273.1    rna-XM_022514057.1    rna-XM_023250688.2    rna-XM_036099871.1, rna-XM_036121011.1    rna-XM_039220013.1, rna-XM_039253885.1    rna-XM_032866749.1    rna-XM_046020922.1    rna-XM_032304549.1    rna-XM_059172148.1    rna-XM_059396202.1    rna-XM_045087941.1    mNeoVis.2.1.1.chr4_hap1.g00079.t01    rna-XM_044920022.1    rna-XM_004417804.1    rna-XM_042974552.1    rna-XM_008697394.2    rna-XM_027574309.2    mNeoVis.3.1.1.chr6.g00080.t01    rna-gnl|WGS_ZZZZ|egapxtmp_001738-R1    mNeoVis.evergladensis.2.1.chr6.g00083.t01    mNeoVis.vulgivaga.2.1.chr6.g00082.t01      1028    N0.HOG0001026   OG0000088  n94   rna-XM_034647088.1, rna-XM_034667672.1    rna-XM_038527281.1    rna-XM_022492376.1, rna-XM_022492490.1, rna-XM_022492516.1, rna-XM_022492622.1, rna-XM_022499678.1    rna-XM_045042673.1    rna-XM_036079441.1, rna-XM_036082329.1    rna-XM_039221109.1    rna-XM_032849969.1, rna-XM_032849970.1, rna-XM_032850865.1, rna-XM_032857956.1, rna-XM_032883114.1    rna-XM_046019191.1, rna-XM_046021654.1, rna-XM_046027167.1    rna-XM_032304397.1, rna-XM_032307716.1, rna-XM_032313635.1, rna-XM_032355501.1    rna-XM_059142804.1, rna-XM_059167384.1, rna-XM_059170309.1, rna-XM_059170746.1, rna-XM_059172512.1    rna-XM_059386860.1, rna-XM_059392746.1, rna-XM_059395806.1, rna-XM_059397081.1, rna-XM_059412117.1    rna-XM_004739486.3, rna-XM_045066826.1, rna-XM_045067369.1, rna-XM_045083036.1    mNeoVis.2.1.1.chr2_hap1.g00630.t01, mNeoVis.2.1.1.chr3_hap1.g00989.t01, mNeoVis.2.1.1.chr4_hap1.g00599.t01    rna-XM_021694387.1    rna-XM_004417259.1, rna-XM_004417315.1    rna-XM_042964400.1    rna-XM_040620317.1, rna-XM_040636285.1    rna-XM_027591300.2, rna-XM_027616719.2    mNeoVis.3.1.1.chr3.g00978.t01, mNeoVis.3.1.1.chr5.g00626.t01, mNeoVis.3.1.1.chr6.g00615.t01    rna-gnl|WGS_ZZZZ|egapxtmp_001992-R1, rna-gnl|WGS_ZZZZ|egapxtmp_009187-R1, rna-gnl|WGS_ZZZZ|egapxtmp_041228-R1    mNeoVis.evergladensis.2.1.chr3.g00985.t01, mNeoVis.evergladensis.2.1.chr5.g00631.t01, mNeoVis.evergladensis.2.1.chr6.g00604.t01    mNeoVis.vulgivaga.2.1.chr3.g01005.t01, mNeoVis.vulgivaga.2.1.chr5.g00625.t01, mNeoVis.vulgivaga.2.1.chr6.g00603.t01      1029    N0.HOG0001027   OG0000089  n2   rna-XM_002930882.2    rna-XM_038564467.1, rna-XM_038564468.1    rna-XM_022507713.1    rna-XM_011286950.2    rna-XM_036110589.1    rna-XM_039245039.1    rna-XM_032879893.1    rna-XM_046015774.1    rna-XM_032359903.1    rna-XM_059140569.1    rna-XM_059395016.1    rna-XM_013044540.2    mNeoVis.2.1.1.chr7_hap1.g02226.t01    rna-XM_021684874.1    rna-XM_004409321.1    rna-XM_007092140.1    rna-XM_008694291.1    rna-XM_027577846.2    mNeoVis.3.1.1.chr7.g02046.t01      mNeoVis.evergladensis.2.1.chr7.g02174.t01    mNeoVis.vulgivaga.2.1.chr7.g02193.t01      1030    N0.HOG0001028   OG0000089  n23   rna-XM_002928228.2, rna-XM_034644822.1    rna-NM_001389022.1    rna-XM_022507710.1    rna-XM_003993467.3    rna-XM_036110588.1    rna-XM_039245037.1    rna-XM_032879892.1    rna-XM_046015776.1    rna-XM_032359902.1    rna-XM_059158983.1    rna-XM_059387406.1    rna-XM_013048483.1    mNeoVis.2.1.1.chr7_hap1.g02217.t01    rna-XM_021685226.2      rna-XM_007092678.2    rna-XM_008694287.1    rna-XM_027577847.2    mNeoVis.3.1.1.chr7.g02041.t01    rna-gnl|WGS_ZZZZ|egapxtmp_013490-R1    mNeoVis.evergladensis.2.1.chr7.g02166.t01    mNeoVis.vulgivaga.2.1.chr7.g02188.t01      1031    N0.HOG0001029   OG0000089  n42   rna-XM_002928227.4, rna-XM_034652224.1    rna-XM_038564469.1    rna-XM_022507586.1    rna-XM_003993468.4      rna-XM_039245038.1    rna-XM_032879028.1    rna-XM_046017197.1    rna-XM_032360529.1    rna-XM_059158984.1    rna-XM_059387610.1    rna-XM_004777634.2    mNeoVis.2.1.1.chr7_hap1.g02218.t01        rna-XM_007092679.2    rna-XM_008694173.2    rna-XM_027580206.1    mNeoVis.3.1.1.chr7.g02042.t01    rna-gnl|WGS_ZZZZ|egapxtmp_010307-R1    mNeoVis.evergladensis.2.1.chr7.g02167.t01        1032    N0.HOG0001030   OG0000089  n59   rna-XM_002928226.4, rna-XM_034644998.1, rna-XM_034651846.1, rna-XM_034651847.1, rna-XM_034651862.1, rna-XM_034651900.1, rna-XM_034651938.1    rna-NM_001388580.1, rna-NM_001388823.1, rna-NM_001388874.1, rna-XM_038562459.1    rna-XM_022506617.1, rna-XM_022507641.1, rna-XM_022507642.1    rna-XM_045039340.1, rna-XM_045039341.1, rna-XM_045039342.1    rna-XM_036110580.1    rna-XM_039245101.1, rna-XM_039245102.1, rna-XM_039245118.1    rna-XM_032879029.1, rna-XM_032879030.1    rna-XM_046015775.1, rna-XM_046017233.1, rna-XM_046017234.1, rna-XM_046017305.1, rna-XM_046017306.1, rna-XM_046017307.1, rna-XM_046017311.1, rna-XM_046017338.1    rna-XM_032356401.1, rna-XM_032359420.1, rna-XM_032360498.1, rna-XM_032360541.1, rna-XM_032360542.1    rna-XM_059158986.1, rna-XM_059158987.1, rna-XM_059158988.1, rna-XM_059158989.1, rna-XM_059158990.1, rna-XM_059158991.1    rna-XM_059387571.1, rna-XM_059387605.1, rna-XM_059387607.1, rna-XM_059387608.1, rna-XM_059387609.1    rna-XM_004777675.2, rna-XM_004777676.2, rna-XM_004777677.2, rna-XM_004777678.2, rna-XM_004777679.1, rna-XM_004777680.2    mNeoVis.2.1.1.chr7_hap1.g02219.t01, mNeoVis.2.1.1.chr7_hap1.g02220.t01, mNeoVis.2.1.1.chr7_hap1.g02221.t01, mNeoVis.2.1.1.chr7_hap1.g02222.t01, mNeoVis.2.1.1.chr7_hap1.g02223.t01, mNeoVis.2.1.1.chr7_hap1.g02224.t01, mNeoVis.2.1.1.chr7_hap1.g02225.t01    rna-XM_021684873.1    rna-XM_004409320.1    rna-XM_007092141.2, rna-XM_042958741.1, rna-XM_042958742.1    rna-XM_008694175.2, rna-XM_008694288.2, rna-XM_008694289.2, rna-XM_008694290.2, rna-XM_040638729.1, rna-XM_040638730.1, rna-XM_040638764.1    rna-XM_027580208.1    mNeoVis.3.1.1.chr7.g02043.t01, mNeoVis.3.1.1.chr7.g02044.t01, mNeoVis.3.1.1.chr7.g02045.t01    rna-gnl|WGS_ZZZZ|egapxtmp_012653-R1, rna-gnl|WGS_ZZZZ|egapxtmp_012654-R1, rna-gnl|WGS_ZZZZ|egapxtmp_013504-R1, rna-gnl|WGS_ZZZZ|egapxtmp_013568-R1    mNeoVis.evergladensis.2.1.chr7.g02168.t01, mNeoVis.evergladensis.2.1.chr7.g02169.t01, mNeoVis.evergladensis.2.1.chr7.g02170.t01, mNeoVis.evergladensis.2.1.chr7.g02171.t01, mNeoVis.evergladensis.2.1.chr7.g02172.t01, mNeoVis.evergladensis.2.1.chr7.g02173.t01    mNeoVis.vulgivaga.2.1.chr7.g02189.t01, mNeoVis.vulgivaga.2.1.chr7.g02190.t01, mNeoVis.vulgivaga.2.1.chr7.g02191.t01, mNeoVis.vulgivaga.2.1.chr7.g02192.t01      1033    N0.HOG0001031   OG0000090  n1   rna-XM_034644934.1    rna-XM_038563974.1    rna-XM_022506456.1    rna-XM_023240050.2    rna-XM_036101204.1    rna-XM_039244766.1    rna-XM_032879438.1    rna-XM_046015602.1    rna-XM_032357273.1    rna-XM_059146911.1    rna-XM_059401707.1    rna-XM_013044643.2    mNeoVis.2.1.1.chr7_hap1.g02398.t01    rna-XM_044919285.1    rna-XM_004393954.1    rna-XM_042957578.1    rna-XM_040639386.1    rna-XM_027580363.1    mNeoVis.3.1.1.chr7.g02209.t01    rna-gnl|WGS_ZZZZ|egapxtmp_012692-R1    mNeoVis.evergladensis.2.1.chr7.g02344.t01    mNeoVis.vulgivaga.2.1.chr7.g02364.t01      1034    N0.HOG0001032   OG0000090  n22   rna-XM_002922393.4    rna-XM_038542895.1    rna-XM_022508235.1    rna-XM_045049152.1    rna-XM_036118911.1    rna-XM_039235845.1    rna-XM_032838850.1    rna-XM_045982473.1    rna-XM_032317211.1    rna-XM_059145984.1    rna-XM_059412932.1    rna-XM_004761872.3    mNeoVis.2.1.1.chr14_hap1.g00248.t01    rna-XM_021679483.1    rna-XM_004393527.1    rna-XM_042976867.1    rna-XM_040636147.1    rna-XM_027611687.2        mNeoVis.evergladensis.2.1.chr14.g00248.t01    mNeoVis.vulgivaga.2.1.chr14.g00249.t01      1035    N0.HOG0001033   OG0000090  n40   rna-XM_034667628.1    rna-XM_038542896.1    rna-XM_022508247.1    rna-XM_045049254.1    rna-XM_036079120.1    rna-XM_039235795.1    rna-XM_032838811.1    rna-XM_045982292.1    rna-XM_032314615.1, rna-XM_032319136.1    rna-XM_059145392.1    rna-XM_059413940.1    rna-XM_045069643.1, rna-XM_045072313.1    mNeoVis.2.1.1.chr14_hap1.g00249.t01    rna-XM_044912313.1, rna-XM_044916204.1    rna-XM_004393526.1    rna-XM_042976845.1    rna-XM_008695408.2    rna-XM_027611998.2    mNeoVis.3.1.1.chr14.g00252.t01      mNeoVis.evergladensis.2.1.chr14.g00249.t01    mNeoVis.vulgivaga.2.1.chr14.g00250.t01      1036    N0.HOG0001034   OG0000090  n63   rna-XM_019794440.2, rna-XM_034647810.1    rna-XM_038554191.1    rna-XM_022503878.1    rna-XM_019830649.3    rna-XM_036082766.1    rna-XM_039231160.1    rna-XM_032862059.1    rna-XM_046006030.1    rna-XM_032340103.1      rna-XM_059399072.1    rna-XM_004739803.3    mNeoVis.2.1.1.chr1_hap1.g01609.t01    rna-XM_021692053.1    rna-XM_004408129.1    rna-XM_015538911.2    rna-XM_040643420.1    rna-XM_027602634.2    mNeoVis.3.1.1.chr1.g01557.t01    rna-gnl|WGS_ZZZZ|egapxtmp_020846-R1    mNeoVis.evergladensis.2.1.chr1.g01575.t01    mNeoVis.vulgivaga.2.1.chr1.g01572.t01      1037    N0.HOG0001035   OG0000090  n86   rna-XM_034639205.1    rna-XM_038528972.1    rna-XM_022525265.1    rna-XM_045046693.1    rna-XM_036118469.1    rna-XM_039217756.1    rna-XM_032841843.1    rna-XM_045986877.1    rna-XM_032326158.1    rna-XM_059151177.1    rna-XM_059383460.1    rna-XM_045077251.1    mNeoVis.2.1.1.chr7_hap1.g00515.t01    rna-XM_021701121.2    rna-XM_004396411.1    rna-XM_042969369.1    rna-XM_040633939.1    rna-XM_035724899.1    mNeoVis.3.1.1.chr7.g00500.t01      mNeoVis.evergladensis.2.1.chr7.g00516.t01    mNeoVis.vulgivaga.2.1.chr7.g00519.t01      1038    N0.HOG0001036   OG0000090  n107   rna-XM_034663574.1    rna-XM_038533935.1    rna-XM_022523670.1    rna-XM_003993929.5    rna-XM_036106970.1    rna-XM_039241257.1    rna-XM_032841127.1    rna-XM_046028015.1    rna-XM_032313469.1    rna-XM_059170496.1    rna-XM_059395795.1    rna-XM_004776002.3    mNeoVis.2.1.1.chr2_hap1.g00655.t01    rna-XM_021696151.1    rna-XM_004404856.1    rna-XM_007086009.3    rna-XM_008689538.2    rna-XM_027586950.2    mNeoVis.3.1.1.chr5.g00630.t01      mNeoVis.evergladensis.2.1.chr5.g00638.t01    mNeoVis.vulgivaga.2.1.chr5.g00632.t01      1039    N0.HOG0001037   OG0000090  n127   rna-XM_002913197.4    rna-XM_038580161.1    rna-XM_022524305.1    rna-XM_003987244.6    rna-XM_036084628.1    rna-XM_039232743.1    rna-XM_032856734.1    rna-XM_046010379.1    rna-XM_032343433.1    rna-XM_059181272.1    rna-XM_059372616.1    rna-XM_004751191.3    mNeoVis.2.1.1.chr10_hap1.g00701.t01    rna-XM_021695621.1    rna-XM_004400387.1    rna-XM_007081703.3    rna-XM_008702196.2    rna-XM_027572315.2    mNeoVis.3.1.1.chr9.g00661.t01      mNeoVis.evergladensis.2.1.chr8.g00682.t01    mNeoVis.vulgivaga.2.1.chr9.g00683.t01      1040    N0.HOG0001038   OG0000091  n1   rna-XM_034645319.1    rna-XM_038577737.1    rna-XM_022519357.1    rna-XM_003988735.4    rna-XM_036096394.1    rna-XM_039224951.1    rna-XM_032876713.1    rna-XM_046011962.1    rna-XM_032348289.1    rna-XM_059185269.1    rna-XM_059404852.1    rna-XM_004774687.2    mNeoVis.2.1.1.chr9_hap1.g00521.t01    rna-XM_021686664.2    rna-XM_004400790.1    rna-XM_007073099.2    rna-XM_008704607.2    rna-XM_027592376.2    mNeoVis.3.1.1.chr10.g00494.t01    rna-gnl|WGS_ZZZZ|egapxtmp_024678-R1    mNeoVis.evergladensis.2.1.chr10.g00513.t01    mNeoVis.vulgivaga.2.1.chr10.g00504.t01      1041    N0.HOG0001039   OG0000091  n23   rna-XM_002930972.4    rna-XM_038577735.1    rna-XM_022519361.1    rna-XM_003988733.5    rna-XM_036096400.1    rna-XM_039224945.1    rna-XM_032876726.1    rna-XM_046012519.1    rna-XM_032348283.1    rna-XM_059185271.1    rna-XM_059403184.1    rna-XM_045073198.1    mNeoVis.2.1.1.chr9_hap1.g00523.t01    rna-XM_021686735.1    rna-XM_004400795.1    rna-XM_007073054.2    rna-XM_008704605.2    rna-XM_035729240.1    mNeoVis.3.1.1.chr10.g00496.t01    rna-gnl|WGS_ZZZZ|egapxtmp_022626-R1    mNeoVis.evergladensis.2.1.chr10.g00515.t01    mNeoVis.vulgivaga.2.1.chr10.g00507.t01      1042    N0.HOG0001040   OG0000091  n42   rna-XM_034645322.1    rna-NM_001346052.1    rna-XM_022519359.1    rna-XM_003988734.5    rna-XM_036096396.1    rna-XM_039224943.1    rna-XM_032876722.1    rna-XM_046011963.1      rna-XM_059185274.1    rna-XM_059403182.1    rna-XM_004774688.3    mNeoVis.2.1.1.chr9_hap1.g00522.t01    rna-XM_044914835.1    rna-XM_004400791.1    rna-XM_042990824.1    rna-XM_040642504.1    rna-XM_027592390.1    mNeoVis.3.1.1.chr10.g00495.t01      mNeoVis.evergladensis.2.1.chr10.g00514.t01    mNeoVis.vulgivaga.2.1.chr10.g00506.t01      1043    N0.HOG0001041   OG0000091  n62             rna-XM_039241343.1    rna-XM_032876508.1          rna-XM_059404518.1    rna-XM_045073226.1    mNeoVis.2.1.1.chr9_hap1.g00528.t01              mNeoVis.3.1.1.chr10.g00499.t01    rna-gnl|WGS_ZZZZ|egapxtmp_022635-R1    mNeoVis.evergladensis.2.1.chr10.g00520.t01        1044    N0.HOG0001042   OG0000091  n70           rna-XM_036096405.1                      rna-XM_004400893.1        rna-XM_035729249.1              1045    N0.HOG0001043   OG0000091  n73   rna-XM_034651744.1                                  rna-XM_040642501.1                1046    N0.HOG0001044   OG0000091  n76   rna-XM_034645785.1    rna-XM_038576168.1    rna-XM_022519516.1    rna-XM_011283950.4, rna-XM_023257096.2    rna-XM_036096406.1    rna-XM_039224952.1    rna-XM_032876731.1    rna-XM_046012545.1    rna-XM_032348288.1    rna-XM_059183579.1    rna-XM_059404520.1    rna-XM_004774691.2    mNeoVis.2.1.1.chr9_hap1.g00525.t01      rna-XM_004400793.1    rna-XM_042990822.1, rna-XM_042992353.1    rna-XM_040642503.1    rna-XM_027592387.1      rna-gnl|WGS_ZZZZ|egapxtmp_022628-R1    mNeoVis.evergladensis.2.1.chr10.g00517.t01        1047    N0.HOG0001045   OG0000091  n94       rna-XM_022519386.1          rna-XM_032876728.1                                    1048    N0.HOG0001046   OG0000091  n96                     rna-XM_059185279.1    rna-XM_059403177.1    rna-XM_004774694.3                          1049    N0.HOG0001047   OG0000091  n101                           mNeoVis.2.1.1.chr9_hap1.g00529.t01              mNeoVis.3.1.1.chr10.g00500.t01            1050    N0.HOG0001048   OG0000091  n104   rna-XM_034645784.1    rna-XM_038577736.1    rna-XM_022519512.1    rna-XM_003988732.5    rna-XM_036096409.1    rna-XM_039224953.1    rna-XM_032876729.1    rna-XM_046012580.1    rna-XM_032348282.1    rna-XM_059185276.1    rna-XM_059403176.1    rna-XM_004774690.3    mNeoVis.2.1.1.chr9_hap1.g00524.t01      rna-XM_004400794.1    rna-XM_042990823.1    rna-XM_040642502.1    rna-XM_027592393.1      rna-gnl|WGS_ZZZZ|egapxtmp_022631-R1    mNeoVis.evergladensis.2.1.chr10.g00516.t01        1051    N0.HOG0001049   OG0000091  n118   rna-XM_034645786.1    rna-XM_038577731.1    rna-XM_022519508.1    rna-XM_003988731.5    rna-XM_036096397.1    rna-XM_039241380.1    rna-XM_032876727.1    rna-XM_046012513.1    rna-XM_032348281.1    rna-XM_059183580.1    rna-XM_059404850.1    rna-XM_045072991.1    mNeoVis.2.1.1.chr9_hap1.g00526.t01      rna-XM_004400792.1    rna-XM_042990821.1    rna-XM_008704764.2    rna-XM_035729234.1        mNeoVis.evergladensis.2.1.chr10.g00518.t01        1052    N0.HOG0001050   OG0000092  n1   rna-XM_002930756.2    rna-XM_038544015.1    rna-XM_022492246.1    rna-XM_003980607.2    rna-XM_036102007.1    rna-XM_039218805.1    rna-XM_032847586.1    rna-XM_046001115.1    rna-XM_032335041.1    rna-XM_059173398.1    rna-XM_059416419.1    rna-XM_004781506.2    mNeoVis.2.1.1.chr1_hap1.g01015.t01    rna-XM_021689904.1      rna-XM_042958519.1    rna-XM_008712038.1    rna-XM_027624406.1    mNeoVis.3.1.1.chr1.g01016.t01    rna-gnl|WGS_ZZZZ|egapxtmp_003779-R1    mNeoVis.evergladensis.2.1.chr1.g01008.t01    mNeoVis.vulgivaga.2.1.chr1.g01020.t01      1053    N0.HOG0001051   OG0000092  n20   rna-XM_002930755.3    rna-NM_001388806.1    rna-XM_022492240.1    rna-XM_003980610.5    rna-XM_036102017.1, rna-XM_036121604.1    rna-XM_039218799.1    rna-XM_032847577.1    rna-XM_046000436.1    rna-XM_032335162.1    rna-XM_059173508.1    rna-XM_059416736.1    rna-XM_004778437.1, rna-XM_013050494.2    mNeoVis.2.1.1.chr1_hap1.g01005.t01    rna-XM_021689905.1      rna-XM_007084988.1    rna-XM_008712036.2    rna-XM_035727693.1    mNeoVis.3.1.1.chr1.g01005.t01    rna-gnl|WGS_ZZZZ|egapxtmp_020948-R1    mNeoVis.evergladensis.2.1.chr1.g00997.t01    mNeoVis.vulgivaga.2.1.chr1.g01011.t01      1054    N0.HOG0001052   OG0000092  n42   rna-XM_019809347.2    rna-NM_001389036.1    rna-XM_022490608.1, rna-XM_022490609.1, rna-XM_022490610.1, rna-XM_022490611.1, rna-XM_022490612.1, rna-XM_022490667.1    rna-XM_003985644.4, rna-XM_003985649.3, rna-XM_003985651.5, rna-XM_003985652.4, rna-XM_023254639.2, rna-XM_045057031.1    rna-XM_036120487.1, rna-XM_036120537.1, rna-XM_036120546.1, rna-XM_036120556.1    rna-XM_039224771.1, rna-XM_039224937.1, rna-XM_039224938.1, rna-XM_039257547.1, rna-XM_039257592.1, rna-XM_039257593.1    rna-XM_032847837.1, rna-XM_032847838.1, rna-XM_032847839.1, rna-XM_032847840.1, rna-XM_032847841.1, rna-XM_032850826.1    rna-XM_046006122.1, rna-XM_046006128.1, rna-XM_046006514.1, rna-XM_046006548.1, rna-XM_046006730.1, rna-XM_046006913.1    rna-XM_032338386.1, rna-XM_032341398.1, rna-XM_032341399.1, rna-XM_032341400.1, rna-XM_032341401.1, rna-XM_032341402.1    rna-XM_059176415.1, rna-XM_059176416.1, rna-XM_059176417.1, rna-XM_059176418.1, rna-XM_059176419.1, rna-XM_059179380.1    rna-XM_059398965.1, rna-XM_059398966.1, rna-XM_059398967.1, rna-XM_059398968.1, rna-XM_059401512.1    rna-XM_004779989.1, rna-XM_004779990.2, rna-XM_004779991.2, rna-XM_004779992.1, rna-XM_004779993.2, rna-XM_004779994.2    mNeoVis.2.1.1.chr1_hap1.g01035.t01, mNeoVis.2.1.1.chr1_hap1.g01036.t01, mNeoVis.2.1.1.chr1_hap1.g01037.t01, mNeoVis.2.1.1.chr1_hap1.g01038.t01, mNeoVis.2.1.1.chr1_hap1.g01039.t01, mNeoVis.2.1.1.chr1_hap1.g01040.t01    rna-XM_021697253.2, rna-XM_044917359.1, rna-XM_044917360.1    rna-XM_004411303.1, rna-XM_004411304.1, rna-XM_004411305.1    rna-XM_007085933.2, rna-XM_007085934.2, rna-XM_007085937.1, rna-XM_007085938.1, rna-XM_007085940.2, rna-XM_042985250.1    rna-XM_008711567.1, rna-XM_008711568.2, rna-XM_008711569.1, rna-XM_008711570.1, rna-XM_008711571.2, rna-XM_040644889.1    rna-XM_027603237.2, rna-XM_027603359.2, rna-XM_027604126.2    mNeoVis.3.1.1.chr1.g01032.t01, mNeoVis.3.1.1.chr1.g01033.t01, mNeoVis.3.1.1.chr1.g01034.t01    rna-gnl|WGS_ZZZZ|egapxtmp_021097-R1, rna-gnl|WGS_ZZZZ|egapxtmp_031428-R1, rna-gnl|WGS_ZZZZ|egapxtmp_031436-R1, rna-gnl|WGS_ZZZZ|egapxtmp_031455-R1, rna-gnl|WGS_ZZZZ|egapxtmp_031465-R1, rna-gnl|WGS_ZZZZ|egapxtmp_031474-R1    mNeoVis.evergladensis.2.1.chr1.g01027.t01, mNeoVis.evergladensis.2.1.chr1.g01028.t01, mNeoVis.evergladensis.2.1.chr1.g01029.t01, mNeoVis.evergladensis.2.1.chr1.g01030.t01, mNeoVis.evergladensis.2.1.chr1.g01031.t01, mNeoVis.evergladensis.2.1.chr1.g01032.t01    mNeoVis.vulgivaga.2.1.chr1.g01039.t01, mNeoVis.vulgivaga.2.1.chr1.g01040.t01, mNeoVis.vulgivaga.2.1.chr1.g01041.t01, mNeoVis.vulgivaga.2.1.chr1.g01042.t01      1055    N0.HOG0001053   OG0000093  n2   rna-XM_002928213.2    rna-NM_001388798.1    rna-XM_022513354.1    rna-XM_011286415.4    rna-XM_036081753.1, rna-XM_036081756.1    rna-XM_039250269.1    rna-XM_032869026.1    rna-XM_046017464.1    rna-XM_032358778.1    rna-XM_059156111.1    rna-XM_059392524.1    rna-XM_004750247.3    mNeoVis.2.1.1.chr7_hap1.g01311.t01    rna-XM_021696633.2    rna-XM_004417548.1    rna-XM_015538919.2    rna-XM_008689929.2    rna-XM_027577941.1        mNeoVis.evergladensis.2.1.chr7.g01298.t01    mNeoVis.vulgivaga.2.1.chr7.g01311.t01      1056    N0.HOG0001054   OG0000093  n21   rna-XM_002928209.4, rna-XM_002928210.4, rna-XM_002928211.1, rna-XM_034651616.1, rna-XM_034667710.1    rna-NM_001388796.1, rna-NM_001388797.1, rna-XM_038535497.1    rna-XM_022513241.1, rna-XM_022513301.1, rna-XM_022513351.1    rna-XM_003992504.1, rna-XM_011286416.2    rna-XM_036081754.1, rna-XM_036081755.1, rna-XM_036081757.1    rna-XM_039250149.1, rna-XM_039250265.1    rna-XM_032869028.1, rna-XM_032869029.1, rna-XM_032869030.1    rna-XM_046015968.1, rna-XM_046017304.1, rna-XM_046017422.1    rna-XM_032358103.1, rna-XM_032359588.1, rna-XM_032359589.1, rna-XM_032360259.1    rna-XM_059156117.1, rna-XM_059156131.1, rna-XM_059156144.1, rna-XM_059171494.1    rna-XM_059392500.1, rna-XM_059392512.1    rna-XM_004750121.2, rna-XM_004750122.2, rna-XM_004750248.1    mNeoVis.2.1.1.chr7_hap1.g01310.t01, mNeoVis.2.1.1.chr7_hap1.g01312.t01    rna-XM_021696507.1, rna-XM_021696515.1    rna-XM_004417545.1, rna-XM_004417547.1    rna-XM_007074034.2, rna-XM_015538922.2    rna-XM_008689926.2, rna-XM_008689928.2, rna-XM_008690164.2, rna-XM_008690165.2    rna-XM_035722979.1, rna-XM_035722982.1    mNeoVis.3.1.1.chr7.g01233.t01, mNeoVis.3.1.1.chr7.g01234.t01    rna-gnl|WGS_ZZZZ|egapxtmp_010607-R1, rna-gnl|WGS_ZZZZ|egapxtmp_013368-R1, rna-gnl|WGS_ZZZZ|egapxtmp_013372-R1      mNeoVis.vulgivaga.2.1.chr7.g01310.t01, mNeoVis.vulgivaga.2.1.chr7.g01312.t01      1057    N0.HOG0001055   OG0000093  n78   rna-XM_002928214.4    rna-XM_038535568.1    rna-XM_022511926.1, rna-XM_022511927.1, rna-XM_022512361.1, rna-XM_022513358.1    rna-XM_003987439.6, rna-XM_011286455.2, rna-XM_023255575.2, rna-XM_045059662.1    rna-XM_036081751.1, rna-XM_036114627.1, rna-XM_036114628.1    rna-XM_039245245.1, rna-XM_039245247.1, rna-XM_039245277.1, rna-XM_039250234.1    rna-XM_032857102.1, rna-XM_032857103.1, rna-XM_032857104.1, rna-XM_032869025.1    rna-XM_046007820.1, rna-XM_046007821.1, rna-XM_046009867.1, rna-XM_046010062.1, rna-XM_046017596.1    rna-XM_032341745.1, rna-XM_032341746.1, rna-XM_032343033.1, rna-XM_032358774.1    rna-XM_059156098.1, rna-XM_059179599.1, rna-XM_059183372.1, rna-XM_059183373.1    rna-XM_059372080.1, rna-XM_059372081.1, rna-XM_059372082.1, rna-XM_059392536.1    rna-XM_004750120.3, rna-XM_004758793.3, rna-XM_045067422.1, rna-XM_045067526.1    mNeoVis.2.1.1.chr10_hap1.g00903.t01, mNeoVis.2.1.1.chr10_hap1.g00904.t01, mNeoVis.2.1.1.chr7_hap1.g01308.t01    rna-XM_021695583.1, rna-XM_021695698.1, rna-XM_021696632.1    rna-XM_004402115.1    rna-XM_007091330.2, rna-XM_007091331.2, rna-XM_007091396.2, rna-XM_015538819.2    rna-XM_008689930.1, rna-XM_008710451.2, rna-XM_008711137.1, rna-XM_008711140.2    rna-XM_027572864.2, rna-XM_027577942.2    mNeoVis.3.1.1.chr7.g01232.t01, mNeoVis.3.1.1.chr9.g00831.t01, mNeoVis.3.1.1.chr9.g00832.t01    rna-gnl|WGS_ZZZZ|egapxtmp_013347-R2, rna-gnl|WGS_ZZZZ|egapxtmp_015531-R1, rna-gnl|WGS_ZZZZ|egapxtmp_015533-R1    mNeoVis.evergladensis.2.1.chr7.g01296.t01, mNeoVis.evergladensis.2.1.chr8.g00873.t01    mNeoVis.vulgivaga.2.1.chr7.g01309.t01, mNeoVis.vulgivaga.2.1.chr9.g00860.t01, mNeoVis.vulgivaga.2.1.chr9.g00861.t01      1058    N0.HOG0001056   OG0000094  n1   rna-XM_002928449.4    rna-XM_038542552.1    rna-XM_022508473.1    rna-XM_006942693.4    rna-XM_036080536.1    rna-XM_039225370.1    rna-XM_032862994.1    rna-XM_045982807.1    rna-XM_032317171.1    rna-XM_059145903.1    rna-XM_059412439.1    rna-XM_004756698.3    mNeoVis.2.1.1.chr14_hap1.g00037.t01    rna-XM_021682637.1    rna-XM_004416476.1    rna-XM_042976678.1    rna-XM_008696131.2    rna-XM_027612271.1    mNeoVis.3.1.1.chr14.g00034.t01    rna-gnl|WGS_ZZZZ|egapxtmp_032790-R1    mNeoVis.evergladensis.2.1.chr14.g00037.t01    mNeoVis.vulgivaga.2.1.chr14.g00039.t01      1059    N0.HOG0001057   OG0000094  n24   rna-XM_019809211.2    rna-XM_038542559.1    rna-XM_022508173.1    rna-XM_019821608.3    rna-XM_036080544.1    rna-XM_039221688.1    rna-XM_032863002.1    rna-XM_045983188.1    rna-XM_032317474.1    rna-XM_059146772.1    rna-XM_059412431.1    rna-XM_004756691.3    mNeoVis.2.1.1.chr14_hap1.g00036.t01    rna-XM_021682639.1    rna-XM_004408900.1    rna-XM_015540488.2    rna-XM_008696087.2    rna-XM_027612275.2    mNeoVis.3.1.1.chr14.g00032.t01    rna-gnl|WGS_ZZZZ|egapxtmp_032791-R1    mNeoVis.evergladensis.2.1.chr14.g00035.t01    mNeoVis.vulgivaga.2.1.chr14.g00037.t01      1060    N0.HOG0001058   OG0000094  n45   rna-XM_034665596.1    rna-XM_038586442.1    rna-XM_022508980.1      rna-XM_036097600.1    rna-XM_039222226.1    rna-XM_032872978.1    rna-XM_045983410.1    rna-XM_032318345.1    rna-XM_059145398.1    rna-XM_059413954.1    rna-XM_013055549.2    mNeoVis.2.1.1.chr14_hap1.g00452.t01        rna-XM_015535685.1        mNeoVis.3.1.1.chr14.g00448.t01    rna-gnl|WGS_ZZZZ|egapxtmp_031931-R1    mNeoVis.evergladensis.2.1.chr14.g00449.t01    mNeoVis.vulgivaga.2.1.chr14.g00451.t01      1061    N0.HOG0001059   OG0000094  n60   rna-XM_034654196.1    rna-XM_038561965.1    rna-XM_022526125.1    rna-XM_023259065.2    rna-XM_036089200.1    rna-XM_039224071.1    rna-XM_032848122.1    rna-XM_046007962.1    rna-XM_032303330.1    rna-XM_059136597.1    rna-XM_059375939.1    rna-XM_004779605.3    mNeoVis.2.1.1.chr2_hap1.g00867.t01    rna-XM_021688097.1    rna-XM_004417450.1, rna-XM_004417624.1    rna-XM_007092578.2    rna-XM_040626844.1    rna-XM_035727130.1    mNeoVis.3.1.1.chr5.g00802.t01    rna-gnl|WGS_ZZZZ|egapxtmp_008233-R3    mNeoVis.evergladensis.2.1.chr5.g00820.t01    mNeoVis.vulgivaga.2.1.chr5.g00811.t01      1062    N0.HOG0001060   OG0000094  n82   rna-XM_019809209.1    rna-NM_001003061.1    rna-XM_022508416.1    rna-XM_003999187.5    rna-XM_036080518.1    rna-XM_039225374.1    rna-XM_032862999.1    rna-XM_045983190.1    rna-XM_032317175.1    rna-XM_059145300.1    rna-XM_059412438.1    rna-XM_004756696.3      rna-XM_021682874.1    rna-XM_004416480.1    rna-XM_042976683.1    rna-XM_008696105.2    rna-XM_027612289.2    mNeoVis.3.1.1.chr14.g00033.t01    rna-gnl|WGS_ZZZZ|egapxtmp_032784-R2    mNeoVis.evergladensis.2.1.chr14.g00036.t01    mNeoVis.vulgivaga.2.1.chr14.g00038.t01      1063    N0.HOG0001061   OG0000094  n103   rna-XM_019809196.2    rna-XM_038542561.1    rna-XM_022508428.1    rna-XM_003999184.5    rna-XM_036080519.1    rna-XM_039221686.1    rna-XM_032863004.1    rna-XM_045983193.1    rna-XM_032317470.1    rna-XM_059146778.1    rna-XM_059412433.1    rna-XM_004756694.3    mNeoVis.2.1.1.chr14_hap1.g00035.t01    rna-XM_021682640.1      rna-XM_007088721.3    rna-XM_008696054.2    rna-XM_027611908.1    mNeoVis.3.1.1.chr14.g00031.t01    rna-gnl|WGS_ZZZZ|egapxtmp_032785-R1    mNeoVis.evergladensis.2.1.chr14.g00034.t01    mNeoVis.vulgivaga.2.1.chr14.g00036.t01      1064    N0.HOG0001062   OG0000094  n122   rna-XM_034667726.1    rna-NM_001003060.2    rna-XM_022508493.1    rna-XM_003999196.3    rna-XM_036080523.1    rna-XM_039221695.1    rna-XM_032863003.1    rna-XM_045983191.1    rna-XM_032317477.1    rna-XM_059146773.1    rna-XM_059412427.1    rna-XM_004756690.3    mNeoVis.2.1.1.chr14_hap1.g00034.t01    rna-XM_021682641.1    rna-XM_004408899.1    rna-XM_007088731.3    rna-XM_008696043.1    rna-XM_027612284.2    mNeoVis.3.1.1.chr14.g00030.t01    rna-gnl|WGS_ZZZZ|egapxtmp_032640-R1    mNeoVis.evergladensis.2.1.chr14.g00033.t01    mNeoVis.vulgivaga.2.1.chr14.g00035.t01      1065    N0.HOG0001063   OG0000095  n1   rna-XM_002930724.4, rna-XM_034652910.1, rna-XM_034663726.1    rna-NM_001388817.1    rna-XM_022501913.1    rna-XM_011288457.3, rna-XM_045043969.1    rna-XM_036107851.1    rna-XM_039239042.1    rna-XM_032850788.1, rna-XM_032866357.1    rna-XM_046021807.1, rna-XM_046021809.1    rna-XM_032308517.1, rna-XM_032308518.1, rna-XM_032308519.1    rna-XM_059143085.1, rna-XM_059143086.1    rna-XM_059387260.1, rna-XM_059412151.1    rna-XM_013050791.2, rna-XM_013058814.2, rna-XM_045082912.1    mNeoVis.2.1.1.chr13_hap1.g00265.t01, mNeoVis.2.1.1.chr13_hap1.g00266.t01, mNeoVis.2.1.1.chr13_hap1.g00267.t01    rna-XM_021691517.1    rna-XM_004392620.1    rna-XM_015543036.2, rna-XM_015543037.2, rna-XM_042964031.1    rna-XM_008698796.2, rna-XM_008698797.2    rna-XM_027616377.1    mNeoVis.3.1.1.chr13.g00211.t01    rna-gnl|WGS_ZZZZ|egapxtmp_030991-R1    mNeoVis.evergladensis.2.1.chr13.g00258.t01    mNeoVis.vulgivaga.2.1.chr13.g00255.t01      1066    N0.HOG0001064   OG0000095  n38   rna-XM_002915958.4, rna-XM_002915959.2, rna-XM_002930725.3, rna-XM_002930728.3, rna-XM_019793500.2, rna-XM_034653272.1    rna-NM_001389082.1, rna-XM_038546067.1, rna-XM_038546179.1    rna-XM_022492680.1, rna-XM_022501450.1, rna-XM_022501698.1, rna-XM_022501735.1, rna-XM_022501737.1, rna-XM_022501765.1    rna-XM_003995832.3, rna-XM_003995835.4, rna-XM_003995846.4, rna-XM_011288459.3, rna-XM_045043131.1, rna-XM_045043133.1, rna-XM_045043181.1, rna-XM_045043272.1, rna-XM_045043959.1, rna-XM_045043965.1, rna-XM_045043967.1    rna-XM_036107845.1, rna-XM_036107849.1, rna-XM_036107850.1, rna-XM_036107906.1    rna-XM_039224355.1, rna-XM_039224357.1, rna-XM_039224358.1, rna-XM_039224359.1, rna-XM_039225113.1, rna-XM_039239033.1, rna-XM_039239043.1, rna-XM_039239044.1, rna-XM_039239064.1    rna-XM_032866351.1, rna-XM_032866352.1, rna-XM_032866356.1, rna-XM_032866366.1, rna-XM_032866370.1    rna-XM_046021810.1, rna-XM_046021820.1, rna-XM_046021821.1, rna-XM_046023560.1, rna-XM_046023613.1, rna-XM_046023620.1, rna-XM_046023652.1, rna-XM_046023962.1    rna-XM_032308521.1, rna-XM_032308527.1, rna-XM_032308528.1, rna-XM_032308633.1    rna-XM_059142843.1, rna-XM_059143087.1, rna-XM_059143095.1, rna-XM_059143096.1    rna-XM_059412152.1, rna-XM_059412213.1    rna-XM_004782113.1, rna-XM_004782570.2, rna-XM_013047353.2, rna-XM_045082909.1, rna-XM_045082910.1    mNeoVis.2.1.1.chr13_hap1.g00268.t01, mNeoVis.2.1.1.chr13_hap1.g00282.t01, mNeoVis.2.1.1.chr13_hap1.g00283.t01    rna-XM_021691582.1, rna-XM_021691593.2, rna-XM_021691595.1    rna-XM_004392621.1, rna-XM_004392626.1, rna-XM_004392648.1    rna-XM_007094369.2, rna-XM_007094511.2, rna-XM_007094525.2, rna-XM_007094526.2, rna-XM_015543039.2, rna-XM_042964062.1, rna-XM_042964575.1    rna-XM_008698798.2, rna-XM_008698805.2, rna-XM_008698927.1, rna-XM_008698932.2, rna-XM_008698942.2, rna-XM_008698943.2, rna-XM_008698944.1, rna-XM_040644525.1    rna-XM_027614694.1, rna-XM_027614695.1, rna-XM_035723668.1, rna-XM_035723736.1    mNeoVis.3.1.1.chr13.g00212.t01, mNeoVis.3.1.1.chr13.g00224.t01, mNeoVis.3.1.1.chr13.g00225.t01, mNeoVis.3.1.1.chr13.g00226.t01    rna-gnl|WGS_ZZZZ|egapxtmp_030161-R1, rna-gnl|WGS_ZZZZ|egapxtmp_030992-R1    mNeoVis.evergladensis.2.1.chr13.g00260.t01, mNeoVis.evergladensis.2.1.chr13.g00272.t01, mNeoVis.evergladensis.2.1.chr13.g00273.t01, mNeoVis.evergladensis.2.1.chr13.g00274.t01    mNeoVis.vulgivaga.2.1.chr13.g00256.t01, mNeoVis.vulgivaga.2.1.chr13.g00269.t01, mNeoVis.vulgivaga.2.1.chr13.g00270.t01      1067    N0.HOG0001065   OG0000096  n1   rna-XM_019793613.2, rna-XM_034650688.1    rna-NM_001388771.1, rna-NM_001388965.1, rna-NM_001388997.1, rna-NM_001389070.1, rna-XM_038582476.1    rna-XM_022508748.1, rna-XM_022509145.1, rna-XM_022509301.1    rna-XM_019811147.2    rna-XM_036070307.1    rna-XM_039225334.1    rna-XM_032852643.1, rna-XM_032882267.1, rna-XM_032882268.1    rna-XM_046002693.1, rna-XM_046003302.1, rna-XM_046003409.1    rna-XM_032301908.1, rna-XM_032301955.1, rna-XM_032301964.1    rna-XM_059165255.1, rna-XM_059165260.1, rna-XM_059165261.1    rna-XM_059388154.1, rna-XM_059388159.1, rna-XM_059388160.1    rna-XM_004771989.2, rna-XM_004772994.2, rna-XM_004772995.2    mNeoVis.2.1.1.chr5_hap1.g01663.t01, mNeoVis.2.1.1.chr5_hap1.g01670.t01, mNeoVis.2.1.1.chr5_hap1.g01671.t01    rna-XM_021687952.1    rna-XM_004391676.1    rna-XM_015543714.2    rna-XM_008689683.2, rna-XM_008689685.2, rna-XM_040621705.1      mNeoVis.3.1.1.chr2.g01585.t01    rna-gnl|WGS_ZZZZ|egapxtmp_037317-R1    mNeoVis.evergladensis.2.1.chr2.g01650.t01    mNeoVis.vulgivaga.2.1.chr2.g01670.t01, mNeoVis.vulgivaga.2.1.chr2.g01675.t01, mNeoVis.vulgivaga.2.1.chr2.g01676.t01      1068    N0.HOG0001066   OG0000096  n46   rna-XM_011232699.2    rna-NM_001389132.1    rna-XM_022502126.1    rna-XM_003991559.4    rna-XM_036070318.1    rna-XM_039221239.1    rna-XM_032852626.1    rna-XM_046003685.1    rna-XM_032344602.1    rna-XM_059165073.1    rna-XM_059388369.1    rna-XM_004771991.1    mNeoVis.2.1.1.chr5_hap1.g01660.t01        rna-XM_007095941.2    rna-XM_008689700.2, rna-XM_008702038.2    rna-XM_027581795.1    mNeoVis.3.1.1.chr2.g01578.t01    rna-gnl|WGS_ZZZZ|egapxtmp_035671-R1    mNeoVis.evergladensis.2.1.chr2.g01647.t01    mNeoVis.vulgivaga.2.1.chr2.g01667.t01      1069    N0.HOG0001067   OG0000096  n64   rna-XM_002931023.3    rna-NM_001388994.1    rna-XM_022508688.1    rna-XM_011285984.2      rna-XM_039225333.1    rna-XM_032852625.1    rna-XM_046003361.1    rna-XM_032344614.1, rna-XM_032344625.1    rna-XM_059165075.1    rna-XM_059387366.1    rna-XM_004771990.2    mNeoVis.2.1.1.chr5_hap1.g01662.t01        rna-XM_007095940.2    rna-XM_008689688.1, rna-XM_008689701.1      mNeoVis.3.1.1.chr2.g01580.t01    rna-gnl|WGS_ZZZZ|egapxtmp_037307-R1    mNeoVis.evergladensis.2.1.chr2.g01649.t01    mNeoVis.vulgivaga.2.1.chr2.g01669.t01      1070    N0.HOG0001068   OG0000096  n82   rna-XM_011217954.3      rna-XM_022508656.1, rna-XM_022509177.1          rna-XM_032852641.1, rna-XM_032852642.1, rna-XM_032852644.1    rna-XM_046003028.1, rna-XM_046003286.1, rna-XM_046003317.1, rna-XM_046003320.1, rna-XM_046003322.1, rna-XM_046003323.1, rna-XM_046003338.1, rna-XM_046003339.1, rna-XM_046003340.1, rna-XM_046003343.1, rna-XM_046003345.1, rna-XM_046003394.1    rna-XM_032301896.1, rna-XM_032301934.1, rna-XM_032351423.1, rna-XM_032351431.1, rna-XM_032351442.1    rna-XM_059165077.1, rna-XM_059165254.1, rna-XM_059165257.1, rna-XM_059165258.1    rna-XM_059387402.1, rna-XM_059388156.1, rna-XM_059388157.1, rna-XM_059388158.1, rna-XM_059388370.1    rna-XM_004771900.3, rna-XM_004772991.2, rna-XM_004773025.3, rna-XM_045088054.1    mNeoVis.2.1.1.chr5_hap1.g01661.t01, mNeoVis.2.1.1.chr5_hap1.g01665.t01, mNeoVis.2.1.1.chr5_hap1.g01666.t01, mNeoVis.2.1.1.chr5_hap1.g01667.t01, mNeoVis.2.1.1.chr5_hap1.g01668.t01          rna-XM_008689697.2, rna-XM_008689699.2, rna-XM_008689703.2, rna-XM_040621716.1      mNeoVis.3.1.1.chr2.g01579.t01, mNeoVis.3.1.1.chr2.g01582.t01, mNeoVis.3.1.1.chr2.g01583.t01    rna-gnl|WGS_ZZZZ|egapxtmp_037308-R1, rna-gnl|WGS_ZZZZ|egapxtmp_037310-R1, rna-gnl|WGS_ZZZZ|egapxtmp_037311-R1, rna-gnl|WGS_ZZZZ|egapxtmp_037312-R1, rna-gnl|WGS_ZZZZ|egapxtmp_037316-R1    mNeoVis.evergladensis.2.1.chr2.g01648.t01, mNeoVis.evergladensis.2.1.chr2.g01652.t01    mNeoVis.vulgivaga.2.1.chr2.g01668.t01, mNeoVis.vulgivaga.2.1.chr2.g01672.t01, mNeoVis.vulgivaga.2.1.chr2.g01673.t01      1071    N0.HOG0001069   OG0000097  n3   rna-XM_034643051.1    rna-XM_038544058.1        rna-XM_036077180.1, rna-XM_036113075.1, rna-XM_036115816.1    rna-XM_039248589.1                mNeoVis.2.1.1.chr12_hap1.g00459.t01, mNeoVis.2.1.1.chr4_hap1.g01010.t01    rna-XM_044911921.1, rna-XM_044913295.1, rna-XM_044914903.1, rna-XM_044920628.1    rna-XM_004413199.1    rna-XM_042963981.1, rna-XM_042993713.1      rna-XM_035726340.1, rna-XM_035727786.1    mNeoVis.3.1.1.chr12.g00420.t01, mNeoVis.3.1.1.chr2.g01039.t01, mNeoVis.3.1.1.chr6.g01027.t01, mNeoVis.3.1.1.chr8.g00699.t01    rna-gnl|WGS_ZZZZ|egapxtmp_002456-R1    mNeoVis.evergladensis.2.1.chr12.g00415.t01, mNeoVis.evergladensis.2.1.chr6.g01014.t01    mNeoVis.vulgivaga.2.1.chr6.g01006.t01      1072    N0.HOG0001070   OG0000097  n27     rna-XM_038533688.1        rna-XM_036070950.1      rna-XM_032864562.1    rna-XM_046012687.1    rna-XM_032347025.1    rna-XM_059186338.1    rna-XM_059404699.1      mNeoVis.2.1.1.chr7_hap1.g01191.t01, mNeoVis.2.1.1.chr9_hap1.g00931.t01    rna-XM_044915738.1, rna-XM_044918405.1        rna-XM_040625847.1    rna-XM_035723177.1    mNeoVis.3.1.1.chr10.g00884.t01, mNeoVis.3.1.1.chr7.g01115.t01    rna-gnl|WGS_ZZZZ|egapxtmp_010243-R1, rna-gnl|WGS_ZZZZ|egapxtmp_021705-R1    mNeoVis.evergladensis.2.1.chr10.g00916.t01, mNeoVis.evergladensis.2.1.chr7.g01180.t01    mNeoVis.vulgivaga.2.1.chr10.g00904.t01      1073    N0.HOG0001071   OG0000097  n43     rna-XM_038529824.1              rna-XM_045986912.1, rna-XM_046012859.1    rna-XM_032335863.1, rna-XM_032338359.1, rna-XM_032357876.1    rna-XM_059176378.1    rna-XM_059399047.1      mNeoVis.2.1.1.chr1_hap1.g01562.t01            rna-XM_035722381.1    mNeoVis.3.1.1.chr1.g01515.t01, mNeoVis.3.1.1.chr2.g01139.t01    rna-gnl|WGS_ZZZZ|egapxtmp_023599-R1, rna-gnl|WGS_ZZZZ|egapxtmp_035567-R1    mNeoVis.evergladensis.2.1.chr1.g01529.t01    mNeoVis.vulgivaga.2.1.chr1.g01525.t01      1074    N0.HOG0001072   OG0000097  n55           rna-XM_036072769.1        rna-XM_046002879.1      rna-XM_059172636.1          rna-XM_044913139.1      rna-XM_042987746.1, rna-XM_042989264.1      rna-XM_035726768.1, rna-XM_035727604.1, rna-XM_035729358.1              1075    N0.HOG0001073   OG0000097  n63   rna-XM_034651201.1        rna-XM_045042041.1    rna-XM_036093279.1, rna-XM_036101470.1, rna-XM_036102995.1, rna-XM_036104736.1, rna-XM_036104866.1    rna-XM_039226492.1      rna-XM_045995523.1        rna-XM_059395791.1    rna-XM_045087244.1    mNeoVis.2.1.1.chr5_hap1.g01130.t01    rna-XM_044911497.1, rna-XM_044912597.1, rna-XM_044913171.1, rna-XM_044916987.1    rna-XM_004395649.1, rna-XM_004398781.1, rna-XM_004416866.1    rna-XM_042956961.1    rna-XM_040645497.1    rna-XM_035723780.1, rna-XM_035726279.1, rna-XM_035727052.1, rna-XM_035727319.1, rna-XM_035728235.1        mNeoVis.evergladensis.2.1.chr2.g01120.t01    mNeoVis.vulgivaga.2.1.chr2.g00896.t01, mNeoVis.vulgivaga.2.1.chr2.g01126.t01      1076    N0.HOG0001074   OG0000097  n91   rna-NM_001304930.1, rna-XM_034640664.1, rna-XM_034651293.1    rna-NM_001252169.1    rna-XM_022498234.1    rna-XM_003982504.5    rna-XM_036073596.1    rna-XM_039220940.1, rna-XM_039231333.1, rna-XM_039247142.1    rna-XM_032841663.1, rna-XM_032843416.1, rna-XM_032850844.1, rna-XM_032850845.1    rna-XM_045990821.1, rna-XM_046012721.1, rna-XM_046020524.1, rna-XM_046027356.1    rna-XM_032337550.1    rna-XM_059161810.1    rna-XM_059390372.1    rna-XM_013054801.2    mNeoVis.2.1.1.chr1_hap1.g00601.t01, mNeoVis.2.1.1.chr5_hap1.g00894.t01, mNeoVis.2.1.1.chr5_hap1.g00958.t01    rna-XM_021695169.1, rna-XM_044912587.1    rna-XM_004399979.1, rna-XM_004401493.1, rna-XM_004411344.1    rna-XM_007075436.3    rna-XM_008697854.2    rna-XM_027584165.1, rna-XM_035721845.1, rna-XM_035723733.1, rna-XM_035725627.1, rna-XM_035727970.1    mNeoVis.3.1.1.chr1.g00618.t01, mNeoVis.3.1.1.chr2.g00806.t01    rna-gnl|WGS_ZZZZ|egapxtmp_035569-R1, rna-gnl|WGS_ZZZZ|egapxtmp_037037-R1    mNeoVis.evergladensis.2.1.chr1.g00606.t01, mNeoVis.evergladensis.2.1.chr2.g00894.t01    mNeoVis.vulgivaga.2.1.chr1.g00615.t01, mNeoVis.vulgivaga.2.1.chr2.g00959.t01      1077    N0.HOG0001075   OG0000098  n1   rna-XM_002930581.3, rna-XM_034652629.1, rna-XM_034653289.1    rna-XM_038568117.1    rna-XM_022522045.1    rna-XM_011286644.2      rna-XM_039224513.1    rna-XM_032853660.1    rna-XM_046016780.1    rna-XM_032360266.1    rna-XM_059157802.1    rna-XM_059389689.1    rna-XM_004779042.2, rna-XM_045088013.1    mNeoVis.2.1.1.chr7_hap1.g01714.t01      rna-XM_004414462.1    rna-XM_007083264.1    rna-XM_008709563.2    rna-XM_027580317.1    mNeoVis.3.1.1.chr7.g01598.t01    rna-gnl|WGS_ZZZZ|egapxtmp_012830-R1    mNeoVis.evergladensis.2.1.chr7.g01683.t01    mNeoVis.vulgivaga.2.1.chr7.g01699.t01      1078    N0.HOG0001076   OG0000098  n24   rna-XM_019794055.2      rna-XM_022521699.1        rna-XM_039224503.1      rna-XM_046015882.1                  rna-XM_007083263.1    rna-XM_008709564.1    rna-XM_027580285.1              1079    N0.HOG0001077   OG0000098  n31   rna-XM_011217419.3    rna-XM_038568119.1          rna-XM_039224499.1                        rna-XM_008709627.2    rna-XM_035722971.1              1080    N0.HOG0001078   OG0000098  n36   rna-XM_002930582.2    rna-NM_001083631.1      rna-XM_011286570.4          rna-XM_046017104.1    rna-XM_032360306.1    rna-XM_059157809.1    rna-XM_059389677.1    rna-XM_045078262.1    mNeoVis.2.1.1.chr7_hap1.g01715.t01          rna-XM_008709625.2          mNeoVis.evergladensis.2.1.chr7.g01685.t01    mNeoVis.vulgivaga.2.1.chr7.g01700.t01      1081    N0.HOG0001079   OG0000098  n47   rna-XM_002930587.4, rna-XM_034651443.1    rna-NM_001388716.1    rna-XM_022522039.1    rna-XM_003992909.6      rna-XM_039224512.1    rna-XM_032852715.1    rna-XM_046017108.1    rna-XM_032359784.1    rna-XM_059157822.1    rna-XM_059389666.1    rna-XM_004779041.2, rna-XM_045078263.1    mNeoVis.2.1.1.chr7_hap1.g01716.t01      rna-XM_004414466.1    rna-XM_007083133.1    rna-XM_008709624.1    rna-XM_027580278.1      rna-gnl|WGS_ZZZZ|egapxtmp_012786-R1    mNeoVis.evergladensis.2.1.chr7.g01684.t01        1082    N0.HOG0001080   OG0000098  n68     rna-XM_038568175.1    rna-XM_022521701.1    rna-XM_003992875.5    rna-XM_036076014.1    rna-XM_039224515.1    rna-XM_032850891.1    rna-XM_046017061.1      rna-XM_059182469.1    rna-XM_059393782.1    rna-XM_004779049.3    mNeoVis.2.1.1.chr7_hap1.g01704.t01    rna-XM_021685640.1      rna-XM_007092199.1    rna-XM_008709634.1    rna-XM_027580533.1      rna-gnl|WGS_ZZZZ|egapxtmp_012531-R1    mNeoVis.evergladensis.2.1.chr7.g01675.t01    mNeoVis.vulgivaga.2.1.chr7.g01691.t01      1083    N0.HOG0001081   OG0000098  n85   rna-XM_002926087.4    rna-XM_038568116.1    rna-XM_022522086.1    rna-XM_003992903.2      rna-XM_039224505.1    rna-XM_032853656.1    rna-XM_046017109.1    rna-XM_032360369.1    rna-XM_059157756.1    rna-XM_059389733.1    rna-XM_004778989.2    mNeoVis.2.1.1.chr7_hap1.g01710.t01        rna-XM_007083258.1    rna-XM_008709565.2    rna-XM_035722998.1      rna-gnl|WGS_ZZZZ|egapxtmp_012790-R1    mNeoVis.evergladensis.2.1.chr7.g01680.t01    mNeoVis.vulgivaga.2.1.chr7.g01697.t01      1084    N0.HOG0001082   OG0000098  n99   rna-XM_034667616.1    rna-NM_001388992.1    rna-XM_022522085.1    rna-XM_011286571.2        rna-XM_032853657.1    rna-XM_046015883.1    rna-XM_032360359.1          mNeoVis.2.1.1.chr7_hap1.g01711.t01        rna-XM_007083132.1    rna-XM_008709631.1        rna-gnl|WGS_ZZZZ|egapxtmp_012880-R1    mNeoVis.evergladensis.2.1.chr7.g01679.t01    mNeoVis.vulgivaga.2.1.chr7.g01698.t01      1085    N0.HOG0001083   OG0000098  n111   rna-XM_034667730.1    rna-NM_001388714.1    rna-XM_022521700.1, rna-XM_022522096.1    rna-XM_003992876.4      rna-XM_039224506.1    rna-XM_032853654.1, rna-XM_032853655.1    rna-XM_046017110.1    rna-XM_032359499.1, rna-XM_032360372.1    rna-XM_059157742.1    rna-XM_059389746.1, rna-XM_059389794.1    rna-XM_004778990.2, rna-XM_013049226.2    mNeoVis.2.1.1.chr7_hap1.g01706.t01, mNeoVis.2.1.1.chr7_hap1.g01709.t01      rna-XM_004414464.1    rna-XM_007083131.1    rna-XM_008709566.1    rna-XM_027580505.1    mNeoVis.3.1.1.chr7.g01595.t01    rna-gnl|WGS_ZZZZ|egapxtmp_012792-R1, rna-gnl|WGS_ZZZZ|egapxtmp_012796-R1    mNeoVis.evergladensis.2.1.chr7.g01677.t01, mNeoVis.evergladensis.2.1.chr7.g01678.t01    mNeoVis.vulgivaga.2.1.chr7.g01693.t01, mNeoVis.vulgivaga.2.1.chr7.g01696.t01      1086    N0.HOG0001084   OG0000099  n1   rna-XM_034658832.1    rna-XM_038565537.1    rna-XM_022498477.1    rna-XM_023250056.2    rna-XM_036073653.1    rna-XM_039254953.1    rna-XM_032850385.1, rna-XM_032850388.1    rna-XM_045991763.1    rna-XM_032316063.1    rna-XM_059161846.1    rna-XM_059390396.1    rna-XM_013054910.2    mNeoVis.2.1.1.chr5_hap1.g00910.t01    rna-XM_021695124.1    rna-XM_004395671.1    rna-XM_042979586.1    rna-XM_040628409.1    rna-XM_027587412.2    mNeoVis.3.1.1.chr2.g00823.t01    rna-gnl|WGS_ZZZZ|egapxtmp_035978-R4    mNeoVis.evergladensis.2.1.chr2.g00910.t01    mNeoVis.vulgivaga.2.1.chr2.g00914.t01      1087    N0.HOG0001085   OG0000099  n19   rna-XM_034658773.1, rna-XM_034663769.1    rna-XM_038565636.1    rna-XM_022498306.1    rna-XM_023250021.2    rna-XM_036079882.1    rna-XM_039231408.1    rna-XM_032841507.1    rna-XM_045992023.1    rna-XM_032327230.1    rna-XM_059161739.1    rna-XM_059390335.1    rna-XM_045063314.1    mNeoVis.2.1.1.chr5_hap1.g00878.t01    rna-XM_021695019.1    rna-XM_004392675.1    rna-XM_007075461.3    rna-XM_008697885.2    rna-XM_027586460.2    mNeoVis.3.1.1.chr2.g00791.t01    rna-gnl|WGS_ZZZZ|egapxtmp_037033-R2    mNeoVis.evergladensis.2.1.chr2.g00879.t01    mNeoVis.vulgivaga.2.1.chr2.g00881.t01      1088    N0.HOG0001086   OG0000099  n30   rna-XM_034649988.1, rna-XM_034653515.1    rna-XM_038588661.1    rna-XM_022492760.1    rna-XM_023249498.2, rna-XM_045050887.1, rna-XM_045050892.1, rna-XM_045051697.1    rna-XM_036074456.1    rna-XM_039219272.1    rna-XM_032846386.1, rna-XM_032846387.1, rna-XM_032851161.1    rna-XM_045995485.1    rna-XM_032329745.1, rna-XM_032330734.1    rna-XM_059158700.1    rna-XM_059387452.1, rna-XM_059387465.1    rna-XM_045068075.1, rna-XM_045068218.1    mNeoVis.2.1.1.chr12_hap1.g00026.t01, mNeoVis.2.1.1.chr12_hap1.g00062.t01    rna-XM_021682851.1, rna-XM_044912444.1    rna-XM_004409164.1    rna-XM_042973989.1, rna-XM_042973992.1, rna-XM_042974803.1    rna-XM_040627001.1    rna-XM_035725748.1    mNeoVis.3.1.1.chr12.g00053.t01    rna-gnl|WGS_ZZZZ|egapxtmp_029229-R1    mNeoVis.evergladensis.2.1.chr12.g00052.t01    mNeoVis.vulgivaga.2.1.chr12.g00054.t01      1089    N0.HOG0001087   OG0000099  n66   rna-XM_002929093.4    rna-XM_038565638.1    rna-XM_022498202.1    rna-XM_003982487.6    rna-XM_036079875.1    rna-XM_039231416.1    rna-XM_032841502.1    rna-XM_045991487.1    rna-XM_032327206.1    rna-XM_059161737.1    rna-XM_059390333.1    rna-XM_004738533.3    mNeoVis.2.1.1.chr5_hap1.g00877.t01    rna-XM_021694958.1    rna-XM_004392676.1    rna-XM_007075526.3    rna-XM_040628449.1    rna-XM_027586058.2    mNeoVis.3.1.1.chr2.g00790.t01      mNeoVis.evergladensis.2.1.chr2.g00878.t01    mNeoVis.vulgivaga.2.1.chr2.g00880.t01      1090    N0.HOG0001088   OG0000099  n86   rna-XM_019793751.2    rna-XM_038576397.1    rna-XM_022522294.1    rna-XM_045061234.1    rna-XM_036092417.1    rna-XM_039236364.1    rna-XM_032874915.1    rna-XM_046010666.1    rna-XM_032349167.1    rna-XM_059184466.1    rna-XM_059404960.1    rna-XM_013046491.2    mNeoVis.2.1.1.chr9_hap1.g00209.t01    rna-XM_021690558.1    rna-XM_004411943.1    rna-XM_042991540.1    rna-XM_008704846.2    rna-XM_027594643.2    mNeoVis.3.1.1.chr10.g00199.t01      mNeoVis.evergladensis.2.1.chr10.g00205.t01    mNeoVis.vulgivaga.2.1.chr10.g00202.t01      1091    N0.HOG0001089   OG0000099  n106   rna-XM_034645916.1    rna-NM_001003322.3    rna-XM_022522285.1    rna-XM_045061238.1    rna-XM_036092413.1    rna-XM_039236289.1    rna-XM_032874895.1    rna-XM_046013044.1    rna-XM_032349163.1    rna-XM_059184476.1    rna-XM_059403840.1    rna-XM_045071617.1    mNeoVis.2.1.1.chr9_hap1.g00210.t01    rna-XM_021690488.1    rna-XM_004411944.1    rna-XM_007092839.3    rna-XM_040642022.1    rna-XM_027594636.2    mNeoVis.3.1.1.chr10.g00200.t01      mNeoVis.evergladensis.2.1.chr10.g00206.t01    mNeoVis.vulgivaga.2.1.chr10.g00203.t01      1092    N0.HOG0001090   OG0000100  n3         rna-NM_001305026.1, rna-XM_023250215.2, rna-XM_045052203.1      id-LOC120225672, id-LOC120227278, id-LOC120227279, rna-XM_039225573.1                      id-LOC102953986, id-LOC102963058, id-LOC102963604                  1093    N0.HOG0001091   OG0000100  n13   id-LOC105234544, id-LOC109488455, id-LOC117795072, rna-XM_034665944.1    rna-XM_038562531.1        id-LOC118551592      id-LOC116876850      id-LOC116568784    id-LOC131830166    rna-XM_059397080.1    id-LOC101684077      id-LOC110574272        id-LOC103674605, id-LOC103674607, id-LOC103674700    id-LOC118356144      id-egapxtmp_004081          1094    N0.HOG0001092   OG0000100  n32   id-LOC109488456    id-LOC100683694                                id-LOC103674604                1095    N0.HOG0001093   OG0000100  n35   id-LOC105234752, id-LOC105234754    id-LOC119863964, id-LOC119864164    id-LOC111155279, rna-XM_022515349.1          id-LOC116876853, id-LOC116876953, id-LOC116876955, id-LOC116877878    id-LOC123951548, id-LOC123951549, id-LOC123951551, id-LOC123951552, id-LOC123951553, id-LOC123952176, id-LOC123952177, id-LOC123952178, id-LOC123952180, id-LOC123952181, id-LOC123952182, id-LOC123952184    id-LOC116568647, id-LOC116568649, id-LOC116568650, id-LOC116568651    id-LOC131829216, rna-XM_059170753.1    id-LOC132009053, id-LOC132015129, id-LOC132016057, id-LOC132016346, id-LOC132016347    id-LOC101683812, id-LOC101691347, id-LOC101692183, id-LOC123394826, rna-XM_045089638.1        id-LOC101370678, id-LOC101370970, id-LOC101371863      id-LOC103674598, id-LOC103674600    id-LOC113911689, id-LOC113931732      id-egapxtmp_004080, id-egapxtmp_004082, id-egapxtmp_004083, id-egapxtmp_004084, id-egapxtmp_004085, id-egapxtmp_004087    mNeoVis.evergladensis.2.1.chr6.g00602.t01        1096    N0.HOG0001094   OG0000100  n86   rna-XM_034665897.1    id-LOC100856335, id-LOC119864100, id-LOC119864163, id-LOC119877224, id-LOC119878508, id-LOC609263, rna-XM_038562675.1    id-LOC111155232            id-LOC123952179                    id-LOC103674603, id-LOC103674702                1097    N0.HOG0001095   OG0000100  n97     id-LOC102152421, id-LOC119877225          rna-XM_039223855.1    id-LOC116876846      id-LOC116568781      id-LOC132016343    id-LOC106006555          id-LOC102963333                  1098    N0.HOG0001096   OG0000100  n105         rna-XM_045052201.1      id-LOC120227618                      id-LOC107178973                  1099    N0.HOG0001097   OG0000100  n108                                 id-LOC102951021, id-LOC102963898                  1100    N0.HOG0001098   OG0000100  n113   id-LOC105234961, id-LOC105240638                                  id-LOC103674609                1101    N0.HOG0001099   OG0000100  n117       id-LOC111155234          rna-XM_032873430.1      rna-XM_032304197.1    rna-XM_059170754.1      rna-XM_045089639.1                          1102    N0.HOG0001100   OG0000100  n125     id-LOC119864167                    id-LOC132027493                            1103    N0.HOG0001101   OG0000100  n126     id-LOC119864166            id-LOC116876848      id-LOC116568783        id-LOC101684369                          1104    N0.HOG0001102   OG0000100  n129           rna-XM_036084033.1                    rna-XM_021682921.1    id-LOC101371268                    1105    N0.HOG0001103   OG0000100  n131       id-LOC111155322      id-LOC118551588      rna-XM_032873432.1      rna-XM_032304424.1    id-LOC131829222    id-LOC132016344    rna-XM_013059022.2    mNeoVis.2.1.1.chr4_hap1.g00597.t01    id-LOC110574271            mNeoVis.3.1.1.chr6.g00609.t01    rna-gnl|WGS_ZZZZ|egapxtmp_004086-R1    mNeoVis.evergladensis.2.1.chr6.g00601.t01    mNeoVis.vulgivaga.2.1.chr6.g00601.t01      1106    N0.HOG0001104   OG0000100  n140             id-LOC120227116, id-LOC120227508                                      1107    N0.HOG0001105   OG0000101  n1   rna-XM_002920345.3    rna-XM_038530204.1    rna-XM_022520744.1    rna-XM_006933209.4    rna-XM_036086162.1    rna-XM_039240375.1    rna-XM_032881660.1    rna-XM_046013563.1    rna-XM_032349847.1    rna-XM_059183714.1    rna-XM_059404003.1    rna-XM_004757924.3    mNeoVis.2.1.1.chr9_hap1.g00022.t01    rna-XM_021696811.2    rna-XM_004411820.1    rna-XM_007081170.3    rna-XM_008705401.2    rna-XM_027595474.1    mNeoVis.3.1.1.chr10.g00021.t01    rna-gnl|WGS_ZZZZ|egapxtmp_022919-R1    mNeoVis.evergladensis.2.1.chr10.g00024.t01    mNeoVis.vulgivaga.2.1.chr10.g00022.t01      1108    N0.HOG0001106   OG0000101  n19   rna-XM_002920344.3    rna-XM_038529611.1    rna-XM_022520722.1    rna-XM_011283583.4    rna-XM_036086171.1    rna-XM_039240356.1    rna-XM_032881592.1    rna-XM_046013036.1    rna-XM_032349848.1    rna-XM_059187554.1    rna-XM_059404981.1    rna-XM_013059125.2    mNeoVis.2.1.1.chr9_hap1.g00021.t01    rna-XM_021696763.2      rna-XM_015537140.2    rna-XM_008705400.2    rna-XM_027595438.2    mNeoVis.3.1.1.chr10.g00020.t01      mNeoVis.evergladensis.2.1.chr10.g00023.t01        1109    N0.HOG0001107   OG0000101  n36   rna-XM_002920781.4, rna-XM_002923050.4, rna-XM_011216806.3, rna-XM_034652206.1    rna-XM_038528500.1, rna-XM_038545548.1, rna-XM_038551229.1    rna-XM_022505084.1, rna-XM_022516253.1, rna-XM_022523860.1    rna-XM_003983965.6, rna-XM_023245109.2, rna-XM_023255948.2, rna-XM_045050629.1    rna-XM_036064445.1, rna-XM_036067271.1, rna-XM_036072327.1, rna-XM_036080833.1, rna-XM_036087381.1, rna-XM_036088887.1, rna-XM_036090380.1, rna-XM_036090855.1, rna-XM_036099241.1, rna-XM_036106706.1, rna-XM_036107932.1, rna-XM_036119967.1, rna-XM_036121652.1    rna-XM_039232540.1, rna-XM_039245925.1, rna-XM_039247430.1    rna-XM_032839012.1, rna-XM_032840304.1, rna-XM_032878625.1    rna-XM_045981186.1, rna-XM_045987323.1, rna-XM_045997098.1, rna-XM_046007970.1, rna-XM_046018224.1, rna-XM_046020866.1    rna-XM_032324222.1, rna-XM_032336180.1, rna-XM_032344685.1, rna-XM_032351841.1    rna-XM_059134543.1, rna-XM_059151829.1, rna-XM_059182515.1    rna-XM_059373530.1, rna-XM_059382064.1, rna-XM_059395790.1, rna-XM_059406249.1    rna-XM_004739064.2, rna-XM_004753012.3, rna-XM_004767522.3    mNeoVis.2.1.1.chr10_hap1.g00253.t01, mNeoVis.2.1.1.chr11_hap1.g00607.t01, mNeoVis.2.1.1.chr2_hap1.g01303.t01, mNeoVis.2.1.1.chr7_hap1.g00990.t01    rna-XM_021679818.1, rna-XM_021681345.1, rna-XM_021703176.2, rna-XM_044913615.1, rna-XM_044913640.1, rna-XM_044915221.1, rna-XM_044918942.1    rna-XM_004394469.1, rna-XM_004400947.1, rna-XM_004401169.1, rna-XM_004401539.1, rna-XM_004403384.1, rna-XM_004410852.1    rna-XM_007084706.3, rna-XM_015538385.2, rna-XM_015540097.2, rna-XM_042969237.1, rna-XM_042990362.1    rna-XM_008698739.2, rna-XM_008699841.2, rna-XM_040634200.1    rna-XM_027569245.2, rna-XM_027578889.1, rna-XM_027600431.2, rna-XM_027605884.2, rna-XM_027606846.2, rna-XM_027618998.1, rna-XM_027621942.1, rna-XM_035721810.1, rna-XM_035723695.1, rna-XM_035726845.1    mNeoVis.3.1.1.chr11.g00569.t01, mNeoVis.3.1.1.chr5.g01228.t01, mNeoVis.3.1.1.chr7.g00929.t01, mNeoVis.3.1.1.chr9.g00223.t01    rna-gnl|WGS_ZZZZ|egapxtmp_006481-R1, rna-gnl|WGS_ZZZZ|egapxtmp_006483-R1, rna-gnl|WGS_ZZZZ|egapxtmp_027804-R1    mNeoVis.evergladensis.2.1.chr11.g00595.t01, mNeoVis.evergladensis.2.1.chr5.g01245.t01, mNeoVis.evergladensis.2.1.chr7.g00983.t01, mNeoVis.evergladensis.2.1.chr8.g00238.t01    mNeoVis.vulgivaga.2.1.chr11.g00603.t01, mNeoVis.vulgivaga.2.1.chr5.g01240.t01, mNeoVis.vulgivaga.2.1.chr7.g00989.t01, mNeoVis.vulgivaga.2.1.chr9.g00239.t01      1110    N0.HOG0001108   OG0000102  n3     rna-XM_038568166.1      rna-XM_045038127.1      rna-XM_039224657.1      rna-XM_046017936.1      rna-XM_059157438.1    rna-XM_059390019.1      mNeoVis.2.1.1.chr7_hap1.g01677.t01              mNeoVis.3.1.1.chr7.g01572.t01    rna-gnl|WGS_ZZZZ|egapxtmp_012804-R1    mNeoVis.evergladensis.2.1.chr7.g01650.t01    mNeoVis.vulgivaga.2.1.chr7.g01667.t01      1111    N0.HOG0001109   OG0000102  n12   rna-XM_011217985.2    rna-XM_038569360.1        rna-XM_036081405.1          rna-XM_032359770.1    rna-XM_059181290.1    rna-XM_059395132.1    rna-XM_045071358.1    mNeoVis.2.1.1.chr7_hap1.g01661.t01          rna-XM_008709657.1      mNeoVis.3.1.1.chr7.g01559.t01      mNeoVis.evergladensis.2.1.chr7.g01634.t01    mNeoVis.vulgivaga.2.1.chr7.g01650.t01      1112    N0.HOG0001110   OG0000102  n23   rna-XM_002926080.3    rna-NM_001388700.1    rna-XM_022521709.1      rna-XM_036087714.1    rna-XM_039224659.1    rna-XM_032852704.1    rna-XM_046017787.1    rna-XM_032359772.1    rna-XM_059157424.1    rna-XM_059390035.1    rna-XM_004782550.2          rna-XM_015542092.2    rna-XM_008709604.2      mNeoVis.3.1.1.chr7.g01571.t01    rna-gnl|WGS_ZZZZ|egapxtmp_012848-R1    mNeoVis.evergladensis.2.1.chr7.g01649.t01    mNeoVis.vulgivaga.2.1.chr7.g01666.t01      1113    N0.HOG0001111   OG0000102  n40   rna-XM_002926102.1    rna-NM_001388703.1      rna-XM_003992888.2        rna-XM_032853639.1    rna-XM_046016949.1    rna-XM_032359773.1    rna-XM_059157474.1    rna-XM_059389990.1    rna-XM_004779059.2    mNeoVis.2.1.1.chr7_hap1.g01680.t01        rna-XM_007092220.2    rna-XM_008709647.1        rna-gnl|WGS_ZZZZ|egapxtmp_012782-R1    mNeoVis.evergladensis.2.1.chr7.g01652.t01    mNeoVis.vulgivaga.2.1.chr7.g01669.t01      1114    N0.HOG0001112   OG0000102  n56   rna-XM_002926078.1      rna-XM_022522071.1    rna-XM_003992863.3      rna-XM_039223760.1    rna-XM_032853640.1    rna-XM_046016747.1    rna-XM_032359668.1    rna-XM_059157486.1    rna-XM_059389980.1    rna-XM_004779016.2    mNeoVis.2.1.1.chr7_hap1.g01678.t01        rna-XM_015542098.2    rna-XM_008709600.1          mNeoVis.evergladensis.2.1.chr7.g01651.t01    mNeoVis.vulgivaga.2.1.chr7.g01670.t01      1115    N0.HOG0001113   OG0000102  n71   rna-XM_011233445.1    rna-NM_001388705.1    rna-XM_022522077.1    rna-XM_003992892.2      rna-XM_039223792.1    rna-XM_032853644.1    rna-XM_046016970.1    rna-XM_032359660.1    rna-XM_059157547.1    rna-XM_059394768.1    rna-XM_004779058.3          rna-XM_007092241.1    rna-XM_008709646.1                1116    N0.HOG0001114   OG0000102  n83   rna-XM_002926076.1, rna-XM_002926077.1    rna-NM_001388956.1    rna-XM_022522074.1, rna-XM_022522078.1    rna-XM_003992864.2, rna-XM_003992891.2      rna-XM_039223761.1, rna-XM_039223785.1    rna-XM_032853642.1    rna-XM_046016754.1, rna-XM_046016967.1    rna-XM_032359626.1, rna-XM_032359655.1    rna-XM_059157523.1, rna-XM_059157534.1    rna-XM_059389954.1, rna-XM_059389969.1    rna-XM_004779012.1, rna-XM_004779013.2    mNeoVis.2.1.1.chr7_hap1.g01683.t01, mNeoVis.2.1.1.chr7_hap1.g01684.t01    rna-XM_021685683.1      rna-XM_007092216.1, rna-XM_007092217.2    rna-XM_008709597.2, rna-XM_008709598.1      mNeoVis.3.1.1.chr7.g01575.t01, mNeoVis.3.1.1.chr7.g01576.t01    rna-gnl|WGS_ZZZZ|egapxtmp_012523-R1, rna-gnl|WGS_ZZZZ|egapxtmp_012825-R1    mNeoVis.evergladensis.2.1.chr7.g01654.t01, mNeoVis.evergladensis.2.1.chr7.g01655.t01    mNeoVis.vulgivaga.2.1.chr7.g01672.t01, mNeoVis.vulgivaga.2.1.chr7.g01673.t01      1117    N0.HOG0001115   OG0000102  n114   rna-XM_011233429.2    rna-XM_038568169.1    rna-XM_022522046.1    rna-XM_003992889.3, rna-XM_011286649.2      rna-XM_039223784.1    rna-XM_032853641.1    rna-XM_046016630.1, rna-XM_046016703.1    rna-XM_032359595.1    rna-XM_059157498.1, rna-XM_059157510.1    rna-XM_059387444.1, rna-XM_059394887.1    rna-XM_004779014.1, rna-XM_004779015.2    mNeoVis.2.1.1.chr7_hap1.g01681.t01, mNeoVis.2.1.1.chr7_hap1.g01682.t01        rna-XM_007092218.2, rna-XM_007092219.2    rna-XM_008709599.1      mNeoVis.3.1.1.chr7.g01574.t01    rna-gnl|WGS_ZZZZ|egapxtmp_012846-R1    mNeoVis.evergladensis.2.1.chr7.g01653.t01    mNeoVis.vulgivaga.2.1.chr7.g01671.t01      1118    N0.HOG0001116   OG0000103  n1   rna-XM_011227054.3, rna-XM_034638066.1, rna-XM_034652157.1, rna-XM_034660110.1    rna-XM_038525552.1, rna-XM_038565384.1    rna-XM_022492255.1, rna-XM_022492256.1, rna-XM_022492739.1, rna-XM_022525009.1, rna-XM_022525592.1    rna-XM_019816358.3, rna-XM_045043587.1, rna-XM_045045722.1, rna-XM_045058467.1    rna-XM_036079195.1, rna-XM_036108522.1, rna-XM_036111010.1, rna-XM_036120129.1, rna-XM_036121997.1    rna-XM_039231952.1, rna-XM_039231953.1, rna-XM_039231956.1, rna-XM_039257541.1, rna-XM_039257582.1    rna-XM_032847795.1, rna-XM_032848331.1, rna-XM_032850825.1, rna-XM_032850963.1, rna-XM_032850964.1, rna-XM_032850965.1, rna-XM_032883459.1    rna-XM_045984898.1, rna-XM_045986803.1, rna-XM_046004875.1, rna-XM_046023202.1    rna-XM_032324150.1, rna-XM_032326060.1, rna-XM_032326084.1, rna-XM_032338232.1    rna-XM_059142763.1, rna-XM_059150684.1, rna-XM_059151209.1, rna-XM_059151221.1, rna-XM_059176233.1    rna-XM_059383359.1, rna-XM_059399677.1, rna-XM_059412223.1, rna-XM_059416531.1    rna-XM_004828401.3, rna-XM_013052792.2, rna-XM_045076996.1, rna-XM_045077278.1, rna-XM_045081724.1    mNeoVis.2.1.1.chr13_hap1.g00342.t01, mNeoVis.2.1.1.chr1_hap1.g01053.t01, mNeoVis.2.1.1.chr7_hap1.g00468.t01    rna-XM_021697185.1, rna-XM_021700821.2, rna-XM_044911346.1, rna-XM_044917241.1      rna-XM_042968074.1, rna-XM_042969974.1, rna-XM_042970015.1, rna-XM_042977251.1, rna-XM_042985496.1, rna-XM_042999286.1    rna-XM_040632781.1, rna-XM_040632878.1, rna-XM_040633557.1, rna-XM_040645164.1    rna-XM_035724052.1, rna-XM_035725015.1    mNeoVis.3.1.1.chr1.g01047.t01, mNeoVis.3.1.1.chr1.g01048.t01, mNeoVis.3.1.1.chr1.g01072.t01, mNeoVis.3.1.1.chr7.g00455.t01, mNeoVis.3.1.1.chr7.g00733.t01, mNeoVis.3.1.1.chr8.g00892.t01    rna-gnl|WGS_ZZZZ|egapxtmp_011766-R1, rna-gnl|WGS_ZZZZ|egapxtmp_029882-R1, rna-gnl|WGS_ZZZZ|egapxtmp_031484-R2, rna-gnl|WGS_ZZZZ|egapxtmp_033908-R1    mNeoVis.evergladensis.2.1.chr1.g01045.t01, mNeoVis.evergladensis.2.1.chr1.g01046.t01, mNeoVis.evergladensis.2.1.chr13.g00332.t01, mNeoVis.evergladensis.2.1.chr13.g00398.t01, mNeoVis.evergladensis.2.1.chr7.g00476.t01, mNeoVis.evergladensis.2.1.chr7.g00488.t01, mNeoVis.evergladensis.2.1.chr7.g00664.t01, mNeoVis.evergladensis.2.1.chr7.g00764.t01    mNeoVis.vulgivaga.2.1.chr1.g01061.t01, mNeoVis.vulgivaga.2.1.chr4.g01095.t01, mNeoVis.vulgivaga.2.1.chr7.g00475.t01, mNeoVis.vulgivaga.2.1.chr7.g00491.t01      1119    N0.HOG0001117   OG0000103  n89     rna-XM_038527557.1                          rna-XM_044911299.1                      1120    N0.HOG0001118   OG0000103  n90   rna-XM_011217823.3, rna-XM_034639125.1, rna-XM_034639397.1    rna-XM_038527954.1    rna-XM_022492097.1, rna-XM_022492253.1, rna-XM_022523766.1    rna-XM_023246258.2, rna-XM_045042475.1    rna-XM_036104091.1      rna-XM_032839048.1, rna-XM_032851126.1, rna-XM_032851184.1, rna-XM_032851196.1    rna-XM_045986828.1, rna-XM_045989562.1    rna-XM_032313758.1, rna-XM_032313759.1, rna-XM_032324058.1    rna-XM_059152425.1    rna-XM_059381967.1, rna-XM_059383370.1, rna-XM_059387504.1, rna-XM_059387519.1, rna-XM_059387526.1, rna-XM_059387594.1    rna-XM_013050684.2, rna-XM_045073828.1, rna-XM_045077356.1, rna-XM_045087806.1      rna-XM_044912520.1    rna-XM_004405696.1, rna-XM_004415493.1    rna-XM_042970051.1    rna-XM_008710954.1, rna-XM_040632937.1    rna-XM_027618315.1    mNeoVis.3.1.1.chr12.g00574.t01, mNeoVis.3.1.1.chr3.g00682.t01, mNeoVis.3.1.1.chr7.g00469.t01, mNeoVis.3.1.1.chr7.g00743.t01    rna-gnl|WGS_ZZZZ|egapxtmp_010584-R1, rna-gnl|WGS_ZZZZ|egapxtmp_010921-R1, rna-gnl|WGS_ZZZZ|egapxtmp_028882-R1    mNeoVis.evergladensis.2.1.chr7.g00791.t01    mNeoVis.vulgivaga.2.1.chr7.g00797.t01      1121    N0.HOG0001119   OG0000104  n1   rna-XM_011221732.3    rna-XM_038583093.1    rna-XM_022501084.1, rna-XM_022515092.1    rna-XM_045045522.1, rna-XM_045062035.1    rna-XM_036069445.1, rna-XM_036072551.1, rna-XM_036083614.1, rna-XM_036094606.1, rna-XM_036102668.1, rna-XM_036117152.1, rna-XM_036121643.1    rna-XM_039217819.1, rna-XM_039220704.1, rna-XM_039226728.1, rna-XM_039228121.1, rna-XM_039229884.1, rna-XM_039232019.1, rna-XM_039238209.1, rna-XM_039238866.1, rna-XM_039239617.1, rna-XM_039245363.1, rna-XM_039249736.1, rna-XM_039250109.1, rna-XM_039250436.1, rna-XM_039253033.1, rna-XM_039256220.1    rna-XM_032848960.1, rna-XM_032869434.1, rna-XM_032872142.1, rna-XM_032875102.1    rna-XM_045978242.1, rna-XM_046002434.1    rna-XM_032317872.1, rna-XM_032330092.1, rna-XM_032336098.1, rna-XM_032338606.1, rna-XM_032345619.1    rna-XM_059162978.1, rna-XM_059183991.1, rna-XM_059187458.1    rna-XM_059391143.1, rna-XM_059392816.1    rna-XM_004745391.3, rna-XM_045066467.1, rna-XM_045069606.1, rna-XM_045073217.1    mNeoVis.2.1.1.chr15_hap1.g00621.t01    rna-XM_021692427.1, rna-XM_044911825.1, rna-XM_044918098.1    rna-XM_004391619.1, rna-XM_004391907.1, rna-XM_004400569.1, rna-XM_004400770.1    rna-XM_015544811.1, rna-XM_042957110.1, rna-XM_042969105.1, rna-XM_042992683.1    rna-XM_008708413.2    rna-XM_027587185.2, rna-XM_027601472.1, rna-XM_035721871.1, rna-XM_035722286.1, rna-XM_035722437.1, rna-XM_035723895.1, rna-XM_035726494.1, rna-XM_035727370.1, rna-XM_035727797.1, rna-XM_035727856.1        mNeoVis.evergladensis.2.1.chr15.g00443.t01        1122    N0.HOG0001120   OG0000104  n39   rna-XM_034661025.1    rna-XM_038571346.1      rna-XM_003991729.5, rna-XM_011280867.3, rna-XM_045033444.1, rna-XM_045034127.1    rna-XM_036080835.1, rna-XM_036087063.1, rna-XM_036092668.1    rna-XM_039216502.1, rna-XM_039216732.1, rna-XM_039219684.1, rna-XM_039220171.1, rna-XM_039228078.1, rna-XM_039245604.1, rna-XM_039251369.1, rna-XM_039254543.1, rna-XM_039255142.1    rna-XM_032837452.1, rna-XM_032880939.1    rna-XM_046008912.1    rna-XM_032315481.1, rna-XM_032341332.1, rna-XM_032341497.1, rna-XM_032345586.1    rna-XM_059176331.1    rna-XM_059399215.1    rna-XM_045067714.1    mNeoVis.2.1.1.chr10_hap1.g00497.t01, mNeoVis.2.1.1.chr13_hap1.g00511.t01, mNeoVis.2.1.1.chr2_hap1.g00819.t01, mNeoVis.2.1.1.chr6_hap1.g00660.t01, mNeoVis.2.1.1.chr7_hap1.g00959.t01, mNeoVis.2.1.1.chr9_hap1.g01099.t01    rna-XM_044913830.1, rna-XM_044918891.1    rna-XM_004393782.1    rna-XM_042993985.1      rna-XM_035722203.1, rna-XM_035722438.1, rna-XM_035724509.1, rna-XM_035724510.1, rna-XM_035726878.1, rna-XM_035727231.1, rna-XM_035727763.1, rna-XM_035728196.1, rna-XM_035728524.1, rna-XM_035728946.1, rna-XM_035728966.1, rna-XM_035729011.1    mNeoVis.3.1.1.chr2.g01143.t01, mNeoVis.3.1.1.chr4.g00668.t01, mNeoVis.3.1.1.chr5.g00766.t01, mNeoVis.3.1.1.chr9.g00459.t01    rna-gnl|WGS_ZZZZ|egapxtmp_006209-R1, rna-gnl|WGS_ZZZZ|egapxtmp_008189-R1, rna-gnl|WGS_ZZZZ|egapxtmp_012349-R1, rna-gnl|WGS_ZZZZ|egapxtmp_014728-R1, rna-gnl|WGS_ZZZZ|egapxtmp_036194-R1    mNeoVis.evergladensis.2.1.chr10.g01088.t01, mNeoVis.evergladensis.2.1.chr13.g00492.t01, mNeoVis.evergladensis.2.1.chr5.g00776.t01, mNeoVis.evergladensis.2.1.chr7.g00952.t01, mNeoVis.evergladensis.2.1.chr8.g00477.t01    mNeoVis.vulgivaga.2.1.chr10.g01078.t01, mNeoVis.vulgivaga.2.1.chr13.g00497.t01, mNeoVis.vulgivaga.2.1.chr2.g01071.t01, mNeoVis.vulgivaga.2.1.chr5.g00769.t01, mNeoVis.vulgivaga.2.1.chr7.g00958.t01, mNeoVis.vulgivaga.2.1.chr9.g00480.t01      1123    N0.HOG0001121   OG0000105  n1   rna-XM_034640621.1    rna-XM_038571808.1    rna-XM_022493970.1    rna-XM_045053803.1    rna-XM_036093280.1    rna-XM_039237673.1    rna-XM_032859025.1    rna-XM_045980649.1    rna-XM_032351209.1    rna-XM_059133741.1    rna-XM_059386455.1, rna-XM_059406673.1    rna-XM_004772926.3    mNeoVis.2.1.1.chr11_hap1.g00394.t01    rna-XM_044918664.1    rna-XM_004398138.1    rna-XM_042980757.1    rna-XM_040638084.1    rna-XM_027621254.2    mNeoVis.3.1.1.chr11.g00360.t01    rna-gnl|WGS_ZZZZ|egapxtmp_025176-R1    mNeoVis.evergladensis.2.1.chr11.g00390.t01    mNeoVis.vulgivaga.2.1.chr11.g00395.t01      1124    N0.HOG0001122   OG0000105  n23       rna-XM_022492569.1, rna-XM_022492570.1, rna-XM_022494723.1          rna-XM_032859022.1, rna-XM_032859023.1    rna-XM_045980934.1, rna-XM_045981627.1    rna-XM_032351169.1, rna-XM_032351206.1, rna-XM_032351208.1    rna-XM_059133261.1, rna-XM_059133714.1, rna-XM_059133722.1, rna-XM_059133732.1, rna-XM_059133735.1    rna-XM_059386456.1, rna-XM_059406674.1, rna-XM_059406678.1, rna-XM_059406680.1, rna-XM_059407929.1    rna-XM_004772929.3, rna-XM_045079459.1, rna-XM_045079696.1, rna-XM_045079859.1, rna-XM_045080085.1, rna-XM_045083994.1    mNeoVis.2.1.1.chr11_hap1.g00389.t01, mNeoVis.2.1.1.chr11_hap1.g00390.t01, mNeoVis.2.1.1.chr11_hap1.g00391.t01, mNeoVis.2.1.1.chr11_hap1.g00392.t01              mNeoVis.3.1.1.chr11.g00355.t01, mNeoVis.3.1.1.chr11.g00356.t01, mNeoVis.3.1.1.chr11.g00357.t01    rna-gnl|WGS_ZZZZ|egapxtmp_025011-R1    mNeoVis.evergladensis.2.1.chr11.g00384.t01, mNeoVis.evergladensis.2.1.chr11.g00385.t01, mNeoVis.evergladensis.2.1.chr11.g00386.t01, mNeoVis.evergladensis.2.1.chr11.g00387.t01, mNeoVis.evergladensis.2.1.chr11.g00388.t01    mNeoVis.vulgivaga.2.1.chr11.g00389.t01, mNeoVis.vulgivaga.2.1.chr11.g00390.t01, mNeoVis.vulgivaga.2.1.chr11.g00391.t01, mNeoVis.vulgivaga.2.1.chr11.g00392.t01, mNeoVis.vulgivaga.2.1.chr11.g00393.t01      1125    N0.HOG0001123   OG0000105  n67   rna-XM_034650500.1          rna-XM_036093290.1, rna-XM_036093299.1                    rna-XM_044919031.1    rna-XM_004398216.1, rna-XM_004417652.1      rna-XM_008703283.2                1126    N0.HOG0001124   OG0000105  n74     rna-XM_038571814.1, rna-XM_038571819.1      rna-XM_023251382.2, rna-XM_045053800.1, rna-XM_045053801.1      rna-XM_039237698.1, rna-XM_039237704.1, rna-XM_039237705.1, rna-XM_039237794.1, rna-XM_039237816.1                      rna-XM_042980740.1, rna-XM_042980750.1                  1127    N0.HOG0001125   OG0000105  n87   rna-XM_019793828.2    rna-XM_038571820.1    rna-XM_022494721.1              rna-XM_032351200.1      rna-XM_059406682.1    rna-XM_004746765.2    mNeoVis.2.1.1.chr11_hap1.g00388.t01      rna-XM_004416483.1      rna-XM_040638663.1    rna-XM_027623806.2    mNeoVis.3.1.1.chr11.g00353.t01    rna-gnl|WGS_ZZZZ|egapxtmp_025009-R1    mNeoVis.evergladensis.2.1.chr11.g00383.t01    mNeoVis.vulgivaga.2.1.chr11.g00388.t01      1128    N0.HOG0001126   OG0000105  n102   rna-XM_011218021.3, rna-XM_034640216.1    rna-XM_038571822.1    rna-XM_022494397.1, rna-XM_022494523.1    rna-XM_045053797.1    rna-XM_036093198.1    rna-XM_039237774.1    rna-XM_032859018.1, rna-XM_032859021.1    rna-XM_045980560.1, rna-XM_045980942.1    rna-XM_032351197.1, rna-XM_032351202.1    rna-XM_059132967.1, rna-XM_059133711.1    rna-XM_059406683.1, rna-XM_059407673.1    rna-XM_013053450.2, rna-XM_045083995.1    mNeoVis.2.1.1.chr11_hap1.g00386.t01, mNeoVis.2.1.1.chr11_hap1.g00387.t01    rna-XM_021689329.1    rna-XM_004416482.1    rna-XM_042980743.1      rna-XM_027623812.2      rna-gnl|WGS_ZZZZ|egapxtmp_025012-R8    mNeoVis.evergladensis.2.1.chr11.g00382.t01    mNeoVis.vulgivaga.2.1.chr11.g00386.t01, mNeoVis.vulgivaga.2.1.chr11.g00387.t01      1129    N0.HOG0001127   OG0000105  n131   rna-XM_034641525.1    rna-XM_038571827.1        rna-XM_036082399.1                    rna-XM_021689307.1    rna-XM_004393179.1      rna-XM_008703278.2    rna-XM_027624255.2, rna-XM_035729029.1              1130    N0.HOG0001128   OG0000105  n138         rna-XM_045053802.1      rna-XM_039237815.1                  rna-XM_044919030.1    rna-XM_004416826.1                    1131    N0.HOG0001129   OG0000106  n2   rna-XM_002924184.4    rna-NM_001388568.1    rna-XM_022510107.1, rna-XM_022510433.1    rna-XM_003983180.4    rna-XM_036087249.1, rna-XM_036087253.1    rna-XM_039243069.1    rna-XM_032866279.1, rna-XM_032866943.1    rna-XM_046021218.1    rna-XM_032305818.1, rna-XM_032305819.1    rna-XM_059172698.1    rna-XM_059395577.1    rna-XM_013045637.1    mNeoVis.2.1.1.chr4_hap1.g00099.t01    rna-XM_044920019.1    rna-XM_004414594.1    rna-XM_007090895.1    rna-XM_040643874.1    rna-XM_027574703.2, rna-XM_027574957.2    mNeoVis.3.1.1.chr6.g00099.t01    rna-gnl|WGS_ZZZZ|egapxtmp_002856-R1, rna-gnl|WGS_ZZZZ|egapxtmp_003135-R1    mNeoVis.evergladensis.2.1.chr6.g00100.t01, mNeoVis.evergladensis.2.1.chr6.g00101.t01        1132    N0.HOG0001130   OG0000106  n31   rna-XM_002921819.4    rna-NM_001388869.1    rna-XM_022510105.1    rna-XM_003983183.2    rna-XM_036087246.1    rna-XM_039243070.1    rna-XM_032866254.1    rna-XM_046021291.1    rna-XM_032306037.1    rna-XM_059172704.1      rna-XM_004772100.1    mNeoVis.2.1.1.chr4_hap1.g00092.t01      rna-XM_004409611.1    rna-XM_007090897.1    rna-XM_040643870.1    rna-XM_027574271.2    mNeoVis.3.1.1.chr6.g00093.t01    rna-gnl|WGS_ZZZZ|egapxtmp_003136-R1    mNeoVis.evergladensis.2.1.chr6.g00095.t01    mNeoVis.vulgivaga.2.1.chr6.g00092.t01      1133    N0.HOG0001131   OG0000106  n47   rna-XM_002921830.2    rna-NM_001388868.1    rna-XM_022510399.1    rna-XM_003983174.2      rna-XM_039243071.1    rna-XM_032865123.1    rna-XM_046021290.1    rna-XM_032306025.1    rna-XM_059172705.1    rna-XM_059395573.1    rna-XM_004772101.2    mNeoVis.2.1.1.chr4_hap1.g00091.t01      rna-XM_004409610.1    rna-XM_007090898.1    rna-XM_040643869.1    rna-XM_027574270.2    mNeoVis.3.1.1.chr6.g00092.t01    rna-gnl|WGS_ZZZZ|egapxtmp_003134-R1    mNeoVis.evergladensis.2.1.chr6.g00094.t01    mNeoVis.vulgivaga.2.1.chr6.g00091.t01      1134    N0.HOG0001132   OG0000106  n66   rna-XM_002921820.4    rna-XM_038559212.1    rna-XM_022510398.1    rna-XM_003983173.2    rna-XM_036087237.1    rna-XM_039243062.1    rna-XM_032865136.1    rna-XM_046021269.1    rna-XM_032305824.1    rna-XM_059172703.1    rna-XM_059395574.1    rna-XM_004772078.1    mNeoVis.2.1.1.chr4_hap1.g00093.t01    rna-XM_021692975.1    rna-XM_004409612.1    rna-XM_007090853.2    rna-XM_008697285.2    rna-XM_027574702.2      rna-gnl|WGS_ZZZZ|egapxtmp_003131-R1      mNeoVis.vulgivaga.2.1.chr6.g00093.t01      1135    N0.HOG0001133   OG0000106  n87   rna-XM_002924173.4, rna-XM_019804790.2    rna-XM_038559213.1      rna-XM_023250678.2, rna-XM_045041951.1      rna-XM_039243047.1, rna-XM_039243048.1    rna-XM_032865153.1, rna-XM_032865159.1    rna-XM_046021311.1, rna-XM_046021320.1    rna-XM_032305823.1, rna-XM_032305992.1    rna-XM_059172699.1, rna-XM_059172700.1    rna-XM_059387261.1, rna-XM_059395576.1    rna-XM_004772075.1, rna-XM_004772076.1    mNeoVis.2.1.1.chr4_hap1.g00095.t01, mNeoVis.2.1.1.chr4_hap1.g00096.t01        rna-XM_007090850.2, rna-XM_007090852.1    rna-XM_008697287.2, rna-XM_040643868.1      mNeoVis.3.1.1.chr6.g00095.t01    rna-gnl|WGS_ZZZZ|egapxtmp_003112-R1    mNeoVis.evergladensis.2.1.chr6.g00097.t01    mNeoVis.vulgivaga.2.1.chr6.g00095.t01, mNeoVis.vulgivaga.2.1.chr6.g00096.t01      1136    N0.HOG0001134   OG0000106  n116     rna-NM_001389074.1              rna-XM_046020235.1                                  1137    N0.HOG0001135   OG0000106  n117   rna-XM_002921821.4    rna-NM_001017516.1    rna-XM_022510368.1    rna-XM_045052923.1    rna-XM_036087236.1    rna-XM_039243080.1    rna-XM_032865143.1      rna-XM_032305822.1    rna-XM_059172701.1      rna-XM_004772077.2    mNeoVis.2.1.1.chr4_hap1.g00094.t01    rna-XM_021692979.1, rna-XM_044920020.1    rna-XM_004409605.1    rna-XM_007090851.2    rna-XM_008697286.2    rna-XM_027574268.2    mNeoVis.3.1.1.chr6.g00094.t01    rna-gnl|WGS_ZZZZ|egapxtmp_003102-R1    mNeoVis.evergladensis.2.1.chr6.g00096.t01    mNeoVis.vulgivaga.2.1.chr6.g00094.t01      1138    N0.HOG0001136   OG0000107  n1   rna-XM_002923464.4    rna-XM_038539159.1    rna-XM_022501524.1    rna-XM_003998539.6    rna-XM_036072418.1    rna-XM_039243514.1    rna-XM_032872468.1    rna-XM_045993733.1    rna-XM_032327321.1    rna-XM_059154588.1    rna-XM_059414654.1    rna-XM_013061034.2    mNeoVis.2.1.1.chr15_hap1.g00718.t01    rna-XM_021700489.2    rna-XM_004399078.1    rna-XM_042970822.1    rna-XM_040634906.1    rna-XM_027611704.2    mNeoVis.3.1.1.chr15.g00495.t01    rna-gnl|WGS_ZZZZ|egapxtmp_033662-R1    mNeoVis.evergladensis.2.1.chr15.g00540.t01, mNeoVis.evergladensis.2.1.chr15.g00541.t01    mNeoVis.vulgivaga.2.1.chr15.g00553.t01      1139    N0.HOG0001137   OG0000107  n7   rna-XM_011220238.3    rna-XM_038575750.1    rna-XM_022522868.1    rna-XM_023241551.2    rna-XM_036074625.1    rna-XM_039225912.1    rna-XM_032883227.1    rna-XM_046025586.1    rna-XM_032310317.1    rna-XM_059139754.1    rna-XM_059408918.1    rna-XM_004766358.3    mNeoVis.2.1.1.chr3_hap1.g00392.t01    rna-XM_021703618.2    rna-XM_004400006.1    rna-XM_042962318.1    rna-XM_040619772.1    rna-XM_027576682.2    mNeoVis.3.1.1.chr3.g00384.t01    rna-gnl|WGS_ZZZZ|egapxtmp_041076-R1    mNeoVis.evergladensis.2.1.chr3.g00399.t01    mNeoVis.vulgivaga.2.1.chr3.g00393.t01      1140    N0.HOG0001138   OG0000107  n12   rna-XM_034641934.1    rna-XM_038572424.1    rna-XM_022493884.1    rna-XM_045053573.1    rna-XM_036072057.1, rna-XM_036111184.1    rna-XM_039248469.1    rna-XM_032838639.1    rna-XM_045980912.1    rna-XM_032350566.1    rna-XM_059133066.1    rna-XM_059387491.1, rna-XM_059407081.1    rna-XM_045082166.1, rna-XM_045087794.1    mNeoVis.2.1.1.chr11_hap1.g00203.t01    rna-XM_021688888.2, rna-XM_044916679.1    rna-XM_004414105.1    rna-XM_042980320.1    rna-XM_040638382.1    rna-XM_027621565.2, rna-XM_035727681.1    mNeoVis.3.1.1.chr11.g00172.t01    rna-gnl|WGS_ZZZZ|egapxtmp_027414-R1    mNeoVis.evergladensis.2.1.chr11.g00200.t01    mNeoVis.vulgivaga.2.1.chr11.g00203.t01      1141    N0.HOG0001139   OG0000107  n25                                     rna-XM_035727980.1    mNeoVis.3.1.1.chr11.g01116.t01            1142    N0.HOG0001140   OG0000107  n27     rna-XM_038535519.1                        mNeoVis.2.1.1.chr13_hap1.g00343.t01      rna-XM_004405720.1          mNeoVis.3.1.1.chr13.g00285.t01    rna-gnl|WGS_ZZZZ|egapxtmp_031284-R1      mNeoVis.vulgivaga.2.1.chr11.g01169.t01, mNeoVis.vulgivaga.2.1.chr13.g00331.t01      1143    N0.HOG0001141   OG0000107  n32   rna-XM_034659925.1    rna-XM_038560751.1    rna-XM_022513528.1    rna-XM_003984508.6    rna-XM_036097545.1    rna-XM_039231054.1    rna-XM_032866825.1    rna-XM_045979881.1    rna-XM_032353364.1    rna-XM_059132684.1    rna-XM_059405132.1    rna-XM_004745775.3    mNeoVis.2.1.1.chr11_hap1.g01173.t01    rna-XM_021697743.1    rna-XM_004401798.1    rna-XM_042982522.1    rna-XM_040622816.1    rna-XM_027621464.2      rna-gnl|WGS_ZZZZ|egapxtmp_024979-R1          1144    N0.HOG0001142   OG0000107  n36   rna-XM_011237078.3    rna-XM_038555215.1    rna-XM_022502235.1    rna-XM_006943327.5    rna-XM_036072723.1    rna-XM_039233782.1    rna-XM_032858579.1    rna-XM_046005479.1    rna-XM_032315551.1    rna-XM_059165905.1    rna-XM_059394935.1    rna-XM_004768885.3      rna-XM_021688701.2    rna-XM_004402344.1    rna-XM_042972625.1    rna-XM_040625717.1    rna-XM_027571855.2    mNeoVis.3.1.1.chr6.g00925.t01            1145    N0.HOG0001143   OG0000107  n37     rna-XM_038559174.1, rna-XM_038562614.1, rna-XM_038579241.1        rna-XM_036101002.1          rna-XM_032333312.1    rna-XM_059140201.1, rna-XM_059140202.1, rna-XM_059140203.1, rna-XM_059140228.1          rna-XM_044917327.1    rna-XM_004406798.1      rna-XM_040629655.1                1146    N0.HOG0001144   OG0000107  n47                           mNeoVis.2.1.1.chr6_hap1.g00913.t01                rna-gnl|WGS_ZZZZ|egapxtmp_001256-R1    mNeoVis.evergladensis.2.1.chr6.g00918.t01    mNeoVis.vulgivaga.2.1.chr4.g00884.t01      1147    N0.HOG0001145   OG0000107  n49         rna-XM_045042315.1                          rna-XM_015536806.2      rna-XM_035728015.1      rna-gnl|WGS_ZZZZ|egapxtmp_004982-R1          1148    N0.HOG0001146   OG0000108  n1   rna-XM_019806504.2    rna-NM_001007142.2    rna-XM_022526394.1, rna-XM_022526408.1    rna-XM_003998628.5    rna-XM_036115356.1    rna-XM_039238069.1    rna-XM_032842703.1    rna-XM_045993187.1    rna-XM_032326705.1    rna-XM_059155185.1    rna-XM_059414797.1    rna-XM_004773798.3    mNeoVis.2.1.1.chr15_hap1.g00003.t01    rna-XM_021700612.1    rna-XM_004415857.1    rna-XM_007094187.3    rna-XM_040635026.1    rna-XM_035722224.1    mNeoVis.3.1.1.chr15.g00003.t01    rna-gnl|WGS_ZZZZ|egapxtmp_034019-R1    mNeoVis.evergladensis.2.1.chr15.g00003.t01    mNeoVis.vulgivaga.2.1.chr15.g00003.t01      1149    N0.HOG0001147   OG0000108  n23   rna-XM_034670235.1    rna-XM_038540013.1    rna-XM_022501867.1    rna-XM_003998755.5    rna-XM_036112499.1    rna-XM_039234408.1    rna-XM_032872870.1    rna-XM_045993082.1    rna-XM_032326922.1    rna-XM_059155366.1    rna-XM_059415162.1    rna-XM_004778344.3    mNeoVis.2.1.1.chr15_hap1.g00161.t01    rna-XM_021686859.1    rna-XM_004398791.1    rna-XM_007084087.3    rna-XM_008697149.2    rna-XM_027613261.1    mNeoVis.3.1.1.chr15.g00140.t01    rna-gnl|WGS_ZZZZ|egapxtmp_033548-R1    mNeoVis.evergladensis.2.1.chr15.g00147.t01    mNeoVis.vulgivaga.2.1.chr15.g00150.t01      1150    N0.HOG0001148   OG0000108  n45   rna-XM_002927839.4    rna-XM_038558107.1    rna-XM_022496803.1    rna-XM_003990064.6    rna-XM_036106597.1    rna-XM_039224652.1    rna-XM_032846666.1    rna-XM_046016798.1    rna-XM_032302925.1    rna-XM_059136949.1    rna-XM_059374663.1    rna-XM_004740356.3    mNeoVis.2.1.1.chr2_hap1.g01315.t01    rna-XM_044914503.1    rna-XM_004413157.1    rna-XM_042996088.1    rna-XM_008699232.1    rna-XM_027612612.2    mNeoVis.3.1.1.chr5.g01240.t01    rna-gnl|WGS_ZZZZ|egapxtmp_005854-R1    mNeoVis.evergladensis.2.1.chr5.g01256.t01    mNeoVis.vulgivaga.2.1.chr5.g01252.t01      1151    N0.HOG0001149   OG0000108  n66   rna-XM_034647288.1, rna-XM_034647289.1    rna-XM_038537072.1    rna-XM_022491432.1    rna-XM_003996333.3    rna-XM_036073456.1    rna-XM_039227363.1    rna-XM_032841315.1    rna-XM_045983734.1    rna-XM_032319583.1    rna-XM_059147184.1    rna-XM_059380416.1    rna-XM_045083662.1    mNeoVis.2.1.1.chr8_hap1.g00879.t01    rna-XM_021694485.1    rna-XM_004405131.1    rna-XM_042965926.1    rna-XM_040628340.1, rna-XM_040628341.1, rna-XM_040630131.1    rna-XM_027625020.2    mNeoVis.3.1.1.chr8.g00826.t01    rna-gnl|WGS_ZZZZ|egapxtmp_019251-R1    mNeoVis.evergladensis.2.1.chr9.g00867.t01    mNeoVis.vulgivaga.2.1.chr8.g00869.t01      1152    N0.HOG0001150   OG0000108  n91     rna-XM_038574509.1      rna-XM_045042328.1    rna-XM_036077737.1              rna-XM_059409913.1            rna-XM_007092298.2      rna-XM_035723990.1              1153    N0.HOG0001151   OG0000108  n97   rna-XM_034638771.1    rna-XM_038574527.1    rna-XM_022522878.1    rna-XM_003994827.4    rna-XM_036074523.1    rna-XM_039225900.1    rna-XM_032883254.1    rna-XM_046025494.1    rna-XM_032311155.1    rna-XM_059140326.1    rna-XM_059409911.1    rna-XM_004766355.2    mNeoVis.2.1.1.chr3_hap1.g00390.t01    rna-XM_044921034.1      rna-XM_042961976.1    rna-XM_008710581.2    rna-XM_027577003.2    mNeoVis.3.1.1.chr3.g00382.t01    rna-gnl|WGS_ZZZZ|egapxtmp_040894-R1    mNeoVis.evergladensis.2.1.chr3.g00397.t01    mNeoVis.vulgivaga.2.1.chr3.g00391.t01      1154    N0.HOG0001152   OG0000108  n116   rna-XM_011220146.3    rna-NM_001007141.1    rna-XM_022522847.1    rna-NM_001195405.1    rna-XM_036074626.1    rna-XM_039225885.1    rna-XM_032883228.1    rna-XM_046024603.1    rna-XM_032311020.1    rna-XM_059140096.1    rna-XM_059409910.1    rna-XM_045068227.1    mNeoVis.2.1.1.chr3_hap1.g00391.t01    rna-XM_044921247.1    rna-XM_004400004.1    rna-XM_007082583.3    rna-XM_008710584.2    rna-XM_027577412.2    mNeoVis.3.1.1.chr3.g00383.t01    rna-gnl|WGS_ZZZZ|egapxtmp_041925-R1    mNeoVis.evergladensis.2.1.chr3.g00398.t01    mNeoVis.vulgivaga.2.1.chr3.g00392.t01      1155    N0.HOG0001153   OG0000109  n2   rna-XM_002926104.1, rna-XM_011233447.3, rna-XM_011233448.3    rna-NM_001388702.1, rna-XM_038568167.1    rna-XM_022521708.1    rna-XM_003992862.2, rna-XM_003992887.5, rna-XM_006937035.5    rna-XM_036087713.1, rna-XM_036087719.1    rna-XM_039223789.1, rna-XM_039224673.1, rna-XM_039224675.1    rna-XM_032853638.1    rna-XM_046015899.1, rna-XM_046017262.1    rna-XM_032356446.1, rna-XM_032356898.1    rna-XM_059157451.1, rna-XM_059157464.1    rna-XM_059390005.1    rna-XM_004779017.2, rna-XM_004779060.3    mNeoVis.2.1.1.chr7_hap1.g01679.t01    rna-XM_021685632.2      rna-XM_007092221.1, rna-XM_007092223.2, rna-XM_007092242.2    rna-XM_008709602.2, rna-XM_008709648.2, rna-XM_008709649.2    rna-XM_027577904.1    mNeoVis.3.1.1.chr7.g01569.t01, mNeoVis.3.1.1.chr7.g01573.t01    rna-gnl|WGS_ZZZZ|egapxtmp_012522-R1, rna-gnl|WGS_ZZZZ|egapxtmp_012805-R1      mNeoVis.vulgivaga.2.1.chr7.g01668.t01      1156    N0.HOG0001154   OG0000109  n40   rna-XM_034650074.1, rna-XM_034650121.1    rna-XM_038569353.1    rna-XM_022521729.1    rna-XM_003992836.2    rna-XM_036114426.1    rna-XM_039223164.1    rna-XM_032853566.1    rna-XM_046015921.1    rna-XM_032359732.1, rna-XM_032359735.1    rna-XM_059178327.1, rna-XM_059178591.1    rna-XM_059387367.1    rna-XM_004768238.3, rna-XM_004782228.2    mNeoVis.2.1.1.chr7_hap1.g01623.t01        rna-XM_007089405.1    rna-XM_008710701.1    rna-XM_027581101.1    mNeoVis.3.1.1.chr7.g01527.t01      mNeoVis.evergladensis.2.1.chr7.g01594.t01        1157    N0.HOG0001155   OG0000109  n61   rna-XM_002931012.2, rna-XM_034650088.1    rna-NM_001388847.1    rna-XM_022522013.1    rna-XM_003992799.3    rna-XM_036114416.1    rna-XM_039223173.1    rna-XM_032853602.1    rna-XM_046016300.1    rna-XM_032359341.1, rna-XM_032359935.1    rna-XM_059156735.1, rna-XM_059156810.1    rna-XM_059387332.1    rna-XM_045071470.1, rna-XM_045071484.1    mNeoVis.2.1.1.chr7_hap1.g01622.t01        rna-XM_007089347.1    rna-XM_008710700.1      mNeoVis.3.1.1.chr7.g01526.t01    rna-gnl|WGS_ZZZZ|egapxtmp_012832-R1    mNeoVis.evergladensis.2.1.chr7.g01593.t01    mNeoVis.vulgivaga.2.1.chr7.g01612.t01      1158    N0.HOG0001156   OG0000109  n81   rna-XM_011217515.3, rna-XM_034667606.1    rna-XM_038569355.1    rna-XM_022521716.1    rna-XM_045038090.1    rna-XM_036081419.1      rna-XM_032852686.1    rna-XM_046015911.1    rna-XM_032359754.1    rna-XM_059179593.1    rna-XM_059396207.1    rna-XM_004768259.1    mNeoVis.2.1.1.chr7_hap1.g01644.t01    rna-XM_021685623.1    rna-XM_004407573.1      rna-XM_008710761.2      mNeoVis.3.1.1.chr7.g01546.t01    rna-gnl|WGS_ZZZZ|egapxtmp_012776-R2      mNeoVis.vulgivaga.2.1.chr7.g01634.t01      1159    N0.HOG0001157   OG0000109  n99         rna-XM_011286663.2, rna-XM_023238990.2      rna-XM_039225376.1, rna-XM_039225381.1                      rna-XM_007089394.2, rna-XM_007089395.3                  1160    N0.HOG0001158   OG0000109  n105   rna-XM_034650119.1, rna-XM_034650183.1    rna-NM_001389067.1    rna-XM_022522098.1, rna-XM_022522194.1      rna-XM_036081404.1, rna-XM_036081420.1      rna-XM_032853618.1    rna-XM_046017541.1, rna-XM_046017680.1    rna-XM_032356455.1, rna-XM_032359755.1    rna-XM_059157051.1, rna-XM_059157066.1    rna-XM_059390350.1, rna-XM_059390365.1    rna-XM_004768258.1, rna-XM_004768260.2    mNeoVis.2.1.1.chr7_hap1.g01643.t01    rna-XM_044919657.1        rna-XM_008710724.2, rna-XM_008710725.1      mNeoVis.3.1.1.chr7.g01545.t01    rna-gnl|WGS_ZZZZ|egapxtmp_012910-R1    mNeoVis.evergladensis.2.1.chr7.g01617.t01, mNeoVis.evergladensis.2.1.chr7.g01619.t01    mNeoVis.vulgivaga.2.1.chr7.g01633.t01      1161    N0.HOG0001159   OG0000109  n131     rna-NM_001389073.1                        mNeoVis.2.1.1.chr7_hap1.g01645.t01                    mNeoVis.vulgivaga.2.1.chr7.g01635.t01      1162    N0.HOG0001160   OG0000110  n0   rna-XM_034670299.1    rna-XM_038540172.1, rna-XM_038540188.1      rna-XM_023246914.2, rna-XM_045038555.1, rna-XM_045038560.1, rna-XM_045038562.1, rna-XM_045038570.1, rna-XM_045043097.1, rna-XM_045048163.1, rna-XM_045051391.1, rna-XM_045051398.1, rna-XM_045051406.1, rna-XM_045051426.1, rna-XM_045051427.1, rna-XM_045051477.1, rna-XM_045051536.1, rna-XM_045053147.1, rna-XM_045053166.1, rna-XM_045053173.1, rna-XM_045053178.1, rna-XM_045053183.1, rna-XM_045060648.1    rna-XM_036064712.1, rna-XM_036067282.1, rna-XM_036073025.1, rna-XM_036079184.1, rna-XM_036080820.1, rna-XM_036112053.1, rna-XM_036115234.1, rna-XM_036123648.1    rna-XM_039225448.1    rna-XM_032836156.1, rna-XM_032842698.1, rna-XM_032844027.1, rna-XM_032850857.1    rna-XM_045992717.1, rna-XM_045992718.1, rna-XM_045992722.1, rna-XM_045992724.1, rna-XM_045992727.1, rna-XM_045992728.1, rna-XM_045992734.1, rna-XM_045992738.1, rna-XM_045992739.1, rna-XM_045992741.1, rna-XM_045992951.1, rna-XM_045992952.1, rna-XM_045992953.1, rna-XM_045992955.1, rna-XM_045992956.1, rna-XM_045992957.1, rna-XM_045993872.1, rna-XM_045993874.1, rna-XM_045994187.1, rna-XM_045996859.1, rna-XM_045996893.1    rna-XM_032327485.1, rna-XM_032327489.1, rna-XM_032327497.1, rna-XM_032327652.1, rna-XM_032328622.1, rna-XM_032328623.1, rna-XM_032328642.1    rna-XM_059154637.1, rna-XM_059155327.1, rna-XM_059155328.1, rna-XM_059155514.1, rna-XM_059155537.1, rna-XM_059155539.1, rna-XM_059155541.1, rna-XM_059155716.1, rna-XM_059155746.1    rna-XM_059387198.1, rna-XM_059387530.1, rna-XM_059387588.1    rna-XM_045071908.1, rna-XM_045071913.1, rna-XM_045071916.1, rna-XM_045072060.1, rna-XM_045072735.1, rna-XM_045075206.1, rna-XM_045075212.1, rna-XM_045075222.1, rna-XM_045075223.1, rna-XM_045078485.1, rna-XM_045078488.1, rna-XM_045078947.1, rna-XM_045078948.1, rna-XM_045082928.1, rna-XM_045082929.1, rna-XM_045087979.1    mNeoVis.2.1.1.chr10_hap1.g01108.t01, mNeoVis.2.1.1.chr10_hap1.g01109.t01, mNeoVis.2.1.1.chr15_hap1.g00179.t01, mNeoVis.2.1.1.chr15_hap1.g00180.t01, mNeoVis.2.1.1.chr15_hap1.g00181.t01, mNeoVis.2.1.1.chr15_hap1.g00183.t01, mNeoVis.2.1.1.chr15_hap1.g00531.t01, mNeoVis.2.1.1.chr3_hap1.g00703.t01, mNeoVis.2.1.1.chr3_hap1.g00705.t01, mNeoVis.2.1.1.chr3_hap1.g00707.t01, mNeoVis.2.1.1.chr5_hap1.g01025.t01, mNeoVis.2.1.1.chr6_hap1.g00887.t01, mNeoVis.2.1.1.chr6_hap1.g00890.t01, mNeoVis.2.1.1.chr6_hap1.g00891.t01, mNeoVis.2.1.1.chr6_hap1.g00893.t01, mNeoVis.2.1.1.chr6_hap1.g00896.t01, mNeoVis.2.1.1.chr6_hap1.g00897.t01, mNeoVis.2.1.1.chr6_hap1.g00899.t01, mNeoVis.2.1.1.chr6_hap1.g00901.t01, mNeoVis.2.1.1.chr6_hap1.g00904.t01, mNeoVis.2.1.1.chr6_hap1.g00908.t01, mNeoVis.2.1.1.chr6_hap1.g00910.t01    rna-XM_044912257.1      rna-XM_042972055.1, rna-XM_042977125.1, rna-XM_042979400.1, rna-XM_042979406.1, rna-XM_042979408.1    rna-XM_008696971.2, rna-XM_040635253.1    rna-XM_035722089.1, rna-XM_035722216.1    mNeoVis.3.1.1.chr15.g00156.t01, mNeoVis.3.1.1.chr15.g00158.t01, mNeoVis.3.1.1.chr15.g00353.t01    rna-gnl|WGS_ZZZZ|egapxtmp_033469-R1, rna-gnl|WGS_ZZZZ|egapxtmp_033629-R1, rna-gnl|WGS_ZZZZ|egapxtmp_036086-R1, rna-gnl|WGS_ZZZZ|egapxtmp_038272-R1    mNeoVis.evergladensis.2.1.chr15.g00396.t01, mNeoVis.evergladensis.2.1.chr3.g00707.t01, mNeoVis.evergladensis.2.1.chr8.g01085.t01    mNeoVis.vulgivaga.2.1.chr15.g00163.t01, mNeoVis.vulgivaga.2.1.chr15.g00403.t01, mNeoVis.vulgivaga.2.1.chr2.g01022.t01      1163    N0.HOG0001161   OG0000111  n1   rna-NM_001304847.1    rna-NM_001252044.1, rna-XM_038575231.1, rna-XM_038585098.1    rna-XM_022494951.1, rna-XM_022522560.1    rna-XM_003980267.6, rna-XM_011286063.4    rna-XM_036077028.1, rna-XM_036093422.1, rna-XM_036109391.1, rna-XM_036118398.1    rna-XM_039227570.1, rna-XM_039255572.1    rna-XM_032838168.1, rna-XM_032850846.1, rna-XM_032857654.1    rna-XM_045978537.1, rna-XM_045998223.1, rna-XM_046003494.1    rna-XM_032315322.1, rna-XM_032351447.1    rna-XM_059144037.1    rna-XM_059378143.1, rna-XM_059388568.1    rna-XM_004782063.3    mNeoVis.2.1.1.chr1_hap1.g01061.t01, mNeoVis.2.1.1.chr1_hap1.g02023.t01, mNeoVis.2.1.1.chr1_hap1.g02128.t01, mNeoVis.2.1.1.chr8_hap1.g01227.t01, mNeoVis.2.1.1.chr9_hap1.g00769.t01    rna-XM_021678335.1, rna-XM_021698959.1, rna-XM_044913914.1    rna-XM_004393722.1, rna-XM_004408359.1, rna-XM_004409591.1, rna-XM_004413669.1, rna-XM_004415448.1    rna-XM_007074350.3, rna-XM_042963353.1    rna-XM_008692172.2    rna-XM_035723439.1, rna-XM_035728657.1, rna-XM_035728734.1    mNeoVis.3.1.1.chr1.g01054.t01, mNeoVis.3.1.1.chr1.g01977.t01, mNeoVis.3.1.1.chr10.g00736.t01, mNeoVis.3.1.1.chr8.g01122.t01    rna-gnl|WGS_ZZZZ|egapxtmp_014205-R1, rna-gnl|WGS_ZZZZ|egapxtmp_014509-R1, rna-gnl|WGS_ZZZZ|egapxtmp_020045-R1, rna-gnl|WGS_ZZZZ|egapxtmp_021442-R1, rna-gnl|WGS_ZZZZ|egapxtmp_024740-R1    mNeoVis.evergladensis.2.1.chr1.g01052.t01, mNeoVis.evergladensis.2.1.chr1.g01989.t01, mNeoVis.evergladensis.2.1.chr10.g00760.t01, mNeoVis.evergladensis.2.1.chr9.g01180.t01    mNeoVis.vulgivaga.2.1.chr10.g00748.t01, mNeoVis.vulgivaga.2.1.chr8.g01182.t01      1164    N0.HOG0001162   OG0000111  n35                             rna-XM_044913146.1          rna-XM_035722104.1              1165    N0.HOG0001163   OG0000111  n39     rna-XM_038562365.1                              rna-XM_042962897.1, rna-XM_042995080.1      rna-XM_035722326.1              1166    N0.HOG0001164   OG0000111  n43         rna-XM_045034766.1      rna-XM_039237139.1    rna-XM_032855391.1, rna-XM_032871303.1, rna-XM_032877043.1    rna-XM_045996857.1, rna-XM_046005947.1, rna-XM_046008603.1              rna-XM_044913299.1    rna-XM_004396922.1, rna-XM_004410250.1    rna-XM_042997088.1      rna-XM_035723290.1          mNeoVis.vulgivaga.2.1.chr4.g00992.t01      1167    N0.HOG0001165   OG0000111  n61                           mNeoVis.2.1.1.chr2_hap1.g01183.t01                rna-gnl|WGS_ZZZZ|egapxtmp_009085-R1    mNeoVis.evergladensis.2.1.chr5.g01126.t01    mNeoVis.vulgivaga.2.1.chr5.g01112.t01      1168    N0.HOG0001166   OG0000111  n70         rna-XM_045043723.1    rna-XM_036078791.1, rna-XM_036096494.1, rna-XM_036102950.1        rna-XM_046009080.1, rna-XM_046023642.1              rna-XM_044918897.1, rna-XM_044919138.1      rna-XM_042964619.1      rna-XM_035728461.1              1169    N0.HOG0001167   OG0000111  n80           rna-XM_036106704.1                    rna-XM_044917723.1                      1170    N0.HOG0001168   OG0000111  n83     rna-XM_038541795.1        rna-XM_036074020.1                    rna-XM_044920724.1                      1171    N0.HOG0001169   OG0000111  n85   rna-XM_034644346.1, rna-XM_034660148.1    rna-XM_038535845.1                          rna-XM_044916032.1      rna-XM_042963493.1    rna-XM_040621939.1, rna-XM_040642618.1                1172    N0.HOG0001170   OG0000111  n91         rna-XM_045039873.1                          rna-XM_042961041.1    rna-XM_008689880.2                1173    N0.HOG0001171   OG0000111  n93             rna-XM_039247352.1, rna-XM_039256375.1                                      1174    N0.HOG0001172   OG0000111  n94             rna-XM_039239012.1                    rna-XM_004404317.1    rna-XM_042973242.1                  1175    N0.HOG0001173   OG0000111  n96   rna-XM_034651454.1                              rna-XM_004415198.1        rna-XM_027568165.2              1176    N0.HOG0001174   OG0000111  n98         rna-XM_045055069.1      rna-XM_039227054.1      rna-XM_045996960.1, rna-XM_046002719.1, rna-XM_046023164.1, rna-XM_046023524.1        rna-XM_059412331.1        rna-XM_021703115.2, rna-XM_044913375.1, rna-XM_044913693.1      rna-XM_042979657.1      rna-XM_035724220.1              1177    N0.HOG0001175   OG0000111  n109           rna-XM_036111609.1                            rna-XM_035725323.1              1178    N0.HOG0001176   OG0000112  n1   rna-XM_034653156.1, rna-XM_034660207.1, rna-XM_034661690.1    rna-NM_001020810.1    rna-XM_022510930.1, rna-XM_022510973.1      rna-XM_036072466.1    rna-XM_039225792.1    rna-XM_032847170.1, rna-XM_032847260.1    rna-XM_046006703.1, rna-XM_046006712.1      rna-XM_059178978.1    rna-XM_059400558.1    rna-XM_045065179.1, rna-XM_045065228.1    mNeoVis.2.1.1.chr1_hap1.g01288.t01, mNeoVis.2.1.1.chr1_hap1.g01319.t01, mNeoVis.2.1.1.chr1_hap1.g01695.t01    rna-XM_021684701.1, rna-XM_021697221.2    rna-XM_004409075.1    rna-XM_042985880.1    rna-XM_008700933.2    rna-XM_035728503.1    mNeoVis.3.1.1.chr1.g01242.t01, mNeoVis.3.1.1.chr1.g01269.t01, mNeoVis.3.1.1.chr1.g01650.t01    rna-gnl|WGS_ZZZZ|egapxtmp_010411-R1, rna-gnl|WGS_ZZZZ|egapxtmp_025570-R1, rna-gnl|WGS_ZZZZ|egapxtmp_033750-R1    mNeoVis.evergladensis.2.1.chr1.g01260.t01, mNeoVis.evergladensis.2.1.chr1.g01290.t01, mNeoVis.evergladensis.2.1.chr1.g01662.t01    mNeoVis.vulgivaga.2.1.chr1.g01259.t01      1179    N0.HOG0001177   OG0000112  n34   rna-NM_001304845.1, rna-XM_034653258.1, rna-XM_034653412.1, rna-XM_034660459.1, rna-XM_034660460.1, rna-XM_034660461.1    rna-NM_001014378.1, rna-NM_001014767.1, rna-XM_038553482.1    rna-XM_022510596.1, rna-XM_022510597.1, rna-XM_022510599.1    rna-NM_001048161.1, rna-XM_011282114.4, rna-XM_023249590.2, rna-XM_045057242.1, rna-XM_045057243.1, rna-XM_045057249.1, rna-XM_045058647.1    rna-XM_036072464.1, rna-XM_036103869.1, rna-XM_036108170.1, rna-XM_036120368.1    rna-XM_039219501.1, rna-XM_039219562.1, rna-XM_039219587.1, rna-XM_039220578.1    rna-XM_032847174.1, rna-XM_032847262.1, rna-XM_032847263.1, rna-XM_032847264.1, rna-XM_032847321.1, rna-XM_032847325.1, rna-XM_032847332.1, rna-XM_032847334.1    rna-XM_046006101.1, rna-XM_046006707.1, rna-XM_046006710.1, rna-XM_046006711.1, rna-XM_046006719.1    rna-XM_032338085.1, rna-XM_032338123.1, rna-XM_032338126.1, rna-XM_032338151.1, rna-XM_032338395.1    rna-XM_059176409.1, rna-XM_059178969.1, rna-XM_059178972.1    rna-XM_059387443.1, rna-XM_059399795.1, rna-XM_059399796.1, rna-XM_059399798.1, rna-XM_059401521.1, rna-XM_059401523.1    rna-XM_045065165.1, rna-XM_045065168.1, rna-XM_045065174.1, rna-XM_045065176.1, rna-XM_045065224.1, rna-XM_045065225.1, rna-XM_045087959.1    mNeoVis.2.1.1.chr1_hap1.g01322.t01, mNeoVis.2.1.1.chr1_hap1.g01323.t01, mNeoVis.2.1.1.chr1_hap1.g01324.t01, mNeoVis.2.1.1.chr1_hap1.g01325.t01, mNeoVis.2.1.1.chr1_hap1.g01326.t01    rna-XM_044912439.1    rna-XM_004415284.1, rna-XM_004417055.1    rna-XM_007090571.3, rna-XM_042985929.1, rna-XM_042985931.1, rna-XM_042985932.1, rna-XM_042985933.1, rna-XM_042985939.1, rna-XM_042985941.1    rna-XM_040627474.1, rna-XM_040627478.1, rna-XM_040637049.1, rna-XM_040644868.1    rna-XM_027603913.2, rna-XM_027603919.2, rna-XM_027604091.2, rna-XM_035728706.1, rna-XM_035728708.1, rna-XM_035728709.1, rna-XM_035728710.1, rna-XM_035728711.1    mNeoVis.3.1.1.chr1.g01270.t01, mNeoVis.3.1.1.chr1.g01273.t01, mNeoVis.3.1.1.chr1.g01274.t01, mNeoVis.3.1.1.chr1.g01275.t01    rna-gnl|WGS_ZZZZ|egapxtmp_010492-R1, rna-gnl|WGS_ZZZZ|egapxtmp_010502-R1, rna-gnl|WGS_ZZZZ|egapxtmp_025426-R1    mNeoVis.evergladensis.2.1.chr1.g01291.t01, mNeoVis.evergladensis.2.1.chr1.g01292.t01, mNeoVis.evergladensis.2.1.chr1.g01293.t01    mNeoVis.vulgivaga.2.1.chr1.g01289.t01, mNeoVis.vulgivaga.2.1.chr1.g01292.t01, mNeoVis.vulgivaga.2.1.chr1.g01293.t01, mNeoVis.vulgivaga.2.1.chr1.g01294.t01      1180    N0.HOG0001178   OG0000113  n1   rna-XM_019794315.2    rna-XM_038576060.1    rna-XM_022499911.1    rna-XM_019814247.2    rna-XM_036079333.1    rna-XM_039223330.1    rna-XM_032883045.1    rna-XM_046027639.1    rna-XM_032311732.1, rna-XM_032313377.1    rna-XM_059172470.1    rna-XM_059395801.1    rna-XM_013050134.2    mNeoVis.2.1.1.chr2_hap1.g00598.t01    rna-XM_021696169.2    rna-XM_004409690.1    rna-XM_007079263.2    rna-XM_008689599.1    rna-XM_027596654.2    mNeoVis.3.1.1.chr5.g00593.t01    rna-gnl|WGS_ZZZZ|egapxtmp_009475-R2    mNeoVis.evergladensis.2.1.chr5.g00601.t01    mNeoVis.vulgivaga.2.1.chr5.g00595.t01      1181    N0.HOG0001179   OG0000113  n23   rna-XM_034663564.1    rna-XM_038576051.1    rna-XM_022499984.1    rna-XM_023240310.2    rna-XM_036079330.1    rna-XM_039223338.1    rna-XM_032883035.1    rna-XM_046027689.1    rna-XM_032313453.1    rna-XM_059170294.1, rna-XM_059172484.1    rna-XM_059387268.1    rna-XM_045081459.1, rna-XM_045087079.1    mNeoVis.2.1.1.chr2_hap1.g00593.t01    rna-XM_021696195.1    rna-XM_004409694.1    rna-XM_042960354.1    rna-XM_008689596.2    rna-XM_027596356.1    mNeoVis.3.1.1.chr5.g00588.t01    rna-gnl|WGS_ZZZZ|egapxtmp_009482-R1    mNeoVis.evergladensis.2.1.chr5.g00598.t01    mNeoVis.vulgivaga.2.1.chr5.g00592.t01      1182    N0.HOG0001180   OG0000113  n49   rna-XM_019794268.2    rna-XM_038576064.1    rna-XM_022499941.1    rna-XM_019814244.3    rna-XM_036079327.1      rna-XM_032883047.1    rna-XM_046027688.1    rna-XM_032313380.1    rna-XM_059172483.1    rna-XM_059397640.1    rna-XM_004780778.3, rna-XM_045087080.1    mNeoVis.2.1.1.chr2_hap1.g00600.t01    rna-XM_021696129.1    rna-XM_004409689.1    rna-XM_007079264.3    rna-XM_008689600.1    rna-XM_027596320.2    mNeoVis.3.1.1.chr5.g00594.t01    rna-gnl|WGS_ZZZZ|egapxtmp_009474-R3    mNeoVis.evergladensis.2.1.chr5.g00602.t01    mNeoVis.vulgivaga.2.1.chr5.g00597.t01      1183    N0.HOG0001181   OG0000113  n69   rna-XM_019794269.2    rna-XM_038576063.1    rna-XM_022499939.1    rna-XM_003993868.5    rna-XM_036079338.1    rna-XM_039223324.1    rna-XM_032883049.1    rna-XM_046027824.1    rna-XM_032313375.1    rna-XM_059172471.1    rna-XM_059397388.1    rna-XM_004780775.3    mNeoVis.2.1.1.chr2_hap1.g00601.t01    rna-XM_021696158.1    rna-XM_004409688.1    rna-XM_007079265.3    rna-XM_008689667.1    rna-XM_027596406.2    mNeoVis.3.1.1.chr5.g00595.t01    rna-gnl|WGS_ZZZZ|egapxtmp_009472-R1    mNeoVis.evergladensis.2.1.chr5.g00603.t01    mNeoVis.vulgivaga.2.1.chr5.g00598.t01      1184    N0.HOG0001182   OG0000113  n89   rna-XM_034662403.1    rna-XM_038576080.1    rna-XM_022499551.1    rna-XM_003993874.5    rna-XM_036079359.1    rna-XM_039222021.1    rna-XM_032883060.1    rna-XM_046027635.1    rna-XM_032312368.1    rna-XM_059172466.1    rna-XM_059397400.1    rna-XM_045073844.1    mNeoVis.2.1.1.chr2_hap1.g00607.t01    rna-XM_021696075.1    rna-XM_004412996.1    rna-XM_042959894.1    rna-XM_040624746.1    rna-XM_035724334.1    mNeoVis.3.1.1.chr5.g00601.t01    rna-gnl|WGS_ZZZZ|egapxtmp_009258-R1    mNeoVis.evergladensis.2.1.chr5.g00609.t01    mNeoVis.vulgivaga.2.1.chr5.g00604.t01      1185    N0.HOG0001183   OG0000113  n109   rna-XM_002914386.2    rna-NM_001003209.1    rna-XM_022500058.1    rna-XM_003993865.4    rna-XM_036079332.1    rna-XM_039223342.1    rna-XM_032883036.1    rna-XM_046026282.1    rna-XM_032313433.1    rna-XM_059170298.1, rna-XM_059172468.1      rna-XM_045073849.1, rna-XM_045081457.1    mNeoVis.2.1.1.chr2_hap1.g00594.t01    rna-XM_021696175.2    rna-XM_004409691.1    rna-XM_007079262.1    rna-XM_008689597.2    rna-XM_027596364.1    mNeoVis.3.1.1.chr5.g00589.t01    rna-gnl|WGS_ZZZZ|egapxtmp_009477-R7    mNeoVis.evergladensis.2.1.chr5.g00599.t01    mNeoVis.vulgivaga.2.1.chr5.g00593.t01      1186    N0.HOG0001184   OG0000114  n3   rna-XM_011221745.3    rna-XM_038566990.1    rna-XM_022497804.1    rna-XM_003982018.6    rna-XM_036095400.1    rna-XM_039243298.1    rna-XM_032867528.1    rna-XM_045991040.1    rna-XM_032314462.1    rna-XM_059160337.1    rna-XM_059389288.1    rna-XM_004761098.3    mNeoVis.2.1.1.chr10_hap1.g01115.t01, mNeoVis.2.1.1.chr5_hap1.g00451.t01    rna-XM_021683286.1    rna-XM_004404573.1    rna-XM_042978467.1    rna-XM_040629123.1, rna-XM_040629125.1    rna-XM_027583295.2    mNeoVis.3.1.1.chr2.g00374.t01, mNeoVis.3.1.1.chr9.g01042.t01, mNeoVis.3.1.1.chr9.g01043.t01    rna-gnl|WGS_ZZZZ|egapxtmp_015428-R1, rna-gnl|WGS_ZZZZ|egapxtmp_034903-R1    mNeoVis.evergladensis.2.1.chr2.g00445.t01, mNeoVis.evergladensis.2.1.chr8.g01090.t01    mNeoVis.vulgivaga.2.1.chr2.g00447.t01, mNeoVis.vulgivaga.2.1.chr9.g01070.t01      1187    N0.HOG0001185   OG0000114  n21   rna-XM_002918878.4, rna-XM_034650814.1, rna-XM_034651328.1    rna-XM_038552778.1    rna-XM_022500572.1    rna-XM_023242428.2    rna-XM_036081279.1    rna-XM_039218595.1    rna-XM_032860516.1    rna-XM_046023078.1    rna-XM_032307408.1    rna-XM_059142679.1    rna-XM_059411391.1    rna-XM_004747532.3    mNeoVis.2.1.1.chr13_hap1.g00500.t01    rna-XM_021691173.1    rna-XM_004392255.1    rna-XM_015535011.2    rna-XM_040631579.1    rna-XM_027614490.2    mNeoVis.3.1.1.chr13.g00435.t01    rna-gnl|WGS_ZZZZ|egapxtmp_029741-R2    mNeoVis.evergladensis.2.1.chr13.g00481.t01    mNeoVis.vulgivaga.2.1.chr13.g00482.t01      1188    N0.HOG0001186   OG0000114  n31   rna-XM_034652078.1, rna-XM_034652638.1, rna-XM_034660876.1    rna-XM_038580628.1    rna-XM_022511894.1    rna-XM_045059202.1    rna-XM_036081128.1    rna-XM_039220016.1    rna-XM_032864798.1    rna-XM_046010399.1    rna-XM_032342775.1    rna-XM_059180717.1    rna-XM_059372205.1    rna-XM_013059566.2    mNeoVis.2.1.1.chr10_hap1.g00977.t01, mNeoVis.2.1.1.chr14_hap1.g00273.t01    rna-XM_044917962.1    rna-XM_004411799.1    rna-XM_042988716.1    rna-XM_040620556.1    rna-XM_027569443.2    mNeoVis.3.1.1.chr14.g00276.t01    rna-gnl|WGS_ZZZZ|egapxtmp_033108-R1    mNeoVis.evergladensis.2.1.chr14.g00273.t01, mNeoVis.evergladensis.2.1.chr8.g00947.t01    mNeoVis.vulgivaga.2.1.chr14.g00275.t01, mNeoVis.vulgivaga.2.1.chr9.g00933.t01      1189    N0.HOG0001187   OG0000114  n52           rna-XM_036086826.1        rna-XM_045983293.1              rna-XM_044915842.1        rna-XM_008697022.2    rna-XM_027610229.2              1190    N0.HOG0001188   OG0000114  n56   rna-XM_019803058.2        rna-XM_019822522.3        rna-XM_032845207.1      rna-XM_032318280.1    rna-XM_059145748.1      rna-XM_045086881.1    mNeoVis.2.1.1.chr14_hap1.g00682.t01      rna-XM_004403002.1    rna-XM_007087937.3        mNeoVis.3.1.1.chr14.g00652.t01      mNeoVis.evergladensis.2.1.chr14.g00680.t01    mNeoVis.vulgivaga.2.1.chr14.g00680.t01      1191    N0.HOG0001189   OG0000114  n68   rna-XM_034667562.1    rna-XM_038535476.1    rna-XM_022490712.1    rna-XM_003999790.5, rna-XM_045055859.1    rna-XM_036086844.1    rna-XM_039218944.1    rna-XM_032845219.1      rna-XM_032318288.1    rna-XM_059146514.1    rna-XM_059413862.1    rna-XM_045086865.1      rna-XM_021681870.1    rna-XM_004403000.1    rna-XM_015536734.2, rna-XM_042976826.1    rna-XM_008696620.2    rna-XM_027613190.2        mNeoVis.evergladensis.2.1.chr8.g00948.t01        1192    N0.HOG0001190   OG0000114  n86     rna-XM_038586542.1        rna-XM_036078202.1                    rna-XM_044921237.1                      1193    N0.HOG0001191   OG0000114  n88   rna-XM_034652062.1      rna-XM_022491016.1      rna-XM_036100644.1      rna-XM_032852635.1      rna-XM_032312971.1      rna-XM_059413866.1                rna-XM_035727068.1    mNeoVis.3.1.1.chr9.g00907.t01    rna-gnl|WGS_ZZZZ|egapxtmp_016699-R3, rna-gnl|WGS_ZZZZ|egapxtmp_032545-R2          1194    N0.HOG0001192   OG0000114  n97   rna-XM_034652092.1, rna-XM_034652188.1    rna-NM_001252211.1          rna-XM_039218983.1      rna-XM_045982145.1                    rna-XM_040635700.1                1195    N0.HOG0001193   OG0000115  n1   rna-XM_034646423.1    rna-XM_038577708.1    rna-XM_022519534.1    rna-XM_019834949.3    rna-XM_036098861.1    rna-XM_039241403.1    rna-XM_032876753.1    rna-XM_046012942.1    rna-XM_032348235.1    rna-XM_059185303.1    rna-XM_059403148.1    rna-XM_004774717.3    mNeoVis.2.1.1.chr9_hap1.g00543.t01    rna-XM_044915201.1    rna-XM_004400810.1    rna-XM_042992322.1    rna-XM_040642469.1    rna-XM_027593334.2    mNeoVis.3.1.1.chr10.g00512.t01    rna-gnl|WGS_ZZZZ|egapxtmp_023718-R1    mNeoVis.evergladensis.2.1.chr10.g00536.t01    mNeoVis.vulgivaga.2.1.chr10.g00522.t01      1196    N0.HOG0001194   OG0000115  n25   rna-XM_034655625.1    rna-XM_038530325.1    rna-XM_022493315.1    rna-XM_045059964.1    rna-XM_036123912.1    rna-XM_039252709.1    rna-XM_032847484.1    rna-XM_045999406.1    rna-XM_032336497.1    rna-XM_059174765.1    rna-XM_059417727.1    rna-XM_013052425.2    mNeoVis.2.1.1.chr1_hap1.g00489.t01    rna-XM_021701839.1    rna-XM_004397791.1    rna-XM_007077889.3    rna-XM_040622702.1    rna-XM_027605674.1    mNeoVis.3.1.1.chr1.g00487.t01    rna-gnl|WGS_ZZZZ|egapxtmp_031091-R1    mNeoVis.evergladensis.2.1.chr1.g00487.t01    mNeoVis.vulgivaga.2.1.chr1.g00497.t01      1197    N0.HOG0001195   OG0000115  n46   rna-XM_019799963.2    rna-XM_038556290.1    rna-XM_022505690.1    rna-XM_019829396.3    rna-XM_036108816.1    rna-XM_039251773.1    rna-XM_032843239.1    rna-XM_045987257.1    rna-XM_032334536.1    rna-XM_059159531.1    rna-XM_059385089.1    rna-XM_045079852.1    mNeoVis.2.1.1.chr6_hap1.g00094.t01    rna-XM_021702448.1    rna-XM_004392904.1    rna-XM_042984402.1    rna-XM_008693197.2    rna-XM_027598153.2    mNeoVis.3.1.1.chr4.g00092.t01    rna-gnl|WGS_ZZZZ|egapxtmp_044310-R1    mNeoVis.evergladensis.2.1.chr4.g00093.t01    mNeoVis.vulgivaga.2.1.chr4.g00092.t01      1198    N0.HOG0001196   OG0000115  n66   rna-XM_034659387.1, rna-XM_034670970.1, rna-XM_034670972.1, rna-XM_034670973.1    rna-XM_038561722.1    rna-XM_022504307.1    rna-XM_019827439.3    rna-XM_036087601.1    rna-XM_039240142.1    rna-XM_032880778.1    rna-XM_045979541.1    rna-XM_032352498.1    rna-XM_059188097.1    rna-XM_059405800.1    rna-XM_045079036.1    mNeoVis.2.1.1.chr11_hap1.g00817.t01    rna-XM_044918882.1    rna-XM_004398712.1    rna-XM_042981759.1    rna-XM_040623308.1    rna-XM_027622887.2    mNeoVis.3.1.1.chr11.g00780.t01    rna-gnl|WGS_ZZZZ|egapxtmp_025303-R1    mNeoVis.evergladensis.2.1.chr11.g00801.t01    mNeoVis.vulgivaga.2.1.chr11.g00821.t01      1199    N0.HOG0001197   OG0000115  n90   rna-XM_034662094.1    rna-XM_038570656.1    rna-XM_022515250.1    rna-XM_045037714.1    rna-XM_036091418.1    rna-XM_039242714.1    rna-XM_032861006.1    rna-XM_046001499.1    rna-XM_032324384.1    rna-XM_059162430.1    rna-XM_059390812.1    rna-XM_045088993.1    mNeoVis.2.1.1.chr5_hap1.g01075.t01    rna-XM_021678816.1    rna-XM_004411783.1    rna-XM_042956780.1    rna-XM_040636884.1    rna-XM_027587930.2    mNeoVis.3.1.1.chr2.g00982.t01    rna-gnl|WGS_ZZZZ|egapxtmp_034308-R1    mNeoVis.evergladensis.2.1.chr2.g01065.t01    mNeoVis.vulgivaga.2.1.chr2.g01070.t01      1200    N0.HOG0001198   OG0000115  n107   rna-XM_034654639.1    rna-XM_038584654.1    rna-XM_022503036.1    rna-XM_019838161.3    rna-XM_036116753.1    rna-XM_039253448.1    rna-XM_032880286.1    rna-XM_046019392.1    rna-XM_032355736.1    rna-XM_059167171.1    rna-XM_059394074.1    rna-XM_004743943.3    mNeoVis.2.1.1.chr3_hap1.g00942.t01    rna-XM_044913311.1    rna-XM_004410187.1    rna-XM_042997242.1    rna-XM_040644559.1    rna-XM_027590050.2    mNeoVis.3.1.1.chr3.g00932.t01    rna-gnl|WGS_ZZZZ|egapxtmp_039148-R1    mNeoVis.evergladensis.2.1.chr3.g00937.t01    mNeoVis.vulgivaga.2.1.chr3.g00956.t01      1201    N0.HOG0001199   OG0000116  n1   rna-XM_002924997.2, rna-XM_002925001.2, rna-XM_034651771.1, rna-XM_034666476.1    rna-NM_001388729.1    rna-XM_022522125.1, rna-XM_022522127.1    rna-XM_003992948.3, rna-XM_003992951.4    rna-XM_036095918.1    rna-XM_039218870.1, rna-XM_039218871.1    rna-XM_032853676.1, rna-XM_032853678.1    rna-XM_046017303.1    rna-XM_032360410.1, rna-XM_032360517.1    rna-XM_059158002.1, rna-XM_059158035.1    rna-XM_059389434.1, rna-XM_059389472.1    rna-XM_004781199.2, rna-XM_045078270.1, rna-XM_045078272.1    mNeoVis.2.1.1.chr7_hap1.g01734.t01, mNeoVis.2.1.1.chr7_hap1.g01737.t01    rna-XM_021685693.1      rna-XM_007083137.2, rna-XM_015537938.2, rna-XM_042959103.1, rna-XM_042959118.1    rna-XM_008710986.1    rna-XM_027580746.1    mNeoVis.3.1.1.chr7.g01612.t01, mNeoVis.3.1.1.chr7.g01615.t01    rna-gnl|WGS_ZZZZ|egapxtmp_012879-R1    mNeoVis.evergladensis.2.1.chr7.g01702.t01    mNeoVis.vulgivaga.2.1.chr7.g01714.t01      1202    N0.HOG0001200   OG0000116  n39   rna-XM_002924985.4      rna-XM_022522151.1        rna-XM_039218865.1    rna-XM_032853679.1    rna-XM_046017294.1    rna-XM_032360516.1    rna-XM_059158049.1    rna-XM_059389420.1    rna-XM_004781202.1    mNeoVis.2.1.1.chr7_hap1.g01738.t01          rna-XM_008710985.1      mNeoVis.3.1.1.chr7.g01616.t01    rna-gnl|WGS_ZZZZ|egapxtmp_012791-R1    mNeoVis.evergladensis.2.1.chr7.g01703.t01    mNeoVis.vulgivaga.2.1.chr7.g01717.t01      1203    N0.HOG0001201   OG0000116  n52   rna-XM_002924998.4        rna-XM_011286696.2    rna-XM_036095957.1    rna-XM_039218869.1                  rna-XM_021685642.2      rna-XM_042959161.1    rna-XM_008711004.1    rna-XM_027580748.1              1204    N0.HOG0001202   OG0000116  n60   rna-XM_011218129.3, rna-XM_034650117.1, rna-XM_034666107.1, rna-XM_034666171.1    rna-NM_001017522.1, rna-NM_001389014.1, rna-XM_038568857.1    rna-XM_022522146.1    rna-XM_019812942.2, rna-XM_023239722.2, rna-XM_045038130.1, rna-XM_045038131.1      rna-XM_039224504.1, rna-XM_039224517.1, rna-XM_039224520.1    rna-XM_032853659.1    rna-XM_046017418.1    rna-XM_032356336.1    rna-XM_059157766.1    rna-XM_059389721.1    rna-XM_004779043.3, rna-XM_045088012.1    mNeoVis.2.1.1.chr7_hap1.g01712.t01      rna-XM_004414463.1    rna-XM_007083260.1, rna-XM_007083261.2, rna-XM_007083262.1, rna-XM_042959274.1    rna-XM_008709628.2, rna-XM_008709629.2, rna-XM_008709630.2      mNeoVis.3.1.1.chr7.g01596.t01    rna-gnl|WGS_ZZZZ|egapxtmp_012795-R1    mNeoVis.evergladensis.2.1.chr7.g01681.t01    mNeoVis.vulgivaga.2.1.chr7.g01695.t01      1205    N0.HOG0001203   OG0000116  n92     rna-NM_001388989.1    rna-XM_022492608.1, rna-XM_022522172.1    rna-XM_003992902.4, rna-XM_011279814.3      rna-XM_039224507.1    rna-XM_032850890.1, rna-XM_032852713.1    rna-XM_046015884.1, rna-XM_046015886.1, rna-XM_046015887.1, rna-XM_046017558.1, rna-XM_046017565.1    rna-XM_032359778.1, rna-XM_032359779.1, rna-XM_032359781.1    rna-XM_059157683.1, rna-XM_059182598.1, rna-XM_059182846.1    rna-XM_059393253.1, rna-XM_059393523.1, rna-XM_059393659.1    rna-XM_004779046.2, rna-XM_004779048.2    mNeoVis.2.1.1.chr7_hap1.g01705.t01, mNeoVis.2.1.1.chr7_hap1.g01707.t01, mNeoVis.2.1.1.chr7_hap1.g01708.t01        rna-XM_007092232.2, rna-XM_015537923.2, rna-XM_042959611.1    rna-XM_008709632.1, rna-XM_008709633.2    rna-XM_027577899.2    mNeoVis.3.1.1.chr7.g01594.t01    rna-gnl|WGS_ZZZZ|egapxtmp_012809-R1    mNeoVis.evergladensis.2.1.chr7.g01676.t01    mNeoVis.vulgivaga.2.1.chr7.g01692.t01, mNeoVis.vulgivaga.2.1.chr7.g01694.t01      1206    N0.HOG0001204   OG0000117  n1   rna-XM_019808319.2    rna-XM_038540141.1    rna-XM_022500938.1    rna-XM_019820825.3    rna-XM_036087380.1    rna-XM_039233669.1    rna-XM_032843948.1    rna-XM_045992949.1    rna-XM_032327122.1    rna-XM_059154472.1    rna-XM_059415167.1    rna-XM_045072730.1    mNeoVis.2.1.1.chr15_hap1.g00172.t01    rna-XM_021692238.1    rna-XM_004415756.1    rna-XM_007084114.3    rna-XM_040635247.1    rna-XM_027612190.2    mNeoVis.3.1.1.chr15.g00148.t01    rna-gnl|WGS_ZZZZ|egapxtmp_034138-R3    mNeoVis.evergladensis.2.1.chr15.g00155.t01    mNeoVis.vulgivaga.2.1.chr15.g00158.t01      1207    N0.HOG0001205   OG0000117  n24   rna-XM_019799401.2    rna-XM_038566258.1    rna-XM_022497930.1    rna-XM_045051061.1    rna-XM_036102512.1    rna-XM_039243684.1    rna-XM_032868239.1    rna-XM_045991948.1    rna-XM_032321474.1    rna-XM_059161082.1    rna-XM_059389879.1    rna-XM_045079199.1    mNeoVis.2.1.1.chr5_hap1.g00716.t01    rna-XM_021694986.1    rna-XM_004411523.1    rna-XM_042979084.1    rna-XM_040628716.1    rna-XM_027587893.1    mNeoVis.3.1.1.chr2.g00628.t01    rna-gnl|WGS_ZZZZ|egapxtmp_035427-R1    mNeoVis.evergladensis.2.1.chr2.g00716.t01    mNeoVis.vulgivaga.2.1.chr2.g00719.t01      1208    N0.HOG0001206   OG0000117  n46   rna-XM_034667630.1    rna-XM_038542857.1    rna-XM_022508439.1    rna-XM_045049335.1    rna-XM_036068919.1, rna-XM_036120908.1    rna-XM_039235797.1    rna-XM_032838780.1    rna-XM_045982293.1    rna-XM_032319061.1    rna-XM_059145622.1    rna-XM_059414046.1    rna-XM_045069521.1    mNeoVis.2.1.1.chr14_hap1.g00238.t01    rna-XM_044912108.1, rna-XM_044916203.1    rna-XM_004393557.1    rna-XM_042976694.1    rna-XM_040636128.1    rna-XM_027613953.2    mNeoVis.3.1.1.chr14.g00240.t01    rna-gnl|WGS_ZZZZ|egapxtmp_031864-R1    mNeoVis.evergladensis.2.1.chr14.g00240.t01    mNeoVis.vulgivaga.2.1.chr14.g00241.t01      1209    N0.HOG0001207   OG0000117  n69   rna-XM_034659452.1    rna-XM_038561598.1    rna-XM_022504259.1    rna-XM_045054471.1    rna-XM_036117335.1    rna-XM_039217619.1    rna-XM_032880676.1    rna-XM_045978683.1    rna-XM_032352621.1    rna-XM_059188231.1    rna-XM_059405712.1    rna-XM_004742302.3    mNeoVis.2.1.1.chr11_hap1.g00856.t01    rna-XM_021697409.1    rna-XM_004415552.1    rna-XM_042981846.1    rna-XM_008700061.2    rna-XM_027623437.2    mNeoVis.3.1.1.chr11.g00815.t01    rna-gnl|WGS_ZZZZ|egapxtmp_027540-R2    mNeoVis.evergladensis.2.1.chr11.g00839.t01    mNeoVis.vulgivaga.2.1.chr11.g00860.t01      1210    N0.HOG0001208   OG0000117  n91   rna-XM_034658502.1    rna-XM_038566103.1    rna-XM_022497828.1    rna-XM_045051717.1    rna-XM_036102442.1    rna-XM_039242492.1    rna-XM_032868373.1    rna-XM_045992256.1    rna-XM_032323076.1    rna-XM_059161254.1    rna-XM_059387847.1    rna-XM_013053167.2    mNeoVis.2.1.1.chr5_hap1.g00757.t01    rna-XM_044916801.1      rna-XM_042979210.1    rna-XM_040628655.1    rna-XM_027587796.2    mNeoVis.3.1.1.chr2.g00666.t01    rna-gnl|WGS_ZZZZ|egapxtmp_036270-R1    mNeoVis.evergladensis.2.1.chr2.g00758.t01    mNeoVis.vulgivaga.2.1.chr2.g00763.t01      1211    N0.HOG0001209   OG0000117  n111   rna-XM_034642450.1    rna-XM_038585215.1    rna-XM_022517313.1    rna-XM_011285443.4    rna-XM_036089415.1, rna-XM_036097299.1    rna-XM_039248955.1    rna-XM_032860188.1    rna-XM_046018191.1    rna-XM_032354245.1    rna-XM_059168545.1    rna-XM_059393094.1    rna-XM_045078496.1    mNeoVis.2.1.1.chr3_hap1.g01325.t01    rna-XM_021703031.2    rna-XM_004393519.1    rna-XM_042994953.1    rna-XM_040636571.1    rna-XM_027588498.2    mNeoVis.3.1.1.chr3.g01302.t01    rna-gnl|WGS_ZZZZ|egapxtmp_042743-R1    mNeoVis.evergladensis.2.1.chr3.g01324.t01    mNeoVis.vulgivaga.2.1.chr3.g01345.t01      1212    N0.HOG0001210   OG0000118  n1   rna-XM_034642590.1    rna-XM_038573998.1    rna-XM_022495232.1    rna-XM_019838774.3    rna-XM_036073797.1    rna-XM_039257611.1    rna-XM_032857373.1    rna-XM_046019417.1    rna-XM_032355022.1    rna-XM_059167847.1    rna-XM_059393619.1    rna-XM_013047894.2    mNeoVis.2.1.1.chr3_hap1.g01123.t01    rna-XM_021682956.1    rna-XM_004409761.1    rna-XM_042995517.1    rna-XM_008690566.2    rna-XM_027590008.2    mNeoVis.3.1.1.chr3.g01112.t01    rna-gnl|WGS_ZZZZ|egapxtmp_042043-R1    mNeoVis.evergladensis.2.1.chr3.g01123.t01    mNeoVis.vulgivaga.2.1.chr3.g01138.t01      1213    N0.HOG0001211   OG0000118  n24   rna-XM_034655281.1    rna-NM_001031818.1    rna-XM_022496451.1    rna-XM_045034560.1    rna-XM_036073732.1    rna-XM_039257625.1    rna-XM_032857371.1    rna-XM_046019313.1    rna-XM_032355016.1    rna-XM_059167837.1    rna-XM_059393614.1    rna-XM_045078801.1    mNeoVis.2.1.1.chr3_hap1.g01125.t01    rna-XM_021682955.1    rna-XM_004409763.1    rna-XM_042995498.1    rna-XM_040639708.1    rna-XM_027589925.2    mNeoVis.3.1.1.chr3.g01113.t01    rna-gnl|WGS_ZZZZ|egapxtmp_042044-R1    mNeoVis.evergladensis.2.1.chr3.g01124.t01    mNeoVis.vulgivaga.2.1.chr3.g01139.t01      1214    N0.HOG0001212   OG0000118  n46   rna-XM_034665466.1    rna-XM_038570040.1    rna-XM_022502888.1    rna-XM_023253164.2    rna-XM_036091057.1    rna-XM_039226664.1    rna-XM_032881048.1    rna-XM_045977871.1    rna-XM_032315040.1    rna-XM_059143851.1    rna-XM_059377881.1    rna-XM_045081318.1    mNeoVis.2.1.1.chr8_hap1.g01509.t01    rna-XM_021696009.1    rna-XM_004409597.1    rna-XM_042980099.1    rna-XM_040624556.1    rna-XM_035726796.1    mNeoVis.3.1.1.chr8.g01406.t01, mNeoVis.3.1.1.chr8.g01407.t01    rna-gnl|WGS_ZZZZ|egapxtmp_017689-R1    mNeoVis.evergladensis.2.1.chr9.g01458.t01    mNeoVis.vulgivaga.2.1.chr8.g01461.t01      1215    N0.HOG0001213   OG0000118  n68   rna-XM_034650007.1    rna-XM_038588046.1    rna-XM_022524861.1    rna-XM_011279393.2    rna-XM_036106897.1    rna-XM_039233538.1    rna-XM_032837168.1    rna-XM_045996295.1    rna-XM_032331816.1    rna-XM_059156530.1    rna-XM_059385604.1    rna-XM_045068260.1    mNeoVis.2.1.1.chr12_hap1.g00295.t01    rna-XM_044911855.1    rna-XM_004414417.1    rna-XM_042974020.1    rna-XM_040627574.1    rna-XM_027609383.1    mNeoVis.3.1.1.chr12.g00266.t01    rna-gnl|WGS_ZZZZ|egapxtmp_029486-R1    mNeoVis.evergladensis.2.1.chr12.g00263.t01    mNeoVis.vulgivaga.2.1.chr12.g00270.t01      1216    N0.HOG0001214   OG0000118  n90   rna-XM_034665468.1    rna-XM_038570042.1    rna-XM_022502638.1    rna-XM_023253172.2    rna-XM_036123776.1    rna-XM_039226667.1    rna-XM_032881049.1    rna-XM_045977872.1    rna-XM_032315042.1    rna-XM_059143853.1    rna-XM_059377883.1    rna-XM_045081320.1    mNeoVis.2.1.1.chr8_hap1.g01508.t01    rna-XM_021695995.2    rna-XM_004409590.1    rna-XM_042980109.1    rna-XM_040624541.1    rna-XM_027590257.2    mNeoVis.3.1.1.chr8.g01405.t01    rna-gnl|WGS_ZZZZ|egapxtmp_019582-R1    mNeoVis.evergladensis.2.1.chr9.g01457.t01    mNeoVis.vulgivaga.2.1.chr8.g01460.t01      1217    N0.HOG0001215   OG0000118  n111   rna-XM_002925583.4    rna-NM_001002979.1    rna-XM_022524848.1    rna-XM_023249434.2    rna-XM_036112455.1    rna-XM_039233532.1    rna-XM_032837167.1    rna-XM_045995411.1    rna-XM_032331819.1    rna-XM_059158581.1    rna-XM_059385657.1    rna-XM_004769031.3    mNeoVis.2.1.1.chr12_hap1.g00294.t01    rna-XM_021678553.2    rna-XM_004399766.1    rna-XM_042974017.1    rna-XM_040627518.1    rna-XM_027609389.1    mNeoVis.3.1.1.chr12.g00262.t01    rna-gnl|WGS_ZZZZ|egapxtmp_029487-R1    mNeoVis.evergladensis.2.1.chr12.g00262.t01    mNeoVis.vulgivaga.2.1.chr12.g00269.t01      1218    N0.HOG0001216   OG0000119  n3     rna-XM_038557530.1    rna-XM_022491966.1, rna-XM_022492472.1    rna-XM_003980584.2    rna-XM_036100234.1    rna-XM_039222536.1      rna-XM_045999663.1                rna-XM_004406930.1    rna-XM_007092967.2    rna-XM_008709889.2                1219    N0.HOG0001217   OG0000119  n12     rna-XM_038557531.1    rna-XM_022491626.1      rna-XM_036100217.1    rna-XM_039225430.1    rna-XM_032851163.1      rna-XM_032335003.1    rna-XM_059173417.1    rna-XM_059416445.1    rna-XM_004781735.1    mNeoVis.2.1.1.chr1_hap1.g00981.t01, mNeoVis.2.1.1.chr1_hap1.g00982.t01    rna-XM_021690945.1    rna-XM_004406931.1      rna-XM_008709888.1                1220    N0.HOG0001218   OG0000119  n25   rna-XM_002926905.1, rna-XM_011217033.2    rna-NM_001389148.1    rna-XM_022491627.1, rna-XM_022492478.1    rna-XM_003980570.2, rna-XM_011283154.2, rna-XM_019833844.1      rna-XM_039218785.1, rna-XM_039222539.1, rna-XM_039222541.1    rna-XM_032847582.1, rna-XM_032848096.1, rna-XM_032848504.1    rna-XM_045998699.1, rna-XM_045999717.1, rna-XM_046001122.1    rna-XM_032334926.1, rna-XM_032334983.1, rna-XM_032335005.1, rna-XM_032335047.1    rna-XM_059173383.1, rna-XM_059173386.1, rna-XM_059173503.1, rna-XM_059173513.1    rna-XM_059416407.1, rna-XM_059416410.1, rna-XM_059416726.1, rna-XM_059416745.1    rna-XM_004780237.3, rna-XM_004781737.1, rna-XM_004782325.2, rna-XM_013050520.2    mNeoVis.2.1.1.chr1_hap1.g00929.t01, mNeoVis.2.1.1.chr1_hap1.g00979.t01, mNeoVis.2.1.1.chr1_hap1.g01006.t01        rna-XM_007088188.1, rna-XM_007092960.2, rna-XM_007092969.1    rna-XM_008683749.2, rna-XM_008709869.2, rna-XM_008709891.2, rna-XM_008710076.2, rna-XM_008710106.2, rna-XM_040622740.1    rna-XM_027604474.1, rna-XM_027604484.1    mNeoVis.3.1.1.chr1.g00943.t01, mNeoVis.3.1.1.chr1.g00986.t01, mNeoVis.3.1.1.chr1.g01006.t01    rna-gnl|WGS_ZZZZ|egapxtmp_020958-R1, rna-gnl|WGS_ZZZZ|egapxtmp_021061-R1    mNeoVis.evergladensis.2.1.chr1.g00924.t01, mNeoVis.evergladensis.2.1.chr1.g00974.t01, mNeoVis.evergladensis.2.1.chr1.g00998.t01    mNeoVis.vulgivaga.2.1.chr1.g00937.t01, mNeoVis.vulgivaga.2.1.chr1.g00987.t01, mNeoVis.vulgivaga.2.1.chr1.g01012.t01      1221    N0.HOG0001219   OG0000119  n81     rna-NM_001389000.1    rna-XM_022491625.1        rna-XM_039225549.1    rna-XM_032844374.1, rna-XM_032849957.1    rna-XM_045999718.1, rna-XM_046001107.1    rna-XM_032335002.1    rna-XM_059173416.1    rna-XM_059416446.1    rna-XM_004781734.1    mNeoVis.2.1.1.chr1_hap1.g00983.t01      rna-XM_004406932.1      rna-XM_008709887.2    rna-XM_027604475.1    mNeoVis.3.1.1.chr1.g00988.t01    rna-gnl|WGS_ZZZZ|egapxtmp_019703-R1    mNeoVis.evergladensis.2.1.chr1.g00976.t01    mNeoVis.vulgivaga.2.1.chr1.g00988.t01      1222    N0.HOG0001220   OG0000119  n99     rna-XM_038557529.1    rna-XM_022491622.1    rna-XM_011283147.3    rna-XM_036100219.1    rna-XM_039222537.1, rna-XM_039222538.1    rna-XM_032850410.1    rna-XM_045996807.1, rna-XM_045996813.1, rna-XM_045996936.1, rna-XM_045997025.1, rna-XM_045997030.1, rna-XM_046001100.1, rna-XM_046001108.1    rna-XM_032335004.1    rna-XM_059158833.1, rna-XM_059173411.1, rna-XM_059173418.1    rna-XM_059416444.1    rna-XM_004781736.1    mNeoVis.2.1.1.chr1_hap1.g00980.t01    rna-XM_021690943.2    rna-XM_004406929.1    rna-XM_007092968.1    rna-XM_008709868.2, rna-XM_008709890.1, rna-XM_008710077.2, rna-XM_040621943.1      mNeoVis.3.1.1.chr1.g00987.t01    rna-gnl|WGS_ZZZZ|egapxtmp_021563-R1    mNeoVis.evergladensis.2.1.chr1.g00975.t01        1223    N0.HOG0001221   OG0000120  n1   rna-XM_034638257.1    rna-NM_001003342.1    rna-XM_022525141.1    rna-XM_045046841.1    rna-XM_036067884.1    rna-XM_039249588.1    rna-XM_032842052.1    rna-XM_045988668.1    rna-XM_032323605.1    rna-XM_059153019.1    rna-XM_059383703.1    rna-XM_045075940.1    mNeoVis.2.1.1.chr7_hap1.g00434.t01    rna-XM_021700795.2    rna-XM_004417088.1    rna-XM_042968888.1    rna-XM_040634385.1    rna-XM_027617396.1    mNeoVis.3.1.1.chr7.g00421.t01    rna-gnl|WGS_ZZZZ|egapxtmp_012219-R1    mNeoVis.evergladensis.2.1.chr7.g00438.t01    mNeoVis.vulgivaga.2.1.chr7.g00436.t01      1224   N0.HOG0001222  OG0000120  n23   rna-XM_034664929.1    rna-XM_038573552.1    rna-XM_022522577.1    rna-XM_045037131.1    rna-XM_036097131.1    rna-XM_039256937.1    rna-XM_032838242.1    rna-XM_045978493.1    rna-XM_032315213.1    rna-XM_059144546.1    rna-XM_059377687.1    rna-XM_004778223.3    mNeoVis.2.1.1.chr8_hap1.g01156.t01    rna-XM_021678182.2    rna-XM_004401374.1    rna-XM_042985700.1    rna-XM_040620881.1    rna-XM_035726841.1, rna-XM_035729023.1    mNeoVis.3.1.1.chr8.g01071.t01    rna-gnl|WGS_ZZZZ|egapxtmp_019875-R1    mNeoVis.evergladensis.2.1.chr9.g01128.t01    mNeoVis.vulgivaga.2.1.chr8.g01127.t01      1225   N0.HOG0001223  OG0000120  n46   rna-XM_034667296.1    rna-XM_038586359.1    rna-XM_022490725.1    rna-XM_023247846.2    rna-XM_036091443.1    rna-XM_039246428.1    rna-XM_032845612.1    rna-XM_045982322.1    rna-XM_032318338.1    rna-XM_059145861.1    rna-XM_059413532.1    rna-XM_004775837.2    mNeoVis.2.1.1.chr14_hap1.g00526.t01    rna-XM_021681913.2    rna-XM_004407862.1    rna-XM_042975542.1    rna-XM_040634481.1    rna-XM_027612787.1    mNeoVis.3.1.1.chr14.g00521.t01    rna-gnl|WGS_ZZZZ|egapxtmp_032178-R1    mNeoVis.evergladensis.2.1.chr14.g00525.t01    mNeoVis.vulgivaga.2.1.chr14.g00523.t01      1226    N0.HOG0001224   OG0000120  n69   rna-XM_019804554.2, rna-XM_034667301.1    rna-XM_038528773.1    rna-XM_022490724.1, rna-XM_022491763.1    rna-XM_019822324.3, rna-XM_045047210.1    rna-XM_036091445.1, rna-XM_036103267.1    rna-XM_039246431.1, rna-XM_039249910.1    rna-XM_032845613.1, rna-XM_032849601.1    rna-XM_045982320.1, rna-XM_045989050.1    rna-XM_032318340.1, rna-XM_032323108.1    rna-XM_059145862.1, rna-XM_059150592.1    rna-XM_059381671.1, rna-XM_059413531.1    rna-XM_004775836.3, rna-XM_045077228.1    mNeoVis.2.1.1.chr14_hap1.g00527.t01, mNeoVis.2.1.1.chr7_hap1.g01094.t01    rna-XM_021692257.1, rna-XM_044916278.1    rna-XM_004407861.1, rna-XM_004408162.1    rna-XM_042969815.1, rna-XM_042975545.1    rna-XM_040634419.1, rna-XM_040634480.1    rna-XM_027620030.2, rna-XM_035722283.1    mNeoVis.3.1.1.chr14.g00522.t01, mNeoVis.3.1.1.chr7.g01022.t01      mNeoVis.evergladensis.2.1.chr14.g00526.t01, mNeoVis.evergladensis.2.1.chr7.g01085.t01    mNeoVis.vulgivaga.2.1.chr14.g00524.t01, mNeoVis.vulgivaga.2.1.chr7.g01097.t01      1227    N0.HOG0001225   OG0000120  n96   rna-XM_019804162.2    rna-NM_001389224.1    rna-XM_022496884.1    rna-XM_011285086.4    rna-XM_036113355.1    rna-XM_039242723.1    rna-XM_032881343.1    rna-XM_046026893.1    rna-XM_032361022.1    rna-XM_059135103.1    rna-XM_059375844.1    rna-XM_045081865.1    mNeoVis.2.1.1.chr2_hap1.g00901.t01    rna-XM_021690233.2    rna-XM_004416508.1    rna-XM_042993813.1    rna-XM_040626867.1    rna-XM_035727346.1    mNeoVis.3.1.1.chr5.g00834.t01      mNeoVis.evergladensis.2.1.chr5.g00856.t01    mNeoVis.vulgivaga.2.1.chr5.g00845.t01      1228    N0.HOG0001226   OG0000121  n1   rna-XM_034645977.1    rna-NM_001002932.1    rna-XM_022522452.1    rna-XM_045060755.1    rna-XM_036117632.1    rna-XM_039218403.1    rna-XM_032874808.1    rna-XM_046011417.1    rna-XM_032349034.1    rna-XM_059184573.1    rna-XM_059403765.1    rna-XM_004766734.3    mNeoVis.2.1.1.chr9_hap1.g00242.t01    rna-XM_044915225.1    rna-XM_004413880.1    rna-XM_007089645.3    rna-XM_040642061.1    rna-XM_027595203.2    mNeoVis.3.1.1.chr10.g00234.t01    rna-gnl|WGS_ZZZZ|egapxtmp_024615-R1    mNeoVis.evergladensis.2.1.chr10.g00240.t01    mNeoVis.vulgivaga.2.1.chr10.g00236.t01      1229   N0.HOG0001227  OG0000121  n24   rna-XM_034646029.1    rna-XM_038577308.1, rna-XM_038577313.1    rna-XM_022519637.1    rna-XM_045062664.1    rna-XM_036086737.1    rna-XM_039223568.1, rna-XM_039225640.1    rna-XM_032854660.1, rna-XM_032876522.1    rna-XM_046011813.1, rna-XM_046011814.1    rna-XM_032347777.1    rna-XM_059187502.1, rna-XM_059187504.1      rna-XM_045072972.1, rna-XM_045073242.1    mNeoVis.2.1.1.chr9_hap1.g00668.t01, mNeoVis.2.1.1.chr9_hap1.g00669.t01      rna-XM_004402277.1    rna-XM_042993686.1    rna-XM_040641940.1    rna-XM_035729324.1    mNeoVis.3.1.1.chr10.g00633.t01    rna-gnl|WGS_ZZZZ|egapxtmp_024719-R1    mNeoVis.evergladensis.2.1.chr10.g00664.t01    mNeoVis.vulgivaga.2.1.chr10.g00647.t01      1230    N0.HOG0001228   OG0000121  n50   rna-XM_034644860.1    rna-XM_038564485.1, rna-XM_038564486.1    rna-XM_022506559.1        rna-XM_039221591.1, rna-XM_039225011.1    rna-XM_032879098.1    rna-XM_046014606.1, rna-XM_046014607.1, rna-XM_046016340.1    rna-XM_032359926.1    rna-XM_059143666.1      rna-XM_045089727.1    mNeoVis.2.1.1.chr7_hap1.g02647.t01      rna-XM_004403824.1    rna-XM_042959649.1        mNeoVis.3.1.1.chr7.g02297.t01    rna-gnl|WGS_ZZZZ|egapxtmp_012733-R1    mNeoVis.evergladensis.2.1.chr7.g02459.t01    mNeoVis.vulgivaga.2.1.chr7.g02481.t01      1231    N0.HOG0001229   OG0000121  n71   rna-XM_034644859.1    rna-XM_038563537.1    rna-XM_022506560.1    rna-XM_045038110.1    rna-XM_036083750.1    rna-XM_039225012.1    rna-XM_032879097.1    rna-XM_046014987.1    rna-XM_032358329.1    rna-XM_059151754.1    rna-XM_059398249.1    rna-XM_045066792.1    mNeoVis.2.1.1.chr7_hap1.g02646.t01    rna-XM_044919533.1    rna-XM_004403825.1    rna-XM_042959647.1    rna-XM_040638798.1    rna-XM_035723154.1      rna-gnl|WGS_ZZZZ|egapxtmp_012732-R1    mNeoVis.evergladensis.2.1.chr7.g02458.t01    mNeoVis.vulgivaga.2.1.chr7.g02480.t01      1232    N0.HOG0001230   OG0000121  n92   rna-XM_034645861.1    rna-XM_038563538.1      rna-XM_045039740.1    rna-XM_036083630.1, rna-XM_036085807.1    rna-XM_039225013.1    rna-XM_032879792.1    rna-XM_046014610.1    rna-XM_032358322.1    rna-XM_059151769.1    rna-XM_059386835.1    rna-XM_045066788.1    mNeoVis.2.1.1.chr7_hap1.g02645.t01        rna-XM_042959646.1    rna-XM_040639685.1    rna-XM_035723153.1          mNeoVis.vulgivaga.2.1.chr7.g02479.t01      1233    N0.HOG0001231   OG0000121  n108   rna-XM_034644858.1, rna-XM_034653153.1    rna-XM_038564487.1    rna-XM_022506565.1    rna-XM_045039739.1    rna-XM_036070745.1, rna-XM_036085808.1    rna-XM_039224105.1, rna-XM_039225014.1    rna-XM_032879096.1    rna-XM_046014619.1    rna-XM_032358328.1    rna-XM_059151719.1      rna-XM_045066780.1    mNeoVis.2.1.1.chr7_hap1.g02644.t01      rna-XM_004403826.1    rna-XM_042959645.1    rna-XM_040638797.1    rna-XM_035723152.1      rna-gnl|WGS_ZZZZ|egapxtmp_012731-R1    mNeoVis.evergladensis.2.1.chr7.g02457.t01    mNeoVis.vulgivaga.2.1.chr7.g02478.t01      1234   N0.HOG0001232  OG0000122  n1   rna-XM_011234811.3    rna-XM_038586386.1    rna-XM_022490811.1    rna-XM_019822340.3, rna-XM_045048638.1    rna-XM_036069767.1    rna-XM_039246435.1    rna-XM_032845649.1    rna-XM_045983046.1    rna-XM_032318407.1    rna-XM_059146128.1    rna-XM_059414147.1    rna-XM_013047454.2    mNeoVis.2.1.1.chr14_hap1.g00546.t01    rna-XM_021681928.2    rna-XM_004407922.1    rna-XM_042975723.1    rna-XM_008700716.2    rna-XM_027611949.2    mNeoVis.3.1.1.chr14.g00540.t01    rna-gnl|WGS_ZZZZ|egapxtmp_033039-R1    mNeoVis.evergladensis.2.1.chr14.g00545.t01    mNeoVis.vulgivaga.2.1.chr14.g00544.t01      1235   N0.HOG0001233  OG0000122  n22     rna-NM_001388778.1    rna-XM_022490812.1    rna-XM_003999605.4    rna-XM_036069832.1    rna-XM_039246417.1    rna-XM_032845659.1    rna-XM_045983457.1    rna-XM_032319359.1    rna-XM_059146129.1    rna-XM_059414144.1    rna-XM_004775818.1    mNeoVis.2.1.1.chr14_hap1.g00544.t01    rna-XM_021681926.1      rna-XM_015538464.2    rna-XM_008700777.2      mNeoVis.3.1.1.chr14.g00537.t01    rna-gnl|WGS_ZZZZ|egapxtmp_033059-R1    mNeoVis.evergladensis.2.1.chr14.g00542.t01    mNeoVis.vulgivaga.2.1.chr14.g00541.t01      1236    N0.HOG0001234   OG0000122  n39   rna-XM_011237480.3, rna-XM_011237481.2, rna-XM_034650114.1    rna-NM_001388780.1    rna-XM_022490739.1    rna-XM_045048637.1    rna-XM_036069772.1    rna-XM_039246518.1    rna-XM_032845721.1    rna-XM_045983332.1    rna-XM_032317572.1          mNeoVis.2.1.1.chr14_hap1.g00542.t01    rna-XM_021681836.2      rna-XM_042975560.1    rna-XM_008700713.2    rna-XM_027611948.1, rna-XM_035722021.1    mNeoVis.3.1.1.chr14.g00533.t01    rna-gnl|WGS_ZZZZ|egapxtmp_033055-R1    mNeoVis.evergladensis.2.1.chr14.g00540.t01    mNeoVis.vulgivaga.2.1.chr14.g00539.t01      1237    N0.HOG0001235   OG0000122  n59   rna-XM_002927275.4, rna-XM_011234815.2, rna-XM_034667322.1    rna-NM_001389196.1, rna-XM_038586387.1    rna-XM_022490813.1, rna-XM_022490814.1, rna-XM_022490818.1, rna-XM_022491169.1    rna-XM_003999604.5, rna-XM_003999606.4, rna-XM_045048479.1    rna-XM_036069770.1, rna-XM_036069773.1, rna-XM_036069774.1, rna-XM_036069833.1    rna-XM_039246436.1, rna-XM_039246437.1, rna-XM_039246515.1    rna-XM_032845626.1, rna-XM_032845662.1, rna-XM_032845719.1    rna-XM_045983331.1, rna-XM_045983448.1, rna-XM_045983481.1, rna-XM_045983508.1    rna-XM_032316981.1, rna-XM_032317573.1, rna-XM_032318408.1, rna-XM_032319217.1    rna-XM_059146131.1, rna-XM_059146623.1, rna-XM_059146882.1    rna-XM_059414146.1, rna-XM_059414150.1, rna-XM_059414216.1    rna-XM_004775964.2, rna-XM_004775966.2, rna-XM_045086616.1    mNeoVis.2.1.1.chr14_hap1.g00543.t01, mNeoVis.2.1.1.chr14_hap1.g00545.t01, mNeoVis.2.1.1.chr14_hap1.g00548.t01    rna-XM_021681838.1, rna-XM_021681927.1, rna-XM_021681929.1      rna-XM_007084303.2, rna-XM_042975558.1, rna-XM_042975559.1    rna-XM_008700715.2, rna-XM_040634575.1    rna-XM_027612863.1, rna-XM_027613060.2    mNeoVis.3.1.1.chr14.g00536.t01, mNeoVis.3.1.1.chr14.g00538.t01, mNeoVis.3.1.1.chr14.g00541.t01    rna-gnl|WGS_ZZZZ|egapxtmp_033053-R3, rna-gnl|WGS_ZZZZ|egapxtmp_033054-R1, rna-gnl|WGS_ZZZZ|egapxtmp_033061-R1    mNeoVis.evergladensis.2.1.chr14.g00541.t01, mNeoVis.evergladensis.2.1.chr14.g00543.t01, mNeoVis.evergladensis.2.1.chr14.g00544.t01, mNeoVis.evergladensis.2.1.chr14.g00547.t01    mNeoVis.vulgivaga.2.1.chr14.g00540.t01, mNeoVis.vulgivaga.2.1.chr14.g00542.t01, mNeoVis.vulgivaga.2.1.chr14.g00543.t01, mNeoVis.vulgivaga.2.1.chr14.g00546.t01      1238    N0.HOG0001236   OG0000123  n2     rna-XM_038581136.1, rna-XM_038581137.1, rna-XM_038581138.1, rna-XM_038581139.1, rna-XM_038581140.1, rna-XM_038581141.1, rna-XM_038581142.1, rna-XM_038581143.1    rna-XM_022492756.1, rna-XM_022515895.1, rna-XM_022515896.1      rna-XM_036118549.1, rna-XM_036118550.1, rna-XM_036118551.1      rna-XM_032865557.1, rna-XM_032865560.1, rna-XM_032865561.1, rna-XM_032865562.1    rna-XM_046003710.1, rna-XM_046003711.1    rna-XM_032302636.1, rna-XM_032302659.1, rna-XM_032302705.1, rna-XM_032302715.1, rna-XM_032302728.1, rna-XM_032302742.1    rna-XM_059165100.1, rna-XM_059165101.1, rna-XM_059165102.1, rna-XM_059165103.1, rna-XM_059165104.1, rna-XM_059165106.1    rna-XM_059388392.1, rna-XM_059388393.1, rna-XM_059388394.1    rna-XM_004757559.2, rna-XM_013058884.2, rna-XM_013058885.2, rna-XM_013058888.1, rna-XM_013058889.1    mNeoVis.2.1.1.chr5_hap1.g01772.t01, mNeoVis.2.1.1.chr5_hap1.g01774.t01, mNeoVis.2.1.1.chr5_hap1.g01776.t01            rna-XM_027581785.1, rna-XM_035724975.1, rna-XM_035727990.1, rna-XM_035728165.1, rna-XM_035728273.1, rna-XM_035728381.1    mNeoVis.3.1.1.chr2.g01682.t01, mNeoVis.3.1.1.chr2.g01685.t01    rna-gnl|WGS_ZZZZ|egapxtmp_034589-R1, rna-gnl|WGS_ZZZZ|egapxtmp_034592-R1, rna-gnl|WGS_ZZZZ|egapxtmp_034595-R1, rna-gnl|WGS_ZZZZ|egapxtmp_034603-R1    mNeoVis.evergladensis.2.1.chr2.g01757.t01, mNeoVis.evergladensis.2.1.chr2.g01760.t01    mNeoVis.vulgivaga.2.1.chr2.g01772.t01, mNeoVis.vulgivaga.2.1.chr2.g01775.t01      1239   N0.HOG0001237  OG0000123  n54   rna-XM_034661549.1, rna-XM_034661675.1, rna-XM_034661722.1    rna-XM_038581732.1      rna-XM_045036417.1, rna-XM_045036418.1, rna-XM_045036419.1, rna-XM_045036420.1, rna-XM_045036421.1, rna-XM_045036422.1, rna-XM_045036423.1, rna-XM_045036424.1, rna-XM_045036425.1, rna-XM_045036426.1, rna-XM_045036427.1, rna-XM_045036428.1, rna-XM_045036429.1, rna-XM_045036430.1, rna-XM_045036431.1    rna-XM_036118552.1    rna-XM_039228520.1, rna-XM_039229600.1, rna-XM_039229612.1, rna-XM_039229626.1, rna-XM_039229640.1, rna-XM_039229651.1    rna-XM_032865558.1, rna-XM_032865559.1    rna-XM_046003708.1, rna-XM_046003712.1, rna-XM_046003713.1            mNeoVis.2.1.1.chr5_hap1.g01773.t01, mNeoVis.2.1.1.chr5_hap1.g01775.t01    rna-XM_044919015.1      rna-XM_042998294.1, rna-XM_042998295.1, rna-XM_042998296.1, rna-XM_042998297.1, rna-XM_042998298.1, rna-XM_042998300.1, rna-XM_042998301.1, rna-XM_042998302.1, rna-XM_042998303.1, rna-XM_042998304.1, rna-XM_042998305.1, rna-XM_042998306.1, rna-XM_042998307.1, rna-XM_042998308.1, rna-XM_042998309.1, rna-XM_042998542.1, rna-XM_042998603.1, rna-XM_042998799.1, rna-XM_042998825.1    rna-XM_040621686.1, rna-XM_040621689.1, rna-XM_040621690.1, rna-XM_040621691.1, rna-XM_040621692.1, rna-XM_040621693.1    rna-XM_027581771.2    mNeoVis.3.1.1.chr2.g01681.t01, mNeoVis.3.1.1.chr2.g01683.t01, mNeoVis.3.1.1.chr2.g01684.t01    rna-gnl|WGS_ZZZZ|egapxtmp_034588-R1, rna-gnl|WGS_ZZZZ|egapxtmp_034596-R1    mNeoVis.evergladensis.2.1.chr2.g01758.t01, mNeoVis.evergladensis.2.1.chr2.g01759.t01    mNeoVis.vulgivaga.2.1.chr2.g01773.t01, mNeoVis.vulgivaga.2.1.chr2.g01774.t01      1240    N0.HOG0001238   OG0000124  n1   rna-XM_034671993.1    rna-XM_038533538.1, rna-XM_038581644.1    rna-XM_022523470.1    rna-XM_045056492.1    rna-XM_036099862.1      rna-XM_032853759.1    rna-XM_045998274.1    rna-XM_032334038.1    rna-XM_059164147.1    rna-XM_059395899.1    rna-XM_045076273.1    mNeoVis.2.1.1.chr6_hap1.g00254.t01    rna-XM_021679421.1    rna-XM_004402877.1    rna-XM_042984818.1    rna-XM_040625314.1    rna-XM_027598905.1    mNeoVis.3.1.1.chr4.g00253.t01    rna-gnl|WGS_ZZZZ|egapxtmp_001292-R1    mNeoVis.evergladensis.2.1.chr4.g00260.t01    mNeoVis.vulgivaga.2.1.chr4.g00259.t01      1241    N0.HOG0001239   OG0000124  n16   rna-XM_019799498.2      rna-XM_022516202.1    rna-XM_023238639.2    rna-XM_036112731.1    rna-XM_039257113.1    rna-XM_032865252.1, rna-XM_032865510.1    rna-XM_045999262.1, rna-XM_046001707.1, rna-XM_046018905.1    rna-XM_032322716.1, rna-XM_032346572.1, rna-XM_032357110.1    rna-XM_059164929.1    rna-XM_059392503.1    rna-XM_045084343.1    mNeoVis.2.1.1.chr5_hap1.g01908.t01    rna-XM_021688695.1, rna-XM_044913286.1, rna-XM_044921820.1    rna-XM_004392932.1, rna-XM_004405799.1, rna-XM_004410647.1    rna-XM_042998678.1    rna-XM_040621816.1    rna-XM_027582869.2, rna-XM_027602053.2, rna-XM_027624556.2, rna-XM_035726248.1    mNeoVis.3.1.1.chr2.g01796.t01, mNeoVis.3.1.1.chr8.g00123.t01      mNeoVis.evergladensis.2.1.chr2.g01887.t01    mNeoVis.vulgivaga.2.1.chr2.g01900.t01, mNeoVis.vulgivaga.2.1.chr8.g00154.t01      1242    N0.HOG0001240   OG0000124  n43           rna-XM_036067065.1, rna-XM_036091383.1                    rna-XM_021678828.1          rna-XM_027574535.1              1243    N0.HOG0001241   OG0000124  n47                           mNeoVis.2.1.1.chr3_hap1.g01098.t01    rna-XM_044916894.1            mNeoVis.3.1.1.chr3.g01086.t01      mNeoVis.evergladensis.2.1.chr3.g01097.t01    mNeoVis.vulgivaga.2.1.chr3.g01113.t01      1244    N0.HOG0001242   OG0000124  n53     rna-NM_001252289.1        rna-XM_036088716.1    rna-XM_039226265.1    rna-XM_032854132.1    rna-XM_045986313.1    rna-XM_032331231.1    rna-XM_059170596.1    rna-XM_059378900.1      mNeoVis.2.1.1.chr8_hap1.g00154.t01      rna-XM_004394348.1      rna-XM_040640248.1        rna-gnl|WGS_ZZZZ|egapxtmp_020272-R1    mNeoVis.evergladensis.2.1.chr9.g00151.t01        1245    N0.HOG0001243   OG0000124  n54   rna-XM_019800307.2, rna-XM_019803114.2, rna-XM_034648255.1    rna-XM_038541765.1      rna-XM_003997214.6, rna-XM_045050717.1    rna-XM_036103438.1, rna-XM_036108381.1, rna-XM_036111786.1, rna-XM_036113514.1, rna-XM_036121311.1    rna-XM_039233484.1, rna-XM_039236687.1    rna-XM_032855549.1, rna-XM_032870646.1    rna-XM_046016800.1    rna-XM_032335883.1, rna-XM_032357959.1    rna-XM_059148669.1, rna-XM_059175322.1    rna-XM_059416654.1    rna-XM_045085931.1    mNeoVis.2.1.1.chr1_hap1.g00696.t01    rna-XM_021688287.1, rna-XM_021696483.1, rna-XM_021698489.2, rna-XM_044912259.1, rna-XM_044912686.1, rna-XM_044919456.1, rna-XM_044919457.1    rna-XM_004393641.1, rna-XM_004406692.1, rna-XM_004406731.1, rna-XM_004411779.1    rna-XM_007091245.3, rna-XM_042974631.1    rna-XM_008684893.1, rna-XM_040644918.1    rna-XM_027580787.1, rna-XM_027581739.2, rna-XM_035723752.1, rna-XM_035727415.1, rna-XM_035728331.1    mNeoVis.3.1.1.chr1.g00394.t01, mNeoVis.3.1.1.chr1.g00715.t01    rna-gnl|WGS_ZZZZ|egapxtmp_020946-R1, rna-gnl|WGS_ZZZZ|egapxtmp_031825-R1    mNeoVis.evergladensis.2.1.chr1.g00703.t01    mNeoVis.vulgivaga.2.1.chr1.g00716.t01      1246    N0.HOG0001244   OG0000125  n3                                   rna-XM_008684240.2, rna-XM_040620958.1                1247    N0.HOG0001245   OG0000125  n5   rna-XM_002931322.2    rna-NM_001388961.1      rna-XM_003987356.2    rna-XM_036082599.1    rna-XM_039218364.1    rna-XM_032857070.1    rna-XM_046010219.1                  rna-XM_007087492.2    rna-XM_008684239.2, rna-XM_008710153.2                1248    N0.HOG0001246   OG0000125  n14   rna-XM_034652662.1    rna-NM_001388966.1    rna-XM_022491259.1, rna-XM_022521777.1    rna-XM_011283342.3, rna-XM_045059601.1    rna-XM_036082595.1, rna-XM_036082597.1    rna-XM_039218356.1, rna-XM_039218362.1    rna-XM_032857071.1, rna-XM_032857073.1    rna-XM_046009662.1, rna-XM_046009673.1    rna-XM_032341775.1, rna-XM_032341778.1    rna-XM_059179636.1, rna-XM_059179637.1    rna-XM_059371140.1    rna-XM_004782098.2, rna-XM_004782101.2    mNeoVis.2.1.1.chr10_hap1.g00805.t01, mNeoVis.2.1.1.chr10_hap1.g00808.t01      rna-XM_004417620.1    rna-XM_007087491.1, rna-XM_015539951.1    rna-XM_040626191.1    rna-XM_027571592.2    mNeoVis.3.1.1.chr9.g00760.t01    rna-gnl|WGS_ZZZZ|egapxtmp_016926-R1, rna-gnl|WGS_ZZZZ|egapxtmp_017365-R1    mNeoVis.evergladensis.2.1.chr8.g00785.t01        1249    N0.HOG0001247   OG0000125  n45   rna-XM_011218364.3, rna-XM_034651700.1    rna-NM_001388767.1    rna-XM_022490688.1    rna-XM_003987315.1    rna-XM_036082601.1    rna-XM_039218365.1    rna-XM_032857068.1    rna-XM_046009588.1    rna-XM_032341781.1    rna-XM_059179638.1    rna-XM_059371142.1    rna-XM_004782103.2    mNeoVis.2.1.1.chr10_hap1.g00803.t01    rna-XM_021695724.1    rna-XM_004414392.1    rna-XM_007087494.1      rna-XM_027571594.1      rna-gnl|WGS_ZZZZ|egapxtmp_017332-R1    mNeoVis.evergladensis.2.1.chr8.g00779.t01, mNeoVis.evergladensis.2.1.chr8.g00783.t01    mNeoVis.vulgivaga.2.1.chr9.g00781.t01      1250    N0.HOG0001248   OG0000125  n66     rna-XM_038579867.1    rna-XM_022490683.1, rna-XM_022521936.1, rna-XM_022521947.1    rna-XM_003987316.4, rna-XM_023256171.2    rna-XM_036082600.1    rna-XM_039218328.1, rna-XM_039218329.1    rna-XM_032857065.1, rna-XM_032857067.1, rna-XM_032857069.1    rna-XM_046007845.1, rna-XM_046007846.1, rna-XM_046007847.1, rna-XM_046007850.1    rna-XM_032341780.1, rna-XM_032341783.1, rna-XM_032341784.1, rna-XM_032341789.1, rna-XM_032341791.1    rna-XM_059183400.1, rna-XM_059183401.1, rna-XM_059183402.1, rna-XM_059183404.1, rna-XM_059183405.1, rna-XM_059183406.1    rna-XM_059372104.1, rna-XM_059372105.1, rna-XM_059372106.1, rna-XM_059372107.1, rna-XM_059372108.1, rna-XM_059372109.1    rna-XM_004782102.1, rna-XM_004782267.2, rna-XM_004782419.1, rna-XM_004782423.2, rna-XM_004782482.1, rna-XM_045067933.1, rna-XM_045076709.1    mNeoVis.2.1.1.chr10_hap1.g00796.t01, mNeoVis.2.1.1.chr10_hap1.g00799.t01, mNeoVis.2.1.1.chr10_hap1.g00804.t01    rna-XM_021695723.1, rna-XM_021695725.1    rna-XM_004414391.1    rna-XM_007087493.2, rna-XM_007087495.2    rna-XM_008710151.2    rna-XM_027570325.2, rna-XM_027570327.1    mNeoVis.3.1.1.chr9.g00755.t01, mNeoVis.3.1.1.chr9.g00756.t01, mNeoVis.3.1.1.chr9.g00758.t01    rna-gnl|WGS_ZZZZ|egapxtmp_016917-R1, rna-gnl|WGS_ZZZZ|egapxtmp_016919-R1, rna-gnl|WGS_ZZZZ|egapxtmp_016923-R1    mNeoVis.evergladensis.2.1.chr8.g00780.t01, mNeoVis.evergladensis.2.1.chr8.g00784.t01    mNeoVis.vulgivaga.2.1.chr9.g00777.t01, mNeoVis.vulgivaga.2.1.chr9.g00782.t01      1251    N0.HOG0001249   OG0000126  n1   rna-XM_034640436.1    rna-XM_038546685.1    rna-XM_022519847.1    rna-XM_019818205.2    rna-XM_036079020.1    rna-XM_039229809.1    rna-XM_032871079.1    rna-XM_045986515.1    rna-XM_032319547.1    rna-XM_059147738.1    rna-XM_059378959.1    rna-XM_045085971.1    mNeoVis.2.1.1.chr8_hap1.g00189.t01    rna-XM_021678472.1    rna-XM_004401848.1    rna-XM_042967755.1    rna-XM_040640303.1    rna-XM_027626406.1    mNeoVis.3.1.1.chr8.g00149.t01    rna-gnl|WGS_ZZZZ|egapxtmp_020252-R2    mNeoVis.evergladensis.2.1.chr9.g00181.t01    mNeoVis.vulgivaga.2.1.chr8.g00185.t01      1252    N0.HOG0001250   OG0000126  n23   rna-XM_034640292.1    rna-XM_038546666.1    rna-XM_022523205.1, rna-XM_022523207.1    rna-XM_023244065.2    rna-XM_036078997.1    rna-XM_039226521.1    rna-XM_032870697.1    rna-XM_045986282.1    rna-XM_032323013.1    rna-XM_059148754.1    rna-XM_059381127.1    rna-XM_013054454.2    mNeoVis.2.1.1.chr8_hap1.g00176.t01    rna-XM_021678383.1    rna-XM_004394404.1    rna-XM_015541659.2    rna-XM_040640289.1    rna-XM_027626525.2    mNeoVis.3.1.1.chr8.g00145.t01      mNeoVis.evergladensis.2.1.chr9.g00173.t01    mNeoVis.vulgivaga.2.1.chr8.g00176.t01      1253    N0.HOG0001251   OG0000126  n44   rna-XM_002931051.4, rna-XM_034650782.1    rna-XM_038546678.1    rna-XM_022519832.1, rna-XM_022519833.1, rna-XM_022519834.1    rna-XM_045044701.1    rna-XM_036079002.1, rna-XM_036079004.1, rna-XM_036079012.1, rna-XM_036079015.1    rna-XM_039229813.1    rna-XM_032850831.1, rna-XM_032870684.1, rna-XM_032870687.1, rna-XM_032870698.1    rna-XM_045984874.1, rna-XM_045984877.1, rna-XM_045984879.1, rna-XM_045984881.1, rna-XM_045984883.1, rna-XM_045984885.1, rna-XM_045984887.1, rna-XM_045984890.1, rna-XM_045985572.1, rna-XM_045985849.1, rna-XM_045986281.1    rna-XM_032319992.1, rna-XM_032319997.1, rna-XM_032320001.1, rna-XM_032320004.1, rna-XM_032320005.1    rna-XM_059147438.1, rna-XM_059147440.1, rna-XM_059147450.1    rna-XM_059379372.1, rna-XM_059380465.1, rna-XM_059380469.1, rna-XM_059380470.1    rna-XM_004749006.3, rna-XM_004749008.3, rna-XM_004749013.3, rna-XM_004749017.3, rna-XM_045085953.1, rna-XM_045085966.1    mNeoVis.2.1.1.chr8_hap1.g00180.t01, mNeoVis.2.1.1.chr8_hap1.g00181.t01, mNeoVis.2.1.1.chr8_hap1.g00183.t01, mNeoVis.2.1.1.chr8_hap1.g00186.t01, mNeoVis.2.1.1.chr8_hap1.g00187.t01    rna-XM_021678473.1, rna-XM_044912162.1, rna-XM_044912163.1, rna-XM_044912277.1, rna-XM_044912279.1    rna-XM_004394405.1, rna-XM_004394406.1, rna-XM_004394407.1, rna-XM_004394408.1    rna-XM_042966721.1    rna-XM_008693957.2, rna-XM_040640299.1    rna-XM_027624519.2, rna-XM_035724378.1    mNeoVis.3.1.1.chr8.g00147.t01    rna-gnl|WGS_ZZZZ|egapxtmp_019341-R1, rna-gnl|WGS_ZZZZ|egapxtmp_019342-R1, rna-gnl|WGS_ZZZZ|egapxtmp_020256-R1    mNeoVis.evergladensis.2.1.chr9.g00177.t01, mNeoVis.evergladensis.2.1.chr9.g00178.t01, mNeoVis.evergladensis.2.1.chr9.g00179.t01    mNeoVis.vulgivaga.2.1.chr8.g00179.t01, mNeoVis.vulgivaga.2.1.chr8.g00181.t01, mNeoVis.vulgivaga.2.1.chr8.g00183.t01      1254    N0.HOG0001252   OG0000126  n115     rna-XM_038546090.1      rna-XM_006940555.4      rna-XM_039229750.1      rna-XM_045984889.1    rna-XM_032320006.1    rna-XM_059147455.1      rna-XM_045085952.1    mNeoVis.2.1.1.chr8_hap1.g00182.t01        rna-XM_007091221.3              mNeoVis.vulgivaga.2.1.chr8.g00180.t01      1255    N0.HOG0001253   OG0000127  n1   rna-XM_034657896.1    rna-NM_001048110.1    rna-XM_022527023.1    rna-XM_023246285.2    rna-XM_036066524.1    rna-XM_039252287.1    rna-XM_032835910.1    rna-XM_045992207.1    rna-XM_032313516.1    rna-XM_059160222.1    rna-XM_059389219.1    rna-XM_045082982.1    mNeoVis.2.1.1.chr5_hap1.g00370.t01    rna-XM_021705572.1    rna-XM_004399013.1    rna-XM_042978386.1    rna-XM_040629166.1    rna-XM_027582305.2    mNeoVis.3.1.1.chr2.g00303.t01    rna-gnl|WGS_ZZZZ|egapxtmp_034864-R1    mNeoVis.evergladensis.2.1.chr2.g00371.t01    mNeoVis.vulgivaga.2.1.chr2.g00366.t01      1256    N0.HOG0001254   OG0000127  n23         rna-XM_045047173.1      rna-XM_039252350.1                      rna-XM_042978416.1                  1257    N0.HOG0001255   OG0000127  n26   rna-XM_011218247.3, rna-XM_034653260.1, rna-XM_034653340.1, rna-XM_034657920.1, rna-XM_034657921.1    rna-NM_001038667.1, rna-XM_038565312.1    rna-XM_022527025.1      rna-XM_036066553.1, rna-XM_036091761.1      rna-XM_032835912.1    rna-XM_045989925.1    rna-XM_032313709.1    rna-XM_059160249.1    rna-XM_059389222.1    rna-XM_013054060.2    mNeoVis.2.1.1.chr5_hap1.g00381.t01    rna-XM_044912216.1, rna-XM_044913010.1    rna-XM_004416771.1, rna-XM_004416772.1      rna-XM_008711694.2, rna-XM_040629161.1    rna-XM_027587391.1    mNeoVis.3.1.1.chr2.g00314.t01    rna-gnl|WGS_ZZZZ|egapxtmp_034860-R1    mNeoVis.evergladensis.2.1.chr2.g00382.t01    mNeoVis.vulgivaga.2.1.chr2.g00377.t01      1258    N0.HOG0001256   OG0000127  n55   rna-XM_034657167.1      rna-XM_022492799.1, rna-XM_022526938.1    rna-XM_045047218.1    rna-XM_036091757.1, rna-XM_036091759.1    rna-XM_039252368.1    rna-XM_032850966.1, rna-XM_032883483.1, rna-XM_032883484.1      rna-XM_032313655.1    rna-XM_059160264.1    rna-XM_059389245.1    rna-XM_045082782.1      rna-XM_044912997.1      rna-XM_042978419.1    rna-XM_008711688.2    rna-XM_027586083.1    mNeoVis.3.1.1.chr2.g00317.t01    rna-gnl|WGS_ZZZZ|egapxtmp_034766-R1    mNeoVis.evergladensis.2.1.chr2.g00385.t01    mNeoVis.vulgivaga.2.1.chr2.g00380.t01      1259    N0.HOG0001257   OG0000127  n77   rna-XM_034650944.1, rna-XM_034657165.1    rna-NM_001038666.1    rna-XM_022526927.1    rna-XM_023246420.2    rna-XM_036066531.1    rna-XM_039252353.1    rna-XM_032835913.1    rna-XM_045992189.1    rna-XM_032313561.1    rna-XM_059160251.1    rna-XM_059389226.1    rna-XM_004748211.3    mNeoVis.2.1.1.chr5_hap1.g00379.t01    rna-XM_021705345.1    rna-XM_004399015.1    rna-XM_042978406.1    rna-XM_040629168.1    rna-XM_027582309.2    mNeoVis.3.1.1.chr2.g00311.t01, mNeoVis.3.1.1.chr2.g00312.t01    rna-gnl|WGS_ZZZZ|egapxtmp_034916-R1    mNeoVis.evergladensis.2.1.chr2.g00380.t01    mNeoVis.vulgivaga.2.1.chr2.g00375.t01      1260    N0.HOG0001258   OG0000127  n99   rna-XM_034653036.1, rna-XM_034653280.1    rna-XM_038565376.1                                              1261    N0.HOG0001259   OG0000127  n101   rna-XM_034650742.1    rna-XM_038567030.1    rna-XM_022492710.1, rna-XM_022526936.1, rna-XM_022526937.1    rna-XM_023246468.2    rna-XM_036091753.1    rna-XM_039252367.1    rna-XM_032835930.1, rna-XM_032835931.1    rna-XM_045992187.1, rna-XM_045992188.1    rna-XM_032313526.1    rna-XM_059160261.1    rna-XM_059389242.1    rna-XM_004748203.3    mNeoVis.2.1.1.chr5_hap1.g00383.t01    rna-XM_044913002.1      rna-XM_007098127.3    rna-XM_008711693.2    rna-XM_027585170.2    mNeoVis.3.1.1.chr2.g00316.t01    rna-gnl|WGS_ZZZZ|egapxtmp_034765-R1    mNeoVis.evergladensis.2.1.chr2.g00384.t01    mNeoVis.vulgivaga.2.1.chr2.g00379.t01      1262    N0.HOG0001260   OG0000128  n1   rna-XM_002912252.4, rna-XM_019797986.2    rna-NM_001003150.1    rna-XM_022518791.1    rna-XM_003998055.6, rna-XM_019819767.2    rna-XM_036068945.1    rna-XM_039232640.1, rna-XM_039232647.1, rna-XM_039232675.1    rna-XM_032854634.1    rna-XM_045988004.1, rna-XM_046007851.1    rna-XM_032325127.1    rna-XM_059151171.1, rna-XM_059153143.1    rna-XM_059381661.1      mNeoVis.2.1.1.chr7_hap1.g00310.t01    rna-XM_021704552.1    rna-XM_004395807.1    rna-XM_007090188.3, rna-XM_042968842.1    rna-XM_040633326.1, rna-XM_040633327.1    rna-XM_027619083.2, rna-XM_035725308.1    mNeoVis.3.1.1.chr7.g00295.t01, mNeoVis.3.1.1.chr7.g00296.t01    rna-gnl|WGS_ZZZZ|egapxtmp_009606-R1, rna-gnl|WGS_ZZZZ|egapxtmp_013271-R1    mNeoVis.evergladensis.2.1.chr7.g00310.t01, mNeoVis.evergladensis.2.1.chr7.g00311.t01    mNeoVis.vulgivaga.2.1.chr7.g00313.t01, mNeoVis.vulgivaga.2.1.chr7.g00314.t01      1263    N0.HOG0001261   OG0000128  n22     rna-NM_001003149.1        rna-XM_036068947.1                    rna-XM_021704540.2    rna-XM_004395805.1        rna-XM_027618841.2, rna-XM_035722942.1              1264    N0.HOG0001262   OG0000128  n30         rna-XM_023245328.2, rna-XM_045046514.1                          rna-XM_042968867.1, rna-XM_042968874.1                  1265    N0.HOG0001263   OG0000128  n34       rna-XM_022518745.1      rna-XM_036073292.1      rna-XM_032854433.1    rna-XM_045988080.1, rna-XM_045988082.1, rna-XM_046024647.1    rna-XM_032325831.1    rna-XM_059153144.1    rna-XM_059383753.1    rna-XM_013052135.2    mNeoVis.2.1.1.chr7_hap1.g00308.t01    rna-XM_021689827.2, rna-XM_044912458.1    rna-XM_004396289.1        rna-XM_035725068.1, rna-XM_035725069.1, rna-XM_035725070.1, rna-XM_035727282.1, rna-XM_035728140.1    mNeoVis.3.1.1.chr7.g00293.t01    rna-gnl|WGS_ZZZZ|egapxtmp_013270-R1    mNeoVis.evergladensis.2.1.chr7.g00308.t01    mNeoVis.vulgivaga.2.1.chr7.g00311.t01      1266    N0.HOG0001264   OG0000128  n50     rna-NM_001003173.1, rna-XM_038531005.1, rna-XM_038544030.1    rna-XM_022518748.1          rna-XM_032854432.1    rna-XM_045988081.1    rna-XM_032318173.1, rna-XM_032325830.1    rna-XM_059153024.1, rna-XM_059161375.1, rna-XM_059183794.1    rna-XM_059383752.1, rna-XM_059387865.1, rna-XM_059404988.1    rna-XM_004744215.3, rna-XM_045087400.1, rna-XM_045088556.1    mNeoVis.2.1.1.chr7_hap1.g00309.t01    rna-XM_021682439.1          rna-XM_035725071.1    mNeoVis.3.1.1.chr7.g00294.t01    rna-gnl|WGS_ZZZZ|egapxtmp_013269-R1    mNeoVis.evergladensis.2.1.chr7.g00309.t01    mNeoVis.vulgivaga.2.1.chr7.g00312.t01      1267    N0.HOG0001265   OG0000128  n66   rna-XM_011222327.1    rna-XM_038532119.1    rna-XM_022525993.1                              rna-XM_040621950.1    rna-XM_035725064.1              1268    N0.HOG0001266   OG0000128  n70   rna-XM_002912251.4, rna-XM_002912253.4    rna-XM_038560607.1    rna-XM_022518749.1        rna-XM_039249175.1    rna-XM_032854434.1      rna-XM_032325832.1, rna-XM_032339689.1    rna-XM_059151974.1, rna-XM_059177724.1    rna-XM_059381179.1, rna-XM_059399126.1    rna-XM_004744216.3, rna-XM_045071278.1    mNeoVis.2.1.1.chr1_hap1.g01737.t01, mNeoVis.2.1.1.chr7_hap1.g00307.t01          rna-XM_008702614.2      mNeoVis.3.1.1.chr1.g01692.t01    rna-gnl|WGS_ZZZZ|egapxtmp_002172-R1, rna-gnl|WGS_ZZZZ|egapxtmp_013273-R1    mNeoVis.evergladensis.2.1.chr1.g01705.t01, mNeoVis.evergladensis.2.1.chr7.g00307.t01    mNeoVis.vulgivaga.2.1.chr1.g01702.t01, mNeoVis.vulgivaga.2.1.chr7.g00310.t01      1269    N0.HOG0001267   OG0000128  n88           rna-XM_036068948.1                          rna-XM_008702617.2                1270    N0.HOG0001268   OG0000129  n1   rna-XM_034644823.1    rna-NM_001388577.1    rna-XM_022507156.1    rna-XM_003993410.3    rna-XM_036110586.1    rna-XM_039245077.1    rna-XM_032879035.1    rna-XM_046016644.1    rna-XM_032359326.1    rna-XM_059143237.1    rna-XM_059387352.1    rna-XM_004770472.1    mNeoVis.2.1.1.chr7_hap1.g02231.t01    rna-XM_021684877.1    rna-XM_004409303.1    rna-XM_007092138.2    rna-XM_008694293.2    rna-XM_027580215.1    mNeoVis.3.1.1.chr7.g02050.t01    rna-gnl|WGS_ZZZZ|egapxtmp_010316-R1    mNeoVis.evergladensis.2.1.chr7.g02179.t01    mNeoVis.vulgivaga.2.1.chr7.g02197.t01      1271    N0.HOG0001269   OG0000129  n24   rna-XM_002930877.1    rna-NM_001388578.1    rna-XM_022507222.1    rna-XM_003993409.3    rna-XM_036110584.1    rna-XM_039245083.1    rna-XM_032879034.1    rna-XM_046016722.1    rna-XM_032359329.1    rna-XM_059143258.1    rna-XM_059387408.1    rna-XM_004770473.1    mNeoVis.2.1.1.chr7_hap1.g02230.t01    rna-XM_021684876.1    rna-XM_004409302.1    rna-XM_007092043.2    rna-XM_008694178.2    rna-XM_027580216.1    mNeoVis.3.1.1.chr7.g02049.t01    rna-gnl|WGS_ZZZZ|egapxtmp_013518-R1    mNeoVis.evergladensis.2.1.chr7.g02178.t01    mNeoVis.vulgivaga.2.1.chr7.g02196.t01      1272    N0.HOG0001270   OG0000129  n44   rna-XM_002930878.3    rna-NM_001389154.1    rna-XM_022507226.1    rna-XM_003993472.2    rna-XM_036110579.1    rna-XM_039245081.1    rna-XM_032879032.1    rna-XM_046016723.1    rna-XM_032359324.1    rna-XM_059143208.1    rna-XM_059387469.1    rna-XM_004770475.1    mNeoVis.2.1.1.chr7_hap1.g02228.t01    rna-XM_021684875.1    rna-XM_004409301.1    rna-XM_007092044.2    rna-XM_008694177.2    rna-XM_027580210.1      rna-gnl|WGS_ZZZZ|egapxtmp_013493-R1    mNeoVis.evergladensis.2.1.chr7.g02176.t01    mNeoVis.vulgivaga.2.1.chr7.g02195.t01      1273    N0.HOG0001271   OG0000129  n64   rna-XM_034646554.1    rna-NM_001388576.1    rna-XM_022507146.1    rna-XM_003993411.2    rna-XM_036110587.1    rna-XM_039245076.1    rna-XM_032879036.1    rna-XM_046016713.1    rna-XM_032359327.1    rna-XM_059143250.1    rna-XM_059387287.1    rna-XM_004770441.2    mNeoVis.2.1.1.chr7_hap1.g02232.t01    rna-XM_021684878.1    rna-XM_004409304.1    rna-XM_007092042.2    rna-XM_008694179.1    rna-XM_027580214.1    mNeoVis.3.1.1.chr7.g02051.t01    rna-gnl|WGS_ZZZZ|egapxtmp_012656-R1    mNeoVis.evergladensis.2.1.chr7.g02180.t01    mNeoVis.vulgivaga.2.1.chr7.g02198.t01      1274   N0.HOG0001272  OG0000129  n82   rna-XM_034646605.1    rna-NM_001388873.1    rna-XM_022507217.1    rna-XM_003993412.2    rna-XM_036110585.1    rna-XM_039245082.1    rna-XM_032879037.1    rna-XM_046016711.1    rna-XM_032359325.1    rna-XM_059143223.1    rna-XM_059387220.1    rna-XM_013044709.2    mNeoVis.2.1.1.chr7_hap1.g02233.t01    rna-XM_021685230.2    rna-XM_004409305.1    rna-XM_007092041.2    rna-XM_008694180.1    rna-XM_027580211.1          mNeoVis.vulgivaga.2.1.chr7.g02199.t01      1275   N0.HOG0001273  OG0000129  n99   rna-XM_002930879.2    rna-XM_038563167.1    rna-XM_022507158.1    rna-XM_003993473.2    rna-XM_036110583.1    rna-XM_039245040.1    rna-XM_032879894.1    rna-XM_046015772.1    rna-XM_032359906.1    rna-XM_059140827.1    rna-XM_059397386.1    rna-XM_004770474.1    mNeoVis.2.1.1.chr7_hap1.g02229.t01    rna-XM_021685229.1    rna-XM_004409322.1    rna-XM_007092139.2    rna-XM_008694292.2    rna-XM_027580217.2    mNeoVis.3.1.1.chr7.g02048.t01    rna-gnl|WGS_ZZZZ|egapxtmp_010306-R1    mNeoVis.evergladensis.2.1.chr7.g02177.t01        1276    N0.HOG0001274   OG0000130  n1   rna-XM_011236399.3    rna-XM_038584342.1    rna-XM_022527128.1    rna-XM_003985721.3    rna-XM_036112322.1    rna-XM_039257273.1    rna-XM_032861712.1    rna-XM_046006261.1    rna-XM_032338240.1    rna-XM_059176247.1    rna-XM_059399603.1    rna-XM_013049110.2    mNeoVis.2.1.1.chr1_hap1.g01143.t01    rna-XM_021696843.2    rna-XM_004406518.1    rna-XM_007074490.3    rna-XM_008692959.2    rna-XM_027602229.2    mNeoVis.3.1.1.chr1.g01103.t01      mNeoVis.evergladensis.2.1.chr1.g01113.t01    mNeoVis.vulgivaga.2.1.chr1.g01114.t01      1277    N0.HOG0001275   OG0000130  n22   rna-XM_011236409.3    rna-XM_038584341.1      rna-XM_045058493.1    rna-XM_036112321.1    rna-XM_039257271.1                  rna-XM_044917581.1    rna-XM_004406513.1    rna-XM_042985524.1    rna-XM_040645167.1    rna-XM_027602227.2              1278    N0.HOG0001276   OG0000130  n33               rna-XM_032861709.1    rna-XM_046006254.1    rna-XM_032338248.1    rna-XM_059176246.1    rna-XM_059399601.1    rna-XM_013049109.2                    mNeoVis.evergladensis.2.1.chr1.g01115.t01        1279   N0.HOG0001277  OG0000130  n38   rna-XM_002927084.4    rna-XM_038584431.1    rna-XM_022492747.1    rna-XM_003985684.3    rna-XM_036121558.1    rna-XM_039223542.1    rna-XM_032861706.1    rna-XM_046006255.1    rna-XM_032338241.1    rna-XM_059176239.1      rna-XM_004781526.3    mNeoVis.2.1.1.chr1_hap1.g01081.t01    rna-XM_021687640.2    rna-XM_004413630.1    rna-XM_007074476.3    rna-XM_040645165.1    rna-XM_035728414.1    mNeoVis.3.1.1.chr1.g01067.t01    rna-gnl|WGS_ZZZZ|egapxtmp_019447-R1    mNeoVis.evergladensis.2.1.chr1.g01065.t01    mNeoVis.vulgivaga.2.1.chr1.g01083.t01      1280    N0.HOG0001278   OG0000130  n56   rna-XM_019809269.2    rna-XM_038584338.1      rna-XM_003985709.6, rna-XM_045050003.1    rna-XM_036112314.1    rna-XM_039257267.1      rna-XM_046006256.1    rna-XM_032338242.1    rna-XM_059176242.1      rna-XM_045082251.1    mNeoVis.2.1.1.chr1_hap1.g01122.t01      rna-XM_004413654.1    rna-XM_042985521.1    rna-XM_040645166.1    rna-XM_027581661.2    mNeoVis.3.1.1.chr1.g01092.t01      mNeoVis.evergladensis.2.1.chr1.g01095.t01        1281    N0.HOG0001279   OG0000130  n71                       rna-XM_059387584.1                  mNeoVis.3.1.1.chr1.g01097.t01        mNeoVis.vulgivaga.2.1.chr1.g01111.t01      1282    N0.HOG0001280   OG0000130  n75   rna-XM_002928550.4    rna-XM_038584339.1, rna-XM_038584487.1        rna-XM_036112319.1, rna-XM_036112320.1        rna-XM_046006259.1, rna-XM_046006262.1    rna-XM_032338246.1, rna-XM_032338396.1    rna-XM_059176245.1, rna-XM_059176250.1      rna-XM_045082252.1, rna-XM_045082254.1    mNeoVis.2.1.1.chr1_hap1.g01130.t01, mNeoVis.2.1.1.chr1_hap1.g01135.t01      rna-XM_004406523.1, rna-XM_004413656.1      rna-XM_040645159.1    rna-XM_027602102.2      rna-gnl|WGS_ZZZZ|egapxtmp_010118-R1    mNeoVis.evergladensis.2.1.chr1.g01106.t01    mNeoVis.vulgivaga.2.1.chr1.g01107.t01      1283    N0.HOG0001281   OG0000130  n95   rna-XM_002928526.4    rna-XM_038584337.1      rna-XM_003985718.5, rna-XM_045058491.1    rna-XM_036112315.1    rna-XM_039257268.1, rna-XM_039257269.1, rna-XM_039257275.1    rna-XM_032861707.1    rna-XM_046006257.1    rna-XM_032338244.1    rna-XM_059176243.1      rna-XM_004778776.3    mNeoVis.2.1.1.chr1_hap1.g01139.t01    rna-XM_044912199.1    rna-XM_004406521.1    rna-XM_007074491.3, rna-XM_042985532.1    rna-XM_008693087.2    rna-XM_035728415.1        mNeoVis.evergladensis.2.1.chr1.g01109.t01        1284    N0.HOG0001282   OG0000130  n113       rna-XM_022527126.1                                  mNeoVis.3.1.1.chr1.g01105.t01            1285    N0.HOG0001283   OG0000131  n1   rna-XM_034649850.1, rna-XM_034650227.1    rna-XM_038557808.1    rna-XM_022496264.1    rna-XM_003989886.5    rna-XM_036102790.1    rna-XM_039224582.1    rna-XM_032877363.1    rna-XM_046012298.1    rna-XM_032301722.1    rna-XM_059136385.1    rna-XM_059377381.1    rna-XM_045064268.1    mNeoVis.2.1.1.chr2_hap1.g01462.t01    rna-XM_044912424.1, rna-XM_044912485.1, rna-XM_044913983.1    rna-XM_004396945.1, rna-XM_004402615.1    rna-XM_007077409.2    rna-XM_008694354.2    rna-XM_027611206.2, rna-XM_035725660.1    mNeoVis.3.1.1.chr5.g01382.t01    rna-gnl|WGS_ZZZZ|egapxtmp_009562-R1    mNeoVis.evergladensis.2.1.chr5.g01400.t01    mNeoVis.vulgivaga.2.1.chr5.g01400.t01      1286    N0.HOG0001284   OG0000131  n20   rna-XM_019809838.2    rna-XM_038587828.1    rna-XM_022521671.1    rna-XM_023249244.2    rna-XM_036069936.1    rna-XM_039224342.1    rna-XM_032881810.1    rna-XM_045995224.1    rna-XM_032331449.1    rna-XM_059158129.1    rna-XM_059386162.1    rna-XM_004782203.3    mNeoVis.2.1.1.chr12_hap1.g00407.t01    rna-XM_021693035.1    rna-XM_004414665.1    rna-XM_007085588.3    rna-XM_008700633.2    rna-XM_027607546.2    mNeoVis.3.1.1.chr12.g00366.t01    rna-gnl|WGS_ZZZZ|egapxtmp_029350-R2    mNeoVis.evergladensis.2.1.chr12.g00362.t01    mNeoVis.vulgivaga.2.1.chr12.g00371.t01      1287    N0.HOG0001285   OG0000131  n37   rna-XM_011223150.3    rna-XM_038564629.1    rna-XM_022506251.1    rna-XM_023252811.2    rna-XM_036087688.1    rna-XM_039216730.1    rna-XM_032857653.1    rna-XM_045997437.1    rna-XM_032333420.1    rna-XM_059168673.1    rna-XM_059395534.1    rna-XM_013053897.2    mNeoVis.2.1.1.chr6_hap1.g00424.t01    rna-XM_021679347.1      rna-XM_007091803.3    rna-XM_008685399.1    rna-XM_027597413.2    mNeoVis.3.1.1.chr4.g00425.t01    rna-gnl|WGS_ZZZZ|egapxtmp_000966-R1    mNeoVis.evergladensis.2.1.chr4.g00431.t01    mNeoVis.vulgivaga.2.1.chr4.g00434.t01      1288    N0.HOG0001286   OG0000131  n53   rna-XM_011236261.2, rna-XM_034651779.1, rna-XM_034653535.1, rna-XM_034668382.1    rna-XM_038555212.1    rna-XM_022502016.1    rna-XM_023248198.2    rna-XM_036072726.1, rna-XM_036079316.1    rna-XM_039222483.1, rna-XM_039233781.1    rna-XM_032850984.1, rna-XM_032858580.1    rna-XM_045992396.1, rna-XM_046010968.1    rna-XM_032315926.1    rna-XM_059165909.1    rna-XM_059394934.1    rna-XM_045077993.1, rna-XM_045088673.1    mNeoVis.2.1.1.chr4_hap1.g00912.t01    rna-XM_021688699.1, rna-XM_021692635.1, rna-XM_044916095.1    rna-XM_004402346.1    rna-XM_042972982.1    rna-XM_008700638.2, rna-XM_040626135.1    rna-XM_027606046.2, rna-XM_035725481.1    mNeoVis.3.1.1.chr6.g00923.t01    rna-gnl|WGS_ZZZZ|egapxtmp_002768-R2    mNeoVis.evergladensis.2.1.chr6.g00916.t01    mNeoVis.vulgivaga.2.1.chr6.g00911.t01      1289    N0.HOG0001287   OG0000131  n73   rna-XM_034641930.1    rna-XM_038572427.1    rna-XM_022494145.1    rna-XM_045053565.1    rna-XM_036072049.1    rna-XM_039248497.1    rna-XM_032838641.1    rna-XM_045981242.1    rna-XM_032350564.1    rna-XM_059133068.1    rna-XM_059407080.1    rna-XM_004746374.3    mNeoVis.2.1.1.chr11_hap1.g00202.t01    rna-XM_021689294.2    rna-XM_004414106.1    rna-XM_042980319.1    rna-XM_040638383.1    rna-XM_027624262.2    mNeoVis.3.1.1.chr11.g00171.t01    rna-gnl|WGS_ZZZZ|egapxtmp_026813-R1    mNeoVis.evergladensis.2.1.chr11.g00199.t01    mNeoVis.vulgivaga.2.1.chr11.g00202.t01      1290    N0.HOG0001288   OG0000132  n1   rna-XM_002926190.3    rna-XM_038569300.1    rna-XM_022507293.1    rna-XM_006937163.5    rna-XM_036098491.1    rna-XM_039223042.1    rna-XM_032843425.1    rna-XM_046014186.1    rna-XM_032359705.1    rna-XM_059138646.1    rna-XM_059406676.1    rna-XM_004755898.3    mNeoVis.2.1.1.chr7_hap1.g01910.t01    rna-XM_021684710.1    rna-XM_004412889.1    rna-XM_007099418.3    rna-XM_008686943.2    rna-XM_027581490.2    mNeoVis.3.1.1.chr7.g01780.t01    rna-gnl|WGS_ZZZZ|egapxtmp_014022-R1    mNeoVis.evergladensis.2.1.chr7.g01872.t01    mNeoVis.vulgivaga.2.1.chr7.g01889.t01      1291    N0.HOG0001289   OG0000132  n19   rna-XM_011236154.3    rna-NM_001003329.1    rna-XM_022496961.1    rna-XM_045033270.1    rna-XM_036076408.1    rna-XM_039251077.1    rna-XM_032881493.1    rna-XM_045978139.1    rna-XM_032303102.1    rna-XM_059136917.1    rna-XM_059375701.1    rna-XM_045081927.1    mNeoVis.2.1.1.chr2_hap1.g00964.t01    rna-XM_021690143.1    rna-XM_004407154.1    rna-XM_015534958.2    rna-XM_040626919.1    rna-XM_027621336.2    mNeoVis.3.1.1.chr5.g00897.t01      mNeoVis.evergladensis.2.1.chr5.g00917.t01    mNeoVis.vulgivaga.2.1.chr5.g00905.t01      1292    N0.HOG0001290   OG0000132  n30   rna-XM_002920437.4    rna-NM_001313824.1    rna-XM_022522394.1    rna-XM_019834392.3    rna-XM_036094746.1    rna-XM_039255765.1    rna-XM_032874812.1    rna-XM_046013766.1    rna-XM_032349039.1    rna-XM_059184568.1    rna-XM_059403766.1    rna-XM_004781513.3    mNeoVis.2.1.1.chr9_hap1.g00239.t01    rna-XM_021690657.1    rna-XM_004413371.1    rna-XM_007089648.3    rna-XM_008683578.1    rna-XM_027594581.2    mNeoVis.3.1.1.chr10.g00231.t01      mNeoVis.evergladensis.2.1.chr10.g00236.t01    mNeoVis.vulgivaga.2.1.chr10.g00233.t01      1293    N0.HOG0001291   OG0000132  n41   rna-XM_002928302.3    rna-XM_038541029.1    rna-XM_022496347.1    rna-XM_019839096.3    rna-XM_036076402.1    rna-XM_039251114.1    rna-XM_032881488.1    rna-XM_046010494.1    rna-XM_032301646.1    rna-XM_059135894.1    rna-XM_059375702.1    rna-XM_004769886.2    mNeoVis.2.1.1.chr2_hap1.g00963.t01    rna-XM_021690128.1    rna-XM_004407153.1    rna-XM_042998157.1    rna-XM_008704229.2    rna-XM_027617014.2    mNeoVis.3.1.1.chr5.g00896.t01    rna-gnl|WGS_ZZZZ|egapxtmp_005650-R1    mNeoVis.evergladensis.2.1.chr5.g00916.t01    mNeoVis.vulgivaga.2.1.chr5.g00904.t01      1294    N0.HOG0001292   OG0000132  n64   rna-XM_011226481.3    rna-XM_038576550.1    rna-XM_022522384.1    rna-XM_019834391.3    rna-XM_036068341.1    rna-XM_039255767.1    rna-XM_032874814.1    rna-XM_046013292.1    rna-XM_032349041.1    rna-XM_059184567.1    rna-XM_059403772.1    rna-XM_004781508.3    mNeoVis.2.1.1.chr9_hap1.g00238.t01    rna-XM_021690670.2    rna-XM_004417261.1    rna-XM_042993746.1    rna-XM_008709341.2    rna-XM_027595330.2    mNeoVis.3.1.1.chr10.g00230.t01      mNeoVis.evergladensis.2.1.chr10.g00235.t01        1295    N0.HOG0001293   OG0000132  n76   rna-XM_002920436.4    rna-NM_001006645.2    rna-XM_022522387.1    rna-XM_006933430.5    rna-XM_036094747.1    rna-XM_039255766.1    rna-XM_032874811.1    rna-XM_046010810.1    rna-XM_032349038.1    rna-XM_059184569.1    rna-XM_059404947.1    rna-NM_001310202.1      rna-XM_021690641.1    rna-XM_004413372.1    rna-XM_042991620.1    rna-XM_040641904.1    rna-XM_027594580.2      rna-gnl|WGS_ZZZZ|egapxtmp_024461-R1    mNeoVis.evergladensis.2.1.chr10.g00237.t01        1296    N0.HOG0001294   OG0000133  n2         rna-XM_023249129.2            rna-XM_032331447.1    rna-XM_059158279.1    rna-XM_059387501.1    rna-XM_045068169.1    mNeoVis.2.1.1.chr12_hap1.g00077.t01    rna-XM_021689081.1      rna-XM_042974916.1    rna-XM_040627116.1        rna-gnl|WGS_ZZZZ|egapxtmp_028642-R1    mNeoVis.evergladensis.2.1.chr12.g00064.t01    mNeoVis.vulgivaga.2.1.chr12.g00067.t01      1297    N0.HOG0001295   OG0000133  n14   rna-XM_034661228.1    rna-XM_038573764.1    rna-XM_022516108.1    rna-XM_019828405.2    rna-XM_036106178.1    rna-XM_039250441.1    rna-XM_032855629.1    rna-XM_045986683.1    rna-XM_032332374.1    rna-XM_059176329.1    rna-XM_059372964.1    rna-XM_013047169.2    mNeoVis.2.1.1.chr6_hap1.g00742.t01    rna-XM_021698973.1    rna-XM_004417761.1    rna-XM_007093922.3    rna-XM_008702049.1    rna-XM_035726358.1      rna-gnl|WGS_ZZZZ|egapxtmp_043873-R1          1298    N0.HOG0001296   OG0000133  n33   rna-XM_002914683.4    rna-XM_038545228.1    rna-XM_022505316.1    rna-XM_003987811.5    rna-XM_036117835.1    rna-XM_039232192.1    rna-XM_032837397.1    rna-XM_046007049.1    rna-XM_032344997.1    rna-XM_059182863.1    rna-XM_059373790.1    rna-XM_004739311.3      rna-XM_021679782.1    rna-XM_004399386.1    rna-XM_007095086.3    rna-XM_008699936.2    rna-XM_027570716.2              1299    N0.HOG0001297   OG0000133  n44     rna-XM_038545980.1    rna-XM_022505340.1    rna-XM_023255976.2    rna-XM_036071578.1    rna-XM_039217280.1    rna-XM_032837395.1    rna-XM_046009909.1    rna-XM_032345101.1    rna-XM_059182957.1    rna-XM_059373853.1    rna-XM_004754794.2      rna-XM_021679653.1    rna-XM_004417740.1    rna-XM_015543436.2    rna-XM_040641504.1                1300    N0.HOG0001298   OG0000133  n59   rna-XM_034649790.1    rna-XM_038586652.1    rna-XM_022491281.1    rna-XM_023249750.2, rna-XM_045050947.1, rna-XM_045051670.1, rna-XM_045051673.1, rna-XM_045051680.1, rna-XM_045051682.1    rna-XM_036082035.1      rna-XM_032850992.1    rna-XM_045995494.1, rna-XM_045995503.1    rna-XM_032331277.1, rna-XM_032331311.1    rna-XM_059158059.1, rna-XM_059158132.1    rna-XM_059387583.1    rna-XM_045068077.1, rna-XM_045068078.1    mNeoVis.2.1.1.chr12_hap1.g00075.t01, mNeoVis.2.1.1.chr12_hap1.g00076.t01      rna-XM_004417387.1    rna-XM_042974314.1, rna-XM_042974317.1    rna-XM_008703045.2    rna-XM_035725596.1, rna-XM_035725598.1, rna-XM_035725616.1              1301    N0.HOG0001299   OG0000133  n82   rna-XM_002929088.4, rna-XM_034650052.1    rna-XM_038588637.1, rna-XM_038588853.1    rna-XM_022491283.1    rna-XM_045050399.1, rna-XM_045050400.1, rna-XM_045050402.1, rna-XM_045050985.1, rna-XM_045051648.1, rna-XM_045051661.1    rna-XM_036106653.1    rna-XM_039219270.1, rna-XM_039220687.1    rna-XM_032846381.1    rna-XM_045994845.1    rna-XM_032329743.1    rna-XM_059156858.1    rna-XM_059385921.1    rna-XM_045068007.1    mNeoVis.2.1.1.chr12_hap1.g00071.t01    rna-XM_044911905.1    rna-XM_004409191.1    rna-XM_042973984.1, rna-XM_042975067.1, rna-XM_042975136.1    rna-XM_040633509.1    rna-XM_035725561.1, rna-XM_035725651.1    mNeoVis.3.1.1.chr12.g00061.t01    rna-gnl|WGS_ZZZZ|egapxtmp_029112-R1    mNeoVis.evergladensis.2.1.chr12.g00061.t01    mNeoVis.vulgivaga.2.1.chr12.g00063.t01      1302    N0.HOG0001300   OG0000134  n1   rna-XM_002928206.2    rna-NM_001389098.1    rna-XM_022513348.1    rna-XM_003992506.5    rna-XM_036081758.1    rna-XM_039250272.1    rna-XM_032868998.1    rna-XM_046017060.1    rna-XM_032358106.1    rna-XM_059171858.1    rna-XM_059404242.1    rna-XM_004750251.3    mNeoVis.2.1.1.chr7_hap1.g01314.t01    rna-XM_044919599.1      rna-XM_007074035.2    rna-XM_008690159.2    rna-XM_027577940.1    mNeoVis.3.1.1.chr7.g01236.t01    rna-gnl|WGS_ZZZZ|egapxtmp_013353-R1    mNeoVis.evergladensis.2.1.chr7.g01300.t01    mNeoVis.vulgivaga.2.1.chr7.g01314.t01      1303    N0.HOG0001301   OG0000134  n21   rna-XM_002928205.4    rna-XM_038535567.1    rna-XM_022513323.1    rna-XM_045039079.1    rna-XM_036081759.1    rna-XM_039250233.1    rna-XM_032868802.1    rna-XM_046017566.1    rna-XM_032360518.1    rna-XM_059156154.1    rna-XM_059392490.1    rna-XM_004750252.3      rna-XM_044919641.1      rna-XM_015538823.2    rna-XM_008683638.1    rna-XM_027581151.1              1304    N0.HOG0001302   OG0000134  n37   rna-XM_002928191.1    rna-XM_038535496.1    rna-XM_022513290.1      rna-XM_036113943.1    rna-XM_039250232.1    rna-XM_032869031.1    rna-XM_046017239.1      rna-XM_059156179.1    rna-XM_059392463.1    rna-XM_004750140.1, rna-XM_045085346.1    mNeoVis.2.1.1.chr7_hap1.g01319.t01        rna-XM_015538827.1    rna-XM_008683624.1    rna-XM_027580643.2    mNeoVis.3.1.1.chr7.g01238.t01    rna-gnl|WGS_ZZZZ|egapxtmp_013365-R1    mNeoVis.evergladensis.2.1.chr7.g01305.t01    mNeoVis.vulgivaga.2.1.chr7.g01320.t01      1305    N0.HOG0001303   OG0000134  n58   rna-XM_002928190.4    rna-XM_038535494.1      rna-XM_003992486.2    rna-XM_036113944.1    rna-XM_039250262.1                    rna-XM_004413690.1    rna-XM_007074038.2    rna-XM_008683622.2    rna-XM_027580801.1              1306    N0.HOG0001304   OG0000134  n66   rna-XM_002928192.2    rna-NM_001388793.1    rna-XM_022513288.1, rna-XM_022513296.1    rna-XM_003992509.2    rna-XM_036113950.1    rna-XM_039250259.1    rna-XM_032868999.1, rna-XM_032869000.1    rna-XM_046017198.1, rna-XM_046017302.1    rna-XM_032358107.1, rna-XM_032358113.1    rna-XM_059156167.1, rna-XM_059172339.1    rna-XM_059392475.1, rna-XM_059404014.1    rna-XM_004750253.1, rna-XM_004750256.2    mNeoVis.2.1.1.chr7_hap1.g01318.t01, mNeoVis.2.1.1.chr7_hap1.g01322.t01      rna-XM_004417291.1    rna-XM_007074076.2    rna-XM_008683625.2    rna-XM_027580701.1      rna-gnl|WGS_ZZZZ|egapxtmp_012434-R1, rna-gnl|WGS_ZZZZ|egapxtmp_013366-R1    mNeoVis.evergladensis.2.1.chr7.g01304.t01    mNeoVis.vulgivaga.2.1.chr7.g01319.t01      1307    N0.HOG0001305   OG0000134  n96   rna-XM_002928207.2    rna-NM_001388794.1    rna-XM_022513349.1          rna-XM_032868996.1    rna-XM_046017559.1    rna-XM_032358105.1    rna-XM_059171740.1    rna-XM_059404367.1    rna-XM_004750250.3            rna-XM_008690160.2                1308    N0.HOG0001306   OG0000134  n105   rna-XM_002928208.4    rna-NM_001388795.1    rna-XM_022513350.1    rna-XM_003992505.1      rna-XM_039250271.1    rna-XM_032868995.1    rna-XM_046015967.1    rna-XM_032358104.1    rna-XM_059171615.1    rna-XM_059404460.1    rna-XM_004750249.1    mNeoVis.2.1.1.chr7_hap1.g01313.t01      rna-XM_004417546.1    rna-XM_007074075.3    rna-XM_008690161.1      mNeoVis.3.1.1.chr7.g01235.t01    rna-gnl|WGS_ZZZZ|egapxtmp_013358-R1    mNeoVis.evergladensis.2.1.chr7.g01299.t01    mNeoVis.vulgivaga.2.1.chr7.g01313.t01      1309    N0.HOG0001307   OG0000135  n1   rna-XM_019798410.2    rna-NM_001110799.1    rna-XM_022504630.1    rna-XM_003991081.4    rna-XM_036070885.1    rna-XM_039240655.1    rna-XM_032860273.1    rna-XM_046019426.1    rna-XM_032355381.1    rna-XM_059168994.1    rna-XM_059392656.1    rna-XM_004764271.1    mNeoVis.2.1.1.chr3_hap1.g01234.t01    rna-XM_021702953.2    rna-XM_004405391.1    rna-XM_042997535.1    rna-XM_008687438.2    rna-XM_027590298.1    mNeoVis.3.1.1.chr3.g01213.t01    rna-gnl|WGS_ZZZZ|egapxtmp_040948-R1    mNeoVis.evergladensis.2.1.chr3.g01233.t01    mNeoVis.vulgivaga.2.1.chr3.g01247.t01      1310    N0.HOG0001308   OG0000135  n21   rna-XM_011223470.2, rna-XM_011223472.2    rna-NM_001113170.1    rna-XM_022496680.1, rna-XM_022504802.1    rna-XM_003991080.3, rna-XM_003991082.5    rna-XM_036070882.1, rna-XM_036070886.1    rna-XM_039240653.1, rna-XM_039240658.1    rna-XM_032860272.1, rna-XM_032860276.1    rna-XM_046019421.1, rna-XM_046019428.1    rna-XM_032355382.1, rna-XM_032355383.1    rna-XM_059168993.1, rna-XM_059168996.1    rna-XM_059392654.1, rna-XM_059392659.1    rna-XM_004764273.2, rna-XM_045078938.1    mNeoVis.2.1.1.chr3_hap1.g01233.t01, mNeoVis.2.1.1.chr3_hap1.g01235.t01    rna-XM_021702982.1, rna-XM_044912366.1, rna-XM_044913390.1    rna-XM_004405390.1, rna-XM_004405392.1    rna-XM_007089931.2, rna-XM_042997536.1    rna-XM_008687436.1, rna-XM_040636525.1    rna-XM_027589605.2, rna-XM_027590367.1      rna-gnl|WGS_ZZZZ|egapxtmp_040946-R1, rna-gnl|WGS_ZZZZ|egapxtmp_040950-R1    mNeoVis.evergladensis.2.1.chr3.g01231.t01, mNeoVis.evergladensis.2.1.chr3.g01234.t01    mNeoVis.vulgivaga.2.1.chr3.g01245.t01, mNeoVis.vulgivaga.2.1.chr3.g01248.t01      1311    N0.HOG0001309   OG0000135  n62                               rna-XM_004405394.1        rna-XM_027589607.2              1312    N0.HOG0001310   OG0000135  n64   rna-XM_002918093.3, rna-XM_034644512.1    rna-XM_038585109.1, rna-XM_038585581.1    rna-XM_022496862.1, rna-XM_022500155.1    rna-XM_003991078.4, rna-XM_003991087.4    rna-XM_036070901.1, rna-XM_036098830.1    rna-XM_039240656.1, rna-XM_039249985.1    rna-XM_032860270.1, rna-XM_032860279.1    rna-XM_046019474.1, rna-XM_046019540.1    rna-XM_032354121.1, rna-XM_032354611.1    rna-XM_059168990.1, rna-XM_059168999.1    rna-XM_059392652.1, rna-XM_059392663.1    rna-XM_004767058.2, rna-XM_013062267.2    mNeoVis.2.1.1.chr3_hap1.g01228.t01, mNeoVis.2.1.1.chr3_hap1.g01237.t01    rna-XM_021703014.1, rna-XM_021703215.1    rna-XM_004405384.1    rna-XM_042997533.1, rna-XM_042997539.1    rna-XM_008687450.2, rna-XM_008687557.1    rna-XM_027590932.2      rna-gnl|WGS_ZZZZ|egapxtmp_040683-R1, rna-gnl|WGS_ZZZZ|egapxtmp_040949-R1    mNeoVis.evergladensis.2.1.chr3.g01226.t01    mNeoVis.vulgivaga.2.1.chr3.g01250.t01      1313    N0.HOG0001311   OG0000135  n101   rna-XM_034644487.1    rna-NM_001113169.1    rna-XM_022504978.1    rna-XM_003991079.4    rna-XM_036070881.1    rna-XM_039240660.1    rna-XM_032860278.1    rna-XM_046019479.1    rna-XM_032354122.1    rna-XM_059168998.1    rna-XM_059392653.1    rna-XM_004764268.1    mNeoVis.2.1.1.chr3_hap1.g01236.t01      rna-XM_004405395.1    rna-XM_007089933.2    rna-XM_008687435.2    rna-XM_027590436.1    mNeoVis.3.1.1.chr3.g01214.t01      mNeoVis.evergladensis.2.1.chr3.g01235.t01    mNeoVis.vulgivaga.2.1.chr3.g01249.t01      1314    N0.HOG0001312   OG0000136  n1   rna-XM_034649056.1, rna-XM_034650053.1, rna-XM_034653538.1, rna-XM_034653541.1, rna-XM_034653580.1    rna-XM_038586650.1    rna-XM_022491280.1, rna-XM_022491332.1, rna-XM_022491405.1, rna-XM_022491406.1      rna-XM_036073698.1, rna-XM_036074796.1, rna-XM_036080416.1, rna-XM_036100876.1, rna-XM_036100877.1, rna-XM_036100948.1, rna-XM_036100960.1, rna-XM_036100973.1, rna-XM_036112092.1, rna-XM_036122706.1, rna-XM_036122708.1      rna-XM_032845280.1, rna-XM_032845298.1, rna-XM_032851054.1    rna-XM_045994652.1, rna-XM_045994654.1, rna-XM_045994656.1, rna-XM_045994658.1, rna-XM_045996048.1, rna-XM_045996085.1    rna-XM_032329735.1, rna-XM_032329738.1, rna-XM_032329739.1, rna-XM_032329741.1, rna-XM_032330810.1, rna-XM_032331632.1    rna-XM_059157485.1, rna-XM_059157535.1, rna-XM_059158459.1, rna-XM_059158461.1, rna-XM_059158504.1    rna-XM_059385924.1, rna-XM_059385935.1, rna-XM_059387543.1    rna-XM_045068035.1, rna-XM_045068036.1, rna-XM_045068070.1, rna-XM_045068086.1, rna-XM_045068096.1, rna-XM_045068193.1, rna-XM_045068208.1    mNeoVis.2.1.1.chr12_hap1.g00081.t01, mNeoVis.2.1.1.chr12_hap1.g00085.t01, mNeoVis.2.1.1.chr12_hap1.g00100.t01, mNeoVis.2.1.1.chr12_hap1.g00101.t01, mNeoVis.2.1.1.chr12_hap1.g00103.t01, mNeoVis.2.1.1.chr12_hap1.g00106.t01, mNeoVis.2.1.1.chr12_hap1.g00107.t01    rna-XM_044912200.1    rna-XM_004412686.1, rna-XM_004416270.1, rna-XM_004416992.1, rna-XM_004417040.1      rna-XM_008690230.2, rna-XM_008690238.2, rna-XM_008690259.2, rna-XM_040634694.1, rna-XM_040635956.1    rna-XM_027575132.2, rna-XM_035722241.1, rna-XM_035725375.1, rna-XM_035725395.1, rna-XM_035725410.1, rna-XM_035725415.1    mNeoVis.3.1.1.chr12.g00094.t01    rna-gnl|WGS_ZZZZ|egapxtmp_028520-R1    mNeoVis.evergladensis.2.1.chr12.g00068.t01    mNeoVis.vulgivaga.2.1.chr12.g00071.t01      1315    N0.HOG0001313   OG0000136  n76     rna-XM_038586645.1, rna-XM_038586646.1, rna-XM_038586647.1, rna-XM_038588504.1, rna-XM_038588505.1, rna-XM_038588506.1, rna-XM_038588525.1, rna-XM_038588584.1, rna-XM_038588850.1, rna-XM_038589589.1, rna-XM_038589614.1      rna-XM_023249541.2, rna-XM_045050360.1, rna-XM_045050398.1, rna-XM_045050617.1, rna-XM_045050618.1, rna-XM_045050910.1, rna-XM_045050911.1, rna-XM_045050912.1, rna-XM_045050921.1, rna-XM_045051104.1    rna-XM_036074795.1    rna-XM_039220688.1, rna-XM_039225460.1, rna-XM_039249884.1, rna-XM_039249885.1, rna-XM_039249886.1                  rna-XM_044911910.1    rna-XM_004407117.1, rna-XM_004412687.1, rna-XM_004414236.1, rna-XM_004414237.1, rna-XM_004416269.1    rna-XM_042974565.1, rna-XM_042975190.1, rna-XM_042975196.1, rna-XM_042975197.1, rna-XM_042975271.1, rna-XM_042977164.1, rna-XM_042977166.1, rna-XM_042977169.1      rna-XM_027607645.2, rna-XM_035725727.1, rna-XM_035725728.1, rna-XM_035727744.1              1316    N0.HOG0001314   OG0000137  n1   rna-XM_034639362.1, rna-XM_034650440.1    rna-XM_038559694.1    rna-XM_022513926.1    rna-XM_045052983.1    rna-XM_036122651.1    rna-XM_039253935.1    rna-XM_032840233.1      rna-XM_032305807.1    rna-XM_059172565.1    rna-XM_059395975.1    rna-XM_045086262.1    mNeoVis.2.1.1.chr4_hap1.g00054.t01    rna-XM_021692910.1    rna-XM_004408878.1    rna-XM_042975079.1    rna-XM_040643818.1    rna-XM_035723206.1    mNeoVis.3.1.1.chr6.g00057.t01    rna-gnl|WGS_ZZZZ|egapxtmp_003028-R1    mNeoVis.evergladensis.2.1.chr6.g00059.t01    mNeoVis.vulgivaga.2.1.chr6.g00060.t01      1317    N0.HOG0001315   OG0000137  n23                       rna-XM_059395974.1    rna-XM_045086261.1                          1318    N0.HOG0001316   OG0000137  n27   rna-XM_034651301.1    rna-XM_038559698.1      rna-XM_023249556.2, rna-XM_045052649.1, rna-XM_045053007.1    rna-XM_036082324.1, rna-XM_036107744.1, rna-XM_036107747.1, rna-XM_036115233.1    rna-XM_039230694.1, rna-XM_039230764.1, rna-XM_039253875.1        rna-XM_032305806.1    rna-XM_059172566.1          rna-XM_044920157.1      rna-XM_042975117.1, rna-XM_042975140.1, rna-XM_042975205.1, rna-XM_042977230.1    rna-XM_040641230.1    rna-XM_035723323.1    mNeoVis.3.1.1.chr6.g00056.t01    rna-gnl|WGS_ZZZZ|egapxtmp_003027-R1    mNeoVis.evergladensis.2.1.chr6.g00058.t01        1319    N0.HOG0001317   OG0000137  n51   rna-XM_034650665.1, rna-XM_034652734.1    rna-XM_038559707.1      rna-XM_003983236.5    rna-XM_036075783.1    rna-XM_039230781.1                  rna-XM_021692845.1    rna-XM_004408871.1    rna-XM_042975260.1    rna-XM_008707809.2, rna-XM_008709359.2, rna-XM_040643812.1                1320    N0.HOG0001318   OG0000137  n62   rna-XM_034639424.1, rna-XM_034649171.1    rna-XM_038559189.1, rna-XM_038559705.1      rna-XM_006929498.5    rna-XM_036102140.1, rna-XM_036107742.1    rna-XM_039230695.1                  rna-XM_021692906.1    rna-XM_004408802.1    rna-XM_007087509.3    rna-XM_040643813.1    rna-XM_035723324.1              1321    N0.HOG0001319   OG0000137  n74   rna-XM_034650292.1, rna-XM_034650713.1, rna-XM_034651197.1    rna-XM_038559701.1    rna-XM_022513927.1    rna-XM_006929496.5    rna-XM_036075846.1    rna-XM_039230791.1    rna-XM_032840231.1              mNeoVis.2.1.1.chr4_hap1.g00053.t01    rna-XM_021692844.1    rna-XM_004408879.1    rna-XM_042975224.1    rna-XM_040643814.1      mNeoVis.3.1.1.chr6.g00055.t01    rna-gnl|WGS_ZZZZ|egapxtmp_003026-R1    mNeoVis.evergladensis.2.1.chr6.g00057.t01    mNeoVis.vulgivaga.2.1.chr6.g00059.t01      1322    N0.HOG0001320   OG0000137  n91   rna-XM_034639395.1    rna-XM_038559697.1      rna-XM_011279623.4, rna-XM_045052984.1    rna-XM_036075745.1    rna-XM_039225665.1, rna-XM_039230768.1                  rna-XM_044919909.1    rna-XM_004408880.1    rna-XM_007085941.3, rna-XM_015541430.2    rna-XM_008709401.1                1323    N0.HOG0001321   OG0000137  n102   rna-XM_034639434.1, rna-XM_034639446.1    rna-XM_038559700.1      rna-XM_045052985.1, rna-XM_045052999.1, rna-XM_045053008.1, rna-XM_045053014.1, rna-XM_045053016.1    rna-XM_036075753.1, rna-XM_036082321.1    rna-XM_039230769.1                  rna-XM_021692938.1      rna-XM_042975176.1, rna-XM_042975195.1, rna-XM_042975203.1, rna-XM_042975228.1, rna-XM_042975279.1, rna-XM_042977228.1    rna-XM_008688624.2, rna-XM_008697253.2                1324   N0.HOG0001322  OG0000138  n1   rna-XM_002915055.4    rna-XM_038525968.1, rna-XM_038548428.1, rna-XM_038586830.1    rna-XM_022491724.1, rna-XM_022518316.1    rna-XM_011291607.2    rna-XM_036065579.1, rna-XM_036076640.1, rna-XM_036099491.1    rna-XM_039234268.1    rna-XM_032852277.1, rna-XM_032880195.1    rna-XM_045996180.1, rna-XM_045996644.1, rna-XM_046012252.1    rna-XM_032329760.1, rna-XM_032331954.1    rna-XM_059156413.1, rna-XM_059156443.1, rna-XM_059171453.1, rna-XM_059179693.1    rna-XM_059385147.1, rna-XM_059386445.1    rna-XM_013059598.2, rna-XM_013062137.2, rna-XM_045087479.1    mNeoVis.2.1.1.chr12_hap1.g00801.t01    rna-XM_021682361.1, rna-XM_044916973.1    rna-XM_004403937.1, rna-XM_004409216.1, rna-XM_004412026.1    rna-XM_042973762.1    rna-XM_040634475.1    rna-XM_027606625.1, rna-XM_027615482.2, rna-XM_027617997.2, rna-XM_035725770.1, rna-XM_035725923.1, rna-XM_035726763.1    mNeoVis.3.1.1.chr12.g00731.t01      mNeoVis.evergladensis.2.1.chr12.g00728.t01    mNeoVis.vulgivaga.2.1.chr12.g00758.t01      1325   N0.HOG0001323  OG0000138  n7                 rna-XM_046014706.1      rna-XM_059172776.1    rna-XM_059394590.1    rna-XM_045079363.1                          1326    N0.HOG0001324   OG0000138  n12         rna-XM_045043071.1, rna-XM_045053818.1, rna-XM_045056974.1    rna-XM_036067260.1, rna-XM_036084729.1, rna-XM_036108809.1    rna-XM_039219927.1        rna-XM_032312904.1    rna-XM_059152471.1    rna-XM_059381994.1    rna-XM_045077358.1      rna-XM_044916286.1, rna-XM_044916932.1, rna-XM_044922206.1    rna-XM_004396765.1, rna-XM_004409148.1    rna-XM_042963882.1, rna-XM_042982716.1    rna-XM_008712277.2    rna-XM_035721940.1, rna-XM_035723393.1, rna-XM_035725300.1              1327    N0.HOG0001325   OG0000138  n34     rna-XM_038565618.1                            rna-XM_004415026.1        rna-XM_035725822.1              1328    N0.HOG0001326   OG0000138  n36     rna-XM_038578051.1                rna-XM_032313374.1, rna-XM_032332898.1    rna-XM_059165938.1    rna-XM_059392925.1    rna-XM_045077984.1              rna-XM_035724087.1, rna-XM_035725530.1, rna-XM_035728177.1              1329   N0.HOG0001327  OG0000138  n44     rna-XM_038573683.1, rna-XM_038575797.1                    rna-XM_059391897.1                            1330    N0.HOG0001328   OG0000138  n47     rna-XM_038559455.1                rna-XM_032361612.1    rna-XM_059165050.1      rna-XM_045088126.1        rna-XM_004411541.1        rna-XM_035725560.1      rna-gnl|WGS_ZZZZ|egapxtmp_035600-R1          1331    N0.HOG0001329   OG0000138  n56     rna-XM_038538182.1        rna-XM_036084952.1        rna-XM_045989234.1    rna-XM_032325082.1    rna-XM_059153525.1    rna-XM_059382133.1    rna-XM_045076355.1    mNeoVis.2.1.1.chr7_hap1.g00088.t01    rna-XM_044922070.1            mNeoVis.3.1.1.chr7.g00073.t01    rna-gnl|WGS_ZZZZ|egapxtmp_010563-R1    mNeoVis.evergladensis.2.1.chr7.g00087.t01    mNeoVis.vulgivaga.2.1.chr7.g00087.t01      1332    N0.HOG0001330   OG0000138  n64     rna-XM_038530011.1        rna-XM_036097057.1        rna-XM_046020695.1              rna-XM_044914996.1                      1333    N0.HOG0001331   OG0000138  n67   rna-XM_034651637.1        rna-XM_045050233.1    rna-XM_036103183.1        rna-XM_046000567.1      rna-XM_059170067.1    rna-XM_059397608.1    rna-XM_045083446.1      rna-XM_044916880.1    rna-XM_004411834.1    rna-XM_042972864.1                  1334   N0.HOG0001332  OG0000139  n1   rna-XM_002930862.2    rna-NM_001388803.1    rna-XM_022513332.1    rna-XM_003992483.3    rna-XM_036081745.1    rna-XM_039250239.1    rna-XM_032869018.1    rna-XM_046017788.1    rna-XM_032357030.1    rna-XM_059156022.1    rna-XM_059392591.1    rna-XM_004750114.1    mNeoVis.2.1.1.chr7_hap1.g01300.t01    rna-XM_044919639.1    rna-XM_004413544.1    rna-XM_007095495.2    rna-XM_008689934.2    rna-XM_027581378.1    mNeoVis.3.1.1.chr7.g01225.t01    rna-gnl|WGS_ZZZZ|egapxtmp_013836-R1    mNeoVis.evergladensis.2.1.chr7.g01289.t01    mNeoVis.vulgivaga.2.1.chr7.g01301.t01      1335   N0.HOG0001333  OG0000139  n20   rna-XM_002926909.1, rna-XM_011234376.2      rna-XM_022491648.1    rna-XM_003980588.2, rna-XM_023240355.2, rna-XM_045061005.1    rna-XM_036079674.1, rna-XM_036079684.1    rna-XM_039222542.1, rna-XM_039222543.1, rna-XM_039222544.1    rna-XM_032848502.1, rna-XM_032848503.1    rna-XM_045999714.1, rna-XM_045999715.1    rna-XM_032335007.1, rna-XM_032335008.1    rna-XM_059173419.1, rna-XM_059173420.1    rna-XM_059416441.1, rna-XM_059416443.1    rna-XM_004782018.2, rna-XM_045086940.1    mNeoVis.2.1.1.chr1_hap1.g00977.t01, mNeoVis.2.1.1.chr1_hap1.g00978.t01      rna-XM_004406164.1    rna-XM_007092970.1, rna-XM_007092971.2, rna-XM_042999028.1    rna-XM_008709873.1, rna-XM_008709894.2, rna-XM_008709896.2, rna-XM_008709897.1, rna-XM_008709898.1, rna-XM_008709899.2, rna-XM_008709902.1, rna-XM_008709903.2, rna-XM_008709904.2, rna-XM_008709906.2, rna-XM_040622725.1      mNeoVis.3.1.1.chr1.g00984.t01, mNeoVis.3.1.1.chr1.g00985.t01    rna-gnl|WGS_ZZZZ|egapxtmp_019102-R1    mNeoVis.evergladensis.2.1.chr1.g00972.t01    mNeoVis.vulgivaga.2.1.chr1.g00986.t01      1336    N0.HOG0001334   OG0000139  n65   rna-XM_002928218.2, rna-XM_034665653.1    rna-NM_001388800.1    rna-XM_022513325.1    rna-XM_003992497.4    rna-XM_036081699.1    rna-XM_039250221.1    rna-XM_032868992.1    rna-XM_046016749.1    rna-XM_032358086.1    rna-XM_059156055.1    rna-XM_059392588.1    rna-XM_004750246.1    mNeoVis.2.1.1.chr7_hap1.g01303.t01        rna-XM_042959184.1    rna-XM_008690169.2    rna-XM_035722861.1    mNeoVis.3.1.1.chr7.g01226.t01    rna-gnl|WGS_ZZZZ|egapxtmp_013835-R1    mNeoVis.evergladensis.2.1.chr7.g01291.t01    mNeoVis.vulgivaga.2.1.chr7.g01303.t01      1337    N0.HOG0001335   OG0000139  n83   rna-XM_011217739.3, rna-XM_019793656.2    rna-NM_001388801.1, rna-NM_001388802.1    rna-XM_022492704.1    rna-XM_003992484.3    rna-XM_036081746.1, rna-XM_036081747.1    rna-XM_039250237.1, rna-XM_039250263.1    rna-XM_032869019.1, rna-XM_032869021.1    rna-XM_046016748.1, rna-XM_046017232.1    rna-XM_032356724.1, rna-XM_032356888.1    rna-XM_059156034.1, rna-XM_059156043.1    rna-XM_059387427.1    rna-XM_004750115.2, rna-XM_045080532.1    mNeoVis.2.1.1.chr7_hap1.g01301.t01, mNeoVis.2.1.1.chr7_hap1.g01302.t01    rna-XM_021689437.2    rna-XM_004413545.1, rna-XM_004413548.1    rna-XM_007074031.2, rna-XM_007095494.2    rna-XM_008689933.2, rna-XM_008690170.2    rna-XM_035723104.1, rna-XM_035723105.1        mNeoVis.evergladensis.2.1.chr7.g01290.t01    mNeoVis.vulgivaga.2.1.chr7.g01302.t01      1338    N0.HOG0001336   OG0000140  n1   rna-XM_011234358.2    rna-NM_001388929.1    rna-XM_022506642.1    rna-XM_003993350.4    rna-XM_036107501.1    rna-XM_039224051.1    rna-XM_032846544.1    rna-XM_046016192.1    rna-XM_032360017.1    rna-XM_059158947.1    rna-XM_059387869.1    rna-XM_004777563.2    mNeoVis.2.1.1.chr7_hap1.g02154.t01    rna-XM_021684836.1      rna-XM_007088785.1    rna-XM_008694254.1    rna-XM_027580396.1    mNeoVis.3.1.1.chr7.g01981.t01    rna-gnl|WGS_ZZZZ|egapxtmp_013484-R1    mNeoVis.evergladensis.2.1.chr7.g02101.t01    mNeoVis.vulgivaga.2.1.chr7.g02121.t01      1339   N0.HOG0001337  OG0000140  n22   rna-XM_034644816.1    rna-XM_038563255.1    rna-XM_022506644.1    rna-XM_045039299.1      rna-XM_039224039.1      rna-XM_046015790.1    rna-XM_032359877.1    rna-XM_059138976.1    rna-XM_059414681.1    rna-XM_004777640.1          rna-XM_015540527.2    rna-XM_008694253.1                1340    N0.HOG0001338   OG0000140  n34   rna-XM_002926865.2    rna-NM_001389029.1    rna-XM_022507175.1    rna-XM_003993329.3      rna-XM_039224006.1    rna-XM_032846556.1    rna-XM_046015804.1    rna-XM_032359860.1    rna-XM_059137414.1    rna-XM_059380392.1    rna-XM_004781691.2      rna-XM_021685203.2      rna-XM_007088805.2    rna-XM_008694226.1            mNeoVis.vulgivaga.2.1.chr7.g02089.t01      1341    N0.HOG0001339   OG0000140  n50   rna-XM_011234340.2    rna-NM_001389182.1      rna-XM_006937307.2      rna-XM_039223516.1      rna-XM_046017195.1                  rna-XM_007088778.2    rna-XM_008694097.1                1342    N0.HOG0001340   OG0000140  n56   rna-XM_002926852.4    rna-NM_001389184.1      rna-XM_011286833.4    rna-XM_036092778.1, rna-XM_036092789.1    rna-XM_039223517.1    rna-XM_032846493.1    rna-XM_046015800.1    rna-XM_032359864.1    rna-XM_059158910.1      rna-XM_013050581.2    mNeoVis.2.1.1.chr7_hap1.g02125.t01    rna-XM_021685207.1      rna-XM_015540525.2    rna-XM_008694232.2    rna-XM_027577864.2    mNeoVis.3.1.1.chr7.g01958.t01      mNeoVis.evergladensis.2.1.chr7.g02071.t01        1343    N0.HOG0001341   OG0000140  n73   rna-XM_002926873.2, rna-XM_011234347.3    rna-XM_038564477.1, rna-XM_038564478.1    rna-XM_022507433.1    rna-XM_011286832.2, rna-XM_011286879.2      rna-XM_039223490.1    rna-XM_032846486.1, rna-XM_032846551.1    rna-XM_046015798.1, rna-XM_046016769.1, rna-XM_046017069.1    rna-XM_032358924.1, rna-XM_032359865.1, rna-XM_032360059.1, rna-XM_032360264.1    rna-XM_059137963.1, rna-XM_059138332.1, rna-XM_059158912.1, rna-XM_059158913.1    rna-XM_059387599.1    rna-XM_004781668.2, rna-XM_004781669.2, rna-XM_004781670.3, rna-XM_004781682.2    mNeoVis.2.1.1.chr7_hap1.g02126.t01, mNeoVis.2.1.1.chr7_hap1.g02127.t01, mNeoVis.2.1.1.chr7_hap1.g02128.t01    rna-XM_044919560.1, rna-XM_044919566.1    rna-XM_004411989.1, rna-XM_004411990.1    rna-XM_007088813.2, rna-XM_015540522.2    rna-XM_008694096.1, rna-XM_008694233.2, rna-XM_008694234.2, rna-XM_008694235.2    rna-XM_027577863.2, rna-XM_027581069.2    mNeoVis.3.1.1.chr7.g01959.t01    rna-gnl|WGS_ZZZZ|egapxtmp_010752-R1, rna-gnl|WGS_ZZZZ|egapxtmp_013475-R1    mNeoVis.evergladensis.2.1.chr7.g02072.t01, mNeoVis.evergladensis.2.1.chr7.g02073.t01, mNeoVis.evergladensis.2.1.chr7.g02074.t01    mNeoVis.vulgivaga.2.1.chr7.g02097.t01      1344    N0.HOG0001342   OG0000141  n1   rna-XM_034653420.1, rna-XM_034660919.1    rna-NM_001287089.1    rna-XM_022512164.1    rna-XM_006932501.5    rna-XM_036107025.1    rna-XM_039230656.1    rna-XM_032864853.1    rna-XM_046010353.1    rna-XM_032341452.1    rna-XM_059180623.1    rna-XM_059372011.1    rna-XM_045067405.1    mNeoVis.2.1.1.chr10_hap1.g01001.t01    rna-XM_021693859.1    rna-XM_004406845.1    rna-XM_042988765.1    rna-XM_040620594.1    rna-XM_027569385.2    mNeoVis.3.1.1.chr9.g00931.t01    rna-gnl|WGS_ZZZZ|egapxtmp_016594-R2    mNeoVis.evergladensis.2.1.chr8.g00972.t01    mNeoVis.vulgivaga.2.1.chr9.g00959.t01      1345    N0.HOG0001343   OG0000141  n20   rna-XM_034647339.1    rna-NM_001286977.1    rna-XM_022512598.1    rna-XM_003995407.4    rna-XM_036077746.1    rna-XM_039241592.1    rna-XM_032866634.1    rna-XM_046023790.1    rna-XM_032306349.1, rna-XM_032320368.1    rna-XM_059143478.1    rna-XM_059411857.1    rna-XM_004769202.3    mNeoVis.2.1.1.chr13_hap1.g00735.t01    rna-XM_021694448.2    rna-XM_004416603.1    rna-XM_007077497.3    rna-XM_008691031.2, rna-XM_040620131.1, rna-XM_040620239.1    rna-XM_027616564.1    mNeoVis.3.1.1.chr13.g00652.t01    rna-gnl|WGS_ZZZZ|egapxtmp_030210-R1    mNeoVis.evergladensis.2.1.chr13.g00700.t01    mNeoVis.vulgivaga.2.1.chr13.g00698.t01      1346    N0.HOG0001344   OG0000141  n44   rna-XM_034667951.1    rna-XM_038532642.1    rna-XM_022511078.1    rna-XM_023254989.2    rna-XM_036092897.1    rna-XM_039235080.1    rna-XM_032835996.1    rna-XM_046007384.1    rna-XM_032342212.1    rna-XM_059180163.1    rna-XM_059371673.1    rna-XM_004759840.3    mNeoVis.2.1.1.chr10_hap1.g01148.t01    rna-XM_021680501.2    rna-XM_004405529.1    rna-XM_042988213.1    rna-XM_040642692.1    rna-XM_027570182.2    mNeoVis.3.1.1.chr9.g01083.t01    rna-gnl|WGS_ZZZZ|egapxtmp_017024-R1    mNeoVis.evergladensis.2.1.chr8.g01127.t01    mNeoVis.vulgivaga.2.1.chr9.g01108.t01      1347    N0.HOG0001345   OG0000141  n63   rna-XM_002914936.4    rna-XM_038552887.1    rna-XM_022501779.1    rna-XM_019816209.3    rna-XM_036100531.1    rna-XM_039234833.1    rna-XM_032860653.1    rna-XM_046023812.1    rna-XM_032308686.1    rna-XM_059142866.1    rna-XM_059411253.1    rna-XM_004747655.3    mNeoVis.2.1.1.chr13_hap1.g00453.t01    rna-XM_021691566.2    rna-XM_004392315.1    rna-XM_042964604.1    rna-XM_008707855.2    rna-XM_027616626.2    mNeoVis.3.1.1.chr13.g00391.t01    rna-gnl|WGS_ZZZZ|egapxtmp_030181-R7    mNeoVis.evergladensis.2.1.chr13.g00435.t01    mNeoVis.vulgivaga.2.1.chr13.g00438.t01      1348    N0.HOG0001346   OG0000141  n84   rna-XM_002913054.4    rna-XM_038575112.1    rna-XM_022517094.1    rna-XM_003994673.6    rna-XM_036076617.1    rna-XM_039242977.1    rna-XM_032876256.1    rna-XM_046024453.1    rna-XM_032310772.1    rna-XM_059140235.1    rna-XM_059408437.1    rna-XM_004753243.2    mNeoVis.2.1.1.chr3_hap1.g00578.t01    rna-XM_021698544.1    rna-XM_004395838.1, rna-XM_004417418.1    rna-XM_042961693.1    rna-XM_008688888.2    rna-XM_035721882.1, rna-XM_035723901.1, rna-XM_035724018.1, rna-XM_035726480.1, rna-XM_035728918.1, rna-XM_035729291.1    mNeoVis.3.1.1.chr3.g00570.t01    rna-gnl|WGS_ZZZZ|egapxtmp_041566-R2    mNeoVis.evergladensis.2.1.chr3.g00586.t01    mNeoVis.vulgivaga.2.1.chr3.g00589.t01      1349    N0.HOG0001347   OG0000142  n1   rna-XM_011218403.2, rna-XM_034666139.1, rna-XM_034667585.1    rna-XM_038568268.1, rna-XM_038568273.1    rna-XM_022513915.1, rna-XM_022514151.1    rna-XM_023239324.2, rna-XM_023239325.2    rna-XM_036074965.1, rna-XM_036074973.1    rna-XM_039254992.1, rna-XM_039254999.1    rna-XM_032853543.1    rna-XM_046015950.1, rna-XM_046017501.1    rna-XM_032359708.1, rna-XM_032359709.1    rna-XM_059174175.1, rna-XM_059186153.1    rna-XM_059401610.1, rna-XM_059401636.1    rna-XM_045087161.1, rna-XM_045087172.1    mNeoVis.2.1.1.chr7_hap1.g01426.t01, mNeoVis.2.1.1.chr7_hap1.g01427.t01    rna-XM_021687549.2, rna-XM_044919779.1    rna-XM_004412437.1, rna-XM_004412438.1    rna-XM_042958222.1, rna-XM_042959469.1    rna-XM_008688422.2, rna-XM_040630635.1    rna-XM_035722921.1, rna-XM_035723119.1    mNeoVis.3.1.1.chr7.g01335.t01    rna-gnl|WGS_ZZZZ|egapxtmp_014263-R1    mNeoVis.evergladensis.2.1.chr7.g01408.t01    mNeoVis.vulgivaga.2.1.chr7.g01421.t01      1350    N0.HOG0001348   OG0000142  n40             rna-XM_039254991.1                      rna-XM_042959021.1                  1351    N0.HOG0001349   OG0000142  n43   rna-XM_011232752.3    rna-XM_038558352.1    rna-XM_022496406.1    rna-XM_003989992.4    rna-XM_036086076.1    rna-XM_039238632.1    rna-XM_032877537.1    rna-XM_046006376.1    rna-XM_032303232.1    rna-XM_059136927.1    rna-XM_059374446.1    rna-XM_004740565.3    mNeoVis.2.1.1.chr2_hap1.g01388.t01    rna-XM_021684255.2    rna-XM_004400654.1    rna-XM_042996876.1    rna-XM_040627408.1    rna-XM_027614330.1    mNeoVis.3.1.1.chr5.g01314.t01    rna-gnl|WGS_ZZZZ|egapxtmp_007073-R1, rna-gnl|WGS_ZZZZ|egapxtmp_008292-R1    mNeoVis.evergladensis.2.1.chr5.g01330.t01    mNeoVis.vulgivaga.2.1.chr5.g01328.t01      1352    N0.HOG0001350   OG0000142  n63   rna-XM_011232486.3    rna-XM_038552329.1    rna-XM_022517552.1    rna-XM_045042970.1    rna-XM_036065799.1, rna-XM_036070789.1    rna-XM_039217112.1, rna-XM_039223641.1    rna-XM_032845561.1, rna-XM_032845569.1    rna-XM_046023271.1    rna-XM_032308067.1    rna-XM_059142109.1    rna-XM_059411674.1    rna-XM_045069836.1    mNeoVis.2.1.1.chr13_hap1.g00641.t01    rna-XM_044920760.1    rna-XM_004393520.1    rna-XM_042965101.1    rna-XM_040631434.1    rna-XM_027615394.2    mNeoVis.3.1.1.chr13.g00568.t01    rna-gnl|WGS_ZZZZ|egapxtmp_031188-R1, rna-gnl|WGS_ZZZZ|egapxtmp_031189-R1    mNeoVis.evergladensis.2.1.chr13.g00615.t01    mNeoVis.vulgivaga.2.1.chr13.g00618.t01      1353    N0.HOG0001351   OG0000142  n88   rna-XM_034657572.1, rna-XM_034657575.1    rna-XM_038567694.1    rna-XM_022525606.1    rna-XM_045044228.1    rna-XM_036111879.1    rna-XM_039247594.1    rna-XM_032883664.1    rna-XM_045979957.1, rna-XM_045990415.1    rna-XM_032307085.1    rna-XM_059159572.1    rna-XM_059388723.1    rna-XM_045086050.1    mNeoVis.2.1.1.chr5_hap1.g00138.t01, mNeoVis.2.1.1.chr5_hap1.g01072.t01    rna-XM_044916750.1    rna-XM_004395309.1    rna-XM_042977864.1    rna-XM_040629528.1    rna-XM_027584577.2    mNeoVis.3.1.1.chr2.g00979.t01    rna-gnl|WGS_ZZZZ|egapxtmp_036303-R2, rna-gnl|WGS_ZZZZ|egapxtmp_038572-R1    mNeoVis.evergladensis.2.1.chr2.g00137.t01, mNeoVis.evergladensis.2.1.chr2.g01063.t01    mNeoVis.vulgivaga.2.1.chr2.g00133.t01, mNeoVis.vulgivaga.2.1.chr2.g01067.t01      1354    N0.HOG0001352   OG0000143  n4   rna-XM_002930339.2, rna-XM_002930341.2    rna-NM_001388854.1      rna-XM_003988831.2      rna-XM_039216977.1, rna-XM_039217023.1      rna-XM_046013647.1, rna-XM_046013686.1    rna-XM_032345704.1    rna-XM_059185070.1, rna-XM_059185072.1    rna-XM_059387242.1    rna-XM_004782524.1, rna-XM_045073467.1    mNeoVis.2.1.1.chr9_hap1.g00448.t01        rna-XM_042990850.1    rna-XM_040641934.1, rna-XM_040641962.1        rna-gnl|WGS_ZZZZ|egapxtmp_024661-R1          1355    N0.HOG0001353   OG0000143  n21   rna-XM_011217113.2    rna-NM_001388756.1    rna-XM_022519768.1    rna-XM_003988828.2      rna-XM_039216969.1    rna-XM_032876530.1, rna-XM_032876533.1    rna-XM_046013736.1    rna-XM_032345714.1    rna-XM_059183560.1    rna-XM_059404885.1    rna-XM_004774514.1    mNeoVis.2.1.1.chr9_hap1.g00451.t01        rna-XM_007081438.2    rna-XM_008711596.2      mNeoVis.3.1.1.chr10.g00426.t01    rna-gnl|WGS_ZZZZ|egapxtmp_024662-R1    mNeoVis.evergladensis.2.1.chr10.g00445.t01    mNeoVis.vulgivaga.2.1.chr10.g00434.t01      1356    N0.HOG0001354   OG0000143  n36   rna-XM_002930338.2, rna-XM_011217112.3    rna-NM_001389077.1    rna-XM_022519765.1, rna-XM_022519773.1    rna-XM_003988830.5, rna-XM_006933786.4, rna-XM_006933788.4, rna-XM_023257798.2, rna-XM_045062670.1    rna-XM_036072863.1    rna-XM_039216976.1, rna-XM_039217025.1    rna-XM_032876528.1, rna-XM_032876532.1    rna-XM_046013530.1, rna-XM_046013631.1, rna-XM_046013632.1, rna-XM_046013651.1, rna-XM_046013652.1, rna-XM_046013750.1    rna-XM_032345644.1, rna-XM_032348506.1    rna-XM_059183558.1      rna-XM_045073223.1, rna-XM_045073455.1      rna-XM_021686743.1      rna-XM_007081439.2, rna-XM_007081563.2, rna-XM_007081564.2, rna-XM_007081565.2, rna-XM_042993695.1    rna-XM_008711593.2, rna-XM_008711595.2      mNeoVis.3.1.1.chr10.g00425.t01      mNeoVis.evergladensis.2.1.chr10.g00444.t01, mNeoVis.evergladensis.2.1.chr3.g00396.t01    mNeoVis.vulgivaga.2.1.chr10.g00433.t01      1357    N0.HOG0001355   OG0000143  n72   rna-XM_002930337.4    rna-NM_001017533.1, rna-NM_001388755.1    rna-XM_022519757.1    rna-XM_003988836.5, rna-XM_006933785.4, rna-XM_011283984.3    rna-XM_036072832.1, rna-XM_036072839.1    rna-XM_039216971.1, rna-XM_039216972.1    rna-XM_032876527.1, rna-XM_032876531.1    rna-XM_046011829.1, rna-XM_046011830.1, rna-XM_046013752.1    rna-XM_032345643.1, rna-XM_032347858.1, rna-XM_032347859.1    rna-XM_059183556.1, rna-XM_059183559.1, rna-XM_059187484.1    rna-XM_059387524.1, rna-XM_059404524.1    rna-XM_004774741.3, rna-XM_004782163.3, rna-XM_045073439.1    mNeoVis.2.1.1.chr9_hap1.g00446.t01, mNeoVis.2.1.1.chr9_hap1.g00449.t01, mNeoVis.2.1.1.chr9_hap1.g00450.t01    rna-XM_021686801.1, rna-XM_021686802.1    rna-XM_004416009.1, rna-XM_004416010.1    rna-XM_007081440.3, rna-XM_007081562.2, rna-XM_007081567.2    rna-XM_008711604.2, rna-XM_008711606.2    rna-XM_027595175.1      rna-gnl|WGS_ZZZZ|egapxtmp_022983-R1    mNeoVis.evergladensis.2.1.chr10.g00442.t01        1358    N0.HOG0001356   OG0000144  n1   rna-XM_034652064.1    rna-NM_001206933.1    rna-XM_022496960.1    rna-XM_023258650.2    rna-XM_036093460.1    rna-XM_039225525.1    rna-XM_032846631.1    rna-XM_046013110.1    rna-XM_032360890.1    rna-XM_059137072.1    rna-XM_059374647.1    rna-XM_004740369.3    mNeoVis.2.1.1.chr2_hap1.g01323.t01    rna-XM_021683881.1    rna-XM_004413166.1    rna-XM_042997019.1    rna-XM_008699240.2    rna-XM_027575740.2    mNeoVis.3.1.1.chr5.g01247.t01    rna-gnl|WGS_ZZZZ|egapxtmp_008290-R1    mNeoVis.evergladensis.2.1.chr5.g01264.t01    mNeoVis.vulgivaga.2.1.chr5.g01260.t01      1359    N0.HOG0001357   OG0000144  n20   rna-XM_002919479.4    rna-XM_038558134.1    rna-XM_022496795.1    rna-XM_011284813.4      rna-XM_039224526.1    rna-XM_032877696.1    rna-XM_046028281.1    rna-XM_032360891.1    rna-XM_059138403.1    rna-XM_059374646.1    rna-XM_045064476.1    mNeoVis.2.1.1.chr2_hap1.g01327.t01    rna-XM_044914024.1    rna-XM_004413169.1    rna-XM_015535333.2    rna-XM_008699242.2    rna-XM_027623304.1    mNeoVis.3.1.1.chr5.g01252.t01    rna-gnl|WGS_ZZZZ|egapxtmp_008398-R1    mNeoVis.evergladensis.2.1.chr5.g01270.t01    mNeoVis.vulgivaga.2.1.chr5.g01267.t01      1360    N0.HOG0001358   OG0000144  n38   rna-XM_002931373.4, rna-XM_034652079.1, rna-XM_034655292.1, rna-XM_034655331.1    rna-NM_001048017.1, rna-NM_001048025.1, rna-NM_001048034.1    rna-XM_022496104.1, rna-XM_022496107.1, rna-XM_022496359.1, rna-XM_022496600.1, rna-XM_022496613.1    rna-XM_023249631.2, rna-XM_023258648.2, rna-XM_045035760.1, rna-XM_045035774.1    rna-XM_036081584.1, rna-XM_036093466.1, rna-XM_036093467.1, rna-XM_036110154.1    rna-XM_039224522.1, rna-XM_039225706.1, rna-XM_039225770.1, rna-XM_039225795.1    rna-XM_032846630.1, rna-XM_032851142.1, rna-XM_032851147.1, rna-XM_032851179.1, rna-XM_032851182.1, rna-XM_032877697.1    rna-XM_046000701.1, rna-XM_046000719.1, rna-XM_046028103.1    rna-XM_032303735.1, rna-XM_032360892.1, rna-XM_032360895.1    rna-XM_059137457.1, rna-XM_059138399.1, rna-XM_059138401.1, rna-XM_059138509.1, rna-XM_059138510.1    rna-XM_059374645.1, rna-XM_059377080.1    rna-XM_004740371.2, rna-XM_004740375.2, rna-XM_004740378.3, rna-XM_045063428.1    mNeoVis.2.1.1.chr2_hap1.g01324.t01, mNeoVis.2.1.1.chr2_hap1.g01325.t01, mNeoVis.2.1.1.chr2_hap1.g01326.t01    rna-XM_044912172.1    rna-XM_004413168.1    rna-XM_007077244.3, rna-XM_015535334.2, rna-XM_042997021.1    rna-XM_008699239.2    rna-XM_035727046.1, rna-XM_035727244.1    mNeoVis.3.1.1.chr5.g01248.t01, mNeoVis.3.1.1.chr5.g01249.t01, mNeoVis.3.1.1.chr5.g01250.t01, mNeoVis.3.1.1.chr5.g01251.t01    rna-gnl|WGS_ZZZZ|egapxtmp_009344-R2, rna-gnl|WGS_ZZZZ|egapxtmp_009345-R1, rna-gnl|WGS_ZZZZ|egapxtmp_009346-R1    mNeoVis.evergladensis.2.1.chr5.g01265.t01, mNeoVis.evergladensis.2.1.chr5.g01266.t01, mNeoVis.evergladensis.2.1.chr5.g01267.t01, mNeoVis.evergladensis.2.1.chr5.g01268.t01, mNeoVis.evergladensis.2.1.chr5.g01269.t01    mNeoVis.vulgivaga.2.1.chr5.g01261.t01, mNeoVis.vulgivaga.2.1.chr5.g01262.t01, mNeoVis.vulgivaga.2.1.chr5.g01263.t01, mNeoVis.vulgivaga.2.1.chr5.g01264.t01, mNeoVis.vulgivaga.2.1.chr5.g01265.t01, mNeoVis.vulgivaga.2.1.chr5.g01266.t01      1361    N0.HOG0001359   OG0000145  n1   rna-XM_019806489.2    rna-XM_038562871.1    rna-XM_022501363.1    rna-XM_045047899.1    rna-XM_036115336.1    rna-XM_039238108.1    rna-XM_032872341.1    rna-XM_045993586.1    rna-XM_032328067.1    rna-XM_059155175.1    rna-XM_059415938.1    rna-XM_045075221.1    mNeoVis.2.1.1.chr15_hap1.g00787.t01    rna-XM_021700440.2    rna-XM_004417703.1    rna-XM_007079104.2    rna-XM_040635015.1    rna-XM_027612394.1    mNeoVis.3.1.1.chr15.g00565.t01    rna-gnl|WGS_ZZZZ|egapxtmp_034164-R2    mNeoVis.evergladensis.2.1.chr15.g00608.t01    mNeoVis.vulgivaga.2.1.chr15.g00621.t01      1362    N0.HOG0001360   OG0000145  n23   rna-XM_034655998.1    rna-XM_038552138.1    rna-XM_022493241.1    rna-XM_011282055.4    rna-XM_036106412.1    rna-XM_039221368.1    rna-XM_032837779.1    rna-XM_045999445.1    rna-XM_032336756.1    rna-XM_059174549.1    rna-XM_059417544.1    rna-XM_004744996.3    mNeoVis.2.1.1.chr1_hap1.g00400.t01    rna-XM_021702133.1    rna-XM_004408260.1    rna-XM_042983714.1    rna-XM_040622462.1    rna-XM_027604509.2    mNeoVis.3.1.1.chr1.g00399.t01      mNeoVis.evergladensis.2.1.chr1.g00399.t01    mNeoVis.vulgivaga.2.1.chr1.g00404.t01      1363    N0.HOG0001361   OG0000145  n40   rna-XM_034649928.1    rna-XM_038588182.1    rna-XM_022518488.1    rna-XM_019824216.3    rna-XM_036087314.1    rna-XM_039219832.1    rna-XM_032882281.1    rna-XM_045994950.1    rna-XM_032329576.1    rna-XM_059156664.1    rna-XM_059385804.1    rna-XM_013061231.2    mNeoVis.2.1.1.chr12_hap1.g00245.t01    rna-XM_044911986.1    rna-XM_004405217.1    rna-XM_042973727.1    rna-XM_040628599.1    rna-XM_027608049.1    mNeoVis.3.1.1.chr12.g00212.t01    rna-gnl|WGS_ZZZZ|egapxtmp_028076-R1    mNeoVis.evergladensis.2.1.chr12.g00214.t01    mNeoVis.vulgivaga.2.1.chr12.g00219.t01      1364    N0.HOG0001362   OG0000145  n60   rna-XM_002929790.4    rna-XM_038582359.1    rna-XM_022505859.1    rna-XM_011281732.3    rna-XM_036097821.1    rna-XM_039230955.1    rna-XM_032835833.1    rna-XM_045997518.1    rna-XM_032334663.1    rna-XM_059158183.1    rna-XM_059385479.1    rna-XM_045079910.1    mNeoVis.2.1.1.chr6_hap1.g00047.t01    rna-XM_021702259.1    rna-XM_004406320.1    rna-XM_042984338.1    rna-XM_040624857.1    rna-XM_027598442.2    mNeoVis.3.1.1.chr4.g00043.t01    rna-gnl|WGS_ZZZZ|egapxtmp_042943-R2    mNeoVis.evergladensis.2.1.chr4.g00049.t01    mNeoVis.vulgivaga.2.1.chr4.g00044.t01      1365    N0.HOG0001363   OG0000145  n79   rna-XM_034658903.1    rna-XM_038550827.1    rna-XM_022516353.1    rna-XM_019826929.3    rna-XM_036110331.1    rna-XM_039224321.1    rna-XM_032842397.1, rna-XM_032852848.1    rna-XM_045981813.1, rna-XM_046020867.1    rna-XM_032351625.1    rna-XM_059134264.1    rna-XM_059406400.1    rna-XM_045077592.1    mNeoVis.2.1.1.chr11_hap1.g00555.t01, mNeoVis.2.1.1.chr13_hap1.g00407.t01    rna-XM_044918487.1    rna-XM_004398406.1    rna-XM_042981034.1    rna-XM_040623686.1    rna-XM_027613606.2, rna-XM_027621617.2    mNeoVis.3.1.1.chr11.g00515.t01, mNeoVis.3.1.1.chr13.g00344.t01    rna-gnl|WGS_ZZZZ|egapxtmp_024924-R5, rna-gnl|WGS_ZZZZ|egapxtmp_030732-R1    mNeoVis.evergladensis.2.1.chr11.g00541.t01, mNeoVis.evergladensis.2.1.chr11.g00542.t01    mNeoVis.vulgivaga.2.1.chr11.g00549.t01, mNeoVis.vulgivaga.2.1.chr13.g00393.t01      1366    N0.HOG0001364   OG0000146  n1   rna-XM_002928776.4    rna-XM_038546999.1    rna-XM_022519929.1    rna-XM_045044384.1    rna-XM_036103123.1    rna-XM_039250670.1    rna-XM_032870953.1    rna-XM_045983739.1    rna-XM_032321467.1    rna-XM_059149054.1    rna-XM_059379237.1    rna-XM_004772447.3    mNeoVis.2.1.1.chr8_hap1.g00284.t01    rna-XM_021678385.1    rna-XM_004411103.1    rna-XM_042966736.1    rna-XM_008693571.2    rna-XM_027625140.1    mNeoVis.3.1.1.chr8.g00252.t01    rna-gnl|WGS_ZZZZ|egapxtmp_020439-R1    mNeoVis.evergladensis.2.1.chr9.g00282.t01    mNeoVis.vulgivaga.2.1.chr8.g00288.t01      1367    N0.HOG0001365   OG0000146  n7   rna-XM_034652323.1, rna-XM_034655757.1    rna-NM_001251927.2    rna-XM_022491649.1    rna-XM_003980625.5    rna-XM_036124267.1    rna-XM_039220250.1    rna-XM_032848509.1    rna-XM_046000890.1    rna-XM_032335124.1    rna-XM_059176104.1    rna-XM_059416296.1    rna-XM_004780601.3    mNeoVis.2.1.1.chr1_hap1.g00954.t01    rna-XM_021688654.2, rna-XM_044915763.1    rna-XM_004407063.1    rna-XM_042981322.1    rna-XM_040622755.1    rna-XM_027624395.2, rna-XM_035721843.1, rna-XM_035729136.1    mNeoVis.3.1.1.chr1.g00966.t01      mNeoVis.evergladensis.2.1.chr1.g00949.t01    mNeoVis.vulgivaga.2.1.chr1.g00960.t01      1368    N0.HOG0001366   OG0000146  n14   rna-XM_002927194.4      rna-XM_022519469.1    rna-XM_006933660.5    rna-XM_036121138.1    rna-XM_039221055.1, rna-XM_039248945.1    rna-XM_032876895.1    rna-XM_046011231.1    rna-XM_032348065.1    rna-XM_059185490.1    rna-XM_059403014.1    rna-XM_004778902.3    mNeoVis.2.1.1.chr9_hap1.g00598.t01    rna-XM_021704783.2    rna-XM_004400873.1    rna-XM_042992202.1    rna-XM_008704531.2, rna-XM_040625226.1    rna-XM_027593047.2    mNeoVis.3.1.1.chr10.g00566.t01    rna-gnl|WGS_ZZZZ|egapxtmp_024477-R2    mNeoVis.evergladensis.2.1.chr10.g00592.t01    mNeoVis.vulgivaga.2.1.chr10.g00578.t01      1369    N0.HOG0001367   OG0000146  n20   rna-XM_002913417.3    rna-XM_038556625.1    rna-XM_022510223.1    rna-XM_003982365.6, rna-XM_019825068.3    rna-XM_036101552.1, rna-XM_036107930.1    rna-XM_039242444.1, rna-XM_039243466.1    rna-XM_032869942.1    rna-XM_046020794.1    rna-XM_032305082.1    rna-XM_059171768.1    rna-XM_059396468.1    rna-XM_004741839.3    mNeoVis.2.1.1.chr4_hap1.g00242.t01    rna-XM_021689732.2, rna-XM_021699433.1    rna-XM_004398041.1    rna-XM_042971522.1, rna-XM_042979224.1    rna-XM_040644050.1    rna-XM_027574060.2, rna-XM_027611810.2    mNeoVis.3.1.1.chr6.g00241.t01      mNeoVis.evergladensis.2.1.chr6.g00241.t01    mNeoVis.vulgivaga.2.1.chr6.g00242.t01      1370    N0.HOG0001368   OG0000146  n28   rna-XM_011231373.3    rna-XM_038566102.1    rna-XM_022498540.1          rna-XM_032868379.1    rna-XM_045991678.1    rna-XM_032323082.1    rna-XM_059161260.1    rna-XM_059390010.1    rna-XM_004738239.3      rna-XM_021695225.1    rna-XM_004412370.1      rna-XM_008703067.2    rna-XM_027585053.2        mNeoVis.evergladensis.2.1.chr2.g00760.t01        1371    N0.HOG0001369   OG0000146  n30                           mNeoVis.2.1.1.chr5_hap1.g00759.t01            rna-XM_035722435.1    mNeoVis.3.1.1.chr2.g00668.t01    rna-gnl|WGS_ZZZZ|egapxtmp_035442-R1      mNeoVis.vulgivaga.2.1.chr2.g00765.t01      1372    N0.HOG0001370   OG0000147  n1   rna-XM_034651204.1, rna-XM_034668875.1    rna-XM_038533054.1    rna-XM_022511229.1    rna-XM_045058696.1    rna-XM_036067623.1    rna-XM_039257144.1    rna-XM_032836824.1    rna-XM_046008330.1    rna-XM_032341855.1    rna-XM_059179768.1    rna-XM_059371377.1    rna-XM_004781710.2    mNeoVis.2.1.1.chr10_hap1.g01283.t01    rna-XM_021680703.1    rna-XM_004403600.1, rna-XM_004417814.1    rna-XM_042987840.1    rna-XM_040642916.1    rna-XM_027569292.2    mNeoVis.3.1.1.chr13.g00494.t01, mNeoVis.3.1.1.chr9.g01216.t01    rna-gnl|WGS_ZZZZ|egapxtmp_002945-R1    mNeoVis.evergladensis.2.1.chr6.g00789.t01, mNeoVis.evergladensis.2.1.chr8.g01261.t01    mNeoVis.vulgivaga.2.1.chr9.g01246.t01      1373    N0.HOG0001371   OG0000147  n28                     rna-XM_059161456.1              rna-XM_042986817.1                  1374   N0.HOG0001372  OG0000147  n30           rna-XM_036120366.1    rna-XM_039247276.1                mNeoVis.2.1.1.chr4_hap1.g00784.t01              mNeoVis.3.1.1.chr5.g01211.t01    rna-gnl|WGS_ZZZZ|egapxtmp_005863-R1    mNeoVis.evergladensis.2.1.chr13.g00536.t01    mNeoVis.vulgivaga.2.1.chr12.g00655.t01      1375   N0.HOG0001373  OG0000147  n38     rna-XM_038541793.1          rna-XM_039223791.1                mNeoVis.2.1.1.chr2_hap1.g00108.t01                        1376    N0.HOG0001374   OG0000147  n42     rna-XM_038579652.1      rna-XM_023243391.2, rna-XM_045034450.1, rna-XM_045038987.1, rna-XM_045055131.1    rna-XM_036079566.1    rna-XM_039247057.1, rna-XM_039256454.1    rna-XM_032853027.1, rna-XM_032880524.1    rna-XM_045994768.1, rna-XM_045997556.1, rna-XM_046010035.1, rna-XM_046020576.1    rna-XM_032307686.1, rna-XM_032336149.1, rna-XM_032344421.1    rna-XM_059166284.1, rna-XM_059178479.1, rna-XM_059180303.1    rna-XM_059376616.1, rna-XM_059395763.1, rna-XM_059399015.1, rna-XM_059418494.1    rna-XM_045074780.1, rna-XM_045075662.1    mNeoVis.2.1.1.chr1_hap1.g01956.t01, mNeoVis.2.1.1.chr5_hap1.g01589.t01    rna-XM_021695114.2    rna-XM_004395696.1, rna-XM_004398335.1, rna-XM_004398365.1    rna-XM_042958331.1, rna-XM_042996260.1    rna-XM_040627942.1    rna-XM_035722475.1, rna-XM_035726481.1, rna-XM_035727369.1, rna-XM_035727883.1      rna-gnl|WGS_ZZZZ|egapxtmp_038477-R1    mNeoVis.evergladensis.2.1.chr2.g01576.t01    mNeoVis.vulgivaga.2.1.chr12.g00149.t01, mNeoVis.vulgivaga.2.1.chr2.g01590.t01, mNeoVis.vulgivaga.2.1.chr6.g00463.t01      1377    N0.HOG0001375   OG0000147  n83   rna-XM_034664016.1    rna-NM_001252143.1, rna-XM_038551085.1, rna-XM_038552596.1    rna-XM_022501823.1    rna-XM_019816079.3    rna-XM_036065050.1      rna-XM_032841625.1, rna-XM_032860361.1    rna-XM_045979411.1, rna-XM_046023401.1    rna-XM_032308310.1    rna-XM_059137350.1, rna-XM_059170909.1    rna-XM_059411509.1    rna-XM_045079689.1    mNeoVis.2.1.1.chr13_hap1.g00553.t01, mNeoVis.2.1.1.chr13_hap1.g00790.t01    rna-XM_021691145.2      rna-XM_042965191.1, rna-XM_042981041.1    rna-XM_040631504.1    rna-XM_035723585.1      rna-gnl|WGS_ZZZZ|egapxtmp_031649-R1      mNeoVis.vulgivaga.2.1.chr13.g00540.t01      1378    N0.HOG0001376   OG0000147  n84         rna-XM_045050689.1        rna-XM_032860787.1, rna-XM_032869484.1        rna-XM_059179102.1        mNeoVis.2.1.1.chr12_hap1.g00702.t01                rna-gnl|WGS_ZZZZ|egapxtmp_028574-R1          1379   N0.HOG0001377  OG0000148  n1   rna-XM_034667380.1    rna-XM_038542911.1    rna-XM_022490772.1    rna-XM_045048498.1    rna-XM_036069823.1    rna-XM_039218481.1    rna-XM_032845708.1    rna-XM_045983221.1    rna-XM_032318041.1    rna-XM_059145605.1    rna-XM_059413587.1    rna-XM_045086664.1    mNeoVis.2.1.1.chr14_hap1.g00589.t01    rna-XM_044916010.1    rna-XM_004415958.1    rna-XM_042975564.1    rna-XM_040635857.1    rna-XM_027610895.2    mNeoVis.3.1.1.chr14.g00575.t01    rna-gnl|WGS_ZZZZ|egapxtmp_031960-R1    mNeoVis.evergladensis.2.1.chr14.g00587.t01    mNeoVis.vulgivaga.2.1.chr14.g00583.t01      1380    N0.HOG0001378   OG0000148  n23   rna-XM_019810202.2    rna-XM_038542899.1    rna-XM_022490729.1      rna-XM_036069846.1        rna-XM_045981948.1      rna-XM_059145412.1      rna-XM_045086536.1, rna-XM_045086645.1    mNeoVis.2.1.1.chr14_hap1.g00585.t01    rna-XM_044912284.1        rna-XM_008700756.1          mNeoVis.evergladensis.2.1.chr14.g00583.t01        1381    N0.HOG0001379   OG0000148  n36   rna-XM_019810192.2    rna-XM_038542904.1    rna-XM_022490698.1    rna-XM_019822360.3    rna-XM_036069807.1    rna-XM_039218493.1    rna-XM_032845646.1    rna-XM_045983673.1    rna-XM_032317713.1    rna-XM_059145120.1    rna-XM_059413577.1    rna-XM_004775776.3    mNeoVis.2.1.1.chr14_hap1.g00587.t01    rna-XM_021681854.1    rna-XM_004415956.1    rna-XM_042976002.1    rna-XM_040635864.1    rna-XM_027610891.2    mNeoVis.3.1.1.chr1.g02110.t01      mNeoVis.evergladensis.2.1.chr14.g00585.t01    mNeoVis.vulgivaga.2.1.chr14.g00581.t01      1382    N0.HOG0001380   OG0000148  n57   rna-XM_034667362.1    rna-XM_038541803.1      rna-XM_023247510.2    rna-XM_036069808.1    rna-XM_039218469.1    rna-XM_032845634.1    rna-XM_045982842.1    rna-XM_032317691.1    rna-XM_059145113.1    rna-XM_059413571.1    rna-XM_045086646.1      rna-XM_021681856.1    rna-XM_004415959.1    rna-XM_007084284.3    rna-XM_040635863.1    rna-XM_027610885.2    mNeoVis.3.1.1.chr14.g00574.t01    rna-gnl|WGS_ZZZZ|egapxtmp_033080-R1    mNeoVis.evergladensis.2.1.chr14.g00586.t01        1383    N0.HOG0001381   OG0000148  n75   rna-XM_011237545.3, rna-XM_034652902.1    rna-XM_038542902.1    rna-XM_022490696.1    rna-XM_019822375.3      rna-XM_039218492.1    rna-XM_032845636.1    rna-XM_045982849.1    rna-XM_032317702.1    rna-XM_059145115.1    rna-XM_059413573.1    rna-XM_045086651.1    mNeoVis.2.1.1.chr14_hap1.g00588.t01    rna-XM_044916012.1    rna-XM_004415960.1      rna-XM_008700763.2    rna-XM_035722020.1    mNeoVis.3.1.1.chr14.g00573.t01    rna-gnl|WGS_ZZZZ|egapxtmp_031959-R1      mNeoVis.vulgivaga.2.1.chr14.g00582.t01      1384    N0.HOG0001382   OG0000148  n94   rna-XM_034667375.1    rna-XM_038542907.1    rna-XM_022490764.1    rna-XM_023247237.2    rna-XM_036069813.1    rna-XM_039218500.1    rna-XM_032845705.1    rna-XM_045983217.1    rna-XM_032318034.1    rna-XM_059145602.1    rna-XM_059413586.1    rna-XM_045086659.1    mNeoVis.2.1.1.chr14_hap1.g00590.t01        rna-XM_042975563.1    rna-XM_040635862.1    rna-XM_027610881.2    mNeoVis.3.1.1.chr14.g00576.t01, mNeoVis.3.1.1.chr7.g00930.t01    rna-gnl|WGS_ZZZZ|egapxtmp_031962-R1    mNeoVis.evergladensis.2.1.chr14.g00588.t01    mNeoVis.vulgivaga.2.1.chr14.g00584.t01      1385    N0.HOG0001383   OG0000149  n1   rna-XM_034649354.1    rna-XM_038586734.1    rna-XM_022518391.1    rna-XM_023249531.2    rna-XM_036097789.1    rna-XM_039245319.1    rna-XM_032880240.1    rna-XM_045995046.1, rna-XM_045996613.1    rna-XM_032330593.1    rna-XM_059158526.1    rna-XM_059385691.1    rna-XM_004768299.3    mNeoVis.2.1.1.chr12_hap1.g00832.t01    rna-XM_021684776.1    rna-XM_004399987.1    rna-XM_042974422.1    rna-XM_008694316.2    rna-XM_027607786.2    mNeoVis.3.1.1.chr12.g00762.t01    rna-gnl|WGS_ZZZZ|egapxtmp_029183-R1    mNeoVis.evergladensis.2.1.chr12.g00760.t01    mNeoVis.vulgivaga.2.1.chr12.g00788.t01      1386    N0.HOG0001384   OG0000149  n25   rna-XM_034649881.1    rna-XM_038586735.1    rna-XM_022518352.1    rna-XM_019823520.3    rna-XM_036097775.1    rna-XM_039245283.1    rna-XM_032880127.1    rna-XM_045995596.1, rna-XM_045996696.1    rna-XM_032331547.1    rna-XM_059158484.1    rna-XM_059385312.1    rna-XM_004768298.2    mNeoVis.2.1.1.chr12_hap1.g00829.t01    rna-XM_021697713.1    rna-XM_004400003.1    rna-XM_007091022.2    rna-XM_008694320.2    rna-XM_027608961.2    mNeoVis.3.1.1.chr12.g00759.t01    rna-gnl|WGS_ZZZZ|egapxtmp_028407-R1    mNeoVis.evergladensis.2.1.chr12.g00757.t01    mNeoVis.vulgivaga.2.1.chr12.g00785.t01      1387    N0.HOG0001385   OG0000149  n48   rna-XM_034649880.1    rna-NM_001048123.1    rna-XM_022518427.1    rna-XM_023249256.2    rna-XM_036097802.1    rna-XM_039245284.1    rna-XM_032880241.1    rna-XM_045995042.1, rna-XM_045996609.1    rna-XM_032331544.1    rna-XM_059158482.1    rna-XM_059385313.1    rna-XM_013064399.2    mNeoVis.2.1.1.chr12_hap1.g00830.t01    rna-XM_021677829.1    rna-XM_004400002.1    rna-XM_007091014.3    rna-XM_008694341.2    rna-XM_027608688.2    mNeoVis.3.1.1.chr12.g00760.t01    rna-gnl|WGS_ZZZZ|egapxtmp_028406-R1    mNeoVis.evergladensis.2.1.chr12.g00758.t01    mNeoVis.vulgivaga.2.1.chr12.g00786.t01      1388    N0.HOG0001386   OG0000149  n70   rna-XM_034649457.1    rna-XM_038586766.1    rna-XM_022518368.1    rna-XM_004000254.5    rna-XM_036087974.1    rna-XM_039238554.1    rna-XM_032880230.1    rna-XM_045996298.1, rna-XM_045996745.1    rna-XM_032329501.1    rna-XM_059158704.1    rna-XM_059385846.1    rna-XM_004768287.3    mNeoVis.2.1.1.chr12_hap1.g00823.t01    rna-XM_021678538.1    rna-XM_004414787.1    rna-XM_042973771.1    rna-XM_008694325.2    rna-XM_027607863.2    mNeoVis.3.1.1.chr12.g00753.t01    rna-gnl|WGS_ZZZZ|egapxtmp_028709-R1    mNeoVis.evergladensis.2.1.chr12.g00751.t01    mNeoVis.vulgivaga.2.1.chr12.g00778.t01      1389    N0.HOG0001387   OG0000149  n92   rna-XM_034649360.1    rna-NM_001048122.1    rna-XM_022518393.1    rna-XM_023249528.2    rna-XM_036097794.1    rna-XM_039245285.1    rna-XM_032880270.1    rna-XM_045995043.1, rna-XM_045996611.1    rna-XM_032330590.1    rna-XM_059158524.1    rna-XM_059385688.1    rna-XM_045089263.1    mNeoVis.2.1.1.chr12_hap1.g00831.t01    rna-XM_021698731.1    rna-XM_004400001.1    rna-XM_007091013.2    rna-XM_040626350.1    rna-XM_035725356.1    mNeoVis.3.1.1.chr12.g00761.t01    rna-gnl|WGS_ZZZZ|egapxtmp_029181-R1    mNeoVis.evergladensis.2.1.chr12.g00759.t01    mNeoVis.vulgivaga.2.1.chr12.g00787.t01      1390    N0.HOG0001388   OG0000150  n1   rna-XM_011229231.2    rna-NM_001388602.1    rna-XM_022507450.1    rna-XM_003993358.1    rna-XM_036107492.1    rna-XM_039224053.1    rna-XM_032846488.1    rna-XM_046016796.1    rna-XM_032359090.1    rna-XM_059158957.1    rna-XM_059387844.1    rna-XM_004777564.1    mNeoVis.2.1.1.chr7_hap1.g02162.t01    rna-XM_044919498.1      rna-XM_042958408.1    rna-XM_008694126.1    rna-XM_027580538.2    mNeoVis.3.1.1.chr7.g01987.t01    rna-gnl|WGS_ZZZZ|egapxtmp_013497-R1    mNeoVis.evergladensis.2.1.chr7.g02110.t01    mNeoVis.vulgivaga.2.1.chr7.g02130.t01      1391    N0.HOG0001389   OG0000150  n21   rna-XM_002922682.2    rna-NM_001388603.1    rna-XM_022507411.1    rna-XM_003993357.1    rna-XM_036107493.1    rna-XM_039224036.1    rna-XM_032846540.1    rna-XM_046016889.1    rna-XM_032359883.1    rna-XM_059158956.1    rna-XM_059410097.1    rna-XM_004777647.1    mNeoVis.2.1.1.chr7_hap1.g02161.t01    rna-XM_021684839.1      rna-XM_015540514.2    rna-XM_008694125.1      mNeoVis.3.1.1.chr7.g01986.t01    rna-gnl|WGS_ZZZZ|egapxtmp_013502-R1    mNeoVis.evergladensis.2.1.chr7.g02108.t01    mNeoVis.vulgivaga.2.1.chr7.g02129.t01      1392    N0.HOG0001390   OG0000150  n40   rna-XM_002922683.4, rna-XM_002922684.4    rna-NM_001389117.1    rna-XM_022506639.1    rna-XM_011286872.3, rna-XM_023239741.2      rna-XM_039225620.1    rna-XM_032846537.1, rna-XM_032846538.1    rna-XM_046015787.1, rna-XM_046016998.1, rna-XM_046017030.1, rna-XM_046017330.1    rna-XM_032359885.1, rna-XM_032359886.1, rna-XM_032359888.1    rna-XM_059158959.1, rna-XM_059158960.1    rna-XM_059387825.1, rna-XM_059406952.1    rna-XM_004777649.2, rna-XM_004777650.2    mNeoVis.2.1.1.chr7_hap1.g02164.t01    rna-XM_021685214.2      rna-XM_007088837.2, rna-XM_007088838.1    rna-XM_008694127.2, rna-XM_008694261.2    rna-XM_027580566.1    mNeoVis.3.1.1.chr7.g01990.t01, mNeoVis.3.1.1.chr7.g01991.t01    rna-gnl|WGS_ZZZZ|egapxtmp_012638-R1, rna-gnl|WGS_ZZZZ|egapxtmp_013533-R1    mNeoVis.evergladensis.2.1.chr7.g02112.t01    mNeoVis.vulgivaga.2.1.chr7.g02133.t01, mNeoVis.vulgivaga.2.1.chr7.g02134.t01      1393    N0.HOG0001391   OG0000150  n75   rna-XM_011229253.2    rna-NM_001388601.1    rna-XM_022506640.1, rna-XM_022507545.1    rna-XM_011286870.4, rna-XM_045039304.1    rna-XM_036107498.1    rna-XM_039225619.1, rna-XM_039225712.1, rna-XM_039225790.1    rna-XM_032846485.1, rna-XM_032846539.1    rna-XM_046015788.1, rna-XM_046016868.1    rna-XM_032359089.1, rna-XM_032359884.1    rna-XM_059139320.1, rna-XM_059158958.1    rna-XM_059387837.1, rna-XM_059408956.1    rna-XM_004777648.2, rna-XM_045068439.1    mNeoVis.2.1.1.chr7_hap1.g02160.t01, mNeoVis.2.1.1.chr7_hap1.g02163.t01    rna-XM_021685213.1      rna-XM_007088839.1, rna-XM_007088840.1    rna-XM_008694259.2, rna-XM_008694260.2    rna-XM_027580536.1    mNeoVis.3.1.1.chr7.g01988.t01, mNeoVis.3.1.1.chr7.g01989.t01    rna-gnl|WGS_ZZZZ|egapxtmp_012636-R1, rna-gnl|WGS_ZZZZ|egapxtmp_012637-R1    mNeoVis.evergladensis.2.1.chr7.g02109.t01, mNeoVis.evergladensis.2.1.chr7.g02111.t01    mNeoVis.vulgivaga.2.1.chr7.g02131.t01, mNeoVis.vulgivaga.2.1.chr7.g02132.t01      1394    N0.HOG0001392   OG0000151  n1   rna-XM_002925622.4    rna-XM_038554232.1    rna-XM_022503934.1    rna-XM_011282370.4    rna-XM_036115011.1    rna-XM_039231210.1    rna-XM_032861946.1, rna-XM_032862020.1    rna-XM_046004416.1    rna-XM_032340153.1    rna-XM_059178146.1    rna-XM_059400335.1, rna-XM_059400337.1    rna-XM_004739775.3    mNeoVis.2.1.1.chr1_hap1.g01595.t01    rna-XM_021692062.2    rna-XM_004408431.1    rna-XM_042986489.1    rna-XM_008691844.2, rna-XM_040643553.1    rna-XM_027602434.2    mNeoVis.3.1.1.chr1.g01545.t01    rna-gnl|WGS_ZZZZ|egapxtmp_027475-R1    mNeoVis.evergladensis.2.1.chr1.g01562.t01    mNeoVis.vulgivaga.2.1.chr1.g01559.t01      1395    N0.HOG0001393   OG0000151  n25   rna-XM_002930018.4    rna-NM_001252138.1, rna-XM_038560140.1    rna-XM_022510870.1    rna-XM_003985936.5    rna-XM_036108278.1    rna-XM_039219594.1    rna-XM_032847143.1    rna-XM_046006350.1    rna-XM_032340861.1    rna-XM_059178893.1    rna-XM_059399864.1    rna-XM_004780957.3    mNeoVis.2.1.1.chr1_hap1.g01352.t01    rna-XM_021697119.2    rna-XM_004415248.1    rna-XM_007098779.3    rna-XM_008711884.2    rna-XM_027603870.1, rna-XM_035725273.1, rna-XM_035728876.1    mNeoVis.3.1.1.chr1.g01301.t01      mNeoVis.evergladensis.2.1.chr1.g01319.t01    mNeoVis.vulgivaga.2.1.chr1.g01320.t01      1396    N0.HOG0001394   OG0000151  n37   rna-XM_034646826.1    rna-XM_038531560.1    rna-XM_022514631.1    rna-XM_023258315.2    rna-XM_036088994.1    rna-XM_039231251.1    rna-XM_032863791.1    rna-XM_046018531.1    rna-XM_032360599.1    rna-XM_059138570.1    rna-XM_059376686.1    rna-XM_004741149.3    mNeoVis.2.1.1.chr2_hap1.g01666.t01    rna-XM_044914438.1    rna-XM_004394739.1    rna-XM_042995062.1    rna-XM_040619001.1    rna-XM_027577692.2    mNeoVis.3.1.1.chr5.g01589.t01    rna-gnl|WGS_ZZZZ|egapxtmp_009584-R1    mNeoVis.evergladensis.2.1.chr5.g01607.t01    mNeoVis.vulgivaga.2.1.chr5.g01609.t01      1397    N0.HOG0001395   OG0000151  n45   rna-XM_002930770.4    rna-NM_001195154.1    rna-XM_022491328.1    rna-XM_019824524.3    rna-XM_036119674.1    rna-XM_039222675.1    rna-XM_032846348.1    rna-XM_045995322.1    rna-XM_032330394.1    rna-XM_059157481.1    rna-XM_059384881.1    rna-XM_045068094.1    mNeoVis.2.1.1.chr12_hap1.g00006.t01    rna-XM_021682636.1    rna-XM_004416918.1    rna-XM_007088957.3    rna-XM_008686767.2    rna-XM_027609343.1      rna-gnl|WGS_ZZZZ|egapxtmp_028982-R1    mNeoVis.evergladensis.2.1.chr12.g00004.t01    mNeoVis.vulgivaga.2.1.chr12.g00004.t01      1398    N0.HOG0001396   OG0000151  n61   rna-XM_034655872.1    rna-XM_038581423.1    rna-XM_022516185.1    rna-XM_023238767.2    rna-XM_036101887.1    rna-XM_039228938.1    rna-XM_032865374.1    rna-XM_046002977.1    rna-XM_032354363.1    rna-XM_059164668.1    rna-XM_059392305.1    rna-XM_004757538.3    mNeoVis.2.1.1.chr5_hap1.g01815.t01    rna-XM_021677975.1    rna-XM_004409626.1    rna-XM_042956873.1    rna-XM_008701865.2    rna-XM_027585143.2    mNeoVis.3.1.1.chr2.g01721.t01    rna-gnl|WGS_ZZZZ|egapxtmp_037488-R1    mNeoVis.evergladensis.2.1.chr2.g01801.t01    mNeoVis.vulgivaga.2.1.chr2.g01815.t01      1399    N0.HOG0001397   OG0000152  n3     rna-XM_038581129.1    rna-XM_022515951.1        rna-XM_039229574.1                        rna-XM_040621697.1                1400    N0.HOG0001398   OG0000152  n8     rna-XM_038581133.1      rna-XM_023238574.2    rna-XM_036118540.1      rna-XM_032865565.1    rna-XM_046001807.1              rna-XM_044913622.1      rna-XM_042956339.1      rna-XM_027582299.1              1401    N0.HOG0001399   OG0000152  n15     rna-XM_038581131.1    rna-XM_022515913.1, rna-XM_022515917.1    rna-XM_023238824.2          rna-XM_046001844.1    rna-XM_032302627.1      rna-XM_059388207.1    rna-XM_045085198.1          rna-XM_042998966.1        mNeoVis.3.1.1.chr2.g01676.t01            1402    N0.HOG0001400   OG0000152  n30                                           mNeoVis.evergladensis.2.1.chr2.g01754.t01    mNeoVis.vulgivaga.2.1.chr2.g01767.t01      1403    N0.HOG0001401   OG0000152  n33     rna-XM_038581135.1              rna-XM_046001819.1    rna-XM_032303182.1          mNeoVis.2.1.1.chr5_hap1.g01771.t01                rna-gnl|WGS_ZZZZ|egapxtmp_034602-R1    mNeoVis.evergladensis.2.1.chr2.g01755.t01    mNeoVis.vulgivaga.2.1.chr2.g01770.t01      1404    N0.HOG0001402   OG0000152  n38   rna-XM_034661836.1    rna-XM_038581134.1    rna-XM_022515914.1          rna-XM_032865564.1    rna-XM_046002014.1        rna-XM_059388208.1    rna-XM_045085197.1    mNeoVis.2.1.1.chr5_hap1.g01770.t01          rna-XM_008701814.1        rna-gnl|WGS_ZZZZ|egapxtmp_034598-R1    mNeoVis.evergladensis.2.1.chr2.g01753.t01    mNeoVis.vulgivaga.2.1.chr2.g01769.t01      1405    N0.HOG0001403   OG0000152  n47   rna-XM_034661792.1                                    rna-XM_027581716.2              1406    N0.HOG0001404   OG0000152  n48   rna-XM_034655632.1, rna-XM_034661905.1, rna-XM_034661990.1, rna-XM_034662066.1    rna-XM_038581128.1    rna-XM_022515903.1, rna-XM_022515905.1    rna-XM_011285738.4, rna-XM_011285739.3, rna-XM_023260848.2, rna-XM_023260849.2, rna-XM_023260850.2    rna-XM_036101102.1, rna-XM_036118555.1, rna-XM_036118557.1    rna-XM_039228268.1, rna-XM_039228286.1    rna-XM_032865528.1, rna-XM_032865566.1, rna-XM_032865567.1, rna-XM_032865568.1    rna-XM_046001552.1    rna-XM_032302576.1, rna-XM_032302585.1, rna-XM_032302593.1, rna-XM_032302607.1    rna-XM_059165291.1, rna-XM_059165292.1    rna-XM_059388203.1, rna-XM_059388204.1, rna-XM_059388205.1, rna-XM_059388206.1    rna-XM_004757489.2, rna-XM_004757554.1, rna-XM_045085199.1, rna-XM_045085200.1    mNeoVis.2.1.1.chr5_hap1.g01763.t01, mNeoVis.2.1.1.chr5_hap1.g01764.t01    rna-XM_021678014.1, rna-XM_044913623.1, rna-XM_044913627.1, rna-XM_044919021.1      rna-XM_007075068.2, rna-XM_015543474.2, rna-XM_042956446.1, rna-XM_042956447.1, rna-XM_042998317.1    rna-XM_040621664.1, rna-XM_040621698.1, rna-XM_040621699.1, rna-XM_040621700.1    rna-XM_027587924.1, rna-XM_035724758.1    mNeoVis.3.1.1.chr2.g01670.t01, mNeoVis.3.1.1.chr2.g01671.t01    rna-gnl|WGS_ZZZZ|egapxtmp_034599-R1, rna-gnl|WGS_ZZZZ|egapxtmp_034600-R1    mNeoVis.evergladensis.2.1.chr2.g01745.t01, mNeoVis.evergladensis.2.1.chr2.g01746.t01, mNeoVis.evergladensis.2.1.chr2.g01747.t01    mNeoVis.vulgivaga.2.1.chr2.g01761.t01, mNeoVis.vulgivaga.2.1.chr2.g01762.t01, mNeoVis.vulgivaga.2.1.chr2.g01763.t01      1407    N0.HOG0001405   OG0000153  n2   rna-XM_034666467.1    rna-NM_001388709.1      rna-XM_045038832.1    rna-XM_036076015.1    rna-XM_039224511.1      rna-XM_046017095.1, rna-XM_046017112.1    rna-XM_032360275.1      rna-XM_059387269.1, rna-XM_059389849.1    rna-XM_004779051.2, rna-XM_004782413.2    mNeoVis.2.1.1.chr7_hap1.g01700.t01        rna-XM_015542099.2    rna-XM_008709570.1        rna-gnl|WGS_ZZZZ|egapxtmp_012810-R1    mNeoVis.evergladensis.2.1.chr7.g01671.t01        1408    N0.HOG0001406   OG0000153  n17   rna-XM_002926097.2    rna-NM_001388707.1    rna-XM_022522047.1    rna-XM_003992895.5      rna-XM_039223787.1    rna-XM_032853650.1    rna-XM_046017458.1    rna-XM_032359663.1    rna-XM_059157610.1    rna-XM_059389885.1    rna-XM_013049231.1    mNeoVis.2.1.1.chr7_hap1.g01695.t01        rna-XM_015542088.1    rna-XM_008709640.2      mNeoVis.3.1.1.chr7.g01583.t01    rna-gnl|WGS_ZZZZ|egapxtmp_012811-R1    mNeoVis.evergladensis.2.1.chr7.g01665.t01    mNeoVis.vulgivaga.2.1.chr7.g01684.t01      1409    N0.HOG0001407   OG0000153  n34   rna-XM_011233415.1    rna-XM_038568856.1    rna-XM_022522105.1    rna-XM_023238997.2    rna-XM_036075928.1    rna-XM_039224516.1    rna-XM_032853652.1    rna-XM_046015889.1, rna-XM_046015890.1, rna-XM_046016979.1    rna-XM_032360370.1    rna-XM_059157623.1    rna-XM_059389874.1    rna-XM_013049228.2    mNeoVis.2.1.1.chr7_hap1.g01699.t01        rna-XM_007092235.2    rna-XM_008709571.1        rna-gnl|WGS_ZZZZ|egapxtmp_012835-R1    mNeoVis.evergladensis.2.1.chr7.g01670.t01        1410    N0.HOG0001408   OG0000153  n52   rna-XM_002926100.4    rna-NM_001388704.1, rna-NM_001388981.1, rna-XM_038568170.1    rna-XM_022521980.1    rna-XM_045038125.1      rna-XM_039223777.1, rna-XM_039223779.1    rna-XM_032852706.1    rna-XM_046016462.1    rna-XM_032359199.1    rna-XM_059157591.1      rna-XM_004779007.2    mNeoVis.2.1.1.chr7_hap1.g01693.t01        rna-XM_007092210.2    rna-XM_008709590.2        rna-gnl|WGS_ZZZZ|egapxtmp_012817-R1    mNeoVis.evergladensis.2.1.chr7.g01663.t01    mNeoVis.vulgivaga.2.1.chr7.g01681.t01      1411    N0.HOG0001409   OG0000153  n70   rna-XM_002926072.4, rna-XM_034667722.1    rna-NM_001388706.1, rna-XM_038568171.1    rna-XM_022521706.1    rna-XM_045038126.1      rna-XM_039223780.1, rna-XM_039223782.1    rna-XM_032852707.1    rna-XM_046015893.1    rna-XM_032359775.1    rna-XM_059181991.1    rna-XM_059389916.1    rna-XM_013049232.1          rna-XM_007092209.2    rna-XM_008709589.2, rna-XM_008709642.2            mNeoVis.vulgivaga.2.1.chr7.g01683.t01      1412    N0.HOG0001410   OG0000153  n87   rna-XM_034653264.1    rna-NM_001388899.1    rna-XM_022521989.1    rna-XM_003992868.3    rna-XM_036087690.1    rna-XM_039223774.1    rna-XM_032853649.1    rna-XM_046016480.1    rna-XM_032359424.1    rna-XM_059157602.1    rna-XM_059389900.1    rna-XM_004779055.3    mNeoVis.2.1.1.chr7_hap1.g01694.t01    rna-XM_021685686.1    rna-XM_004407566.1    rna-XM_007092208.2    rna-XM_040630671.1, rna-XM_040631307.1    rna-XM_027580136.1    mNeoVis.3.1.1.chr7.g01582.t01    rna-gnl|WGS_ZZZZ|egapxtmp_012526-R1      mNeoVis.vulgivaga.2.1.chr7.g01682.t01      1413    N0.HOG0001411   OG0000154  n1   rna-XM_034665560.1    rna-XM_038552727.1    rna-XM_022501375.1    rna-XM_006939264.4    rna-XM_036081297.1    rna-XM_039218563.1    rna-XM_032860499.1    rna-XM_046021836.1    rna-XM_032308023.1    rna-XM_059142437.1    rna-XM_059410122.1    rna-XM_004747511.3    mNeoVis.2.1.1.chr13_hap1.g00509.t01    rna-XM_021691636.2    rna-XM_004392352.1    rna-XM_007076509.3    rna-XM_040632328.1    rna-XM_027614402.2    mNeoVis.3.1.1.chr13.g00444.t01, mNeoVis.3.1.1.chr13.g00445.t01    rna-gnl|WGS_ZZZZ|egapxtmp_030957-R1    mNeoVis.evergladensis.2.1.chr13.g00490.t01    mNeoVis.vulgivaga.2.1.chr13.g00491.t01, mNeoVis.vulgivaga.2.1.chr13.g00493.t01      1414    N0.HOG0001412   OG0000154  n26   rna-XM_034660003.1    rna-XM_038567944.1    rna-XM_022525417.1    rna-XM_023243212.2    rna-XM_036069989.1    rna-XM_039248152.1    rna-XM_032847094.1    rna-XM_045991380.1    rna-XM_032303135.1    rna-XM_059159294.1    rna-XM_059392608.1    rna-XM_045088410.1    mNeoVis.2.1.1.chr5_hap1.g00041.t01    rna-XM_021705533.1    rna-XM_004395486.1    rna-XM_042977599.1    rna-XM_040628303.1    rna-XM_027582221.2    mNeoVis.3.1.1.chr2.g00023.t01    rna-gnl|WGS_ZZZZ|egapxtmp_036490-R1    mNeoVis.evergladensis.2.1.chr2.g00041.t01    mNeoVis.vulgivaga.2.1.chr2.g00041.t01      1415    N0.HOG0001413   OG0000154  n47   rna-XM_002920263.4    rna-XM_038525923.1    rna-XM_022513713.1    rna-XM_011288155.4    rna-XM_036097509.1    rna-XM_039230508.1    rna-XM_032882416.1    rna-XM_046024392.1    rna-XM_032309129.1    rna-XM_059140999.1    rna-XM_059409737.1    rna-XM_004745551.3    mNeoVis.2.1.1.chr3_hap1.g00077.t01    rna-XM_021686141.2    rna-XM_004409812.1    rna-XM_042961341.1    rna-XM_008705311.2    rna-XM_027576119.1    mNeoVis.3.1.1.chr3.g00077.t01    rna-gnl|WGS_ZZZZ|egapxtmp_042112-R2    mNeoVis.evergladensis.2.1.chr3.g00081.t01    mNeoVis.vulgivaga.2.1.chr3.g00076.t01      1416    N0.HOG0001414   OG0000154  n68   rna-XM_034667535.1    rna-XM_038543229.1    rna-XM_022490743.1    rna-XM_011291237.3    rna-XM_036086854.1    rna-XM_039218930.1    rna-XM_032844981.1    rna-XM_045982672.1    rna-XM_032319229.1    rna-XM_059145451.1    rna-XM_059413851.1    rna-XM_045086851.1    mNeoVis.2.1.1.chr14_hap1.g00687.t01    rna-XM_021682172.2    rna-XM_004402989.1    rna-XM_042975974.1    rna-XM_040635715.1    rna-XM_027612325.2    mNeoVis.3.1.1.chr14.g00657.t01      mNeoVis.evergladensis.2.1.chr14.g00686.t01    mNeoVis.vulgivaga.2.1.chr14.g00686.t01      1417    N0.HOG0001415   OG0000154  n88   rna-XM_034660997.1    rna-XM_038580388.1    rna-XM_022524918.1    rna-XM_045059333.1    rna-XM_036101919.1    rna-XM_039237152.1    rna-XM_032856521.1    rna-XM_046009963.1    rna-XM_032343682.1    rna-XM_059181509.1    rna-XM_059372800.1    rna-XM_004755458.3    mNeoVis.2.1.1.chr10_hap1.g00624.t01    rna-XM_021693958.1    rna-XM_004396579.1    rna-XM_042988887.1    rna-XM_008705952.2    rna-XM_027572558.2    mNeoVis.3.1.1.chr9.g00581.t01    rna-gnl|WGS_ZZZZ|egapxtmp_016870-R1, rna-gnl|WGS_ZZZZ|egapxtmp_016871-R1    mNeoVis.evergladensis.2.1.chr8.g00605.t01    mNeoVis.vulgivaga.2.1.chr9.g00605.t01, mNeoVis.vulgivaga.2.1.chr9.g00606.t01      1418    N0.HOG0001416   OG0000155  n1   rna-XM_034650083.1, rna-XM_034650103.1, rna-XM_034650104.1, rna-XM_034650185.1, rna-XM_034666461.1    rna-NM_001389003.1    rna-XM_022521724.1, rna-XM_022521726.1, rna-XM_022521727.1    rna-XM_011286667.3, rna-XM_011286668.4    rna-XM_036081415.1, rna-XM_036114429.1    rna-XM_039223182.1    rna-XM_032852671.1, rna-XM_032852672.1    rna-XM_046015916.1, rna-XM_046015919.1    rna-XM_032359741.1    rna-XM_059178841.1    rna-XM_059397022.1, rna-XM_059397265.1    rna-XM_004768243.2, rna-XM_013064264.2    mNeoVis.2.1.1.chr7_hap1.g01629.t01    rna-XM_021685670.1      rna-XM_007089404.1, rna-XM_015540745.2    rna-XM_008710703.1, rna-XM_008710748.1, rna-XM_008710750.2    rna-XM_035723127.1    mNeoVis.3.1.1.chr7.g01533.t01    rna-gnl|WGS_ZZZZ|egapxtmp_012851-R1    mNeoVis.evergladensis.2.1.chr7.g01600.t01    mNeoVis.vulgivaga.2.1.chr7.g01620.t01      1419    N0.HOG0001417   OG0000155  n36   rna-XM_002931011.2      rna-XM_022522123.1      rna-XM_036114414.1    rna-XM_039223166.1    rna-XM_032853601.1    rna-XM_046017320.1    rna-XM_032356276.1    rna-XM_059178456.1    rna-XM_059390719.1    rna-XM_004782222.1    mNeoVis.2.1.1.chr7_hap1.g01621.t01    rna-XM_021685666.1      rna-XM_015540753.1    rna-XM_008710747.1      mNeoVis.3.1.1.chr7.g01525.t01    rna-gnl|WGS_ZZZZ|egapxtmp_012849-R1    mNeoVis.evergladensis.2.1.chr7.g01592.t01    mNeoVis.vulgivaga.2.1.chr7.g01611.t01      1420    N0.HOG0001418   OG0000155  n51   rna-XM_002931013.4    rna-NM_001388845.1    rna-XM_022521728.1    rna-XM_003992800.5    rna-XM_036114428.1    rna-XM_039223180.1, rna-XM_039223185.1    rna-XM_032852670.1    rna-XM_046015920.1    rna-XM_032359737.1    rna-XM_059178721.1    rna-XM_059397380.1    rna-XM_004768240.1          rna-XM_007089346.3    rna-XM_008710702.2, rna-XM_008710707.2    rna-XM_027577915.2, rna-XM_035723128.1              1421    N0.HOG0001419   OG0000155  n69   rna-XM_002928570.2    rna-XM_038568844.1      rna-XM_003992795.2    rna-XM_036114422.1    rna-XM_039223190.1, rna-XM_039225566.1    rna-XM_032853594.1    rna-XM_046017609.1    rna-XM_032356339.1    rna-XM_059156787.1    rna-XM_059390815.1    rna-XM_004782225.2    mNeoVis.2.1.1.chr7_hap1.g01614.t01        rna-XM_007089408.1    rna-XM_008710694.2      mNeoVis.3.1.1.chr7.g01520.t01    rna-gnl|WGS_ZZZZ|egapxtmp_012838-R1    mNeoVis.evergladensis.2.1.chr7.g01586.t01    mNeoVis.vulgivaga.2.1.chr7.g01604.t01      1422    N0.HOG0001420   OG0000155  n87   rna-XM_002931213.4    rna-NM_001389114.1    rna-XM_022522159.1    rna-XM_003992798.3    rna-XM_036114413.1    rna-XM_039223181.1    rna-XM_032853600.1    rna-XM_046017381.1    rna-XM_032356257.1    rna-XM_059156806.1    rna-XM_059387546.1, rna-XM_059390733.1    rna-XM_013050828.1    mNeoVis.2.1.1.chr7_hap1.g01620.t01    rna-XM_021685619.1      rna-XM_007089348.2    rna-XM_008710699.1    rna-XM_027580952.1    mNeoVis.3.1.1.chr7.g01524.t01    rna-gnl|WGS_ZZZZ|egapxtmp_012884-R1    mNeoVis.evergladensis.2.1.chr7.g01591.t01    mNeoVis.vulgivaga.2.1.chr7.g01610.t01      1423    N0.HOG0001421   OG0000156  n1   rna-XM_002930210.4    rna-XM_038526806.1    rna-XM_022491226.1    rna-XM_011288293.3    rna-XM_036115812.1    rna-XM_039245662.1    rna-XM_032840477.1    rna-XM_045995524.1, rna-XM_046023956.1    rna-XM_032308202.1    rna-XM_059142612.1    rna-XM_059411690.1    rna-XM_004777423.3    mNeoVis.2.1.1.chr13_hap1.g00653.t01    rna-XM_021677897.1    rna-XM_004416500.1    rna-XM_007089015.3    rna-XM_040620137.1    rna-XM_027616741.2    mNeoVis.3.1.1.chr13.g00578.t01    rna-gnl|WGS_ZZZZ|egapxtmp_031359-R1    mNeoVis.evergladensis.2.1.chr13.g00626.t01    mNeoVis.vulgivaga.2.1.chr13.g00629.t01      1424    N0.HOG0001422   OG0000156  n22   rna-XM_011226331.3      rna-XM_022491238.1      rna-XM_036084772.1, rna-XM_036086530.1      rna-XM_032840472.1    rna-XM_046022879.1    rna-XM_032307575.1    rna-XM_059142794.1    rna-XM_059412313.1    rna-XM_013050869.2    mNeoVis.2.1.1.chr13_hap1.g00670.t01    rna-XM_021693498.2    rna-XM_004407457.1      rna-XM_040620160.1    rna-XM_027615669.2    mNeoVis.3.1.1.chr13.g00592.t01    rna-gnl|WGS_ZZZZ|egapxtmp_030811-R1    mNeoVis.evergladensis.2.1.chr13.g00643.t01    mNeoVis.vulgivaga.2.1.chr13.g00645.t01      1425    N0.HOG0001423   OG0000156  n41   rna-XM_034646739.1        rna-XM_023242238.2, rna-XM_045042711.1, rna-XM_045043284.1    rna-XM_036084776.1, rna-XM_036089475.1    rna-XM_039222399.1, rna-XM_039222403.1, rna-XM_039222407.1, rna-XM_039222409.1                  rna-XM_044912306.1    rna-XM_004410997.1    rna-XM_007089000.3, rna-XM_015540621.2, rna-XM_015541574.1, rna-XM_042963911.1      rna-XM_027616245.2, rna-XM_027616250.2              1426    N0.HOG0001424   OG0000156  n59   rna-XM_034647078.1    rna-XM_038525613.1    rna-XM_022491245.1    rna-XM_023249690.2, rna-XM_023249692.2    rna-XM_036089471.1    rna-XM_039222398.1, rna-XM_039222406.1    rna-XM_032840454.1, rna-XM_032840471.1    rna-XM_046023244.1, rna-XM_046023345.1    rna-XM_032307576.1, rna-XM_032307577.1, rna-XM_032307578.1, rna-XM_032307579.1, rna-XM_032307617.1    rna-XM_059141241.1, rna-XM_059142030.1, rna-XM_059142497.1    rna-XM_059387365.1, rna-XM_059387544.1, rna-XM_059387545.1    rna-XM_045086098.1, rna-XM_045086147.1    mNeoVis.2.1.1.chr13_hap1.g00669.t01, mNeoVis.2.1.1.chr13_hap1.g00671.t01    rna-XM_044912305.1    rna-XM_004410998.1    rna-XM_007089001.3, rna-XM_042963966.1    rna-XM_040620161.1    rna-XM_035723666.1      rna-gnl|WGS_ZZZZ|egapxtmp_029969-R1    mNeoVis.evergladensis.2.1.chr13.g00642.t01        1427    N0.HOG0001425   OG0000156  n93     rna-XM_038525504.1    rna-XM_022512730.1    rna-XM_023242239.2, rna-XM_045043357.1    rna-XM_036089473.1    rna-XM_039222405.1, rna-XM_039224698.1                mNeoVis.2.1.1.chr13_hap1.g00672.t01        rna-XM_042963927.1, rna-XM_042964141.1        mNeoVis.3.1.1.chr13.g00595.t01    rna-gnl|WGS_ZZZZ|egapxtmp_029971-R1    mNeoVis.evergladensis.2.1.chr13.g00644.t01    mNeoVis.vulgivaga.2.1.chr13.g00647.t01      1428    N0.HOG0001426   OG0000156  n103   rna-XM_034652783.1        rna-XM_003995465.6      rna-XM_039224701.1                      rna-XM_007089005.3                  1429    N0.HOG0001427   OG0000157  n0   id-LOC105235011, id-LOC117797294, id-LOC117797296, id-LOC117797712, id-LOC117797713, id-LOC117797714, id-LOC117800371, id-LOC117800376    id-LOC102156901    id-LOC111139339, id-LOC111139392, id-LOC111139464, id-LOC111139507, id-LOC111139529, id-LOC111139550, id-LOC111139578    id-LOC101081388, id-LOC101083182    id-LOC118522990, id-LOC118522996, id-LOC118550120, id-LOC118550121, id-LOC118550133    id-LOC120240459, id-LOC120240460, id-LOC120240461, id-LOC120240464, id-LOC120240474    id-LOC116867079, id-LOC116867080, id-LOC116867082, id-LOC116867087, id-LOC116867363, id-LOC116867366, id-LOC116867374, id-LOC116867378, id-LOC116867384, id-LOC116867386    id-LOC123943596, id-LOC123943598, id-LOC123943604, id-LOC123943606, id-LOC123943612, id-LOC123943616, id-LOC123943619    id-LOC116591092, id-LOC116591105, id-LOC116591111, id-LOC116591113, id-LOC116591116    id-LOC131836309, id-LOC131836311, id-LOC131836313, id-LOC131836317, id-LOC131836330, id-LOC131836334, id-LOC131837199, id-LOC131837201    id-LOC131998743, id-LOC131999390, id-LOC131999397, id-LOC131999399, id-LOC131999774, id-LOC131999775, id-LOC131999783, id-LOC132008865    id-LOC101673081, id-LOC101674908, id-LOC101682384, id-LOC101683268, id-LOC101689035, id-LOC101690454, id-LOC123388329, id-LOC123391339, id-LOC123391340, id-LOC123391343      id-LOC123325716    id-LOC101363625, id-LOC101364502, id-LOC101366298, id-LOC101368051, id-LOC101368624, id-LOC101373069, id-LOC101373920, rna-XM_004402168.1    id-LOC102965132, id-LOC102966293, id-LOC102968010, id-LOC107180491    id-LOC103680323, id-LOC103680418, id-LOC103680422, id-LOC103680426, id-LOC103680428, id-LOC103680948, id-LOC103681517, id-LOC103681901, id-LOC103681906, id-LOC103681910, id-LOC103682248, id-LOC103682252, id-LOC103682255, id-LOC121100922, id-LOC121100927, rna-XM_040637047.1    id-LOC113908993, id-LOC113908994, id-LOC113908995, id-LOC113909249, id-LOC113909257, id-LOC113909260, id-LOC113909261, id-LOC113909262              1430    N0.HOG0001428   OG0000158  n2   rna-XM_034665597.1    rna-NM_001388925.1    rna-XM_022522072.1, rna-XM_022522076.1    rna-XM_003992963.2    rna-XM_036089257.1    rna-XM_039224075.1    rna-XM_032851033.1, rna-XM_032851034.1, rna-XM_032851066.1, rna-XM_032851067.1, rna-XM_032852723.1    rna-XM_046016259.1, rna-XM_046016790.1, rna-XM_046017682.1    rna-XM_032359963.1, rna-XM_032360324.1    rna-XM_059158230.1, rna-XM_059183815.1    rna-XM_059389216.1, rna-XM_059392323.1    rna-XM_004780186.1, rna-XM_004780187.2    mNeoVis.2.1.1.chr7_hap1.g01770.t01    rna-XM_021685725.2    rna-XM_004394536.1    rna-XM_007083280.2    rna-XM_008683826.2, rna-XM_008710991.2    rna-XM_027580640.2      rna-gnl|WGS_ZZZZ|egapxtmp_010079-R1    mNeoVis.evergladensis.2.1.chr7.g01731.t01    mNeoVis.vulgivaga.2.1.chr7.g01747.t01      1431    N0.HOG0001429   OG0000158  n34   rna-XM_002924991.2          rna-XM_036095922.1                      rna-XM_004394616.1      rna-XM_008710996.1    rna-XM_027577897.1              1432    N0.HOG0001430   OG0000158  n39                 rna-XM_046017007.1    rna-XM_032359794.1    rna-XM_059158116.1    rna-XM_059389317.1    rna-XM_013050983.2    mNeoVis.2.1.1.chr7_hap1.g01760.t01                rna-gnl|WGS_ZZZZ|egapxtmp_010082-R1      mNeoVis.vulgivaga.2.1.chr7.g01738.t01      1433    N0.HOG0001431   OG0000158  n47         rna-XM_023239003.2      rna-XM_039224082.1                      rna-XM_007083164.3                  1434    N0.HOG0001432   OG0000158  n51   rna-XM_002924990.1    rna-NM_001389100.1      rna-XM_003992958.4    rna-XM_036095963.1    rna-XM_039224081.1      rna-XM_046016999.1    rna-XM_032360276.1    rna-XM_059158124.1, rna-XM_059158146.1    rna-XM_059389277.1, rna-XM_059389304.1    rna-XM_045065883.1, rna-XM_045078275.1, rna-XM_045078277.1          rna-XM_007083277.2    rna-XM_008710963.2                1435    N0.HOG0001433   OG0000158  n66   rna-XM_002924967.2    rna-NM_001389097.1    rna-XM_022522156.1    rna-XM_045038881.1    rna-XM_036095966.1    rna-XM_039224083.1    rna-XM_032853683.1    rna-XM_046017029.1                  rna-XM_015537939.1    rna-XM_008710965.2                1436    N0.HOG0001434   OG0000158  n75     rna-NM_001017517.1, rna-NM_001388743.1, rna-NM_001388744.1, rna-NM_001388745.1, rna-NM_001389115.1, rna-XM_038568925.1    rna-XM_022522040.1, rna-XM_022522069.1    rna-XM_045038885.1    rna-XM_036089273.1      rna-XM_032850794.1, rna-XM_032850795.1, rna-XM_032853691.1    rna-XM_046015872.1, rna-XM_046016649.1    rna-XM_032359318.1, rna-XM_032359799.1    rna-XM_059158240.1, rna-XM_059158247.1    rna-XM_059389203.1    rna-XM_004780179.3, rna-XM_004780185.3    mNeoVis.2.1.1.chr7_hap1.g01769.t01, mNeoVis.2.1.1.chr7_hap1.g01771.t01    rna-XM_021685726.1    rna-XM_004394538.1    rna-XM_007083281.2    rna-XM_008683827.2, rna-XM_008683831.2, rna-XM_008710957.2    rna-XM_027580617.1      rna-gnl|WGS_ZZZZ|egapxtmp_010080-R1, rna-gnl|WGS_ZZZZ|egapxtmp_010083-R1    mNeoVis.evergladensis.2.1.chr7.g01730.t01, mNeoVis.evergladensis.2.1.chr7.g01732.t01    mNeoVis.vulgivaga.2.1.chr7.g01746.t01      1437    N0.HOG0001435   OG0000159  n1   rna-XM_034645671.1    rna-XM_038576782.1    rna-XM_022494971.1      rna-XM_036121270.1      rna-XM_032854986.1    rna-XM_046011793.1, rna-XM_046012700.1    rna-XM_032347278.1, rna-XM_032347287.1, rna-XM_032347754.1    rna-XM_059186100.1, rna-XM_059186115.1, rna-XM_059187524.1    rna-XM_059404470.1, rna-XM_059404738.1    rna-XM_004779753.3, rna-XM_013050907.2, rna-XM_045073465.1    mNeoVis.2.1.1.chr9_hap1.g00834.t01, mNeoVis.2.1.1.chr9_hap1.g00835.t01, mNeoVis.2.1.1.chr9_hap1.g00838.t01    rna-XM_044915219.1        rna-XM_008697442.2, rna-XM_008697444.2    rna-XM_027593954.1    mNeoVis.3.1.1.chr10.g00795.t01    rna-gnl|WGS_ZZZZ|egapxtmp_023581-R1, rna-gnl|WGS_ZZZZ|egapxtmp_024750-R1, rna-gnl|WGS_ZZZZ|egapxtmp_024754-R1    mNeoVis.evergladensis.2.1.chr10.g00822.t01, mNeoVis.evergladensis.2.1.chr10.g00823.t01, mNeoVis.evergladensis.2.1.chr10.g00826.t01    mNeoVis.vulgivaga.2.1.chr10.g00812.t01, mNeoVis.vulgivaga.2.1.chr10.g00813.t01, mNeoVis.vulgivaga.2.1.chr10.g00814.t01      1438    N0.HOG0001436   OG0000159  n35   rna-XM_002928725.2, rna-XM_034645560.1, rna-XM_034645612.1, rna-XM_034645622.1    rna-XM_038576128.1, rna-XM_038576784.1    rna-XM_022494938.1, rna-XM_022494939.1, rna-XM_022494940.1, rna-XM_022494941.1, rna-XM_022494969.1, rna-XM_022494970.1    rna-XM_019834512.2, rna-XM_045061363.1, rna-XM_045061368.1, rna-XM_045061369.1, rna-XM_045061371.1, rna-XM_045061372.1, rna-XM_045062657.1    rna-XM_036100109.1, rna-XM_036100111.1    rna-XM_039222314.1, rna-XM_039222315.1, rna-XM_039222317.1, rna-XM_039222323.1    rna-XM_032854984.1, rna-XM_032854985.1, rna-XM_032854987.1, rna-XM_032854988.1, rna-XM_032854989.1, rna-XM_032855381.1    rna-XM_046012466.1, rna-XM_046012633.1, rna-XM_046012637.1    rna-XM_032347271.1, rna-XM_032347276.1, rna-XM_032347279.1, rna-XM_032347755.1    rna-XM_059186104.1, rna-XM_059186108.1, rna-XM_059186109.1, rna-XM_059186113.1, rna-XM_059186117.1, rna-XM_059187523.1    rna-XM_059404471.1, rna-XM_059404739.1, rna-XM_059404740.1, rna-XM_059404741.1    rna-XM_004779752.3, rna-XM_004779754.3, rna-XM_045073066.1, rna-XM_045073074.1, rna-XM_045073104.1    mNeoVis.2.1.1.chr9_hap1.g00837.t01    rna-XM_021690444.1    rna-XM_004417046.1, rna-XM_004417607.1    rna-XM_042991755.1, rna-XM_042991758.1, rna-XM_042991759.1, rna-XM_042993676.1, rna-XM_042993677.1, rna-XM_042993679.1, rna-XM_042993680.1    rna-XM_008697437.2, rna-XM_040642605.1, rna-XM_040642606.1    rna-XM_027593953.1, rna-XM_027594635.1    mNeoVis.3.1.1.chr10.g00796.t01, mNeoVis.3.1.1.chr10.g00797.t01    rna-gnl|WGS_ZZZZ|egapxtmp_024751-R1, rna-gnl|WGS_ZZZZ|egapxtmp_024752-R1    mNeoVis.evergladensis.2.1.chr10.g00824.t01, mNeoVis.evergladensis.2.1.chr10.g00825.t01, mNeoVis.evergladensis.2.1.chr10.g00827.t01    mNeoVis.vulgivaga.2.1.chr10.g00815.t01, mNeoVis.vulgivaga.2.1.chr10.g00816.t01      1439    N0.HOG0001437   OG0000160  n1     rna-XM_038526998.1    rna-XM_022512537.1, rna-XM_022512552.1    rna-XM_045042625.1, rna-XM_045042626.1    rna-XM_036075421.1    rna-XM_039221101.1    rna-XM_032850098.1, rna-XM_032850178.1, rna-XM_032866706.1    rna-XM_046021860.1      rna-XM_059142841.1      rna-XM_004778660.3    mNeoVis.2.1.1.chr13_hap1.g00695.t01, mNeoVis.2.1.1.chr13_hap1.g00708.t01    rna-XM_021677939.2, rna-XM_044912198.1        rna-XM_008705522.2    rna-XM_035723451.1    mNeoVis.3.1.1.chr13.g00621.t01, mNeoVis.3.1.1.chr13.g00622.t01, mNeoVis.3.1.1.chr13.g00627.t01    rna-gnl|WGS_ZZZZ|egapxtmp_029991-R1, rna-gnl|WGS_ZZZZ|egapxtmp_030519-R1, rna-gnl|WGS_ZZZZ|egapxtmp_031025-R1    mNeoVis.evergladensis.2.1.chr13.g00671.t01, mNeoVis.evergladensis.2.1.chr13.g00675.t01    mNeoVis.vulgivaga.2.1.chr13.g00672.t01, mNeoVis.vulgivaga.2.1.chr13.g00673.t01      1440    N0.HOG0001438   OG0000160  n30   rna-XM_034651952.1, rna-XM_034652086.1    rna-XM_038527005.1    rna-XM_022510956.1    rna-XM_045042617.1, rna-XM_045042622.1, rna-XM_045042624.1    rna-XM_036082882.1, rna-XM_036095858.1    rna-XM_039257203.1    rna-XM_032849881.1    rna-XM_046006088.1, rna-XM_046006089.1, rna-XM_046006091.1, rna-XM_046021861.1    rna-XM_032306504.1, rna-XM_032306506.1, rna-XM_032306508.1, rna-XM_032306509.1, rna-XM_032306510.1, rna-XM_032341355.1, rna-XM_032341357.1, rna-XM_032341358.1    rna-XM_059141243.1, rna-XM_059141244.1, rna-XM_059141246.1, rna-XM_059141247.1, rna-XM_059142508.1, rna-XM_059142628.1, rna-XM_059142972.1, rna-XM_059143273.1, rna-XM_059143375.1, rna-XM_059179338.1, rna-XM_059179339.1, rna-XM_059179340.1    rna-XM_059386865.1, rna-XM_059387560.1    rna-XM_013050638.2, rna-XM_045065079.1, rna-XM_045070399.1, rna-XM_045070400.1, rna-XM_045074853.1, rna-XM_045074854.1, rna-XM_045085385.1, rna-XM_045087511.1    mNeoVis.2.1.1.chr13_hap1.g00696.t01, mNeoVis.2.1.1.chr13_hap1.g00700.t01, mNeoVis.2.1.1.chr13_hap1.g00701.t01, mNeoVis.2.1.1.chr13_hap1.g00702.t01, mNeoVis.2.1.1.chr13_hap1.g00703.t01, mNeoVis.2.1.1.chr13_hap1.g00706.t01, mNeoVis.2.1.1.chr1_hap1.g01480.t01, mNeoVis.2.1.1.chr1_hap1.g01481.t01, mNeoVis.2.1.1.chr1_hap1.g01482.t01, mNeoVis.2.1.1.chr1_hap1.g01484.t01    rna-XM_021677938.1, rna-XM_044912197.1    rna-XM_004415470.1    rna-XM_042963963.1, rna-XM_042965055.1, rna-XM_042965059.1    rna-XM_008705524.2    rna-XM_035723424.1, rna-XM_035723800.1    mNeoVis.3.1.1.chr13.g00617.t01, mNeoVis.3.1.1.chr13.g00619.t01, mNeoVis.3.1.1.chr13.g00620.t01, mNeoVis.3.1.1.chr13.g00623.t01, mNeoVis.3.1.1.chr13.g00625.t01, mNeoVis.3.1.1.chr13.g00626.t01    rna-gnl|WGS_ZZZZ|egapxtmp_029986-R1, rna-gnl|WGS_ZZZZ|egapxtmp_030518-R1, rna-gnl|WGS_ZZZZ|egapxtmp_030522-R1, rna-gnl|WGS_ZZZZ|egapxtmp_031030-R1    mNeoVis.evergladensis.2.1.chr13.g00665.t01, mNeoVis.evergladensis.2.1.chr13.g00669.t01, mNeoVis.evergladensis.2.1.chr13.g00670.t01, mNeoVis.evergladensis.2.1.chr13.g00672.t01, mNeoVis.evergladensis.2.1.chr13.g00674.t01    mNeoVis.vulgivaga.2.1.chr13.g00670.t01, mNeoVis.vulgivaga.2.1.chr13.g00671.t01, mNeoVis.vulgivaga.2.1.chr13.g00674.t01, mNeoVis.vulgivaga.2.1.chr13.g00675.t01      1441    N0.HOG0001439   OG0000161  n1   rna-XM_011226315.3    rna-NM_001006951.1    rna-XM_022519548.1    rna-XM_045061679.1    rna-XM_036099333.1    rna-XM_039243135.1    rna-XM_032876934.1    rna-XM_046012421.1    rna-XM_032348009.1    rna-XM_059187500.1    rna-XM_059404501.1    rna-XM_004765626.3    mNeoVis.2.1.1.chr9_hap1.g00633.t01    rna-XM_021704764.1    rna-XM_004395960.1    rna-XM_042992175.1    rna-XM_040642382.1    rna-XM_027593381.2    mNeoVis.3.1.1.chr10.g00598.t01    rna-gnl|WGS_ZZZZ|egapxtmp_024702-R1    mNeoVis.evergladensis.2.1.chr10.g00627.t01    mNeoVis.vulgivaga.2.1.chr10.g00610.t01      1442    N0.HOG0001440   OG0000161  n22   rna-XM_011224217.3    rna-XM_038585075.1    rna-XM_022504510.1    rna-XM_003990961.4    rna-XM_036119965.1    rna-XM_039246568.1    rna-XM_032860256.1    rna-XM_046019055.1    rna-XM_032354138.1    rna-XM_059168631.1    rna-XM_059393024.1    rna-XM_004763363.2    mNeoVis.2.1.1.chr3_hap1.g01362.t01    rna-XM_021702871.2    rna-XM_004392391.1    rna-XM_007094663.3    rna-XM_040643007.1    rna-XM_027590457.2    mNeoVis.3.1.1.chr3.g01337.t01    rna-gnl|WGS_ZZZZ|egapxtmp_041314-R1    mNeoVis.evergladensis.2.1.chr3.g01364.t01    mNeoVis.vulgivaga.2.1.chr3.g01383.t01      1443    N0.HOG0001441   OG0000161  n44   rna-XM_002918229.4    rna-NM_001003187.1    rna-XM_022503091.1    rna-XM_003982764.6    rna-XM_036123357.1    rna-XM_039236240.1    rna-XM_032874973.1    rna-XM_046020796.1    rna-XM_032305376.1    rna-XM_059171147.1    rna-XM_059396839.1    rna-XM_004767862.3    mNeoVis.2.1.1.chr4_hap1.g00480.t01    rna-XM_021689553.2    rna-XM_004394078.1    rna-XM_007076695.3    rna-XM_008686254.2    rna-XM_027572996.2    mNeoVis.3.1.1.chr6.g00483.t01    rna-gnl|WGS_ZZZZ|egapxtmp_001822-R2    mNeoVis.evergladensis.2.1.chr6.g00487.t01    mNeoVis.vulgivaga.2.1.chr6.g00485.t01      1444    N0.HOG0001442   OG0000161  n64   rna-XM_034640604.1    rna-NM_001003090.1    rna-XM_022509224.1    rna-XM_003996699.5    rna-XM_036094674.1    rna-XM_039227823.1    rna-XM_032872121.1    rna-XM_045985138.1    rna-XM_032320704.1    rna-XM_059148627.1    rna-XM_059379981.1    rna-XM_004764821.3    mNeoVis.2.1.1.chr8_hap1.g00572.t01    rna-XM_021704328.2    rna-XM_004395150.1    rna-XM_042966748.1    rna-XM_040640811.1    rna-XM_027567559.2    mNeoVis.3.1.1.chr8.g00527.t01    rna-gnl|WGS_ZZZZ|egapxtmp_017438-R1    mNeoVis.evergladensis.2.1.chr9.g00562.t01    mNeoVis.vulgivaga.2.1.chr8.g00569.t01      1445    N0.HOG0001443   OG0000161  n81   rna-XM_011224218.3    rna-XM_038585076.1    rna-XM_022494132.1    rna-XM_003990962.5    rna-XM_036119963.1    rna-XM_039223887.1    rna-XM_032860255.1    rna-XM_046019102.1    rna-XM_032354139.1    rna-XM_059168629.1    rna-XM_059393025.1    rna-XM_004763364.3    mNeoVis.2.1.1.chr3_hap1.g01361.t01    rna-XM_021703353.2    rna-XM_004392392.1    rna-XM_007094662.3    rna-XM_008687250.2    rna-XM_027590566.2    mNeoVis.3.1.1.chr3.g01335.t01, mNeoVis.3.1.1.chr3.g01336.t01    rna-gnl|WGS_ZZZZ|egapxtmp_041313-R1    mNeoVis.evergladensis.2.1.chr3.g01363.t01    mNeoVis.vulgivaga.2.1.chr3.g01381.t01, mNeoVis.vulgivaga.2.1.chr3.g01382.t01      1446    N0.HOG0001444   OG0000162  n2   rna-XM_002931164.4, rna-XM_002931307.1, rna-XM_011217934.2, rna-XM_019793765.2, rna-XM_034655715.1    rna-NM_001360012.1, rna-XM_038581160.1, rna-XM_038581161.1      rna-NM_001167474.1, rna-XM_045058639.1      rna-XM_039225771.1      rna-XM_045999664.1    rna-XM_032335021.1    rna-XM_059173431.1    rna-XM_059416427.1    rna-XM_045086937.1    mNeoVis.2.1.1.chr1_hap1.g00904.t01        rna-XM_042987630.1    rna-XM_008709550.1, rna-XM_008709551.1, rna-XM_008709553.1, rna-XM_008709555.1, rna-XM_008710131.1, rna-XM_008710140.2, rna-XM_008710141.2, rna-XM_008710142.2, rna-XM_008710143.1, rna-XM_008710145.2, rna-XM_008710146.2, rna-XM_040621940.1, rna-XM_040622716.1, rna-XM_040622717.1      mNeoVis.3.1.1.chr1.g00926.t01    rna-gnl|WGS_ZZZZ|egapxtmp_038816-R1    mNeoVis.evergladensis.2.1.chr1.g00907.t01        1447    N0.HOG0001445   OG0000162  n36   rna-XM_011217213.3    rna-XM_038565256.1    rna-XM_022491975.1, rna-XM_022491976.1    rna-NM_001167475.1      rna-XM_039218781.1, rna-XM_039218782.1, rna-XM_039218783.1, rna-XM_039225654.1, rna-XM_039225656.1    rna-XM_032848097.1, rna-XM_032848098.1      rna-XM_032335201.1, rna-XM_032337897.1, rna-XM_032341128.1    rna-XM_059173380.1, rna-XM_059173381.1, rna-XM_059173426.1    rna-XM_059416405.1, rna-XM_059416406.1, rna-XM_059416433.1    rna-XM_013049797.2, rna-XM_013049800.1, rna-XM_045086939.1    mNeoVis.2.1.1.chr1_hap1.g00928.t01    rna-XM_044912696.1      rna-XM_015540096.2, rna-XM_042977172.1, rna-XM_042977173.1, rna-XM_042977174.1    rna-XM_008683754.1, rna-XM_008697973.1, rna-XM_008708277.2    rna-XM_027581886.1    mNeoVis.3.1.1.chr1.g00942.t01    rna-gnl|WGS_ZZZZ|egapxtmp_039114-R1    mNeoVis.evergladensis.2.1.chr1.g00923.t01    mNeoVis.vulgivaga.2.1.chr1.g00936.t01      1448    N0.HOG0001446   OG0000162  n72     rna-XM_038584402.1    rna-XM_022490620.1    rna-NM_001167486.1, rna-NM_001167487.1, rna-NM_001167488.1, rna-XM_045058490.1    rna-XM_036112168.1, rna-XM_036112326.1    rna-XM_039224914.1    rna-XM_032861772.1, rna-XM_032861773.1    rna-XM_046006115.1    rna-XM_032341388.1, rna-XM_032341389.1    rna-XM_059179371.1, rna-XM_059179372.1    rna-XM_059401503.1, rna-XM_059401504.1    rna-XM_004778785.3, rna-XM_004778786.1    mNeoVis.2.1.1.chr1_hap1.g01103.t01, mNeoVis.2.1.1.chr1_hap1.g01104.t01    rna-XM_021696841.1, rna-XM_021697244.1      rna-XM_015540739.2    rna-XM_008693093.2, rna-XM_008693095.1, rna-XM_008693096.2    rna-XM_035728354.1, rna-XM_035728411.1, rna-XM_035728412.1    mNeoVis.3.1.1.chr1.g01080.t01, mNeoVis.3.1.1.chr1.g01081.t01    rna-gnl|WGS_ZZZZ|egapxtmp_021274-R1, rna-gnl|WGS_ZZZZ|egapxtmp_021280-R1    mNeoVis.evergladensis.2.1.chr1.g01083.t01, mNeoVis.evergladensis.2.1.chr1.g01084.t01    mNeoVis.vulgivaga.2.1.chr1.g01097.t01, mNeoVis.vulgivaga.2.1.chr1.g01098.t01      1449    N0.HOG0001447   OG0000163  n1   rna-XM_002917593.4, rna-XM_034642201.1, rna-XM_034643164.1, rna-XM_034643165.1    rna-XM_038541812.1, rna-XM_038543505.1    rna-XM_022496017.1    rna-XM_006938853.5, rna-XM_045041277.1    rna-XM_036089346.1, rna-XM_036089360.1, rna-XM_036089363.1    rna-XM_039238877.1, rna-XM_039238878.1, rna-XM_039238880.1    rna-XM_032859611.1, rna-XM_032859612.1, rna-XM_032859614.1    rna-XM_046024851.1, rna-XM_046024854.1, rna-XM_046024855.1    rna-XM_032309453.1, rna-XM_032310940.1    rna-XM_059139357.1, rna-XM_059139358.1, rna-XM_059139361.1    rna-XM_059409468.1, rna-XM_059409470.1, rna-XM_059409471.1    rna-XM_004752305.3, rna-XM_045065770.1    mNeoVis.2.1.1.chr3_hap1.g00162.t01, mNeoVis.2.1.1.chr3_hap1.g00163.t01, mNeoVis.2.1.1.chr3_hap1.g00165.t01    rna-XM_044921294.1    rna-XM_004404381.1, rna-XM_004404383.1, rna-XM_004404404.1    rna-XM_007090741.3, rna-XM_042961572.1    rna-XM_008691128.2, rna-XM_040632605.1    rna-XM_027577161.2, rna-XM_027577171.2    mNeoVis.3.1.1.chr3.g00164.t01, mNeoVis.3.1.1.chr3.g00165.t01    rna-gnl|WGS_ZZZZ|egapxtmp_040732-R2, rna-gnl|WGS_ZZZZ|egapxtmp_040733-R1, rna-gnl|WGS_ZZZZ|egapxtmp_040734-R2    mNeoVis.evergladensis.2.1.chr3.g00168.t01, mNeoVis.evergladensis.2.1.chr3.g00169.t01, mNeoVis.evergladensis.2.1.chr3.g00171.t01    mNeoVis.vulgivaga.2.1.chr3.g00165.t01, mNeoVis.vulgivaga.2.1.chr3.g00166.t01, mNeoVis.vulgivaga.2.1.chr3.g00167.t01      1450    N0.HOG0001448   OG0000163  n49   rna-XM_019793032.2, rna-XM_034660066.1    rna-XM_038583983.1      rna-XM_019830086.3, rna-XM_023254674.2    rna-XM_036089240.1, rna-XM_036089246.1    rna-XM_039257564.1, rna-XM_039257572.1    rna-XM_032847812.1, rna-XM_032847834.1    rna-XM_046004750.1, rna-XM_046004767.1    rna-XM_032338163.1, rna-XM_032338192.1    rna-XM_059176177.1, rna-XM_059176199.1    rna-XM_059399635.1    rna-XM_045073966.1    mNeoVis.2.1.1.chr1_hap1.g01046.t01, mNeoVis.2.1.1.chr1_hap1.g01047.t01    rna-XM_021697095.2, rna-XM_021697189.1    rna-XM_004416709.1, rna-XM_004416710.1    rna-XM_015542286.2, rna-XM_042985427.1    rna-XM_008711511.2, rna-XM_008711516.2    rna-XM_027602162.2, rna-XM_027602202.2    mNeoVis.3.1.1.chr1.g01041.t01    rna-gnl|WGS_ZZZZ|egapxtmp_031246-R1, rna-gnl|WGS_ZZZZ|egapxtmp_031345-R3    mNeoVis.evergladensis.2.1.chr1.g01038.t01, mNeoVis.evergladensis.2.1.chr1.g01039.t01    mNeoVis.vulgivaga.2.1.chr1.g01049.t01, mNeoVis.vulgivaga.2.1.chr1.g01050.t01      1451    N0.HOG0001449   OG0000163  n86   rna-XM_034660082.1    rna-XM_038584442.1      rna-XM_006931350.5      rna-XM_039257587.1      rna-XM_046004868.1    rna-XM_032338178.1    rna-XM_059176217.1    rna-XM_059399680.1      mNeoVis.2.1.1.chr1_hap1.g01056.t01    rna-XM_044917240.1    rna-XM_004416705.1    rna-XM_042985478.1    rna-XM_040645140.1    rna-XM_027602181.2    mNeoVis.3.1.1.chr1.g01051.t01    rna-gnl|WGS_ZZZZ|egapxtmp_031264-R5    mNeoVis.evergladensis.2.1.chr1.g01049.t01    mNeoVis.vulgivaga.2.1.chr1.g01064.t01      1452    N0.HOG0001450   OG0000164  n1   rna-XM_034648060.1    rna-XM_038554102.1    rna-XM_022503975.1    rna-XM_003986149.6    rna-XM_036124052.1    rna-XM_039231156.1    rna-XM_032862140.1    rna-XM_046004512.1    rna-XM_032340007.1    rna-XM_059178017.1    rna-XM_059400432.1    rna-XM_013062136.2    mNeoVis.2.1.1.chr1_hap1.g01640.t01    rna-XM_021690382.2, rna-XM_021692116.2    rna-XM_004408086.1    rna-XM_042986384.1    rna-XM_040643495.1    rna-XM_027601070.2    mNeoVis.3.1.1.chr1.g01590.t01    rna-gnl|WGS_ZZZZ|egapxtmp_022154-R1    mNeoVis.evergladensis.2.1.chr1.g01606.t01    mNeoVis.vulgivaga.2.1.chr1.g01603.t01      1453    N0.HOG0001451   OG0000164  n19   rna-XM_011217898.3    rna-XM_038544899.1    rna-XM_022523477.1    rna-XM_011283256.4    rna-XM_036067735.1    rna-XM_039242110.1    rna-XM_032875239.1    rna-XM_046008672.1    rna-XM_032344429.1    rna-XM_059182271.1    rna-XM_059373343.1    rna-XM_004738897.3    mNeoVis.2.1.1.chr10_hap1.g00328.t01    rna-XM_021701542.2    rna-XM_004409354.1    rna-XM_042989856.1    rna-XM_008701085.2    rna-XM_027570005.2    mNeoVis.3.1.1.chr9.g00298.t01    rna-gnl|WGS_ZZZZ|egapxtmp_016014-R1    mNeoVis.evergladensis.2.1.chr8.g00309.t01    mNeoVis.vulgivaga.2.1.chr9.g00310.t01      1454    N0.HOG0001452   OG0000164  n26   rna-XM_019796911.2    rna-XM_038564026.1    rna-XM_022507285.1    rna-XM_045039519.1    rna-XM_036101164.1    rna-XM_039244827.1    rna-XM_032879412.1    rna-XM_046016330.1    rna-XM_032357242.1    rna-XM_059146617.1    rna-XM_059401984.1    rna-XM_004770108.3    mNeoVis.2.1.1.chr7_hap1.g02380.t01    rna-XM_021685014.2    rna-XM_004393969.1    rna-XM_015541990.2    rna-XM_008708982.2    rna-XM_027579242.2    mNeoVis.3.1.1.chr7.g02191.t01      mNeoVis.evergladensis.2.1.chr7.g02326.t01    mNeoVis.vulgivaga.2.1.chr7.g02345.t01      1455    N0.HOG0001453   OG0000164  n38   rna-XM_011222266.3    rna-XM_038542077.1    rna-XM_022508187.1    rna-XM_003999320.6    rna-XM_036101756.1    rna-XM_039230095.1    rna-XM_032862799.1    rna-XM_045982101.1    rna-XM_032318520.1    rna-XM_059145301.1    rna-XM_059412624.1    rna-XM_045084707.1    mNeoVis.2.1.1.chr14_hap1.g00106.t01    rna-XM_021682750.2    rna-XM_004414487.1    rna-XM_007096166.2    rna-XM_008701172.2    rna-XM_027613639.2    mNeoVis.3.1.1.chr14.g00100.t01    rna-gnl|WGS_ZZZZ|egapxtmp_031697-R1    mNeoVis.evergladensis.2.1.chr14.g00105.t01    mNeoVis.vulgivaga.2.1.chr14.g00106.t01      1456    N0.HOG0001454   OG0000164  n50   rna-XM_002926735.4    rna-XM_038545745.1    rna-XM_022505353.1    rna-XM_023256045.2    rna-XM_036086240.1, rna-XM_036120794.1    rna-XM_039251165.1    rna-XM_032839577.1    rna-XM_046009354.1    rna-XM_032345305.1    rna-XM_059183132.1    rna-XM_059373995.1    rna-XM_045073823.1    mNeoVis.2.1.1.chr10_hap1.g00068.t01    rna-XM_021679849.2    rna-XM_004395558.1    rna-XM_042990563.1    rna-XM_040641390.1    rna-XM_027568707.2    mNeoVis.3.1.1.chr9.g00047.t01    rna-gnl|WGS_ZZZZ|egapxtmp_014328-R11    mNeoVis.evergladensis.2.1.chr8.g00067.t01    mNeoVis.vulgivaga.2.1.chr9.g00067.t01      1457    N0.HOG0001455   OG0000165  n1   rna-XM_034670228.1    rna-XM_038540002.1    rna-XM_022526692.1    rna-XM_023247121.2    rna-XM_036076600.1    rna-XM_039234381.1    rna-XM_032842665.1    rna-XM_045993653.1    rna-XM_032328420.1    rna-XM_059155177.1    rna-XM_059415948.1    rna-XM_004774044.2    mNeoVis.2.1.1.chr15_hap1.g00109.t01    rna-XM_021700565.1    rna-XM_004397133.1    rna-XM_042971107.1    rna-XM_008685352.2    rna-XM_027612307.2    mNeoVis.3.1.1.chr15.g00102.t01    rna-gnl|WGS_ZZZZ|egapxtmp_033541-R1    mNeoVis.evergladensis.2.1.chr15.g00110.t01    mNeoVis.vulgivaga.2.1.chr15.g00113.t01      1458    N0.HOG0001456   OG0000165  n24   rna-XM_034670623.1    rna-XM_038538886.1    rna-XM_022516957.1    rna-XM_003998949.5    rna-XM_036083290.1    rna-XM_039246661.1    rna-XM_032846009.1    rna-XM_045992896.1    rna-XM_032326828.1    rna-XM_059154959.1    rna-XM_059415684.1    rna-XM_004750991.2    mNeoVis.2.1.1.chr15_hap1.g00372.t01, mNeoVis.2.1.1.chr15_hap1.g00409.t01, mNeoVis.2.1.1.chr15_hap1.g00446.t01    rna-XM_021698158.1    rna-XM_004403145.1    rna-XM_042971618.1    rna-XM_040635542.1    rna-XM_027612861.1              1459    N0.HOG0001457   OG0000165  n43   rna-XM_034651651.1    rna-XM_038536942.1    rna-XM_022524822.1    rna-XM_023243537.2    rna-XM_036094737.1    rna-XM_039227356.1    rna-XM_032841304.1    rna-XM_045983872.1    rna-XM_032321787.1    rna-XM_059149342.1    rna-XM_059378587.1    rna-XM_045085390.1    mNeoVis.2.1.1.chr8_hap1.g00971.t01    rna-XM_021694810.1    rna-XM_004417328.1    rna-XM_042965685.1    rna-XM_040630055.1    rna-XM_027624585.2      rna-gnl|WGS_ZZZZ|egapxtmp_017890-R2    mNeoVis.evergladensis.2.1.chr9.g00944.t01    mNeoVis.vulgivaga.2.1.chr8.g00942.t01      1460    N0.HOG0001458   OG0000165  n63   rna-XM_034646947.1, rna-XM_034647203.1    rna-XM_038536946.1    rna-XM_022521087.1    rna-XM_011288909.4    rna-XM_036094738.1    rna-XM_039227365.1    rna-XM_032841271.1    rna-XM_045986617.1    rna-XM_032321534.1    rna-XM_059149080.1    rna-XM_059379143.1    rna-XM_004760137.3    mNeoVis.2.1.1.chr8_hap1.g00969.t01    rna-XM_021694523.2    rna-XM_004415872.1    rna-XM_042966693.1    rna-XM_040630058.1    rna-XM_027568205.2    mNeoVis.3.1.1.chr8.g00901.t01, mNeoVis.3.1.1.chr8.g00903.t01    rna-gnl|WGS_ZZZZ|egapxtmp_017542-R1    mNeoVis.evergladensis.2.1.chr9.g00942.t01    mNeoVis.vulgivaga.2.1.chr8.g00940.t01      1461    N0.HOG0001459   OG0000165  n85   rna-XM_034670262.1    rna-XM_038540049.1    rna-XM_022526426.1, rna-XM_022526694.1    rna-XM_003998814.6    rna-XM_036097343.1    rna-XM_039234438.1    rna-XM_032843988.1    rna-XM_045992494.1    rna-XM_032328549.1    rna-XM_059154834.1    rna-XM_059415114.1    rna-XM_013049729.2    mNeoVis.2.1.1.chr15_hap1.g00139.t01    rna-XM_021686822.1    rna-XM_004398812.1    rna-XM_007084163.3    rna-XM_040635221.1    rna-XM_027610706.2    mNeoVis.3.1.1.chr15.g00117.t01    rna-gnl|WGS_ZZZZ|egapxtmp_033217-R3    mNeoVis.evergladensis.2.1.chr15.g00125.t01, mNeoVis.evergladensis.2.1.chr15.g00126.t01    mNeoVis.vulgivaga.2.1.chr15.g00129.t01      1462    N0.HOG0001460   OG0000166  n1   rna-XM_011220649.3    rna-XM_038565825.1    rna-XM_022498236.1    rna-XM_023249978.2    rna-XM_036079841.1    rna-XM_039219877.1    rna-XM_032867430.1    rna-XM_045991202.1    rna-XM_032325481.1    rna-XM_059161537.1    rna-XM_059390186.1    rna-XM_004738393.2    mNeoVis.2.1.1.chr5_hap1.g00827.t01    rna-XM_021695279.2    rna-XM_004413174.1    rna-XM_007076625.3    rna-XM_008703154.2    rna-XM_027587787.1    mNeoVis.3.1.1.chr2.g00738.t01    rna-gnl|WGS_ZZZZ|egapxtmp_038606-R1    mNeoVis.evergladensis.2.1.chr2.g00828.t01    mNeoVis.vulgivaga.2.1.chr2.g00831.t01      1463    N0.HOG0001461   OG0000166  n24   rna-XM_034658663.1    rna-XM_038565816.1    rna-XM_022498106.1    rna-XM_045051879.1    rna-XM_036066178.1    rna-XM_039240289.1    rna-XM_032867432.1    rna-XM_045992257.1    rna-XM_032325501.1    rna-XM_059161541.1    rna-XM_059390189.1    rna-XM_004738394.3    mNeoVis.2.1.1.chr5_hap1.g00828.t01    rna-XM_021694959.1    rna-XM_004399442.1    rna-XM_042979430.1    rna-XM_040628130.1    rna-XM_027586926.2    mNeoVis.3.1.1.chr2.g00739.t01    rna-gnl|WGS_ZZZZ|egapxtmp_034282-R1    mNeoVis.evergladensis.2.1.chr2.g00830.t01    mNeoVis.vulgivaga.2.1.chr2.g00832.t01      1464    N0.HOG0001462   OG0000166  n44   rna-XM_034645472.1    rna-XM_038577318.1    rna-XM_022519374.1    rna-XM_011283870.3    rna-XM_036080144.1    rna-XM_039249355.1    rna-XM_032877020.1    rna-XM_046011118.1    rna-XM_032347814.1    rna-XM_059185639.1    rna-XM_059402902.1    rna-XM_004765541.3    mNeoVis.2.1.1.chr9_hap1.g00667.t01    rna-XM_044915209.1    rna-XM_004402271.1    rna-XM_042990807.1    rna-XM_008704710.1    rna-XM_027593329.2    mNeoVis.3.1.1.chr10.g00632.t01    rna-gnl|WGS_ZZZZ|egapxtmp_021687-R1    mNeoVis.evergladensis.2.1.chr10.g00663.t01    mNeoVis.vulgivaga.2.1.chr10.g00646.t01      1465    N0.HOG0001463   OG0000166  n63   rna-XM_034666129.1    rna-XM_038568215.1    rna-XM_022513913.1    rna-XM_045038579.1    rna-XM_036081938.1, rna-XM_036087775.1    rna-XM_039221110.1    rna-XM_032852733.1    rna-XM_046016150.1    rna-XM_032358816.1    rna-XM_059185809.1    rna-XM_059391830.1    rna-XM_045080322.1    mNeoVis.2.1.1.chr7_hap1.g01415.t01    rna-XM_044919780.1    rna-XM_004413742.1    rna-XM_042959596.1    rna-XM_008688424.2    rna-XM_027581317.2    mNeoVis.3.1.1.chr7.g01325.t01    rna-gnl|WGS_ZZZZ|egapxtmp_013700-R1    mNeoVis.evergladensis.2.1.chr7.g01398.t01    mNeoVis.vulgivaga.2.1.chr7.g01410.t01      1466    N0.HOG0001464   OG0000166  n85   rna-XM_034658650.1    rna-XM_038565832.1    rna-XM_022498228.1    rna-XM_023249967.2    rna-XM_036081027.1    rna-XM_039230558.1    rna-XM_032867417.1    rna-XM_045992376.1    rna-XM_032325229.1    rna-XM_059161515.1    rna-XM_059390166.1    rna-XM_045063248.1    mNeoVis.2.1.1.chr5_hap1.g00824.t01    rna-XM_021695008.1    rna-XM_004409343.1    rna-XM_042977355.1    rna-XM_008703149.2    rna-XM_027585015.2    mNeoVis.3.1.1.chr2.g00735.t01    rna-gnl|WGS_ZZZZ|egapxtmp_035459-R1    mNeoVis.evergladensis.2.1.chr2.g00825.t01    mNeoVis.vulgivaga.2.1.chr2.g00828.t01      1467    N0.HOG0001465   OG0000167  n1   rna-XM_002926769.3    rna-XM_038581728.1, rna-XM_038581729.1    rna-XM_022515950.1, rna-XM_022516207.1    rna-XM_045036454.1    rna-XM_036101099.1, rna-XM_036101100.1    rna-XM_039229379.1    rna-XM_032865530.1, rna-XM_032865570.1    rna-XM_046003627.1, rna-XM_046003701.1    rna-XM_032302543.1, rna-XM_032302551.1    rna-XM_059165094.1, rna-XM_059165288.1    rna-XM_059388200.1, rna-XM_059388385.1    rna-XM_004757487.3    mNeoVis.2.1.1.chr5_hap1.g01759.t01, mNeoVis.2.1.1.chr5_hap1.g01760.t01    rna-XM_044913643.1, rna-XM_044913653.1    rna-XM_004406302.1, rna-XM_004406303.1    rna-XM_007075065.2    rna-XM_008701810.2, rna-XM_008701941.2    rna-XM_035723067.1, rna-XM_035723216.1    mNeoVis.3.1.1.chr2.g01667.t01    rna-gnl|WGS_ZZZZ|egapxtmp_038021-R1, rna-gnl|WGS_ZZZZ|egapxtmp_038023-R1    mNeoVis.evergladensis.2.1.chr2.g01741.t01, mNeoVis.evergladensis.2.1.chr2.g01742.t01    mNeoVis.vulgivaga.2.1.chr2.g01757.t01, mNeoVis.vulgivaga.2.1.chr2.g01758.t01      1468    N0.HOG0001466   OG0000167  n35   rna-XM_002926760.4, rna-XM_002926761.4, rna-XM_011234236.2, rna-XM_019807532.2    rna-XM_038581123.1, rna-XM_038581124.1, rna-XM_038581126.1    rna-XM_022516152.1, rna-XM_022516153.1, rna-XM_022516218.1, rna-XM_022516224.1    rna-XM_003991505.5, rna-XM_003991507.6, rna-XM_003991508.6, rna-XM_003991509.6, rna-XM_023260851.2    rna-XM_036101095.1, rna-XM_036101096.1, rna-XM_036101097.1    rna-XM_039229822.1, rna-XM_039229848.1, rna-XM_039229856.1, rna-XM_039229867.1, rna-XM_039229877.1, rna-XM_039229887.1    rna-XM_032865531.1, rna-XM_032865571.1, rna-XM_032865572.1, rna-XM_032865573.1    rna-XM_046003142.1, rna-XM_046003873.1    rna-XM_032302511.1, rna-XM_032302523.1, rna-XM_032302533.1, rna-XM_032345581.1    rna-XM_059165093.1, rna-XM_059165284.1, rna-XM_059165286.1, rna-XM_059165287.1    rna-XM_059388197.1, rna-XM_059388198.1, rna-XM_059388199.1    rna-XM_004757485.3, rna-XM_004757486.3, rna-XM_004757551.3    mNeoVis.2.1.1.chr5_hap1.g01757.t01, mNeoVis.2.1.1.chr5_hap1.g01758.t01    rna-XM_021678016.1, rna-XM_021678017.1, rna-XM_021678019.1    rna-XM_004406313.1, rna-XM_004406314.1    rna-XM_007075052.2, rna-XM_007075053.3, rna-XM_007075064.3, rna-XM_007075066.2, rna-XM_042956287.1, rna-XM_042956772.1    rna-XM_008701807.2, rna-XM_008701808.1, rna-XM_008701809.2    rna-XM_027581777.2, rna-XM_027586529.1, rna-XM_027586683.1, rna-XM_035722103.1    mNeoVis.3.1.1.chr2.g01665.t01, mNeoVis.3.1.1.chr2.g01666.t01    rna-gnl|WGS_ZZZZ|egapxtmp_035698-R1, rna-gnl|WGS_ZZZZ|egapxtmp_038020-R1    mNeoVis.evergladensis.2.1.chr2.g01739.t01, mNeoVis.evergladensis.2.1.chr2.g01740.t01    mNeoVis.vulgivaga.2.1.chr2.g01755.t01, mNeoVis.vulgivaga.2.1.chr2.g01756.t01      1469    N0.HOG0001467   OG0000168  n1   rna-XM_002922354.4    rna-XM_038585641.1    rna-XM_022501224.1    rna-XM_003991103.5    rna-XM_036098769.1    rna-XM_039249982.1    rna-XM_032859898.1    rna-XM_046018357.1    rna-XM_032354669.1    rna-XM_059168156.1    rna-XM_059392665.1    rna-XM_004767021.3    mNeoVis.2.1.1.chr3_hap1.g01222.t01    rna-XM_021702703.1    rna-XM_004405369.1    rna-XM_007086082.3    rna-XM_008687463.2    rna-XM_027588205.2    mNeoVis.3.1.1.chr3.g01203.t01    rna-gnl|WGS_ZZZZ|egapxtmp_040813-R1    mNeoVis.evergladensis.2.1.chr3.g01219.t01    mNeoVis.vulgivaga.2.1.chr3.g01234.t01      1470    N0.HOG0001468   OG0000168  n22   rna-XM_002915980.4    rna-XM_038549663.1    rna-XM_022523122.1    rna-XM_006933807.5    rna-XM_036077438.1    rna-XM_039216846.1    rna-XM_032874448.1    rna-XM_046014007.1    rna-XM_032348604.1    rna-XM_059185005.1    rna-XM_059403395.1    rna-XM_004773092.3    mNeoVis.2.1.1.chr9_hap1.g00384.t01    rna-XM_021687176.2    rna-XM_004401658.1    rna-XM_007081490.3    rna-XM_040637125.1    rna-XM_027592540.1    mNeoVis.3.1.1.chr10.g00376.t01      mNeoVis.evergladensis.2.1.chr10.g00381.t01    mNeoVis.vulgivaga.2.1.chr10.g00378.t01      1471    N0.HOG0001469   OG0000168  n38   rna-XM_002915981.4, rna-XM_034651362.1    rna-XM_038549665.1    rna-XM_022523123.1    rna-XM_003988881.4    rna-XM_036077440.1    rna-XM_039216848.1    rna-XM_032874450.1    rna-XM_046014011.1    rna-XM_032348606.1    rna-XM_059185006.1    rna-XM_059403398.1    rna-XM_013046176.2    mNeoVis.2.1.1.chr9_hap1.g00383.t01    rna-XM_021687178.1    rna-XM_004401659.1    rna-XM_007081491.3    rna-XM_008712291.2    rna-XM_027592542.2    mNeoVis.3.1.1.chr10.g00375.t01      mNeoVis.evergladensis.2.1.chr10.g00380.t01    mNeoVis.vulgivaga.2.1.chr10.g00377.t01      1472    N0.HOG0001470   OG0000168  n46   rna-XM_034652005.1    rna-XM_038566641.1    rna-XM_022497662.1    rna-XM_006928689.5    rna-XM_036107690.1    rna-XM_039228301.1    rna-XM_032868525.1    rna-XM_045991970.1    rna-XM_032317815.1    rna-XM_059160718.1    rna-XM_059389580.1    rna-XM_004760785.2    mNeoVis.2.1.1.chr5_hap1.g00593.t01    rna-XM_021687071.2    rna-XM_004399161.1    rna-XM_007088426.3    rna-XM_040628156.1    rna-XM_027585828.2    mNeoVis.3.1.1.chr2.g00514.t01    rna-gnl|WGS_ZZZZ|egapxtmp_038593-R1    mNeoVis.evergladensis.2.1.chr2.g00589.t01    mNeoVis.vulgivaga.2.1.chr2.g00591.t01      1473    N0.HOG0001471   OG0000168  n62   rna-XM_034639740.1    rna-XM_038547021.1    rna-XM_022519995.1    rna-XM_045045135.1    rna-XM_036121182.1    rna-XM_039250650.1    rna-XM_032870928.1    rna-XM_045985811.1    rna-XM_032320176.1    rna-XM_059147704.1    rna-XM_059380789.1    rna-XM_013045964.2    mNeoVis.2.1.1.chr8_hap1.g00278.t01    rna-XM_021678390.2    rna-XM_004411087.1    rna-XM_007088234.3    rna-XM_008693578.2    rna-XM_027625172.1    mNeoVis.3.1.1.chr8.g00245.t01    rna-gnl|WGS_ZZZZ|egapxtmp_019644-R4    mNeoVis.evergladensis.2.1.chr9.g00276.t01    mNeoVis.vulgivaga.2.1.chr8.g00282.t01      1474    N0.HOG0001472   OG0000169  n1   rna-XM_002930051.2    rna-NM_001388616.1    rna-XM_022507177.1    rna-XM_003993322.5    rna-XM_036104528.1    rna-XM_039224012.1    rna-XM_032846560.1    rna-XM_046016886.1    rna-XM_032359856.1    rna-XM_059137083.1    rna-XM_059382757.1    rna-XM_004781695.1    mNeoVis.2.1.1.chr7_hap1.g02110.t01    rna-XM_021685199.2    rna-XM_004411999.1    rna-XM_007088802.2    rna-XM_008694224.1    rna-XM_027577868.2    mNeoVis.3.1.1.chr7.g01947.t01    rna-gnl|WGS_ZZZZ|egapxtmp_012621-R1    mNeoVis.evergladensis.2.1.chr7.g02058.t01    mNeoVis.vulgivaga.2.1.chr7.g02082.t01      1475    N0.HOG0001473   OG0000169  n23   rna-XM_034645355.1    rna-NM_001013420.2, rna-NM_001389206.1    rna-XM_022506392.1    rna-XM_003993271.2    rna-XM_036104526.1    rna-XM_039223999.1    rna-XM_032846571.1    rna-XM_046015805.1, rna-XM_046017946.1    rna-XM_032359083.1    rna-XM_059158901.1    rna-XM_059388069.1    rna-XM_004781674.2      rna-XM_021685201.1      rna-XM_007088771.2    rna-XM_008694086.2      mNeoVis.3.1.1.chr7.g01949.t01        mNeoVis.vulgivaga.2.1.chr7.g02084.t01      1476    N0.HOG0001474   OG0000169  n42   rna-XM_034644584.1    rna-NM_001389008.1    rna-XM_022507509.1    rna-XM_011286835.3    rna-XM_036104519.1    rna-XM_039224020.1      rna-XM_046016827.1    rna-XM_032359085.1    rna-XM_059158900.1    rna-XM_059388073.1    rna-XM_004781675.2    mNeoVis.2.1.1.chr7_hap1.g02114.t01    rna-XM_021685200.2      rna-XM_007088770.2    rna-XM_008694087.1    rna-XM_027581065.2      rna-gnl|WGS_ZZZZ|egapxtmp_012622-R1    mNeoVis.evergladensis.2.1.chr7.g02061.t01        1477    N0.HOG0001475   OG0000169  n59   rna-XM_002931289.4, rna-XM_034650430.1, rna-XM_034651772.1    rna-NM_001389065.1    rna-XM_022507518.1    rna-XM_045038152.1    rna-XM_036104527.1    rna-XM_039224013.1    rna-XM_032846559.1    rna-XM_046017158.1    rna-XM_032359857.1    rna-XM_059158898.1    rna-XM_059388085.1    rna-XM_004781694.3    mNeoVis.2.1.1.chr7_hap1.g02112.t01      rna-XM_004411998.1    rna-XM_007088803.3    rna-XM_008694084.2        rna-gnl|WGS_ZZZZ|egapxtmp_013532-R1    mNeoVis.evergladensis.2.1.chr7.g02059.t01    mNeoVis.vulgivaga.2.1.chr7.g02083.t01      1478    N0.HOG0001476   OG0000169  n78   rna-XM_034644814.1    rna-NM_001389208.1, rna-XM_038562468.1    rna-XM_022507504.1, rna-XM_022507520.1    rna-XM_011286836.1, rna-XM_045038103.1    rna-XM_036104520.1, rna-XM_036104523.1      rna-XM_032846515.1, rna-XM_032846517.1    rna-XM_046017247.1, rna-XM_046017329.1, rna-XM_046017337.1    rna-XM_032359028.1    rna-XM_059158899.1    rna-XM_059388076.1    rna-XM_004781676.2    mNeoVis.2.1.1.chr7_hap1.g02111.t01, mNeoVis.2.1.1.chr7_hap1.g02113.t01    rna-XM_021684830.1    rna-XM_004411976.1, rna-XM_004411977.1    rna-XM_015540524.2    rna-XM_008694083.2, rna-XM_008694085.2    rna-XM_027580964.1      rna-gnl|WGS_ZZZZ|egapxtmp_010276-R1    mNeoVis.evergladensis.2.1.chr7.g02060.t01        1479    N0.HOG0001477   OG0000170  n1   rna-XM_019807462.2    rna-XM_038585689.1    rna-XM_022492619.1, rna-XM_022493730.1, rna-XM_022496124.1    rna-XM_004001367.6    rna-XM_036071804.1, rna-XM_036085337.1    rna-XM_039233078.1    rna-XM_032847457.1    rna-XM_045984619.1, rna-XM_045997259.1, rna-XM_046012157.1, rna-XM_046012970.1, rna-XM_046018031.1, rna-XM_046020707.1, rna-XM_046022083.1    rna-XM_032302293.1, rna-XM_032319025.1, rna-XM_032337671.1, rna-XM_032340952.1, rna-XM_032350297.1, rna-XM_032354743.1, rna-XM_032358788.1    rna-XM_059137241.1, rna-XM_059168097.1, rna-XM_059172043.1, rna-XM_059173441.1, rna-XM_059175314.1    rna-XM_059393443.1, rna-XM_059401606.1, rna-XM_059412456.1, rna-XM_059416479.1    rna-XM_004762714.3, rna-XM_013055653.2, rna-XM_013056771.2, rna-XM_045079627.1, rna-XM_045084733.1    mNeoVis.2.1.1.chr3_hap1.g01199.t01, mNeoVis.2.1.1.chr6_hap1.g00112.t01, mNeoVis.2.1.1.chr9_hap1.g00739.t01    rna-XM_021701498.2, rna-XM_021702813.1, rna-XM_044912168.1    rna-XM_004395710.1, rna-XM_004395835.1, rna-XM_004401226.1    rna-XM_007086112.2    rna-XM_008690483.2    rna-XM_027572796.2, rna-XM_027574416.2, rna-XM_027587474.2, rna-XM_027587485.2, rna-XM_027590599.2, rna-XM_027591183.1, rna-XM_027606388.2    mNeoVis.3.1.1.chr1.g00090.t01, mNeoVis.3.1.1.chr10.g00706.t01, mNeoVis.3.1.1.chr3.g01186.t01, mNeoVis.3.1.1.chr4.g00107.t01    rna-gnl|WGS_ZZZZ|egapxtmp_022162-R1, rna-gnl|WGS_ZZZZ|egapxtmp_041801-R1, rna-gnl|WGS_ZZZZ|egapxtmp_044466-R1    mNeoVis.evergladensis.2.1.chr10.g00105.t01, mNeoVis.evergladensis.2.1.chr3.g01201.t01, mNeoVis.evergladensis.2.1.chr4.g00113.t01    mNeoVis.vulgivaga.2.1.chr1.g00084.t01, mNeoVis.vulgivaga.2.1.chr10.g00720.t01, mNeoVis.vulgivaga.2.1.chr3.g01216.t01, mNeoVis.vulgivaga.2.1.chr4.g00108.t01, mNeoVis.vulgivaga.2.1.chr5.g01302.t01      1480    N0.HOG0001478   OG0000170  n44     rna-XM_038538479.1, rna-XM_038552585.1    rna-XM_022495970.1, rna-XM_022508566.1, rna-XM_022516321.1      rna-XM_036100186.1      rna-XM_032862998.1, rna-XM_032868515.1    rna-XM_045995722.1, rna-XM_046022220.1    rna-XM_032306021.1, rna-XM_032325254.1, rna-XM_032354066.1    rna-XM_059135893.1, rna-XM_059161516.1, rna-XM_059171546.1, rna-XM_059179111.1    rna-XM_059387875.1, rna-XM_059388938.1, rna-XM_059397514.1    rna-XM_013053909.2, rna-XM_013055813.2, rna-XM_013061471.2    mNeoVis.2.1.1.chr4_hap1.g00334.t01, mNeoVis.2.1.1.chr9_hap1.g00412.t01    rna-XM_021678517.1, rna-XM_021687617.2, rna-XM_021704291.2          rna-XM_027589645.2, rna-XM_027593710.2, rna-XM_027622879.1    mNeoVis.3.1.1.chr10.g00404.t01    rna-gnl|WGS_ZZZZ|egapxtmp_002924-R1, rna-gnl|WGS_ZZZZ|egapxtmp_022986-R1    mNeoVis.evergladensis.2.1.chr10.g00410.t01, mNeoVis.evergladensis.2.1.chr6.g00336.t01    mNeoVis.vulgivaga.2.1.chr10.g00405.t01, mNeoVis.vulgivaga.2.1.chr6.g00330.t01      1481    N0.HOG0001479   OG0000171  n2   rna-XM_034669567.1    rna-XM_038540168.1    rna-XM_022500866.1    rna-XM_003998788.5    rna-XM_036087822.1    rna-XM_039233671.1    rna-XM_032843969.1    rna-XM_045992950.1    rna-XM_032327494.1, rna-XM_032327498.1    rna-XM_059155772.1    rna-XM_059416062.1    rna-XM_045071912.1, rna-XM_045072156.1, rna-XM_045072741.1    mNeoVis.2.1.1.chr15_hap1.g00177.t01    rna-XM_044915082.1    rna-XM_004415762.1    rna-XM_007084168.2      rna-XM_027612160.1        mNeoVis.evergladensis.2.1.chr15.g00162.t01        1482    N0.HOG0001480   OG0000171  n23   rna-XM_019808303.2    rna-XM_038540171.1    rna-XM_022500407.1, rna-XM_022516636.1    rna-XM_045047436.1    rna-XM_036087550.1    rna-XM_039233662.1    rna-XM_032843967.1    rna-XM_045994264.1    rna-XM_032327477.1    rna-XM_059155743.1    rna-XM_059415189.1    rna-XM_045072736.1      rna-XM_021692226.1    rna-XM_004415759.1    rna-XM_042971718.1    rna-XM_008709371.2    rna-XM_027612158.1      rna-gnl|WGS_ZZZZ|egapxtmp_033624-R1    mNeoVis.evergladensis.2.1.chr15.g00160.t01    mNeoVis.vulgivaga.2.1.chr15.g00161.t01      1483    N0.HOG0001481   OG0000171  n43   rna-XM_002929075.4    rna-XM_038540204.1    rna-XM_022516536.1    rna-XM_045048158.1    rna-XM_036080825.1    rna-XM_039223813.1, rna-XM_039225704.1    rna-XM_032836170.1    rna-XM_045994050.1    rna-XM_032327475.1    rna-XM_059154670.1    rna-XM_059415200.1    rna-XM_004750271.3    mNeoVis.2.1.1.chr15_hap1.g00188.t01    rna-XM_044915081.1    rna-XM_004417841.1    rna-XM_042971881.1    rna-XM_008696967.2    rna-XM_027612694.2    mNeoVis.3.1.1.chr15.g00164.t01    rna-gnl|WGS_ZZZZ|egapxtmp_033623-R1    mNeoVis.evergladensis.2.1.chr15.g00166.t01    mNeoVis.vulgivaga.2.1.chr15.g00168.t01      1484    N0.HOG0001482   OG0000171  n66         rna-XM_019820891.3      rna-XM_039223809.1, rna-XM_039225282.1, rna-XM_039225284.1, rna-XM_039225449.1, rna-XM_039225657.1                      rna-XM_015539872.2                  1485    N0.HOG0001483   OG0000171  n73   rna-XM_002929071.4    rna-XM_038540193.1    rna-XM_022516722.1      rna-XM_036080826.1      rna-XM_032836163.1    rna-XM_045993902.1    rna-XM_032327481.1    rna-XM_059154930.1    rna-XM_059415197.1    rna-XM_004750269.3    mNeoVis.2.1.1.chr15_hap1.g00187.t01    rna-XM_044912244.1    rna-XM_004417394.1      rna-XM_008696970.2    rna-XM_027611874.2    mNeoVis.3.1.1.chr15.g00160.t01    rna-gnl|WGS_ZZZZ|egapxtmp_033635-R2    mNeoVis.evergladensis.2.1.chr15.g00165.t01    mNeoVis.vulgivaga.2.1.chr15.g00167.t01      1486    N0.HOG0001484   OG0000171  n91   rna-XM_034670298.1    rna-XM_038541720.1    rna-XM_022516746.1      rna-XM_036080824.1      rna-XM_032836162.1    rna-XM_045993875.1    rna-XM_032327483.1    rna-XM_059155100.1    rna-XM_059415193.1    rna-XM_004750268.3    mNeoVis.2.1.1.chr15_hap1.g00186.t01      rna-XM_004417396.1      rna-XM_040635254.1    rna-XM_027612690.1    mNeoVis.3.1.1.chr15.g00159.t01    rna-gnl|WGS_ZZZZ|egapxtmp_033625-R1    mNeoVis.evergladensis.2.1.chr15.g00164.t01    mNeoVis.vulgivaga.2.1.chr15.g00166.t01      1487    N0.HOG0001485   OG0000172  n2     rna-XM_038557547.1    rna-XM_022497400.1    rna-XM_023258581.2      rna-XM_039238726.1      rna-XM_046028322.1    rna-XM_032303549.1    rna-XM_059137754.1    rna-XM_059376453.1    rna-XM_004741539.2    mNeoVis.2.1.1.chr2_hap1.g01405.t01        rna-XM_007077351.1        mNeoVis.3.1.1.chr5.g01331.t01    rna-gnl|WGS_ZZZZ|egapxtmp_008841-R1    mNeoVis.evergladensis.2.1.chr5.g01347.t01    mNeoVis.vulgivaga.2.1.chr5.g01346.t01      1488    N0.HOG0001486   OG0000172  n16   rna-XM_002930965.1    rna-NM_001388549.1    rna-XM_022497377.1    rna-XM_019836826.2    rna-XM_036073430.1    rna-XM_039238723.1    rna-XM_032877035.1    rna-XM_046027320.1    rna-XM_032360611.1    rna-XM_059137913.1    rna-XM_059376452.1    rna-XM_004740609.2    mNeoVis.2.1.1.chr2_hap1.g01407.t01    rna-XM_021683794.1    rna-XM_004400674.1    rna-XM_007077439.1    rna-XM_008699337.1    rna-XM_027572143.1    mNeoVis.3.1.1.chr5.g01333.t01    rna-gnl|WGS_ZZZZ|egapxtmp_008843-R1      mNeoVis.vulgivaga.2.1.chr5.g01348.t01      1489    N0.HOG0001487   OG0000172  n38   rna-XM_002929412.4    rna-NM_001388550.1    rna-XM_022497370.1    rna-XM_045035654.1      rna-XM_039238720.1    rna-XM_032877033.1    rna-XM_046026116.1    rna-XM_032303203.1    rna-XM_059137832.1    rna-XM_059376449.1    rna-XM_004740610.1    mNeoVis.2.1.1.chr2_hap1.g01409.t01    rna-XM_021683793.1      rna-XM_007077440.2    rna-XM_008699433.1    rna-XM_035727321.1    mNeoVis.3.1.1.chr5.g01335.t01    rna-gnl|WGS_ZZZZ|egapxtmp_008842-R1    mNeoVis.evergladensis.2.1.chr5.g01349.t01    mNeoVis.vulgivaga.2.1.chr5.g01349.t01      1490    N0.HOG0001488   OG0000172  n56   rna-XM_002930966.4, rna-XM_034654320.1    rna-NM_001389144.1    rna-XM_022496077.1, rna-XM_022497365.1    rna-XM_003989967.3      rna-XM_039238722.1, rna-XM_039238725.1    rna-XM_032877034.1, rna-XM_032877702.1    rna-XM_045978631.1, rna-XM_046026013.1    rna-XM_032303204.1, rna-XM_032303739.1    rna-XM_059137088.1, rna-XM_059138507.1    rna-XM_059376450.1, rna-XM_059377075.1    rna-XM_004741540.1    mNeoVis.2.1.1.chr2_hap1.g01406.t01, mNeoVis.2.1.1.chr2_hap1.g01408.t01    rna-XM_021683795.1    rna-XM_004400686.1    rna-XM_007077352.2    rna-XM_008699334.2, rna-XM_008699335.2, rna-XM_008699336.2, rna-XM_008699432.2    rna-XM_027616870.1    mNeoVis.3.1.1.chr5.g01332.t01, mNeoVis.3.1.1.chr5.g01334.t01    rna-gnl|WGS_ZZZZ|egapxtmp_005880-R1    mNeoVis.evergladensis.2.1.chr5.g01348.t01, mNeoVis.evergladensis.2.1.chr5.g01350.t01    mNeoVis.vulgivaga.2.1.chr5.g01347.t01      1491    N0.HOG0001489   OG0000172  n90   rna-XM_002925572.3      rna-XM_022497399.1    rna-XM_003989986.3      rna-XM_039238736.1    rna-XM_032877036.1    rna-XM_045978278.1    rna-XM_032360693.1    rna-XM_059137966.1    rna-XM_059376454.1    rna-XM_004740608.2    mNeoVis.2.1.1.chr2_hap1.g01404.t01      rna-XM_004400685.1    rna-XM_007077438.2    rna-XM_008699430.2    rna-XM_027616893.2    mNeoVis.3.1.1.chr5.g01330.t01      mNeoVis.evergladensis.2.1.chr5.g01346.t01    mNeoVis.vulgivaga.2.1.chr5.g01345.t01      1492    N0.HOG0001490   OG0000173  n1   rna-XM_034641250.1    rna-NM_001346066.1    rna-XM_022509508.1    rna-XM_006940332.5    rna-XM_036081475.1    rna-XM_039223295.1    rna-XM_032871590.1    rna-XM_045985456.1    rna-XM_032321934.1    rna-XM_059148591.1    rna-XM_059378398.1    rna-XM_004764582.3    mNeoVis.2.1.1.chr8_hap1.g00454.t01    rna-XM_021704118.1    rna-XM_004395005.1    rna-XM_007085820.2    rna-XM_008689577.2    rna-XM_027624665.1    mNeoVis.3.1.1.chr8.g00410.t01    rna-gnl|WGS_ZZZZ|egapxtmp_019799-R1    mNeoVis.evergladensis.2.1.chr9.g00445.t01    mNeoVis.vulgivaga.2.1.chr8.g00450.t01      1493    N0.HOG0001491   OG0000173  n25   rna-XM_019805634.2    rna-XM_038546135.1    rna-XM_022509261.1    rna-XM_003996835.6    rna-XM_036081473.1    rna-XM_039227780.1    rna-XM_032871583.1    rna-XM_045985571.1    rna-XM_032322347.1    rna-XM_059148128.1    rna-XM_059380922.1    rna-XM_004764584.3    mNeoVis.2.1.1.chr8_hap1.g00456.t01    rna-XM_021704154.1    rna-XM_004417723.1    rna-XM_042966287.1    rna-XM_040640653.1    rna-XM_027568480.2    mNeoVis.3.1.1.chr8.g00412.t01    rna-gnl|WGS_ZZZZ|egapxtmp_019793-R1    mNeoVis.evergladensis.2.1.chr9.g00447.t01    mNeoVis.vulgivaga.2.1.chr8.g00452.t01      1494    N0.HOG0001492   OG0000173  n44   rna-XM_034641315.1    rna-NM_001346072.1    rna-XM_022509448.1    rna-XM_045044175.1    rna-XM_036081476.1    rna-XM_039223300.1    rna-XM_032871597.1    rna-XM_045985113.1    rna-XM_032321933.1    rna-XM_059148327.1    rna-XM_059380974.1    rna-XM_004764581.3    mNeoVis.2.1.1.chr8_hap1.g00452.t01      rna-XM_004395003.1    rna-XM_042966828.1    rna-XM_040639887.1    rna-XM_035724484.1    mNeoVis.3.1.1.chr8.g00408.t01    rna-gnl|WGS_ZZZZ|egapxtmp_019798-R1    mNeoVis.evergladensis.2.1.chr9.g00443.t01    mNeoVis.vulgivaga.2.1.chr8.g00448.t01      1495    N0.HOG0001493   OG0000173  n62   rna-XM_019805635.2    rna-NM_001346065.1    rna-XM_022509394.1    rna-XM_011289152.4    rna-XM_036081477.1    rna-XM_039227968.1    rna-XM_032871592.1    rna-XM_045985548.1    rna-XM_032321078.1    rna-XM_059148339.1    rna-XM_059380969.1    rna-XM_004764583.2    mNeoVis.2.1.1.chr8_hap1.g00455.t01    rna-XM_021704068.2    rna-XM_004395006.1    rna-XM_042966288.1    rna-XM_040640650.1    rna-XM_035724656.1    mNeoVis.3.1.1.chr8.g00411.t01    rna-gnl|WGS_ZZZZ|egapxtmp_019819-R1    mNeoVis.evergladensis.2.1.chr9.g00446.t01    mNeoVis.vulgivaga.2.1.chr8.g00451.t01      1496    N0.HOG0001494   OG0000173  n78   rna-XM_002924942.4    rna-NM_001346069.1    rna-XM_022509416.1    rna-XM_003996838.5    rna-XM_036081478.1    rna-XM_039223301.1    rna-XM_032871591.1    rna-XM_045985111.1    rna-XM_032321935.1    rna-XM_059149301.1    rna-XM_059379082.1    rna-XM_004764580.3    mNeoVis.2.1.1.chr8_hap1.g00453.t01    rna-XM_021704077.1    rna-XM_004395004.1    rna-XM_007085819.3    rna-XM_040640652.1    rna-XM_035724651.1    mNeoVis.3.1.1.chr8.g00409.t01    rna-gnl|WGS_ZZZZ|egapxtmp_019801-R1    mNeoVis.evergladensis.2.1.chr9.g00444.t01    mNeoVis.vulgivaga.2.1.chr8.g00449.t01      1497    N0.HOG0001495   OG0000174  n1   rna-XM_034638670.1    rna-XM_038525663.1    rna-XM_022491791.1    rna-XM_019819461.3    rna-XM_036103261.1    rna-XM_039247555.1    rna-XM_032849624.1    rna-XM_045989049.1    rna-XM_032324895.1    rna-XM_059153353.1    rna-XM_059382466.1    rna-XM_045077219.1    mNeoVis.2.1.1.chr7_hap1.g01088.t01    rna-XM_021692251.1    rna-XM_004408170.1    rna-XM_042969459.1    rna-XM_008709698.2    rna-XM_027617132.2    mNeoVis.3.1.1.chr7.g01017.t01      mNeoVis.evergladensis.2.1.chr7.g01079.t01    mNeoVis.vulgivaga.2.1.chr7.g01091.t01      1498    N0.HOG0001496   OG0000174  n22   rna-XM_034650274.1, rna-XM_034650343.1, rna-XM_034650379.1, rna-XM_034672380.1, rna-XM_034672383.1                                  rna-XM_008684400.2, rna-XM_040634035.1, rna-XM_040634065.1, rna-XM_040634068.1, rna-XM_040634069.1, rna-XM_040634073.1, rna-XM_040634076.1                1499    N0.HOG0001497   OG0000174  n35   rna-XM_034639007.1    rna-NM_001097557.1    rna-XM_022525020.1    rna-XM_011289901.4      rna-XM_039248280.1    rna-XM_032841853.1, rna-XM_032851160.1    rna-XM_045988110.1    rna-XM_032326540.1    rna-XM_059151987.1    rna-XM_059383645.1    rna-XM_045076846.1    mNeoVis.2.1.1.chr7_hap1.g00597.t01    rna-XM_044911371.1    rna-XM_004408188.1    rna-XM_007086642.2      rna-XM_027617440.1    mNeoVis.3.1.1.chr7.g00578.t01    rna-gnl|WGS_ZZZZ|egapxtmp_011570-R3    mNeoVis.evergladensis.2.1.chr7.g00599.t01    mNeoVis.vulgivaga.2.1.chr7.g00602.t01      1500    N0.HOG0001498   OG0000174  n54     rna-NM_001097546.1, rna-NM_001097552.1, rna-NM_001113459.1, rna-XM_038525553.1      rna-XM_045046148.1, rna-XM_045046150.1      rna-XM_039246392.1, rna-XM_039246415.1                      rna-XM_042968720.1, rna-XM_042968725.1                  1501    N0.HOG0001499   OG0000174  n63   rna-XM_034650286.1      rna-XM_022491784.1      rna-XM_036066772.1, rna-XM_036083161.1, rna-XM_036103240.1, rna-XM_036103253.1, rna-XM_036103259.1, rna-XM_036103340.1, rna-XM_036103341.1, rna-XM_036103347.1, rna-XM_036115895.1    rna-XM_039248212.1    rna-XM_032851086.1    rna-XM_045988046.1, rna-XM_045988047.1, rna-XM_045988048.1, rna-XM_045988053.1    rna-XM_032324908.1    rna-XM_059151793.1, rna-XM_059151794.1, rna-XM_059151802.1, rna-XM_059151806.1    rna-XM_059382207.1, rna-XM_059383826.1    rna-XM_013064087.2, rna-XM_045077220.1    mNeoVis.2.1.1.chr7_hap1.g01084.t01, mNeoVis.2.1.1.chr7_hap1.g01085.t01, mNeoVis.2.1.1.chr7_hap1.g01086.t01    rna-XM_044911370.1, rna-XM_044911437.1, rna-XM_044912260.1, rna-XM_044912337.1    rna-XM_004417053.1        rna-XM_027619914.2    mNeoVis.3.1.1.chr7.g01013.t01, mNeoVis.3.1.1.chr7.g01015.t01      mNeoVis.evergladensis.2.1.chr7.g01075.t01, mNeoVis.evergladensis.2.1.chr7.g01076.t01, mNeoVis.evergladensis.2.1.chr7.g01077.t01    mNeoVis.vulgivaga.2.1.chr7.g01087.t01, mNeoVis.vulgivaga.2.1.chr7.g01088.t01, mNeoVis.vulgivaga.2.1.chr7.g01089.t01      1502    N0.HOG0001500   OG0000175  n1   rna-XM_034646133.1    rna-XM_038564234.1    rna-XM_022507627.1    rna-XM_003993528.6    rna-XM_036122158.1    rna-XM_039257692.1    rna-XM_032879278.1    rna-XM_046014375.1    rna-XM_032357082.1    rna-XM_059144889.1    rna-XM_059403572.1    rna-XM_004770262.3    mNeoVis.2.1.1.chr7_hap1.g02324.t01    rna-XM_021685492.2    rna-XM_004413783.1    rna-XM_042957765.1    rna-XM_040639257.1    rna-XM_027578478.1    mNeoVis.3.1.1.chr7.g02140.t01    rna-gnl|WGS_ZZZZ|egapxtmp_011384-R1    mNeoVis.evergladensis.2.1.chr7.g02271.t01    mNeoVis.vulgivaga.2.1.chr7.g02288.t01      1503    N0.HOG0001501   OG0000175  n23   rna-XM_034650057.1, rna-XM_034653521.1, rna-XM_034653583.1    rna-XM_038587943.1, rna-XM_038587948.1, rna-XM_038589687.1    rna-XM_022492317.1    rna-XM_045051675.1    rna-XM_036085631.1, rna-XM_036102631.1    rna-XM_039225826.1    rna-XM_032849674.1    rna-XM_045994650.1, rna-XM_045996341.1    rna-XM_032331545.1    rna-XM_059158378.1    rna-XM_059385202.1    rna-XM_045068329.1    mNeoVis.2.1.1.chr12_hap1.g00352.t01, mNeoVis.2.1.1.chr12_hap1.g00354.t01    rna-XM_021692174.1, rna-XM_044911593.1      rna-XM_042973962.1    rna-XM_040619946.1, rna-XM_040628748.1    rna-XM_027608531.1, rna-XM_035725368.1, rna-XM_035725618.1              1504    N0.HOG0001502   OG0000175  n51   rna-XM_011216822.2    rna-XM_038588819.1    rna-XM_022492309.1    rna-XM_023248970.1    rna-XM_036084053.1          rna-XM_032329710.1                rna-XM_042973961.1              mNeoVis.vulgivaga.2.1.chr12.g00322.t01      1505    N0.HOG0001503   OG0000175  n58   rna-XM_034653558.1    rna-XM_038564537.1, rna-XM_038587892.1, rna-XM_038587917.1    rna-XM_022492322.1, rna-XM_022492323.1    rna-XM_011291825.4, rna-XM_045050388.1    rna-XM_036079122.1, rna-XM_036109510.1    rna-XM_039224098.1, rna-XM_039224262.1    rna-XM_032881818.1, rna-XM_032881822.1    rna-XM_045995527.1, rna-XM_045995949.1, rna-XM_045995950.1    rna-XM_032330620.1, rna-XM_032331583.1, rna-XM_032331655.1    rna-XM_059157362.1, rna-XM_059157376.1, rna-XM_059158373.1    rna-XM_059384651.1, rna-XM_059384737.1, rna-XM_059416554.1    rna-XM_004777128.3, rna-XM_045085342.1, rna-XM_045085343.1    mNeoVis.2.1.1.chr12_hap1.g00360.t01, mNeoVis.2.1.1.chr12_hap1.g00361.t01, mNeoVis.2.1.1.chr12_hap1.g00367.t01    rna-XM_021697485.1, rna-XM_021697914.1    rna-XM_004414938.1    rna-XM_042974234.1, rna-XM_042974238.1    rna-XM_008703974.2, rna-XM_040619816.1, rna-XM_040629773.1    rna-XM_035725637.1, rna-XM_035725640.1, rna-XM_035725647.1    mNeoVis.3.1.1.chr12.g00330.t01    rna-gnl|WGS_ZZZZ|egapxtmp_027986-R1    mNeoVis.evergladensis.2.1.chr12.g00318.t01, mNeoVis.evergladensis.2.1.chr12.g00324.t01    mNeoVis.vulgivaga.2.1.chr12.g00328.t01      1506    N0.HOG0001504   OG0000176  n1   rna-XM_002929382.4    rna-XM_038550864.1    rna-XM_022516341.1    rna-XM_003983869.6    rna-XM_036087224.1    rna-XM_039222195.1    rna-XM_032842416.1    rna-XM_045981654.1    rna-XM_032351652.1    rna-XM_059134299.1    rna-XM_059407871.1    rna-XM_004764349.3    mNeoVis.2.1.1.chr11_hap1.g00563.t01    rna-XM_021680274.1    rna-XM_004414459.1    rna-XM_007097759.3    rna-XM_008699073.2    rna-XM_027622118.2    mNeoVis.3.1.1.chr11.g00523.t01    rna-gnl|WGS_ZZZZ|egapxtmp_027450-R2    mNeoVis.evergladensis.2.1.chr11.g00550.t01    mNeoVis.vulgivaga.2.1.chr11.g00558.t01      1507    N0.HOG0001505   OG0000176  n20   rna-XM_002930286.4    rna-XM_038550862.1    rna-XM_022516270.1    rna-XM_011281074.4    rna-XM_036120462.1    rna-XM_039225755.1    rna-XM_032842417.1    rna-XM_045981067.1    rna-XM_032351655.1    rna-XM_059134301.1    rna-XM_059407870.1    rna-XM_045077607.1    mNeoVis.2.1.1.chr11_hap1.g00564.t01    rna-XM_021680275.1    rna-XM_004414460.1    rna-XM_042979753.1    rna-XM_008699075.2    rna-XM_027622119.1    mNeoVis.3.1.1.chr11.g00524.t01    rna-gnl|WGS_ZZZZ|egapxtmp_027449-R2    mNeoVis.evergladensis.2.1.chr11.g00551.t01    mNeoVis.vulgivaga.2.1.chr11.g00559.t01      1508    N0.HOG0001506   OG0000176  n42   rna-XM_002929461.4    rna-NM_001195835.1    rna-XM_022506124.1    rna-XM_045056263.1    rna-XM_036076112.1    rna-XM_039251765.1    rna-XM_032845783.1    rna-XM_045995602.1    rna-XM_032334482.1    rna-XM_059159924.1    rna-XM_059396652.1    rna-XM_045079834.1    mNeoVis.2.1.1.chr6_hap1.g00117.t01    rna-XM_021702403.1    rna-XM_004392889.1    rna-XM_042982737.1    rna-XM_008683684.2    rna-XM_027599339.2    mNeoVis.3.1.1.chr4.g00112.t01    rna-gnl|WGS_ZZZZ|egapxtmp_001513-R3    mNeoVis.evergladensis.2.1.chr4.g00118.t01    mNeoVis.vulgivaga.2.1.chr4.g00112.t01      1509    N0.HOG0001507   OG0000176  n62   rna-XM_011237499.3    rna-XM_038556239.1    rna-XM_022506029.1    rna-XM_045056260.1      rna-XM_039251783.1    rna-XM_032845591.1    rna-XM_045997161.1    rna-XM_032334485.1    rna-XM_059159902.1    rna-XM_059384942.1    rna-XM_004766331.3    mNeoVis.2.1.1.chr6_hap1.g00116.t01        rna-XM_007086913.3    rna-XM_008683686.2      mNeoVis.3.1.1.chr4.g00111.t01    rna-gnl|WGS_ZZZZ|egapxtmp_001515-R1    mNeoVis.evergladensis.2.1.chr4.g00117.t01    mNeoVis.vulgivaga.2.1.chr4.g00111.t01      1510    N0.HOG0001508   OG0000176  n77   rna-XM_034658913.1    rna-XM_038550865.1    rna-XM_022516369.1    rna-XM_019826951.3    rna-XM_036120449.1    rna-XM_039225352.1, rna-XM_039225538.1, rna-XM_039225639.1    rna-XM_032842428.1    rna-XM_045981069.1    rna-XM_032351656.1    rna-XM_059133201.1    rna-XM_059407868.1    rna-XM_045077608.1    mNeoVis.2.1.1.chr11_hap1.g00565.t01      rna-XM_004414296.1    rna-XM_007097761.3    rna-XM_040622914.1    rna-XM_027622796.1    mNeoVis.3.1.1.chr11.g00525.t01    rna-gnl|WGS_ZZZZ|egapxtmp_027448-R1    mNeoVis.evergladensis.2.1.chr11.g00552.t01    mNeoVis.vulgivaga.2.1.chr11.g00560.t01      1511    N0.HOG0001509   OG0000177  n3   rna-XM_034641186.1, rna-XM_034650471.1, rna-XM_034650833.1    rna-XM_038538960.1, rna-XM_038541148.1, rna-XM_038555965.1, rna-XM_038555967.1, rna-XM_038555970.1, rna-XM_038582547.1, rna-XM_038584498.1, rna-XM_038584697.1, rna-XM_038588798.1      rna-XM_045053965.1, rna-XM_045053966.1                          rna-XM_042977076.1, rna-XM_042977078.1, rna-XM_042977082.1, rna-XM_042977084.1, rna-XM_042977086.1, rna-XM_042977090.1, rna-XM_042977093.1, rna-XM_042977095.1, rna-XM_042977098.1, rna-XM_042977100.1, rna-XM_042977103.1, rna-XM_042981028.1, rna-XM_042981030.1                  1512    N0.HOG0001510   OG0000177  n18   rna-XM_034650473.1, rna-XM_034650475.1    rna-XM_038530877.1, rna-XM_038543941.1    rna-XM_022517569.1      rna-XM_036067670.1, rna-XM_036070340.1, rna-XM_036086352.1, rna-XM_036118831.1      rna-XM_032850378.1, rna-XM_032850379.1    rna-XM_045981672.1, rna-XM_045981675.1, rna-XM_045981677.1, rna-XM_045981682.1    rna-XM_032302352.1, rna-XM_032302353.1, rna-XM_032302357.1, rna-XM_032302943.1, rna-XM_032302945.1, rna-XM_032302947.1, rna-XM_032314803.1    rna-XM_059134349.1    rna-XM_059387218.1    rna-XM_045081793.1, rna-XM_045081797.1, rna-XM_045081802.1, rna-XM_045081804.1, rna-XM_045081806.1, rna-XM_045081809.1, rna-XM_045081978.1, rna-XM_045081979.1, rna-XM_045081980.1, rna-XM_045081982.1, rna-XM_045081983.1, rna-XM_045081984.1, rna-XM_045083409.1    mNeoVis.2.1.1.chr11_hap1.g00546.t01, mNeoVis.2.1.1.chr3_hap1.g00470.t01      rna-XM_004417264.1    rna-XM_042989206.1    rna-XM_040620926.1, rna-XM_040637930.1    rna-XM_027623317.2    mNeoVis.3.1.1.chr1.g00998.t01, mNeoVis.3.1.1.chr11.g00505.t01, mNeoVis.3.1.1.chr5.g00682.t01    rna-gnl|WGS_ZZZZ|egapxtmp_027282-R1, rna-gnl|WGS_ZZZZ|egapxtmp_037265-R1    mNeoVis.evergladensis.2.1.chr1.g00987.t01, mNeoVis.evergladensis.2.1.chr1.g01009.t01, mNeoVis.evergladensis.2.1.chr11.g00532.t01    mNeoVis.vulgivaga.2.1.chr1.g01000.t01, mNeoVis.vulgivaga.2.1.chr11.g00540.t01      1513    N0.HOG0001511   OG0000177  n69   rna-XM_034650193.1, rna-XM_034653313.1, rna-XM_034664754.1    rna-XM_038548877.1    rna-XM_022500557.1    rna-XM_019816714.2    rna-XM_036107955.1    rna-XM_039229591.1    rna-XM_032866163.1    rna-XM_046023312.1    rna-XM_032306200.1    rna-XM_059143400.1    rna-XM_059410429.1    rna-XM_045082662.1    mNeoVis.2.1.1.chr13_hap1.g00119.t01    rna-XM_021691361.1    rna-XM_004392426.1    rna-XM_042963725.1    rna-XM_040632123.1    rna-XM_027616294.2    mNeoVis.3.1.1.chr13.g00063.t01    rna-gnl|WGS_ZZZZ|egapxtmp_031064-R1    mNeoVis.evergladensis.2.1.chr13.g00116.t01    mNeoVis.vulgivaga.2.1.chr13.g00113.t01      1514    N0.HOG0001512   OG0000178  n1   rna-XM_002922000.4, rna-XM_034660632.1    rna-NM_001313800.1    rna-XM_022523005.1    rna-XM_003987612.5    rna-XM_036076258.1    rna-XM_039217341.1, rna-XM_039221815.1, rna-XM_039233036.1, rna-XM_039246992.1    rna-XM_032871186.1    rna-XM_046007597.1    rna-XM_032344155.1    rna-XM_059182029.1, rna-XM_059186736.1    rna-XM_059373147.1, rna-XM_059392910.1, rna-XM_059404627.1, rna-XM_059407581.1    rna-XM_004738727.3    mNeoVis.2.1.1.chr10_hap1.g00431.t01, mNeoVis.2.1.1.chr12_hap1.g00374.t01, mNeoVis.2.1.1.chr8_hap1.g01016.t01    rna-XM_021701454.1    rna-XM_004408013.1    rna-XM_042984147.1, rna-XM_042989638.1    rna-XM_008693515.2, rna-XM_008701715.2    rna-XM_027568987.1, rna-XM_035721977.1, rna-XM_035724680.1, rna-XM_035726144.1, rna-XM_035728510.1    mNeoVis.3.1.1.chr12.g00338.t01, mNeoVis.3.1.1.chr9.g00397.t01    rna-gnl|WGS_ZZZZ|egapxtmp_028676-R1    mNeoVis.evergladensis.2.1.chr12.g00332.t01, mNeoVis.evergladensis.2.1.chr8.g00411.t01    mNeoVis.vulgivaga.2.1.chr9.g00413.t01      1515    N0.HOG0001513   OG0000178  n5           rna-XM_036099649.1                            rna-XM_027594726.1              1516    N0.HOG0001514   OG0000178  n10           rna-XM_036067140.1    rna-XM_039224099.1              rna-XM_004833882.2    mNeoVis.2.1.1.chr9_hap1.g00665.t01    rna-XM_021695869.2    rna-XM_004417743.1        rna-XM_027607978.1, rna-XM_035728810.1      rna-gnl|WGS_ZZZZ|egapxtmp_018507-R1    mNeoVis.evergladensis.2.1.chr10.g00660.t01    mNeoVis.vulgivaga.2.1.chr10.g00644.t01      1517    N0.HOG0001515   OG0000178  n20   rna-XM_034651166.1                          mNeoVis.2.1.1.chr2_hap1.g01468.t01                        1518    N0.HOG0001516   OG0000178  n23     rna-XM_038565252.1    rna-XM_022497978.1, rna-XM_022521645.1    rna-XM_011288275.3    rna-XM_036067730.1, rna-XM_036093051.1, rna-XM_036114544.1    rna-XM_039217201.1, rna-XM_039255385.1, rna-XM_039256599.1    rna-XM_032881815.1    rna-XM_045999594.1    rna-XM_032329709.1    rna-XM_059156818.1      rna-XM_045085345.1      rna-XM_021690663.1, rna-XM_021701466.2    rna-XM_004396619.1, rna-XM_004408421.1, rna-XM_004415855.1    rna-XM_015535453.2      rna-XM_027567588.2, rna-XM_027572905.1, rna-XM_035722436.1, rna-XM_035728958.1      rna-gnl|WGS_ZZZZ|egapxtmp_020154-R1    mNeoVis.evergladensis.2.1.chr9.g00990.t01    mNeoVis.vulgivaga.2.1.chr8.g00991.t01      1519    N0.HOG0001517   OG0000178  n50     rna-XM_038561853.1      rna-XM_011291081.4    rna-XM_036072676.1    rna-XM_039222902.1, rna-XM_039231810.1                  rna-XM_021681524.1, rna-XM_021688539.1    rna-XM_004405671.1, rna-XM_004411925.1    rna-XM_015545072.2      rna-XM_027608505.1    mNeoVis.3.1.1.chr8.g00947.t01        mNeoVis.vulgivaga.2.1.chr12.g00339.t01      1520    N0.HOG0001518   OG0000178  n62         rna-XM_023249779.2      rna-XM_039218089.1, rna-XM_039242002.1              rna-XM_004833869.2        rna-XM_004410008.1    rna-XM_007094082.2      rna-XM_027575238.1              1521    N0.HOG0001519   OG0000179  n2   rna-XM_034665605.1    rna-NM_001388730.1, rna-NM_001388908.1    rna-XM_022521691.1, rna-XM_022521692.1    rna-XM_045038840.1, rna-XM_045038848.1    rna-XM_036095956.1    rna-XM_039218866.1, rna-XM_039218868.1    rna-XM_032852721.1, rna-XM_032853675.1    rna-XM_046017292.1, rna-XM_046017328.1    rna-XM_032359791.1, rna-XM_032360484.1    rna-XM_059157991.1, rna-XM_059158013.1    rna-XM_059389458.1, rna-XM_059389485.1    rna-XM_004781203.2, rna-XM_045078269.1, rna-XM_045078271.1    mNeoVis.2.1.1.chr7_hap1.g01733.t01, mNeoVis.2.1.1.chr7_hap1.g01735.t01    rna-XM_021685692.1      rna-XM_007083272.3, rna-XM_042959035.1    rna-XM_008711005.2    rna-XM_027581401.1      rna-gnl|WGS_ZZZZ|egapxtmp_012900-R1    mNeoVis.evergladensis.2.1.chr7.g01701.t01    mNeoVis.vulgivaga.2.1.chr7.g01713.t01      1522    N0.HOG0001520   OG0000179  n34   rna-XM_002925000.2    rna-XM_038568180.1    rna-XM_022521779.1      rna-XM_036095926.1    rna-XM_039218826.1    rna-XM_032853677.1    rna-XM_046014598.1    rna-XM_032358897.1    rna-XM_059158022.1    rna-XM_059389445.1    rna-XM_004781198.1          rna-XM_007083273.3    rna-XM_008710987.2      mNeoVis.3.1.1.chr7.g01613.t01    rna-gnl|WGS_ZZZZ|egapxtmp_012906-R1      mNeoVis.vulgivaga.2.1.chr7.g01715.t01      1523    N0.HOG0001521   OG0000179  n49   rna-XM_002931295.4, rna-XM_034666459.1, rna-XM_034666460.1    rna-NM_001388979.1, rna-XM_038568843.1    rna-XM_022522122.1, rna-XM_022522147.1    rna-XM_003992794.4, rna-XM_006937026.5, rna-XM_045038816.1    rna-XM_036114421.1    rna-XM_039225845.1    rna-XM_032850792.1, rna-XM_032853592.1, rna-XM_032853593.1    rna-XM_046017000.1, rna-XM_046017116.1, rna-XM_046017146.1, rna-XM_046017148.1    rna-XM_032360373.1, rna-XM_032360460.1, rna-XM_032360461.1, rna-XM_032360462.1    rna-XM_059156745.1, rna-XM_059156760.1, rna-XM_059156784.1    rna-XM_059390844.1, rna-XM_059390858.1    rna-XM_004782219.2, rna-XM_004782221.2, rna-XM_004782226.2, rna-XM_004782227.2, rna-XM_045071509.1    mNeoVis.2.1.1.chr7_hap1.g01607.t01, mNeoVis.2.1.1.chr7_hap1.g01608.t01, mNeoVis.2.1.1.chr7_hap1.g01609.t01, mNeoVis.2.1.1.chr7_hap1.g01612.t01    rna-XM_021685659.2    rna-XM_004407568.1    rna-XM_007089409.2, rna-XM_007089410.2, rna-XM_042959058.1    rna-XM_008710692.1, rna-XM_008710737.2, rna-XM_008710739.2, rna-XM_008710741.2    rna-XM_027580951.1    mNeoVis.3.1.1.chr7.g01517.t01    rna-gnl|WGS_ZZZZ|egapxtmp_012508-R1, rna-gnl|WGS_ZZZZ|egapxtmp_012870-R1, rna-gnl|WGS_ZZZZ|egapxtmp_012875-R2    mNeoVis.evergladensis.2.1.chr7.g01582.t01, mNeoVis.evergladensis.2.1.chr7.g01583.t01, mNeoVis.evergladensis.2.1.chr7.g01584.t01    mNeoVis.vulgivaga.2.1.chr7.g01601.t01, mNeoVis.vulgivaga.2.1.chr7.g01603.t01      1524    N0.HOG0001522   OG0000180  n0   rna-XM_011235221.3      rna-XM_022523660.1    rna-XM_003993932.5    rna-XM_036106965.1    rna-XM_039241244.1    rna-XM_032841117.1    rna-XM_046026895.1    rna-XM_032311729.1, rna-XM_032312576.1, rna-XM_032332009.1, rna-XM_032340908.1    rna-XM_059137762.1, rna-XM_059137876.1, rna-XM_059137884.1, rna-XM_059137959.1, rna-XM_059137980.1, rna-XM_059144063.1, rna-XM_059145432.1, rna-XM_059147585.1, rna-XM_059148081.1, rna-XM_059148084.1, rna-XM_059148543.1, rna-XM_059148579.1, rna-XM_059148590.1, rna-XM_059148609.1, rna-XM_059148626.1, rna-XM_059148648.1, rna-XM_059149234.1, rna-XM_059149361.1, rna-XM_059149363.1, rna-XM_059151594.1, rna-XM_059151923.1, rna-XM_059155865.1, rna-XM_059155870.1, rna-XM_059155919.1, rna-XM_059156489.1, rna-XM_059156490.1, rna-XM_059156524.1, rna-XM_059156526.1, rna-XM_059156799.1, rna-XM_059156863.1, rna-XM_059156865.1, rna-XM_059157557.1, rna-XM_059157560.1, rna-XM_059157605.1, rna-XM_059157647.1, rna-XM_059157650.1, rna-XM_059157721.1, rna-XM_059157959.1, rna-XM_059158023.1, rna-XM_059158082.1, rna-XM_059158087.1, rna-XM_059158423.1, rna-XM_059158431.1, rna-XM_059158435.1, rna-XM_059158438.1, rna-XM_059158439.1, rna-XM_059158440.1, rna-XM_059158443.1, rna-XM_059158445.1, rna-XM_059170501.1, rna-XM_059170506.1, rna-XM_059178952.1, rna-XM_059178971.1, rna-XM_059179024.1, rna-XM_059180077.1, rna-XM_059180078.1, rna-XM_059180080.1, rna-XM_059180087.1, rna-XM_059180089.1, rna-XM_059180091.1    rna-XM_059387339.1, rna-XM_059387474.1, rna-XM_059387487.1, rna-XM_059397349.1    rna-XM_045067115.1, rna-XM_045067199.1, rna-XM_045067361.1, rna-XM_045067385.1, rna-XM_045069859.1, rna-XM_045069864.1, rna-XM_045069869.1, rna-XM_045069918.1, rna-XM_045072794.1, rna-XM_045073808.1, rna-XM_045079341.1, rna-XM_045081109.1, rna-XM_045083152.1, rna-XM_045086234.1, rna-XM_045087223.1, rna-XM_045088048.1    mNeoVis.2.1.1.chr2_hap1.g00653.t01, mNeoVis.2.1.1.chr2_hap1.g00719.t01, mNeoVis.2.1.1.chr7_hap1.g01110.t01    rna-XM_021696059.2    rna-XM_004404854.1    rna-XM_042960136.1    rna-XM_008689535.2    rna-XM_027595315.2    mNeoVis.3.1.1.chr5.g00629.t01    rna-gnl|WGS_ZZZZ|egapxtmp_008780-R1, rna-gnl|WGS_ZZZZ|egapxtmp_009495-R1    mNeoVis.evergladensis.2.1.chr5.g00636.t01    mNeoVis.vulgivaga.2.1.chr5.g00630.t01, mNeoVis.vulgivaga.2.1.chr5.g00669.t01      1525    N0.HOG0001523   OG0000181  n1         rna-XM_011291198.4, rna-XM_045048597.1, rna-XM_045048600.1, rna-XM_045048613.1      rna-XM_039218483.1, rna-XM_039218484.1, rna-XM_039225261.1, rna-XM_039225274.1, rna-XM_039225452.1, rna-XM_039225456.1, rna-XM_039225598.1                      rna-XM_042975839.1, rna-XM_042975843.1, rna-XM_042975845.1, rna-XM_042975955.1                  1526    N0.HOG0001524   OG0000181  n16   rna-XM_034650094.1, rna-XM_034650096.1, rna-XM_034650106.1, rna-XM_034667391.1, rna-XM_034667394.1    rna-XM_038541778.1, rna-XM_038541804.1, rna-XM_038541805.1, rna-XM_038541806.1, rna-XM_038542914.1, rna-XM_038542925.1, rna-XM_038542932.1    rna-XM_022490721.1, rna-XM_022490722.1, rna-XM_022490723.1, rna-XM_022490727.1, rna-XM_022490728.1, rna-XM_022492632.1      rna-XM_036069847.1, rna-XM_036118233.1, rna-XM_036118234.1      rna-XM_032845630.1, rna-XM_032845709.1, rna-XM_032851164.1    rna-XM_045981994.1, rna-XM_045981998.1, rna-XM_045982001.1, rna-XM_045982007.1, rna-XM_045982008.1, rna-XM_045982009.1, rna-XM_045982010.1    rna-XM_032318084.1, rna-XM_032318112.1, rna-XM_032318142.1, rna-XM_032318150.1    rna-XM_059146233.1, rna-XM_059146240.1, rna-XM_059146249.1, rna-XM_059146256.1, rna-XM_059146260.1, rna-XM_059146267.1    rna-XM_059413589.1, rna-XM_059413590.1, rna-XM_059413596.1, rna-XM_059413969.1, rna-XM_059414184.1    rna-XM_004778446.3, rna-XM_045086665.1, rna-XM_045086673.1, rna-XM_045086674.1, rna-XM_045086676.1, rna-XM_045086679.1, rna-XM_045086694.1    mNeoVis.2.1.1.chr14_hap1.g00591.t01, mNeoVis.2.1.1.chr14_hap1.g00592.t01, mNeoVis.2.1.1.chr14_hap1.g00594.t01, mNeoVis.2.1.1.chr14_hap1.g00595.t01    rna-XM_044912194.1        rna-XM_008696528.2, rna-XM_008700772.2, rna-XM_040635844.1, rna-XM_040635848.1, rna-XM_040635849.1, rna-XM_040635851.1, rna-XM_040635855.1    rna-XM_035722145.1, rna-XM_035722150.1, rna-XM_035722393.1, rna-XM_035722395.1        mNeoVis.evergladensis.2.1.chr14.g00589.t01, mNeoVis.evergladensis.2.1.chr14.g00590.t01, mNeoVis.evergladensis.2.1.chr14.g00592.t01    mNeoVis.vulgivaga.2.1.chr14.g00585.t01, mNeoVis.vulgivaga.2.1.chr14.g00586.t01, mNeoVis.vulgivaga.2.1.chr14.g00588.t01      1527    N0.HOG0001525   OG0000181  n90   rna-XM_034667385.1    rna-XM_038542915.1    rna-XM_022491005.1          rna-XM_032845619.1    rna-XM_045982005.1    rna-XM_032318133.1    rna-XM_059146262.1    rna-XM_059413592.1    rna-XM_013048998.2    mNeoVis.2.1.1.chr14_hap1.g00593.t01          rna-XM_040635845.1      mNeoVis.3.1.1.chr14.g00577.t01      mNeoVis.evergladensis.2.1.chr14.g00591.t01    mNeoVis.vulgivaga.2.1.chr14.g00587.t01      1528    N0.HOG0001526   OG0000182  n1   rna-NM_001304867.1    rna-NM_001251956.1    rna-XM_022501591.1    rna-XM_003995898.4, rna-XM_045037703.1    rna-XM_036076202.1, rna-XM_036085613.1, rna-XM_036092420.1, rna-XM_036098252.1, rna-XM_036099266.1, rna-XM_036106822.1, rna-XM_036108741.1, rna-XM_036112887.1, rna-XM_036119442.1, rna-XM_036122877.1    rna-XM_039217107.1, rna-XM_039221943.1, rna-XM_039229583.1, rna-XM_039245382.1    rna-XM_032865914.1    rna-XM_046022898.1    rna-XM_032308408.1    rna-XM_059141376.1, rna-XM_059179687.1    rna-XM_059410782.1    rna-XM_004757271.3    mNeoVis.2.1.1.chr13_hap1.g00217.t01    rna-XM_021691727.1, rna-XM_021696152.2, rna-XM_044917307.1, rna-XM_044919017.1, rna-XM_044919771.1, rna-XM_044920942.1, rna-XM_044921349.1    rna-XM_004392635.1, rna-XM_004392869.1, rna-XM_004393192.1, rna-XM_004396172.1, rna-XM_004398448.1, rna-XM_004400456.1, rna-XM_004402308.1, rna-XM_004404765.1, rna-XM_004406167.1, rna-XM_004411610.1, rna-XM_004414882.1, rna-XM_004416045.1, rna-XM_004416926.1, rna-XM_004417490.1, rna-XM_004417565.1, rna-XM_004417609.1    rna-XM_042963334.1    rna-XM_008698738.2, rna-XM_040625570.1    rna-XM_027614325.2, rna-XM_035721974.1, rna-XM_035723071.1, rna-XM_035723116.1, rna-XM_035723210.1, rna-XM_035723702.1, rna-XM_035723754.1, rna-XM_035723904.1, rna-XM_035724039.1, rna-XM_035724752.1, rna-XM_035725262.1, rna-XM_035726176.1, rna-XM_035726426.1, rna-XM_035726736.1, rna-XM_035727400.1, rna-XM_035727502.1, rna-XM_035727601.1, rna-XM_035728523.1, rna-XM_035728588.1, rna-XM_035728618.1, rna-XM_035728924.1, rna-XM_035729028.1, rna-XM_035729031.1    mNeoVis.3.1.1.chr13.g00168.t01      mNeoVis.evergladensis.2.1.chr13.g00215.t01    mNeoVis.vulgivaga.2.1.chr13.g00212.t01      1529    N0.HOG0001527   OG0000182  n64   rna-XM_034651667.1, rna-XM_034668308.1    rna-XM_038574504.1, rna-XM_038576084.1      rna-XM_045035062.1, rna-XM_045053332.1    rna-XM_036065074.1, rna-XM_036114487.1, rna-XM_036116163.1    rna-XM_039216593.1        rna-XM_032337892.1          mNeoVis.2.1.1.chr10_hap1.g00468.t01    rna-XM_044918580.1, rna-XM_044920583.1    rna-XM_004405959.1    rna-XM_042976106.1    rna-XM_040621365.1, rna-XM_040631400.1, rna-XM_040638645.1    rna-XM_027616329.1, rna-XM_035727907.1    mNeoVis.3.1.1.chr9.g00434.t01    rna-gnl|WGS_ZZZZ|egapxtmp_015500-R1    mNeoVis.evergladensis.2.1.chr8.g00447.t01    mNeoVis.vulgivaga.2.1.chr9.g00451.t01      1530    N0.HOG0001528   OG0000183  n2   rna-XM_002927263.1    rna-NM_001388896.1    rna-XM_022491172.1    rna-XM_011291102.2      rna-XM_039246514.1    rna-XM_032845673.1    rna-XM_045983476.1    rna-XM_032319331.1    rna-XM_059146729.1    rna-XM_059414157.1    rna-XM_013047458.1    mNeoVis.2.1.1.chr14_hap1.g00559.t01    rna-XM_021681937.2      rna-XM_015538461.2    rna-XM_008700800.1    rna-XM_035722549.1    mNeoVis.3.1.1.chr14.g00551.t01    rna-gnl|WGS_ZZZZ|egapxtmp_033043-R1    mNeoVis.evergladensis.2.1.chr14.g00559.t01    mNeoVis.vulgivaga.2.1.chr14.g00557.t01      1531    N0.HOG0001529   OG0000183  n18   rna-XM_011234800.3    rna-XM_038585937.1, rna-XM_038589677.1    rna-XM_022491090.1    rna-XM_023249729.2    rna-XM_036069837.1    rna-XM_039246425.1    rna-XM_032845718.1    rna-XM_045983324.1    rna-XM_032317574.1    rna-XM_059145405.1    rna-XM_059413960.1    rna-XM_004775960.1    mNeoVis.2.1.1.chr14_hap1.g00556.t01    rna-XM_021681841.1      rna-XM_007084294.2    rna-XM_008700733.2      mNeoVis.3.1.1.chr14.g00547.t01    rna-gnl|WGS_ZZZZ|egapxtmp_033044-R1    mNeoVis.evergladensis.2.1.chr14.g00555.t01    mNeoVis.vulgivaga.2.1.chr14.g00553.t01      1532    N0.HOG0001530   OG0000183  n40     rna-XM_038585939.1      rna-XM_045048639.1    rna-XM_036069838.1    rna-XM_039246521.1                      rna-XM_042975725.1    rna-XM_040634572.1                1533    N0.HOG0001531   OG0000183  n48   rna-XM_034667677.1    rna-XM_038585938.1    rna-XM_022490737.1          rna-XM_032845716.1    rna-XM_045983320.1    rna-XM_032317576.1    rna-XM_059145407.1    rna-XM_059413962.1    rna-XM_004775958.2      rna-XM_044916271.1    rna-XM_004415687.1      rna-XM_040634476.1    rna-XM_027611952.1    mNeoVis.3.1.1.chr14.g00549.t01    rna-gnl|WGS_ZZZZ|egapxtmp_033052-R1    mNeoVis.evergladensis.2.1.chr14.g00557.t01    mNeoVis.vulgivaga.2.1.chr14.g00555.t01      1534    N0.HOG0001532   OG0000183  n63   rna-XM_034667676.1    rna-NM_001388891.1, rna-XM_038589676.1    rna-XM_022490738.1          rna-XM_032845717.1      rna-XM_032317575.1    rna-XM_059145406.1    rna-XM_059413961.1    rna-XM_004775959.1            rna-XM_008700797.1    rna-XM_027611951.2    mNeoVis.3.1.1.chr14.g00548.t01    rna-gnl|WGS_ZZZZ|egapxtmp_031951-R1    mNeoVis.evergladensis.2.1.chr14.g00556.t01    mNeoVis.vulgivaga.2.1.chr14.g00554.t01      1535    N0.HOG0001533   OG0000183  n77         rna-XM_045048395.1      rna-XM_039246426.1                      rna-XM_007084293.2                  1536    N0.HOG0001534   OG0000183  n79   rna-XM_002927264.2    rna-NM_001388894.1    rna-XM_022491159.1    rna-XM_006943103.3    rna-XM_036069839.1    rna-XM_039246519.1    rna-XM_032845715.1    rna-XM_045983455.1    rna-XM_032317577.1    rna-XM_059146419.1    rna-XM_059414156.1    rna-XM_004775957.2    mNeoVis.2.1.1.chr14_hap1.g00558.t01    rna-XM_021681936.1      rna-XM_007084291.2    rna-XM_008700799.2    rna-XM_027611953.2    mNeoVis.3.1.1.chr14.g00550.t01    rna-gnl|WGS_ZZZZ|egapxtmp_033047-R1    mNeoVis.evergladensis.2.1.chr14.g00558.t01    mNeoVis.vulgivaga.2.1.chr14.g00556.t01      1537    N0.HOG0001535   OG0000184  n1   rna-XM_002922190.4    rna-XM_038564702.1    rna-XM_022518272.1    rna-XM_004001095.6    rna-XM_036079415.1    rna-XM_039256266.1    rna-XM_032858019.1    rna-XM_045990544.1    rna-XM_032333379.1    rna-XM_059168910.1    rna-XM_059395502.1    rna-XM_004747937.3    mNeoVis.2.1.1.chr6_hap1.g00439.t01    rna-XM_021681468.1    rna-XM_004393607.1    rna-XM_007091812.2    rna-XM_008684396.2    rna-XM_027600690.2    mNeoVis.3.1.1.chr4.g00439.t01    rna-gnl|WGS_ZZZZ|egapxtmp_042998-R1    mNeoVis.evergladensis.2.1.chr4.g00446.t01    mNeoVis.vulgivaga.2.1.chr4.g00448.t01      1538    N0.HOG0001536   OG0000184  n21   rna-XM_019801319.2    rna-XM_038586713.1    rna-XM_022518364.1    rna-XM_023248888.2    rna-XM_036119275.1    rna-XM_039245327.1    rna-XM_032880253.1    rna-XM_045996139.1, rna-XM_045996729.1    rna-XM_032331740.1    rna-XM_059157330.1    rna-XM_059385880.1    rna-XM_045089248.1    mNeoVis.2.1.1.chr12_hap1.g00842.t01    rna-XM_044911639.1    rna-XM_004407840.1    rna-XM_042974595.1    rna-XM_040635105.1    rna-XM_027609054.2, rna-XM_027613785.2    mNeoVis.3.1.1.chr12.g00771.t01    rna-gnl|WGS_ZZZZ|egapxtmp_029464-R1    mNeoVis.evergladensis.2.1.chr12.g00770.t01    mNeoVis.vulgivaga.2.1.chr12.g00799.t01      1539    N0.HOG0001537   OG0000184  n36   rna-XM_034647604.1, rna-XM_034651687.1    rna-XM_038560400.1    rna-XM_022498689.1    rna-XM_023252323.2    rna-XM_036070611.1    rna-XM_039253865.1    rna-XM_032869900.1    rna-XM_045998457.1    rna-XM_032328883.1    rna-XM_059155852.1    rna-XM_059384138.1    rna-XM_004757734.3    mNeoVis.2.1.1.chr6_hap1.g01068.t01    rna-XM_021694230.2    rna-XM_004412733.1    rna-XM_042982980.1    rna-XM_040643391.1    rna-XM_027599077.2    mNeoVis.3.1.1.chr4.g01023.t01      mNeoVis.evergladensis.2.1.chr4.g01010.t01    mNeoVis.vulgivaga.2.1.chr4.g01037.t01      1540    N0.HOG0001538   OG0000184  n53       rna-XM_022518518.1          rna-XM_032882270.1    rna-XM_045996259.1                                  1541    N0.HOG0001539   OG0000184  n55                           mNeoVis.2.1.1.chr13_hap1.g00610.t01                rna-gnl|WGS_ZZZZ|egapxtmp_030191-R1      mNeoVis.vulgivaga.2.1.chr13.g00586.t01      1542    N0.HOG0001540   OG0000184  n60                   rna-XM_032330248.1    rna-XM_059156875.1    rna-XM_059386012.1    rna-XM_045068292.1    mNeoVis.2.1.1.chr12_hap1.g00249.t01                rna-gnl|WGS_ZZZZ|egapxtmp_028917-R1    mNeoVis.evergladensis.2.1.chr12.g00218.t01        1543    N0.HOG0001541   OG0000184  n63     rna-XM_038535556.1                          rna-XM_044912795.1          rna-XM_035726373.1    mNeoVis.3.1.1.chr13.g00534.t01        mNeoVis.vulgivaga.2.1.chr12.g00223.t01      1544    N0.HOG0001542   OG0000184  n67     rna-XM_038588176.1      rna-XM_011279441.3, rna-XM_045047343.1    rna-XM_036092565.1    rna-XM_039219823.1                  rna-XM_044911565.1, rna-XM_044913609.1    rna-XM_004405238.1, rna-XM_004416975.1    rna-XM_015539890.2, rna-XM_042975017.1    rna-XM_040628665.1    rna-XM_027609418.2              1545    N0.HOG0001543   OG0000185  n6     rna-XM_038557537.1                rna-XM_032353737.1      rna-XM_059409291.1    rna-XM_004742782.3, rna-XM_004748594.3              rna-XM_027589463.2, rna-XM_035724634.1        mNeoVis.evergladensis.2.1.chr11.g01104.t01        1546    N0.HOG0001544   OG0000185  n15     rna-XM_038534075.1, rna-XM_038570450.1            rna-XM_032860947.1          rna-XM_059404670.1      mNeoVis.2.1.1.chr6_hap1.g00672.t01    rna-XM_021686231.1            mNeoVis.3.1.1.chr4.g00681.t01    rna-gnl|WGS_ZZZZ|egapxtmp_027640-R1    mNeoVis.evergladensis.2.1.chr1.g00926.t01, mNeoVis.evergladensis.2.1.chr10.g00990.t01    mNeoVis.vulgivaga.2.1.chr4.g00693.t01      1547    N0.HOG0001545   OG0000185  n31     rna-XM_038548096.1                                  rna-XM_027616116.2              1548    N0.HOG0001546   OG0000185  n32     rna-XM_038568458.1                          rna-XM_021703485.1          rna-XM_027577038.1              1549    N0.HOG0001547   OG0000185  n34     rna-XM_038577895.1      rna-XM_045042721.1    rna-XM_036078433.1, rna-XM_036094585.1        rna-XM_045979581.1    rna-XM_032346850.1    rna-XM_059183638.1    rna-XM_059394966.1, rna-XM_059407878.1    rna-XM_013064760.2      rna-XM_021687099.1, rna-XM_021687269.2, rna-XM_021703619.1    rna-XM_004404771.1    rna-XM_042958052.1, rna-XM_042963494.1      rna-XM_035728362.1              1550    N0.HOG0001548   OG0000185  n52   rna-XM_034644424.1    rna-NM_001252131.2, rna-XM_038573760.1    rna-XM_022495277.1    rna-XM_023257300.2    rna-XM_036077454.1    rna-XM_039236607.1    rna-XM_032874218.1    rna-XM_046013190.1    rna-XM_032331340.1, rna-XM_032346780.1    rna-XM_059153023.1, rna-XM_059186517.1    rna-XM_059402355.1    rna-XM_004748550.3      rna-XM_021687520.1    rna-XM_004397471.1, rna-XM_004400282.1    rna-XM_042992889.1    rna-XM_040637316.1    rna-XM_035722137.1, rna-XM_035726302.1, rna-XM_035729226.1              1551    N0.HOG0001549   OG0000185  n64     rna-XM_038579698.1            rna-XM_032852331.1                        rna-XM_027616597.2              1552    N0.HOG0001550   OG0000185  n66     rna-XM_038550985.1, rna-XM_038582498.1        rna-XM_036079636.1      rna-XM_032865168.1      rna-XM_032352464.1          mNeoVis.2.1.1.chr9_hap1.g01003.t01    rna-XM_021695847.1          rna-XM_035728332.1      rna-gnl|WGS_ZZZZ|egapxtmp_024519-R2          1553    N0.HOG0001551   OG0000185  n74           rna-XM_036102179.1      rna-XM_032868016.1    rna-XM_045995521.1      rna-XM_059157195.1, rna-XM_059158364.1    rna-XM_059387289.1    rna-XM_013050793.2              rna-XM_027585708.2              1554    N0.HOG0001552   OG0000186  n1   rna-XM_034647369.1, rna-XM_034653821.1    rna-NM_001388571.1    rna-XM_022510415.1    rna-XM_045052874.1    rna-XM_036086593.1    rna-XM_039234072.1, rna-XM_039234073.1    rna-XM_032866448.1    rna-XM_046021436.1, rna-XM_046021505.1    rna-XM_032305835.1, rna-XM_032305956.1    rna-XM_059172682.1, rna-XM_059172683.1    rna-XM_059395591.1    rna-XM_004741608.1    mNeoVis.2.1.1.chr4_hap1.g00132.t01, mNeoVis.2.1.1.chr4_hap1.g00133.t01    rna-XM_021692878.1      rna-XM_007090826.1    rna-XM_040643913.1, rna-XM_040643914.1      mNeoVis.3.1.1.chr6.g00131.t01, mNeoVis.3.1.1.chr6.g00132.t01    rna-gnl|WGS_ZZZZ|egapxtmp_002782-R1, rna-gnl|WGS_ZZZZ|egapxtmp_002783-R1    mNeoVis.evergladensis.2.1.chr6.g00134.t01, mNeoVis.evergladensis.2.1.chr6.g00135.t01    mNeoVis.vulgivaga.2.1.chr6.g00132.t01, mNeoVis.vulgivaga.2.1.chr6.g00133.t01      1555    N0.HOG0001553   OG0000186  n32   rna-XM_011228038.3    rna-NM_001388858.1    rna-XM_022510257.1    rna-XM_045052873.1    rna-XM_036086594.1    rna-XM_039234075.1    rna-XM_032866422.1    rna-XM_046020240.1    rna-XM_032305833.1    rna-XM_059172555.1    rna-XM_059395985.1    rna-XM_004741984.3    mNeoVis.2.1.1.chr4_hap1.g00130.t01        rna-XM_007090825.2    rna-XM_040643912.1      mNeoVis.3.1.1.chr6.g00130.t01    rna-gnl|WGS_ZZZZ|egapxtmp_004367-R1    mNeoVis.evergladensis.2.1.chr6.g00132.t01    mNeoVis.vulgivaga.2.1.chr6.g00130.t01      1556    N0.HOG0001554   OG0000186  n50   rna-XM_002921671.2    rna-NM_001389135.1    rna-XM_022510256.1    rna-XM_003983116.5        rna-XM_032866434.1    rna-XM_046021493.1    rna-XM_032305834.1    rna-XM_059172684.1    rna-XM_059395590.1    rna-XM_004741985.1    mNeoVis.2.1.1.chr4_hap1.g00131.t01        rna-XM_042973914.1    rna-XM_040643910.1          mNeoVis.evergladensis.2.1.chr6.g00133.t01    mNeoVis.vulgivaga.2.1.chr6.g00131.t01      1557    N0.HOG0001555   OG0000186  n63   rna-XM_002924162.2    rna-NM_001389059.1      rna-XM_003983163.6, rna-XM_045052916.1, rna-XM_045052917.1    rna-XM_036095791.1    rna-XM_039243019.1, rna-XM_039243021.1, rna-XM_039243022.1    rna-XM_032865183.1    rna-XM_046021512.1, rna-XM_046021513.1, rna-XM_046021521.1    rna-XM_032305319.1, rna-XM_032305320.1    rna-XM_059172556.1, rna-XM_059172694.1    rna-XM_059395581.1    rna-XM_004781415.2, rna-XM_004781416.2    mNeoVis.2.1.1.chr4_hap1.g00112.t01, mNeoVis.2.1.1.chr4_hap1.g00113.t01      rna-XM_004414573.1, rna-XM_004414574.1    rna-XM_007090839.2, rna-XM_007090840.2, rna-XM_042977431.1    rna-XM_008697310.2, rna-XM_008697311.2    rna-XM_027573584.1, rna-XM_035723265.1    mNeoVis.3.1.1.chr6.g00112.t01    rna-gnl|WGS_ZZZZ|egapxtmp_003544-R1, rna-gnl|WGS_ZZZZ|egapxtmp_003545-R1    mNeoVis.evergladensis.2.1.chr6.g00114.t01, mNeoVis.evergladensis.2.1.chr6.g00115.t01    mNeoVis.vulgivaga.2.1.chr6.g00111.t01, mNeoVis.vulgivaga.2.1.chr6.g00112.t01      1558    N0.HOG0001556   OG0000187  n1     rna-XM_038553645.1    rna-XM_022492387.1    rna-XM_045051504.1, rna-XM_045051513.1, rna-XM_045051515.1, rna-XM_045051519.1, rna-XM_045057323.1, rna-XM_045061216.1    rna-XM_036064628.1, rna-XM_036114612.1    rna-XM_039219620.1        rna-XM_032335023.1    rna-XM_059176044.1      rna-XM_045069238.1    mNeoVis.2.1.1.chr1_hap1.g00992.t01        rna-XM_015542396.2, rna-XM_042986047.1    rna-XM_040622728.1, rna-XM_040622729.1    rna-XM_027620381.1, rna-XM_035727775.1    mNeoVis.3.1.1.chr1.g00997.t01    rna-gnl|WGS_ZZZZ|egapxtmp_020782-R1    mNeoVis.evergladensis.2.1.chr1.g00986.t01    mNeoVis.vulgivaga.2.1.chr1.g00999.t01      1559    N0.HOG0001557   OG0000187  n25   rna-XM_034653409.1, rna-XM_034660501.1    rna-XM_038553314.1    rna-XM_022492662.1, rna-XM_022492787.1, rna-XM_022510779.1, rna-XM_022510925.1, rna-XM_022510938.1, rna-XM_022510940.1      rna-XM_036108172.1, rna-XM_036108174.1      rna-XM_032849685.1, rna-XM_032849692.1, rna-XM_032849693.1    rna-XM_046004418.1, rna-XM_046004420.1, rna-XM_046004426.1    rna-XM_032340746.1, rna-XM_032340747.1, rna-XM_032340748.1, rna-XM_032340750.1, rna-XM_032340751.1, rna-XM_032341368.1    rna-XM_059178766.1, rna-XM_059178769.1, rna-XM_059178778.1, rna-XM_059178783.1, rna-XM_059178784.1    rna-XM_059387250.1, rna-XM_059398991.1, rna-XM_059399951.1, rna-XM_059399952.1, rna-XM_059401525.1, rna-XM_059401526.1    rna-XM_013050877.2, rna-XM_013050892.2, rna-XM_045065116.1, rna-XM_045065119.1, rna-XM_045065121.1, rna-XM_045074645.1, rna-XM_045084933.1    mNeoVis.2.1.1.chr1_hap1.g01385.t01, mNeoVis.2.1.1.chr1_hap1.g01387.t01, mNeoVis.2.1.1.chr1_hap1.g01388.t01, mNeoVis.2.1.1.chr1_hap1.g01389.t01, mNeoVis.2.1.1.chr1_hap1.g01390.t01, mNeoVis.2.1.1.chr1_hap1.g01391.t01    rna-XM_021697208.1, rna-XM_044917230.1    rna-XM_004417272.1, rna-XM_004417382.1, rna-XM_004417383.1      rna-XM_008711842.2, rna-XM_040644615.1, rna-XM_040644616.1, rna-XM_040644617.1, rna-XM_040644791.1    rna-XM_035728589.1    mNeoVis.3.1.1.chr1.g01332.t01, mNeoVis.3.1.1.chr1.g01334.t01, mNeoVis.3.1.1.chr1.g01335.t01, mNeoVis.3.1.1.chr1.g01336.t01, mNeoVis.3.1.1.chr1.g01337.t01    rna-gnl|WGS_ZZZZ|egapxtmp_021698-R1, rna-gnl|WGS_ZZZZ|egapxtmp_021707-R1, rna-gnl|WGS_ZZZZ|egapxtmp_026572-R1, rna-gnl|WGS_ZZZZ|egapxtmp_026610-R1, rna-gnl|WGS_ZZZZ|egapxtmp_026643-R1    mNeoVis.evergladensis.2.1.chr1.g01350.t01, mNeoVis.evergladensis.2.1.chr1.g01352.t01, mNeoVis.evergladensis.2.1.chr1.g01353.t01, mNeoVis.evergladensis.2.1.chr1.g01354.t01, mNeoVis.evergladensis.2.1.chr1.g01355.t01    mNeoVis.vulgivaga.2.1.chr1.g01352.t01, mNeoVis.vulgivaga.2.1.chr1.g01354.t01, mNeoVis.vulgivaga.2.1.chr1.g01355.t01, mNeoVis.vulgivaga.2.1.chr1.g01356.t01      1560    N0.HOG0001558   OG0000188  n3   rna-XM_034651877.1    rna-XM_038586566.1      rna-XM_011279388.4    rna-XM_036069030.1    rna-XM_039229641.1, rna-XM_039256076.1    rna-XM_032866014.1    rna-XM_046021963.1    rna-XM_032307631.1    rna-XM_059141670.1    rna-XM_059410621.1    rna-XM_045082549.1    mNeoVis.2.1.1.chr11_hap1.g00705.t01, mNeoVis.2.1.1.chr13_hap1.g00181.t01    rna-XM_044911654.1    rna-XM_004393559.1    rna-XM_042963791.1, rna-XM_042974816.1      rna-XM_027615413.2, rna-XM_027621914.2, rna-XM_035725713.1, rna-XM_035728587.1    mNeoVis.3.1.1.chr11.g00667.t01, mNeoVis.3.1.1.chr13.g00132.t01        mNeoVis.vulgivaga.2.1.chr11.g00705.t01      1561    N0.HOG0001559   OG0000188  n18   rna-XM_011219729.3    rna-XM_038549156.1    rna-XM_022492685.1                                rna-XM_035728334.1              1562    N0.HOG0001560   OG0000188  n21       rna-XM_022526430.1              rna-XM_032359827.1    rna-XM_059147429.1, rna-XM_059158776.1    rna-XM_059378588.1, rna-XM_059386848.1, rna-XM_059387046.1, rna-XM_059387587.1    rna-XM_004760523.3, rna-XM_013050569.2                mNeoVis.3.1.1.chr12.g00646.t01    rna-gnl|WGS_ZZZZ|egapxtmp_031092-R1          1563    N0.HOG0001561   OG0000188  n32   rna-XM_034652327.1    rna-NM_001003031.2    rna-XM_022492683.1, rna-XM_022498087.1, rna-XM_022509856.1, rna-XM_022522386.1, rna-XM_022524567.1    rna-XM_011283669.4, rna-XM_045042812.1    rna-XM_036099196.1      rna-XM_032839972.1, rna-XM_032848663.1, rna-XM_032850145.1, rna-XM_032854058.1, rna-XM_032857984.1, rna-XM_032861924.1, rna-XM_032868082.1, rna-XM_032871852.1    rna-XM_046000428.1, rna-XM_046009217.1, rna-XM_046010897.1    rna-XM_032314856.1, rna-XM_032316003.1, rna-XM_032322053.1    rna-XM_059142740.1, rna-XM_059156800.1, rna-XM_059162439.1, rna-XM_059163184.1    rna-XM_059379073.1, rna-XM_059387958.1, rna-XM_059388032.1, rna-XM_059392842.1    rna-XM_013057175.2, rna-XM_045070054.1, rna-XM_045070339.1, rna-XM_045088719.1    mNeoVis.2.1.1.chr1_hap1.g01312.t01, mNeoVis.2.1.1.chr1_hap1.g01608.t01, mNeoVis.2.1.1.chr3_hap1.g00801.t01, mNeoVis.2.1.1.chr3_hap1.g01272.t01, mNeoVis.2.1.1.chr5_hap1.g00753.t01, mNeoVis.2.1.1.chr5_hap1.g00829.t01    rna-XM_021693541.1, rna-XM_021694606.2, rna-XM_044920758.1, rna-XM_044920835.1    rna-XM_004392507.1      rna-XM_008686838.2, rna-XM_040632004.1    rna-XM_027582681.2    mNeoVis.3.1.1.chr2.g00740.t01    rna-gnl|WGS_ZZZZ|egapxtmp_035462-R1    mNeoVis.evergladensis.2.1.chr13.g00179.t01, mNeoVis.evergladensis.2.1.chr2.g00829.t01, mNeoVis.evergladensis.2.1.chr7.g01442.t01    mNeoVis.vulgivaga.2.1.chr1.g01282.t01, mNeoVis.vulgivaga.2.1.chr11.g00556.t01, mNeoVis.vulgivaga.2.1.chr13.g00176.t01, mNeoVis.vulgivaga.2.1.chr2.g00759.t01, mNeoVis.vulgivaga.2.1.chr2.g00833.t01, mNeoVis.vulgivaga.2.1.chr7.g01456.t01      1564    N0.HOG0001562   OG0000189  n1   rna-XM_034665613.1    rna-NM_001388833.1    rna-XM_022522187.1    rna-XM_003992807.2        rna-XM_032853620.1    rna-XM_046017540.1    rna-XM_032356602.1    rna-XM_059157097.1    rna-XM_059390311.1    rna-XM_004768266.1    mNeoVis.2.1.1.chr7_hap1.g01650.t01    rna-XM_021685675.1    rna-XM_004407577.1    rna-XM_007089388.1    rna-XM_008710767.2    rna-XM_027581044.2    mNeoVis.3.1.1.chr7.g01553.t01    rna-gnl|WGS_ZZZZ|egapxtmp_012873-R1    mNeoVis.evergladensis.2.1.chr7.g01618.t01    mNeoVis.vulgivaga.2.1.chr7.g01642.t01      1565    N0.HOG0001563   OG0000189  n20   rna-XM_034665585.1    rna-NM_001389123.1    rna-XM_022521711.1    rna-XM_023239716.2    rna-XM_036081424.1    rna-XM_039224664.1    rna-XM_032852700.1    rna-XM_046017518.1    rna-XM_032359771.1    rna-XM_059181417.1    rna-XM_059390175.1    rna-XM_013049233.2    mNeoVis.2.1.1.chr7_hap1.g01667.t01    rna-XM_021685628.2      rna-XM_007092245.1        mNeoVis.3.1.1.chr7.g01562.t01    rna-gnl|WGS_ZZZZ|egapxtmp_012827-R1    mNeoVis.evergladensis.2.1.chr7.g01638.t01    mNeoVis.vulgivaga.2.1.chr7.g01655.t01      1566    N0.HOG0001564   OG0000189  n39     rna-XM_038569358.1, rna-XM_038569359.1    rna-XM_022522165.1        rna-XM_039225649.1, rna-XM_039225836.1    rna-XM_032852696.1    rna-XM_046015905.1                    rna-XM_008710272.2                1567    N0.HOG0001565   OG0000189  n46   rna-XM_002931089.2, rna-XM_034650087.1    rna-NM_001388696.1      rna-XM_011286656.3, rna-XM_023239714.2      rna-XM_039225652.1    rna-XM_032852698.1    rna-XM_046016720.1    rna-XM_032359765.1, rna-XM_032359766.1    rna-XM_059157151.1, rna-XM_059157163.1, rna-XM_059157176.1    rna-XM_059390267.1    rna-XM_045071505.1, rna-XM_045071527.1    mNeoVis.2.1.1.chr7_hap1.g01653.t01, mNeoVis.2.1.1.chr7_hap1.g01654.t01        rna-XM_007089339.3, rna-XM_007089379.3    rna-XM_008710271.1    rna-XM_027577908.1      rna-gnl|WGS_ZZZZ|egapxtmp_012881-R2    mNeoVis.evergladensis.2.1.chr7.g01629.t01        1568    N0.HOG0001566   OG0000189  n69   rna-XM_034650110.1    rna-NM_001388836.1    rna-XM_022521714.1    rna-XM_003992850.4, rna-XM_045038117.1      rna-XM_039225730.1, rna-XM_039225802.1    rna-XM_032852689.1, rna-XM_032852694.1    rna-XM_046015901.1, rna-XM_046015904.1    rna-XM_032359757.1, rna-XM_032359762.1    rna-XM_059179727.1, rna-XM_059180284.1    rna-XM_059395563.1, rna-XM_059395969.1    rna-XM_004768262.1, rna-XM_004768267.1    mNeoVis.2.1.1.chr7_hap1.g01647.t01    rna-XM_021685625.1    rna-XM_004407575.1    rna-XM_007089387.2, rna-XM_007089392.3    rna-XM_008710763.2, rna-XM_008710768.2      mNeoVis.3.1.1.chr7.g01548.t01    rna-gnl|WGS_ZZZZ|egapxtmp_012517-R1, rna-gnl|WGS_ZZZZ|egapxtmp_012521-R1    mNeoVis.evergladensis.2.1.chr7.g01621.t01, mNeoVis.evergladensis.2.1.chr7.g01625.t01        1569    N0.HOG0001567   OG0000190  n2     rna-XM_038586664.1, rna-XM_038589326.1    rna-XM_022491408.1    rna-XM_023249158.2, rna-XM_045051103.1    rna-XM_036094206.1, rna-XM_036110898.1    rna-XM_039224788.1    rna-XM_032846385.1, rna-XM_032846390.1    rna-XM_045994590.1, rna-XM_045994591.1, rna-XM_045995473.1    rna-XM_032330920.1, rna-XM_032331613.1    rna-XM_059158012.1, rna-XM_059158278.1    rna-XM_059384711.1, rna-XM_059385524.1    rna-XM_045068136.1, rna-XM_045068143.1    mNeoVis.2.1.1.chr12_hap1.g00020.t01, mNeoVis.2.1.1.chr12_hap1.g00033.t01    rna-XM_021692601.2, rna-XM_044917934.1    rna-XM_004414523.1    rna-XM_042974493.1, rna-XM_042975083.1, rna-XM_042975126.1    rna-XM_040639339.1    rna-XM_027607849.1, rna-XM_035728658.1    mNeoVis.3.1.1.chr12.g00024.t01        mNeoVis.vulgivaga.2.1.chr12.g00017.t01      1570    N0.HOG0001568   OG0000190  n36     rna-XM_038586667.1, rna-XM_038589357.1    rna-XM_022491289.1    rna-XM_045050403.1, rna-XM_045050405.1, rna-XM_045050406.1, rna-XM_045050407.1, rna-XM_045050408.1, rna-XM_045051687.1, rna-XM_045051688.1    rna-XM_036077757.1, rna-XM_036077762.1    rna-XM_039225833.1        rna-XM_032329748.1, rna-XM_032329749.1    rna-XM_059156860.1    rna-XM_059385910.1    rna-XM_004782552.2    mNeoVis.2.1.1.chr12_hap1.g00019.t01, mNeoVis.2.1.1.chr12_hap1.g00021.t01      rna-XM_004400286.1    rna-XM_042974718.1, rna-XM_042975080.1, rna-XM_042975108.1, rna-XM_042975123.1, rna-XM_042975128.1      rna-XM_027594799.1, rna-XM_027607846.1, rna-XM_035725763.1              1571    N0.HOG0001569   OG0000190  n59     rna-XM_038586666.1    rna-XM_022491265.1    rna-XM_045050886.1, rna-XM_045050888.1, rna-XM_045050893.1, rna-XM_045050898.1, rna-XM_045050903.1, rna-XM_045050908.1, rna-XM_045051058.1, rna-XM_045051078.1, rna-XM_045051082.1, rna-XM_045051691.1, rna-XM_045051692.1, rna-XM_045051695.1, rna-XM_045051696.1      rna-XM_039225807.1    rna-XM_032846415.1    rna-XM_045994961.1    rna-XM_032329468.1    rna-XM_059158643.1      rna-XM_004779310.3    mNeoVis.2.1.1.chr12_hap1.g00022.t01        rna-XM_007096639.2, rna-XM_042974478.1, rna-XM_042974483.1, rna-XM_042974509.1, rna-XM_042974510.1, rna-XM_042974721.1, rna-XM_042974728.1, rna-XM_042975111.1      rna-XM_027607852.1, rna-XM_035725840.1, rna-XM_035725841.1, rna-XM_035725842.1, rna-XM_035725843.1, rna-XM_035725844.1      rna-gnl|WGS_ZZZZ|egapxtmp_028311-R1    mNeoVis.evergladensis.2.1.chr12.g00016.t01        1572    N0.HOG0001570   OG0000191  n2   rna-XM_034670575.1    rna-XM_038540710.1    rna-XM_022517060.1    rna-XM_045047651.1    rna-XM_036083169.1    rna-XM_039246764.1    rna-XM_032846064.1      rna-XM_032327069.1    rna-XM_059155523.1    rna-XM_059415897.1    rna-XM_045071938.1    mNeoVis.2.1.1.chr15_hap1.g00346.t01    rna-XM_044915254.1      rna-XM_042972322.1    rna-XM_040634450.1      mNeoVis.3.1.1.chr15.g00312.t01    rna-gnl|WGS_ZZZZ|egapxtmp_033992-R1    mNeoVis.evergladensis.2.1.chr15.g00324.t01    mNeoVis.vulgivaga.2.1.chr15.g00331.t01      1573    N0.HOG0001571   OG0000191  n20   rna-XM_034670576.1    rna-XM_038540712.1    rna-XM_022516568.1    rna-XM_023247118.2    rna-XM_036083212.1    rna-XM_039246814.1    rna-XM_032846070.1    rna-XM_045992578.1    rna-XM_032327079.1    rna-XM_059153880.1    rna-XM_059415624.1    rna-XM_045072220.1    mNeoVis.2.1.1.chr15_hap1.g00347.t01, mNeoVis.2.1.1.chr15_hap1.g00384.t01, mNeoVis.2.1.1.chr15_hap1.g00421.t01    rna-XM_021697998.2    rna-XM_004403173.1    rna-XM_042972327.1    rna-XM_040635498.1    rna-XM_027610262.2      rna-gnl|WGS_ZZZZ|egapxtmp_033375-R2          1574    N0.HOG0001572   OG0000191  n41   rna-XM_034669564.1    rna-XM_038540129.1    rna-XM_022501838.1      rna-XM_036113189.1    rna-XM_039234412.1    rna-XM_032850808.1, rna-XM_032872325.1    rna-XM_045992947.1    rna-XM_032328502.1    rna-XM_059155049.1    rna-XM_059416057.1    rna-XM_045072708.1    mNeoVis.2.1.1.chr15_hap1.g00165.t01    rna-XM_044915268.1    rna-XM_004398832.1      rna-XM_008697189.2    rna-XM_035722222.1    mNeoVis.3.1.1.chr15.g00141.t01    rna-gnl|WGS_ZZZZ|egapxtmp_033816-R1    mNeoVis.evergladensis.2.1.chr15.g00148.t01    mNeoVis.vulgivaga.2.1.chr15.g00151.t01      1575    N0.HOG0001573   OG0000191  n62         rna-XM_045047740.1      rna-XM_039234460.1                      rna-XM_042971603.1                  1576    N0.HOG0001574   OG0000191  n65   rna-XM_034669566.1    rna-XM_038540113.1    rna-XM_022501616.1      rna-XM_036113190.1      rna-XM_032844014.1    rna-XM_045992948.1    rna-XM_032328561.1    rna-XM_059155338.1    rna-XM_059416059.1    rna-XM_045072686.1    mNeoVis.2.1.1.chr15_hap1.g00167.t01    rna-XM_044915266.1    rna-XM_004415763.1      rna-XM_040636564.1    rna-XM_027612191.2    mNeoVis.3.1.1.chr15.g00143.t01    rna-gnl|WGS_ZZZZ|egapxtmp_033817-R1    mNeoVis.evergladensis.2.1.chr15.g00150.t01    mNeoVis.vulgivaga.2.1.chr15.g00153.t01      1577    N0.HOG0001575   OG0000191  n83   rna-XM_034669565.1    rna-XM_038540101.1    rna-XM_022501678.1      rna-XM_036113188.1      rna-XM_032843986.1    rna-XM_045993739.1    rna-XM_032328554.1    rna-XM_059155003.1    rna-XM_059416058.1    rna-XM_004780131.2    mNeoVis.2.1.1.chr15_hap1.g00166.t01    rna-XM_044915267.1        rna-XM_040636563.1    rna-XM_035722218.1, rna-XM_035722351.1    mNeoVis.3.1.1.chr15.g00142.t01    rna-gnl|WGS_ZZZZ|egapxtmp_033950-R1    mNeoVis.evergladensis.2.1.chr15.g00149.t01    mNeoVis.vulgivaga.2.1.chr15.g00152.t01      1578    N0.HOG0001576   OG0000192  n2   rna-XM_011234660.3      rna-XM_022490573.1    rna-XM_006931364.3    rna-XM_036074502.1    rna-XM_039223553.1    rna-XM_032861776.1    rna-XM_046005950.1    rna-XM_032338290.1    rna-XM_059176291.1    rna-XM_059399690.1      mNeoVis.2.1.1.chr1_hap1.g01071.t01    rna-XM_021697181.2      rna-XM_007074481.2    rna-XM_008693027.2      mNeoVis.3.1.1.chr1.g01060.t01        mNeoVis.vulgivaga.2.1.chr1.g01075.t01      1579    N0.HOG0001577   OG0000192  n17   rna-XM_002927095.2    rna-XM_038583966.1    rna-XM_022527116.1        rna-XM_039223549.1    rna-XM_032861775.1    rna-XM_046005965.1    rna-XM_032341393.1    rna-XM_059176222.1    rna-XM_059399671.1      mNeoVis.2.1.1.chr1_hap1.g01074.t01          rna-XM_008693100.1      mNeoVis.3.1.1.chr1.g01063.t01      mNeoVis.evergladensis.2.1.chr1.g01059.t01    mNeoVis.vulgivaga.2.1.chr1.g01078.t01      1580    N0.HOG0001578   OG0000192  n29   rna-XM_011217611.1    rna-XM_038583967.1    rna-XM_022492241.1    rna-XM_003980608.3, rna-XM_003980609.2, rna-XM_003985755.2      rna-XM_039218786.1, rna-XM_039223535.1      rna-XM_046001114.1, rna-XM_046006205.1    rna-XM_032335040.1, rna-XM_032338230.1    rna-XM_059173399.1, rna-XM_059173403.1, rna-XM_059176231.1    rna-XM_059399675.1, rna-XM_059416420.1, rna-XM_059416760.1    rna-XM_004781505.1    mNeoVis.2.1.1.chr1_hap1.g01004.t01, mNeoVis.2.1.1.chr1_hap1.g01072.t01        rna-XM_007074480.2, rna-XM_007084989.2, rna-XM_015538813.2    rna-XM_008693026.1, rna-XM_008712037.2      mNeoVis.3.1.1.chr1.g01004.t01, mNeoVis.3.1.1.chr1.g01061.t01    rna-gnl|WGS_ZZZZ|egapxtmp_014494-R1, rna-gnl|WGS_ZZZZ|egapxtmp_020935-R1    mNeoVis.evergladensis.2.1.chr1.g00996.t01, mNeoVis.evergladensis.2.1.chr1.g01057.t01    mNeoVis.vulgivaga.2.1.chr1.g01010.t01, mNeoVis.vulgivaga.2.1.chr1.g01076.t01      1581    N0.HOG0001579   OG0000192  n58   rna-XM_002926903.2, rna-XM_011234384.2    rna-NM_001388984.1    rna-XM_022491660.1, rna-XM_022491679.1    rna-XM_003980573.2, rna-XM_003980593.3    rna-XM_036081038.1    rna-XM_039225179.1    rna-XM_032848484.1    rna-XM_045999699.1, rna-XM_046000637.1, rna-XM_046000820.1    rna-XM_032334913.1, rna-XM_032335010.1    rna-XM_059173423.1, rna-XM_059176070.1    rna-XM_059416438.1, rna-XM_059416743.1    rna-XM_004782009.2, rna-XM_004782326.1    mNeoVis.2.1.1.chr1_hap1.g00972.t01, mNeoVis.2.1.1.chr1_hap1.g00973.t01    rna-XM_021690931.1    rna-XM_004406161.1    rna-XM_007092945.1, rna-XM_007092975.2    rna-XM_008709907.2, rna-XM_040622683.1    rna-XM_027606535.1    mNeoVis.3.1.1.chr1.g00980.t01    rna-gnl|WGS_ZZZZ|egapxtmp_018917-R1    mNeoVis.evergladensis.2.1.chr1.g00967.t01, mNeoVis.evergladensis.2.1.chr1.g00968.t01    mNeoVis.vulgivaga.2.1.chr1.g00980.t01, mNeoVis.vulgivaga.2.1.chr1.g00981.t01      1582    N0.HOG0001580   OG0000193  n1   rna-XM_002920906.4    rna-XM_038529385.1    rna-XM_022525377.1    rna-XM_045047117.1    rna-XM_036108520.1      rna-XM_032841933.1    rna-XM_045989322.1    rna-XM_032324285.1    rna-XM_059152864.1    rna-XM_059381617.1    rna-XM_004765020.3    mNeoVis.2.1.1.chr7_hap1.g00408.t01      rna-XM_004417118.1    rna-XM_007096750.3    rna-XM_008712074.2    rna-XM_035724893.1    mNeoVis.3.1.1.chr7.g00393.t01    rna-gnl|WGS_ZZZZ|egapxtmp_012210-R1    mNeoVis.evergladensis.2.1.chr7.g00411.t01    mNeoVis.vulgivaga.2.1.chr7.g00411.t01      1583    N0.HOG0001581   OG0000193  n20   rna-XM_011231661.3, rna-XM_034639497.1, rna-XM_034639527.1, rna-XM_034639534.1    rna-XM_038527584.1    rna-XM_022492069.1, rna-XM_022492085.1, rna-XM_022492607.1    rna-XM_006940780.5, rna-XM_019819067.3, rna-XM_045046741.1, rna-XM_045046751.1    rna-XM_036091887.1, rna-XM_036091894.1, rna-XM_036091900.1, rna-XM_036091934.1    rna-XM_039219010.1, rna-XM_039219037.1, rna-XM_039219048.1    rna-XM_032847637.1, rna-XM_032847699.1, rna-XM_032847715.1, rna-XM_032851077.1, rna-XM_032851140.1, rna-XM_032851152.1, rna-XM_032851153.1, rna-XM_032851154.1, rna-XM_032851155.1, rna-XM_032851159.1, rna-XM_032851205.1    rna-XM_045987760.1, rna-XM_045987762.1, rna-XM_045989336.1, rna-XM_045989585.1    rna-XM_032324634.1, rna-XM_032324641.1, rna-XM_032324660.1, rna-XM_032324809.1    rna-XM_059152308.1, rna-XM_059152342.1, rna-XM_059152428.1, rna-XM_059152472.1    rna-XM_059381970.1, rna-XM_059383171.1, rna-XM_059383183.1, rna-XM_059383192.1    rna-XM_004776867.3, rna-XM_004776895.3, rna-XM_013048059.2, rna-XM_013048064.2    mNeoVis.2.1.1.chr7_hap1.g00407.t01, mNeoVis.2.1.1.chr7_hap1.g00647.t01, mNeoVis.2.1.1.chr7_hap1.g00648.t01, mNeoVis.2.1.1.chr7_hap1.g00650.t01    rna-XM_021682272.1, rna-XM_021682338.2, rna-XM_044911296.1    rna-XM_004410912.1    rna-XM_007089533.3, rna-XM_042969525.1, rna-XM_042970645.1, rna-XM_042970663.1    rna-XM_008706538.2, rna-XM_008706539.2, rna-XM_040633738.1, rna-XM_040633782.1    rna-XM_027618138.2, rna-XM_027620321.2, rna-XM_035724954.1, rna-XM_035724963.1    mNeoVis.3.1.1.chr7.g00609.t01, mNeoVis.3.1.1.chr7.g00614.t01    rna-gnl|WGS_ZZZZ|egapxtmp_011651-R1, rna-gnl|WGS_ZZZZ|egapxtmp_012980-R1, rna-gnl|WGS_ZZZZ|egapxtmp_012983-R1    mNeoVis.evergladensis.2.1.chr7.g00644.t01, mNeoVis.evergladensis.2.1.chr7.g00645.t01, mNeoVis.evergladensis.2.1.chr7.g00647.t01    mNeoVis.vulgivaga.2.1.chr7.g00646.t01, mNeoVis.vulgivaga.2.1.chr7.g00647.t01      1584    N0.HOG0001582   OG0000194  n1   rna-XM_034640149.1    rna-XM_038547075.1    rna-XM_022519951.1    rna-XM_019818001.2    rna-XM_036116816.1    rna-XM_039220098.1    rna-XM_032871013.1    rna-XM_045986526.1    rna-XM_032320606.1    rna-XM_059149067.1    rna-XM_059378719.1    rna-XM_004772496.3    mNeoVis.2.1.1.chr8_hap1.g00306.t01    rna-XM_044921875.1    rna-XM_004407241.1    rna-XM_042967337.1    rna-XM_040639864.1    rna-XM_027567785.2    mNeoVis.3.1.1.chr8.g00275.t01    rna-gnl|WGS_ZZZZ|egapxtmp_019424-R1    mNeoVis.evergladensis.2.1.chr9.g00305.t01    mNeoVis.vulgivaga.2.1.chr8.g00312.t01      1585    N0.HOG0001583   OG0000194  n21   rna-XM_002918630.4, rna-XM_034651823.1    rna-NM_001287023.1, rna-XM_038573316.1    rna-XM_022515654.1    rna-XM_003988131.4    rna-XM_036097060.1    rna-XM_039245877.1    rna-XM_032880898.1    rna-XM_045994776.1, rna-XM_046003669.1, rna-XM_046013582.1    rna-XM_032349515.1    rna-XM_059184032.1    rna-XM_059404239.1    rna-XM_004757574.3    mNeoVis.2.1.1.chr6_hap1.g00277.t01, mNeoVis.2.1.1.chr9_hap1.g00116.t01    rna-XM_021696765.1    rna-XM_004392950.1, rna-XM_004409536.1    rna-XM_042991170.1    rna-XM_008692083.2    rna-XM_027593295.1, rna-XM_035721868.1    mNeoVis.3.1.1.chr10.g00113.t01, mNeoVis.3.1.1.chr4.g00274.t01    rna-gnl|WGS_ZZZZ|egapxtmp_000707-R1, rna-gnl|WGS_ZZZZ|egapxtmp_023208-R1    mNeoVis.evergladensis.2.1.chr10.g00116.t01, mNeoVis.evergladensis.2.1.chr4.g00284.t01    mNeoVis.vulgivaga.2.1.chr10.g00113.t01, mNeoVis.vulgivaga.2.1.chr4.g00283.t01      1586    N0.HOG0001584   OG0000194  n53   rna-XM_011231793.3    rna-NM_001012394.1    rna-XM_022512645.1    rna-XM_003991178.5    rna-XM_036082965.1    rna-XM_039216698.1    rna-XM_032857284.1    rna-XM_046019645.1    rna-XM_032354894.1    rna-XM_059167963.1    rna-XM_059393540.1    rna-XM_004762851.3    mNeoVis.2.1.1.chr3_hap1.g01158.t01    rna-XM_021702795.2    rna-XM_004402753.1    rna-XM_007086222.2    rna-XM_008690580.2    rna-XM_027589123.2    mNeoVis.3.1.1.chr3.g01144.t01      mNeoVis.evergladensis.2.1.chr3.g01158.t01    mNeoVis.vulgivaga.2.1.chr3.g01173.t01      1587    N0.HOG0001585   OG0000194  n69   rna-XM_011228858.3    rna-XM_038577548.1    rna-XM_022519656.1    rna-XM_011283902.4    rna-XM_036121118.1    rna-XM_039241388.1    rna-XM_032876872.1    rna-XM_046010609.1    rna-XM_032348096.1    rna-XM_059185458.1    rna-XM_059404840.1    rna-XM_013049139.2    mNeoVis.2.1.1.chr9_hap1.g00586.t01    rna-XM_044914855.1    rna-XM_004400857.1    rna-XM_042992208.1    rna-XM_040642406.1    rna-XM_027593644.2    mNeoVis.3.1.1.chr10.g00556.t01      mNeoVis.evergladensis.2.1.chr10.g00579.t01    mNeoVis.vulgivaga.2.1.chr10.g00567.t01      1588    N0.HOG0001586   OG0000195  n0   rna-XM_002926094.4, rna-XM_002928571.2, rna-XM_002928582.1, rna-XM_034650092.1, rna-XM_034653229.1, rna-XM_034667604.1    rna-XM_038569345.1, rna-XM_038569346.1    rna-XM_022521681.1, rna-XM_022521703.1, rna-XM_022521733.1, rna-XM_022522162.1, rna-XM_022522193.1    rna-XM_003992821.2, rna-XM_019813037.2, rna-XM_023238989.2, rna-XM_045038091.1    rna-XM_036074408.1    rna-XM_039223764.1, rna-XM_039252056.1, rna-XM_039252057.1, rna-XM_039252064.1    rna-XM_032852709.1, rna-XM_032853576.1, rna-XM_032853577.1, rna-XM_032853578.1, rna-XM_032853579.1, rna-XM_032853580.1    rna-XM_046014262.1, rna-XM_046015891.1, rna-XM_046016033.1, rna-XM_046016034.1, rna-XM_046017580.1, rna-XM_046017916.1, rna-XM_046017935.1    rna-XM_032356475.1, rna-XM_032356546.1, rna-XM_032359721.1, rna-XM_032359722.1, rna-XM_032360390.1, rna-XM_032360512.1    rna-XM_059134489.1, rna-XM_059156587.1, rna-XM_059156600.1, rna-XM_059176833.1, rna-XM_059177078.1, rna-XM_059177204.1, rna-XM_059177323.1    rna-XM_059391038.1, rna-XM_059391049.1, rna-XM_059394040.1, rna-XM_059398468.1, rna-XM_059398603.1, rna-XM_059398725.1, rna-XM_059398924.1    rna-XM_004768194.2, rna-XM_004768223.2, rna-XM_004768224.2, rna-XM_004768225.1, rna-XM_004768226.1, rna-XM_004779053.2, rna-XM_013064261.2    mNeoVis.2.1.1.chr7_hap1.g01581.t01, mNeoVis.2.1.1.chr7_hap1.g01582.t01, mNeoVis.2.1.1.chr7_hap1.g01583.t01, mNeoVis.2.1.1.chr7_hap1.g01584.t01, mNeoVis.2.1.1.chr7_hap1.g01585.t01    rna-XM_021704696.1      rna-XM_007089360.1, rna-XM_007089361.2, rna-XM_007089421.1, rna-XM_015542087.1    rna-XM_008707263.1, rna-XM_008707400.1, rna-XM_008707401.1, rna-XM_008707402.1, rna-XM_008707403.2, rna-XM_008707405.1      mNeoVis.3.1.1.chr7.g01492.t01, mNeoVis.3.1.1.chr7.g01493.t01, mNeoVis.3.1.1.chr7.g01494.t01, mNeoVis.3.1.1.chr7.g01495.t01, mNeoVis.3.1.1.chr7.g01586.t01    rna-gnl|WGS_ZZZZ|egapxtmp_012502-R1, rna-gnl|WGS_ZZZZ|egapxtmp_012823-R1, rna-gnl|WGS_ZZZZ|egapxtmp_012824-R1, rna-gnl|WGS_ZZZZ|egapxtmp_012828-R1, rna-gnl|WGS_ZZZZ|egapxtmp_012890-R1    mNeoVis.evergladensis.2.1.chr7.g01561.t01, mNeoVis.evergladensis.2.1.chr7.g01562.t01, mNeoVis.evergladensis.2.1.chr7.g01563.t01, mNeoVis.evergladensis.2.1.chr7.g01564.t01, mNeoVis.evergladensis.2.1.chr7.g01668.t01    mNeoVis.vulgivaga.2.1.chr7.g01579.t01, mNeoVis.vulgivaga.2.1.chr7.g01580.t01, mNeoVis.vulgivaga.2.1.chr7.g01581.t01, mNeoVis.vulgivaga.2.1.chr7.g01582.t01, mNeoVis.vulgivaga.2.1.chr7.g01583.t01      1589    N0.HOG0001587   OG0000196  n1   rna-XM_011222974.3    rna-XM_038541449.1    rna-XM_022496561.1    rna-XM_003990287.4    rna-XM_036099448.1    rna-XM_039216639.1    rna-XM_032867065.1    rna-XM_046028327.1    rna-XM_032303278.1    rna-XM_059136605.1    rna-XM_059375315.1    rna-XM_004752644.3    mNeoVis.2.1.1.chr2_hap1.g01107.t01    rna-XM_021689998.1    rna-XM_004408946.1    rna-XM_007079154.3    rna-XM_008690625.2    rna-XM_027574602.2    mNeoVis.3.1.1.chr5.g01030.t01    rna-gnl|WGS_ZZZZ|egapxtmp_009521-R1    mNeoVis.evergladensis.2.1.chr5.g01050.t01    mNeoVis.vulgivaga.2.1.chr5.g01041.t01      1590    N0.HOG0001588   OG0000196  n23   rna-XM_002930364.4    rna-XM_038541450.1      rna-XM_019837391.2      rna-XM_039216628.1                    rna-XM_004408966.1    rna-XM_007079146.3    rna-XM_008690621.2    rna-XM_027593524.2              1591    N0.HOG0001589   OG0000196  n30   rna-XM_002930367.4, rna-XM_034653832.1, rna-XM_034653842.1, rna-XM_034653843.1    rna-XM_038541447.1    rna-XM_022496405.1, rna-XM_022496635.1, rna-XM_022496935.1    rna-XM_003990288.5, rna-XM_003990323.6, rna-XM_011284884.3    rna-XM_036118578.1    rna-XM_039216631.1, rna-XM_039216635.1, rna-XM_039216636.1, rna-XM_039221836.1    rna-XM_032867067.1, rna-XM_032867069.1, rna-XM_032867268.1, rna-XM_032867269.1    rna-XM_045979780.1, rna-XM_046007874.1, rna-XM_046014089.1, rna-XM_046025035.1    rna-XM_032302012.1, rna-XM_032303272.1, rna-XM_032303721.1    rna-XM_059136269.1, rna-XM_059136270.1, rna-XM_059138015.1, rna-XM_059138019.1    rna-XM_059375317.1, rna-XM_059375319.1, rna-XM_059375323.1, rna-XM_059377056.1    rna-XM_004752637.3, rna-XM_004752638.3, rna-XM_045063436.1, rna-XM_045064818.1    mNeoVis.2.1.1.chr2_hap1.g01101.t01, mNeoVis.2.1.1.chr2_hap1.g01102.t01, mNeoVis.2.1.1.chr2_hap1.g01105.t01, mNeoVis.2.1.1.chr2_hap1.g01106.t01    rna-XM_021690308.1      rna-XM_007073769.3, rna-XM_007079153.3, rna-XM_042997789.1    rna-XM_008690616.2, rna-XM_008690622.2, rna-XM_040626586.1    rna-XM_027593387.2, rna-XM_035727055.1    mNeoVis.3.1.1.chr5.g01026.t01, mNeoVis.3.1.1.chr5.g01027.t01, mNeoVis.3.1.1.chr5.g01028.t01, mNeoVis.3.1.1.chr5.g01029.t01    rna-gnl|WGS_ZZZZ|egapxtmp_007005-R1, rna-gnl|WGS_ZZZZ|egapxtmp_009522-R1, rna-gnl|WGS_ZZZZ|egapxtmp_009523-R1, rna-gnl|WGS_ZZZZ|egapxtmp_009524-R1    mNeoVis.evergladensis.2.1.chr5.g01046.t01, mNeoVis.evergladensis.2.1.chr5.g01047.t01, mNeoVis.evergladensis.2.1.chr5.g01048.t01, mNeoVis.evergladensis.2.1.chr5.g01049.t01    mNeoVis.vulgivaga.2.1.chr5.g01037.t01, mNeoVis.vulgivaga.2.1.chr5.g01038.t01, mNeoVis.vulgivaga.2.1.chr5.g01039.t01, mNeoVis.vulgivaga.2.1.chr5.g01040.t01      1592    N0.HOG0001590   OG0000197  n1   rna-XM_011233432.2    rna-XM_038568847.1    rna-XM_022521774.1    rna-XM_003992858.3    rna-XM_036081413.1    rna-XM_039224668.1    rna-XM_032853630.1    rna-XM_046015658.1    rna-XM_032357426.1    rna-XM_059134350.1    rna-XM_059390162.1    rna-XM_004779023.1    mNeoVis.2.1.1.chr7_hap1.g01670.t01    rna-XM_021685680.1    rna-XM_004407561.1    rna-XM_007092228.3    rna-XM_040630710.1    rna-XM_027579485.1    mNeoVis.3.1.1.chr7.g01563.t01      mNeoVis.evergladensis.2.1.chr7.g01641.t01    mNeoVis.vulgivaga.2.1.chr7.g01658.t01      1593    N0.HOG0001591   OG0000197  n20   rna-XM_002926084.4    rna-XM_038568848.1    rna-XM_022521775.1    rna-XM_003992859.3      rna-XM_039224666.1    rna-XM_032853631.1    rna-XM_046015651.1    rna-XM_032359497.1    rna-XM_059134354.1    rna-XM_059390149.1    rna-XM_004779022.2      rna-XM_021685681.1    rna-XM_004407562.1    rna-XM_007092227.3    rna-XM_008709609.2    rna-XM_027579484.1    mNeoVis.3.1.1.chr7.g01564.t01    rna-gnl|WGS_ZZZZ|egapxtmp_012898-R1    mNeoVis.evergladensis.2.1.chr7.g01642.t01    mNeoVis.vulgivaga.2.1.chr7.g01659.t01      1594    N0.HOG0001592   OG0000197  n37   rna-NM_001304883.1, rna-NM_001304885.1    rna-NM_001270883.1, rna-NM_001270884.1, rna-XM_038568846.1    rna-XM_022521772.1    rna-XM_003992857.4, rna-XM_003992882.5    rna-XM_036081410.1, rna-XM_036081411.1, rna-XM_036081412.1    rna-XM_039224665.1, rna-XM_039224676.1, rna-XM_039224677.1    rna-XM_032851177.1, rna-XM_032851178.1, rna-XM_032853273.1    rna-XM_046016642.1, rna-XM_046016643.1, rna-XM_046017762.1    rna-XM_032357422.1, rna-XM_032357423.1, rna-XM_032357424.1, rna-XM_032357425.1, rna-XM_032359451.1    rna-XM_059134333.1, rna-XM_059134341.1, rna-XM_059134346.1    rna-XM_059409589.1, rna-XM_059409601.1    rna-XM_004779025.3, rna-XM_004779026.2, rna-XM_045071441.1    mNeoVis.2.1.1.chr7_hap1.g01668.t01, mNeoVis.2.1.1.chr7_hap1.g01669.t01    rna-XM_044912552.1, rna-XM_044912553.1, rna-XM_044919260.1    rna-XM_004407558.1, rna-XM_004407559.1, rna-XM_004407560.1, rna-XM_004417700.1    rna-XM_007092229.3, rna-XM_007092230.3    rna-XM_008709612.2    rna-XM_027579486.1, rna-XM_027579488.2, rna-XM_035722873.1      rna-gnl|WGS_ZZZZ|egapxtmp_012895-R1, rna-gnl|WGS_ZZZZ|egapxtmp_012896-R1, rna-gnl|WGS_ZZZZ|egapxtmp_012897-R1    mNeoVis.evergladensis.2.1.chr7.g01639.t01, mNeoVis.evergladensis.2.1.chr7.g01640.t01    mNeoVis.vulgivaga.2.1.chr7.g01656.t01, mNeoVis.vulgivaga.2.1.chr7.g01657.t01      1595    N0.HOG0001593   OG0000198  n0   id-LOC105234963, id-LOC109488640, id-LOC117797252, rna-XM_034648724.1    id-LOC100683976, id-LOC111097255, id-LOC119875826, id-LOC119875828, id-LOC119875832, id-LOC119879435, id-LOC607750    id-LOC111139360, id-LOC111139454, id-LOC111139485, id-LOC111139497, id-LOC111139522, id-LOC111139544, id-LOC111140234    id-LOC101081143, id-LOC101101319, id-LOC105260658    id-LOC118522989, id-LOC118522992, id-LOC118550132    id-LOC120240427, id-LOC120240456    id-LOC116867364, id-LOC116867369, id-LOC116867372, id-LOC116867375, id-LOC116867379, id-LOC116867385    id-LOC123943597, id-LOC123943600, id-LOC123943611, id-LOC123943618, id-LOC123944446, id-LOC123944843    id-LOC116590166, id-LOC116590169, id-LOC116590174, id-LOC116591099, id-LOC116591100, id-LOC116591101, id-LOC116591102, id-LOC116591109, id-LOC116591115    id-LOC131836314, id-LOC131836315, id-LOC131836322, id-LOC131836326, id-LOC131836327, id-LOC131837185, id-LOC131837198    id-LOC131998742, id-LOC131998746, id-LOC131998747, id-LOC131999374, id-LOC131999379, id-LOC131999386, id-LOC131999389, id-LOC131999394, id-LOC131999401, rna-XM_059372129.1, rna-XM_059372132.1    id-LOC101681815, id-LOC101683568, id-LOC101685606, id-LOC101686463, id-LOC101687609, id-LOC101689914, id-LOC101694496, id-LOC106006084, id-LOC123386896, id-LOC123391344        id-LOC101369822, id-LOC101370707, id-LOC101371892, id-LOC101374211, id-LOC101375079, rna-XM_004402167.1, rna-XM_004402169.1    id-LOC102952952, id-LOC102965722, id-LOC102966591, id-LOC102967437, id-LOC122239502    id-LOC103680322, id-LOC103682249, id-LOC103682259, id-LOC103682262, id-LOC121100924, id-LOC121100925    id-LOC113908996, id-LOC113908997, id-LOC113909256, id-LOC118357071              1596    N0.HOG0001594   OG0000199  n0   rna-XM_034669288.1    rna-XM_038525484.1, rna-XM_038525588.1, rna-XM_038525590.1    rna-XM_022503508.1, rna-XM_022504137.1, rna-XM_022504138.1, rna-XM_022520151.1    rna-XM_019831384.3, rna-XM_045058196.1    rna-XM_036075019.1, rna-XM_036075021.1, rna-XM_036075024.1    rna-XM_039240974.1, rna-XM_039240978.1    rna-XM_032850270.1, rna-XM_032850272.1, rna-XM_032850967.1, rna-XM_032851014.1, rna-XM_032851015.1, rna-XM_032851073.1, rna-XM_032851129.1, rna-XM_032873000.1, rna-XM_032873002.1    rna-XM_046004069.1, rna-XM_046004359.1, rna-XM_046005942.1, rna-XM_046006038.1, rna-XM_046006059.1, rna-XM_046006476.1    rna-XM_032338314.1, rna-XM_032338758.1, rna-XM_032338759.1, rna-XM_032338762.1, rna-XM_032338764.1, rna-XM_032339256.1, rna-XM_032339258.1, rna-XM_032341319.1, rna-XM_032341322.1, rna-XM_032341324.1, rna-XM_032341334.1    rna-XM_059176315.1, rna-XM_059176316.1, rna-XM_059176786.1, rna-XM_059177262.1, rna-XM_059179263.1    rna-XM_059399202.1, rna-XM_059401252.1, rna-XM_059401253.1, rna-XM_059401256.1    rna-XM_004753555.3, rna-XM_004753556.3, rna-XM_013056267.2, rna-XM_013056731.2, rna-XM_045080190.1, rna-XM_045080273.1    mNeoVis.2.1.1.chr1_hap1.g01881.t01, mNeoVis.2.1.1.chr1_hap1.g02060.t01, mNeoVis.2.1.1.chr1_hap1.g02062.t01, mNeoVis.2.1.1.chr1_hap1.g02063.t01, mNeoVis.2.1.1.chr1_hap1.g02065.t01, mNeoVis.2.1.1.chr1_hap1.g02066.t01, mNeoVis.2.1.1.chr1_hap1.g02067.t01    rna-XM_021689528.2, rna-XM_044917518.1      rna-XM_042987311.1, rna-XM_042987313.1    rna-XM_008695004.2, rna-XM_008709388.1, rna-XM_040627954.1    rna-XM_027603374.2, rna-XM_035728715.1    mNeoVis.3.1.1.chr1.g01835.t01, mNeoVis.3.1.1.chr1.g02014.t01, mNeoVis.3.1.1.chr1.g02015.t01, mNeoVis.3.1.1.chr1.g02016.t01, mNeoVis.3.1.1.chr1.g02017.t01    rna-gnl|WGS_ZZZZ|egapxtmp_022709-R1, rna-gnl|WGS_ZZZZ|egapxtmp_025166-R1, rna-gnl|WGS_ZZZZ|egapxtmp_029791-R1, rna-gnl|WGS_ZZZZ|egapxtmp_035112-R1, rna-gnl|WGS_ZZZZ|egapxtmp_036383-R1    mNeoVis.evergladensis.2.1.chr1.g01847.t01, mNeoVis.evergladensis.2.1.chr1.g02027.t01, mNeoVis.evergladensis.2.1.chr1.g02028.t01, mNeoVis.evergladensis.2.1.chr1.g02030.t01, mNeoVis.evergladensis.2.1.chr1.g02031.t01, mNeoVis.evergladensis.2.1.chr1.g02034.t01, mNeoVis.evergladensis.2.1.chr1.g02035.t01    mNeoVis.vulgivaga.2.1.chr1.g01844.t01, mNeoVis.vulgivaga.2.1.chr1.g01845.t01, mNeoVis.vulgivaga.2.1.chr1.g02021.t01, mNeoVis.vulgivaga.2.1.chr1.g02022.t01, mNeoVis.vulgivaga.2.1.chr1.g02024.t01, mNeoVis.vulgivaga.2.1.chr1.g02025.t01, mNeoVis.vulgivaga.2.1.chr1.g02027.t01, mNeoVis.vulgivaga.2.1.chr1.g02028.t01      1597    N0.HOG0001595   OG0000200  n1   rna-XM_034654082.1    rna-XM_038581715.1    rna-XM_022516242.1    rna-XM_023238577.2    rna-XM_036081655.1    rna-XM_039222149.1    rna-XM_032865534.1    rna-XM_046002497.1    rna-XM_032358089.1    rna-XM_059165006.1    rna-XM_059392585.1    rna-XM_004780261.3    mNeoVis.2.1.1.chr5_hap1.g01930.t01    rna-XM_021679212.2    rna-XM_004416199.1    rna-XM_042956564.1    rna-XM_008710157.2    rna-XM_027585030.1    mNeoVis.3.1.1.chr2.g01817.t01    rna-gnl|WGS_ZZZZ|egapxtmp_037494-R2    mNeoVis.evergladensis.2.1.chr2.g01908.t01    mNeoVis.vulgivaga.2.1.chr2.g01925.t01      1598    N0.HOG0001596   OG0000200  n16     rna-XM_038562418.1      rna-XM_045033538.1    rna-XM_036106068.1      rna-XM_032843663.1, rna-XM_032851211.1, rna-XM_032851219.1      rna-XM_032302635.1    rna-XM_059138546.1    rna-XM_059387435.1    rna-XM_045083055.1    mNeoVis.2.1.1.chr2_hap1.g00753.t01      rna-XM_004417434.1      rna-XM_040626711.1      mNeoVis.3.1.1.chr5.g00701.t01    rna-gnl|WGS_ZZZZ|egapxtmp_007492-R1    mNeoVis.evergladensis.2.1.chr5.g00709.t01    mNeoVis.vulgivaga.2.1.chr5.g00703.t01      1599    N0.HOG0001597   OG0000200  n32   rna-XM_034641900.1, rna-XM_034641904.1    rna-XM_038562285.1    rna-XM_022526116.1, rna-XM_022526117.1    rna-XM_045033628.1, rna-XM_045033629.1, rna-XM_045033630.1    rna-XM_036106062.1, rna-XM_036106063.1, rna-XM_036106071.1    rna-XM_039246237.1    rna-XM_032843689.1, rna-XM_032843696.1    rna-XM_046016866.1    rna-XM_032302339.1, rna-XM_032302342.1    rna-XM_059138545.1, rna-XM_059138547.1    rna-XM_059377124.1    rna-XM_045083053.1      rna-XM_044914625.1    rna-XM_004416681.1    rna-XM_042993875.1, rna-XM_042993876.1    rna-XM_040626710.1      mNeoVis.3.1.1.chr5.g00699.t01, mNeoVis.3.1.1.chr5.g00700.t01    rna-gnl|WGS_ZZZZ|egapxtmp_006931-R1, rna-gnl|WGS_ZZZZ|egapxtmp_006932-R1    mNeoVis.evergladensis.2.1.chr5.g00708.t01    mNeoVis.vulgivaga.2.1.chr5.g00700.t01, mNeoVis.vulgivaga.2.1.chr5.g00701.t01, mNeoVis.vulgivaga.2.1.chr5.g00702.t01, mNeoVis.vulgivaga.2.1.chr5.g00704.t01      1600    N0.HOG0001598   OG0000200  n66   rna-XM_034654299.1    rna-XM_038562283.1    rna-XM_022492727.1    rna-XM_045033533.1    rna-XM_036106072.1, rna-XM_036118795.1    rna-XM_039225824.1    rna-XM_032843571.1    rna-XM_046017106.1    rna-XM_032361624.1    rna-XM_059138550.1    rna-XM_059377122.1, rna-XM_059387570.1    rna-XM_045083056.1    mNeoVis.2.1.1.chr2_hap1.g00756.t01    rna-XM_044914348.1    rna-XM_004404290.1    rna-XM_042996673.1    rna-XM_040626732.1    rna-XM_035727245.1    mNeoVis.3.1.1.chr5.g00703.t01    rna-gnl|WGS_ZZZZ|egapxtmp_006545-R1    mNeoVis.evergladensis.2.1.chr5.g00712.t01        1601    N0.HOG0001599   OG0000201  n1   rna-XM_034648528.1    rna-XM_038558427.1    rna-XM_022512039.1    rna-XM_006932699.5    rna-XM_036117957.1    rna-XM_039245221.1    rna-XM_032856985.1    rna-XM_046009030.1    rna-XM_032343102.1    rna-XM_059180999.1    rna-XM_059372410.1    rna-XM_004758548.3    mNeoVis.2.1.1.chr10_hap1.g00882.t01, mNeoVis.2.1.1.chr10_hap1.g00887.t01    rna-XM_021695462.1    rna-XM_004402137.1    rna-XM_007091348.3    rna-XM_040620690.1    rna-XM_027570508.2    mNeoVis.3.1.1.chr9.g00814.t01    rna-gnl|WGS_ZZZZ|egapxtmp_016240-R1    mNeoVis.evergladensis.2.1.chr8.g00856.t01    mNeoVis.vulgivaga.2.1.chr9.g00843.t01      1602    N0.HOG0001600   OG0000201  n20   rna-XM_034639030.1, rna-XM_034652129.1    rna-XM_038530897.1    rna-XM_022518660.1    rna-XM_011290141.3    rna-XM_036124293.1    rna-XM_039232603.1    rna-XM_032854329.1    rna-XM_045988166.1    rna-XM_032325893.1    rna-XM_059152698.1    rna-XM_059382847.1    rna-XM_004744282.2    mNeoVis.2.1.1.chr7_hap1.g00273.t01    rna-XM_021704263.1    rna-XM_004395759.1    rna-XM_042968752.1    rna-XM_040633381.1    rna-XM_027618778.2    mNeoVis.3.1.1.chr7.g00257.t01    rna-gnl|WGS_ZZZZ|egapxtmp_009656-R1    mNeoVis.evergladensis.2.1.chr7.g00273.t01    mNeoVis.vulgivaga.2.1.chr7.g00276.t01      1603    N0.HOG0001601   OG0000201  n37   rna-XM_034640525.1    rna-XM_038572223.1    rna-XM_022494480.1    rna-XM_019826549.3    rna-XM_036108072.1    rna-XM_039227138.1    rna-XM_032845815.1    rna-XM_045981719.1, rna-XM_046013558.1    rna-XM_032350759.1    rna-XM_059133293.1    rna-XM_059406955.1    rna-XM_045084167.1    mNeoVis.2.1.1.chr11_hap1.g00262.t01, mNeoVis.2.1.1.chr1_hap1.g01498.t01    rna-XM_044918794.1    rna-XM_004393151.1    rna-XM_042980456.1    rna-XM_040638277.1    rna-XM_027622719.2    mNeoVis.3.1.1.chr1.g01441.t01, mNeoVis.3.1.1.chr11.g00233.t01    rna-gnl|WGS_ZZZZ|egapxtmp_011238-R1, rna-gnl|WGS_ZZZZ|egapxtmp_026640-R1    mNeoVis.evergladensis.2.1.chr1.g01460.t01, mNeoVis.evergladensis.2.1.chr11.g00260.t01    mNeoVis.vulgivaga.2.1.chr1.g01461.t01, mNeoVis.vulgivaga.2.1.chr11.g00261.t01      1604    N0.HOG0001602   OG0000201  n56   rna-XM_034663411.1    rna-NM_001284434.1    rna-XM_022502437.1    rna-XM_023248618.2    rna-XM_036091136.1    rna-XM_039240191.1    rna-XM_032873493.1    rna-XM_045979334.1    rna-XM_032315612.1    rna-XM_059165568.1    rna-XM_059395202.1    rna-XM_004743589.3    mNeoVis.2.1.1.chr4_hap1.g01029.t01    rna-XM_021688816.2    rna-XM_004413994.1    rna-XM_042973616.1    rna-XM_040625791.1    rna-XM_027597692.1    mNeoVis.3.1.1.chr6.g01048.t01    rna-gnl|WGS_ZZZZ|egapxtmp_004714-R2    mNeoVis.evergladensis.2.1.chr6.g01034.t01    mNeoVis.vulgivaga.2.1.chr6.g01026.t01      1605    N0.HOG0001603   OG0000202  n0   rna-XM_034652271.1, rna-XM_034653537.1          rna-XM_036066378.1, rna-XM_036068830.1, rna-XM_036068832.1, rna-XM_036075386.1, rna-XM_036076983.1, rna-XM_036077334.1, rna-XM_036077335.1, rna-XM_036081910.1, rna-XM_036081912.1, rna-XM_036085157.1, rna-XM_036088498.1, rna-XM_036089750.1, rna-XM_036094102.1, rna-XM_036099823.1, rna-XM_036101749.1, rna-XM_036102392.1, rna-XM_036102949.1, rna-XM_036111820.1, rna-XM_036112988.1, rna-XM_036116778.1, rna-XM_036123843.1, rna-XM_036124496.1, rna-XM_036124497.1        rna-XM_045996591.1, rna-XM_045996771.1, rna-XM_045996772.1, rna-XM_045996773.1, rna-XM_045996775.1, rna-XM_045996776.1, rna-XM_045996777.1, rna-XM_045996778.1, rna-XM_045996779.1, rna-XM_045996780.1, rna-XM_045996781.1, rna-XM_045996782.1, rna-XM_045996783.1, rna-XM_045996784.1, rna-XM_045996786.1, rna-XM_045996787.1, rna-XM_045996788.1, rna-XM_045996789.1, rna-XM_045996790.1, rna-XM_045996791.1, rna-XM_045996795.1, rna-XM_045996796.1    rna-XM_032331905.1, rna-XM_032331960.1, rna-XM_032331961.1    rna-XM_059158779.1, rna-XM_059158780.1, rna-XM_059158781.1, rna-XM_059158782.1, rna-XM_059158783.1, rna-XM_059158784.1, rna-XM_059158788.1, rna-XM_059158789.1, rna-XM_059158792.1, rna-XM_059158793.1, rna-XM_059158795.1, rna-XM_059158796.1, rna-XM_059158797.1, rna-XM_059158798.1, rna-XM_059158803.1, rna-XM_059158805.1, rna-XM_059158806.1, rna-XM_059158807.1, rna-XM_059158808.1, rna-XM_059158809.1, rna-XM_059158811.1, rna-XM_059158812.1, rna-XM_059158814.1, rna-XM_059158815.1, rna-XM_059158817.1, rna-XM_059158819.1, rna-XM_059158822.1, rna-XM_059158823.1    rna-XM_059387292.1    rna-XM_045072826.1, rna-XM_045074380.1, rna-XM_045087311.1, rna-XM_045087314.1, rna-XM_045087315.1, rna-XM_045087316.1, rna-XM_045087320.1, rna-XM_045087321.1      rna-XM_044912423.1, rna-XM_044912481.1, rna-XM_044912493.1, rna-XM_044912500.1, rna-XM_044912550.1    rna-XM_004413626.1      rna-XM_040636116.1    rna-XM_035725885.1, rna-XM_035725976.1              1606    N0.HOG0001604   OG0000203  n2   rna-XM_002926082.2      rna-XM_022522135.1          rna-XM_032853633.1    rna-XM_046016819.1    rna-XM_032359656.1    rna-XM_059157361.1    rna-XM_059390123.1    rna-XM_004779064.2    mNeoVis.2.1.1.chr7_hap1.g01672.t01          rna-XM_008709655.1        rna-gnl|WGS_ZZZZ|egapxtmp_012843-R1    mNeoVis.evergladensis.2.1.chr7.g01644.t01    mNeoVis.vulgivaga.2.1.chr7.g01661.t01      1607    N0.HOG0001605   OG0000203  n15   rna-XM_011233431.2    rna-NM_001388963.1      rna-XM_011286650.2    rna-XM_036087718.1    rna-XM_039224672.1    rna-XM_032853634.1    rna-XM_046016803.1    rna-XM_032359602.1    rna-XM_059157375.1    rna-XM_059390100.1    rna-XM_004779020.1    mNeoVis.2.1.1.chr7_hap1.g01673.t01    rna-XM_021685630.1      rna-XM_007092225.1    rna-XM_008709654.2    rna-XM_027580688.1    mNeoVis.3.1.1.chr7.g01566.t01    rna-gnl|WGS_ZZZZ|egapxtmp_012885-R1    mNeoVis.evergladensis.2.1.chr7.g01645.t01    mNeoVis.vulgivaga.2.1.chr7.g01662.t01      1608    N0.HOG0001606   OG0000203  n34   rna-XM_002926107.1    rna-NM_001389190.1      rna-XM_023239717.2      rna-XM_039224662.1    rna-XM_032852701.1    rna-XM_046016791.1    rna-XM_032358840.1    rna-XM_059157395.1    rna-XM_059390088.1    rna-XM_004779063.1    mNeoVis.2.1.1.chr7_hap1.g01674.t01          rna-XM_008709652.2      mNeoVis.3.1.1.chr7.g01567.t01    rna-gnl|WGS_ZZZZ|egapxtmp_012844-R1    mNeoVis.evergladensis.2.1.chr7.g01646.t01    mNeoVis.vulgivaga.2.1.chr7.g01663.t01      1609    N0.HOG0001607   OG0000203  n48         rna-XM_003992861.2      rna-XM_039224669.1                      rna-XM_042959160.1                  1610    N0.HOG0001608   OG0000203  n51   rna-XM_002926105.4    rna-NM_001388931.1                                rna-XM_008709650.2                1611    N0.HOG0001609   OG0000203  n53   rna-XM_002926106.4, rna-XM_034667610.1    rna-XM_038568164.1        rna-XM_036087717.1      rna-XM_032853635.1    rna-XM_046016802.1    rna-XM_032358841.1    rna-XM_059157405.1    rna-XM_059390076.1    rna-XM_004779019.1    mNeoVis.2.1.1.chr7_hap1.g01675.t01          rna-XM_008709651.1      mNeoVis.3.1.1.chr7.g01568.t01    rna-gnl|WGS_ZZZZ|egapxtmp_012883-R1    mNeoVis.evergladensis.2.1.chr7.g01647.t01    mNeoVis.vulgivaga.2.1.chr7.g01664.t01      1612    N0.HOG0001610   OG0000203  n68   rna-XM_002926083.3      rna-XM_022522136.1      rna-XM_036087692.1      rna-XM_032853632.1    rna-XM_046016768.1    rna-XM_032359604.1    rna-XM_059157351.1    rna-XM_059390135.1    rna-XM_004779021.1    mNeoVis.2.1.1.chr7_hap1.g01671.t01    rna-XM_021685682.1        rna-XM_008709607.1    rna-XM_027580691.1    mNeoVis.3.1.1.chr7.g01565.t01    rna-gnl|WGS_ZZZZ|egapxtmp_012860-R1    mNeoVis.evergladensis.2.1.chr7.g01643.t01    mNeoVis.vulgivaga.2.1.chr7.g01660.t01      1613    N0.HOG0001611   OG0000203  n85     rna-XM_038568163.1      rna-XM_011286652.2      rna-XM_039224663.1                      rna-XM_015542097.2                  1614    N0.HOG0001612   OG0000203  n88     rna-NM_001388697.1      rna-XM_003992860.2      rna-XM_039224670.1                      rna-XM_007092226.2                  1615    N0.HOG0001613   OG0000204  n1   rna-XM_002922707.4    rna-XM_038563232.1    rna-XM_022507554.1      rna-XM_036076556.1    rna-XM_039245007.1    rna-XM_032846445.1    rna-XM_046017688.1    rna-XM_032358701.1    rna-XM_059142716.1    rna-XM_059405170.1    rna-XM_004777586.2    mNeoVis.2.1.1.chr7_hap1.g02178.t01    rna-XM_021684852.1    rna-XM_004409271.1    rna-XM_015542256.2    rna-XM_040639128.1    rna-XM_027581313.2    mNeoVis.3.1.1.chr7.g02006.t01      mNeoVis.evergladensis.2.1.chr7.g02127.t01    mNeoVis.vulgivaga.2.1.chr7.g02149.t01      1616    N0.HOG0001614   OG0000204  n22     rna-XM_038570233.1                            rna-XM_004400511.1        rna-XM_027576155.2              1617    N0.HOG0001615   OG0000204  n24   rna-XM_002926289.4    rna-XM_038575984.1    rna-XM_022495774.1, rna-XM_022504646.1, rna-XM_022508103.1    rna-XM_019814934.3    rna-XM_036076451.1    rna-XM_039222468.1, rna-XM_039239305.1, rna-XM_039242863.1    rna-XM_032873610.1, rna-XM_032875973.1    rna-XM_045978267.1, rna-XM_046009602.1, rna-XM_046023724.1, rna-XM_046025680.1    rna-XM_032305736.1, rna-XM_032309816.1    rna-XM_059138799.1    rna-XM_059395911.1, rna-XM_059404680.1, rna-XM_059409003.1    rna-XM_045065269.1, rna-XM_045069427.1    mNeoVis.2.1.1.chr3_hap1.g00344.t01    rna-XM_021687930.2, rna-XM_021699452.1, rna-XM_044921235.1    rna-XM_004412456.1, rna-XM_004415998.1      rna-XM_040619546.1    rna-XM_027576172.2, rna-XM_027577314.2, rna-XM_027577635.2, rna-XM_027599945.2, rna-XM_027619728.1, rna-XM_035727311.1    mNeoVis.3.1.1.chr10.g01201.t01, mNeoVis.3.1.1.chr3.g00337.t01    rna-gnl|WGS_ZZZZ|egapxtmp_022119-R1, rna-gnl|WGS_ZZZZ|egapxtmp_041900-R1    mNeoVis.evergladensis.2.1.chr10.g01240.t01, mNeoVis.evergladensis.2.1.chr3.g00352.t01    mNeoVis.vulgivaga.2.1.chr3.g00347.t01      1618    N0.HOG0001616   OG0000204  n47                 rna-XM_045989805.1              rna-XM_044916507.1    rna-XM_004410474.1                    1619    N0.HOG0001617   OG0000204  n49           rna-XM_036078724.1                    rna-XM_021687010.2, rna-XM_021689422.2, rna-XM_044912030.1, rna-XM_044912214.1, rna-XM_044912345.1, rna-XM_044912353.1, rna-XM_044912384.1, rna-XM_044912389.1, rna-XM_044912420.1, rna-XM_044912425.1, rna-XM_044912434.1, rna-XM_044912442.1, rna-XM_044912449.1, rna-XM_044912450.1, rna-XM_044912457.1, rna-XM_044912467.1, rna-XM_044912494.1, rna-XM_044912498.1, rna-XM_044912499.1, rna-XM_044912521.1, rna-XM_044912525.1, rna-XM_044912528.1, rna-XM_044912542.1, rna-XM_044912543.1        rna-XM_008689381.1                1620    N0.HOG0001618   OG0000205  n0     rna-XM_038561084.1, rna-XM_038561105.1, rna-XM_038589132.1, rna-XM_038589268.1, rna-XM_038589329.1, rna-XM_038589331.1, rna-XM_038589349.1, rna-XM_038589351.1, rna-XM_038589390.1, rna-XM_038589433.1    rna-XM_022491514.1, rna-XM_022491530.1, rna-XM_022492805.1    rna-XM_019827751.3    rna-XM_036090166.1      rna-XM_032851042.1, rna-XM_032851226.1, rna-XM_032851231.1    rna-XM_045978645.1, rna-XM_045979027.1, rna-XM_045979391.1, rna-XM_045980515.1, rna-XM_045980516.1, rna-XM_045980518.1, rna-XM_045980520.1, rna-XM_045980521.1, rna-XM_045980522.1, rna-XM_045980523.1, rna-XM_045980524.1, rna-XM_045980526.1, rna-XM_045980527.1, rna-XM_045980528.1, rna-XM_045980530.1, rna-XM_045980531.1, rna-XM_045980532.1, rna-XM_045980533.1, rna-XM_045980534.1, rna-XM_045980536.1, rna-XM_045980537.1, rna-XM_045980540.1, rna-XM_045980541.1, rna-XM_045980542.1, rna-XM_045980543.1, rna-XM_045980544.1, rna-XM_045980545.1, rna-XM_045980546.1, rna-XM_045980548.1    rna-XM_032352954.1, rna-XM_032353027.1, rna-XM_032353084.1, rna-XM_032353708.1    rna-XM_059132384.1, rna-XM_059134372.1, rna-XM_059134374.1, rna-XM_059134377.1, rna-XM_059134382.1, rna-XM_059188595.1, rna-XM_059188706.1, rna-XM_059188762.1    rna-XM_059387520.1, rna-XM_059405407.1, rna-XM_059405408.1, rna-XM_059405409.1, rna-XM_059407614.1, rna-XM_059407619.1    rna-XM_004765483.3, rna-XM_004782476.3, rna-XM_045073871.1, rna-XM_045073872.1, rna-XM_045073873.1, rna-XM_045073874.1, rna-XM_045073899.1, rna-XM_045073901.1, rna-XM_045079332.1, rna-XM_045079336.1, rna-XM_045088015.1    mNeoVis.2.1.1.chr11_hap1.g00987.t01, mNeoVis.2.1.1.chr11_hap1.g00988.t01, mNeoVis.2.1.1.chr11_hap1.g01006.t01, mNeoVis.2.1.1.chr11_hap1.g01046.t01, mNeoVis.2.1.1.chr11_hap1.g01047.t01, mNeoVis.2.1.1.chr11_hap1.g01048.t01, mNeoVis.2.1.1.chr11_hap1.g01081.t01    rna-XM_044918863.1      rna-XM_042982275.1      rna-XM_027621981.2    mNeoVis.3.1.1.chr11.g00940.t01, mNeoVis.3.1.1.chr11.g00954.t01, mNeoVis.3.1.1.chr11.g00955.t01, mNeoVis.3.1.1.chr11.g00996.t01    rna-gnl|WGS_ZZZZ|egapxtmp_025341-R1, rna-gnl|WGS_ZZZZ|egapxtmp_025361-R1    mNeoVis.evergladensis.2.1.chr11.g01023.t01    mNeoVis.vulgivaga.2.1.chr11.g00989.t01, mNeoVis.vulgivaga.2.1.chr11.g01045.t01, mNeoVis.vulgivaga.2.1.chr11.g01046.t01      1621    N0.HOG0001619   OG0000206  n1   rna-XM_034646697.1    rna-XM_038527093.1    rna-XM_022512593.1    rna-XM_045042771.1    rna-XM_036117912.1    rna-XM_039219178.1    rna-XM_032866600.1    rna-XM_046022643.1    rna-XM_032308281.1    rna-XM_059143480.1    rna-XM_059411872.1    rna-XM_045065091.1    mNeoVis.2.1.1.chr13_hap1.g00747.t01    rna-XM_044920577.1    rna-XM_004397550.1    rna-XM_042963470.1    rna-XM_040620250.1    rna-XM_027615317.2    mNeoVis.3.1.1.chr13.g00663.t01    rna-gnl|WGS_ZZZZ|egapxtmp_030001-R1, rna-gnl|WGS_ZZZZ|egapxtmp_031464-R1    mNeoVis.evergladensis.2.1.chr13.g00712.t01    mNeoVis.vulgivaga.2.1.chr13.g00709.t01      1622    N0.HOG0001620   OG0000206  n24   rna-XM_019804740.1    rna-XM_038533683.1    rna-XM_022511028.1    rna-XM_019831960.3    rna-XM_036076846.1, rna-XM_036096907.1    rna-XM_039217193.1    rna-XM_032835937.1    rna-XM_046007799.1    rna-XM_032342257.1    rna-XM_059183364.1    rna-XM_059371236.1    rna-XM_045070417.1    mNeoVis.2.1.1.chr10_hap1.g01132.t01    rna-XM_044918300.1    rna-XM_004409745.1    rna-XM_042988262.1    rna-XM_040642667.1    rna-XM_027570304.2    mNeoVis.3.1.1.chr9.g01066.t01, mNeoVis.3.1.1.chr9.g01067.t01    rna-gnl|WGS_ZZZZ|egapxtmp_014649-R1    mNeoVis.evergladensis.2.1.chr8.g01111.t01    mNeoVis.vulgivaga.2.1.chr9.g01089.t01, mNeoVis.vulgivaga.2.1.chr9.g01091.t01      1623    N0.HOG0001621   OG0000206  n49   rna-XM_034646980.1    rna-XM_038527064.1    rna-XM_022512777.1    rna-XM_045042481.1    rna-XM_036070462.1    rna-XM_039223140.1    rna-XM_032866640.1    rna-XM_046022196.1    rna-XM_032307749.1    rna-XM_059142327.1    rna-XM_059411850.1    rna-XM_045085383.1    mNeoVis.2.1.1.chr13_hap1.g00731.t01    rna-XM_021694318.1    rna-XM_004396731.1    rna-XM_042963460.1    rna-XM_040620130.1    rna-XM_027616433.2    mNeoVis.3.1.1.chr13.g00647.t01    rna-gnl|WGS_ZZZZ|egapxtmp_030202-R1    mNeoVis.evergladensis.2.1.chr13.g00695.t01    mNeoVis.vulgivaga.2.1.chr13.g00693.t01      1624    N0.HOG0001622   OG0000206  n70   rna-XM_011226462.3, rna-XM_034653418.1    rna-XM_038580408.1    rna-XM_022520679.1    rna-XM_011282861.4    rna-XM_036101931.1    rna-XM_039237105.1    rna-XM_032856497.1    rna-XM_046007338.1    rna-XM_032343696.1    rna-XM_059181532.1    rna-XM_059372816.1    rna-XM_045067914.1    mNeoVis.2.1.1.chr10_hap1.g00617.t01    rna-XM_044918200.1    rna-XM_004396553.1    rna-XM_042988877.1    rna-XM_008705960.2    rna-XM_027568726.1    mNeoVis.3.1.1.chr9.g00574.t01    rna-gnl|WGS_ZZZZ|egapxtmp_016576-R1, rna-gnl|WGS_ZZZZ|egapxtmp_016866-R1    mNeoVis.evergladensis.2.1.chr8.g00599.t01    mNeoVis.vulgivaga.2.1.chr9.g00597.t01      1625    N0.HOG0001623   OG0000207  n1   rna-XM_002930591.4    rna-XM_038586513.1    rna-XM_022491641.1    rna-XM_006943778.5    rna-XM_036123769.1    rna-XM_039222566.1    rna-XM_032848760.1    rna-XM_045994501.1    rna-XM_032331519.1    rna-XM_059158315.1    rna-XM_059384920.1    rna-XM_004782115.3    mNeoVis.2.1.1.chr12_hap1.g00566.t01    rna-XM_021678627.1    rna-XM_004411654.1    rna-XM_015542822.2    rna-XM_008684009.2    rna-XM_027608430.2    mNeoVis.3.1.1.chr12.g00521.t01    rna-gnl|WGS_ZZZZ|egapxtmp_029503-R1    mNeoVis.evergladensis.2.1.chr12.g00508.t01    mNeoVis.vulgivaga.2.1.chr12.g00522.t01      1626    N0.HOG0001624   OG0000207  n14   rna-XM_002919610.4, rna-XM_034649025.1    rna-XM_038527329.1    rna-XM_022512813.1    rna-XM_023243216.2    rna-XM_036070080.1    rna-XM_039241531.1    rna-XM_032866423.1    rna-XM_046021967.1    rna-XM_032308653.1    rna-XM_059142996.1    rna-XM_059412080.1    rna-XM_045083029.1    mNeoVis.2.1.1.chr13_hap1.g00820.t01    rna-XM_021694452.2    rna-XM_004409855.1    rna-XM_042965000.1    rna-XM_008690888.2, rna-XM_040623114.1    rna-XM_027614515.2, rna-XM_035728928.1    mNeoVis.3.1.1.chr13.g00740.t01, mNeoVis.3.1.1.chr5.g01253.t01      mNeoVis.evergladensis.2.1.chr13.g00782.t01    mNeoVis.vulgivaga.2.1.chr13.g00780.t01      1627    N0.HOG0001625   OG0000207  n30   rna-XM_011218141.3, rna-XM_034652511.1, rna-XM_034652517.1    rna-XM_038587563.1, rna-XM_038587569.1    rna-XM_022492249.1, rna-XM_022492717.1    rna-XM_011291764.3, rna-XM_045050754.1    rna-XM_036082883.1, rna-XM_036114302.1, rna-XM_036122750.1, rna-XM_036122755.1    rna-XM_039222802.1, rna-XM_039222806.1, rna-XM_039222811.1, rna-XM_039222817.1    rna-XM_032849246.1, rna-XM_032851109.1    rna-XM_045995454.1, rna-XM_045995458.1, rna-XM_045996131.1    rna-XM_032330832.1, rna-XM_032330837.1, rna-XM_032331415.1      rna-XM_059386310.1, rna-XM_059387382.1    rna-XM_004781657.3, rna-XM_004781659.2, rna-XM_013050578.2    mNeoVis.2.1.1.chr12_hap1.g00560.t01, mNeoVis.2.1.1.chr12_hap1.g00561.t01    rna-XM_021692605.1, rna-XM_021692618.1    rna-XM_004413078.1, rna-XM_004414136.1, rna-XM_004414137.1    rna-XM_042974558.1, rna-XM_042975120.1    rna-XM_008684238.1, rna-XM_008711932.2    rna-XM_027608556.2, rna-XM_035725805.1, rna-XM_035725807.1    mNeoVis.3.1.1.chr12.g00517.t01    rna-gnl|WGS_ZZZZ|egapxtmp_029616-R1    mNeoVis.evergladensis.2.1.chr12.g00504.t01    mNeoVis.vulgivaga.2.1.chr12.g00518.t01      1628    N0.HOG0001626   OG0000208  n1   rna-XM_034639832.1    rna-XM_038572891.1    rna-XM_022494823.1    rna-XM_023250951.2    rna-XM_036085301.1    rna-XM_039236946.1    rna-XM_032878772.1    rna-XM_045980271.1    rna-XM_032353614.1    rna-XM_059133999.1    rna-XM_059407439.1    rna-XM_045070692.1    mNeoVis.2.1.1.chr11_hap1.g00023.t01    rna-XM_021688039.1    rna-XM_004410114.1    rna-XM_042979910.1    rna-XM_040638610.1    rna-XM_027620936.2    mNeoVis.3.1.1.chr11.g00013.t01    rna-gnl|WGS_ZZZZ|egapxtmp_034317-R1    mNeoVis.evergladensis.2.1.chr11.g00023.t01    mNeoVis.vulgivaga.2.1.chr11.g00022.t01      1629    N0.HOG0001627   OG0000208  n24   rna-XM_011233556.3    rna-XM_038565200.1    rna-XM_022525290.1    rna-XM_003990761.6, rna-XM_045054468.1    rna-XM_036074687.1    rna-XM_039223895.1    rna-XM_032857793.1    rna-XM_045996018.1, rna-XM_046018604.1    rna-XM_032331429.1, rna-XM_032355570.1    rna-XM_059146343.1, rna-XM_059165735.1, rna-XM_059167343.1    rna-XM_059392951.1, rna-XM_059393959.1    rna-XM_004743857.3, rna-XM_045087023.1, rna-XM_045089513.1    mNeoVis.2.1.1.chr3_hap1.g00976.t01    rna-XM_021702767.2    rna-XM_004415325.1    rna-XM_042994439.1    rna-XM_040644981.1    rna-XM_027588567.2    mNeoVis.3.1.1.chr3.g00966.t01, mNeoVis.3.1.1.chr6.g00317.t01    rna-gnl|WGS_ZZZZ|egapxtmp_041651-R1    mNeoVis.evergladensis.2.1.chr3.g00971.t01        1630    N0.HOG0001628   OG0000208  n37   rna-XM_011234459.3    rna-XM_038529960.1    rna-XM_022515835.1    rna-XM_006933289.5    rna-XM_036109802.1    rna-XM_039245874.1    rna-XM_032880868.1    rna-XM_046012796.1    rna-XM_032349556.1    rna-XM_059183998.1    rna-XM_059404222.1    rna-XM_004757587.3    mNeoVis.2.1.1.chr9_hap1.g00107.t01    rna-XM_021696769.1    rna-XM_004392970.1    rna-XM_007081273.3    rna-XM_008692060.2    rna-XM_027593724.1    mNeoVis.3.1.1.chr10.g00103.t01    rna-gnl|WGS_ZZZZ|egapxtmp_024071-R1    mNeoVis.evergladensis.2.1.chr10.g00107.t01    mNeoVis.vulgivaga.2.1.chr10.g00103.t01      1631    N0.HOG0001629   OG0000208  n44   rna-XM_034640465.1    rna-XM_038547453.1    rna-XM_022509765.1    rna-XM_023244155.2    rna-XM_036112014.1    rna-XM_039227881.1    rna-XM_032871470.1    rna-XM_045983843.1    rna-XM_032321553.1    rna-XM_059149278.1    rna-XM_059379405.1    rna-XM_004764681.3    mNeoVis.2.1.1.chr8_hap1.g00496.t01    rna-XM_021704355.1    rna-XM_004395064.1    rna-XM_042967465.1    rna-XM_040640706.1    rna-XM_027626057.2    mNeoVis.3.1.1.chr8.g00452.t01      mNeoVis.evergladensis.2.1.chr9.g00487.t01    mNeoVis.vulgivaga.2.1.chr8.g00493.t01      1632    N0.HOG0001630   OG0000209  n1   rna-XM_034647191.1    rna-XM_038536617.1    rna-XM_022524539.1    rna-XM_019817072.3    rna-XM_036066257.1    rna-XM_039231601.1    rna-XM_032839991.1    rna-XM_045984150.1    rna-XM_032322427.1    rna-XM_059147655.1    rna-XM_059378709.1    rna-XM_045083251.1    mNeoVis.2.1.1.chr8_hap1.g01080.t01    rna-XM_021694502.2    rna-XM_004398542.1    rna-XM_042965826.1    rna-XM_040629919.1    rna-XM_027625328.1    mNeoVis.3.1.1.chr8.g01003.t01    rna-gnl|WGS_ZZZZ|egapxtmp_017897-R1    mNeoVis.evergladensis.2.1.chr9.g01055.t01    mNeoVis.vulgivaga.2.1.chr8.g01055.t01      1633    N0.HOG0001631   OG0000209  n23   rna-XM_019798334.2    rna-XM_038560602.1, rna-XM_038587632.1    rna-XM_022524433.1    rna-XM_011291795.4    rna-XM_036066127.1, rna-XM_036111422.1    rna-XM_039253330.1    rna-XM_032847770.1    rna-XM_045995323.1    rna-XM_032330263.1    rna-XM_059157126.1    rna-XM_059386002.1    rna-XM_004777197.3    mNeoVis.2.1.1.chr12_hap1.g00495.t01    rna-XM_044911783.1    rna-XM_004396613.1    rna-XM_042974485.1    rna-XM_040630762.1    rna-XM_027608791.2, rna-XM_035722552.1    mNeoVis.3.1.1.chr12.g00450.t01    rna-gnl|WGS_ZZZZ|egapxtmp_029265-R3    mNeoVis.evergladensis.2.1.chr12.g00443.t01    mNeoVis.vulgivaga.2.1.chr12.g00456.t01      1634    N0.HOG0001632   OG0000209  n41   rna-XM_034666214.1    rna-XM_038568453.1    rna-XM_022513930.1    rna-XM_045039809.1    rna-XM_036082205.1, rna-XM_036099712.1    rna-XM_039231805.1    rna-XM_032852923.1    rna-XM_046015636.1    rna-XM_032356653.1    rna-XM_059187674.1    rna-XM_059411951.1    rna-XM_004763483.3    mNeoVis.2.1.1.chr7_hap1.g01479.t01    rna-XM_021686447.2    rna-XM_004402391.1, rna-XM_004415764.1    rna-XM_007078972.3, rna-XM_042959603.1    rna-XM_008704379.2    rna-XM_027578649.2    mNeoVis.3.1.1.chr7.g01386.t01, mNeoVis.3.1.1.chr7.g01387.t01    rna-gnl|WGS_ZZZZ|egapxtmp_013405-R15    mNeoVis.evergladensis.2.1.chr7.g01460.t01    mNeoVis.vulgivaga.2.1.chr7.g01475.t01      1635    N0.HOG0001633   OG0000209  n65   rna-XM_034660398.1    rna-XM_038583189.1    rna-XM_022515064.1    rna-XM_019839943.3    rna-XM_036108018.1    rna-XM_039250838.1    rna-XM_032869311.1    rna-XM_046001679.1    rna-XM_032337041.1    rna-XM_059163069.1    rna-XM_059391228.1    rna-XM_045088103.1    mNeoVis.2.1.1.chr5_hap1.g01255.t01    rna-XM_021692589.1    rna-XM_004391872.1    rna-XM_042956289.1    rna-XM_040621402.1    rna-XM_027583079.1    mNeoVis.3.1.1.chr2.g01173.t01    rna-gnl|WGS_ZZZZ|egapxtmp_037263-R1    mNeoVis.evergladensis.2.1.chr2.g01243.t01    mNeoVis.vulgivaga.2.1.chr2.g01248.t01      1636    N0.HOG0001634   OG0000210  n2   rna-XM_034656008.1      rna-XM_022493612.1      rna-XM_036088937.1, rna-XM_036089250.1      rna-XM_032837550.1    rna-XM_045982040.1, rna-XM_046000652.1    rna-XM_032336725.1    rna-XM_059182055.1    rna-XM_059371279.1, rna-XM_059417568.1    rna-XM_045069321.1    mNeoVis.2.1.1.chr2_hap1.g00959.t01    rna-XM_021705845.1    rna-XM_004405446.1      rna-XM_040622452.1    rna-XM_027605263.2, rna-XM_027613208.2, rna-XM_035728586.1    mNeoVis.3.1.1.chr1.g00409.t01, mNeoVis.3.1.1.chr5.g00892.t01    rna-gnl|WGS_ZZZZ|egapxtmp_002375-R1, rna-gnl|WGS_ZZZZ|egapxtmp_007979-R1, rna-gnl|WGS_ZZZZ|egapxtmp_014821-R1    mNeoVis.evergladensis.2.1.chr1.g00411.t01, mNeoVis.evergladensis.2.1.chr6.g00449.t01    mNeoVis.vulgivaga.2.1.chr1.g00416.t01      1637    N0.HOG0001635   OG0000210  n24             rna-XM_039242290.1                      rna-XM_042983906.1                  1638    N0.HOG0001636   OG0000210  n26     rna-XM_038536346.1      rna-XM_045039343.1                                          1639    N0.HOG0001637   OG0000210  n27     rna-XM_038552160.1      rna-XM_023255158.2                                          1640    N0.HOG0001638   OG0000210  n29   rna-XM_002921973.4    rna-XM_038534443.1, rna-XM_038583232.1    rna-XM_022500333.1    rna-XM_019813333.3    rna-XM_036095296.1    rna-XM_039235101.1, rna-XM_039236922.1    rna-XM_032839508.1    rna-XM_045979592.1, rna-XM_046027040.1    rna-XM_032312540.1    rna-XM_059169819.1    rna-XM_059397872.1    rna-XM_004769345.3    mNeoVis.2.1.1.chr2_hap1.g00452.t01    rna-XM_021700204.1, rna-XM_044914750.1    rna-XM_004407053.1    rna-XM_042961011.1    rna-XM_040624576.1    rna-XM_027595980.2    mNeoVis.3.1.1.chr5.g00443.t01    rna-gnl|WGS_ZZZZ|egapxtmp_009449-R1    mNeoVis.evergladensis.2.1.chr5.g00454.t01    mNeoVis.vulgivaga.2.1.chr5.g00452.t01      1641    N0.HOG0001639   OG0000210  n47   rna-XM_019798705.2    rna-NM_001251965.2    rna-XM_022498715.1    rna-XM_023252565.2    rna-XM_036069869.1, rna-XM_036102323.1    rna-XM_039235610.1    rna-XM_032837984.1    rna-XM_045998104.1    rna-XM_032329060.1, rna-XM_032338231.1    rna-XM_059156069.1, rna-XM_059176232.1    rna-XM_059384513.1, rna-XM_059399676.1    rna-XM_004775706.3, rna-XM_045073977.1    mNeoVis.2.1.1.chr6_hap1.g00930.t01    rna-XM_044912307.1    rna-XM_004407454.1, rna-XM_004408493.1, rna-XM_004414081.1, rna-XM_004415732.1, rna-XM_004416978.1    rna-XM_042983365.1    rna-XM_040643216.1    rna-XM_027593772.2, rna-XM_027600139.2    mNeoVis.3.1.1.chr4.g00891.t01    rna-gnl|WGS_ZZZZ|egapxtmp_001369-R2    mNeoVis.evergladensis.2.1.chr4.g00882.t01    mNeoVis.vulgivaga.2.1.chr4.g00902.t01      1642    N0.HOG0001640   OG0000211  n1   rna-XM_011237265.3    rna-XM_038541163.1    rna-XM_022497221.1    rna-XM_003990389.6    rna-XM_036109943.1    rna-XM_039254974.1    rna-XM_032867264.1    rna-XM_046018199.1    rna-XM_032301618.1    rna-XM_059136090.1    rna-XM_059375597.1    rna-XM_004777508.3    mNeoVis.2.1.1.chr2_hap1.g01018.t01    rna-XM_021689996.2    rna-XM_004405560.1    rna-XM_042998092.1    rna-XM_008704212.2    rna-XM_027615418.2    mNeoVis.3.1.1.chr5.g00945.t01    rna-gnl|WGS_ZZZZ|egapxtmp_009195-R2    mNeoVis.evergladensis.2.1.chr5.g00965.t01    mNeoVis.vulgivaga.2.1.chr5.g00953.t01      1643    N0.HOG0001641   OG0000211  n25   rna-XM_019809937.2    rna-XM_038541153.1      rna-XM_045033236.1    rna-XM_036109804.1                    rna-XM_021690295.1      rna-XM_042996572.1    rna-XM_008704215.1    rna-XM_027601331.1              1644    N0.HOG0001642   OG0000211  n32     rna-XM_038541159.1    rna-XM_022496298.1    rna-XM_011285036.4    rna-XM_036109962.1    rna-XM_039254970.1    rna-XM_032867271.1      rna-XM_032302514.1    rna-XM_059136343.1    rna-XM_059377107.1    rna-XM_045081988.1    mNeoVis.2.1.1.chr2_hap1.g01017.t01    rna-XM_021690163.1      rna-XM_007090057.2    rna-XM_008704214.2    rna-XM_035727057.1, rna-XM_035728056.1    mNeoVis.3.1.1.chr5.g00944.t01    rna-gnl|WGS_ZZZZ|egapxtmp_006980-R1    mNeoVis.evergladensis.2.1.chr5.g00964.t01    mNeoVis.vulgivaga.2.1.chr5.g00952.t01      1645    N0.HOG0001643   OG0000211  n52   rna-XM_034664594.1    rna-XM_038549222.1    rna-XM_022501743.1    rna-XM_023242759.2    rna-XM_036086146.1    rna-XM_039229644.1    rna-XM_032865954.1    rna-XM_046022337.1    rna-XM_032307149.1    rna-XM_059141905.1    rna-XM_059410706.1    rna-XM_045082458.1    mNeoVis.2.1.1.chr13_hap1.g00198.t01    rna-XM_021691772.2    rna-XM_004392538.1    rna-XM_042963890.1    rna-XM_040631959.1    rna-XM_027616252.2    mNeoVis.3.1.1.chr13.g00149.t01    rna-gnl|WGS_ZZZZ|egapxtmp_030036-R1    mNeoVis.evergladensis.2.1.chr13.g00196.t01    mNeoVis.vulgivaga.2.1.chr13.g00193.t01      1646    N0.HOG0001644   OG0000211  n70   rna-XM_034657622.1    rna-XM_038567555.1    rna-XM_022525811.1    rna-XM_023244217.2    rna-XM_036120653.1    rna-XM_039221351.1    rna-XM_032883756.1    rna-XM_045992075.1    rna-XM_032308069.1    rna-XM_059159663.1    rna-XM_059388794.1    rna-XM_045085541.1    mNeoVis.2.1.1.chr5_hap1.g00168.t01    rna-XM_044918435.1    rna-XM_004395282.1    rna-XM_042977946.1    rna-XM_040629484.1    rna-XM_027585243.2    mNeoVis.3.1.1.chr2.g00117.t01    rna-gnl|WGS_ZZZZ|egapxtmp_034855-R1    mNeoVis.evergladensis.2.1.chr2.g00164.t01    mNeoVis.vulgivaga.2.1.chr2.g00163.t01      1647    N0.HOG0001645   OG0000212  n1   rna-XM_034639995.1, rna-XM_034652010.1    rna-XM_038559527.1    rna-XM_022505891.1, rna-XM_022518216.1    rna-XM_045050552.1, rna-XM_045052089.1    rna-XM_036070679.1, rna-XM_036073235.1, rna-XM_036085208.1, rna-XM_036099060.1, rna-XM_036099446.1, rna-XM_036102587.1, rna-XM_036116241.1    rna-XM_039216837.1, rna-XM_039243013.1, rna-XM_039257514.1    rna-XM_032851812.1, rna-XM_032875030.1    rna-XM_046020915.1, rna-XM_046021659.1    rna-XM_032304384.1    rna-XM_059170667.1    rna-XM_059397141.1    rna-XM_045089585.1    mNeoVis.2.1.1.chr4_hap1.g00627.t01    rna-XM_021678392.1, rna-XM_044913303.1, rna-XM_044913922.1, rna-XM_044918272.1, rna-XM_044920204.1    rna-XM_004393656.1, rna-XM_004398788.1, rna-XM_004414065.1, rna-XM_004415001.1    rna-XM_042962959.1, rna-XM_042974616.1    rna-XM_040644573.1    rna-XM_035723075.1, rna-XM_035723402.1, rna-XM_035724828.1, rna-XM_035726183.1, rna-XM_035726710.1, rna-XM_035726775.1, rna-XM_035728956.1, rna-XM_035729217.1        mNeoVis.evergladensis.2.1.chr6.g00636.t01    mNeoVis.vulgivaga.2.1.chr6.g00633.t01      1648    N0.HOG0001646   OG0000212  n49   rna-XM_002915995.4, rna-XM_034651267.1    rna-NM_001252258.1    rna-XM_022493182.1, rna-XM_022522938.1, rna-XM_022522964.1    rna-XM_045040191.1, rna-XM_045061952.1    rna-XM_036116915.1    rna-XM_039216836.1    rna-XM_032874502.1    rna-XM_046011418.1, rna-XM_046011423.1    rna-XM_032348696.1    rna-XM_059184707.1, rna-XM_059184903.1    rna-XM_059403471.1    rna-XM_004773154.3    mNeoVis.2.1.1.chr11_hap1.g00466.t01, mNeoVis.2.1.1.chr3_hap1.g00748.t01, mNeoVis.2.1.1.chr3_hap1.g00829.t01, mNeoVis.2.1.1.chr9_hap1.g00362.t01    rna-XM_044915382.1, rna-XM_044920092.1    rna-XM_004401767.1    rna-XM_007081515.3, rna-XM_042960771.1, rna-XM_042992558.1    rna-XM_040637146.1, rna-XM_040637147.1    rna-XM_027592249.1, rna-XM_027592475.2, rna-XM_035723193.1, rna-XM_035726922.1, rna-XM_035727075.1, rna-XM_035727398.1    mNeoVis.3.1.1.chr10.g00354.t01    rna-gnl|WGS_ZZZZ|egapxtmp_024643-R1, rna-gnl|WGS_ZZZZ|egapxtmp_027081-R1, rna-gnl|WGS_ZZZZ|egapxtmp_041190-R1    mNeoVis.evergladensis.2.1.chr10.g00359.t01    mNeoVis.vulgivaga.2.1.chr10.g00356.t01, mNeoVis.vulgivaga.2.1.chr3.g00836.t01      1649    N0.HOG0001647   OG0000213  n1   rna-XM_034649026.1    rna-XM_038587590.1    rna-XM_022491591.1, rna-XM_022491592.1    rna-XM_004000574.5    rna-XM_036076393.1, rna-XM_036107491.1    rna-XM_039223452.1    rna-XM_032846902.1    rna-XM_045995746.1    rna-XM_032329699.1, rna-XM_032331479.1    rna-XM_059156801.1, rna-XM_059158072.1    rna-XM_059385240.1    rna-XM_045081218.1, rna-XM_045081221.1    mNeoVis.2.1.1.chr12_hap1.g00520.t01, mNeoVis.2.1.1.chr12_hap1.g00521.t01    rna-XM_044911677.1    rna-XM_004404839.1    rna-XM_042974171.1    rna-XM_008710665.2    rna-XM_027609509.1    mNeoVis.3.1.1.chr12.g00474.t01    rna-gnl|WGS_ZZZZ|egapxtmp_028182-R1    mNeoVis.evergladensis.2.1.chr12.g00467.t01    mNeoVis.vulgivaga.2.1.chr12.g00480.t01      1650    N0.HOG0001648   OG0000213  n29   rna-XM_034667587.1    rna-XM_038568301.1    rna-XM_022514128.1    rna-XM_003992613.6    rna-XM_036075008.1    rna-XM_039255008.1    rna-XM_032852790.1    rna-XM_046017259.1    rna-XM_032358246.1    rna-XM_059186348.1    rna-XM_059412792.1    rna-XM_004754506.3    mNeoVis.2.1.1.chr7_hap1.g01438.t01    rna-XM_021679082.1    rna-XM_004410752.1    rna-XM_007079029.3    rna-XM_008688360.1    rna-XM_027580503.2    mNeoVis.3.1.1.chr7.g01345.t01    rna-gnl|WGS_ZZZZ|egapxtmp_013216-R2    mNeoVis.evergladensis.2.1.chr7.g01420.t01    mNeoVis.vulgivaga.2.1.chr7.g01432.t01      1651    N0.HOG0001649   OG0000213  n51   rna-XM_002927043.4    rna-XM_038570192.1    rna-XM_022509037.1    rna-XM_019810577.3    rna-XM_036115640.1    rna-XM_039241037.1    rna-XM_032852660.1    rna-XM_046003438.1    rna-XM_032301496.1    rna-XM_059163807.1    rna-XM_059388091.1    rna-XM_004755505.3    mNeoVis.2.1.1.chr5_hap1.g01500.t01    rna-XM_021685950.1    rna-XM_004405802.1    rna-XM_042998355.1    rna-XM_008707324.2    rna-XM_027581862.1    mNeoVis.3.1.1.chr2.g01421.t01    rna-gnl|WGS_ZZZZ|egapxtmp_034343-R1    mNeoVis.evergladensis.2.1.chr2.g01487.t01    mNeoVis.vulgivaga.2.1.chr2.g01501.t01      1652    N0.HOG0001650   OG0000213  n72   rna-XM_034661847.1    rna-XM_038571174.1    rna-XM_022503821.1    rna-XM_011286166.4    rna-XM_036115616.1    rna-XM_039241027.1    rna-XM_032852450.1    rna-XM_046001709.1    rna-XM_032345950.1    rna-XM_059163808.1    rna-XM_059391759.1    rna-XM_004755496.3    mNeoVis.2.1.1.chr5_hap1.g01501.t01    rna-XM_021685984.1    rna-XM_004405803.1    rna-XM_042998529.1    rna-XM_008707355.2    rna-XM_027586992.2    mNeoVis.3.1.1.chr2.g01422.t01    rna-gnl|WGS_ZZZZ|egapxtmp_034342-R1    mNeoVis.evergladensis.2.1.chr2.g01488.t01    mNeoVis.vulgivaga.2.1.chr2.g01502.t01      1653    N0.HOG0001651   OG0000214  n1   rna-XM_002930742.4, rna-XM_034667681.1    rna-NM_001388902.1    rna-XM_022492579.1    rna-XM_003999612.2    rna-XM_036069855.1    rna-XM_039218509.1    rna-XM_032845666.1    rna-XM_045983344.1, rna-XM_045983365.1    rna-XM_032319346.1    rna-XM_059146637.1    rna-XM_059414164.1    rna-XM_004775805.2    mNeoVis.2.1.1.chr14_hap1.g00570.t01    rna-XM_021681943.1    rna-XM_004415676.1    rna-XM_007084272.2    rna-XM_008700739.1    rna-XM_027613420.1    mNeoVis.3.1.1.chr14.g00557.t01    rna-gnl|WGS_ZZZZ|egapxtmp_033048-R1    mNeoVis.evergladensis.2.1.chr14.g00568.t01    mNeoVis.vulgivaga.2.1.chr14.g00566.t01      1654    N0.HOG0001652   OG0000214  n23   rna-XM_002930745.3    rna-NM_001388903.1    rna-XM_022491210.1    rna-XM_045048642.1    rna-XM_036069858.1    rna-XM_039218480.1    rna-XM_032845710.1    rna-XM_045983569.1    rna-XM_032317583.1    rna-XM_059147019.1    rna-XM_059414165.1    rna-XM_004775950.2    mNeoVis.2.1.1.chr14_hap1.g00572.t01      rna-XM_004415675.1      rna-XM_008700805.2    rna-XM_027613964.1    mNeoVis.3.1.1.chr14.g00559.t01    rna-gnl|WGS_ZZZZ|egapxtmp_031955-R1    mNeoVis.evergladensis.2.1.chr14.g00570.t01    mNeoVis.vulgivaga.2.1.chr14.g00568.t01      1655    N0.HOG0001653   OG0000214  n42   rna-XM_002927257.4, rna-XM_019807955.1    rna-NM_001388898.1    rna-XM_022490733.1, rna-XM_022491173.1, rna-XM_022492577.1, rna-XM_022492578.1, rna-XM_022492788.1    rna-XM_011291149.1, rna-XM_045048641.1    rna-XM_036069842.1, rna-XM_036069844.1    rna-XM_039218478.1, rna-XM_039218479.1    rna-XM_032845697.1, rna-XM_032845713.1    rna-XM_045983318.1, rna-XM_045983319.1, rna-XM_045983464.1, rna-XM_045983474.1    rna-XM_032317259.1, rna-XM_032317374.1, rna-XM_032317580.1, rna-XM_032317581.1    rna-XM_059145410.1, rna-XM_059146758.1, rna-XM_059146823.1    rna-XM_059413965.1, rna-XM_059414162.1    rna-XM_004775806.2, rna-XM_013047450.1, rna-XM_045086545.1    mNeoVis.2.1.1.chr14_hap1.g00565.t01, mNeoVis.2.1.1.chr14_hap1.g00566.t01, mNeoVis.2.1.1.chr14_hap1.g00568.t01    rna-XM_021681846.1, rna-XM_021681847.1    rna-XM_004415684.1, rna-XM_004415685.1    rna-XM_015538449.2    rna-XM_008700802.1, rna-XM_008700803.1    rna-XM_027611958.1, rna-XM_027613765.2    mNeoVis.3.1.1.chr14.g00556.t01    rna-gnl|WGS_ZZZZ|egapxtmp_031953-R1    mNeoVis.evergladensis.2.1.chr14.g00565.t01, mNeoVis.evergladensis.2.1.chr14.g00566.t01    mNeoVis.vulgivaga.2.1.chr14.g00563.t01, mNeoVis.vulgivaga.2.1.chr14.g00564.t01      1656    N0.HOG0001654   OG0000215  n1   rna-XM_002922679.1    rna-NM_001388605.1    rna-XM_022507547.1    rna-XM_003993355.1    rna-XM_036107495.1    rna-XM_039224050.1    rna-XM_032846492.1    rna-XM_046017209.1    rna-XM_032360377.1, rna-XM_032360432.1    rna-XM_059158951.1, rna-XM_059158953.1    rna-XM_059412375.1    rna-XM_004777644.2    mNeoVis.2.1.1.chr7_hap1.g02158.t01    rna-XM_021684838.1      rna-XM_015540520.2    rna-XM_008694256.1    rna-XM_027580749.2    mNeoVis.3.1.1.chr7.g01985.t01    rna-gnl|WGS_ZZZZ|egapxtmp_013520-R1    mNeoVis.evergladensis.2.1.chr7.g02105.t01    mNeoVis.vulgivaga.2.1.chr7.g02126.t01      1657    N0.HOG0001655   OG0000215  n24   rna-XM_002922678.4, rna-XM_034644898.1, rna-XM_034651891.1, rna-XM_034651918.1    rna-NM_001388606.1, rna-XM_038562463.1    rna-XM_022507575.1    rna-XM_023239021.2      rna-XM_039224052.1    rna-XM_032846516.1    rna-XM_046016359.1, rna-XM_046016361.1, rna-XM_046017217.1    rna-XM_032360333.1, rna-XM_032360515.1    rna-XM_059158949.1, rna-XM_059158950.1, rna-XM_059158952.1    rna-XM_059387457.1, rna-XM_059387585.1    rna-XM_045068424.1, rna-XM_045068431.1    mNeoVis.2.1.1.chr7_hap1.g02157.t01        rna-XM_015540521.2    rna-XM_008694122.2, rna-XM_008694123.1, rna-XM_008694124.1    rna-XM_027580747.1    mNeoVis.3.1.1.chr7.g01984.t01    rna-gnl|WGS_ZZZZ|egapxtmp_010279-R1    mNeoVis.evergladensis.2.1.chr7.g02104.t01    mNeoVis.vulgivaga.2.1.chr7.g02125.t01      1658    N0.HOG0001656   OG0000215  n57   rna-XM_002922681.3    rna-NM_001388884.1    rna-XM_022506641.1    rna-XM_006937320.1    rna-XM_036107499.1    rna-XM_039224049.1    rna-XM_032846541.1    rna-XM_046017181.1    rna-XM_032359882.1    rna-XM_059139193.1    rna-XM_059411305.1    rna-XM_004777646.2      rna-XM_021685212.1      rna-XM_007088835.2    rna-XM_008694258.1    rna-XM_027577859.1        mNeoVis.evergladensis.2.1.chr7.g02107.t01    mNeoVis.vulgivaga.2.1.chr7.g02128.t01      1659    N0.HOG0001657   OG0000215  n74   rna-XM_011229251.2    rna-NM_001388604.1    rna-XM_022507687.1    rna-XM_011286874.4    rna-XM_036107494.1    rna-XM_039224055.1    rna-XM_032846542.1    rna-XM_046017266.1    rna-XM_032359881.1    rna-XM_059158955.1    rna-XM_059387853.1    rna-XM_004777645.3    mNeoVis.2.1.1.chr7_hap1.g02159.t01      rna-XM_004411982.1    rna-XM_007088834.1    rna-XM_008694257.2    rna-XM_027581024.1      rna-gnl|WGS_ZZZZ|egapxtmp_012635-R1    mNeoVis.evergladensis.2.1.chr7.g02106.t01    mNeoVis.vulgivaga.2.1.chr7.g02127.t01      1660    N0.HOG0001658   OG0000216  n1   rna-XM_002926231.4    rna-XM_038564343.1    rna-XM_022501826.1, rna-XM_022506767.1    rna-XM_045039426.1    rna-XM_036122259.1    rna-XM_039253577.1    rna-XM_032879205.1    rna-XM_046014173.1    rna-XM_032356959.1    rna-XM_059144273.1    rna-XM_059404210.1    rna-XM_004770323.3    mNeoVis.2.1.1.chr7_hap1.g02293.t01    rna-XM_021684911.1    rna-XM_004410050.1    rna-XM_042958879.1    rna-XM_040639207.1    rna-XM_027580702.1    mNeoVis.3.1.1.chr7.g02111.t01    rna-gnl|WGS_ZZZZ|egapxtmp_009967-R1    mNeoVis.evergladensis.2.1.chr7.g02240.t01    mNeoVis.vulgivaga.2.1.chr7.g02258.t01      1661    N0.HOG0001659   OG0000216  n18   rna-XM_002926221.4    rna-XM_038564342.1    rna-XM_022506706.1    rna-XM_023239925.2    rna-XM_036122281.1    rna-XM_039253499.1    rna-XM_032879200.1    rna-XM_046014487.1    rna-XM_032356956.1    rna-XM_059144185.1    rna-XM_059404235.1    rna-XM_045063347.1    mNeoVis.2.1.1.chr7_hap1.g02292.t01    rna-XM_021685470.2    rna-XM_004410051.1    rna-XM_042959591.1    rna-XM_040639205.1    rna-XM_027580434.2      rna-gnl|WGS_ZZZZ|egapxtmp_012671-R1    mNeoVis.evergladensis.2.1.chr7.g02239.t01    mNeoVis.vulgivaga.2.1.chr7.g02257.t01      1662    N0.HOG0001660   OG0000216  n38   rna-XM_002922685.4, rna-XM_011233618.3    rna-XM_038564347.1, rna-XM_038564353.1, rna-XM_038564354.1    rna-XM_022506638.1, rna-XM_022506707.1    rna-XM_003993434.6, rna-XM_045039301.1, rna-XM_045039306.1, rna-XM_045039308.1    rna-XM_036076612.1, rna-XM_036122284.1    rna-XM_039225588.1, rna-XM_039225621.1, rna-XM_039225636.1, rna-XM_039253495.1    rna-XM_032846461.1, rna-XM_032879204.1    rna-XM_046014488.1, rna-XM_046016881.1    rna-XM_032356961.1, rna-XM_032358460.1    rna-XM_059144243.1, rna-XM_059158961.1    rna-XM_059404284.1, rna-XM_059405657.1    rna-XM_004770328.3, rna-XM_004777565.3    mNeoVis.2.1.1.chr7_hap1.g02165.t01, mNeoVis.2.1.1.chr7_hap1.g02291.t01    rna-XM_021685215.1, rna-XM_021685472.2    rna-XM_004409307.1, rna-XM_004410052.1    rna-XM_007088836.3, rna-XM_007092652.3, rna-XM_042959592.1    rna-XM_008694128.1, rna-XM_040638767.1    rna-XM_027577857.2, rna-XM_027580436.2    mNeoVis.3.1.1.chr7.g01992.t01, mNeoVis.3.1.1.chr7.g02109.t01    rna-gnl|WGS_ZZZZ|egapxtmp_009968-R4, rna-gnl|WGS_ZZZZ|egapxtmp_010334-R1    mNeoVis.evergladensis.2.1.chr7.g02113.t01, mNeoVis.evergladensis.2.1.chr7.g02238.t01    mNeoVis.vulgivaga.2.1.chr7.g02135.t01, mNeoVis.vulgivaga.2.1.chr7.g02256.t01      1663    N0.HOG0001661   OG0000217  n1   rna-XM_034640580.1    rna-NM_001024638.1    rna-XM_022509212.1    rna-XM_023243660.2      rna-XM_039227824.1    rna-XM_032872039.1    rna-XM_045985721.1, rna-XM_045991357.1    rna-XM_032316667.1, rna-XM_032322949.1, rna-XM_032360502.1    rna-XM_059150545.1, rna-XM_059165129.1    rna-XM_059378825.1, rna-XM_059387576.1    rna-XM_004764855.3, rna-XM_045088851.1    mNeoVis.2.1.1.chr8_hap1.g00593.t01    rna-XM_021704011.1    rna-XM_004395210.1    rna-XM_042966752.1    rna-XM_040640846.1    rna-XM_027626319.2    mNeoVis.3.1.1.chr8.g00548.t01    rna-gnl|WGS_ZZZZ|egapxtmp_019671-R1    mNeoVis.evergladensis.2.1.chr9.g00584.t01    mNeoVis.vulgivaga.2.1.chr8.g00591.t01      1664    N0.HOG0001662   OG0000217  n20   rna-XM_034651855.1    rna-XM_038538939.1      rna-XM_023246416.2    rna-XM_036083337.1    rna-XM_039246689.1    rna-XM_032846111.1    rna-XM_045993320.1    rna-XM_032327286.1    rna-XM_059155422.1    rna-XM_059415712.1    rna-XM_045072194.1    mNeoVis.2.1.1.chr15_hap1.g00381.t01, mNeoVis.2.1.1.chr15_hap1.g00455.t01    rna-XM_021698090.1    rna-XM_004403133.1    rna-XM_042970952.1    rna-XM_040635554.1    rna-XM_027614138.2              1665    N0.HOG0001663   OG0000217  n37         rna-XM_006940203.4                          rna-XM_042966753.1                  1666    N0.HOG0001664   OG0000217  n39       rna-XM_022509213.1          rna-XM_032872041.1    rna-XM_045985722.1, rna-XM_046004533.1    rna-XM_032322947.1, rna-XM_032325818.1    rna-XM_059150547.1, rna-XM_059153186.1    rna-XM_059381896.1    rna-XM_004764854.3, rna-XM_045076022.1    mNeoVis.2.1.1.chr7_hap1.g00357.t01, mNeoVis.2.1.1.chr8_hap1.g00592.t01              mNeoVis.3.1.1.chr7.g00345.t01, mNeoVis.3.1.1.chr8.g00547.t01    rna-gnl|WGS_ZZZZ|egapxtmp_019670-R1    mNeoVis.evergladensis.2.1.chr7.g00360.t01, mNeoVis.evergladensis.2.1.chr9.g00583.t01    mNeoVis.vulgivaga.2.1.chr7.g00362.t01, mNeoVis.vulgivaga.2.1.chr8.g00590.t01      1667    N0.HOG0001665   OG0000217  n51   rna-XM_034653474.1, rna-XM_034663576.1    rna-XM_038529558.1, rna-XM_038539023.1, rna-XM_038544918.1, rna-XM_038568260.1    rna-XM_022506063.1          rna-XM_032864738.1    rna-XM_045998093.1                                  1668    N0.HOG0001666   OG0000217  n59   rna-XM_011233315.3    rna-NM_001024637.1        rna-XM_036108484.1          rna-XM_032334010.1    rna-XM_059164475.1    rna-XM_059395869.1    rna-XM_045076149.1    mNeoVis.2.1.1.chr6_hap1.g00263.t01    rna-XM_021704008.1        rna-XM_008687852.2    rna-XM_027626320.1    mNeoVis.3.1.1.chr4.g00262.t01    rna-gnl|WGS_ZZZZ|egapxtmp_001535-R1    mNeoVis.evergladensis.2.1.chr4.g00270.t01    mNeoVis.vulgivaga.2.1.chr4.g00269.t01      1669    N0.HOG0001667   OG0000218  n1   rna-XM_034657739.1    rna-XM_038567358.1    rna-XM_022526905.1    rna-XM_045045327.1    rna-XM_036119642.1    rna-XM_039250988.1    rna-XM_032883936.1    rna-XM_045991414.1    rna-XM_032310546.1    rna-XM_059159903.1    rna-XM_059388955.1    rna-XM_004748484.2    mNeoVis.2.1.1.chr5_hap1.g00265.t01    rna-XM_044918087.1    rna-XM_004398843.1    rna-XM_042973974.1    rna-XM_008708137.2    rna-XM_027586299.1    mNeoVis.3.1.1.chr2.g00207.t01    rna-gnl|WGS_ZZZZ|egapxtmp_034652-R1    mNeoVis.evergladensis.2.1.chr2.g00263.t01    mNeoVis.vulgivaga.2.1.chr2.g00258.t01      1670    N0.HOG0001668   OG0000218  n23   rna-XM_034660529.1    rna-XM_038553692.1    rna-XM_022510584.1    rna-XM_045057375.1    rna-XM_036072332.1    rna-XM_039241825.1    rna-XM_032850584.1, rna-XM_032850594.1    rna-XM_046004542.1    rna-XM_032340680.1    rna-XM_059178692.1    rna-XM_059399984.1    rna-XM_045074680.1    mNeoVis.2.1.1.chr1_hap1.g01418.t01    rna-XM_021684541.1    rna-XM_004407687.1    rna-XM_007093234.3    rna-XM_040644776.1    rna-XM_027603567.2    mNeoVis.3.1.1.chr1.g01365.t01    rna-gnl|WGS_ZZZZ|egapxtmp_021748-R1    mNeoVis.evergladensis.2.1.chr1.g01380.t01    mNeoVis.vulgivaga.2.1.chr1.g01384.t01      1671    N0.HOG0001669   OG0000218  n46   rna-XM_034663748.1, rna-XM_034663749.1    rna-XM_038548863.1    rna-XM_022500736.1    rna-XM_023242951.2    rna-XM_036108705.1    rna-XM_039255152.1    rna-XM_032866237.1    rna-XM_046022302.1    rna-XM_032306372.1    rna-XM_059143471.1    rna-XM_059410379.1    rna-XM_045082847.1    mNeoVis.2.1.1.chr13_hap1.g00082.t01    rna-XM_044912078.1, rna-XM_044912079.1, rna-XM_044920661.1    rna-XM_004411394.1    rna-XM_042965314.1    rna-XM_040631357.1    rna-XM_035723591.1    mNeoVis.3.1.1.chr13.g00045.t01    rna-gnl|WGS_ZZZZ|egapxtmp_031056-R1    mNeoVis.evergladensis.2.1.chr13.g00083.t01    mNeoVis.vulgivaga.2.1.chr13.g00080.t01      1672    N0.HOG0001670   OG0000218  n70   rna-XM_011224596.3    rna-XM_038541181.1    rna-XM_022496424.1    rna-XM_023258940.2    rna-XM_036112142.1    rna-XM_039216657.1    rna-XM_032867236.1    rna-XM_046022249.1    rna-XM_032302425.1    rna-XM_059137189.1    rna-XM_059377355.1    rna-XM_004775159.3    mNeoVis.2.1.1.chr2_hap1.g01029.t01    rna-XM_021690257.1    rna-XM_004401456.1    rna-XM_042998028.1    rna-XM_008697639.2    rna-XM_027571529.2    mNeoVis.3.1.1.chr5.g00955.t01    rna-gnl|WGS_ZZZZ|egapxtmp_009520-R1    mNeoVis.evergladensis.2.1.chr5.g00975.t01    mNeoVis.vulgivaga.2.1.chr5.g00964.t01      1673    N0.HOG0001671   OG0000219  n1   rna-XM_034658681.1    rna-XM_038565780.1    rna-XM_022498291.1    rna-XM_011280176.4    rna-XM_036099245.1, rna-XM_036101751.1    rna-XM_039243211.1    rna-XM_032867455.1    rna-XM_045989695.1    rna-XM_032325977.1    rna-XM_059161568.1    rna-XM_059390206.1    rna-XM_045063251.1    mNeoVis.2.1.1.chr5_hap1.g00842.t01    rna-XM_021694901.1    rna-XM_004412197.1    rna-XM_042979452.1    rna-XM_008697935.2    rna-XM_027586605.2    mNeoVis.3.1.1.chr2.g00752.t01    rna-gnl|WGS_ZZZZ|egapxtmp_035469-R1, rna-gnl|WGS_ZZZZ|egapxtmp_037206-R1    mNeoVis.evergladensis.2.1.chr2.g00844.t01    mNeoVis.vulgivaga.2.1.chr2.g00845.t01      1674    N0.HOG0001672   OG0000219  n17   rna-XM_034655851.1    rna-XM_038551774.1    rna-XM_022521222.1    rna-XM_023254137.2    rna-XM_036080124.1    rna-XM_039252854.1    rna-XM_032846865.1    rna-XM_045999123.1    rna-XM_032335326.1    rna-XM_059175873.1    rna-XM_059416713.1    rna-XM_004759086.3    mNeoVis.2.1.1.chr1_hap1.g00880.t01    rna-XM_021690935.1    rna-XM_004412679.1    rna-XM_042958659.1    rna-XM_040622611.1    rna-XM_027604733.1    mNeoVis.3.1.1.chr1.g00902.t01      mNeoVis.evergladensis.2.1.chr1.g00883.t01    mNeoVis.vulgivaga.2.1.chr1.g00894.t01      1675    N0.HOG0001673   OG0000219  n37   rna-XM_034660566.1    rna-XM_038553771.1    rna-XM_022510812.1    rna-XM_023253859.2    rna-XM_036115773.1    rna-XM_039241816.1    rna-XM_032849150.1    rna-XM_046006405.1    rna-XM_032340602.1    rna-XM_059178602.1    rna-XM_059400048.1    rna-XM_045074721.1    mNeoVis.2.1.1.chr1_hap1.g01445.t01    rna-XM_021684584.1    rna-XM_004407720.1    rna-XM_042986143.1    rna-XM_040644746.1    rna-XM_027604169.2    mNeoVis.3.1.1.chr1.g01391.t01, mNeoVis.3.1.1.chr1.g01392.t01    rna-gnl|WGS_ZZZZ|egapxtmp_021781-R1    mNeoVis.evergladensis.2.1.chr1.g01407.t01    mNeoVis.vulgivaga.2.1.chr1.g01411.t01      1676    N0.HOG0001674   OG0000219  n58   rna-XM_002913413.4    rna-XM_038556627.1    rna-XM_022510280.1    rna-XM_045052711.1    rna-XM_036095992.1, rna-XM_036123636.1    rna-XM_039243493.1    rna-XM_032870003.1    rna-XM_046019994.1    rna-XM_032305558.1    rna-XM_059171757.1    rna-XM_059396474.1    rna-XM_004741849.3    mNeoVis.2.1.1.chr4_hap1.g00245.t01    rna-XM_021699435.1    rna-XM_004398046.1    rna-XM_042971356.1    rna-XM_008705831.2, rna-XM_040644057.1    rna-XM_027573922.1    mNeoVis.3.1.1.chr6.g00244.t01    rna-gnl|WGS_ZZZZ|egapxtmp_003933-R1, rna-gnl|WGS_ZZZZ|egapxtmp_003934-R1    mNeoVis.evergladensis.2.1.chr6.g00244.t01    mNeoVis.vulgivaga.2.1.chr6.g00245.t01      1677    N0.HOG0001675   OG0000220  n1   rna-XM_002923596.4    rna-XM_038534961.1    rna-XM_022493249.1    rna-XM_003981365.5    rna-XM_036114252.1    rna-XM_039224724.1    rna-XM_032851618.1    rna-XM_046000553.1    rna-XM_032337241.1    rna-XM_059174100.1    rna-XM_059417205.1    rna-XM_004737770.3    mNeoVis.2.1.1.chr1_hap1.g00229.t01    rna-XM_021697251.1    rna-XM_004409709.1    rna-XM_015535468.2    rna-XM_008692468.1    rna-XM_035727550.1    mNeoVis.3.1.1.chr1.g00229.t01    rna-gnl|WGS_ZZZZ|egapxtmp_018003-R1    mNeoVis.evergladensis.2.1.chr1.g00226.t01    mNeoVis.vulgivaga.2.1.chr1.g00227.t01      1678    N0.HOG0001676   OG0000220  n15   rna-XM_034661467.1    rna-XM_038559096.1    rna-XM_022512987.1    rna-XM_011281602.4    rna-XM_036088961.1    rna-XM_039221125.1, rna-XM_039256866.1    rna-XM_032863056.1    rna-XM_045997209.1    rna-XM_032332693.1    rna-XM_059173994.1    rna-XM_059374664.1    rna-XM_004772044.3    mNeoVis.2.1.1.chr6_hap1.g00649.t01    rna-XM_021699026.1    rna-XM_004394234.1    rna-XM_007084208.3    rna-XM_008685537.1    rna-XM_027598314.2    mNeoVis.3.1.1.chr4.g00656.t01    rna-gnl|WGS_ZZZZ|egapxtmp_001579-R2    mNeoVis.evergladensis.2.1.chr4.g00657.t01    mNeoVis.vulgivaga.2.1.chr4.g00667.t01      1679    N0.HOG0001677   OG0000220  n32   rna-XM_034649234.1    rna-XM_038588235.1    rna-XM_022518525.1    rna-XM_019824236.2    rna-XM_036094379.1    rna-XM_039238823.1    rna-XM_032882315.1    rna-XM_045995279.1    rna-XM_032330365.1    rna-XM_059156924.1    rna-XM_059384587.1    rna-XM_013061246.2    mNeoVis.2.1.1.chr12_hap1.g00220.t01    rna-XM_021686020.1    rna-XM_004400187.1    rna-XM_042974909.1    rna-XM_040628391.1    rna-XM_027607886.2    mNeoVis.3.1.1.chr12.g00189.t01, mNeoVis.3.1.1.chr12.g00190.t01    rna-gnl|WGS_ZZZZ|egapxtmp_029135-R1    mNeoVis.evergladensis.2.1.chr12.g00192.t01    mNeoVis.vulgivaga.2.1.chr12.g00196.t01      1680    N0.HOG0001678   OG0000220  n47   rna-XM_034665671.1    rna-XM_038536334.1, rna-XM_038589465.1, rna-XM_038589466.1, rna-XM_038589549.1    rna-XM_022512979.1    rna-XM_045039797.1    rna-XM_036103773.1    rna-XM_039250622.1    rna-XM_032838408.1    rna-XM_046014154.1    rna-XM_032357837.1    rna-XM_059179531.1    rna-XM_059418401.1    rna-XM_004764438.3    mNeoVis.2.1.1.chr7_hap1.g01146.t01    rna-XM_021696567.1    rna-XM_004412767.1    rna-XM_042958444.1    rna-XM_008690134.1    rna-XM_027578964.2    mNeoVis.3.1.1.chr7.g01069.t01    rna-gnl|WGS_ZZZZ|egapxtmp_011342-R5    mNeoVis.evergladensis.2.1.chr7.g01133.t01    mNeoVis.vulgivaga.2.1.chr7.g01145.t01      1681    N0.HOG0001679   OG0000221  n1   rna-XM_034667401.1    rna-XM_038542972.1    rna-XM_022491171.1    rna-XM_023247315.2    rna-XM_036118262.1    rna-XM_039251308.1    rna-XM_032845048.1    rna-XM_045983206.1    rna-XM_032318078.1    rna-XM_059146083.1    rna-XM_059413617.1    rna-XM_045086705.1    mNeoVis.2.1.1.chr14_hap1.g00600.t01    rna-XM_044916008.1    rna-XM_004402887.1    rna-XM_042976099.1    rna-XM_040635836.1    rna-XM_027613635.2    mNeoVis.3.1.1.chr14.g00581.t01    rna-gnl|WGS_ZZZZ|egapxtmp_031963-R1    mNeoVis.evergladensis.2.1.chr14.g00597.t01    mNeoVis.vulgivaga.2.1.chr14.g00593.t01      1682    N0.HOG0001680   OG0000221  n24     rna-XM_038542978.1    rna-XM_022490805.1    rna-XM_023247543.2    rna-XM_036118268.1    rna-XM_039251312.1    rna-XM_032845067.1      rna-XM_032318083.1    rna-XM_059146377.1    rna-XM_059413623.1    rna-XM_045086710.1    mNeoVis.2.1.1.chr14_hap1.g00601.t01    rna-XM_044912071.1, rna-XM_044912072.1    rna-XM_004402889.1    rna-XM_042976291.1    rna-XM_040634474.1    rna-XM_027612047.2    mNeoVis.3.1.1.chr14.g00582.t01, mNeoVis.3.1.1.chr8.g00443.t01    rna-gnl|WGS_ZZZZ|egapxtmp_031964-R1    mNeoVis.evergladensis.2.1.chr14.g00598.t01    mNeoVis.vulgivaga.2.1.chr14.g00594.t01      1683    N0.HOG0001681   OG0000221  n46   rna-XM_019794985.2    rna-XM_038526948.1    rna-XM_022512746.1    rna-XM_023242218.2    rna-XM_036113467.1    rna-XM_039222730.1, rna-XM_039222733.1, rna-XM_039222734.1    rna-XM_032840563.1    rna-XM_046023045.1    rna-XM_032307203.1    rna-XM_059142000.1    rna-XM_059411758.1    rna-XM_045086077.1    mNeoVis.2.1.1.chr13_hap1.g00683.t01    rna-XM_021677950.2    rna-XM_004407588.1    rna-XM_042963894.1    rna-XM_040620196.1    rna-XM_027614950.2    mNeoVis.3.1.1.chr13.g00605.t01    rna-gnl|WGS_ZZZZ|egapxtmp_031363-R1    mNeoVis.evergladensis.2.1.chr13.g00654.t01    mNeoVis.vulgivaga.2.1.chr13.g00656.t01      1684    N0.HOG0001682   OG0000221  n66   rna-XM_019799995.2    rna-XM_038532784.1    rna-XM_022511090.1    rna-XM_045058825.1    rna-XM_036116308.1    rna-XM_039217000.1, rna-XM_039217027.1    rna-XM_032836989.1    rna-XM_046008586.1    rna-XM_032342082.1    rna-XM_059180038.1    rna-XM_059371609.1    rna-XM_045067153.1    mNeoVis.2.1.1.chr10_hap1.g01187.t01    rna-XM_021680526.1    rna-XM_004398317.1    rna-XM_042988103.1    rna-XM_040642772.1    rna-XM_027570490.2    mNeoVis.3.1.1.chr9.g01118.t01    rna-gnl|WGS_ZZZZ|egapxtmp_017039-R1    mNeoVis.evergladensis.2.1.chr8.g01163.t01    mNeoVis.vulgivaga.2.1.chr9.g01146.t01      1685    N0.HOG0001683   OG0000222  n1   rna-XM_034668916.1    rna-XM_038530577.1      rna-XM_045038369.1    rna-XM_036072784.1, rna-XM_036073962.1, rna-XM_036074242.1, rna-XM_036074698.1, rna-XM_036080596.1, rna-XM_036083959.1, rna-XM_036084362.1, rna-XM_036092405.1, rna-XM_036105930.1, rna-XM_036113945.1, rna-XM_036117178.1          rna-XM_032310043.1, rna-XM_032312015.1    rna-XM_059170194.1, rna-XM_059177443.1, rna-XM_059180008.1    rna-XM_059395818.1, rna-XM_059399188.1, rna-XM_059418468.1    rna-XM_045067185.1, rna-XM_045069907.1      rna-XM_044912570.1, rna-XM_044913436.1, rna-XM_044914013.1, rna-XM_044916218.1, rna-XM_044916924.1, rna-XM_044917479.1, rna-XM_044917767.1, rna-XM_044918155.1, rna-XM_044918848.1, rna-XM_044919526.1, rna-XM_044920107.1, rna-XM_044921401.1, rna-XM_044921676.1    rna-XM_004391700.1, rna-XM_004395518.1, rna-XM_004405960.1, rna-XM_004413611.1, rna-XM_004414398.1    rna-XM_042989269.1      rna-XM_035722464.1, rna-XM_035723491.1, rna-XM_035723750.1, rna-XM_035726223.1, rna-XM_035726760.1, rna-XM_035728939.1, rna-XM_035729051.1, rna-XM_035729071.1          mNeoVis.vulgivaga.2.1.chr9.g01039.t01      1686    N0.HOG0001684   OG0000222  n53   rna-NM_001304898.1    rna-NM_001252203.1, rna-XM_038565022.1, rna-XM_038586535.1    rna-XM_022516396.1    rna-XM_045054064.1    rna-XM_036087947.1    rna-XM_039253125.1    rna-XM_032840351.1    rna-XM_045981734.1    rna-XM_032351784.1    rna-XM_059134488.1    rna-XM_059376240.1    rna-XM_004753039.3, rna-XM_045066312.1    mNeoVis.2.1.1.chr11_hap1.g00597.t01    rna-XM_021680358.1, rna-XM_021694346.1    rna-XM_004401567.1, rna-XM_004402396.1    rna-XM_007097783.3    rna-XM_008699749.2    rna-XM_027623179.2, rna-XM_035726140.1, rna-XM_035727691.1, rna-XM_035728112.1    mNeoVis.3.1.1.chr11.g00558.t01    rna-gnl|WGS_ZZZZ|egapxtmp_014742-R1    mNeoVis.evergladensis.2.1.chr11.g00584.t01, mNeoVis.evergladensis.2.1.chr8.g01058.t01    mNeoVis.vulgivaga.2.1.chr11.g00593.t01      1687    N0.HOG0001685   OG0000222  n54     rna-XM_038582343.1          rna-XM_039257242.1      rna-XM_046003278.1        rna-XM_059407853.1            rna-XM_042989266.1    rna-XM_040630557.1    rna-XM_035722865.1, rna-XM_035723201.1, rna-XM_035724293.1, rna-XM_035726284.1              1688    N0.HOG0001686   OG0000223  n1   rna-XM_002917747.4    rna-XM_038587436.1    rna-XM_022507823.1    rna-XM_023249415.1    rna-XM_036082417.1    rna-XM_039218256.1    rna-XM_032842879.1    rna-XM_045995148.1    rna-XM_032330093.1    rna-XM_059156876.1    rna-XM_059385627.1    rna-XM_004755090.1    mNeoVis.2.1.1.chr12_hap1.g00594.t01    rna-XM_021678783.1    rna-XM_004396817.1    rna-XM_042973929.1    rna-XM_040631640.1    rna-XM_027608743.1    mNeoVis.3.1.1.chr12.g00541.t01    rna-gnl|WGS_ZZZZ|egapxtmp_028211-R1    mNeoVis.evergladensis.2.1.chr12.g00531.t01    mNeoVis.vulgivaga.2.1.chr12.g00545.t01      1689    N0.HOG0001687   OG0000223  n24   rna-XM_019794519.2    rna-XM_038541920.1    rna-XM_022508677.1    rna-NM_001038605.2    rna-XM_036086684.1    rna-XM_039227512.1    rna-XM_032878228.1    rna-XM_045982783.1    rna-XM_032318911.1    rna-XM_059146380.1    rna-XM_059414074.1    rna-XM_004756256.3    mNeoVis.2.1.1.chr14_hap1.g00315.t01    rna-XM_021682480.1    rna-XM_004392733.1    rna-XM_042976391.1    rna-XM_008685179.2    rna-XM_027611501.2    mNeoVis.3.1.1.chr14.g00314.t01    rna-gnl|WGS_ZZZZ|egapxtmp_033022-R1    mNeoVis.evergladensis.2.1.chr14.g00316.t01    mNeoVis.vulgivaga.2.1.chr14.g00316.t01      1690    N0.HOG0001688   OG0000223  n45   rna-XM_019793444.2    rna-XM_038576272.1    rna-XM_022522233.1    rna-XM_045061160.1    rna-XM_036095077.1, rna-XM_036101410.1    rna-XM_039250015.1    rna-XM_032844463.1    rna-XM_046012080.1    rna-XM_032348855.1, rna-XM_032349969.1    rna-XM_059184355.1, rna-XM_059187467.1    rna-XM_059403901.1, rna-XM_059404543.1    rna-XM_045071752.1, rna-XM_045071878.1    mNeoVis.2.1.1.chr9_hap1.g00193.t01    rna-XM_021690718.2    rna-XM_004406879.1    rna-XM_042991494.1    rna-XM_040641983.1    rna-XM_027592946.2    mNeoVis.3.1.1.chr10.g00185.t01    rna-gnl|WGS_ZZZZ|egapxtmp_024605-R1    mNeoVis.evergladensis.2.1.chr10.g00188.t01    mNeoVis.vulgivaga.2.1.chr10.g00185.t01      1691    N0.HOG0001689   OG0000223  n71   rna-XM_034658480.1    rna-XM_038566148.1    rna-XM_022497560.1    rna-XM_045051390.1    rna-XM_036102518.1    rna-XM_039243604.1    rna-XM_032868310.1    rna-XM_045990264.1    rna-XM_032322272.1    rna-XM_059161179.1    rna-XM_059389949.1    rna-XM_045078797.1    mNeoVis.2.1.1.chr5_hap1.g00740.t01    rna-XM_021694968.1    rna-XM_004411446.1    rna-XM_042977073.1    rna-XM_040628679.1    rna-XM_035727534.1    mNeoVis.3.1.1.chr2.g00650.t01    rna-gnl|WGS_ZZZZ|egapxtmp_035433-R1    mNeoVis.evergladensis.2.1.chr2.g00742.t01    mNeoVis.vulgivaga.2.1.chr2.g00746.t01      1692    N0.HOG0001690   OG0000224  n1   rna-XM_019810249.2    rna-XM_038531341.1    rna-XM_022496278.1    rna-XM_045035400.1    rna-XM_036115402.1    rna-XM_039232699.1    rna-XM_032863982.1    rna-XM_046012571.1    rna-XM_032360832.1    rna-XM_059136122.1    rna-XM_059376920.1    rna-XM_045064074.1    mNeoVis.2.1.1.chr2_hap1.g01583.t01    rna-XM_044913935.1    rna-XM_004394630.1    rna-XM_042996063.1    rna-XM_008694493.2    rna-XM_027569837.2    mNeoVis.3.1.1.chr5.g01505.t01    rna-gnl|WGS_ZZZZ|egapxtmp_007118-R1    mNeoVis.evergladensis.2.1.chr5.g01522.t01    mNeoVis.vulgivaga.2.1.chr5.g01524.t01      1693    N0.HOG0001691   OG0000224  n25   rna-XM_034669107.1    rna-XM_038526752.1    rna-XM_022503785.1    rna-XM_023254330.2    rna-XM_036105406.1    rna-XM_039256923.1    rna-XM_032862503.1    rna-XM_046004500.1    rna-XM_032339563.1    rna-XM_059177599.1    rna-XM_059400697.1    rna-XM_004740175.3    mNeoVis.2.1.1.chr1_hap1.g01775.t01    rna-XM_044917510.1    rna-XM_004413170.1    rna-XM_042987143.1    rna-XM_008691238.2    rna-XM_027601758.2    mNeoVis.3.1.1.chr1.g01727.t01    rna-gnl|WGS_ZZZZ|egapxtmp_013219-R1    mNeoVis.evergladensis.2.1.chr1.g01742.t01    mNeoVis.vulgivaga.2.1.chr1.g01739.t01      1694    N0.HOG0001692   OG0000224  n47   rna-XM_034641590.1    rna-XM_038572340.1    rna-XM_022494012.1    rna-XM_023251232.2    rna-XM_036081752.1    rna-XM_039227066.1    rna-XM_032838508.1    rna-XM_045981306.1    rna-XM_032350627.1    rna-XM_059133123.1    rna-XM_059407017.1    rna-XM_045082202.1    mNeoVis.2.1.1.chr11_hap1.g00224.t01    rna-XM_044919044.1    rna-XM_004417013.1    rna-XM_042980385.1    rna-XM_040638322.1    rna-XM_027622214.1    mNeoVis.3.1.1.chr1.g00079.t01, mNeoVis.3.1.1.chr11.g00195.t01    rna-gnl|WGS_ZZZZ|egapxtmp_023983-R1, rna-gnl|WGS_ZZZZ|egapxtmp_026821-R1    mNeoVis.evergladensis.2.1.chr11.g00222.t01    mNeoVis.vulgivaga.2.1.chr11.g00224.t01      1695    N0.HOG0001693   OG0000224  n69   rna-XM_034642280.1    rna-XM_038543828.1    rna-XM_022495810.1    rna-XM_019814992.3    rna-XM_036100796.1    rna-XM_039255234.1    rna-XM_032875724.1    rna-XM_046025007.1    rna-XM_032310610.1    rna-XM_059139376.1    rna-XM_059409212.1    rna-XM_045065752.1    mNeoVis.2.1.1.chr3_hap1.g00254.t01    rna-XM_044921209.1    rna-XM_004405342.1    rna-XM_042962258.1    rna-XM_040632384.1    rna-XM_027575435.2    mNeoVis.3.1.1.chr3.g00256.t01    rna-gnl|WGS_ZZZZ|egapxtmp_041049-R1    mNeoVis.evergladensis.2.1.chr3.g00264.t01    mNeoVis.vulgivaga.2.1.chr3.g00255.t01      1696    N0.HOG0001694   OG0000225  n1     rna-NM_001252170.1    rna-XM_022517547.1    rna-XM_045054074.1    rna-XM_036091736.1    rna-XM_039244425.1    rna-XM_032880514.1    rna-XM_046023806.1    rna-XM_032308922.1    rna-XM_059143125.1    rna-XM_059411579.1    rna-XM_045079662.1    mNeoVis.2.1.1.chr13_hap1.g00613.t01    rna-XM_021682457.2, rna-XM_044914042.1    rna-XM_004405664.1    rna-XM_042998457.1    rna-XM_008689807.2    rna-XM_027616653.1      rna-gnl|WGS_ZZZZ|egapxtmp_030972-R1      mNeoVis.vulgivaga.2.1.chr13.g00589.t01      1697    N0.HOG0001695   OG0000225  n9                 rna-XM_046002619.1      rna-XM_059164265.1, rna-XM_059164406.1    rna-XM_059388141.1, rna-XM_059388167.1    rna-XM_045088245.1, rna-XM_045088288.1              rna-XM_035726940.1, rna-XM_035729137.1              1698    N0.HOG0001696   OG0000225  n18                     rna-XM_059159713.1                  rna-XM_035723641.1              1699    N0.HOG0001697   OG0000225  n20                                     rna-XM_035721892.1, rna-XM_035728164.1              1700    N0.HOG0001698   OG0000225  n24     rna-XM_038561616.1    rna-XM_022505644.1    rna-XM_045037333.1    rna-XM_036085621.1, rna-XM_036087273.1, rna-XM_036091342.1      rna-XM_032840917.1, rna-XM_032856938.1      rna-XM_032354346.1            rna-XM_021692228.1      rna-XM_042987729.1, rna-XM_042997608.1    rna-XM_040631334.1, rna-XM_040635246.1    rna-XM_035726897.1, rna-XM_035729172.1    mNeoVis.3.1.1.chr13.g00537.t01            1701    N0.HOG0001699   OG0000225  n40     rna-XM_038583287.1        rna-XM_036073068.1                    rna-XM_044919183.1          rna-XM_035723370.1              1702    N0.HOG0001700   OG0000225  n46                                     rna-XM_035722505.1, rna-XM_035722812.1, rna-XM_035723285.1              1703    N0.HOG0001701   OG0000225  n54     rna-XM_038536292.1, rna-XM_038538803.1      rna-XM_003995497.4      rna-XM_039224412.1      rna-XM_046005547.1    rna-XM_032322705.1              rna-XM_004403987.1              mNeoVis.evergladensis.2.1.chr13.g00589.t01        1704    N0.HOG0001702   OG0000225  n61     rna-XM_038587311.1        rna-XM_036102027.1    rna-XM_039256903.1          rna-XM_059148665.1, rna-XM_059162172.1    rna-XM_059380481.1    rna-XM_045085929.1          rna-XM_007099113.3    rna-XM_040623428.1    rna-XM_035726898.1, rna-XM_035728551.1          mNeoVis.vulgivaga.2.1.chr6.g00031.t01      1705    N0.HOG0001703   OG0000225  n72                               rna-XM_004405433.1        rna-XM_035726216.1              1706    N0.HOG0001704   OG0000225  n73           rna-XM_036116048.1        rna-XM_046027570.1                      rna-XM_035726853.1, rna-XM_035728286.1        mNeoVis.evergladensis.2.1.chr4.g00149.t01, mNeoVis.evergladensis.2.1.chr6.g00026.t01        1707    N0.HOG0001705   OG0000225  n78                         rna-XM_045089081.1              rna-XM_035721992.1              1708    N0.HOG0001706   OG0000226  n1   rna-XM_034661231.1    rna-XM_038573766.1    rna-XM_022513979.1    rna-XM_011281440.4    rna-XM_036106184.1    rna-XM_039250424.1    rna-XM_032849437.1, rna-XM_032855791.1    rna-XM_045991553.1    rna-XM_032332214.1    rna-XM_059177834.1    rna-XM_059372035.1    rna-XM_013061798.2    mNeoVis.2.1.1.chr6_hap1.g00808.t01    rna-XM_044912845.1    rna-XM_004412986.1    rna-XM_042983184.1    rna-XM_008702052.2    rna-XM_027598419.1    mNeoVis.3.1.1.chr4.g00818.t01    rna-gnl|WGS_ZZZZ|egapxtmp_000618-R1    mNeoVis.evergladensis.2.1.chr4.g00810.t01    mNeoVis.vulgivaga.2.1.chr4.g00830.t01      1709    N0.HOG0001707   OG0000226  n13   rna-XM_002924149.4    rna-XM_038533057.1    rna-XM_022506114.1    rna-XM_023253331.2    rna-XM_036104661.1    rna-XM_039232158.1    rna-XM_032855388.1    rna-XM_045997863.1    rna-XM_032333777.1    rna-XM_059165694.1    rna-XM_059380499.1    rna-XM_004770526.3    mNeoVis.2.1.1.chr6_hap1.g00328.t01    rna-XM_021677609.1    rna-XM_004416863.1    rna-XM_042984800.1    rna-XM_008685817.2    rna-XM_027600554.2    mNeoVis.3.1.1.chr4.g00329.t01    rna-gnl|WGS_ZZZZ|egapxtmp_042988-R1    mNeoVis.evergladensis.2.1.chr4.g00334.t01    mNeoVis.vulgivaga.2.1.chr4.g00331.t01      1710    N0.HOG0001708   OG0000226  n24   rna-XM_034656220.1    rna-XM_038530528.1    rna-XM_022518090.1    rna-XM_019833906.3    rna-XM_036070010.1, rna-XM_036100442.1    rna-XM_039256697.1    rna-XM_032850790.1    rna-XM_046000714.1    rna-XM_032336274.1    rna-XM_059174966.1    rna-XM_059417851.1    rna-XM_004744681.3, rna-XM_045073794.1    mNeoVis.2.1.1.chr1_hap1.g00557.t01    rna-XM_044917125.1    rna-XM_004414632.1    rna-XM_042986515.1    rna-XM_040622265.1    rna-XM_027605525.2    mNeoVis.3.1.1.chr1.g00572.t01    rna-gnl|WGS_ZZZZ|egapxtmp_026048-R1    mNeoVis.evergladensis.2.1.chr1.g00563.t01    mNeoVis.vulgivaga.2.1.chr1.g00573.t01      1711    N0.HOG0001709   OG0000226  n39   rna-XM_034663452.1    rna-XM_038579147.1    rna-XM_022499470.1    rna-XM_045040866.1    rna-XM_036070117.1    rna-XM_039252407.1    rna-XM_032882060.1    rna-XM_045985817.1, rna-XM_046027911.1    rna-XM_032312720.1    rna-XM_059172807.1    rna-XM_059398840.1    rna-XM_045068516.1    mNeoVis.2.1.1.chr2_hap1.g00035.t01    rna-XM_021703757.1    rna-XM_004402228.1    rna-XM_042961127.1    rna-XM_040624104.1    rna-XM_027586830.2    mNeoVis.3.1.1.chr5.g00028.t01    rna-gnl|WGS_ZZZZ|egapxtmp_008432-R1    mNeoVis.evergladensis.2.1.chr5.g00035.t01    mNeoVis.vulgivaga.2.1.chr5.g00033.t01      1712    N0.HOG0001710   OG0000227  n1   rna-XM_002927024.4    rna-NM_001252418.1, rna-XM_038577489.1    rna-XM_022518845.1    rna-XM_045062429.1    rna-XM_036072609.1, rna-XM_036084035.1, rna-XM_036100869.1, rna-XM_036101400.1, rna-XM_036103408.1, rna-XM_036105997.1    rna-XM_039221383.1, rna-XM_039235510.1    rna-XM_032840821.1, rna-XM_032850786.1, rna-XM_032856901.1, rna-XM_032857629.1, rna-XM_032867119.1, rna-XM_032871304.1, rna-XM_032874373.1, rna-XM_032882923.1    rna-XM_046003882.1, rna-XM_046012890.1, rna-XM_046023257.1, rna-XM_046027286.1    rna-XM_032320227.1, rna-XM_032349024.1    rna-XM_059169446.1, rna-XM_059170585.1, rna-XM_059171286.1, rna-XM_059176062.1, rna-XM_059182146.1    rna-XM_059371283.1    rna-XM_013051696.2, rna-XM_045069658.1    mNeoVis.2.1.1.chr10_hap1.g00385.t01    rna-XM_021679527.1, rna-XM_021687710.1, rna-XM_044916884.1, rna-XM_044919934.1    rna-XM_004397431.1, rna-XM_004399823.1    rna-XM_007079438.3, rna-XM_042992820.1    rna-XM_008702962.2    rna-XM_027591962.2, rna-XM_035726160.1, rna-XM_035726364.1    mNeoVis.3.1.1.chr10.g01151.t01    rna-gnl|WGS_ZZZZ|egapxtmp_014981-R1    mNeoVis.evergladensis.2.1.chr8.g00367.t01    mNeoVis.vulgivaga.2.1.chr9.g00369.t01      1713    N0.HOG0001711   OG0000227  n48     rna-XM_038544821.1            rna-XM_032843457.1, rna-XM_032844090.1, rna-XM_032873595.1    rna-XM_045994760.1    rna-XM_032305881.1, rna-XM_032346168.1, rna-XM_032351651.1, rna-XM_032353775.1    rna-XM_059137449.1, rna-XM_059140182.1, rna-XM_059155313.1, rna-XM_059163254.1, rna-XM_059164556.1, rna-XM_059166456.1, rna-XM_059166681.1, rna-XM_059187078.1, rna-XM_059188677.1    rna-XM_059371205.1, rna-XM_059386563.1, rna-XM_059391374.1, rna-XM_059393848.1, rna-XM_059401938.1, rna-XM_059407879.1, rna-XM_059416680.1    rna-XM_045078470.1, rna-XM_045083799.1, rna-XM_045087158.1    mNeoVis.2.1.1.chr12_hap1.g00393.t01, mNeoVis.2.1.1.chr12_hap1.g00402.t01, mNeoVis.2.1.1.chr5_hap1.g00834.t01, mNeoVis.2.1.1.chr7_hap1.g00493.t01, mNeoVis.2.1.1.chr9_hap1.g01201.t01      rna-XM_004403611.1          mNeoVis.3.1.1.chr11.g00354.t01, mNeoVis.3.1.1.chr6.g00298.t01    rna-gnl|WGS_ZZZZ|egapxtmp_024402-R1, rna-gnl|WGS_ZZZZ|egapxtmp_043894-R1    mNeoVis.evergladensis.2.1.chr10.g01188.t01    mNeoVis.vulgivaga.2.1.chr10.g01176.t01      1714    N0.HOG0001712   OG0000228  n3     rna-NM_001388852.1    rna-XM_022500150.1      rna-XM_036094096.1    rna-XM_039252390.1    rna-XM_032882188.1    rna-XM_046027904.1    rna-XM_032313824.1    rna-XM_059172571.1    rna-XM_059395966.1    rna-XM_004782370.2                          1715    N0.HOG0001713   OG0000228  n12   rna-XM_002931116.4    rna-NM_001388765.1    rna-XM_022492591.1, rna-XM_022492759.1    rna-XM_045040336.1      rna-XM_039233647.1    rna-XM_032851125.1, rna-XM_032877700.1    rna-XM_045996945.1, rna-XM_046028066.1, rna-XM_046028090.1, rna-XM_046028091.1, rna-XM_046028092.1    rna-XM_032312168.1, rna-XM_032312912.1    rna-XM_059172606.1, rna-XM_059172608.1, rna-XM_059172612.1    rna-XM_059387467.1, rna-XM_059387559.1, rna-XM_059395886.1    rna-XM_004765882.3, rna-XM_013050933.2, rna-XM_045068495.1    mNeoVis.2.1.1.chr2_hap1.g00370.t01, mNeoVis.2.1.1.chr2_hap1.g00373.t01    rna-XM_021700161.1    rna-XM_004408620.1    rna-XM_015536603.2    rna-XM_008696186.2    rna-XM_027586865.2    mNeoVis.3.1.1.chr5.g00358.t01    rna-gnl|WGS_ZZZZ|egapxtmp_009439-R1    mNeoVis.evergladensis.2.1.chr5.g00372.t01    mNeoVis.vulgivaga.2.1.chr5.g00370.t01      1716    N0.HOG0001714   OG0000228  n45   rna-XM_019794057.2, rna-XM_034655706.1, rna-XM_034663546.1    rna-XM_038556359.1    rna-XM_022492383.1, rna-XM_022499664.1    rna-XM_003980599.1, rna-XM_003994554.4    rna-XM_036109026.1    rna-XM_039222993.1, rna-XM_039252387.1    rna-XM_032844360.1, rna-XM_032847590.1, rna-XM_032882187.1    rna-XM_045998931.1, rna-XM_046026618.1    rna-XM_032312212.1, rna-XM_032334980.1, rna-XM_032334981.1    rna-XM_059172432.1, rna-XM_059173511.1, rna-XM_059173517.1    rna-XM_059397801.1, rna-XM_059416739.1, rna-XM_059416747.1    rna-XM_004781393.1, rna-XM_045068865.1    mNeoVis.2.1.1.chr2_hap1.g00005.t01    rna-XM_021689901.1    rna-XM_004402393.1    rna-XM_007079751.1, rna-XM_007084981.1    rna-XM_008687284.1, rna-XM_008710083.2    rna-XM_027587779.1, rna-XM_027604485.1    mNeoVis.3.1.1.chr1.g01009.t01, mNeoVis.3.1.1.chr5.g00004.t01, mNeoVis.3.1.1.chr5.g00005.t01    rna-gnl|WGS_ZZZZ|egapxtmp_008878-R1, rna-gnl|WGS_ZZZZ|egapxtmp_020999-R1    mNeoVis.evergladensis.2.1.chr1.g01001.t01, mNeoVis.evergladensis.2.1.chr5.g00003.t01, mNeoVis.evergladensis.2.1.chr5.g00004.t01    mNeoVis.vulgivaga.2.1.chr1.g01015.t01, mNeoVis.vulgivaga.2.1.chr5.g00003.t01      1717    N0.HOG0001715   OG0000229  n1   rna-XM_002918140.4    rna-XM_038535035.1    rna-XM_022493634.1    rna-XM_023260438.2    rna-XM_036113565.1    rna-XM_039239465.1    rna-XM_032851558.1    rna-XM_045999539.1    rna-XM_032337312.1    rna-XM_059174052.1    rna-XM_059417172.1    rna-XM_045079577.1    mNeoVis.2.1.1.chr1_hap1.g00206.t01    rna-XM_021700902.1    rna-XM_004402820.1    rna-XM_042994260.1    rna-XM_040630401.1    rna-XM_027604782.2    mNeoVis.3.1.1.chr1.g00207.t01    rna-gnl|WGS_ZZZZ|egapxtmp_015276-R1    mNeoVis.evergladensis.2.1.chr1.g00201.t01    mNeoVis.vulgivaga.2.1.chr1.g00204.t01      1718    N0.HOG0001716   OG0000229  n16   rna-XM_002920394.4    rna-XM_038534375.1    rna-XM_022499819.1    rna-XM_023240569.2    rna-XM_036110269.1    rna-XM_039235189.1    rna-XM_032840203.1    rna-XM_046028051.1    rna-XM_032313197.1    rna-XM_059169860.1    rna-XM_059397816.1    rna-XM_004769415.3    mNeoVis.2.1.1.chr2_hap1.g00462.t01    rna-XM_021699571.1    rna-XM_004407024.1    rna-XM_042960050.1    rna-XM_008696056.2    rna-XM_027588901.2    mNeoVis.3.1.1.chr5.g00455.t01      mNeoVis.evergladensis.2.1.chr5.g00465.t01    mNeoVis.vulgivaga.2.1.chr5.g00464.t01      1719    N0.HOG0001717   OG0000229  n33   rna-XM_034671694.1    rna-XM_038581779.1    rna-XM_022517412.1    rna-XM_045055930.1    rna-XM_036120990.1    rna-XM_039223306.1, rna-XM_039223352.1    rna-XM_032858895.1    rna-XM_045997495.1    rna-XM_032333263.1    rna-XM_059169785.1    rna-XM_059395250.1    rna-XM_004748022.2, rna-XM_045085569.1    mNeoVis.2.1.1.chr6_hap1.g00484.t01    rna-XM_021684468.1    rna-XM_004409258.1    rna-XM_042984004.1    rna-XM_008689458.1    rna-XM_027598851.1    mNeoVis.3.1.1.chr4.g00487.t01, mNeoVis.3.1.1.chr4.g00488.t01    rna-gnl|WGS_ZZZZ|egapxtmp_044428-R1    mNeoVis.evergladensis.2.1.chr4.g00489.t01    mNeoVis.vulgivaga.2.1.chr4.g00497.t01      1720    N0.HOG0001718   OG0000229  n57   rna-XM_034671406.1    rna-XM_038581773.1    rna-XM_022517076.1    rna-XM_019829011.3    rna-XM_036091794.1    rna-XM_039224531.1, rna-XM_039253807.1    rna-XM_032858829.1    rna-XM_045986881.1    rna-XM_032333268.1    rna-XM_059169713.1    rna-XM_059377783.1    rna-XM_004748013.3    mNeoVis.2.1.1.chr6_hap1.g00481.t01    rna-XM_021684465.1    rna-XM_004401604.1    rna-XM_007087621.3    rna-XM_008689460.2    rna-XM_027597405.1    mNeoVis.3.1.1.chr4.g00484.t01    rna-gnl|WGS_ZZZZ|egapxtmp_044586-R1    mNeoVis.evergladensis.2.1.chr4.g00487.t01    mNeoVis.vulgivaga.2.1.chr4.g00495.t01      1721    N0.HOG0001719   OG0000230  n1   rna-XM_019797143.2    rna-XM_038552537.1    rna-XM_022522484.1    rna-XM_045043753.1    rna-XM_036082669.1    rna-XM_039219929.1    rna-XM_032860297.1    rna-XM_046021694.1    rna-XM_032308345.1    rna-XM_059142846.1    rna-XM_059411547.1    rna-XM_045079675.1    mNeoVis.2.1.1.chr13_hap1.g00580.t01    rna-XM_021681019.1    rna-XM_004405438.1    rna-XM_042964162.1    rna-XM_040631480.1    rna-XM_027616907.1    mNeoVis.3.1.1.chr13.g00517.t01    rna-gnl|WGS_ZZZZ|egapxtmp_030365-R1    mNeoVis.evergladensis.2.1.chr13.g00564.t01    mNeoVis.vulgivaga.2.1.chr13.g00566.t01      1722    N0.HOG0001720   OG0000230  n14   rna-XM_002921911.4    rna-XM_038546823.1, rna-XM_038567486.1    rna-XM_022525488.1    rna-XM_023244561.2    rna-XM_036066037.1, rna-XM_036107380.1    rna-XM_039251557.1    rna-XM_032883822.1    rna-XM_045991989.1    rna-XM_032309104.1    rna-XM_059159772.1    rna-XM_059388855.1    rna-XM_045085058.1    mNeoVis.2.1.1.chr5_hap1.g00203.t01    rna-XM_044915702.1    rna-XM_004412148.1    rna-XM_042978011.1    rna-XM_040629426.1    rna-XM_027582679.1    mNeoVis.3.1.1.chr2.g00153.t01    rna-gnl|WGS_ZZZZ|egapxtmp_034635-R1    mNeoVis.evergladensis.2.1.chr2.g00200.t01    mNeoVis.vulgivaga.2.1.chr2.g00199.t01      1723    N0.HOG0001721   OG0000230  n22   rna-XM_034657817.1    rna-XM_038567204.1    rna-XM_022526954.1, rna-XM_022526970.1    rna-XM_023245678.2    rna-XM_036088886.1    rna-XM_039250985.1    rna-XM_032884085.1    rna-XM_045992124.1    rna-XM_032311837.1    rna-XM_059160057.1    rna-XM_059389080.1    rna-XM_004748380.2    mNeoVis.2.1.1.chr5_hap1.g00313.t01    rna-XM_021705297.1    rna-XM_004398909.1    rna-XM_042977296.1    rna-XM_040628269.1    rna-XM_027587088.2    mNeoVis.3.1.1.chr2.g00251.t01    rna-gnl|WGS_ZZZZ|egapxtmp_037741-R2    mNeoVis.evergladensis.2.1.chr2.g00312.t01    mNeoVis.vulgivaga.2.1.chr2.g00308.t01      1724   N0.HOG0001722  OG0000230  n32   rna-XM_019802062.2    rna-XM_038558078.1    rna-XM_022496700.1    rna-XM_019836975.3    rna-XM_036105521.1, rna-XM_036123046.1    rna-XM_039245403.1    rna-XM_032846651.1    rna-XM_045979520.1    rna-XM_032302766.1    rna-XM_059137440.1    rna-XM_059374676.1    rna-XM_004774502.3    mNeoVis.2.1.1.chr2_hap1.g01311.t01    rna-XM_021683896.1    rna-XM_004413208.1    rna-XM_042997090.1    rna-XM_008699229.2    rna-XM_027582945.2    mNeoVis.3.1.1.chr5.g01236.t01    rna-gnl|WGS_ZZZZ|egapxtmp_006286-R2    mNeoVis.evergladensis.2.1.chr5.g01252.t01    mNeoVis.vulgivaga.2.1.chr5.g01248.t01      1725   N0.HOG0001723  OG0000231  n1   rna-XM_011218027.2    rna-NM_001388814.1, rna-XM_038546157.1    rna-XM_022509827.1, rna-XM_022521046.1    rna-XM_023243940.2    rna-XM_036065002.1, rna-XM_036118499.1    rna-XM_039253153.1    rna-XM_032846729.1, rna-XM_032849010.1    rna-XM_045985920.1    rna-XM_032319726.1, rna-XM_032321710.1    rna-XM_059149919.1    rna-XM_059379786.1    rna-XM_004782336.2    mNeoVis.2.1.1.chr8_hap1.g00916.t01, mNeoVis.2.1.1.chr8_hap1.g00921.t01    rna-XM_021704435.2      rna-XM_007076231.3    rna-XM_008695528.2    rna-XM_027567881.1    mNeoVis.3.1.1.chr8.g00811.t01    rna-gnl|WGS_ZZZZ|egapxtmp_018992-R1    mNeoVis.evergladensis.2.1.chr9.g00853.t01    mNeoVis.vulgivaga.2.1.chr8.g00855.t01      1726    N0.HOG0001724   OG0000231  n28   rna-XM_011217730.3        rna-XM_003996427.3      rna-XM_039253145.1                      rna-XM_042966850.1    rna-XM_008695260.2                1727    N0.HOG0001725   OG0000231  n34   rna-XM_002930852.2    rna-NM_001388813.1      rna-XM_003996401.2      rna-XM_039253134.1      rna-XM_045985942.1    rna-XM_032322472.1    rna-XM_059148847.1    rna-XM_059378577.1    rna-XM_004781935.3    mNeoVis.2.1.1.chr8_hap1.g00864.t01        rna-XM_007076145.2    rna-XM_008695529.2, rna-XM_040639840.1    rna-XM_027624687.1    mNeoVis.3.1.1.chr8.g00810.t01      mNeoVis.evergladensis.2.1.chr9.g00852.t01    mNeoVis.vulgivaga.2.1.chr8.g00854.t01      1728    N0.HOG0001726   OG0000231  n48   rna-XM_002930851.3    rna-NM_001388812.1    rna-XM_022509791.1    rna-XM_003996402.2    rna-XM_036076647.1    rna-XM_039253761.1    rna-XM_032846721.1    rna-XM_045985948.1    rna-XM_032321476.1    rna-XM_059149056.1    rna-XM_059378983.1    rna-XM_004781934.2    mNeoVis.2.1.1.chr8_hap1.g00863.t01    rna-XM_021704434.2      rna-XM_007076144.1    rna-XM_008695261.2    rna-XM_027567829.2    mNeoVis.3.1.1.chr8.g00809.t01    rna-gnl|WGS_ZZZZ|egapxtmp_019056-R1    mNeoVis.evergladensis.2.1.chr9.g00851.t01    mNeoVis.vulgivaga.2.1.chr8.g00853.t01      1729   N0.HOG0001727  OG0000231  n65   rna-XM_002930850.3    rna-NM_001388811.1    rna-XM_022509792.1    rna-XM_006939962.4    rna-XM_036076648.1    rna-XM_039253755.1    rna-XM_032846722.1    rna-XM_045986045.1, rna-XM_045986246.1    rna-XM_032321449.1    rna-XM_059149019.1    rna-XM_059378984.1    rna-XM_013050704.2    mNeoVis.2.1.1.chr8_hap1.g00862.t01    rna-XM_021704307.1      rna-XM_007076143.1    rna-XM_040639910.1    rna-XM_027567823.1    mNeoVis.3.1.1.chr8.g00808.t01    rna-gnl|WGS_ZZZZ|egapxtmp_018145-R1    mNeoVis.evergladensis.2.1.chr9.g00850.t01    mNeoVis.vulgivaga.2.1.chr8.g00852.t01      1730    N0.HOG0001728   OG0000232  n1   rna-XM_019793773.2    rna-XM_038568118.1, rna-XM_038584862.1    rna-XM_022498419.1    rna-XM_003990864.6    rna-XM_036105519.1    rna-XM_039239179.1    rna-XM_032880464.1    rna-XM_046019046.1    rna-XM_032355967.1    rna-XM_059167032.1    rna-XM_059394170.1    rna-XM_004744080.3    mNeoVis.2.1.1.chr3_hap1.g00884.t01    rna-XM_021702736.1    rna-XM_004407643.1    rna-XM_042994710.1    rna-XM_040643885.1    rna-XM_027588726.2    mNeoVis.3.1.1.chr3.g00867.t01    rna-gnl|WGS_ZZZZ|egapxtmp_042695-R1    mNeoVis.evergladensis.2.1.chr3.g00879.t01    mNeoVis.vulgivaga.2.1.chr3.g00894.t01      1731    N0.HOG0001729   OG0000232  n18   rna-XM_034653048.1, rna-XM_034667952.1    rna-XM_038556875.1    rna-XM_022503186.1    rna-XM_003982778.6    rna-XM_036115684.1    rna-XM_039236226.1    rna-XM_032874712.1    rna-XM_046020763.1    rna-XM_032304877.1    rna-XM_059171172.1    rna-XM_059396818.1    rna-XM_004767845.3    mNeoVis.2.1.1.chr4_hap1.g00472.t01    rna-XM_021689775.1    rna-XM_004394066.1    rna-XM_007076688.3    rna-XM_008686270.2    rna-XM_027573484.2    mNeoVis.3.1.1.chr6.g00473.t01      mNeoVis.evergladensis.2.1.chr6.g00479.t01    mNeoVis.vulgivaga.2.1.chr6.g00475.t01      1732    N0.HOG0001730   OG0000232  n35   rna-XM_019804337.2    rna-XM_038547635.1    rna-XM_022509303.1    rna-XM_003996734.4    rna-XM_036094751.1    rna-XM_039227942.1    rna-XM_032872128.1    rna-XM_045984187.1    rna-XM_032319636.1    rna-XM_059149129.1    rna-XM_059380110.1    rna-XM_004764804.2    mNeoVis.2.1.1.chr8_hap1.g00567.t01    rna-XM_021703985.1    rna-XM_004395146.1    rna-XM_042965558.1    rna-XM_040640807.1    rna-XM_027625175.2    mNeoVis.3.1.1.chr8.g00522.t01    rna-gnl|WGS_ZZZZ|egapxtmp_017719-R1    mNeoVis.evergladensis.2.1.chr9.g00557.t01    mNeoVis.vulgivaga.2.1.chr8.g00564.t01      1733    N0.HOG0001731   OG0000232  n51   rna-XM_011234129.3    rna-XM_038587102.1    rna-XM_022507907.1    rna-XM_045050452.1    rna-XM_036096517.1    rna-XM_039257243.1    rna-XM_032842294.1    rna-XM_045994471.1, rna-XM_046016446.1    rna-XM_032329859.1    rna-XM_059156712.1    rna-XM_059385829.1    rna-XM_004769153.3    mNeoVis.2.1.1.chr12_hap1.g00729.t01    rna-XM_021705650.1, rna-XM_044916971.1    rna-XM_004417019.1    rna-XM_042975332.1    rna-XM_040633390.1    rna-XM_027609454.2    mNeoVis.3.1.1.chr12.g00664.t01    rna-gnl|WGS_ZZZZ|egapxtmp_028254-R1    mNeoVis.evergladensis.2.1.chr12.g00661.t01    mNeoVis.vulgivaga.2.1.chr12.g00686.t01      1734   N0.HOG0001732  OG0000233  n0   rna-XM_034650327.1, rna-XM_034653291.1, rna-XM_034665991.1, rna-XM_034670764.1    rna-XM_038546171.1    rna-XM_022506146.1    rna-XM_045036481.1, rna-XM_045036483.1, rna-XM_045036485.1, rna-XM_045043269.1, rna-XM_045043686.1, rna-XM_045051585.1    rna-XM_036084681.1, rna-XM_036084691.1, rna-XM_036094319.1, rna-XM_036095236.1, rna-XM_036110369.1        rna-XM_046011856.1    rna-XM_032308426.1, rna-XM_032316026.1, rna-XM_032328621.1, rna-XM_032338397.1, rna-XM_032347892.1, rna-XM_032349389.1, rna-XM_032359930.1    rna-XM_059168404.1, rna-XM_059184158.1, rna-XM_059184165.1, rna-XM_059187452.1, rna-XM_059187617.1      rna-XM_045069129.1, rna-XM_045069176.1, rna-XM_045069178.1, rna-XM_045069188.1, rna-XM_045069191.1    mNeoVis.2.1.1.chr11_hap1.g00001.t01, mNeoVis.2.1.1.chr11_hap1.g00896.t01, mNeoVis.2.1.1.chr11_hap1.g00897.t01, mNeoVis.2.1.1.chr11_hap1.g00898.t01, mNeoVis.2.1.1.chr11_hap1.g00899.t01, mNeoVis.2.1.1.chr11_hap1.g01206.t01, mNeoVis.2.1.1.chr13_hap1.g00115.t01, mNeoVis.2.1.1.chr13_hap1.g00836.t01, mNeoVis.2.1.1.chr15_hap1.g00162.t01, mNeoVis.2.1.1.chr15_hap1.g00163.t01, mNeoVis.2.1.1.chr15_hap1.g00164.t01, mNeoVis.2.1.1.chr1_hap1.g00001.t01, mNeoVis.2.1.1.chr1_hap1.g02166.t01, mNeoVis.2.1.1.chr2_hap1.g00001.t01, mNeoVis.2.1.1.chr3_hap1.g01374.t01, mNeoVis.2.1.1.chr5_hap1.g01932.t01, mNeoVis.2.1.1.chr6_hap1.g00001.t01, mNeoVis.2.1.1.chr6_hap1.g00043.t01, mNeoVis.2.1.1.chr8_hap1.g00001.t01, mNeoVis.2.1.1.chr9_hap1.g01329.t01        rna-XM_042963511.1, rna-XM_042964233.1, rna-XM_042964234.1, rna-XM_042964236.1, rna-XM_042964744.1, rna-XM_042977132.1, rna-XM_042977133.1, rna-XM_042977137.1, rna-XM_042977138.1, rna-XM_042977175.1      rna-XM_027613264.2, rna-XM_027615986.2, rna-XM_027624421.2, rna-XM_035721898.1, rna-XM_035722542.1, rna-XM_035723274.1, rna-XM_035723488.1, rna-XM_035723975.1, rna-XM_035724139.1, rna-XM_035724432.1, rna-XM_035724668.1, rna-XM_035725964.1, rna-XM_035725978.1, rna-XM_035725981.1, rna-XM_035726001.1, rna-XM_035726076.1, rna-XM_035726292.1, rna-XM_035727331.1, rna-XM_035727541.1, rna-XM_035727553.1, rna-XM_035728080.1, rna-XM_035728544.1        mNeoVis.evergladensis.2.1.chr15.g00610.t01, mNeoVis.evergladensis.2.1.chr4.g00046.t01, mNeoVis.evergladensis.2.1.chr4.g00809.t01    mNeoVis.vulgivaga.2.1.chr15.g00624.t01      1735   N0.HOG0001733  OG0000234  n0   rna-XM_002923022.3, rna-XM_002923023.2, rna-XM_002923031.3, rna-XM_002923032.2, rna-XM_002923033.4, rna-XM_011229642.2, rna-XM_011229644.3    rna-NM_001197146.1, rna-XM_038589004.1    rna-XM_022503500.1, rna-XM_022504052.1, rna-XM_022504053.1, rna-XM_022504054.1    rna-XM_003986562.3, rna-XM_003986563.3, rna-XM_003986564.3, rna-XM_003986565.6, rna-XM_003986587.2, rna-XM_023254740.2    rna-XM_036120860.1, rna-XM_036120861.1    rna-XM_039240408.1, rna-XM_039240441.1, rna-XM_039240442.1, rna-XM_039240443.1, rna-XM_039240444.1, rna-XM_039240446.1    rna-XM_032843495.1, rna-XM_032843496.1, rna-XM_032843497.1, rna-XM_032843512.1    rna-XM_046005752.1, rna-XM_046006375.1, rna-XM_046006380.1, rna-XM_046006390.1, rna-XM_046006395.1    rna-XM_032338337.1, rna-XM_032338338.1, rna-XM_032338339.1, rna-XM_032338340.1, rna-XM_032339495.1    rna-XM_059176343.1, rna-XM_059176345.1, rna-XM_059176347.1, rna-XM_059176348.1, rna-XM_059176349.1    rna-XM_059399158.1, rna-XM_059399159.1, rna-XM_059399160.1, rna-XM_059399162.1    rna-XM_004740211.1, rna-XM_004740212.1, rna-XM_004740213.1, rna-XM_004740347.1    mNeoVis.2.1.1.chr1_hap1.g01797.t01, mNeoVis.2.1.1.chr1_hap1.g01798.t01, mNeoVis.2.1.1.chr1_hap1.g01799.t01, mNeoVis.2.1.1.chr1_hap1.g01801.t01    rna-XM_021693530.2, rna-XM_021693534.1, rna-XM_021693629.1    rna-XM_004417788.1, rna-XM_004417792.1, rna-XM_004417793.1, rna-XM_004417821.1    rna-XM_007078616.3, rna-XM_007078617.3, rna-XM_007078618.3, rna-XM_007078765.2, rna-XM_042987166.1    rna-XM_008691087.1, rna-XM_008691089.2, rna-XM_008691090.2, rna-XM_008691091.2, rna-XM_008691092.2, rna-XM_008691231.2, rna-XM_008691232.2    rna-XM_027603297.1, rna-XM_027604031.1    mNeoVis.3.1.1.chr1.g01755.t01, mNeoVis.3.1.1.chr1.g01756.t01    rna-gnl|WGS_ZZZZ|egapxtmp_005612-R1, rna-gnl|WGS_ZZZZ|egapxtmp_022565-R1, rna-gnl|WGS_ZZZZ|egapxtmp_022574-R1, rna-gnl|WGS_ZZZZ|egapxtmp_022585-R1    mNeoVis.evergladensis.2.1.chr1.g01763.t01, mNeoVis.evergladensis.2.1.chr1.g01764.t01, mNeoVis.evergladensis.2.1.chr1.g01765.t01    mNeoVis.vulgivaga.2.1.chr1.g01760.t01, mNeoVis.vulgivaga.2.1.chr1.g01761.t01, mNeoVis.vulgivaga.2.1.chr10.g00911.t01      1736    N0.HOG0001734   OG0000235  n1   rna-XM_002928067.4    rna-XM_038530114.1    rna-XM_022515707.1    rna-XM_045060865.1    rna-XM_036068428.1    rna-XM_039224763.1    rna-XM_032837893.1    rna-XM_046002446.1, rna-XM_046013033.1    rna-XM_032349415.1    rna-XM_059184137.1    rna-XM_059404325.1    rna-XM_004774892.3    mNeoVis.2.1.1.chr9_hap1.g00153.t01    rna-XM_021696802.2    rna-XM_004409126.1    rna-XM_042991048.1    rna-XM_040645414.1    rna-XM_027592067.2    mNeoVis.3.1.1.chr10.g00144.t01    rna-gnl|WGS_ZZZZ|egapxtmp_023986-R1    mNeoVis.evergladensis.2.1.chr10.g00147.t01    mNeoVis.vulgivaga.2.1.chr10.g00145.t01      1737    N0.HOG0001735   OG0000235  n17   rna-XM_034649642.1    rna-XM_038586658.1    rna-XM_022491363.1    rna-XM_045051381.1    rna-XM_036118573.1    rna-XM_039219239.1    rna-XM_032846372.1    rna-XM_045994494.1    rna-XM_032330737.1    rna-XM_059158292.1    rna-XM_059385766.1    rna-XM_004779320.3    mNeoVis.2.1.1.chr12_hap1.g00063.t01    rna-XM_044911786.1    rna-XM_004409162.1    rna-XM_042974309.1    rna-XM_040630181.1, rna-XM_040639640.1, rna-XM_040639766.1    rna-XM_027608445.1    mNeoVis.3.1.1.chr12.g00054.t01    rna-gnl|WGS_ZZZZ|egapxtmp_028905-R1    mNeoVis.evergladensis.2.1.chr12.g00053.t01    mNeoVis.vulgivaga.2.1.chr12.g00055.t01      1738    N0.HOG0001736   OG0000235  n39   rna-XM_034658730.1    rna-XM_038565744.1    rna-XM_022497886.1    rna-XM_045051936.1    rna-XM_036079814.1    rna-XM_039231325.1    rna-XM_032841525.1    rna-XM_045990126.1    rna-XM_032326432.1    rna-XM_059161629.1    rna-XM_059390253.1    rna-XM_004738469.3    mNeoVis.2.1.1.chr5_hap1.g00857.t01    rna-XM_044916496.1    rna-XM_004392704.1    rna-XM_042979487.1    rna-XM_040628517.1    rna-XM_027586751.2    mNeoVis.3.1.1.chr2.g00769.t01    rna-gnl|WGS_ZZZZ|egapxtmp_037618-R1    mNeoVis.evergladensis.2.1.chr2.g00858.t01    mNeoVis.vulgivaga.2.1.chr2.g00859.t01      1739   N0.HOG0001737  OG0000235  n55   rna-XM_019802624.2    rna-XM_038542032.1    rna-XM_022526744.1    rna-XM_011290967.4    rna-XM_036114186.1    rna-XM_039230109.1    rna-XM_032878122.1    rna-XM_045983375.1    rna-XM_032318410.1    rna-XM_059147070.1    rna-XM_059412975.1    rna-XM_013047211.2    mNeoVis.2.1.1.chr14_hap1.g00267.t01    rna-XM_021682522.1    rna-XM_004408766.1    rna-XM_042975970.1    rna-XM_040636387.1    rna-XM_027609718.1    mNeoVis.3.1.1.chr14.g00268.t01    rna-gnl|WGS_ZZZZ|egapxtmp_031731-R1    mNeoVis.evergladensis.2.1.chr14.g00267.t01    mNeoVis.vulgivaga.2.1.chr14.g00268.t01      1740    N0.HOG0001738   OG0000236  n1   rna-XM_019810190.1    rna-XM_038558772.1    rna-XM_022494904.1    rna-XM_019835379.2    rna-XM_036086049.1    rna-XM_039245597.1    rna-XM_032874321.1    rna-XM_046012101.1    rna-XM_032346678.1    rna-XM_059186625.1    rna-XM_059404155.1    rna-XM_004742937.2    mNeoVis.2.1.1.chr9_hap1.g01035.t01    rna-XM_021687646.1      rna-XM_007085654.2    rna-XM_008687762.1    rna-XM_027592208.1    mNeoVis.3.1.1.chr10.g00996.t01    rna-gnl|WGS_ZZZZ|egapxtmp_023492-R1    mNeoVis.evergladensis.2.1.chr10.g01023.t01    mNeoVis.vulgivaga.2.1.chr10.g01016.t01      1741    N0.HOG0001739   OG0000236  n22   rna-XM_034653326.1    rna-XM_038556456.1    rna-XM_022491659.1    rna-XM_045056483.1    rna-XM_036079676.1      rna-XM_032848479.1    rna-XM_045999701.1    rna-XM_032335053.1    rna-XM_059176063.1    rna-XM_059416440.1    rna-XM_004782011.3    mNeoVis.2.1.1.chr1_hap1.g00976.t01    rna-XM_021690942.1      rna-XM_042958431.1    rna-XM_008709905.2      mNeoVis.3.1.1.chr1.g00983.t01    rna-gnl|WGS_ZZZZ|egapxtmp_018993-R1    mNeoVis.evergladensis.2.1.chr1.g00971.t01    mNeoVis.vulgivaga.2.1.chr1.g00984.t01, mNeoVis.vulgivaga.2.1.chr1.g00985.t01      1742    N0.HOG0001740   OG0000236  n41   rna-XM_011222271.3    rna-XM_038541742.1    rna-XM_022508127.1    rna-XM_003999318.5    rna-XM_036109757.1      rna-XM_032850972.1    rna-XM_045982072.1, rna-XM_045982301.1, rna-XM_045982302.1, rna-XM_045982303.1, rna-XM_045982304.1, rna-XM_045982998.1, rna-XM_045983000.1, rna-XM_045983007.1, rna-XM_045983039.1, rna-XM_045983050.1, rna-XM_045983690.1, rna-XM_045996961.1, rna-XM_045996962.1, rna-XM_045996963.1, rna-XM_045996964.1, rna-XM_045996965.1    rna-XM_032319193.1, rna-XM_032347856.1    rna-XM_059145271.1, rna-XM_059145272.1, rna-XM_059146407.1, rna-XM_059146408.1, rna-XM_059146409.1, rna-XM_059146410.1, rna-XM_059146411.1, rna-XM_059146412.1, rna-XM_059146414.1, rna-XM_059146415.1, rna-XM_059146908.1, rna-XM_059187485.1    rna-XM_059387461.1    rna-XM_045073222.1, rna-XM_045084708.1, rna-XM_045084709.1, rna-XM_045084737.1    mNeoVis.2.1.1.chr14_hap1.g00110.t01, mNeoVis.2.1.1.chr14_hap1.g00111.t01    rna-XM_021682545.1    rna-XM_004414483.1    rna-XM_042975932.1    rna-XM_008701166.2    rna-XM_027612431.2    mNeoVis.3.1.1.chr14.g00104.t01            1743    N0.HOG0001741   OG0000237  n1   rna-XM_034649083.1    rna-XM_038557920.1    rna-XM_022497318.1    rna-XM_019836650.3    rna-XM_036108129.1    rna-XM_039251695.1    rna-XM_032877230.1    rna-XM_046003341.1    rna-XM_032361418.1    rna-XM_059137227.1    rna-XM_059377244.1    rna-XM_013046035.2    mNeoVis.2.1.1.chr2_hap1.g01505.t01    rna-XM_044914477.1    rna-XM_004402671.1    rna-XM_042996472.1    rna-XM_040619252.1    rna-XM_027605974.2    mNeoVis.3.1.1.chr5.g01427.t01    rna-gnl|WGS_ZZZZ|egapxtmp_005553-R4    mNeoVis.evergladensis.2.1.chr5.g01442.t01    mNeoVis.vulgivaga.2.1.chr5.g01443.t01      1744    N0.HOG0001742   OG0000237  n13   rna-XM_019797391.2    rna-XM_038557915.1    rna-XM_022496240.1    rna-XM_003989853.6    rna-XM_036108125.1    rna-XM_039251692.1    rna-XM_032877236.1    rna-XM_046003044.1    rna-XM_032360803.1    rna-XM_059137231.1    rna-XM_059377247.1    rna-XM_004740797.3    mNeoVis.2.1.1.chr2_hap1.g01504.t01    rna-XM_021683695.2    rna-XM_004402673.1    rna-XM_042996471.1    rna-XM_008694400.2    rna-XM_027605945.2    mNeoVis.3.1.1.chr5.g01426.t01    rna-gnl|WGS_ZZZZ|egapxtmp_005550-R1    mNeoVis.evergladensis.2.1.chr5.g01441.t01    mNeoVis.vulgivaga.2.1.chr5.g01442.t01      1745    N0.HOG0001743   OG0000237  n16   rna-XM_034649064.1    rna-XM_038557924.1    rna-XM_022496987.1    rna-XM_023258472.2    rna-XM_036108141.1    rna-XM_039251685.1    rna-XM_032877226.1    rna-XM_046010351.1    rna-XM_032361328.1    rna-XM_059135439.1    rna-XM_059377243.1    rna-XM_045064187.1    mNeoVis.2.1.1.chr2_hap1.g01506.t01    rna-XM_021684204.1    rna-XM_004402676.1    rna-XM_042996470.1    rna-XM_040619250.1    rna-XM_027604977.2    mNeoVis.3.1.1.chr5.g01428.t01    rna-gnl|WGS_ZZZZ|egapxtmp_005554-R1    mNeoVis.evergladensis.2.1.chr5.g01443.t01    mNeoVis.vulgivaga.2.1.chr5.g01444.t01      1746    N0.HOG0001744   OG0000237  n29   rna-XM_019801180.2    rna-XM_038555605.1    rna-XM_022502255.1, rna-XM_022514915.1    rna-XM_019823195.3    rna-XM_036067159.1, rna-XM_036109417.1    rna-XM_039232982.1    rna-XM_032873473.1    rna-XM_046001845.1    rna-XM_032316688.1    rna-XM_059165544.1    rna-XM_059395214.1    rna-XM_045078166.1    mNeoVis.2.1.1.chr4_hap1.g01043.t01    rna-XM_021688819.1, rna-XM_021703835.1    rna-XM_004405570.1    rna-XM_042973408.1    rna-XM_040626152.1    rna-XM_027598826.2    mNeoVis.3.1.1.chr6.g01063.t01    rna-gnl|WGS_ZZZZ|egapxtmp_002528-R2    mNeoVis.evergladensis.2.1.chr6.g01048.t01    mNeoVis.vulgivaga.2.1.chr6.g01040.t01      1747    N0.HOG0001745   OG0000238  n1   rna-XM_019808049.2    rna-XM_038539910.1    rna-XM_022526649.1    rna-XM_023246655.2    rna-XM_036091214.1    rna-XM_039247835.1    rna-XM_032842578.1    rna-XM_045994149.1    rna-XM_032328692.1    rna-XM_059155688.1    rna-XM_059414995.1    rna-XM_045072613.1    mNeoVis.2.1.1.chr15_hap1.g00080.t01    rna-XM_021700470.1    rna-XM_004397171.1    rna-XM_007093462.3    rna-XM_040635125.1    rna-XM_027614053.2    mNeoVis.3.1.1.chr15.g00074.t01      mNeoVis.evergladensis.2.1.chr15.g00081.t01    mNeoVis.vulgivaga.2.1.chr15.g00082.t01      1748    N0.HOG0001746   OG0000238  n21   rna-XM_034640817.1    rna-XM_038548370.1    rna-XM_022509385.1    rna-XM_045044916.1    rna-XM_036094641.1    rna-XM_039253447.1    rna-XM_032848437.1    rna-XM_045985769.1    rna-XM_032321386.1    rna-XM_059148688.1    rna-XM_059380837.1    rna-XM_004746931.3    mNeoVis.2.1.1.chr8_hap1.g00796.t01    rna-XM_044921578.1    rna-XM_004399693.1    rna-XM_042966626.1    rna-XM_040641084.1    rna-XM_027568041.2        mNeoVis.evergladensis.2.1.chr9.g00784.t01    mNeoVis.vulgivaga.2.1.chr8.g00787.t01      1749    N0.HOG0001747   OG0000238  n35   rna-XM_019801860.2    rna-XM_038563751.1    rna-XM_022507249.1    rna-XM_023240114.2    rna-XM_036067549.1    rna-XM_039235468.1    rna-XM_032879640.1    rna-XM_046015593.1    rna-XM_032357630.1    rna-XM_059149340.1    rna-XM_059399909.1    rna-XM_004759517.2    mNeoVis.2.1.1.chr7_hap1.g02520.t01, mNeoVis.2.1.1.chr7_hap1.g02536.t01, mNeoVis.2.1.1.chr7_hap1.g02564.t01    rna-XM_021685121.1    rna-XM_004393839.1    rna-XM_042959181.1    rna-XM_040639514.1    rna-XM_027581038.2              1750    N0.HOG0001748   OG0000238  n53   rna-XM_034638726.1    rna-XM_038575432.1    rna-XM_022517398.1    rna-XM_023241509.2    rna-XM_036085938.1    rna-XM_039243361.1    rna-XM_032847370.1    rna-XM_046024701.1    rna-XM_032309607.1, rna-XM_032316684.1    rna-XM_059139187.1, rna-XM_059165747.1    rna-XM_059392947.1, rna-XM_059408619.1    rna-XM_004779086.3, rna-XM_045078227.1    mNeoVis.2.1.1.chr3_hap1.g00491.t01, mNeoVis.2.1.1.chr4_hap1.g00953.t01    rna-XM_021703633.1    rna-XM_004416163.1    rna-XM_042961597.1    rna-XM_040619820.1    rna-XM_027576157.2    mNeoVis.3.1.1.chr3.g00482.t01, mNeoVis.3.1.1.chr6.g00966.t01    rna-gnl|WGS_ZZZZ|egapxtmp_003111-R1, rna-gnl|WGS_ZZZZ|egapxtmp_041939-R2    mNeoVis.evergladensis.2.1.chr3.g00498.t01, mNeoVis.evergladensis.2.1.chr6.g00959.t01    mNeoVis.vulgivaga.2.1.chr3.g00501.t01      1751    N0.HOG0001749   OG0000239  n3   rna-NM_001304921.1    rna-NM_001024636.2, rna-XM_038539447.1, rna-XM_038570180.1    rna-XM_022523873.1    rna-NM_001048150.1, rna-XM_045056971.1      rna-XM_039245579.1, rna-XM_039250111.1    rna-XM_032839224.1, rna-XM_032856939.1    rna-XM_045989032.1, rna-XM_046005541.1    rna-XM_032325963.1, rna-XM_032360604.1    rna-XM_059151378.1, rna-XM_059175082.1    rna-XM_059383756.1    rna-XM_004767363.3    mNeoVis.2.1.1.chr14_hap1.g00421.t01, mNeoVis.2.1.1.chr7_hap1.g00918.t01      rna-XM_004415086.1    rna-XM_007074150.2, rna-XM_042982715.1    rna-XM_008684525.2    rna-XM_035724689.1, rna-XM_035724952.1, rna-XM_035728609.1    mNeoVis.3.1.1.chr14.g00419.t01, mNeoVis.3.1.1.chr7.g00858.t01    rna-gnl|WGS_ZZZZ|egapxtmp_011334-R1, rna-gnl|WGS_ZZZZ|egapxtmp_032144-R1    mNeoVis.evergladensis.2.1.chr14.g00419.t01, mNeoVis.evergladensis.2.1.chr7.g00911.t01    mNeoVis.vulgivaga.2.1.chr14.g00421.t01, mNeoVis.vulgivaga.2.1.chr7.g00916.t01      1752    N0.HOG0001750   OG0000239  n28   rna-XM_034638695.1    rna-XM_038528905.1      rna-XM_045046177.1    rna-XM_036094965.1, rna-XM_036124545.1    rna-XM_039250120.1    rna-XM_032841823.1    rna-XM_045989402.1    rna-XM_032325742.1, rna-XM_032341341.1    rna-XM_059152085.1, rna-XM_059153316.1    rna-XM_059383978.1    rna-XM_045076808.1    mNeoVis.2.1.1.chr7_hap1.g00551.t01    rna-XM_021681255.1, rna-XM_044914980.1, rna-XM_044916848.1, rna-XM_044917546.1, rna-XM_044922295.1      rna-XM_042969595.1    rna-XM_040633968.1    rna-XM_035725343.1    mNeoVis.3.1.1.chr7.g00534.t01      mNeoVis.evergladensis.2.1.chr7.g00552.t01    mNeoVis.vulgivaga.2.1.chr7.g00554.t01      1753    N0.HOG0001751   OG0000239  n51                             rna-XM_044920142.1          rna-XM_035724257.1              1754    N0.HOG0001752   OG0000239  n52   rna-XM_034657272.1    rna-XM_038587169.1      rna-XM_023249429.2, rna-XM_045058635.1    rna-XM_036082200.1, rna-XM_036100235.1    rna-XM_039237397.1, rna-XM_039256725.1    rna-XM_032883164.1    rna-XM_045996065.1    rna-XM_032331779.1    rna-XM_059158420.1    rna-XM_059385048.1    rna-XM_045089458.1    mNeoVis.2.1.1.chr12_hap1.g00675.t01    rna-XM_021691763.1, rna-XM_044911797.1, rna-XM_044916682.1    rna-XM_004409807.1    rna-XM_042987618.1    rna-XM_040625156.1, rna-XM_040631327.1    rna-XM_035725621.1    mNeoVis.3.1.1.chr12.g00624.t01    rna-gnl|WGS_ZZZZ|egapxtmp_029510-R1    mNeoVis.evergladensis.2.1.chr12.g00613.t01    mNeoVis.vulgivaga.2.1.chr12.g00633.t01      1755    N0.HOG0001753   OG0000240  n1   rna-XM_034639001.1    rna-XM_038528595.1    rna-XM_022524078.1    rna-XM_023246016.2    rna-XM_036091534.1    rna-XM_039248654.1    rna-XM_032838948.1    rna-XM_045987632.1    rna-XM_032325102.1    rna-XM_059152225.1    rna-XM_059381203.1    rna-XM_004767609.3    mNeoVis.2.1.1.chr7_hap1.g01028.t01    rna-XM_021681332.1    rna-XM_004400984.1    rna-XM_042968605.1    rna-XM_040634152.1    rna-XM_027617379.2    mNeoVis.3.1.1.chr7.g00966.t01    rna-gnl|WGS_ZZZZ|egapxtmp_010950-R1    mNeoVis.evergladensis.2.1.chr7.g01021.t01    mNeoVis.vulgivaga.2.1.chr7.g01029.t01      1756    N0.HOG0001754   OG0000240  n22   rna-XM_034645950.1    rna-XM_038564155.1    rna-XM_022506538.1    rna-XM_006937449.5    rna-XM_036100406.1    rna-XM_039257726.1    rna-XM_032879312.1    rna-XM_046017876.1    rna-XM_032357142.1    rna-XM_059145344.1    rna-XM_059403094.1    rna-XM_004770215.3    mNeoVis.2.1.1.chr7_hap1.g02343.t01    rna-XM_021685498.2    rna-XM_004394029.1    rna-XM_042958559.1    rna-XM_008708945.2    rna-XM_027573451.1, rna-XM_027579260.1    mNeoVis.3.1.1.chr7.g02157.t01    rna-gnl|WGS_ZZZZ|egapxtmp_013073-R1    mNeoVis.evergladensis.2.1.chr7.g02289.t01    mNeoVis.vulgivaga.2.1.chr7.g02307.t01      1757    N0.HOG0001755   OG0000240  n41   rna-XM_034667205.1    rna-XM_038586176.1    rna-XM_022508659.1, rna-XM_022508695.1    rna-XM_023247971.2    rna-XM_036097643.1    rna-XM_039250604.1    rna-XM_032872954.1    rna-XM_045982876.1    rna-XM_032318182.1, rna-XM_032329879.1    rna-XM_059145835.1    rna-XM_059413369.1    rna-XM_045081166.1    mNeoVis.2.1.1.chr14_hap1.g00438.t01    rna-XM_044915894.1    rna-XM_004415144.1    rna-XM_042975820.1    rna-XM_040636008.1    rna-XM_027611597.1    mNeoVis.3.1.1.chr14.g00436.t01    rna-gnl|WGS_ZZZZ|egapxtmp_031797-R1    mNeoVis.evergladensis.2.1.chr14.g00436.t01    mNeoVis.vulgivaga.2.1.chr14.g00438.t01      1758    N0.HOG0001756   OG0000240  n63   rna-XM_034642939.1    rna-XM_038545817.1    rna-XM_022505043.1    rna-XM_023256094.2    rna-XM_036099518.1    rna-XM_039255811.1    rna-XM_032839625.1    rna-XM_046010402.1    rna-XM_032345408.1    rna-XM_059183232.1    rna-XM_059374060.1    rna-XM_045074209.1    mNeoVis.2.1.1.chr10_hap1.g00047.t01    rna-XM_021679863.2    rna-XM_004395581.1    rna-XM_042990647.1    rna-XM_008706111.2    rna-XM_027570019.2    mNeoVis.3.1.1.chr9.g00027.t01    rna-gnl|WGS_ZZZZ|egapxtmp_014554-R1    mNeoVis.evergladensis.2.1.chr8.g00046.t01    mNeoVis.vulgivaga.2.1.chr9.g00046.t01      1759    N0.HOG0001757   OG0000241  n1   rna-XM_034657461.1    rna-XM_038567854.1    rna-XM_022525447.1    rna-XM_023243595.2    rna-XM_036116126.1    rna-XM_039247545.1    rna-XM_032883512.1    rna-XM_045989834.1    rna-XM_032305653.1    rna-XM_059159395.1    rna-XM_059388601.1    rna-XM_045086926.1    mNeoVis.2.1.1.chr5_hap1.g00072.t01    rna-XM_044918596.1    rna-XM_004395369.1    rna-XM_042977716.1    rna-XM_040629630.1    rna-XM_027583528.1    mNeoVis.3.1.1.chr2.g00038.t01    rna-gnl|WGS_ZZZZ|egapxtmp_034392-R1    mNeoVis.evergladensis.2.1.chr2.g00073.t01    mNeoVis.vulgivaga.2.1.chr2.g00071.t01      1760    N0.HOG0001758   OG0000241  n23   rna-XM_034660739.1    rna-XM_038580847.1    rna-XM_022511983.1    rna-XM_023255107.2    rna-XM_036113609.1    rna-XM_039237826.1    rna-XM_032864930.1    rna-XM_046007999.1    rna-XM_032342580.1    rna-XM_059180529.1    rna-XM_059371936.1    rna-XM_004758325.3    mNeoVis.2.1.1.chr10_hap1.g01030.t01    rna-XM_044918250.1    rna-XM_004415877.1    rna-XM_042988495.1    rna-XM_040643044.1    rna-XM_027569247.2    mNeoVis.3.1.1.chr9.g00963.t01    rna-gnl|WGS_ZZZZ|egapxtmp_014535-R1    mNeoVis.evergladensis.2.1.chr8.g01003.t01    mNeoVis.vulgivaga.2.1.chr9.g00987.t01      1761    N0.HOG0001759   OG0000241  n39   rna-XM_002927980.4    rna-XM_038527021.1    rna-XM_022512511.1    rna-XM_023243373.2    rna-XM_036095832.1    rna-XM_039257632.1    rna-XM_032866689.1    rna-XM_046022604.1    rna-XM_032306460.1, rna-XM_032337649.1    rna-XM_059141111.1    rna-XM_059411799.1    rna-XM_045085350.1    mNeoVis.2.1.1.chr13_hap1.g00714.t01    rna-XM_044920675.1    rna-XM_004396706.1    rna-XM_042964050.1    rna-XM_040620214.1    rna-XM_027615221.1    mNeoVis.3.1.1.chr13.g00630.t01    rna-gnl|WGS_ZZZZ|egapxtmp_031541-R1, rna-gnl|WGS_ZZZZ|egapxtmp_031542-R1    mNeoVis.evergladensis.2.1.chr13.g00677.t01    mNeoVis.vulgivaga.2.1.chr13.g00678.t01      1762    N0.HOG0001760   OG0000241  n56   rna-XM_034647083.1    rna-XM_038527009.1    rna-XM_022512682.1    rna-XM_011288270.4    rna-XM_036113960.1    rna-XM_039224058.1    rna-XM_032866702.1    rna-XM_046022190.1    rna-XM_032307669.1    rna-XM_059143458.1    rna-XM_059411795.1    rna-XM_004778656.3    mNeoVis.2.1.1.chr13_hap1.g00709.t01    rna-XM_044920519.1    rna-XM_004408422.1    rna-XM_042963720.1    rna-XM_040620210.1    rna-XM_027615297.1    mNeoVis.3.1.1.chr13.g00624.t01    rna-gnl|WGS_ZZZZ|egapxtmp_030523-R1    mNeoVis.evergladensis.2.1.chr13.g00673.t01    mNeoVis.vulgivaga.2.1.chr13.g00676.t01      1763    N0.HOG0001761   OG0000242  n1   rna-XM_034641602.1    rna-XM_038571787.1    rna-XM_022494584.1    rna-XM_023251386.2    rna-XM_036093158.1    rna-XM_039224940.1    rna-XM_032859038.1    rna-XM_045981871.1    rna-XM_032351233.1    rna-XM_059133766.1    rna-XM_059386484.1, rna-XM_059406659.1    rna-XM_045086054.1    mNeoVis.2.1.1.chr11_hap1.g00403.t01    rna-XM_021689088.1    rna-XM_004398147.1    rna-XM_042980768.1    rna-XM_040638074.1    rna-XM_027622006.2    mNeoVis.3.1.1.chr11.g00369.t01    rna-gnl|WGS_ZZZZ|egapxtmp_026057-R1    mNeoVis.evergladensis.2.1.chr11.g00399.t01    mNeoVis.vulgivaga.2.1.chr11.g00404.t01      1764    N0.HOG0001762   OG0000242  n23   rna-XM_002914426.4    rna-XM_038573773.1    rna-XM_022515354.1    rna-XM_023252458.2    rna-XM_036079082.1    rna-XM_039250430.1    rna-XM_032855789.1    rna-XM_045997172.1    rna-XM_032332212.1    rna-XM_059177820.1    rna-XM_059393196.1    rna-XM_004763239.3    mNeoVis.2.1.1.chr6_hap1.g00809.t01    rna-XM_021698818.1    rna-XM_004412984.1    rna-XM_042983190.1    rna-XM_040626486.1    rna-XM_027600719.2    mNeoVis.3.1.1.chr4.g00819.t01    rna-gnl|WGS_ZZZZ|egapxtmp_000860-R1    mNeoVis.evergladensis.2.1.chr4.g00811.t01    mNeoVis.vulgivaga.2.1.chr4.g00831.t01      1765    N0.HOG0001763   OG0000242  n39   rna-XM_011219330.3    rna-XM_038534887.1    rna-XM_022493681.1    rna-XM_045036218.1    rna-XM_036103648.1    rna-XM_039245352.1    rna-XM_032851682.1    rna-XM_046001039.1    rna-XM_032337165.1    rna-XM_059174172.1    rna-XM_059417262.1    rna-XM_004737712.2    mNeoVis.2.1.1.chr1_hap1.g00258.t01    rna-XM_021697770.1    rna-XM_004404788.1    rna-XM_042993469.1    rna-XM_040630353.1    rna-XM_027605968.1    mNeoVis.3.1.1.chr1.g00259.t01    rna-gnl|WGS_ZZZZ|egapxtmp_007466-R1    mNeoVis.evergladensis.2.1.chr1.g00256.t01    mNeoVis.vulgivaga.2.1.chr1.g00261.t01      1766    N0.HOG0001764   OG0000242  n52   rna-XM_011229350.3, rna-XM_034652194.1    rna-XM_038579120.1    rna-XM_022500096.1    rna-XM_023241102.2    rna-XM_036120720.1    rna-XM_039245745.1    rna-XM_032882065.1    rna-XM_046027146.1    rna-XM_032312992.1    rna-XM_059172814.1    rna-XM_059398823.1    rna-XM_045068738.1    mNeoVis.2.1.1.chr2_hap1.g00040.t01    rna-XM_044916622.1    rna-XM_004412839.1    rna-XM_042960580.1    rna-XM_040624505.1    rna-XM_027596731.2    mNeoVis.3.1.1.chr5.g00034.t01    rna-gnl|WGS_ZZZZ|egapxtmp_005899-R1    mNeoVis.evergladensis.2.1.chr5.g00040.t01    mNeoVis.vulgivaga.2.1.chr5.g00037.t01      1767    N0.HOG0001765   OG0000243  n5           rna-XM_036099979.1    rna-XM_039220158.1      rna-XM_045995511.1              rna-XM_044913585.1                      1768    N0.HOG0001766   OG0000243  n9                                     rna-XM_035724247.1, rna-XM_035724971.1              1769    N0.HOG0001767   OG0000243  n13                         rna-XM_004750562.3        rna-XM_004395980.1                    1770    N0.HOG0001768   OG0000243  n15     rna-NM_001002937.2    rna-XM_022513207.1      rna-XM_036099670.1      rna-XM_032880001.1    rna-XM_045978241.1    rna-XM_032315428.1    rna-XM_059144737.1    rna-XM_059377873.1, rna-XM_059384559.1    rna-XM_045069656.1    mNeoVis.2.1.1.chr8_hap1.g01255.t01    rna-XM_021678344.2        rna-XM_008692194.2    rna-XM_027590207.1        mNeoVis.evergladensis.2.1.chr9.g01205.t01    mNeoVis.vulgivaga.2.1.chr8.g01205.t01      1771    N0.HOG0001769   OG0000243  n16   rna-XM_034649906.1, rna-XM_034652057.1    rna-XM_038586471.1    rna-XM_022492734.1    rna-XM_003980278.6, rna-XM_045036439.1, rna-XM_045061751.1    rna-XM_036077738.1, rna-XM_036087516.1, rna-XM_036099238.1, rna-XM_036118366.1    rna-XM_039255139.1    rna-XM_032851396.1    rna-XM_045997305.1, rna-XM_046000000.1      rna-XM_059154198.1    rna-XM_059404965.1, rna-XM_059415986.1      mNeoVis.2.1.1.chr2_hap1.g01184.t01    rna-XM_044918837.1    rna-XM_004406635.1    rna-XM_042993505.1, rna-XM_042996539.1      rna-XM_035723755.1, rna-XM_035725425.1      rna-gnl|WGS_ZZZZ|egapxtmp_009206-R1    mNeoVis.evergladensis.2.1.chr9.g01204.t01        1772    N0.HOG0001770   OG0000243  n40                                       mNeoVis.3.1.1.chr1.g00839.t01    rna-gnl|WGS_ZZZZ|egapxtmp_023458-R1, rna-gnl|WGS_ZZZZ|egapxtmp_035504-R1      mNeoVis.vulgivaga.2.1.chr1.g00835.t01      1773    N0.HOG0001771   OG0000243  n43   rna-XM_034652528.1                rna-XM_045997728.1                                  1774   N0.HOG0001772  OG0000243  n44   rna-XM_002912667.4    rna-XM_038573603.1    rna-XM_022514580.1    rna-XM_003984858.6, rna-XM_045055103.1    rna-XM_036075529.1    rna-XM_039243236.1    rna-XM_032855772.1    rna-XM_045988454.1    rna-XM_032332226.1    rna-XM_059177638.1    rna-XM_059372092.1    rna-XM_004766103.2    mNeoVis.2.1.1.chr6_hap1.g00800.t01, mNeoVis.2.1.1.chr7_hap1.g00001.t01    rna-XM_021698813.1    rna-XM_004395236.1, rna-XM_004398088.1    rna-XM_042979839.1, rna-XM_042983614.1    rna-XM_008705051.2    rna-XM_035725562.1, rna-XM_035726249.1, rna-XM_035726743.1    mNeoVis.3.1.1.chr7.g00001.t01    rna-gnl|WGS_ZZZZ|egapxtmp_042915-R1    mNeoVis.evergladensis.2.1.chr4.g00802.t01, mNeoVis.evergladensis.2.1.chr7.g00001.t01    mNeoVis.vulgivaga.2.1.chr4.g00822.t01, mNeoVis.vulgivaga.2.1.chr7.g00001.t01      1775   N0.HOG0001773  OG0000244  n1   rna-XM_019802748.2    rna-XM_038563631.1    rna-XM_022506567.1    rna-XM_045039683.1    rna-XM_036077842.1, rna-XM_036099585.1    rna-XM_039235388.1    rna-XM_032879731.1    rna-XM_046014591.1    rna-XM_032360195.1    rna-XM_059150570.1    rna-XM_059399135.1    rna-XM_045066326.1    mNeoVis.2.1.1.chr7_hap1.g02603.t01    rna-XM_044919746.1    rna-XM_004393794.1    rna-XM_042957596.1    rna-XM_040639595.1    rna-XM_027579922.2    mNeoVis.3.1.1.chr7.g02265.t01, mNeoVis.3.1.1.chr8.g00080.t01    rna-gnl|WGS_ZZZZ|egapxtmp_012713-R1    mNeoVis.evergladensis.2.1.chr7.g02410.t01    mNeoVis.vulgivaga.2.1.chr7.g02432.t01      1776    N0.HOG0001774   OG0000244  n25   rna-XM_034672774.1    rna-XM_038528293.1    rna-XM_022523746.1    rna-XM_023245985.2    rna-XM_036068285.1    rna-XM_039249597.1    rna-XM_032839292.1    rna-XM_045989308.1    rna-XM_032326016.1    rna-XM_059152777.1    rna-XM_059383303.1    rna-XM_045077044.1    mNeoVis.2.1.1.chr7_hap1.g00909.t01    rna-XM_044911396.1    rna-XM_004415076.1      rna-XM_040633071.1, rna-XM_040634433.1    rna-XM_027619595.2    mNeoVis.3.1.1.chr7.g00850.t01    rna-gnl|WGS_ZZZZ|egapxtmp_013655-R1    mNeoVis.evergladensis.2.1.chr7.g00902.t01    mNeoVis.vulgivaga.2.1.chr7.g00906.t01      1777    N0.HOG0001775   OG0000244  n43   rna-XM_034667661.1    rna-XM_038541825.1    rna-XM_022508964.1    rna-XM_023247297.2    rna-XM_036079572.1    rna-XM_039227597.1    rna-XM_032878306.1    rna-XM_045982983.1    rna-XM_032317880.1    rna-XM_059146145.1    rna-XM_059413134.1    rna-XM_045081024.1    mNeoVis.2.1.1.chr14_hap1.g00350.t01    rna-XM_044916324.1    rna-XM_004392782.1    rna-XM_042976353.1    rna-XM_040636480.1    rna-XM_027612697.2    mNeoVis.3.1.1.chr14.g00348.t01    rna-gnl|WGS_ZZZZ|egapxtmp_031900-R1    mNeoVis.evergladensis.2.1.chr14.g00350.t01    mNeoVis.vulgivaga.2.1.chr14.g00351.t01      1778    N0.HOG0001776   OG0000244  n64   rna-XM_034643745.1    rna-XM_038558539.1    rna-XM_022495008.1    rna-XM_019835275.3    rna-XM_036099031.1    rna-XM_039235562.1    rna-XM_032854666.1    rna-XM_046013266.1    rna-XM_032346888.1    rna-XM_059186425.1    rna-XM_059402392.1    rna-XM_045082940.1    mNeoVis.2.1.1.chr9_hap1.g00964.t01    rna-XM_044914752.1    rna-XM_004407320.1    rna-XM_042992819.1    rna-XM_008710340.2    rna-XM_027592669.2    mNeoVis.3.1.1.chr10.g00919.t01    rna-gnl|WGS_ZZZZ|egapxtmp_023786-R2    mNeoVis.evergladensis.2.1.chr10.g00951.t01    mNeoVis.vulgivaga.2.1.chr10.g00941.t01      1779   N0.HOG0001777  OG0000245  n1   rna-XM_002931009.4    rna-XM_038568182.1    rna-XM_022522034.1    rna-XM_003992945.3    rna-XM_036089256.1    rna-XM_039224077.1    rna-XM_032853690.1    rna-XM_046016258.1    rna-XM_032360323.1    rna-XM_059158224.1    rna-XM_059389230.1    rna-XM_004780180.2    mNeoVis.2.1.1.chr7_hap1.g01768.t01    rna-XM_021685724.1    rna-XM_004394535.1    rna-XM_007083168.2    rna-XM_008710958.1    rna-XM_027580558.2    mNeoVis.3.1.1.chr7.g01642.t01    rna-gnl|WGS_ZZZZ|egapxtmp_010076-R1    mNeoVis.evergladensis.2.1.chr7.g01729.t01    mNeoVis.vulgivaga.2.1.chr7.g01745.t01      1780    N0.HOG0001778   OG0000245  n20   rna-XM_011218119.2    rna-NM_001389139.1      rna-XM_003988897.2    rna-XM_036111278.1, rna-XM_036115190.1    rna-XM_039216973.1, rna-XM_039216974.1    rna-XM_032876505.1    rna-XM_046011831.1    rna-XM_032347861.1    rna-XM_059187483.1    rna-XM_059404525.1    rna-XM_004782165.1    mNeoVis.2.1.1.chr9_hap1.g00443.t01    rna-XM_021686746.1      rna-XM_015537225.2    rna-XM_008711503.1, rna-XM_008711602.1    rna-XM_035729313.1    mNeoVis.3.1.1.chr10.g00422.t01    rna-gnl|WGS_ZZZZ|egapxtmp_024660-R1    mNeoVis.evergladensis.2.1.chr10.g00439.t01    mNeoVis.vulgivaga.2.1.chr10.g00429.t01      1781    N0.HOG0001779   OG0000245  n42   rna-XM_002931008.4, rna-XM_002931010.4, rna-XM_034667724.1    rna-NM_001388739.1, rna-NM_001388741.1, rna-NM_001388742.1, rna-XM_038568181.1    rna-XM_022522027.1, rna-XM_022522033.1    rna-XM_023239004.2, rna-XM_045038133.1, rna-XM_045038134.1    rna-XM_036089253.1, rna-XM_036095960.1, rna-XM_036095965.1    rna-XM_039224078.1, rna-XM_039224080.1    rna-XM_032853688.1, rna-XM_032853689.1    rna-XM_046016673.1, rna-XM_046016789.1    rna-XM_032359376.1, rna-XM_032359480.1, rna-XM_032360321.1    rna-XM_059158186.1, rna-XM_059158199.1, rna-XM_059158208.1      rna-XM_045065880.1, rna-XM_045065882.1    mNeoVis.2.1.1.chr7_hap1.g01766.t01, mNeoVis.2.1.1.chr7_hap1.g01767.t01    rna-XM_021685723.2, rna-XM_044912374.1, rna-XM_044912395.1    rna-XM_004394534.1    rna-XM_007083166.2, rna-XM_007083167.2    rna-XM_008710959.2, rna-XM_008710960.2    rna-XM_027580556.1    mNeoVis.3.1.1.chr7.g01641.t01    rna-gnl|WGS_ZZZZ|egapxtmp_010078-R1    mNeoVis.evergladensis.2.1.chr7.g01728.t01    mNeoVis.vulgivaga.2.1.chr7.g01743.t01, mNeoVis.vulgivaga.2.1.chr7.g01744.t01      1782    N0.HOG0001780   OG0000246  n1   rna-XM_034637908.1    rna-NM_001197119.1    rna-XM_022517341.1    rna-XM_023241276.2    rna-XM_036107603.1    rna-XM_039249567.1    rna-XM_032846891.1, rna-XM_032846897.1    rna-XM_046026085.1    rna-XM_032310375.1    rna-XM_059139894.1    rna-XM_059408191.1    rna-XM_045073630.1    mNeoVis.2.1.1.chr3_hap1.g00663.t01    rna-XM_021698576.1    rna-XM_004415296.1    rna-XM_042962803.1    rna-XM_040619422.1    rna-XM_027575182.2    mNeoVis.3.1.1.chr3.g00655.t01    rna-gnl|WGS_ZZZZ|egapxtmp_040076-R1    mNeoVis.evergladensis.2.1.chr3.g00671.t01    mNeoVis.vulgivaga.2.1.chr3.g00674.t01, mNeoVis.vulgivaga.2.1.chr3.g00675.t01      1783    N0.HOG0001781   OG0000246  n25   rna-XM_034637910.1, rna-XM_034637932.1    rna-XM_038574739.1    rna-XM_022517096.1    rna-XM_023241277.2, rna-XM_023242150.2    rna-XM_036081006.1, rna-XM_036082893.1, rna-XM_036103179.1    rna-XM_039216658.1, rna-XM_039236821.1    rna-XM_032876106.1, rna-XM_032876500.1    rna-XM_046025794.1    rna-XM_032309480.1, rna-XM_032310581.1    rna-XM_059138991.1, rna-XM_059139801.1    rna-XM_059408192.1, rna-XM_059409893.1    rna-XM_013056400.2, rna-XM_045073646.1    mNeoVis.2.1.1.chr3_hap1.g00660.t01, mNeoVis.2.1.1.chr3_hap1.g00662.t01    rna-XM_021698575.1, rna-XM_021698640.2    rna-XM_004395951.1, rna-XM_004417598.1    rna-XM_015543562.2, rna-XM_042961913.1    rna-XM_008688953.2, rna-XM_040619421.1    rna-XM_027576954.2, rna-XM_035724011.1    mNeoVis.3.1.1.chr3.g00652.t01    rna-gnl|WGS_ZZZZ|egapxtmp_041144-R1, rna-gnl|WGS_ZZZZ|egapxtmp_041146-R1    mNeoVis.evergladensis.2.1.chr3.g00668.t01, mNeoVis.evergladensis.2.1.chr3.g00670.t01    mNeoVis.vulgivaga.2.1.chr3.g00672.t01      1784    N0.HOG0001782   OG0000246  n64   rna-XM_034639208.1    rna-XM_038574733.1    rna-XM_022517432.1    rna-XM_023241271.2    rna-XM_036108159.1    rna-XM_039249585.1    rna-XM_032846888.1    rna-XM_046024139.1    rna-XM_032310915.1    rna-XM_059140271.1    rna-XM_059409890.1, rna-XM_059409929.1    rna-XM_045073706.1    mNeoVis.2.1.1.chr3_hap1.g00666.t01    rna-XM_021698657.1    rna-XM_004415295.1    rna-XM_015541026.2, rna-XM_042962801.1    rna-XM_040620044.1    rna-XM_027576431.2    mNeoVis.3.1.1.chr3.g00657.t01    rna-gnl|WGS_ZZZZ|egapxtmp_041147-R1, rna-gnl|WGS_ZZZZ|egapxtmp_041983-R1    mNeoVis.evergladensis.2.1.chr3.g00674.t01    mNeoVis.vulgivaga.2.1.chr3.g00678.t01      1785    N0.HOG0001783   OG0000247  n1   rna-XM_034669529.1    rna-XM_038539345.1, rna-XM_038539350.1, rna-XM_038589238.1      rna-NM_001246271.1, rna-NM_001246278.2      rna-XM_039223729.1, rna-XM_039225508.1, rna-XM_039226068.1      rna-XM_045992644.1, rna-XM_045996922.1    rna-XM_032328294.1    rna-XM_059153852.1    rna-XM_059387288.1, rna-XM_059387357.1    rna-XM_045075070.1    mNeoVis.2.1.1.chr15_hap1.g00655.t01        rna-XM_007081062.3, rna-XM_007081063.3, rna-XM_015537097.2    rna-XM_008698387.2      mNeoVis.3.1.1.chr15.g00439.t01    rna-gnl|WGS_ZZZZ|egapxtmp_034049-R1    mNeoVis.evergladensis.2.1.chr15.g00477.t01    mNeoVis.vulgivaga.2.1.chr15.g00487.t01      1786    N0.HOG0001784   OG0000247  n25   rna-XM_034650322.1, rna-XM_034650408.1, rna-XM_034650548.1, rna-XM_034651423.1, rna-XM_034651487.1    rna-NM_001003338.1, rna-NM_001003340.1, rna-XM_038539354.1, rna-XM_038589237.1, rna-XM_038589240.1, rna-XM_038589742.1, rna-XM_038589743.1    rna-XM_022500609.1, rna-XM_022500610.1, rna-XM_022500615.1      rna-XM_036064786.1, rna-XM_036078731.1, rna-XM_036078737.1      rna-XM_032872312.1, rna-XM_032872624.1, rna-XM_032872634.1    rna-XM_045992626.1, rna-XM_045992642.1, rna-XM_045992643.1, rna-XM_045992661.1, rna-XM_045996919.1, rna-XM_045996921.1, rna-XM_045996924.1, rna-XM_045996959.1    rna-XM_032328297.1, rna-XM_032328302.1    rna-XM_059153854.1, rna-XM_059153858.1, rna-XM_059153861.1    rna-XM_059414480.1, rna-XM_059414500.1    rna-XM_045075072.1, rna-XM_045075076.1, rna-XM_045075078.1    mNeoVis.2.1.1.chr15_hap1.g00653.t01, mNeoVis.2.1.1.chr15_hap1.g00654.t01, mNeoVis.2.1.1.chr15_hap1.g00656.t01    rna-XM_021687656.1, rna-XM_044915282.1, rna-XM_044915283.1, rna-XM_044915717.1    rna-XM_004414732.1, rna-XM_004417518.1      rna-XM_008698381.2, rna-XM_008698388.2    rna-XM_035722051.1, rna-XM_035722052.1, rna-XM_035722053.1, rna-XM_035722054.1    mNeoVis.3.1.1.chr15.g00435.t01, mNeoVis.3.1.1.chr15.g00436.t01, mNeoVis.3.1.1.chr15.g00440.t01    rna-gnl|WGS_ZZZZ|egapxtmp_033907-R1, rna-gnl|WGS_ZZZZ|egapxtmp_034048-R4, rna-gnl|WGS_ZZZZ|egapxtmp_034050-R3    mNeoVis.evergladensis.2.1.chr15.g00475.t01, mNeoVis.evergladensis.2.1.chr15.g00478.t01    mNeoVis.vulgivaga.2.1.chr15.g00484.t01, mNeoVis.vulgivaga.2.1.chr15.g00485.t01, mNeoVis.vulgivaga.2.1.chr15.g00488.t01      1787    N0.HOG0001785   OG0000248  n1   rna-XM_002930697.4, rna-XM_002930698.2, rna-XM_002930703.4    rna-XM_038568159.1    rna-XM_022521683.1, rna-XM_022521722.1, rna-XM_022522211.1    rna-XM_003992844.4, rna-XM_045038819.1, rna-XM_045038822.1, rna-XM_045038823.1      rna-XM_039223189.1, rna-XM_039225377.1, rna-XM_039225674.1    rna-XM_032852683.1, rna-XM_032853615.1, rna-XM_032853616.1    rna-XM_046015912.1, rna-XM_046017615.1, rna-XM_046017620.1    rna-XM_032356673.1, rna-XM_032356828.1, rna-XM_032359750.1    rna-XM_059156956.1, rna-XM_059156995.1, rna-XM_059157009.1    rna-XM_059390403.1, rna-XM_059390417.1, rna-XM_059390463.1    rna-XM_004768251.2, rna-XM_004768254.2, rna-XM_004768255.2    mNeoVis.2.1.1.chr7_hap1.g01610.t01, mNeoVis.2.1.1.chr7_hap1.g01639.t01, mNeoVis.2.1.1.chr7_hap1.g01640.t01    rna-XM_021685622.1      rna-XM_007089341.1, rna-XM_007089343.1, rna-XM_007089396.2, rna-XM_007089398.1    rna-XM_008710717.1, rna-XM_008710721.2      mNeoVis.3.1.1.chr7.g01541.t01, mNeoVis.3.1.1.chr7.g01543.t01    rna-gnl|WGS_ZZZZ|egapxtmp_012829-R3, rna-gnl|WGS_ZZZZ|egapxtmp_012872-R1    mNeoVis.evergladensis.2.1.chr7.g01609.t01, mNeoVis.evergladensis.2.1.chr7.g01613.t01, mNeoVis.evergladensis.2.1.chr7.g01614.t01    mNeoVis.vulgivaga.2.1.chr7.g01627.t01, mNeoVis.vulgivaga.2.1.chr7.g01629.t01, mNeoVis.vulgivaga.2.1.chr7.g01630.t01      1788    N0.HOG0001786   OG0000248  n52   rna-XM_011217513.2    rna-XM_038569354.1    rna-XM_022521718.1, rna-XM_022521723.1    rna-XM_006937028.4      rna-XM_039223162.1    rna-XM_032852678.1, rna-XM_032852685.1    rna-XM_046015909.1, rna-XM_046015913.1    rna-XM_032359743.1    rna-XM_059179393.1    rna-XM_059396662.1    rna-XM_013064266.2          rna-XM_042959609.1    rna-XM_008710755.2, rna-XM_008710760.2      mNeoVis.3.1.1.chr7.g01535.t01    rna-gnl|WGS_ZZZZ|egapxtmp_012784-R1      mNeoVis.vulgivaga.2.1.chr7.g01624.t01      1789    N0.HOG0001787   OG0000248  n71     rna-NM_001389045.1    rna-XM_022521721.1    rna-XM_045038821.1      rna-XM_039223158.1    rna-XM_032852682.1    rna-XM_046017466.1    rna-XM_032359748.1    rna-XM_059156942.1    rna-XM_059390476.1    rna-XM_004768250.1        rna-XM_004407572.1    rna-XM_015540743.2    rna-XM_040630665.1      mNeoVis.3.1.1.chr7.g01540.t01    rna-gnl|WGS_ZZZZ|egapxtmp_010636-R1    mNeoVis.evergladensis.2.1.chr7.g01608.t01    mNeoVis.vulgivaga.2.1.chr7.g01626.t01      1790    N0.HOG0001788   OG0000249  n1   rna-XM_002916155.4, rna-XM_034651215.1, rna-XM_034671590.1    rna-NM_001252225.1, rna-XM_038572542.1    rna-XM_022521807.1    rna-XM_003992976.5    rna-XM_036073625.1, rna-XM_036088513.1, rna-XM_036121418.1    rna-XM_039244613.1, rna-XM_039248701.1    rna-XM_032853411.1    rna-XM_046017618.1    rna-XM_032357982.1    rna-XM_059140198.1    rna-XM_059408824.1    rna-XM_013055797.2    mNeoVis.2.1.1.chr4_hap1.g00020.t01, mNeoVis.2.1.1.chr7_hap1.g01796.t01    rna-XM_021685748.2, rna-XM_044915086.1, rna-XM_044921037.1    rna-XM_004394553.1, rna-XM_004405517.1, rna-XM_004411713.1    rna-NM_001290589.1    rna-XM_008683804.2    rna-XM_027579029.2, rna-XM_035723292.1, rna-XM_035723606.1, rna-XM_035724547.1, rna-XM_035726058.1, rna-XM_035726739.1, rna-XM_035728009.1, rna-XM_035728585.1    mNeoVis.3.1.1.chr4.g00498.t01, mNeoVis.3.1.1.chr6.g00021.t01    rna-gnl|WGS_ZZZZ|egapxtmp_004127-R1    mNeoVis.evergladensis.2.1.chr6.g00020.t01    mNeoVis.vulgivaga.2.1.chr6.g00022.t01, mNeoVis.vulgivaga.2.1.chr7.g01775.t01      1791    N0.HOG0001789   OG0000249  n26                           mNeoVis.2.1.1.chr6_hap1.g00493.t01                rna-gnl|WGS_ZZZZ|egapxtmp_043855-R1    mNeoVis.evergladensis.2.1.chr4.g00499.t01        1792    N0.HOG0001790   OG0000249  n29     rna-XM_038544313.1, rna-XM_038544822.1                                  rna-XM_035722454.1              1793    N0.HOG0001791   OG0000249  n31     rna-XM_038543979.1                                  rna-XM_035727769.1              1794    N0.HOG0001792   OG0000249  n32                                     rna-XM_035724768.1        mNeoVis.evergladensis.2.1.chr7.g01757.t01        1795    N0.HOG0001793   OG0000249  n37                             rna-XM_021685475.1        rna-XM_040626793.1    rna-XM_035725299.1              1796    N0.HOG0001794   OG0000249  n42                                     rna-XM_035722465.1, rna-XM_035729354.1              1797    N0.HOG0001795   OG0000249  n45     rna-XM_038547749.1                  rna-XM_059187465.1    rna-XM_059404545.1        rna-XM_044915890.1        rna-XM_040621740.1    rna-XM_035723674.1      rna-gnl|WGS_ZZZZ|egapxtmp_012547-R1          1798    N0.HOG0001796   OG0000249  n52           rna-XM_036078639.1    rna-XM_039244264.1                      rna-XM_042989362.1      rna-XM_035721877.1              1799    N0.HOG0001797   OG0000249  n57                   rna-XM_032354523.1    rna-XM_059168353.1    rna-XM_059392639.1    rna-XM_045078637.1                          1800    N0.HOG0001798   OG0000249  n63   rna-XM_034649509.1    rna-XM_038569706.1                                              1801    N0.HOG0001799   OG0000249  n64     rna-XM_038569220.1    rna-XM_022498944.1              rna-XM_032328902.1                                1802    N0.HOG0001800   OG0000249  n66                       rna-XM_059412195.1    rna-XM_045082409.1                          1803    N0.HOG0001801   OG0000250  n1   rna-XM_034666841.1    rna-XM_038542200.1    rna-XM_022508204.1    rna-XM_045048892.1    rna-XM_036101502.1    rna-XM_039249319.1    rna-XM_032862704.1    rna-XM_045982495.1    rna-XM_032317766.1    rna-XM_059145253.1    rna-XM_059412718.1    rna-XM_045074304.1    mNeoVis.2.1.1.chr14_hap1.g00142.t01    rna-XM_021682567.1    rna-XM_004396889.1    rna-XM_042976069.1    rna-XM_040634502.1    rna-XM_035722083.1    mNeoVis.3.1.1.chr14.g00138.t01, mNeoVis.3.1.1.chr14.g00139.t01    rna-gnl|WGS_ZZZZ|egapxtmp_032739-R1    mNeoVis.evergladensis.2.1.chr14.g00137.t01    mNeoVis.vulgivaga.2.1.chr14.g00140.t01      1804    N0.HOG0001802   OG0000250  n24   rna-XM_034663768.1    rna-XM_038548580.1    rna-XM_022500758.1, rna-XM_022500760.1    rna-XM_023243153.2    rna-XM_036064991.1    rna-XM_039220724.1    rna-XM_032865594.1    rna-XM_045983149.1, rna-XM_045996803.1, rna-XM_045997001.1, rna-XM_045997011.1, rna-XM_046021348.1, rna-XM_046022820.1    rna-XM_032307656.1    rna-XM_059142283.1    rna-XM_059387303.1, rna-XM_059412121.1    rna-XM_045082838.1      rna-XM_021691077.1    rna-XM_004411308.1, rna-XM_004411309.1, rna-XM_004413623.1, rna-XM_004415903.1, rna-XM_004416362.1, rna-XM_004416569.1, rna-XM_004416865.1, rna-XM_004417022.1, rna-XM_004417047.1, rna-XM_004417153.1    rna-XM_042963523.1    rna-XM_040632311.1    rna-XM_035722240.1, rna-XM_035722809.1, rna-XM_035722949.1, rna-XM_035723273.1, rna-XM_035723545.1, rna-XM_035724105.1, rna-XM_035724264.1, rna-XM_035724667.1, rna-XM_035725983.1, rna-XM_035726449.1, rna-XM_035727686.1, rna-XM_035728825.1, rna-XM_035729292.1              1805    N0.HOG0001803   OG0000250  n68   rna-XM_034657160.1    rna-XM_038567087.1    rna-XM_022527035.1    rna-XM_045046579.1    rna-XM_036110776.1    rna-XM_039252248.1    rna-XM_032835843.1    rna-XM_045990599.1    rna-XM_032312778.1    rna-XM_059160168.1    rna-XM_059389169.1    rna-XM_045083255.1    mNeoVis.2.1.1.chr5_hap1.g00355.t01    rna-XM_044913037.1    rna-XM_004398962.1    rna-XM_042976796.1    rna-XM_040629220.1    rna-XM_027586358.2    mNeoVis.3.1.1.chr2.g00291.t01    rna-gnl|WGS_ZZZZ|egapxtmp_034737-R1    mNeoVis.evergladensis.2.1.chr2.g00356.t01    mNeoVis.vulgivaga.2.1.chr2.g00351.t01      1806    N0.HOG0001804   OG0000251  n1   rna-XM_034647268.1    rna-XM_038536676.1    rna-XM_022524695.1    rna-XM_023244702.2    rna-XM_036066293.1    rna-XM_039231500.1    rna-XM_032839898.1    rna-XM_045986205.1    rna-XM_032321421.1    rna-XM_059147973.1    rna-XM_059380342.1    rna-XM_045083274.1    mNeoVis.2.1.1.chr8_hap1.g01060.t01    rna-XM_044921536.1    rna-XM_004398518.1    rna-XM_042966207.1    rna-XM_040628211.1    rna-XM_027568634.2    mNeoVis.3.1.1.chr8.g00986.t01    rna-gnl|WGS_ZZZZ|egapxtmp_017567-R1    mNeoVis.evergladensis.2.1.chr9.g01035.t01    mNeoVis.vulgivaga.2.1.chr8.g01035.t01      1807    N0.HOG0001805   OG0000251  n19   rna-XM_034662710.1    rna-XM_038583931.1    rna-XM_022503350.1    rna-XM_019840094.3    rna-XM_036082224.1    rna-XM_039218377.1    rna-XM_032842375.1    rna-XM_046001280.1    rna-XM_032340946.1    rna-XM_059163446.1    rna-XM_059391526.1    rna-XM_013058327.2    mNeoVis.2.1.1.chr5_hap1.g01380.t01    rna-XM_021702594.1    rna-XM_004393257.1    rna-XM_042999129.1    rna-XM_040621590.1    rna-XM_027586035.2    mNeoVis.3.1.1.chr2.g01304.t01    rna-gnl|WGS_ZZZZ|egapxtmp_035982-R5    mNeoVis.evergladensis.2.1.chr2.g01366.t01    mNeoVis.vulgivaga.2.1.chr2.g01375.t01      1808    N0.HOG0001806   OG0000251  n35   rna-XM_034648895.1    rna-XM_038587664.1    rna-XM_022524462.1    rna-XM_019823941.2    rna-XM_036119551.1    rna-XM_039253336.1    rna-XM_032850529.1, rna-XM_032850532.1    rna-XM_045996348.1    rna-XM_032330947.1    rna-XM_059156611.1    rna-XM_059384677.1    rna-XM_004777217.3    mNeoVis.2.1.1.chr12_hap1.g00485.t01    rna-XM_021680444.2    rna-XM_004396601.1    rna-XM_015540666.2    rna-XM_040630824.1    rna-XM_027609288.1    mNeoVis.3.1.1.chr12.g00442.t01    rna-gnl|WGS_ZZZZ|egapxtmp_027993-R1    mNeoVis.evergladensis.2.1.chr12.g00436.t01    mNeoVis.vulgivaga.2.1.chr12.g00447.t01      1809    N0.HOG0001807   OG0000251  n52   rna-XM_034649473.1    rna-XM_038586740.1    rna-XM_022518433.1    rna-XM_019823521.3    rna-XM_036068458.1    rna-XM_039238547.1    rna-XM_032880232.1    rna-XM_045994380.1, rna-XM_045996734.1    rna-XM_032331040.1    rna-XM_059158040.1    rna-XM_059384861.1    rna-XM_013064453.2    mNeoVis.2.1.1.chr12_hap1.g00825.t01    rna-XM_044911672.1    rna-XM_004417375.1    rna-XM_015541539.2    rna-XM_040638489.1    rna-XM_027609210.2    mNeoVis.3.1.1.chr12.g00755.t01    rna-gnl|WGS_ZZZZ|egapxtmp_029286-R1    mNeoVis.evergladensis.2.1.chr12.g00753.t01    mNeoVis.vulgivaga.2.1.chr12.g00781.t01      1810    N0.HOG0001808   OG0000252  n1   rna-XM_002923478.4    rna-NM_001003088.1    rna-XM_022501625.1    rna-XM_011290351.4    rna-XM_036112580.1    rna-XM_039243523.1    rna-XM_032872429.1    rna-XM_045993900.1    rna-XM_032328365.1    rna-XM_059154997.1    rna-XM_059414700.1    rna-XM_004761671.3    mNeoVis.2.1.1.chr15_hap1.g00745.t01    rna-XM_021700542.1    rna-XM_004399106.1    rna-XM_042971569.1    rna-XM_040634528.1    rna-XM_027612050.2    mNeoVis.3.1.1.chr15.g00523.t01    rna-gnl|WGS_ZZZZ|egapxtmp_034070-R1    mNeoVis.evergladensis.2.1.chr15.g00567.t01    mNeoVis.vulgivaga.2.1.chr15.g00579.t01      1811    N0.HOG0001809   OG0000252  n19   rna-XM_002923479.4    rna-XM_038539095.1    rna-XM_022501219.1    rna-XM_006942058.3    rna-XM_036112581.1    rna-XM_039243476.1, rna-XM_039255203.1    rna-XM_032872428.1    rna-XM_045993901.1    rna-XM_032328121.1    rna-XM_059154842.1    rna-XM_059414703.1    rna-XM_004761672.3    mNeoVis.2.1.1.chr15_hap1.g00747.t01    rna-XM_021700525.2    rna-XM_004399107.1    rna-XM_015538082.2    rna-XM_040634946.1    rna-XM_027612052.2    mNeoVis.3.1.1.chr15.g00524.t01      mNeoVis.evergladensis.2.1.chr15.g00569.t01    mNeoVis.vulgivaga.2.1.chr15.g00580.t01      1812    N0.HOG0001810   OG0000252  n35   rna-XM_034670530.1    rna-XM_038540635.1    rna-XM_022516904.1    rna-XM_045047569.1    rna-XM_036089949.1    rna-XM_039246808.1    rna-XM_032850304.1, rna-XM_032850328.1    rna-XM_045993996.1    rna-XM_032328509.1    rna-XM_059155031.1    rna-XM_059415562.1    rna-XM_004750608.3    mNeoVis.2.1.1.chr15_hap1.g00323.t01    rna-XM_021698267.1    rna-XM_004403194.1    rna-XM_042972168.1    rna-XM_008696773.1    rna-XM_027612688.1    mNeoVis.3.1.1.chr15.g00294.t01      mNeoVis.evergladensis.2.1.chr15.g00300.t01    mNeoVis.vulgivaga.2.1.chr15.g00305.t01      1813    N0.HOG0001811   OG0000252  n51   rna-XM_034648661.1, rna-XM_034670529.1    rna-XM_038540634.1    rna-XM_022516901.1    rna-XM_045047570.1    rna-XM_036089930.1    rna-XM_039242383.1, rna-XM_039246647.1    rna-XM_032850303.1, rna-XM_032850326.1    rna-XM_045994138.1    rna-XM_032327759.1    rna-XM_059154614.1    rna-XM_059415559.1    rna-XM_004750607.3    mNeoVis.2.1.1.chr15_hap1.g00322.t01    rna-XM_021698289.1    rna-XM_004403195.1    rna-XM_042970889.1    rna-XM_008696774.1    rna-XM_027612735.1    mNeoVis.3.1.1.chr15.g00293.t01      mNeoVis.evergladensis.2.1.chr15.g00299.t01    mNeoVis.vulgivaga.2.1.chr15.g00304.t01      1814    N0.HOG0001812   OG0000253  n1   rna-XM_011217345.3    rna-XM_038567183.1    rna-XM_022526870.1    rna-XM_019817923.3    rna-XM_036110874.1    rna-XM_039250963.1    rna-XM_032884101.1    rna-XM_045990704.1    rna-XM_032312005.1    rna-XM_059160074.1    rna-XM_059387723.1    rna-XM_013054124.2    mNeoVis.2.1.1.chr5_hap1.g00321.t01    rna-XM_044913069.1    rna-XM_004399007.1    rna-XM_042978245.1    rna-XM_008711750.2    rna-XM_027587211.2    mNeoVis.3.1.1.chr2.g00259.t01    rna-gnl|WGS_ZZZZ|egapxtmp_037176-R1    mNeoVis.evergladensis.2.1.chr2.g00320.t01    mNeoVis.vulgivaga.2.1.chr2.g00317.t01      1815    N0.HOG0001813   OG0000253  n23   rna-XM_019801541.2    rna-XM_038567179.1    rna-XM_022526874.1    rna-XM_011289265.4    rna-XM_036110860.1    rna-XM_039252230.1    rna-XM_032884104.1    rna-XM_045989757.1      rna-XM_059168625.1    rna-XM_059389097.1        rna-XM_021705010.2, rna-XM_044913052.1, rna-XM_044913063.1    rna-XM_004398918.1, rna-XM_004399008.1      rna-XM_040629276.1    rna-XM_027586456.2              1816    N0.HOG0001814   OG0000253  n41   rna-XM_034657828.1    rna-XM_038567881.1        rna-XM_036100967.1        rna-XM_045990463.1      rna-XM_059159390.1    rna-XM_059388592.1        rna-XM_044913136.1        rna-XM_040629654.1    rna-XM_035723137.1              1817    N0.HOG0001815   OG0000253  n49   rna-XM_019793332.2    rna-XM_038567186.1    rna-XM_022526876.1    rna-XM_045045888.1    rna-XM_036088885.1    rna-XM_039250966.1    rna-XM_032884100.1    rna-XM_045990705.1    rna-XM_032312036.1    rna-XM_059160075.1    rna-XM_059389096.1    rna-XM_013054125.2    mNeoVis.2.1.1.chr5_hap1.g00319.t01    rna-XM_044917944.1      rna-XM_042978251.1    rna-XM_040629277.1    rna-XM_035723753.1    mNeoVis.3.1.1.chr2.g00257.t01      mNeoVis.evergladensis.2.1.chr2.g00318.t01    mNeoVis.vulgivaga.2.1.chr2.g00315.t01      1818    N0.HOG0001816   OG0000253  n68   rna-XM_034657834.1    rna-XM_038567167.1    rna-XM_022526852.1    rna-XM_045046149.1    rna-XM_036110876.1    rna-XM_039252244.1    rna-XM_032884106.1    rna-XM_045992359.1    rna-XM_032312058.1    rna-XM_059160076.1    rna-XM_059389099.1    rna-XM_045083617.1    mNeoVis.2.1.1.chr5_hap1.g00322.t01    rna-XM_044917927.1    rna-XM_004398920.1    rna-XM_042976790.1    rna-XM_040629273.1    rna-XM_027585555.2    mNeoVis.3.1.1.chr2.g00260.t01    rna-gnl|WGS_ZZZZ|egapxtmp_037178-R1    mNeoVis.evergladensis.2.1.chr2.g00321.t01    mNeoVis.vulgivaga.2.1.chr2.g00314.t01      1819    N0.HOG0001817   OG0000254  n1   rna-XM_034666127.1    rna-XM_038568209.1    rna-XM_022514037.1    rna-XM_003992599.6    rna-XM_036081768.1    rna-XM_039221800.1    rna-XM_032852729.1    rna-XM_046017387.1    rna-XM_032360563.1    rna-XM_059185780.1    rna-XM_059413285.1    rna-XM_004780869.3    mNeoVis.2.1.1.chr7_hap1.g01414.t01    rna-XM_021679062.2    rna-XM_004409736.1    rna-XM_007089137.3    rna-XM_008688390.2    rna-XM_035723017.1    mNeoVis.3.1.1.chr7.g01324.t01    rna-gnl|WGS_ZZZZ|egapxtmp_009776-R2    mNeoVis.evergladensis.2.1.chr7.g01397.t01    mNeoVis.vulgivaga.2.1.chr7.g01409.t01      1820    N0.HOG0001818   OG0000254  n22   rna-XM_034650021.1    rna-XM_038587598.1    rna-XM_022524453.1    rna-XM_023249184.2    rna-XM_036066928.1, rna-XM_036071417.1      rna-XM_032845576.1    rna-XM_045995231.1    rna-XM_032329487.1    rna-XM_059158453.1    rna-XM_059385802.1    rna-XM_045071280.1    mNeoVis.2.1.1.chr12_hap1.g00515.t01    rna-XM_021679709.2    rna-XM_004396638.1    rna-XM_042974015.1    rna-XM_040630677.1    rna-XM_027608512.1    mNeoVis.3.1.1.chr12.g00470.t01    rna-gnl|WGS_ZZZZ|egapxtmp_027953-R1    mNeoVis.evergladensis.2.1.chr12.g00463.t01    mNeoVis.vulgivaga.2.1.chr12.g00476.t01      1821    N0.HOG0001819   OG0000254  n44   rna-XM_034661541.1    rna-XM_038558960.1    rna-XM_022513786.1    rna-XM_003984958.5    rna-XM_036117141.1    rna-XM_039253774.1    rna-XM_032855552.1    rna-XM_045997932.1    rna-XM_032330629.1    rna-XM_059175783.1    rna-XM_059373466.1    rna-XM_045078333.1    mNeoVis.2.1.1.chr6_hap1.g00715.t01    rna-XM_021698766.2    rna-XM_004404755.1    rna-XM_042983811.1    rna-XM_008685462.2    rna-XM_027599802.2    mNeoVis.3.1.1.chr4.g00725.t01    rna-gnl|WGS_ZZZZ|egapxtmp_000826-R1    mNeoVis.evergladensis.2.1.chr4.g00722.t01    mNeoVis.vulgivaga.2.1.chr4.g00739.t01      1822    N0.HOG0001820   OG0000254  n63   rna-XM_019795170.2    rna-XM_038530484.1    rna-XM_022518047.1    rna-XM_011282717.4    rna-XM_036099122.1, rna-XM_036101517.1    rna-XM_039236147.1    rna-XM_032844264.1    rna-XM_045998907.1    rna-XM_032336318.1    rna-XM_059174922.1    rna-XM_059417816.1    rna-XM_045075346.1    mNeoVis.2.1.1.chr1_hap1.g00542.t01    rna-XM_021702215.1    rna-XM_004413506.1    rna-XM_042986218.1    rna-XM_040622289.1    rna-XM_035727445.1    mNeoVis.3.1.1.chr1.g00556.t01    rna-gnl|WGS_ZZZZ|egapxtmp_032682-R1    mNeoVis.evergladensis.2.1.chr1.g00549.t01    mNeoVis.vulgivaga.2.1.chr1.g00558.t01      1823    N0.HOG0001821   OG0000255  n1   rna-XM_002922588.4    rna-XM_038587994.1    rna-XM_022524878.1    rna-XM_045050706.1    rna-XM_036088286.1    rna-XM_039233476.1    rna-XM_032837076.1    rna-XM_045996262.1    rna-XM_032330774.1    rna-XM_059157831.1    rna-XM_059385376.1    rna-XM_004768999.3    mNeoVis.2.1.1.chr12_hap1.g00311.t01    rna-XM_021678170.1    rna-XM_004415208.1    rna-XM_007093775.3    rna-XM_008697233.2    rna-XM_027609465.1    mNeoVis.3.1.1.chr12.g00283.t01    rna-gnl|WGS_ZZZZ|egapxtmp_028496-R1    mNeoVis.evergladensis.2.1.chr12.g00279.t01    mNeoVis.vulgivaga.2.1.chr12.g00286.t01      1824    N0.HOG0001822   OG0000255  n23   rna-XM_034657266.1    rna-XM_038550990.1    rna-XM_022516391.1    rna-XM_023251572.2    rna-XM_036087938.1    rna-XM_039253121.1    rna-XM_032840334.1    rna-XM_045981725.1    rna-XM_032351773.1    rna-XM_059134477.1    rna-XM_059406271.1    rna-XM_045077690.1    mNeoVis.2.1.1.chr11_hap1.g00596.t01    rna-XM_044918840.1    rna-XM_004402395.1    rna-XM_042981164.1    rna-XM_040623594.1    rna-XM_027623182.2    mNeoVis.3.1.1.chr11.g00557.t01    rna-gnl|WGS_ZZZZ|egapxtmp_025229-R1    mNeoVis.evergladensis.2.1.chr11.g00583.t01    mNeoVis.vulgivaga.2.1.chr11.g00592.t01      1825    N0.HOG0001823   OG0000255  n45   rna-XM_019801701.2    rna-XM_038551737.1    rna-XM_022521133.1    rna-XM_023254017.2    rna-XM_036071193.1    rna-XM_039252824.1    rna-XM_032846870.1    rna-XM_045999224.1    rna-XM_032335284.1    rna-XM_059175925.1    rna-XM_059416169.1    rna-XM_045069354.1    mNeoVis.2.1.1.chr1_hap1.g00894.t01    rna-XM_021689942.1    rna-XM_004412667.1    rna-XM_042981959.1    rna-XM_040622632.1    rna-XM_027605553.2    mNeoVis.3.1.1.chr1.g00916.t01    rna-gnl|WGS_ZZZZ|egapxtmp_027208-R1, rna-gnl|WGS_ZZZZ|egapxtmp_038687-R1    mNeoVis.evergladensis.2.1.chr1.g00897.t01    mNeoVis.vulgivaga.2.1.chr1.g00908.t01      1826    N0.HOG0001824   OG0000255  n67   rna-XM_034661737.1    rna-XM_038564570.1    rna-XM_022505709.1    rna-XM_023252784.2    rna-XM_036078930.1    rna-XM_039249155.1    rna-XM_032857195.1    rna-XM_045996808.1    rna-XM_032333469.1    rna-XM_059168266.1    rna-XM_059378601.1    rna-XM_004768462.2    mNeoVis.2.1.1.chr6_hap1.g00405.t01    rna-XM_021679352.2    rna-XM_004405946.1    rna-XM_042983856.1    rna-XM_040626332.1    rna-XM_027598309.2    mNeoVis.3.1.1.chr4.g00407.t01    rna-gnl|WGS_ZZZZ|egapxtmp_000753-R1    mNeoVis.evergladensis.2.1.chr4.g00413.t01    mNeoVis.vulgivaga.2.1.chr4.g00415.t01      1827    N0.HOG0001825   OG0000256  n1   rna-XM_034661738.1    rna-NM_001301862.1    rna-XM_022505989.1    rna-XM_011281688.4    rna-XM_036112412.1    rna-XM_039222286.1    rna-XM_032857571.1    rna-XM_045997678.1    rna-XM_032333426.1    rna-XM_059168662.1    rna-XM_059395542.1    rna-XM_045069780.1    mNeoVis.2.1.1.chr6_hap1.g00421.t01    rna-XM_044912617.1    rna-XM_004405924.1    rna-XM_042983896.1    rna-XM_008685403.2    rna-XM_027599476.2    mNeoVis.3.1.1.chr4.g00422.t01    rna-gnl|WGS_ZZZZ|egapxtmp_000756-R1    mNeoVis.evergladensis.2.1.chr4.g00428.t01    mNeoVis.vulgivaga.2.1.chr4.g00431.t01      1828    N0.HOG0001826   OG0000256  n22   rna-XM_034639045.1    rna-NM_001284482.1    rna-XM_022511795.1    rna-XM_011290283.4    rna-XM_036100920.1    rna-XM_039226260.1    rna-XM_032853791.1    rna-XM_045988633.1    rna-XM_032324965.1    rna-XM_059152531.1    rna-XM_059381565.1    rna-XM_045076412.1    mNeoVis.2.1.1.chr7_hap1.g00045.t01    rna-XM_044922180.1    rna-XM_004392109.1    rna-XM_042969717.1    rna-XM_040633200.1    rna-XM_027619554.2    mNeoVis.3.1.1.chr7.g00036.t01    rna-gnl|WGS_ZZZZ|egapxtmp_010779-R1    mNeoVis.evergladensis.2.1.chr7.g00047.t01    mNeoVis.vulgivaga.2.1.chr7.g00048.t01      1829    N0.HOG0001827   OG0000256  n44   rna-XM_002929956.4    rna-XM_038529452.1    rna-XM_022525292.1    rna-XM_003997949.6    rna-XM_036118410.1    rna-XM_039231965.1    rna-XM_032841998.1    rna-XM_045989015.1    rna-XM_032325130.1    rna-XM_059152487.1    rna-XM_059383652.1    rna-XM_004765209.3    mNeoVis.2.1.1.chr7_hap1.g00392.t01    rna-XM_021701067.1    rna-XM_004405357.1    rna-XM_042970158.1    rna-XM_040634370.1    rna-XM_027619249.1    mNeoVis.3.1.1.chr7.g00378.t01    rna-gnl|WGS_ZZZZ|egapxtmp_014104-R1    mNeoVis.evergladensis.2.1.chr7.g00396.t01    mNeoVis.vulgivaga.2.1.chr7.g00396.t01      1830    N0.HOG0001828   OG0000256  n64   rna-XM_002926011.4    rna-NM_001287169.1    rna-XM_022523041.1    rna-XM_006932735.5    rna-XM_036114821.1    rna-XM_039245168.1    rna-XM_032856374.1    rna-XM_046007221.1    rna-XM_032343828.1    rna-XM_059181653.1    rna-XM_059372923.1    rna-XM_004755327.3    mNeoVis.2.1.1.chr10_hap1.g00568.t01    rna-XM_021695410.2    rna-XM_004402088.1    rna-XM_042989398.1    rna-XM_008709407.1, rna-XM_008710543.2    rna-XM_027569131.2    mNeoVis.3.1.1.chr9.g00523.t01    rna-gnl|WGS_ZZZZ|egapxtmp_016856-R1    mNeoVis.evergladensis.2.1.chr8.g00551.t01    mNeoVis.vulgivaga.2.1.chr9.g00553.t01      1831    N0.HOG0001829   OG0000257  n1   rna-XM_034666952.1, rna-XM_034666956.1    rna-XM_038541998.1    rna-XM_022508678.1    rna-XM_045049264.1    rna-XM_036118200.1    rna-XM_039252672.1    rna-XM_032878162.1    rna-XM_045982406.1    rna-XM_032316985.1    rna-XM_059145020.1    rna-XM_059413015.1    rna-XM_004756182.3    mNeoVis.2.1.1.chr14_hap1.g00280.t01    rna-XM_021682512.1    rna-XM_004415531.1    rna-XM_015543808.2    rna-XM_008685073.2    rna-XM_027610458.2    mNeoVis.3.1.1.chr14.g00282.t01    rna-gnl|WGS_ZZZZ|egapxtmp_032416-R3    mNeoVis.evergladensis.2.1.chr14.g00283.t01    mNeoVis.vulgivaga.2.1.chr14.g00281.t01      1832    N0.HOG0001830   OG0000257  n23   rna-XM_019800849.2    rna-XM_038530182.1    rna-XM_022520811.1    rna-XM_019833901.3    rna-XM_036090045.1    rna-XM_039240362.1    rna-XM_032881641.1    rna-XM_046010951.1    rna-XM_032349809.1    rna-XM_059183751.1    rna-XM_059404033.1    rna-XM_045072357.1    mNeoVis.2.1.1.chr9_hap1.g00032.t01    rna-XM_044915549.1    rna-XM_004411809.1    rna-XM_042990975.1    rna-XM_040645448.1    rna-XM_027594036.2    mNeoVis.3.1.1.chr10.g00031.t01      mNeoVis.evergladensis.2.1.chr10.g00035.t01    mNeoVis.vulgivaga.2.1.chr10.g00032.t01      1833    N0.HOG0001831   OG0000257  n44   rna-XM_019806736.2    rna-XM_038566538.1    rna-XM_022498078.1    rna-XM_045049837.1    rna-XM_036079458.1    rna-XM_039228289.1    rna-XM_032867963.1    rna-XM_045991138.1    rna-XM_032318854.1    rna-XM_059160822.1    rna-XM_059389652.1    rna-XM_045080314.1    mNeoVis.2.1.1.chr5_hap1.g00619.t01    rna-XM_021687039.1    rna-XM_004399194.1    rna-XM_042978858.1    rna-XM_008703818.2    rna-XM_027586484.1    mNeoVis.3.1.1.chr2.g00540.t01    rna-gnl|WGS_ZZZZ|egapxtmp_037749-R1    mNeoVis.evergladensis.2.1.chr2.g00616.t01    mNeoVis.vulgivaga.2.1.chr2.g00617.t01      1834    N0.HOG0001832   OG0000257  n64   rna-XM_034649492.1    rna-XM_038587526.1    rna-XM_022492184.1    rna-XM_004000551.5    rna-XM_036106954.1, rna-XM_036114519.1    rna-XM_039224820.1    rna-XM_032849928.1    rna-XM_045995343.1    rna-XM_032330799.1    rna-XM_059157592.1    rna-XM_059386115.1    rna-XM_004782130.3    mNeoVis.2.1.1.chr12_hap1.g00549.t01    rna-XM_021700194.1    rna-XM_004414685.1    rna-XM_007094064.3    rna-XM_008711679.2    rna-XM_027608467.2    mNeoVis.3.1.1.chr12.g00507.t01    rna-gnl|WGS_ZZZZ|egapxtmp_029615-R2    mNeoVis.evergladensis.2.1.chr12.g00494.t01    mNeoVis.vulgivaga.2.1.chr12.g00507.t01      1835    N0.HOG0001833   OG0000258  n1   rna-XM_034639013.1    rna-XM_038528850.1    rna-XM_022525019.1    rna-XM_023246182.2    rna-XM_036094986.1    rna-XM_039248427.1    rna-XM_032842170.1    rna-XM_045987767.1    rna-XM_032324364.1    rna-XM_059153190.1    rna-XM_059382623.1    rna-XM_045076823.1    mNeoVis.2.1.1.chr7_hap1.g00568.t01    rna-XM_044911372.1    rna-XM_004396348.1    rna-XM_042969534.1    rna-XM_040633991.1    rna-XM_027619090.1    mNeoVis.3.1.1.chr7.g00551.t01    rna-gnl|WGS_ZZZZ|egapxtmp_012251-R1    mNeoVis.evergladensis.2.1.chr7.g00570.t01    mNeoVis.vulgivaga.2.1.chr7.g00572.t01      1836    N0.HOG0001834   OG0000258  n23   rna-XM_034642336.1    rna-NM_001220481.1    rna-XM_022524907.1    rna-XM_023255769.2    rna-XM_036120214.1    rna-XM_039242145.1    rna-XM_032875187.1    rna-XM_046007936.1    rna-XM_032344489.1    rna-XM_059182339.1    rna-XM_059373394.1    rna-XM_004738933.3    mNeoVis.2.1.1.chr10_hap1.g00312.t01    rna-XM_021701335.2    rna-XM_004409368.1    rna-XM_042989898.1    rna-XM_040641794.1    rna-XM_027568865.2    mNeoVis.3.1.1.chr9.g00285.t01    rna-gnl|WGS_ZZZZ|egapxtmp_016796-R1    mNeoVis.evergladensis.2.1.chr8.g00296.t01    mNeoVis.vulgivaga.2.1.chr9.g00296.t01      1837    N0.HOG0001835   OG0000258  n45   rna-XM_034645430.1    rna-XM_038563796.1    rna-XM_022506851.1    rna-XM_019812716.3    rna-XM_036121482.1    rna-XM_039244758.1    rna-XM_032879577.1    rna-XM_046015617.1    rna-XM_032357559.1    rna-XM_059148589.1    rna-XM_059400441.1    rna-XM_013059915.2    mNeoVis.2.1.1.chr7_hap1.g02489.t01    rna-XM_021685571.2    rna-XM_004393862.1    rna-XM_042958040.1    rna-XM_040639474.1    rna-XM_035723003.1    mNeoVis.3.1.1.chr7.g02236.t01    rna-gnl|WGS_ZZZZ|egapxtmp_012701-R1    mNeoVis.evergladensis.2.1.chr7.g02378.t01    mNeoVis.vulgivaga.2.1.chr7.g02397.t01      1838    N0.HOG0001836   OG0000258  n66   rna-XM_002914046.4    rna-XM_038551274.1    rna-XM_022504166.1    rna-XM_023251677.2    rna-XM_036111377.1    rna-XM_039249201.1    rna-XM_032877774.1    rna-XM_045979662.1    rna-XM_032352123.1    rna-XM_059187669.1    rna-XM_059406073.1    rna-XM_004775675.3    mNeoVis.2.1.1.chr11_hap1.g00693.t01    rna-XM_044918587.1    rna-XM_004403662.1    rna-XM_042981398.1    rna-XM_040623478.1    rna-XM_027622169.2    mNeoVis.3.1.1.chr11.g00654.t01, mNeoVis.3.1.1.chr11.g00655.t01    rna-gnl|WGS_ZZZZ|egapxtmp_025264-R1    mNeoVis.evergladensis.2.1.chr11.g00677.t01    mNeoVis.vulgivaga.2.1.chr11.g00693.t01      1839    N0.HOG0001837   OG0000259  n1   rna-XM_034649058.1    rna-XM_038588255.1    rna-XM_022518548.1    rna-XM_011279453.3    rna-XM_036111183.1    rna-XM_039238819.1    rna-XM_032882345.1    rna-XM_045995540.1    rna-XM_032329990.1    rna-XM_059156964.1    rna-XM_059385607.1    rna-XM_045068252.1    mNeoVis.2.1.1.chr12_hap1.g00213.t01    rna-XM_021685960.1    rna-XM_004400190.1    rna-XM_042974325.1    rna-XM_040628223.1    rna-XM_027608817.2    mNeoVis.3.1.1.chr12.g00182.t01    rna-gnl|WGS_ZZZZ|egapxtmp_029000-R1    mNeoVis.evergladensis.2.1.chr12.g00186.t01    mNeoVis.vulgivaga.2.1.chr12.g00190.t01      1840    N0.HOG0001838   OG0000259  n24   rna-XM_034655661.1    rna-XM_038534806.1    rna-XM_022493193.1    rna-XM_045040320.1    rna-XM_036106732.1    rna-XM_039247655.1    rna-XM_032851741.1    rna-XM_046000940.1    rna-XM_032337070.1    rna-XM_059174241.1    rna-XM_059416332.1    rna-XM_045075621.1    mNeoVis.2.1.1.chr1_hap1.g00284.t01    rna-XM_044917119.1    rna-XM_004392032.1    rna-XM_042992777.1    rna-XM_040630317.1    rna-XM_027606095.2    mNeoVis.3.1.1.chr1.g00288.t01    rna-gnl|WGS_ZZZZ|egapxtmp_007253-R1    mNeoVis.evergladensis.2.1.chr1.g00283.t01    mNeoVis.vulgivaga.2.1.chr1.g00287.t01      1841    N0.HOG0001839   OG0000259  n45   rna-XM_034647494.1    rna-XM_038560460.1    rna-XM_022498770.1    rna-XM_045056763.1    rna-XM_036087409.1    rna-XM_039250689.1    rna-XM_032869853.1    rna-XM_045997092.1    rna-XM_032328838.1    rna-XM_059156020.1    rna-XM_059384375.1    rna-XM_013058964.2    mNeoVis.2.1.1.chr6_hap1.g01086.t01    rna-XM_021694125.1    rna-XM_004415376.1    rna-XM_042982909.1    rna-XM_040643363.1    rna-XM_027598498.2    mNeoVis.3.1.1.chr4.g01040.t01    rna-gnl|WGS_ZZZZ|egapxtmp_000914-R1    mNeoVis.evergladensis.2.1.chr4.g01026.t01    mNeoVis.vulgivaga.2.1.chr4.g01053.t01      1842    N0.HOG0001840   OG0000259  n65   rna-XM_034666254.1    rna-XM_038568485.1    rna-XM_022514097.1    rna-XM_023238928.2    rna-XM_036121066.1    rna-XM_039228710.1    rna-XM_032852968.1    rna-XM_046016487.1    rna-XM_032357365.1    rna-XM_059187848.1    rna-XM_059411810.1    rna-XM_013061902.2    mNeoVis.2.1.1.chr7_hap1.g01485.t01    rna-XM_021704752.2    rna-XM_004391833.1    rna-XM_042957496.1    rna-XM_040630912.1    rna-XM_035722936.1    mNeoVis.3.1.1.chr7.g01393.t01    rna-gnl|WGS_ZZZZ|egapxtmp_011363-R2    mNeoVis.evergladensis.2.1.chr7.g01466.t01    mNeoVis.vulgivaga.2.1.chr7.g01483.t01      1843    N0.HOG0001841   OG0000260  n1   rna-XM_034657257.1    rna-XM_038565449.1    rna-XM_022521359.1    rna-XM_003982562.6    rna-XM_036078523.1    rna-XM_039248624.1    rna-XM_032842870.1    rna-XM_045990673.1    rna-XM_032339163.1    rna-XM_059161929.1    rna-XM_059390460.1    rna-XM_045063397.1    mNeoVis.2.1.1.chr5_hap1.g00948.t01    rna-XM_021694917.1    rna-XM_004395717.1    rna-XM_042962373.1    rna-XM_040628359.1    rna-XM_027584334.2    mNeoVis.3.1.1.chr2.g00863.t01    rna-gnl|WGS_ZZZZ|egapxtmp_035491-R1    mNeoVis.evergladensis.2.1.chr2.g00945.t01    mNeoVis.vulgivaga.2.1.chr2.g00949.t01      1844    N0.HOG0001842   OG0000260  n23   rna-XM_034649273.1    rna-XM_038588724.1    rna-XM_022491273.1    rna-XM_045050648.1    rna-XM_036077760.1    rna-XM_039224810.1    rna-XM_032846331.1    rna-XM_045996059.1    rna-XM_032330841.1    rna-XM_059157662.1    rna-XM_059385913.1    rna-XM_004780736.3    mNeoVis.2.1.1.chr12_hap1.g00034.t01    rna-XM_044911896.1    rna-XM_004414522.1    rna-XM_042974811.1    rna-XM_040626535.1, rna-XM_040626743.1    rna-XM_035725834.1    mNeoVis.3.1.1.chr12.g00025.t01    rna-gnl|WGS_ZZZZ|egapxtmp_029103-R1    mNeoVis.evergladensis.2.1.chr12.g00024.t01    mNeoVis.vulgivaga.2.1.chr12.g00026.t01      1845    N0.HOG0001843   OG0000260  n46   rna-XM_011233642.3    rna-XM_038542031.1    rna-XM_022526720.1    rna-XM_003999337.5    rna-XM_036094629.1    rna-XM_039230123.1    rna-XM_032878124.1    rna-XM_045982465.1    rna-XM_032318661.1    rna-XM_059146058.1    rna-XM_059412985.1    rna-XM_004775277.2    mNeoVis.2.1.1.chr14_hap1.g00268.t01    rna-XM_021682521.2    rna-XM_004415540.1    rna-XM_042975512.1    rna-XM_008701248.2    rna-XM_035722087.1    mNeoVis.3.1.1.chr14.g00269.t01    rna-gnl|WGS_ZZZZ|egapxtmp_031869-R1    mNeoVis.evergladensis.2.1.chr14.g00268.t01    mNeoVis.vulgivaga.2.1.chr14.g00269.t01      1846    N0.HOG0001844   OG0000260  n66   rna-XM_034672299.1    rna-XM_038556478.1    rna-XM_022510219.1    rna-XM_045052763.1    rna-XM_036078053.1    rna-XM_039243563.1    rna-XM_032869059.1    rna-XM_046020616.1    rna-XM_032305147.1    rna-XM_059171885.1    rna-XM_059396393.1    rna-XM_045062995.1    mNeoVis.2.1.1.chr4_hap1.g00197.t01    rna-XM_021692787.1    rna-XM_004397980.1    rna-XM_042972611.1    rna-XM_008712055.2    rna-XM_027573479.2    mNeoVis.3.1.1.chr6.g00198.t01    rna-gnl|WGS_ZZZZ|egapxtmp_003820-R1    mNeoVis.evergladensis.2.1.chr6.g00196.t01    mNeoVis.vulgivaga.2.1.chr6.g00197.t01      1847    N0.HOG0001845   OG0000261  n1   rna-XM_002921465.4    rna-NM_001284460.1    rna-XM_022504639.1    rna-XM_003984153.5    rna-XM_036087587.1    rna-XM_039240181.1    rna-XM_032880731.1    rna-XM_045978846.1    rna-XM_032352572.1    rna-XM_059188153.1    rna-XM_059405763.1    rna-XM_004742262.2    mNeoVis.2.1.1.chr11_hap1.g00835.t01    rna-XM_021699100.1    rna-XM_004398738.1    rna-XM_007083482.3    rna-XM_008700037.2    rna-XM_035728736.1    mNeoVis.3.1.1.chr11.g00797.t01    rna-gnl|WGS_ZZZZ|egapxtmp_027338-R1    mNeoVis.evergladensis.2.1.chr11.g00818.t01    mNeoVis.vulgivaga.2.1.chr11.g00838.t01      1848    N0.HOG0001846   OG0000261  n20   rna-XM_019796749.2    rna-XM_038533159.1    rna-XM_022506126.1    rna-XM_023253440.2    rna-XM_036085633.1, rna-XM_036102449.1    rna-XM_039232092.1    rna-XM_032855704.1    rna-XM_045986227.1    rna-XM_032333713.1    rna-XM_059166138.1    rna-XM_059380117.1    rna-XM_004772220.3    mNeoVis.2.1.1.chr6_hap1.g00345.t01    rna-XM_021677673.2, rna-XM_021687610.2    rna-XM_004396187.1    rna-XM_042985056.1    rna-XM_008706338.2    rna-XM_027599929.2    mNeoVis.3.1.1.chr4.g00346.t01    rna-gnl|WGS_ZZZZ|egapxtmp_000730-R1    mNeoVis.evergladensis.2.1.chr4.g00351.t01    mNeoVis.vulgivaga.2.1.chr4.g00348.t01      1849    N0.HOG0001847   OG0000261  n44   rna-XM_034663385.1    rna-XM_038577982.1    rna-XM_022499332.1    rna-XM_023241037.1    rna-XM_036070235.1    rna-XM_039250449.1    rna-XM_032882150.1    rna-XM_046027899.1    rna-XM_032312585.1    rna-XM_059172954.1    rna-XM_059398746.1    rna-XM_045068501.1    mNeoVis.2.1.1.chr2_hap1.g00082.t01    rna-XM_021703829.2    rna-XM_004404000.1    rna-XM_042960478.1    rna-XM_040624481.1    rna-XM_027596451.2    mNeoVis.3.1.1.chr5.g00076.t01      mNeoVis.evergladensis.2.1.chr5.g00083.t01    mNeoVis.vulgivaga.2.1.chr5.g00079.t01      1850    N0.HOG0001848   OG0000261  n62   rna-XM_034647779.1    rna-XM_038560023.1    rna-XM_022498894.1    rna-XM_023252584.2    rna-XM_036118928.1    rna-XM_039242559.1    rna-XM_032838046.1    rna-XM_045997446.1    rna-XM_032329021.1    rna-XM_059156326.1    rna-XM_059384125.1    rna-XM_045070986.1    mNeoVis.2.1.1.chr6_hap1.g00956.t01    rna-XM_021681586.2    rna-XM_004394455.1    rna-XM_042983418.1    rna-XM_040643232.1    rna-XM_027600383.2    mNeoVis.3.1.1.chr4.g00917.t01    rna-gnl|WGS_ZZZZ|egapxtmp_001007-R1    mNeoVis.evergladensis.2.1.chr4.g00907.t01    mNeoVis.vulgivaga.2.1.chr4.g00928.t01      1851    N0.HOG0001849   OG0000262  n1   rna-XM_002912448.4    rna-XM_038583418.1    rna-XM_022505283.1    rna-XM_023260374.2    rna-XM_036115913.1    rna-XM_039254332.1    rna-XM_032875397.1    rna-XM_046001418.1    rna-XM_032340303.1    rna-XM_059163410.1    rna-XM_059391498.1    rna-XM_004756629.3    mNeoVis.2.1.1.chr5_hap1.g01367.t01    rna-XM_021692456.2    rna-XM_004398356.1    rna-XM_042956938.1    rna-XM_040621577.1    rna-XM_027585909.2    mNeoVis.3.1.1.chr2.g01290.t01    rna-gnl|WGS_ZZZZ|egapxtmp_035052-R1    mNeoVis.evergladensis.2.1.chr2.g01354.t01    mNeoVis.vulgivaga.2.1.chr2.g01363.t01      1852    N0.HOG0001850   OG0000262  n11   rna-XM_002927300.4    rna-XM_038539243.1    rna-XM_022500555.1    rna-XM_003998493.6    rna-XM_036073365.1    rna-XM_039221917.1    rna-XM_032872543.1    rna-XM_045994269.1    rna-XM_032328216.1    rna-XM_059153951.1    rna-XM_059414571.1    rna-XM_004761548.3    mNeoVis.2.1.1.chr15_hap1.g00678.t01    rna-XM_021680066.1    rna-XM_004399039.1    rna-XM_042971181.1    rna-XM_040634868.1    rna-XM_027614141.2, rna-XM_035725619.1, rna-XM_035727840.1    mNeoVis.3.1.1.chr15.g00459.t01      mNeoVis.evergladensis.2.1.chr15.g00500.t01    mNeoVis.vulgivaga.2.1.chr15.g00509.t01      1853    N0.HOG0001851   OG0000262  n16   rna-XM_011231236.3    rna-NM_001003236.1    rna-XM_022514285.1    rna-XM_023257953.2    rna-XM_036089665.1    rna-XM_039248413.1    rna-XM_032863174.1    rna-XM_046006895.1    rna-XM_032361066.1    rna-XM_059135018.1    rna-XM_059376720.1    rna-XM_045063556.1    mNeoVis.2.1.1.chr2_hap1.g01910.t01    rna-XM_021683996.2    rna-XM_004407369.1    rna-XM_042994946.1    rna-XM_040645653.1    rna-XM_027625850.1    mNeoVis.3.1.1.chr5.g01822.t01    rna-gnl|WGS_ZZZZ|egapxtmp_005571-R2    mNeoVis.evergladensis.2.1.chr5.g01848.t01    mNeoVis.vulgivaga.2.1.chr5.g01851.t01      1854    N0.HOG0001852   OG0000262  n18   rna-XM_002922232.4    rna-NM_001003004.1    rna-XM_022522371.1    rna-XM_003988332.6    rna-XM_036069330.1    rna-XM_039217530.1    rna-XM_032874718.1    rna-XM_046011401.1    rna-XM_032348950.1    rna-XM_059184670.1    rna-XM_059403683.1    rna-XM_004766655.3    mNeoVis.2.1.1.chr9_hap1.g00269.t01    rna-XM_021690783.2    rna-XM_004413920.1    rna-XM_007089597.3    rna-XM_008683533.2    rna-XM_027591819.2    mNeoVis.3.1.1.chr10.g00260.t01    rna-gnl|WGS_ZZZZ|egapxtmp_023356-R1    mNeoVis.evergladensis.2.1.chr10.g00268.t01    mNeoVis.vulgivaga.2.1.chr10.g00263.t01      1855    N0.HOG0001853   OG0000263  n1   rna-XM_034668845.1    rna-XM_038557033.1    rna-XM_022503097.1    rna-XM_019825556.2    rna-XM_036113160.1    rna-XM_039220715.1    rna-XM_032873686.1    rna-XM_046020993.1    rna-XM_032304492.1    rna-XM_059171292.1    rna-XM_059396742.1    rna-XM_045062750.1    mNeoVis.2.1.1.chr4_hap1.g00429.t01    rna-XM_021689746.2    rna-XM_004400208.1    rna-XM_042967858.1    rna-XM_040644215.1    rna-XM_027573169.2    mNeoVis.3.1.1.chr6.g00428.t01    rna-gnl|WGS_ZZZZ|egapxtmp_002726-R5    mNeoVis.evergladensis.2.1.chr6.g00433.t01    mNeoVis.vulgivaga.2.1.chr6.g00427.t01      1856    N0.HOG0001854   OG0000263  n24   rna-XM_034655536.1    rna-XM_038574376.1    rna-XM_022517842.1    rna-XM_023259937.2    rna-XM_036072923.1    rna-XM_039228574.1    rna-XM_032857621.1    rna-XM_046018152.1    rna-XM_032355368.1    rna-XM_059167526.1    rna-XM_059393836.1    rna-XM_004743734.3    mNeoVis.2.1.1.chr3_hap1.g01026.t01    rna-XM_044913776.1    rna-XM_004410446.1    rna-XM_042995872.1    rna-XM_040638418.1    rna-XM_027588585.2    mNeoVis.3.1.1.chr3.g01013.t01    rna-gnl|WGS_ZZZZ|egapxtmp_041237-R1    mNeoVis.evergladensis.2.1.chr3.g01024.t01    mNeoVis.vulgivaga.2.1.chr3.g01041.t01      1857    N0.HOG0001855   OG0000263  n45   rna-XM_034646532.1    rna-XM_038577457.1    rna-XM_022519478.1    rna-XM_011283885.3    rna-XM_036099344.1    rna-XM_039243090.1    rna-XM_032876942.1    rna-XM_046010925.1    rna-XM_032347983.1    rna-XM_059185542.1    rna-XM_059402975.1    rna-XM_004765614.3    mNeoVis.2.1.1.chr9_hap1.g00636.t01    rna-XM_021704876.2    rna-XM_004395963.1    rna-XM_042992157.1    rna-XM_040642376.1    rna-XM_027592832.2    mNeoVis.3.1.1.chr10.g00601.t01, mNeoVis.3.1.1.chr10.g00602.t01    rna-gnl|WGS_ZZZZ|egapxtmp_024703-R1    mNeoVis.evergladensis.2.1.chr10.g00630.t01    mNeoVis.vulgivaga.2.1.chr10.g00614.t01      1858    N0.HOG0001856   OG0000263  n67   rna-XM_034639790.1    rna-XM_038547138.1    rna-XM_022519808.1    rna-XM_023244460.2    rna-XM_036071273.1    rna-XM_039220901.1    rna-XM_032849069.1    rna-XM_045984396.1    rna-XM_032322991.1    rna-XM_059147529.1    rna-XM_059379888.1    rna-XM_045080758.1    mNeoVis.2.1.1.chr8_hap1.g00329.t01    rna-XM_021681774.1    rna-XM_004407273.1    rna-XM_042965657.1    rna-XM_040640540.1    rna-XM_027625795.1    mNeoVis.3.1.1.chr8.g00295.t01    rna-gnl|WGS_ZZZZ|egapxtmp_020117-R1    mNeoVis.evergladensis.2.1.chr9.g00328.t01    mNeoVis.vulgivaga.2.1.chr8.g00336.t01      1859    N0.HOG0001857   OG0000264  n1   rna-XM_034656680.1    rna-XM_038534674.1    rna-XM_022493679.1    rna-XM_019838343.3    rna-XM_036084850.1    rna-XM_039239773.1    rna-XM_032851885.1    rna-XM_045999398.1    rna-XM_032336918.1    rna-XM_059174390.1    rna-XM_059417427.1    rna-XM_045075575.1    mNeoVis.2.1.1.chr1_hap1.g00343.t01    rna-XM_021698695.1    rna-XM_004401429.1    rna-XM_015539032.2    rna-XM_008692334.2    rna-XM_027606665.2    mNeoVis.3.1.1.chr1.g00347.t01    rna-gnl|WGS_ZZZZ|egapxtmp_028790-R1    mNeoVis.evergladensis.2.1.chr1.g00341.t01    mNeoVis.vulgivaga.2.1.chr1.g00347.t01      1860    N0.HOG0001858   OG0000264  n23   rna-XM_002921461.4    rna-XM_038561644.1    rna-XM_022504365.1    rna-XM_045054458.1    rna-XM_036099425.1    rna-XM_039240129.1    rna-XM_032880704.1    rna-XM_045979838.1    rna-XM_032352577.1    rna-XM_059188165.1    rna-XM_059405736.1    rna-XM_004742274.3    mNeoVis.2.1.1.chr11_hap1.g00840.t01    rna-XM_021699257.1    rna-XM_004398745.1    rna-XM_042981816.1    rna-XM_008700043.2    rna-XM_027621289.2    mNeoVis.3.1.1.chr11.g00802.t01    rna-gnl|WGS_ZZZZ|egapxtmp_026083-R1    mNeoVis.evergladensis.2.1.chr11.g00823.t01    mNeoVis.vulgivaga.2.1.chr11.g00843.t01      1861    N0.HOG0001859   OG0000264  n43   rna-XM_011234262.3, rna-XM_034653451.1    rna-XM_038534165.1    rna-XM_022499266.1    rna-XM_006937834.5    rna-XM_036113802.1    rna-XM_039233252.1    rna-XM_032841458.1    rna-XM_046026755.1    rna-XM_032312018.1    rna-XM_059170062.1    rna-XM_059397611.1    rna-XM_013044273.2    mNeoVis.2.1.1.chr2_hap1.g00522.t01    rna-XM_021700191.1    rna-XM_004393679.1    rna-XM_007095743.2    rna-XM_040624044.1    rna-XM_027596272.2    mNeoVis.3.1.1.chr5.g00516.t01    rna-gnl|WGS_ZZZZ|egapxtmp_008465-R1    mNeoVis.evergladensis.2.1.chr5.g00523.t01    mNeoVis.vulgivaga.2.1.chr5.g00523.t01      1862    N0.HOG0001860   OG0000264  n64   rna-XM_002913727.4    rna-XM_038533722.1    rna-XM_022499684.1    rna-XM_003994002.5    rna-XM_036113810.1    rna-XM_039233271.1    rna-XM_032841473.1    rna-XM_046027243.1    rna-XM_032313596.1    rna-XM_059170065.1    rna-XM_059395828.1    rna-XM_004769545.3    mNeoVis.2.1.1.chr2_hap1.g00524.t01    rna-XM_021699731.1    rna-XM_004393680.1    rna-XM_007095746.3    rna-XM_008695968.1    rna-XM_027597127.1    mNeoVis.3.1.1.chr5.g00518.t01    rna-gnl|WGS_ZZZZ|egapxtmp_008466-R1    mNeoVis.evergladensis.2.1.chr5.g00525.t01    mNeoVis.vulgivaga.2.1.chr5.g00525.t01      1863    N0.HOG0001861   OG0000265  n1   rna-XM_034656660.1    rna-XM_038534629.1    rna-XM_022493003.1    rna-XM_023259108.2    rna-XM_036084879.1    rna-XM_039239725.1    rna-XM_032851934.1    rna-XM_045999764.1    rna-XM_032336867.1    rna-XM_059174443.1    rna-XM_059417467.1    rna-XM_045075563.1    mNeoVis.2.1.1.chr1_hap1.g00357.t01    rna-XM_044916810.1    rna-XM_004401407.1    rna-XM_042991564.1    rna-XM_040630233.1    rna-XM_027605610.1    mNeoVis.3.1.1.chr1.g00359.t01    rna-gnl|WGS_ZZZZ|egapxtmp_029085-R1    mNeoVis.evergladensis.2.1.chr1.g00355.t01    mNeoVis.vulgivaga.2.1.chr1.g00363.t01      1864    N0.HOG0001862   OG0000265  n20   rna-XM_011221440.3    rna-XM_038533188.1    rna-XM_022505796.1    rna-XM_019829825.3    rna-XM_036085659.1    rna-XM_039232054.1    rna-XM_032855952.1    rna-XM_045993415.1    rna-XM_032333653.1    rna-XM_059166575.1    rna-XM_059379932.1    rna-XM_013045752.2    mNeoVis.2.1.1.chr6_hap1.g00357.t01    rna-XM_044913031.1    rna-XM_004396252.1    rna-XM_042985077.1    rna-XM_040625150.1    rna-XM_027598978.2    mNeoVis.3.1.1.chr4.g00362.t01, mNeoVis.3.1.1.chr4.g00363.t01    rna-gnl|WGS_ZZZZ|egapxtmp_001429-R1    mNeoVis.evergladensis.2.1.chr4.g00365.t01    mNeoVis.vulgivaga.2.1.chr4.g00360.t01, mNeoVis.vulgivaga.2.1.chr4.g00361.t01      1865    N0.HOG0001863   OG0000265  n44   rna-XM_034665534.1    rna-XM_038570132.1    rna-XM_022503001.1    rna-XM_003980540.5    rna-XM_036068850.1    rna-XM_039221632.1    rna-XM_032880979.1    rna-XM_045978476.1    rna-XM_032314248.1    rna-XM_059144497.1    rna-XM_059378061.1    rna-XM_004745983.3    mNeoVis.2.1.1.chr8_hap1.g01538.t01    rna-XM_021695966.1    rna-XM_004412264.1    rna-XM_042958281.1    rna-XM_008684928.2    rna-XM_027590419.2    mNeoVis.3.1.1.chr8.g01432.t01      mNeoVis.evergladensis.2.1.chr9.g01485.t01    mNeoVis.vulgivaga.2.1.chr8.g01490.t01      1866    N0.HOG0001864   OG0000265  n64   rna-XM_011219972.3    rna-XM_038578947.1    rna-XM_022499248.1    rna-XM_003994462.6    rna-XM_036123833.1    rna-XM_039237364.1    rna-XM_032878681.1    rna-XM_046027594.1    rna-XM_032312593.1    rna-XM_059173046.1    rna-XM_059398697.1    rna-XM_004751835.3    mNeoVis.2.1.1.chr2_hap1.g00099.t01    rna-XM_044916610.1    rna-XM_004404029.1    rna-XM_042960773.1    rna-XM_008685100.2    rna-XM_027587046.1    mNeoVis.3.1.1.chr5.g00095.t01    rna-gnl|WGS_ZZZZ|egapxtmp_005979-R1    mNeoVis.evergladensis.2.1.chr5.g00104.t01    mNeoVis.vulgivaga.2.1.chr5.g00101.t01      1867    N0.HOG0001865   OG0000266  n1   rna-XM_034670738.1      rna-XM_022516696.1    rna-XM_011290755.4    rna-XM_036083563.1    rna-XM_039223411.1    rna-XM_032846039.1      rna-XM_032327656.1    rna-XM_059155507.1      rna-XM_045072048.1    mNeoVis.2.1.1.chr15_hap1.g00517.t01    rna-XM_021698220.1    rna-XM_004403073.1    rna-XM_042971647.1    rna-XM_040634468.1, rna-XM_040635648.1    rna-XM_027609738.1    mNeoVis.3.1.1.chr15.g00346.t01    rna-gnl|WGS_ZZZZ|egapxtmp_026746-R1    mNeoVis.evergladensis.2.1.chr15.g00383.t01    mNeoVis.vulgivaga.2.1.chr15.g00392.t01      1868    N0.HOG0001866   OG0000266  n21   rna-XM_002920155.4, rna-XM_034651908.1, rna-XM_034670739.1    rna-NM_001270885.1, rna-NM_001270886.1    rna-XM_022516624.1    rna-XM_003998972.5, rna-XM_023246781.2    rna-XM_036083560.1, rna-XM_036083562.1    rna-XM_039223408.1, rna-XM_039223409.1    rna-XM_032846038.1    rna-XM_045993573.1, rna-XM_045993584.1    rna-XM_032327659.1, rna-XM_032327660.1    rna-XM_059155510.1, rna-XM_059155511.1      rna-XM_004750932.2, rna-XM_045072050.1    mNeoVis.2.1.1.chr15_hap1.g00518.t01, mNeoVis.2.1.1.chr15_hap1.g00519.t01      rna-XM_004403070.1, rna-XM_004417685.1    rna-XM_007093647.3, rna-XM_042971463.1    rna-XM_008696690.2    rna-XM_027613422.2, rna-XM_027613605.2        mNeoVis.evergladensis.2.1.chr15.g00384.t01, mNeoVis.evergladensis.2.1.chr15.g00385.t01    mNeoVis.vulgivaga.2.1.chr15.g00393.t01      1869    N0.HOG0001867   OG0000266  n52   rna-XM_002920153.4, rna-XM_034669550.1    rna-XM_038538893.1, rna-XM_038540974.1    rna-XM_022492595.1, rna-XM_022516774.1    rna-XM_023246784.2, rna-XM_023246785.2    rna-XM_036083564.1    rna-XM_039223410.1, rna-XM_039223426.1    rna-XM_032846013.1    rna-XM_045993616.1, rna-XM_045993658.1    rna-XM_032327661.1, rna-XM_032327663.1    rna-XM_059154697.1, rna-XM_059155512.1    rna-XM_059415861.1, rna-XM_059416015.1    rna-XM_004750934.3, rna-XM_045072051.1    mNeoVis.2.1.1.chr15_hap1.g00521.t01, mNeoVis.2.1.1.chr15_hap1.g00522.t01    rna-XM_021697978.1, rna-XM_021698235.2    rna-XM_004403071.1    rna-XM_042972590.1, rna-XM_042972591.1    rna-XM_008697030.2, rna-XM_040635649.1    rna-XM_027613424.1, rna-XM_035722389.1      rna-gnl|WGS_ZZZZ|egapxtmp_033714-R1    mNeoVis.evergladensis.2.1.chr15.g00388.t01    mNeoVis.vulgivaga.2.1.chr15.g00395.t01      1870    N0.HOG0001868   OG0000267  n1   rna-XM_034656299.1    rna-XM_038529583.1    rna-XM_022517982.1    rna-XM_023257117.2    rna-XM_036106779.1    rna-XM_039217708.1    rna-XM_032849379.1    rna-XM_045999065.1    rna-XM_032336080.1    rna-XM_059175136.1    rna-XM_059417990.1    rna-XM_045075265.1    mNeoVis.2.1.1.chr1_hap1.g00625.t01    rna-XM_044916860.1    rna-XM_004402259.1    rna-XM_042987724.1    rna-XM_040622219.1    rna-XM_027604891.2    mNeoVis.3.1.1.chr1.g00644.t01    rna-gnl|WGS_ZZZZ|egapxtmp_033961-R1    mNeoVis.evergladensis.2.1.chr1.g00631.t01    mNeoVis.vulgivaga.2.1.chr1.g00641.t01      1871    N0.HOG0001869   OG0000267  n23   rna-XM_011222140.3    rna-XM_038566842.1    rna-XM_022497664.1    rna-XM_019819687.3    rna-XM_036093881.1    rna-XM_039243320.1    rna-XM_032867652.1    rna-XM_045990082.1    rna-XM_032315784.1    rna-XM_059160493.1    rna-XM_059389403.1    rna-XM_013060576.2    mNeoVis.2.1.1.chr5_hap1.g00506.t01    rna-XM_044921952.1    rna-XM_004404627.1    rna-XM_042978572.1    rna-XM_040628254.1    rna-XM_035728760.1    mNeoVis.3.1.1.chr2.g00429.t01    rna-gnl|WGS_ZZZZ|egapxtmp_034931-R1    mNeoVis.evergladensis.2.1.chr2.g00501.t01    mNeoVis.vulgivaga.2.1.chr2.g00503.t01      1872    N0.HOG0001870   OG0000267  n45   rna-XM_034657853.1    rna-XM_038567132.1    rna-XM_022526901.1    rna-XM_023245932.2    rna-XM_036110820.1    rna-XM_039252249.1    rna-XM_032884137.1    rna-XM_045991936.1    rna-XM_032312371.1    rna-XM_059160114.1    rna-XM_059389128.1    rna-XM_004748341.2    mNeoVis.2.1.1.chr5_hap1.g00339.t01    rna-XM_021704963.2    rna-XM_004398940.1    rna-XM_042978282.1    rna-XM_040629243.1    rna-XM_027587546.2    mNeoVis.3.1.1.chr2.g00277.t01    rna-gnl|WGS_ZZZZ|egapxtmp_038314-R1    mNeoVis.evergladensis.2.1.chr2.g00338.t01, mNeoVis.evergladensis.2.1.chr2.g00339.t01    mNeoVis.vulgivaga.2.1.chr2.g00335.t01      1873    N0.HOG0001871   OG0000267  n66   rna-XM_034651722.1    rna-XM_038558182.1    rna-XM_022496734.1    rna-XM_023258635.2    rna-XM_036100069.1    rna-XM_039254813.1    rna-XM_032877648.1    rna-XM_046010751.1    rna-XM_032361575.1    rna-XM_059137764.1    rna-XM_059374594.1    rna-XM_013044326.2    mNeoVis.2.1.1.chr2_hap1.g01347.t01    rna-XM_044914016.1    rna-XM_004400589.1    rna-XM_042996974.1    rna-XM_040627348.1    rna-XM_027616549.2    mNeoVis.3.1.1.chr5.g01274.t01    rna-gnl|WGS_ZZZZ|egapxtmp_005698-R1    mNeoVis.evergladensis.2.1.chr5.g01290.t01    mNeoVis.vulgivaga.2.1.chr5.g01287.t01      1874    N0.HOG0001872   OG0000268  n1   rna-XM_034656261.1    rna-XM_038530650.1    rna-XM_022518023.1    rna-XM_023256805.2    rna-XM_036077326.1    rna-XM_039243827.1    rna-XM_032850058.1    rna-XM_045999294.1    rna-XM_032334874.1    rna-XM_059175042.1    rna-XM_059417914.1    rna-XM_004744605.3    mNeoVis.2.1.1.chr1_hap1.g00594.t01    rna-XM_021702161.1    rna-XM_004408737.1    rna-XM_042987253.1    rna-XM_040622246.1    rna-XM_027605098.2    mNeoVis.3.1.1.chr1.g00610.t01    rna-gnl|WGS_ZZZZ|egapxtmp_011485-R9    mNeoVis.evergladensis.2.1.chr1.g00599.t01    mNeoVis.vulgivaga.2.1.chr1.g00608.t01      1875    N0.HOG0001873   OG0000268  n20   rna-XM_011222201.3    rna-XM_038566835.1    rna-XM_022497671.1    rna-XM_019819750.3    rna-XM_036093965.1    rna-XM_039243276.1    rna-XM_032867678.1    rna-XM_045991104.1    rna-XM_032315970.1    rna-XM_059160521.1    rna-XM_059389427.1    rna-XM_045081583.1    mNeoVis.2.1.1.chr5_hap1.g00511.t01    rna-XM_021683114.1    rna-XM_004404633.1    rna-XM_042978594.1    rna-XM_008703946.2    rna-XM_027587374.2    mNeoVis.3.1.1.chr2.g00434.t01      mNeoVis.evergladensis.2.1.chr2.g00506.t01    mNeoVis.vulgivaga.2.1.chr2.g00508.t01      1876    N0.HOG0001874   OG0000268  n40   rna-XM_034657764.1    rna-XM_038567316.1    rna-XM_022527006.1    rna-XM_011289052.4    rna-XM_036109264.1    rna-XM_039250976.1    rna-XM_032883977.1    rna-XM_045990974.1    rna-XM_032310948.1    rna-XM_059159947.1    rna-XM_059388992.1    rna-XM_013054154.2    mNeoVis.2.1.1.chr5_hap1.g00285.t01    rna-XM_021705046.1    rna-XM_004399002.1    rna-XM_042978142.1    rna-XM_040629355.1, rna-XM_040629360.1    rna-XM_035724909.1    mNeoVis.3.1.1.chr2.g00225.t01    rna-gnl|WGS_ZZZZ|egapxtmp_034857-R1    mNeoVis.evergladensis.2.1.chr2.g00283.t01    mNeoVis.vulgivaga.2.1.chr2.g00278.t01      1877    N0.HOG0001875   OG0000268  n61   rna-XM_019804216.2    rna-XM_038537222.1    rna-XM_022496458.1    rna-XM_045035976.1    rna-XM_036077702.1    rna-XM_039233423.1    rna-XM_032870016.1    rna-XM_046017685.1    rna-XM_032302838.1    rna-XM_059135175.1    rna-XM_059375023.1    rna-XM_045064680.1    mNeoVis.2.1.1.chr2_hap1.g01209.t01    rna-XM_021680003.1    rna-XM_004411860.1    rna-XM_042997497.1    rna-XM_008690765.2, rna-XM_008709425.2    rna-XM_027619904.2    mNeoVis.3.1.1.chr5.g01125.t01    rna-gnl|WGS_ZZZZ|egapxtmp_005660-R1    mNeoVis.evergladensis.2.1.chr5.g01151.t01    mNeoVis.vulgivaga.2.1.chr5.g01138.t01      1878    N0.HOG0001876   OG0000269  n1   rna-XM_034638340.1    rna-XM_038559358.1    rna-XM_022510273.1    rna-XM_045052642.1    rna-XM_036086601.1    rna-XM_039234017.1    rna-XM_032867306.1    rna-XM_046020164.1    rna-XM_032304957.1    rna-XM_059172069.1    rna-XM_059395984.1    rna-XM_045063084.1    mNeoVis.2.1.1.chr4_hap1.g00126.t01    rna-XM_021692830.2    rna-XM_004416554.1    rna-XM_042973964.1    rna-XM_040643909.1    rna-XM_035723415.1    mNeoVis.3.1.1.chr6.g00126.t01    rna-gnl|WGS_ZZZZ|egapxtmp_004366-R1    mNeoVis.evergladensis.2.1.chr6.g00128.t01    mNeoVis.vulgivaga.2.1.chr6.g00126.t01      1879    N0.HOG0001877   OG0000269  n24   rna-XM_034640095.1, rna-XM_034653173.1    rna-XM_038546375.1    rna-XM_022523559.1    rna-XM_006940651.5    rna-XM_036067381.1    rna-XM_039226443.1    rna-XM_032870438.1    rna-XM_045985067.1    rna-XM_032320432.1    rna-XM_059147327.1    rna-XM_059379818.1    rna-XM_004748755.2    mNeoVis.2.1.1.chr8_hap1.g00078.t01    rna-XM_021680932.2    rna-XM_004404511.1    rna-XM_042966434.1    rna-XM_040640079.1    rna-XM_027625593.2    mNeoVis.3.1.1.chr8.g00054.t01      mNeoVis.evergladensis.2.1.chr9.g00075.t01    mNeoVis.vulgivaga.2.1.chr8.g00076.t01, mNeoVis.vulgivaga.2.1.chr8.g00077.t01      1880    N0.HOG0001878   OG0000269  n46   rna-XM_011224093.3    rna-XM_038583278.1    rna-XM_022506428.1    rna-XM_045036311.1    rna-XM_036083581.1    rna-XM_039255023.1    rna-XM_032836045.1    rna-XM_046002560.1    rna-XM_032361662.1    rna-XM_059163586.1    rna-XM_059388067.1    rna-XM_004774414.2    mNeoVis.2.1.1.chr5_hap1.g01422.t01    rna-XM_021702632.1    rna-XM_004393197.1    rna-XM_007086237.2    rna-XM_040636582.1    rna-XM_027584972.2    mNeoVis.3.1.1.chr2.g01344.t01    rna-gnl|WGS_ZZZZ|egapxtmp_034722-R1    mNeoVis.evergladensis.2.1.chr2.g01407.t01    mNeoVis.vulgivaga.2.1.chr2.g01416.t01      1881    N0.HOG0001879   OG0000269  n67   rna-XM_002921683.4    rna-XM_038559361.1    rna-XM_022510248.1    rna-XM_045052868.1    rna-XM_036086589.1    rna-XM_039234021.1    rna-XM_032867280.1    rna-XM_046020171.1    rna-XM_032304954.1    rna-XM_059172072.1    rna-XM_059396245.1    rna-XM_045063081.1    mNeoVis.2.1.1.chr4_hap1.g00127.t01    rna-XM_021692831.1    rna-XM_004416552.1    rna-XM_042973967.1    rna-XM_008697325.2    rna-XM_027575000.2    mNeoVis.3.1.1.chr6.g00127.t01    rna-gnl|WGS_ZZZZ|egapxtmp_001906-R1    mNeoVis.evergladensis.2.1.chr6.g00129.t01    mNeoVis.vulgivaga.2.1.chr6.g00127.t01      1882    N0.HOG0001880   OG0000270  n1   rna-XM_002914169.4    rna-XM_038586228.1    rna-XM_022508988.1    rna-XM_006942988.5    rna-XM_036065660.1    rna-XM_039222847.1    rna-XM_032872992.1    rna-XM_045982817.1    rna-XM_032319341.1    rna-XM_059145472.1    rna-XM_059413407.1    rna-XM_004751432.3    mNeoVis.2.1.1.chr14_hap1.g00463.t01    rna-XM_021698415.1    rna-XM_004403470.1    rna-XM_042976717.1    rna-XM_040635989.1    rna-XM_027610702.2    mNeoVis.3.1.1.chr14.g00458.t01    rna-gnl|WGS_ZZZZ|egapxtmp_032900-R1    mNeoVis.evergladensis.2.1.chr14.g00459.t01    mNeoVis.vulgivaga.2.1.chr14.g00461.t01      1883    N0.HOG0001881   OG0000270  n9   rna-XM_019796489.2    rna-XM_038549344.1    rna-XM_022501070.1    rna-XM_023242717.2    rna-XM_036112903.1    rna-XM_039221080.1, rna-XM_039238983.1    rna-XM_032865873.1    rna-XM_046022724.1    rna-XM_032306234.1    rna-XM_059141024.1    rna-XM_059410798.1    rna-XM_004757308.3    mNeoVis.2.1.1.chr13_hap1.g00227.t01    rna-XM_021691236.1    rna-XM_004392583.1    rna-XM_042965228.1    rna-XM_040631898.1    rna-XM_027616686.1    mNeoVis.3.1.1.chr13.g00178.t01    rna-gnl|WGS_ZZZZ|egapxtmp_031323-R1    mNeoVis.evergladensis.2.1.chr13.g00225.t01    mNeoVis.vulgivaga.2.1.chr13.g00222.t01      1884    N0.HOG0001882   OG0000270  n16   rna-XM_019797700.2    rna-XM_038566717.1    rna-XM_022498379.1    rna-XM_003982174.5    rna-XM_036093978.1    rna-XM_039228203.1    rna-XM_032867800.1    rna-XM_045992227.1    rna-XM_032316978.1    rna-XM_059160644.1    rna-XM_059389523.1    rna-XM_045081014.1    mNeoVis.2.1.1.chr5_hap1.g00555.t01    rna-XM_044917358.1    rna-XM_004404689.1    rna-XM_042978704.1    rna-XM_040628946.1    rna-XM_027584215.2    mNeoVis.3.1.1.chr2.g00476.t01      mNeoVis.evergladensis.2.1.chr2.g00549.t01    mNeoVis.vulgivaga.2.1.chr2.g00552.t01      1885    N0.HOG0001883   OG0000270  n35   rna-XM_002929647.4    rna-XM_038553615.1    rna-XM_022510554.1    rna-XM_003985970.6    rna-XM_036108183.1    rna-XM_039219609.1    rna-XM_032850415.1, rna-XM_032850438.1    rna-XM_046005565.1    rna-XM_032340774.1    rna-XM_059178798.1    rna-XM_059399945.1    rna-XM_004780880.2    mNeoVis.2.1.1.chr1_hap1.g01382.t01    rna-XM_021697040.2    rna-XM_004415281.1    rna-XM_007093270.2    rna-XM_008711848.2    rna-XM_027601535.2    mNeoVis.3.1.1.chr1.g01329.t01      mNeoVis.evergladensis.2.1.chr1.g01347.t01    mNeoVis.vulgivaga.2.1.chr1.g01349.t01      1886    N0.HOG0001884   OG0000271  n1   rna-XM_011217300.3    rna-XM_038577894.1    rna-XM_022522310.1    rna-XM_006933413.5    rna-XM_036082072.1    rna-XM_039236332.1    rna-XM_032874341.1    rna-XM_046012239.1    rna-XM_032349112.1    rna-XM_059184511.1    rna-XM_059403799.1    rna-XM_045071689.1    mNeoVis.2.1.1.chr9_hap1.g00221.t01    rna-XM_021690797.1    rna-XM_004411952.1    rna-XM_042991577.1    rna-XM_040642037.1    rna-XM_027595157.2    mNeoVis.3.1.1.chr10.g00212.t01    rna-gnl|WGS_ZZZZ|egapxtmp_024074-R1    mNeoVis.evergladensis.2.1.chr10.g00218.t01    mNeoVis.vulgivaga.2.1.chr10.g00215.t01      1887    N0.HOG0001885   OG0000271  n23   rna-XM_034645738.1    rna-XM_038569059.1    rna-XM_022521893.1    rna-XM_011286720.4    rna-XM_036087863.1    rna-XM_039240892.1    rna-XM_032853505.1    rna-XM_046015377.1    rna-XM_032358729.1    rna-XM_059136498.1    rna-XM_059408043.1    rna-XM_045065803.1    mNeoVis.2.1.1.chr7_hap1.g01827.t01    rna-XM_021685311.2    rna-XM_004394601.1    rna-XM_042959701.1    rna-XM_040638827.1    rna-XM_027578584.2    mNeoVis.3.1.1.chr7.g01694.t01    rna-gnl|WGS_ZZZZ|egapxtmp_012559-R1, rna-gnl|WGS_ZZZZ|egapxtmp_012560-R1    mNeoVis.evergladensis.2.1.chr7.g01789.t01    mNeoVis.vulgivaga.2.1.chr7.g01807.t01      1888    N0.HOG0001886   OG0000271  n38   rna-XM_002917850.4    rna-XM_038528249.1    rna-XM_022523797.1    rna-XM_006941008.4    rna-XM_036068275.1    rna-XM_039246853.1    rna-XM_032839475.1    rna-XM_045989082.1    rna-XM_032324537.1    rna-XM_059151650.1    rna-XM_059383224.1    rna-XM_004767334.3    mNeoVis.2.1.1.chr7_hap1.g00902.t01    rna-XM_021681267.2    rna-XM_004409864.1    rna-XM_042968851.1    rna-XM_040634289.1    rna-XM_027618824.1    mNeoVis.3.1.1.chr7.g00843.t01      mNeoVis.evergladensis.2.1.chr7.g00895.t01    mNeoVis.vulgivaga.2.1.chr7.g00899.t01      1889    N0.HOG0001887   OG0000271  n51   rna-XM_034660595.1    rna-XM_038553836.1    rna-XM_022510950.1    rna-XM_045058649.1    rna-XM_036096614.1    rna-XM_039235315.1    rna-XM_032864311.1    rna-XM_046006431.1    rna-XM_032340532.1    rna-XM_059178519.1    rna-XM_059401532.1    rna-XM_045074761.1    mNeoVis.2.1.1.chr1_hap1.g01464.t01    rna-XM_021684664.2    rna-XM_004407764.1    rna-XM_042987593.1    rna-XM_040644610.1    rna-XM_035728342.1    mNeoVis.3.1.1.chr1.g01412.t01    rna-gnl|WGS_ZZZZ|egapxtmp_012303-R1    mNeoVis.evergladensis.2.1.chr1.g01430.t01    mNeoVis.vulgivaga.2.1.chr1.g01431.t01      1890    N0.HOG0001888   OG0000272  n1   rna-XM_034660620.1    rna-NM_001003206.1    rna-XM_022510556.1    rna-XM_003986062.4    rna-XM_036096954.1    rna-XM_039235271.1    rna-XM_032864338.1    rna-XM_046006498.1    rna-XM_032340506.1    rna-XM_059178498.1    rna-XM_059400127.1    rna-XM_004770713.3    mNeoVis.2.1.1.chr1_hap1.g01473.t01    rna-XM_021684676.2    rna-XM_004409796.1    rna-XM_042986207.1    rna-XM_040644699.1    rna-XM_027601582.2    mNeoVis.3.1.1.chr1.g01421.t01    rna-gnl|WGS_ZZZZ|egapxtmp_021812-R1    mNeoVis.evergladensis.2.1.chr1.g01440.t01    mNeoVis.vulgivaga.2.1.chr1.g01441.t01      1891    N0.HOG0001889   OG0000272  n12   rna-XM_034643700.1    rna-XM_038550136.1    rna-XM_022519201.1    rna-XM_023257660.2    rna-XM_036113777.1    rna-XM_039237464.1    rna-XM_032882712.1    rna-XM_046011065.1    rna-XM_032345814.1    rna-XM_059187371.1    rna-XM_059401699.1    rna-XM_004775383.3    mNeoVis.2.1.1.chr9_hap1.g01307.t01    rna-XM_021687830.1    rna-XM_004409663.1    rna-XM_042993610.1    rna-XM_040637759.1    rna-XM_027595708.1    mNeoVis.3.1.1.chr10.g01245.t01    rna-gnl|WGS_ZZZZ|egapxtmp_022363-R1    mNeoVis.evergladensis.2.1.chr10.g01294.t01    mNeoVis.vulgivaga.2.1.chr10.g01284.t01      1892    N0.HOG0001890   OG0000272  n30   rna-XM_034644266.1    rna-XM_038550132.1    rna-XM_022519210.1    rna-XM_023257659.2    rna-XM_036113768.1    rna-XM_039236659.1    rna-XM_032882709.1    rna-XM_046011063.1    rna-XM_032345807.1    rna-XM_059187366.1    rna-XM_059401709.1    rna-XM_045087577.1    mNeoVis.2.1.1.chr9_hap1.g01306.t01    rna-XM_021687828.1    rna-XM_004409664.1    rna-XM_042993609.1    rna-XM_040637758.1    rna-XM_027593561.1    mNeoVis.3.1.1.chr10.g01244.t01    rna-gnl|WGS_ZZZZ|egapxtmp_022364-R1    mNeoVis.evergladensis.2.1.chr10.g01293.t01    mNeoVis.vulgivaga.2.1.chr10.g01283.t01      1893    N0.HOG0001891   OG0000272  n49   rna-XM_002914294.3    rna-XM_038553858.1    rna-XM_022510720.1    rna-XM_023253909.2    rna-XM_036096919.1    rna-XM_039235308.1    rna-XM_032864335.1    rna-XM_046006423.1    rna-XM_032340504.1    rna-XM_059178485.1    rna-XM_059400130.1    rna-XM_004770716.2    mNeoVis.2.1.1.chr1_hap1.g01474.t01    rna-XM_021684606.2    rna-XM_004409794.1    rna-XM_042986214.1    rna-XM_040644698.1    rna-XM_027603370.2    mNeoVis.3.1.1.chr1.g01422.t01      mNeoVis.evergladensis.2.1.chr1.g01441.t01    mNeoVis.vulgivaga.2.1.chr1.g01442.t01      1894    N0.HOG0001892   OG0000273  n1   rna-XM_019794883.2    rna-XM_038552903.1    rna-XM_022500877.1    rna-XM_023242477.2    rna-XM_036100535.1    rna-XM_039234761.1    rna-XM_032860663.1    rna-XM_046023323.1    rna-XM_032306625.1    rna-XM_059141481.1    rna-XM_059411236.1    rna-XM_045081609.1    mNeoVis.2.1.1.chr13_hap1.g00448.t01    rna-XM_021691172.2    rna-XM_004392320.1    rna-XM_042963930.1    rna-XM_040631643.1    rna-XM_027614459.2    mNeoVis.3.1.1.chr13.g00386.t01    rna-gnl|WGS_ZZZZ|egapxtmp_029910-R1    mNeoVis.evergladensis.2.1.chr13.g00430.t01    mNeoVis.vulgivaga.2.1.chr13.g00433.t01      1895    N0.HOG0001893   OG0000273  n18   rna-XM_011225134.3    rna-XM_038562218.1    rna-XM_022526128.1    rna-XM_006935111.4    rna-XM_036118709.1    rna-XM_039246359.1    rna-XM_032843551.1    rna-XM_046013614.1    rna-XM_032302139.1    rna-XM_059135390.1    rna-XM_059374515.1    rna-XM_004776742.3    mNeoVis.2.1.1.chr2_hap1.g00783.t01    rna-XM_021688020.1    rna-XM_004404246.1    rna-XM_007088105.2    rna-XM_040626767.1    rna-XM_027613419.1    mNeoVis.3.1.1.chr5.g00729.t01      mNeoVis.evergladensis.2.1.chr5.g00739.t01    mNeoVis.vulgivaga.2.1.chr5.g00733.t01      1896    N0.HOG0001894   OG0000273  n34   rna-XM_011226470.3    rna-XM_038580428.1    rna-XM_022525033.1    rna-XM_023255360.2    rna-XM_036116632.1    rna-XM_039237143.1    rna-XM_032856474.1    rna-XM_046008786.1, rna-XM_046016548.1    rna-XM_032343731.1    rna-XM_059181553.1    rna-XM_059372829.1    rna-XM_004755423.3    mNeoVis.2.1.1.chr10_hap1.g00606.t01    rna-XM_021693989.1    rna-XM_004396565.1    rna-XM_042988844.1    rna-XM_040620634.1    rna-XM_027569006.2    mNeoVis.3.1.1.chr9.g00563.t01    rna-gnl|WGS_ZZZZ|egapxtmp_016316-R1    mNeoVis.evergladensis.2.1.chr8.g00588.t01    mNeoVis.vulgivaga.2.1.chr9.g00588.t01      1897    N0.HOG0001895   OG0000273  n51   rna-XM_011226473.3    rna-XM_038580430.1    rna-XM_022523020.1    rna-XM_045059302.1    rna-XM_036116629.1    rna-XM_039237135.1    rna-XM_032856469.1    rna-XM_046009140.1    rna-XM_032343730.1    rna-XM_059181559.1    rna-XM_059372834.1    rna-XM_004755422.3    mNeoVis.2.1.1.chr10_hap1.g00605.t01    rna-XM_044918151.1    rna-XM_004396568.1    rna-XM_007081854.3    rna-XM_008705973.2    rna-XM_027571650.2    mNeoVis.3.1.1.chr9.g00562.t01    rna-gnl|WGS_ZZZZ|egapxtmp_016320-R1    mNeoVis.evergladensis.2.1.chr8.g00587.t01    mNeoVis.vulgivaga.2.1.chr9.g00587.t01      1898    N0.HOG0001896   OG0000274  n1   rna-XM_034641039.1    rna-XM_038572493.1    rna-XM_022493909.1    rna-XM_023251177.2    rna-XM_036075392.1    rna-XM_039249449.1    rna-XM_032838556.1    rna-XM_045981545.1    rna-XM_032350483.1    rna-XM_059133391.1    rna-XM_059407131.1    rna-XM_045082142.1    mNeoVis.2.1.1.chr11_hap1.g00165.t01    rna-XM_021689204.1    rna-XM_004405259.1    rna-XM_007077788.3    rna-XM_040638406.1    rna-XM_035728880.1    mNeoVis.3.1.1.chr11.g00137.t01    rna-gnl|WGS_ZZZZ|egapxtmp_026800-R1    mNeoVis.evergladensis.2.1.chr11.g00162.t01    mNeoVis.vulgivaga.2.1.chr11.g00164.t01      1899    N0.HOG0001897   OG0000274  n21   rna-XM_034637793.1    rna-XM_038583361.1    rna-XM_022512431.1    rna-XM_023239871.2    rna-XM_036113658.1    rna-XM_039238010.1    rna-XM_032861829.1    rna-XM_045998840.1    rna-XM_032337848.1    rna-XM_059173568.1    rna-XM_059416815.1    rna-XM_045089247.1    mNeoVis.2.1.1.chr1_hap1.g00016.t01    rna-XM_044917016.1    rna-XM_004411626.1    rna-XM_042998090.1    rna-XM_040630580.1    rna-XM_027605540.2      rna-gnl|WGS_ZZZZ|egapxtmp_027658-R1    mNeoVis.evergladensis.2.1.chr1.g00015.t01    mNeoVis.vulgivaga.2.1.chr1.g00015.t01      1900    N0.HOG0001898   OG0000274  n42   rna-XM_034639210.1    rna-NM_001010952.1    rna-XM_022511839.1    rna-XM_003998143.4    rna-XM_036114987.1    rna-XM_039223524.1    rna-XM_032854142.1    rna-XM_045987447.1    rna-XM_032326350.1    rna-XM_059153280.1    rna-XM_059382709.1    rna-XM_004744478.3    mNeoVis.2.1.1.chr7_hap1.g00204.t01    rna-XM_044922288.1    rna-XM_004393355.1    rna-XM_042970753.1    rna-XM_040633471.1    rna-XM_027619219.1    mNeoVis.3.1.1.chr7.g00187.t01    rna-gnl|WGS_ZZZZ|egapxtmp_010445-R1    mNeoVis.evergladensis.2.1.chr7.g00203.t01    mNeoVis.vulgivaga.2.1.chr7.g00204.t01      1901    N0.HOG0001899   OG0000274  n60   rna-XM_034661217.1    rna-XM_038579902.1    rna-XM_022520521.1    rna-XM_003987308.6    rna-XM_036082572.1    rna-XM_039218321.1    rna-XM_032856928.1    rna-XM_046007954.1    rna-XM_032343170.1    rna-XM_059181072.1    rna-XM_059372445.1    rna-XM_004751016.3    mNeoVis.2.1.1.chr10_hap1.g00786.t01    rna-XM_021695773.1    rna-XM_004414381.1    rna-XM_042989272.1    rna-XM_008702292.2, rna-XM_008709392.2    rna-XM_027569102.2    mNeoVis.3.1.1.chr9.g00746.t01    rna-gnl|WGS_ZZZZ|egapxtmp_016592-R1    mNeoVis.evergladensis.2.1.chr8.g00767.t01    mNeoVis.vulgivaga.2.1.chr9.g00767.t01      1902    N0.HOG0001900   OG0000275  n1   rna-XM_011226890.2    rna-XM_038535459.1    rna-XM_022493512.1    rna-XM_003981371.5    rna-XM_036096118.1    rna-XM_039239496.1    rna-XM_032851614.1    rna-XM_046000255.1    rna-XM_032337249.1    rna-XM_059173345.1    rna-XM_059416329.1    rna-XM_045079434.1    mNeoVis.2.1.1.chr1_hap1.g00226.t01    rna-XM_021697229.1    rna-XM_004402841.1    rna-XM_007093428.2    rna-XM_040630616.1    rna-XM_027606407.1    mNeoVis.3.1.1.chr1.g00226.t01    rna-gnl|WGS_ZZZZ|egapxtmp_027436-R1    mNeoVis.evergladensis.2.1.chr1.g00223.t01    mNeoVis.vulgivaga.2.1.chr1.g00225.t01      1903    N0.HOG0001901   OG0000275  n19   rna-XM_002922632.3    rna-XM_038573623.1    rna-XM_022517247.1    rna-XM_023252660.2    rna-XM_036092327.1    rna-XM_039243238.1    rna-XM_032855758.1    rna-XM_045997855.1    rna-XM_032332237.1    rna-XM_059177450.1    rna-XM_059393318.1    rna-XM_013063301.2    mNeoVis.2.1.1.chr6_hap1.g00792.t01    rna-XM_021698908.1    rna-XM_004395248.1    rna-XM_007097140.3    rna-XM_008705064.2    rna-XM_027598923.2    mNeoVis.3.1.1.chr4.g00801.t01    rna-gnl|WGS_ZZZZ|egapxtmp_044456-R1    mNeoVis.evergladensis.2.1.chr4.g00795.t01    mNeoVis.vulgivaga.2.1.chr4.g00814.t01      1904    N0.HOG0001902   OG0000275  n39   rna-XM_002925346.3    rna-XM_038586857.1    rna-XM_022518436.1    rna-XM_004000283.5    rna-XM_036071492.1    rna-XM_039233918.1    rna-XM_032880155.1    rna-XM_045995593.1    rna-XM_032331174.1    rna-XM_059158385.1    rna-XM_059384685.1    rna-XM_004758872.2    mNeoVis.2.1.1.chr12_hap1.g00792.t01    rna-XM_021682428.1    rna-XM_004410422.1    rna-XM_007090072.3    rna-XM_008696382.1    rna-XM_027608190.1    mNeoVis.3.1.1.chr12.g00723.t01    rna-gnl|WGS_ZZZZ|egapxtmp_029641-R1    mNeoVis.evergladensis.2.1.chr12.g00719.t01    mNeoVis.vulgivaga.2.1.chr12.g00748.t01      1905    N0.HOG0001903   OG0000275  n53   rna-XM_002925922.3    rna-XM_038587971.1    rna-XM_022524890.1    rna-XM_004000744.5    rna-XM_036067716.1    rna-XM_039223741.1    rna-XM_032849750.1    rna-XM_045995785.1    rna-XM_032331657.1    rna-XM_059158403.1    rna-XM_059384643.1    rna-XM_004778678.2    mNeoVis.2.1.1.chr12_hap1.g00339.t01    rna-XM_021691337.1    rna-XM_004415775.1    rna-XM_007086431.3    rna-XM_040645299.1    rna-XM_027608439.1    mNeoVis.3.1.1.chr12.g00301.t01    rna-gnl|WGS_ZZZZ|egapxtmp_029010-R1    mNeoVis.evergladensis.2.1.chr12.g00297.t01    mNeoVis.vulgivaga.2.1.chr12.g00305.t01      1906    N0.HOG0001904   OG0000276  n1   rna-XM_002915455.4    rna-XM_038558862.1    rna-XM_022495192.1    rna-XM_019835418.3    rna-XM_036104211.1    rna-XM_039252483.1    rna-XM_032844331.1    rna-XM_046012589.1    rna-XM_032346577.1    rna-XM_059186686.1    rna-XM_059402224.1    rna-XM_004742996.3    mNeoVis.2.1.1.chr9_hap1.g01061.t01    rna-XM_021687577.1    rna-XM_004413185.1    rna-XM_007085673.3    rna-XM_008705690.2    rna-XM_027593701.2    mNeoVis.3.1.1.chr10.g01021.t01    rna-gnl|WGS_ZZZZ|egapxtmp_024800-R1    mNeoVis.evergladensis.2.1.chr10.g01051.t01    mNeoVis.vulgivaga.2.1.chr10.g01042.t01      1907    N0.HOG0001905   OG0000276  n21   rna-XM_034646222.1    rna-NM_001002966.1    rna-XM_022507303.1    rna-XM_003993778.4    rna-XM_036074735.1    rna-XM_039225072.1    rna-XM_032879091.1    rna-XM_046016420.1    rna-XM_032358573.1    rna-XM_059151173.1    rna-XM_059398707.1    rna-XM_004759661.3    mNeoVis.2.1.1.chr7_hap1.g02626.t01    rna-XM_044919484.1    rna-XM_004403804.1    rna-XM_042959222.1    rna-XM_040638794.1    rna-XM_027580256.2    mNeoVis.3.1.1.chr7.g02282.t01      mNeoVis.evergladensis.2.1.chr7.g02433.t01    mNeoVis.vulgivaga.2.1.chr7.g02456.t01      1908    N0.HOG0001906   OG0000276  n41   rna-XM_019809468.2    rna-NM_001197191.1    rna-XM_022507390.1    rna-XM_023239630.2    rna-XM_036116315.1    rna-XM_039236762.1    rna-XM_032881113.1    rna-XM_046016059.1    rna-XM_032358524.1    rna-XM_059137688.1    rna-XM_059391684.1    rna-XM_045065878.1    mNeoVis.2.1.1.chr7_hap1.g01859.t01    rna-XM_021685833.1    rna-XM_004412465.1    rna-XM_007098430.3    rna-XM_008708933.2    rna-XM_027580181.1, rna-XM_035724331.1    mNeoVis.3.1.1.chr7.g01728.t01    rna-gnl|WGS_ZZZZ|egapxtmp_011088-R1    mNeoVis.evergladensis.2.1.chr7.g01822.t01    mNeoVis.vulgivaga.2.1.chr7.g01839.t01      1909    N0.HOG0001907   OG0000276  n59   rna-XM_019795719.2    rna-NM_001197120.1    rna-XM_022495179.1    rna-XM_003989052.4    rna-XM_036075662.1    rna-XM_039221426.1    rna-XM_032854782.1    rna-XM_046012910.1    rna-XM_032347012.1    rna-XM_059186343.1    rna-XM_059404696.1    rna-XM_004752418.3    mNeoVis.2.1.1.chr9_hap1.g00934.t01    rna-XM_021678577.1    rna-XM_004406014.1    rna-XM_015537566.2    rna-XM_008684308.2    rna-XM_027594361.1    mNeoVis.3.1.1.chr10.g00887.t01    rna-gnl|WGS_ZZZZ|egapxtmp_024769-R1    mNeoVis.evergladensis.2.1.chr10.g00920.t01    mNeoVis.vulgivaga.2.1.chr10.g00908.t01      1910    N0.HOG0001908   OG0000277  n1   rna-XM_034667076.1    rna-XM_038585953.1    rna-XM_022508985.1    rna-XM_003999388.6    rna-XM_036079557.1    rna-XM_039227537.1    rna-XM_032878329.1    rna-XM_045982284.1    rna-XM_032319296.1    rna-XM_059146978.1    rna-XM_059413152.1    rna-XM_013058162.2    mNeoVis.2.1.1.chr14_hap1.g00360.t01    rna-XM_021686531.2    rna-XM_004392795.1    rna-XM_042976840.1    rna-XM_040636487.1    rna-XM_027611043.1    mNeoVis.3.1.1.chr14.g00359.t01    rna-gnl|WGS_ZZZZ|egapxtmp_031903-R1    mNeoVis.evergladensis.2.1.chr14.g00360.t01    mNeoVis.vulgivaga.2.1.chr14.g00361.t01      1911    N0.HOG0001909   OG0000277  n22   rna-XM_034644142.1    rna-NM_001197084.2    rna-XM_022495296.1    rna-XM_006933920.4    rna-XM_036097284.1    rna-XM_039240225.1    rna-XM_032874201.1    rna-XM_046013876.1    rna-XM_032346795.1    rna-XM_059186494.1    rna-XM_059402363.1    rna-XM_013054206.2    mNeoVis.2.1.1.chr9_hap1.g00988.t01    rna-XM_044915360.1    rna-XM_004414839.1    rna-XM_042992881.1    rna-XM_040637306.1    rna-XM_027591799.2    mNeoVis.3.1.1.chr10.g00947.t01    rna-gnl|WGS_ZZZZ|egapxtmp_023091-R2    mNeoVis.evergladensis.2.1.chr10.g00975.t01    mNeoVis.vulgivaga.2.1.chr10.g00967.t01      1912    N0.HOG0001910   OG0000277  n39   rna-XM_034658770.1    rna-XM_038565653.1    rna-XM_022498136.1    rna-XM_045051952.1    rna-XM_036079871.1    rna-XM_039231384.1    rna-XM_032841666.1    rna-XM_045990373.1    rna-XM_032327044.1    rna-XM_059161728.1    rna-XM_059390321.1    rna-XM_045063309.1    mNeoVis.2.1.1.chr5_hap1.g00876.t01    rna-XM_044916427.1    rna-XM_004392677.1    rna-XM_042979511.1    rna-XM_040628450.1    rna-XM_027582262.2    mNeoVis.3.1.1.chr2.g00789.t01    rna-gnl|WGS_ZZZZ|egapxtmp_038149-R1    mNeoVis.evergladensis.2.1.chr2.g00877.t01    mNeoVis.vulgivaga.2.1.chr2.g00879.t01      1913    N0.HOG0001911   OG0000277  n60   rna-XM_034649277.1    rna-XM_038588649.1    rna-XM_022491314.1    rna-XM_023248933.2    rna-XM_036094256.1    rna-XM_039219251.1    rna-XM_032846307.1    rna-XM_045995792.1    rna-XM_032329473.1    rna-XM_059156628.1    rna-XM_059385667.1    rna-XM_045068003.1    mNeoVis.2.1.1.chr12_hap1.g00066.t01    rna-XM_044911643.1    rna-XM_004409155.1    rna-XM_042974204.1    rna-XM_040639912.1    rna-XM_027609151.1    mNeoVis.3.1.1.chr12.g00057.t01    rna-gnl|WGS_ZZZZ|egapxtmp_029326-R7    mNeoVis.evergladensis.2.1.chr12.g00056.t01    mNeoVis.vulgivaga.2.1.chr12.g00058.t01      1914    N0.HOG0001912   OG0000278  n1   rna-XM_034667955.1    rna-XM_038532633.1    rna-XM_022511116.1    rna-XM_023254992.2    rna-XM_036092874.1    rna-XM_039235060.1    rna-XM_032835983.1    rna-XM_046008469.1    rna-XM_032342223.1    rna-XM_059180173.1    rna-XM_059371682.1    rna-XM_045070491.1    mNeoVis.2.1.1.chr10_hap1.g01143.t01    rna-XM_021680644.2    rna-XM_004405544.1    rna-XM_042988221.1    rna-XM_040642686.1    rna-XM_027570880.2    mNeoVis.3.1.1.chr14.g00524.t01, mNeoVis.3.1.1.chr9.g01078.t01    rna-gnl|WGS_ZZZZ|egapxtmp_017022-R1    mNeoVis.evergladensis.2.1.chr8.g01122.t01    mNeoVis.vulgivaga.2.1.chr9.g01103.t01      1915    N0.HOG0001913   OG0000278  n25   rna-XM_034660000.1    rna-XM_038567979.1    rna-XM_022525786.1    rna-XM_023242932.2    rna-XM_036100691.1    rna-XM_039248115.1    rna-XM_032846978.1    rna-XM_045990832.1    rna-XM_032305186.1    rna-XM_059159241.1    rna-XM_059388483.1    rna-XM_004778054.2    mNeoVis.2.1.1.chr5_hap1.g00032.t01    rna-XM_044913174.1    rna-XM_004395492.1    rna-XM_042977568.1    rna-XM_040629740.1, rna-XM_040629776.1    rna-XM_027587435.2    mNeoVis.3.1.1.chr2.g00017.t01      mNeoVis.evergladensis.2.1.chr2.g00032.t01    mNeoVis.vulgivaga.2.1.chr2.g00032.t01      1916    N0.HOG0001914   OG0000278  n46   rna-XM_034667858.1    rna-XM_038532884.1    rna-XM_022511072.1    rna-XM_023254894.2    rna-XM_036095657.1    rna-XM_039257596.1    rna-XM_032836973.1    rna-XM_046010429.1    rna-XM_032345513.1    rna-XM_059179954.1    rna-XM_059371534.1    rna-XM_004763700.3    mNeoVis.2.1.1.chr10_hap1.g01225.t01    rna-XM_021680675.1    rna-XM_004398274.1    rna-XM_042988009.1    rna-XM_040642828.1    rna-XM_027570172.2    mNeoVis.3.1.1.chr9.g01155.t01      mNeoVis.evergladensis.2.1.chr8.g01202.t01    mNeoVis.vulgivaga.2.1.chr9.g01187.t01      1917    N0.HOG0001915   OG0000278  n65   rna-XM_011231366.3    rna-XM_038527060.1    rna-XM_022512624.1    rna-XM_023243354.2    rna-XM_036103195.1    rna-XM_039242170.1    rna-XM_032866664.1    rna-XM_046022867.1    rna-XM_032308750.1    rna-XM_059143274.1    rna-XM_059411830.1    rna-XM_045085372.1    mNeoVis.2.1.1.chr13_hap1.g00726.t01    rna-XM_044920816.1    rna-XM_004396723.1    rna-XM_042963381.1    rna-XM_008691034.2    rna-XM_027616045.2    mNeoVis.3.1.1.chr13.g00642.t01    rna-gnl|WGS_ZZZZ|egapxtmp_031456-R1    mNeoVis.evergladensis.2.1.chr13.g00690.t01    mNeoVis.vulgivaga.2.1.chr13.g00688.t01      1918    N0.HOG0001916   OG0000279  n5     rna-XM_038569303.1      rna-XM_023239867.2    rna-XM_036110595.1    rna-XM_039245024.1    rna-XM_032879112.1      rna-XM_032356801.1    rna-XM_059143332.1    rna-XM_059404798.1    rna-XM_013044614.2    mNeoVis.2.1.1.chr7_hap1.g02239.t01        rna-XM_042957398.1      rna-XM_027578459.1    mNeoVis.3.1.1.chr7.g02059.t01    rna-gnl|WGS_ZZZZ|egapxtmp_013106-R5    mNeoVis.evergladensis.2.1.chr7.g02187.t01    mNeoVis.vulgivaga.2.1.chr7.g02206.t01      1919    N0.HOG0001917   OG0000279  n19     rna-XM_038589120.1              rna-XM_046014326.1              rna-XM_044919311.1    rna-XM_004417539.1        rna-XM_027579849.1              1920    N0.HOG0001918   OG0000279  n25   rna-XM_034670092.1    rna-XM_038539780.1    rna-XM_022526631.1    rna-XM_023247009.2    rna-XM_036118945.1    rna-XM_039247956.1    rna-XM_032842678.1    rna-XM_045992930.1    rna-XM_032327331.1    rna-XM_059154379.1    rna-XM_059414868.1    rna-XM_045072578.1    mNeoVis.2.1.1.chr15_hap1.g00023.t01    rna-XM_021700541.2    rna-XM_004397244.1    rna-XM_015542082.2    rna-XM_040635041.1    rna-XM_027612852.2    mNeoVis.3.1.1.chr15.g00021.t01      mNeoVis.evergladensis.2.1.chr15.g00024.t01    mNeoVis.vulgivaga.2.1.chr15.g00025.t01      1921    N0.HOG0001919   OG0000279  n29   rna-XM_034669991.1    rna-XM_038539100.1    rna-XM_022500478.1    rna-XM_023247027.2    rna-XM_036112569.1    rna-XM_039243469.1    rna-XM_032872433.1    rna-XM_045994304.1    rna-XM_032327670.1    rna-XM_059154693.1    rna-XM_059414695.1    rna-XM_045074956.1    mNeoVis.2.1.1.chr15_hap1.g00742.t01    rna-XM_044915138.1    rna-XM_004399104.1    rna-XM_042972149.1    rna-XM_040634944.1    rna-XM_027613537.2    mNeoVis.3.1.1.chr15.g00520.t01      mNeoVis.evergladensis.2.1.chr15.g00564.t01    mNeoVis.vulgivaga.2.1.chr15.g00576.t01      1922    N0.HOG0001920   OG0000279  n40   rna-XM_034672841.1    rna-XM_038574713.1    rna-XM_022517294.1    rna-XM_045041704.1    rna-XM_036106511.1    rna-XM_039249580.1    rna-XM_032846876.1    rna-XM_046025537.1    rna-XM_032310114.1    rna-XM_059139369.1    rna-XM_059408174.1    rna-XM_045087634.1    mNeoVis.2.1.1.chr3_hap1.g00670.t01    rna-XM_044921010.1    rna-XM_004415454.1    rna-XM_042961658.1    rna-XM_008688785.2    rna-XM_027575838.1    mNeoVis.3.1.1.chr3.g00662.t01    rna-gnl|WGS_ZZZZ|egapxtmp_041759-R1    mNeoVis.evergladensis.2.1.chr3.g00678.t01    mNeoVis.vulgivaga.2.1.chr3.g00682.t01      1923    N0.HOG0001921   OG0000280  n1   rna-XM_034641728.1    rna-XM_038547274.1    rna-XM_022509191.1    rna-XM_019817908.2    rna-XM_036095451.1    rna-XM_039256794.1    rna-XM_032871671.1    rna-XM_045986530.1    rna-XM_032321174.1    rna-XM_059149808.1    rna-XM_059379449.1    rna-XM_013045938.2    mNeoVis.2.1.1.chr8_hap1.g00380.t01    rna-XM_044921731.1    rna-XM_004410539.1    rna-XM_042967416.1    rna-XM_040640597.1    rna-XM_027567771.2    mNeoVis.3.1.1.chr8.g00346.t01    rna-gnl|WGS_ZZZZ|egapxtmp_019477-R3    mNeoVis.evergladensis.2.1.chr9.g00378.t01    mNeoVis.vulgivaga.2.1.chr8.g00385.t01      1924    N0.HOG0001922   OG0000280  n19   rna-XM_034672885.1    rna-XM_038559431.1    rna-XM_022510244.1    rna-XM_019825880.3    rna-XM_036106912.1    rna-XM_039234026.1    rna-XM_032868274.1    rna-XM_046021529.1    rna-XM_032304686.1    rna-XM_059171972.1    rna-XM_059396322.1    rna-XM_004741680.3    mNeoVis.2.1.1.chr4_hap1.g00167.t01    rna-XM_021692893.1    rna-XM_004416688.1    rna-XM_042973280.1    rna-XM_008697374.2    rna-XM_027574776.1    mNeoVis.3.1.1.chr6.g00166.t01    rna-gnl|WGS_ZZZZ|egapxtmp_004026-R5    mNeoVis.evergladensis.2.1.chr6.g00168.t01    mNeoVis.vulgivaga.2.1.chr6.g00167.t01      1925    N0.HOG0001923   OG0000280  n41   rna-XM_034644872.1    rna-XM_038563709.1    rna-XM_022507392.1    rna-XM_023240141.2    rna-XM_036069902.1    rna-XM_039235467.1    rna-XM_032879665.1    rna-XM_046014718.1    rna-XM_032357670.1    rna-XM_059149644.1    rna-XM_059399683.1    rna-XM_004759550.2    mNeoVis.2.1.1.chr7_hap1.g02502.t01, mNeoVis.2.1.1.chr7_hap1.g02554.t01, mNeoVis.2.1.1.chr7_hap1.g02582.t01    rna-XM_021685145.1    rna-XM_004393818.1    rna-XM_042959574.1    rna-XM_040639542.1    rna-XM_035723111.1, rna-XM_035724239.1    mNeoVis.3.1.1.chr1.g00861.t01            1926    N0.HOG0001924   OG0000280  n62   rna-XM_011226562.3    rna-XM_038574805.1    rna-XM_022517274.1    rna-XM_003994607.6    rna-XM_036080740.1    rna-XM_039234513.1    rna-XM_032876458.1    rna-XM_046024942.1    rna-XM_032309795.1    rna-XM_059139303.1    rna-XM_059408251.1    rna-XM_004753093.3    mNeoVis.2.1.1.chr3_hap1.g00650.t01    rna-XM_021698490.1    rna-XM_004395923.1    rna-XM_042961518.1    rna-XM_040620021.1    rna-XM_027576038.1    mNeoVis.3.1.1.chr3.g00641.t01    rna-gnl|WGS_ZZZZ|egapxtmp_041140-R1    mNeoVis.evergladensis.2.1.chr3.g00658.t01    mNeoVis.vulgivaga.2.1.chr3.g00662.t01      1927    N0.HOG0001925   OG0000281  n1   rna-XM_002917818.4    rna-XM_038528082.1    rna-XM_022523818.1    rna-XM_019818567.3    rna-XM_036104016.1    rna-XM_039245051.1    rna-XM_032839340.1    rna-XM_045988245.1    rna-XM_032324388.1    rna-XM_059151045.1    rna-XM_059382119.1    rna-XM_045077402.1    mNeoVis.2.1.1.chr7_hap1.g00850.t01    rna-XM_021681389.1    rna-XM_004409936.1    rna-XM_042970347.1    rna-XM_040632935.1    rna-XM_027617291.2    mNeoVis.3.1.1.chr7.g00792.t01    rna-gnl|WGS_ZZZZ|egapxtmp_012327-R1    mNeoVis.evergladensis.2.1.chr7.g00841.t01    mNeoVis.vulgivaga.2.1.chr7.g00845.t01      1928    N0.HOG0001926   OG0000281  n20   rna-XM_034667621.1    rna-XM_038568931.1    rna-XM_022522154.1    rna-XM_023239527.2    rna-XM_036089265.1    rna-XM_039222435.1    rna-XM_032853372.1    rna-XM_046017567.1    rna-XM_032359108.1    rna-XM_059134964.1    rna-XM_059409112.1    rna-XM_004780172.3    mNeoVis.2.1.1.chr7_hap1.g01776.t01    rna-XM_021685650.1    rna-XM_004394621.1    rna-XM_042959357.1    rna-XM_008683847.2    rna-XM_027577800.1    mNeoVis.3.1.1.chr7.g01647.t01    rna-gnl|WGS_ZZZZ|egapxtmp_012542-R1    mNeoVis.evergladensis.2.1.chr7.g01737.t01    mNeoVis.vulgivaga.2.1.chr7.g01753.t01      1929    N0.HOG0001927   OG0000281  n40   rna-XM_034654138.1    rna-XM_038561826.1    rna-XM_022497321.1    rna-XM_019837655.3    rna-XM_036088444.1    rna-XM_039249107.1    rna-XM_032881388.1    rna-XM_046026776.1    rna-XM_032302961.1    rna-XM_059137542.1    rna-XM_059374331.1    rna-XM_004769856.3    mNeoVis.2.1.1.chr2_hap1.g00919.t01    rna-XM_021690206.1    rna-XM_004401292.1    rna-XM_015534941.2    rna-XM_040626673.1    rna-XM_027581178.2    mNeoVis.3.1.1.chr5.g00851.t01    rna-gnl|WGS_ZZZZ|egapxtmp_006963-R1    mNeoVis.evergladensis.2.1.chr5.g00874.t01    mNeoVis.vulgivaga.2.1.chr5.g00863.t01      1930    N0.HOG0001928   OG0000281  n58   rna-XM_002916584.3    rna-XM_038577247.1    rna-XM_022524956.1    rna-XM_045061645.1    rna-XM_036091828.1    rna-XM_039222553.1    rna-XM_032855340.1    rna-XM_046012658.1    rna-XM_032347720.1    rna-XM_059185663.1    rna-XM_059404800.1    rna-XM_004771824.3    mNeoVis.2.1.1.chr9_hap1.g00680.t01    rna-XM_021690387.1    rna-XM_004415639.1    rna-XM_042992093.1    rna-XM_008704447.1    rna-XM_027595639.1    mNeoVis.3.1.1.chr10.g00645.t01    rna-gnl|WGS_ZZZZ|egapxtmp_024723-R1    mNeoVis.evergladensis.2.1.chr10.g00675.t01    mNeoVis.vulgivaga.2.1.chr10.g00658.t01      1931    N0.HOG0001929   OG0000282  n1   rna-XM_002928596.4    rna-XM_038533826.1    rna-XM_022499152.1    rna-XM_006937742.5    rna-XM_036074097.1    rna-XM_039247189.1    rna-XM_032883102.1    rna-XM_046027879.1    rna-XM_032312976.1    rna-XM_059170330.1    rna-XM_059397373.1    rna-XM_004774788.3    mNeoVis.2.1.1.chr2_hap1.g00623.t01    rna-XM_021696114.1    rna-XM_004413821.1    rna-XM_042961193.1    rna-XM_040623802.1    rna-XM_027595095.2    mNeoVis.3.1.1.chr5.g00621.t01    rna-gnl|WGS_ZZZZ|egapxtmp_006905-R1    mNeoVis.evergladensis.2.1.chr5.g00626.t01    mNeoVis.vulgivaga.2.1.chr5.g00620.t01      1932    N0.HOG0001930   OG0000282  n18   rna-XM_034645217.1    rna-XM_038563816.1    rna-XM_022506854.1    rna-XM_003993681.5    rna-XM_036121502.1    rna-XM_039244675.1    rna-XM_032879564.1    rna-XM_046016351.1    rna-XM_032359661.1    rna-XM_059148396.1    rna-XM_059382717.1    rna-XM_004759439.3    mNeoVis.2.1.1.chr7_hap1.g02483.t01    rna-XM_021685091.1    rna-XM_004393874.1    rna-XM_042959637.1    rna-XM_040638745.1    rna-XM_027579547.1    mNeoVis.3.1.1.chr7.g02230.t01    rna-gnl|WGS_ZZZZ|egapxtmp_012698-R1    mNeoVis.evergladensis.2.1.chr7.g02371.t01    mNeoVis.vulgivaga.2.1.chr7.g02391.t01      1933    N0.HOG0001931   OG0000282  n38   rna-XM_034642784.1    rna-XM_038545049.1    rna-XM_022505293.1    rna-XM_003987769.6    rna-XM_036117433.1    rna-XM_039240707.1    rna-XM_032875104.1    rna-XM_046008205.1    rna-XM_032344588.1    rna-XM_059182422.1    rna-XM_059373467.1    rna-XM_004739003.3    mNeoVis.2.1.1.chr10_hap1.g00285.t01    rna-XM_044918077.1    rna-XM_004397596.1    rna-XM_042989968.1    rna-XM_040641736.1    rna-XM_027571051.2    mNeoVis.3.1.1.chr9.g00258.t01    rna-gnl|WGS_ZZZZ|egapxtmp_016789-R1    mNeoVis.evergladensis.2.1.chr8.g00270.t01    mNeoVis.vulgivaga.2.1.chr9.g00269.t01      1934    N0.HOG0001932   OG0000282  n52   rna-XM_002925838.4    rna-XM_038528968.1    rna-XM_022525038.1    rna-XM_019819583.3    rna-XM_036118460.1    rna-XM_039217755.1    rna-XM_032841910.1    rna-XM_045986876.1    rna-XM_032325361.1    rna-XM_059153488.1    rna-XM_059381416.1    rna-XM_045076778.1    mNeoVis.2.1.1.chr7_hap1.g00514.t01    rna-XM_044911479.1    rna-XM_004396412.1    rna-XM_042970130.1    rna-XM_040633938.1    rna-XM_027617467.2    mNeoVis.3.1.1.chr7.g00499.t01    rna-gnl|WGS_ZZZZ|egapxtmp_011844-R2    mNeoVis.evergladensis.2.1.chr7.g00515.t01    mNeoVis.vulgivaga.2.1.chr7.g00518.t01      1935    N0.HOG0001933   OG0000283  n1   rna-XM_002912683.4    rna-XM_038535624.1    rna-XM_022513176.1    rna-XM_003992579.5    rna-XM_036086287.1    rna-XM_039240269.1    rna-XM_032868943.1    rna-XM_046017465.1    rna-XM_032358234.1    rna-XM_059185144.1    rna-XM_059413694.1    rna-XM_004763929.3    mNeoVis.2.1.1.chr7_hap1.g01396.t01    rna-XM_021696439.2    rna-XM_004412032.1    rna-XM_007082678.2    rna-XM_040631019.1    rna-XM_027579342.2    mNeoVis.3.1.1.chr7.g01308.t01    rna-gnl|WGS_ZZZZ|egapxtmp_011786-R2    mNeoVis.evergladensis.2.1.chr7.g01380.t01    mNeoVis.vulgivaga.2.1.chr7.g01391.t01      1936    N0.HOG0001934   OG0000283  n24   rna-XM_002928738.4    rna-XM_038586287.1    rna-XM_022490998.1    rna-XM_003999565.6    rna-XM_036101431.1    rna-XM_039246499.1    rna-XM_032849511.1    rna-XM_045982706.1    rna-XM_032317964.1    rna-XM_059145066.1    rna-XM_059413459.1    rna-XM_045086564.1    mNeoVis.2.1.1.chr14_hap1.g00494.t01    rna-XM_021681882.1    rna-XM_004407910.1    rna-XM_042976569.1    rna-XM_008709363.2, rna-XM_040634482.1    rna-XM_027610896.2    mNeoVis.3.1.1.chr14.g00488.t01      mNeoVis.evergladensis.2.1.chr14.g00492.t01    mNeoVis.vulgivaga.2.1.chr14.g00493.t01      1937    N0.HOG0001935   OG0000283  n45   rna-XM_034656486.1    rna-XM_038581727.1    rna-XM_022509618.1    rna-XM_019839482.3    rna-XM_036120240.1    rna-XM_039237285.1    rna-XM_032865488.1    rna-XM_046003171.1    rna-XM_032353286.1    rna-XM_059164530.1    rna-XM_059392202.1    rna-XM_004757448.3    mNeoVis.2.1.1.chr5_hap1.g01738.t01    rna-XM_021678039.1    rna-XM_004413667.1    rna-XM_007075058.3    rna-XM_008701934.2    rna-XM_027585231.2    mNeoVis.3.1.1.chr2.g01646.t01    rna-gnl|WGS_ZZZZ|egapxtmp_035691-R1    mNeoVis.evergladensis.2.1.chr2.g01718.t01    mNeoVis.vulgivaga.2.1.chr2.g01734.t01      1938    N0.HOG0001936   OG0000283  n66   rna-XM_034665811.1    rna-XM_038535561.1    rna-XM_022513333.1    rna-XM_019812925.3    rna-XM_036086285.1    rna-XM_039240258.1    rna-XM_032868942.1    rna-XM_046014200.1    rna-XM_032358163.1    rna-XM_059185134.1    rna-XM_059413717.1    rna-XM_004763928.3    mNeoVis.2.1.1.chr7_hap1.g01395.t01    rna-XM_021696488.2    rna-XM_004412033.1    rna-XM_042958707.1    rna-XM_040630642.1    rna-XM_027579343.1    mNeoVis.3.1.1.chr7.g01307.t01    rna-gnl|WGS_ZZZZ|egapxtmp_013968-R1    mNeoVis.evergladensis.2.1.chr7.g01379.t01    mNeoVis.vulgivaga.2.1.chr7.g01390.t01      1939    N0.HOG0001937   OG0000284  n1   rna-XM_034647302.1    rna-XM_038531449.1    rna-XM_022514261.1    rna-XM_019836447.3    rna-XM_036089390.1    rna-XM_039231207.1    rna-XM_032863842.1    rna-XM_046007013.1    rna-XM_032301428.1    rna-XM_059135616.1    rna-XM_059376760.1    rna-XM_004741096.3    mNeoVis.2.1.1.chr2_hap1.g01633.t01    rna-XM_021684141.1    rna-XM_004394700.1    rna-XM_015535710.2    rna-XM_040619061.1    rna-XM_027616353.2    mNeoVis.3.1.1.chr5.g01555.t01    rna-gnl|WGS_ZZZZ|egapxtmp_007125-R1    mNeoVis.evergladensis.2.1.chr5.g01573.t01    mNeoVis.vulgivaga.2.1.chr5.g01575.t01      1940    N0.HOG0001938   OG0000284  n19   rna-XM_019799101.2    rna-XM_038526526.1    rna-XM_022520947.1    rna-XM_045058346.1    rna-XM_036122902.1    rna-XM_039257159.1    rna-XM_032837181.1    rna-XM_046005543.1    rna-XM_032338508.1    rna-XM_059176553.1    rna-XM_059401412.1    rna-XM_045080056.1    mNeoVis.2.1.1.chr1_hap1.g02141.t01    rna-XM_044917246.1    rna-XM_004401546.1    rna-XM_042987516.1    rna-XM_008688469.2    rna-XM_027601223.2    mNeoVis.3.1.1.chr1.g02094.t01    rna-gnl|WGS_ZZZZ|egapxtmp_023325-R1    mNeoVis.evergladensis.2.1.chr1.g02110.t01    mNeoVis.vulgivaga.2.1.chr1.g02101.t01      1941    N0.HOG0001939   OG0000284  n41   rna-XM_034649862.1    rna-XM_038587035.1    rna-XM_022507919.1    rna-XM_004000334.6    rna-XM_036071087.1    rna-XM_039247388.1    rna-XM_032840814.1    rna-XM_045995202.1    rna-XM_032329459.1    rna-XM_059158694.1    rna-XM_059385860.1    rna-XM_045089390.1    mNeoVis.2.1.1.chr12_hap1.g00749.t01    rna-XM_021682226.2    rna-XM_004401524.1      rna-XM_040633566.1    rna-XM_027607855.2    mNeoVis.3.1.1.chr12.g00682.t01    rna-gnl|WGS_ZZZZ|egapxtmp_028399-R1    mNeoVis.evergladensis.2.1.chr12.g00678.t01    mNeoVis.vulgivaga.2.1.chr12.g00704.t01      1942    N0.HOG0001940   OG0000284  n51   rna-XM_034649643.1    rna-XM_038587788.1    rna-XM_022491688.1    rna-XM_023249024.2    rna-XM_036067731.1    rna-XM_039223354.1    rna-XM_032840613.1    rna-XM_045995522.1    rna-XM_032330485.1    rna-XM_059157144.1    rna-XM_059385145.1    rna-XM_004776042.3    mNeoVis.2.1.1.chr12_hap1.g00419.t01    rna-XM_021690974.1    rna-XM_004413568.1    rna-XM_007087880.3, rna-XM_042974970.1    rna-XM_040630311.1    rna-XM_035725759.1    mNeoVis.3.1.1.chr12.g00378.t01    rna-gnl|WGS_ZZZZ|egapxtmp_029559-R3    mNeoVis.evergladensis.2.1.chr12.g00374.t01    mNeoVis.vulgivaga.2.1.chr12.g00383.t01      1943    N0.HOG0001941   OG0000285  n1   rna-XM_034657475.1    rna-XM_038567850.1    rna-XM_022525528.1    rna-XM_023243647.2    rna-XM_036116115.1    rna-XM_039247616.1    rna-XM_032883524.1    rna-XM_045991604.1    rna-XM_032305868.1    rna-XM_059159435.1    rna-XM_059388615.1    rna-XM_045086810.1    mNeoVis.2.1.1.chr5_hap1.g00088.t01    rna-XM_044918567.1    rna-XM_004395361.1    rna-XM_042977729.1    rna-XM_040629622.1    rna-XM_027586553.2    mNeoVis.3.1.1.chr2.g00043.t01    rna-gnl|WGS_ZZZZ|egapxtmp_034399-R1    mNeoVis.evergladensis.2.1.chr2.g00085.t01    mNeoVis.vulgivaga.2.1.chr2.g00082.t01      1944    N0.HOG0001942   OG0000285  n15   rna-XM_019800165.2    rna-XM_038580894.1    rna-XM_022512001.1    rna-XM_023255088.2    rna-XM_036095193.1    rna-XM_039237867.1    rna-XM_032864985.1    rna-XM_046009774.1    rna-XM_032342534.1    rna-XM_059180461.1    rna-XM_059371893.1    rna-XM_004758357.3    mNeoVis.2.1.1.chr10_hap1.g01039.t01    rna-XM_021693813.1    rna-XM_004415340.1    rna-XM_042988452.1    rna-XM_040643025.1    rna-XM_027571872.1    mNeoVis.3.1.1.chr9.g00972.t01    rna-gnl|WGS_ZZZZ|egapxtmp_016707-R1    mNeoVis.evergladensis.2.1.chr8.g01012.t01    mNeoVis.vulgivaga.2.1.chr9.g00996.t01      1945    N0.HOG0001943   OG0000285  n25   rna-XM_034654282.1    rna-XM_038562256.1    rna-XM_022526076.1    rna-XM_006935142.5    rna-XM_036118751.1    rna-XM_039246328.1    rna-XM_032843572.1    rna-XM_046020624.1    rna-XM_032302701.1    rna-XM_059135200.1    rna-XM_059376161.1    rna-XM_013048018.2    mNeoVis.2.1.1.chr2_hap1.g00770.t01    rna-XM_021688007.1    rna-XM_004404273.1    rna-XM_042994125.1    rna-XM_040626617.1    rna-XM_027625067.2    mNeoVis.3.1.1.chr5.g00716.t01      mNeoVis.evergladensis.2.1.chr5.g00726.t01    mNeoVis.vulgivaga.2.1.chr5.g00720.t01      1946    N0.HOG0001944   OG0000285  n36   rna-XM_034642654.1    rna-XM_038543416.1    rna-XM_022495493.1    rna-XM_023241897.2    rna-XM_036069461.1    rna-XM_039238902.1    rna-XM_032859519.1    rna-XM_046025022.1    rna-XM_032310472.1    rna-XM_059139814.1    rna-XM_059409508.1    rna-XM_045065530.1    mNeoVis.2.1.1.chr3_hap1.g00149.t01    rna-XM_044920953.1    rna-XM_004417467.1    rna-XM_042961614.1    rna-XM_040632565.1    rna-XM_027576177.2    mNeoVis.3.1.1.chr3.g00151.t01    rna-gnl|WGS_ZZZZ|egapxtmp_039444-R1    mNeoVis.evergladensis.2.1.chr3.g00154.t01    mNeoVis.vulgivaga.2.1.chr3.g00152.t01      1947    N0.HOG0001945   OG0000286  n1   rna-XM_002912702.4    rna-NM_001003152.1    rna-XM_022498218.1    rna-XM_003982019.6    rna-XM_036095329.1    rna-XM_039243352.1    rna-XM_032867531.1    rna-XM_045991801.1    rna-XM_032314472.1    rna-XM_059160339.1    rna-XM_059389292.1    rna-XM_004761096.3    mNeoVis.2.1.1.chr5_hap1.g00452.t01    rna-XM_021683164.2    rna-XM_004404576.1    rna-XM_042978474.1    rna-XM_008706396.2    rna-XM_027586888.2    mNeoVis.3.1.1.chr2.g00375.t01    rna-gnl|WGS_ZZZZ|egapxtmp_034904-R1    mNeoVis.evergladensis.2.1.chr2.g00446.t01    mNeoVis.vulgivaga.2.1.chr2.g00448.t01      1948    N0.HOG0001946   OG0000286  n11   rna-XM_002914231.4    rna-XM_038580633.1    rna-XM_022512105.1      rna-XM_036081154.1    rna-XM_039219369.1    rna-XM_032864774.1    rna-XM_046008429.1    rna-XM_032342788.1    rna-XM_059180722.1    rna-XM_059372209.1    rna-XM_045067527.1    mNeoVis.2.1.1.chr10_hap1.g00973.t01    rna-XM_021693700.2    rna-XM_004411792.1    rna-XM_042988705.1    rna-XM_008708690.2    rna-XM_027570412.1    mNeoVis.3.1.1.chr9.g00904.t01    rna-gnl|WGS_ZZZZ|egapxtmp_016979-R1    mNeoVis.evergladensis.2.1.chr8.g00943.t01    mNeoVis.vulgivaga.2.1.chr9.g00929.t01      1949    N0.HOG0001947   OG0000286  n19   rna-XM_002922498.4    rna-XM_038543279.1    rna-XM_022490969.1    rna-XM_006943173.5, rna-XM_045049090.1    rna-XM_036086813.1    rna-XM_039218927.1, rna-XM_039218949.1    rna-XM_032845076.1    rna-XM_045982543.1    rna-XM_032317657.1    rna-XM_059145758.1    rna-XM_059413891.1    rna-XM_045086901.1    mNeoVis.2.1.1.chr14_hap1.g00678.t01    rna-XM_044916236.1    rna-XM_004403019.1    rna-XM_042976960.1, rna-XM_042976961.1    rna-XM_008696628.2    rna-XM_035722562.1    mNeoVis.3.1.1.chr14.g00648.t01    rna-gnl|WGS_ZZZZ|egapxtmp_029687-R1    mNeoVis.evergladensis.2.1.chr14.g00676.t01    mNeoVis.vulgivaga.2.1.chr14.g00676.t01      1950    N0.HOG0001948   OG0000286  n41   rna-XM_002922497.4    rna-NM_001313859.1, rna-XM_038536231.1    rna-XM_022491024.1          rna-XM_032845138.1    rna-XM_045982332.1    rna-XM_032317403.1    rna-XM_059145725.1    rna-XM_059413872.1    rna-XM_045086874.1    mNeoVis.2.1.1.chr14_hap1.g00679.t01      rna-XM_004403016.1, rna-XM_004404896.1      rna-XM_040635692.1    rna-XM_027569173.2, rna-XM_027611975.2    mNeoVis.3.1.1.chr14.g00649.t01      mNeoVis.evergladensis.2.1.chr14.g00677.t01    mNeoVis.vulgivaga.2.1.chr14.g00677.t01      1951    N0.HOG0001949   OG0000287  n1   rna-XM_011232902.3    rna-XM_038530547.1    rna-XM_022518073.1    rna-XM_023256432.2    rna-XM_036065730.1    rna-XM_039219188.1    rna-XM_032850701.1    rna-XM_046000202.1    rna-XM_032336255.1    rna-XM_059174977.1    rna-XM_059417860.1    rna-XM_004744676.3    mNeoVis.2.1.1.chr1_hap1.g00559.t01    rna-XM_021701782.1    rna-XM_004414630.1    rna-XM_007079954.2    rna-XM_040622263.1    rna-XM_027606201.2    mNeoVis.3.1.1.chr1.g00574.t01    rna-gnl|WGS_ZZZZ|egapxtmp_010102-R1    mNeoVis.evergladensis.2.1.chr1.g00565.t01    mNeoVis.vulgivaga.2.1.chr1.g00575.t01      1952    N0.HOG0001950   OG0000287  n17   rna-XM_034672142.1    rna-XM_038533406.1    rna-XM_022505707.1    rna-XM_023253323.2    rna-XM_036104675.1    rna-XM_039232030.1    rna-XM_032855291.1    rna-XM_045998427.1    rna-XM_032333797.1    rna-XM_059165580.1    rna-XM_059380607.1    rna-XM_045075501.1    mNeoVis.2.1.1.chr6_hap1.g00323.t01    rna-XM_021677665.2    rna-XM_004413735.1    rna-XM_042984969.1    rna-XM_040625448.1    rna-XM_027600380.2    mNeoVis.3.1.1.chr4.g00323.t01      mNeoVis.evergladensis.2.1.chr4.g00328.t01    mNeoVis.vulgivaga.2.1.chr4.g00326.t01      1953    N0.HOG0001951   OG0000287  n31   rna-XM_019794340.2    rna-XM_038573761.1    rna-XM_022513121.1    rna-XM_011281434.4    rna-XM_036106175.1    rna-XM_039250438.1    rna-XM_032851058.1, rna-XM_032855631.1    rna-XM_045994851.1    rna-XM_032332371.1    rna-XM_059176324.1    rna-XM_059372919.1    rna-XM_004775083.3    mNeoVis.2.1.1.chr6_hap1.g00743.t01    rna-XM_044912564.1    rna-XM_004412990.1    rna-XM_042983180.1    rna-XM_008702048.2    rna-XM_027598882.2    mNeoVis.3.1.1.chr4.g00752.t01    rna-gnl|WGS_ZZZZ|egapxtmp_000613-R1    mNeoVis.evergladensis.2.1.chr4.g00747.t01    mNeoVis.vulgivaga.2.1.chr4.g00766.t01      1954    N0.HOG0001952   OG0000287  n48   rna-XM_002915801.4    rna-XM_038579170.1    rna-XM_022499670.1    rna-XM_045040876.1    rna-XM_036070115.1    rna-XM_039252378.1    rna-XM_032882057.1    rna-XM_046027181.1    rna-XM_032312639.1    rna-XM_059172782.1    rna-XM_059398866.1    rna-XM_004751703.2    mNeoVis.2.1.1.chr2_hap1.g00032.t01    rna-XM_021703809.1    rna-XM_004402219.1    rna-XM_042960929.1    rna-XM_040624293.1    rna-XM_027596144.1      rna-gnl|WGS_ZZZZ|egapxtmp_028280-R1    mNeoVis.evergladensis.2.1.chr5.g00032.t01    mNeoVis.vulgivaga.2.1.chr5.g00030.t01      1955    N0.HOG0001953   OG0000288  n1   rna-XM_002931209.4    rna-XM_038545976.1    rna-XM_022505596.1    rna-XM_006932956.5    rna-XM_036107267.1    rna-XM_039251847.1    rna-XM_032837455.1    rna-XM_046010357.1    rna-XM_032345090.1    rna-XM_059182942.1    rna-XM_059371337.1    rna-XM_004828346.2    mNeoVis.2.1.1.chr10_hap1.g00138.t01    rna-XM_021679903.1    rna-XM_004416348.1    rna-XM_007095130.3    rna-XM_008700191.2    rna-XM_027568894.1    mNeoVis.3.1.1.chr9.g00113.t01    rna-gnl|WGS_ZZZZ|egapxtmp_017071-R1    mNeoVis.evergladensis.2.1.chr8.g00134.t01    mNeoVis.vulgivaga.2.1.chr9.g00133.t01      1956    N0.HOG0001954   OG0000288  n24   rna-XM_002930892.2    rna-XM_038573637.1    rna-XM_022511178.1    rna-XM_003984851.5    rna-XM_036092446.1      rna-XM_032855920.1    rna-XM_045992819.1    rna-XM_032332242.1    rna-XM_059155527.1    rna-XM_059409093.1    rna-XM_004766085.2    mNeoVis.2.1.1.chr6_hap1.g00789.t01    rna-XM_021698749.1    rna-XM_004395266.1    rna-XM_015544209.2    rna-XM_040625647.1    rna-XM_027599820.2    mNeoVis.3.1.1.chr4.g00797.t01    rna-gnl|WGS_ZZZZ|egapxtmp_001477-R1    mNeoVis.evergladensis.2.1.chr4.g00792.t01    mNeoVis.vulgivaga.2.1.chr4.g00811.t01      1957    N0.HOG0001955   OG0000288  n44   rna-XM_011229173.3    rna-XM_038573631.1    rna-XM_022511933.1    rna-XM_006930770.5    rna-XM_036092447.1    rna-XM_039243222.1    rna-XM_032851103.1, rna-XM_032851104.1    rna-XM_045997427.1    rna-XM_032332240.1    rna-XM_059155536.1    rna-XM_059408957.1    rna-XM_045083045.1    mNeoVis.2.1.1.chr6_hap1.g00791.t01    rna-XM_021698748.2      rna-XM_042983595.1    rna-XM_008705066.1    rna-XM_027599185.1    mNeoVis.3.1.1.chr4.g00799.t01    rna-gnl|WGS_ZZZZ|egapxtmp_001476-R1    mNeoVis.evergladensis.2.1.chr4.g00794.t01    mNeoVis.vulgivaga.2.1.chr4.g00813.t01      1958    N0.HOG0001956   OG0000288  n65   rna-XM_002931292.4    rna-XM_038545099.1    rna-XM_022505559.1    rna-XM_003987789.6    rna-XM_036107268.1    rna-XM_039251838.1    rna-XM_032837458.1    rna-XM_046010375.1    rna-XM_032345093.1    rna-XM_059182943.1    rna-XM_059372159.1    rna-XM_004754801.2    mNeoVis.2.1.1.chr10_hap1.g00137.t01    rna-XM_021679538.1    rna-XM_004416347.1    rna-XM_007095327.3    rna-XM_008700193.1    rna-XM_027568895.2    mNeoVis.3.1.1.chr9.g00112.t01    rna-gnl|WGS_ZZZZ|egapxtmp_017072-R2    mNeoVis.evergladensis.2.1.chr8.g00133.t01    mNeoVis.vulgivaga.2.1.chr9.g00132.t01      1959    N0.HOG0001957   OG0000289  n1   rna-XM_011223856.3    rna-XM_038546429.1    rna-XM_022523611.1    rna-XM_045045292.1    rna-XM_036069592.1    rna-XM_039226576.1    rna-XM_032870477.1    rna-XM_045984214.1    rna-XM_032321959.1    rna-XM_059149389.1    rna-XM_059378393.1    rna-XM_013054355.2    mNeoVis.2.1.1.chr8_hap1.g00094.t01    rna-XM_021680940.1    rna-XM_004416253.1    rna-XM_042967612.1    rna-XM_040640102.1    rna-XM_027625015.2    mNeoVis.3.1.1.chr8.g00069.t01    rna-gnl|WGS_ZZZZ|egapxtmp_020099-R1    mNeoVis.evergladensis.2.1.chr9.g00091.t01    mNeoVis.vulgivaga.2.1.chr8.g00094.t01      1960    N0.HOG0001958   OG0000289  n21   rna-XM_034638522.1    rna-XM_038528368.1    rna-XM_022524092.1    rna-XM_023245252.2    rna-XM_036098681.1, rna-XM_036099564.1    rna-XM_039248037.1    rna-XM_032838855.1    rna-XM_045989139.1    rna-XM_032323856.1    rna-XM_059152131.1    rna-XM_059381443.1    rna-XM_004767430.3    mNeoVis.2.1.1.chr7_hap1.g00942.t01    rna-XM_021681228.1, rna-XM_021702971.1    rna-XM_004415112.1    rna-XM_042969414.1    rna-XM_040634255.1    rna-XM_035725058.1    mNeoVis.3.1.1.chr7.g00882.t01      mNeoVis.evergladensis.2.1.chr7.g00935.t01    mNeoVis.vulgivaga.2.1.chr7.g00941.t01      1961    N0.HOG0001959   OG0000289  n42   rna-XM_019796678.2    rna-XM_038539483.1    rna-XM_022501004.1    rna-XM_023246645.2    rna-XM_036123605.1    rna-XM_039226044.1    rna-XM_032872728.1    rna-XM_045993023.1    rna-XM_032328743.1    rna-XM_059155279.1    rna-XM_059414391.1    rna-XM_045075125.1    mNeoVis.2.1.1.chr15_hap1.g00619.t01    rna-XM_021680235.2    rna-XM_004394163.1    rna-XM_042971930.1    rna-XM_040636543.1    rna-XM_027611391.2        mNeoVis.evergladensis.2.1.chr15.g00441.t01    mNeoVis.vulgivaga.2.1.chr15.g00449.t01      1962    N0.HOG0001960   OG0000289  n59   rna-XM_034643487.1    rna-XM_038550659.1    rna-XM_022518993.1    rna-XM_045062326.1    rna-XM_036108706.1    rna-XM_039246179.1    rna-XM_032836597.1    rna-XM_046013334.1    rna-XM_032346369.1    rna-XM_059186898.1    rna-XM_059402074.1    rna-XM_045083903.1    mNeoVis.2.1.1.chr9_hap1.g01145.t01    rna-XM_044915816.1    rna-XM_004410802.1    rna-XM_042993204.1    rna-XM_008705579.2    rna-XM_027595096.2    mNeoVis.3.1.1.chr10.g01100.t01      mNeoVis.evergladensis.2.1.chr10.g01133.t01    mNeoVis.vulgivaga.2.1.chr10.g01125.t01      1963    N0.HOG0001961   OG0000290  n1   rna-XM_002912520.4    rna-XM_038534838.1    rna-XM_022493395.1    rna-XM_006927959.5    rna-XM_036081034.1    rna-XM_039219358.1    rna-XM_032851732.1    rna-XM_046001082.1    rna-XM_032337110.1    rna-XM_059174228.1    rna-XM_059417304.1    rna-XM_004737672.3    mNeoVis.2.1.1.chr1_hap1.g00274.t01    rna-XM_021698020.1    rna-XM_004392045.1    rna-XM_015535483.2    rna-XM_008692409.1    rna-XM_027606857.2    mNeoVis.3.1.1.chr1.g00277.t01    rna-gnl|WGS_ZZZZ|egapxtmp_018720-R3    mNeoVis.evergladensis.2.1.chr1.g00272.t01    mNeoVis.vulgivaga.2.1.chr1.g00278.t01      1964    N0.HOG0001962   OG0000290  n14   rna-XM_019802399.2    rna-XM_038532575.1    rna-XM_022525997.1    rna-XM_019831994.3    rna-XM_036083577.1    rna-XM_039218224.1    rna-XM_032843104.1    rna-XM_046008807.1    rna-XM_032342300.1    rna-XM_059180243.1    rna-XM_059418490.1    rna-XM_004779161.3    mNeoVis.2.1.1.chr10_hap1.g01120.t01    rna-XM_021680472.2    rna-XM_004404722.1    rna-XM_015545123.2    rna-XM_008706320.2    rna-XM_027570991.1    mNeoVis.3.1.1.chr9.g01050.t01    rna-gnl|WGS_ZZZZ|egapxtmp_014401-R3    mNeoVis.evergladensis.2.1.chr8.g01098.t01    mNeoVis.vulgivaga.2.1.chr9.g01074.t01      1965    N0.HOG0001963   OG0000290  n23   rna-XM_002927241.4    rna-XM_038588582.1    rna-XM_022491349.1    rna-XM_004000982.6      rna-XM_039220703.1    rna-XM_032845269.1    rna-XM_045996524.1    rna-XM_032330179.1    rna-XM_059158036.1    rna-XM_059385929.1    rna-XM_004776653.2    mNeoVis.2.1.1.chr12_hap1.g00083.t01    rna-XM_021683536.1    rna-XM_004412688.1    rna-XM_042973981.1    rna-XM_008685641.2    rna-XM_027607297.1    mNeoVis.3.1.1.chr12.g00069.t01    rna-gnl|WGS_ZZZZ|egapxtmp_029246-R1    mNeoVis.evergladensis.2.1.chr12.g00070.t01    mNeoVis.vulgivaga.2.1.chr12.g00074.t01      1966    N0.HOG0001964   OG0000290  n40   rna-XM_002917015.4    rna-XM_038555989.1    rna-XM_022506033.1    rna-XM_011281893.4    rna-XM_036072263.1    rna-XM_039218183.1    rna-XM_032852458.1    rna-XM_045984539.1    rna-XM_032334214.1    rna-XM_059162281.1    rna-XM_059383399.1    rna-XM_013062707.2    mNeoVis.2.1.1.chr6_hap1.g00200.t01    rna-XM_044912723.1    rna-XM_004396310.1    rna-XM_015534764.2    rna-XM_040624855.1    rna-XM_027599558.2    mNeoVis.3.1.1.chr4.g00195.t01    rna-gnl|WGS_ZZZZ|egapxtmp_000562-R1    mNeoVis.evergladensis.2.1.chr4.g00205.t01    mNeoVis.vulgivaga.2.1.chr4.g00200.t01      1967    N0.HOG0001965   OG0000291  n0   rna-NM_001304901.1, rna-XM_002923230.4, rna-XM_011237248.3    rna-XM_038555090.1    rna-XM_022495324.1, rna-XM_022495784.1, rna-XM_022496093.1, rna-XM_022499743.1, rna-XM_022502142.1, rna-XM_022518284.1    rna-XM_006943321.5    rna-XM_036081441.1, rna-XM_036086509.1, rna-XM_036107507.1, rna-XM_036112110.1, rna-XM_036124399.1    rna-XM_039219337.1, rna-XM_039222941.1, rna-XM_039237093.1, rna-XM_039242233.1, rna-XM_039255475.1    rna-XM_032858631.1, rna-XM_032864264.1, rna-XM_032864554.1, rna-XM_032874139.1, rna-XM_032874140.1    rna-XM_045978490.1, rna-XM_045987329.1, rna-XM_045993842.1    rna-XM_032303738.1, rna-XM_032307020.1, rna-XM_032316517.1    rna-XM_059132669.1, rna-XM_059146937.1, rna-XM_059157690.1, rna-XM_059165955.1    rna-XM_059385514.1, rna-XM_059392919.1, rna-XM_059413067.1    rna-XM_004768917.3, rna-XM_013048181.2, rna-XM_013058122.2    mNeoVis.2.1.1.chr12_hap1.g00394.t01, mNeoVis.2.1.1.chr2_hap1.g00428.t01, mNeoVis.2.1.1.chr4_hap1.g00892.t01, mNeoVis.2.1.1.chr8_hap1.g00184.t01, mNeoVis.2.1.1.chr9_hap1.g00201.t01, mNeoVis.2.1.1.chr9_hap1.g00970.t01    rna-XM_021680430.1, rna-XM_021681145.1, rna-XM_021684438.1, rna-XM_021688261.1, rna-XM_021693493.1, rna-XM_021698969.1    rna-XM_004400679.1, rna-XM_004402367.1    rna-XM_007073294.3, rna-XM_015537968.2, rna-XM_015539917.2, rna-XM_015540678.2, rna-XM_015544884.1    rna-XM_008689196.2    rna-XM_027570029.1, rna-XM_027605267.2, rna-XM_035723957.1, rna-XM_035724262.1, rna-XM_035727045.1, rna-XM_035728167.1    mNeoVis.3.1.1.chr10.g00927.t01, mNeoVis.3.1.1.chr12.g00356.t01, mNeoVis.3.1.1.chr5.g00419.t01, mNeoVis.3.1.1.chr6.g00904.t01    rna-gnl|WGS_ZZZZ|egapxtmp_003500-R1, rna-gnl|WGS_ZZZZ|egapxtmp_007468-R1, rna-gnl|WGS_ZZZZ|egapxtmp_019340-R1, rna-gnl|WGS_ZZZZ|egapxtmp_023076-R1, rna-gnl|WGS_ZZZZ|egapxtmp_028804-R1    mNeoVis.evergladensis.2.1.chr10.g00197.t01, mNeoVis.evergladensis.2.1.chr10.g00956.t01, mNeoVis.evergladensis.2.1.chr12.g00351.t01, mNeoVis.evergladensis.2.1.chr5.g00427.t01, mNeoVis.evergladensis.2.1.chr6.g00898.t01    mNeoVis.vulgivaga.2.1.chr10.g00194.t01, mNeoVis.vulgivaga.2.1.chr10.g00949.t01, mNeoVis.vulgivaga.2.1.chr12.g00358.t01, mNeoVis.vulgivaga.2.1.chr5.g00427.t01, mNeoVis.vulgivaga.2.1.chr6.g00891.t01      1968    N0.HOG0001966   OG0000292  n1   rna-XM_034672537.1    rna-NM_001048109.1    rna-XM_022525277.1    rna-XM_019819629.3    rna-XM_036069016.1    rna-XM_039231929.1    rna-XM_032841956.1    rna-XM_045987903.1    rna-XM_032325854.1    rna-XM_059151001.1    rna-XM_059381408.1    rna-XM_045075995.1    mNeoVis.2.1.1.chr7_hap1.g00384.t01    rna-XM_021701085.1    rna-XM_004405352.1    rna-XM_042969189.1    rna-XM_040633697.1    rna-XM_027617828.2    mNeoVis.3.1.1.chr7.g00371.t01    rna-gnl|WGS_ZZZZ|egapxtmp_011555-R1    mNeoVis.evergladensis.2.1.chr7.g00389.t01    mNeoVis.vulgivaga.2.1.chr7.g00388.t01      1969    N0.HOG0001967   OG0000292  n21   rna-XM_002915004.4    rna-XM_038560867.1    rna-XM_022513654.1    rna-XM_003984477.6    rna-XM_036112716.1    rna-XM_039220199.1    rna-XM_032866909.1    rna-XM_045980051.1    rna-XM_032353210.1    rna-XM_059132555.1    rna-XM_059407555.1    rna-XM_004745876.3    mNeoVis.2.1.1.chr11_hap1.g01128.t01    rna-XM_021697689.1      rna-XM_007080721.3    rna-XM_008700385.2      mNeoVis.3.1.1.chr11.g01069.t01    rna-gnl|WGS_ZZZZ|egapxtmp_026237-R1    mNeoVis.evergladensis.2.1.chr11.g01097.t01    mNeoVis.vulgivaga.2.1.chr11.g01125.t01      1970    N0.HOG0001968   OG0000292  n38   rna-XM_034672638.1    rna-XM_038538518.1    rna-XM_022511661.1    rna-XM_023245051.2    rna-XM_036109179.1    rna-XM_039222401.1    rna-XM_032854119.1    rna-XM_045988965.1    rna-XM_032325210.1    rna-XM_059153058.1    rna-XM_059383561.1    rna-XM_045076211.1    mNeoVis.2.1.1.chr7_hap1.g00195.t01    rna-XM_021705737.2    rna-XM_004393345.1    rna-XM_042968700.1    rna-XM_040633498.1    rna-XM_027618590.2    mNeoVis.3.1.1.chr7.g00178.t01    rna-gnl|WGS_ZZZZ|egapxtmp_012164-R1    mNeoVis.evergladensis.2.1.chr7.g00194.t01    mNeoVis.vulgivaga.2.1.chr7.g00195.t01      1971    N0.HOG0001969   OG0000292  n57   rna-XM_034648605.1    rna-XM_038544154.1    rna-XM_022520441.1    rna-XM_045059672.1    rna-XM_036114798.1    rna-XM_039245191.1    rna-XM_032856412.1    rna-XM_046008305.1    rna-XM_032343795.1    rna-XM_059181623.1    rna-XM_059372875.1    rna-XM_013057655.2    mNeoVis.2.1.1.chr10_hap1.g00585.t01    rna-XM_044917851.1    rna-XM_004402112.1    rna-XM_007097951.3    rna-XM_040620709.1    rna-XM_027570433.2    mNeoVis.3.1.1.chr9.g00540.t01, mNeoVis.3.1.1.chr9.g00541.t01    rna-gnl|WGS_ZZZZ|egapxtmp_016689-R1    mNeoVis.evergladensis.2.1.chr8.g00569.t01    mNeoVis.vulgivaga.2.1.chr9.g00570.t01      1972    N0.HOG0001970   OG0000293  n1   rna-XM_034656678.1    rna-XM_038534668.1    rna-XM_022493099.1    rna-XM_045034946.1    rna-XM_036084894.1    rna-XM_039239756.1    rna-XM_032851903.1    rna-XM_046000640.1    rna-XM_032336909.1    rna-XM_059174407.1    rna-XM_059417444.1    rna-XM_013057661.2    mNeoVis.2.1.1.chr1_hap1.g00347.t01    rna-XM_021698742.2    rna-XM_004401423.1    rna-XM_042991687.1    rna-XM_008692330.2    rna-XM_027605506.1    mNeoVis.3.1.1.chr4.g00388.t01    rna-gnl|WGS_ZZZZ|egapxtmp_021560-R1    mNeoVis.evergladensis.2.1.chr1.g00345.t01    mNeoVis.vulgivaga.2.1.chr1.g00351.t01      1973    N0.HOG0001971   OG0000293  n23   rna-XM_034672218.1    rna-XM_038533270.1    rna-XM_022506076.1    rna-XM_023253489.2    rna-XM_036070184.1    rna-XM_039232123.1    rna-XM_032856411.1    rna-XM_045994710.1    rna-XM_032333572.1    rna-XM_059167116.1    rna-XM_059379394.1    rna-XM_045074658.1    mNeoVis.2.1.1.chr6_hap1.g00375.t01    rna-XM_044913042.1    rna-XM_004396217.1    rna-XM_042985147.1    rna-XM_040625496.1    rna-XM_027599597.2      rna-gnl|WGS_ZZZZ|egapxtmp_000740-R1    mNeoVis.evergladensis.2.1.chr4.g00382.t01    mNeoVis.vulgivaga.2.1.chr4.g00381.t01      1974    N0.HOG0001972   OG0000293  n44   rna-XM_034647457.1    rna-XM_038560058.1    rna-XM_022498707.1    rna-XM_011281520.4    rna-XM_036109882.1    rna-XM_039242529.1    rna-XM_032838086.1    rna-XM_045992145.1    rna-XM_032328750.1    rna-XM_059156341.1    rna-XM_059384025.1    rna-XM_013059267.2    mNeoVis.2.1.1.chr6_hap1.g00959.t01    rna-XM_044912831.1    rna-XM_004394430.1    rna-XM_042983435.1    rna-XM_040643242.1    rna-XM_027597974.2    mNeoVis.3.1.1.chr4.g00920.t01      mNeoVis.evergladensis.2.1.chr4.g00910.t01    mNeoVis.vulgivaga.2.1.chr4.g00931.t01      1975    N0.HOG0001973   OG0000293  n58   rna-XM_034663629.1    rna-NM_001003336.1    rna-XM_022499149.1, rna-XM_022499650.1    rna-XM_019813918.3    rna-XM_036110999.1    rna-XM_039250471.1    rna-XM_032882175.1    rna-XM_046026648.1    rna-XM_032314041.1    rna-XM_059172998.1    rna-XM_059398717.1    rna-XM_004751803.3    mNeoVis.2.1.1.chr2_hap1.g00089.t01    rna-XM_021703819.2    rna-XM_004404016.1    rna-XM_042960103.1    rna-XM_008685213.1    rna-XM_027593735.1    mNeoVis.3.1.1.chr5.g00083.t01    rna-gnl|WGS_ZZZZ|egapxtmp_005919-R1    mNeoVis.evergladensis.2.1.chr5.g00092.t01    mNeoVis.vulgivaga.2.1.chr5.g00088.t01      1976    N0.HOG0001974   OG0000294  n1   rna-XM_011236049.3    rna-NM_001197190.1    rna-XM_022499344.1    rna-XM_003994346.6    rna-XM_036086262.1    rna-XM_039243950.1    rna-XM_032881723.1    rna-XM_046027883.1    rna-XM_032311845.1    rna-XM_059169115.1    rna-XM_059398400.1    rna-XM_004749420.3    mNeoVis.2.1.1.chr2_hap1.g00218.t01    rna-XM_021699464.1    rna-XM_004401953.1    rna-XM_007080196.2    rna-XM_040624653.1    rna-XM_027593673.2    mNeoVis.3.1.1.chr5.g00209.t01    rna-gnl|WGS_ZZZZ|egapxtmp_006133-R1    mNeoVis.evergladensis.2.1.chr5.g00222.t01    mNeoVis.vulgivaga.2.1.chr5.g00219.t01      1977    N0.HOG0001975   OG0000294  n18   rna-XM_034672026.1    rna-XM_038533576.1    rna-XM_022506024.1    rna-XM_019829694.2        rna-XM_032854145.1    rna-XM_045998522.1    rna-XM_032333977.1      rna-XM_059381728.1    rna-XM_004762691.3    mNeoVis.2.1.1.chr6_hap1.g00274.t01    rna-XM_021679401.1          rna-XM_027600587.2      rna-gnl|WGS_ZZZZ|egapxtmp_044161-R5    mNeoVis.evergladensis.2.1.chr4.g00281.t01    mNeoVis.vulgivaga.2.1.chr4.g00280.t01      1978    N0.HOG0001976   OG0000294  n26   rna-XM_034656744.1    rna-XM_038534779.1    rna-XM_022493254.1    rna-XM_045035708.1    rna-XM_036079455.1, rna-XM_036120817.1    rna-XM_039216594.1, rna-XM_039235721.1    rna-XM_032851782.1    rna-XM_046000394.1    rna-XM_032337033.1    rna-XM_059164741.1, rna-XM_059174279.1    rna-XM_059417342.1    rna-XM_004737617.3    mNeoVis.2.1.1.chr1_hap1.g00297.t01    rna-XM_021698208.1    rna-XM_004392014.1, rna-XM_004415163.1    rna-XM_042982751.1, rna-XM_042992492.1    rna-XM_008692387.2, rna-XM_040625201.1    rna-XM_027606966.2    mNeoVis.3.1.1.chr1.g00302.t01, mNeoVis.3.1.1.chr4.g00271.t01    rna-gnl|WGS_ZZZZ|egapxtmp_020625-R1    mNeoVis.evergladensis.2.1.chr1.g00297.t01    mNeoVis.vulgivaga.2.1.chr1.g00301.t01      1979    N0.HOG0001977   OG0000294  n45   rna-XM_034659581.1    rna-XM_038561397.1    rna-XM_022520465.1    rna-XM_003984274.6    rna-XM_036074874.1    rna-XM_039231741.1    rna-XM_032843865.1    rna-XM_045980024.1    rna-XM_032352809.1    rna-XM_059188432.1    rna-XM_059405554.1    rna-XM_004770942.3    mNeoVis.2.1.1.chr11_hap1.g00938.t01    rna-XM_021697498.2    rna-XM_004399843.1    rna-XM_015537627.2    rna-XM_040623166.1    rna-XM_027623931.2    mNeoVis.3.1.1.chr11.g00892.t01      mNeoVis.evergladensis.2.1.chr11.g00916.t01    mNeoVis.vulgivaga.2.1.chr11.g00940.t01      1980    N0.HOG0001978   OG0000295  n1   rna-XM_002924018.4    rna-XM_038531832.1    rna-XM_022514561.1    rna-XM_003989525.3    rna-XM_036066977.1    rna-XM_039229057.1    rna-XM_032863560.1    rna-XM_045982105.1    rna-XM_032302662.1    rna-XM_059137267.1    rna-XM_059376395.1    rna-XM_045063840.1    mNeoVis.2.1.1.chr2_hap1.g01759.t01    rna-XM_021683496.1    rna-XM_004397079.1    rna-XM_042996200.1    rna-XM_008694677.2    rna-XM_027616018.2    mNeoVis.3.1.1.chr5.g01685.t01    rna-gnl|WGS_ZZZZ|egapxtmp_007164-R1    mNeoVis.evergladensis.2.1.chr5.g01699.t01    mNeoVis.vulgivaga.2.1.chr5.g01700.t01      1981    N0.HOG0001979   OG0000295  n23   rna-XM_002924019.4    rna-XM_038531831.1    rna-XM_022514662.1    rna-XM_011284582.4    rna-XM_036066975.1    rna-XM_039228932.1    rna-XM_032863561.1    rna-XM_045982093.1    rna-XM_032301911.1    rna-XM_059137268.1    rna-XM_059374862.1    rna-XM_004741360.2    mNeoVis.2.1.1.chr2_hap1.g01758.t01    rna-XM_021683497.1    rna-XM_004397078.1    rna-XM_042996654.1    rna-XM_008694676.2    rna-XM_035727368.1    mNeoVis.3.1.1.chr5.g01684.t01    rna-gnl|WGS_ZZZZ|egapxtmp_005870-R1    mNeoVis.evergladensis.2.1.chr5.g01698.t01    mNeoVis.vulgivaga.2.1.chr5.g01699.t01      1982    N0.HOG0001980   OG0000295  n46   rna-XM_034671264.1    rna-XM_038531830.1    rna-XM_022514674.1    rna-XM_003989527.5    rna-XM_036066973.1    rna-XM_039229074.1    rna-XM_032863562.1    rna-XM_046015815.1    rna-XM_032302852.1    rna-XM_059137240.1    rna-XM_059376396.1    rna-XM_004741359.3    mNeoVis.2.1.1.chr2_hap1.g01757.t01    rna-XM_021683498.1    rna-XM_004397077.1    rna-XM_042996520.1    rna-XM_008694674.2    rna-XM_027583505.2    mNeoVis.3.1.1.chr5.g01683.t01      mNeoVis.evergladensis.2.1.chr5.g01697.t01    mNeoVis.vulgivaga.2.1.chr5.g01698.t01      1983    N0.HOG0001981   OG0000295  n66   rna-XM_019804628.2    rna-XM_038531833.1    rna-XM_022514634.1    rna-XM_003989524.6    rna-XM_036066980.1    rna-XM_039228969.1    rna-XM_032863559.1    rna-XM_046026061.1    rna-XM_032302050.1    rna-XM_059136646.1    rna-XM_059376393.1    rna-XM_004741362.3    mNeoVis.2.1.1.chr2_hap1.g01760.t01    rna-XM_021684383.2    rna-XM_004397080.1    rna-XM_042994947.1    rna-XM_008694904.2    rna-XM_027625645.2    mNeoVis.3.1.1.chr5.g01686.t01      mNeoVis.evergladensis.2.1.chr5.g01700.t01    mNeoVis.vulgivaga.2.1.chr5.g01701.t01      1984    N0.HOG0001982   OG0000296  n1   rna-XM_034672245.1    rna-NM_001003286.1    rna-XM_022506334.1    rna-XM_045056696.1    rna-XM_036086535.1    rna-XM_039220406.1    rna-XM_032853384.1    rna-XM_045997696.1    rna-XM_032333546.1    rna-XM_059154254.1    rna-XM_059371453.1    rna-XM_045074519.1    mNeoVis.2.1.1.chr6_hap1.g00385.t01    rna-XM_021677654.1    rna-XM_004396230.1    rna-XM_007093186.3    rna-XM_008705635.2    rna-XM_027599692.1    mNeoVis.3.1.1.chr4.g00391.t01    rna-gnl|WGS_ZZZZ|egapxtmp_001620-R1    mNeoVis.evergladensis.2.1.chr4.g00392.t01    mNeoVis.vulgivaga.2.1.chr4.g00392.t01      1985    N0.HOG0001983   OG0000296  n20   rna-XM_034655874.1    rna-XM_038551821.1    rna-XM_022521242.1    rna-XM_003980660.6    rna-XM_036066087.1    rna-XM_039254623.1    rna-XM_032842281.1    rna-XM_046001038.1    rna-XM_032335375.1    rna-XM_059175848.1    rna-XM_059416119.1    rna-XM_004759051.3    mNeoVis.2.1.1.chr1_hap1.g00864.t01    rna-XM_021701730.1    rna-XM_004414332.1    rna-XM_007083128.3    rna-XM_040622571.1    rna-XM_035727633.1    mNeoVis.3.1.1.chr1.g00886.t01      mNeoVis.evergladensis.2.1.chr1.g00867.t01    mNeoVis.vulgivaga.2.1.chr1.g00880.t01      1986    N0.HOG0001984   OG0000296  n35   rna-XM_002924063.4    rna-XM_038551822.1    rna-XM_022521252.1    rna-XM_003980699.6    rna-XM_036066097.1    rna-XM_039254625.1    rna-XM_032842282.1    rna-XM_046001030.1    rna-XM_032335376.1    rna-XM_059175846.1    rna-XM_059416117.1    rna-XM_004759052.3    mNeoVis.2.1.1.chr1_hap1.g00863.t01    rna-XM_021701956.1    rna-XM_004414333.1    rna-XM_007083129.3    rna-XM_040622572.1    rna-XM_027605397.2    mNeoVis.3.1.1.chr1.g00887.t01    rna-gnl|WGS_ZZZZ|egapxtmp_015391-R1    mNeoVis.evergladensis.2.1.chr1.g00866.t01    mNeoVis.vulgivaga.2.1.chr1.g00881.t01      1987    N0.HOG0001985   OG0000296  n55   rna-XM_019804645.2, rna-XM_034655724.1    rna-XM_038559556.1    rna-XM_022518235.1    rna-XM_019825201.3    rna-XM_036085235.1    rna-XM_039243009.1    rna-XM_032875052.1    rna-XM_046020950.1    rna-XM_032305512.1    rna-XM_059170698.1    rna-XM_059397128.1    rna-XM_013059761.2    mNeoVis.2.1.1.chr4_hap1.g00617.t01    rna-XM_021703695.1    rna-XM_004414051.1    rna-XM_042962858.1    rna-XM_040644468.1    rna-XM_027573794.2    mNeoVis.3.1.1.chr6.g00632.t01      mNeoVis.evergladensis.2.1.chr6.g00626.t01    mNeoVis.vulgivaga.2.1.chr6.g00623.t01      1988    N0.HOG0001986   OG0000297  n0   rna-XM_019799530.2    rna-XM_038525511.1    rna-XM_022492377.1, rna-XM_022492378.1, rna-XM_022492672.1, rna-XM_022522269.1    rna-XM_023243265.2    rna-XM_036092645.1    rna-XM_039221107.1, rna-XM_039221108.1    rna-XM_032849971.1, rna-XM_032850297.1, rna-XM_032850540.1, rna-XM_032851071.1    rna-XM_046020071.1, rna-XM_046021907.1, rna-XM_046027741.1    rna-XM_032305459.1, rna-XM_032305982.1, rna-XM_032307798.1, rna-XM_032308953.1, rna-XM_032312750.1, rna-XM_032313472.1, rna-XM_032313761.1, rna-XM_032325231.1, rna-XM_032325236.1    rna-XM_059142476.1, rna-XM_059170297.1, rna-XM_059170370.1, rna-XM_059170500.1, rna-XM_059170505.1, rna-XM_059172506.1, rna-XM_059172507.1, rna-XM_059172517.1, rna-XM_059172646.1, rna-XM_059172647.1    rna-XM_059397587.1    rna-XM_004739714.2, rna-XM_004766590.3, rna-XM_045067372.1, rna-XM_045083038.1, rna-XM_045087211.1, rna-XM_045087298.1, rna-XM_045087300.1, rna-XM_045089635.1    mNeoVis.2.1.1.chr1_hap1.g00948.t01, mNeoVis.2.1.1.chr2_hap1.g00595.t01, mNeoVis.2.1.1.chr2_hap1.g00631.t01, mNeoVis.2.1.1.chr2_hap1.g00636.t01, mNeoVis.2.1.1.chr2_hap1.g00637.t01, mNeoVis.2.1.1.chr2_hap1.g00638.t01, mNeoVis.2.1.1.chr2_hap1.g00639.t01, mNeoVis.2.1.1.chr2_hap1.g00640.t01, mNeoVis.2.1.1.chr2_hap1.g00641.t01, mNeoVis.2.1.1.chr2_hap1.g00642.t01, mNeoVis.2.1.1.chr2_hap1.g00647.t01, mNeoVis.2.1.1.chr2_hap1.g00648.t01, mNeoVis.2.1.1.chr2_hap1.g00649.t01, mNeoVis.2.1.1.chr2_hap1.g00652.t01, mNeoVis.2.1.1.chr2_hap1.g00711.t01, mNeoVis.2.1.1.chr2_hap1.g00712.t01, mNeoVis.2.1.1.chr2_hap1.g00713.t01, mNeoVis.2.1.1.chr2_hap1.g00716.t01, mNeoVis.2.1.1.chr2_hap1.g00717.t01, mNeoVis.2.1.1.chr2_hap1.g00718.t01, mNeoVis.2.1.1.chr4_hap1.g00600.t01    rna-XM_021694373.1    rna-XM_004417314.1    rna-XM_042964998.1    rna-XM_040620136.1    rna-XM_035723485.1    mNeoVis.3.1.1.chr1.g00960.t01, mNeoVis.3.1.1.chr10.g00302.t01, mNeoVis.3.1.1.chr6.g00616.t01, mNeoVis.3.1.1.chr6.g00617.t01    rna-gnl|WGS_ZZZZ|egapxtmp_005052-R1, rna-gnl|WGS_ZZZZ|egapxtmp_043068-R1    mNeoVis.evergladensis.2.1.chr1.g00943.t01, mNeoVis.evergladensis.2.1.chr5.g00632.t01, mNeoVis.evergladensis.2.1.chr6.g00605.t01, mNeoVis.evergladensis.2.1.chr6.g00608.t01, mNeoVis.evergladensis.2.1.chr6.g00609.t01    mNeoVis.vulgivaga.2.1.chr1.g00954.t01, mNeoVis.vulgivaga.2.1.chr5.g00626.t01, mNeoVis.vulgivaga.2.1.chr6.g00604.t01, mNeoVis.vulgivaga.2.1.chr6.g00606.t01, mNeoVis.vulgivaga.2.1.chr6.g00607.t01      1989    N0.HOG0001987   OG0000298  n0         rna-XM_019824831.2, rna-XM_023249249.2, rna-XM_045033233.1, rna-XM_045037327.1, rna-XM_045040652.1, rna-XM_045046257.1, rna-XM_045050953.1, rna-XM_045052616.1, rna-XM_045052942.1, rna-XM_045055863.1, rna-XM_045056278.1, rna-XM_045056581.1, rna-XM_045057087.1, rna-XM_045057201.1, rna-XM_045061635.1    rna-XM_036065830.1, rna-XM_036067353.1, rna-XM_036069242.1, rna-XM_036069442.1, rna-XM_036073070.1, rna-XM_036073933.1, rna-XM_036075522.1, rna-XM_036079084.1, rna-XM_036081026.1, rna-XM_036081923.1, rna-XM_036083128.1, rna-XM_036084149.1, rna-XM_036085899.1, rna-XM_036092616.1, rna-XM_036095809.1, rna-XM_036101044.1, rna-XM_036106166.1, rna-XM_036108334.1, rna-XM_036110946.1, rna-XM_036111125.1, rna-XM_036116372.1, rna-XM_036116594.1, rna-XM_036120527.1    rna-XM_039220842.1, rna-XM_039221658.1, rna-XM_039225483.1, rna-XM_039240295.1      rna-XM_045992721.1, rna-XM_045994681.1, rna-XM_045998249.1, rna-XM_046004230.1, rna-XM_046010150.1, rna-XM_046015533.1, rna-XM_046027321.1    rna-XM_032337445.1, rna-XM_032337917.1, rna-XM_032343165.1    rna-XM_059134326.1, rna-XM_059149360.1, rna-XM_059156808.1, rna-XM_059160260.1, rna-XM_059167468.1, rna-XM_059176384.1, rna-XM_059178786.1        mNeoVis.2.1.1.chr12_hap1.g00519.t01    rna-XM_021683653.2, rna-XM_044912309.1, rna-XM_044912409.1, rna-XM_044912533.1      rna-XM_042970072.1, rna-XM_042975135.1, rna-XM_042984244.1, rna-XM_042988903.1, rna-XM_042997795.1    rna-XM_040645070.1    rna-XM_027588038.2, rna-XM_027595168.2, rna-XM_027599582.2, rna-XM_027602277.2, rna-XM_027607112.1, rna-XM_027607549.2, rna-XM_027609408.1, rna-XM_027614665.1, rna-XM_035723079.1, rna-XM_035724387.1, rna-XM_035726410.1, rna-XM_035726474.1, rna-XM_035728304.1, rna-XM_035728472.1    mNeoVis.3.1.1.chr12.g00473.t01    rna-gnl|WGS_ZZZZ|egapxtmp_028181-R1    mNeoVis.evergladensis.2.1.chr12.g00466.t01        1990    N0.HOG0001988   OG0000299  n2   rna-XM_034660293.1    rna-XM_038532071.1        rna-XM_036113954.1, rna-XM_036121167.1    rna-XM_039234273.1    rna-XM_032852978.1      rna-XM_032316013.1            rna-XM_044920327.1    rna-XM_004405822.1      rna-XM_040625600.1, rna-XM_040645083.1    rna-XM_035721929.1, rna-XM_035722494.1              1991    N0.HOG0001989   OG0000299  n14   rna-XM_034661696.1, rna-XM_034665581.1    rna-XM_038565107.1      rna-XM_045057070.1, rna-XM_045062154.1    rna-XM_036096091.1, rna-XM_036097190.1    rna-XM_039217389.1, rna-XM_039229939.1      rna-XM_045984079.1    rna-XM_032320098.1    rna-XM_059147416.1    rna-XM_059380130.1    rna-XM_045070262.1    mNeoVis.2.1.1.chr8_hap1.g00799.t01      rna-XM_004399698.1, rna-XM_004401371.1    rna-XM_042960265.1, rna-XM_042985395.1, rna-XM_042992890.1    rna-XM_040639803.1    rna-XM_035724403.1    mNeoVis.3.1.1.chr6.g00974.t01, mNeoVis.3.1.1.chr8.g00749.t01    rna-gnl|WGS_ZZZZ|egapxtmp_017501-R1    mNeoVis.evergladensis.2.1.chr9.g00786.t01    mNeoVis.vulgivaga.2.1.chr8.g00792.t01      1992    N0.HOG0001990   OG0000299  n39   rna-XM_034651368.1, rna-XM_034665042.1    rna-XM_038559540.1    rna-XM_022518206.1    rna-NM_001009370.1, rna-XM_011285920.4    rna-XM_036071064.1, rna-XM_036085218.1, rna-XM_036106285.1    rna-XM_039232191.1, rna-XM_039243005.1, rna-XM_039251286.1, rna-XM_039251939.1    rna-XM_032868853.1, rna-XM_032875042.1, rna-XM_032876441.1    rna-XM_046020738.1    rna-XM_032304477.1    rna-XM_059134679.1, rna-XM_059140241.1, rna-XM_059170685.1    rna-XM_059397133.1, rna-XM_059407819.1    rna-XM_045077771.1, rna-XM_045089593.1    mNeoVis.2.1.1.chr3_hap1.g00985.t01, mNeoVis.2.1.1.chr4_hap1.g00620.t01    rna-XM_044914595.1, rna-XM_044919908.1    rna-XM_004392776.1, rna-XM_004406612.1, rna-XM_004414057.1    rna-XM_042977371.1    rna-XM_008688478.2, rna-XM_040638738.1, rna-XM_040644472.1    rna-XM_035723339.1, rna-XM_035728166.1, rna-XM_035728380.1    mNeoVis.3.1.1.chr3.g00975.t01, mNeoVis.3.1.1.chr6.g00634.t01    rna-gnl|WGS_ZZZZ|egapxtmp_041227-R1    mNeoVis.evergladensis.2.1.chr3.g00982.t01, mNeoVis.evergladensis.2.1.chr6.g00629.t01    mNeoVis.vulgivaga.2.1.chr3.g01000.t01, mNeoVis.vulgivaga.2.1.chr6.g00626.t01      1993    N0.HOG0001991   OG0000300  n1   rna-XM_002917884.4    rna-XM_038528421.1    rna-XM_022523896.1    rna-XM_003997613.6    rna-XM_036114094.1    rna-XM_039249939.1    rna-XM_032839109.1    rna-XM_045989210.1    rna-XM_032324925.1    rna-XM_059151395.1    rna-XM_059381488.1    rna-XM_004767464.3    mNeoVis.2.1.1.chr7_hap1.g00964.t01    rna-XM_021681206.2    rna-XM_004400918.1    rna-XM_007074109.3    rna-XM_040634428.1    rna-XM_027618111.2    mNeoVis.3.1.1.chr7.g00903.t01    rna-gnl|WGS_ZZZZ|egapxtmp_012352-R1    mNeoVis.evergladensis.2.1.chr7.g00957.t01    mNeoVis.vulgivaga.2.1.chr7.g00962.t01      1994    N0.HOG0001992   OG0000300  n18   rna-XM_011234499.3    rna-XM_038561200.1    rna-XM_022513509.1    rna-XM_003984350.5    rna-XM_036119135.1    rna-XM_039224228.1    rna-XM_032881834.1    rna-XM_045979538.1    rna-XM_032353051.1    rna-XM_059188694.1    rna-XM_059405381.1    rna-XM_004777918.2    mNeoVis.2.1.1.chr11_hap1.g01016.t01    rna-XM_021697504.1    rna-XM_004409579.1    rna-XM_007088841.3    rna-XM_008700558.2    rna-XM_027623048.2    mNeoVis.3.1.1.chr11.g00967.t01      mNeoVis.evergladensis.2.1.chr11.g00991.t01    mNeoVis.vulgivaga.2.1.chr11.g01014.t01      1995    N0.HOG0001993   OG0000300  n30   rna-XM_011221616.3    rna-XM_038564057.1    rna-XM_022506515.1    rna-XM_003993572.6    rna-XM_036101162.1    rna-XM_039244765.1    rna-XM_032879409.1    rna-XM_046014516.1    rna-XM_032357241.1    rna-XM_059146578.1    rna-XM_059402013.1    rna-XM_045064591.1    mNeoVis.2.1.1.chr7_hap1.g02378.t01    rna-XM_044919638.1    rna-XM_004393972.1    rna-XM_042957641.1    rna-XM_040639334.1    rna-XM_027578400.2    mNeoVis.3.1.1.chr7.g02189.t01      mNeoVis.evergladensis.2.1.chr7.g02324.t01    mNeoVis.vulgivaga.2.1.chr7.g02343.t01      1996    N0.HOG0001994   OG0000300  n48   rna-XM_002913202.4    rna-XM_038580145.1    rna-XM_022491063.1    rna-XM_003987252.6    rna-XM_036084596.1    rna-XM_039232879.1    rna-XM_032856759.1    rna-XM_046009630.1    rna-XM_032343399.1    rna-XM_059181253.1    rna-XM_059372603.1    rna-XM_004751173.3    mNeoVis.2.1.1.chr10_hap1.g00708.t01    rna-XM_021695627.1    rna-XM_004400398.1    rna-XM_007081786.2    rna-XM_040620493.1    rna-XM_027570598.2    mNeoVis.3.1.1.chr9.g00668.t01      mNeoVis.evergladensis.2.1.chr8.g00689.t01    mNeoVis.vulgivaga.2.1.chr9.g00690.t01      1997    N0.HOG0001995   OG0000301  n1   rna-XM_034654569.1    rna-XM_038565201.1    rna-XM_022494058.1    rna-XM_045033810.1    rna-XM_036074678.1    rna-XM_039223897.1    rna-XM_032857798.1    rna-XM_046019260.1    rna-XM_032355576.1    rna-XM_059167339.1    rna-XM_059393960.1    rna-XM_045066743.1    mNeoVis.2.1.1.chr3_hap1.g00975.t01    rna-XM_021702867.1    rna-XM_004415321.1    rna-XM_042994442.1    rna-XM_040644965.1    rna-XM_027590445.2    mNeoVis.3.1.1.chr3.g00965.t01    rna-gnl|WGS_ZZZZ|egapxtmp_038953-R2    mNeoVis.evergladensis.2.1.chr3.g00970.t01    mNeoVis.vulgivaga.2.1.chr3.g00989.t01      1998    N0.HOG0001996   OG0000301  n17   rna-XM_034645057.1    rna-XM_038577512.1    rna-XM_022519463.1    rna-XM_003988622.5    rna-XM_036121146.1    rna-XM_039221083.1    rna-XM_032876906.1    rna-XM_046012962.1    rna-XM_032348057.1    rna-XM_059185497.1    rna-XM_059403010.1    rna-XM_004778907.3    mNeoVis.2.1.1.chr9_hap1.g00603.t01    rna-XM_021704920.2    rna-XM_004400878.1    rna-XM_007072962.2    rna-XM_040642395.1    rna-XM_027593507.1    mNeoVis.3.1.1.chr10.g00571.t01      mNeoVis.evergladensis.2.1.chr10.g00597.t01    mNeoVis.vulgivaga.2.1.chr10.g00583.t01      1999    N0.HOG0001997   OG0000301  n27   rna-XM_034640461.1    rna-XM_038547449.1    rna-XM_022509777.1    rna-XM_019817811.3    rna-XM_036111983.1    rna-XM_039227730.1    rna-XM_032871471.1    rna-XM_045986522.1    rna-XM_032321775.1    rna-XM_059149105.1    rna-XM_059378431.1    rna-XM_004764677.3    mNeoVis.2.1.1.chr8_hap1.g00495.t01    rna-XM_021704321.1    rna-XM_004395062.1    rna-XM_042967313.1    rna-XM_040640703.1    rna-XM_027626052.2    mNeoVis.3.1.1.chr8.g00451.t01      mNeoVis.evergladensis.2.1.chr9.g00486.t01    mNeoVis.vulgivaga.2.1.chr8.g00492.t01      2000    N0.HOG0001998   OG0000301  n44   rna-XM_034643922.1    rna-XM_038529965.1    rna-XM_022515676.1    rna-XM_023256405.2    rna-XM_036109799.1    rna-XM_039245863.1    rna-XM_032880871.1    rna-XM_046011561.1    rna-XM_032349539.1    rna-XM_059184001.1    rna-XM_059404224.1    rna-XM_045085007.1    mNeoVis.2.1.1.chr9_hap1.g00110.t01    rna-XM_021696790.1    rna-XM_004392959.1    rna-XM_042991177.1    rna-XM_008692055.2    rna-XM_027594125.2    mNeoVis.3.1.1.chr10.g00106.t01    rna-gnl|WGS_ZZZZ|egapxtmp_024068-R1    mNeoVis.evergladensis.2.1.chr10.g00110.t01    mNeoVis.vulgivaga.2.1.chr10.g00106.t01      2001    N0.HOG0001999   OG0000302  n1   rna-XM_019796172.2    rna-XM_038549701.1    rna-XM_022522954.1    rna-XM_011284101.4    rna-XM_036114437.1, rna-XM_036114439.1, rna-XM_036117083.1    rna-XM_039216803.1    rna-XM_032874491.1    rna-XM_046012831.1    rna-XM_032348684.1    rna-XM_059184918.1    rna-XM_059403455.1    rna-XM_045071854.1    mNeoVis.2.1.1.chr9_hap1.g00366.t01    rna-XM_021687262.2    rna-XM_004401689.1    rna-XM_007081509.2    rna-XM_040637139.1    rna-XM_027593137.1    mNeoVis.3.1.1.chr10.g00358.t01    rna-gnl|WGS_ZZZZ|egapxtmp_023545-R2    mNeoVis.evergladensis.2.1.chr10.g00363.t01    mNeoVis.vulgivaga.2.1.chr10.g00360.t01      2002    N0.HOG0002000   OG0000302  n24   rna-XM_034654609.1    rna-XM_038584637.1    rna-XM_022497462.1, rna-XM_022526542.1    rna-XM_045033860.1    rna-XM_036071010.1    rna-XM_039237556.1    rna-XM_032857928.1    rna-XM_046018664.1    rna-XM_032355715.1    rna-XM_059167194.1    rna-XM_059394056.1    rna-XM_013051988.2    mNeoVis.2.1.1.chr3_hap1.g00948.t01    rna-XM_044913661.1    rna-XM_004394832.1    rna-XM_042994546.1    rna-XM_040644589.1    rna-XM_027588516.2    mNeoVis.3.1.1.chr3.g00937.t01    rna-gnl|WGS_ZZZZ|egapxtmp_038808-R1    mNeoVis.evergladensis.2.1.chr3.g00942.t01    mNeoVis.vulgivaga.2.1.chr3.g00962.t01      2003    N0.HOG0002001   OG0000302  n39   rna-XM_034661831.1, rna-XM_034662106.1    rna-XM_038570628.1    rna-XM_022492451.1, rna-XM_022492454.1, rna-XM_022514916.1    rna-XM_023260637.2    rna-XM_036093021.1    rna-XM_039222358.1, rna-XM_039242720.1    rna-XM_032860907.1, rna-XM_032861014.1    rna-XM_046002641.1, rna-XM_046003638.1    rna-XM_032359695.1, rna-XM_032359768.1    rna-XM_059162401.1, rna-XM_059165033.1    rna-XM_059388320.1, rna-XM_059390793.1    rna-XM_013057186.2, rna-XM_045088141.1    mNeoVis.2.1.1.chr5_hap1.g01066.t01, mNeoVis.2.1.1.chr5_hap1.g01067.t01    rna-XM_021678924.1    rna-XM_004413753.1, rna-XM_004417280.1    rna-XM_015544414.2    rna-XM_008709549.2, rna-XM_040636890.1    rna-XM_027581874.2, rna-XM_027584287.2    mNeoVis.3.1.1.chr2.g00975.t01    rna-gnl|WGS_ZZZZ|egapxtmp_035525-R1, rna-gnl|WGS_ZZZZ|egapxtmp_038484-R1    mNeoVis.evergladensis.2.1.chr2.g01058.t01, mNeoVis.evergladensis.2.1.chr2.g01059.t01    mNeoVis.vulgivaga.2.1.chr2.g01062.t01, mNeoVis.vulgivaga.2.1.chr2.g01063.t01      2004    N0.HOG0002002   OG0000303  n1   rna-XM_034656487.1    rna-XM_038532339.1    rna-XM_022504963.1    rna-XM_023258405.2    rna-XM_036094054.1    rna-XM_039239397.1    rna-XM_032846469.1    rna-XM_046000575.1    rna-XM_032335755.1    rna-XM_059175465.1    rna-XM_059418233.1    rna-XM_045074409.1    mNeoVis.2.1.1.chr1_hap1.g00743.t01    rna-XM_044916852.1    rna-XM_004414676.1    rna-XM_042959593.1    rna-XM_040622094.1    rna-XM_027605205.2    mNeoVis.3.1.1.chr1.g00766.t01, mNeoVis.3.1.1.chr1.g00767.t01    rna-gnl|WGS_ZZZZ|egapxtmp_009013-R1    mNeoVis.evergladensis.2.1.chr1.g00751.t01    mNeoVis.vulgivaga.2.1.chr1.g00765.t01      2005    N0.HOG0002003   OG0000303  n13   rna-XM_019804770.2    rna-XM_038532726.1    rna-XM_022511129.1    rna-XM_023254958.2    rna-XM_036082887.1    rna-XM_039248232.1    rna-XM_032850603.1, rna-XM_032850606.1    rna-XM_046008953.1    rna-XM_032342152.1    rna-XM_059180105.1    rna-XM_059371634.1    rna-XM_045070727.1    mNeoVis.2.1.1.chr10_hap1.g01165.t01    rna-XM_021680512.2    rna-XM_004405778.1    rna-XM_042988141.1    rna-XM_040642731.1    rna-XM_027570440.2    mNeoVis.3.1.1.chr9.g01097.t01      mNeoVis.evergladensis.2.1.chr8.g01143.t01    mNeoVis.vulgivaga.2.1.chr9.g01123.t01      2006    N0.HOG0002004   OG0000303  n14   rna-XM_034649980.1    rna-XM_038588108.1      rna-XM_045051084.1        rna-XM_032837111.1    rna-XM_045995395.1    rna-XM_032331542.1    rna-XM_059158480.1    rna-XM_059385895.1    rna-XM_045068346.1    mNeoVis.2.1.1.chr12_hap1.g00270.t01    rna-XM_044911687.1      rna-XM_042974344.1    rna-XM_040628943.1, rna-XM_040628955.1    rna-XM_035725850.1    mNeoVis.3.1.1.chr12.g00238.t01    rna-gnl|WGS_ZZZZ|egapxtmp_029661-R1    mNeoVis.evergladensis.2.1.chr12.g00239.t01    mNeoVis.vulgivaga.2.1.chr12.g00246.t01      2007    N0.HOG0002005   OG0000303  n30   rna-XM_034658011.1    rna-XM_038566927.1    rna-XM_022498266.1    rna-XM_023247004.2    rna-XM_036095331.1    rna-XM_039243432.1    rna-XM_032867582.1    rna-XM_045989727.1    rna-XM_032315014.1    rna-XM_059160432.1    rna-XM_059389350.1    rna-XM_045082037.1    mNeoVis.2.1.1.chr5_hap1.g00474.t01    rna-XM_021683147.1    rna-XM_004404599.1    rna-XM_042978519.1    rna-XM_040629079.1    rna-XM_027587668.1    mNeoVis.3.1.1.chr2.g00398.t01    rna-gnl|WGS_ZZZZ|egapxtmp_017029-R1    mNeoVis.evergladensis.2.1.chr2.g00469.t01    mNeoVis.vulgivaga.2.1.chr2.g00471.t01      2008    N0.HOG0002006   OG0000304  n1   rna-XM_002921127.4    rna-XM_038551805.1    rna-XM_022521157.1    rna-XM_019830843.3    rna-XM_036096459.1    rna-XM_039252822.1    rna-XM_032846856.1    rna-XM_045999482.1    rna-XM_032335344.1    rna-XM_059175867.1    rna-XM_059416136.1    rna-XM_004759029.3    mNeoVis.2.1.1.chr1_hap1.g00872.t01    rna-XM_021690964.1    rna-XM_004416412.1    rna-XM_042982295.1    rna-XM_008709495.2    rna-XM_027605733.1    mNeoVis.3.1.1.chr1.g00895.t01    rna-gnl|WGS_ZZZZ|egapxtmp_021300-R1    mNeoVis.evergladensis.2.1.chr1.g00875.t01    mNeoVis.vulgivaga.2.1.chr1.g00889.t01      2009    N0.HOG0002007   OG0000304  n23   rna-XM_034668053.1    rna-XM_038580987.1    rna-XM_022512055.1    rna-XM_003986949.5    rna-XM_036117746.1    rna-XM_039238232.1    rna-XM_032865048.1    rna-XM_046008247.1    rna-XM_032342454.1    rna-XM_059180385.1    rna-XM_059371832.1    rna-XM_004758423.3    mNeoVis.2.1.1.chr10_hap1.g01068.t01    rna-XM_021693885.2    rna-XM_004407800.1    rna-XM_042988398.1    rna-XM_040642997.1    rna-XM_027572754.1    mNeoVis.3.1.1.chr9.g01000.t01      mNeoVis.evergladensis.2.1.chr8.g01044.t01    mNeoVis.vulgivaga.2.1.chr9.g01025.t01      2010    N0.HOG0002008   OG0000304  n39   rna-XM_034667494.1    rna-XM_038543119.1    rna-XM_022490959.1    rna-XM_006944267.4    rna-XM_036124400.1    rna-XM_039251434.1    rna-XM_032845045.1    rna-XM_045982622.1    rna-XM_032318226.1    rna-XM_059146435.1    rna-XM_059413770.1    rna-XM_004778578.3    mNeoVis.2.1.1.chr14_hap1.g00646.t01    rna-XM_044916259.1    rna-XM_004402946.1    rna-XM_042976857.1    rna-XM_040635759.1    rna-XM_027611977.2    mNeoVis.3.1.1.chr14.g00621.t01      mNeoVis.evergladensis.2.1.chr14.g00644.t01    mNeoVis.vulgivaga.2.1.chr14.g00642.t01      2011    N0.HOG0002009   OG0000304  n59   rna-XM_002915842.4    rna-XM_038585307.1    rna-XM_022496058.1    rna-XM_019838441.3    rna-XM_036098539.1    rna-XM_039217814.1    rna-XM_032860106.1    rna-XM_046019272.1    rna-XM_032354375.1    rna-XM_059168457.1    rna-XM_059393156.1    rna-XM_045078536.1    mNeoVis.2.1.1.chr3_hap1.g01294.t01    rna-XM_021703251.2    rna-XM_004393476.1    rna-XM_007090013.2    rna-XM_008687337.2    rna-XM_027589398.2    mNeoVis.3.1.1.chr3.g01271.t01    rna-gnl|WGS_ZZZZ|egapxtmp_038989-R1    mNeoVis.evergladensis.2.1.chr3.g01293.t01    mNeoVis.vulgivaga.2.1.chr3.g01313.t01      2012    N0.HOG0002010   OG0000305  n1   rna-XM_034647526.1    rna-XM_038560123.1    rna-XM_022498941.1    rna-XM_023252618.2    rna-XM_036081555.1    rna-XM_039220338.1    rna-XM_032838130.1    rna-XM_045985538.1    rna-XM_032329085.1    rna-XM_059156223.1    rna-XM_059384170.1    rna-XM_013059289.2    mNeoVis.2.1.1.chr6_hap1.g00981.t01    rna-XM_021681506.1    rna-XM_004411924.1    rna-XM_042983506.1    rna-XM_040643172.1    rna-XM_027600316.2    mNeoVis.3.1.1.chr4.g00942.t01    rna-gnl|WGS_ZZZZ|egapxtmp_000075-R1    mNeoVis.evergladensis.2.1.chr4.g00931.t01    mNeoVis.vulgivaga.2.1.chr4.g00954.t01      2013    N0.HOG0002011   OG0000305  n24   rna-XM_034656684.1    rna-XM_038534675.1    rna-XM_022493514.1    rna-XM_011285307.3    rna-XM_036084931.1    rna-XM_039239772.1    rna-XM_032851888.1    rna-XM_045999399.1    rna-XM_032336924.1    rna-XM_059174394.1    rna-XM_059417432.1    rna-XM_004737514.3    mNeoVis.2.1.1.chr1_hap1.g00342.t01    rna-XM_021698682.1    rna-XM_004401428.1    rna-XM_042991743.1    rna-XM_040630609.1    rna-XM_027606667.2      rna-gnl|WGS_ZZZZ|egapxtmp_028678-R1    mNeoVis.evergladensis.2.1.chr1.g00340.t01    mNeoVis.vulgivaga.2.1.chr1.g00346.t01      2014    N0.HOG0002012   OG0000305  n43   rna-XM_019795351.2    rna-XM_038582098.1    rna-XM_022515897.1    rna-XM_019829187.3      rna-XM_039241298.1    rna-XM_032861390.1    rna-XM_045998229.1    rna-XM_032332944.1    rna-XM_059172547.1    rna-XM_059376235.1    rna-XM_004756033.2    mNeoVis.2.1.1.chr6_hap1.g00574.t01    rna-XM_021680394.1    rna-XM_004391492.1    rna-XM_015542240.2    rna-XM_008707176.2    rna-XM_027599020.1    mNeoVis.3.1.1.chr4.g00579.t01    rna-gnl|WGS_ZZZZ|egapxtmp_000310-R1    mNeoVis.evergladensis.2.1.chr4.g00578.t01    mNeoVis.vulgivaga.2.1.chr4.g00588.t01      2015    N0.HOG0002013   OG0000305  n62   rna-XM_019807732.2    rna-XM_038534250.1    rna-XM_022499906.1    rna-XM_006937850.5    rna-XM_036083937.1    rna-XM_039255547.1    rna-XM_032841073.1    rna-XM_046027401.1    rna-XM_032312544.1    rna-XM_059170002.1    rna-XM_059397660.1    rna-XM_004769499.2    mNeoVis.2.1.1.chr2_hap1.g00500.t01    rna-XM_021699568.2    rna-XM_004393659.1    rna-XM_042960636.1    rna-XM_040623782.1    rna-XM_027596516.2    mNeoVis.3.1.1.chr5.g00493.t01    rna-gnl|WGS_ZZZZ|egapxtmp_006868-R1    mNeoVis.evergladensis.2.1.chr5.g00501.t01    mNeoVis.vulgivaga.2.1.chr5.g00501.t01      2016    N0.HOG0002014   OG0000306  n1   rna-XM_019803409.2    rna-XM_038581789.1    rna-XM_022512551.1    rna-XM_019829018.3    rna-XM_036079687.1    rna-XM_039220002.1    rna-XM_032859036.1    rna-XM_045991861.1    rna-XM_032333238.1    rna-XM_059170007.1    rna-XM_059377541.1    rna-XM_013053952.2    mNeoVis.2.1.1.chr6_hap1.g00488.t01    rna-XM_044912890.1    rna-XM_004406836.1, rna-XM_004409259.1    rna-XM_015540216.2    rna-XM_008689455.2    rna-XM_027600484.2    mNeoVis.3.1.1.chr4.g00493.t01    rna-gnl|WGS_ZZZZ|egapxtmp_000578-R1    mNeoVis.evergladensis.2.1.chr4.g00494.t01    mNeoVis.vulgivaga.2.1.chr4.g00502.t01      2017    N0.HOG0002015   OG0000306  n18   rna-XM_034656874.1    rna-XM_038535043.1    rna-XM_022493053.1    rna-XM_011286079.4    rna-XM_036113576.1    rna-XM_039239418.1    rna-XM_032851539.1    rna-XM_045999567.1    rna-XM_032337330.1    rna-XM_059174033.1    rna-XM_059417157.1    rna-XM_045079564.1    mNeoVis.2.1.1.chr1_hap1.g00201.t01    rna-XM_021697078.2    rna-XM_004402812.1    rna-XM_042994349.1    rna-XM_008692512.2    rna-XM_027605583.2    mNeoVis.3.1.1.chr1.g00202.t01      mNeoVis.evergladensis.2.1.chr1.g00196.t01    mNeoVis.vulgivaga.2.1.chr1.g00199.t01      2018    N0.HOG0002016   OG0000306  n25   rna-XM_034662704.1    rna-XM_038534384.1    rna-XM_022500313.1    rna-XM_023240571.2    rna-XM_036110272.1    rna-XM_039235117.1    rna-XM_032839800.1    rna-XM_046026464.1    rna-XM_032313202.1    rna-XM_059169864.1    rna-XM_059397828.1    rna-XM_004769375.3    mNeoVis.2.1.1.chr2_hap1.g00461.t01    rna-XM_021699837.1    rna-XM_004407025.1    rna-XM_042960914.1    rna-XM_040623986.1    rna-XM_027591866.1    mNeoVis.3.1.1.chr5.g00454.t01      mNeoVis.evergladensis.2.1.chr5.g00464.t01    mNeoVis.vulgivaga.2.1.chr5.g00463.t01      2019    N0.HOG0002017   OG0000306  n43   rna-XM_034664942.1    rna-XM_038559562.1    rna-XM_022518167.1    rna-XM_045052077.1    rna-XM_036085265.1    rna-XM_039242970.1    rna-XM_032875055.1    rna-XM_046020294.1    rna-XM_032305404.1    rna-XM_059170706.1    rna-XM_059397110.1    rna-XM_004739466.2    mNeoVis.2.1.1.chr4_hap1.g00613.t01    rna-XM_021703733.1    rna-XM_004414037.1    rna-XM_042962816.1    rna-XM_040644465.1    rna-XM_027574153.2      rna-gnl|WGS_ZZZZ|egapxtmp_004467-R1    mNeoVis.evergladensis.2.1.chr6.g00622.t01    mNeoVis.vulgivaga.2.1.chr6.g00619.t01      2020    N0.HOG0002018   OG0000307  n0   rna-XM_002924190.4, rna-XM_034654274.1, rna-XM_034655630.1    rna-XM_038527682.1, rna-XM_038530451.1, rna-XM_038552090.1, rna-XM_038579321.1, rna-XM_038586175.1    rna-XM_022501158.1    rna-XM_019820444.2, rna-XM_045055056.1, rna-XM_045056233.1    rna-XM_036064851.1, rna-XM_036066132.1, rna-XM_036068773.1, rna-XM_036080834.1, rna-XM_036087850.1, rna-XM_036115809.1, rna-XM_036118361.1    rna-XM_039225327.1, rna-XM_039226003.1, rna-XM_039249086.1, rna-XM_039256079.1    rna-XM_032865146.1, rna-XM_032867030.1, rna-XM_032872656.1    rna-XM_045985854.1, rna-XM_045993237.1, rna-XM_046007367.1    rna-XM_032318787.1, rna-XM_032328486.1, rna-XM_032353202.1    rna-XM_059134224.1, rna-XM_059155358.1    rna-XM_059407564.1, rna-XM_059414449.1    rna-XM_045073811.1, rna-XM_045075094.1    mNeoVis.2.1.1.chr11_hap1.g01120.t01, mNeoVis.2.1.1.chr15_hap1.g00646.t01    rna-XM_021679303.1, rna-XM_021679344.1, rna-XM_021685925.2, rna-XM_021687458.2, rna-XM_021689420.1, rna-XM_021702025.1, rna-XM_021702659.1, rna-XM_044912603.1, rna-XM_044915721.1, rna-XM_044920757.1    rna-XM_004393212.1, rna-XM_004393616.1, rna-XM_004416339.1, rna-XM_004416502.1, rna-XM_004416506.1, rna-XM_004417644.1    rna-XM_007084936.3, rna-XM_042959033.1, rna-XM_042972118.1, rna-XM_042982735.1    rna-XM_008691685.1, rna-XM_008698361.2, rna-XM_040620568.1, rna-XM_040624815.1, rna-XM_040625728.1    rna-XM_027586929.1, rna-XM_027606431.1, rna-XM_027607215.2, rna-XM_027611001.2, rna-XM_027616803.1, rna-XM_035722521.1, rna-XM_035723378.1, rna-XM_035723387.1, rna-XM_035724385.1, rna-XM_035724410.1, rna-XM_035725935.1, rna-XM_035729174.1    mNeoVis.3.1.1.chr11.g01062.t01, mNeoVis.3.1.1.chr15.g00427.t01    rna-gnl|WGS_ZZZZ|egapxtmp_026613-R1    mNeoVis.evergladensis.2.1.chr11.g01089.t01, mNeoVis.evergladensis.2.1.chr15.g00468.t01    mNeoVis.vulgivaga.2.1.chr11.g01115.t01, mNeoVis.vulgivaga.2.1.chr11.g01116.t01, mNeoVis.vulgivaga.2.1.chr15.g00475.t01, mNeoVis.vulgivaga.2.1.chr15.g00476.t01      2021    N0.HOG0002019   OG0000308  n1   rna-XM_034653844.1    rna-XM_038541444.1    rna-XM_022496717.1    rna-XM_023258884.2    rna-XM_036118589.1    rna-XM_039221829.1    rna-XM_032867071.1    rna-XM_046010462.1    rna-XM_032301511.1    rna-XM_059135775.1    rna-XM_059375325.1    rna-XM_045064823.1    mNeoVis.2.1.1.chr2_hap1.g01100.t01    rna-XM_021690199.2    rna-XM_004417325.1    rna-XM_042997803.1    rna-XM_008690613.2    rna-XM_027592734.2    mNeoVis.3.1.1.chr5.g01025.t01    rna-gnl|WGS_ZZZZ|egapxtmp_005511-R2    mNeoVis.evergladensis.2.1.chr5.g01044.t01    mNeoVis.vulgivaga.2.1.chr5.g01035.t01      2022    N0.HOG0002020   OG0000308  n20   rna-XM_002915194.4    rna-XM_038549108.1    rna-XM_022501900.1    rna-XM_019816655.3    rna-XM_036069125.1    rna-XM_039229647.1    rna-XM_032866035.1    rna-XM_046021738.1    rna-XM_032306261.1    rna-XM_059143097.1    rna-XM_059410594.1    rna-XM_004757153.3    mNeoVis.2.1.1.chr13_hap1.g00173.t01    rna-XM_021691474.2    rna-XM_004392499.1    rna-XM_042963577.1    rna-XM_040632029.1    rna-XM_027615032.2      rna-gnl|WGS_ZZZZ|egapxtmp_030030-R1    mNeoVis.evergladensis.2.1.chr13.g00171.t01    mNeoVis.vulgivaga.2.1.chr13.g00169.t01      2023    N0.HOG0002021   OG0000308  n36   rna-XM_034653852.1    rna-XM_038541439.1    rna-XM_022497413.1    rna-XM_023258891.2    rna-XM_036118587.1    rna-XM_039221837.1    rna-XM_032867073.1    rna-XM_046023090.1    rna-XM_032361345.1    rna-XM_059137320.1    rna-XM_059375343.1    rna-XM_004752632.3    mNeoVis.2.1.1.chr2_hap1.g01099.t01    rna-XM_021690147.1    rna-XM_004417326.1    rna-XM_007073762.3    rna-XM_040627061.1    rna-XM_027592524.2    mNeoVis.3.1.1.chr5.g01024.t01    rna-gnl|WGS_ZZZZ|egapxtmp_005513-R1    mNeoVis.evergladensis.2.1.chr5.g01043.t01    mNeoVis.vulgivaga.2.1.chr5.g01034.t01      2024   N0.HOG0002022  OG0000308  n52   rna-XM_034657675.1    rna-XM_038567476.1    rna-XM_022525870.1    rna-XM_045045044.1    rna-XM_036066007.1    rna-XM_039251648.1    rna-XM_032883837.1    rna-XM_045991046.1    rna-XM_032309295.1    rna-XM_059159782.1    rna-XM_059388861.1    rna-XM_004771690.3    mNeoVis.2.1.1.chr5_hap1.g00209.t01    rna-XM_021705367.2    rna-XM_004412154.1    rna-XM_042978020.1    rna-XM_040629423.1    rna-XM_027585279.2    mNeoVis.3.1.1.chr2.g00158.t01      mNeoVis.evergladensis.2.1.chr2.g00206.t01    mNeoVis.vulgivaga.2.1.chr2.g00205.t01      2025   N0.HOG0002023  OG0000309  n1   rna-XM_002921354.4    rna-XM_038567241.1    rna-XM_022526918.1    rna-XM_019817741.3    rna-XM_036075448.1    rna-XM_039251063.1    rna-XM_032884054.1    rna-XM_045992063.1    rna-XM_032311661.1    rna-XM_059160014.1    rna-XM_059389053.1    rna-XM_004748400.3    mNeoVis.2.1.1.chr5_hap1.g00305.t01    rna-XM_021705129.1    rna-XM_004398902.1    rna-XM_042978209.1    rna-XM_040629300.1    rna-XM_027586130.2    mNeoVis.3.1.1.chr2.g00241.t01    rna-gnl|WGS_ZZZZ|egapxtmp_035252-R2    mNeoVis.evergladensis.2.1.chr2.g00303.t01    mNeoVis.vulgivaga.2.1.chr2.g00299.t01      2026    N0.HOG0002024   OG0000309  n18   rna-XM_002912726.4    rna-XM_038566840.1    rna-XM_022497689.1    rna-XM_003982073.6    rna-XM_036093972.1    rna-XM_039243404.1    rna-XM_032867675.1    rna-XM_045991875.1    rna-XM_032316022.1    rna-XM_059160516.1    rna-XM_059389424.1    rna-XM_004760956.3    mNeoVis.2.1.1.chr5_hap1.g00510.t01    rna-XM_021683260.2    rna-XM_004404632.1    rna-XM_007088651.3    rna-XM_040629021.1    rna-XM_027586293.2    mNeoVis.3.1.1.chr2.g00433.t01      mNeoVis.evergladensis.2.1.chr2.g00505.t01    mNeoVis.vulgivaga.2.1.chr2.g00507.t01      2027    N0.HOG0002025   OG0000309  n21   rna-XM_019798663.2    rna-XM_038530648.1    rna-XM_022518064.1    rna-XM_045061019.1      rna-XM_039243831.1    rna-XM_032850167.1    rna-XM_046000833.1    rna-XM_032336181.1    rna-XM_059175040.1    rna-XM_059417912.1    rna-XM_004744619.3    mNeoVis.2.1.1.chr1_hap1.g00590.t01    rna-XM_021701998.1    rna-XM_004408736.1    rna-XM_042959193.1    rna-XM_008691395.2    rna-XM_027606044.2        mNeoVis.evergladensis.2.1.chr1.g00598.t01    mNeoVis.vulgivaga.2.1.chr1.g00607.t01      2028    N0.HOG0002026   OG0000309  n34   rna-XM_002915722.4    rna-XM_038558047.1    rna-XM_022497205.1    rna-XM_019837002.3    rna-XM_036108407.1, rna-XM_036121334.1    rna-XM_039237964.1    rna-XM_032844150.1    rna-XM_046014567.1    rna-XM_032302878.1    rna-XM_059137066.1    rna-XM_059374723.1    rna-XM_004774473.3    mNeoVis.2.1.1.chr2_hap1.g01298.t01    rna-XM_021683902.1, rna-XM_044913364.1    rna-XM_004400311.1    rna-XM_042997154.1    rna-XM_040627316.1    rna-XM_027624191.2    mNeoVis.3.1.1.chr5.g01223.t01    rna-gnl|WGS_ZZZZ|egapxtmp_006324-R2    mNeoVis.evergladensis.2.1.chr5.g01241.t01    mNeoVis.vulgivaga.2.1.chr5.g01235.t01      2029   N0.HOG0002027  OG0000310  n1   rna-XM_034672135.1    rna-XM_038556514.1    rna-XM_022510311.1    rna-XM_045052755.1    rna-XM_036078425.1    rna-XM_039242593.1    rna-XM_032869218.1    rna-XM_046021154.1    rna-XM_032305428.1    rna-XM_059171869.1    rna-XM_059395616.1    rna-XM_013050476.2    mNeoVis.2.1.1.chr4_hap1.g00207.t01    rna-XM_021699320.1    rna-XM_004397991.1    rna-XM_042972400.1    rna-XM_008705882.2    rna-XM_027573940.1    mNeoVis.3.1.1.chr6.g00207.t01    rna-gnl|WGS_ZZZZ|egapxtmp_003918-R1    mNeoVis.evergladensis.2.1.chr6.g00206.t01    mNeoVis.vulgivaga.2.1.chr6.g00207.t01      2030    N0.HOG0002028   OG0000310  n23   rna-XM_002913440.4    rna-XM_038556509.1    rna-XM_022510090.1    rna-XM_003983055.6    rna-XM_036078399.1    rna-XM_039243065.1      rna-XM_046021152.1    rna-XM_032305427.1    rna-XM_059172674.1    rna-XM_059395613.1      mNeoVis.2.1.1.chr4_hap1.g00206.t01    rna-XM_021699322.1    rna-XM_004397989.1    rna-XM_007078092.3    rna-XM_040644542.1    rna-XM_027574738.1    mNeoVis.3.1.1.chr6.g00206.t01    rna-gnl|WGS_ZZZZ|egapxtmp_004384-R1    mNeoVis.evergladensis.2.1.chr6.g00205.t01    mNeoVis.vulgivaga.2.1.chr6.g00206.t01      2031    N0.HOG0002029   OG0000310  n43   rna-XM_002913437.4    rna-XM_038556524.1    rna-XM_022510199.1    rna-XM_003983050.6    rna-XM_036078488.1    rna-XM_039242581.1    rna-XM_032865355.1    rna-XM_046021103.1    rna-XM_032305403.1    rna-XM_059171865.1    rna-XM_059395618.1    rna-XM_004741758.3    mNeoVis.2.1.1.chr4_hap1.g00209.t01    rna-XM_021699373.1    rna-XM_004397995.1    rna-XM_007078087.3    rna-XM_040644543.1    rna-XM_027573207.2    mNeoVis.3.1.1.chr6.g00209.t01    rna-gnl|WGS_ZZZZ|egapxtmp_003920-R1    mNeoVis.evergladensis.2.1.chr6.g00208.t01    mNeoVis.vulgivaga.2.1.chr6.g00209.t01      2032    N0.HOG0002030   OG0000310  n61   rna-XM_011230386.3    rna-XM_038556519.1    rna-XM_022510302.1    rna-XM_003983051.5    rna-XM_036078452.1    rna-XM_039243054.1    rna-XM_032865341.1    rna-XM_046021250.1    rna-XM_032305600.1    rna-XM_059171867.1    rna-XM_059395617.1    rna-XM_004741755.2    mNeoVis.2.1.1.chr4_hap1.g00208.t01    rna-XM_021699331.1    rna-XM_004397993.1    rna-XM_007078094.3    rna-XM_008705881.1    rna-XM_027573971.2    mNeoVis.3.1.1.chr6.g00208.t01    rna-gnl|WGS_ZZZZ|egapxtmp_003917-R1    mNeoVis.evergladensis.2.1.chr6.g00207.t01    mNeoVis.vulgivaga.2.1.chr6.g00208.t01      2033    N0.HOG0002031   OG0000311  n1   rna-XM_034644201.1    rna-XM_038550027.1    rna-XM_022495223.1    rna-XM_023257247.2    rna-XM_036096913.1    rna-XM_039249907.1    rna-XM_032854763.1    rna-XM_046013903.1    rna-XM_032346991.1    rna-XM_059186349.1    rna-XM_059404693.1    rna-XM_004752406.2    mNeoVis.2.1.1.chr9_hap1.g00939.t01    rna-XM_021678566.1    rna-XM_004404816.1    rna-XM_042992764.1    rna-XM_040637266.1    rna-XM_027594673.1    mNeoVis.3.1.1.chr10.g00892.t01    rna-gnl|WGS_ZZZZ|egapxtmp_023255-R1    mNeoVis.evergladensis.2.1.chr10.g00924.t01    mNeoVis.vulgivaga.2.1.chr10.g00914.t01      2034   N0.HOG0002032  OG0000311  n19   rna-XM_002928762.4    rna-NM_001003113.1    rna-XM_022507334.1    rna-XM_003993030.6    rna-XM_036116316.1    rna-XM_039236720.1    rna-XM_032881150.1    rna-XM_046016022.1    rna-XM_032356890.1    rna-XM_059137634.1    rna-XM_059407579.1    rna-XM_004752079.3    mNeoVis.2.1.1.chr7_hap1.g01857.t01    rna-XM_021685831.1    rna-XM_004412467.1    rna-XM_007098468.3    rna-XM_008708932.2    rna-XM_027578808.2    mNeoVis.3.1.1.chr7.g01725.t01      mNeoVis.evergladensis.2.1.chr7.g01820.t01    mNeoVis.vulgivaga.2.1.chr7.g01837.t01      2035   N0.HOG0002033  OG0000311  n26     rna-XM_038528112.1    rna-XM_022524044.1    rna-XM_023245069.2    rna-XM_036068138.1    rna-XM_039244302.1    rna-XM_032838895.1    rna-XM_045988211.1    rna-XM_032325779.1    rna-XM_059151026.1    rna-XM_059382798.1    rna-XM_013063837.2    mNeoVis.2.1.1.chr7_hap1.g00862.t01    rna-XM_044911354.1    rna-XM_004409919.1    rna-XM_042970339.1    rna-XM_040634344.1    rna-XM_035724866.1    mNeoVis.3.1.1.chr7.g00804.t01      mNeoVis.evergladensis.2.1.chr7.g00854.t01    mNeoVis.vulgivaga.2.1.chr7.g00860.t01      2036    N0.HOG0002034   OG0000311  n45   rna-XM_034654056.1    rna-XM_038541036.1    rna-XM_022497125.1    rna-XM_006934983.5    rna-XM_036076422.1    rna-XM_039251068.1    rna-XM_032881503.1    rna-XM_046025694.1    rna-XM_032302537.1    rna-XM_059136532.1    rna-XM_059375679.1    rna-XM_004769730.3    mNeoVis.2.1.1.chr2_hap1.g00970.t01    rna-XM_044914578.1    rna-XM_004407162.1    rna-XM_042998144.1    rna-XM_040626929.1    rna-XM_027575835.2    mNeoVis.3.1.1.chr5.g00903.t01    rna-gnl|WGS_ZZZZ|egapxtmp_009063-R2    mNeoVis.evergladensis.2.1.chr5.g00924.t01    mNeoVis.vulgivaga.2.1.chr5.g00911.t01      2037    N0.HOG0002035   OG0000312  n1   rna-XM_034648400.1, rna-XM_034651737.1, rna-XM_034654619.1    rna-XM_038581150.1, rna-XM_038581633.1, rna-XM_038581634.1    rna-XM_022492675.1, rna-XM_022515964.1    rna-XM_003991381.6, rna-XM_011285802.4, rna-XM_045036513.1    rna-XM_036112719.1    rna-XM_039257132.1    rna-XM_032865251.1, rna-XM_032865540.1    rna-XM_046001293.1, rna-XM_046002685.1, rna-XM_046002686.1    rna-XM_032355675.1, rna-XM_032357063.1    rna-XM_059164922.1, rna-XM_059165304.1, rna-XM_059165317.1    rna-XM_059388244.1, rna-XM_059388397.1    rna-XM_004762271.3, rna-XM_013061260.2, rna-XM_013061279.2    mNeoVis.2.1.1.chr5_hap1.g01855.t01, mNeoVis.2.1.1.chr5_hap1.g01904.t01    rna-XM_021679198.1    rna-XM_004406590.1    rna-XM_015540875.2, rna-XM_042956386.1, rna-XM_042956585.1    rna-XM_008710177.2, rna-XM_040621650.1, rna-XM_040621651.1    rna-XM_027582895.2, rna-XM_027586458.1      rna-gnl|WGS_ZZZZ|egapxtmp_038699-R1          2038    N0.HOG0002036   OG0000312  n39   rna-XM_034651736.1    rna-XM_038581635.1    rna-XM_022516204.1    rna-XM_011285801.3, rna-XM_023260843.2    rna-XM_036097264.1, rna-XM_036112720.1, rna-XM_036112732.1      rna-XM_032865250.1, rna-XM_032865254.1, rna-XM_032865542.1    rna-XM_046002683.1, rna-XM_046002687.1, rna-XM_046003822.1    rna-XM_032302951.1, rna-XM_032302959.1, rna-XM_032346558.1, rna-XM_032355666.1, rna-XM_032357077.1    rna-XM_059164923.1, rna-XM_059164924.1, rna-XM_059165301.1, rna-XM_059165302.1, rna-XM_059165303.1    rna-XM_059387483.1, rna-XM_059388236.1    rna-XM_004762270.3, rna-XM_045084357.1, rna-XM_045084555.1, rna-XM_045084556.1    mNeoVis.2.1.1.chr5_hap1.g01854.t01, mNeoVis.2.1.1.chr5_hap1.g01903.t01, mNeoVis.2.1.1.chr5_hap1.g01905.t01    rna-XM_021679199.1    rna-XM_004406555.1    rna-XM_042956387.1, rna-XM_042998873.1    rna-XM_008709326.2, rna-XM_040621649.1    rna-XM_027586124.2, rna-XM_035728731.1, rna-XM_035728732.1        mNeoVis.evergladensis.2.1.chr2.g01883.t01    mNeoVis.vulgivaga.2.1.chr2.g01897.t01      2039   N0.HOG0002037  OG0000313  n3                     rna-XM_059150688.1    rna-XM_059386793.1                            2040    N0.HOG0002038   OG0000313  n6                           mNeoVis.2.1.1.chr1_hap1.g00985.t01              mNeoVis.3.1.1.chr1.g00990.t01    rna-gnl|WGS_ZZZZ|egapxtmp_020118-R1    mNeoVis.evergladensis.2.1.chr1.g00979.t01    mNeoVis.vulgivaga.2.1.chr1.g00991.t01, mNeoVis.vulgivaga.2.1.chr1.g00992.t01      2041    N0.HOG0002039   OG0000313  n12   rna-XM_034645624.1    rna-XM_038530865.1, rna-XM_038533676.1, rna-XM_038535451.1, rna-XM_038551595.1, rna-XM_038555111.1, rna-XM_038555919.1, rna-XM_038555921.1, rna-XM_038557990.1, rna-XM_038557991.1, rna-XM_038560640.1, rna-XM_038562925.1, rna-XM_038564540.1, rna-XM_038568132.1, rna-XM_038571387.1, rna-XM_038585080.1                      rna-XM_045066399.1                      mNeoVis.vulgivaga.2.1.chr7.g02439.t01      2042    N0.HOG0002040   OG0000313  n29   rna-XM_034644853.1    rna-XM_038535558.1, rna-XM_038546048.1            rna-XM_032851202.1, rna-XM_032851204.1              mNeoVis.2.1.1.chr7_hap1.g00720.t01, mNeoVis.2.1.1.chr7_hap1.g00724.t01, mNeoVis.2.1.1.chr7_hap1.g00727.t01            rna-XM_027579541.2, rna-XM_027579796.2    mNeoVis.3.1.1.chr7.g00679.t01    rna-gnl|WGS_ZZZZ|egapxtmp_014176-R1    mNeoVis.evergladensis.2.1.chr7.g00678.t01, mNeoVis.evergladensis.2.1.chr7.g00723.t01, mNeoVis.evergladensis.2.1.chr7.g02486.t01    mNeoVis.vulgivaga.2.1.chr7.g00718.t01      2043    N0.HOG0002041   OG0000313  n43                           mNeoVis.2.1.1.chr7_hap1.g00722.t01, mNeoVis.2.1.1.chr7_hap1.g00725.t01              mNeoVis.3.1.1.chr7.g00685.t01        mNeoVis.vulgivaga.2.1.chr7.g00719.t01      2044    N0.HOG0002042   OG0000313  n46           rna-XM_036083649.1, rna-XM_036104349.1                                        2045    N0.HOG0002043   OG0000313  n47     rna-XM_038568097.1, rna-XM_038569363.1, rna-XM_038584525.1, rna-XM_038585121.1    rna-XM_022492602.1      rna-XM_036083650.1      rna-XM_032867014.1, rna-XM_032879102.1, rna-XM_032879918.1      rna-XM_032326291.1, rna-XM_032360296.1, rna-XM_032360297.1    rna-XM_059152483.1      rna-XM_045077462.1, rna-XM_045077463.1, rna-XM_045089739.1    mNeoVis.2.1.1.chr11_hap1.g01200.t01, mNeoVis.2.1.1.chr7_hap1.g00679.t01, mNeoVis.2.1.1.chr7_hap1.g00721.t01, mNeoVis.2.1.1.chr7_hap1.g00723.t01, mNeoVis.2.1.1.chr7_hap1.g00726.t01, mNeoVis.2.1.1.chr7_hap1.g00732.t01, mNeoVis.2.1.1.chr7_hap1.g02610.t01, mNeoVis.2.1.1.chr7_hap1.g02673.t01              mNeoVis.3.1.1.chr7.g00644.t01, mNeoVis.3.1.1.chr7.g00645.t01, mNeoVis.3.1.1.chr7.g00684.t01, mNeoVis.3.1.1.chr7.g02274.t01    rna-gnl|WGS_ZZZZ|egapxtmp_010437-R1, rna-gnl|WGS_ZZZZ|egapxtmp_025401-R1    mNeoVis.evergladensis.2.1.chr11.g01173.t01, mNeoVis.evergladensis.2.1.chr7.g02485.t01    mNeoVis.vulgivaga.2.1.chr11.g01199.t01, mNeoVis.vulgivaga.2.1.chr7.g00675.t01, mNeoVis.vulgivaga.2.1.chr7.g00724.t01, mNeoVis.vulgivaga.2.1.chr7.g02440.t01, mNeoVis.vulgivaga.2.1.chr7.g02507.t01      2046    N0.HOG0002044   OG0000314  n1   rna-XM_002919635.4    rna-XM_038535146.1    rna-XM_022493555.1    rna-XM_011286847.4    rna-XM_036111774.1    rna-XM_039234146.1    rna-XM_032851441.1    rna-XM_046000839.1    rna-XM_032337446.1    rna-XM_059173942.1    rna-XM_059417097.1    rna-XM_004737910.3    mNeoVis.2.1.1.chr1_hap1.g00156.t01    rna-XM_021695347.1    rna-XM_004410969.1, rna-XM_004415854.1    rna-XM_042960321.1    rna-XM_040630450.1    rna-XM_027604233.2, rna-XM_035729328.1    mNeoVis.3.1.1.chr1.g00159.t01    rna-gnl|WGS_ZZZZ|egapxtmp_021575-R1    mNeoVis.evergladensis.2.1.chr1.g00153.t01    mNeoVis.vulgivaga.2.1.chr1.g00154.t01      2047    N0.HOG0002045   OG0000314  n25   rna-XM_034661692.1    rna-XM_038579892.1    rna-XM_022512230.1    rna-XM_023255072.2    rna-XM_036065820.1    rna-XM_039237885.1    rna-XM_032865007.1    rna-XM_046009092.1    rna-XM_032342517.1    rna-XM_059180443.1    rna-XM_059371875.1    rna-XM_045070662.1    mNeoVis.2.1.1.chr10_hap1.g01050.t01    rna-XM_021694006.2    rna-XM_004417189.1    rna-XM_042988433.1    rna-XM_040643016.1    rna-XM_027572906.2    mNeoVis.3.1.1.chr9.g00982.t01    rna-gnl|WGS_ZZZZ|egapxtmp_016500-R4    mNeoVis.evergladensis.2.1.chr8.g01023.t01    mNeoVis.vulgivaga.2.1.chr9.g01006.t01      2048    N0.HOG0002046   OG0000314  n48   rna-XM_034659995.1    rna-XM_038565398.1      rna-XM_023242496.2    rna-XM_036100723.1    rna-XM_039248096.1    rna-XM_032847088.1    rna-XM_045990504.1    rna-XM_032304161.1    rna-XM_059159132.1    rna-XM_059392604.1    rna-XM_045088239.1      rna-XM_044913197.1    rna-XM_004395503.1    rna-XM_042977475.1    rna-XM_040628314.1    rna-XM_027583453.2              2049    N0.HOG0002047   OG0000314  n63   rna-XM_034667487.1    rna-XM_038543111.1    rna-XM_022491030.1    rna-XM_023247384.2    rna-XM_036124389.1    rna-XM_039251402.1    rna-XM_032845233.1    rna-XM_045982616.1    rna-XM_032318220.1    rna-XM_059146424.1    rna-XM_059413767.1    rna-XM_004778579.3    mNeoVis.2.1.1.chr14_hap1.g00645.t01    rna-XM_021681979.1    rna-XM_004402945.1    rna-XM_042976862.1    rna-XM_040635765.1    rna-XM_027610479.1    mNeoVis.3.1.1.chr14.g00620.t01      mNeoVis.evergladensis.2.1.chr14.g00643.t01    mNeoVis.vulgivaga.2.1.chr14.g00641.t01      2050    N0.HOG0002048   OG0000315  n1   rna-XM_019794508.2    rna-XM_038541912.1    rna-XM_022508831.1    rna-XM_006942848.4    rna-XM_036086703.1    rna-XM_039227548.1    rna-XM_032878243.1    rna-XM_045983169.1    rna-XM_032318937.1    rna-XM_059146555.1    rna-XM_059414076.1    rna-XM_013058139.1    mNeoVis.2.1.1.chr14_hap1.g00322.t01    rna-XM_021682475.2    rna-XM_004392749.1    rna-XM_007086449.2    rna-XM_008685110.1    rna-XM_027609900.2    mNeoVis.3.1.1.chr14.g00320.t01    rna-gnl|WGS_ZZZZ|egapxtmp_031891-R1    mNeoVis.evergladensis.2.1.chr14.g00322.t01    mNeoVis.vulgivaga.2.1.chr14.g00323.t01      2051    N0.HOG0002049   OG0000315  n22   rna-XM_034645859.1    rna-XM_038562489.1    rna-XM_022506572.1    rna-XM_011287069.3    rna-XM_036074792.1    rna-XM_039223688.1    rna-XM_032879787.1    rna-XM_046016265.1    rna-XM_032359143.1        rna-XM_004759693.3      rna-XM_044919351.1    rna-XM_004403793.1    rna-XM_042958644.1    rna-XM_040639665.1    rna-XM_027580372.1              2052    N0.HOG0002050   OG0000315  n35   rna-XM_034645860.1    rna-XM_038563560.1    rna-XM_022507685.1    rna-XM_023240201.2    rna-XM_036074732.1    rna-XM_039223685.1    rna-XM_032879923.1    rna-XM_046016263.1    rna-XM_032359142.1        rna-XM_045066658.1    mNeoVis.2.1.1.chr7_hap1.g02633.t01, mNeoVis.2.1.1.chr7_hap1.g02634.t01    rna-XM_044919350.1    rna-XM_004403828.1    rna-XM_042958296.1    rna-XM_040638755.1, rna-XM_040638795.1    rna-XM_035723151.1    mNeoVis.3.1.1.chr7.g02287.t01, mNeoVis.3.1.1.chr7.g02288.t01    rna-gnl|WGS_ZZZZ|egapxtmp_011259-R1    mNeoVis.evergladensis.2.1.chr7.g02441.t01, mNeoVis.evergladensis.2.1.chr7.g02442.t01    mNeoVis.vulgivaga.2.1.chr7.g02463.t01, mNeoVis.vulgivaga.2.1.chr7.g02464.t01      2053    N0.HOG0002051   OG0000315  n56   rna-XM_011233781.3    rna-NM_001003041.1    rna-XM_022492447.1    rna-NM_001009237.1    rna-XM_036096905.1    rna-XM_039244243.1    rna-XM_032848699.1    rna-XM_045987116.1    rna-XM_032325416.1    rna-XM_059153503.1    rna-XM_059382723.1    rna-XM_013048160.2    mNeoVis.2.1.1.chr7_hap1.g00715.t01    rna-XM_021682363.1    rna-XM_004405030.1    rna-XM_042968809.1        mNeoVis.3.1.1.chr7.g00674.t01      mNeoVis.evergladensis.2.1.chr7.g00713.t01    mNeoVis.vulgivaga.2.1.chr7.g00713.t01      2054    N0.HOG0002052   OG0000316  n3     rna-XM_038586570.1    rna-XM_022524847.1      rna-XM_036095684.1, rna-XM_036096103.1, rna-XM_036118875.1, rna-XM_036118876.1      rna-XM_032846200.1      rna-XM_032331480.1, rna-XM_032331481.1, rna-XM_032331509.1, rna-XM_032331510.1, rna-XM_032331600.1    rna-XM_059157469.1, rna-XM_059158125.1, rna-XM_059158126.1, rna-XM_059158127.1      rna-XM_045068290.1    mNeoVis.2.1.1.chr12_hap1.g00331.t01    rna-XM_021687691.1, rna-XM_044912234.1    rna-XM_004406377.1, rna-XM_004406378.1, rna-XM_004406380.1, rna-XM_004406381.1, rna-XM_004406382.1, rna-XM_004406383.1, rna-XM_004412215.1, rna-XM_004412216.1, rna-XM_004412217.1, rna-XM_004414183.1, rna-XM_004415788.1, rna-XM_004415935.1, rna-XM_004415936.1, rna-XM_004415937.1, rna-XM_004417336.1, rna-XM_004417337.1        rna-XM_027607617.1, rna-XM_027609246.1, rna-XM_035725663.1, rna-XM_035725854.1    mNeoVis.3.1.1.chr12.g00298.t01        mNeoVis.vulgivaga.2.1.chr12.g00302.t01      2055    N0.HOG0002053   OG0000316  n44   rna-XM_002925934.2, rna-XM_034649033.1, rna-XM_034652923.1    rna-XM_038587973.1, rna-XM_038589581.1    rna-XM_022492639.1, rna-XM_022524869.1      rna-XM_036096096.1, rna-XM_036096104.1, rna-XM_036118874.1, rna-XM_036118877.1      rna-XM_032846203.1, rna-XM_032851021.1, rna-XM_032851095.1      rna-XM_032329713.1, rna-XM_032329714.1          mNeoVis.2.1.1.chr12_hap1.g00329.t01, mNeoVis.2.1.1.chr12_hap1.g00330.t01, mNeoVis.2.1.1.chr12_hap1.g00332.t01, mNeoVis.2.1.1.chr12_hap1.g00333.t01, mNeoVis.2.1.1.chr12_hap1.g00334.t01, mNeoVis.2.1.1.chr12_hap1.g00336.t01    rna-XM_021677997.1    rna-XM_004406375.1, rna-XM_004406376.1, rna-XM_004406379.1, rna-XM_004406384.1, rna-XM_004412218.1, rna-XM_004412219.1, rna-XM_004412425.1, rna-XM_004417237.1, rna-XM_004417462.1      rna-XM_040628263.1, rna-XM_040628345.1, rna-XM_040629552.1, rna-XM_040629564.1    rna-XM_035725434.1, rna-XM_035725851.1, rna-XM_035725852.1, rna-XM_035725853.1              2056    N0.HOG0002054   OG0000317  n1   rna-XM_011233979.2    rna-XM_038540680.1    rna-XM_022516640.1    rna-XM_045047958.1    rna-XM_036089421.1    rna-XM_039246844.1    rna-XM_032846043.1    rna-XM_045993963.1    rna-XM_032328286.1    rna-XM_059155080.1    rna-XM_059415597.1    rna-XM_045071952.1    mNeoVis.2.1.1.chr15_hap1.g00333.t01    rna-XM_044912315.1    rna-XM_004403289.1    rna-XM_042971816.1    rna-XM_040634547.1    rna-XM_027612594.1    mNeoVis.3.1.1.chr15.g00302.t01    rna-gnl|WGS_ZZZZ|egapxtmp_033991-R1    mNeoVis.evergladensis.2.1.chr15.g00310.t01    mNeoVis.vulgivaga.2.1.chr15.g00316.t01      2057    N0.HOG0002055   OG0000317  n21   rna-XM_011231732.3    rna-XM_038539745.1    rna-XM_022526573.1    rna-XM_003998697.3    rna-XM_036118971.1    rna-XM_039247805.1    rna-XM_032842714.1    rna-XM_045993340.1    rna-XM_032328430.1    rna-XM_059154893.1    rna-XM_059414837.1    rna-XM_004773821.3    mNeoVis.2.1.1.chr15_hap1.g00015.t01    rna-XM_044915688.1    rna-XM_004397275.1    rna-XM_015542071.2    rna-XM_040634535.1      mNeoVis.3.1.1.chr15.g00015.t01      mNeoVis.evergladensis.2.1.chr15.g00016.t01    mNeoVis.vulgivaga.2.1.chr15.g00017.t01      2058    N0.HOG0002056   OG0000317  n40   rna-XM_034669583.1    rna-XM_038540675.1    rna-XM_022516638.1    rna-XM_011290768.4    rna-XM_036089375.1    rna-XM_039246762.1    rna-XM_032846045.1    rna-XM_045993958.1    rna-XM_032328280.1    rna-XM_059155077.1    rna-XM_059415600.1    rna-XM_004750651.3      rna-XM_044912302.1    rna-XM_004403288.1    rna-XM_015545031.2    rna-XM_040635484.1    rna-XM_027612019.2    mNeoVis.3.1.1.chr15.g00303.t01      mNeoVis.evergladensis.2.1.chr15.g00311.t01    mNeoVis.vulgivaga.2.1.chr15.g00317.t01      2059    N0.HOG0002057   OG0000317  n59   rna-XM_034670079.1    rna-XM_038541715.1    rna-XM_022526568.1    rna-XM_045048282.1    rna-XM_036118940.1    rna-XM_039247863.1    rna-XM_032842715.1    rna-XM_045993645.1    rna-XM_032328421.1    rna-XM_059154878.1    rna-XM_059414826.1    rna-XM_045072631.1    mNeoVis.2.1.1.chr15_hap1.g00014.t01    rna-XM_021688076.2    rna-XM_004397255.1    rna-XM_042971464.1    rna-XM_040634531.1    rna-XM_035722230.1    mNeoVis.3.1.1.chr15.g00014.t01    rna-gnl|WGS_ZZZZ|egapxtmp_033909-R1    mNeoVis.evergladensis.2.1.chr15.g00015.t01    mNeoVis.vulgivaga.2.1.chr15.g00016.t01      2060    N0.HOG0002058   OG0000318  n0   rna-XM_002930722.4, rna-XM_002930726.2, rna-XM_002931318.2, rna-XM_011217582.2    rna-NM_001388816.1    rna-XM_022501563.1, rna-XM_022501914.1, rna-XM_022501920.1, rna-XM_022501921.1    rna-XM_003995848.3, rna-XM_004001198.4, rna-XM_006939443.4, rna-XM_019815553.3, rna-XM_045042722.1, rna-XM_045042723.1    rna-XM_036107739.1, rna-XM_036107910.1, rna-XM_036107911.1, rna-XM_036107912.1    rna-XM_039239028.1, rna-XM_039239029.1, rna-XM_039239030.1, rna-XM_039239031.1    rna-XM_032850789.1, rna-XM_032866358.1, rna-XM_032866359.1, rna-XM_032866371.1    rna-XM_046021804.1, rna-XM_046021805.1, rna-XM_046021806.1, rna-XM_046023459.1, rna-XM_046023517.1    rna-XM_032308514.1, rna-XM_032308515.1, rna-XM_032308516.1, rna-XM_032308848.1    rna-XM_059142587.1, rna-XM_059143080.1, rna-XM_059143081.1, rna-XM_059143083.1, rna-XM_059143084.1    rna-XM_059387373.1, rna-XM_059412149.1, rna-XM_059412150.1    rna-XM_004757433.1, rna-XM_004757434.1, rna-XM_004757435.2, rna-XM_045082913.1    mNeoVis.2.1.1.chr13_hap1.g00258.t01, mNeoVis.2.1.1.chr13_hap1.g00259.t01, mNeoVis.2.1.1.chr13_hap1.g00260.t01, mNeoVis.2.1.1.chr13_hap1.g00261.t01, mNeoVis.2.1.1.chr13_hap1.g00262.t01, mNeoVis.2.1.1.chr13_hap1.g00263.t01, mNeoVis.2.1.1.chr13_hap1.g00264.t01    rna-XM_021691579.1, rna-XM_021691580.1    rna-XM_004392619.1, rna-XM_004392641.1, rna-XM_004392642.1    rna-XM_007094527.2, rna-XM_007094528.2, rna-XM_007094532.2, rna-XM_042963498.1, rna-XM_042963499.1, rna-XM_042963500.1    rna-XM_008698923.1, rna-XM_008698924.2, rna-XM_008698925.2, rna-XM_008698926.2    rna-XM_027614696.1, rna-XM_027615828.1, rna-XM_027615829.1    mNeoVis.3.1.1.chr13.g00208.t01, mNeoVis.3.1.1.chr13.g00209.t01, mNeoVis.3.1.1.chr13.g00210.t01    rna-gnl|WGS_ZZZZ|egapxtmp_030164-R1, rna-gnl|WGS_ZZZZ|egapxtmp_030311-R1    mNeoVis.evergladensis.2.1.chr13.g00256.t01, mNeoVis.evergladensis.2.1.chr13.g00257.t01, mNeoVis.evergladensis.2.1.chr13.g00259.t01    mNeoVis.vulgivaga.2.1.chr13.g00252.t01, mNeoVis.vulgivaga.2.1.chr13.g00253.t01, mNeoVis.vulgivaga.2.1.chr13.g00254.t01      2061    N0.HOG0002059   OG0000319  n1   rna-XM_034654698.1    rna-NM_001143932.1    rna-XM_022495988.1    rna-NM_001287551.1    rna-XM_036072015.1    rna-XM_039248815.1    rna-XM_032880385.1    rna-XM_046019400.1    rna-XM_032355860.1    rna-XM_059167108.1    rna-XM_059394120.1    rna-XM_004744016.2    mNeoVis.2.1.1.chr3_hap1.g00914.t01    rna-XM_021702735.2    rna-XM_004396908.1    rna-XM_007074605.3    rna-XM_040636677.1    rna-XM_027590501.2    mNeoVis.3.1.1.chr3.g00897.t01    rna-gnl|WGS_ZZZZ|egapxtmp_040420-R1    mNeoVis.evergladensis.2.1.chr3.g00908.t01    mNeoVis.vulgivaga.2.1.chr3.g00924.t01      2062    N0.HOG0002060   OG0000319  n23   rna-XM_034653248.1    rna-XM_038557071.1    rna-XM_022503327.1    rna-XM_045052460.1    rna-XM_036113142.1    rna-XM_039238854.1    rna-XM_032865778.1    rna-XM_046020966.1    rna-XM_032304029.1    rna-XM_059171324.1    rna-XM_059395700.1    rna-XM_045089731.1    mNeoVis.2.1.1.chr4_hap1.g00422.t01    rna-XM_021689644.1    rna-XM_004416848.1    rna-XM_007089072.2    rna-XM_008686372.1    rna-XM_027573670.2    mNeoVis.3.1.1.chr6.g00422.t01    rna-gnl|WGS_ZZZZ|egapxtmp_002372-R3    mNeoVis.evergladensis.2.1.chr6.g00426.t01    mNeoVis.vulgivaga.2.1.chr6.g00420.t01      2063    N0.HOG0002061   OG0000319  n45   rna-XM_034667274.1    rna-XM_038556684.1    rna-XM_022515381.1    rna-XM_023250316.2    rna-XM_036080859.1    rna-XM_039219983.1    rna-XM_032876031.1      rna-XM_032304513.1    rna-XM_059171002.1    rna-XM_059396927.1    rna-XM_045089759.1    mNeoVis.2.1.1.chr4_hap1.g00522.t01    rna-XM_021689832.1    rna-XM_004411007.1        rna-XM_027573568.2    mNeoVis.3.1.1.chr6.g00529.t01    rna-gnl|WGS_ZZZZ|egapxtmp_004438-R1    mNeoVis.evergladensis.2.1.chr6.g00527.t01    mNeoVis.vulgivaga.2.1.chr6.g00530.t01      2064    N0.HOG0002062   OG0000319  n63   rna-XM_019809991.2    rna-NM_001003215.2    rna-XM_022515357.1    rna-NM_001171064.2    rna-XM_036080413.1    rna-XM_039219982.1      rna-XM_046020209.1    rna-XM_032304512.1    rna-XM_059171010.1    rna-XM_059396925.1    rna-XM_045089761.1    mNeoVis.2.1.1.chr4_hap1.g00521.t01    rna-XM_021689632.1    rna-XM_004411008.1    rna-XM_042965581.1    rna-XM_008686201.2    rna-XM_027573453.1    mNeoVis.3.1.1.chr6.g00528.t01    rna-gnl|WGS_ZZZZ|egapxtmp_004882-R1    mNeoVis.evergladensis.2.1.chr6.g00526.t01    mNeoVis.vulgivaga.2.1.chr6.g00529.t01      2065    N0.HOG0002063   OG0000320  n1   rna-XM_019807760.2    rna-XM_038561202.1    rna-XM_022513491.1    rna-XM_045053317.1, rna-XM_045054719.1    rna-XM_036119133.1    rna-XM_039225548.1, rna-XM_039225595.1, rna-XM_039225738.1    rna-XM_032881835.1    rna-XM_045979359.1    rna-XM_032353048.1    rna-XM_059188692.1    rna-XM_059405382.1    rna-XM_004777899.3    mNeoVis.2.1.1.chr11_hap1.g01015.t01    rna-XM_021697708.2    rna-XM_004409580.1    rna-XM_042982217.1, rna-XM_042982219.1    rna-XM_008700260.2    rna-XM_027620817.2    mNeoVis.3.1.1.chr11.g00966.t01    rna-gnl|WGS_ZZZZ|egapxtmp_025347-R1    mNeoVis.evergladensis.2.1.chr11.g00990.t01    mNeoVis.vulgivaga.2.1.chr11.g01013.t01      2066    N0.HOG0002064   OG0000320  n27   rna-XM_034654925.1      rna-XM_022512769.1      rna-XM_036098492.1      rna-XM_032860125.1    rna-XM_046019078.1    rna-XM_032354359.1    rna-XM_059168474.1    rna-XM_059393151.1    rna-XM_045078530.1    mNeoVis.2.1.1.chr3_hap1.g01300.t01    rna-XM_044913618.1    rna-XM_004393479.1    rna-XM_042993908.1    rna-XM_040642184.1    rna-XM_027589610.2    mNeoVis.3.1.1.chr3.g01276.t01    rna-gnl|WGS_ZZZZ|egapxtmp_041295-R1    mNeoVis.evergladensis.2.1.chr3.g01299.t01    mNeoVis.vulgivaga.2.1.chr3.g01319.t01      2067    N0.HOG0002065   OG0000320  n46         rna-XM_045033200.1      rna-XM_039217804.1                      rna-XM_015541013.2                  2068    N0.HOG0002066   OG0000320  n49   rna-XM_034641094.1    rna-NM_001044749.1    rna-XM_022493391.1      rna-XM_036098514.1      rna-XM_032860120.1    rna-XM_046018192.1    rna-XM_032354360.1    rna-XM_059168472.1    rna-XM_059392634.1    rna-XM_004781999.2    mNeoVis.2.1.1.chr3_hap1.g01298.t01    rna-XM_021702938.1        rna-XM_008687327.1    rna-XM_027591222.2    mNeoVis.3.1.1.chr3.g01274.t01    rna-gnl|WGS_ZZZZ|egapxtmp_042217-R1    mNeoVis.evergladensis.2.1.chr3.g01297.t01    mNeoVis.vulgivaga.2.1.chr3.g01317.t01      2069    N0.HOG0002067   OG0000320  n66   rna-XM_002915840.4    rna-NM_001048132.1    rna-XM_022496084.1      rna-XM_036098620.1      rna-XM_032860124.1    rna-XM_046019211.1    rna-XM_032354367.1    rna-XM_059168893.1    rna-XM_059393153.1    rna-XM_045078535.1    mNeoVis.2.1.1.chr3_hap1.g01297.t01    rna-XM_021703015.2        rna-XM_040636568.1    rna-XM_035726724.1    mNeoVis.3.1.1.chr3.g01273.t01    rna-gnl|WGS_ZZZZ|egapxtmp_042216-R1    mNeoVis.evergladensis.2.1.chr3.g01296.t01    mNeoVis.vulgivaga.2.1.chr3.g01316.t01      2070    N0.HOG0002068   OG0000321  n0   rna-XM_002919832.4    rna-XM_038557265.1    rna-XM_022523377.1    rna-XM_003997138.6    rna-XM_036065482.1, rna-XM_036088714.1, rna-XM_036091839.1, rna-XM_036097230.1, rna-XM_036105229.1, rna-XM_036113140.1, rna-XM_036123953.1    rna-XM_039226317.1    rna-XM_032843828.1, rna-XM_032844351.1, rna-XM_032849813.1, rna-XM_032849838.1, rna-XM_032849854.1, rna-XM_032853911.1, rna-XM_032870664.1, rna-XM_032881650.1, rna-XM_032883169.1, rna-XM_032883240.1    rna-XM_045979570.1, rna-XM_045979700.1, rna-XM_045985734.1, rna-XM_045986554.1, rna-XM_046003248.1, rna-XM_046006043.1    rna-XM_032314534.1, rna-XM_032315890.1, rna-XM_032322714.1, rna-XM_032329637.1, rna-XM_032331452.1, rna-XM_032352052.1    rna-XM_059147955.1, rna-XM_059181078.1    rna-XM_059372449.1, rna-XM_059379911.1, rna-XM_059395515.1    rna-XM_004748970.3, rna-XM_045066419.1, rna-XM_045083449.1, rna-XM_045083965.1    mNeoVis.2.1.1.chr1_hap1.g00812.t01, mNeoVis.2.1.1.chr8_hap1.g00161.t01, mNeoVis.2.1.1.chr9_hap1.g01110.t01    rna-XM_021681014.1, rna-XM_044915701.1, rna-XM_044918177.1    rna-XM_004394356.1, rna-XM_004401553.1, rna-XM_004407111.1, rna-XM_004408454.1, rna-XM_004409198.1    rna-XM_007091236.3    rna-XM_040640253.1    rna-XM_027625616.2, rna-XM_035721854.1, rna-XM_035724033.1, rna-XM_035724116.1, rna-XM_035724325.1, rna-XM_035724863.1, rna-XM_035725820.1, rna-XM_035726012.1, rna-XM_035726938.1, rna-XM_035727136.1, rna-XM_035727736.1, rna-XM_035728236.1, rna-XM_035728750.1, rna-XM_035728763.1, rna-XM_035729228.1    mNeoVis.3.1.1.chr1.g00757.t01, mNeoVis.3.1.1.chr8.g00130.t01, mNeoVis.3.1.1.chr9.g00257.t01    rna-gnl|WGS_ZZZZ|egapxtmp_020255-R2, rna-gnl|WGS_ZZZZ|egapxtmp_023268-R1, rna-gnl|WGS_ZZZZ|egapxtmp_023544-R1    mNeoVis.evergladensis.2.1.chr1.g00819.t01, mNeoVis.evergladensis.2.1.chr10.g01100.t01, mNeoVis.evergladensis.2.1.chr9.g00158.t01    mNeoVis.vulgivaga.2.1.chr1.g00834.t01, mNeoVis.vulgivaga.2.1.chr10.g01089.t01, mNeoVis.vulgivaga.2.1.chr8.g00161.t01      2071    N0.HOG0002069   OG0000322  n1   rna-XM_002931356.4, rna-XM_034653305.1    rna-NM_001003193.1    rna-XM_022501713.1    rna-XM_023242939.2    rna-XM_036108086.1    rna-XM_039229735.1    rna-XM_032866171.1    rna-XM_046022907.1    rna-XM_032306218.1    rna-XM_059143411.1    rna-XM_059410423.1    rna-XM_013058590.2    mNeoVis.2.1.1.chr13_hap1.g00118.t01    rna-XM_021691521.2    rna-XM_004392425.1    rna-XM_042964726.1    rna-XM_040632127.1    rna-XM_027616967.2        mNeoVis.evergladensis.2.1.chr13.g00115.t01    mNeoVis.vulgivaga.2.1.chr13.g00112.t01      2072    N0.HOG0002070   OG0000322  n22   rna-XM_011218383.1, rna-XM_019801402.2    rna-XM_038529397.1          rna-XM_039225701.1, rna-XM_039225702.1          rna-XM_059153287.1                rna-XM_040633033.1, rna-XM_040633034.1                2073    N0.HOG0002071   OG0000322  n29     rna-NM_001284436.1, rna-XM_038533793.1, rna-XM_038548818.1    rna-XM_022501440.1, rna-XM_022501441.1    rna-XM_045042762.1, rna-XM_045043266.1    rna-XM_036064899.1, rna-XM_036078423.1, rna-XM_036100647.1, rna-XM_036104392.1, rna-XM_036108149.1, rna-XM_036110155.1, rna-XM_036110370.1, rna-XM_036112051.1, rna-XM_036112052.1    rna-XM_039225842.1    rna-XM_032859381.1, rna-XM_032866173.1    rna-XM_046022900.1, rna-XM_046022916.1    rna-XM_032307303.1    rna-XM_059141572.1    rna-XM_059410421.1      mNeoVis.2.1.1.chr13_hap1.g00117.t01    rna-XM_021691553.1, rna-XM_044912398.1, rna-XM_044915247.1    rna-XM_004394981.1, rna-XM_004403260.1, rna-XM_004415196.1, rna-XM_004415582.1, rna-XM_004415692.1, rna-XM_004416423.1    rna-XM_042963385.1, rna-XM_042977131.1, rna-XM_042977134.1, rna-XM_042977136.1    rna-XM_008709455.2, rna-XM_040632129.1, rna-XM_040645500.1    rna-XM_035722333.1, rna-XM_035723713.1, rna-XM_035723743.1, rna-XM_035723784.1, rna-XM_035724431.1, rna-XM_035727088.1, rna-XM_035728081.1, rna-XM_035729346.1    mNeoVis.3.1.1.chr11.g01146.t01, mNeoVis.3.1.1.chr13.g00061.t01    rna-gnl|WGS_ZZZZ|egapxtmp_031242-R1    mNeoVis.evergladensis.2.1.chr13.g00114.t01    mNeoVis.vulgivaga.2.1.chr13.g00111.t01      2074   N0.HOG0002072  OG0000323  n1   rna-XM_002921201.4    rna-XM_038582058.1    rna-XM_022511321.1    rna-XM_003985150.5    rna-XM_036106581.1    rna-XM_039254245.1    rna-XM_032861160.1    rna-XM_045994634.1    rna-XM_032332971.1    rna-XM_059172465.1    rna-XM_059376347.1    rna-XM_004748191.3    mNeoVis.2.1.1.chr6_hap1.g00566.t01    rna-XM_021680385.1    rna-XM_004391483.1    rna-XM_007080973.3    rna-XM_008707190.2    rna-XM_027599819.1    mNeoVis.3.1.1.chr4.g00572.t01    rna-gnl|WGS_ZZZZ|egapxtmp_000301-R1    mNeoVis.evergladensis.2.1.chr4.g00570.t01    mNeoVis.vulgivaga.2.1.chr4.g00580.t01      2075   N0.HOG0002073  OG0000323  n22     rna-XM_038582458.1                            rna-XM_004417843.1                    2076    N0.HOG0002074   OG0000323  n23   rna-XM_002928003.4    rna-XM_038582055.1    rna-XM_022515242.1    rna-XM_003985148.6    rna-XM_036102699.1    rna-XM_039254247.1    rna-XM_032861146.1    rna-XM_045998197.1    rna-XM_032332975.1    rna-XM_059172430.1    rna-XM_059376358.1    rna-XM_004748187.3    mNeoVis.2.1.1.chr6_hap1.g00563.t01    rna-XM_021680383.1    rna-XM_004391592.1    rna-XM_007080970.3    rna-XM_008707193.2    rna-XM_027600749.2    mNeoVis.3.1.1.chr4.g00569.t01    rna-gnl|WGS_ZZZZ|egapxtmp_000304-R1    mNeoVis.evergladensis.2.1.chr4.g00567.t01    mNeoVis.vulgivaga.2.1.chr4.g00577.t01      2077    N0.HOG0002075   OG0000323  n42           rna-XM_036102678.1      rna-XM_032857919.1    rna-XM_045998594.1    rna-XM_032332974.1    rna-XM_059154891.1    rna-XM_059394684.1    rna-XM_004748190.1    mNeoVis.2.1.1.chr6_hap1.g00565.t01    rna-XM_021680404.1    rna-XM_004391481.1      rna-XM_008707191.1    rna-XM_027599210.1      rna-gnl|WGS_ZZZZ|egapxtmp_000793-R1    mNeoVis.evergladensis.2.1.chr4.g00569.t01    mNeoVis.vulgivaga.2.1.chr4.g00579.t01      2078    N0.HOG0002076   OG0000323  n56   rna-XM_002921200.4    rna-NM_001252155.1    rna-XM_022513033.1    rna-XM_003985151.3    rna-XM_036106579.1    rna-XM_039232161.1, rna-XM_039254234.1    rna-XM_032861174.1    rna-XM_045995089.1    rna-XM_032332973.1    rna-XM_059172486.1    rna-XM_059376330.1    rna-XM_013054058.2    mNeoVis.2.1.1.chr6_hap1.g00567.t01    rna-XM_021680386.2    rna-XM_004391484.1    rna-XM_007080974.3    rna-XM_008707189.2    rna-XM_027599817.1    mNeoVis.3.1.1.chr4.g00571.t01    rna-gnl|WGS_ZZZZ|egapxtmp_000303-R1    mNeoVis.evergladensis.2.1.chr4.g00571.t01        2079   N0.HOG0002077  OG0000324  n1   rna-XM_002918432.4    rna-XM_038538548.1    rna-XM_022511518.1    rna-XM_006941636.4    rna-XM_036115000.1    rna-XM_039223562.1    rna-XM_032854148.1    rna-XM_045987853.1    rna-XM_032326358.1    rna-XM_059153286.1    rna-XM_059381564.1    rna-XM_004744468.3    mNeoVis.2.1.1.chr7_hap1.g00207.t01    rna-XM_021703524.1    rna-XM_004393363.1    rna-XM_042970759.1    rna-XM_040633478.1    rna-XM_027619225.2      rna-gnl|WGS_ZZZZ|egapxtmp_010896-R1    mNeoVis.evergladensis.2.1.chr7.g00206.t01    mNeoVis.vulgivaga.2.1.chr7.g00207.t01      2080    N0.HOG0002078   OG0000324  n19   rna-XM_034672683.1      rna-XM_022511866.1    rna-XM_023246162.2    rna-XM_036088034.1    rna-XM_039256425.1    rna-XM_032853708.1    rna-XM_045986720.1    rna-XM_032324249.1    rna-XM_059153352.1    rna-XM_059387440.1    rna-XM_045076316.1    mNeoVis.2.1.1.chr7_hap1.g00124.t01    rna-XM_021705399.1    rna-XM_004417717.1    rna-XM_042968085.1    rna-XM_008689765.2      mNeoVis.3.1.1.chr7.g00107.t01      mNeoVis.evergladensis.2.1.chr7.g00126.t01    mNeoVis.vulgivaga.2.1.chr7.g00124.t01      2081    N0.HOG0002079   OG0000324  n38   rna-XM_019808444.2    rna-XM_038538311.1    rna-XM_022511641.1    rna-XM_023245390.2    rna-XM_036088033.1    rna-XM_039256431.1    rna-XM_032854619.1    rna-XM_045986722.1    rna-XM_032323107.1    rna-XM_059153354.1      rna-XM_045076310.1      rna-XM_044922147.1    rna-XM_004413072.1    rna-XM_042968873.1      rna-XM_027619551.1          mNeoVis.vulgivaga.2.1.chr7.g00126.t01      2082    N0.HOG0002080   OG0000324  n53     rna-NM_001197055.1    rna-XM_022511438.1    rna-XM_011290248.3    rna-XM_036088103.1    rna-XM_039256437.1    rna-XM_032854618.1    rna-XM_045988466.1    rna-XM_032325734.1, rna-XM_032325736.1    rna-XM_059151873.1, rna-XM_059152099.1, rna-XM_059152129.1, rna-XM_059152234.1    rna-XM_059387341.1, rna-XM_059387407.1    rna-XM_004753990.3, rna-XM_045076311.1, rna-XM_045076312.1    mNeoVis.2.1.1.chr7_hap1.g00125.t01, mNeoVis.2.1.1.chr7_hap1.g00126.t01    rna-XM_021705707.1    rna-XM_004413059.1    rna-XM_042969428.1    rna-XM_040633025.1    rna-XM_027618876.1          mNeoVis.vulgivaga.2.1.chr7.g00125.t01      2083    N0.HOG0002081   OG0000325  n1   rna-XM_034647080.1    rna-NM_001003249.1    rna-XM_022512555.1    rna-XM_023243369.2    rna-XM_036103226.1    rna-XM_039242176.1    rna-XM_032866683.1    rna-XM_046022777.1    rna-XM_032306308.1    rna-XM_059142054.1    rna-XM_059411816.1    rna-XM_045085365.1    mNeoVis.2.1.1.chr13_hap1.g00718.t01    rna-XM_021689365.2    rna-XM_004396715.1    rna-XM_042965339.1    rna-XM_040620227.1    rna-XM_027616175.2    mNeoVis.3.1.1.chr13.g00633.t01    rna-gnl|WGS_ZZZZ|egapxtmp_030891-R1    mNeoVis.evergladensis.2.1.chr13.g00681.t01    mNeoVis.vulgivaga.2.1.chr13.g00681.t01      2084    N0.HOG0002082   OG0000325  n8   rna-XM_002923556.4, rna-XM_034653008.1    rna-NM_001195842.1    rna-XM_022525875.1    rna-XM_023243621.2    rna-XM_036116124.1    rna-XM_039247632.1    rna-XM_032883518.1    rna-XM_045990058.1    rna-XM_032305792.1    rna-XM_059159421.1    rna-XM_059388607.1    rna-XM_045086908.1      rna-XM_021705614.2    rna-XM_004395367.1    rna-XM_042977718.1    rna-XM_040629629.1    rna-XM_027585856.2              2085    N0.HOG0002083   OG0000325  n18   rna-XM_002925446.4    rna-XM_038527036.1    rna-XM_022512556.1    rna-XM_003995421.6    rna-XM_036103224.1    rna-XM_039242164.1    rna-XM_032866723.1      rna-XM_032306309.1    rna-XM_059142061.1    rna-XM_059411817.1    rna-XM_004772134.3    mNeoVis.2.1.1.chr13_hap1.g00720.t01    rna-XM_021689376.1    rna-XM_004396716.1    rna-XM_042964070.1    rna-XM_040620228.1    rna-XM_027616830.2    mNeoVis.3.1.1.chr13.g00634.t01      mNeoVis.evergladensis.2.1.chr13.g00683.t01    mNeoVis.vulgivaga.2.1.chr13.g00682.t01      2086    N0.HOG0002084   OG0000325  n34   rna-XM_034657469.1    rna-XM_038567853.1    rna-XM_022525646.1    rna-XM_023243623.2    rna-XM_036116123.1    rna-XM_039247550.1    rna-XM_032883517.1    rna-XM_045991953.1    rna-XM_032305815.1    rna-XM_059159422.1    rna-XM_059388608.1    rna-XM_004771348.3    mNeoVis.2.1.1.chr5_hap1.g00085.t01    rna-XM_021705143.1    rna-XM_004395366.1    rna-XM_042977719.1    rna-XM_040629628.1    rna-XM_027587800.2    mNeoVis.3.1.1.chr2.g00042.t01    rna-gnl|WGS_ZZZZ|egapxtmp_034398-R1    mNeoVis.evergladensis.2.1.chr2.g00082.t01    mNeoVis.vulgivaga.2.1.chr2.g00079.t01      2087    N0.HOG0002085   OG0000326  n1   rna-XM_019805516.2    rna-XM_038576799.1    rna-XM_022494946.1    rna-XM_006933518.5    rna-XM_036076220.1    rna-XM_039222100.1    rna-XM_032855002.1    rna-XM_046013834.1, rna-XM_046014027.1    rna-XM_032347318.1    rna-XM_059186085.1    rna-XM_059402611.1    rna-XM_045073020.1    mNeoVis.2.1.1.chr9_hap1.g00819.t01    rna-XM_021690586.1    rna-XM_004411059.1    rna-XM_007076320.3    rna-XM_008706985.2    rna-XM_027594882.2    mNeoVis.3.1.1.chr10.g00783.t01    rna-gnl|WGS_ZZZZ|egapxtmp_023643-R1    mNeoVis.evergladensis.2.1.chr10.g00809.t01    mNeoVis.vulgivaga.2.1.chr10.g00797.t01      2088    N0.HOG0002086   OG0000326  n21   rna-XM_034649887.1    rna-XM_038588382.1, rna-XM_038588394.1    rna-XM_022526339.1, rna-XM_022526362.1    rna-XM_045050876.1, rna-XM_045050977.1    rna-XM_036080115.1, rna-XM_036083767.1, rna-XM_036096125.1    rna-XM_039225501.1, rna-XM_039225547.1, rna-XM_039239894.1    rna-XM_032849308.1, rna-XM_032849311.1    rna-XM_045994614.1, rna-XM_045994619.1    rna-XM_032331222.1, rna-XM_032331384.1    rna-XM_059158061.1, rna-XM_059158173.1    rna-XM_059385504.1, rna-XM_059386120.1    rna-XM_045068146.1, rna-XM_045068148.1    mNeoVis.2.1.1.chr12_hap1.g00159.t01, mNeoVis.2.1.1.chr12_hap1.g00163.t01      rna-XM_004416303.1    rna-XM_042974183.1, rna-XM_042975064.1    rna-XM_040627559.1, rna-XM_040641224.1, rna-XM_040641249.1    rna-XM_027607634.2, rna-XM_027609682.2, rna-XM_035725849.1    mNeoVis.3.1.1.chr12.g00127.t01    rna-gnl|WGS_ZZZZ|egapxtmp_028087-R1      mNeoVis.vulgivaga.2.1.chr12.g00136.t01      2089    N0.HOG0002087   OG0000326  n50   rna-XM_011221328.3    rna-XM_038541713.1      rna-XM_006944025.5    rna-XM_036102992.1      rna-XM_032872314.1      rna-XM_032328142.1      rna-XM_059414389.1      mNeoVis.2.1.1.chr3_hap1.g00389.t01    rna-XM_044912529.1    rna-XM_004394213.1, rna-XM_004394961.1    rna-XM_015542640.2        mNeoVis.3.1.1.chr15.g00401.t01, mNeoVis.3.1.1.chr3.g00381.t01    rna-gnl|WGS_ZZZZ|egapxtmp_042433-R1    mNeoVis.evergladensis.2.1.chr15.g00440.t01, mNeoVis.evergladensis.2.1.chr3.g00395.t01        2090    N0.HOG0002088   OG0000327  n1   rna-XM_034640309.1, rna-XM_034653104.1    rna-XM_038546661.1    rna-XM_022523468.1    rna-XM_019818212.3    rna-XM_036088610.1    rna-XM_039226392.1    rna-XM_032870686.1    rna-XM_045985309.1    rna-XM_032323011.1    rna-XM_059148757.1    rna-XM_059378985.1    rna-XM_004748999.3    mNeoVis.2.1.1.chr8_hap1.g00175.t01    rna-XM_021678476.1    rna-XM_004394403.1    rna-XM_042965498.1    rna-XM_040640271.1    rna-XM_027625883.2    mNeoVis.3.1.1.chr8.g00143.t01    rna-gnl|WGS_ZZZZ|egapxtmp_020262-R1    mNeoVis.evergladensis.2.1.chr9.g00172.t01    mNeoVis.vulgivaga.2.1.chr8.g00175.t01      2091    N0.HOG0002089   OG0000327  n24   rna-XM_034640645.1    rna-XM_038546679.1    rna-XM_022519835.1    rna-XM_006940554.3    rna-XM_036079019.1    rna-XM_039229751.1    rna-XM_032871051.1    rna-XM_045985850.1    rna-XM_032322265.1    rna-XM_059147452.1    rna-XM_059378606.1    rna-XM_004749019.3    mNeoVis.2.1.1.chr8_hap1.g00188.t01      rna-XM_004401847.1    rna-XM_007091220.3    rna-XM_008693718.2    rna-XM_035724713.1    mNeoVis.3.1.1.chr8.g00148.t01    rna-gnl|WGS_ZZZZ|egapxtmp_020269-R1    mNeoVis.evergladensis.2.1.chr9.g00180.t01    mNeoVis.vulgivaga.2.1.chr8.g00184.t01      2092    N0.HOG0002090   OG0000327  n45     rna-XM_038546676.1          rna-XM_039229831.1                                      2093    N0.HOG0002091   OG0000327  n46   rna-XM_019793805.2, rna-XM_034640307.1    rna-XM_038546086.1, rna-XM_038546087.1    rna-XM_022523508.1    rna-XM_006940557.4, rna-XM_011289341.3    rna-XM_036079001.1    rna-XM_039226266.1, rna-XM_039226523.1      rna-XM_045983789.1, rna-XM_045983790.1, rna-XM_045983962.1, rna-XM_045983963.1    rna-XM_032323034.1, rna-XM_032323035.1    rna-XM_059149312.1, rna-XM_059149314.1    rna-XM_059378434.1    rna-XM_004749005.3, rna-XM_045085950.1    mNeoVis.2.1.1.chr8_hap1.g00178.t01, mNeoVis.2.1.1.chr8_hap1.g00179.t01    rna-XM_021678474.1    rna-XM_004394377.1    rna-XM_007091222.2, rna-XM_007091290.3    rna-XM_008693725.2, rna-XM_040640296.1    rna-XM_027567484.1    mNeoVis.3.1.1.chr8.g00146.t01    rna-gnl|WGS_ZZZZ|egapxtmp_020266-R1, rna-gnl|WGS_ZZZZ|egapxtmp_020267-R1    mNeoVis.evergladensis.2.1.chr9.g00175.t01, mNeoVis.evergladensis.2.1.chr9.g00176.t01    mNeoVis.vulgivaga.2.1.chr8.g00178.t01      2094    N0.HOG0002092   OG0000328  n1   rna-XM_019805519.2, rna-XM_034667651.1    rna-XM_038541796.1, rna-XM_038541797.1    rna-XM_022508452.1    rna-XM_019821711.3    rna-XM_036070946.1    rna-XM_039257196.1    rna-XM_032862607.1    rna-XM_045983150.1            mNeoVis.2.1.1.chr14_hap1.g00181.t01    rna-XM_021682595.1      rna-XM_015543866.2    rna-XM_008697696.2    rna-XM_027612346.1    mNeoVis.3.1.1.chr14.g00179.t01    rna-gnl|WGS_ZZZZ|egapxtmp_032748-R1    mNeoVis.evergladensis.2.1.chr14.g00179.t01    mNeoVis.vulgivaga.2.1.chr14.g00182.t01      2095    N0.HOG0002093   OG0000328  n17   rna-XM_034650064.1, rna-XM_034650082.1, rna-XM_034667166.1    rna-XM_038586088.1    rna-XM_022508240.1, rna-XM_022508653.1, rna-XM_022508655.1    rna-XM_045048525.1, rna-XM_045048530.1, rna-XM_045048630.1    rna-XM_036084492.1, rna-XM_036084528.1    rna-XM_039217448.1, rna-XM_039225386.1, rna-XM_039225387.1, rna-XM_039225653.1, rna-XM_039225666.1, rna-XM_039225834.1, rna-XM_039227428.1    rna-XM_032851096.1, rna-XM_032851136.1, rna-XM_032872879.1, rna-XM_032872880.1, rna-XM_032878497.1    rna-XM_045982288.1, rna-XM_045982451.1, rna-XM_045983285.1, rna-XM_045983286.1    rna-XM_032317896.1, rna-XM_032317900.1, rna-XM_032317905.1, rna-XM_032317908.1    rna-XM_059145354.1, rna-XM_059145356.1, rna-XM_059145366.1, rna-XM_059145369.1    rna-XM_059413296.1, rna-XM_059413298.1, rna-XM_059413306.1    rna-XM_004756469.3, rna-XM_004756472.2, rna-XM_004756477.3, rna-XM_013058263.2    mNeoVis.2.1.1.chr14_hap1.g00404.t01, mNeoVis.2.1.1.chr14_hap1.g00405.t01    rna-XM_044916048.1, rna-XM_044916050.1    rna-XM_004392866.1, rna-XM_004404844.1    rna-XM_042975713.1, rna-XM_042976088.1    rna-XM_040636538.1    rna-XM_035722196.1, rna-XM_035722257.1    mNeoVis.3.1.1.chr14.g00403.t01, mNeoVis.3.1.1.chr14.g00405.t01    rna-gnl|WGS_ZZZZ|egapxtmp_031795-R1, rna-gnl|WGS_ZZZZ|egapxtmp_031914-R1    mNeoVis.evergladensis.2.1.chr14.g00404.t01, mNeoVis.evergladensis.2.1.chr14.g00406.t01    mNeoVis.vulgivaga.2.1.chr14.g00405.t01, mNeoVis.vulgivaga.2.1.chr14.g00406.t01      2096    N0.HOG0002094   OG0000329  n1   rna-XM_002916903.4    rna-XM_038579672.1    rna-XM_022502368.1    rna-XM_003999953.6    rna-XM_036072659.1    rna-XM_039223468.1    rna-XM_032869087.1    rna-XM_046019334.1    rna-XM_032316230.1    rna-XM_059166080.1    rna-XM_059387508.1, rna-XM_059394822.1    rna-XM_004766969.3    mNeoVis.2.1.1.chr4_hap1.g00843.t01    rna-XM_021688210.2    rna-XM_004393762.1    rna-XM_042973018.1    rna-XM_008689116.2    rna-XM_027617193.2    mNeoVis.3.1.1.chr6.g00852.t01    rna-gnl|WGS_ZZZZ|egapxtmp_003988-R1    mNeoVis.evergladensis.2.1.chr6.g00847.t01    mNeoVis.vulgivaga.2.1.chr6.g00842.t01      2097    N0.HOG0002095   OG0000329  n21   rna-XM_002912257.4    rna-XM_038530985.1    rna-XM_022518670.1    rna-XM_023245147.2    rna-XM_036068964.1    rna-XM_039232666.1    rna-XM_032854409.1    rna-XM_045989164.1    rna-XM_032325189.1    rna-XM_059153365.1    rna-XM_059383873.1    rna-XM_045076107.1    mNeoVis.2.1.1.chr7_hap1.g00302.t01    rna-XM_021705807.2    rna-XM_004395794.1    rna-XM_042968735.1    rna-XM_008702611.2    rna-XM_027619119.2, rna-XM_035727444.1    mNeoVis.3.1.1.chr7.g00288.t01    rna-gnl|WGS_ZZZZ|egapxtmp_010450-R1    mNeoVis.evergladensis.2.1.chr7.g00302.t01    mNeoVis.vulgivaga.2.1.chr7.g00305.t01      2098    N0.HOG0002096   OG0000329  n42   rna-XM_019796611.2, rna-XM_019796614.2    rna-NM_001171748.1, rna-NM_001171749.1    rna-XM_022494474.1, rna-XM_022494477.1    rna-XM_023251288.2    rna-XM_036084378.1    rna-XM_039227090.1    rna-XM_032853496.1, rna-XM_032858766.1    rna-XM_045981585.1    rna-XM_032350813.1    rna-XM_059133367.1    rna-XM_059406896.1    rna-XM_004746533.3, rna-XM_045074272.1, rna-XM_045084129.1    mNeoVis.2.1.1.chr11_hap1.g00278.t01    rna-XM_021688999.1    rna-XM_004393115.1, rna-XM_004393194.1    rna-XM_042980510.1    rna-XM_040638240.1, rna-XM_040638242.1    rna-XM_027623386.1, rna-XM_035723977.1, rna-XM_035727674.1    mNeoVis.3.1.1.chr11.g00248.t01, mNeoVis.3.1.1.chr11.g00249.t01    rna-gnl|WGS_ZZZZ|egapxtmp_025155-R1, rna-gnl|WGS_ZZZZ|egapxtmp_025520-R1    mNeoVis.evergladensis.2.1.chr11.g00275.t01    mNeoVis.vulgivaga.2.1.chr11.g00276.t01, mNeoVis.vulgivaga.2.1.chr11.g00277.t01      2099    N0.HOG0002097   OG0000330  n1   rna-XM_002923047.4, rna-XM_034648483.1    rna-XM_038525651.1    rna-XM_022522157.1, rna-XM_022524195.1    rna-XM_023255687.2    rna-XM_036090341.1, rna-XM_036109211.1    rna-XM_039256121.1    rna-XM_032839299.1    rna-XM_045987324.1, rna-XM_046009990.1    rna-XM_032326427.1, rna-XM_032344224.1    rna-XM_059153400.1, rna-XM_059182088.1    rna-XM_059373198.1, rna-XM_059382813.1    rna-XM_045077468.1    mNeoVis.2.1.1.chr10_hap1.g00406.t01, mNeoVis.2.1.1.chr7_hap1.g00988.t01    rna-XM_021681181.1, rna-XM_021701433.1    rna-XM_004400946.1, rna-XM_004408049.1    rna-XM_042969713.1, rna-XM_042989697.1    rna-XM_008701539.2, rna-XM_040633054.1    rna-XM_027571988.1, rna-XM_027620152.1    mNeoVis.3.1.1.chr7.g00927.t01, mNeoVis.3.1.1.chr9.g00374.t01      mNeoVis.evergladensis.2.1.chr7.g00981.t01, mNeoVis.evergladensis.2.1.chr8.g00388.t01    mNeoVis.vulgivaga.2.1.chr7.g00987.t01, mNeoVis.vulgivaga.2.1.chr9.g00390.t01      2100    N0.HOG0002098   OG0000330  n14   rna-XM_019806097.2    rna-XM_038564264.1    rna-XM_022506667.1    rna-XM_045038375.1, rna-XM_045039461.1    rna-XM_036122213.1    rna-XM_039257668.1    rna-XM_032879264.1    rna-XM_046015042.1    rna-XM_032357420.1    rna-XM_059144791.1    rna-XM_059403784.1      mNeoVis.2.1.1.chr7_hap1.g02316.t01    rna-XM_021684934.1    rna-XM_004413776.1    rna-XM_042957923.1, rna-XM_042959296.1    rna-XM_008707516.2    rna-XM_027578722.2    mNeoVis.3.1.1.chr7.g02132.t01      mNeoVis.evergladensis.2.1.chr7.g02263.t01    mNeoVis.vulgivaga.2.1.chr7.g02280.t01      2101    N0.HOG0002099   OG0000330  n17   rna-XM_034662481.1    rna-XM_038533847.1    rna-XM_022523649.1    rna-XM_019814292.3    rna-XM_036109755.1    rna-XM_039247193.1    rna-XM_032849791.1    rna-XM_046026145.1    rna-XM_032312160.1    rna-XM_059170400.1    rna-XM_059397276.1    rna-XM_013048576.2    mNeoVis.2.1.1.chr2_hap1.g00691.t01    rna-XM_021696198.1    rna-XM_004413832.1    rna-XM_007086040.3    rna-XM_008689568.1    rna-XM_027597194.2    mNeoVis.3.1.1.chr5.g00664.t01    rna-gnl|WGS_ZZZZ|egapxtmp_007380-R1    mNeoVis.evergladensis.2.1.chr5.g00673.t01    mNeoVis.vulgivaga.2.1.chr5.g00664.t01      2102    N0.HOG0002100   OG0000331  n1   rna-XM_034654394.1    rna-XM_038564915.1    rna-XM_022498289.1    rna-XM_023259241.2    rna-XM_036099019.1    rna-XM_039234906.1    rna-XM_032840916.1    rna-XM_046019519.1    rna-XM_032353901.1    rna-XM_059166610.1    rna-XM_059394475.1    rna-XM_004759777.3    mNeoVis.2.1.1.chr3_hap1.g00741.t01    rna-XM_021695872.1    rna-XM_004404732.1    rna-XM_042994229.1    rna-XM_040636877.1    rna-XM_027589892.1    mNeoVis.3.1.1.chr3.g00725.t01    rna-gnl|WGS_ZZZZ|egapxtmp_040911-R1    mNeoVis.evergladensis.2.1.chr3.g00741.t01    mNeoVis.vulgivaga.2.1.chr3.g00748.t01      2103    N0.HOG0002101   OG0000331  n23   rna-XM_034641389.1    rna-XM_038547442.1    rna-XM_022509493.1    rna-XM_019817814.3    rna-XM_036111982.1    rna-XM_039227784.1                  rna-XM_021704373.1    rna-XM_004395219.1    rna-XM_015539143.2      rna-XM_035724415.1              2104    N0.HOG0002102   OG0000331  n33   rna-XM_011227879.3    rna-XM_038577680.1    rna-XM_022519640.1    rna-XM_006933695.5    rna-XM_036098885.1    rna-XM_039241376.1    rna-XM_032876786.1    rna-XM_046012914.1    rna-XM_032348189.1    rna-XM_059185330.1    rna-XM_059403130.1    rna-XM_045072957.1    mNeoVis.2.1.1.chr9_hap1.g00550.t01    rna-XM_044914842.1    rna-XM_004400896.1    rna-XM_007073027.2    rna-XM_008704583.2    rna-XM_027594543.1    mNeoVis.3.1.1.chr10.g00519.t01    rna-gnl|WGS_ZZZZ|egapxtmp_022644-R1    mNeoVis.evergladensis.2.1.chr10.g00543.t01    mNeoVis.vulgivaga.2.1.chr10.g00529.t01      2105    N0.HOG0002103   OG0000331  n46   rna-XM_011224718.3    rna-XM_038570312.1    rna-XM_022521604.1    rna-XM_023260794.2    rna-XM_036085781.1    rna-XM_039253742.1    rna-XM_032861315.1    rna-XM_046001456.1    rna-XM_032302793.1    rna-XM_059162204.1    rna-XM_059390634.1    rna-XM_004774185.3    mNeoVis.2.1.1.chr5_hap1.g00972.t01, mNeoVis.2.1.1.chr8_hap1.g01164.t01    rna-XM_021677575.1    rna-XM_004408971.1    rna-XM_042956349.1    rna-XM_040637001.1    rna-XM_027587502.2    mNeoVis.3.1.1.chr2.g00890.t01, mNeoVis.3.1.1.chr8.g01079.t01    rna-gnl|WGS_ZZZZ|egapxtmp_019276-R1, rna-gnl|WGS_ZZZZ|egapxtmp_035499-R1    mNeoVis.evergladensis.2.1.chr2.g00968.t01, mNeoVis.evergladensis.2.1.chr9.g01136.t01    mNeoVis.vulgivaga.2.1.chr2.g00974.t01, mNeoVis.vulgivaga.2.1.chr8.g01136.t01      2106    N0.HOG0002104   OG0000332  n1   rna-XM_002930414.3    rna-XM_038569333.1    rna-XM_022514185.1    rna-XM_045038589.1    rna-XM_036075003.1    rna-XM_039254982.1    rna-XM_032853571.1    rna-XM_046017111.1    rna-XM_032359937.1    rna-XM_059186331.1    rna-XM_059412828.1    rna-XM_004754509.3    mNeoVis.2.1.1.chr7_hap1.g01436.t01    rna-XM_021679101.1    rna-XM_004410756.1    rna-XM_042958583.1    rna-XM_040630688.1    rna-XM_027580382.2    mNeoVis.3.1.1.chr7.g01343.t01, mNeoVis.3.1.1.chr8.g00668.t01      mNeoVis.evergladensis.2.1.chr7.g01418.t01    mNeoVis.vulgivaga.2.1.chr7.g01430.t01      2107    N0.HOG0002105   OG0000332  n22   rna-XM_034666428.1    rna-XM_038568124.1, rna-XM_038568768.1    rna-XM_022521926.1    rna-XM_003992769.6    rna-XM_036119436.1    rna-XM_039239595.1, rna-XM_039252043.1    rna-XM_032853203.1    rna-XM_046017003.1    rna-XM_032359555.1    rna-XM_059133652.1    rna-XM_059391170.1    rna-XM_013064361.2    mNeoVis.2.1.1.chr7_hap1.g01560.t01    rna-XM_021704630.1    rna-XM_004391736.1    rna-XM_015537017.2    rna-XM_040630810.1    rna-XM_027580875.1    mNeoVis.3.1.1.chr7.g01471.t01    rna-gnl|WGS_ZZZZ|egapxtmp_010256-R1      mNeoVis.vulgivaga.2.1.chr7.g01558.t01      2108    N0.HOG0002106   OG0000332  n44   rna-XM_034665603.1    rna-XM_038568764.1, rna-XM_038568793.1    rna-XM_022521925.1, rna-XM_022521933.1    rna-XM_023239466.2, rna-XM_045038786.1    rna-XM_036119435.1, rna-XM_036119441.1    rna-XM_039252045.1, rna-XM_039252047.1    rna-XM_032853197.1, rna-XM_032853556.1    rna-XM_046016090.1, rna-XM_046017001.1    rna-XM_032357488.1    rna-XM_059133633.1    rna-XM_059410099.1    rna-XM_004768141.3, rna-XM_004768220.2    mNeoVis.2.1.1.chr7_hap1.g01559.t01, mNeoVis.2.1.1.chr7_hap1.g01561.t01    rna-XM_021704629.1, rna-XM_021704650.2    rna-XM_004391737.1    rna-XM_007080893.3, rna-XM_007080895.3    rna-XM_008707305.2, rna-XM_008707413.2    rna-XM_027580877.2    mNeoVis.3.1.1.chr7.g01470.t01      mNeoVis.evergladensis.2.1.chr7.g01540.t01, mNeoVis.evergladensis.2.1.chr7.g01541.t01    mNeoVis.vulgivaga.2.1.chr7.g01557.t01, mNeoVis.vulgivaga.2.1.chr7.g01559.t01      2109    N0.HOG0002107   OG0000333  n2   rna-XM_034657373.1    rna-XM_038568002.1    rna-XM_022525750.1    rna-XM_045042770.1    rna-XM_036100679.1    rna-XM_039248072.1    rna-XM_032847006.1    rna-XM_045991301.1    rna-XM_032305065.1    rna-XM_059159214.1    rna-XM_059388475.1    rna-XM_045087831.1    mNeoVis.2.1.1.chr5_hap1.g00025.t01    rna-XM_021705364.1    rna-XM_004395413.1    rna-XM_042977544.1    rna-XM_040629679.1    rna-XM_027582073.1    mNeoVis.3.1.1.chr2.g00014.t01      mNeoVis.evergladensis.2.1.chr2.g00025.t01    mNeoVis.vulgivaga.2.1.chr2.g00025.t01      2110    N0.HOG0002108   OG0000333  n22   rna-XM_011218329.2    rna-XM_038588873.1, rna-XM_038589232.1, rna-XM_038589383.1, rna-XM_038589399.1, rna-XM_038589438.1, rna-XM_038589443.1, rna-XM_038589509.1, rna-XM_038589559.1, rna-XM_038589643.1    rna-XM_022492522.1    rna-XM_045050969.1, rna-XM_045051662.1        rna-XM_032850583.1    rna-XM_045996181.1, rna-XM_045996356.1    rna-XM_032329742.1, rna-XM_032331803.1    rna-XM_059157630.1, rna-XM_059157740.1    rna-XM_059385922.1    rna-XM_045068129.1, rna-XM_045068153.1    mNeoVis.2.1.1.chr12_hap1.g00074.t01    rna-XM_021692636.1, rna-XM_044912136.1    rna-XM_004417389.1    rna-XM_042975055.1, rna-XM_042975103.1    rna-XM_008709271.1    rna-XM_035725737.1, rna-XM_035725804.1    mNeoVis.3.1.1.chr12.g00064.t01    rna-gnl|WGS_ZZZZ|egapxtmp_027977-R1, rna-gnl|WGS_ZZZZ|egapxtmp_029113-R1    mNeoVis.evergladensis.2.1.chr12.g00063.t01    mNeoVis.vulgivaga.2.1.chr12.g00066.t01      2111    N0.HOG0002109   OG0000333  n56   rna-XM_034649514.1    rna-XM_038587979.1    rna-XM_022524909.1    rna-XM_023248910.2    rna-XM_036117283.1    rna-XM_039225220.1    rna-XM_032846179.1    rna-XM_045994523.1    rna-XM_032330307.1    rna-XM_059157965.1    rna-XM_059386236.1    rna-XM_013049668.2    mNeoVis.2.1.1.chr12_hap1.g00316.t01    rna-XM_021678081.1    rna-XM_004413869.1    rna-XM_042975020.1    rna-XM_008697225.2    rna-XM_027607511.2    mNeoVis.3.1.1.chr12.g00286.t01    rna-gnl|WGS_ZZZZ|egapxtmp_028129-R1    mNeoVis.evergladensis.2.1.chr12.g00282.t01    mNeoVis.vulgivaga.2.1.chr12.g00289.t01      2112    N0.HOG0002110   OG0000334  n1   rna-NM_001304877.1    rna-NM_001252147.1    rna-XM_022516663.1    rna-XM_003998831.4    rna-XM_036090486.1, rna-XM_036097659.1, rna-XM_036098954.1    rna-XM_039233187.1, rna-XM_039241640.1, rna-XM_039255896.1    rna-XM_032836218.1    rna-XM_045994346.1, rna-XM_046006027.1, rna-XM_046024518.1    rna-XM_032327717.1, rna-XM_032340138.1    rna-XM_059154283.1    rna-XM_059415240.1    rna-XM_004750316.3    mNeoVis.2.1.1.chr15_hap1.g00206.t01    rna-XM_021698350.1, rna-XM_044913811.1, rna-XM_044915898.1, rna-XM_044919523.1    rna-XM_004410314.1    rna-XM_007087318.3, rna-XM_015538703.2    rna-XM_040635269.1    rna-XM_027600352.1, rna-XM_027613375.1, rna-XM_035724318.1, rna-XM_035727008.1, rna-XM_035727141.1    mNeoVis.3.1.1.chr15.g00179.t01    rna-gnl|WGS_ZZZZ|egapxtmp_034205-R1    mNeoVis.evergladensis.2.1.chr15.g00185.t01    mNeoVis.vulgivaga.2.1.chr15.g00187.t01      2113    N0.HOG0002111   OG0000334  n14     rna-XM_038550379.1      rna-XM_045044676.1              rna-XM_059172667.1    rna-XM_059395649.1    rna-XM_045062892.1              rna-XM_035727749.1              2114    N0.HOG0002112   OG0000334  n19     rna-XM_038552666.1        rna-XM_036084467.1, rna-XM_036108431.1      rna-XM_032836713.1    rna-XM_046013537.1, rna-XM_046013959.1, rna-XM_046017380.1    rna-XM_032333092.1, rna-XM_032340918.1, rna-XM_032346550.1    rna-XM_059145665.1, rna-XM_059160625.1    rna-XM_059416250.1    rna-XM_045084851.1, rna-XM_045085571.1    mNeoVis.2.1.1.chr6_hap1.g00526.t01, mNeoVis.2.1.1.chr9_hap1.g01074.t01    rna-XM_021693029.1, rna-XM_044912648.1, rna-XM_044913378.1, rna-XM_044914508.1, rna-XM_044921195.1    rna-XM_004391805.1, rna-XM_004393321.1, rna-XM_004416933.1    rna-XM_042976480.1      rna-XM_035726919.1, rna-XM_035727726.1    mNeoVis.3.1.1.chr10.g01035.t01, mNeoVis.3.1.1.chr4.g00531.t01    rna-gnl|WGS_ZZZZ|egapxtmp_000783-R1, rna-gnl|WGS_ZZZZ|egapxtmp_023330-R1    mNeoVis.evergladensis.2.1.chr10.g01065.t01, mNeoVis.evergladensis.2.1.chr3.g01281.t01, mNeoVis.evergladensis.2.1.chr4.g00531.t01    mNeoVis.vulgivaga.2.1.chr10.g01055.t01, mNeoVis.vulgivaga.2.1.chr4.g00540.t01      2115    N0.HOG0002113   OG0000335  n2   rna-XM_034660570.1    rna-NM_001252389.1    rna-XM_022510699.1      rna-XM_036115797.1    rna-XM_039241809.1    rna-XM_032850490.1, rna-XM_032850496.1    rna-XM_046004427.1    rna-XM_032340591.1    rna-XM_059178595.1    rna-XM_059400054.1    rna-XM_004770653.3    mNeoVis.2.1.1.chr1_hap1.g01448.t01    rna-XM_021684655.2      rna-XM_042986150.1    rna-XM_040644740.1    rna-XM_027603477.2    mNeoVis.3.1.1.chr1.g01395.t01      mNeoVis.evergladensis.2.1.chr1.g01410.t01    mNeoVis.vulgivaga.2.1.chr1.g01414.t01      2116    N0.HOG0002114   OG0000335  n13   rna-XM_002931377.4    rna-XM_038587470.1, rna-XM_038587472.1, rna-XM_038589541.1, rna-XM_038589572.1    rna-XM_022508037.1    rna-XM_006943744.4, rna-XM_045050834.1    rna-XM_036112666.1, rna-XM_036118378.1    rna-XM_039225082.1, rna-XM_039225083.1, rna-XM_039225722.1    rna-XM_032842990.1    rna-XM_045995417.1, rna-XM_045995571.1    rna-XM_032331507.1, rna-XM_032331521.1    rna-XM_059157436.1, rna-XM_059158375.1    rna-XM_059387604.1    rna-XM_004755128.3, rna-XM_004755130.3    mNeoVis.2.1.1.chr12_hap1.g00578.t01, mNeoVis.2.1.1.chr12_hap1.g00581.t01    rna-XM_021678699.1, rna-XM_021678705.1    rna-XM_004416520.1, rna-XM_004416521.1    rna-XM_042974833.1, rna-XM_042974941.1    rna-XM_008709340.2    rna-XM_035725355.1, rna-XM_035725667.1      rna-gnl|WGS_ZZZZ|egapxtmp_029266-R1          2117    N0.HOG0002115   OG0000335  n36   rna-XM_034644422.1    rna-XM_038558639.1    rna-XM_022495267.1    rna-XM_006933953.5    rna-XM_036077434.1    rna-XM_039236603.1    rna-XM_032874215.1    rna-XM_046011454.1    rna-XM_032346778.1    rna-XM_059155207.1, rna-XM_059172345.1, rna-XM_059186511.1    rna-XM_059402356.1    rna-XM_004748548.3    mNeoVis.2.1.1.chr9_hap1.g01002.t01    rna-XM_044915356.1    rna-XM_004400283.1    rna-XM_042992888.1    rna-XM_040637314.1    rna-XM_027593650.2    mNeoVis.3.1.1.chr10.g00961.t01    rna-gnl|WGS_ZZZZ|egapxtmp_021731-R1    mNeoVis.evergladensis.2.1.chr10.g00989.t01    mNeoVis.vulgivaga.2.1.chr10.g00981.t01      2118    N0.HOG0002116   OG0000336  n0   rna-XM_002931201.3, rna-XM_034652676.1, rna-XM_034652763.1, rna-XM_034652764.1      rna-XM_022491027.1, rna-XM_022491462.1, rna-XM_022492673.1, rna-XM_022492833.1    rna-XM_003987357.3, rna-XM_003987363.5, rna-XM_023256172.2    rna-XM_036076878.1, rna-XM_036082557.1, rna-XM_036082594.1, rna-XM_036082596.1    rna-XM_039218330.1, rna-XM_039218348.1, rna-XM_039225476.1    rna-XM_032857135.1, rna-XM_032857136.1, rna-XM_032857139.1, rna-XM_032857144.1    rna-XM_046007093.1, rna-XM_046007233.1, rna-XM_046007344.1, rna-XM_046007843.1, rna-XM_046009578.1, rna-XM_046010102.1    rna-XM_032341683.1, rna-XM_032341774.1, rna-XM_032341776.1, rna-XM_032341777.1    rna-XM_059179627.1, rna-XM_059179671.1, rna-XM_059183396.1, rna-XM_059183398.1, rna-XM_059183399.1    rna-XM_059371129.1, rna-XM_059371138.1, rna-XM_059372102.1, rna-XM_059372103.1, rna-XM_059387384.1    rna-XM_004781930.3, rna-XM_004782097.2, rna-XM_004782099.1, rna-XM_004782100.2, rna-XM_013050784.2    mNeoVis.2.1.1.chr10_hap1.g00806.t01, mNeoVis.2.1.1.chr10_hap1.g00807.t01, mNeoVis.2.1.1.chr10_hap1.g00810.t01, mNeoVis.2.1.1.chr10_hap1.g00820.t01    rna-XM_021695516.1, rna-XM_021695542.1, rna-XM_044918283.1    rna-XM_004413512.1, rna-XM_004417619.1    rna-XM_007087460.1, rna-XM_007087490.2, rna-XM_015539947.2    rna-XM_008684022.2, rna-XM_008684026.2    rna-XM_027571495.2, rna-XM_027571631.2, rna-XM_027572117.1    mNeoVis.3.1.1.chr9.g00759.t01, mNeoVis.3.1.1.chr9.g00761.t01, mNeoVis.3.1.1.chr9.g00763.t01, mNeoVis.3.1.1.chr9.g00770.t01    rna-gnl|WGS_ZZZZ|egapxtmp_017333-R1, rna-gnl|WGS_ZZZZ|egapxtmp_017339-R1, rna-gnl|WGS_ZZZZ|egapxtmp_017346-R1, rna-gnl|WGS_ZZZZ|egapxtmp_017348-R1    mNeoVis.evergladensis.2.1.chr8.g00786.t01, mNeoVis.evergladensis.2.1.chr8.g00788.t01, mNeoVis.evergladensis.2.1.chr8.g00798.t01    mNeoVis.vulgivaga.2.1.chr9.g00784.t01, mNeoVis.vulgivaga.2.1.chr9.g00785.t01, mNeoVis.vulgivaga.2.1.chr9.g00787.t01, mNeoVis.vulgivaga.2.1.chr9.g00796.t01      2119    N0.HOG0002117   OG0000337  n1   rna-XM_019808212.2    rna-XM_038558831.1    rna-XM_022492520.1, rna-XM_022495037.1    rna-XM_023257371.2    rna-XM_036080804.1, rna-XM_036082017.1, rna-XM_036082022.1, rna-XM_036082025.1, rna-XM_036096686.1, rna-XM_036105923.1, rna-XM_036107291.1, rna-XM_036111197.1    rna-XM_039250241.1, rna-XM_039252504.1    rna-XM_032844298.1, rna-XM_032850799.1, rna-XM_032850800.1, rna-XM_032878075.1    rna-XM_045996270.1, rna-XM_045996283.1, rna-XM_045996288.1, rna-XM_046011475.1    rna-XM_032331617.1, rna-XM_032331637.1, rna-XM_032331744.1, rna-XM_032346603.1    rna-XM_059155790.1, rna-XM_059186666.1    rna-XM_059392862.1, rna-XM_059402247.1    rna-XM_004742980.3, rna-XM_004782374.2, rna-XM_045068076.1    mNeoVis.2.1.1.chr5_hap1.g00493.t01, mNeoVis.2.1.1.chr9_hap1.g01052.t01    rna-XM_021682411.2, rna-XM_021687561.1, rna-XM_021692643.2, rna-XM_044912161.1    rna-XM_004417300.1    rna-XM_042993020.1    rna-XM_008705707.2, rna-XM_008709237.2    rna-XM_027593758.1    mNeoVis.3.1.1.chr10.g01012.t01, mNeoVis.3.1.1.chr2.g00417.t01    rna-gnl|WGS_ZZZZ|egapxtmp_022103-R1    mNeoVis.evergladensis.2.1.chr10.g01042.t01, mNeoVis.evergladensis.2.1.chr2.g00488.t01    mNeoVis.vulgivaga.2.1.chr10.g01032.t01, mNeoVis.vulgivaga.2.1.chr2.g00490.t01      2120    N0.HOG0002118   OG0000337  n47     rna-XM_038588870.1, rna-XM_038589139.1, rna-XM_038589405.1, rna-XM_038589439.1, rna-XM_038589515.1    rna-XM_022492521.1    rna-XM_023249246.2, rna-XM_023249731.2, rna-XM_045051669.1, rna-XM_045051678.1    rna-XM_036082021.1, rna-XM_036105925.1      rna-XM_032850796.1    rna-XM_045996395.1, rna-XM_045996402.1, rna-XM_045996596.1    rna-XM_032331693.1    rna-XM_059156857.1    rna-XM_059387319.1    rna-XM_045068190.1    mNeoVis.2.1.1.chr12_hap1.g00073.t01    rna-XM_044911906.1      rna-XM_015543525.2, rna-XM_042974366.1    rna-XM_040634444.1      mNeoVis.3.1.1.chr12.g00063.t01        mNeoVis.vulgivaga.2.1.chr12.g00065.t01      2121    N0.HOG0002119   OG0000338  n0   rna-XM_034651558.1, rna-XM_034652040.1, rna-XM_034652382.1    rna-NM_001252165.1    rna-XM_022496317.1    rna-XM_003990035.5    rna-XM_036068593.1, rna-XM_036075754.1, rna-XM_036079229.1, rna-XM_036080913.1, rna-XM_036097014.1, rna-XM_036112663.1, rna-XM_036113092.1    rna-XM_039254781.1    rna-XM_032877631.1    rna-XM_045994836.1, rna-XM_046007871.1, rna-XM_046009501.1, rna-XM_046016719.1, rna-XM_046017164.1    rna-XM_032302394.1, rna-XM_032320629.1, rna-XM_032354374.1, rna-XM_032361160.1    rna-XM_059135757.1, rna-XM_059166112.1    rna-XM_059374579.1, rna-XM_059392907.1, rna-XM_059394734.1    rna-XM_013063756.1, rna-XM_045064443.1, rna-XM_045067086.1    mNeoVis.2.1.1.chr10_hap1.g01021.t01, mNeoVis.2.1.1.chr1_hap1.g00120.t01, mNeoVis.2.1.1.chr1_hap1.g01860.t01, mNeoVis.2.1.1.chr2_hap1.g01354.t01, mNeoVis.2.1.1.chr4_hap1.g00955.t01, mNeoVis.2.1.1.chr9_hap1.g00101.t01, mNeoVis.2.1.1.chr9_hap1.g01038.t01    rna-XM_021683771.1, rna-XM_044913185.1, rna-XM_044914015.1, rna-XM_044914281.1, rna-XM_044916950.1, rna-XM_044920418.1, rna-XM_044921366.1    rna-XM_004400598.1, rna-XM_004411218.1    rna-XM_007077274.3, rna-XM_007080334.3    rna-XM_008699284.2    rna-XM_027578668.2, rna-XM_035722328.1, rna-XM_035722332.1, rna-XM_035723299.1, rna-XM_035723332.1, rna-XM_035723961.1, rna-XM_035725254.1, rna-XM_035726924.1, rna-XM_035727211.1, rna-XM_035727247.1, rna-XM_035727750.1, rna-XM_035727965.1, rna-XM_035729353.1    mNeoVis.3.1.1.chr5.g01281.t01, mNeoVis.3.1.1.chr7.g00008.t01, mNeoVis.3.1.1.chr9.g00953.t01    rna-gnl|WGS_ZZZZ|egapxtmp_004525-R1, rna-gnl|WGS_ZZZZ|egapxtmp_007409-R1, rna-gnl|WGS_ZZZZ|egapxtmp_020652-R1, rna-gnl|WGS_ZZZZ|egapxtmp_023501-R1    mNeoVis.evergladensis.2.1.chr10.g01027.t01, mNeoVis.evergladensis.2.1.chr5.g01297.t01, mNeoVis.evergladensis.2.1.chr6.g00961.t01    mNeoVis.vulgivaga.2.1.chr1.g01822.t01, mNeoVis.vulgivaga.2.1.chr2.g01727.t01, mNeoVis.vulgivaga.2.1.chr5.g01294.t01, mNeoVis.vulgivaga.2.1.chr7.g00007.t01, mNeoVis.vulgivaga.2.1.chr9.g00979.t01      2122    N0.HOG0002120   OG0000339  n1   rna-XM_034654275.1    rna-XM_038581674.1    rna-XM_022516071.1    rna-XM_045036486.1    rna-XM_036099479.1    rna-XM_039257116.1    rna-XM_032865191.1    rna-XM_046002832.1    rna-XM_032357481.1    rna-XM_059164957.1    rna-XM_059392530.1    rna-XM_045084301.1    mNeoVis.2.1.1.chr5_hap1.g01916.t01    rna-XM_021679127.1    rna-XM_004410640.1    rna-XM_042957291.1    rna-XM_040621781.1    rna-XM_035726636.1    mNeoVis.3.1.1.chr2.g01802.t01    rna-gnl|WGS_ZZZZ|egapxtmp_036407-R2    mNeoVis.evergladensis.2.1.chr2.g01895.t01    mNeoVis.vulgivaga.2.1.chr2.g01911.t01      2123    N0.HOG0002121   OG0000339  n21   rna-XM_034659304.1    rna-XM_038549549.1, rna-XM_038586487.1    rna-XM_022504345.1    rna-XM_011281113.3    rna-XM_036088263.1    rna-XM_039239581.1    rna-XM_032873717.1    rna-XM_045978758.1    rna-XM_032341736.1, rna-XM_032352356.1    rna-XM_059187905.1    rna-XM_059405921.1    rna-XM_004742131.2    mNeoVis.2.1.1.chr11_hap1.g00773.t01    rna-XM_021699043.1    rna-XM_004398641.1    rna-XM_007083401.2    rna-XM_008699957.2    rna-XM_027620707.1    mNeoVis.3.1.1.chr11.g00737.t01      mNeoVis.evergladensis.2.1.chr11.g00755.t01    mNeoVis.vulgivaga.2.1.chr11.g00775.t01      2124   N0.HOG0002122  OG0000339  n28   rna-XM_034644791.1    rna-XM_038575761.1    rna-XM_022519609.1    rna-XM_019835000.2    rna-XM_036124123.1      rna-XM_032876633.1          rna-XM_059385414.1, rna-XM_059403255.1      mNeoVis.2.1.1.chr11_hap1.g01136.t01, mNeoVis.2.1.1.chr9_hap1.g00484.t01    rna-XM_044914337.1, rna-XM_044915392.1    rna-XM_004406213.1    rna-XM_042992402.1      rna-XM_035727050.1    mNeoVis.3.1.1.chr10.g00457.t01, mNeoVis.3.1.1.chr11.g01078.t01    rna-gnl|WGS_ZZZZ|egapxtmp_022997-R14, rna-gnl|WGS_ZZZZ|egapxtmp_027368-R1    mNeoVis.evergladensis.2.1.chr10.g00477.t01, mNeoVis.evergladensis.2.1.chr11.g01106.t01    mNeoVis.vulgivaga.2.1.chr10.g00466.t01, mNeoVis.vulgivaga.2.1.chr11.g01133.t01      2125   N0.HOG0002123  OG0000339  n47     rna-XM_038577795.1          rna-XM_039255051.1    rna-XM_032842074.1    rna-XM_046012433.1    rna-XM_032348400.1    rna-XM_059185164.1      rna-XM_045073294.1            rna-XM_040642533.1    rna-XM_027592155.2              2126    N0.HOG0002124   OG0000339  n49       rna-XM_022502854.1              rna-XM_032305990.1                                2127    N0.HOG0002125   OG0000340  n1   rna-XM_034645805.1    rna-XM_038562457.1    rna-XM_022507719.1    rna-XM_023239866.2    rna-XM_036110603.1    rna-XM_039245013.1    rna-XM_032879896.1    rna-XM_046015770.1    rna-XM_032359909.1    rna-XM_059141132.1    rna-XM_059390500.1    rna-XM_004770469.1    mNeoVis.2.1.1.chr7_hap1.g02238.t01    rna-XM_021685234.2    rna-XM_004417622.1    rna-XM_007092037.3    rna-XM_040639143.1    rna-XM_027581150.1    mNeoVis.3.1.1.chr7.g02057.t01    rna-gnl|WGS_ZZZZ|egapxtmp_013508-R1    mNeoVis.evergladensis.2.1.chr7.g02185.t01    mNeoVis.vulgivaga.2.1.chr7.g02205.t01      2128    N0.HOG0002126   OG0000340  n20   rna-XM_034645810.1    rna-XM_038562458.1    rna-XM_022507712.1    rna-XM_023239026.2    rna-XM_036110594.1    rna-XM_039245042.1      rna-XM_046015771.1    rna-XM_032359908.1    rna-XM_059141010.1    rna-XM_059391799.1    rna-XM_004770470.1      rna-XM_021684881.2      rna-XM_015541989.2    rna-XM_008694188.1    rna-XM_027581132.1          mNeoVis.vulgivaga.2.1.chr7.g02204.t01      2129   N0.HOG0002127  OG0000340  n35   rna-XM_034645811.1    rna-NM_001388574.1    rna-XM_022507744.1    rna-XM_045039346.1    rna-XM_036110611.1    rna-XM_039245015.1    rna-XM_032879038.1    rna-XM_046014291.1    rna-XM_032357031.1    rna-XM_059158994.1    rna-XM_059387040.1    rna-XM_004770437.1    mNeoVis.2.1.1.chr7_hap1.g02237.t01    rna-XM_021684880.1    rna-XM_004417621.1    rna-XM_007092038.2    rna-XM_008694184.2    rna-XM_027581155.1    mNeoVis.3.1.1.chr7.g02056.t01    rna-gnl|WGS_ZZZZ|egapxtmp_013558-R1    mNeoVis.evergladensis.2.1.chr7.g02184.t01    mNeoVis.vulgivaga.2.1.chr7.g02203.t01      2130    N0.HOG0002128   OG0000340  n53   rna-XM_002929662.4    rna-NM_001388575.1    rna-XM_022507438.1      rna-XM_036110555.1      rna-XM_032879895.1    rna-XM_046017686.1    rna-XM_032359907.1    rna-XM_059143310.1    rna-XM_059392764.1    rna-XM_004770438.3    mNeoVis.2.1.1.chr7_hap1.g02236.t01    rna-XM_021685233.1    rna-XM_004409324.1      rna-XM_008694183.1    rna-XM_027579358.2    mNeoVis.3.1.1.chr7.g02055.t01    rna-gnl|WGS_ZZZZ|egapxtmp_013487-R1    mNeoVis.evergladensis.2.1.chr7.g02183.t01    mNeoVis.vulgivaga.2.1.chr7.g02202.t01      2131    N0.HOG0002129   OG0000341  n3       rna-XM_022515195.1            rna-XM_046001852.1      rna-XM_059162488.1    rna-XM_059390857.1      mNeoVis.2.1.1.chr5_hap1.g01087.t01              mNeoVis.3.1.1.chr2.g00995.t01        mNeoVis.vulgivaga.2.1.chr2.g01084.t01      2132    N0.HOG0002130   OG0000341  n7     rna-XM_038559858.1        rna-XM_036111349.1      rna-XM_032860964.1      rna-XM_032329736.1        rna-XM_045088965.1        rna-XM_004404351.1        rna-XM_027600056.1, rna-XM_035724271.1              2133    N0.HOG0002131   OG0000341  n13   rna-XM_011229533.3    rna-XM_038570712.1      rna-XM_023260604.2      rna-XM_039254592.1                  rna-XM_021678914.2    rna-XM_004415586.1    rna-XM_042957131.1    rna-XM_040636866.1    rna-XM_027582469.2              2134   N0.HOG0002132  OG0000341  n15   rna-XM_034662067.1    rna-XM_038570719.1    rna-XM_022515002.1      rna-XM_036080339.1, rna-XM_036111438.1    rna-XM_039254577.1    rna-XM_032860953.1    rna-XM_046003183.1, rna-XM_046022971.1    rna-XM_032330531.1    rna-XM_059162497.1    rna-XM_059390860.1    rna-XM_045088962.1    mNeoVis.2.1.1.chr5_hap1.g01088.t01    rna-XM_044920568.1    rna-XM_004415583.1    rna-XM_042956335.1    rna-XM_040636863.1    rna-XM_027586093.1, rna-XM_035722801.1    mNeoVis.3.1.1.chr2.g00996.t01    rna-gnl|WGS_ZZZZ|egapxtmp_037245-R1    mNeoVis.evergladensis.2.1.chr2.g01078.t01    mNeoVis.vulgivaga.2.1.chr2.g01085.t01      2135   N0.HOG0002133  OG0000341  n31   rna-XM_034654813.1    rna-XM_038585017.1    rna-XM_022494756.1    rna-XM_023259524.2    rna-XM_036069545.1, rna-XM_036070133.1, rna-XM_036079912.1    rna-XM_039239643.1, rna-XM_039254350.1    rna-XM_032856048.1, rna-XM_032864541.1    rna-XM_046019848.1, rna-XM_046027305.1    rna-XM_032356113.1    rna-XM_059166870.1    rna-XM_059394277.1, rna-XM_059395486.1    rna-XM_045066649.1    mNeoVis.2.1.1.chr3_hap1.g00828.t01    rna-XM_021702810.2, rna-XM_044922262.1    rna-XM_004403744.1    rna-XM_042994819.1    rna-XM_040643272.1    rna-XM_027591006.2, rna-XM_035726993.1, rna-XM_035729154.1    mNeoVis.3.1.1.chr3.g00808.t01    rna-gnl|WGS_ZZZZ|egapxtmp_040166-R2    mNeoVis.evergladensis.2.1.chr3.g00821.t01    mNeoVis.vulgivaga.2.1.chr3.g00835.t01      2136    N0.HOG0002134   OG0000342  n3     rna-XM_038582854.1                          rna-XM_044921634.1    rna-XM_004395045.1        rna-XM_027594367.1              2137    N0.HOG0002135   OG0000342  n6   rna-XM_034654224.1    rna-XM_038573057.1        rna-XM_036073066.1, rna-XM_036109831.1      rna-XM_032841446.1    rna-XM_045998137.1, rna-XM_046000074.1    rna-XM_032303034.1, rna-XM_032330595.1, rna-XM_032333748.1    rna-XM_059136058.1    rna-XM_059376583.1      mNeoVis.2.1.1.chr2_hap1.g01307.t01    rna-XM_044917377.1, rna-XM_044922265.1    rna-XM_004412422.1        rna-XM_027574420.1, rna-XM_027600918.2    mNeoVis.3.1.1.chr12.g00109.t01, mNeoVis.3.1.1.chr3.g00944.t01, mNeoVis.3.1.1.chr5.g01232.t01    rna-gnl|WGS_ZZZZ|egapxtmp_006394-R1, rna-gnl|WGS_ZZZZ|egapxtmp_029060-R1      mNeoVis.vulgivaga.2.1.chr5.g01244.t01      2138    N0.HOG0002136   OG0000342  n31           rna-XM_036091689.1              rna-XM_059404967.1                            2139   N0.HOG0002137  OG0000342  n34   rna-NM_001304860.1    rna-XM_038559013.1    rna-XM_022518125.1    rna-XM_003984944.4    rna-XM_036089202.1    rna-XM_039234371.1    rna-XM_032855536.1    rna-XM_045995151.1    rna-XM_032332508.1    rna-XM_059175606.1      rna-XM_004758048.3    mNeoVis.2.1.1.chr6_hap1.g00709.t01    rna-XM_021699007.1    rna-XM_004402419.1    rna-XM_007084173.3    rna-XM_008685470.2    rna-XM_027598773.1      rna-gnl|WGS_ZZZZ|egapxtmp_044440-R1    mNeoVis.evergladensis.2.1.chr4.g00716.t01    mNeoVis.vulgivaga.2.1.chr4.g00732.t01      2140    N0.HOG0002138   OG0000342  n35   rna-XM_034652117.1    rna-XM_038545873.1, rna-XM_038586490.1      rna-XM_045059072.1    rna-XM_036110760.1, rna-XM_036117105.1      rna-XM_032852854.1        rna-XM_059165985.1    rna-XM_059373651.1, rna-XM_059395701.1    rna-XM_045075331.1    mNeoVis.2.1.1.chr5_hap1.g01338.t01      rna-XM_004410993.1        rna-XM_035726593.1, rna-XM_035726789.1, rna-XM_035728982.1    mNeoVis.3.1.1.chr4.g00719.t01    rna-gnl|WGS_ZZZZ|egapxtmp_038434-R1, rna-gnl|WGS_ZZZZ|egapxtmp_041041-R1    mNeoVis.evergladensis.2.1.chr2.g01324.t01, mNeoVis.evergladensis.2.1.chr3.g00234.t01    mNeoVis.vulgivaga.2.1.chr2.g01332.t01      2141    N0.HOG0002139   OG0000342  n56     rna-XM_038570186.1        rna-XM_036102024.1    rna-XM_039225546.1                  rna-XM_044918798.1                      2142    N0.HOG0002140   OG0000343  n1   rna-XM_011230559.3    rna-XM_038564731.1    rna-XM_022516570.1    rna-XM_019829000.3    rna-XM_036069426.1    rna-XM_039241603.1    rna-XM_032858250.1    rna-XM_045997558.1    rna-XM_032333347.1    rna-XM_059169017.1    rna-XM_059395460.1    rna-XM_004747966.2    mNeoVis.2.1.1.chr6_hap1.g00450.t01    rna-XM_021681478.1    rna-XM_004401577.1    rna-XM_007087588.3    rna-XM_008689491.2    rna-XM_027599071.2    mNeoVis.3.1.1.chr4.g00452.t01    rna-gnl|WGS_ZZZZ|egapxtmp_043467-R1    mNeoVis.evergladensis.2.1.chr4.g00456.t01    mNeoVis.vulgivaga.2.1.chr4.g00462.t01      2143    N0.HOG0002141   OG0000343  n23   rna-XM_002922620.3    rna-XM_038543700.1    rna-XM_022495871.1    rna-XM_003995056.5    rna-XM_036082049.1    rna-XM_039218222.1    rna-XM_032859836.1    rna-XM_046025840.1    rna-XM_032310229.1    rna-XM_059140341.1    rna-XM_059410030.1    rna-XM_004742806.3    mNeoVis.2.1.1.chr3_hap1.g00224.t01    rna-XM_021686238.1    rna-XM_004411204.1    rna-XM_007075942.2    rna-XM_008690510.2    rna-XM_027576525.1    mNeoVis.3.1.1.chr3.g00226.t01    rna-gnl|WGS_ZZZZ|egapxtmp_040745-R1    mNeoVis.evergladensis.2.1.chr3.g00235.t01    mNeoVis.vulgivaga.2.1.chr3.g00227.t01      2144    N0.HOG0002142   OG0000343  n36   rna-XM_011216812.3    rna-XM_038586651.1    rna-XM_022491293.1    rna-XM_004000983.5    rna-XM_036088404.1, rna-XM_036120835.1    rna-XM_039220701.1    rna-XM_032845247.1, rna-XM_032878071.1    rna-XM_045994623.1    rna-XM_032330589.1    rna-XM_059157519.1    rna-XM_059386246.1    rna-XM_004776658.3    mNeoVis.2.1.1.chr10_hap1.g00979.t01, mNeoVis.2.1.1.chr12_hap1.g00080.t01, mNeoVis.2.1.1.chr9_hap1.g00313.t01    rna-XM_021683547.1, rna-XM_044918852.1    rna-XM_004397538.1    rna-XM_007088978.3    rna-XM_040627189.1    rna-XM_027574312.2, rna-XM_027608259.2, rna-XM_035723703.1    mNeoVis.3.1.1.chr12.g00067.t01, mNeoVis.3.1.1.chr9.g00910.t01    rna-gnl|WGS_ZZZZ|egapxtmp_015740-R1, rna-gnl|WGS_ZZZZ|egapxtmp_028009-R1    mNeoVis.evergladensis.2.1.chr10.g00310.t01, mNeoVis.evergladensis.2.1.chr12.g00067.t01, mNeoVis.evergladensis.2.1.chr8.g00950.t01    mNeoVis.vulgivaga.2.1.chr12.g00070.t01, mNeoVis.vulgivaga.2.1.chr9.g00935.t01      2145    N0.HOG0002143   OG0000344  n1   rna-XM_002915618.2    rna-NM_001388877.1    rna-XM_022509712.1    rna-XM_003996640.4    rna-XM_036084601.1    rna-XM_039241170.1    rna-XM_032871764.1    rna-XM_045985902.1    rna-XM_032321331.1    rna-XM_059149221.1    rna-XM_059379085.1    rna-XM_004747323.2    mNeoVis.2.1.1.chr8_hap1.g00626.t01      rna-XM_004397627.1    rna-XM_007086711.1    rna-XM_008695511.1    rna-XM_027624670.2    mNeoVis.3.1.1.chr8.g00578.t01    rna-gnl|WGS_ZZZZ|egapxtmp_017727-R1    mNeoVis.evergladensis.2.1.chr9.g00617.t01    mNeoVis.vulgivaga.2.1.chr8.g00623.t01      2146    N0.HOG0002144   OG0000344  n21   rna-XM_002929402.2, rna-XM_011220260.2, rna-XM_034639720.1, rna-XM_034650748.1    rna-NM_001388809.1, rna-NM_001388810.1, rna-XM_038546150.1    rna-XM_022509717.1, rna-XM_022509723.1, rna-XM_022509739.1    rna-XM_003996638.4, rna-XM_045044722.1    rna-XM_036082468.1, rna-XM_036083582.1, rna-XM_036096265.1    rna-XM_039241176.1, rna-XM_039241177.1    rna-XM_032871763.1, rna-XM_032871803.1    rna-XM_045985903.1, rna-XM_045985957.1, rna-XM_045986001.1    rna-XM_032321330.1, rna-XM_032322231.1, rna-XM_032322277.1    rna-XM_059149096.1, rna-XM_059149170.1, rna-XM_059149222.1    rna-XM_059379087.1, rna-XM_059379125.1    rna-XM_004747322.3, rna-XM_004747399.3, rna-XM_004747400.3    mNeoVis.2.1.1.chr8_hap1.g00627.t01, mNeoVis.2.1.1.chr8_hap1.g00628.t01, mNeoVis.2.1.1.chr8_hap1.g00629.t01    rna-XM_044921825.1    rna-XM_004397701.1, rna-XM_004417689.1    rna-XM_007086712.3, rna-XM_007086713.3    rna-XM_008695510.2, rna-XM_008695612.1, rna-XM_008695613.2, rna-XM_008695614.2    rna-XM_027567677.2, rna-XM_035724412.1, rna-XM_035724413.1, rna-XM_035724709.1    mNeoVis.3.1.1.chr7.g02052.t01, mNeoVis.3.1.1.chr8.g00579.t01, mNeoVis.3.1.1.chr8.g00580.t01    rna-gnl|WGS_ZZZZ|egapxtmp_017726-R1    mNeoVis.evergladensis.2.1.chr9.g00618.t01, mNeoVis.evergladensis.2.1.chr9.g00619.t01    mNeoVis.vulgivaga.2.1.chr8.g00624.t01, mNeoVis.vulgivaga.2.1.chr8.g00625.t01      2147    N0.HOG0002145   OG0000345  n1   rna-XM_034667660.1    rna-XM_038541856.1    rna-XM_022508684.1    rna-XM_023247926.2    rna-XM_036079614.1, rna-XM_036092962.1    rna-XM_039227457.1    rna-XM_032878288.1    rna-XM_045982822.1    rna-XM_032317308.1    rna-XM_059147051.1    rna-XM_059413120.1    rna-XM_004756318.3    mNeoVis.2.1.1.chr14_hap1.g00340.t01    rna-XM_021682708.2    rna-XM_004392773.1    rna-XM_042976879.1    rna-XM_040636465.1    rna-XM_027610957.2    mNeoVis.3.1.1.chr14.g00339.t01    rna-gnl|WGS_ZZZZ|egapxtmp_031898-R1    mNeoVis.evergladensis.2.1.chr14.g00341.t01    mNeoVis.vulgivaga.2.1.chr14.g00342.t01      2148    N0.HOG0002146   OG0000345  n23   rna-XM_002914106.4    rna-XM_038576395.1    rna-XM_022522216.1    rna-XM_006933402.5    rna-XM_036092409.1    rna-XM_039236294.1    rna-XM_032874917.1    rna-XM_046011476.1    rna-XM_032349171.1    rna-XM_059184463.1    rna-XM_059403849.1    rna-XM_013046431.2    mNeoVis.2.1.1.chr9_hap1.g00208.t01    rna-XM_021690804.2    rna-XM_004411942.1    rna-XM_042991539.1    rna-XM_008704845.2    rna-XM_027592596.2    mNeoVis.3.1.1.chr10.g00198.t01    rna-gnl|WGS_ZZZZ|egapxtmp_023432-R1    mNeoVis.evergladensis.2.1.chr10.g00204.t01    mNeoVis.vulgivaga.2.1.chr10.g00201.t01      2149    N0.HOG0002147   OG0000345  n43   rna-XM_011234257.3    rna-NM_001048032.1    rna-XM_022492193.1, rna-XM_022492358.1    rna-XM_023249335.2    rna-XM_036102777.1, rna-XM_036114902.1    rna-XM_039220988.1    rna-XM_032847968.1    rna-XM_045994940.1, rna-XM_045996660.1    rna-XM_032329679.1, rna-XM_032331916.1    rna-XM_059158392.1    rna-XM_059385713.1, rna-XM_059386417.1    rna-XM_004781211.3, rna-XM_045087438.1    mNeoVis.2.1.1.chr12_hap1.g00533.t01    rna-XM_021702688.2, rna-XM_021703000.2    rna-XM_004406957.1, rna-XM_004416058.1    rna-XM_042973807.1    rna-XM_040631376.1, rna-XM_040631386.1, rna-XM_040636012.1    rna-XM_027607892.2, rna-XM_035725916.1    mNeoVis.3.1.1.chr12.g00484.t01    rna-gnl|WGS_ZZZZ|egapxtmp_028379-R1    mNeoVis.evergladensis.2.1.chr12.g00477.t01    mNeoVis.vulgivaga.2.1.chr12.g00489.t01      2150    N0.HOG0002148   OG0000346  n1   rna-XM_011231800.3    rna-XM_038570961.1    rna-XM_022514974.1    rna-XM_023260539.2    rna-XM_036080958.1, rna-XM_036120530.1    rna-XM_039222707.1    rna-XM_032869634.1    rna-XM_046002311.1    rna-XM_032333876.1    rna-XM_059162687.1    rna-XM_059390976.1    rna-XM_013061415.2    mNeoVis.2.1.1.chr5_hap1.g01156.t01    rna-XM_021678881.2    rna-XM_004414861.1    rna-XM_042998577.1    rna-XM_040636781.1    rna-XM_027584268.2    mNeoVis.3.1.1.chr2.g01067.t01    rna-gnl|WGS_ZZZZ|egapxtmp_037065-R1    mNeoVis.evergladensis.2.1.chr2.g01146.t01    mNeoVis.vulgivaga.2.1.chr2.g01153.t01      2151    N0.HOG0002149   OG0000346  n22   rna-XM_034648821.1, rna-XM_034649878.1    rna-XM_038586701.1    rna-XM_022518317.1    rna-XM_045051187.1    rna-XM_036088917.1, rna-XM_036096464.1    rna-XM_039245305.1, rna-XM_039245306.1    rna-XM_032880260.1    rna-XM_045995152.1, rna-XM_045996625.1    rna-XM_032331702.1    rna-XM_059157744.1    rna-XM_059385881.1, rna-XM_059387305.1, rna-XM_059387314.1    rna-XM_045089233.1    mNeoVis.2.1.1.chr12_hap1.g00850.t01    rna-XM_021690237.2, rna-XM_044911642.1, rna-XM_044912505.1    rna-XM_004399992.1, rna-XM_004399993.1    rna-XM_042974463.1, rna-XM_042975207.1    rna-XM_040635171.1, rna-XM_040638732.1    rna-XM_035725502.1    mNeoVis.3.1.1.chr12.g00779.t01    rna-gnl|WGS_ZZZZ|egapxtmp_028284-R1    mNeoVis.evergladensis.2.1.chr12.g00778.t01    mNeoVis.vulgivaga.2.1.chr12.g00807.t01      2152    N0.HOG0002150   OG0000346  n54   rna-XM_002922296.4    rna-XM_038588251.1    rna-XM_022518498.1    rna-XM_011279456.4      rna-XM_039238809.1    rna-XM_032882337.1    rna-XM_045994926.1    rna-XM_032329452.1    rna-XM_059158632.1    rna-XM_059385182.1    rna-XM_004762025.2    mNeoVis.2.1.1.chr12_hap1.g00216.t01    rna-XM_021686638.1    rna-XM_004400193.1    rna-XM_042973900.1    rna-XM_040628264.1    rna-XM_035725493.1    mNeoVis.3.1.1.chr12.g00186.t01    rna-gnl|WGS_ZZZZ|egapxtmp_029533-R8    mNeoVis.evergladensis.2.1.chr12.g00189.t01    mNeoVis.vulgivaga.2.1.chr12.g00193.t01      2153    N0.HOG0002151   OG0000347  n1   rna-XM_019793421.2, rna-XM_034655267.1    rna-XM_038539213.1    rna-XM_022491638.1, rna-XM_022501515.1    rna-XM_003998527.4, rna-XM_011285654.3    rna-XM_036078817.1, rna-XM_036098711.1    rna-XM_039236836.1, rna-XM_039243461.1    rna-XM_032857340.1, rna-XM_032872524.1    rna-XM_045994203.1, rna-XM_046018808.1    rna-XM_032327234.1, rna-XM_032354972.1    rna-XM_059155686.1, rna-XM_059167901.1    rna-XM_059393586.1, rna-XM_059414602.1    rna-XM_004761583.3, rna-XM_004762908.3    mNeoVis.2.1.1.chr15_hap1.g00693.t01, mNeoVis.2.1.1.chr3_hap1.g01136.t01    rna-XM_021682993.1, rna-XM_021692439.1    rna-XM_004399053.1, rna-XM_004407471.1    rna-XM_042971386.1, rna-XM_042995461.1    rna-XM_040634884.1, rna-XM_040639845.1    rna-XM_027589695.2, rna-XM_027612418.2    mNeoVis.3.1.1.chr15.g00474.t01, mNeoVis.3.1.1.chr3.g01124.t01    rna-gnl|WGS_ZZZZ|egapxtmp_034163-R1, rna-gnl|WGS_ZZZZ|egapxtmp_042210-R1    mNeoVis.evergladensis.2.1.chr15.g00515.t01, mNeoVis.evergladensis.2.1.chr3.g01135.t01    mNeoVis.vulgivaga.2.1.chr3.g01150.t01      2154    N0.HOG0002152   OG0000347  n23   rna-XM_034649638.1    rna-XM_038586889.1    rna-XM_022491719.1, rna-XM_022518333.1    rna-XM_023248821.2    rna-XM_036080337.1    rna-XM_039233939.1    rna-XM_032840864.1, rna-XM_032843483.1    rna-XM_045995169.1, rna-XM_045996751.1    rna-XM_032330079.1, rna-XM_032331967.1, rna-XM_032347109.1    rna-XM_059158599.1    rna-XM_059384583.1    rna-XM_045089324.1    mNeoVis.2.1.1.chr12_hap1.g00778.t01, mNeoVis.2.1.1.chr2_hap1.g00627.t01    rna-XM_044911972.1    rna-XM_004412030.1, rna-XM_004413876.1, rna-XM_004415695.1    rna-XM_042973711.1    rna-XM_008683968.2, rna-XM_040634059.1    rna-XM_027608721.2, rna-XM_035725874.1    mNeoVis.3.1.1.chr12.g00710.t01, mNeoVis.3.1.1.chr5.g00625.t01    rna-gnl|WGS_ZZZZ|egapxtmp_006365-R1, rna-gnl|WGS_ZZZZ|egapxtmp_029514-R1    mNeoVis.evergladensis.2.1.chr12.g00705.t01, mNeoVis.evergladensis.2.1.chr5.g00630.t01    mNeoVis.vulgivaga.2.1.chr12.g00734.t01, mNeoVis.vulgivaga.2.1.chr5.g00624.t01      2155    N0.HOG0002153   OG0000348  n4   rna-XM_034655695.1, rna-XM_034663272.1    rna-XM_038532782.1    rna-XM_022492964.1, rna-XM_022493398.1      rna-XM_036068457.1, rna-XM_036111186.1    rna-XM_039244265.1    rna-XM_032851977.1, rna-XM_032851997.1    rna-XM_046000297.1              rna-XM_044916934.1        rna-XM_008692758.1    rna-XM_035727734.1              2156    N0.HOG0002154   OG0000348  n17     rna-XM_038536230.1        rna-XM_036091048.1    rna-XM_039221200.1                  rna-XM_021686367.2          rna-XM_035727618.1              2157    N0.HOG0002155   OG0000348  n21   rna-XM_011232680.3, rna-XM_034651532.1    rna-NM_001252042.1    rna-XM_022494047.1, rna-XM_022524424.1    rna-XM_004000627.5    rna-XM_036087371.1, rna-XM_036087998.1    rna-XM_039219869.1, rna-XM_039234683.1    rna-XM_032843152.1, rna-XM_032848812.1, rna-XM_032852137.1    rna-XM_045996370.1, rna-XM_045996579.1, rna-XM_045998098.1, rna-XM_045998267.1    rna-XM_032329477.1, rna-XM_032330898.1, rna-XM_032333320.1, rna-XM_032342458.1    rna-XM_059162487.1, rna-XM_059162630.1, rna-XM_059170870.1, rna-XM_059173679.1, rna-XM_059180403.1    rna-XM_059371839.1, rna-XM_059377322.1, rna-XM_059384944.1, rna-XM_059387962.1, rna-XM_059392594.1, rna-XM_059404110.1    rna-XM_004743022.3, rna-XM_004777255.3, rna-XM_013061410.2, rna-XM_045070442.1, rna-XM_045085542.1, rna-XM_045088967.1    mNeoVis.2.1.1.chr12_hap1.g00467.t01, mNeoVis.2.1.1.chr4_hap1.g00906.t01    rna-XM_021680682.1    rna-XM_004396441.1, rna-XM_004413673.1    rna-XM_042974900.1    rna-XM_008709723.2    rna-XM_027608570.1, rna-XM_027622062.2, rna-XM_027624198.2    mNeoVis.3.1.1.chr2.g00991.t01, mNeoVis.3.1.1.chr5.g00669.t01    rna-gnl|WGS_ZZZZ|egapxtmp_004549-R1, rna-gnl|WGS_ZZZZ|egapxtmp_028866-R1, rna-gnl|WGS_ZZZZ|egapxtmp_036182-R1    mNeoVis.evergladensis.2.1.chr12.g00420.t01, mNeoVis.evergladensis.2.1.chr2.g01073.t01    mNeoVis.vulgivaga.2.1.chr12.g00427.t01, mNeoVis.vulgivaga.2.1.chr6.g00904.t01      2158    N0.HOG0002156   OG0000349  n1   rna-XM_019809893.2    rna-XM_038533965.1, rna-XM_038551623.1    rna-XM_022499766.1    rna-XM_023240453.2    rna-XM_036078149.1    rna-XM_039225304.1    rna-XM_032842748.1    rna-XM_046027739.1    rna-XM_032313338.1    rna-XM_059170219.1    rna-XM_059395812.1    rna-XM_013050091.2    mNeoVis.2.1.1.chr2_hap1.g00572.t01    rna-XM_021699577.2    rna-XM_004396103.1    rna-XM_042960471.1    rna-XM_040623931.1    rna-XM_027594203.2    mNeoVis.3.1.1.chr5.g00566.t01    rna-gnl|WGS_ZZZZ|egapxtmp_005465-R1    mNeoVis.evergladensis.2.1.chr5.g00574.t01    mNeoVis.vulgivaga.2.1.chr5.g00571.t01      2159    N0.HOG0002157   OG0000349  n14               rna-XM_032846972.1                        rna-XM_035721883.1              2160    N0.HOG0002158   OG0000349  n16         rna-XM_045059811.1                  rna-XM_045066268.1    mNeoVis.2.1.1.chr1_hap1.g00479.t01            rna-XM_027605266.2    mNeoVis.3.1.1.chr1.g00477.t01    rna-gnl|WGS_ZZZZ|egapxtmp_033188-R1    mNeoVis.evergladensis.2.1.chr1.g00477.t01    mNeoVis.vulgivaga.2.1.chr1.g00487.t01      2161    N0.HOG0002159   OG0000349  n23   rna-XM_034656105.1    rna-XM_038530299.1    rna-XM_022493089.1      rna-XM_036088874.1    rna-XM_039252739.1    rna-XM_032847461.1    rna-XM_045999599.1    rna-XM_032336549.1    rna-XM_059174752.1    rna-XM_059417710.1    rna-XM_004744824.3      rna-XM_021701769.1    rna-XM_004397805.1      rna-XM_040622371.1                2162    N0.HOG0002160   OG0000349  n24   rna-XM_034671438.1    rna-XM_038582013.1    rna-XM_022517996.1    rna-XM_006930940.4    rna-XM_036070637.1, rna-XM_036095782.1    rna-XM_039220816.1    rna-XM_032860741.1    rna-XM_045997189.1    rna-XM_032333039.1    rna-XM_059171916.1    rna-XM_059376698.1    rna-XM_013054038.2    mNeoVis.2.1.1.chr6_hap1.g00546.t01    rna-XM_021680415.2, rna-XM_021685301.2, rna-XM_021687947.1    rna-XM_004400778.1, rna-XM_004401857.1    rna-XM_042984117.1, rna-XM_042985228.1    rna-XM_040624982.1    rna-XM_027598537.2, rna-XM_035728594.1, rna-XM_035728938.1    mNeoVis.3.1.1.chr4.g00551.t01    rna-gnl|WGS_ZZZZ|egapxtmp_000221-R1, rna-gnl|WGS_ZZZZ|egapxtmp_043488-R1    mNeoVis.evergladensis.2.1.chr4.g00551.t01    mNeoVis.vulgivaga.2.1.chr4.g00560.t01, mNeoVis.vulgivaga.2.1.chr4.g00631.t01      2163    N0.HOG0002161   OG0000350  n1   rna-XM_034663169.1    rna-XM_038562393.1    rna-XM_022513421.1    rna-XM_045041775.1    rna-XM_036086322.1          rna-XM_032353716.1    rna-XM_059134387.1    rna-XM_059407603.1    rna-XM_004745966.1    mNeoVis.2.1.1.chr11_hap1.g01141.t01    rna-XM_044916584.1      rna-XM_042977386.1    rna-XM_040624270.1    rna-XM_035724203.1    mNeoVis.3.1.1.chr11.g01082.t01    rna-gnl|WGS_ZZZZ|egapxtmp_025382-R1    mNeoVis.evergladensis.2.1.chr11.g01110.t01    mNeoVis.vulgivaga.2.1.chr11.g01138.t01      2164    N0.HOG0002162   OG0000350  n16   rna-XM_034642292.1        rna-XM_045049545.1    rna-XM_036118966.1                        rna-XM_042973577.1    rna-XM_040625485.1                2165    N0.HOG0002163   OG0000350  n23                             rna-XM_044912618.1    rna-XM_004401094.1            rna-gnl|WGS_ZZZZ|egapxtmp_041298-R1          2166    N0.HOG0002164   OG0000350  n26   rna-XM_034651342.1        rna-XM_011286713.4                                          2167    N0.HOG0002165   OG0000350  n28                     rna-XM_059155360.1    rna-XM_059387176.1    rna-XM_045078332.1                          2168    N0.HOG0002166   OG0000350  n30     rna-XM_038544880.1    rna-XM_022497140.1      rna-XM_036079236.1                    rna-XM_021678043.2, rna-XM_021678273.1, rna-XM_021696516.2, rna-XM_021702936.2, rna-XM_044915423.1    rna-XM_004393499.1        rna-XM_035722134.1, rna-XM_035726720.1, rna-XM_035727732.1, rna-XM_035728624.1, rna-XM_035729216.1    mNeoVis.3.1.1.chr3.g01291.t01            2169    N0.HOG0002167   OG0000350  n32     rna-XM_038551672.1        rna-XM_036118608.1                    rna-XM_044918743.1          rna-XM_035728995.1              2170    N0.HOG0002168   OG0000350  n35     rna-XM_038540320.1        rna-XM_036076876.1, rna-XM_036083870.1, rna-XM_036087334.1, rna-XM_036119506.1    rna-XM_039225741.1                  rna-XM_044914649.1, rna-XM_044914901.1, rna-XM_044918231.1, rna-XM_044918853.1, rna-XM_044919634.1    rna-XM_004391841.1, rna-XM_004393738.1, rna-XM_004403291.1, rna-XM_004408433.1    rna-XM_042993704.1      rna-XM_035722207.1, rna-XM_035722818.1, rna-XM_035723541.1, rna-XM_035723658.1, rna-XM_035724095.1, rna-XM_035727564.1, rna-XM_035728996.1, rna-XM_035729325.1              2171    N0.HOG0002169   OG0000350  n58                               rna-XM_004397247.1, rna-XM_004413789.1        rna-XM_035722229.1              2172    N0.HOG0002170   OG0000351  n1   rna-XM_002916419.3    rna-NM_001252166.1    rna-XM_022493844.1    rna-XM_003983555.4    rna-XM_036064603.1    rna-XM_039227134.1    rna-XM_032845745.1    rna-XM_045980390.1    rna-XM_032350776.1    rna-XM_059133314.1    rna-XM_059406950.1    rna-XM_004746496.3    mNeoVis.2.1.1.chr11_hap1.g00266.t01    rna-XM_021689351.2    rna-XM_004393144.1    rna-XM_042980470.1    rna-XM_040638263.1    rna-XM_027620702.1    mNeoVis.3.1.1.chr11.g00237.t01      mNeoVis.evergladensis.2.1.chr11.g00264.t01    mNeoVis.vulgivaga.2.1.chr11.g00265.t01      2173    N0.HOG0002171   OG0000351  n4         rna-XM_003995041.5      rna-XM_039236652.1, rna-XM_039243196.1    rna-XM_032875677.1      rna-XM_032309706.1                    rna-XM_035727288.1              2174   N0.HOG0002172  OG0000351  n11   rna-XM_019808246.2      rna-XM_022495803.1      rna-XM_036122339.1        rna-XM_046024835.1      rna-XM_059138711.1    rna-XM_059409289.1    rna-XM_004742780.3    mNeoVis.2.1.1.chr3_hap1.g00238.t01    rna-XM_021686041.1    rna-XM_004412346.1    rna-XM_007089490.3    rna-XM_008690304.2    rna-XM_027575856.1    mNeoVis.3.1.1.chr3.g00240.t01    rna-gnl|WGS_ZZZZ|egapxtmp_042788-R1    mNeoVis.evergladensis.2.1.chr3.g00249.t01    mNeoVis.vulgivaga.2.1.chr3.g00239.t01      2175   N0.HOG0002173  OG0000351  n16   rna-XM_034642232.1    rna-XM_038543728.1    rna-XM_022495804.1      rna-XM_036122341.1      rna-XM_032855463.1, rna-XM_032875679.1      rna-XM_032310304.1, rna-XM_032335557.1    rna-XM_059138713.1, rna-XM_059162318.1    rna-XM_059387929.1, rna-XM_059399058.1, rna-XM_059409290.1    rna-XM_004739723.3, rna-XM_004742779.2, rna-XM_045063392.1    mNeoVis.2.1.1.chr3_hap1.g00239.t01    rna-XM_021686039.2, rna-XM_044918538.1    rna-XM_004412349.1      rna-XM_040632691.1    rna-XM_027575858.1, rna-XM_035724537.1, rna-XM_035725922.1, rna-XM_035725927.1    mNeoVis.3.1.1.chr3.g00241.t01    rna-gnl|WGS_ZZZZ|egapxtmp_038840-R1    mNeoVis.evergladensis.2.1.chr3.g00250.t01    mNeoVis.vulgivaga.2.1.chr3.g00240.t01      2176    N0.HOG0002174   OG0000351  n28     rna-XM_038543725.1      rna-XM_011288088.3      rna-XM_039243201.1                      rna-XM_042962775.1                  2177    N0.HOG0002175   OG0000352  n1   rna-XM_034672212.1    rna-XM_038583520.1    rna-XM_022508588.1    rna-XM_045037323.1    rna-XM_036122535.1    rna-XM_039241720.1    rna-XM_032875484.1    rna-XM_046003386.1    rna-XM_032339236.1    rna-XM_059163330.1    rna-XM_059391430.1    rna-XM_045088683.1    mNeoVis.2.1.1.chr5_hap1.g01340.t01    rna-XM_021692633.1      rna-XM_007083636.1    rna-XM_040621517.1    rna-XM_027585088.1    mNeoVis.3.1.1.chr2.g01257.t01    rna-gnl|WGS_ZZZZ|egapxtmp_038292-R1    mNeoVis.evergladensis.2.1.chr2.g01326.t01    mNeoVis.vulgivaga.2.1.chr2.g01334.t01      2178    N0.HOG0002176   OG0000352  n23   rna-XM_002915691.2    rna-XM_038583518.1    rna-XM_022508599.1    rna-XM_003991865.3    rna-XM_036122492.1    rna-XM_039241681.1    rna-XM_032875486.1    rna-XM_046003390.1    rna-XM_032339227.1    rna-XM_059163326.1    rna-XM_059392595.1    rna-XM_004745119.3      rna-XM_021692518.2    rna-XM_004406788.1    rna-XM_007083633.2    rna-XM_008705258.1                2179   N0.HOG0002177  OG0000352  n38   rna-XM_034672293.1    rna-XM_038583263.1    rna-XM_022508431.1    rna-XM_045037201.1    rna-XM_036122435.1    rna-XM_039241719.1    rna-XM_032875483.1    rna-XM_046003393.1    rna-XM_032339192.1    rna-XM_059163328.1    rna-XM_059388044.1    rna-XM_045088682.1    mNeoVis.2.1.1.chr5_hap1.g01342.t01      rna-XM_004406807.1    rna-XM_015535497.1    rna-XM_008705257.2    rna-XM_035724269.1    mNeoVis.3.1.1.chr2.g01259.t01    rna-gnl|WGS_ZZZZ|egapxtmp_038293-R1    mNeoVis.evergladensis.2.1.chr2.g01328.t01    mNeoVis.vulgivaga.2.1.chr2.g01336.t01      2180    N0.HOG0002178   OG0000352  n58   rna-XM_002915695.4    rna-XM_038583264.1    rna-XM_022508563.1    rna-XM_003991866.3    rna-XM_036122491.1    rna-XM_039241721.1    rna-XM_032850935.1    rna-XM_046003392.1    rna-XM_032339173.1    rna-XM_059163329.1    rna-XM_059388043.1    rna-XM_004745113.3    mNeoVis.2.1.1.chr5_hap1.g01341.t01    rna-XM_021692530.2      rna-XM_007077549.2    rna-XM_040621736.1    rna-XM_035724266.1    mNeoVis.3.1.1.chr2.g01258.t01      mNeoVis.evergladensis.2.1.chr2.g01327.t01    mNeoVis.vulgivaga.2.1.chr2.g01335.t01      2181    N0.HOG0002179   OG0000353  n1   rna-XM_002928368.4    rna-NM_001388683.1    rna-XM_022525423.1    rna-XM_003981789.2    rna-XM_036066094.1    rna-XM_039251576.1    rna-XM_032883465.1    rna-XM_045989672.1    rna-XM_032308475.1    rna-XM_059159715.1    rna-XM_059387693.1    rna-XM_013045223.2    mNeoVis.2.1.1.chr5_hap1.g00183.t01    rna-XM_021705503.1      rna-XM_007090685.2    rna-XM_008711297.2    rna-XM_027582172.1    mNeoVis.3.1.1.chr2.g00133.t01    rna-gnl|WGS_ZZZZ|egapxtmp_037734-R1    mNeoVis.evergladensis.2.1.chr2.g00180.t01    mNeoVis.vulgivaga.2.1.chr2.g00179.t01      2182    N0.HOG0002180   OG0000353  n22   rna-XM_034655710.1    rna-NM_001389191.1, rna-XM_038588893.1, rna-XM_038588895.1, rna-XM_038588910.1, rna-XM_038588914.1, rna-XM_038589212.1, rna-XM_038589430.1, rna-XM_038589657.1    rna-XM_022491973.1    rna-XM_023239879.2    rna-XM_036071423.1    rna-XM_039220839.1    rna-XM_032848095.1    rna-XM_045996820.1, rna-XM_045996842.1, rna-XM_045996877.1, rna-XM_045996978.1, rna-XM_045997034.1, rna-XM_045999730.1, rna-XM_045999759.1, rna-XM_045999770.1, rna-XM_045999774.1, rna-XM_046001104.1    rna-XM_032337895.1    rna-XM_059173385.1    rna-XM_059416409.1    rna-XM_013050515.2      rna-XM_021690949.2    rna-XM_004403495.1    rna-XM_007092961.2    rna-XM_008683750.2    rna-XM_027604481.1    mNeoVis.3.1.1.chr1.g00950.t01    rna-gnl|WGS_ZZZZ|egapxtmp_039500-R1    mNeoVis.evergladensis.2.1.chr1.g00933.t01    mNeoVis.vulgivaga.2.1.chr1.g00944.t01      2183    N0.HOG0002181   OG0000353  n49   rna-XM_002930718.1    rna-XM_038565302.1    rna-XM_022497964.1    rna-XM_003982048.3    rna-XM_036089056.1    rna-XM_039243288.1    rna-XM_032868485.1    rna-XM_045989953.1    rna-XM_032303847.1    rna-XM_059159103.1    rna-XM_059388267.1    rna-XM_004761184.1    mNeoVis.2.1.1.chr5_hap1.g00448.t01        rna-XM_007088624.2    rna-XM_008706431.1    rna-XM_035726741.1    mNeoVis.3.1.1.chr2.g00373.t01    rna-gnl|WGS_ZZZZ|egapxtmp_038072-R1    mNeoVis.evergladensis.2.1.chr2.g00443.t01    mNeoVis.vulgivaga.2.1.chr2.g00445.t01      2184    N0.HOG0002182   OG0000354  n1   rna-XM_002929814.4    rna-XM_038554325.1    rna-XM_022496949.1, rna-XM_022512301.1    rna-XM_003987059.6, rna-XM_011290956.3, rna-XM_045037671.1    rna-XM_036075073.1, rna-XM_036081132.1, rna-XM_036112019.1    rna-XM_039218621.1    rna-XM_032864775.1, rna-XM_032881411.1    rna-XM_046010297.1, rna-XM_046012641.1    rna-XM_032342776.1    rna-XM_059137262.1, rna-XM_059180721.1    rna-XM_059372208.1    rna-XM_004758755.3, rna-XM_045081887.1    mNeoVis.2.1.1.chr10_hap1.g00974.t01    rna-XM_021682020.1, rna-XM_021693877.2, rna-XM_044912539.1    rna-XM_004411794.1    rna-XM_042957117.1, rna-XM_042988702.1    rna-XM_040620541.1, rna-XM_040626885.1, rna-XM_040632994.1    rna-XM_027572560.1, rna-XM_027600071.2    mNeoVis.3.1.1.chr9.g00905.t01    rna-gnl|WGS_ZZZZ|egapxtmp_016246-R1    mNeoVis.evergladensis.2.1.chr8.g00944.t01    mNeoVis.vulgivaga.2.1.chr9.g00930.t01      2185    N0.HOG0002183   OG0000354  n21   rna-XM_002922496.4    rna-NM_001252286.1    rna-XM_022491026.1    rna-XM_023248096.2    rna-XM_036086846.1      rna-XM_032845241.1    rna-XM_045982333.1    rna-XM_032317405.1    rna-XM_059145761.1    rna-XM_059413876.1    rna-XM_045086888.1    mNeoVis.2.1.1.chr14_hap1.g00680.t01      rna-XM_004403015.1      rna-XM_040635695.1    rna-XM_027612419.2, rna-XM_027612527.1    mNeoVis.3.1.1.chr14.g00650.t01      mNeoVis.evergladensis.2.1.chr14.g00678.t01    mNeoVis.vulgivaga.2.1.chr14.g00678.t01      2186    N0.HOG0002184   OG0000354  n22     rna-XM_038586493.1      rna-XM_019814310.3, rna-XM_019821401.3, rna-XM_023253214.2    rna-XM_036066625.1    rna-XM_039245335.1      rna-XM_046021057.1                rna-XM_004416741.1    rna-XM_015535495.2      rna-XM_027614399.2              2187    N0.HOG0002185   OG0000354  n31             rna-XM_039218928.1                          rna-XM_027607034.2              2188    N0.HOG0002186   OG0000354  n32     rna-XM_038562455.1, rna-XM_038589597.1        rna-XM_036088497.1, rna-XM_036110320.1                    rna-XM_021693617.1    rna-XM_004407015.1      rna-XM_040624429.1    rna-XM_027589726.2, rna-XM_027593621.1              2189    N0.HOG0002187   OG0000355  n1   rna-XM_011223132.3    rna-XM_038587248.1    rna-XM_022508057.1    rna-XM_019823356.3    rna-XM_036089383.1    rna-XM_039216790.1    rna-XM_032842903.1    rna-XM_045996092.1    rna-XM_032330455.1    rna-XM_059157209.1    rna-XM_059384622.1    rna-XM_045068405.1    mNeoVis.2.1.1.chr12_hap1.g00638.t01    rna-XM_021679419.2    rna-XM_004396756.1    rna-XM_007081605.3    rna-XM_040632332.1    rna-XM_035725793.1    mNeoVis.3.1.1.chr12.g00582.t01    rna-gnl|WGS_ZZZZ|egapxtmp_028218-R1    mNeoVis.evergladensis.2.1.chr12.g00576.t01    mNeoVis.vulgivaga.2.1.chr12.g00592.t01      2190    N0.HOG0002188   OG0000355  n24   rna-XM_011235635.3    rna-XM_038527930.1    rna-XM_022524110.1    rna-XM_006940922.5    rna-XM_036104060.1    rna-XM_039249116.1    rna-XM_032839406.1    rna-XM_045988212.1    rna-XM_032324011.1    rna-XM_059151041.1    rna-XM_059382645.1    rna-XM_013064051.2    mNeoVis.2.1.1.chr7_hap1.g00812.t01    rna-XM_021681387.1    rna-XM_004415491.1    rna-XM_007074218.3    rna-XM_008684741.2    rna-XM_027618297.2        mNeoVis.evergladensis.2.1.chr7.g00801.t01        2191    N0.HOG0002189   OG0000355  n42   rna-XM_034649372.1    rna-XM_038587320.1    rna-XM_022508079.1    rna-XM_011291715.4    rna-XM_036111838.1    rna-XM_039225035.1    rna-XM_032842872.1    rna-XM_045996122.1    rna-XM_032330322.1    rna-XM_059157214.1    rna-XM_059385085.1    rna-XM_004754995.3    mNeoVis.2.1.1.chr12_hap1.g00630.t01    rna-XM_044911613.1    rna-XM_004396761.1    rna-XM_042974460.1    rna-XM_040632351.1    rna-XM_035725566.1    mNeoVis.3.1.1.chr12.g00575.t01      mNeoVis.evergladensis.2.1.chr12.g00568.t01    mNeoVis.vulgivaga.2.1.chr12.g00584.t01      2192    N0.HOG0002190   OG0000355  n61   rna-XM_019793385.2    rna-XM_038529029.1    rna-XM_022525372.1    rna-XM_045047009.1      rna-XM_039217741.1    rna-XM_032842065.1    rna-XM_045987961.1    rna-XM_032325598.1    rna-XM_059153715.1    rna-XM_059382415.1    rna-XM_045075900.1    mNeoVis.2.1.1.chr7_hap1.g00491.t01              mNeoVis.3.1.1.chr7.g00473.t01    rna-gnl|WGS_ZZZZ|egapxtmp_012053-R1    mNeoVis.evergladensis.2.1.chr7.g00494.t01        2193    N0.HOG0002191   OG0000356  n1   rna-XM_002916525.4, rna-XM_019805233.1    rna-XM_038553080.1    rna-XM_022501211.1, rna-XM_022516988.1    rna-XM_023242558.2    rna-XM_036090883.1, rna-XM_036123561.1    rna-XM_039250050.1    rna-XM_032860829.1    rna-XM_046023496.1    rna-XM_032307563.1    rna-XM_059141711.1    rna-XM_059411095.1    rna-XM_004747780.3    mNeoVis.2.1.1.chr13_hap1.g00365.t01    rna-XM_021691166.2, rna-XM_044917194.1, rna-XM_044919052.1    rna-XM_004396991.1, rna-XM_004413608.1    rna-XM_015537544.2, rna-XM_042964277.1    rna-XM_040631695.1, rna-XM_040644787.1    rna-XM_027601615.2, rna-XM_027616677.2    mNeoVis.3.1.1.chr13.g00308.t01    rna-gnl|WGS_ZZZZ|egapxtmp_031507-R1    mNeoVis.evergladensis.2.1.chr13.g00353.t01    mNeoVis.vulgivaga.2.1.chr13.g00354.t01      2194    N0.HOG0002192   OG0000356  n27   rna-XM_019796725.2, rna-XM_034651584.1    rna-XM_038553081.1    rna-XM_022500409.1    rna-XM_019816309.3    rna-XM_036090885.1    rna-XM_039240887.1    rna-XM_032860884.1    rna-XM_046023351.1    rna-XM_032308745.1    rna-XM_059143165.1    rna-XM_059412231.1    rna-XM_004747783.2    mNeoVis.2.1.1.chr13_hap1.g00364.t01    rna-XM_044920664.1    rna-XM_004397017.1    rna-XM_007094279.3    rna-XM_040631342.1    rna-XM_027616668.1    mNeoVis.3.1.1.chr13.g00307.t01    rna-gnl|WGS_ZZZZ|egapxtmp_030575-R1    mNeoVis.evergladensis.2.1.chr13.g00352.t01    mNeoVis.vulgivaga.2.1.chr13.g00353.t01      2195    N0.HOG0002193   OG0000356  n46   rna-XM_019802274.2    rna-XM_038543868.1    rna-XM_022495657.1    rna-XM_003995017.5    rna-XM_036092473.1    rna-XM_039219091.1    rna-XM_032875763.1    rna-XM_046025731.1    rna-XM_032311265.1    rna-XM_059140524.1    rna-XM_059408057.1    rna-XM_004742703.3    mNeoVis.2.1.1.chr3_hap1.g00264.t01    rna-XM_021689479.2    rna-XM_004416111.1    rna-XM_042962583.1    rna-XM_008692538.2    rna-XM_027576701.2    mNeoVis.3.1.1.chr3.g00267.t01    rna-gnl|WGS_ZZZZ|egapxtmp_041454-R1    mNeoVis.evergladensis.2.1.chr3.g00273.t01    mNeoVis.vulgivaga.2.1.chr3.g00266.t01      2196    N0.HOG0002194   OG0000357  n2                           mNeoVis.2.1.1.chr9_hap1.g00317.t01                  mNeoVis.evergladensis.2.1.chr1.g01789.t01        2197    N0.HOG0002195   OG0000357  n4   rna-NM_001304869.1, rna-NM_001304869.1-2, rna-XM_011218040.3    rna-XM_038546178.1        rna-XM_036075801.1, rna-XM_036105515.1    rna-XM_039223464.1, rna-XM_039247091.1    rna-XM_032875896.1    rna-XM_046004213.1, rna-XM_046009001.1        rna-XM_059418489.1        rna-XM_021704337.2, rna-XM_044913300.1    rna-XM_004391834.1, rna-XM_004397315.1, rna-XM_004408380.1        rna-XM_027571109.2, rna-XM_035725587.1    mNeoVis.3.1.1.chr2.g00937.t01            2198    N0.HOG0002196   OG0000357  n23   rna-XM_034652551.1    rna-XM_038571129.1    rna-XM_022520820.1      rna-XM_036078728.1, rna-XM_036090969.1, rna-XM_036092448.1, rna-XM_036095272.1, rna-XM_036118638.1    rna-XM_039237192.1        rna-XM_032329690.1    rna-XM_059156804.1    rna-XM_059384676.1    rna-XM_045068419.1      rna-XM_021680407.2, rna-XM_044914878.1    rna-XM_004409205.1      rna-XM_040641273.1    rna-XM_035726352.1, rna-XM_035726836.1              2199    N0.HOG0002197   OG0000357  n43     rna-NM_001252230.1    rna-XM_022521330.1    rna-XM_003992219.5    rna-XM_036076823.1        rna-XM_046003794.1    rna-XM_032335970.1, rna-XM_032346520.1    rna-XM_059162015.1    rna-XM_059390507.1    rna-XM_004759305.3    mNeoVis.2.1.1.chr5_hap1.g01018.t01    rna-XM_021677549.2, rna-XM_021694259.2      rna-XM_007081014.2    rna-XM_008710003.2    rna-XM_035722025.1, rna-XM_035722098.1, rna-XM_035726139.1      rna-gnl|WGS_ZZZZ|egapxtmp_034305-R1    mNeoVis.evergladensis.2.1.chr2.g01014.t01    mNeoVis.vulgivaga.2.1.chr2.g01019.t01      2200    N0.HOG0002198   OG0000357  n45           rna-XM_036095178.1                      rna-XM_004411061.1        rna-XM_027571532.2              2201    N0.HOG0002199   OG0000357  n48     rna-XM_038579301.1              rna-XM_045997377.1    rna-XM_032347729.1      rna-XM_059404657.1    rna-XM_045084847.1        rna-XM_004412117.1                    2202    N0.HOG0002200   OG0000357  n53     rna-XM_038544994.1                                  rna-XM_027589536.2, rna-XM_027594096.2, rna-XM_027602843.2, rna-XM_035727649.1              2203    N0.HOG0002201   OG0000358  n2   rna-XM_002915497.3    rna-XM_038559714.1    rna-XM_022514152.1    rna-XM_019826009.3    rna-XM_036102065.1    rna-XM_039230662.1    rna-XM_032840200.1    rna-XM_046021200.1    rna-XM_032305728.1    rna-XM_059172240.1    rna-XM_059395560.1    rna-XM_013063179.2    mNeoVis.2.1.1.chr4_hap1.g00050.t01    rna-XM_021692962.1    rna-XM_004408803.1    rna-XM_042977434.1    rna-XM_008688655.1    rna-XM_027573093.1      rna-gnl|WGS_ZZZZ|egapxtmp_003023-R1    mNeoVis.evergladensis.2.1.chr6.g00054.t01    mNeoVis.vulgivaga.2.1.chr6.g00056.t01      2204   N0.HOG0002202  OG0000358  n22     rna-XM_038547224.1    rna-XM_022509233.1    rna-XM_003996911.5    rna-XM_036095525.1    rna-XM_039225753.1, rna-XM_039225757.1    rna-XM_032871721.1      rna-XM_032320253.1    rna-XM_059148568.1    rna-XM_059380087.1    rna-XM_004772641.3    mNeoVis.2.1.1.chr8_hap1.g00359.t01, mNeoVis.2.1.1.chr8_hap1.g00360.t01    rna-XM_021681789.2    rna-XM_004410582.1    rna-XM_015540029.2    rna-XM_008701317.2    rna-XM_027625792.2    mNeoVis.3.1.1.chr8.g00326.t01      mNeoVis.evergladensis.2.1.chr9.g00358.t01    mNeoVis.vulgivaga.2.1.chr8.g00364.t01      2205   N0.HOG0002203  OG0000358  n47   rna-XM_034639883.1, rna-XM_034639884.1    rna-XM_038546018.1, rna-XM_038547221.1    rna-XM_022509334.1    rna-XM_011289125.4    rna-XM_036095522.1, rna-XM_036095524.1    rna-XM_039224296.1    rna-XM_032871726.1    rna-XM_045984966.1      rna-XM_059149063.1    rna-XM_059380085.1        rna-XM_021681667.2, rna-XM_021681788.2, rna-XM_044921835.1    rna-XM_004410579.1    rna-XM_042967099.1    rna-XM_008701361.2    rna-XM_035724393.1, rna-XM_035724394.1    mNeoVis.3.1.1.chr8.g00325.t01      mNeoVis.evergladensis.2.1.chr9.g00357.t01    mNeoVis.vulgivaga.2.1.chr8.g00363.t01      2206    N0.HOG0002204   OG0000358  n70     rna-XM_038547229.1    rna-XM_022509504.1          rna-XM_032871725.1        rna-XM_059149152.1    rna-XM_059380084.1    rna-XM_045080730.1                          2207    N0.HOG0002205   OG0000358  n75                                           mNeoVis.evergladensis.2.1.chr9.g00356.t01    mNeoVis.vulgivaga.2.1.chr8.g00362.t01      2208    N0.HOG0002206   OG0000359  n1   rna-XM_034641182.1    rna-XM_038546754.1    rna-XM_022519928.1    rna-XM_045044291.1    rna-XM_036075602.1    rna-XM_039229786.1    rna-XM_032870754.1    rna-XM_045984737.1    rna-XM_032321809.1    rna-XM_059147860.1    rna-XM_059380260.1    rna-XM_004749078.3    mNeoVis.2.1.1.chr8_hap1.g00216.t01    rna-XM_021678446.2    rna-XM_004405753.1    rna-XM_042966660.1    rna-XM_040640322.1    rna-XM_027626115.2    mNeoVis.3.1.1.chr8.g00174.t01    rna-gnl|WGS_ZZZZ|egapxtmp_019209-R2    mNeoVis.evergladensis.2.1.chr9.g00206.t01    mNeoVis.vulgivaga.2.1.chr8.g00212.t01      2209   N0.HOG0002207  OG0000359  n23   rna-XM_034640714.1        rna-XM_045044431.1    rna-XM_036075569.1    rna-XM_039229795.1              rna-XM_045080651.1      rna-XM_044921848.1    rna-XM_004405774.1    rna-XM_007073441.3    rna-XM_040641210.1    rna-XM_035724424.1              2210    N0.HOG0002208   OG0000359  n33   rna-XM_011232843.3    rna-XM_038546766.1    rna-XM_022519946.1    rna-XM_045044130.1    rna-XM_036075568.1    rna-XM_039229814.1    rna-XM_032870758.1    rna-XM_045984943.1    rna-XM_032320132.1    rna-XM_059147526.1    rna-XM_059380070.1    rna-XM_045080950.1    mNeoVis.2.1.1.chr8_hap1.g00218.t01    rna-XM_044921643.1, rna-XM_044921846.1    rna-XM_004405776.1    rna-XM_042966546.1    rna-XM_008693677.2    rna-XM_027625912.2    mNeoVis.3.1.1.chr8.g00176.t01    rna-gnl|WGS_ZZZZ|egapxtmp_019213-R2    mNeoVis.evergladensis.2.1.chr9.g00208.t01    mNeoVis.vulgivaga.2.1.chr8.g00214.t01      2211    N0.HOG0002209   OG0000359  n55   rna-XM_034641179.1    rna-XM_038546751.1    rna-XM_022519849.1    rna-XM_003997105.5    rna-XM_036075592.1    rna-XM_039229787.1    rna-XM_032870756.1    rna-XM_045985943.1    rna-XM_032321798.1    rna-XM_059147856.1    rna-XM_059378412.1    rna-XM_045080951.1    mNeoVis.2.1.1.chr8_hap1.g00217.t01    rna-XM_044921847.1    rna-XM_004405775.1    rna-XM_007073434.3    rna-XM_008693680.1    rna-XM_027626111.2    mNeoVis.3.1.1.chr8.g00175.t01    rna-gnl|WGS_ZZZZ|egapxtmp_019212-R2    mNeoVis.evergladensis.2.1.chr9.g00207.t01    mNeoVis.vulgivaga.2.1.chr8.g00213.t01      2212    N0.HOG0002210   OG0000360  n1   rna-XM_034643000.1    rna-XM_038545226.1    rna-XM_022505533.1    rna-XM_045060197.1, rna-XM_045060198.1, rna-XM_045060199.1, rna-XM_045060200.1    rna-XM_036117833.1    rna-XM_039232302.1    rna-XM_032837412.1    rna-XM_046009925.1    rna-XM_032344998.1, rna-XM_032345007.1    rna-XM_059182864.1, rna-XM_059182873.1    rna-XM_059373794.1    rna-XM_045072611.1    mNeoVis.2.1.1.chr10_hap1.g00159.t01, mNeoVis.2.1.1.chr10_hap1.g00160.t01    rna-XM_021679548.2    rna-XM_004399388.1    rna-XM_042987767.1    rna-XM_040641675.1    rna-XM_027571512.2    mNeoVis.3.1.1.chr9.g00132.t01      mNeoVis.evergladensis.2.1.chr8.g00153.t01    mNeoVis.vulgivaga.2.1.chr9.g00153.t01      2213    N0.HOG0002211   OG0000360  n28   rna-XM_034643002.1    rna-XM_038545223.1    rna-XM_022492740.1    rna-XM_006932977.5      rna-XM_039232292.1    rna-XM_032837392.1    rna-XM_046009924.1    rna-XM_032345009.1    rna-XM_059182874.1, rna-XM_059182875.1, rna-XM_059183438.1, rna-XM_059183439.1, rna-XM_059183440.1, rna-XM_059183441.1    rna-XM_059387351.1    rna-XM_045072614.1, rna-XM_045074608.1    mNeoVis.2.1.1.chr10_hap1.g00158.t01    rna-XM_021679547.2    rna-XM_004399391.1    rna-XM_042987766.1, rna-XM_042990071.1      rna-XM_027571514.2      rna-gnl|WGS_ZZZZ|egapxtmp_015990-R1      mNeoVis.vulgivaga.2.1.chr9.g00152.t01      2214   N0.HOG0002212  OG0000360  n50   rna-XM_011219025.3    rna-XM_038545220.1      rna-XM_003987826.5    rna-XM_036117869.1    rna-XM_039232370.1    rna-XM_032837535.1    rna-XM_046008612.1    rna-XM_032345008.1, rna-XM_032345013.1    rna-XM_059156496.1, rna-XM_059182879.1    rna-XM_059372182.1, rna-XM_059396993.1    rna-XM_045072637.1, rna-XM_045079929.1    mNeoVis.2.1.1.chr10_hap1.g00157.t01    rna-XM_044912321.1    rna-XM_004399427.1    rna-XM_042990070.1    rna-XM_040641677.1    rna-XM_027569874.1    mNeoVis.3.1.1.chr9.g00131.t01    rna-gnl|WGS_ZZZZ|egapxtmp_015679-R1    mNeoVis.evergladensis.2.1.chr8.g00152.t01    mNeoVis.vulgivaga.2.1.chr9.g00151.t01      2215   N0.HOG0002213  OG0000361  n0     rna-XM_038581519.1    rna-XM_022516004.1    rna-XM_023238731.2, rna-XM_045036438.1, rna-XM_045048037.1, rna-XM_045057765.1    rna-XM_036072647.1, rna-XM_036076210.1, rna-XM_036077527.1, rna-XM_036093494.1, rna-XM_036108437.1, rna-XM_036115256.1, rna-XM_036119637.1, rna-XM_036123443.1    rna-XM_039242721.1, rna-XM_039256115.1, rna-XM_039257515.1    rna-XM_032861218.1, rna-XM_032865313.1    rna-XM_045989841.1, rna-XM_045991238.1, rna-XM_045994788.1, rna-XM_046003716.1, rna-XM_046023881.1    rna-XM_032344304.1, rna-XM_032346234.1    rna-XM_059165312.1, rna-XM_059171272.1    rna-XM_059388396.1, rna-XM_059392599.1    rna-XM_013050051.2, rna-XM_045084491.1    mNeoVis.2.1.1.chr12_hap1.g00647.t01, mNeoVis.2.1.1.chr2_hap1.g00585.t01, mNeoVis.2.1.1.chr5_hap1.g01840.t01, mNeoVis.2.1.1.chr7_hap1.g01107.t01    rna-XM_021699432.2, rna-XM_044912629.1, rna-XM_044912709.1, rna-XM_044915342.1, rna-XM_044915698.1, rna-XM_044916038.1, rna-XM_044918211.1, rna-XM_044918867.1    rna-XM_004401099.1, rna-XM_004409742.1, rna-XM_004411362.1    rna-XM_042962734.1, rna-XM_042963483.1, rna-XM_042971566.1, rna-XM_042998446.1    rna-XM_040621893.1, rna-XM_040636572.1    rna-XM_027587377.2, rna-XM_035723365.1, rna-XM_035724859.1, rna-XM_035725577.1, rna-XM_035727367.1, rna-XM_035727401.1, rna-XM_035727835.1, rna-XM_035728430.1, rna-XM_035728713.1, rna-XM_035728827.1, rna-XM_035729125.1, rna-XM_035729212.1, rna-XM_035729357.1    mNeoVis.3.1.1.chr2.g01742.t01    rna-gnl|WGS_ZZZZ|egapxtmp_005749-R1, rna-gnl|WGS_ZZZZ|egapxtmp_010230-R1, rna-gnl|WGS_ZZZZ|egapxtmp_036400-R1    mNeoVis.evergladensis.2.1.chr2.g01823.t01, mNeoVis.evergladensis.2.1.chr3.g00705.t01, mNeoVis.evergladensis.2.1.chr5.g00590.t01    mNeoVis.vulgivaga.2.1.chr12.g00602.t01, mNeoVis.vulgivaga.2.1.chr2.g01838.t01, mNeoVis.vulgivaga.2.1.chr5.g00584.t01      2216    N0.HOG0002214   OG0000362  n1   rna-XM_002930325.4    rna-XM_038571838.1    rna-XM_022494566.1    rna-XM_045053793.1    rna-XM_036121757.1    rna-XM_039237787.1    rna-XM_032859014.1    rna-XM_045981116.1    rna-XM_032351159.1    rna-XM_059133700.1    rna-XM_059406695.1    rna-XM_013053445.2    mNeoVis.2.1.1.chr11_hap1.g00382.t01    rna-XM_021689033.2    rna-XM_004392992.1    rna-XM_042980731.1    rna-XM_008703264.2    rna-XM_035728909.1    mNeoVis.3.1.1.chr11.g00349.t01      mNeoVis.evergladensis.2.1.chr11.g00378.t01    mNeoVis.vulgivaga.2.1.chr11.g00382.t01      2217    N0.HOG0002215   OG0000362  n19   rna-XM_002918335.4, rna-XM_034660193.1    rna-XM_038565270.1, rna-XM_038579369.1    rna-XM_022504842.1, rna-XM_022515557.1    rna-XM_006943225.4    rna-XM_036082295.1, rna-XM_036087316.1, rna-XM_036088294.1, rna-XM_036100809.1    rna-XM_039221102.1, rna-XM_039225103.1    rna-XM_032870821.1, rna-XM_032873231.1, rna-XM_032875431.1, rna-XM_032882600.1    rna-XM_046010716.1, rna-XM_046018848.1, rna-XM_046027445.1    rna-XM_032316927.1, rna-XM_032317910.1, rna-XM_032324149.1    rna-XM_059134308.1, rna-XM_059166397.1, rna-XM_059170991.1, rna-XM_059171202.1    rna-XM_059394613.1, rna-XM_059395090.1, rna-XM_059395761.1, rna-XM_059413368.1    rna-XM_045075764.1, rna-XM_045089424.1, rna-XM_045089755.1    mNeoVis.2.1.1.chr11_hap1.g00486.t01, mNeoVis.2.1.1.chr4_hap1.g00728.t01, mNeoVis.2.1.1.chr8_hap1.g00655.t01    rna-XM_021688155.2, rna-XM_021697065.1, rna-XM_044911805.1    rna-XM_004398768.1    rna-XM_042973380.1    rna-XM_040625522.1    rna-XM_027595174.2, rna-XM_027612075.2    mNeoVis.3.1.1.chr2.g01280.t01, mNeoVis.3.1.1.chr6.g00736.t01    rna-gnl|WGS_ZZZZ|egapxtmp_004513-R1, rna-gnl|WGS_ZZZZ|egapxtmp_019110-R1, rna-gnl|WGS_ZZZZ|egapxtmp_035856-R1    mNeoVis.evergladensis.2.1.chr2.g01343.t01, mNeoVis.evergladensis.2.1.chr6.g00735.t01, mNeoVis.evergladensis.2.1.chr6.g00736.t01    mNeoVis.vulgivaga.2.1.chr2.g01353.t01, mNeoVis.vulgivaga.2.1.chr6.g00732.t01      2218    N0.HOG0002216   OG0000363  n1   rna-XM_034667620.1    rna-XM_038568930.1    rna-XM_022521676.1    rna-XM_011286755.4    rna-XM_036089272.1    rna-XM_039225172.1    rna-XM_032853369.1    rna-XM_046017717.1    rna-XM_032358476.1    rna-XM_059134894.1    rna-XM_059389142.1    rna-XM_045065868.1    mNeoVis.2.1.1.chr7_hap1.g01775.t01    rna-XM_021685729.1    rna-XM_004394540.1    rna-XM_042958826.1    rna-XM_040630678.1    rna-XM_027580095.2    mNeoVis.3.1.1.chr7.g01646.t01    rna-gnl|WGS_ZZZZ|egapxtmp_013873-R1    mNeoVis.evergladensis.2.1.chr7.g01736.t01    mNeoVis.vulgivaga.2.1.chr7.g01752.t01      2219   N0.HOG0002217  OG0000363  n23           rna-XM_036072373.1                                rna-gnl|WGS_ZZZZ|egapxtmp_029201-R1          2220    N0.HOG0002218   OG0000363  n25       rna-XM_022492509.1            rna-XM_045996706.1    rna-XM_032331903.1    rna-XM_059158787.1    rna-XM_059387286.1    rna-XM_045087319.1, rna-XM_045087327.1      rna-XM_021690224.2, rna-XM_021690464.1    rna-XM_004414298.1, rna-XM_004416013.1      rna-XM_008684014.2    rna-XM_027569084.2, rna-XM_027576663.2, rna-XM_035725859.1              2221    N0.HOG0002219   OG0000363  n40   rna-XM_002919915.4    rna-XM_038588429.1    rna-XM_022491609.1    rna-XM_011279490.3    rna-XM_036103437.1, rna-XM_036115876.1    rna-XM_039217137.1    rna-XM_032841382.1    rna-XM_045995643.1    rna-XM_032309374.1, rna-XM_032330218.1    rna-XM_059157584.1, rna-XM_059171247.1    rna-XM_059384605.1    rna-XM_004773650.3, rna-XM_013051704.2    mNeoVis.2.1.1.chr12_hap1.g00144.t01    rna-XM_021685051.1    rna-XM_004410476.1, rna-XM_004415381.1    rna-XM_042974419.1    rna-XM_008701544.2    rna-XM_035725602.1    mNeoVis.3.1.1.chr12.g00113.t01      mNeoVis.evergladensis.2.1.chr12.g00120.t01    mNeoVis.vulgivaga.2.1.chr12.g00120.t01      2222   N0.HOG0002220  OG0000363  n49   rna-XM_034652490.1    rna-XM_038545158.1                  rna-XM_059158202.1    rna-XM_059386163.1    rna-XM_013049659.2    mNeoVis.2.1.1.chr12_hap1.g00447.t01            rna-XM_027572942.2    mNeoVis.3.1.1.chr12.g00409.t01    rna-gnl|WGS_ZZZZ|egapxtmp_029424-R1    mNeoVis.evergladensis.2.1.chr12.g00404.t01    mNeoVis.vulgivaga.2.1.chr12.g00412.t01      2223   N0.HOG0002221  OG0000364  n1   rna-XM_034652926.1    rna-XM_038554192.1, rna-XM_038588777.1    rna-XM_022503516.1    rna-XM_023254019.2    rna-XM_036082753.1    rna-XM_039231126.1    rna-XM_032850151.1, rna-XM_032862050.1, rna-XM_032871302.1    rna-XM_046005570.1    rna-XM_032334437.1, rna-XM_032340124.1, rna-XM_032350242.1, rna-XM_032353446.1    rna-XM_059178104.1    rna-XM_059400355.1    rna-XM_004739794.3    mNeoVis.2.1.1.chr1_hap1.g01605.t01    rna-XM_021691868.2    rna-XM_004408131.1    rna-XM_007085124.3    rna-XM_008691859.2, rna-XM_040639804.1    rna-XM_027601720.2, rna-XM_035728672.1, rna-XM_035729042.1    mNeoVis.3.1.1.chr1.g01555.t01    rna-gnl|WGS_ZZZZ|egapxtmp_000796-R1    mNeoVis.evergladensis.2.1.chr1.g01572.t01, mNeoVis.evergladensis.2.1.chr4.g00253.t01    mNeoVis.vulgivaga.2.1.chr1.g01569.t01      2224   N0.HOG0002222  OG0000364  n22                   rna-XM_032345323.1                        rna-gnl|WGS_ZZZZ|egapxtmp_005760-R1          2225   N0.HOG0002223  OG0000364  n24   rna-XM_034647892.1, rna-XM_034667189.1    rna-XM_038551649.1        rna-XM_036123528.1                    rna-XM_044914237.1                  mNeoVis.vulgivaga.2.1.chr4.g00585.t01      2226   N0.HOG0002224  OG0000364  n29   rna-XM_034642571.1    rna-XM_038545759.1    rna-XM_022505137.1    rna-XM_023256047.2    rna-XM_036086255.1    rna-XM_039251186.1    rna-XM_032839544.1    rna-XM_045995997.1, rna-XM_046002303.1, rna-XM_046008978.1    rna-XM_032311033.1    rna-XM_059167925.1, rna-XM_059175733.1, rna-XM_059183146.1    rna-XM_059374009.1, rna-XM_059393571.1, rna-XM_059410041.1    rna-XM_045073887.1, rna-XM_045078781.1    mNeoVis.2.1.1.chr10_hap1.g00065.t01, mNeoVis.2.1.1.chr7_hap1.g01208.t01    rna-XM_044917983.1    rna-XM_004417839.1    rna-XM_042990574.1    rna-XM_008702568.2, rna-XM_040641401.1    rna-XM_027591119.2, rna-XM_035728107.1    mNeoVis.3.1.1.chr14.g00404.t01, mNeoVis.3.1.1.chr9.g00044.t01    rna-gnl|WGS_ZZZZ|egapxtmp_015606-R1, rna-gnl|WGS_ZZZZ|egapxtmp_033028-R1    mNeoVis.evergladensis.2.1.chr14.g00405.t01, mNeoVis.evergladensis.2.1.chr8.g00064.t01    mNeoVis.vulgivaga.2.1.chr9.g00064.t01      2227   N0.HOG0002225  OG0000365  n1   rna-XM_002925213.4    rna-XM_038579474.1    rna-XM_022492471.1, rna-XM_022517855.1    rna-XM_023248571.2    rna-XM_036064630.1    rna-XM_039216921.1    rna-XM_032848425.1, rna-XM_032850926.1, rna-XM_032851166.1    rna-XM_045979874.1, rna-XM_045979916.1, rna-XM_045979925.1    rna-XM_032316150.1    rna-XM_059166267.1    rna-XM_059387175.1, rna-XM_059395513.1    rna-XM_045063230.1    mNeoVis.2.1.1.chr4_hap1.g00772.t01    rna-XM_044914225.1    rna-XM_004413206.1    rna-XM_042973055.1    rna-XM_040625832.1    rna-XM_035727132.1    mNeoVis.3.1.1.chr6.g00776.t01, mNeoVis.3.1.1.chr6.g00777.t01    rna-gnl|WGS_ZZZZ|egapxtmp_004965-R1    mNeoVis.evergladensis.2.1.chr6.g00776.t01    mNeoVis.vulgivaga.2.1.chr6.g00770.t01      2228   N0.HOG0002226  OG0000365  n29   rna-XM_034639807.1    rna-XM_038572997.1    rna-XM_022493862.1    rna-XM_023251134.2    rna-XM_036099072.1    rna-XM_039230026.1    rna-XM_032879024.1    rna-XM_045980873.1, rna-XM_045996816.1, rna-XM_045996818.1, rna-XM_045996824.1, rna-XM_045996832.1, rna-XM_045996849.1, rna-XM_045996889.1, rna-XM_045996891.1, rna-XM_045996918.1, rna-XM_045996928.1, rna-XM_046010605.1, rna-XM_046011283.1    rna-XM_032316126.1, rna-XM_032350412.1    rna-XM_059132900.1, rna-XM_059168848.1, rna-XM_059168850.1    rna-XM_059386867.1, rna-XM_059407188.1    rna-XM_004746244.3, rna-XM_045063237.1, rna-XM_045063240.1, rna-XM_045075833.1, rna-XM_045081209.1    mNeoVis.2.1.1.chr11_hap1.g00149.t01, mNeoVis.2.1.1.chr4_hap1.g00768.t01, mNeoVis.2.1.1.chr4_hap1.g00771.t01    rna-XM_044919013.1    rna-XM_004412533.1    rna-XM_042980207.1    rna-XM_040638439.1    rna-XM_027621888.2    mNeoVis.3.1.1.chr11.g00120.t01, mNeoVis.3.1.1.chr6.g00774.t01    rna-gnl|WGS_ZZZZ|egapxtmp_002934-R1, rna-gnl|WGS_ZZZZ|egapxtmp_026797-R1    mNeoVis.evergladensis.2.1.chr11.g00145.t01, mNeoVis.evergladensis.2.1.chr6.g00774.t01, mNeoVis.evergladensis.2.1.chr6.g00775.t01    mNeoVis.vulgivaga.2.1.chr11.g00148.t01      2229   N0.HOG0002227  OG0000366  n1   rna-XM_034668518.1    rna-XM_038555366.1    rna-XM_022502159.1    rna-XM_011291447.4    rna-XM_036101514.1, rna-XM_036106140.1    rna-XM_039219338.1    rna-XM_032858445.1    rna-XM_046025880.1    rna-XM_032316319.1    rna-XM_059165746.1    rna-XM_059395523.1    rna-XM_004743466.3    mNeoVis.2.1.1.chr4_hap1.g00954.t01    rna-XM_021688377.1    rna-XM_004400753.1, rna-XM_004406459.1    rna-XM_042973239.1    rna-XM_008689272.2    rna-XM_027574666.2    mNeoVis.3.1.1.chr6.g00971.t01, mNeoVis.3.1.1.chr6.g00972.t01    rna-gnl|WGS_ZZZZ|egapxtmp_004559-R1    mNeoVis.evergladensis.2.1.chr6.g00960.t01    mNeoVis.vulgivaga.2.1.chr6.g00953.t01      2230   N0.HOG0002228  OG0000366  n27                       rna-XM_059395524.1                  mNeoVis.3.1.1.chr6.g00973.t01            2231   N0.HOG0002229  OG0000366  n29   rna-XM_034647758.1    rna-XM_038560546.1    rna-XM_022498737.1    rna-XM_045056760.1    rna-XM_036074224.1, rna-XM_036121171.1    rna-XM_039222836.1, rna-XM_039255776.1    rna-XM_032869815.1    rna-XM_045997494.1    rna-XM_032328946.1    rna-XM_059156007.1    rna-XM_059384481.1    rna-XM_045086171.1    mNeoVis.2.1.1.chr6_hap1.g01117.t01      rna-XM_004401153.1    rna-XM_042982757.1    rna-XM_040643176.1    rna-XM_027598604.2    mNeoVis.3.1.1.chr4.g01065.t01    rna-gnl|WGS_ZZZZ|egapxtmp_001046-R2    mNeoVis.evergladensis.2.1.chr4.g01051.t01    mNeoVis.vulgivaga.2.1.chr4.g01084.t01      2232   N0.HOG0002230  OG0000366  n51   rna-XM_034645185.1    rna-XM_038557633.1    rna-XM_022496499.1    rna-XM_023258437.2    rna-XM_036065276.1    rna-XM_039231736.1    rna-XM_032877161.1    rna-XM_045979530.1, rna-XM_045979636.1, rna-XM_045980312.1    rna-XM_032302436.1    rna-XM_059136925.1    rna-XM_059377195.1    rna-XM_045064161.1    mNeoVis.2.1.1.chr2_hap1.g01526.t01    rna-XM_044913827.1    rna-XM_004406457.1, rna-XM_004417125.1    rna-XM_042993830.1    rna-XM_040619227.1    rna-XM_027615123.2    mNeoVis.3.1.1.chr5.g01448.t01    rna-gnl|WGS_ZZZZ|egapxtmp_007099-R1    mNeoVis.evergladensis.2.1.chr5.g01464.t01    mNeoVis.vulgivaga.2.1.chr5.g01465.t01      2233   N0.HOG0002231  OG0000367  n1   rna-XM_019807444.2    rna-XM_038550375.1    rna-XM_022519167.1    rna-XM_045062453.1    rna-XM_036098262.1, rna-XM_036098288.1    rna-XM_039236931.1    rna-XM_032882865.1    rna-XM_046011772.1, rna-XM_046013719.1    rna-XM_032346074.1    rna-XM_059187131.1, rna-XM_059187133.1    rna-XM_059401889.1    rna-XM_045083770.1    mNeoVis.2.1.1.chr9_hap1.g01228.t01    rna-XM_021687747.1, rna-XM_044915768.1    rna-XM_004397403.1    rna-XM_042993396.1    rna-XM_040637615.1, rna-XM_040637616.1    rna-XM_027594954.2    mNeoVis.3.1.1.chr10.g01178.t01    rna-gnl|WGS_ZZZZ|egapxtmp_024156-R1    mNeoVis.evergladensis.2.1.chr10.g01216.t01    mNeoVis.vulgivaga.2.1.chr10.g01204.t01      2234   N0.HOG0002232  OG0000367  n28   rna-XM_034670424.1    rna-XM_038540489.1    rna-XM_022516881.1    rna-XM_019820989.3    rna-XM_036091444.1    rna-XM_039239231.1    rna-XM_032843785.1    rna-XM_045994097.1    rna-XM_032327450.1    rna-XM_059154115.1    rna-XM_059415417.1    rna-XM_004750445.3    mNeoVis.2.1.1.chr15_hap1.g00276.t01    rna-XM_021698014.1    rna-XM_004403247.1    rna-XM_042971580.1    rna-XM_008696852.2    rna-XM_027612534.1    mNeoVis.3.1.1.chr15.g00253.t01    rna-gnl|WGS_ZZZZ|egapxtmp_033373-R1    mNeoVis.evergladensis.2.1.chr15.g00253.t01    mNeoVis.vulgivaga.2.1.chr15.g00257.t01      2235   N0.HOG0002233  OG0000367  n48   rna-XM_034640103.1    rna-XM_038546489.1    rna-XM_022523410.1    rna-XM_045045287.1    rna-XM_036110441.1    rna-XM_039226600.1    rna-XM_032870506.1    rna-XM_045985287.1    rna-XM_032320129.1    rna-XM_059150300.1    rna-XM_059378414.1    rna-XM_004748817.3    mNeoVis.2.1.1.chr8_hap1.g00108.t01    rna-XM_021680960.1    rna-XM_004394277.1    rna-XM_042967572.1    rna-XM_040640141.1    rna-XM_027619262.2, rna-XM_035723675.1, rna-XM_035724513.1, rna-XM_035726054.1, rna-XM_035728629.1    mNeoVis.3.1.1.chr8.g00081.t01    rna-gnl|WGS_ZZZZ|egapxtmp_020236-R1    mNeoVis.evergladensis.2.1.chr9.g00105.t01    mNeoVis.vulgivaga.2.1.chr8.g00108.t01      2236   N0.HOG0002234  OG0000368  n1   rna-XM_011222406.3    rna-XM_038578149.1    rna-XM_022499602.1    rna-XM_006937998.5    rna-XM_036084690.1    rna-XM_039257740.1    rna-XM_032836910.1    rna-XM_046027230.1    rna-XM_032314072.1    rna-XM_059169544.1    rna-XM_059398086.1    rna-XM_004749694.2    mNeoVis.2.1.1.chr2_hap1.g00343.t01    rna-XM_021699854.1    rna-XM_004410241.1    rna-XM_042960976.1    rna-XM_040624082.1    rna-XM_027596868.2    mNeoVis.3.1.1.chr5.g00333.t01    rna-gnl|WGS_ZZZZ|egapxtmp_006838-R1    mNeoVis.evergladensis.2.1.chr5.g00345.t01    mNeoVis.vulgivaga.2.1.chr5.g00346.t01      2237   N0.HOG0002235  OG0000368  n18   rna-XM_019794275.2, rna-XM_019794276.2    rna-XM_038533778.1, rna-XM_038533779.1    rna-XM_022500175.1    rna-XM_003993890.6, rna-XM_006937727.5, rna-XM_045040958.1    rna-XM_036079372.1, rna-XM_036079374.1, rna-XM_036079375.1    rna-XM_039222006.1, rna-XM_039222016.1    rna-XM_032850819.1, rna-XM_032850821.1, rna-XM_032883081.1, rna-XM_032883085.1    rna-XM_046027165.1, rna-XM_046027845.1, rna-XM_046027847.1    rna-XM_032313771.1, rna-XM_032313970.1, rna-XM_032313971.1    rna-XM_059172455.1, rna-XM_059172456.1    rna-XM_059397402.1, rna-XM_059397403.1    rna-XM_004780757.3, rna-XM_004780758.3    mNeoVis.2.1.1.chr2_hap1.g00609.t01    rna-XM_021696204.2, rna-XM_021696206.1    rna-XM_004412998.1, rna-XM_004413005.1    rna-XM_007079274.3, rna-XM_015536272.2, rna-XM_015536273.2    rna-XM_008689616.2, rna-XM_040623790.1    rna-XM_035724320.1, rna-XM_035724332.1, rna-XM_035724333.1    mNeoVis.3.1.1.chr5.g00604.t01, mNeoVis.3.1.1.chr5.g00606.t01, mNeoVis.3.1.1.chr5.g00607.t01    rna-gnl|WGS_ZZZZ|egapxtmp_006902-R1, rna-gnl|WGS_ZZZZ|egapxtmp_008774-R1, rna-gnl|WGS_ZZZZ|egapxtmp_008776-R1    mNeoVis.evergladensis.2.1.chr5.g00612.t01, mNeoVis.evergladensis.2.1.chr5.g00613.t01, mNeoVis.evergladensis.2.1.chr5.g00614.t01    mNeoVis.vulgivaga.2.1.chr5.g00606.t01, mNeoVis.vulgivaga.2.1.chr5.g00608.t01      2238   N0.HOG0002236  OG0000369  n0   rna-XM_034651285.1    rna-XM_038545997.1, rna-XM_038545998.1    rna-XM_022492347.1, rna-XM_022492348.1, rna-XM_022492349.1    rna-XM_045060649.1, rna-XM_045060713.1, rna-XM_045060714.1, rna-XM_045060715.1, rna-XM_045060716.1, rna-XM_045060717.1, rna-XM_045060718.1, rna-XM_045060719.1, rna-XM_045060720.1    rna-XM_036073683.1, rna-XM_036073684.1    rna-XM_039224173.1    rna-XM_032850642.1, rna-XM_032851017.1, rna-XM_032851018.1    rna-XM_046007429.1, rna-XM_046007431.1, rna-XM_046007914.1    rna-XM_032331952.1, rna-XM_032341731.1, rna-XM_032341837.1, rna-XM_032345471.1, rna-XM_032345472.1    rna-XM_059179722.1, rna-XM_059179723.1, rna-XM_059179724.1, rna-XM_059183454.1    rna-XM_059372177.1, rna-XM_059372178.1, rna-XM_059372179.1, rna-XM_059372181.1    rna-XM_004781973.2, rna-XM_004781974.2, rna-XM_004781976.3, rna-XM_013050719.2, rna-XM_045071907.1    mNeoVis.2.1.1.chr10_hap1.g00003.t01, mNeoVis.2.1.1.chr10_hap1.g00004.t01, mNeoVis.2.1.1.chr10_hap1.g00005.t01, mNeoVis.2.1.1.chr10_hap1.g00006.t01    rna-XM_021689545.1, rna-XM_044918027.1    rna-XM_004416827.1, rna-XM_004417695.1    rna-XM_042989289.1, rna-XM_042990760.1, rna-XM_042990761.1, rna-XM_042990762.1, rna-XM_042990763.1    rna-XM_008683969.2, rna-XM_008683971.2, rna-XM_008683972.2    rna-XM_027571493.1, rna-XM_035728153.1    mNeoVis.3.1.1.chr9.g00001.t01, mNeoVis.3.1.1.chr9.g00002.t01    rna-gnl|WGS_ZZZZ|egapxtmp_014902-R1, rna-gnl|WGS_ZZZZ|egapxtmp_015435-R1, rna-gnl|WGS_ZZZZ|egapxtmp_016427-R1    mNeoVis.evergladensis.2.1.chr8.g00001.t01, mNeoVis.evergladensis.2.1.chr8.g00002.t01, mNeoVis.evergladensis.2.1.chr8.g00003.t01, mNeoVis.evergladensis.2.1.chr8.g00004.t01    mNeoVis.vulgivaga.2.1.chr9.g00001.t01, mNeoVis.vulgivaga.2.1.chr9.g00002.t01, mNeoVis.vulgivaga.2.1.chr9.g00003.t01, mNeoVis.vulgivaga.2.1.chr9.g00004.t01, mNeoVis.vulgivaga.2.1.chr9.g00005.t01, mNeoVis.vulgivaga.2.1.chr9.g00006.t01      2239   N0.HOG0002237  OG0000370  n0   rna-XM_011234734.3, rna-XM_034653363.1    rna-XM_038561085.1    rna-XM_022520386.1    rna-XM_003984381.6    rna-XM_036090196.1    rna-XM_039238491.1    rna-XM_032849555.1, rna-XM_032850920.1, rna-XM_032850921.1, rna-XM_032850999.1, rna-XM_032882027.1    rna-XM_045978919.1, rna-XM_045979025.1, rna-XM_045981688.1    rna-XM_032353038.1, rna-XM_032353064.1, rna-XM_032353082.1, rna-XM_032353712.1    rna-XM_059132385.1, rna-XM_059132388.1, rna-XM_059132398.1, rna-XM_059132404.1, rna-XM_059132412.1, rna-XM_059132413.1, rna-XM_059132418.1, rna-XM_059132419.1, rna-XM_059132421.1, rna-XM_059132422.1, rna-XM_059134371.1, rna-XM_059134383.1, rna-XM_059188765.1    rna-XM_059387239.1, rna-XM_059387240.1, rna-XM_059387503.1, rna-XM_059407613.1, rna-XM_059407617.1    rna-XM_045073908.1, rna-XM_045079267.1, rna-XM_045079268.1, rna-XM_045079287.1, rna-XM_045087492.1, rna-XM_045087493.1, rna-XM_045087494.1, rna-XM_045087497.1, rna-XM_045088014.1, rna-XM_045088016.1, rna-XM_045088017.1, rna-XM_045088019.1, rna-XM_045088037.1, rna-XM_045088040.1    mNeoVis.2.1.1.chr11_hap1.g00989.t01, mNeoVis.2.1.1.chr11_hap1.g01005.t01, mNeoVis.2.1.1.chr11_hap1.g01022.t01, mNeoVis.2.1.1.chr11_hap1.g01051.t01, mNeoVis.2.1.1.chr11_hap1.g01080.t01, mNeoVis.2.1.1.chr11_hap1.g01082.t01, mNeoVis.2.1.1.chr11_hap1.g01083.t01    rna-XM_021697304.2    rna-XM_004394863.1    rna-XM_007088883.3    rna-XM_008700306.2    rna-XM_027621238.2    mNeoVis.3.1.1.chr11.g00974.t01, mNeoVis.3.1.1.chr11.g00999.t01    rna-gnl|WGS_ZZZZ|egapxtmp_025352-R1, rna-gnl|WGS_ZZZZ|egapxtmp_025363-R1    mNeoVis.evergladensis.2.1.chr11.g00967.t01, mNeoVis.evergladensis.2.1.chr11.g00997.t01, mNeoVis.evergladensis.2.1.chr11.g00998.t01, mNeoVis.evergladensis.2.1.chr11.g01022.t01, mNeoVis.evergladensis.2.1.chr6.g00294.t01    mNeoVis.vulgivaga.2.1.chr11.g01004.t01, mNeoVis.vulgivaga.2.1.chr11.g01048.t01, mNeoVis.vulgivaga.2.1.chr11.g01077.t01      2240   N0.HOG0002238  OG0000371  n1   rna-XM_034650332.1, rna-XM_034660513.1    rna-NM_001014381.1    rna-XM_022510505.1, rna-XM_022510867.1      rna-XM_036118532.1, rna-XM_036118535.1    rna-XM_039241862.1    rna-XM_032849270.1    rna-XM_046005053.1, rna-XM_046005244.1, rna-XM_046005270.1    rna-XM_032340721.1, rna-XM_032340722.1    rna-XM_059178723.1, rna-XM_059178725.1    rna-XM_059399963.1, rna-XM_059399964.1    rna-XM_004782396.3, rna-XM_004782469.3, rna-XM_045072314.1, rna-XM_045074664.1    mNeoVis.2.1.1.chr1_hap1.g01400.t01, mNeoVis.2.1.1.chr1_hap1.g01403.t01    rna-XM_021684532.2    rna-XM_004410634.1, rna-XM_004417613.1      rna-XM_008693470.2    rna-XM_027601665.2    mNeoVis.3.1.1.chr1.g01350.t01    rna-gnl|WGS_ZZZZ|egapxtmp_006966-R1, rna-gnl|WGS_ZZZZ|egapxtmp_007007-R1, rna-gnl|WGS_ZZZZ|egapxtmp_021739-R1    mNeoVis.evergladensis.2.1.chr1.g01365.t01, mNeoVis.evergladensis.2.1.chr1.g01369.t01    mNeoVis.vulgivaga.2.1.chr1.g01369.t01, mNeoVis.vulgivaga.2.1.chr1.g01372.t01      2241   N0.HOG0002239  OG0000371  n37   rna-XM_034660514.1, rna-XM_034660515.1    rna-NM_001014768.1    rna-XM_022510546.1, rna-XM_022510709.1    rna-NM_001305030.1, rna-XM_003985987.6, rna-XM_023249583.2    rna-XM_036095319.1    rna-XM_039241794.1, rna-XM_039241795.1    rna-XM_032849266.1    rna-XM_046005207.1, rna-XM_046005208.1    rna-XM_032340728.1    rna-XM_059178734.1    rna-XM_059399959.1    rna-XM_004782005.3, rna-XM_045074281.1, rna-XM_045074661.1    mNeoVis.2.1.1.chr1_hap1.g01401.t01    rna-XM_044917229.1, rna-XM_044917627.1    rna-XM_004407680.1, rna-XM_004417611.1    rna-XM_007093259.3, rna-XM_007093264.2, rna-XM_042986094.1    rna-XM_008693469.2, rna-XM_008693472.2    rna-XM_027602130.2, rna-XM_035728484.1    mNeoVis.3.1.1.chr1.g01348.t01    rna-gnl|WGS_ZZZZ|egapxtmp_007023-R1    mNeoVis.evergladensis.2.1.chr1.g01367.t01    mNeoVis.vulgivaga.2.1.chr1.g01366.t01, mNeoVis.vulgivaga.2.1.chr1.g01367.t01      2242    N0.HOG0002240   OG0000372  n1   rna-XM_019802107.2    rna-XM_038577662.1    rna-XM_022519594.1    rna-XM_023257069.2    rna-XM_036098911.1    rna-XM_039241363.1    rna-XM_032876797.1    rna-XM_046013869.1    rna-XM_032348208.1    rna-XM_059185352.1    rna-XM_059403112.1    rna-XM_004782070.3    mNeoVis.2.1.1.chr9_hap1.g00554.t01    rna-XM_021704912.1    rna-XM_004400825.1    rna-XM_042992295.1    rna-XM_040642453.1    rna-XM_027595376.2    mNeoVis.3.1.1.chr10.g00524.t01    rna-gnl|WGS_ZZZZ|egapxtmp_022057-R4    mNeoVis.evergladensis.2.1.chr10.g00547.t01    mNeoVis.vulgivaga.2.1.chr10.g00533.t01      2243    N0.HOG0002241   OG0000372  n20   rna-XM_034644787.1          rna-XM_036096367.1    rna-XM_039241398.1        rna-XM_032347853.1    rna-XM_059187489.1    rna-XM_059404517.1      mNeoVis.2.1.1.chr9_hap1.g00531.t01        rna-XM_042993691.1        mNeoVis.3.1.1.chr10.g00502.t01    rna-gnl|WGS_ZZZZ|egapxtmp_024680-R1    mNeoVis.evergladensis.2.1.chr10.g00523.t01    mNeoVis.vulgivaga.2.1.chr10.g00510.t01      2244   N0.HOG0002242  OG0000372  n30       rna-XM_022519593.1          rna-XM_032876796.1    rna-XM_046013868.1    rna-XM_032348207.1    rna-XM_059185354.1    rna-XM_059403111.1    rna-XM_004782069.3                mNeoVis.3.1.1.chr10.g00525.t01        mNeoVis.vulgivaga.2.1.chr10.g00534.t01      2245   N0.HOG0002243  OG0000372  n39         rna-XM_003988690.6    rna-XM_036098922.1    rna-XM_039241362.1                  rna-XM_021704911.1    rna-XM_004400826.1        rna-XM_027593567.2              2246    N0.HOG0002244   OG0000372  n45   rna-XM_002921571.4    rna-XM_038577661.1                                rna-XM_008704577.2                2247    N0.HOG0002245   OG0000372  n47   rna-XM_002930313.4    rna-XM_038549579.1    rna-XM_022523130.1    rna-XM_003988851.5    rna-XM_036086568.1    rna-XM_039216896.1    rna-XM_032874386.1    rna-XM_046012574.1    rna-XM_032348531.1    rna-XM_059185066.1    rna-XM_059403345.1    rna-XM_004773038.2    mNeoVis.2.1.1.chr9_hap1.g00407.t01    rna-XM_021687159.1    rna-XM_004401765.1    rna-XM_042992482.1    rna-XM_008711969.2    rna-XM_027594934.2    mNeoVis.3.1.1.chr10.g00399.t01    rna-gnl|WGS_ZZZZ|egapxtmp_022605-R1    mNeoVis.evergladensis.2.1.chr10.g00404.t01    mNeoVis.vulgivaga.2.1.chr10.g00400.t01      2248    N0.HOG0002246   OG0000373  n1   rna-XM_002918191.4    rna-XM_038525619.1      rna-XM_019822043.3    rna-XM_036079549.1    rna-XM_039217605.1, rna-XM_039227444.1      rna-XM_045983403.1      rna-XM_059146902.1    rna-XM_059413169.1    rna-XM_004756375.3    mNeoVis.2.1.1.chr11_hap1.g01099.t01, mNeoVis.2.1.1.chr14_hap1.g00364.t01    rna-XM_021686544.2, rna-XM_021700552.1    rna-XM_004392805.1, rna-XM_004400308.1    rna-XM_007086520.3    rna-XM_008685152.2    rna-XM_035728174.1    mNeoVis.3.1.1.chr11.g01040.t01, mNeoVis.3.1.1.chr14.g00363.t01    rna-gnl|WGS_ZZZZ|egapxtmp_026612-R1    mNeoVis.evergladensis.2.1.chr11.g01066.t01, mNeoVis.evergladensis.2.1.chr14.g00364.t01    mNeoVis.vulgivaga.2.1.chr11.g01094.t01, mNeoVis.vulgivaga.2.1.chr14.g00365.t01      2249   N0.HOG0002247  OG0000373  n11   rna-XM_002927155.4    rna-XM_038585970.1    rna-XM_022491644.1, rna-XM_022508737.1      rna-XM_036072501.1, rna-XM_036084474.1, rna-XM_036088935.1, rna-XM_036089984.1, rna-XM_036094131.1, rna-XM_036099710.1, rna-XM_036106543.1, rna-XM_036113334.1, rna-XM_036120583.1      rna-XM_032877150.1, rna-XM_032878348.1, rna-XM_032882659.1    rna-XM_045980734.1, rna-XM_045996493.1, rna-XM_045997758.1, rna-XM_046020025.1    rna-XM_032316758.1, rna-XM_032319288.1, rna-XM_032331115.1    rna-XM_059168921.1    rna-XM_059392861.1    rna-XM_045075807.1    mNeoVis.2.1.1.chr4_hap1.g00741.t01    rna-XM_021699282.1, rna-XM_044913799.1, rna-XM_044914221.1, rna-XM_044914769.1, rna-XM_044916556.1, rna-XM_044917502.1, rna-XM_044919643.1, rna-XM_044920952.1    rna-XM_004398787.1, rna-XM_004403912.1, rna-XM_004415622.1, rna-XM_004416938.1        rna-XM_027595186.1, rna-XM_027595301.2, rna-XM_027606115.1, rna-XM_027612116.1, rna-XM_035727328.1      rna-gnl|WGS_ZZZZ|egapxtmp_002423-R1, rna-gnl|WGS_ZZZZ|egapxtmp_032080-R1    mNeoVis.evergladensis.2.1.chr6.g00750.t01    mNeoVis.vulgivaga.2.1.chr6.g00746.t01      2250    N0.HOG0002248   OG0000374  n1   rna-XM_002931044.2    rna-NM_001389104.1, rna-NM_001389112.1, rna-NM_001389147.1, rna-XM_038589548.1    rna-XM_022507196.1      rna-XM_036073550.1      rna-XM_032850677.1, rna-XM_032850810.1    rna-XM_046014281.1, rna-XM_046014290.1, rna-XM_046015826.1, rna-XM_046015827.1, rna-XM_046015831.1    rna-XM_032359837.1, rna-XM_032359838.1, rna-XM_032359841.1, rna-XM_032360078.1    rna-XM_059134802.1, rna-XM_059158872.1    rna-XM_059387535.1, rna-XM_059388263.1    rna-XM_004782274.2    mNeoVis.2.1.1.chr7_hap1.g02079.t01    rna-XM_021685191.2    rna-XM_004412008.1      rna-XM_008694202.1    rna-XM_027581073.1    mNeoVis.3.1.1.chr7.g01930.t01, mNeoVis.3.1.1.chr7.g01932.t01    rna-gnl|WGS_ZZZZ|egapxtmp_010708-R1, rna-gnl|WGS_ZZZZ|egapxtmp_010714-R1, rna-gnl|WGS_ZZZZ|egapxtmp_010729-R1    mNeoVis.evergladensis.2.1.chr7.g02034.t01, mNeoVis.evergladensis.2.1.chr7.g02035.t01    mNeoVis.vulgivaga.2.1.chr7.g02052.t01      2251    N0.HOG0002249   OG0000374  n36   rna-XM_002931301.4, rna-XM_034651800.1    rna-NM_001389099.1    rna-XM_022492748.1, rna-XM_022507188.1, rna-XM_022507190.1    rna-XM_003993309.3, rna-XM_011286839.2      rna-XM_039224974.1    rna-XM_032846568.1    rna-XM_046015814.1, rna-XM_046015818.1, rna-XM_046015819.1, rna-XM_046015820.1    rna-XM_032359844.1    rna-XM_059135856.1, rna-XM_059135983.1    rna-XM_059387805.1, rna-XM_059387887.1, rna-XM_059388051.1    rna-XM_004782173.2, rna-XM_004782178.1, rna-XM_004782180.1    mNeoVis.2.1.1.chr7_hap1.g02085.t01, mNeoVis.2.1.1.chr7_hap1.g02087.t01, mNeoVis.2.1.1.chr7_hap1.g02096.t01        rna-XM_007088788.2, rna-XM_007095372.2, rna-XM_042958199.1    rna-XM_008694065.2, rna-XM_040638711.1      mNeoVis.3.1.1.chr7.g01936.t01, mNeoVis.3.1.1.chr7.g01939.t01    rna-gnl|WGS_ZZZZ|egapxtmp_010747-R1    mNeoVis.evergladensis.2.1.chr7.g02040.t01, mNeoVis.evergladensis.2.1.chr7.g02046.t01    mNeoVis.vulgivaga.2.1.chr7.g02059.t01, mNeoVis.vulgivaga.2.1.chr7.g02068.t01      2252    N0.HOG0002250   OG0000375  n1   rna-XM_034669549.1    rna-XM_038539734.1    rna-XM_022526652.1    rna-XM_006942119.4    rna-XM_036118982.1    rna-XM_039247854.1    rna-XM_032842575.1    rna-XM_045994364.1    rna-XM_032327893.1    rna-XM_059154719.1    rna-XM_059414822.1    rna-XM_045072499.1    mNeoVis.2.1.1.chr15_hap1.g00010.t01    rna-XM_044915116.1    rna-XM_004416985.1    rna-XM_042971172.1    rna-XM_008685340.2    rna-XM_027613154.2    mNeoVis.3.1.1.chr15.g00010.t01    rna-gnl|WGS_ZZZZ|egapxtmp_033795-R1    mNeoVis.evergladensis.2.1.chr15.g00010.t01    mNeoVis.vulgivaga.2.1.chr15.g00010.t01      2253    N0.HOG0002251   OG0000375  n23   rna-XM_002924749.4    rna-XM_038539733.1    rna-XM_022526415.1    rna-XM_006942113.4    rna-XM_036118992.1    rna-XM_039247885.1    rna-XM_032842758.1    rna-XM_045994214.1    rna-XM_032327765.1    rna-XM_059154601.1    rna-XM_059414817.1    rna-XM_045072492.1    mNeoVis.2.1.1.chr15_hap1.g00009.t01    rna-XM_021700345.1    rna-XM_004416984.1    rna-XM_042971232.1, rna-XM_042971625.1    rna-XM_008685342.2    rna-XM_027610679.2    mNeoVis.3.1.1.chr15.g00009.t01      mNeoVis.evergladensis.2.1.chr15.g00009.t01    mNeoVis.vulgivaga.2.1.chr15.g00009.t01      2254   N0.HOG0002252  OG0000375  n44   rna-XM_034652525.1, rna-XM_034669548.1    rna-NM_001110802.1, rna-XM_038539730.1    rna-XM_022526417.1    rna-XM_023246699.2, rna-XM_023246717.2    rna-XM_036115339.1    rna-XM_039238049.1    rna-XM_032842625.1, rna-XM_032842642.1    rna-XM_045993497.1, rna-XM_045993505.1    rna-XM_032327903.1    rna-XM_059154867.1    rna-XM_059416047.1    rna-XM_045072510.1    mNeoVis.2.1.1.chr15_hap1.g00007.t01    rna-XM_021700385.1    rna-XM_004415862.1    rna-XM_042972110.1, rna-XM_042972158.1    rna-XM_040635028.1    rna-XM_035722379.1, rna-XM_035722480.1    mNeoVis.3.1.1.chr15.g00007.t01      mNeoVis.evergladensis.2.1.chr15.g00007.t01    mNeoVis.vulgivaga.2.1.chr15.g00007.t01      2255   N0.HOG0002253  OG0000376  n2   rna-XM_034641585.1    rna-XM_038572463.1    rna-XM_022494137.1    rna-XM_023251195.2      rna-XM_039249430.1          rna-XM_059133543.1    rna-XM_059407700.1    rna-XM_004746778.3    mNeoVis.2.1.1.chr11_hap1.g00183.t01    rna-XM_021689288.1    rna-XM_004417421.1    rna-XM_042980294.1        mNeoVis.3.1.1.chr11.g00152.t01    rna-gnl|WGS_ZZZZ|egapxtmp_026806-R1    mNeoVis.evergladensis.2.1.chr11.g00180.t01    mNeoVis.vulgivaga.2.1.chr11.g00182.t01      2256    N0.HOG0002254   OG0000376  n15   rna-XM_011217997.1    rna-XM_038572454.1    rna-XM_022494138.1    rna-XM_045053548.1    rna-XM_036103860.1    rna-XM_039249380.1    rna-XM_032838628.1    rna-XM_045980428.1, rna-XM_045980655.1    rna-XM_032352172.1    rna-XM_059133548.1    rna-XM_059407699.1    rna-XM_045082217.1      rna-XM_044918944.1      rna-XM_042980296.1      rna-XM_027623889.1              2257    N0.HOG0002255   OG0000376  n31     rna-NM_001003165.1, rna-XM_038571378.1        rna-XM_036103861.1, rna-XM_036103866.1    rna-XM_039249426.1        rna-XM_032350107.1    rna-XM_059133549.1      rna-XM_013053151.2    mNeoVis.2.1.1.chr11_hap1.g00184.t01    rna-XM_044919045.1          rna-XM_027623888.2    mNeoVis.3.1.1.chr11.g00153.t01    rna-gnl|WGS_ZZZZ|egapxtmp_025609-R1    mNeoVis.evergladensis.2.1.chr11.g00181.t01    mNeoVis.vulgivaga.2.1.chr11.g00183.t01      2258    N0.HOG0002256   OG0000376  n46     rna-XM_038572466.1    rna-XM_022494136.1    rna-XM_023251198.2      rna-XM_039249432.1    rna-XM_032838626.1    rna-XM_045980581.1      rna-XM_059133542.1    rna-XM_059407701.1    rna-XM_004746777.2    mNeoVis.2.1.1.chr11_hap1.g00182.t01        rna-XM_007077768.2    rna-XM_008698227.1    rna-XM_027623892.1        mNeoVis.evergladensis.2.1.chr11.g00179.t01    mNeoVis.vulgivaga.2.1.chr11.g00181.t01      2259   N0.HOG0002257  OG0000376  n59   rna-XM_034650507.1    rna-XM_038571379.1    rna-XM_022493833.1      rna-XM_036103865.1    rna-XM_039249379.1    rna-XM_032851020.1    rna-XM_045980580.1        rna-XM_059407697.1        rna-XM_044919046.1    rna-XM_004417485.1      rna-XM_040638680.1    rna-XM_027623890.1              2260    N0.HOG0002258   OG0000377  n0   rna-XM_002924628.4    rna-XM_038546845.1    rna-XM_022519900.1    rna-XM_045044649.1    rna-XM_036116165.1, rna-XM_036120492.1    rna-XM_039236012.1, rna-XM_039241997.1    rna-XM_032859195.1, rna-XM_032859414.1, rna-XM_032870819.1    rna-XM_045979417.1, rna-XM_045986654.1, rna-XM_045993781.1, rna-XM_045995541.1, rna-XM_046000064.1, rna-XM_046002975.1, rna-XM_046005783.1, rna-XM_046006974.1, rna-XM_046012471.1    rna-XM_032308302.1, rna-XM_032311695.1, rna-XM_032321667.1, rna-XM_032344477.1, rna-XM_032345572.1    rna-XM_059149111.1, rna-XM_059173675.1, rna-XM_059179179.1, rna-XM_059181067.1    rna-XM_059395864.1, rna-XM_059401617.1, rna-XM_059416465.1    rna-XM_045062915.1, rna-XM_045067016.1, rna-XM_045080913.1, rna-XM_045082029.1, rna-XM_045087208.1    mNeoVis.2.1.1.chr10_hap1.g01114.t01, mNeoVis.2.1.1.chr3_hap1.g00288.t01, mNeoVis.2.1.1.chr8_hap1.g00241.t01, mNeoVis.2.1.1.chr9_hap1.g00900.t01    rna-XM_021678416.2, rna-XM_044913332.1, rna-XM_044913338.1, rna-XM_044918828.1    rna-XM_004405897.1    rna-XM_042966848.1    rna-XM_008693635.2, rna-XM_040643408.1    rna-XM_035724727.1, rna-XM_035724835.1, rna-XM_035725432.1, rna-XM_035728123.1, rna-XM_035728261.1, rna-XM_035728878.1    mNeoVis.3.1.1.chr1.g01454.t01, mNeoVis.3.1.1.chr8.g00198.t01    rna-gnl|WGS_ZZZZ|egapxtmp_016180-R1, rna-gnl|WGS_ZZZZ|egapxtmp_017846-R1, rna-gnl|WGS_ZZZZ|egapxtmp_028131-R1, rna-gnl|WGS_ZZZZ|egapxtmp_033802-R1, rna-gnl|WGS_ZZZZ|egapxtmp_035324-R1, rna-gnl|WGS_ZZZZ|egapxtmp_043050-R1    mNeoVis.evergladensis.2.1.chr12.g00513.t01, mNeoVis.evergladensis.2.1.chr2.g01271.t01, mNeoVis.evergladensis.2.1.chr2.g01411.t01, mNeoVis.evergladensis.2.1.chr9.g00231.t01    mNeoVis.vulgivaga.2.1.chr1.g01472.t01, mNeoVis.vulgivaga.2.1.chr12.g00527.t01, mNeoVis.vulgivaga.2.1.chr2.g01422.t01, mNeoVis.vulgivaga.2.1.chr4.g00692.t01, mNeoVis.vulgivaga.2.1.chr8.g00239.t01      2261    N0.HOG0002259   OG0000378  n2     rna-XM_038588771.1, rna-XM_038588772.1, rna-XM_038589389.1      rna-XM_045051098.1, rna-XM_045051099.1    rna-XM_036086496.1, rna-XM_036088514.1    rna-XM_039257236.1    rna-XM_032842331.1, rna-XM_032842333.1    rna-XM_045994727.1, rna-XM_045994728.1    rna-XM_032331584.1, rna-XM_032331585.1    rna-XM_059158522.1, rna-XM_059158523.1    rna-XM_059385294.1, rna-XM_059385296.1    rna-XM_004782560.2, rna-XM_045089205.1    mNeoVis.2.1.1.chr12_hap1.g00730.t01, mNeoVis.2.1.1.chr12_hap1.g00731.t01    rna-XM_044911625.1, rna-XM_044911626.1    rna-XM_004410042.1    rna-XM_042973933.1, rna-XM_042973934.1    rna-XM_040636964.1    rna-XM_035725457.1, rna-XM_035725458.1      rna-gnl|WGS_ZZZZ|egapxtmp_028255-R1          2262    N0.HOG0002260   OG0000378  n32   rna-XM_034649874.1    rna-XM_038587124.1      rna-XM_045050817.1    rna-XM_036091829.1    rna-XM_039255352.1    rna-XM_032844354.1        rna-XM_059158476.1    rna-XM_059385906.1    rna-XM_045089201.1    mNeoVis.2.1.1.chr12_hap1.g00715.t01    rna-XM_044911892.1    rna-XM_004411842.1    rna-XM_042975202.1    rna-XM_040626449.1    rna-XM_035725720.1    mNeoVis.3.1.1.chr12.g00653.t01    rna-gnl|WGS_ZZZZ|egapxtmp_029087-R1    mNeoVis.evergladensis.2.1.chr12.g00646.t01    mNeoVis.vulgivaga.2.1.chr12.g00668.t01      2263    N0.HOG0002261   OG0000378  n50   rna-XM_034665078.1    rna-XM_038573181.1    rna-XM_022520670.1    rna-XM_006927235.3    rna-XM_036066907.1    rna-XM_039253616.1    rna-XM_032880092.1    rna-XM_045978245.1    rna-XM_032305303.1, rna-XM_032314342.1    rna-XM_059143691.1    rna-XM_059378105.1    rna-XM_004770760.3    mNeoVis.2.1.1.chr8_hap1.g01288.t01    rna-XM_044913567.1    rna-XM_004399903.1    rna-XM_042957914.1    rna-XM_040622129.1    rna-XM_027590945.2    mNeoVis.3.1.1.chr8.g01184.t01    rna-gnl|WGS_ZZZZ|egapxtmp_019699-R1    mNeoVis.evergladensis.2.1.chr9.g01236.t01    mNeoVis.vulgivaga.2.1.chr8.g01238.t01      2264   N0.HOG0002262  OG0000379  n1   rna-XM_002912810.4    rna-XM_038566799.1    rna-XM_022498298.1    rna-XM_023247604.2    rna-XM_036093811.1    rna-XM_039228259.1    rna-XM_032867714.1    rna-XM_045992320.1    rna-XM_032316462.1    rna-XM_059160576.1    rna-XM_059389470.1    rna-XM_045081402.1    mNeoVis.2.1.1.chr5_hap1.g00528.t01    rna-XM_021683106.1    rna-XM_004404720.1    rna-XM_042978637.1    rna-XM_040629001.1    rna-XM_027584845.2    mNeoVis.3.1.1.chr2.g00450.t01    rna-gnl|WGS_ZZZZ|egapxtmp_036564-R1    mNeoVis.evergladensis.2.1.chr2.g00522.t01    mNeoVis.vulgivaga.2.1.chr2.g00524.t01      2265   N0.HOG0002263  OG0000379  n19   rna-XM_002918534.4    rna-XM_038552526.1    rna-XM_022517592.1    rna-XM_045042845.1    rna-XM_036082649.1    rna-XM_039221264.1    rna-XM_032880499.1    rna-XM_045981718.1, rna-XM_046023706.1    rna-XM_032306986.1    rna-XM_059142265.1    rna-XM_059411562.1    rna-XM_004761816.3    mNeoVis.2.1.1.chr13_hap1.g00597.t01    rna-XM_021697284.1    rna-XM_004412912.1    rna-XM_042963641.1    rna-XM_008689793.2    rna-XM_027616089.2    mNeoVis.3.1.1.chr13.g00527.t01    rna-gnl|WGS_ZZZZ|egapxtmp_030738-R1    mNeoVis.evergladensis.2.1.chr13.g00574.t01    mNeoVis.vulgivaga.2.1.chr13.g00574.t01      2266    N0.HOG0002264   OG0000379  n34   rna-XM_034648762.1    rna-XM_038588136.1    rna-XM_022518507.1    rna-XM_004000821.6    rna-XM_036083590.1    rna-XM_039244402.1    rna-XM_032844757.1    rna-XM_045996358.1    rna-XM_032316776.1, rna-XM_032316778.1, rna-XM_032316789.1, rna-XM_032330765.1    rna-XM_059157695.1, rna-XM_059166281.1    rna-XM_059385191.1    rna-XM_045063235.1, rna-XM_045068271.1, rna-XM_045075687.1    mNeoVis.2.1.1.chr12_hap1.g00260.t01    rna-XM_021686331.1    rna-XM_004414299.1    rna-XM_007091736.3    rna-XM_008706952.2    rna-XM_027609344.2    mNeoVis.3.1.1.chr12.g00227.t01    rna-gnl|WGS_ZZZZ|egapxtmp_029592-R5    mNeoVis.evergladensis.2.1.chr12.g00228.t01    mNeoVis.vulgivaga.2.1.chr12.g00235.t01      2267    N0.HOG0002265   OG0000380  n1   rna-XM_034643547.1    rna-XM_038557079.1    rna-XM_022503160.1    rna-XM_045052469.1    rna-XM_036113133.1    rna-XM_039238848.1    rna-XM_032873525.1    rna-XM_046021406.1    rna-XM_032305983.1    rna-XM_059171331.1    rna-XM_059397525.1    rna-XM_013051673.2    mNeoVis.2.1.1.chr4_hap1.g00418.t01    rna-XM_021689842.2    rna-XM_004397377.1    rna-XM_042967928.1    rna-XM_040644201.1    rna-XM_027574450.2    mNeoVis.3.1.1.chr6.g00417.t01    rna-gnl|WGS_ZZZZ|egapxtmp_003555-R1    mNeoVis.evergladensis.2.1.chr6.g00421.t01    mNeoVis.vulgivaga.2.1.chr6.g00415.t01      2268    N0.HOG0002266   OG0000380  n23   rna-XM_034647281.1    rna-XM_038536892.1    rna-XM_022524635.1    rna-XM_023244531.2    rna-XM_036105275.1, rna-XM_036106911.1    rna-XM_039231630.1    rna-XM_032840101.1    rna-XM_045984493.1    rna-XM_032322973.1    rna-XM_059150223.1    rna-XM_059380821.1    rna-XM_045083401.1    mNeoVis.2.1.1.chr8_hap1.g00977.t01    rna-XM_021694498.1    rna-XM_004416565.1, rna-XM_004417503.1, rna-XM_004417610.1    rna-XM_042967844.1    rna-XM_040630049.1    rna-XM_035724677.1, rna-XM_035724711.1    mNeoVis.3.1.1.chr8.g00910.t01    rna-gnl|WGS_ZZZZ|egapxtmp_017546-R1    mNeoVis.evergladensis.2.1.chr9.g00950.t01, mNeoVis.evergladensis.2.1.chr9.g00951.t01    mNeoVis.vulgivaga.2.1.chr8.g00948.t01, mNeoVis.vulgivaga.2.1.chr8.g00949.t01, mNeoVis.vulgivaga.2.1.chr8.g00951.t01      2269   N0.HOG0002267  OG0000380  n51   rna-XM_034641543.1    rna-XM_038546433.1    rna-XM_022523225.1    rna-XM_023244151.2    rna-XM_036069589.1    rna-XM_039226469.1    rna-XM_032870483.1    rna-XM_045984411.1    rna-XM_032321862.1    rna-XM_059149924.1    rna-XM_059379091.1    rna-XM_013054360.2    mNeoVis.2.1.1.chr8_hap1.g00095.t01    rna-XM_021680944.1    rna-XM_004416243.1    rna-XM_042966512.1    rna-XM_040640104.1    rna-XM_035724561.1    mNeoVis.3.1.1.chr8.g00070.t01    rna-gnl|WGS_ZZZZ|egapxtmp_017842-R1    mNeoVis.evergladensis.2.1.chr9.g00092.t01    mNeoVis.vulgivaga.2.1.chr8.g00095.t01      2270    N0.HOG0002268   OG0000381  n1                           mNeoVis.2.1.1.chr2_hap1.g00760.t01    rna-XM_044916986.1            mNeoVis.3.1.1.chr5.g00706.t01    rna-gnl|WGS_ZZZZ|egapxtmp_007712-R1    mNeoVis.evergladensis.2.1.chr5.g00716.t01    mNeoVis.vulgivaga.2.1.chr5.g00710.t01      2271    N0.HOG0002269   OG0000381  n6     rna-XM_038582721.1        rna-XM_036097117.1, rna-XM_036118787.1      rna-XM_032867458.1      rna-XM_032326034.1, rna-XM_032333451.1    rna-XM_059161570.1    rna-XM_059390208.1    rna-XM_045063252.1      rna-XM_021687995.1, rna-XM_044912581.1    rna-XM_004393449.1, rna-XM_004395233.1        rna-XM_035723312.1, rna-XM_035726391.1, rna-XM_035726871.1, rna-XM_035729088.1              2272   N0.HOG0002270  OG0000381  n21   rna-XM_034654309.1    rna-XM_038562280.1    rna-XM_022526133.1    rna-XM_023259176.2    rna-XM_036108376.1    rna-XM_039246343.1    rna-XM_032843529.1, rna-XM_032877953.1    rna-XM_046002700.1    rna-XM_032307950.1, rna-XM_032330709.1, rna-XM_032361632.1    rna-XM_059132954.1, rna-XM_059135232.1, rna-XM_059142052.1, rna-XM_059168923.1, rna-XM_059186083.1    rna-XM_059376179.1, rna-XM_059392850.1, rna-XM_059404751.1, rna-XM_059407512.1    rna-XM_013053223.2, rna-XM_045075809.1, rna-XM_045081653.1, rna-XM_045083057.1    mNeoVis.2.1.1.chr11_hap1.g00154.t01, mNeoVis.2.1.1.chr12_hap1.g00821.t01, mNeoVis.2.1.1.chr5_hap1.g01780.t01      rna-XM_004404283.1, rna-XM_004417297.1    rna-XM_042994143.1    rna-XM_040626734.1, rna-XM_040629059.1    rna-XM_027583243.2, rna-XM_027590242.2, rna-XM_035724252.1, rna-XM_035725002.1, rna-XM_035726437.1, rna-XM_035726918.1, rna-XM_035728898.1    mNeoVis.3.1.1.chr11.g00126.t01    rna-gnl|WGS_ZZZZ|egapxtmp_026977-R1, rna-gnl|WGS_ZZZZ|egapxtmp_028486-R1, rna-gnl|WGS_ZZZZ|egapxtmp_035217-R1    mNeoVis.evergladensis.2.1.chr11.g00151.t01, mNeoVis.evergladensis.2.1.chr12.g00749.t01, mNeoVis.evergladensis.2.1.chr2.g01764.t01    mNeoVis.vulgivaga.2.1.chr11.g00153.t01, mNeoVis.vulgivaga.2.1.chr12.g00776.t01, mNeoVis.vulgivaga.2.1.chr2.g01779.t01      2273   N0.HOG0002271  OG0000382  n0   rna-XM_002924951.4, rna-XM_002924952.4, rna-XM_034639730.1    rna-XM_038546129.1, rna-XM_038546130.1, rna-XM_038546131.1, rna-XM_038546132.1    rna-XM_022509173.1, rna-XM_022509178.1, rna-XM_022509179.1    rna-XM_003996929.4, rna-XM_003996930.6, rna-XM_019817869.3, rna-XM_023244788.2    rna-XM_036068533.1, rna-XM_036068534.1, rna-XM_036068535.1, rna-XM_036068540.1    rna-XM_039223268.1, rna-XM_039223269.1, rna-XM_039223270.1, rna-XM_039223294.1    rna-XM_032850712.1, rna-XM_032850713.1, rna-XM_032850714.1, rna-XM_032850729.1, rna-XM_032850730.1, rna-XM_032850731.1    rna-XM_045985540.1, rna-XM_045985730.1, rna-XM_045986573.1    rna-XM_032321535.1, rna-XM_032322481.1, rna-XM_032322482.1, rna-XM_032322781.1    rna-XM_059149663.1, rna-XM_059149664.1, rna-XM_059149665.1    rna-XM_059378551.1    rna-XM_004764569.3, rna-XM_004764570.3, rna-XM_045080657.1, rna-XM_045080969.1    mNeoVis.2.1.1.chr8_hap1.g00437.t01, mNeoVis.2.1.1.chr8_hap1.g00438.t01, mNeoVis.2.1.1.chr8_hap1.g00439.t01, mNeoVis.2.1.1.chr8_hap1.g00440.t01    rna-XM_021704359.2    rna-XM_004394994.1, rna-XM_004394995.1, rna-XM_004394996.1    rna-XM_042965638.1, rna-XM_042965639.1, rna-XM_042965640.1    rna-XM_008689728.2, rna-XM_008689737.1, rna-XM_008689957.2    rna-XM_035724695.1, rna-XM_035724740.1, rna-XM_035724769.1, rna-XM_035724770.1    mNeoVis.3.1.1.chr8.g00394.t01, mNeoVis.3.1.1.chr8.g00395.t01, mNeoVis.3.1.1.chr8.g00396.t01    rna-gnl|WGS_ZZZZ|egapxtmp_019105-R1, rna-gnl|WGS_ZZZZ|egapxtmp_019812-R1, rna-gnl|WGS_ZZZZ|egapxtmp_019824-R1    mNeoVis.evergladensis.2.1.chr9.g00427.t01, mNeoVis.evergladensis.2.1.chr9.g00428.t01, mNeoVis.evergladensis.2.1.chr9.g00429.t01, mNeoVis.evergladensis.2.1.chr9.g00430.t01    mNeoVis.vulgivaga.2.1.chr8.g00435.t01, mNeoVis.vulgivaga.2.1.chr8.g00436.t01      2274   N0.HOG0002272  OG0000383  n1   rna-XM_019795557.2    rna-XM_038559755.1, rna-XM_038563685.1, rna-XM_038563905.1, rna-XM_038588822.1      rna-XM_019826036.3    rna-XM_036102099.1    rna-XM_039230758.1    rna-XM_032840242.1    rna-XM_045986718.1, rna-XM_046021323.1    rna-XM_032303890.1, rna-XM_032342729.1    rna-XM_059172277.1    rna-XM_059396096.1    rna-XM_004758783.3    mNeoVis.2.1.1.chr4_hap1.g00036.t01    rna-XM_021692801.2      rna-XM_042975561.1    rna-XM_008688605.2    rna-XM_027573954.2    mNeoVis.3.1.1.chr1.g00973.t01, mNeoVis.3.1.1.chr6.g00039.t01    rna-gnl|WGS_ZZZZ|egapxtmp_021264-R1, rna-gnl|WGS_ZZZZ|egapxtmp_021316-R1    mNeoVis.evergladensis.2.1.chr6.g00038.t01    mNeoVis.vulgivaga.2.1.chr1.g00973.t01, mNeoVis.vulgivaga.2.1.chr1.g01112.t01, mNeoVis.vulgivaga.2.1.chr6.g00042.t01      2275   N0.HOG0002273  OG0000383  n28   rna-XM_002915831.4, rna-XM_019795911.2    rna-XM_038584310.1, rna-XM_038584316.1    rna-XM_022490560.1    rna-XM_003985736.5, rna-XM_003985737.5    rna-XM_036112343.1, rna-XM_036112344.1    rna-XM_039257325.1, rna-XM_039257326.1    rna-XM_032861727.1, rna-XM_032861731.1    rna-XM_046005945.1, rna-XM_046006776.1    rna-XM_032338385.1, rna-XM_032341280.1    rna-XM_059176413.1, rna-XM_059176414.1    rna-XM_059398951.1, rna-XM_059398952.1    rna-XM_004778755.3, rna-XM_045082303.1    mNeoVis.2.1.1.chr1_hap1.g01158.t01, mNeoVis.2.1.1.chr1_hap1.g01160.t01    rna-XM_021693624.1, rna-XM_021693631.1    rna-XM_004406497.1, rna-XM_004406498.1    rna-XM_007074428.3, rna-XM_007074429.3    rna-XM_008692943.1, rna-XM_008692945.2    rna-XM_027602244.2, rna-XM_035728680.1      rna-gnl|WGS_ZZZZ|egapxtmp_018239-R6, rna-gnl|WGS_ZZZZ|egapxtmp_018273-R1    mNeoVis.evergladensis.2.1.chr1.g01072.t01, mNeoVis.evergladensis.2.1.chr1.g01104.t01, mNeoVis.evergladensis.2.1.chr1.g01131.t01, mNeoVis.evergladensis.2.1.chr1.g01133.t01, mNeoVis.evergladensis.2.1.chr5.g00786.t01    mNeoVis.vulgivaga.2.1.chr1.g01131.t01      2276   N0.HOG0002274  OG0000384  n1   rna-XM_034660544.1, rna-XM_034661684.1    rna-XM_038553724.1    rna-XM_022510917.1    rna-XM_045057393.1    rna-XM_036115292.1    rna-XM_039241842.1    rna-XM_032849157.1    rna-XM_046005674.1    rna-XM_032340625.1    rna-XM_059178625.1    rna-XM_059400026.1    rna-XM_045074700.1    mNeoVis.2.1.1.chr1_hap1.g01436.t01    rna-XM_021684573.1    rna-XM_004407712.1    rna-XM_042986129.1    rna-XM_040644751.1    rna-XM_027601077.1    mNeoVis.3.1.1.chr1.g01381.t01, mNeoVis.3.1.1.chr1.g01382.t01    rna-gnl|WGS_ZZZZ|egapxtmp_021766-R1    mNeoVis.evergladensis.2.1.chr1.g01398.t01    mNeoVis.vulgivaga.2.1.chr1.g01402.t01      2277   N0.HOG0002275  OG0000384  n22   rna-XM_034664445.1    rna-XM_038549475.1    rna-XM_022501452.1    rna-XM_023242672.2    rna-XM_036107798.1    rna-XM_039234632.1    rna-XM_032865768.1    rna-XM_046022289.1    rna-XM_032308603.1    rna-XM_059142882.1    rna-XM_059410882.1    rna-XM_045082344.1    mNeoVis.2.1.1.chr13_hap1.g00291.t01    rna-XM_021691785.2    rna-XM_004411565.1    rna-XM_042963800.1    rna-XM_040631828.1    rna-XM_027615200.2    mNeoVis.3.1.1.chr13.g00234.t01    rna-gnl|WGS_ZZZZ|egapxtmp_031115-R1    mNeoVis.evergladensis.2.1.chr13.g00282.t01    mNeoVis.vulgivaga.2.1.chr13.g00278.t01      2278   N0.HOG0002276  OG0000384  n39   rna-XM_011220939.3    rna-XM_038542400.1    rna-XM_022508613.1    rna-XM_045048587.1    rna-XM_036095381.1, rna-XM_036120909.1    rna-XM_039255386.1    rna-XM_032838785.1, rna-XM_032862569.1    rna-XM_045982544.1    rna-XM_032317288.1    rna-XM_059145570.1    rna-XM_059412827.1    rna-XM_004761972.3    mNeoVis.2.1.1.chr14_hap1.g00194.t01    rna-XM_021682604.2    rna-XM_004398123.1    rna-XM_015543797.2    rna-XM_008696743.2    rna-XM_027610620.2    mNeoVis.3.1.1.chr14.g00193.t01    rna-gnl|WGS_ZZZZ|egapxtmp_032687-R1    mNeoVis.evergladensis.2.1.chr14.g00193.t01, mNeoVis.evergladensis.2.1.chr14.g00194.t01    mNeoVis.vulgivaga.2.1.chr14.g00197.t01, mNeoVis.vulgivaga.2.1.chr14.g00198.t01      2279   N0.HOG0002277  OG0000385  n1   rna-XM_034651967.1    rna-XM_038579852.1        rna-XM_036075865.1, rna-XM_036082725.1, rna-XM_036083178.1, rna-XM_036088737.1, rna-XM_036096765.1, rna-XM_036100894.1, rna-XM_036104505.1, rna-XM_036109099.1, rna-XM_036116658.1        rna-XM_045995798.1, rna-XM_045996764.1, rna-XM_045997997.1, rna-XM_046002703.1, rna-XM_046015949.1, rna-XM_046019241.1, rna-XM_046021847.1      rna-XM_059183824.1    rna-XM_059404993.1      mNeoVis.2.1.1.chr1_hap1.g01195.t01, mNeoVis.2.1.1.chr3_hap1.g00804.t01    rna-XM_044912679.1, rna-XM_044916052.1, rna-XM_044916797.1, rna-XM_044917025.1, rna-XM_044918287.1    rna-XM_004410947.1, rna-XM_004416014.1, rna-XM_004417008.1        rna-XM_027570135.2, rna-XM_035722506.1, rna-XM_035723073.1, rna-XM_035725205.1, rna-XM_035726235.1, rna-XM_035726656.1, rna-XM_035727071.1, rna-XM_035727772.1, rna-XM_035728968.1, rna-XM_035729043.1    mNeoVis.3.1.1.chr1.g01733.t01, mNeoVis.3.1.1.chr3.g00787.t01    rna-gnl|WGS_ZZZZ|egapxtmp_040422-R1, rna-gnl|WGS_ZZZZ|egapxtmp_042761-R1    mNeoVis.evergladensis.2.1.chr1.g01169.t01    mNeoVis.vulgivaga.2.1.chr1.g01169.t01, mNeoVis.vulgivaga.2.1.chr1.g01740.t01, mNeoVis.vulgivaga.2.1.chr3.g00811.t01      2280   N0.HOG0002278  OG0000385  n48   rna-NM_001304849.1    rna-NM_001252145.1    rna-XM_022510627.1    rna-XM_003986099.6    rna-XM_036096660.1    rna-XM_039235328.1    rna-XM_032864310.1    rna-XM_046006432.1    rna-XM_032340533.1    rna-XM_059178531.1    rna-XM_059400102.1    rna-XM_013049357.2, rna-XM_045074765.1    mNeoVis.2.1.1.chr1_hap1.g01463.t01    rna-XM_021684663.1    rna-XM_004407748.1    rna-XM_007085277.3    rna-XM_040644711.1    rna-XM_027602091.1, rna-XM_035723243.1    mNeoVis.3.1.1.chr1.g01411.t01    rna-gnl|WGS_ZZZZ|egapxtmp_012958-R1    mNeoVis.evergladensis.2.1.chr1.g01429.t01    mNeoVis.vulgivaga.2.1.chr1.g01430.t01      2281   N0.HOG0002279  OG0000386  n1   rna-XM_034658038.1    rna-XM_038566890.1    rna-XM_022498299.1    rna-XM_023247146.2    rna-XM_036118826.1    rna-XM_039243366.1    rna-XM_032867620.1    rna-XM_045991645.1    rna-XM_032315301.1    rna-XM_059160459.1    rna-XM_059389381.1    rna-XM_004761008.3    mNeoVis.2.1.1.chr5_hap1.g00494.t01    rna-XM_021683126.1    rna-XM_004404717.1    rna-XM_042977060.1    rna-XM_040628168.1    rna-XM_027586530.2    mNeoVis.3.1.1.chr2.g00418.t01    rna-gnl|WGS_ZZZZ|egapxtmp_035380-R1    mNeoVis.evergladensis.2.1.chr2.g00489.t01    mNeoVis.vulgivaga.2.1.chr2.g00491.t01      2282    N0.HOG0002280   OG0000386  n22   rna-XM_034664078.1      rna-XM_022501228.1    rna-XM_045042885.1    rna-XM_036081330.1          rna-XM_032308022.1        rna-XM_045081554.1        rna-XM_004392243.1    rna-XM_042965021.1        mNeoVis.3.1.1.chr13.g00451.t01, mNeoVis.3.1.1.chr13.g00452.t01    rna-gnl|WGS_ZZZZ|egapxtmp_031007-R1      mNeoVis.vulgivaga.2.1.chr13.g00495.t01, mNeoVis.vulgivaga.2.1.chr13.g00496.t01      2283    N0.HOG0002281   OG0000386  n35     rna-XM_038552716.1          rna-XM_039218602.1    rna-XM_032860471.1    rna-XM_046023713.1      rna-XM_059142420.1    rna-XM_059411406.1      mNeoVis.2.1.1.chr13_hap1.g00510.t01    rna-XM_044920652.1        rna-XM_040631563.1    rna-XM_027614547.2    mNeoVis.3.1.1.chr13.g00446.t01    rna-gnl|WGS_ZZZZ|egapxtmp_030106-R1    mNeoVis.evergladensis.2.1.chr13.g00491.t01    mNeoVis.vulgivaga.2.1.chr13.g00494.t01      2284   N0.HOG0002282  OG0000386  n48   rna-XM_034661708.1    rna-XM_038580464.1    rna-XM_022512170.1    rna-XM_045059288.1    rna-XM_036083872.1    rna-XM_039254229.1    rna-XM_032864917.1    rna-XM_046009970.1    rna-XM_032342611.1    rna-XM_059180554.1    rna-XM_059371954.1    rna-XM_045073496.1    mNeoVis.2.1.1.chr10_hap1.g01023.t01    rna-XM_044917815.1    rna-XM_004403963.1    rna-XM_042988813.1    rna-XM_008705990.2    rna-XM_027572007.2    mNeoVis.3.1.1.chr9.g00955.t01    rna-gnl|WGS_ZZZZ|egapxtmp_016705-R2    mNeoVis.evergladensis.2.1.chr8.g00994.t01, mNeoVis.evergladensis.2.1.chr8.g00995.t01    mNeoVis.vulgivaga.2.1.chr9.g00980.t01      2285   N0.HOG0002283  OG0000387  n1   rna-XM_034653454.1, rna-XM_034663566.1    rna-XM_038533814.1    rna-XM_022499183.1    rna-XM_023240358.2    rna-XM_036089262.1    rna-XM_039254352.1    rna-XM_032883087.1    rna-XM_046026812.1    rna-XM_032312090.1    rna-XM_059170336.1    rna-XM_059397367.1    rna-XM_045087357.1    mNeoVis.2.1.1.chr2_hap1.g00620.t01    rna-XM_021696142.1    rna-XM_004411338.1, rna-XM_004416580.1, rna-XM_004416581.1    rna-XM_042960054.1    rna-XM_008689669.1    rna-XM_027588196.2    mNeoVis.3.1.1.chr5.g00617.t01    rna-gnl|WGS_ZZZZ|egapxtmp_006167-R1    mNeoVis.evergladensis.2.1.chr5.g00624.t01    mNeoVis.vulgivaga.2.1.chr5.g00617.t01      2286    N0.HOG0002284   OG0000387  n26   rna-XM_034653440.1, rna-XM_034661206.1    rna-XM_038579910.1    rna-XM_022522093.1    rna-XM_045059588.1    rna-XM_036082604.1    rna-XM_039247258.1    rna-XM_032856920.1    rna-XM_046008318.1    rna-XM_032343182.1    rna-XM_059183407.1    rna-XM_059372112.1    rna-XM_004751026.3    mNeoVis.2.1.1.chr10_hap1.g00780.t01    rna-XM_021695474.2    rna-XM_004409035.1    rna-XM_042989263.1    rna-XM_040620426.1    rna-XM_027569112.2    mNeoVis.3.1.1.chr9.g00740.t01    rna-gnl|WGS_ZZZZ|egapxtmp_016358-R1    mNeoVis.evergladensis.2.1.chr8.g00761.t01    mNeoVis.vulgivaga.2.1.chr9.g00761.t01      2287    N0.HOG0002285   OG0000387  n48   rna-XM_034639016.1, rna-XM_034650410.1    rna-XM_038528976.1    rna-XM_022524996.1    rna-XM_023245508.2    rna-XM_036118459.1    rna-XM_039217730.1    rna-XM_032841976.1    rna-XM_045987235.1    rna-XM_032325359.1    rna-XM_059152534.1    rna-XM_059382868.1    rna-XM_045076770.1    mNeoVis.2.1.1.chr7_hap1.g00511.t01    rna-XM_021700963.2    rna-XM_004396495.1    rna-XM_042970107.1    rna-XM_040633935.1    rna-XM_027617543.2    mNeoVis.3.1.1.chr1.g02105.t01, mNeoVis.3.1.1.chr7.g00496.t01    rna-gnl|WGS_ZZZZ|egapxtmp_012234-R1    mNeoVis.evergladensis.2.1.chr7.g00512.t01    mNeoVis.vulgivaga.2.1.chr7.g00515.t01      2288    N0.HOG0002286   OG0000388  n1   rna-XM_002917019.4    rna-XM_038556017.1    rna-XM_022505679.1    rna-XM_003985402.6    rna-XM_036096666.1    rna-XM_039241109.1    rna-XM_032852364.1    rna-XM_045997778.1    rna-XM_032334224.1    rna-XM_059162069.1    rna-XM_059383410.1    rna-XM_013062691.2    mNeoVis.2.1.1.chr6_hap1.g00194.t01    rna-XM_021701654.1    rna-XM_004396302.1    rna-XM_007075911.3    rna-XM_008701451.2    rna-XM_035726135.1    mNeoVis.3.1.1.chr4.g00188.t01, mNeoVis.3.1.1.chr4.g00189.t01, mNeoVis.3.1.1.chr4.g00190.t01    rna-gnl|WGS_ZZZZ|egapxtmp_000681-R1, rna-gnl|WGS_ZZZZ|egapxtmp_042975-R1    mNeoVis.evergladensis.2.1.chr4.g00198.t01, mNeoVis.evergladensis.2.1.chr4.g00199.t01    mNeoVis.vulgivaga.2.1.chr4.g00194.t01      2289   N0.HOG0002287  OG0000388  n27   rna-XM_019795040.2    rna-XM_038534847.1    rna-XM_022493759.1    rna-XM_045035992.1    rna-XM_036078843.1    rna-XM_039245355.1    rna-XM_032851724.1    rna-XM_045999267.1    rna-XM_032337123.1    rna-XM_059174217.1    rna-XM_059417295.1    rna-XM_004737682.3    mNeoVis.2.1.1.chr1_hap1.g00271.t01    rna-XM_044917124.1    rna-XM_004404767.1    rna-XM_042992945.1    rna-XM_008692415.1    rna-XM_027606547.2    mNeoVis.3.1.1.chr1.g00274.t01    rna-gnl|WGS_ZZZZ|egapxtmp_027514-R1    mNeoVis.evergladensis.2.1.chr1.g00268.t01, mNeoVis.evergladensis.2.1.chr1.g00269.t01    mNeoVis.vulgivaga.2.1.chr1.g00273.t01, mNeoVis.vulgivaga.2.1.chr1.g00274.t01      2290    N0.HOG0002288   OG0000388  n50   rna-XM_034668907.1    rna-XM_038533681.1    rna-XM_022525987.1    rna-XM_023255028.2    rna-XM_036096087.1    rna-XM_039218643.1    rna-XM_032843114.1    rna-XM_046008749.1    rna-XM_032342290.1    rna-XM_059180232.1    rna-XM_059371731.1    rna-XM_045070732.1    mNeoVis.2.1.1.chr10_hap1.g01122.t01    rna-XM_021680630.1    rna-XM_004411784.1    rna-XM_042988299.1    rna-XM_040642656.1    rna-XM_027571792.2    mNeoVis.3.1.1.chr9.g01052.t01    rna-gnl|WGS_ZZZZ|egapxtmp_014539-R1    mNeoVis.evergladensis.2.1.chr8.g01101.t01    mNeoVis.vulgivaga.2.1.chr9.g01077.t01      2291    N0.HOG0002289   OG0000389  n1   rna-XM_034638166.1, rna-XM_034654852.1    rna-XM_038585146.1    rna-XM_022497864.1    rna-XM_045034133.1    rna-XM_036075492.1    rna-XM_039234699.1    rna-XM_032860248.1    rna-XM_046018891.1    rna-XM_032354165.1    rna-XM_059168620.1    rna-XM_059393046.1    rna-XM_004763372.3    mNeoVis.2.1.1.chr3_hap1.g01353.t01    rna-XM_021702835.1    rna-XM_004392402.1    rna-XM_007094679.3    rna-XM_008687258.2    rna-XM_027588746.2    mNeoVis.3.1.1.chr3.g01328.t01    rna-gnl|WGS_ZZZZ|egapxtmp_041525-R1    mNeoVis.evergladensis.2.1.chr3.g01355.t01    mNeoVis.vulgivaga.2.1.chr3.g01374.t01      2292    N0.HOG0002290   OG0000389  n11   rna-XM_011237374.3    rna-XM_038547407.1    rna-XM_022510009.1    rna-XM_003996790.6    rna-XM_036079973.1    rna-XM_039227971.1    rna-XM_032871525.1    rna-XM_045986637.1    rna-XM_032321293.1    rna-XM_059148916.1    rna-XM_059379196.1    rna-XM_004764629.3    mNeoVis.2.1.1.chr8_hap1.g00477.t01    rna-XM_021704095.2    rna-XM_004395036.1    rna-XM_042965392.1    rna-XM_040640680.1    rna-XM_027568120.2      rna-gnl|WGS_ZZZZ|egapxtmp_019896-R2    mNeoVis.evergladensis.2.1.chr9.g00468.t01    mNeoVis.vulgivaga.2.1.chr8.g00473.t01      2293    N0.HOG0002291   OG0000389  n15     rna-XM_038576749.1    rna-XM_022523059.1    rna-XM_003988859.5, rna-XM_045059360.1    rna-XM_036086586.1    rna-XM_039216820.1    rna-XM_032874399.1    rna-XM_046011531.1    rna-XM_032348548.1    rna-XM_059185057.1    rna-XM_059403352.1    rna-XM_004773047.3    mNeoVis.2.1.1.chr9_hap1.g00401.t01    rna-XM_021687167.1    rna-XM_004399902.1, rna-XM_004401633.1    rna-XM_007081467.3, rna-XM_042989215.1    rna-XM_008712022.2, rna-XM_040633391.1    rna-XM_027592357.2, rna-XM_035727240.1    mNeoVis.3.1.1.chr1.g02043.t01, mNeoVis.3.1.1.chr10.g00392.t01    rna-gnl|WGS_ZZZZ|egapxtmp_024346-R1    mNeoVis.evergladensis.2.1.chr10.g00398.t01, mNeoVis.evergladensis.2.1.chr8.g01084.t01    mNeoVis.vulgivaga.2.1.chr10.g00394.t01      2294   N0.HOG0002292  OG0000390  n2   rna-XM_011231678.3    rna-XM_038527582.1    rna-XM_022492076.1    rna-XM_045046780.1    rna-XM_036091929.1    rna-XM_039219028.1    rna-XM_032847679.1    rna-XM_045987935.1    rna-XM_032324766.1    rna-XM_059152399.1    rna-XM_059381955.1    rna-XM_004776891.3    mNeoVis.2.1.1.chr7_hap1.g00643.t01    rna-XM_044911297.1      rna-XM_042970686.1    rna-XM_008706526.2, rna-XM_008709409.2    rna-XM_027618144.2    mNeoVis.3.1.1.chr7.g00612.t01    rna-gnl|WGS_ZZZZ|egapxtmp_012262-R1    mNeoVis.evergladensis.2.1.chr7.g00640.t01    mNeoVis.vulgivaga.2.1.chr7.g00643.t01      2295   N0.HOG0002293  OG0000390  n23   rna-XM_034639394.1    rna-XM_038525518.1    rna-XM_022492268.1    rna-XM_019818989.3    rna-XM_036088413.1      rna-XM_032847686.1    rna-XM_045989570.1      rna-XM_059152367.1    rna-XM_059381938.1    rna-XM_004780499.3    mNeoVis.2.1.1.chr7_hap1.g00624.t01    rna-XM_021691821.2    rna-XM_004405695.1, rna-XM_004405698.1    rna-XM_042969900.1    rna-XM_008706493.2    rna-XM_035725178.1        mNeoVis.evergladensis.2.1.chr7.g00626.t01        2296    N0.HOG0002294   OG0000390  n41   rna-XM_019805397.2    rna-XM_038527572.1    rna-XM_022492077.1        rna-XM_039219020.1    rna-XM_032847595.1        rna-XM_059152424.1      rna-XM_013048119.2      rna-XM_044911298.1    rna-XM_004410924.1    rna-XM_042970684.1    rna-XM_040633756.1                2297    N0.HOG0002295   OG0000390  n51   rna-XM_034639499.1    rna-XM_038527575.1    rna-XM_022492072.1    rna-XM_045046779.1    rna-XM_036091952.1    rna-XM_039219026.1    rna-XM_032847658.1    rna-XM_045987927.1    rna-XM_032324686.1    rna-XM_059152359.1    rna-XM_059381918.1    rna-XM_013048091.2    mNeoVis.2.1.1.chr7_hap1.g00645.t01    rna-XM_021682340.2    rna-XM_004410923.1    rna-XM_042970676.1    rna-XM_040633740.1    rna-XM_027618142.2      rna-gnl|WGS_ZZZZ|egapxtmp_009696-R1    mNeoVis.evergladensis.2.1.chr7.g00642.t01    mNeoVis.vulgivaga.2.1.chr7.g00644.t01      2298    N0.HOG0002296   OG0000391  n1   rna-XM_034638090.1    rna-XM_038527899.1    rna-XM_022491918.1    rna-XM_023246002.2    rna-XM_036096794.1    rna-XM_039244228.1    rna-XM_032848715.1    rna-XM_045987165.1    rna-XM_032326574.1    rna-XM_059153189.1    rna-XM_059381546.1    rna-XM_004777016.3    mNeoVis.2.1.1.chr7_hap1.g00705.t01    rna-XM_044922021.1    rna-XM_004405078.1    rna-XM_042969967.1    rna-XM_040633807.1    rna-XM_027618516.1    mNeoVis.3.1.1.chr7.g00665.t01      mNeoVis.evergladensis.2.1.chr7.g00703.t01    mNeoVis.vulgivaga.2.1.chr7.g00703.t01      2299   N0.HOG0002297  OG0000391  n19         rna-XM_003997899.6, rna-XM_023249716.2                          rna-XM_007096778.3, rna-XM_015545024.2                  2300    N0.HOG0002298   OG0000391  n25     rna-XM_038529617.1        rna-XM_036067912.1                      rna-XM_004405321.1                    2301    N0.HOG0002299   OG0000391  n30   rna-XM_019797276.1    rna-XM_038584252.1    rna-XM_022515636.1, rna-XM_022525127.1      rna-XM_036070132.1      rna-XM_032846584.1      rna-XM_032316340.1, rna-XM_032323783.1    rna-XM_059152705.1    rna-XM_059377396.1, rna-XM_059382043.1, rna-XM_059395505.1    rna-XM_004765091.3    mNeoVis.2.1.1.chr9_hap1.g00854.t01    rna-XM_021693544.1, rna-XM_021702820.1        rna-XM_040627577.1    rna-XM_027591244.1    mNeoVis.3.1.1.chr6.g00715.t01    rna-gnl|WGS_ZZZZ|egapxtmp_022525-R1    mNeoVis.evergladensis.2.1.chr10.g00843.t01    mNeoVis.vulgivaga.2.1.chr10.g00832.t01, mNeoVis.vulgivaga.2.1.chr6.g00710.t01      2302    N0.HOG0002300   OG0000391  n46     rna-XM_038529320.1, rna-XM_038576134.1        rna-XM_036072002.1        rna-XM_045978691.1              rna-XM_021700906.1          rna-XM_027588725.1, rna-XM_027605116.2, rna-XM_027619089.1              2303    N0.HOG0002301   OG0000391  n53   rna-XM_002920875.4            rna-XM_039249587.1    rna-XM_032841881.1    rna-XM_045988909.1                rna-XM_004404429.1      rna-XM_008712113.2    rna-XM_027614098.1, rna-XM_027617982.1, rna-XM_035722099.1              2304   N0.HOG0002302  OG0000392  n1   rna-XM_034650016.1    rna-XM_038586903.1    rna-XM_022518365.1    rna-XM_011291619.2    rna-XM_036092964.1    rna-XM_039233861.1    rna-XM_032840826.1    rna-XM_045995544.1    rna-XM_032329572.1, rna-XM_032349647.1    rna-XM_059158648.1    rna-XM_059384807.1    rna-XM_045089337.1    mNeoVis.2.1.1.chr12_hap1.g00770.t01    rna-XM_044911835.1    rna-XM_004413150.1    rna-XM_042973752.1    rna-XM_040633937.1    rna-XM_027607569.1, rna-XM_035722412.1    mNeoVis.3.1.1.chr12.g00701.t01    rna-gnl|WGS_ZZZZ|egapxtmp_029275-R4    mNeoVis.evergladensis.2.1.chr12.g00698.t01    mNeoVis.vulgivaga.2.1.chr12.g00725.t01      2305   N0.HOG0002303  OG0000392  n21   rna-XM_019802455.2    rna-XM_038531208.1    rna-XM_022497032.1, rna-XM_022520751.1    rna-XM_003989824.6    rna-XM_036065200.1, rna-XM_036072148.1, rna-XM_036102765.1    rna-XM_039232231.1    rna-XM_032877098.1, rna-XM_032881276.1, rna-XM_032881597.1    rna-XM_045979612.1, rna-XM_045981634.1, rna-XM_045998700.1, rna-XM_045998878.1, rna-XM_046000281.1, rna-XM_046009058.1, rna-XM_046016135.1, rna-XM_046016401.1, rna-XM_046016978.1    rna-XM_032339610.1, rna-XM_032360789.1    rna-XM_059138109.1, rna-XM_059163955.1    rna-XM_059377060.1, rna-XM_059388105.1    rna-XM_013046045.2, rna-XM_045064129.1, rna-XM_045088433.1    mNeoVis.2.1.1.chr2_hap1.g01543.t01    rna-XM_021683661.2, rna-XM_021686415.1, rna-XM_044911766.1, rna-XM_044915373.1    rna-XM_004406454.1, rna-XM_004416912.1    rna-XM_042996324.1    rna-XM_040619193.1    rna-XM_027567555.2, rna-XM_027605828.2, rna-XM_035721786.1, rna-XM_035725448.1, rna-XM_035729341.1    mNeoVis.3.1.1.chr5.g01467.t01    rna-gnl|WGS_ZZZZ|egapxtmp_006595-R1    mNeoVis.evergladensis.2.1.chr5.g01481.t01    mNeoVis.vulgivaga.2.1.chr5.g01485.t01      2306    N0.HOG0002304   OG0000393  n1   rna-XM_034660383.1    rna-XM_038587862.1    rna-XM_022521654.1    rna-XM_019824066.3    rna-XM_036077337.1    rna-XM_039218090.1    rna-XM_032881755.1    rna-XM_045994464.1    rna-XM_032330904.1    rna-XM_059158014.1    rna-XM_059386215.1    rna-XM_045085303.1    mNeoVis.2.1.1.chr12_hap1.g00378.t01    rna-XM_044911695.1    rna-XM_004417556.1    rna-XM_015542894.2    rna-XM_040630042.1    rna-XM_027609084.1    mNeoVis.3.1.1.chr12.g00342.t01    rna-gnl|WGS_ZZZZ|egapxtmp_028138-R1    mNeoVis.evergladensis.2.1.chr12.g00336.t01    mNeoVis.vulgivaga.2.1.chr12.g00343.t01      2307    N0.HOG0002305   OG0000393  n23   rna-XM_034649821.1    rna-NM_001003343.1    rna-XM_022508030.1    rna-XM_045050787.1    rna-XM_036076649.1, rna-XM_036077838.1, rna-XM_036095621.1    rna-XM_039248767.1, rna-XM_039253866.1    rna-XM_032883122.1    rna-XM_045994538.1    rna-XM_032330374.1    rna-XM_059157897.1    rna-XM_059385278.1    rna-XM_045089434.1    mNeoVis.2.1.1.chr12_hap1.g00703.t01    rna-XM_021677947.1    rna-XM_004416747.1, rna-XM_004417391.1    rna-XM_042974140.1    rna-XM_040633145.1    rna-XM_027607979.2    mNeoVis.3.1.1.chr12.g00643.t01    rna-gnl|WGS_ZZZZ|egapxtmp_029570-R1    mNeoVis.evergladensis.2.1.chr12.g00633.t01    mNeoVis.vulgivaga.2.1.chr12.g00656.t01      2308    N0.HOG0002306   OG0000393  n48   rna-XM_034669247.1, rna-XM_034670789.1    rna-NM_001012395.1    rna-XM_022503757.1    rna-XM_045058162.1    rna-XM_036096786.1, rna-XM_036112463.1    rna-XM_039230876.1    rna-XM_032873099.1    rna-XM_046006017.1    rna-XM_032338007.1    rna-XM_059177334.1    rna-XM_059400878.1    rna-XM_004752756.3    mNeoVis.2.1.1.chr1_hap1.g01859.t01    rna-XM_044917201.1    rna-XM_004397039.1    rna-XM_042987266.1    rna-XM_040627923.1    rna-XM_027601860.2    mNeoVis.3.1.1.chr1.g01812.t01    rna-gnl|WGS_ZZZZ|egapxtmp_022691-R1    mNeoVis.evergladensis.2.1.chr1.g01826.t01    mNeoVis.vulgivaga.2.1.chr1.g01821.t01      2309   N0.HOG0002307  OG0000394  n1   rna-XM_002928765.4, rna-XM_034645542.1    rna-XM_038569193.1, rna-XM_038569194.1, rna-XM_038569195.1, rna-XM_038569196.1      rna-XM_045039038.1, rna-XM_045039042.1    rna-XM_036107301.1, rna-XM_036107304.1, rna-XM_036107308.1, rna-XM_036112154.1, rna-XM_036112156.1    rna-XM_039236738.1, rna-XM_039236749.1    rna-XM_032881162.1    rna-XM_046014240.1      rna-XM_059137732.1    rna-XM_059407517.1    rna-XM_004752088.3    mNeoVis.2.1.1.chr7_hap1.g01862.t01      rna-XM_004412464.1, rna-XM_004415668.1, rna-XM_004417499.1    rna-XM_007098431.3, rna-XM_007098434.3    rna-XM_008708926.2, rna-XM_040638876.1    rna-XM_027580577.2, rna-XM_027581055.2, rna-XM_035723089.1, rna-XM_035723090.1      rna-gnl|WGS_ZZZZ|egapxtmp_012573-R1    mNeoVis.evergladensis.2.1.chr7.g01825.t01    mNeoVis.vulgivaga.2.1.chr7.g01841.t01      2310    N0.HOG0002308   OG0000394  n35   rna-XM_034651894.1    rna-NM_001003050.1, rna-NM_001313872.1    rna-XM_022506629.1, rna-XM_022506958.1    rna-XM_023249659.2, rna-XM_045038040.1      rna-XM_039236736.1, rna-XM_039236751.1    rna-XM_032851156.1, rna-XM_032881164.1, rna-XM_032881249.1    rna-XM_046014242.1, rna-XM_046016275.1    rna-XM_032358259.1, rna-XM_032359355.1    rna-XM_059137715.1, rna-XM_059137747.1    rna-XM_059407520.1, rna-XM_059407521.1    rna-XM_004752086.3, rna-XM_004752090.3    mNeoVis.2.1.1.chr7_hap1.g01861.t01    rna-XM_044919325.1, rna-XM_044919877.1    rna-XM_004415671.1      rna-XM_008686930.2, rna-XM_008686954.1    rna-XM_027581040.2, rna-XM_027614264.1, rna-XM_027614266.1, rna-XM_027617011.2, rna-XM_035723091.1, rna-XM_035723092.1    mNeoVis.3.1.1.chr7.g01730.t01    rna-gnl|WGS_ZZZZ|egapxtmp_010854-R1    mNeoVis.evergladensis.2.1.chr7.g01824.t01        2311    N0.HOG0002309   OG0000395  n3       rna-XM_022513923.1            rna-XM_045978240.1              rna-XM_044911822.1          rna-XM_035728694.1    mNeoVis.3.1.1.chr11.g00279.t01            2312    N0.HOG0002310   OG0000395  n9   rna-XM_011234062.3    rna-XM_038570517.1    rna-XM_022521548.1    rna-XM_023260700.2    rna-XM_036111611.1, rna-XM_036123005.1    rna-XM_039237195.1    rna-XM_032861129.1    rna-XM_046003529.1    rna-XM_032344929.1    rna-XM_059161991.1    rna-XM_059390487.1    rna-XM_004780245.3      rna-XM_021705848.1, rna-XM_044912523.1, rna-XM_044916566.1    rna-XM_004412109.1    rna-XM_042957166.1    rna-XM_040636934.1    rna-XM_027587555.2, rna-XM_035728598.1    mNeoVis.3.1.1.chr2.g00942.t01        mNeoVis.vulgivaga.2.1.chr2.g01025.t01      2313    N0.HOG0002311   OG0000395  n15     rna-XM_038581943.1, rna-XM_038588923.1      rna-XM_045051175.1        rna-XM_032839431.1    rna-XM_046027697.1    rna-XM_032314128.1    rna-XM_059136222.1, rna-XM_059172426.1    rna-XM_059375986.1, rna-XM_059395954.1    rna-XM_045064939.1, rna-XM_045083582.1        rna-XM_004410283.1    rna-XM_007086427.3      rna-XM_035728643.1              2314   N0.HOG0002312  OG0000395  n29   rna-XM_034652516.1                rna-XM_046024142.1                                  2315   N0.HOG0002313  OG0000395  n30   rna-NM_001304841.1    rna-NM_001286937.1    rna-XM_022509147.1    rna-XM_003996863.6    rna-XM_036066712.1, rna-XM_036069124.1, rna-XM_036073687.1    rna-XM_039256811.1    rna-XM_032871621.1, rna-XM_032876353.1    rna-XM_045983743.1    rna-XM_032321470.1, rna-XM_032322857.1    rna-XM_059149442.1    rna-XM_059378735.1    rna-XM_004772741.3    mNeoVis.2.1.1.chr8_hap1.g00407.t01    rna-XM_044921722.1    rna-XM_004410502.1, rna-XM_004416911.1    rna-XM_042966907.1    rna-XM_008701269.2    rna-XM_027626064.2, rna-XM_035726157.1    mNeoVis.3.1.1.chr8.g00371.t01        mNeoVis.vulgivaga.2.1.chr8.g00411.t01      2316    N0.HOG0002314   OG0000396  n1   rna-XM_002917287.4    rna-XM_038578118.1    rna-XM_022499387.1    rna-XM_003994192.6    rna-XM_036087071.1, rna-XM_036106387.1    rna-XM_039257752.1    rna-XM_032837202.1    rna-XM_046027378.1    rna-XM_032313827.1    rna-XM_059162095.1, rna-XM_059169550.1    rna-XM_059398039.1    rna-XM_004749714.3    mNeoVis.2.1.1.chr2_hap1.g00350.t01    rna-XM_021700073.1    rna-XM_004410231.1    rna-XM_042961215.1    rna-XM_040624088.1    rna-XM_035724214.1    mNeoVis.3.1.1.chr5.g00343.t01    rna-gnl|WGS_ZZZZ|egapxtmp_008083-R1    mNeoVis.evergladensis.2.1.chr5.g00351.t01    mNeoVis.vulgivaga.2.1.chr5.g00352.t01      2317    N0.HOG0002315   OG0000396  n6   rna-XM_011237456.3    rna-XM_038587872.1    rna-XM_022521621.1    rna-XM_006943867.5    rna-XM_036067137.1, rna-XM_036079392.1    rna-XM_039218088.1    rna-XM_032881744.1    rna-XM_045996290.1    rna-XM_032330498.1    rna-XM_059157812.1    rna-XM_059385150.1    rna-XM_045085309.1    mNeoVis.2.1.1.chr12_hap1.g00372.t01    rna-XM_021696057.2, rna-XM_044916730.1    rna-XM_004412676.1, rna-XM_004417552.1    rna-XM_007094084.3    rna-XM_040629949.1    rna-XM_027607975.2    mNeoVis.3.1.1.chr1.g00908.t01, mNeoVis.3.1.1.chr12.g00308.t01, mNeoVis.3.1.1.chr12.g00336.t01    rna-gnl|WGS_ZZZZ|egapxtmp_028675-R2    mNeoVis.evergladensis.2.1.chr12.g00330.t01    mNeoVis.vulgivaga.2.1.chr12.g00336.t01      2318    N0.HOG0002316   OG0000396  n19   rna-XM_034655843.1    rna-XM_038551767.1, rna-XM_038585110.1    rna-XM_022521142.1    rna-XM_006927511.3    rna-XM_036097104.1    rna-XM_039252810.1    rna-XM_032843417.1, rna-XM_032846800.1    rna-XM_045999943.1    rna-XM_032335312.1    rna-XM_059175894.1    rna-XM_059416156.1    rna-XM_004759018.3    mNeoVis.2.1.1.chr1_hap1.g00886.t01      rna-XM_004410902.1    rna-XM_042982135.1    rna-XM_040622619.1    rna-XM_027607090.1        mNeoVis.evergladensis.2.1.chr1.g00889.t01    mNeoVis.vulgivaga.2.1.chr1.g00900.t01      2319   N0.HOG0002317  OG0000397  n1   rna-XM_034650463.1                                  rna-XM_040640456.1                2320    N0.HOG0002318   OG0000397  n4           rna-XM_036103177.1, rna-XM_036108438.1, rna-XM_036121575.1, rna-XM_036121637.1, rna-XM_036121646.1, rna-XM_036121692.1, rna-XM_036121765.1                    rna-XM_044912475.1    rna-XM_004416947.1        rna-XM_035724420.1, rna-XM_035724421.1              2321    N0.HOG0002319   OG0000397  n15     rna-XM_038547033.1                                  rna-XM_035724418.1, rna-XM_035724422.1, rna-XM_035724663.1              2322   N0.HOG0002320  OG0000397  n18     rna-XM_038589373.1, rna-XM_038589378.1        rna-XM_036108439.1, rna-XM_036121589.1, rna-XM_036121629.1                    rna-XM_044912526.1, rna-XM_044912540.1          rna-XM_035724458.1, rna-XM_035724471.1, rna-XM_035724833.1              2323   N0.HOG0002321  OG0000397  n27     rna-XM_038547022.1, rna-XM_038547034.1, rna-XM_038556883.1, rna-XM_038589363.1, rna-XM_038589371.1, rna-XM_038589374.1, rna-XM_038589375.1, rna-XM_038589377.1, rna-XM_038589479.1, rna-XM_038589585.1    rna-XM_022519908.1    rna-XM_045044297.1, rna-XM_045044337.1, rna-XM_045044342.1, rna-XM_045045022.1, rna-XM_045045023.1, rna-XM_045045024.1, rna-XM_045045025.1, rna-XM_045051601.1      rna-XM_039225318.1, rna-XM_039225819.1, rna-XM_039241128.1    rna-XM_032870966.1, rna-XM_032871039.1    rna-XM_045986275.1    rna-XM_032320811.1    rna-XM_059149058.1    rna-XM_059380215.1    rna-XM_045080821.1    mNeoVis.2.1.1.chr8_hap1.g00292.t01        rna-XM_042966759.1, rna-XM_042966771.1, rna-XM_042966772.1, rna-XM_042967923.1, rna-XM_042975437.1, rna-XM_042975438.1, rna-XM_042975456.1, rna-XM_042977147.1, rna-XM_042977148.1, rna-XM_042977158.1      rna-XM_027624355.2    mNeoVis.3.1.1.chr8.g00261.t01    rna-gnl|WGS_ZZZZ|egapxtmp_020110-R1    mNeoVis.evergladensis.2.1.chr9.g00291.t01    mNeoVis.vulgivaga.2.1.chr8.g00297.t01      2324   N0.HOG0002322  OG0000398  n1   rna-XM_034642838.1    rna-XM_038575832.1    rna-XM_022495875.1    rna-XM_003994978.5    rna-XM_036076496.1    rna-XM_039242927.1    rna-XM_032876000.1    rna-XM_046025984.1    rna-XM_032310911.1    rna-XM_059140181.1    rna-XM_059408985.1    rna-XM_004742506.3    mNeoVis.2.1.1.chr3_hap1.g00359.t01    rna-XM_021687924.1    rna-XM_004400523.1    rna-XM_042963007.1    rna-XM_040619529.1    rna-XM_027577266.1      rna-gnl|WGS_ZZZZ|egapxtmp_042426-R1    mNeoVis.evergladensis.2.1.chr3.g00368.t01    mNeoVis.vulgivaga.2.1.chr3.g00362.t01      2325   N0.HOG0002323  OG0000398  n20   rna-XM_002925797.4    rna-XM_038538984.1    rna-XM_022501871.1, rna-XM_022501947.1, rna-XM_022514029.1    rna-XM_003998609.5    rna-XM_036102424.1, rna-XM_036115290.1    rna-XM_039222664.1, rna-XM_039238048.1    rna-XM_032846513.1, rna-XM_032872355.1    rna-XM_045982991.1, rna-XM_045992423.1, rna-XM_046012939.1, rna-XM_046024857.1    rna-XM_032328116.1, rna-XM_032339074.1, rna-XM_032342451.1, rna-XM_032346195.1    rna-XM_059154864.1    rna-XM_059414778.1    rna-XM_045074888.1, rna-XM_045081610.1    mNeoVis.2.1.1.chr15_hap1.g00779.t01    rna-XM_044914640.1, rna-XM_044915120.1, rna-XM_044918189.1    rna-XM_004396264.1, rna-XM_004405422.1, rna-XM_004413037.1    rna-XM_007079094.3    rna-XM_040635009.1    rna-XM_027613426.2, rna-XM_035722466.1, rna-XM_035722585.1, rna-XM_035723194.1, rna-XM_035723971.1, rna-XM_035724309.1, rna-XM_035724462.1, rna-XM_035725744.1, rna-XM_035726413.1, rna-XM_035726725.1, rna-XM_035727243.1, rna-XM_035727278.1, rna-XM_035727310.1, rna-XM_035728287.1, rna-XM_035728920.1    mNeoVis.3.1.1.chr15.g00557.t01      mNeoVis.evergladensis.2.1.chr15.g00601.t01    mNeoVis.vulgivaga.2.1.chr15.g00613.t01      2326   N0.HOG0002324  OG0000399  n1   rna-XM_034653101.1    rna-XM_038583011.1    rna-XM_022514956.1    rna-XM_019839839.2    rna-XM_036093687.1    rna-XM_039240608.1    rna-XM_032869527.1    rna-XM_046002409.1    rna-XM_032334962.1    rna-XM_059162836.1    rna-XM_059391059.1    rna-XM_004745464.3    mNeoVis.2.1.1.chr5_hap1.g01200.t01    rna-XM_021692709.2    rna-XM_004391946.1    rna-XM_042956389.1    rna-XM_040621322.1    rna-XM_027583536.2    mNeoVis.3.1.1.chr2.g01112.t01    rna-gnl|WGS_ZZZZ|egapxtmp_035795-R1    mNeoVis.evergladensis.2.1.chr2.g01188.t01    mNeoVis.vulgivaga.2.1.chr2.g01194.t01      2327   N0.HOG0002325  OG0000399  n11   rna-XM_034669031.1    rna-XM_038526635.1    rna-XM_022504061.1    rna-XM_023254279.2    rna-XM_036082265.1    rna-XM_039223752.1    rna-XM_032862373.1    rna-XM_046006905.1    rna-XM_032339721.1    rna-XM_059177751.1    rna-XM_059400603.1    rna-XM_013063023.2    mNeoVis.2.1.1.chr1_hap1.g01722.t01    rna-XM_021693444.1    rna-XM_004397946.1    rna-XM_007078691.3    rna-XM_008691179.2    rna-XM_027603495.2    mNeoVis.3.1.1.chr1.g01677.t01    rna-gnl|WGS_ZZZZ|egapxtmp_040010-R1    mNeoVis.evergladensis.2.1.chr1.g01691.t01    mNeoVis.vulgivaga.2.1.chr1.g01687.t01      2328   N0.HOG0002326  OG0000399  n26   rna-XM_034648312.1    rna-XM_038531256.1    rna-XM_022496793.1    rna-XM_003989809.6    rna-XM_036081831.1    rna-XM_039231134.1    rna-XM_032877055.1    rna-XM_046000282.1    rna-XM_032361257.1    rna-XM_059136401.1    rna-XM_059377010.1, rna-XM_059384509.1    rna-XM_045064112.1, rna-XM_045073777.1    mNeoVis.2.1.1.chr2_hap1.g01562.t01, mNeoVis.2.1.1.chr7_hap1.g01112.t01    rna-XM_021684182.1    rna-XM_004406451.1    rna-XM_007092405.3    rna-XM_008694476.2    rna-XM_027598208.2    mNeoVis.3.1.1.chr5.g01483.t01, mNeoVis.3.1.1.chr7.g01035.t01    rna-gnl|WGS_ZZZZ|egapxtmp_007774-R1    mNeoVis.evergladensis.2.1.chr5.g01500.t01    mNeoVis.vulgivaga.2.1.chr5.g01504.t01, mNeoVis.vulgivaga.2.1.chr7.g01111.t01      2329   N0.HOG0002327  OG0000400  n1   rna-XM_034653948.1    rna-XM_038541286.1    rna-XM_022496445.1    rna-XM_011284861.4    rna-XM_036122690.1    rna-XM_039228452.1, rna-XM_039254497.1    rna-XM_032867183.1, rna-XM_032872361.1    rna-XM_046005546.1, rna-XM_046019425.1    rna-XM_032360598.1    rna-XM_059137746.1    rna-XM_059375459.1    rna-XM_004752561.3    mNeoVis.2.1.1.chr2_hap1.g01062.t01    rna-XM_021690327.1    rna-XM_004402492.1    rna-XM_042997941.1    rna-XM_040627013.1, rna-XM_040635004.1    rna-XM_027611154.2    mNeoVis.3.1.1.chr5.g00988.t01    rna-gnl|WGS_ZZZZ|egapxtmp_006249-R1    mNeoVis.evergladensis.2.1.chr5.g01006.t01    mNeoVis.vulgivaga.2.1.chr5.g00996.t01      2330   N0.HOG0002328  OG0000400  n17   rna-XM_002930829.4    rna-XM_038567193.1    rna-XM_022526944.1    rna-XM_023245709.2    rna-XM_036088912.1    rna-XM_039250994.1    rna-XM_032884093.1    rna-XM_045992395.1    rna-XM_032311923.1    rna-XM_059160066.1    rna-XM_059387722.1    rna-NM_001310211.1    mNeoVis.2.1.1.chr5_hap1.g00316.t01    rna-XM_021705225.2    rna-XM_004396836.1, rna-XM_004398914.1    rna-XM_042978241.1    rna-XM_040628185.1    rna-XM_035722678.1, rna-XM_035722822.1, rna-XM_035725717.1    mNeoVis.3.1.1.chr2.g00254.t01    rna-gnl|WGS_ZZZZ|egapxtmp_034701-R1    mNeoVis.evergladensis.2.1.chr2.g00315.t01    mNeoVis.vulgivaga.2.1.chr2.g00311.t01      2331   N0.HOG0002329  OG0000400  n36   rna-XM_034657362.1    rna-XM_038568039.1    rna-XM_022525736.1    rna-XM_045042592.1    rna-XM_036096915.1, rna-XM_036100773.1    rna-XM_039248194.1    rna-XM_032847099.1    rna-XM_045992069.1    rna-XM_032304801.1    rna-XM_059159196.1    rna-XM_059388456.1    rna-XM_045088055.1    mNeoVis.2.1.1.chr5_hap1.g00012.t01    rna-XM_021705137.2    rna-XM_004395431.1    rna-XM_042977514.1    rna-XM_040629727.1    rna-XM_027587582.1        mNeoVis.evergladensis.2.1.chr2.g00012.t01    mNeoVis.vulgivaga.2.1.chr2.g00012.t01      2332   N0.HOG0002330  OG0000401  n1   rna-XM_011219432.3    rna-NM_001287052.1    rna-XM_022513498.1    rna-XM_003984470.4    rna-XM_036112888.1    rna-XM_039220197.1    rna-XM_032866912.1    rna-XM_045979927.1    rna-XM_032353209.1    rna-XM_059132554.1    rna-XM_059405200.1    rna-XM_004745878.3    mNeoVis.2.1.1.chr11_hap1.g01126.t01    rna-XM_021697397.1    rna-XM_004405518.1    rna-XM_042982437.1    rna-XM_040622931.1    rna-XM_027623444.2    mNeoVis.3.1.1.chr1.g02095.t01    rna-gnl|WGS_ZZZZ|egapxtmp_027550-R1    mNeoVis.evergladensis.2.1.chr11.g01095.t01    mNeoVis.vulgivaga.2.1.chr11.g01122.t01, mNeoVis.vulgivaga.2.1.chr11.g01123.t01      2333   N0.HOG0002331  OG0000401  n5   rna-XM_034654109.1    rna-XM_038562364.1    rna-XM_022496469.1    rna-XM_011285072.4    rna-XM_036122593.1    rna-XM_039254923.1    rna-XM_032881424.1    rna-XM_046005549.1    rna-XM_032360733.1    rna-XM_059137726.1    rna-XM_059375761.1    rna-XM_004769801.3    mNeoVis.2.1.1.chr2_hap1.g00936.t01    rna-XM_021690068.1    rna-XM_004401312.1    rna-XM_042998209.1    rna-XM_040626895.1    rna-XM_027597268.2    mNeoVis.3.1.1.chr5.g00867.t01      mNeoVis.evergladensis.2.1.chr5.g00890.t01    mNeoVis.vulgivaga.2.1.chr5.g00878.t01      2334   N0.HOG0002332  OG0000401  n7   rna-XM_002920545.4    rna-NM_001003273.3    rna-XM_022498093.1    rna-XM_023248788.2    rna-XM_036100573.1    rna-XM_039228392.1, rna-XM_039257644.1    rna-XM_032868052.1    rna-XM_045992012.1    rna-XM_032319734.1    rna-XM_059140323.1, rna-XM_059160898.1    rna-XM_059389726.1    rna-XM_045079927.1    mNeoVis.2.1.1.chr5_hap1.g00648.t01    rna-XM_021686899.2, rna-XM_044912764.1    rna-XM_004399236.1, rna-XM_004411849.1    rna-XM_042978943.1    rna-XM_040628815.1    rna-XM_027582346.2, rna-XM_027608315.2    mNeoVis.3.1.1.chr2.g00567.t01, mNeoVis.3.1.1.chr2.g00568.t01      mNeoVis.evergladensis.2.1.chr2.g00645.t01    mNeoVis.vulgivaga.2.1.chr2.g00646.t01      2335   N0.HOG0002333  OG0000402  n2   rna-XM_002925899.4    rna-XM_038567887.1    rna-XM_022525596.1    rna-XM_023243428.2    rna-XM_036100997.1    rna-XM_039248014.1    rna-XM_032847058.1      rna-XM_032359067.1    rna-XM_059159392.1    rna-XM_059388578.1    rna-XM_045087166.1    mNeoVis.2.1.1.chr5_hap1.g00067.t01    rna-XM_044918628.1    rna-XM_004395377.1    rna-XM_042977687.1    rna-XM_040629665.1    rna-XM_027582598.2    mNeoVis.3.1.1.chr2.g00036.t01      mNeoVis.evergladensis.2.1.chr2.g00067.t01    mNeoVis.vulgivaga.2.1.chr2.g00067.t01      2336   N0.HOG0002334  OG0000402  n8     rna-XM_038541700.1    rna-XM_022496324.1    rna-XM_045035996.1    rna-XM_036103740.1, rna-XM_036103850.1    rna-XM_039217401.1    rna-XM_032844300.1, rna-XM_032870073.1    rna-XM_046021610.1    rna-XM_032346635.1    rna-XM_059183647.1    rna-XM_059375082.1, rna-XM_059404643.1    rna-XM_045064710.1, rna-XM_045084852.1    mNeoVis.2.1.1.chr2_hap1.g01197.t01, mNeoVis.2.1.1.chr9_hap1.g01044.t01    rna-XM_021680025.1, rna-XM_044918961.1    rna-XM_004399933.1, rna-XM_004415173.1    rna-XM_007089813.3    rna-XM_040627146.1      mNeoVis.3.1.1.chr10.g01004.t01, mNeoVis.3.1.1.chr5.g01113.t01    rna-gnl|WGS_ZZZZ|egapxtmp_022179-R1    mNeoVis.evergladensis.2.1.chr10.g01033.t01, mNeoVis.evergladensis.2.1.chr5.g01139.t01    mNeoVis.vulgivaga.2.1.chr10.g01024.t01, mNeoVis.vulgivaga.2.1.chr5.g01126.t01      2337   N0.HOG0002335  OG0000402  n24   rna-XM_034657845.1    rna-XM_038567164.1    rna-XM_022526841.1    rna-XM_006928464.5    rna-XM_036110899.1    rna-XM_039252296.1    rna-XM_032884118.1    rna-XM_045990594.1    rna-XM_032312263.1    rna-XM_059160098.1    rna-XM_059389110.1    rna-XM_045083601.1    mNeoVis.2.1.1.chr5_hap1.g00327.t01    rna-XM_021705020.2    rna-XM_004398926.1    rna-XM_042978256.1    rna-XM_040629266.1    rna-XM_027584332.1    mNeoVis.3.1.1.chr2.g00265.t01      mNeoVis.evergladensis.2.1.chr2.g00326.t01    mNeoVis.vulgivaga.2.1.chr2.g00322.t01, mNeoVis.vulgivaga.2.1.chr2.g00323.t01      2338   N0.HOG0002336  OG0000403  n1   rna-XM_034660408.1    rna-XM_038584022.1    rna-XM_022517720.1    rna-XM_006931498.5    rna-XM_036087295.1    rna-XM_039246538.1    rna-XM_032861457.1    rna-XM_046005854.1    rna-XM_032340959.1    rna-XM_059179120.1    rna-XM_059399339.1    rna-XM_045066976.1    mNeoVis.2.1.1.chr1_hap1.g01273.t01    rna-XM_021696975.1    rna-XM_004408410.1    rna-XM_042985863.1    rna-XM_008692869.2    rna-XM_027601750.1    mNeoVis.3.1.1.chr1.g01229.t01    rna-gnl|WGS_ZZZZ|egapxtmp_043419-R4    mNeoVis.evergladensis.2.1.chr1.g01245.t01    mNeoVis.vulgivaga.2.1.chr1.g01245.t01      2339   N0.HOG0002337  OG0000403  n21   rna-XM_011216767.3    rna-XM_038584027.1    rna-XM_022517728.1    rna-XM_006931495.5    rna-XM_036087300.1    rna-XM_039246545.1    rna-XM_032861464.1    rna-XM_046006551.1    rna-XM_032330085.1, rna-XM_032340963.1    rna-XM_059133860.1, rna-XM_059179116.1    rna-XM_059373703.1, rna-XM_059399346.1    rna-XM_004753617.3, rna-XM_045069057.1    mNeoVis.2.1.1.chr1_hap1.g01272.t01    rna-XM_021696938.1    rna-XM_004408407.1    rna-XM_015534916.2    rna-XM_040644999.1    rna-XM_027602763.1    mNeoVis.3.1.1.chr1.g01228.t01    rna-gnl|WGS_ZZZZ|egapxtmp_043455-R1    mNeoVis.evergladensis.2.1.chr1.g01244.t01    mNeoVis.vulgivaga.2.1.chr1.g01244.t01, mNeoVis.vulgivaga.2.1.chr4.g00250.t01      2340   N0.HOG0002338  OG0000403  n46   rna-XM_019807101.2    rna-XM_038525838.1    rna-XM_022513733.1    rna-XM_045041078.1    rna-XM_036067640.1    rna-XM_039251905.1    rna-XM_032872263.1    rna-XM_046024053.1    rna-XM_032310553.1    rna-XM_059139338.1    rna-XM_059410090.1    rna-XM_045065706.1    mNeoVis.2.1.1.chr3_hap1.g00041.t01    rna-XM_021686154.2    rna-XM_004398070.1    rna-XM_042962442.1    rna-XM_040632447.1    rna-XM_027576080.2    mNeoVis.3.1.1.chr3.g00044.t01    rna-gnl|WGS_ZZZZ|egapxtmp_040706-R3    mNeoVis.evergladensis.2.1.chr3.g00047.t01    mNeoVis.vulgivaga.2.1.chr3.g00041.t01      2341   N0.HOG0002339  OG0000404  n1   rna-XM_034651985.1, rna-XM_034652202.1, rna-XM_034653377.1    rna-XM_038562623.1    rna-XM_022518253.1    rna-XM_006929086.4    rna-XM_036122804.1    rna-XM_039242949.1    rna-XM_032874987.1    rna-XM_046020187.1    rna-XM_032304894.1    rna-XM_059170603.1    rna-XM_059397207.1    rna-XM_004739376.3    mNeoVis.2.1.1.chr4_hap1.g00659.t01    rna-XM_021703751.2    rna-XM_004416233.1    rna-XM_042963486.1    rna-XM_040644520.1    rna-XM_027573139.2    mNeoVis.3.1.1.chr6.g00670.t01    rna-gnl|WGS_ZZZZ|egapxtmp_004219-R1    mNeoVis.evergladensis.2.1.chr6.g00666.t01    mNeoVis.vulgivaga.2.1.chr6.g00663.t01      2342    N0.HOG0002340   OG0000404  n24   rna-XM_019802140.2    rna-XM_038585647.1    rna-XM_022490816.1    rna-XM_019838610.3    rna-XM_036089853.1, rna-XM_036123355.1    rna-XM_039252967.1    rna-XM_032859886.1    rna-XM_046019455.1    rna-XM_032354680.1    rna-XM_059168147.1    rna-XM_059395472.1    rna-XM_004767010.3    mNeoVis.2.1.1.chr3_hap1.g01219.t01    rna-XM_021702992.1    rna-XM_004401230.1, rna-XM_004405409.1    rna-XM_042993912.1    rna-XM_008687562.2    rna-XM_027590421.2    mNeoVis.3.1.1.chr3.g01200.t01    rna-gnl|WGS_ZZZZ|egapxtmp_039007-R3    mNeoVis.evergladensis.2.1.chr3.g01216.t01    mNeoVis.vulgivaga.2.1.chr3.g01231.t01      2343    N0.HOG0002341   OG0000404  n45   rna-XM_002915974.4    rna-XM_038549633.1    rna-XM_022523079.1    rna-NM_001319274.1    rna-XM_036077423.1    rna-XM_039216851.1    rna-XM_032874433.1    rna-XM_046011712.1    rna-XM_032348591.1    rna-XM_059185011.1    rna-XM_059403382.1    rna-XM_004773086.3    mNeoVis.2.1.1.chr9_hap1.g00388.t01    rna-XM_021687175.1    rna-XM_004401653.1    rna-XM_042992519.1    rna-XM_008684345.2, rna-XM_040637123.1    rna-XM_027591980.2    mNeoVis.3.1.1.chr10.g00380.t01    rna-gnl|WGS_ZZZZ|egapxtmp_024465-R1    mNeoVis.evergladensis.2.1.chr10.g00385.t01    mNeoVis.vulgivaga.2.1.chr10.g00382.t01      2344   N0.HOG0002342  OG0000405  n1   rna-XM_002923650.4    rna-NM_001284474.1    rna-XM_022511620.1    rna-XM_023245698.2    rna-XM_036070803.1    rna-XM_039222183.1    rna-XM_032854054.1    rna-XM_045988596.1    rna-XM_032323083.1    rna-XM_059150800.1    rna-XM_059383233.1    rna-XM_045076244.1    mNeoVis.2.1.1.chr7_hap1.g00173.t01    rna-XM_021703263.1    rna-XM_004393312.1    rna-XM_042969661.1    rna-XM_040633530.1    rna-XM_027619043.2    mNeoVis.3.1.1.chr7.g00155.t01    rna-gnl|WGS_ZZZZ|egapxtmp_011536-R1    mNeoVis.evergladensis.2.1.chr7.g00173.t01    mNeoVis.vulgivaga.2.1.chr7.g00173.t01      2345   N0.HOG0002343  OG0000405  n21   rna-XM_034641270.1    rna-XM_038572048.1    rna-XM_022493949.1    rna-XM_003983597.6    rna-XM_036092250.1    rna-XM_039248704.1    rna-XM_032858844.1    rna-XM_045980760.1    rna-XM_032350907.1    rna-XM_059133471.1    rna-XM_059406828.1    rna-XM_045084101.1    mNeoVis.2.1.1.chr11_hap1.g00303.t01    rna-XM_021689029.1    rna-XM_004393077.1    rna-XM_042980570.1    rna-XM_040638196.1    rna-XM_027622084.1    mNeoVis.3.1.1.chr11.g00273.t01    rna-gnl|WGS_ZZZZ|egapxtmp_026987-R1    mNeoVis.evergladensis.2.1.chr11.g00300.t01    mNeoVis.vulgivaga.2.1.chr11.g00301.t01      2346    N0.HOG0002344   OG0000405  n42   rna-XM_034668266.1    rna-XM_038579667.1    rna-XM_022502089.1    rna-XM_011291318.3    rna-XM_036072660.1, rna-XM_036080504.1    rna-XM_039223456.1    rna-XM_032869088.1    rna-XM_046005073.1    rna-XM_032314884.1, rna-XM_032316196.1    rna-XM_059166083.1    rna-XM_059378287.1, rna-XM_059394814.1    rna-XM_045063146.1, rna-XM_045069623.1, rna-XM_045072312.1    mNeoVis.2.1.1.chr4_hap1.g00841.t01    rna-XM_021688666.2    rna-XM_004393760.1    rna-XM_042972645.1    rna-XM_040625785.1    rna-XM_027581555.2    mNeoVis.3.1.1.chr6.g00849.t01    rna-gnl|WGS_ZZZZ|egapxtmp_003989-R2    mNeoVis.evergladensis.2.1.chr6.g00844.t01    mNeoVis.vulgivaga.2.1.chr6.g00839.t01      2347    N0.HOG0002345   OG0000406  n1   rna-XM_002927910.4    rna-XM_038527144.1    rna-XM_022512564.1    rna-XM_003995370.6    rna-XM_036100433.1    rna-XM_039249526.1    rna-XM_032866569.1    rna-XM_046022090.1    rna-XM_032306745.1    rna-XM_059141855.1    rna-XM_059411900.1    rna-XM_013044158.2    mNeoVis.2.1.1.chr13_hap1.g00760.t01    rna-XM_021694308.1    rna-XM_004397527.1    rna-XM_042964806.1    rna-XM_040620277.1    rna-XM_027615117.2    mNeoVis.3.1.1.chr13.g00677.t01      mNeoVis.evergladensis.2.1.chr13.g00724.t01    mNeoVis.vulgivaga.2.1.chr13.g00722.t01      2348    N0.HOG0002346   OG0000406  n16   rna-XM_034649459.1    rna-XM_038588356.1    rna-XM_022526358.1    rna-XM_045050334.1    rna-XM_036109672.1    rna-XM_039239875.1    rna-XM_032849299.1    rna-XM_045996510.1    rna-XM_032331075.1, rna-XM_032341803.1    rna-XM_059157767.1, rna-XM_059183413.1    rna-XM_059385446.1    rna-XM_004778403.3    mNeoVis.2.1.1.chr10_hap1.g00649.t01, mNeoVis.2.1.1.chr12_hap1.g00167.t01    rna-XM_044911920.1      rna-XM_042975285.1    rna-XM_040627514.1    rna-XM_035725564.1    mNeoVis.3.1.1.chr12.g00133.t01, mNeoVis.3.1.1.chr12.g00134.t01, mNeoVis.3.1.1.chr9.g00608.t01    rna-gnl|WGS_ZZZZ|egapxtmp_016216-R1, rna-gnl|WGS_ZZZZ|egapxtmp_028996-R1    mNeoVis.evergladensis.2.1.chr12.g00140.t01, mNeoVis.evergladensis.2.1.chr8.g00631.t01    mNeoVis.vulgivaga.2.1.chr12.g00142.t01, mNeoVis.vulgivaga.2.1.chr9.g00632.t01      2349   N0.HOG0002347  OG0000406  n43   rna-XM_011224356.3    rna-NM_001313831.1    rna-XM_022526312.1    rna-XM_006944018.4      rna-XM_039239861.1    rna-XM_032849286.1    rna-XM_045995634.1    rna-XM_032330466.1    rna-XM_059157301.1    rna-XM_059384629.1    rna-XM_013048920.2    mNeoVis.2.1.1.chr12_hap1.g00168.t01    rna-XM_044911529.1    rna-XM_004401073.1    rna-XM_042975291.1    rna-XM_040627507.1    rna-XM_027608901.2    mNeoVis.3.1.1.chr12.g00135.t01    rna-gnl|WGS_ZZZZ|egapxtmp_029403-R1    mNeoVis.evergladensis.2.1.chr12.g00141.t01    mNeoVis.vulgivaga.2.1.chr12.g00143.t01      2350    N0.HOG0002348   OG0000407  n1   rna-XM_002922227.3    rna-XM_038576640.1    rna-XM_022522282.1    rna-XM_011283710.4    rna-XM_036069325.1    rna-XM_039217564.1    rna-XM_032874709.1    rna-XM_046013541.1    rna-XM_032348943.1    rna-XM_059184666.1    rna-XM_059403667.1    rna-XM_045071832.1    mNeoVis.2.1.1.chr9_hap1.g00275.t01    rna-XM_044914942.1    rna-XM_004413927.1    rna-XM_042991689.1    rna-XM_040642115.1    rna-XM_027594995.2    mNeoVis.3.1.1.chr7.g00053.t01    rna-gnl|WGS_ZZZZ|egapxtmp_021647-R2    mNeoVis.evergladensis.2.1.chr10.g00274.t01    mNeoVis.vulgivaga.2.1.chr10.g00270.t01      2351    N0.HOG0002349   OG0000407  n15   rna-XM_011237023.3, rna-XM_034652967.1    rna-XM_038536537.1    rna-XM_022524528.1    rna-XM_003996145.5    rna-XM_036078301.1    rna-XM_039254532.1    rna-XM_032839917.1    rna-XM_045985643.1    rna-XM_032320893.1    rna-XM_059148268.1    rna-XM_059379672.1    rna-XM_004760451.3    mNeoVis.2.1.1.chr8_hap1.g01109.t01    rna-XM_021694736.1    rna-XM_004398582.1    rna-XM_042965796.1    rna-XM_040629875.1    rna-XM_035724635.1        mNeoVis.evergladensis.2.1.chr9.g01087.t01    mNeoVis.vulgivaga.2.1.chr8.g01086.t01      2352    N0.HOG0002350   OG0000407  n33   rna-XM_019799082.2, rna-XM_034651418.1, rna-XM_034651802.1, rna-XM_034652860.1    rna-XM_038537883.1    rna-XM_022514267.1    rna-XM_003989469.5    rna-XM_036090537.1    rna-XM_039234952.1    rna-XM_032863072.1, rna-XM_032863326.1    rna-XM_046016165.1    rna-XM_032301880.1    rna-XM_059136615.1    rna-XM_059374878.1    rna-XM_045063686.1    mNeoVis.2.1.1.chr2_hap1.g01847.t01    rna-XM_044914379.1    rna-XM_004397560.1    rna-XM_042994384.1    rna-XM_040645743.1    rna-XM_027614080.2, rna-XM_035722537.1, rna-XM_035724119.1    mNeoVis.3.1.1.chr5.g01774.t01    rna-gnl|WGS_ZZZZ|egapxtmp_009593-R2    mNeoVis.evergladensis.2.1.chr5.g01785.t01    mNeoVis.vulgivaga.2.1.chr5.g01790.t01      2353    N0.HOG0002351   OG0000408  n1   rna-XM_034646073.1, rna-XM_034646076.1    rna-XM_038577621.1    rna-XM_022519394.1    rna-XM_003988680.6    rna-XM_036112975.1    rna-XM_039241381.1    rna-XM_032876826.1    rna-XM_046013204.1    rna-XM_032348149.1    rna-XM_059185393.1    rna-XM_059403077.1    rna-XM_004778840.3    mNeoVis.2.1.1.chr9_hap1.g00567.t01    rna-XM_021704927.1    rna-XM_004400837.1    rna-XM_042992249.1    rna-XM_040642431.1    rna-XM_027592620.2    mNeoVis.3.1.1.chr10.g00536.t01    rna-gnl|WGS_ZZZZ|egapxtmp_024088-R1    mNeoVis.evergladensis.2.1.chr10.g00560.t01    mNeoVis.vulgivaga.2.1.chr10.g00548.t01      2354   N0.HOG0002352  OG0000408  n6   rna-XM_034639705.1    rna-XM_038559752.1    rna-XM_022514016.1    rna-XM_019826029.3    rna-XM_036102097.1    rna-XM_039230724.1    rna-XM_032840280.1    rna-XM_046020508.1    rna-XM_032304485.1    rna-XM_059172280.1    rna-XM_059396087.1    rna-XM_045086368.1    mNeoVis.2.1.1.chr4_hap1.g00033.t01    rna-XM_044919987.1    rna-XM_004408823.1, rna-XM_004408844.1    rna-XM_042975580.1    rna-XM_040643792.1    rna-XM_027573519.2    mNeoVis.3.1.1.chr6.g00037.t01    rna-gnl|WGS_ZZZZ|egapxtmp_002309-R1    mNeoVis.evergladensis.2.1.chr6.g00035.t01    mNeoVis.vulgivaga.2.1.chr6.g00039.t01      2355   N0.HOG0002353  OG0000408  n18     rna-XM_038525480.1                      rna-XM_013052457.2                mNeoVis.3.1.1.chr8.g00214.t01    rna-gnl|WGS_ZZZZ|egapxtmp_019779-R1          2356    N0.HOG0002354   OG0000408  n21   rna-XM_034640996.1    rna-XM_038546900.1    rna-XM_022519828.1    rna-XM_011289308.3    rna-XM_036116178.1    rna-XM_039235915.1    rna-XM_032870861.1    rna-XM_045984854.1    rna-XM_032323058.1    rna-XM_059150065.1    rna-XM_059380502.1    rna-XM_045080898.1    mNeoVis.2.1.1.chr8_hap1.g00257.t01    rna-XM_021678500.1    rna-XM_004405865.1    rna-XM_042966919.1    rna-XM_040640412.1    rna-XM_027605242.2, rna-XM_035724742.1        mNeoVis.evergladensis.2.1.chr9.g00246.t01    mNeoVis.vulgivaga.2.1.chr8.g00254.t01      2357    N0.HOG0002355   OG0000409  n1   rna-XM_034656084.1    rna-XM_038552316.1    rna-XM_022493010.1    rna-XM_011282359.4    rna-XM_036084224.1, rna-XM_036088848.1    rna-XM_039224866.1    rna-XM_032847546.1, rna-XM_032850572.1, rna-XM_032850573.1    rna-XM_045998668.1    rna-XM_032336573.1    rna-XM_059174693.1    rna-XM_059417668.1    rna-XM_013052450.2    mNeoVis.2.1.1.chr1_hap1.g00466.t01    rna-XM_021701882.1    rna-XM_004397823.1    rna-XM_042984805.1    rna-XM_040622383.1    rna-XM_027606834.1    mNeoVis.3.1.1.chr1.g00464.t01    rna-gnl|WGS_ZZZZ|egapxtmp_030719-R1    mNeoVis.evergladensis.2.1.chr1.g00464.t01    mNeoVis.vulgivaga.2.1.chr1.g00472.t01, mNeoVis.vulgivaga.2.1.chr1.g00473.t01, mNeoVis.vulgivaga.2.1.chr1.g00474.t01      2358    N0.HOG0002356   OG0000409  n25   rna-XM_034657281.1, rna-XM_034659447.1    rna-XM_038561604.1    rna-XM_022504361.1    rna-XM_023251808.2    rna-XM_036087001.1    rna-XM_039238115.1    rna-XM_032880681.1    rna-XM_045979218.1    rna-XM_032352608.1    rna-XM_059188197.1    rna-XM_059405714.1    rna-XM_004742295.3    mNeoVis.2.1.1.chr11_hap1.g00851.t01    rna-XM_044919115.1    rna-XM_004411583.1    rna-XM_042981836.1    rna-XM_040623260.1    rna-XM_027622199.2      rna-gnl|WGS_ZZZZ|egapxtmp_027659-R9    mNeoVis.evergladensis.2.1.chr11.g00834.t01    mNeoVis.vulgivaga.2.1.chr11.g00856.t01      2359   N0.HOG0002357  OG0000409  n37   rna-XM_034662590.1    rna-XM_038534069.1    rna-XM_022499291.1    rna-XM_045040980.1    rna-XM_036110933.1    rna-XM_039233287.1    rna-XM_032842041.1    rna-XM_046026581.1    rna-XM_032312382.1    rna-XM_059170124.1    rna-XM_059397521.1    rna-XM_045069885.1    mNeoVis.2.1.1.chr2_hap1.g00545.t01    rna-XM_021700133.1    rna-XM_004393721.1    rna-XM_042960559.1    rna-XM_008695933.2    rna-XM_027595012.1    mNeoVis.3.1.1.chr5.g00540.t01    rna-gnl|WGS_ZZZZ|egapxtmp_009015-R2    mNeoVis.evergladensis.2.1.chr5.g00547.t01    mNeoVis.vulgivaga.2.1.chr5.g00546.t01      2360    N0.HOG0002358   OG0000410  n1   rna-XM_034671732.1    rna-XM_038556238.1    rna-XM_022505991.1    rna-XM_045056264.1    rna-XM_036076110.1    rna-XM_039251792.1    rna-XM_032845892.1    rna-XM_045996930.1    rna-XM_032334475.1    rna-XM_059159945.1    rna-XM_059384871.1    rna-XM_045079833.1    mNeoVis.2.1.1.chr6_hap1.g00118.t01    rna-XM_044912712.1    rna-XM_004392885.1    rna-XM_042982738.1    rna-XM_040624862.1    rna-XM_027600724.2    mNeoVis.3.1.1.chr4.g00113.t01    rna-gnl|WGS_ZZZZ|egapxtmp_001516-R1    mNeoVis.evergladensis.2.1.chr4.g00119.t01    mNeoVis.vulgivaga.2.1.chr4.g00113.t01      2361    N0.HOG0002359   OG0000410  n21   rna-XM_002930857.4    rna-XM_038556234.1      rna-XM_003985311.5                          rna-XM_007086904.3    rna-XM_040625430.1                2362    N0.HOG0002360   OG0000410  n27     rna-XM_038555084.1        rna-XM_036109114.1                    rna-XM_021702275.2                      2363    N0.HOG0002361   OG0000410  n29   rna-XM_034650561.1    rna-NM_001003381.1, rna-XM_038556224.1, rna-XM_038556225.1, rna-XM_038556226.1, rna-XM_038556227.1, rna-XM_038556228.1    rna-XM_022505818.1, rna-XM_022505848.1      rna-XM_036109097.1      rna-XM_032846554.1    rna-XM_045993074.1, rna-XM_045993104.1, rna-XM_045993118.1, rna-XM_045993141.1    rna-XM_032334471.1, rna-XM_032334472.1, rna-XM_032334474.1    rna-XM_059160003.1, rna-XM_059160026.1, rna-XM_059160041.1    rna-XM_059384846.1, rna-XM_059384858.1    rna-XM_045079832.1, rna-XM_045086256.1, rna-XM_045086257.1    mNeoVis.2.1.1.chr6_hap1.g00119.t01, mNeoVis.2.1.1.chr6_hap1.g00120.t01    rna-XM_021702443.2    rna-XM_004392882.1      rna-XM_008683652.2, rna-XM_040625251.1, rna-XM_040625491.1    rna-XM_027600475.1    mNeoVis.3.1.1.chr4.g00114.t01      mNeoVis.evergladensis.2.1.chr4.g00120.t01, mNeoVis.evergladensis.2.1.chr4.g00121.t01    mNeoVis.vulgivaga.2.1.chr4.g00114.t01, mNeoVis.vulgivaga.2.1.chr4.g00115.t01      2364   N0.HOG0002362  OG0000411  n1   rna-XM_034663775.1    rna-XM_038573445.1    rna-XM_022522597.1    rna-XM_045043129.1    rna-XM_036106757.1    rna-XM_039255593.1    rna-XM_032838221.1    rna-XM_045978322.1    rna-XM_032314760.1    rna-XM_059144204.1    rna-XM_059377600.1    rna-XM_045069586.1    mNeoVis.2.1.1.chr8_hap1.g01213.t01    rna-XM_021678237.1    rna-XM_004417294.1, rna-XM_004417494.1    rna-XM_042984757.1    rna-XM_040621665.1    rna-XM_027591112.2    mNeoVis.3.1.1.chr8.g01109.t01, mNeoVis.3.1.1.chr8.g01110.t01    rna-gnl|WGS_ZZZZ|egapxtmp_017597-R1, rna-gnl|WGS_ZZZZ|egapxtmp_020386-R2    mNeoVis.evergladensis.2.1.chr9.g01170.t01, mNeoVis.evergladensis.2.1.chr9.g01171.t01    mNeoVis.vulgivaga.2.1.chr8.g01168.t01, mNeoVis.vulgivaga.2.1.chr8.g01169.t01      2365   N0.HOG0002363  OG0000411  n26   rna-XM_034663776.1    rna-XM_038573428.1    rna-XM_022522564.1    rna-XM_019821051.3    rna-XM_036101368.1    rna-XM_039254618.1      rna-XM_045978516.1    rna-XM_032314708.1    rna-XM_059143871.1    rna-XM_059378234.1    rna-XM_045069633.1    mNeoVis.2.1.1.chr8_hap1.g01215.t01    rna-XM_021678187.2    rna-XM_004416854.1    rna-XM_042985568.1    rna-XM_008692279.2    rna-XM_027589587.2    mNeoVis.3.1.1.chr8.g01112.t01    rna-gnl|WGS_ZZZZ|egapxtmp_018158-R1    mNeoVis.evergladensis.2.1.chr9.g01173.t01    mNeoVis.vulgivaga.2.1.chr8.g01172.t01, mNeoVis.vulgivaga.2.1.chr8.g01173.t01      2366    N0.HOG0002364   OG0000411  n47   rna-XM_019801970.2    rna-XM_038555959.1    rna-XM_022506005.1    rna-XM_045056433.1    rna-XM_036076974.1    rna-XM_039226991.1    rna-XM_032852621.1    rna-XM_045988323.1    rna-XM_032334198.1    rna-XM_059162405.1    rna-XM_059383312.1    rna-XM_004764082.3    mNeoVis.2.1.1.chr6_hap1.g00207.t01    rna-XM_044912996.1    rna-XM_004396120.1    rna-XM_042984609.1    rna-XM_040624838.1    rna-XM_027597680.2    mNeoVis.3.1.1.chr4.g00203.t01    rna-gnl|WGS_ZZZZ|egapxtmp_000688-R1    mNeoVis.evergladensis.2.1.chr4.g00212.t01    mNeoVis.vulgivaga.2.1.chr4.g00208.t01      2367    N0.HOG0002365   OG0000412  n1   rna-XM_034655704.1    rna-XM_038581445.1    rna-XM_022516179.1    rna-XM_003991442.6    rna-XM_036075283.1    rna-XM_039228483.1    rna-XM_032865363.1    rna-XM_046001524.1    rna-XM_032354595.1    rna-XM_059164685.1    rna-XM_059392320.1    rna-XM_004762133.3    mNeoVis.2.1.1.chr5_hap1.g01821.t01    rna-XM_021677971.2    rna-XM_004409617.1    rna-XM_042998497.1    rna-XM_008701886.2    rna-XM_027587525.2    mNeoVis.3.1.1.chr2.g01725.t01      mNeoVis.evergladensis.2.1.chr2.g01805.t01    mNeoVis.vulgivaga.2.1.chr2.g01819.t01      2368    N0.HOG0002366   OG0000412  n20   rna-XM_002917066.4, rna-XM_034655708.1    rna-XM_038581443.1, rna-XM_038581444.1, rna-XM_038581734.1    rna-XM_022516194.1    rna-XM_003991445.4, rna-XM_003991446.5, rna-XM_011285830.4    rna-XM_036103986.1    rna-XM_039228971.1, rna-XM_039229209.1, rna-XM_039229261.1    rna-XM_032865364.1    rna-XM_046001519.1    rna-XM_032354580.1    rna-XM_059164684.1    rna-XM_059392319.1    rna-XM_045084528.1    mNeoVis.2.1.1.chr5_hap1.g01820.t01      rna-XM_004409625.1    rna-XM_007090487.3, rna-XM_042957203.1    rna-XM_008701882.1, rna-XM_008701883.2, rna-XM_008701954.2, rna-XM_008701956.2    rna-XM_035722590.1    mNeoVis.3.1.1.chr2.g01724.t01    rna-gnl|WGS_ZZZZ|egapxtmp_037884-R1      mNeoVis.vulgivaga.2.1.chr2.g01818.t01      2369   N0.HOG0002367  OG0000412  n49   rna-XM_011222090.3    rna-XM_038581442.1    rna-XM_022516016.1    rna-XM_011285768.4    rna-XM_036103984.1    rna-XM_039229230.1    rna-XM_032865365.1    rna-XM_046003599.1    rna-XM_032354574.1    rna-XM_059164681.1    rna-XM_059392318.1    rna-XM_004762132.2    mNeoVis.2.1.1.chr5_hap1.g01818.t01, mNeoVis.2.1.1.chr5_hap1.g01819.t01    rna-XM_021677973.2, rna-XM_044913449.1      rna-XM_007090451.3    rna-XM_008701884.2          mNeoVis.evergladensis.2.1.chr2.g01804.t01        2370    N0.HOG0002368   OG0000413  n1   rna-XM_034661597.1    rna-XM_038583623.1    rna-XM_022506726.1    rna-XM_019840016.3    rna-XM_036084706.1    rna-XM_039241723.1    rna-XM_032875575.1    rna-XM_046002613.1    rna-XM_032338344.1    rna-XM_059163212.1    rna-XM_059391340.1    rna-XM_013052595.2    mNeoVis.2.1.1.chr5_hap1.g01312.t01    rna-XM_021692336.2    rna-XM_004406734.1    rna-XM_042998598.1    rna-XM_040621481.1    rna-XM_027587359.2    mNeoVis.3.1.1.chr2.g01232.t01    rna-gnl|WGS_ZZZZ|egapxtmp_035585-R1    mNeoVis.evergladensis.2.1.chr2.g01299.t01    mNeoVis.vulgivaga.2.1.chr2.g01306.t01      2371    N0.HOG0002369   OG0000413  n20   rna-XM_002916827.4    rna-XM_038547606.1    rna-XM_022509641.1    rna-XM_003996719.5    rna-XM_036095282.1    rna-XM_039227920.1    rna-XM_032871771.1    rna-XM_045985797.1    rna-XM_032321679.1    rna-XM_059148174.1    rna-XM_059378721.1    rna-XM_045070165.1    mNeoVis.2.1.1.chr1_hap1.g00543.t01    rna-XM_021704247.1    rna-XM_004395121.1    rna-XM_007085717.2    rna-XM_040640765.1    rna-XM_035724690.1, rna-XM_035728308.1    mNeoVis.3.1.1.chr1.g00557.t01, mNeoVis.3.1.1.chr8.g00504.t01    rna-gnl|WGS_ZZZZ|egapxtmp_010522-R1    mNeoVis.evergladensis.2.1.chr1.g00550.t01    mNeoVis.vulgivaga.2.1.chr1.g00559.t01      2372   N0.HOG0002370  OG0000413  n35   rna-XM_002921331.4    rna-XM_038557101.1    rna-XM_022503170.1    rna-XM_003982846.5    rna-XM_036120865.1    rna-XM_039222511.1    rna-XM_032873271.1    rna-XM_046021074.1    rna-XM_032303932.1    rna-XM_059171355.1    rna-XM_059397520.1    rna-XM_004743336.3    mNeoVis.2.1.1.chr4_hap1.g00405.t01    rna-XM_021689680.1    rna-XM_004397332.1    rna-XM_007089055.3    rna-XM_008686388.2    rna-XM_027573663.2    mNeoVis.3.1.1.chr6.g00405.t01    rna-gnl|WGS_ZZZZ|egapxtmp_004420-R1    mNeoVis.evergladensis.2.1.chr6.g00408.t01    mNeoVis.vulgivaga.2.1.chr6.g00404.t01      2373   N0.HOG0002371  OG0000414  n1     rna-XM_038573563.1      rna-XM_045055057.1    rna-XM_036099864.1, rna-XM_036117505.1          rna-XM_032331543.1    rna-XM_059158481.1, rna-XM_059182226.1    rna-XM_059385894.1, rna-XM_059392627.1    rna-XM_045068074.1, rna-XM_045080476.1    mNeoVis.2.1.1.chr12_hap1.g00178.t01    rna-XM_044911498.1, rna-XM_044913108.1    rna-XM_004402875.1, rna-XM_004408898.1, rna-XM_004411537.1    rna-XM_042998000.1    rna-XM_040625504.1    rna-XM_035726416.1, rna-XM_035727018.1, rna-XM_035727909.1    mNeoVis.3.1.1.chr12.g00148.t01    rna-gnl|WGS_ZZZZ|egapxtmp_029062-R1, rna-gnl|WGS_ZZZZ|egapxtmp_029063-R1    mNeoVis.evergladensis.2.1.chr12.g00151.t01    mNeoVis.vulgivaga.2.1.chr12.g00154.t01      2374   N0.HOG0002372  OG0000414  n26     rna-XM_038581778.1    rna-XM_022504841.1, rna-XM_022510631.1      rna-XM_036086051.1    rna-XM_039222854.1    rna-XM_032847071.1, rna-XM_032861092.1    rna-XM_045999866.1, rna-XM_046025362.1    rna-XM_032335808.1, rna-XM_032344350.1, rna-XM_032360431.1    rna-XM_059141413.1, rna-XM_059175420.1    rna-XM_059412250.1, rna-XM_059418198.1    rna-XM_045074410.1, rna-XM_045081639.1, rna-XM_045088056.1    mNeoVis.2.1.1.chr1_hap1.g00726.t01    rna-XM_044914732.1      rna-XM_007079774.3, rna-XM_007097416.2    rna-XM_040643731.1      mNeoVis.3.1.1.chr1.g00749.t01    rna-gnl|WGS_ZZZZ|egapxtmp_033447-R1    mNeoVis.evergladensis.2.1.chr1.g00736.t01, mNeoVis.evergladensis.2.1.chr3.g01232.t01    mNeoVis.vulgivaga.2.1.chr1.g00748.t01, mNeoVis.vulgivaga.2.1.chr3.g01246.t01      2375   N0.HOG0002373  OG0000414  n44   rna-XM_002913621.4    rna-XM_038532400.1                            rna-XM_004403132.1, rna-XM_004411196.1, rna-XM_004411554.1      rna-XM_008700813.2                2376   N0.HOG0002374  OG0000414  n46     rna-XM_038580318.1        rna-XM_036096629.1                    rna-XM_021699517.1                      2377   N0.HOG0002375  OG0000414  n48                                     rna-XM_027585843.2, rna-XM_027607053.2, rna-XM_027623077.1, rna-XM_035727360.1              2378   N0.HOG0002376  OG0000415  n1   rna-XM_002927439.4    rna-NM_001003317.2    rna-XM_022515121.1    rna-XM_003992157.6    rna-XM_036099169.1    rna-XM_039219716.1    rna-XM_032860936.1    rna-XM_046002706.1    rna-XM_032332053.1    rna-XM_059162513.1    rna-XM_059390864.1    rna-XM_004779891.3    mNeoVis.2.1.1.chr5_hap1.g01095.t01    rna-XM_021678913.1    rna-XM_004397915.1    rna-XM_042956976.1    rna-XM_040636858.1    rna-XM_027583247.1    mNeoVis.3.1.1.chr2.g01001.t01    rna-gnl|WGS_ZZZZ|egapxtmp_037778-R1    mNeoVis.evergladensis.2.1.chr2.g01085.t01    mNeoVis.vulgivaga.2.1.chr2.g01092.t01      2379   N0.HOG0002377  OG0000415  n4   rna-XM_019796144.2    rna-XM_038549619.1    rna-XM_022497245.1, rna-XM_022522985.1    rna-XM_003988870.6    rna-XM_036086614.1    rna-XM_039216900.1    rna-XM_032870108.1, rna-XM_032874415.1    rna-XM_046011586.1    rna-XM_032348563.1    rna-XM_059143479.1, rna-XM_059185042.1    rna-XM_059403370.1    rna-XM_004773063.3    mNeoVis.2.1.1.chr9_hap1.g00392.t01    rna-XM_021687171.2    rna-XM_004401645.1    rna-XM_007081477.3    rna-XM_008712124.2    rna-XM_027592279.2    mNeoVis.3.1.1.chr10.g00384.t01    rna-gnl|WGS_ZZZZ|egapxtmp_024201-R1    mNeoVis.evergladensis.2.1.chr10.g00389.t01    mNeoVis.vulgivaga.2.1.chr10.g00386.t01      2380   N0.HOG0002378  OG0000415  n10   rna-XM_019802714.2    rna-NM_001003261.2    rna-XM_022509830.1    rna-XM_003996878.6    rna-XM_036069183.1    rna-XM_039243087.1, rna-XM_039256812.1    rna-XM_032871655.1    rna-XM_045985931.1    rna-XM_032320558.1    rna-XM_059148749.1    rna-XM_059379163.1    rna-XM_045080703.1    mNeoVis.2.1.1.chr8_hap1.g00388.t01    rna-XM_021681697.2    rna-XM_004410525.1    rna-XM_007087658.3    rna-XM_008701284.2    rna-XM_027568473.2    mNeoVis.3.1.1.chr8.g00352.t01    rna-gnl|WGS_ZZZZ|egapxtmp_020006-R2    mNeoVis.evergladensis.2.1.chr9.g00386.t01    mNeoVis.vulgivaga.2.1.chr8.g00393.t01      2381   N0.HOG0002379  OG0000416  n0   rna-XM_002921986.4, rna-XM_034650980.1, rna-XM_034659446.1, rna-XM_034665595.1    rna-XM_038544653.1    rna-XM_022524872.1    rna-XM_003987610.4, rna-XM_045055290.1    rna-XM_036076088.1, rna-XM_036107065.1, rna-XM_036120617.1, rna-XM_036122306.1    rna-XM_039217332.1    rna-XM_032871191.1    rna-XM_045990414.1, rna-XM_045997797.1, rna-XM_046007264.1    rna-XM_032337230.1, rna-XM_032344156.1    rna-XM_059133259.1, rna-XM_059171330.1, rna-XM_059174111.1, rna-XM_059182016.1    rna-XM_059371265.1, rna-XM_059395698.1, rna-XM_059407511.1, rna-XM_059417214.1    rna-XM_004738723.3, rna-XM_045062770.1, rna-XM_045079596.1, rna-XM_045084213.1    mNeoVis.2.1.1.chr10_hap1.g00433.t01, mNeoVis.2.1.1.chr1_hap1.g00235.t01, mNeoVis.2.1.1.chr1_hap1.g00281.t01, mNeoVis.2.1.1.chr4_hap1.g00404.t01    rna-XM_021699629.1, rna-XM_021701479.1, rna-XM_044915008.1, rna-XM_044916543.1, rna-XM_044918216.1, rna-XM_044918606.1    rna-XM_004394115.1, rna-XM_004408010.1    rna-XM_007097545.1, rna-XM_042960474.1    rna-XM_008693567.2, rna-XM_040630686.1, rna-XM_040633032.1    rna-XM_027571069.2, rna-XM_027586748.2, rna-XM_035723368.1, rna-XM_035726006.1, rna-XM_035727299.1, rna-XM_035727788.1, rna-XM_035728095.1, rna-XM_035728664.1    mNeoVis.3.1.1.chr1.g00235.t01, mNeoVis.3.1.1.chr1.g00284.t01, mNeoVis.3.1.1.chr6.g00404.t01    rna-gnl|WGS_ZZZZ|egapxtmp_003058-R1, rna-gnl|WGS_ZZZZ|egapxtmp_012491-R1, rna-gnl|WGS_ZZZZ|egapxtmp_036848-R1    mNeoVis.evergladensis.2.1.chr1.g00232.t01, mNeoVis.evergladensis.2.1.chr1.g00280.t01, mNeoVis.evergladensis.2.1.chr6.g00407.t01, mNeoVis.evergladensis.2.1.chr8.g00413.t01    mNeoVis.vulgivaga.2.1.chr1.g00236.t01, mNeoVis.vulgivaga.2.1.chr1.g00284.t01, mNeoVis.vulgivaga.2.1.chr6.g00403.t01, mNeoVis.vulgivaga.2.1.chr9.g00415.t01      2382    N0.HOG0002380   OG0000417  n1   rna-XM_034640863.1    rna-XM_038548576.1    rna-XM_022521018.1, rna-XM_022521370.1    rna-XM_011289055.3    rna-XM_036077298.1    rna-XM_039223014.1    rna-XM_032842774.1, rna-XM_032848975.1    rna-XM_045985896.1    rna-XM_032319644.1    rna-XM_059150181.1    rna-XM_059379878.1    rna-XM_045085448.1    mNeoVis.2.1.1.chr8_hap1.g00943.t01    rna-XM_044921914.1    rna-XM_004415963.1    rna-XM_042965724.1    rna-XM_040641182.1    rna-XM_027625549.2    mNeoVis.3.1.1.chr8.g00875.t01    rna-gnl|WGS_ZZZZ|egapxtmp_020559-R1    mNeoVis.evergladensis.2.1.chr9.g00917.t01    mNeoVis.vulgivaga.2.1.chr8.g00917.t01      2383    N0.HOG0002381   OG0000417  n25   rna-XM_019800487.2    rna-NM_001003214.1    rna-XM_022517141.1    rna-NM_001009216.1    rna-XM_036089203.1, rna-XM_036103439.1    rna-XM_039245169.1    rna-XM_032876319.1    rna-XM_046024660.1    rna-XM_032311489.1    rna-XM_059140138.1    rna-XM_059408392.1    rna-XM_004753221.3    mNeoVis.2.1.1.chr3_hap1.g00597.t01    rna-XM_044921312.1    rna-XM_004395860.1    rna-XM_042962471.1    rna-XM_040619944.1    rna-XM_027576175.1    mNeoVis.3.1.1.chr3.g00589.t01    rna-gnl|WGS_ZZZZ|egapxtmp_041746-R1    mNeoVis.evergladensis.2.1.chr3.g00605.t01    mNeoVis.vulgivaga.2.1.chr3.g00608.t01      2384   N0.HOG0002382  OG0000417  n46   rna-XM_002927328.4, rna-XM_034652711.1    rna-XM_038539954.1    rna-XM_022526587.1    rna-XM_003998734.6    rna-XM_036066527.1    rna-XM_039247886.1    rna-XM_032842731.1    rna-XM_045994351.1    rna-XM_032326785.1    rna-XM_059154093.1    rna-XM_059415040.1    rna-XM_013046579.2    mNeoVis.2.1.1.chr15_hap1.g00093.t01    rna-XM_044915650.1    rna-XM_004397146.1    rna-XM_042971020.1    rna-XM_008685219.2    rna-XM_027612919.1    mNeoVis.3.1.1.chr15.g00087.t01    rna-gnl|WGS_ZZZZ|egapxtmp_033445-R1    mNeoVis.evergladensis.2.1.chr15.g00094.t01    mNeoVis.vulgivaga.2.1.chr15.g00096.t01      2385   N0.HOG0002383  OG0000418  n1   rna-XM_034638602.1    rna-XM_038528435.1    rna-XM_022524045.1    rna-XM_023245973.2    rna-XM_036114070.1    rna-XM_039244491.1    rna-XM_032839204.1    rna-XM_045989414.1    rna-XM_032325219.1    rna-XM_059152665.1    rna-XM_059381651.1    rna-XM_004767473.3    mNeoVis.2.1.1.chr7_hap1.g00969.t01    rna-XM_021681200.2    rna-XM_004400922.1    rna-XM_042969547.1    rna-XM_040633056.1    rna-XM_027620292.2    mNeoVis.3.1.1.chr7.g00907.t01    rna-gnl|WGS_ZZZZ|egapxtmp_011574-R2    mNeoVis.evergladensis.2.1.chr7.g00962.t01    mNeoVis.vulgivaga.2.1.chr7.g00967.t01      2386    N0.HOG0002384   OG0000418  n22   rna-XM_011222069.3    rna-XM_038545083.1    rna-XM_022505330.1    rna-XM_045060130.1    rna-XM_036107253.1    rna-XM_039223377.1    rna-XM_032850231.1, rna-XM_032850236.1, rna-XM_032878680.1    rna-XM_046007068.1    rna-XM_032344617.1    rna-XM_059182454.1    rna-XM_059373490.1    rna-XM_045071036.1    mNeoVis.2.1.1.chr10_hap1.g00273.t01    rna-XM_021679535.1    rna-XM_004397613.1    rna-XM_042990008.1    rna-XM_008703142.2    rna-XM_027568794.1    mNeoVis.3.1.1.chr9.g00243.t01    rna-gnl|WGS_ZZZZ|egapxtmp_014576-R1    mNeoVis.evergladensis.2.1.chr8.g00258.t01    mNeoVis.vulgivaga.2.1.chr9.g00257.t01      2387    N0.HOG0002385   OG0000418  n44   rna-XM_034653367.1, rna-XM_034659600.1    rna-NM_001304962.1    rna-XM_022520438.1    rna-NM_001009848.1    rna-XM_036065649.1, rna-XM_036077344.1    rna-XM_039222495.1    rna-XM_032881994.1    rna-XM_045979200.1    rna-XM_032352861.1    rna-XM_059188487.1    rna-XM_059405523.1    rna-XM_013044992.2    mNeoVis.2.1.1.chr11_hap1.g00958.t01    rna-XM_044919007.1    rna-XM_004399818.1    rna-XM_042982106.1    rna-XM_008700195.2    rna-XM_027621061.2    mNeoVis.3.1.1.chr11.g00912.t01    rna-gnl|WGS_ZZZZ|egapxtmp_025565-R5    mNeoVis.evergladensis.2.1.chr11.g00936.t01    mNeoVis.vulgivaga.2.1.chr11.g00960.t01      2388    N0.HOG0002386   OG0000419  n1   rna-XM_002920750.4    rna-XM_038534974.1    rna-XM_022493437.1    rna-XM_006927970.5    rna-XM_036087664.1    rna-XM_039239503.1    rna-XM_032851606.1    rna-XM_046000247.1    rna-XM_032337256.1    rna-XM_059174084.1    rna-XM_059417194.1    rna-XM_004737782.3    mNeoVis.2.1.1.chr1_hap1.g00222.t01    rna-XM_021701017.1    rna-XM_004402836.1    rna-XM_042993913.1    rna-XM_008692484.2    rna-XM_027605684.2    mNeoVis.3.1.1.chr1.g00222.t01    rna-gnl|WGS_ZZZZ|egapxtmp_021496-R1    mNeoVis.evergladensis.2.1.chr1.g00219.t01    mNeoVis.vulgivaga.2.1.chr1.g00221.t01      2389   N0.HOG0002387  OG0000419  n23   rna-XM_034666164.1    rna-XM_038568332.1    rna-XM_022514093.1    rna-XM_019811653.3    rna-XM_036111825.1    rna-XM_039222360.1    rna-XM_032852829.1    rna-XM_046016531.1    rna-XM_032359434.1    rna-XM_059186746.1    rna-XM_059412525.1    rna-XM_004763573.3    mNeoVis.2.1.1.chr7_hap1.g01448.t01    rna-XM_044919382.1    rna-XM_004410744.1    rna-XM_042959599.1, rna-XM_042959600.1    rna-XM_040631005.1    rna-XM_035722989.1    mNeoVis.3.1.1.chr7.g01356.t01    rna-gnl|WGS_ZZZZ|egapxtmp_013849-R1    mNeoVis.evergladensis.2.1.chr7.g01431.t01    mNeoVis.vulgivaga.2.1.chr7.g01443.t01      2390    N0.HOG0002388   OG0000419  n45   rna-XM_011226817.3    rna-XM_038560321.1    rna-XM_022498823.1    rna-XM_045055240.1    rna-XM_036112387.1    rna-XM_039241195.1    rna-XM_032838150.1    rna-XM_045997973.1    rna-XM_032329212.1    rna-XM_059156316.1    rna-XM_059384252.1    rna-XM_045073780.1, rna-XM_045073791.1, rna-XM_045073793.1    mNeoVis.2.1.1.chr6_hap1.g01003.t01    rna-XM_021694126.1    rna-XM_004407940.1    rna-XM_042956795.1, rna-XM_042983010.1    rna-XM_008710459.2    rna-XM_027599211.2    mNeoVis.3.1.1.chr4.g00960.t01    rna-gnl|WGS_ZZZZ|egapxtmp_000341-R1    mNeoVis.evergladensis.2.1.chr4.g00948.t01    mNeoVis.vulgivaga.2.1.chr4.g00972.t01      2391    N0.HOG0002389   OG0000420  n2   rna-XM_034650113.1    rna-NM_001205191.1    rna-XM_022522131.1      rna-XM_036075999.1    rna-XM_039223772.1    rna-XM_032853279.1    rna-XM_046017449.1    rna-XM_032360383.1    rna-XM_059134363.1    rna-XM_059409552.1    rna-XM_004778997.3      rna-XM_021685638.1    rna-XM_004407582.1        rna-XM_035722655.1    mNeoVis.3.1.1.chr7.g01587.t01      mNeoVis.evergladensis.2.1.chr7.g01666.t01        2392    N0.HOG0002390   OG0000420  n17   rna-XM_011233425.3    rna-NM_001205190.2    rna-XM_022522133.1    rna-XM_003992869.5    rna-XM_036075993.1    rna-XM_039223769.1    rna-XM_032853280.1    rna-XM_046017450.1    rna-XM_032360389.1    rna-XM_059134415.1    rna-XM_059409475.1    rna-XM_004779000.3    mNeoVis.2.1.1.chr7_hap1.g01697.t01    rna-XM_021685292.1    rna-XM_004407567.1    rna-XM_042959564.1    rna-XM_040631310.1    rna-XM_027580390.2    mNeoVis.3.1.1.chr7.g01585.t01    rna-gnl|WGS_ZZZZ|egapxtmp_012529-R1    mNeoVis.evergladensis.2.1.chr7.g01667.t01        2393    N0.HOG0002391   OG0000420  n37   rna-XM_034667615.1    rna-XM_038568855.1    rna-XM_022522130.1    rna-XM_019811931.2, rna-XM_045038022.1    rna-XM_036075989.1, rna-XM_036076006.1    rna-XM_039223788.1    rna-XM_032853289.1, rna-XM_032853293.1    rna-XM_046017447.1, rna-XM_046017455.1    rna-XM_032360381.1, rna-XM_032360394.1    rna-XM_059182225.1    rna-XM_059409464.1, rna-XM_059409567.1    rna-XM_004778994.3, rna-XM_004778999.3    mNeoVis.2.1.1.chr7_hap1.g01698.t01    rna-XM_021685293.1, rna-XM_021685294.1    rna-XM_004407583.1    rna-XM_007092206.3, rna-XM_015542100.2    rna-XM_008709572.2, rna-XM_008709581.2    rna-XM_027580389.2, rna-XM_035722823.1    mNeoVis.3.1.1.chr7.g01588.t01    rna-gnl|WGS_ZZZZ|egapxtmp_012771-R1    mNeoVis.evergladensis.2.1.chr7.g01669.t01    mNeoVis.vulgivaga.2.1.chr7.g01686.t01      2394   N0.HOG0002392  OG0000421  n2   rna-XM_034644841.1    rna-XM_038563848.1    rna-XM_022507246.1    rna-XM_023240082.2    rna-XM_036121535.1    rna-XM_039244860.1    rna-XM_032879535.1    rna-XM_046017694.1    rna-XM_032357505.1    rna-XM_059148079.1    rna-XM_059401021.1    rna-XM_004769961.3    mNeoVis.2.1.1.chr7_hap1.g02469.t01    rna-XM_021685071.2    rna-XM_004393895.1    rna-XM_042958278.1    rna-XM_008709038.2    rna-XM_027580929.1        mNeoVis.evergladensis.2.1.chr7.g02357.t01    mNeoVis.vulgivaga.2.1.chr7.g02377.t01      2395   N0.HOG0002393  OG0000421  n6   rna-XM_034666595.1    rna-XM_038542871.1    rna-XM_022508571.1        rna-XM_039235775.1    rna-XM_032838808.1    rna-XM_045982951.1    rna-XM_032319105.1    rna-XM_059146632.1    rna-XM_059414048.1    rna-XM_004761890.3    mNeoVis.2.1.1.chr14_hap1.g00240.t01    rna-XM_021679500.1    rna-XM_004393536.1      rna-XM_040636133.1    rna-XM_027611539.1    mNeoVis.3.1.1.chr14.g00241.t01    rna-gnl|WGS_ZZZZ|egapxtmp_031865-R1    mNeoVis.evergladensis.2.1.chr14.g00241.t01    mNeoVis.vulgivaga.2.1.chr14.g00242.t01      2396    N0.HOG0002394   OG0000421  n16           rna-XM_036095794.1                      rna-XM_004391591.1                    2397    N0.HOG0002395   OG0000421  n18     rna-XM_038588919.1, rna-XM_038589653.1, rna-XM_038589661.1, rna-XM_038589692.1    rna-XM_022492813.1      rna-XM_036104544.1    rna-XM_039223060.1    rna-XM_032875352.1                rna-XM_044917836.1                      2398    N0.HOG0002396   OG0000421  n23   rna-XM_002919646.4    rna-XM_038544756.1      rna-XM_003987648.6      rna-XM_039240452.1      rna-XM_046009798.1    rna-XM_032344266.1    rna-XM_059182136.1    rna-XM_059373239.1    rna-XM_004738813.3    mNeoVis.2.1.1.chr10_hap1.g00389.t01    rna-XM_044918839.1    rna-XM_004393074.1    rna-XM_042989724.1      rna-XM_027571957.1    mNeoVis.3.1.1.chr9.g00359.t01    rna-gnl|WGS_ZZZZ|egapxtmp_014594-R2    mNeoVis.evergladensis.2.1.chr8.g00371.t01    mNeoVis.vulgivaga.2.1.chr9.g00373.t01      2399   N0.HOG0002397  OG0000422  n1   rna-XM_002915371.4    rna-XM_038556382.1    rna-XM_022503380.1    rna-XM_045052457.1    rna-XM_036120261.1    rna-XM_039223477.1    rna-XM_032865838.1    rna-XM_046021258.1    rna-XM_032305809.1    rna-XM_059172655.1    rna-XM_059395709.1    rna-XM_013051500.2    mNeoVis.2.1.1.chr4_hap1.g00430.t01    rna-XM_021689707.1    rna-XM_004414402.1    rna-XM_015540637.2    rna-XM_008686365.1    rna-XM_027573177.1, rna-XM_027608835.1    mNeoVis.3.1.1.chr6.g00429.t01    rna-gnl|WGS_ZZZZ|egapxtmp_003754-R1    mNeoVis.evergladensis.2.1.chr6.g00434.t01    mNeoVis.vulgivaga.2.1.chr6.g00429.t01      2400    N0.HOG0002398   OG0000422  n21       rna-XM_022524891.1                      mNeoVis.2.1.1.chr14_hap1.g00198.t01              mNeoVis.3.1.1.chr14.g00196.t01      mNeoVis.evergladensis.2.1.chr14.g00198.t01    mNeoVis.vulgivaga.2.1.chr14.g00202.t01      2401    N0.HOG0002399   OG0000422  n28   rna-XM_019795045.2    rna-XM_038586823.1    rna-XM_022518416.1    rna-XM_006943557.5      rna-XM_039234244.1    rna-XM_032880198.1    rna-XM_045995314.1    rna-XM_032330283.1    rna-XM_059157309.1    rna-XM_059384848.1    rna-XM_045089290.1    mNeoVis.2.1.1.chr12_hap1.g00804.t01    rna-XM_044911967.1    rna-XM_004409220.1    rna-XM_042974167.1    rna-XM_040634507.1    rna-XM_027608091.1    mNeoVis.3.1.1.chr12.g00734.t01    rna-gnl|WGS_ZZZZ|egapxtmp_027970-R1    mNeoVis.evergladensis.2.1.chr12.g00731.t01    mNeoVis.vulgivaga.2.1.chr12.g00761.t01      2402    N0.HOG0002400   OG0000422  n39     rna-XM_038569043.1            rna-XM_032837021.1                rna-XM_044911506.1              rna-gnl|WGS_ZZZZ|egapxtmp_029006-R2    mNeoVis.evergladensis.2.1.chr12.g00271.t01    mNeoVis.vulgivaga.2.1.chr12.g00278.t01      2403    N0.HOG0002401   OG0000422  n44   rna-XM_002922593.4    rna-XM_038588013.1      rna-XM_004000771.5    rna-XM_036101398.1    rna-XM_039233530.1      rna-XM_045996401.1    rna-XM_032330571.1    rna-XM_059157692.1    rna-XM_059384998.1    rna-XM_004769011.3    mNeoVis.2.1.1.chr12_hap1.g00303.t01        rna-XM_007090638.3    rna-XM_040629390.1      mNeoVis.3.1.1.chr12.g00275.t01            2404    N0.HOG0002402   OG0000423  n1   rna-XM_034656164.1    rna-XM_038530398.1    rna-XM_022518002.1    rna-XM_023256080.2    rna-XM_036072241.1    rna-XM_039236213.1    rna-XM_032849948.1    rna-XM_045999121.1    rna-XM_032336375.1    rna-XM_059174874.1    rna-XM_059417781.1    rna-XM_004744762.3    mNeoVis.2.1.1.chr1_hap1.g00527.t01, mNeoVis.2.1.1.chr1_hap1.g00528.t01    rna-XM_044916728.1    rna-XM_004397747.1    rna-XM_007077911.2    rna-XM_040622307.1    rna-XM_027606347.2    mNeoVis.3.1.1.chr1.g00542.t01    rna-gnl|WGS_ZZZZ|egapxtmp_032596-R1    mNeoVis.evergladensis.2.1.chr1.g00534.t01    mNeoVis.vulgivaga.2.1.chr1.g00543.t01      2405    N0.HOG0002403   OG0000423  n23   rna-XM_034649855.1    rna-XM_038587833.1    rna-XM_022521608.1    rna-XM_019824053.3      rna-XM_039223066.1, rna-XM_039223208.1    rna-XM_032881797.1    rna-XM_045995095.1    rna-XM_032331686.1    rna-XM_059156884.1    rna-XM_059385748.1    rna-XM_013047532.2    mNeoVis.2.1.1.chr12_hap1.g00398.t01    rna-XM_044911596.1    rna-XM_004412220.1    rna-XM_042974379.1    rna-XM_040630183.1    rna-XM_035725382.1    mNeoVis.3.1.1.chr12.g00359.t01    rna-gnl|WGS_ZZZZ|egapxtmp_029019-R1    mNeoVis.evergladensis.2.1.chr12.g00355.t01    mNeoVis.vulgivaga.2.1.chr12.g00362.t01, mNeoVis.vulgivaga.2.1.chr12.g00364.t01      2406    N0.HOG0002404   OG0000423  n46   rna-XM_019796631.2    rna-XM_038569685.1    rna-XM_022502823.1    rna-XM_006927330.5    rna-XM_036085425.1    rna-XM_039247287.1    rna-XM_032836998.1    rna-XM_046028128.1    rna-XM_032314557.1    rna-XM_059144199.1    rna-XM_059377539.1    rna-XM_013047223.2, rna-XM_045086153.1    mNeoVis.2.1.1.chr8_hap1.g01402.t01    rna-XM_021697835.2    rna-XM_004393646.1    rna-XM_015536788.2    rna-XM_040623355.1    rna-XM_027591357.2    mNeoVis.3.1.1.chr8.g01297.t01    rna-gnl|WGS_ZZZZ|egapxtmp_017643-R1    mNeoVis.evergladensis.2.1.chr9.g01346.t01    mNeoVis.vulgivaga.2.1.chr8.g01354.t01      2407    N0.HOG0002405   OG0000424  n1   rna-XM_034672400.1    rna-XM_038583477.1    rna-XM_022504421.1    rna-XM_023260368.2    rna-XM_036095300.1    rna-XM_039242339.1    rna-XM_032875436.1    rna-XM_046001867.1    rna-XM_032335411.1, rna-XM_032339824.1    rna-XM_059163376.1    rna-XM_059391466.1    rna-XM_045088659.1    mNeoVis.2.1.1.chr5_hap1.g01354.t01    rna-XM_021692582.1    rna-XM_004406804.1    rna-XM_042956602.1    rna-XM_040621552.1    rna-XM_027582777.1    mNeoVis.3.1.1.chr2.g01273.t01, mNeoVis.3.1.1.chr2.g01274.t01    rna-gnl|WGS_ZZZZ|egapxtmp_035592-R1    mNeoVis.evergladensis.2.1.chr2.g01339.t01    mNeoVis.vulgivaga.2.1.chr2.g01350.t01      2408    N0.HOG0002406   OG0000424  n23   rna-XM_034649970.1    rna-XM_038588444.1    rna-XM_022521287.1    rna-XM_019824382.3    rna-XM_036081783.1    rna-XM_039219991.1    rna-XM_032841419.1    rna-XM_045995007.1    rna-XM_032329561.1    rna-XM_059156475.1    rna-XM_059384661.1    rna-XM_004773628.3    mNeoVis.2.1.1.chr12_hap1.g00133.t01    rna-XM_021684582.1    rna-XM_004405909.1    rna-XM_042974882.1    rna-XM_040627861.1    rna-XM_027607943.1    mNeoVis.3.1.1.chr12.g00104.t01    rna-gnl|WGS_ZZZZ|egapxtmp_028065-R1    mNeoVis.evergladensis.2.1.chr12.g00110.t01    mNeoVis.vulgivaga.2.1.chr12.g00111.t01      2409    N0.HOG0002407   OG0000424  n43   rna-XM_034665503.1    rna-XM_038570080.1    rna-XM_022502742.1    rna-XM_023253346.2    rna-XM_036068107.1    rna-XM_039226631.1    rna-XM_032881024.1    rna-XM_045977936.1    rna-XM_032314940.1    rna-XM_059144408.1    rna-XM_059377892.1    rna-XM_013053042.2    mNeoVis.2.1.1.chr8_hap1.g01524.t01    rna-XM_044913327.1    rna-XM_004412248.1    rna-XM_042980389.1    rna-XM_040620383.1    rna-XM_027589024.2    mNeoVis.3.1.1.chr8.g01420.t01    rna-gnl|WGS_ZZZZ|egapxtmp_017693-R1    mNeoVis.evergladensis.2.1.chr9.g01471.t01    mNeoVis.vulgivaga.2.1.chr8.g01475.t01      2410    N0.HOG0002408   OG0000425  n1   rna-XM_002916927.4    rna-XM_038534725.1    rna-XM_022493581.1    rna-XM_019838717.3    rna-XM_036068860.1, rna-XM_036120349.1    rna-XM_039222216.1, rna-XM_039235764.1    rna-XM_032851837.1    rna-XM_045999280.1    rna-XM_032336994.1    rna-XM_059174331.1    rna-XM_059417380.1    rna-XM_004737580.3    mNeoVis.2.1.1.chr1_hap1.g00321.t01    rna-XM_044917108.1    rna-XM_004391985.1    rna-XM_042992077.1    rna-XM_040630613.1    rna-XM_027604980.2    mNeoVis.3.1.1.chr1.g00325.t01    rna-gnl|WGS_ZZZZ|egapxtmp_004777-R1    mNeoVis.evergladensis.2.1.chr1.g00319.t01    mNeoVis.vulgivaga.2.1.chr1.g00324.t01      2411    N0.HOG0002409   OG0000425  n18   rna-XM_034653453.1, rna-XM_034662947.1, rna-XM_034662954.1    rna-XM_038578187.1    rna-XM_022499781.1    rna-XM_023240724.2    rna-XM_036079070.1    rna-XM_039230189.1    rna-XM_032836415.1    rna-XM_046026522.1    rna-XM_032313576.1    rna-XM_059169454.1    rna-XM_059398129.1    rna-XM_004749661.3    mNeoVis.2.1.1.chr2_hap1.g00330.t01    rna-XM_021699786.1    rna-XM_004410980.1    rna-XM_042961001.1    rna-XM_040624688.1    rna-XM_027596641.2    mNeoVis.3.1.1.chr5.g00320.t01    rna-gnl|WGS_ZZZZ|egapxtmp_008451-R1    mNeoVis.evergladensis.2.1.chr5.g00332.t01    mNeoVis.vulgivaga.2.1.chr5.g00333.t01      2412    N0.HOG0002410   OG0000425  n40   rna-XM_034672103.1    rna-XM_038533350.1    rna-XM_022505938.1    rna-XM_045056569.1    rna-XM_036096546.1    rna-XM_039241567.1    rna-XM_032854796.1    rna-XM_045994755.1    rna-XM_032333864.1    rna-XM_059165210.1    rna-XM_059381132.1    rna-XM_045075685.1    mNeoVis.2.1.1.chr6_hap1.g00297.t01    rna-XM_021679433.2    rna-XM_004410886.1    rna-XM_042984909.1    rna-XM_040625069.1    rna-XM_027598238.2    mNeoVis.3.1.1.chr4.g00296.t01    rna-gnl|WGS_ZZZZ|egapxtmp_001201-R1    mNeoVis.evergladensis.2.1.chr4.g00304.t01    mNeoVis.vulgivaga.2.1.chr4.g00302.t01      2413    N0.HOG0002411   OG0000426  n1   rna-XM_034663392.1    rna-XM_038579021.1    rna-XM_022499853.1    rna-XM_023241051.2    rna-XM_036067145.1    rna-XM_039237960.1    rna-XM_032882129.1    rna-XM_046026166.1    rna-XM_032312218.1    rna-XM_059172912.1    rna-XM_059398759.1    rna-XM_004751756.3    mNeoVis.2.1.1.chr2_hap1.g00068.t01    rna-XM_044916228.1    rna-XM_004416397.1    rna-XM_042960604.1    rna-XM_040624398.1    rna-XM_027596158.2    mNeoVis.3.1.1.chr5.g00062.t01    rna-gnl|WGS_ZZZZ|egapxtmp_005913-R1    mNeoVis.evergladensis.2.1.chr5.g00069.t01    mNeoVis.vulgivaga.2.1.chr5.g00065.t01      2414    N0.HOG0002412   OG0000426  n22   rna-XM_002918780.4    rna-XM_038571778.1    rna-XM_022494491.1    rna-XM_003983725.5    rna-XM_036093166.1    rna-XM_039224846.1    rna-XM_032859042.1    rna-XM_045981187.1    rna-XM_032351235.1    rna-XM_059133771.1    rna-XM_059386510.1, rna-XM_059406658.1    rna-XM_045086052.1    mNeoVis.2.1.1.chr11_hap1.g00404.t01    rna-XM_021689146.1    rna-XM_004398148.1    rna-XM_042980772.1    rna-XM_040638072.1    rna-XM_027622703.2    mNeoVis.3.1.1.chr11.g00370.t01      mNeoVis.evergladensis.2.1.chr11.g00400.t01    mNeoVis.vulgivaga.2.1.chr11.g00405.t01      2415    N0.HOG0002413   OG0000426  n40   rna-XM_034639755.1, rna-XM_034650213.1, rna-XM_034650909.1    rna-XM_038559575.1    rna-XM_022518217.1, rna-XM_022518288.1    rna-XM_023250109.2    rna-XM_036085277.1    rna-XM_039242982.1    rna-XM_032875063.1    rna-XM_046021221.1    rna-XM_032304148.1    rna-XM_059170717.1    rna-XM_059397103.1    rna-XM_004739710.3    mNeoVis.2.1.1.chr4_hap1.g00608.t01    rna-XM_021693136.2, rna-XM_044920203.1    rna-XM_004414031.1    rna-XM_042962758.1    rna-XM_040644445.1    rna-XM_027574814.2    mNeoVis.3.1.1.chr6.g00625.t01    rna-gnl|WGS_ZZZZ|egapxtmp_002812-R1    mNeoVis.evergladensis.2.1.chr6.g00617.t01    mNeoVis.vulgivaga.2.1.chr6.g00614.t01      2416    N0.HOG0002414   OG0000427  n1   rna-XM_034670130.1    rna-XM_038539815.1    rna-XM_022526504.1    rna-XM_045048309.1    rna-XM_036099798.1    rna-XM_039247716.1    rna-XM_032842644.1    rna-XM_045992939.1    rna-XM_032327851.1    rna-XM_059154131.1    rna-XM_059414919.1    rna-XM_004773889.3    mNeoVis.2.1.1.chr15_hap1.g00047.t01    rna-XM_021700401.2    rna-XM_004397218.1    rna-XM_042971094.1    rna-XM_040636557.1    rna-XM_027611010.2    mNeoVis.3.1.1.chr15.g00044.t01    rna-gnl|WGS_ZZZZ|egapxtmp_033515-R1    mNeoVis.evergladensis.2.1.chr15.g00047.t01    mNeoVis.vulgivaga.2.1.chr15.g00049.t01      2417    N0.HOG0002415   OG0000427  n21   rna-XM_034642541.1    rna-XM_038575894.1    rna-XM_022495632.1    rna-XM_023241666.2, rna-XM_045041668.1    rna-XM_036104239.1    rna-XM_039242871.1    rna-XM_032875893.1    rna-XM_046025688.1    rna-XM_032310463.1    rna-XM_059139932.1    rna-XM_059409065.1    rna-XM_004742603.3    mNeoVis.2.1.1.chr3_hap1.g00305.t01    rna-XM_021690886.2    rna-XM_004414916.1    rna-XM_042962498.1, rna-XM_042962530.1    rna-XM_040619581.1    rna-XM_027577419.2        mNeoVis.evergladensis.2.1.chr3.g00314.t01    mNeoVis.vulgivaga.2.1.chr3.g00308.t01      2418    N0.HOG0002416   OG0000427  n32   rna-XM_034639927.1    rna-XM_038547897.1    rna-XM_022509295.1, rna-XM_022509472.1    rna-XM_019817610.3    rna-XM_036082740.1    rna-XM_039247327.1    rna-XM_032871943.1    rna-XM_045984124.1    rna-XM_032319832.1    rna-XM_059147370.1, rna-XM_059147408.1    rna-XM_059379381.1    rna-XM_004747285.3    mNeoVis.2.1.1.chr8_hap1.g00639.t01    rna-XM_044921734.1    rna-XM_004397641.1, rna-XM_004397702.1    rna-XM_007086722.3    rna-XM_008695503.2, rna-XM_008695606.2    rna-XM_027567500.2    mNeoVis.3.1.1.chr8.g00590.t01    rna-gnl|WGS_ZZZZ|egapxtmp_019228-R1    mNeoVis.evergladensis.2.1.chr9.g00629.t01    mNeoVis.vulgivaga.2.1.chr8.g00635.t01      2419    N0.HOG0002417   OG0000428  n1   rna-XM_034653623.1    rna-XM_038537234.1    rna-XM_022496942.1    rna-XM_023258782.2    rna-XM_036110032.1    rna-XM_039233451.1    rna-XM_032844851.1, rna-XM_032876980.1    rna-XM_046019523.1    rna-XM_032303780.1    rna-XM_059135424.1    rna-XM_059374992.1    rna-XM_004762942.3    mNeoVis.2.1.1.chr2_hap1.g01213.t01    rna-XM_021679964.2    rna-XM_004411852.1    rna-XM_042997476.1    rna-XM_040627192.1    rna-XM_027611016.2    mNeoVis.3.1.1.chr5.g01129.t01    rna-gnl|WGS_ZZZZ|egapxtmp_005673-R1    mNeoVis.evergladensis.2.1.chr5.g01155.t01    mNeoVis.vulgivaga.2.1.chr5.g01143.t01      2420    N0.HOG0002418   OG0000428  n21   rna-XM_034655892.1    rna-XM_038551840.1    rna-XM_022521128.1    rna-XM_045062602.1    rna-XM_036078986.1                    rna-XM_044917048.1    rna-XM_004402598.1    rna-XM_042958739.1    rna-XM_040622566.1    rna-XM_035727511.1    mNeoVis.3.1.1.chr1.g00877.t01    rna-gnl|WGS_ZZZZ|egapxtmp_037933-R1    mNeoVis.evergladensis.2.1.chr1.g00857.t01    mNeoVis.vulgivaga.2.1.chr1.g00872.t01      2421    N0.HOG0002419   OG0000428  n34   rna-XM_034647108.1    rna-XM_038527254.1, rna-XM_038544374.1, rna-XM_038546987.1, rna-XM_038547032.1, rna-XM_038549494.1, rna-XM_038576146.1, rna-XM_038589365.1, rna-XM_038589372.1, rna-XM_038589376.1, rna-XM_038589485.1    rna-XM_022512657.1    rna-XM_019815521.3    rna-XM_036101251.1    rna-XM_039230809.1    rna-XM_032866515.1    rna-XM_046022521.1    rna-XM_032307548.1    rna-XM_059141071.1    rna-XM_059411983.1    rna-XM_004766180.3    mNeoVis.2.1.1.chr13_hap1.g00781.t01    rna-XM_021694397.2    rna-XM_004417764.1    rna-XM_042964132.1    rna-XM_008690919.2, rna-XM_040631542.1    rna-XM_027599160.2, rna-XM_027614830.2    mNeoVis.3.1.1.chr13.g00699.t01    rna-gnl|WGS_ZZZZ|egapxtmp_031367-R1    mNeoVis.evergladensis.2.1.chr13.g00747.t01    mNeoVis.vulgivaga.2.1.chr13.g00743.t01      2422    N0.HOG0002420   OG0000429  n1   rna-XM_034647178.1    rna-XM_038527147.1    rna-XM_022512673.1    rna-XM_045043274.1    rna-XM_036090844.1    rna-XM_039249540.1    rna-XM_032866560.1    rna-XM_046021705.1    rna-XM_032308620.1    rna-XM_059142619.1    rna-XM_059411918.1    rna-XM_045065054.1    mNeoVis.2.1.1.chr13_hap1.g00765.t01    rna-XM_021694440.1    rna-XM_004397518.1    rna-XM_042964328.1    rna-XM_040620284.1    rna-XM_027614627.2    mNeoVis.3.1.1.chr13.g00681.t01    rna-gnl|WGS_ZZZZ|egapxtmp_030781-R1    mNeoVis.evergladensis.2.1.chr13.g00730.t01    mNeoVis.vulgivaga.2.1.chr13.g00727.t01      2423    N0.HOG0002421   OG0000429  n23   rna-XM_034653591.1    rna-XM_038537305.1    rna-XM_022497505.1    rna-XM_045035942.1    rna-XM_036091359.1, rna-XM_036091793.1, rna-XM_036109047.1    rna-XM_039255285.1    rna-XM_032844877.1    rna-XM_046006260.1    rna-XM_032303467.1    rna-XM_059137856.1    rna-XM_059374949.1    rna-XM_045064638.1    mNeoVis.2.1.1.chr2_hap1.g01225.t01    rna-XM_044914530.1    rna-XM_004411284.1    rna-XM_042997426.1    rna-XM_040627213.1    rna-XM_027620886.2    mNeoVis.3.1.1.chr5.g01143.t01    rna-gnl|WGS_ZZZZ|egapxtmp_008826-R1    mNeoVis.evergladensis.2.1.chr5.g01169.t01    mNeoVis.vulgivaga.2.1.chr5.g01159.t01      2424   N0.HOG0002422  OG0000429  n46   rna-XM_034657801.1    rna-XM_038567244.1    rna-XM_022526757.1    rna-XM_045045731.1    rna-XM_036075292.1    rna-XM_039250982.1    rna-XM_032884048.1    rna-XM_045991112.1    rna-XM_032311556.1    rna-XM_059160006.1    rna-XM_059389041.1    rna-XM_045083811.1    mNeoVis.2.1.1.chr5_hap1.g00302.t01    rna-XM_021705108.2    rna-XM_004398890.1    rna-XM_042978208.1    rna-XM_040628272.1    rna-XM_027587631.2    mNeoVis.3.1.1.chr2.g00238.t01    rna-gnl|WGS_ZZZZ|egapxtmp_034669-R1    mNeoVis.evergladensis.2.1.chr2.g00300.t01    mNeoVis.vulgivaga.2.1.chr2.g00296.t01      2425   N0.HOG0002423  OG0000430  n1   rna-XM_034665569.1    rna-XM_038552999.1    rna-XM_022500621.1    rna-XM_045042502.1    rna-XM_036111443.1    rna-XM_039240113.1    rna-XM_032860758.1    rna-XM_046022411.1    rna-XM_032308438.1    rna-XM_059143293.1    rna-XM_059411157.1    rna-XM_013053829.2    mNeoVis.2.1.1.chr13_hap1.g00389.t01    rna-XM_021691219.2    rna-XM_004409735.1    rna-XM_042965354.1    rna-XM_040631674.1    rna-XM_035723567.1    mNeoVis.3.1.1.chr13.g00333.t01    rna-gnl|WGS_ZZZZ|egapxtmp_029897-R1    mNeoVis.evergladensis.2.1.chr13.g00378.t01    mNeoVis.vulgivaga.2.1.chr13.g00378.t01      2426    N0.HOG0002424   OG0000430  n23   rna-XM_034657364.1    rna-XM_038565395.1    rna-XM_022525533.1    rna-XM_045042071.1    rna-XM_036100724.1    rna-XM_039248161.1    rna-XM_032847100.1    rna-XM_045990151.1    rna-XM_032304602.1    rna-XM_059159175.1    rna-XM_059388444.1    rna-XM_045087963.1    mNeoVis.2.1.1.chr5_hap1.g00013.t01    rna-XM_021705543.1    rna-XM_004395495.1      rna-XM_040628311.1, rna-XM_040629757.1, rna-XM_040629758.1    rna-XM_027587777.2    mNeoVis.3.1.1.chr2.g00007.t01    rna-gnl|WGS_ZZZZ|egapxtmp_038715-R1    mNeoVis.evergladensis.2.1.chr2.g00013.t01    mNeoVis.vulgivaga.2.1.chr2.g00013.t01      2427    N0.HOG0002425   OG0000430  n45   rna-XM_019798266.2    rna-NM_001003360.2    rna-XM_022496358.1    rna-XM_019837464.3    rna-XM_036094473.1    rna-XM_039236476.1    rna-XM_032867270.1    rna-XM_045993755.1, rna-XM_046025848.1    rna-XM_032302587.1    rna-XM_059136745.1    rna-XM_059375441.1    rna-XM_004752571.3    mNeoVis.2.1.1.chr2_hap1.g01067.t01    rna-XM_021690258.1    rna-XM_004402504.1    rna-XM_042971563.1, rna-XM_042997926.1    rna-XM_040627018.1    rna-XM_027603436.2    mNeoVis.3.1.1.chr5.g00993.t01    rna-gnl|WGS_ZZZZ|egapxtmp_002440-R1, rna-gnl|WGS_ZZZZ|egapxtmp_008517-R2    mNeoVis.evergladensis.2.1.chr5.g01011.t01    mNeoVis.vulgivaga.2.1.chr5.g01001.t01      2428    N0.HOG0002426   OG0000431  n1   rna-XM_002915125.4    rna-XM_038541246.1    rna-XM_022496076.1    rna-XM_006934931.5    rna-XM_036109367.1    rna-XM_039228442.1    rna-XM_032867194.1    rna-XM_046005612.1    rna-XM_032361055.1    rna-XM_059135197.1    rna-XM_059375473.1    rna-XM_004752541.3    mNeoVis.2.1.1.chr2_hap1.g01057.t01    rna-XM_021689948.2    rna-XM_004409208.1    rna-XM_042997961.1    rna-XM_040626999.1    rna-XM_027598559.2    mNeoVis.3.1.1.chr5.g00984.t01    rna-gnl|WGS_ZZZZ|egapxtmp_008513-R6    mNeoVis.evergladensis.2.1.chr5.g01001.t01    mNeoVis.vulgivaga.2.1.chr5.g00991.t01      2429   N0.HOG0002427  OG0000431  n12         rna-XM_045042445.1      rna-XM_039248038.1        rna-XM_032304372.1    rna-XM_059159152.1    rna-XM_059388426.1    rna-XM_004778121.3    mNeoVis.2.1.1.chr5_hap1.g00001.t01        rna-XM_042977489.1        mNeoVis.3.1.1.chr2.g00001.t01      mNeoVis.evergladensis.2.1.chr2.g00001.t01    mNeoVis.vulgivaga.2.1.chr2.g00001.t01      2430    N0.HOG0002428   OG0000431  n20     rna-XM_038579629.1    rna-XM_022525476.1      rna-XM_036100752.1        rna-XM_045990895.1              rna-XM_021705615.1    rna-XM_004395439.1        rna-XM_027584834.1, rna-XM_035728007.1              2431    N0.HOG0002429   OG0000431  n26   rna-XM_034659997.1    rna-XM_038568050.1            rna-XM_032846924.1    rna-XM_046018140.1                                  2432    N0.HOG0002430   OG0000431  n29   rna-XM_034664281.1    rna-XM_038553111.1    rna-XM_022501117.1    rna-XM_023242569.2    rna-XM_036090931.1    rna-XM_039240496.1    rna-XM_032851180.1, rna-XM_032865602.1    rna-XM_046023752.1    rna-XM_032308468.1    rna-XM_059143259.1    rna-XM_059411050.1    rna-XM_045081709.1    mNeoVis.2.1.1.chr13_hap1.g00350.t01    rna-XM_021691711.2    rna-XM_004397013.1    rna-XM_042964615.1    rna-XM_040631719.1    rna-XM_027614598.2    mNeoVis.3.1.1.chr13.g00293.t01    rna-gnl|WGS_ZZZZ|egapxtmp_030495-R1    mNeoVis.evergladensis.2.1.chr13.g00339.t01    mNeoVis.vulgivaga.2.1.chr13.g00338.t01      2433    N0.HOG0002431   OG0000432  n4   rna-XM_034640938.1          rna-XM_036116146.1, rna-XM_036116231.1, rna-XM_036121179.1                    rna-XM_021678370.1, rna-XM_021678393.1    rna-XM_004405896.1, rna-XM_004411110.1      rna-XM_008693581.2, rna-XM_008693929.1    rna-XM_027568521.2, rna-XM_027624497.2              2434   N0.HOG0002432  OG0000432  n16         rna-XM_045044749.1                          rna-XM_007088236.3, rna-XM_042965462.1, rna-XM_042966011.1                  2435   N0.HOG0002433  OG0000432  n20                     rna-XM_059149641.1                    mNeoVis.3.1.1.chr8.g00240.t01, mNeoVis.3.1.1.chr8.g00241.t01      mNeoVis.evergladensis.2.1.chr9.g00272.t01    mNeoVis.vulgivaga.2.1.chr8.g00278.t01      2436    N0.HOG0002434   OG0000432  n27     rna-XM_038546960.1          rna-XM_039250638.1                                      2437    N0.HOG0002435   OG0000432  n28     rna-XM_038549497.1          rna-XM_039250636.1                            mNeoVis.3.1.1.chr8.g00238.t01            2438    N0.HOG0002436   OG0000432  n30     rna-XM_038546985.1    rna-XM_022520111.1, rna-XM_022520115.1          rna-XM_032870911.1, rna-XM_032871037.1, rna-XM_032871038.1    rna-XM_045985397.1, rna-XM_045986277.1    rna-XM_032320145.1, rna-XM_032320157.1    rna-XM_059149515.1, rna-XM_059149640.1, rna-XM_059149705.1    rna-XM_059379097.1, rna-XM_059379546.1    rna-XM_004749234.3, rna-XM_013054250.2, rna-XM_045073825.1, rna-XM_045080864.1    mNeoVis.2.1.1.chr8_hap1.g00276.t01        rna-XM_042965463.1        mNeoVis.3.1.1.chr8.g00235.t01, mNeoVis.3.1.1.chr8.g00236.t01, mNeoVis.3.1.1.chr8.g00237.t01, mNeoVis.3.1.1.chr8.g00239.t01, mNeoVis.3.1.1.chr8.g00243.t01    rna-gnl|WGS_ZZZZ|egapxtmp_018875-R1, rna-gnl|WGS_ZZZZ|egapxtmp_018877-R2, rna-gnl|WGS_ZZZZ|egapxtmp_019366-R1    mNeoVis.evergladensis.2.1.chr9.g00267.t01, mNeoVis.evergladensis.2.1.chr9.g00268.t01, mNeoVis.evergladensis.2.1.chr9.g00269.t01, mNeoVis.evergladensis.2.1.chr9.g00270.t01, mNeoVis.evergladensis.2.1.chr9.g00271.t01, mNeoVis.evergladensis.2.1.chr9.g00274.t01    mNeoVis.vulgivaga.2.1.chr8.g00274.t01, mNeoVis.vulgivaga.2.1.chr8.g00275.t01, mNeoVis.vulgivaga.2.1.chr8.g00276.t01, mNeoVis.vulgivaga.2.1.chr8.g00277.t01, mNeoVis.vulgivaga.2.1.chr8.g00280.t01      2439   N0.HOG0002437  OG0000433  n2   rna-XM_034657670.1    rna-XM_038567499.1    rna-XM_022525857.1    rna-XM_003981782.6    rna-XM_036066093.1    rna-XM_039251622.1    rna-XM_032883810.1    rna-XM_045991947.1    rna-XM_032308956.1    rna-XM_059159756.1    rna-XM_059388837.1    rna-XM_004771657.3    mNeoVis.2.1.1.chr5_hap1.g00193.t01    rna-XM_021705584.2    rna-XM_004412138.1    rna-XM_042978000.1    rna-XM_040628286.1    rna-XM_027582334.2    mNeoVis.3.1.1.chr2.g00143.t01    rna-gnl|WGS_ZZZZ|egapxtmp_034460-R1    mNeoVis.evergladensis.2.1.chr2.g00190.t01    mNeoVis.vulgivaga.2.1.chr2.g00188.t01      2440    N0.HOG0002438   OG0000433  n11   rna-XM_034667465.1    rna-NM_001313865.1    rna-XM_022490994.1    rna-XM_003999627.6    rna-XM_036067900.1    rna-XM_039251294.1    rna-XM_032845012.1    rna-XM_045982167.1    rna-XM_032317778.1    rna-XM_059145342.1    rna-XM_059413692.1    rna-XM_004778542.3    mNeoVis.2.1.1.chr14_hap1.g00630.t01    rna-XM_021681969.1    rna-XM_004402928.1    rna-XM_042975903.1    rna-XM_008696562.2    rna-XM_035722417.1    mNeoVis.3.1.1.chr14.g00609.t01    rna-gnl|WGS_ZZZZ|egapxtmp_031972-R1    mNeoVis.evergladensis.2.1.chr14.g00627.t01    mNeoVis.vulgivaga.2.1.chr14.g00624.t01      2441    N0.HOG0002439   OG0000433  n30   rna-XM_002914258.4    rna-NM_001003276.3    rna-XM_022504893.1    rna-XM_003987022.6    rna-XM_036087119.1    rna-XM_039218671.1    rna-XM_032864661.1    rna-XM_046007988.1    rna-XM_032331328.1    rna-XM_059180867.1    rna-XM_059372303.1    rna-XM_004758646.3    mNeoVis.2.1.1.chr10_hap1.g00936.t01    rna-XM_021693806.2    rna-XM_004414526.1    rna-XM_007084524.3    rna-XM_008708567.2    rna-XM_027569519.1    mNeoVis.3.1.1.chr9.g00866.t01    rna-gnl|WGS_ZZZZ|egapxtmp_017198-R1    mNeoVis.evergladensis.2.1.chr8.g00906.t01    mNeoVis.vulgivaga.2.1.chr9.g00893.t01      2442    N0.HOG0002440   OG0000434  n1   rna-XM_034661717.1    rna-XM_038580039.1    rna-XM_022526632.1    rna-XM_023255495.2    rna-XM_036110665.1    rna-XM_039232853.1    rna-XM_032856852.1    rna-XM_046009409.1    rna-XM_032343272.1    rna-XM_059181147.1    rna-XM_059372511.1    rna-XM_004751091.3    mNeoVis.2.1.1.chr10_hap1.g00746.t01    rna-XM_021695393.1    rna-XM_004400455.1    rna-XM_042989159.1    rna-XM_040620457.1, rna-XM_040620463.1    rna-XM_035728113.1    mNeoVis.3.1.1.chr9.g00705.t01, mNeoVis.3.1.1.chr9.g00706.t01    rna-gnl|WGS_ZZZZ|egapxtmp_016909-R1    mNeoVis.evergladensis.2.1.chr8.g00727.t01    mNeoVis.vulgivaga.2.1.chr9.g00727.t01      2443    N0.HOG0002441   OG0000434  n24   rna-XM_034640285.1    rna-XM_038571644.1    rna-XM_022493977.1    rna-XM_023251450.2    rna-XM_036122370.1    rna-XM_039224551.1    rna-XM_032859164.1    rna-XM_045980928.1    rna-XM_032351387.1    rna-XM_059133978.1    rna-XM_059386583.1, rna-XM_059406554.1    rna-XM_045086010.1    mNeoVis.2.1.1.chr11_hap1.g00460.t01    rna-XM_021688914.1    rna-XM_004414792.1    rna-XM_042980903.1    rna-XM_040637987.1    rna-XM_027621223.2    mNeoVis.3.1.1.chr11.g00423.t01    rna-gnl|WGS_ZZZZ|egapxtmp_025188-R1    mNeoVis.evergladensis.2.1.chr11.g00452.t01    mNeoVis.vulgivaga.2.1.chr11.g00460.t01      2444    N0.HOG0002442   OG0000434  n43   rna-XM_019796927.2    rna-XM_038564128.1    rna-XM_022507314.1    rna-XM_045039495.1    rna-XM_036100365.1    rna-XM_039244868.1    rna-XM_032879351.1    rna-XM_046014652.1    rna-XM_032357183.1    rna-XM_059145768.1    rna-XM_059402728.1    rna-XM_045064213.1    mNeoVis.2.1.1.chr7_hap1.g02359.t01    rna-XM_021684986.2    rna-XM_004394006.1    rna-XM_042958509.1    rna-XM_040639301.1    rna-XM_027581117.2    mNeoVis.3.1.1.chr7.g02171.t01    rna-gnl|WGS_ZZZZ|egapxtmp_013058-R1    mNeoVis.evergladensis.2.1.chr7.g02305.t01    mNeoVis.vulgivaga.2.1.chr7.g02323.t01      2445    N0.HOG0002443   OG0000435  n1   rna-XM_002922967.4    rna-XM_038579989.1    rna-XM_022522558.1    rna-XM_023255512.2    rna-XM_036072546.1    rna-XM_039217966.1    rna-XM_032856884.1    rna-XM_046008451.1    rna-XM_032343226.1    rna-XM_059181113.1    rna-XM_059372474.1    rna-XM_045067842.1    mNeoVis.2.1.1.chr10_hap1.g00765.t01    rna-XM_021695675.2    rna-XM_004409638.1    rna-XM_042989190.1    rna-XM_040620440.1    rna-XM_027570456.1    mNeoVis.3.1.1.chr9.g00723.t01, mNeoVis.3.1.1.chr9.g00724.t01    rna-gnl|WGS_ZZZZ|egapxtmp_015861-R1    mNeoVis.evergladensis.2.1.chr8.g00746.t01    mNeoVis.vulgivaga.2.1.chr9.g00746.t01      2446    N0.HOG0002444   OG0000435  n10   rna-XM_034659232.1    rna-XM_038551459.1    rna-XM_022504371.1    rna-XM_019827300.3    rna-XM_036085922.1    rna-XM_039250903.1    rna-XM_032873763.1    rna-XM_045979502.1    rna-XM_032352294.1    rna-XM_059187841.1    rna-XM_059405957.1    rna-XM_045077917.1    mNeoVis.2.1.1.chr11_hap1.g00748.t01    rna-XM_021699080.2    rna-XM_004397491.1    rna-XM_042981564.1    rna-XM_040623398.1    rna-XM_035728929.1    mNeoVis.3.1.1.chr11.g00710.t01    rna-gnl|WGS_ZZZZ|egapxtmp_025286-R1    mNeoVis.evergladensis.2.1.chr11.g00730.t01    mNeoVis.vulgivaga.2.1.chr11.g00749.t01      2447    N0.HOG0002445   OG0000435  n21   rna-XM_034638989.1    rna-XM_038528440.1    rna-XM_022524026.1    rna-XM_023245975.2    rna-XM_036114075.1    rna-XM_039244427.1    rna-XM_032839258.1    rna-XM_045989363.1    rna-XM_032325220.1    rna-XM_059152904.1    rna-XM_059382243.1, rna-XM_059395819.1    rna-XM_004767478.2, rna-XM_045069898.1    mNeoVis.2.1.1.chr7_hap1.g00970.t01    rna-XM_021681199.2    rna-XM_004400924.1    rna-XM_042968372.1    rna-XM_040634221.1    rna-XM_027620172.1    mNeoVis.3.1.1.chr7.g00909.t01    rna-gnl|WGS_ZZZZ|egapxtmp_012355-R1    mNeoVis.evergladensis.2.1.chr7.g00963.t01    mNeoVis.vulgivaga.2.1.chr7.g00968.t01      2448    N0.HOG0002446   OG0000436  n1   rna-XM_034656750.1, rna-XM_034656751.1    rna-XM_038534784.1    rna-XM_022493104.1    rna-XM_003981290.4    rna-XM_036079461.1    rna-XM_039235715.1    rna-XM_032851775.1    rna-XM_046000057.1    rna-XM_032337047.1    rna-XM_059174267.1    rna-XM_059416333.1    rna-XM_004737633.3    mNeoVis.2.1.1.chr1_hap1.g00292.t01    rna-XM_021698161.2    rna-XM_004392021.1    rna-XM_042992600.1    rna-XM_040630307.1    rna-XM_027605817.2    mNeoVis.3.1.1.chr1.g00297.t01    rna-gnl|WGS_ZZZZ|egapxtmp_027890-R1    mNeoVis.evergladensis.2.1.chr1.g00291.t01    mNeoVis.vulgivaga.2.1.chr1.g00295.t01      2449    N0.HOG0002447   OG0000436  n24   rna-XM_034661244.1    rna-XM_038573777.1    rna-XM_022512914.1    rna-XM_023252463.2    rna-XM_036078988.1, rna-XM_036101094.1, rna-XM_036116490.1    rna-XM_039250418.1    rna-XM_032855782.1    rna-XM_045997731.1    rna-XM_032332193.1    rna-XM_059177750.1    rna-XM_059372023.1    rna-XM_004763249.2    mNeoVis.2.1.1.chr6_hap1.g00813.t01    rna-XM_044912561.1    rna-XM_004414134.1    rna-XM_042983204.1    rna-XM_040626490.1    rna-XM_027597925.2    mNeoVis.3.1.1.chr4.g00823.t01    rna-gnl|WGS_ZZZZ|egapxtmp_000864-R1    mNeoVis.evergladensis.2.1.chr4.g00815.t01    mNeoVis.vulgivaga.2.1.chr4.g00835.t01      2450    N0.HOG0002448   OG0000436  n47   rna-XM_034663641.1    rna-XM_038579104.1    rna-XM_022499695.1    rna-XM_011287620.4    rna-XM_036101940.1    rna-XM_039250757.1    rna-XM_032882087.1    rna-XM_046026751.1    rna-XM_032314015.1    rna-XM_059172848.1    rna-XM_059398805.1    rna-XM_045068856.1    mNeoVis.2.1.1.chr2_hap1.g00048.t01    rna-XM_044916230.1    rna-XM_004404353.1    rna-XM_042961087.1    rna-XM_040623969.1    rna-XM_027591584.2    mNeoVis.3.1.1.chr5.g00041.t01    rna-gnl|WGS_ZZZZ|egapxtmp_005904-R1    mNeoVis.evergladensis.2.1.chr5.g00048.t01    mNeoVis.vulgivaga.2.1.chr5.g00045.t01      2451    N0.HOG0002449   OG0000437  n1   rna-XM_002918964.4    rna-XM_038566169.1    rna-XM_022497584.1    rna-XM_019822871.2    rna-XM_036102555.1    rna-XM_039243644.1    rna-XM_032868305.1    rna-XM_045990937.1    rna-XM_032322220.1    rna-XM_059161176.1    rna-XM_059389946.1    rna-XM_004738190.3    mNeoVis.2.1.1.chr5_hap1.g00738.t01, mNeoVis.2.1.1.chr7_hap1.g01905.t01    rna-XM_021695165.2    rna-XM_004411453.1    rna-XM_042979161.1    rna-XM_040628685.1    rna-XM_027582515.2    mNeoVis.3.1.1.chr7.g01774.t01    rna-gnl|WGS_ZZZZ|egapxtmp_013017-R1, rna-gnl|WGS_ZZZZ|egapxtmp_037919-R1    mNeoVis.evergladensis.2.1.chr2.g00740.t01, mNeoVis.evergladensis.2.1.chr7.g01867.t01    mNeoVis.vulgivaga.2.1.chr2.g00743.t01, mNeoVis.vulgivaga.2.1.chr7.g01884.t01      2452    N0.HOG0002450   OG0000437  n25   rna-XM_002927811.3, rna-XM_011236732.3    rna-XM_038588703.1    rna-XM_022491347.1, rna-XM_022518709.1    rna-XM_003984891.5, rna-XM_023249077.2    rna-XM_036094708.1, rna-XM_036111210.1    rna-XM_039219230.1, rna-XM_039244546.1    rna-XM_032846359.1, rna-XM_032875573.1    rna-XM_045994519.1, rna-XM_045998199.1    rna-XM_032330941.1, rna-XM_032331020.1    rna-XM_059155230.1, rna-XM_059157388.1    rna-XM_059385797.1, rna-XM_059394010.1    rna-XM_004780708.3, rna-XM_013047327.2    mNeoVis.2.1.1.chr12_hap1.g00043.t01, mNeoVis.2.1.1.chr6_hap1.g00664.t01    rna-XM_021698956.1, rna-XM_044911776.1    rna-XM_004409189.1, rna-XM_004412401.1    rna-XM_007084228.3, rna-XM_042974758.1    rna-XM_008685595.2, rna-XM_040626852.1    rna-XM_027598178.2, rna-XM_027607317.2    mNeoVis.3.1.1.chr12.g00034.t01, mNeoVis.3.1.1.chr4.g00673.t01    rna-gnl|WGS_ZZZZ|egapxtmp_029396-R2, rna-gnl|WGS_ZZZZ|egapxtmp_043044-R1    mNeoVis.evergladensis.2.1.chr12.g00033.t01, mNeoVis.evergladensis.2.1.chr4.g00672.t01    mNeoVis.vulgivaga.2.1.chr12.g00035.t01, mNeoVis.vulgivaga.2.1.chr4.g00683.t01      2453    N0.HOG0002451   OG0000438  n1   rna-XM_002930909.4    rna-NM_001388768.1    rna-XM_022492350.1    rna-XM_003987317.3, rna-XM_003987319.3    rna-XM_036082607.1    rna-XM_039218359.1, rna-XM_039221977.1    rna-XM_032857140.1    rna-XM_046009800.1    rna-XM_032341689.1    rna-XM_059179632.1    rna-XM_059371135.1    rna-XM_004781927.1    mNeoVis.2.1.1.chr10_hap1.g00813.t01    rna-XM_021695577.1    rna-XM_004413510.1    rna-XM_007087459.1, rna-XM_007087461.2    rna-XM_008684021.1, rna-XM_008684023.1    rna-XM_035728192.1    mNeoVis.3.1.1.chr9.g00765.t01    rna-gnl|WGS_ZZZZ|egapxtmp_016928-R1    mNeoVis.evergladensis.2.1.chr8.g00791.t01    mNeoVis.vulgivaga.2.1.chr9.g00790.t01      2454    N0.HOG0002452   OG0000438  n24     rna-NM_001388552.1    rna-XM_022491585.1    rna-XM_011283095.2    rna-XM_036076881.1    rna-XM_039221956.1    rna-XM_032857153.1    rna-XM_046009766.1    rna-XM_032341673.1    rna-XM_059179614.1    rna-XM_059371114.1, rna-XM_059371115.1    rna-XM_013050612.1    mNeoVis.2.1.1.chr10_hap1.g00835.t01    rna-XM_021695578.1      rna-XM_042989295.1    rna-XM_008710337.2, rna-XM_008710434.2    rna-XM_035728197.1      rna-gnl|WGS_ZZZZ|egapxtmp_017347-R1    mNeoVis.evergladensis.2.1.chr8.g00809.t01        2455    N0.HOG0002453   OG0000438  n41   rna-XM_034653411.1, rna-XM_034653437.1    rna-NM_001388551.1, rna-XM_038589595.1    rna-XM_022491377.1    rna-XM_003987371.4    rna-XM_036076882.1    rna-XM_039221958.1    rna-XM_032857151.1    rna-XM_046009764.1    rna-XM_032341674.1    rna-XM_059179615.1    rna-XM_059371117.1    rna-XM_004781738.2    mNeoVis.2.1.1.chr10_hap1.g00834.t01    rna-XM_021695570.1    rna-XM_004415199.1    rna-XM_007091389.1    rna-XM_008710435.2    rna-XM_027570338.2      rna-gnl|WGS_ZZZZ|egapxtmp_017344-R1    mNeoVis.evergladensis.2.1.chr8.g00808.t01    mNeoVis.vulgivaga.2.1.chr9.g00805.t01      2456    N0.HOG0002454   OG0000439  n1   rna-XM_019794050.2, rna-XM_034638885.1    rna-XM_038538348.1    rna-XM_022511572.1    rna-XM_019820034.3    rna-XM_036088061.1    rna-XM_039223001.1    rna-XM_032853967.1    rna-XM_045989100.1    rna-XM_032326226.1    rna-XM_059153312.1    rna-XM_059381500.1    rna-XM_004754013.3    mNeoVis.2.1.1.chr7_hap1.g00141.t01    rna-XM_044911492.1    rna-XM_004393274.1    rna-XM_007074885.3    rna-XM_008702349.2    rna-XM_035725072.1    mNeoVis.3.1.1.chr7.g00123.t01    rna-gnl|WGS_ZZZZ|egapxtmp_009998-R1    mNeoVis.evergladensis.2.1.chr7.g00141.t01    mNeoVis.vulgivaga.2.1.chr7.g00142.t01      2457    N0.HOG0002455   OG0000439  n23   rna-XM_019807194.2    rna-XM_038535710.1    rna-XM_022513046.1    rna-XM_023239279.2    rna-XM_036099247.1    rna-XM_039254855.1    rna-XM_032868866.1    rna-XM_046016129.1    rna-XM_032358546.1    rna-XM_059184115.1    rna-XM_059392104.1    rna-XM_004750201.3    mNeoVis.2.1.1.chr7_hap1.g01370.t01    rna-XM_021696321.1    rna-XM_004408794.1    rna-XM_042957756.1    rna-XM_040631063.1    rna-XM_027580815.2    mNeoVis.3.1.1.chr7.g01280.t01    rna-gnl|WGS_ZZZZ|egapxtmp_013010-R1    mNeoVis.evergladensis.2.1.chr7.g01353.t01    mNeoVis.vulgivaga.2.1.chr7.g01364.t01      2458    N0.HOG0002456   OG0000439  n43   rna-XM_034638886.1    rna-XM_038538353.1    rna-XM_022511564.1    rna-XM_003998235.5    rna-XM_036088065.1    rna-XM_039223033.1    rna-XM_032853971.1, rna-XM_032881796.1    rna-XM_045988990.1    rna-XM_032326231.1    rna-XM_059152164.1    rna-XM_059383780.1    rna-XM_013057024.2    mNeoVis.2.1.1.chr7_hap1.g00142.t01    rna-XM_021703053.1    rna-XM_004393277.1    rna-XM_007074889.3    rna-XM_040633572.1    rna-XM_035725099.1    mNeoVis.3.1.1.chr7.g00124.t01, mNeoVis.3.1.1.chr7.g00125.t01    rna-gnl|WGS_ZZZZ|egapxtmp_009999-R1    mNeoVis.evergladensis.2.1.chr7.g00142.t01    mNeoVis.vulgivaga.2.1.chr7.g00143.t01      2459    N0.HOG0002457   OG0000440  n1   rna-XM_002917214.3    rna-XM_038532518.1    rna-XM_022504892.1    rna-XM_003981081.6    rna-XM_036065564.1    rna-XM_039245561.1    rna-XM_032848182.1    rna-XM_046001090.1    rna-XM_032335926.1    rna-XM_059175276.1    rna-XM_059418097.1    rna-XM_013063701.2    mNeoVis.2.1.1.chr1_hap1.g00674.t01    rna-XM_021700088.1    rna-XM_004413385.1    rna-XM_007079832.3    rna-XM_008691273.2    rna-XM_027606820.2    mNeoVis.3.1.1.chr1.g00693.t01    rna-gnl|WGS_ZZZZ|egapxtmp_034479-R1, rna-gnl|WGS_ZZZZ|egapxtmp_034564-R1    mNeoVis.evergladensis.2.1.chr1.g00681.t01    mNeoVis.vulgivaga.2.1.chr1.g00694.t01      2460    N0.HOG0002458   OG0000440  n24   rna-XM_034642864.1    rna-XM_038562080.1    rna-XM_022526241.1    rna-XM_003990559.6    rna-XM_036098379.1    rna-XM_039246192.1    rna-XM_032843568.1    rna-XM_045981012.1    rna-XM_032303381.1    rna-XM_059136555.1    rna-XM_059376009.1    rna-XM_004776827.3    mNeoVis.2.1.1.chr2_hap1.g00823.t01    rna-XM_044914137.1    rna-XM_004404181.1    rna-XM_042993992.1    rna-XM_008695742.2    rna-XM_027607068.2    mNeoVis.3.1.1.chr5.g00770.t01    rna-gnl|WGS_ZZZZ|egapxtmp_005490-R1    mNeoVis.evergladensis.2.1.chr5.g00780.t01    mNeoVis.vulgivaga.2.1.chr5.g00773.t01      2461    N0.HOG0002459   OG0000440  n41   rna-XM_034667911.1    rna-XM_038532758.1    rna-XM_022511064.1    rna-XM_019831918.3    rna-XM_036096134.1    rna-XM_039245763.1    rna-XM_032842336.1    rna-XM_046010100.1    rna-XM_032342104.1    rna-XM_059180062.1    rna-XM_059371624.1    rna-XM_045067211.1    mNeoVis.2.1.1.chr10_hap1.g01177.t01    rna-XM_021680521.1    rna-XM_004408221.1    rna-XM_007078246.3    rna-XM_008687651.2    rna-XM_027572485.2    mNeoVis.3.1.1.chr9.g01111.t01    rna-gnl|WGS_ZZZZ|egapxtmp_014880-R2    mNeoVis.evergladensis.2.1.chr8.g01155.t01    mNeoVis.vulgivaga.2.1.chr9.g01133.t01      2462    N0.HOG0002460   OG0000441  n1   rna-XM_034648181.1    rna-XM_038531318.1    rna-XM_022496521.1    rna-XM_023258399.2    rna-XM_036098436.1    rna-XM_039230108.1    rna-XM_032864007.1    rna-XM_045995205.1    rna-XM_032302847.1    rna-XM_059137416.1    rna-XM_059376955.1    rna-XM_004740953.3    mNeoVis.2.1.1.chr2_hap1.g01577.t01    rna-XM_021683634.2    rna-XM_004406448.1    rna-XM_042996165.1    rna-XM_040619159.1    rna-XM_027614255.2    mNeoVis.3.1.1.chr5.g01499.t01    rna-gnl|WGS_ZZZZ|egapxtmp_006302-R1    mNeoVis.evergladensis.2.1.chr5.g01516.t01    mNeoVis.vulgivaga.2.1.chr5.g01519.t01      2463    N0.HOG0002461   OG0000441  n9   rna-XM_034669080.1    rna-XM_038526700.1    rna-XM_022503870.1    rna-XM_045057984.1    rna-XM_036108594.1    rna-XM_039256203.1    rna-XM_032862439.1    rna-XM_046006829.1    rna-XM_032339650.1    rna-XM_059177666.1    rna-XM_059400652.1    rna-XM_013063305.2    mNeoVis.2.1.1.chr1_hap1.g01754.t01    rna-XM_021693163.1    rna-XM_004403626.1    rna-XM_007078666.3    rna-XM_008691247.2    rna-XM_027603723.2    mNeoVis.3.1.1.chr1.g01707.t01    rna-gnl|WGS_ZZZZ|egapxtmp_020599-R4    mNeoVis.evergladensis.2.1.chr1.g01722.t01    mNeoVis.vulgivaga.2.1.chr1.g01718.t01      2464    N0.HOG0002462   OG0000441  n23   rna-XM_034641563.1    rna-XM_038572831.1              rna-XM_045980596.1              rna-XM_021677830.1                      2465    N0.HOG0002463   OG0000441  n26   rna-XM_034668114.1    rna-XM_038579381.1    rna-XM_022515541.1    rna-XM_023248646.2      rna-XM_039254597.1    rna-XM_032882626.1    rna-XM_045979715.1    rna-XM_032316115.1    rna-XM_059168851.1    rna-XM_059395511.1    rna-XM_013046102.2    mNeoVis.2.1.1.chr4_hap1.g00736.t01    rna-XM_021688357.1    rna-XM_004398789.1    rna-XM_042972876.1    rna-XM_040625948.1    rna-XM_027572733.2    mNeoVis.3.1.1.chr6.g00745.t01    rna-gnl|WGS_ZZZZ|egapxtmp_002161-R1    mNeoVis.evergladensis.2.1.chr6.g00745.t01    mNeoVis.vulgivaga.2.1.chr6.g00742.t01      2466    N0.HOG0002464   OG0000442  n1   rna-XM_034668143.1    rna-NM_001190428.1    rna-XM_022515527.1    rna-XM_011291369.3    rna-XM_036077150.1, rna-XM_036102406.1    rna-XM_039246981.1    rna-XM_032847908.1    rna-XM_046020112.1    rna-XM_032315597.1    rna-XM_059157935.1, rna-XM_059166309.1    rna-XM_059394668.1    rna-XM_045075712.1    mNeoVis.2.1.1.chr4_hap1.g00758.t01    rna-XM_021688617.2    rna-XM_004402285.1    rna-XM_042973362.1    rna-XM_040625653.1    rna-XM_027617079.2    mNeoVis.3.1.1.chr6.g00764.t01    rna-gnl|WGS_ZZZZ|egapxtmp_004908-R3    mNeoVis.evergladensis.2.1.chr6.g00765.t01    mNeoVis.vulgivaga.2.1.chr6.g00762.t01      2467    N0.HOG0002465   OG0000442  n16   rna-XM_002922291.4    rna-XM_038526060.1    rna-XM_022503858.1    rna-XM_019831272.3    rna-XM_036097438.1    rna-XM_039242043.1    rna-XM_032844788.1    rna-XM_046005794.1    rna-XM_032339454.1    rna-XM_059177498.1    rna-XM_059400758.1    rna-XM_004740240.3    mNeoVis.2.1.1.chr1_hap1.g01813.t01    rna-XM_044917491.1    rna-XM_004401178.1    rna-XM_042987179.1    rna-XM_040627815.1    rna-XM_027598750.2, rna-XM_035728555.1    mNeoVis.3.1.1.chr1.g01766.t01    rna-gnl|WGS_ZZZZ|egapxtmp_022595-R1    mNeoVis.evergladensis.2.1.chr1.g01777.t01    mNeoVis.vulgivaga.2.1.chr1.g01773.t01      2468    N0.HOG0002466   OG0000442  n36   rna-XM_034641589.1    rna-XM_038572353.1    rna-XM_022494360.1    rna-XM_045053601.1    rna-XM_036072514.1, rna-XM_036096075.1    rna-XM_039248484.1    rna-XM_032838576.1    rna-XM_045980842.1    rna-XM_032350622.1    rna-XM_059133108.1    rna-XM_059407039.1    rna-XM_004746414.3    mNeoVis.2.1.1.chr11_hap1.g00221.t01    rna-XM_021688978.1    rna-XM_004417143.1    rna-XM_042980377.1    rna-XM_008698091.2    rna-XM_027623956.1    mNeoVis.3.1.1.chr11.g00191.t01      mNeoVis.evergladensis.2.1.chr11.g00219.t01    mNeoVis.vulgivaga.2.1.chr11.g00221.t01      2469    N0.HOG0002467   OG0000443  n1   rna-XM_002916890.4    rna-XM_038579662.1    rna-XM_022502155.1, rna-XM_022508967.1      rna-XM_036080492.1      rna-XM_032851266.1, rna-XM_032855078.1, rna-XM_032872883.1    rna-XM_045997743.1, rna-XM_045998240.1, rna-XM_046020960.1, rna-XM_046025821.1    rna-XM_032318475.1, rna-XM_032353773.1    rna-XM_059133137.1, rna-XM_059146750.1, rna-XM_059170128.1    rna-XM_059395822.1, rna-XM_059407891.1, rna-XM_059414118.1    rna-XM_013055548.2, rna-XM_045069888.1, rna-XM_045078482.1    mNeoVis.2.1.1.chr4_hap1.g00832.t01    rna-XM_021688203.1, rna-XM_021696269.1, rna-XM_044921215.1    rna-XM_004393753.1, rna-XM_004401772.1        rna-XM_027571846.2, rna-XM_027586790.2, rna-XM_027600312.1    mNeoVis.3.1.1.chr6.g00841.t01    rna-gnl|WGS_ZZZZ|egapxtmp_003979-R1          2470    N0.HOG0002468   OG0000443  n27   rna-XM_019801782.2    rna-XM_038583261.1    rna-XM_022510571.1    rna-XM_003999941.5    rna-XM_036088206.1, rna-XM_036122719.1    rna-XM_039242320.1, rna-XM_039253481.1    rna-XM_032869102.1    rna-XM_046004004.1    rna-XM_032316361.1, rna-XM_032361534.1    rna-XM_059165223.1, rna-XM_059166102.1    rna-XM_059388046.1, rna-XM_059394808.1    rna-XM_013063769.2    mNeoVis.2.1.1.chr5_hap1.g01356.t01    rna-XM_021695121.1, rna-XM_021696544.1, rna-XM_021702027.1    rna-XM_004417655.1    rna-XM_007084314.3, rna-XM_015543292.2    rna-XM_008689101.2, rna-XM_008710062.1    rna-XM_027581023.2, rna-XM_027587258.1, rna-XM_027599780.2    mNeoVis.3.1.1.chr2.g01278.t01    rna-gnl|WGS_ZZZZ|egapxtmp_038299-R1    mNeoVis.evergladensis.2.1.chr2.g01341.t01, mNeoVis.evergladensis.2.1.chr6.g00835.t01    mNeoVis.vulgivaga.2.1.chr2.g01351.t01, mNeoVis.vulgivaga.2.1.chr6.g00831.t01      2471    N0.HOG0002469   OG0000444  n1   rna-XM_002912309.4    rna-XM_038548342.1    rna-XM_022510019.1    rna-XM_045044975.1    rna-XM_036111869.1    rna-XM_039251701.1    rna-XM_032848449.1    rna-XM_045984362.1    rna-XM_032321230.1    rna-XM_059148069.1    rna-XM_059380827.1    rna-XM_045069950.1    mNeoVis.2.1.1.chr8_hap1.g00791.t01    rna-XM_021704581.2    rna-XM_004399685.1    rna-XM_007076068.3    rna-XM_040641074.1    rna-XM_027568207.2    mNeoVis.3.1.1.chr8.g00742.t01    rna-gnl|WGS_ZZZZ|egapxtmp_017736-R1    mNeoVis.evergladensis.2.1.chr9.g00779.t01    mNeoVis.vulgivaga.2.1.chr8.g00782.t01      2472    N0.HOG0002470   OG0000444  n10   rna-XM_034637788.1    rna-XM_038575265.1    rna-XM_022517211.1    rna-XM_023241471.2    rna-XM_036083260.1    rna-XM_039226949.1    rna-XM_032876176.1    rna-XM_046024808.1    rna-XM_032309141.1    rna-XM_059140871.1    rna-XM_059408535.1    rna-XM_004753331.3    mNeoVis.2.1.1.chr3_hap1.g00532.t01    rna-XM_021703617.2    rna-XM_004396684.1    rna-XM_042962001.1    rna-XM_040619864.1    rna-XM_027576560.2    mNeoVis.3.1.1.chr3.g00521.t01, mNeoVis.3.1.1.chr3.g00523.t01    rna-gnl|WGS_ZZZZ|egapxtmp_041949-R3    mNeoVis.evergladensis.2.1.chr3.g00541.t01    mNeoVis.vulgivaga.2.1.chr3.g00542.t01      2473    N0.HOG0002471   OG0000444  n32   rna-XM_019808033.2    rna-XM_038539859.1    rna-XM_022526495.1    rna-XM_006942168.5    rna-XM_036091225.1    rna-XM_039247789.1    rna-XM_032842524.1    rna-XM_045994321.1    rna-XM_032327820.1    rna-XM_059155552.1    rna-XM_059414955.1    rna-XM_013046519.2    mNeoVis.2.1.1.chr15_hap1.g00067.t01    rna-XM_021700354.1    rna-XM_004397189.1    rna-XM_042972536.1    rna-XM_008685259.2    rna-XM_027613919.1    mNeoVis.3.1.1.chr15.g00062.t01    rna-gnl|WGS_ZZZZ|egapxtmp_033509-R1, rna-gnl|WGS_ZZZZ|egapxtmp_033593-R1    mNeoVis.evergladensis.2.1.chr15.g00067.t01, mNeoVis.evergladensis.2.1.chr15.g00068.t01    mNeoVis.vulgivaga.2.1.chr15.g00069.t01      2474   N0.HOG0002472  OG0000445  n1   rna-XM_034672372.1    rna-XM_038527804.1    rna-XM_022491939.1    rna-XM_006940894.5    rna-XM_036065430.1    rna-XM_039244253.1    rna-XM_032848620.1    rna-XM_045988535.1    rna-XM_032323590.1    rna-XM_059153613.1    rna-XM_059381654.1    rna-XM_013049590.2    mNeoVis.2.1.1.chr7_hap1.g00764.t01    rna-XM_044911339.1    rna-XM_004409541.1    rna-XM_042969553.1    rna-XM_040633896.1    rna-XM_027618299.2    mNeoVis.3.1.1.chr7.g00714.t01    rna-gnl|WGS_ZZZZ|egapxtmp_012291-R1    mNeoVis.evergladensis.2.1.chr7.g00752.t01    mNeoVis.vulgivaga.2.1.chr7.g00754.t01      2475   N0.HOG0002473  OG0000445  n14   rna-XM_034670243.1    rna-XM_038540027.1    rna-XM_022526437.1    rna-XM_023246794.2    rna-XM_036111159.1    rna-XM_039234397.1    rna-XM_032843946.1    rna-XM_045993020.1    rna-XM_032326948.1    rna-XM_059154819.1    rna-XM_059415140.1    rna-XM_045072661.1    mNeoVis.2.1.1.chr15_hap1.g00155.t01    rna-XM_021686834.1    rna-XM_004398799.1    rna-XM_042971550.1    rna-XM_040635210.1    rna-XM_027611156.2    mNeoVis.3.1.1.chr15.g00132.t01, mNeoVis.3.1.1.chr15.g00133.t01, mNeoVis.3.1.1.chr15.g00134.t01    rna-gnl|WGS_ZZZZ|egapxtmp_033947-R1    mNeoVis.evergladensis.2.1.chr15.g00140.t01    mNeoVis.vulgivaga.2.1.chr15.g00144.t01      2476    N0.HOG0002474   OG0000445  n26   rna-XM_002921719.4    rna-XM_038546804.1    rna-XM_022519916.1    rna-XM_023244468.2    rna-XM_036106825.1    rna-XM_039235921.1    rna-XM_032870794.1    rna-XM_045986064.1    rna-XM_032320329.1    rna-XM_059148178.1    rna-XM_059379464.1    rna-XM_045080929.1    mNeoVis.2.1.1.chr8_hap1.g00231.t01    rna-XM_021678425.1    rna-XM_004415769.1    rna-XM_042967753.1    rna-XM_040640368.1    rna-XM_027625640.2    mNeoVis.3.1.1.chr8.g00189.t01    rna-gnl|WGS_ZZZZ|egapxtmp_018576-R1    mNeoVis.evergladensis.2.1.chr9.g00221.t01    mNeoVis.vulgivaga.2.1.chr8.g00227.t01      2477    N0.HOG0002475   OG0000446  n1   rna-XM_034644835.1, rna-XM_034650451.1    rna-XM_038564062.1    rna-XM_022506760.1    rna-XM_011287057.4    rna-XM_036101142.1    rna-XM_039244789.1    rna-XM_032879398.1    rna-XM_046016581.1    rna-XM_032357235.1    rna-XM_059146318.1    rna-XM_059402119.1    rna-XM_045064548.1    mNeoVis.2.1.1.chr7_hap1.g02375.t01    rna-XM_044912379.1, rna-XM_044912380.1, rna-XM_044919558.1    rna-XM_004393976.1    rna-XM_042958229.1    rna-XM_040639311.1    rna-XM_035723035.1    mNeoVis.3.1.1.chr7.g02186.t01    rna-gnl|WGS_ZZZZ|egapxtmp_014319-R1    mNeoVis.evergladensis.2.1.chr7.g02321.t01    mNeoVis.vulgivaga.2.1.chr7.g02340.t01      2478    N0.HOG0002476   OG0000446  n26   rna-XM_034659618.1    rna-XM_038561329.1    rna-XM_022520408.1    rna-XM_023251934.2    rna-XM_036067875.1    rna-XM_039251257.1    rna-XM_032881980.1    rna-XM_045979484.1    rna-XM_032352887.1    rna-XM_059188514.1    rna-XM_059405517.1    rna-XM_045079190.1    mNeoVis.2.1.1.chr11_hap1.g00963.t01    rna-XM_021697772.1    rna-XM_004410006.1    rna-XM_042982110.1    rna-XM_040623136.1    rna-XM_027621088.2    mNeoVis.3.1.1.chr11.g00916.t01    rna-gnl|WGS_ZZZZ|egapxtmp_027862-R1    mNeoVis.evergladensis.2.1.chr11.g00942.t01    mNeoVis.vulgivaga.2.1.chr11.g00965.t01      2479   N0.HOG0002477  OG0000446  n46   rna-XM_034648593.1    rna-XM_038544670.1    rna-XM_022491566.1    rna-XM_045059911.1    rna-XM_036110758.1    rna-XM_039217336.1    rna-XM_032871150.1    rna-XM_046009653.1    rna-XM_032344182.1    rna-XM_059182048.1    rna-XM_059373163.1    rna-XM_045069292.1    mNeoVis.2.1.1.chr10_hap1.g00416.t01    rna-XM_044918069.1    rna-XM_004408032.1    rna-XM_042989675.1    rna-XM_040620878.1    rna-XM_027569988.1    mNeoVis.3.1.1.chr9.g00383.t01    rna-gnl|WGS_ZZZZ|egapxtmp_016205-R1    mNeoVis.evergladensis.2.1.chr8.g00397.t01    mNeoVis.vulgivaga.2.1.chr9.g00399.t01      2480    N0.HOG0002478   OG0000447  n1   rna-XM_002921364.3    rna-XM_038567277.1    rna-XM_022526844.1    rna-XM_023245382.2    rna-XM_036096945.1    rna-XM_039250939.1    rna-XM_032884011.1    rna-XM_045991269.1    rna-XM_032311218.1    rna-XM_059159976.1    rna-XM_059389020.1    rna-XM_045083992.1, rna-XM_045087515.1    mNeoVis.2.1.1.chr5_hap1.g00292.t01    rna-XM_044914664.1    rna-XM_004398879.1    rna-XM_042978167.1    rna-XM_040629323.1    rna-XM_027587279.2    mNeoVis.3.1.1.chr2.g00229.t01    rna-gnl|WGS_ZZZZ|egapxtmp_034658-R1    mNeoVis.evergladensis.2.1.chr2.g00290.t01    mNeoVis.vulgivaga.2.1.chr2.g00286.t01      2481    N0.HOG0002479   OG0000447  n23   rna-XM_034663734.1    rna-NM_001131049.1    rna-XM_022500810.1    rna-XM_023242776.2    rna-XM_036117921.1    rna-XM_039229537.1    rna-XM_032865968.1    rna-XM_046022574.1    rna-XM_032308970.1    rna-XM_059142930.1    rna-XM_059410689.1    rna-XM_045082472.1    mNeoVis.2.1.1.chr13_hap1.g00193.t01    rna-XM_021691695.2    rna-XM_004392530.1    rna-XM_042964228.1    rna-XM_040631975.1    rna-XM_027615690.2    mNeoVis.3.1.1.chr13.g00145.t01    rna-gnl|WGS_ZZZZ|egapxtmp_031097-R1    mNeoVis.evergladensis.2.1.chr13.g00191.t01    mNeoVis.vulgivaga.2.1.chr13.g00188.t01      2482    N0.HOG0002480   OG0000447  n44   rna-XM_034666734.1    rna-XM_038542508.1    rna-XM_022508131.1    rna-XM_045049387.1    rna-XM_036074172.1, rna-XM_036115393.1    rna-XM_039222176.1, rna-XM_039222417.1    rna-XM_032862977.1    rna-XM_045982933.1    rna-XM_032317383.1    rna-XM_059147002.1    rna-XM_059412461.1    rna-XM_045084619.1    mNeoVis.2.1.1.chr14_hap1.g00042.t01    rna-XM_021682629.1    rna-XM_004412443.1    rna-XM_042975633.1    rna-XM_040636212.1    rna-XM_027613356.2    mNeoVis.3.1.1.chr14.g00039.t01    rna-gnl|WGS_ZZZZ|egapxtmp_032914-R1    mNeoVis.evergladensis.2.1.chr14.g00043.t01    mNeoVis.vulgivaga.2.1.chr14.g00044.t01      2483    N0.HOG0002481   OG0000448  n1   rna-XM_034658052.1    rna-XM_038566861.1    rna-XM_022498413.1    rna-XM_045048232.1    rna-XM_036093949.1    rna-XM_039243264.1    rna-XM_032867648.1    rna-XM_045989761.1    rna-XM_032315542.1    rna-XM_059160486.1    rna-XM_059389398.1    rna-XM_045081719.1    mNeoVis.2.1.1.chr5_hap1.g00503.t01    rna-XM_044917473.1    rna-XM_004404623.1    rna-XM_042977329.1    rna-XM_040629030.1    rna-XM_027585425.1    mNeoVis.3.1.1.chr2.g00427.t01    rna-gnl|WGS_ZZZZ|egapxtmp_035382-R1    mNeoVis.evergladensis.2.1.chr2.g00498.t01    mNeoVis.vulgivaga.2.1.chr2.g00500.t01      2484    N0.HOG0002482   OG0000448  n22   rna-XM_019803046.2    rna-XM_038543211.1    rna-XM_022490943.1    rna-XM_023248059.2    rna-XM_036086757.1    rna-XM_039251430.1    rna-XM_032845124.1    rna-XM_045982229.1    rna-XM_032318168.1    rna-XM_059145512.1    rna-XM_059413841.1    rna-XM_045086841.1    mNeoVis.2.1.1.chr14_hap1.g00693.t01    rna-XM_021682010.1    rna-XM_004402986.1    rna-XM_042975676.1    rna-XM_040635725.1    rna-XM_027613344.2    mNeoVis.3.1.1.chr14.g00663.t01    rna-gnl|WGS_ZZZZ|egapxtmp_031755-R1    mNeoVis.evergladensis.2.1.chr14.g00693.t01    mNeoVis.vulgivaga.2.1.chr14.g00692.t01      2485    N0.HOG0002483   OG0000448  n44   rna-XM_034655880.1    rna-XM_038551826.1    rna-XM_022521232.1    rna-XM_019830963.3    rna-XM_036067068.1, rna-XM_036083803.1    rna-XM_039254648.1    rna-XM_032842238.1    rna-XM_046000439.1    rna-XM_032335384.1    rna-XM_059175837.1    rna-XM_059416113.1    rna-XM_004759056.3    mNeoVis.2.1.1.chr1_hap1.g00858.t01    rna-XM_021701941.2    rna-XM_004416598.1    rna-XM_042958702.1    rna-XM_040622570.1    rna-XM_027605754.2    mNeoVis.3.1.1.chr1.g00882.t01    rna-gnl|WGS_ZZZZ|egapxtmp_033348-R2, rna-gnl|WGS_ZZZZ|egapxtmp_038010-R1    mNeoVis.evergladensis.2.1.chr1.g00862.t01    mNeoVis.vulgivaga.2.1.chr1.g00877.t01      2486    N0.HOG0002484   OG0000449  n1         rna-XM_019824099.3, rna-XM_023249181.2, rna-XM_023249506.2      rna-XM_039223739.1, rna-XM_039224096.1, rna-XM_039224190.1                      rna-XM_007091723.3, rna-XM_042974248.1, rna-XM_042974352.1                  2487    N0.HOG0002485   OG0000449  n10   rna-XM_034649800.1, rna-XM_034653560.1    rna-XM_038587909.1, rna-XM_038587958.1    rna-XM_022492328.1, rna-XM_022524894.1      rna-XM_036109513.1, rna-XM_036117158.1      rna-XM_032849751.1, rna-XM_032881710.1    rna-XM_045995407.1, rna-XM_045995836.1    rna-XM_032330678.1, rna-XM_032331701.1, rna-XM_032349721.1    rna-XM_059158206.1, rna-XM_059158475.1    rna-XM_059386317.1, rna-XM_059387234.1, rna-XM_059387236.1, rna-XM_059404998.1    rna-XM_004757665.3, rna-XM_004777120.3, rna-XM_004778674.3    mNeoVis.2.1.1.chr12_hap1.g00342.t01, mNeoVis.2.1.1.chr12_hap1.g00365.t01    rna-XM_021691283.1, rna-XM_021697734.1    rna-XM_004415784.1, rna-XM_004417529.1      rna-XM_008683643.2, rna-XM_040629292.1, rna-XM_040645368.1    rna-XM_027607293.2, rna-XM_027608521.2    mNeoVis.3.1.1.chr12.g00304.t01, mNeoVis.3.1.1.chr12.g00328.t01    rna-gnl|WGS_ZZZZ|egapxtmp_029423-R1, rna-gnl|WGS_ZZZZ|egapxtmp_029600-R1    mNeoVis.evergladensis.2.1.chr12.g00300.t01, mNeoVis.evergladensis.2.1.chr12.g00322.t01    mNeoVis.vulgivaga.2.1.chr12.g00307.t01      2488    N0.HOG0002486   OG0000449  n49   rna-XM_034649595.1    rna-XM_038543875.1, rna-XM_038587952.1    rna-XM_022492372.1      rna-XM_036099068.1      rna-XM_032849673.1    rna-XM_045995566.1    rna-XM_032331523.1    rna-XM_059158407.1    rna-XM_059384734.1    rna-XM_013050613.2        rna-XM_004416285.1      rna-XM_008703778.1    rna-XM_027608310.2    mNeoVis.3.1.1.chr12.g00314.t01    rna-gnl|WGS_ZZZZ|egapxtmp_028082-R1    mNeoVis.evergladensis.2.1.chr12.g00308.t01    mNeoVis.vulgivaga.2.1.chr12.g00315.t01      2489    N0.HOG0002487   OG0000450  n1   rna-XM_002919914.3    rna-XM_038588431.1    rna-XM_022491617.1    rna-XM_023249131.2    rna-XM_036120157.1    rna-XM_039218378.1    rna-XM_032841416.1    rna-XM_045995644.1    rna-XM_032331451.1    rna-XM_059158368.1    rna-XM_059385542.1    rna-XM_004773649.3    mNeoVis.2.1.1.chr12_hap1.g00141.t01    rna-XM_021684641.1    rna-XM_004416925.1    rna-XM_042974556.1    rna-XM_008701558.2, rna-XM_040629782.1, rna-XM_040630485.1    rna-XM_027608321.1    mNeoVis.3.1.1.chr12.g00110.t01    rna-gnl|WGS_ZZZZ|egapxtmp_028072-R1    mNeoVis.evergladensis.2.1.chr12.g00117.t01    mNeoVis.vulgivaga.2.1.chr12.g00117.t01      2490    N0.HOG0002488   OG0000450  n21   rna-XM_034661857.1    rna-XM_038570191.1    rna-XM_022506800.1    rna-XM_003992031.5    rna-XM_036111789.1    rna-XM_039222250.1    rna-XM_032852433.1    rna-XM_046003618.1    rna-XM_032346328.1    rna-XM_059163832.1    rna-XM_059391778.1    rna-XM_004754447.3    mNeoVis.2.1.1.chr5_hap1.g01510.t01    rna-XM_021685992.1    rna-XM_004405815.1    rna-XM_015539346.2    rna-XM_008711073.2    rna-XM_027585209.2    mNeoVis.3.1.1.chr2.g01430.t01    rna-gnl|WGS_ZZZZ|egapxtmp_036806-R1    mNeoVis.evergladensis.2.1.chr2.g01496.t01    mNeoVis.vulgivaga.2.1.chr2.g01510.t01      2491    N0.HOG0002489   OG0000450  n28   rna-XM_011219283.2    rna-XM_038569387.1    rna-XM_022502986.1    rna-XM_023252910.2    rna-XM_036090468.1    rna-XM_039254279.1    rna-XM_032850703.1, rna-XM_032850704.1    rna-XM_046028211.1    rna-XM_032315101.1    rna-XM_059144514.1    rna-XM_059377525.1    rna-XM_045081361.1    mNeoVis.2.1.1.chr8_hap1.g01480.t01    rna-XM_021696013.1    rna-XM_004404932.1    rna-XM_042958060.1    rna-XM_008685045.2    rna-XM_027590058.2    mNeoVis.3.1.1.chr8.g01376.t01    rna-gnl|WGS_ZZZZ|egapxtmp_017773-R1    mNeoVis.evergladensis.2.1.chr9.g01429.t01    mNeoVis.vulgivaga.2.1.chr8.g01433.t01      2492    N0.HOG0002490   OG0000451  n1   rna-XM_019804353.2    rna-XM_038583717.1    rna-XM_022515224.1    rna-XM_023260301.2, rna-XM_045037126.1, rna-XM_045051532.1    rna-XM_036077503.1    rna-XM_039230393.1    rna-XM_032849968.1, rna-XM_032850094.1    rna-XM_046003818.1    rna-XM_032337736.1    rna-XM_059165212.1    rna-XM_059388018.1    rna-XM_004745278.2    mNeoVis.2.1.1.chr5_hap1.g01273.t01    rna-XM_021692578.1    rna-XM_004391855.1    rna-XM_042956329.1, rna-XM_042977110.1    rna-XM_040621712.1    rna-XM_027586206.2    mNeoVis.3.1.1.chr2.g01195.t01    rna-gnl|WGS_ZZZZ|egapxtmp_037079-R5    mNeoVis.evergladensis.2.1.chr2.g01260.t01    mNeoVis.vulgivaga.2.1.chr2.g01266.t01      2493    N0.HOG0002491   OG0000451  n21   rna-XM_034644830.1, rna-XM_034644831.1    rna-XM_038562512.1, rna-XM_038564190.1    rna-XM_022506441.1      rna-XM_036100426.1    rna-XM_039257658.1    rna-XM_032879908.1    rna-XM_046017505.1    rna-XM_032358591.1    rna-XM_059145240.1    rna-XM_059383999.1    rna-XM_004770225.3    mNeoVis.2.1.1.chr7_hap1.g02336.t01, mNeoVis.2.1.1.chr7_hap1.g02337.t01    rna-XM_021684960.1    rna-XM_004394038.1    rna-XM_042958482.1    rna-XM_008708937.2, rna-XM_040639276.1    rna-XM_035722669.1        mNeoVis.evergladensis.2.1.chr7.g02283.t01    mNeoVis.vulgivaga.2.1.chr7.g02300.t01      2494    N0.HOG0002492   OG0000451  n43   rna-XM_034651807.1, rna-XM_034651821.1    rna-XM_038564195.1    rna-XM_022507106.1    rna-XM_006937445.3    rna-XM_036122135.1, rna-XM_036122149.1    rna-XM_039257711.1    rna-XM_032879068.1    rna-XM_046014338.1    rna-XM_032358827.1    rna-XM_059145195.1    rna-XM_059403278.1    rna-XM_045063967.1      rna-XM_044912335.1    rna-XM_004394065.1, rna-XM_004413805.1    rna-XM_015542009.2    rna-XM_008683959.2    rna-XM_035723107.1              2495    N0.HOG0002493   OG0000452  n1   rna-XM_002928709.3, rna-XM_002931139.4    rna-NM_001003339.1    rna-XM_022499543.1, rna-XM_022500292.1    rna-NM_001048010.1, rna-XM_019813568.3, rna-XM_045039855.1    rna-XM_036094092.1, rna-XM_036111713.1    rna-XM_039230216.1, rna-XM_039230218.1, rna-XM_039252439.1    rna-XM_032835939.1, rna-XM_032882034.1    rna-XM_046027750.1, rna-XM_046027813.1    rna-XM_032312651.1, rna-XM_032313320.1    rna-XM_059169401.1, rna-XM_059172717.1    rna-XM_059398167.1, rna-XM_059398920.1    rna-XM_004749633.2, rna-XM_045068886.1    mNeoVis.2.1.1.chr2_hap1.g00004.t01, mNeoVis.2.1.1.chr2_hap1.g00305.t01    rna-XM_021697277.1, rna-XM_021699824.1, rna-XM_044917663.1    rna-XM_004394968.1, rna-XM_004412554.1    rna-XM_007079713.3, rna-XM_015536660.2    rna-XM_008686141.1    rna-XM_027595941.1, rna-XM_027596150.1    mNeoVis.3.1.1.chr5.g00003.t01, mNeoVis.3.1.1.chr5.g00298.t01    rna-gnl|WGS_ZZZZ|egapxtmp_006828-R1    mNeoVis.evergladensis.2.1.chr5.g00002.t01, mNeoVis.evergladensis.2.1.chr5.g00310.t01    mNeoVis.vulgivaga.2.1.chr5.g00002.t01, mNeoVis.vulgivaga.2.1.chr5.g00311.t01      2496    N0.HOG0002494   OG0000452  n41   rna-XM_034662973.1    rna-NM_001197044.1    rna-XM_022499564.1    rna-XM_019813570.3    rna-XM_036111709.1    rna-XM_039230221.1    rna-XM_032835932.1    rna-XM_045996984.1, rna-XM_046027649.1, rna-XM_046027685.1    rna-XM_032313316.1    rna-XM_059169399.1    rna-XM_059398171.1    rna-XM_004749629.2    mNeoVis.2.1.1.chr2_hap1.g00304.t01    rna-XM_021699489.1    rna-XM_004416880.1    rna-XM_015536661.2    rna-XM_008699299.2, rna-XM_040623769.1    rna-XM_027588917.1    mNeoVis.3.1.1.chr5.g00297.t01    rna-gnl|WGS_ZZZZ|egapxtmp_007866-R1    mNeoVis.evergladensis.2.1.chr5.g00309.t01    mNeoVis.vulgivaga.2.1.chr5.g00310.t01      2497    N0.HOG0002495   OG0000453  n1   rna-XM_034670980.1    rna-XM_038533096.1    rna-XM_022506082.1    rna-XM_023253426.2    rna-XM_036085527.1    rna-XM_039232113.1    rna-XM_032855524.1    rna-XM_045998356.1    rna-XM_032333741.1    rna-XM_059165935.1    rna-XM_059380313.1    rna-XM_045075320.1    mNeoVis.2.1.1.chr6_hap1.g00341.t01    rna-XM_044913028.1    rna-XM_004396247.1    rna-XM_042985024.1    rna-XM_040625236.1    rna-XM_027597353.2    mNeoVis.3.1.1.chr4.g00342.t01    rna-gnl|WGS_ZZZZ|egapxtmp_000727-R1    mNeoVis.evergladensis.2.1.chr4.g00347.t01    mNeoVis.vulgivaga.2.1.chr4.g00344.t01      2498    N0.HOG0002496   OG0000453  n22   rna-XM_034663620.1    rna-XM_038578740.1    rna-XM_022499799.1    rna-XM_023240898.2    rna-XM_036083129.1, rna-XM_036118383.1    rna-XM_039218779.1    rna-XM_032880553.1    rna-XM_046028096.1    rna-XM_032313389.1    rna-XM_059173261.1    rna-XM_059398521.1    rna-XM_004749297.3    mNeoVis.2.1.1.chr2_hap1.g00170.t01    rna-XM_021699769.1    rna-XM_004416773.1    rna-XM_042960443.1    rna-XM_008686652.2    rna-XM_027597074.2    mNeoVis.3.1.1.chr5.g00161.t01    rna-gnl|WGS_ZZZZ|egapxtmp_006125-R1    mNeoVis.evergladensis.2.1.chr5.g00173.t01    mNeoVis.vulgivaga.2.1.chr5.g00170.t01      2499    N0.HOG0002497   OG0000453  n43   rna-XM_034663251.1    rna-XM_038578743.1    rna-XM_022499251.1    rna-XM_045040656.1    rna-XM_036100150.1    rna-XM_039217305.1    rna-XM_032880513.1    rna-XM_046027663.1    rna-XM_032312688.1    rna-XM_059173260.1    rna-XM_059398526.1    rna-XM_045068616.1    mNeoVis.2.1.1.chr2_hap1.g00168.t01    rna-XM_021699529.1    rna-XM_004416789.1, rna-XM_004417492.1    rna-XM_042960644.1    rna-XM_040624056.1    rna-XM_027596987.2    mNeoVis.3.1.1.chr5.g00157.t01, mNeoVis.3.1.1.chr5.g00158.t01    rna-gnl|WGS_ZZZZ|egapxtmp_009141-R1    mNeoVis.evergladensis.2.1.chr5.g00170.t01    mNeoVis.vulgivaga.2.1.chr5.g00167.t01      2500    N0.HOG0002498   OG0000454  n2   rna-XM_034648542.1    rna-XM_038544355.1    rna-XM_022524102.1    rna-XM_006932792.4    rna-XM_036114759.1    rna-XM_039254671.1    rna-XM_032856226.1    rna-XM_046010303.1    rna-XM_032343975.1    rna-XM_059181808.1    rna-XM_059371234.1    rna-XM_045067827.1    mNeoVis.2.1.1.chr10_hap1.g00515.t01    rna-XM_021695495.1    rna-XM_004402016.1    rna-XM_007098066.3    rna-XM_008708819.2    rna-XM_027572147.2    mNeoVis.3.1.1.chr9.g00475.t01      mNeoVis.evergladensis.2.1.chr8.g00496.t01        2501    N0.HOG0002499   OG0000454  n21   rna-XM_034648630.1    rna-XM_038544354.1    rna-XM_022492021.1, rna-XM_022526203.1    rna-XM_003987569.4, rna-XM_006932791.4    rna-XM_036114756.1, rna-XM_036114761.1    rna-XM_039254669.1, rna-XM_039254678.1    rna-XM_032857052.1    rna-XM_046010312.1    rna-XM_032343974.1    rna-XM_059179683.1    rna-XM_059371232.1    rna-XM_004755171.3      rna-XM_021695688.1, rna-XM_021695689.1    rna-XM_004402151.1, rna-XM_004402152.1    rna-XM_007098065.3, rna-XM_007098104.2    rna-XM_008708815.2    rna-XM_027570352.1, rna-XM_027572148.1      rna-gnl|WGS_ZZZZ|egapxtmp_016850-R1    mNeoVis.evergladensis.2.1.chr8.g00495.t01    mNeoVis.vulgivaga.2.1.chr9.g00496.t01      2502    N0.HOG0002500   OG0000454  n48   rna-XM_002921032.4    rna-XM_038544357.1    rna-XM_022524843.1    rna-XM_006932823.4    rna-XM_036114758.1    rna-XM_039254679.1    rna-XM_032856225.1    rna-XM_046008913.1    rna-XM_032343978.1    rna-XM_059181811.1    rna-XM_059371235.1    rna-XM_004755165.3    mNeoVis.2.1.1.chr10_hap1.g00514.t01    rna-XM_021695430.1    rna-XM_004402015.1    rna-XM_007098105.3    rna-XM_008708817.2    rna-XM_027572146.1      rna-gnl|WGS_ZZZZ|egapxtmp_017287-R1    mNeoVis.evergladensis.2.1.chr8.g00494.t01    mNeoVis.vulgivaga.2.1.chr9.g00495.t01      2503    N0.HOG0002501   OG0000455  n1   rna-XM_034649429.1    rna-XM_038557875.1    rna-XM_022496723.1    rna-XM_003989866.6    rna-XM_036119078.1    rna-XM_039244493.1    rna-XM_032877281.1    rna-XM_046028203.1    rna-XM_032302638.1    rna-XM_059137080.1    rna-XM_059377288.1    rna-XM_004740761.3    mNeoVis.2.1.1.chr2_hap1.g01490.t01    rna-XM_044912077.1, rna-XM_044912186.1    rna-XM_004402652.1    rna-XM_042996562.1    rna-XM_040619278.1    rna-XM_027580703.2    mNeoVis.3.1.1.chr5.g01412.t01    rna-gnl|WGS_ZZZZ|egapxtmp_008409-R1    mNeoVis.evergladensis.2.1.chr5.g01427.t01    mNeoVis.vulgivaga.2.1.chr5.g01428.t01      2504    N0.HOG0002502   OG0000455  n19   rna-XM_034670902.1    rna-XM_038554811.1    rna-XM_022520192.1    rna-XM_011282447.4    rna-XM_036078060.1    rna-XM_039244665.1    rna-XM_032858207.1    rna-XM_046006541.1    rna-XM_032338955.1    rna-XM_059176999.1    rna-XM_059401114.1    rna-XM_004767094.2    mNeoVis.2.1.1.chr1_hap1.g01985.t01    rna-XM_021693614.1    rna-XM_004408661.1    rna-XM_042986881.1    rna-XM_008704998.1    rna-XM_027602788.2    mNeoVis.3.1.1.chr1.g01940.t01    rna-gnl|WGS_ZZZZ|egapxtmp_028529-R1    mNeoVis.evergladensis.2.1.chr1.g01947.t01    mNeoVis.vulgivaga.2.1.chr1.g01946.t01      2505    N0.HOG0002503   OG0000455  n32   rna-XM_002924137.2    rna-XM_038581312.1    rna-XM_022516054.1    rna-XM_019839466.3    rna-XM_036111232.1    rna-XM_039219386.1    rna-XM_032865451.1    rna-XM_046002597.1    rna-XM_032353659.1    rna-XM_059164562.1    rna-XM_059392238.1    rna-XM_045085170.1    mNeoVis.2.1.1.chr5_hap1.g01750.t01    rna-XM_044913681.1    rna-XM_004394378.1    rna-XM_042956726.1    rna-XM_040621050.1    rna-XM_027584982.2    mNeoVis.3.1.1.chr2.g01659.t01    rna-gnl|WGS_ZZZZ|egapxtmp_036832-R2    mNeoVis.evergladensis.2.1.chr2.g01732.t01    mNeoVis.vulgivaga.2.1.chr2.g01748.t01      2506    N0.HOG0002504   OG0000456  n0   rna-XM_034659656.1    rna-XM_038561280.1, rna-XM_038564712.1, rna-XM_038584720.1    rna-XM_022514064.1, rna-XM_022520362.1    rna-XM_019827684.3    rna-XM_036070416.1, rna-XM_036074699.1, rna-XM_036098672.1, rna-XM_036104626.1    rna-XM_039241935.1, rna-XM_039247316.1, rna-XM_039257212.1    rna-XM_032839398.1, rna-XM_032843694.1, rna-XM_032844810.1, rna-XM_032859601.1, rna-XM_032877899.1, rna-XM_032878022.1, rna-XM_032881913.1    rna-XM_045978542.1, rna-XM_045979513.1, rna-XM_045989194.1, rna-XM_046019178.1, rna-XM_046025763.1, rna-XM_046026212.1    rna-XM_032352964.1    rna-XM_059173445.1, rna-XM_059188591.1    rna-XM_059405460.1, rna-XM_059416507.1    rna-XM_013062957.2, rna-XM_045079630.1    mNeoVis.2.1.1.chr11_hap1.g00985.t01, mNeoVis.2.1.1.chr1_hap1.g00161.t01, mNeoVis.2.1.1.chr1_hap1.g01247.t01, mNeoVis.2.1.1.chr8_hap1.g01375.t01    rna-XM_021689573.1, rna-XM_021693325.2, rna-XM_021693643.1, rna-XM_021697415.1, rna-XM_021697692.1      rna-XM_042982162.1    rna-XM_040621756.1, rna-XM_040623109.1    rna-XM_027575061.1, rna-XM_027577934.2, rna-XM_027587523.1, rna-XM_027604144.2, rna-XM_027608374.1, rna-XM_027622067.1, rna-XM_027623920.1    mNeoVis.3.1.1.chr1.g00165.t01, mNeoVis.3.1.1.chr1.g01205.t01, mNeoVis.3.1.1.chr11.g00938.t01, mNeoVis.3.1.1.chr8.g01271.t01    rna-gnl|WGS_ZZZZ|egapxtmp_002759-R1, rna-gnl|WGS_ZZZZ|egapxtmp_020486-R1, rna-gnl|WGS_ZZZZ|egapxtmp_024621-R1, rna-gnl|WGS_ZZZZ|egapxtmp_027353-R2    mNeoVis.evergladensis.2.1.chr1.g00157.t01, mNeoVis.evergladensis.2.1.chr1.g01221.t01, mNeoVis.evergladensis.2.1.chr11.g00964.t01, mNeoVis.evergladensis.2.1.chr9.g01321.t01    mNeoVis.vulgivaga.2.1.chr1.g00158.t01, mNeoVis.vulgivaga.2.1.chr1.g01223.t01, mNeoVis.vulgivaga.2.1.chr11.g00986.t01, mNeoVis.vulgivaga.2.1.chr8.g01327.t01      2507    N0.HOG0002505   OG0000457  n1   rna-XM_034660525.1    rna-NM_001048087.1    rna-XM_022510575.1    rna-XM_023253811.2    rna-XM_036072356.1    rna-XM_039241766.1    rna-XM_032849258.1    rna-XM_046005058.1    rna-XM_032340706.1    rna-XM_059178704.1    rna-XM_059399979.1    rna-XM_004770579.3    mNeoVis.2.1.1.chr1_hap1.g01414.t01    rna-XM_021684647.2, rna-XM_044914355.1    rna-XM_004407674.1    rna-XM_007093239.3    rna-XM_040644777.1    rna-XM_027601634.2    mNeoVis.3.1.1.chr1.g01360.t01    rna-gnl|WGS_ZZZZ|egapxtmp_021317-R1    mNeoVis.evergladensis.2.1.chr1.g01376.t01    mNeoVis.vulgivaga.2.1.chr1.g01380.t01      2508    N0.HOG0002506   OG0000457  n22   rna-XM_034664762.1    rna-XM_038548842.1    rna-XM_022501246.1    rna-XM_019816736.3    rna-XM_036108525.1    rna-XM_039255153.1    rna-XM_032866248.1    rna-XM_046022979.1    rna-XM_032307832.1    rna-XM_059141214.1    rna-XM_059410381.1    rna-XM_045082871.1    mNeoVis.2.1.1.chr13_hap1.g00077.t01    rna-XM_021691255.1    rna-XM_004411391.1    rna-XM_042965206.1    rna-XM_040632154.1    rna-XM_027615477.2    mNeoVis.3.1.1.chr13.g00042.t01    rna-gnl|WGS_ZZZZ|egapxtmp_029775-R1    mNeoVis.evergladensis.2.1.chr13.g00078.t01    mNeoVis.vulgivaga.2.1.chr13.g00076.t01      2509    N0.HOG0002507   OG0000457  n41   rna-XM_034657934.1    rna-XM_038567019.1    rna-XM_022498614.1    rna-XM_023246610.2    rna-XM_036089133.1    rna-XM_039243377.1    rna-XM_032867484.1    rna-XM_045990523.1    rna-XM_032302431.1, rna-XM_032313966.1    rna-XM_059160287.1    rna-XM_059389262.1    rna-XM_045082570.1    mNeoVis.2.1.1.chr5_hap1.g00425.t01    rna-XM_044914057.1    rna-XM_004408368.1    rna-XM_042978438.1    rna-XM_040629146.1    rna-XM_035726073.1    mNeoVis.3.1.1.chr2.g00355.t01    rna-gnl|WGS_ZZZZ|egapxtmp_035367-R1    mNeoVis.evergladensis.2.1.chr2.g00423.t01    mNeoVis.vulgivaga.2.1.chr2.g00424.t01      2510    N0.HOG0002508   OG0000458  n1   rna-XM_002929624.4    rna-XM_038577248.1    rna-XM_022524935.1    rna-XM_003988551.6    rna-XM_036082452.1    rna-XM_039224596.1    rna-XM_032855344.1    rna-XM_046013749.1    rna-XM_032347726.1    rna-XM_059185658.1    rna-XM_059404497.1    rna-XM_004771828.3    mNeoVis.2.1.1.chr9_hap1.g00677.t01    rna-XM_021690383.1    rna-XM_004402260.1    rna-XM_007090800.3    rna-XM_008704450.2    rna-XM_027594830.2    mNeoVis.3.1.1.chr10.g00642.t01    rna-gnl|WGS_ZZZZ|egapxtmp_024212-R3    mNeoVis.evergladensis.2.1.chr10.g00672.t01    mNeoVis.vulgivaga.2.1.chr10.g00655.t01      2511    N0.HOG0002509   OG0000458  n17   rna-XM_002925036.4    rna-XM_038565742.1    rna-XM_022498337.1    rna-XM_011280193.3    rna-XM_036079799.1    rna-XM_039231323.1    rna-XM_032841540.1    rna-XM_045990147.1    rna-XM_032326403.1    rna-XM_059161614.1    rna-XM_059390243.1    rna-XM_004738460.3    mNeoVis.2.1.1.chr5_hap1.g00854.t01    rna-XM_021695190.1    rna-XM_004392708.1    rna-XM_042979485.1    rna-XM_008697919.2    rna-XM_027584936.1    mNeoVis.3.1.1.chr2.g00766.t01    rna-gnl|WGS_ZZZZ|egapxtmp_037032-R1    mNeoVis.evergladensis.2.1.chr2.g00855.t01    mNeoVis.vulgivaga.2.1.chr2.g00856.t01      2512    N0.HOG0002510   OG0000458  n35   rna-XM_034660641.1    rna-XM_038553892.1    rna-XM_022503884.1    rna-XM_023253925.2    rna-XM_036097283.1    rna-XM_039235331.1    rna-XM_032862351.1    rna-XM_046006700.1    rna-XM_032339757.1    rna-XM_059177787.1    rna-XM_059399110.1    rna-XM_013062943.2    mNeoVis.2.1.1.chr1_hap1.g01707.t01    rna-XM_044917427.1    rna-XM_004409787.1    rna-XM_042986224.1    rna-XM_040644674.1    rna-XM_027603969.2    mNeoVis.3.1.1.chr1.g01662.t01, mNeoVis.3.1.1.chr4.g00020.t01    rna-gnl|WGS_ZZZZ|egapxtmp_022362-R1    mNeoVis.evergladensis.2.1.chr1.g01674.t01    mNeoVis.vulgivaga.2.1.chr1.g01670.t01, mNeoVis.vulgivaga.2.1.chr1.g01671.t01      2513    N0.HOG0002511   OG0000459  n1   rna-XM_002914752.4    rna-XM_038573339.1    rna-XM_022520630.1    rna-XM_023250512.2    rna-XM_036070484.1    rna-XM_039221123.1    rna-XM_032880006.1    rna-XM_045977998.1    rna-XM_032314645.1    rna-XM_059144142.1    rna-XM_059377434.1    rna-XM_004774803.2    mNeoVis.2.1.1.chr8_hap1.g01263.t01    rna-XM_021678271.1    rna-XM_004406639.1    rna-XM_007074330.3    rna-XM_040621933.1    rna-XM_027590366.2    mNeoVis.3.1.1.chr8.g01155.t01    rna-gnl|WGS_ZZZZ|egapxtmp_019546-R1    mNeoVis.evergladensis.2.1.chr9.g01211.t01    mNeoVis.vulgivaga.2.1.chr8.g01212.t01      2514    N0.HOG0002512   OG0000459  n21   rna-XM_002912908.4    rna-NM_001048016.1    rna-XM_022493015.1    rna-XM_003980762.5    rna-XM_036070329.1    rna-XM_039221373.1, rna-XM_039252594.1    rna-XM_032837758.1    rna-XM_046001204.1    rna-XM_032336729.1    rna-XM_059174571.1    rna-XM_059387301.1, rna-XM_059417565.1    rna-XM_004744982.3    mNeoVis.2.1.1.chr1_hap1.g00409.t01    rna-XM_021702053.1    rna-XM_004408272.1    rna-XM_042983846.1    rna-XM_008691617.2    rna-XM_027605117.2    mNeoVis.3.1.1.chr1.g00406.t01    rna-gnl|WGS_ZZZZ|egapxtmp_038608-R1    mNeoVis.evergladensis.2.1.chr1.g00407.t01    mNeoVis.vulgivaga.2.1.chr1.g00413.t01      2515    N0.HOG0002513   OG0000459  n43   rna-XM_019802739.2    rna-XM_038564700.1    rna-XM_022515544.1    rna-XM_004001096.6    rna-XM_036079418.1    rna-XM_039256232.1    rna-XM_032857985.1    rna-XM_045990530.1    rna-XM_032333368.1    rna-XM_059168894.1    rna-XM_059378215.1    rna-XM_004747935.3    mNeoVis.2.1.1.chr6_hap1.g00437.t01    rna-XM_021681467.1    rna-XM_004393606.1    rna-XM_007091811.2    rna-XM_040626293.1    rna-XM_027598608.1    mNeoVis.3.1.1.chr4.g00438.t01    rna-gnl|WGS_ZZZZ|egapxtmp_043832-R1    mNeoVis.evergladensis.2.1.chr4.g00445.t01    mNeoVis.vulgivaga.2.1.chr4.g00447.t01      2516    N0.HOG0002514   OG0000460  n1   rna-XM_034665629.1    rna-XM_038536382.1    rna-XM_022513178.1    rna-XM_023239035.2    rna-XM_036115890.1    rna-XM_039216535.1    rna-XM_032844086.1    rna-XM_046014309.1    rna-XM_032358686.1    rna-XM_059179084.1    rna-XM_059371246.1    rna-XM_045077476.1    mNeoVis.2.1.1.chr7_hap1.g01170.t01    rna-XM_044919428.1    rna-XM_004415193.1    rna-XM_042958350.1    rna-XM_040630660.1    rna-XM_027579667.2    mNeoVis.3.1.1.chr7.g01094.t01    rna-gnl|WGS_ZZZZ|egapxtmp_012994-R2    mNeoVis.evergladensis.2.1.chr7.g01159.t01    mNeoVis.vulgivaga.2.1.chr7.g01171.t01      2517    N0.HOG0002515   OG0000460  n23   rna-XM_034661662.1    rna-XM_038564748.1    rna-XM_022518516.1    rna-XM_023252836.2    rna-XM_036069402.1, rna-XM_036099191.1    rna-XM_039241613.1    rna-XM_032858375.1    rna-XM_045997185.1    rna-XM_032333321.1    rna-XM_059169058.1    rna-XM_059378001.1    rna-XM_004747976.3    mNeoVis.2.1.1.chr6_hap1.g00455.t01    rna-XM_021681482.1    rna-XM_004401582.1    rna-XM_042982686.1    rna-XM_008689484.2    rna-XM_027597702.1    mNeoVis.3.1.1.chr4.g00457.t01    rna-gnl|WGS_ZZZZ|egapxtmp_001316-R1    mNeoVis.evergladensis.2.1.chr4.g00461.t01    mNeoVis.vulgivaga.2.1.chr4.g00467.t01      2518    N0.HOG0002516   OG0000460  n44   rna-XM_034655736.1    rna-XM_038551660.1    rna-XM_022493385.1    rna-XM_006927698.4    rna-XM_036114313.1    rna-XM_039242302.1    rna-XM_032837670.1    rna-XM_046000290.1    rna-XM_032336641.1    rna-XM_059174643.1    rna-XM_059416602.1    rna-XM_045075655.1    mNeoVis.2.1.1.chr1_hap1.g00443.t01    rna-XM_021702093.1    rna-XM_004405477.1    rna-XM_042956330.1    rna-XM_008691582.2    rna-XM_027606215.2    mNeoVis.3.1.1.chr1.g00439.t01, mNeoVis.3.1.1.chr1.g00440.t01    rna-gnl|WGS_ZZZZ|egapxtmp_030272-R1    mNeoVis.evergladensis.2.1.chr1.g00440.t01    mNeoVis.vulgivaga.2.1.chr1.g00446.t01      2519    N0.HOG0002517   OG0000461  n1   rna-XM_034645522.1    rna-XM_038577652.1    rna-XM_022519419.1    rna-XM_045061777.1    rna-XM_036098923.1    rna-XM_039241344.1    rna-XM_032876808.1    rna-XM_046011907.1    rna-XM_032348169.1    rna-XM_059185365.1    rna-XM_059403106.1    rna-XM_045073200.1    mNeoVis.2.1.1.chr9_hap1.g00559.t01    rna-XM_044915421.1    rna-XM_004400829.1    rna-XM_042992273.1    rna-XM_040642443.1    rna-XM_027592568.1    mNeoVis.3.1.1.chr10.g00529.t01    rna-gnl|WGS_ZZZZ|egapxtmp_023003-R1    mNeoVis.evergladensis.2.1.chr10.g00552.t01    mNeoVis.vulgivaga.2.1.chr10.g00539.t01      2520    N0.HOG0002518   OG0000461  n19   rna-XM_034644335.1, rna-XM_034653195.1    rna-XM_038530232.1    rna-XM_022520749.1    rna-XM_023256254.2    rna-XM_036097398.1    rna-XM_039241633.1    rna-XM_032881692.1    rna-XM_046011076.1    rna-XM_032349894.1    rna-XM_059183671.1    rna-XM_059403972.1    rna-XM_045072331.1    mNeoVis.2.1.1.chr9_hap1.g00002.t01    rna-XM_021690907.1    rna-XM_004412214.1    rna-XM_042990907.1    rna-XM_040645478.1    rna-XM_027593628.2    mNeoVis.3.1.1.chr10.g00002.t01    rna-gnl|WGS_ZZZZ|egapxtmp_023269-R1    mNeoVis.evergladensis.2.1.chr10.g00002.t01    mNeoVis.vulgivaga.2.1.chr10.g00002.t01      2521    N0.HOG0002519   OG0000461  n40   rna-XM_034654059.1, rna-XM_034654093.1    rna-XM_038581712.1    rna-XM_022515900.1    rna-XM_023238578.2    rna-XM_036081656.1    rna-XM_039222142.1    rna-XM_032865173.1    rna-XM_046002483.1    rna-XM_032358012.1    rna-XM_059164996.1    rna-XM_059392575.1    rna-XM_045084272.1    mNeoVis.2.1.1.chr5_hap1.g01929.t01    rna-XM_044919232.1    rna-XM_004416200.1    rna-XM_042956560.1    rna-XM_040621761.1    rna-XM_027582434.2    mNeoVis.3.1.1.chr2.g01816.t01    rna-gnl|WGS_ZZZZ|egapxtmp_035751-R1    mNeoVis.evergladensis.2.1.chr2.g01907.t01    mNeoVis.vulgivaga.2.1.chr2.g01924.t01      2522    N0.HOG0002520   OG0000462  n0   rna-XM_034644134.1, rna-XM_034651688.1, rna-XM_034651712.1    rna-NM_001253720.1, rna-NM_001253721.1, rna-XM_038530092.1, rna-XM_038530094.1, rna-XM_038530105.1, rna-XM_038589022.1, rna-XM_038589023.1, rna-XM_038589025.1, rna-XM_038589027.1, rna-XM_038589125.1, rna-XM_038589126.1, rna-XM_038589172.1, rna-XM_038589173.1, rna-XM_038589461.1, rna-XM_038589462.1, rna-XM_038589629.1, rna-XM_038589656.1    rna-XM_022492609.1, rna-XM_022515686.1    rna-XM_006933252.5    rna-XM_036068446.1, rna-XM_036097629.1    rna-XM_039240042.1    rna-XM_032850771.1    rna-XM_046010792.1, rna-XM_046011860.1, rna-XM_046011862.1, rna-XM_046011865.1, rna-XM_046011866.1, rna-XM_046012335.1, rna-XM_046012338.1, rna-XM_046012363.1, rna-XM_046012364.1, rna-XM_046012527.1, rna-XM_046012579.1, rna-XM_046013649.1    rna-XM_032349429.1, rna-XM_032349430.1, rna-XM_032349433.1    rna-XM_059184121.1, rna-XM_059184122.1, rna-XM_059184123.1    rna-XM_059387355.1, rna-XM_059387464.1, rna-XM_059405021.1    rna-XM_045069121.1, rna-XM_045069122.1    mNeoVis.2.1.1.chr9_hap1.g00147.t01, mNeoVis.2.1.1.chr9_hap1.g00148.t01    rna-XM_044915006.1    rna-XM_004408358.1, rna-XM_004409122.1    rna-XM_007081224.3    rna-XM_008711978.1, rna-XM_040645408.1    rna-XM_027594452.2, rna-XM_027611939.2    mNeoVis.3.1.1.chr10.g00138.t01, mNeoVis.3.1.1.chr10.g00139.t01    rna-gnl|WGS_ZZZZ|egapxtmp_023209-R1, rna-gnl|WGS_ZZZZ|egapxtmp_024324-R2    mNeoVis.evergladensis.2.1.chr10.g00140.t01, mNeoVis.evergladensis.2.1.chr10.g00141.t01    mNeoVis.vulgivaga.2.1.chr10.g00139.t01, mNeoVis.vulgivaga.2.1.chr10.g00140.t01      2523    N0.HOG0002521   OG0000463  n1   rna-XM_034640027.1    rna-XM_038547466.1    rna-XM_022509376.1, rna-XM_022509557.1    rna-XM_045044685.1    rna-XM_036091639.1    rna-XM_039227662.1    rna-XM_032871463.1    rna-XM_045984383.1    rna-XM_032320852.1    rna-XM_059147671.1    rna-XM_059379550.1    rna-XM_045070344.1    mNeoVis.2.1.1.chr8_hap1.g00503.t01    rna-XM_044921467.1    rna-XM_004395072.1    rna-XM_042965684.1    rna-XM_040640718.1    rna-XM_027625231.2    mNeoVis.3.1.1.chr8.g00459.t01    rna-gnl|WGS_ZZZZ|egapxtmp_017427-R1    mNeoVis.evergladensis.2.1.chr9.g00494.t01    mNeoVis.vulgivaga.2.1.chr8.g00500.t01      2524   N0.HOG0002522  OG0000463  n12   rna-XM_002913152.4    rna-XM_038529823.1    rna-XM_022515695.1    rna-XM_003988149.5    rna-XM_036092628.1    rna-XM_039233047.1    rna-XM_032880827.1    rna-XM_046012352.1    rna-XM_032349607.1    rna-XM_059183941.1    rna-XM_059404184.1    rna-XM_045084981.1    mNeoVis.2.1.1.chr9_hap1.g00091.t01    rna-XM_021696687.2    rna-XM_004392982.1    rna-XM_042991212.1    rna-XM_040645330.1    rna-XM_027592629.2    mNeoVis.3.1.1.chr10.g00088.t01    rna-gnl|WGS_ZZZZ|egapxtmp_021755-R1    mNeoVis.evergladensis.2.1.chr10.g00092.t01    mNeoVis.vulgivaga.2.1.chr10.g00085.t01      2525   N0.HOG0002523  OG0000463  n26   rna-XM_011220773.3    rna-XM_038549794.1    rna-XM_022522997.1    rna-XM_003988950.6    rna-XM_036077954.1, rna-XM_036100184.1    rna-XM_039256172.1    rna-XM_032874588.1    rna-XM_046010995.1    rna-XM_032348790.1    rna-XM_059184818.1    rna-XM_059403556.1    rna-XM_004773239.3    mNeoVis.2.1.1.chr9_hap1.g00335.t01    rna-XM_021687288.1    rna-XM_004401733.1    rna-XM_042992621.1    rna-XM_008684358.2    rna-XM_027594517.2    mNeoVis.3.1.1.chr10.g00326.t01    rna-gnl|WGS_ZZZZ|egapxtmp_022941-R1    mNeoVis.evergladensis.2.1.chr10.g00332.t01    mNeoVis.vulgivaga.2.1.chr10.g00328.t01      2526    N0.HOG0002524   OG0000464  n1   rna-XM_034645910.1    rna-XM_038569273.1    rna-XM_022506952.1    rna-XM_023239658.2    rna-XM_036100910.1    rna-XM_039218371.1    rna-XM_032881226.1    rna-XM_046014190.1    rna-XM_032359616.1    rna-XM_059138525.1    rna-XM_059406869.1    rna-XM_004755876.3    mNeoVis.2.1.1.chr7_hap1.g01896.t01    rna-XM_021685869.1    rna-XM_004416870.1    rna-XM_007098476.3    rna-XM_040638905.1    rna-XM_035722983.1    mNeoVis.3.1.1.chr7.g01765.t01    rna-gnl|WGS_ZZZZ|egapxtmp_014307-R1    mNeoVis.evergladensis.2.1.chr7.g01858.t01    mNeoVis.vulgivaga.2.1.chr7.g01875.t01      2527    N0.HOG0002525   OG0000464  n21   rna-XM_034645216.1    rna-XM_038562912.1    rna-XM_022506555.1    rna-XM_019812902.3    rna-XM_036098217.1    rna-XM_039221527.1    rna-XM_032879866.1    rna-XM_046015099.1    rna-XM_032358492.1    rna-XM_059152828.1    rna-XM_059397565.1    rna-XM_045067143.1    mNeoVis.2.1.1.chr7_hap1.g02677.t01    rna-XM_021692207.1    rna-XM_004403817.1    rna-XM_042958426.1    rna-XM_040639757.1    rna-XM_027579954.1    mNeoVis.3.1.1.chr7.g02314.t01      mNeoVis.evergladensis.2.1.chr7.g02490.t01    mNeoVis.vulgivaga.2.1.chr7.g02511.t01      2528    N0.HOG0002526   OG0000464  n41   rna-XM_034644141.1    rna-XM_038558799.1    rna-XM_022495363.1    rna-XM_045062224.1    rna-XM_036086024.1, rna-XM_036120166.1, rna-XM_036123353.1    rna-XM_039245598.1    rna-XM_032844283.1    rna-XM_046014083.1    rna-XM_032346621.1    rna-XM_059186643.1    rna-XM_059402257.1    rna-XM_004742962.3    mNeoVis.2.1.1.chr9_hap1.g01045.t01    rna-XM_044914740.1    rna-XM_004403854.1    rna-XM_042993008.1    rna-XM_040637073.1    rna-XM_027593728.1    mNeoVis.3.1.1.chr10.g01005.t01    rna-gnl|WGS_ZZZZ|egapxtmp_022708-R2    mNeoVis.evergladensis.2.1.chr10.g01034.t01, mNeoVis.evergladensis.2.1.chr10.g01035.t01    mNeoVis.vulgivaga.2.1.chr10.g01025.t01      2529   N0.HOG0002527  OG0000465  n1   rna-XM_019794488.2    rna-XM_038550665.1    rna-XM_022519321.1    rna-XM_045062320.1    rna-XM_036108714.1    rna-XM_039246082.1    rna-XM_032836587.1    rna-XM_046013831.1    rna-XM_032346390.1    rna-XM_059186881.1    rna-XM_059402084.1    rna-XM_045083912.1    mNeoVis.2.1.1.chr9_hap1.g01139.t01    rna-XM_044915817.1    rna-XM_004410823.1    rna-XM_042993188.1    rna-XM_040637493.1    rna-XM_027594655.1    mNeoVis.3.1.1.chr10.g01094.t01    rna-gnl|WGS_ZZZZ|egapxtmp_023125-R1    mNeoVis.evergladensis.2.1.chr10.g01127.t01    mNeoVis.vulgivaga.2.1.chr10.g01119.t01      2530    N0.HOG0002528   OG0000465  n22     rna-XM_038540684.1    rna-XM_022516628.1    rna-XM_023246972.2    rna-XM_036083210.1    rna-XM_039246855.1    rna-XM_032846127.1    rna-XM_045992585.1    rna-XM_032327585.1    rna-XM_059155083.1    rna-XM_059415605.1    rna-XM_004750659.3    mNeoVis.2.1.1.chr15_hap1.g00338.t01    rna-XM_021698198.1    rna-XM_004403185.1    rna-XM_042971667.1    rna-XM_040635488.1    rna-XM_027613076.2      rna-gnl|WGS_ZZZZ|egapxtmp_033468-R1    mNeoVis.evergladensis.2.1.chr15.g00316.t01    mNeoVis.vulgivaga.2.1.chr15.g00323.t01      2531    N0.HOG0002529   OG0000465  n35   rna-XM_011225800.3    rna-XM_038546639.1    rna-XM_022523507.1      rna-XM_036088705.1    rna-XM_039226316.1    rna-XM_032870661.1    rna-XM_045986551.1    rna-XM_032322704.1    rna-XM_059147952.1    rna-XM_059379907.1    rna-XM_004749282.3    mNeoVis.2.1.1.chr6_hap1.g01125.t01, mNeoVis.2.1.1.chr8_hap1.g00160.t01    rna-XM_021681013.1    rna-XM_004394355.1    rna-XM_007091305.2    rna-XM_040640251.1    rna-XM_027625615.2    mNeoVis.3.1.1.chr4.g01072.t01, mNeoVis.3.1.1.chr8.g00129.t01    rna-gnl|WGS_ZZZZ|egapxtmp_001159-R1, rna-gnl|WGS_ZZZZ|egapxtmp_020257-R1    mNeoVis.evergladensis.2.1.chr4.g01057.t01, mNeoVis.evergladensis.2.1.chr9.g00157.t01    mNeoVis.vulgivaga.2.1.chr4.g01091.t01, mNeoVis.vulgivaga.2.1.chr8.g00160.t01      2532    N0.HOG0002530   OG0000466  n1   rna-XM_011231049.3    rna-NM_001003120.3    rna-XM_022520036.1    rna-XM_023243719.2    rna-XM_036083783.1    rna-XM_039229747.1    rna-XM_032870738.1    rna-XM_045986454.1    rna-XM_032320763.1    rna-XM_059148608.1    rna-XM_059379255.1    rna-XM_013054487.2    mNeoVis.2.1.1.chr8_hap1.g00210.t01    rna-XM_021678450.1    rna-XM_004405749.1    rna-XM_042966900.1    rna-XM_008693693.2    rna-XM_027626617.2    mNeoVis.3.1.1.chr8.g00170.t01      mNeoVis.evergladensis.2.1.chr9.g00201.t01    mNeoVis.vulgivaga.2.1.chr8.g00206.t01      2533    N0.HOG0002531   OG0000466  n16   rna-XM_011218833.3    rna-XM_038550581.1    rna-XM_022519265.1    rna-XM_003989260.6    rna-XM_036102245.1    rna-XM_039246150.1    rna-XM_032836715.1    rna-XM_046014032.1    rna-XM_032346280.1    rna-XM_059186976.1    rna-XM_059404598.1    rna-XM_045083864.1    mNeoVis.2.1.1.chr9_hap1.g01163.t01    rna-XM_021687668.1    rna-XM_004410770.1    rna-XM_042993267.1    rna-XM_040637064.1, rna-XM_040637535.1    rna-XM_027595494.2    mNeoVis.3.1.1.chr10.g01116.t01      mNeoVis.evergladensis.2.1.chr10.g01151.t01    mNeoVis.vulgivaga.2.1.chr10.g01143.t01      2534   N0.HOG0002532  OG0000466  n31   rna-XM_034646685.1    rna-NM_001242714.1    rna-XM_022491457.1    rna-XM_023243493.2    rna-XM_036101860.1    rna-XM_039227366.1    rna-XM_032848992.1, rna-XM_032850396.1, rna-XM_032851005.1, rna-XM_032853569.1    rna-XM_045984316.1      rna-XM_059148860.1    rna-XM_059380614.1    rna-XM_004776465.3    mNeoVis.2.1.1.chr8_hap1.g00865.t01, mNeoVis.2.1.1.chr8_hap1.g00907.t01, mNeoVis.2.1.1.chr8_hap1.g00908.t01    rna-XM_021694790.1    rna-XM_004405173.1    rna-XM_042966095.1    rna-XM_008685959.2, rna-XM_040630179.1    rna-XM_027624477.2    mNeoVis.3.1.1.chr8.g00812.t01      mNeoVis.evergladensis.2.1.chr9.g00895.t01        2535   N0.HOG0002533  OG0000467  n1   rna-XM_034656482.1    rna-XM_038532352.1    rna-XM_022504991.1    rna-XM_019836585.3    rna-XM_036072307.1    rna-XM_039219377.1    rna-XM_032846863.1    rna-XM_046000028.1    rna-XM_032335795.1    rna-XM_059175452.1    rna-XM_059418220.1    rna-XM_045074479.1    mNeoVis.2.1.1.chr1_hap1.g00732.t01    rna-XM_044916994.1    rna-XM_004403306.1    rna-XM_042989769.1    rna-XM_008700829.2    rna-XM_027604537.2    mNeoVis.3.1.1.chr1.g00754.t01    rna-gnl|WGS_ZZZZ|egapxtmp_035547-R1    mNeoVis.evergladensis.2.1.chr1.g00742.t01    mNeoVis.vulgivaga.2.1.chr1.g00753.t01      2536    N0.HOG0002534   OG0000467  n9   rna-XM_034667439.1    rna-XM_038543013.1    rna-XM_022490762.1    rna-XM_023247965.2    rna-XM_036118274.1    rna-XM_039251337.1    rna-XM_032845220.1    rna-XM_045982169.1    rna-XM_032318102.1    rna-XM_059145273.1    rna-XM_059413657.1    rna-XM_045086731.1    mNeoVis.2.1.1.chr14_hap1.g00617.t01    rna-XM_021682089.1    rna-XM_004402906.1    rna-XM_042975574.1    rna-XM_040635811.1    rna-XM_027611523.2    mNeoVis.3.1.1.chr14.g00598.t01      mNeoVis.evergladensis.2.1.chr14.g00614.t01    mNeoVis.vulgivaga.2.1.chr14.g00610.t01      2537    N0.HOG0002535   OG0000467  n29   rna-XM_034667933.1    rna-XM_038532680.1    rna-XM_022511039.1    rna-XM_023254974.2    rna-XM_036082305.1    rna-XM_039235051.1    rna-XM_032836016.1    rna-XM_046007644.1    rna-XM_032342133.1, rna-XM_032342150.1, rna-XM_032342182.1    rna-XM_059180093.1, rna-XM_059180145.1    rna-XM_059387347.1    rna-XM_045070582.1    mNeoVis.2.1.1.chr10_hap1.g01154.t01    rna-XM_044918230.1    rna-XM_004405535.1    rna-XM_042988173.1    rna-XM_040642702.1    rna-XM_027569116.2    mNeoVis.3.1.1.chr9.g01089.t01    rna-gnl|WGS_ZZZZ|egapxtmp_017124-R7    mNeoVis.evergladensis.2.1.chr8.g01133.t01    mNeoVis.vulgivaga.2.1.chr9.g01114.t01      2538    N0.HOG0002536   OG0000468  n1   rna-XM_034657503.1    rna-XM_038567799.1    rna-XM_022525938.1    rna-XM_023243812.2    rna-XM_036116078.1    rna-XM_039247493.1    rna-XM_032883559.1    rna-XM_045990473.1    rna-XM_032306142.1    rna-XM_059159469.1    rna-XM_059388649.1    rna-XM_004771405.3    mNeoVis.2.1.1.chr5_hap1.g00102.t01    rna-XM_021705200.1, rna-XM_044912224.1    rna-XM_004395343.1    rna-XM_042977764.1    rna-XM_040629590.1, rna-XM_040629591.1    rna-XM_027587904.2    mNeoVis.3.1.1.chr2.g00057.t01    rna-gnl|WGS_ZZZZ|egapxtmp_034404-R1    mNeoVis.evergladensis.2.1.chr2.g00100.t01    mNeoVis.vulgivaga.2.1.chr2.g00097.t01      2539   N0.HOG0002537  OG0000468  n25   rna-XM_034667971.1    rna-XM_038532619.1    rna-XM_022511172.1    rna-XM_045058911.1    rna-XM_036106233.1    rna-XM_039217195.1    rna-XM_032835963.1    rna-XM_046009954.1    rna-XM_032342243.1    rna-XM_059180195.1    rna-XM_059371702.1    rna-XM_045070615.1    mNeoVis.2.1.1.chr10_hap1.g01137.t01    rna-XM_021680490.2    rna-XM_004409751.1    rna-XM_042988249.1    rna-XM_040642672.1    rna-XM_027572490.1    mNeoVis.3.1.1.chr9.g01073.t01    rna-gnl|WGS_ZZZZ|egapxtmp_016603-R12    mNeoVis.evergladensis.2.1.chr8.g01116.t01    mNeoVis.vulgivaga.2.1.chr9.g01096.t01      2540    N0.HOG0002538   OG0000468  n46   rna-XM_034647205.1    rna-XM_038527133.1    rna-XM_022512776.1    rna-XM_023243334.2    rna-XM_036117890.1    rna-XM_039249534.1    rna-XM_032866582.1    rna-XM_046023677.1    rna-XM_032308664.1    rna-XM_059141276.1    rna-XM_059411894.1    rna-XM_045065075.1    mNeoVis.2.1.1.chr13_hap1.g00754.t01    rna-XM_021694416.1    rna-XM_004397535.1    rna-XM_042965241.1    rna-XM_040620261.1    rna-XM_027615657.2    mNeoVis.3.1.1.chr13.g00671.t01    rna-gnl|WGS_ZZZZ|egapxtmp_030003-R1    mNeoVis.evergladensis.2.1.chr13.g00718.t01    mNeoVis.vulgivaga.2.1.chr13.g00716.t01      2541    N0.HOG0002539   OG0000469  n1   rna-XM_034666149.1    rna-XM_038568289.1    rna-XM_022514000.1    rna-XM_019811619.2    rna-XM_036074966.1    rna-XM_039255007.1    rna-XM_032852774.1    rna-XM_046015612.1    rna-XM_032359433.1    rna-XM_059186213.1    rna-XM_059391777.1    rna-XM_013057288.2    mNeoVis.2.1.1.chr7_hap1.g01430.t01    rna-XM_021679075.2    rna-XM_004412430.1    rna-XM_042959100.1    rna-XM_008688368.2    rna-XM_027580118.1    mNeoVis.3.1.1.chr7.g01338.t01    rna-gnl|WGS_ZZZZ|egapxtmp_014266-R1    mNeoVis.evergladensis.2.1.chr7.g01412.t01    mNeoVis.vulgivaga.2.1.chr7.g01424.t01      2542    N0.HOG0002540   OG0000469  n22   rna-XM_034642185.1    rna-XM_038575345.1, rna-XM_038575361.1, rna-XM_038575456.1, rna-XM_038576091.1    rna-XM_022522792.1    rna-XM_045042081.1    rna-XM_036105212.1    rna-XM_039225607.1    rna-XM_032883238.1    rna-XM_046025551.1    rna-XM_032309471.1    rna-XM_059138987.1    rna-XM_059409905.1    rna-XM_045081527.1    mNeoVis.2.1.1.chr3_hap1.g00469.t01    rna-XM_021703471.1    rna-XM_004412462.1    rna-XM_042961555.1      rna-XM_035723979.1    mNeoVis.3.1.1.chr3.g00461.t01    rna-gnl|WGS_ZZZZ|egapxtmp_041092-R1    mNeoVis.evergladensis.2.1.chr3.g00477.t01    mNeoVis.vulgivaga.2.1.chr3.g00480.t01      2543    N0.HOG0002541   OG0000469  n45   rna-XM_019807885.2    rna-XM_038574729.1    rna-XM_022517389.1    rna-XM_019814415.3    rna-XM_036094837.1    rna-XM_039249569.1    rna-XM_032846881.1    rna-XM_046025592.1    rna-XM_032311032.1    rna-XM_059140274.1    rna-XM_059409926.1    rna-XM_004778210.3    mNeoVis.2.1.1.chr3_hap1.g00668.t01    rna-XM_021698637.1    rna-XM_004415452.1    rna-XM_042963111.1    rna-XM_008688788.1    rna-XM_027577541.2    mNeoVis.3.1.1.chr3.g00660.t01    rna-gnl|WGS_ZZZZ|egapxtmp_041148-R1    mNeoVis.evergladensis.2.1.chr3.g00676.t01    mNeoVis.vulgivaga.2.1.chr3.g00680.t01      2544    N0.HOG0002542   OG0000470  n1   rna-XM_034661871.1    rna-NM_001128100.1    rna-XM_022503597.1    rna-XM_045037202.1    rna-XM_036122454.1    rna-XM_039241661.1    rna-XM_032875474.1    rna-XM_046003380.1    rna-XM_032339260.1    rna-XM_059163332.1    rna-XM_059391434.1    rna-XM_004745106.3    mNeoVis.2.1.1.chr5_hap1.g01343.t01    rna-XM_021692586.2    rna-XM_004406789.1    rna-XM_007077551.3    rna-XM_008705260.2    rna-XM_027587196.2    mNeoVis.3.1.1.chr2.g01261.t01, mNeoVis.3.1.1.chr2.g01262.t01    rna-gnl|WGS_ZZZZ|egapxtmp_035044-R1, rna-gnl|WGS_ZZZZ|egapxtmp_035589-R1    mNeoVis.evergladensis.2.1.chr2.g01329.t01    mNeoVis.vulgivaga.2.1.chr2.g01337.t01      2545    N0.HOG0002543   OG0000470  n25   rna-XM_034639357.1    rna-XM_038531122.1    rna-XM_022518750.1    rna-XM_019819692.3    rna-XM_036124326.1    rna-XM_039241814.1    rna-XM_032854553.1    rna-XM_045988497.1    rna-XM_032326256.1    rna-XM_059153000.1    rna-XM_059383869.1    rna-XM_004744119.2    mNeoVis.2.1.1.chr7_hap1.g00351.t01    rna-XM_044922053.1    rna-XM_004417048.1    rna-XM_042969680.1    rna-XM_040633258.1    rna-XM_027617235.2    mNeoVis.3.1.1.chr7.g00338.t01    rna-gnl|WGS_ZZZZ|egapxtmp_013138-R5    mNeoVis.evergladensis.2.1.chr7.g00353.t01        2546    N0.HOG0002544   OG0000470  n44   rna-XM_034638965.1    rna-XM_038531102.1    rna-XM_022518638.1    rna-XM_003998010.6    rna-XM_036124324.1    rna-XM_039242111.1    rna-XM_032854551.1    rna-XM_045988125.1    rna-XM_032326510.1    rna-XM_059151922.1    rna-XM_059381174.1    rna-XM_004744123.3    mNeoVis.2.1.1.chr7_hap1.g00350.t01    rna-XM_021705834.1    rna-XM_004417049.1    rna-XM_042968080.1, rna-XM_042970102.1    rna-XM_008702776.2    rna-XM_027617233.2    mNeoVis.3.1.1.chr7.g00337.t01    rna-gnl|WGS_ZZZZ|egapxtmp_013140-R1    mNeoVis.evergladensis.2.1.chr7.g00352.t01        2547    N0.HOG0002545   OG0000471  n1   rna-XM_034657482.1    rna-XM_038567825.1    rna-XM_022525740.1    rna-XM_045043479.1    rna-XM_036116095.1    rna-XM_039247539.1    rna-XM_032883535.1    rna-XM_045989651.1    rna-XM_032305958.1    rna-XM_059159440.1    rna-XM_059388624.1    rna-XM_045086753.1    mNeoVis.2.1.1.chr5_hap1.g00089.t01    rna-XM_021705411.2    rna-XM_004395355.1    rna-XM_042977731.1    rna-XM_040629616.1    rna-XM_027582935.2    mNeoVis.3.1.1.chr2.g00044.t01      mNeoVis.evergladensis.2.1.chr2.g00086.t01    mNeoVis.vulgivaga.2.1.chr2.g00083.t01      2548    N0.HOG0002546   OG0000471  n18   rna-XM_034653513.1    rna-XM_038537346.1    rna-XM_022496466.1    rna-XM_045035930.1    rna-XM_036070039.1    rna-XM_039238382.1    rna-XM_032839785.1    rna-XM_046007054.1    rna-XM_032303339.1    rna-XM_059138244.1    rna-XM_059374929.1    rna-XM_045064620.1    mNeoVis.2.1.1.chr2_hap1.g01232.t01    rna-XM_021679991.2    rna-XM_004411272.1    rna-XM_042997401.1    rna-XM_040627223.1    rna-XM_027597911.2    mNeoVis.3.1.1.chr5.g01151.t01    rna-gnl|WGS_ZZZZ|egapxtmp_009207-R2    mNeoVis.evergladensis.2.1.chr5.g01176.t01    mNeoVis.vulgivaga.2.1.chr5.g01166.t01      2549    N0.HOG0002547   OG0000471  n37   rna-XM_034653098.1, rna-XM_034657861.1    rna-XM_038567101.1    rna-XM_022526776.1    rna-XM_019818198.3    rna-XM_036074203.1, rna-XM_036110792.1    rna-XM_039252259.1    rna-XM_032884163.1    rna-XM_045990004.1    rna-XM_032312672.1    rna-XM_059160152.1    rna-XM_059389147.1    rna-XM_045083394.1    mNeoVis.2.1.1.chr5_hap1.g00349.t01    rna-XM_044913186.1    rna-XM_004398951.1    rna-XM_042978307.1    rna-XM_040627221.1, rna-XM_040629231.1    rna-XM_035727864.1    mNeoVis.3.1.1.chr2.g00285.t01      mNeoVis.evergladensis.2.1.chr2.g00349.t01, mNeoVis.evergladensis.2.1.chr2.g00350.t01    mNeoVis.vulgivaga.2.1.chr2.g00345.t01      2550    N0.HOG0002548   OG0000472  n1   rna-XM_011221882.3    rna-XM_038535588.1    rna-XM_022513331.1    rna-XM_023239701.2    rna-XM_036088117.1    rna-XM_039253783.1    rna-XM_032868980.1    rna-XM_046016473.1    rna-XM_032359045.1    rna-XM_059185654.1    rna-XM_059402417.1    rna-XM_013062172.2    mNeoVis.2.1.1.chr7_hap1.g01409.t01    rna-XM_021696239.1    rna-XM_004416082.1    rna-XM_042959595.1    rna-XM_008688409.2    rna-XM_027580242.1    mNeoVis.3.1.1.chr7.g01319.t01    rna-gnl|WGS_ZZZZ|egapxtmp_012455-R1    mNeoVis.evergladensis.2.1.chr7.g01392.t01    mNeoVis.vulgivaga.2.1.chr7.g01402.t01      2551    N0.HOG0002549   OG0000472  n22   rna-XM_034666119.1    rna-XM_038535560.1    rna-XM_022513115.1    rna-XM_019813033.3    rna-XM_036088125.1    rna-XM_039253791.1    rna-XM_032868982.1    rna-XM_046016668.1    rna-XM_032359040.1    rna-XM_059185700.1    rna-XM_059402294.1    rna-XM_013062132.2    mNeoVis.2.1.1.chr7_hap1.g01410.t01    rna-XM_044919781.1    rna-XM_004414354.1    rna-XM_042958481.1    rna-XM_040630641.1    rna-XM_027580167.2    mNeoVis.3.1.1.chr7.g01320.t01    rna-gnl|WGS_ZZZZ|egapxtmp_013693-R1    mNeoVis.evergladensis.2.1.chr7.g01393.t01    mNeoVis.vulgivaga.2.1.chr7.g01404.t01      2552    N0.HOG0002550   OG0000472  n43   rna-XM_011221878.3, rna-XM_034650091.1    rna-XM_038535577.1    rna-XM_022513116.1    rna-XM_045038551.1    rna-XM_036088114.1    rna-XM_039253782.1    rna-XM_032868983.1    rna-XM_046014105.1    rna-XM_032358281.1    rna-XM_059185713.1    rna-XM_059413315.1    rna-XM_045080324.1    mNeoVis.2.1.1.chr7_hap1.g01411.t01    rna-XM_021696241.1    rna-XM_004411370.1, rna-XM_004414351.1    rna-XM_007082660.3    rna-XM_008688341.2    rna-XM_027579892.2    mNeoVis.3.1.1.chr7.g01321.t01    rna-gnl|WGS_ZZZZ|egapxtmp_013695-R1    mNeoVis.evergladensis.2.1.chr7.g01394.t01    mNeoVis.vulgivaga.2.1.chr7.g01405.t01      2553    N0.HOG0002551   OG0000473  n1   rna-XM_011232247.3    rna-XM_038533472.1    rna-XM_022506215.1    rna-XM_045056460.1    rna-XM_036112298.1    rna-XM_039227000.1    rna-XM_032853272.1    rna-XM_045998075.1    rna-XM_032334104.1    rna-XM_059163380.1    rna-XM_059382595.1    rna-XM_045076582.1    mNeoVis.2.1.1.chr6_hap1.g00232.t01    rna-XM_021701567.1    rna-XM_004396162.1    rna-XM_015545115.2    rna-XM_040625227.1    rna-XM_027599089.2    mNeoVis.3.1.1.chr4.g00230.t01    rna-gnl|WGS_ZZZZ|egapxtmp_042985-R1    mNeoVis.evergladensis.2.1.chr4.g00239.t01    mNeoVis.vulgivaga.2.1.chr4.g00235.t01      2554    N0.HOG0002552   OG0000473  n19   rna-XM_002928403.4    rna-XM_038587824.1    rna-XM_022521668.1      rna-XM_036092468.1, rna-XM_036105309.1    rna-XM_039221751.1    rna-XM_032881811.1, rna-XM_032881817.1    rna-XM_045994793.1, rna-XM_045995409.1    rna-XM_032331382.1    rna-XM_059157916.1    rna-XM_059385213.1    rna-XM_045089522.1    mNeoVis.2.1.1.chr12_hap1.g00409.t01    rna-XM_021692332.1    rna-XM_004416763.1    rna-XM_007085586.2    rna-XM_040630221.1    rna-XM_027608379.2    mNeoVis.3.1.1.chr12.g00368.t01    rna-gnl|WGS_ZZZZ|egapxtmp_029349-R1    mNeoVis.evergladensis.2.1.chr12.g00364.t01    mNeoVis.vulgivaga.2.1.chr12.g00373.t01      2555    N0.HOG0002553   OG0000473  n42   rna-XM_034663802.1    rna-XM_038569718.1    rna-XM_022502960.1    rna-XM_011288605.4    rna-XM_036121357.1    rna-XM_039244387.1    rna-XM_032841065.1    rna-XM_045978113.1    rna-XM_032315139.1    rna-XM_059144513.1    rna-XM_059377441.1    rna-XM_045080638.1    mNeoVis.2.1.1.chr8_hap1.g01408.t01    rna-XM_021697925.1    rna-XM_004407339.1    rna-XM_042957762.1    rna-XM_008686634.2    rna-XM_027590019.2    mNeoVis.3.1.1.chr8.g01306.t01    rna-gnl|WGS_ZZZZ|egapxtmp_020066-R1    mNeoVis.evergladensis.2.1.chr9.g01354.t01    mNeoVis.vulgivaga.2.1.chr8.g01361.t01      2556    N0.HOG0002554   OG0000474  n1   rna-XM_034647122.1, rna-XM_034649386.1    rna-XM_038536550.1    rna-XM_022524532.1    rna-XM_023243426.2    rna-XM_036078291.1, rna-XM_036099059.1    rna-XM_039254542.1    rna-XM_032840108.1    rna-XM_045985648.1    rna-XM_032320905.1    rna-XM_059148289.1    rna-XM_059379696.1    rna-XM_045083208.1    mNeoVis.2.1.1.chr4_hap1.g00073.t01, mNeoVis.2.1.1.chr8_hap1.g01108.t01    rna-XM_021694735.2    rna-XM_004398584.1, rna-XM_004401830.1, rna-XM_004406534.1    rna-XM_042965806.1    rna-XM_040629881.1    rna-XM_027567696.2, rna-XM_035727027.1, rna-XM_035728016.1, rna-XM_035728278.1    mNeoVis.3.1.1.chr8.g01029.t01      mNeoVis.evergladensis.2.1.chr9.g01086.t01    mNeoVis.vulgivaga.2.1.chr6.g00077.t01, mNeoVis.vulgivaga.2.1.chr8.g01085.t01      2557    N0.HOG0002555   OG0000474  n16   rna-XM_034661835.1    rna-XM_038571200.1    rna-XM_022509524.1    rna-XM_023260463.2    rna-XM_036095592.1    rna-XM_039241040.1    rna-XM_032852476.1    rna-XM_046003401.1    rna-XM_032345717.1    rna-XM_059163790.1    rna-XM_059391752.1    rna-XM_004755524.2      rna-XM_021685946.2    rna-XM_004405018.1    rna-XM_042999146.1    rna-XM_040636683.1    rna-XM_027587716.2              2558    N0.HOG0002556   OG0000474  n22   rna-XM_034656667.1    rna-XM_038534642.1    rna-XM_022493762.1    rna-XM_003981209.5    rna-XM_036084855.1    rna-XM_039239817.1    rna-XM_032851265.1    rna-XM_045998939.1    rna-XM_032336886.1    rna-XM_059173464.1    rna-XM_059416570.1    rna-XM_004737472.3    mNeoVis.2.1.1.chr1_hap1.g00355.t01    rna-XM_021698795.1    rna-XM_004401411.1    rna-XM_007085535.3    rna-XM_008692314.2    rna-XM_027605900.1      rna-gnl|WGS_ZZZZ|egapxtmp_039836-R1    mNeoVis.evergladensis.2.1.chr1.g00353.t01    mNeoVis.vulgivaga.2.1.chr1.g00361.t01      2559    N0.HOG0002557   OG0000475  n1   rna-XM_002918733.4    rna-XM_038588473.1    rna-XM_022521294.1    rna-XM_004000949.6    rna-XM_036101272.1, rna-XM_036104533.1    rna-XM_039256018.1    rna-XM_032846612.1    rna-XM_045995330.1    rna-XM_032329909.1    rna-XM_059156975.1    rna-XM_059386195.1    rna-XM_013048662.2    mNeoVis.2.1.1.chr12_hap1.g00114.t01    rna-XM_021682534.2    rna-XM_004413877.1    rna-XM_042973825.1    rna-XM_040627976.1    rna-XM_027608227.2    mNeoVis.3.1.1.chr12.g00089.t01    rna-gnl|WGS_ZZZZ|egapxtmp_028489-R1    mNeoVis.evergladensis.2.1.chr12.g00089.t01    mNeoVis.vulgivaga.2.1.chr12.g00093.t01      2560    N0.HOG0002558   OG0000475  n22   rna-XM_019792907.2    rna-XM_038583462.1    rna-XM_022503509.1    rna-XM_045037225.1    rna-XM_036095302.1    rna-XM_039221192.1    rna-XM_032875422.1    rna-XM_046001674.1    rna-XM_032339896.1    rna-XM_059163381.1    rna-XM_059391471.1    rna-XM_004745072.3    mNeoVis.2.1.1.chr5_hap1.g01359.t01    rna-XM_021692624.2    rna-XM_004398337.1    rna-XM_007094569.3    rna-XM_008705290.2    rna-XM_027583863.1      rna-gnl|WGS_ZZZZ|egapxtmp_035596-R1    mNeoVis.evergladensis.2.1.chr2.g01345.t01, mNeoVis.evergladensis.2.1.chr2.g01346.t01    mNeoVis.vulgivaga.2.1.chr2.g01355.t01      2561    N0.HOG0002559   OG0000475  n38   rna-XM_034646975.1    rna-XM_038536709.1    rna-XM_022524721.1    rna-XM_023243850.2, rna-XM_045049771.1    rna-XM_036066347.1    rna-XM_039231474.1    rna-XM_032839849.1    rna-XM_045985786.1    rna-XM_032322643.1    rna-XM_059150473.1    rna-XM_059378524.1    rna-XM_004760268.2    mNeoVis.2.1.1.chr8_hap1.g01043.t01    rna-XM_021694500.1    rna-XM_004398495.1    rna-XM_042967624.1    rna-XM_040629972.1    rna-XM_027568500.2    mNeoVis.3.1.1.chr8.g00972.t01    rna-gnl|WGS_ZZZZ|egapxtmp_017563-R1    mNeoVis.evergladensis.2.1.chr9.g01017.t01    mNeoVis.vulgivaga.2.1.chr8.g01017.t01      2562    N0.HOG0002560   OG0000476  n1   rna-XM_034641605.1    rna-XM_038571702.1    rna-XM_022493829.1    rna-XM_045053837.1    rna-XM_036093410.1    rna-XM_039224102.1    rna-XM_032859432.1    rna-XM_045980337.1    rna-XM_032353671.1    rna-XM_059134343.1    rna-XM_059386584.1, rna-XM_059406604.1    rna-XM_045086057.1    mNeoVis.2.1.1.chr11_hap1.g00434.t01    rna-XM_044919029.1    rna-XM_004398190.1    rna-XM_007073271.2    rna-XM_040638023.1    rna-XM_027624003.1    mNeoVis.3.1.1.chr11.g00394.t01    rna-gnl|WGS_ZZZZ|egapxtmp_026649-R1    mNeoVis.evergladensis.2.1.chr11.g00425.t01, mNeoVis.evergladensis.2.1.chr11.g00426.t01    mNeoVis.vulgivaga.2.1.chr11.g00433.t01      2563    N0.HOG0002561   OG0000476  n24   rna-XM_002918611.4    rna-XM_038534797.1    rna-XM_022493482.1    rna-XM_023259796.2    rna-XM_036106745.1    rna-XM_039247665.1    rna-XM_032851767.1    rna-XM_045999826.1    rna-XM_032334905.1    rna-XM_059174256.1    rna-XM_059417326.1    rna-XM_004737645.2    mNeoVis.2.1.1.chr1_hap1.g00288.t01    rna-XM_044917115.1    rna-XM_004392028.1    rna-XM_042992729.1    rna-XM_008692399.2    rna-XM_027606991.2    mNeoVis.3.1.1.chr1.g00293.t01    rna-gnl|WGS_ZZZZ|egapxtmp_019930-R1    mNeoVis.evergladensis.2.1.chr1.g00287.t01    mNeoVis.vulgivaga.2.1.chr1.g00291.t01      2564    N0.HOG0002562   OG0000476  n33   rna-XM_034662933.1    rna-XM_038578167.1    rna-XM_022499606.1    rna-XM_023240708.2    rna-XM_036084692.1    rna-XM_039257754.1    rna-XM_032836809.1    rna-XM_046027124.1    rna-XM_032312131.1    rna-XM_059169502.1    rna-XM_059398094.1    rna-XM_004749685.3    mNeoVis.2.1.1.chr2_hap1.g00341.t01    rna-XM_021699586.2    rna-XM_004410225.1    rna-XM_042960895.1    rna-XM_040624126.1    rna-XM_027595840.2    mNeoVis.3.1.1.chr5.g00331.t01    rna-gnl|WGS_ZZZZ|egapxtmp_006520-R1    mNeoVis.evergladensis.2.1.chr5.g00342.t01    mNeoVis.vulgivaga.2.1.chr5.g00344.t01      2565    N0.HOG0002563   OG0000477  n1   rna-XM_034656228.1    rna-XM_038530554.1    rna-XM_022518097.1    rna-XM_045060664.1    rna-XM_036065724.1    rna-XM_039219194.1    rna-XM_032850643.1    rna-XM_045999746.1    rna-XM_032336247.1    rna-XM_059174980.1    rna-XM_059417862.1    rna-XM_004744669.3    mNeoVis.2.1.1.chr1_hap1.g00560.t01    rna-XM_021701889.1    rna-XM_004414626.1    rna-XM_042986671.1    rna-XM_040622262.1    rna-XM_027606588.2    mNeoVis.3.1.1.chr1.g00575.t01    rna-gnl|WGS_ZZZZ|egapxtmp_032400-R8    mNeoVis.evergladensis.2.1.chr1.g00566.t01    mNeoVis.vulgivaga.2.1.chr1.g00576.t01      2566    N0.HOG0002564   OG0000477  n18   rna-XM_034672150.1    rna-XM_038533413.1    rna-XM_022506020.1    rna-XM_045056614.1    rna-XM_036104667.1    rna-XM_039232028.1    rna-XM_032855324.1    rna-XM_045998595.1    rna-XM_032333784.1    rna-XM_059165614.1    rna-XM_059380551.1    rna-XM_045075476.1    mNeoVis.2.1.1.chr6_hap1.g00325.t01    rna-XM_021677604.1    rna-XM_004416859.1    rna-XM_042984799.1, rna-XM_042984971.1    rna-XM_040625109.1      mNeoVis.3.1.1.chr4.g00325.t01, mNeoVis.3.1.1.chr4.g00326.t01    rna-gnl|WGS_ZZZZ|egapxtmp_000721-R1    mNeoVis.evergladensis.2.1.chr4.g00330.t01, mNeoVis.evergladensis.2.1.chr4.g00331.t01    mNeoVis.vulgivaga.2.1.chr4.g00328.t01      2567    N0.HOG0002565   OG0000477  n41   rna-XM_034663457.1    rna-XM_038579155.1    rna-XM_022499723.1    rna-XM_023241120.2    rna-XM_036070087.1    rna-XM_039252402.1    rna-XM_032882058.1    rna-XM_046027703.1    rna-XM_032313109.1    rna-XM_059172793.1    rna-XM_059398850.1    rna-XM_045068758.1    mNeoVis.2.1.1.chr2_hap1.g00033.t01    rna-XM_044916231.1    rna-XM_004402220.1    rna-XM_042959864.1    rna-XM_040624572.1    rna-XM_027597093.2    mNeoVis.3.1.1.chr5.g00025.t01    rna-gnl|WGS_ZZZZ|egapxtmp_005898-R1    mNeoVis.evergladensis.2.1.chr5.g00033.t01    mNeoVis.vulgivaga.2.1.chr5.g00031.t01      2568    N0.HOG0002566   OG0000478  n1   rna-XM_034655406.1    rna-XM_038574217.1    rna-XM_022498064.1    rna-XM_045034632.1    rna-XM_036109561.1, rna-XM_036109577.1    rna-XM_039234463.1    rna-XM_032857512.1    rna-XM_046019449.1    rna-XM_032355191.1    rna-XM_059167692.1    rna-XM_059393734.1    rna-XM_004773501.3    mNeoVis.2.1.1.chr3_hap1.g01065.t01    rna-XM_044913771.1    rna-XM_004396061.1, rna-XM_004396086.1    rna-XM_042995667.1    rna-XM_040638927.1    rna-XM_027591226.1    mNeoVis.3.1.1.chr3.g01057.t01    rna-gnl|WGS_ZZZZ|egapxtmp_040263-R1    mNeoVis.evergladensis.2.1.chr3.g01065.t01    mNeoVis.vulgivaga.2.1.chr3.g01082.t01      2569    N0.HOG0002567   OG0000478  n25   rna-XM_034671458.1    rna-XM_038531862.1    rna-XM_022514655.1    rna-XM_045035144.1    rna-XM_036067010.1    rna-XM_039229041.1    rna-XM_032863520.1    rna-XM_046016952.1    rna-XM_032302076.1    rna-XM_059136021.1    rna-XM_059376318.1    rna-XM_045063816.1    mNeoVis.2.1.1.chr2_hap1.g01770.t01    rna-XM_021684056.1    rna-XM_004397126.1    rna-XM_042997050.1    rna-XM_040645831.1    rna-XM_027623311.1    mNeoVis.3.1.1.chr5.g01695.t01    rna-gnl|WGS_ZZZZ|egapxtmp_007169-R1    mNeoVis.evergladensis.2.1.chr5.g01710.t01    mNeoVis.vulgivaga.2.1.chr5.g01714.t01      2570    N0.HOG0002568   OG0000478  n45   rna-XM_002921824.4    rna-XM_038559281.1    rna-XM_022510332.1    rna-XM_045052645.1    rna-XM_036087227.1    rna-XM_039243024.1    rna-XM_032866833.1    rna-XM_046021016.1    rna-XM_032304785.1    rna-XM_059172144.1    rna-XM_059396204.1    rna-XM_045063113.1    mNeoVis.2.1.1.chr4_hap1.g00083.t01    rna-XM_044920102.1    rna-XM_004409600.1    rna-XM_042974437.1    rna-XM_040644587.1    rna-XM_027573178.2    mNeoVis.3.1.1.chr6.g00084.t01    rna-gnl|WGS_ZZZZ|egapxtmp_003078-R1    mNeoVis.evergladensis.2.1.chr6.g00087.t01    mNeoVis.vulgivaga.2.1.chr6.g00085.t01      2571    N0.HOG0002569   OG0000479  n1   rna-XM_019804772.2    rna-XM_038559354.1    rna-XM_022510193.1    rna-XM_003983123.4    rna-XM_036086605.1    rna-XM_039243039.1    rna-XM_032867259.1    rna-XM_046020405.1    rna-XM_032306129.1    rna-XM_059172083.1    rna-XM_059395833.1    rna-XM_004781428.3    mNeoVis.2.1.1.chr4_hap1.g00119.t01    rna-XM_021693011.1    rna-XM_004414568.1    rna-XM_007090834.2    rna-XM_040643907.1    rna-XM_027573040.1    mNeoVis.3.1.1.chr6.g00120.t01      mNeoVis.evergladensis.2.1.chr6.g00121.t01    mNeoVis.vulgivaga.2.1.chr6.g00118.t01      2572    N0.HOG0002570   OG0000479  n20   rna-XM_034651935.1, rna-XM_034651936.1    rna-XM_038559355.1    rna-XM_022492657.1, rna-XM_022492666.1, rna-XM_022510201.1    rna-XM_003983122.4    rna-XM_036086595.1, rna-XM_036086603.1    rna-XM_039234067.1    rna-XM_032865211.1, rna-XM_032865223.1    rna-XM_046020409.1, rna-XM_046020410.1, rna-XM_046020411.1, rna-XM_046020412.1    rna-XM_032306127.1, rna-XM_032306130.1, rna-XM_032306131.1    rna-XM_059172081.1, rna-XM_059172690.1, rna-XM_059172692.1    rna-XM_059395584.1, rna-XM_059395585.1, rna-XM_059395587.1    rna-XM_004781430.3, rna-XM_045063086.1, rna-XM_045063087.1    mNeoVis.2.1.1.chr4_hap1.g00120.t01, mNeoVis.2.1.1.chr4_hap1.g00121.t01, mNeoVis.2.1.1.chr4_hap1.g00122.t01    rna-XM_044912237.1, rna-XM_044912238.1    rna-XM_004416546.1    rna-XM_007090833.2    rna-XM_008697319.1, rna-XM_008697320.2    rna-XM_027573515.1    mNeoVis.3.1.1.chr6.g00121.t01, mNeoVis.3.1.1.chr6.g00122.t01    rna-gnl|WGS_ZZZZ|egapxtmp_002858-R1, rna-gnl|WGS_ZZZZ|egapxtmp_004720-R2    mNeoVis.evergladensis.2.1.chr6.g00122.t01, mNeoVis.evergladensis.2.1.chr6.g00123.t01    mNeoVis.vulgivaga.2.1.chr6.g00119.t01, mNeoVis.vulgivaga.2.1.chr6.g00120.t01, mNeoVis.vulgivaga.2.1.chr6.g00121.t01      2573    N0.HOG0002571   OG0000480  n1   rna-XM_002929575.2    rna-XM_038553297.1    rna-XM_022501609.1    rna-XM_003995717.2    rna-XM_036111430.1    rna-XM_039220410.1    rna-XM_032860895.1    rna-XM_046021830.1    rna-XM_032306484.1    rna-XM_059141222.1    rna-XM_059410116.1    rna-XM_004747892.2    mNeoVis.2.1.1.chr13_hap1.g00405.t01    rna-XM_021691619.2    rna-XM_004416758.1    rna-XM_007094470.1    rna-XM_008708026.1    rna-XM_027614670.1    mNeoVis.3.1.1.chr13.g00343.t01    rna-gnl|WGS_ZZZZ|egapxtmp_031000-R1    mNeoVis.evergladensis.2.1.chr13.g00389.t01    mNeoVis.vulgivaga.2.1.chr13.g00391.t01      2574   N0.HOG0002572  OG0000480  n23   rna-XM_019796723.2, rna-XM_034664256.1    rna-NM_001389130.1    rna-XM_022501540.1    rna-XM_003995753.5    rna-XM_036090959.1    rna-XM_039240549.1    rna-XM_032860887.1    rna-XM_046023685.1    rna-XM_032306781.1    rna-XM_059141421.1    rna-XM_059412227.1    rna-XM_004747904.3    mNeoVis.2.1.1.chr13_hap1.g00360.t01    rna-XM_021691685.1, rna-XM_044912281.1      rna-XM_007094285.3    rna-XM_040631704.1    rna-XM_027616495.1    mNeoVis.3.1.1.chr13.g00303.t01    rna-gnl|WGS_ZZZZ|egapxtmp_030801-R1    mNeoVis.evergladensis.2.1.chr13.g00348.t01    mNeoVis.vulgivaga.2.1.chr13.g00349.t01      2575   N0.HOG0002573  OG0000480  n44   rna-XM_002929572.2    rna-NM_001017515.1    rna-XM_022501617.1    rna-XM_045042509.1, rna-XM_045042510.1, rna-XM_045042514.1    rna-XM_036111446.1    rna-XM_039220412.1, rna-XM_039220421.1    rna-XM_032860899.1    rna-XM_046021827.1, rna-XM_046021828.1    rna-XM_032306479.1          mNeoVis.2.1.1.chr13_hap1.g00399.t01    rna-XM_021691617.1    rna-XM_004416761.1    rna-XM_015543001.2, rna-XM_015543002.1    rna-XM_008708028.2    rna-XM_027615814.1      rna-gnl|WGS_ZZZZ|egapxtmp_030088-R1    mNeoVis.evergladensis.2.1.chr13.g00384.t01    mNeoVis.vulgivaga.2.1.chr13.g00386.t01      2576    N0.HOG0002574   OG0000481  n1   rna-XM_034653455.1    rna-XM_038533940.1    rna-XM_022523617.1, rna-XM_022523619.1    rna-XM_023240434.2    rna-XM_036106959.1    rna-XM_039241238.1    rna-XM_032841093.1    rna-XM_046026557.1    rna-XM_032311941.1    rna-XM_059170535.1    rna-XM_059397251.1    rna-XM_045087094.1    mNeoVis.2.1.1.chr2_hap1.g00721.t01    rna-XM_021696126.1    rna-XM_004404852.1    rna-XM_042960401.1    rna-XM_040624115.1    rna-XM_027595935.2    mNeoVis.3.1.1.chr5.g00671.t01    rna-gnl|WGS_ZZZZ|egapxtmp_006921-R1    mNeoVis.evergladensis.2.1.chr5.g00680.t01    mNeoVis.vulgivaga.2.1.chr5.g00671.t01      2577    N0.HOG0002575   OG0000481  n22   rna-XM_034657296.1    rna-XM_038561198.1    rna-XM_022513416.1, rna-XM_022513524.1    rna-XM_019827708.3    rna-XM_036119136.1    rna-XM_039238458.1    rna-XM_032881832.1    rna-XM_045979482.1    rna-XM_032353052.1    rna-XM_059188697.1    rna-XM_059407590.1    rna-XM_004777897.3    mNeoVis.2.1.1.chr11_hap1.g01018.t01    rna-XM_021697574.1    rna-XM_004409576.1    rna-XM_042982220.1    rna-XM_040623060.1    rna-XM_027622059.2    mNeoVis.3.1.1.chr11.g00969.t01    rna-gnl|WGS_ZZZZ|egapxtmp_025349-R1    mNeoVis.evergladensis.2.1.chr11.g00993.t01    mNeoVis.vulgivaga.2.1.chr11.g01016.t01      2578    N0.HOG0002576   OG0000481  n42   rna-XM_002917025.4    rna-XM_038545060.1    rna-XM_022505302.1    rna-XM_003987773.6    rna-XM_036107230.1    rna-XM_039240749.1    rna-XM_032849393.1    rna-XM_046009896.1    rna-XM_032344596.1    rna-XM_059182431.1    rna-XM_059373473.1    rna-XM_004739009.3    mNeoVis.2.1.1.chr10_hap1.g00282.t01    rna-XM_021679524.1    rna-XM_004397601.1    rna-XM_042989986.1    rna-XM_040641728.1    rna-XM_027570846.2    mNeoVis.3.1.1.chr9.g00254.t01    rna-gnl|WGS_ZZZZ|egapxtmp_016788-R1    mNeoVis.evergladensis.2.1.chr8.g00267.t01    mNeoVis.vulgivaga.2.1.chr9.g00266.t01      2579   N0.HOG0002577  OG0000482  n1   rna-XM_002927655.4    rna-NM_001205115.1    rna-XM_022515590.1    rna-XM_045049643.1, rna-XM_045050373.1    rna-XM_036094049.1    rna-XM_039223657.1    rna-XM_032841192.1    rna-XM_046000271.1, rna-XM_046010466.1    rna-XM_032315628.1    rna-XM_059166485.1    rna-XM_059394564.1    rna-XM_045082041.1    mNeoVis.2.1.1.chr4_hap1.g00691.t01    rna-XM_021688149.1    rna-XM_004395744.1    rna-XM_007084798.3, rna-XM_042963593.1    rna-XM_040625853.1    rna-XM_027603945.1    mNeoVis.3.1.1.chr6.g00696.t01    rna-gnl|WGS_ZZZZ|egapxtmp_002466-R1    mNeoVis.evergladensis.2.1.chr6.g00695.t01    mNeoVis.vulgivaga.2.1.chr6.g00690.t01      2580    N0.HOG0002578   OG0000482  n12   rna-XM_011225015.2    rna-NM_001205096.1    rna-XM_022493938.1, rna-XM_022517236.1    rna-XM_045056957.1, rna-XM_045059327.1    rna-XM_036064666.1    rna-XM_039229991.1, rna-XM_039231955.1    rna-XM_032876337.1, rna-XM_032877833.1, rna-XM_032878989.1    rna-XM_045980602.1    rna-XM_032350371.1    rna-XM_059132856.1, rna-XM_059165196.1, rna-XM_059165197.1    rna-XM_059388319.1, rna-XM_059407219.1    rna-XM_004746213.3, rna-XM_045088142.1, rna-XM_045088143.1    mNeoVis.2.1.1.chr11_hap1.g00131.t01, mNeoVis.2.1.1.chr1_hap1.g02033.t01    rna-XM_021685892.2    rna-XM_004394056.1, rna-XM_004403961.1    rna-XM_042963646.1, rna-XM_042972663.1, rna-XM_042980170.1, rna-XM_042996216.1    rna-XM_008698016.2    rna-XM_027606759.2, rna-XM_027610879.2, rna-XM_027621508.2, rna-XM_035728383.1    mNeoVis.3.1.1.chr1.g01987.t01, mNeoVis.3.1.1.chr3.g00154.t01    rna-gnl|WGS_ZZZZ|egapxtmp_002515-R1, rna-gnl|WGS_ZZZZ|egapxtmp_026974-R1    mNeoVis.evergladensis.2.1.chr1.g02000.t01, mNeoVis.evergladensis.2.1.chr11.g00128.t01    mNeoVis.vulgivaga.2.1.chr1.g01993.t01      2581    N0.HOG0002579   OG0000483  n1   rna-XM_019800455.2    rna-XM_038565787.1    rna-XM_022498062.1    rna-XM_023249983.2    rna-XM_036075554.1    rna-XM_039240299.1    rna-XM_032867448.1    rna-XM_045991794.1    rna-XM_032325752.1    rna-XM_059161557.1    rna-XM_059390201.1    rna-XM_013054016.2    mNeoVis.2.1.1.chr5_hap1.g00837.t01    rna-XM_021695118.2    rna-XM_004399434.1    rna-XM_007076615.3    rna-XM_008709432.2, rna-XM_040628531.1    rna-XM_027586400.2    mNeoVis.3.1.1.chr2.g00747.t01    rna-gnl|WGS_ZZZZ|egapxtmp_035464-R1    mNeoVis.evergladensis.2.1.chr2.g00838.t01    mNeoVis.vulgivaga.2.1.chr2.g00840.t01      2582    N0.HOG0002580   OG0000483  n24   rna-XM_034653422.1, rna-XM_034660564.1    rna-XM_038553756.1    rna-XM_022510983.1    rna-XM_023253857.2    rna-XM_036115683.1    rna-XM_039241803.1    rna-XM_032849136.1    rna-XM_046004189.1    rna-XM_032340606.1    rna-XM_059178608.1    rna-XM_059400042.1    rna-XM_045074716.1    mNeoVis.2.1.1.chr1_hap1.g01440.t01    rna-XM_044917632.1    rna-XM_004407716.1    rna-XM_042986136.1    rna-XM_040644612.1    rna-XM_035728560.1    mNeoVis.3.1.1.chr1.g01386.t01      mNeoVis.evergladensis.2.1.chr1.g01402.t01    mNeoVis.vulgivaga.2.1.chr1.g01406.t01      2583    N0.HOG0002581   OG0000483  n45   rna-XM_034646080.1    rna-XM_038577122.1    rna-XM_022522705.1    rna-XM_019834731.3    rna-XM_036080192.1    rna-XM_039240789.1    rna-XM_032855215.1    rna-XM_046011092.1    rna-XM_032347608.1    rna-XM_059185826.1    rna-XM_059402745.1    rna-XM_004776400.3    mNeoVis.2.1.1.chr9_hap1.g00719.t01    rna-XM_044915443.1    rna-XM_004415811.1    rna-XM_042991993.1    rna-XM_040642268.1    rna-XM_027593395.2    mNeoVis.3.1.1.chr10.g00688.t01    rna-gnl|WGS_ZZZZ|egapxtmp_024731-R1    mNeoVis.evergladensis.2.1.chr10.g00712.t01    mNeoVis.vulgivaga.2.1.chr10.g00702.t01      2584    N0.HOG0002582   OG0000484  n1   rna-XM_011230582.3    rna-XM_038580442.1    rna-XM_022525016.1    rna-XM_019832406.3    rna-XM_036116646.1    rna-XM_039237163.1    rna-XM_032856457.1    rna-XM_046009077.1    rna-XM_032343752.1    rna-XM_059181572.1    rna-XM_059372842.1    rna-XM_004755408.3    mNeoVis.2.1.1.chr10_hap1.g00599.t01    rna-XM_021693732.1    rna-XM_004396574.1    rna-XM_042988833.1    rna-XM_040620422.1    rna-XM_027572805.2    mNeoVis.3.1.1.chr9.g00556.t01    rna-gnl|WGS_ZZZZ|egapxtmp_016864-R1    mNeoVis.evergladensis.2.1.chr8.g00581.t01    mNeoVis.vulgivaga.2.1.chr9.g00581.t01      2585    N0.HOG0002583   OG0000484  n23   rna-XM_034642874.1    rna-XM_038543976.1    rna-XM_022495717.1    rna-XM_045042264.1    rna-XM_036078211.1    rna-XM_039228822.1    rna-XM_032850479.1, rna-XM_032882532.1    rna-XM_046024282.1    rna-XM_032310011.1    rna-XM_059138812.1    rna-XM_059409632.1    rna-XM_045065613.1    mNeoVis.2.1.1.chr3_hap1.g00110.t01    rna-XM_044920870.1    rna-XM_004410471.1    rna-XM_042961931.1    rna-XM_040632520.1    rna-XM_027575563.2    mNeoVis.3.1.1.chr3.g00114.t01    rna-gnl|WGS_ZZZZ|egapxtmp_041013-R1    mNeoVis.evergladensis.2.1.chr3.g00114.t01    mNeoVis.vulgivaga.2.1.chr3.g00114.t01      2586    N0.HOG0002584   OG0000484  n45   rna-XM_034660960.1    rna-XM_038580443.1    rna-XM_022492892.1    rna-XM_023255339.2    rna-XM_036116649.1    rna-XM_039237099.1    rna-XM_032856449.1    rna-XM_046008072.1    rna-XM_032343753.1    rna-XM_059181577.1    rna-XM_059372844.1    rna-XM_045067906.1    mNeoVis.2.1.1.chr10_hap1.g00598.t01    rna-XM_044918191.1    rna-XM_004396575.1    rna-XM_042988826.1    rna-XM_040620421.1    rna-XM_027570252.2    mNeoVis.3.1.1.chr9.g00555.t01    rna-gnl|WGS_ZZZZ|egapxtmp_016863-R1    mNeoVis.evergladensis.2.1.chr8.g00580.t01    mNeoVis.vulgivaga.2.1.chr9.g00580.t01      2587    N0.HOG0002585   OG0000485  n1   rna-XM_034663052.1    rna-XM_038578385.1    rna-XM_022499930.1    rna-XM_023240792.2    rna-XM_036111621.1    rna-XM_039230278.1    rna-XM_032883365.1    rna-XM_046027312.1    rna-XM_032313404.1    rna-XM_059169311.1    rna-XM_059395909.1    rna-XM_045068820.1    mNeoVis.2.1.1.chr2_hap1.g00291.t01    rna-XM_021699869.1    rna-XM_004409484.1    rna-XM_042960931.1    rna-XM_040624553.1    rna-XM_027597114.2    mNeoVis.3.1.1.chr5.g00285.t01    rna-gnl|WGS_ZZZZ|egapxtmp_006824-R1, rna-gnl|WGS_ZZZZ|egapxtmp_008449-R1    mNeoVis.evergladensis.2.1.chr5.g00296.t01    mNeoVis.vulgivaga.2.1.chr5.g00296.t01      2588    N0.HOG0002586   OG0000485  n24   rna-XM_019794734.2    rna-XM_038573862.1    rna-XM_022517772.1    rna-XM_045055405.1    rna-XM_036075172.1    rna-XM_039243741.1    rna-XM_032855860.1    rna-XM_045992022.1    rna-XM_032332113.1    rna-XM_059178536.1    rna-XM_059371423.1    rna-XM_004763336.2    mNeoVis.2.1.1.chr6_hap1.g00841.t01    rna-XM_021698765.2    rna-XM_004402191.1    rna-XM_042983290.1    rna-XM_040626544.1    rna-XM_027598416.2    mNeoVis.3.1.1.chr4.g00852.t01    rna-gnl|WGS_ZZZZ|egapxtmp_000872-R1    mNeoVis.evergladensis.2.1.chr4.g00843.t01    mNeoVis.vulgivaga.2.1.chr4.g00863.t01      2589    N0.HOG0002587   OG0000485  n45   rna-XM_034661408.1    rna-XM_038559153.1    rna-XM_022516745.1    rna-XM_023252694.2    rna-XM_036112138.1    rna-XM_039249957.1    rna-XM_032855437.1    rna-XM_045998280.1    rna-XM_032332599.1    rna-XM_059174740.1    rna-XM_059374212.1    rna-XM_004779539.3    mNeoVis.2.1.1.chr6_hap1.g00682.t01    rna-XM_021698858.1    rna-XM_004392671.1    rna-XM_042984746.1    rna-XM_040625543.1    rna-XM_035726294.1    mNeoVis.3.1.1.chr4.g00689.t01    rna-gnl|WGS_ZZZZ|egapxtmp_043501-R1    mNeoVis.evergladensis.2.1.chr4.g00688.t01    mNeoVis.vulgivaga.2.1.chr4.g00703.t01      2590    N0.HOG0002588   OG0000486  n1   rna-XM_002924688.1, rna-XM_034655684.1    rna-XM_038529616.1    rna-XM_022518164.1    rna-XM_003980945.5    rna-XM_036070018.1    rna-XM_039256681.1    rna-XM_032844171.1    rna-XM_046001193.1    rna-XM_032334953.1    rna-XM_059174960.1    rna-XM_059417844.1    rna-XM_004744697.2    mNeoVis.2.1.1.chr1_hap1.g00554.t01    rna-XM_021702028.1    rna-XM_004413481.1    rna-XM_007079960.2    rna-XM_008691434.1    rna-XM_027607084.2    mNeoVis.3.1.1.chr1.g00569.t01    rna-gnl|WGS_ZZZZ|egapxtmp_017709-R1    mNeoVis.evergladensis.2.1.chr1.g00560.t01    mNeoVis.vulgivaga.2.1.chr1.g00570.t01      2591    N0.HOG0002589   OG0000486  n9   rna-XM_002916241.2    rna-XM_038557609.1    rna-XM_022514640.1    rna-XM_011281629.4    rna-XM_036077766.1    rna-XM_039254288.1    rna-XM_032855911.1    rna-XM_045998510.1    rna-XM_032330885.1    rna-XM_059155383.1    rna-XM_059393754.1    rna-XM_004758019.2    mNeoVis.2.1.1.chr6_hap1.g00721.t01    rna-XM_021679305.1    rna-XM_004411755.1    rna-XM_042982678.1    rna-XM_040626040.1    rna-XM_027600434.1    mNeoVis.3.1.1.chr4.g00732.t01    rna-gnl|WGS_ZZZZ|egapxtmp_044243-R1    mNeoVis.evergladensis.2.1.chr4.g00728.t01    mNeoVis.vulgivaga.2.1.chr4.g00746.t01      2592    N0.HOG0002590   OG0000486  n21   rna-XM_034663805.1    rna-XM_038569375.1    rna-XM_022503022.1    rna-XM_023252424.2    rna-XM_036066932.1    rna-XM_039217787.1    rna-XM_032864258.1    rna-XM_045978374.1    rna-XM_032315239.1    rna-XM_059144218.1    rna-XM_059377838.1    rna-XM_045070894.1    mNeoVis.2.1.1.chr8_hap1.g01434.t01    rna-XM_044913457.1    rna-XM_004410620.1    rna-XM_042957854.1    rna-XM_040623731.1    rna-XM_027590385.2    mNeoVis.3.1.1.chr8.g01331.t01    rna-gnl|WGS_ZZZZ|egapxtmp_020490-R1    mNeoVis.evergladensis.2.1.chr9.g01379.t01    mNeoVis.vulgivaga.2.1.chr8.g01388.t01      2593    N0.HOG0002591   OG0000487  n1   rna-XM_011220726.3    rna-NM_001313795.1    rna-XM_022523105.1    rna-XM_003988894.5    rna-XM_036117109.1    rna-XM_039216854.1    rna-XM_032874484.1    rna-XM_046012244.1    rna-XM_032348691.1    rna-XM_059184934.1    rna-XM_059403447.1    rna-XM_004773142.2    mNeoVis.2.1.1.chr9_hap1.g00369.t01    rna-XM_021687258.1    rna-XM_004401686.1    rna-XM_007081508.3    rna-XM_040637135.1    rna-XM_027594444.1    mNeoVis.3.1.1.chr10.g00361.t01    rna-gnl|WGS_ZZZZ|egapxtmp_024647-R1    mNeoVis.evergladensis.2.1.chr10.g00366.t01    mNeoVis.vulgivaga.2.1.chr10.g00363.t01      2594    N0.HOG0002592   OG0000487  n17   rna-XM_034646292.1    rna-NM_001313833.1    rna-XM_022519624.1    rna-XM_023257024.2    rna-XM_036112965.1    rna-XM_039241427.1    rna-XM_032876838.1    rna-XM_046010651.1    rna-XM_032348141.1    rna-XM_059185414.1    rna-XM_059403061.1    rna-XM_004778844.2    mNeoVis.2.1.1.chr9_hap1.g00572.t01    rna-XM_021704838.1    rna-XM_004400842.1    rna-XM_042990815.1    rna-XM_040641918.1, rna-XM_040641946.1    rna-XM_027595720.1    mNeoVis.3.1.1.chr10.g00542.t01    rna-gnl|WGS_ZZZZ|egapxtmp_023521-R1    mNeoVis.evergladensis.2.1.chr10.g00565.t01    mNeoVis.vulgivaga.2.1.chr10.g00553.t01      2595    N0.HOG0002593   OG0000487  n33   rna-XM_034646390.1    rna-XM_038576165.1    rna-XM_022519623.1    rna-XM_003988711.6    rna-XM_036112982.1    rna-XM_039241402.1    rna-XM_032876837.1    rna-XM_046012784.1    rna-XM_032348142.1    rna-XM_059185413.1    rna-XM_059403062.1    rna-XM_004778924.2    mNeoVis.2.1.1.chr9_hap1.g00571.t01    rna-XM_021704790.2    rna-XM_004400901.1    rna-XM_007072994.3    rna-XM_008704750.2    rna-XM_027595355.1    mNeoVis.3.1.1.chr10.g00540.t01    rna-gnl|WGS_ZZZZ|egapxtmp_023520-R1    mNeoVis.evergladensis.2.1.chr10.g00564.t01    mNeoVis.vulgivaga.2.1.chr10.g00552.t01      2596    N0.HOG0002594   OG0000488  n1   rna-XM_034644806.1    rna-NM_001048106.1    rna-XM_022506643.1    rna-XM_019812396.2    rna-XM_036105069.1    rna-XM_039226906.1    rna-XM_032844560.1    rna-XM_046015851.1    rna-XM_032360144.1    rna-XM_059141936.1    rna-XM_059386733.1, rna-XM_059387044.1    rna-XM_013061711.2    mNeoVis.2.1.1.chr7_hap1.g02004.t01    rna-XM_021684802.1    rna-XM_004399524.1    rna-XM_042958215.1    rna-XM_040639074.1    rna-XM_027581074.1    mNeoVis.3.1.1.chr7.g01871.t01    rna-gnl|WGS_ZZZZ|egapxtmp_012601-R1    mNeoVis.evergladensis.2.1.chr7.g01968.t01    mNeoVis.vulgivaga.2.1.chr7.g01985.t01      2597    N0.HOG0002595   OG0000488  n23   rna-XM_002917819.4    rna-XM_038528087.1    rna-XM_022523814.1    rna-XM_006940954.5    rna-XM_036068100.1    rna-XM_039248836.1    rna-XM_032838911.1    rna-XM_045988202.1    rna-XM_032325774.1    rna-XM_059151018.1    rna-XM_059383673.1    rna-XM_013063922.2    mNeoVis.2.1.1.chr7_hap1.g00857.t01    rna-XM_021681095.2    rna-XM_004409924.1    rna-XM_042970054.1    rna-XM_008684576.2    rna-XM_027617276.2    mNeoVis.3.1.1.chr7.g00799.t01    rna-gnl|WGS_ZZZZ|egapxtmp_012328-R1    mNeoVis.evergladensis.2.1.chr7.g00849.t01    mNeoVis.vulgivaga.2.1.chr7.g00853.t01      2598    N0.HOG0002596   OG0000488  n44   rna-XM_034643674.1    rna-XM_038558817.1    rna-XM_022495362.1    rna-XM_023257370.2    rna-XM_036107384.1    rna-XM_039252509.1    rna-XM_032844278.1    rna-XM_046011783.1    rna-XM_032346604.1    rna-XM_059186658.1    rna-XM_059404642.1    rna-XM_013051516.2    mNeoVis.2.1.1.chr9_hap1.g01050.t01    rna-XM_021687690.1    rna-XM_004417303.1    rna-XM_042993709.1    rna-XM_008705772.2    rna-XM_027594316.2    mNeoVis.3.1.1.chr10.g01010.t01    rna-gnl|WGS_ZZZZ|egapxtmp_023594-R2    mNeoVis.evergladensis.2.1.chr10.g01040.t01    mNeoVis.vulgivaga.2.1.chr10.g01030.t01      2599    N0.HOG0002597   OG0000489  n1   rna-XM_019802056.2    rna-XM_038550747.1    rna-XM_022518974.1    rna-XM_023257413.2    rna-XM_036093503.1    rna-XM_039230444.1    rna-XM_032836530.1    rna-XM_046011478.1    rna-XM_032346464.1    rna-XM_059186781.1    rna-XM_059402159.1    rna-XM_004743091.3    mNeoVis.2.1.1.chr9_hap1.g01105.t01    rna-XM_021687385.2    rna-XM_004414186.1    rna-XM_042993111.1    rna-XM_008705634.1    rna-XM_027595572.2    mNeoVis.3.1.1.chr10.g01062.t01    rna-gnl|WGS_ZZZZ|egapxtmp_024546-R1    mNeoVis.evergladensis.2.1.chr10.g01095.t01    mNeoVis.vulgivaga.2.1.chr10.g01085.t01      2600    N0.HOG0002598   OG0000489  n21   rna-XM_034657253.1    rna-XM_038565619.1    rna-XM_022498269.1    rna-XM_023250026.2    rna-XM_036073560.1    rna-XM_039231379.1    rna-XM_032841678.1    rna-XM_045991996.1    rna-XM_032327577.1    rna-XM_059161765.1    rna-XM_059390346.1    rna-XM_013052398.2    mNeoVis.2.1.1.chr5_hap1.g00885.t01    rna-XM_044916385.1    rna-XM_004395723.1    rna-XM_042979524.1    rna-XM_040628439.1    rna-XM_027583091.2    mNeoVis.3.1.1.chr2.g00796.t01    rna-gnl|WGS_ZZZZ|egapxtmp_037218-R1    mNeoVis.evergladensis.2.1.chr2.g00885.t01    mNeoVis.vulgivaga.2.1.chr2.g00887.t01      2601    N0.HOG0002599   OG0000489  n39   rna-XM_002917774.4    rna-XM_038587312.1    rna-XM_022508075.1    rna-XM_019823364.3    rna-XM_036064895.1, rna-XM_036096662.1    rna-XM_039216754.1    rna-XM_032843020.1    rna-XM_045995099.1    rna-XM_032330002.1    rna-XM_059156895.1    rna-XM_059385699.1    rna-XM_004754979.3    mNeoVis.2.1.1.chr12_hap1.g00635.t01    rna-XM_021679489.2    rna-XM_004396752.1    rna-XM_042974094.1    rna-XM_040632344.1    rna-XM_027607353.2    mNeoVis.3.1.1.chr12.g00579.t01    rna-gnl|WGS_ZZZZ|egapxtmp_029439-R1    mNeoVis.evergladensis.2.1.chr12.g00573.t01    mNeoVis.vulgivaga.2.1.chr12.g00589.t01      2602    N0.HOG0002600   OG0000490  n1   rna-XM_034670386.1    rna-XM_038540435.1    rna-XM_022516602.1    rna-XM_045047971.1    rna-XM_036092298.1    rna-XM_039218654.1    rna-XM_032843792.1    rna-XM_045993965.1    rna-XM_032326741.1    rna-XM_059154415.1    rna-XM_059415356.1    rna-XM_013055176.2    mNeoVis.2.1.1.chr15_hap1.g00263.t01    rna-XM_021698134.1    rna-XM_004413374.1    rna-XM_042971113.1    rna-XM_040635351.1    rna-XM_027611422.2    mNeoVis.3.1.1.chr15.g00241.t01    rna-gnl|WGS_ZZZZ|egapxtmp_033835-R18    mNeoVis.evergladensis.2.1.chr15.g00241.t01    mNeoVis.vulgivaga.2.1.chr15.g00244.t01      2603    N0.HOG0002601   OG0000490  n22   rna-XM_034640550.1    rna-XM_038546392.1    rna-XM_022523212.1, rna-XM_022523393.1    rna-XM_023244202.2    rna-XM_036084525.1    rna-XM_039226511.1    rna-XM_032870449.1    rna-XM_045986467.1    rna-XM_032320540.1    rna-XM_059147628.1    rna-XM_059379591.1    rna-XM_045085780.1    mNeoVis.2.1.1.chr8_hap1.g00087.t01    rna-XM_021680810.1    rna-XM_004416250.1    rna-XM_042967864.1    rna-XM_040640089.1    rna-XM_027567993.2    mNeoVis.3.1.1.chr8.g00062.t01    rna-gnl|WGS_ZZZZ|egapxtmp_019320-R1    mNeoVis.evergladensis.2.1.chr9.g00084.t01    mNeoVis.vulgivaga.2.1.chr8.g00087.t01      2604    N0.HOG0002602   OG0000490  n41   rna-XM_034644166.1    rna-XM_038550702.1    rna-XM_022518998.1    rna-XM_023257426.2    rna-XM_036100164.1    rna-XM_039230419.1    rna-XM_032836550.1    rna-XM_046012661.1    rna-XM_032346415.1    rna-XM_059186827.1    rna-XM_059402107.1    rna-XM_045083930.1    mNeoVis.2.1.1.chr9_hap1.g01122.t01    rna-XM_044915336.1    rna-XM_004416094.1    rna-XM_042993152.1    rna-XM_040637443.1    rna-XM_027591858.2    mNeoVis.3.1.1.chr10.g01077.t01    rna-gnl|WGS_ZZZZ|egapxtmp_022720-R1    mNeoVis.evergladensis.2.1.chr10.g01112.t01    mNeoVis.vulgivaga.2.1.chr10.g01101.t01      2605    N0.HOG0002603   OG0000491  n1   rna-XM_034643694.1, rna-XM_034653091.1    rna-XM_038551560.1    rna-XM_022519170.1    rna-XM_011284343.4    rna-XM_036072825.1    rna-XM_039237380.1    rna-XM_032882764.1    rna-XM_046010582.1    rna-XM_032345900.1    rna-XM_059187287.1    rna-XM_059401768.1    rna-XM_004775466.3    mNeoVis.2.1.1.chr9_hap1.g01286.t01    rna-XM_044915313.1    rna-XM_004415831.1    rna-XM_042993550.1    rna-XM_040637722.1    rna-XM_035729132.1    mNeoVis.3.1.1.chr10.g01232.t01    rna-gnl|WGS_ZZZZ|egapxtmp_024858-R1    mNeoVis.evergladensis.2.1.chr10.g01271.t01    mNeoVis.vulgivaga.2.1.chr10.g01260.t01      2606    N0.HOG0002604   OG0000491  n24   rna-XM_034654027.1    rna-XM_038541086.1    rna-XM_022496435.1    rna-XM_011285046.4    rna-XM_036109808.1    rna-XM_039251115.1    rna-XM_032881555.1    rna-XM_046015704.1    rna-XM_032361356.1    rna-XM_059135785.1    rna-XM_059375633.1    rna-XM_045081948.1    mNeoVis.2.1.1.chr2_hap1.g00993.t01    rna-XM_021689982.2    rna-XM_004407194.1    rna-XM_042998123.1    rna-XM_040626944.1    rna-XM_027622982.2    mNeoVis.3.1.1.chr5.g00926.t01    rna-gnl|WGS_ZZZZ|egapxtmp_008360-R1    mNeoVis.evergladensis.2.1.chr5.g00946.t01    mNeoVis.vulgivaga.2.1.chr5.g00933.t01      2607    N0.HOG0002605   OG0000491  n45   rna-XM_034658269.1    rna-XM_038566534.1    rna-XM_022497754.1    rna-XM_006928732.5    rna-XM_036079548.1    rna-XM_039228115.1    rna-XM_032867970.1    rna-XM_045990493.1    rna-XM_032318906.1    rna-XM_059160827.1    rna-XM_059389663.1    rna-XM_045080243.1    mNeoVis.2.1.1.chr5_hap1.g00625.t01    rna-XM_021686936.1    rna-XM_004399202.1    rna-XM_042978870.1    rna-XM_040628853.1    rna-XM_027584969.2    mNeoVis.3.1.1.chr2.g00545.t01    rna-gnl|WGS_ZZZZ|egapxtmp_037751-R2    mNeoVis.evergladensis.2.1.chr2.g00622.t01    mNeoVis.vulgivaga.2.1.chr2.g00623.t01      2608    N0.HOG0002606   OG0000492  n1   rna-XM_034643641.1    rna-XM_038550111.1    rna-XM_022518851.1    rna-XM_023257665.2    rna-XM_036113781.1    rna-XM_039234774.1    rna-XM_032882691.1    rna-XM_046010725.1    rna-XM_032345782.1    rna-XM_059187391.1    rna-XM_059401682.1    rna-XM_045087556.1    mNeoVis.2.1.1.chr9_hap1.g01310.t01    rna-XM_044915303.1    rna-XM_004409675.1    rna-XM_042993625.1    rna-XM_040637766.1    rna-XM_027595619.2    mNeoVis.3.1.1.chr10.g01247.t01    rna-gnl|WGS_ZZZZ|egapxtmp_024273-R1    mNeoVis.evergladensis.2.1.chr10.g01297.t01    mNeoVis.vulgivaga.2.1.chr10.g01288.t01      2609    N0.HOG0002607   OG0000492  n24   rna-XM_034672408.1    rna-XM_038527770.1    rna-XM_022491921.1    rna-XM_045046408.1    rna-XM_036096882.1    rna-XM_039244099.1    rna-XM_032848599.1    rna-XM_045987070.1    rna-XM_032323874.1    rna-XM_059151808.1    rna-XM_059383313.1    rna-XM_045076924.1    mNeoVis.2.1.1.chr7_hap1.g00710.t01    rna-XM_044921993.1    rna-XM_004405034.1    rna-XM_042967999.1    rna-XM_040633818.1    rna-XM_027618267.1    mNeoVis.3.1.1.chr7.g00670.t01    rna-gnl|WGS_ZZZZ|egapxtmp_014180-R1    mNeoVis.evergladensis.2.1.chr7.g00708.t01    mNeoVis.vulgivaga.2.1.chr7.g00708.t01      2610    N0.HOG0002608   OG0000492  n44   rna-XM_034644845.1    rna-XM_038563686.1    rna-XM_022507091.1    rna-XM_045039655.1    rna-XM_036105407.1    rna-XM_039235405.1    rna-XM_032879679.1    rna-XM_046014883.1    rna-XM_032357780.1    rna-XM_059149836.1    rna-XM_059399447.1    rna-XM_045066044.1    mNeoVis.2.1.1.chr7_hap1.g02586.t01    rna-XM_044919467.1    rna-XM_004393812.1    rna-XM_042957935.1    rna-XM_040639557.1    rna-XM_027577819.2    mNeoVis.3.1.1.chr7.g02250.t01    rna-gnl|WGS_ZZZZ|egapxtmp_012705-R1    mNeoVis.evergladensis.2.1.chr7.g02392.t01    mNeoVis.vulgivaga.2.1.chr7.g02412.t01      2611    N0.HOG0002609   OG0000493  n1   rna-XM_034637994.1    rna-XM_038528204.1    rna-XM_022524065.1    rna-XM_019819201.3    rna-XM_036068198.1    rna-XM_039250264.1    rna-XM_032839056.1    rna-XM_045989361.1    rna-XM_032325006.1    rna-XM_059150808.1    rna-XM_059381496.1    rna-XM_004767279.3    mNeoVis.2.1.1.chr7_hap1.g00880.t01    rna-XM_021681299.1    rna-XM_004409895.1    rna-XM_042969543.1    rna-XM_040634316.1    rna-XM_027620143.2    mNeoVis.3.1.1.chr7.g00821.t01    rna-gnl|WGS_ZZZZ|egapxtmp_012333-R1    mNeoVis.evergladensis.2.1.chr7.g00874.t01    mNeoVis.vulgivaga.2.1.chr7.g00878.t01      2612    N0.HOG0002610   OG0000493  n22   rna-XM_034643911.1    rna-XM_038550078.1    rna-XM_022518924.1    rna-XM_003989427.6    rna-XM_036093538.1    rna-XM_039236533.1    rna-XM_032882671.1    rna-XM_046012286.1    rna-XM_032345744.1    rna-XM_059187412.1    rna-XM_059401658.1    rna-XM_004775345.3    mNeoVis.2.1.1.chr9_hap1.g01322.t01    rna-XM_021687843.2    rna-XM_004409651.1    rna-XM_007094016.3    rna-XM_040637785.1    rna-XM_027594298.1    mNeoVis.3.1.1.chr10.g01260.t01      mNeoVis.evergladensis.2.1.chr10.g01309.t01    mNeoVis.vulgivaga.2.1.chr10.g01301.t01      2613    N0.HOG0002611   OG0000493  n42   rna-XM_019809980.2    rna-NM_001286860.1, rna-XM_038589301.1, rna-XM_038589391.1, rna-XM_038589435.1    rna-XM_022506406.1    rna-XM_023240171.2    rna-XM_036073424.1    rna-XM_039235421.1    rna-XM_032879696.1    rna-XM_046014730.1    rna-XM_032357830.1    rna-XM_059150018.1    rna-XM_059399414.1    rna-XM_045066141.1    mNeoVis.2.1.1.chr7_hap1.g02589.t01    rna-XM_044919727.1    rna-XM_004393806.1    rna-XM_042957705.1    rna-XM_040639573.1    rna-XM_027578165.1    mNeoVis.3.1.1.chr7.g02253.t01      mNeoVis.evergladensis.2.1.chr7.g02395.t01    mNeoVis.vulgivaga.2.1.chr7.g02416.t01      2614    N0.HOG0002612   OG0000494  n1   rna-XM_002912353.3    rna-XM_038548134.1    rna-XM_022509525.1    rna-XM_023243509.2    rna-XM_036122794.1    rna-XM_039253921.1    rna-XM_032843251.1    rna-XM_045983837.1    rna-XM_032321661.1    rna-XM_059149050.1    rna-XM_059379024.1    rna-XM_013053586.2    mNeoVis.2.1.1.chr8_hap1.g00724.t01    rna-XM_021704196.1    rna-XM_004417196.1    rna-XM_042967847.1    rna-XM_040640998.1    rna-XM_027567842.2    mNeoVis.3.1.1.chr8.g00676.t01    rna-gnl|WGS_ZZZZ|egapxtmp_017479-R1    mNeoVis.evergladensis.2.1.chr9.g00712.t01    mNeoVis.vulgivaga.2.1.chr8.g00716.t01      2615    N0.HOG0002613   OG0000494  n16   rna-XM_034653125.1    rna-XM_038559720.1    rna-XM_022513853.1    rna-XM_011280641.3    rna-XM_036102131.1    rna-XM_039230773.1    rna-XM_032840196.1    rna-XM_046021158.1    rna-XM_032305074.1    rna-XM_059172248.1    rna-XM_059396112.1    rna-XM_004765937.3    mNeoVis.2.1.1.chr4_hap1.g00045.t01    rna-XM_021693043.1    rna-XM_004408809.1    rna-XM_042977435.1    rna-XM_040644583.1    rna-XM_035723315.1    mNeoVis.3.1.1.chr6.g00048.t01    rna-gnl|WGS_ZZZZ|egapxtmp_002534-R1    mNeoVis.evergladensis.2.1.chr6.g00047.t01    mNeoVis.vulgivaga.2.1.chr6.g00051.t01      2616    N0.HOG0002614   OG0000494  n35   rna-XM_011228879.3, rna-XM_034652894.1    rna-XM_038577619.1    rna-XM_022519560.1    rna-XM_019834899.3    rna-XM_036112968.1    rna-XM_039241443.1    rna-XM_032876827.1    rna-XM_046013203.1    rna-XM_032348153.1    rna-XM_059185391.1    rna-XM_059403076.1    rna-XM_045073470.1    mNeoVis.2.1.1.chr9_hap1.g00568.t01    rna-XM_021704848.2    rna-XM_004400839.1    rna-XM_007073000.3    rna-XM_008704561.2    rna-XM_027592607.2    mNeoVis.3.1.1.chr10.g00537.t01    rna-gnl|WGS_ZZZZ|egapxtmp_024087-R6    mNeoVis.evergladensis.2.1.chr10.g00561.t01    mNeoVis.vulgivaga.2.1.chr10.g00549.t01      2617    N0.HOG0002615   OG0000495  n1   rna-XM_002920932.4    rna-XM_038529214.1    rna-XM_022525202.1    rna-XM_019818828.2    rna-XM_036068093.1    rna-XM_039249654.1    rna-XM_032842166.1, rna-XM_032877008.1    rna-XM_045988911.1    rna-XM_032323770.1    rna-XM_059151986.1    rna-XM_059383631.1    rna-XM_004765160.3    mNeoVis.2.1.1.chr3_hap1.g00603.t01, mNeoVis.2.1.1.chr7_hap1.g00465.t01    rna-XM_021700907.1    rna-XM_004396448.1    rna-XM_007096817.3    rna-XM_008712182.2    rna-XM_027617985.1, rna-XM_035723613.1, rna-XM_035725277.1    mNeoVis.3.1.1.chr3.g00595.t01    rna-gnl|WGS_ZZZZ|egapxtmp_039791-R1    mNeoVis.evergladensis.2.1.chr3.g00611.t01, mNeoVis.evergladensis.2.1.chr7.g00473.t01    mNeoVis.vulgivaga.2.1.chr3.g00614.t01, mNeoVis.vulgivaga.2.1.chr7.g00473.t01      2618    N0.HOG0002616   OG0000495  n22     rna-XM_038544741.1, rna-XM_038588934.1          rna-XM_039219018.1, rna-XM_039236331.1, rna-XM_039245796.1    rna-XM_032843079.1    rna-XM_045995674.1, rna-XM_046006811.1    rna-XM_032330351.1    rna-XM_059157208.1    rna-XM_059385589.1    rna-XM_045068374.1    mNeoVis.2.1.1.chr12_hap1.g00605.t01      rna-XM_004406615.1, rna-XM_004412909.1        rna-XM_027580109.2, rna-XM_027603579.1, rna-XM_035722604.1      rna-gnl|WGS_ZZZZ|egapxtmp_020654-R1, rna-gnl|WGS_ZZZZ|egapxtmp_029080-R1    mNeoVis.evergladensis.2.1.chr12.g00542.t01    mNeoVis.vulgivaga.2.1.chr12.g00557.t01      2619    N0.HOG0002617   OG0000495  n40   rna-XM_019806204.2    rna-XM_038554546.1    rna-XM_022503843.1          rna-XM_032873168.1    rna-XM_046004464.1    rna-XM_032320261.1, rna-XM_032339254.1    rna-XM_059177261.1    rna-XM_059400940.1    rna-XM_004752784.3    mNeoVis.2.1.1.chr1_hap1.g01882.t01    rna-XM_021693571.2, rna-XM_044921197.1      rna-XM_007090121.3    rna-XM_008684910.2          mNeoVis.evergladensis.2.1.chr1.g01849.t01    mNeoVis.vulgivaga.2.1.chr1.g01846.t01      2620    N0.HOG0002618   OG0000496  n1   rna-XM_034648785.1    rna-XM_038587780.1    rna-XM_022524309.1    rna-XM_045051271.1    rna-XM_036099591.1    rna-XM_039221266.1    rna-XM_032840635.1    rna-XM_045995124.1    rna-XM_032329762.1    rna-XM_059156492.1    rna-XM_059385220.1    rna-XM_013050048.2    mNeoVis.2.1.1.chr12_hap1.g00425.t01    rna-XM_021689738.1    rna-XM_004407332.1    rna-XM_042975141.1    rna-XM_040630327.1    rna-XM_027608726.2    mNeoVis.3.1.1.chr12.g00384.t01    rna-gnl|WGS_ZZZZ|egapxtmp_028157-R1    mNeoVis.evergladensis.2.1.chr12.g00380.t01    mNeoVis.vulgivaga.2.1.chr12.g00389.t01      2621    N0.HOG0002619   OG0000496  n22   rna-XM_034654200.1    rna-XM_038581506.1    rna-XM_022515908.1    rna-XM_023238732.2    rna-XM_036072653.1    rna-XM_039228911.1    rna-XM_032865315.1    rna-XM_046003479.1    rna-XM_032355130.1    rna-XM_059164738.1    rna-XM_059392357.1    rna-XM_045084493.1    mNeoVis.2.1.1.chr5_hap1.g01839.t01    rna-XM_021679271.2    rna-XM_004411363.1    rna-XM_042956480.1    rna-XM_040621894.1    rna-XM_027587774.1    mNeoVis.3.1.1.chr2.g01741.t01    rna-gnl|WGS_ZZZZ|egapxtmp_035715-R1    mNeoVis.evergladensis.2.1.chr2.g01822.t01    mNeoVis.vulgivaga.2.1.chr2.g01837.t01      2622    N0.HOG0002620   OG0000496  n43   rna-XM_034648088.1    rna-XM_038554572.1    rna-XM_022504076.1    rna-XM_023254153.2    rna-XM_036082210.1    rna-XM_039216422.1    rna-XM_032873199.1    rna-XM_046004559.1    rna-XM_032339209.1    rna-XM_059177235.1    rna-XM_059400964.1    rna-XM_045066232.1    mNeoVis.2.1.1.chr1_hap1.g01894.t01    rna-XM_044917665.1    rna-XM_004399926.1    rna-XM_042986700.1    rna-XM_040643689.1    rna-XM_027603244.2    mNeoVis.3.1.1.chr1.g01847.t01    rna-gnl|WGS_ZZZZ|egapxtmp_005856-R1    mNeoVis.evergladensis.2.1.chr1.g01861.t01    mNeoVis.vulgivaga.2.1.chr1.g01859.t01      2623    N0.HOG0002621   OG0000497  n1   rna-XM_034670952.1    rna-XM_038556011.1    rna-XM_022505961.1    rna-XM_011281892.4    rna-XM_036071963.1    rna-XM_039241103.1    rna-XM_032852416.1    rna-XM_045997484.1    rna-XM_032334220.1    rna-XM_059162170.1    rna-XM_059396136.1    rna-XM_004764920.3    mNeoVis.2.1.1.chr6_hap1.g00196.t01    rna-XM_021701656.1    rna-XM_004396304.1    rna-XM_015534761.2    rna-XM_008701446.2    rna-XM_027599170.2    mNeoVis.3.1.1.chr4.g00192.t01    rna-gnl|WGS_ZZZZ|egapxtmp_000682-R1    mNeoVis.evergladensis.2.1.chr4.g00201.t01    mNeoVis.vulgivaga.2.1.chr4.g00196.t01      2624   N0.HOG0002622  OG0000497  n11   rna-XM_034656770.1    rna-XM_038534843.1    rna-XM_022493475.1    rna-XM_011285651.4    rna-XM_036081030.1    rna-XM_039219362.1    rna-XM_032851728.1    rna-XM_046000987.1    rna-XM_032337114.1    rna-XM_059174221.1    rna-XM_059417298.1    rna-XM_013064899.2    mNeoVis.2.1.1.chr1_hap1.g00272.t01    rna-XM_021697985.1    rna-XM_004392048.1    rna-XM_042959971.1    rna-XM_008692411.2    rna-XM_027606576.1    mNeoVis.3.1.1.chr1.g00275.t01    rna-gnl|WGS_ZZZZ|egapxtmp_039670-R2    mNeoVis.evergladensis.2.1.chr1.g00270.t01    mNeoVis.vulgivaga.2.1.chr1.g00275.t01      2625   N0.HOG0002623  OG0000497  n21   rna-XM_034668908.1    rna-XM_038532578.1    rna-XM_022525990.1    rna-XM_019831992.3    rna-XM_036083608.1    rna-XM_039224345.1    rna-XM_032843092.1    rna-XM_046007017.1    rna-XM_032342296.1    rna-XM_059163796.1, rna-XM_059180239.1    rna-XM_059371734.1    rna-XM_004779156.3    mNeoVis.2.1.1.chr10_hap1.g01121.t01    rna-XM_021680475.1    rna-XM_004414355.1    rna-XM_015545130.2    rna-XM_040642655.1    rna-XM_027572184.2    mNeoVis.3.1.1.chr9.g01051.t01    rna-gnl|WGS_ZZZZ|egapxtmp_017122-R1    mNeoVis.evergladensis.2.1.chr8.g01099.t01    mNeoVis.vulgivaga.2.1.chr9.g01075.t01      2626    N0.HOG0002624   OG0000498  n1   rna-XM_034639753.1    rna-XM_038571376.1    rna-XM_022493778.1    rna-XM_023251210.2    rna-XM_036096080.1    rna-XM_039248516.1    rna-XM_032838563.1    rna-XM_045981775.1    rna-XM_032350567.1    rna-XM_059133566.1    rna-XM_059407075.1    rna-XM_045082167.1    mNeoVis.2.1.1.chr11_hap1.g00205.t01    rna-XM_021689235.1    rna-XM_004414103.1    rna-XM_042980321.1    rna-XM_040638378.1    rna-XM_027623061.1    mNeoVis.3.1.1.chr11.g00174.t01    rna-gnl|WGS_ZZZZ|egapxtmp_026814-R1    mNeoVis.evergladensis.2.1.chr11.g00202.t01    mNeoVis.vulgivaga.2.1.chr11.g00205.t01      2627    N0.HOG0002625   OG0000498  n21   rna-XM_034668526.1    rna-XM_038555603.1    rna-XM_022502551.1    rna-XM_011291471.4    rna-XM_036078730.1, rna-XM_036116422.1    rna-XM_039232954.1    rna-XM_032873480.1    rna-XM_045978348.1    rna-XM_032316440.1    rna-XM_059165550.1    rna-XM_059395211.1    rna-XM_013051782.2    mNeoVis.2.1.1.chr4_hap1.g01038.t01    rna-XM_021688637.1    rna-XM_004405567.1    rna-XM_042973084.1    rna-XM_040625535.1    rna-XM_027574989.2    mNeoVis.3.1.1.chr6.g01059.t01    rna-gnl|WGS_ZZZZ|egapxtmp_004582-R1    mNeoVis.evergladensis.2.1.chr6.g01044.t01    mNeoVis.vulgivaga.2.1.chr6.g01036.t01      2628    N0.HOG0002626   OG0000498  n42   rna-XM_034659777.1    rna-XM_038561008.1    rna-XM_022513653.1    rna-XM_003984414.6    rna-XM_036090189.1    rna-XM_039238315.1    rna-XM_032867008.1    rna-XM_045979704.1    rna-XM_032353086.1    rna-XM_059132429.1    rna-XM_059405293.1    rna-XM_045079288.1    mNeoVis.2.1.1.chr11_hap1.g01086.t01    rna-XM_021697670.1    rna-XM_004394897.1    rna-XM_042982327.1    rna-XM_040623000.1    rna-XM_027621906.2    mNeoVis.3.1.1.chr11.g01026.t01    rna-gnl|WGS_ZZZZ|egapxtmp_025370-R1    mNeoVis.evergladensis.2.1.chr11.g01053.t01    mNeoVis.vulgivaga.2.1.chr11.g01080.t01      2629   N0.HOG0002627  OG0000499  n1   rna-XM_002913726.4    rna-NM_001012617.1    rna-XM_022500235.1    rna-XM_045040068.1    rna-XM_036113803.1    rna-XM_039233254.1    rna-XM_032843100.1    rna-XM_046026758.1    rna-XM_032312019.1    rna-XM_059170063.1    rna-XM_059397670.1    rna-XM_004769538.3    mNeoVis.2.1.1.chr2_hap1.g00521.t01    rna-XM_021700046.1    rna-XM_004393678.1    rna-XM_007095745.2    rna-XM_040624046.1    rna-XM_027596274.2    mNeoVis.3.1.1.chr5.g00515.t01    rna-gnl|WGS_ZZZZ|egapxtmp_006043-R1    mNeoVis.evergladensis.2.1.chr5.g00522.t01    mNeoVis.vulgivaga.2.1.chr5.g00522.t01      2630    N0.HOG0002628   OG0000499  n23   rna-XM_034659427.1    rna-NM_001012619.1    rna-XM_022504588.1    rna-XM_003984160.5    rna-XM_036099428.1    rna-XM_039240100.1    rna-XM_032880697.1    rna-XM_045978592.1    rna-XM_032352583.1    rna-XM_059188169.1    rna-XM_059405735.1    rna-XM_004742277.2    mNeoVis.2.1.1.chr11_hap1.g00842.t01    rna-XM_021699208.1    rna-XM_004398747.1    rna-XM_007083489.3    rna-XM_008700044.1    rna-XM_027624112.2    mNeoVis.3.1.1.chr11.g00804.t01    rna-gnl|WGS_ZZZZ|egapxtmp_027856-R1    mNeoVis.evergladensis.2.1.chr11.g00825.t01    mNeoVis.vulgivaga.2.1.chr11.g00845.t01      2631    N0.HOG0002629   OG0000499  n39   rna-XM_034653509.1, rna-XM_034671726.1    rna-NM_001097541.1    rna-XM_022518154.1    rna-XM_003985120.5    rna-XM_036113512.1    rna-XM_039223755.1    rna-XM_032860664.1    rna-XM_045997972.1    rna-XM_032333048.1    rna-XM_059171769.1    rna-XM_059394873.1    rna-XM_004748139.2    mNeoVis.2.1.1.chr6_hap1.g00539.t01    rna-XM_021680369.1    rna-XM_004401872.1    rna-XM_007080980.3    rna-XM_008689504.2    rna-XM_027599647.1    mNeoVis.3.1.1.chr4.g00544.t01    rna-gnl|WGS_ZZZZ|egapxtmp_043476-R1    mNeoVis.evergladensis.2.1.chr4.g00544.t01    mNeoVis.vulgivaga.2.1.chr4.g00553.t01      2632    N0.HOG0002630   OG0000500  n1   rna-XM_002928032.4    rna-XM_038569046.1    rna-XM_022521983.1    rna-XM_003993000.5    rna-XM_036087929.1    rna-XM_039240915.1    rna-XM_032853479.1    rna-XM_046017675.1    rna-XM_032358168.1    rna-XM_059136258.1    rna-XM_059408147.1    rna-XM_004751990.3    mNeoVis.2.1.1.chr7_hap1.g01821.t01    rna-XM_021685303.2    rna-XM_004394589.1    rna-XM_042957905.1    rna-XM_040638810.1    rna-XM_027580482.2    mNeoVis.3.1.1.chr7.g01688.t01    rna-gnl|WGS_ZZZZ|egapxtmp_013034-R1    mNeoVis.evergladensis.2.1.chr7.g01783.t01    mNeoVis.vulgivaga.2.1.chr7.g01801.t01      2633    N0.HOG0002631   OG0000500  n20   rna-XM_034670022.1    rna-XM_038539024.1    rna-XM_022501704.1    rna-XM_011290337.3    rna-XM_036074021.1, rna-XM_036085715.1    rna-XM_039238102.1    rna-XM_032872378.1    rna-XM_045993859.1    rna-XM_032328265.1    rna-XM_059154780.1    rna-XM_059414740.1    rna-XM_004761697.3    mNeoVis.2.1.1.chr15_hap1.g00764.t01    rna-XM_021700553.2    rna-XM_004406476.1    rna-XM_042971396.1    rna-XM_040634984.1    rna-XM_027613677.2    mNeoVis.3.1.1.chr15.g00543.t01    rna-gnl|WGS_ZZZZ|egapxtmp_034130-R1    mNeoVis.evergladensis.2.1.chr15.g00585.t01    mNeoVis.vulgivaga.2.1.chr15.g00597.t01      2634   N0.HOG0002632  OG0000500  n38   rna-XM_034672864.1    rna-XM_038574675.1    rna-XM_022492170.1    rna-XM_023241248.2    rna-XM_036108753.1    rna-XM_039221303.1    rna-XM_032849203.1    rna-XM_046025832.1    rna-XM_032311543.1    rna-XM_059139977.1    rna-XM_059408144.1    rna-XM_004779569.3    mNeoVis.2.1.1.chr3_hap1.g00681.t01    rna-XM_044921044.1    rna-XM_004416895.1    rna-XM_042962361.1    rna-XM_040619425.1, rna-XM_040643414.1    rna-XM_035724027.1    mNeoVis.3.1.1.chr3.g00671.t01      mNeoVis.evergladensis.2.1.chr3.g00689.t01    mNeoVis.vulgivaga.2.1.chr3.g00693.t01      2635   N0.HOG0002633  OG0000501  n1   rna-XM_034641847.1    rna-NM_001313848.1    rna-XM_022509598.1    rna-XM_045044087.1    rna-XM_036122558.1    rna-XM_039253833.1    rna-XM_032845440.1    rna-XM_045983750.1    rna-XM_032322279.1    rna-XM_059149013.1    rna-XM_059378744.1    rna-XM_045069936.1    mNeoVis.2.1.1.chr8_hap1.g00753.t01    rna-XM_021703959.2    rna-XM_004399635.1    rna-XM_042965681.1    rna-XM_040641031.1    rna-XM_027567746.2    mNeoVis.3.1.1.chr8.g00703.t01    rna-gnl|WGS_ZZZZ|egapxtmp_017490-R1    mNeoVis.evergladensis.2.1.chr9.g00740.t01    mNeoVis.vulgivaga.2.1.chr8.g00744.t01      2636    N0.HOG0002634   OG0000501  n23   rna-XM_019795567.2    rna-NM_001003158.2    rna-XM_022513958.1    rna-XM_003983218.6    rna-XM_036102119.1    rna-XM_039230741.1    rna-XM_032840153.1    rna-XM_046021147.1    rna-XM_032304659.1    rna-XM_059172241.1    rna-XM_059396116.1    rna-XM_004765923.3    mNeoVis.2.1.1.chr4_hap1.g00048.t01    rna-XM_021692988.2    rna-XM_004408806.1    rna-XM_042975385.1    rna-XM_040643807.1    rna-XM_027574572.2    mNeoVis.3.1.1.chr5.g01206.t01, mNeoVis.3.1.1.chr6.g00051.t01    rna-gnl|WGS_ZZZZ|egapxtmp_003020-R1    mNeoVis.evergladensis.2.1.chr6.g00050.t01    mNeoVis.vulgivaga.2.1.chr6.g00054.t01      2637    N0.HOG0002635   OG0000501  n45   rna-XM_034638647.1    rna-NM_001195145.1    rna-XM_022517288.1    rna-XM_011287802.4    rna-XM_036115842.1    rna-XM_039233421.1    rna-XM_032876188.1    rna-XM_046025871.1    rna-XM_032309123.1    rna-XM_059140981.1    rna-XM_059408524.1    rna-XM_013056573.2    mNeoVis.2.1.1.chr3_hap1.g00539.t01    rna-XM_021698514.1    rna-XM_004396691.1    rna-XM_042962140.1    rna-XM_040619415.1    rna-XM_027577102.2    mNeoVis.3.1.1.chr3.g00529.t01    rna-gnl|WGS_ZZZZ|egapxtmp_038876-R1    mNeoVis.evergladensis.2.1.chr3.g00547.t01    mNeoVis.vulgivaga.2.1.chr3.g00550.t01      2638    N0.HOG0002636   OG0000502  n1   rna-XM_034664713.1    rna-XM_038571212.1    rna-XM_022509227.1    rna-XM_019811114.3    rna-XM_036095578.1    rna-XM_039241022.1    rna-XM_032852494.1    rna-XM_046003938.1    rna-XM_032345548.1    rna-XM_059163764.1    rna-XM_059391732.1    rna-XM_004755546.3    mNeoVis.2.1.1.chr5_hap1.g01484.t01    rna-XM_021685971.1    rna-XM_004405011.1    rna-XM_042956451.1    rna-XM_008710983.2    rna-XM_027585608.2    mNeoVis.3.1.1.chr2.g01405.t01    rna-gnl|WGS_ZZZZ|egapxtmp_036358-R1    mNeoVis.evergladensis.2.1.chr2.g01473.t01    mNeoVis.vulgivaga.2.1.chr2.g01484.t01      2639   N0.HOG0002637  OG0000502  n8   rna-XM_002928474.4, rna-XM_019795048.1, rna-XM_034653044.1    rna-XM_038531109.1    rna-XM_022518409.1, rna-XM_022518643.1    rna-XM_003998013.6, rna-XM_023249182.2    rna-XM_036072285.1, rna-XM_036124332.1    rna-XM_039233928.1, rna-XM_039242158.1    rna-XM_032854545.1, rna-XM_032880202.1    rna-XM_045988130.1, rna-XM_045995847.1    rna-XM_032326515.1, rna-XM_032329887.1    rna-XM_059153392.1, rna-XM_059157260.1    rna-XM_059383466.1, rna-XM_059385588.1    rna-XM_004744126.3, rna-XM_004758836.3    mNeoVis.2.1.1.chr12_hap1.g00807.t01, mNeoVis.2.1.1.chr7_hap1.g00348.t01    rna-XM_021684029.1, rna-XM_021705831.1    rna-XM_004409224.1, rna-XM_004414639.1    rna-XM_007092265.3, rna-XM_007096700.3    rna-XM_008696369.2, rna-XM_008702658.2    rna-XM_027609163.2, rna-XM_027618650.2    mNeoVis.3.1.1.chr12.g00736.t01, mNeoVis.3.1.1.chr12.g00737.t01, mNeoVis.3.1.1.chr7.g00335.t01    rna-gnl|WGS_ZZZZ|egapxtmp_013141-R1, rna-gnl|WGS_ZZZZ|egapxtmp_028638-R1    mNeoVis.evergladensis.2.1.chr12.g00733.t01, mNeoVis.evergladensis.2.1.chr7.g00350.t01    mNeoVis.vulgivaga.2.1.chr12.g00763.t01, mNeoVis.vulgivaga.2.1.chr7.g00352.t01      2640    N0.HOG0002638   OG0000503  n1   rna-XM_034640499.1    rna-NM_001110771.1    rna-XM_022509611.1    rna-XM_045044407.1    rna-XM_036111867.1    rna-XM_039253617.1    rna-XM_032848453.1    rna-XM_045986174.1    rna-XM_032319676.1    rna-XM_059147279.1    rna-XM_059378840.1    rna-XM_013053501.2    mNeoVis.2.1.1.chr8_hap1.g00801.t01    rna-XM_044921813.1    rna-XM_004399702.1    rna-XM_007076086.2    rna-XM_040641089.1    rna-XM_027568529.2    mNeoVis.3.1.1.chr8.g00752.t01    rna-gnl|WGS_ZZZZ|egapxtmp_018290-R1    mNeoVis.evergladensis.2.1.chr9.g00789.t01    mNeoVis.vulgivaga.2.1.chr8.g00794.t01      2641    N0.HOG0002639   OG0000503  n21   rna-XM_019799247.2    rna-NM_001136500.1    rna-XM_022512471.1    rna-XM_045036648.1    rna-XM_036113650.1    rna-XM_039238042.1    rna-XM_032861932.1    rna-XM_046000738.1    rna-XM_032337833.1    rna-XM_059173583.1    rna-XM_059416454.1    rna-XM_004768387.3    mNeoVis.2.1.1.chr1_hap1.g00021.t01    rna-XM_044917014.1    rna-XM_004411632.1    rna-XM_042957153.1    rna-XM_040630576.1    rna-XM_027606423.2    mNeoVis.3.1.1.chr1.g00019.t01    rna-gnl|WGS_ZZZZ|egapxtmp_001789-R1    mNeoVis.evergladensis.2.1.chr1.g00019.t01    mNeoVis.vulgivaga.2.1.chr1.g00020.t01      2642    N0.HOG0002640   OG0000503  n39   rna-XM_034638961.1    rna-NM_001136502.1    rna-XM_022518751.1    rna-XM_045045425.1    rna-XM_036088792.1    rna-XM_039242003.1    rna-XM_032854447.1    rna-XM_045988597.1    rna-XM_032324927.1    rna-XM_059153704.1    rna-XM_059381481.1    rna-XM_013052128.2    mNeoVis.2.1.1.chr7_hap1.g00319.t01    rna-XM_021704620.1    rna-XM_004395817.1    rna-XM_042969816.1    rna-XM_040633010.1    rna-XM_027620271.1    mNeoVis.3.1.1.chr7.g00305.t01    rna-gnl|WGS_ZZZZ|egapxtmp_012190-R1    mNeoVis.evergladensis.2.1.chr7.g00319.t01    mNeoVis.vulgivaga.2.1.chr7.g00323.t01      2643    N0.HOG0002641   OG0000504  n1   rna-XM_019805031.2    rna-XM_038537639.1    rna-XM_022514528.1    rna-XM_023257949.2    rna-XM_036089658.1    rna-XM_039248347.1    rna-XM_032863169.1    rna-XM_046010618.1    rna-XM_032361060.1    rna-XM_059135007.1    rna-XM_059376232.1    rna-XM_045063545.1    mNeoVis.2.1.1.chr2_hap1.g01912.t01    rna-XM_021683990.1    rna-XM_004407371.1    rna-XM_042995458.1    rna-XM_040645644.1    rna-XM_027606723.2    mNeoVis.3.1.1.chr5.g01824.t01    rna-gnl|WGS_ZZZZ|egapxtmp_008336-R1    mNeoVis.evergladensis.2.1.chr5.g01850.t01    mNeoVis.vulgivaga.2.1.chr5.g01853.t01      2644    N0.HOG0002642   OG0000504  n24   rna-XM_002926797.4    rna-XM_038538001.1    rna-XM_022511377.1    rna-XM_003998365.5    rna-XM_036119762.1    rna-XM_039226321.1    rna-XM_032853743.1    rna-XM_045987557.1, rna-XM_046010583.1    rna-XM_032326503.1    rna-XM_059152822.1    rna-XM_059382383.1    rna-XM_045076490.1    mNeoVis.2.1.1.chr7_hap1.g00017.t01    rna-XM_021705654.2    rna-XM_004392132.1    rna-XM_042970538.1    rna-XM_040633127.1    rna-XM_027617117.2    mNeoVis.3.1.1.chr3.g01152.t01, mNeoVis.3.1.1.chr7.g00017.t01    rna-gnl|WGS_ZZZZ|egapxtmp_013804-R1    mNeoVis.evergladensis.2.1.chr7.g00018.t01    mNeoVis.vulgivaga.2.1.chr7.g00018.t01      2645    N0.HOG0002643   OG0000504  n46   rna-XM_034671044.1    rna-XM_038537630.1      rna-XM_004001136.6    rna-XM_036089636.1      rna-XM_032863166.1      rna-XM_032361235.1    rna-XM_059135100.1    rna-XM_059376227.1    rna-XM_045063537.1    mNeoVis.2.1.1.chr2_hap1.g01913.t01    rna-XM_021683353.1    rna-XM_004407372.1    rna-XM_042996149.1    rna-XM_040645641.1    rna-XM_027612756.2    mNeoVis.3.1.1.chr5.g01825.t01    rna-gnl|WGS_ZZZZ|egapxtmp_007211-R1    mNeoVis.evergladensis.2.1.chr5.g01851.t01    mNeoVis.vulgivaga.2.1.chr5.g01854.t01      2646    N0.HOG0002644   OG0000504  n64     rna-XM_038588781.1          rna-XM_039248329.1                                      2647    N0.HOG0002645   OG0000505  n1   rna-XM_034667715.1    rna-XM_038535565.1, rna-XM_038535790.1    rna-XM_022513339.1    rna-XM_045038479.1    rna-XM_036123279.1    rna-XM_039250223.1    rna-XM_032868821.1    rna-XM_046016051.1    rna-XM_032357963.1    rna-XM_059183790.1    rna-XM_059392182.1    rna-XM_045080387.1    mNeoVis.2.1.1.chr7_hap1.g01349.t01    rna-XM_021696226.1    rna-XM_004413711.1    rna-XM_042959273.1    rna-XM_040630692.1    rna-XM_027581451.2    mNeoVis.3.1.1.chr7.g01259.t01    rna-gnl|WGS_ZZZZ|egapxtmp_011063-R1    mNeoVis.evergladensis.2.1.chr7.g01332.t01    mNeoVis.vulgivaga.2.1.chr7.g01345.t01      2648    N0.HOG0002646   OG0000505  n24   rna-XM_034656798.1    rna-XM_038581202.1    rna-XM_022504872.1    rna-XM_019839525.2    rna-XM_036096495.1    rna-XM_039237972.1    rna-XM_032878031.1    rna-XM_046002525.1    rna-XM_032352334.1    rna-XM_059164447.1    rna-XM_059392151.1    rna-XM_045088210.1    mNeoVis.2.1.1.chr5_hap1.g01706.t01    rna-XM_044919552.1    rna-XM_004416843.1    rna-XM_042998321.1    rna-XM_040621098.1    rna-XM_035727373.1    mNeoVis.3.1.1.chr2.g01618.t01    rna-gnl|WGS_ZZZZ|egapxtmp_037468-R1    mNeoVis.evergladensis.2.1.chr2.g01688.t01    mNeoVis.vulgivaga.2.1.chr2.g01703.t01      2649    N0.HOG0002647   OG0000505  n45   rna-XM_034656877.1    rna-XM_038581193.1    rna-XM_022504882.1    rna-XM_045036746.1    rna-XM_036121221.1    rna-XM_039237973.1    rna-XM_032878057.1    rna-XM_046003038.1    rna-XM_032352245.1    rna-XM_059164441.1    rna-XM_059392146.1    rna-XM_004770827.3    mNeoVis.2.1.1.chr5_hap1.g01705.t01    rna-XM_044919565.1    rna-XM_004412193.1    rna-XM_042956242.1    rna-XM_008701732.2    rna-XM_027585188.2    mNeoVis.3.1.1.chr2.g01617.t01    rna-gnl|WGS_ZZZZ|egapxtmp_036058-R1    mNeoVis.evergladensis.2.1.chr2.g01687.t01    mNeoVis.vulgivaga.2.1.chr2.g01702.t01      2650    N0.HOG0002648   OG0000506  n1   rna-XM_002926710.3    rna-XM_038586477.1    rna-XM_022507809.1    rna-XM_045051003.1    rna-XM_036082395.1    rna-XM_039223092.1    rna-XM_032842304.1    rna-XM_045996182.1    rna-XM_032330359.1    rna-XM_059157600.1    rna-XM_059385289.1    rna-XM_045089209.1    mNeoVis.2.1.1.chr12_hap1.g00719.t01    rna-XM_021677710.1    rna-XM_004415593.1    rna-XM_042975114.1    rna-XM_008696448.2, rna-XM_040636570.1    rna-XM_035725599.1    mNeoVis.3.1.1.chr12.g00657.t01    rna-gnl|WGS_ZZZZ|egapxtmp_028435-R1    mNeoVis.evergladensis.2.1.chr12.g00651.t01    mNeoVis.vulgivaga.2.1.chr12.g00674.t01      2651    N0.HOG0002649   OG0000506  n24   rna-XM_011226933.3    rna-XM_038545575.1    rna-XM_022505055.1    rna-XM_011283449.3    rna-XM_036097515.1    rna-XM_039232487.1    rna-XM_032878644.1    rna-XM_046007997.1    rna-XM_032344656.1    rna-XM_059182493.1    rna-XM_059373507.1    rna-XM_004739051.3    mNeoVis.2.1.1.chr10_hap1.g00260.t01    rna-XM_021679820.1    rna-XM_004394475.1    rna-XM_042990387.1    rna-XM_040641520.1    rna-XM_027569487.2    mNeoVis.3.1.1.chr9.g00230.t01    rna-gnl|WGS_ZZZZ|egapxtmp_016783-R1    mNeoVis.evergladensis.2.1.chr8.g00245.t01    mNeoVis.vulgivaga.2.1.chr9.g00245.t01      2652    N0.HOG0002650   OG0000506  n41   rna-XM_034659157.1    rna-XM_038551297.1    rna-XM_022504192.1    rna-XM_003984012.6    rna-XM_036069886.1    rna-XM_039218156.1    rna-XM_032877734.1    rna-XM_045979762.1    rna-XM_032352165.1    rna-XM_059187705.1    rna-XM_059406046.1    rna-XM_004780298.3    mNeoVis.2.1.1.chr11_hap1.g00702.t01    rna-XM_044918996.1    rna-XM_004403644.1    rna-XM_007075387.3    rna-XM_040623467.1    rna-XM_027620725.2    mNeoVis.3.1.1.chr11.g00664.t01    rna-gnl|WGS_ZZZZ|egapxtmp_027016-R1    mNeoVis.evergladensis.2.1.chr11.g00686.t01    mNeoVis.vulgivaga.2.1.chr11.g00702.t01      2653    N0.HOG0002651   OG0000507  n1   rna-XM_034661809.1    rna-XM_038571123.1    rna-XM_022504104.1    rna-XM_003992039.6    rna-XM_036088812.1    rna-XM_039222819.1    rna-XM_032852421.1    rna-XM_046003499.1    rna-XM_032346639.1    rna-XM_059163845.1    rna-XM_059391790.1    rna-XM_004754462.3    mNeoVis.2.1.1.chr5_hap1.g01518.t01    rna-XM_021678860.1    rna-XM_004397309.1    rna-XM_007086333.3    rna-XM_008707662.2    rna-XM_027584623.2    mNeoVis.3.1.1.chr2.g01437.t01    rna-gnl|WGS_ZZZZ|egapxtmp_035081-R1    mNeoVis.evergladensis.2.1.chr2.g01504.t01    mNeoVis.vulgivaga.2.1.chr2.g01518.t01      2654    N0.HOG0002652   OG0000507  n21   rna-XM_034649423.1    rna-XM_038587258.1    rna-XM_022507797.1    rna-XM_023249454.2    rna-XM_036090706.1    rna-XM_039216779.1    rna-XM_032842965.1    rna-XM_045995561.1    rna-XM_032329784.1    rna-XM_059158749.1    rna-XM_059384789.1    rna-XM_045068421.1    mNeoVis.2.1.1.chr12_hap1.g00648.t01    rna-XM_044911617.1    rna-XM_004417033.1    rna-XM_042973842.1    rna-XM_040632600.1, rna-XM_040634311.1    rna-XM_027607350.2    mNeoVis.3.1.1.chr12.g00593.t01    rna-gnl|WGS_ZZZZ|egapxtmp_029507-R1    mNeoVis.evergladensis.2.1.chr12.g00585.t01    mNeoVis.vulgivaga.2.1.chr12.g00603.t01      2655    N0.HOG0002653   OG0000507  n40   rna-XM_034648921.1    rna-XM_038588401.1    rna-XM_022491613.1    rna-XM_019824357.2    rna-XM_036098309.1    rna-XM_039217140.1    rna-XM_032841452.1    rna-XM_045994975.1    rna-XM_032330338.1    rna-XM_059157277.1    rna-XM_059384809.1    rna-XM_045068027.1    mNeoVis.2.1.1.chr12_hap1.g00153.t01    rna-XM_021684716.1    rna-XM_004410484.1    rna-XM_042974150.1    rna-XM_040627674.1    rna-XM_027608666.2    mNeoVis.3.1.1.chr12.g00122.t01    rna-gnl|WGS_ZZZZ|egapxtmp_027935-R1    mNeoVis.evergladensis.2.1.chr12.g00129.t01    mNeoVis.vulgivaga.2.1.chr12.g00130.t01      2656    N0.HOG0002654   OG0000508  n0   rna-XM_019801391.2, rna-XM_034649038.1, rna-XM_034649992.1, rna-XM_034651555.1    rna-XM_038536999.1    rna-XM_022491747.1, rna-XM_022507953.1    rna-XM_023243704.2, rna-XM_023249250.2, rna-XM_023249273.2, rna-XM_045051074.1    rna-XM_036074633.1, rna-XM_036090165.1, rna-XM_036094630.1, rna-XM_036112668.1    rna-XM_039224909.1, rna-XM_039224910.1, rna-XM_039227375.1    rna-XM_032841299.1, rna-XM_032842989.1, rna-XM_032849730.1, rna-XM_032849731.1    rna-XM_045984670.1, rna-XM_045995766.1, rna-XM_045996014.1    rna-XM_032321683.1, rna-XM_032329694.1, rna-XM_032331517.1, rna-XM_032331546.1    rna-XM_059148739.1, rna-XM_059156791.1, rna-XM_059157412.1, rna-XM_059158419.1    rna-XM_059379351.1, rna-XM_059384857.1, rna-XM_059384921.1, rna-XM_059385254.1    rna-XM_004779890.2, rna-XM_045068412.1, rna-XM_045068413.1, rna-XM_045068418.1    mNeoVis.2.1.1.chr12_hap1.g00575.t01, mNeoVis.2.1.1.chr12_hap1.g00576.t01, mNeoVis.2.1.1.chr8_hap1.g00954.t01    rna-XM_044911612.1, rna-XM_044911837.1, rna-XM_044911839.1, rna-XM_044921802.1    rna-XM_004417720.1    rna-XM_042967401.1, rna-XM_042974563.1    rna-XM_040630086.1, rna-XM_040631531.1, rna-XM_040631537.1, rna-XM_040631556.1    rna-XM_027607590.2, rna-XM_027625511.2, rna-XM_035725666.1    mNeoVis.3.1.1.chr8.g00887.t01    rna-gnl|WGS_ZZZZ|egapxtmp_019682-R1, rna-gnl|WGS_ZZZZ|egapxtmp_028209-R1, rna-gnl|WGS_ZZZZ|egapxtmp_028932-R1    mNeoVis.evergladensis.2.1.chr12.g00517.t01, mNeoVis.evergladensis.2.1.chr9.g00927.t01    mNeoVis.vulgivaga.2.1.chr12.g00530.t01, mNeoVis.vulgivaga.2.1.chr12.g00532.t01, mNeoVis.vulgivaga.2.1.chr8.g00927.t01      2657    N0.HOG0002655   OG0000509  n1   rna-XM_019798992.2    rna-XM_038573959.1    rna-XM_022499619.1, rna-XM_022500044.1    rna-XM_019838741.3    rna-XM_036094744.1    rna-XM_039236822.1    rna-XM_032857349.1    rna-XM_046018392.1    rna-XM_032354981.1    rna-XM_059167882.1    rna-XM_059393593.1    rna-XM_004762924.3    mNeoVis.2.1.1.chr3_hap1.g01129.t01    rna-XM_044913434.1    rna-XM_004407462.1    rna-XM_042995482.1    rna-XM_040639806.1    rna-XM_035726538.1    mNeoVis.3.1.1.chr3.g01117.t01    rna-gnl|WGS_ZZZZ|egapxtmp_041506-R1    mNeoVis.evergladensis.2.1.chr3.g01128.t01    mNeoVis.vulgivaga.2.1.chr3.g01143.t01      2658    N0.HOG0002656   OG0000509  n24   rna-XM_034649105.1    rna-XM_038588149.1    rna-XM_022518519.1    rna-XM_023249251.2    rna-XM_036094670.1    rna-XM_039244393.1    rna-XM_032844782.1    rna-XM_045995226.1    rna-XM_032331694.1    rna-XM_059158508.1    rna-XM_059385729.1    rna-XM_045068322.1    mNeoVis.2.1.1.chr12_hap1.g00258.t01    rna-XM_044911764.1    rna-XM_004405234.1    rna-XM_042973744.1    rna-XM_040628795.1    rna-XM_035725507.1    mNeoVis.3.1.1.chr12.g00225.t01    rna-gnl|WGS_ZZZZ|egapxtmp_028354-R1    mNeoVis.evergladensis.2.1.chr12.g00226.t01    mNeoVis.vulgivaga.2.1.chr12.g00232.t01      2659    N0.HOG0002657   OG0000509  n45   rna-XM_034665400.1    rna-XM_038569905.1    rna-XM_022502987.1    rna-XM_019828858.3    rna-XM_036120931.1    rna-XM_039252510.1    rna-XM_032864085.1    rna-XM_046028133.1    rna-XM_032314496.1    rna-XM_059144085.1    rna-XM_059377648.1    rna-XM_004746097.3    mNeoVis.2.1.1.chr8_hap1.g01473.t01    rna-XM_044913528.1    rna-XM_004404940.1    rna-XM_042979606.1    rna-XM_040624036.1    rna-XM_027590614.2    mNeoVis.3.1.1.chr8.g01368.t01    rna-gnl|WGS_ZZZZ|egapxtmp_020421-R5    mNeoVis.evergladensis.2.1.chr9.g01421.t01    mNeoVis.vulgivaga.2.1.chr8.g01426.t01      2660    N0.HOG0002658   OG0000510  n1   rna-XM_034645105.1    rna-XM_038576569.1    rna-XM_022522230.1    rna-XM_045061292.1    rna-XM_036117655.1    rna-XM_039218390.1    rna-XM_032874794.1    rna-XM_046010693.1    rna-XM_032349027.1    rna-XM_059184597.1    rna-XM_059403747.1    rna-XM_045071543.1    mNeoVis.2.1.1.chr9_hap1.g00249.t01    rna-XM_021690579.1    rna-XM_004413888.1    rna-XM_042991646.1    rna-XM_008683567.2    rna-XM_027593898.2    mNeoVis.3.1.1.chr10.g00241.t01      mNeoVis.evergladensis.2.1.chr10.g00247.t01    mNeoVis.vulgivaga.2.1.chr10.g00242.t01      2661    N0.HOG0002659   OG0000510  n22     rna-XM_038536804.1    rna-XM_022524626.1    rna-XM_023244701.2      rna-XM_039231593.1    rna-XM_032839866.1    rna-XM_045986496.1    rna-XM_032319988.1    rna-XM_059150446.1    rna-XM_059378796.1    rna-XM_045083349.1    mNeoVis.2.1.1.chr8_hap1.g01020.t01    rna-XM_044921796.1    rna-XM_004398470.1    rna-XM_042965966.1    rna-XM_040629990.1    rna-XM_027624769.2    mNeoVis.3.1.1.chr8.g00951.t01    rna-gnl|WGS_ZZZZ|egapxtmp_018006-R1    mNeoVis.evergladensis.2.1.chr9.g00994.t01    mNeoVis.vulgivaga.2.1.chr8.g00995.t01      2662    N0.HOG0002660   OG0000510  n25   rna-XM_034671172.1    rna-XM_038537837.1    rna-XM_022514792.1    rna-XM_023258054.2    rna-XM_036064583.1, rna-XM_036072385.1, rna-XM_036102029.1    rna-XM_039234936.1    rna-XM_032863307.1    rna-XM_045978586.1    rna-XM_032302027.1    rna-XM_059136833.1    rna-XM_059374987.1    rna-XM_045063682.1    mNeoVis.2.1.1.chr2_hap1.g01856.t01    rna-XM_021683403.1, rna-XM_044915745.1    rna-XM_004411022.1    rna-XM_042994240.1    rna-XM_040645730.1    rna-XM_027571044.2, rna-XM_035724864.1, rna-XM_035727857.1      rna-gnl|WGS_ZZZZ|egapxtmp_009119-R1    mNeoVis.evergladensis.2.1.chr5.g01794.t01    mNeoVis.vulgivaga.2.1.chr5.g01798.t01      2663    N0.HOG0002661   OG0000511  n1   rna-XM_002922217.4    rna-XM_038576666.1    rna-XM_022522426.1    rna-XM_023256740.2    rna-XM_036116279.1    rna-XM_039217509.1    rna-XM_032874694.1    rna-XM_046013447.1    rna-XM_032348905.1    rna-XM_059184695.1    rna-XM_059403648.1    rna-XM_004766623.3    mNeoVis.2.1.1.chr9_hap1.g00286.t01    rna-XM_044914937.1    rna-XM_004407623.1    rna-XM_042991696.1    rna-XM_008683514.2    rna-XM_027595799.2    mNeoVis.3.1.1.chr10.g00276.t01    rna-gnl|WGS_ZZZZ|egapxtmp_021649-R1    mNeoVis.evergladensis.2.1.chr10.g00285.t01    mNeoVis.vulgivaga.2.1.chr10.g00281.t01      2664    N0.HOG0002662   OG0000511  n22   rna-XM_034661817.1    rna-XM_038571050.1    rna-XM_022514927.1    rna-XM_019810639.3    rna-XM_036105942.1    rna-XM_039248556.1    rna-XM_032869258.1    rna-XM_046002347.1    rna-XM_032334718.1    rna-XM_059162790.1    rna-XM_059391040.1    rna-XM_045088847.1    mNeoVis.2.1.1.chr5_hap1.g01183.t01    rna-XM_044912336.1    rna-XM_004397312.1, rna-XM_004417606.1    rna-XM_042998727.1    rna-XM_008711208.1    rna-XM_027584719.2    mNeoVis.3.1.1.chr2.g01091.t01    rna-gnl|WGS_ZZZZ|egapxtmp_035558-R1    mNeoVis.evergladensis.2.1.chr2.g01171.t01    mNeoVis.vulgivaga.2.1.chr2.g01177.t01      2665    N0.HOG0002663   OG0000511  n44   rna-XM_034671179.1    rna-XM_038537920.1    rna-XM_022514399.1    rna-XM_023258083.2    rna-XM_036090513.1    rna-XM_039234948.1    rna-XM_032863349.1    rna-XM_046011895.1    rna-XM_032303020.1    rna-XM_059135470.1    rna-XM_059375106.1    rna-XM_045063701.1    mNeoVis.2.1.1.chr2_hap1.g01838.t01    rna-XM_044914380.1    rna-XM_004397622.1    rna-XM_042996586.1    rna-XM_040645538.1    rna-XM_027599017.2    mNeoVis.3.1.1.chr5.g01764.t01    rna-gnl|WGS_ZZZZ|egapxtmp_007187-R1    mNeoVis.evergladensis.2.1.chr5.g01776.t01    mNeoVis.vulgivaga.2.1.chr5.g01781.t01      2666    N0.HOG0002664   OG0000512  n1   rna-XM_002923226.4    rna-XM_038565189.1    rna-XM_022495811.1    rna-XM_003990756.6    rna-XM_036111208.1, rna-XM_036121234.1    rna-XM_039223566.1    rna-XM_032857749.1    rna-XM_046019868.1    rna-XM_032355514.1    rna-XM_059167379.1    rna-XM_059393950.1    rna-XM_004743824.3    mNeoVis.2.1.1.chr3_hap1.g00984.t01    rna-XM_021702804.1    rna-XM_004401528.1    rna-XM_007078538.3    rna-XM_008686971.2    rna-XM_027590201.2    mNeoVis.3.1.1.chr3.g00974.t01    rna-gnl|WGS_ZZZZ|egapxtmp_042708-R1    mNeoVis.evergladensis.2.1.chr3.g00981.t01    mNeoVis.vulgivaga.2.1.chr3.g00999.t01      2667    N0.HOG0002665   OG0000512  n10   rna-XM_034641052.1    rna-XM_038547210.1    rna-XM_022509713.1    rna-XM_023243773.2    rna-XM_036095542.1    rna-XM_039224273.1    rna-XM_032871739.1    rna-XM_045983923.1    rna-XM_032321953.1    rna-XM_059149343.1    rna-XM_059379311.1    rna-XM_045080744.1    mNeoVis.2.1.1.chr8_hap1.g00350.t01    rna-XM_021681787.2    rna-XM_004410592.1    rna-XM_007087724.2    rna-XM_008701364.2    rna-XM_027568579.2    mNeoVis.3.1.1.chr8.g00317.t01      mNeoVis.evergladensis.2.1.chr9.g00348.t01    mNeoVis.vulgivaga.2.1.chr8.g00355.t01      2668    N0.HOG0002666   OG0000512  n20   rna-XM_011234752.3    rna-XM_038577515.1    rna-XM_022519579.1    rna-XM_003988627.5    rna-XM_036121151.1    rna-XM_039221047.1    rna-XM_032876898.1    rna-XM_046010750.1    rna-XM_032348062.1    rna-XM_059185494.1    rna-XM_059404831.1    rna-XM_004778905.3    mNeoVis.2.1.1.chr9_hap1.g00601.t01    rna-XM_021704890.1    rna-XM_004400876.1    rna-XM_015540308.2    rna-XM_008704527.2    rna-XM_027595252.2, rna-XM_035723757.1    mNeoVis.3.1.1.chr10.g00569.t01    rna-gnl|WGS_ZZZZ|egapxtmp_024009-R1    mNeoVis.evergladensis.2.1.chr10.g00595.t01    mNeoVis.vulgivaga.2.1.chr10.g00581.t01      2669    N0.HOG0002667   OG0000513  n1   rna-XM_034659106.1    rna-XM_038551252.1    rna-XM_022504343.1    rna-XM_019827164.3    rna-XM_036109967.1    rna-XM_039220336.1, rna-XM_039223099.1    rna-XM_032877795.1    rna-XM_045979150.1    rna-XM_032352071.1    rna-XM_059134773.1    rna-XM_059406087.1    rna-XM_004752832.2    mNeoVis.2.1.1.chr11_hap1.g00683.t01    rna-XM_021701263.1    rna-XM_004405650.1    rna-XM_042981355.1    rna-XM_040623491.1    rna-XM_027622286.2    mNeoVis.3.1.1.chr11.g00644.t01    rna-gnl|WGS_ZZZZ|egapxtmp_027460-R1    mNeoVis.evergladensis.2.1.chr11.g00667.t01    mNeoVis.vulgivaga.2.1.chr11.g00680.t01      2670    N0.HOG0002668   OG0000513  n21   rna-XM_034645151.1    rna-XM_038564086.1    rna-XM_022507049.1    rna-XM_023240000.1    rna-XM_036101114.1    rna-XM_039244800.1    rna-XM_032879373.1    rna-XM_046015147.1    rna-XM_032357695.1    rna-XM_059146057.1    rna-XM_059402227.1    rna-XM_045064346.1    mNeoVis.2.1.1.chr7_hap1.g02371.t01    rna-XM_021685004.1    rna-XM_004393984.1    rna-XM_042958364.1    rna-XM_040639309.1    rna-XM_027580338.2    mNeoVis.3.1.1.chr7.g02182.t01      mNeoVis.evergladensis.2.1.chr7.g02317.t01    mNeoVis.vulgivaga.2.1.chr7.g02335.t01      2671    N0.HOG0002669   OG0000513  n41   rna-XM_034643175.1    rna-XM_038545435.1    rna-XM_022505141.1    rna-XM_019833384.3    rna-XM_036084340.1    rna-XM_039221796.1    rna-XM_032878508.1    rna-XM_046009302.1    rna-XM_032341500.1    rna-XM_059182641.1    rna-XM_059373596.1    rna-XM_045074585.1    mNeoVis.2.1.1.chr10_hap1.g00227.t01    rna-XM_021679606.2    rna-XM_004399418.1    rna-XM_042990246.1    rna-XM_040641594.1    rna-XM_035728020.1    mNeoVis.3.1.1.chr9.g00195.t01    rna-gnl|WGS_ZZZZ|egapxtmp_014563-R1    mNeoVis.evergladensis.2.1.chr8.g00211.t01    mNeoVis.vulgivaga.2.1.chr9.g00214.t01      2672    N0.HOG0002670   OG0000514  n1   rna-XM_002926269.4    rna-XM_038551430.1    rna-XM_022504460.1    rna-XM_023251712.2    rna-XM_036107080.1    rna-XM_039243158.1    rna-XM_032873786.1    rna-XM_045979621.1    rna-XM_032352266.1    rna-XM_059187822.1    rna-XM_059405972.1    rna-XM_004742065.3    mNeoVis.2.1.1.chr11_hap1.g00737.t01    rna-XM_044918537.1    rna-XM_004397477.1    rna-XM_042981531.1    rna-XM_040623415.1    rna-XM_027623047.1    mNeoVis.3.1.1.chr11.g00700.t01    rna-gnl|WGS_ZZZZ|egapxtmp_025786-R1    mNeoVis.evergladensis.2.1.chr11.g00719.t01    mNeoVis.vulgivaga.2.1.chr11.g00739.t01      2673    N0.HOG0002671   OG0000514  n6   rna-XM_034648627.1    rna-XM_038544799.1    rna-XM_022492592.1    rna-XM_023255714.2    rna-XM_036108995.1    rna-XM_039255137.1    rna-XM_032875318.1    rna-XM_046009516.1    rna-XM_032344301.1    rna-XM_059182173.1    rna-XM_059373271.1    rna-XM_045069790.1    mNeoVis.2.1.1.chr10_hap1.g00374.t01    rna-XM_021701376.2    rna-XM_004404898.1    rna-XM_042989775.1    rna-XM_040641869.1    rna-XM_027571670.2    mNeoVis.3.1.1.chr9.g00345.t01    rna-gnl|WGS_ZZZZ|egapxtmp_014591-R1    mNeoVis.evergladensis.2.1.chr8.g00356.t01    mNeoVis.vulgivaga.2.1.chr9.g00357.t01      2674   N0.HOG0002672  OG0000514  n22   rna-XM_002927799.4, rna-XM_034644842.1    rna-XM_038563830.1    rna-XM_022507068.1    rna-XM_023240084.2    rna-XM_036121516.1    rna-XM_039244671.1    rna-XM_032879556.1    rna-XM_046015359.1    rna-XM_032357524.1    rna-XM_059148346.1    rna-XM_059400835.1    rna-XM_004759428.3    mNeoVis.2.1.1.chr7_hap1.g02479.t01    rna-XM_021685087.2    rna-XM_004393881.1    rna-XM_042959369.1    rna-XM_040639458.1    rna-XM_027579451.2    mNeoVis.3.1.1.chr7.g02226.t01    rna-gnl|WGS_ZZZZ|egapxtmp_009981-R1    mNeoVis.evergladensis.2.1.chr7.g02367.t01    mNeoVis.vulgivaga.2.1.chr7.g02387.t01      2675   N0.HOG0002673  OG0000515  n1   rna-XM_034662635.1    rna-XM_038534220.1    rna-XM_022500254.1    rna-XM_045040106.1    rna-XM_036083945.1    rna-XM_039233303.1    rna-XM_032841150.1    rna-XM_046026583.1    rna-XM_032312193.1    rna-XM_059170022.1    rna-XM_059397652.1    rna-XM_004769511.3    mNeoVis.2.1.1.chr2_hap1.g00504.t01    rna-XM_044916488.1    rna-XM_004393661.1    rna-XM_042960498.1    rna-XM_040624771.1    rna-XM_027596864.2    mNeoVis.3.1.1.chr5.g00498.t01    rna-gnl|WGS_ZZZZ|egapxtmp_006871-R1    mNeoVis.evergladensis.2.1.chr5.g00505.t01    mNeoVis.vulgivaga.2.1.chr5.g00505.t01      2676    N0.HOG0002674   OG0000515  n23   rna-XM_034655721.1    rna-XM_038551648.1    rna-XM_022521186.1    rna-XM_023254116.2    rna-XM_036080139.1    rna-XM_039252863.1    rna-XM_032846790.1    rna-XM_046000432.1    rna-XM_032335321.1    rna-XM_059175886.1    rna-XM_059416151.1    rna-XM_013059699.2    mNeoVis.2.1.1.chr1_hap1.g00883.t01    rna-XM_044916657.1    rna-XM_004412678.1    rna-XM_042999277.1    rna-XM_040622615.1    rna-XM_027604731.2    mNeoVis.3.1.1.chr1.g00904.t01    rna-gnl|WGS_ZZZZ|egapxtmp_038532-R1    mNeoVis.evergladensis.2.1.chr1.g00887.t01    mNeoVis.vulgivaga.2.1.chr1.g00898.t01      2677    N0.HOG0002675   OG0000515  n45   rna-XM_034671705.1    rna-XM_038556297.1    rna-XM_022506007.1    rna-XM_011281725.4    rna-XM_036078046.1    rna-XM_039230951.1    rna-XM_032842910.1    rna-XM_045992326.1    rna-XM_032334543.1    rna-XM_059159482.1    rna-XM_059385162.1    rna-XM_045079854.1    mNeoVis.2.1.1.chr6_hap1.g00091.t01    rna-XM_021702380.1    rna-XM_004392908.1    rna-XM_042984395.1    rna-XM_008693199.2    rna-XM_027600232.2    mNeoVis.3.1.1.chr4.g00088.t01    rna-gnl|WGS_ZZZZ|egapxtmp_044021-R1    mNeoVis.evergladensis.2.1.chr4.g00090.t01    mNeoVis.vulgivaga.2.1.chr4.g00089.t01      2678    N0.HOG0002676   OG0000516  n1   rna-XM_034663127.1    rna-NM_001003081.1    rna-XM_022499481.1    rna-XM_045040545.1    rna-XM_036116950.1    rna-XM_039249245.1    rna-XM_032882459.1    rna-XM_046026178.1    rna-XM_032313456.1    rna-XM_059169222.1    rna-XM_059398333.1    rna-XM_004749506.2    mNeoVis.2.1.1.chr2_hap1.g00252.t01    rna-XM_021699979.2    rna-XM_004401905.1    rna-XM_015536686.2    rna-XM_040623765.1    rna-XM_027587451.2    mNeoVis.3.1.1.chr5.g00245.t01    rna-gnl|WGS_ZZZZ|egapxtmp_006812-R1    mNeoVis.evergladensis.2.1.chr5.g00256.t01    mNeoVis.vulgivaga.2.1.chr5.g00254.t01      2679   N0.HOG0002677  OG0000516  n23   rna-XM_034639977.1    rna-XM_038547687.1    rna-XM_022509600.1    rna-XM_045044363.1    rna-XM_036076302.1    rna-XM_039227876.1    rna-XM_032872060.1    rna-XM_045984783.1    rna-XM_032319888.1    rna-XM_059150386.1    rna-XM_059378542.1    rna-XM_045070331.1    mNeoVis.2.1.1.chr8_hap1.g00585.t01    rna-XM_021704040.1    rna-XM_004395231.1    rna-XM_042965451.1    rna-XM_040640835.1    rna-XM_027625248.2    mNeoVis.3.1.1.chr6.g00582.t01, mNeoVis.3.1.1.chr8.g00540.t01    rna-gnl|WGS_ZZZZ|egapxtmp_017442-R1    mNeoVis.evergladensis.2.1.chr9.g00575.t01    mNeoVis.vulgivaga.2.1.chr8.g00583.t01      2680    N0.HOG0002678   OG0000516  n45   rna-XM_019807751.2    rna-NM_001002971.1    rna-XM_022516989.1    rna-XM_023246871.2    rna-XM_036115484.1    rna-XM_039233599.1    rna-XM_032836225.1    rna-XM_045993322.1    rna-XM_032326777.1    rna-XM_059153924.1    rna-XM_059415249.1    rna-XM_045072205.1    mNeoVis.2.1.1.chr15_hap1.g00211.t01    rna-XM_044915634.1    rna-XM_004408941.1    rna-XM_042971135.1    rna-XM_040635271.1    rna-XM_035722135.1    mNeoVis.3.1.1.chr15.g00185.t01    rna-gnl|WGS_ZZZZ|egapxtmp_033956-R1    mNeoVis.evergladensis.2.1.chr15.g00190.t01    mNeoVis.vulgivaga.2.1.chr15.g00191.t01      2681    N0.HOG0002679   OG0000517  n1   rna-XM_019804047.2    rna-XM_038539160.1    rna-XM_022501992.1    rna-XM_023246643.2    rna-XM_036072414.1    rna-XM_039243518.1    rna-XM_032872469.1    rna-XM_045994206.1    rna-XM_032326897.1    rna-XM_059155183.1    rna-XM_059415931.1    rna-XM_045074983.1    mNeoVis.2.1.1.chr15_hap1.g00716.t01    rna-XM_021700590.1    rna-XM_004399077.1    rna-XM_042970816.1    rna-XM_040634905.1    rna-XM_027612092.1    mNeoVis.3.1.1.chr15.g00494.t01      mNeoVis.evergladensis.2.1.chr15.g00538.t01    mNeoVis.vulgivaga.2.1.chr15.g00552.t01      2682    N0.HOG0002680   OG0000517  n22         rna-XM_045039986.1                          rna-XM_042960069.1                  2683    N0.HOG0002681   OG0000517  n24   rna-XM_034663632.1    rna-XM_038579001.1      rna-XM_045040780.1    rna-XM_036070200.1    rna-XM_039237952.1, rna-XM_039237953.1, rna-XM_039250443.1    rna-XM_032882148.1    rna-XM_046026304.1    rna-XM_032313979.1    rna-XM_059172937.1      rna-XM_045068963.1    mNeoVis.2.1.1.chr2_hap1.g00081.t01    rna-XM_021703830.2    rna-XM_004417815.1    rna-XM_042961225.1    rna-XM_040624307.1    rna-XM_035724237.1    mNeoVis.3.1.1.chr5.g00075.t01    rna-gnl|WGS_ZZZZ|egapxtmp_005916-R1    mNeoVis.evergladensis.2.1.chr5.g00082.t01    mNeoVis.vulgivaga.2.1.chr5.g00078.t01      2684    N0.HOG0002682   OG0000517  n45   rna-XM_011218244.3, rna-XM_034663633.1    rna-XM_038577983.1    rna-XM_022499653.1    rna-XM_045039987.1    rna-XM_036070226.1, rna-XM_036070267.1    rna-XM_039237951.1    rna-XM_032882195.1    rna-XM_046026610.1    rna-XM_032313978.1    rna-XM_059172439.1    rna-XM_059397731.1    rna-XM_045068873.1    mNeoVis.2.1.1.chr2_hap1.g00079.t01    rna-XM_044916227.1        rna-XM_040624305.1    rna-XM_035724137.1    mNeoVis.3.1.1.chr5.g00073.t01    rna-gnl|WGS_ZZZZ|egapxtmp_005915-R1    mNeoVis.evergladensis.2.1.chr5.g00080.t01    mNeoVis.vulgivaga.2.1.chr5.g00076.t01      2685    N0.HOG0002683   OG0000518  n1   rna-XM_034671063.1    rna-XM_038537602.1    rna-XM_022514356.1    rna-XM_011284401.3    rna-XM_036089672.1    rna-XM_039248333.1    rna-XM_032863128.1    rna-XM_046002104.1    rna-XM_032361700.1    rna-XM_059136245.1    rna-XM_059376510.1    rna-XM_045063505.1    mNeoVis.2.1.1.chr2_hap1.g01927.t01    rna-XM_021683982.2    rna-XM_004407395.1    rna-XM_042996380.1    rna-XM_040645609.1    rna-XM_027612140.2    mNeoVis.3.1.1.chr5.g01832.t01    rna-gnl|WGS_ZZZZ|egapxtmp_008736-R1    mNeoVis.evergladensis.2.1.chr5.g01866.t01    mNeoVis.vulgivaga.2.1.chr5.g01868.t01      2686    N0.HOG0002684   OG0000518  n20   rna-XM_019795793.2    rna-XM_038583521.1    rna-XM_022505691.1, rna-XM_022506296.1    rna-XM_023260347.2    rna-XM_036122475.1    rna-XM_039241695.1    rna-XM_032875489.1    rna-XM_046002637.1    rna-XM_032339160.1    rna-XM_059163317.1    rna-XM_059391428.1    rna-XM_004745114.2    mNeoVis.2.1.1.chr5_hap1.g01337.t01    rna-XM_021692483.2    rna-XM_004406784.1    rna-XM_042956252.1    rna-XM_040621516.1    rna-XM_027586204.2    mNeoVis.3.1.1.chr2.g01255.t01    rna-gnl|WGS_ZZZZ|egapxtmp_034538-R1    mNeoVis.evergladensis.2.1.chr2.g01323.t01    mNeoVis.vulgivaga.2.1.chr2.g01331.t01      2687    N0.HOG0002685   OG0000518  n38   rna-XM_034647180.1    rna-XM_038536628.1    rna-XM_022524606.1    rna-XM_003996159.5    rna-XM_036066267.1    rna-XM_039231605.1    rna-XM_032839993.1    rna-XM_045984154.1    rna-XM_032322439.1    rna-XM_059147659.1    rna-XM_059378713.1    rna-XM_004760353.3    mNeoVis.2.1.1.chr8_hap1.g01078.t01    rna-XM_044921939.1    rna-XM_004398540.1    rna-XM_042965840.1    rna-XM_040629926.1, rna-XM_040629928.1    rna-XM_027625325.1    mNeoVis.3.1.1.chr8.g01000.t01      mNeoVis.evergladensis.2.1.chr9.g01053.t01    mNeoVis.vulgivaga.2.1.chr8.g01053.t01      2688    N0.HOG0002686   OG0000519  n1   rna-XM_034672184.1    rna-XM_038533182.1    rna-XM_022506298.1    rna-XM_045056939.1    rna-XM_036085700.1    rna-XM_039232122.1    rna-XM_032855844.1    rna-XM_045985460.1    rna-XM_032333694.1    rna-XM_059166277.1    rna-XM_059380026.1    rna-XM_004772234.3    mNeoVis.2.1.1.chr6_hap1.g00353.t01    rna-XM_021677624.1    rna-XM_004396195.1    rna-XM_042982761.1    rna-XM_040625383.1    rna-XM_027600763.2    mNeoVis.3.1.1.chr4.g00356.t01    rna-gnl|WGS_ZZZZ|egapxtmp_044578-R1    mNeoVis.evergladensis.2.1.chr4.g00360.t01    mNeoVis.vulgivaga.2.1.chr4.g00355.t01      2689    N0.HOG0002687   OG0000519  n23   rna-XM_034656650.1    rna-XM_038534641.1    rna-XM_022493524.1    rna-XM_045054223.1    rna-XM_036084852.1    rna-XM_039239812.1    rna-XM_032851930.1    rna-XM_046000047.1    rna-XM_032336883.1    rna-XM_059174436.1    rna-XM_059417466.1    rna-XM_004737470.3    mNeoVis.2.1.1.chr1_hap1.g00356.t01    rna-XM_021698807.1    rna-XM_004401410.1    rna-XM_042991596.1    rna-XM_040630237.1, rna-XM_040630607.1    rna-XM_027605899.1    mNeoVis.3.1.1.chr1.g00358.t01    rna-gnl|WGS_ZZZZ|egapxtmp_028997-R1    mNeoVis.evergladensis.2.1.chr1.g00354.t01    mNeoVis.vulgivaga.2.1.chr1.g00362.t01      2690    N0.HOG0002688   OG0000519  n45   rna-XM_034672186.1    rna-NM_001031815.2    rna-XM_022506195.1    rna-XM_011281954.4    rna-XM_036085699.1    rna-XM_039232108.1    rna-XM_032855866.1    rna-XM_045987005.1    rna-XM_032333687.1    rna-XM_059166298.1    rna-XM_059380005.1    rna-XM_045075136.1      rna-XM_021677677.1, rna-XM_021692940.1    rna-XM_004396196.1    rna-XM_042984804.1    rna-XM_040624930.1    rna-XM_035726114.1    mNeoVis.3.1.1.chr4.g00357.t01    rna-gnl|WGS_ZZZZ|egapxtmp_000732-R1    mNeoVis.evergladensis.2.1.chr4.g00361.t01    mNeoVis.vulgivaga.2.1.chr4.g00356.t01      2691    N0.HOG0002689   OG0000520  n1   rna-XM_002912326.4, rna-XM_034650752.1    rna-NM_001286945.1    rna-XM_022509462.1    rna-XM_003996488.6    rna-XM_036122049.1    rna-XM_039252473.1    rna-XM_032845386.1    rna-XM_045986317.1    rna-XM_032320303.1    rna-XM_059148798.1    rna-XM_059379083.1    rna-XM_004746996.3    mNeoVis.2.1.1.chr8_hap1.g00769.t01    rna-XM_044921849.1    rna-XM_004399654.1    rna-XM_007076050.3    rna-XM_008695346.2    rna-XM_027625983.2    mNeoVis.3.1.1.chr8.g00719.t01    rna-gnl|WGS_ZZZZ|egapxtmp_019915-R2    mNeoVis.evergladensis.2.1.chr9.g00756.t01    mNeoVis.vulgivaga.2.1.chr8.g00760.t01      2692    N0.HOG0002690   OG0000520  n18   rna-XM_011233874.3    rna-XM_038552971.1    rna-XM_022500884.1    rna-XM_006939334.4    rna-XM_036065901.1    rna-XM_039225558.1    rna-XM_032860732.1    rna-XM_046023864.1    rna-XM_032306440.1    rna-XM_059142732.1    rna-XM_059411180.1    rna-XM_004747710.3    mNeoVis.2.1.1.chr13_hap1.g00414.t01    rna-XM_021691434.1      rna-XM_007089262.3    rna-XM_008707894.1    rna-XM_027616608.1    mNeoVis.3.1.1.chr13.g00354.t01    rna-gnl|WGS_ZZZZ|egapxtmp_031163-R1    mNeoVis.evergladensis.2.1.chr13.g00397.t01    mNeoVis.vulgivaga.2.1.chr13.g00401.t01      2693    N0.HOG0002691   OG0000520  n37   rna-XM_034670157.1    rna-XM_038539886.1    rna-XM_022526386.1    rna-XM_003998684.5    rna-XM_036091199.1    rna-XM_039247824.1    rna-XM_032842517.1    rna-XM_045993868.1    rna-XM_032327771.1    rna-XM_059155691.1    rna-XM_059414977.1    rna-XM_004773940.2    mNeoVis.2.1.1.chr15_hap1.g00074.t01    rna-XM_044915675.1    rna-XM_004397178.1    rna-XM_042971190.1    rna-XM_008685248.2    rna-XM_027611463.1    mNeoVis.3.1.1.chr15.g00068.t01      mNeoVis.evergladensis.2.1.chr15.g00075.t01    mNeoVis.vulgivaga.2.1.chr15.g00076.t01      2694    N0.HOG0002692   OG0000520  n56   rna-XM_034652872.1    rna-XM_038557542.1                                              2695    N0.HOG0002693   OG0000521  n1   rna-XM_019797269.2    rna-XM_038545075.1    rna-XM_022505377.1    rna-XM_003987779.4    rna-XM_036107249.1    rna-XM_039240704.1    rna-XM_032849387.1    rna-XM_046008108.1    rna-XM_032344609.1    rna-XM_059182444.1    rna-XM_059373481.1    rna-XM_013057507.2    mNeoVis.2.1.1.chr10_hap1.g00276.t01    rna-XM_044917917.1    rna-XM_004393622.1, rna-XM_004397607.1    rna-XM_007095145.3    rna-XM_040641722.1    rna-XM_027571426.1    mNeoVis.3.1.1.chr9.g00246.t01    rna-gnl|WGS_ZZZZ|egapxtmp_016747-R1    mNeoVis.evergladensis.2.1.chr8.g00261.t01    mNeoVis.vulgivaga.2.1.chr9.g00260.t01      2696    N0.HOG0002694   OG0000521  n8   rna-XM_011222580.3    rna-XM_038572342.1    rna-XM_022494400.1    rna-XM_023251231.2    rna-XM_036096079.1    rna-XM_039248458.1    rna-XM_032838491.1    rna-XM_045981360.1    rna-XM_032350625.1    rna-XM_059133119.1    rna-XM_059407030.1    rna-XM_004746416.3    mNeoVis.2.1.1.chr11_hap1.g00223.t01    rna-XM_021689279.1    rna-XM_004417144.1    rna-XM_042980381.1    rna-XM_040638324.1    rna-XM_027622599.2    mNeoVis.3.1.1.chr11.g00193.t01    rna-gnl|WGS_ZZZZ|egapxtmp_026820-R1    mNeoVis.evergladensis.2.1.chr11.g00221.t01    mNeoVis.vulgivaga.2.1.chr11.g00223.t01      2697    N0.HOG0002695   OG0000521  n15   rna-XM_002929558.4    rna-XM_038531347.1    rna-XM_022496166.1    rna-XM_003989779.5    rna-XM_036115443.1    rna-XM_039231166.1    rna-XM_032863977.1    rna-XM_046002819.1    rna-XM_032360865.1    rna-XM_059138307.1    rna-XM_059376931.1    rna-XM_004740978.3    mNeoVis.2.1.1.chr2_hap1.g01585.t01    rna-XM_044914447.1    rna-XM_004394636.1    rna-XM_042996043.1    rna-XM_040619139.1    rna-XM_027570844.2    mNeoVis.3.1.1.chr5.g01507.t01    rna-gnl|WGS_ZZZZ|egapxtmp_005383-R1    mNeoVis.evergladensis.2.1.chr5.g01524.t01    mNeoVis.vulgivaga.2.1.chr5.g01526.t01      2698    N0.HOG0002696   OG0000522  n1   rna-XM_019795857.2    rna-XM_038537451.1    rna-XM_022497428.1    rna-XM_019837059.3    rna-XM_036121445.1    rna-XM_039238461.1    rna-XM_032844112.1    rna-XM_046021880.1    rna-XM_032303680.1    rna-XM_059137424.1    rna-XM_059374833.1    rna-XM_045064567.1    mNeoVis.2.1.1.chr2_hap1.g01261.t01    rna-XM_021684332.2    rna-XM_004400357.1    rna-XM_042997282.1    rna-XM_040627266.1    rna-XM_035727081.1    mNeoVis.3.1.1.chr5.g01183.t01    rna-gnl|WGS_ZZZZ|egapxtmp_009092-R1    mNeoVis.evergladensis.2.1.chr5.g01205.t01    mNeoVis.vulgivaga.2.1.chr5.g01198.t01      2699    N0.HOG0002697   OG0000522  n13   rna-XM_034657198.1    rna-XM_038566810.1    rna-XM_022497677.1    rna-XM_045048524.1    rna-XM_036093917.1    rna-XM_039228098.1    rna-XM_032867693.1    rna-XM_045992245.1    rna-XM_032316185.1    rna-XM_059160543.1    rna-XM_059389442.1    rna-XM_045081504.1    mNeoVis.2.1.1.chr5_hap1.g00519.t01    rna-XM_021683256.2    rna-XM_004404639.1    rna-XM_042978602.1    rna-XM_040629013.1    rna-XM_027586007.2      rna-gnl|WGS_ZZZZ|egapxtmp_037011-R1    mNeoVis.evergladensis.2.1.chr2.g00513.t01    mNeoVis.vulgivaga.2.1.chr2.g00515.t01      2700    N0.HOG0002698   OG0000522  n32   rna-XM_011220987.3    rna-XM_038532429.1    rna-XM_022504775.1    rna-XM_045033195.1    rna-XM_036067208.1, rna-XM_036113341.1    rna-XM_039220108.1    rna-XM_032847362.1    rna-XM_046000541.1    rna-XM_032334847.1    rna-XM_059175380.1    rna-XM_059418171.1    rna-XM_004759144.3, rna-XM_004782540.3    mNeoVis.2.1.1.chr1_hap1.g00713.t01    rna-XM_044916974.1    rna-XM_004405611.1    rna-XM_007079789.3    rna-XM_008711041.2    rna-XM_027604661.2      rna-gnl|WGS_ZZZZ|egapxtmp_041643-R1    mNeoVis.evergladensis.2.1.chr1.g00722.t01    mNeoVis.vulgivaga.2.1.chr1.g00733.t01      2701    N0.HOG0002699   OG0000523  n1   rna-XM_002923361.4    rna-XM_038535322.1    rna-XM_022493566.1    rna-XM_019811923.3    rna-XM_036107412.1    rna-XM_039250523.1, rna-XM_039250530.1    rna-XM_032851321.1    rna-XM_045998683.1    rna-XM_032337602.1    rna-XM_059173795.1    rna-XM_059416998.1    rna-XM_004738018.3    mNeoVis.2.1.1.chr1_hap1.g00107.t01    rna-XM_044916961.1    rna-XM_004403946.1    rna-XM_042960499.1, rna-XM_042996530.1    rna-XM_008692641.2, rna-XM_040630207.1    rna-XM_027605177.2    mNeoVis.3.1.1.chr1.g00110.t01    rna-gnl|WGS_ZZZZ|egapxtmp_027588-R1    mNeoVis.evergladensis.2.1.chr1.g00105.t01    mNeoVis.vulgivaga.2.1.chr1.g00103.t01      2702    N0.HOG0002700   OG0000523  n25   rna-XM_034657016.1    rna-XM_038533756.1, rna-XM_038535324.1    rna-XM_022493556.1, rna-XM_022493619.1    rna-XM_019811937.3    rna-XM_036107414.1    rna-XM_039248263.1    rna-XM_032851323.1, rna-XM_032851325.1    rna-XM_046001075.1    rna-XM_032334971.1        rna-XM_013045377.2    mNeoVis.2.1.1.chr1_hap1.g00106.t01    rna-XM_044916680.1    rna-XM_004403948.1    rna-XM_042996558.1    rna-XM_008692644.2, rna-XM_040630208.1    rna-XM_035727720.1    mNeoVis.3.1.1.chr1.g00108.t01, mNeoVis.3.1.1.chr1.g00109.t01    rna-gnl|WGS_ZZZZ|egapxtmp_024516-R1, rna-gnl|WGS_ZZZZ|egapxtmp_024595-R1    mNeoVis.evergladensis.2.1.chr1.g00103.t01        2703    N0.HOG0002701   OG0000523  n48   rna-XM_034657015.1    rna-XM_038535318.1      rna-XM_011286527.4, rna-XM_045038118.1    rna-XM_036107413.1    rna-XM_039248268.1, rna-XM_039248272.1      rna-XM_046001077.1    rna-XM_032337606.1    rna-XM_059173337.1    rna-XM_059416490.1    rna-XM_004738121.2        rna-XM_004403947.1    rna-XM_015544392.2, rna-XM_042996546.1    rna-XM_008692643.1    rna-XM_035727821.1              2704   N0.HOG0002702  OG0000524  n1   rna-XM_034644711.1    rna-XM_038563392.1    rna-XM_022506748.1    rna-XM_019812392.2    rna-XM_036105102.1    rna-XM_039226754.1    rna-XM_032844539.1    rna-XM_046014710.1    rna-XM_032360163.1    rna-XM_059141900.1    rna-XM_059386731.1, rna-XM_059387042.1    rna-XM_045065942.1    mNeoVis.2.1.1.chr7_hap1.g01999.t01    rna-XM_021684795.2    rna-XM_004399539.1    rna-XM_042959156.1    rna-XM_040639048.1    rna-XM_027578272.2    mNeoVis.3.1.1.chr7.g01866.t01    rna-gnl|WGS_ZZZZ|egapxtmp_009959-R1    mNeoVis.evergladensis.2.1.chr7.g01963.t01    mNeoVis.vulgivaga.2.1.chr7.g01980.t01      2705   N0.HOG0002703  OG0000524  n21   rna-XM_034660569.1    rna-XM_038553792.1    rna-XM_022510911.1    rna-XM_023253869.2    rna-XM_036072167.1    rna-XM_039241777.1    rna-XM_032864266.1    rna-XM_046006449.1    rna-XM_032340586.1    rna-XM_059178588.1    rna-XM_059400062.1    rna-XM_004770656.2    mNeoVis.2.1.1.chr1_hap1.g01450.t01    rna-XM_021684588.1    rna-XM_004407727.1    rna-XM_042986153.1    rna-XM_040644738.1    rna-XM_027603175.2    mNeoVis.3.1.1.chr1.g01397.t01      mNeoVis.evergladensis.2.1.chr1.g01412.t01    mNeoVis.vulgivaga.2.1.chr1.g01416.t01      2706    N0.HOG0002704   OG0000524  n36   rna-XM_034644176.1    rna-XM_038550305.1    rna-XM_022519093.1    rna-XM_019835675.3    rna-XM_036088352.1    rna-XM_039235052.1    rna-XM_032882835.1    rna-XM_046010531.1    rna-XM_032346033.1    rna-XM_059187175.1    rna-XM_059401852.1    rna-XM_045083749.1    mNeoVis.2.1.1.chr9_hap1.g01244.t01    rna-XM_044915314.1    rna-XM_004397389.1    rna-XM_007079520.3    rna-XM_040637660.1    rna-XM_027593245.1    mNeoVis.3.1.1.chr10.g01194.t01    rna-gnl|WGS_ZZZZ|egapxtmp_023800-R3    mNeoVis.evergladensis.2.1.chr10.g01233.t01    mNeoVis.vulgivaga.2.1.chr10.g01220.t01      2707    N0.HOG0002705   OG0000525  n1   rna-XM_034660585.1    rna-XM_038553817.1    rna-XM_022510780.1    rna-XM_019831576.3    rna-XM_036096202.1    rna-XM_039241783.1    rna-XM_032864289.1    rna-XM_046006725.1    rna-XM_032340544.1    rna-XM_059178539.1    rna-XM_059400080.1    rna-XM_045074744.1    mNeoVis.2.1.1.chr1_hap1.g01458.t01    rna-XM_044917443.1    rna-XM_004407740.1    rna-XM_042986181.1    rna-XM_040644728.1    rna-XM_027604083.2    mNeoVis.3.1.1.chr1.g01405.t01    rna-gnl|WGS_ZZZZ|egapxtmp_021807-R1    mNeoVis.evergladensis.2.1.chr1.g01423.t01    mNeoVis.vulgivaga.2.1.chr1.g01424.t01      2708    N0.HOG0002706   OG0000525  n21   rna-XM_011230029.3    rna-XM_038550290.1    rna-XM_022519283.1    rna-XM_023257570.2    rna-XM_036088362.1    rna-XM_039234658.1    rna-XM_032882823.1    rna-XM_046013125.1    rna-XM_032346018.1    rna-XM_059187196.1    rna-XM_059404574.1    rna-XM_045083742.1    mNeoVis.2.1.1.chr9_hap1.g01250.t01    rna-XM_044915773.1    rna-XM_004397384.1    rna-XM_042993464.1    rna-XM_040637674.1    rna-XM_027594583.2    mNeoVis.3.1.1.chr10.g01200.t01    rna-gnl|WGS_ZZZZ|egapxtmp_022739-R1    mNeoVis.evergladensis.2.1.chr10.g01239.t01    mNeoVis.vulgivaga.2.1.chr10.g01227.t01      2709   N0.HOG0002707  OG0000525  n42   rna-XM_034666532.1    rna-XM_038569006.1    rna-XM_022521767.1    rna-XM_023239543.2    rna-XM_036121386.1    rna-XM_039244582.1    rna-XM_032853430.1    rna-XM_046015738.1    rna-XM_032357953.1    rna-XM_059135552.1    rna-XM_059408632.1    rna-XM_045065847.1    mNeoVis.2.1.1.chr7_hap1.g01803.t01    rna-XM_021685760.1    rna-XM_004394561.1    rna-XM_042958347.1    rna-XM_040631316.1    rna-XM_027579591.2    mNeoVis.3.1.1.chr7.g01672.t01    rna-gnl|WGS_ZZZZ|egapxtmp_009794-R1    mNeoVis.evergladensis.2.1.chr7.g01764.t01    mNeoVis.vulgivaga.2.1.chr7.g01782.t01      2710    N0.HOG0002708   OG0000526  n1   rna-XM_002921554.4    rna-XM_038535131.1    rna-XM_022492960.1    rna-XM_045037296.1    rna-XM_036114460.1    rna-XM_039256206.1    rna-XM_032851453.1    rna-XM_045998899.1    rna-XM_032337429.1    rna-XM_059173951.1    rna-XM_059417104.1    rna-XM_004737905.3    mNeoVis.2.1.1.chr1_hap1.g00162.t01    rna-XM_021695399.1    rna-XM_004410964.1    rna-XM_015535080.2    rna-XM_040630445.1    rna-XM_027606606.1    mNeoVis.3.1.1.chr1.g00167.t01    rna-gnl|WGS_ZZZZ|egapxtmp_025879-R1    mNeoVis.evergladensis.2.1.chr1.g00158.t01    mNeoVis.vulgivaga.2.1.chr1.g00160.t01      2711    N0.HOG0002709   OG0000526  n23   rna-XM_011231480.3    rna-XM_038562033.1    rna-XM_022526135.1    rna-XM_003990541.6    rna-XM_036112996.1    rna-XM_039217848.1    rna-XM_032848153.1    rna-XM_046026073.1    rna-XM_032361317.1    rna-XM_059134866.1    rna-XM_059375987.1    rna-XM_004779649.3    mNeoVis.2.1.1.chr2_hap1.g00849.t01    rna-XM_021688081.1    rna-XM_004398385.1    rna-XM_007092558.3    rna-XM_008707753.1    rna-XM_027579851.2    mNeoVis.3.1.1.chr5.g00786.t01    rna-gnl|WGS_ZZZZ|egapxtmp_006947-R1    mNeoVis.evergladensis.2.1.chr5.g00803.t01    mNeoVis.vulgivaga.2.1.chr5.g00794.t01      2712    N0.HOG0002710   OG0000526  n44   rna-XM_034660769.1    rna-XM_038580816.1    rna-XM_022511924.1    rna-XM_023255135.2    rna-XM_036087083.1    rna-XM_039218561.1    rna-XM_032864605.1    rna-XM_046010344.1    rna-XM_032343008.1    rna-XM_059180927.1    rna-XM_059372356.1    rna-XM_045067443.1    mNeoVis.2.1.1.chr10_hap1.g00915.t01    rna-XM_021693715.1    rna-XM_004412236.1    rna-XM_042988522.1    rna-XM_040643059.1    rna-XM_027570941.2    mNeoVis.3.1.1.chr9.g00843.t01    rna-gnl|WGS_ZZZZ|egapxtmp_016961-R1    mNeoVis.evergladensis.2.1.chr8.g00885.t01    mNeoVis.vulgivaga.2.1.chr9.g00872.t01      2713    N0.HOG0002711   OG0000527  n1   rna-XM_034664023.1    rna-XM_038552610.1    rna-XM_022501298.1    rna-XM_011288588.4    rna-XM_036069563.1    rna-XM_039247027.1    rna-XM_032860389.1    rna-XM_046022874.1    rna-XM_032307048.1    rna-XM_059142718.1    rna-XM_059411490.1    rna-XM_004765332.3    mNeoVis.2.1.1.chr13_hap1.g00545.t01    rna-XM_021691262.1    rna-XM_004392189.1    rna-XM_007076570.2    rna-XM_040631530.1    rna-XM_027616904.2    mNeoVis.3.1.1.chr13.g00486.t01    rna-gnl|WGS_ZZZZ|egapxtmp_031429-R1    mNeoVis.evergladensis.2.1.chr13.g00528.t01    mNeoVis.vulgivaga.2.1.chr13.g00531.t01      2714   N0.HOG0002712  OG0000527  n18   rna-XM_002920668.4    rna-XM_038580537.1    rna-XM_022512255.1    rna-XM_003987104.6    rna-XM_036107027.1    rna-XM_039230649.1    rna-XM_032864852.1    rna-XM_046007167.1    rna-XM_032342675.1    rna-XM_059180624.1    rna-XM_059372012.1    rna-XM_004777288.3    mNeoVis.2.1.1.chr10_hap1.g01000.t01    rna-XM_021693876.2    rna-XM_004406847.1    rna-XM_007084459.3    rna-XM_008706025.2    rna-XM_027569387.2    mNeoVis.3.1.1.chr9.g00930.t01    rna-gnl|WGS_ZZZZ|egapxtmp_016986-R1    mNeoVis.evergladensis.2.1.chr8.g00971.t01    mNeoVis.vulgivaga.2.1.chr9.g00958.t01      2715   N0.HOG0002713  OG0000527  n36   rna-XM_034667541.1    rna-XM_038543234.1    rna-XM_022490713.1    rna-XM_019822497.3    rna-XM_036086868.1    rna-XM_039218935.1    rna-XM_032845153.1    rna-XM_045982675.1    rna-XM_032318487.1    rna-XM_059146300.1    rna-XM_059414190.1    rna-XM_013049932.2    mNeoVis.2.1.1.chr14_hap1.g00686.t01    rna-XM_021681869.1    rna-XM_004403057.1    rna-XM_042975978.1    rna-XM_008696607.2    rna-XM_027612331.2    mNeoVis.3.1.1.chr14.g00656.t01    rna-gnl|WGS_ZZZZ|egapxtmp_032479-R1    mNeoVis.evergladensis.2.1.chr14.g00684.t01    mNeoVis.vulgivaga.2.1.chr14.g00684.t01      2716    N0.HOG0002714   OG0000528  n1   rna-XM_011218992.3    rna-NM_001003268.1    rna-XM_022513673.1    rna-XM_019815315.3    rna-XM_036080728.1    rna-XM_039256089.1    rna-XM_032882433.1    rna-XM_046024539.1    rna-XM_032311422.1    rna-XM_059139402.1    rna-XM_059409713.1    rna-XM_045065657.1    mNeoVis.2.1.1.chr3_hap1.g00085.t01    rna-XM_044921065.1    rna-XM_004405827.1    rna-XM_042963032.1    rna-XM_040632482.1    rna-XM_027576822.2    mNeoVis.3.1.1.chr3.g00086.t01    rna-gnl|WGS_ZZZZ|egapxtmp_041429-R2    mNeoVis.evergladensis.2.1.chr3.g00088.t01    mNeoVis.vulgivaga.2.1.chr3.g00085.t01      2717    N0.HOG0002715   OG0000528  n15   rna-XM_019801206.2    rna-XM_038580520.1    rna-XM_022511943.1    rna-XM_045059253.1    rna-XM_036107004.1    rna-XM_039230611.1    rna-XM_032864863.1    rna-XM_046007718.1    rna-XM_032341444.1    rna-XM_059180606.1    rna-XM_059371992.1    rna-XM_004777303.3    mNeoVis.2.1.1.chr10_hap1.g01005.t01    rna-XM_021693892.1    rna-XM_004406837.1    rna-XM_042988778.1    rna-XM_040620598.1    rna-XM_027569322.2    mNeoVis.3.1.1.chr9.g00936.t01    rna-gnl|WGS_ZZZZ|egapxtmp_016484-R2    mNeoVis.evergladensis.2.1.chr8.g00976.t01    mNeoVis.vulgivaga.2.1.chr9.g00963.t01      2718    N0.HOG0002716   OG0000528  n31   rna-XM_034657392.1    rna-XM_038567945.1    rna-XM_022525553.1    rna-XM_023243101.2    rna-XM_036075733.1    rna-XM_039248127.1    rna-XM_032846943.1    rna-XM_045990743.1    rna-XM_032358361.1    rna-XM_059159271.1    rna-XM_059388515.1    rna-XM_045087496.1    mNeoVis.2.1.1.chr5_hap1.g00039.t01    rna-XM_021705060.2    rna-XM_004395402.1    rna-XM_042977586.1    rna-XM_040629681.1    rna-XM_027583340.1    mNeoVis.3.1.1.chr2.g00021.t01    rna-gnl|WGS_ZZZZ|egapxtmp_034421-R1    mNeoVis.evergladensis.2.1.chr2.g00039.t01    mNeoVis.vulgivaga.2.1.chr2.g00039.t01      2719   N0.HOG0002717  OG0000529  n1   rna-XM_034657671.1    rna-XM_038567487.1    rna-XM_022525653.1    rna-XM_045045016.1    rna-XM_036066022.1    rna-XM_039251578.1    rna-XM_032883468.1    rna-XM_045991463.1    rna-XM_032309031.1    rna-XM_059159767.1    rna-XM_059388849.1    rna-XM_045085108.1    mNeoVis.2.1.1.chr5_hap1.g00201.t01    rna-XM_044913115.1    rna-XM_004412189.1    rna-XM_042963462.1    rna-XM_040628285.1    rna-XM_035723607.1    mNeoVis.3.1.1.chr2.g00151.t01    rna-gnl|WGS_ZZZZ|egapxtmp_034634-R1    mNeoVis.evergladensis.2.1.chr2.g00198.t01    mNeoVis.vulgivaga.2.1.chr2.g00197.t01      2720    N0.HOG0002718   OG0000529  n23   rna-XM_002917548.4    rna-NM_001287085.1    rna-XM_022520875.1    rna-XM_023255387.2    rna-XM_036075961.1    rna-XM_039218035.1    rna-XM_032856557.1    rna-XM_046008182.1    rna-XM_032343645.1    rna-XM_059181480.1    rna-XM_059372774.1    rna-XM_045067863.1    mNeoVis.2.1.1.chr10_hap1.g00634.t01    rna-XM_021693975.2    rna-XM_004396527.1    rna-XM_042988929.1    rna-XM_008705933.2    rna-XM_027569939.2    mNeoVis.3.1.1.chr9.g00594.t01    rna-gnl|WGS_ZZZZ|egapxtmp_016872-R1    mNeoVis.evergladensis.2.1.chr8.g00615.t01    mNeoVis.vulgivaga.2.1.chr9.g00618.t01      2721    N0.HOG0002719   OG0000529  n44   rna-XM_034655954.1    rna-XM_038552058.1    rna-XM_022493138.1    rna-XM_003980732.5    rna-XM_036072328.1    rna-XM_039236691.1    rna-XM_032851970.1    rna-XM_046000771.1    rna-XM_032336836.1    rna-XM_059174478.1    rna-XM_059417490.1    rna-XM_004737427.3    mNeoVis.2.1.1.chr1_hap1.g00373.t01    rna-XM_021701751.1    rna-XM_004413259.1    rna-XM_007087821.2    rna-XM_040622515.1    rna-XM_027606724.2    mNeoVis.3.1.1.chr1.g00372.t01    rna-gnl|WGS_ZZZZ|egapxtmp_034406-R1    mNeoVis.evergladensis.2.1.chr1.g00370.t01    mNeoVis.vulgivaga.2.1.chr1.g00378.t01      2722   N0.HOG0002720  OG0000530  n1   rna-XM_011217768.3    rna-XM_038530995.1    rna-XM_022518705.1    rna-XM_023245212.2    rna-XM_036068951.1    rna-XM_039232583.1    rna-XM_032854422.1    rna-XM_045989131.1    rna-XM_032323982.1    rna-XM_059152746.1    rna-XM_059383439.1    rna-XM_004744220.2    mNeoVis.2.1.1.chr7_hap1.g00305.t01    rna-XM_021704504.1    rna-XM_004395799.1    rna-XM_007090194.3    rna-XM_040633330.1    rna-XM_035724948.1    mNeoVis.3.1.1.chr7.g00291.t01    rna-gnl|WGS_ZZZZ|egapxtmp_013137-R2    mNeoVis.evergladensis.2.1.chr7.g00305.t01    mNeoVis.vulgivaga.2.1.chr7.g00308.t01      2723   N0.HOG0002721  OG0000530  n23   rna-XM_034655729.1    rna-XM_038552056.1    rna-XM_022493158.1    rna-XM_011283447.3    rna-XM_036072324.1    rna-XM_039236698.1    rna-XM_032851968.1    rna-XM_045999901.1    rna-XM_032336837.1    rna-XM_059174472.1    rna-XM_059417486.1    rna-NM_001310166.1    mNeoVis.2.1.1.chr1_hap1.g00370.t01    rna-XM_021702155.2    rna-XM_004413260.1    rna-XM_015540046.2    rna-XM_008699034.2    rna-XM_027604681.2    mNeoVis.3.1.1.chr1.g00371.t01    rna-gnl|WGS_ZZZZ|egapxtmp_034420-R1    mNeoVis.evergladensis.2.1.chr1.g00367.t01    mNeoVis.vulgivaga.2.1.chr1.g00375.t01      2724   N0.HOG0002722  OG0000530  n42   rna-XM_002917549.3    rna-XM_038580355.1    rna-XM_022524246.1    rna-XM_003987165.3    rna-XM_036075954.1    rna-XM_039218038.1    rna-XM_032856562.1    rna-XM_046009199.1    rna-XM_032343634.1    rna-XM_059181472.1    rna-XM_059372768.1    rna-NM_001310193.1    mNeoVis.2.1.1.chr10_hap1.g00637.t01    rna-XM_044917786.1    rna-XM_004396523.1    rna-XM_007081816.3    rna-XM_008705930.2    rna-XM_027570290.2    mNeoVis.3.1.1.chr9.g00596.t01    rna-gnl|WGS_ZZZZ|egapxtmp_016329-R1    mNeoVis.evergladensis.2.1.chr8.g00618.t01    mNeoVis.vulgivaga.2.1.chr9.g00621.t01      2725   N0.HOG0002723  OG0000531  n1   rna-XM_002916969.4    rna-XM_038578475.1    rna-XM_022499777.1    rna-XM_023240813.2    rna-XM_036080663.1    rna-XM_039221206.1    rna-XM_032882769.1    rna-XM_046026183.1    rna-XM_032312077.1    rna-XM_059169255.1    rna-XM_059398313.1    rna-XM_013054658.2    mNeoVis.2.1.1.chr2_hap1.g00264.t01    rna-XM_021699634.1    rna-XM_004415745.1    rna-XM_007080157.3    rna-XM_040624695.1    rna-XM_027586786.2    mNeoVis.3.1.1.chr5.g00257.t01    rna-gnl|WGS_ZZZZ|egapxtmp_006817-R1    mNeoVis.evergladensis.2.1.chr5.g00268.t01    mNeoVis.vulgivaga.2.1.chr5.g00266.t01      2726   N0.HOG0002724  OG0000531  n23   rna-XM_019801992.2    rna-XM_038561650.1    rna-XM_022504633.1    rna-XM_019827460.3    rna-XM_036087576.1    rna-XM_039240174.1    rna-XM_032880718.1    rna-XM_045978837.1    rna-XM_032352539.1    rna-XM_059188127.1    rna-XM_059405737.1    rna-XM_013051176.2    mNeoVis.2.1.1.chr11_hap1.g00836.t01    rna-XM_021699193.2    rna-XM_004398762.1    rna-XM_042981797.1    rna-XM_008700034.2    rna-XM_027620490.2    mNeoVis.3.1.1.chr11.g00798.t01    rna-gnl|WGS_ZZZZ|egapxtmp_027337-R1    mNeoVis.evergladensis.2.1.chr11.g00819.t01    mNeoVis.vulgivaga.2.1.chr11.g00839.t01      2727   N0.HOG0002725  OG0000531  n44   rna-XM_034661260.1    rna-XM_038573799.1    rna-XM_022514934.1    rna-XM_045055385.1    rna-XM_036075095.1    rna-XM_039243769.1    rna-XM_032855810.1    rna-XM_045998134.1    rna-XM_032332167.1    rna-XM_059178117.1    rna-XM_059371842.1    rna-XM_004763265.3    mNeoVis.2.1.1.chr6_hap1.g00825.t01    rna-XM_044912559.1    rna-XM_004402211.1    rna-XM_042983236.1    rna-XM_008702068.2    rna-XM_035726206.1    mNeoVis.3.1.1.chr4.g00836.t01    rna-gnl|WGS_ZZZZ|egapxtmp_000868-R1    mNeoVis.evergladensis.2.1.chr4.g00827.t01    mNeoVis.vulgivaga.2.1.chr4.g00847.t01      2728   N0.HOG0002726  OG0000532  n1   rna-XM_011223578.3    rna-NM_001308463.1    rna-XM_022514384.1    rna-XM_003984867.6    rna-XM_036085545.1    rna-XM_039219208.1    rna-XM_032855475.1    rna-XM_045995026.1    rna-XM_032332568.1    rna-XM_059175071.1    rna-XM_059374113.1    rna-XM_004758101.3    mNeoVis.2.1.1.chr6_hap1.g00692.t01    rna-XM_021699008.2    rna-XM_004392655.1    rna-XM_042983637.1    rna-XM_008685632.2    rna-XM_027598101.2    mNeoVis.3.1.1.chr4.g00702.t01    rna-gnl|WGS_ZZZZ|egapxtmp_043504-R1    mNeoVis.evergladensis.2.1.chr4.g00699.t01    mNeoVis.vulgivaga.2.1.chr4.g00714.t01      2729   N0.HOG0002727  OG0000532  n14   rna-XM_034649394.1    rna-XM_038586800.1    rna-XM_022518413.1    rna-XM_019823537.3    rna-XM_036068843.1    rna-XM_039234250.1    rna-XM_032880216.1    rna-XM_045995057.1    rna-XM_032330617.1    rna-XM_059157310.1    rna-XM_059385775.1    rna-XM_004758812.3    mNeoVis.2.1.1.chr12_hap1.g00813.t01    rna-XM_021684124.1    rna-XM_004409250.1    rna-XM_042974508.1    rna-XM_040634595.1    rna-XM_027608056.2    mNeoVis.3.1.1.chr12.g00744.t01    rna-gnl|WGS_ZZZZ|egapxtmp_029180-R1    mNeoVis.evergladensis.2.1.chr12.g00740.t01    mNeoVis.vulgivaga.2.1.chr12.g00769.t01      2730   N0.HOG0002728  OG0000532  n34   rna-XM_002917742.4    rna-XM_038587444.1    rna-XM_022507882.1    rna-XM_011291722.4    rna-XM_036103903.1    rna-XM_039221177.1    rna-XM_032843044.1    rna-XM_045995030.1    rna-XM_032329511.1    rna-XM_059156426.1    rna-XM_059385861.1    rna-XM_004755112.3    mNeoVis.2.1.1.chr12_hap1.g00588.t01    rna-XM_021678741.1    rna-XM_004396829.1    rna-XM_042974411.1    rna-XM_008690162.2    rna-XM_027608571.2    mNeoVis.3.1.1.chr12.g00535.t01    rna-gnl|WGS_ZZZZ|egapxtmp_028510-R1    mNeoVis.evergladensis.2.1.chr12.g00525.t01    mNeoVis.vulgivaga.2.1.chr12.g00539.t01      2731   N0.HOG0002729  OG0000533  n1   rna-XM_011221023.3    rna-XM_038558911.1    rna-XM_022513604.1    rna-XM_023252779.2    rna-XM_036077773.1    rna-XM_039254315.1    rna-XM_032855598.1    rna-XM_045997910.1    rna-XM_032332407.1    rna-XM_059176135.1    rna-XM_059373253.1    rna-XM_004758000.3    mNeoVis.2.1.1.chr6_hap1.g00729.t01    rna-XM_021679358.1    rna-XM_004411761.1    rna-XM_042983833.1    rna-XM_040626343.1    rna-XM_027598110.2    mNeoVis.3.1.1.chr4.g00739.t01    rna-gnl|WGS_ZZZZ|egapxtmp_000992-R1    mNeoVis.evergladensis.2.1.chr4.g00736.t01    mNeoVis.vulgivaga.2.1.chr4.g00754.t01      2732   N0.HOG0002730  OG0000533  n5   rna-XM_002912928.4    rna-XM_038552236.1    rna-XM_022493329.1    rna-XM_006927618.5    rna-XM_036114320.1    rna-XM_039242273.1    rna-XM_032837759.1    rna-XM_045999865.1    rna-XM_032336642.1    rna-XM_059174641.1    rna-XM_059417621.1    rna-XM_004744911.3    mNeoVis.2.1.1.chr1_hap1.g00441.t01    rna-XM_021701887.1    rna-XM_004405475.1    rna-XM_042984410.1    rna-XM_040622411.1    rna-XM_035727634.1    mNeoVis.3.1.1.chr1.g00436.t01    rna-gnl|WGS_ZZZZ|egapxtmp_032609-R1    mNeoVis.evergladensis.2.1.chr1.g00438.t01    mNeoVis.vulgivaga.2.1.chr1.g00444.t01      2733   N0.HOG0002731  OG0000533  n11   rna-XM_011217477.3    rna-XM_038573204.1    rna-XM_022520640.1    rna-XM_023250901.2    rna-XM_036090993.1    rna-XM_039253651.1    rna-XM_032880082.1    rna-XM_045978142.1    rna-XM_032314649.1    rna-XM_059143936.1    rna-XM_059377821.1    rna-XM_045069536.1    mNeoVis.2.1.1.chr8_hap1.g01285.t01    rna-XM_021678299.1    rna-XM_004399908.1    rna-XM_042957568.1    rna-XM_008692237.2    rna-XM_027589555.2    mNeoVis.3.1.1.chr8.g01181.t01    rna-gnl|WGS_ZZZZ|egapxtmp_020056-R1    mNeoVis.evergladensis.2.1.chr9.g01233.t01    mNeoVis.vulgivaga.2.1.chr8.g01235.t01      2734   N0.HOG0002732  OG0000534  n1   rna-XM_034644089.1    rna-XM_038530215.1    rna-XM_022520719.1    rna-XM_045060809.1    rna-XM_036088808.1    rna-XM_039240369.1    rna-XM_032881677.1    rna-XM_046012169.1    rna-XM_032349868.1    rna-XM_059183690.1    rna-XM_059403986.1    rna-XM_004757906.2    mNeoVis.2.1.1.chr9_hap1.g00012.t01    rna-XM_021690903.1    rna-XM_004409345.1    rna-XM_042990929.1    rna-XM_040645468.1    rna-XM_035721885.1    mNeoVis.3.1.1.chr10.g00010.t01    rna-gnl|WGS_ZZZZ|egapxtmp_023345-R1    mNeoVis.evergladensis.2.1.chr10.g00014.t01    mNeoVis.vulgivaga.2.1.chr10.g00013.t01      2735   N0.HOG0002733  OG0000534  n23   rna-XM_034654791.1    rna-XM_038581620.1    rna-XM_022515926.1    rna-XM_023238651.2    rna-XM_036097234.1    rna-XM_039245433.1    rna-XM_032865215.1    rna-XM_046002435.1    rna-XM_032356663.1    rna-XM_059164893.1    rna-XM_059392474.1    rna-XM_045084361.1    mNeoVis.2.1.1.chr5_hap1.g01892.t01    rna-XM_044919313.1    rna-XM_004410657.1    rna-XM_042998843.1    rna-XM_040621826.1    rna-XM_027586378.1        mNeoVis.evergladensis.2.1.chr2.g01872.t01    mNeoVis.vulgivaga.2.1.chr2.g01887.t01      2736   N0.HOG0002734  OG0000534  n41   rna-XM_019800807.2    rna-XM_038577468.1, rna-XM_038581745.1    rna-XM_022519705.1    rna-XM_019834856.2    rna-XM_036099319.1    rna-XM_039243116.1    rna-XM_032870086.1, rna-XM_032876921.1    rna-XM_046013950.1    rna-XM_032348013.1    rna-XM_059185522.1    rna-XM_059402988.1    rna-XM_013063007.2    mNeoVis.2.1.1.chr9_hap1.g00631.t01    rna-XM_021704784.1    rna-XM_004395955.1    rna-XM_015539474.2    rna-XM_040642385.1    rna-XM_027592757.1    mNeoVis.3.1.1.chr10.g00596.t01    rna-gnl|WGS_ZZZZ|egapxtmp_021678-R2    mNeoVis.evergladensis.2.1.chr10.g00625.t01    mNeoVis.vulgivaga.2.1.chr10.g00608.t01      2737   N0.HOG0002735  OG0000535  n1   rna-XM_034643655.1    rna-XM_038529873.1    rna-XM_022515766.1    rna-XM_003988210.6    rna-XM_036099595.1    rna-XM_039248658.1    rna-XM_032837906.1    rna-XM_046011711.1    rna-XM_032349343.1    rna-XM_059184213.1    rna-XM_059404366.1    rna-XM_004777869.3    mNeoVis.2.1.1.chr9_hap1.g00166.t01    rna-XM_021686504.1    rna-XM_004401819.1    rna-XM_007092757.2    rna-XM_008683948.2    rna-XM_027592258.2    mNeoVis.3.1.1.chr10.g00157.t01    rna-gnl|WGS_ZZZZ|egapxtmp_022586-R1    mNeoVis.evergladensis.2.1.chr10.g00161.t01    mNeoVis.vulgivaga.2.1.chr10.g00157.t01      2738   N0.HOG0002736  OG0000535  n12   rna-XM_011230488.3    rna-XM_038565167.1    rna-XM_022494987.1    rna-XM_011285287.4    rna-XM_036092566.1    rna-XM_039244578.1    rna-XM_032856096.1    rna-XM_046019619.1    rna-XM_032356183.1    rna-XM_059166812.1    rna-XM_059394320.1    rna-XM_004757858.2    mNeoVis.2.1.1.chr3_hap1.g00806.t01    rna-XM_021703005.1    rna-XM_004401538.1    rna-XM_007078543.3    rna-XM_040645119.1    rna-XM_027589084.2    mNeoVis.3.1.1.chr3.g00789.t01    rna-gnl|WGS_ZZZZ|egapxtmp_041599-R1    mNeoVis.evergladensis.2.1.chr3.g00800.t01    mNeoVis.vulgivaga.2.1.chr3.g00813.t01      2739   N0.HOG0002737  OG0000535  n27   rna-XM_019796218.2    rna-XM_038549790.1    rna-XM_022522990.1    rna-XM_011284094.4    rna-XM_036077952.1    rna-XM_039256158.1    rna-XM_032874578.1    rna-XM_046012619.1    rna-XM_032348777.1    rna-XM_059184832.1    rna-XM_059403538.1    rna-XM_004773229.3    mNeoVis.2.1.1.chr9_hap1.g00336.t01    rna-XM_021687287.1    rna-XM_004401731.1    rna-XM_007074981.3    rna-XM_040637176.1    rna-XM_027595259.2    mNeoVis.3.1.1.chr10.g00327.t01    rna-gnl|WGS_ZZZZ|egapxtmp_021667-R1    mNeoVis.evergladensis.2.1.chr10.g00333.t01    mNeoVis.vulgivaga.2.1.chr10.g00329.t01      2740   N0.HOG0002738  OG0000536  n1   rna-XM_019796140.2    rna-XM_038549603.1    rna-XM_022522970.1    rna-XM_019835058.3    rna-XM_036086618.1    rna-XM_039216840.1    rna-XM_032874403.1    rna-XM_046011574.1    rna-XM_032348551.1    rna-XM_059185027.1    rna-XM_059403362.1    rna-XM_004773057.3    mNeoVis.2.1.1.chr9_hap1.g00395.t01    rna-XM_044915389.1    rna-XM_004401638.1    rna-XM_007081471.3    rna-XM_040637116.1    rna-XM_027592645.2    mNeoVis.3.1.1.chr10.g00387.t01    rna-gnl|WGS_ZZZZ|egapxtmp_023546-R1    mNeoVis.evergladensis.2.1.chr10.g00392.t01    mNeoVis.vulgivaga.2.1.chr10.g00389.t01      2741   N0.HOG0002739  OG0000536  n23   rna-XM_034668553.1    rna-XM_038556977.1    rna-XM_022503288.1    rna-XM_045052433.1    rna-XM_036106040.1    rna-XM_039221452.1    rna-XM_032873966.1    rna-XM_046021573.1    rna-XM_032305329.1    rna-XM_059171260.1    rna-XM_059396759.1    rna-XM_004743406.3    mNeoVis.2.1.1.chr4_hap1.g00444.t01    rna-XM_021689585.1    rna-XM_004400194.1    rna-XM_042967486.1    rna-XM_040644224.1    rna-XM_027574755.2    mNeoVis.3.1.1.chr6.g00443.t01    rna-gnl|WGS_ZZZZ|egapxtmp_002376-R1    mNeoVis.evergladensis.2.1.chr6.g00450.t01    mNeoVis.vulgivaga.2.1.chr6.g00443.t01      2742    N0.HOG0002740   OG0000536  n40   rna-XM_002915692.4    rna-XM_038583634.1    rna-XM_022505659.1    rna-XM_023260324.2    rna-XM_036066662.1    rna-XM_039241717.1    rna-XM_032875583.1    rna-XM_046003528.1    rna-XM_032338301.1    rna-XM_059163206.1    rna-XM_059391333.1    rna-XM_004745197.3    mNeoVis.2.1.1.chr5_hap1.g01309.t01    rna-XM_021692349.1    rna-XM_004413403.1    rna-XM_042956236.1    rna-XM_040621478.1    rna-XM_027586641.2    mNeoVis.3.1.1.chr2.g01229.t01    rna-gnl|WGS_ZZZZ|egapxtmp_037445-R1    mNeoVis.evergladensis.2.1.chr2.g01296.t01    mNeoVis.vulgivaga.2.1.chr2.g01303.t01      2743    N0.HOG0002741   OG0000537  n1   rna-XM_019795559.2    rna-XM_038559731.1    rna-XM_022513846.1    rna-XM_023250737.2    rna-XM_036102106.1    rna-XM_039230700.1    rna-XM_032840143.1    rna-XM_046020431.1    rna-XM_032303871.1    rna-XM_059172253.1    rna-XM_059396099.1    rna-XM_045086374.1    mNeoVis.2.1.1.chr4_hap1.g00040.t01    rna-XM_044920262.1    rna-XM_004408818.1    rna-XM_042975467.1    rna-XM_040643795.1    rna-XM_027574373.1    mNeoVis.3.1.1.chr6.g00043.t01    rna-gnl|WGS_ZZZZ|egapxtmp_003018-R1    mNeoVis.evergladensis.2.1.chr6.g00042.t01    mNeoVis.vulgivaga.2.1.chr6.g00046.t01      2744   N0.HOG0002742  OG0000537  n23   rna-XM_034655451.1    rna-XM_038574267.1    rna-XM_022519024.1    rna-XM_023259835.2    rna-XM_036091840.1    rna-XM_039228587.1    rna-XM_032857548.1    rna-XM_046018359.1    rna-XM_032355251.1    rna-XM_059167635.1    rna-XM_059393772.1    rna-XM_004777552.3    mNeoVis.2.1.1.chr3_hap1.g01050.t01    rna-XM_021679032.1    rna-XM_004417398.1    rna-XM_042995745.1    rna-XM_040638721.1    rna-XM_027588838.2    mNeoVis.3.1.1.chr3.g01038.t01    rna-gnl|WGS_ZZZZ|egapxtmp_038956-R1    mNeoVis.evergladensis.2.1.chr3.g01049.t01    mNeoVis.vulgivaga.2.1.chr3.g01067.t01      2745   N0.HOG0002743  OG0000537  n44   rna-XM_034644311.1    rna-XM_038549814.1    rna-XM_022523047.1    rna-XM_011284083.4    rna-XM_036077970.1    rna-XM_039256190.1    rna-XM_032874596.1    rna-XM_046011491.1    rna-XM_032348799.1    rna-XM_059184793.1    rna-XM_059403567.1    rna-XM_004773264.2    mNeoVis.2.1.1.chr9_hap1.g00329.t01    rna-XM_044914778.1    rna-XM_004401750.1    rna-XM_042992634.1    rna-XM_040637191.1    rna-XM_027593003.2    mNeoVis.3.1.1.chr10.g00319.t01    rna-gnl|WGS_ZZZZ|egapxtmp_022601-R1    mNeoVis.evergladensis.2.1.chr10.g00326.t01    mNeoVis.vulgivaga.2.1.chr10.g00322.t01      2746    N0.HOG0002744   OG0000538  n1   rna-XM_034658695.1    rna-XM_038565766.1    rna-XM_022498254.1    rna-XM_019824937.3    rna-XM_036096153.1    rna-XM_039231356.1    rna-XM_032841604.1    rna-XM_045991582.1    rna-XM_032326246.1    rna-XM_059161597.1    rna-XM_059390212.1    rna-XM_004738433.3    mNeoVis.2.1.1.chr5_hap1.g00849.t01    rna-XM_021695153.1    rna-XM_004392717.1    rna-XM_042979459.1    rna-XM_040628128.1    rna-XM_027584523.2    mNeoVis.3.1.1.chr2.g00760.t01    rna-gnl|WGS_ZZZZ|egapxtmp_034291-R3    mNeoVis.evergladensis.2.1.chr2.g00850.t01    mNeoVis.vulgivaga.2.1.chr2.g00851.t01      2747    N0.HOG0002745   OG0000538  n14   rna-XM_034644551.1    rna-XM_038549883.1    rna-XM_022522307.1    rna-XM_011284066.4    rna-XM_036070968.1    rna-XM_039234288.1    rna-XM_032874653.1    rna-XM_046012397.1    rna-XM_032348867.1    rna-XM_059184738.1    rna-XM_059403611.1    rna-XM_004766597.3    mNeoVis.2.1.1.chr9_hap1.g00305.t01    rna-XM_021678634.1    rna-XM_004413807.1    rna-XM_007082210.2    rna-XM_008684244.2    rna-XM_027593051.2    mNeoVis.3.1.1.chr10.g00294.t01    rna-gnl|WGS_ZZZZ|egapxtmp_024334-R3    mNeoVis.evergladensis.2.1.chr10.g00303.t01    mNeoVis.vulgivaga.2.1.chr10.g00297.t01      2748    N0.HOG0002746   OG0000538  n33   rna-XM_034667155.1    rna-XM_038586064.1    rna-XM_022508663.1    rna-XM_006942921.5    rna-XM_036077188.1    rna-XM_039227559.1    rna-XM_032878469.1    rna-XM_045982173.1    rna-XM_032318979.1    rna-XM_059146580.1    rna-XM_059413272.1    rna-XM_004756459.3    mNeoVis.2.1.1.chr14_hap1.g00400.t01    rna-XM_021686555.2    rna-XM_004392843.1    rna-XM_015535288.2    rna-XM_008699550.2    rna-XM_027610047.1    mNeoVis.3.1.1.chr14.g00399.t01    rna-gnl|WGS_ZZZZ|egapxtmp_031913-R1    mNeoVis.evergladensis.2.1.chr14.g00400.t01    mNeoVis.vulgivaga.2.1.chr14.g00401.t01      2749   N0.HOG0002747  OG0000539  n1   rna-XM_002919247.4    rna-XM_038558705.1    rna-XM_022495026.1    rna-XM_023257340.2    rna-XM_036085856.1    rna-XM_039236560.1    rna-XM_032874278.1    rna-XM_046010835.1    rna-XM_032346706.1    rna-XM_059186575.1    rna-XM_059402310.1    rna-XM_004748492.3    mNeoVis.2.1.1.chr9_hap1.g01025.t01    rna-XM_044914734.1    rna-XM_004400255.1    rna-XM_042992950.1    rna-XM_040637343.1    rna-XM_027591965.2    mNeoVis.3.1.1.chr10.g00986.t01    rna-gnl|WGS_ZZZZ|egapxtmp_024256-R2    mNeoVis.evergladensis.2.1.chr10.g01012.t01    mNeoVis.vulgivaga.2.1.chr10.g01006.t01      2750    N0.HOG0002748   OG0000539  n9   rna-XM_034667124.1    rna-XM_038586030.1    rna-XM_022508787.1    rna-XM_019822119.2    rna-XM_036077735.1    rna-XM_039227617.1    rna-XM_032878440.1    rna-XM_045982486.1    rna-XM_032318268.1    rna-XM_059145773.1    rna-XM_059413244.1    rna-XM_045081097.1    mNeoVis.2.1.1.chr14_hap1.g00385.t01    rna-XM_021686617.1    rna-XM_004392827.1    rna-XM_042976665.1    rna-XM_040634517.1    rna-XM_027611694.2    mNeoVis.3.1.1.chr14.g00384.t01      mNeoVis.evergladensis.2.1.chr14.g00385.t01    mNeoVis.vulgivaga.2.1.chr14.g00386.t01      2751    N0.HOG0002749   OG0000539  n29   rna-XM_019798121.2    rna-XM_038587280.1    rna-XM_022508084.1    rna-XM_011291660.3    rna-XM_036117295.1    rna-XM_039216775.1    rna-XM_032842873.1    rna-XM_045995300.1    rna-XM_032331568.1    rna-XM_059156616.1    rna-XM_059385417.1    rna-XM_045068423.1    mNeoVis.2.1.1.chr12_hap1.g00640.t01    rna-XM_021679110.1    rna-XM_004396746.1    rna-XM_007081603.3    rna-XM_040632492.1    rna-XM_027607359.1    mNeoVis.3.1.1.chr12.g00585.t01, mNeoVis.3.1.1.chr12.g00586.t01    rna-gnl|WGS_ZZZZ|egapxtmp_029271-R1    mNeoVis.evergladensis.2.1.chr12.g00578.t01    mNeoVis.vulgivaga.2.1.chr12.g00595.t01      2752    N0.HOG0002750   OG0000540  n1   rna-XM_002914059.4    rna-NM_001168414.1    rna-XM_022507241.1    rna-XM_011286746.4    rna-XM_036117203.1    rna-XM_039253284.1    rna-XM_032881219.1    rna-XM_046016266.1    rna-XM_032359479.1    rna-XM_059138432.1    rna-XM_059391313.1    rna-XM_004752155.3    mNeoVis.2.1.1.chr7_hap1.g01889.t01    rna-XM_021685861.1    rna-XM_004409202.1    rna-XM_042958097.1    rna-XM_008686898.1    rna-XM_027577817.2    mNeoVis.3.1.1.chr7.g01758.t01    rna-gnl|WGS_ZZZZ|egapxtmp_011158-R1    mNeoVis.evergladensis.2.1.chr7.g01851.t01    mNeoVis.vulgivaga.2.1.chr7.g01868.t01      2753    N0.HOG0002751   OG0000540  n15   rna-XM_002917845.4    rna-XM_038525643.1    rna-XM_022523852.1    rna-XM_023245257.2    rna-XM_036068250.1    rna-XM_039250389.1    rna-XM_032839225.1    rna-XM_045986858.1    rna-XM_032323238.1    rna-XM_059153717.1    rna-XM_059382230.1    rna-XM_004767315.2    mNeoVis.2.1.1.chr7_hap1.g00897.t01    rna-XM_021681275.1    rna-XM_004409876.1    rna-XM_007074168.3    rna-XM_008684677.2    rna-XM_027618972.2    mNeoVis.3.1.1.chr7.g00838.t01    rna-gnl|WGS_ZZZZ|egapxtmp_013653-R1    mNeoVis.evergladensis.2.1.chr7.g00890.t01    mNeoVis.vulgivaga.2.1.chr7.g00894.t01      2754   N0.HOG0002752  OG0000540  n27   rna-XM_002928487.4    rna-XM_038558784.1    rna-XM_022495252.1    rna-XM_003989158.5    rna-XM_036086030.1    rna-XM_039245592.1    rna-XM_032844325.1    rna-XM_046010760.1    rna-XM_032346640.1    rna-XM_059183645.1    rna-XM_059404648.1    rna-XM_004742953.3    mNeoVis.2.1.1.chr9_hap1.g01041.t01    rna-XM_021687550.2    rna-XM_004403859.1    rna-XM_007085659.3    rna-XM_008705721.2    rna-XM_027595599.2    mNeoVis.3.1.1.chr10.g01001.t01    rna-gnl|WGS_ZZZZ|egapxtmp_021735-R1    mNeoVis.evergladensis.2.1.chr10.g01030.t01    mNeoVis.vulgivaga.2.1.chr10.g01021.t01      2755   N0.HOG0002753  OG0000541  n1   rna-XM_034669782.1    rna-XM_038539488.1    rna-XM_022500976.1    rna-XM_011290425.4    rna-XM_036103001.1    rna-XM_039226088.1    rna-XM_032872732.1    rna-XM_045993574.1    rna-XM_032328339.1    rna-XM_059154721.1    rna-XM_059414382.1    rna-XM_045075130.1    mNeoVis.2.1.1.chr15_hap1.g00615.t01    rna-XM_021680241.1    rna-XM_004394158.1    rna-XM_042971332.1    rna-XM_040634763.1    rna-XM_027611397.2    mNeoVis.3.1.1.chr15.g00398.t01    rna-gnl|WGS_ZZZZ|egapxtmp_034223-R1    mNeoVis.evergladensis.2.1.chr15.g00437.t01    mNeoVis.vulgivaga.2.1.chr15.g00445.t01      2756    N0.HOG0002754   OG0000541  n5   rna-XM_002917875.4, rna-XM_034643685.1    rna-XM_038528376.1    rna-XM_022519277.1, rna-XM_022524138.1    rna-XM_003997697.6, rna-XM_019835585.3    rna-XM_036099615.1, rna-XM_036102254.1    rna-XM_039234239.1, rna-XM_039246125.1    rna-XM_032836681.1, rna-XM_032839018.1    rna-XM_045989223.1, rna-XM_046014060.1    rna-XM_032325101.1, rna-XM_032346277.1    rna-XM_059152230.1, rna-XM_059186982.1    rna-XM_059381394.1, rna-XM_059402002.1    rna-XM_004767436.3, rna-XM_045083863.1    mNeoVis.2.1.1.chr7_hap1.g00946.t01, mNeoVis.2.1.1.chr9_hap1.g01164.t01    rna-XM_021681225.1, rna-XM_021687421.1    rna-XM_004407140.1, rna-XM_004410769.1, rna-XM_004415116.1    rna-NM_001320169.1, rna-XM_042969777.1    rna-XM_040634252.1, rna-XM_040637540.1    rna-XM_027581851.1, rna-XM_027595032.2, rna-XM_027619838.2, rna-XM_035726361.1    mNeoVis.3.1.1.chr10.g01117.t01, mNeoVis.3.1.1.chr7.g00886.t01      mNeoVis.evergladensis.2.1.chr10.g01152.t01, mNeoVis.evergladensis.2.1.chr7.g00939.t01    mNeoVis.vulgivaga.2.1.chr10.g01144.t01, mNeoVis.vulgivaga.2.1.chr7.g00945.t01      2757    N0.HOG0002755   OG0000542  n1   rna-XM_002924207.4    rna-XM_038539453.1    rna-XM_022501365.1    rna-XM_019820247.3    rna-XM_036064891.1    rna-XM_039225965.1    rna-XM_032872680.1    rna-XM_045994349.1    rna-XM_032328329.1    rna-XM_059154973.1    rna-XM_059415919.1    rna-XM_004775031.2    mNeoVis.2.1.1.chr15_hap1.g00634.t01    rna-XM_021680109.1    rna-XM_004394178.1    rna-XM_042972072.1    rna-XM_040634522.1    rna-XM_027614194.2    mNeoVis.3.1.1.chr15.g00415.t01    rna-gnl|WGS_ZZZZ|egapxtmp_033304-R1    mNeoVis.evergladensis.2.1.chr15.g00456.t01    mNeoVis.vulgivaga.2.1.chr15.g00463.t01      2758    N0.HOG0002756   OG0000542  n20   rna-XM_034641546.1    rna-XM_038546357.1    rna-XM_022523576.1, rna-XM_022523577.1    rna-XM_023244271.2    rna-XM_036067403.1    rna-XM_039226527.1    rna-XM_032870428.1    rna-XM_045984031.1    rna-XM_032321258.1    rna-XM_059148849.1    rna-XM_059378619.1    rna-XM_045085739.1    mNeoVis.2.1.1.chr8_hap1.g00072.t01    rna-XM_044921732.1    rna-XM_004404507.1    rna-XM_042967190.1    rna-XM_040639852.1    rna-XM_027568024.2    mNeoVis.3.1.1.chr8.g00048.t01      mNeoVis.evergladensis.2.1.chr9.g00069.t01    mNeoVis.vulgivaga.2.1.chr8.g00069.t01      2759   N0.HOG0002757  OG0000542  n39   rna-XM_019794475.2    rna-XM_038551569.1    rna-XM_022518889.1    rna-XM_045062382.1    rna-XM_036078617.1    rna-XM_039246091.1    rna-XM_032836694.1    rna-XM_046011597.1    rna-XM_032346254.1    rna-XM_059187003.1    rna-XM_059404442.1    rna-XM_013051636.2    mNeoVis.2.1.1.chr9_hap1.g01174.t01    rna-XM_021687719.2    rna-XM_004410815.1    rna-XM_042993719.1    rna-XM_040637087.1    rna-XM_027594327.2    mNeoVis.3.1.1.chr10.g01128.t01    rna-gnl|WGS_ZZZZ|egapxtmp_024827-R1    mNeoVis.evergladensis.2.1.chr10.g01163.t01    mNeoVis.vulgivaga.2.1.chr10.g01154.t01      2760    N0.HOG0002758   OG0000543  n1   rna-XM_034643687.1    rna-XM_038551567.1    rna-XM_022519008.1    rna-XM_045062392.1    rna-XM_036078583.1    rna-XM_039236135.1    rna-XM_032882957.1    rna-XM_046012718.1    rna-XM_032346215.1    rna-XM_059187032.1    rna-XM_059401965.1    rna-XM_045083830.1    mNeoVis.2.1.1.chr9_hap1.g01190.t01    rna-XM_044915784.1    rna-XM_004397467.1    rna-XM_042993317.1    rna-XM_040637569.1    rna-XM_035721872.1    mNeoVis.3.1.1.chr10.g01140.t01    rna-gnl|WGS_ZZZZ|egapxtmp_024834-R1    mNeoVis.evergladensis.2.1.chr10.g01178.t01    mNeoVis.vulgivaga.2.1.chr10.g01166.t01      2761    N0.HOG0002759   OG0000543  n23   rna-XM_034669596.1    rna-XM_038540857.1    rna-XM_022516542.1    rna-XM_023246207.2    rna-XM_036083381.1    rna-XM_039246746.1    rna-XM_032846005.1    rna-XM_045993374.1    rna-XM_032327700.1    rna-XM_059154536.1    rna-XM_059415754.1    rna-XM_045071962.1    mNeoVis.2.1.1.chr15_hap1.g00476.t01    rna-XM_044915568.1    rna-XM_004403111.1    rna-XM_042971567.1    rna-XM_040635586.1    rna-XM_027612986.2    mNeoVis.3.1.1.chr15.g00323.t01    rna-gnl|WGS_ZZZZ|egapxtmp_034112-R1    mNeoVis.evergladensis.2.1.chr15.g00343.t01    mNeoVis.vulgivaga.2.1.chr15.g00351.t01      2762    N0.HOG0002760   OG0000543  n44   rna-XM_034640751.1    rna-XM_038547659.1    rna-XM_022509041.1    rna-XM_023244165.2    rna-XM_036076266.1    rna-XM_039227831.1    rna-XM_032872067.1    rna-XM_045985211.1    rna-XM_032320462.1    rna-XM_059148513.1    rna-XM_059379235.1    rna-XM_045070093.1    mNeoVis.2.1.1.chr8_hap1.g00584.t01    rna-XM_021703916.2    rna-XM_004395166.1    rna-XM_042966370.1    rna-XM_040640834.1    rna-XM_027625233.2    mNeoVis.3.1.1.chr8.g00539.t01    rna-gnl|WGS_ZZZZ|egapxtmp_017441-R1    mNeoVis.evergladensis.2.1.chr9.g00574.t01    mNeoVis.vulgivaga.2.1.chr8.g00582.t01      2763    N0.HOG0002761   OG0000544  n1   rna-XM_034646834.1    rna-XM_038531559.1    rna-XM_022514308.1    rna-XM_023258316.2    rna-XM_036089049.1    rna-XM_039230253.1    rna-XM_032863792.1    rna-XM_046016696.1    rna-XM_032361785.1    rna-XM_059135863.1    rna-XM_059376350.1    rna-XM_045063977.1    mNeoVis.2.1.1.chr2_hap1.g01665.t01    rna-XM_044913905.1    rna-XM_004394738.1    rna-XM_042995302.1    rna-XM_040619004.1    rna-XM_027569163.2    mNeoVis.3.1.1.chr5.g01588.t01    rna-gnl|WGS_ZZZZ|egapxtmp_005568-R1    mNeoVis.evergladensis.2.1.chr5.g01606.t01    mNeoVis.vulgivaga.2.1.chr5.g01608.t01      2764   N0.HOG0002762  OG0000544  n23   rna-XM_034655847.1    rna-XM_038581435.1    rna-XM_022516231.1    rna-XM_023238765.2    rna-XM_036101888.1    rna-XM_039228411.1    rna-XM_032865370.1    rna-XM_046002999.1    rna-XM_032354396.1    rna-XM_059164670.1    rna-XM_059392308.1    rna-XM_004757540.3    mNeoVis.2.1.1.chr5_hap1.g01816.t01    rna-XM_044919441.1    rna-XM_004409619.1    rna-XM_042956909.1    rna-XM_040620989.1    rna-XM_027583257.2    mNeoVis.3.1.1.chr2.g01722.t01    rna-gnl|WGS_ZZZZ|egapxtmp_037711-R2    mNeoVis.evergladensis.2.1.chr2.g01802.t01    mNeoVis.vulgivaga.2.1.chr2.g01816.t01      2765   N0.HOG0002763  OG0000544  n38   rna-XM_034648166.1    rna-XM_038554213.1    rna-XM_022503664.1    rna-XM_045058566.1    rna-XM_036115291.1    rna-XM_039231165.1    rna-XM_032862021.1    rna-XM_046005129.1    rna-XM_032340139.1    rna-XM_059178130.1    rna-XM_059400338.1    rna-XM_045071039.1    mNeoVis.2.1.1.chr1_hap1.g01596.t01    rna-XM_021691894.2    rna-XM_004408428.1    rna-XM_042986479.1    rna-XM_040643544.1    rna-XM_027601296.2    mNeoVis.3.1.1.chr1.g01546.t01    rna-gnl|WGS_ZZZZ|egapxtmp_012085-R1    mNeoVis.evergladensis.2.1.chr1.g01563.t01    mNeoVis.vulgivaga.2.1.chr1.g01560.t01      2766    N0.HOG0002764   OG0000545  n1   rna-XM_002915544.4    rna-NM_001197052.1    rna-XM_022494473.1    rna-XM_011280760.4    rna-XM_036109668.1    rna-XM_039236906.1    rna-XM_032878918.1    rna-XM_045981850.1    rna-XM_032350267.1    rna-XM_059133183.1    rna-XM_059407281.1    rna-XM_045070638.1, rna-XM_045082044.1    mNeoVis.2.1.1.chr11_hap1.g00096.t01    rna-XM_021677894.1    rna-XM_004416538.1    rna-XM_042980100.1    rna-XM_040638496.1    rna-XM_027624209.2    mNeoVis.3.1.1.chr11.g00068.t01    rna-gnl|WGS_ZZZZ|egapxtmp_027395-R1    mNeoVis.evergladensis.2.1.chr11.g00094.t01    mNeoVis.vulgivaga.2.1.chr11.g00094.t01      2767    N0.HOG0002765   OG0000545  n18   rna-XM_002918922.4    rna-XM_038554362.1    rna-XM_022510845.1    rna-XM_003986274.5    rna-XM_036120597.1    rna-XM_039244297.1    rna-XM_032864502.1    rna-XM_046004088.1    rna-XM_032340298.1    rna-XM_059178289.1    rna-XM_059400224.1    rna-XM_004772404.3    mNeoVis.2.1.1.chr1_hap1.g01530.t01    rna-XM_021692027.2    rna-XM_004402457.1    rna-XM_007095542.3    rna-XM_008691760.1    rna-XM_035728572.1    mNeoVis.3.1.1.chr1.g01482.t01    rna-gnl|WGS_ZZZZ|egapxtmp_019658-R1    mNeoVis.evergladensis.2.1.chr1.g01498.t01    mNeoVis.vulgivaga.2.1.chr1.g01492.t01      2768    N0.HOG0002766   OG0000545  n36     rna-XM_038584127.1    rna-XM_022517690.1    rna-XM_023253693.2    rna-XM_036120641.1    rna-XM_039254415.1    rna-XM_032861532.1    rna-XM_046004657.1    rna-XM_032341059.1    rna-XM_059179214.1    rna-XM_059399404.1    rna-XM_004753751.3    mNeoVis.2.1.1.chr1_hap1.g01240.t01    rna-XM_021697031.1    rna-XM_004405638.1    rna-XM_042985756.1    rna-XM_040645033.1    rna-XM_027601685.1    mNeoVis.3.1.1.chr1.g01198.t01    rna-gnl|WGS_ZZZZ|egapxtmp_004173-R1    mNeoVis.evergladensis.2.1.chr1.g01213.t01    mNeoVis.vulgivaga.2.1.chr1.g01216.t01      2769   N0.HOG0002767  OG0000546  n1   rna-XM_034648304.1    rna-XM_038531267.1    rna-XM_022497132.1    rna-XM_045035471.1    rna-XM_036081840.1    rna-XM_039230723.1    rna-XM_032877050.1    rna-XM_046009481.1    rna-XM_032361269.1    rna-XM_059135551.1    rna-XM_059376995.1    rna-XM_045064104.1    mNeoVis.2.1.1.chr2_hap1.g01565.t01    rna-XM_021684180.2    rna-XM_004406401.1    rna-XM_042996272.1    rna-XM_040619171.1    rna-XM_027607456.2    mNeoVis.3.1.1.chr5.g01486.t01    rna-gnl|WGS_ZZZZ|egapxtmp_009104-R1    mNeoVis.evergladensis.2.1.chr5.g01503.t01    mNeoVis.vulgivaga.2.1.chr5.g01507.t01      2770    N0.HOG0002768   OG0000546  n7   rna-XM_034667594.1                                  rna-XM_008704423.2                2771    N0.HOG0002769   OG0000546  n10     rna-XM_038554427.1      rna-XM_045057702.1              rna-XM_059178352.1    rna-XM_059400188.1            rna-XM_042986608.1                  2772   N0.HOG0002770  OG0000546  n14     rna-XM_038568366.1                        mNeoVis.2.1.1.chr1_hap1.g01514.t01                rna-gnl|WGS_ZZZZ|egapxtmp_039247-R3          2773   N0.HOG0002771  OG0000546  n16       rna-XM_022514165.1      rna-XM_036082148.1      rna-XM_032853572.1    rna-XM_046017218.1    rna-XM_032360033.1    rna-XM_059187156.1    rna-XM_059391562.1    rna-XM_045071471.1    mNeoVis.2.1.1.chr7_hap1.g01468.t01    rna-XM_021686465.1    rna-XM_004417231.1    rna-XM_042959287.1        mNeoVis.3.1.1.chr7.g01374.t01    rna-gnl|WGS_ZZZZ|egapxtmp_010998-R1    mNeoVis.evergladensis.2.1.chr7.g01449.t01    mNeoVis.vulgivaga.2.1.chr7.g01462.t01      2774   N0.HOG0002772  OG0000546  n29   rna-XM_019806668.2    rna-XM_038576970.1    rna-XM_022510721.1      rna-XM_036102743.1    rna-XM_039223082.1    rna-XM_032864454.1    rna-XM_046005548.1    rna-XM_032340364.1        rna-XM_004775650.3    mNeoVis.2.1.1.chr8_hap1.g00674.t01    rna-XM_021691953.2    rna-XM_004417604.1      rna-XM_040643599.1    rna-XM_027601766.1    mNeoVis.3.1.1.chr8.g00623.t01      mNeoVis.evergladensis.2.1.chr1.g01480.t01    mNeoVis.vulgivaga.2.1.chr1.g01475.t01      2775   N0.HOG0002773  OG0000547  n1   rna-XM_034648079.1    rna-XM_038554438.1    rna-XM_022510749.1    rna-XM_011282404.4    rna-XM_036072668.1    rna-XM_039245799.1    rna-XM_032864447.1    rna-XM_046006639.1    rna-XM_032340373.1    rna-XM_059178363.1    rna-XM_059400179.1    rna-XM_045069381.1    mNeoVis.2.1.1.chr1_hap1.g01508.t01    rna-XM_021691966.2    rna-XM_004399970.1    rna-XM_042986617.1    rna-XM_040643606.1    rna-XM_027603095.2    mNeoVis.3.1.1.chr1.g01452.t01    rna-gnl|WGS_ZZZZ|egapxtmp_015188-R2    mNeoVis.evergladensis.2.1.chr1.g01472.t01    mNeoVis.vulgivaga.2.1.chr1.g01470.t01      2776   N0.HOG0002774  OG0000547  n22   rna-XM_034668700.1, rna-XM_034668701.1    rna-XM_038555664.1    rna-XM_022502194.1    rna-XM_045049947.1    rna-XM_036112774.1    rna-XM_039232996.1    rna-XM_032873457.1    rna-XM_046014090.1    rna-XM_032316350.1    rna-XM_059168689.1    rna-XM_059395241.1    rna-XM_045078201.1    mNeoVis.2.1.1.chr4_hap1.g01052.t01    rna-XM_044914309.1    rna-XM_004405579.1    rna-XM_042973196.1    rna-XM_040625641.1    rna-XM_027573057.2      rna-gnl|WGS_ZZZZ|egapxtmp_004587-R1    mNeoVis.evergladensis.2.1.chr6.g01057.t01    mNeoVis.vulgivaga.2.1.chr6.g01050.t01      2777   N0.HOG0002775  OG0000547  n43   rna-XM_034648282.1    rna-XM_038531286.1    rna-XM_022496376.1    rna-XM_006934526.5    rna-XM_036098634.1    rna-XM_039232784.1    rna-XM_032850541.1, rna-XM_032850543.1    rna-XM_046021676.1    rna-XM_032302563.1    rna-XM_059136986.1    rna-XM_059376981.1    rna-XM_004740940.3    mNeoVis.2.1.1.chr2_hap1.g01567.t01    rna-XM_044913942.1    rna-XM_004406400.1    rna-XM_042996738.1    rna-XM_040619384.1    rna-XM_027582017.1        mNeoVis.evergladensis.2.1.chr5.g01506.t01    mNeoVis.vulgivaga.2.1.chr5.g01509.t01      2778   N0.HOG0002776  OG0000548  n1   rna-XM_019807682.2    rna-XM_038543008.1    rna-XM_022491067.1    rna-XM_006943075.5    rna-XM_036118241.1    rna-XM_039251396.1    rna-XM_032845107.1    rna-XM_045982426.1    rna-XM_032318067.1    rna-XM_059146102.1    rna-XM_059413656.1    rna-XM_045086730.1    mNeoVis.2.1.1.chr14_hap1.g00616.t01    rna-XM_044916268.1    rna-XM_004403047.1    rna-XM_042976368.1    rna-XM_040635812.1    rna-XM_027614208.2    mNeoVis.3.1.1.chr14.g00597.t01    rna-gnl|WGS_ZZZZ|egapxtmp_032823-R1    mNeoVis.evergladensis.2.1.chr14.g00613.t01    mNeoVis.vulgivaga.2.1.chr14.g00609.t01      2779   N0.HOG0002777  OG0000548  n23   rna-XM_034656385.1    rna-XM_038532512.1    rna-XM_022504674.1    rna-XM_011284171.4    rna-XM_036072249.1    rna-XM_039245540.1    rna-XM_032848125.1    rna-XM_045999265.1    rna-XM_032335919.1    rna-XM_059175277.1    rna-XM_059418098.1    rna-XM_004766846.3    mNeoVis.2.1.1.chr1_hap1.g00675.t01    rna-XM_044916696.1    rna-XM_004413394.1    rna-XM_042988703.1    rna-XM_040622170.1    rna-XM_027604836.2    mNeoVis.3.1.1.chr1.g00695.t01    rna-gnl|WGS_ZZZZ|egapxtmp_021521-R3    mNeoVis.evergladensis.2.1.chr1.g00683.t01    mNeoVis.vulgivaga.2.1.chr1.g00695.t01      2780   N0.HOG0002778  OG0000548  n44   rna-XM_019800245.2    rna-XM_038532895.1    rna-XM_022511279.1    rna-XM_023254889.2    rna-XM_036082898.1    rna-XM_039256899.1    rna-XM_032836822.1    rna-XM_046007996.1    rna-XM_032342036.1    rna-XM_059179946.1    rna-XM_059371530.1    rna-XM_045067235.1    mNeoVis.2.1.1.chr10_hap1.g01229.t01    rna-XM_021680679.1    rna-XM_004398272.1    rna-XM_042988003.1    rna-XM_040642830.1    rna-XM_027569682.2    mNeoVis.3.1.1.chr9.g01158.t01    rna-gnl|WGS_ZZZZ|egapxtmp_014414-R1    mNeoVis.evergladensis.2.1.chr8.g01205.t01    mNeoVis.vulgivaga.2.1.chr9.g01190.t01      2781   N0.HOG0002779  OG0000549  n1   rna-XM_034667925.1    rna-XM_038532717.1    rna-XM_022511183.1    rna-XM_023254966.2    rna-XM_036077486.1    rna-XM_039235040.1    rna-XM_032836038.1    rna-XM_046008028.1    rna-XM_032342166.1    rna-XM_059180118.1    rna-XM_059371653.1    rna-XM_004759809.2    mNeoVis.2.1.1.chr10_hap1.g01159.t01    rna-XM_021680652.2    rna-XM_004405547.1    rna-XM_042988145.1    rna-XM_040642718.1    rna-XM_035727932.1    mNeoVis.3.1.1.chr9.g01093.t01    rna-gnl|WGS_ZZZZ|egapxtmp_017028-R1    mNeoVis.evergladensis.2.1.chr8.g01138.t01    mNeoVis.vulgivaga.2.1.chr9.g01118.t01      2782    N0.HOG0002780   OG0000549  n23   rna-XM_034667412.1    rna-XM_038542974.1    rna-XM_022490804.1    rna-XM_023247539.2    rna-XM_036118311.1    rna-XM_039251304.1    rna-XM_032845066.1    rna-XM_045983483.1    rna-XM_032318075.1    rna-XM_059146376.1    rna-XM_059413620.1    rna-XM_045086711.1    mNeoVis.2.1.1.chr14_hap1.g00602.t01    rna-XM_021681951.2, rna-XM_044912233.1    rna-XM_004402888.1    rna-XM_042976292.1    rna-XM_040635833.1    rna-XM_027612046.1    mNeoVis.3.1.1.chr14.g00583.t01      mNeoVis.evergladensis.2.1.chr14.g00599.t01    mNeoVis.vulgivaga.2.1.chr14.g00595.t01      2783    N0.HOG0002781   OG0000549  n44   rna-XM_034657716.1    rna-XM_038567406.1    rna-XM_022525765.1    rna-XM_023244792.2    rna-XM_036096481.1    rna-XM_039251635.1    rna-XM_032883877.1    rna-XM_045989626.1    rna-XM_032309594.1    rna-XM_059159852.1    rna-XM_059388898.1    rna-XM_045084533.1    mNeoVis.2.1.1.chr5_hap1.g00229.t01    rna-XM_044915489.1    rna-XM_004412177.1    rna-XM_042965366.1    rna-XM_040629381.1    rna-XM_027582996.2    mNeoVis.3.1.1.chr2.g00175.t01    rna-gnl|WGS_ZZZZ|egapxtmp_034642-R1    mNeoVis.evergladensis.2.1.chr2.g00225.t01    mNeoVis.vulgivaga.2.1.chr2.g00225.t01      2784   N0.HOG0002782  OG0000550  n1   rna-XM_034667987.1    rna-XM_038532572.1    rna-XM_022525999.1    rna-XM_019831997.3    rna-XM_036101822.1    rna-XM_039218219.1    rna-XM_032843107.1    rna-XM_046006975.1    rna-XM_032342305.1    rna-XM_059180244.1    rna-XM_059371735.1    rna-XM_045070423.1    mNeoVis.2.1.1.chr10_hap1.g01118.t01    rna-XM_044917764.1    rna-XM_004404723.1    rna-XM_042988303.1    rna-XM_040642653.1    rna-XM_027571970.1    mNeoVis.3.1.1.chr9.g01046.t01    rna-gnl|WGS_ZZZZ|egapxtmp_014402-R2    mNeoVis.evergladensis.2.1.chr8.g01094.t01    mNeoVis.vulgivaga.2.1.chr9.g01073.t01      2785   N0.HOG0002783  OG0000550  n17   rna-XM_034656768.1    rna-XM_038534835.1    rna-XM_022493342.1    rna-XM_003981341.6    rna-XM_036081035.1    rna-XM_039219360.1    rna-XM_032851734.1    rna-XM_046001222.1    rna-XM_032337107.1    rna-XM_059174229.1    rna-XM_059417306.1    rna-XM_004737670.3    mNeoVis.2.1.1.chr1_hap1.g00275.t01    rna-XM_021698030.1    rna-XM_004392043.1    rna-XM_042992923.1    rna-XM_040630323.1    rna-XM_027607054.2    mNeoVis.3.1.1.chr1.g00278.t01    rna-gnl|WGS_ZZZZ|egapxtmp_002104-R1    mNeoVis.evergladensis.2.1.chr1.g00273.t01    mNeoVis.vulgivaga.2.1.chr1.g00279.t01      2786    N0.HOG0002784   OG0000550  n28   rna-XM_002921754.4    rna-XM_038555986.1    rna-XM_022506133.1    rna-XM_003985410.6    rna-XM_036072268.1    rna-XM_039218182.1    rna-XM_032852471.1    rna-XM_045998096.1    rna-XM_032334211.1    rna-XM_059162296.1    rna-XM_059383365.1    rna-XM_004764906.3    mNeoVis.2.1.1.chr6_hap1.g00201.t01    rna-XM_021701659.1    rna-XM_004417060.1    rna-XM_007075917.3    rna-XM_008701439.1    rna-XM_027600580.2    mNeoVis.3.1.1.chr4.g00196.t01    rna-gnl|WGS_ZZZZ|egapxtmp_000397-R3    mNeoVis.evergladensis.2.1.chr4.g00206.t01    mNeoVis.vulgivaga.2.1.chr4.g00201.t01      2787    N0.HOG0002785   OG0000551  n1   rna-XM_011235510.3    rna-XM_038558464.1    rna-XM_022511972.1    rna-XM_011283119.4    rna-XM_036117922.1    rna-XM_039245155.1    rna-XM_032857021.1    rna-XM_046008156.1    rna-XM_032343041.1    rna-XM_059180949.1    rna-XM_059372384.1    rna-XM_013059458.2    mNeoVis.2.1.1.chr10_hap1.g00898.t01    rna-XM_021695777.2    rna-XM_004402122.1    rna-XM_007091336.3    rna-XM_040620676.1    rna-XM_027572808.1    mNeoVis.3.1.1.chr9.g00826.t01    rna-gnl|WGS_ZZZZ|egapxtmp_015539-R1    mNeoVis.evergladensis.2.1.chr8.g00868.t01    mNeoVis.vulgivaga.2.1.chr9.g00855.t01      2788    N0.HOG0002786   OG0000551  n23   rna-XM_011220466.3    rna-XM_038531048.1    rna-XM_022518623.1    rna-XM_045046396.1    rna-XM_036081670.1    rna-XM_039241337.1    rna-XM_032854469.1    rna-XM_045987027.1    rna-XM_032325864.1    rna-XM_059152717.1    rna-XM_059381331.1    rna-XM_045076062.1    mNeoVis.2.1.1.chr7_hap1.g00330.t01    rna-XM_044922304.1    rna-XM_004395829.1    rna-XM_042970467.1    rna-XM_040633303.1    rna-XM_027617213.2    mNeoVis.3.1.1.chr7.g00317.t01    rna-gnl|WGS_ZZZZ|egapxtmp_009886-R1    mNeoVis.evergladensis.2.1.chr7.g00331.t01    mNeoVis.vulgivaga.2.1.chr7.g00334.t01      2789   N0.HOG0002787  OG0000551  n44   rna-XM_011220367.3    rna-XM_038579376.1    rna-XM_022515513.1    rna-XM_045050025.1    rna-XM_036099459.1    rna-XM_039249132.1    rna-XM_032882612.1    rna-XM_046003377.1    rna-XM_032316781.1    rna-XM_059166372.1    rna-XM_059394627.1    rna-XM_045075751.1    mNeoVis.2.1.1.chr4_hap1.g00733.t01    rna-XM_021688597.2    rna-XM_004398778.1    rna-XM_042972706.1    rna-XM_040626012.1    rna-XM_027597269.2    mNeoVis.3.1.1.chr6.g00742.t01    rna-gnl|WGS_ZZZZ|egapxtmp_003496-R1    mNeoVis.evergladensis.2.1.chr6.g00742.t01    mNeoVis.vulgivaga.2.1.chr6.g00738.t01      2790    N0.HOG0002788   OG0000552  n1   rna-XM_034668123.1    rna-XM_038579389.1    rna-XM_022515577.1    rna-XM_011291374.4    rna-XM_036088919.1    rna-XM_039254598.1    rna-XM_032882634.1    rna-XM_046011138.1    rna-XM_032315724.1    rna-XM_059166352.1    rna-XM_059394647.1    rna-XM_004772946.2    mNeoVis.2.1.1.chr4_hap1.g00740.t01    rna-XM_021688606.2    rna-XM_004398784.1    rna-XM_042972691.1    rna-XM_040625658.1    rna-XM_035727122.1    mNeoVis.3.1.1.chr6.g00749.t01    rna-gnl|WGS_ZZZZ|egapxtmp_004102-R1    mNeoVis.evergladensis.2.1.chr6.g00749.t01    mNeoVis.vulgivaga.2.1.chr6.g00745.t01      2791    N0.HOG0002789   OG0000552  n15   rna-XM_034647895.1    rna-XM_038531353.1    rna-XM_022497113.1    rna-XM_023258384.2    rna-XM_036123445.1    rna-XM_039232729.1    rna-XM_032863952.1    rna-XM_046022301.1    rna-XM_032301530.1    rna-XM_059136214.1    rna-XM_059376888.1    rna-XM_004741004.3    mNeoVis.2.1.1.chr2_hap1.g01589.t01    rna-XM_044913931.1    rna-XM_004394641.1    rna-XM_042996017.1    rna-XM_040619136.1    rna-XM_027623097.2    mNeoVis.3.1.1.chr5.g01510.t01    rna-gnl|WGS_ZZZZ|egapxtmp_009582-R1    mNeoVis.evergladensis.2.1.chr5.g01528.t01    mNeoVis.vulgivaga.2.1.chr5.g01531.t01      2792    N0.HOG0002790   OG0000552  n27   rna-XM_034640155.1    rna-XM_038572828.1    rna-XM_022494426.1    rna-XM_023251009.2    rna-XM_036084699.1    rna-XM_039237046.1    rna-XM_032878812.1    rna-XM_045981038.1    rna-XM_032353542.1    rna-XM_059132846.1    rna-XM_059407388.1    rna-XM_045070488.1    mNeoVis.2.1.1.chr11_hap1.g00040.t01    rna-XM_044918638.1    rna-XM_004410129.1    rna-XM_042979953.1    rna-XM_040638587.1    rna-XM_027622919.2    mNeoVis.3.1.1.chr11.g00020.t01    rna-gnl|WGS_ZZZZ|egapxtmp_027220-R1    mNeoVis.evergladensis.2.1.chr11.g00041.t01    mNeoVis.vulgivaga.2.1.chr11.g00039.t01      2793    N0.HOG0002791   OG0000553  n1   rna-XM_034641397.1    rna-XM_038572509.1    rna-XM_022494259.1    rna-XM_023251154.2    rna-XM_036069693.1    rna-XM_039249361.1    rna-XM_032838579.1    rna-XM_045981144.1    rna-XM_032350455.1    rna-XM_059132949.1    rna-XM_059407158.1    rna-XM_045082132.1    mNeoVis.2.1.1.chr11_hap1.g00153.t01    rna-XM_021689061.1    rna-XM_004405272.1    rna-XM_042980247.1    rna-XM_040638424.1    rna-XM_027622030.2    mNeoVis.3.1.1.chr11.g00125.t01    rna-gnl|WGS_ZZZZ|egapxtmp_026798-R1    mNeoVis.evergladensis.2.1.chr11.g00150.t01    mNeoVis.vulgivaga.2.1.chr11.g00152.t01      2794   N0.HOG0002792  OG0000553  n23   rna-XM_034668177.1    rna-XM_038579501.1    rna-XM_022517868.1    rna-XM_045050234.1    rna-XM_036109411.1    rna-XM_039238391.1    rna-XM_032869202.1    rna-XM_046013512.1    rna-XM_032316539.1    rna-XM_059166236.1    rna-XM_059394705.1    rna-XM_013061397.2    mNeoVis.2.1.1.chr4_hap1.g00780.t01    rna-XM_021688168.2    rna-XM_004408705.1    rna-XM_007084782.3    rna-XM_008710789.2    rna-XM_027582479.1    mNeoVis.3.1.1.chr6.g00790.t01    rna-gnl|WGS_ZZZZ|egapxtmp_003964-R2    mNeoVis.evergladensis.2.1.chr6.g00785.t01    mNeoVis.vulgivaga.2.1.chr6.g00779.t01      2795   N0.HOG0002793  OG0000553  n44   rna-XM_011228766.3    rna-XM_038526049.1    rna-XM_022503825.1    rna-XM_006932029.5    rna-XM_036097415.1    rna-XM_039242044.1    rna-XM_032843485.1    rna-XM_046006790.1    rna-XM_032339472.1    rna-XM_059177515.1    rna-XM_059401572.1    rna-XM_045066338.1    mNeoVis.2.1.1.chr1_hap1.g01809.t01    rna-XM_021693330.1    rna-XM_004401201.1    rna-XM_042987169.1    rna-XM_040627489.1    rna-XM_027603546.2    mNeoVis.3.1.1.chr1.g01762.t01    rna-gnl|WGS_ZZZZ|egapxtmp_016258-R8    mNeoVis.evergladensis.2.1.chr1.g01773.t01    mNeoVis.vulgivaga.2.1.chr1.g01768.t01      2796    N0.HOG0002794   OG0000554  n1   rna-XM_011231299.3    rna-XM_038533670.1    rna-XM_022505894.1, rna-XM_022505915.1    rna-NM_001009874.3    rna-XM_036094317.1    rna-XM_039220373.1    rna-XM_032853519.1    rna-XM_045997397.1    rna-XM_032329817.1    rna-XM_059155382.1    rna-XM_059371348.1    rna-XM_004772382.3    mNeoVis.2.1.1.chr6_hap1.g00396.t01    rna-XM_044913066.1    rna-XM_004396238.1    rna-XM_042984806.1    rna-XM_040625091.1    rna-XM_027598861.1    mNeoVis.3.1.1.chr4.g00399.t01    rna-gnl|WGS_ZZZZ|egapxtmp_000749-R1    mNeoVis.evergladensis.2.1.chr4.g00404.t01    mNeoVis.vulgivaga.2.1.chr4.g00405.t01      2797    N0.HOG0002795   OG0000554  n22   rna-XM_019800430.1    rna-NM_001003069.1    rna-XM_022517080.1    rna-XM_003994663.6    rna-XM_036080593.1    rna-XM_039255309.1    rna-XM_032876296.1    rna-XM_046025229.1    rna-XM_032311036.1    rna-XM_059140418.1    rna-XM_059409942.1    rna-XM_004753241.3    mNeoVis.2.1.1.chr3_hap1.g00585.t01    rna-XM_021698519.1    rna-XM_004395844.1    rna-XM_007079605.3    rna-XM_008688989.2    rna-XM_027576615.1    mNeoVis.3.1.1.chr3.g00577.t01    rna-gnl|WGS_ZZZZ|egapxtmp_040644-R1    mNeoVis.evergladensis.2.1.chr3.g00593.t01    mNeoVis.vulgivaga.2.1.chr3.g00596.t01      2798    N0.HOG0002796   OG0000554  n38   rna-XM_011233080.3    rna-XM_038538848.1    rna-XM_022501592.1    rna-XM_045047906.1    rna-XM_036075608.1    rna-XM_039243527.1    rna-XM_032872333.1    rna-XM_045993736.1    rna-XM_032328136.1    rna-XM_059154770.1    rna-XM_059415929.1    rna-XM_004761599.3    mNeoVis.2.1.1.chr15_hap1.g00703.t01    rna-XM_021700567.1    rna-XM_004399062.1    rna-XM_042971843.1    rna-XM_040634525.1    rna-XM_027610527.1    mNeoVis.3.1.1.chr15.g00484.t01    rna-gnl|WGS_ZZZZ|egapxtmp_034088-R1    mNeoVis.evergladensis.2.1.chr15.g00525.t01        2799   N0.HOG0002797  OG0000555  n1   rna-XM_034638122.1, rna-XM_034653314.1    rna-XM_038575365.1    rna-XM_022517139.1    rna-XM_023241539.2    rna-XM_036081011.1    rna-XM_039241180.1    rna-XM_032847429.1    rna-XM_046024464.1    rna-XM_032311177.1    rna-XM_059139007.1    rna-XM_059408674.1    rna-XM_004779152.3    mNeoVis.2.1.1.chr3_hap1.g00472.t01    rna-XM_021703447.1    rna-XM_004416192.1    rna-XM_007092348.2    rna-XM_040619778.1    rna-XM_027576245.1    mNeoVis.3.1.1.chr3.g00463.t01    rna-gnl|WGS_ZZZZ|egapxtmp_042668-R2    mNeoVis.evergladensis.2.1.chr3.g00479.t01    mNeoVis.vulgivaga.2.1.chr3.g00482.t01      2800    N0.HOG0002798   OG0000555  n23   rna-XM_034640815.1    rna-XM_038548364.1    rna-XM_022509665.1    rna-XM_006940036.5    rna-XM_036094648.1    rna-XM_039253418.1    rna-XM_032848433.1    rna-XM_045985760.1    rna-XM_032321383.1    rna-XM_059148684.1    rna-XM_059380834.1    rna-XM_045070112.1    mNeoVis.2.1.1.chr8_hap1.g00795.t01    rna-XM_021704306.1    rna-XM_004399692.1    rna-XM_042966625.1    rna-XM_040641083.1    rna-XM_027568040.2    mNeoVis.3.1.1.chr8.g00746.t01      mNeoVis.evergladensis.2.1.chr9.g00783.t01    mNeoVis.vulgivaga.2.1.chr8.g00786.t01      2801    N0.HOG0002799   OG0000555  n42   rna-XM_034646462.1    rna-XM_038563768.1    rna-XM_022507451.1    rna-XM_019812751.3    rna-XM_036067529.1    rna-XM_039235529.1    rna-XM_032879604.1    rna-XM_046017079.1    rna-XM_032358067.1    rna-XM_059148833.1    rna-XM_059400248.1    rna-XM_004759711.3    mNeoVis.2.1.1.chr7_hap1.g02499.t01    rna-XM_021685102.1    rna-XM_004393850.1    rna-XM_042959011.1    rna-XM_040639495.1    rna-XM_027578172.2    mNeoVis.3.1.1.chr7.g02243.t01    rna-gnl|WGS_ZZZZ|egapxtmp_011748-R1    mNeoVis.evergladensis.2.1.chr7.g02388.t01    mNeoVis.vulgivaga.2.1.chr7.g02407.t01      2802    N0.HOG0002800   OG0000556  n1   rna-XM_034638063.1    rna-XM_038527676.1    rna-XM_022492098.1    rna-XM_003997378.3      rna-XM_039244031.1    rna-XM_032851040.1    rna-XM_045986825.1    rna-XM_032326143.1    rna-XM_059151195.1      rna-XM_004777072.1    mNeoVis.2.1.1.chr7_hap1.g00659.t01    rna-XM_021682348.2    rna-XM_004410930.1    rna-XM_007080628.2    rna-XM_008706576.2    rna-XM_027619320.1    mNeoVis.3.1.1.chr7.g00622.t01    rna-gnl|WGS_ZZZZ|egapxtmp_011481-R1    mNeoVis.evergladensis.2.1.chr7.g00655.t01    mNeoVis.vulgivaga.2.1.chr7.g00658.t01      2803    N0.HOG0002801   OG0000556  n19   rna-XM_034639535.1    rna-XM_038527673.1    rna-XM_022492099.1    rna-XM_045046802.1      rna-XM_039244055.1    rna-XM_032851041.1    rna-XM_045987603.1    rna-XM_032324821.1    rna-XM_059151194.1      rna-XM_004777071.3    mNeoVis.2.1.1.chr7_hap1.g00658.t01    rna-XM_021682285.1    rna-XM_004410931.1    rna-XM_007080627.2      rna-XM_027619319.2    mNeoVis.3.1.1.chr7.g00621.t01    rna-gnl|WGS_ZZZZ|egapxtmp_011120-R1    mNeoVis.evergladensis.2.1.chr7.g00654.t01    mNeoVis.vulgivaga.2.1.chr7.g00657.t01      2804    N0.HOG0002802   OG0000556  n35   rna-XM_034639539.1    rna-NM_001389179.1, rna-XM_038527677.1      rna-XM_023246084.2      rna-XM_039244032.1      rna-XM_045987604.1, rna-XM_045987605.1    rna-XM_032324815.1, rna-XM_032324822.1    rna-XM_059151196.1, rna-XM_059152443.1    rna-XM_059387346.1, rna-XM_059387387.1    rna-XM_004776954.1, rna-XM_004777073.2    mNeoVis.2.1.1.chr7_hap1.g00657.t01, mNeoVis.2.1.1.chr7_hap1.g00660.t01        rna-XM_007080602.2    rna-XM_040633765.1, rna-XM_040633766.1    rna-XM_027617535.2, rna-XM_035724940.1    mNeoVis.3.1.1.chr7.g00623.t01    rna-gnl|WGS_ZZZZ|egapxtmp_011652-R1    mNeoVis.evergladensis.2.1.chr7.g00656.t01    mNeoVis.vulgivaga.2.1.chr7.g00656.t01, mNeoVis.vulgivaga.2.1.chr7.g00659.t01      2805    N0.HOG0002803   OG0000557  n1   rna-XM_002914832.4    rna-XM_038561275.1    rna-XM_022520358.1    rna-XM_023251955.2    rna-XM_036070403.1    rna-XM_039257218.1    rna-XM_032881908.1    rna-XM_045979509.1    rna-XM_032352957.1    rna-XM_059188586.1    rna-XM_059405456.1    rna-XM_045079231.1    mNeoVis.2.1.1.chr11_hap1.g00982.t01    rna-XM_021697416.2    rna-XM_004409985.1    rna-XM_042982152.1    rna-XM_040623103.1    rna-XM_027620980.2    mNeoVis.3.1.1.chr11.g00935.t01    rna-gnl|WGS_ZZZZ|egapxtmp_025808-R1    mNeoVis.evergladensis.2.1.chr11.g00961.t01    mNeoVis.vulgivaga.2.1.chr11.g00983.t01      2806    N0.HOG0002804   OG0000557  n22   rna-XM_034648652.1    rna-XM_038544379.1    rna-XM_022525090.1    rna-XM_023255617.2    rna-XM_036083611.1    rna-XM_039256378.1    rna-XM_032856215.1    rna-XM_046009920.1    rna-XM_032343992.1    rna-XM_059181820.1    rna-XM_059373029.1    rna-XM_004755158.3    mNeoVis.2.1.1.chr10_hap1.g00504.t01    rna-XM_021695290.1    rna-XM_004399945.1    rna-XM_042989494.1    rna-XM_040620799.1    rna-XM_027569852.2    mNeoVis.3.1.1.chr9.g00467.t01    rna-gnl|WGS_ZZZZ|egapxtmp_014848-R1    mNeoVis.evergladensis.2.1.chr8.g00484.t01    mNeoVis.vulgivaga.2.1.chr9.g00487.t01      2807    N0.HOG0002805   OG0000557  n41   rna-XM_034638992.1    rna-XM_038528489.1    rna-XM_022523805.1    rna-XM_011289800.4    rna-XM_036114004.1    rna-XM_039249822.1    rna-XM_032839399.1    rna-XM_045987915.1    rna-XM_032326165.1    rna-XM_059152949.1    rna-XM_059383741.1    rna-XM_004767517.3    mNeoVis.2.1.1.chr7_hap1.g00986.t01    rna-XM_021681349.2    rna-XM_004400943.1    rna-XM_042968821.1    rna-XM_040634203.1    rna-XM_027619137.2    mNeoVis.3.1.1.chr7.g00925.t01    rna-gnl|WGS_ZZZZ|egapxtmp_011593-R1    mNeoVis.evergladensis.2.1.chr7.g00979.t01    mNeoVis.vulgivaga.2.1.chr7.g00985.t01      2808    N0.HOG0002806   OG0000558  n1   rna-XM_011235551.3    rna-XM_038561347.1    rna-XM_022520570.1    rna-XM_019827633.3    rna-XM_036074889.1    rna-XM_039231681.1    rna-XM_032882015.1    rna-XM_045979105.1    rna-XM_032352852.1    rna-XM_059188469.1    rna-XM_059405537.1    rna-XM_004770918.3    mNeoVis.2.1.1.chr11_hap1.g00955.t01    rna-XM_021697806.2    rna-XM_004399825.1    rna-XM_042982093.1    rna-XM_008700190.2    rna-XM_027621109.2    mNeoVis.3.1.1.chr11.g00909.t01    rna-gnl|WGS_ZZZZ|egapxtmp_025332-R1    mNeoVis.evergladensis.2.1.chr11.g00933.t01    mNeoVis.vulgivaga.2.1.chr11.g00957.t01      2809    N0.HOG0002807   OG0000558  n23   rna-XM_019809255.2    rna-XM_038528561.1    rna-XM_022523995.1    rna-XM_023245803.2    rna-XM_036091494.1    rna-XM_039247800.1    rna-XM_032838971.1    rna-XM_045988644.1    rna-XM_032325302.1    rna-XM_059152478.1    rna-XM_059383693.1    rna-XM_004767575.3    mNeoVis.2.1.1.chr7_hap1.g01015.t01    rna-XM_044911446.1    rna-XM_004400967.1    rna-XM_042968741.1    rna-XM_040634170.1    rna-XM_027618713.2    mNeoVis.3.1.1.chr7.g00953.t01    rna-gnl|WGS_ZZZZ|egapxtmp_012364-R1    mNeoVis.evergladensis.2.1.chr7.g01008.t01    mNeoVis.vulgivaga.2.1.chr7.g01016.t01      2810    N0.HOG0002808   OG0000558  n44   rna-XM_034642088.1    rna-XM_038545703.1    rna-XM_022505152.1    rna-XM_023256205.2    rna-XM_036065379.1    rna-XM_039252959.1    rna-XM_032839534.1    rna-XM_046007633.1    rna-XM_032345244.1    rna-XM_059183089.1    rna-XM_059373956.1    rna-XM_045073589.1    mNeoVis.2.1.1.chr10_hap1.g00084.t01    rna-XM_044918353.1    rna-XM_004395543.1    rna-XM_042990512.1    rna-XM_040641440.1    rna-XM_027571727.2    mNeoVis.3.1.1.chr9.g00059.t01    rna-gnl|WGS_ZZZZ|egapxtmp_016623-R1    mNeoVis.evergladensis.2.1.chr8.g00082.t01    mNeoVis.vulgivaga.2.1.chr9.g00081.t01      2811    N0.HOG0002809   OG0000559  n1   rna-XM_034638822.1    rna-XM_038525656.1    rna-XM_022523734.1    rna-XM_045046997.1    rna-XM_036091501.1    rna-XM_039247176.1    rna-XM_032839435.1    rna-XM_045987652.1    rna-XM_032323339.1    rna-XM_059153475.1    rna-XM_059382473.1    rna-XM_045077153.1    mNeoVis.2.1.1.chr7_hap1.g01019.t01    rna-XM_021681069.2    rna-XM_004401033.1    rna-XM_042970602.1    rna-XM_008684610.2    rna-XM_027619493.2    mNeoVis.3.1.1.chr7.g00957.t01    rna-gnl|WGS_ZZZZ|egapxtmp_010937-R1    mNeoVis.evergladensis.2.1.chr7.g01012.t01    mNeoVis.vulgivaga.2.1.chr7.g01020.t01      2812    N0.HOG0002810   OG0000559  n23   rna-XM_011225515.3    rna-XM_038544861.1    rna-XM_022527120.1    rna-XM_006932902.5    rna-XM_036099590.1    rna-XM_039240760.1    rna-XM_032875264.1    rna-XM_046007353.1    rna-XM_032344378.1    rna-XM_059182242.1    rna-XM_059373327.1    rna-XM_004738876.3    mNeoVis.2.1.1.chr10_hap1.g00346.t01    rna-XM_021701434.2    rna-XM_004415711.1    rna-XM_042989823.1    rna-XM_040641817.1    rna-XM_027569076.1    mNeoVis.3.1.1.chr9.g00316.t01    rna-gnl|WGS_ZZZZ|egapxtmp_016017-R1    mNeoVis.evergladensis.2.1.chr8.g00328.t01    mNeoVis.vulgivaga.2.1.chr9.g00329.t01      2813    N0.HOG0002811   OG0000559  n39   rna-XM_002912403.4    rna-XM_038551177.1    rna-XM_022516313.1    rna-XM_003983935.5    rna-XM_036087391.1    rna-XM_039232386.1    rna-XM_032877966.1    rna-XM_045981803.1    rna-XM_032351878.1    rna-XM_059134596.1    rna-XM_059406220.1    rna-XM_004752977.3    mNeoVis.2.1.1.chr11_hap1.g00625.t01    rna-XM_021701150.1    rna-XM_004400167.1    rna-XM_042981296.1    rna-XM_040623518.1    rna-XM_027622723.1    mNeoVis.3.1.1.chr11.g00587.t01    rna-gnl|WGS_ZZZZ|egapxtmp_025040-R1    mNeoVis.evergladensis.2.1.chr11.g00613.t01    mNeoVis.vulgivaga.2.1.chr11.g00622.t01      2814    N0.HOG0002812   OG0000560  n1   rna-XM_034638745.1    rna-XM_038529007.1    rna-XM_022525240.1    rna-XM_019819593.3    rna-XM_036118431.1    rna-XM_039217718.1    rna-XM_032841934.1    rna-XM_045986901.1    rna-XM_032325933.1    rna-XM_059153010.1    rna-XM_059382330.1    rna-XM_004776250.3    mNeoVis.2.1.1.chr7_hap1.g00499.t01    rna-XM_021701032.2    rna-XM_004396426.1    rna-XM_042968822.1    rna-XM_040633910.1    rna-XM_027618209.2    mNeoVis.3.1.1.chr7.g00484.t01    rna-gnl|WGS_ZZZZ|egapxtmp_013147-R2    mNeoVis.evergladensis.2.1.chr7.g00500.t01    mNeoVis.vulgivaga.2.1.chr7.g00503.t01      2815    N0.HOG0002813   OG0000560  n12   rna-XM_034645292.1    rna-XM_038563968.1    rna-XM_022506873.1    rna-XM_006937501.3, rna-XM_045036436.1      rna-XM_039244846.1    rna-XM_032879446.1    rna-XM_046016018.1    rna-XM_032357282.1    rna-XM_059146997.1    rna-XM_059401645.1    rna-XM_004770066.3    mNeoVis.2.1.1.chr7_hap1.g02403.t01    rna-XM_044919201.1    rna-XM_004393949.1    rna-XM_042958337.1    rna-XM_040639391.1    rna-XM_027580849.2    mNeoVis.3.1.1.chr7.g02213.t01    rna-gnl|WGS_ZZZZ|egapxtmp_012694-R1    mNeoVis.evergladensis.2.1.chr7.g02349.t01    mNeoVis.vulgivaga.2.1.chr7.g02369.t01      2816    N0.HOG0002814   OG0000560  n22   rna-XM_034642251.1    rna-XM_038545159.1    rna-XM_022505262.1    rna-XM_045060171.1    rna-XM_036099730.1    rna-XM_039219937.1    rna-XM_032837279.1    rna-XM_046009580.1    rna-XM_032345048.1    rna-XM_059182916.1    rna-XM_059373821.1    rna-XM_013057477.2    mNeoVis.2.1.1.chr10_hap1.g00148.t01    rna-XM_021679542.1    rna-XM_004399429.1    rna-XM_042990038.1    rna-XM_008703117.2    rna-XM_027569208.1    mNeoVis.3.1.1.chr9.g00122.t01    rna-gnl|WGS_ZZZZ|egapxtmp_015984-R1    mNeoVis.evergladensis.2.1.chr8.g00142.t01    mNeoVis.vulgivaga.2.1.chr9.g00142.t01      2817    N0.HOG0002815   OG0000561  n1   rna-XM_002918415.4    rna-XM_038538668.1    rna-XM_022511671.1    rna-XM_006941610.4    rna-XM_036115074.1    rna-XM_039222626.1    rna-XM_032854206.1    rna-XM_045987914.1    rna-XM_032324346.1    rna-XM_059151791.1    rna-XM_059381768.1    rna-XM_004744388.3    mNeoVis.2.1.1.chr7_hap1.g00227.t01    rna-XM_021705763.2    rna-XM_004393387.1    rna-XM_007090283.3    rna-XM_008702469.2    rna-XM_027618580.1    mNeoVis.3.1.1.chr7.g00209.t01    rna-gnl|WGS_ZZZZ|egapxtmp_010905-R1    mNeoVis.evergladensis.2.1.chr7.g00226.t01    mNeoVis.vulgivaga.2.1.chr7.g00228.t01      2818    N0.HOG0002816   OG0000561  n15   rna-XM_019808558.2    rna-XM_038558431.1    rna-XM_022512261.1    rna-XM_045059650.1    rna-XM_036117971.1    rna-XM_039245230.1    rna-XM_032857169.1    rna-XM_046009917.1    rna-XM_032341654.1    rna-XM_059179602.1    rna-XM_059418545.1    rna-XM_004758550.2    mNeoVis.2.1.1.chr10_hap1.g00888.t01    rna-XM_021695561.2    rna-XM_004402136.1    rna-XM_042989345.1    rna-XM_040620693.1    rna-XM_027570520.2    mNeoVis.3.1.1.chr9.g00815.t01      mNeoVis.evergladensis.2.1.chr8.g00858.t01    mNeoVis.vulgivaga.2.1.chr9.g00844.t01      2819    N0.HOG0002817   OG0000561  n34   rna-XM_002926635.4    rna-XM_038583380.1    rna-XM_022512487.1    rna-XM_045039511.1    rna-XM_036078471.1    rna-XM_039238035.1    rna-XM_032861821.1    rna-XM_046000805.1    rna-XM_032337866.1    rna-XM_059173558.1    rna-XM_059416809.1    rna-XM_045089406.1    mNeoVis.2.1.1.chr1_hap1.g00011.t01    rna-XM_021687087.1    rna-XM_004411623.1    rna-XM_042977194.1, rna-XM_042998180.1    rna-XM_040630591.1    rna-XM_027606905.2    mNeoVis.3.1.1.chr1.g00013.t01    rna-gnl|WGS_ZZZZ|egapxtmp_021157-R1    mNeoVis.evergladensis.2.1.chr1.g00010.t01    mNeoVis.vulgivaga.2.1.chr1.g00010.t01      2820    N0.HOG0002818   OG0000562  n1   rna-XM_002912736.4    rna-XM_038566792.1    rna-XM_022498067.1    rna-XM_003982085.5    rna-XM_036093832.1    rna-XM_039227977.1    rna-XM_032867728.1    rna-XM_045991214.1    rna-XM_032316385.1    rna-XM_059160566.1    rna-XM_059387796.1    rna-XM_004760918.3    mNeoVis.2.1.1.chr5_hap1.g00530.t01    rna-XM_021683104.1    rna-XM_004404659.1    rna-XM_042978650.1    rna-XM_040628994.1    rna-XM_027584714.2    mNeoVis.3.1.1.chr2.g00452.t01    rna-gnl|WGS_ZZZZ|egapxtmp_035238-R1    mNeoVis.evergladensis.2.1.chr2.g00524.t01    mNeoVis.vulgivaga.2.1.chr2.g00527.t01      2821    N0.HOG0002819   OG0000562  n22   rna-XM_034650112.1, rna-XM_034667506.1    rna-XM_038543125.1    rna-XM_022491112.1    rna-XM_023247381.2    rna-XM_036124407.1    rna-XM_039251385.1    rna-XM_032845061.1    rna-XM_045982608.1    rna-XM_032317150.1    rna-XM_059146420.1    rna-XM_059413780.1    rna-XM_004778591.3    mNeoVis.2.1.1.chr14_hap1.g00650.t01    rna-XM_021681989.2    rna-XM_004402956.1    rna-XM_042976847.1    rna-XM_008696583.1    rna-XM_027611624.2    mNeoVis.3.1.1.chr14.g00626.t01      mNeoVis.evergladensis.2.1.chr14.g00648.t01    mNeoVis.vulgivaga.2.1.chr14.g00648.t01      2822    N0.HOG0002820   OG0000562  n43   rna-XM_002916215.4    rna-XM_038532462.1    rna-XM_022504752.1    rna-XM_003981108.6    rna-XM_036098188.1    rna-XM_039247477.1    rna-XM_032847723.1    rna-XM_046001211.1    rna-XM_032335865.1    rna-XM_059175338.1    rna-XM_059418142.1    rna-XM_004766802.3    mNeoVis.2.1.1.chr1_hap1.g00701.t01    rna-XM_021699801.1    rna-XM_004405600.1    rna-XM_007079980.2    rna-XM_040622141.1    rna-XM_027607118.1    mNeoVis.3.1.1.chr1.g00721.t01    rna-gnl|WGS_ZZZZ|egapxtmp_035106-R1    mNeoVis.evergladensis.2.1.chr1.g00709.t01    mNeoVis.vulgivaga.2.1.chr1.g00722.t01      2823    N0.HOG0002821   OG0000563  n1   rna-XM_034649986.1    rna-XM_038587581.1    rna-XM_022491580.1    rna-XM_004000571.6    rna-XM_036072147.1    rna-XM_039222261.1    rna-XM_032846916.1, rna-XM_032877139.1    rna-XM_045996175.1    rna-XM_032329644.1    rna-XM_059156627.1    rna-XM_059384660.1    rna-XM_004780009.3    mNeoVis.2.1.1.chr12_hap1.g00523.t01    rna-XM_021680558.2    rna-XM_004417411.1    rna-XM_007094653.3    rna-XM_008710661.2    rna-XM_027608080.1    mNeoVis.3.1.1.chr12.g00476.t01    rna-gnl|WGS_ZZZZ|egapxtmp_028183-R1    mNeoVis.evergladensis.2.1.chr12.g00469.t01    mNeoVis.vulgivaga.2.1.chr12.g00482.t01      2824   N0.HOG0002822  OG0000563  n16   rna-XM_019797680.2    rna-XM_038536238.1    rna-XM_022513172.1    rna-XM_023239076.2    rna-XM_036106888.1    rna-XM_039229141.1    rna-XM_032838364.1    rna-XM_046016101.1    rna-XM_032356357.1    rna-XM_059179918.1    rna-XM_059417782.1    rna-XM_045077544.1    mNeoVis.2.1.1.chr7_hap1.g01122.t01    rna-XM_044919756.1    rna-XM_004401065.1    rna-XM_042957703.1    rna-XM_040631244.1    rna-XM_027578046.1      rna-gnl|WGS_ZZZZ|egapxtmp_011862-R1    mNeoVis.evergladensis.2.1.chr7.g01108.t01    mNeoVis.vulgivaga.2.1.chr7.g01120.t01      2825   N0.HOG0002823  OG0000563  n32   rna-XM_011219561.3    rna-NM_001286989.1    rna-XM_022520942.1    rna-XM_023254543.2    rna-XM_036110078.1    rna-XM_039252073.1    rna-XM_032836475.1    rna-XM_046006437.1    rna-XM_032338593.1    rna-XM_059176639.1    rna-XM_059401362.1    rna-XM_045080099.1    mNeoVis.2.1.1.chr1_hap1.g02107.t01    rna-XM_021693661.2    rna-XM_004416816.1    rna-XM_042987445.1    rna-XM_040628015.1    rna-XM_027601356.2    mNeoVis.3.1.1.chr1.g02063.t01    rna-gnl|WGS_ZZZZ|egapxtmp_023215-R1    mNeoVis.evergladensis.2.1.chr1.g02077.t01    mNeoVis.vulgivaga.2.1.chr1.g02071.t01      2826    N0.HOG0002824   OG0000564  n1   rna-XM_034648811.1    rna-XM_038588540.1    rna-XM_022491372.1    rna-XM_023249466.2    rna-XM_036102384.1    rna-XM_039249890.1    rna-XM_032845289.1    rna-XM_045995088.1    rna-XM_032329808.1    rna-XM_059156455.1    rna-XM_059385074.1    rna-XM_004776624.3    mNeoVis.2.1.1.chr12_hap1.g00095.t01    rna-XM_021683399.2    rna-XM_004407115.1    rna-XM_042975268.1    rna-XM_040628113.1    rna-XM_027608330.1    mNeoVis.3.1.1.chr12.g00080.t01    rna-gnl|WGS_ZZZZ|egapxtmp_028991-R1    mNeoVis.evergladensis.2.1.chr12.g00081.t01    mNeoVis.vulgivaga.2.1.chr12.g00085.t01      2827    N0.HOG0002825   OG0000564  n21   rna-XM_002925711.4    rna-XM_038568229.1    rna-XM_022514135.1    rna-XM_011286508.3      rna-XM_039220322.1    rna-XM_032852743.1    rna-XM_046014969.1    rna-XM_032360397.1    rna-XM_059185906.1    rna-XM_059413188.1    rna-XM_004754545.3    mNeoVis.2.1.1.chr7_hap1.g01420.t01    rna-XM_021679109.2    rna-XM_004398083.1    rna-XM_007079043.3    rna-XM_040630977.1    rna-XM_027579543.2    mNeoVis.3.1.1.chr7.g01329.t01    rna-gnl|WGS_ZZZZ|egapxtmp_010990-R1    mNeoVis.evergladensis.2.1.chr7.g01402.t01    mNeoVis.vulgivaga.2.1.chr7.g01415.t01      2828    N0.HOG0002826   OG0000564  n36   rna-XM_019805245.2    rna-XM_038588545.1    rna-XM_022491268.1    rna-XM_023249233.2    rna-XM_036102300.1    rna-XM_039249875.1    rna-XM_032845288.1    rna-XM_045995255.1    rna-XM_032331049.1    rna-XM_059156542.1    rna-XM_059386137.1    rna-XM_045068170.1    mNeoVis.2.1.1.chr12_hap1.g00094.t01    rna-XM_021683645.1    rna-XM_004407114.1    rna-XM_042975266.1    rna-XM_040628092.1    rna-XM_027607952.1, rna-XM_035724906.1    mNeoVis.3.1.1.chr12.g00079.t01    rna-gnl|WGS_ZZZZ|egapxtmp_028989-R1    mNeoVis.evergladensis.2.1.chr12.g00080.t01    mNeoVis.vulgivaga.2.1.chr12.g00084.t01      2829   N0.HOG0002827  OG0000565  n1   rna-XM_002916748.4    rna-XM_038583165.1    rna-XM_022514960.1    rna-XM_003991756.6    rna-XM_036108626.1    rna-XM_039250886.1    rna-XM_032869347.1    rna-XM_046001668.1    rna-XM_032336833.1    rna-XM_059163040.1    rna-XM_059391214.1    rna-XM_004745330.3    mNeoVis.2.1.1.chr5_hap1.g01249.t01    rna-XM_021692344.1    rna-XM_004391974.1    rna-XM_007090962.3    rna-XM_040621394.1    rna-XM_027587697.1    mNeoVis.3.1.1.chr2.g01167.t01    rna-gnl|WGS_ZZZZ|egapxtmp_038198-R1    mNeoVis.evergladensis.2.1.chr2.g01237.t01    mNeoVis.vulgivaga.2.1.chr2.g01242.t01      2830    N0.HOG0002828   OG0000565  n13   rna-XM_034666289.1    rna-XM_038568521.1    rna-XM_022514006.1    rna-XM_023239404.2    rna-XM_036073007.1    rna-XM_039228722.1    rna-XM_032852997.1    rna-XM_046016053.1    rna-XM_032356795.1    rna-XM_059188195.1    rna-XM_059411533.1    rna-XM_013050253.2    mNeoVis.2.1.1.chr7_hap1.g01499.t01    rna-XM_021704727.1    rna-XM_004391818.1    rna-XM_042959496.1    rna-XM_040630897.1    rna-XM_027580607.2    mNeoVis.3.1.1.chr7.g01408.t01    rna-gnl|WGS_ZZZZ|egapxtmp_011907-R1    mNeoVis.evergladensis.2.1.chr7.g01481.t01    mNeoVis.vulgivaga.2.1.chr7.g01498.t01      2831    N0.HOG0002829   OG0000565  n23   rna-XM_034648868.1    rna-XM_038588082.1    rna-XM_022524359.1    rna-XM_023249300.2    rna-XM_036074519.1    rna-XM_039247416.1    rna-XM_032837026.1    rna-XM_045995443.1    rna-XM_032330607.1    rna-XM_059157369.1    rna-XM_059384573.1    rna-XM_045068266.1    mNeoVis.2.1.1.chr12_hap1.g00283.t01    rna-XM_044911971.1    rna-XM_004399749.1    rna-XM_042974188.1    rna-XM_040629053.1    rna-XM_035725555.1    mNeoVis.3.1.1.chr12.g00251.t01    rna-gnl|WGS_ZZZZ|egapxtmp_028125-R1    mNeoVis.evergladensis.2.1.chr12.g00251.t01    mNeoVis.vulgivaga.2.1.chr12.g00258.t01      2832    N0.HOG0002830   OG0000566  n1   rna-XM_011230841.1    rna-XM_038586579.1    rna-XM_022524394.1    rna-XM_006943916.4    rna-XM_036078947.1    rna-XM_039233529.1    rna-XM_032837144.1    rna-XM_045994808.1    rna-XM_032329717.1    rna-XM_059156827.1    rna-XM_059385195.1    rna-XM_045068288.1    mNeoVis.2.1.1.chr12_hap1.g00292.t01    rna-XM_021678327.1    rna-XM_004399761.1    rna-XM_007091050.2    rna-XM_008709304.2    rna-XM_027608799.1    mNeoVis.3.1.1.chr12.g00260.t01    rna-gnl|WGS_ZZZZ|egapxtmp_028126-R1    mNeoVis.evergladensis.2.1.chr12.g00260.t01    mNeoVis.vulgivaga.2.1.chr12.g00267.t01      2833    N0.HOG0002831   OG0000566  n23   rna-XM_011219816.3    rna-XM_038569340.1    rna-XM_022521745.1    rna-XM_045038087.1    rna-XM_036082858.1    rna-XM_039228707.1    rna-XM_032853552.1    rna-XM_046015935.1    rna-XM_032359718.1    rna-XM_059176133.1    rna-XM_059399387.1    rna-XM_004768213.3    mNeoVis.2.1.1.chr7_hap1.g01507.t01    rna-XM_021704684.1    rna-XM_004391844.1    rna-XM_042959606.1    rna-XM_040630876.1    rna-XM_027577923.2    mNeoVis.3.1.1.chr7.g01416.t01    rna-gnl|WGS_ZZZZ|egapxtmp_012479-R1    mNeoVis.evergladensis.2.1.chr7.g01489.t01    mNeoVis.vulgivaga.2.1.chr7.g01505.t01      2834   N0.HOG0002832  OG0000566  n44   rna-XM_034647239.1    rna-NM_001003207.1    rna-XM_022524711.1    rna-XM_023243814.2    rna-XM_036094517.1    rna-XM_039231454.1    rna-XM_032839852.1    rna-XM_045986440.1    rna-XM_032322473.1    rna-XM_059147432.1    rna-XM_059378595.1    rna-XM_004760239.3    mNeoVis.2.1.1.chr8_hap1.g01026.t01    rna-XM_021694807.1    rna-XM_004398477.1    rna-XM_042967406.1    rna-XM_040628219.1    rna-XM_027624608.2    mNeoVis.3.1.1.chr8.g00956.t01    rna-gnl|WGS_ZZZZ|egapxtmp_017561-R1    mNeoVis.evergladensis.2.1.chr9.g01000.t01    mNeoVis.vulgivaga.2.1.chr8.g01001.t01      2835   N0.HOG0002833  OG0000567  n1   rna-XM_011232992.3    rna-XM_038542788.1    rna-XM_022508346.1    rna-XM_045049424.1    rna-XM_036080882.1    rna-XM_039235873.1    rna-XM_032862838.1    rna-XM_045983594.1    rna-XM_032317316.1    rna-XM_059145519.1    rna-XM_059412594.1    rna-XM_004756819.3    mNeoVis.2.1.1.chr14_hap1.g00096.t01    rna-XM_044915969.1    rna-XM_004393563.1    rna-XM_042976800.1    rna-XM_040636100.1, rna-XM_040636101.1    rna-XM_027612772.2    mNeoVis.3.1.1.chr14.g00090.t01    rna-gnl|WGS_ZZZZ|egapxtmp_031981-R1    mNeoVis.evergladensis.2.1.chr14.g00095.t01    mNeoVis.vulgivaga.2.1.chr14.g00095.t01      2836    N0.HOG0002834   OG0000567  n23   rna-XM_034645574.1    rna-XM_038564059.1    rna-XM_022506695.1    rna-XM_023240022.2    rna-XM_036101135.1    rna-XM_039244837.1    rna-XM_032879399.1    rna-XM_046016001.1    rna-XM_032357492.1    rna-XM_059146536.1    rna-XM_059402028.1    rna-XM_045064565.1    mNeoVis.2.1.1.chr7_hap1.g02377.t01    rna-XM_021685523.2    rna-XM_004393974.1    rna-XM_042958332.1    rna-XM_040638775.1    rna-XM_027580912.1    mNeoVis.3.1.1.chr7.g02188.t01      mNeoVis.evergladensis.2.1.chr7.g02323.t01    mNeoVis.vulgivaga.2.1.chr7.g02342.t01      2837    N0.HOG0002835   OG0000567  n43   rna-XM_034642068.1    rna-XM_038545798.1    rna-XM_022505164.1    rna-XM_023256085.2    rna-XM_036099507.1    rna-XM_039251183.1    rna-XM_032839704.1    rna-XM_046008756.1    rna-XM_032345368.1    rna-XM_059183202.1    rna-XM_059374046.1    rna-XM_045074019.1    mNeoVis.2.1.1.chr10_hap1.g00055.t01    rna-XM_044918120.1    rna-XM_004395577.1    rna-XM_042990640.1    rna-XM_040641386.1    rna-XM_027569275.2    mNeoVis.3.1.1.chr9.g00033.t01    rna-gnl|WGS_ZZZZ|egapxtmp_015586-R1    mNeoVis.evergladensis.2.1.chr8.g00054.t01    mNeoVis.vulgivaga.2.1.chr9.g00054.t01      2838    N0.HOG0002836   OG0000568  n1   rna-XM_034664395.1    rna-XM_038553236.1    rna-XM_022501493.1    rna-XM_011288472.4    rna-XM_036090482.1    rna-XM_039234611.1    rna-XM_032865723.1    rna-XM_046021693.1    rna-XM_032308161.1    rna-XM_059142617.1    rna-XM_059410930.1    rna-XM_004775642.3    mNeoVis.2.1.1.chr13_hap1.g00307.t01    rna-XM_021691503.1    rna-XM_004411399.1    rna-XM_042964823.1    rna-XM_008698837.2    rna-XM_027616802.2    mNeoVis.3.1.1.chr13.g00250.t01    rna-gnl|WGS_ZZZZ|egapxtmp_031267-R1    mNeoVis.evergladensis.2.1.chr13.g00298.t01    mNeoVis.vulgivaga.2.1.chr13.g00294.t01      2839   N0.HOG0002837  OG0000568  n16   rna-XM_034667184.1    rna-XM_038586119.1    rna-XM_022508861.1    rna-XM_019822169.2    rna-XM_036112049.1    rna-XM_039221314.1    rna-XM_032872908.1    rna-XM_045983654.1    rna-XM_032318837.1    rna-XM_059146396.1    rna-XM_059413324.1    rna-XM_004751486.3    mNeoVis.2.1.1.chr14_hap1.g00422.t01    rna-XM_021686643.2    rna-XM_004409740.1    rna-XM_042975995.1    rna-XM_008685663.1    rna-XM_027613990.2    mNeoVis.3.1.1.chr14.g00420.t01    rna-gnl|WGS_ZZZZ|egapxtmp_032589-R2    mNeoVis.evergladensis.2.1.chr14.g00420.t01    mNeoVis.vulgivaga.2.1.chr14.g00422.t01      2840    N0.HOG0002838   OG0000568  n35   rna-XM_011220937.3    rna-XM_038542418.1    rna-XM_022508265.1    rna-XM_045048928.1    rna-XM_036066944.1    rna-XM_039225140.1    rna-XM_032838741.1    rna-XM_045981943.1    rna-XM_032318909.1    rna-XM_059146404.1    rna-XM_059412843.1    rna-XM_004761967.3    mNeoVis.2.1.1.chr14_hap1.g00199.t01    rna-XM_021682606.1    rna-XM_004398128.1    rna-XM_042976370.1    rna-XM_040634596.1    rna-XM_027612318.2    mNeoVis.3.1.1.chr14.g00197.t01    rna-gnl|WGS_ZZZZ|egapxtmp_032054-R1    mNeoVis.evergladensis.2.1.chr14.g00199.t01    mNeoVis.vulgivaga.2.1.chr14.g00203.t01      2841    N0.HOG0002839   OG0000569  n1   rna-XM_011231948.3    rna-XM_038578793.1    rna-XM_022499245.1    rna-XM_045040672.1    rna-XM_036076697.1    rna-XM_039237643.1    rna-XM_032880116.1    rna-XM_046026866.1    rna-XM_032311865.1    rna-XM_059173196.1    rna-XM_059398576.1    rna-XM_004763846.3    mNeoVis.2.1.1.chr2_hap1.g00148.t01    rna-XM_021700105.2    rna-XM_004412774.1    rna-XM_007087176.3    rna-XM_008684563.2    rna-XM_027596340.1    mNeoVis.3.1.1.chr5.g00141.t01    rna-gnl|WGS_ZZZZ|egapxtmp_006794-R1    mNeoVis.evergladensis.2.1.chr5.g00148.t01    mNeoVis.vulgivaga.2.1.chr5.g00150.t01      2842    N0.HOG0002840   OG0000569  n23   rna-XM_019798955.2    rna-XM_038552101.1    rna-XM_022493145.1    rna-XM_023254930.2    rna-XM_036106453.1    rna-XM_039252570.1    rna-XM_032837558.1    rna-XM_046000494.1    rna-XM_032336800.1    rna-XM_059174504.1    rna-XM_059417516.1    rna-XM_045075539.1    mNeoVis.2.1.1.chr1_hap1.g00386.t01    rna-XM_021702203.1    rna-XM_004408239.1    rna-XM_007087779.3    rna-XM_008698965.2    rna-XM_027605677.2    mNeoVis.3.1.1.chr1.g00384.t01    rna-gnl|WGS_ZZZZ|egapxtmp_021512-R2    mNeoVis.evergladensis.2.1.chr1.g00383.t01    mNeoVis.vulgivaga.2.1.chr1.g00390.t01      2843    N0.HOG0002841   OG0000569  n44   rna-XM_034647513.1    rna-XM_038560406.1    rna-XM_022498902.1    rna-XM_023252322.2    rna-XM_036070618.1    rna-XM_039253845.1    rna-XM_032869897.1    rna-XM_045993805.1    rna-XM_032328802.1    rna-XM_059156408.1    rna-XM_059384350.1    rna-XM_045069462.1    mNeoVis.2.1.1.chr6_hap1.g01070.t01    rna-XM_021694263.1    rna-XM_004412760.1    rna-XM_042982974.1    rna-XM_040643409.1    rna-XM_027597897.1    mNeoVis.3.1.1.chr4.g01025.t01    rna-gnl|WGS_ZZZZ|egapxtmp_000011-R1    mNeoVis.evergladensis.2.1.chr4.g01012.t01    mNeoVis.vulgivaga.2.1.chr4.g01039.t01      2844    N0.HOG0002842   OG0000570  n1   rna-XM_002913478.4    rna-NM_001100291.1    rna-XM_022507774.1    rna-XM_004000394.5    rna-XM_036081569.1    rna-XM_039255842.1    rna-XM_032883141.1    rna-XM_045996246.1    rna-XM_032330286.1    rna-XM_059157445.1    rna-XM_059384811.1    rna-XM_045089449.1    mNeoVis.2.1.1.chr12_hap1.g00682.t01    rna-XM_021681634.1    rna-XM_004399711.1    rna-XM_042974012.1    rna-XM_008696457.2    rna-XM_027607454.2    mNeoVis.3.1.1.chr12.g00630.t01    rna-gnl|WGS_ZZZZ|egapxtmp_029028-R1    mNeoVis.evergladensis.2.1.chr12.g00619.t01    mNeoVis.vulgivaga.2.1.chr12.g00639.t01      2845    N0.HOG0002843   OG0000570  n19   rna-XM_034660533.1    rna-XM_038587853.1    rna-XM_022521657.1    rna-XM_004000704.6    rna-XM_036071224.1    rna-XM_039218095.1    rna-XM_032881773.1    rna-XM_045995740.1    rna-XM_032330130.1    rna-XM_059156569.1    rna-XM_059384699.1    rna-XM_045085298.1    mNeoVis.2.1.1.chr12_hap1.g00384.t01    rna-XM_044911651.1    rna-XM_004415445.1    rna-XM_042974732.1    rna-XM_040620577.1    rna-XM_027608599.2    mNeoVis.3.1.1.chr12.g00348.t01    rna-gnl|WGS_ZZZZ|egapxtmp_028461-R3    mNeoVis.evergladensis.2.1.chr12.g00342.t01    mNeoVis.vulgivaga.2.1.chr12.g00349.t01      2846    N0.HOG0002844   OG0000570  n38   rna-XM_002927151.2    rna-NM_001101833.1    rna-XM_022520977.1    rna-XM_023254521.2    rna-XM_036112084.1    rna-XM_039234124.1    rna-XM_032836484.1    rna-XM_046006516.1    rna-XM_032338641.1    rna-XM_059179262.1    rna-XM_059399290.1    rna-XM_045080272.1    mNeoVis.2.1.1.chr1_hap1.g02094.t01    rna-XM_021693497.1    rna-XM_004416597.1    rna-XM_042985362.1    rna-XM_008688526.1    rna-XM_027603243.1    mNeoVis.3.1.1.chr1.g02047.t01    rna-gnl|WGS_ZZZZ|egapxtmp_000767-R1    mNeoVis.evergladensis.2.1.chr1.g02063.t01    mNeoVis.vulgivaga.2.1.chr1.g02055.t01      2847    N0.HOG0002845   OG0000571  n1   rna-XM_034649192.1    rna-XM_038587382.1    rna-XM_022507959.1    rna-XM_023249138.2    rna-XM_036095747.1    rna-XM_039218243.1    rna-XM_032842950.1    rna-XM_045995037.1    rna-XM_032330347.1    rna-XM_059157205.1    rna-XM_059385154.1    rna-XM_004755056.3    mNeoVis.2.1.1.chr12_hap1.g00606.t01    rna-XM_021678837.1    rna-XM_004396797.1    rna-XM_042974319.1    rna-XM_040631808.1    rna-XM_027608550.2    mNeoVis.3.1.1.chr12.g00552.t01    rna-gnl|WGS_ZZZZ|egapxtmp_028513-R1    mNeoVis.evergladensis.2.1.chr12.g00543.t01    mNeoVis.vulgivaga.2.1.chr12.g00558.t01      2848    N0.HOG0002846   OG0000571  n16   rna-XM_034654085.1    rna-XM_038562369.1    rna-XM_022497024.1    rna-XM_023258982.2    rna-XM_036122695.1    rna-XM_039254930.1    rna-XM_032881445.1    rna-XM_046026937.1    rna-XM_032302410.1    rna-XM_059136811.1    rna-XM_059375741.1    rna-XM_045081791.1    mNeoVis.2.1.1.chr2_hap1.g00943.t01    rna-XM_021690096.1    rna-XM_004401322.1    rna-XM_042998192.1    rna-XM_008704078.2    rna-XM_027615773.1    mNeoVis.3.1.1.chr5.g00874.t01      mNeoVis.evergladensis.2.1.chr5.g00896.t01    mNeoVis.vulgivaga.2.1.chr5.g00885.t01      2849    N0.HOG0002847   OG0000571  n29   rna-XM_034671074.1    rna-XM_038556397.1, rna-XM_038557473.1    rna-XM_022510274.1    rna-XM_011280521.3    rna-XM_036092984.1    rna-XM_039255016.1    rna-XM_032870642.1    rna-XM_046021625.1    rna-XM_032305064.1    rna-XM_059171679.1    rna-XM_059396517.1    rna-NM_001310177.1    mNeoVis.2.1.1.chr4_hap1.g00276.t01    rna-XM_021699359.1    rna-XM_004404406.1    rna-XM_042977411.1    rna-XM_008705802.2    rna-XM_027574815.2    mNeoVis.3.1.1.chr6.g00276.t01    rna-gnl|WGS_ZZZZ|egapxtmp_003189-R1    mNeoVis.evergladensis.2.1.chr6.g00277.t01    mNeoVis.vulgivaga.2.1.chr6.g00273.t01      2850    N0.HOG0002848   OG0000572  n1   rna-XM_011216839.3    rna-XM_038588122.1    rna-XM_022524358.1    rna-XM_006943944.4    rna-XM_036071848.1    rna-XM_039256077.1    rna-XM_032837036.1    rna-XM_045996256.1    rna-XM_032329856.1    rna-XM_059156700.1    rna-XM_059386340.1    rna-XM_004777538.3    mNeoVis.2.1.1.chr12_hap1.g00265.t01    rna-XM_044911958.1    rna-XM_004412011.1    rna-XM_042974026.1    rna-XM_040628910.1    rna-XM_027609094.1    mNeoVis.3.1.1.chr12.g00234.t01    rna-gnl|WGS_ZZZZ|egapxtmp_028670-R1    mNeoVis.evergladensis.2.1.chr12.g00234.t01    mNeoVis.vulgivaga.2.1.chr12.g00241.t01      2851    N0.HOG0002849   OG0000572  n14   rna-XM_011219397.3    rna-XM_038569538.1    rna-XM_022503060.1    rna-XM_003980396.6    rna-XM_036078540.1    rna-XM_039254371.1    rna-XM_032874009.1    rna-XM_045977992.1    rna-XM_032315429.1    rna-XM_059144808.1    rna-XM_059378039.1    rna-XM_013059807.2    mNeoVis.2.1.1.chr8_hap1.g01354.t01    rna-XM_044913183.1    rna-XM_004408581.1    rna-XM_042972366.1    rna-XM_008706839.2    rna-XM_035726654.1    mNeoVis.3.1.1.chr8.g01249.t01    rna-gnl|WGS_ZZZZ|egapxtmp_019713-R1    mNeoVis.evergladensis.2.1.chr9.g01300.t01    mNeoVis.vulgivaga.2.1.chr8.g01304.t01      2852    N0.HOG0002850   OG0000572  n33   rna-XM_034661814.1    rna-XM_038571108.1    rna-XM_022507847.1    rna-XM_045037491.1    rna-XM_036096337.1    rna-XM_039219318.1    rna-XM_032852406.1    rna-XM_046002107.1    rna-XM_032346866.1    rna-XM_059163869.1    rna-XM_059391809.1    rna-XM_045088478.1    mNeoVis.2.1.1.chr5_hap1.g01524.t01    rna-XM_021678729.1    rna-XM_004397302.1    rna-XM_042999038.1    rna-XM_008707709.1    rna-XM_027584210.2    mNeoVis.3.1.1.chr2.g01442.t01    rna-gnl|WGS_ZZZZ|egapxtmp_037860-R1    mNeoVis.evergladensis.2.1.chr2.g01510.t01    mNeoVis.vulgivaga.2.1.chr2.g01524.t01      2853    N0.HOG0002851   OG0000573  n1   rna-XM_034664297.1    rna-XM_038571239.1    rna-XM_022503016.1    rna-XM_019810547.3    rna-XM_036072154.1    rna-XM_039221408.1    rna-XM_032852517.1    rna-XM_046002500.1    rna-XM_032344932.1    rna-XM_059163721.1    rna-XM_059391718.1    rna-XM_045088529.1    mNeoVis.2.1.1.chr5_hap1.g01472.t01    rna-XM_044920224.1    rna-XM_004404995.1    rna-XM_015539324.2    rna-XM_040636651.1    rna-XM_027584005.2    mNeoVis.3.1.1.chr2.g01392.t01    rna-gnl|WGS_ZZZZ|egapxtmp_038700-R2    mNeoVis.evergladensis.2.1.chr2.g01461.t01    mNeoVis.vulgivaga.2.1.chr2.g01471.t01      2854    N0.HOG0002852   OG0000573  n11   rna-XM_019794792.2    rna-XM_038569886.1    rna-XM_022502871.1    rna-XM_011280623.4    rna-XM_036068291.1    rna-XM_039217422.1    rna-XM_032864130.1    rna-XM_045978428.1    rna-XM_032314981.1    rna-XM_059144488.1    rna-XM_059377941.1    rna-XM_004746120.3    mNeoVis.2.1.1.chr8_hap1.g01467.t01    rna-XM_044913532.1    rna-XM_004408716.1    rna-XM_042979456.1    rna-XM_040623938.1    rna-XM_027589043.2    mNeoVis.3.1.1.chr8.g01362.t01    rna-gnl|WGS_ZZZZ|egapxtmp_018089-R7    mNeoVis.evergladensis.2.1.chr9.g01415.t01    mNeoVis.vulgivaga.2.1.chr8.g01419.t01      2855    N0.HOG0002853   OG0000573  n26   rna-XM_019803451.2    rna-XM_038588317.1    rna-XM_022526346.1    rna-XM_023249052.2    rna-XM_036100309.1    rna-XM_039239857.1    rna-XM_032843352.1    rna-XM_045995270.1    rna-XM_032330747.1    rna-XM_059157472.1    rna-XM_059384657.1    rna-XM_045068098.1    mNeoVis.2.1.1.chr12_hap1.g00179.t01    rna-XM_044911965.1    rna-XM_004401086.1    rna-XM_042973893.1    rna-XM_040627442.1    rna-XM_027608781.2    mNeoVis.3.1.1.chr12.g00149.t01    rna-gnl|WGS_ZZZZ|egapxtmp_029588-R1    mNeoVis.evergladensis.2.1.chr12.g00152.t01    mNeoVis.vulgivaga.2.1.chr12.g00155.t01      2856    N0.HOG0002854   OG0000574  n1   rna-XM_011231492.2    rna-XM_038588852.1    rna-XM_022491373.1    rna-XM_011279517.4    rna-XM_036086724.1    rna-XM_039249866.1    rna-XM_032845271.1    rna-XM_045994843.1    rna-XM_032331320.1    rna-XM_059158092.1    rna-XM_059386226.1    rna-XM_004776651.3    mNeoVis.2.1.1.chr12_hap1.g00086.t01    rna-XM_021683492.1    rna-XM_004407107.1    rna-XM_015543250.2    rna-XM_008690249.1    rna-XM_027609309.1    mNeoVis.3.1.1.chr12.g00071.t01    rna-gnl|WGS_ZZZZ|egapxtmp_028822-R1    mNeoVis.evergladensis.2.1.chr12.g00072.t01    mNeoVis.vulgivaga.2.1.chr12.g00076.t01      2857    N0.HOG0002855   OG0000574  n22   rna-XM_002919972.4    rna-NM_001301112.1    rna-XM_022516403.1    rna-XM_045054104.1    rna-XM_036110721.1    rna-XM_039232378.1    rna-XM_032877712.1    rna-XM_045980186.1    rna-XM_032351984.1    rna-XM_059134685.1    rna-XM_059407816.1    rna-XM_004752902.3    mNeoVis.2.1.1.chr11_hap1.g00655.t01    rna-XM_021701243.1    rna-XM_004400140.1    rna-XM_042981226.1    rna-XM_008699792.1    rna-XM_027620450.1    mNeoVis.3.1.1.chr11.g00614.t01    rna-gnl|WGS_ZZZZ|egapxtmp_026219-R1    mNeoVis.evergladensis.2.1.chr11.g00641.t01    mNeoVis.vulgivaga.2.1.chr11.g00649.t01      2858    N0.HOG0002856   OG0000574  n42   rna-XM_011222034.3    rna-NM_001003222.1    rna-XM_022506174.1    rna-XM_006931133.4    rna-XM_036096664.1    rna-XM_039241099.1    rna-XM_032852289.1    rna-XM_045997482.1    rna-XM_032334239.1    rna-XM_059161982.1    rna-XM_059383519.1    rna-XM_045079932.1    mNeoVis.2.1.1.chr6_hap1.g00191.t01    rna-XM_021701666.1    rna-XM_004396298.1    rna-XM_007075906.2    rna-XM_008701454.2    rna-XM_027599894.2    mNeoVis.3.1.1.chr4.g00185.t01    rna-gnl|WGS_ZZZZ|egapxtmp_044130-R1    mNeoVis.evergladensis.2.1.chr4.g00195.t01    mNeoVis.vulgivaga.2.1.chr4.g00191.t01      2859    N0.HOG0002857   OG0000575  n1   rna-XM_034647085.1    rna-NM_001159327.1    rna-XM_022524638.1    rna-XM_023244732.2    rna-XM_036066286.1    rna-XM_039231543.1    rna-XM_032839904.1    rna-XM_045983705.1    rna-XM_032322068.1    rna-XM_059149341.1    rna-XM_059379035.1    rna-XM_004760345.3    mNeoVis.2.1.1.chr8_hap1.g01072.t01    rna-XM_021694489.2    rna-XM_004398533.1    rna-XM_042966138.1    rna-XM_040629937.1    rna-XM_027625938.2    mNeoVis.3.1.1.chr8.g00994.t01      mNeoVis.evergladensis.2.1.chr9.g01047.t01    mNeoVis.vulgivaga.2.1.chr8.g01047.t01      2860    N0.HOG0002858   OG0000575  n21   rna-XM_011237436.3    rna-NM_001159326.1    rna-XM_022497010.1    rna-XM_003989965.6    rna-XM_036073453.1    rna-XM_039238712.1    rna-XM_032877497.1    rna-XM_046020835.1    rna-XM_032332007.1    rna-XM_059136473.1    rna-XM_059374407.1    rna-XM_045064347.1    mNeoVis.2.1.1.chr2_hap1.g01412.t01    rna-XM_021684246.2    rna-XM_004406649.1    rna-XM_042996839.1    rna-XM_008699338.2    rna-XM_027608422.2    mNeoVis.3.1.1.chr5.g01338.t01      mNeoVis.evergladensis.2.1.chr5.g01353.t01    mNeoVis.vulgivaga.2.1.chr5.g01352.t01      2861    N0.HOG0002859   OG0000575  n39   rna-XM_002929153.4    rna-XM_038576696.1    rna-XM_022522292.1    rna-XM_019834462.3    rna-XM_036116583.1    rna-XM_039220642.1    rna-XM_032874661.1    rna-XM_046012995.1    rna-XM_032348875.1    rna-XM_059184732.1    rna-XM_059403616.1    rna-XM_004766603.3    mNeoVis.2.1.1.chr9_hap1.g00297.t01, mNeoVis.2.1.1.chr9_hap1.g00770.t01    rna-XM_021690811.1    rna-XM_004406902.1, rna-XM_004407631.1, rna-XM_004417352.1    rna-XM_042991719.1    rna-XM_008683490.2    rna-XM_035721798.1, rna-XM_035721823.1, rna-XM_035721931.1, rna-XM_035729150.1              2862    N0.HOG0002860   OG0000576  n1   rna-XM_011227859.3    rna-XM_038577719.1    rna-XM_022519502.1    rna-XM_045061835.1    rna-XM_036077875.1    rna-XM_039241466.1    rna-XM_032876738.1    rna-XM_046011720.1    rna-XM_032348265.1    rna-XM_059185295.1    rna-XM_059403164.1    rna-XM_045073434.1    mNeoVis.2.1.1.chr9_hap1.g00537.t01    rna-XM_044915405.1    rna-XM_004400803.1    rna-XM_042992340.1    rna-XM_040642483.1    rna-XM_027592881.2    mNeoVis.3.1.1.chr10.g00508.t01    rna-gnl|WGS_ZZZZ|egapxtmp_024207-R1    mNeoVis.evergladensis.2.1.chr10.g00530.t01    mNeoVis.vulgivaga.2.1.chr10.g00516.t01      2863    N0.HOG0002861   OG0000576  n15   rna-XM_034665564.1    rna-XM_038552924.1    rna-XM_022501448.1    rna-XM_023242499.2    rna-XM_036074146.1    rna-XM_039234814.1    rna-XM_032860698.1    rna-XM_046023220.1    rna-XM_032307023.1    rna-XM_059141809.1    rna-XM_059411204.1    rna-XM_045081623.1    mNeoVis.2.1.1.chr13_hap1.g00431.t01    rna-XM_021691624.2    rna-XM_004392336.1    rna-XM_042965347.1    rna-XM_040631660.1    rna-XM_027615585.2    mNeoVis.3.1.1.chr13.g00369.t01    rna-gnl|WGS_ZZZZ|egapxtmp_031294-R1    mNeoVis.evergladensis.2.1.chr13.g00412.t01    mNeoVis.vulgivaga.2.1.chr13.g00416.t01      2864    N0.HOG0002862   OG0000576  n31   rna-XM_011230540.3    rna-XM_038584545.1    rna-XM_022496540.1    rna-XM_023259333.2    rna-XM_036092204.1    rna-XM_039237543.1    rna-XM_032857853.1    rna-XM_046019692.1    rna-XM_032355636.1    rna-XM_059167284.1    rna-XM_059393999.1    rna-XM_004743893.3    mNeoVis.2.1.1.chr3_hap1.g00962.t01    rna-XM_021703040.1    rna-XM_004394815.1    rna-XM_042994473.1    rna-XM_040644727.1    rna-XM_027590824.1    mNeoVis.3.1.1.chr3.g00952.t01    rna-gnl|WGS_ZZZZ|egapxtmp_042704-R6    mNeoVis.evergladensis.2.1.chr3.g00955.t01    mNeoVis.vulgivaga.2.1.chr3.g00975.t01      2865    N0.HOG0002863   OG0000577  n1   rna-XM_034654587.1    rna-XM_038584533.1    rna-XM_022525491.1    rna-XM_006935294.5    rna-XM_036092211.1    rna-XM_039253011.1    rna-XM_032857840.1    rna-XM_046019563.1    rna-XM_032355618.1    rna-XM_059167296.1    rna-XM_059393991.1    rna-XM_013051942.2    mNeoVis.2.1.1.chr3_hap1.g00967.t01    rna-XM_021702924.1    rna-XM_004394805.1    rna-XM_007078494.3    rna-XM_040644769.1    rna-XM_035726900.1    mNeoVis.3.1.1.chr3.g00957.t01    rna-gnl|WGS_ZZZZ|egapxtmp_042072-R1    mNeoVis.evergladensis.2.1.chr3.g00960.t01    mNeoVis.vulgivaga.2.1.chr3.g00980.t01      2866    N0.HOG0002864   OG0000577  n14   rna-XM_034664192.1    rna-XM_038552931.1    rna-XM_022501602.1    rna-XM_023242504.2    rna-XM_036074141.1    rna-XM_039256568.1    rna-XM_032860703.1    rna-XM_046023928.1    rna-XM_032307954.1    rna-XM_059142627.1    rna-XM_059411200.1    rna-XM_045081626.1    mNeoVis.2.1.1.chr13_hap1.g00427.t01    rna-XM_021691497.1    rna-XM_004392339.1    rna-XM_042963880.1    rna-XM_040631663.1    rna-XM_027615335.2    mNeoVis.3.1.1.chr13.g00365.t01    rna-gnl|WGS_ZZZZ|egapxtmp_030101-R1    mNeoVis.evergladensis.2.1.chr13.g00409.t01    mNeoVis.vulgivaga.2.1.chr13.g00412.t01      2867    N0.HOG0002865   OG0000577  n30   rna-XM_034645345.1    rna-NM_001003227.1    rna-XM_022519589.1    rna-XM_006933714.3    rna-XM_036096377.1    rna-XM_039241418.1    rna-XM_032876737.1    rna-XM_046013167.1    rna-XM_032348270.1    rna-XM_059185292.1    rna-XM_059403169.1    rna-XM_045072951.1    mNeoVis.2.1.1.chr9_hap1.g00535.t01    rna-XM_021704824.1    rna-XM_004400800.1    rna-XM_042992343.1    rna-XM_040642490.1    rna-XM_027595425.2    mNeoVis.3.1.1.chr10.g00506.t01      mNeoVis.evergladensis.2.1.chr10.g00527.t01, mNeoVis.evergladensis.2.1.chr10.g00528.t01    mNeoVis.vulgivaga.2.1.chr10.g00514.t01      2868    N0.HOG0002866   OG0000578  n1   rna-XM_019804012.2    rna-XM_038577869.1    rna-XM_022519559.1    rna-XM_011283982.4    rna-XM_036124251.1    rna-XM_039216996.1    rna-XM_032876562.1    rna-XM_046012041.1    rna-XM_032348485.1    rna-XM_059185078.1    rna-XM_059403330.1    rna-XM_045073036.1    mNeoVis.2.1.1.chr9_hap1.g00456.t01    rna-XM_021686797.1    rna-XM_004406264.1    rna-XM_042992457.1    rna-XM_008704694.2    rna-XM_027593067.2    mNeoVis.3.1.1.chr10.g00431.t01    rna-gnl|WGS_ZZZZ|egapxtmp_022607-R1    mNeoVis.evergladensis.2.1.chr10.g00450.t01    mNeoVis.vulgivaga.2.1.chr10.g00439.t01      2869    N0.HOG0002867   OG0000578  n18   rna-XM_034654817.1    rna-NM_001003091.1    rna-XM_022496393.1    rna-XM_011285382.4    rna-XM_036080150.1    rna-XM_039239650.1    rna-XM_032856066.1    rna-XM_046019690.1    rna-XM_032356140.1    rna-XM_059166853.1    rna-XM_059394291.1    rna-XM_045066781.1    mNeoVis.2.1.1.chr3_hap1.g00820.t01    rna-XM_021702875.1    rna-XM_004392369.1    rna-XM_042994850.1    rna-XM_008687226.1    rna-XM_027591025.2    mNeoVis.3.1.1.chr3.g00801.t01    rna-gnl|WGS_ZZZZ|egapxtmp_041608-R3    mNeoVis.evergladensis.2.1.chr3.g00814.t01    mNeoVis.vulgivaga.2.1.chr3.g00827.t01      2870    N0.HOG0002868   OG0000578  n38   rna-XM_034670080.1    rna-XM_038557256.1    rna-XM_022503266.1    rna-XM_019825660.2    rna-XM_036098663.1    rna-XM_039251155.1    rna-XM_032872082.1    rna-XM_046020562.1    rna-XM_032304733.1    rna-XM_059171502.1    rna-XM_059396607.1    rna-XM_045062853.1    mNeoVis.2.1.1.chr4_hap1.g00347.t01    rna-XM_021689572.2    rna-XM_004414998.1    rna-XM_007090530.3    rna-XM_008686465.2    rna-XM_027575051.2    mNeoVis.3.1.1.chr6.g00349.t01    rna-gnl|WGS_ZZZZ|egapxtmp_003839-R1    mNeoVis.evergladensis.2.1.chr6.g00351.t01    mNeoVis.vulgivaga.2.1.chr6.g00344.t01      2871    N0.HOG0002869   OG0000579  n1   rna-XM_011222755.2    rna-XM_038563067.1    rna-XM_022506903.1    rna-XM_023239680.2    rna-XM_036107269.1    rna-XM_039233331.1    rna-XM_032841022.1    rna-XM_046015638.1    rna-XM_032356373.1    rna-XM_059139211.1    rna-XM_059406222.1    rna-XM_045066058.1    mNeoVis.2.1.1.chr7_hap1.g01930.t01    rna-XM_021685344.2    rna-XM_004406098.1    rna-XM_042959171.1    rna-XM_040638926.1    rna-XM_027578391.2    mNeoVis.3.1.1.chr7.g01802.t01    rna-gnl|WGS_ZZZZ|egapxtmp_013880-R1    mNeoVis.evergladensis.2.1.chr7.g01894.t01    mNeoVis.vulgivaga.2.1.chr7.g01910.t01      2872    N0.HOG0002870   OG0000579  n23   rna-XM_034637906.1    rna-NM_001014769.1    rna-XM_022510082.1    rna-XM_023250618.2    rna-XM_036119981.1    rna-XM_039233986.1    rna-XM_032867811.1    rna-XM_046020149.1    rna-XM_032305957.1    rna-XM_059172009.1    rna-XM_059396286.1    rna-XM_045063046.1    mNeoVis.2.1.1.chr4_hap1.g00155.t01    rna-XM_044920289.1    rna-XM_004414156.1    rna-XM_042973497.1    rna-XM_040643945.1    rna-XM_035723207.1    mNeoVis.3.1.1.chr6.g00155.t01    rna-gnl|WGS_ZZZZ|egapxtmp_004371-R1    mNeoVis.evergladensis.2.1.chr6.g00157.t01    mNeoVis.vulgivaga.2.1.chr6.g00154.t01      2873    N0.HOG0002871   OG0000579  n44   rna-XM_011223666.3    rna-XM_038561792.1    rna-XM_022496217.1    rna-XM_011285077.4    rna-XM_036088455.1    rna-XM_039249069.1    rna-XM_032881392.1    rna-XM_046009280.1    rna-XM_032360591.1    rna-XM_059138319.1    rna-XM_059375803.1    rna-XM_045081884.1    mNeoVis.2.1.1.chr2_hap1.g00921.t01    rna-XM_044914588.1    rna-XM_004401294.1    rna-XM_015534944.2    rna-XM_040626721.1    rna-XM_027606217.2    mNeoVis.3.1.1.chr5.g00853.t01    rna-gnl|WGS_ZZZZ|egapxtmp_006227-R7    mNeoVis.evergladensis.2.1.chr5.g00876.t01    mNeoVis.vulgivaga.2.1.chr5.g00865.t01      2874   N0.HOG0002872  OG0000580  n1   rna-XM_002921995.4    rna-XM_038544675.1    rna-XM_022524144.1    rna-XM_003987628.6    rna-XM_036110770.1    rna-XM_039217337.1    rna-XM_032871157.1    rna-XM_046009657.1    rna-XM_032344185.1    rna-XM_059182052.1    rna-XM_059373160.1    rna-XM_004738749.2    mNeoVis.2.1.1.chr10_hap1.g00415.t01    rna-XM_044918068.1    rna-XM_004408033.1    rna-XM_007097557.3    rna-XM_008693194.2    rna-XM_027572370.1    mNeoVis.3.1.1.chr9.g00382.t01    rna-gnl|WGS_ZZZZ|egapxtmp_016206-R1    mNeoVis.evergladensis.2.1.chr8.g00396.t01    mNeoVis.vulgivaga.2.1.chr9.g00398.t01      2875   N0.HOG0002873  OG0000580  n19   rna-XM_034659649.1    rna-XM_038561296.1    rna-XM_022520451.1    rna-XM_023251943.2    rna-XM_036070376.1    rna-XM_039251235.1    rna-XM_032881935.1    rna-XM_045979191.1    rna-XM_032352936.1    rna-XM_059188561.1    rna-XM_059405475.1    rna-XM_004765501.3    mNeoVis.2.1.1.chr11_hap1.g00976.t01    rna-XM_021697323.2    rna-XM_004410015.1    rna-XM_042982139.1    rna-XM_040623125.1    rna-XM_027621637.1    mNeoVis.3.1.1.chr11.g00928.t01    rna-gnl|WGS_ZZZZ|egapxtmp_026923-R3    mNeoVis.evergladensis.2.1.chr11.g00955.t01    mNeoVis.vulgivaga.2.1.chr11.g00976.t01      2876    N0.HOG0002874   OG0000580  n40   rna-XM_002916677.4    rna-XM_038564180.1    rna-XM_022506435.1    rna-XM_023239979.2    rna-XM_036100421.1    rna-XM_039257673.1    rna-XM_032879303.1    rna-XM_046016060.1    rna-XM_032357116.1    rna-XM_059145215.1    rna-XM_059403241.1    rna-XM_004770224.3    mNeoVis.2.1.1.chr7_hap1.g02338.t01    rna-XM_021685495.2    rna-XM_004394037.1    rna-XM_042958140.1    rna-XM_040639279.1    rna-XM_027578741.2    mNeoVis.3.1.1.chr7.g02152.t01    rna-gnl|WGS_ZZZZ|egapxtmp_013060-R3    mNeoVis.evergladensis.2.1.chr7.g02284.t01    mNeoVis.vulgivaga.2.1.chr7.g02301.t01      2877    N0.HOG0002875   OG0000581  n1   rna-XM_034648393.1    rna-XM_038544738.1    rna-XM_022522851.1    rna-XM_023255692.2    rna-XM_036098413.1    rna-XM_039240453.1    rna-XM_032871083.1    rna-XM_046009405.1    rna-XM_032344253.1    rna-XM_059182117.1    rna-XM_059373224.1    rna-XM_013056286.2    mNeoVis.2.1.1.chr10_hap1.g00394.t01    rna-XM_044918402.1    rna-XM_004415825.1    rna-XM_042989706.1    rna-XM_040641844.1    rna-XM_027570452.2    mNeoVis.3.1.1.chr9.g00364.t01    rna-gnl|WGS_ZZZZ|egapxtmp_017257-R1    mNeoVis.evergladensis.2.1.chr8.g00376.t01    mNeoVis.vulgivaga.2.1.chr9.g00378.t01      2878    N0.HOG0002876   OG0000581  n18   rna-XM_034646583.1    rna-XM_038564140.1    rna-XM_022506622.1    rna-XM_003993551.6    rna-XM_036100369.1    rna-XM_039244917.1    rna-XM_032879332.1    rna-XM_046015048.1    rna-XM_032357161.1    rna-XM_059145556.1    rna-XM_059402857.1    rna-XM_004770181.3    mNeoVis.2.1.1.chr7_hap1.g02351.t01    rna-XM_021684967.2    rna-XM_004394017.1    rna-XM_042958160.1    rna-XM_040639294.1    rna-XM_027580976.1    mNeoVis.3.1.1.chr7.g02164.t01      mNeoVis.evergladensis.2.1.chr7.g02297.t01    mNeoVis.vulgivaga.2.1.chr7.g02315.t01      2879   N0.HOG0002877  OG0000581  n38   rna-XM_034641057.1    rna-XM_038571649.1    rna-XM_022494520.1    rna-XM_045053862.1    rna-XM_036095823.1    rna-XM_039224884.1    rna-XM_032859159.1    rna-XM_045981854.1    rna-XM_032351382.1    rna-XM_059133917.1    rna-XM_059386525.1, rna-XM_059406562.1    rna-XM_004751536.3    mNeoVis.2.1.1.chr11_hap1.g00454.t01    rna-XM_021689173.1    rna-XM_004398215.1    rna-XM_007073250.3    rna-XM_008703333.2    rna-XM_027623781.1    mNeoVis.3.1.1.chr11.g00418.t01    rna-gnl|WGS_ZZZZ|egapxtmp_026656-R1    mNeoVis.evergladensis.2.1.chr11.g00446.t01    mNeoVis.vulgivaga.2.1.chr11.g00454.t01      2880    N0.HOG0002878   OG0000582  n1   rna-XM_034644766.1    rna-XM_038564077.1    rna-XM_022506520.1    rna-XM_003993567.6    rna-XM_036101141.1    rna-XM_039244893.1    rna-XM_032879387.1    rna-XM_046014175.1    rna-XM_032357214.1    rna-XM_059146307.1    rna-XM_059402215.1    rna-XM_004770132.3    mNeoVis.2.1.1.chr7_hap1.g02373.t01    rna-XM_021685011.1    rna-XM_004393978.1    rna-XM_042957846.1    rna-XM_040639310.1    rna-XM_027578950.1    mNeoVis.3.1.1.chr7.g02184.t01    rna-gnl|WGS_ZZZZ|egapxtmp_013090-R1    mNeoVis.evergladensis.2.1.chr7.g02319.t01    mNeoVis.vulgivaga.2.1.chr7.g02338.t01      2881    N0.HOG0002879   OG0000582  n21   rna-XM_034662353.1    rna-XM_038570227.1    rna-XM_022517548.1    rna-XM_006930010.5    rna-XM_036110019.1    rna-XM_039224427.1    rna-XM_032849408.1    rna-XM_045978851.1    rna-XM_032351602.1    rna-XM_059134249.1    rna-XM_059406411.1    rna-XM_004781236.3    mNeoVis.2.1.1.chr11_hap1.g00551.t01    rna-XM_044919133.1    rna-XM_004398410.1    rna-XM_042981019.1    rna-XM_040620949.1    rna-XM_027622799.2    mNeoVis.3.1.1.chr11.g00510.t01    rna-gnl|WGS_ZZZZ|egapxtmp_026997-R1    mNeoVis.evergladensis.2.1.chr11.g00537.t01    mNeoVis.vulgivaga.2.1.chr11.g00545.t01      2882    N0.HOG0002880   OG0000582  n39   rna-XM_011228429.3    rna-NM_001287162.1    rna-XM_022521833.1    rna-XM_023255684.2    rna-XM_036110982.1    rna-XM_039256113.1    rna-XM_032871135.1    rna-XM_046008691.1    rna-XM_032344184.1    rna-XM_059182070.1    rna-XM_059373175.1    rna-XM_004738765.3    mNeoVis.2.1.1.chr10_hap1.g00411.t01    rna-XM_021701347.1    rna-XM_004408041.1    rna-XM_042989680.1    rna-XM_040620897.1    rna-XM_027572369.2    mNeoVis.3.1.1.chr9.g00378.t01    rna-gnl|WGS_ZZZZ|egapxtmp_016812-R1    mNeoVis.evergladensis.2.1.chr8.g00392.t01    mNeoVis.vulgivaga.2.1.chr9.g00394.t01      2883    N0.HOG0002881   OG0000583  n1   rna-XM_002912764.4    rna-NM_001287076.1    rna-XM_022498279.1    rna-XM_011291127.3    rna-XM_036107657.1    rna-XM_039228234.1    rna-XM_032867852.1    rna-XM_045992358.1    rna-XM_032317684.1    rna-XM_059160702.1    rna-XM_059389566.1    rna-XM_045080747.1    mNeoVis.2.1.1.chr5_hap1.g00582.t01    rna-XM_021683046.1    rna-XM_004399155.1    rna-XM_042978769.1    rna-XM_008703877.2    rna-XM_027587678.2    mNeoVis.3.1.1.chr2.g00502.t01    rna-gnl|WGS_ZZZZ|egapxtmp_035897-R2    mNeoVis.evergladensis.2.1.chr2.g00577.t01    mNeoVis.vulgivaga.2.1.chr2.g00579.t01      2884    N0.HOG0002882   OG0000583  n22   rna-XM_011230832.3    rna-XM_038570924.1    rna-XM_022514902.1    rna-XM_019810683.2    rna-XM_036077090.1    rna-XM_039243839.1    rna-XM_032869670.1    rna-XM_046002913.1    rna-XM_032333507.1    rna-XM_059162647.1    rna-XM_059390946.1    rna-XM_045088893.1    mNeoVis.2.1.1.chr5_hap1.g01142.t01    rna-XM_021678891.1    rna-XM_004412077.1    rna-XM_042998359.1    rna-XM_040636803.1    rna-XM_027583617.2    mNeoVis.3.1.1.chr2.g01051.t01    rna-gnl|WGS_ZZZZ|egapxtmp_035549-R1    mNeoVis.evergladensis.2.1.chr2.g01132.t01    mNeoVis.vulgivaga.2.1.chr2.g01138.t01      2885    N0.HOG0002883   OG0000583  n43   rna-XM_034661978.1    rna-XM_038570921.1    rna-XM_022515107.1    rna-XM_006936404.5    rna-XM_036077097.1    rna-XM_039243838.1    rna-XM_032869671.1    rna-XM_046003613.1    rna-XM_032333462.1    rna-XM_059162644.1    rna-XM_059390942.1    rna-XM_045088894.1    mNeoVis.2.1.1.chr5_hap1.g01141.t01    rna-XM_021678776.1    rna-XM_004412076.1    rna-XM_007087396.3    rna-XM_008708505.2    rna-XM_027583615.2    mNeoVis.3.1.1.chr2.g01050.t01    rna-gnl|WGS_ZZZZ|egapxtmp_038515-R1    mNeoVis.evergladensis.2.1.chr2.g01131.t01    mNeoVis.vulgivaga.2.1.chr2.g01137.t01      2886    N0.HOG0002884   OG0000584  n1   rna-XM_034662839.1    rna-XM_038578014.1    rna-XM_022499410.1    rna-XM_023240656.2    rna-XM_036103162.1    rna-XM_039233686.1    rna-XM_032837989.1    rna-XM_046027000.1    rna-XM_032311961.1    rna-XM_059169666.1    rna-XM_059397974.1    rna-XM_045068974.1    mNeoVis.2.1.1.chr2_hap1.g00395.t01    rna-XM_021700252.2    rna-XM_004408586.1    rna-XM_042961236.1    rna-XM_040623831.1    rna-XM_027594728.1    mNeoVis.3.1.1.chr5.g00381.t01    rna-gnl|WGS_ZZZZ|egapxtmp_006525-R6    mNeoVis.evergladensis.2.1.chr5.g00394.t01    mNeoVis.vulgivaga.2.1.chr5.g00394.t01      2887    N0.HOG0002885   OG0000584  n8   rna-XM_019806256.2    rna-XM_038582223.1    rna-XM_022513374.1    rna-XM_045056130.1    rna-XM_036095665.1    rna-XM_039250559.1    rna-XM_032862324.1    rna-XM_045997787.1    rna-XM_032332788.1    rna-XM_059173364.1    rna-XM_059375288.1    rna-XM_045079391.1    mNeoVis.2.1.1.chr6_hap1.g00617.t01    rna-XM_044912681.1    rna-XM_004391548.1    rna-XM_042984242.1    rna-XM_008706167.2    rna-XM_027600541.2    mNeoVis.3.1.1.chr4.g00623.t01    rna-gnl|WGS_ZZZZ|egapxtmp_043132-R2    mNeoVis.evergladensis.2.1.chr4.g00622.t01    mNeoVis.vulgivaga.2.1.chr4.g00632.t01      2888    N0.HOG0002886   OG0000584  n20   rna-XM_002927784.4    rna-XM_038551716.1    rna-XM_022521200.1    rna-XM_023253986.2    rna-XM_036069310.1    rna-XM_039252828.1    rna-XM_032846837.1    rna-XM_045999394.1    rna-XM_032335270.1    rna-XM_059175947.1    rna-XM_059416195.1    rna-XM_004758990.3    mNeoVis.2.1.1.chr1_hap1.g00897.t01    rna-XM_044916749.1    rna-XM_004412664.1    rna-XM_042981890.1    rna-XM_040622640.1    rna-XM_027604800.2    mNeoVis.3.1.1.chr1.g00919.t01    rna-gnl|WGS_ZZZZ|egapxtmp_038258-R2    mNeoVis.evergladensis.2.1.chr1.g00900.t01    mNeoVis.vulgivaga.2.1.chr1.g00912.t01      2889    N0.HOG0002887   OG0000585  n1   rna-XM_002916098.4    rna-XM_038578213.1    rna-XM_022499523.1    rna-XM_045040420.1    rna-XM_036072486.1    rna-XM_039230252.1    rna-XM_032836198.1    rna-XM_046026442.1    rna-XM_032313394.1    rna-XM_059169420.1    rna-XM_059398151.1    rna-XM_004749643.3    mNeoVis.2.1.1.chr2_hap1.g00315.t01    rna-XM_021699800.2    rna-XM_004412567.1    rna-XM_042960763.1    rna-XM_008696253.2    rna-XM_027587160.2    mNeoVis.3.1.1.chr5.g00306.t01    rna-gnl|WGS_ZZZZ|egapxtmp_008061-R1    mNeoVis.evergladensis.2.1.chr5.g00319.t01    mNeoVis.vulgivaga.2.1.chr5.g00320.t01      2890    N0.HOG0002888   OG0000585  n23   rna-XM_002924420.4    rna-NM_001002934.2    rna-XM_022505981.1    rna-XM_003985610.4    rna-XM_036094301.1    rna-XM_039220409.1    rna-XM_032869725.1    rna-XM_045996885.1    rna-XM_032331652.1    rna-XM_059154493.1    rna-XM_059395602.1    rna-XM_004772362.3    mNeoVis.2.1.1.chr6_hap1.g00398.t01    rna-XM_021677659.1    rna-XM_004396240.1    rna-XM_042982769.1    rna-XM_008705527.2    rna-XM_027598852.2    mNeoVis.3.1.1.chr4.g00401.t01    rna-gnl|WGS_ZZZZ|egapxtmp_000750-R1    mNeoVis.evergladensis.2.1.chr4.g00406.t01    mNeoVis.vulgivaga.2.1.chr4.g00406.t01      2891    N0.HOG0002889   OG0000585  n43   rna-XM_034656887.1    rna-NM_001003073.1    rna-XM_022493244.1    rna-XM_045036991.1    rna-XM_036080653.1    rna-XM_039239468.1    rna-XM_032851520.1    rna-XM_046000840.1    rna-XM_032337350.1    rna-XM_059174014.1    rna-XM_059416522.1    rna-XM_004737847.3    mNeoVis.2.1.1.chr1_hap1.g00194.t01    rna-XM_021697017.1    rna-XM_004402803.1    rna-XM_007093376.3    rna-XM_040630419.1    rna-XM_027606333.1    mNeoVis.3.1.1.chr1.g00195.t01    rna-gnl|WGS_ZZZZ|egapxtmp_023430-R1    mNeoVis.evergladensis.2.1.chr1.g00189.t01    mNeoVis.vulgivaga.2.1.chr1.g00192.t01      2892    N0.HOG0002890   OG0000586  n1   rna-XM_011222001.3    rna-XM_038578405.1    rna-XM_022499320.1    rna-XM_019813631.3    rna-XM_036111605.1    rna-XM_039230187.1    rna-XM_032883205.1    rna-XM_046027075.1    rna-XM_032313090.1    rna-XM_059169308.1    rna-XM_059398257.1    rna-XM_004749578.3    mNeoVis.2.1.1.chr2_hap1.g00286.t01    rna-XM_021699806.1    rna-XM_004409478.1    rna-XM_042960064.1    rna-XM_040624555.1    rna-XM_035724321.1    mNeoVis.3.1.1.chr5.g00279.t01    rna-gnl|WGS_ZZZZ|egapxtmp_008051-R1    mNeoVis.evergladensis.2.1.chr5.g00291.t01    mNeoVis.vulgivaga.2.1.chr5.g00290.t01      2893    N0.HOG0002891   OG0000586  n23   rna-XM_034656754.1    rna-XM_038535455.1    rna-XM_022493179.1    rna-XM_023259788.2    rna-XM_036106743.1    rna-XM_039247657.1    rna-XM_032851770.1    rna-XM_045999729.1    rna-XM_032334904.1    rna-XM_059174255.1    rna-XM_059417324.1    rna-XM_004737643.2    mNeoVis.2.1.1.chr1_hap1.g00289.t01    rna-XM_044917114.1    rna-XM_004392025.1    rna-XM_042992684.1    rna-XM_040630315.1    rna-XM_027606988.2    mNeoVis.3.1.1.chr1.g00294.t01    rna-gnl|WGS_ZZZZ|egapxtmp_027812-R1    mNeoVis.evergladensis.2.1.chr1.g00288.t01    mNeoVis.vulgivaga.2.1.chr1.g00292.t01      2894    N0.HOG0002892   OG0000586  n44   rna-XM_019796533.2    rna-XM_038533590.1    rna-XM_022505792.1    rna-XM_023253260.2    rna-XM_036120769.1    rna-XM_039216599.1    rna-XM_032854235.1    rna-XM_045997781.1    rna-XM_032333963.1    rna-XM_059164780.1    rna-XM_059381673.1    rna-XM_013061510.2    mNeoVis.2.1.1.chr6_hap1.g00276.t01    rna-XM_021679395.2    rna-XM_004415159.1    rna-XM_007093741.2    rna-XM_040625100.1    rna-XM_027599650.2    mNeoVis.3.1.1.chr4.g00273.t01    rna-gnl|WGS_ZZZZ|egapxtmp_000962-R1    mNeoVis.evergladensis.2.1.chr4.g00283.t01    mNeoVis.vulgivaga.2.1.chr4.g00282.t01      2895    N0.HOG0002893   OG0000587  n1   rna-XM_034663279.1    rna-XM_038577959.1    rna-XM_022499504.1    rna-NM_001039955.1    rna-XM_036099054.1    rna-XM_039250722.1    rna-XM_032880190.1    rna-XM_046028060.1    rna-XM_032312586.1    rna-XM_059173219.1    rna-XM_059398566.1    rna-XM_045068707.1    mNeoVis.2.1.1.chr2_hap1.g00153.t01    rna-XM_021699900.2    rna-XM_004411589.1    rna-XM_042961040.1    rna-XM_040624821.1    rna-XM_027597135.2      rna-gnl|WGS_ZZZZ|egapxtmp_005589-R1    mNeoVis.evergladensis.2.1.chr5.g00153.t01    mNeoVis.vulgivaga.2.1.chr5.g00154.t01      2896    N0.HOG0002894   OG0000587  n21   rna-XM_034659579.1    rna-XM_038560594.1    rna-XM_022520500.1    rna-XM_006930331.4    rna-XM_036074877.1    rna-XM_039231783.1    rna-XM_032843907.1    rna-XM_045978634.1    rna-XM_032352825.1    rna-XM_059188443.1    rna-XM_059405551.1    rna-XM_004770934.3    mNeoVis.2.1.1.chr11_hap1.g00941.t01    rna-XM_021697667.1    rna-XM_004399838.1    rna-XM_042979789.1    rna-XM_040623163.1, rna-XM_040623165.1    rna-XM_027622732.2    mNeoVis.3.1.1.chr11.g00895.t01      mNeoVis.evergladensis.2.1.chr11.g00919.t01    mNeoVis.vulgivaga.2.1.chr11.g00943.t01      2897    N0.HOG0002895   OG0000587  n40   rna-XM_034670985.1    rna-XM_038533165.1    rna-XM_022506300.1    rna-XM_023253342.2    rna-XM_036085637.1    rna-XM_039232091.1    rna-XM_032855800.1    rna-XM_045997165.1    rna-XM_032333700.1    rna-XM_059166259.1    rna-XM_059395672.1    rna-XM_045075177.1    mNeoVis.2.1.1.chr6_hap1.g00351.t01    rna-XM_021677676.1    rna-XM_004396193.1    rna-XM_042982759.1    rna-XM_040624887.1, rna-XM_040624929.1    rna-XM_027599118.2      rna-gnl|WGS_ZZZZ|egapxtmp_043272-R1    mNeoVis.evergladensis.2.1.chr4.g00357.t01    mNeoVis.vulgivaga.2.1.chr4.g00353.t01      2898    N0.HOG0002896   OG0000588  n1   rna-XM_002918684.4    rna-XM_038537884.1    rna-XM_022514633.1    rna-XM_023258070.2    rna-XM_036090488.1    rna-XM_039234926.1    rna-XM_032863327.1    rna-XM_046018347.1    rna-XM_032302558.1    rna-XM_059137128.1    rna-XM_059374893.1    rna-XM_004768615.3    mNeoVis.2.1.1.chr2_hap1.g01845.t01    rna-XM_021684017.1    rna-XM_004397621.1    rna-XM_042996493.1    rna-XM_040645746.1    rna-XM_035726946.1    mNeoVis.3.1.1.chr5.g01772.t01    rna-gnl|WGS_ZZZZ|egapxtmp_007190-R1    mNeoVis.evergladensis.2.1.chr5.g01783.t01    mNeoVis.vulgivaga.2.1.chr5.g01788.t01      2899    N0.HOG0002897   OG0000588  n21   rna-XM_019805772.2    rna-XM_038575818.1    rna-XM_022495911.1    rna-XM_045041682.1    rna-XM_036076497.1    rna-XM_039242857.1    rna-XM_032876013.1    rna-XM_046025635.1    rna-XM_032310925.1    rna-XM_059140785.1    rna-XM_059408965.1    rna-XM_004742487.3    mNeoVis.2.1.1.chr3_hap1.g00363.t01    rna-XM_021687896.1    rna-XM_004400529.1    rna-XM_042962381.1    rna-XM_040619517.1    rna-XM_027577269.2    mNeoVis.3.1.1.chr3.g00355.t01    rna-gnl|WGS_ZZZZ|egapxtmp_041068-R1    mNeoVis.evergladensis.2.1.chr3.g00371.t01    mNeoVis.vulgivaga.2.1.chr3.g00366.t01      2900    N0.HOG0002898   OG0000588  n42   rna-XM_034672122.1    rna-NM_001130835.2    rna-XM_022506017.1    rna-XM_019829750.3    rna-XM_036110535.1    rna-XM_039248988.1    rna-XM_032855044.1    rna-XM_045997128.1    rna-XM_032333828.1    rna-XM_059165331.1    rna-XM_059380851.1    rna-XM_013044748.2    mNeoVis.2.1.1.chr6_hap1.g00309.t01    rna-XM_021677592.1    rna-XM_004415890.1    rna-XM_042984935.1    rna-XM_008685761.2    rna-XM_027598278.2    mNeoVis.3.1.1.chr4.g00309.t01    rna-gnl|WGS_ZZZZ|egapxtmp_044179-R1    mNeoVis.evergladensis.2.1.chr4.g00315.t01    mNeoVis.vulgivaga.2.1.chr4.g00312.t01      2901    N0.HOG0002899   OG0000589  n1   rna-XM_019808121.2    rna-XM_038537949.1    rna-XM_022514364.1    rna-XM_023258093.2    rna-XM_036123468.1    rna-XM_039234996.1    rna-XM_032863361.1    rna-XM_045998371.1    rna-XM_032301545.1    rna-XM_059135590.1    rna-XM_059375327.1    rna-XM_045063717.1    mNeoVis.2.1.1.chr2_hap1.g01831.t01    rna-XM_044914382.1    rna-XM_004411129.1    rna-XM_042997511.1    rna-XM_008694741.2    rna-XM_035726999.1    mNeoVis.3.1.1.chr5.g01756.t01    rna-gnl|WGS_ZZZZ|egapxtmp_009111-R2    mNeoVis.evergladensis.2.1.chr5.g01770.t01    mNeoVis.vulgivaga.2.1.chr5.g01774.t01      2902    N0.HOG0002900   OG0000589  n24   rna-XM_034647116.1    rna-XM_038536511.1    rna-XM_022524698.1    rna-XM_023243539.2    rna-XM_036078304.1    rna-XM_039254531.1    rna-XM_032839846.1    rna-XM_045985301.1    rna-XM_032320889.1    rna-XM_059148264.1    rna-XM_059380231.1    rna-XM_045083201.1    mNeoVis.2.1.1.chr8_hap1.g01113.t01    rna-XM_021694877.1    rna-XM_004398590.1    rna-XM_042967133.1    rna-XM_040629864.1    rna-XM_035724639.1    mNeoVis.3.1.1.chr8.g01032.t01      mNeoVis.evergladensis.2.1.chr9.g01091.t01    mNeoVis.vulgivaga.2.1.chr8.g01090.t01      2903    N0.HOG0002901   OG0000589  n44   rna-XM_034655549.1    rna-NM_001168473.2    rna-XM_022517434.1    rna-XM_023259973.2    rna-XM_036117566.1    rna-XM_039228553.1    rna-XM_032857674.1    rna-XM_046018520.1    rna-XM_032355408.1    rna-XM_059167471.1    rna-XM_059393862.1    rna-XM_013051867.2    mNeoVis.2.1.1.chr3_hap1.g01012.t01    rna-XM_044913468.1    rna-XM_004412695.1    rna-XM_042995912.1    rna-XM_040638149.1    rna-XM_027590345.2    mNeoVis.3.1.1.chr3.g01002.t01    rna-gnl|WGS_ZZZZ|egapxtmp_041233-R1    mNeoVis.evergladensis.2.1.chr3.g01009.t01    mNeoVis.vulgivaga.2.1.chr3.g01028.t01      2904    N0.HOG0002902   OG0000590  n1   rna-XM_002927156.4    rna-XM_038551335.1    rna-XM_022504553.1    rna-XM_011281009.4    rna-XM_036077009.1    rna-XM_039240325.1    rna-XM_032873865.1    rna-XM_045978789.1    rna-XM_032352212.1    rna-XM_059187749.1    rna-XM_059406020.1    rna-XM_004742010.3    mNeoVis.2.1.1.chr11_hap1.g00712.t01    rna-XM_021699054.2    rna-XM_004397713.1    rna-XM_007075373.3    rna-XM_040623446.1    rna-XM_027621858.1    mNeoVis.3.1.1.chr11.g00675.t01    rna-gnl|WGS_ZZZZ|egapxtmp_027023-R1    mNeoVis.evergladensis.2.1.chr11.g00696.t01    mNeoVis.vulgivaga.2.1.chr11.g00712.t01      2905    N0.HOG0002903   OG0000590  n24   rna-XM_011221305.3    rna-XM_038539578.1    rna-XM_022501634.1    rna-XM_011290434.4    rna-XM_036116561.1    rna-XM_039226157.1    rna-XM_032872770.1    rna-XM_045993991.1    rna-XM_032326844.1    rna-XM_059154028.1    rna-XM_059414332.1    rna-XM_013047109.2    mNeoVis.2.1.1.chr15_hap1.g00598.t01    rna-XM_021680132.1      rna-XM_015537116.2    rna-XM_040634723.1    rna-XM_027610869.2    mNeoVis.3.1.1.chr15.g00381.t01    rna-gnl|WGS_ZZZZ|egapxtmp_034017-R1    mNeoVis.evergladensis.2.1.chr15.g00420.t01    mNeoVis.vulgivaga.2.1.chr15.g00427.t01      2906    N0.HOG0002904   OG0000590  n43   rna-XM_011234825.3    rna-XM_038545674.1      rna-XM_019833613.3    rna-XM_036104697.1                    rna-XM_021679831.2    rna-XM_004416486.1    rna-XM_042990488.1    rna-XM_040641455.1    rna-XM_027570220.2              2907    N0.HOG0002905   OG0000590  n51       rna-XM_022505458.1          rna-XM_032837314.1    rna-XM_046009281.1    rna-XM_032345209.1    rna-XM_059183059.1    rna-XM_059373928.1    rna-XM_004754724.2    mNeoVis.2.1.1.chr10_hap1.g00095.t01            rna-XM_035724036.1    mNeoVis.3.1.1.chr9.g00069.t01    rna-gnl|WGS_ZZZZ|egapxtmp_015614-R1    mNeoVis.evergladensis.2.1.chr8.g00090.t01    mNeoVis.vulgivaga.2.1.chr9.g00090.t01      2908    N0.HOG0002906   OG0000591  n1   rna-XM_011236941.2    rna-XM_038536399.1    rna-XM_022524759.1    rna-XM_006939708.3    rna-XM_036078394.1    rna-XM_039254521.1    rna-XM_032840026.1    rna-XM_045985949.1    rna-XM_032322723.1    rna-XM_059149494.1    rna-XM_059380025.1    rna-XM_013060383.2    mNeoVis.2.1.1.chr8_hap1.g01141.t01    rna-XM_044921788.1    rna-XM_004406137.1    rna-XM_042967040.1    rna-XM_040629834.1    rna-XM_027568213.2    mNeoVis.3.1.1.chr8.g01059.t01    rna-gnl|WGS_ZZZZ|egapxtmp_020373-R1    mNeoVis.evergladensis.2.1.chr9.g01118.t01    mNeoVis.vulgivaga.2.1.chr8.g01116.t01      2909    N0.HOG0002907   OG0000591  n23   rna-XM_034647261.1    rna-XM_038536408.1    rna-XM_022524536.1    rna-XM_023244680.2    rna-XM_036078389.1, rna-XM_036098452.1    rna-XM_039254491.1    rna-XM_032840024.1    rna-XM_045986358.1    rna-XM_032322720.1    rna-XM_059149490.1    rna-XM_059380021.1    rna-XM_045083158.1    mNeoVis.2.1.1.chr8_hap1.g01140.t01    rna-XM_021694557.2    rna-XM_004406132.1    rna-XM_042966284.1    rna-XM_040629840.1    rna-XM_027625708.1    mNeoVis.3.1.1.chr8.g01058.t01    rna-gnl|WGS_ZZZZ|egapxtmp_020371-R1    mNeoVis.evergladensis.2.1.chr9.g01117.t01    mNeoVis.vulgivaga.2.1.chr8.g01115.t01      2910    N0.HOG0002908   OG0000591  n45   rna-XM_034670205.1    rna-XM_038539969.1    rna-XM_022526459.1    rna-XM_045047644.1    rna-XM_036066675.1    rna-XM_039247868.1    rna-XM_032842586.1    rna-XM_045994021.1    rna-XM_032327118.1    rna-XM_059154571.1    rna-XM_059415058.1    rna-XM_004773997.2    mNeoVis.2.1.1.chr15_hap1.g00097.t01    rna-XM_021700458.1    rna-XM_004397141.1    rna-XM_042971515.1    rna-XM_040635167.1    rna-XM_027611618.1    mNeoVis.3.1.1.chr15.g00092.t01      mNeoVis.evergladensis.2.1.chr15.g00098.t01    mNeoVis.vulgivaga.2.1.chr15.g00101.t01      2911    N0.HOG0002909   OG0000592  n1   rna-XM_002920035.3    rna-XM_038583249.1    rna-XM_022509744.1    rna-XM_006935978.4    rna-XM_036070319.1    rna-XM_039221983.1    rna-XM_032882250.1    rna-XM_046003633.1    rna-XM_032351470.1    rna-XM_059165269.1    rna-XM_059388166.1    rna-XM_004773003.3    mNeoVis.2.1.1.chr5_hap1.g01679.t01    rna-XM_021687950.1    rna-XM_004391677.1    rna-XM_042956257.1    rna-XM_008689678.1    rna-XM_027586613.1    mNeoVis.3.1.1.chr2.g01591.t01    rna-gnl|WGS_ZZZZ|egapxtmp_035202-R2    mNeoVis.evergladensis.2.1.chr2.g01662.t01    mNeoVis.vulgivaga.2.1.chr2.g01679.t01      2912    N0.HOG0002910   OG0000592  n22   rna-XM_034652823.1, rna-XM_034670813.1    rna-XM_038553358.1    rna-XM_022521102.1    rna-XM_003986368.4    rna-XM_036065177.1    rna-XM_039232185.1    rna-XM_032858367.1    rna-XM_046004878.1    rna-XM_032338324.1    rna-XM_059176327.1    rna-XM_059399242.1    rna-XM_004766917.3    mNeoVis.2.1.1.chr1_hap1.g01946.t01    rna-XM_021693454.1    rna-XM_004404297.1    rna-XM_042986799.1    rna-XM_008687538.1    rna-XM_027603019.2      rna-gnl|WGS_ZZZZ|egapxtmp_022852-R1    mNeoVis.evergladensis.2.1.chr1.g01912.t01    mNeoVis.vulgivaga.2.1.chr1.g01909.t01      2913    N0.HOG0002911   OG0000592  n42   rna-XM_002922666.4    rna-XM_038554722.1    rna-XM_022521086.1    rna-XM_003986370.5    rna-XM_036065163.1    rna-XM_039232176.1    rna-XM_032858103.1    rna-XM_046004323.1    rna-XM_032338322.1    rna-XM_059176326.1    rna-XM_059399243.1    rna-XM_004766916.3    mNeoVis.2.1.1.chr1_hap1.g01947.t01    rna-XM_021693422.1    rna-XM_004404299.1    rna-XM_007076927.3    rna-XM_008687544.2    rna-XM_027603018.2    mNeoVis.3.1.1.chr1.g01895.t01    rna-gnl|WGS_ZZZZ|egapxtmp_016596-R1    mNeoVis.evergladensis.2.1.chr1.g01913.t01    mNeoVis.vulgivaga.2.1.chr1.g01910.t01      2914    N0.HOG0002912   OG0000593  n1   rna-XM_034650823.1    rna-XM_038558355.1    rna-XM_022497499.1    rna-XM_045035680.1    rna-XM_036086084.1    rna-XM_039238634.1    rna-XM_032877524.1    rna-XM_046010924.1    rna-XM_032303624.1    rna-XM_059137968.1    rna-XM_059374433.1    rna-XM_045064357.1    mNeoVis.2.1.1.chr2_hap1.g01389.t01    rna-XM_021684250.1    rna-XM_004400656.1    rna-XM_042996871.1    rna-XM_040627409.1    rna-XM_027609217.2    mNeoVis.3.1.1.chr5.g01315.t01    rna-gnl|WGS_ZZZZ|egapxtmp_007915-R1    mNeoVis.evergladensis.2.1.chr5.g01331.t01    mNeoVis.vulgivaga.2.1.chr5.g01329.t01      2915    N0.HOG0002913   OG0000593  n23   rna-XM_034660015.1    rna-XM_038567676.1    rna-XM_022525462.1    rna-XM_023250840.2    rna-XM_036111892.1    rna-XM_039247556.1    rna-XM_032883676.1    rna-XM_045990103.1    rna-XM_032307167.1    rna-XM_059159586.1    rna-XM_059388742.1    rna-XM_045086013.1    mNeoVis.2.1.1.chr5_hap1.g00139.t01    rna-XM_044916652.1    rna-XM_004395308.1    rna-XM_042977875.1    rna-XM_040629520.1    rna-XM_035727229.1    mNeoVis.3.1.1.chr2.g00091.t01    rna-gnl|WGS_ZZZZ|egapxtmp_034414-R1    mNeoVis.evergladensis.2.1.chr2.g00138.t01    mNeoVis.vulgivaga.2.1.chr2.g00134.t01      2916    N0.HOG0002914   OG0000593  n44   rna-XM_034663855.1    rna-XM_038552342.1    rna-XM_022517585.1    rna-XM_019815913.3    rna-XM_036082477.1    rna-XM_039252611.1    rna-XM_032845508.1    rna-XM_046022091.1    rna-XM_032307451.1    rna-XM_059143037.1    rna-XM_059411633.1    rna-XM_045069786.1    mNeoVis.2.1.1.chr13_hap1.g00639.t01    rna-XM_044920761.1    rna-XM_004405919.1    rna-XM_042963696.1    rna-XM_040632317.1    rna-XM_027615436.2    mNeoVis.3.1.1.chr13.g00566.t01    rna-gnl|WGS_ZZZZ|egapxtmp_031443-R3    mNeoVis.evergladensis.2.1.chr13.g00613.t01    mNeoVis.vulgivaga.2.1.chr13.g00616.t01      2917    N0.HOG0002915   OG0000594  n1   rna-XM_034663750.1    rna-XM_038548841.1    rna-XM_022501979.1    rna-XM_045043733.1    rna-XM_036108482.1    rna-XM_039255164.1    rna-XM_032866244.1    rna-XM_046022975.1    rna-XM_032307816.1    rna-XM_059142138.1    rna-XM_059410390.1    rna-XM_045082864.1    mNeoVis.2.1.1.chr13_hap1.g00075.t01    rna-XM_044920438.1    rna-XM_004411388.1    rna-XM_042964032.1    rna-XM_040632153.1    rna-XM_027615347.2    mNeoVis.3.1.1.chr13.g00040.t01    rna-gnl|WGS_ZZZZ|egapxtmp_031055-R1    mNeoVis.evergladensis.2.1.chr13.g00076.t01    mNeoVis.vulgivaga.2.1.chr13.g00074.t01      2918    N0.HOG0002916   OG0000594  n22   rna-XM_034657627.1    rna-XM_038567529.1    rna-XM_022525439.1    rna-XM_006928302.4    rna-XM_036066103.1    rna-XM_039225019.1    rna-XM_032883774.1    rna-XM_045992133.1    rna-XM_032308315.1    rna-XM_059159697.1    rna-XM_059388814.1    rna-XM_004771616.3    mNeoVis.2.1.1.chr5_hap1.g00179.t01    rna-XM_021705580.1    rna-XM_004395270.1    rna-XM_042977968.1    rna-XM_008711294.2    rna-XM_027585291.2    mNeoVis.3.1.1.chr2.g00127.t01    rna-gnl|WGS_ZZZZ|egapxtmp_034631-R1    mNeoVis.evergladensis.2.1.chr2.g00175.t01    mNeoVis.vulgivaga.2.1.chr2.g00174.t01      2919    N0.HOG0002917   OG0000594  n41   rna-XM_002928305.4    rna-XM_038541164.1    rna-XM_022496164.1    rna-XM_011285035.4    rna-XM_036089450.1    rna-XM_039254969.1    rna-XM_032867257.1    rna-XM_046003440.1    rna-XM_032302316.1    rna-XM_059137588.1    rna-XM_059375593.1    rna-XM_045081970.1    mNeoVis.2.1.1.chr2_hap1.g01020.t01    rna-XM_021690322.1    rna-XM_004405562.1    rna-XM_007090051.2    rna-XM_008704211.2    rna-XM_027620157.2    mNeoVis.3.1.1.chr5.g00947.t01    rna-gnl|WGS_ZZZZ|egapxtmp_005324-R2    mNeoVis.evergladensis.2.1.chr5.g00967.t01    mNeoVis.vulgivaga.2.1.chr5.g00956.t01      2920    N0.HOG0002918   OG0000595  n1   rna-XM_034652921.1, rna-XM_034653237.1    rna-XM_038561960.1    rna-XM_022526120.1    rna-XM_045033375.1, rna-XM_045033384.1    rna-XM_036090103.1      rna-XM_032848189.1, rna-XM_032850927.1    rna-XM_046022588.1    rna-XM_032302416.1, rna-XM_032302737.1, rna-XM_032302741.1, rna-XM_032302743.1, rna-XM_032302745.1, rna-XM_032302747.1, rna-XM_032302748.1    rna-XM_059137874.1      rna-XM_004782494.3, rna-XM_045081985.1, rna-XM_045081987.1    mNeoVis.2.1.1.chr2_hap1.g00869.t01, mNeoVis.2.1.1.chr2_hap1.g00871.t01, mNeoVis.2.1.1.chr2_hap1.g01008.t01, mNeoVis.2.1.1.chr2_hap1.g01009.t01, mNeoVis.2.1.1.chr2_hap1.g01010.t01, mNeoVis.2.1.1.chr2_hap1.g01011.t01, mNeoVis.2.1.1.chr2_hap1.g01012.t01    rna-XM_044914346.1      rna-XM_042993920.1, rna-XM_042993922.1    rna-XM_040626581.1, rna-XM_040626716.1, rna-XM_040626850.1    rna-XM_035725994.1    mNeoVis.3.1.1.chr5.g00804.t01, mNeoVis.3.1.1.chr5.g00806.t01, mNeoVis.3.1.1.chr5.g00938.t01    rna-gnl|WGS_ZZZZ|egapxtmp_006550-R1, rna-gnl|WGS_ZZZZ|egapxtmp_008252-R1    mNeoVis.evergladensis.2.1.chr5.g00822.t01    mNeoVis.vulgivaga.2.1.chr5.g00813.t01, mNeoVis.vulgivaga.2.1.chr5.g00947.t01, mNeoVis.vulgivaga.2.1.chr5.g00948.t01      2921    N0.HOG0002919   OG0000595  n41   rna-XM_034641839.1    rna-XM_038562062.1    rna-XM_022526121.1    rna-XM_019837757.3    rna-XM_036098403.1      rna-XM_032843665.1    rna-XM_046018190.1    rna-XM_032360606.1    rna-XM_059137998.1    rna-XM_059377119.1    rna-XM_013048001.2    mNeoVis.2.1.1.chr2_hap1.g00839.t01    rna-XM_021687978.2    rna-XM_004404286.1    rna-XM_042993970.1    rna-XM_008695758.2    rna-XM_027608924.2    mNeoVis.3.1.1.chr5.g00777.t01    rna-gnl|WGS_ZZZZ|egapxtmp_006204-R1    mNeoVis.evergladensis.2.1.chr5.g00793.t01    mNeoVis.vulgivaga.2.1.chr5.g00783.t01, mNeoVis.vulgivaga.2.1.chr5.g00784.t01      2922    N0.HOG0002920   OG0000596  n1   rna-XM_034654840.1    rna-XM_038558248.1    rna-XM_022496715.1    rna-XM_019836903.3    rna-XM_036080597.1    rna-XM_039238630.1    rna-XM_032877615.1    rna-XM_046023252.1    rna-XM_032361603.1    rna-XM_059135338.1    rna-XM_059374562.1    rna-XM_004741536.3    mNeoVis.2.1.1.chr2_hap1.g01362.t01    rna-XM_021683842.1    rna-XM_004400608.1    rna-XM_007077431.3    rna-XM_040627363.1    rna-XM_027573834.2    mNeoVis.3.1.1.chr5.g01289.t01    rna-gnl|WGS_ZZZZ|egapxtmp_007061-R1    mNeoVis.evergladensis.2.1.chr5.g01305.t01    mNeoVis.vulgivaga.2.1.chr5.g01301.t01      2923    N0.HOG0002921   OG0000596  n21   rna-XM_034657871.1    rna-XM_038567080.1    rna-XM_022526956.1    rna-XM_023246142.2    rna-XM_036066461.1    rna-XM_039252335.1    rna-XM_032835870.1    rna-XM_045991961.1    rna-XM_032313071.1    rna-XM_059160176.1    rna-XM_059389185.1    rna-XM_004748273.3    mNeoVis.2.1.1.chr5_hap1.g00359.t01    rna-XM_021705588.2    rna-XM_004398969.1    rna-XM_042978336.1    rna-XM_040629212.1    rna-XM_027585150.2    mNeoVis.3.1.1.chr2.g00293.t01      mNeoVis.evergladensis.2.1.chr2.g00360.t01    mNeoVis.vulgivaga.2.1.chr2.g00355.t01      2924   N0.HOG0002922  OG0000596  n39   rna-XM_034655634.1    rna-XM_038530694.1    rna-XM_022518037.1    rna-XM_023256902.2    rna-XM_036070961.1    rna-XM_039255588.1    rna-XM_032849910.1    rna-XM_045999662.1    rna-XM_032336162.1    rna-XM_059175070.1    rna-XM_059417938.1    rna-XM_045075298.1    mNeoVis.2.1.1.chr1_hap1.g00600.t01    rna-XM_044916658.1    rna-XM_004416018.1    rna-XM_042987450.1    rna-XM_008691674.1    rna-XM_027604835.2    mNeoVis.3.1.1.chr1.g00617.t01    rna-gnl|WGS_ZZZZ|egapxtmp_038638-R1    mNeoVis.evergladensis.2.1.chr1.g00605.t01    mNeoVis.vulgivaga.2.1.chr1.g00614.t01      2925   N0.HOG0002923  OG0000597  n1   rna-XM_034653807.1    rna-XM_038541520.1    rna-XM_022496291.1, rna-XM_022496606.1    rna-XM_019837340.2    rna-XM_036108857.1    rna-XM_039246913.1    rna-XM_032870269.1    rna-XM_046011747.1    rna-XM_032301960.1    rna-XM_059135703.1    rna-XM_059375284.1    rna-XM_045064793.1    mNeoVis.2.1.1.chr2_hap1.g01133.t01    rna-XM_021690103.1    rna-XM_004413455.1    rna-XM_042997706.1    rna-XM_040627087.1    rna-XM_035726937.1    mNeoVis.3.1.1.chr5.g01052.t01    rna-gnl|WGS_ZZZZ|egapxtmp_009202-R1    mNeoVis.evergladensis.2.1.chr5.g01075.t01    mNeoVis.vulgivaga.2.1.chr5.g01065.t01      2926    N0.HOG0002924   OG0000597  n24   rna-XM_034657703.1    rna-XM_038567419.1    rna-XM_022525859.1    rna-XM_006928355.5    rna-XM_036065960.1    rna-XM_039251581.1    rna-XM_032883873.1    rna-XM_045989856.1    rna-XM_032309798.1    rna-XM_059159836.1    rna-XM_059388897.1    rna-XM_045084790.1    mNeoVis.2.1.1.chr5_hap1.g00224.t01    rna-XM_021704987.2    rna-XM_004412176.1    rna-XM_042978061.1    rna-XM_040629402.1    rna-XM_027583011.2    mNeoVis.3.1.1.chr2.g00171.t01      mNeoVis.evergladensis.2.1.chr2.g00220.t01    mNeoVis.vulgivaga.2.1.chr2.g00220.t01      2927    N0.HOG0002925   OG0000597  n40   rna-XM_002919391.4    rna-XM_038541512.1    rna-XM_022497002.1    rna-XM_045036117.1    rna-XM_036108856.1    rna-XM_039246930.1    rna-XM_032870275.1    rna-XM_046014831.1    rna-XM_032302795.1    rna-XM_059135701.1    rna-XM_059376974.1    rna-XM_045064798.1    mNeoVis.2.1.1.chr2_hap1.g01132.t01    rna-XM_044914058.1    rna-XM_004413453.1    rna-XM_007079174.3    rna-XM_008690651.2    rna-XM_027604146.1    mNeoVis.3.1.1.chr5.g01051.t01    rna-gnl|WGS_ZZZZ|egapxtmp_009200-R1    mNeoVis.evergladensis.2.1.chr5.g01074.t01    mNeoVis.vulgivaga.2.1.chr5.g01064.t01      2928    N0.HOG0002926   OG0000598  n1   rna-XM_002912864.4    rna-XM_038557135.1    rna-XM_022503374.1    rna-XM_006929256.5    rna-XM_036097210.1    rna-XM_039221791.1    rna-XM_032872996.1    rna-XM_046020812.1    rna-XM_032304466.1    rna-XM_059171380.1    rna-XM_059396712.1    rna-XM_004743308.3    mNeoVis.2.1.1.chr4_hap1.g00396.t01    rna-XM_021689656.1    rna-XM_004397345.1    rna-XM_042968434.1    rna-XM_040644174.1    rna-XM_027573545.2    mNeoVis.3.1.1.chr6.g00396.t01    rna-gnl|WGS_ZZZZ|egapxtmp_002365-R1    mNeoVis.evergladensis.2.1.chr6.g00399.t01    mNeoVis.vulgivaga.2.1.chr6.g00395.t01      2929   N0.HOG0002927  OG0000598  n21   rna-XM_034654792.1    rna-XM_038584953.1    rna-XM_022505318.1    rna-XM_023259488.2    rna-XM_036099143.1    rna-XM_039222479.1    rna-XM_032855984.1    rna-XM_046019015.1    rna-XM_032356047.1    rna-XM_059166931.1    rna-XM_059394230.1    rna-XM_004768981.2    mNeoVis.2.1.1.chr3_hap1.g00841.t01    rna-XM_021702724.2    rna-XM_004403732.1    rna-XM_042994792.1    rna-XM_008687184.2    rna-XM_027587966.2    mNeoVis.3.1.1.chr3.g00824.t01    rna-gnl|WGS_ZZZZ|egapxtmp_041195-R1    mNeoVis.evergladensis.2.1.chr3.g00838.t01    mNeoVis.vulgivaga.2.1.chr3.g00849.t01      2930    N0.HOG0002928   OG0000598  n39   rna-XM_002916810.4    rna-XM_038547521.1    rna-XM_022509793.1    rna-XM_045044620.1    rna-XM_036119688.1    rna-XM_039227931.1    rna-XM_032871403.1    rna-XM_045986193.1    rna-XM_032321519.1    rna-XM_059148872.1    rna-XM_059378752.1    rna-XM_004764709.3    mNeoVis.2.1.1.chr8_hap1.g00519.t01    rna-XM_044921527.1    rna-XM_004395087.1    rna-XM_042967069.1    rna-XM_040640735.1    rna-XM_027625896.2    mNeoVis.3.1.1.chr8.g00475.t01    rna-gnl|WGS_ZZZZ|egapxtmp_017431-R1    mNeoVis.evergladensis.2.1.chr9.g00510.t01    mNeoVis.vulgivaga.2.1.chr8.g00516.t01      2931    N0.HOG0002929   OG0000599  n0   rna-XM_002931024.4, rna-XM_002931025.4, rna-XM_002931222.4    rna-NM_001256454.1    rna-XM_022509407.1, rna-XM_022509897.1, rna-XM_022509971.1    rna-XM_003996404.2, rna-XM_003996405.2, rna-XM_003996406.2    rna-XM_036076641.1, rna-XM_036076646.1, rna-XM_036096073.1, rna-XM_036114217.1    rna-XM_039254067.1, rna-XM_039254196.1, rna-XM_039254211.1, rna-XM_039254220.1    rna-XM_032846727.1, rna-XM_032846766.1    rna-XM_045986235.1, rna-XM_045986245.1, rna-XM_045986380.1, rna-XM_045986406.1, rna-XM_045986418.1    rna-XM_032320652.1, rna-XM_032321711.1, rna-XM_032321817.1    rna-XM_059149582.1, rna-XM_059149678.1, rna-XM_059149679.1    rna-XM_059378576.1, rna-XM_059378581.1, rna-XM_059379612.1    rna-XM_004782400.1, rna-XM_013053455.2, rna-XM_013053476.2, rna-XM_045070287.1, rna-XM_045085472.1    mNeoVis.2.1.1.chr8_hap1.g00856.t01, mNeoVis.2.1.1.chr8_hap1.g00857.t01, mNeoVis.2.1.1.chr8_hap1.g00858.t01, mNeoVis.2.1.1.chr8_hap1.g00859.t01    rna-XM_021687125.2, rna-XM_021704396.1, rna-XM_021704443.2    rna-XM_004412129.1, rna-XM_004412131.1    rna-XM_007076139.2, rna-XM_007076140.2, rna-XM_007076141.2    rna-XM_008695264.2, rna-XM_008695265.2, rna-XM_008695266.2, rna-XM_008695531.2    rna-XM_027567946.2, rna-XM_027567962.1, rna-XM_027620383.1    mNeoVis.3.1.1.chr8.g00804.t01, mNeoVis.3.1.1.chr8.g00805.t01    rna-gnl|WGS_ZZZZ|egapxtmp_018661-R1, rna-gnl|WGS_ZZZZ|egapxtmp_018662-R1    mNeoVis.evergladensis.2.1.chr9.g00846.t01, mNeoVis.evergladensis.2.1.chr9.g00847.t01    mNeoVis.vulgivaga.2.1.chr8.g00848.t01, mNeoVis.vulgivaga.2.1.chr8.g00849.t01      2932    N0.HOG0002930   OG0000600  n1   rna-XM_019807529.2    rna-XM_038581130.1      rna-XM_045036452.1      rna-XM_039229587.1    rna-XM_032865522.1    rna-XM_046003707.1    rna-XM_032345596.1    rna-XM_059165096.1    rna-XM_059388387.1    rna-XM_004757556.3    mNeoVis.2.1.1.chr5_hap1.g01765.t01        rna-XM_042998315.1      rna-XM_027581773.2, rna-XM_035722473.1, rna-XM_035722485.1    mNeoVis.3.1.1.chr2.g01673.t01      mNeoVis.evergladensis.2.1.chr2.g01748.t01    mNeoVis.vulgivaga.2.1.chr2.g01764.t01      2933    N0.HOG0002931   OG0000600  n16   rna-XM_034655389.1, rna-XM_034655577.1    rna-XM_038581132.1, rna-XM_038581730.1, rna-XM_038581731.1    rna-XM_022515952.1, rna-XM_022515953.1, rna-XM_022515954.1, rna-XM_022515956.1    rna-XM_006935954.3    rna-XM_036118542.1, rna-XM_036118543.1      rna-XM_032865523.1, rna-XM_032865524.1, rna-XM_032865525.1, rna-XM_032865527.1    rna-XM_046003704.1, rna-XM_046003706.1    rna-XM_032302616.1, rna-XM_032345602.1, rna-XM_032345611.1, rna-XM_032345614.1    rna-XM_059165098.1, rna-XM_059165099.1    rna-XM_059388388.1, rna-XM_059388389.1, rna-XM_059388390.1    rna-XM_004757557.1, rna-XM_004757558.1, rna-XM_045085189.1, rna-XM_045085190.1    mNeoVis.2.1.1.chr5_hap1.g01766.t01, mNeoVis.2.1.1.chr5_hap1.g01768.t01    rna-XM_044919016.1        rna-XM_008701942.2, rna-XM_008701943.2, rna-XM_040621663.1, rna-XM_040621695.1      mNeoVis.3.1.1.chr2.g01674.t01, mNeoVis.3.1.1.chr2.g01675.t01    rna-gnl|WGS_ZZZZ|egapxtmp_034591-R1, rna-gnl|WGS_ZZZZ|egapxtmp_034597-R1, rna-gnl|WGS_ZZZZ|egapxtmp_035699-R1    mNeoVis.evergladensis.2.1.chr2.g01749.t01, mNeoVis.evergladensis.2.1.chr2.g01750.t01, mNeoVis.evergladensis.2.1.chr2.g01751.t01    mNeoVis.vulgivaga.2.1.chr2.g01765.t01, mNeoVis.vulgivaga.2.1.chr2.g01766.t01      2934   N0.HOG0002932  OG0000601  n1   rna-XM_034656545.1, rna-XM_034656548.1    rna-XM_038532268.1    rna-XM_022505004.1    rna-XM_023258660.2    rna-XM_036105314.1    rna-XM_039239383.1    rna-XM_032845992.1    rna-XM_045999842.1    rna-XM_032335684.1    rna-XM_059175518.1    rna-XM_059418265.1    rna-XM_004769945.3    mNeoVis.2.1.1.chr1_hap1.g00761.t01    rna-XM_021701690.1    rna-XM_004406604.1    rna-XM_042990525.1    rna-XM_040622052.1    rna-XM_027605318.2    mNeoVis.3.1.1.chr1.g00786.t01    rna-gnl|WGS_ZZZZ|egapxtmp_036078-R1    mNeoVis.evergladensis.2.1.chr1.g00769.t01    mNeoVis.vulgivaga.2.1.chr1.g00783.t01      2935   N0.HOG0002933  OG0000601  n24   rna-XM_034656549.1    rna-XM_038532266.1    rna-XM_022504871.1      rna-XM_036105312.1    rna-XM_039239353.1    rna-XM_032845967.1    rna-XM_046000909.1    rna-XM_032335683.1    rna-XM_059175540.1    rna-XM_059418279.1    rna-XM_004769949.3    mNeoVis.2.1.1.chr1_hap1.g00762.t01    rna-XM_021699218.1    rna-XM_004406605.1    rna-XM_042956647.1    rna-XM_040622048.1    rna-XM_027606703.2    mNeoVis.3.1.1.chr1.g00788.t01    rna-gnl|WGS_ZZZZ|egapxtmp_043127-R1    mNeoVis.evergladensis.2.1.chr1.g00770.t01    mNeoVis.vulgivaga.2.1.chr1.g00785.t01      2936    N0.HOG0002934   OG0000601  n44   rna-XM_011225481.3    rna-XM_038532256.1    rna-XM_022504721.1    rna-XM_045033957.1    rna-XM_036113416.1    rna-XM_039239367.1    rna-XM_032845937.1    rna-XM_045999457.1    rna-XM_032335681.1    rna-XM_059175541.1    rna-XM_059418282.1    rna-XM_004769950.3    mNeoVis.2.1.1.chr1_hap1.g00763.t01    rna-XM_021701642.1    rna-XM_004406606.1    rna-XM_042990660.1    rna-XM_008700890.2    rna-XM_027606697.2    mNeoVis.3.1.1.chr1.g00789.t01    rna-gnl|WGS_ZZZZ|egapxtmp_043245-R1    mNeoVis.evergladensis.2.1.chr1.g00771.t01    mNeoVis.vulgivaga.2.1.chr1.g00786.t01      2937    N0.HOG0002935   OG0000602  n1   rna-XM_002928643.2    rna-NM_001003213.1    rna-XM_022527130.1    rna-XM_045058476.1    rna-XM_036089247.1    rna-XM_039257585.1    rna-XM_032847824.1    rna-XM_046004785.1    rna-XM_032338243.1    rna-XM_059176307.1    rna-XM_059399682.1    rna-XM_004779971.3    mNeoVis.2.1.1.chr1_hap1.g01044.t01    rna-XM_021697100.1    rna-XM_004411295.1    rna-XM_007092719.3    rna-XM_008711508.1    rna-XM_027602230.1    mNeoVis.3.1.1.chr1.g01039.t01    rna-gnl|WGS_ZZZZ|egapxtmp_021128-R1    mNeoVis.evergladensis.2.1.chr1.g01036.t01    mNeoVis.vulgivaga.2.1.chr1.g01047.t01      2938    N0.HOG0002936   OG0000602  n23   rna-XM_002928644.3    rna-NM_001256320.1    rna-XM_022527129.1    rna-XM_003985655.6    rna-XM_036089248.1    rna-XM_039257584.1    rna-XM_032847823.1    rna-XM_046004784.1    rna-XM_032338245.1    rna-XM_059176244.1    rna-XM_059399681.1    rna-XM_004779972.2    mNeoVis.2.1.1.chr1_hap1.g01045.t01    rna-XM_021697101.1    rna-XM_004411297.1    rna-XM_042985448.1    rna-XM_040645172.1    rna-XM_027602225.2    mNeoVis.3.1.1.chr1.g01040.t01    rna-gnl|WGS_ZZZZ|egapxtmp_031582-R1    mNeoVis.evergladensis.2.1.chr1.g01037.t01    mNeoVis.vulgivaga.2.1.chr1.g01048.t01      2939   N0.HOG0002937  OG0000602  n42   rna-XM_002920757.4    rna-NM_001164454.1    rna-XM_022493103.1    rna-XM_003981387.6    rna-XM_036090652.1    rna-XM_039239511.1    rna-XM_032851590.1    rna-XM_046000538.1    rna-XM_032337281.1    rna-XM_059174075.1    rna-XM_059417187.1    rna-XM_004737800.3    mNeoVis.2.1.1.chr1_hap1.g00214.t01    rna-XM_044917141.1    rna-XM_004402830.1    rna-XM_007093409.3    rna-XM_040630394.1    rna-XM_027605857.1    mNeoVis.3.1.1.chr1.g00215.t01      mNeoVis.evergladensis.2.1.chr1.g00210.t01    mNeoVis.vulgivaga.2.1.chr1.g00213.t01      2940    N0.HOG0002938   OG0000603  n1   rna-XM_002915830.4    rna-XM_038584317.1    rna-XM_022527090.1    rna-XM_023253587.2    rna-XM_036112280.1    rna-XM_039257327.1    rna-XM_032861679.1    rna-XM_046005004.1    rna-XM_032341269.1    rna-XM_059179506.1    rna-XM_059398954.1    rna-XM_004778756.2    mNeoVis.2.1.1.chr1_hap1.g01157.t01    rna-XM_021697061.1    rna-XM_004406530.1    rna-XM_042985404.1    rna-XM_008693080.1    rna-XM_027602198.2    mNeoVis.3.1.1.chr1.g01116.t01    rna-gnl|WGS_ZZZZ|egapxtmp_018428-R1    mNeoVis.evergladensis.2.1.chr1.g01130.t01    mNeoVis.vulgivaga.2.1.chr1.g01128.t01      2941    N0.HOG0002939   OG0000603  n23   rna-XM_002915828.3    rna-XM_038584327.1    rna-XM_022527089.1    rna-XM_019830130.3    rna-XM_036112166.1    rna-XM_039257334.1    rna-XM_032861770.1    rna-XM_046005002.1    rna-XM_032341383.1    rna-XM_059179369.1    rna-XM_059398956.1    rna-XM_045082304.1    mNeoVis.2.1.1.chr1_hap1.g01155.t01      rna-XM_004406500.1    rna-XM_015540480.2    rna-XM_008693082.1    rna-XM_027602178.1    mNeoVis.3.1.1.chr1.g01114.t01    rna-gnl|WGS_ZZZZ|egapxtmp_021345-R1    mNeoVis.evergladensis.2.1.chr1.g01128.t01    mNeoVis.vulgivaga.2.1.chr1.g01126.t01      2942    N0.HOG0002940   OG0000603  n43   rna-XM_034660233.1    rna-XM_038584319.1    rna-XM_022527085.1    rna-XM_023253585.2    rna-XM_036112287.1    rna-XM_039257322.1    rna-XM_032861680.1    rna-XM_046005596.1    rna-XM_032341282.1    rna-XM_059179519.1    rna-XM_059398957.1    rna-XM_013049104.2    mNeoVis.2.1.1.chr1_hap1.g01154.t01    rna-XM_021697016.1    rna-XM_004406501.1    rna-XM_007074433.3    rna-XM_008692946.1    rna-XM_027602186.1    mNeoVis.3.1.1.chr1.g01113.t01    rna-gnl|WGS_ZZZZ|egapxtmp_018059-R3    mNeoVis.evergladensis.2.1.chr1.g01127.t01    mNeoVis.vulgivaga.2.1.chr1.g01125.t01      2943    N0.HOG0002941   OG0000604  n1   rna-XM_011218571.3    rna-XM_038553705.1    rna-XM_022510587.1    rna-XM_023253789.2    rna-XM_036072340.1    rna-XM_039241753.1    rna-XM_032850587.1, rna-XM_032850598.1    rna-XM_046004548.1    rna-XM_032340687.1    rna-XM_059178681.1    rna-XM_059399993.1    rna-XM_004770595.3    mNeoVis.2.1.1.chr1_hap1.g01419.t01    rna-XM_044917610.1    rna-XM_004407690.1    rna-XM_042986104.1    rna-XM_040644771.1    rna-XM_027601289.1    mNeoVis.3.1.1.chr1.g01366.t01      mNeoVis.evergladensis.2.1.chr1.g01381.t01    mNeoVis.vulgivaga.2.1.chr1.g01385.t01      2944    N0.HOG0002942   OG0000604  n22   rna-XM_002914172.4    rna-XM_038586222.1    rna-XM_022508723.1    rna-XM_003999535.6    rna-XM_036113586.1    rna-XM_039221247.1    rna-XM_032872988.1    rna-XM_045983631.1    rna-XM_032318468.1    rna-XM_059146736.1    rna-XM_059413396.1    rna-XM_013055544.2    mNeoVis.2.1.1.chr14_hap1.g00460.t01    rna-XM_021698429.1    rna-XM_004408340.1    rna-XM_042976383.1    rna-XM_040635992.1    rna-XM_027613416.2    mNeoVis.3.1.1.chr14.g00456.t01    rna-gnl|WGS_ZZZZ|egapxtmp_032595-R1    mNeoVis.evergladensis.2.1.chr14.g00457.t01    mNeoVis.vulgivaga.2.1.chr14.g00459.t01      2945    N0.HOG0002943   OG0000604  n41   rna-XM_034664761.1    rna-XM_038548860.1    rna-XM_022501410.1    rna-XM_006939617.5    rna-XM_036108654.1    rna-XM_039255157.1    rna-XM_032866238.1    rna-XM_046022598.1    rna-XM_032308990.1    rna-XM_059143581.1    rna-XM_059410359.1    rna-XM_013058548.2    mNeoVis.2.1.1.chr13_hap1.g00079.t01    rna-XM_044920391.1    rna-XM_004411393.1    rna-XM_042964700.1    rna-XM_040632159.1    rna-XM_027616268.2    mNeoVis.3.1.1.chr13.g00043.t01      mNeoVis.evergladensis.2.1.chr13.g00080.t01    mNeoVis.vulgivaga.2.1.chr13.g00078.t01      2946    N0.HOG0002944   OG0000605  n1   rna-XM_034671449.1    rna-NM_001197096.1    rna-XM_022514218.1    rna-XM_003985136.5    rna-XM_036095790.1    rna-XM_039223803.1    rna-XM_032860901.1    rna-XM_045992690.1    rna-XM_032333006.1    rna-XM_059172149.1    rna-XM_059376531.1    rna-XM_004748167.3    mNeoVis.2.1.1.chr6_hap1.g00551.t01    rna-XM_044912904.1    rna-XM_004391464.1    rna-XM_042984125.1    rna-XM_040625167.1    rna-XM_027598213.2    mNeoVis.3.1.1.chr4.g00557.t01    rna-gnl|WGS_ZZZZ|egapxtmp_000790-R1    mNeoVis.evergladensis.2.1.chr4.g00556.t01    mNeoVis.vulgivaga.2.1.chr4.g00565.t01      2947    N0.HOG0002945   OG0000605  n14   rna-XM_034662693.1    rna-XM_038534358.1    rna-XM_022499269.1    rna-XM_023240548.2    rna-XM_036110315.1    rna-XM_039235171.1    rna-XM_032840448.1    rna-XM_046027340.1    rna-XM_032311759.1    rna-XM_059169926.1    rna-XM_059397791.1    rna-XM_045083481.1    mNeoVis.2.1.1.chr2_hap1.g00471.t01    rna-XM_021700269.1    rna-XM_004407011.1      rna-XM_040624610.1    rna-XM_027588618.2    mNeoVis.3.1.1.chr5.g00463.t01    rna-gnl|WGS_ZZZZ|egapxtmp_005617-R1    mNeoVis.evergladensis.2.1.chr5.g00474.t01    mNeoVis.vulgivaga.2.1.chr5.g00473.t01      2948    N0.HOG0002946   OG0000605  n25   rna-XM_034661279.1    rna-XM_038573848.1    rna-XM_022513570.1    rna-XM_023252503.2    rna-XM_036075144.1    rna-XM_039243813.1    rna-XM_032855846.1    rna-XM_045996785.1    rna-XM_032332141.1    rna-XM_059178419.1    rna-XM_059371515.1    rna-XM_004763293.3    mNeoVis.2.1.1.chr6_hap1.g00836.t01    rna-XM_044912840.1    rna-XM_004402199.1    rna-XM_042983277.1    rna-XM_040626539.1    rna-XM_027597464.2    mNeoVis.3.1.1.chr4.g00847.t01    rna-gnl|WGS_ZZZZ|egapxtmp_001000-R1    mNeoVis.evergladensis.2.1.chr4.g00838.t01    mNeoVis.vulgivaga.2.1.chr4.g00858.t01      2949    N0.HOG0002947   OG0000606  n1   rna-XM_002921943.4    rna-XM_038538927.1    rna-XM_022496754.1    rna-XM_023257813.2    rna-XM_036103729.1    rna-XM_039233473.1    rna-XM_032870072.1    rna-XM_045986119.1    rna-XM_032302502.1    rna-XM_059137218.1    rna-XM_059375079.1    rna-XM_004762981.3    mNeoVis.2.1.1.chr2_hap1.g01199.t01    rna-XM_021679981.1    rna-XM_004415174.1    rna-XM_042997542.1    rna-XM_040627153.1    rna-XM_027571068.1    mNeoVis.3.1.1.chr5.g01115.t01      mNeoVis.evergladensis.2.1.chr5.g01141.t01    mNeoVis.vulgivaga.2.1.chr5.g01128.t01      2950    N0.HOG0002948   OG0000606  n10   rna-XM_034647166.1    rna-XM_038525621.1    rna-XM_022512768.1    rna-XM_023243239.2    rna-XM_036075051.1    rna-XM_039241475.1    rna-XM_032866416.1    rna-XM_046023076.1    rna-XM_032308422.1    rna-XM_059142867.1    rna-XM_059412339.1    rna-XM_004778803.3    mNeoVis.2.1.1.chr13_hap1.g00828.t01    rna-XM_021694337.1    rna-XM_004409847.1    rna-XM_042964445.1    rna-XM_040620337.1, rna-XM_040620339.1, rna-XM_040620391.1    rna-XM_027614587.2      rna-gnl|WGS_ZZZZ|egapxtmp_031629-R1    mNeoVis.evergladensis.2.1.chr13.g00790.t01    mNeoVis.vulgivaga.2.1.chr13.g00787.t01      2951    N0.HOG0002949   OG0000606  n21   rna-XM_002925898.4    rna-XM_038565343.1    rna-XM_022525747.1    rna-XM_003981616.6    rna-XM_036100994.1    rna-XM_039248008.1    rna-XM_032847056.1    rna-XM_045991233.1    rna-XM_032359042.1    rna-XM_059159366.1    rna-XM_059387671.1    rna-XM_004777977.2    mNeoVis.2.1.1.chr5_hap1.g00066.t01    rna-XM_044918629.1    rna-XM_004395378.1    rna-XM_042977250.1    rna-XM_008711216.2    rna-XM_027582594.2    mNeoVis.3.1.1.chr2.g00035.t01      mNeoVis.evergladensis.2.1.chr2.g00066.t01    mNeoVis.vulgivaga.2.1.chr2.g00066.t01      2952    N0.HOG0002950   OG0000607  n1   rna-XM_002913156.4, rna-XM_019796483.2    rna-XM_038529771.1    rna-XM_022515815.1    rna-XM_006933313.5, rna-XM_019816516.3    rna-XM_036072705.1, rna-XM_036112922.1    rna-XM_039233054.1, rna-XM_039238980.1    rna-XM_032865851.1, rna-XM_032880805.1    rna-XM_046011050.1, rna-XM_046023086.1    rna-XM_032308729.1, rna-XM_032349651.1    rna-XM_059141629.1, rna-XM_059183905.1    rna-XM_059404162.1, rna-XM_059410817.1    rna-XM_004757327.3, rna-XM_004757623.3    mNeoVis.2.1.1.chr13_hap1.g00230.t01, mNeoVis.2.1.1.chr9_hap1.g00082.t01    rna-XM_044914985.1, rna-XM_044920593.1    rna-XM_004392592.1, rna-XM_004403981.1    rna-XM_042965239.1, rna-XM_042991242.1    rna-XM_008698771.2, rna-XM_040645322.1    rna-XM_027595384.1, rna-XM_035723461.1    mNeoVis.3.1.1.chr10.g00078.t01, mNeoVis.3.1.1.chr13.g00181.t01    rna-gnl|WGS_ZZZZ|egapxtmp_029723-R1    mNeoVis.evergladensis.2.1.chr10.g00084.t01, mNeoVis.evergladensis.2.1.chr13.g00228.t01    mNeoVis.vulgivaga.2.1.chr10.g00077.t01, mNeoVis.vulgivaga.2.1.chr13.g00225.t01      2953    N0.HOG0002951   OG0000607  n34   rna-XM_034649916.1    rna-NM_001003357.2    rna-XM_022526309.1    rna-XM_023248905.2    rna-XM_036085830.1, rna-XM_036112038.1    rna-XM_039239860.1    rna-XM_032843345.1    rna-XM_045996285.1    rna-XM_032330824.1    rna-XM_059157660.1    rna-XM_059385450.1    rna-XM_045068095.1    mNeoVis.2.1.1.chr12_hap1.g00170.t01    rna-XM_021684341.1    rna-XM_004401077.1    rna-XM_042973725.1    rna-XM_040627500.1    rna-XM_027608382.2, rna-XM_035728584.1    mNeoVis.3.1.1.chr12.g00137.t01    rna-gnl|WGS_ZZZZ|egapxtmp_029529-R1    mNeoVis.evergladensis.2.1.chr12.g00144.t01    mNeoVis.vulgivaga.2.1.chr12.g00145.t01      2954    N0.HOG0002952   OG0000608  n1   rna-XM_002914532.4    rna-XM_038550661.1    rna-XM_022518914.1    rna-XM_023257477.2    rna-XM_036108707.1    rna-XM_039246168.1    rna-XM_032836594.1    rna-XM_046012624.1    rna-XM_032346372.1    rna-XM_059186902.1    rna-XM_059402075.1    rna-XM_004743172.3    mNeoVis.2.1.1.chr9_hap1.g01144.t01    rna-XM_021687635.1    rna-XM_004410803.1    rna-XM_042993200.1    rna-XM_008705580.2    rna-XM_027594893.2      rna-gnl|WGS_ZZZZ|egapxtmp_024398-R1    mNeoVis.evergladensis.2.1.chr10.g01132.t01    mNeoVis.vulgivaga.2.1.chr10.g01124.t01      2955    N0.HOG0002953   OG0000608  n9   rna-XM_011224660.3    rna-XM_038546276.1    rna-XM_022523279.1    rna-XM_023244353.2    rna-XM_036093590.1, rna-XM_036099009.1    rna-XM_039226424.1    rna-XM_032870355.1    rna-XM_045985632.1    rna-XM_032322174.1    rna-XM_059149855.1        mNeoVis.2.1.1.chr8_hap1.g00031.t01    rna-XM_021680711.2    rna-XM_004404466.1    rna-XM_042966139.1    rna-XM_040639846.1    rna-XM_027568134.2        mNeoVis.evergladensis.2.1.chr9.g00029.t01    mNeoVis.vulgivaga.2.1.chr8.g00028.t01      2956    N0.HOG0002954   OG0000608  n16   rna-XM_034669785.1    rna-NM_001003274.2    rna-XM_022500681.1    rna-XM_023246216.2    rna-XM_036103004.1    rna-XM_039226032.1    rna-XM_032872729.1    rna-XM_045994013.1    rna-XM_032328337.1    rna-XM_059155203.1    rna-XM_059414386.1    rna-XM_045075128.1    mNeoVis.2.1.1.chr15_hap1.g00617.t01    rna-XM_021680236.1, rna-XM_044920411.1    rna-XM_004394160.1, rna-XM_004416902.1    rna-XM_042971949.1    rna-XM_040634767.1    rna-XM_027611399.2, rna-XM_035726750.1    mNeoVis.3.1.1.chr15.g00400.t01    rna-gnl|WGS_ZZZZ|egapxtmp_033244-R1    mNeoVis.evergladensis.2.1.chr15.g00439.t01    mNeoVis.vulgivaga.2.1.chr15.g00447.t01      2957    N0.HOG0002955   OG0000609  n1   rna-XM_019796341.2    rna-XM_038568947.1    rna-XM_022522056.1    rna-XM_023239530.2    rna-XM_036089224.1    rna-XM_039222430.1    rna-XM_032853386.1    rna-XM_046016304.1    rna-XM_032356274.1    rna-XM_059135098.1    rna-XM_059409059.1    rna-XM_004780170.3    mNeoVis.2.1.1.chr7_hap1.g01779.t01    rna-XM_021685731.1    rna-XM_004394543.1    rna-XM_042959178.1    rna-XM_040630744.1    rna-XM_027579021.2    mNeoVis.3.1.1.chr7.g01650.t01      mNeoVis.evergladensis.2.1.chr7.g01740.t01    mNeoVis.vulgivaga.2.1.chr7.g01756.t01      2958    N0.HOG0002956   OG0000609  n22   rna-XM_002925431.4    rna-NM_001160425.1    rna-XM_022508721.1    rna-XM_003999379.5    rna-XM_036079607.1    rna-XM_039227630.1    rna-XM_032878302.1    rna-XM_045983576.1    rna-XM_032318663.1    rna-XM_059146375.1    rna-XM_059414080.1    rna-XM_004756324.1    mNeoVis.2.1.1.chr14_hap1.g00347.t01    rna-XM_021686578.1    rna-XM_004392780.1    rna-XM_007086473.2    rna-XM_008685135.2    rna-XM_027614027.2    mNeoVis.3.1.1.chr14.g00345.t01    rna-gnl|WGS_ZZZZ|egapxtmp_032201-R1    mNeoVis.evergladensis.2.1.chr14.g00347.t01    mNeoVis.vulgivaga.2.1.chr14.g00348.t01      2959    N0.HOG0002957   OG0000609  n41   rna-XM_034644019.1    rna-NM_001048084.1    rna-XM_022518880.1    rna-XM_045062469.1    rna-XM_036088364.1    rna-XM_039236219.1    rna-XM_032882849.1    rna-XM_046011649.1    rna-XM_032346062.1, rna-XM_032359072.1    rna-XM_059187167.1    rna-XM_059401867.1    rna-XM_045083759.1    mNeoVis.2.1.1.chr9_hap1.g01241.t01    rna-XM_021687779.1    rna-XM_004397392.1    rna-XM_042993433.1    rna-XM_008702920.2    rna-XM_027594797.2    mNeoVis.3.1.1.chr10.g01192.t01      mNeoVis.evergladensis.2.1.chr10.g01230.t01    mNeoVis.vulgivaga.2.1.chr10.g01217.t01      2960    N0.HOG0002958   OG0000610  n1   rna-XM_034649931.1    rna-XM_038588662.1    rna-XM_022491285.1    rna-XM_023249474.2    rna-XM_036120480.1    rna-XM_039219277.1    rna-XM_032846342.1    rna-XM_045995959.1    rna-XM_032330900.1    rna-XM_059157685.1    rna-XM_059385918.1    rna-XM_045067978.1    mNeoVis.2.1.1.chr12_hap1.g00059.t01    rna-XM_044912328.1    rna-XM_004409167.1    rna-XM_042974840.1    rna-XM_040629481.1    rna-XM_027607714.1    mNeoVis.3.1.1.chr12.g00050.t01    rna-gnl|WGS_ZZZZ|egapxtmp_029108-R1    mNeoVis.evergladensis.2.1.chr12.g00049.t01    mNeoVis.vulgivaga.2.1.chr12.g00051.t01      2961    N0.HOG0002959   OG0000610  n23   rna-XM_034643795.1    rna-XM_038550108.1    rna-XM_022518895.1    rna-XM_045062572.1    rna-XM_036113772.1    rna-XM_039235535.1    rna-XM_032882699.1    rna-XM_046011427.1    rna-XM_032345772.1    rna-XM_059187372.1    rna-XM_059401688.1    rna-XM_004775375.3    mNeoVis.2.1.1.chr9_hap1.g01308.t01    rna-XM_044915307.1    rna-XM_004409662.1    rna-XM_042993617.1    rna-XM_040637760.1    rna-XM_027592961.1      rna-gnl|WGS_ZZZZ|egapxtmp_023515-R1    mNeoVis.evergladensis.2.1.chr10.g01295.t01    mNeoVis.vulgivaga.2.1.chr10.g01285.t01      2962    N0.HOG0002960   OG0000610  n43   rna-XM_034658246.1    rna-XM_038566553.1    rna-XM_022497740.1    rna-XM_023248461.2    rna-XM_036079275.1    rna-XM_039228175.1    rna-XM_032867942.1    rna-XM_045990433.1    rna-XM_032318609.1    rna-XM_059160796.1    rna-XM_059389631.1    rna-XM_013060491.2    mNeoVis.2.1.1.chr5_hap1.g00613.t01    rna-XM_021686944.1    rna-XM_004399188.1    rna-XM_042978843.1    rna-XM_040628870.1    rna-XM_027584555.2    mNeoVis.3.1.1.chr2.g00535.t01    rna-gnl|WGS_ZZZZ|egapxtmp_035411-R1    mNeoVis.evergladensis.2.1.chr2.g00610.t01    mNeoVis.vulgivaga.2.1.chr2.g00611.t01      2963    N0.HOG0002961   OG0000611  n1   rna-XM_002915435.4    rna-NM_001110768.1    rna-XM_022510875.1    rna-XM_006931794.4    rna-XM_036094455.1    rna-XM_039222909.1    rna-XM_032864572.1    rna-XM_046006820.1    rna-XM_032340222.1    rna-XM_059178212.1    rna-XM_059399053.1    rna-XM_004771896.2    mNeoVis.2.1.1.chr1_hap1.g01569.t01    rna-XM_021691933.1    rna-XM_004414234.1    rna-XM_007095504.3    rna-XM_008691811.2    rna-XM_027603091.2    mNeoVis.3.1.1.chr1.g01522.t01    rna-gnl|WGS_ZZZZ|egapxtmp_036796-R1    mNeoVis.evergladensis.2.1.chr1.g01536.t01    mNeoVis.vulgivaga.2.1.chr1.g01532.t01      2964    N0.HOG0002962   OG0000611  n22   rna-XM_011225409.3    rna-NM_001048022.1    rna-XM_022511159.1    rna-XM_003986810.6    rna-XM_036069896.1    rna-XM_039216788.1    rna-XM_032836862.1    rna-XM_046007231.1    rna-XM_032345546.1    rna-XM_059180019.1    rna-XM_059371591.1    rna-XM_004763755.3    mNeoVis.2.1.1.chr10_hap1.g01201.t01    rna-XM_021680665.2    rna-XM_004398305.1    rna-XM_042987660.1    rna-XM_040642790.1    rna-XM_027568767.2    mNeoVis.3.1.1.chr9.g01131.t01    rna-gnl|WGS_ZZZZ|egapxtmp_017044-R1    mNeoVis.evergladensis.2.1.chr8.g01177.t01    mNeoVis.vulgivaga.2.1.chr9.g01160.t01      2965    N0.HOG0002963   OG0000611  n42   rna-XM_002926944.4    rna-NM_001003017.2    rna-XM_022491041.1    rna-XM_023247170.2    rna-XM_036070215.1    rna-XM_039251341.1    rna-XM_032844999.1    rna-XM_045982812.1    rna-XM_032318122.1    rna-XM_059145580.1    rna-XM_059413662.1    rna-XM_004778501.2    mNeoVis.2.1.1.chr14_hap1.g00618.t01    rna-XM_021681967.1    rna-XM_004402909.1    rna-XM_042975962.1    rna-XM_040635809.1    rna-XM_027610824.2    mNeoVis.3.1.1.chr14.g00599.t01      mNeoVis.evergladensis.2.1.chr14.g00615.t01    mNeoVis.vulgivaga.2.1.chr14.g00611.t01      2966    N0.HOG0002964   OG0000612  n1   rna-XM_034648055.1    rna-XM_038554596.1    rna-XM_022503904.1    rna-XM_019831599.3    rna-XM_036069427.1    rna-XM_039216420.1    rna-XM_032873210.1    rna-XM_046005221.1    rna-XM_032339208.1    rna-XM_059177228.1    rna-XM_059400973.1    rna-XM_004752803.2    mNeoVis.2.1.1.chr1_hap1.g01897.t01    rna-XM_021693598.1    rna-XM_004399934.1    rna-XM_042987611.1    rna-XM_040643694.1    rna-XM_027604051.2    mNeoVis.3.1.1.chr1.g01850.t01    rna-gnl|WGS_ZZZZ|egapxtmp_033827-R4    mNeoVis.evergladensis.2.1.chr1.g01864.t01    mNeoVis.vulgivaga.2.1.chr1.g01863.t01      2967    N0.HOG0002965   OG0000612  n13   rna-XM_002930275.4    rna-XM_038587784.1    rna-XM_022524328.1    rna-XM_011291813.4      rna-XM_039225471.1    rna-XM_032840610.1    rna-XM_045994467.1    rna-XM_032330999.1    rna-XM_059157931.1    rna-XM_059386320.1    rna-XM_013050085.2    mNeoVis.2.1.1.chr12_hap1.g00423.t01    rna-XM_021689990.1    rna-XM_004407336.1    rna-XM_007087561.3    rna-XM_008700613.2    rna-XM_027609474.2    mNeoVis.3.1.1.chr12.g00383.t01    rna-gnl|WGS_ZZZZ|egapxtmp_028463-R1    mNeoVis.evergladensis.2.1.chr12.g00378.t01    mNeoVis.vulgivaga.2.1.chr12.g00387.t01      2968    N0.HOG0002966   OG0000612  n24   rna-XM_034654132.1    rna-XM_038581514.1    rna-XM_022515959.1    rna-XM_045036603.1    rna-XM_036102973.1    rna-XM_039228191.1    rna-XM_032865519.1    rna-XM_046001612.1    rna-XM_032355270.1    rna-XM_059164747.1    rna-XM_059392361.1    rna-XM_004762183.3    mNeoVis.2.1.1.chr5_hap1.g01843.t01    rna-XM_044919374.1    rna-XM_004411369.1    rna-XM_007090474.3    rna-XM_008701921.2    rna-XM_035727181.1    mNeoVis.3.1.1.chr2.g01745.t01    rna-gnl|WGS_ZZZZ|egapxtmp_036404-R1    mNeoVis.evergladensis.2.1.chr2.g01826.t01    mNeoVis.vulgivaga.2.1.chr2.g01841.t01      2969    N0.HOG0002967   OG0000613  n1   rna-XM_034660019.1    rna-XM_038567591.1    rna-XM_022525851.1    rna-XM_019816243.3    rna-XM_036120682.1    rna-XM_039221342.1    rna-XM_032883732.1    rna-XM_045990206.1    rna-XM_032307841.1    rna-XM_059159647.1    rna-XM_059387631.1    rna-XM_004771562.2    mNeoVis.2.1.1.chr5_hap1.g00158.t01    rna-XM_021705351.1    rna-XM_004395466.1    rna-XM_042977923.1    rna-XM_008711443.2    rna-XM_027585145.1    mNeoVis.3.1.1.chr2.g00106.t01    rna-gnl|WGS_ZZZZ|egapxtmp_036721-R1    mNeoVis.evergladensis.2.1.chr2.g00154.t01    mNeoVis.vulgivaga.2.1.chr2.g00152.t01      2970    N0.HOG0002968   OG0000613  n23   rna-XM_034660020.1    rna-XM_038567604.1    rna-XM_022525429.1    rna-XM_045044510.1    rna-XM_036120716.1    rna-XM_039221360.1    rna-XM_032883735.1    rna-XM_045991449.1    rna-XM_032307873.1    rna-XM_059159650.1    rna-XM_059387685.1    rna-XM_004771564.3    mNeoVis.2.1.1.chr5_hap1.g00159.t01    rna-XM_044913122.1    rna-XM_004395465.1    rna-XM_042977931.1    rna-XM_008711444.2    rna-XM_035722138.1    mNeoVis.3.1.1.chr2.g00107.t01      mNeoVis.evergladensis.2.1.chr2.g00155.t01    mNeoVis.vulgivaga.2.1.chr2.g00153.t01      2971    N0.HOG0002969   OG0000613  n43   rna-XM_034667755.1    rna-XM_038533044.1    rna-XM_022511037.1    rna-XM_019831747.3    rna-XM_036067597.1    rna-XM_039216807.1    rna-XM_032836914.1    rna-XM_046009912.1    rna-XM_032341858.1    rna-XM_059179771.1    rna-XM_059371379.1    rna-XM_045067140.1    mNeoVis.2.1.1.chr10_hap1.g01282.t01    rna-XM_044918458.1    rna-XM_004403614.1    rna-XM_042987844.1    rna-XM_040642915.1    rna-XM_027569398.2    mNeoVis.3.1.1.chr9.g01215.t01    rna-gnl|WGS_ZZZZ|egapxtmp_014416-R1    mNeoVis.evergladensis.2.1.chr8.g01260.t01    mNeoVis.vulgivaga.2.1.chr9.g01245.t01      2972    N0.HOG0002970   OG0000614  n1   rna-XM_034641438.1    rna-XM_038572019.1    rna-XM_022494316.1    rna-XM_003983612.5    rna-XM_036079487.1    rna-XM_039248734.1    rna-XM_032858877.1    rna-XM_045981221.1    rna-XM_032350950.1    rna-XM_059133523.1    rna-XM_059406802.1    rna-XM_013053399.2    mNeoVis.2.1.1.chr11_hap1.g00315.t01    rna-XM_021689038.2    rna-XM_004393060.1    rna-XM_042980600.1    rna-XM_040638174.1    rna-XM_027624292.2    mNeoVis.3.1.1.chr11.g00286.t01    rna-gnl|WGS_ZZZZ|egapxtmp_027396-R1    mNeoVis.evergladensis.2.1.chr11.g00312.t01    mNeoVis.vulgivaga.2.1.chr11.g00313.t01      2973    N0.HOG0002971   OG0000614  n22   rna-XM_019796850.2    rna-XM_038579733.1    rna-XM_022502151.1    rna-XM_023248593.2    rna-XM_036081518.1    rna-XM_039240568.1    rna-XM_032858741.1    rna-XM_046023573.1    rna-XM_032316914.1    rna-XM_059166052.1    rna-XM_059394840.1    rna-XM_004768592.3    mNeoVis.2.1.1.chr4_hap1.g00860.t01    rna-XM_021688233.1    rna-XM_004414312.1    rna-XM_042972825.1    rna-XM_008689135.2    rna-XM_027572929.2    mNeoVis.3.1.1.chr6.g00871.t01    rna-gnl|WGS_ZZZZ|egapxtmp_002111-R1    mNeoVis.evergladensis.2.1.chr6.g00866.t01    mNeoVis.vulgivaga.2.1.chr6.g00860.t01      2974   N0.HOG0002972  OG0000614  n36   rna-XM_034672621.1    rna-XM_038538042.1    rna-XM_022511686.1    rna-XM_023244909.2    rna-XM_036119705.1    rna-XM_039227712.1    rna-XM_032853764.1    rna-XM_045987341.1    rna-XM_032326115.1    rna-XM_059150735.1    rna-XM_059383362.1    rna-XM_004753806.3    mNeoVis.2.1.1.chr7_hap1.g00028.t01    rna-XM_021702349.1    rna-XM_004392163.1    rna-XM_042968762.1    rna-XM_040633163.1    rna-XM_035725215.1      rna-gnl|WGS_ZZZZ|egapxtmp_013125-R1    mNeoVis.evergladensis.2.1.chr7.g00029.t01    mNeoVis.vulgivaga.2.1.chr7.g00028.t01      2975   N0.HOG0002973  OG0000615  n1   rna-XM_034647292.1    rna-XM_038536438.1    rna-XM_022524735.1    rna-XM_023243754.2    rna-XM_036078352.1    rna-XM_039254538.1    rna-XM_032839841.1    rna-XM_045986463.1    rna-XM_032321660.1    rna-XM_059149038.1    rna-XM_059380918.1    rna-XM_045083172.1    mNeoVis.2.1.1.chr8_hap1.g01128.t01    rna-XM_044921791.1    rna-XM_004398604.1    rna-XM_042967179.1    rna-XM_040628316.1    rna-XM_027568447.2    mNeoVis.3.1.1.chr8.g01046.t01    rna-gnl|WGS_ZZZZ|egapxtmp_017579-R1    mNeoVis.evergladensis.2.1.chr9.g01105.t01    mNeoVis.vulgivaga.2.1.chr8.g01104.t01      2976    N0.HOG0002974   OG0000615  n24   rna-XM_002921283.4    rna-XM_038563738.1    rna-XM_022507651.1    rna-XM_023240127.2    rna-XM_036067559.1    rna-XM_039235375.1    rna-XM_032879641.1    rna-XM_046016208.1    rna-XM_032357645.1    rna-XM_059149438.1    rna-XM_059400236.1    rna-XM_045065867.1    mNeoVis.2.1.1.chr7_hap1.g02515.t01, mNeoVis.2.1.1.chr7_hap1.g02541.t01, mNeoVis.2.1.1.chr7_hap1.g02569.t01    rna-XM_021685127.1    rna-XM_004393832.1    rna-XM_042957656.1    rna-XM_040639529.1    rna-XM_027579103.2              2977    N0.HOG0002975   OG0000615  n42   rna-XM_034638678.1    rna-XM_038527020.1, rna-XM_038574860.1    rna-XM_022517171.1    rna-XM_023241340.2    rna-XM_036080180.1    rna-XM_039256977.1    rna-XM_032876403.1    rna-XM_046024888.1    rna-XM_032310645.1    rna-XM_059140001.1    rna-XM_059408286.1    rna-XM_004753141.3    mNeoVis.2.1.1.chr3_hap1.g00636.t01    rna-XM_044921298.1    rna-XM_004395908.1    rna-XM_042961785.1    rna-XM_040619997.1    rna-XM_027576740.2      rna-gnl|WGS_ZZZZ|egapxtmp_042487-R1    mNeoVis.evergladensis.2.1.chr3.g00644.t01    mNeoVis.vulgivaga.2.1.chr3.g00648.t01      2978    N0.HOG0002976   OG0000616  n2   rna-XM_002921278.4    rna-NM_001003064.2    rna-XM_022506886.1    rna-XM_023240118.2    rna-XM_036067544.1    rna-XM_039235463.1    rna-XM_032879617.1    rna-XM_046014529.1    rna-XM_032357639.1    rna-XM_059148945.1    rna-XM_059400100.1    rna-XM_004759505.2    mNeoVis.2.1.1.chr7_hap1.g02525.t01, mNeoVis.2.1.1.chr7_hap1.g02531.t01, mNeoVis.2.1.1.chr7_hap1.g02559.t01    rna-XM_021685113.1    rna-XM_004393844.1    rna-XM_042957967.1    rna-XM_040639510.1    rna-XM_027580316.1              2979   N0.HOG0002977  OG0000616  n4   rna-XM_034659708.1    rna-NM_001003034.1    rna-XM_022491524.1    rna-XM_006930392.5    rna-XM_036113240.1    rna-XM_039238422.1    rna-XM_032845862.1    rna-XM_045979452.1      rna-XM_059188729.1    rna-XM_059405367.1    rna-XM_004777483.3    mNeoVis.2.1.1.chr11_hap1.g01034.t01    rna-XM_021697810.1    rna-XM_004394842.1    rna-XM_042982255.1    rna-XM_040623046.1    rna-XM_027622078.2    mNeoVis.3.1.1.chr11.g00985.t01    rna-gnl|WGS_ZZZZ|egapxtmp_027051-R1    mNeoVis.evergladensis.2.1.chr11.g01011.t01    mNeoVis.vulgivaga.2.1.chr11.g01033.t01      2980    N0.HOG0002978   OG0000616  n8   rna-XM_034637888.1    rna-NM_001003033.1    rna-XM_022499208.1, rna-XM_022517201.1    rna-XM_003994662.6    rna-XM_036080034.1    rna-XM_039243478.1    rna-XM_032842734.1, rna-XM_032876303.1    rna-XM_046025236.1    rna-XM_032310010.1    rna-XM_059139393.1    rna-XM_059408408.1    rna-XM_004753239.3    mNeoVis.2.1.1.chr3_hap1.g00587.t01    rna-XM_021698679.2    rna-XM_004395846.1    rna-XM_007079658.3    rna-XM_040619930.1    rna-XM_027576609.1    mNeoVis.3.1.1.chr3.g00579.t01    rna-gnl|WGS_ZZZZ|egapxtmp_041127-R1    mNeoVis.evergladensis.2.1.chr3.g00595.t01    mNeoVis.vulgivaga.2.1.chr3.g00598.t01      2981    N0.HOG0002979   OG0000617  n1   rna-XM_034637939.1    rna-XM_038528327.1    rna-XM_022524040.1    rna-XM_003997529.4    rna-XM_036124517.1    rna-XM_039249571.1    rna-XM_032839268.1    rna-XM_045989054.1    rna-XM_032324256.1    rna-XM_059153747.1    rna-XM_059382154.1    rna-XM_004767391.3    mNeoVis.2.1.1.chr7_hap1.g00928.t01    rna-XM_021681241.1    rna-XM_004415097.1    rna-XM_042969168.1    rna-XM_008684514.2    rna-XM_027620338.2    mNeoVis.3.1.1.chr7.g00869.t01      mNeoVis.evergladensis.2.1.chr7.g00921.t01    mNeoVis.vulgivaga.2.1.chr7.g00926.t01      2982    N0.HOG0002980   OG0000617  n20   rna-XM_011223250.3    rna-XM_038528331.1    rna-XM_022524023.1    rna-XM_045047041.1    rna-XM_036099444.1, rna-XM_036124513.1    rna-XM_039245466.1    rna-XM_032839226.1    rna-XM_045988187.1    rna-XM_032323296.1    rna-XM_059153346.1    rna-XM_059381228.1    rna-XM_004767409.3    mNeoVis.2.1.1.chr7_hap1.g00933.t01    rna-XM_044911361.1    rna-XM_004415101.1    rna-XM_042969337.1, rna-XM_042969341.1    rna-XM_008684666.2, rna-XM_040639816.1    rna-XM_035724840.1    mNeoVis.3.1.1.chr7.g00874.t01    rna-gnl|WGS_ZZZZ|egapxtmp_013820-R1    mNeoVis.evergladensis.2.1.chr7.g00926.t01    mNeoVis.vulgivaga.2.1.chr7.g00931.t01      2983    N0.HOG0002981   OG0000617  n44   rna-XM_034638537.1    rna-XM_038528347.1    rna-XM_022524025.1      rna-XM_036124511.1    rna-XM_039245412.1    rna-XM_032839392.1    rna-XM_045987372.1    rna-XM_032323299.1    rna-XM_059153755.1    rna-XM_059383267.1    rna-XM_004767408.3    mNeoVis.2.1.1.chr7_hap1.g00932.t01    rna-XM_021681236.2    rna-XM_004415100.1      rna-XM_008684667.2    rna-XM_035724845.1    mNeoVis.3.1.1.chr7.g00873.t01      mNeoVis.evergladensis.2.1.chr7.g00925.t01    mNeoVis.vulgivaga.2.1.chr7.g00930.t01      2984    N0.HOG0002982   OG0000618  n1   rna-XM_011224044.3    rna-XM_038544400.1    rna-XM_022520611.1    rna-XM_023255634.2    rna-XM_036093119.1    rna-XM_039247116.1    rna-XM_032848913.1    rna-XM_046008902.1    rna-XM_032341481.1    rna-XM_059181847.1    rna-XM_059373046.1    rna-XM_004755147.3    mNeoVis.2.1.1.chr10_hap1.g00500.t01    rna-XM_021695779.2    rna-XM_004399950.1    rna-XM_042989521.1    rna-XM_040620802.1    rna-XM_027569650.2    mNeoVis.3.1.1.chr9.g00463.t01    rna-gnl|WGS_ZZZZ|egapxtmp_016841-R1    mNeoVis.evergladensis.2.1.chr8.g00480.t01    mNeoVis.vulgivaga.2.1.chr9.g00483.t01      2985    N0.HOG0002983   OG0000618  n22   rna-XM_034638218.1    rna-XM_038528491.1    rna-XM_022523807.1    rna-XM_023245655.2    rna-XM_036114009.1    rna-XM_039245947.1    rna-XM_032839175.1    rna-XM_045989174.1    rna-XM_032326168.1    rna-XM_059152952.1    rna-XM_059383744.1    rna-XM_045077134.1    mNeoVis.2.1.1.chr7_hap1.g00985.t01    rna-XM_021681346.1    rna-XM_004400941.1    rna-XM_042968105.1    rna-XM_040634427.1    rna-XM_027617548.2    mNeoVis.3.1.1.chr7.g00924.t01      mNeoVis.evergladensis.2.1.chr7.g00978.t01    mNeoVis.vulgivaga.2.1.chr7.g00984.t01      2986    N0.HOG0002984   OG0000618  n42   rna-XM_011219234.2    rna-XM_038561265.1    rna-XM_022520522.1    rna-XM_023251959.2    rna-XM_036121513.1    rna-XM_039257205.1    rna-XM_032881870.1    rna-XM_045978985.1    rna-XM_032353005.1    rna-XM_059188645.1    rna-XM_059405421.1    rna-XM_004765442.3    mNeoVis.2.1.1.chr11_hap1.g01000.t01    rna-XM_044918761.1    rna-XM_004409979.1    rna-XM_042982169.1    rna-XM_040623097.1    rna-XM_027624105.2    mNeoVis.3.1.1.chr11.g00950.t01    rna-gnl|WGS_ZZZZ|egapxtmp_027481-R1    mNeoVis.evergladensis.2.1.chr11.g00978.t01    mNeoVis.vulgivaga.2.1.chr11.g00999.t01      2987    N0.HOG0002985   OG0000619  n1   rna-XM_002928500.3    rna-XM_038528528.1    rna-XM_022524210.1    rna-XM_023245573.2    rna-XM_036091586.1    rna-XM_039245027.1    rna-XM_032839306.1    rna-XM_045987728.1    rna-XM_032326396.1    rna-XM_059153259.1    rna-XM_059383125.1    rna-XM_004767553.3    mNeoVis.2.1.1.chr7_hap1.g01004.t01    rna-XM_021681169.1    rna-XM_004400958.1    rna-XM_042969383.1    rna-XM_040633052.1    rna-XM_027619175.1    mNeoVis.3.1.1.chr7.g00943.t01    rna-gnl|WGS_ZZZZ|egapxtmp_012360-R1    mNeoVis.evergladensis.2.1.chr7.g00997.t01    mNeoVis.vulgivaga.2.1.chr7.g01003.t01      2988    N0.HOG0002986   OG0000619  n22   rna-XM_019806468.2    rna-XM_038544550.1    rna-XM_022492849.1    rna-XM_045059834.1    rna-XM_036103875.1    rna-XM_039249561.1    rna-XM_032871239.1    rna-XM_046009944.1    rna-XM_032344068.1    rna-XM_059181946.1    rna-XM_059373108.1    rna-XM_004773313.3    mNeoVis.2.1.1.chr10_hap1.g00462.t01    rna-XM_021701459.2    rna-XM_004399785.1    rna-XM_042989581.1    rna-XM_040620910.1    rna-XM_027568846.2    mNeoVis.3.1.1.chr9.g00428.t01    rna-gnl|WGS_ZZZZ|egapxtmp_014599-R1    mNeoVis.evergladensis.2.1.chr8.g00441.t01    mNeoVis.vulgivaga.2.1.chr9.g00444.t01      2989    N0.HOG0002987   OG0000619  n39   rna-XM_002924124.4    rna-XM_038571420.1    rna-XM_022494667.1    rna-XM_003983832.5    rna-XM_036069601.1    rna-XM_039223900.1    rna-XM_032859315.1    rna-XM_045981074.1    rna-XM_032351566.1    rna-XM_059134202.1    rna-XM_059406440.1    rna-XM_004754117.3    mNeoVis.2.1.1.chr11_hap1.g00523.t01    rna-XM_021700723.1    rna-XM_004415130.1    rna-XM_042979747.1    rna-XM_040637816.1    rna-XM_027622404.2    mNeoVis.3.1.1.chr11.g00485.t01      mNeoVis.evergladensis.2.1.chr11.g00511.t01    mNeoVis.vulgivaga.2.1.chr11.g00520.t01      2990    N0.HOG0002988   OG0000620  n1   rna-XM_002912975.3, rna-XM_034642221.1    rna-XM_038572605.1    rna-XM_022494700.1, rna-XM_022513700.1    rna-XM_023242134.2, rna-XM_023251103.2    rna-XM_036064689.1, rna-XM_036074507.1    rna-XM_039230042.1, rna-XM_039245821.1    rna-XM_032872152.1, rna-XM_032878966.1    rna-XM_045980287.1, rna-XM_046025134.1    rna-XM_032310850.1, rna-XM_032350344.1    rna-XM_059133740.1, rna-XM_059138617.1    rna-XM_059407231.1, rna-XM_059409867.1    rna-XM_045082078.1, rna-XM_045085275.1    mNeoVis.2.1.1.chr11_hap1.g00121.t01, mNeoVis.2.1.1.chr3_hap1.g00008.t01    rna-XM_021685911.1, rna-XM_021686032.1    rna-XM_004412504.1, rna-XM_004416593.1    rna-XM_042962738.1, rna-XM_042980153.1    rna-XM_040632347.1, rna-XM_040637848.1    rna-XM_027600804.2, rna-XM_027624157.2    mNeoVis.3.1.1.chr11.g00092.t01, mNeoVis.3.1.1.chr3.g00007.t01    rna-gnl|WGS_ZZZZ|egapxtmp_041678-R1    mNeoVis.evergladensis.2.1.chr11.g00118.t01, mNeoVis.evergladensis.2.1.chr3.g00009.t01    mNeoVis.vulgivaga.2.1.chr11.g00120.t01, mNeoVis.vulgivaga.2.1.chr11.g00121.t01, mNeoVis.vulgivaga.2.1.chr3.g00007.t01      2991    N0.HOG0002989   OG0000620  n39   rna-XM_034639607.1    rna-XM_038538145.1    rna-XM_022511835.1    rna-XM_019820145.3    rna-XM_036104930.1    rna-XM_039227271.1    rna-XM_032854615.1    rna-XM_045988074.1    rna-XM_032325003.1    rna-XM_059153033.1    rna-XM_059383982.1    rna-XM_004753896.2    mNeoVis.2.1.1.chr7_hap1.g00075.t01    rna-XM_021702578.1    rna-XM_004392151.1    rna-XM_042970131.1    rna-XM_008689789.2    rna-XM_027619814.2    mNeoVis.3.1.1.chr7.g00063.t01    rna-gnl|WGS_ZZZZ|egapxtmp_010791-R1    mNeoVis.evergladensis.2.1.chr7.g00076.t01    mNeoVis.vulgivaga.2.1.chr7.g00076.t01      2992    N0.HOG0002990   OG0000621  n1   rna-XM_019799152.2    rna-XM_038586686.1, rna-XM_038588495.1, rna-XM_038588526.1    rna-XM_022491333.1    rna-XM_023249079.2, rna-XM_045050359.1    rna-XM_036100875.1      rna-XM_032845296.1    rna-XM_045994418.1, rna-XM_045996147.1    rna-XM_032331626.1, rna-XM_032331629.1    rna-XM_059157536.1    rna-XM_059386094.1    rna-XM_045068069.1, rna-XM_045068085.1    mNeoVis.2.1.1.chr12_hap1.g00099.t01, mNeoVis.2.1.1.chr12_hap1.g00108.t01    rna-XM_021682459.1    rna-XM_004417487.1    rna-XM_007074505.2, rna-XM_042974740.1, rna-XM_042975269.1    rna-XM_040628186.1    rna-XM_035725408.1, rna-XM_035725541.1              2993    N0.HOG0002991   OG0000621  n26   rna-XM_034649057.1    rna-XM_038588510.1, rna-XM_038588851.1    rna-XM_022491279.1, rna-XM_022492360.1    rna-XM_045050397.1    rna-XM_036065831.1, rna-XM_036080420.1    rna-XM_039225459.1    rna-XM_032845297.1    rna-XM_045995639.1, rna-XM_045995951.1, rna-XM_045996578.1    rna-XM_032329737.1, rna-XM_032331625.1, rna-XM_032331957.1    rna-XM_059157493.1, rna-XM_059158384.1    rna-XM_059384740.1, rna-XM_059386434.1    rna-XM_045068080.1, rna-XM_045068081.1, rna-XM_045087434.1    mNeoVis.2.1.1.chr12_hap1.g00102.t01, mNeoVis.2.1.1.chr12_hap1.g00105.t01    rna-XM_021683363.1, rna-XM_021702683.1    rna-XM_004407832.1, rna-XM_004410286.1, rna-XM_004412406.1, rna-XM_004417350.1      rna-XM_008699042.1, rna-XM_040635958.1    rna-XM_035725925.1, rna-XM_035725926.1, rna-XM_035725928.1, rna-XM_035725929.1, rna-XM_035727779.1              2994    N0.HOG0002992   OG0000622  n1   rna-XM_034649027.1    rna-XM_038588803.1    rna-XM_022524477.1    rna-XM_019823921.3    rna-XM_036065885.1    rna-XM_039253367.1    rna-XM_032847757.1    rna-XM_045996025.1    rna-XM_032330679.1    rna-XM_059156802.1    rna-XM_059387403.1    rna-XM_045072769.1    mNeoVis.2.1.1.chr12_hap1.g00499.t01    rna-XM_021680303.1    rna-XM_004396621.1    rna-XM_042973951.1    rna-XM_040622856.1    rna-XM_027608281.1    mNeoVis.3.1.1.chr12.g00454.t01    rna-gnl|WGS_ZZZZ|egapxtmp_028175-R1    mNeoVis.evergladensis.2.1.chr12.g00447.t01    mNeoVis.vulgivaga.2.1.chr12.g00460.t01      2995    N0.HOG0002993   OG0000622  n23   rna-XM_034647412.1    rna-XM_038536584.1    rna-XM_022524703.1    rna-XM_023243688.2    rna-XM_036066190.1    rna-XM_039224881.1    rna-XM_032839996.1    rna-XM_045986108.1    rna-XM_032321312.1    rna-XM_059147688.1    rna-XM_059378449.1    rna-XM_004760398.3    mNeoVis.2.1.1.chr8_hap1.g01094.t01    rna-XM_021694560.2    rna-XM_004398561.1    rna-XM_042966551.1    rna-XM_040629892.1    rna-XM_027625051.1    mNeoVis.3.1.1.chr8.g01016.t01    rna-gnl|WGS_ZZZZ|egapxtmp_017910-R1    mNeoVis.evergladensis.2.1.chr9.g01070.t01    mNeoVis.vulgivaga.2.1.chr8.g01070.t01      2996    N0.HOG0002994   OG0000622  n42   rna-XM_019795365.2    rna-XM_038568609.1    rna-XM_022521927.1    rna-XM_006936919.5    rna-XM_036082806.1    rna-XM_039228667.1    rna-XM_032853046.1    rna-XM_046017434.1    rna-XM_032356407.1    rna-XM_059132366.1    rna-XM_059410950.1    rna-XM_045071504.1    mNeoVis.2.1.1.chr7_hap1.g01521.t01    rna-XM_021704613.1    rna-XM_004391793.1    rna-XM_042959223.1    rna-XM_040630850.1    rna-XM_027578754.2    mNeoVis.3.1.1.chr7.g01432.t01      mNeoVis.evergladensis.2.1.chr7.g01502.t01    mNeoVis.vulgivaga.2.1.chr7.g01518.t01      2997    N0.HOG0002995   OG0000623  n1   rna-XM_002915009.4    rna-XM_038526976.1    rna-XM_022512655.1    rna-XM_011288266.3    rna-XM_036090578.1    rna-XM_039250406.1    rna-XM_032840560.1    rna-XM_046022744.1    rna-XM_032306892.1    rna-XM_059141355.1    rna-XM_059411788.1    rna-XM_004774076.3    mNeoVis.2.1.1.chr13_hap1.g00692.t01    rna-XM_021677961.1    rna-XM_004407606.1    rna-XM_042965243.1    rna-XM_040620205.1    rna-XM_027615574.2    mNeoVis.3.1.1.chr13.g00613.t01    rna-gnl|WGS_ZZZZ|egapxtmp_031020-R1    mNeoVis.evergladensis.2.1.chr13.g00662.t01    mNeoVis.vulgivaga.2.1.chr13.g00665.t01      2998    N0.HOG0002996   OG0000623  n20   rna-XM_034667687.1    rna-XM_038543059.1    rna-XM_022490757.1    rna-XM_045048522.1    rna-XM_036067819.1    rna-XM_039251293.1    rna-XM_032845011.1    rna-XM_045982165.1    rna-XM_032317776.1    rna-XM_059145339.1    rna-XM_059413689.1    rna-XM_045086752.1    mNeoVis.2.1.1.chr14_hap1.g00632.t01    rna-XM_021682113.2    rna-XM_004402931.1    rna-XM_042975899.1    rna-XM_040635787.1, rna-XM_040635789.1    rna-XM_027610821.2        mNeoVis.evergladensis.2.1.chr14.g00629.t01    mNeoVis.vulgivaga.2.1.chr14.g00626.t01      2999    N0.HOG0002997   OG0000623  n40   rna-XM_002931232.4    rna-XM_038587545.1    rna-XM_022492178.1    rna-XM_004000561.6    rna-XM_036068776.1    rna-XM_039222815.1    rna-XM_032848967.1    rna-XM_045996406.1    rna-XM_032330812.1    rna-XM_059157834.1    rna-XM_059386470.1    rna-XM_004782353.3    mNeoVis.2.1.1.chr12_hap1.g00557.t01    rna-XM_044911761.1    rna-XM_004413188.1    rna-XM_007093168.2    rna-XM_008703185.2    rna-XM_027608153.2    mNeoVis.3.1.1.chr12.g00515.t01    rna-gnl|WGS_ZZZZ|egapxtmp_027994-R1    mNeoVis.evergladensis.2.1.chr12.g00502.t01    mNeoVis.vulgivaga.2.1.chr12.g00515.t01      3000    N0.HOG0002998   OG0000624  n1   rna-XM_002912498.4    rna-XM_038534931.1    rna-XM_022493516.1    rna-XM_019839908.3    rna-XM_036096205.1    rna-XM_039249450.1    rna-XM_032851650.1    rna-XM_046000930.1    rna-XM_032337201.1    rna-XM_059174135.1    rna-XM_059416542.1    rna-XM_004737737.3    mNeoVis.2.1.1.chr1_hap1.g00242.t01    rna-XM_021697353.2    rna-XM_004416578.1    rna-XM_042993762.1    rna-XM_008692457.2    rna-XM_027606658.2    mNeoVis.3.1.1.chr1.g00242.t01    rna-gnl|WGS_ZZZZ|egapxtmp_027209-R1    mNeoVis.evergladensis.2.1.chr1.g00239.t01    mNeoVis.vulgivaga.2.1.chr1.g00244.t01      3001    N0.HOG0002999   OG0000624  n18   rna-XM_034647674.1    rna-XM_038559195.1    rna-XM_022498868.1    rna-XM_019828324.3    rna-XM_036068584.1    rna-XM_039221153.1    rna-XM_032843404.1    rna-XM_045998548.1    rna-XM_032329030.1    rna-XM_059156376.1    rna-XM_059384510.1      mNeoVis.2.1.1.chr6_hap1.g01028.t01    rna-XM_021694220.1    rna-XM_004417368.1    rna-XM_042982634.1    rna-XM_008710886.2    rna-XM_027600618.2    mNeoVis.3.1.1.chr4.g00987.t01    rna-gnl|WGS_ZZZZ|egapxtmp_001268-R1    mNeoVis.evergladensis.2.1.chr4.g00971.t01    mNeoVis.vulgivaga.2.1.chr4.g00998.t01      3002    N0.HOG0003000   OG0000624  n38   rna-XM_011226784.3    rna-NM_001014276.2    rna-XM_022522602.1    rna-XM_023249893.2    rna-XM_036074309.1    rna-XM_039255345.1    rna-XM_032838196.1    rna-XM_046028349.1    rna-XM_032314593.1    rna-XM_059144615.1    rna-XM_059378288.1    rna-XM_004774340.3    mNeoVis.2.1.1.chr8_hap1.g01181.t01    rna-XM_021678228.2    rna-XM_004409517.1    rna-XM_042981327.1    rna-XM_008692138.2    rna-XM_027591103.2    mNeoVis.3.1.1.chr8.g01096.t01    rna-gnl|WGS_ZZZZ|egapxtmp_020381-R1    mNeoVis.evergladensis.2.1.chr9.g01153.t01        3003    N0.HOG0003001   OG0000625  n1   rna-XM_034649904.1    rna-XM_038586989.1    rna-XM_022507830.1    rna-XM_045050412.1    rna-XM_036086120.1    rna-XM_039247384.1    rna-XM_032840732.1    rna-XM_045994893.1    rna-XM_032330215.1    rna-XM_059157232.1    rna-XM_059385745.1    rna-XM_013059635.2    mNeoVis.2.1.1.chr12_hap1.g00754.t01    rna-XM_021683941.2    rna-XM_004401514.1    rna-XM_042975280.1    rna-XM_040633703.1    rna-XM_035725496.1    mNeoVis.3.1.1.chr12.g00687.t01    rna-gnl|WGS_ZZZZ|egapxtmp_027966-R1    mNeoVis.evergladensis.2.1.chr12.g00683.t01    mNeoVis.vulgivaga.2.1.chr12.g00709.t01      3004    N0.HOG0003002   OG0000625  n23   rna-XM_034645075.1    rna-XM_038529592.1    rna-XM_022497497.1    rna-XM_011284512.4    rna-XM_036116431.1    rna-XM_039231568.1    rna-XM_032863882.1    rna-XM_045980563.1    rna-XM_032302465.1    rna-XM_059136326.1    rna-XM_059376809.1    rna-XM_013047166.2    mNeoVis.2.1.1.chr2_hap1.g01617.t01    rna-XM_044913823.1    rna-XM_004394786.1    rna-XM_042996159.1    rna-XM_040645561.1    rna-XM_035727041.1    mNeoVis.3.1.1.chr5.g01538.t01      mNeoVis.evergladensis.2.1.chr5.g01556.t01    mNeoVis.vulgivaga.2.1.chr5.g01559.t01      3005    N0.HOG0003003   OG0000625  n43   rna-XM_034669190.1    rna-XM_038526115.1    rna-XM_022503520.1    rna-XM_023254424.2    rna-XM_036097478.1    rna-XM_039219698.1    rna-XM_032844826.1    rna-XM_046005093.1    rna-XM_032339409.1    rna-XM_059177456.1    rna-XM_059400786.1    rna-XM_004740273.3    mNeoVis.2.1.1.chr1_hap1.g01823.t01    rna-XM_021693127.2    rna-XM_004410252.1    rna-XM_042987196.1    rna-XM_008691227.2    rna-XM_027602930.1    mNeoVis.3.1.1.chr1.g01776.t01    rna-gnl|WGS_ZZZZ|egapxtmp_040439-R1    mNeoVis.evergladensis.2.1.chr1.g01788.t01    mNeoVis.vulgivaga.2.1.chr1.g01783.t01      3006    N0.HOG0003004   OG0000626  n1   rna-XM_034649942.1    rna-XM_038588273.1    rna-XM_022518573.1    rna-XM_011279473.4    rna-XM_036101622.1    rna-XM_039250809.1    rna-XM_032882365.1    rna-XM_045996050.1    rna-XM_032330282.1    rna-XM_059156453.1    rna-XM_059386240.1    rna-XM_013050011.2    mNeoVis.2.1.1.chr12_hap1.g00202.t01    rna-XM_021685796.1    rna-XM_004400226.1    rna-XM_042973792.1    rna-XM_040627241.1    rna-XM_027607907.1    mNeoVis.3.1.1.chr12.g00170.t01    rna-gnl|WGS_ZZZZ|egapxtmp_028998-R1    mNeoVis.evergladensis.2.1.chr12.g00174.t01    mNeoVis.vulgivaga.2.1.chr12.g00178.t01      3007    N0.HOG0003005   OG0000626  n13   rna-XM_034669376.1    rna-XM_038526424.1    rna-XM_022520889.1    rna-XM_045058291.1    rna-XM_036084192.1    rna-XM_039252072.1    rna-XM_032836436.1    rna-XM_046005508.1    rna-XM_032338621.1    rna-XM_059176669.1    rna-XM_059401334.1    rna-XM_045080124.1    mNeoVis.2.1.1.chr1_hap1.g02098.t01    rna-XM_021693291.1    rna-XM_004403891.1      rna-XM_040627999.1    rna-XM_027603528.2    mNeoVis.3.1.1.chr1.g02053.t01    rna-gnl|WGS_ZZZZ|egapxtmp_023159-R1    mNeoVis.evergladensis.2.1.chr1.g02068.t01    mNeoVis.vulgivaga.2.1.chr1.g02061.t01      3008    N0.HOG0003006   OG0000626  n33   rna-XM_011229408.3    rna-XM_038531439.1    rna-XM_022514652.1    rna-XM_045035359.1    rna-XM_036116443.1    rna-XM_039231762.1    rna-XM_032863854.1    rna-XM_046003002.1    rna-XM_032361442.1    rna-XM_059135960.1    rna-XM_059376768.1    rna-XM_045064015.1    mNeoVis.2.1.1.chr2_hap1.g01630.t01    rna-XM_044913822.1    rna-XM_004394787.1    rna-XM_042995677.1    rna-XM_040619071.1    rna-XM_027598680.1    mNeoVis.3.1.1.chr5.g01550.t01    rna-gnl|WGS_ZZZZ|egapxtmp_007810-R1    mNeoVis.evergladensis.2.1.chr5.g01570.t01    mNeoVis.vulgivaga.2.1.chr5.g01572.t01      3009    N0.HOG0003007   OG0000627  n1   rna-XM_019803449.2    rna-XM_038588310.1    rna-XM_022526351.1    rna-XM_004000900.5    rna-XM_036123053.1    rna-XM_039239889.1    rna-XM_032843381.1    rna-XM_045996517.1    rna-XM_032330297.1    rna-XM_059157252.1    rna-XM_059386218.1    rna-XM_004778720.3    mNeoVis.2.1.1.chr12_hap1.g00183.t01    rna-XM_021684217.2    rna-XM_004401090.1    rna-XM_042973816.1    rna-XM_040627401.1    rna-XM_027609351.2    mNeoVis.3.1.1.chr12.g00152.t01    rna-gnl|WGS_ZZZZ|egapxtmp_027937-R1    mNeoVis.evergladensis.2.1.chr12.g00156.t01    mNeoVis.vulgivaga.2.1.chr12.g00158.t01      3010    N0.HOG0003008   OG0000627  n10   rna-XM_034665395.1    rna-XM_038569902.1    rna-XM_022502785.1    rna-XM_003980489.6    rna-XM_036119606.1    rna-XM_039252534.1    rna-XM_032864120.1    rna-XM_045978475.1    rna-XM_032315066.1    rna-XM_059144406.1    rna-XM_059378046.1    rna-XM_004746113.3    mNeoVis.2.1.1.chr8_hap1.g01471.t01    rna-XM_044913530.1    rna-XM_004404944.1    rna-XM_007098395.3    rna-XM_040624008.1    rna-XM_027589282.2    mNeoVis.3.1.1.chr8.g01366.t01    rna-gnl|WGS_ZZZZ|egapxtmp_017684-R1    mNeoVis.evergladensis.2.1.chr9.g01419.t01    mNeoVis.vulgivaga.2.1.chr8.g01424.t01      3011    N0.HOG0003009   OG0000627  n25   rna-XM_034655588.1    rna-NM_001286859.1    rna-XM_022491619.1    rna-XM_023260011.2    rna-XM_036081864.1    rna-XM_039228538.1    rna-XM_032857717.1    rna-XM_046019907.1    rna-XM_032355475.1    rna-XM_059167420.1    rna-XM_059393934.1    rna-XM_013051897.2    mNeoVis.2.1.1.chr3_hap1.g01000.t01    rna-XM_021679053.1    rna-XM_004412702.1    rna-XM_042995938.1    rna-XM_040638021.1    rna-XM_027588252.2    mNeoVis.3.1.1.chr3.g00989.t01      mNeoVis.evergladensis.2.1.chr3.g00996.t01    mNeoVis.vulgivaga.2.1.chr3.g01016.t01      3012    N0.HOG0003010   OG0000628  n1   rna-XM_034649591.1    rna-XM_038588435.1    rna-XM_022521315.1    rna-XM_045051000.1    rna-XM_036078804.1    rna-XM_039220955.1    rna-XM_032841401.1    rna-XM_045996541.1    rna-XM_032329640.1    rna-XM_059157055.1    rna-XM_059386042.1    rna-XM_045068230.1    mNeoVis.2.1.1.chr12_hap1.g00136.t01    rna-XM_044911590.1    rna-XM_004405912.1    rna-XM_042974636.1    rna-XM_040627830.1    rna-XM_027609278.2    mNeoVis.3.1.1.chr12.g00107.t01    rna-gnl|WGS_ZZZZ|egapxtmp_029527-R2    mNeoVis.evergladensis.2.1.chr12.g00113.t01    mNeoVis.vulgivaga.2.1.chr12.g00114.t01      3013    N0.HOG0003011   OG0000628  n8   rna-XM_002927247.4    rna-XM_038583704.1    rna-XM_022507198.1    rna-XM_003991788.6    rna-XM_036109236.1    rna-XM_039221705.1    rna-XM_032875651.1    rna-XM_046003433.1    rna-XM_032337776.1    rna-XM_059163129.1    rna-XM_059391273.1    rna-XM_004745272.3    mNeoVis.2.1.1.chr5_hap1.g01275.t01    rna-XM_021692471.1    rna-XM_004391851.1    rna-XM_042999107.1    rna-XM_008705152.2    rna-XM_027587164.2      rna-gnl|WGS_ZZZZ|egapxtmp_037082-R1    mNeoVis.evergladensis.2.1.chr2.g01262.t01    mNeoVis.vulgivaga.2.1.chr2.g01268.t01      3014    N0.HOG0003012   OG0000628  n11   rna-XM_034665441.1    rna-XM_038569988.1    rna-XM_022502930.1    rna-XM_019829208.3    rna-XM_036090392.1    rna-XM_039222032.1    rna-XM_032881086.1    rna-XM_045978511.1    rna-XM_032315400.1    rna-XM_059143847.1    rna-XM_059377578.1    rna-XM_004746069.3    mNeoVis.2.1.1.chr8_hap1.g01486.t01    rna-XM_021695973.1    rna-XM_004404919.1    rna-XM_042979778.1    rna-XM_008684988.1    rna-XM_027591057.2    mNeoVis.3.1.1.chr8.g01385.t01    rna-gnl|WGS_ZZZZ|egapxtmp_019309-R2    mNeoVis.evergladensis.2.1.chr9.g01436.t01    mNeoVis.vulgivaga.2.1.chr8.g01440.t01      3015    N0.HOG0003013   OG0000629  n1   rna-XM_019804143.2    rna-XM_038556574.1    rna-XM_022510184.1    rna-XM_045052722.1    rna-XM_036089285.1    rna-XM_039242669.1    rna-XM_032869591.1    rna-XM_046021478.1    rna-XM_032304729.1    rna-XM_059171805.1    rna-XM_059396444.1    rna-XM_004741803.3    mNeoVis.2.1.1.chr4_hap1.g00227.t01    rna-XM_021699407.2    rna-XM_004398017.1    rna-XM_042971780.1    rna-XM_040644024.1    rna-XM_027574643.2    mNeoVis.3.1.1.chr6.g00226.t01    rna-gnl|WGS_ZZZZ|egapxtmp_031073-R1    mNeoVis.evergladensis.2.1.chr6.g00226.t01    mNeoVis.vulgivaga.2.1.chr6.g00227.t01      3016    N0.HOG0003014   OG0000629  n23   rna-XM_034649967.1    rna-XM_038588707.1      rna-XM_023249411.2    rna-XM_036065458.1    rna-XM_039225582.1    rna-XM_032846378.1    rna-XM_045994702.1    rna-XM_032331782.1    rna-XM_059156521.1    rna-XM_059384970.1    rna-XM_013050098.2    mNeoVis.2.1.1.chr12_hap1.g00042.t01    rna-XM_044911792.1    rna-XM_004414495.1    rna-XM_042973882.1    rna-XM_040626822.1    rna-XM_027607986.1    mNeoVis.3.1.1.chr12.g00033.t01    rna-gnl|WGS_ZZZZ|egapxtmp_029104-R1    mNeoVis.evergladensis.2.1.chr12.g00032.t01    mNeoVis.vulgivaga.2.1.chr12.g00034.t01      3017    N0.HOG0003015   OG0000629  n43   rna-XM_002913523.4    rna-XM_038566049.1    rna-XM_022498584.1    rna-XM_019823423.3    rna-XM_036071705.1    rna-XM_039242456.1    rna-XM_032868420.1    rna-XM_045990981.1    rna-XM_032323456.1    rna-XM_059161308.1    rna-XM_059390036.1    rna-XM_004738244.3    mNeoVis.2.1.1.chr5_hap1.g00762.t01    rna-XM_021694902.2    rna-XM_004412376.1    rna-XM_015538545.2    rna-XM_008703081.2    rna-XM_027582672.2    mNeoVis.3.1.1.chr2.g00672.t01    rna-gnl|WGS_ZZZZ|egapxtmp_038370-R1    mNeoVis.evergladensis.2.1.chr2.g00763.t01    mNeoVis.vulgivaga.2.1.chr2.g00768.t01      3018    N0.HOG0003016   OG0000630  n2   rna-XM_034651136.1    rna-NM_001003190.1      rna-XM_019815490.3    rna-XM_036123545.1    rna-XM_039219666.1, rna-XM_039219679.1    rna-XM_032866389.1      rna-XM_032307884.1    rna-XM_059142569.1, rna-XM_059142570.1    rna-XM_059412144.1, rna-XM_059412147.1    rna-XM_004756978.3    mNeoVis.2.1.1.chr13_hap1.g00089.t01    rna-XM_021691050.1    rna-XM_004410739.1    rna-XM_042964811.1        mNeoVis.3.1.1.chr13.g00048.t01    rna-gnl|WGS_ZZZZ|egapxtmp_031057-R1    mNeoVis.evergladensis.2.1.chr13.g00089.t01    mNeoVis.vulgivaga.2.1.chr13.g00087.t01      3019    N0.HOG0003017   OG0000630  n22   rna-XM_019805330.2    rna-NM_001003189.2      rna-XM_045043438.1    rna-XM_036113103.1, rna-XM_036113106.1    rna-XM_039219660.1                  rna-XM_021691054.1, rna-XM_021691055.1    rna-XM_004410733.1      rna-XM_040631364.1, rna-XM_040631365.1    rna-XM_027614713.2, rna-XM_027616028.1              3020    N0.HOG0003018   OG0000630  n36   rna-XM_034664800.1    rna-XM_038548815.1      rna-XM_023243389.2    rna-XM_036113107.1    rna-XM_039219661.1                  rna-XM_044920442.1    rna-XM_004410734.1    rna-XM_042964269.1    rna-XM_040631363.1    rna-XM_035723811.1              3021    N0.HOG0003019   OG0000630  n45   rna-XM_019806630.2    rna-NM_001284461.1    rna-XM_022501815.1    rna-XM_019815497.3      rna-XM_039240800.1    rna-XM_032865628.1    rna-XM_046023203.1    rna-XM_032308091.1    rna-XM_059142765.1    rna-XM_059412224.1    rna-XM_004747830.2    mNeoVis.2.1.1.chr13_hap1.g00344.t01    rna-XM_021691710.1    rna-XM_004396973.1    rna-XM_015543007.2    rna-XM_008707964.2      mNeoVis.3.1.1.chr13.g00286.t01    rna-gnl|WGS_ZZZZ|egapxtmp_030798-R1    mNeoVis.evergladensis.2.1.chr13.g00333.t01    mNeoVis.vulgivaga.2.1.chr13.g00330.t01      3022    N0.HOG0003020   OG0000631  n1   rna-XM_034670259.1    rna-NM_001313864.1    rna-XM_022526633.1    rna-XM_011290618.4    rna-XM_036105236.1    rna-XM_039234383.1    rna-XM_032843977.1    rna-XM_045992415.1    rna-XM_032328088.1    rna-XM_059154713.1    rna-XM_059415120.1    rna-XM_045072676.1    mNeoVis.2.1.1.chr15_hap1.g00143.t01    rna-XM_021686845.1    rna-XM_004398810.1    rna-XM_042971109.1    rna-XM_008697165.1    rna-XM_035722515.1    mNeoVis.3.1.1.chr15.g00120.t01    rna-gnl|WGS_ZZZZ|egapxtmp_033616-R1    mNeoVis.evergladensis.2.1.chr15.g00129.t01    mNeoVis.vulgivaga.2.1.chr15.g00132.t01      3023    N0.HOG0003021   OG0000631  n23   rna-XM_034671010.1    rna-XM_038537582.1    rna-XM_022514851.1    rna-XM_045034910.1    rna-XM_036089739.1    rna-XM_039248313.1    rna-XM_032863118.1    rna-XM_046006127.1    rna-XM_032361522.1    rna-XM_059136249.1    rna-XM_059375119.1    rna-XM_045063500.1    mNeoVis.2.1.1.chr2_hap1.g01933.t01    rna-XM_021684351.1    rna-XM_004407429.1    rna-XM_042994295.1    rna-XM_040645522.1    rna-XM_027615009.2    mNeoVis.3.1.1.chr5.g01836.t01      mNeoVis.evergladensis.2.1.chr5.g01872.t01    mNeoVis.vulgivaga.2.1.chr5.g01874.t01      3024   N0.HOG0003022  OG0000631  n43   rna-XM_034645097.1    rna-XM_038576561.1    rna-XM_022522378.1    rna-XM_045061290.1    rna-XM_036117634.1    rna-XM_039218396.1    rna-XM_032874797.1    rna-XM_046010683.1    rna-XM_032349020.1    rna-XM_059184584.1    rna-XM_059403753.1    rna-XM_013063583.2    mNeoVis.2.1.1.chr9_hap1.g00247.t01    rna-XM_021690807.1    rna-XM_004413936.1    rna-XM_042991642.1    rna-XM_040642067.1    rna-XM_027593765.2    mNeoVis.3.1.1.chr10.g00239.t01    rna-gnl|WGS_ZZZZ|egapxtmp_024616-R1    mNeoVis.evergladensis.2.1.chr10.g00245.t01    mNeoVis.vulgivaga.2.1.chr10.g00240.t01      3025   N0.HOG0003023  OG0000632  n2   rna-XM_034646552.1    rna-XM_038577092.1    rna-XM_022522716.1    rna-XM_045061560.1    rna-XM_036109646.1    rna-XM_039221710.1    rna-XM_032855235.1    rna-XM_046011300.1    rna-XM_032347641.1    rna-XM_059185794.1    rna-XM_059402781.1    rna-XM_045073368.1    mNeoVis.2.1.1.chr9_hap1.g00714.t01    rna-XM_044915445.1    rna-XM_004412407.1    rna-XM_042991969.1    rna-XM_040642260.1    rna-XM_027593037.2      rna-gnl|WGS_ZZZZ|egapxtmp_024012-R5      mNeoVis.vulgivaga.2.1.chr10.g00697.t01      3026    N0.HOG0003024   OG0000632  n8   rna-XM_002928613.4    rna-NM_001287065.1    rna-XM_022496841.1    rna-XM_019837665.2    rna-XM_036088484.1    rna-XM_039249082.1    rna-XM_032881380.1    rna-XM_046012049.1    rna-XM_032302661.1    rna-XM_059138074.1    rna-XM_059375811.1    rna-XM_004769870.3    mNeoVis.2.1.1.chr2_hap1.g00914.t01    rna-XM_021690020.1    rna-XM_004401284.1    rna-XM_007076341.3    rna-XM_008697746.2    rna-XM_027610029.2    mNeoVis.3.1.1.chr5.g00846.t01    rna-gnl|WGS_ZZZZ|egapxtmp_008664-R1    mNeoVis.evergladensis.2.1.chr5.g00869.t01    mNeoVis.vulgivaga.2.1.chr5.g00858.t01      3027    N0.HOG0003025   OG0000632  n15   rna-XM_019810185.2    rna-NM_001287069.1    rna-XM_022492599.1    rna-XM_023240249.2    rna-XM_036083660.1    rna-XM_039221471.1    rna-XM_032879861.1    rna-XM_046016085.1    rna-XM_032358454.1    rna-XM_059152704.1    rna-XM_059397635.1, rna-XM_059416459.1    rna-XM_045067090.1    mNeoVis.2.1.1.chr7_hap1.g02672.t01    rna-XM_044919209.1    rna-XM_004403761.1    rna-XM_042959122.1    rna-XM_008697503.1, rna-XM_040639746.1    rna-XM_027580477.1      rna-gnl|WGS_ZZZZ|egapxtmp_009848-R2    mNeoVis.evergladensis.2.1.chr7.g02484.t01    mNeoVis.vulgivaga.2.1.chr7.g02506.t01      3028    N0.HOG0003026   OG0000633  n1   rna-XM_034641225.1    rna-XM_038547035.1    rna-XM_022519864.1    rna-XM_023244520.2    rna-XM_036096647.1    rna-XM_039220045.1    rna-XM_032870980.1    rna-XM_045983716.1    rna-XM_032320821.1    rna-XM_059150512.1    rna-XM_059380796.1    rna-XM_004772469.3    mNeoVis.2.1.1.chr8_hap1.g00297.t01    rna-XM_044921592.1    rna-XM_004407234.1    rna-XM_042967008.1    rna-XM_040640471.1    rna-XM_027625034.2    mNeoVis.3.1.1.chr8.g00266.t01    rna-gnl|WGS_ZZZZ|egapxtmp_019421-R1    mNeoVis.evergladensis.2.1.chr9.g00296.t01    mNeoVis.vulgivaga.2.1.chr8.g00303.t01      3029   N0.HOG0003027  OG0000633  n23   rna-XM_034646507.1    rna-XM_038577587.1    rna-XM_022519407.1    rna-XM_011283910.4    rna-XM_036121086.1    rna-XM_039241421.1    rna-XM_032876850.1    rna-XM_046011500.1    rna-XM_032348111.1    rna-XM_059185426.1    rna-XM_059403041.1    rna-XM_004778861.3    mNeoVis.2.1.1.chr9_hap1.g00577.t01    rna-XM_021704791.2    rna-XM_004400848.1    rna-XM_007072987.3    rna-XM_040642419.1    rna-XM_027592127.2    mNeoVis.3.1.1.chr10.g00547.t01      mNeoVis.evergladensis.2.1.chr10.g00570.t01    mNeoVis.vulgivaga.2.1.chr10.g00558.t01      3030    N0.HOG0003028   OG0000633  n41   rna-XM_034654575.1    rna-XM_038565210.1    rna-XM_022520953.1    rna-XM_045033814.1    rna-XM_036085436.1    rna-XM_039219404.1    rna-XM_032857803.1    rna-XM_046019155.1    rna-XM_032355571.1    rna-XM_059167306.1    rna-XM_059393967.1    rna-XM_004743860.3    mNeoVis.2.1.1.chr3_hap1.g00973.t01    rna-XM_044913290.1    rna-XM_004394833.1    rna-XM_042994446.1    rna-XM_040644919.1    rna-XM_027589168.2    mNeoVis.3.1.1.chr3.g00963.t01    rna-gnl|WGS_ZZZZ|egapxtmp_041225-R1    mNeoVis.evergladensis.2.1.chr3.g00968.t01    mNeoVis.vulgivaga.2.1.chr3.g00987.t01      3031    N0.HOG0003029   OG0000634  n1   rna-XM_034639893.1    rna-NM_001012645.1    rna-XM_022509229.1    rna-XM_019817836.3    rna-XM_036080006.1    rna-XM_039227711.1    rna-XM_032871548.1    rna-XM_045985039.1    rna-XM_032322487.1    rna-XM_059149562.1    rna-XM_059380407.1    rna-XM_004764594.3    mNeoVis.2.1.1.chr8_hap1.g00464.t01    rna-XM_044921934.1    rna-XM_004395013.1    rna-XM_042966344.1    rna-XM_040640661.1    rna-XM_027625534.2    mNeoVis.3.1.1.chr8.g00419.t01      mNeoVis.evergladensis.2.1.chr9.g00455.t01    mNeoVis.vulgivaga.2.1.chr8.g00460.t01      3032    N0.HOG0003030   OG0000634  n12   rna-XM_034662078.1    rna-XM_038570687.1    rna-XM_022515245.1    rna-XM_023260626.2    rna-XM_036072138.1    rna-XM_039251935.1    rna-XM_032860980.1    rna-XM_046001560.1    rna-XM_032327731.1    rna-XM_059162464.1    rna-XM_059390835.1    rna-XM_004754347.3    mNeoVis.2.1.1.chr5_hap1.g01082.t01    rna-XM_021678917.2    rna-XM_004416473.1    rna-XM_007097468.3    rna-XM_008709331.2, rna-XM_008711574.2    rna-XM_035724196.1    mNeoVis.3.1.1.chr2.g00989.t01    rna-gnl|WGS_ZZZZ|egapxtmp_035531-R1    mNeoVis.evergladensis.2.1.chr2.g01071.t01    mNeoVis.vulgivaga.2.1.chr2.g01078.t01      3033    N0.HOG0003031   OG0000634  n30   rna-XM_002923276.4    rna-XM_038577764.1    rna-XM_022519739.1    rna-XM_023257108.2    rna-XM_036124083.1    rna-XM_039255120.1    rna-XM_032876676.1    rna-XM_046013659.1    rna-XM_032348354.1    rna-XM_059185212.1    rna-XM_059403222.1    rna-XM_004774635.3    mNeoVis.2.1.1.chr9_hap1.g00495.t01    rna-XM_021686696.2    rna-XM_004406231.1    rna-XM_007081369.3    rna-XM_008704631.2    rna-XM_027591689.1    mNeoVis.3.1.1.chr10.g00468.t01      mNeoVis.evergladensis.2.1.chr10.g00488.t01    mNeoVis.vulgivaga.2.1.chr10.g00477.t01      3034   N0.HOG0003032  OG0000635  n1   rna-XM_034645882.1    rna-XM_038568197.1    rna-XM_022506615.1    rna-XM_006937350.4      rna-XM_039245084.1    rna-XM_032879043.1    rna-XM_046016474.1    rna-XM_032359473.1    rna-XM_059143443.1    rna-XM_059388186.1    rna-XM_004770424.3    mNeoVis.2.1.1.chr7_hap1.g02246.t01        rna-XM_015542035.2    rna-XM_008689021.2      mNeoVis.3.1.1.chr7.g02066.t01    rna-gnl|WGS_ZZZZ|egapxtmp_010737-R1    mNeoVis.evergladensis.2.1.chr7.g02194.t01    mNeoVis.vulgivaga.2.1.chr7.g02212.t01      3035   N0.HOG0003033  OG0000635  n19   rna-XM_034646230.1    rna-XM_038568200.1    rna-XM_022506614.1        rna-XM_039253607.1    rna-XM_032879900.1      rna-XM_032360307.1    rna-XM_059158998.1      rna-XM_004770417.3    mNeoVis.2.1.1.chr7_hap1.g02248.t01        rna-XM_007092027.2    rna-XM_040639166.1      mNeoVis.3.1.1.chr7.g02068.t01    rna-gnl|WGS_ZZZZ|egapxtmp_012662-R1    mNeoVis.evergladensis.2.1.chr7.g02196.t01    mNeoVis.vulgivaga.2.1.chr7.g02215.t01      3036    N0.HOG0003034   OG0000635  n32   rna-XM_034645729.1    rna-XM_038569320.1    rna-XM_022506462.1    rna-XM_045039354.1      rna-XM_039245098.1    rna-XM_032879048.1      rna-XM_032359322.1    rna-XM_059143491.1      rna-XM_013044704.2          rna-XM_007092028.3    rna-XM_008689025.2                3037    N0.HOG0003035   OG0000635  n42   rna-XM_019808336.2    rna-XM_038568206.1      rna-XM_045039361.1    rna-XM_036110659.1    rna-XM_039253569.1    rna-XM_032879051.1    rna-XM_046016471.1    rna-XM_032359259.1    rna-XM_059159002.1    rna-XM_059386192.1    rna-XM_004770412.3    mNeoVis.2.1.1.chr7_hap1.g02251.t01    rna-XM_021685239.1    rna-XM_004417253.1    rna-XM_042958558.1    rna-XM_008707454.2    rna-XM_035723142.1    mNeoVis.3.1.1.chr7.g02071.t01    rna-gnl|WGS_ZZZZ|egapxtmp_012663-R1    mNeoVis.evergladensis.2.1.chr7.g02199.t01    mNeoVis.vulgivaga.2.1.chr7.g02218.t01      3038    N0.HOG0003036   OG0000636  n1   rna-XM_034646400.1    rna-XM_038564288.1    rna-XM_022506401.1    rna-XM_023239949.2    rna-XM_036122225.1    rna-XM_039257688.1    rna-XM_032879237.1    rna-XM_046014496.1    rna-XM_032357050.1    rna-XM_059144510.1    rna-XM_059404022.1    rna-XM_045063508.1    mNeoVis.2.1.1.chr7_hap1.g02304.t01    rna-XM_021684923.1    rna-XM_004413757.1    rna-XM_042957675.1    rna-XM_040639237.1    rna-XM_027579413.2    mNeoVis.3.1.1.chr7.g02121.t01    rna-gnl|WGS_ZZZZ|egapxtmp_012677-R1    mNeoVis.evergladensis.2.1.chr7.g02251.t01    mNeoVis.vulgivaga.2.1.chr7.g02268.t01      3039   N0.HOG0003037  OG0000636  n13   rna-XM_034659594.1    rna-XM_038561356.1    rna-XM_022520432.1    rna-XM_023251915.2    rna-XM_036065126.1    rna-XM_039231778.1    rna-XM_032843831.1    rna-XM_045979575.1    rna-XM_032352837.1    rna-XM_059188455.1    rna-XM_059405539.1    rna-XM_045079165.1    mNeoVis.2.1.1.chr11_hap1.g00952.t01    rna-XM_044918686.1    rna-XM_004399875.1    rna-XM_042982070.1    rna-XM_040623148.1    rna-XM_027621295.2    mNeoVis.3.1.1.chr11.g00906.t01    rna-gnl|WGS_ZZZZ|egapxtmp_027672-R1    mNeoVis.evergladensis.2.1.chr11.g00930.t01    mNeoVis.vulgivaga.2.1.chr11.g00954.t01      3040    N0.HOG0003038   OG0000636  n34   rna-XM_034643029.1    rna-XM_038545953.1    rna-XM_022505106.1    rna-XM_023256149.2    rna-XM_036100073.1    rna-XM_039223940.1    rna-XM_032839670.1    rna-XM_046007695.1    rna-XM_032341581.1    rna-XM_059183349.1    rna-XM_059374187.1    rna-XM_013050753.2    mNeoVis.2.1.1.chr10_hap1.g00011.t01    rna-XM_021679892.2    rna-XM_004395633.1    rna-XM_042990749.1    rna-XM_040641294.1    rna-XM_027571153.1      rna-gnl|WGS_ZZZZ|egapxtmp_016617-R1    mNeoVis.evergladensis.2.1.chr8.g00011.t01    mNeoVis.vulgivaga.2.1.chr9.g00012.t01      3041    N0.HOG0003039   OG0000637  n1   rna-XM_002917530.3    rna-XM_038578899.1    rna-XM_022499833.1    rna-XM_003994441.4    rna-XM_036096313.1    rna-XM_039237342.1    rna-XM_032877249.1    rna-XM_046027960.1    rna-XM_032313294.1    rna-XM_059173080.1    rna-XM_059395934.1    rna-XM_004751858.2    mNeoVis.2.1.1.chr2_hap1.g00121.t01    rna-XM_021699853.1    rna-XM_004413852.1    rna-XM_042960503.1    rna-XM_040624438.1    rna-XM_027595953.1    mNeoVis.3.1.1.chr5.g00115.t01    rna-gnl|WGS_ZZZZ|egapxtmp_009394-R1    mNeoVis.evergladensis.2.1.chr5.g00124.t01    mNeoVis.vulgivaga.2.1.chr5.g00121.t01      3042    N0.HOG0003040   OG0000637  n23   rna-XM_011222805.3    rna-NM_001003319.2    rna-XM_022499552.1    rna-XM_006938208.3    rna-XM_036096288.1    rna-XM_039237347.1    rna-XM_032877261.1    rna-XM_046026264.1    rna-XM_032313293.1    rna-XM_059172585.1    rna-XM_059395933.1    rna-XM_013055771.2    mNeoVis.2.1.1.chr2_hap1.g00122.t01    rna-XM_021699885.1    rna-XM_004413851.1    rna-XM_007084847.1    rna-XM_040624489.1    rna-XM_027591554.2    mNeoVis.3.1.1.chr5.g00116.t01      mNeoVis.evergladensis.2.1.chr5.g00125.t01    mNeoVis.vulgivaga.2.1.chr5.g00122.t01      3043    N0.HOG0003041   OG0000637  n40   rna-XM_011222804.3    rna-XM_038578898.1    rna-XM_022499562.1    rna-XM_006938207.2    rna-XM_036096286.1    rna-XM_039237334.1    rna-XM_032877274.1    rna-XM_046027429.1    rna-XM_032313355.1    rna-XM_059173081.1    rna-XM_059395931.1    rna-XM_004751861.2    mNeoVis.2.1.1.chr2_hap1.g00123.t01    rna-XM_021700128.1    rna-XM_004413850.1    rna-XM_015538763.2    rna-XM_040624632.1    rna-XM_027595919.1    mNeoVis.3.1.1.chr5.g00117.t01    rna-gnl|WGS_ZZZZ|egapxtmp_009392-R1    mNeoVis.evergladensis.2.1.chr5.g00126.t01    mNeoVis.vulgivaga.2.1.chr5.g00123.t01      3044    N0.HOG0003042   OG0000638  n1   rna-XM_034670976.1    rna-XM_038533402.1    rna-XM_022506161.1    rna-XM_045056598.1    rna-XM_036085907.1    rna-XM_039232111.1    rna-XM_032855239.1    rna-XM_045986344.1    rna-XM_032333810.1    rna-XM_059165449.1    rna-XM_059380690.1    rna-XM_004770541.3    mNeoVis.2.1.1.chr6_hap1.g00319.t01    rna-XM_021677602.1    rna-XM_004413737.1    rna-XM_042984951.1    rna-XM_008685823.2    rna-XM_027597375.2    mNeoVis.3.1.1.chr4.g00319.t01    rna-gnl|WGS_ZZZZ|egapxtmp_000715-R1    mNeoVis.evergladensis.2.1.chr4.g00325.t01    mNeoVis.vulgivaga.2.1.chr4.g00323.t01      3045    N0.HOG0003043   OG0000638  n23   rna-XM_034663464.1    rna-XM_038579172.1    rna-XM_022499938.1    rna-XM_045040894.1      rna-XM_039252434.1    rna-XM_032882055.1    rna-XM_046027044.1    rna-XM_032312635.1    rna-XM_059172785.1    rna-XM_059398867.1    rna-XM_045068593.1    mNeoVis.2.1.1.chr2_hap1.g00031.t01    rna-XM_021703799.1    rna-XM_004402218.1    rna-XM_042959873.1    rna-XM_040624502.1    rna-XM_027596714.2    mNeoVis.3.1.1.chr5.g00023.t01    rna-gnl|WGS_ZZZZ|egapxtmp_005897-R1    mNeoVis.evergladensis.2.1.chr5.g00031.t01    mNeoVis.vulgivaga.2.1.chr5.g00029.t01      3046    N0.HOG0003044   OG0000638  n42   rna-XM_011232903.3    rna-XM_038530544.1    rna-XM_022518140.1    rna-XM_006927743.5    rna-XM_036065733.1    rna-XM_039219197.1    rna-XM_032850718.1    rna-XM_045998648.1    rna-XM_032336257.1    rna-XM_059174974.1    rna-XM_059417858.1    rna-XM_013052335.2    mNeoVis.2.1.1.chr1_hap1.g00558.t01    rna-XM_021701957.1    rna-XM_004414638.1    rna-XM_042986632.1    rna-XM_008691687.2    rna-XM_027605843.2    mNeoVis.3.1.1.chr1.g00573.t01    rna-gnl|WGS_ZZZZ|egapxtmp_026056-R2    mNeoVis.evergladensis.2.1.chr1.g00564.t01    mNeoVis.vulgivaga.2.1.chr1.g00574.t01      3047    N0.HOG0003045   OG0000639  n1   rna-XM_034671743.1    rna-XM_038556190.1    rna-XM_022505656.1    rna-XM_045056289.1    rna-XM_036075848.1    rna-XM_039217419.1    rna-XM_032849431.1    rna-XM_045993784.1    rna-XM_032334415.1    rna-XM_059160314.1    rna-XM_059384586.1    rna-XM_045079821.1    mNeoVis.2.1.1.chr6_hap1.g00136.t01    rna-XM_044912942.1    rna-XM_004394948.1    rna-XM_042984447.1    rna-XM_040624907.1    rna-XM_035726125.1    mNeoVis.3.1.1.chr4.g00131.t01    rna-gnl|WGS_ZZZZ|egapxtmp_000948-R17    mNeoVis.evergladensis.2.1.chr4.g00140.t01    mNeoVis.vulgivaga.2.1.chr4.g00133.t01      3048    N0.HOG0003046   OG0000639  n22   rna-XM_034657893.1    rna-XM_038567055.1    rna-XM_022526990.1    rna-XM_023246270.2    rna-XM_036066520.1    rna-XM_039252322.1    rna-XM_032835896.1    rna-XM_045989891.1    rna-XM_032313423.1    rna-XM_059160215.1    rna-XM_059389209.1    rna-XM_004748235.2    mNeoVis.2.1.1.chr5_hap1.g00369.t01    rna-XM_044911267.1    rna-XM_004398983.1    rna-XM_042978378.1    rna-XM_040629193.1    rna-XM_027582890.2    mNeoVis.3.1.1.chr2.g00302.t01      mNeoVis.evergladensis.2.1.chr2.g00370.t01    mNeoVis.vulgivaga.2.1.chr2.g00365.t01      3049    N0.HOG0003047   OG0000639  n40   rna-XM_034653773.1    rna-NM_001190488.1    rna-XM_022496875.1    rna-XM_045036065.1    rna-XM_036076672.1    rna-XM_039218413.1    rna-XM_032870191.1    rna-XM_046010332.1    rna-XM_032360948.1    rna-XM_059136196.1    rna-XM_059375220.1    rna-XM_045064782.1    mNeoVis.2.1.1.chr2_hap1.g01149.t01    rna-XM_021690193.1    rna-XM_004411114.1    rna-XM_042997664.1    rna-XM_040627104.1    rna-XM_027575091.2    mNeoVis.3.1.1.chr5.g01068.t01    rna-gnl|WGS_ZZZZ|egapxtmp_009299-R1    mNeoVis.evergladensis.2.1.chr5.g01091.t01    mNeoVis.vulgivaga.2.1.chr5.g01081.t01      3050    N0.HOG0003048   OG0000640  n1   rna-XM_034671956.1    rna-NM_001173127.1    rna-XM_022506295.1    rna-XM_045056469.1    rna-XM_036112416.1    rna-XM_039227037.1    rna-XM_032853444.1    rna-XM_045998221.1    rna-XM_032334087.1    rna-XM_059163505.1    rna-XM_059395979.1    rna-XM_013062218.2    mNeoVis.2.1.1.chr6_hap1.g00237.t01    rna-XM_021701665.1    rna-XM_004396166.1    rna-XM_042984723.1    rna-XM_040624977.1    rna-XM_027598950.1    mNeoVis.3.1.1.chr4.g00236.t01    rna-gnl|WGS_ZZZZ|egapxtmp_001186-R1    mNeoVis.evergladensis.2.1.chr4.g00244.t01    mNeoVis.vulgivaga.2.1.chr4.g00241.t01      3051    N0.HOG0003049   OG0000640  n23   rna-XM_019808911.2    rna-NM_001146143.1    rna-XM_022505615.1    rna-XM_003985440.4    rna-XM_036112508.1    rna-XM_039226970.1    rna-XM_032853370.1    rna-XM_045997906.1    rna-XM_032334094.1    rna-XM_059163534.1    rna-XM_059382486.1    rna-XM_045076557.1    mNeoVis.2.1.1.chr6_hap1.g00236.t01    rna-XM_021701636.1    rna-XM_004396165.1    rna-XM_007098862.3      rna-XM_035726261.1    mNeoVis.3.1.1.chr4.g00234.t01    rna-gnl|WGS_ZZZZ|egapxtmp_001188-R1      mNeoVis.vulgivaga.2.1.chr4.g00240.t01      3052    N0.HOG0003050   OG0000640  n42   rna-XM_019808910.2    rna-XM_038533484.1    rna-XM_022505613.1    rna-XM_003985467.5    rna-XM_036112533.1    rna-XM_039226969.1    rna-XM_032853345.1    rna-XM_045997901.1    rna-XM_032334077.1    rna-XM_059163423.1    rna-XM_059382413.1    rna-XM_013062213.2    mNeoVis.2.1.1.chr6_hap1.g00235.t01    rna-XM_021701660.1    rna-XM_004396174.1    rna-XM_007098861.3    rna-XM_040624979.1    rna-XM_035726265.1    mNeoVis.3.1.1.chr4.g00233.t01    rna-gnl|WGS_ZZZZ|egapxtmp_001187-R1    mNeoVis.evergladensis.2.1.chr4.g00242.t01, mNeoVis.evergladensis.2.1.chr4.g00243.t01    mNeoVis.vulgivaga.2.1.chr4.g00239.t01      3053    N0.HOG0003051   OG0000641  n1   rna-XM_034648683.1    rna-XM_038546024.1    rna-XM_022509498.1    rna-XM_006940397.4    rna-XM_036069111.1    rna-XM_039256739.1    rna-XM_032871813.1    rna-XM_045986263.1    rna-XM_032319484.1    rna-XM_059149657.1    rna-XM_059380225.1    rna-XM_004772774.3    mNeoVis.2.1.1.chr8_hap1.g00420.t01    rna-XM_044921778.1    rna-XM_004410601.1    rna-XM_015540007.2    rna-XM_008701258.1    rna-XM_027626089.2    mNeoVis.3.1.1.chr8.g00384.t01    rna-gnl|WGS_ZZZZ|egapxtmp_019795-R2    mNeoVis.evergladensis.2.1.chr9.g00417.t01    mNeoVis.vulgivaga.2.1.chr8.g00425.t01      3054    N0.HOG0003052   OG0000641  n22   rna-XM_034648397.1, rna-XM_034648399.1, rna-XM_034652424.1    rna-XM_038546023.1, rna-XM_038546103.1    rna-XM_022509141.1, rna-XM_022509499.1    rna-XM_006940341.5, rna-XM_011289139.4    rna-XM_036069112.1, rna-XM_036069198.1    rna-XM_039256740.1, rna-XM_039256742.1    rna-XM_032871605.1, rna-XM_032871606.1    rna-XM_045985358.1, rna-XM_045986264.1    rna-XM_032319485.1, rna-XM_032321714.1    rna-XM_059148189.1, rna-XM_059149656.1    rna-XM_059380224.1, rna-XM_059380865.1    rna-XM_004772752.3, rna-XM_004772773.3    mNeoVis.2.1.1.chr8_hap1.g00418.t01, mNeoVis.2.1.1.chr8_hap1.g00419.t01    rna-XM_021681721.1, rna-XM_021681748.1    rna-XM_004410494.1    rna-XM_007087636.2, rna-XM_007087639.2    rna-XM_008701259.2, rna-XM_008701261.2    rna-XM_035724619.1, rna-XM_035724625.1    mNeoVis.3.1.1.chr8.g00382.t01, mNeoVis.3.1.1.chr8.g00383.t01    rna-gnl|WGS_ZZZZ|egapxtmp_019810-R1    mNeoVis.evergladensis.2.1.chr9.g00415.t01, mNeoVis.evergladensis.2.1.chr9.g00416.t01    mNeoVis.vulgivaga.2.1.chr8.g00423.t01, mNeoVis.vulgivaga.2.1.chr8.g00424.t01      3055    N0.HOG0003053   OG0000642  n1   rna-XM_034647287.1    rna-XM_038537058.1    rna-XM_022491463.1    rna-XM_006939892.4    rna-XM_036073481.1    rna-XM_039227350.1    rna-XM_032841259.1    rna-XM_045985357.1    rna-XM_032321095.1    rna-XM_059148632.1    rna-XM_059380770.1    rna-XM_004776562.3    mNeoVis.2.1.1.chr8_hap1.g00874.t01    rna-XM_021694740.1    rna-XM_004405123.1    rna-XM_042965403.1    rna-XM_040630121.1, rna-XM_040630122.1    rna-XM_027626465.1    mNeoVis.3.1.1.chr8.g00820.t01      mNeoVis.evergladensis.2.1.chr9.g00862.t01    mNeoVis.vulgivaga.2.1.chr8.g00864.t01      3056    N0.HOG0003054   OG0000642  n20         rna-XM_003989748.5                          rna-XM_007078163.2                  3057    N0.HOG0003055   OG0000642  n22   rna-XM_034653492.1, rna-XM_034653493.1    rna-XM_038548865.1    rna-XM_022500368.1, rna-XM_022501399.1      rna-XM_036108733.1    rna-XM_039255150.1    rna-XM_032851030.1    rna-XM_046023122.1, rna-XM_046023204.1    rna-XM_032306374.1, rna-XM_032308045.1    rna-XM_059143473.1, rna-XM_059143475.1    rna-XM_059410357.1    rna-XM_045082888.1, rna-XM_045082893.1    mNeoVis.2.1.1.chr13_hap1.g00083.t01, mNeoVis.2.1.1.chr13_hap1.g00084.t01    rna-XM_021691547.1    rna-XM_004417763.1      rna-XM_008709446.2, rna-XM_040631358.1    rna-XM_035723738.1    mNeoVis.3.1.1.chr13.g00046.t01    rna-gnl|WGS_ZZZZ|egapxtmp_030706-R1    mNeoVis.evergladensis.2.1.chr13.g00084.t01    mNeoVis.vulgivaga.2.1.chr13.g00081.t01      3058    N0.HOG0003056   OG0000642  n49   rna-XM_034645088.1    rna-XM_038529591.1    rna-XM_022497498.1      rna-XM_036116432.1    rna-XM_039230855.1    rna-XM_032864037.1      rna-XM_032303750.1          mNeoVis.2.1.1.chr2_hap1.g01616.t01    rna-XM_021684408.1        rna-XM_040645562.1      mNeoVis.3.1.1.chr5.g01537.t01      mNeoVis.evergladensis.2.1.chr5.g01555.t01    mNeoVis.vulgivaga.2.1.chr5.g01558.t01      3059    N0.HOG0003057   OG0000643  n0   rna-XM_011230274.3, rna-XM_019804125.2, rna-XM_034653378.1    rna-XM_038567872.1    rna-XM_022492640.1, rna-XM_022525580.1    rna-XM_006928243.5, rna-XM_006928244.5, rna-XM_019815693.3, rna-XM_019826123.3, rna-XM_045043285.1    rna-XM_036100949.1, rna-XM_036100950.1, rna-XM_036116142.1    rna-XM_039247551.1, rna-XM_039247552.1, rna-XM_039247553.1, rna-XM_039247629.1, rna-XM_039247970.1, rna-XM_039247977.1, rna-XM_039248059.1    rna-XM_032883509.1    rna-XM_045990440.1, rna-XM_045990445.1, rna-XM_045991421.1    rna-XM_032303212.1, rna-XM_032303226.1, rna-XM_032305539.1, rna-XM_032305570.1, rna-XM_032305592.1    rna-XM_059159367.1, rna-XM_059159371.1, rna-XM_059159372.1, rna-XM_059159378.1    rna-XM_059387308.1, rna-XM_059387318.1, rna-XM_059388591.1, rna-XM_059388593.1    rna-XM_004771335.3, rna-XM_004777961.3, rna-XM_045087000.1, rna-XM_045087012.1    mNeoVis.2.1.1.chr5_hap1.g00075.t01, mNeoVis.2.1.1.chr5_hap1.g00077.t01, mNeoVis.2.1.1.chr5_hap1.g00078.t01, mNeoVis.2.1.1.chr5_hap1.g00079.t01, mNeoVis.2.1.1.chr5_hap1.g00080.t01    rna-XM_044913133.1    rna-XM_004395479.1, rna-XM_004395480.1, rna-XM_004417727.1    rna-XM_007075242.3, rna-XM_015543923.2, rna-XM_015543929.2, rna-XM_042977704.1, rna-XM_042977713.1    rna-XM_008711237.2, rna-XM_040629648.1    rna-XM_027582543.2, rna-XM_027582563.2, rna-XM_035726537.1      rna-gnl|WGS_ZZZZ|egapxtmp_036237-R1, rna-gnl|WGS_ZZZZ|egapxtmp_037374-R1    mNeoVis.evergladensis.2.1.chr2.g00077.t01    mNeoVis.vulgivaga.2.1.chr2.g00073.t01      3060    N0.HOG0003058   OG0000644  n1   rna-XM_034654971.1    rna-XM_038585390.1    rna-XM_022493971.1    rna-XM_019839139.3    rna-XM_036098598.1    rna-XM_039257414.1    rna-XM_032860064.1    rna-XM_046019746.1    rna-XM_032354451.1    rna-XM_059168392.1    rna-XM_059393217.1    rna-XM_004764173.3    mNeoVis.2.1.1.chr3_hap1.g01277.t01    rna-XM_021702845.1    rna-XM_004393512.1    rna-XM_042995079.1    rna-XM_008687547.2    rna-XM_027590853.2    mNeoVis.3.1.1.chr3.g01254.t01    rna-gnl|WGS_ZZZZ|egapxtmp_042736-R1    mNeoVis.evergladensis.2.1.chr3.g01275.t01    mNeoVis.vulgivaga.2.1.chr3.g01293.t01      3061    N0.HOG0003059   OG0000644  n19   rna-XM_034641363.1    rna-XM_038546095.1    rna-XM_022520022.1    rna-XM_011289270.3    rna-XM_036092706.1    rna-XM_039220024.1    rna-XM_032850726.1, rna-XM_032850727.1    rna-XM_045985412.1    rna-XM_032322823.1    rna-XM_059149238.1    rna-XM_059380880.1    rna-XM_004772766.3    mNeoVis.2.1.1.chr8_hap1.g00316.t01    rna-XM_021681627.1    rna-XM_004407252.1    rna-XM_007093851.2    rna-XM_008693904.2    rna-XM_027626252.2    mNeoVis.3.1.1.chr8.g00284.t01      mNeoVis.evergladensis.2.1.chr9.g00315.t01    mNeoVis.vulgivaga.2.1.chr8.g00323.t01      3062    N0.HOG0003060   OG0000644  n26   rna-XM_002927753.4    rna-XM_038556765.1    rna-XM_022503315.1    rna-XM_003982740.6    rna-XM_036071489.1    rna-XM_039250039.1    rna-XM_032875502.1    rna-XM_046020426.1    rna-XM_032304895.1    rna-XM_059171076.1    rna-XM_059396883.1    rna-XM_004776459.3    mNeoVis.2.1.1.chr4_hap1.g00504.t01    rna-XM_021689613.1    rna-XM_004394105.1    rna-XM_042966117.1    rna-XM_008686304.2    rna-XM_027574428.2      rna-gnl|WGS_ZZZZ|egapxtmp_004778-R1    mNeoVis.evergladensis.2.1.chr6.g00508.t01    mNeoVis.vulgivaga.2.1.chr6.g00509.t01      3063    N0.HOG0003061   OG0000645  n1   rna-XM_002915492.4    rna-XM_038559728.1    rna-XM_022513844.1    rna-XM_023250732.2    rna-XM_036102104.1    rna-XM_039230696.1    rna-XM_032840187.1    rna-XM_046021053.1    rna-XM_032304043.1    rna-XM_059172249.1    rna-XM_059396102.1    rna-XM_004765951.3    mNeoVis.2.1.1.chr4_hap1.g00043.t01    rna-XM_021692995.1    rna-XM_004408812.1    rna-XM_042975455.1    rna-XM_040643798.1    rna-XM_027573809.2    mNeoVis.3.1.1.chr6.g00046.t01    rna-gnl|WGS_ZZZZ|egapxtmp_003019-R1    mNeoVis.evergladensis.2.1.chr6.g00045.t01    mNeoVis.vulgivaga.2.1.chr6.g00049.t01      3064    N0.HOG0003062   OG0000645  n21   rna-XM_019800201.2    rna-NM_001048031.1    rna-XM_022520021.1    rna-XM_019817978.3    rna-XM_036071252.1    rna-XM_039220927.1    rna-XM_032849047.1    rna-XM_045986269.1    rna-XM_032321832.1    rna-XM_059149458.1    rna-XM_059378405.1    rna-XM_004772543.3    mNeoVis.2.1.1.chr8_hap1.g00323.t01    rna-XM_021681636.1    rna-XM_004407263.1    rna-XM_042967106.1    rna-XM_040640530.1    rna-XM_027624655.2      rna-gnl|WGS_ZZZZ|egapxtmp_019427-R1    mNeoVis.evergladensis.2.1.chr9.g00322.t01    mNeoVis.vulgivaga.2.1.chr8.g00330.t01      3065    N0.HOG0003063   OG0000645  n40   rna-XM_034655253.1    rna-NM_001289433.1    rna-XM_022494442.1    rna-XM_045034528.1    rna-XM_036082902.1    rna-XM_039216692.1    rna-XM_032857315.1    rna-XM_046019777.1    rna-XM_032354932.1    rna-XM_059167927.1    rna-XM_059393567.1    rna-XM_045078780.1    mNeoVis.2.1.1.chr3_hap1.g01147.t01    rna-XM_044913427.1    rna-XM_004402775.1    rna-XM_042995406.1    rna-XM_040639984.1    rna-XM_035726695.1    mNeoVis.3.1.1.chr3.g01134.t01    rna-gnl|WGS_ZZZZ|egapxtmp_042334-R1    mNeoVis.evergladensis.2.1.chr3.g01147.t01    mNeoVis.vulgivaga.2.1.chr3.g01162.t01      3066    N0.HOG0003064   OG0000646  n1   rna-XM_034657960.1    rna-XM_038567008.1    rna-XM_022497645.1    rna-XM_019818723.3    rna-XM_036089089.1    rna-XM_039243374.1    rna-XM_032867504.1    rna-XM_045992301.1    rna-XM_032314206.1    rna-XM_059160316.1    rna-XM_059389278.1    rna-XM_004761123.3    mNeoVis.2.1.1.chr5_hap1.g00432.t01    rna-XM_021683172.1    rna-XM_004408378.1    rna-XM_042978453.1    rna-XM_008706435.2    rna-XM_027585544.1    mNeoVis.3.1.1.chr2.g00360.t01    rna-gnl|WGS_ZZZZ|egapxtmp_034917-R4    mNeoVis.evergladensis.2.1.chr2.g00429.t01    mNeoVis.vulgivaga.2.1.chr2.g00431.t01      3067    N0.HOG0003065   OG0000646  n23   rna-XM_019793754.2          rna-XM_036089076.1                      rna-XM_004408393.1      rna-XM_008706417.2    rna-XM_027586090.1              3068    N0.HOG0003066   OG0000646  n28     rna-XM_038566999.1      rna-XM_045047544.1, rna-XM_045047564.1, rna-XM_045047577.1, rna-XM_045047580.1, rna-XM_045047591.1                          rna-XM_042978455.1                  3069    N0.HOG0003067   OG0000646  n34   rna-XM_034657966.1    rna-XM_038566998.1    rna-XM_022497731.1, rna-XM_022497733.1    rna-XM_045047599.1    rna-XM_036089153.1    rna-XM_039225709.1, rna-XM_039243370.1    rna-XM_032867506.1, rna-XM_032867513.1    rna-XM_045990038.1, rna-XM_045990041.1    rna-XM_032314159.1, rna-XM_032314227.1    rna-XM_059160317.1, rna-XM_059160321.1    rna-XM_059389280.1, rna-XM_059389286.1    rna-XM_004761113.2, rna-XM_045082388.1    mNeoVis.2.1.1.chr5_hap1.g00433.t01    rna-XM_021683170.1    rna-XM_004408379.1    rna-XM_042978456.1    rna-XM_040629131.1    rna-XM_027586087.2    mNeoVis.3.1.1.chr2.g00361.t01, mNeoVis.3.1.1.chr2.g00362.t01    rna-gnl|WGS_ZZZZ|egapxtmp_034918-R2    mNeoVis.evergladensis.2.1.chr2.g00430.t01    mNeoVis.vulgivaga.2.1.chr2.g00432.t01      3070    N0.HOG0003068   OG0000647  n1   rna-XM_002926661.4    rna-NM_001313860.1    rna-XM_022495835.1    rna-XM_019815118.2    rna-XM_036096702.1    rna-XM_039254452.1    rna-XM_032859702.1    rna-XM_046025358.1    rna-XM_032309189.1    rna-XM_059138997.1    rna-XM_059410044.1    rna-XM_013051477.2    mNeoVis.2.1.1.chr3_hap1.g00184.t01    rna-XM_021686077.2    rna-XM_004412931.1    rna-XM_015542481.2    rna-XM_008690930.2    rna-XM_035723870.1    mNeoVis.3.1.1.chr3.g00185.t01    rna-gnl|WGS_ZZZZ|egapxtmp_042389-R1    mNeoVis.evergladensis.2.1.chr3.g00193.t01    mNeoVis.vulgivaga.2.1.chr3.g00186.t01      3071    N0.HOG0003069   OG0000647  n22   rna-XM_034652451.1    rna-XM_038543611.1    rna-XM_022496001.1    rna-XM_006938833.4    rna-XM_036096707.1, rna-XM_036111048.1    rna-XM_039254440.1    rna-XM_032859709.1    rna-XM_046025311.1    rna-XM_032311058.1    rna-XM_059140268.1    rna-XM_059410042.1    rna-XM_013051475.2      rna-XM_021686075.2    rna-XM_004412937.1    rna-XM_042961568.1    rna-XM_040632618.1    rna-XM_027577660.2    mNeoVis.3.1.1.chr3.g00186.t01    rna-gnl|WGS_ZZZZ|egapxtmp_042390-R6    mNeoVis.evergladensis.2.1.chr3.g00195.t01        3072    N0.HOG0003070   OG0000647  n42   rna-XM_034652452.1    rna-XM_038543607.1    rna-XM_022495698.1    rna-XM_011288060.4    rna-XM_036096703.1    rna-XM_039254438.1    rna-XM_032859704.1    rna-XM_046025527.1    rna-XM_032310750.1    rna-XM_059139810.1    rna-XM_059409402.1    rna-XM_045065482.1    mNeoVis.2.1.1.chr3_hap1.g00185.t01    rna-XM_021686276.2    rna-XM_004412933.1    rna-XM_042961975.1    rna-XM_040632617.1    rna-XM_027577030.2    mNeoVis.3.1.1.chr3.g00187.t01    rna-gnl|WGS_ZZZZ|egapxtmp_039392-R1    mNeoVis.evergladensis.2.1.chr3.g00194.t01    mNeoVis.vulgivaga.2.1.chr3.g00187.t01      3073    N0.HOG0003071   OG0000648  n1   rna-XM_034654042.1, rna-XM_034654043.1    rna-XM_038541061.1, rna-XM_038565269.1    rna-XM_022496847.1    rna-XM_045033608.1    rna-XM_036109868.1    rna-XM_039251131.1    rna-XM_032881274.1    rna-XM_046001015.1, rna-XM_046001117.1    rna-XM_032302287.1, rna-XM_032302288.1    rna-XM_059136362.1, rna-XM_059136367.1    rna-XM_059375665.1, rna-XM_059375668.1    rna-XM_045081939.1, rna-XM_045081940.1    mNeoVis.2.1.1.chr2_hap1.g00979.t01, mNeoVis.2.1.1.chr2_hap1.g00980.t01    rna-XM_021689999.1    rna-XM_004417775.1    rna-XM_042998136.1    rna-XM_008704023.2, rna-XM_008704027.2    rna-XM_027621856.1    mNeoVis.3.1.1.chr5.g00913.t01      mNeoVis.evergladensis.2.1.chr5.g00933.t01    mNeoVis.vulgivaga.2.1.chr5.g00920.t01      3074   N0.HOG0003072  OG0000648  n30   rna-XM_011224982.3, rna-XM_034641640.1, rna-XM_034651431.1, rna-XM_034652227.1, rna-XM_034654037.1, rna-XM_034654040.1    rna-XM_038541060.1    rna-XM_022496519.1    rna-XM_003990413.6    rna-XM_036082624.1, rna-XM_036109867.1    rna-XM_039251075.1    rna-XM_032851195.1, rna-XM_032881520.1    rna-XM_045979212.1, rna-XM_046000897.1    rna-XM_032303165.1    rna-XM_059137594.1    rna-XM_059374237.1    rna-XM_004769717.3    mNeoVis.2.1.1.chr2_hap1.g00978.t01    rna-XM_021690047.1    rna-XM_004407173.1, rna-XM_004417335.1    rna-XM_007076413.3    rna-XM_008704029.2    rna-XM_027578167.2, rna-XM_027599715.2, rna-XM_027619940.2    mNeoVis.3.1.1.chr5.g00911.t01, mNeoVis.3.1.1.chr5.g00912.t01    rna-gnl|WGS_ZZZZ|egapxtmp_007391-R1    mNeoVis.evergladensis.2.1.chr5.g00932.t01    mNeoVis.vulgivaga.2.1.chr5.g00919.t01      3075   N0.HOG0003073  OG0000649  n1   rna-XM_034655935.1    rna-XM_038551979.1    rna-XM_022504896.1    rna-XM_045058295.1    rna-XM_036118798.1    rna-XM_039253424.1    rna-XM_032844385.1    rna-XM_046000035.1    rna-XM_032335489.1    rna-XM_059175731.1    rna-XM_059418417.1    rna-XM_045081421.1    mNeoVis.2.1.1.chr1_hap1.g00815.t01    rna-XM_044917123.1    rna-XM_004402565.1    rna-XM_042982865.1    rna-XM_040622527.1    rna-XM_027605455.2    mNeoVis.3.1.1.chr1.g00842.t01    rna-gnl|WGS_ZZZZ|egapxtmp_032959-R1    mNeoVis.evergladensis.2.1.chr1.g00822.t01    mNeoVis.vulgivaga.2.1.chr1.g00838.t01      3076    N0.HOG0003074   OG0000649  n10   rna-XM_034657412.1    rna-XM_038567926.1    rna-XM_022525489.1    rna-XM_023243302.2    rna-XM_036069974.1    rna-XM_039248032.1    rna-XM_032846919.1    rna-XM_045990495.1    rna-XM_032358692.1    rna-XM_059159312.1    rna-XM_059388542.1    rna-XM_045087404.1      rna-XM_021705602.2    rna-XM_004395398.1    rna-XM_042977617.1    rna-XM_040629722.1    rna-XM_027585690.2    mNeoVis.3.1.1.chr2.g00026.t01    rna-gnl|WGS_ZZZZ|egapxtmp_034432-R1      mNeoVis.vulgivaga.2.1.chr2.g00047.t01      3077    N0.HOG0003075   OG0000649  n28   rna-XM_034660837.1    rna-XM_038580664.1    rna-XM_022512045.1    rna-XM_045059174.1    rna-XM_036083832.1    rna-XM_039219050.1    rna-XM_032864726.1    rna-XM_046008576.1    rna-XM_032342837.1    rna-XM_059180767.1    rna-XM_059372231.1    rna-XM_045067413.1    mNeoVis.2.1.1.chr10_hap1.g00961.t01    rna-XM_044918446.1    rna-XM_004409418.1    rna-XM_042988654.1    rna-XM_008708611.2    rna-XM_027569965.1    mNeoVis.3.1.1.chr9.g00891.t01    rna-gnl|WGS_ZZZZ|egapxtmp_015870-R11    mNeoVis.evergladensis.2.1.chr8.g00931.t01    mNeoVis.vulgivaga.2.1.chr9.g00918.t01      3078    N0.HOG0003076   OG0000650  n1   rna-XM_011226764.3    rna-XM_038558499.1    rna-XM_022494921.1    rna-XM_023257263.2    rna-XM_036099826.1    rna-XM_039249913.1    rna-XM_032854726.1    rna-XM_046012050.1    rna-XM_032346937.1    rna-XM_059186380.1    rna-XM_059402420.1    rna-XM_045069086.1    mNeoVis.2.1.1.chr9_hap1.g00948.t01    rna-XM_044915365.1    rna-XM_004404830.1    rna-XM_042992785.1    rna-XM_040637280.1    rna-XM_027591882.2    mNeoVis.3.1.1.chr10.g00902.t01    rna-gnl|WGS_ZZZZ|egapxtmp_024373-R2    mNeoVis.evergladensis.2.1.chr10.g00933.t01    mNeoVis.vulgivaga.2.1.chr10.g00923.t01      3079   N0.HOG0003077  OG0000650  n21   rna-XM_034645288.1    rna-XM_038563612.1    rna-XM_022507643.1    rna-XM_019812838.3    rna-XM_036067772.1    rna-XM_039235453.1    rna-XM_032879746.1    rna-XM_046016687.1    rna-XM_032358439.1    rna-XM_059150716.1    rna-XM_059398935.1    rna-XM_045066409.1    mNeoVis.2.1.1.chr7_hap1.g02612.t01    rna-XM_044919540.1    rna-XM_004393789.1    rna-XM_042958646.1    rna-XM_040639616.1    rna-XM_027579872.2        mNeoVis.evergladensis.2.1.chr7.g02418.t01    mNeoVis.vulgivaga.2.1.chr7.g02442.t01      3080    N0.HOG0003078   OG0000650  n40   rna-XM_034646232.1    rna-XM_038563608.1    rna-XM_022507349.1    rna-XM_045039704.1    rna-XM_036067780.1    rna-XM_039235495.1    rna-XM_032879086.1, rna-XM_032879919.1    rna-XM_046016109.1    rna-XM_032358139.1    rna-XM_059150759.1    rna-XM_059398934.1    rna-XM_045066418.1    mNeoVis.2.1.1.chr7_hap1.g02613.t01    rna-XM_021685160.2      rna-XM_042958211.1    rna-XM_040638790.1, rna-XM_040639619.1    rna-XM_027581098.2        mNeoVis.evergladensis.2.1.chr7.g02419.t01    mNeoVis.vulgivaga.2.1.chr7.g02443.t01      3081    N0.HOG0003079   OG0000651  n1   rna-XM_034648130.1    rna-NM_001314095.1    rna-XM_022510648.1    rna-XM_011282429.4    rna-XM_036092711.1, rna-XM_036092722.1    rna-XM_039222946.1    rna-XM_032864530.1    rna-XM_046005029.1    rna-XM_032340256.1    rna-XM_059178258.1    rna-XM_059387306.1, rna-XM_059401547.1    rna-XM_004771867.3    mNeoVis.2.1.1.chr1_hap1.g01548.t01    rna-XM_021691962.1    rna-XM_004410693.1, rna-XM_004410701.1    rna-XM_007095518.3, rna-XM_015543560.2    rna-XM_008691781.2, rna-XM_008691784.2    rna-XM_027602855.1, rna-XM_035728382.1    mNeoVis.3.1.1.chr1.g01501.t01    rna-gnl|WGS_ZZZZ|egapxtmp_025919-R1    mNeoVis.evergladensis.2.1.chr1.g01516.t01    mNeoVis.vulgivaga.2.1.chr1.g01511.t01      3082    N0.HOG0003080   OG0000651  n28     rna-XM_038554317.1    rna-XM_022510738.1    rna-XM_023254061.2      rna-XM_039222949.1    rna-XM_032864528.1    rna-XM_046004722.1    rna-XM_032340259.1    rna-XM_059178259.1    rna-XM_059400245.1    rna-XM_045071162.1    mNeoVis.2.1.1.chr1_hap1.g01547.t01        rna-XM_007095519.3    rna-XM_008691782.1    rna-XM_027601259.1    mNeoVis.3.1.1.chr1.g01500.t01    rna-gnl|WGS_ZZZZ|egapxtmp_025906-R1    mNeoVis.evergladensis.2.1.chr1.g01515.t01    mNeoVis.vulgivaga.2.1.chr1.g01510.t01      3083    N0.HOG0003081   OG0000651  n42     rna-XM_038554310.1    rna-XM_022510837.1        rna-XM_039222957.1, rna-XM_039225570.1    rna-XM_032864555.1    rna-XM_046004948.1    rna-XM_032340255.1    rna-XM_059178256.1    rna-XM_059401548.1      mNeoVis.2.1.1.chr1_hap1.g01550.t01      rna-XM_004410702.1    rna-XM_042986557.1    rna-XM_008691787.2    rna-XM_027602683.1    mNeoVis.3.1.1.chr1.g01503.t01      mNeoVis.evergladensis.2.1.chr1.g01517.t01    mNeoVis.vulgivaga.2.1.chr1.g01512.t01      3084    N0.HOG0003082   OG0000652  n1   rna-XM_019794673.2    rna-XM_038527086.1    rna-XM_022512661.1    rna-XM_023243342.2    rna-XM_036080031.1    rna-XM_039241591.1    rna-XM_032866625.1    rna-XM_046022603.1    rna-XM_032306759.1    rna-XM_059141018.1    rna-XM_059411864.1    rna-XM_045065097.1    mNeoVis.2.1.1.chr13_hap1.g00743.t01    rna-XM_021694311.1    rna-XM_004397544.1    rna-XM_042963843.1    rna-XM_040620243.1    rna-XM_027615775.2    mNeoVis.3.1.1.chr13.g00659.t01    rna-gnl|WGS_ZZZZ|egapxtmp_031576-R1    mNeoVis.evergladensis.2.1.chr13.g00708.t01    mNeoVis.vulgivaga.2.1.chr13.g00705.t01      3085    N0.HOG0003083   OG0000652  n11   rna-XM_034657411.1    rna-XM_038567939.1    rna-XM_022525797.1    rna-XM_003981612.6    rna-XM_036069987.1    rna-XM_039248099.1    rna-XM_032847087.1    rna-XM_045990988.1    rna-XM_032358802.1    rna-XM_059159308.1    rna-XM_059388537.1    rna-XM_045087435.1    mNeoVis.2.1.1.chr5_hap1.g00044.t01    rna-XM_021705165.1    rna-XM_004395400.1    rna-XM_042977612.1    rna-XM_040629728.1    rna-XM_027587442.1        mNeoVis.evergladensis.2.1.chr2.g00044.t01    mNeoVis.vulgivaga.2.1.chr2.g00044.t01      3086    N0.HOG0003084   OG0000652  n25   rna-XM_034668880.1    rna-XM_038532997.1    rna-XM_022511089.1    rna-XM_023254806.2    rna-XM_036079161.1    rna-XM_039217253.1    rna-XM_032836870.1    rna-XM_046008408.1    rna-XM_032341928.1    rna-XM_059179830.1    rna-XM_059371435.1    rna-XM_013062019.2    mNeoVis.2.1.1.chr10_hap1.g01264.t01    rna-XM_021680461.1    rna-XM_004403571.1    rna-XM_042987905.1    rna-XM_040642887.1    rna-XM_027571379.2      rna-gnl|WGS_ZZZZ|egapxtmp_015602-R1    mNeoVis.evergladensis.2.1.chr8.g01241.t01    mNeoVis.vulgivaga.2.1.chr9.g01224.t01      3087    N0.HOG0003085   OG0000653  n2   rna-NM_001304917.1    rna-XM_038579346.1    rna-XM_022515570.1    rna-XM_003999819.6    rna-XM_036106087.1, rna-XM_036112589.1    rna-XM_039220286.1    rna-XM_032882583.1    rna-XM_045988149.1, rna-XM_046015821.1, rna-XM_046019338.1    rna-XM_032316496.1    rna-XM_059166421.1    rna-XM_059394599.1, rna-XM_059404641.1    rna-XM_004768802.1, rna-XM_045084781.1    mNeoVis.2.1.1.chr4_hap1.g00717.t01    rna-XM_021688198.1, rna-XM_021693374.1, rna-XM_021697740.2, rna-XM_021702952.1    rna-XM_004401426.1, rna-XM_004411123.1, rna-XM_004414440.1    rna-NM_001290591.1    rna-XM_008706704.2    rna-XM_027613934.2      rna-gnl|WGS_ZZZZ|egapxtmp_002644-R1    mNeoVis.evergladensis.2.1.chr6.g00723.t01        3088    N0.HOG0003086   OG0000653  n4     rna-XM_038569665.1                  rna-XM_059183648.1                              3089    N0.HOG0003087   OG0000653  n5     rna-XM_038579882.1        rna-XM_036076217.1            rna-XM_059152521.1          rna-XM_044912715.1, rna-XM_044912722.1, rna-XM_044914925.1          rna-XM_035726158.1              3090    N0.HOG0003088   OG0000653  n11   rna-XM_034648676.1    rna-XM_038559003.1      rna-XM_019825346.2, rna-XM_045035338.1, rna-XM_045038431.1    rna-XM_036084276.1, rna-XM_036089212.1    rna-XM_039247695.1          rna-XM_059136039.1    rna-XM_059383555.1        rna-XM_044916765.1    rna-XM_004396146.1, rna-XM_004412114.1    rna-XM_015535706.2, rna-XM_015541603.2, rna-XM_042996199.1    rna-XM_040642159.1, rna-XM_040644091.1    rna-XM_035722446.1, rna-XM_035727605.1    mNeoVis.3.1.1.chr12.g00228.t01        mNeoVis.vulgivaga.2.1.chr6.g00718.t01      3091    N0.HOG0003089   OG0000654  n1   rna-XM_034668398.1    rna-XM_038555233.1    rna-XM_022502339.1    rna-XM_019822967.2    rna-XM_036080157.1    rna-XM_039233846.1    rna-XM_032858567.1    rna-XM_046016599.1    rna-XM_032316832.1    rna-XM_059165892.1    rna-XM_059394947.1    rna-XM_013064753.2    mNeoVis.2.1.1.chr4_hap1.g00919.t01    rna-XM_044914269.1    rna-XM_004402339.1    rna-XM_042973308.1    rna-XM_008689216.2    rna-XM_027622109.2    mNeoVis.3.1.1.chr6.g00930.t01    rna-gnl|WGS_ZZZZ|egapxtmp_004699-R2    mNeoVis.evergladensis.2.1.chr6.g00923.t01    mNeoVis.vulgivaga.2.1.chr6.g00919.t01      3092    N0.HOG0003090   OG0000654  n5   rna-XM_019802443.2    rna-XM_038531183.1    rna-XM_022497084.1    rna-XM_006934543.5    rna-XM_036065234.1    rna-XM_039233773.1    rna-XM_032877128.1    rna-XM_046022497.1    rna-XM_032303004.1    rna-XM_059136761.1    rna-XM_059377147.1    rna-XM_045064150.1    mNeoVis.2.1.1.chr2_hap1.g01537.t01    rna-XM_021683667.1    rna-XM_004406434.1    rna-XM_042996215.1    rna-XM_008694451.2    rna-XM_027581398.1    mNeoVis.3.1.1.chr5.g01461.t01      mNeoVis.evergladensis.2.1.chr5.g01475.t01    mNeoVis.vulgivaga.2.1.chr5.g01479.t01      3093    N0.HOG0003091   OG0000654  n8   rna-XM_002921512.4    rna-XM_038560767.1    rna-XM_022513497.1    rna-XM_003984505.5    rna-XM_036097554.1    rna-XM_039231077.1    rna-XM_032866839.1    rna-XM_045979393.1    rna-XM_032353332.1    rna-XM_059132651.1    rna-XM_059405148.1    rna-XM_004745795.3    mNeoVis.2.1.1.chr11_hap1.g01167.t01    rna-XM_021697376.1    rna-XM_004401792.1    rna-XM_042982511.1    rna-XM_040622803.1    rna-XM_027623960.2    mNeoVis.3.1.1.chr11.g01108.t01      mNeoVis.evergladensis.2.1.chr11.g01136.t01    mNeoVis.vulgivaga.2.1.chr11.g01163.t01      3094    N0.HOG0003092   OG0000655  n1   rna-XM_011220176.3    rna-XM_038575637.1    rna-XM_022522783.1    rna-XM_003994790.6    rna-XM_036123221.1    rna-XM_039225909.1    rna-XM_032883409.1    rna-XM_046024286.1    rna-XM_032309321.1    rna-XM_059140552.1    rna-XM_059408822.1    rna-XM_004766438.3    mNeoVis.2.1.1.chr3_hap1.g00418.t01    rna-XM_021703622.1    rna-XM_004400046.1    rna-XM_042962647.1    rna-XM_040619740.1    rna-XM_027576338.2    mNeoVis.3.1.1.chr3.g00410.t01    rna-gnl|WGS_ZZZZ|egapxtmp_042253-R1    mNeoVis.evergladensis.2.1.chr3.g00425.t01    mNeoVis.vulgivaga.2.1.chr3.g00422.t01      3095    N0.HOG0003093   OG0000655  n19   rna-XM_019795713.2    rna-XM_038575646.1    rna-XM_022522841.1    rna-XM_019814777.3    rna-XM_036098484.1    rna-XM_039225923.1    rna-XM_032883412.1    rna-XM_046025474.1    rna-XM_032309325.1    rna-XM_059140554.1    rna-XM_059408825.1    rna-XM_004766432.3      rna-XM_021703605.2    rna-XM_004400041.1    rna-XM_042962716.1    rna-XM_008710614.2    rna-XM_027577641.2    mNeoVis.3.1.1.chr3.g00407.t01    rna-gnl|WGS_ZZZZ|egapxtmp_041081-R1, rna-gnl|WGS_ZZZZ|egapxtmp_042664-R1      mNeoVis.vulgivaga.2.1.chr3.g00419.t01      3096    N0.HOG0003094   OG0000655  n39   rna-XM_002915564.4    rna-XM_038574526.1    rna-XM_022522880.1    rna-XM_003994793.4    rna-XM_036098486.1    rna-XM_039225924.1    rna-XM_032883411.1    rna-XM_046024288.1    rna-XM_032309324.1    rna-XM_059140553.1    rna-XM_059408823.1    rna-XM_004766433.3    mNeoVis.2.1.1.chr3_hap1.g00416.t01    rna-XM_021703616.1    rna-XM_004400043.1    rna-XM_007082551.3    rna-XM_008710615.1    rna-XM_027577580.2    mNeoVis.3.1.1.chr3.g00408.t01      mNeoVis.evergladensis.2.1.chr3.g00423.t01    mNeoVis.vulgivaga.2.1.chr3.g00420.t01      3097    N0.HOG0003095   OG0000656  n1   rna-XM_034638103.1    rna-XM_038527819.1    rna-XM_022492428.1    rna-XM_023246043.2    rna-XM_036092083.1    rna-XM_039244191.1    rna-XM_032848635.1    rna-XM_045989485.1    rna-XM_032324475.1    rna-XM_059153617.1    rna-XM_059381592.1    rna-XM_004779810.2    mNeoVis.2.1.1.chr7_hap1.g00739.t01    rna-XM_021682252.1, rna-XM_044912221.1    rna-XM_004415864.1    rna-XM_042968252.1    rna-XM_040633859.1    rna-XM_027618404.2    mNeoVis.3.1.1.chr7.g00691.t01    rna-gnl|WGS_ZZZZ|egapxtmp_012282-R1    mNeoVis.evergladensis.2.1.chr7.g00729.t01    mNeoVis.vulgivaga.2.1.chr7.g00731.t01      3098    N0.HOG0003096   OG0000656  n23   rna-XM_034669718.1    rna-XM_038539618.1    rna-XM_022500644.1    rna-XM_023247112.2    rna-XM_036101739.1    rna-XM_039226013.1    rna-XM_032872790.1    rna-XM_045992795.1    rna-XM_032326847.1    rna-XM_059154029.1    rna-XM_059414310.1    rna-XM_045075171.1    mNeoVis.2.1.1.chr15_hap1.g00577.t01, mNeoVis.2.1.1.chr15_hap1.g00587.t01    rna-XM_021680139.2    rna-XM_004394135.1    rna-XM_042971620.1    rna-XM_040634697.1    rna-XM_027613121.2              3099    N0.HOG0003097   OG0000656  n40   rna-XM_034640717.1    rna-XM_038546708.1    rna-XM_022520078.1    rna-XM_003997129.5    rna-XM_036079040.1    rna-XM_039229801.1    rna-XM_032870711.1    rna-XM_045984556.1    rna-XM_032319537.1    rna-XM_059149055.1    rna-XM_059378451.1    rna-XM_004749029.3    mNeoVis.2.1.1.chr8_hap1.g00195.t01    rna-XM_021678464.2    rna-XM_004401835.1    rna-XM_042967005.1    rna-XM_040640305.1    rna-XM_027568203.2    mNeoVis.3.1.1.chr8.g00155.t01    rna-gnl|WGS_ZZZZ|egapxtmp_019344-R1    mNeoVis.evergladensis.2.1.chr9.g00187.t01    mNeoVis.vulgivaga.2.1.chr8.g00191.t01      3100    N0.HOG0003098   OG0000657  n1   rna-XM_011223567.3    rna-XM_038542729.1    rna-XM_022508564.1    rna-XM_045049316.1    rna-XM_036097011.1    rna-XM_039237499.1    rna-XM_032862896.1    rna-XM_045982267.1    rna-XM_032317283.1    rna-XM_059146697.1    rna-XM_059412531.1    rna-XM_045084743.1    mNeoVis.2.1.1.chr14_hap1.g00076.t01    rna-XM_044916426.1    rna-XM_004400461.1    rna-XM_042976947.1    rna-XM_040636074.1    rna-XM_027610408.2    mNeoVis.3.1.1.chr14.g00072.t01    rna-gnl|WGS_ZZZZ|egapxtmp_032926-R2    mNeoVis.evergladensis.2.1.chr14.g00077.t01    mNeoVis.vulgivaga.2.1.chr14.g00076.t01      3101    N0.HOG0003099   OG0000657  n13   rna-XM_034638446.1    rna-NM_001012340.1    rna-XM_022524988.1    rna-XM_045046067.1    rna-XM_036095050.1    rna-XM_039251411.1    rna-XM_032841964.1    rna-XM_045987543.1    rna-XM_032323412.1    rna-XM_059150919.1    rna-XM_059383080.1    rna-XM_045076813.1    mNeoVis.2.1.1.chr7_hap1.g00560.t01    rna-XM_021701010.2    rna-XM_004396362.1    rna-XM_042970090.1    rna-XM_040633979.1    rna-XM_027617591.2    mNeoVis.3.1.1.chr7.g00543.t01    rna-gnl|WGS_ZZZZ|egapxtmp_013302-R2    mNeoVis.evergladensis.2.1.chr7.g00561.t01    mNeoVis.vulgivaga.2.1.chr7.g00563.t01      3102    N0.HOG0003100   OG0000657  n33   rna-XM_034642003.1    rna-XM_038545911.1    rna-XM_022505561.1    rna-NM_001322435.1    rna-XM_036113820.1    rna-XM_039255817.1    rna-XM_032839543.1    rna-XM_046007917.1    rna-XM_032341596.1    rna-XM_059183306.1    rna-XM_059374136.1    rna-XM_045074538.1    mNeoVis.2.1.1.chr10_hap1.g00028.t01    rna-XM_021679884.2    rna-XM_004395604.1    rna-XM_042990712.1    rna-XM_040641319.1    rna-XM_027571789.2        mNeoVis.evergladensis.2.1.chr8.g00026.t01    mNeoVis.vulgivaga.2.1.chr9.g00027.t01      3103    N0.HOG0003101   OG0000658  n1   rna-XM_034642678.1    rna-NM_001287156.1    rna-XM_022495596.1    rna-XM_023241848.2    rna-XM_036072061.1    rna-XM_039219201.1    rna-XM_032859711.1    rna-XM_046025814.1    rna-XM_032309845.1    rna-XM_059139331.1    rna-XM_059409398.1    rna-XM_004742872.3    mNeoVis.2.1.1.chr3_hap1.g00187.t01    rna-XM_021686273.2    rna-XM_004413395.1    rna-XM_042962397.1    rna-XM_040632382.1    rna-XM_027576021.2    mNeoVis.3.1.1.chr3.g00189.t01    rna-gnl|WGS_ZZZZ|egapxtmp_039075-R1    mNeoVis.evergladensis.2.1.chr3.g00197.t01    mNeoVis.vulgivaga.2.1.chr3.g00189.t01      3104    N0.HOG0003102   OG0000658  n20   rna-XM_034639051.1    rna-XM_038538035.1    rna-XM_022511797.1    rna-XM_023245523.2    rna-XM_036119807.1    rna-XM_039223619.1    rna-XM_032853758.1    rna-XM_045989404.1    rna-XM_032325041.1    rna-XM_059150729.1    rna-XM_059383913.1    rna-XM_045076456.1    mNeoVis.2.1.1.chr7_hap1.g00026.t01    rna-XM_021702339.1    rna-XM_004392125.1    rna-XM_042970269.1    rna-XM_040633005.1    rna-XM_027617063.2        mNeoVis.evergladensis.2.1.chr7.g00027.t01    mNeoVis.vulgivaga.2.1.chr7.g00026.t01      3105    N0.HOG0003103   OG0000658  n39   rna-XM_034641573.1    rna-XM_038572777.1    rna-XM_022494091.1    rna-XM_023251048.2    rna-XM_036105351.1    rna-XM_039237054.1    rna-XM_032878710.1    rna-XM_045981757.1    rna-XM_032350165.1    rna-XM_059133210.1    rna-XM_059407342.1    rna-XM_045070576.1    mNeoVis.2.1.1.chr11_hap1.g00065.t01    rna-XM_044918535.1    rna-XM_004415151.1    rna-XM_042980003.1    rna-XM_040638543.1    rna-XM_027622753.2    mNeoVis.3.1.1.chr11.g00039.t01    rna-gnl|WGS_ZZZZ|egapxtmp_025493-R1    mNeoVis.evergladensis.2.1.chr11.g00065.t01    mNeoVis.vulgivaga.2.1.chr11.g00065.t01      3106    N0.HOG0003104   OG0000659  n1   rna-XM_011227289.3    rna-XM_038544021.1    rna-XM_022524504.1    rna-XM_006932755.5    rna-XM_036114641.1    rna-XM_039254741.1    rna-XM_032856330.1    rna-XM_046008772.1    rna-XM_032343877.1    rna-XM_059181704.1    rna-XM_059371224.1    rna-XM_004755282.3    mNeoVis.2.1.1.chr10_hap1.g00548.t01    rna-XM_021695417.2    rna-XM_004402065.1    rna-XM_042987742.1    rna-XM_040620904.1    rna-XM_027570958.1    mNeoVis.3.1.1.chr9.g00506.t01    rna-gnl|WGS_ZZZZ|egapxtmp_014380-R1    mNeoVis.evergladensis.2.1.chr8.g00530.t01    mNeoVis.vulgivaga.2.1.chr9.g00530.t01      3107    N0.HOG0003105   OG0000659  n18   rna-XM_002922652.4    rna-XM_038570851.1    rna-XM_022515133.1    rna-XM_045037596.1    rna-XM_036106774.1    rna-XM_039242236.1    rna-XM_032869701.1    rna-XM_046001314.1    rna-XM_032333091.1    rna-XM_059162603.1    rna-XM_059387975.1    rna-XM_004762506.3    mNeoVis.2.1.1.chr5_hap1.g01129.t01    rna-XM_021678784.1    rna-XM_004415628.1    rna-XM_042956642.1    rna-XM_008708689.1    rna-XM_027586333.2    mNeoVis.3.1.1.chr2.g01038.t01    rna-gnl|WGS_ZZZZ|egapxtmp_037051-R1    mNeoVis.evergladensis.2.1.chr2.g01119.t01    mNeoVis.vulgivaga.2.1.chr2.g01125.t01      3108    N0.HOG0003106   OG0000659  n37   rna-XM_034638611.1    rna-XM_038538014.1    rna-XM_022511637.1      rna-XM_036119779.1    rna-XM_039226812.1    rna-XM_032853751.1    rna-XM_045989498.1    rna-XM_032324566.1    rna-XM_059152578.1    rna-XM_059381456.1    rna-XM_045076469.1    mNeoVis.2.1.1.chr7_hap1.g00021.t01    rna-XM_021705057.1    rna-XM_004392166.1    rna-XM_042968749.1    rna-XM_040633004.1    rna-XM_027618845.1    mNeoVis.3.1.1.chr7.g00018.t01      mNeoVis.evergladensis.2.1.chr7.g00022.t01    mNeoVis.vulgivaga.2.1.chr7.g00021.t01      3109    N0.HOG0003107   OG0000660  n1   rna-XM_034647224.1    rna-XM_038527186.1    rna-XM_022512762.1    rna-XM_045043101.1    rna-XM_036076099.1    rna-XM_039230858.1    rna-XM_032866529.1    rna-XM_046022097.1, rna-XM_046023089.1    rna-XM_032308672.1    rna-XM_059142824.1    rna-XM_059411945.1    rna-XM_004766195.3    mNeoVis.2.1.1.chr13_hap1.g00775.t01    rna-XM_044920625.1    rna-XM_004394522.1    rna-XM_042963651.1    rna-XM_040620294.1    rna-XM_035723664.1    mNeoVis.3.1.1.chr13.g00693.t01    rna-gnl|WGS_ZZZZ|egapxtmp_031473-R15    mNeoVis.evergladensis.2.1.chr13.g00741.t01    mNeoVis.vulgivaga.2.1.chr13.g00737.t01      3110    N0.HOG0003108   OG0000660  n23   rna-XM_034652098.1, rna-XM_034657162.1    rna-XM_038567062.1    rna-XM_022526782.1    rna-XM_045046809.1    rna-XM_036066497.1    rna-XM_039252314.1    rna-XM_032835881.1    rna-XM_045990934.1    rna-XM_032313251.1    rna-XM_059160198.1    rna-XM_059389197.1    rna-XM_045083090.1      rna-XM_044917766.1    rna-XM_004398973.1    rna-XM_042978362.1    rna-XM_040628265.1, rna-XM_040629205.1    rna-XM_027583985.1              3111    N0.HOG0003109   OG0000660  n41   rna-XM_034657606.1    rna-XM_038567609.1    rna-XM_022525485.1    rna-XM_023250874.2    rna-XM_036120674.1    rna-XM_039221337.1    rna-XM_032883726.1    rna-XM_045990186.1    rna-XM_032307678.1    rna-XM_059159641.1    rna-XM_059388781.1    rna-XM_004771558.3    mNeoVis.2.1.1.chr5_hap1.g00157.t01    rna-XM_021705569.2    rna-XM_004395289.1    rna-XM_042977914.1    rna-XM_040629496.1    rna-XM_027583635.2    mNeoVis.3.1.1.chr2.g00105.t01    rna-gnl|WGS_ZZZZ|egapxtmp_038095-R4    mNeoVis.evergladensis.2.1.chr2.g00153.t01    mNeoVis.vulgivaga.2.1.chr2.g00151.t01      3112    N0.HOG0003110   OG0000661  n1   rna-XM_034655948.1    rna-XM_038552012.1    rna-XM_022493529.1    rna-XM_019835509.3    rna-XM_036070350.1    rna-XM_039236695.1    rna-XM_032851952.1    rna-XM_046000831.1    rna-XM_032336856.1    rna-XM_059174457.1    rna-XM_059417483.1    rna-XM_004737438.3    mNeoVis.2.1.1.chr1_hap1.g00362.t01    rna-XM_021701767.1    rna-XM_004413270.1    rna-XM_042983005.1    rna-XM_040622522.1    rna-XM_027606705.2    mNeoVis.3.1.1.chr1.g00364.t01    rna-gnl|WGS_ZZZZ|egapxtmp_008988-R1    mNeoVis.evergladensis.2.1.chr1.g00360.t01    mNeoVis.vulgivaga.2.1.chr1.g00368.t01      3113    N0.HOG0003111   OG0000661  n20   rna-XM_034667463.1    rna-NM_001287151.1    rna-XM_022490938.1    rna-XM_023247978.2    rna-XM_036068168.1    rna-XM_039251446.1    rna-XM_032845023.1    rna-XM_045982649.1    rna-XM_032317167.1    rna-XM_059146988.1    rna-XM_059413687.1    rna-XM_004778539.3    mNeoVis.2.1.1.chr14_hap1.g00628.t01    rna-XM_044915866.1    rna-XM_004402924.1    rna-XM_042975770.1    rna-XM_040635791.1    rna-XM_027610820.2    mNeoVis.3.1.1.chr14.g00608.t01    rna-gnl|WGS_ZZZZ|egapxtmp_031970-R1    mNeoVis.evergladensis.2.1.chr14.g00625.t01    mNeoVis.vulgivaga.2.1.chr14.g00622.t01      3114    N0.HOG0003112   OG0000661  n39   rna-XM_034657441.1    rna-XM_038567889.1    rna-XM_022525861.1    rna-XM_023243403.2    rna-XM_036100990.1    rna-XM_039247994.1    rna-XM_032846928.1    rna-XM_045990028.1    rna-XM_032358518.1    rna-XM_059159363.1    rna-XM_059388575.1    rna-XM_004778156.2      rna-XM_044918635.1    rna-XM_004395379.1    rna-XM_042977683.1    rna-XM_040629698.1    rna-XM_027582555.2      rna-gnl|WGS_ZZZZ|egapxtmp_034316-R1    mNeoVis.evergladensis.2.1.chr2.g00065.t01    mNeoVis.vulgivaga.2.1.chr2.g00065.t01      3115    N0.HOG0003113   OG0000662  n1   rna-XM_034647273.1    rna-XM_038536742.1    rna-XM_022524579.1    rna-XM_023243920.2    rna-XM_036094519.1    rna-XM_039231637.1    rna-XM_032840072.1    rna-XM_045984219.1    rna-XM_032322516.1    rna-XM_059147195.1    rna-XM_059379856.1    rna-XM_045083311.1    mNeoVis.2.1.1.chr8_hap1.g01031.t01    rna-XM_044921861.1    rna-XM_004398623.1    rna-XM_042967544.1    rna-XM_040629979.1    rna-XM_027625738.2    mNeoVis.3.1.1.chr8.g00961.t01      mNeoVis.evergladensis.2.1.chr9.g01005.t01    mNeoVis.vulgivaga.2.1.chr8.g01006.t01      3116    N0.HOG0003114   OG0000662  n22   rna-XM_011228723.3    rna-XM_038576575.1    rna-XM_022522217.1    rna-XM_023256718.2    rna-XM_036117661.1    rna-XM_039217518.1    rna-XM_032874747.1    rna-XM_046011259.1    rna-XM_032348988.1    rna-XM_059184600.1    rna-XM_059403716.1    rna-XM_013063550.2    mNeoVis.2.1.1.chr9_hap1.g00256.t01    rna-XM_021690768.2    rna-XM_004413897.1    rna-XM_042991657.1    rna-XM_040642081.1    rna-XM_035729218.1    mNeoVis.3.1.1.chr10.g00249.t01    rna-gnl|WGS_ZZZZ|egapxtmp_024078-R1    mNeoVis.evergladensis.2.1.chr10.g00254.t01    mNeoVis.vulgivaga.2.1.chr10.g00250.t01      3117    N0.HOG0003115   OG0000662  n42   rna-XM_034671443.1    rna-XM_038537763.1    rna-XM_022514624.1    rna-XM_045034994.1    rna-XM_036064502.1    rna-XM_039253242.1    rna-XM_032863257.1    rna-XM_046017918.1    rna-XM_032302454.1    rna-XM_059135116.1    rna-XM_059374472.1    rna-XM_045063640.1    mNeoVis.2.1.1.chr2_hap1.g01875.t01    rna-XM_044913810.1    rna-XM_004411045.1    rna-XM_042996981.1    rna-XM_040645694.1    rna-XM_027572268.2    mNeoVis.3.1.1.chr5.g01796.t01      mNeoVis.evergladensis.2.1.chr5.g01813.t01    mNeoVis.vulgivaga.2.1.chr5.g01816.t01      3118    N0.HOG0003116   OG0000663  n1   rna-XM_034645728.1    rna-XM_038576747.1    rna-XM_022495007.1    rna-XM_019834473.3    rna-XM_036067278.1, rna-XM_036092642.1    rna-XM_039224416.1    rna-XM_032854939.1    rna-XM_046013391.1, rna-XM_046013446.1    rna-XM_032347205.1    rna-XM_059186178.1, rna-XM_059186209.1    rna-XM_059402573.1, rna-XM_059404466.1    rna-XM_013049561.1, rna-XM_045073219.1    mNeoVis.2.1.1.chr9_hap1.g00865.t01, mNeoVis.2.1.1.chr9_hap1.g00868.t01    rna-XM_021690432.1    rna-XM_004410890.1    rna-XM_015542908.2    rna-XM_040642140.1    rna-XM_027593964.2    mNeoVis.3.1.1.chr10.g00822.t01    rna-gnl|WGS_ZZZZ|egapxtmp_023243-R1    mNeoVis.evergladensis.2.1.chr10.g00853.t01        3119    N0.HOG0003117   OG0000663  n28   rna-XM_034645726.1, rna-XM_034650230.1    rna-XM_038577901.1      rna-XM_045061347.1    rna-XM_036112936.1    rna-XM_039224417.1      rna-XM_046011791.1              rna-XM_044915222.1    rna-XM_004410904.1    rna-XM_015542907.2    rna-XM_008697488.2                3120    N0.HOG0003118   OG0000663  n38   rna-XM_034645615.1    rna-XM_038576746.1    rna-XM_022495190.1    rna-XM_006933497.4    rna-XM_036112935.1, rna-XM_036112937.1    rna-XM_039224415.1    rna-XM_032854938.1    rna-XM_046010673.1    rna-XM_032347186.1    rna-XM_059186180.1, rna-XM_059186183.1    rna-XM_059386471.1, rna-XM_059402571.1    rna-XM_045073424.1, rna-XM_045073425.1    mNeoVis.2.1.1.chr9_hap1.g00870.t01    rna-XM_021690766.2    rna-XM_004410891.1    rna-XM_007094105.3    rna-XM_008697489.2    rna-XM_027594069.2    mNeoVis.3.1.1.chr10.g00823.t01    rna-gnl|WGS_ZZZZ|egapxtmp_024110-R1    mNeoVis.evergladensis.2.1.chr10.g00854.t01    mNeoVis.vulgivaga.2.1.chr10.g00844.t01      3121    N0.HOG0003119   OG0000664  n0   rna-XM_002927630.4, rna-XM_034644824.1, rna-XM_034646603.1    rna-XM_038564503.1    rna-XM_022506448.1, rna-XM_022506608.1, rna-XM_022507689.1    rna-XM_003993486.5, rna-XM_003993487.5, rna-XM_045038104.1    rna-XM_036110554.1, rna-XM_036110624.1, rna-XM_036110660.1    rna-XM_039253537.1, rna-XM_039253542.1    rna-XM_032879059.1, rna-XM_032879060.1, rna-XM_032879901.1    rna-XM_046016290.1, rna-XM_046016699.1, rna-XM_046017605.1    rna-XM_032359460.1, rna-XM_032359910.1, rna-XM_032360048.1    rna-XM_059143599.1, rna-XM_059159010.1, rna-XM_059159011.1    rna-XM_059385940.1, rna-XM_059386061.1, rna-XM_059400152.1    rna-XM_004770390.2, rna-XM_004770465.2, rna-XM_045068764.1    mNeoVis.2.1.1.chr7_hap1.g02256.t01, mNeoVis.2.1.1.chr7_hap1.g02260.t01, mNeoVis.2.1.1.chr7_hap1.g02261.t01    rna-XM_021684889.1, rna-XM_021684890.1, rna-XM_021685244.1    rna-XM_004410082.1, rna-XM_004410083.1, rna-XM_004417257.1    rna-XM_007092129.2, rna-XM_007092130.2, rna-XM_042958338.1    rna-XM_008707458.1, rna-XM_008707462.1, rna-XM_008707463.2, rna-XM_040639167.1    rna-XM_027577842.2, rna-XM_027580549.2, rna-XM_027580582.1    mNeoVis.3.1.1.chr7.g02076.t01, mNeoVis.3.1.1.chr7.g02081.t01, mNeoVis.3.1.1.chr7.g02082.t01    rna-gnl|WGS_ZZZZ|egapxtmp_010761-R1, rna-gnl|WGS_ZZZZ|egapxtmp_011183-R1, rna-gnl|WGS_ZZZZ|egapxtmp_012665-R1    mNeoVis.evergladensis.2.1.chr7.g02204.t01, mNeoVis.evergladensis.2.1.chr7.g02208.t01, mNeoVis.evergladensis.2.1.chr7.g02209.t01    mNeoVis.vulgivaga.2.1.chr7.g02223.t01, mNeoVis.vulgivaga.2.1.chr7.g02226.t01, mNeoVis.vulgivaga.2.1.chr7.g02227.t01      3122    N0.HOG0003120   OG0000665  n1   rna-XM_019806087.1    rna-XM_038564214.1    rna-XM_022507389.1    rna-XM_019812564.3    rna-XM_036122150.1    rna-XM_039257691.1    rna-XM_032879292.1    rna-XM_046014331.1    rna-XM_032359782.1    rna-XM_059144989.1    rna-XM_059403470.1    rna-XM_004770244.2    mNeoVis.2.1.1.chr7_hap1.g02330.t01    rna-XM_021684950.1    rna-XM_004413792.1    rna-XM_042958719.1    rna-XM_040638771.1    rna-XM_027580005.1    mNeoVis.3.1.1.chr7.g02146.t01    rna-gnl|WGS_ZZZZ|egapxtmp_013577-R1    mNeoVis.evergladensis.2.1.chr7.g02277.t01    mNeoVis.vulgivaga.2.1.chr7.g02294.t01      3123    N0.HOG0003121   OG0000665  n21   rna-XM_002916700.4    rna-XM_038563972.1    rna-XM_022506590.1      rna-XM_036119245.1      rna-XM_032879070.1    rna-XM_046014284.1    rna-XM_032360051.1    rna-XM_059146940.1, rna-XM_059159022.1    rna-XM_059386977.1    rna-XM_004770071.3    mNeoVis.2.1.1.chr7_hap1.g02400.t01    rna-XM_021685260.1    rna-XM_004394057.1      rna-XM_008708998.2, rna-XM_040639390.1    rna-XM_027579664.1    mNeoVis.3.1.1.chr7.g02210.t01    rna-gnl|WGS_ZZZZ|egapxtmp_012693-R1    mNeoVis.evergladensis.2.1.chr7.g02346.t01    mNeoVis.vulgivaga.2.1.chr7.g02366.t01      3124   N0.HOG0003122  OG0000665  n41     rna-XM_038563976.1    rna-XM_022507593.1    rna-XM_003993589.4    rna-XM_036113700.1    rna-XM_039244685.1    rna-XM_032879441.1    rna-XM_046014404.1    rna-XM_032357276.1    rna-XM_059146929.1    rna-XM_059383518.1    rna-XM_004770072.3    mNeoVis.2.1.1.chr7_hap1.g02399.t01    rna-XM_021685031.1    rna-XM_004393953.1    rna-XM_042957579.1    rna-XM_040639388.1    rna-XM_027581208.1      rna-gnl|WGS_ZZZZ|egapxtmp_013736-R3    mNeoVis.evergladensis.2.1.chr7.g02345.t01    mNeoVis.vulgivaga.2.1.chr7.g02365.t01      3125   N0.HOG0003123  OG0000666  n1   rna-XM_002925076.4    rna-XM_038571525.1    rna-XM_022494213.1    rna-XM_011280964.3    rna-XM_036120498.1    rna-XM_039256523.1    rna-XM_032859233.1    rna-XM_045980697.1    rna-XM_032351473.1    rna-XM_059134110.1    rna-XM_059406498.1    rna-XM_004754180.2    mNeoVis.2.1.1.chr11_hap1.g00494.t01    rna-XM_021700681.2    rna-XM_004404158.1    rna-XM_042980949.1    rna-XM_040637956.1    rna-XM_027612201.2, rna-XM_027620911.2    mNeoVis.3.1.1.chr11.g00457.t01    rna-gnl|WGS_ZZZZ|egapxtmp_027801-R1    mNeoVis.evergladensis.2.1.chr11.g00484.t01    mNeoVis.vulgivaga.2.1.chr11.g00491.t01      3126    N0.HOG0003124   OG0000666  n19   rna-XM_002916633.4    rna-XM_038563891.1    rna-XM_022506540.1    rna-XM_003993723.5    rna-XM_036077605.1    rna-XM_039244888.1    rna-XM_032879487.1    rna-XM_046015003.1    rna-XM_032357380.1    rna-XM_059147499.1    rna-XM_059401482.1    rna-XM_004770004.3    mNeoVis.2.1.1.chr7_hap1.g02419.t01, mNeoVis.2.1.1.chr7_hap1.g02448.t01    rna-XM_021685052.1    rna-XM_004393925.1    rna-XM_042958667.1    rna-XM_008709009.2    rna-XM_027578015.1              3127    N0.HOG0003125   OG0000666  n37   rna-XM_019803324.2    rna-XM_038544473.1    rna-XM_022524957.1    rna-XM_023255655.2    rna-XM_036075480.1    rna-XM_039256978.1    rna-XM_032871295.1    rna-XM_046008841.1    rna-XM_032344014.1    rna-XM_059183422.1    rna-XM_059373069.1    rna-XM_045067637.1    mNeoVis.2.1.1.chr10_hap1.g00484.t01    rna-XM_021695553.1    rna-XM_004399817.1    rna-XM_042989552.1    rna-XM_008694245.2    rna-XM_035728234.1    mNeoVis.3.1.1.chr9.g00449.t01    rna-gnl|WGS_ZZZZ|egapxtmp_014610-R1    mNeoVis.evergladensis.2.1.chr8.g00463.t01    mNeoVis.vulgivaga.2.1.chr9.g00466.t01      3128    N0.HOG0003126   OG0000667  n2   rna-XM_019805094.2    rna-XM_038536655.1    rna-XM_022524518.1    rna-XM_023244544.2    rna-XM_036066298.1    rna-XM_039231514.1    rna-XM_032840063.1    rna-XM_045985272.1    rna-XM_032322566.1    rna-XM_059147975.1    rna-XM_059379498.1    rna-XM_004760325.3    mNeoVis.2.1.1.chr8_hap1.g01066.t01    rna-XM_021694488.2    rna-XM_004398524.1    rna-XM_042965779.1    rna-XM_040629942.1    rna-XM_027567534.1      rna-gnl|WGS_ZZZZ|egapxtmp_017568-R1    mNeoVis.evergladensis.2.1.chr9.g01041.t01    mNeoVis.vulgivaga.2.1.chr8.g01041.t01      3129   N0.HOG0003127  OG0000667  n22   rna-XM_034672024.1    rna-XM_038537592.1    rna-XM_022514704.1    rna-XM_045034906.1    rna-XM_036089667.1    rna-XM_039248324.1    rna-XM_032863119.1    rna-XM_045997521.1    rna-XM_032361520.1    rna-XM_059136241.1    rna-XM_059376504.1    rna-XM_045063497.1    mNeoVis.2.1.1.chr2_hap1.g01932.t01    rna-XM_044913841.1    rna-XM_004407400.1    rna-XM_042994290.1    rna-XM_040645596.1    rna-XM_027611257.2    mNeoVis.3.1.1.chr5.g01835.t01      mNeoVis.evergladensis.2.1.chr5.g01871.t01    mNeoVis.vulgivaga.2.1.chr5.g01873.t01      3130    N0.HOG0003128   OG0000667  n42   rna-XM_034660734.1    rna-XM_038583224.1    rna-XM_022514935.1    rna-XM_023260282.2    rna-XM_036114406.1    rna-XM_039230372.1    rna-XM_032869293.1    rna-XM_046002023.1    rna-XM_032337278.1    rna-XM_059163093.1    rna-XM_059391248.1    rna-XM_013052619.2    mNeoVis.2.1.1.chr5_hap1.g01260.t01    rna-XM_044919883.1    rna-XM_004391867.1    rna-XM_042999275.1    rna-XM_040621408.1    rna-XM_027582803.2    mNeoVis.3.1.1.chr2.g01180.t01    rna-gnl|WGS_ZZZZ|egapxtmp_038654-R1    mNeoVis.evergladensis.2.1.chr2.g01248.t01    mNeoVis.vulgivaga.2.1.chr2.g01253.t01      3131    N0.HOG0003129   OG0000668  n1   rna-XM_002921159.4    rna-XM_038536788.1    rna-XM_022524808.1    rna-XM_003996187.5    rna-XM_036094566.1    rna-XM_039231538.1    rna-XM_032839921.1    rna-XM_045986640.1    rna-XM_032321792.1    rna-XM_059147928.1    rna-XM_059379905.1    rna-XM_004760246.3    mNeoVis.2.1.1.chr8_hap1.g01029.t01    rna-XM_021694754.1    rna-XM_004398482.1    rna-XM_007091647.2    rna-XM_040629981.1    rna-XM_027626216.2    mNeoVis.3.1.1.chr8.g00959.t01      mNeoVis.evergladensis.2.1.chr9.g01003.t01    mNeoVis.vulgivaga.2.1.chr8.g01004.t01      3132    N0.HOG0003130   OG0000668  n19   rna-XM_034664047.1    rna-XM_038571281.1    rna-XM_022509326.1    rna-XM_045037401.1    rna-XM_036122376.1    rna-XM_039236026.1    rna-XM_032852575.1    rna-XM_046001738.1    rna-XM_032344007.1    rna-XM_059163683.1    rna-XM_059391681.1    rna-XM_004755616.3    mNeoVis.2.1.1.chr5_hap1.g01456.t01    rna-XM_021702559.2    rna-XM_004412872.1    rna-XM_007086274.2    rna-XM_008706834.2    rna-XM_027587679.2    mNeoVis.3.1.1.chr2.g01376.t01    rna-gnl|WGS_ZZZZ|egapxtmp_034333-R1    mNeoVis.evergladensis.2.1.chr2.g01446.t01    mNeoVis.vulgivaga.2.1.chr2.g01457.t01      3133    N0.HOG0003131   OG0000668  n38   rna-XM_034671122.1    rna-XM_038537772.1    rna-XM_022514612.1    rna-XM_019836063.3    rna-XM_036064511.1    rna-XM_039253262.1    rna-XM_032863254.1    rna-XM_046013672.1    rna-XM_032303799.1    rna-XM_059137165.1    rna-XM_059377266.1    rna-XM_004768698.3    mNeoVis.2.1.1.chr2_hap1.g01876.t01    rna-XM_044913856.1    rna-XM_004411046.1    rna-XM_042996987.1    rna-XM_040645693.1    rna-XM_027605054.2    mNeoVis.3.1.1.chr5.g01797.t01      mNeoVis.evergladensis.2.1.chr5.g01814.t01    mNeoVis.vulgivaga.2.1.chr5.g01817.t01      3134   N0.HOG0003132  OG0000669  n1   rna-XM_002914905.4    rna-XM_038551517.1    rna-XM_022504451.1    rna-XM_003984090.6    rna-XM_036088285.1    rna-XM_039239594.1    rna-XM_032873719.1    rna-XM_045979070.1    rna-XM_032352353.1    rna-XM_059187904.1    rna-XM_059405924.1    rna-XM_004742128.3    mNeoVis.2.1.1.chr11_hap1.g00771.t01    rna-XM_021699078.2    rna-XM_004398639.1    rna-XM_042981645.1    rna-XM_040623348.1    rna-XM_027622565.2    mNeoVis.3.1.1.chr11.g00735.t01    rna-gnl|WGS_ZZZZ|egapxtmp_026414-R1    mNeoVis.evergladensis.2.1.chr11.g00753.t01    mNeoVis.vulgivaga.2.1.chr11.g00773.t01      3135   N0.HOG0003133  OG0000669  n13   rna-XM_011221473.3    rna-XM_038533258.1    rna-XM_022505922.1    rna-XM_023253533.2    rna-XM_036067292.1    rna-XM_039232117.1    rna-XM_032856261.1    rna-XM_045996768.1    rna-XM_032333606.1    rna-XM_059166860.1    rna-XM_059379547.1    rna-XM_045074764.1    mNeoVis.2.1.1.chr6_hap1.g00367.t01    rna-XM_044913040.1    rna-XM_004396211.1    rna-XM_042985127.1    rna-XM_040625248.1    rna-XM_027599809.2    mNeoVis.3.1.1.chr4.g00374.t01      mNeoVis.evergladensis.2.1.chr4.g00375.t01    mNeoVis.vulgivaga.2.1.chr4.g00372.t01      3136    N0.HOG0003134   OG0000669  n30   rna-XM_011235974.3    rna-XM_038534654.1    rna-XM_022493074.1    rna-XM_003981248.6    rna-XM_036084870.1    rna-XM_039239713.1    rna-XM_032851917.1    rna-XM_046000500.1    rna-XM_032336903.1    rna-XM_059174418.1    rna-XM_059417453.1    rna-XM_004737493.3    mNeoVis.2.1.1.chr1_hap1.g00351.t01    rna-XM_021698755.1    rna-XM_004401415.1    rna-XM_007085525.3    rna-XM_040630249.1    rna-XM_027606141.1    mNeoVis.3.1.1.chr1.g00354.t01      mNeoVis.evergladensis.2.1.chr1.g00349.t01    mNeoVis.vulgivaga.2.1.chr1.g00357.t01      3137    N0.HOG0003135   OG0000670  n1   rna-XM_019801364.2    rna-XM_038583888.1    rna-XM_022503698.1    rna-XM_019840116.3    rna-XM_036075618.1    rna-XM_039236515.1    rna-XM_032847923.1    rna-XM_046001979.1    rna-XM_032341356.1    rna-XM_059163479.1    rna-XM_059391550.1    rna-XM_004756575.3    mNeoVis.2.1.1.chr5_hap1.g01390.t01    rna-XM_021702658.2    rna-XM_004393236.1    rna-XM_042998969.1    rna-XM_040621605.1    rna-XM_027583486.2    mNeoVis.3.1.1.chr2.g01314.t01    rna-gnl|WGS_ZZZZ|egapxtmp_035602-R1    mNeoVis.evergladensis.2.1.chr2.g01376.t01    mNeoVis.vulgivaga.2.1.chr2.g01386.t01      3138    N0.HOG0003136   OG0000670  n21   rna-XM_034647266.1    rna-XM_038536552.1    rna-XM_022524559.1    rna-XM_045045203.1      rna-XM_039224883.1    rna-XM_032840046.1    rna-XM_045985247.1    rna-XM_032319862.1    rna-XM_059150155.1    rna-XM_059380923.1    rna-XM_045083209.1    mNeoVis.2.1.1.chr8_hap1.g01104.t01    rna-XM_044921395.1    rna-XM_004398575.1    rna-XM_042967770.1    rna-XM_040628318.1    rna-XM_035724599.1    mNeoVis.3.1.1.chr8.g01025.t01      mNeoVis.evergladensis.2.1.chr9.g01080.t01    mNeoVis.vulgivaga.2.1.chr8.g01080.t01      3139   N0.HOG0003137  OG0000670  n40   rna-XM_034658959.1    rna-XM_038550931.1    rna-XM_022516444.1    rna-XM_045053998.1    rna-XM_036087789.1    rna-XM_039253053.1    rna-XM_032840353.1    rna-XM_045981418.1    rna-XM_032351735.1    rna-XM_059134413.1    rna-XM_059406288.1    rna-XM_045077635.1    mNeoVis.2.1.1.chr11_hap1.g00591.t01    rna-XM_021680340.2    rna-XM_004402401.1    rna-XM_042981097.1    rna-XM_040623599.1    rna-XM_027622421.2    mNeoVis.3.1.1.chr11.g00552.t01    rna-gnl|WGS_ZZZZ|egapxtmp_027501-R1    mNeoVis.evergladensis.2.1.chr11.g00578.t01    mNeoVis.vulgivaga.2.1.chr11.g00587.t01      3140    N0.HOG0003138   OG0000671  n1   rna-XM_034647202.1    rna-XM_038535523.1    rna-XM_022524692.1    rna-XM_023243919.2    rna-XM_036094518.1    rna-XM_039231594.1    rna-XM_032839851.1    rna-XM_045986042.1    rna-XM_032321245.1    rna-XM_059148290.1    rna-XM_059378472.1    rna-XM_004760236.3    mNeoVis.2.1.1.chr8_hap1.g01025.t01    rna-XM_021694799.2    rna-XM_004398474.1    rna-XM_042967225.1    rna-XM_040629986.1    rna-XM_027567621.2    mNeoVis.3.1.1.chr8.g00955.t01      mNeoVis.evergladensis.2.1.chr9.g00999.t01    mNeoVis.vulgivaga.2.1.chr8.g01000.t01      3141    N0.HOG0003139   OG0000671  n22   rna-XM_011227413.3    rna-XM_038536792.1    rna-XM_022524678.1    rna-XM_019817110.3    rna-XM_036094531.1    rna-XM_039231562.1    rna-XM_032839942.1    rna-XM_045986590.1    rna-XM_032322054.1    rna-XM_059149304.1    rna-XM_059379051.1    rna-XM_004760233.3    mNeoVis.2.1.1.chr8_hap1.g01023.t01    rna-XM_044921541.1    rna-XM_004398622.1    rna-XM_042967204.1    rna-XM_040628222.1    rna-XM_027626039.2    mNeoVis.3.1.1.chr8.g00953.t01      mNeoVis.evergladensis.2.1.chr9.g00997.t01    mNeoVis.vulgivaga.2.1.chr8.g00998.t01      3142    N0.HOG0003140   OG0000671  n41   rna-XM_034647274.1    rna-XM_038535524.1    rna-XM_022524756.1    rna-XM_003996189.3    rna-XM_036094489.1    rna-XM_039231451.1    rna-XM_032839848.1    rna-XM_045985627.1    rna-XM_032322405.1    rna-XM_059149682.1    rna-XM_059378594.1    rna-XM_004760235.2    mNeoVis.2.1.1.chr8_hap1.g01024.t01    rna-XM_021694684.2    rna-XM_004398473.1    rna-XM_042967224.1    rna-XM_040628220.1    rna-XM_027567680.1    mNeoVis.3.1.1.chr8.g00954.t01    rna-gnl|WGS_ZZZZ|egapxtmp_017560-R1    mNeoVis.evergladensis.2.1.chr9.g00998.t01    mNeoVis.vulgivaga.2.1.chr8.g00999.t01      3143    N0.HOG0003141   OG0000672  n1   rna-XM_019801383.2    rna-XM_038537039.1    rna-XM_022491484.1    rna-XM_045044156.1    rna-XM_036073463.1    rna-XM_039227406.1    rna-XM_032841292.1    rna-XM_045986395.1    rna-XM_032320531.1    rna-XM_059147761.1    rna-XM_059379565.1    rna-XM_004776575.3    mNeoVis.2.1.1.chr8_hap1.g00868.t01    rna-XM_044921803.1    rna-XM_004405120.1    rna-XM_007080787.3    rna-XM_040628338.1    rna-XM_027624586.2    mNeoVis.3.1.1.chr8.g00814.t01    rna-gnl|WGS_ZZZZ|egapxtmp_017520-R1    mNeoVis.evergladensis.2.1.chr9.g00856.t01    mNeoVis.vulgivaga.2.1.chr8.g00858.t01      3144    N0.HOG0003142   OG0000672  n22   rna-XM_011227022.3, rna-XM_034647284.1    rna-XM_038537050.1    rna-XM_022491483.1, rna-XM_022491509.1    rna-XM_003996362.6, rna-XM_045044350.1    rna-XM_036073455.1, rna-XM_036073472.1    rna-XM_039227343.1, rna-XM_039227358.1    rna-XM_032841321.1, rna-XM_032841323.1    rna-XM_045984872.1    rna-XM_032322057.1, rna-XM_032322060.1    rna-XM_059149345.1, rna-XM_059149346.1    rna-XM_059378669.1, rna-XM_059381121.1    rna-XM_004776566.3, rna-XM_004776570.3    mNeoVis.2.1.1.chr8_hap1.g00871.t01, mNeoVis.2.1.1.chr8_hap1.g00872.t01    rna-XM_021694514.1, rna-XM_044921804.1    rna-XM_004405175.1, rna-XM_004405176.1    rna-XM_042965888.1, rna-XM_042965890.1    rna-XM_040628339.1, rna-XM_040630118.1    rna-XM_027624695.1, rna-XM_027626435.2    mNeoVis.3.1.1.chr8.g00816.t01, mNeoVis.3.1.1.chr8.g00818.t01    rna-gnl|WGS_ZZZZ|egapxtmp_020555-R1, rna-gnl|WGS_ZZZZ|egapxtmp_020556-R1    mNeoVis.evergladensis.2.1.chr9.g00859.t01, mNeoVis.evergladensis.2.1.chr9.g00860.t01    mNeoVis.vulgivaga.2.1.chr8.g00861.t01, mNeoVis.vulgivaga.2.1.chr8.g00862.t01      3145    N0.HOG0003143   OG0000673  n1   rna-XM_034657736.1    rna-XM_038567364.1    rna-XM_022526913.1    rna-XM_023245077.2    rna-XM_036119645.1    rna-XM_039250934.1    rna-XM_032883934.1    rna-XM_045989708.1    rna-XM_032310506.1    rna-XM_059159900.1    rna-XM_059388953.1    rna-XM_045084331.1    mNeoVis.2.1.1.chr5_hap1.g00264.t01    rna-XM_044918095.1    rna-XM_004398842.1    rna-XM_042978112.1    rna-XM_040629377.1    rna-XM_035728404.1    mNeoVis.3.1.1.chr2.g00206.t01      mNeoVis.evergladensis.2.1.chr2.g00262.t01    mNeoVis.vulgivaga.2.1.chr2.g00257.t01      3146    N0.HOG0003144   OG0000673  n17   rna-XM_034664759.1    rna-XM_038548866.1    rna-XM_022500367.1    rna-XM_023242949.2    rna-XM_036108740.1    rna-XM_039255182.1    rna-XM_032866233.1    rna-XM_046023391.1    rna-XM_032307947.1    rna-XM_059142418.1    rna-XM_059410355.1    rna-XM_045082873.1    mNeoVis.2.1.1.chr13_hap1.g00085.t01    rna-XM_021691206.2    rna-XM_004410723.1    rna-XM_042964837.1    rna-XM_040632171.1    rna-XM_035723807.1    mNeoVis.3.1.1.chr13.g00047.t01      mNeoVis.evergladensis.2.1.chr13.g00085.t01    mNeoVis.vulgivaga.2.1.chr13.g00083.t01      3147    N0.HOG0003145   OG0000673  n32   rna-XM_011224601.3    rna-XM_038541186.1    rna-XM_022496677.1    rna-XM_003990375.6    rna-XM_036103977.1    rna-XM_039223358.1    rna-XM_032867233.1    rna-XM_045980322.1    rna-XM_032302689.1    rna-XM_059137171.1    rna-XM_059375563.1    rna-XM_004775167.3    mNeoVis.2.1.1.chr2_hap1.g01030.t01    rna-XM_021690120.1    rna-XM_004401460.1    rna-XM_042996553.1    rna-XM_008697643.2    rna-XM_027579489.2    mNeoVis.3.1.1.chr5.g00956.t01    rna-gnl|WGS_ZZZZ|egapxtmp_006246-R1    mNeoVis.evergladensis.2.1.chr5.g00976.t01    mNeoVis.vulgivaga.2.1.chr5.g00966.t01      3148    N0.HOG0003146   OG0000674  n1   rna-XM_002925567.4    rna-XM_038553278.1, rna-XM_038558374.1    rna-XM_022496609.1    rna-XM_003989976.5    rna-XM_036086065.1    rna-XM_039238665.1    rna-XM_032877520.1    rna-XM_046004987.1    rna-XM_032303601.1    rna-XM_059135949.1    rna-XM_059376462.1    rna-XM_045064356.1    mNeoVis.2.1.1.chr2_hap1.g01392.t01    rna-XM_021683807.1    rna-XM_004400661.1    rna-XM_007077332.2    rna-XM_008699326.2    rna-XM_027603826.2    mNeoVis.3.1.1.chr5.g01318.t01    rna-gnl|WGS_ZZZZ|egapxtmp_007414-R1    mNeoVis.evergladensis.2.1.chr5.g01334.t01    mNeoVis.vulgivaga.2.1.chr5.g01332.t01      3149    N0.HOG0003147   OG0000674  n22   rna-XM_019806296.2    rna-XM_038558378.1    rna-XM_022496449.1    rna-XM_003989973.5    rna-XM_036086069.1    rna-XM_039238670.1    rna-XM_032877518.1    rna-XM_046004949.1    rna-XM_032303600.1    rna-XM_059135946.1    rna-XM_059374425.1    rna-XM_004740585.2      rna-XM_021683805.2    rna-XM_004400663.1    rna-XM_042996855.1    rna-XM_040627413.1    rna-XM_027614033.2    mNeoVis.3.1.1.chr5.g01319.t01        mNeoVis.vulgivaga.2.1.chr5.g01333.t01      3150    N0.HOG0003148   OG0000674  n37   rna-XM_034656264.1    rna-XM_038530678.1    rna-XM_022518144.1    rna-XM_023256840.2    rna-XM_036101088.1    rna-XM_039243834.1    rna-XM_032850017.1    rna-XM_046000711.1    rna-XM_032336174.1    rna-XM_059175062.1    rna-XM_059417930.1    rna-XM_004744602.2    mNeoVis.2.1.1.chr1_hap1.g00596.t01    rna-XM_021701861.1    rna-XM_004408748.1    rna-XM_042987350.1    rna-XM_008691391.2    rna-XM_027605605.2    mNeoVis.3.1.1.chr1.g00612.t01    rna-gnl|WGS_ZZZZ|egapxtmp_011456-R1    mNeoVis.evergladensis.2.1.chr1.g00601.t01    mNeoVis.vulgivaga.2.1.chr1.g00610.t01      3151    N0.HOG0003149   OG0000675  n1   rna-XM_011228456.3    rna-XM_038546919.1    rna-XM_022520042.1    rna-XM_019818085.3    rna-XM_036116201.1    rna-XM_039235990.1    rna-XM_032870876.1    rna-XM_045985624.1    rna-XM_032322763.1    rna-XM_059149685.1    rna-XM_059380194.1    rna-XM_013054254.2    mNeoVis.2.1.1.chr8_hap1.g00267.t01    rna-XM_021678398.1    rna-XM_004405882.1    rna-XM_042967525.1    rna-XM_008693589.2    rna-XM_027568611.2    mNeoVis.3.1.1.chr8.g00224.t01      mNeoVis.evergladensis.2.1.chr9.g00256.t01    mNeoVis.vulgivaga.2.1.chr8.g00264.t01      3152    N0.HOG0003150   OG0000675  n21   rna-XM_034639517.1    rna-NM_001003145.1    rna-XM_022514033.1    rna-XM_023250727.2    rna-XM_036102121.1    rna-XM_039230787.1    rna-XM_032840181.1    rna-XM_046021287.1    rna-XM_032304661.1    rna-XM_059172242.1    rna-XM_059396119.1    rna-XM_004765930.3    mNeoVis.2.1.1.chr4_hap1.g00049.t01    rna-XM_044919918.1    rna-XM_004408804.1    rna-XM_042975344.1    rna-XM_040643809.1    rna-XM_027574575.2    mNeoVis.3.1.1.chr6.g00052.t01      mNeoVis.evergladensis.2.1.chr6.g00052.t01    mNeoVis.vulgivaga.2.1.chr6.g00055.t01      3153    N0.HOG0003151   OG0000675  n41   rna-XM_034654650.1    rna-XM_038584668.1    rna-XM_022496465.1    rna-XM_023259392.2    rna-XM_036068803.1    rna-XM_039253434.1    rna-XM_032880306.1    rna-XM_046018071.1    rna-XM_032355757.1    rna-XM_059167165.1    rna-XM_059392755.1    rna-XM_004743962.3    mNeoVis.2.1.1.chr3_hap1.g00936.t01    rna-XM_021702769.1    rna-XM_004410197.1    rna-XM_042994571.1    rna-XM_040644531.1    rna-XM_027591037.2    mNeoVis.3.1.1.chr3.g00926.t01    rna-gnl|WGS_ZZZZ|egapxtmp_038799-R1    mNeoVis.evergladensis.2.1.chr3.g00931.t01    mNeoVis.vulgivaga.2.1.chr3.g00949.t01      3154    N0.HOG0003152   OG0000676  n1   rna-XM_002929399.4    rna-XM_038547866.1    rna-XM_022509529.1    rna-XM_023244022.2    rna-XM_036092977.1    rna-XM_039247360.1    rna-XM_032871968.1    rna-XM_045985465.1    rna-XM_032321838.1    rna-XM_059149333.1    rna-XM_059379308.1, rna-XM_059404957.1    rna-XM_004747319.3    mNeoVis.2.1.1.chr8_hap1.g00632.t01    rna-XM_021704237.1    rna-XM_004397630.1    rna-XM_042965906.1    rna-XM_040639898.1    rna-XM_035724675.1    mNeoVis.3.1.1.chr8.g00583.t01    rna-gnl|WGS_ZZZZ|egapxtmp_017455-R1    mNeoVis.evergladensis.2.1.chr9.g00622.t01    mNeoVis.vulgivaga.2.1.chr8.g00628.t01      3155    N0.HOG0003153   OG0000676  n22   rna-XM_002929408.4    rna-XM_038547859.1    rna-XM_022509359.1      rna-XM_036094093.1      rna-XM_032871971.1    rna-XM_045986059.1    rna-XM_032321685.1    rna-XM_059149081.1    rna-XM_059378528.1    rna-XM_004747320.3    mNeoVis.2.1.1.chr8_hap1.g00630.t01    rna-XM_021704281.1    rna-XM_004397628.1    rna-XM_042965449.1    rna-XM_040639826.1    rna-XM_035724804.1    mNeoVis.3.1.1.chr8.g00581.t01    rna-gnl|WGS_ZZZZ|egapxtmp_017454-R1    mNeoVis.evergladensis.2.1.chr9.g00620.t01    mNeoVis.vulgivaga.2.1.chr8.g00626.t01      3156    N0.HOG0003154   OG0000676  n41   rna-XM_019810086.2    rna-XM_038547871.1    rna-XM_022509714.1    rna-XM_019816875.3    rna-XM_036091943.1    rna-XM_039247351.1    rna-XM_032871967.1    rna-XM_045985693.1    rna-XM_032321974.1    rna-XM_059149511.1    rna-XM_059379209.1    rna-XM_004747318.2    mNeoVis.2.1.1.chr8_hap1.g00633.t01    rna-XM_044921545.1    rna-XM_004397632.1    rna-XM_042965910.1    rna-XM_040640894.1    rna-XM_027567492.1    mNeoVis.3.1.1.chr8.g00584.t01      mNeoVis.evergladensis.2.1.chr9.g00623.t01    mNeoVis.vulgivaga.2.1.chr8.g00629.t01      3157    N0.HOG0003155   OG0000677  n0   rna-XM_019805654.2, rna-XM_034640681.1, rna-XM_034640682.1    rna-XM_038547363.1, rna-XM_038547364.1, rna-XM_038547365.1, rna-XM_038547366.1    rna-XM_022509175.1, rna-XM_022509496.1    rna-XM_045044176.1, rna-XM_045044474.1, rna-XM_045044475.1, rna-XM_045044476.1    rna-XM_036068532.1, rna-XM_036068536.1, rna-XM_036068537.1, rna-XM_036068538.1    rna-XM_039223291.1    rna-XM_032850711.1, rna-XM_032850715.1, rna-XM_032850733.1, rna-XM_032850734.1    rna-XM_045985535.1, rna-XM_045985536.1, rna-XM_045985537.1, rna-XM_045985539.1    rna-XM_032319480.1, rna-XM_032322776.1, rna-XM_032322777.1    rna-XM_059148677.1, rna-XM_059148678.1, rna-XM_059148679.1, rna-XM_059149795.1    rna-XM_059378736.1, rna-XM_059387482.1    rna-XM_013062501.2, rna-XM_045080654.1, rna-XM_045080656.1    mNeoVis.2.1.1.chr8_hap1.g00441.t01, mNeoVis.2.1.1.chr8_hap1.g00442.t01    rna-XM_044912509.1, rna-XM_044921834.1    rna-XM_004417669.1    rna-XM_007085828.2, rna-XM_015539135.2, rna-XM_042965456.1    rna-XM_008689927.2, rna-XM_008689938.2, rna-XM_040639885.1, rna-XM_040640645.1    rna-XM_027568053.1, rna-XM_027568054.1, rna-XM_027568055.1, rna-XM_027568057.1    mNeoVis.3.1.1.chr8.g00397.t01, mNeoVis.3.1.1.chr8.g00398.t01    rna-gnl|WGS_ZZZZ|egapxtmp_019107-R1, rna-gnl|WGS_ZZZZ|egapxtmp_019811-R1, rna-gnl|WGS_ZZZZ|egapxtmp_019823-R1    mNeoVis.evergladensis.2.1.chr9.g00431.t01, mNeoVis.evergladensis.2.1.chr9.g00432.t01    mNeoVis.vulgivaga.2.1.chr8.g00437.t01, mNeoVis.vulgivaga.2.1.chr8.g00438.t01, mNeoVis.vulgivaga.2.1.chr8.g00439.t01      3158    N0.HOG0003156   OG0000678  n1   rna-XM_034641282.1    rna-XM_038572789.1    rna-XM_022494146.1    rna-XM_011280785.4    rna-XM_036098175.1, rna-XM_036105373.1    rna-XM_039236962.1    rna-XM_032878851.1    rna-XM_045980612.1    rna-XM_032350144.1    rna-XM_059132607.1    rna-XM_059407345.1    rna-XM_045070853.1    mNeoVis.2.1.1.chr11_hap1.g00058.t01    rna-XM_021677867.1, rna-XM_021686066.2    rna-XM_004410163.1, rna-XM_004413717.1    rna-XM_042979979.1    rna-XM_008688179.2    rna-XM_027576566.2    mNeoVis.3.1.1.chr11.g00033.t01    rna-gnl|WGS_ZZZZ|egapxtmp_027125-R1    mNeoVis.evergladensis.2.1.chr11.g00058.t01    mNeoVis.vulgivaga.2.1.chr11.g00057.t01      3159    N0.HOG0003157   OG0000678  n27   rna-XM_034643239.1    rna-XM_038543635.1    rna-XM_022495736.1    rna-XM_045041822.1      rna-XM_039217802.1    rna-XM_032859742.1    rna-XM_046024703.1    rna-XM_032309390.1    rna-XM_059140420.1    rna-XM_059409375.1    rna-XM_045065459.1    mNeoVis.2.1.1.chr3_hap1.g00201.t01        rna-XM_007075995.2    rna-XM_040632633.1      mNeoVis.3.1.1.chr3.g00205.t01    rna-gnl|WGS_ZZZZ|egapxtmp_042130-R1    mNeoVis.evergladensis.2.1.chr3.g00212.t01    mNeoVis.vulgivaga.2.1.chr3.g00204.t01      3160    N0.HOG0003158   OG0000678  n44   rna-XM_002925734.4    rna-XM_038572796.1    rna-XM_022494634.1    rna-XM_023251035.2    rna-XM_036105378.1    rna-XM_039237012.1    rna-XM_032878858.1    rna-XM_045980321.1    rna-XM_032350141.1    rna-XM_059134746.1    rna-XM_059407713.1    rna-XM_045070730.1    mNeoVis.2.1.1.chr11_hap1.g00057.t01    rna-XM_021677864.1      rna-XM_042979984.1    rna-XM_040638559.1    rna-XM_027623923.2    mNeoVis.3.1.1.chr2.g00857.t01      mNeoVis.evergladensis.2.1.chr11.g00057.t01    mNeoVis.vulgivaga.2.1.chr11.g00056.t01      3161    N0.HOG0003159   OG0000679  n1     rna-XM_038549530.1      rna-XM_019826438.3, rna-XM_045044507.1    rna-XM_036077501.1, rna-XM_036107259.1        rna-XM_046015941.1      rna-XM_059168243.1, rna-XM_059177563.1    rna-XM_059393330.1, rna-XM_059399155.1    rna-XM_045066221.1, rna-XM_045078663.1    mNeoVis.2.1.1.chr11_hap1.g00146.t01, mNeoVis.2.1.1.chr5_hap1.g01013.t01    rna-XM_044920660.1    rna-XM_004404146.1    rna-XM_042967383.1      rna-XM_027573399.1, rna-XM_027606948.2, rna-XM_035728227.1, rna-XM_035728464.1    mNeoVis.3.1.1.chr11.g00117.t01, mNeoVis.3.1.1.chr11.g00148.t01, mNeoVis.3.1.1.chr2.g00931.t01, mNeoVis.3.1.1.chr4.g00298.t01    rna-gnl|WGS_ZZZZ|egapxtmp_025115-R1, rna-gnl|WGS_ZZZZ|egapxtmp_038385-R1, rna-gnl|WGS_ZZZZ|egapxtmp_043818-R1    mNeoVis.evergladensis.2.1.chr2.g01009.t01, mNeoVis.evergladensis.2.1.chr4.g00306.t01    mNeoVis.vulgivaga.2.1.chr11.g00146.t01, mNeoVis.vulgivaga.2.1.chr2.g01014.t01      3162    N0.HOG0003160   OG0000679  n29   rna-XM_011235853.3    rna-XM_038572472.1    rna-XM_022492351.1, rna-XM_022494348.1      rna-XM_036081053.1, rna-XM_036103857.1    rna-XM_039249365.1    rna-XM_032838664.1    rna-XM_045981252.1, rna-XM_046003355.1, rna-XM_046009783.1    rna-XM_032350522.1    rna-XM_059133014.1    rna-XM_059407993.1    rna-XM_004746329.3    mNeoVis.2.1.1.chr11_hap1.g00178.t01    rna-XM_021689160.1    rna-XM_004405244.1, rna-XM_004414553.1    rna-XM_042979705.1    rna-XM_040637843.1    rna-XM_027603330.1, rna-XM_027603521.1, rna-XM_035723019.1, rna-XM_035723407.1, rna-XM_035724006.1, rna-XM_035724660.1, rna-XM_035728175.1, rna-XM_035728933.1, rna-XM_035729222.1        mNeoVis.evergladensis.2.1.chr11.g00175.t01    mNeoVis.vulgivaga.2.1.chr11.g00177.t01      3163    N0.HOG0003161   OG0000680  n1   rna-NM_001304870.1    rna-XM_038565536.1    rna-XM_022498422.1    rna-XM_003982524.5    rna-XM_036070419.1, rna-XM_036073662.1    rna-XM_039254943.1    rna-XM_032849446.1    rna-XM_045989832.1    rna-XM_032310140.1, rna-XM_032327411.1    rna-XM_059161857.1    rna-XM_059390400.1    rna-XM_004738598.3, rna-XM_045075558.1    mNeoVis.2.1.1.chr10_hap1.g00324.t01, mNeoVis.2.1.1.chr11_hap1.g00660.t01, mNeoVis.2.1.1.chr2_hap1.g01717.t01, mNeoVis.2.1.1.chr8_hap1.g00423.t01    rna-XM_021694993.1, rna-XM_021697337.1    rna-XM_004391709.1, rna-XM_004395672.1, rna-XM_004412279.1    rna-XM_015545133.2    rna-XM_008697830.2    rna-XM_027582101.2, rna-XM_027589449.2, rna-XM_035729364.1    mNeoVis.3.1.1.chr2.g00825.t01    rna-gnl|WGS_ZZZZ|egapxtmp_007632-R1, rna-gnl|WGS_ZZZZ|egapxtmp_019813-R1, rna-gnl|WGS_ZZZZ|egapxtmp_027294-R1, rna-gnl|WGS_ZZZZ|egapxtmp_037631-R1    mNeoVis.evergladensis.2.1.chr2.g00911.t01, mNeoVis.evergladensis.2.1.chr5.g01656.t01    mNeoVis.vulgivaga.2.1.chr2.g00915.t01, mNeoVis.vulgivaga.2.1.chr5.g01658.t01, mNeoVis.vulgivaga.2.1.chr8.g00428.t01      3164    N0.HOG0003162   OG0000680  n12     rna-XM_038558200.1        rna-XM_036080879.1                                        3165    N0.HOG0003163   OG0000680  n13     rna-XM_038569123.1        rna-XM_036069577.1, rna-XM_036097442.1, rna-XM_036119321.1                    rna-XM_044911622.1          rna-XM_035723108.1, rna-XM_035725456.1, rna-XM_035725504.1, rna-XM_035729057.1              3166    N0.HOG0003164   OG0000680  n21     rna-XM_038573465.1          rna-XM_039248807.1    rna-XM_032866553.1, rna-XM_032872958.1    rna-XM_045978388.1      rna-XM_059144183.1          rna-XM_044918658.1          rna-XM_027597112.2, rna-XM_035724056.1, rna-XM_035729020.1    mNeoVis.3.1.1.chr5.g01641.t01            3167    N0.HOG0003165   OG0000681  n1   rna-XM_011220261.3    rna-XM_038547852.1    rna-XM_022509421.1    rna-XM_045044851.1    rna-XM_036105809.1    rna-XM_039241157.1    rna-XM_032871974.1    rna-XM_045985575.1    rna-XM_032320644.1    rna-XM_059148067.1    rna-XM_059380758.1    rna-XM_004747325.3    mNeoVis.2.1.1.chr8_hap1.g00624.t01    rna-XM_021704526.2    rna-XM_004405200.1    rna-XM_042967438.1    rna-XM_040640892.1    rna-XM_027625059.2    mNeoVis.3.1.1.chr8.g00577.t01      mNeoVis.evergladensis.2.1.chr9.g00615.t01    mNeoVis.vulgivaga.2.1.chr8.g00622.t01      3168    N0.HOG0003166   OG0000681  n4   rna-XM_002914437.3, rna-XM_011217191.3, rna-XM_034651331.1    rna-XM_038575328.1    rna-XM_022498074.1, rna-XM_022508803.1, rna-XM_022517144.1, rna-XM_022517226.1    rna-XM_019814621.3    rna-XM_036068804.1, rna-XM_036069064.1, rna-XM_036081995.1    rna-XM_039220540.1    rna-XM_032855444.1, rna-XM_032876147.1    rna-XM_045982036.1, rna-XM_045997054.1, rna-XM_046024082.1    rna-XM_032309139.1, rna-XM_032338378.1    rna-XM_059140676.1    rna-XM_059408584.1    rna-XM_004753366.3    mNeoVis.2.1.1.chr3_hap1.g00511.t01, mNeoVis.2.1.1.chr9_hap1.g00011.t01    rna-XM_021685921.1, rna-XM_021692796.1, rna-XM_021694218.1, rna-XM_021703603.2    rna-XM_004396664.1, rna-XM_004413204.1    rna-XM_042962739.1    rna-XM_008705141.1, rna-XM_040619846.1    rna-XM_027573887.2, rna-XM_027576509.2    mNeoVis.3.1.1.chr3.g00147.t01, mNeoVis.3.1.1.chr3.g00503.t01    rna-gnl|WGS_ZZZZ|egapxtmp_023344-R1    mNeoVis.evergladensis.2.1.chr10.g00013.t01, mNeoVis.evergladensis.2.1.chr3.g00520.t01    mNeoVis.vulgivaga.2.1.chr3.g00522.t01      3169    N0.HOG0003167   OG0000682  n1   rna-XM_002928027.3    rna-NM_001077439.1    rna-XM_022492029.1    rna-XM_003988173.6    rna-XM_036092301.1    rna-XM_039220509.1, rna-XM_039221774.1    rna-XM_032848953.1    rna-XM_046010596.1    rna-XM_032349663.1    rna-XM_059183891.1    rna-XM_059404154.1    rna-XM_004757636.3    mNeoVis.2.1.1.chr9_hap1.g00076.t01    rna-XM_021687153.1    rna-XM_004403976.1    rna-XM_007081308.3    rna-XM_008709679.1    rna-XM_027595472.2    mNeoVis.3.1.1.chr10.g00072.t01    rna-gnl|WGS_ZZZZ|egapxtmp_022566-R2    mNeoVis.evergladensis.2.1.chr10.g00074.t01    mNeoVis.vulgivaga.2.1.chr10.g00070.t01      3170    N0.HOG0003168   OG0000682  n20   rna-XM_034642210.1    rna-XM_038529561.1    rna-XM_022513787.1      rna-XM_036083766.1    rna-XM_039230532.1    rna-XM_032872173.1    rna-XM_046024366.1    rna-XM_032311266.1    rna-XM_059139634.1    rna-XM_059410073.1    rna-XM_013052718.2    mNeoVis.2.1.1.chr3_hap1.g00062.t01        rna-XM_015536799.1    rna-XM_008705411.2      mNeoVis.3.1.1.chr3.g00063.t01    rna-gnl|WGS_ZZZZ|egapxtmp_040998-R1    mNeoVis.evergladensis.2.1.chr3.g00065.t01    mNeoVis.vulgivaga.2.1.chr3.g00059.t01      3171    N0.HOG0003169   OG0000682  n36   rna-XM_011230649.3    rna-NM_001097543.1    rna-XM_022507120.1    rna-XM_019835945.3    rna-XM_036067303.1    rna-XM_039220477.1    rna-XM_032880431.1    rna-XM_046017983.1    rna-XM_032355914.1    rna-XM_059167064.1    rna-XM_059394145.1    rna-XM_004744054.3    mNeoVis.2.1.1.chr3_hap1.g00897.t01    rna-XM_021702751.1    rna-XM_004407667.1    rna-XM_007074588.3    rna-XM_008687105.1    rna-XM_027589840.2    mNeoVis.3.1.1.chr3.g00881.t01    rna-gnl|WGS_ZZZZ|egapxtmp_041208-R1    mNeoVis.evergladensis.2.1.chr3.g00891.t01    mNeoVis.vulgivaga.2.1.chr3.g00908.t01      3172    N0.HOG0003170   OG0000683  n0   rna-XM_002916117.4, rna-XM_034644345.1, rna-XM_034669462.1    rna-XM_038555092.1    rna-XM_022503120.1    rna-XM_011280439.4, rna-XM_045036442.1    rna-XM_036082223.1, rna-XM_036099980.1    rna-XM_039217440.1, rna-XM_039229872.1, rna-XM_039237410.1    rna-XM_032866919.1, rna-XM_032870113.1, rna-XM_032872169.1, rna-XM_032872906.1    rna-XM_046020984.1    rna-XM_032305651.1, rna-XM_032311779.1    rna-XM_059171399.1, rna-XM_059174467.1, rna-XM_059177123.1, rna-XM_059187735.1    rna-XM_059396706.1, rna-XM_059399240.1, rna-XM_059416574.1    rna-XM_004742416.2, rna-XM_045062803.1, rna-XM_045075554.1, rna-XM_045075852.1, rna-XM_045080195.1    mNeoVis.2.1.1.chr12_hap1.g00271.t01, mNeoVis.2.1.1.chr1_hap1.g00140.t01, mNeoVis.2.1.1.chr2_hap1.g00465.t01, mNeoVis.2.1.1.chr4_hap1.g00390.t01    rna-XM_021689810.2, rna-XM_021692909.1, rna-XM_044916093.1, rna-XM_044916222.1, rna-XM_044917245.1, rna-XM_044921022.1    rna-XM_004397354.1, rna-XM_004410400.1    rna-XM_007094557.2, rna-XM_015535282.2, rna-XM_042977288.1    rna-XM_008686422.2    rna-XM_027573622.2, rna-XM_027584046.2, rna-XM_035721916.1, rna-XM_035723465.1, rna-XM_035723771.1, rna-XM_035723922.1, rna-XM_035726953.1, rna-XM_035729282.1    mNeoVis.3.1.1.chr6.g00390.t01    rna-gnl|WGS_ZZZZ|egapxtmp_002725-R1, rna-gnl|WGS_ZZZZ|egapxtmp_025330-R1    mNeoVis.evergladensis.2.1.chr12.g00240.t01, mNeoVis.evergladensis.2.1.chr5.g00468.t01, mNeoVis.evergladensis.2.1.chr6.g00393.t01    mNeoVis.vulgivaga.2.1.chr5.g00467.t01, mNeoVis.vulgivaga.2.1.chr6.g00389.t01      3173    N0.HOG0003171   OG0000684  n2   rna-XM_002920446.4    rna-XM_038576513.1    rna-XM_022522356.1    rna-XM_003988283.6    rna-XM_036082112.1    rna-XM_039255756.1    rna-XM_032874838.1    rna-XM_046011535.1    rna-XM_032349075.1    rna-XM_059184549.1    rna-XM_059403780.1    rna-XM_004778282.3    mNeoVis.2.1.1.chr9_hap1.g00227.t01    rna-XM_021690518.2    rna-XM_004415976.1    rna-XM_007089662.3    rna-XM_008704875.2    rna-XM_027591956.2    mNeoVis.3.1.1.chr10.g00220.t01      mNeoVis.evergladensis.2.1.chr10.g00224.t01    mNeoVis.vulgivaga.2.1.chr10.g00221.t01      3174   N0.HOG0003172  OG0000684  n7   rna-XM_002914487.4    rna-XM_038554047.1    rna-XM_022504112.1    rna-XM_011282227.4    rna-XM_036102088.1    rna-XM_039242594.1    rna-XM_032862193.1    rna-XM_046004483.1    rna-XM_032339943.1    rna-XM_059177949.1    rna-XM_059400488.1    rna-XM_004739917.3    mNeoVis.2.1.1.chr1_hap1.g01658.t01    rna-XM_021691950.2    rna-XM_004408065.1    rna-XM_007085197.3    rna-XM_040643465.1    rna-XM_027602008.2    mNeoVis.3.1.1.chr1.g01607.t01      mNeoVis.evergladensis.2.1.chr1.g01624.t01    mNeoVis.vulgivaga.2.1.chr1.g01621.t01      3175   N0.HOG0003173  OG0000684  n21   rna-XM_034644999.1    rna-NM_001005757.1    rna-XM_022506549.1    rna-XM_011287102.4    rna-XM_036099568.1    rna-XM_039235370.1    rna-XM_032879711.1    rna-XM_046014964.1    rna-XM_032358060.1    rna-XM_059150341.1    rna-XM_059399195.1    rna-XM_004759617.3    mNeoVis.2.1.1.chr7_hap1.g02596.t01    rna-XM_021685594.1    rna-XM_004393801.1    rna-XM_042959425.1    rna-XM_040639580.1    rna-XM_027578897.2        mNeoVis.evergladensis.2.1.chr7.g02402.t01    mNeoVis.vulgivaga.2.1.chr7.g02425.t01      3176    N0.HOG0003174   OG0000685  n1   rna-XM_034663957.1    rna-XM_038552480.1    rna-XM_022517618.1    rna-XM_045042681.1    rna-XM_036089858.1    rna-XM_039240918.1    rna-XM_032880525.1    rna-XM_046023343.1    rna-XM_032307120.1    rna-XM_059141999.1    rna-XM_059411592.1    rna-XM_045079654.1    mNeoVis.2.1.1.chr13_hap1.g00622.t01    rna-XM_021682418.1    rna-XM_004416743.1    rna-XM_042964872.1    rna-XM_008689815.1    rna-XM_027616871.2    mNeoVis.3.1.1.chr13.g00547.t01    rna-gnl|WGS_ZZZZ|egapxtmp_031434-R1    mNeoVis.evergladensis.2.1.chr13.g00597.t01    mNeoVis.vulgivaga.2.1.chr13.g00598.t01      3177    N0.HOG0003175   OG0000685  n19   rna-XM_034657541.1    rna-XM_038567743.1    rna-XM_022525469.1    rna-XM_023244015.2    rna-XM_036116024.1    rna-XM_039247574.1    rna-XM_032883597.1    rna-XM_045989800.1    rna-XM_032306757.1, rna-XM_032348419.1    rna-XM_059159518.1    rna-XM_059388688.1    rna-XM_004771449.3    mNeoVis.2.1.1.chr5_hap1.g00123.t01    rna-XM_021705334.2    rna-XM_004395325.1    rna-XM_042977797.1    rna-XM_040629569.1    rna-XM_027584397.2        mNeoVis.evergladensis.2.1.chr2.g00122.t01    mNeoVis.vulgivaga.2.1.chr2.g00118.t01      3178    N0.HOG0003176   OG0000685  n32   rna-XM_034667806.1    rna-XM_038532974.1    rna-XM_022511203.1    rna-XM_023254841.2    rna-XM_036072907.1    rna-XM_039216448.1    rna-XM_032836846.1    rna-XM_046008794.1    rna-XM_032341961.1    rna-XM_059179854.1    rna-XM_059371459.1    rna-XM_004763656.3      rna-XM_021680592.1    rna-XM_004417333.1    rna-XM_042987929.1    rna-XM_040642872.1    rna-XM_027571927.2    mNeoVis.3.1.1.chr9.g01181.t01    rna-gnl|WGS_ZZZZ|egapxtmp_017059-R1    mNeoVis.evergladensis.2.1.chr8.g01228.t01        3179   N0.HOG0003177  OG0000686  n1   rna-XM_034637947.1    rna-XM_038538229.1    rna-XM_022511869.1    rna-XM_023245367.2    rna-XM_036084973.1    rna-XM_039227204.1    rna-XM_032854607.1    rna-XM_045988938.1    rna-XM_032326530.1    rna-XM_059153112.1    rna-XM_059381634.1    rna-XM_004754070.3    mNeoVis.2.1.1.chr7_hap1.g00105.t01    rna-XM_044922212.1    rna-XM_004392062.1    rna-XM_042969660.1    rna-XM_040633628.1    rna-XM_027618626.2    mNeoVis.3.1.1.chr7.g00089.t01      mNeoVis.evergladensis.2.1.chr7.g00106.t01    mNeoVis.vulgivaga.2.1.chr7.g00103.t01      3180    N0.HOG0003178   OG0000686  n22   rna-XM_019798909.2    rna-XM_038571375.1    rna-XM_022494087.1    rna-XM_023251254.2    rna-XM_036121385.1    rna-XM_039227189.1    rna-XM_032838542.1    rna-XM_045981615.1    rna-XM_032350670.1    rna-XM_059133228.1    rna-XM_059406999.1    rna-XM_013053312.2    mNeoVis.2.1.1.chr11_hap1.g00231.t01    rna-XM_021688954.2    rna-XM_004411970.1    rna-XM_042980400.1    rna-XM_040638310.1    rna-XM_027623613.1      rna-gnl|WGS_ZZZZ|egapxtmp_026849-R1    mNeoVis.evergladensis.2.1.chr11.g00230.t01    mNeoVis.vulgivaga.2.1.chr11.g00231.t01      3181    N0.HOG0003179   OG0000686  n30   rna-XM_034668533.1    rna-XM_038555104.1    rna-XM_022502399.1    rna-XM_023248573.2    rna-XM_036105730.1    rna-XM_039254008.1    rna-XM_032848293.1    rna-XM_046021178.1    rna-XM_032316834.1    rna-XM_059168784.1    rna-XM_059392984.1    rna-XM_045074204.1    mNeoVis.2.1.1.chr4_hap1.g01078.t01    rna-XM_021688614.1    rna-XM_004400697.1    rna-XM_042973295.1    rna-XM_040625905.1    rna-XM_027623087.2    mNeoVis.3.1.1.chr6.g01077.t01      mNeoVis.evergladensis.2.1.chr6.g01079.t01    mNeoVis.vulgivaga.2.1.chr6.g01072.t01      3182    N0.HOG0003180   OG0000687  n1   rna-XM_002927358.4    rna-XM_038539905.1    rna-XM_022526387.1    rna-XM_003998688.6    rna-XM_036091206.1    rna-XM_039247867.1    rna-XM_032842521.1    rna-XM_045994072.1    rna-XM_032328704.1    rna-XM_059155697.1    rna-XM_059414980.1    rna-XM_045072617.1    mNeoVis.2.1.1.chr15_hap1.g00075.t01    rna-XM_021700315.1    rna-XM_004397184.1    rna-XM_007093466.3    rna-XM_040635120.1, rna-XM_040643166.1    rna-XM_027613833.2    mNeoVis.3.1.1.chr15.g00069.t01      mNeoVis.evergladensis.2.1.chr15.g00076.t01    mNeoVis.vulgivaga.2.1.chr15.g00077.t01      3183    N0.HOG0003181   OG0000687  n6   rna-XM_002912913.4, rna-XM_034653201.1    rna-NM_001003063.1    rna-XM_022493349.1    rna-XM_003980768.6    rna-XM_036089054.1    rna-XM_039242276.1    rna-XM_032837775.1    rna-XM_046000142.1            mNeoVis.2.1.1.chr1_hap1.g00415.t01    rna-XM_021702225.2    rna-XM_004405449.1    rna-XM_042983946.1    rna-XM_040622443.1    rna-XM_027606243.1, rna-XM_035725557.1    mNeoVis.3.1.1.chr1.g00412.t01        mNeoVis.vulgivaga.2.1.chr1.g00419.t01      3184    N0.HOG0003182   OG0000687  n9   rna-XM_019801339.2, rna-XM_034653097.1    rna-XM_038560171.1    rna-XM_022498869.1    rna-XM_006930608.4    rna-XM_036107084.1    rna-XM_039221099.1    rna-XM_032838115.1    rna-XM_045997968.1    rna-XM_032329407.1    rna-XM_059156219.1    rna-XM_059384121.1    rna-XM_045073776.1    mNeoVis.2.1.1.chr6_hap1.g00997.t01    rna-XM_021694118.2    rna-XM_004408280.1    rna-XM_042983137.1    rna-XM_040643283.1    rna-XM_027599658.2    mNeoVis.3.1.1.chr4.g00954.t01    rna-gnl|WGS_ZZZZ|egapxtmp_000894-R1    mNeoVis.evergladensis.2.1.chr4.g00942.t01    mNeoVis.vulgivaga.2.1.chr4.g00966.t01      3185    N0.HOG0003183   OG0000688  n1   rna-XM_034655619.1    rna-XM_038581478.1    rna-XM_022516141.1    rna-XM_003991434.6    rna-XM_036109247.1    rna-XM_039229763.1    rna-XM_032865328.1    rna-XM_046003617.1    rna-XM_032354960.1    rna-XM_059164724.1    rna-XM_059392349.1    rna-XM_004762159.3    mNeoVis.2.1.1.chr5_hap1.g01834.t01    rna-XM_021677963.1    rna-XM_004414138.1    rna-XM_015541195.2    rna-XM_008701902.2    rna-XM_027586904.2    mNeoVis.3.1.1.chr2.g01736.t01    rna-gnl|WGS_ZZZZ|egapxtmp_034612-R2    mNeoVis.evergladensis.2.1.chr2.g01817.t01    mNeoVis.vulgivaga.2.1.chr2.g01832.t01      3186    N0.HOG0003184   OG0000688  n10   rna-XM_034651156.1    rna-XM_038586375.1    rna-XM_022491125.1    rna-XM_045048457.1    rna-XM_036091463.1    rna-XM_039246416.1    rna-XM_032845653.1    rna-XM_045982092.1    rna-XM_032318585.1    rna-XM_059147158.1    rna-XM_059413538.1    rna-XM_004775969.3    mNeoVis.2.1.1.chr14_hap1.g00529.t01    rna-XM_021681915.2    rna-XM_004407859.1    rna-XM_042976224.1    rna-XM_040635906.1    rna-XM_027612462.2        mNeoVis.evergladensis.2.1.chr14.g00528.t01    mNeoVis.vulgivaga.2.1.chr14.g00526.t01      3187    N0.HOG0003185   OG0000688  n21   rna-XM_019796888.2    rna-XM_038535660.1    rna-XM_022513267.1    rna-XM_006936792.4    rna-XM_036086103.1    rna-XM_039240263.1    rna-XM_032868910.1    rna-XM_046014184.1    rna-XM_032359464.1    rna-XM_059184814.1    rna-XM_059392050.1    rna-XM_004763902.3    mNeoVis.2.1.1.chr7_hap1.g01385.t01    rna-XM_021696501.1    rna-XM_004412047.1    rna-XM_015537718.2    rna-XM_008688309.2    rna-XM_027580570.2    mNeoVis.3.1.1.chr7.g01295.t01      mNeoVis.evergladensis.2.1.chr7.g01368.t01    mNeoVis.vulgivaga.2.1.chr7.g01380.t01      3188    N0.HOG0003186   OG0000689  n1   rna-XM_002928760.4    rna-XM_038586536.1    rna-XM_022524343.1    rna-XM_004000660.5    rna-XM_036110545.1    rna-XM_039222718.1    rna-XM_032840625.1    rna-XM_045996409.1    rna-XM_032331149.1    rna-XM_059157659.1    rna-XM_059385687.1    rna-XM_004777113.3    mNeoVis.2.1.1.chr12_hap1.g00442.t01    rna-XM_021687942.1    rna-XM_004415149.1    rna-XM_042975118.1    rna-XM_040630467.1    rna-XM_027608873.1    mNeoVis.3.1.1.chr12.g00404.t01    rna-gnl|WGS_ZZZZ|egapxtmp_029667-R1    mNeoVis.evergladensis.2.1.chr12.g00399.t01    mNeoVis.vulgivaga.2.1.chr12.g00407.t01      3189    N0.HOG0003187   OG0000689  n10   rna-XM_002928925.4    rna-XM_038573486.1    rna-XM_022522620.1    rna-XM_003980257.6    rna-XM_036065483.1    rna-XM_039255396.1    rna-XM_032838224.1    rna-XM_046028118.1    rna-XM_032315019.1    rna-XM_059144312.1    rna-XM_059377770.1    rna-XM_004774339.3    mNeoVis.2.1.1.chr8_hap1.g01180.t01    rna-XM_021678227.1    rna-XM_004401339.1    rna-XM_007074361.3    rna-XM_008692136.2    rna-XM_027587980.1    mNeoVis.3.1.1.chr8.g01095.t01      mNeoVis.evergladensis.2.1.chr9.g01152.t01    mNeoVis.vulgivaga.2.1.chr8.g01152.t01      3190    N0.HOG0003188   OG0000689  n12   rna-XM_034647697.1    rna-XM_038559235.1    rna-XM_022499013.1    rna-XM_023252220.2    rna-XM_036101487.1    rna-XM_039256276.1    rna-XM_032844942.1    rna-XM_045998451.1    rna-XM_032329331.1    rna-XM_059156300.1    rna-XM_059384310.1      mNeoVis.2.1.1.chr6_hap1.g01023.t01    rna-XM_021694262.1    rna-XM_004409519.1    rna-XM_042982632.1    rna-XM_008710935.2    rna-XM_027600277.1    mNeoVis.3.1.1.chr4.g00983.t01    rna-gnl|WGS_ZZZZ|egapxtmp_044261-R1    mNeoVis.evergladensis.2.1.chr4.g00967.t01        3191    N0.HOG0003189   OG0000690  n1   rna-XM_034647254.1    rna-XM_038537137.1    rna-XM_022491448.1    rna-XM_023244640.2    rna-XM_036101863.1    rna-XM_039227288.1    rna-XM_032841224.1    rna-XM_045984319.1    rna-XM_032319677.1    rna-XM_059147604.1    rna-XM_059380283.1    rna-XM_013047853.2    mNeoVis.2.1.1.chr8_hap1.g00906.t01    rna-XM_021694663.2    rna-XM_004405168.1    rna-XM_042966324.1    rna-XM_040630174.1    rna-XM_027624800.2    mNeoVis.3.1.1.chr8.g00853.t01    rna-gnl|WGS_ZZZZ|egapxtmp_017529-R1    mNeoVis.evergladensis.2.1.chr9.g00894.t01    mNeoVis.vulgivaga.2.1.chr8.g00897.t01      3192    N0.HOG0003190   OG0000690  n18   rna-XM_019793908.2    rna-XM_038586511.1      rna-XM_045050949.1    rna-XM_036105442.1    rna-XM_039225663.1    rna-XM_032850721.1    rna-XM_045994583.1    rna-XM_032331381.1    rna-XM_059158018.1      rna-XM_013050794.2    mNeoVis.2.1.1.chr12_hap1.g00553.t01    rna-XM_021700883.1    rna-XM_004416833.1    rna-XM_015544191.2    rna-XM_008703188.2    rna-XM_027608408.2      rna-gnl|WGS_ZZZZ|egapxtmp_029612-R2    mNeoVis.evergladensis.2.1.chr12.g00498.t01    mNeoVis.vulgivaga.2.1.chr12.g00511.t01      3193    N0.HOG0003191   OG0000690  n34   rna-XM_034649017.1    rna-XM_038588789.1    rna-XM_022507951.1    rna-XM_045050376.1    rna-XM_036071617.1    rna-XM_039221172.1    rna-XM_032842986.1    rna-XM_045994757.1    rna-XM_032329692.1    rna-XM_059156790.1    rna-XM_059385255.1    rna-XM_004755111.2    mNeoVis.2.1.1.chr12_hap1.g00589.t01    rna-XM_021679298.1    rna-XM_004396842.1    rna-XM_042973943.1    rna-XM_040633649.1    rna-XM_027607588.2    mNeoVis.3.1.1.chr12.g00536.t01    rna-gnl|WGS_ZZZZ|egapxtmp_028293-R1    mNeoVis.evergladensis.2.1.chr12.g00526.t01    mNeoVis.vulgivaga.2.1.chr12.g00540.t01      3194    N0.HOG0003192   OG0000691  n1   rna-XM_002927188.4    rna-XM_038577537.1    rna-XM_022519772.1    rna-XM_003988641.6    rna-XM_036121128.1    rna-XM_039221085.1    rna-XM_032876884.1    rna-XM_046010503.1    rna-XM_032348080.1    rna-XM_059185470.1    rna-XM_059404839.1    rna-XM_045073180.1    mNeoVis.2.1.1.chr9_hap1.g00591.t01    rna-XM_021704889.1    rna-XM_004400864.1    rna-XM_042992206.1    rna-XM_040642408.1    rna-XM_027593410.2    mNeoVis.3.1.1.chr10.g00559.t01      mNeoVis.evergladensis.2.1.chr10.g00585.t01    mNeoVis.vulgivaga.2.1.chr10.g00571.t01      3195    N0.HOG0003193   OG0000691  n11   rna-XM_002913694.4    rna-XM_038585773.1    rna-XM_022500123.1    rna-XM_019839142.3    rna-XM_036083044.1    rna-XM_039233166.1    rna-XM_032857232.1    rna-XM_046019490.1    rna-XM_032354837.1    rna-XM_059168008.1    rna-XM_059393500.1    rna-XM_004762933.3    mNeoVis.2.1.1.chr3_hap1.g01165.t01    rna-XM_021703159.1    rna-XM_004402732.1    rna-XM_042997613.1    rna-XM_040637237.1    rna-XM_027588047.2    mNeoVis.3.1.1.chr3.g01153.t01      mNeoVis.evergladensis.2.1.chr3.g01167.t01    mNeoVis.vulgivaga.2.1.chr3.g01182.t01      3196    N0.HOG0003194   OG0000691  n29   rna-XM_034650041.1    rna-XM_038559814.1    rna-XM_022514177.1    rna-XM_045052658.1    rna-XM_036069604.1    rna-XM_039230668.1    rna-XM_032840275.1    rna-XM_046019990.1    rna-XM_032305348.1    rna-XM_059172368.1    rna-XM_059396017.1    rna-XM_045086293.1    mNeoVis.2.1.1.chr4_hap1.g00012.t01    rna-XM_021689395.1    rna-XM_004403479.1    rna-XM_042977233.1    rna-XM_008688638.2    rna-XM_027572985.1      rna-gnl|WGS_ZZZZ|egapxtmp_003449-R1    mNeoVis.evergladensis.2.1.chr6.g00012.t01    mNeoVis.vulgivaga.2.1.chr6.g00014.t01      3197    N0.HOG0003195   OG0000692  n3     rna-XM_038565266.1    rna-XM_022519553.1    rna-XM_045045200.1    rna-XM_036065884.1, rna-XM_036102268.1      rna-XM_032838619.1, rna-XM_032866922.1, rna-XM_032876607.1, rna-XM_032883917.1    rna-XM_045977931.1, rna-XM_046011816.1, rna-XM_046013829.1    rna-XM_032304811.1, rna-XM_032321185.1, rna-XM_032330186.1, rna-XM_032348424.1, rna-XM_032358725.1    rna-XM_059134014.1, rna-XM_059157043.1, rna-XM_059185142.1    rna-XM_059388179.1, rna-XM_059403287.1, rna-XM_059407910.1    rna-XM_045068898.1, rna-XM_045073102.1, rna-XM_045086004.1, rna-XM_045088188.1    mNeoVis.2.1.1.chr12_hap1.g00775.t01, mNeoVis.2.1.1.chr5_hap1.g01291.t01, mNeoVis.2.1.1.chr9_hap1.g00479.t01    rna-XM_044913355.1, rna-XM_044920185.1    rna-XM_004397977.1    rna-XM_042977235.1, rna-XM_042993891.1      rna-XM_035723388.1, rna-XM_035727562.1    mNeoVis.3.1.1.chr10.g00453.t01, mNeoVis.3.1.1.chr12.g00707.t01    rna-gnl|WGS_ZZZZ|egapxtmp_023552-R1, rna-gnl|WGS_ZZZZ|egapxtmp_038213-R1    mNeoVis.evergladensis.2.1.chr10.g00473.t01, mNeoVis.evergladensis.2.1.chr2.g01278.t01    mNeoVis.vulgivaga.2.1.chr7.g01497.t01, mNeoVis.vulgivaga.2.1.chr7.g01793.t01      3198    N0.HOG0003196   OG0000692  n43     rna-XM_038542983.1, rna-XM_038547727.1, rna-XM_038577821.1      rna-XM_045061874.1    rna-XM_036124175.1    rna-XM_039255119.1                  rna-XM_044914824.1    rna-XM_004406204.1, rna-XM_004406359.1    rna-XM_015538892.2, rna-XM_042963333.1, rna-XM_042992426.1      rna-XM_027593222.1, rna-XM_027620327.2, rna-XM_035726088.1, rna-XM_035726282.1              3199    N0.HOG0003197   OG0000693  n1     rna-XM_038562448.1, rna-XM_038562450.1, rna-XM_038562905.1    rna-XM_022507323.1    rna-XM_003993844.6    rna-XM_036098245.1    rna-XM_039221534.1      rna-XM_046015723.1, rna-XM_046015733.1    rna-XM_032358521.1    rna-XM_059152984.1        mNeoVis.2.1.1.chr7_hap1.g02682.t01      rna-XM_004403815.1, rna-XM_004403915.1    rna-XM_007094211.3    rna-XM_008699591.2, rna-XM_008699593.2    rna-XM_035722926.1, rna-XM_035722929.1, rna-XM_035723015.1, rna-XM_035723376.1, rna-XM_035725437.1, rna-XM_035725636.1, rna-XM_035726606.1, rna-XM_035726748.1, rna-XM_035728703.1, rna-XM_035728978.1        mNeoVis.evergladensis.2.1.chr7.g02495.t01    mNeoVis.vulgivaga.2.1.chr7.g02515.t01      3200    N0.HOG0003198   OG0000693  n27   rna-XM_002931254.4, rna-XM_034644985.1    rna-XM_038562447.1, rna-XM_038562449.1    rna-XM_022506907.1    rna-XM_003993816.6, rna-XM_006937693.5    rna-XM_036098241.1, rna-XM_036098243.1    rna-XM_039221533.1, rna-XM_039221535.1    rna-XM_032855500.1, rna-XM_032879871.1    rna-XM_046015729.1, rna-XM_046015732.1    rna-XM_032358517.1, rna-XM_032358522.1    rna-XM_059152962.1, rna-XM_059152972.1    rna-XM_059387497.1, rna-XM_059387531.1    rna-XM_045067221.1, rna-XM_045067236.1    mNeoVis.2.1.1.chr7_hap1.g02681.t01, mNeoVis.2.1.1.chr7_hap1.g02683.t01    rna-XM_021699189.1, rna-XM_021703572.1    rna-XM_004403814.1    rna-XM_007094212.3, rna-XM_007094213.3    rna-XM_008699589.2, rna-XM_008699592.2    rna-XM_035722927.1    mNeoVis.3.1.1.chr7.g02318.t01            3201    N0.HOG0003199   OG0000694  n1   rna-XM_034669534.1    rna-XM_038539208.1    rna-XM_022501519.1    rna-XM_045047337.1    rna-XM_036078820.1    rna-XM_039243536.1    rna-XM_032872516.1    rna-XM_045993265.1    rna-XM_032327760.1    rna-XM_059153904.1    rna-XM_059414606.1    rna-XM_045075024.1    mNeoVis.2.1.1.chr15_hap1.g00697.t01    rna-XM_044915154.1    rna-XM_004399055.1    rna-XM_042971626.1    rna-XM_040634885.1    rna-XM_027613347.2    mNeoVis.3.1.1.chr15.g00478.t01      mNeoVis.evergladensis.2.1.chr15.g00519.t01    mNeoVis.vulgivaga.2.1.chr15.g00530.t01      3202    N0.HOG0003200   OG0000694  n20   rna-XM_002924869.4, rna-XM_034671211.1    rna-NM_001284442.1    rna-XM_022505145.1, rna-XM_022514340.1    rna-XM_003989499.4, rna-XM_045037478.1    rna-XM_036069555.1, rna-XM_036078142.1    rna-XM_039216490.1, rna-XM_039224840.1    rna-XM_032852425.1, rna-XM_032863428.1    rna-XM_046000403.1, rna-XM_046002161.1    rna-XM_032301721.1, rna-XM_032346492.1    rna-XM_059135455.1, rna-XM_059163837.1    rna-XM_059375602.1, rna-XM_059391784.1    rna-XM_004754459.3, rna-XM_045063753.1    mNeoVis.2.1.1.chr2_hap1.g01804.t01, mNeoVis.2.1.1.chr5_hap1.g01515.t01    rna-XM_021683447.1, rna-XM_021696812.1    rna-XM_004411162.1, rna-XM_004417564.1    rna-XM_042956431.1, rna-XM_042995730.1    rna-XM_008711108.2, rna-XM_040645785.1    rna-XM_027570409.2, rna-XM_027586476.1    mNeoVis.3.1.1.chr2.g01435.t01, mNeoVis.3.1.1.chr5.g01729.t01    rna-gnl|WGS_ZZZZ|egapxtmp_034559-R3    mNeoVis.evergladensis.2.1.chr2.g01502.t01, mNeoVis.evergladensis.2.1.chr5.g01743.t01    mNeoVis.vulgivaga.2.1.chr2.g01515.t01, mNeoVis.vulgivaga.2.1.chr5.g01747.t01      3203    N0.HOG0003201   OG0000695  n1   rna-XM_034647125.1    rna-XM_038536612.1    rna-XM_022524714.1    rna-XM_045044714.1      rna-XM_039231638.1    rna-XM_032839936.1    rna-XM_045985365.1    rna-XM_032321251.1    rna-XM_059148318.1    rna-XM_059379144.1    rna-XM_045083243.1    mNeoVis.2.1.1.chr8_hap1.g01082.t01    rna-XM_021694702.1    rna-XM_004398549.1    rna-XM_042966335.1    rna-XM_040629912.1    rna-XM_027626377.1    mNeoVis.3.1.1.chr8.g01006.t01    rna-gnl|WGS_ZZZZ|egapxtmp_017571-R1    mNeoVis.evergladensis.2.1.chr9.g01058.t01    mNeoVis.vulgivaga.2.1.chr8.g01058.t01      3204   N0.HOG0003202  OG0000695  n22   rna-XM_034647312.1    rna-XM_038536638.1    rna-XM_022524715.1    rna-XM_023244113.2    rna-XM_036066274.1      rna-XM_032839938.1    rna-XM_045986113.1    rna-XM_032322312.1    rna-XM_059149624.1    rna-XM_059378634.1      mNeoVis.2.1.1.chr8_hap1.g01081.t01    rna-XM_044921506.1    rna-XM_004398626.1    rna-XM_042966147.1    rna-XM_040628210.1, rna-XM_040628322.1      mNeoVis.3.1.1.chr8.g01004.t01    rna-gnl|WGS_ZZZZ|egapxtmp_017911-R1    mNeoVis.evergladensis.2.1.chr9.g01056.t01    mNeoVis.vulgivaga.2.1.chr8.g01056.t01      3205   N0.HOG0003203  OG0000695  n41   rna-XM_034647210.1    rna-XM_038536615.1    rna-XM_022524569.1    rna-XM_023243845.2    rna-XM_036066253.1    rna-XM_039231639.1    rna-XM_032839939.1    rna-XM_045986111.1    rna-XM_032321246.1    rna-XM_059148315.1    rna-XM_059378637.1    rna-XM_045083246.1      rna-XM_021694722.1    rna-XM_004398547.1    rna-XM_042966336.1    rna-XM_040629915.1    rna-XM_027568549.1    mNeoVis.3.1.1.chr7.g01466.t01, mNeoVis.3.1.1.chr8.g01005.t01    rna-gnl|WGS_ZZZZ|egapxtmp_017915-R1    mNeoVis.evergladensis.2.1.chr9.g01057.t01    mNeoVis.vulgivaga.2.1.chr8.g01057.t01      3206    N0.HOG0003204   OG0000696  n2   rna-XM_019799374.2    rna-XM_038556863.1    rna-XM_022503271.1    rna-XM_003982776.6    rna-XM_036115685.1    rna-XM_039236272.1    rna-XM_032874752.1    rna-XM_046020132.1    rna-XM_032304634.1    rna-XM_059171166.1    rna-XM_059396820.1    rna-XM_004767850.3    mNeoVis.2.1.1.chr4_hap1.g00474.t01    rna-XM_021689729.2    rna-XM_004394070.1    rna-XM_007076690.3    rna-XM_008686265.2    rna-XM_035723369.1    mNeoVis.3.1.1.chr6.g00476.t01    rna-gnl|WGS_ZZZZ|egapxtmp_003640-R1    mNeoVis.evergladensis.2.1.chr6.g00482.t01        3207    N0.HOG0003205   OG0000696  n22   rna-XM_034667914.1    rna-NM_001003205.2    rna-XM_022503210.1    rna-XM_023250366.2    rna-XM_036115697.1    rna-XM_039236241.1    rna-XM_032874738.1    rna-XM_046020131.1    rna-XM_032304631.1    rna-XM_059171169.1    rna-XM_059396819.1    rna-XM_004767847.2      rna-XM_021689574.1    rna-XM_004394068.1    rna-XM_042966767.1    rna-XM_040644259.1    rna-XM_027574662.2    mNeoVis.3.1.1.chr6.g00475.t01    rna-gnl|WGS_ZZZZ|egapxtmp_003641-R1      mNeoVis.vulgivaga.2.1.chr6.g00477.t01      3208    N0.HOG0003206   OG0000696  n40   rna-XM_002918938.4    rna-XM_038556862.1    rna-XM_022503325.1    rna-XM_003982775.4    rna-XM_036115695.1    rna-XM_039236227.1    rna-XM_032874765.1    rna-XM_046021419.1    rna-XM_032304633.1    rna-XM_059171164.1    rna-XM_059395736.1    rna-XM_004767851.3    mNeoVis.2.1.1.chr4_hap1.g00475.t01    rna-XM_021689518.1    rna-XM_004394071.1    rna-XM_007076691.3    rna-XM_008686263.2    rna-XM_027573461.1    mNeoVis.3.1.1.chr6.g00477.t01    rna-gnl|WGS_ZZZZ|egapxtmp_003639-R1    mNeoVis.evergladensis.2.1.chr6.g00483.t01    mNeoVis.vulgivaga.2.1.chr6.g00478.t01      3209   N0.HOG0003207  OG0000697  n1   rna-XM_019792841.2    rna-NM_001014376.1, rna-XM_038562808.1    rna-XM_022510744.1    rna-XM_003985913.6    rna-XM_036108327.1    rna-XM_039219579.1    rna-XM_032847314.1    rna-XM_046006485.1    rna-XM_032338143.1    rna-XM_059178962.1    rna-XM_059399808.1    rna-XM_004781007.3    mNeoVis.2.1.1.chr1_hap1.g01329.t01    rna-XM_021697263.1    rna-XM_004415217.1    rna-XM_042985961.1    rna-XM_040644864.1    rna-XM_027603909.1    mNeoVis.3.1.1.chr1.g01278.t01      mNeoVis.evergladensis.2.1.chr1.g01296.t01    mNeoVis.vulgivaga.2.1.chr1.g01297.t01      3210    N0.HOG0003208   OG0000697  n20   rna-XM_002915775.4    rna-XM_038553221.1    rna-XM_022495806.1, rna-XM_022501627.1    rna-XM_003995777.6    rna-XM_036065828.1, rna-XM_036099392.1, rna-XM_036121900.1    rna-XM_039240102.1    rna-XM_032865695.1    rna-XM_046023461.1    rna-XM_032307609.1    rna-XM_059141454.1    rna-XM_059410964.1    rna-XM_004774053.3    mNeoVis.2.1.1.chr13_hap1.g00319.t01, mNeoVis.2.1.1.chr5_hap1.g00825.t01    rna-XM_021691808.1, rna-XM_044914270.1, rna-XM_044918425.1    rna-XM_004396947.1, rna-XM_004401873.1, rna-XM_004411111.1, rna-XM_004412772.1, rna-XM_004416722.1, rna-XM_004416745.1    rna-XM_007094327.3    rna-XM_008698849.2    rna-XM_027616243.2, rna-XM_027619719.2, rna-XM_035724664.1, rna-XM_035725746.1    mNeoVis.3.1.1.chr13.g00263.t01, mNeoVis.3.1.1.chr2.g00736.t01    rna-gnl|WGS_ZZZZ|egapxtmp_029727-R1, rna-gnl|WGS_ZZZZ|egapxtmp_029728-R1    mNeoVis.evergladensis.2.1.chr13.g00310.t01, mNeoVis.evergladensis.2.1.chr2.g00826.t01    mNeoVis.vulgivaga.2.1.chr13.g00307.t01, mNeoVis.vulgivaga.2.1.chr2.g00829.t01      3211    N0.HOG0003209   OG0000698  n1   rna-XM_002922023.4    rna-XM_038536353.1    rna-XM_022513028.1    rna-XM_003992305.5    rna-XM_036068514.1    rna-XM_039216555.1    rna-XM_032844089.1    rna-XM_046017796.1    rna-XM_032358217.1    rna-XM_059179313.1    rna-XM_059418539.1    rna-XM_045077498.1    mNeoVis.2.1.1.chr7_hap1.g01159.t01    rna-XM_021696352.1    rna-XM_004415990.1    rna-XM_007075966.3    rna-XM_008690148.2    rna-XM_027580714.2    mNeoVis.3.1.1.chr7.g01082.t01      mNeoVis.evergladensis.2.1.chr7.g01147.t01    mNeoVis.vulgivaga.2.1.chr7.g01158.t01      3212    N0.HOG0003210   OG0000698  n22   rna-XM_034652786.1, rna-XM_034652919.1                                                3213    N0.HOG0003211   OG0000698  n24   rna-XM_002922024.4    rna-XM_038538798.1    rna-XM_022513112.1      rna-XM_036068517.1    rna-XM_039216550.1    rna-XM_032844070.1      rna-XM_032358213.1    rna-XM_059179289.1    rna-XM_059418514.1    rna-XM_013062429.2    mNeoVis.2.1.1.chr7_hap1.g01158.t01    rna-XM_021696596.1      rna-XM_015534785.2    rna-XM_008690147.2      mNeoVis.3.1.1.chr7.g01081.t01    rna-gnl|WGS_ZZZZ|egapxtmp_013672-R1    mNeoVis.evergladensis.2.1.chr7.g01146.t01    mNeoVis.vulgivaga.2.1.chr7.g01156.t01, mNeoVis.vulgivaga.2.1.chr7.g01157.t01      3214   N0.HOG0003212  OG0000698  n42   rna-XM_034665647.1    rna-XM_038536352.1    rna-XM_022512989.1    rna-XM_003992316.4    rna-XM_036068518.1    rna-XM_039221143.1    rna-XM_032844061.1    rna-XM_046017795.1    rna-XM_032358220.1    rna-XM_059179321.1    rna-XM_059418512.1    rna-XM_004764453.3    mNeoVis.2.1.1.chr7_hap1.g01157.t01    rna-XM_021696411.1      rna-XM_007075965.3    rna-XM_008690145.2    rna-XM_027581505.2    mNeoVis.3.1.1.chr7.g01079.t01      mNeoVis.evergladensis.2.1.chr7.g01144.t01    mNeoVis.vulgivaga.2.1.chr7.g01155.t01      3215   N0.HOG0003213  OG0000699  n1   rna-XM_002914534.4    rna-XM_038550652.1    rna-XM_022518994.1    rna-XM_045062329.1    rna-XM_036102149.1    rna-XM_039246126.1    rna-XM_032836607.1    rna-XM_046013005.1    rna-XM_032346355.1    rna-XM_059186901.1    rna-XM_059402068.1    rna-XM_045083901.1    mNeoVis.2.1.1.chr9_hap1.g01147.t01    rna-XM_021687639.1    rna-XM_004410799.1    rna-XM_042993210.1    rna-XM_040637492.1    rna-XM_027594879.2    mNeoVis.3.1.1.chr10.g01101.t01      mNeoVis.evergladensis.2.1.chr10.g01135.t01    mNeoVis.vulgivaga.2.1.chr10.g01127.t01      3216    N0.HOG0003214   OG0000699  n20   rna-XM_011224635.2    rna-XM_038546272.1    rna-XM_022523240.1    rna-XM_006940701.5    rna-XM_036093451.1    rna-XM_039226420.1    rna-XM_032870348.1    rna-XM_045984694.1    rna-XM_032322170.1    rna-XM_059147901.1    rna-XM_059378853.1    rna-XM_013054279.2    mNeoVis.2.1.1.chr8_hap1.g00028.t01    rna-XM_021680710.2    rna-XM_004404464.1    rna-XM_042966569.1    rna-XM_040639969.1    rna-XM_027626022.1      rna-gnl|WGS_ZZZZ|egapxtmp_017965-R1    mNeoVis.evergladensis.2.1.chr9.g00026.t01    mNeoVis.vulgivaga.2.1.chr8.g00025.t01      3217    N0.HOG0003215   OG0000699  n40   rna-XM_034669713.1    rna-XM_038539620.1    rna-XM_022501638.1    rna-XM_023246563.2    rna-XM_036101695.1    r
