## Supplementary material for "Mink by mink: stitching together signatures of subspecies adaptation through a pangenome of threatened mustelids": Table S14: Table S14.html

  
        A  B  C  D  E  F  G  H  I  J       1   125260621:   mNeoVis.vulgivaga.1.1.chr8.g00505.t01#4    mNeoVis.evergladensis.1.1.chr8.g00506.t01#3    mNeoVis.evergladensis.2.1.chr9.g00510.t01#2    mNeoVis.vulgivaga.2.1.chr8.g00516.t01#1   mNeoVis.2.1.1.chr8_hap1.g00519.t01#5       2   125260622:   mNeoVis.vulgivaga.1.1.chr2.g00147.t01#4    mNeoVis.evergladensis.1.1.chr2.g00145.t01#3    mNeoVis.evergladensis.2.1.chr2.g00146.t01#2    mNeoVis.3.1.1.chr2.g00098.t01#6    mNeoVis.vulgivaga.2.1.chr2.g00143.t01#1   mNeoVis.2.1.1.chr5_hap1.g00148.t01#5       3   125260623:   mNeoVis.3.1.1.chr15.g00559.t01#6    mNeoVis.evergladensis.1.1.chr15.g00008.t01#3    mNeoVis.vulgivaga.2.1.chr15.g00616.t01#1    mNeoVis.vulgivaga.1.1.chr15.g00606.t01#4    mNeoVis.2.1.1.chr15_hap1.g00781.t01#5   mNeoVis.evergladensis.2.1.chr15.g00603.t01#2       4   125260624:   mNeoVis.vulgivaga.1.1.chr8.g00506.t01#4    mNeoVis.evergladensis.1.1.chr8.g00507.t01#3    mNeoVis.vulgivaga.2.1.chr8.g00517.t01#1    mNeoVis.3.1.1.chr8.g00476.t01#6    mNeoVis.2.1.1.chr8_hap1.g00520.t01#5   mNeoVis.evergladensis.2.1.chr9.g00511.t01#2       5   125260625:   mNeoVis.evergladensis.1.1.chr2.g00147.t01#3    mNeoVis.evergladensis.1.1.chr2.g00146.t01#3    mNeoVis.3.1.1.chr2.g00099.t01#6    mNeoVis.2.1.1.chr5_hap1.g00149.t01#5    mNeoVis.vulgivaga.1.1.chr2.g00148.t01#4    mNeoVis.vulgivaga.2.1.chr2.g00144.t01#1   mNeoVis.evergladensis.2.1.chr2.g00147.t01#2     6   125260626:   mNeoVis.evergladensis.2.1.chr9.g00512.t01#2    mNeoVis.2.1.1.chr8_hap1.g00521.t01#5    mNeoVis.vulgivaga.1.1.chr8.g00507.t01#4    mNeoVis.evergladensis.1.1.chr8.g00508.t01#3    mNeoVis.vulgivaga.2.1.chr8.g00518.t01#1   mNeoVis.3.1.1.chr8.g00477.t01#6       7   125260627:   mNeoVis.2.1.1.chr5_hap1.g00150.t01#5    mNeoVis.3.1.1.chr2.g00100.t01#6    mNeoVis.vulgivaga.1.1.chr2.g00149.t01#4    mNeoVis.evergladensis.1.1.chr2.g00148.t01#3    mNeoVis.vulgivaga.2.1.chr2.g00145.t01#1   mNeoVis.evergladensis.2.1.chr2.g00148.t01#2       8   125260628:   mNeoVis.evergladensis.1.1.chr8.g00509.t01#3    mNeoVis.vulgivaga.1.1.chr8.g00508.t01#4    mNeoVis.3.1.1.chr8.g00478.t01#6    mNeoVis.vulgivaga.2.1.chr8.g00519.t01#1    mNeoVis.evergladensis.2.1.chr9.g00513.t01#2   mNeoVis.2.1.1.chr8_hap1.g00522.t01#5       9   125260629:  mNeoVis.vulgivaga.2.1.chr2.g00146.t01#1               10   125260630:   mNeoVis.evergladensis.1.1.chr8.g00510.t01#3    mNeoVis.vulgivaga.1.1.chr8.g00509.t01#4    mNeoVis.3.1.1.chr8.g00479.t01#6    mNeoVis.vulgivaga.2.1.chr8.g00520.t01#1    mNeoVis.2.1.1.chr8_hap1.g00523.t01#5   mNeoVis.evergladensis.2.1.chr9.g00514.t01#2       11   125260631:   mNeoVis.evergladensis.2.1.chr9.g00515.t01#2    mNeoVis.evergladensis.1.1.chr8.g00511.t01#3    mNeoVis.3.1.1.chr8.g00480.t01#6    mNeoVis.2.1.1.chr8_hap1.g00524.t01#5    mNeoVis.vulgivaga.2.1.chr8.g00521.t01#1   mNeoVis.vulgivaga.1.1.chr8.g00510.t01#4       12   125260632:   mNeoVis.evergladensis.1.1.chr7.g00597.t01#3    mNeoVis.3.1.1.chr7.g00570.t01#6    mNeoVis.vulgivaga.2.1.chr7.g00592.t01#1    mNeoVis.2.1.1.chr7_hap1.g00587.t01#5    mNeoVis.vulgivaga.1.1.chr7.g00596.t01#4   mNeoVis.evergladensis.2.1.chr7.g00589.t01#2       13   125260633:   mNeoVis.vulgivaga.2.1.chr2.g00147.t01#1    mNeoVis.evergladensis.1.1.chr2.g00150.t01#3    mNeoVis.vulgivaga.1.1.chr2.g00151.t01#4    mNeoVis.2.1.1.chr5_hap1.g00154.t01#5    mNeoVis.evergladensis.2.1.chr2.g00150.t01#2   mNeoVis.3.1.1.chr2.g00102.t01#6       14   125260634:   mNeoVis.vulgivaga.2.1.chr2.g00148.t01#1    mNeoVis.evergladensis.1.1.chr2.g00151.t01#3    mNeoVis.evergladensis.2.1.chr2.g00151.t01#2    mNeoVis.3.1.1.chr2.g00103.t01#6    mNeoVis.vulgivaga.1.1.chr2.g00152.t01#4   mNeoVis.2.1.1.chr5_hap1.g00155.t01#5       15   125260635:   mNeoVis.vulgivaga.2.1.chr8.g00522.t01#1    mNeoVis.3.1.1.chr8.g00481.t01#6    mNeoVis.evergladensis.1.1.chr8.g00512.t01#3    mNeoVis.2.1.1.chr8_hap1.g00525.t01#5    mNeoVis.vulgivaga.1.1.chr8.g00511.t01#4   mNeoVis.evergladensis.2.1.chr9.g00516.t01#2       16   125260636:   mNeoVis.evergladensis.1.1.chr2.g00152.t01#3    mNeoVis.vulgivaga.2.1.chr2.g00150.t01#1    mNeoVis.2.1.1.chr5_hap1.g00156.t01#5    mNeoVis.vulgivaga.1.1.chr2.g00153.t01#4    mNeoVis.evergladensis.2.1.chr2.g00152.t01#2    mNeoVis.3.1.1.chr2.g00104.t01#6   mNeoVis.vulgivaga.2.1.chr2.g00149.t01#1     17   125260637:   mNeoVis.vulgivaga.2.1.chr8.g00523.t01#1    mNeoVis.3.1.1.chr8.g00482.t01#6    mNeoVis.evergladensis.1.1.chr8.g00513.t01#3    mNeoVis.2.1.1.chr8_hap1.g00526.t01#5    mNeoVis.evergladensis.2.1.chr9.g00517.t01#2   mNeoVis.vulgivaga.1.1.chr8.g00512.t01#4       18   125260638:   mNeoVis.3.1.1.chr8.g00483.t01#6    mNeoVis.2.1.1.chr8_hap1.g00527.t01#5    mNeoVis.evergladensis.2.1.chr9.g00518.t01#2    mNeoVis.vulgivaga.1.1.chr8.g00513.t01#4    mNeoVis.vulgivaga.2.1.chr8.g00524.t01#1   mNeoVis.evergladensis.1.1.chr8.g00514.t01#3       19   125260639:   mNeoVis.evergladensis.2.1.chr2.g00153.t01#2    mNeoVis.2.1.1.chr5_hap1.g00157.t01#5    mNeoVis.vulgivaga.1.1.chr2.g00154.t01#4    mNeoVis.evergladensis.1.1.chr2.g00153.t01#3   mNeoVis.vulgivaga.2.1.chr2.g00151.t01#1       20   125260640:   mNeoVis.2.1.1.chr9_hap1.g00463.t01#5    mNeoVis.vulgivaga.1.1.chr10.g00437.t01#4    mNeoVis.3.1.1.chr10.g00439.t01#6    mNeoVis.evergladensis.1.1.chr10.g00462.t01#3    mNeoVis.evergladensis.2.1.chr10.g00457.t01#2   mNeoVis.vulgivaga.2.1.chr10.g00446.t01#1       21   125260641:   mNeoVis.vulgivaga.1.1.chr8.g00514.t01#4    mNeoVis.evergladensis.1.1.chr8.g00515.t01#3   mNeoVis.vulgivaga.2.1.chr8.g00525.t01#1           22   125260642:   mNeoVis.evergladensis.2.1.chr2.g00154.t01#2    mNeoVis.vulgivaga.1.1.chr2.g00155.t01#4    mNeoVis.2.1.1.chr5_hap1.g00158.t01#5    mNeoVis.evergladensis.1.1.chr2.g00154.t01#3    mNeoVis.3.1.1.chr2.g00106.t01#6   mNeoVis.vulgivaga.2.1.chr2.g00152.t01#1       23   125260643:  mNeoVis.vulgivaga.2.1.chr8.g00526.t01#1               24   125260644:   mNeoVis.3.1.1.chr8.g00485.t01#6    mNeoVis.evergladensis.1.1.chr8.g00516.t01#3    mNeoVis.2.1.1.chr8_hap1.g00529.t01#5    mNeoVis.vulgivaga.1.1.chr8.g00515.t01#4    mNeoVis.vulgivaga.2.1.chr8.g00527.t01#1   mNeoVis.evergladensis.2.1.chr9.g00520.t01#2       25   125260645:   mNeoVis.vulgivaga.2.1.chr8.g00528.t01#1    mNeoVis.evergladensis.2.1.chr9.g00521.t01#2    mNeoVis.2.1.1.chr8_hap1.g00530.t01#5    mNeoVis.3.1.1.chr8.g00486.t01#6    mNeoVis.vulgivaga.1.1.chr8.g00516.t01#4   mNeoVis.evergladensis.1.1.chr8.g00517.t01#3       26   125260646:   mNeoVis.vulgivaga.1.1.chr2.g00156.t01#4    mNeoVis.evergladensis.1.1.chr2.g00155.t01#3    mNeoVis.2.1.1.chr5_hap1.g00159.t01#5    mNeoVis.evergladensis.2.1.chr2.g00155.t01#2    mNeoVis.3.1.1.chr2.g00107.t01#6   mNeoVis.vulgivaga.2.1.chr2.g00153.t01#1       27   125260647:   mNeoVis.evergladensis.2.1.chr9.g00522.t01#2    mNeoVis.vulgivaga.2.1.chr8.g00529.t01#1    mNeoVis.vulgivaga.1.1.chr8.g00517.t01#4    mNeoVis.3.1.1.chr8.g00487.t01#6    mNeoVis.evergladensis.1.1.chr8.g00518.t01#3   mNeoVis.2.1.1.chr8_hap1.g00531.t01#5       28   125260648:   mNeoVis.vulgivaga.2.1.chr8.g00530.t01#1    mNeoVis.evergladensis.2.1.chr9.g00523.t01#2    mNeoVis.vulgivaga.1.1.chr8.g00518.t01#4    mNeoVis.3.1.1.chr8.g00488.t01#6    mNeoVis.2.1.1.chr8_hap1.g00532.t01#5   mNeoVis.evergladensis.1.1.chr8.g00519.t01#3       29   125260649:   mNeoVis.2.1.1.chr15_hap1.g00782.t01#5    mNeoVis.evergladensis.2.1.chr15.g00604.t01#2    mNeoVis.evergladensis.1.1.chr15.g00007.t01#3    mNeoVis.vulgivaga.2.1.chr15.g00617.t01#1    mNeoVis.vulgivaga.1.1.chr15.g00607.t01#4   mNeoVis.3.1.1.chr15.g00560.t01#6       30   125260650:   mNeoVis.2.1.1.chr8_hap1.g00717.t01#5    mNeoVis.vulgivaga.2.1.chr8.g00711.t01#1    mNeoVis.evergladensis.2.1.chr9.g00705.t01#2    mNeoVis.vulgivaga.1.1.chr8.g00695.t01#4    mNeoVis.evergladensis.1.1.chr8.g00698.t01#3   mNeoVis.3.1.1.chr8.g00667.t01#6       31   125260651:   mNeoVis.2.1.1.chr5_hap1.g00059.t01#5    mNeoVis.evergladensis.1.1.chr2.g00060.t01#3    mNeoVis.vulgivaga.1.1.chr2.g00060.t01#4    mNeoVis.vulgivaga.2.1.chr2.g00059.t01#1   mNeoVis.evergladensis.2.1.chr2.g00059.t01#2       32   125260652:   mNeoVis.evergladensis.2.1.chr9.g00524.t01#2    mNeoVis.evergladensis.1.1.chr8.g00520.t01#3    mNeoVis.vulgivaga.2.1.chr8.g00531.t01#1    mNeoVis.vulgivaga.1.1.chr8.g00519.t01#4    mNeoVis.3.1.1.chr8.g00489.t01#6   mNeoVis.2.1.1.chr8_hap1.g00533.t01#5       33   125260653:   mNeoVis.vulgivaga.2.1.chr2.g00060.t01#1    mNeoVis.evergladensis.1.1.chr2.g00061.t01#3    mNeoVis.2.1.1.chr5_hap1.g00060.t01#5    mNeoVis.evergladensis.2.1.chr2.g00060.t01#2   mNeoVis.vulgivaga.1.1.chr2.g00061.t01#4       34   125260654:   mNeoVis.vulgivaga.1.1.chr2.g00062.t01#4    mNeoVis.evergladensis.1.1.chr2.g00062.t01#3    mNeoVis.vulgivaga.2.1.chr2.g00061.t01#1   mNeoVis.2.1.1.chr5_hap1.g00061.t01#5         35   125260655:   mNeoVis.2.1.1.chr8_hap1.g00534.t01#5    mNeoVis.vulgivaga.1.1.chr8.g00520.t01#4    mNeoVis.vulgivaga.2.1.chr8.g00532.t01#1    mNeoVis.evergladensis.2.1.chr9.g00525.t01#2   mNeoVis.evergladensis.1.1.chr8.g00521.t01#3       36   125260656:   mNeoVis.vulgivaga.1.1.chr8.g00521.t01#4    mNeoVis.2.1.1.chr8_hap1.g00535.t01#5    mNeoVis.evergladensis.1.1.chr8.g00522.t01#3    mNeoVis.3.1.1.chr8.g00490.t01#6    mNeoVis.evergladensis.2.1.chr9.g00526.t01#2   mNeoVis.vulgivaga.2.1.chr8.g00533.t01#1       37   125260657:   mNeoVis.vulgivaga.2.1.chr2.g00062.t01#1    mNeoVis.evergladensis.1.1.chr2.g00063.t01#3    mNeoVis.evergladensis.2.1.chr2.g00062.t01#2    mNeoVis.2.1.1.chr5_hap1.g00062.t01#5   mNeoVis.vulgivaga.1.1.chr2.g00063.t01#4       38   125260658:   mNeoVis.vulgivaga.2.1.chr8.g00534.t01#1    mNeoVis.2.1.1.chr8_hap1.g00536.t01#5    mNeoVis.vulgivaga.1.1.chr8.g00522.t01#4    mNeoVis.evergladensis.1.1.chr8.g00523.t01#3    mNeoVis.3.1.1.chr8.g00491.t01#6   mNeoVis.evergladensis.2.1.chr9.g00527.t01#2       39   125260659:   mNeoVis.vulgivaga.1.1.chr8.g00523.t01#4    mNeoVis.evergladensis.1.1.chr8.g00524.t01#3    mNeoVis.evergladensis.2.1.chr9.g00528.t01#2    mNeoVis.2.1.1.chr8_hap1.g00537.t01#5    mNeoVis.3.1.1.chr8.g00492.t01#6   mNeoVis.vulgivaga.2.1.chr8.g00535.t01#1       40   125260660:   mNeoVis.vulgivaga.2.1.chr8.g00536.t01#1    mNeoVis.evergladensis.1.1.chr8.g00525.t01#3    mNeoVis.vulgivaga.1.1.chr8.g00524.t01#4    mNeoVis.2.1.1.chr8_hap1.g00538.t01#5    mNeoVis.3.1.1.chr8.g00493.t01#6   mNeoVis.evergladensis.2.1.chr9.g00529.t01#2       41   125260661:   mNeoVis.vulgivaga.2.1.chr2.g00063.t01#1    mNeoVis.evergladensis.1.1.chr2.g00064.t01#3    mNeoVis.evergladensis.2.1.chr2.g00063.t01#2    mNeoVis.2.1.1.chr5_hap1.g00063.t01#5   mNeoVis.vulgivaga.1.1.chr2.g00064.t01#4       42   125260662:   mNeoVis.evergladensis.1.1.chr8.g00526.t01#3    mNeoVis.evergladensis.2.1.chr9.g00530.t01#2    mNeoVis.vulgivaga.1.1.chr8.g00525.t01#4    mNeoVis.3.1.1.chr8.g00494.t01#6    mNeoVis.vulgivaga.2.1.chr8.g00537.t01#1   mNeoVis.2.1.1.chr8_hap1.g00539.t01#5       43   125260663:   mNeoVis.2.1.1.chr8_hap1.g00540.t01#5    mNeoVis.vulgivaga.2.1.chr8.g00538.t01#1    mNeoVis.evergladensis.2.1.chr9.g00531.t01#2    mNeoVis.evergladensis.1.1.chr8.g00527.t01#3   mNeoVis.vulgivaga.1.1.chr8.g00526.t01#4       44   125260664:   mNeoVis.3.1.1.chr8.g00495.t01#6    mNeoVis.vulgivaga.1.1.chr8.g00527.t01#4    mNeoVis.2.1.1.chr8_hap1.g00541.t01#5    mNeoVis.vulgivaga.2.1.chr8.g00539.t01#1   mNeoVis.evergladensis.2.1.chr9.g00532.t01#2       45   125260665:   mNeoVis.3.1.1.chr8.g00497.t01#6    mNeoVis.vulgivaga.2.1.chr8.g00540.t01#1    mNeoVis.evergladensis.2.1.chr9.g00533.t01#2    mNeoVis.2.1.1.chr8_hap1.g00543.t01#5    mNeoVis.evergladensis.1.1.chr8.g00528.t01#3   mNeoVis.vulgivaga.1.1.chr8.g00528.t01#4       46   125260666:   mNeoVis.2.1.1.chr8_hap1.g00544.t01#5    mNeoVis.vulgivaga.1.1.chr8.g00529.t01#4    mNeoVis.evergladensis.1.1.chr8.g00529.t01#3    mNeoVis.3.1.1.chr8.g00498.t01#6    mNeoVis.vulgivaga.2.1.chr8.g00541.t01#1   mNeoVis.evergladensis.2.1.chr9.g00534.t01#2       47   125260667:   mNeoVis.2.1.1.chr5_hap1.g00064.t01#5    mNeoVis.vulgivaga.1.1.chr2.g00065.t01#4    mNeoVis.evergladensis.2.1.chr2.g00064.t01#2    mNeoVis.vulgivaga.2.1.chr2.g00064.t01#1   mNeoVis.evergladensis.1.1.chr2.g00065.t01#3       48   125260668:   mNeoVis.evergladensis.1.1.chr2.g00066.t01#3   mNeoVis.vulgivaga.2.1.chr2.g00065.t01#1             49   125260669:   mNeoVis.vulgivaga.1.1.chr8.g00530.t01#4    mNeoVis.3.1.1.chr8.g00499.t01#6    mNeoVis.evergladensis.1.1.chr8.g00530.t01#3    mNeoVis.2.1.1.chr8_hap1.g00545.t01#5    mNeoVis.vulgivaga.2.1.chr8.g00542.t01#1   mNeoVis.evergladensis.2.1.chr9.g00535.t01#2       50   125260670:   mNeoVis.2.1.1.chr5_hap1.g00066.t01#5    mNeoVis.evergladensis.1.1.chr2.g00067.t01#3    mNeoVis.vulgivaga.2.1.chr2.g00066.t01#1    mNeoVis.evergladensis.2.1.chr2.g00066.t01#2    mNeoVis.vulgivaga.1.1.chr2.g00067.t01#4   mNeoVis.3.1.1.chr2.g00035.t01#6       51   125260671:   mNeoVis.2.1.1.chr2_hap1.g01197.t01#5    mNeoVis.evergladensis.2.1.chr5.g01139.t01#2    mNeoVis.evergladensis.1.1.chr5.g01134.t01#3    mNeoVis.3.1.1.chr5.g01113.t01#6    mNeoVis.evergladensis.1.1.chr10.g01033.t01#3    mNeoVis.evergladensis.2.1.chr10.g01033.t01#2    mNeoVis.vulgivaga.1.1.chr10.g01012.t01#4    mNeoVis.vulgivaga.2.1.chr10.g01024.t01#1    mNeoVis.vulgivaga.2.1.chr5.g01126.t01#1      52   125260672:   mNeoVis.evergladensis.2.1.chr2.g00067.t01#2    mNeoVis.3.1.1.chr2.g00036.t01#6    mNeoVis.vulgivaga.2.1.chr2.g00067.t01#1    mNeoVis.vulgivaga.1.1.chr2.g00068.t01#4   mNeoVis.2.1.1.chr5_hap1.g00067.t01#5       53   125260673:   mNeoVis.vulgivaga.2.1.chr8.g00543.t01#1    mNeoVis.2.1.1.chr8_hap1.g00546.t01#5    mNeoVis.evergladensis.1.1.chr8.g00531.t01#3    mNeoVis.vulgivaga.1.1.chr8.g00531.t01#4    mNeoVis.3.1.1.chr8.g00500.t01#6   mNeoVis.evergladensis.2.1.chr9.g00536.t01#2       54   125260674:   mNeoVis.vulgivaga.2.1.chr2.g00068.t01#1    mNeoVis.evergladensis.1.1.chr2.g00069.t01#3    mNeoVis.2.1.1.chr5_hap1.g00068.t01#5    mNeoVis.evergladensis.2.1.chr2.g00068.t01#2    mNeoVis.vulgivaga.1.1.chr2.g00069.t01#4   mNeoVis.3.1.1.chr2.g00867.t01#6       55   125260675:   mNeoVis.vulgivaga.2.1.chr13.g00788.t01#1    mNeoVis.vulgivaga.1.1.chr13.g00781.t01#4    mNeoVis.3.1.1.chr13.g00749.t01#6    mNeoVis.evergladensis.1.1.chr13.g00785.t01#3    mNeoVis.evergladensis.2.1.chr13.g00791.t01#2   mNeoVis.2.1.1.chr13_hap1.g00829.t01#5       56   125260676:   mNeoVis.evergladensis.1.1.chr2.g00070.t01#3    mNeoVis.evergladensis.2.1.chr2.g00069.t01#2    mNeoVis.2.1.1.chr5_hap1.g00069.t01#5    mNeoVis.vulgivaga.1.1.chr2.g00070.t01#4   mNeoVis.vulgivaga.2.1.chr2.g00069.t01#1       57   125260677:   mNeoVis.evergladensis.1.1.chr8.g00532.t01#3    mNeoVis.evergladensis.2.1.chr9.g00537.t01#2    mNeoVis.3.1.1.chr8.g00501.t01#6    mNeoVis.vulgivaga.2.1.chr8.g00544.t01#1    mNeoVis.2.1.1.chr8_hap1.g00547.t01#5   mNeoVis.vulgivaga.1.1.chr8.g00532.t01#4       58   125260678:   mNeoVis.2.1.1.chr5_hap1.g00070.t01#5    mNeoVis.3.1.1.chr2.g00037.t01#6    mNeoVis.vulgivaga.1.1.chr2.g00071.t01#4    mNeoVis.evergladensis.1.1.chr2.g00071.t01#3    mNeoVis.vulgivaga.2.1.chr2.g00070.t01#1   mNeoVis.evergladensis.2.1.chr2.g00070.t01#2       59   125260679:   mNeoVis.evergladensis.1.1.chr2.g00074.t01#3    mNeoVis.vulgivaga.2.1.chr2.g00071.t01#1    mNeoVis.evergladensis.2.1.chr2.g00073.t01#2    mNeoVis.2.1.1.chr5_hap1.g00072.t01#5    mNeoVis.vulgivaga.1.1.chr2.g00073.t01#4   mNeoVis.3.1.1.chr2.g00038.t01#6       60   125260680:   mNeoVis.evergladensis.1.1.chr8.g00533.t01#3    mNeoVis.3.1.1.chr8.g00502.t01#6    mNeoVis.evergladensis.2.1.chr9.g00538.t01#2    mNeoVis.2.1.1.chr8_hap1.g00548.t01#5    mNeoVis.vulgivaga.2.1.chr8.g00545.t01#1   mNeoVis.vulgivaga.1.1.chr8.g00533.t01#4       61   125260681:   mNeoVis.evergladensis.1.1.chr8.g00534.t01#3    mNeoVis.2.1.1.chr8_hap1.g00549.t01#5    mNeoVis.evergladensis.2.1.chr9.g00539.t01#2    mNeoVis.vulgivaga.2.1.chr8.g00546.t01#1    mNeoVis.vulgivaga.1.1.chr8.g00534.t01#4   mNeoVis.3.1.1.chr8.g00503.t01#6       62   125260682:   mNeoVis.evergladensis.1.1.chr7.g00598.t01#3    mNeoVis.vulgivaga.2.1.chr7.g00593.t01#1    mNeoVis.vulgivaga.1.1.chr7.g00597.t01#4    mNeoVis.2.1.1.chr7_hap1.g00588.t01#5    mNeoVis.evergladensis.2.1.chr7.g00590.t01#2   mNeoVis.3.1.1.chr7.g00571.t01#6       63   125260683:   mNeoVis.evergladensis.2.1.chr9.g00540.t01#2    mNeoVis.evergladensis.1.1.chr8.g00535.t01#3    mNeoVis.vulgivaga.2.1.chr8.g00547.t01#1    mNeoVis.3.1.1.chr8.g00504.t01#6    mNeoVis.2.1.1.chr8_hap1.g00550.t01#5   mNeoVis.vulgivaga.1.1.chr8.g00535.t01#4       64   125260684:   mNeoVis.evergladensis.1.1.chr1.g00550.t01#3    mNeoVis.3.1.1.chr1.g00557.t01#6    mNeoVis.evergladensis.2.1.chr1.g00550.t01#2    mNeoVis.2.1.1.chr1_hap1.g00543.t01#5    mNeoVis.vulgivaga.2.1.chr1.g00559.t01#1   mNeoVis.vulgivaga.1.1.chr1.g00554.t01#4       65   125260685:   mNeoVis.2.1.1.chr5_hap1.g00073.t01#5    mNeoVis.vulgivaga.2.1.chr2.g00072.t01#1    mNeoVis.evergladensis.2.1.chr2.g00074.t01#2    mNeoVis.3.1.1.chr2.g00039.t01#6    mNeoVis.vulgivaga.1.1.chr2.g00074.t01#4   mNeoVis.evergladensis.1.1.chr2.g00075.t01#3       66   125260686:   mNeoVis.evergladensis.1.1.chr8.g00536.t01#3    mNeoVis.3.1.1.chr8.g00505.t01#6    mNeoVis.2.1.1.chr8_hap1.g00551.t01#5    mNeoVis.evergladensis.2.1.chr9.g00541.t01#2    mNeoVis.vulgivaga.1.1.chr8.g00536.t01#4   mNeoVis.vulgivaga.2.1.chr8.g00548.t01#1       67   125260687:   mNeoVis.vulgivaga.1.1.chr2.g00075.t01#4    mNeoVis.vulgivaga.2.1.chr2.g00073.t01#1   mNeoVis.2.1.1.chr5_hap1.g00075.t01#5           68   125260688:   mNeoVis.2.1.1.chr5_hap1.g00076.t01#5    mNeoVis.evergladensis.1.1.chr2.g00076.t01#3    mNeoVis.evergladensis.2.1.chr2.g00076.t01#2    mNeoVis.vulgivaga.1.1.chr2.g00076.t01#4   mNeoVis.vulgivaga.2.1.chr2.g00074.t01#1       69   125260689:   mNeoVis.vulgivaga.1.1.chr8.g00537.t01#4    mNeoVis.vulgivaga.2.1.chr8.g00549.t01#1    mNeoVis.3.1.1.chr8.g00506.t01#6    mNeoVis.evergladensis.1.1.chr8.g00537.t01#3    mNeoVis.2.1.1.chr8_hap1.g00552.t01#5   mNeoVis.evergladensis.2.1.chr9.g00542.t01#2       70   125260690:   mNeoVis.evergladensis.2.1.chr9.g00543.t01#2    mNeoVis.vulgivaga.2.1.chr8.g00550.t01#1    mNeoVis.3.1.1.chr8.g00507.t01#6    mNeoVis.evergladensis.1.1.chr8.g00538.t01#3    mNeoVis.2.1.1.chr8_hap1.g00553.t01#5   mNeoVis.vulgivaga.1.1.chr8.g00538.t01#4       71   125260691:   mNeoVis.vulgivaga.2.1.chr2.g00075.t01#1    mNeoVis.evergladensis.2.1.chr2.g00078.t01#2    mNeoVis.2.1.1.chr5_hap1.g00081.t01#5   mNeoVis.evergladensis.1.1.chr2.g00077.t01#3       72   125260692:  mNeoVis.vulgivaga.2.1.chr2.g00076.t01#1               73   125260693:   mNeoVis.2.1.1.chr5_hap1.g00083.t01#5    mNeoVis.evergladensis.2.1.chr2.g00080.t01#2    mNeoVis.3.1.1.chr2.g00040.t01#6    mNeoVis.evergladensis.1.1.chr2.g00079.t01#3    mNeoVis.vulgivaga.2.1.chr2.g00077.t01#1   mNeoVis.vulgivaga.1.1.chr2.g00079.t01#4       74   125260694:   mNeoVis.evergladensis.2.1.chr2.g00081.t01#2    mNeoVis.2.1.1.chr5_hap1.g00084.t01#5    mNeoVis.vulgivaga.2.1.chr2.g00078.t01#1    mNeoVis.evergladensis.1.1.chr2.g00080.t01#3   mNeoVis.vulgivaga.1.1.chr2.g00080.t01#4       75   125260695:   mNeoVis.evergladensis.2.1.chr2.g00071.t01#2   mNeoVis.3.1.1.chr2.g00041.t01#6             76   125260696:   mNeoVis.evergladensis.2.1.chr2.g00082.t01#2    mNeoVis.vulgivaga.1.1.chr2.g00081.t01#4    mNeoVis.vulgivaga.2.1.chr2.g00079.t01#1    mNeoVis.3.1.1.chr2.g00042.t01#6    mNeoVis.evergladensis.1.1.chr2.g00081.t01#3   mNeoVis.2.1.1.chr5_hap1.g00085.t01#5       77   125260697:   mNeoVis.vulgivaga.2.1.chr8.g00551.t01#1    mNeoVis.2.1.1.chr8_hap1.g00554.t01#5    mNeoVis.evergladensis.1.1.chr8.g00539.t01#3    mNeoVis.evergladensis.2.1.chr9.g00544.t01#2    mNeoVis.vulgivaga.1.1.chr8.g00539.t01#4   mNeoVis.3.1.1.chr8.g00508.t01#6       78   125260698:   mNeoVis.3.1.1.chr8.g00509.t01#6    mNeoVis.evergladensis.1.1.chr8.g00540.t01#3    mNeoVis.evergladensis.2.1.chr9.g00545.t01#2    mNeoVis.2.1.1.chr8_hap1.g00555.t01#5    mNeoVis.vulgivaga.1.1.chr8.g00540.t01#4   mNeoVis.vulgivaga.2.1.chr8.g00552.t01#1       79   125260699:   mNeoVis.3.1.1.chr8.g00510.t01#6    mNeoVis.evergladensis.1.1.chr8.g00541.t01#3    mNeoVis.2.1.1.chr8_hap1.g00556.t01#5    mNeoVis.vulgivaga.2.1.chr8.g00553.t01#1    mNeoVis.evergladensis.2.1.chr9.g00546.t01#2   mNeoVis.vulgivaga.1.1.chr8.g00541.t01#4       80   125260700:   mNeoVis.evergladensis.1.1.chr8.g00542.t01#3    mNeoVis.3.1.1.chr8.g00511.t01#6    mNeoVis.evergladensis.2.1.chr9.g00547.t01#2    mNeoVis.2.1.1.chr8_hap1.g00557.t01#5    mNeoVis.vulgivaga.1.1.chr8.g00542.t01#4   mNeoVis.vulgivaga.2.1.chr8.g00554.t01#1       81   125260701:   mNeoVis.vulgivaga.1.1.chr8.g00543.t01#4    mNeoVis.evergladensis.1.1.chr8.g00543.t01#3    mNeoVis.3.1.1.chr8.g00513.t01#6    mNeoVis.evergladensis.2.1.chr9.g00548.t01#2    mNeoVis.2.1.1.chr8_hap1.g00558.t01#5   mNeoVis.vulgivaga.2.1.chr8.g00555.t01#1       82   125260702:   mNeoVis.evergladensis.2.1.chr9.g00549.t01#2    mNeoVis.evergladensis.1.1.chr8.g00544.t01#3    mNeoVis.3.1.1.chr8.g00514.t01#6    mNeoVis.vulgivaga.2.1.chr8.g00556.t01#1    mNeoVis.2.1.1.chr8_hap1.g00559.t01#5   mNeoVis.vulgivaga.1.1.chr8.g00544.t01#4       83   125260703:   mNeoVis.2.1.1.chr3_hap1.g00787.t01#5    mNeoVis.evergladensis.1.1.chr3.g00788.t01#3    mNeoVis.3.1.1.chr3.g00773.t01#6    mNeoVis.vulgivaga.1.1.chr3.g00781.t01#4    mNeoVis.evergladensis.2.1.chr3.g00784.t01#2   mNeoVis.vulgivaga.2.1.chr3.g00792.t01#1       84   125260704:   mNeoVis.vulgivaga.1.1.chr15.g00609.t01#4    mNeoVis.3.1.1.chr15.g00561.t01#6    mNeoVis.evergladensis.1.1.chr15.g00006.t01#3    mNeoVis.2.1.1.chr15_hap1.g00784.t01#5    mNeoVis.vulgivaga.2.1.chr15.g00618.t01#1   mNeoVis.evergladensis.2.1.chr15.g00605.t01#2       85   125260705:   mNeoVis.vulgivaga.1.1.chr2.g00082.t01#4    mNeoVis.vulgivaga.2.1.chr2.g00080.t01#1    mNeoVis.evergladensis.1.1.chr2.g00082.t01#3    mNeoVis.evergladensis.2.1.chr2.g00083.t01#2   mNeoVis.2.1.1.chr5_hap1.g00086.t01#5       86   125260706:   mNeoVis.2.1.1.chr5_hap1.g00087.t01#5    mNeoVis.vulgivaga.2.1.chr2.g00081.t01#1    mNeoVis.vulgivaga.1.1.chr2.g00083.t01#4    mNeoVis.evergladensis.1.1.chr2.g00083.t01#3   mNeoVis.evergladensis.2.1.chr2.g00084.t01#2       87   125260707:   mNeoVis.evergladensis.2.1.chr2.g00085.t01#2    mNeoVis.2.1.1.chr5_hap1.g00088.t01#5    mNeoVis.vulgivaga.2.1.chr2.g00082.t01#1    mNeoVis.evergladensis.1.1.chr2.g00084.t01#3    mNeoVis.vulgivaga.1.1.chr2.g00084.t01#4   mNeoVis.3.1.1.chr2.g00043.t01#6       88   125260708:   mNeoVis.2.1.1.chr8_hap1.g00560.t01#5    mNeoVis.3.1.1.chr8.g00515.t01#6    mNeoVis.vulgivaga.1.1.chr8.g00545.t01#4    mNeoVis.evergladensis.2.1.chr9.g00550.t01#2    mNeoVis.evergladensis.1.1.chr8.g00545.t01#3   mNeoVis.vulgivaga.2.1.chr8.g00557.t01#1       89   125260709:   mNeoVis.evergladensis.2.1.chr9.g00551.t01#2    mNeoVis.3.1.1.chr8.g00516.t01#6    mNeoVis.evergladensis.1.1.chr8.g00546.t01#3    mNeoVis.vulgivaga.1.1.chr8.g00546.t01#4    mNeoVis.vulgivaga.2.1.chr8.g00558.t01#1   mNeoVis.2.1.1.chr8_hap1.g00561.t01#5       90   125260710:   mNeoVis.3.1.1.chr8.g00517.t01#6    mNeoVis.evergladensis.1.1.chr8.g00547.t01#3    mNeoVis.vulgivaga.1.1.chr8.g00547.t01#4    mNeoVis.evergladensis.2.1.chr9.g00552.t01#2    mNeoVis.vulgivaga.2.1.chr8.g00559.t01#1   mNeoVis.2.1.1.chr8_hap1.g00562.t01#5       91   125260711:   mNeoVis.2.1.1.chr13_hap1.g00634.t01#5    mNeoVis.evergladensis.1.1.chr13.g00607.t01#3    mNeoVis.evergladensis.2.1.chr13.g00609.t01#2    mNeoVis.3.1.1.chr13.g00560.t01#6    mNeoVis.vulgivaga.2.1.chr13.g00611.t01#1   mNeoVis.vulgivaga.1.1.chr13.g00594.t01#4       92   125260712:   mNeoVis.3.1.1.chr5.g01151.t01#6    mNeoVis.2.1.1.chr2_hap1.g01232.t01#5    mNeoVis.vulgivaga.2.1.chr5.g01166.t01#1    mNeoVis.vulgivaga.1.1.chr5.g01137.t01#4    mNeoVis.evergladensis.2.1.chr5.g01176.t01#2   mNeoVis.evergladensis.1.1.chr5.g01171.t01#3       93   125260713:   mNeoVis.vulgivaga.1.1.chr2.g00086.t01#4    mNeoVis.evergladensis.2.1.chr2.g00086.t01#2    mNeoVis.vulgivaga.2.1.chr2.g00083.t01#1    mNeoVis.evergladensis.1.1.chr2.g00085.t01#3   mNeoVis.2.1.1.chr5_hap1.g00089.t01#5       94   125260714:   mNeoVis.vulgivaga.2.1.chr2.g00345.t01#1    mNeoVis.vulgivaga.1.1.chr2.g00344.t01#4    mNeoVis.evergladensis.2.1.chr2.g00350.t01#2    mNeoVis.2.1.1.chr5_hap1.g00349.t01#5   mNeoVis.evergladensis.1.1.chr2.g00347.t01#3       95   125260715:   mNeoVis.evergladensis.2.1.chr2.g00349.t01#2    mNeoVis.vulgivaga.1.1.chr2.g00085.t01#4    mNeoVis.3.1.1.chr2.g00044.t01#6   mNeoVis.3.1.1.chr2.g00285.t01#6         96   125260716:   mNeoVis.evergladensis.1.1.chr8.g00548.t01#3    mNeoVis.vulgivaga.2.1.chr8.g00560.t01#1    mNeoVis.vulgivaga.1.1.chr8.g00548.t01#4    mNeoVis.evergladensis.2.1.chr9.g00553.t01#2    mNeoVis.3.1.1.chr8.g00518.t01#6   mNeoVis.2.1.1.chr8_hap1.g00563.t01#5       97   125260717:   mNeoVis.evergladensis.2.1.chr7.g00591.t01#2    mNeoVis.evergladensis.1.1.chr7.g00599.t01#3    mNeoVis.vulgivaga.1.1.chr7.g00598.t01#4    mNeoVis.2.1.1.chr7_hap1.g00589.t01#5   mNeoVis.vulgivaga.2.1.chr7.g00594.t01#1       98   125260718:   mNeoVis.evergladensis.2.1.chr2.g00087.t01#2    mNeoVis.vulgivaga.1.1.chr2.g00087.t01#4    mNeoVis.2.1.1.chr5_hap1.g00090.t01#5    mNeoVis.vulgivaga.2.1.chr2.g00084.t01#1    mNeoVis.3.1.1.chr2.g00045.t01#6   mNeoVis.evergladensis.1.1.chr2.g00087.t01#3       99   125260719:  mNeoVis.vulgivaga.2.1.chr8.g00561.t01#1               100   125260720:   mNeoVis.evergladensis.2.1.chr2.g00088.t01#2    mNeoVis.vulgivaga.1.1.chr2.g00088.t01#4    mNeoVis.3.1.1.chr2.g00046.t01#6    mNeoVis.evergladensis.1.1.chr2.g00088.t01#3    mNeoVis.vulgivaga.2.1.chr2.g00085.t01#1   mNeoVis.2.1.1.chr5_hap1.g00091.t01#5       101   125260721:   mNeoVis.vulgivaga.1.1.chr8.g00550.t01#4    mNeoVis.evergladensis.2.1.chr9.g00555.t01#2    mNeoVis.evergladensis.1.1.chr8.g00550.t01#3    mNeoVis.2.1.1.chr8_hap1.g00565.t01#5   mNeoVis.vulgivaga.2.1.chr8.g00562.t01#1       102   125260722:   mNeoVis.vulgivaga.1.1.chr2.g00089.t01#4    mNeoVis.vulgivaga.2.1.chr2.g00086.t01#1    mNeoVis.2.1.1.chr5_hap1.g00092.t01#5    mNeoVis.3.1.1.chr2.g00047.t01#6    mNeoVis.evergladensis.2.1.chr2.g00089.t01#2   mNeoVis.evergladensis.1.1.chr2.g00089.t01#3       103   125260723:   mNeoVis.evergladensis.1.1.chr2.g00090.t01#3    mNeoVis.vulgivaga.1.1.chr2.g00090.t01#4    mNeoVis.vulgivaga.2.1.chr2.g00087.t01#1    mNeoVis.3.1.1.chr2.g00048.t01#6    mNeoVis.evergladensis.2.1.chr2.g00090.t01#2   mNeoVis.2.1.1.chr5_hap1.g00093.t01#5       104   125260724:   mNeoVis.evergladensis.1.1.chr8.g00551.t01#3    mNeoVis.2.1.1.chr8_hap1.g00566.t01#5    mNeoVis.vulgivaga.2.1.chr8.g00563.t01#1    mNeoVis.3.1.1.chr8.g00521.t01#6    mNeoVis.vulgivaga.1.1.chr8.g00551.t01#4   mNeoVis.evergladensis.2.1.chr9.g00556.t01#2       105   125260725:   mNeoVis.evergladensis.1.1.chr2.g00091.t01#3    mNeoVis.evergladensis.2.1.chr2.g00091.t01#2    mNeoVis.3.1.1.chr2.g00049.t01#6    mNeoVis.vulgivaga.2.1.chr2.g00088.t01#1    mNeoVis.vulgivaga.1.1.chr2.g00091.t01#4   mNeoVis.2.1.1.chr5_hap1.g00094.t01#5       106   125260726:   mNeoVis.2.1.1.chr5_hap1.g00095.t01#5    mNeoVis.evergladensis.1.1.chr2.g00092.t01#3    mNeoVis.vulgivaga.1.1.chr2.g00092.t01#4    mNeoVis.vulgivaga.2.1.chr2.g00089.t01#1    mNeoVis.evergladensis.2.1.chr2.g00092.t01#2   mNeoVis.3.1.1.chr2.g00050.t01#6       107   125260727:   mNeoVis.evergladensis.2.1.chr9.g00557.t01#2    mNeoVis.vulgivaga.1.1.chr8.g00552.t01#4    mNeoVis.evergladensis.1.1.chr8.g00552.t01#3    mNeoVis.3.1.1.chr8.g00522.t01#6    mNeoVis.2.1.1.chr8_hap1.g00567.t01#5   mNeoVis.vulgivaga.2.1.chr8.g00564.t01#1       108   125260728:   mNeoVis.evergladensis.1.1.chr2.g00093.t01#3    mNeoVis.vulgivaga.2.1.chr2.g00090.t01#1    mNeoVis.3.1.1.chr2.g00051.t01#6    mNeoVis.evergladensis.2.1.chr2.g00093.t01#2    mNeoVis.vulgivaga.1.1.chr2.g00093.t01#4   mNeoVis.2.1.1.chr5_hap1.g00096.t01#5       109   125260729:   mNeoVis.3.1.1.chr15.g00562.t01#6    mNeoVis.evergladensis.1.1.chr15.g00005.t01#3    mNeoVis.vulgivaga.2.1.chr15.g00619.t01#1    mNeoVis.2.1.1.chr15_hap1.g00785.t01#5    mNeoVis.evergladensis.2.1.chr15.g00606.t01#2   mNeoVis.vulgivaga.1.1.chr15.g00610.t01#4       110   125260730:   mNeoVis.vulgivaga.1.1.chr2.g00094.t01#4    mNeoVis.vulgivaga.2.1.chr2.g00091.t01#1    mNeoVis.evergladensis.2.1.chr2.g00094.t01#2    mNeoVis.evergladensis.1.1.chr2.g00094.t01#3   mNeoVis.3.1.1.chr2.g00052.t01#6       111   125260731:   mNeoVis.evergladensis.1.1.chr8.g00553.t01#3    mNeoVis.3.1.1.chr8.g00523.t01#6    mNeoVis.2.1.1.chr8_hap1.g00568.t01#5    mNeoVis.vulgivaga.2.1.chr8.g00565.t01#1    mNeoVis.evergladensis.2.1.chr9.g00558.t01#2   mNeoVis.vulgivaga.1.1.chr8.g00553.t01#4       112   125260732:   mNeoVis.vulgivaga.2.1.chr2.g00092.t01#1    mNeoVis.evergladensis.2.1.chr2.g00095.t01#2   mNeoVis.3.1.1.chr2.g00053.t01#6           113   125260733:   mNeoVis.vulgivaga.1.1.chr2.g00095.t01#4    mNeoVis.2.1.1.chr5_hap1.g00098.t01#5    mNeoVis.evergladensis.2.1.chr2.g00096.t01#2    mNeoVis.evergladensis.1.1.chr2.g00095.t01#3   mNeoVis.vulgivaga.2.1.chr2.g00093.t01#1       114   125260734:   mNeoVis.vulgivaga.1.1.chr8.g00554.t01#4    mNeoVis.evergladensis.1.1.chr8.g00554.t01#3    mNeoVis.vulgivaga.2.1.chr8.g00566.t01#1    mNeoVis.3.1.1.chr8.g00524.t01#6    mNeoVis.2.1.1.chr8_hap1.g00569.t01#5   mNeoVis.evergladensis.2.1.chr9.g00559.t01#2       115   125260735:   mNeoVis.evergladensis.2.1.chr2.g00097.t01#2    mNeoVis.2.1.1.chr5_hap1.g00099.t01#5    mNeoVis.evergladensis.1.1.chr2.g00096.t01#3    mNeoVis.vulgivaga.2.1.chr2.g00094.t01#1   mNeoVis.vulgivaga.1.1.chr2.g00096.t01#4       116   125260736:   mNeoVis.evergladensis.2.1.chr7.g00593.t01#2    mNeoVis.vulgivaga.1.1.chr7.g00600.t01#4    mNeoVis.evergladensis.1.1.chr7.g00600.t01#3    mNeoVis.3.1.1.chr7.g00573.t01#6    mNeoVis.2.1.1.chr7_hap1.g00591.t01#5   mNeoVis.vulgivaga.2.1.chr7.g00595.t01#1       117   125260737:   mNeoVis.3.1.1.chr2.g00055.t01#6    mNeoVis.vulgivaga.2.1.chr2.g00095.t01#1    mNeoVis.evergladensis.1.1.chr2.g00097.t01#3    mNeoVis.evergladensis.2.1.chr2.g00098.t01#2    mNeoVis.vulgivaga.1.1.chr2.g00097.t01#4   mNeoVis.2.1.1.chr5_hap1.g00100.t01#5       118   125260738:   mNeoVis.evergladensis.2.1.chr9.g00560.t01#2    mNeoVis.3.1.1.chr8.g00525.t01#6    mNeoVis.2.1.1.chr8_hap1.g00570.t01#5    mNeoVis.vulgivaga.1.1.chr8.g00555.t01#4    mNeoVis.vulgivaga.2.1.chr8.g00567.t01#1   mNeoVis.evergladensis.1.1.chr8.g00555.t01#3       119   125260739:   mNeoVis.evergladensis.1.1.chr8.g00556.t01#3    mNeoVis.vulgivaga.1.1.chr8.g00556.t01#4    mNeoVis.3.1.1.chr8.g00526.t01#6    mNeoVis.2.1.1.chr8_hap1.g00571.t01#5    mNeoVis.vulgivaga.2.1.chr8.g00568.t01#1   mNeoVis.evergladensis.2.1.chr9.g00561.t01#2       120   125260740:  mNeoVis.vulgivaga.2.1.chr8.g00569.t01#1               121   125260741:   mNeoVis.vulgivaga.1.1.chr2.g00098.t01#4    mNeoVis.2.1.1.chr5_hap1.g00101.t01#5    mNeoVis.evergladensis.2.1.chr2.g00099.t01#2    mNeoVis.evergladensis.1.1.chr2.g00098.t01#3    mNeoVis.vulgivaga.2.1.chr2.g00096.t01#1   mNeoVis.3.1.1.chr2.g00056.t01#6       122   125260742:   mNeoVis.vulgivaga.1.1.chr8.g00456.t01#4    mNeoVis.vulgivaga.2.1.chr8.g00466.t01#1    mNeoVis.2.1.1.chr8_hap1.g00470.t01#5    mNeoVis.evergladensis.2.1.chr9.g00461.t01#2    mNeoVis.3.1.1.chr8.g00425.t01#6   mNeoVis.evergladensis.1.1.chr8.g00456.t01#3       123   125260743:   mNeoVis.vulgivaga.1.1.chr2.g00099.t01#4    mNeoVis.2.1.1.chr5_hap1.g00102.t01#5    mNeoVis.3.1.1.chr2.g00057.t01#6    mNeoVis.vulgivaga.2.1.chr2.g00097.t01#1    mNeoVis.evergladensis.2.1.chr2.g00100.t01#2   mNeoVis.evergladensis.1.1.chr2.g00099.t01#3       124   125260744:   mNeoVis.evergladensis.2.1.chr9.g00462.t01#2    mNeoVis.vulgivaga.1.1.chr8.g00457.t01#4    mNeoVis.3.1.1.chr8.g00426.t01#6    mNeoVis.2.1.1.chr8_hap1.g00471.t01#5    mNeoVis.evergladensis.1.1.chr8.g00457.t01#3   mNeoVis.vulgivaga.2.1.chr8.g00467.t01#1       125   125260745:   mNeoVis.2.1.1.chr5_hap1.g01083.t01#5    mNeoVis.evergladensis.2.1.chr2.g01072.t01#2    mNeoVis.3.1.1.chr2.g00990.t01#6    mNeoVis.vulgivaga.2.1.chr2.g01080.t01#1    mNeoVis.vulgivaga.1.1.chr2.g01075.t01#4   mNeoVis.evergladensis.1.1.chr2.g01074.t01#3       126   125260746:   mNeoVis.vulgivaga.2.1.chr2.g00098.t01#1    mNeoVis.3.1.1.chr2.g00058.t01#6    mNeoVis.evergladensis.1.1.chr2.g00100.t01#3    mNeoVis.2.1.1.chr5_hap1.g00103.t01#5    mNeoVis.vulgivaga.1.1.chr2.g00100.t01#4   mNeoVis.evergladensis.2.1.chr2.g00101.t01#2       127   125260747:   mNeoVis.vulgivaga.2.1.chr8.g00468.t01#1    mNeoVis.evergladensis.1.1.chr8.g00458.t01#3    mNeoVis.2.1.1.chr8_hap1.g00472.t01#5    mNeoVis.3.1.1.chr8.g00427.t01#6    mNeoVis.vulgivaga.1.1.chr8.g00458.t01#4   mNeoVis.evergladensis.2.1.chr9.g00463.t01#2       128   125260748:   mNeoVis.evergladensis.2.1.chr2.g00102.t01#2    mNeoVis.2.1.1.chr5_hap1.g00104.t01#5    mNeoVis.3.1.1.chr2.g00059.t01#6    mNeoVis.evergladensis.1.1.chr2.g00101.t01#3    mNeoVis.vulgivaga.1.1.chr2.g00101.t01#4   mNeoVis.vulgivaga.2.1.chr2.g00099.t01#1       129   125260749:   mNeoVis.3.1.1.chr2.g00060.t01#6    mNeoVis.2.1.1.chr5_hap1.g00105.t01#5    mNeoVis.vulgivaga.1.1.chr2.g00102.t01#4    mNeoVis.evergladensis.2.1.chr2.g00103.t01#2    mNeoVis.vulgivaga.2.1.chr2.g00100.t01#1   mNeoVis.evergladensis.1.1.chr2.g00102.t01#3       130   125260750:   mNeoVis.3.1.1.chr7.g00572.t01#6    mNeoVis.2.1.1.chr7_hap1.g00590.t01#5    mNeoVis.evergladensis.1.1.chr7.g00601.t01#3    mNeoVis.vulgivaga.1.1.chr7.g00599.t01#4    mNeoVis.evergladensis.2.1.chr7.g00592.t01#2   mNeoVis.vulgivaga.2.1.chr7.g00596.t01#1       131   125260751:   mNeoVis.2.1.1.chr5_hap1.g00106.t01#5    mNeoVis.vulgivaga.1.1.chr2.g00103.t01#4    mNeoVis.vulgivaga.2.1.chr2.g00101.t01#1    mNeoVis.evergladensis.2.1.chr2.g00104.t01#2   mNeoVis.evergladensis.1.1.chr2.g00103.t01#3       132   125260752:   mNeoVis.vulgivaga.2.1.chr8.g00469.t01#1    mNeoVis.vulgivaga.1.1.chr8.g00459.t01#4    mNeoVis.2.1.1.chr8_hap1.g00473.t01#5    mNeoVis.evergladensis.2.1.chr9.g00464.t01#2    mNeoVis.evergladensis.1.1.chr8.g00459.t01#3   mNeoVis.3.1.1.chr8.g00428.t01#6       133   125260753:   mNeoVis.3.1.1.chr8.g00429.t01#6    mNeoVis.vulgivaga.2.1.chr8.g00470.t01#1    mNeoVis.evergladensis.2.1.chr9.g00465.t01#2    mNeoVis.evergladensis.1.1.chr8.g00460.t01#3    mNeoVis.vulgivaga.1.1.chr8.g00460.t01#4   mNeoVis.2.1.1.chr8_hap1.g00474.t01#5       134   125260754:   mNeoVis.vulgivaga.1.1.chr2.g00104.t01#4    mNeoVis.evergladensis.1.1.chr2.g00104.t01#3    mNeoVis.2.1.1.chr5_hap1.g00107.t01#5    mNeoVis.3.1.1.chr2.g00062.t01#6    mNeoVis.vulgivaga.2.1.chr2.g00102.t01#1   mNeoVis.evergladensis.2.1.chr2.g00105.t01#2       135   125260755:   mNeoVis.vulgivaga.2.1.chr7.g00597.t01#1    mNeoVis.evergladensis.1.1.chr7.g00602.t01#3    mNeoVis.evergladensis.2.1.chr7.g00594.t01#2    mNeoVis.2.1.1.chr7_hap1.g00592.t01#5    mNeoVis.vulgivaga.1.1.chr7.g00601.t01#4   mNeoVis.3.1.1.chr7.g00574.t01#6       136   125260756:   mNeoVis.vulgivaga.1.1.chr15.g00611.t01#4    mNeoVis.evergladensis.2.1.chr15.g00607.t01#2    mNeoVis.evergladensis.1.1.chr15.g00004.t01#3    mNeoVis.vulgivaga.2.1.chr15.g00620.t01#1    mNeoVis.3.1.1.chr15.g00563.t01#6   mNeoVis.2.1.1.chr15_hap1.g00786.t01#5       137   125260757:   mNeoVis.evergladensis.2.1.chr2.g00106.t01#2    mNeoVis.vulgivaga.2.1.chr2.g00103.t01#1    mNeoVis.2.1.1.chr5_hap1.g00108.t01#5    mNeoVis.evergladensis.1.1.chr2.g00105.t01#3   mNeoVis.vulgivaga.1.1.chr2.g00105.t01#4       138   125260758:   mNeoVis.2.1.1.chr8_hap1.g00475.t01#5    mNeoVis.vulgivaga.2.1.chr8.g00471.t01#1    mNeoVis.3.1.1.chr8.g00430.t01#6    mNeoVis.evergladensis.2.1.chr9.g00466.t01#2    mNeoVis.vulgivaga.1.1.chr8.g00461.t01#4   mNeoVis.evergladensis.1.1.chr8.g00461.t01#3       139   125260759:   mNeoVis.vulgivaga.1.1.chr8.g00462.t01#4    mNeoVis.evergladensis.1.1.chr8.g00462.t01#3    mNeoVis.vulgivaga.2.1.chr8.g00472.t01#1    mNeoVis.2.1.1.chr8_hap1.g00476.t01#5   mNeoVis.evergladensis.2.1.chr9.g00467.t01#2       140   125260760:   mNeoVis.vulgivaga.1.1.chr10.g00390.t01#4    mNeoVis.evergladensis.2.1.chr8.g01084.t01#2    mNeoVis.3.1.1.chr10.g00392.t01#6    mNeoVis.evergladensis.2.1.chr10.g00398.t01#2    mNeoVis.vulgivaga.2.1.chr10.g00394.t01#1    mNeoVis.2.1.1.chr9_hap1.g00401.t01#5   mNeoVis.evergladensis.1.1.chr10.g00405.t01#3     141   125260761:   mNeoVis.vulgivaga.2.1.chr3.g01374.t01#1    mNeoVis.evergladensis.1.1.chr3.g01356.t01#3    mNeoVis.evergladensis.2.1.chr3.g01355.t01#2    mNeoVis.vulgivaga.1.1.chr3.g01348.t01#4    mNeoVis.2.1.1.chr3_hap1.g01353.t01#5   mNeoVis.3.1.1.chr3.g01328.t01#6       142   125260762:   mNeoVis.evergladensis.2.1.chr9.g00468.t01#2    mNeoVis.2.1.1.chr8_hap1.g00477.t01#5    mNeoVis.vulgivaga.2.1.chr8.g00473.t01#1    mNeoVis.evergladensis.1.1.chr8.g00463.t01#3   mNeoVis.vulgivaga.1.1.chr8.g00463.t01#4       143   125260763:  mNeoVis.3.1.1.chr1.g02043.t01#6               144   125260764:   mNeoVis.vulgivaga.1.1.chr2.g00106.t01#4    mNeoVis.2.1.1.chr5_hap1.g00109.t01#5    mNeoVis.evergladensis.1.1.chr2.g00106.t01#3    mNeoVis.3.1.1.chr2.g00063.t01#6    mNeoVis.evergladensis.2.1.chr2.g00107.t01#2   mNeoVis.vulgivaga.2.1.chr2.g00104.t01#1       145   125260765:   mNeoVis.vulgivaga.2.1.chr8.g00474.t01#1    mNeoVis.evergladensis.2.1.chr9.g00469.t01#2    mNeoVis.2.1.1.chr8_hap1.g00478.t01#5   mNeoVis.3.1.1.chr8.g00431.t01#6         146   125260766:   mNeoVis.vulgivaga.1.1.chr8.g00464.t01#4    mNeoVis.2.1.1.chr8_hap1.g00479.t01#5    mNeoVis.3.1.1.chr8.g00432.t01#6    mNeoVis.vulgivaga.2.1.chr8.g00475.t01#1    mNeoVis.evergladensis.1.1.chr8.g00464.t01#3   mNeoVis.evergladensis.2.1.chr9.g00470.t01#2       147   125260767:   mNeoVis.vulgivaga.2.1.chr2.g00105.t01#1    mNeoVis.vulgivaga.1.1.chr2.g00107.t01#4    mNeoVis.evergladensis.1.1.chr2.g00107.t01#3    mNeoVis.2.1.1.chr5_hap1.g00110.t01#5   mNeoVis.evergladensis.2.1.chr2.g00108.t01#2       148   125260768:   mNeoVis.evergladensis.2.1.chr9.g00471.t01#2    mNeoVis.vulgivaga.1.1.chr8.g00465.t01#4    mNeoVis.vulgivaga.2.1.chr8.g00476.t01#1    mNeoVis.2.1.1.chr8_hap1.g00480.t01#5    mNeoVis.3.1.1.chr8.g00433.t01#6   mNeoVis.evergladensis.1.1.chr8.g00465.t01#3       149   125260769:   mNeoVis.2.1.1.chr15_hap1.g00787.t01#5    mNeoVis.evergladensis.1.1.chr15.g00002.t01#3    mNeoVis.3.1.1.chr15.g00565.t01#6    mNeoVis.evergladensis.2.1.chr15.g00608.t01#2    mNeoVis.vulgivaga.1.1.chr15.g00612.t01#4   mNeoVis.vulgivaga.2.1.chr15.g00621.t01#1       150   125260770:   mNeoVis.evergladensis.2.1.chr2.g00109.t01#2    mNeoVis.3.1.1.chr2.g00064.t01#6    mNeoVis.vulgivaga.1.1.chr2.g00108.t01#4    mNeoVis.2.1.1.chr5_hap1.g00111.t01#5    mNeoVis.evergladensis.1.1.chr2.g00108.t01#3   mNeoVis.vulgivaga.2.1.chr2.g00106.t01#1       151   125260771:   mNeoVis.vulgivaga.2.1.chr9.g00888.t01#1    mNeoVis.3.1.1.chr9.g00862.t01#6    mNeoVis.vulgivaga.1.1.chr9.g00900.t01#4    mNeoVis.evergladensis.1.1.chr9.g00896.t01#3    mNeoVis.evergladensis.2.1.chr8.g00901.t01#2   mNeoVis.2.1.1.chr10_hap1.g00931.t01#5       152   125260772:   mNeoVis.vulgivaga.1.1.chr2.g00109.t01#4    mNeoVis.vulgivaga.2.1.chr2.g00107.t01#1    mNeoVis.2.1.1.chr5_hap1.g00112.t01#5    mNeoVis.evergladensis.2.1.chr2.g00110.t01#2   mNeoVis.evergladensis.1.1.chr2.g00109.t01#3       153   125260773:   mNeoVis.evergladensis.2.1.chr9.g00472.t01#2    mNeoVis.vulgivaga.2.1.chr8.g00477.t01#1    mNeoVis.vulgivaga.1.1.chr8.g00466.t01#4    mNeoVis.2.1.1.chr8_hap1.g00481.t01#5    mNeoVis.3.1.1.chr8.g00434.t01#6   mNeoVis.evergladensis.1.1.chr8.g00466.t01#3       154   125260774:   mNeoVis.vulgivaga.2.1.chr8.g00478.t01#1    mNeoVis.vulgivaga.1.1.chr8.g00467.t01#4    mNeoVis.3.1.1.chr8.g00435.t01#6    mNeoVis.2.1.1.chr8_hap1.g00482.t01#5    mNeoVis.evergladensis.1.1.chr8.g00467.t01#3   mNeoVis.evergladensis.2.1.chr9.g00473.t01#2       155   125260775:   mNeoVis.evergladensis.2.1.chr2.g00111.t01#2    mNeoVis.3.1.1.chr2.g00066.t01#6    mNeoVis.vulgivaga.1.1.chr2.g00110.t01#4    mNeoVis.vulgivaga.2.1.chr2.g00108.t01#1    mNeoVis.2.1.1.chr5_hap1.g00113.t01#5   mNeoVis.evergladensis.1.1.chr2.g00110.t01#3       156   125260776:   mNeoVis.vulgivaga.1.1.chr8.g00468.t01#4    mNeoVis.3.1.1.chr8.g00436.t01#6    mNeoVis.evergladensis.1.1.chr8.g00468.t01#3    mNeoVis.2.1.1.chr8_hap1.g00483.t01#5    mNeoVis.evergladensis.2.1.chr9.g00474.t01#2   mNeoVis.vulgivaga.2.1.chr8.g00479.t01#1       157   125260777:   mNeoVis.evergladensis.1.1.chr2.g00111.t01#3    mNeoVis.vulgivaga.2.1.chr2.g00109.t01#1    mNeoVis.2.1.1.chr5_hap1.g00114.t01#5    mNeoVis.evergladensis.2.1.chr2.g00112.t01#2    mNeoVis.3.1.1.chr2.g00067.t01#6   mNeoVis.vulgivaga.1.1.chr2.g00111.t01#4       158   125260778:   mNeoVis.evergladensis.2.1.chr2.g00113.t01#2    mNeoVis.vulgivaga.1.1.chr2.g00112.t01#4    mNeoVis.2.1.1.chr5_hap1.g00115.t01#5    mNeoVis.evergladensis.1.1.chr2.g00112.t01#3   mNeoVis.vulgivaga.2.1.chr2.g00110.t01#1       159   125260779:   mNeoVis.evergladensis.2.1.chr7.g00595.t01#2    mNeoVis.vulgivaga.2.1.chr7.g00598.t01#1    mNeoVis.evergladensis.1.1.chr7.g00603.t01#3    mNeoVis.2.1.1.chr7_hap1.g00593.t01#5    mNeoVis.3.1.1.chr7.g00575.t01#6   mNeoVis.vulgivaga.1.1.chr7.g00602.t01#4       160   125260780:   mNeoVis.3.1.1.chr8.g00438.t01#6    mNeoVis.evergladensis.2.1.chr9.g00475.t01#2    mNeoVis.vulgivaga.2.1.chr8.g00480.t01#1    mNeoVis.vulgivaga.1.1.chr8.g00469.t01#4    mNeoVis.evergladensis.1.1.chr8.g00469.t01#3   mNeoVis.2.1.1.chr8_hap1.g00484.t01#5       161   125260781:   mNeoVis.evergladensis.1.1.chr8.g00470.t01#3    mNeoVis.evergladensis.2.1.chr9.g00476.t01#2    mNeoVis.vulgivaga.2.1.chr8.g00481.t01#1    mNeoVis.vulgivaga.1.1.chr8.g00470.t01#4    mNeoVis.2.1.1.chr8_hap1.g00485.t01#5   mNeoVis.3.1.1.chr8.g00439.t01#6       162   125260782:   mNeoVis.evergladensis.1.1.chr2.g00014.t01#3    mNeoVis.vulgivaga.1.1.chr2.g00014.t01#4    mNeoVis.evergladensis.2.1.chr2.g00014.t01#2    mNeoVis.vulgivaga.2.1.chr2.g00014.t01#1   mNeoVis.2.1.1.chr5_hap1.g00014.t01#5       163   125260783:   mNeoVis.vulgivaga.2.1.chr8.g00482.t01#1    mNeoVis.2.1.1.chr8_hap1.g00486.t01#5    mNeoVis.vulgivaga.1.1.chr8.g00471.t01#4    mNeoVis.3.1.1.chr8.g00440.t01#6    mNeoVis.evergladensis.2.1.chr9.g00477.t01#2   mNeoVis.evergladensis.1.1.chr8.g00471.t01#3       164   125260784:   mNeoVis.vulgivaga.2.1.chr2.g00015.t01#1    mNeoVis.vulgivaga.1.1.chr2.g00015.t01#4    mNeoVis.3.1.1.chr4.g00693.t01#6    mNeoVis.evergladensis.2.1.chr2.g00015.t01#2    mNeoVis.2.1.1.chr5_hap1.g00015.t01#5   mNeoVis.evergladensis.1.1.chr2.g00015.t01#3       165   125260785:   mNeoVis.3.1.1.chr8.g00441.t01#6    mNeoVis.evergladensis.2.1.chr9.g00478.t01#2    mNeoVis.vulgivaga.2.1.chr8.g00483.t01#1    mNeoVis.vulgivaga.1.1.chr8.g00472.t01#4    mNeoVis.evergladensis.1.1.chr8.g00472.t01#3   mNeoVis.2.1.1.chr8_hap1.g00487.t01#5       166   125260786:   mNeoVis.3.1.1.chr8.g00442.t01#6    mNeoVis.vulgivaga.2.1.chr8.g00484.t01#1    mNeoVis.vulgivaga.1.1.chr8.g00473.t01#4    mNeoVis.evergladensis.2.1.chr9.g00479.t01#2    mNeoVis.2.1.1.chr8_hap1.g00488.t01#5   mNeoVis.evergladensis.1.1.chr8.g00473.t01#3       167   125260787:   mNeoVis.vulgivaga.2.1.chr8.g00485.t01#1    mNeoVis.vulgivaga.1.1.chr8.g00474.t01#4    mNeoVis.3.1.1.chr8.g00444.t01#6    mNeoVis.2.1.1.chr8_hap1.g00489.t01#5    mNeoVis.evergladensis.2.1.chr9.g00480.t01#2   mNeoVis.evergladensis.1.1.chr8.g00474.t01#3       168   125260788:   mNeoVis.vulgivaga.1.1.chr2.g00016.t01#4    mNeoVis.evergladensis.1.1.chr7.g01876.t01#3    mNeoVis.vulgivaga.2.1.chr2.g00016.t01#1    mNeoVis.2.1.1.chr5_hap1.g00016.t01#5    mNeoVis.evergladensis.1.1.chr2.g00016.t01#3   mNeoVis.evergladensis.2.1.chr2.g00016.t01#2       169   125260789:   mNeoVis.evergladensis.2.1.chr2.g00017.t01#2    mNeoVis.vulgivaga.1.1.chr2.g00017.t01#4    mNeoVis.vulgivaga.2.1.chr2.g00017.t01#1    mNeoVis.2.1.1.chr5_hap1.g00017.t01#5   mNeoVis.evergladensis.1.1.chr2.g00017.t01#3       170   125260790:   mNeoVis.vulgivaga.2.1.chr8.g00486.t01#1    mNeoVis.vulgivaga.1.1.chr8.g00475.t01#4    mNeoVis.2.1.1.chr8_hap1.g00490.t01#5    mNeoVis.3.1.1.chr8.g00446.t01#6    mNeoVis.evergladensis.2.1.chr9.g00481.t01#2   mNeoVis.evergladensis.1.1.chr8.g00476.t01#3       171   125260791:   mNeoVis.vulgivaga.2.1.chr8.g00487.t01#1    mNeoVis.vulgivaga.1.1.chr8.g00476.t01#4    mNeoVis.2.1.1.chr8_hap1.g00491.t01#5    mNeoVis.3.1.1.chr8.g00447.t01#6    mNeoVis.evergladensis.2.1.chr9.g00482.t01#2   mNeoVis.evergladensis.1.1.chr8.g00478.t01#3       172   125260792:   mNeoVis.evergladensis.2.1.chr2.g00018.t01#2    mNeoVis.evergladensis.1.1.chr2.g00018.t01#3    mNeoVis.2.1.1.chr5_hap1.g00018.t01#5    mNeoVis.vulgivaga.1.1.chr2.g00019.t01#4   mNeoVis.vulgivaga.2.1.chr2.g00018.t01#1       173   125260793:   mNeoVis.vulgivaga.1.1.chr8.g00477.t01#4    mNeoVis.3.1.1.chr8.g00448.t01#6    mNeoVis.evergladensis.2.1.chr9.g00483.t01#2    mNeoVis.2.1.1.chr8_hap1.g00492.t01#5    mNeoVis.evergladensis.1.1.chr8.g00479.t01#3   mNeoVis.vulgivaga.2.1.chr8.g00488.t01#1       174   125260794:   mNeoVis.vulgivaga.2.1.chr2.g00975.t01#1    mNeoVis.2.1.1.chr5_hap1.g00973.t01#5    mNeoVis.evergladensis.2.1.chr2.g00969.t01#2    mNeoVis.vulgivaga.1.1.chr2.g00972.t01#4    mNeoVis.evergladensis.1.1.chr2.g00968.t01#3   mNeoVis.3.1.1.chr2.g00891.t01#6       175   125260795:   mNeoVis.evergladensis.1.1.chr8.g00480.t01#3    mNeoVis.vulgivaga.2.1.chr8.g00489.t01#1   mNeoVis.vulgivaga.1.1.chr8.g00478.t01#4           176   125260796:   mNeoVis.vulgivaga.1.1.chr2.g00020.t01#4    mNeoVis.3.1.1.chr2.g00010.t01#6    mNeoVis.evergladensis.2.1.chr2.g00019.t01#2    mNeoVis.evergladensis.1.1.chr2.g00019.t01#3    mNeoVis.2.1.1.chr5_hap1.g00019.t01#5   mNeoVis.vulgivaga.2.1.chr2.g00019.t01#1       177   125260797:   mNeoVis.vulgivaga.1.1.chr8.g00479.t01#4    mNeoVis.3.1.1.chr8.g00449.t01#6    mNeoVis.evergladensis.1.1.chr8.g00481.t01#3    mNeoVis.2.1.1.chr8_hap1.g00493.t01#5    mNeoVis.vulgivaga.2.1.chr8.g00490.t01#1   mNeoVis.evergladensis.2.1.chr9.g00484.t01#2       178   125260798:   mNeoVis.evergladensis.1.1.chr2.g00020.t01#3    mNeoVis.evergladensis.2.1.chr2.g00020.t01#2    mNeoVis.vulgivaga.2.1.chr2.g00020.t01#1    mNeoVis.vulgivaga.1.1.chr2.g00021.t01#4   mNeoVis.2.1.1.chr5_hap1.g00020.t01#5       179   125260799:   mNeoVis.vulgivaga.1.1.chr2.g00022.t01#4    mNeoVis.2.1.1.chr5_hap1.g00021.t01#5    mNeoVis.vulgivaga.2.1.chr2.g00021.t01#1    mNeoVis.evergladensis.2.1.chr2.g00021.t01#2    mNeoVis.evergladensis.1.1.chr2.g00021.t01#3   mNeoVis.3.1.1.chr2.g00012.t01#6       180   125260800:   mNeoVis.3.1.1.chr8.g00450.t01#6    mNeoVis.2.1.1.chr8_hap1.g00494.t01#5    mNeoVis.vulgivaga.1.1.chr8.g00480.t01#4    mNeoVis.evergladensis.1.1.chr8.g00482.t01#3    mNeoVis.vulgivaga.2.1.chr8.g00491.t01#1   mNeoVis.evergladensis.2.1.chr9.g00485.t01#2       181   125260801:   mNeoVis.evergladensis.1.1.chr1.g01430.t01#3    mNeoVis.3.1.1.chr1.g01409.t01#6    mNeoVis.evergladensis.2.1.chr1.g01427.t01#2    mNeoVis.vulgivaga.1.1.chr1.g01429.t01#4   mNeoVis.vulgivaga.2.1.chr1.g01428.t01#1       182   125260802:   mNeoVis.2.1.1.chr8_hap1.g00495.t01#5    mNeoVis.3.1.1.chr8.g00451.t01#6    mNeoVis.evergladensis.2.1.chr9.g00486.t01#2    mNeoVis.vulgivaga.1.1.chr8.g00481.t01#4    mNeoVis.evergladensis.1.1.chr8.g00483.t01#3   mNeoVis.vulgivaga.2.1.chr8.g00492.t01#1       183   125260803:   mNeoVis.vulgivaga.1.1.chr2.g00023.t01#4   mNeoVis.vulgivaga.2.1.chr2.g00022.t01#1             184   125260804:   mNeoVis.evergladensis.2.1.chr2.g00023.t01#2    mNeoVis.vulgivaga.1.1.chr2.g00024.t01#4    mNeoVis.2.1.1.chr5_hap1.g00023.t01#5    mNeoVis.vulgivaga.2.1.chr2.g00023.t01#1   mNeoVis.evergladensis.1.1.chr2.g00024.t01#3       185   125260805:   mNeoVis.vulgivaga.2.1.chr2.g00024.t01#1    mNeoVis.2.1.1.chr5_hap1.g00024.t01#5   mNeoVis.vulgivaga.1.1.chr2.g00025.t01#4           186   125260806:   mNeoVis.vulgivaga.1.1.chr2.g00026.t01#4    mNeoVis.evergladensis.2.1.chr2.g00025.t01#2    mNeoVis.3.1.1.chr2.g00014.t01#6    mNeoVis.vulgivaga.2.1.chr2.g00025.t01#1    mNeoVis.2.1.1.chr5_hap1.g00025.t01#5   mNeoVis.evergladensis.1.1.chr2.g00026.t01#3       187   125260807:   mNeoVis.vulgivaga.2.1.chr8.g00493.t01#1    mNeoVis.3.1.1.chr8.g00452.t01#6    mNeoVis.2.1.1.chr8_hap1.g00496.t01#5    mNeoVis.evergladensis.1.1.chr8.g00484.t01#3    mNeoVis.vulgivaga.1.1.chr8.g00482.t01#4   mNeoVis.evergladensis.2.1.chr9.g00487.t01#2       188   125260808:   mNeoVis.evergladensis.2.1.chr10.g00107.t01#2    mNeoVis.vulgivaga.2.1.chr10.g00103.t01#1    mNeoVis.vulgivaga.1.1.chr10.g00100.t01#4    mNeoVis.evergladensis.1.1.chr10.g00110.t01#3    mNeoVis.3.1.1.chr10.g00103.t01#6   mNeoVis.2.1.1.chr9_hap1.g00107.t01#5       189   125260809:   mNeoVis.vulgivaga.1.1.chr3.g00978.t01#4    mNeoVis.vulgivaga.2.1.chr3.g00990.t01#1    mNeoVis.evergladensis.1.1.chr3.g00984.t01#3    mNeoVis.3.1.1.chr3.g00966.t01#6    mNeoVis.evergladensis.2.1.chr3.g00971.t01#2   mNeoVis.2.1.1.chr3_hap1.g00976.t01#5       190   125260810:   mNeoVis.2.1.1.chr8_hap1.g00497.t01#5    mNeoVis.3.1.1.chr8.g00453.t01#6    mNeoVis.evergladensis.1.1.chr8.g00485.t01#3    mNeoVis.vulgivaga.1.1.chr8.g00483.t01#4    mNeoVis.evergladensis.2.1.chr9.g00488.t01#2   mNeoVis.vulgivaga.2.1.chr8.g00494.t01#1       191   125260811:   mNeoVis.vulgivaga.2.1.chr2.g00026.t01#1   mNeoVis.vulgivaga.1.1.chr2.g00027.t01#4             192   125260812:   mNeoVis.evergladensis.1.1.chr8.g00486.t01#3    mNeoVis.vulgivaga.2.1.chr8.g00495.t01#1    mNeoVis.3.1.1.chr8.g00454.t01#6    mNeoVis.2.1.1.chr8_hap1.g00498.t01#5    mNeoVis.vulgivaga.1.1.chr8.g00484.t01#4   mNeoVis.evergladensis.2.1.chr9.g00489.t01#2       193   125260813:   mNeoVis.evergladensis.2.1.chr15.g00609.t01#2    mNeoVis.vulgivaga.1.1.chr15.g00613.t01#4    mNeoVis.evergladensis.1.1.chr15.g00003.t01#3    mNeoVis.2.1.1.chr15_hap1.g00788.t01#5   mNeoVis.vulgivaga.2.1.chr15.g00622.t01#1       194   125260814:   mNeoVis.evergladensis.1.1.chr2.g00028.t01#3    mNeoVis.evergladensis.2.1.chr2.g00027.t01#2    mNeoVis.vulgivaga.2.1.chr2.g00027.t01#1    mNeoVis.vulgivaga.1.1.chr2.g00028.t01#4   mNeoVis.2.1.1.chr5_hap1.g00027.t01#5       195   125260815:   mNeoVis.evergladensis.1.1.chr7.g00604.t01#3    mNeoVis.2.1.1.chr7_hap1.g00594.t01#5    mNeoVis.evergladensis.2.1.chr7.g00596.t01#2    mNeoVis.vulgivaga.1.1.chr7.g00603.t01#4   mNeoVis.vulgivaga.2.1.chr7.g00599.t01#1       196   125260816:   mNeoVis.vulgivaga.1.1.chr2.g00029.t01#4    mNeoVis.3.1.1.chr6.g00531.t01#6    mNeoVis.vulgivaga.2.1.chr2.g00028.t01#1    mNeoVis.evergladensis.1.1.chr2.g00029.t01#3    mNeoVis.evergladensis.2.1.chr2.g00028.t01#2   mNeoVis.2.1.1.chr5_hap1.g00028.t01#5       197   125260817:   mNeoVis.vulgivaga.1.1.chr8.g00485.t01#4    mNeoVis.evergladensis.2.1.chr9.g00490.t01#2    mNeoVis.vulgivaga.2.1.chr8.g00496.t01#1    mNeoVis.3.1.1.chr8.g00455.t01#6   mNeoVis.2.1.1.chr8_hap1.g00499.t01#5       198   125260818:   mNeoVis.vulgivaga.1.1.chr8.g00486.t01#4    mNeoVis.evergladensis.1.1.chr8.g00488.t01#3    mNeoVis.3.1.1.chr8.g00456.t01#6    mNeoVis.vulgivaga.2.1.chr8.g00497.t01#1    mNeoVis.2.1.1.chr8_hap1.g00500.t01#5   mNeoVis.evergladensis.2.1.chr9.g00491.t01#2       199   125260819:   mNeoVis.evergladensis.2.1.chr2.g00029.t01#2    mNeoVis.2.1.1.chr5_hap1.g00029.t01#5    mNeoVis.vulgivaga.2.1.chr2.g00029.t01#1    mNeoVis.evergladensis.1.1.chr2.g00030.t01#3    mNeoVis.3.1.1.chr2.g00016.t01#6   mNeoVis.vulgivaga.1.1.chr2.g00030.t01#4       200   125260820:   mNeoVis.3.1.1.chr8.g00457.t01#6    mNeoVis.vulgivaga.2.1.chr8.g00498.t01#1    mNeoVis.2.1.1.chr8_hap1.g00501.t01#5    mNeoVis.evergladensis.1.1.chr8.g00489.t01#3    mNeoVis.vulgivaga.1.1.chr8.g00487.t01#4   mNeoVis.evergladensis.2.1.chr9.g00492.t01#2       201   125260821:  mNeoVis.vulgivaga.2.1.chr15.g00623.t01#1               202   125260822:   mNeoVis.evergladensis.2.1.chr7.g00597.t01#2    mNeoVis.vulgivaga.1.1.chr7.g00604.t01#4    mNeoVis.evergladensis.1.1.chr7.g00605.t01#3    mNeoVis.2.1.1.chr7_hap1.g00595.t01#5   mNeoVis.vulgivaga.2.1.chr7.g00600.t01#1       203   125260823:   mNeoVis.vulgivaga.2.1.chr2.g00030.t01#1    mNeoVis.vulgivaga.1.1.chr2.g00031.t01#4    mNeoVis.evergladensis.1.1.chr2.g00031.t01#3    mNeoVis.evergladensis.2.1.chr2.g00030.t01#2   mNeoVis.2.1.1.chr5_hap1.g00030.t01#5       204   125260824:   mNeoVis.evergladensis.2.1.chr9.g00493.t01#2    mNeoVis.3.1.1.chr8.g00458.t01#6    mNeoVis.2.1.1.chr8_hap1.g00502.t01#5    mNeoVis.evergladensis.1.1.chr8.g00490.t01#3    mNeoVis.vulgivaga.2.1.chr8.g00499.t01#1   mNeoVis.vulgivaga.1.1.chr8.g00488.t01#4       205   125260825:   mNeoVis.2.1.1.chr8_hap1.g00503.t01#5    mNeoVis.evergladensis.2.1.chr9.g00494.t01#2    mNeoVis.evergladensis.1.1.chr8.g00491.t01#3    mNeoVis.vulgivaga.2.1.chr8.g00500.t01#1    mNeoVis.3.1.1.chr8.g00459.t01#6   mNeoVis.vulgivaga.1.1.chr8.g00489.t01#4       206   125260826:   mNeoVis.evergladensis.2.1.chr10.g00092.t01#2    mNeoVis.vulgivaga.2.1.chr10.g00085.t01#1    mNeoVis.2.1.1.chr9_hap1.g00091.t01#5    mNeoVis.evergladensis.1.1.chr10.g00094.t01#3    mNeoVis.vulgivaga.1.1.chr10.g00085.t01#4   mNeoVis.3.1.1.chr10.g00088.t01#6       207   125260827:   mNeoVis.evergladensis.1.1.chr2.g00032.t01#3    mNeoVis.vulgivaga.1.1.chr2.g00032.t01#4    mNeoVis.vulgivaga.2.1.chr2.g00031.t01#1    mNeoVis.evergladensis.2.1.chr2.g00031.t01#2   mNeoVis.2.1.1.chr5_hap1.g00031.t01#5       208   125260828:  mNeoVis.vulgivaga.2.1.chr2.g00032.t01#1               209   125260829:   mNeoVis.vulgivaga.2.1.chr8.g00501.t01#1    mNeoVis.vulgivaga.1.1.chr8.g00490.t01#4    mNeoVis.evergladensis.2.1.chr9.g00495.t01#2    mNeoVis.evergladensis.1.1.chr8.g00492.t01#3    mNeoVis.3.1.1.chr8.g00460.t01#6   mNeoVis.2.1.1.chr8_hap1.g00504.t01#5       210   125260830:   mNeoVis.vulgivaga.2.1.chr8.g00502.t01#1    mNeoVis.3.1.1.chr8.g00461.t01#6    mNeoVis.vulgivaga.1.1.chr8.g00491.t01#4    mNeoVis.evergladensis.1.1.chr8.g00493.t01#3    mNeoVis.evergladensis.2.1.chr9.g00496.t01#2   mNeoVis.2.1.1.chr8_hap1.g00505.t01#5       211   125260831:   mNeoVis.evergladensis.2.1.chr9.g00497.t01#2    mNeoVis.vulgivaga.1.1.chr8.g00492.t01#4    mNeoVis.2.1.1.chr8_hap1.g00506.t01#5    mNeoVis.vulgivaga.2.1.chr8.g00503.t01#1    mNeoVis.3.1.1.chr8.g00462.t01#6   mNeoVis.evergladensis.1.1.chr8.g00494.t01#3       212   125260832:  mNeoVis.vulgivaga.2.1.chr2.g00033.t01#1               213   125260833:   mNeoVis.vulgivaga.2.1.chr2.g00034.t01#1    mNeoVis.evergladensis.1.1.chr2.g00035.t01#3    mNeoVis.vulgivaga.1.1.chr2.g00035.t01#4    mNeoVis.2.1.1.chr5_hap1.g00034.t01#5   mNeoVis.evergladensis.2.1.chr2.g00034.t01#2       214   125260834:   mNeoVis.3.1.1.chr8.g00463.t01#6    mNeoVis.evergladensis.1.1.chr8.g00495.t01#3    mNeoVis.evergladensis.2.1.chr9.g00498.t01#2    mNeoVis.vulgivaga.2.1.chr8.g00504.t01#1    mNeoVis.vulgivaga.1.1.chr8.g00493.t01#4   mNeoVis.2.1.1.chr8_hap1.g00507.t01#5       215   125260835:   mNeoVis.vulgivaga.1.1.chr2.g00036.t01#4    mNeoVis.vulgivaga.2.1.chr2.g00035.t01#1    mNeoVis.evergladensis.1.1.chr2.g00036.t01#3    mNeoVis.2.1.1.chr5_hap1.g00035.t01#5   mNeoVis.evergladensis.2.1.chr2.g00035.t01#2       216   125260836:   mNeoVis.vulgivaga.2.1.chr8.g00505.t01#1    mNeoVis.evergladensis.1.1.chr8.g00496.t01#3    mNeoVis.2.1.1.chr8_hap1.g00508.t01#5    mNeoVis.3.1.1.chr8.g00464.t01#6    mNeoVis.evergladensis.2.1.chr9.g00499.t01#2   mNeoVis.vulgivaga.1.1.chr8.g00494.t01#4       217   125260837:   mNeoVis.evergladensis.2.1.chr15.g00610.t01#2    mNeoVis.vulgivaga.2.1.chr15.g00624.t01#1    mNeoVis.vulgivaga.1.1.chr15.g00614.t01#4   mNeoVis.evergladensis.1.1.chr15.g00001.t01#3       218   125260838:   mNeoVis.3.1.1.chr8.g00465.t01#6    mNeoVis.vulgivaga.2.1.chr8.g00506.t01#1    mNeoVis.evergladensis.2.1.chr9.g00500.t01#2    mNeoVis.evergladensis.1.1.chr8.g00497.t01#3    mNeoVis.vulgivaga.1.1.chr8.g00495.t01#4   mNeoVis.2.1.1.chr8_hap1.g00509.t01#5       219   125260839:   mNeoVis.2.1.1.chr7_hap1.g00596.t01#5    mNeoVis.vulgivaga.1.1.chr7.g00605.t01#4    mNeoVis.3.1.1.chr7.g00576.t01#6    mNeoVis.vulgivaga.2.1.chr7.g00601.t01#1    mNeoVis.evergladensis.1.1.chr7.g00606.t01#3   mNeoVis.evergladensis.2.1.chr7.g00598.t01#2       220   125260840:   mNeoVis.evergladensis.1.1.chr2.g00037.t01#3    mNeoVis.vulgivaga.1.1.chr2.g00037.t01#4    mNeoVis.2.1.1.chr5_hap1.g00036.t01#5    mNeoVis.vulgivaga.2.1.chr2.g00036.t01#1    mNeoVis.evergladensis.2.1.chr2.g00036.t01#2   mNeoVis.3.1.1.chr2.g00018.t01#6       221   125260841:   mNeoVis.vulgivaga.2.1.chr2.g00037.t01#1    mNeoVis.3.1.1.chr2.g00019.t01#6    mNeoVis.evergladensis.1.1.chr2.g00038.t01#3    mNeoVis.vulgivaga.1.1.chr2.g00038.t01#4    mNeoVis.evergladensis.2.1.chr2.g00037.t01#2   mNeoVis.2.1.1.chr5_hap1.g00037.t01#5       222   125260842:   mNeoVis.3.1.1.chr2.g00020.t01#6    mNeoVis.2.1.1.chr5_hap1.g00038.t01#5    mNeoVis.vulgivaga.1.1.chr2.g00039.t01#4    mNeoVis.evergladensis.2.1.chr2.g00038.t01#2    mNeoVis.vulgivaga.2.1.chr2.g00038.t01#1   mNeoVis.evergladensis.1.1.chr2.g00039.t01#3       223   125260843:   mNeoVis.vulgivaga.2.1.chr2.g00039.t01#1    mNeoVis.evergladensis.1.1.chr2.g00040.t01#3    mNeoVis.2.1.1.chr5_hap1.g00039.t01#5    mNeoVis.vulgivaga.1.1.chr2.g00040.t01#4    mNeoVis.evergladensis.2.1.chr2.g00039.t01#2   mNeoVis.3.1.1.chr2.g00021.t01#6       224   125260844:   mNeoVis.evergladensis.1.1.chr9.g00973.t01#3    mNeoVis.evergladensis.2.1.chr8.g00976.t01#2    mNeoVis.3.1.1.chr9.g00936.t01#6    mNeoVis.2.1.1.chr10_hap1.g01005.t01#5    mNeoVis.vulgivaga.2.1.chr9.g00963.t01#1   mNeoVis.vulgivaga.1.1.chr9.g00977.t01#4       225   125260845:   mNeoVis.evergladensis.2.1.chr9.g00501.t01#2    mNeoVis.evergladensis.1.1.chr8.g00498.t01#3    mNeoVis.3.1.1.chr8.g00466.t01#6    mNeoVis.vulgivaga.1.1.chr8.g00496.t01#4    mNeoVis.vulgivaga.2.1.chr8.g00507.t01#1   mNeoVis.2.1.1.chr8_hap1.g00510.t01#5       226   125260846:   mNeoVis.evergladensis.2.1.chr2.g00040.t01#2    mNeoVis.vulgivaga.1.1.chr2.g00041.t01#4    mNeoVis.vulgivaga.2.1.chr2.g00040.t01#1    mNeoVis.2.1.1.chr5_hap1.g00040.t01#5   mNeoVis.evergladensis.1.1.chr2.g00041.t01#3       227   125260847:   mNeoVis.vulgivaga.2.1.chr2.g00041.t01#1    mNeoVis.evergladensis.1.1.chr2.g00042.t01#3    mNeoVis.evergladensis.2.1.chr2.g00041.t01#2    mNeoVis.2.1.1.chr5_hap1.g00041.t01#5   mNeoVis.vulgivaga.1.1.chr2.g00042.t01#4       228   125260848:   mNeoVis.3.1.1.chr7.g00578.t01#6    mNeoVis.evergladensis.2.1.chr7.g00599.t01#2    mNeoVis.vulgivaga.2.1.chr7.g00602.t01#1    mNeoVis.evergladensis.1.1.chr7.g00607.t01#3    mNeoVis.2.1.1.chr7_hap1.g00597.t01#5   mNeoVis.vulgivaga.1.1.chr7.g00606.t01#4       229   125260849:   mNeoVis.2.1.1.chr7_hap1.g01084.t01#5    mNeoVis.vulgivaga.2.1.chr7.g01087.t01#1    mNeoVis.vulgivaga.1.1.chr7.g01077.t01#4    mNeoVis.3.1.1.chr7.g01013.t01#6    mNeoVis.evergladensis.2.1.chr7.g01075.t01#2   mNeoVis.evergladensis.1.1.chr7.g01080.t01#3       230   125260850:   mNeoVis.vulgivaga.2.1.chr8.g00508.t01#1    mNeoVis.3.1.1.chr8.g00467.t01#6    mNeoVis.evergladensis.2.1.chr9.g00502.t01#2   mNeoVis.2.1.1.chr8_hap1.g00511.t01#5         231   125260851:   mNeoVis.3.1.1.chr8.g00468.t01#6    mNeoVis.vulgivaga.1.1.chr8.g00497.t01#4    mNeoVis.vulgivaga.2.1.chr8.g00509.t01#1    mNeoVis.evergladensis.1.1.chr8.g00499.t01#3    mNeoVis.evergladensis.2.1.chr9.g00503.t01#2   mNeoVis.2.1.1.chr8_hap1.g00512.t01#5       232   125260852:   mNeoVis.vulgivaga.2.1.chr8.g00510.t01#1    mNeoVis.2.1.1.chr8_hap1.g00513.t01#5    mNeoVis.evergladensis.2.1.chr9.g00504.t01#2    mNeoVis.evergladensis.1.1.chr8.g00500.t01#3   mNeoVis.3.1.1.chr8.g00469.t01#6       233   125260853:   mNeoVis.evergladensis.2.1.chr2.g00042.t01#2    mNeoVis.vulgivaga.1.1.chr2.g00043.t01#4    mNeoVis.3.1.1.chr2.g00024.t01#6    mNeoVis.vulgivaga.2.1.chr2.g00042.t01#1    mNeoVis.evergladensis.1.1.chr2.g00043.t01#3   mNeoVis.2.1.1.chr5_hap1.g00042.t01#5       234   125260854:  mNeoVis.vulgivaga.2.1.chr2.g00043.t01#1               235   125260855:   mNeoVis.2.1.1.chr8_hap1.g00514.t01#5    mNeoVis.3.1.1.chr8.g00470.t01#6    mNeoVis.vulgivaga.2.1.chr8.g00511.t01#1    mNeoVis.evergladensis.1.1.chr8.g00501.t01#3    mNeoVis.evergladensis.2.1.chr9.g00505.t01#2   mNeoVis.vulgivaga.1.1.chr8.g00500.t01#4       236   125260856:   mNeoVis.vulgivaga.1.1.chr13.g00699.t01#4    mNeoVis.3.1.1.chr13.g00659.t01#6    mNeoVis.2.1.1.chr13_hap1.g00743.t01#5    mNeoVis.vulgivaga.2.1.chr13.g00705.t01#1    mNeoVis.evergladensis.1.1.chr13.g00697.t01#3   mNeoVis.evergladensis.2.1.chr13.g00708.t01#2       237   125260857:   mNeoVis.2.1.1.chr5_hap1.g00044.t01#5    mNeoVis.vulgivaga.1.1.chr2.g00045.t01#4    mNeoVis.vulgivaga.2.1.chr2.g00044.t01#1    mNeoVis.evergladensis.1.1.chr2.g00045.t01#3   mNeoVis.evergladensis.2.1.chr2.g00044.t01#2       238   125260858:   mNeoVis.evergladensis.1.1.chr9.g01235.t01#3    mNeoVis.vulgivaga.2.1.chr9.g01224.t01#1    mNeoVis.2.1.1.chr10_hap1.g01264.t01#5    mNeoVis.vulgivaga.1.1.chr9.g01240.t01#4   mNeoVis.evergladensis.2.1.chr8.g01241.t01#2       239   125260859:   mNeoVis.evergladensis.1.1.chr2.g00046.t01#3    mNeoVis.vulgivaga.2.1.chr2.g00045.t01#1    mNeoVis.3.1.1.chr2.g00025.t01#6   mNeoVis.evergladensis.2.1.chr2.g00045.t01#2       240   125260860:   mNeoVis.evergladensis.1.1.chr2.g00047.t01#3    mNeoVis.2.1.1.chr5_hap1.g00045.t01#5    mNeoVis.vulgivaga.2.1.chr2.g00046.t01#1    mNeoVis.evergladensis.2.1.chr2.g00046.t01#2   mNeoVis.vulgivaga.1.1.chr2.g00047.t01#4       241   125260861:   mNeoVis.2.1.1.chr1_hap1.g00815.t01#5    mNeoVis.3.1.1.chr1.g00842.t01#6    mNeoVis.vulgivaga.2.1.chr1.g00838.t01#1    mNeoVis.evergladensis.2.1.chr1.g00822.t01#2    mNeoVis.evergladensis.1.1.chr1.g00834.t01#3   mNeoVis.vulgivaga.1.1.chr1.g00835.t01#4       242   125260862:   mNeoVis.vulgivaga.2.1.chr9.g00918.t01#1    mNeoVis.evergladensis.1.1.chr9.g00927.t01#3    mNeoVis.vulgivaga.1.1.chr9.g00930.t01#4    mNeoVis.3.1.1.chr9.g00891.t01#6    mNeoVis.evergladensis.2.1.chr8.g00931.t01#2   mNeoVis.2.1.1.chr10_hap1.g00961.t01#5       243   125260863:   mNeoVis.vulgivaga.2.1.chr2.g00047.t01#1    mNeoVis.evergladensis.1.1.chr2.g00048.t01#3   mNeoVis.3.1.1.chr2.g00026.t01#6           244   125260864:   mNeoVis.evergladensis.2.1.chr9.g00399.t01#2    mNeoVis.evergladensis.1.1.chr8.g00396.t01#3    mNeoVis.3.1.1.chr8.g00365.t01#6    mNeoVis.vulgivaga.2.1.chr8.g00405.t01#1    mNeoVis.2.1.1.chr8_hap1.g00401.t01#5   mNeoVis.vulgivaga.1.1.chr8.g00392.t01#4       245   125260865:   mNeoVis.vulgivaga.1.1.chr15.g00558.t01#4    mNeoVis.vulgivaga.2.1.chr15.g00567.t01#1    mNeoVis.2.1.1.chr15_hap1.g00733.t01#5   mNeoVis.evergladensis.2.1.chr15.g00555.t01#2       246   125260866:   mNeoVis.vulgivaga.2.1.chr8.g00406.t01#1    mNeoVis.evergladensis.2.1.chr9.g00400.t01#2    mNeoVis.evergladensis.1.1.chr8.g00397.t01#3    mNeoVis.3.1.1.chr8.g00366.t01#6    mNeoVis.2.1.1.chr8_hap1.g00402.t01#5   mNeoVis.vulgivaga.1.1.chr8.g00393.t01#4       247   125260867:   mNeoVis.2.1.1.chr5_hap1.g00048.t01#5   mNeoVis.vulgivaga.2.1.chr2.g00048.t01#1             248   125260868:   mNeoVis.vulgivaga.1.1.chr2.g00049.t01#4   mNeoVis.2.1.1.chr3_hap1.g00212.t01#5             249   125260869:   mNeoVis.2.1.1.chr5_hap1.g00049.t01#5    mNeoVis.vulgivaga.2.1.chr2.g00049.t01#1    mNeoVis.vulgivaga.1.1.chr2.g00050.t01#4    mNeoVis.evergladensis.1.1.chr2.g00050.t01#3   mNeoVis.evergladensis.2.1.chr2.g00049.t01#2       250   125260870:   mNeoVis.vulgivaga.2.1.chr8.g00407.t01#1    mNeoVis.2.1.1.chr8_hap1.g00403.t01#5    mNeoVis.vulgivaga.1.1.chr8.g00394.t01#4    mNeoVis.evergladensis.2.1.chr9.g00401.t01#2    mNeoVis.3.1.1.chr8.g00367.t01#6   mNeoVis.evergladensis.1.1.chr8.g00398.t01#3       251   125260871:   mNeoVis.evergladensis.1.1.chr7.g00608.t01#3    mNeoVis.2.1.1.chr7_hap1.g00598.t01#5    mNeoVis.3.1.1.chr7.g00581.t01#6    mNeoVis.vulgivaga.2.1.chr7.g00603.t01#1    mNeoVis.evergladensis.2.1.chr7.g00600.t01#2   mNeoVis.vulgivaga.1.1.chr7.g00607.t01#4       252   125260872:   mNeoVis.2.1.1.chr8_hap1.g00404.t01#5    mNeoVis.vulgivaga.1.1.chr8.g00395.t01#4    mNeoVis.evergladensis.2.1.chr9.g00402.t01#2    mNeoVis.vulgivaga.2.1.chr8.g00408.t01#1    mNeoVis.3.1.1.chr8.g00368.t01#6   mNeoVis.evergladensis.1.1.chr8.g00399.t01#3       253   125260873:   mNeoVis.3.1.1.chr2.g00028.t01#6    mNeoVis.vulgivaga.2.1.chr2.g00050.t01#1    mNeoVis.vulgivaga.1.1.chr2.g00051.t01#4    mNeoVis.evergladensis.2.1.chr2.g00050.t01#2    mNeoVis.2.1.1.chr5_hap1.g00050.t01#5   mNeoVis.evergladensis.1.1.chr2.g00051.t01#3       254   125260874:   mNeoVis.vulgivaga.1.1.chr13.g00540.t01#4    mNeoVis.evergladensis.2.1.chr13.g00551.t01#2    mNeoVis.evergladensis.1.1.chr13.g00546.t01#3    mNeoVis.vulgivaga.2.1.chr13.g00553.t01#1    mNeoVis.2.1.1.chr13_hap1.g00567.t01#5   mNeoVis.3.1.1.chr13.g00505.t01#6       255   125260875:   mNeoVis.vulgivaga.1.1.chr5.g01242.t01#4    mNeoVis.evergladensis.2.1.chr5.g01274.t01#2    mNeoVis.2.1.1.chr2_hap1.g01331.t01#5    mNeoVis.evergladensis.1.1.chr5.g01274.t01#3    mNeoVis.3.1.1.chr5.g01257.t01#6   mNeoVis.vulgivaga.2.1.chr5.g01271.t01#1       256   125260876:   mNeoVis.vulgivaga.2.1.chr2.g00051.t01#1    mNeoVis.evergladensis.1.1.chr2.g00052.t01#3    mNeoVis.3.1.1.chr2.g00029.t01#6    mNeoVis.2.1.1.chr5_hap1.g00051.t01#5    mNeoVis.evergladensis.2.1.chr2.g00051.t01#2   mNeoVis.vulgivaga.1.1.chr2.g00052.t01#4       257   125260877:   mNeoVis.evergladensis.1.1.chr15.g00057.t01#3    mNeoVis.3.1.1.chr15.g00511.t01#6    mNeoVis.2.1.1.chr15_hap1.g00734.t01#5    mNeoVis.evergladensis.2.1.chr15.g00556.t01#2    mNeoVis.vulgivaga.2.1.chr15.g00568.t01#1   mNeoVis.vulgivaga.1.1.chr15.g00559.t01#4       258   125260878:   mNeoVis.evergladensis.2.1.chr2.g00052.t01#2    mNeoVis.vulgivaga.2.1.chr2.g00052.t01#1    mNeoVis.vulgivaga.1.1.chr2.g00053.t01#4    mNeoVis.2.1.1.chr5_hap1.g00052.t01#5    mNeoVis.3.1.1.chr2.g00030.t01#6   mNeoVis.evergladensis.1.1.chr2.g00053.t01#3       259   125260879:   mNeoVis.evergladensis.2.1.chr9.g00403.t01#2    mNeoVis.vulgivaga.1.1.chr8.g00396.t01#4    mNeoVis.evergladensis.1.1.chr8.g00400.t01#3   mNeoVis.vulgivaga.2.1.chr8.g00409.t01#1         260   125260880:   mNeoVis.vulgivaga.1.1.chr7.g00608.t01#4    mNeoVis.evergladensis.1.1.chr7.g00609.t01#3    mNeoVis.vulgivaga.2.1.chr7.g00604.t01#1    mNeoVis.2.1.1.chr7_hap1.g00599.t01#5   mNeoVis.evergladensis.2.1.chr7.g00601.t01#2       261   125260881:   mNeoVis.vulgivaga.2.1.chr8.g00410.t01#1    mNeoVis.vulgivaga.1.1.chr8.g00397.t01#4    mNeoVis.3.1.1.chr8.g00370.t01#6    mNeoVis.evergladensis.1.1.chr8.g00401.t01#3    mNeoVis.evergladensis.2.1.chr9.g00404.t01#2   mNeoVis.2.1.1.chr8_hap1.g00406.t01#5       262   125260882:   mNeoVis.3.1.1.chr15.g00512.t01#6    mNeoVis.vulgivaga.1.1.chr15.g00560.t01#4    mNeoVis.vulgivaga.2.1.chr15.g00569.t01#1    mNeoVis.evergladensis.1.1.chr15.g00056.t01#3    mNeoVis.2.1.1.chr15_hap1.g00735.t01#5   mNeoVis.evergladensis.2.1.chr15.g00557.t01#2       263   125260883:   mNeoVis.3.1.1.chr2.g00942.t01#6    mNeoVis.vulgivaga.1.1.chr2.g01023.t01#4    mNeoVis.evergladensis.1.1.chr8.g00402.t01#3    mNeoVis.2.1.1.chr8_hap1.g00407.t01#5    mNeoVis.vulgivaga.1.1.chr8.g00398.t01#4    mNeoVis.vulgivaga.2.1.chr8.g00411.t01#1    mNeoVis.vulgivaga.2.1.chr2.g01025.t01#1    mNeoVis.evergladensis.2.1.chr9.g00405.t01#2    mNeoVis.3.1.1.chr8.g00371.t01#6      264   125260884:   mNeoVis.3.1.1.chr6.g00775.t01#6    mNeoVis.vulgivaga.2.1.chr8.g00414.t01#1    mNeoVis.vulgivaga.1.1.chr8.g00401.t01#4    mNeoVis.vulgivaga.1.1.chr8.g00399.t01#4    mNeoVis.vulgivaga.2.1.chr8.g00412.t01#1    mNeoVis.evergladensis.1.1.chr8.g00403.t01#3    mNeoVis.evergladensis.2.1.chr9.g00406.t01#2   mNeoVis.2.1.1.chr8_hap1.g00408.t01#5     265   125260885:   mNeoVis.evergladensis.1.1.chr8.g00404.t01#3    mNeoVis.evergladensis.2.1.chr9.g00407.t01#2    mNeoVis.vulgivaga.2.1.chr8.g00413.t01#1    mNeoVis.2.1.1.chr8_hap1.g00409.t01#5    mNeoVis.vulgivaga.1.1.chr8.g00400.t01#4    mNeoVis.3.1.1.chr8.g00373.t01#6   mNeoVis.3.1.1.chr8.g00372.t01#6       266   125260886:   mNeoVis.vulgivaga.2.1.chr8.g00415.t01#1    mNeoVis.evergladensis.2.1.chr9.g00408.t01#2    mNeoVis.evergladensis.1.1.chr8.g00405.t01#3    mNeoVis.vulgivaga.1.1.chr8.g00402.t01#4    mNeoVis.3.1.1.chr8.g00374.t01#6   mNeoVis.2.1.1.chr8_hap1.g00410.t01#5       267   125260887:   mNeoVis.vulgivaga.2.1.chr2.g00053.t01#1    mNeoVis.evergladensis.2.1.chr2.g00053.t01#2    mNeoVis.vulgivaga.1.1.chr2.g00054.t01#4    mNeoVis.evergladensis.1.1.chr2.g00054.t01#3   mNeoVis.2.1.1.chr5_hap1.g00053.t01#5       268   125260888:   mNeoVis.vulgivaga.2.1.chr2.g00054.t01#1    mNeoVis.evergladensis.2.1.chr2.g00054.t01#2    mNeoVis.2.1.1.chr5_hap1.g00054.t01#5    mNeoVis.vulgivaga.1.1.chr2.g00055.t01#4   mNeoVis.evergladensis.1.1.chr2.g00055.t01#3       269   125260889:   mNeoVis.vulgivaga.2.1.chr8.g00416.t01#1    mNeoVis.vulgivaga.1.1.chr8.g00403.t01#4    mNeoVis.2.1.1.chr8_hap1.g00411.t01#5    mNeoVis.evergladensis.2.1.chr9.g00409.t01#2    mNeoVis.evergladensis.1.1.chr8.g00406.t01#3   mNeoVis.3.1.1.chr8.g00375.t01#6       270   125260890:   mNeoVis.evergladensis.2.1.chr2.g00055.t01#2    mNeoVis.vulgivaga.1.1.chr2.g00056.t01#4    mNeoVis.vulgivaga.2.1.chr2.g00055.t01#1    mNeoVis.2.1.1.chr5_hap1.g00055.t01#5   mNeoVis.evergladensis.1.1.chr2.g00056.t01#3       271   125260891:   mNeoVis.vulgivaga.1.1.chr8.g00404.t01#4    mNeoVis.evergladensis.1.1.chr8.g00407.t01#3    mNeoVis.3.1.1.chr8.g00376.t01#6    mNeoVis.2.1.1.chr8_hap1.g00412.t01#5    mNeoVis.vulgivaga.2.1.chr8.g00417.t01#1   mNeoVis.evergladensis.2.1.chr9.g00410.t01#2       272   125260892:   mNeoVis.vulgivaga.2.1.chr2.g00056.t01#1    mNeoVis.evergladensis.2.1.chr2.g00056.t01#2    mNeoVis.evergladensis.1.1.chr2.g00057.t01#3    mNeoVis.2.1.1.chr5_hap1.g00056.t01#5   mNeoVis.vulgivaga.1.1.chr2.g00057.t01#4       273   125260893:   mNeoVis.vulgivaga.1.1.chr2.g00058.t01#4    mNeoVis.2.1.1.chr5_hap1.g00057.t01#5    mNeoVis.vulgivaga.2.1.chr2.g00057.t01#1    mNeoVis.evergladensis.1.1.chr2.g00058.t01#3   mNeoVis.evergladensis.2.1.chr2.g00057.t01#2       274   125260894:   mNeoVis.vulgivaga.1.1.chr8.g00405.t01#4    mNeoVis.evergladensis.1.1.chr8.g00408.t01#3    mNeoVis.evergladensis.2.1.chr9.g00411.t01#2    mNeoVis.2.1.1.chr8_hap1.g00413.t01#5    mNeoVis.2.1.1.chr9_hap1.g00811.t01#5    mNeoVis.evergladensis.1.1.chr10.g00801.t01#3    mNeoVis.evergladensis.2.1.chr10.g00801.t01#2    mNeoVis.3.1.1.chr10.g00776.t01#6    mNeoVis.vulgivaga.1.1.chr10.g00786.t01#4      275   125260895:   mNeoVis.vulgivaga.1.1.chr8.g00406.t01#4    mNeoVis.evergladensis.1.1.chr8.g00409.t01#3    mNeoVis.evergladensis.2.1.chr9.g00412.t01#2    mNeoVis.3.1.1.chr8.g00378.t01#6    mNeoVis.2.1.1.chr8_hap1.g00414.t01#5   mNeoVis.vulgivaga.2.1.chr8.g00419.t01#1       276   125260896:  mNeoVis.vulgivaga.1.1.chr3.g00401.t01#4               277   125260897:  mNeoVis.vulgivaga.2.1.chr2.g00058.t01#1               278   125260898:   mNeoVis.2.1.1.chr8_hap1.g00415.t01#5    mNeoVis.vulgivaga.2.1.chr8.g00420.t01#1    mNeoVis.3.1.1.chr8.g00379.t01#6    mNeoVis.evergladensis.2.1.chr9.g00413.t01#2    mNeoVis.evergladensis.1.1.chr8.g00410.t01#3   mNeoVis.vulgivaga.1.1.chr8.g00407.t01#4       279   125260899:   mNeoVis.evergladensis.1.1.chr1.g02106.t01#3    mNeoVis.3.1.1.chr1.g02091.t01#6    mNeoVis.vulgivaga.2.1.chr1.g02096.t01#1    mNeoVis.2.1.1.chr1_hap1.g02136.t01#5    mNeoVis.evergladensis.2.1.chr1.g02106.t01#2   mNeoVis.vulgivaga.1.1.chr1.g02110.t01#4       280   125260900:   mNeoVis.3.1.1.chr8.g00380.t01#6    mNeoVis.2.1.1.chr8_hap1.g00416.t01#5    mNeoVis.vulgivaga.2.1.chr8.g00421.t01#1    mNeoVis.evergladensis.1.1.chr8.g00411.t01#3   mNeoVis.vulgivaga.1.1.chr8.g00408.t01#4       281   125260901:   mNeoVis.evergladensis.1.1.chr1.g02107.t01#3    mNeoVis.vulgivaga.1.1.chr1.g02111.t01#4    mNeoVis.vulgivaga.2.1.chr1.g02097.t01#1   mNeoVis.2.1.1.chr1_hap1.g02137.t01#5         282   125260902:   mNeoVis.evergladensis.2.1.chr9.g00414.t01#2    mNeoVis.vulgivaga.2.1.chr8.g00422.t01#1    mNeoVis.3.1.1.chr8.g00381.t01#6    mNeoVis.2.1.1.chr8_hap1.g00417.t01#5    mNeoVis.evergladensis.1.1.chr8.g00412.t01#3   mNeoVis.vulgivaga.1.1.chr8.g00409.t01#4       283   125260903:   mNeoVis.2.1.1.chr1_hap1.g02138.t01#5    mNeoVis.vulgivaga.1.1.chr1.g02112.t01#4    mNeoVis.vulgivaga.2.1.chr1.g02098.t01#1    mNeoVis.evergladensis.1.1.chr1.g02108.t01#3   mNeoVis.evergladensis.2.1.chr1.g02107.t01#2       284   125260904:   mNeoVis.evergladensis.1.1.chr1.g02109.t01#3    mNeoVis.2.1.1.chr1_hap1.g02139.t01#5    mNeoVis.3.1.1.chr1.g02092.t01#6    mNeoVis.evergladensis.2.1.chr1.g02108.t01#2    mNeoVis.vulgivaga.1.1.chr1.g02113.t01#4   mNeoVis.vulgivaga.2.1.chr1.g02099.t01#1       285   125260905:   mNeoVis.2.1.1.chr8_hap1.g00420.t01#5    mNeoVis.3.1.1.chr8.g00384.t01#6    mNeoVis.vulgivaga.1.1.chr8.g00413.t01#4    mNeoVis.evergladensis.1.1.chr8.g00415.t01#3    mNeoVis.evergladensis.2.1.chr9.g00417.t01#2   mNeoVis.vulgivaga.2.1.chr8.g00425.t01#1       286   125260906:   mNeoVis.2.1.1.chr8_hap1.g00418.t01#5    mNeoVis.2.1.1.chr8_hap1.g00419.t01#5    mNeoVis.evergladensis.1.1.chr8.g00413.t01#3    mNeoVis.evergladensis.1.1.chr8.g00414.t01#3    mNeoVis.3.1.1.chr8.g00383.t01#6   mNeoVis.evergladensis.2.1.chr9.g00416.t01#2       287   125260907:   mNeoVis.3.1.1.chr8.g00382.t01#6    mNeoVis.vulgivaga.1.1.chr8.g00410.t01#4    mNeoVis.vulgivaga.2.1.chr8.g00423.t01#1   mNeoVis.evergladensis.2.1.chr9.g00415.t01#2       288   125260908:   mNeoVis.vulgivaga.1.1.chr8.g00412.t01#4   mNeoVis.vulgivaga.2.1.chr8.g00424.t01#1             289   125260909:  mNeoVis.vulgivaga.1.1.chr8.g00411.t01#4               290   125260910:   mNeoVis.vulgivaga.2.1.chr1.g02100.t01#1    mNeoVis.evergladensis.2.1.chr1.g02109.t01#2    mNeoVis.3.1.1.chr1.g02093.t01#6    mNeoVis.vulgivaga.1.1.chr1.g02114.t01#4    mNeoVis.evergladensis.1.1.chr1.g02110.t01#3   mNeoVis.2.1.1.chr1_hap1.g02140.t01#5       291   125260911:   mNeoVis.2.1.1.chr1_hap1.g02141.t01#5    mNeoVis.vulgivaga.1.1.chr1.g02115.t01#4    mNeoVis.evergladensis.1.1.chr1.g02111.t01#3    mNeoVis.vulgivaga.2.1.chr1.g02101.t01#1    mNeoVis.3.1.1.chr1.g02094.t01#6   mNeoVis.evergladensis.2.1.chr1.g02110.t01#2       292   125260912:   mNeoVis.3.1.1.chr5.g01555.t01#6    mNeoVis.2.1.1.chr2_hap1.g01633.t01#5    mNeoVis.vulgivaga.1.1.chr5.g01551.t01#4    mNeoVis.evergladensis.2.1.chr5.g01573.t01#2    mNeoVis.evergladensis.1.1.chr5.g01578.t01#3   mNeoVis.vulgivaga.2.1.chr5.g01575.t01#1       293   125260913:   mNeoVis.evergladensis.2.1.chr12.g00678.t01#2    mNeoVis.evergladensis.1.1.chr12.g00686.t01#3    mNeoVis.3.1.1.chr12.g00682.t01#6    mNeoVis.vulgivaga.2.1.chr12.g00704.t01#1    mNeoVis.vulgivaga.1.1.chr12.g00699.t01#4   mNeoVis.2.1.1.chr12_hap1.g00749.t01#5       294   125260914:   mNeoVis.evergladensis.2.1.chr9.g00418.t01#2    mNeoVis.vulgivaga.2.1.chr8.g00426.t01#1    mNeoVis.2.1.1.chr8_hap1.g00421.t01#5    mNeoVis.vulgivaga.1.1.chr8.g00414.t01#4    mNeoVis.3.1.1.chr8.g00385.t01#6    mNeoVis.evergladensis.1.1.chr8.g00416.t01#3    mNeoVis.evergladensis.2.1.chr9.g00420.t01#2   mNeoVis.evergladensis.1.1.chr8.g00417.t01#3     295   125260915:   mNeoVis.2.1.1.chr8_hap1.g00422.t01#5    mNeoVis.vulgivaga.2.1.chr8.g00429.t01#1    mNeoVis.vulgivaga.1.1.chr8.g00415.t01#4    mNeoVis.3.1.1.chr8.g00386.t01#6    mNeoVis.vulgivaga.2.1.chr8.g00427.t01#1    mNeoVis.2.1.1.chr8_hap1.g00424.t01#5   mNeoVis.evergladensis.2.1.chr9.g00419.t01#2     296   125260916:   mNeoVis.vulgivaga.1.1.chr8.g00417.t01#4   mNeoVis.vulgivaga.1.1.chr8.g00416.t01#4             297   125260917:   mNeoVis.evergladensis.1.1.chr7.g00611.t01#3    mNeoVis.2.1.1.chr7_hap1.g00600.t01#5    mNeoVis.evergladensis.2.1.chr7.g00602.t01#2    mNeoVis.vulgivaga.1.1.chr7.g00609.t01#4    mNeoVis.vulgivaga.2.1.chr7.g00605.t01#1   mNeoVis.3.1.1.chr7.g00583.t01#6       298   125260918:   mNeoVis.evergladensis.1.1.chr1.g02112.t01#3    mNeoVis.vulgivaga.2.1.chr1.g02102.t01#1    mNeoVis.2.1.1.chr1_hap1.g02142.t01#5    mNeoVis.vulgivaga.1.1.chr1.g02116.t01#4    mNeoVis.evergladensis.2.1.chr1.g02111.t01#2   mNeoVis.3.1.1.chr1.g02096.t01#6       299   125260919:   mNeoVis.3.1.1.chr1.g02097.t01#6    mNeoVis.evergladensis.1.1.chr1.g02113.t01#3    mNeoVis.2.1.1.chr1_hap1.g02143.t01#5    mNeoVis.evergladensis.2.1.chr1.g02112.t01#2   mNeoVis.vulgivaga.2.1.chr1.g02103.t01#1       300   125260920:   mNeoVis.2.1.1.chr8_hap1.g00423.t01#5    mNeoVis.2.1.1.chr10_hap1.g00324.t01#5    mNeoVis.vulgivaga.2.1.chr2.g00915.t01#1    mNeoVis.evergladensis.1.1.chr11.g00648.t01#3    mNeoVis.vulgivaga.1.1.chr5.g01636.t01#4    mNeoVis.2.1.1.chr2_hap1.g01717.t01#5    mNeoVis.vulgivaga.2.1.chr5.g01658.t01#1    mNeoVis.evergladensis.2.1.chr2.g00911.t01#2    mNeoVis.evergladensis.2.1.chr5.g01656.t01#2      301   125260921:   mNeoVis.vulgivaga.1.1.chr1.g01999.t01#4    mNeoVis.evergladensis.2.1.chr1.g01997.t01#2    mNeoVis.2.1.1.chr1_hap1.g02030.t01#5    mNeoVis.evergladensis.1.1.chr1.g01995.t01#3    mNeoVis.vulgivaga.2.1.chr1.g01990.t01#1   mNeoVis.3.1.1.chr1.g01984.t01#6       302   125260922:   mNeoVis.vulgivaga.2.1.chr1.g02104.t01#1    mNeoVis.evergladensis.2.1.chr1.g02113.t01#2    mNeoVis.vulgivaga.1.1.chr1.g02118.t01#4    mNeoVis.3.1.1.chr1.g02098.t01#6    mNeoVis.evergladensis.1.1.chr1.g02114.t01#3   mNeoVis.2.1.1.chr1_hap1.g02144.t01#5       303   125260923:   mNeoVis.vulgivaga.2.1.chr8.g00430.t01#1    mNeoVis.evergladensis.2.1.chr9.g00421.t01#2    mNeoVis.evergladensis.1.1.chr8.g00418.t01#3    mNeoVis.2.1.1.chr8_hap1.g00425.t01#5    mNeoVis.3.1.1.chr8.g00388.t01#6   mNeoVis.vulgivaga.1.1.chr8.g00418.t01#4       304   125260924:   mNeoVis.vulgivaga.1.1.chr1.g02119.t01#4    mNeoVis.2.1.1.chr1_hap1.g02145.t01#5    mNeoVis.vulgivaga.2.1.chr1.g02105.t01#1    mNeoVis.evergladensis.1.1.chr1.g02115.t01#3    mNeoVis.3.1.1.chr1.g02099.t01#6   mNeoVis.evergladensis.2.1.chr1.g02114.t01#2       305   125260925:  mNeoVis.vulgivaga.2.1.chr1.g02106.t01#1               306   125260926:   mNeoVis.vulgivaga.2.1.chr8.g00431.t01#1    mNeoVis.evergladensis.1.1.chr8.g00419.t01#3    mNeoVis.3.1.1.chr8.g00389.t01#6    mNeoVis.evergladensis.2.1.chr9.g00422.t01#2    mNeoVis.2.1.1.chr8_hap1.g00426.t01#5   mNeoVis.vulgivaga.1.1.chr8.g00419.t01#4       307   125260927:   mNeoVis.vulgivaga.1.1.chr8.g00420.t01#4    mNeoVis.3.1.1.chr8.g00390.t01#6    mNeoVis.evergladensis.2.1.chr9.g00423.t01#2    mNeoVis.2.1.1.chr8_hap1.g00427.t01#5    mNeoVis.evergladensis.1.1.chr8.g00420.t01#3   mNeoVis.vulgivaga.2.1.chr8.g00432.t01#1       308   125260928:   mNeoVis.evergladensis.2.1.chr9.g00425.t01#2    mNeoVis.evergladensis.1.1.chr8.g00421.t01#3    mNeoVis.vulgivaga.1.1.chr8.g00421.t01#4   mNeoVis.3.1.1.chr8.g00391.t01#6         309   125260929:   mNeoVis.vulgivaga.2.1.chr8.g00434.t01#1    mNeoVis.vulgivaga.1.1.chr8.g00422.t01#4    mNeoVis.evergladensis.2.1.chr9.g00426.t01#2   mNeoVis.evergladensis.1.1.chr8.g00422.t01#3       310   125260930:  mNeoVis.vulgivaga.2.1.chr8.g00433.t01#1               311   125260931:   mNeoVis.evergladensis.1.1.chr7.g00610.t01#3    mNeoVis.vulgivaga.2.1.chr7.g00606.t01#1    mNeoVis.3.1.1.chr7.g00584.t01#6    mNeoVis.3.1.1.chr7.g00579.t01#6    mNeoVis.2.1.1.chr7_hap1.g00601.t01#5    mNeoVis.evergladensis.2.1.chr7.g00603.t01#2   mNeoVis.vulgivaga.1.1.chr7.g00610.t01#4     312   125260932:   mNeoVis.3.1.1.chr1.g02101.t01#6    mNeoVis.evergladensis.1.1.chr1.g02118.t01#3    mNeoVis.evergladensis.2.1.chr1.g02116.t01#2    mNeoVis.vulgivaga.1.1.chr1.g02121.t01#4    mNeoVis.vulgivaga.2.1.chr1.g02107.t01#1   mNeoVis.2.1.1.chr1_hap1.g02147.t01#5       313   125260933:   mNeoVis.evergladensis.1.1.chr1.g02119.t01#3    mNeoVis.3.1.1.chr1.g02102.t01#6    mNeoVis.vulgivaga.1.1.chr1.g02122.t01#4    mNeoVis.vulgivaga.2.1.chr1.g02108.t01#1    mNeoVis.2.1.1.chr1_hap1.g02148.t01#5   mNeoVis.evergladensis.2.1.chr1.g02117.t01#2       314   125260934:   mNeoVis.vulgivaga.1.1.chr8.g00423.t01#4    mNeoVis.evergladensis.1.1.chr8.g00423.t01#3    mNeoVis.3.1.1.chr8.g00394.t01#6    mNeoVis.evergladensis.2.1.chr9.g00427.t01#2    mNeoVis.2.1.1.chr8_hap1.g00437.t01#5   mNeoVis.vulgivaga.2.1.chr8.g00435.t01#1       315   125260935:   mNeoVis.2.1.1.chr8_hap1.g00440.t01#5    mNeoVis.evergladensis.1.1.chr8.g00425.t01#3    mNeoVis.vulgivaga.1.1.chr8.g00426.t01#4    mNeoVis.evergladensis.2.1.chr9.g00430.t01#2    mNeoVis.vulgivaga.2.1.chr8.g00436.t01#1   mNeoVis.2.1.1.chr8_hap1.g00438.t01#5       316   125260936:   mNeoVis.2.1.1.chr8_hap1.g00439.t01#5    mNeoVis.vulgivaga.1.1.chr8.g00425.t01#4    mNeoVis.evergladensis.1.1.chr8.g00424.t01#3    mNeoVis.3.1.1.chr8.g00396.t01#6   mNeoVis.evergladensis.2.1.chr9.g00429.t01#2       317   125260937:  mNeoVis.evergladensis.2.1.chr9.g00428.t01#2             318   125260938:   mNeoVis.vulgivaga.2.1.chr8.g00439.t01#1    mNeoVis.evergladensis.1.1.chr8.g00428.t01#3    mNeoVis.evergladensis.2.1.chr9.g00432.t01#2    mNeoVis.2.1.1.chr8_hap1.g00442.t01#5    mNeoVis.vulgivaga.1.1.chr8.g00428.t01#4    mNeoVis.2.1.1.chr8_hap1.g00441.t01#5    mNeoVis.evergladensis.1.1.chr8.g00427.t01#3    mNeoVis.vulgivaga.2.1.chr8.g00438.t01#1    mNeoVis.vulgivaga.1.1.chr8.g00427.t01#4      319   125260939:   mNeoVis.vulgivaga.2.1.chr8.g00437.t01#1    mNeoVis.evergladensis.1.1.chr8.g00426.t01#3    mNeoVis.3.1.1.chr8.g00398.t01#6   mNeoVis.evergladensis.2.1.chr9.g00431.t01#2       320   125260940:   mNeoVis.vulgivaga.1.1.chr8.g00430.t01#4    mNeoVis.3.1.1.chr8.g00401.t01#6    mNeoVis.3.1.1.chr8.g00400.t01#6    mNeoVis.evergladensis.2.1.chr9.g00436.t01#2    mNeoVis.2.1.1.chr8_hap1.g00445.t01#5    mNeoVis.evergladensis.1.1.chr8.g00430.t01#3    mNeoVis.evergladensis.2.1.chr9.g00435.t01#2    mNeoVis.vulgivaga.1.1.chr8.g00431.t01#4    mNeoVis.2.1.1.chr8_hap1.g00444.t01#5      321   125260941:   mNeoVis.vulgivaga.2.1.chr8.g00440.t01#1    mNeoVis.2.1.1.chr8_hap1.g00443.t01#5    mNeoVis.evergladensis.2.1.chr9.g00433.t01#2    mNeoVis.evergladensis.1.1.chr8.g00429.t01#3    mNeoVis.vulgivaga.1.1.chr8.g00429.t01#4   mNeoVis.3.1.1.chr8.g00399.t01#6       322   125260942:   mNeoVis.3.1.1.chr1.g02103.t01#6    mNeoVis.evergladensis.1.1.chr1.g02120.t01#3    mNeoVis.2.1.1.chr1_hap1.g02149.t01#5   mNeoVis.vulgivaga.2.1.chr1.g02109.t01#1         323   125260943:   mNeoVis.vulgivaga.1.1.chr1.g02123.t01#4    mNeoVis.evergladensis.2.1.chr1.g02118.t01#2    mNeoVis.3.1.1.chr1.g02104.t01#6    mNeoVis.evergladensis.1.1.chr1.g02121.t01#3    mNeoVis.2.1.1.chr1_hap1.g02150.t01#5   mNeoVis.vulgivaga.2.1.chr1.g02110.t01#1       324   125260944:   mNeoVis.evergladensis.1.1.chr8.g00432.t01#3    mNeoVis.evergladensis.2.1.chr9.g00437.t01#2    mNeoVis.2.1.1.chr8_hap1.g00446.t01#5    mNeoVis.vulgivaga.1.1.chr8.g00432.t01#4    mNeoVis.vulgivaga.2.1.chr8.g00442.t01#1   mNeoVis.3.1.1.chr8.g00402.t01#6       325   125260945:   mNeoVis.evergladensis.1.1.chr8.g00433.t01#3    mNeoVis.evergladensis.2.1.chr9.g00438.t01#2    mNeoVis.2.1.1.chr8_hap1.g00447.t01#5    mNeoVis.vulgivaga.1.1.chr8.g00433.t01#4    mNeoVis.vulgivaga.2.1.chr8.g00443.t01#1   mNeoVis.3.1.1.chr8.g00403.t01#6       326   125260946:   mNeoVis.vulgivaga.1.1.chr8.g00434.t01#4    mNeoVis.evergladensis.1.1.chr8.g00434.t01#3    mNeoVis.evergladensis.2.1.chr9.g00439.t01#2    mNeoVis.2.1.1.chr8_hap1.g00448.t01#5    mNeoVis.vulgivaga.2.1.chr8.g00444.t01#1   mNeoVis.3.1.1.chr8.g00404.t01#6       327   125260947:   mNeoVis.2.1.1.chr8_hap1.g00449.t01#5    mNeoVis.evergladensis.1.1.chr8.g00435.t01#3    mNeoVis.evergladensis.2.1.chr9.g00440.t01#2    mNeoVis.vulgivaga.2.1.chr8.g00445.t01#1    mNeoVis.3.1.1.chr8.g00405.t01#6   mNeoVis.vulgivaga.1.1.chr8.g00435.t01#4       328   125260948:   mNeoVis.evergladensis.2.1.chr7.g00604.t01#2    mNeoVis.vulgivaga.1.1.chr7.g00611.t01#4    mNeoVis.vulgivaga.2.1.chr7.g00607.t01#1    mNeoVis.evergladensis.1.1.chr7.g00612.t01#3    mNeoVis.2.1.1.chr7_hap1.g00602.t01#5    mNeoVis.vulgivaga.2.1.chr7.g00608.t01#1   mNeoVis.3.1.1.chr7.g00585.t01#6       329   125260949:  mNeoVis.vulgivaga.2.1.chr1.g02111.t01#1               330   125260950:   mNeoVis.3.1.1.chr1.g02106.t01#6    mNeoVis.2.1.1.chr1_hap1.g02151.t01#5    mNeoVis.evergladensis.1.1.chr1.g02122.t01#3    mNeoVis.vulgivaga.2.1.chr1.g02112.t01#1   mNeoVis.evergladensis.2.1.chr1.g02119.t01#2       331   125260951:   mNeoVis.2.1.1.chr8_hap1.g00450.t01#5    mNeoVis.vulgivaga.2.1.chr8.g00446.t01#1    mNeoVis.evergladensis.1.1.chr8.g00436.t01#3    mNeoVis.3.1.1.chr8.g00406.t01#6    mNeoVis.vulgivaga.1.1.chr8.g00436.t01#4   mNeoVis.evergladensis.2.1.chr9.g00441.t01#2       332   125260952:   mNeoVis.vulgivaga.2.1.chr8.g00447.t01#1    mNeoVis.3.1.1.chr8.g00407.t01#6    mNeoVis.evergladensis.1.1.chr8.g00437.t01#3    mNeoVis.vulgivaga.1.1.chr8.g00437.t01#4    mNeoVis.evergladensis.2.1.chr9.g00442.t01#2   mNeoVis.2.1.1.chr8_hap1.g00451.t01#5       333   125260953:   mNeoVis.vulgivaga.2.1.chr15.g00570.t01#1    mNeoVis.3.1.1.chr15.g00513.t01#6    mNeoVis.2.1.1.chr15_hap1.g00736.t01#5    mNeoVis.evergladensis.2.1.chr15.g00558.t01#2    mNeoVis.evergladensis.1.1.chr15.g00055.t01#3   mNeoVis.vulgivaga.1.1.chr15.g00561.t01#4       334   125260954:   mNeoVis.vulgivaga.2.1.chr1.g02113.t01#1   mNeoVis.3.1.1.chr1.g02107.t01#6             335   125260955:   mNeoVis.3.1.1.chr8.g00408.t01#6    mNeoVis.vulgivaga.2.1.chr8.g00448.t01#1    mNeoVis.vulgivaga.1.1.chr8.g00438.t01#4    mNeoVis.evergladensis.1.1.chr8.g00438.t01#3    mNeoVis.evergladensis.2.1.chr9.g00443.t01#2   mNeoVis.2.1.1.chr8_hap1.g00452.t01#5       336   125260956:   mNeoVis.vulgivaga.2.1.chr8.g00449.t01#1    mNeoVis.vulgivaga.1.1.chr8.g00439.t01#4    mNeoVis.2.1.1.chr8_hap1.g00453.t01#5    mNeoVis.evergladensis.2.1.chr9.g00444.t01#2    mNeoVis.evergladensis.1.1.chr8.g00439.t01#3   mNeoVis.3.1.1.chr8.g00409.t01#6       337   125260957:   mNeoVis.evergladensis.1.1.chr8.g00441.t01#3    mNeoVis.vulgivaga.1.1.chr8.g00441.t01#4    mNeoVis.vulgivaga.2.1.chr8.g00451.t01#1    mNeoVis.2.1.1.chr8_hap1.g00455.t01#5    mNeoVis.3.1.1.chr8.g00411.t01#6   mNeoVis.evergladensis.2.1.chr9.g00446.t01#2       338   125260958:   mNeoVis.evergladensis.1.1.chr1.g02123.t01#3    mNeoVis.evergladensis.2.1.chr1.g02120.t01#2    mNeoVis.3.1.1.chr1.g00007.t01#6    mNeoVis.2.1.1.chr1_hap1.g02152.t01#5    mNeoVis.vulgivaga.2.1.chr1.g02114.t01#1   mNeoVis.vulgivaga.1.1.chr1.g02125.t01#4       339   125260959:   mNeoVis.2.1.1.chr1_hap1.g02153.t01#5    mNeoVis.evergladensis.1.1.chr1.g02124.t01#3    mNeoVis.vulgivaga.2.1.chr1.g02115.t01#1    mNeoVis.evergladensis.2.1.chr1.g02121.t01#2    mNeoVis.vulgivaga.1.1.chr1.g02126.t01#4   mNeoVis.3.1.1.chr1.g02108.t01#6       340   125260960:   mNeoVis.vulgivaga.2.1.chr8.g00450.t01#1    mNeoVis.evergladensis.2.1.chr9.g00445.t01#2    mNeoVis.vulgivaga.1.1.chr8.g00440.t01#4    mNeoVis.2.1.1.chr8_hap1.g00454.t01#5    mNeoVis.evergladensis.1.1.chr8.g00440.t01#3   mNeoVis.3.1.1.chr8.g00410.t01#6       341   125260961:   mNeoVis.evergladensis.1.1.chr1.g02125.t01#3    mNeoVis.2.1.1.chr1_hap1.g02154.t01#5    mNeoVis.evergladensis.2.1.chr1.g02122.t01#2   mNeoVis.vulgivaga.2.1.chr1.g02116.t01#1         342   125260962:   mNeoVis.vulgivaga.1.1.chr8.g00442.t01#4    mNeoVis.evergladensis.1.1.chr8.g00442.t01#3    mNeoVis.2.1.1.chr8_hap1.g00456.t01#5    mNeoVis.vulgivaga.2.1.chr8.g00452.t01#1    mNeoVis.3.1.1.chr8.g00412.t01#6   mNeoVis.evergladensis.2.1.chr9.g00447.t01#2       343   125260963:   mNeoVis.2.1.1.chr15_hap1.g00737.t01#5    mNeoVis.3.1.1.chr15.g00514.t01#6    mNeoVis.vulgivaga.2.1.chr15.g00571.t01#1    mNeoVis.evergladensis.2.1.chr15.g00559.t01#2    mNeoVis.evergladensis.1.1.chr15.g00054.t01#3    mNeoVis.vulgivaga.1.1.chr15.g00562.t01#4   mNeoVis.3.1.1.chr8.g00437.t01#6       344   125260964:   mNeoVis.2.1.1.chr7_hap1.g00603.t01#5    mNeoVis.vulgivaga.2.1.chr7.g00609.t01#1    mNeoVis.vulgivaga.1.1.chr7.g00612.t01#4    mNeoVis.evergladensis.1.1.chr7.g00613.t01#3    mNeoVis.3.1.1.chr7.g00586.t01#6   mNeoVis.evergladensis.2.1.chr7.g00605.t01#2       345   125260965:   mNeoVis.evergladensis.2.1.chr1.g02123.t01#2    mNeoVis.vulgivaga.2.1.chr1.g02117.t01#1    mNeoVis.2.1.1.chr1_hap1.g02155.t01#5   mNeoVis.vulgivaga.1.1.chr1.g02128.t01#4         346   125260966:   mNeoVis.evergladensis.2.1.chr9.g00448.t01#2    mNeoVis.2.1.1.chr8_hap1.g00457.t01#5    mNeoVis.3.1.1.chr8.g00413.t01#6    mNeoVis.evergladensis.1.1.chr8.g00443.t01#3    mNeoVis.vulgivaga.2.1.chr8.g00453.t01#1   mNeoVis.vulgivaga.1.1.chr8.g00443.t01#4       347   125260967:   mNeoVis.vulgivaga.1.1.chr8.g00444.t01#4    mNeoVis.3.1.1.chr11.g00645.t01#6    mNeoVis.2.1.1.chr8_hap1.g00458.t01#5    mNeoVis.vulgivaga.2.1.chr8.g00454.t01#1    mNeoVis.evergladensis.2.1.chr9.g00449.t01#2   mNeoVis.vulgivaga.2.1.chr11.g00682.t01#1       348   125260968:   mNeoVis.evergladensis.1.1.chr8.g00444.t01#3   mNeoVis.3.1.1.chr8.g00414.t01#6             349   125260969:   mNeoVis.evergladensis.2.1.chr9.g00450.t01#2    mNeoVis.evergladensis.1.1.chr8.g00445.t01#3    mNeoVis.vulgivaga.1.1.chr8.g00445.t01#4    mNeoVis.2.1.1.chr8_hap1.g00459.t01#5   mNeoVis.vulgivaga.2.1.chr8.g00455.t01#1       350   125260970:   mNeoVis.evergladensis.1.1.chr1.g02127.t01#3    mNeoVis.vulgivaga.2.1.chr1.g02118.t01#1    mNeoVis.evergladensis.2.1.chr1.g02124.t01#2    mNeoVis.3.1.1.chr1.g02114.t01#6    mNeoVis.vulgivaga.1.1.chr1.g02129.t01#4   mNeoVis.2.1.1.chr1_hap1.g02156.t01#5       351   125260971:   mNeoVis.3.1.1.chr8.g00415.t01#6    mNeoVis.vulgivaga.1.1.chr8.g00446.t01#4    mNeoVis.2.1.1.chr8_hap1.g00460.t01#5    mNeoVis.evergladensis.2.1.chr9.g00451.t01#2    mNeoVis.evergladensis.1.1.chr8.g00446.t01#3   mNeoVis.vulgivaga.2.1.chr8.g00456.t01#1       352   125260972:   mNeoVis.2.1.1.chr1_hap1.g02157.t01#5    mNeoVis.evergladensis.2.1.chr1.g02125.t01#2    mNeoVis.vulgivaga.2.1.chr1.g02119.t01#1    mNeoVis.3.1.1.chr1.g02115.t01#6    mNeoVis.vulgivaga.1.1.chr1.g02130.t01#4   mNeoVis.evergladensis.1.1.chr1.g02128.t01#3       353   125260973:   mNeoVis.3.1.1.chr8.g00416.t01#6    mNeoVis.vulgivaga.1.1.chr8.g00447.t01#4    mNeoVis.evergladensis.1.1.chr8.g00447.t01#3    mNeoVis.2.1.1.chr8_hap1.g00461.t01#5    mNeoVis.evergladensis.2.1.chr9.g00452.t01#2   mNeoVis.vulgivaga.2.1.chr8.g00457.t01#1       354   125260974:   mNeoVis.vulgivaga.2.1.chr1.g02120.t01#1    mNeoVis.vulgivaga.1.1.chr1.g02131.t01#4    mNeoVis.3.1.1.chr1.g02116.t01#6    mNeoVis.evergladensis.1.1.chr1.g02129.t01#3   mNeoVis.2.1.1.chr1_hap1.g02158.t01#5       355   125260975:   mNeoVis.evergladensis.1.1.chr1.g02130.t01#3    mNeoVis.evergladensis.2.1.chr1.g02127.t01#2    mNeoVis.2.1.1.chr1_hap1.g02159.t01#5    mNeoVis.vulgivaga.2.1.chr1.g02121.t01#1    mNeoVis.3.1.1.chr1.g02117.t01#6   mNeoVis.vulgivaga.1.1.chr1.g02132.t01#4       356   125260976:   mNeoVis.vulgivaga.2.1.chr8.g00458.t01#1    mNeoVis.evergladensis.1.1.chr8.g00448.t01#3    mNeoVis.2.1.1.chr8_hap1.g00462.t01#5    mNeoVis.evergladensis.2.1.chr9.g00453.t01#2    mNeoVis.vulgivaga.1.1.chr8.g00448.t01#4   mNeoVis.3.1.1.chr8.g00417.t01#6       357   125260977:   mNeoVis.2.1.1.chr7_hap1.g00604.t01#5    mNeoVis.3.1.1.chr7.g00587.t01#6    mNeoVis.vulgivaga.1.1.chr7.g00613.t01#4    mNeoVis.evergladensis.1.1.chr7.g00614.t01#3    mNeoVis.vulgivaga.2.1.chr7.g00610.t01#1   mNeoVis.evergladensis.2.1.chr7.g00606.t01#2       358   125260978:   mNeoVis.vulgivaga.1.1.chr1.g02133.t01#4    mNeoVis.vulgivaga.2.1.chr1.g02122.t01#1    mNeoVis.evergladensis.2.1.chr1.g02128.t01#2    mNeoVis.2.1.1.chr1_hap1.g02160.t01#5   mNeoVis.evergladensis.1.1.chr1.g02131.t01#3       359   125260979:   mNeoVis.3.1.1.chr1.g02118.t01#6    mNeoVis.2.1.1.chr1_hap1.g02161.t01#5    mNeoVis.evergladensis.2.1.chr1.g02129.t01#2    mNeoVis.evergladensis.1.1.chr1.g02132.t01#3    mNeoVis.vulgivaga.2.1.chr1.g02123.t01#1   mNeoVis.vulgivaga.1.1.chr1.g02134.t01#4       360   125260980:   mNeoVis.2.1.1.chr15_hap1.g00738.t01#5    mNeoVis.vulgivaga.2.1.chr15.g00572.t01#1    mNeoVis.3.1.1.chr15.g00515.t01#6    mNeoVis.evergladensis.1.1.chr15.g00053.t01#3    mNeoVis.3.1.1.chr10.g01189.t01#6    mNeoVis.evergladensis.2.1.chr15.g00560.t01#2   mNeoVis.vulgivaga.1.1.chr15.g00563.t01#4     361   125260981:   mNeoVis.vulgivaga.2.1.chr1.g02124.t01#1    mNeoVis.vulgivaga.1.1.chr1.g02135.t01#4    mNeoVis.evergladensis.2.1.chr1.g02130.t01#2    mNeoVis.2.1.1.chr1_hap1.g02162.t01#5   mNeoVis.evergladensis.1.1.chr1.g02133.t01#3       362   125260982:   mNeoVis.vulgivaga.1.1.chr8.g00449.t01#4    mNeoVis.vulgivaga.2.1.chr8.g00459.t01#1    mNeoVis.3.1.1.chr8.g00418.t01#6    mNeoVis.2.1.1.chr8_hap1.g00463.t01#5    mNeoVis.evergladensis.2.1.chr9.g00454.t01#2   mNeoVis.evergladensis.1.1.chr8.g00449.t01#3       363   125260983:   mNeoVis.3.1.1.chr8.g00419.t01#6    mNeoVis.vulgivaga.1.1.chr8.g00450.t01#4    mNeoVis.evergladensis.1.1.chr8.g00450.t01#3    mNeoVis.vulgivaga.2.1.chr8.g00460.t01#1    mNeoVis.evergladensis.2.1.chr9.g00455.t01#2   mNeoVis.2.1.1.chr8_hap1.g00464.t01#5       364   125260984:   mNeoVis.evergladensis.2.1.chr1.g02131.t01#2    mNeoVis.2.1.1.chr1_hap1.g02163.t01#5    mNeoVis.3.1.1.chr1.g02120.t01#6    mNeoVis.evergladensis.1.1.chr1.g02134.t01#3    mNeoVis.vulgivaga.1.1.chr1.g02136.t01#4   mNeoVis.vulgivaga.2.1.chr1.g02125.t01#1       365   125260985:   mNeoVis.2.1.1.chr8_hap1.g00465.t01#5    mNeoVis.3.1.1.chr8.g00420.t01#6    mNeoVis.evergladensis.1.1.chr8.g00451.t01#3    mNeoVis.evergladensis.2.1.chr9.g00456.t01#2    mNeoVis.vulgivaga.1.1.chr8.g00451.t01#4   mNeoVis.vulgivaga.2.1.chr8.g00461.t01#1       366   125260986:   mNeoVis.3.1.1.chr1.g02121.t01#6    mNeoVis.evergladensis.2.1.chr1.g02132.t01#2    mNeoVis.vulgivaga.1.1.chr1.g02137.t01#4    mNeoVis.evergladensis.1.1.chr1.g02135.t01#3    mNeoVis.2.1.1.chr1_hap1.g02164.t01#5   mNeoVis.vulgivaga.2.1.chr1.g02126.t01#1       367   125260987:   mNeoVis.vulgivaga.1.1.chr8.g00452.t01#4    mNeoVis.vulgivaga.2.1.chr8.g00462.t01#1    mNeoVis.3.1.1.chr8.g00421.t01#6    mNeoVis.2.1.1.chr8_hap1.g00466.t01#5   mNeoVis.evergladensis.1.1.chr8.g00452.t01#3       368   125260988:   mNeoVis.vulgivaga.2.1.chr1.g02127.t01#1    mNeoVis.vulgivaga.1.1.chr1.g02138.t01#4    mNeoVis.evergladensis.1.1.chr1.g02136.t01#3    mNeoVis.3.1.1.chr1.g02122.t01#6    mNeoVis.2.1.1.chr1_hap1.g02165.t01#5   mNeoVis.evergladensis.2.1.chr1.g02133.t01#2       369   125260989:   mNeoVis.evergladensis.2.1.chr7.g00607.t01#2    mNeoVis.2.1.1.chr7_hap1.g00605.t01#5    mNeoVis.3.1.1.chr7.g00588.t01#6    mNeoVis.vulgivaga.1.1.chr7.g00614.t01#4    mNeoVis.vulgivaga.2.1.chr7.g00611.t01#1   mNeoVis.evergladensis.1.1.chr7.g00615.t01#3       370   125260990:   mNeoVis.vulgivaga.1.1.chr2.g00001.t01#4    mNeoVis.2.1.1.chr5_hap1.g00001.t01#5    mNeoVis.evergladensis.1.1.chr2.g00001.t01#3    mNeoVis.3.1.1.chr2.g00001.t01#6    mNeoVis.vulgivaga.2.1.chr2.g00001.t01#1   mNeoVis.evergladensis.2.1.chr2.g00001.t01#2       371   125260991:   mNeoVis.vulgivaga.2.1.chr8.g00463.t01#1    mNeoVis.evergladensis.2.1.chr9.g00458.t01#2    mNeoVis.vulgivaga.1.1.chr8.g00453.t01#4    mNeoVis.2.1.1.chr8_hap1.g00467.t01#5    mNeoVis.3.1.1.chr8.g00422.t01#6   mNeoVis.evergladensis.1.1.chr8.g00453.t01#3       372   125260992:   mNeoVis.vulgivaga.1.1.chr2.g00002.t01#4    mNeoVis.2.1.1.chr5_hap1.g00002.t01#5    mNeoVis.evergladensis.1.1.chr2.g00002.t01#3    mNeoVis.vulgivaga.2.1.chr2.g00002.t01#1    mNeoVis.3.1.1.chr2.g00002.t01#6   mNeoVis.evergladensis.2.1.chr2.g00002.t01#2       373   125260993:   mNeoVis.vulgivaga.2.1.chr8.g00464.t01#1    mNeoVis.vulgivaga.1.1.chr8.g00454.t01#4    mNeoVis.evergladensis.2.1.chr9.g00459.t01#2    mNeoVis.2.1.1.chr8_hap1.g00468.t01#5    mNeoVis.3.1.1.chr8.g00423.t01#6   mNeoVis.evergladensis.1.1.chr8.g00454.t01#3       374   125260994:   mNeoVis.evergladensis.1.1.chr2.g00003.t01#3    mNeoVis.2.1.1.chr5_hap1.g00003.t01#5    mNeoVis.evergladensis.2.1.chr2.g00003.t01#2    mNeoVis.vulgivaga.1.1.chr2.g00003.t01#4   mNeoVis.vulgivaga.2.1.chr2.g00003.t01#1       375   125260995:   mNeoVis.vulgivaga.2.1.chr2.g00004.t01#1   mNeoVis.evergladensis.2.1.chr2.g00004.t01#2           376   125260996:   mNeoVis.vulgivaga.2.1.chr8.g00465.t01#1    mNeoVis.evergladensis.1.1.chr8.g00455.t01#3    mNeoVis.vulgivaga.1.1.chr8.g00455.t01#4    mNeoVis.3.1.1.chr8.g00424.t01#6    mNeoVis.2.1.1.chr8_hap1.g00469.t01#5   mNeoVis.evergladensis.2.1.chr9.g00460.t01#2       377   125260997:   mNeoVis.vulgivaga.1.1.chr2.g00005.t01#4    mNeoVis.vulgivaga.2.1.chr2.g00005.t01#1    mNeoVis.evergladensis.1.1.chr2.g00005.t01#3    mNeoVis.2.1.1.chr5_hap1.g00005.t01#5   mNeoVis.evergladensis.2.1.chr2.g00005.t01#2       378   125260998:   mNeoVis.3.1.1.chr8.g00319.t01#6    mNeoVis.vulgivaga.2.1.chr8.g00357.t01#1    mNeoVis.evergladensis.1.1.chr8.g00351.t01#3   mNeoVis.evergladensis.2.1.chr9.g00350.t01#2       379   125260999:   mNeoVis.vulgivaga.1.1.chr2.g00006.t01#4    mNeoVis.vulgivaga.2.1.chr2.g00006.t01#1    mNeoVis.evergladensis.1.1.chr2.g00006.t01#3    mNeoVis.evergladensis.2.1.chr2.g00006.t01#2   mNeoVis.2.1.1.chr5_hap1.g00006.t01#5       380   125261000:   mNeoVis.3.1.1.chr8.g00320.t01#6    mNeoVis.2.1.1.chr8_hap1.g00353.t01#5    mNeoVis.evergladensis.1.1.chr8.g00352.t01#3    mNeoVis.evergladensis.2.1.chr9.g00351.t01#2    mNeoVis.vulgivaga.1.1.chr8.g00345.t01#4   mNeoVis.vulgivaga.2.1.chr8.g00358.t01#1       381   125261001:   mNeoVis.evergladensis.1.1.chr8.g00353.t01#3    mNeoVis.vulgivaga.1.1.chr8.g00346.t01#4    mNeoVis.vulgivaga.2.1.chr8.g00359.t01#1    mNeoVis.evergladensis.2.1.chr9.g00352.t01#2    mNeoVis.2.1.1.chr8_hap1.g00354.t01#5   mNeoVis.3.1.1.chr8.g00321.t01#6       382   125261002:   mNeoVis.vulgivaga.2.1.chr2.g00007.t01#1    mNeoVis.2.1.1.chr5_hap1.g00007.t01#5    mNeoVis.evergladensis.2.1.chr2.g00007.t01#2    mNeoVis.vulgivaga.1.1.chr2.g00007.t01#4   mNeoVis.evergladensis.1.1.chr2.g00007.t01#3       383   125261003:   mNeoVis.vulgivaga.1.1.chr8.g00348.t01#4    mNeoVis.3.1.1.chr8.g00323.t01#6    mNeoVis.evergladensis.2.1.chr9.g00354.t01#2    mNeoVis.vulgivaga.2.1.chr8.g00360.t01#1   mNeoVis.2.1.1.chr8_hap1.g00356.t01#5       384   125261004:   mNeoVis.3.1.1.chr8.g00322.t01#6    mNeoVis.vulgivaga.1.1.chr8.g00347.t01#4    mNeoVis.evergladensis.2.1.chr9.g00353.t01#2   mNeoVis.2.1.1.chr8_hap1.g00355.t01#5         385   125261005:  mNeoVis.evergladensis.1.1.chr8.g00354.t01#3             386   125261006:   mNeoVis.vulgivaga.1.1.chr2.g00008.t01#4    mNeoVis.evergladensis.1.1.chr2.g00008.t01#3    mNeoVis.evergladensis.2.1.chr2.g00008.t01#2    mNeoVis.2.1.1.chr5_hap1.g00008.t01#5   mNeoVis.vulgivaga.2.1.chr2.g00008.t01#1       387   125261007:   mNeoVis.vulgivaga.2.1.chr2.g00009.t01#1    mNeoVis.2.1.1.chr5_hap1.g00009.t01#5    mNeoVis.evergladensis.1.1.chr2.g00009.t01#3    mNeoVis.vulgivaga.1.1.chr2.g00009.t01#4   mNeoVis.3.1.1.chr2.g00004.t01#6       388   125261008:   mNeoVis.vulgivaga.2.1.chr2.g00010.t01#1    mNeoVis.2.1.1.chr5_hap1.g00010.t01#5    mNeoVis.evergladensis.2.1.chr2.g00010.t01#2    mNeoVis.evergladensis.1.1.chr2.g00010.t01#3   mNeoVis.vulgivaga.1.1.chr2.g00010.t01#4       389   125261009:   mNeoVis.evergladensis.2.1.chr2.g00011.t01#2    mNeoVis.3.1.1.chr2.g00006.t01#6    mNeoVis.vulgivaga.1.1.chr2.g00011.t01#4    mNeoVis.evergladensis.1.1.chr2.g00011.t01#3    mNeoVis.2.1.1.chr5_hap1.g00011.t01#5   mNeoVis.vulgivaga.2.1.chr2.g00011.t01#1       390   125261010:   mNeoVis.2.1.1.chr8_hap1.g00357.t01#5    mNeoVis.3.1.1.chr8.g00324.t01#6    mNeoVis.evergladensis.1.1.chr8.g00355.t01#3    mNeoVis.evergladensis.2.1.chr9.g00355.t01#2    mNeoVis.vulgivaga.1.1.chr8.g00349.t01#4   mNeoVis.vulgivaga.2.1.chr8.g00361.t01#1       391   125261011:   mNeoVis.evergladensis.1.1.chr8.g00356.t01#3    mNeoVis.evergladensis.2.1.chr9.g00357.t01#2    mNeoVis.2.1.1.chr8_hap1.g00358.t01#5    mNeoVis.3.1.1.chr8.g00325.t01#6    mNeoVis.vulgivaga.2.1.chr8.g00363.t01#1   mNeoVis.vulgivaga.1.1.chr8.g00350.t01#4       392   125261012:   mNeoVis.vulgivaga.2.1.chr8.g00364.t01#1    mNeoVis.evergladensis.2.1.chr9.g00358.t01#2    mNeoVis.3.1.1.chr8.g00326.t01#6   mNeoVis.2.1.1.chr8_hap1.g00360.t01#5         393   125261013:   mNeoVis.vulgivaga.1.1.chr8.g00351.t01#4    mNeoVis.vulgivaga.2.1.chr8.g00362.t01#1   mNeoVis.evergladensis.2.1.chr9.g00356.t01#2         394   125261014:   mNeoVis.2.1.1.chr5_hap1.g00012.t01#5    mNeoVis.evergladensis.1.1.chr2.g00012.t01#3    mNeoVis.evergladensis.2.1.chr2.g00012.t01#2    mNeoVis.vulgivaga.2.1.chr2.g00012.t01#1   mNeoVis.vulgivaga.1.1.chr2.g00012.t01#4       395   125261015:   mNeoVis.evergladensis.2.1.chr2.g00013.t01#2    mNeoVis.2.1.1.chr5_hap1.g00013.t01#5    mNeoVis.evergladensis.1.1.chr2.g00013.t01#3    mNeoVis.vulgivaga.2.1.chr2.g00013.t01#1   mNeoVis.vulgivaga.1.1.chr2.g00013.t01#4       396   125261016:   mNeoVis.vulgivaga.2.1.chr8.g00365.t01#1    mNeoVis.evergladensis.2.1.chr9.g00359.t01#2    mNeoVis.vulgivaga.1.1.chr8.g00352.t01#4    mNeoVis.2.1.1.chr8_hap1.g00361.t01#5    mNeoVis.evergladensis.1.1.chr8.g00358.t01#3    mNeoVis.3.1.1.chr8.g00327.t01#6   mNeoVis.vulgivaga.2.1.chr8.g00366.t01#1     397   125261017:   mNeoVis.vulgivaga.2.1.chr15.g00573.t01#1    mNeoVis.2.1.1.chr15_hap1.g00739.t01#5    mNeoVis.vulgivaga.1.1.chr15.g00564.t01#4    mNeoVis.evergladensis.2.1.chr15.g00561.t01#2    mNeoVis.evergladensis.1.1.chr15.g00052.t01#3   mNeoVis.3.1.1.chr15.g00517.t01#6       398   125261018:   mNeoVis.2.1.1.chr8_hap1.g00362.t01#5    mNeoVis.evergladensis.1.1.chr8.g00359.t01#3    mNeoVis.vulgivaga.1.1.chr8.g00353.t01#4    mNeoVis.3.1.1.chr8.g00328.t01#6    mNeoVis.vulgivaga.2.1.chr8.g00367.t01#1   mNeoVis.evergladensis.2.1.chr9.g00360.t01#2       399   125261019:   mNeoVis.evergladensis.2.1.chr9.g00361.t01#2    mNeoVis.2.1.1.chr8_hap1.g00363.t01#5    mNeoVis.vulgivaga.1.1.chr8.g00354.t01#4    mNeoVis.vulgivaga.2.1.chr8.g00368.t01#1    mNeoVis.3.1.1.chr8.g00329.t01#6   mNeoVis.evergladensis.1.1.chr8.g00360.t01#3       400   125261020:   mNeoVis.evergladensis.2.1.chr9.g00362.t01#2    mNeoVis.2.1.1.chr8_hap1.g00364.t01#5    mNeoVis.vulgivaga.2.1.chr8.g00369.t01#1    mNeoVis.vulgivaga.1.1.chr8.g00355.t01#4   mNeoVis.3.1.1.chr8.g00330.t01#6       401   125261021:   mNeoVis.2.1.1.chr8_hap1.g00365.t01#5    mNeoVis.vulgivaga.1.1.chr8.g00356.t01#4    mNeoVis.3.1.1.chr8.g00331.t01#6    mNeoVis.vulgivaga.2.1.chr8.g00370.t01#1    mNeoVis.evergladensis.2.1.chr9.g00363.t01#2   mNeoVis.evergladensis.1.1.chr8.g00361.t01#3       402   125261022:   mNeoVis.evergladensis.1.1.chr8.g00362.t01#3    mNeoVis.vulgivaga.2.1.chr8.g00371.t01#1    mNeoVis.2.1.1.chr8_hap1.g00366.t01#5    mNeoVis.3.1.1.chr8.g00332.t01#6    mNeoVis.vulgivaga.1.1.chr8.g00357.t01#4   mNeoVis.evergladensis.2.1.chr9.g00364.t01#2       403   125261023:   mNeoVis.evergladensis.2.1.chr9.g00365.t01#2    mNeoVis.evergladensis.1.1.chr8.g00363.t01#3    mNeoVis.vulgivaga.2.1.chr8.g00372.t01#1    mNeoVis.3.1.1.chr8.g00333.t01#6    mNeoVis.2.1.1.chr8_hap1.g00367.t01#5   mNeoVis.vulgivaga.1.1.chr8.g00358.t01#4       404   125261024:   mNeoVis.vulgivaga.1.1.chr1.g02070.t01#4    mNeoVis.evergladensis.1.1.chr1.g02067.t01#3    mNeoVis.2.1.1.chr1_hap1.g02096.t01#5    mNeoVis.3.1.1.chr1.g02051.t01#6    mNeoVis.vulgivaga.2.1.chr1.g02059.t01#1   mNeoVis.evergladensis.2.1.chr1.g02066.t01#2       405   125261025:   mNeoVis.2.1.1.chr8_hap1.g00368.t01#5    mNeoVis.evergladensis.2.1.chr9.g00366.t01#2    mNeoVis.vulgivaga.1.1.chr8.g00359.t01#4    mNeoVis.vulgivaga.2.1.chr8.g00373.t01#1    mNeoVis.3.1.1.chr8.g00334.t01#6   mNeoVis.evergladensis.1.1.chr8.g00364.t01#3       406   125261026:   mNeoVis.vulgivaga.1.1.chr4.g00251.t01#4    mNeoVis.vulgivaga.2.1.chr4.g00255.t01#1    mNeoVis.evergladensis.1.1.chr4.g00252.t01#3    mNeoVis.2.1.1.chr7_hap1.g01865.t01#5    mNeoVis.3.1.1.chr9.g00937.t01#6    mNeoVis.vulgivaga.2.1.chr2.g00886.t01#1    mNeoVis.2.1.1.chr5_hap1.g00884.t01#5    mNeoVis.3.1.1.chr7.g01733.t01#6    mNeoVis.vulgivaga.1.1.chr7.g01835.t01#4      407   125261027:   mNeoVis.evergladensis.1.1.chr10.g00692.t01#3    mNeoVis.3.1.1.chr10.g00664.t01#6    mNeoVis.evergladensis.1.1.chr10.g00068.t01#3    mNeoVis.evergladensis.2.1.chr10.g00072.t01#2    mNeoVis.2.1.1.chr9_hap1.g00074.t01#5    mNeoVis.vulgivaga.1.1.chr6.g00418.t01#4    mNeoVis.2.1.1.chr4_hap1.g00412.t01#5    mNeoVis.evergladensis.2.1.chr6.g00415.t01#2    mNeoVis.vulgivaga.2.1.chr13.g00396.t01#1      408   125261028:   mNeoVis.evergladensis.1.1.chr1.g02068.t01#3    mNeoVis.vulgivaga.2.1.chr1.g02060.t01#1    mNeoVis.vulgivaga.1.1.chr1.g02071.t01#4    mNeoVis.evergladensis.2.1.chr1.g02067.t01#2    mNeoVis.2.1.1.chr1_hap1.g02097.t01#5   mNeoVis.3.1.1.chr1.g02052.t01#6       409   125261029:   mNeoVis.evergladensis.2.1.chr7.g01828.t01#2    mNeoVis.3.1.1.chr7.g01734.t01#6    mNeoVis.evergladensis.1.1.chr7.g01825.t01#3    mNeoVis.vulgivaga.1.1.chr7.g01836.t01#4   mNeoVis.2.1.1.chr7_hap1.g01866.t01#5       410   125261030:   mNeoVis.vulgivaga.2.1.chr15.g00574.t01#1    mNeoVis.vulgivaga.1.1.chr15.g00565.t01#4    mNeoVis.evergladensis.1.1.chr15.g00051.t01#3    mNeoVis.2.1.1.chr15_hap1.g00740.t01#5    mNeoVis.evergladensis.2.1.chr15.g00562.t01#2   mNeoVis.3.1.1.chr15.g00518.t01#6       411   125261031:   mNeoVis.3.1.1.chr1.g02053.t01#6    mNeoVis.vulgivaga.2.1.chr1.g02061.t01#1    mNeoVis.vulgivaga.1.1.chr1.g02072.t01#4   mNeoVis.2.1.1.chr1_hap1.g02098.t01#5         412   125261032:   mNeoVis.vulgivaga.2.1.chr1.g02062.t01#1    mNeoVis.evergladensis.2.1.chr1.g02069.t01#2    mNeoVis.vulgivaga.1.1.chr1.g02073.t01#4    mNeoVis.2.1.1.chr1_hap1.g02099.t01#5    mNeoVis.evergladensis.1.1.chr1.g02070.t01#3   mNeoVis.3.1.1.chr1.g02054.t01#6       413   125261033:   mNeoVis.vulgivaga.2.1.chr8.g00374.t01#1    mNeoVis.evergladensis.2.1.chr9.g00367.t01#2    mNeoVis.3.1.1.chr8.g00335.t01#6    mNeoVis.evergladensis.1.1.chr8.g00365.t01#3    mNeoVis.2.1.1.chr8_hap1.g00369.t01#5   mNeoVis.vulgivaga.1.1.chr8.g00360.t01#4       414   125261034:   mNeoVis.2.1.1.chr5_hap1.g01378.t01#5    mNeoVis.evergladensis.2.1.chr15.g00563.t01#2    mNeoVis.2.1.1.chr15_hap1.g00741.t01#5    mNeoVis.vulgivaga.2.1.chr15.g00575.t01#1    mNeoVis.evergladensis.1.1.chr2.g01375.t01#3    mNeoVis.vulgivaga.1.1.chr2.g01379.t01#4    mNeoVis.vulgivaga.2.1.chr2.g01374.t01#1    mNeoVis.evergladensis.2.1.chr2.g01365.t01#2    mNeoVis.vulgivaga.1.1.chr15.g00566.t01#4      415   125261035:   mNeoVis.vulgivaga.1.1.chr1.g02074.t01#4    mNeoVis.evergladensis.1.1.chr1.g02071.t01#3    mNeoVis.2.1.1.chr1_hap1.g02100.t01#5    mNeoVis.3.1.1.chr1.g02055.t01#6    mNeoVis.vulgivaga.2.1.chr1.g02063.t01#1   mNeoVis.evergladensis.2.1.chr1.g02070.t01#2       416   125261036:   mNeoVis.evergladensis.1.1.chr8.g00366.t01#3    mNeoVis.2.1.1.chr8_hap1.g00370.t01#5    mNeoVis.vulgivaga.1.1.chr8.g00361.t01#4    mNeoVis.evergladensis.2.1.chr9.g00368.t01#2    mNeoVis.3.1.1.chr8.g00336.t01#6   mNeoVis.vulgivaga.2.1.chr8.g00375.t01#1       417   125261037:   mNeoVis.vulgivaga.2.1.chr8.g00376.t01#1    mNeoVis.3.1.1.chr8.g00337.t01#6    mNeoVis.vulgivaga.1.1.chr8.g00362.t01#4    mNeoVis.evergladensis.2.1.chr9.g00369.t01#2    mNeoVis.2.1.1.chr8_hap1.g00371.t01#5   mNeoVis.evergladensis.1.1.chr8.g00367.t01#3       418   125261038:   mNeoVis.2.1.1.chr8_hap1.g00372.t01#5    mNeoVis.evergladensis.1.1.chr8.g00369.t01#3    mNeoVis.evergladensis.2.1.chr9.g00370.t01#2    mNeoVis.2.1.1.chr8_hap1.g00373.t01#5   mNeoVis.vulgivaga.2.1.chr8.g00378.t01#1       419   125261039:   mNeoVis.vulgivaga.2.1.chr8.g00377.t01#1    mNeoVis.3.1.1.chr8.g00338.t01#6   mNeoVis.vulgivaga.1.1.chr8.g00364.t01#4           420   125261040:   mNeoVis.evergladensis.2.1.chr9.g00371.t01#2    mNeoVis.3.1.1.chr8.g00339.t01#6   mNeoVis.vulgivaga.1.1.chr8.g00363.t01#4           421   125261041:  mNeoVis.evergladensis.1.1.chr8.g00368.t01#3             422   125261042:   mNeoVis.3.1.1.chr1.g02056.t01#6    mNeoVis.evergladensis.2.1.chr1.g02071.t01#2    mNeoVis.vulgivaga.2.1.chr1.g02064.t01#1    mNeoVis.vulgivaga.1.1.chr1.g02075.t01#4    mNeoVis.2.1.1.chr1_hap1.g02101.t01#5   mNeoVis.evergladensis.1.1.chr1.g02072.t01#3       423   125261043:   mNeoVis.vulgivaga.2.1.chr8.g00379.t01#1    mNeoVis.evergladensis.1.1.chr8.g00370.t01#3    mNeoVis.2.1.1.chr8_hap1.g00374.t01#5    mNeoVis.vulgivaga.1.1.chr8.g00365.t01#4    mNeoVis.evergladensis.2.1.chr9.g00372.t01#2   mNeoVis.3.1.1.chr8.g00340.t01#6       424   125261044:   mNeoVis.2.1.1.chr1_hap1.g02102.t01#5    mNeoVis.3.1.1.chr1.g02057.t01#6    mNeoVis.evergladensis.2.1.chr1.g02072.t01#2    mNeoVis.evergladensis.1.1.chr1.g02073.t01#3    mNeoVis.vulgivaga.2.1.chr1.g02065.t01#1   mNeoVis.vulgivaga.1.1.chr1.g02076.t01#4       425   125261045:   mNeoVis.2.1.1.chr8_hap1.g00375.t01#5    mNeoVis.vulgivaga.2.1.chr8.g00380.t01#1    mNeoVis.vulgivaga.1.1.chr8.g00366.t01#4    mNeoVis.3.1.1.chr8.g00341.t01#6    mNeoVis.evergladensis.1.1.chr8.g00371.t01#3   mNeoVis.evergladensis.2.1.chr9.g00373.t01#2       426   125261046:   mNeoVis.vulgivaga.2.1.chr1.g02066.t01#1   mNeoVis.evergladensis.1.1.chr1.g02074.t01#3           427   125261047:   mNeoVis.evergladensis.1.1.chr1.g02075.t01#3    mNeoVis.vulgivaga.1.1.chr1.g02078.t01#4    mNeoVis.evergladensis.2.1.chr1.g02073.t01#2    mNeoVis.3.1.1.chr1.g02059.t01#6    mNeoVis.vulgivaga.2.1.chr1.g02067.t01#1   mNeoVis.2.1.1.chr1_hap1.g02103.t01#5       428   125261048:   mNeoVis.vulgivaga.1.1.chr8.g00367.t01#4    mNeoVis.evergladensis.2.1.chr9.g00374.t01#2    mNeoVis.evergladensis.1.1.chr8.g00372.t01#3    mNeoVis.3.1.1.chr8.g00342.t01#6    mNeoVis.2.1.1.chr8_hap1.g00376.t01#5   mNeoVis.vulgivaga.2.1.chr8.g00381.t01#1       429   125261049:   mNeoVis.vulgivaga.1.1.chr8.g00368.t01#4    mNeoVis.evergladensis.2.1.chr9.g00375.t01#2    mNeoVis.3.1.1.chr8.g00343.t01#6    mNeoVis.vulgivaga.2.1.chr8.g00382.t01#1    mNeoVis.evergladensis.1.1.chr8.g00373.t01#3   mNeoVis.2.1.1.chr8_hap1.g00377.t01#5       430   125261050:   mNeoVis.vulgivaga.1.1.chr1.g02079.t01#4    mNeoVis.evergladensis.1.1.chr1.g02076.t01#3    mNeoVis.2.1.1.chr1_hap1.g02104.t01#5    mNeoVis.evergladensis.2.1.chr1.g02074.t01#2    mNeoVis.3.1.1.chr1.g02060.t01#6   mNeoVis.vulgivaga.2.1.chr1.g02068.t01#1       431   125261051:   mNeoVis.3.1.1.chr15.g00520.t01#6    mNeoVis.vulgivaga.1.1.chr15.g00567.t01#4    mNeoVis.vulgivaga.2.1.chr15.g00576.t01#1    mNeoVis.evergladensis.1.1.chr15.g00049.t01#3    mNeoVis.2.1.1.chr15_hap1.g00742.t01#5   mNeoVis.evergladensis.2.1.chr15.g00564.t01#2       432   125261052:   mNeoVis.vulgivaga.1.1.chr15.g00024.t01#4    mNeoVis.2.1.1.chr15_hap1.g00023.t01#5    mNeoVis.3.1.1.chr15.g00021.t01#6    mNeoVis.evergladensis.1.1.chr15.g00591.t01#3    mNeoVis.vulgivaga.2.1.chr15.g00025.t01#1   mNeoVis.evergladensis.2.1.chr15.g00024.t01#2       433   125261053:   mNeoVis.3.1.1.chr8.g00344.t01#6    mNeoVis.evergladensis.1.1.chr8.g00374.t01#3    mNeoVis.vulgivaga.2.1.chr8.g00383.t01#1    mNeoVis.evergladensis.2.1.chr9.g00376.t01#2    mNeoVis.2.1.1.chr8_hap1.g00378.t01#5   mNeoVis.vulgivaga.1.1.chr8.g00369.t01#4       434   125261054:   mNeoVis.evergladensis.2.1.chr9.g00377.t01#2    mNeoVis.vulgivaga.1.1.chr8.g00370.t01#4    mNeoVis.3.1.1.chr8.g00345.t01#6    mNeoVis.vulgivaga.2.1.chr8.g00384.t01#1    mNeoVis.evergladensis.1.1.chr8.g00375.t01#3   mNeoVis.2.1.1.chr8_hap1.g00379.t01#5       435   125261055:   mNeoVis.vulgivaga.1.1.chr1.g02080.t01#4    mNeoVis.3.1.1.chr1.g02061.t01#6    mNeoVis.vulgivaga.2.1.chr1.g02069.t01#1    mNeoVis.2.1.1.chr1_hap1.g02105.t01#5    mNeoVis.evergladensis.2.1.chr1.g02075.t01#2   mNeoVis.evergladensis.1.1.chr1.g02077.t01#3       436   125261056:   mNeoVis.evergladensis.1.1.chr8.g00376.t01#3    mNeoVis.vulgivaga.1.1.chr8.g00371.t01#4    mNeoVis.3.1.1.chr8.g00346.t01#6    mNeoVis.vulgivaga.2.1.chr8.g00385.t01#1    mNeoVis.2.1.1.chr8_hap1.g00380.t01#5   mNeoVis.evergladensis.2.1.chr9.g00378.t01#2       437   125261057:   mNeoVis.3.1.1.chr1.g02062.t01#6    mNeoVis.evergladensis.1.1.chr1.g02078.t01#3    mNeoVis.evergladensis.2.1.chr1.g02076.t01#2    mNeoVis.2.1.1.chr1_hap1.g02106.t01#5    mNeoVis.vulgivaga.2.1.chr1.g02070.t01#1   mNeoVis.vulgivaga.1.1.chr1.g02081.t01#4       438   125261058:   mNeoVis.vulgivaga.2.1.chr1.g02071.t01#1    mNeoVis.evergladensis.1.1.chr1.g02079.t01#3    mNeoVis.3.1.1.chr1.g02063.t01#6    mNeoVis.evergladensis.2.1.chr1.g02077.t01#2    mNeoVis.2.1.1.chr1_hap1.g02107.t01#5   mNeoVis.vulgivaga.1.1.chr1.g02082.t01#4       439   125261059:   mNeoVis.vulgivaga.2.1.chr7.g01120.t01#1    mNeoVis.evergladensis.2.1.chr7.g01108.t01#2    mNeoVis.evergladensis.1.1.chr7.g01112.t01#3    mNeoVis.3.1.1.chr7.g01045.t01#6    mNeoVis.vulgivaga.1.1.chr7.g01108.t01#4   mNeoVis.2.1.1.chr7_hap1.g01122.t01#5       440   125261060:   mNeoVis.vulgivaga.1.1.chr12.g00479.t01#4    mNeoVis.3.1.1.chr12.g00476.t01#6    mNeoVis.2.1.1.chr12_hap1.g00523.t01#5    mNeoVis.evergladensis.1.1.chr12.g00477.t01#3    mNeoVis.evergladensis.2.1.chr12.g00469.t01#2   mNeoVis.vulgivaga.2.1.chr12.g00482.t01#1       441   125261061:   mNeoVis.2.1.1.chr1_hap1.g02108.t01#5    mNeoVis.vulgivaga.2.1.chr1.g02072.t01#1    mNeoVis.evergladensis.1.1.chr1.g02080.t01#3    mNeoVis.vulgivaga.1.1.chr1.g02083.t01#4    mNeoVis.evergladensis.2.1.chr1.g02078.t01#2   mNeoVis.3.1.1.chr1.g02064.t01#6       442   125261062:   mNeoVis.2.1.1.chr8_hap1.g00381.t01#5    mNeoVis.vulgivaga.2.1.chr8.g00386.t01#1    mNeoVis.vulgivaga.1.1.chr8.g00372.t01#4    mNeoVis.3.1.1.chr8.g00347.t01#6    mNeoVis.evergladensis.1.1.chr8.g00377.t01#3   mNeoVis.evergladensis.2.1.chr9.g00379.t01#2       443   125261063:   mNeoVis.evergladensis.1.1.chr8.g00378.t01#3    mNeoVis.3.1.1.chr8.g00348.t01#6    mNeoVis.evergladensis.2.1.chr9.g00380.t01#2    mNeoVis.vulgivaga.2.1.chr8.g00387.t01#1    mNeoVis.2.1.1.chr8_hap1.g00382.t01#5   mNeoVis.vulgivaga.1.1.chr8.g00373.t01#4       444   125261064:   mNeoVis.evergladensis.2.1.chr1.g02079.t01#2    mNeoVis.evergladensis.1.1.chr1.g02081.t01#3    mNeoVis.3.1.1.chr1.g02065.t01#6    mNeoVis.2.1.1.chr1_hap1.g02109.t01#5    mNeoVis.vulgivaga.2.1.chr1.g02073.t01#1    mNeoVis.evergladensis.2.1.chr1.g02080.t01#2   mNeoVis.vulgivaga.1.1.chr1.g02084.t01#4     445   125261065:   mNeoVis.3.1.1.chr8.g00349.t01#6    mNeoVis.evergladensis.1.1.chr8.g00379.t01#3    mNeoVis.2.1.1.chr8_hap1.g00383.t01#5    mNeoVis.evergladensis.2.1.chr9.g00381.t01#2    mNeoVis.vulgivaga.1.1.chr8.g00374.t01#4   mNeoVis.vulgivaga.2.1.chr8.g00388.t01#1       446   125261066:   mNeoVis.vulgivaga.2.1.chr8.g00389.t01#1    mNeoVis.evergladensis.1.1.chr8.g00380.t01#3   mNeoVis.vulgivaga.1.1.chr8.g00375.t01#4           447   125261067:   mNeoVis.evergladensis.2.1.chr9.g00382.t01#2   mNeoVis.3.1.1.chr8.g00350.t01#6             448   125261068:  mNeoVis.2.1.1.chr8_hap1.g00384.t01#5               449   125261069:   mNeoVis.vulgivaga.1.1.chr1.g02085.t01#4    mNeoVis.evergladensis.2.1.chr1.g02081.t01#2    mNeoVis.3.1.1.chr1.g02066.t01#6    mNeoVis.evergladensis.1.1.chr1.g02082.t01#3    mNeoVis.vulgivaga.2.1.chr1.g02074.t01#1   mNeoVis.2.1.1.chr1_hap1.g02110.t01#5       450   125261070:   mNeoVis.2.1.1.chr15_hap1.g00743.t01#5    mNeoVis.evergladensis.2.1.chr15.g00565.t01#2    mNeoVis.vulgivaga.1.1.chr15.g00568.t01#4    mNeoVis.vulgivaga.2.1.chr15.g00577.t01#1    mNeoVis.3.1.1.chr15.g00521.t01#6   mNeoVis.evergladensis.1.1.chr15.g00048.t01#3       451   125261071:   mNeoVis.evergladensis.1.1.chr1.g02084.t01#3    mNeoVis.evergladensis.2.1.chr1.g02083.t01#2    mNeoVis.vulgivaga.2.1.chr1.g02075.t01#1    mNeoVis.vulgivaga.1.1.chr1.g02087.t01#4    mNeoVis.2.1.1.chr1_hap1.g02113.t01#5   mNeoVis.3.1.1.chr1.g02068.t01#6       452   125261072:   mNeoVis.vulgivaga.1.1.chr1.g02088.t01#4    mNeoVis.vulgivaga.1.1.chr1.g01257.t01#4    mNeoVis.evergladensis.1.1.chr1.g02085.t01#3    mNeoVis.2.1.1.chr1_hap1.g02114.t01#5    mNeoVis.evergladensis.2.1.chr1.g02084.t01#2   mNeoVis.vulgivaga.2.1.chr1.g02076.t01#1       453   125261073:   mNeoVis.2.1.1.chr1_hap1.g01765.t01#5    mNeoVis.evergladensis.1.1.chr1.g01731.t01#3    mNeoVis.3.1.1.chr1.g01719.t01#6    mNeoVis.vulgivaga.1.1.chr1.g01735.t01#4    mNeoVis.vulgivaga.2.1.chr1.g01729.t01#1   mNeoVis.evergladensis.2.1.chr1.g01732.t01#2       454   125261074:   mNeoVis.evergladensis.2.1.chr9.g00383.t01#2    mNeoVis.2.1.1.chr8_hap1.g00385.t01#5    mNeoVis.vulgivaga.1.1.chr8.g00376.t01#4    mNeoVis.vulgivaga.2.1.chr8.g00390.t01#1   mNeoVis.evergladensis.1.1.chr8.g00381.t01#3       455   125261075:   mNeoVis.3.1.1.chr1.g02070.t01#6    mNeoVis.evergladensis.1.1.chr1.g02086.t01#3    mNeoVis.2.1.1.chr1_hap1.g02115.t01#5    mNeoVis.evergladensis.2.1.chr1.g02085.t01#2    mNeoVis.vulgivaga.2.1.chr1.g02077.t01#1   mNeoVis.vulgivaga.1.1.chr1.g02089.t01#4       456   125261076:   mNeoVis.vulgivaga.1.1.chr8.g00377.t01#4    mNeoVis.vulgivaga.2.1.chr8.g00391.t01#1    mNeoVis.evergladensis.2.1.chr9.g00384.t01#2    mNeoVis.2.1.1.chr8_hap1.g00386.t01#5   mNeoVis.evergladensis.1.1.chr8.g00382.t01#3       457   125261077:   mNeoVis.evergladensis.1.1.chr1.g02087.t01#3    mNeoVis.vulgivaga.1.1.chr1.g02090.t01#4    mNeoVis.vulgivaga.2.1.chr1.g02078.t01#1    mNeoVis.2.1.1.chr1_hap1.g02116.t01#5    mNeoVis.3.1.1.chr1.g02072.t01#6   mNeoVis.evergladensis.2.1.chr1.g02086.t01#2       458   125261078:   mNeoVis.2.1.1.chr8_hap1.g00387.t01#5    mNeoVis.evergladensis.1.1.chr8.g00383.t01#3    mNeoVis.evergladensis.2.1.chr9.g00385.t01#2    mNeoVis.vulgivaga.2.1.chr8.g00392.t01#1    mNeoVis.3.1.1.chr8.g00351.t01#6   mNeoVis.vulgivaga.1.1.chr8.g00378.t01#4       459   125261079:   mNeoVis.2.1.1.chr1_hap1.g02117.t01#5    mNeoVis.3.1.1.chr1.g02073.t01#6    mNeoVis.vulgivaga.2.1.chr1.g02079.t01#1    mNeoVis.evergladensis.1.1.chr1.g02088.t01#3    mNeoVis.vulgivaga.1.1.chr1.g02091.t01#4   mNeoVis.evergladensis.2.1.chr1.g02087.t01#2       460   125261080:   mNeoVis.3.1.1.chr15.g00522.t01#6    mNeoVis.vulgivaga.1.1.chr15.g00569.t01#4    mNeoVis.vulgivaga.2.1.chr15.g00578.t01#1    mNeoVis.2.1.1.chr15_hap1.g00744.t01#5    mNeoVis.evergladensis.2.1.chr15.g00566.t01#2   mNeoVis.evergladensis.1.1.chr15.g00047.t01#3       461   125261081:   mNeoVis.2.1.1.chr8_hap1.g00388.t01#5    mNeoVis.vulgivaga.2.1.chr8.g00393.t01#1    mNeoVis.evergladensis.1.1.chr8.g00384.t01#3    mNeoVis.3.1.1.chr8.g00352.t01#6    mNeoVis.evergladensis.2.1.chr9.g00386.t01#2   mNeoVis.vulgivaga.1.1.chr8.g00379.t01#4       462   125261082:   mNeoVis.evergladensis.2.1.chr10.g00389.t01#2    mNeoVis.3.1.1.chr10.g00384.t01#6    mNeoVis.vulgivaga.1.1.chr10.g00381.t01#4    mNeoVis.vulgivaga.2.1.chr10.g00386.t01#1    mNeoVis.2.1.1.chr9_hap1.g00392.t01#5   mNeoVis.evergladensis.1.1.chr10.g00394.t01#3       463   125261083:   mNeoVis.2.1.1.chr5_hap1.g01095.t01#5    mNeoVis.evergladensis.2.1.chr2.g01085.t01#2    mNeoVis.vulgivaga.1.1.chr2.g01089.t01#4    mNeoVis.evergladensis.1.1.chr2.g01088.t01#3    mNeoVis.3.1.1.chr2.g01001.t01#6   mNeoVis.vulgivaga.2.1.chr2.g01092.t01#1       464   125261084:   mNeoVis.evergladensis.2.1.chr1.g02088.t01#2    mNeoVis.3.1.1.chr1.g02074.t01#6    mNeoVis.vulgivaga.1.1.chr1.g02092.t01#4    mNeoVis.evergladensis.1.1.chr1.g02089.t01#3    mNeoVis.2.1.1.chr1_hap1.g02118.t01#5   mNeoVis.vulgivaga.2.1.chr1.g02080.t01#1       465   125261085:   mNeoVis.vulgivaga.2.1.chr8.g00394.t01#1    mNeoVis.evergladensis.2.1.chr9.g00387.t01#2    mNeoVis.evergladensis.1.1.chr8.g00385.t01#3    mNeoVis.2.1.1.chr8_hap1.g00389.t01#5    mNeoVis.3.1.1.chr8.g00353.t01#6   mNeoVis.vulgivaga.1.1.chr8.g00380.t01#4       466   125261086:   mNeoVis.vulgivaga.2.1.chr8.g00395.t01#1    mNeoVis.evergladensis.2.1.chr9.g00388.t01#2    mNeoVis.2.1.1.chr8_hap1.g00390.t01#5    mNeoVis.3.1.1.chr8.g00354.t01#6    mNeoVis.evergladensis.1.1.chr8.g00386.t01#3   mNeoVis.vulgivaga.1.1.chr8.g00381.t01#4       467   125261087:   mNeoVis.vulgivaga.2.1.chr1.g02081.t01#1    mNeoVis.evergladensis.2.1.chr1.g02089.t01#2    mNeoVis.evergladensis.1.1.chr1.g02090.t01#3    mNeoVis.2.1.1.chr1_hap1.g02119.t01#5    mNeoVis.3.1.1.chr1.g02075.t01#6   mNeoVis.vulgivaga.1.1.chr1.g02093.t01#4       468   125261088:   mNeoVis.evergladensis.1.1.chr1.g02091.t01#3    mNeoVis.vulgivaga.1.1.chr1.g02094.t01#4    mNeoVis.3.1.1.chr1.g02076.t01#6    mNeoVis.vulgivaga.2.1.chr1.g02082.t01#1    mNeoVis.evergladensis.2.1.chr1.g02090.t01#2   mNeoVis.2.1.1.chr1_hap1.g02120.t01#5       469   125261089:   mNeoVis.3.1.1.chr8.g00355.t01#6    mNeoVis.vulgivaga.1.1.chr8.g00382.t01#4    mNeoVis.2.1.1.chr8_hap1.g00391.t01#5    mNeoVis.evergladensis.2.1.chr9.g00389.t01#2    mNeoVis.vulgivaga.2.1.chr8.g00396.t01#1   mNeoVis.evergladensis.1.1.chr8.g00387.t01#3       470   125261090:   mNeoVis.2.1.1.chr8_hap1.g00392.t01#5    mNeoVis.evergladensis.2.1.chr9.g00390.t01#2    mNeoVis.3.1.1.chr8.g00357.t01#6    mNeoVis.vulgivaga.1.1.chr8.g00384.t01#4    mNeoVis.evergladensis.1.1.chr8.g00388.t01#3    mNeoVis.3.1.1.chr8.g00356.t01#6    mNeoVis.vulgivaga.1.1.chr8.g00383.t01#4    mNeoVis.vulgivaga.2.1.chr8.g00397.t01#1    mNeoVis.evergladensis.2.1.chr9.g00391.t01#2      471   125261091:   mNeoVis.2.1.1.chr8_hap1.g00394.t01#5    mNeoVis.evergladensis.2.1.chr9.g00392.t01#2    mNeoVis.vulgivaga.2.1.chr8.g00398.t01#1    mNeoVis.vulgivaga.1.1.chr8.g00385.t01#4   mNeoVis.evergladensis.1.1.chr8.g00389.t01#3       472   125261092:   mNeoVis.vulgivaga.1.1.chr8.g00386.t01#4    mNeoVis.3.1.1.chr8.g00359.t01#6    mNeoVis.vulgivaga.2.1.chr8.g00399.t01#1    mNeoVis.evergladensis.1.1.chr8.g00390.t01#3    mNeoVis.evergladensis.2.1.chr9.g00393.t01#2   mNeoVis.2.1.1.chr8_hap1.g00395.t01#5       473   125261093:   mNeoVis.vulgivaga.1.1.chr15.g00570.t01#4    mNeoVis.3.1.1.chr15.g00523.t01#6    mNeoVis.2.1.1.chr15_hap1.g00745.t01#5    mNeoVis.evergladensis.1.1.chr15.g00046.t01#3    mNeoVis.vulgivaga.2.1.chr15.g00579.t01#1   mNeoVis.evergladensis.2.1.chr15.g00567.t01#2       474   125261094:   mNeoVis.evergladensis.1.1.chr1.g02092.t01#3    mNeoVis.3.1.1.chr1.g02077.t01#6    mNeoVis.evergladensis.2.1.chr1.g02091.t01#2    mNeoVis.vulgivaga.2.1.chr1.g02083.t01#1    mNeoVis.2.1.1.chr1_hap1.g02121.t01#5   mNeoVis.vulgivaga.1.1.chr1.g02095.t01#4       475   125261095:   mNeoVis.vulgivaga.1.1.chr8.g00387.t01#4    mNeoVis.3.1.1.chr8.g00360.t01#6    mNeoVis.vulgivaga.2.1.chr8.g00400.t01#1    mNeoVis.evergladensis.1.1.chr8.g00391.t01#3    mNeoVis.evergladensis.2.1.chr9.g00394.t01#2   mNeoVis.2.1.1.chr8_hap1.g00396.t01#5       476   125261096:   mNeoVis.evergladensis.2.1.chr1.g02092.t01#2    mNeoVis.2.1.1.chr1_hap1.g02122.t01#5    mNeoVis.vulgivaga.1.1.chr1.g02096.t01#4    mNeoVis.evergladensis.1.1.chr1.g02093.t01#3    mNeoVis.3.1.1.chr1.g02078.t01#6   mNeoVis.vulgivaga.2.1.chr1.g02084.t01#1       477   125261097:   mNeoVis.evergladensis.1.1.chr1.g02094.t01#3    mNeoVis.vulgivaga.2.1.chr1.g02085.t01#1    mNeoVis.3.1.1.chr1.g02079.t01#6    mNeoVis.vulgivaga.1.1.chr1.g02097.t01#4    mNeoVis.evergladensis.2.1.chr1.g02093.t01#2   mNeoVis.2.1.1.chr1_hap1.g02123.t01#5       478   125261098:   mNeoVis.evergladensis.2.1.chr1.g02094.t01#2    mNeoVis.vulgivaga.1.1.chr1.g02098.t01#4    mNeoVis.evergladensis.1.1.chr1.g02095.t01#3    mNeoVis.2.1.1.chr1_hap1.g02124.t01#5    mNeoVis.vulgivaga.2.1.chr1.g02086.t01#1   mNeoVis.3.1.1.chr1.g02080.t01#6       479   125261099:   mNeoVis.2.1.1.chr8_hap1.g00397.t01#5    mNeoVis.evergladensis.1.1.chr8.g00392.t01#3    mNeoVis.evergladensis.2.1.chr9.g00395.t01#2    mNeoVis.vulgivaga.1.1.chr8.g00388.t01#4    mNeoVis.vulgivaga.2.1.chr8.g00401.t01#1   mNeoVis.3.1.1.chr8.g00361.t01#6       480   125261100:   mNeoVis.vulgivaga.1.1.chr8.g00389.t01#4    mNeoVis.3.1.1.chr8.g00362.t01#6    mNeoVis.vulgivaga.2.1.chr8.g00402.t01#1    mNeoVis.2.1.1.chr8_hap1.g00398.t01#5    mNeoVis.evergladensis.1.1.chr8.g00393.t01#3   mNeoVis.evergladensis.2.1.chr9.g00396.t01#2       481   125261101:   mNeoVis.2.1.1.chr15_hap1.g00747.t01#5    mNeoVis.evergladensis.1.1.chr15.g00044.t01#3    mNeoVis.evergladensis.2.1.chr15.g00569.t01#2    mNeoVis.vulgivaga.1.1.chr15.g00571.t01#4    mNeoVis.3.1.1.chr15.g00524.t01#6   mNeoVis.vulgivaga.2.1.chr15.g00580.t01#1       482   125261102:   mNeoVis.3.1.1.chr1.g02081.t01#6    mNeoVis.vulgivaga.1.1.chr1.g02099.t01#4    mNeoVis.evergladensis.2.1.chr1.g02095.t01#2    mNeoVis.vulgivaga.2.1.chr1.g02087.t01#1    mNeoVis.evergladensis.1.1.chr1.g02096.t01#3   mNeoVis.2.1.1.chr1_hap1.g02125.t01#5       483   125261103:   mNeoVis.3.1.1.chr8.g00363.t01#6    mNeoVis.vulgivaga.1.1.chr8.g00390.t01#4    mNeoVis.vulgivaga.2.1.chr8.g00403.t01#1    mNeoVis.evergladensis.2.1.chr9.g00397.t01#2    mNeoVis.2.1.1.chr8_hap1.g00399.t01#5   mNeoVis.evergladensis.1.1.chr8.g00394.t01#3       484   125261104:   mNeoVis.evergladensis.1.1.chr8.g00395.t01#3    mNeoVis.evergladensis.2.1.chr9.g00398.t01#2    mNeoVis.3.1.1.chr8.g00364.t01#6    mNeoVis.2.1.1.chr8_hap1.g00400.t01#5    mNeoVis.vulgivaga.1.1.chr8.g00391.t01#4   mNeoVis.vulgivaga.2.1.chr8.g00404.t01#1       485   125261105:   mNeoVis.evergladensis.1.1.chr15.g00043.t01#3    mNeoVis.2.1.1.chr15_hap1.g00748.t01#5    mNeoVis.3.1.1.chr15.g00525.t01#6    mNeoVis.evergladensis.2.1.chr15.g00570.t01#2    mNeoVis.vulgivaga.2.1.chr15.g00581.t01#1   mNeoVis.vulgivaga.1.1.chr15.g00572.t01#4       486   125261106:   mNeoVis.vulgivaga.2.1.chr8.g00313.t01#1    mNeoVis.vulgivaga.1.1.chr8.g00300.t01#4    mNeoVis.evergladensis.1.1.chr8.g00306.t01#3    mNeoVis.evergladensis.2.1.chr9.g00306.t01#2   mNeoVis.2.1.1.chr8_hap1.g00307.t01#5       487   125261107:   mNeoVis.3.1.1.chr8.g00276.t01#6    mNeoVis.evergladensis.1.1.chr8.g00307.t01#3    mNeoVis.vulgivaga.1.1.chr8.g00301.t01#4    mNeoVis.evergladensis.2.1.chr9.g00307.t01#2    mNeoVis.vulgivaga.2.1.chr8.g00314.t01#1   mNeoVis.2.1.1.chr8_hap1.g00308.t01#5       488   125261108:   mNeoVis.3.1.1.chr1.g02082.t01#6    mNeoVis.evergladensis.2.1.chr1.g02096.t01#2    mNeoVis.evergladensis.1.1.chr1.g02097.t01#3    mNeoVis.2.1.1.chr1_hap1.g02126.t01#5    mNeoVis.vulgivaga.1.1.chr1.g02100.t01#4   mNeoVis.vulgivaga.2.1.chr1.g02088.t01#1       489   125261109:   mNeoVis.2.1.1.chr8_hap1.g00309.t01#5    mNeoVis.vulgivaga.1.1.chr8.g00302.t01#4    mNeoVis.evergladensis.1.1.chr8.g00308.t01#3    mNeoVis.3.1.1.chr8.g00277.t01#6    mNeoVis.evergladensis.2.1.chr9.g00308.t01#2   mNeoVis.vulgivaga.2.1.chr8.g00315.t01#1       490   125261110:   mNeoVis.evergladensis.2.1.chr1.g02097.t01#2    mNeoVis.3.1.1.chr1.g02083.t01#6    mNeoVis.evergladensis.1.1.chr1.g02098.t01#3    mNeoVis.2.1.1.chr1_hap1.g02127.t01#5    mNeoVis.vulgivaga.2.1.chr1.g02089.t01#1   mNeoVis.vulgivaga.1.1.chr1.g02101.t01#4       491   125261111:   mNeoVis.2.1.1.chr12_hap1.g00539.t01#5    mNeoVis.vulgivaga.1.1.chr12.g00396.t01#4    mNeoVis.3.1.1.chr1.g01952.t01#6    mNeoVis.2.1.1.chr6_hap1.g00469.t01#5    mNeoVis.evergladensis.2.1.chr1.g01962.t01#2    mNeoVis.evergladensis.1.1.chr1.g01963.t01#3    mNeoVis.2.1.1.chr1_hap1.g01998.t01#5    mNeoVis.evergladensis.2.1.chr12.g00385.t01#2    mNeoVis.vulgivaga.2.1.chr1.g01961.t01#1      492   125261112:   mNeoVis.evergladensis.1.1.chr1.g02099.t01#3    mNeoVis.vulgivaga.2.1.chr1.g02090.t01#1    mNeoVis.2.1.1.chr1_hap1.g02129.t01#5    mNeoVis.3.1.1.chr1.g02084.t01#6   mNeoVis.vulgivaga.1.1.chr1.g02103.t01#4       493   125261113:   mNeoVis.3.1.1.chr1.g02085.t01#6    mNeoVis.evergladensis.1.1.chr1.g02100.t01#3    mNeoVis.evergladensis.2.1.chr1.g02098.t01#2    mNeoVis.vulgivaga.2.1.chr1.g02091.t01#1    mNeoVis.evergladensis.2.1.chr1.g02099.t01#2   mNeoVis.2.1.1.chr1_hap1.g02130.t01#5       494   125261114:   mNeoVis.evergladensis.2.1.chr9.g00309.t01#2    mNeoVis.3.1.1.chr8.g00278.t01#6    mNeoVis.evergladensis.1.1.chr8.g00309.t01#3    mNeoVis.2.1.1.chr8_hap1.g00310.t01#5    mNeoVis.vulgivaga.2.1.chr8.g00316.t01#1   mNeoVis.vulgivaga.1.1.chr8.g00303.t01#4       495   125261115:   mNeoVis.evergladensis.2.1.chr9.g00310.t01#2    mNeoVis.vulgivaga.2.1.chr8.g00317.t01#1    mNeoVis.vulgivaga.1.1.chr8.g00304.t01#4    mNeoVis.3.1.1.chr8.g00279.t01#6    mNeoVis.evergladensis.1.1.chr8.g00310.t01#3   mNeoVis.2.1.1.chr8_hap1.g00311.t01#5       496   125261116:   mNeoVis.evergladensis.1.1.chr8.g00311.t01#3    mNeoVis.2.1.1.chr8_hap1.g00312.t01#5    mNeoVis.vulgivaga.1.1.chr8.g00305.t01#4    mNeoVis.evergladensis.2.1.chr9.g00311.t01#2    mNeoVis.vulgivaga.2.1.chr8.g00318.t01#1   mNeoVis.3.1.1.chr8.g00280.t01#6       497   125261117:   mNeoVis.3.1.1.chr1.g02086.t01#6    mNeoVis.evergladensis.2.1.chr1.g02100.t01#2    mNeoVis.vulgivaga.2.1.chr1.g02092.t01#1   mNeoVis.2.1.1.chr1_hap1.g02131.t01#5         498   125261118:   mNeoVis.2.1.1.chr1_hap1.g02133.t01#5    mNeoVis.3.1.1.chr1.g02088.t01#6    mNeoVis.vulgivaga.1.1.chr1.g02107.t01#4    mNeoVis.evergladensis.1.1.chr1.g02101.t01#3    mNeoVis.evergladensis.2.1.chr1.g02103.t01#2   mNeoVis.vulgivaga.2.1.chr1.g02093.t01#1       499   125261119:   mNeoVis.evergladensis.1.1.chr1.g02103.t01#3    mNeoVis.2.1.1.chr1_hap1.g02134.t01#5    mNeoVis.vulgivaga.2.1.chr1.g02094.t01#1    mNeoVis.3.1.1.chr1.g02089.t01#6    mNeoVis.vulgivaga.1.1.chr1.g02108.t01#4   mNeoVis.evergladensis.2.1.chr1.g02104.t01#2       500   125261120:  mNeoVis.vulgivaga.2.1.chr8.g00319.t01#1               501   125261121:   mNeoVis.vulgivaga.1.1.chr1.g02109.t01#4    mNeoVis.evergladensis.2.1.chr1.g02105.t01#2    mNeoVis.evergladensis.1.1.chr1.g02104.t01#3    mNeoVis.vulgivaga.2.1.chr1.g02095.t01#1   mNeoVis.2.1.1.chr1_hap1.g02135.t01#5       502   125261122:   mNeoVis.2.1.1.chr15_hap1.g00749.t01#5    mNeoVis.vulgivaga.2.1.chr15.g00582.t01#1    mNeoVis.evergladensis.2.1.chr15.g00571.t01#2    mNeoVis.evergladensis.1.1.chr15.g00042.t01#3    mNeoVis.vulgivaga.1.1.chr15.g00573.t01#4   mNeoVis.3.1.1.chr15.g00526.t01#6       503   125261123:   mNeoVis.vulgivaga.1.1.chr8.g00306.t01#4    mNeoVis.evergladensis.1.1.chr8.g00312.t01#3    mNeoVis.3.1.1.chr8.g00281.t01#6    mNeoVis.vulgivaga.2.1.chr8.g00320.t01#1    mNeoVis.evergladensis.2.1.chr9.g00312.t01#2   mNeoVis.2.1.1.chr8_hap1.g00313.t01#5       504   125261124:   mNeoVis.vulgivaga.1.1.chr8.g00307.t01#4    mNeoVis.evergladensis.1.1.chr8.g00313.t01#3    mNeoVis.vulgivaga.2.1.chr8.g00321.t01#1    mNeoVis.2.1.1.chr8_hap1.g00314.t01#5    mNeoVis.3.1.1.chr8.g00282.t01#6   mNeoVis.evergladensis.2.1.chr9.g00313.t01#2       505   125335273:   mNeoVis.3.1.1.chr1.g02008.t01#6    mNeoVis.evergladensis.2.1.chr1.g02020.t01#2    mNeoVis.2.1.1.chr1_hap1.g02053.t01#5    mNeoVis.vulgivaga.1.1.chr1.g02023.t01#4    mNeoVis.evergladensis.1.1.chr1.g02020.t01#3   mNeoVis.vulgivaga.2.1.chr1.g02013.t01#1       506   125335274:   mNeoVis.2.1.1.chr8_hap1.g00315.t01#5    mNeoVis.evergladensis.1.1.chr8.g00314.t01#3    mNeoVis.3.1.1.chr8.g00283.t01#6    mNeoVis.vulgivaga.2.1.chr8.g00322.t01#1    mNeoVis.vulgivaga.1.1.chr8.g00308.t01#4   mNeoVis.evergladensis.2.1.chr9.g00314.t01#2       507   125335275:   mNeoVis.2.1.1.chr1_hap1.g02054.t01#5    mNeoVis.vulgivaga.1.1.chr1.g02024.t01#4    mNeoVis.evergladensis.2.1.chr1.g02021.t01#2    mNeoVis.evergladensis.1.1.chr1.g02021.t01#3    mNeoVis.vulgivaga.2.1.chr1.g02014.t01#1   mNeoVis.3.1.1.chr1.g02009.t01#6       508   125261675:   mNeoVis.evergladensis.1.1.chr1.g02022.t01#3    mNeoVis.3.1.1.chr1.g02010.t01#6    mNeoVis.vulgivaga.2.1.chr1.g02015.t01#1    mNeoVis.vulgivaga.1.1.chr1.g02025.t01#4    mNeoVis.evergladensis.2.1.chr1.g02022.t01#2   mNeoVis.2.1.1.chr1_hap1.g02055.t01#5       509   125261676:   mNeoVis.evergladensis.2.1.chr9.g00315.t01#2    mNeoVis.3.1.1.chr8.g00284.t01#6    mNeoVis.2.1.1.chr8_hap1.g00316.t01#5    mNeoVis.vulgivaga.1.1.chr8.g00309.t01#4    mNeoVis.vulgivaga.2.1.chr8.g00323.t01#1   mNeoVis.evergladensis.1.1.chr8.g00315.t01#3       510   125413060:   mNeoVis.evergladensis.2.1.chr3.g01275.t01#2    mNeoVis.evergladensis.1.1.chr3.g01283.t01#3    mNeoVis.2.1.1.chr3_hap1.g01277.t01#5    mNeoVis.3.1.1.chr3.g01254.t01#6    mNeoVis.vulgivaga.2.1.chr3.g01293.t01#1   mNeoVis.vulgivaga.1.1.chr3.g01269.t01#4       511   125413061:  mNeoVis.vulgivaga.2.1.chr1.g02016.t01#1               512   125413062:   mNeoVis.vulgivaga.2.1.chr8.g00324.t01#1    mNeoVis.evergladensis.2.1.chr9.g00316.t01#2    mNeoVis.evergladensis.1.1.chr8.g00316.t01#3    mNeoVis.2.1.1.chr8_hap1.g00317.t01#5    mNeoVis.vulgivaga.1.1.chr8.g00310.t01#4   mNeoVis.3.1.1.chr8.g00285.t01#6       513   125413063:   mNeoVis.evergladensis.1.1.chr15.g00040.t01#3    mNeoVis.vulgivaga.1.1.chr15.g00574.t01#4    mNeoVis.3.1.1.chr15.g00528.t01#6    mNeoVis.2.1.1.chr15_hap1.g00751.t01#5    mNeoVis.vulgivaga.2.1.chr15.g00583.t01#1   mNeoVis.evergladensis.2.1.chr15.g00572.t01#2       514   125413064:   mNeoVis.vulgivaga.2.1.chr1.g02017.t01#1    mNeoVis.evergladensis.2.1.chr1.g02023.t01#2    mNeoVis.3.1.1.chr1.g02011.t01#6    mNeoVis.evergladensis.1.1.chr1.g02024.t01#3    mNeoVis.vulgivaga.1.1.chr1.g02026.t01#4    mNeoVis.2.1.1.chr1_hap1.g02056.t01#5   mNeoVis.evergladensis.1.1.chr1.g02023.t01#3     515   125413065:   mNeoVis.vulgivaga.2.1.chr8.g00325.t01#1    mNeoVis.2.1.1.chr8_hap1.g00318.t01#5    mNeoVis.vulgivaga.1.1.chr8.g00311.t01#4    mNeoVis.evergladensis.1.1.chr8.g00317.t01#3   mNeoVis.evergladensis.2.1.chr9.g00317.t01#2       516   125413066:   mNeoVis.3.1.1.chr1.g02012.t01#6    mNeoVis.vulgivaga.1.1.chr1.g02027.t01#4    mNeoVis.vulgivaga.2.1.chr1.g02018.t01#1    mNeoVis.2.1.1.chr1_hap1.g02057.t01#5    mNeoVis.evergladensis.2.1.chr1.g02024.t01#2   mNeoVis.evergladensis.1.1.chr1.g02025.t01#3       517   125384910:   mNeoVis.vulgivaga.2.1.chr1.g02019.t01#1    mNeoVis.evergladensis.2.1.chr1.g02025.t01#2    mNeoVis.evergladensis.1.1.chr1.g02026.t01#3    mNeoVis.vulgivaga.1.1.chr1.g02028.t01#4   mNeoVis.2.1.1.chr1_hap1.g02058.t01#5       518   125384911:   mNeoVis.vulgivaga.2.1.chr1.g02020.t01#1    mNeoVis.vulgivaga.1.1.chr1.g02029.t01#4    mNeoVis.2.1.1.chr1_hap1.g02059.t01#5    mNeoVis.3.1.1.chr1.g02013.t01#6    mNeoVis.evergladensis.1.1.chr1.g02027.t01#3   mNeoVis.evergladensis.2.1.chr1.g02026.t01#2       519   125384912:   mNeoVis.2.1.1.chr1_hap1.g02063.t01#5    mNeoVis.vulgivaga.1.1.chr1.g02033.t01#4    mNeoVis.evergladensis.1.1.chr1.g02032.t01#3    mNeoVis.evergladensis.2.1.chr1.g02031.t01#2    mNeoVis.3.1.1.chr1.g02016.t01#6   mNeoVis.vulgivaga.2.1.chr1.g02025.t01#1       520   125359047:   mNeoVis.vulgivaga.2.1.chr1.g02021.t01#1    mNeoVis.evergladensis.1.1.chr1.g02028.t01#3    mNeoVis.evergladensis.2.1.chr1.g02027.t01#2   mNeoVis.vulgivaga.1.1.chr1.g02030.t01#4         521   125359048:   mNeoVis.2.1.1.chr1_hap1.g02066.t01#5    mNeoVis.vulgivaga.2.1.chr1.g02022.t01#1    mNeoVis.vulgivaga.1.1.chr1.g02037.t01#4   mNeoVis.2.1.1.chr1_hap1.g02060.t01#5         522   125359049:   mNeoVis.3.1.1.chr1.g02014.t01#6   mNeoVis.evergladensis.1.1.chr1.g02029.t01#3           523   125359050:  mNeoVis.evergladensis.2.1.chr1.g02028.t01#2             524   125359051:  mNeoVis.vulgivaga.2.1.chr1.g02023.t01#1               525   125359052:   mNeoVis.evergladensis.2.1.chr9.g00318.t01#2    mNeoVis.3.1.1.chr8.g00287.t01#6    mNeoVis.2.1.1.chr8_hap1.g00319.t01#5    mNeoVis.evergladensis.1.1.chr8.g00318.t01#3    mNeoVis.vulgivaga.1.1.chr8.g00312.t01#4   mNeoVis.vulgivaga.2.1.chr8.g00326.t01#1       526   125359053:   mNeoVis.vulgivaga.2.1.chr1.g02024.t01#1    mNeoVis.evergladensis.1.1.chr1.g02031.t01#3    mNeoVis.2.1.1.chr1_hap1.g02062.t01#5    mNeoVis.vulgivaga.1.1.chr1.g02032.t01#4    mNeoVis.evergladensis.2.1.chr1.g02030.t01#2   mNeoVis.3.1.1.chr1.g02015.t01#6       527   125359054:   mNeoVis.vulgivaga.2.1.chr8.g00327.t01#1    mNeoVis.evergladensis.2.1.chr9.g00319.t01#2    mNeoVis.3.1.1.chr8.g00288.t01#6    mNeoVis.2.1.1.chr8_hap1.g00320.t01#5    mNeoVis.vulgivaga.1.1.chr8.g00313.t01#4   mNeoVis.evergladensis.1.1.chr8.g00319.t01#3       528   125359055:   mNeoVis.vulgivaga.1.1.chr1.g02035.t01#4    mNeoVis.vulgivaga.2.1.chr1.g02026.t01#1    mNeoVis.evergladensis.1.1.chr1.g02033.t01#3   mNeoVis.evergladensis.2.1.chr1.g02033.t01#2       529   125359056:   mNeoVis.evergladensis.2.1.chr1.g02032.t01#2   mNeoVis.vulgivaga.1.1.chr1.g02034.t01#4             530   125359057:  mNeoVis.2.1.1.chr1_hap1.g02064.t01#5               531   125359058:   mNeoVis.evergladensis.2.1.chr1.g02034.t01#2    mNeoVis.3.1.1.chr1.g02017.t01#6    mNeoVis.vulgivaga.2.1.chr1.g02027.t01#1    mNeoVis.2.1.1.chr1_hap1.g02065.t01#5   mNeoVis.vulgivaga.1.1.chr1.g02036.t01#4       532   125359059:   mNeoVis.vulgivaga.1.1.chr1.g02038.t01#4    mNeoVis.evergladensis.2.1.chr1.g02035.t01#2    mNeoVis.vulgivaga.2.1.chr1.g02028.t01#1    mNeoVis.2.1.1.chr1_hap1.g02067.t01#5   mNeoVis.evergladensis.1.1.chr1.g02034.t01#3       533   125359060:   mNeoVis.vulgivaga.1.1.chr8.g00314.t01#4    mNeoVis.vulgivaga.2.1.chr8.g00328.t01#1    mNeoVis.evergladensis.1.1.chr8.g00320.t01#3    mNeoVis.evergladensis.2.1.chr9.g00320.t01#2   mNeoVis.2.1.1.chr8_hap1.g00321.t01#5       534   125359061:   mNeoVis.evergladensis.1.1.chr1.g02035.t01#3    mNeoVis.2.1.1.chr1_hap1.g02068.t01#5    mNeoVis.evergladensis.2.1.chr1.g02036.t01#2    mNeoVis.3.1.1.chr1.g02018.t01#6    mNeoVis.vulgivaga.2.1.chr1.g02029.t01#1   mNeoVis.vulgivaga.1.1.chr1.g02039.t01#4       535   125359062:   mNeoVis.vulgivaga.1.1.chr1.g02040.t01#4    mNeoVis.3.1.1.chr1.g02019.t01#6    mNeoVis.vulgivaga.2.1.chr1.g02030.t01#1    mNeoVis.2.1.1.chr1_hap1.g02069.t01#5    mNeoVis.evergladensis.1.1.chr1.g02036.t01#3   mNeoVis.evergladensis.2.1.chr1.g02037.t01#2       536   125314002:   mNeoVis.evergladensis.1.1.chr8.g00321.t01#3    mNeoVis.2.1.1.chr8_hap1.g00322.t01#5    mNeoVis.evergladensis.2.1.chr9.g00321.t01#2    mNeoVis.vulgivaga.1.1.chr8.g00315.t01#4    mNeoVis.vulgivaga.2.1.chr8.g00329.t01#1   mNeoVis.3.1.1.chr8.g00289.t01#6       537   125314003:   mNeoVis.vulgivaga.1.1.chr1.g02041.t01#4    mNeoVis.evergladensis.2.1.chr1.g02039.t01#2    mNeoVis.vulgivaga.2.1.chr1.g02031.t01#1    mNeoVis.2.1.1.chr1_hap1.g02071.t01#5    mNeoVis.3.1.1.chr1.g02021.t01#6   mNeoVis.evergladensis.1.1.chr1.g02038.t01#3       538   125314004:   mNeoVis.vulgivaga.1.1.chr8.g00316.t01#4    mNeoVis.evergladensis.2.1.chr9.g00322.t01#2    mNeoVis.evergladensis.1.1.chr8.g00322.t01#3    mNeoVis.2.1.1.chr8_hap1.g00323.t01#5   mNeoVis.vulgivaga.2.1.chr8.g00330.t01#1       539   125314005:   mNeoVis.vulgivaga.1.1.chr7.g00520.t01#4    mNeoVis.evergladensis.2.1.chr7.g00513.t01#2    mNeoVis.vulgivaga.2.1.chr7.g00516.t01#1    mNeoVis.2.1.1.chr7_hap1.g00512.t01#5    mNeoVis.3.1.1.chr7.g00497.t01#6   mNeoVis.evergladensis.1.1.chr7.g00522.t01#3       540   125314006:   mNeoVis.3.1.1.chr1.g02023.t01#6    mNeoVis.2.1.1.chr1_hap1.g02072.t01#5    mNeoVis.vulgivaga.2.1.chr1.g02032.t01#1    mNeoVis.evergladensis.1.1.chr1.g02039.t01#3    mNeoVis.evergladensis.1.1.chr1.g02040.t01#3    mNeoVis.evergladensis.2.1.chr1.g02040.t01#2   mNeoVis.vulgivaga.1.1.chr1.g02044.t01#4     541   125314007:   mNeoVis.3.1.1.chr9.g00292.t01#6    mNeoVis.2.1.1.chr10_hap1.g00319.t01#5    mNeoVis.vulgivaga.2.1.chr9.g00304.t01#1    mNeoVis.vulgivaga.1.1.chr9.g00304.t01#4    mNeoVis.evergladensis.1.1.chr9.g00307.t01#3   mNeoVis.evergladensis.2.1.chr8.g00303.t01#2       542   125314008:   mNeoVis.3.1.1.chr1.g02022.t01#6   mNeoVis.vulgivaga.1.1.chr1.g02042.t01#4             543   125314009:  mNeoVis.vulgivaga.1.1.chr1.g02043.t01#4               544   125314010:   mNeoVis.evergladensis.2.1.chr9.g00323.t01#2    mNeoVis.evergladensis.1.1.chr8.g00323.t01#3    mNeoVis.2.1.1.chr8_hap1.g00324.t01#5    mNeoVis.3.1.1.chr8.g00290.t01#6    mNeoVis.vulgivaga.1.1.chr8.g00317.t01#4   mNeoVis.vulgivaga.2.1.chr8.g00331.t01#1       545   125314011:   mNeoVis.vulgivaga.2.1.chr1.g02033.t01#1    mNeoVis.evergladensis.2.1.chr1.g02042.t01#2    mNeoVis.2.1.1.chr1_hap1.g02073.t01#5    mNeoVis.vulgivaga.1.1.chr1.g02045.t01#4    mNeoVis.evergladensis.1.1.chr1.g02041.t01#3   mNeoVis.3.1.1.chr1.g02024.t01#6       546   125293065:   mNeoVis.vulgivaga.1.1.chr1.g02046.t01#4    mNeoVis.evergladensis.2.1.chr1.g02043.t01#2    mNeoVis.3.1.1.chr1.g02025.t01#6    mNeoVis.vulgivaga.2.1.chr1.g02034.t01#1    mNeoVis.2.1.1.chr1_hap1.g02074.t01#5   mNeoVis.evergladensis.1.1.chr1.g02042.t01#3       547   125293066:   mNeoVis.vulgivaga.2.1.chr8.g00332.t01#1    mNeoVis.evergladensis.2.1.chr9.g00324.t01#2    mNeoVis.2.1.1.chr8_hap1.g00325.t01#5    mNeoVis.vulgivaga.1.1.chr8.g00318.t01#4    mNeoVis.3.1.1.chr8.g00291.t01#6   mNeoVis.evergladensis.1.1.chr8.g00324.t01#3       548   125293067:   mNeoVis.2.1.1.chr1_hap1.g02075.t01#5    mNeoVis.3.1.1.chr1.g02026.t01#6    mNeoVis.evergladensis.1.1.chr1.g02043.t01#3    mNeoVis.vulgivaga.2.1.chr1.g02035.t01#1    mNeoVis.vulgivaga.1.1.chr1.g02047.t01#4   mNeoVis.evergladensis.2.1.chr1.g02044.t01#2       549   125293068:   mNeoVis.3.1.1.chr8.g00292.t01#6    mNeoVis.vulgivaga.1.1.chr8.g00319.t01#4    mNeoVis.vulgivaga.2.1.chr8.g00333.t01#1    mNeoVis.evergladensis.2.1.chr9.g00325.t01#2    mNeoVis.2.1.1.chr8_hap1.g00326.t01#5   mNeoVis.evergladensis.1.1.chr8.g00325.t01#3       550   125293069:   mNeoVis.vulgivaga.2.1.chr1.g02036.t01#1    mNeoVis.vulgivaga.1.1.chr1.g02048.t01#4    mNeoVis.evergladensis.2.1.chr1.g02045.t01#2    mNeoVis.2.1.1.chr1_hap1.g02076.t01#5    mNeoVis.3.1.1.chr1.g02027.t01#6   mNeoVis.evergladensis.1.1.chr1.g02044.t01#3       551   125293070:   mNeoVis.vulgivaga.2.1.chr1.g02037.t01#1    mNeoVis.evergladensis.1.1.chr1.g02045.t01#3    mNeoVis.3.1.1.chr1.g02029.t01#6    mNeoVis.2.1.1.chr1_hap1.g02077.t01#5    mNeoVis.evergladensis.2.1.chr1.g02046.t01#2   mNeoVis.vulgivaga.1.1.chr1.g02049.t01#4       552   125293071:   mNeoVis.2.1.1.chr8_hap1.g00327.t01#5    mNeoVis.evergladensis.1.1.chr8.g00326.t01#3    mNeoVis.evergladensis.2.1.chr9.g00326.t01#2    mNeoVis.vulgivaga.2.1.chr8.g00334.t01#1    mNeoVis.vulgivaga.1.1.chr8.g00320.t01#4   mNeoVis.3.1.1.chr8.g00293.t01#6       553   125261677:   mNeoVis.evergladensis.1.1.chr8.g00327.t01#3    mNeoVis.evergladensis.2.1.chr9.g00327.t01#2    mNeoVis.vulgivaga.2.1.chr8.g00335.t01#1    mNeoVis.2.1.1.chr8_hap1.g00328.t01#5    mNeoVis.3.1.1.chr8.g00294.t01#6   mNeoVis.vulgivaga.1.1.chr8.g00321.t01#4       554   125261678:  mNeoVis.vulgivaga.2.1.chr1.g02038.t01#1               555   125261679:   mNeoVis.3.1.1.chr8.g00295.t01#6    mNeoVis.vulgivaga.1.1.chr8.g00322.t01#4    mNeoVis.evergladensis.1.1.chr8.g00328.t01#3    mNeoVis.evergladensis.2.1.chr9.g00328.t01#2    mNeoVis.vulgivaga.2.1.chr8.g00336.t01#1   mNeoVis.2.1.1.chr8_hap1.g00329.t01#5       556   125261680:   mNeoVis.vulgivaga.1.1.chr1.g02051.t01#4    mNeoVis.3.1.1.chr1.g02030.t01#6    mNeoVis.2.1.1.chr1_hap1.g02078.t01#5    mNeoVis.evergladensis.1.1.chr1.g02046.t01#3    mNeoVis.vulgivaga.1.1.chr1.g02050.t01#4    mNeoVis.vulgivaga.2.1.chr1.g02039.t01#1   mNeoVis.evergladensis.2.1.chr1.g02047.t01#2     557   125261681:   mNeoVis.evergladensis.1.1.chr1.g02047.t01#3    mNeoVis.vulgivaga.2.1.chr1.g02040.t01#1    mNeoVis.3.1.1.chr1.g02031.t01#6    mNeoVis.2.1.1.chr1_hap1.g02079.t01#5    mNeoVis.vulgivaga.1.1.chr1.g02052.t01#4   mNeoVis.evergladensis.2.1.chr1.g02048.t01#2       558   125261682:   mNeoVis.evergladensis.2.1.chr9.g00329.t01#2    mNeoVis.vulgivaga.1.1.chr8.g00323.t01#4    mNeoVis.vulgivaga.2.1.chr8.g00337.t01#1    mNeoVis.2.1.1.chr8_hap1.g00330.t01#5    mNeoVis.evergladensis.1.1.chr8.g00329.t01#3   mNeoVis.3.1.1.chr8.g00296.t01#6       559   125261683:   mNeoVis.evergladensis.2.1.chr9.g00330.t01#2    mNeoVis.vulgivaga.2.1.chr8.g01357.t01#1    mNeoVis.vulgivaga.1.1.chr8.g00324.t01#4    mNeoVis.vulgivaga.1.1.chr8.g01339.t01#4    mNeoVis.vulgivaga.2.1.chr8.g00338.t01#1    mNeoVis.3.1.1.chr8.g01301.t01#6    mNeoVis.evergladensis.1.1.chr8.g00330.t01#3    mNeoVis.3.1.1.chr8.g00297.t01#6    mNeoVis.evergladensis.2.1.chr9.g01350.t01#2      560   125261684:   mNeoVis.3.1.1.chr8.g00298.t01#6    mNeoVis.2.1.1.chr8_hap1.g00332.t01#5    mNeoVis.evergladensis.2.1.chr9.g00331.t01#2    mNeoVis.vulgivaga.1.1.chr8.g00325.t01#4    mNeoVis.evergladensis.1.1.chr8.g00331.t01#3   mNeoVis.vulgivaga.2.1.chr8.g00339.t01#1       561   125261685:   mNeoVis.vulgivaga.2.1.chr8.g00340.t01#1    mNeoVis.evergladensis.1.1.chr8.g00332.t01#3    mNeoVis.2.1.1.chr8_hap1.g00333.t01#5    mNeoVis.evergladensis.2.1.chr9.g00332.t01#2    mNeoVis.3.1.1.chr8.g00299.t01#6   mNeoVis.vulgivaga.1.1.chr8.g00326.t01#4       562   125261686:   mNeoVis.evergladensis.1.1.chr8.g00333.t01#3    mNeoVis.vulgivaga.1.1.chr8.g00327.t01#4    mNeoVis.2.1.1.chr8_hap1.g00334.t01#5    mNeoVis.evergladensis.2.1.chr9.g00333.t01#2    mNeoVis.3.1.1.chr8.g00300.t01#6   mNeoVis.vulgivaga.2.1.chr8.g00341.t01#1       563   125261687:   mNeoVis.2.1.1.chr8_hap1.g00336.t01#5    mNeoVis.evergladensis.2.1.chr9.g00335.t01#2    mNeoVis.evergladensis.1.1.chr8.g00335.t01#3    mNeoVis.3.1.1.chr8.g00302.t01#6    mNeoVis.vulgivaga.2.1.chr8.g00342.t01#1   mNeoVis.vulgivaga.1.1.chr8.g00329.t01#4       564   125261688:   mNeoVis.vulgivaga.1.1.chr8.g00330.t01#4    mNeoVis.3.1.1.chr8.g00303.t01#6    mNeoVis.evergladensis.2.1.chr9.g00336.t01#2    mNeoVis.2.1.1.chr8_hap1.g00337.t01#5    mNeoVis.vulgivaga.2.1.chr8.g00343.t01#1   mNeoVis.evergladensis.1.1.chr8.g00336.t01#3       565   125261689:   mNeoVis.vulgivaga.1.1.chr8.g00331.t01#4    mNeoVis.vulgivaga.2.1.chr8.g00344.t01#1    mNeoVis.evergladensis.1.1.chr8.g00337.t01#3    mNeoVis.2.1.1.chr8_hap1.g00338.t01#5    mNeoVis.3.1.1.chr8.g00304.t01#6   mNeoVis.evergladensis.2.1.chr9.g00337.t01#2       566   125261690:   mNeoVis.3.1.1.chr8.g00305.t01#6    mNeoVis.evergladensis.2.1.chr9.g00338.t01#2    mNeoVis.vulgivaga.2.1.chr8.g00345.t01#1    mNeoVis.evergladensis.1.1.chr8.g00338.t01#3    mNeoVis.vulgivaga.1.1.chr8.g00332.t01#4   mNeoVis.2.1.1.chr8_hap1.g00339.t01#5       567   125261691:   mNeoVis.evergladensis.2.1.chr9.g00138.t01#2    mNeoVis.3.1.1.chr8.g00111.t01#6    mNeoVis.vulgivaga.1.1.chr8.g00135.t01#4    mNeoVis.evergladensis.1.1.chr8.g00136.t01#3    mNeoVis.2.1.1.chr8_hap1.g00141.t01#5   mNeoVis.vulgivaga.2.1.chr8.g00141.t01#1       568   125261692:   mNeoVis.evergladensis.1.1.chr8.g00137.t01#3    mNeoVis.vulgivaga.1.1.chr8.g00136.t01#4    mNeoVis.evergladensis.2.1.chr9.g00139.t01#2    mNeoVis.vulgivaga.2.1.chr8.g00142.t01#1   mNeoVis.2.1.1.chr8_hap1.g00142.t01#5       569   125261693:   mNeoVis.vulgivaga.1.1.chr15.g00575.t01#4    mNeoVis.evergladensis.1.1.chr15.g00039.t01#3    mNeoVis.vulgivaga.2.1.chr15.g00584.t01#1    mNeoVis.2.1.1.chr15_hap1.g00752.t01#5    mNeoVis.evergladensis.2.1.chr15.g00573.t01#2   mNeoVis.3.1.1.chr15.g00529.t01#6       570   125261694:   mNeoVis.3.1.1.chr8.g00112.t01#6    mNeoVis.evergladensis.1.1.chr8.g00138.t01#3    mNeoVis.vulgivaga.1.1.chr8.g00137.t01#4    mNeoVis.evergladensis.2.1.chr9.g00140.t01#2    mNeoVis.vulgivaga.2.1.chr8.g00143.t01#1   mNeoVis.2.1.1.chr8_hap1.g00143.t01#5       571   125261695:   mNeoVis.evergladensis.2.1.chr9.g00141.t01#2    mNeoVis.vulgivaga.2.1.chr8.g00144.t01#1    mNeoVis.vulgivaga.1.1.chr8.g00138.t01#4    mNeoVis.2.1.1.chr8_hap1.g00144.t01#5   mNeoVis.evergladensis.1.1.chr8.g00139.t01#3       572   125261696:   mNeoVis.evergladensis.1.1.chr1.g02049.t01#3    mNeoVis.2.1.1.chr1_hap1.g02080.t01#5    mNeoVis.evergladensis.2.1.chr1.g02049.t01#2    mNeoVis.vulgivaga.2.1.chr1.g02041.t01#1    mNeoVis.vulgivaga.1.1.chr1.g02053.t01#4   mNeoVis.3.1.1.chr1.g02032.t01#6       573   125261697:   mNeoVis.vulgivaga.1.1.chr1.g02054.t01#4    mNeoVis.2.1.1.chr1_hap1.g02081.t01#5    mNeoVis.vulgivaga.2.1.chr1.g02042.t01#1    mNeoVis.3.1.1.chr1.g02033.t01#6    mNeoVis.evergladensis.1.1.chr1.g02050.t01#3   mNeoVis.evergladensis.2.1.chr1.g02050.t01#2       574   125261698:   mNeoVis.evergladensis.2.1.chr1.g02051.t01#2    mNeoVis.evergladensis.1.1.chr1.g02051.t01#3    mNeoVis.3.1.1.chr1.g02034.t01#6    mNeoVis.vulgivaga.1.1.chr1.g02055.t01#4    mNeoVis.vulgivaga.2.1.chr1.g02043.t01#1   mNeoVis.2.1.1.chr1_hap1.g02082.t01#5       575   125261699:   mNeoVis.3.1.1.chr1.g02035.t01#6    mNeoVis.evergladensis.2.1.chr1.g02052.t01#2    mNeoVis.evergladensis.1.1.chr1.g02052.t01#3    mNeoVis.vulgivaga.2.1.chr1.g02044.t01#1    mNeoVis.2.1.1.chr1_hap1.g02083.t01#5   mNeoVis.vulgivaga.1.1.chr1.g02056.t01#4       576   125261700:   mNeoVis.evergladensis.2.1.chr1.g02053.t01#2    mNeoVis.3.1.1.chr1.g02036.t01#6    mNeoVis.evergladensis.1.1.chr1.g02053.t01#3    mNeoVis.2.1.1.chr1_hap1.g02084.t01#5    mNeoVis.vulgivaga.2.1.chr1.g02045.t01#1   mNeoVis.vulgivaga.1.1.chr1.g02057.t01#4       577   125261701:   mNeoVis.evergladensis.2.1.chr1.g02054.t01#2    mNeoVis.2.1.1.chr1_hap1.g02085.t01#5    mNeoVis.3.1.1.chr1.g02037.t01#6    mNeoVis.vulgivaga.2.1.chr1.g02046.t01#1   mNeoVis.evergladensis.1.1.chr1.g02054.t01#3       578   125261702:   mNeoVis.vulgivaga.1.1.chr7.g00521.t01#4    mNeoVis.evergladensis.2.1.chr7.g00514.t01#2    mNeoVis.vulgivaga.2.1.chr7.g00517.t01#1    mNeoVis.3.1.1.chr7.g00498.t01#6    mNeoVis.2.1.1.chr7_hap1.g00513.t01#5   mNeoVis.evergladensis.1.1.chr7.g00523.t01#3       579   125261703:   mNeoVis.evergladensis.1.1.chr1.g02055.t01#3    mNeoVis.vulgivaga.2.1.chr1.g02047.t01#1    mNeoVis.2.1.1.chr1_hap1.g02086.t01#5    mNeoVis.3.1.1.chr1.g02038.t01#6    mNeoVis.vulgivaga.1.1.chr1.g02059.t01#4   mNeoVis.evergladensis.2.1.chr1.g02055.t01#2       580   125261704:   mNeoVis.3.1.1.chr1.g02039.t01#6    mNeoVis.vulgivaga.1.1.chr1.g02060.t01#4    mNeoVis.vulgivaga.2.1.chr1.g02048.t01#1    mNeoVis.evergladensis.1.1.chr1.g02056.t01#3    mNeoVis.evergladensis.2.1.chr1.g02056.t01#2   mNeoVis.2.1.1.chr1_hap1.g02087.t01#5       581   125261705:   mNeoVis.vulgivaga.2.1.chr8.g00069.t01#1    mNeoVis.3.1.1.chr8.g00048.t01#6    mNeoVis.evergladensis.1.1.chr8.g00068.t01#3    mNeoVis.2.1.1.chr8_hap1.g00072.t01#5    mNeoVis.evergladensis.2.1.chr9.g00069.t01#2   mNeoVis.vulgivaga.1.1.chr8.g00068.t01#4       582   125261706:   mNeoVis.vulgivaga.2.1.chr1.g02049.t01#1    mNeoVis.evergladensis.2.1.chr1.g02058.t01#2    mNeoVis.vulgivaga.1.1.chr1.g02061.t01#4   mNeoVis.2.1.1.chr1_hap1.g02089.t01#5         583   125261707:   mNeoVis.vulgivaga.2.1.chr1.g02050.t01#1    mNeoVis.2.1.1.chr1_hap1.g02090.t01#5    mNeoVis.evergladensis.2.1.chr1.g02059.t01#2    mNeoVis.3.1.1.chr1.g02041.t01#6    mNeoVis.evergladensis.1.1.chr1.g02059.t01#3   mNeoVis.vulgivaga.1.1.chr1.g02062.t01#4       584   125261708:   mNeoVis.vulgivaga.1.1.chr8.g00067.t01#4    mNeoVis.evergladensis.1.1.chr8.g00067.t01#3    mNeoVis.vulgivaga.2.1.chr8.g00070.t01#1    mNeoVis.evergladensis.2.1.chr9.g00068.t01#2   mNeoVis.2.1.1.chr8_hap1.g00071.t01#5       585   125261709:   mNeoVis.vulgivaga.1.1.chr8.g00069.t01#4    mNeoVis.evergladensis.2.1.chr9.g00070.t01#2    mNeoVis.evergladensis.1.1.chr8.g00069.t01#3    mNeoVis.vulgivaga.2.1.chr8.g00071.t01#1    mNeoVis.3.1.1.chr8.g00049.t01#6   mNeoVis.2.1.1.chr8_hap1.g00073.t01#5       586   125261710:  mNeoVis.vulgivaga.2.1.chr1.g02051.t01#1               587   125261711:   mNeoVis.vulgivaga.2.1.chr1.g02052.t01#1    mNeoVis.evergladensis.1.1.chr1.g02060.t01#3    mNeoVis.2.1.1.chr1_hap1.g02091.t01#5    mNeoVis.3.1.1.chr1.g02044.t01#6    mNeoVis.evergladensis.2.1.chr1.g02060.t01#2   mNeoVis.vulgivaga.1.1.chr1.g02063.t01#4       588   125261712:   mNeoVis.evergladensis.2.1.chr7.g00515.t01#2    mNeoVis.vulgivaga.2.1.chr7.g00518.t01#1    mNeoVis.3.1.1.chr7.g00499.t01#6    mNeoVis.evergladensis.1.1.chr7.g00524.t01#3    mNeoVis.2.1.1.chr7_hap1.g00514.t01#5   mNeoVis.vulgivaga.1.1.chr7.g00522.t01#4       589   125261713:   mNeoVis.evergladensis.1.1.chr7.g02369.t01#3    mNeoVis.vulgivaga.2.1.chr7.g02391.t01#1    mNeoVis.vulgivaga.1.1.chr7.g02375.t01#4    mNeoVis.3.1.1.chr7.g02230.t01#6    mNeoVis.2.1.1.chr7_hap1.g02483.t01#5   mNeoVis.evergladensis.2.1.chr7.g02371.t01#2       590   125261714:   mNeoVis.vulgivaga.1.1.chr5.g00603.t01#4    mNeoVis.3.1.1.chr5.g00621.t01#6    mNeoVis.evergladensis.2.1.chr5.g00626.t01#2    mNeoVis.2.1.1.chr2_hap1.g00623.t01#5    mNeoVis.vulgivaga.2.1.chr5.g00620.t01#1   mNeoVis.evergladensis.1.1.chr5.g00619.t01#3       591   125261715:   mNeoVis.3.1.1.chr9.g00258.t01#6    mNeoVis.2.1.1.chr10_hap1.g00285.t01#5    mNeoVis.vulgivaga.2.1.chr9.g00269.t01#1   mNeoVis.evergladensis.2.1.chr8.g00270.t01#2       592   125261716:  mNeoVis.evergladensis.1.1.chr9.g00273.t01#3             593   125261717:  mNeoVis.vulgivaga.1.1.chr9.g00270.t01#4               594   125261718:   mNeoVis.vulgivaga.2.1.chr1.g02053.t01#1    mNeoVis.vulgivaga.1.1.chr1.g02064.t01#4    mNeoVis.evergladensis.1.1.chr1.g02061.t01#3   mNeoVis.3.1.1.chr1.g02045.t01#6         595   125261719:   mNeoVis.evergladensis.1.1.chr15.g00038.t01#3    mNeoVis.vulgivaga.1.1.chr15.g00576.t01#4    mNeoVis.vulgivaga.2.1.chr15.g00585.t01#1    mNeoVis.2.1.1.chr15_hap1.g00753.t01#5    mNeoVis.evergladensis.2.1.chr15.g00574.t01#2   mNeoVis.3.1.1.chr15.g00530.t01#6       596   125261720:   mNeoVis.2.1.1.chr1_hap1.g02093.t01#5    mNeoVis.vulgivaga.1.1.chr1.g02065.t01#4    mNeoVis.vulgivaga.2.1.chr1.g02054.t01#1    mNeoVis.evergladensis.1.1.chr1.g02062.t01#3    mNeoVis.evergladensis.2.1.chr1.g02062.t01#2   mNeoVis.3.1.1.chr1.g02046.t01#6       597   125261721:   mNeoVis.evergladensis.1.1.chr1.g02063.t01#3    mNeoVis.3.1.1.chr1.g02047.t01#6    mNeoVis.2.1.1.chr1_hap1.g02094.t01#5    mNeoVis.vulgivaga.1.1.chr1.g02066.t01#4    mNeoVis.evergladensis.2.1.chr1.g02063.t01#2   mNeoVis.vulgivaga.2.1.chr1.g02055.t01#1       598   125261722:   mNeoVis.2.1.1.chr8_hap1.g00074.t01#5    mNeoVis.evergladensis.1.1.chr8.g00070.t01#3    mNeoVis.vulgivaga.2.1.chr8.g00072.t01#1    mNeoVis.3.1.1.chr8.g00050.t01#6    mNeoVis.vulgivaga.1.1.chr8.g00070.t01#4   mNeoVis.evergladensis.2.1.chr9.g00071.t01#2       599   125261723:   mNeoVis.vulgivaga.1.1.chr10.g00278.t01#4    mNeoVis.3.1.1.chr9.g00221.t01#6    mNeoVis.vulgivaga.1.1.chr9.g00236.t01#4    mNeoVis.evergladensis.1.1.chr9.g00240.t01#3    mNeoVis.vulgivaga.2.1.chr9.g00237.t01#1    mNeoVis.evergladensis.2.1.chr8.g00236.t01#2    mNeoVis.2.1.1.chr10_hap1.g00250.t01#5   mNeoVis.2.1.1.chr2_hap1.g00621.t01#5     600   125261724:   mNeoVis.evergladensis.2.1.chr1.g02064.t01#2    mNeoVis.3.1.1.chr1.g02048.t01#6    mNeoVis.vulgivaga.1.1.chr1.g02067.t01#4    mNeoVis.evergladensis.1.1.chr1.g02064.t01#3   mNeoVis.vulgivaga.2.1.chr1.g02056.t01#1       601   125261725:  mNeoVis.vulgivaga.2.1.chr1.g02057.t01#1               602   125261726:   mNeoVis.2.1.1.chr15_hap1.g00754.t01#5    mNeoVis.evergladensis.2.1.chr15.g00575.t01#2    mNeoVis.evergladensis.1.1.chr15.g00037.t01#3    mNeoVis.vulgivaga.2.1.chr15.g00586.t01#1   mNeoVis.vulgivaga.1.1.chr15.g00577.t01#4       603   125261727:   mNeoVis.vulgivaga.1.1.chr1.g02069.t01#4    mNeoVis.2.1.1.chr1_hap1.g02095.t01#5    mNeoVis.evergladensis.1.1.chr1.g02066.t01#3    mNeoVis.3.1.1.chr1.g02050.t01#6   mNeoVis.vulgivaga.2.1.chr1.g02058.t01#1       604   125261728:   mNeoVis.vulgivaga.2.1.chr8.g00073.t01#1    mNeoVis.vulgivaga.1.1.chr8.g00071.t01#4    mNeoVis.evergladensis.1.1.chr8.g00071.t01#3    mNeoVis.2.1.1.chr8_hap1.g00075.t01#5    mNeoVis.evergladensis.2.1.chr9.g00072.t01#2   mNeoVis.3.1.1.chr8.g00051.t01#6       605   125261729:   mNeoVis.2.1.1.chr1_hap1.g02009.t01#5    mNeoVis.evergladensis.2.1.chr1.g01975.t01#2    mNeoVis.3.1.1.chr1.g01964.t01#6    mNeoVis.vulgivaga.2.1.chr1.g01972.t01#1    mNeoVis.evergladensis.1.1.chr1.g01975.t01#3   mNeoVis.vulgivaga.1.1.chr1.g01981.t01#4       606   125261730:   mNeoVis.3.1.1.chr1.g01965.t01#6    mNeoVis.evergladensis.2.1.chr1.g01976.t01#2    mNeoVis.vulgivaga.2.1.chr1.g01973.t01#1    mNeoVis.evergladensis.1.1.chr1.g01976.t01#3   mNeoVis.2.1.1.chr1_hap1.g02010.t01#5       607   125261731:   mNeoVis.vulgivaga.1.1.chr8.g00072.t01#4    mNeoVis.vulgivaga.2.1.chr8.g00074.t01#1    mNeoVis.evergladensis.2.1.chr9.g00073.t01#2    mNeoVis.3.1.1.chr8.g00052.t01#6    mNeoVis.evergladensis.1.1.chr8.g00072.t01#3   mNeoVis.2.1.1.chr8_hap1.g00076.t01#5       608   125261732:   mNeoVis.vulgivaga.1.1.chr1.g01982.t01#4    mNeoVis.2.1.1.chr1_hap1.g02011.t01#5    mNeoVis.evergladensis.1.1.chr1.g01977.t01#3    mNeoVis.vulgivaga.2.1.chr1.g01974.t01#1    mNeoVis.evergladensis.2.1.chr1.g01977.t01#2   mNeoVis.3.1.1.chr1.g01966.t01#6       609   125261733:   mNeoVis.evergladensis.2.1.chr9.g00339.t01#2    mNeoVis.evergladensis.1.1.chr8.g00339.t01#3    mNeoVis.3.1.1.chr8.g00306.t01#6    mNeoVis.vulgivaga.2.1.chr8.g00346.t01#1    mNeoVis.2.1.1.chr8_hap1.g00340.t01#5   mNeoVis.vulgivaga.1.1.chr8.g00333.t01#4       610   125261734:   mNeoVis.vulgivaga.1.1.chr8.g00334.t01#4    mNeoVis.2.1.1.chr8_hap1.g00341.t01#5    mNeoVis.evergladensis.1.1.chr8.g00340.t01#3    mNeoVis.3.1.1.chr8.g00307.t01#6    mNeoVis.evergladensis.2.1.chr9.g00340.t01#2   mNeoVis.vulgivaga.2.1.chr8.g00347.t01#1       611   125261735:   mNeoVis.3.1.1.chr1.g01967.t01#6    mNeoVis.evergladensis.1.1.chr1.g01978.t01#3    mNeoVis.evergladensis.2.1.chr1.g01978.t01#2    mNeoVis.vulgivaga.1.1.chr1.g01983.t01#4    mNeoVis.2.1.1.chr1_hap1.g02012.t01#5   mNeoVis.vulgivaga.2.1.chr1.g01975.t01#1       612   125261736:   mNeoVis.3.1.1.chr8.g00308.t01#6    mNeoVis.evergladensis.2.1.chr9.g00341.t01#2    mNeoVis.2.1.1.chr8_hap1.g00342.t01#5    mNeoVis.vulgivaga.2.1.chr8.g00348.t01#1    mNeoVis.evergladensis.1.1.chr8.g00341.t01#3   mNeoVis.vulgivaga.1.1.chr8.g00335.t01#4       613   125261737:   mNeoVis.2.1.1.chr8_hap1.g00343.t01#5    mNeoVis.vulgivaga.2.1.chr8.g00349.t01#1    mNeoVis.evergladensis.1.1.chr8.g00342.t01#3    mNeoVis.vulgivaga.1.1.chr8.g00336.t01#4    mNeoVis.3.1.1.chr8.g00309.t01#6   mNeoVis.evergladensis.2.1.chr9.g00342.t01#2       614   125261738:   mNeoVis.2.1.1.chr1_hap1.g02013.t01#5    mNeoVis.evergladensis.2.1.chr1.g01979.t01#2    mNeoVis.evergladensis.1.1.chr1.g01979.t01#3    mNeoVis.3.1.1.chr1.g01968.t01#6    mNeoVis.vulgivaga.2.1.chr1.g01976.t01#1   mNeoVis.vulgivaga.1.1.chr1.g01984.t01#4       615   125261739:   mNeoVis.evergladensis.1.1.chr1.g01980.t01#3    mNeoVis.vulgivaga.1.1.chr1.g01985.t01#4    mNeoVis.3.1.1.chr1.g01969.t01#6    mNeoVis.2.1.1.chr1_hap1.g02015.t01#5    mNeoVis.evergladensis.2.1.chr1.g01981.t01#2   mNeoVis.vulgivaga.2.1.chr1.g01977.t01#1       616   125261740:   mNeoVis.vulgivaga.2.1.chr8.g00350.t01#1    mNeoVis.3.1.1.chr8.g00310.t01#6    mNeoVis.vulgivaga.1.1.chr8.g00337.t01#4    mNeoVis.2.1.1.chr8_hap1.g00344.t01#5    mNeoVis.evergladensis.2.1.chr9.g00343.t01#2   mNeoVis.evergladensis.1.1.chr8.g00343.t01#3       617   125261741:   mNeoVis.evergladensis.2.1.chr9.g00074.t01#2    mNeoVis.vulgivaga.2.1.chr8.g00075.t01#1    mNeoVis.evergladensis.1.1.chr8.g00073.t01#3    mNeoVis.vulgivaga.1.1.chr8.g00073.t01#4    mNeoVis.3.1.1.chr8.g00053.t01#6   mNeoVis.2.1.1.chr8_hap1.g00077.t01#5       618   125261742:   mNeoVis.3.1.1.chr15.g00531.t01#6    mNeoVis.2.1.1.chr15_hap1.g00755.t01#5    mNeoVis.evergladensis.2.1.chr15.g00576.t01#2    mNeoVis.evergladensis.1.1.chr15.g00036.t01#3    mNeoVis.vulgivaga.2.1.chr15.g00587.t01#1   mNeoVis.vulgivaga.1.1.chr15.g00578.t01#4       619   125261743:   mNeoVis.vulgivaga.1.1.chr7.g00523.t01#4    mNeoVis.vulgivaga.2.1.chr7.g00519.t01#1    mNeoVis.evergladensis.2.1.chr7.g00516.t01#2    mNeoVis.2.1.1.chr7_hap1.g00515.t01#5   mNeoVis.evergladensis.1.1.chr7.g00525.t01#3       620   125261744:   mNeoVis.evergladensis.2.1.chr1.g01982.t01#2    mNeoVis.evergladensis.1.1.chr1.g01981.t01#3    mNeoVis.vulgivaga.2.1.chr1.g01978.t01#1    mNeoVis.vulgivaga.1.1.chr1.g01986.t01#4    mNeoVis.2.1.1.chr1_hap1.g02016.t01#5   mNeoVis.3.1.1.chr1.g01970.t01#6       621   125261745:   mNeoVis.evergladensis.2.1.chr9.g00344.t01#2    mNeoVis.vulgivaga.1.1.chr8.g00338.t01#4    mNeoVis.2.1.1.chr8_hap1.g00345.t01#5    mNeoVis.vulgivaga.2.1.chr8.g00351.t01#1   mNeoVis.evergladensis.1.1.chr8.g00345.t01#3       622   125261746:   mNeoVis.vulgivaga.1.1.chr1.g01987.t01#4    mNeoVis.evergladensis.1.1.chr1.g01982.t01#3    mNeoVis.evergladensis.2.1.chr1.g01983.t01#2    mNeoVis.2.1.1.chr1_hap1.g02017.t01#5    mNeoVis.vulgivaga.2.1.chr1.g01979.t01#1   mNeoVis.3.1.1.chr1.g01971.t01#6       623   125261747:   mNeoVis.vulgivaga.1.1.chr8.g00339.t01#4    mNeoVis.evergladensis.1.1.chr8.g00346.t01#3    mNeoVis.2.1.1.chr8_hap1.g00347.t01#5    mNeoVis.evergladensis.2.1.chr9.g00345.t01#2   mNeoVis.vulgivaga.2.1.chr8.g00352.t01#1       624   125261748:   mNeoVis.evergladensis.2.1.chr1.g01984.t01#2    mNeoVis.vulgivaga.1.1.chr1.g01988.t01#4    mNeoVis.2.1.1.chr1_hap1.g02018.t01#5    mNeoVis.3.1.1.chr1.g01973.t01#6    mNeoVis.evergladensis.1.1.chr1.g01983.t01#3   mNeoVis.vulgivaga.2.1.chr1.g01980.t01#1       625   125261749:   mNeoVis.3.1.1.chr8.g00313.t01#6    mNeoVis.2.1.1.chr8_hap1.g00348.t01#5    mNeoVis.evergladensis.1.1.chr8.g00347.t01#3    mNeoVis.vulgivaga.2.1.chr8.g00353.t01#1    mNeoVis.vulgivaga.1.1.chr8.g00340.t01#4   mNeoVis.evergladensis.2.1.chr9.g00346.t01#2       626   125261750:   mNeoVis.2.1.1.chr1_hap1.g02019.t01#5    mNeoVis.evergladensis.1.1.chr1.g01984.t01#3    mNeoVis.vulgivaga.1.1.chr1.g01989.t01#4    mNeoVis.3.1.1.chr1.g01974.t01#6    mNeoVis.evergladensis.2.1.chr1.g01985.t01#2   mNeoVis.vulgivaga.2.1.chr1.g01981.t01#1       627   125261751:   mNeoVis.vulgivaga.1.1.chr8.g00341.t01#4    mNeoVis.evergladensis.2.1.chr9.g00347.t01#2    mNeoVis.3.1.1.chr8.g00314.t01#6    mNeoVis.2.1.1.chr8_hap1.g00349.t01#5    mNeoVis.vulgivaga.2.1.chr8.g00354.t01#1   mNeoVis.evergladensis.1.1.chr8.g00348.t01#3       628   125261752:   mNeoVis.3.1.1.chr1.g01975.t01#6    mNeoVis.evergladensis.1.1.chr1.g01985.t01#3    mNeoVis.vulgivaga.2.1.chr1.g01982.t01#1    mNeoVis.evergladensis.2.1.chr1.g01986.t01#2    mNeoVis.2.1.1.chr1_hap1.g02020.t01#5   mNeoVis.vulgivaga.1.1.chr1.g01990.t01#4       629   125261753:   mNeoVis.3.1.1.chr1.g01976.t01#6    mNeoVis.vulgivaga.1.1.chr1.g01991.t01#4    mNeoVis.vulgivaga.2.1.chr1.g01983.t01#1    mNeoVis.evergladensis.1.1.chr1.g01987.t01#3    mNeoVis.evergladensis.2.1.chr1.g01988.t01#2   mNeoVis.2.1.1.chr1_hap1.g02022.t01#5       630   125261754:   mNeoVis.vulgivaga.1.1.chr8.g00342.t01#4    mNeoVis.3.1.1.chr8.g00317.t01#6    mNeoVis.evergladensis.2.1.chr9.g00348.t01#2    mNeoVis.vulgivaga.2.1.chr8.g00355.t01#1    mNeoVis.2.1.1.chr8_hap1.g00350.t01#5   mNeoVis.evergladensis.1.1.chr8.g00349.t01#3       631   125261755:   mNeoVis.2.1.1.chr8_hap1.g00351.t01#5    mNeoVis.3.1.1.chr8.g00318.t01#6    mNeoVis.vulgivaga.1.1.chr8.g00343.t01#4    mNeoVis.evergladensis.2.1.chr9.g00349.t01#2    mNeoVis.evergladensis.1.1.chr8.g00350.t01#3   mNeoVis.vulgivaga.2.1.chr8.g00356.t01#1       632   125261756:   mNeoVis.vulgivaga.1.1.chr15.g00579.t01#4    mNeoVis.2.1.1.chr15_hap1.g00756.t01#5    mNeoVis.3.1.1.chr15.g00532.t01#6    mNeoVis.evergladensis.2.1.chr15.g00577.t01#2    mNeoVis.evergladensis.1.1.chr15.g00035.t01#3   mNeoVis.vulgivaga.2.1.chr15.g00588.t01#1       633   125261757:   mNeoVis.2.1.1.chr1_hap1.g02024.t01#5    mNeoVis.3.1.1.chr1.g01978.t01#6    mNeoVis.evergladensis.1.1.chr1.g01989.t01#3    mNeoVis.evergladensis.2.1.chr1.g01990.t01#2    mNeoVis.vulgivaga.2.1.chr1.g01984.t01#1   mNeoVis.vulgivaga.1.1.chr1.g01993.t01#4       634   125261758:   mNeoVis.vulgivaga.2.1.chr8.g00276.t01#1    mNeoVis.3.1.1.chr8.g00237.t01#6   mNeoVis.evergladensis.2.1.chr9.g00270.t01#2         635   125261759:   mNeoVis.2.1.1.chr1_hap1.g02025.t01#5    mNeoVis.3.1.1.chr1.g01979.t01#6    mNeoVis.evergladensis.1.1.chr1.g01990.t01#3    mNeoVis.vulgivaga.2.1.chr1.g01985.t01#1    mNeoVis.evergladensis.2.1.chr1.g01991.t01#2   mNeoVis.vulgivaga.1.1.chr1.g01994.t01#4       636   125261760:   mNeoVis.vulgivaga.1.1.chr1.g01995.t01#4    mNeoVis.2.1.1.chr1_hap1.g02026.t01#5    mNeoVis.3.1.1.chr1.g01980.t01#6    mNeoVis.evergladensis.2.1.chr1.g01992.t01#2    mNeoVis.vulgivaga.2.1.chr1.g01986.t01#1   mNeoVis.evergladensis.1.1.chr1.g01991.t01#3       637   125261761:   mNeoVis.evergladensis.2.1.chr1.g01993.t01#2    mNeoVis.2.1.1.chr1_hap1.g02027.t01#5    mNeoVis.vulgivaga.2.1.chr1.g01987.t01#1    mNeoVis.3.1.1.chr1.g01981.t01#6    mNeoVis.vulgivaga.1.1.chr1.g01996.t01#4   mNeoVis.evergladensis.1.1.chr1.g01992.t01#3       638   125261762:   mNeoVis.vulgivaga.2.1.chr1.g01988.t01#1    mNeoVis.evergladensis.2.1.chr1.g01995.t01#2    mNeoVis.vulgivaga.1.1.chr1.g01997.t01#4    mNeoVis.3.1.1.chr1.g01982.t01#6   mNeoVis.2.1.1.chr1_hap1.g02028.t01#5       639   125261763:   mNeoVis.evergladensis.2.1.chr1.g01994.t01#2   mNeoVis.evergladensis.1.1.chr1.g01993.t01#3           640   125261764:  mNeoVis.vulgivaga.2.1.chr8.g00277.t01#1               641   125261765:   mNeoVis.vulgivaga.1.1.chr1.g01998.t01#4    mNeoVis.evergladensis.2.1.chr1.g01996.t01#2    mNeoVis.vulgivaga.2.1.chr1.g01989.t01#1    mNeoVis.3.1.1.chr1.g01983.t01#6    mNeoVis.evergladensis.1.1.chr1.g01994.t01#3   mNeoVis.2.1.1.chr1_hap1.g02029.t01#5       642   125261766:   mNeoVis.evergladensis.2.1.chr1.g01998.t01#2    mNeoVis.3.1.1.chr1.g01985.t01#6    mNeoVis.evergladensis.1.1.chr1.g01996.t01#3    mNeoVis.vulgivaga.1.1.chr1.g02000.t01#4    mNeoVis.vulgivaga.2.1.chr1.g01991.t01#1   mNeoVis.2.1.1.chr1_hap1.g02031.t01#5       643   125261767:   mNeoVis.2.1.1.chr8_hap1.g00078.t01#5    mNeoVis.evergladensis.2.1.chr9.g00075.t01#2    mNeoVis.vulgivaga.2.1.chr8.g00076.t01#1    mNeoVis.vulgivaga.1.1.chr8.g00074.t01#4   mNeoVis.evergladensis.1.1.chr8.g00074.t01#3       644   125261768:   mNeoVis.vulgivaga.2.1.chr8.g00077.t01#1   mNeoVis.3.1.1.chr8.g00054.t01#6             645   125261769:   mNeoVis.3.1.1.chr8.g01115.t01#6    mNeoVis.evergladensis.1.1.chr8.g01169.t01#3    mNeoVis.vulgivaga.2.1.chr7.g01716.t01#1    mNeoVis.vulgivaga.1.1.chr7.g01710.t01#4    mNeoVis.2.1.1.chr7_hap1.g01736.t01#5    mNeoVis.evergladensis.1.1.chr7.g01699.t01#3    mNeoVis.evergladensis.2.1.chr9.g01175.t01#2    mNeoVis.2.1.1.chr8_hap1.g01217.t01#5    mNeoVis.3.1.1.chr7.g01614.t01#6      646   125261770:   mNeoVis.vulgivaga.2.1.chr1.g01992.t01#1    mNeoVis.evergladensis.2.1.chr1.g01999.t01#2    mNeoVis.evergladensis.1.1.chr1.g01997.t01#3    mNeoVis.2.1.1.chr1_hap1.g02032.t01#5   mNeoVis.vulgivaga.1.1.chr1.g02001.t01#4       647   125261771:   mNeoVis.3.1.1.chr1.g01986.t01#6   mNeoVis.vulgivaga.1.1.chr7.g01709.t01#4             648   125261772:   mNeoVis.vulgivaga.2.1.chr8.g00078.t01#1    mNeoVis.evergladensis.2.1.chr9.g00076.t01#2    mNeoVis.2.1.1.chr8_hap1.g00079.t01#5    mNeoVis.evergladensis.1.1.chr8.g00075.t01#3   mNeoVis.vulgivaga.1.1.chr8.g00076.t01#4       649   125261773:   mNeoVis.vulgivaga.2.1.chr6.g00690.t01#1    mNeoVis.2.1.1.chr12_hap1.g00222.t01#5    mNeoVis.vulgivaga.2.1.chr12.g00197.t01#1    mNeoVis.vulgivaga.1.1.chr12.g00201.t01#4    mNeoVis.evergladensis.2.1.chr6.g00695.t01#2    mNeoVis.3.1.1.chr6.g00696.t01#6    mNeoVis.evergladensis.1.1.chr6.g00695.t01#3    mNeoVis.2.1.1.chr4_hap1.g00691.t01#5    mNeoVis.vulgivaga.1.1.chr6.g00691.t01#4      650   125261774:   mNeoVis.3.1.1.chr3.g00154.t01#6    mNeoVis.vulgivaga.1.1.chr3.g00157.t01#4    mNeoVis.evergladensis.2.1.chr1.g02000.t01#2    mNeoVis.evergladensis.1.1.chr1.g01998.t01#3    mNeoVis.vulgivaga.2.1.chr1.g01993.t01#1    mNeoVis.2.1.1.chr1_hap1.g02033.t01#5    mNeoVis.vulgivaga.1.1.chr11.g00125.t01#4    mNeoVis.evergladensis.2.1.chr11.g00128.t01#2    mNeoVis.2.1.1.chr11_hap1.g00131.t01#5      651   125261775:   mNeoVis.evergladensis.2.1.chr11.g00127.t01#2    mNeoVis.vulgivaga.2.1.chr11.g00130.t01#1    mNeoVis.3.1.1.chr11.g00101.t01#6    mNeoVis.2.1.1.chr11_hap1.g00130.t01#5   mNeoVis.vulgivaga.1.1.chr11.g00124.t01#4       652   125261776:   mNeoVis.3.1.1.chr11.g00102.t01#6   mNeoVis.vulgivaga.2.1.chr11.g00131.t01#1           653   125261777:  mNeoVis.evergladensis.1.1.chr11.g00129.t01#3             654   125261778:   mNeoVis.evergladensis.1.1.chr1.g01999.t01#3    mNeoVis.vulgivaga.2.1.chr1.g01994.t01#1    mNeoVis.3.1.1.chr1.g01988.t01#6    mNeoVis.2.1.1.chr1_hap1.g02034.t01#5    mNeoVis.evergladensis.2.1.chr1.g02001.t01#2   mNeoVis.vulgivaga.1.1.chr1.g02003.t01#4       655   125261779:   mNeoVis.vulgivaga.1.1.chr1.g02004.t01#4    mNeoVis.2.1.1.chr1_hap1.g02035.t01#5    mNeoVis.evergladensis.2.1.chr1.g02002.t01#2    mNeoVis.vulgivaga.2.1.chr1.g01995.t01#1    mNeoVis.3.1.1.chr1.g01989.t01#6   mNeoVis.evergladensis.1.1.chr1.g02000.t01#3       656   125261780:   mNeoVis.2.1.1.chr8_hap1.g00080.t01#5    mNeoVis.evergladensis.1.1.chr8.g00076.t01#3    mNeoVis.vulgivaga.1.1.chr8.g00077.t01#4    mNeoVis.evergladensis.2.1.chr9.g00077.t01#2   mNeoVis.vulgivaga.2.1.chr8.g00079.t01#1       657   125261781:   mNeoVis.vulgivaga.1.1.chr1.g02005.t01#4    mNeoVis.2.1.1.chr1_hap1.g02036.t01#5    mNeoVis.evergladensis.2.1.chr1.g02003.t01#2    mNeoVis.vulgivaga.2.1.chr1.g01996.t01#1    mNeoVis.3.1.1.chr1.g01990.t01#6   mNeoVis.evergladensis.1.1.chr1.g02001.t01#3       658   125261782:  mNeoVis.vulgivaga.2.1.chr8.g00080.t01#1               659   125261783:   mNeoVis.evergladensis.1.1.chr8.g00077.t01#3    mNeoVis.vulgivaga.1.1.chr8.g00078.t01#4    mNeoVis.2.1.1.chr8_hap1.g00081.t01#5    mNeoVis.vulgivaga.2.1.chr8.g00081.t01#1   mNeoVis.evergladensis.2.1.chr9.g00078.t01#2       660   125261784:   mNeoVis.3.1.1.chr1.g01991.t01#6    mNeoVis.2.1.1.chr1_hap1.g02037.t01#5    mNeoVis.vulgivaga.2.1.chr1.g01997.t01#1    mNeoVis.evergladensis.2.1.chr1.g02004.t01#2    mNeoVis.vulgivaga.1.1.chr1.g02006.t01#4   mNeoVis.evergladensis.1.1.chr1.g02002.t01#3       661   125261785:   mNeoVis.vulgivaga.2.1.chr8.g00082.t01#1    mNeoVis.vulgivaga.1.1.chr8.g00079.t01#4    mNeoVis.evergladensis.2.1.chr9.g00079.t01#2    mNeoVis.2.1.1.chr8_hap1.g00082.t01#5    mNeoVis.3.1.1.chr8.g00057.t01#6   mNeoVis.evergladensis.1.1.chr8.g00078.t01#3       662   125261786:   mNeoVis.vulgivaga.1.1.chr15.g00184.t01#4    mNeoVis.evergladensis.1.1.chr15.g00426.t01#3    mNeoVis.vulgivaga.2.1.chr15.g00188.t01#1    mNeoVis.2.1.1.chr15_hap1.g00208.t01#5    mNeoVis.3.1.1.chr15.g00181.t01#6   mNeoVis.evergladensis.2.1.chr15.g00187.t01#2       663   125261787:   mNeoVis.vulgivaga.2.1.chr1.g01998.t01#1    mNeoVis.3.1.1.chr1.g01992.t01#6    mNeoVis.2.1.1.chr1_hap1.g02038.t01#5    mNeoVis.evergladensis.1.1.chr1.g02003.t01#3    mNeoVis.vulgivaga.1.1.chr1.g02007.t01#4   mNeoVis.evergladensis.2.1.chr1.g02005.t01#2       664   125261788:   mNeoVis.vulgivaga.2.1.chr8.g00083.t01#1    mNeoVis.evergladensis.2.1.chr9.g00080.t01#2    mNeoVis.vulgivaga.1.1.chr8.g00080.t01#4    mNeoVis.evergladensis.1.1.chr8.g00079.t01#3   mNeoVis.2.1.1.chr8_hap1.g00083.t01#5       665   125261789:   mNeoVis.evergladensis.1.1.chr1.g02004.t01#3    mNeoVis.vulgivaga.1.1.chr1.g02008.t01#4    mNeoVis.vulgivaga.2.1.chr1.g01999.t01#1    mNeoVis.3.1.1.chr1.g01993.t01#6    mNeoVis.evergladensis.2.1.chr1.g02006.t01#2   mNeoVis.2.1.1.chr1_hap1.g02039.t01#5       666   125261790:   mNeoVis.evergladensis.2.1.chr1.g00584.t01#2    mNeoVis.2.1.1.chr1_hap1.g00579.t01#5    mNeoVis.evergladensis.2.1.chr11.g00268.t01#2    mNeoVis.evergladensis.1.1.chr11.g00270.t01#3    mNeoVis.2.1.1.chr1_hap1.g00345.t01#5    mNeoVis.evergladensis.2.1.chr13.g00721.t01#2    mNeoVis.evergladensis.2.1.chr2.g01081.t01#2    mNeoVis.evergladensis.1.1.chr1.g00588.t01#3    mNeoVis.2.1.1.chr11_hap1.g00271.t01#5      667   125261791:   mNeoVis.evergladensis.1.1.chr11.g00767.t01#3    mNeoVis.3.1.1.chr11.g00741.t01#6    mNeoVis.vulgivaga.2.1.chr11.g00779.t01#1    mNeoVis.vulgivaga.1.1.chr11.g00759.t01#4    mNeoVis.vulgivaga.1.1.chr1.g00343.t01#4    mNeoVis.vulgivaga.1.1.chr13.g00714.t01#4    mNeoVis.3.1.1.chr5.g01498.t01#6    mNeoVis.vulgivaga.1.1.chr11.g00265.t01#4    mNeoVis.2.1.1.chr11_hap1.g00777.t01#5      668   125261792:   mNeoVis.evergladensis.1.1.chr11.g00720.t01#3    mNeoVis.evergladensis.2.1.chr11.g00713.t01#2    mNeoVis.3.1.1.chr11.g00693.t01#6    mNeoVis.vulgivaga.2.1.chr11.g00730.t01#1    mNeoVis.2.1.1.chr11_hap1.g00730.t01#5    mNeoVis.3.1.1.chr11.g00692.t01#6   mNeoVis.vulgivaga.1.1.chr11.g00713.t01#4     669   125261793:   mNeoVis.3.1.1.chr11.g01068.t01#6    mNeoVis.evergladensis.1.1.chr11.g01113.t01#3    mNeoVis.evergladensis.2.1.chr11.g01096.t01#2    mNeoVis.vulgivaga.1.1.chr11.g01095.t01#4    mNeoVis.2.1.1.chr11_hap1.g01127.t01#5   mNeoVis.vulgivaga.2.1.chr11.g01124.t01#1       670   125261794:   mNeoVis.evergladensis.2.1.chr3.g00201.t01#2    mNeoVis.2.1.1.chr3_hap1.g00190.t01#5    mNeoVis.3.1.1.chr3.g00193.t01#6    mNeoVis.evergladensis.1.1.chr3.g00197.t01#3    mNeoVis.vulgivaga.1.1.chr3.g00195.t01#4   mNeoVis.vulgivaga.2.1.chr3.g00193.t01#1       671   125261795:   mNeoVis.vulgivaga.2.1.chr8.g00084.t01#1    mNeoVis.vulgivaga.2.1.chr11.g00269.t01#1   mNeoVis.vulgivaga.2.1.chr3.g01004.t01#1           672   125261796:   mNeoVis.evergladensis.1.1.chr8.g00080.t01#3   mNeoVis.3.1.1.chr8.g00059.t01#6             673   125261797:   mNeoVis.evergladensis.1.1.chr8.g00081.t01#3    mNeoVis.vulgivaga.1.1.chr8.g00082.t01#4    mNeoVis.2.1.1.chr8_hap1.g00085.t01#5    mNeoVis.evergladensis.2.1.chr9.g00082.t01#2   mNeoVis.vulgivaga.2.1.chr8.g00085.t01#1       674   125261798:   mNeoVis.2.1.1.chr8_hap1.g00086.t01#5    mNeoVis.evergladensis.2.1.chr9.g00083.t01#2    mNeoVis.3.1.1.chr8.g00061.t01#6    mNeoVis.vulgivaga.2.1.chr8.g00086.t01#1   mNeoVis.evergladensis.1.1.chr8.g00082.t01#3       675   125261799:   mNeoVis.vulgivaga.2.1.chr8.g00087.t01#1    mNeoVis.vulgivaga.1.1.chr8.g00083.t01#4    mNeoVis.evergladensis.2.1.chr9.g00084.t01#2    mNeoVis.3.1.1.chr8.g00062.t01#6    mNeoVis.2.1.1.chr8_hap1.g00087.t01#5   mNeoVis.evergladensis.1.1.chr8.g00083.t01#3       676   125261800:   mNeoVis.vulgivaga.2.1.chr1.g02000.t01#1    mNeoVis.evergladensis.2.1.chr1.g02007.t01#2    mNeoVis.evergladensis.1.1.chr1.g02006.t01#3    mNeoVis.3.1.1.chr1.g01994.t01#6    mNeoVis.vulgivaga.1.1.chr1.g02009.t01#4   mNeoVis.2.1.1.chr1_hap1.g02040.t01#5       677   125261801:   mNeoVis.vulgivaga.2.1.chr8.g00088.t01#1    mNeoVis.2.1.1.chr8_hap1.g00088.t01#5    mNeoVis.evergladensis.1.1.chr8.g00084.t01#3    mNeoVis.3.1.1.chr8.g00063.t01#6    mNeoVis.evergladensis.2.1.chr9.g00085.t01#2   mNeoVis.vulgivaga.1.1.chr8.g00084.t01#4       678   125261802:   mNeoVis.vulgivaga.1.1.chr1.g02010.t01#4    mNeoVis.2.1.1.chr1_hap1.g02041.t01#5    mNeoVis.evergladensis.1.1.chr1.g02007.t01#3    mNeoVis.evergladensis.2.1.chr1.g02008.t01#2    mNeoVis.vulgivaga.2.1.chr1.g02001.t01#1   mNeoVis.3.1.1.chr1.g01995.t01#6       679   125261803:   mNeoVis.2.1.1.chr7_hap1.g00516.t01#5    mNeoVis.evergladensis.2.1.chr7.g00518.t01#2    mNeoVis.evergladensis.1.1.chr7.g00526.t01#3    mNeoVis.2.1.1.chr7_hap1.g00517.t01#5    mNeoVis.evergladensis.2.1.chr7.g00517.t01#2   mNeoVis.vulgivaga.1.1.chr7.g00525.t01#4       680   125261804:   mNeoVis.vulgivaga.1.1.chr7.g00524.t01#4    mNeoVis.vulgivaga.2.1.chr7.g00520.t01#1   mNeoVis.3.1.1.chr7.g00501.t01#6           681   125261805:   mNeoVis.evergladensis.1.1.chr8.g00085.t01#3    mNeoVis.vulgivaga.2.1.chr8.g00089.t01#1    mNeoVis.3.1.1.chr8.g00064.t01#6    mNeoVis.2.1.1.chr8_hap1.g00089.t01#5    mNeoVis.vulgivaga.1.1.chr8.g00085.t01#4   mNeoVis.evergladensis.2.1.chr9.g00086.t01#2       682   125261806:   mNeoVis.vulgivaga.1.1.chr1.g02012.t01#4    mNeoVis.vulgivaga.2.1.chr1.g02002.t01#1   mNeoVis.3.1.1.chr1.g01996.t01#6           683   125261807:   mNeoVis.evergladensis.2.1.chr1.g02009.t01#2    mNeoVis.evergladensis.1.1.chr1.g02009.t01#3    mNeoVis.3.1.1.chr1.g01997.t01#6    mNeoVis.vulgivaga.2.1.chr1.g02003.t01#1    mNeoVis.vulgivaga.1.1.chr1.g02013.t01#4   mNeoVis.2.1.1.chr1_hap1.g02042.t01#5       684   125261808:   mNeoVis.evergladensis.1.1.chr8.g00086.t01#3    mNeoVis.evergladensis.1.1.chr8.g00087.t01#3    mNeoVis.vulgivaga.1.1.chr8.g00086.t01#4    mNeoVis.3.1.1.chr8.g00066.t01#6    mNeoVis.2.1.1.chr8_hap1.g00090.t01#5    mNeoVis.vulgivaga.2.1.chr8.g00090.t01#1    mNeoVis.evergladensis.2.1.chr9.g00087.t01#2   mNeoVis.3.1.1.chr8.g00065.t01#6     685   125261809:   mNeoVis.3.1.1.chr1.g01998.t01#6    mNeoVis.vulgivaga.2.1.chr1.g02004.t01#1    mNeoVis.evergladensis.1.1.chr1.g02010.t01#3    mNeoVis.2.1.1.chr1_hap1.g02043.t01#5    mNeoVis.vulgivaga.1.1.chr1.g02014.t01#4   mNeoVis.evergladensis.2.1.chr1.g02010.t01#2       686   125261810:   mNeoVis.3.1.1.chr8.g00067.t01#6    mNeoVis.evergladensis.2.1.chr9.g00089.t01#2    mNeoVis.evergladensis.1.1.chr8.g00089.t01#3    mNeoVis.2.1.1.chr8_hap1.g00092.t01#5   mNeoVis.vulgivaga.2.1.chr8.g00091.t01#1       687   125261811:   mNeoVis.vulgivaga.2.1.chr8.g00092.t01#1   mNeoVis.evergladensis.1.1.chr8.g00090.t01#3           688   125261812:   mNeoVis.vulgivaga.2.1.chr1.g02005.t01#1    mNeoVis.evergladensis.2.1.chr1.g02011.t01#2    mNeoVis.2.1.1.chr1_hap1.g02044.t01#5    mNeoVis.evergladensis.1.1.chr1.g02011.t01#3   mNeoVis.3.1.1.chr1.g01999.t01#6       689   125261813:   mNeoVis.evergladensis.1.1.chr8.g00091.t01#3    mNeoVis.2.1.1.chr8_hap1.g00093.t01#5    mNeoVis.vulgivaga.1.1.chr8.g00088.t01#4    mNeoVis.vulgivaga.2.1.chr8.g00093.t01#1    mNeoVis.evergladensis.2.1.chr9.g00090.t01#2   mNeoVis.3.1.1.chr8.g00068.t01#6       690   125261814:   mNeoVis.evergladensis.1.1.chr1.g02012.t01#3    mNeoVis.3.1.1.chr1.g02000.t01#6    mNeoVis.evergladensis.2.1.chr1.g02012.t01#2    mNeoVis.vulgivaga.1.1.chr1.g02016.t01#4    mNeoVis.vulgivaga.2.1.chr1.g02006.t01#1   mNeoVis.2.1.1.chr1_hap1.g02045.t01#5       691   125261815:   mNeoVis.3.1.1.chr2.g01281.t01#6    mNeoVis.vulgivaga.1.1.chr10.g00997.t01#4    mNeoVis.3.1.1.chr7.g00502.t01#6    mNeoVis.vulgivaga.1.1.chr7.g00526.t01#4    mNeoVis.vulgivaga.2.1.chr2.g01354.t01#1    mNeoVis.2.1.1.chr9_hap1.g01028.t01#5    mNeoVis.2.1.1.chr5_hap1.g01358.t01#5    mNeoVis.evergladensis.1.1.chr10.g01018.t01#3    mNeoVis.evergladensis.1.1.chr7.g00527.t01#3      692   125261816:   mNeoVis.vulgivaga.2.1.chr8.g00094.t01#1    mNeoVis.evergladensis.1.1.chr8.g00092.t01#3    mNeoVis.3.1.1.chr8.g00069.t01#6    mNeoVis.2.1.1.chr8_hap1.g00094.t01#5    mNeoVis.vulgivaga.1.1.chr8.g00089.t01#4   mNeoVis.evergladensis.2.1.chr9.g00091.t01#2       693   125261817:   mNeoVis.evergladensis.2.1.chr7.g00935.t01#2    mNeoVis.vulgivaga.2.1.chr7.g00941.t01#1    mNeoVis.evergladensis.1.1.chr7.g00941.t01#3    mNeoVis.3.1.1.chr7.g00882.t01#6    mNeoVis.vulgivaga.1.1.chr7.g00939.t01#4   mNeoVis.2.1.1.chr7_hap1.g00942.t01#5       694   125261818:   mNeoVis.evergladensis.1.1.chr10.g01128.t01#3    mNeoVis.vulgivaga.2.1.chr10.g01125.t01#1    mNeoVis.2.1.1.chr9_hap1.g01145.t01#5    mNeoVis.vulgivaga.1.1.chr10.g01109.t01#4    mNeoVis.3.1.1.chr10.g01100.t01#6   mNeoVis.evergladensis.2.1.chr10.g01133.t01#2       695   125261819:   mNeoVis.vulgivaga.1.1.chr15.g00442.t01#4    mNeoVis.evergladensis.1.1.chr15.g00173.t01#3    mNeoVis.evergladensis.2.1.chr15.g00441.t01#2    mNeoVis.vulgivaga.2.1.chr15.g00449.t01#1   mNeoVis.2.1.1.chr15_hap1.g00619.t01#5       696   125261820:   mNeoVis.evergladensis.1.1.chr7.g00528.t01#3    mNeoVis.3.1.1.chr7.g00503.t01#6    mNeoVis.2.1.1.chr7_hap1.g00519.t01#5    mNeoVis.evergladensis.2.1.chr7.g00519.t01#2    mNeoVis.vulgivaga.2.1.chr7.g00522.t01#1   mNeoVis.vulgivaga.1.1.chr7.g00527.t01#4       697   125261821:   mNeoVis.evergladensis.1.1.chr8.g00093.t01#3    mNeoVis.evergladensis.2.1.chr9.g00092.t01#2    mNeoVis.vulgivaga.2.1.chr8.g00095.t01#1    mNeoVis.2.1.1.chr8_hap1.g00095.t01#5   mNeoVis.vulgivaga.1.1.chr8.g00090.t01#4       698   125261822:   mNeoVis.vulgivaga.2.1.chr1.g02007.t01#1    mNeoVis.evergladensis.1.1.chr1.g02013.t01#3    mNeoVis.3.1.1.chr1.g02001.t01#6    mNeoVis.2.1.1.chr1_hap1.g02046.t01#5    mNeoVis.vulgivaga.1.1.chr1.g02017.t01#4   mNeoVis.evergladensis.2.1.chr1.g02013.t01#2       699   125261823:   mNeoVis.vulgivaga.2.1.chr1.g02008.t01#1    mNeoVis.evergladensis.1.1.chr1.g02014.t01#3    mNeoVis.evergladensis.2.1.chr1.g02014.t01#2    mNeoVis.3.1.1.chr1.g02002.t01#6    mNeoVis.2.1.1.chr1_hap1.g02047.t01#5   mNeoVis.vulgivaga.1.1.chr1.g02018.t01#4       700   125261824:   mNeoVis.3.1.1.chr15.g00533.t01#6    mNeoVis.evergladensis.1.1.chr15.g00034.t01#3    mNeoVis.vulgivaga.1.1.chr15.g00580.t01#4    mNeoVis.2.1.1.chr15_hap1.g00757.t01#5    mNeoVis.evergladensis.2.1.chr15.g00578.t01#2   mNeoVis.vulgivaga.2.1.chr15.g00589.t01#1       701   125261825:   mNeoVis.evergladensis.2.1.chr7.g01182.t01#2    mNeoVis.evergladensis.2.1.chr1.g02015.t01#2    mNeoVis.evergladensis.2.1.chr5.g01490.t01#2    mNeoVis.vulgivaga.1.1.chr1.g02019.t01#4    mNeoVis.2.1.1.chr7_hap1.g01193.t01#5    mNeoVis.vulgivaga.2.1.chr5.g01494.t01#1    mNeoVis.evergladensis.1.1.chr1.g02015.t01#3    mNeoVis.vulgivaga.1.1.chr5.g01469.t01#4    mNeoVis.2.1.1.chr2_hap1.g01552.t01#5      702   125261826:   mNeoVis.vulgivaga.1.1.chr1.g02020.t01#4    mNeoVis.2.1.1.chr1_hap1.g02049.t01#5    mNeoVis.evergladensis.1.1.chr1.g02016.t01#3    mNeoVis.evergladensis.2.1.chr1.g02016.t01#2    mNeoVis.vulgivaga.2.1.chr1.g02010.t01#1   mNeoVis.3.1.1.chr1.g02005.t01#6       703   125261827:   mNeoVis.evergladensis.2.1.chr1.g02018.t01#2    mNeoVis.vulgivaga.2.1.chr1.g02011.t01#1    mNeoVis.evergladensis.1.1.chr1.g02018.t01#3    mNeoVis.2.1.1.chr1_hap1.g02051.t01#5    mNeoVis.3.1.1.chr1.g02006.t01#6   mNeoVis.vulgivaga.1.1.chr1.g02021.t01#4       704   125261828:   mNeoVis.3.1.1.chr1.g02007.t01#6    mNeoVis.evergladensis.1.1.chr1.g02019.t01#3    mNeoVis.vulgivaga.2.1.chr1.g02012.t01#1    mNeoVis.2.1.1.chr1_hap1.g02052.t01#5    mNeoVis.vulgivaga.1.1.chr1.g02022.t01#4   mNeoVis.evergladensis.2.1.chr1.g02019.t01#2       705   125261829:   mNeoVis.evergladensis.2.1.chr1.g01924.t01#2    mNeoVis.vulgivaga.1.1.chr1.g01933.t01#4    mNeoVis.vulgivaga.2.1.chr1.g01923.t01#1    mNeoVis.3.1.1.chr1.g01914.t01#6    mNeoVis.2.1.1.chr1_hap1.g01962.t01#5   mNeoVis.evergladensis.1.1.chr1.g01924.t01#3       706   125261830:   mNeoVis.3.1.1.chr1.g01915.t01#6    mNeoVis.vulgivaga.1.1.chr3.g00235.t01#4    mNeoVis.vulgivaga.1.1.chr1.g00408.t01#4    mNeoVis.evergladensis.2.1.chr1.g00413.t01#2    mNeoVis.3.1.1.chr3.g00233.t01#6    mNeoVis.3.1.1.chr1.g00411.t01#6    mNeoVis.vulgivaga.2.1.chr1.g01924.t01#1    mNeoVis.vulgivaga.1.1.chr6.g00936.t01#4    mNeoVis.evergladensis.1.1.chr6.g00942.t01#3      707   125261831:   mNeoVis.3.1.1.chr1.g00992.t01#6    mNeoVis.evergladensis.1.1.chr1.g00985.t01#3    mNeoVis.2.1.1.chr1_hap1.g00988.t01#5    mNeoVis.vulgivaga.1.1.chr1.g00994.t01#4    mNeoVis.evergladensis.2.1.chr1.g00982.t01#2    mNeoVis.vulgivaga.2.1.chr1.g00996.t01#1   mNeoVis.vulgivaga.2.1.chr1.g00995.t01#1     708   125261832:   mNeoVis.vulgivaga.2.1.chr1.g01925.t01#1    mNeoVis.evergladensis.1.1.chr1.g01925.t01#3    mNeoVis.2.1.1.chr1_hap1.g01963.t01#5    mNeoVis.evergladensis.2.1.chr1.g01925.t01#2    mNeoVis.3.1.1.chr1.g01917.t01#6   mNeoVis.vulgivaga.1.1.chr1.g01934.t01#4       709   125261833:   mNeoVis.vulgivaga.1.1.chr1.g01935.t01#4    mNeoVis.3.1.1.chr1.g01918.t01#6    mNeoVis.vulgivaga.2.1.chr1.g01926.t01#1    mNeoVis.2.1.1.chr1_hap1.g01964.t01#5    mNeoVis.evergladensis.2.1.chr1.g01926.t01#2   mNeoVis.evergladensis.1.1.chr1.g01926.t01#3       710   125261834:   mNeoVis.evergladensis.2.1.chr1.g01927.t01#2    mNeoVis.2.1.1.chr1_hap1.g01965.t01#5    mNeoVis.evergladensis.1.1.chr1.g01927.t01#3    mNeoVis.vulgivaga.2.1.chr1.g01927.t01#1   mNeoVis.vulgivaga.1.1.chr1.g01936.t01#4       711   125261835:   mNeoVis.vulgivaga.2.1.chr8.g00096.t01#1    mNeoVis.evergladensis.1.1.chr8.g00094.t01#3    mNeoVis.vulgivaga.1.1.chr8.g00091.t01#4    mNeoVis.2.1.1.chr8_hap1.g00096.t01#5    mNeoVis.3.1.1.chr8.g00071.t01#6   mNeoVis.evergladensis.2.1.chr9.g00093.t01#2       712   125261836:   mNeoVis.2.1.1.chr1_hap1.g01966.t01#5    mNeoVis.evergladensis.1.1.chr1.g01928.t01#3    mNeoVis.vulgivaga.2.1.chr1.g01928.t01#1    mNeoVis.3.1.1.chr1.g01920.t01#6    mNeoVis.vulgivaga.1.1.chr1.g01937.t01#4   mNeoVis.evergladensis.2.1.chr1.g01928.t01#2       713   125261837:   mNeoVis.evergladensis.1.1.chr1.g01929.t01#3    mNeoVis.evergladensis.2.1.chr1.g01929.t01#2    mNeoVis.vulgivaga.1.1.chr1.g01938.t01#4    mNeoVis.2.1.1.chr1_hap1.g01967.t01#5    mNeoVis.3.1.1.chr1.g01921.t01#6   mNeoVis.vulgivaga.2.1.chr1.g01929.t01#1       714   125261838:   mNeoVis.2.1.1.chr1_hap1.g01968.t01#5    mNeoVis.3.1.1.chr1.g01922.t01#6    mNeoVis.vulgivaga.2.1.chr1.g01930.t01#1    mNeoVis.evergladensis.1.1.chr1.g01930.t01#3    mNeoVis.evergladensis.2.1.chr1.g01930.t01#2   mNeoVis.vulgivaga.1.1.chr1.g01939.t01#4       715   125261839:   mNeoVis.2.1.1.chr8_hap1.g00097.t01#5    mNeoVis.vulgivaga.1.1.chr8.g00092.t01#4    mNeoVis.evergladensis.1.1.chr8.g00095.t01#3    mNeoVis.evergladensis.2.1.chr9.g00094.t01#2   mNeoVis.vulgivaga.2.1.chr8.g00097.t01#1       716   125261840:   mNeoVis.evergladensis.1.1.chr8.g00096.t01#3    mNeoVis.evergladensis.2.1.chr9.g00095.t01#2    mNeoVis.vulgivaga.2.1.chr8.g00098.t01#1    mNeoVis.2.1.1.chr8_hap1.g00098.t01#5   mNeoVis.vulgivaga.1.1.chr8.g00093.t01#4       717   125261841:   mNeoVis.evergladensis.1.1.chr8.g00097.t01#3    mNeoVis.evergladensis.2.1.chr9.g00096.t01#2    mNeoVis.vulgivaga.2.1.chr8.g00099.t01#1    mNeoVis.2.1.1.chr8_hap1.g00099.t01#5   mNeoVis.vulgivaga.1.1.chr8.g00094.t01#4       718   125261842:   mNeoVis.evergladensis.1.1.chr1.g01931.t01#3    mNeoVis.evergladensis.2.1.chr1.g01931.t01#2    mNeoVis.2.1.1.chr1_hap1.g01969.t01#5    mNeoVis.vulgivaga.1.1.chr1.g01940.t01#4    mNeoVis.vulgivaga.2.1.chr1.g01931.t01#1   mNeoVis.3.1.1.chr1.g01923.t01#6       719   125261843:   mNeoVis.3.1.1.chr8.g00072.t01#6    mNeoVis.vulgivaga.2.1.chr8.g00100.t01#1    mNeoVis.2.1.1.chr8_hap1.g00100.t01#5    mNeoVis.vulgivaga.1.1.chr8.g00095.t01#4    mNeoVis.evergladensis.2.1.chr9.g00097.t01#2   mNeoVis.evergladensis.1.1.chr8.g00098.t01#3       720   125261844:   mNeoVis.evergladensis.1.1.chr8.g00099.t01#3    mNeoVis.2.1.1.chr8_hap1.g00101.t01#5    mNeoVis.vulgivaga.1.1.chr8.g00096.t01#4    mNeoVis.3.1.1.chr8.g00073.t01#6    mNeoVis.vulgivaga.2.1.chr8.g00101.t01#1   mNeoVis.evergladensis.2.1.chr9.g00098.t01#2       721   125261845:   mNeoVis.evergladensis.1.1.chr7.g00529.t01#3    mNeoVis.evergladensis.2.1.chr7.g00520.t01#2    mNeoVis.2.1.1.chr7_hap1.g00520.t01#5    mNeoVis.2.1.1.chr2_hap1.g01431.t01#5    mNeoVis.vulgivaga.2.1.chr7.g00523.t01#1    mNeoVis.evergladensis.1.1.chr13.g00669.t01#3    mNeoVis.vulgivaga.1.1.chr7.g00528.t01#4   mNeoVis.3.1.1.chr7.g00504.t01#6     722   125261846:   mNeoVis.vulgivaga.1.1.chr8.g00097.t01#4    mNeoVis.vulgivaga.2.1.chr8.g00102.t01#1    mNeoVis.3.1.1.chr8.g00074.t01#6    mNeoVis.evergladensis.2.1.chr9.g00099.t01#2    mNeoVis.evergladensis.1.1.chr8.g00100.t01#3   mNeoVis.2.1.1.chr8_hap1.g00102.t01#5       723   125261847:   mNeoVis.vulgivaga.2.1.chr1.g01932.t01#1    mNeoVis.evergladensis.2.1.chr1.g01932.t01#2   mNeoVis.evergladensis.1.1.chr1.g01932.t01#3         724   125261848:   mNeoVis.2.1.1.chr8_hap1.g00103.t01#5    mNeoVis.vulgivaga.1.1.chr8.g00098.t01#4    mNeoVis.3.1.1.chr8.g00075.t01#6    mNeoVis.evergladensis.1.1.chr8.g00101.t01#3    mNeoVis.vulgivaga.2.1.chr8.g00103.t01#1   mNeoVis.evergladensis.2.1.chr9.g00100.t01#2       725   125261849:   mNeoVis.vulgivaga.2.1.chr8.g00104.t01#1    mNeoVis.evergladensis.1.1.chr8.g00102.t01#3    mNeoVis.3.1.1.chr8.g00076.t01#6    mNeoVis.2.1.1.chr8_hap1.g00104.t01#5    mNeoVis.vulgivaga.1.1.chr8.g00099.t01#4   mNeoVis.evergladensis.2.1.chr9.g00101.t01#2       726   125261850:   mNeoVis.vulgivaga.2.1.chr1.g01933.t01#1    mNeoVis.evergladensis.1.1.chr1.g01933.t01#3    mNeoVis.evergladensis.2.1.chr1.g01933.t01#2    mNeoVis.2.1.1.chr1_hap1.g01971.t01#5   mNeoVis.vulgivaga.1.1.chr1.g01942.t01#4       727   125261851:   mNeoVis.vulgivaga.1.1.chr1.g01943.t01#4    mNeoVis.evergladensis.2.1.chr1.g01934.t01#2    mNeoVis.2.1.1.chr1_hap1.g01972.t01#5    mNeoVis.3.1.1.chr1.g01926.t01#6    mNeoVis.evergladensis.1.1.chr1.g01934.t01#3   mNeoVis.vulgivaga.2.1.chr1.g01934.t01#1       728   125261852:   mNeoVis.evergladensis.2.1.chr11.g00567.t01#2    mNeoVis.evergladensis.1.1.chr11.g00571.t01#3    mNeoVis.vulgivaga.1.1.chr11.g00561.t01#4    mNeoVis.2.1.1.chr11_hap1.g00580.t01#5   mNeoVis.vulgivaga.2.1.chr11.g00575.t01#1       729   125261853:   mNeoVis.evergladensis.1.1.chr1.g01935.t01#3    mNeoVis.3.1.1.chr1.g01927.t01#6    mNeoVis.vulgivaga.1.1.chr1.g01944.t01#4    mNeoVis.2.1.1.chr1_hap1.g01973.t01#5    mNeoVis.vulgivaga.2.1.chr1.g01935.t01#1   mNeoVis.evergladensis.2.1.chr1.g01935.t01#2       730   125261854:   mNeoVis.vulgivaga.2.1.chr1.g01936.t01#1    mNeoVis.evergladensis.2.1.chr1.g01936.t01#2    mNeoVis.2.1.1.chr1_hap1.g01974.t01#5    mNeoVis.evergladensis.1.1.chr1.g01936.t01#3   mNeoVis.vulgivaga.1.1.chr1.g01945.t01#4       731   125261855:   mNeoVis.evergladensis.1.1.chr1.g01937.t01#3    mNeoVis.vulgivaga.1.1.chr1.g01946.t01#4    mNeoVis.2.1.1.chr1_hap1.g01975.t01#5    mNeoVis.vulgivaga.2.1.chr1.g01937.t01#1    mNeoVis.evergladensis.2.1.chr1.g01937.t01#2   mNeoVis.3.1.1.chr1.g01930.t01#6       732   125261856:   mNeoVis.evergladensis.1.1.chr1.g01938.t01#3    mNeoVis.evergladensis.2.1.chr1.g01938.t01#2    mNeoVis.evergladensis.1.1.chr1.g01939.t01#3    mNeoVis.vulgivaga.2.1.chr1.g01938.t01#1    mNeoVis.2.1.1.chr1_hap1.g01976.t01#5    mNeoVis.3.1.1.chr1.g01931.t01#6   mNeoVis.vulgivaga.1.1.chr1.g01947.t01#4     733   125261857:   mNeoVis.2.1.1.chr8_hap1.g00026.t01#5    mNeoVis.vulgivaga.1.1.chr8.g00022.t01#4    mNeoVis.evergladensis.1.1.chr8.g00022.t01#3    mNeoVis.vulgivaga.2.1.chr8.g00023.t01#1   mNeoVis.evergladensis.2.1.chr9.g00024.t01#2       734   125261858:   mNeoVis.evergladensis.1.1.chr8.g00023.t01#3    mNeoVis.evergladensis.2.1.chr9.g00025.t01#2    mNeoVis.2.1.1.chr8_hap1.g00027.t01#5    mNeoVis.vulgivaga.1.1.chr8.g00023.t01#4   mNeoVis.vulgivaga.2.1.chr8.g00024.t01#1       735   125261859:   mNeoVis.evergladensis.1.1.chr1.g01940.t01#3    mNeoVis.evergladensis.2.1.chr1.g01939.t01#2    mNeoVis.vulgivaga.1.1.chr1.g01948.t01#4    mNeoVis.2.1.1.chr1_hap1.g01977.t01#5   mNeoVis.vulgivaga.2.1.chr1.g01939.t01#1       736   125261860:   mNeoVis.3.1.1.chr1.g01932.t01#6   mNeoVis.vulgivaga.1.1.chr1.g01949.t01#4             737   125261861:   mNeoVis.evergladensis.2.1.chr9.g00026.t01#2    mNeoVis.vulgivaga.2.1.chr8.g00025.t01#1    mNeoVis.vulgivaga.1.1.chr8.g00024.t01#4    mNeoVis.2.1.1.chr8_hap1.g00028.t01#5   mNeoVis.evergladensis.1.1.chr8.g00024.t01#3       738   125261862:   mNeoVis.evergladensis.2.1.chr1.g01940.t01#2    mNeoVis.vulgivaga.1.1.chr1.g01950.t01#4    mNeoVis.2.1.1.chr1_hap1.g01978.t01#5    mNeoVis.evergladensis.1.1.chr1.g01941.t01#3   mNeoVis.vulgivaga.2.1.chr1.g01940.t01#1       739   125261863:   mNeoVis.2.1.1.chr1_hap1.g01979.t01#5    mNeoVis.evergladensis.2.1.chr1.g01941.t01#2    mNeoVis.evergladensis.1.1.chr1.g01942.t01#3    mNeoVis.vulgivaga.2.1.chr1.g01941.t01#1    mNeoVis.vulgivaga.1.1.chr1.g01951.t01#4   mNeoVis.3.1.1.chr1.g01934.t01#6       740   125261864:   mNeoVis.evergladensis.1.1.chr8.g00025.t01#3    mNeoVis.vulgivaga.2.1.chr8.g00026.t01#1    mNeoVis.2.1.1.chr8_hap1.g00029.t01#5    mNeoVis.vulgivaga.1.1.chr8.g00025.t01#4   mNeoVis.evergladensis.2.1.chr9.g00028.t01#2       741   125261865:   mNeoVis.evergladensis.1.1.chr8.g00026.t01#3    mNeoVis.vulgivaga.1.1.chr8.g00026.t01#4    mNeoVis.2.1.1.chr8_hap1.g00030.t01#5    mNeoVis.evergladensis.2.1.chr9.g00027.t01#2   mNeoVis.vulgivaga.2.1.chr8.g00027.t01#1       742   125261866:   mNeoVis.3.1.1.chr1.g01935.t01#6    mNeoVis.evergladensis.2.1.chr1.g01942.t01#2    mNeoVis.vulgivaga.1.1.chr1.g01952.t01#4    mNeoVis.vulgivaga.2.1.chr1.g01942.t01#1    mNeoVis.evergladensis.1.1.chr1.g01943.t01#3   mNeoVis.2.1.1.chr1_hap1.g01980.t01#5       743   125261867:   mNeoVis.2.1.1.chr7_hap1.g00521.t01#5    mNeoVis.evergladensis.2.1.chr7.g00521.t01#2    mNeoVis.vulgivaga.1.1.chr7.g00529.t01#4    mNeoVis.vulgivaga.2.1.chr7.g00524.t01#1    mNeoVis.evergladensis.1.1.chr7.g00530.t01#3   mNeoVis.3.1.1.chr7.g00505.t01#6       744   125261868:   mNeoVis.vulgivaga.2.1.chr1.g01943.t01#1    mNeoVis.3.1.1.chr1.g01936.t01#6    mNeoVis.vulgivaga.1.1.chr1.g01954.t01#4    mNeoVis.evergladensis.2.1.chr1.g01944.t01#2    mNeoVis.evergladensis.1.1.chr1.g01945.t01#3   mNeoVis.2.1.1.chr1_hap1.g01982.t01#5       745   125261869:   mNeoVis.evergladensis.1.1.chr1.g01946.t01#3    mNeoVis.vulgivaga.2.1.chr1.g01944.t01#1    mNeoVis.2.1.1.chr1_hap1.g01983.t01#5    mNeoVis.evergladensis.2.1.chr1.g01945.t01#2    mNeoVis.3.1.1.chr1.g01937.t01#6   mNeoVis.vulgivaga.1.1.chr1.g01955.t01#4       746   125261870:   mNeoVis.vulgivaga.1.1.chr7.g01569.t01#4    mNeoVis.3.1.1.chr7.g01487.t01#6    mNeoVis.evergladensis.1.1.chr12.g00436.t01#3    mNeoVis.2.1.1.chr7_hap1.g01576.t01#5    mNeoVis.evergladensis.1.1.chr7.g01561.t01#3    mNeoVis.3.1.1.chr7.g01486.t01#6    mNeoVis.vulgivaga.2.1.chr7.g01574.t01#1    mNeoVis.evergladensis.2.1.chr7.g01556.t01#2    mNeoVis.evergladensis.2.1.chr12.g00431.t01#2      747   125261871:   mNeoVis.3.1.1.chr15.g00400.t01#6    mNeoVis.evergladensis.1.1.chr15.g00175.t01#3    mNeoVis.2.1.1.chr15_hap1.g00617.t01#5    mNeoVis.vulgivaga.1.1.chr15.g00440.t01#4    mNeoVis.evergladensis.2.1.chr15.g00439.t01#2   mNeoVis.vulgivaga.2.1.chr15.g00447.t01#1       748   125261872:   mNeoVis.evergladensis.1.1.chr10.g01127.t01#3    mNeoVis.2.1.1.chr8_hap1.g00031.t01#5    mNeoVis.vulgivaga.1.1.chr8.g00027.t01#4    mNeoVis.2.1.1.chr9_hap1.g01144.t01#5    mNeoVis.evergladensis.1.1.chr8.g00027.t01#3    mNeoVis.vulgivaga.1.1.chr10.g01108.t01#4    mNeoVis.vulgivaga.2.1.chr8.g00028.t01#1    mNeoVis.evergladensis.2.1.chr10.g01132.t01#2    mNeoVis.vulgivaga.2.1.chr10.g01124.t01#1      749   125261873:   mNeoVis.vulgivaga.1.1.chr8.g00028.t01#4    mNeoVis.evergladensis.1.1.chr8.g00028.t01#3    mNeoVis.2.1.1.chr8_hap1.g00032.t01#5    mNeoVis.evergladensis.2.1.chr9.g00030.t01#2    mNeoVis.3.1.1.chr8.g00018.t01#6   mNeoVis.vulgivaga.2.1.chr8.g00029.t01#1       750   125261874:   mNeoVis.2.1.1.chr1_hap1.g01984.t01#5    mNeoVis.vulgivaga.1.1.chr1.g01956.t01#4    mNeoVis.3.1.1.chr1.g01938.t01#6    mNeoVis.vulgivaga.2.1.chr1.g01945.t01#1    mNeoVis.evergladensis.2.1.chr1.g01946.t01#2   mNeoVis.evergladensis.1.1.chr1.g01947.t01#3       751   125261875:   mNeoVis.2.1.1.chr8_hap1.g00033.t01#5    mNeoVis.evergladensis.1.1.chr8.g00029.t01#3    mNeoVis.vulgivaga.2.1.chr8.g00030.t01#1    mNeoVis.evergladensis.2.1.chr9.g00031.t01#2   mNeoVis.vulgivaga.1.1.chr8.g00029.t01#4       752   125261876:   mNeoVis.3.1.1.chr8.g00019.t01#6    mNeoVis.vulgivaga.1.1.chr8.g00030.t01#4    mNeoVis.evergladensis.2.1.chr9.g00032.t01#2    mNeoVis.evergladensis.1.1.chr8.g00030.t01#3    mNeoVis.2.1.1.chr8_hap1.g00034.t01#5   mNeoVis.vulgivaga.2.1.chr8.g00031.t01#1       753   125261877:   mNeoVis.2.1.1.chr8_hap1.g00035.t01#5    mNeoVis.evergladensis.2.1.chr9.g00033.t01#2    mNeoVis.evergladensis.1.1.chr8.g00031.t01#3    mNeoVis.evergladensis.1.1.chr8.g00032.t01#3    mNeoVis.vulgivaga.2.1.chr8.g00032.t01#1   mNeoVis.vulgivaga.1.1.chr8.g00031.t01#4       754   125261878:   mNeoVis.vulgivaga.1.1.chr1.g01829.t01#4    mNeoVis.2.1.1.chr1_hap1.g01859.t01#5    mNeoVis.vulgivaga.2.1.chr1.g01821.t01#1    mNeoVis.evergladensis.2.1.chr1.g01826.t01#2    mNeoVis.evergladensis.1.1.chr1.g01825.t01#3   mNeoVis.3.1.1.chr1.g01812.t01#6       755   125261879:   mNeoVis.vulgivaga.1.1.chr8.g00032.t01#4    mNeoVis.evergladensis.1.1.chr8.g00033.t01#3    mNeoVis.vulgivaga.2.1.chr8.g00033.t01#1    mNeoVis.evergladensis.2.1.chr9.g00034.t01#2   mNeoVis.2.1.1.chr8_hap1.g00036.t01#5       756   125261880:   mNeoVis.vulgivaga.2.1.chr7.g00129.t01#1    mNeoVis.3.1.1.chr7.g00110.t01#6    mNeoVis.vulgivaga.1.1.chr7.g00132.t01#4    mNeoVis.evergladensis.2.1.chr7.g00129.t01#2   mNeoVis.2.1.1.chr7_hap1.g00129.t01#5       757   125261881:   mNeoVis.2.1.1.chr14_hap1.g00231.t01#5    mNeoVis.vulgivaga.1.1.chr14.g00230.t01#4    mNeoVis.evergladensis.2.1.chr14.g00233.t01#2    mNeoVis.3.1.1.chr14.g00233.t01#6    mNeoVis.vulgivaga.2.1.chr14.g00234.t01#1   mNeoVis.evergladensis.1.1.chr14.g00239.t01#3       758   125261882:   mNeoVis.vulgivaga.1.1.chr8.g00033.t01#4    mNeoVis.evergladensis.2.1.chr9.g00035.t01#2    mNeoVis.2.1.1.chr8_hap1.g00037.t01#5    mNeoVis.vulgivaga.2.1.chr8.g00034.t01#1   mNeoVis.evergladensis.1.1.chr8.g00034.t01#3       759   125261883:   mNeoVis.3.1.1.chr8.g00021.t01#6    mNeoVis.2.1.1.chr8_hap1.g00038.t01#5    mNeoVis.evergladensis.2.1.chr9.g00036.t01#2    mNeoVis.vulgivaga.2.1.chr8.g00035.t01#1    mNeoVis.evergladensis.1.1.chr8.g00035.t01#3   mNeoVis.vulgivaga.1.1.chr8.g00034.t01#4       760   125261884:   mNeoVis.vulgivaga.1.1.chr8.g00036.t01#4    mNeoVis.2.1.1.chr8_hap1.g00040.t01#5    mNeoVis.vulgivaga.2.1.chr8.g00037.t01#1    mNeoVis.evergladensis.1.1.chr8.g00037.t01#3   mNeoVis.evergladensis.2.1.chr9.g00037.t01#2       761   125261885:   mNeoVis.evergladensis.1.1.chr8.g00036.t01#3    mNeoVis.vulgivaga.1.1.chr8.g00035.t01#4    mNeoVis.vulgivaga.2.1.chr8.g00036.t01#1   mNeoVis.2.1.1.chr8_hap1.g00039.t01#5         762   125261886:   mNeoVis.vulgivaga.2.1.chr8.g00038.t01#1    mNeoVis.vulgivaga.1.1.chr8.g00037.t01#4    mNeoVis.evergladensis.1.1.chr8.g00038.t01#3    mNeoVis.2.1.1.chr8_hap1.g00041.t01#5   mNeoVis.evergladensis.2.1.chr9.g00038.t01#2       763   125261887:   mNeoVis.vulgivaga.1.1.chr8.g00038.t01#4    mNeoVis.evergladensis.2.1.chr9.g00039.t01#2    mNeoVis.2.1.1.chr8_hap1.g00042.t01#5    mNeoVis.vulgivaga.2.1.chr8.g00039.t01#1   mNeoVis.evergladensis.1.1.chr8.g00039.t01#3       764   125261888:   mNeoVis.vulgivaga.2.1.chr8.g00040.t01#1    mNeoVis.evergladensis.1.1.chr8.g00040.t01#3    mNeoVis.vulgivaga.1.1.chr8.g00039.t01#4    mNeoVis.2.1.1.chr8_hap1.g00043.t01#5   mNeoVis.evergladensis.2.1.chr9.g00040.t01#2       765   125261889:   mNeoVis.evergladensis.2.1.chr7.g00130.t01#2    mNeoVis.2.1.1.chr7_hap1.g00130.t01#5    mNeoVis.evergladensis.1.1.chr7.g00131.t01#3    mNeoVis.3.1.1.chr7.g00111.t01#6    mNeoVis.vulgivaga.2.1.chr7.g00130.t01#1   mNeoVis.vulgivaga.1.1.chr7.g00133.t01#4       766   125261890:   mNeoVis.evergladensis.1.1.chr7.g00009.t01#3    mNeoVis.vulgivaga.1.1.chr3.g00694.t01#4    mNeoVis.2.1.1.chr1_hap1.g00120.t01#5    mNeoVis.2.1.1.chr1_hap1.g01860.t01#5    mNeoVis.3.1.1.chr7.g00008.t01#6    mNeoVis.vulgivaga.2.1.chr7.g00007.t01#1    mNeoVis.vulgivaga.2.1.chr9.g00979.t01#1    mNeoVis.2.1.1.chr9_hap1.g00101.t01#5    mNeoVis.2.1.1.chr4_hap1.g00955.t01#5      767   125261891:   mNeoVis.vulgivaga.2.1.chr2.g01727.t01#1    mNeoVis.2.1.1.chr9_hap1.g01038.t01#5    mNeoVis.evergladensis.1.1.chr5.g01298.t01#3    mNeoVis.3.1.1.chr5.g01281.t01#6    mNeoVis.evergladensis.2.1.chr10.g01027.t01#2    mNeoVis.vulgivaga.2.1.chr1.g01822.t01#1   mNeoVis.3.1.1.chr9.g00953.t01#6       768   125261892:   mNeoVis.evergladensis.1.1.chr2.g00333.t01#3    mNeoVis.2.1.1.chr5_hap1.g00335.t01#5    mNeoVis.3.1.1.chr2.g00273.t01#6    mNeoVis.vulgivaga.1.1.chr2.g00330.t01#4    mNeoVis.evergladensis.2.1.chr2.g00334.t01#2   mNeoVis.vulgivaga.2.1.chr2.g00331.t01#1       769   125261893:   mNeoVis.evergladensis.1.1.chr6.g00966.t01#3   mNeoVis.vulgivaga.2.1.chr5.g01294.t01#1             770   125261894:   mNeoVis.2.1.1.chr1_hap1.g01861.t01#5    mNeoVis.3.1.1.chr1.g01814.t01#6    mNeoVis.evergladensis.2.1.chr1.g01827.t01#2    mNeoVis.vulgivaga.1.1.chr1.g01830.t01#4    mNeoVis.3.1.1.chr1.g01813.t01#6    mNeoVis.vulgivaga.2.1.chr1.g01823.t01#1   mNeoVis.evergladensis.1.1.chr1.g01826.t01#3     771   125261895:  mNeoVis.vulgivaga.2.1.chr8.g00041.t01#1               772   125261896:   mNeoVis.vulgivaga.2.1.chr8.g00042.t01#1    mNeoVis.evergladensis.1.1.chr8.g00042.t01#3    mNeoVis.2.1.1.chr8_hap1.g00045.t01#5    mNeoVis.evergladensis.2.1.chr9.g00042.t01#2   mNeoVis.vulgivaga.1.1.chr8.g00041.t01#4       773   125261897:   mNeoVis.evergladensis.2.1.chr1.g01828.t01#2    mNeoVis.3.1.1.chr1.g01815.t01#6    mNeoVis.evergladensis.1.1.chr1.g01827.t01#3    mNeoVis.vulgivaga.1.1.chr1.g01831.t01#4    mNeoVis.2.1.1.chr1_hap1.g01862.t01#5   mNeoVis.vulgivaga.2.1.chr1.g01824.t01#1       774   125261898:   mNeoVis.2.1.1.chr1_hap1.g01863.t01#5    mNeoVis.evergladensis.2.1.chr1.g01829.t01#2    mNeoVis.3.1.1.chr1.g01816.t01#6    mNeoVis.vulgivaga.1.1.chr1.g01832.t01#4    mNeoVis.evergladensis.1.1.chr1.g01828.t01#3   mNeoVis.vulgivaga.2.1.chr1.g01825.t01#1       775   125261899:   mNeoVis.vulgivaga.2.1.chr8.g00043.t01#1    mNeoVis.vulgivaga.1.1.chr8.g00042.t01#4    mNeoVis.evergladensis.1.1.chr8.g00043.t01#3    mNeoVis.evergladensis.2.1.chr9.g00043.t01#2    mNeoVis.2.1.1.chr8_hap1.g00046.t01#5   mNeoVis.3.1.1.chr8.g00023.t01#6       776   125261900:   mNeoVis.evergladensis.1.1.chr8.g00044.t01#3    mNeoVis.2.1.1.chr8_hap1.g00047.t01#5    mNeoVis.3.1.1.chr8.g00024.t01#6    mNeoVis.vulgivaga.2.1.chr8.g00044.t01#1    mNeoVis.vulgivaga.1.1.chr8.g00043.t01#4   mNeoVis.evergladensis.2.1.chr9.g00044.t01#2       777   125261901:   mNeoVis.evergladensis.2.1.chr9.g00045.t01#2    mNeoVis.2.1.1.chr8_hap1.g00048.t01#5    mNeoVis.3.1.1.chr8.g00025.t01#6    mNeoVis.vulgivaga.2.1.chr8.g00045.t01#1    mNeoVis.evergladensis.1.1.chr8.g00045.t01#3   mNeoVis.vulgivaga.1.1.chr8.g00044.t01#4       778   125261902:   mNeoVis.3.1.1.chr15.g00182.t01#6    mNeoVis.evergladensis.1.1.chr15.g00425.t01#3    mNeoVis.2.1.1.chr15_hap1.g00209.t01#5    mNeoVis.vulgivaga.2.1.chr15.g00189.t01#1    mNeoVis.evergladensis.2.1.chr15.g00188.t01#2   mNeoVis.vulgivaga.1.1.chr15.g00185.t01#4       779   125261903:  mNeoVis.vulgivaga.2.1.chr1.g01826.t01#1               780   125261904:   mNeoVis.vulgivaga.2.1.chr8.g00046.t01#1    mNeoVis.vulgivaga.1.1.chr8.g00045.t01#4    mNeoVis.2.1.1.chr8_hap1.g00049.t01#5    mNeoVis.evergladensis.1.1.chr8.g00046.t01#3    mNeoVis.3.1.1.chr8.g00026.t01#6   mNeoVis.evergladensis.2.1.chr9.g00046.t01#2       781   125261905:   mNeoVis.evergladensis.1.1.chr1.g01948.t01#3    mNeoVis.evergladensis.2.1.chr1.g01947.t01#2    mNeoVis.3.1.1.chr1.g01940.t01#6    mNeoVis.vulgivaga.2.1.chr1.g01946.t01#1    mNeoVis.vulgivaga.1.1.chr1.g01957.t01#4   mNeoVis.2.1.1.chr1_hap1.g01985.t01#5       782   125261906:   mNeoVis.evergladensis.1.1.chr5.g01431.t01#3    mNeoVis.2.1.1.chr2_hap1.g01490.t01#5    mNeoVis.3.1.1.chr5.g01412.t01#6    mNeoVis.evergladensis.2.1.chr5.g01427.t01#2    mNeoVis.vulgivaga.2.1.chr5.g01428.t01#1   mNeoVis.vulgivaga.1.1.chr5.g01404.t01#4       783   125261907:   mNeoVis.evergladensis.2.1.chr2.g01732.t01#2    mNeoVis.vulgivaga.1.1.chr2.g01741.t01#4    mNeoVis.3.1.1.chr2.g01659.t01#6    mNeoVis.2.1.1.chr5_hap1.g01750.t01#5    mNeoVis.vulgivaga.2.1.chr2.g01748.t01#1   mNeoVis.evergladensis.1.1.chr2.g01746.t01#3       784   125261908:   mNeoVis.evergladensis.1.1.chr8.g00047.t01#3    mNeoVis.3.1.1.chr8.g00027.t01#6    mNeoVis.vulgivaga.1.1.chr8.g00046.t01#4    mNeoVis.2.1.1.chr8_hap1.g00050.t01#5    mNeoVis.evergladensis.2.1.chr9.g00047.t01#2   mNeoVis.vulgivaga.2.1.chr8.g00047.t01#1       785   125261909:   mNeoVis.2.1.1.chr8_hap1.g00051.t01#5    mNeoVis.vulgivaga.2.1.chr8.g00048.t01#1    mNeoVis.3.1.1.chr8.g00028.t01#6    mNeoVis.evergladensis.2.1.chr9.g00048.t01#2    mNeoVis.vulgivaga.1.1.chr8.g00047.t01#4   mNeoVis.evergladensis.1.1.chr8.g00048.t01#3       786   125261910:   mNeoVis.vulgivaga.2.1.chr3.g00531.t01#1    mNeoVis.3.1.1.chr1.g01941.t01#6    mNeoVis.vulgivaga.1.1.chr1.g01958.t01#4    mNeoVis.3.1.1.chr3.g00512.t01#6    mNeoVis.vulgivaga.1.1.chr3.g00525.t01#4    mNeoVis.evergladensis.2.1.chr3.g00529.t01#2    mNeoVis.evergladensis.1.1.chr3.g00527.t01#3    mNeoVis.vulgivaga.2.1.chr1.g01947.t01#1    mNeoVis.2.1.1.chr3_hap1.g00520.t01#5      787   125261911:   mNeoVis.vulgivaga.2.1.chr1.g01948.t01#1    mNeoVis.evergladensis.1.1.chr1.g01950.t01#3    mNeoVis.2.1.1.chr1_hap1.g01986.t01#5    mNeoVis.vulgivaga.1.1.chr1.g01959.t01#4   mNeoVis.evergladensis.2.1.chr1.g01948.t01#2       788   125261912:   mNeoVis.vulgivaga.1.1.chr8.g00048.t01#4    mNeoVis.2.1.1.chr8_hap1.g00052.t01#5    mNeoVis.3.1.1.chr8.g00029.t01#6    mNeoVis.vulgivaga.2.1.chr8.g00049.t01#1    mNeoVis.evergladensis.2.1.chr9.g00049.t01#2   mNeoVis.evergladensis.1.1.chr8.g00049.t01#3       789   125261913:   mNeoVis.vulgivaga.1.1.chr8.g00049.t01#4    mNeoVis.2.1.1.chr8_hap1.g00053.t01#5    mNeoVis.evergladensis.2.1.chr9.g00050.t01#2    mNeoVis.3.1.1.chr8.g00030.t01#6    mNeoVis.evergladensis.1.1.chr8.g00050.t01#3   mNeoVis.vulgivaga.2.1.chr8.g00050.t01#1       790   125261914:   mNeoVis.2.1.1.chr9_hap1.g00781.t01#5    mNeoVis.vulgivaga.1.1.chr10.g00756.t01#4    mNeoVis.evergladensis.2.1.chr10.g00771.t01#2    mNeoVis.evergladensis.1.1.chr10.g00772.t01#3    mNeoVis.3.1.1.chr10.g00746.t01#6   mNeoVis.vulgivaga.2.1.chr10.g00758.t01#1       791   125261915:   mNeoVis.evergladensis.2.1.chr9.g00051.t01#2    mNeoVis.vulgivaga.1.1.chr8.g00050.t01#4    mNeoVis.2.1.1.chr8_hap1.g00054.t01#5    mNeoVis.vulgivaga.2.1.chr8.g00051.t01#1   mNeoVis.3.1.1.chr8.g00031.t01#6       792   125261916:   mNeoVis.evergladensis.2.1.chr1.g01949.t01#2    mNeoVis.vulgivaga.2.1.chr1.g01949.t01#1    mNeoVis.evergladensis.1.1.chr1.g01951.t01#3    mNeoVis.2.1.1.chr1_hap1.g01987.t01#5    mNeoVis.3.1.1.chr1.g01942.t01#6   mNeoVis.vulgivaga.1.1.chr1.g01960.t01#4       793   125261917:   mNeoVis.3.1.1.chr8.g00032.t01#6    mNeoVis.vulgivaga.2.1.chr8.g00052.t01#1    mNeoVis.evergladensis.1.1.chr8.g00051.t01#3    mNeoVis.vulgivaga.1.1.chr8.g00051.t01#4    mNeoVis.2.1.1.chr8_hap1.g00055.t01#5   mNeoVis.evergladensis.2.1.chr9.g00052.t01#2       794   125261918:   mNeoVis.evergladensis.1.1.chr1.g01952.t01#3    mNeoVis.2.1.1.chr1_hap1.g01988.t01#5    mNeoVis.vulgivaga.2.1.chr1.g01950.t01#1    mNeoVis.3.1.1.chr1.g01943.t01#6    mNeoVis.vulgivaga.1.1.chr1.g01961.t01#4   mNeoVis.evergladensis.2.1.chr1.g01950.t01#2       795   125261919:   mNeoVis.evergladensis.1.1.chr8.g00052.t01#3    mNeoVis.vulgivaga.2.1.chr8.g00053.t01#1    mNeoVis.vulgivaga.1.1.chr8.g00052.t01#4    mNeoVis.2.1.1.chr8_hap1.g00056.t01#5    mNeoVis.3.1.1.chr8.g00033.t01#6   mNeoVis.evergladensis.2.1.chr9.g00053.t01#2       796   125261920:   mNeoVis.3.1.1.chr8.g00036.t01#6    mNeoVis.vulgivaga.2.1.chr8.g00054.t01#1    mNeoVis.evergladensis.2.1.chr9.g00054.t01#2    mNeoVis.vulgivaga.1.1.chr8.g00053.t01#4    mNeoVis.evergladensis.1.1.chr8.g00053.t01#3   mNeoVis.2.1.1.chr8_hap1.g00057.t01#5       797   125261921:   mNeoVis.evergladensis.1.1.chr1.g01953.t01#3    mNeoVis.2.1.1.chr1_hap1.g01989.t01#5    mNeoVis.vulgivaga.2.1.chr1.g01951.t01#1    mNeoVis.3.1.1.chr1.g01944.t01#6    mNeoVis.vulgivaga.1.1.chr1.g01962.t01#4   mNeoVis.evergladensis.2.1.chr1.g01951.t01#2       798   125261922:   mNeoVis.evergladensis.1.1.chr1.g01954.t01#3    mNeoVis.evergladensis.2.1.chr1.g01952.t01#2    mNeoVis.3.1.1.chr1.g01945.t01#6    mNeoVis.vulgivaga.2.1.chr1.g01952.t01#1    mNeoVis.2.1.1.chr1_hap1.g01990.t01#5   mNeoVis.vulgivaga.1.1.chr1.g01963.t01#4       799   125261923:   mNeoVis.2.1.1.chr1_hap1.g01991.t01#5    mNeoVis.3.1.1.chr1.g01946.t01#6    mNeoVis.evergladensis.1.1.chr1.g01955.t01#3    mNeoVis.vulgivaga.1.1.chr1.g01964.t01#4    mNeoVis.evergladensis.2.1.chr1.g01953.t01#2   mNeoVis.vulgivaga.2.1.chr1.g01953.t01#1       800   125261924:   mNeoVis.vulgivaga.1.1.chr8.g00055.t01#4    mNeoVis.2.1.1.chr8_hap1.g00059.t01#5    mNeoVis.vulgivaga.2.1.chr8.g00055.t01#1    mNeoVis.evergladensis.1.1.chr8.g00054.t01#3   mNeoVis.evergladensis.2.1.chr9.g00056.t01#2       801   125261925:   mNeoVis.vulgivaga.2.1.chr1.g01954.t01#1    mNeoVis.2.1.1.chr1_hap1.g01992.t01#5    mNeoVis.evergladensis.1.1.chr1.g01956.t01#3   mNeoVis.evergladensis.2.1.chr1.g01954.t01#2       802   125261926:   mNeoVis.evergladensis.1.1.chr8.g00055.t01#3    mNeoVis.vulgivaga.2.1.chr8.g00056.t01#1    mNeoVis.3.1.1.chr8.g00034.t01#6    mNeoVis.vulgivaga.1.1.chr8.g00054.t01#4    mNeoVis.2.1.1.chr8_hap1.g00058.t01#5   mNeoVis.evergladensis.2.1.chr9.g00055.t01#2       803   125261927:   mNeoVis.evergladensis.1.1.chr1.g01958.t01#3    mNeoVis.vulgivaga.1.1.chr1.g01965.t01#4    mNeoVis.evergladensis.1.1.chr1.g01957.t01#3    mNeoVis.3.1.1.chr1.g01947.t01#6    mNeoVis.vulgivaga.2.1.chr1.g01955.t01#1    mNeoVis.evergladensis.2.1.chr1.g01956.t01#2   mNeoVis.2.1.1.chr1_hap1.g01993.t01#5     804   125261928:   mNeoVis.evergladensis.1.1.chr15.g00191.t01#3    mNeoVis.vulgivaga.2.1.chr5.g00601.t01#1    mNeoVis.2.1.1.chr15_hap1.g00603.t01#5    mNeoVis.vulgivaga.2.1.chr9.g00753.t01#1    mNeoVis.2.1.1.chr11_hap1.g00812.t01#5    mNeoVis.2.1.1.chr8_hap1.g00060.t01#5    mNeoVis.3.1.1.chr9.g00732.t01#6    mNeoVis.evergladensis.1.1.chr11.g00803.t01#3    mNeoVis.3.1.1.chr5.g00598.t01#6      805   125261929:   mNeoVis.vulgivaga.1.1.chr3.g00918.t01#4    mNeoVis.evergladensis.2.1.chr3.g00913.t01#2    mNeoVis.evergladensis.1.1.chr3.g00922.t01#3    mNeoVis.3.1.1.chr3.g00902.t01#6   mNeoVis.vulgivaga.2.1.chr3.g00929.t01#1       806   125261930:   mNeoVis.evergladensis.1.1.chr1.g01959.t01#3    mNeoVis.evergladensis.2.1.chr1.g01957.t01#2    mNeoVis.vulgivaga.2.1.chr1.g01956.t01#1    mNeoVis.vulgivaga.1.1.chr1.g01966.t01#4    mNeoVis.3.1.1.chr1.g01948.t01#6   mNeoVis.2.1.1.chr1_hap1.g01994.t01#5       807   125261931:   mNeoVis.vulgivaga.1.1.chr8.g00057.t01#4    mNeoVis.2.1.1.chr8_hap1.g00061.t01#5    mNeoVis.evergladensis.2.1.chr9.g00058.t01#2    mNeoVis.evergladensis.1.1.chr8.g00057.t01#3   mNeoVis.vulgivaga.2.1.chr8.g00058.t01#1       808   125261932:   mNeoVis.evergladensis.2.1.chr1.g01958.t01#2    mNeoVis.vulgivaga.2.1.chr1.g01957.t01#1   mNeoVis.vulgivaga.1.1.chr1.g01967.t01#4           809   125261933:   mNeoVis.vulgivaga.1.1.chr1.g01968.t01#4    mNeoVis.evergladensis.1.1.chr1.g01960.t01#3    mNeoVis.vulgivaga.2.1.chr1.g01958.t01#1    mNeoVis.3.1.1.chr1.g01949.t01#6    mNeoVis.evergladensis.2.1.chr1.g01959.t01#2   mNeoVis.2.1.1.chr1_hap1.g01995.t01#5       810   125261934:   mNeoVis.vulgivaga.2.1.chr1.g01959.t01#1    mNeoVis.evergladensis.1.1.chr1.g01961.t01#3    mNeoVis.3.1.1.chr1.g01950.t01#6    mNeoVis.evergladensis.2.1.chr1.g01960.t01#2    mNeoVis.2.1.1.chr1_hap1.g01996.t01#5   mNeoVis.vulgivaga.1.1.chr1.g01969.t01#4       811   125261935:   mNeoVis.2.1.1.chr8_hap1.g00062.t01#5    mNeoVis.vulgivaga.2.1.chr8.g00059.t01#1    mNeoVis.evergladensis.1.1.chr8.g00058.t01#3    mNeoVis.vulgivaga.1.1.chr8.g00058.t01#4   mNeoVis.evergladensis.2.1.chr9.g00059.t01#2       812   125261936:   mNeoVis.vulgivaga.2.1.chr1.g01960.t01#1    mNeoVis.vulgivaga.1.1.chr1.g01970.t01#4    mNeoVis.2.1.1.chr1_hap1.g01997.t01#5    mNeoVis.evergladensis.1.1.chr1.g01962.t01#3    mNeoVis.evergladensis.2.1.chr1.g01961.t01#2   mNeoVis.3.1.1.chr1.g01951.t01#6       813   125261937:   mNeoVis.vulgivaga.2.1.chr8.g00060.t01#1    mNeoVis.vulgivaga.1.1.chr8.g00059.t01#4    mNeoVis.evergladensis.1.1.chr8.g00059.t01#3    mNeoVis.2.1.1.chr8_hap1.g00063.t01#5   mNeoVis.evergladensis.2.1.chr9.g00060.t01#2       814   125261938:   mNeoVis.evergladensis.1.1.chr8.g00060.t01#3    mNeoVis.2.1.1.chr8_hap1.g00064.t01#5    mNeoVis.evergladensis.2.1.chr9.g00061.t01#2    mNeoVis.vulgivaga.2.1.chr8.g00061.t01#1   mNeoVis.vulgivaga.1.1.chr8.g00060.t01#4       815   125261939:   mNeoVis.evergladensis.1.1.chr1.g01964.t01#3    mNeoVis.evergladensis.2.1.chr1.g01963.t01#2    mNeoVis.2.1.1.chr1_hap1.g01999.t01#5    mNeoVis.vulgivaga.2.1.chr1.g01962.t01#1    mNeoVis.3.1.1.chr1.g01953.t01#6   mNeoVis.vulgivaga.1.1.chr1.g01971.t01#4       816   125261940:   mNeoVis.vulgivaga.1.1.chr1.g01972.t01#4    mNeoVis.3.1.1.chr1.g01954.t01#6    mNeoVis.vulgivaga.2.1.chr1.g01963.t01#1    mNeoVis.evergladensis.2.1.chr1.g01964.t01#2    mNeoVis.evergladensis.1.1.chr1.g01965.t01#3   mNeoVis.2.1.1.chr1_hap1.g02000.t01#5       817   125261941:   mNeoVis.evergladensis.2.1.chr9.g00062.t01#2    mNeoVis.2.1.1.chr8_hap1.g00065.t01#5    mNeoVis.vulgivaga.2.1.chr8.g00062.t01#1    mNeoVis.evergladensis.1.1.chr8.g00061.t01#3   mNeoVis.vulgivaga.1.1.chr8.g00061.t01#4       818   125261942:   mNeoVis.3.1.1.chr1.g01955.t01#6    mNeoVis.2.1.1.chr1_hap1.g02001.t01#5    mNeoVis.evergladensis.2.1.chr1.g01966.t01#2    mNeoVis.vulgivaga.2.1.chr1.g01964.t01#1    mNeoVis.evergladensis.1.1.chr1.g01966.t01#3    mNeoVis.evergladensis.2.1.chr1.g01965.t01#2   mNeoVis.vulgivaga.1.1.chr1.g01973.t01#4     819   125261943:   mNeoVis.vulgivaga.1.1.chr1.g01974.t01#4    mNeoVis.evergladensis.2.1.chr1.g01967.t01#2    mNeoVis.3.1.1.chr1.g01957.t01#6    mNeoVis.vulgivaga.2.1.chr1.g01965.t01#1    mNeoVis.evergladensis.1.1.chr1.g01967.t01#3    mNeoVis.3.1.1.chr1.g01956.t01#6   mNeoVis.2.1.1.chr1_hap1.g02002.t01#5     820   125261944:   mNeoVis.vulgivaga.1.1.chr1.g01975.t01#4    mNeoVis.vulgivaga.2.1.chr1.g01966.t01#1    mNeoVis.evergladensis.1.1.chr1.g01968.t01#3    mNeoVis.2.1.1.chr1_hap1.g02003.t01#5    mNeoVis.3.1.1.chr1.g01958.t01#6   mNeoVis.evergladensis.2.1.chr1.g01968.t01#2       821   125261945:   mNeoVis.vulgivaga.1.1.chr1.g01976.t01#4    mNeoVis.vulgivaga.2.1.chr1.g01967.t01#1    mNeoVis.evergladensis.2.1.chr1.g01970.t01#2    mNeoVis.evergladensis.1.1.chr1.g01970.t01#3    mNeoVis.3.1.1.chr1.g01959.t01#6   mNeoVis.2.1.1.chr1_hap1.g02004.t01#5       822   125261946:   mNeoVis.2.1.1.chr8_hap1.g00066.t01#5    mNeoVis.evergladensis.1.1.chr8.g00062.t01#3    mNeoVis.vulgivaga.2.1.chr8.g00063.t01#1    mNeoVis.vulgivaga.1.1.chr8.g00062.t01#4   mNeoVis.evergladensis.2.1.chr9.g00063.t01#2       823   125261947:   mNeoVis.evergladensis.2.1.chr9.g00064.t01#2    mNeoVis.vulgivaga.1.1.chr8.g00063.t01#4    mNeoVis.vulgivaga.2.1.chr8.g00064.t01#1    mNeoVis.2.1.1.chr8_hap1.g00067.t01#5   mNeoVis.evergladensis.1.1.chr8.g00063.t01#3       824   125261948:   mNeoVis.evergladensis.1.1.chr7.g00133.t01#3    mNeoVis.vulgivaga.1.1.chr7.g00135.t01#4    mNeoVis.evergladensis.2.1.chr7.g00131.t01#2    mNeoVis.vulgivaga.2.1.chr7.g00131.t01#1    mNeoVis.2.1.1.chr7_hap1.g00131.t01#5   mNeoVis.3.1.1.chr7.g00113.t01#6       825   125261949:   mNeoVis.evergladensis.1.1.chr1.g01971.t01#3    mNeoVis.vulgivaga.2.1.chr1.g01968.t01#1    mNeoVis.2.1.1.chr1_hap1.g02005.t01#5    mNeoVis.evergladensis.2.1.chr1.g01971.t01#2    mNeoVis.3.1.1.chr1.g01960.t01#6   mNeoVis.vulgivaga.1.1.chr1.g01977.t01#4       826   125261950:   mNeoVis.evergladensis.1.1.chr1.g01972.t01#3    mNeoVis.vulgivaga.2.1.chr1.g01969.t01#1    mNeoVis.2.1.1.chr1_hap1.g02006.t01#5    mNeoVis.evergladensis.2.1.chr1.g01972.t01#2    mNeoVis.3.1.1.chr1.g01961.t01#6   mNeoVis.vulgivaga.1.1.chr1.g01978.t01#4       827   125261951:  mNeoVis.vulgivaga.2.1.chr8.g00065.t01#1               828   125261952:   mNeoVis.vulgivaga.1.1.chr8.g00064.t01#4    mNeoVis.evergladensis.2.1.chr9.g00065.t01#2    mNeoVis.2.1.1.chr8_hap1.g00068.t01#5    mNeoVis.evergladensis.1.1.chr8.g00064.t01#3   mNeoVis.vulgivaga.2.1.chr8.g00066.t01#1       829   125261953:   mNeoVis.vulgivaga.1.1.chr1.g01979.t01#4    mNeoVis.vulgivaga.2.1.chr1.g01970.t01#1    mNeoVis.2.1.1.chr1_hap1.g02007.t01#5    mNeoVis.evergladensis.1.1.chr1.g01973.t01#3    mNeoVis.3.1.1.chr1.g01962.t01#6   mNeoVis.evergladensis.2.1.chr1.g01973.t01#2       830   125261954:   mNeoVis.vulgivaga.2.1.chr12.g00452.t01#1    mNeoVis.evergladensis.1.1.chr12.g00446.t01#3    mNeoVis.3.1.1.chr12.g00447.t01#6    mNeoVis.vulgivaga.1.1.chr12.g00449.t01#4    mNeoVis.2.1.1.chr12_hap1.g00490.t01#5   mNeoVis.evergladensis.2.1.chr12.g00441.t01#2       831   125261955:   mNeoVis.evergladensis.2.1.chr1.g01974.t01#2    mNeoVis.2.1.1.chr1_hap1.g02008.t01#5    mNeoVis.evergladensis.1.1.chr1.g01974.t01#3    mNeoVis.vulgivaga.1.1.chr1.g01980.t01#4    mNeoVis.3.1.1.chr1.g01963.t01#6   mNeoVis.vulgivaga.2.1.chr1.g01971.t01#1       832   125261956:   mNeoVis.evergladensis.1.1.chr8.g00065.t01#3    mNeoVis.vulgivaga.1.1.chr8.g00065.t01#4    mNeoVis.vulgivaga.2.1.chr8.g00067.t01#1    mNeoVis.2.1.1.chr8_hap1.g00069.t01#5   mNeoVis.evergladensis.2.1.chr9.g00066.t01#2       833   125261957:   mNeoVis.3.1.1.chr1.g01868.t01#6    mNeoVis.vulgivaga.2.1.chr1.g01882.t01#1    mNeoVis.evergladensis.2.1.chr1.g01885.t01#2    mNeoVis.evergladensis.1.1.chr1.g01886.t01#3    mNeoVis.2.1.1.chr1_hap1.g01917.t01#5   mNeoVis.vulgivaga.1.1.chr1.g01892.t01#4       834   125261958:   mNeoVis.vulgivaga.2.1.chr8.g00068.t01#1    mNeoVis.2.1.1.chr8_hap1.g00070.t01#5    mNeoVis.evergladensis.2.1.chr9.g00067.t01#2   mNeoVis.evergladensis.1.1.chr8.g00066.t01#3       835   125261959:   mNeoVis.evergladensis.1.1.chr1.g01887.t01#3    mNeoVis.2.1.1.chr1_hap1.g01918.t01#5    mNeoVis.vulgivaga.1.1.chr1.g01893.t01#4    mNeoVis.evergladensis.2.1.chr1.g01886.t01#2    mNeoVis.vulgivaga.2.1.chr1.g01883.t01#1   mNeoVis.3.1.1.chr1.g01869.t01#6       836   125261960:   mNeoVis.evergladensis.2.1.chr1.g01887.t01#2    mNeoVis.vulgivaga.2.1.chr1.g01884.t01#1    mNeoVis.3.1.1.chr1.g01870.t01#6    mNeoVis.evergladensis.1.1.chr1.g01888.t01#3    mNeoVis.2.1.1.chr1_hap1.g01919.t01#5   mNeoVis.vulgivaga.1.1.chr1.g01894.t01#4       837   125261961:   mNeoVis.vulgivaga.2.1.chr1.g01885.t01#1   mNeoVis.evergladensis.1.1.chr1.g01889.t01#3           838   125261962:   mNeoVis.evergladensis.1.1.chr1.g01891.t01#3    mNeoVis.vulgivaga.2.1.chr1.g01886.t01#1    mNeoVis.2.1.1.chr1_hap1.g01922.t01#5    mNeoVis.3.1.1.chr1.g01872.t01#6    mNeoVis.vulgivaga.1.1.chr1.g01896.t01#4   mNeoVis.evergladensis.2.1.chr1.g01889.t01#2       839   125261963:   mNeoVis.vulgivaga.2.1.chr7.g02499.t01#1    mNeoVis.evergladensis.1.1.chr7.g02473.t01#3    mNeoVis.2.1.1.chr7_hap1.g02665.t01#5    mNeoVis.evergladensis.2.1.chr7.g02477.t01#2   mNeoVis.vulgivaga.1.1.chr7.g02479.t01#4       840   125261964:   mNeoVis.evergladensis.2.1.chr7.g02478.t01#2    mNeoVis.vulgivaga.1.1.chr7.g02480.t01#4    mNeoVis.vulgivaga.2.1.chr7.g02500.t01#1    mNeoVis.evergladensis.1.1.chr7.g02474.t01#3   mNeoVis.2.1.1.chr7_hap1.g02666.t01#5       841   125261965:   mNeoVis.vulgivaga.1.1.chr7.g02481.t01#4    mNeoVis.vulgivaga.2.1.chr7.g02501.t01#1    mNeoVis.evergladensis.2.1.chr7.g02479.t01#2    mNeoVis.2.1.1.chr7_hap1.g02667.t01#5   mNeoVis.evergladensis.1.1.chr7.g02475.t01#3       842   125261966:   mNeoVis.vulgivaga.2.1.chr7.g02502.t01#1    mNeoVis.evergladensis.1.1.chr7.g02476.t01#3    mNeoVis.2.1.1.chr7_hap1.g02668.t01#5    mNeoVis.evergladensis.2.1.chr7.g02480.t01#2   mNeoVis.vulgivaga.1.1.chr7.g02482.t01#4       843   125261967:   mNeoVis.vulgivaga.1.1.chr1.g01897.t01#4    mNeoVis.evergladensis.2.1.chr1.g01890.t01#2    mNeoVis.3.1.1.chr1.g01873.t01#6    mNeoVis.2.1.1.chr1_hap1.g01923.t01#5    mNeoVis.evergladensis.1.1.chr1.g01892.t01#3   mNeoVis.vulgivaga.2.1.chr1.g01887.t01#1       844   125261968:   mNeoVis.2.1.1.chr7_hap1.g02669.t01#5    mNeoVis.evergladensis.2.1.chr7.g02481.t01#2    mNeoVis.vulgivaga.1.1.chr7.g02483.t01#4    mNeoVis.vulgivaga.2.1.chr7.g02503.t01#1   mNeoVis.evergladensis.1.1.chr7.g02477.t01#3       845   125261969:   mNeoVis.evergladensis.2.1.chr7.g00132.t01#2    mNeoVis.evergladensis.1.1.chr7.g00135.t01#3    mNeoVis.vulgivaga.2.1.chr7.g00132.t01#1    mNeoVis.vulgivaga.1.1.chr7.g00136.t01#4    mNeoVis.2.1.1.chr7_hap1.g00132.t01#5   mNeoVis.3.1.1.chr7.g00114.t01#6       846   125261970:   mNeoVis.evergladensis.1.1.chr15.g00424.t01#3    mNeoVis.2.1.1.chr15_hap1.g00210.t01#5    mNeoVis.vulgivaga.2.1.chr15.g00190.t01#1    mNeoVis.3.1.1.chr15.g00184.t01#6    mNeoVis.vulgivaga.1.1.chr15.g00186.t01#4   mNeoVis.evergladensis.2.1.chr15.g00189.t01#2       847   125261971:   mNeoVis.2.1.1.chr7_hap1.g02670.t01#5    mNeoVis.evergladensis.1.1.chr7.g02478.t01#3    mNeoVis.evergladensis.2.1.chr7.g02482.t01#2    mNeoVis.vulgivaga.1.1.chr7.g02484.t01#4   mNeoVis.vulgivaga.2.1.chr7.g02504.t01#1       848   125261972:   mNeoVis.2.1.1.chr1_hap1.g01864.t01#5    mNeoVis.3.1.1.chr1.g01817.t01#6    mNeoVis.vulgivaga.1.1.chr1.g01833.t01#4    mNeoVis.evergladensis.2.1.chr1.g01830.t01#2    mNeoVis.evergladensis.1.1.chr1.g01829.t01#3   mNeoVis.vulgivaga.2.1.chr1.g01827.t01#1       849   125261973:   mNeoVis.2.1.1.chr1_hap1.g01865.t01#5    mNeoVis.evergladensis.1.1.chr1.g01830.t01#3    mNeoVis.evergladensis.2.1.chr1.g01831.t01#2    mNeoVis.vulgivaga.2.1.chr1.g01828.t01#1   mNeoVis.3.1.1.chr1.g01818.t01#6       850   125261974:   mNeoVis.vulgivaga.2.1.chr7.g02505.t01#1    mNeoVis.2.1.1.chr7_hap1.g02671.t01#5    mNeoVis.evergladensis.1.1.chr7.g02479.t01#3    mNeoVis.vulgivaga.1.1.chr7.g02485.t01#4   mNeoVis.evergladensis.2.1.chr7.g02483.t01#2       851   125261975:   mNeoVis.3.1.1.chr1.g01819.t01#6    mNeoVis.evergladensis.2.1.chr1.g01832.t01#2    mNeoVis.vulgivaga.2.1.chr1.g01829.t01#1    mNeoVis.2.1.1.chr1_hap1.g01866.t01#5    mNeoVis.evergladensis.1.1.chr1.g01831.t01#3   mNeoVis.vulgivaga.1.1.chr1.g01835.t01#4       852   125261976:   mNeoVis.2.1.1.chr2_hap1.g00914.t01#5    mNeoVis.evergladensis.2.1.chr5.g00869.t01#2    mNeoVis.vulgivaga.1.1.chr5.g00838.t01#4    mNeoVis.vulgivaga.2.1.chr10.g00697.t01#1    mNeoVis.2.1.1.chr7_hap1.g02672.t01#5    mNeoVis.vulgivaga.2.1.chr7.g02506.t01#1    mNeoVis.2.1.1.chr9_hap1.g00714.t01#5    mNeoVis.vulgivaga.1.1.chr10.g00692.t01#4    mNeoVis.vulgivaga.2.1.chr5.g00858.t01#1      853   125261977:   mNeoVis.2.1.1.chr11_hap1.g01200.t01#5    mNeoVis.vulgivaga.1.1.chr11.g01170.t01#4    mNeoVis.vulgivaga.2.1.chr7.g02440.t01#1    mNeoVis.vulgivaga.1.1.chr7.g02420.t01#4   mNeoVis.3.1.1.chr7.g02274.t01#6       854   125261978:   mNeoVis.3.1.1.chr7.g00644.t01#6    mNeoVis.vulgivaga.1.1.chr7.g00684.t01#4    mNeoVis.vulgivaga.1.1.chr7.g00731.t01#4    mNeoVis.3.1.1.chr7.g00685.t01#6   mNeoVis.vulgivaga.2.1.chr7.g00719.t01#1       855   125261979:   mNeoVis.2.1.1.chr7_hap1.g00723.t01#5    mNeoVis.2.1.1.chr7_hap1.g00679.t01#5    mNeoVis.2.1.1.chr7_hap1.g00726.t01#5   mNeoVis.2.1.1.chr7_hap1.g00721.t01#5         856   125261980:   mNeoVis.evergladensis.2.1.chr7.g02485.t01#2    mNeoVis.vulgivaga.2.1.chr7.g02507.t01#1   mNeoVis.2.1.1.chr7_hap1.g02673.t01#5           857   125261981:   mNeoVis.vulgivaga.2.1.chr7.g00724.t01#1    mNeoVis.vulgivaga.1.1.chr7.g00730.t01#4   mNeoVis.evergladensis.1.1.chr7.g02482.t01#3         858   125261982:   mNeoVis.evergladensis.2.1.chr11.g01173.t01#2   mNeoVis.vulgivaga.2.1.chr11.g01199.t01#1           859   125261983:   mNeoVis.2.1.1.chr7_hap1.g00732.t01#5   mNeoVis.vulgivaga.2.1.chr7.g02439.t01#1             860   125261984:  mNeoVis.evergladensis.1.1.chr7.g02481.t01#3             861   125261985:  mNeoVis.2.1.1.chr7_hap1.g02610.t01#5               862   125261986:  mNeoVis.3.1.1.chr7.g00684.t01#6               863   125261987:   mNeoVis.evergladensis.2.1.chr8.g00380.t01#2    mNeoVis.vulgivaga.1.1.chr7.g02487.t01#4    mNeoVis.evergladensis.1.1.chr9.g00387.t01#3    mNeoVis.2.1.1.chr10_hap1.g00398.t01#5    mNeoVis.vulgivaga.1.1.chr9.g00379.t01#4    mNeoVis.3.1.1.chr7.g02313.t01#6   mNeoVis.3.1.1.chr9.g00367.t01#6       864   125261988:   mNeoVis.vulgivaga.2.1.chr9.g00382.t01#1    mNeoVis.evergladensis.2.1.chr7.g02487.t01#2    mNeoVis.evergladensis.1.1.chr7.g02483.t01#3    mNeoVis.2.1.1.chr7_hap1.g02674.t01#5   mNeoVis.vulgivaga.2.1.chr7.g02508.t01#1       865   125261989:   mNeoVis.evergladensis.2.1.chr7.g02488.t01#2    mNeoVis.evergladensis.1.1.chr7.g02485.t01#3    mNeoVis.2.1.1.chr7_hap1.g02675.t01#5    mNeoVis.vulgivaga.2.1.chr7.g02509.t01#1    mNeoVis.vulgivaga.1.1.chr7.g02488.t01#4   mNeoVis.evergladensis.1.1.chr7.g02484.t01#3       866   125261990:   mNeoVis.2.1.1.chr7_hap1.g02676.t01#5    mNeoVis.vulgivaga.1.1.chr7.g02489.t01#4    mNeoVis.vulgivaga.2.1.chr7.g02510.t01#1    mNeoVis.evergladensis.1.1.chr7.g02486.t01#3   mNeoVis.evergladensis.2.1.chr7.g02489.t01#2       867   125261991:   mNeoVis.3.1.1.chr7.g02314.t01#6    mNeoVis.evergladensis.1.1.chr7.g02487.t01#3    mNeoVis.evergladensis.2.1.chr7.g02490.t01#2    mNeoVis.2.1.1.chr7_hap1.g02677.t01#5    mNeoVis.vulgivaga.1.1.chr7.g02490.t01#4   mNeoVis.vulgivaga.2.1.chr7.g02511.t01#1       868   125261992:   mNeoVis.evergladensis.2.1.chr1.g01833.t01#2    mNeoVis.evergladensis.1.1.chr1.g01832.t01#3    mNeoVis.2.1.1.chr1_hap1.g01867.t01#5    mNeoVis.vulgivaga.1.1.chr1.g01836.t01#4    mNeoVis.vulgivaga.2.1.chr1.g01830.t01#1   mNeoVis.3.1.1.chr1.g01820.t01#6       869   125261993:   mNeoVis.evergladensis.2.1.chr7.g02491.t01#2    mNeoVis.vulgivaga.1.1.chr7.g02491.t01#4    mNeoVis.evergladensis.1.1.chr7.g02488.t01#3    mNeoVis.2.1.1.chr7_hap1.g02678.t01#5   mNeoVis.vulgivaga.2.1.chr7.g02512.t01#1       870   125261994:   mNeoVis.vulgivaga.2.1.chr1.g01831.t01#1    mNeoVis.evergladensis.2.1.chr1.g01834.t01#2    mNeoVis.2.1.1.chr1_hap1.g01868.t01#5    mNeoVis.3.1.1.chr1.g01821.t01#6    mNeoVis.vulgivaga.1.1.chr1.g01837.t01#4   mNeoVis.evergladensis.1.1.chr1.g01833.t01#3       871   125261995:   mNeoVis.3.1.1.chr1.g01822.t01#6    mNeoVis.vulgivaga.1.1.chr1.g01838.t01#4    mNeoVis.evergladensis.1.1.chr1.g01834.t01#3    mNeoVis.2.1.1.chr1_hap1.g01869.t01#5    mNeoVis.vulgivaga.2.1.chr1.g01832.t01#1   mNeoVis.evergladensis.2.1.chr1.g01835.t01#2       872   125261996:   mNeoVis.evergladensis.2.1.chr7.g02492.t01#2    mNeoVis.vulgivaga.1.1.chr7.g02492.t01#4    mNeoVis.2.1.1.chr7_hap1.g02679.t01#5    mNeoVis.evergladensis.1.1.chr7.g02489.t01#3   mNeoVis.vulgivaga.2.1.chr7.g02513.t01#1       873   125261997:   mNeoVis.vulgivaga.1.1.chr7.g02493.t01#4    mNeoVis.2.1.1.chr7_hap1.g02680.t01#5    mNeoVis.evergladensis.1.1.chr7.g02490.t01#3    mNeoVis.vulgivaga.2.1.chr7.g02514.t01#1   mNeoVis.evergladensis.2.1.chr7.g02493.t01#2       874   125261998:   mNeoVis.2.1.1.chr1_hap1.g01870.t01#5    mNeoVis.evergladensis.2.1.chr1.g01836.t01#2    mNeoVis.vulgivaga.2.1.chr1.g01833.t01#1    mNeoVis.evergladensis.1.1.chr1.g01835.t01#3    mNeoVis.vulgivaga.1.1.chr1.g01839.t01#4   mNeoVis.3.1.1.chr1.g01823.t01#6       875   125261999:   mNeoVis.3.1.1.chr1.g01824.t01#6    mNeoVis.evergladensis.2.1.chr1.g01837.t01#2    mNeoVis.2.1.1.chr1_hap1.g01871.t01#5    mNeoVis.vulgivaga.2.1.chr1.g01834.t01#1    mNeoVis.evergladensis.1.1.chr1.g01836.t01#3   mNeoVis.vulgivaga.1.1.chr1.g01840.t01#4       876   125262000:   mNeoVis.evergladensis.2.1.chr1.g01838.t01#2    mNeoVis.3.1.1.chr1.g01825.t01#6    mNeoVis.vulgivaga.2.1.chr1.g01835.t01#1    mNeoVis.evergladensis.1.1.chr1.g01837.t01#3    mNeoVis.2.1.1.chr1_hap1.g01872.t01#5   mNeoVis.vulgivaga.1.1.chr1.g01841.t01#4       877   125262001:  mNeoVis.vulgivaga.2.1.chr1.g01836.t01#1               878   125262002:   mNeoVis.vulgivaga.1.1.chr1.g01755.t01#4    mNeoVis.vulgivaga.2.1.chr1.g01748.t01#1    mNeoVis.2.1.1.chr1_hap1.g01783.t01#5    mNeoVis.evergladensis.1.1.chr1.g01751.t01#3   mNeoVis.evergladensis.2.1.chr1.g01750.t01#2       879   125262003:   mNeoVis.evergladensis.1.1.chr7.g02492.t01#3    mNeoVis.vulgivaga.2.1.chr7.g02515.t01#1    mNeoVis.vulgivaga.1.1.chr7.g02494.t01#4    mNeoVis.evergladensis.2.1.chr7.g02495.t01#2   mNeoVis.2.1.1.chr7_hap1.g02682.t01#5       880   125262004:   mNeoVis.2.1.1.chr7_hap1.g02683.t01#5    mNeoVis.2.1.1.chr7_hap1.g02681.t01#5   mNeoVis.3.1.1.chr7.g02318.t01#6           881   125262005:   mNeoVis.evergladensis.1.1.chr7.g02494.t01#3    mNeoVis.2.1.1.chr7_hap1.g02685.t01#5    mNeoVis.evergladensis.2.1.chr7.g02496.t01#2    mNeoVis.vulgivaga.2.1.chr7.g02516.t01#1   mNeoVis.vulgivaga.1.1.chr7.g02496.t01#4       882   125262006:   mNeoVis.vulgivaga.2.1.chr7.g00133.t01#1    mNeoVis.2.1.1.chr7_hap1.g00133.t01#5    mNeoVis.evergladensis.1.1.chr7.g00136.t01#3    mNeoVis.3.1.1.chr7.g00115.t01#6    mNeoVis.evergladensis.2.1.chr7.g00133.t01#2   mNeoVis.vulgivaga.1.1.chr7.g00137.t01#4       883   125262007:   mNeoVis.evergladensis.1.1.chr7.g00391.t01#3    mNeoVis.vulgivaga.1.1.chr7.g00391.t01#4    mNeoVis.evergladensis.2.1.chr7.g00384.t01#2    mNeoVis.vulgivaga.2.1.chr7.g00383.t01#1   mNeoVis.2.1.1.chr7_hap1.g00379.t01#5       884   125262008:   mNeoVis.evergladensis.1.1.chr1.g01753.t01#3    mNeoVis.3.1.1.chr1.g01743.t01#6    mNeoVis.2.1.1.chr1_hap1.g01785.t01#5    mNeoVis.vulgivaga.2.1.chr1.g01749.t01#1    mNeoVis.evergladensis.2.1.chr1.g01752.t01#2   mNeoVis.vulgivaga.1.1.chr1.g01757.t01#4       885   125262009:   mNeoVis.evergladensis.2.1.chr7.g02497.t01#2    mNeoVis.3.1.1.chr7.g02319.t01#6    mNeoVis.vulgivaga.2.1.chr7.g02517.t01#1    mNeoVis.2.1.1.chr7_hap1.g02686.t01#5   mNeoVis.evergladensis.1.1.chr7.g02495.t01#3       886   125262010:   mNeoVis.vulgivaga.2.1.chr7.g02518.t01#1    mNeoVis.evergladensis.1.1.chr7.g02496.t01#3   mNeoVis.3.1.1.chr7.g02320.t01#6           887   125262011:   mNeoVis.3.1.1.chr7.g02321.t01#6    mNeoVis.2.1.1.chr7_hap1.g02688.t01#5    mNeoVis.vulgivaga.2.1.chr7.g02519.t01#1    mNeoVis.evergladensis.1.1.chr7.g02497.t01#3    mNeoVis.vulgivaga.1.1.chr7.g02499.t01#4   mNeoVis.evergladensis.2.1.chr7.g02499.t01#2       888   125262012:   mNeoVis.vulgivaga.2.1.chr7.g00134.t01#1    mNeoVis.evergladensis.2.1.chr7.g00134.t01#2    mNeoVis.3.1.1.chr7.g00116.t01#6    mNeoVis.vulgivaga.1.1.chr7.g00138.t01#4    mNeoVis.2.1.1.chr7_hap1.g00134.t01#5   mNeoVis.evergladensis.1.1.chr7.g00137.t01#3       889   125262013:  mNeoVis.vulgivaga.2.1.chr1.g01750.t01#1               890   125262014:   mNeoVis.vulgivaga.1.1.chr1.g01759.t01#4    mNeoVis.evergladensis.2.1.chr1.g01754.t01#2    mNeoVis.evergladensis.1.1.chr1.g01755.t01#3    mNeoVis.3.1.1.chr1.g01745.t01#6    mNeoVis.vulgivaga.2.1.chr1.g01751.t01#1   mNeoVis.2.1.1.chr1_hap1.g01787.t01#5       891   125262015:   mNeoVis.vulgivaga.2.1.chr7.g02520.t01#1    mNeoVis.evergladensis.2.1.chr7.g02500.t01#2    mNeoVis.3.1.1.chr7.g02322.t01#6    mNeoVis.2.1.1.chr7_hap1.g02689.t01#5    mNeoVis.evergladensis.1.1.chr7.g02498.t01#3   mNeoVis.vulgivaga.1.1.chr7.g02500.t01#4       892   125262016:   mNeoVis.vulgivaga.2.1.chr1.g01752.t01#1    mNeoVis.vulgivaga.1.1.chr1.g01761.t01#4    mNeoVis.evergladensis.1.1.chr1.g01757.t01#3    mNeoVis.3.1.1.chr1.g01748.t01#6    mNeoVis.evergladensis.2.1.chr1.g01756.t01#2   mNeoVis.2.1.1.chr1_hap1.g01790.t01#5       893   125262017:   mNeoVis.vulgivaga.2.1.chr1.g01753.t01#1    mNeoVis.evergladensis.1.1.chr1.g01758.t01#3    mNeoVis.evergladensis.2.1.chr1.g01757.t01#2    mNeoVis.2.1.1.chr1_hap1.g01791.t01#5    mNeoVis.3.1.1.chr1.g01749.t01#6   mNeoVis.vulgivaga.1.1.chr1.g01762.t01#4       894   125262018:   mNeoVis.evergladensis.2.1.chr7.g02501.t01#2    mNeoVis.3.1.1.chr7.g02323.t01#6    mNeoVis.2.1.1.chr7_hap1.g02690.t01#5    mNeoVis.evergladensis.1.1.chr7.g02499.t01#3    mNeoVis.vulgivaga.1.1.chr7.g02501.t01#4   mNeoVis.vulgivaga.2.1.chr7.g02521.t01#1       895   125262019:   mNeoVis.evergladensis.2.1.chr7.g02502.t01#2    mNeoVis.vulgivaga.1.1.chr7.g02502.t01#4    mNeoVis.3.1.1.chr7.g02324.t01#6    mNeoVis.2.1.1.chr7_hap1.g02691.t01#5    mNeoVis.evergladensis.1.1.chr7.g02500.t01#3   mNeoVis.vulgivaga.2.1.chr7.g02522.t01#1       896   125262020:   mNeoVis.3.1.1.chr7.g02325.t01#6    mNeoVis.vulgivaga.2.1.chr7.g02523.t01#1    mNeoVis.2.1.1.chr7_hap1.g02692.t01#5    mNeoVis.evergladensis.1.1.chr7.g02501.t01#3    mNeoVis.evergladensis.2.1.chr7.g02503.t01#2   mNeoVis.vulgivaga.1.1.chr7.g02503.t01#4       897   125262021:   mNeoVis.2.1.1.chr7_hap1.g02693.t01#5    mNeoVis.vulgivaga.2.1.chr7.g02524.t01#1    mNeoVis.3.1.1.chr7.g02326.t01#6    mNeoVis.evergladensis.2.1.chr7.g02504.t01#2   mNeoVis.vulgivaga.1.1.chr7.g02504.t01#4       898   125262022:   mNeoVis.3.1.1.chr1.g01750.t01#6    mNeoVis.vulgivaga.2.1.chr1.g01754.t01#1    mNeoVis.vulgivaga.1.1.chr1.g01763.t01#4    mNeoVis.evergladensis.2.1.chr1.g01758.t01#2    mNeoVis.evergladensis.1.1.chr1.g01759.t01#3   mNeoVis.2.1.1.chr1_hap1.g01792.t01#5       899   125262023:   mNeoVis.evergladensis.2.1.chr9.g00001.t01#2    mNeoVis.vulgivaga.1.1.chr8.g00001.t01#4    mNeoVis.vulgivaga.2.1.chr8.g00001.t01#1    mNeoVis.2.1.1.chr8_hap1.g00002.t01#5   mNeoVis.evergladensis.1.1.chr8.g00001.t01#3       900   125262024:   mNeoVis.2.1.1.chr8_hap1.g00003.t01#5   mNeoVis.vulgivaga.2.1.chr8.g00002.t01#1             901   125262025:  mNeoVis.vulgivaga.2.1.chr8.g00003.t01#1               902   125262026:   mNeoVis.vulgivaga.2.1.chr8.g00004.t01#1    mNeoVis.2.1.1.chr8_hap1.g00005.t01#5    mNeoVis.3.1.1.chr8.g00003.t01#6    mNeoVis.evergladensis.2.1.chr9.g00003.t01#2    mNeoVis.vulgivaga.1.1.chr8.g00003.t01#4   mNeoVis.evergladensis.1.1.chr8.g00003.t01#3       903   125262027:   mNeoVis.2.1.1.chr1_hap1.g01793.t01#5    mNeoVis.evergladensis.2.1.chr1.g01759.t01#2    mNeoVis.evergladensis.1.1.chr1.g01760.t01#3    mNeoVis.3.1.1.chr1.g01751.t01#6    mNeoVis.vulgivaga.1.1.chr1.g01764.t01#4   mNeoVis.vulgivaga.2.1.chr1.g01755.t01#1       904   125262028:   mNeoVis.3.1.1.chr1.g01752.t01#6    mNeoVis.vulgivaga.1.1.chr1.g01765.t01#4    mNeoVis.evergladensis.1.1.chr1.g01761.t01#3    mNeoVis.2.1.1.chr1_hap1.g01794.t01#5    mNeoVis.evergladensis.2.1.chr1.g01760.t01#2   mNeoVis.vulgivaga.2.1.chr1.g01756.t01#1       905   125262029:  mNeoVis.vulgivaga.2.1.chr1.g01757.t01#1               906   125262030:   mNeoVis.evergladensis.2.1.chr9.g00004.t01#2    mNeoVis.2.1.1.chr8_hap1.g00006.t01#5    mNeoVis.3.1.1.chr8.g00004.t01#6    mNeoVis.vulgivaga.1.1.chr8.g00004.t01#4    mNeoVis.vulgivaga.2.1.chr8.g00005.t01#1   mNeoVis.evergladensis.1.1.chr8.g00004.t01#3       907   125262031:   mNeoVis.vulgivaga.1.1.chr1.g01766.t01#4    mNeoVis.2.1.1.chr1_hap1.g01795.t01#5    mNeoVis.vulgivaga.2.1.chr1.g01758.t01#1    mNeoVis.evergladensis.1.1.chr1.g01762.t01#3    mNeoVis.3.1.1.chr1.g01753.t01#6   mNeoVis.evergladensis.2.1.chr1.g01761.t01#2       908   125262032:   mNeoVis.2.1.1.chr8_hap1.g00007.t01#5    mNeoVis.3.1.1.chr8.g00006.t01#6    mNeoVis.vulgivaga.1.1.chr8.g00005.t01#4    mNeoVis.evergladensis.1.1.chr8.g00005.t01#3    mNeoVis.evergladensis.2.1.chr9.g00005.t01#2   mNeoVis.vulgivaga.2.1.chr8.g00006.t01#1       909   125262033:   mNeoVis.3.1.1.chr8.g00007.t01#6    mNeoVis.2.1.1.chr8_hap1.g00008.t01#5    mNeoVis.evergladensis.1.1.chr8.g00006.t01#3    mNeoVis.vulgivaga.1.1.chr8.g00006.t01#4    mNeoVis.vulgivaga.2.1.chr8.g00007.t01#1   mNeoVis.evergladensis.2.1.chr9.g00006.t01#2       910   125262034:   mNeoVis.vulgivaga.2.1.chr1.g01759.t01#1    mNeoVis.3.1.1.chr1.g01754.t01#6    mNeoVis.evergladensis.1.1.chr1.g01763.t01#3    mNeoVis.2.1.1.chr1_hap1.g01796.t01#5    mNeoVis.evergladensis.2.1.chr1.g01762.t01#2   mNeoVis.vulgivaga.1.1.chr1.g01767.t01#4       911   125262035:   mNeoVis.evergladensis.2.1.chr2.g00178.t01#2    mNeoVis.vulgivaga.2.1.chr2.g00177.t01#1   mNeoVis.3.1.1.chr2.g00131.t01#6           912   125262036:   mNeoVis.vulgivaga.2.1.chr8.g00008.t01#1    mNeoVis.3.1.1.chr8.g00008.t01#6    mNeoVis.vulgivaga.1.1.chr8.g00007.t01#4    mNeoVis.evergladensis.2.1.chr9.g00008.t01#2    mNeoVis.evergladensis.1.1.chr8.g00008.t01#3   mNeoVis.2.1.1.chr8_hap1.g00010.t01#5       913   125262037:   mNeoVis.vulgivaga.2.1.chr15.g00377.t01#1    mNeoVis.evergladensis.1.1.chr15.g00245.t01#3    mNeoVis.evergladensis.2.1.chr15.g00368.t01#2    mNeoVis.2.1.1.chr15_hap1.g00502.t01#5   mNeoVis.vulgivaga.1.1.chr15.g00369.t01#4       914   125262038:   mNeoVis.evergladensis.2.1.chr1.g01763.t01#2    mNeoVis.2.1.1.chr1_hap1.g01797.t01#5    mNeoVis.vulgivaga.1.1.chr1.g01768.t01#4    mNeoVis.evergladensis.1.1.chr1.g01764.t01#3    mNeoVis.vulgivaga.2.1.chr1.g01760.t01#1   mNeoVis.3.1.1.chr1.g01755.t01#6       915   125262039:   mNeoVis.2.1.1.chr1_hap1.g01799.t01#5    mNeoVis.evergladensis.2.1.chr1.g01765.t01#2    mNeoVis.evergladensis.1.1.chr1.g01766.t01#3    mNeoVis.2.1.1.chr1_hap1.g01801.t01#5   mNeoVis.vulgivaga.2.1.chr10.g00911.t01#1       916   125262040:   mNeoVis.evergladensis.1.1.chr1.g01765.t01#3    mNeoVis.vulgivaga.1.1.chr1.g01770.t01#4    mNeoVis.3.1.1.chr1.g01756.t01#6   mNeoVis.vulgivaga.2.1.chr1.g01761.t01#1         917   125262041:   mNeoVis.evergladensis.1.1.chr1.g01768.t01#3    mNeoVis.2.1.1.chr1_hap1.g01803.t01#5    mNeoVis.vulgivaga.1.1.chr1.g01772.t01#4    mNeoVis.vulgivaga.2.1.chr1.g01762.t01#1   mNeoVis.evergladensis.2.1.chr1.g01767.t01#2       918   125262042:   mNeoVis.evergladensis.2.1.chr9.g00009.t01#2    mNeoVis.2.1.1.chr8_hap1.g00011.t01#5    mNeoVis.vulgivaga.2.1.chr8.g00009.t01#1    mNeoVis.vulgivaga.1.1.chr8.g00008.t01#4    mNeoVis.3.1.1.chr8.g00009.t01#6   mNeoVis.evergladensis.1.1.chr8.g00009.t01#3       919   125262043:   mNeoVis.vulgivaga.1.1.chr1.g01773.t01#4    mNeoVis.vulgivaga.2.1.chr1.g01763.t01#1    mNeoVis.evergladensis.2.1.chr1.g01768.t01#2    mNeoVis.2.1.1.chr1_hap1.g01804.t01#5    mNeoVis.3.1.1.chr1.g01757.t01#6   mNeoVis.evergladensis.1.1.chr1.g01769.t01#3       920   125262044:   mNeoVis.3.1.1.chr1.g01758.t01#6    mNeoVis.2.1.1.chr1_hap1.g01805.t01#5    mNeoVis.evergladensis.2.1.chr1.g01769.t01#2    mNeoVis.vulgivaga.2.1.chr1.g01764.t01#1    mNeoVis.evergladensis.1.1.chr1.g01770.t01#3   mNeoVis.vulgivaga.1.1.chr1.g01774.t01#4       921   125262045:   mNeoVis.evergladensis.1.1.chr1.g01771.t01#3    mNeoVis.evergladensis.2.1.chr1.g01770.t01#2    mNeoVis.3.1.1.chr1.g01759.t01#6    mNeoVis.2.1.1.chr1_hap1.g01806.t01#5    mNeoVis.vulgivaga.2.1.chr1.g01765.t01#1   mNeoVis.vulgivaga.1.1.chr1.g01775.t01#4       922   125262046:   mNeoVis.vulgivaga.1.1.chr13.g00616.t01#4    mNeoVis.evergladensis.1.1.chr13.g00627.t01#3    mNeoVis.2.1.1.chr13_hap1.g00655.t01#5    mNeoVis.evergladensis.2.1.chr13.g00628.t01#2    mNeoVis.3.1.1.chr13.g00580.t01#6   mNeoVis.vulgivaga.2.1.chr13.g00630.t01#1       923   125262047:   mNeoVis.vulgivaga.2.1.chr8.g00010.t01#1   mNeoVis.evergladensis.2.1.chr9.g00011.t01#2           924   125262048:   mNeoVis.2.1.1.chr8_hap1.g00012.t01#5   mNeoVis.vulgivaga.1.1.chr8.g00009.t01#4             925   125262049:  mNeoVis.3.1.1.chr8.g00010.t01#6               926   125262050:   mNeoVis.vulgivaga.1.1.chr8.g00011.t01#4    mNeoVis.evergladensis.2.1.chr9.g00012.t01#2    mNeoVis.evergladensis.1.1.chr8.g00011.t01#3    mNeoVis.vulgivaga.2.1.chr8.g00011.t01#1   mNeoVis.2.1.1.chr8_hap1.g00013.t01#5       927   125262051:   mNeoVis.vulgivaga.2.1.chr1.g01766.t01#1    mNeoVis.3.1.1.chr1.g01760.t01#6    mNeoVis.2.1.1.chr1_hap1.g01807.t01#5    mNeoVis.evergladensis.1.1.chr1.g01772.t01#3    mNeoVis.vulgivaga.1.1.chr1.g01776.t01#4   mNeoVis.evergladensis.2.1.chr1.g01771.t01#2       928   125262052:   mNeoVis.vulgivaga.2.1.chr8.g00012.t01#1    mNeoVis.2.1.1.chr8_hap1.g00014.t01#5    mNeoVis.3.1.1.chr8.g00012.t01#6    mNeoVis.vulgivaga.1.1.chr8.g00012.t01#4    mNeoVis.evergladensis.2.1.chr9.g00013.t01#2   mNeoVis.evergladensis.1.1.chr8.g00012.t01#3       929   125262053:   mNeoVis.evergladensis.2.1.chr1.g01772.t01#2    mNeoVis.vulgivaga.2.1.chr1.g01767.t01#1    mNeoVis.evergladensis.1.1.chr1.g01773.t01#3    mNeoVis.2.1.1.chr1_hap1.g01808.t01#5   mNeoVis.vulgivaga.1.1.chr1.g01777.t01#4       930   125262054:   mNeoVis.vulgivaga.2.1.chr8.g00013.t01#1    mNeoVis.2.1.1.chr8_hap1.g00015.t01#5    mNeoVis.vulgivaga.1.1.chr8.g00013.t01#4    mNeoVis.evergladensis.1.1.chr8.g00013.t01#3   mNeoVis.evergladensis.2.1.chr9.g00014.t01#2       931   125262055:   mNeoVis.vulgivaga.2.1.chr15.g00191.t01#1    mNeoVis.evergladensis.1.1.chr15.g00422.t01#3    mNeoVis.2.1.1.chr15_hap1.g00211.t01#5    mNeoVis.evergladensis.2.1.chr15.g00190.t01#2    mNeoVis.3.1.1.chr15.g00185.t01#6   mNeoVis.vulgivaga.1.1.chr15.g00187.t01#4       932   125262056:   mNeoVis.evergladensis.2.1.chr1.g01773.t01#2    mNeoVis.3.1.1.chr1.g01762.t01#6    mNeoVis.2.1.1.chr1_hap1.g01809.t01#5    mNeoVis.evergladensis.1.1.chr1.g01774.t01#3    mNeoVis.vulgivaga.1.1.chr1.g01778.t01#4   mNeoVis.vulgivaga.2.1.chr1.g01768.t01#1       933   125262057:   mNeoVis.evergladensis.1.1.chr8.g00014.t01#3    mNeoVis.evergladensis.2.1.chr9.g00015.t01#2    mNeoVis.vulgivaga.2.1.chr8.g00014.t01#1    mNeoVis.2.1.1.chr8_hap1.g00016.t01#5   mNeoVis.3.1.1.chr8.g00013.t01#6       934   125262058:   mNeoVis.vulgivaga.1.1.chr1.g01779.t01#4    mNeoVis.evergladensis.2.1.chr1.g01774.t01#2    mNeoVis.evergladensis.1.1.chr1.g01775.t01#3    mNeoVis.vulgivaga.2.1.chr1.g01769.t01#1    mNeoVis.3.1.1.chr1.g01763.t01#6   mNeoVis.2.1.1.chr1_hap1.g01810.t01#5       935   125262059:   mNeoVis.2.1.1.chr1_hap1.g01811.t01#5    mNeoVis.evergladensis.2.1.chr1.g01775.t01#2    mNeoVis.evergladensis.1.1.chr1.g01776.t01#3   mNeoVis.vulgivaga.1.1.chr1.g01780.t01#4         936   125262060:   mNeoVis.evergladensis.2.1.chr12.g00358.t01#2    mNeoVis.2.1.1.chr12_hap1.g00401.t01#5   mNeoVis.evergladensis.1.1.chr12.g00365.t01#3         937   125262061:   mNeoVis.vulgivaga.2.1.chr1.g01770.t01#1   mNeoVis.vulgivaga.1.1.chr12.g00367.t01#4           938   125262062:   mNeoVis.vulgivaga.2.1.chr12.g00366.t01#1   mNeoVis.3.1.1.chr12.g00362.t01#6             939   125262063:   mNeoVis.3.1.1.chr1.g01764.t01#6   mNeoVis.vulgivaga.2.1.chr1.g01771.t01#1             940   125262064:   mNeoVis.vulgivaga.2.1.chr8.g00015.t01#1    mNeoVis.evergladensis.2.1.chr9.g00016.t01#2    mNeoVis.2.1.1.chr8_hap1.g00017.t01#5    mNeoVis.3.1.1.chr8.g00014.t01#6    mNeoVis.evergladensis.1.1.chr8.g00015.t01#3   mNeoVis.vulgivaga.1.1.chr8.g00015.t01#4       941   125262065:   mNeoVis.2.1.1.chr8_hap1.g00019.t01#5    mNeoVis.2.1.1.chr8_hap1.g00018.t01#5   mNeoVis.vulgivaga.2.1.chr8.g00016.t01#1           942   125262066:   mNeoVis.evergladensis.1.1.chr8.g00016.t01#3    mNeoVis.evergladensis.2.1.chr9.g00017.t01#2    mNeoVis.evergladensis.2.1.chr9.g00018.t01#2    mNeoVis.vulgivaga.1.1.chr8.g00016.t01#4    mNeoVis.vulgivaga.2.1.chr8.g00017.t01#1    mNeoVis.3.1.1.chr9.g01028.t01#6   mNeoVis.2.1.1.chr8_hap1.g00020.t01#5     943   125262067:   mNeoVis.2.1.1.chr1_hap1.g01812.t01#5    mNeoVis.3.1.1.chr1.g01765.t01#6    mNeoVis.evergladensis.2.1.chr1.g01776.t01#2    mNeoVis.evergladensis.1.1.chr1.g01777.t01#3    mNeoVis.vulgivaga.1.1.chr1.g01781.t01#4   mNeoVis.vulgivaga.2.1.chr1.g01772.t01#1       944   125262068:   mNeoVis.vulgivaga.2.1.chr8.g00018.t01#1    mNeoVis.vulgivaga.1.1.chr8.g00017.t01#4    mNeoVis.evergladensis.2.1.chr9.g00019.t01#2    mNeoVis.evergladensis.1.1.chr8.g00017.t01#3   mNeoVis.2.1.1.chr8_hap1.g00021.t01#5       945   125262069:   mNeoVis.2.1.1.chr8_hap1.g00022.t01#5    mNeoVis.evergladensis.2.1.chr9.g00020.t01#2    mNeoVis.evergladensis.1.1.chr8.g00018.t01#3    mNeoVis.vulgivaga.2.1.chr8.g00019.t01#1   mNeoVis.vulgivaga.1.1.chr8.g00018.t01#4       946   125262070:   mNeoVis.evergladensis.1.1.chr10.g01146.t01#3   mNeoVis.vulgivaga.2.1.chr10.g01142.t01#1           947   125262071:   mNeoVis.vulgivaga.2.1.chr1.g01773.t01#1    mNeoVis.2.1.1.chr1_hap1.g01813.t01#5    mNeoVis.evergladensis.1.1.chr1.g01778.t01#3    mNeoVis.3.1.1.chr1.g01766.t01#6   mNeoVis.vulgivaga.1.1.chr1.g01782.t01#4       948   125262072:   mNeoVis.evergladensis.1.1.chr8.g00019.t01#3    mNeoVis.vulgivaga.1.1.chr8.g00019.t01#4    mNeoVis.2.1.1.chr8_hap1.g00023.t01#5    mNeoVis.evergladensis.2.1.chr9.g00021.t01#2   mNeoVis.vulgivaga.2.1.chr8.g00020.t01#1       949   125262073:   mNeoVis.evergladensis.2.1.chr1.g01778.t01#2    mNeoVis.3.1.1.chr1.g01768.t01#6    mNeoVis.vulgivaga.2.1.chr1.g01774.t01#1    mNeoVis.vulgivaga.1.1.chr1.g01783.t01#4    mNeoVis.evergladensis.1.1.chr1.g01779.t01#3   mNeoVis.2.1.1.chr1_hap1.g01814.t01#5       950   125262074:  mNeoVis.vulgivaga.2.1.chr8.g00021.t01#1               951   125262075:   mNeoVis.evergladensis.2.1.chr1.g01779.t01#2    mNeoVis.3.1.1.chr1.g01769.t01#6    mNeoVis.evergladensis.1.1.chr1.g01780.t01#3    mNeoVis.vulgivaga.2.1.chr1.g01775.t01#1    mNeoVis.vulgivaga.1.1.chr1.g01784.t01#4   mNeoVis.2.1.1.chr1_hap1.g01815.t01#5       952   125262076:   mNeoVis.vulgivaga.2.1.chr1.g01776.t01#1    mNeoVis.2.1.1.chr1_hap1.g01816.t01#5    mNeoVis.evergladensis.2.1.chr1.g01781.t01#2    mNeoVis.3.1.1.chr1.g01770.t01#6    mNeoVis.vulgivaga.1.1.chr1.g01786.t01#4   mNeoVis.evergladensis.1.1.chr1.g01781.t01#3       953   125262077:   mNeoVis.evergladensis.1.1.chr8.g00021.t01#3    mNeoVis.vulgivaga.2.1.chr8.g00022.t01#1    mNeoVis.evergladensis.2.1.chr9.g00023.t01#2    mNeoVis.2.1.1.chr8_hap1.g00025.t01#5   mNeoVis.vulgivaga.1.1.chr8.g00021.t01#4       954   125262078:   mNeoVis.vulgivaga.2.1.chr7.g00135.t01#1    mNeoVis.vulgivaga.1.1.chr7.g00139.t01#4   mNeoVis.evergladensis.2.1.chr7.g00135.t01#2         955   125262079:   mNeoVis.2.1.1.chr1_hap1.g01817.t01#5    mNeoVis.vulgivaga.1.1.chr1.g01787.t01#4    mNeoVis.evergladensis.2.1.chr1.g01782.t01#2    mNeoVis.vulgivaga.2.1.chr1.g01777.t01#1    mNeoVis.evergladensis.1.1.chr1.g01782.t01#3   mNeoVis.3.1.1.chr1.g01771.t01#6       956   125262080:   mNeoVis.vulgivaga.2.1.chr7.g02461.t01#1    mNeoVis.vulgivaga.1.1.chr7.g02439.t01#4    mNeoVis.3.1.1.chr7.g02286.t01#6    mNeoVis.evergladensis.2.1.chr7.g02439.t01#2    mNeoVis.2.1.1.chr7_hap1.g02631.t01#5   mNeoVis.evergladensis.1.1.chr7.g02436.t01#3       957   125262081:   mNeoVis.3.1.1.chr7.g00117.t01#6    mNeoVis.2.1.1.chr7_hap1.g00135.t01#5    mNeoVis.evergladensis.1.1.chr7.g00138.t01#3    mNeoVis.vulgivaga.1.1.chr7.g00140.t01#4    mNeoVis.vulgivaga.2.1.chr7.g00136.t01#1   mNeoVis.evergladensis.2.1.chr7.g00136.t01#2       958   125262082:   mNeoVis.vulgivaga.2.1.chr1.g01778.t01#1    mNeoVis.vulgivaga.1.1.chr1.g01788.t01#4    mNeoVis.evergladensis.1.1.chr1.g01783.t01#3    mNeoVis.2.1.1.chr1_hap1.g01818.t01#5    mNeoVis.3.1.1.chr1.g01772.t01#6   mNeoVis.evergladensis.2.1.chr1.g01783.t01#2       959   125262083:   mNeoVis.vulgivaga.2.1.chr7.g02462.t01#1    mNeoVis.evergladensis.2.1.chr7.g02440.t01#2    mNeoVis.3.1.1.chr7.g02285.t01#6    mNeoVis.2.1.1.chr7_hap1.g02632.t01#5    mNeoVis.vulgivaga.1.1.chr7.g02440.t01#4   mNeoVis.evergladensis.1.1.chr7.g02435.t01#3       960   125262084:   mNeoVis.3.1.1.chr7.g02287.t01#6    mNeoVis.evergladensis.2.1.chr7.g02441.t01#2    mNeoVis.2.1.1.chr7_hap1.g02633.t01#5    mNeoVis.vulgivaga.1.1.chr7.g02441.t01#4    mNeoVis.vulgivaga.2.1.chr7.g02463.t01#1   mNeoVis.evergladensis.1.1.chr7.g02437.t01#3       961   125262085:   mNeoVis.vulgivaga.2.1.chr7.g02464.t01#1    mNeoVis.3.1.1.chr7.g02288.t01#6    mNeoVis.evergladensis.1.1.chr7.g02438.t01#3   mNeoVis.2.1.1.chr7_hap1.g02634.t01#5         962   125262086:   mNeoVis.vulgivaga.2.1.chr1.g01779.t01#1    mNeoVis.3.1.1.chr1.g01773.t01#6    mNeoVis.2.1.1.chr1_hap1.g01819.t01#5    mNeoVis.vulgivaga.1.1.chr1.g01789.t01#4    mNeoVis.evergladensis.1.1.chr1.g01784.t01#3   mNeoVis.evergladensis.2.1.chr1.g01784.t01#2       963   125262087:   mNeoVis.2.1.1.chr7_hap1.g02635.t01#5    mNeoVis.vulgivaga.2.1.chr7.g02465.t01#1    mNeoVis.evergladensis.2.1.chr7.g02443.t01#2    mNeoVis.evergladensis.1.1.chr7.g02439.t01#3   mNeoVis.vulgivaga.1.1.chr7.g02443.t01#4       964   125262088:   mNeoVis.vulgivaga.2.1.chr1.g01780.t01#1    mNeoVis.2.1.1.chr1_hap1.g01820.t01#5    mNeoVis.evergladensis.1.1.chr1.g01785.t01#3    mNeoVis.vulgivaga.1.1.chr1.g01790.t01#4   mNeoVis.evergladensis.2.1.chr1.g01785.t01#2       965   125262089:   mNeoVis.evergladensis.2.1.chr7.g02444.t01#2    mNeoVis.vulgivaga.1.1.chr7.g02444.t01#4    mNeoVis.vulgivaga.2.1.chr7.g02466.t01#1    mNeoVis.3.1.1.chr7.g02289.t01#6    mNeoVis.evergladensis.1.1.chr7.g02440.t01#3   mNeoVis.2.1.1.chr7_hap1.g02636.t01#5       966   125262090:   mNeoVis.vulgivaga.1.1.chr1.g01791.t01#4    mNeoVis.evergladensis.1.1.chr1.g01786.t01#3    mNeoVis.evergladensis.2.1.chr1.g01786.t01#2    mNeoVis.vulgivaga.2.1.chr1.g01781.t01#1    mNeoVis.3.1.1.chr1.g01775.t01#6   mNeoVis.2.1.1.chr1_hap1.g01821.t01#5       967   125262091:   mNeoVis.vulgivaga.2.1.chr7.g02470.t01#1    mNeoVis.3.1.1.chr7.g02292.t01#6    mNeoVis.evergladensis.1.1.chr7.g02444.t01#3    mNeoVis.vulgivaga.1.1.chr7.g02448.t01#4    mNeoVis.evergladensis.2.1.chr7.g02450.t01#2   mNeoVis.vulgivaga.1.1.chr7.g02450.t01#4       968   125262092:   mNeoVis.vulgivaga.2.1.chr7.g02469.t01#1    mNeoVis.vulgivaga.1.1.chr7.g02446.t01#4    mNeoVis.evergladensis.2.1.chr7.g02448.t01#2    mNeoVis.2.1.1.chr7_hap1.g02639.t01#5   mNeoVis.evergladensis.1.1.chr7.g02442.t01#3       969   125262093:   mNeoVis.evergladensis.2.1.chr7.g02445.t01#2    mNeoVis.vulgivaga.1.1.chr7.g02452.t01#4    mNeoVis.evergladensis.1.1.chr7.g02446.t01#3   mNeoVis.vulgivaga.2.1.chr7.g02473.t01#1         970   125262094:   mNeoVis.vulgivaga.2.1.chr7.g02467.t01#1    mNeoVis.evergladensis.1.1.chr7.g02441.t01#3   mNeoVis.vulgivaga.1.1.chr7.g02445.t01#4           971   125262095:   mNeoVis.evergladensis.2.1.chr7.g02452.t01#2   mNeoVis.vulgivaga.1.1.chr7.g02451.t01#4             972   125262096:   mNeoVis.evergladensis.2.1.chr7.g02449.t01#2   mNeoVis.vulgivaga.1.1.chr7.g02447.t01#4             973   125262097:   mNeoVis.vulgivaga.2.1.chr7.g02472.t01#1   mNeoVis.evergladensis.1.1.chr7.g02443.t01#3           974   125262098:   mNeoVis.evergladensis.2.1.chr7.g02451.t01#2   mNeoVis.evergladensis.1.1.chr7.g02445.t01#3           975   125262099:   mNeoVis.2.1.1.chr7_hap1.g02637.t01#5   mNeoVis.vulgivaga.2.1.chr7.g02471.t01#1             976   125262100:   mNeoVis.3.1.1.chr7.g02291.t01#6   mNeoVis.vulgivaga.2.1.chr7.g02468.t01#1             977   125262101:  mNeoVis.evergladensis.2.1.chr7.g02447.t01#2             978   125262102:   mNeoVis.2.1.1.chr1_hap1.g01822.t01#5    mNeoVis.vulgivaga.2.1.chr1.g01782.t01#1   mNeoVis.evergladensis.2.1.chr1.g01787.t01#2         979   125262103:   mNeoVis.vulgivaga.2.1.chr1.g01783.t01#1    mNeoVis.evergladensis.2.1.chr1.g01788.t01#2    mNeoVis.vulgivaga.1.1.chr1.g01792.t01#4    mNeoVis.2.1.1.chr1_hap1.g01823.t01#5    mNeoVis.evergladensis.1.1.chr1.g01787.t01#3   mNeoVis.3.1.1.chr1.g01776.t01#6       980   125262104:   mNeoVis.2.1.1.chr7_hap1.g02640.t01#5    mNeoVis.vulgivaga.2.1.chr7.g02474.t01#1    mNeoVis.evergladensis.1.1.chr7.g02447.t01#3    mNeoVis.vulgivaga.1.1.chr7.g02453.t01#4    mNeoVis.3.1.1.chr7.g02293.t01#6   mNeoVis.evergladensis.2.1.chr7.g02453.t01#2       981   125262105:   mNeoVis.2.1.1.chr7_hap1.g02641.t01#5    mNeoVis.evergladensis.2.1.chr7.g02454.t01#2    mNeoVis.3.1.1.chr7.g02294.t01#6    mNeoVis.vulgivaga.2.1.chr7.g02475.t01#1   mNeoVis.vulgivaga.1.1.chr7.g02454.t01#4       982   125262106:   mNeoVis.vulgivaga.2.1.chr7.g02476.t01#1    mNeoVis.evergladensis.2.1.chr7.g02455.t01#2    mNeoVis.evergladensis.1.1.chr7.g02449.t01#3    mNeoVis.3.1.1.chr7.g02295.t01#6    mNeoVis.2.1.1.chr7_hap1.g02642.t01#5   mNeoVis.vulgivaga.1.1.chr7.g02455.t01#4       983   125262107:   mNeoVis.vulgivaga.2.1.chr1.g01784.t01#1    mNeoVis.evergladensis.2.1.chr1.g01790.t01#2    mNeoVis.evergladensis.1.1.chr1.g01788.t01#3    mNeoVis.2.1.1.chr1_hap1.g01824.t01#5    mNeoVis.vulgivaga.1.1.chr1.g01793.t01#4   mNeoVis.3.1.1.chr1.g01777.t01#6       984   125262108:   mNeoVis.evergladensis.2.1.chr1.g01791.t01#2    mNeoVis.vulgivaga.2.1.chr1.g01785.t01#1    mNeoVis.3.1.1.chr1.g01778.t01#6    mNeoVis.vulgivaga.1.1.chr1.g01794.t01#4    mNeoVis.evergladensis.1.1.chr1.g01789.t01#3   mNeoVis.2.1.1.chr1_hap1.g01825.t01#5       985   125262109:   mNeoVis.vulgivaga.2.1.chr7.g02477.t01#1    mNeoVis.2.1.1.chr7_hap1.g02643.t01#5    mNeoVis.evergladensis.1.1.chr7.g02450.t01#3    mNeoVis.vulgivaga.1.1.chr7.g02456.t01#4   mNeoVis.evergladensis.2.1.chr7.g02456.t01#2       986   125262110:   mNeoVis.vulgivaga.1.1.chr1.g01795.t01#4    mNeoVis.evergladensis.2.1.chr1.g01792.t01#2    mNeoVis.2.1.1.chr1_hap1.g01826.t01#5    mNeoVis.evergladensis.1.1.chr1.g01790.t01#3    mNeoVis.3.1.1.chr1.g01779.t01#6   mNeoVis.vulgivaga.2.1.chr1.g01786.t01#1       987   125262111:   mNeoVis.vulgivaga.1.1.chr1.g01796.t01#4    mNeoVis.2.1.1.chr1_hap1.g01827.t01#5    mNeoVis.vulgivaga.2.1.chr1.g01787.t01#1    mNeoVis.evergladensis.1.1.chr1.g01791.t01#3    mNeoVis.3.1.1.chr1.g01780.t01#6   mNeoVis.evergladensis.2.1.chr1.g01793.t01#2       988   125262112:   mNeoVis.2.1.1.chr7_hap1.g02644.t01#5    mNeoVis.vulgivaga.2.1.chr7.g02478.t01#1   mNeoVis.evergladensis.2.1.chr7.g02457.t01#2         989   125262113:   mNeoVis.3.1.1.chr1.g01781.t01#6    mNeoVis.evergladensis.1.1.chr1.g01792.t01#3    mNeoVis.vulgivaga.2.1.chr1.g01788.t01#1    mNeoVis.2.1.1.chr1_hap1.g01828.t01#5    mNeoVis.vulgivaga.1.1.chr1.g01797.t01#4   mNeoVis.evergladensis.2.1.chr1.g01794.t01#2       990   125262114:   mNeoVis.vulgivaga.2.1.chr1.g01789.t01#1    mNeoVis.vulgivaga.1.1.chr1.g01798.t01#4    mNeoVis.evergladensis.2.1.chr1.g01795.t01#2    mNeoVis.evergladensis.1.1.chr1.g01793.t01#3   mNeoVis.2.1.1.chr1_hap1.g01829.t01#5       991   125262115:   mNeoVis.2.1.1.chr1_hap1.g01830.t01#5    mNeoVis.evergladensis.2.1.chr1.g01796.t01#2    mNeoVis.vulgivaga.1.1.chr1.g01799.t01#4    mNeoVis.3.1.1.chr1.g01782.t01#6    mNeoVis.evergladensis.1.1.chr1.g01794.t01#3   mNeoVis.vulgivaga.2.1.chr1.g01790.t01#1       992   125262116:   mNeoVis.vulgivaga.1.1.chr1.g01800.t01#4    mNeoVis.evergladensis.2.1.chr1.g01797.t01#2    mNeoVis.2.1.1.chr1_hap1.g01831.t01#5    mNeoVis.3.1.1.chr1.g01783.t01#6    mNeoVis.vulgivaga.2.1.chr1.g01791.t01#1   mNeoVis.evergladensis.1.1.chr1.g01795.t01#3       993   125262117:   mNeoVis.3.1.1.chr1.g01784.t01#6    mNeoVis.evergladensis.1.1.chr1.g01796.t01#3    mNeoVis.vulgivaga.2.1.chr1.g01792.t01#1    mNeoVis.evergladensis.2.1.chr1.g01798.t01#2    mNeoVis.vulgivaga.1.1.chr1.g01801.t01#4   mNeoVis.2.1.1.chr1_hap1.g01832.t01#5       994   125262118:   mNeoVis.vulgivaga.2.1.chr15.g00192.t01#1    mNeoVis.3.1.1.chr15.g00186.t01#6    mNeoVis.2.1.1.chr15_hap1.g00212.t01#5    mNeoVis.evergladensis.1.1.chr15.g00421.t01#3    mNeoVis.vulgivaga.1.1.chr15.g00188.t01#4   mNeoVis.evergladensis.2.1.chr15.g00191.t01#2       995   125262119:   mNeoVis.2.1.1.chr1_hap1.g01737.t01#5    mNeoVis.vulgivaga.1.1.chr7.g00315.t01#4    mNeoVis.vulgivaga.2.1.chr1.g01702.t01#1    mNeoVis.evergladensis.2.1.chr1.g01705.t01#2    mNeoVis.evergladensis.2.1.chr7.g00307.t01#2    mNeoVis.3.1.1.chr7.g00293.t01#6    mNeoVis.vulgivaga.1.1.chr1.g01709.t01#4    mNeoVis.vulgivaga.2.1.chr7.g00310.t01#1    mNeoVis.evergladensis.1.1.chr7.g00315.t01#3      996   125262120:   mNeoVis.vulgivaga.1.1.chr7.g00317.t01#4    mNeoVis.evergladensis.2.1.chr7.g00309.t01#2    mNeoVis.vulgivaga.2.1.chr7.g00312.t01#1    mNeoVis.2.1.1.chr7_hap1.g00308.t01#5    mNeoVis.evergladensis.1.1.chr7.g00317.t01#3    mNeoVis.vulgivaga.1.1.chr7.g00316.t01#4    mNeoVis.evergladensis.2.1.chr7.g00308.t01#2    mNeoVis.3.1.1.chr7.g00294.t01#6    mNeoVis.2.1.1.chr7_hap1.g00309.t01#5      997   125262121:   mNeoVis.2.1.1.chr1_hap1.g01738.t01#5    mNeoVis.vulgivaga.1.1.chr1.g01710.t01#4    mNeoVis.evergladensis.1.1.chr1.g01704.t01#3    mNeoVis.3.1.1.chr1.g01693.t01#6    mNeoVis.vulgivaga.2.1.chr1.g01703.t01#1   mNeoVis.evergladensis.2.1.chr1.g01706.t01#2       998   125262122:   mNeoVis.evergladensis.1.1.chr7.g02452.t01#3    mNeoVis.vulgivaga.2.1.chr7.g02479.t01#1   mNeoVis.vulgivaga.1.1.chr7.g02459.t01#4           999   125262123:   mNeoVis.vulgivaga.2.1.chr7.g00137.t01#1    mNeoVis.3.1.1.chr7.g00118.t01#6    mNeoVis.2.1.1.chr7_hap1.g00136.t01#5   mNeoVis.evergladensis.1.1.chr7.g00139.t01#3       1000   125262124:   mNeoVis.2.1.1.chr1_hap1.g01739.t01#5    mNeoVis.evergladensis.2.1.chr1.g01707.t01#2    mNeoVis.3.1.1.chr1.g01694.t01#6    mNeoVis.vulgivaga.1.1.chr1.g01711.t01#4    mNeoVis.vulgivaga.2.1.chr1.g01704.t01#1   mNeoVis.evergladensis.1.1.chr1.g01705.t01#3       1001   125262125:   mNeoVis.evergladensis.1.1.chr1.g01706.t01#3    mNeoVis.2.1.1.chr1_hap1.g01740.t01#5    mNeoVis.evergladensis.2.1.chr1.g01708.t01#2    mNeoVis.vulgivaga.2.1.chr1.g01705.t01#1    mNeoVis.vulgivaga.1.1.chr1.g01712.t01#4   mNeoVis.3.1.1.chr1.g01695.t01#6       1002   125262126:   mNeoVis.2.1.1.chr15_hap1.g00213.t01#5    mNeoVis.evergladensis.1.1.chr15.g00420.t01#3    mNeoVis.vulgivaga.1.1.chr15.g00189.t01#4    mNeoVis.3.1.1.chr15.g00187.t01#6    mNeoVis.evergladensis.2.1.chr15.g00192.t01#2   mNeoVis.vulgivaga.2.1.chr15.g00193.t01#1       1003   125262127:   mNeoVis.2.1.1.chr8_hap1.g01004.t01#5    mNeoVis.evergladensis.2.1.chr9.g00978.t01#2    mNeoVis.evergladensis.1.1.chr8.g00970.t01#3    mNeoVis.3.1.1.chr8.g00937.t01#6    mNeoVis.vulgivaga.2.1.chr8.g00979.t01#1   mNeoVis.vulgivaga.1.1.chr8.g00964.t01#4       1004   125262128:   mNeoVis.evergladensis.1.1.chr10.g01109.t01#3    mNeoVis.vulgivaga.2.1.chr10.g01105.t01#1    mNeoVis.evergladensis.2.1.chr10.g01114.t01#2    mNeoVis.vulgivaga.1.1.chr10.g01091.t01#4   mNeoVis.2.1.1.chr9_hap1.g01126.t01#5       1005   125262129:   mNeoVis.2.1.1.chr7_hap1.g00137.t01#5    mNeoVis.vulgivaga.1.1.chr7.g00141.t01#4    mNeoVis.evergladensis.2.1.chr7.g00137.t01#2    mNeoVis.evergladensis.1.1.chr7.g00140.t01#3   mNeoVis.vulgivaga.2.1.chr7.g00138.t01#1       1006   125262130:   mNeoVis.2.1.1.chr1_hap1.g01742.t01#5    mNeoVis.vulgivaga.1.1.chr1.g02086.t01#4    mNeoVis.2.1.1.chr9_hap1.g01106.t01#5    mNeoVis.vulgivaga.2.1.chr12.g00217.t01#1    mNeoVis.vulgivaga.1.1.chr10.g00739.t01#4    mNeoVis.3.1.1.chr1.g02067.t01#6    mNeoVis.evergladensis.1.1.chr1.g01667.t01#3    mNeoVis.3.1.1.chr12.g00633.t01#6    mNeoVis.2.1.1.chr12_hap1.g00243.t01#5      1007   125262131:   mNeoVis.evergladensis.1.1.chr11.g00485.t01#3    mNeoVis.3.1.1.chr11.g00453.t01#6    mNeoVis.2.1.1.chr11_hap1.g00491.t01#5    mNeoVis.vulgivaga.1.1.chr11.g00478.t01#4    mNeoVis.vulgivaga.2.1.chr11.g00488.t01#1   mNeoVis.evergladensis.2.1.chr11.g00481.t01#2       1008   125262132:   mNeoVis.3.1.1.chr3.g01250.t01#6    mNeoVis.2.1.1.chr3_hap1.g01273.t01#5    mNeoVis.evergladensis.1.1.chr3.g01279.t01#3    mNeoVis.vulgivaga.2.1.chr3.g01289.t01#1    mNeoVis.evergladensis.2.1.chr3.g01271.t01#2   mNeoVis.evergladensis.1.1.chr3.g01278.t01#3       1009   125262133:   mNeoVis.3.1.1.chr9.g00143.t01#6    mNeoVis.evergladensis.1.1.chr9.g00167.t01#3   mNeoVis.vulgivaga.2.1.chr9.g00164.t01#1           1010   125262134:   mNeoVis.evergladensis.1.1.chr1.g01709.t01#3    mNeoVis.vulgivaga.2.1.chr1.g01707.t01#1    mNeoVis.evergladensis.2.1.chr1.g01711.t01#2    mNeoVis.3.1.1.chr1.g01697.t01#6   mNeoVis.vulgivaga.1.1.chr1.g01713.t01#4       1011   125262135:   mNeoVis.2.1.1.chr1_hap1.g01744.t01#5    mNeoVis.vulgivaga.1.1.chr1.g01714.t01#4    mNeoVis.vulgivaga.2.1.chr1.g01708.t01#1    mNeoVis.3.1.1.chr1.g01698.t01#6    mNeoVis.evergladensis.1.1.chr1.g01710.t01#3   mNeoVis.evergladensis.2.1.chr1.g01712.t01#2       1012   125262136:  mNeoVis.vulgivaga.2.1.chr7.g02480.t01#1               1013   125262137:   mNeoVis.vulgivaga.1.1.chr1.g01715.t01#4    mNeoVis.vulgivaga.2.1.chr1.g01709.t01#1    mNeoVis.evergladensis.1.1.chr1.g01711.t01#3    mNeoVis.evergladensis.2.1.chr1.g01713.t01#2    mNeoVis.2.1.1.chr1_hap1.g01745.t01#5   mNeoVis.3.1.1.chr1.g01699.t01#6       1014   125262138:   mNeoVis.vulgivaga.2.1.chr1.g01710.t01#1    mNeoVis.3.1.1.chr1.g01700.t01#6    mNeoVis.evergladensis.1.1.chr1.g01712.t01#3    mNeoVis.vulgivaga.1.1.chr1.g01716.t01#4    mNeoVis.evergladensis.2.1.chr1.g01714.t01#2   mNeoVis.2.1.1.chr1_hap1.g01746.t01#5       1015   125262139:   mNeoVis.vulgivaga.2.1.chr1.g01711.t01#1    mNeoVis.vulgivaga.1.1.chr1.g01717.t01#4    mNeoVis.2.1.1.chr1_hap1.g01747.t01#5    mNeoVis.evergladensis.1.1.chr1.g01713.t01#3    mNeoVis.3.1.1.chr1.g01701.t01#6   mNeoVis.evergladensis.2.1.chr1.g01715.t01#2       1016   125262140:   mNeoVis.evergladensis.2.1.chr1.g01716.t01#2    mNeoVis.vulgivaga.2.1.chr1.g01712.t01#1   mNeoVis.2.1.1.chr1_hap1.g01748.t01#5           1017   125262141:  mNeoVis.vulgivaga.2.1.chr7.g02481.t01#1               1018   125262142:   mNeoVis.3.1.1.chr1.g01702.t01#6   mNeoVis.vulgivaga.2.1.chr1.g01713.t01#1             1019   125262143:   mNeoVis.evergladensis.2.1.chr1.g01718.t01#2    mNeoVis.vulgivaga.2.1.chr1.g01714.t01#1    mNeoVis.vulgivaga.1.1.chr1.g01719.t01#4    mNeoVis.3.1.1.chr1.g01703.t01#6    mNeoVis.2.1.1.chr1_hap1.g01750.t01#5   mNeoVis.evergladensis.1.1.chr1.g01715.t01#3       1020   125262144:   mNeoVis.vulgivaga.2.1.chr1.g01715.t01#1    mNeoVis.evergladensis.1.1.chr1.g01716.t01#3    mNeoVis.evergladensis.2.1.chr1.g01719.t01#2    mNeoVis.vulgivaga.1.1.chr1.g01720.t01#4    mNeoVis.3.1.1.chr1.g01704.t01#6   mNeoVis.2.1.1.chr1_hap1.g01751.t01#5       1021   125262145:   mNeoVis.evergladensis.2.1.chr1.g01720.t01#2    mNeoVis.2.1.1.chr1_hap1.g01752.t01#5    mNeoVis.3.1.1.chr1.g01705.t01#6    mNeoVis.vulgivaga.1.1.chr1.g01721.t01#4    mNeoVis.evergladensis.1.1.chr1.g01717.t01#3   mNeoVis.vulgivaga.2.1.chr1.g01716.t01#1       1022   125262146:   mNeoVis.evergladensis.2.1.chr6.g00743.t01#2    mNeoVis.vulgivaga.1.1.chr6.g00739.t01#4    mNeoVis.evergladensis.1.1.chr6.g00745.t01#3    mNeoVis.2.1.1.chr4_hap1.g00734.t01#5    mNeoVis.vulgivaga.2.1.chr6.g00740.t01#1   mNeoVis.3.1.1.chr6.g00743.t01#6       1023   125262147:   mNeoVis.2.1.1.chr7_hap1.g00138.t01#5    mNeoVis.evergladensis.2.1.chr7.g00138.t01#2    mNeoVis.vulgivaga.1.1.chr7.g00142.t01#4    mNeoVis.3.1.1.chr7.g00120.t01#6    mNeoVis.evergladensis.1.1.chr7.g00141.t01#3   mNeoVis.vulgivaga.2.1.chr7.g00139.t01#1       1024   125262148:   mNeoVis.evergladensis.1.1.chr1.g01718.t01#3    mNeoVis.vulgivaga.2.1.chr1.g01717.t01#1    mNeoVis.2.1.1.chr1_hap1.g01753.t01#5    mNeoVis.3.1.1.chr1.g01706.t01#6    mNeoVis.vulgivaga.1.1.chr1.g01722.t01#4    mNeoVis.evergladensis.2.1.chr1.g01721.t01#2   mNeoVis.evergladensis.1.1.chr1.g01719.t01#3     1025   125259597:   mNeoVis.evergladensis.1.1.chr7.g00876.t01#3    mNeoVis.vulgivaga.1.1.chr7.g00877.t01#4    mNeoVis.3.1.1.chr7.g00818.t01#6    mNeoVis.vulgivaga.2.1.chr7.g00876.t01#1    mNeoVis.evergladensis.2.1.chr7.g00871.t01#2   mNeoVis.2.1.1.chr7_hap1.g00878.t01#5       1026   125259598:   mNeoVis.2.1.1.chr7_hap1.g02648.t01#5    mNeoVis.evergladensis.1.1.chr7.g02456.t01#3    mNeoVis.vulgivaga.2.1.chr7.g02482.t01#1    mNeoVis.evergladensis.2.1.chr7.g02460.t01#2   mNeoVis.vulgivaga.1.1.chr7.g02462.t01#4       1027   125259599:   mNeoVis.vulgivaga.2.1.chr7.g00140.t01#1    mNeoVis.evergladensis.2.1.chr7.g00139.t01#2    mNeoVis.3.1.1.chr7.g00121.t01#6    mNeoVis.evergladensis.1.1.chr7.g00142.t01#3    mNeoVis.2.1.1.chr7_hap1.g00139.t01#5   mNeoVis.vulgivaga.1.1.chr7.g00143.t01#4       1028   125259600:   mNeoVis.3.1.1.chr7.g00122.t01#6    mNeoVis.vulgivaga.2.1.chr7.g00141.t01#1    mNeoVis.evergladensis.2.1.chr7.g00140.t01#2    mNeoVis.2.1.1.chr7_hap1.g00140.t01#5    mNeoVis.vulgivaga.1.1.chr7.g00144.t01#4   mNeoVis.evergladensis.1.1.chr7.g00143.t01#3       1029   125259601:   mNeoVis.vulgivaga.2.1.chr7.g02483.t01#1    mNeoVis.vulgivaga.1.1.chr7.g02463.t01#4    mNeoVis.evergladensis.1.1.chr7.g02457.t01#3    mNeoVis.2.1.1.chr7_hap1.g02649.t01#5   mNeoVis.evergladensis.2.1.chr7.g02461.t01#2       1030   125259602:   mNeoVis.2.1.1.chr1_hap1.g01754.t01#5    mNeoVis.evergladensis.2.1.chr1.g01722.t01#2    mNeoVis.evergladensis.1.1.chr1.g01720.t01#3    mNeoVis.vulgivaga.1.1.chr1.g01723.t01#4    mNeoVis.vulgivaga.2.1.chr1.g01718.t01#1   mNeoVis.3.1.1.chr1.g01707.t01#6       1031   125259603:   mNeoVis.vulgivaga.2.1.chr7.g02484.t01#1    mNeoVis.vulgivaga.1.1.chr7.g02464.t01#4    mNeoVis.2.1.1.chr7_hap1.g02650.t01#5    mNeoVis.evergladensis.2.1.chr7.g02462.t01#2   mNeoVis.evergladensis.1.1.chr7.g02458.t01#3       1032   125259604:   mNeoVis.evergladensis.1.1.chr1.g01722.t01#3    mNeoVis.vulgivaga.2.1.chr1.g01719.t01#1    mNeoVis.evergladensis.2.1.chr1.g01724.t01#2    mNeoVis.3.1.1.chr1.g01709.t01#6    mNeoVis.vulgivaga.1.1.chr1.g01725.t01#4   mNeoVis.2.1.1.chr1_hap1.g01756.t01#5       1033   125259605:   mNeoVis.vulgivaga.1.1.chr1.g01726.t01#4    mNeoVis.2.1.1.chr1_hap1.g01757.t01#5   mNeoVis.vulgivaga.2.1.chr1.g01720.t01#1           1034   125259606:   mNeoVis.3.1.1.chr1.g01710.t01#6    mNeoVis.evergladensis.1.1.chr1.g01723.t01#3   mNeoVis.evergladensis.2.1.chr1.g01725.t01#2         1035   125259607:   mNeoVis.evergladensis.2.1.chr1.g01726.t01#2    mNeoVis.3.1.1.chr1.g01711.t01#6    mNeoVis.evergladensis.1.1.chr1.g01724.t01#3    mNeoVis.vulgivaga.2.1.chr1.g01721.t01#1    mNeoVis.vulgivaga.1.1.chr1.g01727.t01#4   mNeoVis.2.1.1.chr1_hap1.g01758.t01#5       1036   125259608:   mNeoVis.evergladensis.2.1.chr1.g01727.t01#2    mNeoVis.vulgivaga.1.1.chr1.g01728.t01#4    mNeoVis.2.1.1.chr1_hap1.g01759.t01#5    mNeoVis.evergladensis.1.1.chr1.g01725.t01#3    mNeoVis.3.1.1.chr1.g01712.t01#6   mNeoVis.vulgivaga.2.1.chr1.g01722.t01#1       1037   125259609:   mNeoVis.vulgivaga.1.1.chr1.g01729.t01#4    mNeoVis.2.1.1.chr1_hap1.g01760.t01#5    mNeoVis.3.1.1.chr1.g01713.t01#6    mNeoVis.evergladensis.2.1.chr1.g01728.t01#2    mNeoVis.evergladensis.1.1.chr1.g01726.t01#3   mNeoVis.vulgivaga.2.1.chr1.g01723.t01#1       1038   125259610:   mNeoVis.vulgivaga.1.1.chr1.g01730.t01#4    mNeoVis.2.1.1.chr1_hap1.g01761.t01#5    mNeoVis.evergladensis.1.1.chr1.g01727.t01#3   mNeoVis.vulgivaga.2.1.chr1.g01724.t01#1         1039   125259611:   mNeoVis.vulgivaga.2.1.chr1.g01725.t01#1    mNeoVis.2.1.1.chr1_hap1.g01762.t01#5    mNeoVis.3.1.1.chr1.g01715.t01#6    mNeoVis.evergladensis.1.1.chr1.g01728.t01#3    mNeoVis.vulgivaga.1.1.chr1.g01731.t01#4   mNeoVis.evergladensis.2.1.chr1.g01730.t01#2       1040   125259612:   mNeoVis.evergladensis.2.1.chr7.g02463.t01#2    mNeoVis.vulgivaga.1.1.chr7.g02465.t01#4    mNeoVis.vulgivaga.2.1.chr7.g02485.t01#1    mNeoVis.3.1.1.chr7.g02299.t01#6    mNeoVis.2.1.1.chr7_hap1.g02651.t01#5   mNeoVis.evergladensis.1.1.chr7.g02459.t01#3       1041   125259613:   mNeoVis.vulgivaga.2.1.chr1.g01857.t01#1    mNeoVis.evergladensis.2.1.chr1.g01859.t01#2    mNeoVis.3.1.1.chr1.g01844.t01#6    mNeoVis.evergladensis.1.1.chr1.g01860.t01#3    mNeoVis.vulgivaga.1.1.chr1.g01864.t01#4   mNeoVis.2.1.1.chr1_hap1.g01892.t01#5       1042   125259614:   mNeoVis.vulgivaga.1.1.chr7.g02466.t01#4    mNeoVis.evergladensis.2.1.chr7.g02464.t01#2    mNeoVis.3.1.1.chr7.g02300.t01#6    mNeoVis.2.1.1.chr7_hap1.g02652.t01#5    mNeoVis.vulgivaga.2.1.chr7.g02486.t01#1   mNeoVis.evergladensis.1.1.chr7.g02460.t01#3       1043   125259615:   mNeoVis.vulgivaga.2.1.chr7.g02487.t01#1    mNeoVis.evergladensis.2.1.chr7.g02465.t01#2    mNeoVis.3.1.1.chr7.g02301.t01#6    mNeoVis.2.1.1.chr7_hap1.g02653.t01#5    mNeoVis.evergladensis.1.1.chr7.g02461.t01#3   mNeoVis.vulgivaga.1.1.chr7.g02467.t01#4       1044   125259616:   mNeoVis.evergladensis.1.1.chr7.g02462.t01#3    mNeoVis.vulgivaga.2.1.chr7.g02488.t01#1    mNeoVis.evergladensis.2.1.chr7.g02466.t01#2    mNeoVis.vulgivaga.1.1.chr7.g02468.t01#4    mNeoVis.2.1.1.chr7_hap1.g02654.t01#5   mNeoVis.3.1.1.chr7.g02302.t01#6       1045   125259617:   mNeoVis.evergladensis.1.1.chr7.g02463.t01#3    mNeoVis.vulgivaga.2.1.chr7.g02489.t01#1    mNeoVis.vulgivaga.1.1.chr7.g02469.t01#4    mNeoVis.2.1.1.chr7_hap1.g02655.t01#5    mNeoVis.3.1.1.chr7.g02303.t01#6   mNeoVis.evergladensis.2.1.chr7.g02467.t01#2       1046   125259618:   mNeoVis.evergladensis.2.1.chr7.g02468.t01#2    mNeoVis.vulgivaga.1.1.chr7.g02470.t01#4    mNeoVis.vulgivaga.2.1.chr7.g02490.t01#1    mNeoVis.evergladensis.1.1.chr7.g02464.t01#3    mNeoVis.3.1.1.chr7.g02304.t01#6   mNeoVis.2.1.1.chr7_hap1.g02656.t01#5       1047   125259619:   mNeoVis.evergladensis.2.1.chr7.g02469.t01#2    mNeoVis.vulgivaga.2.1.chr7.g02491.t01#1    mNeoVis.vulgivaga.1.1.chr7.g02471.t01#4    mNeoVis.evergladensis.1.1.chr7.g02465.t01#3    mNeoVis.2.1.1.chr7_hap1.g02657.t01#5   mNeoVis.3.1.1.chr7.g02305.t01#6       1048   125259620:   mNeoVis.vulgivaga.1.1.chr1.g01733.t01#4    mNeoVis.3.1.1.chr1.g01717.t01#6    mNeoVis.vulgivaga.2.1.chr1.g01726.t01#1   mNeoVis.2.1.1.chr1_hap1.g01763.t01#5         1049   125259621:   mNeoVis.evergladensis.1.1.chr1.g01729.t01#3    mNeoVis.vulgivaga.2.1.chr1.g01727.t01#1    mNeoVis.3.1.1.chr1.g01718.t01#6    mNeoVis.vulgivaga.1.1.chr1.g01734.t01#4    mNeoVis.evergladensis.2.1.chr1.g01731.t01#2   mNeoVis.2.1.1.chr1_hap1.g01764.t01#5       1050   125259622:   mNeoVis.evergladensis.1.1.chr7.g02466.t01#3    mNeoVis.evergladensis.2.1.chr7.g02470.t01#2    mNeoVis.3.1.1.chr7.g02306.t01#6    mNeoVis.vulgivaga.1.1.chr7.g02472.t01#4    mNeoVis.2.1.1.chr7_hap1.g02658.t01#5   mNeoVis.vulgivaga.2.1.chr7.g02492.t01#1       1051   125259623:   mNeoVis.2.1.1.chr7_hap1.g02659.t01#5    mNeoVis.vulgivaga.1.1.chr7.g02473.t01#4    mNeoVis.evergladensis.1.1.chr7.g02467.t01#3    mNeoVis.vulgivaga.2.1.chr7.g02493.t01#1   mNeoVis.evergladensis.2.1.chr7.g02471.t01#2       1052   125259624:   mNeoVis.vulgivaga.2.1.chr1.g01728.t01#1   mNeoVis.evergladensis.1.1.chr1.g01730.t01#3           1053   125259625:   mNeoVis.vulgivaga.2.1.chr1.g01730.t01#1    mNeoVis.vulgivaga.1.1.chr1.g01736.t01#4    mNeoVis.2.1.1.chr1_hap1.g01766.t01#5    mNeoVis.evergladensis.1.1.chr1.g01732.t01#3   mNeoVis.evergladensis.2.1.chr1.g01733.t01#2       1054   125259626:   mNeoVis.evergladensis.1.1.chr7.g02468.t01#3    mNeoVis.evergladensis.2.1.chr7.g02472.t01#2   mNeoVis.vulgivaga.2.1.chr7.g02494.t01#1           1055   125259627:   mNeoVis.2.1.1.chr7_hap1.g02660.t01#5    mNeoVis.3.1.1.chr7.g02308.t01#6   mNeoVis.vulgivaga.1.1.chr7.g02474.t01#4           1056   125259628:   mNeoVis.2.1.1.chr1_hap1.g01767.t01#5    mNeoVis.evergladensis.2.1.chr1.g01734.t01#2    mNeoVis.vulgivaga.2.1.chr1.g01731.t01#1    mNeoVis.vulgivaga.1.1.chr1.g01737.t01#4    mNeoVis.evergladensis.1.1.chr1.g01733.t01#3   mNeoVis.3.1.1.chr1.g01720.t01#6       1057   125259629:   mNeoVis.vulgivaga.1.1.chr1.g01739.t01#4    mNeoVis.evergladensis.1.1.chr1.g01734.t01#3    mNeoVis.evergladensis.2.1.chr1.g01735.t01#2    mNeoVis.vulgivaga.2.1.chr1.g01732.t01#1    mNeoVis.2.1.1.chr1_hap1.g01768.t01#5   mNeoVis.3.1.1.chr1.g01721.t01#6       1058   125259630:   mNeoVis.vulgivaga.2.1.chr7.g02495.t01#1    mNeoVis.evergladensis.1.1.chr7.g02469.t01#3    mNeoVis.vulgivaga.1.1.chr7.g02475.t01#4    mNeoVis.2.1.1.chr7_hap1.g02661.t01#5   mNeoVis.evergladensis.2.1.chr7.g02473.t01#2       1059   125259631:   mNeoVis.3.1.1.chr1.g01722.t01#6    mNeoVis.2.1.1.chr1_hap1.g01769.t01#5    mNeoVis.vulgivaga.2.1.chr1.g01733.t01#1    mNeoVis.vulgivaga.1.1.chr1.g01740.t01#4    mNeoVis.evergladensis.1.1.chr1.g01735.t01#3   mNeoVis.evergladensis.2.1.chr1.g01736.t01#2       1060   125259632:   mNeoVis.3.1.1.chr15.g00188.t01#6    mNeoVis.evergladensis.1.1.chr15.g00419.t01#3    mNeoVis.vulgivaga.2.1.chr15.g00194.t01#1    mNeoVis.2.1.1.chr15_hap1.g00214.t01#5    mNeoVis.vulgivaga.1.1.chr15.g00190.t01#4   mNeoVis.evergladensis.2.1.chr15.g00193.t01#2       1061   125259633:   mNeoVis.vulgivaga.1.1.chr1.g01741.t01#4    mNeoVis.2.1.1.chr1_hap1.g01770.t01#5    mNeoVis.vulgivaga.2.1.chr1.g01734.t01#1    mNeoVis.evergladensis.1.1.chr1.g01736.t01#3    mNeoVis.evergladensis.2.1.chr1.g01737.t01#2   mNeoVis.3.1.1.chr1.g01723.t01#6       1062   125259634:   mNeoVis.2.1.1.chr1_hap1.g01771.t01#5    mNeoVis.vulgivaga.2.1.chr1.g01735.t01#1    mNeoVis.vulgivaga.1.1.chr1.g01742.t01#4    mNeoVis.3.1.1.chr1.g01724.t01#6    mNeoVis.evergladensis.1.1.chr1.g01737.t01#3   mNeoVis.evergladensis.2.1.chr1.g01738.t01#2       1063   125259635:   mNeoVis.2.1.1.chr7_hap1.g00142.t01#5    mNeoVis.evergladensis.1.1.chr7.g00145.t01#3    mNeoVis.vulgivaga.1.1.chr7.g00146.t01#4   mNeoVis.evergladensis.2.1.chr7.g00142.t01#2       1064   125259636:   mNeoVis.evergladensis.1.1.chr7.g00144.t01#3    mNeoVis.3.1.1.chr7.g00123.t01#6   mNeoVis.vulgivaga.1.1.chr7.g00145.t01#4           1065   125259637:   mNeoVis.3.1.1.chr7.g00125.t01#6   mNeoVis.vulgivaga.2.1.chr7.g00142.t01#1             1066   125259638:   mNeoVis.evergladensis.2.1.chr7.g00141.t01#2   mNeoVis.2.1.1.chr7_hap1.g00141.t01#5             1067   125259639:   mNeoVis.vulgivaga.2.1.chr7.g00143.t01#1   mNeoVis.3.1.1.chr7.g00124.t01#6             1068   125259640:   mNeoVis.3.1.1.chr1.g01725.t01#6    mNeoVis.vulgivaga.1.1.chr1.g01743.t01#4    mNeoVis.vulgivaga.2.1.chr1.g01736.t01#1    mNeoVis.vulgivaga.2.1.chr2.g00613.t01#1    mNeoVis.3.1.1.chr2.g00537.t01#6    mNeoVis.evergladensis.1.1.chr1.g01738.t01#3    mNeoVis.vulgivaga.1.1.chr2.g00606.t01#4    mNeoVis.evergladensis.2.1.chr1.g01739.t01#2    mNeoVis.evergladensis.1.1.chr2.g00610.t01#3      1069   125259641:   mNeoVis.2.1.1.chr1_hap1.g01773.t01#5    mNeoVis.evergladensis.2.1.chr1.g01740.t01#2    mNeoVis.3.1.1.chr1.g01726.t01#6    mNeoVis.vulgivaga.1.1.chr1.g01744.t01#4    mNeoVis.vulgivaga.2.1.chr1.g01737.t01#1   mNeoVis.evergladensis.1.1.chr1.g01739.t01#3       1070   125259642:   mNeoVis.evergladensis.2.1.chr7.g02474.t01#2   mNeoVis.vulgivaga.2.1.chr7.g02496.t01#1             1071   125259643:   mNeoVis.vulgivaga.1.1.chr7.g02477.t01#4    mNeoVis.vulgivaga.2.1.chr7.g02497.t01#1    mNeoVis.2.1.1.chr7_hap1.g02663.t01#5    mNeoVis.evergladensis.2.1.chr7.g02475.t01#2    mNeoVis.evergladensis.1.1.chr7.g02471.t01#3   mNeoVis.3.1.1.chr7.g02309.t01#6       1072   125259644:   mNeoVis.evergladensis.2.1.chr1.g01741.t01#2    mNeoVis.2.1.1.chr1_hap1.g01774.t01#5    mNeoVis.vulgivaga.2.1.chr1.g01738.t01#1    mNeoVis.vulgivaga.1.1.chr1.g01745.t01#4   mNeoVis.evergladensis.1.1.chr1.g01740.t01#3       1073   125259645:   mNeoVis.2.1.1.chr7_hap1.g02664.t01#5    mNeoVis.evergladensis.2.1.chr7.g02476.t01#2    mNeoVis.vulgivaga.1.1.chr7.g02478.t01#4    mNeoVis.evergladensis.1.1.chr7.g02472.t01#3    mNeoVis.3.1.1.chr7.g02310.t01#6   mNeoVis.vulgivaga.2.1.chr7.g02498.t01#1       1074   125259646:   mNeoVis.vulgivaga.1.1.chr1.g01746.t01#4    mNeoVis.vulgivaga.2.1.chr1.g01739.t01#1    mNeoVis.evergladensis.1.1.chr1.g01741.t01#3    mNeoVis.evergladensis.2.1.chr1.g01742.t01#2   mNeoVis.2.1.1.chr1_hap1.g01775.t01#5       1075   125259647:   mNeoVis.evergladensis.2.1.chr15.g00194.t01#2    mNeoVis.evergladensis.1.1.chr15.g00418.t01#3    mNeoVis.2.1.1.chr15_hap1.g00215.t01#5    mNeoVis.vulgivaga.2.1.chr15.g00195.t01#1    mNeoVis.vulgivaga.1.1.chr15.g00191.t01#4   mNeoVis.3.1.1.chr15.g00189.t01#6       1076   125259648:   mNeoVis.3.1.1.chr7.g02251.t01#6    mNeoVis.2.1.1.chr7_hap1.g02587.t01#5    mNeoVis.vulgivaga.1.1.chr7.g02397.t01#4    mNeoVis.evergladensis.1.1.chr7.g02391.t01#3    mNeoVis.evergladensis.2.1.chr7.g02393.t01#2   mNeoVis.vulgivaga.2.1.chr7.g02413.t01#1       1077   125259649:   mNeoVis.3.1.1.chr1.g01411.t01#6    mNeoVis.evergladensis.2.1.chr1.g01169.t01#2    mNeoVis.evergladensis.1.1.chr1.g01432.t01#3    mNeoVis.vulgivaga.2.1.chr1.g01169.t01#1    mNeoVis.vulgivaga.2.1.chr1.g01430.t01#1    mNeoVis.vulgivaga.1.1.chr1.g01431.t01#4   mNeoVis.2.1.1.chr1_hap1.g01195.t01#5     1078   125259650:   mNeoVis.vulgivaga.1.1.chr1.g01748.t01#4    mNeoVis.2.1.1.chr1_hap1.g01463.t01#5    mNeoVis.3.1.1.chr1.g01733.t01#6    mNeoVis.evergladensis.2.1.chr1.g01429.t01#2   mNeoVis.vulgivaga.2.1.chr1.g01740.t01#1       1079   125259651:   mNeoVis.vulgivaga.2.1.chr7.g02414.t01#1    mNeoVis.3.1.1.chr7.g02252.t01#6    mNeoVis.evergladensis.1.1.chr7.g02392.t01#3    mNeoVis.vulgivaga.1.1.chr7.g02398.t01#4    mNeoVis.2.1.1.chr7_hap1.g02588.t01#5    mNeoVis.vulgivaga.2.1.chr7.g02415.t01#1   mNeoVis.evergladensis.2.1.chr7.g02394.t01#2     1080   125259652:   mNeoVis.2.1.1.chr1_hap1.g01777.t01#5    mNeoVis.evergladensis.1.1.chr1.g01743.t01#3    mNeoVis.3.1.1.chr1.g01735.t01#6    mNeoVis.vulgivaga.1.1.chr1.g01749.t01#4    mNeoVis.vulgivaga.2.1.chr1.g01741.t01#1   mNeoVis.evergladensis.2.1.chr1.g01744.t01#2       1081   125259653:   mNeoVis.2.1.1.chr7_hap1.g02589.t01#5    mNeoVis.vulgivaga.2.1.chr7.g02416.t01#1    mNeoVis.vulgivaga.1.1.chr7.g02399.t01#4    mNeoVis.evergladensis.1.1.chr7.g02393.t01#3    mNeoVis.3.1.1.chr7.g02253.t01#6   mNeoVis.evergladensis.2.1.chr7.g02395.t01#2       1082   125259654:  mNeoVis.vulgivaga.2.1.chr7.g02417.t01#1               1083   125259655:   mNeoVis.evergladensis.1.1.chr7.g02394.t01#3    mNeoVis.3.1.1.chr7.g02254.t01#6   mNeoVis.vulgivaga.2.1.chr7.g02418.t01#1           1084   125259656:   mNeoVis.evergladensis.2.1.chr7.g02396.t01#2   mNeoVis.2.1.1.chr7_hap1.g02590.t01#5             1085   125259657:  mNeoVis.vulgivaga.1.1.chr7.g02400.t01#4               1086   125259658:   mNeoVis.3.1.1.chr7.g02255.t01#6    mNeoVis.evergladensis.1.1.chr7.g02395.t01#3    mNeoVis.vulgivaga.2.1.chr7.g02419.t01#1    mNeoVis.evergladensis.2.1.chr7.g02397.t01#2    mNeoVis.vulgivaga.1.1.chr7.g02401.t01#4   mNeoVis.2.1.1.chr7_hap1.g02591.t01#5       1087   125259659:   mNeoVis.vulgivaga.2.1.chr7.g00144.t01#1    mNeoVis.vulgivaga.1.1.chr7.g00147.t01#4    mNeoVis.evergladensis.2.1.chr7.g00143.t01#2    mNeoVis.2.1.1.chr7_hap1.g00143.t01#5    mNeoVis.evergladensis.1.1.chr7.g00146.t01#3   mNeoVis.3.1.1.chr7.g00126.t01#6       1088   125259660:   mNeoVis.evergladensis.1.1.chr7.g02396.t01#3    mNeoVis.evergladensis.2.1.chr7.g02398.t01#2    mNeoVis.vulgivaga.1.1.chr7.g02402.t01#4    mNeoVis.vulgivaga.2.1.chr7.g02420.t01#1   mNeoVis.2.1.1.chr7_hap1.g02592.t01#5       1089   125259661:  mNeoVis.vulgivaga.2.1.chr1.g01742.t01#1               1090   125259662:   mNeoVis.evergladensis.2.1.chr7.g02399.t01#2    mNeoVis.3.1.1.chr7.g02256.t01#6    mNeoVis.vulgivaga.2.1.chr7.g02421.t01#1    mNeoVis.2.1.1.chr7_hap1.g02593.t01#5    mNeoVis.evergladensis.1.1.chr7.g02397.t01#3   mNeoVis.vulgivaga.1.1.chr7.g02403.t01#4       1091   125259663:   mNeoVis.2.1.1.chr1_hap1.g01778.t01#5    mNeoVis.3.1.1.chr1.g01737.t01#6    mNeoVis.evergladensis.1.1.chr1.g01745.t01#3    mNeoVis.evergladensis.2.1.chr1.g01745.t01#2    mNeoVis.vulgivaga.2.1.chr1.g01743.t01#1   mNeoVis.vulgivaga.1.1.chr1.g01750.t01#4       1092   125259664:   mNeoVis.evergladensis.1.1.chr7.g02398.t01#3    mNeoVis.vulgivaga.2.1.chr7.g02422.t01#1    mNeoVis.vulgivaga.1.1.chr7.g02404.t01#4    mNeoVis.evergladensis.2.1.chr7.g02400.t01#2   mNeoVis.2.1.1.chr7_hap1.g02594.t01#5       1093   125259665:   mNeoVis.2.1.1.chr1_hap1.g01779.t01#5    mNeoVis.evergladensis.1.1.chr1.g01746.t01#3    mNeoVis.3.1.1.chr1.g01738.t01#6    mNeoVis.vulgivaga.2.1.chr1.g01744.t01#1    mNeoVis.vulgivaga.1.1.chr1.g01751.t01#4   mNeoVis.evergladensis.2.1.chr1.g01746.t01#2       1094   125259666:   mNeoVis.evergladensis.2.1.chr7.g00144.t01#2    mNeoVis.vulgivaga.2.1.chr7.g00145.t01#1    mNeoVis.vulgivaga.1.1.chr7.g00148.t01#4    mNeoVis.3.1.1.chr7.g00127.t01#6    mNeoVis.2.1.1.chr7_hap1.g00144.t01#5   mNeoVis.evergladensis.1.1.chr7.g00147.t01#3       1095   125259667:   mNeoVis.3.1.1.chr1.g01739.t01#6    mNeoVis.2.1.1.chr1_hap1.g01780.t01#5    mNeoVis.vulgivaga.2.1.chr1.g01745.t01#1    mNeoVis.vulgivaga.1.1.chr1.g01752.t01#4    mNeoVis.evergladensis.1.1.chr1.g01747.t01#3   mNeoVis.evergladensis.2.1.chr1.g01747.t01#2       1096   125259668:   mNeoVis.evergladensis.1.1.chr15.g00488.t01#3    mNeoVis.3.1.1.chr15.g00119.t01#6    mNeoVis.vulgivaga.2.1.chr15.g00131.t01#1    mNeoVis.2.1.1.chr15_hap1.g00142.t01#5    mNeoVis.evergladensis.2.1.chr15.g00128.t01#2   mNeoVis.vulgivaga.1.1.chr15.g00128.t01#4       1097   125259669:   mNeoVis.evergladensis.2.1.chr7.g02401.t01#2    mNeoVis.evergladensis.1.1.chr7.g02399.t01#3    mNeoVis.2.1.1.chr7_hap1.g02595.t01#5    mNeoVis.3.1.1.chr7.g02258.t01#6    mNeoVis.vulgivaga.1.1.chr7.g02405.t01#4   mNeoVis.vulgivaga.2.1.chr7.g02423.t01#1       1098   125259670:  mNeoVis.vulgivaga.2.1.chr7.g02424.t01#1               1099   125259671:   mNeoVis.evergladensis.2.1.chr7.g02402.t01#2    mNeoVis.vulgivaga.2.1.chr7.g02425.t01#1    mNeoVis.2.1.1.chr7_hap1.g02596.t01#5    mNeoVis.vulgivaga.1.1.chr7.g02406.t01#4   mNeoVis.evergladensis.1.1.chr7.g02400.t01#3       1100   125259672:   mNeoVis.2.1.1.chr1_hap1.g01781.t01#5    mNeoVis.vulgivaga.1.1.chr1.g01753.t01#4    mNeoVis.evergladensis.1.1.chr1.g01748.t01#3    mNeoVis.3.1.1.chr1.g01740.t01#6    mNeoVis.vulgivaga.2.1.chr1.g01746.t01#1   mNeoVis.evergladensis.2.1.chr1.g01748.t01#2       1101   125259673:   mNeoVis.vulgivaga.2.1.chr7.g02426.t01#1    mNeoVis.evergladensis.1.1.chr7.g02401.t01#3   mNeoVis.3.1.1.chr7.g02260.t01#6           1102   125259674:   mNeoVis.evergladensis.2.1.chr7.g02404.t01#2    mNeoVis.vulgivaga.1.1.chr7.g02408.t01#4    mNeoVis.evergladensis.1.1.chr7.g02402.t01#3    mNeoVis.vulgivaga.2.1.chr7.g02427.t01#1    mNeoVis.3.1.1.chr7.g02261.t01#6   mNeoVis.2.1.1.chr7_hap1.g02598.t01#5       1103   125259675:   mNeoVis.evergladensis.1.1.chr15.g00487.t01#3    mNeoVis.3.1.1.chr15.g00120.t01#6    mNeoVis.2.1.1.chr15_hap1.g00143.t01#5    mNeoVis.vulgivaga.1.1.chr15.g00129.t01#4    mNeoVis.evergladensis.2.1.chr15.g00129.t01#2   mNeoVis.vulgivaga.2.1.chr15.g00132.t01#1       1104   125259676:   mNeoVis.3.1.1.chr7.g02262.t01#6    mNeoVis.2.1.1.chr7_hap1.g02599.t01#5    mNeoVis.evergladensis.2.1.chr7.g02405.t01#2    mNeoVis.vulgivaga.2.1.chr7.g02428.t01#1    mNeoVis.vulgivaga.1.1.chr7.g02409.t01#4   mNeoVis.evergladensis.1.1.chr7.g02403.t01#3       1105   125259677:   mNeoVis.evergladensis.2.1.chr1.g01749.t01#2    mNeoVis.vulgivaga.2.1.chr1.g01747.t01#1    mNeoVis.3.1.1.chr1.g01741.t01#6    mNeoVis.evergladensis.1.1.chr1.g01750.t01#3    mNeoVis.2.1.1.chr1_hap1.g01782.t01#5   mNeoVis.vulgivaga.1.1.chr1.g01754.t01#4       1106   125259678:   mNeoVis.vulgivaga.1.1.chr7.g02410.t01#4    mNeoVis.evergladensis.2.1.chr7.g02406.t01#2    mNeoVis.evergladensis.1.1.chr7.g02404.t01#3    mNeoVis.vulgivaga.2.1.chr7.g02429.t01#1   mNeoVis.2.1.1.chr7_hap1.g02600.t01#5       1107   125259679:   mNeoVis.vulgivaga.2.1.chr7.g02430.t01#1    mNeoVis.2.1.1.chr7_hap1.g02601.t01#5    mNeoVis.vulgivaga.1.1.chr7.g02411.t01#4    mNeoVis.evergladensis.2.1.chr7.g02407.t01#2   mNeoVis.evergladensis.1.1.chr7.g02405.t01#3       1108   125259680:   mNeoVis.vulgivaga.2.1.chr7.g00146.t01#1    mNeoVis.2.1.1.chr7_hap1.g00145.t01#5    mNeoVis.evergladensis.2.1.chr7.g00145.t01#2    mNeoVis.vulgivaga.1.1.chr7.g00149.t01#4    mNeoVis.3.1.1.chr7.g00128.t01#6   mNeoVis.evergladensis.1.1.chr7.g00148.t01#3       1109   125259681:   mNeoVis.vulgivaga.2.1.chr15.g00133.t01#1    mNeoVis.evergladensis.2.1.chr15.g00130.t01#2    mNeoVis.vulgivaga.1.1.chr15.g00130.t01#4    mNeoVis.2.1.1.chr15_hap1.g00144.t01#5    mNeoVis.evergladensis.1.1.chr15.g00485.t01#3   mNeoVis.3.1.1.chr15.g00121.t01#6       1110   125259682:   mNeoVis.evergladensis.1.1.chr7.g02406.t01#3    mNeoVis.vulgivaga.2.1.chr7.g02431.t01#1    mNeoVis.evergladensis.2.1.chr7.g02409.t01#2    mNeoVis.3.1.1.chr7.g02264.t01#6    mNeoVis.vulgivaga.1.1.chr7.g02412.t01#4   mNeoVis.2.1.1.chr7_hap1.g02602.t01#5       1111   125259683:   mNeoVis.evergladensis.2.1.chr1.g01655.t01#2    mNeoVis.evergladensis.1.1.chr1.g01655.t01#3    mNeoVis.3.1.1.chr1.g01643.t01#6   mNeoVis.vulgivaga.2.1.chr1.g01653.t01#1         1112   125259684:   mNeoVis.3.1.1.chr7.g02265.t01#6    mNeoVis.vulgivaga.1.1.chr7.g02413.t01#4    mNeoVis.2.1.1.chr7_hap1.g02603.t01#5    mNeoVis.vulgivaga.2.1.chr7.g02432.t01#1    mNeoVis.evergladensis.1.1.chr7.g02407.t01#3   mNeoVis.evergladensis.2.1.chr7.g02410.t01#2       1113   125259685:   mNeoVis.vulgivaga.1.1.chr1.g01659.t01#4    mNeoVis.evergladensis.2.1.chr1.g01657.t01#2    mNeoVis.3.1.1.chr1.g01645.t01#6    mNeoVis.2.1.1.chr1_hap1.g01690.t01#5    mNeoVis.vulgivaga.2.1.chr1.g01654.t01#1   mNeoVis.evergladensis.1.1.chr1.g01657.t01#3       1114   125259686:   mNeoVis.evergladensis.1.1.chr1.g01658.t01#3    mNeoVis.evergladensis.2.1.chr1.g01658.t01#2    mNeoVis.vulgivaga.1.1.chr1.g01660.t01#4    mNeoVis.vulgivaga.2.1.chr1.g01655.t01#1    mNeoVis.3.1.1.chr1.g01646.t01#6   mNeoVis.2.1.1.chr1_hap1.g01691.t01#5       1115   125259687:   mNeoVis.evergladensis.1.1.chr1.g01659.t01#3    mNeoVis.evergladensis.2.1.chr1.g01659.t01#2    mNeoVis.vulgivaga.1.1.chr1.g01661.t01#4    mNeoVis.3.1.1.chr1.g01647.t01#6    mNeoVis.2.1.1.chr1_hap1.g01692.t01#5   mNeoVis.vulgivaga.2.1.chr1.g01656.t01#1       1116   125259688:   mNeoVis.vulgivaga.2.1.chr7.g02433.t01#1    mNeoVis.evergladensis.2.1.chr7.g02411.t01#2    mNeoVis.3.1.1.chr7.g02266.t01#6    mNeoVis.2.1.1.chr7_hap1.g02604.t01#5    mNeoVis.evergladensis.1.1.chr7.g02408.t01#3   mNeoVis.vulgivaga.1.1.chr7.g02414.t01#4       1117   125259689:   mNeoVis.2.1.1.chr1_hap1.g01693.t01#5    mNeoVis.vulgivaga.2.1.chr1.g01657.t01#1    mNeoVis.evergladensis.2.1.chr1.g01660.t01#2    mNeoVis.vulgivaga.1.1.chr1.g01662.t01#4    mNeoVis.evergladensis.1.1.chr1.g01660.t01#3   mNeoVis.3.1.1.chr1.g01648.t01#6       1118   125259690:   mNeoVis.evergladensis.2.1.chr1.g01661.t01#2    mNeoVis.vulgivaga.2.1.chr1.g01658.t01#1    mNeoVis.evergladensis.1.1.chr1.g01661.t01#3    mNeoVis.3.1.1.chr1.g01649.t01#6    mNeoVis.vulgivaga.1.1.chr1.g01663.t01#4   mNeoVis.2.1.1.chr1_hap1.g01694.t01#5       1119   125259691:   mNeoVis.2.1.1.chr7_hap1.g02605.t01#5    mNeoVis.vulgivaga.2.1.chr7.g02434.t01#1    mNeoVis.evergladensis.2.1.chr7.g02412.t01#2   mNeoVis.evergladensis.1.1.chr7.g02409.t01#3       1120   125259692:   mNeoVis.3.1.1.chr1.g01651.t01#6    mNeoVis.2.1.1.chr1_hap1.g01696.t01#5    mNeoVis.evergladensis.2.1.chr1.g01663.t01#2    mNeoVis.vulgivaga.2.1.chr1.g01659.t01#1    mNeoVis.evergladensis.1.1.chr1.g01662.t01#3   mNeoVis.vulgivaga.1.1.chr1.g01665.t01#4       1121   125259693:   mNeoVis.3.1.1.chr7.g02270.t01#6    mNeoVis.2.1.1.chr7_hap1.g02606.t01#5    mNeoVis.evergladensis.1.1.chr7.g02411.t01#3    mNeoVis.vulgivaga.1.1.chr7.g02416.t01#4    mNeoVis.vulgivaga.2.1.chr7.g02435.t01#1   mNeoVis.evergladensis.2.1.chr7.g02413.t01#2       1122   125259694:   mNeoVis.evergladensis.1.1.chr1.g01663.t01#3    mNeoVis.3.1.1.chr1.g01652.t01#6    mNeoVis.2.1.1.chr1_hap1.g01697.t01#5    mNeoVis.evergladensis.2.1.chr1.g01664.t01#2    mNeoVis.vulgivaga.1.1.chr1.g01666.t01#4   mNeoVis.vulgivaga.2.1.chr1.g01660.t01#1       1123   125259695:   mNeoVis.vulgivaga.2.1.chr1.g01661.t01#1    mNeoVis.vulgivaga.1.1.chr1.g01668.t01#4    mNeoVis.evergladensis.2.1.chr1.g01665.t01#2    mNeoVis.2.1.1.chr1_hap1.g01698.t01#5    mNeoVis.3.1.1.chr1.g01653.t01#6   mNeoVis.evergladensis.1.1.chr1.g01664.t01#3       1124   125259696:   mNeoVis.vulgivaga.1.1.chr7.g02417.t01#4    mNeoVis.vulgivaga.2.1.chr7.g02436.t01#1    mNeoVis.evergladensis.1.1.chr7.g02412.t01#3    mNeoVis.2.1.1.chr7_hap1.g02607.t01#5   mNeoVis.evergladensis.2.1.chr7.g02414.t01#2       1125   125259697:   mNeoVis.2.1.1.chr1_hap1.g01699.t01#5    mNeoVis.evergladensis.1.1.chr1.g01665.t01#3    mNeoVis.vulgivaga.1.1.chr1.g01667.t01#4    mNeoVis.vulgivaga.2.1.chr1.g01662.t01#1    mNeoVis.evergladensis.2.1.chr1.g01666.t01#2   mNeoVis.3.1.1.chr1.g01654.t01#6       1126   125259698:   mNeoVis.3.1.1.chr1.g01655.t01#6    mNeoVis.vulgivaga.2.1.chr1.g01663.t01#1    mNeoVis.vulgivaga.1.1.chr1.g01669.t01#4    mNeoVis.2.1.1.chr1_hap1.g01700.t01#5    mNeoVis.evergladensis.2.1.chr1.g01667.t01#2   mNeoVis.evergladensis.1.1.chr1.g01666.t01#3       1127   125259699:   mNeoVis.3.1.1.chr7.g02273.t01#6    mNeoVis.vulgivaga.1.1.chr7.g02418.t01#4    mNeoVis.2.1.1.chr7_hap1.g02608.t01#5    mNeoVis.evergladensis.2.1.chr7.g02415.t01#2    mNeoVis.vulgivaga.2.1.chr7.g02437.t01#1   mNeoVis.evergladensis.1.1.chr7.g02413.t01#3       1128   125259700:   mNeoVis.2.1.1.chr7_hap1.g00728.t01#5    mNeoVis.evergladensis.2.1.chr7.g00719.t01#2    mNeoVis.evergladensis.1.1.chr7.g00728.t01#3   mNeoVis.vulgivaga.2.1.chr7.g00720.t01#1         1129   125259701:   mNeoVis.3.1.1.chr1.g01657.t01#6    mNeoVis.vulgivaga.2.1.chr1.g01665.t01#1    mNeoVis.evergladensis.2.1.chr1.g01669.t01#2    mNeoVis.evergladensis.1.1.chr1.g01668.t01#3    mNeoVis.vulgivaga.1.1.chr1.g01672.t01#4   mNeoVis.2.1.1.chr1_hap1.g01702.t01#5       1130   125259702:  mNeoVis.vulgivaga.2.1.chr7.g02438.t01#1               1131   125259703:   mNeoVis.2.1.1.chr1_hap1.g01703.t01#5    mNeoVis.3.1.1.chr1.g01658.t01#6    mNeoVis.vulgivaga.2.1.chr1.g01666.t01#1    mNeoVis.evergladensis.1.1.chr1.g01669.t01#3    mNeoVis.evergladensis.2.1.chr1.g01670.t01#2   mNeoVis.vulgivaga.1.1.chr1.g01673.t01#4       1132   125259704:   mNeoVis.3.1.1.chr7.g00129.t01#6    mNeoVis.evergladensis.2.1.chr7.g00146.t01#2    mNeoVis.evergladensis.1.1.chr7.g00149.t01#3    mNeoVis.2.1.1.chr7_hap1.g00146.t01#5    mNeoVis.vulgivaga.2.1.chr7.g00147.t01#1   mNeoVis.vulgivaga.1.1.chr7.g00150.t01#4       1133   125259705:   mNeoVis.evergladensis.1.1.chr7.g02415.t01#3    mNeoVis.2.1.1.chr7_hap1.g02611.t01#5    mNeoVis.vulgivaga.1.1.chr7.g02421.t01#4    mNeoVis.vulgivaga.2.1.chr7.g02441.t01#1   mNeoVis.evergladensis.2.1.chr7.g02417.t01#2       1134   125259706:   mNeoVis.evergladensis.1.1.chr7.g02416.t01#3    mNeoVis.evergladensis.2.1.chr7.g02418.t01#2    mNeoVis.2.1.1.chr7_hap1.g02612.t01#5    mNeoVis.vulgivaga.2.1.chr7.g02442.t01#1   mNeoVis.vulgivaga.1.1.chr7.g02422.t01#4       1135   125259707:   mNeoVis.3.1.1.chr15.g00122.t01#6    mNeoVis.2.1.1.chr15_hap1.g00145.t01#5   mNeoVis.vulgivaga.2.1.chr15.g00134.t01#1         1136   125259708:   mNeoVis.2.1.1.chr3_hap1.g00445.t01#5    mNeoVis.evergladensis.2.1.chr3.g00453.t01#2   mNeoVis.3.1.1.chr3.g00438.t01#6           1137   125259709:  mNeoVis.evergladensis.1.1.chr15.g00486.t01#3             1138   125259710:   mNeoVis.vulgivaga.1.1.chr1.g01674.t01#4    mNeoVis.vulgivaga.2.1.chr1.g01667.t01#1    mNeoVis.2.1.1.chr1_hap1.g01704.t01#5    mNeoVis.evergladensis.2.1.chr1.g01671.t01#2    mNeoVis.3.1.1.chr1.g01659.t01#6   mNeoVis.evergladensis.1.1.chr1.g01670.t01#3       1139   125259711:   mNeoVis.vulgivaga.1.1.chr1.g01675.t01#4    mNeoVis.vulgivaga.2.1.chr1.g01668.t01#1    mNeoVis.3.1.1.chr1.g01660.t01#6    mNeoVis.2.1.1.chr1_hap1.g01705.t01#5    mNeoVis.evergladensis.1.1.chr1.g01671.t01#3   mNeoVis.evergladensis.2.1.chr1.g01672.t01#2       1140   125259712:   mNeoVis.vulgivaga.2.1.chr1.g01669.t01#1    mNeoVis.evergladensis.2.1.chr1.g01673.t01#2    mNeoVis.2.1.1.chr1_hap1.g01706.t01#5    mNeoVis.vulgivaga.1.1.chr1.g01676.t01#4    mNeoVis.evergladensis.1.1.chr1.g01672.t01#3   mNeoVis.3.1.1.chr1.g01661.t01#6       1141   125259713:   mNeoVis.evergladensis.1.1.chr1.g01673.t01#3    mNeoVis.vulgivaga.1.1.chr1.g01677.t01#4    mNeoVis.vulgivaga.2.1.chr1.g01670.t01#1    mNeoVis.vulgivaga.1.1.chr1.g01678.t01#4    mNeoVis.vulgivaga.2.1.chr1.g01671.t01#1    mNeoVis.2.1.1.chr1_hap1.g01707.t01#5    mNeoVis.evergladensis.2.1.chr1.g01674.t01#2   mNeoVis.3.1.1.chr1.g01662.t01#6     1142   125259714:   mNeoVis.vulgivaga.2.1.chr2.g00856.t01#1    mNeoVis.evergladensis.1.1.chr2.g00850.t01#3    mNeoVis.3.1.1.chr2.g00766.t01#6    mNeoVis.evergladensis.2.1.chr2.g00855.t01#2    mNeoVis.vulgivaga.1.1.chr2.g00856.t01#4   mNeoVis.2.1.1.chr5_hap1.g00854.t01#5       1143   125259715:   mNeoVis.vulgivaga.1.1.chr10.g00653.t01#4    mNeoVis.evergladensis.1.1.chr10.g00673.t01#3    mNeoVis.2.1.1.chr9_hap1.g00677.t01#5    mNeoVis.vulgivaga.2.1.chr10.g00655.t01#1    mNeoVis.3.1.1.chr10.g00642.t01#6   mNeoVis.evergladensis.2.1.chr10.g00672.t01#2       1144   125259716:  mNeoVis.3.1.1.chr4.g00020.t01#6               1145   125259717:   mNeoVis.vulgivaga.2.1.chr7.g02443.t01#1    mNeoVis.evergladensis.1.1.chr7.g02417.t01#3    mNeoVis.evergladensis.2.1.chr7.g02419.t01#2   mNeoVis.2.1.1.chr7_hap1.g02613.t01#5         1146   125259718:   mNeoVis.3.1.1.chr15.g00123.t01#6    mNeoVis.2.1.1.chr15_hap1.g00146.t01#5    mNeoVis.evergladensis.1.1.chr15.g00484.t01#3    mNeoVis.evergladensis.2.1.chr15.g00131.t01#2    mNeoVis.vulgivaga.1.1.chr15.g00131.t01#4   mNeoVis.vulgivaga.2.1.chr15.g00135.t01#1       1147   125259719:   mNeoVis.vulgivaga.2.1.chr7.g02444.t01#1    mNeoVis.2.1.1.chr7_hap1.g02614.t01#5    mNeoVis.evergladensis.1.1.chr7.g02418.t01#3   mNeoVis.evergladensis.2.1.chr7.g02420.t01#2       1148   125259720:   mNeoVis.vulgivaga.1.1.chr1.g01679.t01#4    mNeoVis.2.1.1.chr1_hap1.g01708.t01#5    mNeoVis.evergladensis.1.1.chr1.g01674.t01#3    mNeoVis.vulgivaga.2.1.chr1.g01672.t01#1    mNeoVis.3.1.1.chr1.g01663.t01#6   mNeoVis.evergladensis.2.1.chr1.g01675.t01#2       1149   125259721:   mNeoVis.evergladensis.1.1.chr1.g01675.t01#3    mNeoVis.evergladensis.2.1.chr1.g01676.t01#2    mNeoVis.vulgivaga.1.1.chr1.g01680.t01#4    mNeoVis.2.1.1.chr1_hap1.g01709.t01#5   mNeoVis.vulgivaga.2.1.chr1.g01673.t01#1       1150   125259722:   mNeoVis.evergladensis.2.1.chr1.g00909.t01#2    mNeoVis.vulgivaga.2.1.chr1.g01674.t01#1    mNeoVis.vulgivaga.1.1.chr1.g01681.t01#4    mNeoVis.3.1.1.chr1.g00928.t01#6    mNeoVis.2.1.1.chr1_hap1.g01710.t01#5    mNeoVis.evergladensis.1.1.chr1.g00917.t01#3    mNeoVis.vulgivaga.2.1.chr1.g00919.t01#1    mNeoVis.3.1.1.chr1.g01664.t01#6    mNeoVis.2.1.1.chr1_hap1.g00905.t01#5      1151   125259723:   mNeoVis.evergladensis.2.1.chr7.g02421.t01#2    mNeoVis.vulgivaga.2.1.chr7.g02445.t01#1    mNeoVis.2.1.1.chr7_hap1.g02615.t01#5    mNeoVis.vulgivaga.1.1.chr7.g02424.t01#4    mNeoVis.3.1.1.chr7.g02275.t01#6   mNeoVis.evergladensis.1.1.chr7.g02419.t01#3       1152   125259724:   mNeoVis.vulgivaga.1.1.chr1.g01682.t01#4    mNeoVis.vulgivaga.2.1.chr1.g01675.t01#1    mNeoVis.3.1.1.chr1.g01665.t01#6    mNeoVis.2.1.1.chr1_hap1.g01711.t01#5    mNeoVis.evergladensis.2.1.chr1.g01678.t01#2   mNeoVis.evergladensis.1.1.chr1.g01677.t01#3       1153   125259725:   mNeoVis.evergladensis.2.1.chr10.g00717.t01#2    mNeoVis.vulgivaga.2.1.chr10.g00707.t01#1    mNeoVis.3.1.1.chr10.g00693.t01#6    mNeoVis.2.1.1.chr9_hap1.g00724.t01#5    mNeoVis.vulgivaga.1.1.chr10.g00702.t01#4   mNeoVis.evergladensis.1.1.chr10.g00720.t01#3       1154   125259726:   mNeoVis.2.1.1.chr7_hap1.g02616.t01#5    mNeoVis.vulgivaga.2.1.chr7.g02446.t01#1   mNeoVis.evergladensis.2.1.chr7.g02422.t01#2         1155   125259727:   mNeoVis.vulgivaga.2.1.chr7.g02447.t01#1    mNeoVis.evergladensis.2.1.chr7.g02423.t01#2    mNeoVis.vulgivaga.1.1.chr7.g02425.t01#4    mNeoVis.2.1.1.chr7_hap1.g02617.t01#5   mNeoVis.evergladensis.1.1.chr7.g02420.t01#3       1156   125259728:   mNeoVis.evergladensis.1.1.chr7.g02421.t01#3    mNeoVis.evergladensis.2.1.chr7.g02424.t01#2    mNeoVis.vulgivaga.2.1.chr7.g02448.t01#1    mNeoVis.vulgivaga.1.1.chr7.g02426.t01#4   mNeoVis.2.1.1.chr7_hap1.g02618.t01#5       1157   125259729:   mNeoVis.evergladensis.1.1.chr1.g01678.t01#3    mNeoVis.2.1.1.chr1_hap1.g01712.t01#5    mNeoVis.3.1.1.chr1.g01666.t01#6    mNeoVis.vulgivaga.1.1.chr1.g01683.t01#4    mNeoVis.vulgivaga.2.1.chr1.g01676.t01#1   mNeoVis.evergladensis.2.1.chr1.g01679.t01#2       1158   125259730:   mNeoVis.2.1.1.chr1_hap1.g01713.t01#5    mNeoVis.vulgivaga.2.1.chr1.g01677.t01#1    mNeoVis.evergladensis.1.1.chr1.g01679.t01#3    mNeoVis.3.1.1.chr1.g01667.t01#6    mNeoVis.vulgivaga.1.1.chr1.g01684.t01#4   mNeoVis.evergladensis.2.1.chr1.g01680.t01#2       1159   125259731:   mNeoVis.2.1.1.chr7_hap1.g02619.t01#5    mNeoVis.vulgivaga.2.1.chr7.g02449.t01#1    mNeoVis.evergladensis.1.1.chr7.g02422.t01#3    mNeoVis.vulgivaga.1.1.chr7.g02427.t01#4    mNeoVis.evergladensis.2.1.chr7.g02425.t01#2   mNeoVis.3.1.1.chr7.g02276.t01#6       1160   125259732:   mNeoVis.3.1.1.chr1.g01668.t01#6    mNeoVis.evergladensis.2.1.chr1.g01681.t01#2    mNeoVis.evergladensis.1.1.chr1.g01680.t01#3    mNeoVis.vulgivaga.2.1.chr1.g01678.t01#1    mNeoVis.vulgivaga.1.1.chr1.g01685.t01#4   mNeoVis.2.1.1.chr1_hap1.g01714.t01#5       1161   125259733:   mNeoVis.vulgivaga.2.1.chr1.g01679.t01#1   mNeoVis.evergladensis.2.1.chr1.g01682.t01#2           1162   125259734:   mNeoVis.evergladensis.1.1.chr7.g02423.t01#3    mNeoVis.vulgivaga.1.1.chr7.g02428.t01#4    mNeoVis.3.1.1.chr7.g02277.t01#6    mNeoVis.evergladensis.2.1.chr7.g02426.t01#2    mNeoVis.2.1.1.chr7_hap1.g02620.t01#5    mNeoVis.evergladensis.1.1.chr7.g02424.t01#3   mNeoVis.vulgivaga.2.1.chr7.g02450.t01#1     1163   125259735:   mNeoVis.vulgivaga.2.1.chr1.g01680.t01#1    mNeoVis.2.1.1.chr1_hap1.g01715.t01#5    mNeoVis.3.1.1.chr1.g01669.t01#6    mNeoVis.evergladensis.1.1.chr1.g01681.t01#3    mNeoVis.vulgivaga.1.1.chr1.g01686.t01#4   mNeoVis.evergladensis.2.1.chr1.g01683.t01#2       1164   125259736:   mNeoVis.vulgivaga.1.1.chr1.g01688.t01#4    mNeoVis.vulgivaga.1.1.chr1.g01689.t01#4    mNeoVis.vulgivaga.2.1.chr1.g01682.t01#1    mNeoVis.3.1.1.chr1.g01672.t01#6    mNeoVis.evergladensis.2.1.chr1.g01686.t01#2    mNeoVis.3.1.1.chr1.g01671.t01#6    mNeoVis.evergladensis.2.1.chr1.g01685.t01#2    mNeoVis.2.1.1.chr1_hap1.g01717.t01#5    mNeoVis.evergladensis.1.1.chr1.g01682.t01#3      1165   125259737:   mNeoVis.vulgivaga.1.1.chr1.g01687.t01#4    mNeoVis.vulgivaga.2.1.chr1.g01681.t01#1    mNeoVis.evergladensis.2.1.chr1.g01684.t01#2   mNeoVis.2.1.1.chr1_hap1.g01716.t01#5         1166   125259738:   mNeoVis.2.1.1.chr1_hap1.g01718.t01#5   mNeoVis.3.1.1.chr1.g01670.t01#6             1167   125259739:   mNeoVis.evergladensis.2.1.chr7.g02428.t01#2    mNeoVis.vulgivaga.1.1.chr7.g02429.t01#4    mNeoVis.vulgivaga.2.1.chr7.g02451.t01#1    mNeoVis.2.1.1.chr7_hap1.g02621.t01#5   mNeoVis.evergladensis.1.1.chr7.g02425.t01#3       1168   125259740:   mNeoVis.vulgivaga.2.1.chr1.g01683.t01#1    mNeoVis.3.1.1.chr1.g01673.t01#6    mNeoVis.evergladensis.2.1.chr1.g01687.t01#2   mNeoVis.evergladensis.1.1.chr1.g01684.t01#3       1169   125259741:   mNeoVis.evergladensis.1.1.chr1.g01685.t01#3    mNeoVis.2.1.1.chr1_hap1.g01719.t01#5    mNeoVis.evergladensis.2.1.chr1.g01688.t01#2    mNeoVis.vulgivaga.2.1.chr1.g01684.t01#1    mNeoVis.3.1.1.chr1.g01674.t01#6   mNeoVis.vulgivaga.1.1.chr1.g01690.t01#4       1170   125259742:   mNeoVis.3.1.1.chr1.g01675.t01#6    mNeoVis.vulgivaga.1.1.chr1.g01691.t01#4    mNeoVis.2.1.1.chr1_hap1.g01720.t01#5    mNeoVis.vulgivaga.2.1.chr1.g01685.t01#1    mNeoVis.evergladensis.1.1.chr1.g01686.t01#3   mNeoVis.evergladensis.2.1.chr1.g01689.t01#2       1171   125259743:   mNeoVis.vulgivaga.1.1.chr1.g01692.t01#4    mNeoVis.2.1.1.chr1_hap1.g01721.t01#5    mNeoVis.3.1.1.chr1.g01676.t01#6    mNeoVis.vulgivaga.2.1.chr1.g01686.t01#1    mNeoVis.evergladensis.2.1.chr1.g01690.t01#2   mNeoVis.evergladensis.1.1.chr1.g01687.t01#3       1172   125259744:   mNeoVis.2.1.1.chr7_hap1.g02622.t01#5    mNeoVis.vulgivaga.1.1.chr7.g02430.t01#4    mNeoVis.evergladensis.1.1.chr7.g02426.t01#3    mNeoVis.3.1.1.chr7.g02278.t01#6    mNeoVis.evergladensis.2.1.chr7.g02429.t01#2   mNeoVis.vulgivaga.2.1.chr7.g02452.t01#1       1173   125259745:   mNeoVis.evergladensis.2.1.chr7.g02430.t01#2    mNeoVis.evergladensis.1.1.chr7.g02427.t01#3    mNeoVis.3.1.1.chr7.g02279.t01#6    mNeoVis.vulgivaga.2.1.chr7.g02453.t01#1   mNeoVis.2.1.1.chr7_hap1.g02623.t01#5       1174   125259746:   mNeoVis.2.1.1.chr7_hap1.g01112.t01#5    mNeoVis.vulgivaga.2.1.chr7.g01111.t01#1    mNeoVis.evergladensis.1.1.chr5.g01507.t01#3    mNeoVis.2.1.1.chr2_hap1.g01562.t01#5    mNeoVis.vulgivaga.2.1.chr5.g01504.t01#1    mNeoVis.vulgivaga.1.1.chr5.g01479.t01#4    mNeoVis.3.1.1.chr5.g01483.t01#6    mNeoVis.evergladensis.2.1.chr5.g01500.t01#2    mNeoVis.3.1.1.chr7.g01035.t01#6      1175   125259747:   mNeoVis.vulgivaga.2.1.chr2.g01194.t01#1    mNeoVis.evergladensis.1.1.chr2.g01191.t01#3    mNeoVis.vulgivaga.1.1.chr2.g01193.t01#4    mNeoVis.3.1.1.chr2.g01112.t01#6    mNeoVis.2.1.1.chr5_hap1.g01200.t01#5   mNeoVis.evergladensis.2.1.chr2.g01188.t01#2       1176   125259748:   mNeoVis.vulgivaga.1.1.chr1.g01694.t01#4    mNeoVis.vulgivaga.1.1.chr1.g01693.t01#4    mNeoVis.3.1.1.chr1.g01677.t01#6    mNeoVis.evergladensis.2.1.chr1.g01691.t01#2    mNeoVis.2.1.1.chr1_hap1.g01722.t01#5   mNeoVis.evergladensis.1.1.chr1.g01688.t01#3       1177   125259749:  mNeoVis.vulgivaga.2.1.chr1.g01687.t01#1               1178   125259750:   mNeoVis.vulgivaga.2.1.chr7.g02454.t01#1    mNeoVis.evergladensis.2.1.chr7.g02431.t01#2    mNeoVis.2.1.1.chr7_hap1.g02624.t01#5    mNeoVis.evergladensis.1.1.chr7.g02428.t01#3   mNeoVis.vulgivaga.1.1.chr7.g02432.t01#4       1179   125259751:   mNeoVis.vulgivaga.1.1.chr1.g01695.t01#4    mNeoVis.evergladensis.1.1.chr1.g01689.t01#3    mNeoVis.3.1.1.chr1.g01678.t01#6    mNeoVis.2.1.1.chr1_hap1.g01723.t01#5    mNeoVis.evergladensis.2.1.chr1.g01692.t01#2   mNeoVis.vulgivaga.2.1.chr1.g01688.t01#1       1180   125259752:   mNeoVis.vulgivaga.2.1.chr1.g01689.t01#1    mNeoVis.3.1.1.chr1.g01679.t01#6    mNeoVis.evergladensis.2.1.chr1.g01693.t01#2    mNeoVis.2.1.1.chr1_hap1.g01724.t01#5    mNeoVis.vulgivaga.1.1.chr1.g01696.t01#4   mNeoVis.evergladensis.1.1.chr1.g01690.t01#3       1181   125259753:   mNeoVis.2.1.1.chr7_hap1.g02625.t01#5    mNeoVis.vulgivaga.1.1.chr7.g02433.t01#4    mNeoVis.vulgivaga.2.1.chr7.g02455.t01#1    mNeoVis.evergladensis.1.1.chr7.g02429.t01#3    mNeoVis.evergladensis.2.1.chr7.g02432.t01#2   mNeoVis.3.1.1.chr7.g02281.t01#6       1182   125259754:   mNeoVis.vulgivaga.2.1.chr7.g02456.t01#1    mNeoVis.evergladensis.1.1.chr7.g02430.t01#3    mNeoVis.2.1.1.chr7_hap1.g02626.t01#5    mNeoVis.vulgivaga.1.1.chr7.g02434.t01#4   mNeoVis.evergladensis.2.1.chr7.g02433.t01#2       1183   125259755:   mNeoVis.evergladensis.1.1.chr15.g00483.t01#3    mNeoVis.2.1.1.chr15_hap1.g00147.t01#5    mNeoVis.evergladensis.2.1.chr15.g00132.t01#2    mNeoVis.vulgivaga.1.1.chr15.g00132.t01#4    mNeoVis.3.1.1.chr15.g00124.t01#6    mNeoVis.3.1.1.chr15.g00125.t01#6   mNeoVis.vulgivaga.2.1.chr15.g00136.t01#1     1184   125259756:   mNeoVis.2.1.1.chr1_hap1.g01725.t01#5    mNeoVis.3.1.1.chr1.g01680.t01#6    mNeoVis.evergladensis.2.1.chr1.g01694.t01#2    mNeoVis.vulgivaga.1.1.chr1.g01697.t01#4    mNeoVis.vulgivaga.2.1.chr1.g01690.t01#1   mNeoVis.evergladensis.1.1.chr1.g01691.t01#3       1185   125259757:   mNeoVis.vulgivaga.1.1.chr7.g02435.t01#4    mNeoVis.2.1.1.chr7_hap1.g02627.t01#5    mNeoVis.evergladensis.2.1.chr7.g02434.t01#2    mNeoVis.vulgivaga.2.1.chr7.g02457.t01#1    mNeoVis.evergladensis.1.1.chr7.g02431.t01#3   mNeoVis.3.1.1.chr7.g02283.t01#6       1186   125259758:   mNeoVis.vulgivaga.1.1.chr7.g02436.t01#4    mNeoVis.evergladensis.1.1.chr7.g02432.t01#3    mNeoVis.3.1.1.chr7.g02284.t01#6    mNeoVis.evergladensis.2.1.chr7.g02435.t01#2    mNeoVis.vulgivaga.2.1.chr7.g02458.t01#1   mNeoVis.2.1.1.chr7_hap1.g02628.t01#5       1187   125259759:   mNeoVis.evergladensis.1.1.chr2.g01434.t01#3    mNeoVis.vulgivaga.1.1.chr2.g01436.t01#4    mNeoVis.2.1.1.chr5_hap1.g01436.t01#5    mNeoVis.evergladensis.2.1.chr2.g01424.t01#2    mNeoVis.vulgivaga.2.1.chr2.g01432.t01#1   mNeoVis.3.1.1.chr2.g01354.t01#6       1188   125259760:   mNeoVis.vulgivaga.1.1.chr7.g02437.t01#4    mNeoVis.2.1.1.chr7_hap1.g02629.t01#5    mNeoVis.evergladensis.2.1.chr7.g02436.t01#2    mNeoVis.vulgivaga.2.1.chr7.g02459.t01#1   mNeoVis.evergladensis.1.1.chr7.g02433.t01#3       1189   125259761:   mNeoVis.vulgivaga.1.1.chr1.g01698.t01#4    mNeoVis.evergladensis.1.1.chr1.g01692.t01#3    mNeoVis.evergladensis.2.1.chr1.g01695.t01#2    mNeoVis.vulgivaga.2.1.chr1.g01691.t01#1    mNeoVis.2.1.1.chr1_hap1.g01726.t01#5   mNeoVis.3.1.1.chr1.g01681.t01#6       1190   125259762:   mNeoVis.2.1.1.chr1_hap1.g01727.t01#5    mNeoVis.vulgivaga.2.1.chr1.g01692.t01#1    mNeoVis.3.1.1.chr1.g01682.t01#6    mNeoVis.evergladensis.1.1.chr1.g01693.t01#3    mNeoVis.vulgivaga.1.1.chr1.g01699.t01#4   mNeoVis.evergladensis.2.1.chr1.g01696.t01#2       1191   125259763:   mNeoVis.evergladensis.2.1.chr7.g02437.t01#2    mNeoVis.vulgivaga.2.1.chr7.g02460.t01#1    mNeoVis.vulgivaga.1.1.chr7.g02438.t01#4    mNeoVis.2.1.1.chr7_hap1.g02630.t01#5   mNeoVis.evergladensis.1.1.chr7.g02434.t01#3       1192   125259764:   mNeoVis.3.1.1.chr7.g02212.t01#6    mNeoVis.2.1.1.chr7_hap1.g02402.t01#5    mNeoVis.vulgivaga.1.1.chr7.g02352.t01#4    mNeoVis.vulgivaga.2.1.chr7.g02368.t01#1    mNeoVis.evergladensis.1.1.chr7.g02344.t01#3   mNeoVis.evergladensis.2.1.chr7.g02348.t01#2       1193   125259765:   mNeoVis.evergladensis.1.1.chr7.g02345.t01#3    mNeoVis.2.1.1.chr7_hap1.g02403.t01#5    mNeoVis.vulgivaga.2.1.chr7.g02369.t01#1    mNeoVis.evergladensis.2.1.chr7.g02349.t01#2    mNeoVis.vulgivaga.1.1.chr7.g02353.t01#4   mNeoVis.3.1.1.chr7.g02213.t01#6       1194   125259766:   mNeoVis.3.1.1.chr7.g00043.t01#6    mNeoVis.vulgivaga.2.1.chr7.g00055.t01#1    mNeoVis.vulgivaga.1.1.chr7.g00056.t01#4    mNeoVis.evergladensis.1.1.chr7.g00057.t01#3    mNeoVis.evergladensis.2.1.chr7.g00055.t01#2   mNeoVis.2.1.1.chr7_hap1.g00054.t01#5       1195   125259767:   mNeoVis.evergladensis.1.1.chr7.g00058.t01#3    mNeoVis.vulgivaga.2.1.chr7.g00056.t01#1    mNeoVis.vulgivaga.1.1.chr7.g00057.t01#4    mNeoVis.2.1.1.chr7_hap1.g00055.t01#5   mNeoVis.evergladensis.2.1.chr7.g00056.t01#2       1196   125259768:   mNeoVis.3.1.1.chr7.g02214.t01#6    mNeoVis.evergladensis.2.1.chr7.g02350.t01#2    mNeoVis.evergladensis.1.1.chr7.g02347.t01#3    mNeoVis.vulgivaga.1.1.chr7.g02354.t01#4    mNeoVis.2.1.1.chr7_hap1.g02404.t01#5    mNeoVis.evergladensis.1.1.chr7.g02346.t01#3   mNeoVis.vulgivaga.2.1.chr7.g02370.t01#1     1197   125259769:   mNeoVis.2.1.1.chr7_hap1.g02405.t01#5    mNeoVis.evergladensis.2.1.chr7.g02351.t01#2    mNeoVis.vulgivaga.2.1.chr7.g02371.t01#1    mNeoVis.3.1.1.chr7.g02215.t01#6    mNeoVis.evergladensis.1.1.chr7.g02348.t01#3   mNeoVis.vulgivaga.1.1.chr7.g02355.t01#4       1198   125259770:   mNeoVis.3.1.1.chr7.g02216.t01#6    mNeoVis.vulgivaga.1.1.chr7.g02356.t01#4    mNeoVis.evergladensis.2.1.chr7.g02352.t01#2    mNeoVis.evergladensis.1.1.chr7.g02349.t01#3    mNeoVis.vulgivaga.2.1.chr7.g02372.t01#1   mNeoVis.2.1.1.chr7_hap1.g02464.t01#5       1199   125259771:   mNeoVis.2.1.1.chr7_hap1.g02465.t01#5    mNeoVis.vulgivaga.2.1.chr7.g02373.t01#1    mNeoVis.vulgivaga.1.1.chr7.g02357.t01#4    mNeoVis.evergladensis.1.1.chr7.g02350.t01#3    mNeoVis.evergladensis.2.1.chr7.g02353.t01#2   mNeoVis.3.1.1.chr7.g02217.t01#6       1200   125259772:   mNeoVis.3.1.1.chr1.g01683.t01#6    mNeoVis.evergladensis.1.1.chr1.g01694.t01#3    mNeoVis.vulgivaga.2.1.chr1.g01693.t01#1    mNeoVis.2.1.1.chr1_hap1.g01728.t01#5    mNeoVis.evergladensis.2.1.chr1.g01697.t01#2   mNeoVis.vulgivaga.1.1.chr1.g01700.t01#4       1201   125259773:   mNeoVis.evergladensis.2.1.chr7.g00057.t01#2    mNeoVis.2.1.1.chr7_hap1.g00056.t01#5    mNeoVis.vulgivaga.2.1.chr7.g00057.t01#1    mNeoVis.3.1.1.chr7.g00044.t01#6    mNeoVis.evergladensis.1.1.chr7.g00059.t01#3   mNeoVis.vulgivaga.1.1.chr7.g00058.t01#4       1202   125259774:   mNeoVis.vulgivaga.1.1.chr1.g01701.t01#4    mNeoVis.vulgivaga.2.1.chr1.g01694.t01#1    mNeoVis.evergladensis.1.1.chr1.g01695.t01#3    mNeoVis.3.1.1.chr1.g01684.t01#6    mNeoVis.2.1.1.chr1_hap1.g01729.t01#5   mNeoVis.evergladensis.2.1.chr1.g01698.t01#2       1203   125259775:   mNeoVis.2.1.1.chr7_hap1.g02466.t01#5    mNeoVis.vulgivaga.2.1.chr7.g02374.t01#1    mNeoVis.vulgivaga.1.1.chr7.g02358.t01#4    mNeoVis.3.1.1.chr7.g02218.t01#6    mNeoVis.evergladensis.2.1.chr7.g02354.t01#2   mNeoVis.evergladensis.1.1.chr7.g02351.t01#3       1204   125259776:   mNeoVis.3.1.1.chr15.g00126.t01#6    mNeoVis.evergladensis.2.1.chr15.g00133.t01#2    mNeoVis.evergladensis.1.1.chr15.g00482.t01#3    mNeoVis.vulgivaga.2.1.chr15.g00137.t01#1    mNeoVis.2.1.1.chr15_hap1.g00148.t01#5   mNeoVis.vulgivaga.1.1.chr15.g00133.t01#4       1205   125259777:   mNeoVis.evergladensis.1.1.chr1.g01696.t01#3    mNeoVis.3.1.1.chr1.g01685.t01#6    mNeoVis.vulgivaga.1.1.chr1.g01702.t01#4    mNeoVis.vulgivaga.2.1.chr1.g01695.t01#1    mNeoVis.2.1.1.chr1_hap1.g01730.t01#5   mNeoVis.evergladensis.2.1.chr1.g01699.t01#2       1206   125259778:   mNeoVis.vulgivaga.1.1.chr1.g01703.t01#4    mNeoVis.evergladensis.1.1.chr1.g01697.t01#3    mNeoVis.3.1.1.chr1.g01686.t01#6    mNeoVis.vulgivaga.2.1.chr1.g01696.t01#1    mNeoVis.evergladensis.2.1.chr1.g01700.t01#2   mNeoVis.2.1.1.chr1_hap1.g01731.t01#5       1207   125259779:   mNeoVis.3.1.1.chr1.g01687.t01#6    mNeoVis.2.1.1.chr1_hap1.g01732.t01#5    mNeoVis.vulgivaga.2.1.chr1.g01697.t01#1   mNeoVis.vulgivaga.1.1.chr1.g01704.t01#4         1208   125259780:   mNeoVis.evergladensis.2.1.chr7.g02355.t01#2    mNeoVis.2.1.1.chr7_hap1.g02467.t01#5    mNeoVis.vulgivaga.2.1.chr7.g02375.t01#1    mNeoVis.vulgivaga.1.1.chr7.g02359.t01#4   mNeoVis.evergladensis.1.1.chr7.g02352.t01#3       1209   125259781:   mNeoVis.evergladensis.2.1.chr1.g01701.t01#2    mNeoVis.3.1.1.chr1.g01688.t01#6    mNeoVis.2.1.1.chr1_hap1.g01733.t01#5    mNeoVis.vulgivaga.2.1.chr1.g01698.t01#1    mNeoVis.vulgivaga.1.1.chr1.g01705.t01#4   mNeoVis.evergladensis.1.1.chr1.g01698.t01#3       1210   125259782:   mNeoVis.evergladensis.2.1.chr1.g01702.t01#2    mNeoVis.2.1.1.chr1_hap1.g01734.t01#5    mNeoVis.vulgivaga.1.1.chr1.g01706.t01#4    mNeoVis.3.1.1.chr1.g01689.t01#6    mNeoVis.evergladensis.1.1.chr1.g01700.t01#3    mNeoVis.vulgivaga.2.1.chr1.g01699.t01#1   mNeoVis.evergladensis.1.1.chr1.g01699.t01#3     1211   125259783:   mNeoVis.3.1.1.chr11.g00706.t01#6    mNeoVis.vulgivaga.1.1.chr11.g00725.t01#4    mNeoVis.evergladensis.2.1.chr11.g00725.t01#2    mNeoVis.2.1.1.chr11_hap1.g00743.t01#5    mNeoVis.vulgivaga.2.1.chr11.g00745.t01#1   mNeoVis.evergladensis.1.1.chr11.g00733.t01#3       1212   125259784:   mNeoVis.evergladensis.2.1.chr7.g02356.t01#2    mNeoVis.2.1.1.chr7_hap1.g02468.t01#5    mNeoVis.vulgivaga.2.1.chr7.g02376.t01#1    mNeoVis.vulgivaga.1.1.chr7.g02360.t01#4   mNeoVis.evergladensis.1.1.chr7.g02353.t01#3       1213   125259785:   mNeoVis.2.1.1.chr1_hap1.g01735.t01#5    mNeoVis.vulgivaga.1.1.chr1.g01707.t01#4    mNeoVis.3.1.1.chr1.g01690.t01#6    mNeoVis.vulgivaga.2.1.chr1.g01700.t01#1    mNeoVis.evergladensis.2.1.chr1.g01703.t01#2   mNeoVis.evergladensis.1.1.chr1.g01701.t01#3       1214   125259786:   mNeoVis.3.1.1.chr14.g00241.t01#6    mNeoVis.vulgivaga.1.1.chr14.g00237.t01#4    mNeoVis.evergladensis.1.1.chr14.g00247.t01#3    mNeoVis.evergladensis.2.1.chr14.g00241.t01#2    mNeoVis.2.1.1.chr14_hap1.g00240.t01#5   mNeoVis.vulgivaga.2.1.chr14.g00242.t01#1       1215   125259787:   mNeoVis.evergladensis.1.1.chr7.g02354.t01#3    mNeoVis.2.1.1.chr7_hap1.g02469.t01#5    mNeoVis.evergladensis.2.1.chr7.g02357.t01#2    mNeoVis.vulgivaga.2.1.chr7.g02377.t01#1   mNeoVis.vulgivaga.1.1.chr7.g02361.t01#4       1216   125259788:   mNeoVis.evergladensis.2.1.chr1.g01704.t01#2    mNeoVis.evergladensis.1.1.chr1.g01702.t01#3    mNeoVis.2.1.1.chr1_hap1.g01736.t01#5    mNeoVis.3.1.1.chr1.g01691.t01#6    mNeoVis.vulgivaga.2.1.chr1.g01701.t01#1   mNeoVis.vulgivaga.1.1.chr1.g01708.t01#4       1217   125259789:   mNeoVis.evergladensis.1.1.chr7.g02355.t01#3    mNeoVis.2.1.1.chr7_hap1.g02470.t01#5    mNeoVis.evergladensis.2.1.chr7.g02358.t01#2    mNeoVis.vulgivaga.2.1.chr7.g02378.t01#1   mNeoVis.vulgivaga.1.1.chr7.g02362.t01#4       1218   125259790:   mNeoVis.evergladensis.2.1.chr1.g01608.t01#2    mNeoVis.2.1.1.chr1_hap1.g01642.t01#5    mNeoVis.3.1.1.chr1.g01592.t01#6    mNeoVis.vulgivaga.2.1.chr1.g01605.t01#1    mNeoVis.vulgivaga.1.1.chr1.g01609.t01#4   mNeoVis.evergladensis.1.1.chr1.g01605.t01#3       1219   125259791:   mNeoVis.2.1.1.chr7_hap1.g02471.t01#5    mNeoVis.evergladensis.2.1.chr7.g02359.t01#2    mNeoVis.evergladensis.1.1.chr7.g02356.t01#3    mNeoVis.vulgivaga.2.1.chr7.g02379.t01#1   mNeoVis.vulgivaga.1.1.chr7.g02363.t01#4       1220   125259792:   mNeoVis.3.1.1.chr7.g01418.t01#6    mNeoVis.vulgivaga.2.1.chr7.g02380.t01#1    mNeoVis.vulgivaga.1.1.chr7.g02364.t01#4    mNeoVis.2.1.1.chr7_hap1.g02472.t01#5    mNeoVis.evergladensis.1.1.chr7.g02357.t01#3   mNeoVis.evergladensis.2.1.chr7.g02360.t01#2       1221   125259793:   mNeoVis.3.1.1.chr1.g01593.t01#6    mNeoVis.2.1.1.chr1_hap1.g01643.t01#5    mNeoVis.vulgivaga.2.1.chr1.g01606.t01#1    mNeoVis.evergladensis.2.1.chr1.g01609.t01#2    mNeoVis.evergladensis.1.1.chr1.g01606.t01#3   mNeoVis.vulgivaga.1.1.chr1.g01610.t01#4       1222   125259794:   mNeoVis.3.1.1.chr1.g01594.t01#6    mNeoVis.2.1.1.chr1_hap1.g01644.t01#5    mNeoVis.vulgivaga.2.1.chr1.g01607.t01#1    mNeoVis.evergladensis.2.1.chr1.g01610.t01#2    mNeoVis.evergladensis.1.1.chr1.g01607.t01#3   mNeoVis.vulgivaga.1.1.chr1.g01611.t01#4       1223   125259795:   mNeoVis.vulgivaga.1.1.chr7.g02365.t01#4    mNeoVis.evergladensis.2.1.chr7.g02361.t01#2    mNeoVis.evergladensis.1.1.chr7.g02358.t01#3    mNeoVis.vulgivaga.2.1.chr7.g02381.t01#1    mNeoVis.3.1.1.chr7.g02220.t01#6    mNeoVis.2.1.1.chr7_hap1.g02473.t01#5   mNeoVis.3.1.1.chr7.g01419.t01#6       1224   125259796:   mNeoVis.evergladensis.2.1.chr15.g00134.t01#2    mNeoVis.vulgivaga.2.1.chr15.g00138.t01#1    mNeoVis.evergladensis.1.1.chr15.g00481.t01#3    mNeoVis.2.1.1.chr15_hap1.g00149.t01#5   mNeoVis.vulgivaga.1.1.chr15.g00134.t01#4       1225   125259797:   mNeoVis.evergladensis.2.1.chr7.g00058.t01#2    mNeoVis.2.1.1.chr7_hap1.g00057.t01#5    mNeoVis.evergladensis.1.1.chr7.g00060.t01#3    mNeoVis.vulgivaga.2.1.chr7.g00058.t01#1    mNeoVis.vulgivaga.1.1.chr7.g00059.t01#4   mNeoVis.3.1.1.chr7.g00045.t01#6       1226   125259798:   mNeoVis.evergladensis.2.1.chr7.g02362.t01#2    mNeoVis.2.1.1.chr7_hap1.g02474.t01#5    mNeoVis.evergladensis.1.1.chr7.g02359.t01#3    mNeoVis.vulgivaga.2.1.chr7.g02382.t01#1    mNeoVis.3.1.1.chr7.g02221.t01#6   mNeoVis.vulgivaga.1.1.chr7.g02366.t01#4       1227   125259799:   mNeoVis.evergladensis.1.1.chr1.g01608.t01#3    mNeoVis.evergladensis.2.1.chr1.g01611.t01#2    mNeoVis.vulgivaga.1.1.chr1.g01612.t01#4    mNeoVis.vulgivaga.2.1.chr1.g01608.t01#1    mNeoVis.3.1.1.chr1.g01595.t01#6   mNeoVis.2.1.1.chr1_hap1.g01645.t01#5       1228   125259800:   mNeoVis.evergladensis.1.1.chr1.g01609.t01#3    mNeoVis.vulgivaga.1.1.chr1.g01613.t01#4    mNeoVis.3.1.1.chr1.g01596.t01#6    mNeoVis.vulgivaga.2.1.chr1.g01609.t01#1    mNeoVis.2.1.1.chr1_hap1.g01646.t01#5   mNeoVis.evergladensis.2.1.chr1.g01612.t01#2       1229   125259801:   mNeoVis.vulgivaga.1.1.chr7.g02367.t01#4    mNeoVis.vulgivaga.2.1.chr7.g02383.t01#1    mNeoVis.2.1.1.chr7_hap1.g02475.t01#5    mNeoVis.evergladensis.2.1.chr7.g02363.t01#2    mNeoVis.evergladensis.1.1.chr7.g02360.t01#3   mNeoVis.3.1.1.chr7.g02222.t01#6       1230   125259802:   mNeoVis.vulgivaga.2.1.chr7.g00059.t01#1    mNeoVis.evergladensis.2.1.chr7.g00059.t01#2    mNeoVis.evergladensis.1.1.chr7.g00061.t01#3   mNeoVis.2.1.1.chr7_hap1.g00058.t01#5         1231   125259803:   mNeoVis.evergladensis.2.1.chr7.g02364.t01#2    mNeoVis.vulgivaga.2.1.chr7.g02384.t01#1    mNeoVis.evergladensis.1.1.chr7.g02361.t01#3    mNeoVis.2.1.1.chr7_hap1.g02476.t01#5    mNeoVis.3.1.1.chr7.g02223.t01#6   mNeoVis.vulgivaga.1.1.chr7.g02368.t01#4       1232   125259804:   mNeoVis.vulgivaga.2.1.chr1.g01610.t01#1    mNeoVis.3.1.1.chr1.g01597.t01#6    mNeoVis.2.1.1.chr1_hap1.g01647.t01#5    mNeoVis.evergladensis.2.1.chr1.g01613.t01#2    mNeoVis.vulgivaga.1.1.chr1.g01614.t01#4   mNeoVis.evergladensis.1.1.chr1.g01610.t01#3       1233   125259805:   mNeoVis.evergladensis.2.1.chr1.g01614.t01#2    mNeoVis.vulgivaga.2.1.chr1.g01611.t01#1    mNeoVis.vulgivaga.1.1.chr1.g01615.t01#4    mNeoVis.3.1.1.chr1.g01598.t01#6    mNeoVis.2.1.1.chr1_hap1.g01648.t01#5   mNeoVis.evergladensis.1.1.chr1.g01611.t01#3       1234   125259806:   mNeoVis.2.1.1.chr1_hap1.g01649.t01#5    mNeoVis.evergladensis.1.1.chr1.g01612.t01#3    mNeoVis.vulgivaga.2.1.chr1.g01612.t01#1    mNeoVis.vulgivaga.1.1.chr1.g01616.t01#4    mNeoVis.3.1.1.chr1.g01599.t01#6   mNeoVis.evergladensis.2.1.chr1.g01615.t01#2       1235   125259807:   mNeoVis.vulgivaga.2.1.chr7.g02385.t01#1    mNeoVis.2.1.1.chr7_hap1.g02477.t01#5    mNeoVis.3.1.1.chr7.g02224.t01#6    mNeoVis.evergladensis.1.1.chr7.g02362.t01#3    mNeoVis.evergladensis.2.1.chr7.g02365.t01#2   mNeoVis.vulgivaga.1.1.chr7.g02369.t01#4       1236   125259808:   mNeoVis.3.1.1.chr15.g00127.t01#6    mNeoVis.2.1.1.chr15_hap1.g00150.t01#5    mNeoVis.vulgivaga.1.1.chr15.g00135.t01#4    mNeoVis.evergladensis.2.1.chr15.g00135.t01#2    mNeoVis.evergladensis.1.1.chr15.g00480.t01#3   mNeoVis.vulgivaga.2.1.chr15.g00139.t01#1       1237   125259809:   mNeoVis.vulgivaga.1.1.chr7.g02370.t01#4    mNeoVis.evergladensis.1.1.chr7.g02363.t01#3    mNeoVis.evergladensis.2.1.chr7.g02366.t01#2    mNeoVis.2.1.1.chr7_hap1.g02478.t01#5    mNeoVis.3.1.1.chr7.g02225.t01#6   mNeoVis.vulgivaga.2.1.chr7.g02386.t01#1       1238   125259810:   mNeoVis.vulgivaga.2.1.chr7.g02387.t01#1    mNeoVis.evergladensis.2.1.chr7.g02367.t01#2    mNeoVis.3.1.1.chr7.g02226.t01#6    mNeoVis.vulgivaga.1.1.chr7.g02371.t01#4    mNeoVis.evergladensis.1.1.chr7.g02364.t01#3   mNeoVis.2.1.1.chr7_hap1.g02479.t01#5       1239   125259811:   mNeoVis.evergladensis.2.1.chr7.g02368.t01#2    mNeoVis.2.1.1.chr7_hap1.g02480.t01#5    mNeoVis.3.1.1.chr7.g02227.t01#6    mNeoVis.vulgivaga.1.1.chr7.g02372.t01#4    mNeoVis.evergladensis.1.1.chr7.g02365.t01#3   mNeoVis.vulgivaga.2.1.chr7.g02388.t01#1       1240   125259812:   mNeoVis.3.1.1.chr7.g02228.t01#6    mNeoVis.vulgivaga.1.1.chr7.g02373.t01#4    mNeoVis.vulgivaga.2.1.chr7.g02389.t01#1    mNeoVis.2.1.1.chr7_hap1.g02481.t01#5    mNeoVis.evergladensis.2.1.chr7.g02369.t01#2   mNeoVis.evergladensis.1.1.chr7.g02367.t01#3       1241   125259813:   mNeoVis.2.1.1.chr1_hap1.g01650.t01#5    mNeoVis.vulgivaga.1.1.chr1.g01617.t01#4    mNeoVis.evergladensis.2.1.chr1.g01616.t01#2    mNeoVis.evergladensis.1.1.chr1.g01613.t01#3    mNeoVis.vulgivaga.2.1.chr1.g01613.t01#1   mNeoVis.3.1.1.chr1.g01600.t01#6       1242   125259814:   mNeoVis.evergladensis.2.1.chr7.g00060.t01#2    mNeoVis.evergladensis.1.1.chr7.g00062.t01#3    mNeoVis.vulgivaga.1.1.chr7.g00060.t01#4    mNeoVis.vulgivaga.2.1.chr7.g00060.t01#1   mNeoVis.2.1.1.chr7_hap1.g00059.t01#5       1243   125259815:   mNeoVis.vulgivaga.1.1.chr1.g01618.t01#4    mNeoVis.evergladensis.1.1.chr2.g00403.t01#3    mNeoVis.3.1.1.chr2.g00335.t01#6    mNeoVis.evergladensis.2.1.chr2.g00402.t01#2    mNeoVis.evergladensis.2.1.chr1.g01617.t01#2    mNeoVis.2.1.1.chr1_hap1.g01651.t01#5    mNeoVis.3.1.1.chr1.g01601.t01#6    mNeoVis.2.1.1.chr5_hap1.g00403.t01#5    mNeoVis.vulgivaga.2.1.chr2.g00402.t01#1      1244   125259816:   mNeoVis.vulgivaga.1.1.chr7.g02374.t01#4    mNeoVis.2.1.1.chr7_hap1.g02482.t01#5    mNeoVis.evergladensis.2.1.chr7.g02370.t01#2    mNeoVis.evergladensis.1.1.chr7.g02368.t01#3   mNeoVis.vulgivaga.2.1.chr7.g02390.t01#1       1245   125259817:   mNeoVis.vulgivaga.1.1.chr1.g01619.t01#4    mNeoVis.vulgivaga.2.1.chr1.g01615.t01#1    mNeoVis.evergladensis.2.1.chr1.g01618.t01#2    mNeoVis.3.1.1.chr1.g01602.t01#6    mNeoVis.2.1.1.chr1_hap1.g01652.t01#5   mNeoVis.evergladensis.1.1.chr1.g01615.t01#3       1246   125259818:   mNeoVis.3.1.1.chr1.g01603.t01#6    mNeoVis.vulgivaga.2.1.chr1.g01616.t01#1    mNeoVis.evergladensis.2.1.chr1.g01619.t01#2    mNeoVis.evergladensis.1.1.chr1.g01616.t01#3    mNeoVis.2.1.1.chr1_hap1.g01653.t01#5   mNeoVis.vulgivaga.1.1.chr1.g01620.t01#4       1247   125259819:   mNeoVis.vulgivaga.2.1.chr1.g01617.t01#1    mNeoVis.vulgivaga.1.1.chr1.g01621.t01#4    mNeoVis.evergladensis.1.1.chr1.g01617.t01#3   mNeoVis.2.1.1.chr1_hap1.g01654.t01#5         1248   125259820:   mNeoVis.2.1.1.chr1_hap1.g01655.t01#5    mNeoVis.vulgivaga.1.1.chr1.g01622.t01#4    mNeoVis.3.1.1.chr1.g01605.t01#6    mNeoVis.evergladensis.1.1.chr1.g01618.t01#3   mNeoVis.vulgivaga.2.1.chr1.g01618.t01#1       1249   125259821:   mNeoVis.vulgivaga.2.1.chr7.g02392.t01#1    mNeoVis.evergladensis.1.1.chr7.g02370.t01#3    mNeoVis.vulgivaga.1.1.chr7.g02376.t01#4    mNeoVis.3.1.1.chr7.g02231.t01#6    mNeoVis.evergladensis.2.1.chr7.g02372.t01#2   mNeoVis.2.1.1.chr7_hap1.g02484.t01#5       1250   125259822:   mNeoVis.vulgivaga.1.1.chr1.g01623.t01#4    mNeoVis.vulgivaga.2.1.chr1.g01619.t01#1    mNeoVis.evergladensis.1.1.chr1.g01619.t01#3    mNeoVis.evergladensis.2.1.chr1.g01622.t01#2    mNeoVis.2.1.1.chr1_hap1.g01656.t01#5   mNeoVis.3.1.1.chr1.g01606.t01#6       1251   125259823:   mNeoVis.2.1.1.chr1_hap1.g01657.t01#5    mNeoVis.evergladensis.1.1.chr1.g01620.t01#3    mNeoVis.vulgivaga.1.1.chr1.g01624.t01#4    mNeoVis.vulgivaga.2.1.chr1.g01620.t01#1    mNeoVis.evergladensis.2.1.chr1.g01623.t01#2   mNeoVis.3.1.1.chr1.g01609.t01#6       1252   125259824:   mNeoVis.vulgivaga.2.1.chr7.g02393.t01#1    mNeoVis.3.1.1.chr7.g02232.t01#6    mNeoVis.evergladensis.1.1.chr7.g02371.t01#3    mNeoVis.evergladensis.2.1.chr7.g02373.t01#2    mNeoVis.vulgivaga.1.1.chr7.g02377.t01#4   mNeoVis.2.1.1.chr7_hap1.g02485.t01#5       1253   125259825:   mNeoVis.vulgivaga.1.1.chr1.g01625.t01#4    mNeoVis.vulgivaga.2.1.chr1.g01621.t01#1    mNeoVis.3.1.1.chr1.g01607.t01#6    mNeoVis.evergladensis.2.1.chr1.g01624.t01#2    mNeoVis.evergladensis.1.1.chr1.g01621.t01#3   mNeoVis.2.1.1.chr1_hap1.g01658.t01#5       1254   125259826:   mNeoVis.vulgivaga.1.1.chr7.g02378.t01#4    mNeoVis.3.1.1.chr7.g02233.t01#6    mNeoVis.2.1.1.chr7_hap1.g02486.t01#5    mNeoVis.vulgivaga.2.1.chr7.g02394.t01#1    mNeoVis.evergladensis.2.1.chr7.g02374.t01#2   mNeoVis.evergladensis.1.1.chr7.g02372.t01#3       1255   125259827:   mNeoVis.vulgivaga.1.1.chr1.g01626.t01#4    mNeoVis.2.1.1.chr1_hap1.g01659.t01#5    mNeoVis.evergladensis.2.1.chr1.g01625.t01#2    mNeoVis.vulgivaga.2.1.chr1.g01622.t01#1    mNeoVis.evergladensis.1.1.chr1.g01622.t01#3   mNeoVis.3.1.1.chr1.g01608.t01#6       1256   125259828:   mNeoVis.vulgivaga.1.1.chr7.g02379.t01#4    mNeoVis.vulgivaga.2.1.chr7.g02395.t01#1    mNeoVis.3.1.1.chr7.g02234.t01#6    mNeoVis.evergladensis.2.1.chr7.g02375.t01#2    mNeoVis.2.1.1.chr7_hap1.g02487.t01#5    mNeoVis.evergladensis.1.1.chr7.g02373.t01#3   mNeoVis.evergladensis.2.1.chr7.g02376.t01#2     1257   125259829:   mNeoVis.evergladensis.2.1.chr1.g01626.t01#2    mNeoVis.vulgivaga.2.1.chr1.g01623.t01#1    mNeoVis.vulgivaga.1.1.chr1.g01627.t01#4    mNeoVis.2.1.1.chr1_hap1.g01660.t01#5    mNeoVis.3.1.1.chr1.g01610.t01#6   mNeoVis.evergladensis.1.1.chr1.g01623.t01#3       1258   125259830:   mNeoVis.evergladensis.1.1.chr7.g02374.t01#3    mNeoVis.2.1.1.chr7_hap1.g02488.t01#5    mNeoVis.3.1.1.chr7.g02235.t01#6    mNeoVis.evergladensis.2.1.chr7.g02377.t01#2    mNeoVis.vulgivaga.1.1.chr7.g02380.t01#4   mNeoVis.vulgivaga.2.1.chr7.g02396.t01#1       1259   125259831:   mNeoVis.vulgivaga.2.1.chr7.g02397.t01#1    mNeoVis.vulgivaga.1.1.chr7.g02381.t01#4    mNeoVis.evergladensis.1.1.chr7.g02375.t01#3    mNeoVis.3.1.1.chr7.g02236.t01#6    mNeoVis.evergladensis.2.1.chr7.g02378.t01#2   mNeoVis.2.1.1.chr7_hap1.g02489.t01#5       1260   125259832:   mNeoVis.vulgivaga.2.1.chr1.g01624.t01#1    mNeoVis.evergladensis.2.1.chr1.g01627.t01#2    mNeoVis.evergladensis.1.1.chr1.g01624.t01#3    mNeoVis.vulgivaga.1.1.chr1.g01628.t01#4    mNeoVis.3.1.1.chr1.g01611.t01#6   mNeoVis.2.1.1.chr1_hap1.g01661.t01#5       1261   125259833:   mNeoVis.vulgivaga.1.1.chr1.g01629.t01#4    mNeoVis.2.1.1.chr1_hap1.g01662.t01#5    mNeoVis.evergladensis.2.1.chr1.g01628.t01#2    mNeoVis.evergladensis.1.1.chr1.g01625.t01#3    mNeoVis.vulgivaga.2.1.chr1.g01625.t01#1   mNeoVis.3.1.1.chr1.g01612.t01#6       1262   125259834:   mNeoVis.3.1.1.chr15.g00128.t01#6    mNeoVis.2.1.1.chr15_hap1.g00151.t01#5    mNeoVis.evergladensis.2.1.chr15.g00136.t01#2    mNeoVis.vulgivaga.1.1.chr15.g00136.t01#4    mNeoVis.vulgivaga.2.1.chr15.g00140.t01#1   mNeoVis.evergladensis.1.1.chr15.g00479.t01#3       1263   125259835:   mNeoVis.vulgivaga.1.1.chr1.g01630.t01#4    mNeoVis.3.1.1.chr1.g01613.t01#6    mNeoVis.vulgivaga.2.1.chr1.g01626.t01#1    mNeoVis.evergladensis.1.1.chr1.g01626.t01#3    mNeoVis.evergladensis.2.1.chr1.g01629.t01#2   mNeoVis.2.1.1.chr1_hap1.g01663.t01#5       1264   125259836:   mNeoVis.evergladensis.2.1.chr1.g01630.t01#2    mNeoVis.vulgivaga.1.1.chr1.g01631.t01#4    mNeoVis.evergladensis.1.1.chr1.g01627.t01#3    mNeoVis.vulgivaga.2.1.chr1.g01627.t01#1    mNeoVis.2.1.1.chr1_hap1.g01664.t01#5   mNeoVis.3.1.1.chr1.g01614.t01#6       1265   125259837:   mNeoVis.3.1.1.chr7.g00048.t01#6    mNeoVis.vulgivaga.2.1.chr7.g00061.t01#1    mNeoVis.evergladensis.2.1.chr7.g00061.t01#2    mNeoVis.2.1.1.chr7_hap1.g00060.t01#5   mNeoVis.evergladensis.1.1.chr7.g00063.t01#3       1266   125259838:   mNeoVis.evergladensis.2.1.chr1.g01631.t01#2    mNeoVis.2.1.1.chr1_hap1.g01665.t01#5    mNeoVis.3.1.1.chr1.g01615.t01#6    mNeoVis.vulgivaga.1.1.chr1.g01632.t01#4    mNeoVis.evergladensis.1.1.chr1.g01628.t01#3   mNeoVis.vulgivaga.2.1.chr1.g01628.t01#1       1267   125259839:   mNeoVis.vulgivaga.2.1.chr7.g02398.t01#1    mNeoVis.evergladensis.1.1.chr7.g02376.t01#3    mNeoVis.3.1.1.chr7.g02237.t01#6    mNeoVis.2.1.1.chr7_hap1.g02490.t01#5    mNeoVis.vulgivaga.1.1.chr7.g02382.t01#4   mNeoVis.evergladensis.2.1.chr7.g02379.t01#2       1268   125259840:   mNeoVis.2.1.1.chr1_hap1.g01666.t01#5    mNeoVis.evergladensis.1.1.chr1.g01629.t01#3    mNeoVis.3.1.1.chr1.g01616.t01#6    mNeoVis.vulgivaga.1.1.chr1.g01633.t01#4    mNeoVis.vulgivaga.2.1.chr1.g01629.t01#1   mNeoVis.evergladensis.2.1.chr1.g01632.t01#2       1269   125259841:   mNeoVis.vulgivaga.1.1.chr1.g01634.t01#4    mNeoVis.evergladensis.1.1.chr1.g01630.t01#3    mNeoVis.evergladensis.2.1.chr1.g01633.t01#2    mNeoVis.vulgivaga.2.1.chr1.g01630.t01#1    mNeoVis.2.1.1.chr1_hap1.g01667.t01#5   mNeoVis.3.1.1.chr1.g01617.t01#6       1270   125259842:   mNeoVis.vulgivaga.2.1.chr7.g02399.t01#1    mNeoVis.evergladensis.2.1.chr7.g02380.t01#2    mNeoVis.3.1.1.chr7.g02238.t01#6    mNeoVis.evergladensis.1.1.chr7.g02377.t01#3   mNeoVis.2.1.1.chr7_hap1.g02491.t01#5       1271   125259843:   mNeoVis.vulgivaga.2.1.chr7.g02400.t01#1    mNeoVis.evergladensis.1.1.chr7.g02378.t01#3    mNeoVis.2.1.1.chr7_hap1.g02492.t01#5    mNeoVis.vulgivaga.1.1.chr7.g02384.t01#4   mNeoVis.evergladensis.2.1.chr7.g02381.t01#2       1272   125259844:   mNeoVis.3.1.1.chr1.g01618.t01#6    mNeoVis.vulgivaga.2.1.chr1.g01631.t01#1   mNeoVis.evergladensis.1.1.chr1.g01632.t01#3         1273   125259845:   mNeoVis.vulgivaga.1.1.chr1.g01636.t01#4    mNeoVis.evergladensis.1.1.chr1.g01633.t01#3    mNeoVis.vulgivaga.2.1.chr1.g01632.t01#1    mNeoVis.evergladensis.2.1.chr1.g01635.t01#2    mNeoVis.3.1.1.chr1.g01619.t01#6   mNeoVis.2.1.1.chr1_hap1.g01669.t01#5       1274   125259846:   mNeoVis.2.1.1.chr15_hap1.g00152.t01#5    mNeoVis.3.1.1.chr15.g00129.t01#6    mNeoVis.vulgivaga.2.1.chr15.g00141.t01#1    mNeoVis.vulgivaga.1.1.chr15.g00137.t01#4    mNeoVis.evergladensis.1.1.chr15.g00478.t01#3   mNeoVis.evergladensis.2.1.chr15.g00137.t01#2       1275   125259847:   mNeoVis.evergladensis.1.1.chr7.g00064.t01#3    mNeoVis.vulgivaga.2.1.chr7.g00062.t01#1    mNeoVis.3.1.1.chr7.g00049.t01#6    mNeoVis.evergladensis.2.1.chr7.g00062.t01#2    mNeoVis.vulgivaga.1.1.chr7.g00062.t01#4   mNeoVis.2.1.1.chr7_hap1.g00061.t01#5       1276   125259848:   mNeoVis.evergladensis.2.1.chr1.g01636.t01#2    mNeoVis.evergladensis.1.1.chr1.g01634.t01#3    mNeoVis.3.1.1.chr1.g01620.t01#6    mNeoVis.vulgivaga.2.1.chr1.g01633.t01#1    mNeoVis.2.1.1.chr1_hap1.g01670.t01#5   mNeoVis.vulgivaga.1.1.chr1.g01637.t01#4       1277   125259849:   mNeoVis.3.1.1.chr1.g01621.t01#6    mNeoVis.2.1.1.chr1_hap1.g01671.t01#5    mNeoVis.vulgivaga.1.1.chr1.g01638.t01#4    mNeoVis.evergladensis.2.1.chr1.g01637.t01#2    mNeoVis.vulgivaga.2.1.chr1.g01634.t01#1   mNeoVis.evergladensis.1.1.chr1.g01635.t01#3       1278   125259850:   mNeoVis.evergladensis.1.1.chr7.g02379.t01#3    mNeoVis.evergladensis.2.1.chr7.g02382.t01#2    mNeoVis.vulgivaga.2.1.chr7.g02401.t01#1    mNeoVis.2.1.1.chr7_hap1.g02493.t01#5   mNeoVis.vulgivaga.1.1.chr7.g02385.t01#4       1279   125259851:   mNeoVis.2.1.1.chr1_hap1.g01672.t01#5    mNeoVis.evergladensis.1.1.chr1.g01636.t01#3    mNeoVis.evergladensis.2.1.chr1.g01638.t01#2    mNeoVis.3.1.1.chr1.g01622.t01#6    mNeoVis.vulgivaga.2.1.chr1.g01635.t01#1   mNeoVis.vulgivaga.1.1.chr1.g01639.t01#4       1280   125259852:   mNeoVis.vulgivaga.2.1.chr7.g02402.t01#1    mNeoVis.3.1.1.chr7.g02239.t01#6    mNeoVis.2.1.1.chr7_hap1.g02494.t01#5    mNeoVis.vulgivaga.1.1.chr7.g02386.t01#4    mNeoVis.evergladensis.2.1.chr7.g02383.t01#2   mNeoVis.evergladensis.1.1.chr7.g02380.t01#3       1281   125259853:   mNeoVis.vulgivaga.1.1.chr7.g02387.t01#4    mNeoVis.2.1.1.chr7_hap1.g02495.t01#5    mNeoVis.evergladensis.2.1.chr7.g02384.t01#2    mNeoVis.evergladensis.1.1.chr7.g02381.t01#3    mNeoVis.3.1.1.chr7.g02240.t01#6   mNeoVis.vulgivaga.2.1.chr7.g02403.t01#1       1282   125259854:   mNeoVis.3.1.1.chr1.g01623.t01#6    mNeoVis.vulgivaga.2.1.chr1.g01636.t01#1    mNeoVis.vulgivaga.1.1.chr1.g01640.t01#4    mNeoVis.evergladensis.1.1.chr1.g01637.t01#3    mNeoVis.evergladensis.2.1.chr1.g01639.t01#2   mNeoVis.2.1.1.chr1_hap1.g01673.t01#5       1283   125259855:   mNeoVis.vulgivaga.2.1.chr7.g02404.t01#1    mNeoVis.2.1.1.chr7_hap1.g02496.t01#5    mNeoVis.evergladensis.2.1.chr7.g02385.t01#2    mNeoVis.3.1.1.chr7.g02241.t01#6    mNeoVis.evergladensis.1.1.chr7.g02382.t01#3   mNeoVis.vulgivaga.1.1.chr7.g02388.t01#4       1284   125259856:   mNeoVis.vulgivaga.1.1.chr7.g00063.t01#4    mNeoVis.3.1.1.chr7.g00050.t01#6    mNeoVis.2.1.1.chr7_hap1.g00062.t01#5    mNeoVis.vulgivaga.2.1.chr7.g00063.t01#1    mNeoVis.evergladensis.1.1.chr7.g00065.t01#3   mNeoVis.evergladensis.2.1.chr7.g00063.t01#2       1285   125259857:   mNeoVis.vulgivaga.1.1.chr1.g01641.t01#4    mNeoVis.3.1.1.chr1.g01624.t01#6    mNeoVis.evergladensis.2.1.chr1.g01640.t01#2    mNeoVis.evergladensis.1.1.chr1.g01638.t01#3    mNeoVis.vulgivaga.2.1.chr1.g01637.t01#1   mNeoVis.2.1.1.chr1_hap1.g01674.t01#5       1286   125259858:   mNeoVis.evergladensis.2.1.chr7.g02386.t01#2    mNeoVis.2.1.1.chr7_hap1.g02497.t01#5    mNeoVis.3.1.1.chr7.g02242.t01#6    mNeoVis.vulgivaga.2.1.chr7.g02405.t01#1    mNeoVis.evergladensis.1.1.chr7.g02383.t01#3   mNeoVis.vulgivaga.1.1.chr7.g02389.t01#4       1287   125259859:   mNeoVis.vulgivaga.1.1.chr1.g01642.t01#4    mNeoVis.evergladensis.2.1.chr1.g01641.t01#2    mNeoVis.evergladensis.1.1.chr1.g01639.t01#3    mNeoVis.3.1.1.chr1.g01625.t01#6    mNeoVis.vulgivaga.2.1.chr1.g01638.t01#1   mNeoVis.2.1.1.chr1_hap1.g01675.t01#5       1288   125259860:   mNeoVis.vulgivaga.1.1.chr3.g00467.t01#4    mNeoVis.evergladensis.1.1.chr3.g00470.t01#3    mNeoVis.3.1.1.chr3.g00454.t01#6    mNeoVis.vulgivaga.2.1.chr3.g00470.t01#1    mNeoVis.evergladensis.2.1.chr3.g00471.t01#2   mNeoVis.2.1.1.chr3_hap1.g00463.t01#5       1289   125259861:   mNeoVis.vulgivaga.2.1.chr7.g02406.t01#1    mNeoVis.vulgivaga.1.1.chr7.g02390.t01#4    mNeoVis.evergladensis.2.1.chr7.g02387.t01#2    mNeoVis.2.1.1.chr7_hap1.g02498.t01#5   mNeoVis.evergladensis.1.1.chr7.g02384.t01#3       1290   125259862:   mNeoVis.evergladensis.2.1.chr7.g00064.t01#2    mNeoVis.vulgivaga.1.1.chr7.g00064.t01#4    mNeoVis.vulgivaga.2.1.chr7.g00064.t01#1    mNeoVis.2.1.1.chr7_hap1.g00063.t01#5   mNeoVis.3.1.1.chr7.g00051.t01#6       1291   125259863:   mNeoVis.vulgivaga.2.1.chr1.g01639.t01#1    mNeoVis.evergladensis.1.1.chr1.g01640.t01#3    mNeoVis.2.1.1.chr1_hap1.g01676.t01#5    mNeoVis.vulgivaga.1.1.chr1.g01643.t01#4    mNeoVis.vulgivaga.1.1.chr1.g01644.t01#4    mNeoVis.3.1.1.chr1.g01626.t01#6   mNeoVis.evergladensis.2.1.chr1.g01642.t01#2     1292   125259864:   mNeoVis.evergladensis.1.1.chr1.g01641.t01#3    mNeoVis.2.1.1.chr1_hap1.g01677.t01#5    mNeoVis.evergladensis.2.1.chr1.g01643.t01#2    mNeoVis.vulgivaga.1.1.chr1.g01645.t01#4    mNeoVis.vulgivaga.2.1.chr1.g01640.t01#1   mNeoVis.3.1.1.chr1.g01627.t01#6       1293   125259865:   mNeoVis.vulgivaga.2.1.chr1.g01641.t01#1    mNeoVis.3.1.1.chr1.g01628.t01#6   mNeoVis.evergladensis.1.1.chr1.g01642.t01#3         1294   125259866:   mNeoVis.evergladensis.2.1.chr7.g02388.t01#2    mNeoVis.3.1.1.chr7.g02243.t01#6    mNeoVis.evergladensis.1.1.chr7.g02385.t01#3    mNeoVis.vulgivaga.2.1.chr7.g02407.t01#1    mNeoVis.vulgivaga.1.1.chr7.g02391.t01#4   mNeoVis.2.1.1.chr7_hap1.g02499.t01#5       1295   125259867:   mNeoVis.evergladensis.1.1.chr1.g01643.t01#3    mNeoVis.vulgivaga.2.1.chr1.g01642.t01#1    mNeoVis.evergladensis.2.1.chr1.g01644.t01#2    mNeoVis.vulgivaga.1.1.chr1.g01646.t01#4    mNeoVis.2.1.1.chr1_hap1.g01678.t01#5   mNeoVis.3.1.1.chr1.g01629.t01#6       1296   125259868:   mNeoVis.vulgivaga.2.1.chr15.g00142.t01#1    mNeoVis.evergladensis.1.1.chr15.g00477.t01#3    mNeoVis.2.1.1.chr15_hap1.g00153.t01#5    mNeoVis.evergladensis.2.1.chr15.g00138.t01#2    mNeoVis.vulgivaga.1.1.chr15.g00138.t01#4   mNeoVis.3.1.1.chr15.g00130.t01#6       1297   125259869:   mNeoVis.vulgivaga.2.1.chr7.g02408.t01#1    mNeoVis.evergladensis.2.1.chr7.g02389.t01#2    mNeoVis.evergladensis.1.1.chr7.g02386.t01#3    mNeoVis.vulgivaga.1.1.chr7.g02392.t01#4    mNeoVis.3.1.1.chr7.g02244.t01#6   mNeoVis.2.1.1.chr7_hap1.g02500.t01#5       1298   125259870:   mNeoVis.3.1.1.chr1.g01630.t01#6    mNeoVis.vulgivaga.1.1.chr1.g01647.t01#4    mNeoVis.evergladensis.2.1.chr1.g01645.t01#2    mNeoVis.vulgivaga.2.1.chr1.g01643.t01#1    mNeoVis.evergladensis.1.1.chr1.g01644.t01#3   mNeoVis.2.1.1.chr1_hap1.g01679.t01#5       1299   125259871:   mNeoVis.evergladensis.2.1.chr1.g01646.t01#2    mNeoVis.3.1.1.chr1.g01631.t01#6    mNeoVis.2.1.1.chr1_hap1.g01680.t01#5    mNeoVis.evergladensis.1.1.chr1.g01645.t01#3    mNeoVis.vulgivaga.1.1.chr1.g01648.t01#4   mNeoVis.vulgivaga.2.1.chr1.g01644.t01#1       1300   125259872:   mNeoVis.evergladensis.1.1.chr1.g01646.t01#3    mNeoVis.vulgivaga.1.1.chr1.g01649.t01#4    mNeoVis.vulgivaga.2.1.chr1.g01645.t01#1    mNeoVis.evergladensis.2.1.chr1.g01647.t01#2    mNeoVis.3.1.1.chr1.g01632.t01#6   mNeoVis.2.1.1.chr1_hap1.g01681.t01#5       1301   125259873:   mNeoVis.vulgivaga.2.1.chr7.g02409.t01#1    mNeoVis.evergladensis.1.1.chr7.g02387.t01#3    mNeoVis.vulgivaga.1.1.chr7.g02393.t01#4   mNeoVis.3.1.1.chr7.g02245.t01#6         1302   125259874:   mNeoVis.evergladensis.1.1.chr7.g02388.t01#3    mNeoVis.2.1.1.chr7_hap1.g02584.t01#5    mNeoVis.vulgivaga.1.1.chr7.g02394.t01#4    mNeoVis.3.1.1.chr7.g02248.t01#6    mNeoVis.evergladensis.2.1.chr7.g02390.t01#2   mNeoVis.vulgivaga.2.1.chr7.g02410.t01#1       1303   125259875:   mNeoVis.evergladensis.1.1.chr7.g02389.t01#3    mNeoVis.vulgivaga.2.1.chr7.g02411.t01#1    mNeoVis.evergladensis.2.1.chr7.g02391.t01#2    mNeoVis.2.1.1.chr7_hap1.g02585.t01#5   mNeoVis.vulgivaga.1.1.chr7.g02395.t01#4       1304   125259876:   mNeoVis.evergladensis.1.1.chr7.g00067.t01#3    mNeoVis.evergladensis.2.1.chr7.g00065.t01#2    mNeoVis.vulgivaga.2.1.chr7.g00065.t01#1   mNeoVis.2.1.1.chr7_hap1.g00064.t01#5         1305   125259877:   mNeoVis.2.1.1.chr1_hap1.g01682.t01#5    mNeoVis.evergladensis.1.1.chr1.g01648.t01#3    mNeoVis.3.1.1.chr1.g01636.t01#6    mNeoVis.evergladensis.2.1.chr1.g01648.t01#2    mNeoVis.vulgivaga.1.1.chr1.g01651.t01#4    mNeoVis.vulgivaga.2.1.chr1.g01646.t01#1   mNeoVis.3.1.1.chr1.g01635.t01#6       1306   125259878:   mNeoVis.3.1.1.chr1.g01633.t01#6    mNeoVis.vulgivaga.1.1.chr1.g01650.t01#4   mNeoVis.evergladensis.1.1.chr1.g01647.t01#3         1307   125259879:   mNeoVis.vulgivaga.2.1.chr7.g02412.t01#1    mNeoVis.3.1.1.chr7.g02250.t01#6    mNeoVis.evergladensis.1.1.chr7.g02390.t01#3    mNeoVis.vulgivaga.1.1.chr7.g02396.t01#4    mNeoVis.2.1.1.chr7_hap1.g02586.t01#5   mNeoVis.evergladensis.2.1.chr7.g02392.t01#2       1308   125259880:   mNeoVis.vulgivaga.1.1.chr7.g02315.t01#4    mNeoVis.3.1.1.chr7.g02178.t01#6    mNeoVis.2.1.1.chr7_hap1.g02365.t01#5    mNeoVis.vulgivaga.2.1.chr7.g02329.t01#1    mNeoVis.evergladensis.2.1.chr7.g02311.t01#2   mNeoVis.evergladensis.1.1.chr7.g02307.t01#3       1309   125259881:   mNeoVis.3.1.1.chr7.g02179.t01#6    mNeoVis.evergladensis.1.1.chr7.g02308.t01#3    mNeoVis.evergladensis.2.1.chr7.g02312.t01#2    mNeoVis.vulgivaga.1.1.chr7.g02316.t01#4    mNeoVis.2.1.1.chr7_hap1.g02366.t01#5   mNeoVis.vulgivaga.2.1.chr7.g02330.t01#1       1310   125259882:   mNeoVis.3.1.1.chr1.g01637.t01#6    mNeoVis.vulgivaga.2.1.chr1.g01647.t01#1    mNeoVis.evergladensis.2.1.chr1.g01649.t01#2    mNeoVis.2.1.1.chr1_hap1.g01683.t01#5    mNeoVis.vulgivaga.1.1.chr1.g01652.t01#4   mNeoVis.evergladensis.1.1.chr1.g01649.t01#3       1311   125259883:   mNeoVis.evergladensis.1.1.chr1.g01650.t01#3    mNeoVis.vulgivaga.2.1.chr1.g01648.t01#1    mNeoVis.vulgivaga.1.1.chr1.g01653.t01#4    mNeoVis.evergladensis.2.1.chr1.g01650.t01#2    mNeoVis.3.1.1.chr1.g01638.t01#6   mNeoVis.2.1.1.chr1_hap1.g01684.t01#5       1312   125259884:   mNeoVis.evergladensis.2.1.chr1.g01651.t01#2    mNeoVis.vulgivaga.1.1.chr1.g01654.t01#4    mNeoVis.evergladensis.1.1.chr1.g01651.t01#3    mNeoVis.3.1.1.chr1.g01639.t01#6    mNeoVis.vulgivaga.2.1.chr1.g01649.t01#1   mNeoVis.2.1.1.chr1_hap1.g01685.t01#5       1313   125259885:   mNeoVis.vulgivaga.2.1.chr7.g02331.t01#1    mNeoVis.2.1.1.chr7_hap1.g02367.t01#5    mNeoVis.evergladensis.1.1.chr7.g02309.t01#3    mNeoVis.vulgivaga.1.1.chr7.g02317.t01#4   mNeoVis.evergladensis.2.1.chr7.g02313.t01#2       1314   125259886:   mNeoVis.vulgivaga.1.1.chr1.g01655.t01#4    mNeoVis.vulgivaga.2.1.chr1.g01650.t01#1    mNeoVis.3.1.1.chr1.g01640.t01#6    mNeoVis.evergladensis.2.1.chr1.g01652.t01#2    mNeoVis.2.1.1.chr1_hap1.g01686.t01#5   mNeoVis.evergladensis.1.1.chr1.g01652.t01#3       1315   125259887:   mNeoVis.evergladensis.1.1.chr1.g01653.t01#3    mNeoVis.3.1.1.chr1.g01641.t01#6    mNeoVis.vulgivaga.1.1.chr1.g01656.t01#4    mNeoVis.evergladensis.2.1.chr1.g01653.t01#2    mNeoVis.2.1.1.chr1_hap1.g01687.t01#5   mNeoVis.vulgivaga.2.1.chr1.g01651.t01#1       1316   125259888:   mNeoVis.vulgivaga.1.1.chr7.g02318.t01#4    mNeoVis.3.1.1.chr7.g02180.t01#6    mNeoVis.vulgivaga.2.1.chr7.g02332.t01#1    mNeoVis.2.1.1.chr7_hap1.g02368.t01#5    mNeoVis.evergladensis.2.1.chr7.g02314.t01#2   mNeoVis.evergladensis.1.1.chr7.g02310.t01#3       1317   125259889:   mNeoVis.vulgivaga.2.1.chr1.g01652.t01#1    mNeoVis.evergladensis.1.1.chr1.g01654.t01#3    mNeoVis.vulgivaga.1.1.chr1.g01657.t01#4    mNeoVis.evergladensis.2.1.chr1.g01654.t01#2    mNeoVis.3.1.1.chr1.g01642.t01#6   mNeoVis.2.1.1.chr1_hap1.g01688.t01#5       1318   125259890:   mNeoVis.evergladensis.2.1.chr7.g02315.t01#2    mNeoVis.2.1.1.chr7_hap1.g02369.t01#5    mNeoVis.vulgivaga.2.1.chr7.g02333.t01#1    mNeoVis.vulgivaga.1.1.chr7.g02319.t01#4    mNeoVis.evergladensis.1.1.chr7.g02311.t01#3   mNeoVis.3.1.1.chr7.g02181.t01#6       1319   125259891:   mNeoVis.vulgivaga.1.1.chr7.g02320.t01#4    mNeoVis.evergladensis.1.1.chr7.g02312.t01#3    mNeoVis.vulgivaga.2.1.chr7.g02334.t01#1    mNeoVis.evergladensis.2.1.chr7.g02316.t01#2   mNeoVis.2.1.1.chr7_hap1.g02370.t01#5       1320   125259892:   mNeoVis.vulgivaga.1.1.chr7.g02321.t01#4    mNeoVis.2.1.1.chr7_hap1.g02371.t01#5    mNeoVis.vulgivaga.2.1.chr7.g02335.t01#1    mNeoVis.evergladensis.1.1.chr7.g02313.t01#3   mNeoVis.evergladensis.2.1.chr7.g02317.t01#2       1321   125259893:   mNeoVis.evergladensis.2.1.chr11.g00667.t01#2    mNeoVis.vulgivaga.1.1.chr11.g00664.t01#4    mNeoVis.3.1.1.chr11.g00644.t01#6   mNeoVis.2.1.1.chr11_hap1.g00683.t01#5         1322   125259894:   mNeoVis.vulgivaga.1.1.chr11.g00663.t01#4   mNeoVis.3.1.1.chr7.g02182.t01#6             1323   125259895:   mNeoVis.vulgivaga.2.1.chr11.g00680.t01#1   mNeoVis.evergladensis.1.1.chr11.g00671.t01#3           1324   125259896:  mNeoVis.vulgivaga.2.1.chr7.g02336.t01#1               1325   125259897:   mNeoVis.vulgivaga.1.1.chr7.g02322.t01#4    mNeoVis.evergladensis.2.1.chr7.g02318.t01#2    mNeoVis.evergladensis.1.1.chr7.g02314.t01#3    mNeoVis.2.1.1.chr7_hap1.g02372.t01#5    mNeoVis.3.1.1.chr7.g02183.t01#6   mNeoVis.vulgivaga.2.1.chr7.g02337.t01#1       1326   125259898:   mNeoVis.vulgivaga.2.1.chr13.g00340.t01#1    mNeoVis.2.1.1.chr13_hap1.g00352.t01#5   mNeoVis.3.1.1.chr13.g00295.t01#6           1327   125259899:   mNeoVis.2.1.1.chr1_hap1.g01601.t01#5    mNeoVis.vulgivaga.1.1.chr1.g01570.t01#4    mNeoVis.3.1.1.chr1.g01551.t01#6    mNeoVis.vulgivaga.1.1.chr1.g01569.t01#4    mNeoVis.evergladensis.2.1.chr1.g01568.t01#2    mNeoVis.evergladensis.1.1.chr1.g01566.t01#3   mNeoVis.vulgivaga.2.1.chr1.g01565.t01#1     1328   125259900:   mNeoVis.vulgivaga.2.1.chr1.g01566.t01#1    mNeoVis.vulgivaga.1.1.chr1.g01571.t01#4    mNeoVis.3.1.1.chr1.g01552.t01#6    mNeoVis.evergladensis.1.1.chr1.g01567.t01#3    mNeoVis.evergladensis.2.1.chr1.g01569.t01#2   mNeoVis.2.1.1.chr1_hap1.g01602.t01#5       1329   125259901:   mNeoVis.vulgivaga.1.1.chr1.g01572.t01#4    mNeoVis.vulgivaga.2.1.chr1.g01567.t01#1    mNeoVis.3.1.1.chr1.g01553.t01#6    mNeoVis.2.1.1.chr1_hap1.g01603.t01#5    mNeoVis.evergladensis.1.1.chr1.g01568.t01#3   mNeoVis.evergladensis.2.1.chr1.g01570.t01#2       1330   125259902:   mNeoVis.2.1.1.chr7_hap1.g02373.t01#5    mNeoVis.evergladensis.1.1.chr7.g02315.t01#3    mNeoVis.vulgivaga.2.1.chr7.g02338.t01#1    mNeoVis.3.1.1.chr7.g02184.t01#6    mNeoVis.evergladensis.2.1.chr7.g02319.t01#2   mNeoVis.vulgivaga.1.1.chr7.g02323.t01#4       1331   125259903:   mNeoVis.2.1.1.chr11_hap1.g00551.t01#5    mNeoVis.evergladensis.1.1.chr11.g00540.t01#3    mNeoVis.evergladensis.2.1.chr11.g00537.t01#2    mNeoVis.vulgivaga.1.1.chr11.g00533.t01#4    mNeoVis.vulgivaga.2.1.chr11.g00545.t01#1   mNeoVis.3.1.1.chr11.g00510.t01#6       1332   125259904:   mNeoVis.3.1.1.chr9.g00378.t01#6    mNeoVis.2.1.1.chr10_hap1.g00411.t01#5    mNeoVis.vulgivaga.2.1.chr9.g00394.t01#1    mNeoVis.vulgivaga.1.1.chr9.g00392.t01#4    mNeoVis.evergladensis.1.1.chr9.g00399.t01#3   mNeoVis.evergladensis.2.1.chr8.g00392.t01#2       1333   125259905:   mNeoVis.2.1.1.chr1_hap1.g01604.t01#5    mNeoVis.3.1.1.chr1.g01554.t01#6    mNeoVis.vulgivaga.2.1.chr1.g01568.t01#1    mNeoVis.evergladensis.1.1.chr1.g01569.t01#3    mNeoVis.evergladensis.2.1.chr1.g01571.t01#2   mNeoVis.vulgivaga.1.1.chr1.g01573.t01#4       1334   125259906:   mNeoVis.vulgivaga.2.1.chr1.g01569.t01#1    mNeoVis.vulgivaga.1.1.chr6.g00819.t01#4    mNeoVis.evergladensis.1.1.chr1.g01570.t01#3    mNeoVis.vulgivaga.1.1.chr1.g01574.t01#4    mNeoVis.evergladensis.1.1.chr6.g00825.t01#3    mNeoVis.3.1.1.chr1.g01555.t01#6    mNeoVis.evergladensis.2.1.chr4.g00253.t01#2    mNeoVis.evergladensis.2.1.chr1.g01572.t01#2    mNeoVis.2.1.1.chr1_hap1.g01605.t01#5      1335   125259907:   mNeoVis.evergladensis.1.1.chr3.g00192.t01#3    mNeoVis.2.1.1.chr10_hap1.g00065.t01#5    mNeoVis.2.1.1.chr7_hap1.g01208.t01#5    mNeoVis.vulgivaga.1.1.chr9.g00058.t01#4    mNeoVis.3.1.1.chr14.g00404.t01#6   mNeoVis.evergladensis.2.1.chr8.g00064.t01#2       1336   125259908:   mNeoVis.evergladensis.1.1.chr5.g00825.t01#3    mNeoVis.evergladensis.1.1.chr4.g00583.t01#3   mNeoVis.vulgivaga.2.1.chr4.g00585.t01#1           1337   125259909:   mNeoVis.evergladensis.1.1.chr9.g00064.t01#3   mNeoVis.vulgivaga.2.1.chr9.g00064.t01#1             1338   125259910:   mNeoVis.evergladensis.2.1.chr1.g01573.t01#2    mNeoVis.2.1.1.chr1_hap1.g01606.t01#5    mNeoVis.vulgivaga.1.1.chr1.g01575.t01#4    mNeoVis.vulgivaga.2.1.chr1.g01570.t01#1    mNeoVis.3.1.1.chr1.g01556.t01#6   mNeoVis.evergladensis.1.1.chr1.g01571.t01#3       1339   125259911:   mNeoVis.vulgivaga.2.1.chr7.g02339.t01#1    mNeoVis.evergladensis.1.1.chr7.g02316.t01#3    mNeoVis.3.1.1.chr7.g02185.t01#6    mNeoVis.2.1.1.chr7_hap1.g02374.t01#5    mNeoVis.vulgivaga.1.1.chr7.g02324.t01#4   mNeoVis.evergladensis.2.1.chr7.g02320.t01#2       1340   125259912:   mNeoVis.2.1.1.chr1_hap1.g01607.t01#5    mNeoVis.evergladensis.1.1.chr1.g01572.t01#3    mNeoVis.vulgivaga.1.1.chr1.g01576.t01#4    mNeoVis.evergladensis.2.1.chr1.g01574.t01#2   mNeoVis.vulgivaga.2.1.chr1.g01571.t01#1       1341   125259913:  mNeoVis.vulgivaga.2.1.chr1.g01572.t01#1               1342   125259914:   mNeoVis.vulgivaga.1.1.chr7.g02325.t01#4    mNeoVis.evergladensis.1.1.chr7.g02317.t01#3    mNeoVis.evergladensis.2.1.chr7.g02321.t01#2    mNeoVis.vulgivaga.2.1.chr7.g02340.t01#1   mNeoVis.2.1.1.chr7_hap1.g02375.t01#5       1343   125259915:   mNeoVis.vulgivaga.2.1.chr1.g01573.t01#1    mNeoVis.vulgivaga.1.1.chr1.g01578.t01#4    mNeoVis.evergladensis.2.1.chr1.g01576.t01#2    mNeoVis.3.1.1.chr1.g01558.t01#6    mNeoVis.2.1.1.chr1_hap1.g01610.t01#5   mNeoVis.evergladensis.1.1.chr1.g01574.t01#3       1344   125259916:   mNeoVis.3.1.1.chr7.g02187.t01#6    mNeoVis.vulgivaga.1.1.chr7.g02326.t01#4    mNeoVis.evergladensis.1.1.chr7.g02318.t01#3    mNeoVis.evergladensis.2.1.chr7.g02322.t01#2    mNeoVis.2.1.1.chr7_hap1.g02376.t01#5   mNeoVis.vulgivaga.2.1.chr7.g02341.t01#1       1345   125259917:   mNeoVis.evergladensis.1.1.chr7.g02319.t01#3    mNeoVis.vulgivaga.2.1.chr7.g02342.t01#1    mNeoVis.evergladensis.2.1.chr7.g02323.t01#2    mNeoVis.2.1.1.chr7_hap1.g02377.t01#5   mNeoVis.vulgivaga.1.1.chr7.g02327.t01#4       1346   125259918:   mNeoVis.vulgivaga.1.1.chr1.g01579.t01#4    mNeoVis.3.1.1.chr1.g01559.t01#6    mNeoVis.2.1.1.chr1_hap1.g01611.t01#5    mNeoVis.evergladensis.1.1.chr1.g01575.t01#3    mNeoVis.evergladensis.2.1.chr1.g01577.t01#2   mNeoVis.vulgivaga.2.1.chr1.g01574.t01#1       1347   125259919:   mNeoVis.2.1.1.chr1_hap1.g01612.t01#5    mNeoVis.evergladensis.1.1.chr1.g01576.t01#3    mNeoVis.vulgivaga.1.1.chr1.g01580.t01#4    mNeoVis.evergladensis.2.1.chr1.g01578.t01#2   mNeoVis.3.1.1.chr1.g01560.t01#6       1348   125259920:   mNeoVis.3.1.1.chr9.g00251.t01#6   mNeoVis.vulgivaga.2.1.chr1.g01575.t01#1             1349   125259921:   mNeoVis.2.1.1.chr15_hap1.g00154.t01#5    mNeoVis.evergladensis.1.1.chr15.g00476.t01#3    mNeoVis.vulgivaga.1.1.chr15.g00139.t01#4    mNeoVis.vulgivaga.2.1.chr15.g00143.t01#1    mNeoVis.evergladensis.2.1.chr15.g00139.t01#2   mNeoVis.3.1.1.chr15.g00131.t01#6       1350   125259922:   mNeoVis.2.1.1.chr7_hap1.g00065.t01#5    mNeoVis.evergladensis.1.1.chr7.g00068.t01#3    mNeoVis.evergladensis.2.1.chr7.g00066.t01#2    mNeoVis.vulgivaga.2.1.chr7.g00066.t01#1   mNeoVis.vulgivaga.1.1.chr7.g00067.t01#4       1351   125259923:   mNeoVis.2.1.1.chr1_hap1.g01613.t01#5    mNeoVis.evergladensis.2.1.chr1.g01579.t01#2    mNeoVis.vulgivaga.1.1.chr1.g01581.t01#4    mNeoVis.vulgivaga.2.1.chr1.g01576.t01#1    mNeoVis.evergladensis.1.1.chr1.g01577.t01#3   mNeoVis.3.1.1.chr1.g01561.t01#6       1352   125259924:   mNeoVis.2.1.1.chr1_hap1.g01614.t01#5    mNeoVis.evergladensis.1.1.chr1.g01578.t01#3    mNeoVis.vulgivaga.2.1.chr1.g01577.t01#1    mNeoVis.evergladensis.2.1.chr1.g01580.t01#2    mNeoVis.vulgivaga.1.1.chr1.g01582.t01#4   mNeoVis.3.1.1.chr1.g01562.t01#6       1353   125259925:   mNeoVis.vulgivaga.1.1.chr1.g01583.t01#4    mNeoVis.3.1.1.chr1.g01563.t01#6    mNeoVis.evergladensis.1.1.chr1.g01579.t01#3    mNeoVis.vulgivaga.2.1.chr1.g01578.t01#1    mNeoVis.evergladensis.2.1.chr1.g01581.t01#2   mNeoVis.2.1.1.chr1_hap1.g01615.t01#5       1354   125259926:   mNeoVis.2.1.1.chr1_hap1.g01616.t01#5    mNeoVis.vulgivaga.2.1.chr1.g01579.t01#1    mNeoVis.vulgivaga.1.1.chr1.g01584.t01#4    mNeoVis.evergladensis.2.1.chr1.g01582.t01#2    mNeoVis.evergladensis.1.1.chr1.g01580.t01#3   mNeoVis.3.1.1.chr1.g01564.t01#6       1355   125259927:   mNeoVis.3.1.1.chr11.g00967.t01#6    mNeoVis.evergladensis.1.1.chr11.g01008.t01#3    mNeoVis.evergladensis.2.1.chr11.g00991.t01#2    mNeoVis.vulgivaga.2.1.chr7.g02343.t01#1    mNeoVis.vulgivaga.1.1.chr11.g00988.t01#4    mNeoVis.vulgivaga.2.1.chr11.g01014.t01#1    mNeoVis.2.1.1.chr7_hap1.g02378.t01#5    mNeoVis.2.1.1.chr11_hap1.g01016.t01#5    mNeoVis.vulgivaga.1.1.chr7.g02328.t01#4      1356   125259928:   mNeoVis.evergladensis.1.1.chr9.g00697.t01#3    mNeoVis.vulgivaga.2.1.chr9.g00690.t01#1    mNeoVis.2.1.1.chr10_hap1.g00708.t01#5    mNeoVis.evergladensis.2.1.chr8.g00689.t01#2    mNeoVis.3.1.1.chr9.g00668.t01#6   mNeoVis.vulgivaga.1.1.chr9.g00698.t01#4       1357   125259929:   mNeoVis.3.1.1.chr7.g02189.t01#6    mNeoVis.evergladensis.1.1.chr7.g02320.t01#3   mNeoVis.evergladensis.2.1.chr7.g02324.t01#2         1358   125259930:   mNeoVis.evergladensis.1.1.chr1.g01581.t01#3    mNeoVis.vulgivaga.1.1.chr1.g01585.t01#4    mNeoVis.3.1.1.chr1.g01565.t01#6    mNeoVis.evergladensis.2.1.chr1.g01583.t01#2    mNeoVis.vulgivaga.2.1.chr1.g01580.t01#1   mNeoVis.2.1.1.chr1_hap1.g01617.t01#5       1359   125259931:   mNeoVis.vulgivaga.2.1.chr7.g02344.t01#1    mNeoVis.vulgivaga.1.1.chr7.g02329.t01#4    mNeoVis.evergladensis.1.1.chr7.g02321.t01#3    mNeoVis.2.1.1.chr7_hap1.g02379.t01#5   mNeoVis.evergladensis.2.1.chr7.g02325.t01#2       1360   125259932:   mNeoVis.vulgivaga.2.1.chr1.g01581.t01#1    mNeoVis.3.1.1.chr1.g01566.t01#6    mNeoVis.2.1.1.chr1_hap1.g01618.t01#5    mNeoVis.vulgivaga.1.1.chr1.g01586.t01#4    mNeoVis.evergladensis.1.1.chr1.g01582.t01#3   mNeoVis.evergladensis.2.1.chr1.g01584.t01#2       1361   125259933:   mNeoVis.evergladensis.1.1.chr15.g00475.t01#3    mNeoVis.2.1.1.chr15_hap1.g00155.t01#5    mNeoVis.3.1.1.chr15.g00133.t01#6    mNeoVis.evergladensis.2.1.chr15.g00140.t01#2    mNeoVis.vulgivaga.2.1.chr15.g00144.t01#1    mNeoVis.vulgivaga.1.1.chr15.g00140.t01#4   mNeoVis.3.1.1.chr15.g00134.t01#6       1362   125259934:   mNeoVis.vulgivaga.1.1.chr8.g00220.t01#4    mNeoVis.2.1.1.chr8_hap1.g00231.t01#5    mNeoVis.evergladensis.2.1.chr9.g00221.t01#2    mNeoVis.vulgivaga.2.1.chr8.g00228.t01#1    mNeoVis.vulgivaga.2.1.chr8.g00227.t01#1    mNeoVis.evergladensis.1.1.chr8.g00221.t01#3   mNeoVis.3.1.1.chr8.g00189.t01#6       1363   125259935:   mNeoVis.evergladensis.2.1.chr7.g00752.t01#2    mNeoVis.2.1.1.chr7_hap1.g00764.t01#5    mNeoVis.vulgivaga.2.1.chr7.g00754.t01#1    mNeoVis.evergladensis.1.1.chr7.g00762.t01#3    mNeoVis.3.1.1.chr7.g00714.t01#6   mNeoVis.vulgivaga.1.1.chr7.g00763.t01#4       1364   125259936:   mNeoVis.2.1.1.chr7_hap1.g00066.t01#5    mNeoVis.evergladensis.2.1.chr7.g00067.t01#2    mNeoVis.evergladensis.1.1.chr7.g00069.t01#3    mNeoVis.vulgivaga.2.1.chr7.g00067.t01#1    mNeoVis.vulgivaga.1.1.chr7.g00068.t01#4   mNeoVis.3.1.1.chr7.g00054.t01#6       1365   125259937:   mNeoVis.3.1.1.chr1.g01567.t01#6    mNeoVis.vulgivaga.2.1.chr1.g01582.t01#1    mNeoVis.evergladensis.1.1.chr1.g01583.t01#3    mNeoVis.2.1.1.chr1_hap1.g01619.t01#5    mNeoVis.vulgivaga.1.1.chr1.g01587.t01#4   mNeoVis.evergladensis.2.1.chr1.g01585.t01#2       1366   125259938:   mNeoVis.vulgivaga.1.1.chr1.g01588.t01#4    mNeoVis.vulgivaga.2.1.chr1.g01583.t01#1    mNeoVis.evergladensis.1.1.chr1.g01584.t01#3    mNeoVis.3.1.1.chr1.g01568.t01#6    mNeoVis.evergladensis.2.1.chr1.g01586.t01#2   mNeoVis.2.1.1.chr1_hap1.g01620.t01#5       1367   125259939:   mNeoVis.vulgivaga.2.1.chr1.g01584.t01#1    mNeoVis.evergladensis.1.1.chr1.g01585.t01#3    mNeoVis.2.1.1.chr1_hap1.g01621.t01#5    mNeoVis.vulgivaga.1.1.chr1.g01589.t01#4    mNeoVis.evergladensis.2.1.chr1.g01587.t01#2   mNeoVis.3.1.1.chr1.g01569.t01#6       1368   125259940:   mNeoVis.vulgivaga.2.1.chr7.g02345.t01#1    mNeoVis.3.1.1.chr7.g02191.t01#6    mNeoVis.2.1.1.chr7_hap1.g02380.t01#5    mNeoVis.evergladensis.2.1.chr7.g02326.t01#2    mNeoVis.vulgivaga.1.1.chr7.g02330.t01#4   mNeoVis.evergladensis.1.1.chr7.g02322.t01#3       1369   125259941:   mNeoVis.3.1.1.chr9.g00298.t01#6    mNeoVis.evergladensis.2.1.chr8.g00309.t01#2    mNeoVis.vulgivaga.1.1.chr9.g00310.t01#4    mNeoVis.evergladensis.1.1.chr9.g00313.t01#3    mNeoVis.2.1.1.chr10_hap1.g00328.t01#5   mNeoVis.vulgivaga.2.1.chr9.g00310.t01#1       1370   125259942:   mNeoVis.evergladensis.1.1.chr14.g00111.t01#3    mNeoVis.vulgivaga.1.1.chr14.g00106.t01#4    mNeoVis.3.1.1.chr14.g00100.t01#6    mNeoVis.evergladensis.2.1.chr14.g00105.t01#2    mNeoVis.2.1.1.chr14_hap1.g00106.t01#5   mNeoVis.vulgivaga.2.1.chr14.g00106.t01#1       1371   125259943:   mNeoVis.evergladensis.1.1.chr7.g02323.t01#3    mNeoVis.evergladensis.2.1.chr7.g02327.t01#2    mNeoVis.2.1.1.chr7_hap1.g02381.t01#5    mNeoVis.3.1.1.chr7.g02192.t01#6    mNeoVis.vulgivaga.2.1.chr7.g02346.t01#1   mNeoVis.vulgivaga.1.1.chr7.g02331.t01#4       1372   125259944:   mNeoVis.vulgivaga.2.1.chr7.g02347.t01#1    mNeoVis.3.1.1.chr7.g02193.t01#6    mNeoVis.2.1.1.chr7_hap1.g02382.t01#5    mNeoVis.vulgivaga.1.1.chr7.g02332.t01#4    mNeoVis.evergladensis.1.1.chr7.g02324.t01#3   mNeoVis.evergladensis.2.1.chr7.g02328.t01#2       1373   125259945:   mNeoVis.vulgivaga.1.1.chr1.g01590.t01#4    mNeoVis.evergladensis.1.1.chr1.g01586.t01#3    mNeoVis.3.1.1.chr1.g01570.t01#6    mNeoVis.evergladensis.2.1.chr1.g01588.t01#2    mNeoVis.2.1.1.chr1_hap1.g01622.t01#5   mNeoVis.vulgivaga.2.1.chr1.g01585.t01#1       1374   125259946:   mNeoVis.evergladensis.1.1.chr7.g02325.t01#3    mNeoVis.evergladensis.2.1.chr7.g02329.t01#2    mNeoVis.3.1.1.chr7.g02194.t01#6    mNeoVis.2.1.1.chr7_hap1.g02383.t01#5    mNeoVis.vulgivaga.2.1.chr7.g02348.t01#1   mNeoVis.vulgivaga.1.1.chr7.g02333.t01#4       1375   125259947:   mNeoVis.vulgivaga.2.1.chr1.g01586.t01#1    mNeoVis.evergladensis.1.1.chr1.g01587.t01#3    mNeoVis.2.1.1.chr1_hap1.g01623.t01#5    mNeoVis.vulgivaga.1.1.chr1.g01591.t01#4    mNeoVis.3.1.1.chr1.g01571.t01#6   mNeoVis.evergladensis.2.1.chr1.g01589.t01#2       1376   125259948:   mNeoVis.evergladensis.1.1.chr7.g02326.t01#3    mNeoVis.2.1.1.chr7_hap1.g02384.t01#5    mNeoVis.3.1.1.chr7.g02195.t01#6    mNeoVis.evergladensis.2.1.chr7.g02330.t01#2    mNeoVis.vulgivaga.2.1.chr7.g02349.t01#1   mNeoVis.vulgivaga.1.1.chr7.g02334.t01#4       1377   125259949:   mNeoVis.vulgivaga.1.1.chr7.g02335.t01#4    mNeoVis.evergladensis.1.1.chr7.g02327.t01#3    mNeoVis.2.1.1.chr7_hap1.g02385.t01#5    mNeoVis.vulgivaga.2.1.chr7.g02350.t01#1    mNeoVis.3.1.1.chr7.g02196.t01#6   mNeoVis.evergladensis.2.1.chr7.g02331.t01#2       1378   125259950:   mNeoVis.vulgivaga.2.1.chr1.g01587.t01#1    mNeoVis.2.1.1.chr1_hap1.g01624.t01#5    mNeoVis.evergladensis.2.1.chr1.g01590.t01#2    mNeoVis.vulgivaga.1.1.chr1.g01592.t01#4    mNeoVis.evergladensis.1.1.chr1.g01588.t01#3   mNeoVis.3.1.1.chr1.g01572.t01#6       1379   125259951:   mNeoVis.vulgivaga.1.1.chr7.g02336.t01#4    mNeoVis.vulgivaga.2.1.chr7.g02351.t01#1    mNeoVis.evergladensis.1.1.chr7.g02328.t01#3    mNeoVis.2.1.1.chr7_hap1.g02386.t01#5    mNeoVis.3.1.1.chr7.g02197.t01#6   mNeoVis.evergladensis.2.1.chr7.g02332.t01#2       1380   125259952:   mNeoVis.2.1.1.chr7_hap1.g02387.t01#5    mNeoVis.vulgivaga.2.1.chr7.g02352.t01#1    mNeoVis.evergladensis.2.1.chr7.g02333.t01#2    mNeoVis.3.1.1.chr7.g02198.t01#6    mNeoVis.vulgivaga.1.1.chr7.g02337.t01#4   mNeoVis.evergladensis.1.1.chr7.g02329.t01#3       1381   125259953:   mNeoVis.evergladensis.1.1.chr2.g00839.t01#3    mNeoVis.2.1.1.chr3_hap1.g01315.t01#5    mNeoVis.vulgivaga.2.1.chr1.g01588.t01#1    mNeoVis.vulgivaga.2.1.chr3.g01334.t01#1    mNeoVis.vulgivaga.1.1.chr2.g00845.t01#4    mNeoVis.2.1.1.chr6_hap1.g00804.t01#5    mNeoVis.vulgivaga.2.1.chr4.g00826.t01#1    mNeoVis.3.1.1.chr4.g00814.t01#6    mNeoVis.2.1.1.chr5_hap1.g00843.t01#5      1382   125259954:   mNeoVis.evergladensis.1.1.chr1.g01589.t01#3    mNeoVis.2.1.1.chr1_hap1.g01626.t01#5    mNeoVis.evergladensis.2.1.chr1.g01592.t01#2    mNeoVis.vulgivaga.1.1.chr1.g01593.t01#4   mNeoVis.vulgivaga.2.1.chr1.g01589.t01#1       1383   125259955:   mNeoVis.3.1.1.chr7.g02199.t01#6    mNeoVis.vulgivaga.1.1.chr7.g02338.t01#4    mNeoVis.evergladensis.1.1.chr7.g02330.t01#3    mNeoVis.evergladensis.2.1.chr7.g02334.t01#2    mNeoVis.vulgivaga.2.1.chr7.g02353.t01#1   mNeoVis.2.1.1.chr7_hap1.g02388.t01#5       1384   125259956:   mNeoVis.vulgivaga.2.1.chr7.g02354.t01#1    mNeoVis.2.1.1.chr7_hap1.g02389.t01#5    mNeoVis.evergladensis.1.1.chr7.g02331.t01#3    mNeoVis.vulgivaga.1.1.chr7.g02339.t01#4    mNeoVis.evergladensis.2.1.chr7.g02335.t01#2   mNeoVis.3.1.1.chr7.g02200.t01#6       1385   125259957:   mNeoVis.evergladensis.2.1.chr1.g01593.t01#2    mNeoVis.vulgivaga.2.1.chr1.g01590.t01#1    mNeoVis.vulgivaga.1.1.chr1.g01594.t01#4    mNeoVis.evergladensis.1.1.chr1.g01590.t01#3    mNeoVis.2.1.1.chr1_hap1.g01627.t01#5   mNeoVis.3.1.1.chr1.g01576.t01#6       1386   125259958:   mNeoVis.vulgivaga.2.1.chr7.g02355.t01#1    mNeoVis.evergladensis.1.1.chr7.g02332.t01#3    mNeoVis.evergladensis.2.1.chr7.g02336.t01#2    mNeoVis.2.1.1.chr7_hap1.g02390.t01#5    mNeoVis.3.1.1.chr7.g02201.t01#6   mNeoVis.vulgivaga.1.1.chr7.g02340.t01#4       1387   125259959:   mNeoVis.3.1.1.chr1.g01577.t01#6    mNeoVis.evergladensis.2.1.chr1.g01594.t01#2    mNeoVis.evergladensis.1.1.chr1.g01591.t01#3    mNeoVis.vulgivaga.2.1.chr1.g01591.t01#1    mNeoVis.2.1.1.chr1_hap1.g01628.t01#5   mNeoVis.vulgivaga.1.1.chr1.g01595.t01#4       1388   125259960:   mNeoVis.3.1.1.chr7.g02202.t01#6    mNeoVis.vulgivaga.1.1.chr7.g02341.t01#4    mNeoVis.evergladensis.2.1.chr7.g02337.t01#2    mNeoVis.vulgivaga.2.1.chr7.g02357.t01#1    mNeoVis.vulgivaga.2.1.chr7.g02356.t01#1    mNeoVis.evergladensis.1.1.chr7.g02333.t01#3   mNeoVis.2.1.1.chr7_hap1.g02391.t01#5     1389   125259961:   mNeoVis.vulgivaga.1.1.chr1.g01596.t01#4    mNeoVis.evergladensis.1.1.chr1.g01592.t01#3    mNeoVis.2.1.1.chr1_hap1.g01629.t01#5    mNeoVis.evergladensis.2.1.chr1.g01595.t01#2   mNeoVis.vulgivaga.2.1.chr1.g01592.t01#1       1390   125259962:   mNeoVis.vulgivaga.2.1.chr7.g02358.t01#1    mNeoVis.evergladensis.2.1.chr7.g02338.t01#2    mNeoVis.2.1.1.chr7_hap1.g02392.t01#5    mNeoVis.evergladensis.1.1.chr7.g02334.t01#3    mNeoVis.vulgivaga.1.1.chr7.g02342.t01#4   mNeoVis.3.1.1.chr7.g02203.t01#6       1391   125259963:   mNeoVis.2.1.1.chr7_hap1.g02393.t01#5    mNeoVis.3.1.1.chr7.g02204.t01#6    mNeoVis.vulgivaga.1.1.chr7.g02343.t01#4    mNeoVis.evergladensis.2.1.chr7.g02339.t01#2    mNeoVis.vulgivaga.2.1.chr7.g02359.t01#1   mNeoVis.evergladensis.1.1.chr7.g02335.t01#3       1392   125259964:   mNeoVis.evergladensis.2.1.chr1.g01596.t01#2    mNeoVis.2.1.1.chr1_hap1.g01630.t01#5    mNeoVis.evergladensis.1.1.chr1.g01593.t01#3    mNeoVis.vulgivaga.2.1.chr1.g01593.t01#1    mNeoVis.vulgivaga.1.1.chr1.g01597.t01#4   mNeoVis.3.1.1.chr1.g01579.t01#6       1393   125259965:   mNeoVis.vulgivaga.2.1.chr1.g01594.t01#1    mNeoVis.evergladensis.2.1.chr1.g01597.t01#2    mNeoVis.evergladensis.1.1.chr1.g01594.t01#3    mNeoVis.vulgivaga.1.1.chr1.g01598.t01#4   mNeoVis.2.1.1.chr1_hap1.g01631.t01#5       1394   125259966:   mNeoVis.3.1.1.chr7.g02205.t01#6    mNeoVis.evergladensis.2.1.chr7.g02340.t01#2    mNeoVis.vulgivaga.2.1.chr7.g02360.t01#1    mNeoVis.evergladensis.1.1.chr7.g02336.t01#3    mNeoVis.vulgivaga.1.1.chr7.g02344.t01#4   mNeoVis.2.1.1.chr7_hap1.g02394.t01#5       1395   125259967:   mNeoVis.2.1.1.chr7_hap1.g02395.t01#5    mNeoVis.vulgivaga.1.1.chr7.g02345.t01#4    mNeoVis.evergladensis.2.1.chr7.g02341.t01#2    mNeoVis.vulgivaga.2.1.chr7.g02361.t01#1    mNeoVis.3.1.1.chr7.g02206.t01#6   mNeoVis.evergladensis.1.1.chr7.g02337.t01#3       1396   125259968:   mNeoVis.evergladensis.1.1.chr7.g02338.t01#3    mNeoVis.evergladensis.2.1.chr7.g02342.t01#2    mNeoVis.2.1.1.chr7_hap1.g02396.t01#5    mNeoVis.vulgivaga.1.1.chr7.g02346.t01#4    mNeoVis.3.1.1.chr7.g02207.t01#6   mNeoVis.vulgivaga.2.1.chr7.g02362.t01#1       1397   125259969:   mNeoVis.evergladensis.1.1.chr7.g02339.t01#3    mNeoVis.3.1.1.chr7.g02208.t01#6    mNeoVis.evergladensis.2.1.chr7.g02343.t01#2    mNeoVis.2.1.1.chr7_hap1.g02397.t01#5    mNeoVis.vulgivaga.1.1.chr7.g02347.t01#4   mNeoVis.vulgivaga.2.1.chr7.g02363.t01#1       1398   125259970:   mNeoVis.3.1.1.chr7.g02209.t01#6    mNeoVis.evergladensis.1.1.chr7.g02340.t01#3    mNeoVis.vulgivaga.2.1.chr7.g02364.t01#1    mNeoVis.2.1.1.chr7_hap1.g02398.t01#5    mNeoVis.vulgivaga.1.1.chr7.g02348.t01#4   mNeoVis.evergladensis.2.1.chr7.g02344.t01#2       1399   125259971:   mNeoVis.3.1.1.chr7.g00055.t01#6    mNeoVis.vulgivaga.2.1.chr7.g00068.t01#1    mNeoVis.2.1.1.chr7_hap1.g00067.t01#5    mNeoVis.evergladensis.2.1.chr7.g00068.t01#2   mNeoVis.evergladensis.1.1.chr7.g00070.t01#3       1400   125259972:   mNeoVis.evergladensis.2.1.chr1.g01598.t01#2    mNeoVis.vulgivaga.2.1.chr1.g01595.t01#1    mNeoVis.3.1.1.chr1.g01582.t01#6    mNeoVis.evergladensis.1.1.chr1.g01595.t01#3    mNeoVis.2.1.1.chr1_hap1.g01632.t01#5   mNeoVis.vulgivaga.1.1.chr1.g01599.t01#4       1401   125259973:   mNeoVis.evergladensis.1.1.chr15.g00474.t01#3    mNeoVis.2.1.1.chr15_hap1.g00156.t01#5    mNeoVis.evergladensis.2.1.chr15.g00141.t01#2    mNeoVis.3.1.1.chr15.g00135.t01#6    mNeoVis.vulgivaga.1.1.chr15.g00141.t01#4   mNeoVis.vulgivaga.2.1.chr15.g00145.t01#1       1402   125259974:   mNeoVis.3.1.1.chr1.g01583.t01#6    mNeoVis.evergladensis.1.1.chr1.g01596.t01#3    mNeoVis.evergladensis.2.1.chr1.g01599.t01#2    mNeoVis.2.1.1.chr1_hap1.g01633.t01#5    mNeoVis.vulgivaga.1.1.chr1.g01600.t01#4   mNeoVis.vulgivaga.2.1.chr1.g01596.t01#1       1403   125259975:   mNeoVis.vulgivaga.2.1.chr7.g02365.t01#1    mNeoVis.evergladensis.1.1.chr7.g02341.t01#3    mNeoVis.evergladensis.2.1.chr7.g02345.t01#2    mNeoVis.2.1.1.chr7_hap1.g02399.t01#5   mNeoVis.vulgivaga.1.1.chr7.g02349.t01#4       1404   125259976:   mNeoVis.vulgivaga.2.1.chr1.g01597.t01#1    mNeoVis.evergladensis.2.1.chr1.g01600.t01#2    mNeoVis.evergladensis.1.1.chr1.g01597.t01#3    mNeoVis.3.1.1.chr1.g01584.t01#6    mNeoVis.2.1.1.chr1_hap1.g01634.t01#5   mNeoVis.vulgivaga.1.1.chr1.g01601.t01#4       1405   125259977:   mNeoVis.2.1.1.chr7_hap1.g02400.t01#5    mNeoVis.vulgivaga.2.1.chr7.g02366.t01#1    mNeoVis.3.1.1.chr7.g02210.t01#6    mNeoVis.vulgivaga.1.1.chr7.g02350.t01#4    mNeoVis.evergladensis.2.1.chr7.g02346.t01#2   mNeoVis.evergladensis.1.1.chr7.g02342.t01#3       1406   125259978:   mNeoVis.vulgivaga.2.1.chr1.g01598.t01#1    mNeoVis.evergladensis.1.1.chr1.g01598.t01#3    mNeoVis.vulgivaga.1.1.chr1.g01602.t01#4    mNeoVis.3.1.1.chr1.g01585.t01#6    mNeoVis.2.1.1.chr1_hap1.g01635.t01#5   mNeoVis.evergladensis.2.1.chr1.g01601.t01#2       1407   125259979:   mNeoVis.3.1.1.chr11.g00530.t01#6    mNeoVis.evergladensis.1.1.chr7.g02343.t01#3    mNeoVis.2.1.1.chr11_hap1.g00569.t01#5    mNeoVis.vulgivaga.2.1.chr11.g00564.t01#1    mNeoVis.vulgivaga.2.1.chr7.g02367.t01#1    mNeoVis.evergladensis.2.1.chr11.g00556.t01#2    mNeoVis.3.1.1.chr7.g02211.t01#6    mNeoVis.vulgivaga.1.1.chr11.g00551.t01#4    mNeoVis.evergladensis.1.1.chr11.g00559.t01#3      1408   125259980:   mNeoVis.3.1.1.chr15.g00136.t01#6    mNeoVis.vulgivaga.2.1.chr15.g00146.t01#1    mNeoVis.2.1.1.chr15_hap1.g00157.t01#5    mNeoVis.vulgivaga.1.1.chr15.g00142.t01#4    mNeoVis.evergladensis.2.1.chr15.g00142.t01#2   mNeoVis.evergladensis.1.1.chr15.g00473.t01#3       1409   125259981:   mNeoVis.evergladensis.2.1.chr1.g01602.t01#2    mNeoVis.evergladensis.1.1.chr1.g01599.t01#3    mNeoVis.3.1.1.chr1.g01586.t01#6    mNeoVis.vulgivaga.1.1.chr1.g01603.t01#4    mNeoVis.2.1.1.chr1_hap1.g01636.t01#5   mNeoVis.vulgivaga.2.1.chr1.g01599.t01#1       1410   125259982:   mNeoVis.2.1.1.chr7_hap1.g02308.t01#5    mNeoVis.evergladensis.1.1.chr7.g02252.t01#3    mNeoVis.vulgivaga.1.1.chr7.g02260.t01#4    mNeoVis.3.1.1.chr7.g02124.t01#6    mNeoVis.vulgivaga.2.1.chr7.g02272.t01#1   mNeoVis.evergladensis.2.1.chr7.g02255.t01#2       1411   125259983:   mNeoVis.vulgivaga.2.1.chr7.g00069.t01#1    mNeoVis.3.1.1.chr7.g00056.t01#6    mNeoVis.2.1.1.chr7_hap1.g00068.t01#5    mNeoVis.vulgivaga.1.1.chr7.g00070.t01#4    mNeoVis.evergladensis.2.1.chr7.g00069.t01#2   mNeoVis.evergladensis.1.1.chr7.g00071.t01#3       1412   125259984:   mNeoVis.evergladensis.2.1.chr1.g01603.t01#2    mNeoVis.evergladensis.1.1.chr1.g01600.t01#3    mNeoVis.vulgivaga.2.1.chr1.g01600.t01#1    mNeoVis.2.1.1.chr1_hap1.g01637.t01#5    mNeoVis.3.1.1.chr1.g01587.t01#6   mNeoVis.vulgivaga.1.1.chr1.g01604.t01#4       1413   125259985:   mNeoVis.evergladensis.2.1.chr7.g02256.t01#2    mNeoVis.evergladensis.1.1.chr7.g02253.t01#3    mNeoVis.vulgivaga.1.1.chr7.g02261.t01#4    mNeoVis.2.1.1.chr7_hap1.g02309.t01#5    mNeoVis.3.1.1.chr7.g02125.t01#6   mNeoVis.vulgivaga.2.1.chr7.g02273.t01#1       1414   125259986:   mNeoVis.evergladensis.2.1.chr1.g01604.t01#2    mNeoVis.vulgivaga.2.1.chr1.g01601.t01#1    mNeoVis.vulgivaga.1.1.chr1.g01605.t01#4    mNeoVis.evergladensis.1.1.chr1.g01601.t01#3    mNeoVis.3.1.1.chr1.g01588.t01#6   mNeoVis.2.1.1.chr1_hap1.g01638.t01#5       1415   125259987:   mNeoVis.evergladensis.1.1.chr7.g02254.t01#3    mNeoVis.3.1.1.chr7.g02126.t01#6    mNeoVis.2.1.1.chr7_hap1.g02310.t01#5    mNeoVis.evergladensis.2.1.chr7.g02257.t01#2    mNeoVis.vulgivaga.2.1.chr7.g02274.t01#1   mNeoVis.vulgivaga.1.1.chr7.g02262.t01#4       1416   125259988:   mNeoVis.vulgivaga.2.1.chr7.g02275.t01#1    mNeoVis.evergladensis.2.1.chr7.g02258.t01#2    mNeoVis.vulgivaga.1.1.chr7.g02263.t01#4    mNeoVis.evergladensis.1.1.chr7.g02255.t01#3    mNeoVis.2.1.1.chr7_hap1.g02311.t01#5   mNeoVis.3.1.1.chr7.g02127.t01#6       1417   125259989:   mNeoVis.3.1.1.chr1.g01589.t01#6    mNeoVis.evergladensis.2.1.chr1.g01605.t01#2    mNeoVis.2.1.1.chr1_hap1.g01639.t01#5    mNeoVis.vulgivaga.1.1.chr1.g01606.t01#4    mNeoVis.vulgivaga.2.1.chr1.g01602.t01#1   mNeoVis.evergladensis.1.1.chr1.g01602.t01#3       1418   125259990:   mNeoVis.evergladensis.2.1.chr8.g00241.t01#2    mNeoVis.evergladensis.1.1.chr9.g00245.t01#3    mNeoVis.3.1.1.chr9.g00226.t01#6   mNeoVis.2.1.1.chr10_hap1.g00256.t01#5         1419   125259991:   mNeoVis.3.1.1.chr7.g02128.t01#6    mNeoVis.2.1.1.chr7_hap1.g02312.t01#5    mNeoVis.evergladensis.2.1.chr7.g02259.t01#2    mNeoVis.vulgivaga.2.1.chr7.g02276.t01#1    mNeoVis.vulgivaga.1.1.chr7.g02264.t01#4   mNeoVis.evergladensis.1.1.chr7.g02256.t01#3       1420   125259992:   mNeoVis.3.1.1.chr7.g02129.t01#6    mNeoVis.2.1.1.chr7_hap1.g02313.t01#5    mNeoVis.evergladensis.2.1.chr7.g02260.t01#2    mNeoVis.vulgivaga.2.1.chr7.g02277.t01#1    mNeoVis.vulgivaga.1.1.chr7.g02265.t01#4   mNeoVis.evergladensis.1.1.chr7.g02257.t01#3       1421   125259993:   mNeoVis.evergladensis.1.1.chr1.g01603.t01#3    mNeoVis.vulgivaga.1.1.chr1.g01607.t01#4    mNeoVis.evergladensis.2.1.chr1.g01606.t01#2    mNeoVis.3.1.1.chr1.g01590.t01#6    mNeoVis.vulgivaga.2.1.chr1.g01603.t01#1   mNeoVis.2.1.1.chr1_hap1.g01640.t01#5       1422   125259994:   mNeoVis.vulgivaga.1.1.chr7.g02266.t01#4    mNeoVis.evergladensis.1.1.chr7.g02258.t01#3    mNeoVis.vulgivaga.2.1.chr7.g02278.t01#1    mNeoVis.2.1.1.chr7_hap1.g02314.t01#5    mNeoVis.evergladensis.2.1.chr7.g02261.t01#2   mNeoVis.3.1.1.chr7.g02130.t01#6       1423   125259995:   mNeoVis.2.1.1.chr7_hap1.g02315.t01#5    mNeoVis.3.1.1.chr7.g02131.t01#6    mNeoVis.vulgivaga.1.1.chr7.g02267.t01#4    mNeoVis.evergladensis.2.1.chr7.g02262.t01#2    mNeoVis.vulgivaga.2.1.chr7.g02279.t01#1   mNeoVis.evergladensis.1.1.chr7.g02259.t01#3       1424   125259996:   mNeoVis.vulgivaga.2.1.chr1.g01604.t01#1    mNeoVis.2.1.1.chr1_hap1.g01641.t01#5    mNeoVis.3.1.1.chr1.g01591.t01#6    mNeoVis.evergladensis.1.1.chr1.g01604.t01#3    mNeoVis.vulgivaga.1.1.chr1.g01608.t01#4   mNeoVis.evergladensis.2.1.chr1.g01607.t01#2       1425   125259997:   mNeoVis.vulgivaga.2.1.chr7.g02280.t01#1    mNeoVis.vulgivaga.1.1.chr7.g02268.t01#4    mNeoVis.evergladensis.1.1.chr7.g02260.t01#3    mNeoVis.3.1.1.chr7.g02132.t01#6    mNeoVis.2.1.1.chr7_hap1.g02316.t01#5   mNeoVis.evergladensis.2.1.chr7.g02263.t01#2       1426   125259998:   mNeoVis.evergladensis.1.1.chr5.g00668.t01#3    mNeoVis.vulgivaga.2.1.chr5.g00664.t01#1    mNeoVis.3.1.1.chr5.g00664.t01#6    mNeoVis.vulgivaga.1.1.chr5.g00648.t01#4    mNeoVis.evergladensis.2.1.chr5.g00673.t01#2   mNeoVis.2.1.1.chr2_hap1.g00691.t01#5       1427   125259999:   mNeoVis.vulgivaga.1.1.chr9.g00387.t01#4    mNeoVis.3.1.1.chr9.g00374.t01#6    mNeoVis.vulgivaga.2.1.chr9.g00390.t01#1    mNeoVis.evergladensis.2.1.chr8.g00388.t01#2   mNeoVis.2.1.1.chr10_hap1.g00406.t01#5       1428   125260000:   mNeoVis.3.1.1.chr7.g02133.t01#6    mNeoVis.2.1.1.chr7_hap1.g02317.t01#5    mNeoVis.evergladensis.2.1.chr7.g02264.t01#2    mNeoVis.vulgivaga.2.1.chr7.g02281.t01#1    mNeoVis.vulgivaga.1.1.chr7.g02269.t01#4   mNeoVis.evergladensis.1.1.chr7.g02261.t01#3       1429   125260001:   mNeoVis.evergladensis.2.1.chr1.g01519.t01#2    mNeoVis.3.1.1.chr1.g01505.t01#6    mNeoVis.evergladensis.1.1.chr1.g01518.t01#3    mNeoVis.2.1.1.chr1_hap1.g01552.t01#5    mNeoVis.vulgivaga.2.1.chr1.g01514.t01#1   mNeoVis.vulgivaga.1.1.chr1.g01521.t01#4       1430   125260002:   mNeoVis.evergladensis.1.1.chr7.g02262.t01#3    mNeoVis.3.1.1.chr7.g02134.t01#6    mNeoVis.evergladensis.2.1.chr7.g02265.t01#2    mNeoVis.2.1.1.chr7_hap1.g02318.t01#5    mNeoVis.vulgivaga.1.1.chr7.g02270.t01#4   mNeoVis.vulgivaga.2.1.chr7.g02282.t01#1       1431   125260003:   mNeoVis.evergladensis.2.1.chr1.g01520.t01#2    mNeoVis.3.1.1.chr1.g01506.t01#6    mNeoVis.vulgivaga.1.1.chr1.g01522.t01#4    mNeoVis.vulgivaga.2.1.chr1.g01515.t01#1    mNeoVis.evergladensis.1.1.chr1.g01519.t01#3   mNeoVis.2.1.1.chr1_hap1.g01553.t01#5       1432   125260004:   mNeoVis.vulgivaga.1.1.chr7.g02271.t01#4    mNeoVis.vulgivaga.2.1.chr7.g02283.t01#1    mNeoVis.evergladensis.1.1.chr7.g02263.t01#3    mNeoVis.3.1.1.chr7.g02135.t01#6    mNeoVis.2.1.1.chr7_hap1.g02319.t01#5   mNeoVis.evergladensis.2.1.chr7.g02266.t01#2       1433   125260005:   mNeoVis.vulgivaga.2.1.chr7.g02284.t01#1    mNeoVis.evergladensis.2.1.chr7.g02267.t01#2    mNeoVis.evergladensis.1.1.chr7.g02264.t01#3    mNeoVis.2.1.1.chr7_hap1.g02320.t01#5    mNeoVis.3.1.1.chr7.g02136.t01#6   mNeoVis.vulgivaga.1.1.chr7.g02272.t01#4       1434   125260006:   mNeoVis.evergladensis.1.1.chr1.g01520.t01#3    mNeoVis.3.1.1.chr1.g01507.t01#6    mNeoVis.vulgivaga.1.1.chr1.g01523.t01#4    mNeoVis.vulgivaga.2.1.chr1.g01516.t01#1    mNeoVis.2.1.1.chr1_hap1.g01554.t01#5   mNeoVis.evergladensis.2.1.chr1.g01521.t01#2       1435   125260007:   mNeoVis.evergladensis.1.1.chr7.g02265.t01#3    mNeoVis.vulgivaga.2.1.chr7.g02285.t01#1    mNeoVis.2.1.1.chr7_hap1.g02321.t01#5    mNeoVis.3.1.1.chr7.g02137.t01#6    mNeoVis.vulgivaga.1.1.chr7.g02273.t01#4   mNeoVis.evergladensis.2.1.chr7.g02268.t01#2       1436   125260008:   mNeoVis.vulgivaga.1.1.chr1.g01524.t01#4    mNeoVis.3.1.1.chr1.g01508.t01#6    mNeoVis.2.1.1.chr1_hap1.g01555.t01#5    mNeoVis.evergladensis.2.1.chr1.g01522.t01#2    mNeoVis.vulgivaga.2.1.chr1.g01517.t01#1   mNeoVis.evergladensis.1.1.chr1.g01521.t01#3       1437   125260009:   mNeoVis.2.1.1.chr1_hap1.g01556.t01#5    mNeoVis.evergladensis.1.1.chr1.g01522.t01#3    mNeoVis.evergladensis.2.1.chr1.g01523.t01#2    mNeoVis.vulgivaga.1.1.chr1.g01525.t01#4   mNeoVis.vulgivaga.2.1.chr1.g01518.t01#1       1438   125260010:   mNeoVis.vulgivaga.1.1.chr1.g01526.t01#4    mNeoVis.vulgivaga.2.1.chr1.g01519.t01#1    mNeoVis.2.1.1.chr1_hap1.g01557.t01#5    mNeoVis.evergladensis.1.1.chr1.g01523.t01#3    mNeoVis.3.1.1.chr1.g01509.t01#6   mNeoVis.evergladensis.2.1.chr1.g01524.t01#2       1439   125260011:   mNeoVis.2.1.1.chr1_hap1.g01558.t01#5    mNeoVis.evergladensis.1.1.chr1.g01524.t01#3    mNeoVis.3.1.1.chr1.g01510.t01#6    mNeoVis.vulgivaga.2.1.chr1.g01520.t01#1    mNeoVis.evergladensis.2.1.chr1.g01525.t01#2   mNeoVis.vulgivaga.1.1.chr1.g01527.t01#4       1440   125260012:   mNeoVis.vulgivaga.1.1.chr7.g02274.t01#4    mNeoVis.2.1.1.chr7_hap1.g02322.t01#5    mNeoVis.evergladensis.1.1.chr7.g02266.t01#3    mNeoVis.vulgivaga.2.1.chr7.g02286.t01#1    mNeoVis.3.1.1.chr7.g02138.t01#6   mNeoVis.evergladensis.2.1.chr7.g02269.t01#2       1441   125260013:   mNeoVis.2.1.1.chr7_hap1.g02323.t01#5    mNeoVis.3.1.1.chr7.g02139.t01#6    mNeoVis.vulgivaga.1.1.chr7.g02275.t01#4    mNeoVis.evergladensis.1.1.chr7.g02267.t01#3    mNeoVis.evergladensis.2.1.chr7.g02270.t01#2   mNeoVis.vulgivaga.2.1.chr7.g02287.t01#1       1442   125260014:   mNeoVis.evergladensis.1.1.chr7.g02268.t01#3    mNeoVis.evergladensis.2.1.chr7.g02271.t01#2    mNeoVis.vulgivaga.2.1.chr7.g02288.t01#1    mNeoVis.3.1.1.chr7.g02140.t01#6    mNeoVis.vulgivaga.1.1.chr7.g02276.t01#4   mNeoVis.2.1.1.chr7_hap1.g02324.t01#5       1443   125260015:   mNeoVis.evergladensis.2.1.chr7.g02272.t01#2    mNeoVis.vulgivaga.1.1.chr7.g02277.t01#4    mNeoVis.2.1.1.chr7_hap1.g02325.t01#5    mNeoVis.vulgivaga.2.1.chr7.g02289.t01#1    mNeoVis.evergladensis.1.1.chr7.g02269.t01#3   mNeoVis.3.1.1.chr7.g02141.t01#6       1444   125260016:   mNeoVis.vulgivaga.2.1.chr7.g02290.t01#1    mNeoVis.evergladensis.2.1.chr7.g02273.t01#2    mNeoVis.3.1.1.chr7.g02142.t01#6    mNeoVis.vulgivaga.1.1.chr7.g02278.t01#4    mNeoVis.2.1.1.chr7_hap1.g02326.t01#5   mNeoVis.evergladensis.1.1.chr7.g02270.t01#3       1445   125260017:   mNeoVis.2.1.1.chr7_hap1.g02327.t01#5    mNeoVis.3.1.1.chr7.g02143.t01#6    mNeoVis.vulgivaga.1.1.chr7.g02279.t01#4    mNeoVis.evergladensis.1.1.chr7.g02271.t01#3    mNeoVis.evergladensis.2.1.chr7.g02274.t01#2   mNeoVis.vulgivaga.2.1.chr7.g02291.t01#1       1446   125260018:   mNeoVis.evergladensis.1.1.chr7.g02272.t01#3    mNeoVis.evergladensis.2.1.chr7.g02275.t01#2    mNeoVis.vulgivaga.2.1.chr7.g02292.t01#1    mNeoVis.vulgivaga.1.1.chr7.g02280.t01#4    mNeoVis.2.1.1.chr7_hap1.g02328.t01#5   mNeoVis.3.1.1.chr7.g02144.t01#6       1447   125260019:   mNeoVis.2.1.1.chr4_hap1.g00757.t01#5    mNeoVis.vulgivaga.1.1.chr6.g00762.t01#4    mNeoVis.evergladensis.1.1.chr6.g00769.t01#3    mNeoVis.vulgivaga.2.1.chr6.g00761.t01#1    mNeoVis.evergladensis.2.1.chr6.g00764.t01#2   mNeoVis.3.1.1.chr6.g00763.t01#6       1448   125260020:   mNeoVis.evergladensis.2.1.chr7.g02276.t01#2    mNeoVis.3.1.1.chr7.g02145.t01#6    mNeoVis.vulgivaga.1.1.chr7.g02281.t01#4    mNeoVis.2.1.1.chr7_hap1.g02329.t01#5    mNeoVis.vulgivaga.2.1.chr7.g02293.t01#1   mNeoVis.evergladensis.1.1.chr7.g02273.t01#3       1449   125260021:   mNeoVis.vulgivaga.2.1.chr7.g02294.t01#1    mNeoVis.2.1.1.chr7_hap1.g02330.t01#5    mNeoVis.evergladensis.1.1.chr7.g02274.t01#3    mNeoVis.evergladensis.2.1.chr7.g02277.t01#2    mNeoVis.vulgivaga.1.1.chr7.g02282.t01#4   mNeoVis.3.1.1.chr7.g02146.t01#6       1450   125260022:   mNeoVis.evergladensis.1.1.chr7.g02275.t01#3    mNeoVis.evergladensis.2.1.chr7.g02278.t01#2    mNeoVis.vulgivaga.1.1.chr7.g02283.t01#4    mNeoVis.vulgivaga.2.1.chr7.g02295.t01#1    mNeoVis.2.1.1.chr7_hap1.g02331.t01#5   mNeoVis.3.1.1.chr7.g02147.t01#6       1451   125260023:   mNeoVis.evergladensis.1.1.chr1.g01525.t01#3    mNeoVis.vulgivaga.2.1.chr1.g01521.t01#1    mNeoVis.3.1.1.chr1.g01511.t01#6   mNeoVis.2.1.1.chr1_hap1.g01559.t01#5         1452   125260024:   mNeoVis.vulgivaga.1.1.chr1.g01529.t01#4    mNeoVis.3.1.1.chr1.g01513.t01#6    mNeoVis.vulgivaga.2.1.chr1.g01522.t01#1    mNeoVis.evergladensis.1.1.chr1.g01526.t01#3    mNeoVis.2.1.1.chr1_hap1.g01560.t01#5   mNeoVis.evergladensis.2.1.chr1.g01527.t01#2       1453   125260025:   mNeoVis.2.1.1.chr7_hap1.g02332.t01#5    mNeoVis.3.1.1.chr7.g02148.t01#6    mNeoVis.evergladensis.2.1.chr7.g02279.t01#2    mNeoVis.evergladensis.1.1.chr7.g02276.t01#3    mNeoVis.vulgivaga.2.1.chr7.g02296.t01#1   mNeoVis.vulgivaga.1.1.chr7.g02284.t01#4       1454   125260026:  mNeoVis.vulgivaga.2.1.chr1.g01523.t01#1               1455   125260027:   mNeoVis.vulgivaga.2.1.chr1.g01524.t01#1    mNeoVis.3.1.1.chr1.g01514.t01#6    mNeoVis.vulgivaga.1.1.chr1.g01530.t01#4    mNeoVis.2.1.1.chr1_hap1.g01561.t01#5    mNeoVis.evergladensis.2.1.chr1.g01528.t01#2   mNeoVis.evergladensis.1.1.chr1.g01527.t01#3       1456   125260028:   mNeoVis.evergladensis.1.1.chr7.g02277.t01#3    mNeoVis.vulgivaga.2.1.chr7.g02297.t01#1    mNeoVis.3.1.1.chr7.g02149.t01#6    mNeoVis.evergladensis.2.1.chr7.g02280.t01#2    mNeoVis.vulgivaga.1.1.chr7.g02285.t01#4   mNeoVis.2.1.1.chr7_hap1.g02333.t01#5       1457   125260029:   mNeoVis.evergladensis.1.1.chr3.g00191.t01#3    mNeoVis.vulgivaga.1.1.chr7.g01181.t01#4    mNeoVis.3.1.1.chr10.g00884.t01#6    mNeoVis.3.1.1.chr2.g00806.t01#6    mNeoVis.3.1.1.chr3.g00188.t01#6    mNeoVis.3.1.1.chr1.g01515.t01#6    mNeoVis.vulgivaga.1.1.chr10.g00898.t01#4    mNeoVis.evergladensis.1.1.chr10.g00916.t01#3    mNeoVis.3.1.1.chr1.g00618.t01#6      1458   125260030:   mNeoVis.3.1.1.chr2.g01139.t01#6    mNeoVis.2.1.1.chr7_hap1.g01191.t01#5    mNeoVis.vulgivaga.2.1.chr1.g01525.t01#1    mNeoVis.vulgivaga.2.1.chr1.g00615.t01#1    mNeoVis.evergladensis.1.1.chr7.g01180.t01#3    mNeoVis.3.1.1.chr7.g01115.t01#6    mNeoVis.2.1.1.chr1_hap1.g01562.t01#5    mNeoVis.vulgivaga.2.1.chr3.g00188.t01#1    mNeoVis.evergladensis.1.1.chr1.g01528.t01#3      1459   125260031:   mNeoVis.evergladensis.2.1.chr2.g00894.t01#2    mNeoVis.evergladensis.2.1.chr3.g00196.t01#2    mNeoVis.evergladensis.2.1.chr1.g01529.t01#2    mNeoVis.evergladensis.2.1.chr1.g00606.t01#2   mNeoVis.evergladensis.2.1.chr2.g01120.t01#2       1460   125260032:   mNeoVis.evergladensis.1.1.chr2.g00891.t01#3    mNeoVis.evergladensis.1.1.chr1.g00611.t01#3   mNeoVis.evergladensis.1.1.chr2.g01123.t01#3         1461   125260033:   mNeoVis.evergladensis.2.1.chr1.g01530.t01#2    mNeoVis.vulgivaga.1.1.chr1.g01532.t01#4    mNeoVis.vulgivaga.2.1.chr1.g01526.t01#1    mNeoVis.evergladensis.1.1.chr1.g01529.t01#3    mNeoVis.2.1.1.chr1_hap1.g01563.t01#5   mNeoVis.3.1.1.chr1.g01516.t01#6       1462   125260034:   mNeoVis.evergladensis.1.1.chr1.g01530.t01#3    mNeoVis.3.1.1.chr1.g01517.t01#6    mNeoVis.vulgivaga.1.1.chr1.g01533.t01#4    mNeoVis.vulgivaga.2.1.chr1.g01527.t01#1    mNeoVis.2.1.1.chr1_hap1.g01564.t01#5   mNeoVis.evergladensis.2.1.chr1.g01531.t01#2       1463   125260035:   mNeoVis.vulgivaga.1.1.chr1.g01534.t01#4    mNeoVis.evergladensis.2.1.chr1.g01532.t01#2    mNeoVis.2.1.1.chr1_hap1.g01565.t01#5    mNeoVis.3.1.1.chr1.g01518.t01#6    mNeoVis.vulgivaga.2.1.chr1.g01528.t01#1   mNeoVis.evergladensis.1.1.chr1.g01531.t01#3       1464   125260036:   mNeoVis.vulgivaga.2.1.chr7.g02298.t01#1    mNeoVis.2.1.1.chr7_hap1.g02334.t01#5    mNeoVis.evergladensis.1.1.chr7.g02278.t01#3    mNeoVis.3.1.1.chr7.g02150.t01#6    mNeoVis.vulgivaga.1.1.chr7.g02286.t01#4   mNeoVis.evergladensis.2.1.chr7.g02281.t01#2       1465   125260037:   mNeoVis.vulgivaga.2.1.chr1.g01529.t01#1    mNeoVis.evergladensis.2.1.chr1.g01533.t01#2    mNeoVis.2.1.1.chr1_hap1.g01566.t01#5    mNeoVis.vulgivaga.1.1.chr1.g01535.t01#4    mNeoVis.3.1.1.chr1.g01519.t01#6   mNeoVis.evergladensis.1.1.chr1.g01532.t01#3       1466   125260038:   mNeoVis.evergladensis.2.1.chr12.g00394.t01#2    mNeoVis.evergladensis.1.1.chr12.g00401.t01#3    mNeoVis.vulgivaga.2.1.chr12.g00402.t01#1    mNeoVis.vulgivaga.1.1.chr12.g00404.t01#4    mNeoVis.3.1.1.chr12.g00398.t01#6   mNeoVis.2.1.1.chr12_hap1.g00437.t01#5       1467   125260039:   mNeoVis.2.1.1.chr1_hap1.g01568.t01#5    mNeoVis.evergladensis.1.1.chr1.g01534.t01#3    mNeoVis.vulgivaga.1.1.chr1.g01537.t01#4    mNeoVis.3.1.1.chr1.g01521.t01#6    mNeoVis.vulgivaga.2.1.chr1.g01531.t01#1   mNeoVis.evergladensis.2.1.chr1.g01535.t01#2       1468   125260040:   mNeoVis.vulgivaga.2.1.chr1.g01530.t01#1    mNeoVis.3.1.1.chr1.g01520.t01#6   mNeoVis.evergladensis.1.1.chr1.g01533.t01#3         1469   125260041:   mNeoVis.evergladensis.2.1.chr1.g01534.t01#2    mNeoVis.2.1.1.chr1_hap1.g01567.t01#5   mNeoVis.vulgivaga.1.1.chr1.g01536.t01#4           1470   125260042:   mNeoVis.vulgivaga.2.1.chr7.g02299.t01#1    mNeoVis.3.1.1.chr7.g02151.t01#6   mNeoVis.evergladensis.1.1.chr7.g02279.t01#3         1471   125260043:   mNeoVis.2.1.1.chr7_hap1.g02336.t01#5   mNeoVis.vulgivaga.2.1.chr7.g02300.t01#1             1472   125260044:   mNeoVis.2.1.1.chr7_hap1.g02337.t01#5   mNeoVis.evergladensis.1.1.chr7.g02280.t01#3           1473   125260045:   mNeoVis.evergladensis.1.1.chr1.g01535.t01#3    mNeoVis.vulgivaga.1.1.chr1.g01538.t01#4    mNeoVis.2.1.1.chr1_hap1.g01569.t01#5    mNeoVis.evergladensis.2.1.chr1.g01536.t01#2    mNeoVis.vulgivaga.2.1.chr1.g01532.t01#1   mNeoVis.3.1.1.chr1.g01522.t01#6       1474   125260046:   mNeoVis.evergladensis.2.1.chr1.g01537.t01#2    mNeoVis.3.1.1.chr1.g01523.t01#6    mNeoVis.vulgivaga.2.1.chr1.g01533.t01#1    mNeoVis.vulgivaga.1.1.chr1.g01539.t01#4   mNeoVis.2.1.1.chr1_hap1.g01570.t01#5       1475   125260047:   mNeoVis.vulgivaga.1.1.chr7.g02289.t01#4    mNeoVis.vulgivaga.2.1.chr7.g02301.t01#1    mNeoVis.evergladensis.1.1.chr7.g02281.t01#3    mNeoVis.evergladensis.2.1.chr7.g02284.t01#2    mNeoVis.2.1.1.chr7_hap1.g02338.t01#5   mNeoVis.3.1.1.chr7.g02152.t01#6       1476   125260048:  mNeoVis.vulgivaga.2.1.chr7.g02302.t01#1               1477   125260049:   mNeoVis.evergladensis.1.1.chr7.g02282.t01#3    mNeoVis.2.1.1.chr7_hap1.g02339.t01#5    mNeoVis.3.1.1.chr7.g02154.t01#6    mNeoVis.evergladensis.2.1.chr7.g02285.t01#2    mNeoVis.vulgivaga.1.1.chr7.g02290.t01#4   mNeoVis.vulgivaga.2.1.chr7.g02303.t01#1       1478   125260050:   mNeoVis.vulgivaga.2.1.chr7.g02304.t01#1    mNeoVis.evergladensis.2.1.chr7.g02286.t01#2    mNeoVis.evergladensis.1.1.chr7.g02283.t01#3    mNeoVis.vulgivaga.1.1.chr7.g02291.t01#4   mNeoVis.2.1.1.chr7_hap1.g02340.t01#5       1479   125260051:   mNeoVis.evergladensis.2.1.chr1.g01538.t01#2    mNeoVis.2.1.1.chr1_hap1.g01571.t01#5    mNeoVis.2.1.1.chr1_hap1.g01574.t01#5    mNeoVis.vulgivaga.2.1.chr1.g01534.t01#1   mNeoVis.evergladensis.1.1.chr1.g01536.t01#3       1480   125260052:   mNeoVis.evergladensis.1.1.chr1.g01537.t01#3    mNeoVis.2.1.1.chr1_hap1.g01573.t01#5    mNeoVis.evergladensis.2.1.chr1.g01541.t01#2    mNeoVis.vulgivaga.2.1.chr1.g01536.t01#1    mNeoVis.vulgivaga.1.1.chr1.g01541.t01#4    mNeoVis.2.1.1.chr1_hap1.g01572.t01#5   mNeoVis.2.1.1.chr1_hap1.g01575.t01#5     1481   125260053:   mNeoVis.vulgivaga.2.1.chr1.g01537.t01#1    mNeoVis.vulgivaga.2.1.chr1.g01535.t01#1   mNeoVis.vulgivaga.1.1.chr1.g01543.t01#4           1482   125260054:   mNeoVis.vulgivaga.1.1.chr1.g01540.t01#4    mNeoVis.3.1.1.chr1.g01526.t01#6   mNeoVis.evergladensis.2.1.chr1.g01539.t01#2         1483   125260055:   mNeoVis.evergladensis.1.1.chr7.g02284.t01#3    mNeoVis.vulgivaga.2.1.chr7.g02305.t01#1    mNeoVis.evergladensis.2.1.chr7.g02287.t01#2    mNeoVis.3.1.1.chr7.g02155.t01#6    mNeoVis.2.1.1.chr7_hap1.g02341.t01#5   mNeoVis.vulgivaga.1.1.chr7.g02292.t01#4       1484   125260056:   mNeoVis.2.1.1.chr1_hap1.g00330.t01#5    mNeoVis.vulgivaga.2.1.chr1.g00334.t01#1    mNeoVis.vulgivaga.1.1.chr1.g00328.t01#4    mNeoVis.3.1.1.chr1.g00334.t01#6    mNeoVis.evergladensis.2.1.chr1.g00328.t01#2   mNeoVis.evergladensis.1.1.chr1.g00337.t01#3       1485   125260057:   mNeoVis.3.1.1.chr7.g02156.t01#6    mNeoVis.2.1.1.chr7_hap1.g02342.t01#5    mNeoVis.evergladensis.1.1.chr7.g02285.t01#3    mNeoVis.vulgivaga.1.1.chr7.g02293.t01#4    mNeoVis.evergladensis.2.1.chr7.g02288.t01#2   mNeoVis.vulgivaga.2.1.chr7.g02306.t01#1       1486   125260058:   mNeoVis.3.1.1.chr7.g02157.t01#6    mNeoVis.2.1.1.chr7_hap1.g02343.t01#5    mNeoVis.evergladensis.2.1.chr7.g02289.t01#2    mNeoVis.evergladensis.1.1.chr7.g02286.t01#3    mNeoVis.vulgivaga.1.1.chr7.g02294.t01#4   mNeoVis.vulgivaga.2.1.chr7.g02307.t01#1       1487   125260059:   mNeoVis.vulgivaga.1.1.chr7.g00071.t01#4    mNeoVis.2.1.1.chr7_hap1.g00069.t01#5    mNeoVis.3.1.1.chr7.g00057.t01#6    mNeoVis.evergladensis.2.1.chr7.g00070.t01#2    mNeoVis.vulgivaga.2.1.chr7.g00070.t01#1   mNeoVis.evergladensis.1.1.chr7.g00072.t01#3       1488   125260060:   mNeoVis.evergladensis.2.1.chr1.g01540.t01#2    mNeoVis.vulgivaga.2.1.chr1.g01538.t01#1    mNeoVis.evergladensis.2.1.chr1.g01542.t01#2    mNeoVis.vulgivaga.1.1.chr1.g01544.t01#4    mNeoVis.3.1.1.chr1.g01525.t01#6    mNeoVis.2.1.1.chr1_hap1.g01576.t01#5   mNeoVis.evergladensis.1.1.chr1.g01539.t01#3     1489   125260061:   mNeoVis.vulgivaga.2.1.chr1.g01539.t01#1    mNeoVis.evergladensis.1.1.chr1.g01540.t01#3    mNeoVis.3.1.1.chr1.g01527.t01#6    mNeoVis.evergladensis.2.1.chr1.g01543.t01#2    mNeoVis.vulgivaga.2.1.chr1.g01540.t01#1    mNeoVis.vulgivaga.1.1.chr1.g01545.t01#4   mNeoVis.2.1.1.chr1_hap1.g01577.t01#5     1490   125260062:   mNeoVis.2.1.1.chr7_hap1.g02344.t01#5    mNeoVis.evergladensis.1.1.chr7.g02287.t01#3    mNeoVis.vulgivaga.2.1.chr7.g02308.t01#1    mNeoVis.vulgivaga.1.1.chr7.g02295.t01#4   mNeoVis.evergladensis.2.1.chr7.g02290.t01#2       1491   125260063:   mNeoVis.3.1.1.chr1.g01528.t01#6    mNeoVis.vulgivaga.1.1.chr1.g01546.t01#4    mNeoVis.evergladensis.1.1.chr1.g01542.t01#3    mNeoVis.evergladensis.2.1.chr1.g01544.t01#2    mNeoVis.2.1.1.chr1_hap1.g01578.t01#5   mNeoVis.vulgivaga.2.1.chr1.g01541.t01#1       1492   125260064:   mNeoVis.evergladensis.2.1.chr7.g02291.t01#2    mNeoVis.vulgivaga.2.1.chr7.g02309.t01#1    mNeoVis.evergladensis.1.1.chr7.g02288.t01#3    mNeoVis.vulgivaga.1.1.chr7.g02296.t01#4    mNeoVis.3.1.1.chr7.g02158.t01#6   mNeoVis.2.1.1.chr7_hap1.g02345.t01#5       1493   125260065:   mNeoVis.2.1.1.chr1_hap1.g01579.t01#5    mNeoVis.evergladensis.2.1.chr1.g01545.t01#2    mNeoVis.vulgivaga.2.1.chr1.g01542.t01#1    mNeoVis.vulgivaga.1.1.chr1.g01547.t01#4    mNeoVis.3.1.1.chr1.g01529.t01#6   mNeoVis.evergladensis.1.1.chr1.g01543.t01#3       1494   125260066:   mNeoVis.2.1.1.chr7_hap1.g02346.t01#5    mNeoVis.evergladensis.2.1.chr7.g02292.t01#2    mNeoVis.vulgivaga.2.1.chr7.g02310.t01#1    mNeoVis.vulgivaga.1.1.chr7.g02297.t01#4   mNeoVis.3.1.1.chr7.g02159.t01#6       1495   125260067:   mNeoVis.evergladensis.1.1.chr7.g02290.t01#3    mNeoVis.vulgivaga.1.1.chr7.g02298.t01#4    mNeoVis.3.1.1.chr7.g02160.t01#6    mNeoVis.2.1.1.chr7_hap1.g02347.t01#5    mNeoVis.evergladensis.2.1.chr7.g02293.t01#2   mNeoVis.vulgivaga.2.1.chr7.g02311.t01#1       1496   125260068:   mNeoVis.2.1.1.chr1_hap1.g01580.t01#5    mNeoVis.vulgivaga.1.1.chr1.g01548.t01#4    mNeoVis.evergladensis.1.1.chr1.g01544.t01#3    mNeoVis.evergladensis.2.1.chr1.g01546.t01#2    mNeoVis.3.1.1.chr1.g01530.t01#6   mNeoVis.vulgivaga.2.1.chr1.g01543.t01#1       1497   125260069:   mNeoVis.vulgivaga.1.1.chr1.g01549.t01#4    mNeoVis.3.1.1.chr1.g01531.t01#6    mNeoVis.vulgivaga.2.1.chr1.g01544.t01#1    mNeoVis.2.1.1.chr1_hap1.g01581.t01#5    mNeoVis.evergladensis.2.1.chr1.g01547.t01#2   mNeoVis.evergladensis.1.1.chr1.g01545.t01#3       1498   125260070:   mNeoVis.evergladensis.1.1.chr7.g02291.t01#3    mNeoVis.vulgivaga.1.1.chr7.g02299.t01#4    mNeoVis.2.1.1.chr7_hap1.g02348.t01#5    mNeoVis.vulgivaga.2.1.chr7.g02312.t01#1   mNeoVis.evergladensis.2.1.chr7.g02294.t01#2       1499   125260071:   mNeoVis.vulgivaga.1.1.chr15.g00143.t01#4    mNeoVis.3.1.1.chr15.g00137.t01#6    mNeoVis.vulgivaga.2.1.chr15.g00147.t01#1    mNeoVis.evergladensis.1.1.chr15.g00472.t01#3    mNeoVis.2.1.1.chr15_hap1.g00158.t01#5   mNeoVis.evergladensis.2.1.chr15.g00143.t01#2       1500   125260072:   mNeoVis.vulgivaga.2.1.chr1.g01545.t01#1    mNeoVis.vulgivaga.1.1.chr1.g01550.t01#4    mNeoVis.2.1.1.chr1_hap1.g01582.t01#5    mNeoVis.evergladensis.2.1.chr1.g01548.t01#2    mNeoVis.evergladensis.1.1.chr1.g01546.t01#3   mNeoVis.3.1.1.chr1.g01532.t01#6       1501   125260073:   mNeoVis.2.1.1.chr7_hap1.g02349.t01#5    mNeoVis.evergladensis.2.1.chr7.g02295.t01#2    mNeoVis.evergladensis.1.1.chr7.g02292.t01#3    mNeoVis.vulgivaga.1.1.chr7.g02300.t01#4    mNeoVis.3.1.1.chr7.g02161.t01#6   mNeoVis.vulgivaga.2.1.chr7.g02313.t01#1       1502   125260074:   mNeoVis.vulgivaga.1.1.chr7.g02301.t01#4    mNeoVis.evergladensis.1.1.chr7.g02293.t01#3    mNeoVis.2.1.1.chr7_hap1.g02350.t01#5    mNeoVis.3.1.1.chr7.g02163.t01#6    mNeoVis.evergladensis.2.1.chr7.g02296.t01#2   mNeoVis.vulgivaga.2.1.chr7.g02314.t01#1       1503   125260075:   mNeoVis.vulgivaga.2.1.chr1.g01546.t01#1    mNeoVis.evergladensis.1.1.chr1.g01547.t01#3    mNeoVis.2.1.1.chr1_hap1.g01583.t01#5    mNeoVis.vulgivaga.1.1.chr1.g01551.t01#4    mNeoVis.evergladensis.2.1.chr1.g01549.t01#2   mNeoVis.3.1.1.chr1.g01533.t01#6       1504   125260076:   mNeoVis.vulgivaga.2.1.chr1.g01547.t01#1    mNeoVis.evergladensis.1.1.chr1.g01548.t01#3    mNeoVis.3.1.1.chr1.g01534.t01#6    mNeoVis.evergladensis.2.1.chr1.g01550.t01#2    mNeoVis.vulgivaga.1.1.chr1.g01552.t01#4   mNeoVis.2.1.1.chr1_hap1.g01584.t01#5       1505   125260077:   mNeoVis.vulgivaga.1.1.chr1.g01553.t01#4    mNeoVis.2.1.1.chr1_hap1.g01585.t01#5    mNeoVis.vulgivaga.2.1.chr1.g01548.t01#1    mNeoVis.evergladensis.1.1.chr1.g01549.t01#3    mNeoVis.3.1.1.chr1.g01535.t01#6   mNeoVis.evergladensis.2.1.chr1.g01551.t01#2       1506   125260078:   mNeoVis.2.1.1.chr7_hap1.g02351.t01#5    mNeoVis.vulgivaga.1.1.chr7.g02302.t01#4    mNeoVis.vulgivaga.2.1.chr7.g02315.t01#1    mNeoVis.evergladensis.1.1.chr7.g02294.t01#3    mNeoVis.evergladensis.2.1.chr7.g02297.t01#2   mNeoVis.3.1.1.chr7.g02164.t01#6       1507   125260079:   mNeoVis.3.1.1.chr7.g02165.t01#6    mNeoVis.2.1.1.chr7_hap1.g02352.t01#5    mNeoVis.evergladensis.1.1.chr7.g02295.t01#3    mNeoVis.vulgivaga.1.1.chr7.g02303.t01#4    mNeoVis.evergladensis.2.1.chr7.g02298.t01#2   mNeoVis.vulgivaga.2.1.chr7.g02316.t01#1       1508   125260080:   mNeoVis.3.1.1.chr7.g02166.t01#6    mNeoVis.vulgivaga.1.1.chr7.g02304.t01#4    mNeoVis.2.1.1.chr7_hap1.g02353.t01#5    mNeoVis.evergladensis.1.1.chr7.g02296.t01#3    mNeoVis.vulgivaga.2.1.chr7.g02317.t01#1   mNeoVis.evergladensis.2.1.chr7.g02299.t01#2       1509   125260081:   mNeoVis.3.1.1.chr7.g02167.t01#6    mNeoVis.evergladensis.2.1.chr7.g02300.t01#2    mNeoVis.evergladensis.1.1.chr7.g02297.t01#3    mNeoVis.2.1.1.chr7_hap1.g02354.t01#5   mNeoVis.vulgivaga.2.1.chr7.g02318.t01#1       1510   125260082:   mNeoVis.3.1.1.chr7.g02168.t01#6    mNeoVis.2.1.1.chr7_hap1.g02355.t01#5    mNeoVis.evergladensis.1.1.chr7.g02298.t01#3    mNeoVis.vulgivaga.2.1.chr7.g02319.t01#1   mNeoVis.evergladensis.2.1.chr7.g02301.t01#2       1511   125260083:   mNeoVis.evergladensis.2.1.chr1.g01552.t01#2    mNeoVis.3.1.1.chr1.g01536.t01#6    mNeoVis.vulgivaga.2.1.chr1.g01549.t01#1    mNeoVis.evergladensis.1.1.chr1.g01550.t01#3    mNeoVis.vulgivaga.1.1.chr1.g01554.t01#4   mNeoVis.2.1.1.chr1_hap1.g01586.t01#5       1512   125260084:   mNeoVis.vulgivaga.1.1.chr7.g02307.t01#4    mNeoVis.vulgivaga.2.1.chr7.g02320.t01#1    mNeoVis.2.1.1.chr7_hap1.g02357.t01#5   mNeoVis.evergladensis.2.1.chr7.g02303.t01#2       1513   125260085:   mNeoVis.evergladensis.2.1.chr1.g01553.t01#2    mNeoVis.3.1.1.chr1.g01537.t01#6    mNeoVis.vulgivaga.2.1.chr1.g01550.t01#1    mNeoVis.evergladensis.1.1.chr1.g01551.t01#3    mNeoVis.2.1.1.chr1_hap1.g01587.t01#5   mNeoVis.vulgivaga.1.1.chr1.g01555.t01#4       1514   125260086:   mNeoVis.vulgivaga.1.1.chr7.g02306.t01#4    mNeoVis.2.1.1.chr7_hap1.g02356.t01#5    mNeoVis.evergladensis.1.1.chr7.g02299.t01#3    mNeoVis.evergladensis.2.1.chr7.g02302.t01#2    mNeoVis.3.1.1.chr7.g02169.t01#6   mNeoVis.vulgivaga.2.1.chr7.g02321.t01#1       1515   125260087:   mNeoVis.3.1.1.chr7.g02170.t01#6    mNeoVis.vulgivaga.2.1.chr7.g02322.t01#1    mNeoVis.2.1.1.chr7_hap1.g02358.t01#5    mNeoVis.evergladensis.1.1.chr7.g02300.t01#3    mNeoVis.evergladensis.2.1.chr7.g02304.t01#2   mNeoVis.vulgivaga.1.1.chr7.g02308.t01#4       1516   125260088:   mNeoVis.evergladensis.1.1.chr7.g02301.t01#3    mNeoVis.vulgivaga.1.1.chr7.g02309.t01#4    mNeoVis.vulgivaga.2.1.chr7.g02323.t01#1    mNeoVis.evergladensis.2.1.chr7.g02305.t01#2   mNeoVis.2.1.1.chr7_hap1.g02359.t01#5       1517   125260089:   mNeoVis.evergladensis.1.1.chr1.g01552.t01#3    mNeoVis.vulgivaga.2.1.chr1.g01551.t01#1    mNeoVis.vulgivaga.1.1.chr1.g01556.t01#4    mNeoVis.2.1.1.chr1_hap1.g01588.t01#5    mNeoVis.evergladensis.2.1.chr1.g01554.t01#2   mNeoVis.3.1.1.chr1.g01538.t01#6       1518   125260090:   mNeoVis.vulgivaga.2.1.chr1.g01552.t01#1    mNeoVis.evergladensis.1.1.chr1.g01553.t01#3    mNeoVis.evergladensis.2.1.chr1.g01555.t01#2    mNeoVis.vulgivaga.1.1.chr1.g01557.t01#4   mNeoVis.2.1.1.chr1_hap1.g01589.t01#5       1519   125260091:   mNeoVis.2.1.1.chr1_hap1.g01590.t01#5    mNeoVis.evergladensis.1.1.chr1.g01554.t01#3    mNeoVis.evergladensis.2.1.chr1.g01556.t01#2    mNeoVis.vulgivaga.2.1.chr1.g01553.t01#1    mNeoVis.vulgivaga.1.1.chr1.g01558.t01#4   mNeoVis.3.1.1.chr1.g01540.t01#6       1520   125260092:   mNeoVis.evergladensis.2.1.chr1.g01557.t01#2    mNeoVis.evergladensis.1.1.chr1.g01555.t01#3    mNeoVis.3.1.1.chr1.g01541.t01#6    mNeoVis.vulgivaga.1.1.chr1.g01559.t01#4    mNeoVis.vulgivaga.2.1.chr1.g01554.t01#1   mNeoVis.2.1.1.chr1_hap1.g01591.t01#5       1521   125260093:   mNeoVis.evergladensis.2.1.chr7.g02306.t01#2    mNeoVis.evergladensis.1.1.chr7.g02302.t01#3    mNeoVis.vulgivaga.2.1.chr7.g02324.t01#1    mNeoVis.vulgivaga.1.1.chr7.g02310.t01#4    mNeoVis.2.1.1.chr7_hap1.g02360.t01#5   mNeoVis.3.1.1.chr7.g02173.t01#6       1522   125260094:   mNeoVis.2.1.1.chr7_hap1.g02361.t01#5    mNeoVis.3.1.1.chr7.g02174.t01#6    mNeoVis.vulgivaga.2.1.chr7.g02325.t01#1    mNeoVis.evergladensis.1.1.chr7.g02303.t01#3    mNeoVis.vulgivaga.1.1.chr7.g02311.t01#4   mNeoVis.evergladensis.2.1.chr7.g02307.t01#2       1523   125260095:   mNeoVis.3.1.1.chr7.g02175.t01#6    mNeoVis.vulgivaga.2.1.chr7.g02326.t01#1    mNeoVis.2.1.1.chr7_hap1.g02362.t01#5    mNeoVis.evergladensis.2.1.chr7.g02308.t01#2    mNeoVis.evergladensis.1.1.chr7.g02304.t01#3   mNeoVis.vulgivaga.1.1.chr7.g02312.t01#4       1524   125260096:   mNeoVis.evergladensis.2.1.chr7.g00071.t01#2    mNeoVis.3.1.1.chr7.g00058.t01#6    mNeoVis.vulgivaga.2.1.chr7.g00071.t01#1    mNeoVis.2.1.1.chr7_hap1.g00070.t01#5   mNeoVis.evergladensis.1.1.chr7.g00073.t01#3       1525   125260097:   mNeoVis.2.1.1.chr1_hap1.g01592.t01#5    mNeoVis.evergladensis.2.1.chr1.g01558.t01#2    mNeoVis.3.1.1.chr1.g01542.t01#6    mNeoVis.vulgivaga.2.1.chr1.g01555.t01#1    mNeoVis.evergladensis.1.1.chr1.g01556.t01#3   mNeoVis.vulgivaga.1.1.chr1.g01560.t01#4       1526   125260098:   mNeoVis.2.1.1.chr4_hap1.g01120.t01#5    mNeoVis.vulgivaga.2.1.chr1.g01556.t01#1   mNeoVis.2.1.1.chr4_hap1.g01135.t01#5           1527   125260099:   mNeoVis.2.1.1.chr7_hap1.g02363.t01#5    mNeoVis.evergladensis.2.1.chr7.g02309.t01#2    mNeoVis.3.1.1.chr7.g02176.t01#6    mNeoVis.evergladensis.1.1.chr7.g02305.t01#3    mNeoVis.vulgivaga.1.1.chr7.g02313.t01#4   mNeoVis.vulgivaga.2.1.chr7.g02327.t01#1       1528   125260100:   mNeoVis.2.1.1.chr1_hap1.g01593.t01#5    mNeoVis.3.1.1.chr1.g01543.t01#6    mNeoVis.vulgivaga.2.1.chr1.g01557.t01#1    mNeoVis.evergladensis.2.1.chr1.g01560.t01#2    mNeoVis.vulgivaga.1.1.chr1.g01561.t01#4    mNeoVis.evergladensis.2.1.chr1.g01559.t01#2   mNeoVis.evergladensis.1.1.chr1.g01557.t01#3     1529   125260101:   mNeoVis.vulgivaga.1.1.chr7.g02314.t01#4    mNeoVis.2.1.1.chr7_hap1.g02364.t01#5    mNeoVis.vulgivaga.2.1.chr7.g02328.t01#1    mNeoVis.3.1.1.chr7.g02177.t01#6    mNeoVis.evergladensis.2.1.chr7.g02310.t01#2   mNeoVis.evergladensis.1.1.chr7.g02306.t01#3       1530   125260102:   mNeoVis.3.1.1.chr1.g01544.t01#6    mNeoVis.vulgivaga.2.1.chr1.g01558.t01#1    mNeoVis.vulgivaga.1.1.chr1.g01562.t01#4    mNeoVis.evergladensis.2.1.chr1.g01561.t01#2    mNeoVis.evergladensis.1.1.chr1.g01558.t01#3   mNeoVis.2.1.1.chr1_hap1.g01594.t01#5       1531   125260103:   mNeoVis.evergladensis.2.1.chr1.g01562.t01#2    mNeoVis.2.1.1.chr1_hap1.g01595.t01#5    mNeoVis.evergladensis.1.1.chr1.g01559.t01#3    mNeoVis.vulgivaga.1.1.chr1.g01563.t01#4    mNeoVis.3.1.1.chr1.g01545.t01#6   mNeoVis.vulgivaga.2.1.chr1.g01559.t01#1       1532   125260104:  mNeoVis.vulgivaga.2.1.chr1.g01560.t01#1               1533   125260105:   mNeoVis.2.1.1.chr7_hap1.g02264.t01#5    mNeoVis.evergladensis.1.1.chr7.g02208.t01#3    mNeoVis.vulgivaga.1.1.chr7.g02215.t01#4    mNeoVis.3.1.1.chr7.g02084.t01#6    mNeoVis.vulgivaga.2.1.chr7.g02229.t01#1   mNeoVis.evergladensis.2.1.chr7.g02211.t01#2       1534   125260106:   mNeoVis.3.1.1.chr7.g02085.t01#6    mNeoVis.vulgivaga.2.1.chr7.g02230.t01#1    mNeoVis.evergladensis.1.1.chr7.g02209.t01#3    mNeoVis.vulgivaga.1.1.chr7.g02216.t01#4    mNeoVis.2.1.1.chr7_hap1.g02265.t01#5   mNeoVis.evergladensis.2.1.chr7.g02212.t01#2       1535   125260107:   mNeoVis.3.1.1.chr1.g01547.t01#6    mNeoVis.vulgivaga.2.1.chr1.g01561.t01#1    mNeoVis.2.1.1.chr1_hap1.g01597.t01#5    mNeoVis.evergladensis.2.1.chr1.g01564.t01#2    mNeoVis.evergladensis.1.1.chr1.g01561.t01#3   mNeoVis.vulgivaga.1.1.chr1.g01565.t01#4       1536   125260108:   mNeoVis.evergladensis.2.1.chr7.g02213.t01#2    mNeoVis.2.1.1.chr7_hap1.g02266.t01#5    mNeoVis.vulgivaga.1.1.chr7.g02217.t01#4    mNeoVis.evergladensis.1.1.chr7.g02210.t01#3    mNeoVis.3.1.1.chr5.g00410.t01#6   mNeoVis.vulgivaga.2.1.chr7.g02231.t01#1       1537   125260109:   mNeoVis.evergladensis.1.1.chr7.g00074.t01#3    mNeoVis.3.1.1.chr7.g00059.t01#6    mNeoVis.evergladensis.2.1.chr7.g00072.t01#2    mNeoVis.vulgivaga.2.1.chr7.g00072.t01#1    mNeoVis.vulgivaga.1.1.chr7.g00073.t01#4   mNeoVis.2.1.1.chr7_hap1.g00071.t01#5       1538   125260110:   mNeoVis.evergladensis.2.1.chr1.g01565.t01#2    mNeoVis.3.1.1.chr1.g01548.t01#6    mNeoVis.2.1.1.chr1_hap1.g01598.t01#5    mNeoVis.evergladensis.1.1.chr1.g01562.t01#3    mNeoVis.vulgivaga.1.1.chr1.g01566.t01#4   mNeoVis.vulgivaga.2.1.chr1.g01562.t01#1       1539   125260111:   mNeoVis.evergladensis.2.1.chr7.g02214.t01#2    mNeoVis.evergladensis.1.1.chr7.g02211.t01#3    mNeoVis.vulgivaga.1.1.chr7.g02218.t01#4    mNeoVis.vulgivaga.2.1.chr7.g02232.t01#1    mNeoVis.3.1.1.chr7.g02086.t01#6   mNeoVis.2.1.1.chr7_hap1.g02267.t01#5       1540   125260112:   mNeoVis.evergladensis.2.1.chr1.g01566.t01#2    mNeoVis.2.1.1.chr1_hap1.g01599.t01#5    mNeoVis.3.1.1.chr1.g01549.t01#6    mNeoVis.vulgivaga.1.1.chr1.g01567.t01#4    mNeoVis.evergladensis.1.1.chr1.g01564.t01#3   mNeoVis.vulgivaga.2.1.chr1.g01563.t01#1       1541   125260113:   mNeoVis.3.1.1.chr1.g01550.t01#6    mNeoVis.evergladensis.1.1.chr1.g01565.t01#3    mNeoVis.evergladensis.2.1.chr1.g01567.t01#2    mNeoVis.2.1.1.chr1_hap1.g01600.t01#5    mNeoVis.vulgivaga.2.1.chr1.g01564.t01#1   mNeoVis.vulgivaga.1.1.chr1.g01568.t01#4       1542   125260114:   mNeoVis.evergladensis.2.1.chr2.g01271.t01#2    mNeoVis.vulgivaga.1.1.chr4.g00688.t01#4    mNeoVis.2.1.1.chr8_hap1.g00241.t01#5    mNeoVis.evergladensis.1.1.chr8.g00232.t01#3    mNeoVis.2.1.1.chr3_hap1.g00288.t01#5    mNeoVis.2.1.1.chr10_hap1.g01114.t01#5    mNeoVis.3.1.1.chr8.g00198.t01#6    mNeoVis.vulgivaga.1.1.chr8.g00230.t01#4    mNeoVis.evergladensis.2.1.chr9.g00231.t01#2      1543   125260115:   mNeoVis.evergladensis.1.1.chr2.g01275.t01#3    mNeoVis.3.1.1.chr1.g01454.t01#6    mNeoVis.vulgivaga.2.1.chr4.g00692.t01#1    mNeoVis.vulgivaga.2.1.chr1.g01472.t01#1    mNeoVis.vulgivaga.2.1.chr2.g01422.t01#1    mNeoVis.vulgivaga.1.1.chr2.g01426.t01#4   mNeoVis.evergladensis.2.1.chr2.g01411.t01#2     1544   125260116:   mNeoVis.vulgivaga.2.1.chr12.g00527.t01#1    mNeoVis.2.1.1.chr9_hap1.g00900.t01#5    mNeoVis.evergladensis.2.1.chr12.g00513.t01#2    mNeoVis.vulgivaga.2.1.chr8.g00239.t01#1   mNeoVis.evergladensis.1.1.chr9.g01079.t01#3       1545   125260117:   mNeoVis.evergladensis.1.1.chr1.g01475.t01#3    mNeoVis.vulgivaga.2.1.chr1.g01473.t01#1   mNeoVis.evergladensis.2.1.chr1.g01474.t01#2         1546   125260118:   mNeoVis.evergladensis.1.1.chr7.g02212.t01#3    mNeoVis.2.1.1.chr7_hap1.g02268.t01#5    mNeoVis.vulgivaga.1.1.chr7.g02219.t01#4    mNeoVis.vulgivaga.2.1.chr7.g02233.t01#1   mNeoVis.evergladensis.2.1.chr7.g02215.t01#2       1547   125260119:   mNeoVis.vulgivaga.2.1.chr7.g02234.t01#1    mNeoVis.evergladensis.1.1.chr7.g02213.t01#3    mNeoVis.evergladensis.2.1.chr7.g02216.t01#2    mNeoVis.3.1.1.chr7.g02088.t01#6    mNeoVis.vulgivaga.1.1.chr7.g02220.t01#4   mNeoVis.2.1.1.chr7_hap1.g02269.t01#5       1548   125260120:   mNeoVis.evergladensis.2.1.chr7.g02217.t01#2    mNeoVis.3.1.1.chr7.g02089.t01#6    mNeoVis.vulgivaga.1.1.chr7.g02221.t01#4    mNeoVis.vulgivaga.2.1.chr7.g02235.t01#1    mNeoVis.2.1.1.chr7_hap1.g02270.t01#5   mNeoVis.evergladensis.1.1.chr7.g02214.t01#3       1549   125260121:   mNeoVis.2.1.1.chr7_hap1.g02271.t01#5    mNeoVis.vulgivaga.1.1.chr7.g02222.t01#4    mNeoVis.3.1.1.chr7.g02090.t01#6    mNeoVis.vulgivaga.2.1.chr7.g02236.t01#1    mNeoVis.evergladensis.2.1.chr7.g02218.t01#2   mNeoVis.evergladensis.1.1.chr7.g02215.t01#3       1550   125260122:   mNeoVis.vulgivaga.1.1.chr7.g00074.t01#4    mNeoVis.vulgivaga.2.1.chr7.g00073.t01#1    mNeoVis.evergladensis.2.1.chr7.g00073.t01#2    mNeoVis.2.1.1.chr7_hap1.g00072.t01#5    mNeoVis.evergladensis.1.1.chr7.g00075.t01#3   mNeoVis.3.1.1.chr7.g00060.t01#6       1551   125260123:   mNeoVis.evergladensis.2.1.chr7.g02219.t01#2    mNeoVis.vulgivaga.1.1.chr7.g02223.t01#4    mNeoVis.2.1.1.chr7_hap1.g02272.t01#5    mNeoVis.vulgivaga.2.1.chr7.g02237.t01#1    mNeoVis.3.1.1.chr7.g02091.t01#6   mNeoVis.evergladensis.1.1.chr7.g02216.t01#3       1552   125260124:   mNeoVis.vulgivaga.2.1.chr7.g02238.t01#1    mNeoVis.2.1.1.chr7_hap1.g02273.t01#5    mNeoVis.3.1.1.chr7.g02092.t01#6    mNeoVis.evergladensis.1.1.chr7.g02217.t01#3    mNeoVis.vulgivaga.1.1.chr7.g02224.t01#4   mNeoVis.evergladensis.2.1.chr7.g02220.t01#2       1553   125260125:   mNeoVis.3.1.1.chr1.g01459.t01#6    mNeoVis.2.1.1.chr1_hap1.g01513.t01#5    mNeoVis.evergladensis.1.1.chr1.g01477.t01#3    mNeoVis.evergladensis.2.1.chr1.g01478.t01#2    mNeoVis.vulgivaga.1.1.chr1.g01480.t01#4   mNeoVis.vulgivaga.2.1.chr1.g01474.t01#1       1554   125260126:   mNeoVis.vulgivaga.1.1.chr7.g02225.t01#4    mNeoVis.3.1.1.chr7.g02093.t01#6    mNeoVis.2.1.1.chr7_hap1.g02274.t01#5    mNeoVis.vulgivaga.2.1.chr7.g02239.t01#1    mNeoVis.evergladensis.2.1.chr7.g02221.t01#2   mNeoVis.evergladensis.1.1.chr7.g02218.t01#3       1555   125260127:   mNeoVis.evergladensis.2.1.chr7.g02222.t01#2    mNeoVis.2.1.1.chr7_hap1.g02275.t01#5    mNeoVis.evergladensis.1.1.chr7.g02219.t01#3    mNeoVis.vulgivaga.2.1.chr7.g02240.t01#1    mNeoVis.vulgivaga.1.1.chr7.g02226.t01#4   mNeoVis.3.1.1.chr7.g02094.t01#6       1556   125260128:   mNeoVis.2.1.1.chr1_hap1.g01514.t01#5    mNeoVis.vulgivaga.2.1.chr7.g01462.t01#1    mNeoVis.3.1.1.chr7.g01374.t01#6    mNeoVis.vulgivaga.2.1.chr1.g01475.t01#1    mNeoVis.3.1.1.chr5.g01486.t01#6    mNeoVis.2.1.1.chr2_hap1.g01565.t01#5    mNeoVis.evergladensis.2.1.chr7.g01449.t01#2    mNeoVis.evergladensis.1.1.chr5.g01510.t01#3    mNeoVis.2.1.1.chr8_hap1.g00674.t01#5      1557   125260129:   mNeoVis.vulgivaga.2.1.chr6.g01045.t01#1    mNeoVis.2.1.1.chr4_hap1.g01048.t01#5    mNeoVis.evergladensis.2.1.chr6.g01053.t01#2    mNeoVis.vulgivaga.1.1.chr6.g01046.t01#4   mNeoVis.evergladensis.1.1.chr6.g01055.t01#3       1558   125260130:   mNeoVis.3.1.1.chr1.g01461.t01#6    mNeoVis.vulgivaga.2.1.chr1.g01476.t01#1   mNeoVis.evergladensis.2.1.chr1.g01481.t01#2         1559   125260131:   mNeoVis.vulgivaga.2.1.chr1.g01477.t01#1    mNeoVis.2.1.1.chr1_hap1.g01516.t01#5    mNeoVis.evergladensis.2.1.chr1.g01482.t01#2    mNeoVis.3.1.1.chr1.g01462.t01#6   mNeoVis.vulgivaga.1.1.chr1.g01483.t01#4       1560   125260132:   mNeoVis.vulgivaga.1.1.chr7.g02228.t01#4    mNeoVis.evergladensis.1.1.chr7.g02220.t01#3    mNeoVis.evergladensis.2.1.chr7.g02223.t01#2    mNeoVis.3.1.1.chr7.g02095.t01#6    mNeoVis.vulgivaga.2.1.chr7.g02241.t01#1   mNeoVis.2.1.1.chr7_hap1.g02276.t01#5       1561   125260133:   mNeoVis.evergladensis.1.1.chr15.g00032.t01#3    mNeoVis.vulgivaga.2.1.chr15.g00590.t01#1    mNeoVis.2.1.1.chr15_hap1.g00758.t01#5    mNeoVis.evergladensis.2.1.chr15.g00579.t01#2    mNeoVis.vulgivaga.1.1.chr15.g00581.t01#4    mNeoVis.evergladensis.1.1.chr15.g00033.t01#3    mNeoVis.3.1.1.chr15.g00534.t01#6   mNeoVis.vulgivaga.1.1.chr15.g00582.t01#4     1562   125260134:   mNeoVis.vulgivaga.2.1.chr1.g01478.t01#1    mNeoVis.evergladensis.2.1.chr1.g01484.t01#2    mNeoVis.vulgivaga.1.1.chr1.g01485.t01#4    mNeoVis.3.1.1.chr1.g01463.t01#6   mNeoVis.2.1.1.chr1_hap1.g01518.t01#5       1563   125260135:  mNeoVis.vulgivaga.2.1.chr1.g01479.t01#1               1564   125260136:   mNeoVis.vulgivaga.2.1.chr7.g02242.t01#1    mNeoVis.2.1.1.chr7_hap1.g02277.t01#5    mNeoVis.vulgivaga.1.1.chr7.g02229.t01#4    mNeoVis.evergladensis.2.1.chr7.g02224.t01#2    mNeoVis.evergladensis.1.1.chr7.g02221.t01#3   mNeoVis.3.1.1.chr7.g02096.t01#6       1565   125260137:   mNeoVis.vulgivaga.1.1.chr1.g01487.t01#4    mNeoVis.2.1.1.chr1_hap1.g01520.t01#5   mNeoVis.vulgivaga.2.1.chr1.g01480.t01#1           1566   125260138:   mNeoVis.3.1.1.chr1.g01466.t01#6    mNeoVis.evergladensis.1.1.chr1.g01485.t01#3    mNeoVis.vulgivaga.2.1.chr1.g01481.t01#1   mNeoVis.2.1.1.chr1_hap1.g01521.t01#5         1567   125260139:   mNeoVis.vulgivaga.2.1.chr1.g01482.t01#1    mNeoVis.3.1.1.chr1.g01467.t01#6    mNeoVis.2.1.1.chr1_hap1.g01522.t01#5    mNeoVis.evergladensis.2.1.chr1.g01487.t01#2    mNeoVis.vulgivaga.1.1.chr1.g01488.t01#4   mNeoVis.evergladensis.1.1.chr1.g01486.t01#3       1568   125260140:   mNeoVis.evergladensis.2.1.chr1.g01488.t01#2    mNeoVis.3.1.1.chr1.g01468.t01#6    mNeoVis.evergladensis.1.1.chr1.g01487.t01#3    mNeoVis.2.1.1.chr1_hap1.g01523.t01#5    mNeoVis.vulgivaga.1.1.chr1.g01489.t01#4   mNeoVis.vulgivaga.2.1.chr1.g01483.t01#1       1569   125260141:   mNeoVis.evergladensis.1.1.chr7.g02222.t01#3    mNeoVis.vulgivaga.1.1.chr7.g02231.t01#4    mNeoVis.vulgivaga.2.1.chr7.g02243.t01#1    mNeoVis.3.1.1.chr7.g02097.t01#6    mNeoVis.vulgivaga.1.1.chr7.g02230.t01#4    mNeoVis.2.1.1.chr7_hap1.g02278.t01#5   mNeoVis.evergladensis.2.1.chr7.g02225.t01#2     1570   125260142:   mNeoVis.3.1.1.chr1.g01469.t01#6    mNeoVis.evergladensis.1.1.chr1.g01488.t01#3    mNeoVis.2.1.1.chr1_hap1.g01524.t01#5    mNeoVis.vulgivaga.2.1.chr1.g01484.t01#1    mNeoVis.evergladensis.2.1.chr1.g01489.t01#2   mNeoVis.vulgivaga.1.1.chr1.g01490.t01#4       1571   125260143:   mNeoVis.evergladensis.2.1.chr1.g01490.t01#2    mNeoVis.vulgivaga.1.1.chr1.g01491.t01#4    mNeoVis.3.1.1.chr1.g01470.t01#6    mNeoVis.evergladensis.1.1.chr1.g01489.t01#3    mNeoVis.2.1.1.chr1_hap1.g01525.t01#5   mNeoVis.vulgivaga.2.1.chr1.g01485.t01#1       1572   125260144:   mNeoVis.evergladensis.1.1.chr7.g02223.t01#3    mNeoVis.vulgivaga.2.1.chr7.g02244.t01#1    mNeoVis.2.1.1.chr7_hap1.g02279.t01#5    mNeoVis.evergladensis.2.1.chr7.g02226.t01#2    mNeoVis.3.1.1.chr7.g02098.t01#6   mNeoVis.vulgivaga.1.1.chr7.g02232.t01#4       1573   125260145:   mNeoVis.evergladensis.2.1.chr1.g01492.t01#2    mNeoVis.vulgivaga.2.1.chr1.g01486.t01#1    mNeoVis.2.1.1.chr1_hap1.g01527.t01#5    mNeoVis.evergladensis.1.1.chr1.g01490.t01#3   mNeoVis.vulgivaga.1.1.chr1.g01494.t01#4       1574   125260146:   mNeoVis.2.1.1.chr7_hap1.g00522.t01#5    mNeoVis.vulgivaga.2.1.chr7.g00525.t01#1    mNeoVis.vulgivaga.1.1.chr7.g00530.t01#4    mNeoVis.evergladensis.2.1.chr7.g00522.t01#2    mNeoVis.3.1.1.chr7.g00506.t01#6   mNeoVis.evergladensis.1.1.chr7.g00531.t01#3       1575   125260147:   mNeoVis.3.1.1.chr1.g01473.t01#6    mNeoVis.vulgivaga.1.1.chr1.g01495.t01#4    mNeoVis.2.1.1.chr1_hap1.g01528.t01#5    mNeoVis.evergladensis.1.1.chr1.g01491.t01#3    mNeoVis.evergladensis.2.1.chr1.g01493.t01#2   mNeoVis.vulgivaga.2.1.chr1.g01487.t01#1       1576   125260148:   mNeoVis.2.1.1.chr7_hap1.g02280.t01#5    mNeoVis.evergladensis.1.1.chr7.g02224.t01#3    mNeoVis.3.1.1.chr7.g02099.t01#6    mNeoVis.vulgivaga.2.1.chr7.g02245.t01#1    mNeoVis.vulgivaga.1.1.chr7.g02233.t01#4   mNeoVis.evergladensis.2.1.chr7.g02227.t01#2       1577   125260149:   mNeoVis.evergladensis.2.1.chr7.g02228.t01#2    mNeoVis.vulgivaga.1.1.chr7.g02234.t01#4    mNeoVis.3.1.1.chr7.g02100.t01#6    mNeoVis.evergladensis.1.1.chr7.g02225.t01#3    mNeoVis.vulgivaga.2.1.chr7.g02246.t01#1   mNeoVis.2.1.1.chr7_hap1.g02281.t01#5       1578   125260150:   mNeoVis.evergladensis.2.1.chr7.g02229.t01#2    mNeoVis.vulgivaga.1.1.chr7.g02235.t01#4    mNeoVis.3.1.1.chr7.g02101.t01#6    mNeoVis.vulgivaga.2.1.chr7.g02247.t01#1    mNeoVis.evergladensis.1.1.chr7.g02226.t01#3   mNeoVis.2.1.1.chr7_hap1.g02282.t01#5       1579   125260151:   mNeoVis.vulgivaga.1.1.chr7.g02236.t01#4    mNeoVis.evergladensis.1.1.chr7.g02227.t01#3    mNeoVis.vulgivaga.2.1.chr7.g02248.t01#1    mNeoVis.3.1.1.chr7.g02102.t01#6    mNeoVis.2.1.1.chr7_hap1.g02283.t01#5   mNeoVis.evergladensis.2.1.chr7.g02230.t01#2       1580   125260152:   mNeoVis.evergladensis.1.1.chr7.g02228.t01#3    mNeoVis.vulgivaga.1.1.chr7.g02237.t01#4    mNeoVis.vulgivaga.2.1.chr7.g02249.t01#1    mNeoVis.3.1.1.chr7.g02103.t01#6    mNeoVis.evergladensis.2.1.chr7.g02231.t01#2   mNeoVis.2.1.1.chr7_hap1.g02284.t01#5       1581   125260153:   mNeoVis.2.1.1.chr7_hap1.g02285.t01#5    mNeoVis.evergladensis.2.1.chr7.g02232.t01#2    mNeoVis.vulgivaga.2.1.chr7.g02250.t01#1    mNeoVis.evergladensis.1.1.chr7.g02229.t01#3    mNeoVis.vulgivaga.1.1.chr7.g02238.t01#4   mNeoVis.3.1.1.chr7.g02104.t01#6       1582   125260154:   mNeoVis.evergladensis.2.1.chr7.g02233.t01#2    mNeoVis.vulgivaga.2.1.chr7.g02251.t01#1    mNeoVis.evergladensis.1.1.chr7.g02230.t01#3    mNeoVis.vulgivaga.1.1.chr7.g02239.t01#4    mNeoVis.3.1.1.chr7.g02105.t01#6   mNeoVis.2.1.1.chr7_hap1.g02286.t01#5       1583   125260155:   mNeoVis.vulgivaga.2.1.chr7.g02252.t01#1    mNeoVis.evergladensis.2.1.chr7.g02234.t01#2    mNeoVis.2.1.1.chr7_hap1.g02287.t01#5    mNeoVis.evergladensis.1.1.chr7.g02231.t01#3   mNeoVis.vulgivaga.1.1.chr7.g02240.t01#4       1584   125260156:   mNeoVis.evergladensis.1.1.chr7.g02232.t01#3    mNeoVis.vulgivaga.1.1.chr7.g02241.t01#4    mNeoVis.evergladensis.2.1.chr7.g02235.t01#2    mNeoVis.3.1.1.chr7.g02106.t01#6    mNeoVis.2.1.1.chr7_hap1.g02288.t01#5   mNeoVis.vulgivaga.2.1.chr7.g02253.t01#1       1585   125260157:   mNeoVis.vulgivaga.2.1.chr1.g01488.t01#1    mNeoVis.3.1.1.chr1.g01474.t01#6    mNeoVis.evergladensis.1.1.chr1.g01493.t01#3   mNeoVis.3.1.1.chr1.g01475.t01#6         1586   125260158:   mNeoVis.evergladensis.2.1.chr1.g01495.t01#2    mNeoVis.vulgivaga.1.1.chr1.g01497.t01#4    mNeoVis.3.1.1.chr1.g01477.t01#6   mNeoVis.vulgivaga.2.1.chr1.g01489.t01#1         1587   125260159:   mNeoVis.3.1.1.chr3.g00197.t01#6    mNeoVis.vulgivaga.1.1.chr3.g00200.t01#4    mNeoVis.evergladensis.2.1.chr3.g00205.t01#2    mNeoVis.vulgivaga.2.1.chr7.g02254.t01#1    mNeoVis.vulgivaga.1.1.chr3.g00199.t01#4    mNeoVis.evergladensis.2.1.chr7.g02236.t01#2    mNeoVis.vulgivaga.2.1.chr3.g00197.t01#1    mNeoVis.vulgivaga.1.1.chr7.g02242.t01#4    mNeoVis.3.1.1.chr7.g02107.t01#6      1588   125260160:   mNeoVis.3.1.1.chr7.g00507.t01#6    mNeoVis.evergladensis.2.1.chr7.g00523.t01#2    mNeoVis.evergladensis.1.1.chr7.g00532.t01#3    mNeoVis.vulgivaga.2.1.chr7.g00526.t01#1    mNeoVis.vulgivaga.1.1.chr7.g00531.t01#4   mNeoVis.2.1.1.chr7_hap1.g00523.t01#5       1589   125260161:   mNeoVis.3.1.1.chr1.g01480.t01#6    mNeoVis.2.1.1.chr1_hap1.g01529.t01#5    mNeoVis.vulgivaga.1.1.chr1.g01499.t01#4    mNeoVis.vulgivaga.2.1.chr1.g01490.t01#1    mNeoVis.evergladensis.2.1.chr1.g01496.t01#2   mNeoVis.evergladensis.1.1.chr1.g01495.t01#3       1590   125260162:   mNeoVis.2.1.1.chr7_hap1.g02290.t01#5    mNeoVis.3.1.1.chr7.g02108.t01#6    mNeoVis.evergladensis.2.1.chr7.g02237.t01#2    mNeoVis.vulgivaga.1.1.chr7.g02243.t01#4    mNeoVis.evergladensis.1.1.chr7.g02234.t01#3   mNeoVis.vulgivaga.2.1.chr7.g02255.t01#1       1591   125260163:   mNeoVis.3.1.1.chr7.g02109.t01#6    mNeoVis.2.1.1.chr7_hap1.g02291.t01#5    mNeoVis.evergladensis.2.1.chr7.g02238.t01#2    mNeoVis.evergladensis.1.1.chr7.g02235.t01#3    mNeoVis.vulgivaga.1.1.chr7.g02244.t01#4   mNeoVis.vulgivaga.2.1.chr7.g02256.t01#1       1592   125260164:   mNeoVis.evergladensis.2.1.chr7.g02239.t01#2    mNeoVis.2.1.1.chr7_hap1.g02292.t01#5    mNeoVis.vulgivaga.2.1.chr7.g02257.t01#1    mNeoVis.evergladensis.1.1.chr7.g02236.t01#3   mNeoVis.vulgivaga.1.1.chr7.g02245.t01#4       1593   125260165:   mNeoVis.vulgivaga.2.1.chr7.g02258.t01#1    mNeoVis.evergladensis.1.1.chr7.g02237.t01#3    mNeoVis.vulgivaga.1.1.chr7.g02246.t01#4    mNeoVis.evergladensis.2.1.chr7.g02240.t01#2    mNeoVis.3.1.1.chr7.g02111.t01#6   mNeoVis.2.1.1.chr7_hap1.g02293.t01#5       1594   125260166:   mNeoVis.2.1.1.chr2_hap1.g00989.t01#5    mNeoVis.vulgivaga.2.1.chr5.g00929.t01#1    mNeoVis.vulgivaga.1.1.chr5.g00910.t01#4    mNeoVis.evergladensis.1.1.chr1.g01497.t01#3    mNeoVis.evergladensis.1.1.chr5.g00933.t01#3    mNeoVis.evergladensis.2.1.chr1.g01497.t01#2    mNeoVis.3.1.1.chr1.g01481.t01#6    mNeoVis.3.1.1.chr5.g00922.t01#6    mNeoVis.vulgivaga.1.1.chr1.g01500.t01#4      1595   125260167:   mNeoVis.evergladensis.1.1.chr1.g01498.t01#3    mNeoVis.vulgivaga.2.1.chr1.g01492.t01#1    mNeoVis.vulgivaga.1.1.chr1.g01501.t01#4    mNeoVis.2.1.1.chr1_hap1.g01530.t01#5    mNeoVis.3.1.1.chr1.g01482.t01#6   mNeoVis.evergladensis.2.1.chr1.g01498.t01#2       1596   125260168:   mNeoVis.evergladensis.1.1.chr1.g01499.t01#3    mNeoVis.vulgivaga.2.1.chr1.g01493.t01#1    mNeoVis.vulgivaga.1.1.chr1.g01502.t01#4    mNeoVis.2.1.1.chr1_hap1.g01531.t01#5    mNeoVis.evergladensis.2.1.chr1.g01499.t01#2   mNeoVis.3.1.1.chr1.g01483.t01#6       1597   125260169:   mNeoVis.2.1.1.chr7_hap1.g02294.t01#5    mNeoVis.evergladensis.2.1.chr7.g02241.t01#2    mNeoVis.3.1.1.chr7.g02112.t01#6    mNeoVis.vulgivaga.2.1.chr7.g02259.t01#1    mNeoVis.evergladensis.1.1.chr7.g02238.t01#3   mNeoVis.vulgivaga.1.1.chr7.g02247.t01#4       1598   125260170:   mNeoVis.vulgivaga.2.1.chr1.g01494.t01#1    mNeoVis.vulgivaga.1.1.chr1.g01503.t01#4    mNeoVis.evergladensis.2.1.chr1.g01500.t01#2    mNeoVis.evergladensis.1.1.chr1.g01500.t01#3    mNeoVis.3.1.1.chr1.g01484.t01#6   mNeoVis.2.1.1.chr1_hap1.g01532.t01#5       1599   125260171:   mNeoVis.vulgivaga.2.1.chr7.g02260.t01#1    mNeoVis.evergladensis.1.1.chr7.g02239.t01#3    mNeoVis.vulgivaga.1.1.chr7.g02248.t01#4    mNeoVis.3.1.1.chr7.g02113.t01#6    mNeoVis.2.1.1.chr7_hap1.g02295.t01#5   mNeoVis.evergladensis.2.1.chr7.g02242.t01#2       1600   125260172:   mNeoVis.vulgivaga.2.1.chr1.g01495.t01#1    mNeoVis.evergladensis.2.1.chr1.g01501.t01#2    mNeoVis.3.1.1.chr1.g01485.t01#6    mNeoVis.evergladensis.1.1.chr1.g01501.t01#3    mNeoVis.vulgivaga.1.1.chr1.g01504.t01#4   mNeoVis.2.1.1.chr1_hap1.g01533.t01#5       1601   125260173:   mNeoVis.evergladensis.2.1.chr5.g01618.t01#2    mNeoVis.evergladensis.1.1.chr5.g01623.t01#3    mNeoVis.vulgivaga.2.1.chr5.g01620.t01#1    mNeoVis.3.1.1.chr5.g01600.t01#6    mNeoVis.2.1.1.chr2_hap1.g01677.t01#5   mNeoVis.vulgivaga.1.1.chr5.g01597.t01#4       1602   125260174:   mNeoVis.evergladensis.2.1.chr1.g01502.t01#2    mNeoVis.vulgivaga.2.1.chr1.g01496.t01#1    mNeoVis.3.1.1.chr1.g01486.t01#6    mNeoVis.evergladensis.1.1.chr1.g01502.t01#3   mNeoVis.2.1.1.chr1_hap1.g01534.t01#5       1603   125260175:   mNeoVis.evergladensis.1.1.chr12.g00628.t01#3    mNeoVis.3.1.1.chr12.g00636.t01#6    mNeoVis.vulgivaga.2.1.chr12.g00642.t01#1    mNeoVis.evergladensis.2.1.chr12.g00622.t01#2    mNeoVis.evergladensis.2.1.chr12.g00625.t01#2    mNeoVis.vulgivaga.1.1.chr12.g00638.t01#4    mNeoVis.3.1.1.chr12.g00637.t01#6    mNeoVis.vulgivaga.2.1.chr12.g00647.t01#1    mNeoVis.evergladensis.1.1.chr12.g00627.t01#3      1604   125260176:   mNeoVis.3.1.1.chr7.g02114.t01#6    mNeoVis.2.1.1.chr7_hap1.g02296.t01#5    mNeoVis.evergladensis.2.1.chr7.g02243.t01#2    mNeoVis.evergladensis.1.1.chr4.g00757.t01#3    mNeoVis.2.1.1.chr6_hap1.g00740.t01#5    mNeoVis.vulgivaga.1.1.chr7.g02249.t01#4    mNeoVis.evergladensis.1.1.chr7.g02240.t01#3   mNeoVis.vulgivaga.2.1.chr7.g02261.t01#1     1605   125260177:   mNeoVis.vulgivaga.2.1.chr12.g00646.t01#1    mNeoVis.evergladensis.1.1.chr12.g00644.t01#3    mNeoVis.vulgivaga.2.1.chr12.g00652.t01#1    mNeoVis.vulgivaga.2.1.chr12.g00644.t01#1    mNeoVis.vulgivaga.2.1.chr12.g00657.t01#1   mNeoVis.vulgivaga.1.1.chr12.g00652.t01#4       1606   125260178:   mNeoVis.3.1.1.chr4.g00428.t01#6    mNeoVis.vulgivaga.2.1.chr4.g00437.t01#1    mNeoVis.evergladensis.2.1.chr4.g00434.t01#2    mNeoVis.evergladensis.1.1.chr4.g00436.t01#3    mNeoVis.vulgivaga.1.1.chr4.g00429.t01#4   mNeoVis.2.1.1.chr6_hap1.g00427.t01#5       1607   125260179:   mNeoVis.evergladensis.1.1.chr1.g00846.t01#3    mNeoVis.evergladensis.2.1.chr1.g00834.t01#2    mNeoVis.vulgivaga.2.1.chr1.g00851.t01#1    mNeoVis.2.1.1.chr1_hap1.g00830.t01#5    mNeoVis.3.1.1.chr1.g00854.t01#6   mNeoVis.vulgivaga.1.1.chr1.g00851.t01#4       1608   125260180:   mNeoVis.evergladensis.1.1.chr8.g01390.t01#3    mNeoVis.vulgivaga.1.1.chr8.g01384.t01#4    mNeoVis.3.1.1.chr8.g01342.t01#6    mNeoVis.2.1.1.chr8_hap1.g01445.t01#5   mNeoVis.evergladensis.2.1.chr9.g01392.t01#2       1609   125260181:   mNeoVis.2.1.1.chr12_hap1.g00691.t01#5    mNeoVis.2.1.1.chr12_hap1.g00704.t01#5    mNeoVis.2.1.1.chr12_hap1.g00690.t01#5    mNeoVis.2.1.1.chr12_hap1.g00686.t01#5   mNeoVis.2.1.1.chr12_hap1.g00685.t01#5       1610   125260182:   mNeoVis.vulgivaga.2.1.chr9.g00379.t01#1    mNeoVis.evergladensis.1.1.chr9.g00384.t01#3   mNeoVis.evergladensis.2.1.chr8.g00377.t01#2         1611   125260183:   mNeoVis.3.1.1.chr12.g00644.t01#6    mNeoVis.evergladensis.2.1.chr12.g00634.t01#2   mNeoVis.2.1.1.chr12_hap1.g00688.t01#5           1612   125260184:   mNeoVis.evergladensis.1.1.chr2.g01723.t01#3   mNeoVis.2.1.1.chr5_hap1.g01729.t01#5             1613   125260185:  mNeoVis.2.1.1.chr10_hap1.g00395.t01#5               1614   125260186:   mNeoVis.evergladensis.2.1.chr1.g01503.t01#2    mNeoVis.2.1.1.chr1_hap1.g01535.t01#5    mNeoVis.vulgivaga.2.1.chr1.g01497.t01#1    mNeoVis.evergladensis.1.1.chr1.g01503.t01#3    mNeoVis.3.1.1.chr1.g01487.t01#6   mNeoVis.vulgivaga.1.1.chr1.g01505.t01#4       1615   125260187:   mNeoVis.vulgivaga.1.1.chr7.g02250.t01#4    mNeoVis.evergladensis.1.1.chr7.g02241.t01#3    mNeoVis.2.1.1.chr7_hap1.g02297.t01#5    mNeoVis.3.1.1.chr7.g02115.t01#6    mNeoVis.evergladensis.2.1.chr7.g02244.t01#2   mNeoVis.vulgivaga.2.1.chr7.g02262.t01#1       1616   125260188:   mNeoVis.vulgivaga.2.1.chr1.g01498.t01#1    mNeoVis.evergladensis.2.1.chr1.g01504.t01#2    mNeoVis.vulgivaga.1.1.chr1.g01506.t01#4    mNeoVis.evergladensis.1.1.chr1.g01504.t01#3    mNeoVis.3.1.1.chr1.g01488.t01#6   mNeoVis.2.1.1.chr1_hap1.g01536.t01#5       1617   125260189:   mNeoVis.evergladensis.1.1.chr1.g01505.t01#3    mNeoVis.evergladensis.2.1.chr1.g01505.t01#2    mNeoVis.2.1.1.chr1_hap1.g01537.t01#5    mNeoVis.3.1.1.chr1.g01489.t01#6    mNeoVis.vulgivaga.2.1.chr1.g01499.t01#1   mNeoVis.vulgivaga.1.1.chr1.g01507.t01#4       1618   125260190:   mNeoVis.vulgivaga.1.1.chr7.g02252.t01#4    mNeoVis.evergladensis.1.1.chr7.g02243.t01#3    mNeoVis.evergladensis.2.1.chr7.g02246.t01#2    mNeoVis.2.1.1.chr7_hap1.g02299.t01#5    mNeoVis.vulgivaga.2.1.chr7.g02263.t01#1   mNeoVis.3.1.1.chr7.g02116.t01#6       1619   125260191:   mNeoVis.2.1.1.chr1_hap1.g01538.t01#5    mNeoVis.3.1.1.chr1.g01491.t01#6    mNeoVis.evergladensis.1.1.chr1.g01506.t01#3    mNeoVis.vulgivaga.2.1.chr1.g01500.t01#1    mNeoVis.vulgivaga.1.1.chr1.g01508.t01#4   mNeoVis.evergladensis.2.1.chr1.g01506.t01#2       1620   125260192:   mNeoVis.evergladensis.1.1.chr7.g02244.t01#3    mNeoVis.evergladensis.2.1.chr7.g02247.t01#2    mNeoVis.2.1.1.chr7_hap1.g02300.t01#5    mNeoVis.vulgivaga.2.1.chr7.g02264.t01#1   mNeoVis.3.1.1.chr7.g02117.t01#6       1621   125260193:   mNeoVis.2.1.1.chr7_hap1.g02301.t01#5    mNeoVis.vulgivaga.1.1.chr7.g02253.t01#4    mNeoVis.vulgivaga.2.1.chr7.g02265.t01#1    mNeoVis.3.1.1.chr7.g02118.t01#6    mNeoVis.evergladensis.2.1.chr7.g02248.t01#2   mNeoVis.evergladensis.1.1.chr7.g02245.t01#3       1622   125260194:   mNeoVis.evergladensis.1.1.chr7.g02246.t01#3    mNeoVis.3.1.1.chr7.g02119.t01#6    mNeoVis.vulgivaga.2.1.chr4.g00950.t01#1    mNeoVis.vulgivaga.1.1.chr7.g02254.t01#4    mNeoVis.vulgivaga.1.1.chr5.g00255.t01#4    mNeoVis.evergladensis.2.1.chr7.g02249.t01#2    mNeoVis.2.1.1.chr7_hap1.g02302.t01#5    mNeoVis.evergladensis.1.1.chr5.g00264.t01#3    mNeoVis.evergladensis.2.1.chr4.g00928.t01#2      1623   125260195:   mNeoVis.vulgivaga.2.1.chr1.g01501.t01#1    mNeoVis.2.1.1.chr1_hap1.g01539.t01#5    mNeoVis.evergladensis.2.1.chr1.g01507.t01#2    mNeoVis.3.1.1.chr1.g01492.t01#6    mNeoVis.vulgivaga.1.1.chr1.g01509.t01#4   mNeoVis.evergladensis.1.1.chr1.g01507.t01#3       1624   125260196:   mNeoVis.vulgivaga.1.1.chr1.g01510.t01#4    mNeoVis.evergladensis.1.1.chr1.g01508.t01#3    mNeoVis.evergladensis.2.1.chr1.g01508.t01#2    mNeoVis.vulgivaga.2.1.chr1.g01502.t01#1    mNeoVis.3.1.1.chr1.g01493.t01#6   mNeoVis.2.1.1.chr1_hap1.g01540.t01#5       1625   125260197:   mNeoVis.evergladensis.2.1.chr7.g02250.t01#2    mNeoVis.evergladensis.1.1.chr7.g02247.t01#3    mNeoVis.3.1.1.chr7.g02120.t01#6    mNeoVis.vulgivaga.1.1.chr7.g02255.t01#4    mNeoVis.2.1.1.chr7_hap1.g02303.t01#5   mNeoVis.vulgivaga.2.1.chr7.g02267.t01#1       1626   125260198:   mNeoVis.vulgivaga.2.1.chr7.g02268.t01#1    mNeoVis.evergladensis.1.1.chr7.g02248.t01#3    mNeoVis.vulgivaga.1.1.chr7.g02256.t01#4    mNeoVis.3.1.1.chr7.g02121.t01#6    mNeoVis.2.1.1.chr7_hap1.g02304.t01#5   mNeoVis.evergladensis.2.1.chr7.g02251.t01#2       1627   125260199:   mNeoVis.vulgivaga.2.1.chr1.g01503.t01#1    mNeoVis.vulgivaga.1.1.chr1.g01511.t01#4    mNeoVis.evergladensis.1.1.chr1.g01509.t01#3    mNeoVis.3.1.1.chr1.g01494.t01#6    mNeoVis.2.1.1.chr1_hap1.g01541.t01#5   mNeoVis.evergladensis.2.1.chr1.g01509.t01#2       1628   125260200:   mNeoVis.evergladensis.1.1.chr1.g01510.t01#3    mNeoVis.evergladensis.2.1.chr1.g01510.t01#2    mNeoVis.2.1.1.chr1_hap1.g01542.t01#5    mNeoVis.vulgivaga.1.1.chr1.g01512.t01#4    mNeoVis.vulgivaga.2.1.chr1.g01504.t01#1   mNeoVis.3.1.1.chr1.g01495.t01#6       1629   125260201:   mNeoVis.vulgivaga.1.1.chr1.g01513.t01#4    mNeoVis.evergladensis.2.1.chr1.g01511.t01#2    mNeoVis.evergladensis.1.1.chr1.g01511.t01#3    mNeoVis.vulgivaga.2.1.chr1.g01505.t01#1    mNeoVis.3.1.1.chr1.g01496.t01#6   mNeoVis.2.1.1.chr1_hap1.g01543.t01#5       1630   125260202:   mNeoVis.2.1.1.chr7_hap1.g02305.t01#5    mNeoVis.evergladensis.2.1.chr7.g02252.t01#2    mNeoVis.vulgivaga.1.1.chr7.g02257.t01#4    mNeoVis.evergladensis.1.1.chr7.g02249.t01#3   mNeoVis.vulgivaga.2.1.chr7.g02269.t01#1       1631   125260203:   mNeoVis.vulgivaga.1.1.chr15.g00583.t01#4    mNeoVis.evergladensis.2.1.chr15.g00580.t01#2    mNeoVis.2.1.1.chr15_hap1.g00759.t01#5    mNeoVis.vulgivaga.2.1.chr15.g00591.t01#1   mNeoVis.evergladensis.1.1.chr15.g00031.t01#3       1632   125260204:   mNeoVis.vulgivaga.2.1.chr1.g01507.t01#1    mNeoVis.evergladensis.2.1.chr1.g01512.t01#2    mNeoVis.evergladensis.1.1.chr1.g01512.t01#3    mNeoVis.3.1.1.chr1.g01497.t01#6    mNeoVis.2.1.1.chr1_hap1.g01544.t01#5    mNeoVis.vulgivaga.2.1.chr1.g01506.t01#1   mNeoVis.vulgivaga.1.1.chr1.g01514.t01#4     1633   125260205:   mNeoVis.evergladensis.2.1.chr1.g01513.t01#2    mNeoVis.vulgivaga.2.1.chr1.g01508.t01#1    mNeoVis.evergladensis.1.1.chr1.g01513.t01#3    mNeoVis.vulgivaga.1.1.chr1.g01515.t01#4    mNeoVis.2.1.1.chr1_hap1.g01545.t01#5   mNeoVis.3.1.1.chr1.g01498.t01#6       1634   125260206:   mNeoVis.3.1.1.chr7.g00508.t01#6    mNeoVis.evergladensis.1.1.chr7.g00533.t01#3    mNeoVis.vulgivaga.2.1.chr7.g00527.t01#1    mNeoVis.evergladensis.2.1.chr7.g00524.t01#2    mNeoVis.2.1.1.chr7_hap1.g00524.t01#5   mNeoVis.vulgivaga.1.1.chr7.g00532.t01#4       1635   125260207:   mNeoVis.evergladensis.2.1.chr1.g01514.t01#2    mNeoVis.vulgivaga.1.1.chr1.g01516.t01#4    mNeoVis.2.1.1.chr1_hap1.g01546.t01#5    mNeoVis.3.1.1.chr1.g01499.t01#6    mNeoVis.evergladensis.1.1.chr1.g01514.t01#3   mNeoVis.vulgivaga.2.1.chr1.g01509.t01#1       1636   125260208:   mNeoVis.3.1.1.chr12.g00380.t01#6    mNeoVis.vulgivaga.1.1.chr12.g00386.t01#4   mNeoVis.evergladensis.1.1.chr12.g00383.t01#3         1637   125260209:   mNeoVis.vulgivaga.2.1.chr1.g01511.t01#1    mNeoVis.evergladensis.2.1.chr1.g01516.t01#2    mNeoVis.3.1.1.chr1.g01501.t01#6    mNeoVis.evergladensis.1.1.chr1.g01515.t01#3    mNeoVis.2.1.1.chr1_hap1.g01548.t01#5   mNeoVis.vulgivaga.1.1.chr1.g01518.t01#4       1638   125260210:   mNeoVis.2.1.1.chr1_hap1.g01547.t01#5    mNeoVis.3.1.1.chr1.g01500.t01#6    mNeoVis.vulgivaga.1.1.chr1.g01517.t01#4    mNeoVis.vulgivaga.2.1.chr1.g01510.t01#1   mNeoVis.evergladensis.2.1.chr1.g01515.t01#2       1639   125260211:   mNeoVis.evergladensis.1.1.chr7.g02250.t01#3    mNeoVis.vulgivaga.2.1.chr7.g02270.t01#1    mNeoVis.2.1.1.chr7_hap1.g02306.t01#5    mNeoVis.evergladensis.2.1.chr7.g02253.t01#2   mNeoVis.vulgivaga.1.1.chr7.g02258.t01#4       1640   125260212:   mNeoVis.vulgivaga.1.1.chr7.g02259.t01#4    mNeoVis.3.1.1.chr7.g02123.t01#6    mNeoVis.evergladensis.1.1.chr7.g02251.t01#3    mNeoVis.vulgivaga.2.1.chr7.g02271.t01#1    mNeoVis.2.1.1.chr7_hap1.g02307.t01#5   mNeoVis.evergladensis.2.1.chr7.g02254.t01#2       1641   125260213:   mNeoVis.evergladensis.2.1.chr7.g02120.t01#2    mNeoVis.evergladensis.1.1.chr7.g02123.t01#3    mNeoVis.vulgivaga.1.1.chr7.g02126.t01#4    mNeoVis.vulgivaga.2.1.chr7.g02142.t01#1   mNeoVis.3.1.1.chr7.g01999.t01#6       1642   125260214:   mNeoVis.3.1.1.chr7.g02000.t01#6    mNeoVis.vulgivaga.2.1.chr7.g02143.t01#1    mNeoVis.evergladensis.1.1.chr7.g02124.t01#3    mNeoVis.2.1.1.chr7_hap1.g02172.t01#5    mNeoVis.evergladensis.2.1.chr7.g02121.t01#2   mNeoVis.vulgivaga.1.1.chr7.g02127.t01#4       1643   125260215:   mNeoVis.3.1.1.chr7.g02001.t01#6    mNeoVis.vulgivaga.2.1.chr7.g02144.t01#1    mNeoVis.evergladensis.2.1.chr7.g02122.t01#2    mNeoVis.evergladensis.1.1.chr7.g02125.t01#3    mNeoVis.2.1.1.chr7_hap1.g02173.t01#5   mNeoVis.vulgivaga.1.1.chr7.g02128.t01#4       1644   125260216:   mNeoVis.3.1.1.chr1.g01503.t01#6    mNeoVis.vulgivaga.2.1.chr1.g01512.t01#1    mNeoVis.evergladensis.2.1.chr1.g01517.t01#2    mNeoVis.evergladensis.1.1.chr1.g01516.t01#3    mNeoVis.2.1.1.chr1_hap1.g01550.t01#5   mNeoVis.vulgivaga.1.1.chr1.g01519.t01#4       1645   125260217:   mNeoVis.2.1.1.chr7_hap1.g02174.t01#5    mNeoVis.evergladensis.2.1.chr7.g02123.t01#2    mNeoVis.vulgivaga.1.1.chr7.g02129.t01#4    mNeoVis.vulgivaga.2.1.chr7.g02145.t01#1   mNeoVis.evergladensis.1.1.chr7.g02126.t01#3       1646   125260218:   mNeoVis.vulgivaga.1.1.chr1.g01520.t01#4    mNeoVis.evergladensis.1.1.chr1.g01517.t01#3    mNeoVis.vulgivaga.2.1.chr1.g01513.t01#1    mNeoVis.3.1.1.chr1.g01504.t01#6    mNeoVis.2.1.1.chr1_hap1.g01551.t01#5   mNeoVis.evergladensis.2.1.chr1.g01518.t01#2       1647   125260219:   mNeoVis.evergladensis.1.1.chr7.g02127.t01#3    mNeoVis.2.1.1.chr7_hap1.g02175.t01#5    mNeoVis.vulgivaga.1.1.chr7.g02130.t01#4    mNeoVis.evergladensis.2.1.chr7.g02124.t01#2    mNeoVis.vulgivaga.2.1.chr7.g02146.t01#1   mNeoVis.3.1.1.chr5.g00435.t01#6       1648   125260220:   mNeoVis.vulgivaga.2.1.chr1.g01441.t01#1    mNeoVis.evergladensis.2.1.chr1.g01440.t01#2    mNeoVis.2.1.1.chr1_hap1.g01473.t01#5    mNeoVis.vulgivaga.1.1.chr1.g01442.t01#4    mNeoVis.3.1.1.chr1.g01421.t01#6   mNeoVis.evergladensis.1.1.chr1.g01443.t01#3       1649   125260221:   mNeoVis.evergladensis.2.1.chr10.g01294.t01#2    mNeoVis.3.1.1.chr10.g01245.t01#6    mNeoVis.evergladensis.1.1.chr10.g01285.t01#3    mNeoVis.vulgivaga.2.1.chr10.g01284.t01#1    mNeoVis.2.1.1.chr9_hap1.g01307.t01#5   mNeoVis.vulgivaga.1.1.chr10.g01271.t01#4       1650   125260222:   mNeoVis.evergladensis.1.1.chr7.g02128.t01#3    mNeoVis.evergladensis.2.1.chr7.g02125.t01#2    mNeoVis.2.1.1.chr7_hap1.g02176.t01#5    mNeoVis.3.1.1.chr7.g02004.t01#6    mNeoVis.vulgivaga.2.1.chr7.g02147.t01#1   mNeoVis.vulgivaga.1.1.chr7.g02131.t01#4       1651   125260223:   mNeoVis.vulgivaga.1.1.chr12.g00112.t01#4    mNeoVis.evergladensis.2.1.chr12.g00107.t01#2   mNeoVis.evergladensis.1.1.chr12.g00106.t01#3         1652   125260224:   mNeoVis.3.1.1.chr7.g02005.t01#6    mNeoVis.evergladensis.2.1.chr7.g02126.t01#2    mNeoVis.vulgivaga.1.1.chr7.g02132.t01#4    mNeoVis.2.1.1.chr7_hap1.g02177.t01#5    mNeoVis.vulgivaga.2.1.chr7.g02148.t01#1   mNeoVis.evergladensis.1.1.chr7.g02129.t01#3       1653   125260225:   mNeoVis.vulgivaga.1.1.chr7.g00533.t01#4    mNeoVis.evergladensis.1.1.chr7.g00534.t01#3    mNeoVis.3.1.1.chr7.g00509.t01#6    mNeoVis.vulgivaga.2.1.chr7.g00528.t01#1    mNeoVis.evergladensis.2.1.chr7.g00525.t01#2   mNeoVis.2.1.1.chr7_hap1.g00525.t01#5       1654   125260226:   mNeoVis.3.1.1.chr1.g01422.t01#6    mNeoVis.2.1.1.chr1_hap1.g01474.t01#5    mNeoVis.evergladensis.1.1.chr1.g01444.t01#3    mNeoVis.vulgivaga.1.1.chr1.g01443.t01#4    mNeoVis.evergladensis.2.1.chr1.g01441.t01#2   mNeoVis.vulgivaga.2.1.chr1.g01442.t01#1       1655   125260227:   mNeoVis.evergladensis.2.1.chr1.g01442.t01#2    mNeoVis.vulgivaga.2.1.chr1.g01443.t01#1    mNeoVis.evergladensis.1.1.chr1.g01445.t01#3    mNeoVis.2.1.1.chr1_hap1.g01475.t01#5    mNeoVis.vulgivaga.1.1.chr1.g01444.t01#4   mNeoVis.3.1.1.chr1.g01423.t01#6       1656   125260228:   mNeoVis.2.1.1.chr7_hap1.g02240.t01#5    mNeoVis.evergladensis.2.1.chr7.g02188.t01#2    mNeoVis.3.1.1.chr7.g02060.t01#6    mNeoVis.evergladensis.1.1.chr7.g02185.t01#3    mNeoVis.vulgivaga.2.1.chr7.g02207.t01#1   mNeoVis.vulgivaga.1.1.chr7.g02192.t01#4       1657   125260229:   mNeoVis.evergladensis.1.1.chr7.g02187.t01#3    mNeoVis.2.1.1.chr7_hap1.g02242.t01#5    mNeoVis.vulgivaga.1.1.chr7.g02194.t01#4    mNeoVis.vulgivaga.2.1.chr7.g02208.t01#1    mNeoVis.evergladensis.2.1.chr7.g02190.t01#2   mNeoVis.3.1.1.chr7.g02062.t01#6       1658   125260230:   mNeoVis.3.1.1.chr15.g00536.t01#6    mNeoVis.evergladensis.2.1.chr15.g00581.t01#2    mNeoVis.2.1.1.chr15_hap1.g00760.t01#5    mNeoVis.vulgivaga.2.1.chr15.g00592.t01#1    mNeoVis.vulgivaga.1.1.chr15.g00584.t01#4   mNeoVis.evergladensis.1.1.chr15.g00030.t01#3       1659   125260231:   mNeoVis.evergladensis.1.1.chr1.g01446.t01#3    mNeoVis.vulgivaga.2.1.chr1.g01444.t01#1    mNeoVis.evergladensis.2.1.chr1.g01443.t01#2    mNeoVis.2.1.1.chr1_hap1.g01476.t01#5    mNeoVis.3.1.1.chr1.g01424.t01#6   mNeoVis.vulgivaga.1.1.chr1.g01445.t01#4       1660   125260232:   mNeoVis.2.1.1.chr7_hap1.g02243.t01#5    mNeoVis.vulgivaga.1.1.chr7.g02195.t01#4    mNeoVis.evergladensis.1.1.chr7.g02188.t01#3    mNeoVis.3.1.1.chr7.g02063.t01#6    mNeoVis.evergladensis.2.1.chr7.g02191.t01#2   mNeoVis.vulgivaga.2.1.chr7.g02209.t01#1       1661   125260233:   mNeoVis.2.1.1.chr1_hap1.g01477.t01#5    mNeoVis.vulgivaga.1.1.chr1.g01446.t01#4    mNeoVis.evergladensis.2.1.chr1.g01444.t01#2    mNeoVis.3.1.1.chr1.g01425.t01#6    mNeoVis.vulgivaga.2.1.chr1.g01445.t01#1   mNeoVis.evergladensis.1.1.chr1.g01447.t01#3       1662   125260234:   mNeoVis.3.1.1.chr7.g02064.t01#6    mNeoVis.vulgivaga.2.1.chr7.g02210.t01#1    mNeoVis.evergladensis.2.1.chr7.g02192.t01#2    mNeoVis.vulgivaga.1.1.chr7.g02196.t01#4    mNeoVis.2.1.1.chr7_hap1.g02244.t01#5   mNeoVis.evergladensis.1.1.chr7.g02189.t01#3       1663   125260235:   mNeoVis.evergladensis.1.1.chr7.g02190.t01#3    mNeoVis.2.1.1.chr7_hap1.g02245.t01#5    mNeoVis.vulgivaga.2.1.chr7.g02211.t01#1    mNeoVis.3.1.1.chr7.g02065.t01#6    mNeoVis.evergladensis.2.1.chr7.g02193.t01#2   mNeoVis.vulgivaga.1.1.chr7.g02197.t01#4       1664   125260236:   mNeoVis.evergladensis.1.1.chr1.g01448.t01#3    mNeoVis.2.1.1.chr1_hap1.g01478.t01#5    mNeoVis.vulgivaga.1.1.chr1.g01447.t01#4    mNeoVis.vulgivaga.2.1.chr1.g01446.t01#1   mNeoVis.3.1.1.chr1.g01426.t01#6       1665   125260237:   mNeoVis.evergladensis.1.1.chr7.g02191.t01#3    mNeoVis.2.1.1.chr7_hap1.g02246.t01#5    mNeoVis.3.1.1.chr7.g02066.t01#6    mNeoVis.vulgivaga.2.1.chr7.g02212.t01#1    mNeoVis.vulgivaga.1.1.chr7.g02198.t01#4   mNeoVis.evergladensis.2.1.chr7.g02194.t01#2       1666   125260238:   mNeoVis.2.1.1.chr7_hap1.g00526.t01#5    mNeoVis.evergladensis.1.1.chr7.g00535.t01#3    mNeoVis.vulgivaga.1.1.chr7.g00534.t01#4    mNeoVis.vulgivaga.2.1.chr7.g00529.t01#1    mNeoVis.evergladensis.2.1.chr7.g00526.t01#2   mNeoVis.3.1.1.chr7.g00510.t01#6       1667   125260239:   mNeoVis.vulgivaga.2.1.chr1.g01447.t01#1    mNeoVis.evergladensis.1.1.chr1.g01449.t01#3    mNeoVis.vulgivaga.1.1.chr1.g01448.t01#4    mNeoVis.2.1.1.chr1_hap1.g01479.t01#5    mNeoVis.evergladensis.2.1.chr1.g01446.t01#2   mNeoVis.3.1.1.chr1.g01427.t01#6       1668   125260240:   mNeoVis.evergladensis.1.1.chr7.g02192.t01#3   mNeoVis.vulgivaga.2.1.chr7.g02213.t01#1             1669   125260241:   mNeoVis.evergladensis.2.1.chr7.g02195.t01#2    mNeoVis.vulgivaga.2.1.chr7.g02214.t01#1    mNeoVis.2.1.1.chr7_hap1.g02247.t01#5    mNeoVis.vulgivaga.1.1.chr7.g02199.t01#4   mNeoVis.3.1.1.chr7.g02067.t01#6       1670   125260242:   mNeoVis.vulgivaga.1.1.chr7.g02200.t01#4    mNeoVis.2.1.1.chr7_hap1.g02248.t01#5    mNeoVis.evergladensis.2.1.chr7.g02196.t01#2    mNeoVis.vulgivaga.2.1.chr7.g02215.t01#1   mNeoVis.3.1.1.chr7.g02068.t01#6       1671   125260243:   mNeoVis.evergladensis.2.1.chr7.g02197.t01#2    mNeoVis.vulgivaga.1.1.chr7.g02201.t01#4    mNeoVis.evergladensis.1.1.chr7.g02194.t01#3    mNeoVis.vulgivaga.2.1.chr7.g02216.t01#1    mNeoVis.2.1.1.chr7_hap1.g02249.t01#5   mNeoVis.3.1.1.chr7.g02069.t01#6       1672   125260244:   mNeoVis.vulgivaga.1.1.chr7.g02202.t01#4    mNeoVis.vulgivaga.2.1.chr7.g02217.t01#1    mNeoVis.evergladensis.2.1.chr7.g02198.t01#2   mNeoVis.2.1.1.chr7_hap1.g02250.t01#5         1673   125260245:   mNeoVis.3.1.1.chr7.g02070.t01#6   mNeoVis.evergladensis.1.1.chr7.g02195.t01#3           1674   125260246:   mNeoVis.vulgivaga.2.1.chr7.g02218.t01#1    mNeoVis.vulgivaga.1.1.chr7.g02203.t01#4    mNeoVis.evergladensis.1.1.chr7.g02196.t01#3    mNeoVis.evergladensis.2.1.chr7.g02199.t01#2    mNeoVis.2.1.1.chr7_hap1.g02251.t01#5   mNeoVis.3.1.1.chr7.g02071.t01#6       1675   125260247:   mNeoVis.3.1.1.chr7.g02072.t01#6    mNeoVis.vulgivaga.1.1.chr7.g02204.t01#4    mNeoVis.vulgivaga.2.1.chr7.g02219.t01#1    mNeoVis.2.1.1.chr7_hap1.g02252.t01#5    mNeoVis.evergladensis.2.1.chr7.g02200.t01#2   mNeoVis.evergladensis.1.1.chr7.g02197.t01#3       1676   125260248:   mNeoVis.vulgivaga.2.1.chr1.g01448.t01#1    mNeoVis.evergladensis.2.1.chr1.g01447.t01#2    mNeoVis.3.1.1.chr1.g01428.t01#6    mNeoVis.2.1.1.chr1_hap1.g01485.t01#5    mNeoVis.evergladensis.1.1.chr1.g01450.t01#3   mNeoVis.vulgivaga.1.1.chr1.g01449.t01#4       1677   125260249:   mNeoVis.vulgivaga.1.1.chr7.g02205.t01#4    mNeoVis.3.1.1.chr7.g02073.t01#6    mNeoVis.evergladensis.2.1.chr7.g02201.t01#2    mNeoVis.vulgivaga.2.1.chr7.g02220.t01#1    mNeoVis.evergladensis.1.1.chr7.g02198.t01#3   mNeoVis.2.1.1.chr7_hap1.g02253.t01#5       1678   125260250:   mNeoVis.vulgivaga.1.1.chr7.g02206.t01#4    mNeoVis.vulgivaga.2.1.chr7.g02221.t01#1    mNeoVis.evergladensis.2.1.chr7.g02202.t01#2    mNeoVis.evergladensis.1.1.chr7.g02199.t01#3   mNeoVis.2.1.1.chr7_hap1.g02254.t01#5       1679   125260251:   mNeoVis.evergladensis.1.1.chr7.g02200.t01#3    mNeoVis.2.1.1.chr7_hap1.g02255.t01#5    mNeoVis.vulgivaga.1.1.chr7.g02207.t01#4    mNeoVis.evergladensis.2.1.chr7.g02203.t01#2    mNeoVis.3.1.1.chr7.g02075.t01#6   mNeoVis.vulgivaga.2.1.chr7.g02222.t01#1       1680   125260252:   mNeoVis.evergladensis.1.1.chr7.g02201.t01#3    mNeoVis.2.1.1.chr7_hap1.g02256.t01#5    mNeoVis.vulgivaga.1.1.chr7.g02208.t01#4    mNeoVis.evergladensis.2.1.chr7.g02204.t01#2    mNeoVis.vulgivaga.2.1.chr7.g02223.t01#1   mNeoVis.3.1.1.chr7.g02076.t01#6       1681   125260253:   mNeoVis.evergladensis.2.1.chr7.g02205.t01#2    mNeoVis.evergladensis.1.1.chr7.g02202.t01#3    mNeoVis.2.1.1.chr7_hap1.g02257.t01#5    mNeoVis.vulgivaga.1.1.chr7.g02209.t01#4    mNeoVis.3.1.1.chr7.g02077.t01#6   mNeoVis.vulgivaga.2.1.chr7.g02224.t01#1       1682   125260254:   mNeoVis.vulgivaga.2.1.chr7.g02225.t01#1    mNeoVis.evergladensis.1.1.chr7.g02203.t01#3    mNeoVis.evergladensis.2.1.chr7.g02206.t01#2    mNeoVis.2.1.1.chr7_hap1.g02258.t01#5    mNeoVis.vulgivaga.1.1.chr7.g02210.t01#4   mNeoVis.3.1.1.chr7.g02078.t01#6       1683   125260255:   mNeoVis.vulgivaga.1.1.chr1.g01450.t01#4    mNeoVis.3.1.1.chr1.g01429.t01#6    mNeoVis.evergladensis.1.1.chr1.g01451.t01#3    mNeoVis.2.1.1.chr1_hap1.g01486.t01#5    mNeoVis.vulgivaga.2.1.chr1.g01449.t01#1   mNeoVis.evergladensis.2.1.chr1.g01448.t01#2       1684   125260256:   mNeoVis.vulgivaga.2.1.chr7.g02226.t01#1    mNeoVis.evergladensis.2.1.chr7.g02208.t01#2    mNeoVis.vulgivaga.1.1.chr7.g02212.t01#4    mNeoVis.evergladensis.1.1.chr7.g02205.t01#3    mNeoVis.3.1.1.chr7.g02081.t01#6   mNeoVis.2.1.1.chr7_hap1.g02260.t01#5       1685   125260257:   mNeoVis.vulgivaga.2.1.chr5.g00489.t01#1    mNeoVis.2.1.1.chr1_hap1.g01487.t01#5    mNeoVis.vulgivaga.1.1.chr5.g00473.t01#4    mNeoVis.evergladensis.2.1.chr1.g01449.t01#2    mNeoVis.vulgivaga.1.1.chr1.g01451.t01#4    mNeoVis.vulgivaga.2.1.chr1.g01450.t01#1    mNeoVis.3.1.1.chr5.g00478.t01#6    mNeoVis.evergladensis.1.1.chr5.g00482.t01#3    mNeoVis.3.1.1.chr1.g01430.t01#6      1686   125260258:   mNeoVis.2.1.1.chr4_hap1.g00682.t01#5    mNeoVis.vulgivaga.2.1.chr5.g00828.t01#1    mNeoVis.3.1.1.chr6.g00689.t01#6    mNeoVis.vulgivaga.1.1.chr12.g00155.t01#4    mNeoVis.evergladensis.1.1.chr11.g00033.t01#3    mNeoVis.2.1.1.chr1_hap1.g01488.t01#5    mNeoVis.evergladensis.1.1.chr2.g01165.t01#3    mNeoVis.2.1.1.chr4_hap1.g00681.t01#5    mNeoVis.vulgivaga.2.1.chr4.g00253.t01#1      1687   125260259:   mNeoVis.3.1.1.chr1.g01431.t01#6    mNeoVis.vulgivaga.2.1.chr8.g00269.t01#1    mNeoVis.evergladensis.2.1.chr1.g01450.t01#2    mNeoVis.3.1.1.chr1.g01453.t01#6    mNeoVis.vulgivaga.2.1.chr3.g00416.t01#1    mNeoVis.vulgivaga.2.1.chr7.g01403.t01#1    mNeoVis.evergladensis.1.1.chr1.g01847.t01#3    mNeoVis.3.1.1.chr1.g00115.t01#6    mNeoVis.vulgivaga.1.1.chr7.g01400.t01#4      1688   125260260:   mNeoVis.3.1.1.chr10.g00296.t01#6   mNeoVis.vulgivaga.2.1.chr10.g00299.t01#1           1689   125260261:   mNeoVis.evergladensis.2.1.chr7.g02209.t01#2    mNeoVis.vulgivaga.2.1.chr7.g02227.t01#1    mNeoVis.3.1.1.chr7.g02082.t01#6    mNeoVis.vulgivaga.1.1.chr7.g02213.t01#4    mNeoVis.evergladensis.1.1.chr7.g02206.t01#3   mNeoVis.2.1.1.chr7_hap1.g02261.t01#5       1690   125260262:   mNeoVis.vulgivaga.2.1.chr7.g02228.t01#1    mNeoVis.evergladensis.2.1.chr7.g02210.t01#2    mNeoVis.2.1.1.chr7_hap1.g02263.t01#5    mNeoVis.vulgivaga.1.1.chr7.g02214.t01#4    mNeoVis.evergladensis.1.1.chr7.g02207.t01#3   mNeoVis.3.1.1.chr7.g02083.t01#6       1691   125260263:   mNeoVis.vulgivaga.1.1.chr1.g01453.t01#4    mNeoVis.evergladensis.2.1.chr1.g01451.t01#2    mNeoVis.3.1.1.chr1.g01432.t01#6    mNeoVis.2.1.1.chr1_hap1.g01489.t01#5    mNeoVis.vulgivaga.2.1.chr1.g01452.t01#1   mNeoVis.evergladensis.1.1.chr1.g01454.t01#3       1692   125260264:   mNeoVis.2.1.1.chr7_hap1.g02178.t01#5    mNeoVis.3.1.1.chr7.g02006.t01#6    mNeoVis.vulgivaga.1.1.chr7.g02133.t01#4    mNeoVis.vulgivaga.2.1.chr7.g02149.t01#1    mNeoVis.evergladensis.2.1.chr7.g02127.t01#2   mNeoVis.evergladensis.1.1.chr7.g02130.t01#3       1693   125260265:   mNeoVis.evergladensis.2.1.chr7.g02128.t01#2    mNeoVis.evergladensis.1.1.chr7.g02131.t01#3    mNeoVis.3.1.1.chr7.g02007.t01#6    mNeoVis.vulgivaga.1.1.chr7.g02134.t01#4    mNeoVis.2.1.1.chr7_hap1.g02179.t01#5   mNeoVis.vulgivaga.2.1.chr7.g02150.t01#1       1694   125260266:   mNeoVis.vulgivaga.1.1.chr3.g00352.t01#4    mNeoVis.2.1.1.chr15_hap1.g00077.t01#5    mNeoVis.3.1.1.chr15.g00071.t01#6    mNeoVis.vulgivaga.2.1.chr3.g00353.t01#1    mNeoVis.evergladensis.2.1.chr3.g00358.t01#2    mNeoVis.vulgivaga.1.1.chr15.g00078.t01#4    mNeoVis.2.1.1.chr8_hap1.g00649.t01#5    mNeoVis.vulgivaga.1.1.chr8.g00628.t01#4    mNeoVis.vulgivaga.2.1.chr8.g00644.t01#1      1695   125260267:   mNeoVis.evergladensis.1.1.chr7.g02132.t01#3    mNeoVis.2.1.1.chr7_hap1.g02180.t01#5    mNeoVis.evergladensis.2.1.chr7.g02129.t01#2    mNeoVis.3.1.1.chr7.g02008.t01#6    mNeoVis.vulgivaga.1.1.chr7.g02135.t01#4   mNeoVis.vulgivaga.2.1.chr7.g02151.t01#1       1696   125260268:   mNeoVis.vulgivaga.2.1.chr1.g01453.t01#1    mNeoVis.3.1.1.chr1.g01433.t01#6    mNeoVis.2.1.1.chr1_hap1.g01490.t01#5    mNeoVis.evergladensis.2.1.chr1.g01452.t01#2    mNeoVis.evergladensis.1.1.chr1.g01455.t01#3   mNeoVis.vulgivaga.1.1.chr1.g01454.t01#4       1697   125260269:   mNeoVis.vulgivaga.2.1.chr7.g02152.t01#1    mNeoVis.evergladensis.1.1.chr7.g02133.t01#3    mNeoVis.3.1.1.chr7.g02009.t01#6    mNeoVis.evergladensis.2.1.chr7.g02130.t01#2    mNeoVis.2.1.1.chr7_hap1.g02181.t01#5    mNeoVis.vulgivaga.1.1.chr7.g02136.t01#4   mNeoVis.evergladensis.1.1.chr7.g02134.t01#3     1698   125260270:   mNeoVis.evergladensis.2.1.chr1.g01453.t01#2    mNeoVis.evergladensis.1.1.chr1.g01456.t01#3    mNeoVis.2.1.1.chr1_hap1.g01491.t01#5    mNeoVis.vulgivaga.2.1.chr1.g01454.t01#1    mNeoVis.vulgivaga.1.1.chr1.g01455.t01#4   mNeoVis.3.1.1.chr1.g01434.t01#6       1699   125260271:   mNeoVis.vulgivaga.1.1.chr12.g00691.t01#4    mNeoVis.2.1.1.chr12_hap1.g00741.t01#5    mNeoVis.evergladensis.1.1.chr12.g00678.t01#3    mNeoVis.vulgivaga.2.1.chr12.g00696.t01#1   mNeoVis.3.1.1.chr12.g00674.t01#6       1700   125260272:   mNeoVis.2.1.1.chr7_hap1.g02182.t01#5    mNeoVis.3.1.1.chr7.g02010.t01#6    mNeoVis.evergladensis.1.1.chr7.g02135.t01#3    mNeoVis.vulgivaga.1.1.chr7.g02137.t01#4    mNeoVis.evergladensis.2.1.chr7.g02131.t01#2   mNeoVis.vulgivaga.2.1.chr7.g02153.t01#1       1701   125260273:   mNeoVis.3.1.1.chr15.g00537.t01#6    mNeoVis.evergladensis.2.1.chr15.g00582.t01#2    mNeoVis.evergladensis.1.1.chr15.g00029.t01#3    mNeoVis.2.1.1.chr15_hap1.g00761.t01#5    mNeoVis.vulgivaga.2.1.chr15.g00593.t01#1   mNeoVis.vulgivaga.1.1.chr15.g00585.t01#4       1702   125260274:   mNeoVis.3.1.1.chr1.g01435.t01#6    mNeoVis.evergladensis.2.1.chr1.g01454.t01#2    mNeoVis.vulgivaga.2.1.chr1.g01455.t01#1    mNeoVis.evergladensis.1.1.chr1.g01457.t01#3    mNeoVis.2.1.1.chr1_hap1.g01492.t01#5   mNeoVis.vulgivaga.1.1.chr1.g01456.t01#4       1703   125260275:   mNeoVis.evergladensis.1.1.chr1.g01458.t01#3    mNeoVis.evergladensis.2.1.chr1.g01455.t01#2    mNeoVis.2.1.1.chr1_hap1.g01493.t01#5    mNeoVis.vulgivaga.2.1.chr1.g01456.t01#1    mNeoVis.3.1.1.chr1.g01436.t01#6   mNeoVis.vulgivaga.1.1.chr1.g01458.t01#4       1704   125260276:   mNeoVis.2.1.1.chr7_hap1.g02183.t01#5    mNeoVis.3.1.1.chr7.g02011.t01#6    mNeoVis.evergladensis.1.1.chr7.g02136.t01#3    mNeoVis.vulgivaga.2.1.chr7.g02154.t01#1    mNeoVis.evergladensis.2.1.chr7.g02132.t01#2   mNeoVis.vulgivaga.1.1.chr7.g02138.t01#4       1705   125260277:   mNeoVis.vulgivaga.2.1.chr1.g01457.t01#1    mNeoVis.2.1.1.chr1_hap1.g01494.t01#5    mNeoVis.3.1.1.chr1.g01437.t01#6    mNeoVis.vulgivaga.1.1.chr1.g01457.t01#4    mNeoVis.evergladensis.2.1.chr1.g01456.t01#2   mNeoVis.evergladensis.1.1.chr1.g01459.t01#3       1706   125260278:   mNeoVis.vulgivaga.2.1.chr7.g02155.t01#1    mNeoVis.vulgivaga.1.1.chr7.g02139.t01#4    mNeoVis.evergladensis.2.1.chr7.g02133.t01#2    mNeoVis.2.1.1.chr7_hap1.g02184.t01#5    mNeoVis.3.1.1.chr7.g02012.t01#6   mNeoVis.evergladensis.1.1.chr7.g02137.t01#3       1707   125260279:   mNeoVis.evergladensis.2.1.chr1.g01457.t01#2    mNeoVis.evergladensis.1.1.chr1.g01460.t01#3    mNeoVis.3.1.1.chr1.g01438.t01#6    mNeoVis.vulgivaga.1.1.chr1.g01459.t01#4    mNeoVis.vulgivaga.2.1.chr1.g01458.t01#1   mNeoVis.2.1.1.chr1_hap1.g01495.t01#5       1708   125260280:   mNeoVis.2.1.1.chr1_hap1.g01496.t01#5    mNeoVis.vulgivaga.2.1.chr1.g01459.t01#1    mNeoVis.evergladensis.1.1.chr1.g01461.t01#3    mNeoVis.evergladensis.2.1.chr1.g01458.t01#2    mNeoVis.3.1.1.chr1.g01439.t01#6   mNeoVis.vulgivaga.1.1.chr1.g01460.t01#4       1709   125260281:   mNeoVis.vulgivaga.1.1.chr7.g02140.t01#4    mNeoVis.evergladensis.1.1.chr7.g02138.t01#3    mNeoVis.vulgivaga.2.1.chr7.g02156.t01#1    mNeoVis.2.1.1.chr7_hap1.g02185.t01#5    mNeoVis.evergladensis.2.1.chr7.g02134.t01#2   mNeoVis.3.1.1.chr7.g02013.t01#6       1710   125260282:   mNeoVis.3.1.1.chr7.g02014.t01#6    mNeoVis.2.1.1.chr7_hap1.g02186.t01#5    mNeoVis.evergladensis.1.1.chr7.g02139.t01#3    mNeoVis.vulgivaga.2.1.chr7.g02157.t01#1    mNeoVis.vulgivaga.1.1.chr7.g02141.t01#4   mNeoVis.evergladensis.2.1.chr7.g02135.t01#2       1711   125260283:   mNeoVis.vulgivaga.2.1.chr7.g02158.t01#1    mNeoVis.evergladensis.2.1.chr7.g02136.t01#2    mNeoVis.evergladensis.1.1.chr7.g02140.t01#3    mNeoVis.2.1.1.chr7_hap1.g02187.t01#5    mNeoVis.3.1.1.chr7.g02015.t01#6   mNeoVis.vulgivaga.1.1.chr7.g02142.t01#4       1712   125260284:   mNeoVis.evergladensis.2.1.chr1.g01459.t01#2    mNeoVis.2.1.1.chr1_hap1.g01497.t01#5    mNeoVis.vulgivaga.2.1.chr1.g01460.t01#1    mNeoVis.evergladensis.1.1.chr1.g01462.t01#3   mNeoVis.vulgivaga.1.1.chr1.g01461.t01#4       1713   125260285:   mNeoVis.vulgivaga.1.1.chr7.g00535.t01#4    mNeoVis.evergladensis.1.1.chr7.g00536.t01#3    mNeoVis.3.1.1.chr7.g00511.t01#6    mNeoVis.vulgivaga.2.1.chr7.g00530.t01#1    mNeoVis.evergladensis.2.1.chr7.g00527.t01#2   mNeoVis.2.1.1.chr7_hap1.g00527.t01#5       1714   125260286:   mNeoVis.3.1.1.chr7.g02016.t01#6    mNeoVis.evergladensis.2.1.chr7.g02137.t01#2    mNeoVis.2.1.1.chr7_hap1.g02188.t01#5    mNeoVis.vulgivaga.1.1.chr7.g02143.t01#4    mNeoVis.vulgivaga.2.1.chr7.g02159.t01#1   mNeoVis.evergladensis.1.1.chr7.g02141.t01#3       1715   125260287:   mNeoVis.vulgivaga.1.1.chr7.g02144.t01#4    mNeoVis.vulgivaga.2.1.chr7.g02160.t01#1    mNeoVis.evergladensis.2.1.chr7.g02138.t01#2    mNeoVis.evergladensis.1.1.chr7.g02142.t01#3   mNeoVis.2.1.1.chr7_hap1.g02189.t01#5       1716   125260288:   mNeoVis.vulgivaga.2.1.chr7.g02161.t01#1    mNeoVis.evergladensis.1.1.chr7.g02143.t01#3    mNeoVis.2.1.1.chr7_hap1.g02190.t01#5    mNeoVis.evergladensis.2.1.chr7.g02139.t01#2   mNeoVis.vulgivaga.1.1.chr7.g02145.t01#4       1717   125260289:   mNeoVis.vulgivaga.2.1.chr7.g02162.t01#1    mNeoVis.3.1.1.chr7.g02017.t01#6    mNeoVis.evergladensis.1.1.chr7.g02144.t01#3    mNeoVis.evergladensis.2.1.chr7.g02140.t01#2    mNeoVis.2.1.1.chr7_hap1.g02191.t01#5   mNeoVis.vulgivaga.1.1.chr7.g02146.t01#4       1718   125260290:   mNeoVis.3.1.1.chr7.g00512.t01#6    mNeoVis.2.1.1.chr7_hap1.g00528.t01#5    mNeoVis.vulgivaga.2.1.chr7.g00531.t01#1    mNeoVis.vulgivaga.1.1.chr7.g00536.t01#4    mNeoVis.evergladensis.1.1.chr7.g00537.t01#3   mNeoVis.evergladensis.2.1.chr7.g00528.t01#2       1719   125260291:   mNeoVis.2.1.1.chr7_hap1.g02192.t01#5    mNeoVis.evergladensis.2.1.chr7.g02141.t01#2    mNeoVis.vulgivaga.2.1.chr7.g02163.t01#1    mNeoVis.vulgivaga.1.1.chr7.g02147.t01#4    mNeoVis.3.1.1.chr7.g02018.t01#6   mNeoVis.evergladensis.1.1.chr7.g02145.t01#3       1720   125260292:   mNeoVis.vulgivaga.1.1.chr1.g01462.t01#4    mNeoVis.evergladensis.2.1.chr1.g01460.t01#2    mNeoVis.3.1.1.chr1.g01441.t01#6    mNeoVis.evergladensis.1.1.chr1.g01463.t01#3    mNeoVis.vulgivaga.2.1.chr1.g01461.t01#1   mNeoVis.2.1.1.chr1_hap1.g01498.t01#5       1721   125260293:   mNeoVis.vulgivaga.2.1.chr11.g00261.t01#1    mNeoVis.2.1.1.chr11_hap1.g00262.t01#5    mNeoVis.vulgivaga.1.1.chr11.g00257.t01#4    mNeoVis.evergladensis.2.1.chr11.g00260.t01#2    mNeoVis.evergladensis.1.1.chr11.g00262.t01#3   mNeoVis.3.1.1.chr11.g00233.t01#6       1722   125260294:   mNeoVis.vulgivaga.1.1.chr7.g02148.t01#4    mNeoVis.evergladensis.1.1.chr7.g02147.t01#3    mNeoVis.vulgivaga.2.1.chr7.g02165.t01#1    mNeoVis.evergladensis.2.1.chr7.g02142.t01#2    mNeoVis.3.1.1.chr7.g02019.t01#6    mNeoVis.vulgivaga.2.1.chr7.g02164.t01#1    mNeoVis.evergladensis.1.1.chr7.g02146.t01#3    mNeoVis.2.1.1.chr7_hap1.g02194.t01#5    mNeoVis.vulgivaga.1.1.chr7.g02149.t01#4      1723   125260295:   mNeoVis.2.1.1.chr1_hap1.g01499.t01#5    mNeoVis.evergladensis.1.1.chr1.g01464.t01#3    mNeoVis.3.1.1.chr1.g01444.t01#6    mNeoVis.evergladensis.2.1.chr1.g01463.t01#2    mNeoVis.vulgivaga.1.1.chr1.g01465.t01#4   mNeoVis.vulgivaga.2.1.chr1.g01462.t01#1       1724   125260296:   mNeoVis.vulgivaga.2.1.chr7.g02166.t01#1    mNeoVis.evergladensis.2.1.chr7.g02144.t01#2    mNeoVis.2.1.1.chr7_hap1.g02195.t01#5    mNeoVis.vulgivaga.1.1.chr7.g02150.t01#4    mNeoVis.3.1.1.chr7.g02020.t01#6   mNeoVis.evergladensis.1.1.chr7.g02148.t01#3       1725   125260297:   mNeoVis.vulgivaga.2.1.chr1.g01463.t01#1    mNeoVis.2.1.1.chr1_hap1.g01500.t01#5    mNeoVis.evergladensis.1.1.chr1.g01465.t01#3    mNeoVis.evergladensis.2.1.chr1.g01464.t01#2    mNeoVis.3.1.1.chr1.g01445.t01#6   mNeoVis.vulgivaga.1.1.chr1.g01466.t01#4       1726   125260298:   mNeoVis.3.1.1.chr7.g02025.t01#6    mNeoVis.vulgivaga.2.1.chr7.g02174.t01#1    mNeoVis.evergladensis.2.1.chr7.g02149.t01#2    mNeoVis.evergladensis.1.1.chr7.g02150.t01#3    mNeoVis.3.1.1.chr7.g02024.t01#6    mNeoVis.2.1.1.chr7_hap1.g02202.t01#5    mNeoVis.evergladensis.2.1.chr7.g02151.t01#2    mNeoVis.2.1.1.chr7_hap1.g02200.t01#5    mNeoVis.vulgivaga.1.1.chr7.g02158.t01#4      1727   125260299:   mNeoVis.2.1.1.chr7_hap1.g02203.t01#5    mNeoVis.evergladensis.2.1.chr7.g02152.t01#2    mNeoVis.vulgivaga.1.1.chr7.g02157.t01#4    mNeoVis.evergladensis.2.1.chr7.g02150.t01#2    mNeoVis.vulgivaga.2.1.chr7.g02175.t01#1    mNeoVis.2.1.1.chr7_hap1.g02201.t01#5    mNeoVis.vulgivaga.1.1.chr7.g02159.t01#4    mNeoVis.vulgivaga.2.1.chr7.g02172.t01#1    mNeoVis.evergladensis.1.1.chr7.g02151.t01#3      1728   125260300:   mNeoVis.evergladensis.2.1.chr7.g02153.t01#2    mNeoVis.vulgivaga.2.1.chr7.g02173.t01#1    mNeoVis.evergladensis.1.1.chr7.g02152.t01#3    mNeoVis.3.1.1.chr7.g02026.t01#6   mNeoVis.2.1.1.chr7_hap1.g02204.t01#5       1729   125260301:   mNeoVis.2.1.1.chr7_hap1.g02196.t01#5    mNeoVis.evergladensis.1.1.chr7.g02149.t01#3   mNeoVis.vulgivaga.2.1.chr7.g02167.t01#1           1730   125260302:   mNeoVis.vulgivaga.1.1.chr7.g02151.t01#4    mNeoVis.evergladensis.2.1.chr7.g02145.t01#2   mNeoVis.evergladensis.2.1.chr7.g02147.t01#2         1731   125260303:   mNeoVis.2.1.1.chr7_hap1.g02199.t01#5    mNeoVis.vulgivaga.2.1.chr7.g02171.t01#1   mNeoVis.vulgivaga.1.1.chr7.g02154.t01#4           1732   125260304:   mNeoVis.vulgivaga.1.1.chr7.g02160.t01#4   mNeoVis.3.1.1.chr7.g02027.t01#6             1733   125260305:  mNeoVis.vulgivaga.2.1.chr7.g02170.t01#1               1734   125260306:  mNeoVis.2.1.1.chr7_hap1.g02198.t01#5               1735   125260307:  mNeoVis.3.1.1.chr7.g02022.t01#6               1736   125260308:  mNeoVis.evergladensis.2.1.chr7.g02148.t01#2             1737   125260309:   mNeoVis.vulgivaga.2.1.chr7.g02169.t01#1    mNeoVis.vulgivaga.1.1.chr7.g02152.t01#4    mNeoVis.vulgivaga.2.1.chr7.g02168.t01#1    mNeoVis.3.1.1.chr7.g02021.t01#6    mNeoVis.evergladensis.2.1.chr7.g02146.t01#2   mNeoVis.2.1.1.chr7_hap1.g02197.t01#5       1738   125260310:   mNeoVis.vulgivaga.2.1.chr1.g01464.t01#1    mNeoVis.evergladensis.1.1.chr1.g01466.t01#3    mNeoVis.vulgivaga.1.1.chr1.g01467.t01#4    mNeoVis.evergladensis.2.1.chr1.g01465.t01#2    mNeoVis.3.1.1.chr1.g01446.t01#6   mNeoVis.2.1.1.chr1_hap1.g01501.t01#5       1739   125260311:   mNeoVis.evergladensis.1.1.chr15.g00028.t01#3    mNeoVis.vulgivaga.1.1.chr15.g00586.t01#4    mNeoVis.vulgivaga.2.1.chr15.g00594.t01#1    mNeoVis.3.1.1.chr15.g00538.t01#6    mNeoVis.evergladensis.2.1.chr15.g00583.t01#2   mNeoVis.2.1.1.chr15_hap1.g00762.t01#5       1740   125260312:   mNeoVis.3.1.1.chr1.g01447.t01#6    mNeoVis.evergladensis.2.1.chr1.g01467.t01#2    mNeoVis.2.1.1.chr1_hap1.g01503.t01#5    mNeoVis.vulgivaga.2.1.chr1.g01465.t01#1    mNeoVis.vulgivaga.1.1.chr1.g01468.t01#4   mNeoVis.evergladensis.1.1.chr1.g01467.t01#3       1741   125260313:   mNeoVis.evergladensis.2.1.chr1.g01468.t01#2    mNeoVis.2.1.1.chr1_hap1.g01504.t01#5    mNeoVis.vulgivaga.1.1.chr1.g01469.t01#4    mNeoVis.evergladensis.1.1.chr1.g01468.t01#3    mNeoVis.vulgivaga.2.1.chr1.g01466.t01#1   mNeoVis.3.1.1.chr1.g01448.t01#6       1742   125260314:   mNeoVis.3.1.1.chr7.g02028.t01#6    mNeoVis.evergladensis.2.1.chr7.g02154.t01#2    mNeoVis.vulgivaga.2.1.chr7.g02176.t01#1    mNeoVis.2.1.1.chr7_hap1.g02205.t01#5   mNeoVis.evergladensis.1.1.chr7.g02154.t01#3       1743   125260315:   mNeoVis.vulgivaga.1.1.chr7.g02161.t01#4    mNeoVis.2.1.1.chr7_hap1.g02206.t01#5    mNeoVis.evergladensis.2.1.chr7.g02155.t01#2    mNeoVis.evergladensis.1.1.chr7.g02155.t01#3    mNeoVis.3.1.1.chr7.g02029.t01#6   mNeoVis.vulgivaga.2.1.chr7.g02177.t01#1       1744   125260316:   mNeoVis.vulgivaga.2.1.chr7.g02178.t01#1    mNeoVis.evergladensis.1.1.chr7.g02156.t01#3    mNeoVis.3.1.1.chr7.g02030.t01#6    mNeoVis.evergladensis.2.1.chr7.g02156.t01#2    mNeoVis.vulgivaga.1.1.chr7.g02162.t01#4   mNeoVis.2.1.1.chr7_hap1.g02207.t01#5       1745   125260317:   mNeoVis.vulgivaga.1.1.chr1.g01470.t01#4    mNeoVis.3.1.1.chr1.g01449.t01#6    mNeoVis.evergladensis.2.1.chr1.g01469.t01#2    mNeoVis.vulgivaga.2.1.chr1.g01467.t01#1    mNeoVis.evergladensis.1.1.chr1.g01469.t01#3   mNeoVis.2.1.1.chr1_hap1.g01505.t01#5       1746   125260318:   mNeoVis.evergladensis.1.1.chr7.g02157.t01#3    mNeoVis.vulgivaga.1.1.chr7.g02163.t01#4    mNeoVis.2.1.1.chr7_hap1.g02208.t01#5    mNeoVis.evergladensis.2.1.chr7.g02157.t01#2    mNeoVis.vulgivaga.2.1.chr7.g02179.t01#1   mNeoVis.3.1.1.chr7.g02031.t01#6       1747   125260319:   mNeoVis.2.1.1.chr7_hap1.g02209.t01#5    mNeoVis.3.1.1.chr7.g02032.t01#6    mNeoVis.evergladensis.2.1.chr7.g02158.t01#2    mNeoVis.vulgivaga.2.1.chr7.g02180.t01#1    mNeoVis.evergladensis.1.1.chr7.g02158.t01#3   mNeoVis.vulgivaga.1.1.chr7.g02164.t01#4       1748   125260320:   mNeoVis.2.1.1.chr7_hap1.g02210.t01#5    mNeoVis.vulgivaga.2.1.chr7.g02181.t01#1    mNeoVis.3.1.1.chr7.g02033.t01#6    mNeoVis.vulgivaga.1.1.chr7.g02165.t01#4    mNeoVis.evergladensis.1.1.chr7.g02159.t01#3   mNeoVis.evergladensis.2.1.chr7.g02159.t01#2       1749   125260321:   mNeoVis.3.1.1.chr7.g00513.t01#6    mNeoVis.vulgivaga.2.1.chr7.g00532.t01#1    mNeoVis.vulgivaga.1.1.chr7.g00537.t01#4    mNeoVis.evergladensis.2.1.chr7.g00529.t01#2    mNeoVis.2.1.1.chr7_hap1.g00529.t01#5   mNeoVis.evergladensis.1.1.chr7.g00538.t01#3       1750   125260322:   mNeoVis.vulgivaga.1.1.chr7.g02166.t01#4    mNeoVis.vulgivaga.2.1.chr7.g02182.t01#1    mNeoVis.evergladensis.2.1.chr7.g02160.t01#2    mNeoVis.2.1.1.chr7_hap1.g02211.t01#5    mNeoVis.3.1.1.chr7.g02034.t01#6   mNeoVis.evergladensis.1.1.chr7.g02160.t01#3       1751   125260323:   mNeoVis.evergladensis.2.1.chr7.g02161.t01#2    mNeoVis.evergladensis.1.1.chr7.g02161.t01#3    mNeoVis.3.1.1.chr7.g02036.t01#6    mNeoVis.vulgivaga.1.1.chr7.g02167.t01#4    mNeoVis.2.1.1.chr7_hap1.g02212.t01#5   mNeoVis.vulgivaga.2.1.chr7.g02183.t01#1       1752   125260324:   mNeoVis.2.1.1.chr7_hap1.g02213.t01#5    mNeoVis.vulgivaga.2.1.chr7.g02184.t01#1    mNeoVis.evergladensis.1.1.chr7.g02162.t01#3    mNeoVis.evergladensis.2.1.chr7.g02162.t01#2    mNeoVis.3.1.1.chr7.g02037.t01#6   mNeoVis.vulgivaga.1.1.chr7.g02168.t01#4       1753   125260325:   mNeoVis.2.1.1.chr1_hap1.g01506.t01#5    mNeoVis.3.1.1.chr1.g01450.t01#6    mNeoVis.vulgivaga.2.1.chr1.g01468.t01#1    mNeoVis.evergladensis.2.1.chr1.g01470.t01#2    mNeoVis.evergladensis.1.1.chr1.g01470.t01#3   mNeoVis.vulgivaga.1.1.chr1.g01471.t01#4       1754   125260326:   mNeoVis.evergladensis.2.1.chr7.g02163.t01#2    mNeoVis.evergladensis.1.1.chr7.g02163.t01#3    mNeoVis.vulgivaga.1.1.chr7.g02169.t01#4    mNeoVis.vulgivaga.2.1.chr7.g02185.t01#1   mNeoVis.2.1.1.chr7_hap1.g02214.t01#5       1755   125260327:   mNeoVis.evergladensis.1.1.chr7.g02164.t01#3    mNeoVis.vulgivaga.1.1.chr7.g02170.t01#4    mNeoVis.evergladensis.2.1.chr7.g02164.t01#2    mNeoVis.3.1.1.chr7.g02039.t01#6    mNeoVis.vulgivaga.2.1.chr7.g02186.t01#1   mNeoVis.2.1.1.chr7_hap1.g02215.t01#5       1756   125260328:   mNeoVis.2.1.1.chr7_hap1.g02216.t01#5    mNeoVis.evergladensis.2.1.chr7.g02165.t01#2    mNeoVis.3.1.1.chr7.g02040.t01#6    mNeoVis.evergladensis.1.1.chr7.g02165.t01#3    mNeoVis.vulgivaga.1.1.chr7.g02171.t01#4   mNeoVis.vulgivaga.2.1.chr7.g02187.t01#1       1757   125260329:   mNeoVis.evergladensis.2.1.chr7.g02172.t01#2    mNeoVis.evergladensis.2.1.chr7.g02169.t01#2    mNeoVis.3.1.1.chr7.g02043.t01#6    mNeoVis.2.1.1.chr7_hap1.g02225.t01#5    mNeoVis.vulgivaga.2.1.chr7.g02191.t01#1    mNeoVis.evergladensis.2.1.chr7.g02171.t01#2    mNeoVis.evergladensis.2.1.chr7.g02170.t01#2    mNeoVis.evergladensis.2.1.chr7.g02168.t01#2    mNeoVis.2.1.1.chr7_hap1.g02220.t01#5      1758   125260330:   mNeoVis.evergladensis.1.1.chr7.g02166.t01#3    mNeoVis.vulgivaga.2.1.chr7.g02188.t01#1    mNeoVis.3.1.1.chr7.g02041.t01#6    mNeoVis.vulgivaga.1.1.chr7.g02172.t01#4    mNeoVis.2.1.1.chr7_hap1.g02217.t01#5   mNeoVis.evergladensis.2.1.chr7.g02166.t01#2       1759   125260331:   mNeoVis.vulgivaga.2.1.chr7.g02189.t01#1    mNeoVis.evergladensis.2.1.chr7.g02167.t01#2    mNeoVis.3.1.1.chr7.g02042.t01#6    mNeoVis.vulgivaga.1.1.chr7.g02173.t01#4    mNeoVis.2.1.1.chr7_hap1.g02218.t01#5   mNeoVis.evergladensis.1.1.chr7.g02167.t01#3       1760   125260332:   mNeoVis.vulgivaga.1.1.chr7.g02178.t01#4    mNeoVis.2.1.1.chr7_hap1.g02226.t01#5    mNeoVis.3.1.1.chr7.g02046.t01#6    mNeoVis.vulgivaga.2.1.chr7.g02193.t01#1   mNeoVis.evergladensis.2.1.chr7.g02174.t01#2       1761   125260333:   mNeoVis.vulgivaga.2.1.chr7.g02194.t01#1    mNeoVis.vulgivaga.1.1.chr7.g02179.t01#4    mNeoVis.evergladensis.1.1.chr7.g02171.t01#3    mNeoVis.2.1.1.chr7_hap1.g02227.t01#5    mNeoVis.3.1.1.chr7.g02047.t01#6   mNeoVis.evergladensis.2.1.chr7.g02175.t01#2       1762   125260334:   mNeoVis.evergladensis.2.1.chr7.g02180.t01#2    mNeoVis.vulgivaga.2.1.chr7.g02198.t01#1    mNeoVis.3.1.1.chr7.g02050.t01#6    mNeoVis.evergladensis.2.1.chr7.g02179.t01#2    mNeoVis.vulgivaga.2.1.chr7.g02197.t01#1    mNeoVis.2.1.1.chr7_hap1.g02233.t01#5    mNeoVis.vulgivaga.2.1.chr7.g02199.t01#1    mNeoVis.evergladensis.1.1.chr7.g02173.t01#3    mNeoVis.2.1.1.chr7_hap1.g02232.t01#5      1763   125260335:   mNeoVis.evergladensis.2.1.chr7.g02178.t01#2    mNeoVis.evergladensis.1.1.chr7.g02174.t01#3    mNeoVis.2.1.1.chr7_hap1.g02230.t01#5    mNeoVis.3.1.1.chr7.g02049.t01#6    mNeoVis.vulgivaga.1.1.chr7.g02181.t01#4   mNeoVis.vulgivaga.2.1.chr7.g02196.t01#1       1764   125260336:   mNeoVis.evergladensis.2.1.chr7.g02176.t01#2    mNeoVis.2.1.1.chr7_hap1.g02228.t01#5    mNeoVis.vulgivaga.2.1.chr7.g02195.t01#1    mNeoVis.vulgivaga.1.1.chr7.g02180.t01#4   mNeoVis.evergladensis.1.1.chr7.g02172.t01#3       1765   125260337:   mNeoVis.evergladensis.1.1.chr7.g02177.t01#3    mNeoVis.3.1.1.chr7.g02052.t01#6   mNeoVis.3.1.1.chr7.g02051.t01#6           1766   125260338:   mNeoVis.evergladensis.1.1.chr7.g00539.t01#3    mNeoVis.evergladensis.2.1.chr7.g00530.t01#2    mNeoVis.3.1.1.chr7.g00514.t01#6    mNeoVis.vulgivaga.2.1.chr7.g00533.t01#1    mNeoVis.vulgivaga.1.1.chr7.g00538.t01#4   mNeoVis.2.1.1.chr7_hap1.g00530.t01#5       1767   125260339:   mNeoVis.2.1.1.chr15_hap1.g00763.t01#5    mNeoVis.3.1.1.chr15.g00540.t01#6    mNeoVis.vulgivaga.2.1.chr15.g00595.t01#1    mNeoVis.evergladensis.1.1.chr15.g00027.t01#3   mNeoVis.evergladensis.2.1.chr15.g00584.t01#2       1768   125260340:   mNeoVis.3.1.1.chr1.g01451.t01#6    mNeoVis.evergladensis.2.1.chr1.g01471.t01#2    mNeoVis.evergladensis.1.1.chr1.g01471.t01#3    mNeoVis.vulgivaga.2.1.chr1.g01469.t01#1    mNeoVis.2.1.1.chr1_hap1.g01507.t01#5   mNeoVis.vulgivaga.1.1.chr1.g01472.t01#4       1769   125260341:   mNeoVis.vulgivaga.2.1.chr7.g02200.t01#1    mNeoVis.3.1.1.chr7.g02053.t01#6    mNeoVis.vulgivaga.1.1.chr7.g02185.t01#4    mNeoVis.2.1.1.chr7_hap1.g02234.t01#5    mNeoVis.evergladensis.2.1.chr7.g02181.t01#2   mNeoVis.evergladensis.1.1.chr7.g02178.t01#3       1770   125260342:   mNeoVis.evergladensis.1.1.chr7.g02179.t01#3    mNeoVis.3.1.1.chr7.g02054.t01#6    mNeoVis.vulgivaga.2.1.chr7.g02201.t01#1    mNeoVis.vulgivaga.1.1.chr7.g02186.t01#4    mNeoVis.evergladensis.2.1.chr7.g02182.t01#2   mNeoVis.2.1.1.chr7_hap1.g02235.t01#5       1771   125260343:   mNeoVis.vulgivaga.1.1.chr1.g01473.t01#4    mNeoVis.3.1.1.chr1.g01452.t01#6    mNeoVis.evergladensis.2.1.chr1.g01472.t01#2    mNeoVis.2.1.1.chr1_hap1.g01508.t01#5    mNeoVis.vulgivaga.2.1.chr1.g01470.t01#1   mNeoVis.evergladensis.1.1.chr1.g01473.t01#3       1772   125260344:   mNeoVis.vulgivaga.2.1.chr7.g02202.t01#1    mNeoVis.2.1.1.chr7_hap1.g02236.t01#5    mNeoVis.evergladensis.2.1.chr7.g02183.t01#2    mNeoVis.3.1.1.chr7.g02055.t01#6    mNeoVis.evergladensis.1.1.chr7.g02180.t01#3   mNeoVis.vulgivaga.1.1.chr7.g02187.t01#4       1773   125260345:   mNeoVis.3.1.1.chr7.g02056.t01#6    mNeoVis.evergladensis.1.1.chr7.g02181.t01#3    mNeoVis.vulgivaga.1.1.chr7.g02188.t01#4    mNeoVis.evergladensis.2.1.chr7.g02184.t01#2    mNeoVis.vulgivaga.2.1.chr7.g02203.t01#1   mNeoVis.2.1.1.chr7_hap1.g02237.t01#5       1774   125260346:   mNeoVis.vulgivaga.1.1.chr7.g02189.t01#4   mNeoVis.vulgivaga.2.1.chr7.g02204.t01#1             1775   125260347:   mNeoVis.vulgivaga.1.1.chr7.g02190.t01#4    mNeoVis.evergladensis.2.1.chr7.g02185.t01#2    mNeoVis.2.1.1.chr7_hap1.g02238.t01#5    mNeoVis.vulgivaga.2.1.chr7.g02205.t01#1    mNeoVis.evergladensis.1.1.chr7.g02182.t01#3   mNeoVis.3.1.1.chr7.g02057.t01#6       1776   125260348:   mNeoVis.vulgivaga.1.1.chr7.g02191.t01#4    mNeoVis.2.1.1.chr7_hap1.g02239.t01#5    mNeoVis.vulgivaga.2.1.chr7.g02206.t01#1    mNeoVis.evergladensis.1.1.chr7.g02184.t01#3   mNeoVis.3.1.1.chr7.g02059.t01#6       1777   125260349:   mNeoVis.vulgivaga.1.1.chr1.g01474.t01#4    mNeoVis.evergladensis.1.1.chr1.g01474.t01#3    mNeoVis.2.1.1.chr1_hap1.g01509.t01#5    mNeoVis.evergladensis.2.1.chr1.g01473.t01#2   mNeoVis.vulgivaga.2.1.chr1.g01471.t01#1       1778   125260350:   mNeoVis.vulgivaga.1.1.chr1.g01394.t01#4    mNeoVis.3.1.1.chr1.g01372.t01#6    mNeoVis.evergladensis.1.1.chr1.g01395.t01#3    mNeoVis.evergladensis.2.1.chr1.g01389.t01#2    mNeoVis.2.1.1.chr1_hap1.g01427.t01#5   mNeoVis.vulgivaga.2.1.chr1.g01393.t01#1       1779   125260351:   mNeoVis.vulgivaga.2.1.chr7.g01999.t01#1    mNeoVis.vulgivaga.1.1.chr7.g01993.t01#4    mNeoVis.3.1.1.chr7.g01883.t01#6    mNeoVis.evergladensis.2.1.chr7.g01981.t01#2   mNeoVis.2.1.1.chr7_hap1.g02017.t01#5       1780   125260352:   mNeoVis.vulgivaga.2.1.chr1.g01394.t01#1    mNeoVis.evergladensis.1.1.chr1.g01396.t01#3    mNeoVis.2.1.1.chr1_hap1.g01428.t01#5    mNeoVis.3.1.1.chr1.g01373.t01#6    mNeoVis.vulgivaga.1.1.chr1.g01395.t01#4   mNeoVis.evergladensis.2.1.chr1.g01390.t01#2       1781   125260353:   mNeoVis.evergladensis.1.1.chr1.g01397.t01#3    mNeoVis.3.1.1.chr1.g01374.t01#6    mNeoVis.vulgivaga.1.1.chr1.g01396.t01#4    mNeoVis.2.1.1.chr1_hap1.g01429.t01#5    mNeoVis.vulgivaga.2.1.chr1.g01395.t01#1   mNeoVis.evergladensis.2.1.chr1.g01391.t01#2       1782   125260354:   mNeoVis.evergladensis.1.1.chr1.g01398.t01#3    mNeoVis.vulgivaga.1.1.chr1.g01397.t01#4    mNeoVis.3.1.1.chr1.g01375.t01#6    mNeoVis.vulgivaga.2.1.chr1.g01396.t01#1    mNeoVis.2.1.1.chr1_hap1.g01430.t01#5   mNeoVis.evergladensis.2.1.chr1.g01392.t01#2       1783   125260355:   mNeoVis.2.1.1.chr7_hap1.g00531.t01#5    mNeoVis.evergladensis.1.1.chr7.g00540.t01#3    mNeoVis.evergladensis.2.1.chr7.g00531.t01#2    mNeoVis.vulgivaga.1.1.chr7.g00539.t01#4    mNeoVis.vulgivaga.2.1.chr7.g00534.t01#1   mNeoVis.3.1.1.chr7.g00515.t01#6       1784   125260356:   mNeoVis.vulgivaga.2.1.chr1.g01397.t01#1    mNeoVis.evergladensis.2.1.chr1.g01393.t01#2    mNeoVis.3.1.1.chr1.g01376.t01#6    mNeoVis.vulgivaga.1.1.chr1.g01398.t01#4    mNeoVis.2.1.1.chr1_hap1.g01431.t01#5   mNeoVis.evergladensis.1.1.chr1.g01399.t01#3       1785   125260357:   mNeoVis.vulgivaga.2.1.chr7.g02000.t01#1   mNeoVis.evergladensis.1.1.chr7.g01988.t01#3           1786   125260358:   mNeoVis.3.1.1.chr1.g01377.t01#6    mNeoVis.evergladensis.2.1.chr1.g01394.t01#2    mNeoVis.vulgivaga.2.1.chr1.g01398.t01#1    mNeoVis.evergladensis.1.1.chr1.g01400.t01#3    mNeoVis.2.1.1.chr1_hap1.g01432.t01#5   mNeoVis.vulgivaga.1.1.chr1.g01399.t01#4       1787   125260359:   mNeoVis.vulgivaga.1.1.chr1.g01400.t01#4    mNeoVis.evergladensis.2.1.chr1.g01395.t01#2    mNeoVis.2.1.1.chr1_hap1.g01433.t01#5    mNeoVis.vulgivaga.2.1.chr1.g01399.t01#1    mNeoVis.evergladensis.1.1.chr1.g01401.t01#3   mNeoVis.3.1.1.chr1.g01378.t01#6       1788   125260360:   mNeoVis.vulgivaga.2.1.chr1.g01400.t01#1    mNeoVis.evergladensis.1.1.chr1.g01402.t01#3    mNeoVis.3.1.1.chr1.g01379.t01#6    mNeoVis.vulgivaga.1.1.chr1.g01401.t01#4    mNeoVis.evergladensis.2.1.chr1.g01396.t01#2   mNeoVis.2.1.1.chr1_hap1.g01434.t01#5       1789   125260361:   mNeoVis.evergladensis.1.1.chr7.g01989.t01#3    mNeoVis.evergladensis.2.1.chr7.g01982.t01#2    mNeoVis.3.1.1.chr7.g01884.t01#6    mNeoVis.2.1.1.chr7_hap1.g02018.t01#5    mNeoVis.vulgivaga.1.1.chr7.g01994.t01#4   mNeoVis.vulgivaga.2.1.chr7.g02001.t01#1       1790   125260362:   mNeoVis.vulgivaga.1.1.chr1.g01402.t01#4    mNeoVis.3.1.1.chr1.g01380.t01#6    mNeoVis.evergladensis.2.1.chr1.g01397.t01#2    mNeoVis.evergladensis.1.1.chr1.g01403.t01#3    mNeoVis.2.1.1.chr1_hap1.g01435.t01#5   mNeoVis.vulgivaga.2.1.chr1.g01401.t01#1       1791   125260363:   mNeoVis.evergladensis.1.1.chr1.g01404.t01#3    mNeoVis.vulgivaga.1.1.chr1.g01403.t01#4    mNeoVis.2.1.1.chr1_hap1.g01436.t01#5    mNeoVis.3.1.1.chr1.g01382.t01#6    mNeoVis.vulgivaga.2.1.chr1.g01402.t01#1   mNeoVis.evergladensis.2.1.chr1.g01398.t01#2       1792   125260364:   mNeoVis.evergladensis.2.1.chr7.g01991.t01#2    mNeoVis.evergladensis.1.1.chr7.g01999.t01#3    mNeoVis.2.1.1.chr7_hap1.g02029.t01#5    mNeoVis.vulgivaga.1.1.chr7.g02006.t01#4    mNeoVis.3.1.1.chr7.g01894.t01#6    mNeoVis.evergladensis.1.1.chr7.g01998.t01#3    mNeoVis.vulgivaga.2.1.chr7.g02011.t01#1    mNeoVis.vulgivaga.2.1.chr7.g02013.t01#1    mNeoVis.2.1.1.chr7_hap1.g02028.t01#5      1793   125260365:   mNeoVis.vulgivaga.1.1.chr7.g02007.t01#4    mNeoVis.vulgivaga.2.1.chr7.g02002.t01#1    mNeoVis.evergladensis.2.1.chr7.g01990.t01#2    mNeoVis.2.1.1.chr7_hap1.g02030.t01#5   mNeoVis.evergladensis.1.1.chr7.g02000.t01#3       1794   125260366:   mNeoVis.evergladensis.1.1.chr7.g02037.t01#3    mNeoVis.evergladensis.2.1.chr7.g02025.t01#2    mNeoVis.evergladensis.1.1.chr7.g02036.t01#3    mNeoVis.3.1.1.chr7.g01923.t01#6    mNeoVis.vulgivaga.1.1.chr7.g02032.t01#4    mNeoVis.vulgivaga.2.1.chr7.g02044.t01#1    mNeoVis.evergladensis.1.1.chr7.g02035.t01#3    mNeoVis.evergladensis.2.1.chr7.g02029.t01#2    mNeoVis.evergladensis.2.1.chr7.g02024.t01#2      1795   125260367:   mNeoVis.evergladensis.1.1.chr7.g02017.t01#3    mNeoVis.2.1.1.chr7_hap1.g02046.t01#5    mNeoVis.evergladensis.2.1.chr7.g02005.t01#2    mNeoVis.vulgivaga.1.1.chr7.g02021.t01#4    mNeoVis.vulgivaga.2.1.chr7.g02028.t01#1    mNeoVis.2.1.1.chr7_hap1.g02052.t01#5    mNeoVis.3.1.1.chr7.g01913.t01#6    mNeoVis.evergladensis.2.1.chr7.g01998.t01#2    mNeoVis.vulgivaga.2.1.chr7.g02021.t01#1      1796   125260368:   mNeoVis.2.1.1.chr7_hap1.g02021.t01#5    mNeoVis.vulgivaga.2.1.chr7.g02004.t01#1    mNeoVis.2.1.1.chr7_hap1.g02020.t01#5    mNeoVis.evergladensis.2.1.chr7.g01984.t01#2    mNeoVis.evergladensis.1.1.chr7.g01991.t01#3    mNeoVis.vulgivaga.2.1.chr7.g02003.t01#1    mNeoVis.vulgivaga.1.1.chr7.g01996.t01#4    mNeoVis.3.1.1.chr7.g01891.t01#6    mNeoVis.3.1.1.chr7.g01886.t01#6      1797   125260369:   mNeoVis.vulgivaga.2.1.chr7.g02034.t01#1    mNeoVis.evergladensis.1.1.chr7.g02024.t01#3    mNeoVis.vulgivaga.1.1.chr7.g02025.t01#4    mNeoVis.2.1.1.chr7_hap1.g02047.t01#5    mNeoVis.3.1.1.chr7.g01907.t01#6    mNeoVis.vulgivaga.1.1.chr7.g02019.t01#4    mNeoVis.evergladensis.1.1.chr7.g02022.t01#3    mNeoVis.evergladensis.2.1.chr7.g02011.t01#2    mNeoVis.evergladensis.1.1.chr7.g02015.t01#3      1798   125260370:   mNeoVis.vulgivaga.1.1.chr7.g02013.t01#4    mNeoVis.vulgivaga.1.1.chr7.g02012.t01#4    mNeoVis.3.1.1.chr7.g01903.t01#6    mNeoVis.evergladensis.2.1.chr7.g02009.t01#2    mNeoVis.vulgivaga.1.1.chr7.g02020.t01#4    mNeoVis.2.1.1.chr7_hap1.g02051.t01#5    mNeoVis.vulgivaga.2.1.chr7.g02030.t01#1    mNeoVis.vulgivaga.2.1.chr7.g02020.t01#1    mNeoVis.evergladensis.1.1.chr7.g02019.t01#3      1799   125260371:   mNeoVis.2.1.1.chr7_hap1.g02063.t01#5    mNeoVis.evergladensis.2.1.chr7.g02004.t01#2    mNeoVis.evergladensis.2.1.chr7.g02019.t01#2    mNeoVis.vulgivaga.1.1.chr7.g02030.t01#4    mNeoVis.2.1.1.chr7_hap1.g02053.t01#5    mNeoVis.evergladensis.1.1.chr7.g02029.t01#3    mNeoVis.vulgivaga.2.1.chr7.g02038.t01#1    mNeoVis.3.1.1.chr7.g01909.t01#6    mNeoVis.vulgivaga.2.1.chr7.g02031.t01#1      1800   125260372:   mNeoVis.evergladensis.2.1.chr7.g02015.t01#2    mNeoVis.vulgivaga.1.1.chr7.g02031.t01#4    mNeoVis.evergladensis.2.1.chr7.g02016.t01#2    mNeoVis.evergladensis.1.1.chr7.g02030.t01#3    mNeoVis.vulgivaga.1.1.chr7.g02027.t01#4    mNeoVis.2.1.1.chr7_hap1.g02060.t01#5    mNeoVis.evergladensis.1.1.chr7.g02028.t01#3    mNeoVis.3.1.1.chr7.g01919.t01#6    mNeoVis.vulgivaga.2.1.chr7.g02036.t01#1      1801   125260373:   mNeoVis.2.1.1.chr7_hap1.g02036.t01#5    mNeoVis.3.1.1.chr7.g01902.t01#6    mNeoVis.evergladensis.1.1.chr7.g02006.t01#3    mNeoVis.vulgivaga.2.1.chr7.g02029.t01#1    mNeoVis.3.1.1.chr7.g01901.t01#6    mNeoVis.evergladensis.1.1.chr7.g02005.t01#3    mNeoVis.vulgivaga.2.1.chr7.g02019.t01#1    mNeoVis.evergladensis.1.1.chr7.g02016.t01#3    mNeoVis.evergladensis.1.1.chr7.g02004.t01#3      1802   125260374:   mNeoVis.2.1.1.chr7_hap1.g02071.t01#5    mNeoVis.2.1.1.chr7_hap1.g02074.t01#5    mNeoVis.2.1.1.chr7_hap1.g02073.t01#5    mNeoVis.2.1.1.chr7_hap1.g02075.t01#5    mNeoVis.2.1.1.chr7_hap1.g02070.t01#5    mNeoVis.2.1.1.chr7_hap1.g02066.t01#5    mNeoVis.2.1.1.chr7_hap1.g02072.t01#5    mNeoVis.2.1.1.chr7_hap1.g02076.t01#5    mNeoVis.2.1.1.chr7_hap1.g02069.t01#5      1803   125260375:   mNeoVis.vulgivaga.1.1.chr7.g02016.t01#4    mNeoVis.evergladensis.1.1.chr7.g02012.t01#3    mNeoVis.evergladensis.2.1.chr7.g02003.t01#2    mNeoVis.vulgivaga.2.1.chr7.g02024.t01#1    mNeoVis.evergladensis.1.1.chr7.g02011.t01#3    mNeoVis.vulgivaga.2.1.chr7.g02025.t01#1    mNeoVis.3.1.1.chr7.g01908.t01#6    mNeoVis.2.1.1.chr7_hap1.g02043.t01#5    mNeoVis.evergladensis.2.1.chr7.g02002.t01#2      1804   125260376:   mNeoVis.2.1.1.chr7_hap1.g02042.t01#5    mNeoVis.evergladensis.2.1.chr7.g02001.t01#2    mNeoVis.3.1.1.chr7.g01906.t01#6    mNeoVis.evergladensis.1.1.chr7.g02010.t01#3    mNeoVis.vulgivaga.1.1.chr7.g02014.t01#4   mNeoVis.vulgivaga.2.1.chr7.g02022.t01#1       1805   125260377:   mNeoVis.evergladensis.2.1.chr7.g02007.t01#2    mNeoVis.3.1.1.chr7.g01914.t01#6    mNeoVis.evergladensis.2.1.chr7.g01999.t01#2    mNeoVis.evergladensis.2.1.chr7.g01995.t01#2   mNeoVis.evergladensis.2.1.chr7.g01996.t01#2       1806   125260378:   mNeoVis.2.1.1.chr7_hap1.g02062.t01#5    mNeoVis.evergladensis.2.1.chr7.g02018.t01#2    mNeoVis.vulgivaga.2.1.chr7.g02035.t01#1    mNeoVis.vulgivaga.1.1.chr7.g02026.t01#4   mNeoVis.evergladensis.1.1.chr7.g02025.t01#3       1807   125260379:   mNeoVis.3.1.1.chr7.g01912.t01#6    mNeoVis.evergladensis.2.1.chr7.g02008.t01#2    mNeoVis.vulgivaga.2.1.chr7.g02027.t01#1   mNeoVis.3.1.1.chr7.g01911.t01#6         1808   125260380:   mNeoVis.vulgivaga.1.1.chr7.g02041.t01#4    mNeoVis.2.1.1.chr7_hap1.g02077.t01#5    mNeoVis.vulgivaga.2.1.chr7.g02049.t01#1   mNeoVis.evergladensis.2.1.chr7.g02032.t01#2       1809   125260381:   mNeoVis.evergladensis.2.1.chr7.g02010.t01#2   mNeoVis.evergladensis.1.1.chr7.g02021.t01#3           1810   125260382:   mNeoVis.evergladensis.2.1.chr7.g02012.t01#2   mNeoVis.evergladensis.2.1.chr7.g02014.t01#2           1811   125260383:   mNeoVis.2.1.1.chr7_hap1.g02054.t01#5   mNeoVis.2.1.1.chr7_hap1.g02057.t01#5             1812   125260384:   mNeoVis.vulgivaga.1.1.chr7.g02023.t01#4   mNeoVis.vulgivaga.2.1.chr7.g02032.t01#1             1813   125260385:  mNeoVis.2.1.1.chr7_hap1.g02040.t01#5               1814   125260386:  mNeoVis.3.1.1.chr7.g01904.t01#6               1815   125260387:  mNeoVis.2.1.1.chr7_hap1.g02049.t01#5               1816   125260388:  mNeoVis.vulgivaga.1.1.chr7.g02018.t01#4               1817   125260389:   mNeoVis.evergladensis.2.1.chr7.g01985.t01#2    mNeoVis.vulgivaga.1.1.chr7.g01998.t01#4    mNeoVis.2.1.1.chr7_hap1.g02022.t01#5    mNeoVis.3.1.1.chr7.g01888.t01#6    mNeoVis.vulgivaga.2.1.chr7.g02005.t01#1   mNeoVis.evergladensis.1.1.chr7.g01992.t01#3       1818   125260390:   mNeoVis.vulgivaga.2.1.chr7.g02006.t01#1    mNeoVis.2.1.1.chr7_hap1.g02023.t01#5    mNeoVis.evergladensis.2.1.chr7.g01986.t01#2    mNeoVis.3.1.1.chr7.g01889.t01#6   mNeoVis.evergladensis.1.1.chr7.g01993.t01#3       1819   125260391:   mNeoVis.vulgivaga.1.1.chr7.g02000.t01#4    mNeoVis.vulgivaga.2.1.chr7.g02007.t01#1    mNeoVis.2.1.1.chr7_hap1.g02024.t01#5    mNeoVis.3.1.1.chr7.g01890.t01#6   mNeoVis.evergladensis.1.1.chr7.g01994.t01#3       1820   125260392:   mNeoVis.vulgivaga.2.1.chr7.g02009.t01#1    mNeoVis.evergladensis.1.1.chr7.g01996.t01#3    mNeoVis.2.1.1.chr7_hap1.g02026.t01#5    mNeoVis.evergladensis.2.1.chr7.g01988.t01#2    mNeoVis.vulgivaga.1.1.chr7.g02002.t01#4   mNeoVis.3.1.1.chr7.g01892.t01#6       1821   125260393:   mNeoVis.vulgivaga.1.1.chr7.g02003.t01#4    mNeoVis.evergladensis.2.1.chr7.g01989.t01#2    mNeoVis.3.1.1.chr7.g01893.t01#6    mNeoVis.vulgivaga.2.1.chr7.g02010.t01#1    mNeoVis.evergladensis.1.1.chr7.g01997.t01#3   mNeoVis.2.1.1.chr7_hap1.g02027.t01#5       1822   125260394:   mNeoVis.3.1.1.chr7.g01922.t01#6    mNeoVis.vulgivaga.2.1.chr7.g02063.t01#1    mNeoVis.2.1.1.chr7_hap1.g02088.t01#5    mNeoVis.vulgivaga.1.1.chr7.g02055.t01#4    mNeoVis.vulgivaga.1.1.chr7.g02033.t01#4    mNeoVis.2.1.1.chr7_hap1.g02034.t01#5    mNeoVis.evergladensis.2.1.chr7.g02042.t01#2    mNeoVis.evergladensis.2.1.chr7.g02023.t01#2    mNeoVis.2.1.1.chr7_hap1.g02093.t01#5      1823   125260395:   mNeoVis.vulgivaga.1.1.chr7.g02010.t01#4    mNeoVis.3.1.1.chr7.g01899.t01#6    mNeoVis.evergladensis.1.1.chr7.g02003.t01#3    mNeoVis.vulgivaga.2.1.chr7.g02016.t01#1    mNeoVis.evergladensis.2.1.chr7.g01994.t01#2   mNeoVis.2.1.1.chr7_hap1.g02035.t01#5       1824   125260396:   mNeoVis.2.1.1.chr7_hap1.g02033.t01#5    mNeoVis.vulgivaga.2.1.chr7.g02014.t01#1    mNeoVis.evergladensis.1.1.chr7.g02001.t01#3    mNeoVis.vulgivaga.1.1.chr7.g02008.t01#4   mNeoVis.3.1.1.chr7.g01897.t01#6       1825   125260397:   mNeoVis.evergladensis.2.1.chr1.g01399.t01#2    mNeoVis.vulgivaga.1.1.chr1.g01404.t01#4    mNeoVis.evergladensis.1.1.chr1.g01405.t01#3    mNeoVis.3.1.1.chr1.g01383.t01#6    mNeoVis.vulgivaga.2.1.chr1.g01403.t01#1   mNeoVis.2.1.1.chr1_hap1.g01437.t01#5       1826   125260398:   mNeoVis.3.1.1.chr1.g01384.t01#6    mNeoVis.vulgivaga.1.1.chr1.g01405.t01#4    mNeoVis.evergladensis.1.1.chr1.g01406.t01#3    mNeoVis.2.1.1.chr1_hap1.g01438.t01#5    mNeoVis.evergladensis.2.1.chr1.g01400.t01#2   mNeoVis.vulgivaga.2.1.chr1.g01404.t01#1       1827   125260399:   mNeoVis.2.1.1.chr1_hap1.g01439.t01#5    mNeoVis.vulgivaga.2.1.chr1.g01405.t01#1    mNeoVis.evergladensis.1.1.chr1.g01407.t01#3    mNeoVis.3.1.1.chr1.g01385.t01#6    mNeoVis.vulgivaga.1.1.chr1.g01406.t01#4   mNeoVis.evergladensis.2.1.chr1.g01401.t01#2       1828   125260400:   mNeoVis.2.1.1.chr1_hap1.g01440.t01#5    mNeoVis.vulgivaga.2.1.chr1.g01406.t01#1    mNeoVis.evergladensis.2.1.chr1.g01402.t01#2    mNeoVis.evergladensis.1.1.chr1.g01408.t01#3    mNeoVis.3.1.1.chr1.g01386.t01#6   mNeoVis.vulgivaga.1.1.chr1.g01407.t01#4       1829   125260401:   mNeoVis.3.1.1.chr7.g01905.t01#6    mNeoVis.evergladensis.1.1.chr7.g02009.t01#3    mNeoVis.vulgivaga.1.1.chr7.g02015.t01#4    mNeoVis.vulgivaga.2.1.chr7.g02023.t01#1    mNeoVis.2.1.1.chr7_hap1.g02041.t01#5   mNeoVis.evergladensis.2.1.chr7.g02000.t01#2       1830   125260402:   mNeoVis.2.1.1.chr1_hap1.g00913.t01#5    mNeoVis.vulgivaga.2.1.chr15.g00596.t01#1    mNeoVis.3.1.1.chr1.g00930.t01#6    mNeoVis.vulgivaga.2.1.chr1.g00922.t01#1    mNeoVis.2.1.1.chr1_hap1.g00908.t01#5   mNeoVis.vulgivaga.1.1.chr1.g00925.t01#4       1831   125260403:   mNeoVis.2.1.1.chr7_hap1.g02055.t01#5    mNeoVis.evergladensis.1.1.chr7.g02023.t01#3    mNeoVis.vulgivaga.1.1.chr7.g02024.t01#4    mNeoVis.vulgivaga.2.1.chr7.g02033.t01#1    mNeoVis.3.1.1.chr7.g01917.t01#6   mNeoVis.evergladensis.2.1.chr7.g02013.t01#2       1832   125260404:   mNeoVis.vulgivaga.1.1.chr1.g01408.t01#4    mNeoVis.3.1.1.chr1.g01387.t01#6    mNeoVis.evergladensis.1.1.chr1.g01409.t01#3    mNeoVis.vulgivaga.2.1.chr1.g01407.t01#1    mNeoVis.2.1.1.chr1_hap1.g01441.t01#5   mNeoVis.evergladensis.2.1.chr1.g01403.t01#2       1833   125260405:   mNeoVis.evergladensis.1.1.chr7.g02031.t01#3    mNeoVis.2.1.1.chr7_hap1.g02059.t01#5    mNeoVis.vulgivaga.1.1.chr7.g02028.t01#4    mNeoVis.evergladensis.2.1.chr7.g02017.t01#2    mNeoVis.3.1.1.chr7.g01921.t01#6    mNeoVis.evergladensis.2.1.chr7.g02021.t01#2    mNeoVis.vulgivaga.2.1.chr7.g02039.t01#1    mNeoVis.evergladensis.1.1.chr7.g02026.t01#3    mNeoVis.2.1.1.chr7_hap1.g02061.t01#5      1834   125260406:   mNeoVis.2.1.1.chr1_hap1.g01442.t01#5    mNeoVis.vulgivaga.2.1.chr1.g01408.t01#1    mNeoVis.evergladensis.2.1.chr1.g01404.t01#2    mNeoVis.3.1.1.chr1.g01388.t01#6    mNeoVis.vulgivaga.1.1.chr1.g01409.t01#4   mNeoVis.evergladensis.1.1.chr1.g01410.t01#3       1835   125260407:   mNeoVis.evergladensis.1.1.chr1.g01411.t01#3    mNeoVis.vulgivaga.2.1.chr1.g01409.t01#1    mNeoVis.3.1.1.chr1.g01389.t01#6    mNeoVis.vulgivaga.1.1.chr1.g01410.t01#4    mNeoVis.evergladensis.2.1.chr1.g01405.t01#2   mNeoVis.2.1.1.chr1_hap1.g01443.t01#5       1836   125260408:   mNeoVis.evergladensis.2.1.chr15.g00585.t01#2    mNeoVis.evergladensis.1.1.chr15.g00025.t01#3    mNeoVis.vulgivaga.2.1.chr15.g00597.t01#1    mNeoVis.2.1.1.chr15_hap1.g00764.t01#5    mNeoVis.3.1.1.chr15.g00543.t01#6   mNeoVis.vulgivaga.1.1.chr15.g00588.t01#4       1837   125260409:   mNeoVis.3.1.1.chr3.g00671.t01#6    mNeoVis.vulgivaga.2.1.chr3.g00693.t01#1    mNeoVis.evergladensis.1.1.chr3.g00689.t01#3   mNeoVis.2.1.1.chr3_hap1.g00681.t01#5         1838   125260410:   mNeoVis.evergladensis.1.1.chr3.g00690.t01#3    mNeoVis.vulgivaga.1.1.chr3.g00684.t01#4   mNeoVis.evergladensis.2.1.chr3.g00689.t01#2         1839   125260411:   mNeoVis.evergladensis.2.1.chr7.g00532.t01#2    mNeoVis.evergladensis.1.1.chr7.g00541.t01#3    mNeoVis.3.1.1.chr7.g00516.t01#6    mNeoVis.2.1.1.chr7_hap1.g00532.t01#5    mNeoVis.vulgivaga.2.1.chr7.g00535.t01#1   mNeoVis.vulgivaga.1.1.chr7.g00540.t01#4       1840   125260412:   mNeoVis.2.1.1.chr1_hap1.g01444.t01#5    mNeoVis.evergladensis.2.1.chr1.g01406.t01#2    mNeoVis.vulgivaga.2.1.chr1.g01410.t01#1    mNeoVis.3.1.1.chr1.g01390.t01#6    mNeoVis.evergladensis.1.1.chr1.g01412.t01#3   mNeoVis.vulgivaga.1.1.chr1.g01411.t01#4       1841   125260413:   mNeoVis.vulgivaga.2.1.chr1.g01411.t01#1    mNeoVis.evergladensis.1.1.chr1.g01413.t01#3    mNeoVis.evergladensis.2.1.chr1.g01407.t01#2    mNeoVis.3.1.1.chr1.g01391.t01#6    mNeoVis.vulgivaga.1.1.chr1.g01412.t01#4    mNeoVis.3.1.1.chr1.g01392.t01#6   mNeoVis.2.1.1.chr1_hap1.g01445.t01#5     1842   125260414:   mNeoVis.2.1.1.chr4_hap1.g00245.t01#5    mNeoVis.evergladensis.1.1.chr6.g00246.t01#3    mNeoVis.3.1.1.chr6.g00244.t01#6    mNeoVis.vulgivaga.2.1.chr6.g00245.t01#1    mNeoVis.evergladensis.2.1.chr6.g00244.t01#2   mNeoVis.vulgivaga.1.1.chr6.g00252.t01#4       1843   125260415:   mNeoVis.vulgivaga.2.1.chr2.g00845.t01#1    mNeoVis.vulgivaga.1.1.chr2.g00844.t01#4    mNeoVis.3.1.1.chr2.g00752.t01#6    mNeoVis.2.1.1.chr5_hap1.g00842.t01#5    mNeoVis.evergladensis.1.1.chr2.g00838.t01#3   mNeoVis.evergladensis.2.1.chr2.g00844.t01#2       1844   125260416:   mNeoVis.3.1.1.chr7.g01931.t01#6    mNeoVis.evergladensis.1.1.chr7.g02048.t01#3    mNeoVis.vulgivaga.1.1.chr7.g02056.t01#4    mNeoVis.vulgivaga.2.1.chr7.g02053.t01#1    mNeoVis.vulgivaga.2.1.chr7.g02069.t01#1    mNeoVis.vulgivaga.2.1.chr7.g02058.t01#1    mNeoVis.evergladensis.2.1.chr7.g02039.t01#2    mNeoVis.evergladensis.2.1.chr7.g02037.t01#2    mNeoVis.evergladensis.1.1.chr7.g02046.t01#3      1845   125260417:   mNeoVis.vulgivaga.2.1.chr7.g02059.t01#1    mNeoVis.2.1.1.chr7_hap1.g02096.t01#5    mNeoVis.vulgivaga.1.1.chr7.g02044.t01#4    mNeoVis.evergladensis.1.1.chr7.g02055.t01#3    mNeoVis.3.1.1.chr7.g01930.t01#6    mNeoVis.3.1.1.chr7.g01936.t01#6    mNeoVis.evergladensis.2.1.chr7.g02046.t01#2    mNeoVis.2.1.1.chr7_hap1.g02087.t01#5    mNeoVis.vulgivaga.2.1.chr7.g02068.t01#1      1846   125260418:   mNeoVis.evergladensis.2.1.chr7.g02049.t01#2    mNeoVis.evergladensis.1.1.chr7.g02047.t01#3    mNeoVis.vulgivaga.2.1.chr7.g02072.t01#1    mNeoVis.vulgivaga.1.1.chr7.g02058.t01#4    mNeoVis.2.1.1.chr7_hap1.g02100.t01#5    mNeoVis.vulgivaga.1.1.chr7.g02057.t01#4    mNeoVis.vulgivaga.2.1.chr7.g02070.t01#1    mNeoVis.evergladensis.2.1.chr7.g02038.t01#2    mNeoVis.evergladensis.1.1.chr7.g02058.t01#3      1847   125260419:   mNeoVis.2.1.1.chr7_hap1.g02089.t01#5    mNeoVis.vulgivaga.2.1.chr7.g02051.t01#1    mNeoVis.2.1.1.chr7_hap1.g02091.t01#5    mNeoVis.vulgivaga.1.1.chr7.g02043.t01#4    mNeoVis.3.1.1.chr7.g01937.t01#6    mNeoVis.evergladensis.1.1.chr7.g02053.t01#3    mNeoVis.evergladensis.2.1.chr7.g02044.t01#2   mNeoVis.vulgivaga.2.1.chr7.g02061.t01#1     1848   125260420:   mNeoVis.evergladensis.1.1.chr7.g02059.t01#3    mNeoVis.vulgivaga.2.1.chr7.g02073.t01#1    mNeoVis.evergladensis.2.1.chr7.g02050.t01#2    mNeoVis.3.1.1.chr7.g01941.t01#6    mNeoVis.vulgivaga.1.1.chr7.g02059.t01#4   mNeoVis.2.1.1.chr7_hap1.g02094.t01#5       1849   125260421:   mNeoVis.3.1.1.chr7.g01942.t01#6    mNeoVis.vulgivaga.1.1.chr7.g02060.t01#4    mNeoVis.2.1.1.chr7_hap1.g02082.t01#5    mNeoVis.vulgivaga.1.1.chr7.g02049.t01#4   mNeoVis.vulgivaga.2.1.chr7.g02055.t01#1       1850   125260422:   mNeoVis.2.1.1.chr7_hap1.g02101.t01#5    mNeoVis.vulgivaga.1.1.chr7.g02061.t01#4    mNeoVis.vulgivaga.2.1.chr7.g02074.t01#1   mNeoVis.3.1.1.chr7.g01943.t01#6         1851   125260423:   mNeoVis.3.1.1.chr7.g01934.t01#6    mNeoVis.evergladensis.1.1.chr7.g02057.t01#3   mNeoVis.evergladensis.2.1.chr7.g02048.t01#2         1852   125260424:  mNeoVis.vulgivaga.1.1.chr7.g02054.t01#4               1853   125260425:   mNeoVis.2.1.1.chr1_hap1.g01446.t01#5    mNeoVis.vulgivaga.1.1.chr1.g01413.t01#4    mNeoVis.evergladensis.2.1.chr1.g01408.t01#2    mNeoVis.3.1.1.chr1.g01393.t01#6    mNeoVis.vulgivaga.2.1.chr1.g01412.t01#1   mNeoVis.evergladensis.1.1.chr1.g01414.t01#3       1854   125260426:   mNeoVis.vulgivaga.2.1.chr1.g01413.t01#1   mNeoVis.3.1.1.chr1.g01394.t01#6             1855   125260427:   mNeoVis.vulgivaga.1.1.chr1.g01415.t01#4    mNeoVis.evergladensis.1.1.chr1.g01416.t01#3    mNeoVis.vulgivaga.2.1.chr1.g01414.t01#1    mNeoVis.3.1.1.chr1.g01395.t01#6    mNeoVis.evergladensis.2.1.chr1.g01410.t01#2   mNeoVis.2.1.1.chr1_hap1.g01448.t01#5       1856   125260428:   mNeoVis.evergladensis.2.1.chr1.g01411.t01#2    mNeoVis.3.1.1.chr1.g01396.t01#6    mNeoVis.vulgivaga.1.1.chr1.g01416.t01#4    mNeoVis.vulgivaga.2.1.chr1.g01415.t01#1    mNeoVis.2.1.1.chr1_hap1.g01449.t01#5   mNeoVis.evergladensis.1.1.chr1.g01417.t01#3       1857   125260429:  mNeoVis.vulgivaga.2.1.chr7.g02062.t01#1               1858   125260430:   mNeoVis.2.1.1.chr1_hap1.g01450.t01#5    mNeoVis.3.1.1.chr1.g01397.t01#6    mNeoVis.vulgivaga.2.1.chr1.g01416.t01#1    mNeoVis.evergladensis.2.1.chr1.g01412.t01#2    mNeoVis.vulgivaga.1.1.chr1.g01417.t01#4   mNeoVis.evergladensis.1.1.chr1.g01418.t01#3       1859   125260431:   mNeoVis.3.1.1.chr1.g01398.t01#6    mNeoVis.evergladensis.1.1.chr1.g01419.t01#3    mNeoVis.2.1.1.chr1_hap1.g01451.t01#5    mNeoVis.vulgivaga.2.1.chr1.g01417.t01#1    mNeoVis.evergladensis.2.1.chr1.g01413.t01#2   mNeoVis.vulgivaga.1.1.chr1.g01418.t01#4       1860   125260432:   mNeoVis.vulgivaga.2.1.chr7.g00536.t01#1    mNeoVis.2.1.1.chr7_hap1.g00533.t01#5    mNeoVis.evergladensis.1.1.chr7.g00542.t01#3    mNeoVis.evergladensis.2.1.chr7.g00533.t01#2   mNeoVis.vulgivaga.1.1.chr7.g00541.t01#4       1861   125260433:   mNeoVis.2.1.1.chr1_hap1.g01452.t01#5    mNeoVis.evergladensis.1.1.chr1.g01420.t01#3    mNeoVis.vulgivaga.2.1.chr1.g01418.t01#1    mNeoVis.vulgivaga.1.1.chr1.g01419.t01#4    mNeoVis.3.1.1.chr1.g01399.t01#6   mNeoVis.evergladensis.2.1.chr1.g01414.t01#2       1862   125260434:   mNeoVis.evergladensis.2.1.chr1.g01415.t01#2    mNeoVis.vulgivaga.2.1.chr1.g01419.t01#1    mNeoVis.evergladensis.1.1.chr4.g01065.t01#3    mNeoVis.3.1.1.chr4.g00270.t01#6    mNeoVis.2.1.1.chr6_hap1.g01099.t01#5    mNeoVis.evergladensis.2.1.chr4.g00280.t01#2   mNeoVis.vulgivaga.1.1.chr4.g00274.t01#4     1863   125260435:   mNeoVis.2.1.1.chr1_hap1.g01453.t01#5    mNeoVis.vulgivaga.1.1.chr1.g01420.t01#4    mNeoVis.3.1.1.chr1.g01400.t01#6   mNeoVis.evergladensis.1.1.chr1.g01421.t01#3       1864   125260436:   mNeoVis.vulgivaga.1.1.chr4.g01058.t01#4   mNeoVis.vulgivaga.2.1.chr4.g01065.t01#1             1865   125260437:   mNeoVis.2.1.1.chr1_hap1.g01454.t01#5    mNeoVis.vulgivaga.2.1.chr1.g01420.t01#1    mNeoVis.vulgivaga.1.1.chr1.g01421.t01#4    mNeoVis.3.1.1.chr1.g01401.t01#6    mNeoVis.evergladensis.2.1.chr1.g01416.t01#2   mNeoVis.evergladensis.1.1.chr1.g01422.t01#3       1866   125260438:   mNeoVis.2.1.1.chr7_hap1.g02102.t01#5    mNeoVis.vulgivaga.1.1.chr7.g02062.t01#4    mNeoVis.vulgivaga.2.1.chr7.g02075.t01#1    mNeoVis.evergladensis.1.1.chr7.g02060.t01#3    mNeoVis.3.1.1.chr7.g01944.t01#6   mNeoVis.evergladensis.2.1.chr7.g02051.t01#2       1867   125260439:   mNeoVis.vulgivaga.2.1.chr7.g02076.t01#1    mNeoVis.vulgivaga.1.1.chr7.g02063.t01#4   mNeoVis.2.1.1.chr7_hap1.g02103.t01#5           1868   125260440:   mNeoVis.evergladensis.2.1.chr7.g02052.t01#2    mNeoVis.evergladensis.1.1.chr7.g02061.t01#3    mNeoVis.evergladensis.2.1.chr7.g02053.t01#2    mNeoVis.2.1.1.chr7_hap1.g02106.t01#5    mNeoVis.vulgivaga.1.1.chr7.g02065.t01#4    mNeoVis.2.1.1.chr7_hap1.g02105.t01#5    mNeoVis.vulgivaga.1.1.chr7.g02064.t01#4    mNeoVis.vulgivaga.2.1.chr7.g02079.t01#1    mNeoVis.vulgivaga.2.1.chr7.g02078.t01#1      1869   125260441:   mNeoVis.evergladensis.2.1.chr7.g02056.t01#2    mNeoVis.evergladensis.1.1.chr7.g02064.t01#3    mNeoVis.vulgivaga.2.1.chr7.g02080.t01#1    mNeoVis.2.1.1.chr7_hap1.g02108.t01#5    mNeoVis.vulgivaga.1.1.chr7.g02067.t01#4    mNeoVis.evergladensis.1.1.chr7.g02063.t01#3    mNeoVis.2.1.1.chr7_hap1.g02107.t01#5   mNeoVis.evergladensis.2.1.chr7.g02055.t01#2     1870   125260442:  mNeoVis.vulgivaga.2.1.chr7.g02081.t01#1               1871   125260443:   mNeoVis.evergladensis.1.1.chr7.g00543.t01#3    mNeoVis.evergladensis.2.1.chr7.g00534.t01#2    mNeoVis.3.1.1.chr7.g00517.t01#6    mNeoVis.vulgivaga.2.1.chr7.g00537.t01#1    mNeoVis.vulgivaga.1.1.chr7.g00542.t01#4   mNeoVis.2.1.1.chr7_hap1.g00534.t01#5       1872   125260444:   mNeoVis.evergladensis.2.1.chr7.g02060.t01#2    mNeoVis.evergladensis.1.1.chr7.g02067.t01#3    mNeoVis.2.1.1.chr7_hap1.g02111.t01#5    mNeoVis.vulgivaga.2.1.chr7.g02082.t01#1    mNeoVis.2.1.1.chr7_hap1.g02110.t01#5    mNeoVis.vulgivaga.1.1.chr7.g02070.t01#4    mNeoVis.2.1.1.chr7_hap1.g02113.t01#5    mNeoVis.evergladensis.1.1.chr7.g02066.t01#3    mNeoVis.vulgivaga.1.1.chr7.g02069.t01#4      1873   125260445:   mNeoVis.2.1.1.chr7_hap1.g02114.t01#5    mNeoVis.vulgivaga.2.1.chr7.g02083.t01#1    mNeoVis.vulgivaga.1.1.chr7.g02072.t01#4    mNeoVis.2.1.1.chr7_hap1.g02112.t01#5    mNeoVis.evergladensis.1.1.chr7.g02068.t01#3    mNeoVis.evergladensis.2.1.chr7.g02059.t01#2   mNeoVis.vulgivaga.1.1.chr7.g02071.t01#4     1874   125260446:   mNeoVis.evergladensis.2.1.chr7.g02061.t01#2   mNeoVis.3.1.1.chr7.g01949.t01#6             1875   125260447:  mNeoVis.3.1.1.chr7.g01948.t01#6               1876   125260448:   mNeoVis.2.1.1.chr1_hap1.g01455.t01#5    mNeoVis.vulgivaga.1.1.chr1.g01422.t01#4    mNeoVis.evergladensis.1.1.chr1.g01423.t01#3    mNeoVis.evergladensis.2.1.chr1.g01417.t01#2    mNeoVis.3.1.1.chr1.g01402.t01#6   mNeoVis.vulgivaga.2.1.chr1.g01421.t01#1       1877   125260449:   mNeoVis.3.1.1.chr15.g00544.t01#6    mNeoVis.vulgivaga.1.1.chr15.g00589.t01#4    mNeoVis.evergladensis.2.1.chr15.g00586.t01#2    mNeoVis.vulgivaga.2.1.chr15.g00598.t01#1   mNeoVis.evergladensis.1.1.chr15.g00026.t01#3       1878   125260450:   mNeoVis.3.1.1.chr1.g01403.t01#6    mNeoVis.vulgivaga.2.1.chr1.g01422.t01#1    mNeoVis.evergladensis.1.1.chr1.g01424.t01#3    mNeoVis.2.1.1.chr1_hap1.g01456.t01#5   mNeoVis.vulgivaga.1.1.chr1.g01423.t01#4       1879   125260451:   mNeoVis.3.1.1.chr7.g01950.t01#6    mNeoVis.2.1.1.chr7_hap1.g02116.t01#5    mNeoVis.vulgivaga.2.1.chr7.g02085.t01#1    mNeoVis.evergladensis.1.1.chr7.g02065.t01#3   mNeoVis.vulgivaga.1.1.chr7.g02074.t01#4       1880   125260452:   mNeoVis.3.1.1.chr7.g01952.t01#6    mNeoVis.vulgivaga.2.1.chr7.g02087.t01#1    mNeoVis.vulgivaga.1.1.chr7.g02075.t01#4    mNeoVis.evergladensis.1.1.chr7.g02071.t01#3   mNeoVis.evergladensis.2.1.chr7.g02063.t01#2       1881   125260453:   mNeoVis.vulgivaga.1.1.chr7.g02078.t01#4    mNeoVis.2.1.1.chr7_hap1.g02119.t01#5    mNeoVis.evergladensis.1.1.chr7.g02073.t01#3    mNeoVis.evergladensis.2.1.chr7.g02065.t01#2   mNeoVis.2.1.1.chr7_hap1.g02118.t01#5       1882   125260454:   mNeoVis.vulgivaga.2.1.chr7.g02086.t01#1    mNeoVis.evergladensis.1.1.chr7.g02070.t01#3   mNeoVis.3.1.1.chr7.g01951.t01#6           1883   125260455:   mNeoVis.evergladensis.1.1.chr7.g02069.t01#3   mNeoVis.vulgivaga.2.1.chr7.g02084.t01#1             1884   125260456:   mNeoVis.vulgivaga.1.1.chr7.g02073.t01#4   mNeoVis.2.1.1.chr7_hap1.g02115.t01#5             1885   125260457:   mNeoVis.3.1.1.chr7.g01954.t01#6   mNeoVis.vulgivaga.1.1.chr7.g02077.t01#4             1886   125260458:  mNeoVis.evergladensis.2.1.chr7.g02062.t01#2             1887   125260459:  mNeoVis.2.1.1.chr7_hap1.g02117.t01#5               1888   125260460:  mNeoVis.evergladensis.2.1.chr7.g02064.t01#2             1889   125260461:  mNeoVis.evergladensis.1.1.chr7.g02072.t01#3             1890   125260462:  mNeoVis.vulgivaga.2.1.chr7.g02090.t01#1               1891   125260463:  mNeoVis.vulgivaga.2.1.chr7.g02088.t01#1               1892   125260464:   mNeoVis.evergladensis.1.1.chr7.g02086.t01#3    mNeoVis.2.1.1.chr7_hap1.g02133.t01#5    mNeoVis.3.1.1.chr7.g01961.t01#6    mNeoVis.evergladensis.2.1.chr7.g02079.t01#2    mNeoVis.vulgivaga.2.1.chr7.g02100.t01#1    mNeoVis.2.1.1.chr7_hap1.g02132.t01#5    mNeoVis.evergladensis.1.1.chr7.g02085.t01#3    mNeoVis.evergladensis.1.1.chr7.g02089.t01#3    mNeoVis.vulgivaga.2.1.chr7.g02102.t01#1      1893   125260465:   mNeoVis.evergladensis.1.1.chr7.g02080.t01#3    mNeoVis.vulgivaga.2.1.chr7.g02089.t01#1    mNeoVis.evergladensis.2.1.chr7.g02071.t01#2    mNeoVis.2.1.1.chr7_hap1.g02125.t01#5    mNeoVis.vulgivaga.1.1.chr7.g02076.t01#4    mNeoVis.evergladensis.2.1.chr7.g02074.t01#2    mNeoVis.3.1.1.chr7.g01958.t01#6   mNeoVis.vulgivaga.1.1.chr7.g02084.t01#4     1894   125260466:   mNeoVis.2.1.1.chr7_hap1.g02127.t01#5    mNeoVis.2.1.1.chr7_hap1.g02126.t01#5    mNeoVis.evergladensis.2.1.chr7.g02073.t01#2    mNeoVis.vulgivaga.2.1.chr7.g02097.t01#1    mNeoVis.3.1.1.chr7.g01959.t01#6    mNeoVis.evergladensis.1.1.chr7.g02082.t01#3    mNeoVis.evergladensis.1.1.chr7.g02081.t01#3   mNeoVis.2.1.1.chr7_hap1.g02128.t01#5     1895   125260467:   mNeoVis.evergladensis.2.1.chr7.g02075.t01#2    mNeoVis.evergladensis.2.1.chr7.g02078.t01#2    mNeoVis.vulgivaga.1.1.chr7.g02085.t01#4    mNeoVis.evergladensis.2.1.chr7.g02080.t01#2    mNeoVis.2.1.1.chr7_hap1.g02129.t01#5    mNeoVis.2.1.1.chr7_hap1.g02134.t01#5   mNeoVis.2.1.1.chr7_hap1.g02131.t01#5     1896   125260468:   mNeoVis.evergladensis.2.1.chr7.g02067.t01#2    mNeoVis.vulgivaga.1.1.chr7.g02080.t01#4    mNeoVis.3.1.1.chr7.g01953.t01#6    mNeoVis.vulgivaga.2.1.chr7.g02092.t01#1    mNeoVis.evergladensis.1.1.chr7.g02075.t01#3   mNeoVis.2.1.1.chr7_hap1.g02121.t01#5       1897   125260469:   mNeoVis.2.1.1.chr7_hap1.g02140.t01#5    mNeoVis.evergladensis.2.1.chr7.g02087.t01#2    mNeoVis.vulgivaga.2.1.chr7.g02109.t01#1    mNeoVis.3.1.1.chr7.g01969.t01#6    mNeoVis.evergladensis.1.1.chr7.g02093.t01#3   mNeoVis.vulgivaga.1.1.chr7.g02094.t01#4       1898   125260470:   mNeoVis.vulgivaga.2.1.chr7.g02106.t01#1    mNeoVis.vulgivaga.1.1.chr7.g02090.t01#4    mNeoVis.evergladensis.2.1.chr7.g02081.t01#2    mNeoVis.vulgivaga.1.1.chr7.g02089.t01#4   mNeoVis.2.1.1.chr7_hap1.g02137.t01#5       1899   125260471:   mNeoVis.vulgivaga.2.1.chr7.g02103.t01#1    mNeoVis.evergladensis.1.1.chr7.g02088.t01#3    mNeoVis.evergladensis.2.1.chr7.g02084.t01#2    mNeoVis.2.1.1.chr7_hap1.g02135.t01#5   mNeoVis.3.1.1.chr7.g01964.t01#6       1900   125260472:   mNeoVis.evergladensis.2.1.chr7.g02057.t01#2    mNeoVis.vulgivaga.2.1.chr7.g02104.t01#1   mNeoVis.vulgivaga.2.1.chr7.g02098.t01#1           1901   125260473:   mNeoVis.3.1.1.chr7.g01966.t01#6    mNeoVis.vulgivaga.1.1.chr7.g02092.t01#4   mNeoVis.evergladensis.2.1.chr7.g02083.t01#2         1902   125260474:  mNeoVis.3.1.1.chr7.g01965.t01#6               1903   125260475:   mNeoVis.2.1.1.chr1_hap1.g01457.t01#5    mNeoVis.evergladensis.1.1.chr1.g01425.t01#3    mNeoVis.vulgivaga.1.1.chr1.g01424.t01#4    mNeoVis.3.1.1.chr1.g01404.t01#6    mNeoVis.vulgivaga.2.1.chr1.g01423.t01#1   mNeoVis.evergladensis.2.1.chr1.g01422.t01#2       1904   125260476:   mNeoVis.vulgivaga.2.1.chr7.g02091.t01#1    mNeoVis.evergladensis.1.1.chr7.g02074.t01#3    mNeoVis.2.1.1.chr7_hap1.g02120.t01#5    mNeoVis.vulgivaga.1.1.chr7.g02079.t01#4   mNeoVis.evergladensis.2.1.chr7.g02066.t01#2       1905   125260477:   mNeoVis.vulgivaga.1.1.chr7.g02081.t01#4    mNeoVis.evergladensis.1.1.chr7.g02076.t01#3    mNeoVis.vulgivaga.2.1.chr7.g02093.t01#1    mNeoVis.2.1.1.chr7_hap1.g02122.t01#5    mNeoVis.3.1.1.chr7.g01955.t01#6   mNeoVis.evergladensis.2.1.chr7.g02068.t01#2       1906   125260478:   mNeoVis.evergladensis.1.1.chr7.g02091.t01#3    mNeoVis.2.1.1.chr7_hap1.g02138.t01#5    mNeoVis.evergladensis.2.1.chr7.g02085.t01#2    mNeoVis.vulgivaga.2.1.chr7.g02107.t01#1   mNeoVis.vulgivaga.1.1.chr7.g02091.t01#4       1907   125260479:   mNeoVis.vulgivaga.2.1.chr7.g02095.t01#1    mNeoVis.evergladensis.1.1.chr7.g02077.t01#3    mNeoVis.evergladensis.2.1.chr7.g02069.t01#2    mNeoVis.vulgivaga.2.1.chr7.g02094.t01#1    mNeoVis.2.1.1.chr7_hap1.g02123.t01#5    mNeoVis.3.1.1.chr7.g01956.t01#6    mNeoVis.vulgivaga.1.1.chr7.g02082.t01#4   mNeoVis.evergladensis.1.1.chr7.g02078.t01#3     1908   125260480:   mNeoVis.evergladensis.2.1.chr1.g01423.t01#2    mNeoVis.evergladensis.1.1.chr1.g01426.t01#3    mNeoVis.vulgivaga.2.1.chr1.g01424.t01#1    mNeoVis.2.1.1.chr1_hap1.g01458.t01#5    mNeoVis.vulgivaga.1.1.chr1.g01425.t01#4   mNeoVis.3.1.1.chr1.g01405.t01#6       1909   125260481:   mNeoVis.evergladensis.2.1.chr7.g02070.t01#2    mNeoVis.3.1.1.chr7.g01957.t01#6    mNeoVis.vulgivaga.1.1.chr7.g02083.t01#4    mNeoVis.vulgivaga.2.1.chr7.g02096.t01#1    mNeoVis.evergladensis.1.1.chr7.g02079.t01#3   mNeoVis.2.1.1.chr7_hap1.g02124.t01#5       1910   125260482:   mNeoVis.3.1.1.chr15.g00546.t01#6    mNeoVis.evergladensis.1.1.chr15.g00023.t01#3    mNeoVis.vulgivaga.2.1.chr15.g00599.t01#1    mNeoVis.2.1.1.chr15_hap1.g00766.t01#5    mNeoVis.vulgivaga.1.1.chr15.g00591.t01#4   mNeoVis.evergladensis.2.1.chr15.g00588.t01#2       1911   125260483:   mNeoVis.vulgivaga.2.1.chr7.g00538.t01#1   mNeoVis.2.1.1.chr7_hap1.g00535.t01#5             1912   125260484:   mNeoVis.evergladensis.1.1.chr7.g00544.t01#3   mNeoVis.evergladensis.2.1.chr7.g00535.t01#2           1913   125260485:   mNeoVis.vulgivaga.1.1.chr1.g01426.t01#4    mNeoVis.vulgivaga.2.1.chr1.g01425.t01#1    mNeoVis.2.1.1.chr1_hap1.g01459.t01#5    mNeoVis.evergladensis.1.1.chr1.g01427.t01#3    mNeoVis.evergladensis.2.1.chr1.g01424.t01#2   mNeoVis.3.1.1.chr1.g01406.t01#6       1914   125260486:   mNeoVis.2.1.1.chr7_hap1.g02136.t01#5    mNeoVis.vulgivaga.2.1.chr7.g02105.t01#1    mNeoVis.evergladensis.2.1.chr7.g02082.t01#2    mNeoVis.evergladensis.1.1.chr7.g02090.t01#3   mNeoVis.vulgivaga.1.1.chr7.g02088.t01#4       1915   125260487:   mNeoVis.vulgivaga.1.1.chr7.g02093.t01#4    mNeoVis.2.1.1.chr7_hap1.g02139.t01#5    mNeoVis.vulgivaga.2.1.chr7.g02108.t01#1    mNeoVis.3.1.1.chr7.g01967.t01#6    mNeoVis.evergladensis.1.1.chr7.g02092.t01#3   mNeoVis.evergladensis.2.1.chr7.g02086.t01#2       1916   125260488:   mNeoVis.3.1.1.chr7.g01968.t01#6    mNeoVis.evergladensis.1.1.chr7.g02094.t01#3    mNeoVis.evergladensis.2.1.chr7.g02088.t01#2    mNeoVis.vulgivaga.1.1.chr7.g02095.t01#4    mNeoVis.2.1.1.chr7_hap1.g02141.t01#5   mNeoVis.vulgivaga.2.1.chr7.g02110.t01#1       1917   125260489:   mNeoVis.evergladensis.2.1.chr7.g02089.t01#2    mNeoVis.evergladensis.1.1.chr7.g02095.t01#3    mNeoVis.vulgivaga.1.1.chr7.g02096.t01#4   mNeoVis.2.1.1.chr7_hap1.g02142.t01#5         1918   125260490:   mNeoVis.evergladensis.2.1.chr1.g01425.t01#2    mNeoVis.vulgivaga.2.1.chr1.g01426.t01#1    mNeoVis.vulgivaga.1.1.chr1.g01427.t01#4    mNeoVis.2.1.1.chr1_hap1.g01460.t01#5    mNeoVis.3.1.1.chr1.g01407.t01#6   mNeoVis.evergladensis.1.1.chr1.g01428.t01#3       1919   125260491:   mNeoVis.vulgivaga.2.1.chr7.g02111.t01#1    mNeoVis.2.1.1.chr7_hap1.g02143.t01#5    mNeoVis.evergladensis.2.1.chr7.g02090.t01#2    mNeoVis.evergladensis.1.1.chr7.g02096.t01#3   mNeoVis.vulgivaga.1.1.chr7.g02097.t01#4       1920   125260492:   mNeoVis.2.1.1.chr7_hap1.g02144.t01#5    mNeoVis.evergladensis.1.1.chr7.g02097.t01#3    mNeoVis.vulgivaga.1.1.chr7.g02098.t01#4    mNeoVis.3.1.1.chr7.g01970.t01#6    mNeoVis.evergladensis.2.1.chr7.g02091.t01#2   mNeoVis.vulgivaga.2.1.chr7.g02112.t01#1       1921   125260493:   mNeoVis.evergladensis.1.1.chr1.g01429.t01#3    mNeoVis.vulgivaga.2.1.chr1.g01427.t01#1    mNeoVis.3.1.1.chr1.g01408.t01#6    mNeoVis.vulgivaga.1.1.chr1.g01428.t01#4    mNeoVis.evergladensis.2.1.chr1.g01426.t01#2   mNeoVis.2.1.1.chr1_hap1.g01461.t01#5       1922   125260494:   mNeoVis.evergladensis.2.1.chr7.g02092.t01#2    mNeoVis.vulgivaga.2.1.chr7.g02113.t01#1    mNeoVis.evergladensis.1.1.chr7.g02098.t01#3    mNeoVis.vulgivaga.1.1.chr7.g02099.t01#4   mNeoVis.3.1.1.chr7.g01971.t01#6       1923   125260495:   mNeoVis.2.1.1.chr7_hap1.g02146.t01#5    mNeoVis.vulgivaga.2.1.chr7.g02114.t01#1    mNeoVis.3.1.1.chr7.g01972.t01#6    mNeoVis.evergladensis.1.1.chr7.g02099.t01#3   mNeoVis.evergladensis.2.1.chr7.g02093.t01#2       1924   125260496:   mNeoVis.3.1.1.chr7.g01976.t01#6    mNeoVis.vulgivaga.1.1.chr7.g02101.t01#4    mNeoVis.2.1.1.chr7_hap1.g02149.t01#5    mNeoVis.evergladensis.2.1.chr7.g02096.t01#2    mNeoVis.vulgivaga.2.1.chr7.g02116.t01#1    mNeoVis.2.1.1.chr7_hap1.g02148.t01#5    mNeoVis.evergladensis.1.1.chr7.g02102.t01#3    mNeoVis.3.1.1.chr7.g01975.t01#6    mNeoVis.vulgivaga.2.1.chr7.g02117.t01#1      1925   125260497:   mNeoVis.3.1.1.chr7.g01973.t01#6    mNeoVis.evergladensis.1.1.chr7.g02100.t01#3    mNeoVis.vulgivaga.1.1.chr7.g02100.t01#4    mNeoVis.evergladensis.2.1.chr7.g02094.t01#2    mNeoVis.2.1.1.chr7_hap1.g02147.t01#5   mNeoVis.vulgivaga.2.1.chr7.g02115.t01#1       1926   125260498:   mNeoVis.evergladensis.2.1.chr1.g01428.t01#2    mNeoVis.vulgivaga.1.1.chr1.g01430.t01#4    mNeoVis.vulgivaga.2.1.chr1.g01429.t01#1    mNeoVis.3.1.1.chr1.g01410.t01#6    mNeoVis.2.1.1.chr1_hap1.g01462.t01#5   mNeoVis.evergladensis.1.1.chr1.g01431.t01#3       1927   125260499:   mNeoVis.2.1.1.chr7_hap1.g02150.t01#5    mNeoVis.evergladensis.1.1.chr7.g02103.t01#3    mNeoVis.evergladensis.2.1.chr7.g02097.t01#2    mNeoVis.vulgivaga.2.1.chr7.g02118.t01#1    mNeoVis.3.1.1.chr7.g01977.t01#6   mNeoVis.vulgivaga.1.1.chr7.g02102.t01#4       1928   125260500:   mNeoVis.vulgivaga.2.1.chr7.g02119.t01#1    mNeoVis.3.1.1.chr7.g01978.t01#6    mNeoVis.3.1.1.chr7.g01979.t01#6    mNeoVis.evergladensis.1.1.chr7.g02104.t01#3    mNeoVis.evergladensis.2.1.chr7.g02098.t01#2    mNeoVis.vulgivaga.1.1.chr7.g02103.t01#4   mNeoVis.2.1.1.chr7_hap1.g02151.t01#5     1929   125260501:   mNeoVis.3.1.1.chr7.g01981.t01#6    mNeoVis.vulgivaga.1.1.chr7.g02107.t01#4    mNeoVis.vulgivaga.2.1.chr7.g02121.t01#1    mNeoVis.evergladensis.2.1.chr7.g02101.t01#2    mNeoVis.evergladensis.1.1.chr7.g02107.t01#3   mNeoVis.2.1.1.chr7_hap1.g02154.t01#5       1930   125260502:   mNeoVis.vulgivaga.1.1.chr7.g02105.t01#4    mNeoVis.evergladensis.1.1.chr7.g02105.t01#3    mNeoVis.evergladensis.2.1.chr7.g02099.t01#2   mNeoVis.vulgivaga.2.1.chr7.g02120.t01#1         1931   125260503:   mNeoVis.vulgivaga.1.1.chr7.g02104.t01#4    mNeoVis.2.1.1.chr7_hap1.g02145.t01#5   mNeoVis.2.1.1.chr7_hap1.g02152.t01#5           1932   125260504:   mNeoVis.2.1.1.chr7_hap1.g02153.t01#5   mNeoVis.evergladensis.2.1.chr7.g02100.t01#2           1933   125260505:   mNeoVis.2.1.1.chr7_hap1.g00537.t01#5    mNeoVis.vulgivaga.2.1.chr7.g00539.t01#1    mNeoVis.evergladensis.1.1.chr7.g00545.t01#3    mNeoVis.evergladensis.2.1.chr7.g00537.t01#2    mNeoVis.3.1.1.chr7.g00519.t01#6   mNeoVis.vulgivaga.1.1.chr7.g00544.t01#4       1934   125260506:   mNeoVis.vulgivaga.2.1.chr7.g02122.t01#1    mNeoVis.3.1.1.chr7.g01980.t01#6    mNeoVis.vulgivaga.1.1.chr7.g02106.t01#4   mNeoVis.evergladensis.1.1.chr7.g02106.t01#3       1935   125260507:   mNeoVis.3.1.1.chr15.g00547.t01#6    mNeoVis.vulgivaga.1.1.chr15.g00592.t01#4    mNeoVis.2.1.1.chr15_hap1.g00767.t01#5    mNeoVis.evergladensis.2.1.chr15.g00590.t01#2    mNeoVis.evergladensis.1.1.chr15.g00022.t01#3    mNeoVis.vulgivaga.2.1.chr15.g00600.t01#1   mNeoVis.evergladensis.1.1.chr15.g00021.t01#3     1936   125260508:   mNeoVis.2.1.1.chr1_hap1.g01464.t01#5    mNeoVis.vulgivaga.2.1.chr1.g01431.t01#1    mNeoVis.evergladensis.1.1.chr1.g01433.t01#3    mNeoVis.vulgivaga.1.1.chr1.g01432.t01#4    mNeoVis.evergladensis.2.1.chr1.g01430.t01#2   mNeoVis.3.1.1.chr1.g01412.t01#6       1937   125260509:   mNeoVis.2.1.1.chr7_hap1.g02155.t01#5    mNeoVis.evergladensis.2.1.chr7.g02102.t01#2    mNeoVis.vulgivaga.2.1.chr7.g02123.t01#1    mNeoVis.evergladensis.1.1.chr7.g02108.t01#3    mNeoVis.3.1.1.chr7.g01982.t01#6   mNeoVis.vulgivaga.1.1.chr7.g02108.t01#4       1938   125260510:   mNeoVis.3.1.1.chr7.g01983.t01#6    mNeoVis.2.1.1.chr7_hap1.g02156.t01#5    mNeoVis.vulgivaga.2.1.chr7.g02124.t01#1    mNeoVis.evergladensis.1.1.chr7.g02109.t01#3    mNeoVis.vulgivaga.1.1.chr7.g02109.t01#4   mNeoVis.evergladensis.2.1.chr7.g02103.t01#2       1939   125260511:   mNeoVis.3.1.1.chr7.g01989.t01#6    mNeoVis.2.1.1.chr7_hap1.g02163.t01#5    mNeoVis.vulgivaga.2.1.chr7.g02132.t01#1    mNeoVis.evergladensis.1.1.chr7.g02116.t01#3    mNeoVis.evergladensis.2.1.chr7.g02109.t01#2    mNeoVis.vulgivaga.1.1.chr7.g02116.t01#4    mNeoVis.evergladensis.1.1.chr7.g02115.t01#3    mNeoVis.3.1.1.chr7.g01988.t01#6    mNeoVis.evergladensis.2.1.chr7.g02111.t01#2      1940   125260512:   mNeoVis.evergladensis.2.1.chr7.g02112.t01#2    mNeoVis.vulgivaga.1.1.chr7.g02117.t01#4    mNeoVis.vulgivaga.2.1.chr7.g02134.t01#1    mNeoVis.3.1.1.chr7.g01990.t01#6    mNeoVis.vulgivaga.2.1.chr7.g02133.t01#1    mNeoVis.3.1.1.chr7.g01991.t01#6    mNeoVis.2.1.1.chr7_hap1.g02164.t01#5   mNeoVis.vulgivaga.1.1.chr7.g02118.t01#4     1941   125260513:   mNeoVis.3.1.1.chr7.g01984.t01#6    mNeoVis.2.1.1.chr7_hap1.g02157.t01#5    mNeoVis.evergladensis.2.1.chr7.g02104.t01#2    mNeoVis.evergladensis.1.1.chr7.g02110.t01#3    mNeoVis.vulgivaga.1.1.chr7.g02110.t01#4   mNeoVis.vulgivaga.2.1.chr7.g02125.t01#1       1942   125260514:   mNeoVis.vulgivaga.2.1.chr7.g02126.t01#1    mNeoVis.evergladensis.2.1.chr7.g02105.t01#2    mNeoVis.2.1.1.chr7_hap1.g02158.t01#5    mNeoVis.3.1.1.chr7.g01985.t01#6    mNeoVis.evergladensis.1.1.chr7.g02111.t01#3   mNeoVis.vulgivaga.1.1.chr7.g02111.t01#4       1943   125260515:   mNeoVis.vulgivaga.1.1.chr7.g02114.t01#4    mNeoVis.evergladensis.2.1.chr7.g02108.t01#2    mNeoVis.evergladensis.1.1.chr7.g02113.t01#3    mNeoVis.vulgivaga.2.1.chr7.g02129.t01#1    mNeoVis.3.1.1.chr7.g01986.t01#6   mNeoVis.2.1.1.chr7_hap1.g02161.t01#5       1944   125260516:   mNeoVis.vulgivaga.2.1.chr7.g02130.t01#1    mNeoVis.2.1.1.chr7_hap1.g02162.t01#5    mNeoVis.evergladensis.2.1.chr7.g02110.t01#2    mNeoVis.vulgivaga.1.1.chr7.g02115.t01#4    mNeoVis.3.1.1.chr7.g01987.t01#6   mNeoVis.evergladensis.1.1.chr7.g02114.t01#3       1945   125260517:   mNeoVis.vulgivaga.1.1.chr7.g02113.t01#4    mNeoVis.vulgivaga.2.1.chr7.g02128.t01#1   mNeoVis.evergladensis.2.1.chr7.g02107.t01#2         1946   125260518:  mNeoVis.2.1.1.chr7_hap1.g02160.t01#5               1947   125260519:   mNeoVis.vulgivaga.2.1.chr7.g02127.t01#1    mNeoVis.3.1.1.chr7.g01960.t01#6    mNeoVis.2.1.1.chr7_hap1.g02159.t01#5    mNeoVis.evergladensis.1.1.chr7.g02112.t01#3    mNeoVis.vulgivaga.1.1.chr7.g02112.t01#4   mNeoVis.evergladensis.2.1.chr7.g02106.t01#2       1948   125260520:   mNeoVis.vulgivaga.2.1.chr1.g01432.t01#1    mNeoVis.2.1.1.chr1_hap1.g01465.t01#5    mNeoVis.evergladensis.2.1.chr1.g01431.t01#2    mNeoVis.evergladensis.1.1.chr1.g01434.t01#3    mNeoVis.vulgivaga.1.1.chr1.g01433.t01#4   mNeoVis.3.1.1.chr1.g01413.t01#6       1949   125260521:   mNeoVis.vulgivaga.1.1.chr1.g01434.t01#4    mNeoVis.3.1.1.chr1.g01414.t01#6    mNeoVis.2.1.1.chr1_hap1.g01466.t01#5    mNeoVis.evergladensis.2.1.chr1.g01432.t01#2    mNeoVis.vulgivaga.2.1.chr1.g01433.t01#1   mNeoVis.evergladensis.1.1.chr1.g01435.t01#3       1950   125260522:   mNeoVis.vulgivaga.2.1.chr7.g02135.t01#1    mNeoVis.vulgivaga.1.1.chr7.g02119.t01#4    mNeoVis.2.1.1.chr7_hap1.g02165.t01#5    mNeoVis.3.1.1.chr7.g01992.t01#6    mNeoVis.evergladensis.2.1.chr7.g02113.t01#2   mNeoVis.evergladensis.1.1.chr7.g02117.t01#3       1951   125260523:   mNeoVis.vulgivaga.2.1.chr1.g01434.t01#1   mNeoVis.vulgivaga.1.1.chr1.g01435.t01#4             1952   125260524:   mNeoVis.vulgivaga.2.1.chr1.g01435.t01#1    mNeoVis.2.1.1.chr1_hap1.g01467.t01#5    mNeoVis.vulgivaga.1.1.chr1.g01436.t01#4    mNeoVis.evergladensis.1.1.chr1.g01437.t01#3    mNeoVis.evergladensis.2.1.chr1.g01433.t01#2   mNeoVis.3.1.1.chr1.g01416.t01#6       1953   125260525:  mNeoVis.vulgivaga.2.1.chr7.g02136.t01#1               1954   125260526:   mNeoVis.vulgivaga.2.1.chr7.g02137.t01#1    mNeoVis.vulgivaga.1.1.chr7.g02121.t01#4    mNeoVis.evergladensis.1.1.chr7.g02118.t01#3    mNeoVis.2.1.1.chr7_hap1.g02166.t01#5    mNeoVis.3.1.1.chr7.g01994.t01#6   mNeoVis.evergladensis.2.1.chr7.g02115.t01#2       1955   125260527:   mNeoVis.vulgivaga.1.1.chr7.g00545.t01#4    mNeoVis.2.1.1.chr7_hap1.g00538.t01#5    mNeoVis.evergladensis.2.1.chr7.g00538.t01#2    mNeoVis.3.1.1.chr7.g00520.t01#6    mNeoVis.vulgivaga.2.1.chr7.g00540.t01#1   mNeoVis.evergladensis.1.1.chr7.g00546.t01#3       1956   125260528:   mNeoVis.evergladensis.2.1.chr7.g02116.t01#2    mNeoVis.vulgivaga.1.1.chr7.g02122.t01#4    mNeoVis.2.1.1.chr7_hap1.g02167.t01#5    mNeoVis.evergladensis.1.1.chr7.g02119.t01#3    mNeoVis.vulgivaga.2.1.chr7.g02138.t01#1   mNeoVis.3.1.1.chr7.g01995.t01#6       1957   125260529:   mNeoVis.vulgivaga.2.1.chr7.g02139.t01#1    mNeoVis.vulgivaga.1.1.chr7.g02123.t01#4    mNeoVis.evergladensis.1.1.chr7.g02120.t01#3    mNeoVis.3.1.1.chr7.g01996.t01#6    mNeoVis.2.1.1.chr7_hap1.g02169.t01#5   mNeoVis.evergladensis.2.1.chr7.g02117.t01#2       1958   125260530:   mNeoVis.vulgivaga.2.1.chr1.g01358.t01#1    mNeoVis.3.1.1.chr1.g01340.t01#6   mNeoVis.evergladensis.2.1.chr1.g01356.t01#2         1959   125260531:   mNeoVis.vulgivaga.1.1.chr1.g01358.t01#4    mNeoVis.evergladensis.1.1.chr1.g01363.t01#3    mNeoVis.evergladensis.2.1.chr1.g01358.t01#2    mNeoVis.vulgivaga.2.1.chr1.g01359.t01#1    mNeoVis.3.1.1.chr1.g01342.t01#6   mNeoVis.2.1.1.chr1_hap1.g01393.t01#5       1960   125260532:  mNeoVis.vulgivaga.2.1.chr1.g01360.t01#1               1961   125260533:   mNeoVis.vulgivaga.2.1.chr1.g01361.t01#1    mNeoVis.evergladensis.2.1.chr1.g01360.t01#2    mNeoVis.2.1.1.chr1_hap1.g01395.t01#5    mNeoVis.3.1.1.chr1.g01344.t01#6    mNeoVis.evergladensis.1.1.chr1.g01364.t01#3   mNeoVis.vulgivaga.1.1.chr1.g01360.t01#4       1962   125260534:   mNeoVis.evergladensis.2.1.chr7.g02118.t01#2    mNeoVis.vulgivaga.2.1.chr7.g02140.t01#1    mNeoVis.2.1.1.chr7_hap1.g02170.t01#5    mNeoVis.evergladensis.1.1.chr7.g02121.t01#3    mNeoVis.vulgivaga.1.1.chr7.g02124.t01#4   mNeoVis.3.1.1.chr7.g01997.t01#6       1963   125260535:   mNeoVis.vulgivaga.2.1.chr7.g02141.t01#1    mNeoVis.2.1.1.chr7_hap1.g02171.t01#5    mNeoVis.3.1.1.chr7.g01998.t01#6    mNeoVis.evergladensis.1.1.chr7.g02122.t01#3    mNeoVis.vulgivaga.1.1.chr7.g02125.t01#4   mNeoVis.evergladensis.2.1.chr7.g02119.t01#2       1964   125260536:   mNeoVis.vulgivaga.1.1.chr1.g01361.t01#4    mNeoVis.evergladensis.1.1.chr1.g01365.t01#3    mNeoVis.2.1.1.chr1_hap1.g01396.t01#5    mNeoVis.3.1.1.chr1.g01345.t01#6    mNeoVis.evergladensis.2.1.chr1.g01361.t01#2   mNeoVis.vulgivaga.2.1.chr1.g01362.t01#1       1965   125260537:   mNeoVis.3.1.1.chr1.g01346.t01#6    mNeoVis.vulgivaga.1.1.chr1.g01363.t01#4    mNeoVis.vulgivaga.2.1.chr1.g01363.t01#1    mNeoVis.evergladensis.2.1.chr1.g01363.t01#2    mNeoVis.evergladensis.1.1.chr1.g01367.t01#3   mNeoVis.2.1.1.chr1_hap1.g01398.t01#5       1966   125260538:   mNeoVis.evergladensis.1.1.chr7.g01951.t01#3    mNeoVis.vulgivaga.2.1.chr7.g01964.t01#1    mNeoVis.evergladensis.2.1.chr7.g01947.t01#2    mNeoVis.3.1.1.chr7.g01851.t01#6    mNeoVis.vulgivaga.1.1.chr7.g01958.t01#4   mNeoVis.2.1.1.chr7_hap1.g01983.t01#5       1967   125260539:   mNeoVis.vulgivaga.2.1.chr7.g00541.t01#1    mNeoVis.3.1.1.chr7.g00521.t01#6    mNeoVis.2.1.1.chr7_hap1.g00539.t01#5    mNeoVis.evergladensis.1.1.chr7.g00547.t01#3    mNeoVis.evergladensis.2.1.chr7.g00539.t01#2   mNeoVis.vulgivaga.1.1.chr7.g00546.t01#4       1968   125260540:   mNeoVis.vulgivaga.1.1.chr9.g00370.t01#4    mNeoVis.vulgivaga.2.1.chr9.g00372.t01#1    mNeoVis.2.1.1.chr10_hap1.g00388.t01#5    mNeoVis.evergladensis.1.1.chr9.g00377.t01#3    mNeoVis.3.1.1.chr9.g00358.t01#6   mNeoVis.evergladensis.2.1.chr8.g00370.t01#2       1969   125260541:   mNeoVis.evergladensis.2.1.chr7.g01948.t01#2    mNeoVis.2.1.1.chr7_hap1.g01984.t01#5    mNeoVis.evergladensis.1.1.chr7.g01952.t01#3    mNeoVis.vulgivaga.2.1.chr7.g01965.t01#1    mNeoVis.vulgivaga.1.1.chr7.g01959.t01#4   mNeoVis.3.1.1.chr7.g01852.t01#6       1970   125260542:   mNeoVis.vulgivaga.1.1.chr1.g01364.t01#4    mNeoVis.3.1.1.chr1.g01347.t01#6    mNeoVis.evergladensis.2.1.chr1.g01364.t01#2    mNeoVis.evergladensis.1.1.chr1.g01368.t01#3    mNeoVis.vulgivaga.2.1.chr1.g01364.t01#1   mNeoVis.2.1.1.chr1_hap1.g01399.t01#5       1971   125260543:   mNeoVis.evergladensis.1.1.chr1.g01369.t01#3    mNeoVis.3.1.1.chr1.g01343.t01#6    mNeoVis.vulgivaga.1.1.chr1.g01365.t01#4   mNeoVis.vulgivaga.2.1.chr1.g01365.t01#1         1972   125260544:  mNeoVis.vulgivaga.2.1.chr1.g01366.t01#1               1973   125260545:   mNeoVis.vulgivaga.2.1.chr1.g01367.t01#1    mNeoVis.vulgivaga.1.1.chr1.g01368.t01#4    mNeoVis.evergladensis.2.1.chr1.g01367.t01#2    mNeoVis.2.1.1.chr1_hap1.g01401.t01#5    mNeoVis.evergladensis.1.1.chr1.g01372.t01#3   mNeoVis.3.1.1.chr1.g01348.t01#6       1974   125260546:   mNeoVis.evergladensis.2.1.chr1.g01365.t01#2    mNeoVis.evergladensis.1.1.chr1.g01370.t01#3    mNeoVis.vulgivaga.1.1.chr1.g01366.t01#4   mNeoVis.2.1.1.chr1_hap1.g01400.t01#5         1975   125260547:   mNeoVis.3.1.1.chr7.g01854.t01#6    mNeoVis.evergladensis.2.1.chr7.g01949.t01#2    mNeoVis.vulgivaga.1.1.chr7.g01960.t01#4    mNeoVis.evergladensis.1.1.chr7.g01953.t01#3    mNeoVis.2.1.1.chr7_hap1.g01985.t01#5   mNeoVis.vulgivaga.2.1.chr7.g01966.t01#1       1976   125260548:   mNeoVis.vulgivaga.1.1.chr1.g01369.t01#4    mNeoVis.evergladensis.1.1.chr5.g00046.t01#3    mNeoVis.evergladensis.2.1.chr5.g00046.t01#2    mNeoVis.2.1.1.chr2_hap1.g00046.t01#5    mNeoVis.vulgivaga.2.1.chr1.g01368.t01#1    mNeoVis.vulgivaga.1.1.chr5.g00044.t01#4    mNeoVis.vulgivaga.2.1.chr5.g00043.t01#1    mNeoVis.2.1.1.chr1_hap1.g01402.t01#5    mNeoVis.3.1.1.chr1.g01349.t01#6      1977   125260549:   mNeoVis.vulgivaga.1.1.chr1.g01372.t01#4    mNeoVis.evergladensis.2.1.chr1.g01368.t01#2    mNeoVis.vulgivaga.2.1.chr1.g01371.t01#1    mNeoVis.2.1.1.chr1_hap1.g01405.t01#5    mNeoVis.evergladensis.1.1.chr1.g01373.t01#3   mNeoVis.3.1.1.chr1.g01352.t01#6       1978   125260550:   mNeoVis.evergladensis.2.1.chr1.g01369.t01#2    mNeoVis.2.1.1.chr1_hap1.g01403.t01#5    mNeoVis.vulgivaga.2.1.chr1.g01372.t01#1    mNeoVis.evergladensis.1.1.chr1.g01374.t01#3    mNeoVis.vulgivaga.2.1.chr1.g01369.t01#1   mNeoVis.vulgivaga.1.1.chr1.g01373.t01#4       1979   125260551:   mNeoVis.evergladensis.1.1.chr7.g01954.t01#3    mNeoVis.vulgivaga.1.1.chr7.g01961.t01#4    mNeoVis.evergladensis.2.1.chr7.g01950.t01#2    mNeoVis.vulgivaga.2.1.chr7.g01967.t01#1   mNeoVis.2.1.1.chr7_hap1.g01986.t01#5       1980   125260552:   mNeoVis.2.1.1.chr1_hap1.g01404.t01#5    mNeoVis.2.1.1.chr10_hap1.g00046.t01#5    mNeoVis.3.1.1.chr1.g01351.t01#6    mNeoVis.vulgivaga.1.1.chr1.g01371.t01#4    mNeoVis.vulgivaga.1.1.chr9.g00039.t01#4    mNeoVis.vulgivaga.2.1.chr9.g00045.t01#1    mNeoVis.3.1.1.chr9.g00026.t01#6    mNeoVis.evergladensis.1.1.chr9.g00045.t01#3    mNeoVis.evergladensis.2.1.chr8.g00045.t01#2      1981   125260553:   mNeoVis.3.1.1.chr7.g00965.t01#6    mNeoVis.vulgivaga.1.1.chr7.g01022.t01#4    mNeoVis.vulgivaga.2.1.chr7.g01028.t01#1    mNeoVis.evergladensis.2.1.chr7.g01020.t01#2    mNeoVis.evergladensis.1.1.chr7.g01026.t01#3   mNeoVis.2.1.1.chr7_hap1.g01027.t01#5       1982   125260554:   mNeoVis.3.1.1.chr1.g01353.t01#6    mNeoVis.vulgivaga.1.1.chr1.g01374.t01#4    mNeoVis.2.1.1.chr1_hap1.g01406.t01#5    mNeoVis.vulgivaga.2.1.chr1.g01373.t01#1   mNeoVis.evergladensis.1.1.chr1.g01375.t01#3       1983   125260555:   mNeoVis.evergladensis.2.1.chr1.g01370.t01#2    mNeoVis.3.1.1.chr1.g01354.t01#6    mNeoVis.vulgivaga.1.1.chr1.g01375.t01#4    mNeoVis.vulgivaga.2.1.chr1.g01374.t01#1    mNeoVis.evergladensis.1.1.chr1.g01376.t01#3   mNeoVis.2.1.1.chr1_hap1.g01407.t01#5       1984   125260556:   mNeoVis.2.1.1.chr7_hap1.g01987.t01#5    mNeoVis.evergladensis.2.1.chr7.g01951.t01#2    mNeoVis.vulgivaga.2.1.chr7.g01968.t01#1    mNeoVis.vulgivaga.1.1.chr7.g01962.t01#4   mNeoVis.evergladensis.1.1.chr7.g01955.t01#3       1985   125260557:   mNeoVis.3.1.1.chr15.g00548.t01#6    mNeoVis.vulgivaga.2.1.chr15.g00601.t01#1    mNeoVis.evergladensis.2.1.chr15.g00591.t01#2    mNeoVis.vulgivaga.1.1.chr15.g00593.t01#4    mNeoVis.2.1.1.chr15_hap1.g00768.t01#5   mNeoVis.evergladensis.1.1.chr15.g00020.t01#3       1986   125260558:   mNeoVis.evergladensis.2.1.chr1.g01371.t01#2    mNeoVis.2.1.1.chr1_hap1.g01408.t01#5    mNeoVis.evergladensis.1.1.chr1.g01377.t01#3    mNeoVis.vulgivaga.1.1.chr1.g01376.t01#4    mNeoVis.vulgivaga.2.1.chr1.g01375.t01#1   mNeoVis.3.1.1.chr1.g01355.t01#6       1987   125260559:   mNeoVis.vulgivaga.2.1.chr7.g01969.t01#1    mNeoVis.evergladensis.1.1.chr7.g01956.t01#3    mNeoVis.2.1.1.chr7_hap1.g01988.t01#5    mNeoVis.vulgivaga.1.1.chr7.g01963.t01#4   mNeoVis.evergladensis.2.1.chr7.g01952.t01#2       1988   125260560:   mNeoVis.vulgivaga.1.1.chr7.g01964.t01#4    mNeoVis.2.1.1.chr7_hap1.g01989.t01#5    mNeoVis.3.1.1.chr7.g01856.t01#6    mNeoVis.evergladensis.2.1.chr7.g01953.t01#2    mNeoVis.vulgivaga.2.1.chr7.g01970.t01#1   mNeoVis.evergladensis.1.1.chr7.g01957.t01#3       1989   125260561:   mNeoVis.vulgivaga.1.1.chr1.g01377.t01#4    mNeoVis.2.1.1.chr1_hap1.g01409.t01#5    mNeoVis.vulgivaga.2.1.chr1.g01376.t01#1    mNeoVis.evergladensis.1.1.chr1.g01378.t01#3    mNeoVis.3.1.1.chr1.g01356.t01#6   mNeoVis.evergladensis.2.1.chr1.g01372.t01#2       1990   125260562:   mNeoVis.3.1.1.chr7.g00522.t01#6    mNeoVis.evergladensis.1.1.chr7.g00548.t01#3    mNeoVis.vulgivaga.2.1.chr7.g00542.t01#1    mNeoVis.vulgivaga.1.1.chr7.g00547.t01#4    mNeoVis.evergladensis.2.1.chr7.g00540.t01#2   mNeoVis.2.1.1.chr7_hap1.g00540.t01#5       1991   125260563:   mNeoVis.3.1.1.chr1.g01357.t01#6    mNeoVis.evergladensis.2.1.chr1.g01373.t01#2    mNeoVis.vulgivaga.1.1.chr1.g01378.t01#4    mNeoVis.vulgivaga.2.1.chr1.g01377.t01#1    mNeoVis.2.1.1.chr1_hap1.g01410.t01#5   mNeoVis.evergladensis.1.1.chr1.g01379.t01#3       1992   125260564:   mNeoVis.vulgivaga.2.1.chr7.g01971.t01#1    mNeoVis.3.1.1.chr7.g01857.t01#6    mNeoVis.evergladensis.1.1.chr7.g01958.t01#3    mNeoVis.vulgivaga.1.1.chr7.g01965.t01#4    mNeoVis.evergladensis.2.1.chr7.g01954.t01#2   mNeoVis.2.1.1.chr7_hap1.g01990.t01#5       1993   125260565:   mNeoVis.vulgivaga.2.1.chr1.g01378.t01#1    mNeoVis.evergladensis.1.1.chr1.g01380.t01#3    mNeoVis.3.1.1.chr1.g01358.t01#6    mNeoVis.evergladensis.2.1.chr1.g01374.t01#2    mNeoVis.2.1.1.chr1_hap1.g01411.t01#5   mNeoVis.vulgivaga.1.1.chr1.g01379.t01#4       1994   125260566:   mNeoVis.3.1.1.chr7.g01858.t01#6    mNeoVis.vulgivaga.1.1.chr7.g01966.t01#4    mNeoVis.evergladensis.1.1.chr7.g01959.t01#3    mNeoVis.2.1.1.chr7_hap1.g01991.t01#5    mNeoVis.evergladensis.2.1.chr7.g01955.t01#2   mNeoVis.vulgivaga.2.1.chr7.g01972.t01#1       1995   125260567:   mNeoVis.evergladensis.1.1.chr1.g01381.t01#3    mNeoVis.3.1.1.chr1.g01359.t01#6    mNeoVis.vulgivaga.1.1.chr1.g01380.t01#4    mNeoVis.2.1.1.chr1_hap1.g01412.t01#5    mNeoVis.vulgivaga.2.1.chr1.g01379.t01#1   mNeoVis.evergladensis.2.1.chr1.g01375.t01#2       1996   125260568:   mNeoVis.vulgivaga.2.1.chr1.g01380.t01#1    mNeoVis.vulgivaga.1.1.chr1.g01381.t01#4    mNeoVis.evergladensis.1.1.chr1.g01382.t01#3    mNeoVis.2.1.1.chr1_hap1.g01414.t01#5    mNeoVis.3.1.1.chr1.g01360.t01#6   mNeoVis.evergladensis.2.1.chr1.g01376.t01#2       1997   125260569:   mNeoVis.vulgivaga.2.1.chr7.g01973.t01#1    mNeoVis.2.1.1.chr7_hap1.g01992.t01#5    mNeoVis.evergladensis.2.1.chr7.g01956.t01#2    mNeoVis.evergladensis.1.1.chr7.g01960.t01#3    mNeoVis.3.1.1.chr7.g01859.t01#6   mNeoVis.vulgivaga.1.1.chr7.g01967.t01#4       1998   125260570:   mNeoVis.evergladensis.2.1.chr1.g01377.t01#2    mNeoVis.2.1.1.chr1_hap1.g01415.t01#5    mNeoVis.3.1.1.chr1.g01361.t01#6    mNeoVis.vulgivaga.1.1.chr1.g01382.t01#4    mNeoVis.evergladensis.1.1.chr1.g01383.t01#3   mNeoVis.vulgivaga.2.1.chr1.g01381.t01#1       1999   125260571:   mNeoVis.2.1.1.chr1_hap1.g01416.t01#5    mNeoVis.vulgivaga.2.1.chr1.g01382.t01#1    mNeoVis.vulgivaga.1.1.chr1.g01383.t01#4    mNeoVis.evergladensis.2.1.chr1.g01378.t01#2    mNeoVis.3.1.1.chr1.g01363.t01#6   mNeoVis.evergladensis.1.1.chr1.g01384.t01#3       2000   125260572:   mNeoVis.vulgivaga.2.1.chr7.g01974.t01#1    mNeoVis.evergladensis.2.1.chr7.g01957.t01#2    mNeoVis.3.1.1.chr7.g01860.t01#6    mNeoVis.vulgivaga.1.1.chr7.g01968.t01#4    mNeoVis.evergladensis.1.1.chr7.g01961.t01#3   mNeoVis.2.1.1.chr7_hap1.g01993.t01#5       2001   125260573:   mNeoVis.vulgivaga.2.1.chr7.g01975.t01#1    mNeoVis.3.1.1.chr7.g01861.t01#6    mNeoVis.evergladensis.1.1.chr7.g01962.t01#3    mNeoVis.vulgivaga.1.1.chr7.g01969.t01#4    mNeoVis.evergladensis.2.1.chr7.g01958.t01#2   mNeoVis.2.1.1.chr7_hap1.g01994.t01#5       2002   125260574:   mNeoVis.vulgivaga.2.1.chr15.g00602.t01#1    mNeoVis.3.1.1.chr15.g00549.t01#6    mNeoVis.evergladensis.2.1.chr15.g00592.t01#2    mNeoVis.vulgivaga.1.1.chr15.g00594.t01#4    mNeoVis.2.1.1.chr15_hap1.g00769.t01#5   mNeoVis.evergladensis.1.1.chr15.g00019.t01#3       2003   125260575:   mNeoVis.evergladensis.2.1.chr1.g01379.t01#2    mNeoVis.evergladensis.1.1.chr1.g01385.t01#3    mNeoVis.3.1.1.chr1.g01364.t01#6    mNeoVis.vulgivaga.1.1.chr1.g01384.t01#4    mNeoVis.vulgivaga.2.1.chr1.g01383.t01#1   mNeoVis.2.1.1.chr1_hap1.g01417.t01#5       2004   125260576:   mNeoVis.evergladensis.1.1.chr1.g01386.t01#3    mNeoVis.3.1.1.chr1.g01365.t01#6    mNeoVis.vulgivaga.1.1.chr1.g01385.t01#4    mNeoVis.2.1.1.chr1_hap1.g01418.t01#5    mNeoVis.evergladensis.2.1.chr1.g01380.t01#2   mNeoVis.vulgivaga.2.1.chr1.g01384.t01#1       2005   125260577:   mNeoVis.vulgivaga.1.1.chr7.g01970.t01#4    mNeoVis.evergladensis.2.1.chr7.g01959.t01#2    mNeoVis.evergladensis.1.1.chr7.g01963.t01#3    mNeoVis.3.1.1.chr7.g01862.t01#6    mNeoVis.2.1.1.chr7_hap1.g01995.t01#5   mNeoVis.vulgivaga.2.1.chr7.g01976.t01#1       2006   125260578:   mNeoVis.2.1.1.chr7_hap1.g01996.t01#5    mNeoVis.vulgivaga.2.1.chr7.g01977.t01#1    mNeoVis.evergladensis.1.1.chr7.g01964.t01#3    mNeoVis.vulgivaga.1.1.chr7.g01971.t01#4    mNeoVis.3.1.1.chr7.g01863.t01#6   mNeoVis.evergladensis.2.1.chr7.g01960.t01#2       2007   125260579:   mNeoVis.2.1.1.chr1_hap1.g01419.t01#5    mNeoVis.vulgivaga.1.1.chr1.g01386.t01#4    mNeoVis.vulgivaga.2.1.chr1.g01385.t01#1    mNeoVis.evergladensis.1.1.chr1.g01387.t01#3    mNeoVis.evergladensis.2.1.chr1.g01381.t01#2   mNeoVis.3.1.1.chr1.g01366.t01#6       2008   125260580:   mNeoVis.vulgivaga.2.1.chr15.g00603.t01#1    mNeoVis.3.1.1.chr15.g00550.t01#6    mNeoVis.evergladensis.2.1.chr15.g00593.t01#2    mNeoVis.vulgivaga.1.1.chr15.g00595.t01#4    mNeoVis.2.1.1.chr15_hap1.g00770.t01#5   mNeoVis.evergladensis.1.1.chr15.g00018.t01#3       2009   125260581:   mNeoVis.evergladensis.2.1.chr7.g00541.t01#2    mNeoVis.evergladensis.1.1.chr7.g00549.t01#3    mNeoVis.2.1.1.chr7_hap1.g00541.t01#5    mNeoVis.3.1.1.chr7.g00523.t01#6    mNeoVis.vulgivaga.2.1.chr7.g00543.t01#1   mNeoVis.vulgivaga.1.1.chr7.g00548.t01#4       2010   125260582:   mNeoVis.evergladensis.2.1.chr1.g01434.t01#2    mNeoVis.vulgivaga.2.1.chr1.g01436.t01#1    mNeoVis.evergladensis.1.1.chr1.g01438.t01#3    mNeoVis.2.1.1.chr1_hap1.g01468.t01#5    mNeoVis.vulgivaga.1.1.chr1.g01437.t01#4   mNeoVis.3.1.1.chr1.g01417.t01#6       2011   125260583:   mNeoVis.evergladensis.2.1.chr1.g01435.t01#2    mNeoVis.vulgivaga.2.1.chr1.g01437.t01#1    mNeoVis.evergladensis.1.1.chr1.g01439.t01#3    mNeoVis.2.1.1.chr1_hap1.g01469.t01#5   mNeoVis.vulgivaga.1.1.chr1.g01438.t01#4       2012   125260584:   mNeoVis.3.1.1.chr1.g01418.t01#6    mNeoVis.evergladensis.1.1.chr1.g01440.t01#3    mNeoVis.evergladensis.2.1.chr1.g01436.t01#2    mNeoVis.2.1.1.chr1_hap1.g01470.t01#5    mNeoVis.vulgivaga.1.1.chr1.g01439.t01#4   mNeoVis.vulgivaga.2.1.chr1.g01438.t01#1       2013   125260585:   mNeoVis.vulgivaga.2.1.chr1.g01439.t01#1    mNeoVis.vulgivaga.1.1.chr1.g01440.t01#4    mNeoVis.evergladensis.2.1.chr1.g01437.t01#2    mNeoVis.3.1.1.chr1.g01419.t01#6    mNeoVis.2.1.1.chr1_hap1.g01471.t01#5   mNeoVis.evergladensis.1.1.chr1.g01441.t01#3       2014   125260586:   mNeoVis.2.1.1.chr7_hap1.g01997.t01#5    mNeoVis.vulgivaga.2.1.chr7.g01978.t01#1    mNeoVis.vulgivaga.1.1.chr7.g01972.t01#4    mNeoVis.3.1.1.chr1.g00031.t01#6    mNeoVis.evergladensis.1.1.chr7.g01965.t01#3   mNeoVis.evergladensis.2.1.chr7.g01961.t01#2       2015   125260587:   mNeoVis.2.1.1.chr7_hap1.g01998.t01#5    mNeoVis.3.1.1.chr7.g01865.t01#6    mNeoVis.evergladensis.2.1.chr7.g01962.t01#2    mNeoVis.vulgivaga.1.1.chr7.g01973.t01#4   mNeoVis.vulgivaga.2.1.chr7.g01979.t01#1       2016   125260588:   mNeoVis.vulgivaga.2.1.chr1.g01440.t01#1    mNeoVis.3.1.1.chr1.g01420.t01#6    mNeoVis.vulgivaga.1.1.chr1.g01441.t01#4    mNeoVis.2.1.1.chr1_hap1.g01472.t01#5    mNeoVis.evergladensis.2.1.chr1.g01439.t01#2   mNeoVis.evergladensis.1.1.chr1.g01442.t01#3       2017   125260589:   mNeoVis.3.1.1.chr1.g01325.t01#6    mNeoVis.2.1.1.chr1_hap1.g01378.t01#5    mNeoVis.evergladensis.2.1.chr1.g01343.t01#2    mNeoVis.vulgivaga.2.1.chr1.g01345.t01#1    mNeoVis.vulgivaga.1.1.chr1.g01345.t01#4   mNeoVis.evergladensis.1.1.chr1.g01347.t01#3       2018   125260590:   mNeoVis.evergladensis.2.1.chr7.g01963.t01#2    mNeoVis.evergladensis.1.1.chr7.g01967.t01#3    mNeoVis.3.1.1.chr7.g01866.t01#6    mNeoVis.vulgivaga.2.1.chr7.g01980.t01#1    mNeoVis.vulgivaga.1.1.chr7.g01974.t01#4   mNeoVis.2.1.1.chr7_hap1.g01999.t01#5       2019   125260591:   mNeoVis.vulgivaga.1.1.chr1.g01346.t01#4    mNeoVis.2.1.1.chr1_hap1.g01379.t01#5    mNeoVis.3.1.1.chr1.g01326.t01#6    mNeoVis.evergladensis.1.1.chr1.g01348.t01#3    mNeoVis.evergladensis.2.1.chr1.g01344.t01#2   mNeoVis.vulgivaga.2.1.chr1.g01346.t01#1       2020   125260592:   mNeoVis.vulgivaga.2.1.chr1.g01347.t01#1    mNeoVis.2.1.1.chr1_hap1.g01380.t01#5    mNeoVis.vulgivaga.1.1.chr1.g01347.t01#4    mNeoVis.3.1.1.chr1.g01327.t01#6    mNeoVis.evergladensis.2.1.chr1.g01345.t01#2   mNeoVis.evergladensis.1.1.chr1.g01349.t01#3       2021   125260593:   mNeoVis.evergladensis.2.1.chr7.g01964.t01#2    mNeoVis.vulgivaga.1.1.chr7.g01975.t01#4    mNeoVis.2.1.1.chr7_hap1.g02000.t01#5    mNeoVis.vulgivaga.2.1.chr7.g01981.t01#1    mNeoVis.evergladensis.1.1.chr7.g01968.t01#3   mNeoVis.3.1.1.chr7.g01867.t01#6       2022   125260594:   mNeoVis.vulgivaga.2.1.chr1.g01348.t01#1    mNeoVis.2.1.1.chr1_hap1.g01381.t01#5    mNeoVis.3.1.1.chr1.g01328.t01#6    mNeoVis.evergladensis.1.1.chr1.g01350.t01#3    mNeoVis.vulgivaga.1.1.chr1.g01348.t01#4   mNeoVis.evergladensis.2.1.chr1.g01346.t01#2       2023   125260595:   mNeoVis.3.1.1.chr7.g01868.t01#6    mNeoVis.evergladensis.1.1.chr7.g01969.t01#3    mNeoVis.evergladensis.2.1.chr7.g01965.t01#2    mNeoVis.vulgivaga.2.1.chr7.g01982.t01#1    mNeoVis.vulgivaga.1.1.chr7.g01976.t01#4   mNeoVis.2.1.1.chr7_hap1.g02001.t01#5       2024   125260596:   mNeoVis.vulgivaga.1.1.chr14.g00454.t01#4    mNeoVis.2.1.1.chr14_hap1.g00463.t01#5    mNeoVis.evergladensis.1.1.chr14.g00461.t01#3    mNeoVis.evergladensis.2.1.chr14.g00460.t01#2    mNeoVis.3.1.1.chr14.g00458.t01#6    mNeoVis.evergladensis.2.1.chr14.g00459.t01#2   mNeoVis.vulgivaga.2.1.chr14.g00461.t01#1     2025   125260597:   mNeoVis.evergladensis.1.1.chr1.g01351.t01#3    mNeoVis.evergladensis.2.1.chr1.g01347.t01#2    mNeoVis.2.1.1.chr1_hap1.g01382.t01#5    mNeoVis.vulgivaga.1.1.chr1.g01349.t01#4    mNeoVis.3.1.1.chr1.g01329.t01#6   mNeoVis.vulgivaga.2.1.chr1.g01349.t01#1       2026   125260598:   mNeoVis.vulgivaga.2.1.chr13.g00222.t01#1    mNeoVis.evergladensis.1.1.chr13.g00224.t01#3    mNeoVis.2.1.1.chr13_hap1.g00227.t01#5    mNeoVis.3.1.1.chr13.g00178.t01#6    mNeoVis.vulgivaga.1.1.chr13.g00219.t01#4   mNeoVis.evergladensis.2.1.chr13.g00225.t01#2       2027   125260599:   mNeoVis.evergladensis.1.1.chr1.g01352.t01#3    mNeoVis.evergladensis.2.1.chr1.g01348.t01#2    mNeoVis.vulgivaga.2.1.chr1.g01350.t01#1    mNeoVis.2.1.1.chr1_hap1.g01383.t01#5    mNeoVis.3.1.1.chr1.g01330.t01#6   mNeoVis.vulgivaga.1.1.chr1.g01350.t01#4       2028   125260600:   mNeoVis.evergladensis.2.1.chr7.g01966.t01#2    mNeoVis.3.1.1.chr7.g01869.t01#6    mNeoVis.vulgivaga.1.1.chr7.g01977.t01#4    mNeoVis.2.1.1.chr7_hap1.g02002.t01#5    mNeoVis.evergladensis.1.1.chr7.g01970.t01#3   mNeoVis.vulgivaga.2.1.chr7.g01983.t01#1       2029   125260601:   mNeoVis.evergladensis.2.1.chr7.g01967.t01#2    mNeoVis.2.1.1.chr7_hap1.g02003.t01#5    mNeoVis.evergladensis.1.1.chr7.g01971.t01#3    mNeoVis.vulgivaga.2.1.chr7.g01984.t01#1    mNeoVis.vulgivaga.1.1.chr7.g01978.t01#4   mNeoVis.3.1.1.chr7.g01870.t01#6       2030   125260602:   mNeoVis.vulgivaga.1.1.chr1.g01351.t01#4    mNeoVis.evergladensis.1.1.chr1.g01353.t01#3    mNeoVis.2.1.1.chr1_hap1.g01384.t01#5    mNeoVis.vulgivaga.2.1.chr1.g01351.t01#1    mNeoVis.3.1.1.chr1.g01331.t01#6   mNeoVis.evergladensis.2.1.chr1.g01349.t01#2       2031   125260603:   mNeoVis.evergladensis.1.1.chr15.g00017.t01#3    mNeoVis.vulgivaga.2.1.chr15.g00604.t01#1    mNeoVis.3.1.1.chr15.g00551.t01#6    mNeoVis.evergladensis.2.1.chr15.g00594.t01#2    mNeoVis.2.1.1.chr15_hap1.g00771.t01#5   mNeoVis.vulgivaga.1.1.chr15.g00596.t01#4       2032   125260604:   mNeoVis.3.1.1.chr1.g01335.t01#6    mNeoVis.3.1.1.chr1.g01338.t01#6    mNeoVis.vulgivaga.1.1.chr1.g01355.t01#4    mNeoVis.evergladensis.1.1.chr1.g01358.t01#3    mNeoVis.vulgivaga.1.1.chr1.g01352.t01#4    mNeoVis.evergladensis.1.1.chr1.g01360.t01#3    mNeoVis.3.1.1.chr1.g01334.t01#6    mNeoVis.evergladensis.1.1.chr1.g01354.t01#3    mNeoVis.2.1.1.chr1_hap1.g01388.t01#5      2033   125260605:   mNeoVis.2.1.1.chr1_hap1.g01390.t01#5    mNeoVis.evergladensis.1.1.chr1.g01359.t01#3    mNeoVis.3.1.1.chr1.g01336.t01#6    mNeoVis.evergladensis.2.1.chr1.g01355.t01#2    mNeoVis.2.1.1.chr1_hap1.g01391.t01#5    mNeoVis.vulgivaga.1.1.chr1.g01354.t01#4    mNeoVis.evergladensis.2.1.chr1.g01354.t01#2    mNeoVis.vulgivaga.2.1.chr1.g01355.t01#1    mNeoVis.vulgivaga.2.1.chr1.g01356.t01#1      2034   125260606:   mNeoVis.evergladensis.2.1.chr6.g00304.t01#2    mNeoVis.3.1.1.chr6.g00302.t01#6    mNeoVis.vulgivaga.1.1.chr3.g00924.t01#4    mNeoVis.vulgivaga.1.1.chr6.g00307.t01#4    mNeoVis.2.1.1.chr11_hap1.g00074.t01#5    mNeoVis.evergladensis.1.1.chr11.g00074.t01#3    mNeoVis.evergladensis.1.1.chr1.g01355.t01#3    mNeoVis.evergladensis.2.1.chr11.g00073.t01#2    mNeoVis.2.1.1.chr4_hap1.g00300.t01#5      2035   125260607:   mNeoVis.evergladensis.2.1.chr1.g01351.t01#2    mNeoVis.3.1.1.chr1.g01333.t01#6    mNeoVis.vulgivaga.2.1.chr1.g01353.t01#1   mNeoVis.2.1.1.chr1_hap1.g01386.t01#5         2036   125260608:   mNeoVis.3.1.1.chr7.g01871.t01#6    mNeoVis.evergladensis.2.1.chr7.g01968.t01#2    mNeoVis.vulgivaga.2.1.chr7.g01985.t01#1    mNeoVis.2.1.1.chr7_hap1.g02004.t01#5    mNeoVis.vulgivaga.1.1.chr7.g01979.t01#4   mNeoVis.evergladensis.1.1.chr7.g01972.t01#3       2037   125260609:   mNeoVis.vulgivaga.1.1.chr15.g00597.t01#4    mNeoVis.evergladensis.1.1.chr15.g00016.t01#3    mNeoVis.2.1.1.chr15_hap1.g00772.t01#5    mNeoVis.evergladensis.2.1.chr15.g00595.t01#2    mNeoVis.vulgivaga.2.1.chr15.g00605.t01#1   mNeoVis.3.1.1.chr15.g00552.t01#6       2038   125260610:   mNeoVis.vulgivaga.2.1.chr7.g00544.t01#1    mNeoVis.evergladensis.2.1.chr7.g00542.t01#2    mNeoVis.2.1.1.chr7_hap1.g00542.t01#5    mNeoVis.evergladensis.1.1.chr7.g00550.t01#3    mNeoVis.3.1.1.chr7.g00524.t01#6   mNeoVis.vulgivaga.1.1.chr7.g00549.t01#4       2039   125260611:   mNeoVis.evergladensis.2.1.chr1.g01357.t01#2    mNeoVis.3.1.1.chr1.g01341.t01#6   mNeoVis.evergladensis.1.1.chr1.g01362.t01#3         2040   125260612:   mNeoVis.3.1.1.chr1.g01339.t01#6   mNeoVis.vulgivaga.2.1.chr1.g01357.t01#1             2041   125260613:  mNeoVis.2.1.1.chr1_hap1.g01392.t01#5               2042   125260614:  mNeoVis.evergladensis.1.1.chr1.g01361.t01#3             2043   125260615:  mNeoVis.vulgivaga.1.1.chr1.g01356.t01#4               2044   125260616:   mNeoVis.3.1.1.chr7.g01872.t01#6    mNeoVis.vulgivaga.1.1.chr7.g01980.t01#4    mNeoVis.evergladensis.1.1.chr7.g01973.t01#3    mNeoVis.evergladensis.2.1.chr7.g01969.t01#2    mNeoVis.2.1.1.chr7_hap1.g02005.t01#5   mNeoVis.vulgivaga.2.1.chr7.g01986.t01#1       2045   125260617:   mNeoVis.evergladensis.2.1.chr1.g01382.t01#2    mNeoVis.evergladensis.1.1.chr1.g01388.t01#3    mNeoVis.3.1.1.chr1.g01367.t01#6    mNeoVis.vulgivaga.1.1.chr1.g01387.t01#4    mNeoVis.2.1.1.chr1_hap1.g01420.t01#5   mNeoVis.vulgivaga.2.1.chr1.g01386.t01#1       2046   125260618:   mNeoVis.vulgivaga.2.1.chr7.g01987.t01#1    mNeoVis.3.1.1.chr7.g01873.t01#6    mNeoVis.vulgivaga.1.1.chr7.g01981.t01#4    mNeoVis.evergladensis.1.1.chr7.g01974.t01#3    mNeoVis.evergladensis.2.1.chr7.g01970.t01#2   mNeoVis.2.1.1.chr7_hap1.g02006.t01#5       2047   125260619:   mNeoVis.evergladensis.2.1.chr1.g01383.t01#2    mNeoVis.evergladensis.1.1.chr1.g01389.t01#3    mNeoVis.vulgivaga.1.1.chr1.g01388.t01#4    mNeoVis.3.1.1.chr1.g01368.t01#6    mNeoVis.vulgivaga.2.1.chr1.g01387.t01#1   mNeoVis.2.1.1.chr1_hap1.g01421.t01#5       2048   125260620:   mNeoVis.3.1.1.chr7.g01874.t01#6    mNeoVis.vulgivaga.1.1.chr7.g01982.t01#4    mNeoVis.evergladensis.2.1.chr7.g01971.t01#2    mNeoVis.evergladensis.1.1.chr7.g01975.t01#3    mNeoVis.vulgivaga.2.1.chr7.g01988.t01#1   mNeoVis.2.1.1.chr7_hap1.g02007.t01#5       2049   125258573:   mNeoVis.vulgivaga.2.1.chr1.g01388.t01#1    mNeoVis.evergladensis.1.1.chr1.g01390.t01#3    mNeoVis.3.1.1.chr1.g01369.t01#6    mNeoVis.vulgivaga.1.1.chr1.g01389.t01#4    mNeoVis.evergladensis.2.1.chr1.g01384.t01#2   mNeoVis.2.1.1.chr1_hap1.g01422.t01#5       2050   125258574:   mNeoVis.evergladensis.1.1.chr1.g01391.t01#3    mNeoVis.evergladensis.2.1.chr1.g01385.t01#2    mNeoVis.3.1.1.chr1.g01370.t01#6    mNeoVis.vulgivaga.2.1.chr1.g01389.t01#1    mNeoVis.vulgivaga.1.1.chr1.g01390.t01#4   mNeoVis.2.1.1.chr1_hap1.g01423.t01#5       2051   125258575:   mNeoVis.3.1.1.chr7.g01875.t01#6    mNeoVis.evergladensis.2.1.chr7.g01972.t01#2    mNeoVis.vulgivaga.1.1.chr7.g01983.t01#4    mNeoVis.2.1.1.chr7_hap1.g02008.t01#5    mNeoVis.vulgivaga.2.1.chr7.g01989.t01#1   mNeoVis.evergladensis.1.1.chr7.g01976.t01#3       2052   125258576:   mNeoVis.2.1.1.chr15_hap1.g00773.t01#5    mNeoVis.3.1.1.chr15.g00553.t01#6    mNeoVis.vulgivaga.1.1.chr15.g00598.t01#4    mNeoVis.evergladensis.2.1.chr15.g00596.t01#2    mNeoVis.vulgivaga.2.1.chr15.g00606.t01#1   mNeoVis.evergladensis.1.1.chr15.g00015.t01#3       2053   125258577:   mNeoVis.3.1.1.chr7.g00525.t01#6    mNeoVis.evergladensis.2.1.chr7.g00543.t01#2    mNeoVis.vulgivaga.1.1.chr7.g00550.t01#4    mNeoVis.vulgivaga.2.1.chr7.g00545.t01#1    mNeoVis.evergladensis.1.1.chr7.g00551.t01#3   mNeoVis.2.1.1.chr7_hap1.g00543.t01#5       2054   125258578:   mNeoVis.vulgivaga.2.1.chr7.g01990.t01#1    mNeoVis.evergladensis.2.1.chr7.g01973.t01#2    mNeoVis.vulgivaga.1.1.chr7.g01984.t01#4    mNeoVis.3.1.1.chr7.g01876.t01#6    mNeoVis.evergladensis.1.1.chr7.g01977.t01#3   mNeoVis.2.1.1.chr7_hap1.g02009.t01#5       2055   125258579:   mNeoVis.vulgivaga.2.1.chr1.g01150.t01#1    mNeoVis.evergladensis.1.1.chr1.g01392.t01#3    mNeoVis.evergladensis.2.1.chr1.g01386.t01#2    mNeoVis.3.1.1.chr1.g01135.t01#6    mNeoVis.vulgivaga.1.1.chr1.g01391.t01#4    mNeoVis.2.1.1.chr1_hap1.g01424.t01#5    mNeoVis.vulgivaga.1.1.chr1.g01147.t01#4    mNeoVis.vulgivaga.2.1.chr1.g01390.t01#1    mNeoVis.evergladensis.1.1.chr1.g01147.t01#3      2056   125258580:   mNeoVis.2.1.1.chr1_hap1.g01425.t01#5    mNeoVis.evergladensis.1.1.chr1.g01393.t01#3    mNeoVis.evergladensis.2.1.chr1.g01387.t01#2    mNeoVis.vulgivaga.1.1.chr1.g01392.t01#4   mNeoVis.vulgivaga.2.1.chr1.g01391.t01#1       2057   125258581:   mNeoVis.vulgivaga.1.1.chr1.g01393.t01#4    mNeoVis.evergladensis.1.1.chr1.g01394.t01#3    mNeoVis.3.1.1.chr1.g01371.t01#6    mNeoVis.vulgivaga.2.1.chr1.g01392.t01#1    mNeoVis.evergladensis.2.1.chr1.g01388.t01#2   mNeoVis.2.1.1.chr1_hap1.g01426.t01#5       2058   125258582:   mNeoVis.3.1.1.chr1.g01287.t01#6    mNeoVis.vulgivaga.2.1.chr1.g01306.t01#1    mNeoVis.vulgivaga.1.1.chr1.g01305.t01#4    mNeoVis.evergladensis.1.1.chr1.g01307.t01#3    mNeoVis.evergladensis.2.1.chr1.g01305.t01#2   mNeoVis.2.1.1.chr1_hap1.g01338.t01#5       2059   125258583:   mNeoVis.3.1.1.chr7.g01879.t01#6    mNeoVis.vulgivaga.1.1.chr7.g01987.t01#4    mNeoVis.vulgivaga.2.1.chr7.g01992.t01#1    mNeoVis.evergladensis.2.1.chr7.g01974.t01#2    mNeoVis.evergladensis.1.1.chr7.g01978.t01#3   mNeoVis.2.1.1.chr7_hap1.g02010.t01#5       2060   125258584:   mNeoVis.vulgivaga.1.1.chr7.g01985.t01#4    mNeoVis.3.1.1.chr7.g01877.t01#6   mNeoVis.vulgivaga.2.1.chr7.g01991.t01#1           2061   125258585:   mNeoVis.vulgivaga.2.1.chr7.g01993.t01#1    mNeoVis.evergladensis.2.1.chr7.g01975.t01#2    mNeoVis.evergladensis.1.1.chr7.g01979.t01#3    mNeoVis.2.1.1.chr7_hap1.g02011.t01#5    mNeoVis.3.1.1.chr7.g01878.t01#6   mNeoVis.vulgivaga.1.1.chr7.g01986.t01#4       2062   125258586:   mNeoVis.vulgivaga.1.1.chr7.g01988.t01#4    mNeoVis.vulgivaga.2.1.chr7.g01994.t01#1    mNeoVis.evergladensis.2.1.chr7.g01976.t01#2    mNeoVis.evergladensis.1.1.chr7.g01980.t01#3   mNeoVis.2.1.1.chr7_hap1.g02012.t01#5       2063   125258587:   mNeoVis.2.1.1.chr7_hap1.g02013.t01#5    mNeoVis.evergladensis.1.1.chr7.g01981.t01#3    mNeoVis.evergladensis.2.1.chr7.g01977.t01#2    mNeoVis.vulgivaga.1.1.chr7.g01989.t01#4   mNeoVis.vulgivaga.2.1.chr7.g01995.t01#1       2064   125258588:   mNeoVis.2.1.1.chr7_hap1.g02014.t01#5    mNeoVis.vulgivaga.2.1.chr7.g01996.t01#1    mNeoVis.3.1.1.chr7.g01880.t01#6    mNeoVis.vulgivaga.1.1.chr7.g01990.t01#4    mNeoVis.evergladensis.2.1.chr7.g01978.t01#2   mNeoVis.evergladensis.1.1.chr7.g01982.t01#3       2065   125258589:   mNeoVis.evergladensis.1.1.chr7.g01983.t01#3    mNeoVis.evergladensis.2.1.chr7.g01979.t01#2    mNeoVis.3.1.1.chr7.g01881.t01#6    mNeoVis.vulgivaga.2.1.chr7.g01997.t01#1    mNeoVis.2.1.1.chr7_hap1.g02015.t01#5   mNeoVis.vulgivaga.1.1.chr7.g01991.t01#4       2066   125258590:   mNeoVis.vulgivaga.2.1.chr7.g00546.t01#1    mNeoVis.evergladensis.1.1.chr7.g00552.t01#3    mNeoVis.evergladensis.2.1.chr7.g00544.t01#2    mNeoVis.2.1.1.chr7_hap1.g00544.t01#5    mNeoVis.3.1.1.chr7.g00526.t01#6   mNeoVis.vulgivaga.1.1.chr7.g00551.t01#4       2067   125258591:   mNeoVis.evergladensis.1.1.chr1.g01308.t01#3    mNeoVis.vulgivaga.1.1.chr1.g01306.t01#4    mNeoVis.2.1.1.chr1_hap1.g01339.t01#5    mNeoVis.evergladensis.2.1.chr1.g01306.t01#2    mNeoVis.3.1.1.chr1.g01288.t01#6   mNeoVis.vulgivaga.2.1.chr1.g01307.t01#1       2068   125258592:   mNeoVis.evergladensis.2.1.chr1.g01307.t01#2    mNeoVis.2.1.1.chr1_hap1.g01340.t01#5    mNeoVis.vulgivaga.1.1.chr1.g01307.t01#4    mNeoVis.3.1.1.chr1.g01289.t01#6    mNeoVis.evergladensis.1.1.chr1.g01309.t01#3    mNeoVis.vulgivaga.1.1.chr1.g01308.t01#4   mNeoVis.vulgivaga.2.1.chr1.g01308.t01#1     2069   125258593:   mNeoVis.vulgivaga.2.1.chr1.g01309.t01#1    mNeoVis.2.1.1.chr1_hap1.g01341.t01#5    mNeoVis.3.1.1.chr1.g01290.t01#6    mNeoVis.vulgivaga.1.1.chr1.g01309.t01#4    mNeoVis.evergladensis.1.1.chr1.g01310.t01#3   mNeoVis.evergladensis.2.1.chr1.g01308.t01#2       2070   125258594:   mNeoVis.3.1.1.chr1.g01291.t01#6    mNeoVis.evergladensis.2.1.chr1.g01174.t01#2    mNeoVis.3.1.1.chr1.g01157.t01#6    mNeoVis.vulgivaga.1.1.chr1.g01310.t01#4    mNeoVis.evergladensis.1.1.chr1.g01311.t01#3    mNeoVis.evergladensis.1.1.chr1.g01172.t01#3    mNeoVis.vulgivaga.1.1.chr1.g01168.t01#4    mNeoVis.evergladensis.2.1.chr1.g01309.t01#2    mNeoVis.vulgivaga.2.1.chr1.g01310.t01#1      2071   125258595:   mNeoVis.2.1.1.chr7_hap1.g02016.t01#5    mNeoVis.3.1.1.chr7.g01882.t01#6    mNeoVis.evergladensis.2.1.chr7.g01980.t01#2    mNeoVis.vulgivaga.1.1.chr7.g01992.t01#4    mNeoVis.vulgivaga.2.1.chr7.g01998.t01#1   mNeoVis.evergladensis.1.1.chr7.g01984.t01#3       2072   125258596:   mNeoVis.vulgivaga.1.1.chr1.g01311.t01#4    mNeoVis.3.1.1.chr1.g01292.t01#6    mNeoVis.evergladensis.1.1.chr1.g01312.t01#3    mNeoVis.evergladensis.2.1.chr1.g01310.t01#2    mNeoVis.2.1.1.chr1_hap1.g01343.t01#5   mNeoVis.vulgivaga.2.1.chr1.g01311.t01#1       2073   125258597:   mNeoVis.evergladensis.2.1.chr1.g01311.t01#2    mNeoVis.vulgivaga.2.1.chr1.g01312.t01#1    mNeoVis.2.1.1.chr1_hap1.g01344.t01#5    mNeoVis.vulgivaga.1.1.chr1.g01312.t01#4    mNeoVis.3.1.1.chr1.g01293.t01#6   mNeoVis.evergladensis.1.1.chr1.g01313.t01#3       2074   125258598:   mNeoVis.evergladensis.2.1.chr7.g01895.t01#2    mNeoVis.2.1.1.chr7_hap1.g01931.t01#5    mNeoVis.vulgivaga.1.1.chr7.g01904.t01#4    mNeoVis.3.1.1.chr7.g01803.t01#6    mNeoVis.vulgivaga.2.1.chr7.g01911.t01#1   mNeoVis.evergladensis.1.1.chr7.g01893.t01#3       2075   125258599:   mNeoVis.vulgivaga.2.1.chr1.g01313.t01#1    mNeoVis.evergladensis.1.1.chr1.g01314.t01#3    mNeoVis.vulgivaga.1.1.chr1.g01313.t01#4    mNeoVis.2.1.1.chr1_hap1.g01345.t01#5    mNeoVis.3.1.1.chr1.g01294.t01#6   mNeoVis.evergladensis.2.1.chr1.g01312.t01#2       2076   125258600:   mNeoVis.evergladensis.1.1.chr7.g01896.t01#3    mNeoVis.3.1.1.chr7.g01804.t01#6    mNeoVis.2.1.1.chr7_hap1.g01932.t01#5    mNeoVis.vulgivaga.2.1.chr7.g01912.t01#1    mNeoVis.evergladensis.2.1.chr7.g01896.t01#2   mNeoVis.vulgivaga.1.1.chr7.g01905.t01#4       2077   125258601:   mNeoVis.vulgivaga.1.1.chr15.g00599.t01#4    mNeoVis.3.1.1.chr15.g00554.t01#6    mNeoVis.evergladensis.2.1.chr15.g00597.t01#2    mNeoVis.vulgivaga.2.1.chr15.g00607.t01#1    mNeoVis.evergladensis.1.1.chr15.g00014.t01#3   mNeoVis.2.1.1.chr15_hap1.g00774.t01#5       2078   125258602:   mNeoVis.vulgivaga.2.1.chr1.g01314.t01#1    mNeoVis.evergladensis.2.1.chr1.g01313.t01#2    mNeoVis.evergladensis.1.1.chr1.g01315.t01#3    mNeoVis.2.1.1.chr1_hap1.g01346.t01#5    mNeoVis.vulgivaga.1.1.chr1.g01314.t01#4   mNeoVis.3.1.1.chr1.g01295.t01#6       2079   125258603:   mNeoVis.evergladensis.2.1.chr7.g01897.t01#2    mNeoVis.2.1.1.chr7_hap1.g01933.t01#5    mNeoVis.vulgivaga.1.1.chr7.g01906.t01#4    mNeoVis.vulgivaga.2.1.chr7.g01913.t01#1    mNeoVis.evergladensis.1.1.chr7.g01897.t01#3   mNeoVis.3.1.1.chr7.g01805.t01#6       2080   125258604:   mNeoVis.evergladensis.1.1.chr1.g01316.t01#3    mNeoVis.vulgivaga.2.1.chr1.g01315.t01#1    mNeoVis.evergladensis.2.1.chr1.g01314.t01#2    mNeoVis.2.1.1.chr1_hap1.g01347.t01#5    mNeoVis.vulgivaga.1.1.chr1.g01315.t01#4   mNeoVis.3.1.1.chr1.g01296.t01#6       2081   125258605:   mNeoVis.evergladensis.1.1.chr1.g01317.t01#3    mNeoVis.vulgivaga.2.1.chr1.g01316.t01#1    mNeoVis.2.1.1.chr1_hap1.g01348.t01#5    mNeoVis.vulgivaga.1.1.chr1.g01316.t01#4    mNeoVis.3.1.1.chr1.g01297.t01#6   mNeoVis.evergladensis.2.1.chr1.g01315.t01#2       2082   125258606:   mNeoVis.evergladensis.2.1.chr1.g01316.t01#2    mNeoVis.evergladensis.1.1.chr1.g01318.t01#3    mNeoVis.vulgivaga.2.1.chr1.g01317.t01#1    mNeoVis.2.1.1.chr1_hap1.g01349.t01#5    mNeoVis.vulgivaga.1.1.chr1.g01317.t01#4   mNeoVis.3.1.1.chr1.g01298.t01#6       2083   125258607:   mNeoVis.vulgivaga.2.1.chr7.g01914.t01#1    mNeoVis.evergladensis.1.1.chr7.g01898.t01#3    mNeoVis.3.1.1.chr7.g01806.t01#6    mNeoVis.2.1.1.chr7_hap1.g01934.t01#5    mNeoVis.evergladensis.2.1.chr7.g01898.t01#2   mNeoVis.vulgivaga.1.1.chr7.g01907.t01#4       2084   125258608:   mNeoVis.evergladensis.1.1.chr1.g01319.t01#3    mNeoVis.2.1.1.chr1_hap1.g01350.t01#5    mNeoVis.vulgivaga.1.1.chr1.g01318.t01#4    mNeoVis.vulgivaga.2.1.chr1.g01318.t01#1    mNeoVis.3.1.1.chr1.g01299.t01#6   mNeoVis.evergladensis.2.1.chr1.g01317.t01#2       2085   125258609:   mNeoVis.vulgivaga.1.1.chr7.g01908.t01#4    mNeoVis.evergladensis.1.1.chr7.g01899.t01#3    mNeoVis.2.1.1.chr7_hap1.g01935.t01#5    mNeoVis.3.1.1.chr7.g01807.t01#6    mNeoVis.vulgivaga.2.1.chr7.g01915.t01#1   mNeoVis.evergladensis.2.1.chr7.g01899.t01#2       2086   125258610:   mNeoVis.evergladensis.2.1.chr1.g01318.t01#2    mNeoVis.vulgivaga.1.1.chr1.g01319.t01#4    mNeoVis.2.1.1.chr1_hap1.g01351.t01#5    mNeoVis.3.1.1.chr1.g01300.t01#6    mNeoVis.vulgivaga.2.1.chr1.g01319.t01#1   mNeoVis.evergladensis.1.1.chr1.g01320.t01#3       2087   125258611:   mNeoVis.vulgivaga.1.1.chr8.g00885.t01#4    mNeoVis.evergladensis.1.1.chr7.g01900.t01#3    mNeoVis.3.1.1.chr7.g01808.t01#6    mNeoVis.2.1.1.chr8_hap1.g00917.t01#5    mNeoVis.2.1.1.chr7_hap1.g01936.t01#5    mNeoVis.vulgivaga.1.1.chr7.g01909.t01#4    mNeoVis.evergladensis.1.1.chr8.g00894.t01#3    mNeoVis.evergladensis.2.1.chr7.g01900.t01#2    mNeoVis.vulgivaga.2.1.chr7.g01916.t01#1      2088   125258612:  mNeoVis.vulgivaga.2.1.chr1.g01320.t01#1               2089   125258613:   mNeoVis.3.1.1.chr1.g01302.t01#6    mNeoVis.evergladensis.2.1.chr1.g01320.t01#2    mNeoVis.vulgivaga.1.1.chr1.g01321.t01#4    mNeoVis.evergladensis.1.1.chr1.g01322.t01#3    mNeoVis.2.1.1.chr1_hap1.g01353.t01#5   mNeoVis.vulgivaga.2.1.chr1.g01321.t01#1       2090   125258614:   mNeoVis.evergladensis.2.1.chr7.g01901.t01#2    mNeoVis.vulgivaga.2.1.chr7.g01917.t01#1    mNeoVis.vulgivaga.1.1.chr7.g01910.t01#4    mNeoVis.3.1.1.chr7.g01809.t01#6    mNeoVis.evergladensis.1.1.chr7.g01901.t01#3   mNeoVis.2.1.1.chr7_hap1.g01937.t01#5       2091   125258615:   mNeoVis.vulgivaga.1.1.chr1.g01322.t01#4    mNeoVis.2.1.1.chr1_hap1.g01354.t01#5    mNeoVis.evergladensis.2.1.chr1.g01321.t01#2    mNeoVis.3.1.1.chr1.g01303.t01#6    mNeoVis.evergladensis.1.1.chr1.g01323.t01#3   mNeoVis.vulgivaga.2.1.chr1.g01322.t01#1       2092   125258616:   mNeoVis.3.1.1.chr1.g01304.t01#6    mNeoVis.evergladensis.1.1.chr1.g01324.t01#3    mNeoVis.evergladensis.2.1.chr1.g01322.t01#2    mNeoVis.vulgivaga.2.1.chr1.g01323.t01#1    mNeoVis.vulgivaga.1.1.chr1.g01323.t01#4   mNeoVis.2.1.1.chr1_hap1.g01355.t01#5       2093   125258617:   mNeoVis.vulgivaga.1.1.chr1.g01324.t01#4    mNeoVis.2.1.1.chr1_hap1.g01356.t01#5    mNeoVis.evergladensis.1.1.chr1.g01325.t01#3    mNeoVis.evergladensis.2.1.chr1.g01323.t01#2   mNeoVis.vulgivaga.2.1.chr1.g01324.t01#1       2094   125258618:   mNeoVis.2.1.1.chr7_hap1.g01938.t01#5    mNeoVis.3.1.1.chr7.g01810.t01#6    mNeoVis.evergladensis.2.1.chr7.g01902.t01#2    mNeoVis.vulgivaga.2.1.chr7.g01918.t01#1    mNeoVis.vulgivaga.1.1.chr7.g01911.t01#4   mNeoVis.evergladensis.1.1.chr7.g01902.t01#3       2095   125258619:   mNeoVis.evergladensis.2.1.chr7.g01903.t01#2    mNeoVis.vulgivaga.2.1.chr7.g01919.t01#1    mNeoVis.3.1.1.chr7.g01811.t01#6    mNeoVis.vulgivaga.1.1.chr7.g01912.t01#4    mNeoVis.evergladensis.1.1.chr7.g01903.t01#3   mNeoVis.2.1.1.chr7_hap1.g01939.t01#5       2096   125258620:   mNeoVis.2.1.1.chr7_hap1.g01940.t01#5    mNeoVis.3.1.1.chr7.g01812.t01#6    mNeoVis.evergladensis.2.1.chr7.g01904.t01#2    mNeoVis.vulgivaga.2.1.chr7.g01920.t01#1    mNeoVis.2.1.1.chr1_hap1.g00636.t01#5    mNeoVis.evergladensis.1.1.chr7.g01904.t01#3   mNeoVis.evergladensis.1.1.chr1.g00647.t01#3     2097   125258621:   mNeoVis.2.1.1.chr12_hap1.g00436.t01#5    mNeoVis.vulgivaga.2.1.chr12.g00401.t01#1    mNeoVis.vulgivaga.1.1.chr12.g00403.t01#4    mNeoVis.evergladensis.2.1.chr12.g00393.t01#2    mNeoVis.3.1.1.chr12.g00397.t01#6   mNeoVis.evergladensis.1.1.chr12.g00400.t01#3       2098   125258622:   mNeoVis.3.1.1.chr7.g01813.t01#6    mNeoVis.vulgivaga.2.1.chr7.g01921.t01#1    mNeoVis.evergladensis.1.1.chr7.g01905.t01#3    mNeoVis.vulgivaga.1.1.chr7.g01913.t01#4    mNeoVis.2.1.1.chr7_hap1.g01941.t01#5   mNeoVis.evergladensis.2.1.chr7.g01905.t01#2       2099   125258623:   mNeoVis.vulgivaga.1.1.chr1.g01325.t01#4    mNeoVis.evergladensis.2.1.chr1.g01324.t01#2    mNeoVis.3.1.1.chr1.g01306.t01#6    mNeoVis.vulgivaga.2.1.chr1.g01325.t01#1    mNeoVis.2.1.1.chr1_hap1.g01357.t01#5   mNeoVis.evergladensis.1.1.chr1.g01326.t01#3       2100   125258624:   mNeoVis.3.1.1.chr7.g01814.t01#6    mNeoVis.evergladensis.2.1.chr7.g01906.t01#2    mNeoVis.vulgivaga.2.1.chr7.g01922.t01#1    mNeoVis.evergladensis.1.1.chr7.g01906.t01#3    mNeoVis.vulgivaga.1.1.chr7.g01914.t01#4   mNeoVis.2.1.1.chr7_hap1.g01942.t01#5       2101   125258625:   mNeoVis.evergladensis.2.1.chr8.g00299.t01#2    mNeoVis.evergladensis.2.1.chr1.g01325.t01#2    mNeoVis.vulgivaga.1.1.chr12.g00650.t01#4    mNeoVis.3.1.1.chr9.g00288.t01#6    mNeoVis.2.1.1.chr10_hap1.g00315.t01#5    mNeoVis.evergladensis.1.1.chr1.g01327.t01#3    mNeoVis.vulgivaga.1.1.chr1.g01326.t01#4    mNeoVis.vulgivaga.2.1.chr1.g01326.t01#1    mNeoVis.2.1.1.chr1_hap1.g01359.t01#5      2102   125258626:   mNeoVis.3.1.1.chr14.g00422.t01#6    mNeoVis.3.1.1.chr1.g00073.t01#6    mNeoVis.vulgivaga.2.1.chr7.g01295.t01#1    mNeoVis.evergladensis.2.1.chr7.g01283.t01#2   mNeoVis.vulgivaga.1.1.chr7.g01285.t01#4       2103   125258627:   mNeoVis.evergladensis.2.1.chr3.g01145.t01#2    mNeoVis.evergladensis.1.1.chr3.g01372.t01#3    mNeoVis.2.1.1.chr7_hap1.g01294.t01#5   mNeoVis.vulgivaga.2.1.chr12.g00654.t01#1       2104   125258628:   mNeoVis.evergladensis.1.1.chr4.g00704.t01#3    mNeoVis.3.1.1.chr7.g01218.t01#6   mNeoVis.vulgivaga.2.1.chr5.g01139.t01#1           2105   125258629:   mNeoVis.vulgivaga.1.1.chr1.g01327.t01#4   mNeoVis.2.1.1.chr10_hap1.g00455.t01#5             2106   125258630:  mNeoVis.vulgivaga.1.1.chr12.g00365.t01#4               2107   125258631:   mNeoVis.2.1.1.chr1_hap1.g01361.t01#5    mNeoVis.evergladensis.1.1.chr1.g01328.t01#3    mNeoVis.vulgivaga.2.1.chr1.g01327.t01#1    mNeoVis.vulgivaga.1.1.chr1.g01328.t01#4    mNeoVis.evergladensis.2.1.chr1.g01326.t01#2   mNeoVis.3.1.1.chr1.g01309.t01#6       2108   125258632:   mNeoVis.evergladensis.2.1.chr1.g01327.t01#2    mNeoVis.2.1.1.chr1_hap1.g01362.t01#5    mNeoVis.vulgivaga.2.1.chr1.g01328.t01#1    mNeoVis.3.1.1.chr1.g01310.t01#6    mNeoVis.vulgivaga.1.1.chr1.g01329.t01#4   mNeoVis.evergladensis.1.1.chr1.g01329.t01#3       2109   125258633:   mNeoVis.vulgivaga.2.1.chr7.g01923.t01#1    mNeoVis.evergladensis.2.1.chr7.g01907.t01#2    mNeoVis.3.1.1.chr7.g01815.t01#6    mNeoVis.evergladensis.1.1.chr7.g01907.t01#3    mNeoVis.vulgivaga.1.1.chr7.g01915.t01#4   mNeoVis.2.1.1.chr7_hap1.g01943.t01#5       2110   125258634:   mNeoVis.evergladensis.1.1.chr7.g01908.t01#3    mNeoVis.evergladensis.2.1.chr7.g01908.t01#2    mNeoVis.vulgivaga.1.1.chr7.g01916.t01#4    mNeoVis.2.1.1.chr7_hap1.g01944.t01#5    mNeoVis.3.1.1.chr7.g01816.t01#6   mNeoVis.vulgivaga.2.1.chr7.g01924.t01#1       2111   125258635:   mNeoVis.2.1.1.chr7_hap1.g00545.t01#5    mNeoVis.vulgivaga.1.1.chr7.g00552.t01#4    mNeoVis.evergladensis.2.1.chr7.g00545.t01#2    mNeoVis.vulgivaga.2.1.chr7.g00547.t01#1    mNeoVis.evergladensis.1.1.chr7.g00553.t01#3   mNeoVis.3.1.1.chr7.g00527.t01#6       2112   125258636:   mNeoVis.evergladensis.1.1.chr1.g01330.t01#3    mNeoVis.2.1.1.chr1_hap1.g01363.t01#5    mNeoVis.vulgivaga.1.1.chr1.g01330.t01#4    mNeoVis.3.1.1.chr1.g01311.t01#6    mNeoVis.vulgivaga.2.1.chr1.g01329.t01#1   mNeoVis.evergladensis.2.1.chr1.g01328.t01#2       2113   125258637:   mNeoVis.2.1.1.chr7_hap1.g01945.t01#5    mNeoVis.evergladensis.2.1.chr7.g01909.t01#2    mNeoVis.vulgivaga.2.1.chr7.g01925.t01#1   mNeoVis.vulgivaga.1.1.chr7.g01917.t01#4         2114   125258638:   mNeoVis.evergladensis.2.1.chr7.g01910.t01#2    mNeoVis.3.1.1.chr7.g01817.t01#6    mNeoVis.vulgivaga.2.1.chr7.g01926.t01#1    mNeoVis.vulgivaga.1.1.chr7.g01918.t01#4    mNeoVis.2.1.1.chr7_hap1.g01946.t01#5   mNeoVis.evergladensis.1.1.chr7.g01909.t01#3       2115   125258639:   mNeoVis.vulgivaga.2.1.chr15.g00608.t01#1    mNeoVis.2.1.1.chr15_hap1.g00775.t01#5    mNeoVis.3.1.1.chr15.g00555.t01#6    mNeoVis.evergladensis.1.1.chr15.g00013.t01#3    mNeoVis.vulgivaga.1.1.chr15.g00600.t01#4    mNeoVis.evergladensis.2.1.chr15.g00598.t01#2   mNeoVis.vulgivaga.2.1.chr15.g00609.t01#1     2116   125258640:   mNeoVis.evergladensis.1.1.chr7.g01910.t01#3    mNeoVis.vulgivaga.2.1.chr7.g01927.t01#1    mNeoVis.vulgivaga.1.1.chr7.g01919.t01#4    mNeoVis.3.1.1.chr7.g01818.t01#6    mNeoVis.2.1.1.chr7_hap1.g01947.t01#5   mNeoVis.evergladensis.2.1.chr7.g01911.t01#2       2117   125258641:   mNeoVis.vulgivaga.1.1.chr1.g01331.t01#4    mNeoVis.3.1.1.chr1.g01312.t01#6    mNeoVis.vulgivaga.2.1.chr1.g01330.t01#1    mNeoVis.2.1.1.chr1_hap1.g01364.t01#5    mNeoVis.evergladensis.1.1.chr1.g01331.t01#3   mNeoVis.evergladensis.2.1.chr1.g01329.t01#2       2118   125258642:   mNeoVis.evergladensis.1.1.chr7.g00554.t01#3    mNeoVis.evergladensis.2.1.chr7.g00546.t01#2    mNeoVis.3.1.1.chr7.g00528.t01#6    mNeoVis.2.1.1.chr7_hap1.g00546.t01#5    mNeoVis.vulgivaga.2.1.chr7.g00548.t01#1   mNeoVis.vulgivaga.1.1.chr7.g00553.t01#4       2119   125258643:   mNeoVis.3.1.1.chr15.g00421.t01#6    mNeoVis.evergladensis.2.1.chr15.g00462.t01#2    mNeoVis.vulgivaga.2.1.chr1.g01331.t01#1    mNeoVis.vulgivaga.2.1.chr15.g00469.t01#1    mNeoVis.2.1.1.chr14_hap1.g00135.t01#5    mNeoVis.3.1.1.chr1.g01313.t01#6    mNeoVis.2.1.1.chr15_hap1.g00640.t01#5    mNeoVis.vulgivaga.1.1.chr14.g00133.t01#4    mNeoVis.evergladensis.2.1.chr14.g00131.t01#2      2120   125258644:   mNeoVis.3.1.1.chr1.g01314.t01#6    mNeoVis.evergladensis.1.1.chr1.g01332.t01#3    mNeoVis.evergladensis.2.1.chr1.g01330.t01#2    mNeoVis.vulgivaga.1.1.chr1.g01332.t01#4    mNeoVis.2.1.1.chr1_hap1.g01365.t01#5   mNeoVis.vulgivaga.2.1.chr1.g01332.t01#1       2121   125258645:   mNeoVis.vulgivaga.2.1.chr7.g01928.t01#1    mNeoVis.evergladensis.1.1.chr7.g01911.t01#3    mNeoVis.evergladensis.2.1.chr7.g01912.t01#2    mNeoVis.2.1.1.chr7_hap1.g01948.t01#5    mNeoVis.3.1.1.chr7.g01819.t01#6   mNeoVis.vulgivaga.1.1.chr7.g01920.t01#4       2122   125258646:   mNeoVis.evergladensis.1.1.chr7.g01912.t01#3    mNeoVis.vulgivaga.2.1.chr7.g01929.t01#1    mNeoVis.2.1.1.chr7_hap1.g01949.t01#5    mNeoVis.vulgivaga.1.1.chr7.g01921.t01#4   mNeoVis.evergladensis.2.1.chr7.g01913.t01#2       2123   125258647:   mNeoVis.vulgivaga.2.1.chr7.g01930.t01#1    mNeoVis.vulgivaga.1.1.chr7.g01922.t01#4    mNeoVis.2.1.1.chr7_hap1.g01950.t01#5    mNeoVis.3.1.1.chr7.g01820.t01#6    mNeoVis.evergladensis.2.1.chr7.g01914.t01#2   mNeoVis.evergladensis.1.1.chr7.g01913.t01#3       2124   125258648:   mNeoVis.evergladensis.2.1.chr1.g01331.t01#2    mNeoVis.2.1.1.chr1_hap1.g01366.t01#5    mNeoVis.vulgivaga.2.1.chr1.g01333.t01#1    mNeoVis.3.1.1.chr1.g01315.t01#6    mNeoVis.vulgivaga.1.1.chr1.g01333.t01#4   mNeoVis.evergladensis.1.1.chr1.g01333.t01#3       2125   125258649:   mNeoVis.vulgivaga.1.1.chr7.g01923.t01#4    mNeoVis.evergladensis.1.1.chr7.g01914.t01#3    mNeoVis.2.1.1.chr7_hap1.g01951.t01#5    mNeoVis.vulgivaga.2.1.chr7.g01931.t01#1    mNeoVis.evergladensis.2.1.chr7.g01915.t01#2   mNeoVis.3.1.1.chr7.g01821.t01#6       2126   125258650:   mNeoVis.2.1.1.chr7_hap1.g01952.t01#5    mNeoVis.vulgivaga.1.1.chr7.g01924.t01#4    mNeoVis.evergladensis.1.1.chr7.g01915.t01#3    mNeoVis.3.1.1.chr7.g01822.t01#6    mNeoVis.evergladensis.2.1.chr7.g01916.t01#2   mNeoVis.vulgivaga.2.1.chr7.g01932.t01#1       2127   125258651:   mNeoVis.vulgivaga.1.1.chr7.g01925.t01#4    mNeoVis.evergladensis.1.1.chr7.g01916.t01#3    mNeoVis.vulgivaga.2.1.chr7.g01933.t01#1    mNeoVis.2.1.1.chr7_hap1.g01953.t01#5    mNeoVis.evergladensis.2.1.chr7.g01917.t01#2   mNeoVis.3.1.1.chr7.g01823.t01#6       2128   125258652:   mNeoVis.vulgivaga.1.1.chr1.g01334.t01#4    mNeoVis.evergladensis.2.1.chr1.g01332.t01#2    mNeoVis.2.1.1.chr1_hap1.g01367.t01#5    mNeoVis.3.1.1.chr1.g01316.t01#6    mNeoVis.vulgivaga.2.1.chr1.g01334.t01#1   mNeoVis.evergladensis.1.1.chr1.g01334.t01#3       2129   125258653:   mNeoVis.evergladensis.1.1.chr7.g01917.t01#3    mNeoVis.3.1.1.chr7.g01824.t01#6    mNeoVis.evergladensis.2.1.chr7.g01918.t01#2    mNeoVis.2.1.1.chr7_hap1.g01954.t01#5    mNeoVis.vulgivaga.1.1.chr7.g01926.t01#4   mNeoVis.vulgivaga.2.1.chr7.g01934.t01#1       2130   125258654:   mNeoVis.2.1.1.chr1_hap1.g01368.t01#5    mNeoVis.vulgivaga.2.1.chr1.g01335.t01#1    mNeoVis.evergladensis.2.1.chr1.g01333.t01#2    mNeoVis.3.1.1.chr1.g01317.t01#6    mNeoVis.evergladensis.1.1.chr1.g01335.t01#3   mNeoVis.vulgivaga.1.1.chr1.g01335.t01#4       2131   125258655:   mNeoVis.vulgivaga.1.1.chr7.g01927.t01#4    mNeoVis.vulgivaga.2.1.chr7.g01935.t01#1    mNeoVis.2.1.1.chr7_hap1.g01955.t01#5    mNeoVis.evergladensis.1.1.chr7.g01918.t01#3    mNeoVis.3.1.1.chr7.g01825.t01#6   mNeoVis.evergladensis.2.1.chr7.g01919.t01#2       2132   125258656:   mNeoVis.evergladensis.1.1.chr7.g01919.t01#3    mNeoVis.3.1.1.chr7.g01826.t01#6    mNeoVis.vulgivaga.1.1.chr7.g01928.t01#4    mNeoVis.vulgivaga.2.1.chr7.g01936.t01#1    mNeoVis.evergladensis.2.1.chr7.g01920.t01#2   mNeoVis.2.1.1.chr7_hap1.g01956.t01#5       2133   125258657:   mNeoVis.evergladensis.1.1.chr7.g01920.t01#3    mNeoVis.2.1.1.chr7_hap1.g01957.t01#5    mNeoVis.vulgivaga.1.1.chr7.g01929.t01#4   mNeoVis.vulgivaga.2.1.chr7.g01937.t01#1         2134   125258658:   mNeoVis.vulgivaga.1.1.chr5.g01015.t01#4    mNeoVis.vulgivaga.2.1.chr5.g01043.t01#1    mNeoVis.evergladensis.1.1.chr5.g01043.t01#3    mNeoVis.evergladensis.2.1.chr5.g01052.t01#2    mNeoVis.3.1.1.chr5.g01032.t01#6   mNeoVis.2.1.1.chr2_hap1.g01109.t01#5       2135   125258659:   mNeoVis.vulgivaga.2.1.chr7.g01938.t01#1    mNeoVis.vulgivaga.1.1.chr7.g01931.t01#4    mNeoVis.2.1.1.chr7_hap1.g01959.t01#5    mNeoVis.evergladensis.1.1.chr7.g01922.t01#3   mNeoVis.3.1.1.chr7.g01827.t01#6       2136   125258660:   mNeoVis.evergladensis.2.1.chr7.g01922.t01#2    mNeoVis.vulgivaga.1.1.chr7.g01932.t01#4    mNeoVis.2.1.1.chr7_hap1.g01960.t01#5    mNeoVis.evergladensis.1.1.chr7.g01923.t01#3    mNeoVis.3.1.1.chr7.g01828.t01#6   mNeoVis.vulgivaga.2.1.chr7.g01939.t01#1       2137   125258661:   mNeoVis.vulgivaga.1.1.chr7.g00554.t01#4    mNeoVis.evergladensis.1.1.chr7.g00555.t01#3    mNeoVis.evergladensis.2.1.chr7.g00547.t01#2    mNeoVis.3.1.1.chr7.g00529.t01#6    mNeoVis.vulgivaga.2.1.chr7.g00549.t01#1   mNeoVis.2.1.1.chr7_hap1.g00547.t01#5       2138   125258662:   mNeoVis.evergladensis.1.1.chr7.g01925.t01#3    mNeoVis.vulgivaga.2.1.chr7.g01940.t01#1    mNeoVis.vulgivaga.1.1.chr7.g01934.t01#4   mNeoVis.evergladensis.2.1.chr7.g01923.t01#2       2139   125258663:   mNeoVis.evergladensis.1.1.chr1.g01337.t01#3    mNeoVis.vulgivaga.2.1.chr1.g01336.t01#1    mNeoVis.vulgivaga.1.1.chr1.g01336.t01#4    mNeoVis.evergladensis.1.1.chr1.g01336.t01#3    mNeoVis.2.1.1.chr1_hap1.g01369.t01#5    mNeoVis.3.1.1.chr1.g01318.t01#6   mNeoVis.evergladensis.2.1.chr1.g01334.t01#2     2140   125258664:   mNeoVis.vulgivaga.1.1.chr1.g01337.t01#4    mNeoVis.evergladensis.1.1.chr1.g01338.t01#3    mNeoVis.2.1.1.chr1_hap1.g01370.t01#5    mNeoVis.vulgivaga.2.1.chr1.g01337.t01#1    mNeoVis.3.1.1.chr1.g01319.t01#6   mNeoVis.evergladensis.2.1.chr1.g01335.t01#2       2141   125258665:   mNeoVis.evergladensis.1.1.chr1.g01339.t01#3    mNeoVis.vulgivaga.1.1.chr1.g01338.t01#4    mNeoVis.2.1.1.chr1_hap1.g01371.t01#5    mNeoVis.evergladensis.2.1.chr1.g01336.t01#2    mNeoVis.3.1.1.chr1.g01320.t01#6   mNeoVis.vulgivaga.2.1.chr1.g01338.t01#1       2142   125258666:   mNeoVis.vulgivaga.2.1.chr7.g01941.t01#1    mNeoVis.evergladensis.1.1.chr7.g01927.t01#3    mNeoVis.vulgivaga.1.1.chr7.g01936.t01#4    mNeoVis.3.1.1.chr7.g01830.t01#6    mNeoVis.evergladensis.2.1.chr7.g01925.t01#2   mNeoVis.2.1.1.chr7_hap1.g01962.t01#5       2143   125258667:   mNeoVis.evergladensis.2.1.chr7.g01926.t01#2    mNeoVis.vulgivaga.2.1.chr7.g01942.t01#1    mNeoVis.evergladensis.1.1.chr7.g01928.t01#3    mNeoVis.3.1.1.chr7.g01831.t01#6    mNeoVis.vulgivaga.1.1.chr7.g01937.t01#4   mNeoVis.2.1.1.chr7_hap1.g01963.t01#5       2144   125258668:  mNeoVis.vulgivaga.2.1.chr7.g01943.t01#1               2145   125258669:   mNeoVis.3.1.1.chr7.g01832.t01#6    mNeoVis.evergladensis.1.1.chr7.g01929.t01#3    mNeoVis.vulgivaga.2.1.chr7.g01944.t01#1    mNeoVis.2.1.1.chr7_hap1.g01964.t01#5    mNeoVis.vulgivaga.1.1.chr7.g01938.t01#4   mNeoVis.evergladensis.2.1.chr7.g01927.t01#2       2146   125258670:   mNeoVis.evergladensis.1.1.chr1.g01340.t01#3    mNeoVis.evergladensis.2.1.chr1.g01337.t01#2    mNeoVis.vulgivaga.1.1.chr1.g01339.t01#4    mNeoVis.2.1.1.chr1_hap1.g01372.t01#5   mNeoVis.vulgivaga.2.1.chr1.g01339.t01#1       2147   125258671:   mNeoVis.evergladensis.2.1.chr7.g01928.t01#2    mNeoVis.vulgivaga.1.1.chr7.g01939.t01#4    mNeoVis.evergladensis.1.1.chr7.g01930.t01#3    mNeoVis.vulgivaga.2.1.chr7.g01945.t01#1    mNeoVis.3.1.1.chr7.g01833.t01#6   mNeoVis.2.1.1.chr7_hap1.g01965.t01#5       2148   125258672:   mNeoVis.2.1.1.chr1_hap1.g01373.t01#5    mNeoVis.evergladensis.2.1.chr1.g01338.t01#2    mNeoVis.vulgivaga.1.1.chr1.g01340.t01#4   mNeoVis.vulgivaga.2.1.chr1.g01340.t01#1         2149   125258673:   mNeoVis.3.1.1.chr7.g01834.t01#6    mNeoVis.evergladensis.1.1.chr7.g01931.t01#3    mNeoVis.vulgivaga.2.1.chr7.g01946.t01#1    mNeoVis.2.1.1.chr7_hap1.g01966.t01#5    mNeoVis.evergladensis.2.1.chr7.g01929.t01#2   mNeoVis.vulgivaga.1.1.chr7.g01940.t01#4       2150   125258674:   mNeoVis.vulgivaga.1.1.chr7.g01941.t01#4    mNeoVis.2.1.1.chr7_hap1.g01967.t01#5    mNeoVis.3.1.1.chr7.g01835.t01#6    mNeoVis.vulgivaga.2.1.chr7.g01947.t01#1    mNeoVis.evergladensis.1.1.chr7.g01932.t01#3   mNeoVis.evergladensis.2.1.chr7.g01930.t01#2       2151   125258675:   mNeoVis.vulgivaga.1.1.chr15.g00601.t01#4    mNeoVis.evergladensis.2.1.chr15.g00599.t01#2    mNeoVis.evergladensis.1.1.chr15.g00012.t01#3    mNeoVis.vulgivaga.2.1.chr15.g00610.t01#1   mNeoVis.2.1.1.chr15_hap1.g00776.t01#5       2152   125258676:   mNeoVis.vulgivaga.1.1.chr7.g01942.t01#4    mNeoVis.evergladensis.2.1.chr7.g01931.t01#2    mNeoVis.vulgivaga.2.1.chr7.g01948.t01#1    mNeoVis.2.1.1.chr7_hap1.g01968.t01#5    mNeoVis.evergladensis.1.1.chr7.g01933.t01#3   mNeoVis.3.1.1.chr7.g01836.t01#6       2153   125258677:   mNeoVis.vulgivaga.1.1.chr1.g01341.t01#4    mNeoVis.3.1.1.chr1.g01322.t01#6    mNeoVis.evergladensis.2.1.chr1.g01339.t01#2    mNeoVis.evergladensis.1.1.chr1.g01343.t01#3    mNeoVis.2.1.1.chr1_hap1.g01374.t01#5   mNeoVis.vulgivaga.2.1.chr1.g01341.t01#1       2154   125258678:   mNeoVis.vulgivaga.2.1.chr7.g01950.t01#1    mNeoVis.vulgivaga.2.1.chr7.g01949.t01#1    mNeoVis.evergladensis.2.1.chr7.g01932.t01#2    mNeoVis.vulgivaga.1.1.chr7.g01943.t01#4    mNeoVis.evergladensis.1.1.chr7.g01935.t01#3    mNeoVis.2.1.1.chr7_hap1.g01969.t01#5    mNeoVis.evergladensis.1.1.chr7.g01934.t01#3    mNeoVis.3.1.1.chr7.g01837.t01#6    mNeoVis.evergladensis.1.1.chr7.g01936.t01#3      2155   125258679:   mNeoVis.evergladensis.1.1.chr1.g01344.t01#3    mNeoVis.2.1.1.chr1_hap1.g01375.t01#5    mNeoVis.evergladensis.2.1.chr1.g01340.t01#2    mNeoVis.vulgivaga.2.1.chr1.g01342.t01#1    mNeoVis.3.1.1.chr1.g01323.t01#6   mNeoVis.vulgivaga.1.1.chr1.g01342.t01#4       2156   125258680:   mNeoVis.vulgivaga.2.1.chr1.g01343.t01#1    mNeoVis.2.1.1.chr1_hap1.g01376.t01#5    mNeoVis.evergladensis.2.1.chr1.g01341.t01#2    mNeoVis.vulgivaga.1.1.chr1.g01343.t01#4   mNeoVis.evergladensis.1.1.chr1.g01345.t01#3       2157   125258681:   mNeoVis.evergladensis.1.1.chr7.g01937.t01#3    mNeoVis.evergladensis.2.1.chr7.g01933.t01#2    mNeoVis.vulgivaga.2.1.chr7.g01951.t01#1    mNeoVis.vulgivaga.1.1.chr7.g01944.t01#4    mNeoVis.3.1.1.chr7.g01838.t01#6   mNeoVis.2.1.1.chr7_hap1.g01970.t01#5       2158   125258682:   mNeoVis.vulgivaga.2.1.chr15.g00611.t01#1    mNeoVis.evergladensis.1.1.chr15.g00011.t01#3    mNeoVis.2.1.1.chr15_hap1.g00778.t01#5    mNeoVis.evergladensis.2.1.chr15.g00600.t01#2    mNeoVis.vulgivaga.2.1.chr15.g00612.t01#1   mNeoVis.vulgivaga.1.1.chr15.g00603.t01#4       2159   125258683:   mNeoVis.vulgivaga.2.1.chr1.g01344.t01#1    mNeoVis.2.1.1.chr1_hap1.g01377.t01#5    mNeoVis.evergladensis.1.1.chr1.g01346.t01#3    mNeoVis.vulgivaga.1.1.chr1.g01344.t01#4    mNeoVis.evergladensis.2.1.chr1.g01342.t01#2   mNeoVis.3.1.1.chr1.g01324.t01#6       2160   125258684:   mNeoVis.evergladensis.2.1.chr15.g00539.t01#2    mNeoVis.vulgivaga.1.1.chr6.g00908.t01#4    mNeoVis.vulgivaga.2.1.chr15.g00551.t01#1    mNeoVis.vulgivaga.1.1.chr7.g01381.t01#4    mNeoVis.evergladensis.1.1.chr8.g00980.t01#3    mNeoVis.evergladensis.1.1.chr7.g01373.t01#3    mNeoVis.2.1.1.chr9_hap1.g00229.t01#5    mNeoVis.vulgivaga.2.1.chr8.g00990.t01#1    mNeoVis.vulgivaga.2.1.chr7.g01386.t01#1      2161   125258685:   mNeoVis.vulgivaga.1.1.chr7.g01945.t01#4    mNeoVis.2.1.1.chr7_hap1.g01971.t01#5    mNeoVis.evergladensis.1.1.chr7.g01938.t01#3    mNeoVis.evergladensis.2.1.chr7.g01934.t01#2    mNeoVis.3.1.1.chr7.g01839.t01#6   mNeoVis.vulgivaga.2.1.chr7.g01952.t01#1       2162   125258686:   mNeoVis.evergladensis.1.1.chr7.g01939.t01#3    mNeoVis.vulgivaga.2.1.chr7.g01953.t01#1    mNeoVis.evergladensis.2.1.chr7.g01935.t01#2    mNeoVis.3.1.1.chr7.g01840.t01#6    mNeoVis.vulgivaga.1.1.chr7.g01946.t01#4   mNeoVis.2.1.1.chr7_hap1.g01972.t01#5       2163   125258687:   mNeoVis.evergladensis.2.1.chr1.g01251.t01#2    mNeoVis.3.1.1.chr1.g01234.t01#6    mNeoVis.vulgivaga.1.1.chr1.g01248.t01#4    mNeoVis.evergladensis.1.1.chr1.g01252.t01#3    mNeoVis.vulgivaga.2.1.chr1.g01250.t01#1   mNeoVis.2.1.1.chr1_hap1.g01278.t01#5       2164   125258688:   mNeoVis.evergladensis.2.1.chr1.g01252.t01#2    mNeoVis.2.1.1.chr1_hap1.g01279.t01#5    mNeoVis.vulgivaga.1.1.chr1.g01249.t01#4    mNeoVis.3.1.1.chr1.g01235.t01#6    mNeoVis.evergladensis.1.1.chr1.g01253.t01#3   mNeoVis.vulgivaga.2.1.chr1.g01251.t01#1       2165   125258689:   mNeoVis.evergladensis.2.1.chr1.g01253.t01#2    mNeoVis.vulgivaga.2.1.chr1.g01252.t01#1    mNeoVis.2.1.1.chr1_hap1.g01280.t01#5    mNeoVis.vulgivaga.1.1.chr1.g01250.t01#4   mNeoVis.evergladensis.1.1.chr1.g01254.t01#3       2166   125258690:   mNeoVis.vulgivaga.1.1.chr1.g01251.t01#4    mNeoVis.2.1.1.chr1_hap1.g01281.t01#5    mNeoVis.evergladensis.1.1.chr1.g01255.t01#3    mNeoVis.vulgivaga.2.1.chr1.g01253.t01#1    mNeoVis.evergladensis.2.1.chr1.g01254.t01#2   mNeoVis.3.1.1.chr1.g01236.t01#6       2167   125258691:   mNeoVis.3.1.1.chr1.g01238.t01#6    mNeoVis.vulgivaga.1.1.chr1.g01252.t01#4    mNeoVis.evergladensis.2.1.chr1.g01256.t01#2    mNeoVis.vulgivaga.2.1.chr1.g01254.t01#1    mNeoVis.evergladensis.1.1.chr1.g01257.t01#3   mNeoVis.2.1.1.chr1_hap1.g01283.t01#5       2168   125258692:   mNeoVis.evergladensis.1.1.chr7.g01940.t01#3    mNeoVis.evergladensis.2.1.chr7.g01936.t01#2    mNeoVis.vulgivaga.2.1.chr7.g01954.t01#1    mNeoVis.2.1.1.chr7_hap1.g01973.t01#5    mNeoVis.vulgivaga.1.1.chr7.g01947.t01#4   mNeoVis.3.1.1.chr7.g01841.t01#6       2169   125258693:   mNeoVis.3.1.1.chr7.g01842.t01#6    mNeoVis.vulgivaga.2.1.chr7.g01955.t01#1    mNeoVis.evergladensis.2.1.chr7.g01937.t01#2    mNeoVis.2.1.1.chr7_hap1.g01974.t01#5    mNeoVis.vulgivaga.1.1.chr7.g01948.t01#4   mNeoVis.evergladensis.1.1.chr7.g01941.t01#3       2170   125258694:   mNeoVis.3.1.1.chr7.g01843.t01#6    mNeoVis.2.1.1.chr7_hap1.g01975.t01#5    mNeoVis.evergladensis.2.1.chr7.g01939.t01#2    mNeoVis.vulgivaga.2.1.chr7.g01956.t01#1    mNeoVis.vulgivaga.1.1.chr7.g01950.t01#4   mNeoVis.evergladensis.1.1.chr7.g01942.t01#3       2171   125258695:   mNeoVis.2.1.1.chr7_hap1.g01976.t01#5    mNeoVis.evergladensis.1.1.chr7.g01944.t01#3    mNeoVis.vulgivaga.2.1.chr7.g01957.t01#1    mNeoVis.3.1.1.chr7.g01844.t01#6    mNeoVis.evergladensis.2.1.chr7.g01940.t01#2   mNeoVis.vulgivaga.1.1.chr7.g01951.t01#4       2172   125258696:   mNeoVis.3.1.1.chr7.g01845.t01#6    mNeoVis.evergladensis.1.1.chr7.g01945.t01#3    mNeoVis.2.1.1.chr7_hap1.g01977.t01#5    mNeoVis.vulgivaga.2.1.chr7.g01958.t01#1    mNeoVis.evergladensis.2.1.chr7.g01941.t01#2   mNeoVis.vulgivaga.1.1.chr7.g01952.t01#4       2173   125258697:   mNeoVis.vulgivaga.1.1.chr15.g00604.t01#4    mNeoVis.vulgivaga.2.1.chr15.g00613.t01#1    mNeoVis.2.1.1.chr15_hap1.g00779.t01#5    mNeoVis.3.1.1.chr15.g00557.t01#6    mNeoVis.evergladensis.1.1.chr15.g00010.t01#3   mNeoVis.evergladensis.2.1.chr15.g00601.t01#2       2174   125258698:   mNeoVis.vulgivaga.2.1.chr1.g01255.t01#1    mNeoVis.evergladensis.1.1.chr1.g01256.t01#3    mNeoVis.2.1.1.chr1_hap1.g01284.t01#5    mNeoVis.vulgivaga.1.1.chr1.g01253.t01#4   mNeoVis.evergladensis.2.1.chr1.g01255.t01#2       2175   125258699:   mNeoVis.evergladensis.2.1.chr7.g01942.t01#2    mNeoVis.evergladensis.1.1.chr7.g01946.t01#3    mNeoVis.vulgivaga.1.1.chr7.g01953.t01#4    mNeoVis.vulgivaga.2.1.chr7.g01959.t01#1    mNeoVis.3.1.1.chr7.g01846.t01#6   mNeoVis.2.1.1.chr7_hap1.g01978.t01#5       2176   125258700:   mNeoVis.3.1.1.chr7.g01847.t01#6    mNeoVis.2.1.1.chr7_hap1.g01979.t01#5    mNeoVis.vulgivaga.2.1.chr7.g01960.t01#1    mNeoVis.evergladensis.1.1.chr7.g01947.t01#3    mNeoVis.evergladensis.2.1.chr7.g01943.t01#2   mNeoVis.vulgivaga.1.1.chr7.g01954.t01#4       2177   125258701:   mNeoVis.2.1.1.chr1_hap1.g01285.t01#5    mNeoVis.vulgivaga.2.1.chr5.g00051.t01#1    mNeoVis.vulgivaga.1.1.chr5.g00052.t01#4    mNeoVis.evergladensis.1.1.chr5.g00054.t01#3    mNeoVis.vulgivaga.1.1.chr1.g01254.t01#4    mNeoVis.3.1.1.chr1.g01239.t01#6    mNeoVis.vulgivaga.2.1.chr1.g01256.t01#1    mNeoVis.evergladensis.2.1.chr5.g00054.t01#2    mNeoVis.evergladensis.2.1.chr1.g01257.t01#2      2178   125258702:   mNeoVis.3.1.1.chr1.g01240.t01#6    mNeoVis.evergladensis.1.1.chr1.g01259.t01#3    mNeoVis.evergladensis.2.1.chr1.g01258.t01#2    mNeoVis.2.1.1.chr1_hap1.g01286.t01#5    mNeoVis.vulgivaga.2.1.chr1.g01257.t01#1   mNeoVis.vulgivaga.1.1.chr1.g01255.t01#4       2179   125258703:   mNeoVis.3.1.1.chr1.g01241.t01#6    mNeoVis.vulgivaga.1.1.chr1.g01256.t01#4    mNeoVis.vulgivaga.2.1.chr1.g01258.t01#1    mNeoVis.evergladensis.2.1.chr1.g01259.t01#2    mNeoVis.evergladensis.1.1.chr1.g01260.t01#3   mNeoVis.2.1.1.chr1_hap1.g01287.t01#5       2180   125258704:   mNeoVis.evergladensis.1.1.chr7.g01948.t01#3    mNeoVis.evergladensis.2.1.chr7.g01944.t01#2    mNeoVis.vulgivaga.2.1.chr7.g01961.t01#1    mNeoVis.3.1.1.chr7.g01848.t01#6    mNeoVis.vulgivaga.1.1.chr7.g01955.t01#4   mNeoVis.2.1.1.chr7_hap1.g01980.t01#5       2181   125258705:   mNeoVis.vulgivaga.2.1.chr15.g00615.t01#1    mNeoVis.3.1.1.chr15.g00558.t01#6    mNeoVis.evergladensis.2.1.chr15.g00602.t01#2    mNeoVis.evergladensis.1.1.chr15.g00009.t01#3    mNeoVis.2.1.1.chr15_hap1.g00780.t01#5    mNeoVis.vulgivaga.1.1.chr15.g00605.t01#4   mNeoVis.vulgivaga.2.1.chr15.g00614.t01#1     2182   125258706:   mNeoVis.evergladensis.2.1.chr1.g01260.t01#2    mNeoVis.3.1.1.chr1.g01242.t01#6    mNeoVis.vulgivaga.1.1.chr1.g01258.t01#4    mNeoVis.vulgivaga.2.1.chr1.g01259.t01#1    mNeoVis.evergladensis.1.1.chr1.g01261.t01#3   mNeoVis.2.1.1.chr1_hap1.g01288.t01#5       2183   125258707:   mNeoVis.2.1.1.chr7_hap1.g01981.t01#5    mNeoVis.evergladensis.1.1.chr7.g01949.t01#3    mNeoVis.vulgivaga.1.1.chr7.g01956.t01#4    mNeoVis.3.1.1.chr7.g01849.t01#6    mNeoVis.vulgivaga.2.1.chr7.g01962.t01#1   mNeoVis.evergladensis.2.1.chr7.g01945.t01#2       2184   125258708:  mNeoVis.vulgivaga.2.1.chr1.g01260.t01#1               2185   125258709:   mNeoVis.vulgivaga.1.1.chr7.g01957.t01#4    mNeoVis.evergladensis.2.1.chr7.g01946.t01#2    mNeoVis.3.1.1.chr7.g01850.t01#6    mNeoVis.2.1.1.chr7_hap1.g01982.t01#5    mNeoVis.vulgivaga.2.1.chr7.g01963.t01#1   mNeoVis.evergladensis.1.1.chr7.g01950.t01#3       2186   125258710:   mNeoVis.vulgivaga.1.1.chr7.g01850.t01#4    mNeoVis.evergladensis.1.1.chr7.g01839.t01#3    mNeoVis.evergladensis.2.1.chr7.g01842.t01#2    mNeoVis.2.1.1.chr7_hap1.g01881.t01#5    mNeoVis.vulgivaga.2.1.chr7.g01860.t01#1   mNeoVis.3.1.1.chr7.g01750.t01#6       2187   125258711:   mNeoVis.vulgivaga.2.1.chr1.g01261.t01#1    mNeoVis.3.1.1.chr1.g01245.t01#6    mNeoVis.2.1.1.chr1_hap1.g01292.t01#5    mNeoVis.evergladensis.2.1.chr1.g01263.t01#2    mNeoVis.evergladensis.1.1.chr1.g01264.t01#3   mNeoVis.vulgivaga.1.1.chr1.g01261.t01#4       2188   125258712:   mNeoVis.evergladensis.2.1.chr1.g01264.t01#2    mNeoVis.vulgivaga.2.1.chr1.g01262.t01#1    mNeoVis.vulgivaga.1.1.chr1.g01262.t01#4    mNeoVis.evergladensis.1.1.chr1.g01265.t01#3    mNeoVis.3.1.1.chr1.g01246.t01#6   mNeoVis.2.1.1.chr1_hap1.g01293.t01#5       2189   125258713:   mNeoVis.vulgivaga.1.1.chr7.g01851.t01#4    mNeoVis.evergladensis.1.1.chr7.g01840.t01#3    mNeoVis.3.1.1.chr7.g01751.t01#6    mNeoVis.evergladensis.2.1.chr7.g01843.t01#2    mNeoVis.2.1.1.chr7_hap1.g01882.t01#5   mNeoVis.vulgivaga.2.1.chr7.g01861.t01#1       2190   125258714:   mNeoVis.evergladensis.2.1.chr1.g01265.t01#2    mNeoVis.vulgivaga.2.1.chr1.g01263.t01#1    mNeoVis.evergladensis.1.1.chr1.g01266.t01#3    mNeoVis.vulgivaga.1.1.chr1.g01263.t01#4    mNeoVis.3.1.1.chr1.g01247.t01#6   mNeoVis.2.1.1.chr1_hap1.g01294.t01#5       2191   125258715:   mNeoVis.vulgivaga.1.1.chr7.g01852.t01#4    mNeoVis.vulgivaga.2.1.chr7.g01862.t01#1    mNeoVis.evergladensis.2.1.chr7.g01844.t01#2    mNeoVis.2.1.1.chr7_hap1.g01883.t01#5    mNeoVis.3.1.1.chr7.g01752.t01#6   mNeoVis.evergladensis.1.1.chr7.g01841.t01#3       2192   125258716:   mNeoVis.2.1.1.chr7_hap1.g00548.t01#5    mNeoVis.vulgivaga.1.1.chr7.g00555.t01#4    mNeoVis.evergladensis.1.1.chr7.g00556.t01#3    mNeoVis.3.1.1.chr7.g00530.t01#6    mNeoVis.vulgivaga.2.1.chr7.g00550.t01#1   mNeoVis.evergladensis.2.1.chr7.g00548.t01#2       2193   125258717:   mNeoVis.vulgivaga.2.1.chr1.g01264.t01#1    mNeoVis.evergladensis.2.1.chr1.g01266.t01#2    mNeoVis.2.1.1.chr1_hap1.g01295.t01#5    mNeoVis.vulgivaga.1.1.chr1.g01264.t01#4    mNeoVis.3.1.1.chr1.g01248.t01#6   mNeoVis.evergladensis.1.1.chr1.g01267.t01#3       2194   125258718:   mNeoVis.vulgivaga.1.1.chr1.g01265.t01#4    mNeoVis.evergladensis.2.1.chr1.g01267.t01#2    mNeoVis.evergladensis.1.1.chr1.g01268.t01#3    mNeoVis.3.1.1.chr1.g01249.t01#6    mNeoVis.vulgivaga.2.1.chr1.g01265.t01#1   mNeoVis.2.1.1.chr1_hap1.g01296.t01#5       2195   125258719:   mNeoVis.vulgivaga.1.1.chr1.g01266.t01#4    mNeoVis.evergladensis.2.1.chr1.g01268.t01#2    mNeoVis.evergladensis.1.1.chr1.g01269.t01#3    mNeoVis.3.1.1.chr1.g01250.t01#6    mNeoVis.vulgivaga.2.1.chr1.g01266.t01#1   mNeoVis.2.1.1.chr1_hap1.g01297.t01#5       2196   125258720:   mNeoVis.3.1.1.chr1.g01251.t01#6    mNeoVis.evergladensis.2.1.chr1.g01269.t01#2    mNeoVis.2.1.1.chr1_hap1.g01298.t01#5    mNeoVis.vulgivaga.2.1.chr1.g01267.t01#1    mNeoVis.vulgivaga.1.1.chr1.g01267.t01#4   mNeoVis.evergladensis.1.1.chr1.g01270.t01#3       2197   125258721:  mNeoVis.vulgivaga.2.1.chr1.g01268.t01#1               2198   125258722:   mNeoVis.evergladensis.2.1.chr7.g01845.t01#2    mNeoVis.2.1.1.chr7_hap1.g01884.t01#5    mNeoVis.vulgivaga.2.1.chr7.g01863.t01#1    mNeoVis.evergladensis.1.1.chr7.g01842.t01#3   mNeoVis.3.1.1.chr7.g01753.t01#6       2199   125258723:   mNeoVis.vulgivaga.1.1.chr8.g00913.t01#4    mNeoVis.3.1.1.chr8.g00888.t01#6    mNeoVis.vulgivaga.2.1.chr9.g00135.t01#1    mNeoVis.vulgivaga.1.1.chr3.g00659.t01#4    mNeoVis.vulgivaga.1.1.chr2.g00514.t01#4    mNeoVis.2.1.1.chr3_hap1.g00656.t01#5    mNeoVis.vulgivaga.2.1.chr3.g00668.t01#1    mNeoVis.evergladensis.1.1.chr2.g00515.t01#3    mNeoVis.evergladensis.2.1.chr3.g00664.t01#2      2200   125258724:   mNeoVis.evergladensis.2.1.chr7.g01846.t01#2    mNeoVis.3.1.1.chr7.g01754.t01#6    mNeoVis.evergladensis.1.1.chr7.g01843.t01#3    mNeoVis.evergladensis.2.1.chr11.g00681.t01#2    mNeoVis.3.1.1.chr11.g00659.t01#6    mNeoVis.vulgivaga.2.1.chr11.g00697.t01#1    mNeoVis.vulgivaga.1.1.chr11.g00679.t01#4    mNeoVis.evergladensis.1.1.chr11.g00688.t01#3    mNeoVis.2.1.1.chr11_hap1.g00697.t01#5      2201   125258725:   mNeoVis.evergladensis.1.1.chr5.g00629.t01#3    mNeoVis.evergladensis.2.1.chr5.g00634.t01#2    mNeoVis.vulgivaga.2.1.chr5.g00629.t01#1    mNeoVis.evergladensis.1.1.chr5.g00628.t01#3    mNeoVis.vulgivaga.2.1.chr2.g00520.t01#1    mNeoVis.3.1.1.chr5.g00628.t01#6   mNeoVis.2.1.1.chr2_hap1.g00651.t01#5     2202   125258726:   mNeoVis.2.1.1.chr1_hap1.g00687.t01#5    mNeoVis.evergladensis.2.1.chr1.g00695.t01#2    mNeoVis.vulgivaga.1.1.chr1.g00699.t01#4    mNeoVis.evergladensis.1.1.chr1.g00699.t01#3    mNeoVis.3.1.1.chr1.g00707.t01#6   mNeoVis.vulgivaga.2.1.chr1.g00707.t01#1       2203   125258727:   mNeoVis.evergladensis.2.1.chr7.g01847.t01#2    mNeoVis.vulgivaga.1.1.chr7.g01857.t01#4    mNeoVis.vulgivaga.1.1.chr7.g01856.t01#4    mNeoVis.vulgivaga.2.1.chr7.g01865.t01#1    mNeoVis.2.1.1.chr7_hap1.g01886.t01#5    mNeoVis.evergladensis.2.1.chr7.g01848.t01#2    mNeoVis.evergladensis.1.1.chr7.g01844.t01#3   mNeoVis.3.1.1.chr7.g01755.t01#6     2204   125258728:   mNeoVis.evergladensis.2.1.chr12.g00047.t01#2    mNeoVis.vulgivaga.1.1.chr12.g00054.t01#4    mNeoVis.3.1.1.chr12.g00048.t01#6    mNeoVis.vulgivaga.2.1.chr12.g00049.t01#1    mNeoVis.evergladensis.1.1.chr12.g00048.t01#3   mNeoVis.2.1.1.chr12_hap1.g00057.t01#5       2205   125258729:   mNeoVis.3.1.1.chr7.g01756.t01#6    mNeoVis.vulgivaga.2.1.chr7.g01866.t01#1   mNeoVis.2.1.1.chr7_hap1.g01887.t01#5           2206   125258730:   mNeoVis.vulgivaga.1.1.chr7.g01859.t01#4    mNeoVis.vulgivaga.2.1.chr7.g01867.t01#1    mNeoVis.2.1.1.chr7_hap1.g01888.t01#5    mNeoVis.3.1.1.chr7.g01757.t01#6    mNeoVis.evergladensis.2.1.chr7.g01850.t01#2   mNeoVis.evergladensis.1.1.chr7.g01846.t01#3       2207   125258731:   mNeoVis.evergladensis.1.1.chr7.g01847.t01#3    mNeoVis.3.1.1.chr7.g01758.t01#6    mNeoVis.2.1.1.chr7_hap1.g01889.t01#5    mNeoVis.evergladensis.2.1.chr7.g01851.t01#2    mNeoVis.vulgivaga.1.1.chr7.g01860.t01#4   mNeoVis.vulgivaga.2.1.chr7.g01868.t01#1       2208   125258732:   mNeoVis.2.1.1.chr7_hap1.g00897.t01#5    mNeoVis.vulgivaga.1.1.chr7.g00895.t01#4    mNeoVis.evergladensis.2.1.chr7.g00890.t01#2    mNeoVis.vulgivaga.2.1.chr7.g00894.t01#1    mNeoVis.evergladensis.1.1.chr7.g00895.t01#3   mNeoVis.3.1.1.chr7.g00838.t01#6       2209   125258733:   mNeoVis.vulgivaga.1.1.chr7.g01861.t01#4    mNeoVis.3.1.1.chr7.g01759.t01#6    mNeoVis.2.1.1.chr7_hap1.g01890.t01#5    mNeoVis.evergladensis.1.1.chr7.g01848.t01#3    mNeoVis.evergladensis.2.1.chr7.g01852.t01#2   mNeoVis.vulgivaga.2.1.chr7.g01869.t01#1       2210   125258734:  mNeoVis.vulgivaga.2.1.chr1.g01269.t01#1               2211   125258735:   mNeoVis.2.1.1.chr1_hap1.g01300.t01#5    mNeoVis.vulgivaga.2.1.chr1.g01270.t01#1    mNeoVis.vulgivaga.1.1.chr1.g01269.t01#4    mNeoVis.evergladensis.2.1.chr1.g01271.t01#2    mNeoVis.evergladensis.1.1.chr1.g01272.t01#3   mNeoVis.3.1.1.chr1.g01253.t01#6       2212   125258736:   mNeoVis.evergladensis.1.1.chr7.g01849.t01#3    mNeoVis.evergladensis.2.1.chr7.g01853.t01#2    mNeoVis.vulgivaga.1.1.chr7.g01862.t01#4    mNeoVis.2.1.1.chr7_hap1.g01891.t01#5    mNeoVis.3.1.1.chr7.g01760.t01#6   mNeoVis.vulgivaga.2.1.chr7.g01870.t01#1       2213   125258737:   mNeoVis.vulgivaga.1.1.chr1.g01270.t01#4    mNeoVis.3.1.1.chr1.g01254.t01#6    mNeoVis.2.1.1.chr1_hap1.g01301.t01#5    mNeoVis.vulgivaga.2.1.chr1.g01271.t01#1    mNeoVis.evergladensis.1.1.chr1.g01273.t01#3   mNeoVis.evergladensis.2.1.chr1.g01272.t01#2       2214   125258738:   mNeoVis.vulgivaga.2.1.chr15.g00514.t01#1    mNeoVis.vulgivaga.1.1.chr15.g00508.t01#4    mNeoVis.evergladensis.2.1.chr15.g00505.t01#2    mNeoVis.3.1.1.chr15.g00464.t01#6    mNeoVis.2.1.1.chr15_hap1.g00683.t01#5   mNeoVis.evergladensis.1.1.chr15.g00108.t01#3       2215   125258739:   mNeoVis.vulgivaga.2.1.chr1.g01272.t01#1    mNeoVis.3.1.1.chr1.g01255.t01#6    mNeoVis.2.1.1.chr1_hap1.g01302.t01#5    mNeoVis.evergladensis.2.1.chr1.g01273.t01#2    mNeoVis.evergladensis.1.1.chr1.g01274.t01#3   mNeoVis.vulgivaga.1.1.chr1.g01271.t01#4       2216   125258740:   mNeoVis.evergladensis.2.1.chr7.g01854.t01#2    mNeoVis.2.1.1.chr7_hap1.g01892.t01#5    mNeoVis.3.1.1.chr7.g01761.t01#6   mNeoVis.vulgivaga.2.1.chr7.g01871.t01#1         2217   125258741:   mNeoVis.vulgivaga.2.1.chr7.g01872.t01#1    mNeoVis.evergladensis.1.1.chr7.g01851.t01#3    mNeoVis.evergladensis.2.1.chr7.g01855.t01#2    mNeoVis.2.1.1.chr7_hap1.g01893.t01#5   mNeoVis.3.1.1.chr7.g01762.t01#6       2218   125258742:   mNeoVis.2.1.1.chr7_hap1.g01894.t01#5    mNeoVis.vulgivaga.2.1.chr7.g01873.t01#1    mNeoVis.evergladensis.1.1.chr7.g01852.t01#3    mNeoVis.evergladensis.2.1.chr7.g01856.t01#2   mNeoVis.vulgivaga.1.1.chr7.g01864.t01#4       2219   125258743:   mNeoVis.evergladensis.1.1.chr7.g01853.t01#3    mNeoVis.evergladensis.2.1.chr7.g01857.t01#2    mNeoVis.vulgivaga.1.1.chr7.g01865.t01#4    mNeoVis.vulgivaga.2.1.chr7.g01874.t01#1   mNeoVis.2.1.1.chr7_hap1.g01895.t01#5       2220   125258744:   mNeoVis.evergladensis.2.1.chr1.g01274.t01#2    mNeoVis.2.1.1.chr5_hap1.g00171.t01#5    mNeoVis.evergladensis.2.1.chr13.g00017.t01#2    mNeoVis.2.1.1.chr1_hap1.g01268.t01#5    mNeoVis.vulgivaga.1.1.chr1.g01237.t01#4    mNeoVis.evergladensis.1.1.chr2.g00169.t01#3    mNeoVis.evergladensis.2.1.chr1.g01239.t01#2    mNeoVis.2.1.1.chr1_hap1.g01267.t01#5    mNeoVis.3.1.1.chr3.g00717.t01#6      2221   125258745:   mNeoVis.3.1.1.chr7.g00010.t01#6    mNeoVis.3.1.1.chr1.g01256.t01#6    mNeoVis.vulgivaga.2.1.chr3.g00277.t01#1    mNeoVis.evergladensis.2.1.chr3.g00284.t01#2    mNeoVis.3.1.1.chr1.g01223.t01#6    mNeoVis.vulgivaga.1.1.chr3.g00277.t01#4    mNeoVis.evergladensis.2.1.chr7.g00010.t01#2    mNeoVis.vulgivaga.1.1.chr2.g00168.t01#4    mNeoVis.vulgivaga.1.1.chr5.g00430.t01#4      2222   125258746:   mNeoVis.evergladensis.1.1.chr3.g00800.t01#3    mNeoVis.vulgivaga.1.1.chr1.g01236.t01#4    mNeoVis.evergladensis.1.1.chr1.g01241.t01#3    mNeoVis.3.1.1.chr3.g00277.t01#6   mNeoVis.evergladensis.2.1.chr1.g01241.t01#2       2223   125258747:   mNeoVis.3.1.1.chr11.g00049.t01#6    mNeoVis.evergladensis.2.1.chr11.g00074.t01#2    mNeoVis.vulgivaga.1.1.chr11.g00072.t01#4    mNeoVis.vulgivaga.2.1.chr11.g00074.t01#1   mNeoVis.2.1.1.chr11_hap1.g00075.t01#5       2224   125258748:   mNeoVis.vulgivaga.1.1.chr1.g01272.t01#4    mNeoVis.vulgivaga.2.1.chr3.g00804.t01#1    mNeoVis.evergladensis.2.1.chr5.g00444.t01#2   mNeoVis.vulgivaga.2.1.chr7.g00009.t01#1         2225   125258749:   mNeoVis.vulgivaga.2.1.chr13.g00016.t01#1    mNeoVis.vulgivaga.2.1.chr1.g01240.t01#1   mNeoVis.vulgivaga.2.1.chr1.g01273.t01#1           2226   125258750:   mNeoVis.evergladensis.2.1.chr1.g01275.t01#2    mNeoVis.vulgivaga.2.1.chr1.g01274.t01#1    mNeoVis.vulgivaga.1.1.chr1.g01273.t01#4    mNeoVis.2.1.1.chr1_hap1.g01304.t01#5    mNeoVis.3.1.1.chr1.g01257.t01#6   mNeoVis.evergladensis.1.1.chr1.g01276.t01#3       2227   125258751:   mNeoVis.2.1.1.chr7_hap1.g01896.t01#5    mNeoVis.vulgivaga.1.1.chr7.g01867.t01#4    mNeoVis.vulgivaga.2.1.chr7.g01875.t01#1    mNeoVis.3.1.1.chr7.g01765.t01#6    mNeoVis.evergladensis.2.1.chr7.g01858.t01#2   mNeoVis.evergladensis.1.1.chr7.g01855.t01#3       2228   125258752:   mNeoVis.evergladensis.1.1.chr1.g01277.t01#3    mNeoVis.2.1.1.chr1_hap1.g01305.t01#5    mNeoVis.vulgivaga.1.1.chr1.g01274.t01#4    mNeoVis.evergladensis.2.1.chr1.g01276.t01#2    mNeoVis.3.1.1.chr1.g01258.t01#6   mNeoVis.vulgivaga.2.1.chr1.g01275.t01#1       2229   125258753:   mNeoVis.vulgivaga.2.1.chr15.g00492.t01#1    mNeoVis.vulgivaga.1.1.chr3.g00118.t01#4    mNeoVis.evergladensis.1.1.chr3.g00115.t01#3    mNeoVis.evergladensis.2.1.chr8.g00470.t01#2    mNeoVis.2.1.1.chr8_hap1.g01401.t01#5    mNeoVis.3.1.1.chr3.g00117.t01#6    mNeoVis.vulgivaga.1.1.chr1.g01275.t01#4    mNeoVis.2.1.1.chr1_hap1.g01306.t01#5    mNeoVis.evergladensis.2.1.chr2.g01639.t01#2      2230   125258754:   mNeoVis.vulgivaga.1.1.chr9.g01116.t01#4   mNeoVis.vulgivaga.2.1.chr9.g01102.t01#1             2231   125258755:   mNeoVis.3.1.1.chr1.g01259.t01#6    mNeoVis.2.1.1.chr1_hap1.g01307.t01#5    mNeoVis.vulgivaga.2.1.chr1.g01277.t01#1    mNeoVis.evergladensis.2.1.chr1.g01278.t01#2    mNeoVis.evergladensis.1.1.chr1.g01279.t01#3   mNeoVis.vulgivaga.1.1.chr1.g01276.t01#4       2232   125258756:   mNeoVis.vulgivaga.2.1.chr1.g01278.t01#1    mNeoVis.evergladensis.1.1.chr1.g01280.t01#3    mNeoVis.3.1.1.chr1.g01260.t01#6    mNeoVis.evergladensis.2.1.chr1.g01279.t01#2    mNeoVis.vulgivaga.1.1.chr1.g01277.t01#4   mNeoVis.2.1.1.chr1_hap1.g01308.t01#5       2233   125258757:   mNeoVis.vulgivaga.2.1.chr7.g01876.t01#1    mNeoVis.3.1.1.chr7.g01766.t01#6    mNeoVis.2.1.1.chr7_hap1.g01897.t01#5    mNeoVis.evergladensis.1.1.chr7.g01856.t01#3    mNeoVis.vulgivaga.1.1.chr7.g01868.t01#4   mNeoVis.evergladensis.2.1.chr7.g01859.t01#2       2234   125258758:   mNeoVis.vulgivaga.2.1.chr7.g01877.t01#1    mNeoVis.vulgivaga.1.1.chr7.g01869.t01#4    mNeoVis.2.1.1.chr7_hap1.g01898.t01#5    mNeoVis.evergladensis.2.1.chr7.g01860.t01#2    mNeoVis.3.1.1.chr7.g01767.t01#6   mNeoVis.evergladensis.1.1.chr7.g01857.t01#3       2235   125258759:   mNeoVis.vulgivaga.2.1.chr1.g01279.t01#1    mNeoVis.evergladensis.2.1.chr1.g01280.t01#2    mNeoVis.vulgivaga.1.1.chr1.g01278.t01#4    mNeoVis.evergladensis.1.1.chr1.g01281.t01#3   mNeoVis.2.1.1.chr1_hap1.g01309.t01#5       2236   125258760:   mNeoVis.vulgivaga.2.1.chr7.g01878.t01#1    mNeoVis.vulgivaga.1.1.chr7.g01870.t01#4    mNeoVis.3.1.1.chr7.g01768.t01#6    mNeoVis.2.1.1.chr7_hap1.g01899.t01#5    mNeoVis.evergladensis.2.1.chr7.g01861.t01#2   mNeoVis.evergladensis.1.1.chr7.g01858.t01#3       2237   125258761:   mNeoVis.vulgivaga.2.1.chr7.g01879.t01#1    mNeoVis.evergladensis.2.1.chr7.g01862.t01#2    mNeoVis.vulgivaga.1.1.chr7.g01871.t01#4    mNeoVis.3.1.1.chr7.g01769.t01#6    mNeoVis.2.1.1.chr7_hap1.g01900.t01#5   mNeoVis.evergladensis.1.1.chr7.g01859.t01#3       2238   125258762:   mNeoVis.evergladensis.1.1.chr7.g00557.t01#3    mNeoVis.3.1.1.chr7.g00531.t01#6    mNeoVis.vulgivaga.1.1.chr7.g00556.t01#4    mNeoVis.vulgivaga.2.1.chr7.g00551.t01#1    mNeoVis.evergladensis.2.1.chr7.g00549.t01#2   mNeoVis.2.1.1.chr7_hap1.g00549.t01#5       2239   125258763:   mNeoVis.evergladensis.2.1.chr7.g01863.t01#2    mNeoVis.vulgivaga.2.1.chr7.g01880.t01#1    mNeoVis.2.1.1.chr7_hap1.g01901.t01#5    mNeoVis.evergladensis.1.1.chr7.g01860.t01#3    mNeoVis.3.1.1.chr7.g01770.t01#6   mNeoVis.vulgivaga.1.1.chr7.g01872.t01#4       2240   125258764:   mNeoVis.vulgivaga.1.1.chr7.g01873.t01#4    mNeoVis.2.1.1.chr7_hap1.g01902.t01#5    mNeoVis.evergladensis.1.1.chr7.g01861.t01#3    mNeoVis.vulgivaga.2.1.chr7.g01881.t01#1    mNeoVis.3.1.1.chr7.g01771.t01#6   mNeoVis.evergladensis.2.1.chr7.g01864.t01#2       2241   125258765:   mNeoVis.vulgivaga.1.1.chr1.g01279.t01#4    mNeoVis.vulgivaga.2.1.chr1.g01280.t01#1    mNeoVis.2.1.1.chr1_hap1.g01310.t01#5    mNeoVis.evergladensis.2.1.chr1.g01281.t01#2   mNeoVis.evergladensis.1.1.chr1.g01282.t01#3       2242   125258766:   mNeoVis.evergladensis.2.1.chr1.g01282.t01#2    mNeoVis.2.1.1.chr1_hap1.g01311.t01#5    mNeoVis.evergladensis.1.1.chr1.g01283.t01#3    mNeoVis.vulgivaga.1.1.chr1.g01280.t01#4   mNeoVis.vulgivaga.2.1.chr1.g01281.t01#1       2243   125258767:   mNeoVis.vulgivaga.2.1.chr2.g00759.t01#1    mNeoVis.2.1.1.chr5_hap1.g00753.t01#5    mNeoVis.2.1.1.chr1_hap1.g01608.t01#5    mNeoVis.evergladensis.2.1.chr13.g00179.t01#2    mNeoVis.evergladensis.1.1.chr1.g01284.t01#3    mNeoVis.2.1.1.chr5_hap1.g00829.t01#5    mNeoVis.2.1.1.chr1_hap1.g01312.t01#5    mNeoVis.2.1.1.chr3_hap1.g01272.t01#5    mNeoVis.2.1.1.chr3_hap1.g00801.t01#5      2244   125258768:   mNeoVis.vulgivaga.2.1.chr11.g00556.t01#1    mNeoVis.vulgivaga.1.1.chr7.g01450.t01#4    mNeoVis.evergladensis.1.1.chr7.g01440.t01#3    mNeoVis.vulgivaga.2.1.chr1.g01282.t01#1    mNeoVis.vulgivaga.2.1.chr7.g01456.t01#1    mNeoVis.evergladensis.2.1.chr7.g01442.t01#2    mNeoVis.evergladensis.2.1.chr2.g00829.t01#2    mNeoVis.3.1.1.chr2.g00740.t01#6    mNeoVis.vulgivaga.2.1.chr2.g00833.t01#1      2245   125258769:   mNeoVis.vulgivaga.1.1.chr13.g00173.t01#4    mNeoVis.evergladensis.1.1.chr13.g00177.t01#3    mNeoVis.2.1.1.chr13_hap1.g00181.t01#5   mNeoVis.3.1.1.chr13.g00132.t01#6         2246   125258770:   mNeoVis.3.1.1.chr11.g00667.t01#6    mNeoVis.2.1.1.chr11_hap1.g00705.t01#5    mNeoVis.evergladensis.1.1.chr11.g00696.t01#3   mNeoVis.vulgivaga.2.1.chr11.g00705.t01#1       2247   125258771:   mNeoVis.3.1.1.chr12.g00646.t01#6   mNeoVis.evergladensis.1.1.chr3.g00803.t01#3           2248   125258772:   mNeoVis.evergladensis.2.1.chr7.g01865.t01#2    mNeoVis.3.1.1.chr7.g01772.t01#6    mNeoVis.vulgivaga.1.1.chr7.g01874.t01#4    mNeoVis.2.1.1.chr7_hap1.g01903.t01#5    mNeoVis.vulgivaga.2.1.chr7.g01882.t01#1   mNeoVis.evergladensis.1.1.chr7.g01862.t01#3       2249   125258773:   mNeoVis.evergladensis.2.1.chr7.g00903.t01#2    mNeoVis.3.1.1.chr7.g00851.t01#6    mNeoVis.evergladensis.1.1.chr7.g00908.t01#3    mNeoVis.2.1.1.chr7_hap1.g00910.t01#5    mNeoVis.vulgivaga.1.1.chr7.g00907.t01#4   mNeoVis.vulgivaga.2.1.chr7.g00907.t01#1       2250   125258774:   mNeoVis.evergladensis.1.1.chr10.g00948.t01#3    mNeoVis.vulgivaga.1.1.chr10.g00928.t01#4    mNeoVis.2.1.1.chr9_hap1.g00961.t01#5    mNeoVis.vulgivaga.2.1.chr10.g00938.t01#1    mNeoVis.evergladensis.2.1.chr10.g00948.t01#2   mNeoVis.3.1.1.chr10.g00916.t01#6       2251   125258775:   mNeoVis.vulgivaga.2.1.chr1.g01283.t01#1   mNeoVis.3.1.1.chr1.g01263.t01#6             2252   125258776:   mNeoVis.evergladensis.1.1.chr1.g01285.t01#3    mNeoVis.2.1.1.chr1_hap1.g01314.t01#5    mNeoVis.vulgivaga.1.1.chr1.g01282.t01#4    mNeoVis.vulgivaga.2.1.chr1.g01284.t01#1    mNeoVis.evergladensis.2.1.chr1.g01284.t01#2   mNeoVis.3.1.1.chr1.g01264.t01#6       2253   125258777:   mNeoVis.vulgivaga.1.1.chr1.g01283.t01#4    mNeoVis.vulgivaga.2.1.chr1.g01285.t01#1    mNeoVis.evergladensis.2.1.chr1.g01285.t01#2    mNeoVis.3.1.1.chr1.g01265.t01#6    mNeoVis.2.1.1.chr1_hap1.g01315.t01#5   mNeoVis.evergladensis.1.1.chr1.g01286.t01#3       2254   125258778:   mNeoVis.evergladensis.1.1.chr7.g01863.t01#3    mNeoVis.3.1.1.chr7.g01773.t01#6    mNeoVis.vulgivaga.2.1.chr7.g01883.t01#1    mNeoVis.vulgivaga.1.1.chr7.g01875.t01#4    mNeoVis.2.1.1.chr7_hap1.g01904.t01#5   mNeoVis.evergladensis.2.1.chr7.g01866.t01#2       2255   125258779:   mNeoVis.vulgivaga.2.1.chr7.g01884.t01#1    mNeoVis.evergladensis.1.1.chr2.g00735.t01#3    mNeoVis.evergladensis.2.1.chr7.g01867.t01#2    mNeoVis.vulgivaga.1.1.chr2.g00733.t01#4    mNeoVis.vulgivaga.1.1.chr7.g01876.t01#4    mNeoVis.evergladensis.1.1.chr7.g01864.t01#3    mNeoVis.vulgivaga.2.1.chr2.g00743.t01#1    mNeoVis.2.1.1.chr7_hap1.g01905.t01#5    mNeoVis.evergladensis.2.1.chr2.g00740.t01#2      2256   125258780:   mNeoVis.evergladensis.1.1.chr7.g01865.t01#3    mNeoVis.vulgivaga.1.1.chr7.g01877.t01#4    mNeoVis.2.1.1.chr7_hap1.g01906.t01#5    mNeoVis.vulgivaga.2.1.chr7.g01885.t01#1    mNeoVis.evergladensis.2.1.chr7.g01868.t01#2   mNeoVis.3.1.1.chr7.g01775.t01#6       2257   125258781:   mNeoVis.vulgivaga.1.1.chr1.g01284.t01#4    mNeoVis.evergladensis.1.1.chr1.g01287.t01#3    mNeoVis.evergladensis.2.1.chr1.g01286.t01#2    mNeoVis.3.1.1.chr1.g01266.t01#6   mNeoVis.2.1.1.chr1_hap1.g01316.t01#5       2258   125258782:   mNeoVis.vulgivaga.2.1.chr1.g01286.t01#1   mNeoVis.evergladensis.2.1.chr1.g01287.t01#2           2259   125258783:   mNeoVis.2.1.1.chr7_hap1.g01907.t01#5    mNeoVis.evergladensis.1.1.chr7.g01866.t01#3    mNeoVis.evergladensis.2.1.chr7.g01869.t01#2    mNeoVis.vulgivaga.1.1.chr7.g01878.t01#4    mNeoVis.vulgivaga.2.1.chr7.g01886.t01#1    mNeoVis.3.1.1.chr7.g01776.t01#6   mNeoVis.evergladensis.1.1.chr7.g01867.t01#3     2260   125258784:   mNeoVis.vulgivaga.1.1.chr1.g01285.t01#4    mNeoVis.evergladensis.2.1.chr1.g01288.t01#2    mNeoVis.3.1.1.chr1.g01267.t01#6    mNeoVis.2.1.1.chr1_hap1.g01317.t01#5    mNeoVis.vulgivaga.2.1.chr1.g01287.t01#1   mNeoVis.evergladensis.1.1.chr1.g01288.t01#3       2261   125258785:   mNeoVis.evergladensis.1.1.chr1.g01289.t01#3    mNeoVis.3.1.1.chr1.g01268.t01#6    mNeoVis.2.1.1.chr1_hap1.g01318.t01#5    mNeoVis.vulgivaga.2.1.chr1.g01288.t01#1    mNeoVis.vulgivaga.1.1.chr1.g01286.t01#4   mNeoVis.evergladensis.2.1.chr1.g01289.t01#2       2262   125258786:   mNeoVis.2.1.1.chr1_hap1.g01323.t01#5    mNeoVis.2.1.1.chr1_hap1.g01324.t01#5    mNeoVis.evergladensis.2.1.chr1.g01292.t01#2    mNeoVis.evergladensis.1.1.chr1.g01294.t01#3    mNeoVis.3.1.1.chr1.g01274.t01#6    mNeoVis.vulgivaga.1.1.chr1.g01290.t01#4   mNeoVis.vulgivaga.2.1.chr1.g01293.t01#1     2263   125258787:   mNeoVis.evergladensis.1.1.chr1.g01290.t01#3    mNeoVis.evergladensis.2.1.chr1.g01290.t01#2    mNeoVis.vulgivaga.1.1.chr1.g01287.t01#4    mNeoVis.3.1.1.chr1.g01269.t01#6   mNeoVis.2.1.1.chr1_hap1.g01319.t01#5       2264   125258788:   mNeoVis.vulgivaga.2.1.chr1.g01292.t01#1    mNeoVis.vulgivaga.1.1.chr1.g01289.t01#4    mNeoVis.evergladensis.1.1.chr1.g01293.t01#3    mNeoVis.3.1.1.chr1.g01273.t01#6   mNeoVis.2.1.1.chr1_hap1.g01322.t01#5       2265   125258789:   mNeoVis.evergladensis.2.1.chr1.g01291.t01#2    mNeoVis.3.1.1.chr1.g01270.t01#6   mNeoVis.vulgivaga.1.1.chr1.g01288.t01#4           2266   125258790:   mNeoVis.evergladensis.1.1.chr1.g01295.t01#3    mNeoVis.2.1.1.chr1_hap1.g01325.t01#5   mNeoVis.vulgivaga.1.1.chr1.g01291.t01#4           2267   125258791:   mNeoVis.evergladensis.1.1.chr1.g01291.t01#3   mNeoVis.vulgivaga.2.1.chr1.g01289.t01#1             2268   125258792:   mNeoVis.evergladensis.2.1.chr7.g01870.t01#2    mNeoVis.evergladensis.1.1.chr7.g01868.t01#3    mNeoVis.vulgivaga.1.1.chr7.g01879.t01#4    mNeoVis.2.1.1.chr7_hap1.g01908.t01#5   mNeoVis.vulgivaga.2.1.chr7.g01887.t01#1       2269   125258793:   mNeoVis.evergladensis.2.1.chr7.g01872.t01#2    mNeoVis.vulgivaga.1.1.chr7.g01882.t01#4    mNeoVis.evergladensis.1.1.chr7.g01870.t01#3    mNeoVis.3.1.1.chr7.g01780.t01#6    mNeoVis.2.1.1.chr7_hap1.g01910.t01#5   mNeoVis.vulgivaga.2.1.chr7.g01889.t01#1       2270   125258794:   mNeoVis.vulgivaga.1.1.chr7.g01883.t01#4    mNeoVis.evergladensis.1.1.chr7.g01871.t01#3    mNeoVis.vulgivaga.2.1.chr7.g01890.t01#1    mNeoVis.3.1.1.chr7.g01781.t01#6    mNeoVis.2.1.1.chr7_hap1.g01911.t01#5   mNeoVis.evergladensis.2.1.chr7.g01873.t01#2       2271   125258795:   mNeoVis.2.1.1.chr7_hap1.g01912.t01#5    mNeoVis.evergladensis.2.1.chr7.g01874.t01#2    mNeoVis.vulgivaga.1.1.chr7.g01884.t01#4    mNeoVis.3.1.1.chr7.g01782.t01#6    mNeoVis.evergladensis.1.1.chr7.g01872.t01#3   mNeoVis.vulgivaga.2.1.chr7.g01891.t01#1       2272   125258796:   mNeoVis.evergladensis.1.1.chr10.g01234.t01#3    mNeoVis.vulgivaga.2.1.chr10.g01229.t01#1    mNeoVis.vulgivaga.2.1.chr10.g01228.t01#1    mNeoVis.3.1.1.chr10.g01202.t01#6    mNeoVis.evergladensis.2.1.chr10.g01241.t01#2    mNeoVis.2.1.1.chr9_hap1.g01251.t01#5   mNeoVis.vulgivaga.1.1.chr10.g01214.t01#4     2273   125258797:   mNeoVis.evergladensis.1.1.chr7.g01873.t01#3    mNeoVis.2.1.1.chr7_hap1.g01913.t01#5    mNeoVis.vulgivaga.2.1.chr7.g01892.t01#1    mNeoVis.vulgivaga.1.1.chr7.g01885.t01#4   mNeoVis.evergladensis.2.1.chr7.g01875.t01#2       2274   125258798:   mNeoVis.evergladensis.2.1.chr4.g00524.t01#2    mNeoVis.3.1.1.chr4.g00521.t01#6    mNeoVis.evergladensis.1.1.chr4.g00529.t01#3    mNeoVis.vulgivaga.2.1.chr1.g01290.t01#1    mNeoVis.vulgivaga.1.1.chr4.g00520.t01#4    mNeoVis.3.1.1.chr1.g01271.t01#6    mNeoVis.2.1.1.chr6_hap1.g00518.t01#5    mNeoVis.2.1.1.chr1_hap1.g01320.t01#5    mNeoVis.vulgivaga.2.1.chr4.g00531.t01#1      2275   125258799:   mNeoVis.evergladensis.1.1.chr14.g00200.t01#3    mNeoVis.2.1.1.chr14_hap1.g00195.t01#5    mNeoVis.vulgivaga.2.1.chr14.g00199.t01#1    mNeoVis.vulgivaga.1.1.chr14.g00194.t01#4   mNeoVis.evergladensis.2.1.chr14.g00195.t01#2       2276   125258800:   mNeoVis.2.1.1.chr1_hap1.g01321.t01#5    mNeoVis.3.1.1.chr1.g01272.t01#6   mNeoVis.vulgivaga.2.1.chr1.g01291.t01#1           2277   125258801:   mNeoVis.evergladensis.1.1.chr7.g01874.t01#3    mNeoVis.vulgivaga.2.1.chr7.g01893.t01#1    mNeoVis.2.1.1.chr7_hap1.g01914.t01#5   mNeoVis.vulgivaga.1.1.chr7.g01886.t01#4         2278   125258802:   mNeoVis.2.1.1.chr15_hap1.g00684.t01#5    mNeoVis.evergladensis.1.1.chr15.g00107.t01#3    mNeoVis.vulgivaga.1.1.chr15.g00509.t01#4    mNeoVis.3.1.1.chr15.g00465.t01#6    mNeoVis.evergladensis.2.1.chr15.g00506.t01#2   mNeoVis.vulgivaga.2.1.chr15.g00515.t01#1       2279   125258803:  mNeoVis.vulgivaga.2.1.chr1.g01294.t01#1               2280   125258804:  mNeoVis.vulgivaga.2.1.chr1.g01295.t01#1               2281   125258805:   mNeoVis.vulgivaga.2.1.chr1.g01296.t01#1    mNeoVis.2.1.1.chr1_hap1.g01328.t01#5    mNeoVis.3.1.1.chr1.g01277.t01#6    mNeoVis.vulgivaga.1.1.chr1.g01295.t01#4    mNeoVis.evergladensis.2.1.chr1.g01295.t01#2   mNeoVis.evergladensis.1.1.chr1.g01297.t01#3       2282   125258806:   mNeoVis.vulgivaga.1.1.chr2.g00366.t01#4    mNeoVis.3.1.1.chr2.g00304.t01#6    mNeoVis.vulgivaga.2.1.chr2.g00367.t01#1    mNeoVis.evergladensis.2.1.chr2.g00372.t01#2    mNeoVis.evergladensis.1.1.chr2.g00370.t01#3   mNeoVis.2.1.1.chr5_hap1.g00371.t01#5       2283   125258807:   mNeoVis.2.1.1.chr1_hap1.g01329.t01#5    mNeoVis.3.1.1.chr1.g01278.t01#6    mNeoVis.evergladensis.2.1.chr1.g01296.t01#2    mNeoVis.vulgivaga.1.1.chr1.g01296.t01#4    mNeoVis.evergladensis.1.1.chr1.g01298.t01#3   mNeoVis.vulgivaga.2.1.chr1.g01297.t01#1       2284   125258808:   mNeoVis.2.1.1.chr7_hap1.g01915.t01#5    mNeoVis.vulgivaga.2.1.chr7.g01894.t01#1    mNeoVis.evergladensis.2.1.chr7.g01879.t01#2    mNeoVis.evergladensis.1.1.chr7.g01875.t01#3    mNeoVis.3.1.1.chr7.g01786.t01#6   mNeoVis.vulgivaga.1.1.chr7.g01887.t01#4       2285   125258809:   mNeoVis.evergladensis.1.1.chr1.g01299.t01#3    mNeoVis.evergladensis.2.1.chr1.g01297.t01#2    mNeoVis.vulgivaga.2.1.chr1.g01298.t01#1    mNeoVis.2.1.1.chr1_hap1.g01330.t01#5    mNeoVis.3.1.1.chr1.g01279.t01#6   mNeoVis.vulgivaga.1.1.chr1.g01297.t01#4       2286   125258810:   mNeoVis.evergladensis.1.1.chr7.g01877.t01#3    mNeoVis.2.1.1.chr7_hap1.g01916.t01#5    mNeoVis.evergladensis.2.1.chr7.g01880.t01#2    mNeoVis.3.1.1.chr1.g00598.t01#6    mNeoVis.vulgivaga.2.1.chr7.g01895.t01#1    mNeoVis.vulgivaga.1.1.chr1.g00596.t01#4    mNeoVis.vulgivaga.2.1.chr1.g00598.t01#1    mNeoVis.3.1.1.chr7.g01787.t01#6    mNeoVis.evergladensis.2.1.chr1.g00589.t01#2      2287   125258811:  mNeoVis.vulgivaga.2.1.chr7.g00552.t01#1               2288   125258812:   mNeoVis.vulgivaga.2.1.chr1.g01299.t01#1    mNeoVis.evergladensis.2.1.chr1.g01298.t01#2    mNeoVis.2.1.1.chr1_hap1.g01331.t01#5    mNeoVis.3.1.1.chr1.g01280.t01#6    mNeoVis.evergladensis.1.1.chr1.g01300.t01#3   mNeoVis.vulgivaga.1.1.chr1.g01298.t01#4       2289   125258813:   mNeoVis.vulgivaga.1.1.chr1.g01299.t01#4    mNeoVis.evergladensis.2.1.chr1.g01299.t01#2    mNeoVis.vulgivaga.2.1.chr1.g01300.t01#1    mNeoVis.2.1.1.chr1_hap1.g01332.t01#5    mNeoVis.3.1.1.chr1.g01281.t01#6   mNeoVis.evergladensis.1.1.chr1.g01301.t01#3       2290   125258814:  mNeoVis.vulgivaga.2.1.chr1.g01301.t01#1               2291   125258815:   mNeoVis.vulgivaga.1.1.chr7.g01889.t01#4    mNeoVis.3.1.1.chr7.g01788.t01#6    mNeoVis.evergladensis.1.1.chr7.g01878.t01#3    mNeoVis.evergladensis.2.1.chr7.g01881.t01#2    mNeoVis.vulgivaga.2.1.chr7.g01896.t01#1   mNeoVis.2.1.1.chr7_hap1.g01917.t01#5       2292   125258816:   mNeoVis.vulgivaga.2.1.chr1.g01302.t01#1    mNeoVis.vulgivaga.1.1.chr1.g01301.t01#4    mNeoVis.3.1.1.chr1.g01284.t01#6    mNeoVis.evergladensis.1.1.chr1.g01303.t01#3    mNeoVis.evergladensis.2.1.chr1.g01301.t01#2   mNeoVis.2.1.1.chr1_hap1.g01334.t01#5       2293   125258817:   mNeoVis.3.1.1.chr1.g01283.t01#6    mNeoVis.vulgivaga.1.1.chr1.g01302.t01#4    mNeoVis.vulgivaga.2.1.chr1.g01303.t01#1    mNeoVis.evergladensis.2.1.chr1.g01302.t01#2    mNeoVis.2.1.1.chr1_hap1.g01335.t01#5   mNeoVis.evergladensis.1.1.chr1.g01304.t01#3       2294   125258818:   mNeoVis.2.1.1.chr1_hap1.g01336.t01#5    mNeoVis.3.1.1.chr1.g01285.t01#6    mNeoVis.evergladensis.1.1.chr1.g01305.t01#3    mNeoVis.evergladensis.2.1.chr1.g01303.t01#2    mNeoVis.vulgivaga.2.1.chr1.g01304.t01#1   mNeoVis.vulgivaga.1.1.chr1.g01303.t01#4       2295   125258819:   mNeoVis.evergladensis.1.1.chr7.g01879.t01#3    mNeoVis.2.1.1.chr7_hap1.g01918.t01#5    mNeoVis.vulgivaga.2.1.chr7.g01897.t01#1    mNeoVis.3.1.1.chr7.g01789.t01#6    mNeoVis.vulgivaga.1.1.chr7.g01890.t01#4   mNeoVis.evergladensis.2.1.chr7.g01882.t01#2       2296   125258820:   mNeoVis.evergladensis.2.1.chr7.g01883.t01#2    mNeoVis.3.1.1.chr1.g00135.t01#6    mNeoVis.vulgivaga.1.1.chr11.g00672.t01#4    mNeoVis.evergladensis.2.1.chr1.g00129.t01#2    mNeoVis.evergladensis.2.1.chr11.g00675.t01#2    mNeoVis.vulgivaga.1.1.chr7.g01891.t01#4    mNeoVis.evergladensis.2.1.chr8.g00331.t01#2    mNeoVis.2.1.1.chr10_hap1.g00349.t01#5    mNeoVis.vulgivaga.2.1.chr3.g01350.t01#1      2297   125258821:  mNeoVis.vulgivaga.2.1.chr1.g00130.t01#1               2298   125258822:   mNeoVis.3.1.1.chr7.g01790.t01#6    mNeoVis.evergladensis.1.1.chr7.g01881.t01#3    mNeoVis.2.1.1.chr7_hap1.g01920.t01#5    mNeoVis.vulgivaga.1.1.chr7.g01893.t01#4    mNeoVis.evergladensis.2.1.chr7.g01884.t01#2   mNeoVis.vulgivaga.2.1.chr7.g01899.t01#1       2299   125258823:   mNeoVis.2.1.1.chr1_hap1.g01337.t01#5    mNeoVis.3.1.1.chr1.g01286.t01#6    mNeoVis.evergladensis.2.1.chr1.g01304.t01#2    mNeoVis.vulgivaga.2.1.chr1.g01305.t01#1    mNeoVis.vulgivaga.1.1.chr1.g01304.t01#4   mNeoVis.evergladensis.1.1.chr1.g01306.t01#3       2300   125258824:   mNeoVis.3.1.1.chr7.g01791.t01#6    mNeoVis.vulgivaga.2.1.chr7.g01900.t01#1    mNeoVis.evergladensis.1.1.chr7.g01882.t01#3    mNeoVis.vulgivaga.1.1.chr7.g01894.t01#4    mNeoVis.evergladensis.2.1.chr7.g01885.t01#2   mNeoVis.2.1.1.chr7_hap1.g01921.t01#5       2301   125258825:   mNeoVis.3.1.1.chr7.g01793.t01#6    mNeoVis.vulgivaga.2.1.chr7.g01901.t01#1    mNeoVis.evergladensis.2.1.chr7.g01886.t01#2    mNeoVis.evergladensis.1.1.chr7.g01883.t01#3    mNeoVis.2.1.1.chr7_hap1.g01922.t01#5   mNeoVis.vulgivaga.1.1.chr7.g01895.t01#4       2302   125258826:   mNeoVis.evergladensis.1.1.chr1.g01193.t01#3    mNeoVis.vulgivaga.1.1.chr1.g01189.t01#4    mNeoVis.vulgivaga.2.1.chr1.g01195.t01#1   mNeoVis.2.1.1.chr1_hap1.g01220.t01#5         2303   125258827:  mNeoVis.vulgivaga.2.1.chr1.g01196.t01#1               2304   125258828:   mNeoVis.evergladensis.2.1.chr7.g01887.t01#2    mNeoVis.evergladensis.1.1.chr7.g01884.t01#3    mNeoVis.3.1.1.chr7.g01794.t01#6    mNeoVis.2.1.1.chr7_hap1.g01923.t01#5    mNeoVis.vulgivaga.2.1.chr7.g01902.t01#1   mNeoVis.vulgivaga.1.1.chr7.g01892.t01#4       2305   125258829:   mNeoVis.2.1.1.chr7_hap1.g01924.t01#5    mNeoVis.3.1.1.chr7.g01796.t01#6    mNeoVis.3.1.1.chr7.g01795.t01#6    mNeoVis.vulgivaga.1.1.chr7.g01896.t01#4    mNeoVis.evergladensis.2.1.chr7.g01888.t01#2    mNeoVis.vulgivaga.2.1.chr7.g01903.t01#1   mNeoVis.evergladensis.1.1.chr7.g01885.t01#3     2306   125258830:   mNeoVis.vulgivaga.1.1.chr1.g01191.t01#4    mNeoVis.3.1.1.chr1.g01180.t01#6    mNeoVis.evergladensis.1.1.chr1.g01195.t01#3    mNeoVis.evergladensis.2.1.chr1.g01195.t01#2    mNeoVis.vulgivaga.2.1.chr1.g01197.t01#1   mNeoVis.2.1.1.chr1_hap1.g01222.t01#5       2307   125258831:   mNeoVis.evergladensis.1.1.chr1.g01196.t01#3    mNeoVis.evergladensis.2.1.chr1.g01196.t01#2    mNeoVis.2.1.1.chr1_hap1.g01223.t01#5    mNeoVis.vulgivaga.2.1.chr1.g01198.t01#1    mNeoVis.vulgivaga.1.1.chr1.g01192.t01#4   mNeoVis.3.1.1.chr1.g01181.t01#6       2308   125258832:   mNeoVis.evergladensis.2.1.chr1.g01197.t01#2    mNeoVis.evergladensis.1.1.chr1.g01197.t01#3    mNeoVis.2.1.1.chr1_hap1.g01224.t01#5    mNeoVis.vulgivaga.2.1.chr1.g01199.t01#1   mNeoVis.vulgivaga.1.1.chr1.g01193.t01#4       2309   125258833:   mNeoVis.vulgivaga.2.1.chr1.g01200.t01#1    mNeoVis.3.1.1.chr1.g01183.t01#6    mNeoVis.evergladensis.1.1.chr1.g01198.t01#3    mNeoVis.2.1.1.chr1_hap1.g01225.t01#5    mNeoVis.vulgivaga.1.1.chr1.g01194.t01#4   mNeoVis.evergladensis.2.1.chr1.g01198.t01#2       2310   125258834:   mNeoVis.vulgivaga.2.1.chr7.g01904.t01#1    mNeoVis.vulgivaga.1.1.chr7.g01897.t01#4    mNeoVis.2.1.1.chr7_hap1.g01925.t01#5    mNeoVis.evergladensis.1.1.chr7.g01886.t01#3   mNeoVis.evergladensis.2.1.chr7.g01889.t01#2       2311   125258835:   mNeoVis.evergladensis.2.1.chr7.g01890.t01#2    mNeoVis.vulgivaga.2.1.chr7.g01905.t01#1    mNeoVis.3.1.1.chr7.g01798.t01#6    mNeoVis.evergladensis.1.1.chr7.g01888.t01#3    mNeoVis.vulgivaga.1.1.chr7.g01898.t01#4   mNeoVis.2.1.1.chr7_hap1.g01926.t01#5       2312   125258836:   mNeoVis.vulgivaga.1.1.chr1.g01195.t01#4    mNeoVis.evergladensis.2.1.chr1.g01199.t01#2    mNeoVis.3.1.1.chr1.g01184.t01#6    mNeoVis.evergladensis.1.1.chr1.g01199.t01#3    mNeoVis.vulgivaga.2.1.chr1.g01201.t01#1   mNeoVis.2.1.1.chr1_hap1.g01226.t01#5       2313   125258837:   mNeoVis.evergladensis.1.1.chr7.g00559.t01#3    mNeoVis.vulgivaga.1.1.chr7.g00558.t01#4    mNeoVis.evergladensis.2.1.chr7.g00551.t01#2    mNeoVis.vulgivaga.2.1.chr7.g00553.t01#1    mNeoVis.2.1.1.chr7_hap1.g00550.t01#5   mNeoVis.3.1.1.chr7.g00533.t01#6       2314   125258838:   mNeoVis.evergladensis.2.1.chr7.g01891.t01#2    mNeoVis.2.1.1.chr7_hap1.g01927.t01#5    mNeoVis.evergladensis.1.1.chr7.g01889.t01#3    mNeoVis.vulgivaga.1.1.chr7.g01899.t01#4    mNeoVis.vulgivaga.2.1.chr7.g01906.t01#1   mNeoVis.3.1.1.chr7.g01799.t01#6       2315   125258839:   mNeoVis.vulgivaga.2.1.chr14.g00421.t01#1    mNeoVis.vulgivaga.1.1.chr7.g00915.t01#4    mNeoVis.evergladensis.2.1.chr14.g00419.t01#2    mNeoVis.evergladensis.1.1.chr14.g00423.t01#3    mNeoVis.vulgivaga.1.1.chr14.g00414.t01#4    mNeoVis.2.1.1.chr14_hap1.g00421.t01#5    mNeoVis.evergladensis.2.1.chr7.g00911.t01#2    mNeoVis.2.1.1.chr7_hap1.g00918.t01#5    mNeoVis.vulgivaga.2.1.chr7.g00916.t01#1      2316   125258840:   mNeoVis.3.1.1.chr7.g00534.t01#6    mNeoVis.2.1.1.chr7_hap1.g00551.t01#5    mNeoVis.evergladensis.2.1.chr7.g00552.t01#2    mNeoVis.vulgivaga.2.1.chr7.g00554.t01#1    mNeoVis.evergladensis.1.1.chr7.g00560.t01#3   mNeoVis.vulgivaga.1.1.chr7.g00559.t01#4       2317   125258841:   mNeoVis.2.1.1.chr1_hap1.g01227.t01#5    mNeoVis.evergladensis.2.1.chr1.g01200.t01#2    mNeoVis.vulgivaga.1.1.chr1.g01196.t01#4    mNeoVis.3.1.1.chr1.g01185.t01#6    mNeoVis.vulgivaga.2.1.chr1.g01202.t01#1   mNeoVis.evergladensis.1.1.chr1.g01200.t01#3       2318   125258842:   mNeoVis.vulgivaga.2.1.chr1.g01203.t01#1    mNeoVis.evergladensis.1.1.chr1.g01201.t01#3    mNeoVis.2.1.1.chr1_hap1.g01228.t01#5    mNeoVis.vulgivaga.1.1.chr1.g01197.t01#4   mNeoVis.evergladensis.2.1.chr1.g01201.t01#2       2319   125258843:   mNeoVis.evergladensis.1.1.chr7.g01890.t01#3    mNeoVis.2.1.1.chr2_hap1.g00547.t01#5    mNeoVis.evergladensis.2.1.chr5.g00549.t01#2    mNeoVis.evergladensis.1.1.chr5.g00542.t01#3    mNeoVis.3.1.1.chr7.g01800.t01#6   mNeoVis.evergladensis.2.1.chr7.g01892.t01#2       2320   125258844:   mNeoVis.vulgivaga.2.1.chr7.g01907.t01#1    mNeoVis.vulgivaga.1.1.chr7.g01900.t01#4   mNeoVis.2.1.1.chr7_hap1.g01928.t01#5           2321   125258845:   mNeoVis.2.1.1.chr7_hap1.g01929.t01#5    mNeoVis.evergladensis.2.1.chr7.g01893.t01#2    mNeoVis.3.1.1.chr7.g01801.t01#6    mNeoVis.vulgivaga.2.1.chr7.g01908.t01#1    mNeoVis.vulgivaga.1.1.chr7.g01901.t01#4    mNeoVis.evergladensis.1.1.chr7.g01891.t01#3   mNeoVis.vulgivaga.2.1.chr7.g01909.t01#1     2322   125258846:   mNeoVis.vulgivaga.2.1.chr15.g00516.t01#1    mNeoVis.vulgivaga.1.1.chr15.g00510.t01#4    mNeoVis.evergladensis.2.1.chr15.g00507.t01#2    mNeoVis.3.1.1.chr15.g00466.t01#6    mNeoVis.evergladensis.1.1.chr15.g00106.t01#3   mNeoVis.2.1.1.chr15_hap1.g00685.t01#5       2323   125258847:   mNeoVis.evergladensis.2.1.chr1.g01202.t01#2    mNeoVis.vulgivaga.1.1.chr1.g01198.t01#4    mNeoVis.evergladensis.1.1.chr1.g01202.t01#3    mNeoVis.3.1.1.chr1.g01186.t01#6    mNeoVis.vulgivaga.2.1.chr1.g01204.t01#1   mNeoVis.2.1.1.chr1_hap1.g01229.t01#5       2324   125258848:   mNeoVis.vulgivaga.2.1.chr7.g00555.t01#1    mNeoVis.3.1.1.chr7.g00535.t01#6    mNeoVis.evergladensis.1.1.chr7.g00561.t01#3    mNeoVis.vulgivaga.1.1.chr7.g00560.t01#4    mNeoVis.evergladensis.2.1.chr7.g00553.t01#2   mNeoVis.2.1.1.chr7_hap1.g00552.t01#5       2325   125258849:   mNeoVis.vulgivaga.1.1.chr1.g01199.t01#4    mNeoVis.3.1.1.chr1.g01187.t01#6    mNeoVis.evergladensis.2.1.chr1.g01203.t01#2    mNeoVis.evergladensis.1.1.chr1.g01203.t01#3    mNeoVis.2.1.1.chr1_hap1.g01230.t01#5   mNeoVis.vulgivaga.2.1.chr1.g01205.t01#1       2326   125258850:   mNeoVis.3.1.1.chr1.g01188.t01#6    mNeoVis.vulgivaga.1.1.chr1.g01200.t01#4    mNeoVis.evergladensis.1.1.chr1.g01204.t01#3    mNeoVis.vulgivaga.2.1.chr1.g01206.t01#1    mNeoVis.2.1.1.chr1_hap1.g01231.t01#5   mNeoVis.evergladensis.2.1.chr1.g01204.t01#2       2327   125258851:   mNeoVis.evergladensis.1.1.chr7.g01892.t01#3    mNeoVis.vulgivaga.2.1.chr7.g01910.t01#1    mNeoVis.2.1.1.chr7_hap1.g01930.t01#5    mNeoVis.3.1.1.chr7.g01802.t01#6    mNeoVis.evergladensis.2.1.chr7.g01894.t01#2   mNeoVis.vulgivaga.1.1.chr7.g01902.t01#4       2328   125258852:   mNeoVis.2.1.1.chr1_hap1.g01232.t01#5    mNeoVis.evergladensis.1.1.chr1.g01205.t01#3    mNeoVis.evergladensis.2.1.chr1.g01205.t01#2    mNeoVis.vulgivaga.1.1.chr1.g01201.t01#4    mNeoVis.vulgivaga.2.1.chr1.g01207.t01#1   mNeoVis.3.1.1.chr1.g01189.t01#6       2329   125258853:   mNeoVis.3.1.1.chr7.g01714.t01#6    mNeoVis.evergladensis.1.1.chr7.g01805.t01#3    mNeoVis.evergladensis.2.1.chr7.g01808.t01#2    mNeoVis.vulgivaga.2.1.chr7.g01826.t01#1    mNeoVis.vulgivaga.1.1.chr7.g01815.t01#4   mNeoVis.2.1.1.chr7_hap1.g01845.t01#5       2330   125258854:   mNeoVis.3.1.1.chr1.g01190.t01#6    mNeoVis.evergladensis.1.1.chr1.g01206.t01#3    mNeoVis.evergladensis.2.1.chr1.g01206.t01#2    mNeoVis.vulgivaga.2.1.chr1.g01208.t01#1    mNeoVis.2.1.1.chr1_hap1.g01233.t01#5   mNeoVis.vulgivaga.1.1.chr1.g01202.t01#4       2331   125258855:   mNeoVis.vulgivaga.1.1.chr1.g01205.t01#4    mNeoVis.vulgivaga.1.1.chr1.g01206.t01#4    mNeoVis.2.1.1.chr1_hap1.g01234.t01#5    mNeoVis.evergladensis.2.1.chr1.g01207.t01#2    mNeoVis.3.1.1.chr1.g01191.t01#6    mNeoVis.evergladensis.1.1.chr1.g01207.t01#3   mNeoVis.vulgivaga.2.1.chr1.g01209.t01#1     2332   125258856:   mNeoVis.vulgivaga.1.1.chr7.g01816.t01#4    mNeoVis.2.1.1.chr7_hap1.g01846.t01#5    mNeoVis.evergladensis.1.1.chr7.g01806.t01#3    mNeoVis.3.1.1.chr7.g01715.t01#6    mNeoVis.vulgivaga.2.1.chr7.g01827.t01#1   mNeoVis.evergladensis.2.1.chr7.g01809.t01#2       2333   125258857:   mNeoVis.2.1.1.chr1_hap1.g01235.t01#5    mNeoVis.vulgivaga.2.1.chr1.g01210.t01#1    mNeoVis.evergladensis.2.1.chr1.g01208.t01#2    mNeoVis.vulgivaga.1.1.chr1.g01207.t01#4    mNeoVis.3.1.1.chr1.g01192.t01#6   mNeoVis.evergladensis.1.1.chr1.g01208.t01#3       2334   125258858:   mNeoVis.vulgivaga.1.1.chr7.g01795.t01#4    mNeoVis.evergladensis.1.1.chr7.g01807.t01#3    mNeoVis.evergladensis.2.1.chr7.g01810.t01#2    mNeoVis.vulgivaga.1.1.chr7.g01817.t01#4    mNeoVis.evergladensis.1.1.chr7.g01784.t01#3    mNeoVis.3.1.1.chr7.g01716.t01#6    mNeoVis.2.1.1.chr7_hap1.g01825.t01#5    mNeoVis.2.1.1.chr7_hap1.g01847.t01#5    mNeoVis.vulgivaga.2.1.chr7.g01805.t01#1      2335   125258859:  mNeoVis.vulgivaga.2.1.chr1.g01211.t01#1               2336   125258860:   mNeoVis.vulgivaga.1.1.chr1.g01208.t01#4    mNeoVis.2.1.1.chr1_hap1.g01236.t01#5    mNeoVis.vulgivaga.2.1.chr1.g01212.t01#1    mNeoVis.evergladensis.1.1.chr1.g01209.t01#3   mNeoVis.evergladensis.2.1.chr1.g01209.t01#2       2337   125258861:   mNeoVis.evergladensis.2.1.chr7.g01811.t01#2    mNeoVis.evergladensis.1.1.chr7.g01808.t01#3    mNeoVis.vulgivaga.1.1.chr7.g01818.t01#4    mNeoVis.2.1.1.chr7_hap1.g01848.t01#5    mNeoVis.3.1.1.chr7.g01717.t01#6   mNeoVis.vulgivaga.2.1.chr7.g01829.t01#1       2338   125258862:   mNeoVis.vulgivaga.2.1.chr1.g01213.t01#1    mNeoVis.evergladensis.2.1.chr1.g01210.t01#2    mNeoVis.vulgivaga.1.1.chr1.g01209.t01#4    mNeoVis.evergladensis.1.1.chr1.g01210.t01#3    mNeoVis.3.1.1.chr1.g01195.t01#6   mNeoVis.2.1.1.chr1_hap1.g01237.t01#5       2339   125258863:   mNeoVis.vulgivaga.1.1.chr1.g01210.t01#4    mNeoVis.3.1.1.chr1.g01196.t01#6    mNeoVis.2.1.1.chr1_hap1.g01238.t01#5    mNeoVis.evergladensis.2.1.chr1.g01211.t01#2    mNeoVis.evergladensis.1.1.chr1.g01211.t01#3   mNeoVis.vulgivaga.2.1.chr1.g01214.t01#1       2340   125258864:   mNeoVis.3.1.1.chr1.g01197.t01#6    mNeoVis.2.1.1.chr1_hap1.g01239.t01#5    mNeoVis.vulgivaga.2.1.chr1.g01215.t01#1    mNeoVis.evergladensis.2.1.chr1.g01212.t01#2    mNeoVis.vulgivaga.1.1.chr1.g01211.t01#4   mNeoVis.evergladensis.1.1.chr1.g01212.t01#3       2341   125258865:   mNeoVis.2.1.1.chr15_hap1.g00686.t01#5    mNeoVis.3.1.1.chr15.g00467.t01#6    mNeoVis.evergladensis.1.1.chr15.g00105.t01#3    mNeoVis.vulgivaga.1.1.chr15.g00511.t01#4    mNeoVis.evergladensis.2.1.chr15.g00508.t01#2   mNeoVis.vulgivaga.2.1.chr15.g00517.t01#1       2342   125258866:   mNeoVis.evergladensis.1.1.chr1.g01213.t01#3    mNeoVis.vulgivaga.1.1.chr1.g01212.t01#4    mNeoVis.vulgivaga.2.1.chr1.g01216.t01#1    mNeoVis.3.1.1.chr1.g01198.t01#6    mNeoVis.evergladensis.2.1.chr1.g01213.t01#2   mNeoVis.2.1.1.chr1_hap1.g01240.t01#5       2343   125258867:   mNeoVis.vulgivaga.1.1.chr7.g01820.t01#4    mNeoVis.evergladensis.1.1.chr7.g01810.t01#3    mNeoVis.2.1.1.chr7_hap1.g01850.t01#5    mNeoVis.vulgivaga.2.1.chr7.g01830.t01#1    mNeoVis.3.1.1.chr7.g01719.t01#6   mNeoVis.evergladensis.2.1.chr7.g01813.t01#2       2344   125258868:   mNeoVis.3.1.1.chr1.g01199.t01#6    mNeoVis.2.1.1.chr1_hap1.g01241.t01#5    mNeoVis.evergladensis.2.1.chr1.g01214.t01#2    mNeoVis.vulgivaga.1.1.chr1.g01213.t01#4    mNeoVis.vulgivaga.2.1.chr1.g01217.t01#1   mNeoVis.evergladensis.1.1.chr1.g01215.t01#3       2345   125258869:   mNeoVis.evergladensis.1.1.chr1.g01216.t01#3    mNeoVis.vulgivaga.1.1.chr1.g01214.t01#4    mNeoVis.3.1.1.chr1.g01200.t01#6    mNeoVis.vulgivaga.2.1.chr1.g01218.t01#1    mNeoVis.2.1.1.chr1_hap1.g01242.t01#5   mNeoVis.evergladensis.2.1.chr1.g01215.t01#2       2346   125258870:   mNeoVis.vulgivaga.1.1.chr7.g01821.t01#4    mNeoVis.evergladensis.1.1.chr7.g01811.t01#3    mNeoVis.vulgivaga.2.1.chr7.g01831.t01#1    mNeoVis.evergladensis.2.1.chr7.g01814.t01#2    mNeoVis.2.1.1.chr7_hap1.g01851.t01#5   mNeoVis.3.1.1.chr7.g01720.t01#6       2347   125258871:   mNeoVis.evergladensis.1.1.chr7.g00562.t01#3    mNeoVis.vulgivaga.1.1.chr7.g00561.t01#4    mNeoVis.evergladensis.2.1.chr7.g00554.t01#2    mNeoVis.2.1.1.chr7_hap1.g00553.t01#5    mNeoVis.vulgivaga.2.1.chr7.g00556.t01#1   mNeoVis.3.1.1.chr7.g00536.t01#6       2348   125258872:   mNeoVis.vulgivaga.1.1.chr1.g00286.t01#4    mNeoVis.evergladensis.2.1.chr1.g00286.t01#2    mNeoVis.evergladensis.1.1.chr1.g00295.t01#3    mNeoVis.3.1.1.chr1.g00292.t01#6    mNeoVis.vulgivaga.2.1.chr1.g00290.t01#1   mNeoVis.2.1.1.chr1_hap1.g00287.t01#5       2349   125258873:   mNeoVis.2.1.1.chr7_hap1.g01852.t01#5    mNeoVis.evergladensis.1.1.chr7.g01812.t01#3    mNeoVis.vulgivaga.2.1.chr7.g01832.t01#1    mNeoVis.vulgivaga.1.1.chr7.g01822.t01#4   mNeoVis.evergladensis.2.1.chr7.g01815.t01#2       2350   125258874:   mNeoVis.evergladensis.1.1.chr7.g01813.t01#3    mNeoVis.3.1.1.chr7.g01721.t01#6    mNeoVis.vulgivaga.2.1.chr7.g01833.t01#1    mNeoVis.vulgivaga.1.1.chr7.g01823.t01#4    mNeoVis.evergladensis.2.1.chr7.g01816.t01#2   mNeoVis.2.1.1.chr7_hap1.g01853.t01#5       2351   125258875:   mNeoVis.vulgivaga.1.1.chr1.g01215.t01#4    mNeoVis.vulgivaga.2.1.chr1.g01219.t01#1    mNeoVis.3.1.1.chr1.g01201.t01#6    mNeoVis.evergladensis.1.1.chr1.g01217.t01#3    mNeoVis.2.1.1.chr1_hap1.g01243.t01#5   mNeoVis.evergladensis.2.1.chr1.g01216.t01#2       2352   125258876:   mNeoVis.evergladensis.1.1.chr1.g01218.t01#3    mNeoVis.2.1.1.chr1_hap1.g01244.t01#5    mNeoVis.evergladensis.2.1.chr1.g01217.t01#2    mNeoVis.vulgivaga.1.1.chr1.g01216.t01#4    mNeoVis.3.1.1.chr1.g01202.t01#6    mNeoVis.evergladensis.2.1.chr1.g01218.t01#2   mNeoVis.vulgivaga.2.1.chr1.g01220.t01#1     2353   125258877:   mNeoVis.vulgivaga.2.1.chr7.g01834.t01#1    mNeoVis.vulgivaga.1.1.chr7.g01824.t01#4    mNeoVis.3.1.1.chr7.g01722.t01#6   mNeoVis.2.1.1.chr7_hap1.g01854.t01#5         2354   125258878:   mNeoVis.3.1.1.chr1.g01203.t01#6    mNeoVis.evergladensis.1.1.chr1.g01219.t01#3    mNeoVis.vulgivaga.2.1.chr1.g01221.t01#1    mNeoVis.evergladensis.2.1.chr1.g01219.t01#2    mNeoVis.2.1.1.chr1_hap1.g01245.t01#5   mNeoVis.vulgivaga.1.1.chr1.g01217.t01#4       2355   125258879:   mNeoVis.evergladensis.2.1.chr1.g01220.t01#2    mNeoVis.2.1.1.chr1_hap1.g01246.t01#5    mNeoVis.vulgivaga.2.1.chr1.g01222.t01#1    mNeoVis.3.1.1.chr1.g01204.t01#6    mNeoVis.evergladensis.1.1.chr1.g01220.t01#3   mNeoVis.vulgivaga.1.1.chr1.g01218.t01#4       2356   125258880:   mNeoVis.vulgivaga.1.1.chr7.g01825.t01#4    mNeoVis.evergladensis.2.1.chr7.g01818.t01#2    mNeoVis.evergladensis.1.1.chr7.g01815.t01#3    mNeoVis.vulgivaga.2.1.chr7.g01835.t01#1   mNeoVis.2.1.1.chr7_hap1.g01855.t01#5       2357   125258881:   mNeoVis.vulgivaga.1.1.chr1.g00157.t01#4    mNeoVis.vulgivaga.1.1.chr8.g01311.t01#4    mNeoVis.evergladensis.2.1.chr9.g01321.t01#2    mNeoVis.evergladensis.1.1.chr1.g01221.t01#3    mNeoVis.vulgivaga.2.1.chr8.g01327.t01#1    mNeoVis.vulgivaga.2.1.chr1.g01223.t01#1    mNeoVis.2.1.1.chr1_hap1.g01247.t01#5    mNeoVis.3.1.1.chr1.g00165.t01#6    mNeoVis.2.1.1.chr1_hap1.g00161.t01#5      2358   125258882:   mNeoVis.3.1.1.chr1.g01206.t01#6    mNeoVis.evergladensis.1.1.chr1.g01222.t01#3    mNeoVis.vulgivaga.2.1.chr1.g01224.t01#1    mNeoVis.evergladensis.2.1.chr1.g01222.t01#2    mNeoVis.vulgivaga.1.1.chr1.g01220.t01#4   mNeoVis.2.1.1.chr1_hap1.g01248.t01#5       2359   125258883:   mNeoVis.vulgivaga.1.1.chr1.g01221.t01#4    mNeoVis.evergladensis.1.1.chr1.g01223.t01#3    mNeoVis.vulgivaga.2.1.chr1.g01225.t01#1    mNeoVis.2.1.1.chr1_hap1.g01249.t01#5    mNeoVis.evergladensis.2.1.chr1.g01223.t01#2   mNeoVis.3.1.1.chr1.g01207.t01#6       2360   125258884:   mNeoVis.evergladensis.1.1.chr1.g01224.t01#3    mNeoVis.vulgivaga.1.1.chr1.g01222.t01#4    mNeoVis.2.1.1.chr1_hap1.g01250.t01#5    mNeoVis.vulgivaga.2.1.chr1.g01226.t01#1    mNeoVis.3.1.1.chr1.g01208.t01#6   mNeoVis.evergladensis.2.1.chr1.g01224.t01#2       2361   125258885:   mNeoVis.2.1.1.chr1_hap1.g01251.t01#5    mNeoVis.evergladensis.1.1.chr1.g01226.t01#3    mNeoVis.3.1.1.chr1.g01209.t01#6    mNeoVis.vulgivaga.2.1.chr1.g01227.t01#1    mNeoVis.evergladensis.2.1.chr1.g01225.t01#2    mNeoVis.evergladensis.1.1.chr1.g01225.t01#3   mNeoVis.vulgivaga.1.1.chr1.g01223.t01#4     2362   125258886:   mNeoVis.2.1.1.chr1_hap1.g01252.t01#5    mNeoVis.evergladensis.2.1.chr1.g01226.t01#2    mNeoVis.evergladensis.1.1.chr1.g01227.t01#3    mNeoVis.vulgivaga.1.1.chr1.g01224.t01#4   mNeoVis.vulgivaga.2.1.chr1.g01228.t01#1       2363   125258887:   mNeoVis.vulgivaga.2.1.chr1.g01229.t01#1    mNeoVis.evergladensis.1.1.chr1.g01228.t01#3    mNeoVis.vulgivaga.1.1.chr1.g01225.t01#4    mNeoVis.2.1.1.chr1_hap1.g01253.t01#5   mNeoVis.evergladensis.2.1.chr1.g01227.t01#2       2364   125258888:   mNeoVis.3.1.1.chr1.g01211.t01#6    mNeoVis.2.1.1.chr1_hap1.g01255.t01#5    mNeoVis.evergladensis.2.1.chr1.g01228.t01#2    mNeoVis.vulgivaga.1.1.chr1.g01226.t01#4    mNeoVis.vulgivaga.2.1.chr1.g01230.t01#1   mNeoVis.evergladensis.1.1.chr1.g01229.t01#3       2365   125258889:   mNeoVis.3.1.1.chr7.g00537.t01#6    mNeoVis.2.1.1.chr7_hap1.g00554.t01#5    mNeoVis.evergladensis.2.1.chr7.g00555.t01#2    mNeoVis.vulgivaga.2.1.chr7.g00557.t01#1    mNeoVis.evergladensis.1.1.chr7.g00563.t01#3   mNeoVis.vulgivaga.1.1.chr7.g00562.t01#4       2366   125258890:   mNeoVis.3.1.1.chr1.g01212.t01#6    mNeoVis.vulgivaga.2.1.chr1.g01231.t01#1    mNeoVis.2.1.1.chr1_hap1.g01256.t01#5    mNeoVis.evergladensis.2.1.chr1.g01229.t01#2    mNeoVis.evergladensis.1.1.chr1.g01230.t01#3   mNeoVis.vulgivaga.1.1.chr1.g01227.t01#4       2367   125258891:   mNeoVis.vulgivaga.2.1.chr7.g01836.t01#1    mNeoVis.evergladensis.1.1.chr7.g01816.t01#3    mNeoVis.evergladensis.2.1.chr7.g01819.t01#2    mNeoVis.vulgivaga.1.1.chr7.g01826.t01#4    mNeoVis.3.1.1.chr7.g01724.t01#6   mNeoVis.2.1.1.chr7_hap1.g01856.t01#5       2368   125258892:   mNeoVis.3.1.1.chr7.g01725.t01#6    mNeoVis.vulgivaga.1.1.chr7.g01827.t01#4    mNeoVis.2.1.1.chr7_hap1.g01857.t01#5    mNeoVis.vulgivaga.2.1.chr7.g01837.t01#1    mNeoVis.evergladensis.1.1.chr7.g01817.t01#3   mNeoVis.evergladensis.2.1.chr7.g01820.t01#2       2369   125258893:   mNeoVis.vulgivaga.2.1.chr1.g01232.t01#1    mNeoVis.evergladensis.2.1.chr1.g01230.t01#2    mNeoVis.3.1.1.chr1.g01213.t01#6    mNeoVis.evergladensis.1.1.chr1.g01231.t01#3    mNeoVis.vulgivaga.1.1.chr1.g01228.t01#4   mNeoVis.2.1.1.chr1_hap1.g01257.t01#5       2370   125258894:   mNeoVis.evergladensis.2.1.chr7.g01821.t01#2    mNeoVis.vulgivaga.1.1.chr7.g01828.t01#4    mNeoVis.3.1.1.chr7.g01727.t01#6    mNeoVis.2.1.1.chr7_hap1.g01858.t01#5    mNeoVis.vulgivaga.2.1.chr7.g01838.t01#1   mNeoVis.evergladensis.1.1.chr7.g01818.t01#3       2371   125258895:   mNeoVis.evergladensis.1.1.chr1.g01233.t01#3    mNeoVis.vulgivaga.1.1.chr1.g01230.t01#4    mNeoVis.evergladensis.2.1.chr1.g01232.t01#2    mNeoVis.3.1.1.chr1.g01215.t01#6    mNeoVis.vulgivaga.2.1.chr1.g01233.t01#1   mNeoVis.2.1.1.chr1_hap1.g01259.t01#5       2372   125258896:   mNeoVis.3.1.1.chr7.g01728.t01#6    mNeoVis.2.1.1.chr7_hap1.g01859.t01#5    mNeoVis.evergladensis.1.1.chr7.g01819.t01#3    mNeoVis.evergladensis.2.1.chr7.g01822.t01#2    mNeoVis.vulgivaga.1.1.chr7.g01829.t01#4   mNeoVis.vulgivaga.2.1.chr7.g01839.t01#1       2373   125258897:   mNeoVis.vulgivaga.2.1.chr3.g01312.t01#1    mNeoVis.evergladensis.1.1.chr3.g01299.t01#3    mNeoVis.3.1.1.chr3.g01270.t01#6    mNeoVis.2.1.1.chr3_hap1.g01293.t01#5    mNeoVis.vulgivaga.1.1.chr15.g00144.t01#4    mNeoVis.vulgivaga.1.1.chr3.g01286.t01#4    mNeoVis.vulgivaga.2.1.chr15.g00148.t01#1    mNeoVis.3.1.1.chr15.g00138.t01#6    mNeoVis.evergladensis.2.1.chr3.g01292.t01#2      2374   125258898:   mNeoVis.2.1.1.chr1_hap1.g01260.t01#5    mNeoVis.vulgivaga.2.1.chr1.g01234.t01#1    mNeoVis.evergladensis.2.1.chr1.g01233.t01#2    mNeoVis.vulgivaga.1.1.chr1.g01231.t01#4    mNeoVis.3.1.1.chr1.g01216.t01#6   mNeoVis.evergladensis.1.1.chr1.g01234.t01#3       2375   125258899:   mNeoVis.evergladensis.2.1.chr7.g01823.t01#2    mNeoVis.vulgivaga.1.1.chr7.g01830.t01#4    mNeoVis.3.1.1.chr7.g01729.t01#6    mNeoVis.2.1.1.chr7_hap1.g01860.t01#5    mNeoVis.evergladensis.1.1.chr7.g01820.t01#3   mNeoVis.vulgivaga.2.1.chr7.g01840.t01#1       2376   125258900:   mNeoVis.evergladensis.2.1.chr1.g01234.t01#2    mNeoVis.2.1.1.chr1_hap1.g01261.t01#5    mNeoVis.3.1.1.chr1.g01217.t01#6   mNeoVis.vulgivaga.2.1.chr1.g01235.t01#1         2377   125258901:   mNeoVis.vulgivaga.1.1.chr7.g01832.t01#4    mNeoVis.vulgivaga.2.1.chr7.g01841.t01#1    mNeoVis.2.1.1.chr7_hap1.g01862.t01#5    mNeoVis.evergladensis.1.1.chr7.g01822.t01#3   mNeoVis.evergladensis.2.1.chr7.g01825.t01#2       2378   125258902:   mNeoVis.2.1.1.chr7_hap1.g01861.t01#5    mNeoVis.3.1.1.chr7.g01730.t01#6   mNeoVis.vulgivaga.1.1.chr7.g01831.t01#4           2379   125258903:   mNeoVis.vulgivaga.1.1.chr12.g00031.t01#4    mNeoVis.evergladensis.1.1.chr12.g00026.t01#3    mNeoVis.vulgivaga.2.1.chr12.g00027.t01#1    mNeoVis.evergladensis.2.1.chr12.g00025.t01#2    mNeoVis.2.1.1.chr12_hap1.g00035.t01#5   mNeoVis.3.1.1.chr12.g00026.t01#6       2380   125258904:   mNeoVis.vulgivaga.2.1.chr1.g01236.t01#1    mNeoVis.evergladensis.2.1.chr1.g01235.t01#2    mNeoVis.2.1.1.chr1_hap1.g01262.t01#5    mNeoVis.3.1.1.chr1.g01218.t01#6   mNeoVis.vulgivaga.1.1.chr1.g01232.t01#4       2381   125258905:   mNeoVis.vulgivaga.1.1.chr1.g01233.t01#4    mNeoVis.2.1.1.chr1_hap1.g01263.t01#5    mNeoVis.3.1.1.chr1.g01219.t01#6    mNeoVis.evergladensis.2.1.chr1.g01236.t01#2    mNeoVis.evergladensis.1.1.chr1.g01237.t01#3   mNeoVis.vulgivaga.2.1.chr1.g01237.t01#1       2382   125258906:   mNeoVis.3.1.1.chr7.g01731.t01#6    mNeoVis.vulgivaga.2.1.chr7.g01842.t01#1    mNeoVis.evergladensis.1.1.chr7.g01823.t01#3    mNeoVis.evergladensis.2.1.chr7.g01826.t01#2    mNeoVis.vulgivaga.1.1.chr7.g01833.t01#4   mNeoVis.2.1.1.chr7_hap1.g01863.t01#5       2383   125258907:   mNeoVis.evergladensis.1.1.chr7.g00076.t01#3    mNeoVis.vulgivaga.2.1.chr7.g00074.t01#1    mNeoVis.vulgivaga.1.1.chr7.g00075.t01#4    mNeoVis.evergladensis.2.1.chr7.g00074.t01#2   mNeoVis.2.1.1.chr7_hap1.g00073.t01#5       2384   125258908:   mNeoVis.evergladensis.2.1.chr15.g00146.t01#2    mNeoVis.evergladensis.2.1.chr4.g00871.t01#2    mNeoVis.3.1.1.chr15.g00139.t01#6    mNeoVis.vulgivaga.2.1.chr4.g00892.t01#1    mNeoVis.2.1.1.chr6_hap1.g00920.t01#5    mNeoVis.2.1.1.chr15_hap1.g00160.t01#5    mNeoVis.evergladensis.1.1.chr15.g00471.t01#3    mNeoVis.evergladensis.1.1.chr4.g00889.t01#3    mNeoVis.3.1.1.chr4.g00880.t01#6      2385   125258909:   mNeoVis.vulgivaga.2.1.chr1.g01238.t01#1    mNeoVis.vulgivaga.1.1.chr1.g01234.t01#4    mNeoVis.evergladensis.2.1.chr1.g01237.t01#2    mNeoVis.3.1.1.chr1.g01221.t01#6    mNeoVis.evergladensis.1.1.chr1.g01239.t01#3   mNeoVis.2.1.1.chr1_hap1.g01265.t01#5       2386   125258910:   mNeoVis.3.1.1.chr1.g01222.t01#6    mNeoVis.evergladensis.2.1.chr1.g01238.t01#2    mNeoVis.vulgivaga.2.1.chr1.g01239.t01#1    mNeoVis.2.1.1.chr1_hap1.g01266.t01#5    mNeoVis.evergladensis.1.1.chr1.g01240.t01#3   mNeoVis.vulgivaga.1.1.chr1.g01235.t01#4       2387   125258911:   mNeoVis.2.1.1.chr1_hap1.g01269.t01#5    mNeoVis.vulgivaga.1.1.chr1.g01238.t01#4    mNeoVis.3.1.1.chr1.g01224.t01#6    mNeoVis.vulgivaga.2.1.chr1.g01241.t01#1    mNeoVis.evergladensis.1.1.chr1.g01243.t01#3   mNeoVis.evergladensis.2.1.chr1.g01240.t01#2       2388   125258912:   mNeoVis.vulgivaga.2.1.chr7.g01843.t01#1    mNeoVis.2.1.1.chr7_hap1.g01864.t01#5    mNeoVis.evergladensis.1.1.chr7.g01824.t01#3    mNeoVis.3.1.1.chr7.g01732.t01#6    mNeoVis.evergladensis.2.1.chr7.g01827.t01#2   mNeoVis.vulgivaga.1.1.chr7.g01834.t01#4       2389   125258913:   mNeoVis.2.1.1.chr1_hap1.g01270.t01#5    mNeoVis.vulgivaga.1.1.chr1.g01239.t01#4    mNeoVis.vulgivaga.2.1.chr1.g01242.t01#1    mNeoVis.evergladensis.2.1.chr1.g01242.t01#2    mNeoVis.3.1.1.chr1.g01225.t01#6   mNeoVis.evergladensis.1.1.chr1.g01244.t01#3       2390   125258914:   mNeoVis.vulgivaga.1.1.chr1.g01240.t01#4    mNeoVis.evergladensis.2.1.chr1.g01243.t01#2    mNeoVis.2.1.1.chr1_hap1.g01271.t01#5    mNeoVis.3.1.1.chr1.g01226.t01#6   mNeoVis.vulgivaga.2.1.chr1.g01243.t01#1       2391   125258915:   mNeoVis.3.1.1.chr7.g01735.t01#6    mNeoVis.evergladensis.1.1.chr7.g01826.t01#3    mNeoVis.vulgivaga.1.1.chr7.g01837.t01#4    mNeoVis.evergladensis.2.1.chr7.g01829.t01#2    mNeoVis.vulgivaga.2.1.chr7.g01845.t01#1   mNeoVis.2.1.1.chr7_hap1.g01867.t01#5       2392   125258916:   mNeoVis.2.1.1.chr2_hap1.g00687.t01#5    mNeoVis.evergladensis.1.1.chr5.g00664.t01#3    mNeoVis.evergladensis.2.1.chr5.g00669.t01#2   mNeoVis.vulgivaga.2.1.chr5.g00661.t01#1         2393   125258917:   mNeoVis.vulgivaga.1.1.chr1.g01242.t01#4    mNeoVis.2.1.1.chr1_hap1.g01272.t01#5    mNeoVis.evergladensis.1.1.chr1.g01246.t01#3    mNeoVis.vulgivaga.2.1.chr4.g00250.t01#1    mNeoVis.3.1.1.chr1.g01228.t01#6    mNeoVis.evergladensis.2.1.chr1.g01244.t01#2   mNeoVis.vulgivaga.2.1.chr1.g01244.t01#1     2394   125258918:   mNeoVis.vulgivaga.1.1.chr1.g01243.t01#4    mNeoVis.evergladensis.1.1.chr1.g01247.t01#3    mNeoVis.2.1.1.chr1_hap1.g01273.t01#5    mNeoVis.3.1.1.chr1.g01229.t01#6    mNeoVis.evergladensis.2.1.chr1.g01245.t01#2   mNeoVis.vulgivaga.2.1.chr1.g01245.t01#1       2395   125258919:   mNeoVis.vulgivaga.1.1.chr7.g01838.t01#4    mNeoVis.2.1.1.chr7_hap1.g01868.t01#5    mNeoVis.evergladensis.2.1.chr7.g01830.t01#2    mNeoVis.vulgivaga.2.1.chr7.g01846.t01#1    mNeoVis.3.1.1.chr7.g01736.t01#6   mNeoVis.evergladensis.1.1.chr7.g01827.t01#3       2396   125258920:   mNeoVis.3.1.1.chr1.g01230.t01#6    mNeoVis.evergladensis.1.1.chr1.g01248.t01#3    mNeoVis.2.1.1.chr1_hap1.g01274.t01#5    mNeoVis.vulgivaga.2.1.chr1.g01246.t01#1    mNeoVis.vulgivaga.1.1.chr1.g01244.t01#4   mNeoVis.evergladensis.2.1.chr1.g01246.t01#2       2397   125258921:   mNeoVis.evergladensis.2.1.chr7.g01831.t01#2    mNeoVis.2.1.1.chr7_hap1.g01870.t01#5    mNeoVis.vulgivaga.1.1.chr7.g01839.t01#4    mNeoVis.evergladensis.1.1.chr7.g01828.t01#3    mNeoVis.vulgivaga.2.1.chr7.g01847.t01#1   mNeoVis.3.1.1.chr7.g01738.t01#6       2398   125258922:   mNeoVis.2.1.1.chr1_hap1.g01275.t01#5    mNeoVis.evergladensis.1.1.chr1.g01249.t01#3    mNeoVis.3.1.1.chr1.g01231.t01#6    mNeoVis.vulgivaga.2.1.chr1.g01247.t01#1    mNeoVis.vulgivaga.1.1.chr1.g01245.t01#4   mNeoVis.evergladensis.2.1.chr1.g01247.t01#2       2399   125258923:   mNeoVis.vulgivaga.2.1.chr7.g01848.t01#1    mNeoVis.evergladensis.2.1.chr7.g01832.t01#2    mNeoVis.vulgivaga.1.1.chr7.g01840.t01#4    mNeoVis.evergladensis.1.1.chr7.g01829.t01#3    mNeoVis.2.1.1.chr7_hap1.g01871.t01#5   mNeoVis.3.1.1.chr7.g01739.t01#6       2400   125258924:   mNeoVis.evergladensis.2.1.chr7.g01833.t01#2    mNeoVis.vulgivaga.1.1.chr7.g01841.t01#4    mNeoVis.evergladensis.1.1.chr7.g01830.t01#3    mNeoVis.2.1.1.chr7_hap1.g01872.t01#5   mNeoVis.vulgivaga.2.1.chr7.g01849.t01#1       2401   125258925:   mNeoVis.2.1.1.chr1_hap1.g01276.t01#5    mNeoVis.evergladensis.1.1.chr1.g01250.t01#3    mNeoVis.3.1.1.chr1.g01232.t01#6    mNeoVis.vulgivaga.1.1.chr1.g01246.t01#4    mNeoVis.vulgivaga.2.1.chr1.g01248.t01#1   mNeoVis.evergladensis.2.1.chr1.g01248.t01#2       2402   125258926:   mNeoVis.3.1.1.chr7.g00347.t01#6    mNeoVis.2.1.1.chr7_hap1.g00359.t01#5    mNeoVis.evergladensis.2.1.chr7.g00362.t01#2    mNeoVis.vulgivaga.1.1.chr7.g00369.t01#4    mNeoVis.evergladensis.1.1.chr7.g00370.t01#3   mNeoVis.vulgivaga.2.1.chr7.g00364.t01#1       2403   125258927:   mNeoVis.vulgivaga.1.1.chr1.g01247.t01#4    mNeoVis.3.1.1.chr1.g01233.t01#6    mNeoVis.vulgivaga.2.1.chr1.g01249.t01#1   mNeoVis.evergladensis.1.1.chr1.g01251.t01#3       2404   125258928:   mNeoVis.evergladensis.1.1.chr15.g00104.t01#3    mNeoVis.vulgivaga.1.1.chr15.g00512.t01#4    mNeoVis.evergladensis.2.1.chr15.g00509.t01#2    mNeoVis.2.1.1.chr15_hap1.g00687.t01#5    mNeoVis.3.1.1.chr15.g00468.t01#6   mNeoVis.vulgivaga.2.1.chr15.g00518.t01#1       2405   125258929:   mNeoVis.3.1.1.chr1.g01124.t01#6    mNeoVis.evergladensis.1.1.chr1.g01136.t01#3    mNeoVis.vulgivaga.1.1.chr1.g00949.t01#4    mNeoVis.vulgivaga.2.1.chr1.g01138.t01#1    mNeoVis.vulgivaga.1.1.chr1.g01137.t01#4    mNeoVis.vulgivaga.2.1.chr1.g00946.t01#1    mNeoVis.3.1.1.chr1.g00952.t01#6    mNeoVis.evergladensis.1.1.chr1.g00941.t01#3    mNeoVis.2.1.1.chr1_hap1.g00939.t01#5      2406   125258930:   mNeoVis.2.1.1.chr7_hap1.g01873.t01#5    mNeoVis.evergladensis.1.1.chr7.g01831.t01#3    mNeoVis.vulgivaga.1.1.chr7.g01842.t01#4    mNeoVis.evergladensis.2.1.chr7.g01834.t01#2    mNeoVis.vulgivaga.2.1.chr7.g01850.t01#1   mNeoVis.3.1.1.chr7.g01741.t01#6       2407   125258931:   mNeoVis.evergladensis.1.1.chr1.g01137.t01#3    mNeoVis.evergladensis.2.1.chr1.g01141.t01#2    mNeoVis.vulgivaga.1.1.chr1.g01138.t01#4    mNeoVis.vulgivaga.2.1.chr1.g01139.t01#1    mNeoVis.3.1.1.chr1.g01125.t01#6   mNeoVis.2.1.1.chr1_hap1.g01167.t01#5       2408   125258932:   mNeoVis.vulgivaga.1.1.chr15.g00513.t01#4    mNeoVis.evergladensis.1.1.chr15.g00103.t01#3    mNeoVis.evergladensis.2.1.chr15.g00510.t01#2    mNeoVis.3.1.1.chr15.g00469.t01#6    mNeoVis.vulgivaga.2.1.chr15.g00519.t01#1   mNeoVis.2.1.1.chr15_hap1.g00688.t01#5       2409   125258933:   mNeoVis.vulgivaga.1.1.chr1.g01139.t01#4    mNeoVis.evergladensis.1.1.chr1.g01138.t01#3    mNeoVis.vulgivaga.2.1.chr1.g01140.t01#1    mNeoVis.2.1.1.chr1_hap1.g01168.t01#5    mNeoVis.3.1.1.chr1.g01126.t01#6   mNeoVis.evergladensis.2.1.chr1.g01142.t01#2       2410   125258934:   mNeoVis.vulgivaga.2.1.chr1.g01141.t01#1    mNeoVis.2.1.1.chr1_hap1.g01169.t01#5    mNeoVis.evergladensis.2.1.chr1.g01143.t01#2    mNeoVis.vulgivaga.1.1.chr1.g01140.t01#4    mNeoVis.evergladensis.1.1.chr1.g01139.t01#3   mNeoVis.3.1.1.chr1.g01127.t01#6       2411   125258935:   mNeoVis.vulgivaga.2.1.chr1.g01142.t01#1   mNeoVis.evergladensis.1.1.chr1.g01140.t01#3           2412   125258936:   mNeoVis.2.1.1.chr7_hap1.g01874.t01#5    mNeoVis.vulgivaga.2.1.chr7.g01851.t01#1    mNeoVis.3.1.1.chr7.g01742.t01#6    mNeoVis.vulgivaga.1.1.chr7.g01843.t01#4    mNeoVis.evergladensis.1.1.chr7.g01832.t01#3   mNeoVis.evergladensis.2.1.chr7.g01835.t01#2       2413   125258937:   mNeoVis.vulgivaga.2.1.chr7.g01853.t01#1    mNeoVis.vulgivaga.1.1.chr7.g01844.t01#4    mNeoVis.evergladensis.1.1.chr7.g01833.t01#3    mNeoVis.evergladensis.2.1.chr7.g01836.t01#2   mNeoVis.2.1.1.chr7_hap1.g01875.t01#5       2414   125258938:   mNeoVis.vulgivaga.2.1.chr7.g01852.t01#1   mNeoVis.3.1.1.chr7.g01743.t01#6             2415   125258939:   mNeoVis.evergladensis.2.1.chr1.g01145.t01#2    mNeoVis.evergladensis.1.1.chr1.g01141.t01#3   mNeoVis.vulgivaga.2.1.chr1.g01143.t01#1           2416   125258940:   mNeoVis.evergladensis.2.1.chr7.g01837.t01#2    mNeoVis.vulgivaga.2.1.chr7.g01854.t01#1    mNeoVis.vulgivaga.1.1.chr7.g01845.t01#4    mNeoVis.3.1.1.chr7.g01744.t01#6    mNeoVis.2.1.1.chr7_hap1.g01876.t01#5   mNeoVis.evergladensis.1.1.chr7.g01834.t01#3       2417   125258941:   mNeoVis.vulgivaga.1.1.chr1.g01142.t01#4    mNeoVis.3.1.1.chr1.g01129.t01#6    mNeoVis.2.1.1.chr1_hap1.g01171.t01#5    mNeoVis.vulgivaga.2.1.chr1.g01144.t01#1    mNeoVis.evergladensis.2.1.chr1.g01146.t01#2   mNeoVis.evergladensis.1.1.chr1.g01142.t01#3       2418   125258942:   mNeoVis.evergladensis.2.1.chr7.g01838.t01#2    mNeoVis.3.1.1.chr7.g01745.t01#6    mNeoVis.vulgivaga.1.1.chr7.g01846.t01#4    mNeoVis.vulgivaga.2.1.chr7.g01855.t01#1    mNeoVis.evergladensis.1.1.chr7.g01835.t01#3   mNeoVis.2.1.1.chr7_hap1.g01877.t01#5       2419   125258943:   mNeoVis.evergladensis.2.1.chr15.g00511.t01#2    mNeoVis.2.1.1.chr15_hap1.g00689.t01#5   mNeoVis.vulgivaga.2.1.chr15.g00520.t01#1         2420   125258944:   mNeoVis.2.1.1.chr1_hap1.g01172.t01#5    mNeoVis.vulgivaga.2.1.chr1.g01145.t01#1    mNeoVis.evergladensis.1.1.chr1.g01143.t01#3    mNeoVis.vulgivaga.1.1.chr1.g01143.t01#4    mNeoVis.evergladensis.2.1.chr1.g01147.t01#2   mNeoVis.3.1.1.chr1.g01130.t01#6       2421   125258945:   mNeoVis.vulgivaga.2.1.chr7.g01856.t01#1   mNeoVis.3.1.1.chr7.g01746.t01#6             2422   125258946:   mNeoVis.2.1.1.chr7_hap1.g01878.t01#5    mNeoVis.evergladensis.1.1.chr7.g01836.t01#3    mNeoVis.vulgivaga.2.1.chr7.g01857.t01#1    mNeoVis.evergladensis.2.1.chr7.g01839.t01#2    mNeoVis.3.1.1.chr7.g01747.t01#6   mNeoVis.vulgivaga.1.1.chr7.g01847.t01#4       2423   125258947:   mNeoVis.vulgivaga.2.1.chr1.g01146.t01#1    mNeoVis.vulgivaga.1.1.chr1.g01144.t01#4    mNeoVis.evergladensis.1.1.chr1.g01144.t01#3    mNeoVis.3.1.1.chr1.g01131.t01#6    mNeoVis.2.1.1.chr1_hap1.g01173.t01#5   mNeoVis.evergladensis.2.1.chr1.g01148.t01#2       2424   125258948:   mNeoVis.evergladensis.2.1.chr1.g01149.t01#2    mNeoVis.vulgivaga.2.1.chr1.g01147.t01#1    mNeoVis.vulgivaga.1.1.chr1.g01145.t01#4    mNeoVis.vulgivaga.2.1.chr1.g01148.t01#1    mNeoVis.evergladensis.1.1.chr1.g01145.t01#3    mNeoVis.3.1.1.chr1.g01132.t01#6   mNeoVis.2.1.1.chr1_hap1.g01174.t01#5     2425   125258949:   mNeoVis.vulgivaga.1.1.chr1.g01146.t01#4    mNeoVis.evergladensis.1.1.chr1.g01146.t01#3    mNeoVis.evergladensis.2.1.chr1.g01150.t01#2    mNeoVis.2.1.1.chr1_hap1.g01175.t01#5   mNeoVis.vulgivaga.2.1.chr1.g01149.t01#1       2426   125258950:   mNeoVis.evergladensis.1.1.chr1.g01148.t01#3    mNeoVis.2.1.1.chr1_hap1.g01177.t01#5    mNeoVis.vulgivaga.1.1.chr1.g01148.t01#4    mNeoVis.evergladensis.2.1.chr1.g01152.t01#2    mNeoVis.vulgivaga.2.1.chr1.g01151.t01#1   mNeoVis.3.1.1.chr1.g01136.t01#6       2427   125258951:   mNeoVis.2.1.1.chr7_hap1.g01879.t01#5    mNeoVis.evergladensis.1.1.chr7.g01837.t01#3    mNeoVis.vulgivaga.2.1.chr7.g01858.t01#1    mNeoVis.evergladensis.2.1.chr7.g01840.t01#2    mNeoVis.3.1.1.chr7.g01748.t01#6   mNeoVis.vulgivaga.1.1.chr7.g01848.t01#4       2428   125258952:   mNeoVis.vulgivaga.1.1.chr7.g01849.t01#4    mNeoVis.evergladensis.2.1.chr7.g01841.t01#2    mNeoVis.2.1.1.chr7_hap1.g01880.t01#5    mNeoVis.vulgivaga.2.1.chr7.g01859.t01#1   mNeoVis.3.1.1.chr7.g01749.t01#6       2429   125258953:   mNeoVis.vulgivaga.1.1.chr1.g01150.t01#4    mNeoVis.evergladensis.2.1.chr1.g01154.t01#2    mNeoVis.vulgivaga.2.1.chr1.g01152.t01#1    mNeoVis.3.1.1.chr1.g01137.t01#6    mNeoVis.2.1.1.chr1_hap1.g01179.t01#5   mNeoVis.evergladensis.1.1.chr1.g01149.t01#3       2430   125258954:   mNeoVis.3.1.1.chr1.g01138.t01#6    mNeoVis.vulgivaga.2.1.chr1.g01153.t01#1   mNeoVis.vulgivaga.1.1.chr1.g01149.t01#4           2431   125258955:   mNeoVis.3.1.1.chr1.g01139.t01#6    mNeoVis.evergladensis.1.1.chr1.g01150.t01#3    mNeoVis.vulgivaga.2.1.chr1.g01154.t01#1    mNeoVis.evergladensis.2.1.chr1.g01156.t01#2    mNeoVis.vulgivaga.1.1.chr1.g01151.t01#4   mNeoVis.2.1.1.chr1_hap1.g01181.t01#5       2432   125258956:   mNeoVis.vulgivaga.2.1.chr7.g00455.t01#1    mNeoVis.vulgivaga.1.1.chr7.g00459.t01#4    mNeoVis.evergladensis.2.1.chr7.g00456.t01#2    mNeoVis.2.1.1.chr7_hap1.g00450.t01#5    mNeoVis.3.1.1.chr7.g00438.t01#6   mNeoVis.evergladensis.1.1.chr7.g00462.t01#3       2433   125258957:   mNeoVis.2.1.1.chr1_hap1.g01183.t01#5    mNeoVis.3.1.1.chr1.g01141.t01#6    mNeoVis.evergladensis.1.1.chr1.g01152.t01#3    mNeoVis.evergladensis.2.1.chr1.g01157.t01#2    mNeoVis.vulgivaga.1.1.chr1.g01153.t01#4   mNeoVis.vulgivaga.2.1.chr1.g01155.t01#1       2434   125258958:   mNeoVis.vulgivaga.1.1.chr7.g01777.t01#4    mNeoVis.3.1.1.chr7.g01676.t01#6    mNeoVis.2.1.1.chr7_hap1.g01807.t01#5    mNeoVis.evergladensis.2.1.chr7.g01769.t01#2    mNeoVis.vulgivaga.2.1.chr7.g01787.t01#1   mNeoVis.evergladensis.1.1.chr7.g01765.t01#3       2435   125258959:   mNeoVis.3.1.1.chr1.g01142.t01#6    mNeoVis.evergladensis.1.1.chr1.g01153.t01#3    mNeoVis.2.1.1.chr1_hap1.g01184.t01#5    mNeoVis.evergladensis.2.1.chr1.g01158.t01#2    mNeoVis.vulgivaga.2.1.chr1.g01156.t01#1   mNeoVis.vulgivaga.1.1.chr1.g01154.t01#4       2436   125258960:   mNeoVis.vulgivaga.2.1.chr1.g01157.t01#1    mNeoVis.evergladensis.1.1.chr1.g01154.t01#3    mNeoVis.3.1.1.chr1.g01143.t01#6    mNeoVis.2.1.1.chr1_hap1.g01185.t01#5    mNeoVis.vulgivaga.1.1.chr1.g01155.t01#4   mNeoVis.evergladensis.2.1.chr1.g01159.t01#2       2437   125258961:   mNeoVis.3.1.1.chr7.g01677.t01#6    mNeoVis.vulgivaga.1.1.chr7.g01778.t01#4    mNeoVis.2.1.1.chr7_hap1.g01808.t01#5    mNeoVis.evergladensis.2.1.chr7.g01770.t01#2    mNeoVis.vulgivaga.2.1.chr7.g01788.t01#1   mNeoVis.evergladensis.1.1.chr7.g01766.t01#3       2438   125258962:   mNeoVis.3.1.1.chr7.g01678.t01#6    mNeoVis.evergladensis.1.1.chr7.g01767.t01#3    mNeoVis.vulgivaga.2.1.chr7.g01789.t01#1    mNeoVis.vulgivaga.1.1.chr7.g01779.t01#4    mNeoVis.evergladensis.2.1.chr7.g01771.t01#2   mNeoVis.2.1.1.chr7_hap1.g01809.t01#5       2439   125258963:   mNeoVis.vulgivaga.2.1.chr15.g00521.t01#1    mNeoVis.evergladensis.1.1.chr15.g00102.t01#3   mNeoVis.3.1.1.chr15.g00470.t01#6           2440   125258964:   mNeoVis.vulgivaga.1.1.chr1.g01156.t01#4    mNeoVis.evergladensis.2.1.chr1.g01160.t01#2    mNeoVis.evergladensis.1.1.chr1.g01155.t01#3    mNeoVis.vulgivaga.2.1.chr1.g01159.t01#1    mNeoVis.vulgivaga.2.1.chr1.g01158.t01#1    mNeoVis.2.1.1.chr1_hap1.g01186.t01#5   mNeoVis.3.1.1.chr1.g01144.t01#6       2441   125258965:  mNeoVis.vulgivaga.2.1.chr1.g01160.t01#1               2442   125258966:   mNeoVis.3.1.1.chr1.g01145.t01#6    mNeoVis.evergladensis.2.1.chr1.g01162.t01#2    mNeoVis.vulgivaga.2.1.chr1.g01161.t01#1    mNeoVis.vulgivaga.1.1.chr1.g01157.t01#4   mNeoVis.2.1.1.chr1_hap1.g01188.t01#5       2443   125258967:   mNeoVis.2.1.1.chr7_hap1.g01810.t01#5    mNeoVis.vulgivaga.1.1.chr7.g01780.t01#4    mNeoVis.vulgivaga.2.1.chr7.g01790.t01#1    mNeoVis.evergladensis.1.1.chr7.g01768.t01#3    mNeoVis.evergladensis.2.1.chr7.g01772.t01#2   mNeoVis.3.1.1.chr7.g01679.t01#6       2444   125258968:   mNeoVis.2.1.1.chr1_hap1.g01189.t01#5    mNeoVis.evergladensis.1.1.chr1.g01157.t01#3    mNeoVis.3.1.1.chr1.g01146.t01#6    mNeoVis.evergladensis.2.1.chr1.g01163.t01#2    mNeoVis.vulgivaga.1.1.chr1.g01158.t01#4   mNeoVis.vulgivaga.2.1.chr1.g01162.t01#1       2445   125258969:   mNeoVis.vulgivaga.2.1.chr1.g01163.t01#1    mNeoVis.evergladensis.1.1.chr1.g01158.t01#3    mNeoVis.3.1.1.chr1.g01147.t01#6   mNeoVis.vulgivaga.1.1.chr1.g01159.t01#4         2446   125258970:   mNeoVis.2.1.1.chr1_hap1.g01190.t01#5    mNeoVis.vulgivaga.1.1.chr1.g01160.t01#4    mNeoVis.evergladensis.2.1.chr1.g01164.t01#2    mNeoVis.vulgivaga.2.1.chr1.g01164.t01#1    mNeoVis.evergladensis.1.1.chr1.g01159.t01#3   mNeoVis.3.1.1.chr1.g01148.t01#6       2447   125258971:   mNeoVis.vulgivaga.2.1.chr15.g00522.t01#1    mNeoVis.3.1.1.chr15.g00471.t01#6    mNeoVis.vulgivaga.2.1.chr15.g00523.t01#1    mNeoVis.evergladensis.2.1.chr15.g00512.t01#2    mNeoVis.evergladensis.1.1.chr15.g00100.t01#3   mNeoVis.2.1.1.chr15_hap1.g00690.t01#5       2448   125258972:   mNeoVis.vulgivaga.1.1.chr15.g00514.t01#4   mNeoVis.evergladensis.1.1.chr15.g00101.t01#3           2449   125258973:   mNeoVis.vulgivaga.2.1.chr1.g01165.t01#1    mNeoVis.evergladensis.1.1.chr1.g01160.t01#3    mNeoVis.3.1.1.chr1.g01149.t01#6    mNeoVis.evergladensis.2.1.chr1.g01165.t01#2    mNeoVis.2.1.1.chr1_hap1.g01191.t01#5   mNeoVis.vulgivaga.1.1.chr1.g01161.t01#4       2450   125258974:   mNeoVis.vulgivaga.2.1.chr7.g01791.t01#1   mNeoVis.evergladensis.2.1.chr7.g01773.t01#2           2451   125258975:   mNeoVis.3.1.1.chr7.g01681.t01#6    mNeoVis.evergladensis.2.1.chr7.g01774.t01#2    mNeoVis.vulgivaga.2.1.chr7.g01792.t01#1    mNeoVis.vulgivaga.1.1.chr7.g01781.t01#4    mNeoVis.evergladensis.1.1.chr7.g01770.t01#3   mNeoVis.2.1.1.chr7_hap1.g01811.t01#5       2452   125258976:   mNeoVis.vulgivaga.1.1.chr12.g00726.t01#4    mNeoVis.evergladensis.1.1.chr2.g01282.t01#3    mNeoVis.vulgivaga.1.1.chr10.g00453.t01#4    mNeoVis.3.1.1.chr12.g00707.t01#6    mNeoVis.2.1.1.chr5_hap1.g01291.t01#5    mNeoVis.2.1.1.chr12_hap1.g00775.t01#5    mNeoVis.evergladensis.1.1.chr4.g00505.t01#3    mNeoVis.evergladensis.2.1.chr2.g01278.t01#2    mNeoVis.vulgivaga.2.1.chr7.g01793.t01#1      2453   125258977:   mNeoVis.evergladensis.2.1.chr1.g01166.t01#2    mNeoVis.vulgivaga.1.1.chr1.g01162.t01#4    mNeoVis.3.1.1.chr1.g01150.t01#6    mNeoVis.2.1.1.chr1_hap1.g01192.t01#5    mNeoVis.evergladensis.1.1.chr1.g01161.t01#3   mNeoVis.vulgivaga.2.1.chr1.g01166.t01#1       2454   125258978:   mNeoVis.2.1.1.chr7_hap1.g01812.t01#5    mNeoVis.evergladensis.2.1.chr7.g01775.t01#2    mNeoVis.vulgivaga.1.1.chr7.g01782.t01#4    mNeoVis.evergladensis.1.1.chr7.g01771.t01#3    mNeoVis.3.1.1.chr7.g01682.t01#6   mNeoVis.vulgivaga.2.1.chr7.g01794.t01#1       2455   125258979:   mNeoVis.evergladensis.1.1.chr7.g00463.t01#3    mNeoVis.3.1.1.chr7.g00439.t01#6   mNeoVis.vulgivaga.2.1.chr7.g00456.t01#1           2456   125258980:   mNeoVis.vulgivaga.2.1.chr1.g01167.t01#1    mNeoVis.evergladensis.2.1.chr1.g01167.t01#2    mNeoVis.3.1.1.chr1.g01151.t01#6    mNeoVis.vulgivaga.1.1.chr1.g01163.t01#4    mNeoVis.2.1.1.chr1_hap1.g01193.t01#5   mNeoVis.evergladensis.1.1.chr1.g01162.t01#3       2457   125258981:   mNeoVis.3.1.1.chr1.g01152.t01#6    mNeoVis.vulgivaga.1.1.chr1.g01164.t01#4    mNeoVis.evergladensis.1.1.chr1.g01166.t01#3    mNeoVis.2.1.1.chr1_hap1.g01194.t01#5    mNeoVis.vulgivaga.2.1.chr1.g01168.t01#1   mNeoVis.evergladensis.2.1.chr1.g01168.t01#2       2458   125258982:   mNeoVis.evergladensis.1.1.chr7.g01772.t01#3    mNeoVis.vulgivaga.1.1.chr7.g01783.t01#4    mNeoVis.2.1.1.chr7_hap1.g01813.t01#5    mNeoVis.evergladensis.2.1.chr7.g01776.t01#2    mNeoVis.vulgivaga.2.1.chr7.g01795.t01#1   mNeoVis.3.1.1.chr7.g01683.t01#6       2459   125258983:   mNeoVis.evergladensis.1.1.chr7.g01774.t01#3    mNeoVis.vulgivaga.2.1.chr7.g01796.t01#1    mNeoVis.2.1.1.chr7_hap1.g01814.t01#5    mNeoVis.vulgivaga.1.1.chr7.g01785.t01#4   mNeoVis.evergladensis.2.1.chr7.g01778.t01#2       2460   125258984:   mNeoVis.evergladensis.1.1.chr7.g01773.t01#3    mNeoVis.vulgivaga.1.1.chr7.g01784.t01#4    mNeoVis.evergladensis.2.1.chr7.g01777.t01#2    mNeoVis.3.1.1.chr7.g01684.t01#6    mNeoVis.vulgivaga.2.1.chr7.g01797.t01#1   mNeoVis.2.1.1.chr7_hap1.g01815.t01#5       2461   125258985:   mNeoVis.vulgivaga.2.1.chr7.g01798.t01#1    mNeoVis.evergladensis.2.1.chr7.g01779.t01#2    mNeoVis.3.1.1.chr7.g01685.t01#6    mNeoVis.evergladensis.1.1.chr7.g01775.t01#3    mNeoVis.vulgivaga.1.1.chr7.g01786.t01#4   mNeoVis.2.1.1.chr7_hap1.g01816.t01#5       2462   125258986:   mNeoVis.2.1.1.chr1_hap1.g01196.t01#5    mNeoVis.evergladensis.2.1.chr1.g01170.t01#2    mNeoVis.vulgivaga.2.1.chr1.g01170.t01#1   mNeoVis.evergladensis.1.1.chr1.g01168.t01#3       2463   125258987:   mNeoVis.evergladensis.1.1.chr1.g01169.t01#3    mNeoVis.3.1.1.chr1.g01154.t01#6    mNeoVis.evergladensis.2.1.chr1.g01171.t01#2    mNeoVis.vulgivaga.2.1.chr1.g01171.t01#1    mNeoVis.2.1.1.chr1_hap1.g01197.t01#5   mNeoVis.vulgivaga.1.1.chr1.g01165.t01#4       2464   125258988:   mNeoVis.vulgivaga.2.1.chr7.g01799.t01#1    mNeoVis.evergladensis.1.1.chr7.g01776.t01#3    mNeoVis.evergladensis.2.1.chr7.g01780.t01#2    mNeoVis.vulgivaga.1.1.chr7.g01787.t01#4    mNeoVis.3.1.1.chr7.g01686.t01#6   mNeoVis.2.1.1.chr7_hap1.g01817.t01#5       2465   125258989:   mNeoVis.evergladensis.2.1.chr7.g00458.t01#2    mNeoVis.vulgivaga.1.1.chr7.g00461.t01#4    mNeoVis.2.1.1.chr7_hap1.g00451.t01#5    mNeoVis.evergladensis.1.1.chr7.g00464.t01#3    mNeoVis.3.1.1.chr7.g00440.t01#6   mNeoVis.vulgivaga.2.1.chr7.g00457.t01#1       2466   125258990:   mNeoVis.vulgivaga.1.1.chr1.g01166.t01#4    mNeoVis.vulgivaga.2.1.chr1.g01172.t01#1    mNeoVis.evergladensis.1.1.chr1.g01170.t01#3    mNeoVis.2.1.1.chr1_hap1.g01198.t01#5    mNeoVis.3.1.1.chr1.g01155.t01#6   mNeoVis.evergladensis.2.1.chr1.g01172.t01#2       2467   125258991:   mNeoVis.2.1.1.chr1_hap1.g01199.t01#5    mNeoVis.evergladensis.1.1.chr1.g01171.t01#3    mNeoVis.3.1.1.chr1.g01156.t01#6    mNeoVis.evergladensis.2.1.chr1.g01173.t01#2    mNeoVis.vulgivaga.2.1.chr1.g01173.t01#1   mNeoVis.vulgivaga.1.1.chr1.g01167.t01#4       2468   125258992:   mNeoVis.2.1.1.chr1_hap1.g01201.t01#5    mNeoVis.evergladensis.1.1.chr1.g01173.t01#3    mNeoVis.vulgivaga.2.1.chr1.g01174.t01#1    mNeoVis.evergladensis.2.1.chr1.g01175.t01#2   mNeoVis.vulgivaga.1.1.chr1.g01169.t01#4       2469   125258993:   mNeoVis.3.1.1.chr7.g01687.t01#6    mNeoVis.vulgivaga.1.1.chr7.g01788.t01#4    mNeoVis.evergladensis.1.1.chr7.g01777.t01#3    mNeoVis.vulgivaga.2.1.chr7.g01800.t01#1    mNeoVis.2.1.1.chr7_hap1.g01818.t01#5   mNeoVis.evergladensis.2.1.chr7.g01781.t01#2       2470   125258994:   mNeoVis.vulgivaga.2.1.chr1.g01175.t01#1    mNeoVis.3.1.1.chr1.g01160.t01#6    mNeoVis.evergladensis.2.1.chr1.g01176.t01#2    mNeoVis.vulgivaga.1.1.chr1.g01170.t01#4    mNeoVis.2.1.1.chr1_hap1.g01202.t01#5   mNeoVis.evergladensis.1.1.chr1.g01174.t01#3       2471   125258995:   mNeoVis.evergladensis.1.1.chr1.g01175.t01#3    mNeoVis.vulgivaga.2.1.chr1.g01176.t01#1    mNeoVis.evergladensis.2.1.chr1.g01177.t01#2    mNeoVis.3.1.1.chr1.g01161.t01#6   mNeoVis.2.1.1.chr1_hap1.g01203.t01#5       2472   125258996:   mNeoVis.3.1.1.chr1.g01162.t01#6    mNeoVis.2.1.1.chr1_hap1.g01204.t01#5    mNeoVis.vulgivaga.1.1.chr1.g01171.t01#4    mNeoVis.evergladensis.1.1.chr1.g01176.t01#3    mNeoVis.vulgivaga.2.1.chr1.g01177.t01#1   mNeoVis.evergladensis.2.1.chr1.g01178.t01#2       2473   125258997:   mNeoVis.vulgivaga.1.1.chr7.g01789.t01#4    mNeoVis.2.1.1.chr7_hap1.g01821.t01#5    mNeoVis.evergladensis.1.1.chr7.g01778.t01#3    mNeoVis.vulgivaga.2.1.chr7.g01801.t01#1    mNeoVis.evergladensis.2.1.chr7.g01783.t01#2   mNeoVis.3.1.1.chr7.g01688.t01#6       2474   125258998:   mNeoVis.evergladensis.2.1.chr1.g01179.t01#2    mNeoVis.vulgivaga.1.1.chr1.g01172.t01#4    mNeoVis.2.1.1.chr1_hap1.g01205.t01#5    mNeoVis.vulgivaga.2.1.chr1.g01178.t01#1    mNeoVis.evergladensis.1.1.chr1.g01177.t01#3   mNeoVis.3.1.1.chr1.g01163.t01#6       2475   125258999:   mNeoVis.evergladensis.1.1.chr9.g00523.t01#3    mNeoVis.2.1.1.chr10_hap1.g00531.t01#5    mNeoVis.3.1.1.chr9.g00490.t01#6    mNeoVis.vulgivaga.2.1.chr9.g00513.t01#1    mNeoVis.evergladensis.2.1.chr8.g00513.t01#2   mNeoVis.vulgivaga.1.1.chr9.g00517.t01#4       2476   125259000:   mNeoVis.vulgivaga.1.1.chr7.g01790.t01#4    mNeoVis.2.1.1.chr7_hap1.g01822.t01#5    mNeoVis.3.1.1.chr7.g01689.t01#6    mNeoVis.evergladensis.2.1.chr7.g01784.t01#2    mNeoVis.evergladensis.1.1.chr7.g01780.t01#3   mNeoVis.vulgivaga.2.1.chr7.g01802.t01#1       2477   125259001:   mNeoVis.3.1.1.chr1.g01164.t01#6    mNeoVis.evergladensis.2.1.chr1.g01180.t01#2    mNeoVis.vulgivaga.2.1.chr1.g01179.t01#1    mNeoVis.evergladensis.1.1.chr1.g01178.t01#3    mNeoVis.2.1.1.chr1_hap1.g01206.t01#5   mNeoVis.vulgivaga.1.1.chr1.g01173.t01#4       2478   125259002:   mNeoVis.vulgivaga.2.1.chr1.g01180.t01#1   mNeoVis.vulgivaga.1.1.chr1.g01174.t01#4             2479   125259003:   mNeoVis.evergladensis.2.1.chr1.g01181.t01#2    mNeoVis.vulgivaga.2.1.chr1.g01181.t01#1    mNeoVis.2.1.1.chr1_hap1.g01207.t01#5    mNeoVis.3.1.1.chr1.g01166.t01#6    mNeoVis.3.1.1.chr1.g01165.t01#6    mNeoVis.vulgivaga.1.1.chr1.g01175.t01#4   mNeoVis.evergladensis.1.1.chr1.g01180.t01#3     2480   125259004:   mNeoVis.evergladensis.1.1.chr1.g01181.t01#3    mNeoVis.evergladensis.2.1.chr1.g01182.t01#2    mNeoVis.2.1.1.chr1_hap1.g01208.t01#5    mNeoVis.vulgivaga.1.1.chr1.g01176.t01#4    mNeoVis.3.1.1.chr1.g01167.t01#6   mNeoVis.vulgivaga.2.1.chr1.g01182.t01#1       2481   125259005:   mNeoVis.3.1.1.chr7.g01690.t01#6    mNeoVis.evergladensis.2.1.chr7.g01785.t01#2    mNeoVis.vulgivaga.1.1.chr7.g01791.t01#4    mNeoVis.evergladensis.1.1.chr7.g01781.t01#3    mNeoVis.2.1.1.chr7_hap1.g01823.t01#5   mNeoVis.vulgivaga.2.1.chr7.g01803.t01#1       2482   125259006:   mNeoVis.vulgivaga.2.1.chr7.g01804.t01#1    mNeoVis.2.1.1.chr7_hap1.g01824.t01#5    mNeoVis.3.1.1.chr7.g01691.t01#6   mNeoVis.evergladensis.2.1.chr7.g01786.t01#2       2483   125259007:   mNeoVis.vulgivaga.1.1.chr1.g01177.t01#4   mNeoVis.vulgivaga.2.1.chr1.g01183.t01#1             2484   125259008:   mNeoVis.2.1.1.chr1_hap1.g01209.t01#5    mNeoVis.evergladensis.1.1.chr1.g01182.t01#3    mNeoVis.3.1.1.chr1.g01168.t01#6    mNeoVis.evergladensis.2.1.chr1.g01183.t01#2    mNeoVis.vulgivaga.2.1.chr1.g01184.t01#1   mNeoVis.vulgivaga.1.1.chr1.g01178.t01#4       2485   125259009:   mNeoVis.evergladensis.1.1.chr1.g01183.t01#3    mNeoVis.vulgivaga.2.1.chr1.g01185.t01#1    mNeoVis.vulgivaga.1.1.chr1.g01179.t01#4    mNeoVis.evergladensis.2.1.chr1.g01184.t01#2    mNeoVis.3.1.1.chr1.g01169.t01#6   mNeoVis.2.1.1.chr1_hap1.g01210.t01#5       2486   125259010:   mNeoVis.3.1.1.chr1.g01170.t01#6    mNeoVis.evergladensis.2.1.chr1.g01185.t01#2    mNeoVis.evergladensis.1.1.chr1.g01184.t01#3    mNeoVis.vulgivaga.2.1.chr1.g01186.t01#1    mNeoVis.vulgivaga.1.1.chr1.g01180.t01#4   mNeoVis.2.1.1.chr1_hap1.g01211.t01#5       2487   125259011:   mNeoVis.3.1.1.chr1.g01171.t01#6    mNeoVis.evergladensis.2.1.chr1.g01186.t01#2    mNeoVis.vulgivaga.1.1.chr1.g01181.t01#4    mNeoVis.2.1.1.chr1_hap1.g01212.t01#5    mNeoVis.evergladensis.1.1.chr1.g01185.t01#3   mNeoVis.vulgivaga.2.1.chr1.g01187.t01#1       2488   125259012:   mNeoVis.evergladensis.1.1.chr7.g01785.t01#3    mNeoVis.2.1.1.chr7_hap1.g01826.t01#5    mNeoVis.vulgivaga.2.1.chr7.g01806.t01#1    mNeoVis.evergladensis.2.1.chr7.g01788.t01#2    mNeoVis.3.1.1.chr7.g01693.t01#6   mNeoVis.vulgivaga.1.1.chr7.g01796.t01#4       2489   125259013:   mNeoVis.evergladensis.2.1.chr10.g01247.t01#2    mNeoVis.2.1.1.chr9_hap1.g01257.t01#5    mNeoVis.vulgivaga.2.1.chr10.g01235.t01#1    mNeoVis.vulgivaga.1.1.chr10.g01220.t01#4    mNeoVis.evergladensis.1.1.chr10.g01240.t01#3   mNeoVis.3.1.1.chr10.g01208.t01#6       2490   125259014:   mNeoVis.evergladensis.2.1.chr1.g01187.t01#2    mNeoVis.vulgivaga.1.1.chr1.g01182.t01#4    mNeoVis.3.1.1.chr1.g01172.t01#6    mNeoVis.2.1.1.chr1_hap1.g01213.t01#5    mNeoVis.evergladensis.1.1.chr1.g01186.t01#3   mNeoVis.vulgivaga.2.1.chr1.g01188.t01#1       2491   125259015:   mNeoVis.3.1.1.chr7.g01694.t01#6    mNeoVis.evergladensis.1.1.chr7.g01786.t01#3    mNeoVis.vulgivaga.2.1.chr7.g01807.t01#1    mNeoVis.2.1.1.chr7_hap1.g01827.t01#5    mNeoVis.evergladensis.2.1.chr7.g01789.t01#2   mNeoVis.vulgivaga.1.1.chr7.g01797.t01#4       2492   125259016:   mNeoVis.3.1.1.chr1.g01173.t01#6    mNeoVis.2.1.1.chr1_hap1.g01214.t01#5    mNeoVis.vulgivaga.1.1.chr1.g01183.t01#4    mNeoVis.evergladensis.1.1.chr1.g01187.t01#3    mNeoVis.evergladensis.2.1.chr1.g01188.t01#2   mNeoVis.vulgivaga.2.1.chr1.g01189.t01#1       2493   125259017:   mNeoVis.evergladensis.1.1.chr1.g01188.t01#3    mNeoVis.vulgivaga.2.1.chr1.g01190.t01#1    mNeoVis.3.1.1.chr1.g01174.t01#6    mNeoVis.2.1.1.chr1_hap1.g01215.t01#5    mNeoVis.vulgivaga.1.1.chr1.g01184.t01#4   mNeoVis.evergladensis.2.1.chr1.g01189.t01#2       2494   125259018:   mNeoVis.vulgivaga.2.1.chr1.g01191.t01#1    mNeoVis.3.1.1.chr1.g01175.t01#6    mNeoVis.evergladensis.1.1.chr1.g01189.t01#3    mNeoVis.evergladensis.2.1.chr1.g01190.t01#2    mNeoVis.vulgivaga.1.1.chr1.g01185.t01#4   mNeoVis.2.1.1.chr1_hap1.g01216.t01#5       2495   125259019:   mNeoVis.evergladensis.2.1.chr7.g01790.t01#2    mNeoVis.vulgivaga.1.1.chr7.g01798.t01#4    mNeoVis.vulgivaga.2.1.chr7.g01808.t01#1    mNeoVis.3.1.1.chr7.g01695.t01#6    mNeoVis.evergladensis.1.1.chr7.g01787.t01#3   mNeoVis.2.1.1.chr7_hap1.g01828.t01#5       2496   125259020:   mNeoVis.vulgivaga.2.1.chr7.g01809.t01#1    mNeoVis.3.1.1.chr7.g01696.t01#6    mNeoVis.evergladensis.1.1.chr7.g01788.t01#3    mNeoVis.2.1.1.chr7_hap1.g01829.t01#5    mNeoVis.evergladensis.2.1.chr7.g01791.t01#2   mNeoVis.vulgivaga.1.1.chr7.g01799.t01#4       2497   125259021:   mNeoVis.2.1.1.chr1_hap1.g01217.t01#5    mNeoVis.vulgivaga.1.1.chr1.g01186.t01#4    mNeoVis.evergladensis.1.1.chr1.g01190.t01#3    mNeoVis.evergladensis.2.1.chr1.g01191.t01#2   mNeoVis.vulgivaga.2.1.chr1.g01192.t01#1       2498   125259022:  mNeoVis.vulgivaga.2.1.chr1.g01193.t01#1               2499   125259023:   mNeoVis.2.1.1.chr7_hap1.g01830.t01#5    mNeoVis.vulgivaga.2.1.chr7.g01810.t01#1    mNeoVis.3.1.1.chr7.g01697.t01#6    mNeoVis.vulgivaga.1.1.chr7.g01800.t01#4    mNeoVis.evergladensis.2.1.chr7.g01792.t01#2   mNeoVis.evergladensis.1.1.chr7.g01789.t01#3       2500   125259024:   mNeoVis.vulgivaga.2.1.chr1.g01194.t01#1    mNeoVis.evergladensis.2.1.chr1.g01193.t01#2    mNeoVis.vulgivaga.1.1.chr1.g01188.t01#4    mNeoVis.3.1.1.chr1.g01178.t01#6    mNeoVis.evergladensis.1.1.chr1.g01192.t01#3   mNeoVis.2.1.1.chr1_hap1.g01219.t01#5       2501   125259025:   mNeoVis.3.1.1.chr7.g01698.t01#6    mNeoVis.2.1.1.chr7_hap1.g01831.t01#5    mNeoVis.vulgivaga.1.1.chr7.g01801.t01#4    mNeoVis.evergladensis.2.1.chr7.g01793.t01#2    mNeoVis.vulgivaga.2.1.chr7.g01811.t01#1   mNeoVis.evergladensis.1.1.chr7.g01790.t01#3       2502   125259026:   mNeoVis.3.1.1.chr15.g00140.t01#6    mNeoVis.evergladensis.1.1.chr15.g00470.t01#3    mNeoVis.vulgivaga.1.1.chr15.g00146.t01#4    mNeoVis.vulgivaga.2.1.chr15.g00150.t01#1    mNeoVis.2.1.1.chr15_hap1.g00161.t01#5   mNeoVis.evergladensis.2.1.chr15.g00147.t01#2       2503   125259027:   mNeoVis.evergladensis.1.1.chr7.g01792.t01#3    mNeoVis.vulgivaga.2.1.chr7.g01812.t01#1    mNeoVis.2.1.1.chr7_hap1.g01833.t01#5    mNeoVis.vulgivaga.1.1.chr7.g01803.t01#4    mNeoVis.3.1.1.chr7.g01700.t01#6   mNeoVis.evergladensis.2.1.chr7.g01795.t01#2       2504   125259028:   mNeoVis.evergladensis.1.1.chr1.g01023.t01#3    mNeoVis.vulgivaga.2.1.chr1.g01032.t01#1    mNeoVis.3.1.1.chr1.g01026.t01#6    mNeoVis.vulgivaga.1.1.chr1.g01032.t01#4    mNeoVis.2.1.1.chr1_hap1.g01027.t01#5   mNeoVis.evergladensis.2.1.chr1.g01021.t01#2       2505   125259029:   mNeoVis.vulgivaga.1.1.chr1.g01033.t01#4    mNeoVis.evergladensis.2.1.chr1.g01022.t01#2    mNeoVis.vulgivaga.2.1.chr1.g01033.t01#1    mNeoVis.evergladensis.1.1.chr1.g01024.t01#3    mNeoVis.3.1.1.chr1.g01027.t01#6   mNeoVis.2.1.1.chr1_hap1.g01028.t01#5       2506   125259030:   mNeoVis.vulgivaga.2.1.chr6.g00608.t01#1    mNeoVis.2.1.1.chr4_hap1.g00602.t01#5    mNeoVis.evergladensis.2.1.chr6.g00610.t01#2    mNeoVis.vulgivaga.1.1.chr6.g00610.t01#4    mNeoVis.3.1.1.chr6.g00619.t01#6   mNeoVis.evergladensis.1.1.chr6.g00613.t01#3       2507   125259031:   mNeoVis.2.1.1.chr1_hap1.g01029.t01#5    mNeoVis.vulgivaga.2.1.chr1.g01034.t01#1    mNeoVis.3.1.1.chr1.g01028.t01#6    mNeoVis.evergladensis.1.1.chr1.g01025.t01#3    mNeoVis.vulgivaga.1.1.chr1.g01034.t01#4   mNeoVis.evergladensis.2.1.chr1.g01023.t01#2       2508   125259032:   mNeoVis.3.1.1.chr4.g01080.t01#6    mNeoVis.evergladensis.2.1.chr4.g01067.t01#2    mNeoVis.vulgivaga.2.1.chr4.g01101.t01#1    mNeoVis.2.1.1.chr6_hap1.g01134.t01#5    mNeoVis.vulgivaga.1.1.chr4.g01091.t01#4   mNeoVis.evergladensis.1.1.chr4.g01099.t01#3       2509   125259033:   mNeoVis.evergladensis.1.1.chr1.g01026.t01#3    mNeoVis.2.1.1.chr1_hap1.g01030.t01#5    mNeoVis.3.1.1.chr1.g01029.t01#6    mNeoVis.vulgivaga.1.1.chr1.g01035.t01#4   mNeoVis.vulgivaga.2.1.chr1.g01035.t01#1       2510   125259034:   mNeoVis.2.1.1.chr1_hap1.g01031.t01#5    mNeoVis.vulgivaga.2.1.chr1.g01036.t01#1    mNeoVis.evergladensis.1.1.chr1.g01027.t01#3   mNeoVis.evergladensis.2.1.chr1.g01024.t01#2       2511   125259035:   mNeoVis.2.1.1.chr1_hap1.g01033.t01#5    mNeoVis.vulgivaga.2.1.chr1.g01037.t01#1    mNeoVis.evergladensis.1.1.chr1.g01028.t01#3    mNeoVis.evergladensis.2.1.chr1.g01025.t01#2    mNeoVis.vulgivaga.1.1.chr1.g01038.t01#4    mNeoVis.3.1.1.chr1.g01030.t01#6    mNeoVis.2.1.1.chr1_hap1.g01032.t01#5   mNeoVis.vulgivaga.1.1.chr1.g01036.t01#4     2512   125259036:   mNeoVis.vulgivaga.2.1.chr1.g01038.t01#1    mNeoVis.evergladensis.2.1.chr1.g01026.t01#2    mNeoVis.3.1.1.chr1.g01031.t01#6    mNeoVis.2.1.1.chr1_hap1.g01034.t01#5    mNeoVis.evergladensis.1.1.chr1.g01030.t01#3   mNeoVis.vulgivaga.1.1.chr1.g01039.t01#4       2513   125259037:   mNeoVis.vulgivaga.2.1.chr1.g01042.t01#1    mNeoVis.2.1.1.chr1_hap1.g01040.t01#5    mNeoVis.3.1.1.chr1.g01033.t01#6    mNeoVis.vulgivaga.2.1.chr1.g01041.t01#1    mNeoVis.evergladensis.1.1.chr1.g01035.t01#3    mNeoVis.evergladensis.2.1.chr1.g01027.t01#2    mNeoVis.3.1.1.chr1.g01034.t01#6    mNeoVis.2.1.1.chr1_hap1.g01038.t01#5    mNeoVis.evergladensis.1.1.chr1.g01034.t01#3      2514   125259038:   mNeoVis.vulgivaga.1.1.chr1.g01037.t01#4   mNeoVis.evergladensis.1.1.chr1.g01029.t01#3           2515   125259039:   mNeoVis.evergladensis.2.1.chr7.g01796.t01#2    mNeoVis.vulgivaga.2.1.chr7.g01813.t01#1    mNeoVis.evergladensis.1.1.chr7.g01793.t01#3   mNeoVis.2.1.1.chr7_hap1.g01834.t01#5         2516   125259040:   mNeoVis.3.1.1.chr7.g01702.t01#6    mNeoVis.2.1.1.chr7_hap1.g01835.t01#5    mNeoVis.vulgivaga.2.1.chr7.g01814.t01#1    mNeoVis.evergladensis.2.1.chr7.g01797.t01#2    mNeoVis.vulgivaga.1.1.chr7.g01804.t01#4   mNeoVis.evergladensis.1.1.chr7.g01794.t01#3       2517   125259041:   mNeoVis.vulgivaga.1.1.chr1.g01043.t01#4    mNeoVis.evergladensis.1.1.chr1.g01037.t01#3    mNeoVis.2.1.1.chr1_hap1.g01041.t01#5    mNeoVis.vulgivaga.2.1.chr1.g01043.t01#1    mNeoVis.evergladensis.2.1.chr1.g01033.t01#2   mNeoVis.3.1.1.chr1.g01035.t01#6       2518   125259042:   mNeoVis.vulgivaga.1.1.chr1.g01044.t01#4    mNeoVis.evergladensis.2.1.chr1.g01034.t01#2    mNeoVis.3.1.1.chr1.g01037.t01#6    mNeoVis.vulgivaga.2.1.chr1.g01044.t01#1    mNeoVis.evergladensis.1.1.chr1.g01038.t01#3   mNeoVis.2.1.1.chr1_hap1.g01042.t01#5       2519   125259043:   mNeoVis.evergladensis.2.1.chr7.g01798.t01#2    mNeoVis.evergladensis.1.1.chr7.g01795.t01#3    mNeoVis.vulgivaga.2.1.chr7.g01815.t01#1   mNeoVis.2.1.1.chr7_hap1.g01836.t01#5         2520   125259044:   mNeoVis.vulgivaga.1.1.chr1.g01045.t01#4    mNeoVis.evergladensis.2.1.chr1.g01035.t01#2    mNeoVis.vulgivaga.2.1.chr1.g01045.t01#1   mNeoVis.3.1.1.chr1.g01038.t01#6         2521   125259045:   mNeoVis.vulgivaga.2.1.chr7.g01816.t01#1    mNeoVis.2.1.1.chr7_hap1.g01837.t01#5    mNeoVis.3.1.1.chr7.g01704.t01#6    mNeoVis.vulgivaga.1.1.chr7.g01806.t01#4    mNeoVis.evergladensis.1.1.chr7.g01796.t01#3   mNeoVis.evergladensis.2.1.chr7.g01799.t01#2       2522   125259046:  mNeoVis.vulgivaga.2.1.chr1.g01046.t01#1               2523   125259047:   mNeoVis.vulgivaga.2.1.chr1.g01047.t01#1    mNeoVis.3.1.1.chr1.g01039.t01#6    mNeoVis.2.1.1.chr1_hap1.g01044.t01#5    mNeoVis.evergladensis.2.1.chr1.g01036.t01#2   mNeoVis.evergladensis.1.1.chr1.g01040.t01#3       2524   125259048:   mNeoVis.evergladensis.1.1.chr1.g01042.t01#3    mNeoVis.vulgivaga.2.1.chr1.g01048.t01#1    mNeoVis.3.1.1.chr1.g01040.t01#6    mNeoVis.2.1.1.chr1_hap1.g01045.t01#5   mNeoVis.evergladensis.2.1.chr1.g01037.t01#2       2525   125259049:   mNeoVis.2.1.1.chr1_hap1.g01046.t01#5    mNeoVis.vulgivaga.2.1.chr1.g01049.t01#1    mNeoVis.evergladensis.2.1.chr1.g01038.t01#2   mNeoVis.vulgivaga.1.1.chr1.g01049.t01#4         2526   125259050:   mNeoVis.evergladensis.1.1.chr7.g01797.t01#3    mNeoVis.vulgivaga.2.1.chr7.g01817.t01#1    mNeoVis.2.1.1.chr7_hap1.g01838.t01#5    mNeoVis.evergladensis.2.1.chr7.g01800.t01#2    mNeoVis.3.1.1.chr7.g01705.t01#6   mNeoVis.vulgivaga.1.1.chr7.g01807.t01#4       2527   125259051:   mNeoVis.evergladensis.2.1.chr1.g01039.t01#2    mNeoVis.vulgivaga.1.1.chr1.g01050.t01#4    mNeoVis.2.1.1.chr1_hap1.g01047.t01#5   mNeoVis.vulgivaga.2.1.chr1.g01050.t01#1         2528   125259052:  mNeoVis.vulgivaga.2.1.chr1.g01051.t01#1               2529   125259053:   mNeoVis.3.1.1.chr7.g01706.t01#6    mNeoVis.vulgivaga.1.1.chr7.g01808.t01#4    mNeoVis.evergladensis.1.1.chr7.g01798.t01#3    mNeoVis.2.1.1.chr7_hap1.g01839.t01#5    mNeoVis.evergladensis.2.1.chr7.g01801.t01#2   mNeoVis.vulgivaga.2.1.chr7.g01818.t01#1       2530   125259054:  mNeoVis.vulgivaga.2.1.chr1.g01052.t01#1               2531   125259055:   mNeoVis.evergladensis.2.1.chr1.g01040.t01#2    mNeoVis.vulgivaga.2.1.chr1.g01053.t01#1    mNeoVis.2.1.1.chr1_hap1.g01048.t01#5    mNeoVis.3.1.1.chr1.g01042.t01#6    mNeoVis.evergladensis.1.1.chr1.g01044.t01#3   mNeoVis.vulgivaga.1.1.chr1.g01051.t01#4       2532   125259056:   mNeoVis.evergladensis.1.1.chr1.g01045.t01#3    mNeoVis.vulgivaga.1.1.chr1.g01052.t01#4    mNeoVis.evergladensis.2.1.chr1.g01041.t01#2    mNeoVis.vulgivaga.2.1.chr1.g01054.t01#1   mNeoVis.2.1.1.chr1_hap1.g01049.t01#5       2533   125259057:   mNeoVis.3.1.1.chr1.g01043.t01#6   mNeoVis.vulgivaga.2.1.chr1.g01055.t01#1             2534   125259058:   mNeoVis.3.1.1.chr7.g01707.t01#6    mNeoVis.vulgivaga.1.1.chr7.g01809.t01#4    mNeoVis.vulgivaga.2.1.chr7.g01819.t01#1    mNeoVis.evergladensis.1.1.chr7.g01799.t01#3    mNeoVis.evergladensis.2.1.chr7.g01802.t01#2   mNeoVis.2.1.1.chr7_hap1.g01840.t01#5       2535   125259059:  mNeoVis.vulgivaga.2.1.chr1.g01056.t01#1               2536   125259060:   mNeoVis.3.1.1.chr1.g01044.t01#6    mNeoVis.vulgivaga.2.1.chr1.g01057.t01#1   mNeoVis.evergladensis.2.1.chr1.g01042.t01#2         2537   125259061:   mNeoVis.evergladensis.1.1.chr1.g01046.t01#3    mNeoVis.vulgivaga.1.1.chr1.g01053.t01#4    mNeoVis.3.1.1.chr1.g01045.t01#6    mNeoVis.evergladensis.2.1.chr1.g01043.t01#2    mNeoVis.2.1.1.chr1_hap1.g01050.t01#5   mNeoVis.vulgivaga.2.1.chr1.g01058.t01#1       2538   125259062:   mNeoVis.2.1.1.chr1_hap1.g01051.t01#5    mNeoVis.vulgivaga.2.1.chr1.g01059.t01#1    mNeoVis.evergladensis.1.1.chr1.g01047.t01#3    mNeoVis.3.1.1.chr1.g01046.t01#6   mNeoVis.evergladensis.2.1.chr1.g01044.t01#2       2539   125259063:   mNeoVis.3.1.1.chr7.g01708.t01#6   mNeoVis.vulgivaga.2.1.chr7.g01820.t01#1             2540   125259064:   mNeoVis.evergladensis.2.1.chr7.g01803.t01#2    mNeoVis.2.1.1.chr7_hap1.g01841.t01#5    mNeoVis.vulgivaga.2.1.chr7.g01821.t01#1   mNeoVis.evergladensis.1.1.chr7.g01800.t01#3       2541   125259065:   mNeoVis.3.1.1.chr7.g01710.t01#6    mNeoVis.vulgivaga.1.1.chr7.g01811.t01#4    mNeoVis.vulgivaga.2.1.chr7.g01822.t01#1    mNeoVis.2.1.1.chr7_hap1.g01842.t01#5   mNeoVis.evergladensis.2.1.chr7.g01804.t01#2       2542   125259066:   mNeoVis.vulgivaga.1.1.chr7.g00076.t01#4    mNeoVis.evergladensis.2.1.chr7.g00075.t01#2    mNeoVis.2.1.1.chr7_hap1.g00074.t01#5    mNeoVis.3.1.1.chr7.g00062.t01#6    mNeoVis.vulgivaga.2.1.chr7.g00075.t01#1   mNeoVis.evergladensis.1.1.chr7.g00077.t01#3       2543   125259067:   mNeoVis.evergladensis.2.1.chr1.g01045.t01#2    mNeoVis.2.1.1.chr1_hap1.g01052.t01#5    mNeoVis.vulgivaga.1.1.chr1.g01054.t01#4    mNeoVis.vulgivaga.2.1.chr1.g01060.t01#1    mNeoVis.evergladensis.1.1.chr1.g01048.t01#3   mNeoVis.3.1.1.chr1.g01047.t01#6       2544   125259068:   mNeoVis.evergladensis.2.1.chr7.g01805.t01#2    mNeoVis.vulgivaga.1.1.chr7.g01812.t01#4    mNeoVis.evergladensis.1.1.chr7.g01802.t01#3    mNeoVis.3.1.1.chr7.g01711.t01#6    mNeoVis.vulgivaga.2.1.chr7.g01823.t01#1   mNeoVis.2.1.1.chr7_hap1.g01843.t01#5       2545   125259069:   mNeoVis.vulgivaga.2.1.chr1.g01061.t01#1    mNeoVis.3.1.1.chr1.g01048.t01#6    mNeoVis.evergladensis.2.1.chr1.g01046.t01#2   mNeoVis.2.1.1.chr1_hap1.g01053.t01#5         2546   125259070:   mNeoVis.evergladensis.1.1.chr1.g01049.t01#3    mNeoVis.evergladensis.2.1.chr1.g01047.t01#2    mNeoVis.vulgivaga.1.1.chr1.g01055.t01#4    mNeoVis.3.1.1.chr1.g01049.t01#6    mNeoVis.vulgivaga.2.1.chr1.g01062.t01#1   mNeoVis.2.1.1.chr1_hap1.g01054.t01#5       2547   125259071:   mNeoVis.2.1.1.chr1_hap1.g01055.t01#5    mNeoVis.3.1.1.chr1.g01050.t01#6    mNeoVis.vulgivaga.2.1.chr1.g01063.t01#1    mNeoVis.vulgivaga.1.1.chr1.g01056.t01#4    mNeoVis.evergladensis.2.1.chr1.g01048.t01#2   mNeoVis.evergladensis.1.1.chr1.g01050.t01#3       2548   125259072:   mNeoVis.3.1.1.chr7.g01712.t01#6    mNeoVis.vulgivaga.1.1.chr7.g01813.t01#4    mNeoVis.vulgivaga.2.1.chr7.g01824.t01#1    mNeoVis.evergladensis.1.1.chr7.g01803.t01#3   mNeoVis.evergladensis.2.1.chr7.g01806.t01#2       2549   125259073:   mNeoVis.3.1.1.chr1.g01051.t01#6    mNeoVis.vulgivaga.2.1.chr1.g01064.t01#1    mNeoVis.2.1.1.chr1_hap1.g01056.t01#5    mNeoVis.evergladensis.1.1.chr1.g01051.t01#3    mNeoVis.evergladensis.2.1.chr1.g01049.t01#2   mNeoVis.vulgivaga.1.1.chr1.g01057.t01#4       2550   125259074:   mNeoVis.vulgivaga.1.1.chr1.g01058.t01#4    mNeoVis.2.1.1.chr1_hap1.g01057.t01#5    mNeoVis.vulgivaga.2.1.chr1.g01065.t01#1   mNeoVis.evergladensis.1.1.chr1.g01052.t01#3       2551   125259075:   mNeoVis.vulgivaga.1.1.chr7.g01814.t01#4    mNeoVis.evergladensis.1.1.chr7.g01804.t01#3    mNeoVis.2.1.1.chr7_hap1.g01844.t01#5    mNeoVis.vulgivaga.2.1.chr7.g01825.t01#1    mNeoVis.3.1.1.chr7.g01713.t01#6   mNeoVis.evergladensis.2.1.chr7.g01807.t01#2       2552   125259076:   mNeoVis.evergladensis.1.1.chr1.g01053.t01#3    mNeoVis.3.1.1.chr1.g01052.t01#6    mNeoVis.vulgivaga.2.1.chr1.g01066.t01#1    mNeoVis.2.1.1.chr1_hap1.g01058.t01#5   mNeoVis.vulgivaga.1.1.chr1.g01059.t01#4       2553   125259077:   mNeoVis.2.1.1.chr1_hap1.g01060.t01#5    mNeoVis.3.1.1.chr1.g01057.t01#6    mNeoVis.vulgivaga.2.1.chr1.g01067.t01#1    mNeoVis.vulgivaga.1.1.chr1.g01063.t01#4    mNeoVis.evergladensis.2.1.chr1.g01050.t01#2    mNeoVis.evergladensis.1.1.chr1.g01059.t01#3    mNeoVis.3.1.1.chr1.g01058.t01#6    mNeoVis.vulgivaga.1.1.chr1.g01064.t01#4    mNeoVis.vulgivaga.2.1.chr1.g01073.t01#1      2554   125259078:   mNeoVis.3.1.1.chr1.g01064.t01#6    mNeoVis.vulgivaga.1.1.chr1.g01069.t01#4    mNeoVis.vulgivaga.2.1.chr1.g01079.t01#1    mNeoVis.evergladensis.2.1.chr1.g01058.t01#2    mNeoVis.vulgivaga.2.1.chr1.g01077.t01#1    mNeoVis.3.1.1.chr1.g01062.t01#6    mNeoVis.evergladensis.2.1.chr1.g01060.t01#2    mNeoVis.evergladensis.1.1.chr1.g01063.t01#3    mNeoVis.evergladensis.1.1.chr1.g01065.t01#3      2555   125259079:   mNeoVis.3.1.1.chr7.g01626.t01#6    mNeoVis.vulgivaga.1.1.chr7.g01721.t01#4    mNeoVis.2.1.1.chr7_hap1.g01748.t01#5    mNeoVis.evergladensis.2.1.chr7.g01713.t01#2    mNeoVis.evergladensis.1.1.chr7.g01710.t01#3   mNeoVis.vulgivaga.2.1.chr7.g01727.t01#1       2556   125259080:   mNeoVis.evergladensis.1.1.chr1.g01060.t01#3    mNeoVis.vulgivaga.2.1.chr1.g01074.t01#1    mNeoVis.2.1.1.chr1_hap1.g01070.t01#5   mNeoVis.vulgivaga.1.1.chr1.g01067.t01#4         2557   125259081:   mNeoVis.vulgivaga.1.1.chr1.g00983.t01#4    mNeoVis.vulgivaga.2.1.chr1.g00981.t01#1    mNeoVis.evergladensis.1.1.chr1.g00973.t01#3    mNeoVis.2.1.1.chr1_hap1.g00972.t01#5    mNeoVis.vulgivaga.2.1.chr1.g00980.t01#1    mNeoVis.3.1.1.chr1.g00980.t01#6    mNeoVis.evergladensis.2.1.chr1.g00968.t01#2    mNeoVis.2.1.1.chr1_hap1.g00973.t01#5    mNeoVis.evergladensis.2.1.chr1.g00967.t01#2      2558   125259082:   mNeoVis.2.1.1.chr1_hap1.g01071.t01#5    mNeoVis.3.1.1.chr1.g01060.t01#6    mNeoVis.vulgivaga.2.1.chr1.g01075.t01#1   mNeoVis.evergladensis.1.1.chr1.g01061.t01#3       2559   125259083:   mNeoVis.vulgivaga.1.1.chr1.g01068.t01#4    mNeoVis.3.1.1.chr1.g01061.t01#6    mNeoVis.evergladensis.2.1.chr1.g01057.t01#2    mNeoVis.2.1.1.chr1_hap1.g01072.t01#5    mNeoVis.vulgivaga.2.1.chr1.g01076.t01#1   mNeoVis.evergladensis.1.1.chr1.g01062.t01#3       2560   125259084:   mNeoVis.evergladensis.2.1.chr1.g01059.t01#2    mNeoVis.3.1.1.chr1.g01063.t01#6    mNeoVis.evergladensis.1.1.chr1.g01064.t01#3    mNeoVis.vulgivaga.2.1.chr1.g01078.t01#1    mNeoVis.vulgivaga.1.1.chr1.g01070.t01#4   mNeoVis.2.1.1.chr1_hap1.g01074.t01#5       2561   125259085:   mNeoVis.vulgivaga.1.1.chr1.g01107.t01#4    mNeoVis.2.1.1.chr1_hap1.g00965.t01#5    mNeoVis.vulgivaga.2.1.chr1.g00973.t01#1    mNeoVis.3.1.1.chr1.g01107.t01#6    mNeoVis.evergladensis.2.1.chr1.g01103.t01#2    mNeoVis.vulgivaga.1.1.chr1.g01089.t01#4    mNeoVis.2.1.1.chr1_hap1.g00966.t01#5    mNeoVis.evergladensis.2.1.chr1.g01131.t01#2    mNeoVis.vulgivaga.2.1.chr1.g00972.t01#1      2562   125259086:   mNeoVis.evergladensis.1.1.chr6.g00037.t01#3    mNeoVis.vulgivaga.1.1.chr6.g00040.t01#4    mNeoVis.vulgivaga.2.1.chr6.g00042.t01#1    mNeoVis.2.1.1.chr4_hap1.g00036.t01#5    mNeoVis.3.1.1.chr6.g00039.t01#6   mNeoVis.evergladensis.2.1.chr6.g00038.t01#2       2563   125259087:   mNeoVis.3.1.1.chr1.g01102.t01#6    mNeoVis.3.1.1.chr1.g01068.t01#6   mNeoVis.vulgivaga.1.1.chr1.g01087.t01#4           2564   125259088:   mNeoVis.vulgivaga.1.1.chr1.g01073.t01#4    mNeoVis.evergladensis.2.1.chr1.g01123.t01#2    mNeoVis.evergladensis.1.1.chr1.g01104.t01#3    mNeoVis.3.1.1.chr1.g01090.t01#6    mNeoVis.3.1.1.chr1.g01109.t01#6    mNeoVis.vulgivaga.1.1.chr1.g01109.t01#4    mNeoVis.2.1.1.chr2_hap1.g00834.t01#5    mNeoVis.2.1.1.chr1_hap1.g01078.t01#5    mNeoVis.2.1.1.chr1_hap1.g01113.t01#5      2565   125259089:   mNeoVis.2.1.1.chr1_hap1.g01146.t01#5    mNeoVis.evergladensis.2.1.chr1.g01124.t01#2    mNeoVis.2.1.1.chr1_hap1.g01088.t01#5    mNeoVis.evergladensis.2.1.chr1.g01064.t01#2    mNeoVis.2.1.1.chr1_hap1.g01123.t01#5    mNeoVis.2.1.1.chr2_hap1.g00835.t01#5    mNeoVis.evergladensis.2.1.chr1.g01092.t01#2    mNeoVis.2.1.1.chr2_hap1.g00833.t01#5    mNeoVis.evergladensis.1.1.chr1.g01070.t01#3      2566   125259090:   mNeoVis.vulgivaga.2.1.chr7.g01265.t01#1    mNeoVis.3.1.1.chr7.g01187.t01#6    mNeoVis.vulgivaga.1.1.chr1.g01118.t01#4    mNeoVis.2.1.1.chr9_hap1.g00789.t01#5    mNeoVis.evergladensis.1.1.chr1.g01072.t01#3    mNeoVis.evergladensis.1.1.chr5.g00775.t01#3    mNeoVis.evergladensis.2.1.chr1.g01096.t01#2    mNeoVis.evergladensis.2.1.chr10.g00779.t01#2    mNeoVis.3.1.1.chr10.g00754.t01#6      2567   125259091:   mNeoVis.evergladensis.1.1.chr1.g01094.t01#3    mNeoVis.evergladensis.1.1.chr5.g00776.t01#3    mNeoVis.2.1.1.chr1_hap1.g01147.t01#5    mNeoVis.2.1.1.chr1_hap1.g01089.t01#5    mNeoVis.vulgivaga.1.1.chr1.g01074.t01#4    mNeoVis.2.1.1.chr1_hap1.g01121.t01#5    mNeoVis.2.1.1.chr1_hap1.g01087.t01#5    mNeoVis.evergladensis.1.1.chr1.g00965.t01#3    mNeoVis.2.1.1.chr1_hap1.g01159.t01#5      2568   125259092:   mNeoVis.evergladensis.2.1.chr14.g00230.t01#2    mNeoVis.evergladensis.1.1.chr8.g00141.t01#3    mNeoVis.vulgivaga.2.1.chr12.g00204.t01#1    mNeoVis.evergladensis.1.1.chr1.g01102.t01#3    mNeoVis.vulgivaga.2.1.chr1.g01118.t01#1    mNeoVis.evergladensis.2.1.chr12.g00199.t01#2    mNeoVis.2.1.1.chr8_hap1.g00146.t01#5    mNeoVis.evergladensis.1.1.chr1.g01068.t01#3    mNeoVis.evergladensis.1.1.chr14.g00236.t01#3      2569   125259093:   mNeoVis.3.1.1.chr1.g01097.t01#6    mNeoVis.vulgivaga.2.1.chr1.g01111.t01#1    mNeoVis.3.1.1.chr1.g01092.t01#6    mNeoVis.vulgivaga.2.1.chr1.g01114.t01#1    mNeoVis.evergladensis.1.1.chr1.g01107.t01#3    mNeoVis.evergladensis.1.1.chr1.g01099.t01#3    mNeoVis.3.1.1.chr1.g01101.t01#6    mNeoVis.evergladensis.2.1.chr1.g01095.t01#2    mNeoVis.2.1.1.chr1_hap1.g01122.t01#5      2570   125259094:   mNeoVis.vulgivaga.1.1.chr1.g01086.t01#4    mNeoVis.vulgivaga.2.1.chr1.g01132.t01#1    mNeoVis.vulgivaga.2.1.chr5.g00775.t01#1    mNeoVis.evergladensis.2.1.chr1.g01116.t01#2    mNeoVis.vulgivaga.2.1.chr10.g00767.t01#1    mNeoVis.evergladensis.1.1.chr1.g01114.t01#3    mNeoVis.vulgivaga.2.1.chr10.g00766.t01#1   mNeoVis.vulgivaga.2.1.chr1.g01084.t01#1     2571   125259095:   mNeoVis.2.1.1.chr1_hap1.g01081.t01#5    mNeoVis.3.1.1.chr1.g01067.t01#6    mNeoVis.vulgivaga.2.1.chr1.g01083.t01#1    mNeoVis.evergladensis.1.1.chr1.g01071.t01#3    mNeoVis.evergladensis.2.1.chr1.g01065.t01#2   mNeoVis.vulgivaga.1.1.chr1.g01075.t01#4       2572   125259096:   mNeoVis.3.1.1.chr1.g01105.t01#6    mNeoVis.vulgivaga.1.1.chr1.g01117.t01#4    mNeoVis.2.1.1.chr1_hap1.g01145.t01#5    mNeoVis.evergladensis.1.1.chr1.g01113.t01#3   mNeoVis.evergladensis.2.1.chr1.g01115.t01#2       2573   125259097:   mNeoVis.vulgivaga.1.1.chr1.g01120.t01#4    mNeoVis.vulgivaga.2.1.chr1.g01117.t01#1    mNeoVis.evergladensis.2.1.chr1.g01118.t01#2    mNeoVis.3.1.1.chr1.g01066.t01#6   mNeoVis.evergladensis.2.1.chr1.g01068.t01#2       2574   125259098:   mNeoVis.2.1.1.chr1_hap1.g01130.t01#5    mNeoVis.vulgivaga.2.1.chr1.g01107.t01#1    mNeoVis.3.1.1.chr1.g01095.t01#6   mNeoVis.evergladensis.1.1.chr1.g01103.t01#3       2575   125259099:   mNeoVis.vulgivaga.1.1.chr1.g01113.t01#4    mNeoVis.2.1.1.chr1_hap1.g01139.t01#5    mNeoVis.evergladensis.2.1.chr1.g01109.t01#2   mNeoVis.evergladensis.1.1.chr1.g01109.t01#3       2576   125259100:   mNeoVis.vulgivaga.1.1.chr1.g01110.t01#4    mNeoVis.2.1.1.chr1_hap1.g01135.t01#5   mNeoVis.evergladensis.2.1.chr1.g01106.t01#2         2577   125259101:   mNeoVis.evergladensis.2.1.chr1.g01061.t01#2   mNeoVis.vulgivaga.1.1.chr1.g01071.t01#4             2578   125259102:   mNeoVis.vulgivaga.1.1.chr5.g00757.t01#4   mNeoVis.3.1.1.chr5.g00773.t01#6             2579   125259103:   mNeoVis.3.1.1.chr1.g01108.t01#6   mNeoVis.vulgivaga.1.1.chr1.g01072.t01#4             2580   125259104:   mNeoVis.vulgivaga.2.1.chr1.g01106.t01#1   mNeoVis.vulgivaga.1.1.chr5.g00762.t01#4             2581   125259105:  mNeoVis.evergladensis.1.1.chr1.g01084.t01#3             2582   125259106:   mNeoVis.evergladensis.1.1.chr7.g01711.t01#3    mNeoVis.vulgivaga.2.1.chr7.g01728.t01#1    mNeoVis.3.1.1.chr7.g01627.t01#6    mNeoVis.evergladensis.2.1.chr7.g01714.t01#2    mNeoVis.2.1.1.chr7_hap1.g01749.t01#5   mNeoVis.vulgivaga.1.1.chr7.g01722.t01#4       2583   125259107:   mNeoVis.3.1.1.chr7.g01628.t01#6    mNeoVis.evergladensis.2.1.chr7.g01715.t01#2    mNeoVis.2.1.1.chr7_hap1.g01750.t01#5    mNeoVis.vulgivaga.1.1.chr7.g01723.t01#4    mNeoVis.evergladensis.1.1.chr7.g01712.t01#3   mNeoVis.vulgivaga.2.1.chr7.g01729.t01#1       2584   125259108:   mNeoVis.vulgivaga.1.1.chr7.g01724.t01#4    mNeoVis.evergladensis.2.1.chr7.g01716.t01#2    mNeoVis.3.1.1.chr7.g01629.t01#6    mNeoVis.vulgivaga.2.1.chr7.g01730.t01#1    mNeoVis.2.1.1.chr7_hap1.g01751.t01#5   mNeoVis.evergladensis.1.1.chr7.g01713.t01#3       2585   125259109:   mNeoVis.evergladensis.1.1.chr7.g01714.t01#3    mNeoVis.vulgivaga.2.1.chr7.g01731.t01#1    mNeoVis.vulgivaga.1.1.chr7.g01725.t01#4    mNeoVis.evergladensis.2.1.chr7.g01717.t01#2    mNeoVis.3.1.1.chr7.g01630.t01#6   mNeoVis.2.1.1.chr7_hap1.g01752.t01#5       2586   125259110:   mNeoVis.2.1.1.chr15_hap1.g00165.t01#5    mNeoVis.vulgivaga.2.1.chr15.g00151.t01#1    mNeoVis.evergladensis.2.1.chr15.g00148.t01#2    mNeoVis.vulgivaga.1.1.chr15.g00147.t01#4    mNeoVis.evergladensis.1.1.chr15.g00465.t01#3   mNeoVis.3.1.1.chr15.g00141.t01#6       2587   125259111:   mNeoVis.3.1.1.chr7.g00063.t01#6    mNeoVis.2.1.1.chr7_hap1.g00075.t01#5    mNeoVis.evergladensis.2.1.chr7.g00076.t01#2    mNeoVis.vulgivaga.2.1.chr7.g00076.t01#1    mNeoVis.evergladensis.1.1.chr7.g00078.t01#3   mNeoVis.vulgivaga.1.1.chr7.g00077.t01#4       2588   125259112:  mNeoVis.vulgivaga.2.1.chr1.g01089.t01#1               2589   125259113:   mNeoVis.2.1.1.chr1_hap1.g01093.t01#5    mNeoVis.vulgivaga.2.1.chr1.g01090.t01#1    mNeoVis.evergladensis.1.1.chr1.g01076.t01#3    mNeoVis.3.1.1.chr1.g01072.t01#6    mNeoVis.vulgivaga.1.1.chr1.g01082.t01#4   mNeoVis.evergladensis.2.1.chr1.g01074.t01#2       2590   125259114:   mNeoVis.vulgivaga.2.1.chr1.g01091.t01#1    mNeoVis.3.1.1.chr1.g01076.t01#6    mNeoVis.evergladensis.2.1.chr1.g01076.t01#2    mNeoVis.2.1.1.chr1_hap1.g01095.t01#5    mNeoVis.evergladensis.1.1.chr1.g01078.t01#3    mNeoVis.vulgivaga.2.1.chr1.g01092.t01#1   mNeoVis.vulgivaga.1.1.chr1.g01083.t01#4     2591   125259115:   mNeoVis.vulgivaga.1.1.chr1.g01084.t01#4    mNeoVis.vulgivaga.2.1.chr1.g01093.t01#1    mNeoVis.evergladensis.1.1.chr1.g01079.t01#3    mNeoVis.evergladensis.2.1.chr1.g01077.t01#2    mNeoVis.3.1.1.chr1.g01077.t01#6   mNeoVis.2.1.1.chr1_hap1.g01096.t01#5       2592   125259116:   mNeoVis.2.1.1.chr14_hap1.g00005.t01#5    mNeoVis.vulgivaga.2.1.chr14.g00005.t01#1    mNeoVis.vulgivaga.1.1.chr14.g00005.t01#4    mNeoVis.evergladensis.1.1.chr14.g00005.t01#3   mNeoVis.evergladensis.2.1.chr14.g00005.t01#2       2593   125259117:   mNeoVis.vulgivaga.2.1.chr1.g01094.t01#1    mNeoVis.evergladensis.1.1.chr1.g01080.t01#3    mNeoVis.2.1.1.chr1_hap1.g01097.t01#5    mNeoVis.evergladensis.2.1.chr1.g01078.t01#2   mNeoVis.vulgivaga.1.1.chr1.g01085.t01#4       2594   125259118:   mNeoVis.2.1.1.chr7_hap1.g00077.t01#5    mNeoVis.evergladensis.1.1.chr7.g00079.t01#3    mNeoVis.evergladensis.2.1.chr7.g00077.t01#2    mNeoVis.3.1.1.chr7.g00064.t01#6    mNeoVis.vulgivaga.2.1.chr7.g00077.t01#1   mNeoVis.vulgivaga.1.1.chr7.g00079.t01#4       2595   125259119:   mNeoVis.vulgivaga.2.1.chr7.g01732.t01#1    mNeoVis.2.1.1.chr7_hap1.g01753.t01#5    mNeoVis.vulgivaga.1.1.chr7.g01726.t01#4    mNeoVis.evergladensis.1.1.chr7.g01715.t01#3    mNeoVis.3.1.1.chr7.g01631.t01#6   mNeoVis.evergladensis.2.1.chr7.g01718.t01#2       2596   125259120:   mNeoVis.vulgivaga.2.1.chr1.g01097.t01#1    mNeoVis.2.1.1.chr1_hap1.g01103.t01#5    mNeoVis.3.1.1.chr1.g01080.t01#6    mNeoVis.evergladensis.1.1.chr1.g01086.t01#3    mNeoVis.vulgivaga.1.1.chr1.g01090.t01#4   mNeoVis.evergladensis.2.1.chr1.g01083.t01#2       2597   125259121:   mNeoVis.evergladensis.1.1.chr1.g01087.t01#3    mNeoVis.2.1.1.chr1_hap1.g01104.t01#5    mNeoVis.evergladensis.2.1.chr1.g01084.t01#2    mNeoVis.vulgivaga.2.1.chr1.g01098.t01#1    mNeoVis.vulgivaga.1.1.chr1.g01091.t01#4   mNeoVis.3.1.1.chr1.g01081.t01#6       2598   125259122:   mNeoVis.2.1.1.chr7_hap1.g01754.t01#5    mNeoVis.vulgivaga.1.1.chr7.g01727.t01#4    mNeoVis.evergladensis.1.1.chr7.g01716.t01#3    mNeoVis.3.1.1.chr7.g01632.t01#6    mNeoVis.vulgivaga.2.1.chr7.g01733.t01#1   mNeoVis.evergladensis.2.1.chr7.g01719.t01#2       2599   125259123:   mNeoVis.vulgivaga.2.1.chr1.g01099.t01#1    mNeoVis.evergladensis.1.1.chr12.g00315.t01#3    mNeoVis.evergladensis.2.1.chr1.g01085.t01#2    mNeoVis.2.1.1.chr12_hap1.g00349.t01#5    mNeoVis.3.1.1.chr1.g01082.t01#6   mNeoVis.2.1.1.chr1_hap1.g01105.t01#5       2600   125259124:   mNeoVis.evergladensis.2.1.chr1.g01086.t01#2    mNeoVis.vulgivaga.2.1.chr1.g01100.t01#1    mNeoVis.3.1.1.chr1.g01084.t01#6    mNeoVis.2.1.1.chr1_hap1.g01108.t01#5    mNeoVis.vulgivaga.1.1.chr1.g01093.t01#4   mNeoVis.evergladensis.1.1.chr1.g01088.t01#3       2601   125259125:   mNeoVis.vulgivaga.1.1.chr5.g01552.t01#4    mNeoVis.3.1.1.chr7.g00627.t01#6    mNeoVis.vulgivaga.2.1.chr7.g00662.t01#1    mNeoVis.vulgivaga.1.1.chr7.g00667.t01#4    mNeoVis.evergladensis.1.1.chr7.g00668.t01#3    mNeoVis.evergladensis.2.1.chr5.g01574.t01#2    mNeoVis.vulgivaga.2.1.chr5.g01576.t01#1    mNeoVis.evergladensis.1.1.chr5.g01579.t01#3    mNeoVis.evergladensis.2.1.chr7.g00659.t01#2      2602   125259126:   mNeoVis.2.1.1.chr1_hap1.g01109.t01#5    mNeoVis.vulgivaga.1.1.chr1.g01094.t01#4    mNeoVis.evergladensis.1.1.chr1.g01089.t01#3    mNeoVis.vulgivaga.2.1.chr1.g01101.t01#1    mNeoVis.evergladensis.2.1.chr1.g01087.t01#2   mNeoVis.3.1.1.chr1.g01085.t01#6       2603   125259127:   mNeoVis.3.1.1.chr10.g00979.t01#6    mNeoVis.vulgivaga.2.1.chr10.g00999.t01#1    mNeoVis.evergladensis.1.1.chr10.g01007.t01#3    mNeoVis.2.1.1.chr9_hap1.g01018.t01#5    mNeoVis.evergladensis.2.1.chr10.g01005.t01#2   mNeoVis.vulgivaga.1.1.chr10.g00987.t01#4       2604   125259128:   mNeoVis.vulgivaga.2.1.chr1.g01102.t01#1    mNeoVis.evergladensis.1.1.chr1.g01090.t01#3    mNeoVis.evergladensis.2.1.chr1.g01088.t01#2    mNeoVis.3.1.1.chr1.g01086.t01#6   mNeoVis.vulgivaga.1.1.chr1.g01095.t01#4       2605   125259129:   mNeoVis.vulgivaga.1.1.chr8.g00842.t01#4    mNeoVis.evergladensis.2.1.chr7.g01720.t01#2    mNeoVis.2.1.1.chr8_hap1.g00870.t01#5    mNeoVis.evergladensis.2.1.chr9.g00858.t01#2    mNeoVis.evergladensis.1.1.chr8.g00851.t01#3    mNeoVis.2.1.1.chr7_hap1.g01755.t01#5    mNeoVis.vulgivaga.1.1.chr7.g01728.t01#4    mNeoVis.3.1.1.chr7.g01633.t01#6    mNeoVis.evergladensis.1.1.chr7.g01717.t01#3      2606   125259130:   mNeoVis.evergladensis.2.1.chr1.g01089.t01#2    mNeoVis.3.1.1.chr1.g01087.t01#6    mNeoVis.vulgivaga.2.1.chr1.g01103.t01#1    mNeoVis.evergladensis.1.1.chr1.g01091.t01#3    mNeoVis.2.1.1.chr1_hap1.g01112.t01#5   mNeoVis.vulgivaga.1.1.chr1.g01096.t01#4       2607   125259131:   mNeoVis.evergladensis.1.1.chr7.g01718.t01#3    mNeoVis.3.1.1.chr7.g01634.t01#6    mNeoVis.evergladensis.2.1.chr7.g01721.t01#2    mNeoVis.2.1.1.chr7_hap1.g01756.t01#5    mNeoVis.vulgivaga.2.1.chr7.g01735.t01#1   mNeoVis.vulgivaga.1.1.chr7.g01729.t01#4       2608   125259132:   mNeoVis.2.1.1.chr7_hap1.g01757.t01#5    mNeoVis.vulgivaga.1.1.chr7.g01730.t01#4    mNeoVis.evergladensis.1.1.chr7.g01719.t01#3    mNeoVis.3.1.1.chr7.g01635.t01#6    mNeoVis.evergladensis.2.1.chr7.g01722.t01#2   mNeoVis.vulgivaga.2.1.chr7.g01736.t01#1       2609   125259133:   mNeoVis.evergladensis.1.1.chr7.g01720.t01#3    mNeoVis.vulgivaga.1.1.chr7.g01731.t01#4    mNeoVis.vulgivaga.2.1.chr7.g01737.t01#1    mNeoVis.2.1.1.chr7_hap1.g01758.t01#5    mNeoVis.3.1.1.chr7.g01636.t01#6   mNeoVis.evergladensis.2.1.chr7.g01723.t01#2       2610   125259134:   mNeoVis.vulgivaga.2.1.chr7.g01746.t01#1    mNeoVis.2.1.1.chr7_hap1.g01767.t01#5    mNeoVis.evergladensis.2.1.chr7.g01732.t01#2    mNeoVis.2.1.1.chr7_hap1.g01771.t01#5    mNeoVis.vulgivaga.1.1.chr7.g01737.t01#4    mNeoVis.evergladensis.2.1.chr7.g01728.t01#2    mNeoVis.vulgivaga.2.1.chr7.g01744.t01#1    mNeoVis.2.1.1.chr7_hap1.g01766.t01#5    mNeoVis.vulgivaga.1.1.chr7.g01740.t01#4      2611   125259135:   mNeoVis.3.1.1.chr7.g01642.t01#6    mNeoVis.evergladensis.2.1.chr7.g01729.t01#2    mNeoVis.evergladensis.1.1.chr7.g01726.t01#3    mNeoVis.vulgivaga.1.1.chr7.g01738.t01#4    mNeoVis.2.1.1.chr7_hap1.g01768.t01#5   mNeoVis.vulgivaga.2.1.chr7.g01745.t01#1       2612   125259136:   mNeoVis.3.1.1.chr7.g01637.t01#6    mNeoVis.vulgivaga.2.1.chr7.g01738.t01#1    mNeoVis.vulgivaga.1.1.chr7.g01732.t01#4   mNeoVis.2.1.1.chr7_hap1.g01760.t01#5         2613   125259137:   mNeoVis.vulgivaga.2.1.chr7.g01743.t01#1   mNeoVis.3.1.1.chr7.g01641.t01#6             2614   125259138:   mNeoVis.vulgivaga.2.1.chr7.g01739.t01#1    mNeoVis.2.1.1.chr7_hap1.g01759.t01#5    mNeoVis.evergladensis.2.1.chr7.g01724.t01#2    mNeoVis.2.1.1.chr7_hap1.g01762.t01#5    mNeoVis.evergladensis.1.1.chr7.g01721.t01#3   mNeoVis.2.1.1.chr7_hap1.g01761.t01#5       2615   125259139:   mNeoVis.vulgivaga.1.1.chr7.g01734.t01#4    mNeoVis.2.1.1.chr7_hap1.g01763.t01#5    mNeoVis.evergladensis.1.1.chr7.g01722.t01#3    mNeoVis.3.1.1.chr7.g01638.t01#6    mNeoVis.vulgivaga.2.1.chr7.g01740.t01#1   mNeoVis.evergladensis.2.1.chr7.g01725.t01#2       2616   125259140:   mNeoVis.evergladensis.1.1.chr7.g01723.t01#3    mNeoVis.3.1.1.chr7.g01639.t01#6    mNeoVis.vulgivaga.1.1.chr7.g01735.t01#4    mNeoVis.evergladensis.2.1.chr7.g01726.t01#2    mNeoVis.2.1.1.chr7_hap1.g01764.t01#5   mNeoVis.vulgivaga.2.1.chr7.g01741.t01#1       2617   125259141:   mNeoVis.vulgivaga.2.1.chr1.g01115.t01#1    mNeoVis.vulgivaga.1.1.chr1.g01116.t01#4    mNeoVis.3.1.1.chr1.g01104.t01#6    mNeoVis.2.1.1.chr1_hap1.g01144.t01#5    mNeoVis.evergladensis.1.1.chr1.g01112.t01#3   mNeoVis.evergladensis.2.1.chr1.g01114.t01#2       2618   125259142:   mNeoVis.evergladensis.2.1.chr7.g01727.t01#2    mNeoVis.3.1.1.chr7.g01640.t01#6    mNeoVis.vulgivaga.1.1.chr7.g01736.t01#4    mNeoVis.2.1.1.chr7_hap1.g01765.t01#5    mNeoVis.vulgivaga.2.1.chr7.g01742.t01#1   mNeoVis.evergladensis.1.1.chr7.g01724.t01#3       2619   125259143:   mNeoVis.evergladensis.1.1.chr7.g01727.t01#3    mNeoVis.evergladensis.2.1.chr7.g01731.t01#2    mNeoVis.2.1.1.chr7_hap1.g01770.t01#5    mNeoVis.vulgivaga.2.1.chr7.g01747.t01#1   mNeoVis.vulgivaga.1.1.chr7.g01739.t01#4       2620   125259144:  mNeoVis.vulgivaga.2.1.chr1.g01120.t01#1               2621   125259145:   mNeoVis.vulgivaga.2.1.chr1.g01123.t01#1    mNeoVis.3.1.1.chr1.g01110.t01#6    mNeoVis.evergladensis.1.1.chr1.g01120.t01#3    mNeoVis.vulgivaga.1.1.chr1.g01123.t01#4   mNeoVis.evergladensis.2.1.chr1.g01125.t01#2       2622   125259146:   mNeoVis.evergladensis.2.1.chr1.g01126.t01#2    mNeoVis.vulgivaga.2.1.chr1.g01124.t01#1    mNeoVis.vulgivaga.1.1.chr1.g01124.t01#4    mNeoVis.2.1.1.chr1_hap1.g01153.t01#5    mNeoVis.3.1.1.chr1.g01112.t01#6   mNeoVis.evergladensis.1.1.chr1.g01121.t01#3       2623   125259147:   mNeoVis.vulgivaga.2.1.chr1.g01125.t01#1    mNeoVis.vulgivaga.1.1.chr1.g01125.t01#4    mNeoVis.2.1.1.chr1_hap1.g01154.t01#5    mNeoVis.3.1.1.chr1.g01113.t01#6    mNeoVis.evergladensis.1.1.chr1.g01122.t01#3   mNeoVis.evergladensis.2.1.chr1.g01127.t01#2       2624   125259148:   mNeoVis.vulgivaga.2.1.chr7.g01748.t01#1    mNeoVis.vulgivaga.1.1.chr7.g01741.t01#4    mNeoVis.evergladensis.1.1.chr7.g01728.t01#3    mNeoVis.evergladensis.2.1.chr7.g01733.t01#2   mNeoVis.3.1.1.chr7.g01643.t01#6       2625   125259149:   mNeoVis.3.1.1.chr7.g01644.t01#6    mNeoVis.vulgivaga.1.1.chr7.g01742.t01#4    mNeoVis.evergladensis.1.1.chr7.g01729.t01#3    mNeoVis.vulgivaga.2.1.chr7.g01749.t01#1   mNeoVis.2.1.1.chr7_hap1.g01773.t01#5       2626   125259150:   mNeoVis.vulgivaga.1.1.chr7.g01743.t01#4    mNeoVis.3.1.1.chr7.g01645.t01#6    mNeoVis.evergladensis.2.1.chr7.g01734.t01#2    mNeoVis.evergladensis.1.1.chr7.g01730.t01#3    mNeoVis.vulgivaga.2.1.chr7.g01750.t01#1   mNeoVis.2.1.1.chr7_hap1.g01772.t01#5       2627   125259151:   mNeoVis.evergladensis.2.1.chr12.g00404.t01#2    mNeoVis.evergladensis.1.1.chr12.g00411.t01#3    mNeoVis.vulgivaga.2.1.chr12.g00412.t01#1    mNeoVis.evergladensis.2.1.chr12.g00120.t01#2    mNeoVis.3.1.1.chr12.g00409.t01#6    mNeoVis.2.1.1.chr12_hap1.g00144.t01#5    mNeoVis.vulgivaga.2.1.chr12.g00120.t01#1    mNeoVis.vulgivaga.1.1.chr12.g00126.t01#4    mNeoVis.evergladensis.1.1.chr12.g00119.t01#3      2628   125259152:   mNeoVis.vulgivaga.2.1.chr7.g01752.t01#1    mNeoVis.vulgivaga.1.1.chr7.g01745.t01#4    mNeoVis.evergladensis.2.1.chr7.g01736.t01#2    mNeoVis.3.1.1.chr7.g01646.t01#6    mNeoVis.2.1.1.chr7_hap1.g01775.t01#5   mNeoVis.evergladensis.1.1.chr7.g01732.t01#3       2629   125259153:   mNeoVis.evergladensis.2.1.chr7.g01737.t01#2    mNeoVis.3.1.1.chr7.g01647.t01#6    mNeoVis.2.1.1.chr7_hap1.g01776.t01#5    mNeoVis.evergladensis.1.1.chr7.g01733.t01#3    mNeoVis.vulgivaga.2.1.chr7.g01753.t01#1   mNeoVis.vulgivaga.1.1.chr7.g01746.t01#4       2630   125259154:   mNeoVis.3.1.1.chr7.g00065.t01#6    mNeoVis.vulgivaga.2.1.chr7.g00078.t01#1    mNeoVis.vulgivaga.1.1.chr7.g00080.t01#4    mNeoVis.2.1.1.chr7_hap1.g00078.t01#5    mNeoVis.evergladensis.2.1.chr7.g00078.t01#2   mNeoVis.evergladensis.1.1.chr7.g00080.t01#3       2631   125259155:   mNeoVis.evergladensis.1.1.chr1.g01123.t01#3    mNeoVis.2.1.1.chr1_hap1.g01155.t01#5    mNeoVis.evergladensis.2.1.chr1.g01128.t01#2    mNeoVis.3.1.1.chr1.g01114.t01#6    mNeoVis.vulgivaga.1.1.chr1.g01126.t01#4   mNeoVis.vulgivaga.2.1.chr1.g01126.t01#1       2632   125259156:   mNeoVis.evergladensis.2.1.chr7.g01738.t01#2    mNeoVis.3.1.1.chr7.g01648.t01#6    mNeoVis.2.1.1.chr7_hap1.g01777.t01#5    mNeoVis.evergladensis.1.1.chr7.g01734.t01#3    mNeoVis.vulgivaga.2.1.chr7.g01754.t01#1   mNeoVis.vulgivaga.1.1.chr7.g01747.t01#4       2633   125259157:   mNeoVis.vulgivaga.1.1.chr7.g01748.t01#4    mNeoVis.evergladensis.2.1.chr7.g01739.t01#2    mNeoVis.vulgivaga.2.1.chr7.g01755.t01#1    mNeoVis.3.1.1.chr7.g01649.t01#6    mNeoVis.2.1.1.chr7_hap1.g01778.t01#5   mNeoVis.evergladensis.1.1.chr7.g01735.t01#3       2634   125259158:   mNeoVis.vulgivaga.1.1.chr1.g01127.t01#4    mNeoVis.evergladensis.2.1.chr1.g01129.t01#2    mNeoVis.vulgivaga.2.1.chr1.g01127.t01#1    mNeoVis.2.1.1.chr1_hap1.g01156.t01#5    mNeoVis.3.1.1.chr1.g01115.t01#6   mNeoVis.evergladensis.1.1.chr1.g01124.t01#3       2635   125259159:  mNeoVis.vulgivaga.2.1.chr1.g01128.t01#1               2636   125259160:   mNeoVis.vulgivaga.2.1.chr7.g01756.t01#1    mNeoVis.evergladensis.2.1.chr7.g01740.t01#2    mNeoVis.vulgivaga.1.1.chr7.g01749.t01#4    mNeoVis.3.1.1.chr7.g01650.t01#6    mNeoVis.2.1.1.chr7_hap1.g01779.t01#5   mNeoVis.evergladensis.1.1.chr7.g01736.t01#3       2637   125259161:   mNeoVis.2.1.1.chr1_hap1.g01162.t01#5    mNeoVis.vulgivaga.1.1.chr1.g01132.t01#4    mNeoVis.3.1.1.chr1.g01119.t01#6    mNeoVis.vulgivaga.2.1.chr1.g01133.t01#1    mNeoVis.evergladensis.2.1.chr1.g01135.t01#2   mNeoVis.evergladensis.1.1.chr1.g01129.t01#3       2638   125259162:   mNeoVis.vulgivaga.2.1.chr1.g01134.t01#1    mNeoVis.2.1.1.chr1_hap1.g01163.t01#5    mNeoVis.evergladensis.2.1.chr1.g01136.t01#2   mNeoVis.3.1.1.chr1.g01120.t01#6         2639   125259163:   mNeoVis.2.1.1.chr7_hap1.g01780.t01#5    mNeoVis.evergladensis.2.1.chr7.g01741.t01#2    mNeoVis.evergladensis.1.1.chr7.g01737.t01#3   mNeoVis.vulgivaga.2.1.chr7.g01757.t01#1         2640   125259164:   mNeoVis.vulgivaga.1.1.chr7.g01750.t01#4    mNeoVis.vulgivaga.2.1.chr7.g01758.t01#1    mNeoVis.vulgivaga.2.1.chr7.g01759.t01#1   mNeoVis.3.1.1.chr7.g01652.t01#6         2641   125259165:   mNeoVis.vulgivaga.1.1.chr7.g01751.t01#4    mNeoVis.2.1.1.chr7_hap1.g01781.t01#5    mNeoVis.evergladensis.2.1.chr7.g01742.t01#2    mNeoVis.vulgivaga.2.1.chr7.g01760.t01#1    mNeoVis.evergladensis.1.1.chr7.g01738.t01#3   mNeoVis.3.1.1.chr7.g01651.t01#6       2642   125259166:   mNeoVis.vulgivaga.2.1.chr7.g01761.t01#1    mNeoVis.evergladensis.1.1.chr7.g01739.t01#3    mNeoVis.evergladensis.2.1.chr7.g01744.t01#2    mNeoVis.2.1.1.chr7_hap1.g01782.t01#5    mNeoVis.vulgivaga.1.1.chr7.g01752.t01#4    mNeoVis.vulgivaga.2.1.chr7.g01762.t01#1    mNeoVis.3.1.1.chr7.g01653.t01#6    mNeoVis.evergladensis.1.1.chr7.g01740.t01#3    mNeoVis.evergladensis.2.1.chr7.g01743.t01#2      2643   125259167:   mNeoVis.3.1.1.chr7.g01654.t01#6    mNeoVis.evergladensis.2.1.chr7.g01745.t01#2    mNeoVis.evergladensis.1.1.chr7.g01741.t01#3    mNeoVis.2.1.1.chr7_hap1.g01783.t01#5    mNeoVis.vulgivaga.2.1.chr7.g01763.t01#1   mNeoVis.vulgivaga.1.1.chr7.g01753.t01#4       2644   125259168:   mNeoVis.evergladensis.1.1.chr7.g00081.t01#3    mNeoVis.evergladensis.2.1.chr7.g00079.t01#2    mNeoVis.3.1.1.chr7.g00066.t01#6    mNeoVis.2.1.1.chr7_hap1.g00079.t01#5    mNeoVis.vulgivaga.2.1.chr7.g00079.t01#1   mNeoVis.vulgivaga.1.1.chr7.g00081.t01#4       2645   125259169:   mNeoVis.2.1.1.chr7_hap1.g01784.t01#5    mNeoVis.evergladensis.1.1.chr7.g01742.t01#3    mNeoVis.vulgivaga.1.1.chr7.g01754.t01#4    mNeoVis.vulgivaga.2.1.chr7.g01764.t01#1   mNeoVis.3.1.1.chr7.g01655.t01#6       2646   125259170:   mNeoVis.2.1.1.chr7_hap1.g01785.t01#5    mNeoVis.evergladensis.1.1.chr7.g01743.t01#3    mNeoVis.vulgivaga.1.1.chr7.g01755.t01#4    mNeoVis.vulgivaga.2.1.chr7.g01766.t01#1    mNeoVis.vulgivaga.2.1.chr7.g01765.t01#1    mNeoVis.3.1.1.chr7.g01656.t01#6    mNeoVis.evergladensis.2.1.chr7.g01746.t01#2    mNeoVis.vulgivaga.1.1.chr7.g01756.t01#4    mNeoVis.2.1.1.chr7_hap1.g01787.t01#5      2647   125259171:   mNeoVis.vulgivaga.1.1.chr7.g01757.t01#4    mNeoVis.evergladensis.1.1.chr7.g01745.t01#3    mNeoVis.2.1.1.chr7_hap1.g01788.t01#5    mNeoVis.evergladensis.2.1.chr7.g01749.t01#2    mNeoVis.vulgivaga.2.1.chr7.g01767.t01#1   mNeoVis.3.1.1.chr7.g01657.t01#6       2648   125259172:   mNeoVis.3.1.1.chr7.g01658.t01#6    mNeoVis.vulgivaga.2.1.chr7.g01768.t01#1    mNeoVis.evergladensis.1.1.chr7.g01746.t01#3    mNeoVis.2.1.1.chr7_hap1.g01789.t01#5    mNeoVis.vulgivaga.1.1.chr7.g01758.t01#4   mNeoVis.evergladensis.2.1.chr7.g01750.t01#2       2649   125259173:  mNeoVis.vulgivaga.2.1.chr7.g01769.t01#1               2650   125259174:  mNeoVis.vulgivaga.2.1.chr1.g01135.t01#1               2651   125259175:   mNeoVis.evergladensis.2.1.chr7.g01752.t01#2    mNeoVis.2.1.1.chr7_hap1.g01791.t01#5    mNeoVis.3.1.1.chr7.g01660.t01#6    mNeoVis.evergladensis.1.1.chr7.g01748.t01#3    mNeoVis.vulgivaga.2.1.chr7.g01770.t01#1   mNeoVis.vulgivaga.1.1.chr7.g01760.t01#4       2652   125259176:   mNeoVis.evergladensis.1.1.chr1.g01134.t01#3    mNeoVis.2.1.1.chr1_hap1.g01164.t01#5    mNeoVis.evergladensis.2.1.chr1.g01138.t01#2    mNeoVis.vulgivaga.1.1.chr1.g01135.t01#4    mNeoVis.3.1.1.chr1.g01122.t01#6   mNeoVis.vulgivaga.2.1.chr1.g01136.t01#1       2653   125259177:   mNeoVis.vulgivaga.1.1.chr7.g01761.t01#4    mNeoVis.vulgivaga.2.1.chr7.g01771.t01#1    mNeoVis.evergladensis.2.1.chr7.g01753.t01#2    mNeoVis.2.1.1.chr7_hap1.g01792.t01#5    mNeoVis.evergladensis.1.1.chr7.g01749.t01#3   mNeoVis.3.1.1.chr7.g01661.t01#6       2654   125259178:   mNeoVis.3.1.1.chr7.g01662.t01#6    mNeoVis.vulgivaga.1.1.chr7.g01762.t01#4    mNeoVis.2.1.1.chr7_hap1.g01793.t01#5    mNeoVis.vulgivaga.2.1.chr7.g01772.t01#1    mNeoVis.evergladensis.2.1.chr7.g01754.t01#2   mNeoVis.evergladensis.1.1.chr7.g01750.t01#3       2655   125259179:   mNeoVis.2.1.1.chr9_hap1.g00772.t01#5    mNeoVis.vulgivaga.2.1.chr10.g00750.t01#1    mNeoVis.evergladensis.1.1.chr10.g00762.t01#3    mNeoVis.3.1.1.chr10.g00738.t01#6    mNeoVis.vulgivaga.1.1.chr10.g00747.t01#4   mNeoVis.evergladensis.2.1.chr10.g00762.t01#2       2656   125259180:   mNeoVis.evergladensis.1.1.chr7.g01751.t01#3    mNeoVis.3.1.1.chr7.g01663.t01#6    mNeoVis.evergladensis.2.1.chr7.g01755.t01#2    mNeoVis.vulgivaga.2.1.chr7.g01773.t01#1    mNeoVis.vulgivaga.1.1.chr7.g01763.t01#4   mNeoVis.2.1.1.chr7_hap1.g01794.t01#5       2657   125259181:   mNeoVis.vulgivaga.1.1.chr15.g00148.t01#4    mNeoVis.3.1.1.chr15.g00142.t01#6    mNeoVis.2.1.1.chr15_hap1.g00166.t01#5    mNeoVis.vulgivaga.2.1.chr15.g00152.t01#1   mNeoVis.evergladensis.2.1.chr15.g00149.t01#2       2658   125259182:   mNeoVis.evergladensis.2.1.chr1.g01139.t01#2    mNeoVis.evergladensis.1.1.chr1.g01135.t01#3    mNeoVis.2.1.1.chr1_hap1.g01165.t01#5    mNeoVis.3.1.1.chr1.g01123.t01#6    mNeoVis.vulgivaga.1.1.chr1.g01136.t01#4   mNeoVis.vulgivaga.2.1.chr1.g01137.t01#1       2659   125259183:   mNeoVis.2.1.1.chr7_hap1.g01795.t01#5    mNeoVis.evergladensis.1.1.chr7.g01752.t01#3    mNeoVis.vulgivaga.2.1.chr7.g01774.t01#1    mNeoVis.evergladensis.2.1.chr7.g01756.t01#2    mNeoVis.3.1.1.chr7.g01664.t01#6   mNeoVis.vulgivaga.1.1.chr7.g01764.t01#4       2660   125259184:   mNeoVis.vulgivaga.1.1.chr1.g00961.t01#4    mNeoVis.2.1.1.chr1_hap1.g00952.t01#5    mNeoVis.evergladensis.1.1.chr1.g00953.t01#3    mNeoVis.evergladensis.2.1.chr1.g00947.t01#2    mNeoVis.vulgivaga.2.1.chr1.g00958.t01#1   mNeoVis.3.1.1.chr1.g00964.t01#6       2661   125259185:   mNeoVis.vulgivaga.2.1.chr1.g00959.t01#1    mNeoVis.3.1.1.chr1.g00965.t01#6    mNeoVis.vulgivaga.1.1.chr1.g00962.t01#4    mNeoVis.evergladensis.2.1.chr1.g00948.t01#2    mNeoVis.2.1.1.chr1_hap1.g00953.t01#5   mNeoVis.evergladensis.1.1.chr1.g00954.t01#3       2662   125259186:   mNeoVis.3.1.1.chr8.g00255.t01#6    mNeoVis.2.1.1.chr8_hap1.g00286.t01#5    mNeoVis.vulgivaga.1.1.chr8.g00279.t01#4    mNeoVis.vulgivaga.2.1.chr8.g00291.t01#1    mNeoVis.evergladensis.1.1.chr8.g00285.t01#3   mNeoVis.evergladensis.2.1.chr9.g00285.t01#2       2663   125259187:   mNeoVis.evergladensis.1.1.chr10.g00594.t01#3    mNeoVis.evergladensis.1.1.chr1.g00955.t01#3    mNeoVis.evergladensis.1.1.chr8.g00282.t01#3    mNeoVis.vulgivaga.2.1.chr8.g00288.t01#1    mNeoVis.3.1.1.chr10.g00566.t01#6    mNeoVis.vulgivaga.2.1.chr1.g00960.t01#1    mNeoVis.2.1.1.chr9_hap1.g00598.t01#5    mNeoVis.evergladensis.2.1.chr9.g00282.t01#2    mNeoVis.vulgivaga.1.1.chr1.g00963.t01#4      2664   125259188:   mNeoVis.evergladensis.1.1.chr6.g00243.t01#3    mNeoVis.2.1.1.chr4_hap1.g00242.t01#5    mNeoVis.vulgivaga.2.1.chr6.g00242.t01#1    mNeoVis.3.1.1.chr6.g00241.t01#6    mNeoVis.vulgivaga.1.1.chr6.g00249.t01#4   mNeoVis.evergladensis.2.1.chr6.g00241.t01#2       2665   125259189:   mNeoVis.3.1.1.chr2.g00668.t01#6    mNeoVis.vulgivaga.1.1.chr2.g00754.t01#4    mNeoVis.vulgivaga.2.1.chr2.g00765.t01#1    mNeoVis.2.1.1.chr5_hap1.g00759.t01#5   mNeoVis.evergladensis.1.1.chr2.g00757.t01#3       2666   125259190:  mNeoVis.evergladensis.2.1.chr2.g00760.t01#2             2667   125259191:   mNeoVis.vulgivaga.2.1.chr1.g00961.t01#1    mNeoVis.2.1.1.chr1_hap1.g00955.t01#5    mNeoVis.vulgivaga.1.1.chr1.g00965.t01#4    mNeoVis.3.1.1.chr1.g00967.t01#6    mNeoVis.evergladensis.2.1.chr1.g00950.t01#2   mNeoVis.evergladensis.1.1.chr1.g00956.t01#3       2668   125259192:   mNeoVis.vulgivaga.1.1.chr4.g00496.t01#4    mNeoVis.vulgivaga.1.1.chr6.g00022.t01#4    mNeoVis.evergladensis.1.1.chr7.g01753.t01#3    mNeoVis.evergladensis.2.1.chr4.g00499.t01#2    mNeoVis.evergladensis.2.1.chr7.g01757.t01#2    mNeoVis.2.1.1.chr4_hap1.g00020.t01#5    mNeoVis.vulgivaga.1.1.chr7.g01765.t01#4    mNeoVis.vulgivaga.2.1.chr7.g01775.t01#1    mNeoVis.evergladensis.2.1.chr6.g00020.t01#2      2669   125259193:   mNeoVis.evergladensis.2.1.chr1.g00951.t01#2    mNeoVis.2.1.1.chr1_hap1.g00956.t01#5    mNeoVis.vulgivaga.2.1.chr1.g00962.t01#1    mNeoVis.evergladensis.1.1.chr1.g00957.t01#3   mNeoVis.vulgivaga.1.1.chr1.g00966.t01#4       2670   125259194:   mNeoVis.vulgivaga.1.1.chr1.g00967.t01#4    mNeoVis.evergladensis.2.1.chr1.g00952.t01#2    mNeoVis.vulgivaga.2.1.chr1.g00963.t01#1    mNeoVis.2.1.1.chr1_hap1.g00957.t01#5   mNeoVis.evergladensis.1.1.chr1.g00958.t01#3       2671   125259195:   mNeoVis.evergladensis.2.1.chr7.g01758.t01#2    mNeoVis.3.1.1.chr7.g01666.t01#6    mNeoVis.vulgivaga.1.1.chr7.g01766.t01#4    mNeoVis.evergladensis.1.1.chr7.g01754.t01#3    mNeoVis.2.1.1.chr7_hap1.g01797.t01#5   mNeoVis.vulgivaga.2.1.chr7.g01776.t01#1       2672   125259196:   mNeoVis.2.1.1.chr1_hap1.g00958.t01#5    mNeoVis.evergladensis.2.1.chr1.g00953.t01#2    mNeoVis.vulgivaga.2.1.chr1.g00964.t01#1    mNeoVis.vulgivaga.1.1.chr1.g00968.t01#4    mNeoVis.evergladensis.1.1.chr1.g00959.t01#3   mNeoVis.3.1.1.chr1.g00968.t01#6       2673   125259197:   mNeoVis.2.1.1.chr1_hap1.g00959.t01#5    mNeoVis.evergladensis.1.1.chr1.g00960.t01#3    mNeoVis.vulgivaga.1.1.chr1.g00969.t01#4    mNeoVis.evergladensis.2.1.chr1.g00954.t01#2    mNeoVis.3.1.1.chr1.g00969.t01#6   mNeoVis.vulgivaga.2.1.chr1.g00965.t01#1       2674   125259198:   mNeoVis.vulgivaga.2.1.chr1.g00966.t01#1    mNeoVis.evergladensis.2.1.chr1.g00955.t01#2    mNeoVis.vulgivaga.1.1.chr1.g00970.t01#4    mNeoVis.evergladensis.1.1.chr1.g00961.t01#3   mNeoVis.2.1.1.chr1_hap1.g00960.t01#5       2675   125259199:   mNeoVis.vulgivaga.2.1.chr7.g01777.t01#1    mNeoVis.evergladensis.2.1.chr7.g01759.t01#2    mNeoVis.3.1.1.chr7.g01667.t01#6    mNeoVis.evergladensis.1.1.chr7.g01755.t01#3    mNeoVis.vulgivaga.1.1.chr7.g01767.t01#4   mNeoVis.2.1.1.chr7_hap1.g01798.t01#5       2676   125259200:   mNeoVis.vulgivaga.2.1.chr7.g01778.t01#1    mNeoVis.vulgivaga.1.1.chr7.g01768.t01#4    mNeoVis.evergladensis.2.1.chr7.g01760.t01#2    mNeoVis.2.1.1.chr7_hap1.g01799.t01#5    mNeoVis.3.1.1.chr7.g01668.t01#6   mNeoVis.evergladensis.1.1.chr7.g01756.t01#3       2677   125259201:   mNeoVis.evergladensis.1.1.chr7.g01757.t01#3    mNeoVis.vulgivaga.1.1.chr7.g01769.t01#4    mNeoVis.2.1.1.chr7_hap1.g01800.t01#5    mNeoVis.3.1.1.chr7.g01669.t01#6    mNeoVis.evergladensis.2.1.chr7.g01761.t01#2   mNeoVis.vulgivaga.2.1.chr7.g01779.t01#1       2678   125259202:   mNeoVis.3.1.1.chr7.g00067.t01#6    mNeoVis.2.1.1.chr7_hap1.g00080.t01#5    mNeoVis.evergladensis.1.1.chr7.g00082.t01#3    mNeoVis.evergladensis.2.1.chr7.g00080.t01#2    mNeoVis.vulgivaga.2.1.chr7.g00080.t01#1   mNeoVis.vulgivaga.1.1.chr7.g00082.t01#4       2679   125259203:   mNeoVis.3.1.1.chr7.g01670.t01#6    mNeoVis.2.1.1.chr7_hap1.g01801.t01#5    mNeoVis.evergladensis.1.1.chr7.g01758.t01#3    mNeoVis.vulgivaga.2.1.chr7.g01780.t01#1    mNeoVis.vulgivaga.1.1.chr7.g01770.t01#4   mNeoVis.evergladensis.2.1.chr7.g01762.t01#2       2680   125259204:   mNeoVis.vulgivaga.1.1.chr7.g01771.t01#4    mNeoVis.evergladensis.2.1.chr7.g01763.t01#2    mNeoVis.3.1.1.chr7.g01671.t01#6    mNeoVis.vulgivaga.2.1.chr7.g01781.t01#1    mNeoVis.2.1.1.chr7_hap1.g01802.t01#5   mNeoVis.evergladensis.1.1.chr7.g01759.t01#3       2681   125259205:   mNeoVis.vulgivaga.1.1.chr7.g01772.t01#4    mNeoVis.evergladensis.2.1.chr7.g01764.t01#2    mNeoVis.vulgivaga.2.1.chr7.g01782.t01#1    mNeoVis.evergladensis.1.1.chr7.g01760.t01#3    mNeoVis.3.1.1.chr7.g01672.t01#6   mNeoVis.2.1.1.chr7_hap1.g01803.t01#5       2682   125259206:   mNeoVis.evergladensis.2.1.chr7.g01765.t01#2    mNeoVis.vulgivaga.2.1.chr7.g01783.t01#1    mNeoVis.evergladensis.1.1.chr7.g01761.t01#3    mNeoVis.3.1.1.chr7.g01673.t01#6    mNeoVis.vulgivaga.1.1.chr7.g01773.t01#4   mNeoVis.2.1.1.chr7_hap1.g01804.t01#5       2683   125259207:   mNeoVis.vulgivaga.2.1.chr15.g00383.t01#1    mNeoVis.evergladensis.2.1.chr15.g00374.t01#2    mNeoVis.evergladensis.1.1.chr15.g00239.t01#3    mNeoVis.2.1.1.chr15_hap1.g00508.t01#5   mNeoVis.3.1.1.chr15.g00338.t01#6       2684   125259208:   mNeoVis.evergladensis.2.1.chr7.g01766.t01#2    mNeoVis.vulgivaga.2.1.chr7.g01784.t01#1    mNeoVis.evergladensis.1.1.chr7.g01762.t01#3   mNeoVis.vulgivaga.1.1.chr7.g01774.t01#4         2685   125259209:   mNeoVis.evergladensis.1.1.chr7.g01763.t01#3    mNeoVis.vulgivaga.1.1.chr7.g01775.t01#4    mNeoVis.evergladensis.2.1.chr7.g01767.t01#2    mNeoVis.3.1.1.chr7.g01674.t01#6    mNeoVis.vulgivaga.2.1.chr7.g01785.t01#1   mNeoVis.2.1.1.chr7_hap1.g01805.t01#5       2686   125259210:   mNeoVis.evergladensis.1.1.chr7.g01764.t01#3    mNeoVis.evergladensis.2.1.chr7.g01768.t01#2    mNeoVis.vulgivaga.1.1.chr7.g01776.t01#4    mNeoVis.3.1.1.chr7.g01675.t01#6    mNeoVis.vulgivaga.2.1.chr7.g01786.t01#1   mNeoVis.2.1.1.chr7_hap1.g01806.t01#5       2687   125259211:   mNeoVis.evergladensis.1.1.chr7.g01566.t01#3    mNeoVis.2.1.1.chr7_hap1.g01582.t01#5    mNeoVis.vulgivaga.1.1.chr7.g01575.t01#4    mNeoVis.vulgivaga.2.1.chr7.g01580.t01#1    mNeoVis.vulgivaga.1.1.chr7.g01574.t01#4    mNeoVis.3.1.1.chr7.g01492.t01#6    mNeoVis.vulgivaga.2.1.chr7.g01581.t01#1    mNeoVis.vulgivaga.1.1.chr7.g01576.t01#4    mNeoVis.evergladensis.1.1.chr7.g01568.t01#3      2688   125259212:   mNeoVis.vulgivaga.1.1.chr7.g01577.t01#4    mNeoVis.evergladensis.1.1.chr7.g01569.t01#3    mNeoVis.2.1.1.chr7_hap1.g01584.t01#5    mNeoVis.evergladensis.2.1.chr7.g01563.t01#2   mNeoVis.vulgivaga.2.1.chr7.g01582.t01#1       2689   125259213:   mNeoVis.vulgivaga.1.1.chr7.g01680.t01#4    mNeoVis.evergladensis.2.1.chr7.g01668.t01#2   mNeoVis.3.1.1.chr7.g01586.t01#6           2690   125259214:   mNeoVis.3.1.1.chr7.g01587.t01#6    mNeoVis.evergladensis.2.1.chr7.g01666.t01#2   mNeoVis.vulgivaga.1.1.chr7.g01678.t01#4           2691   125259215:   mNeoVis.evergladensis.2.1.chr7.g01561.t01#2   mNeoVis.evergladensis.2.1.chr7.g01562.t01#2           2692   125259216:   mNeoVis.2.1.1.chr7_hap1.g01697.t01#5   mNeoVis.evergladensis.1.1.chr7.g01666.t01#3           2693   125259217:  mNeoVis.2.1.1.chr7_hap1.g01581.t01#5               2694   125259218:   mNeoVis.3.1.1.chr7.g01495.t01#6    mNeoVis.2.1.1.chr7_hap1.g01585.t01#5    mNeoVis.evergladensis.2.1.chr7.g01564.t01#2    mNeoVis.vulgivaga.2.1.chr7.g01583.t01#1    mNeoVis.vulgivaga.1.1.chr7.g01578.t01#4   mNeoVis.evergladensis.1.1.chr7.g01570.t01#3       2695   125259219:   mNeoVis.2.1.1.chr7_hap1.g01586.t01#5    mNeoVis.vulgivaga.2.1.chr7.g01584.t01#1    mNeoVis.vulgivaga.1.1.chr7.g01579.t01#4    mNeoVis.evergladensis.1.1.chr7.g01571.t01#3    mNeoVis.3.1.1.chr7.g01496.t01#6   mNeoVis.evergladensis.2.1.chr7.g01565.t01#2       2696   125259220:   mNeoVis.vulgivaga.1.1.chr7.g00083.t01#4    mNeoVis.2.1.1.chr7_hap1.g00081.t01#5    mNeoVis.vulgivaga.2.1.chr7.g00081.t01#1    mNeoVis.evergladensis.2.1.chr7.g00081.t01#2   mNeoVis.evergladensis.1.1.chr7.g00083.t01#3       2697   125259221:   mNeoVis.2.1.1.chr7_hap1.g01587.t01#5    mNeoVis.vulgivaga.2.1.chr7.g01585.t01#1    mNeoVis.evergladensis.2.1.chr7.g01566.t01#2    mNeoVis.evergladensis.1.1.chr7.g01572.t01#3    mNeoVis.vulgivaga.1.1.chr7.g01580.t01#4   mNeoVis.3.1.1.chr7.g01497.t01#6       2698   125259222:  mNeoVis.vulgivaga.2.1.chr1.g00967.t01#1               2699   125259223:   mNeoVis.evergladensis.2.1.chr1.g00956.t01#2    mNeoVis.vulgivaga.1.1.chr1.g00971.t01#4    mNeoVis.vulgivaga.2.1.chr1.g00968.t01#1    mNeoVis.2.1.1.chr1_hap1.g00961.t01#5   mNeoVis.evergladensis.1.1.chr1.g00962.t01#3       2700   125259224:   mNeoVis.evergladensis.2.1.chr7.g01567.t01#2    mNeoVis.vulgivaga.1.1.chr7.g01581.t01#4    mNeoVis.evergladensis.1.1.chr7.g01573.t01#3    mNeoVis.vulgivaga.2.1.chr7.g01586.t01#1    mNeoVis.3.1.1.chr7.g01498.t01#6   mNeoVis.2.1.1.chr7_hap1.g01588.t01#5       2701   125259225:   mNeoVis.evergladensis.1.1.chr7.g01574.t01#3    mNeoVis.vulgivaga.1.1.chr7.g01582.t01#4    mNeoVis.3.1.1.chr7.g01499.t01#6    mNeoVis.evergladensis.2.1.chr7.g01568.t01#2   mNeoVis.2.1.1.chr7_hap1.g01589.t01#5       2702   125259226:   mNeoVis.vulgivaga.2.1.chr7.g01587.t01#1    mNeoVis.2.1.1.chr7_hap1.g01590.t01#5    mNeoVis.evergladensis.2.1.chr7.g01569.t01#2   mNeoVis.vulgivaga.1.1.chr7.g01583.t01#4         2703   125259227:   mNeoVis.evergladensis.2.1.chr1.g00957.t01#2    mNeoVis.2.1.1.chr1_hap1.g00962.t01#5    mNeoVis.vulgivaga.2.1.chr1.g00969.t01#1    mNeoVis.vulgivaga.1.1.chr1.g00972.t01#4   mNeoVis.evergladensis.1.1.chr1.g00963.t01#3       2704   125259228:   mNeoVis.2.1.1.chr7_hap1.g01591.t01#5    mNeoVis.vulgivaga.1.1.chr7.g01585.t01#4    mNeoVis.evergladensis.1.1.chr7.g01577.t01#3    mNeoVis.3.1.1.chr7.g01501.t01#6    mNeoVis.vulgivaga.2.1.chr7.g01589.t01#1   mNeoVis.evergladensis.2.1.chr7.g01571.t01#2       2705   125259229:   mNeoVis.vulgivaga.2.1.chr7.g01588.t01#1    mNeoVis.vulgivaga.1.1.chr7.g01584.t01#4    mNeoVis.3.1.1.chr7.g01500.t01#6    mNeoVis.evergladensis.1.1.chr7.g01576.t01#3   mNeoVis.evergladensis.2.1.chr7.g01570.t01#2       2706   125259230:   mNeoVis.evergladensis.2.1.chr7.g01572.t01#2    mNeoVis.3.1.1.chr7.g01503.t01#6   mNeoVis.2.1.1.chr7_hap1.g01593.t01#5           2707   125259231:   mNeoVis.evergladensis.2.1.chr7.g01573.t01#2    mNeoVis.2.1.1.chr7_hap1.g01594.t01#5    mNeoVis.3.1.1.chr7.g01504.t01#6    mNeoVis.evergladensis.1.1.chr7.g01579.t01#3    mNeoVis.vulgivaga.1.1.chr7.g01586.t01#4   mNeoVis.vulgivaga.2.1.chr7.g01590.t01#1       2708   125259232:   mNeoVis.vulgivaga.2.1.chr7.g01591.t01#1    mNeoVis.evergladensis.2.1.chr7.g01575.t01#2    mNeoVis.2.1.1.chr7_hap1.g01595.t01#5    mNeoVis.evergladensis.1.1.chr7.g01580.t01#3    mNeoVis.3.1.1.chr7.g01506.t01#6   mNeoVis.vulgivaga.1.1.chr7.g01587.t01#4       2709   125259233:   mNeoVis.2.1.1.chr7_hap1.g01596.t01#5    mNeoVis.3.1.1.chr7.g01507.t01#6    mNeoVis.evergladensis.1.1.chr7.g01581.t01#3    mNeoVis.evergladensis.2.1.chr7.g01576.t01#2    mNeoVis.vulgivaga.1.1.chr7.g01588.t01#4   mNeoVis.vulgivaga.2.1.chr7.g01592.t01#1       2710   125259234:   mNeoVis.3.1.1.chr7.g01508.t01#6    mNeoVis.evergladensis.1.1.chr7.g01582.t01#3    mNeoVis.2.1.1.chr7_hap1.g01600.t01#5    mNeoVis.vulgivaga.2.1.chr7.g01594.t01#1    mNeoVis.3.1.1.chr7.g01510.t01#6    mNeoVis.vulgivaga.1.1.chr7.g01589.t01#4    mNeoVis.vulgivaga.2.1.chr7.g01593.t01#1    mNeoVis.evergladensis.2.1.chr7.g01577.t01#2    mNeoVis.2.1.1.chr7_hap1.g01597.t01#5      2711   125259235:   mNeoVis.vulgivaga.1.1.chr7.g01590.t01#4    mNeoVis.2.1.1.chr7_hap1.g01599.t01#5    mNeoVis.evergladensis.2.1.chr7.g01578.t01#2    mNeoVis.3.1.1.chr7.g01509.t01#6   mNeoVis.2.1.1.chr7_hap1.g01598.t01#5       2712   125259236:   mNeoVis.vulgivaga.2.1.chr1.g00975.t01#1    mNeoVis.3.1.1.chr1.g00975.t01#6    mNeoVis.vulgivaga.1.1.chr1.g00977.t01#4    mNeoVis.evergladensis.1.1.chr1.g00968.t01#3    mNeoVis.2.1.1.chr1_hap1.g00967.t01#5   mNeoVis.evergladensis.2.1.chr1.g00962.t01#2       2713   125259237:   mNeoVis.evergladensis.1.1.chr7.g01585.t01#3    mNeoVis.3.1.1.chr7.g01513.t01#6    mNeoVis.vulgivaga.2.1.chr7.g01597.t01#1    mNeoVis.2.1.1.chr7_hap1.g01603.t01#5    mNeoVis.evergladensis.1.1.chr7.g01583.t01#3    mNeoVis.3.1.1.chr7.g01511.t01#6    mNeoVis.vulgivaga.2.1.chr7.g01595.t01#1   mNeoVis.2.1.1.chr7_hap1.g01601.t01#5     2714   125259238:   mNeoVis.vulgivaga.2.1.chr7.g01602.t01#1    mNeoVis.3.1.1.chr7.g01519.t01#6    mNeoVis.vulgivaga.1.1.chr7.g01593.t01#4    mNeoVis.2.1.1.chr7_hap1.g01611.t01#5    mNeoVis.evergladensis.1.1.chr7.g01590.t01#3    mNeoVis.2.1.1.chr7_hap1.g01613.t01#5   mNeoVis.vulgivaga.1.1.chr7.g01601.t01#4     2715   125259239:   mNeoVis.evergladensis.1.1.chr7.g01587.t01#3    mNeoVis.vulgivaga.1.1.chr7.g01597.t01#4    mNeoVis.vulgivaga.2.1.chr7.g01599.t01#1    mNeoVis.2.1.1.chr7_hap1.g01605.t01#5    mNeoVis.3.1.1.chr7.g01515.t01#6   mNeoVis.evergladensis.2.1.chr7.g01581.t01#2       2716   125259240:   mNeoVis.vulgivaga.1.1.chr7.g01598.t01#4    mNeoVis.evergladensis.1.1.chr7.g01588.t01#3    mNeoVis.2.1.1.chr7_hap1.g01606.t01#5    mNeoVis.3.1.1.chr7.g01516.t01#6    mNeoVis.vulgivaga.2.1.chr7.g01600.t01#1   mNeoVis.evergladensis.2.1.chr7.g01579.t01#2       2717   125259241:   mNeoVis.evergladensis.2.1.chr7.g01580.t01#2    mNeoVis.evergladensis.1.1.chr7.g01584.t01#3    mNeoVis.2.1.1.chr7_hap1.g01602.t01#5    mNeoVis.vulgivaga.2.1.chr7.g01596.t01#1    mNeoVis.vulgivaga.1.1.chr7.g01594.t01#4   mNeoVis.3.1.1.chr7.g01512.t01#6       2718   125259242:   mNeoVis.evergladensis.1.1.chr7.g01586.t01#3    mNeoVis.2.1.1.chr7_hap1.g01604.t01#5    mNeoVis.vulgivaga.2.1.chr7.g01598.t01#1    mNeoVis.vulgivaga.1.1.chr7.g01596.t01#4   mNeoVis.3.1.1.chr7.g01514.t01#6       2719   125259243:  mNeoVis.vulgivaga.1.1.chr7.g01592.t01#4               2720   125259244:  mNeoVis.vulgivaga.1.1.chr7.g01595.t01#4               2721   125259245:   mNeoVis.vulgivaga.1.1.chr1.g00978.t01#4    mNeoVis.evergladensis.1.1.chr1.g00969.t01#3    mNeoVis.3.1.1.chr1.g00976.t01#6    mNeoVis.2.1.1.chr1_hap1.g00968.t01#5    mNeoVis.vulgivaga.2.1.chr1.g00976.t01#1   mNeoVis.evergladensis.2.1.chr1.g00963.t01#2       2722   125259246:   mNeoVis.evergladensis.2.1.chr1.g00964.t01#2    mNeoVis.3.1.1.chr1.g00977.t01#6    mNeoVis.2.1.1.chr1_hap1.g00969.t01#5    mNeoVis.evergladensis.1.1.chr1.g00970.t01#3    mNeoVis.vulgivaga.1.1.chr1.g00979.t01#4   mNeoVis.vulgivaga.2.1.chr1.g00977.t01#1       2723   125259247:   mNeoVis.vulgivaga.1.1.chr1.g00980.t01#4    mNeoVis.evergladensis.1.1.chr1.g00971.t01#3    mNeoVis.evergladensis.2.1.chr1.g00965.t01#2    mNeoVis.3.1.1.chr1.g00978.t01#6    mNeoVis.vulgivaga.2.1.chr1.g00978.t01#1   mNeoVis.2.1.1.chr1_hap1.g00970.t01#5       2724   125259248:   mNeoVis.2.1.1.chr7_hap1.g01607.t01#5    mNeoVis.vulgivaga.2.1.chr7.g01601.t01#1    mNeoVis.evergladensis.2.1.chr7.g01582.t01#2    mNeoVis.evergladensis.2.1.chr7.g01583.t01#2    mNeoVis.2.1.1.chr7_hap1.g01609.t01#5    mNeoVis.vulgivaga.1.1.chr7.g01599.t01#4    mNeoVis.evergladensis.1.1.chr7.g01589.t01#3    mNeoVis.3.1.1.chr7.g01517.t01#6    mNeoVis.2.1.1.chr7_hap1.g01608.t01#5      2725   125259249:   mNeoVis.2.1.1.chr7_hap1.g01612.t01#5    mNeoVis.evergladensis.2.1.chr7.g01584.t01#2    mNeoVis.evergladensis.1.1.chr7.g01591.t01#3    mNeoVis.vulgivaga.1.1.chr7.g01602.t01#4   mNeoVis.vulgivaga.2.1.chr7.g01603.t01#1       2726   125259250:   mNeoVis.3.1.1.chr1.g00979.t01#6    mNeoVis.evergladensis.2.1.chr1.g00966.t01#2    mNeoVis.2.1.1.chr1_hap1.g00971.t01#5    mNeoVis.vulgivaga.1.1.chr1.g00981.t01#4    mNeoVis.vulgivaga.2.1.chr1.g00979.t01#1   mNeoVis.evergladensis.1.1.chr1.g00972.t01#3       2727   125259251:   mNeoVis.vulgivaga.2.1.chr1.g00982.t01#1    mNeoVis.evergladensis.2.1.chr1.g00969.t01#2    mNeoVis.vulgivaga.1.1.chr1.g00984.t01#4    mNeoVis.3.1.1.chr1.g00981.t01#6    mNeoVis.evergladensis.1.1.chr1.g00974.t01#3   mNeoVis.2.1.1.chr1_hap1.g00974.t01#5       2728   125259252:   mNeoVis.evergladensis.1.1.chr7.g01592.t01#3    mNeoVis.vulgivaga.1.1.chr7.g01603.t01#4    mNeoVis.evergladensis.2.1.chr7.g01586.t01#2    mNeoVis.3.1.1.chr7.g01520.t01#6    mNeoVis.2.1.1.chr7_hap1.g01614.t01#5   mNeoVis.vulgivaga.2.1.chr7.g01604.t01#1       2729   125259253:   mNeoVis.2.1.1.chr7_hap1.g01620.t01#5    mNeoVis.vulgivaga.2.1.chr7.g01610.t01#1    mNeoVis.3.1.1.chr7.g01524.t01#6    mNeoVis.vulgivaga.1.1.chr7.g01609.t01#4    mNeoVis.evergladensis.2.1.chr7.g01591.t01#2   mNeoVis.evergladensis.1.1.chr7.g01598.t01#3       2730   125259254:   mNeoVis.3.1.1.chr7.g01525.t01#6    mNeoVis.vulgivaga.1.1.chr7.g01610.t01#4    mNeoVis.evergladensis.2.1.chr7.g01592.t01#2    mNeoVis.evergladensis.1.1.chr7.g01599.t01#3    mNeoVis.2.1.1.chr7_hap1.g01621.t01#5   mNeoVis.vulgivaga.2.1.chr7.g01611.t01#1       2731   125259255:   mNeoVis.2.1.1.chr7_hap1.g01615.t01#5    mNeoVis.3.1.1.chr7.g01521.t01#6    mNeoVis.vulgivaga.2.1.chr7.g01605.t01#1    mNeoVis.evergladensis.2.1.chr7.g01587.t01#2    mNeoVis.evergladensis.1.1.chr7.g01593.t01#3   mNeoVis.vulgivaga.1.1.chr7.g01604.t01#4       2732   125259256:   mNeoVis.2.1.1.chr7_hap1.g01616.t01#5    mNeoVis.evergladensis.2.1.chr7.g01588.t01#2    mNeoVis.evergladensis.1.1.chr7.g01594.t01#3    mNeoVis.vulgivaga.2.1.chr7.g01606.t01#1    mNeoVis.vulgivaga.1.1.chr7.g01605.t01#4   mNeoVis.3.1.1.chr7.g01522.t01#6       2733   125259257:   mNeoVis.evergladensis.1.1.chr1.g00975.t01#3    mNeoVis.evergladensis.2.1.chr1.g00970.t01#2    mNeoVis.vulgivaga.1.1.chr1.g00985.t01#4    mNeoVis.3.1.1.chr1.g00982.t01#6    mNeoVis.2.1.1.chr1_hap1.g00975.t01#5   mNeoVis.vulgivaga.2.1.chr1.g00983.t01#1       2734   125259258:   mNeoVis.evergladensis.1.1.chr7.g01595.t01#3    mNeoVis.3.1.1.chr7.g01518.t01#6    mNeoVis.vulgivaga.1.1.chr7.g01606.t01#4    mNeoVis.evergladensis.2.1.chr7.g01589.t01#2    mNeoVis.vulgivaga.2.1.chr7.g01607.t01#1   mNeoVis.2.1.1.chr7_hap1.g01617.t01#5       2735   125259259:   mNeoVis.2.1.1.chr1_hap1.g00976.t01#5    mNeoVis.vulgivaga.1.1.chr1.g00986.t01#4   mNeoVis.evergladensis.2.1.chr1.g00971.t01#2         2736   125259260:   mNeoVis.vulgivaga.2.1.chr1.g00984.t01#1   mNeoVis.evergladensis.1.1.chr1.g00976.t01#3           2737   125259261:  mNeoVis.3.1.1.chr1.g00983.t01#6               2738   125259262:  mNeoVis.vulgivaga.2.1.chr1.g00985.t01#1               2739   125259263:   mNeoVis.vulgivaga.2.1.chr7.g01608.t01#1    mNeoVis.evergladensis.2.1.chr7.g01585.t01#2    mNeoVis.vulgivaga.1.1.chr7.g01607.t01#4    mNeoVis.2.1.1.chr7_hap1.g01618.t01#5   mNeoVis.evergladensis.1.1.chr7.g01596.t01#3       2740   125259264:   mNeoVis.evergladensis.1.1.chr7.g01597.t01#3    mNeoVis.2.1.1.chr7_hap1.g01619.t01#5    mNeoVis.3.1.1.chr7.g01523.t01#6    mNeoVis.vulgivaga.2.1.chr7.g01609.t01#1    mNeoVis.vulgivaga.1.1.chr7.g01608.t01#4   mNeoVis.evergladensis.2.1.chr7.g01590.t01#2       2741   125259265:   mNeoVis.3.1.1.chr1.g00984.t01#6    mNeoVis.evergladensis.2.1.chr1.g00972.t01#2   mNeoVis.vulgivaga.2.1.chr1.g00986.t01#1           2742   125259266:   mNeoVis.2.1.1.chr1_hap1.g00977.t01#5   mNeoVis.evergladensis.1.1.chr1.g00977.t01#3           2743   125259267:   mNeoVis.2.1.1.chr1_hap1.g00978.t01#5   mNeoVis.3.1.1.chr1.g00985.t01#6             2744   125259268:  mNeoVis.vulgivaga.1.1.chr1.g00987.t01#4               2745   125259269:   mNeoVis.evergladensis.1.1.chr7.g01600.t01#3    mNeoVis.evergladensis.2.1.chr7.g01593.t01#2    mNeoVis.2.1.1.chr7_hap1.g01622.t01#5    mNeoVis.vulgivaga.2.1.chr7.g01612.t01#1    mNeoVis.3.1.1.chr7.g01526.t01#6   mNeoVis.vulgivaga.1.1.chr7.g01611.t01#4       2746   125259270:   mNeoVis.evergladensis.1.1.chr7.g01601.t01#3   mNeoVis.vulgivaga.2.1.chr7.g01613.t01#1             2747   125259271:   mNeoVis.vulgivaga.1.1.chr1.g00988.t01#4    mNeoVis.evergladensis.2.1.chr1.g00974.t01#2    mNeoVis.evergladensis.1.1.chr1.g00978.t01#3    mNeoVis.vulgivaga.2.1.chr1.g00987.t01#1    mNeoVis.2.1.1.chr1_hap1.g00979.t01#5   mNeoVis.3.1.1.chr1.g00986.t01#6       2748   125259272:   mNeoVis.vulgivaga.1.1.chr1.g00989.t01#4    mNeoVis.evergladensis.1.1.chr1.g00980.t01#3    mNeoVis.vulgivaga.2.1.chr1.g00988.t01#1    mNeoVis.2.1.1.chr1_hap1.g00983.t01#5    mNeoVis.evergladensis.2.1.chr1.g00976.t01#2   mNeoVis.3.1.1.chr1.g00988.t01#6       2749   125259273:   mNeoVis.vulgivaga.2.1.chr7.g01622.t01#1    mNeoVis.evergladensis.2.1.chr7.g01601.t01#2    mNeoVis.3.1.1.chr7.g01536.t01#6    mNeoVis.evergladensis.2.1.chr7.g01603.t01#2    mNeoVis.vulgivaga.2.1.chr7.g01621.t01#1    mNeoVis.evergladensis.1.1.chr7.g01602.t01#3    mNeoVis.evergladensis.2.1.chr7.g01602.t01#2    mNeoVis.2.1.1.chr7_hap1.g01633.t01#5    mNeoVis.3.1.1.chr7.g01534.t01#6      2750   125259274:   mNeoVis.vulgivaga.2.1.chr7.g01630.t01#1    mNeoVis.evergladensis.2.1.chr7.g01614.t01#2    mNeoVis.3.1.1.chr7.g01541.t01#6    mNeoVis.vulgivaga.2.1.chr7.g01629.t01#1    mNeoVis.vulgivaga.1.1.chr7.g01628.t01#4    mNeoVis.evergladensis.2.1.chr7.g01613.t01#2    mNeoVis.vulgivaga.1.1.chr7.g01627.t01#4    mNeoVis.evergladensis.1.1.chr7.g01619.t01#3    mNeoVis.evergladensis.2.1.chr7.g01609.t01#2      2751   125259275:   mNeoVis.2.1.1.chr7_hap1.g01719.t01#5    mNeoVis.vulgivaga.2.1.chr7.g01702.t01#1    mNeoVis.2.1.1.chr7_hap1.g01718.t01#5    mNeoVis.3.1.1.chr7.g01601.t01#6    mNeoVis.evergladensis.1.1.chr7.g01684.t01#3    mNeoVis.evergladensis.2.1.chr7.g01687.t01#2    mNeoVis.vulgivaga.1.1.chr7.g01697.t01#4    mNeoVis.3.1.1.chr7.g01602.t01#6    mNeoVis.vulgivaga.1.1.chr7.g01696.t01#4      2752   125259276:   mNeoVis.2.1.1.chr7_hap1.g01628.t01#5    mNeoVis.vulgivaga.2.1.chr7.g01619.t01#1    mNeoVis.vulgivaga.1.1.chr7.g01617.t01#4    mNeoVis.2.1.1.chr7_hap1.g01634.t01#5    mNeoVis.evergladensis.2.1.chr7.g01599.t01#2    mNeoVis.evergladensis.1.1.chr7.g01612.t01#3    mNeoVis.3.1.1.chr7.g01532.t01#6   mNeoVis.evergladensis.2.1.chr7.g01605.t01#2     2753   125259277:   mNeoVis.2.1.1.chr7_hap1.g01632.t01#5    mNeoVis.evergladensis.1.1.chr7.g01611.t01#3    mNeoVis.vulgivaga.2.1.chr7.g01623.t01#1    mNeoVis.vulgivaga.1.1.chr7.g01621.t01#4    mNeoVis.2.1.1.chr7_hap1.g01635.t01#5    mNeoVis.2.1.1.chr7_hap1.g01610.t01#5   mNeoVis.evergladensis.2.1.chr7.g01604.t01#2     2754   125259278:   mNeoVis.evergladensis.1.1.chr7.g01606.t01#3    mNeoVis.evergladensis.2.1.chr7.g01606.t01#2    mNeoVis.2.1.1.chr7_hap1.g01627.t01#5    mNeoVis.vulgivaga.1.1.chr7.g01623.t01#4    mNeoVis.vulgivaga.2.1.chr7.g01618.t01#1   mNeoVis.3.1.1.chr7.g01528.t01#6       2755   125259279:   mNeoVis.2.1.1.chr7_hap1.g01636.t01#5    mNeoVis.evergladensis.1.1.chr7.g01609.t01#3    mNeoVis.evergladensis.2.1.chr7.g01607.t01#2    mNeoVis.3.1.1.chr7.g01538.t01#6    mNeoVis.vulgivaga.1.1.chr7.g01624.t01#4   mNeoVis.vulgivaga.2.1.chr7.g01625.t01#1       2756   125259280:   mNeoVis.vulgivaga.2.1.chr7.g01704.t01#1    mNeoVis.2.1.1.chr7_hap1.g01723.t01#5    mNeoVis.evergladensis.2.1.chr7.g01690.t01#2    mNeoVis.3.1.1.chr7.g01605.t01#6    mNeoVis.evergladensis.1.1.chr7.g01686.t01#3   mNeoVis.vulgivaga.1.1.chr7.g01700.t01#4       2757   125259281:   mNeoVis.evergladensis.1.1.chr7.g01685.t01#3    mNeoVis.vulgivaga.2.1.chr7.g01703.t01#1    mNeoVis.vulgivaga.1.1.chr7.g01698.t01#4    mNeoVis.3.1.1.chr7.g01603.t01#6    mNeoVis.2.1.1.chr7_hap1.g01721.t01#5   mNeoVis.evergladensis.2.1.chr7.g01688.t01#2       2758   125259282:   mNeoVis.vulgivaga.1.1.chr7.g01699.t01#4    mNeoVis.3.1.1.chr7.g01604.t01#6    mNeoVis.evergladensis.2.1.chr7.g01689.t01#2    mNeoVis.2.1.1.chr7_hap1.g01722.t01#5   mNeoVis.2.1.1.chr7_hap1.g01720.t01#5       2759   125259283:   mNeoVis.vulgivaga.1.1.chr7.g01626.t01#4    mNeoVis.3.1.1.chr7.g01542.t01#6    mNeoVis.evergladensis.1.1.chr7.g01617.t01#3    mNeoVis.evergladensis.2.1.chr7.g01612.t01#2   mNeoVis.2.1.1.chr7_hap1.g01638.t01#5       2760   125259284:   mNeoVis.vulgivaga.1.1.chr7.g01622.t01#4    mNeoVis.vulgivaga.2.1.chr7.g01624.t01#1    mNeoVis.3.1.1.chr7.g01535.t01#6   mNeoVis.evergladensis.1.1.chr7.g01613.t01#3       2761   125259285:   mNeoVis.evergladensis.1.1.chr7.g01614.t01#3    mNeoVis.evergladensis.2.1.chr7.g01608.t01#2    mNeoVis.vulgivaga.2.1.chr7.g01626.t01#1   mNeoVis.3.1.1.chr7.g01540.t01#6         2762   125259286:   mNeoVis.3.1.1.chr7.g01599.t01#6   mNeoVis.evergladensis.2.1.chr7.g01686.t01#2           2763   125259287:  mNeoVis.2.1.1.chr7_hap1.g01717.t01#5               2764   125259288:  mNeoVis.evergladensis.1.1.chr7.g01683.t01#3             2765   125259289:  mNeoVis.vulgivaga.1.1.chr7.g01695.t01#4               2766   125259290:  mNeoVis.3.1.1.chr7.g01600.t01#6               2767   125259291:  mNeoVis.vulgivaga.2.1.chr7.g01701.t01#1               2768   125259292:   mNeoVis.2.1.1.chr1_hap1.g00984.t01#5    mNeoVis.evergladensis.2.1.chr1.g00977.t01#2    mNeoVis.3.1.1.chr1.g00989.t01#6    mNeoVis.vulgivaga.2.1.chr1.g00990.t01#1    mNeoVis.evergladensis.1.1.chr1.g00981.t01#3    mNeoVis.vulgivaga.2.1.chr1.g00989.t01#1    mNeoVis.evergladensis.2.1.chr1.g00978.t01#2   mNeoVis.vulgivaga.1.1.chr1.g00990.t01#4     2769   125259293:   mNeoVis.evergladensis.2.1.chr1.g00983.t01#2    mNeoVis.3.1.1.chr1.g00993.t01#6    mNeoVis.2.1.1.chr1_hap1.g00989.t01#5    mNeoVis.vulgivaga.2.1.chr1.g00997.t01#1    mNeoVis.vulgivaga.1.1.chr1.g00995.t01#4   mNeoVis.evergladensis.1.1.chr1.g00986.t01#3       2770   125259294:   mNeoVis.vulgivaga.2.1.chr7.g01615.t01#1    mNeoVis.evergladensis.1.1.chr7.g01603.t01#3    mNeoVis.vulgivaga.1.1.chr7.g01614.t01#4   mNeoVis.evergladensis.2.1.chr7.g01596.t01#2       2771   125259295:   mNeoVis.evergladensis.1.1.chr1.g00982.t01#3    mNeoVis.vulgivaga.2.1.chr1.g00992.t01#1    mNeoVis.2.1.1.chr1_hap1.g00985.t01#5    mNeoVis.vulgivaga.1.1.chr1.g00992.t01#4   mNeoVis.3.1.1.chr1.g00990.t01#6       2772   125259296:   mNeoVis.vulgivaga.1.1.chr1.g00991.t01#4   mNeoVis.vulgivaga.2.1.chr1.g00991.t01#1             2773   125259297:   mNeoVis.vulgivaga.1.1.chr7.g01615.t01#4    mNeoVis.3.1.1.chr7.g01530.t01#6    mNeoVis.evergladensis.1.1.chr7.g01604.t01#3    mNeoVis.2.1.1.chr7_hap1.g01625.t01#5    mNeoVis.vulgivaga.2.1.chr7.g01616.t01#1   mNeoVis.evergladensis.2.1.chr7.g01597.t01#2       2774   125259298:   mNeoVis.vulgivaga.1.1.chr1.g00993.t01#4    mNeoVis.2.1.1.chr1_hap1.g00986.t01#5    mNeoVis.vulgivaga.2.1.chr1.g00993.t01#1    mNeoVis.evergladensis.1.1.chr1.g00983.t01#3   mNeoVis.evergladensis.2.1.chr1.g00980.t01#2       2775   125259299:   mNeoVis.2.1.1.chr11_hap1.g00541.t01#5    mNeoVis.3.1.1.chr11.g00502.t01#6    mNeoVis.evergladensis.2.1.chr11.g00529.t01#2    mNeoVis.vulgivaga.2.1.chr11.g00538.t01#1   mNeoVis.evergladensis.1.1.chr11.g00532.t01#3       2776   125259300:   mNeoVis.evergladensis.1.1.chr1.g00984.t01#3    mNeoVis.evergladensis.2.1.chr1.g00981.t01#2    mNeoVis.2.1.1.chr1_hap1.g00987.t01#5   mNeoVis.vulgivaga.2.1.chr1.g00994.t01#1         2777   125259301:   mNeoVis.3.1.1.chr7.g01531.t01#6    mNeoVis.evergladensis.1.1.chr7.g01605.t01#3    mNeoVis.evergladensis.2.1.chr7.g01598.t01#2    mNeoVis.2.1.1.chr7_hap1.g01626.t01#5    mNeoVis.vulgivaga.1.1.chr7.g01616.t01#4   mNeoVis.vulgivaga.2.1.chr7.g01617.t01#1       2778   125259302:   mNeoVis.evergladensis.2.1.chr7.g01600.t01#2    mNeoVis.vulgivaga.1.1.chr7.g01618.t01#4    mNeoVis.evergladensis.1.1.chr7.g01608.t01#3    mNeoVis.2.1.1.chr7_hap1.g01629.t01#5   mNeoVis.vulgivaga.2.1.chr7.g01620.t01#1       2779   125259303:  mNeoVis.vulgivaga.2.1.chr1.g00998.t01#1               2780   125259304:   mNeoVis.2.1.1.chr1_hap1.g00992.t01#5    mNeoVis.vulgivaga.1.1.chr1.g00998.t01#4    mNeoVis.evergladensis.2.1.chr1.g00986.t01#2    mNeoVis.vulgivaga.2.1.chr1.g00999.t01#1    mNeoVis.3.1.1.chr1.g00997.t01#6   mNeoVis.evergladensis.1.1.chr1.g00989.t01#3       2781   125259305:   mNeoVis.evergladensis.2.1.chr7.g01615.t01#2    mNeoVis.vulgivaga.2.1.chr7.g01631.t01#1    mNeoVis.vulgivaga.1.1.chr7.g01629.t01#4    mNeoVis.evergladensis.2.1.chr7.g01610.t01#2    mNeoVis.evergladensis.1.1.chr7.g01616.t01#3   mNeoVis.2.1.1.chr7_hap1.g01641.t01#5       2782   125259306:   mNeoVis.vulgivaga.1.1.chr7.g01625.t01#4    mNeoVis.vulgivaga.2.1.chr7.g01628.t01#1    mNeoVis.evergladensis.2.1.chr7.g01611.t01#2   mNeoVis.2.1.1.chr7_hap1.g01637.t01#5         2783   125259307:   mNeoVis.evergladensis.2.1.chr1.g00987.t01#2   mNeoVis.vulgivaga.2.1.chr1.g01000.t01#1             2784   125259308:   mNeoVis.2.1.1.chr15_hap1.g00167.t01#5    mNeoVis.evergladensis.2.1.chr15.g00150.t01#2    mNeoVis.3.1.1.chr15.g00143.t01#6    mNeoVis.vulgivaga.1.1.chr15.g00149.t01#4    mNeoVis.evergladensis.1.1.chr15.g00464.t01#3   mNeoVis.vulgivaga.2.1.chr15.g00153.t01#1       2785   125259309:   mNeoVis.2.1.1.chr1_hap1.g00946.t01#5    mNeoVis.vulgivaga.1.1.chr1.g00956.t01#4    mNeoVis.3.1.1.chr1.g00958.t01#6    mNeoVis.vulgivaga.2.1.chr1.g01014.t01#1    mNeoVis.3.1.1.chr1.g01008.t01#6    mNeoVis.evergladensis.2.1.chr1.g01000.t01#2    mNeoVis.evergladensis.2.1.chr1.g00941.t01#2    mNeoVis.vulgivaga.2.1.chr1.g00952.t01#1    mNeoVis.evergladensis.1.1.chr1.g00947.t01#3      2786   125259310:   mNeoVis.2.1.1.chr1_hap1.g00997.t01#5    mNeoVis.vulgivaga.1.1.chr1.g01000.t01#4   mNeoVis.vulgivaga.2.1.chr1.g01001.t01#1           2787   125259311:   mNeoVis.vulgivaga.2.1.chr1.g01002.t01#1   mNeoVis.vulgivaga.1.1.chr1.g01001.t01#4             2788   125259312:   mNeoVis.vulgivaga.1.1.chr1.g01002.t01#4    mNeoVis.vulgivaga.2.1.chr1.g01003.t01#1    mNeoVis.2.1.1.chr1_hap1.g00947.t01#5    mNeoVis.vulgivaga.1.1.chr1.g00957.t01#4    mNeoVis.vulgivaga.2.1.chr1.g00953.t01#1    mNeoVis.3.1.1.chr1.g00999.t01#6    mNeoVis.2.1.1.chr1_hap1.g00998.t01#5    mNeoVis.evergladensis.2.1.chr1.g00989.t01#2    mNeoVis.evergladensis.2.1.chr1.g00942.t01#2      2789   125259313:   mNeoVis.2.1.1.chr11_hap1.g00544.t01#5    mNeoVis.evergladensis.2.1.chr11.g00531.t01#2    mNeoVis.evergladensis.1.1.chr11.g00534.t01#3    mNeoVis.vulgivaga.1.1.chr11.g00526.t01#4   mNeoVis.vulgivaga.2.1.chr11.g00539.t01#1       2790   125259314:   mNeoVis.evergladensis.1.1.chr1.g00993.t01#3   mNeoVis.vulgivaga.1.1.chr1.g01004.t01#4             2791   125259315:   mNeoVis.evergladensis.1.1.chr1.g00948.t01#3   mNeoVis.3.1.1.chr1.g00959.t01#6             2792   125259316:   mNeoVis.vulgivaga.2.1.chr7.g01632.t01#1    mNeoVis.vulgivaga.1.1.chr7.g01630.t01#4    mNeoVis.evergladensis.2.1.chr7.g01616.t01#2    mNeoVis.evergladensis.1.1.chr7.g01620.t01#3    mNeoVis.2.1.1.chr7_hap1.g01642.t01#5   mNeoVis.3.1.1.chr7.g01544.t01#6       2793   125259317:   mNeoVis.3.1.1.chr7.g01547.t01#6    mNeoVis.2.1.1.chr7_hap1.g01646.t01#5    mNeoVis.evergladensis.2.1.chr7.g01620.t01#2    mNeoVis.evergladensis.1.1.chr7.g01622.t01#3    mNeoVis.vulgivaga.2.1.chr7.g01636.t01#1   mNeoVis.vulgivaga.1.1.chr7.g01633.t01#4       2794   125259318:   mNeoVis.vulgivaga.2.1.chr7.g01655.t01#1    mNeoVis.3.1.1.chr7.g01562.t01#6    mNeoVis.evergladensis.1.1.chr7.g01638.t01#3    mNeoVis.evergladensis.2.1.chr7.g01638.t01#2    mNeoVis.2.1.1.chr7_hap1.g01667.t01#5   mNeoVis.vulgivaga.1.1.chr7.g01652.t01#4       2795   125259319:   mNeoVis.evergladensis.1.1.chr7.g01623.t01#3    mNeoVis.vulgivaga.1.1.chr7.g01634.t01#4    mNeoVis.vulgivaga.1.1.chr7.g01640.t01#4    mNeoVis.evergladensis.2.1.chr7.g01621.t01#2   mNeoVis.evergladensis.2.1.chr7.g01625.t01#2       2796   125259320:   mNeoVis.2.1.1.chr7_hap1.g01650.t01#5    mNeoVis.vulgivaga.2.1.chr7.g01642.t01#1    mNeoVis.3.1.1.chr7.g01553.t01#6    mNeoVis.vulgivaga.1.1.chr7.g01639.t01#4   mNeoVis.evergladensis.1.1.chr7.g01628.t01#3       2797   125259321:   mNeoVis.vulgivaga.1.1.chr7.g01637.t01#4    mNeoVis.vulgivaga.2.1.chr7.g01640.t01#1    mNeoVis.evergladensis.1.1.chr7.g01626.t01#3   mNeoVis.3.1.1.chr7.g01551.t01#6         2798   125259322:   mNeoVis.vulgivaga.1.1.chr7.g01632.t01#4    mNeoVis.evergladensis.2.1.chr7.g01617.t01#2   mNeoVis.vulgivaga.2.1.chr7.g01633.t01#1           2799   125259323:   mNeoVis.3.1.1.chr7.g01545.t01#6    mNeoVis.2.1.1.chr7_hap1.g01643.t01#5   mNeoVis.evergladensis.2.1.chr7.g01619.t01#2         2800   125259324:   mNeoVis.2.1.1.chr7_hap1.g01645.t01#5   mNeoVis.vulgivaga.2.1.chr7.g01635.t01#1             2801   125259325:   mNeoVis.2.1.1.chr7_hap1.g01654.t01#5   mNeoVis.2.1.1.chr7_hap1.g01653.t01#5             2802   125259326:   mNeoVis.evergladensis.2.1.chr7.g01629.t01#2   mNeoVis.evergladensis.1.1.chr7.g01631.t01#3           2803   125259327:  mNeoVis.evergladensis.2.1.chr7.g01618.t01#2             2804   125259328:   mNeoVis.2.1.1.chr7_hap1.g01644.t01#5    mNeoVis.evergladensis.1.1.chr7.g01621.t01#3    mNeoVis.vulgivaga.2.1.chr7.g01634.t01#1    mNeoVis.3.1.1.chr7.g01546.t01#6   mNeoVis.vulgivaga.1.1.chr7.g01631.t01#4       2805   125259329:   mNeoVis.vulgivaga.1.1.chr1.g01003.t01#4    mNeoVis.evergladensis.1.1.chr1.g00992.t01#3    mNeoVis.evergladensis.2.1.chr1.g00990.t01#2    mNeoVis.vulgivaga.2.1.chr1.g01004.t01#1    mNeoVis.2.1.1.chr1_hap1.g00999.t01#5   mNeoVis.3.1.1.chr1.g01000.t01#6       2806   125259330:   mNeoVis.evergladensis.2.1.chr1.g00991.t01#2   mNeoVis.vulgivaga.2.1.chr1.g01005.t01#1             2807   125259331:   mNeoVis.vulgivaga.2.1.chr7.g01637.t01#1    mNeoVis.evergladensis.1.1.chr7.g01624.t01#3    mNeoVis.3.1.1.chr7.g01549.t01#6    mNeoVis.evergladensis.2.1.chr7.g01622.t01#2    mNeoVis.2.1.1.chr7_hap1.g01647.t01#5   mNeoVis.vulgivaga.1.1.chr7.g01635.t01#4       2808   125259332:  mNeoVis.vulgivaga.2.1.chr7.g01638.t01#1               2809   125259333:   mNeoVis.vulgivaga.1.1.chr1.g01005.t01#4    mNeoVis.evergladensis.2.1.chr1.g00992.t01#2    mNeoVis.2.1.1.chr1_hap1.g01000.t01#5    mNeoVis.vulgivaga.2.1.chr1.g01006.t01#1    mNeoVis.3.1.1.chr1.g01001.t01#6   mNeoVis.evergladensis.1.1.chr1.g00994.t01#3       2810   125259334:   mNeoVis.vulgivaga.2.1.chr1.g01007.t01#1    mNeoVis.3.1.1.chr1.g01002.t01#6    mNeoVis.vulgivaga.1.1.chr1.g01006.t01#4    mNeoVis.evergladensis.1.1.chr1.g00995.t01#3    mNeoVis.2.1.1.chr1_hap1.g01001.t01#5   mNeoVis.evergladensis.2.1.chr1.g00993.t01#2       2811   125259335:   mNeoVis.evergladensis.2.1.chr7.g01623.t01#2    mNeoVis.vulgivaga.2.1.chr7.g01639.t01#1    mNeoVis.vulgivaga.1.1.chr7.g01636.t01#4    mNeoVis.evergladensis.1.1.chr7.g01625.t01#3    mNeoVis.3.1.1.chr7.g01550.t01#6   mNeoVis.2.1.1.chr7_hap1.g01648.t01#5       2812   125259336:   mNeoVis.evergladensis.1.1.chr1.g00996.t01#3    mNeoVis.2.1.1.chr1_hap1.g01003.t01#5    mNeoVis.3.1.1.chr1.g01003.t01#6    mNeoVis.vulgivaga.2.1.chr1.g01009.t01#1    mNeoVis.evergladensis.2.1.chr1.g00994.t01#2    mNeoVis.vulgivaga.1.1.chr1.g01007.t01#4    mNeoVis.evergladensis.1.1.chr1.g00997.t01#3    mNeoVis.vulgivaga.2.1.chr1.g01008.t01#1    mNeoVis.2.1.1.chr1_hap1.g01002.t01#5      2813   125259337:   mNeoVis.evergladensis.1.1.chr1.g00940.t01#3    mNeoVis.evergladensis.2.1.chr1.g00934.t01#2    mNeoVis.vulgivaga.2.1.chr1.g00945.t01#1    mNeoVis.3.1.1.chr1.g00951.t01#6    mNeoVis.2.1.1.chr1_hap1.g00938.t01#5   mNeoVis.vulgivaga.1.1.chr1.g00948.t01#4       2814   125259338:   mNeoVis.2.1.1.chr1_hap1.g01009.t01#5    mNeoVis.vulgivaga.1.1.chr1.g01013.t01#4    mNeoVis.vulgivaga.2.1.chr1.g01016.t01#1    mNeoVis.evergladensis.2.1.chr1.g01002.t01#2    mNeoVis.evergladensis.1.1.chr1.g01003.t01#3   mNeoVis.3.1.1.chr1.g01010.t01#6       2815   125259339:   mNeoVis.3.1.1.chr1.g01011.t01#6    mNeoVis.evergladensis.2.1.chr1.g01003.t01#2    mNeoVis.2.1.1.chr1_hap1.g01010.t01#5    mNeoVis.vulgivaga.1.1.chr1.g01014.t01#4   mNeoVis.evergladensis.1.1.chr1.g01004.t01#3       2816   125259340:   mNeoVis.evergladensis.1.1.chr7.g01633.t01#3    mNeoVis.evergladensis.2.1.chr7.g01632.t01#2    mNeoVis.3.1.1.chr7.g01557.t01#6    mNeoVis.vulgivaga.2.1.chr7.g01649.t01#1    mNeoVis.3.1.1.chr7.g01558.t01#6    mNeoVis.2.1.1.chr7_hap1.g01660.t01#5    mNeoVis.vulgivaga.1.1.chr7.g01650.t01#4    mNeoVis.vulgivaga.2.1.chr7.g01648.t01#1    mNeoVis.2.1.1.chr7_hap1.g01659.t01#5      2817   125259341:   mNeoVis.2.1.1.chr7_hap1.g01651.t01#5    mNeoVis.3.1.1.chr7.g01537.t01#6    mNeoVis.vulgivaga.1.1.chr7.g01641.t01#4    mNeoVis.evergladensis.2.1.chr7.g01626.t01#2    mNeoVis.evergladensis.2.1.chr7.g01627.t01#2    mNeoVis.evergladensis.1.1.chr7.g01629.t01#3    mNeoVis.vulgivaga.2.1.chr7.g01643.t01#1    mNeoVis.vulgivaga.2.1.chr7.g01644.t01#1    mNeoVis.vulgivaga.1.1.chr7.g01642.t01#4      2818   125259342:   mNeoVis.vulgivaga.2.1.chr7.g01652.t01#1    mNeoVis.evergladensis.2.1.chr7.g01635.t01#2    mNeoVis.vulgivaga.1.1.chr7.g01648.t01#4    mNeoVis.3.1.1.chr7.g01560.t01#6    mNeoVis.2.1.1.chr7_hap1.g01664.t01#5    mNeoVis.vulgivaga.2.1.chr7.g01654.t01#1    mNeoVis.evergladensis.1.1.chr7.g01634.t01#3   mNeoVis.2.1.1.chr7_hap1.g01662.t01#5     2819   125259343:   mNeoVis.vulgivaga.2.1.chr7.g01641.t01#1    mNeoVis.vulgivaga.1.1.chr7.g01638.t01#4    mNeoVis.evergladensis.1.1.chr7.g01627.t01#3    mNeoVis.2.1.1.chr7_hap1.g01649.t01#5    mNeoVis.evergladensis.2.1.chr7.g01624.t01#2   mNeoVis.3.1.1.chr7.g01552.t01#6       2820   125259344:   mNeoVis.vulgivaga.2.1.chr7.g01647.t01#1    mNeoVis.evergladensis.1.1.chr7.g01632.t01#3    mNeoVis.2.1.1.chr7_hap1.g01657.t01#5    mNeoVis.2.1.1.chr7_hap1.g01656.t01#5    mNeoVis.vulgivaga.1.1.chr7.g01644.t01#4   mNeoVis.evergladensis.2.1.chr7.g01631.t01#2       2821   125259345:   mNeoVis.3.1.1.chr7.g01559.t01#6    mNeoVis.2.1.1.chr7_hap1.g01661.t01#5    mNeoVis.vulgivaga.2.1.chr7.g01650.t01#1    mNeoVis.evergladensis.2.1.chr7.g01634.t01#2   mNeoVis.vulgivaga.1.1.chr7.g01647.t01#4       2822   125259346:   mNeoVis.2.1.1.chr7_hap1.g01655.t01#5    mNeoVis.3.1.1.chr7.g01554.t01#6   mNeoVis.evergladensis.2.1.chr7.g01630.t01#2         2823   125259347:  mNeoVis.2.1.1.chr7_hap1.g01658.t01#5               2824   125259348:   mNeoVis.vulgivaga.1.1.chr1.g01008.t01#4    mNeoVis.vulgivaga.2.1.chr1.g01010.t01#1    mNeoVis.evergladensis.2.1.chr1.g00996.t01#2    mNeoVis.2.1.1.chr1_hap1.g01004.t01#5    mNeoVis.evergladensis.1.1.chr1.g00998.t01#3   mNeoVis.3.1.1.chr1.g01004.t01#6       2825   125259349:   mNeoVis.vulgivaga.1.1.chr7.g00084.t01#4    mNeoVis.2.1.1.chr7_hap1.g00082.t01#5    mNeoVis.evergladensis.2.1.chr7.g00082.t01#2    mNeoVis.vulgivaga.2.1.chr7.g00082.t01#1   mNeoVis.evergladensis.1.1.chr7.g00084.t01#3       2826   125259350:   mNeoVis.evergladensis.2.1.chr1.g00997.t01#2    mNeoVis.vulgivaga.1.1.chr1.g01009.t01#4    mNeoVis.3.1.1.chr1.g01005.t01#6    mNeoVis.2.1.1.chr1_hap1.g01005.t01#5    mNeoVis.evergladensis.1.1.chr1.g00999.t01#3   mNeoVis.vulgivaga.2.1.chr1.g01011.t01#1       2827   125259351:   mNeoVis.2.1.1.chr1_hap1.g01006.t01#5    mNeoVis.evergladensis.2.1.chr1.g00998.t01#2    mNeoVis.vulgivaga.2.1.chr1.g01012.t01#1    mNeoVis.evergladensis.1.1.chr1.g01000.t01#3    mNeoVis.3.1.1.chr1.g01006.t01#6   mNeoVis.vulgivaga.1.1.chr1.g01010.t01#4       2828   125259352:   mNeoVis.vulgivaga.1.1.chr1.g01011.t01#4    mNeoVis.3.1.1.chr1.g01007.t01#6    mNeoVis.2.1.1.chr1_hap1.g01007.t01#5    mNeoVis.2.1.1.chr11_hap1.g00543.t01#5    mNeoVis.evergladensis.1.1.chr1.g01001.t01#3    mNeoVis.evergladensis.2.1.chr1.g00999.t01#2   mNeoVis.vulgivaga.2.1.chr1.g01013.t01#1     2829   125259353:   mNeoVis.evergladensis.1.1.chr7.g01630.t01#3    mNeoVis.vulgivaga.2.1.chr7.g01645.t01#1    mNeoVis.evergladensis.2.1.chr7.g01628.t01#2   mNeoVis.2.1.1.chr7_hap1.g01652.t01#5         2830   125259354:  mNeoVis.vulgivaga.2.1.chr7.g01646.t01#1               2831   125259355:   mNeoVis.evergladensis.2.1.chr1.g01001.t01#2    mNeoVis.3.1.1.chr1.g01009.t01#6   mNeoVis.vulgivaga.2.1.chr1.g01015.t01#1           2832   125259356:   mNeoVis.evergladensis.1.1.chr1.g01006.t01#3    mNeoVis.evergladensis.2.1.chr1.g01005.t01#2    mNeoVis.3.1.1.chr1.g01013.t01#6    mNeoVis.2.1.1.chr1_hap1.g01012.t01#5    mNeoVis.vulgivaga.2.1.chr1.g01017.t01#1   mNeoVis.vulgivaga.1.1.chr1.g01016.t01#4       2833   125259357:   mNeoVis.vulgivaga.2.1.chr1.g01018.t01#1    mNeoVis.3.1.1.chr1.g01014.t01#6    mNeoVis.vulgivaga.1.1.chr1.g01017.t01#4    mNeoVis.2.1.1.chr1_hap1.g01013.t01#5    mNeoVis.evergladensis.1.1.chr1.g01007.t01#3   mNeoVis.evergladensis.2.1.chr1.g01006.t01#2       2834   125259358:   mNeoVis.2.1.1.chr1_hap1.g01014.t01#5    mNeoVis.vulgivaga.1.1.chr1.g01018.t01#4    mNeoVis.evergladensis.2.1.chr1.g01007.t01#2    mNeoVis.evergladensis.1.1.chr1.g01008.t01#3    mNeoVis.3.1.1.chr1.g01015.t01#6   mNeoVis.vulgivaga.2.1.chr1.g01019.t01#1       2835   125259359:   mNeoVis.evergladensis.2.1.chr7.g01642.t01#2    mNeoVis.vulgivaga.1.1.chr7.g01654.t01#4    mNeoVis.evergladensis.1.1.chr7.g01640.t01#3    mNeoVis.2.1.1.chr7_hap1.g01670.t01#5    mNeoVis.evergladensis.2.1.chr7.g01641.t01#2    mNeoVis.vulgivaga.1.1.chr7.g01655.t01#4    mNeoVis.vulgivaga.2.1.chr7.g01659.t01#1    mNeoVis.vulgivaga.2.1.chr7.g01658.t01#1    mNeoVis.evergladensis.1.1.chr7.g01641.t01#3      2836   125259360:   mNeoVis.2.1.1.chr7_hap1.g01668.t01#5    mNeoVis.evergladensis.2.1.chr7.g01639.t01#2    mNeoVis.vulgivaga.1.1.chr7.g01653.t01#4    mNeoVis.vulgivaga.2.1.chr7.g01656.t01#1   mNeoVis.evergladensis.1.1.chr7.g01639.t01#3       2837   125259361:   mNeoVis.evergladensis.2.1.chr7.g01640.t01#2   mNeoVis.2.1.1.chr7_hap1.g01669.t01#5             2838   125259362:  mNeoVis.vulgivaga.2.1.chr7.g01657.t01#1               2839   125259363:   mNeoVis.evergladensis.1.1.chr1.g01009.t01#3    mNeoVis.vulgivaga.2.1.chr1.g01020.t01#1    mNeoVis.2.1.1.chr1_hap1.g01015.t01#5    mNeoVis.vulgivaga.1.1.chr1.g01019.t01#4    mNeoVis.evergladensis.2.1.chr1.g01008.t01#2   mNeoVis.3.1.1.chr1.g01016.t01#6       2840   125259364:   mNeoVis.evergladensis.2.1.chr1.g01010.t01#2    mNeoVis.3.1.1.chr1.g01017.t01#6    mNeoVis.evergladensis.1.1.chr1.g01011.t01#3    mNeoVis.2.1.1.chr1_hap1.g01016.t01#5    mNeoVis.vulgivaga.1.1.chr1.g01020.t01#4   mNeoVis.vulgivaga.2.1.chr1.g01021.t01#1       2841   125259365:   mNeoVis.evergladensis.1.1.chr7.g01643.t01#3    mNeoVis.vulgivaga.1.1.chr7.g01656.t01#4    mNeoVis.evergladensis.2.1.chr7.g01644.t01#2    mNeoVis.evergladensis.1.1.chr7.g01642.t01#3    mNeoVis.vulgivaga.2.1.chr7.g01660.t01#1    mNeoVis.2.1.1.chr7_hap1.g01672.t01#5    mNeoVis.vulgivaga.2.1.chr7.g01661.t01#1    mNeoVis.2.1.1.chr7_hap1.g01671.t01#5    mNeoVis.3.1.1.chr7.g01565.t01#6      2842   125259366:   mNeoVis.3.1.1.chr7.g01566.t01#6    mNeoVis.2.1.1.chr7_hap1.g01673.t01#5    mNeoVis.vulgivaga.2.1.chr7.g01662.t01#1    mNeoVis.evergladensis.2.1.chr7.g01645.t01#2    mNeoVis.vulgivaga.1.1.chr7.g01657.t01#4   mNeoVis.evergladensis.1.1.chr7.g01644.t01#3       2843   125259367:   mNeoVis.2.1.1.chr7_hap1.g01674.t01#5    mNeoVis.vulgivaga.2.1.chr7.g01663.t01#1    mNeoVis.vulgivaga.1.1.chr7.g01658.t01#4    mNeoVis.evergladensis.1.1.chr7.g01645.t01#3    mNeoVis.evergladensis.2.1.chr7.g01646.t01#2   mNeoVis.3.1.1.chr7.g01567.t01#6       2844   125259368:   mNeoVis.vulgivaga.1.1.chr7.g01659.t01#4    mNeoVis.evergladensis.2.1.chr7.g01647.t01#2    mNeoVis.evergladensis.1.1.chr7.g01646.t01#3    mNeoVis.3.1.1.chr7.g01568.t01#6    mNeoVis.2.1.1.chr7_hap1.g01675.t01#5   mNeoVis.vulgivaga.2.1.chr7.g01664.t01#1       2845   125259369:  mNeoVis.evergladensis.2.1.chr7.g01643.t01#2             2846   125259370:   mNeoVis.3.1.1.chr1.g01018.t01#6    mNeoVis.evergladensis.2.1.chr1.g01011.t01#2    mNeoVis.vulgivaga.2.1.chr1.g01022.t01#1    mNeoVis.2.1.1.chr1_hap1.g01017.t01#5    mNeoVis.evergladensis.1.1.chr1.g01012.t01#3   mNeoVis.vulgivaga.1.1.chr1.g01021.t01#4       2847   125259371:   mNeoVis.evergladensis.1.1.chr7.g01647.t01#3    mNeoVis.evergladensis.2.1.chr7.g01648.t01#2    mNeoVis.vulgivaga.2.1.chr7.g01665.t01#1    mNeoVis.2.1.1.chr7_hap1.g01676.t01#5   mNeoVis.vulgivaga.1.1.chr7.g01660.t01#4       2848   125259372:   mNeoVis.3.1.1.chr7.g01571.t01#6    mNeoVis.evergladensis.2.1.chr7.g01649.t01#2    mNeoVis.evergladensis.1.1.chr7.g01649.t01#3    mNeoVis.vulgivaga.1.1.chr7.g01661.t01#4   mNeoVis.vulgivaga.2.1.chr7.g01666.t01#1       2849   125259373:   mNeoVis.2.1.1.chr7_hap1.g01677.t01#5    mNeoVis.evergladensis.2.1.chr7.g01650.t01#2    mNeoVis.evergladensis.1.1.chr7.g01650.t01#3    mNeoVis.vulgivaga.2.1.chr7.g01667.t01#1   mNeoVis.3.1.1.chr7.g01572.t01#6       2850   125259374:   mNeoVis.3.1.1.chr1.g01019.t01#6    mNeoVis.vulgivaga.2.1.chr1.g01023.t01#1    mNeoVis.evergladensis.2.1.chr1.g01012.t01#2    mNeoVis.2.1.1.chr1_hap1.g01018.t01#5    mNeoVis.evergladensis.1.1.chr1.g01013.t01#3   mNeoVis.vulgivaga.1.1.chr1.g01022.t01#4       2851   125259375:   mNeoVis.vulgivaga.1.1.chr7.g01667.t01#4    mNeoVis.3.1.1.chr7.g01576.t01#6    mNeoVis.evergladensis.2.1.chr7.g01655.t01#2    mNeoVis.vulgivaga.2.1.chr7.g01673.t01#1    mNeoVis.2.1.1.chr7_hap1.g01684.t01#5   mNeoVis.evergladensis.1.1.chr7.g01654.t01#3       2852   125259376:   mNeoVis.vulgivaga.2.1.chr7.g01668.t01#1    mNeoVis.2.1.1.chr7_hap1.g01679.t01#5    mNeoVis.3.1.1.chr7.g01573.t01#6   mNeoVis.vulgivaga.1.1.chr7.g01663.t01#4         2853   125259377:   mNeoVis.2.1.1.chr7_hap1.g01683.t01#5    mNeoVis.3.1.1.chr7.g01575.t01#6    mNeoVis.evergladensis.2.1.chr7.g01654.t01#2   mNeoVis.vulgivaga.2.1.chr7.g01672.t01#1         2854   125259378:   mNeoVis.evergladensis.2.1.chr7.g01651.t01#2    mNeoVis.2.1.1.chr7_hap1.g01678.t01#5   mNeoVis.evergladensis.1.1.chr7.g01651.t01#3         2855   125259379:  mNeoVis.vulgivaga.2.1.chr7.g01670.t01#1               2856   125259380:  mNeoVis.vulgivaga.1.1.chr7.g01665.t01#4               2857   125259381:   mNeoVis.2.1.1.chr7_hap1.g01680.t01#5    mNeoVis.vulgivaga.2.1.chr7.g01669.t01#1    mNeoVis.vulgivaga.1.1.chr7.g01664.t01#4    mNeoVis.evergladensis.1.1.chr7.g01652.t01#3   mNeoVis.evergladensis.2.1.chr7.g01652.t01#2       2858   125259382:   mNeoVis.vulgivaga.2.1.chr1.g01024.t01#1    mNeoVis.evergladensis.1.1.chr1.g01014.t01#3    mNeoVis.2.1.1.chr1_hap1.g01019.t01#5    mNeoVis.vulgivaga.1.1.chr1.g01023.t01#4    mNeoVis.evergladensis.2.1.chr1.g01013.t01#2   mNeoVis.3.1.1.chr1.g01020.t01#6       2859   125259383:   mNeoVis.evergladensis.2.1.chr7.g01653.t01#2    mNeoVis.evergladensis.1.1.chr7.g01653.t01#3    mNeoVis.vulgivaga.1.1.chr7.g01666.t01#4    mNeoVis.2.1.1.chr7_hap1.g01682.t01#5    mNeoVis.3.1.1.chr7.g01574.t01#6    mNeoVis.2.1.1.chr7_hap1.g01681.t01#5   mNeoVis.vulgivaga.2.1.chr7.g01671.t01#1     2860   125259384:   mNeoVis.evergladensis.1.1.chr1.g01015.t01#3    mNeoVis.2.1.1.chr1_hap1.g01020.t01#5    mNeoVis.vulgivaga.2.1.chr1.g01025.t01#1    mNeoVis.vulgivaga.1.1.chr1.g01024.t01#4    mNeoVis.evergladensis.2.1.chr1.g01014.t01#2   mNeoVis.3.1.1.chr1.g01021.t01#6       2861   125259385:   mNeoVis.3.1.1.chr1.g01022.t01#6    mNeoVis.vulgivaga.2.1.chr7.g01140.t01#1    mNeoVis.vulgivaga.1.1.chr7.g01129.t01#4    mNeoVis.evergladensis.2.1.chr7.g01128.t01#2   mNeoVis.2.1.1.chr7_hap1.g01141.t01#5       2862   125259386:   mNeoVis.evergladensis.1.1.chr1.g01016.t01#3    mNeoVis.evergladensis.2.1.chr1.g01015.t01#2    mNeoVis.vulgivaga.1.1.chr1.g01025.t01#4    mNeoVis.2.1.1.chr1_hap1.g01021.t01#5   mNeoVis.vulgivaga.2.1.chr1.g01026.t01#1       2863   125259387:   mNeoVis.evergladensis.2.1.chr1.g01016.t01#2    mNeoVis.vulgivaga.1.1.chr1.g01026.t01#4    mNeoVis.2.1.1.chr1_hap1.g01022.t01#5    mNeoVis.vulgivaga.2.1.chr1.g01027.t01#1   mNeoVis.evergladensis.1.1.chr1.g01017.t01#3       2864   125259388:   mNeoVis.evergladensis.1.1.chr1.g01018.t01#3    mNeoVis.evergladensis.2.1.chr1.g01017.t01#2    mNeoVis.vulgivaga.2.1.chr1.g01028.t01#1    mNeoVis.2.1.1.chr1_hap1.g01023.t01#5   mNeoVis.vulgivaga.1.1.chr1.g01027.t01#4       2865   125259389:   mNeoVis.evergladensis.1.1.chr7.g01657.t01#3    mNeoVis.2.1.1.chr7_hap1.g01687.t01#5    mNeoVis.vulgivaga.2.1.chr7.g01675.t01#1    mNeoVis.vulgivaga.1.1.chr7.g01669.t01#4    mNeoVis.2.1.1.chr7_hap1.g01686.t01#5    mNeoVis.evergladensis.1.1.chr7.g01656.t01#3    mNeoVis.evergladensis.2.1.chr7.g01657.t01#2    mNeoVis.vulgivaga.2.1.chr7.g01674.t01#1    mNeoVis.2.1.1.chr7_hap1.g01685.t01#5      2866   125259390:   mNeoVis.vulgivaga.1.1.chr7.g01670.t01#4    mNeoVis.3.1.1.chr7.g01578.t01#6    mNeoVis.evergladensis.1.1.chr7.g01658.t01#3    mNeoVis.evergladensis.2.1.chr7.g01658.t01#2    mNeoVis.2.1.1.chr7_hap1.g01688.t01#5   mNeoVis.vulgivaga.2.1.chr7.g01676.t01#1       2867   125259391:   mNeoVis.evergladensis.1.1.chr7.g01659.t01#3    mNeoVis.vulgivaga.2.1.chr7.g01677.t01#1    mNeoVis.2.1.1.chr7_hap1.g01689.t01#5    mNeoVis.vulgivaga.1.1.chr7.g01671.t01#4   mNeoVis.evergladensis.2.1.chr7.g01659.t01#2       2868   125259392:   mNeoVis.vulgivaga.2.1.chr7.g01678.t01#1    mNeoVis.evergladensis.1.1.chr7.g01660.t01#3    mNeoVis.3.1.1.chr7.g01579.t01#6    mNeoVis.evergladensis.2.1.chr7.g01660.t01#2    mNeoVis.vulgivaga.1.1.chr7.g01672.t01#4   mNeoVis.2.1.1.chr7_hap1.g01690.t01#5       2869   125259393:   mNeoVis.2.1.1.chr1_hap1.g01024.t01#5    mNeoVis.evergladensis.1.1.chr1.g01019.t01#3    mNeoVis.3.1.1.chr1.g01023.t01#6    mNeoVis.3.1.1.chr13.g00064.t01#6    mNeoVis.evergladensis.2.1.chr1.g01018.t01#2    mNeoVis.vulgivaga.2.1.chr1.g01029.t01#1   mNeoVis.vulgivaga.1.1.chr1.g01028.t01#4     2870   125259394:   mNeoVis.2.1.1.chr7_hap1.g01691.t01#5    mNeoVis.vulgivaga.2.1.chr7.g01679.t01#1    mNeoVis.evergladensis.1.1.chr7.g01655.t01#3    mNeoVis.evergladensis.2.1.chr7.g01661.t01#2    mNeoVis.evergladensis.1.1.chr7.g01661.t01#3    mNeoVis.vulgivaga.1.1.chr7.g01673.t01#4   mNeoVis.3.1.1.chr7.g01580.t01#6       2871   125259395:  mNeoVis.vulgivaga.2.1.chr7.g01680.t01#1               2872   125259396:   mNeoVis.2.1.1.chr1_hap1.g01025.t01#5    mNeoVis.evergladensis.2.1.chr1.g01019.t01#2    mNeoVis.evergladensis.1.1.chr1.g01020.t01#3    mNeoVis.vulgivaga.1.1.chr1.g01029.t01#4   mNeoVis.vulgivaga.2.1.chr1.g01030.t01#1       2873   125259397:   mNeoVis.vulgivaga.2.1.chr7.g01681.t01#1    mNeoVis.evergladensis.1.1.chr7.g01662.t01#3    mNeoVis.vulgivaga.1.1.chr7.g01675.t01#4    mNeoVis.vulgivaga.2.1.chr7.g01683.t01#1    mNeoVis.3.1.1.chr7.g01582.t01#6    mNeoVis.2.1.1.chr7_hap1.g01693.t01#5    mNeoVis.vulgivaga.2.1.chr7.g01682.t01#1    mNeoVis.evergladensis.1.1.chr7.g01663.t01#3    mNeoVis.evergladensis.2.1.chr7.g01663.t01#2      2874   125259398:   mNeoVis.2.1.1.chr7_hap1.g01700.t01#5    mNeoVis.evergladensis.2.1.chr7.g01671.t01#2    mNeoVis.3.1.1.chr7.g01589.t01#6    mNeoVis.vulgivaga.1.1.chr7.g01682.t01#4    mNeoVis.2.1.1.chr7_hap1.g01699.t01#5    mNeoVis.evergladensis.1.1.chr7.g01668.t01#3    mNeoVis.vulgivaga.2.1.chr7.g01687.t01#1   mNeoVis.evergladensis.2.1.chr7.g01670.t01#2     2875   125259399:   mNeoVis.3.1.1.chr7.g01583.t01#6    mNeoVis.2.1.1.chr7_hap1.g01695.t01#5    mNeoVis.vulgivaga.2.1.chr7.g01684.t01#1    mNeoVis.evergladensis.1.1.chr7.g01664.t01#3    mNeoVis.evergladensis.2.1.chr7.g01665.t01#2   mNeoVis.vulgivaga.1.1.chr7.g01677.t01#4       2876   125259400:   mNeoVis.2.1.1.chr7_hap1.g01692.t01#5    mNeoVis.vulgivaga.1.1.chr7.g01674.t01#4   mNeoVis.3.1.1.chr7.g01581.t01#6           2877   125259401:   mNeoVis.vulgivaga.1.1.chr7.g01676.t01#4   mNeoVis.2.1.1.chr7_hap1.g01694.t01#5             2878   125259402:  mNeoVis.evergladensis.2.1.chr7.g01662.t01#2             2879   125259403:   mNeoVis.vulgivaga.2.1.chr7.g01685.t01#1    mNeoVis.2.1.1.chr7_hap1.g01696.t01#5    mNeoVis.evergladensis.2.1.chr7.g01664.t01#2    mNeoVis.evergladensis.1.1.chr7.g01665.t01#3   mNeoVis.3.1.1.chr7.g01584.t01#6       2880   125259404:   mNeoVis.2.1.1.chr1_hap1.g01026.t01#5    mNeoVis.vulgivaga.2.1.chr1.g01031.t01#1    mNeoVis.evergladensis.1.1.chr1.g01021.t01#3    mNeoVis.vulgivaga.1.1.chr1.g01030.t01#4   mNeoVis.evergladensis.2.1.chr1.g01020.t01#2       2881   125259405:  mNeoVis.vulgivaga.2.1.chr7.g01686.t01#1               2882   125259406:   mNeoVis.evergladensis.1.1.chr1.g00884.t01#3    mNeoVis.3.1.1.chr1.g00894.t01#6    mNeoVis.vulgivaga.2.1.chr1.g00888.t01#1    mNeoVis.evergladensis.2.1.chr1.g00874.t01#2    mNeoVis.2.1.1.chr1_hap1.g00871.t01#5   mNeoVis.vulgivaga.1.1.chr1.g00889.t01#4       2883   125259407:   mNeoVis.2.1.1.chr1_hap1.g00872.t01#5    mNeoVis.evergladensis.1.1.chr1.g00885.t01#3    mNeoVis.evergladensis.2.1.chr1.g00875.t01#2    mNeoVis.vulgivaga.1.1.chr1.g00890.t01#4    mNeoVis.3.1.1.chr1.g00895.t01#6   mNeoVis.vulgivaga.2.1.chr1.g00889.t01#1       2884   125259408:   mNeoVis.vulgivaga.2.1.chr3.g01313.t01#1    mNeoVis.evergladensis.1.1.chr3.g01300.t01#3    mNeoVis.2.1.1.chr3_hap1.g01294.t01#5    mNeoVis.vulgivaga.1.1.chr3.g01287.t01#4    mNeoVis.evergladensis.2.1.chr3.g01293.t01#2   mNeoVis.3.1.1.chr3.g01271.t01#6       2885   125259409:   mNeoVis.3.1.1.chr7.g01590.t01#6    mNeoVis.evergladensis.2.1.chr7.g01672.t01#2    mNeoVis.2.1.1.chr7_hap1.g01701.t01#5    mNeoVis.vulgivaga.2.1.chr7.g01688.t01#1    mNeoVis.vulgivaga.1.1.chr7.g01683.t01#4   mNeoVis.evergladensis.1.1.chr7.g01669.t01#3       2886   125259410:   mNeoVis.vulgivaga.1.1.chr7.g01662.t01#4    mNeoVis.vulgivaga.2.1.chr7.g01689.t01#1    mNeoVis.evergladensis.1.1.chr7.g01670.t01#3    mNeoVis.2.1.1.chr7_hap1.g01702.t01#5    mNeoVis.3.1.1.chr7.g01591.t01#6   mNeoVis.evergladensis.2.1.chr7.g01673.t01#2       2887   125259411:   mNeoVis.vulgivaga.2.1.chr7.g01690.t01#1    mNeoVis.evergladensis.2.1.chr7.g01674.t01#2    mNeoVis.evergladensis.1.1.chr7.g01671.t01#3    mNeoVis.2.1.1.chr7_hap1.g01703.t01#5    mNeoVis.vulgivaga.1.1.chr7.g01684.t01#4   mNeoVis.3.1.1.chr7.g01592.t01#6       2888   125259412:   mNeoVis.evergladensis.1.1.chr1.g00887.t01#3    mNeoVis.vulgivaga.1.1.chr1.g00891.t01#4    mNeoVis.vulgivaga.2.1.chr1.g00890.t01#1    mNeoVis.2.1.1.chr1_hap1.g00874.t01#5    mNeoVis.3.1.1.chr1.g00897.t01#6   mNeoVis.evergladensis.2.1.chr1.g00876.t01#2       2889   125259413:   mNeoVis.2.1.1.chr7_hap1.g01704.t01#5    mNeoVis.3.1.1.chr7.g01593.t01#6    mNeoVis.evergladensis.2.1.chr7.g01675.t01#2    mNeoVis.evergladensis.1.1.chr7.g01677.t01#3    mNeoVis.vulgivaga.2.1.chr7.g01698.t01#1    mNeoVis.evergladensis.2.1.chr7.g01680.t01#2    mNeoVis.evergladensis.1.1.chr7.g01676.t01#3    mNeoVis.evergladensis.2.1.chr7.g01679.t01#2    mNeoVis.vulgivaga.1.1.chr7.g01685.t01#4      2890   125259414:   mNeoVis.vulgivaga.2.1.chr7.g01699.t01#1    mNeoVis.2.1.1.chr7_hap1.g01714.t01#5    mNeoVis.evergladensis.2.1.chr7.g01683.t01#2    mNeoVis.vulgivaga.1.1.chr7.g01693.t01#4    mNeoVis.3.1.1.chr7.g01598.t01#6   mNeoVis.evergladensis.1.1.chr7.g01680.t01#3       2891   125259415:   mNeoVis.evergladensis.2.1.chr7.g01684.t01#2    mNeoVis.2.1.1.chr7_hap1.g01715.t01#5    mNeoVis.evergladensis.2.1.chr7.g01685.t01#2    mNeoVis.evergladensis.1.1.chr7.g01681.t01#3   mNeoVis.vulgivaga.2.1.chr7.g01700.t01#1       2892   125259416:   mNeoVis.vulgivaga.1.1.chr7.g01694.t01#4    mNeoVis.evergladensis.1.1.chr7.g01682.t01#3   mNeoVis.2.1.1.chr7_hap1.g01716.t01#5           2893   125259417:  mNeoVis.vulgivaga.2.1.chr7.g01692.t01#1               2894   125259418:   mNeoVis.vulgivaga.1.1.chr1.g00892.t01#4    mNeoVis.evergladensis.2.1.chr1.g00877.t01#2    mNeoVis.vulgivaga.2.1.chr1.g00891.t01#1    mNeoVis.2.1.1.chr1_hap1.g00875.t01#5    mNeoVis.evergladensis.1.1.chr1.g00888.t01#3   mNeoVis.3.1.1.chr1.g00898.t01#6       2895   125259419:   mNeoVis.3.1.1.chr7.g01594.t01#6    mNeoVis.evergladensis.1.1.chr7.g01675.t01#3    mNeoVis.vulgivaga.1.1.chr7.g01687.t01#4    mNeoVis.vulgivaga.2.1.chr7.g01694.t01#1    mNeoVis.2.1.1.chr7_hap1.g01707.t01#5    mNeoVis.evergladensis.2.1.chr7.g01676.t01#2    mNeoVis.2.1.1.chr7_hap1.g01708.t01#5    mNeoVis.evergladensis.1.1.chr7.g01673.t01#3    mNeoVis.2.1.1.chr7_hap1.g01705.t01#5      2896   125259420:   mNeoVis.3.1.1.chr7.g01596.t01#6    mNeoVis.vulgivaga.1.1.chr7.g01691.t01#4    mNeoVis.evergladensis.1.1.chr7.g01678.t01#3    mNeoVis.2.1.1.chr7_hap1.g01712.t01#5    mNeoVis.vulgivaga.2.1.chr7.g01695.t01#1   mNeoVis.evergladensis.2.1.chr7.g01681.t01#2       2897   125259421:   mNeoVis.vulgivaga.1.1.chr1.g00893.t01#4    mNeoVis.evergladensis.2.1.chr1.g00878.t01#2    mNeoVis.2.1.1.chr1_hap1.g00876.t01#5    mNeoVis.evergladensis.1.1.chr1.g00889.t01#3    mNeoVis.3.1.1.chr1.g00899.t01#6   mNeoVis.vulgivaga.2.1.chr1.g00892.t01#1       2898   125259422:   mNeoVis.vulgivaga.2.1.chr7.g00083.t01#1    mNeoVis.vulgivaga.1.1.chr7.g00085.t01#4    mNeoVis.evergladensis.1.1.chr7.g00085.t01#3   mNeoVis.3.1.1.chr7.g00069.t01#6         2899   125259423:   mNeoVis.evergladensis.2.1.chr1.g00882.t01#2    mNeoVis.evergladensis.1.1.chr1.g00891.t01#3    mNeoVis.vulgivaga.1.1.chr1.g00895.t01#4    mNeoVis.2.1.1.chr1_hap1.g00879.t01#5    mNeoVis.3.1.1.chr1.g00901.t01#6   mNeoVis.vulgivaga.2.1.chr1.g00893.t01#1       2900   125259424:   mNeoVis.vulgivaga.1.1.chr1.g00896.t01#4    mNeoVis.2.1.1.chr1_hap1.g00880.t01#5    mNeoVis.vulgivaga.2.1.chr1.g00894.t01#1    mNeoVis.evergladensis.2.1.chr1.g00883.t01#2   mNeoVis.evergladensis.1.1.chr1.g00892.t01#3       2901   125259425:   mNeoVis.3.1.1.chr15.g00144.t01#6    mNeoVis.evergladensis.2.1.chr15.g00151.t01#2    mNeoVis.evergladensis.1.1.chr15.g00463.t01#3    mNeoVis.vulgivaga.1.1.chr15.g00150.t01#4    mNeoVis.vulgivaga.2.1.chr15.g00154.t01#1   mNeoVis.2.1.1.chr15_hap1.g00168.t01#5       2902   125259426:   mNeoVis.evergladensis.1.1.chr1.g00893.t01#3    mNeoVis.evergladensis.2.1.chr1.g00884.t01#2    mNeoVis.vulgivaga.1.1.chr1.g00897.t01#4    mNeoVis.2.1.1.chr1_hap1.g00881.t01#5   mNeoVis.vulgivaga.2.1.chr1.g00895.t01#1       2903   125259427:   mNeoVis.2.1.1.chr7_hap1.g00084.t01#5    mNeoVis.evergladensis.2.1.chr7.g00084.t01#2    mNeoVis.vulgivaga.2.1.chr7.g00084.t01#1    mNeoVis.3.1.1.chr7.g00070.t01#6    mNeoVis.vulgivaga.1.1.chr7.g00086.t01#4   mNeoVis.evergladensis.1.1.chr7.g00086.t01#3       2904   125259428:   mNeoVis.vulgivaga.2.1.chr1.g00896.t01#1    mNeoVis.2.1.1.chr1_hap1.g00882.t01#5    mNeoVis.vulgivaga.2.1.chr1.g00897.t01#1    mNeoVis.evergladensis.1.1.chr1.g00894.t01#3   mNeoVis.evergladensis.2.1.chr1.g00885.t01#2       2905   125259429:   mNeoVis.vulgivaga.1.1.chr1.g00898.t01#4   mNeoVis.3.1.1.chr1.g00903.t01#6             2906   125259430:   mNeoVis.3.1.1.chr7.g01611.t01#6    mNeoVis.vulgivaga.1.1.chr7.g01706.t01#4    mNeoVis.vulgivaga.2.1.chr7.g01712.t01#1    mNeoVis.vulgivaga.1.1.chr7.g01705.t01#4    mNeoVis.evergladensis.2.1.chr7.g01700.t01#2    mNeoVis.vulgivaga.2.1.chr7.g01711.t01#1    mNeoVis.evergladensis.1.1.chr7.g01696.t01#3    mNeoVis.evergladensis.2.1.chr7.g01691.t01#2    mNeoVis.evergladensis.2.1.chr7.g01699.t01#2      2907   125259431:   mNeoVis.evergladensis.2.1.chr7.g01692.t01#2    mNeoVis.evergladensis.1.1.chr7.g01690.t01#3    mNeoVis.vulgivaga.2.1.chr7.g01707.t01#1    mNeoVis.3.1.1.chr7.g01608.t01#6    mNeoVis.2.1.1.chr7_hap1.g01728.t01#5    mNeoVis.vulgivaga.2.1.chr7.g01706.t01#1    mNeoVis.3.1.1.chr7.g01609.t01#6    mNeoVis.evergladensis.1.1.chr7.g01688.t01#3    mNeoVis.evergladensis.2.1.chr7.g01694.t01#2      2908   125259432:   mNeoVis.evergladensis.2.1.chr7.g01693.t01#2    mNeoVis.3.1.1.chr7.g01607.t01#6    mNeoVis.vulgivaga.1.1.chr7.g01702.t01#4    mNeoVis.vulgivaga.2.1.chr7.g01708.t01#1    mNeoVis.2.1.1.chr7_hap1.g01725.t01#5   mNeoVis.evergladensis.1.1.chr7.g01689.t01#3       2909   125259433:  mNeoVis.2.1.1.chr7_hap1.g01727.t01#5               2910   125259434:  mNeoVis.vulgivaga.1.1.chr7.g01701.t01#4               2911   125259435:   mNeoVis.2.1.1.chr1_hap1.g00883.t01#5    mNeoVis.3.1.1.chr1.g00904.t01#6    mNeoVis.vulgivaga.1.1.chr1.g00900.t01#4    mNeoVis.vulgivaga.2.1.chr1.g00898.t01#1   mNeoVis.evergladensis.1.1.chr1.g00895.t01#3       2912   125259436:   mNeoVis.3.1.1.chr7.g01610.t01#6    mNeoVis.2.1.1.chr7_hap1.g01729.t01#5    mNeoVis.evergladensis.2.1.chr7.g01697.t01#2   mNeoVis.vulgivaga.2.1.chr7.g01710.t01#1         2913   125259437:   mNeoVis.2.1.1.chr7_hap1.g01733.t01#5    mNeoVis.3.1.1.chr7.g01613.t01#6    mNeoVis.evergladensis.2.1.chr7.g01701.t01#2    mNeoVis.2.1.1.chr7_hap1.g01735.t01#5    mNeoVis.vulgivaga.2.1.chr7.g01715.t01#1    mNeoVis.vulgivaga.2.1.chr7.g01713.t01#1    mNeoVis.evergladensis.1.1.chr7.g01698.t01#3   mNeoVis.vulgivaga.1.1.chr7.g01708.t01#4     2914   125259438:   mNeoVis.2.1.1.chr15_hap1.g00169.t01#5    mNeoVis.evergladensis.2.1.chr15.g00152.t01#2    mNeoVis.vulgivaga.2.1.chr15.g00155.t01#1    mNeoVis.vulgivaga.1.1.chr15.g00151.t01#4    mNeoVis.evergladensis.1.1.chr15.g00461.t01#3   mNeoVis.3.1.1.chr15.g00145.t01#6       2915   125259439:   mNeoVis.vulgivaga.1.1.chr7.g01711.t01#4    mNeoVis.evergladensis.1.1.chr7.g01700.t01#3    mNeoVis.3.1.1.chr7.g01616.t01#6    mNeoVis.2.1.1.chr7_hap1.g01738.t01#5    mNeoVis.evergladensis.2.1.chr7.g01703.t01#2   mNeoVis.vulgivaga.2.1.chr7.g01717.t01#1       2916   125259440:   mNeoVis.2.1.1.chr7_hap1.g01737.t01#5    mNeoVis.vulgivaga.2.1.chr7.g01714.t01#1    mNeoVis.3.1.1.chr7.g01615.t01#6    mNeoVis.evergladensis.2.1.chr7.g01702.t01#2   mNeoVis.2.1.1.chr7_hap1.g01734.t01#5       2917   125259441:   mNeoVis.vulgivaga.1.1.chr7.g01707.t01#4   mNeoVis.3.1.1.chr7.g01612.t01#6             2918   125259442:  mNeoVis.evergladensis.1.1.chr7.g01697.t01#3             2919   125259443:   mNeoVis.vulgivaga.2.1.chr1.g00899.t01#1    mNeoVis.2.1.1.chr1_hap1.g00885.t01#5    mNeoVis.evergladensis.2.1.chr1.g00888.t01#2    mNeoVis.3.1.1.chr1.g00907.t01#6    mNeoVis.vulgivaga.1.1.chr1.g00902.t01#4   mNeoVis.evergladensis.1.1.chr1.g00896.t01#3       2920   125259444:   mNeoVis.evergladensis.2.1.chr7.g01704.t01#2    mNeoVis.3.1.1.chr7.g01617.t01#6    mNeoVis.vulgivaga.2.1.chr7.g01718.t01#1    mNeoVis.2.1.1.chr7_hap1.g01739.t01#5    mNeoVis.vulgivaga.1.1.chr7.g01712.t01#4   mNeoVis.evergladensis.1.1.chr7.g01701.t01#3       2921   125259445:   mNeoVis.3.1.1.chr7.g01618.t01#6    mNeoVis.vulgivaga.2.1.chr7.g01719.t01#1    mNeoVis.2.1.1.chr7_hap1.g01740.t01#5    mNeoVis.evergladensis.2.1.chr7.g01705.t01#2    mNeoVis.evergladensis.1.1.chr7.g01702.t01#3   mNeoVis.vulgivaga.1.1.chr7.g01713.t01#4       2922   125259446:   mNeoVis.vulgivaga.2.1.chr7.g01720.t01#1    mNeoVis.evergladensis.2.1.chr7.g01706.t01#2    mNeoVis.2.1.1.chr7_hap1.g01741.t01#5    mNeoVis.vulgivaga.1.1.chr7.g01714.t01#4    mNeoVis.evergladensis.1.1.chr7.g01703.t01#3   mNeoVis.3.1.1.chr7.g01619.t01#6       2923   125259447:   mNeoVis.3.1.1.chr1.g00908.t01#6    mNeoVis.vulgivaga.1.1.chr1.g00903.t01#4    mNeoVis.evergladensis.1.1.chr12.g00336.t01#3    mNeoVis.vulgivaga.2.1.chr12.g00336.t01#1    mNeoVis.vulgivaga.1.1.chr12.g00336.t01#4    mNeoVis.evergladensis.2.1.chr12.g00330.t01#2    mNeoVis.evergladensis.1.1.chr1.g00897.t01#3    mNeoVis.vulgivaga.2.1.chr1.g00900.t01#1    mNeoVis.2.1.1.chr12_hap1.g00372.t01#5      2924   125259448:   mNeoVis.vulgivaga.2.1.chr5.g00352.t01#1    mNeoVis.2.1.1.chr2_hap1.g00350.t01#5    mNeoVis.vulgivaga.1.1.chr5.g00344.t01#4    mNeoVis.3.1.1.chr5.g00343.t01#6    mNeoVis.evergladensis.2.1.chr5.g00351.t01#2   mNeoVis.evergladensis.1.1.chr5.g00350.t01#3       2925   125259449:   mNeoVis.evergladensis.1.1.chr7.g00087.t01#3    mNeoVis.3.1.1.chr7.g00071.t01#6    mNeoVis.2.1.1.chr7_hap1.g00085.t01#5    mNeoVis.vulgivaga.2.1.chr7.g00085.t01#1    mNeoVis.vulgivaga.1.1.chr7.g00087.t01#4   mNeoVis.evergladensis.2.1.chr7.g00085.t01#2       2926   125259450:   mNeoVis.vulgivaga.2.1.chr7.g01721.t01#1    mNeoVis.3.1.1.chr7.g01620.t01#6    mNeoVis.evergladensis.2.1.chr7.g01707.t01#2    mNeoVis.2.1.1.chr7_hap1.g01742.t01#5    mNeoVis.vulgivaga.1.1.chr7.g01715.t01#4   mNeoVis.evergladensis.1.1.chr7.g01704.t01#3       2927   125259451:   mNeoVis.3.1.1.chr1.g00909.t01#6    mNeoVis.2.1.1.chr1_hap1.g00887.t01#5    mNeoVis.evergladensis.1.1.chr1.g00898.t01#3    mNeoVis.evergladensis.2.1.chr1.g00890.t01#2    mNeoVis.vulgivaga.1.1.chr1.g00904.t01#4   mNeoVis.vulgivaga.2.1.chr1.g00901.t01#1       2928   125259452:   mNeoVis.vulgivaga.1.1.chr1.g00905.t01#4    mNeoVis.vulgivaga.2.1.chr1.g00902.t01#1    mNeoVis.evergladensis.1.1.chr1.g00899.t01#3    mNeoVis.3.1.1.chr1.g00910.t01#6    mNeoVis.2.1.1.chr1_hap1.g00888.t01#5   mNeoVis.evergladensis.2.1.chr1.g00891.t01#2       2929   125259453:   mNeoVis.2.1.1.chr7_hap1.g01743.t01#5    mNeoVis.vulgivaga.1.1.chr7.g01716.t01#4    mNeoVis.evergladensis.1.1.chr7.g01705.t01#3    mNeoVis.vulgivaga.2.1.chr7.g01722.t01#1    mNeoVis.evergladensis.2.1.chr7.g01708.t01#2   mNeoVis.3.1.1.chr7.g01621.t01#6       2930   125259454:   mNeoVis.2.1.1.chr7_hap1.g01744.t01#5    mNeoVis.evergladensis.2.1.chr7.g01709.t01#2    mNeoVis.vulgivaga.1.1.chr7.g01717.t01#4    mNeoVis.evergladensis.1.1.chr7.g01706.t01#3    mNeoVis.3.1.1.chr7.g01622.t01#6   mNeoVis.vulgivaga.2.1.chr7.g01723.t01#1       2931   125259455:   mNeoVis.vulgivaga.1.1.chr15.g00152.t01#4    mNeoVis.vulgivaga.2.1.chr15.g00156.t01#1    mNeoVis.3.1.1.chr15.g00146.t01#6    mNeoVis.evergladensis.2.1.chr15.g00153.t01#2    mNeoVis.2.1.1.chr15_hap1.g00170.t01#5   mNeoVis.evergladensis.1.1.chr15.g00462.t01#3       2932   125259456:   mNeoVis.evergladensis.1.1.chr6.g00421.t01#3    mNeoVis.3.1.1.chr6.g00420.t01#6    mNeoVis.vulgivaga.2.1.chr6.g00418.t01#1    mNeoVis.2.1.1.chr4_hap1.g00420.t01#5   mNeoVis.evergladensis.2.1.chr6.g00424.t01#2       2933   125259457:   mNeoVis.evergladensis.1.1.chr1.g00900.t01#3    mNeoVis.vulgivaga.1.1.chr1.g00906.t01#4    mNeoVis.3.1.1.chr1.g00911.t01#6    mNeoVis.2.1.1.chr1_hap1.g00889.t01#5    mNeoVis.vulgivaga.2.1.chr1.g00903.t01#1   mNeoVis.evergladensis.2.1.chr1.g00892.t01#2       2934   125259458:   mNeoVis.3.1.1.chr7.g01623.t01#6    mNeoVis.vulgivaga.2.1.chr7.g01724.t01#1    mNeoVis.2.1.1.chr7_hap1.g01745.t01#5    mNeoVis.evergladensis.2.1.chr7.g01710.t01#2    mNeoVis.vulgivaga.1.1.chr7.g01718.t01#4   mNeoVis.evergladensis.1.1.chr7.g01707.t01#3       2935   125259459:   mNeoVis.vulgivaga.1.1.chr7.g01719.t01#4    mNeoVis.2.1.1.chr7_hap1.g01746.t01#5    mNeoVis.evergladensis.1.1.chr7.g01708.t01#3    mNeoVis.3.1.1.chr7.g01624.t01#6    mNeoVis.evergladensis.2.1.chr7.g01711.t01#2   mNeoVis.vulgivaga.2.1.chr7.g01725.t01#1       2936   125259460:   mNeoVis.vulgivaga.1.1.chr1.g00907.t01#4    mNeoVis.2.1.1.chr1_hap1.g00890.t01#5    mNeoVis.evergladensis.1.1.chr1.g00901.t01#3    mNeoVis.evergladensis.2.1.chr1.g00893.t01#2   mNeoVis.vulgivaga.2.1.chr1.g00904.t01#1       2937   125259461:   mNeoVis.2.1.1.chr7_hap1.g01747.t01#5    mNeoVis.vulgivaga.1.1.chr7.g01720.t01#4    mNeoVis.vulgivaga.2.1.chr7.g01726.t01#1    mNeoVis.evergladensis.1.1.chr7.g01709.t01#3    mNeoVis.evergladensis.2.1.chr7.g01712.t01#2   mNeoVis.3.1.1.chr7.g01625.t01#6       2938   125259462:   mNeoVis.2.1.1.chr1_hap1.g00891.t01#5    mNeoVis.evergladensis.2.1.chr1.g00894.t01#2    mNeoVis.evergladensis.1.1.chr1.g00902.t01#3    mNeoVis.vulgivaga.2.1.chr1.g00905.t01#1   mNeoVis.vulgivaga.1.1.chr1.g00908.t01#4       2939   125259463:   mNeoVis.2.1.1.chr7_hap1.g01537.t01#5    mNeoVis.evergladensis.2.1.chr7.g01518.t01#2    mNeoVis.vulgivaga.2.1.chr7.g01535.t01#1    mNeoVis.3.1.1.chr7.g01448.t01#6    mNeoVis.vulgivaga.1.1.chr7.g01531.t01#4   mNeoVis.evergladensis.1.1.chr7.g01521.t01#3       2940   125259464:   mNeoVis.evergladensis.2.1.chr1.g00895.t01#2    mNeoVis.2.1.1.chr1_hap1.g00892.t01#5    mNeoVis.3.1.1.chr1.g00912.t01#6    mNeoVis.vulgivaga.2.1.chr1.g00906.t01#1    mNeoVis.evergladensis.1.1.chr1.g00903.t01#3   mNeoVis.vulgivaga.1.1.chr1.g00909.t01#4       2941   125259465:   mNeoVis.vulgivaga.1.1.chr7.g00090.t01#4    mNeoVis.2.1.1.chr7_hap1.g00087.t01#5    mNeoVis.vulgivaga.2.1.chr7.g00086.t01#1    mNeoVis.vulgivaga.1.1.chr7.g00089.t01#4    mNeoVis.evergladensis.2.1.chr7.g00086.t01#2    mNeoVis.evergladensis.1.1.chr7.g00089.t01#3   mNeoVis.3.1.1.chr7.g00072.t01#6       2942   125259466:   mNeoVis.2.1.1.chr7_hap1.g01538.t01#5    mNeoVis.vulgivaga.2.1.chr7.g01536.t01#1   mNeoVis.3.1.1.chr7.g01449.t01#6           2943   125259467:   mNeoVis.evergladensis.2.1.chr1.g00896.t01#2    mNeoVis.3.1.1.chr1.g00914.t01#6    mNeoVis.vulgivaga.2.1.chr1.g00907.t01#1    mNeoVis.2.1.1.chr1_hap1.g00893.t01#5    mNeoVis.3.1.1.chr1.g00915.t01#6    mNeoVis.vulgivaga.1.1.chr1.g00910.t01#4   mNeoVis.evergladensis.1.1.chr1.g00904.t01#3     2944   125259468:   mNeoVis.2.1.1.chr7_hap1.g01539.t01#5    mNeoVis.vulgivaga.2.1.chr7.g01537.t01#1    mNeoVis.evergladensis.2.1.chr7.g01520.t01#2    mNeoVis.3.1.1.chr7.g01450.t01#6    mNeoVis.vulgivaga.1.1.chr7.g01533.t01#4   mNeoVis.evergladensis.1.1.chr7.g01523.t01#3       2945   125259469:   mNeoVis.2.1.1.chr1_hap1.g00894.t01#5    mNeoVis.3.1.1.chr1.g00916.t01#6    mNeoVis.evergladensis.2.1.chr1.g00897.t01#2    mNeoVis.vulgivaga.2.1.chr1.g00908.t01#1    mNeoVis.vulgivaga.1.1.chr1.g00911.t01#4   mNeoVis.evergladensis.1.1.chr1.g00905.t01#3       2946   125259470:   mNeoVis.vulgivaga.2.1.chr7.g01538.t01#1    mNeoVis.evergladensis.2.1.chr7.g01521.t01#2    mNeoVis.evergladensis.1.1.chr7.g01525.t01#3    mNeoVis.2.1.1.chr7_hap1.g01541.t01#5    mNeoVis.vulgivaga.1.1.chr7.g01534.t01#4   mNeoVis.3.1.1.chr7.g01452.t01#6       2947   125259471:   mNeoVis.vulgivaga.2.1.chr7.g01539.t01#1    mNeoVis.evergladensis.2.1.chr7.g01522.t01#2    mNeoVis.evergladensis.1.1.chr7.g01526.t01#3    mNeoVis.3.1.1.chr7.g01453.t01#6   mNeoVis.2.1.1.chr7_hap1.g01542.t01#5       2948   125259472:   mNeoVis.vulgivaga.1.1.chr1.g00912.t01#4    mNeoVis.2.1.1.chr1_hap1.g00895.t01#5    mNeoVis.vulgivaga.2.1.chr1.g00910.t01#1    mNeoVis.evergladensis.1.1.chr1.g00906.t01#3   mNeoVis.evergladensis.2.1.chr1.g00898.t01#2       2949   125259473:   mNeoVis.3.1.1.chr1.g00917.t01#6   mNeoVis.vulgivaga.2.1.chr1.g00909.t01#1             2950   125259474:   mNeoVis.vulgivaga.2.1.chr7.g01540.t01#1    mNeoVis.evergladensis.1.1.chr7.g01527.t01#3    mNeoVis.2.1.1.chr7_hap1.g01543.t01#5    mNeoVis.evergladensis.2.1.chr7.g01523.t01#2    mNeoVis.3.1.1.chr7.g01454.t01#6   mNeoVis.vulgivaga.1.1.chr7.g01535.t01#4       2951   125259475:   mNeoVis.evergladensis.2.1.chr7.g01524.t01#2    mNeoVis.evergladensis.1.1.chr7.g01528.t01#3    mNeoVis.3.1.1.chr7.g01455.t01#6    mNeoVis.2.1.1.chr7_hap1.g01544.t01#5    mNeoVis.vulgivaga.1.1.chr7.g01536.t01#4   mNeoVis.vulgivaga.2.1.chr7.g01541.t01#1       2952   125259476:   mNeoVis.2.1.1.chr5_hap1.g01144.t01#5    mNeoVis.evergladensis.1.1.chr2.g01137.t01#3    mNeoVis.evergladensis.2.1.chr2.g01134.t01#2    mNeoVis.vulgivaga.2.1.chr2.g01140.t01#1    mNeoVis.3.1.1.chr2.g01053.t01#6   mNeoVis.vulgivaga.1.1.chr2.g01141.t01#4       2953   125259477:   mNeoVis.evergladensis.1.1.chr1.g00907.t01#3    mNeoVis.2.1.1.chr1_hap1.g00896.t01#5    mNeoVis.vulgivaga.1.1.chr1.g00913.t01#4    mNeoVis.vulgivaga.2.1.chr1.g00911.t01#1    mNeoVis.3.1.1.chr1.g00918.t01#6   mNeoVis.evergladensis.2.1.chr1.g00899.t01#2       2954   125259478:   mNeoVis.evergladensis.1.1.chr7.g01529.t01#3    mNeoVis.2.1.1.chr7_hap1.g01545.t01#5    mNeoVis.vulgivaga.2.1.chr7.g01542.t01#1    mNeoVis.vulgivaga.1.1.chr7.g01537.t01#4    mNeoVis.3.1.1.chr7.g01456.t01#6   mNeoVis.evergladensis.2.1.chr7.g01525.t01#2       2955   125259479:   mNeoVis.vulgivaga.1.1.chr4.g00625.t01#4    mNeoVis.evergladensis.1.1.chr4.g00629.t01#3    mNeoVis.vulgivaga.2.1.chr5.g00394.t01#1    mNeoVis.evergladensis.2.1.chr4.g00622.t01#2    mNeoVis.vulgivaga.1.1.chr5.g00381.t01#4    mNeoVis.2.1.1.chr6_hap1.g00617.t01#5    mNeoVis.3.1.1.chr4.g00623.t01#6    mNeoVis.evergladensis.1.1.chr5.g00390.t01#3    mNeoVis.evergladensis.2.1.chr5.g00394.t01#2      2956   125259480:   mNeoVis.evergladensis.2.1.chr1.g00900.t01#2    mNeoVis.vulgivaga.2.1.chr1.g00912.t01#1    mNeoVis.2.1.1.chr1_hap1.g00897.t01#5    mNeoVis.vulgivaga.1.1.chr1.g00914.t01#4    mNeoVis.3.1.1.chr1.g00919.t01#6   mNeoVis.evergladensis.1.1.chr1.g00908.t01#3       2957   125259481:   mNeoVis.vulgivaga.2.1.chr7.g01543.t01#1    mNeoVis.3.1.1.chr7.g01457.t01#6    mNeoVis.vulgivaga.1.1.chr7.g01538.t01#4    mNeoVis.2.1.1.chr7_hap1.g01546.t01#5    mNeoVis.evergladensis.2.1.chr7.g01526.t01#2   mNeoVis.evergladensis.1.1.chr7.g01530.t01#3       2958   125259482:   mNeoVis.vulgivaga.2.1.chr7.g01544.t01#1    mNeoVis.evergladensis.2.1.chr7.g01527.t01#2    mNeoVis.evergladensis.1.1.chr7.g01531.t01#3    mNeoVis.2.1.1.chr7_hap1.g01547.t01#5    mNeoVis.3.1.1.chr7.g01458.t01#6   mNeoVis.vulgivaga.1.1.chr7.g01539.t01#4       2959   125259483:   mNeoVis.evergladensis.2.1.chr1.g00901.t01#2    mNeoVis.evergladensis.1.1.chr1.g00909.t01#3    mNeoVis.2.1.1.chr1_hap1.g00898.t01#5    mNeoVis.3.1.1.chr1.g00920.t01#6    mNeoVis.vulgivaga.2.1.chr1.g00913.t01#1   mNeoVis.vulgivaga.1.1.chr1.g00915.t01#4       2960   125259484:   mNeoVis.2.1.1.chr11_hap1.g00765.t01#5    mNeoVis.vulgivaga.2.1.chr11.g00767.t01#1    mNeoVis.3.1.1.chr11.g00728.t01#6    mNeoVis.vulgivaga.1.1.chr11.g00747.t01#4    mNeoVis.evergladensis.2.1.chr11.g00747.t01#2   mNeoVis.evergladensis.1.1.chr11.g00755.t01#3       2961   125259485:   mNeoVis.evergladensis.2.1.chr7.g01528.t01#2    mNeoVis.evergladensis.1.1.chr7.g01532.t01#3    mNeoVis.2.1.1.chr7_hap1.g01548.t01#5    mNeoVis.3.1.1.chr7.g01460.t01#6    mNeoVis.vulgivaga.1.1.chr7.g01540.t01#4   mNeoVis.vulgivaga.2.1.chr7.g01545.t01#1       2962   125259486:   mNeoVis.3.1.1.chr1.g00921.t01#6    mNeoVis.2.1.1.chr1_hap1.g00899.t01#5    mNeoVis.vulgivaga.1.1.chr1.g00916.t01#4    mNeoVis.evergladensis.2.1.chr1.g00902.t01#2    mNeoVis.vulgivaga.2.1.chr1.g00914.t01#1   mNeoVis.evergladensis.1.1.chr1.g00910.t01#3       2963   125259487:   mNeoVis.evergladensis.1.1.chr4.g00042.t01#3    mNeoVis.evergladensis.2.1.chr4.g00044.t01#2    mNeoVis.vulgivaga.1.1.chr4.g00040.t01#4    mNeoVis.vulgivaga.2.1.chr4.g00040.t01#1    mNeoVis.3.1.1.chr4.g00039.t01#6   mNeoVis.2.1.1.chr6_hap1.g00041.t01#5       2964   125259488:   mNeoVis.3.1.1.chr15.g00147.t01#6    mNeoVis.vulgivaga.2.1.chr15.g00157.t01#1    mNeoVis.evergladensis.1.1.chr15.g00460.t01#3    mNeoVis.vulgivaga.1.1.chr15.g00153.t01#4    mNeoVis.evergladensis.2.1.chr15.g00154.t01#2   mNeoVis.2.1.1.chr15_hap1.g00171.t01#5       2965   125259489:   mNeoVis.2.1.1.chr1_hap1.g00900.t01#5    mNeoVis.vulgivaga.1.1.chr1.g00917.t01#4    mNeoVis.evergladensis.2.1.chr1.g00903.t01#2    mNeoVis.vulgivaga.2.1.chr1.g00915.t01#1    mNeoVis.3.1.1.chr1.g00922.t01#6   mNeoVis.evergladensis.1.1.chr1.g00911.t01#3       2966   125259490:   mNeoVis.vulgivaga.2.1.chr1.g00916.t01#1    mNeoVis.evergladensis.1.1.chr1.g00912.t01#3    mNeoVis.2.1.1.chr1_hap1.g00901.t01#5    mNeoVis.vulgivaga.1.1.chr1.g00918.t01#4   mNeoVis.evergladensis.2.1.chr1.g00904.t01#2       2967   125259491:   mNeoVis.vulgivaga.2.1.chr7.g01546.t01#1    mNeoVis.evergladensis.2.1.chr7.g01529.t01#2    mNeoVis.vulgivaga.1.1.chr7.g01541.t01#4    mNeoVis.2.1.1.chr7_hap1.g01549.t01#5    mNeoVis.3.1.1.chr10.g00541.t01#6   mNeoVis.evergladensis.1.1.chr7.g01533.t01#3       2968   125259492:   mNeoVis.vulgivaga.1.1.chr7.g01542.t01#4    mNeoVis.2.1.1.chr7_hap1.g01550.t01#5    mNeoVis.vulgivaga.2.1.chr7.g01547.t01#1    mNeoVis.3.1.1.chr7.g01461.t01#6    mNeoVis.evergladensis.2.1.chr7.g01530.t01#2   mNeoVis.evergladensis.1.1.chr7.g01534.t01#3       2969   125259493:   mNeoVis.vulgivaga.2.1.chr7.g01548.t01#1    mNeoVis.evergladensis.2.1.chr7.g01531.t01#2    mNeoVis.evergladensis.1.1.chr7.g01535.t01#3    mNeoVis.3.1.1.chr7.g01462.t01#6    mNeoVis.vulgivaga.1.1.chr7.g01543.t01#4   mNeoVis.2.1.1.chr7_hap1.g01551.t01#5       2970   125259494:   mNeoVis.vulgivaga.1.1.chr12.g00303.t01#4    mNeoVis.vulgivaga.2.1.chr12.g00301.t01#1    mNeoVis.3.1.1.chr12.g00297.t01#6    mNeoVis.3.1.1.chr12.g00322.t01#6    mNeoVis.2.1.1.chr12_hap1.g00328.t01#5    mNeoVis.evergladensis.2.1.chr12.g00294.t01#2   mNeoVis.evergladensis.1.1.chr12.g00299.t01#3     2971   125259495:   mNeoVis.2.1.1.chr7_hap1.g00088.t01#5    mNeoVis.vulgivaga.1.1.chr7.g00091.t01#4    mNeoVis.evergladensis.2.1.chr7.g00087.t01#2    mNeoVis.vulgivaga.2.1.chr7.g00087.t01#1    mNeoVis.3.1.1.chr7.g00073.t01#6   mNeoVis.evergladensis.1.1.chr7.g00090.t01#3       2972   125259496:   mNeoVis.vulgivaga.2.1.chr12.g00758.t01#1    mNeoVis.2.1.1.chr12_hap1.g00801.t01#5    mNeoVis.evergladensis.2.1.chr12.g00728.t01#2    mNeoVis.vulgivaga.1.1.chr12.g00753.t01#4    mNeoVis.3.1.1.chr12.g00731.t01#6   mNeoVis.evergladensis.1.1.chr12.g00736.t01#3       2973   125259497:   mNeoVis.3.1.1.chr11.g00817.t01#6    mNeoVis.evergladensis.2.1.chr11.g00841.t01#2    mNeoVis.evergladensis.1.1.chr11.g00849.t01#3    mNeoVis.vulgivaga.2.1.chr11.g00862.t01#1    mNeoVis.vulgivaga.1.1.chr11.g00840.t01#4   mNeoVis.2.1.1.chr11_hap1.g00858.t01#5       2974   125259498:   mNeoVis.evergladensis.1.1.chr1.g00914.t01#3    mNeoVis.vulgivaga.1.1.chr1.g00920.t01#4    mNeoVis.3.1.1.chr1.g00925.t01#6    mNeoVis.evergladensis.2.1.chr1.g00906.t01#2    mNeoVis.vulgivaga.2.1.chr1.g00917.t01#1   mNeoVis.2.1.1.chr1_hap1.g00903.t01#5       2975   125259499:   mNeoVis.2.1.1.chr7_hap1.g01552.t01#5    mNeoVis.vulgivaga.2.1.chr7.g01549.t01#1    mNeoVis.evergladensis.1.1.chr7.g01536.t01#3    mNeoVis.evergladensis.2.1.chr7.g01532.t01#2    mNeoVis.3.1.1.chr7.g01463.t01#6   mNeoVis.vulgivaga.1.1.chr7.g01544.t01#4       2976   125259500:   mNeoVis.3.1.1.chr1.g00927.t01#6    mNeoVis.evergladensis.1.1.chr1.g00916.t01#3    mNeoVis.2.1.1.chr1_hap1.g00907.t01#5    mNeoVis.evergladensis.2.1.chr1.g00908.t01#2    mNeoVis.vulgivaga.1.1.chr1.g00922.t01#4   mNeoVis.vulgivaga.2.1.chr1.g00918.t01#1       2977   125259501:   mNeoVis.2.1.1.chr1_hap1.g00909.t01#5   mNeoVis.2.1.1.chr1_hap1.g00912.t01#5             2978   125259502:   mNeoVis.vulgivaga.1.1.chr1.g00924.t01#4    mNeoVis.vulgivaga.2.1.chr1.g00921.t01#1    mNeoVis.evergladensis.1.1.chr1.g00918.t01#3    mNeoVis.3.1.1.chr1.g00929.t01#6    mNeoVis.evergladensis.2.1.chr1.g00910.t01#2   mNeoVis.vulgivaga.2.1.chr1.g00920.t01#1       2979   125259503:   mNeoVis.evergladensis.1.1.chr7.g00091.t01#3    mNeoVis.vulgivaga.2.1.chr7.g00088.t01#1    mNeoVis.2.1.1.chr7_hap1.g00089.t01#5    mNeoVis.3.1.1.chr7.g00074.t01#6    mNeoVis.vulgivaga.1.1.chr7.g00092.t01#4   mNeoVis.evergladensis.2.1.chr7.g00088.t01#2       2980   125259504:   mNeoVis.vulgivaga.2.1.chr7.g01550.t01#1    mNeoVis.evergladensis.1.1.chr7.g01537.t01#3    mNeoVis.3.1.1.chr7.g01464.t01#6    mNeoVis.2.1.1.chr7_hap1.g01553.t01#5    mNeoVis.vulgivaga.1.1.chr7.g01545.t01#4   mNeoVis.evergladensis.2.1.chr7.g01533.t01#2       2981   125259505:  mNeoVis.vulgivaga.2.1.chr1.g00923.t01#1               2982   125259506:   mNeoVis.evergladensis.1.1.chr7.g01538.t01#3    mNeoVis.vulgivaga.2.1.chr7.g01551.t01#1    mNeoVis.2.1.1.chr7_hap1.g01554.t01#5    mNeoVis.vulgivaga.1.1.chr7.g01546.t01#4   mNeoVis.evergladensis.2.1.chr7.g01534.t01#2       2983   125259507:   mNeoVis.vulgivaga.2.1.chr1.g00924.t01#1    mNeoVis.evergladensis.2.1.chr1.g00912.t01#2    mNeoVis.vulgivaga.1.1.chr1.g00927.t01#4    mNeoVis.2.1.1.chr1_hap1.g00915.t01#5   mNeoVis.evergladensis.1.1.chr1.g00920.t01#3       2984   125259508:   mNeoVis.vulgivaga.1.1.chr15.g00154.t01#4    mNeoVis.evergladensis.1.1.chr15.g00459.t01#3    mNeoVis.vulgivaga.2.1.chr15.g00158.t01#1    mNeoVis.2.1.1.chr15_hap1.g00172.t01#5    mNeoVis.evergladensis.2.1.chr15.g00155.t01#2   mNeoVis.3.1.1.chr15.g00148.t01#6       2985   125259509:   mNeoVis.evergladensis.1.1.chr1.g00921.t01#3    mNeoVis.2.1.1.chr1_hap1.g00916.t01#5    mNeoVis.evergladensis.2.1.chr1.g00913.t01#2    mNeoVis.vulgivaga.2.1.chr1.g00925.t01#1    mNeoVis.vulgivaga.1.1.chr1.g00928.t01#4   mNeoVis.3.1.1.chr1.g00932.t01#6       2986   125259510:   mNeoVis.vulgivaga.1.1.chr1.g00929.t01#4    mNeoVis.evergladensis.1.1.chr1.g00922.t01#3    mNeoVis.2.1.1.chr1_hap1.g00917.t01#5    mNeoVis.vulgivaga.2.1.chr1.g00926.t01#1    mNeoVis.evergladensis.2.1.chr1.g00914.t01#2   mNeoVis.3.1.1.chr1.g00933.t01#6       2987   125259511:   mNeoVis.evergladensis.1.1.chr9.g00744.t01#3    mNeoVis.vulgivaga.1.1.chr9.g00746.t01#4    mNeoVis.3.1.1.chr9.g00713.t01#6    mNeoVis.vulgivaga.2.1.chr1.g00927.t01#1    mNeoVis.vulgivaga.2.1.chr9.g00735.t01#1    mNeoVis.vulgivaga.1.1.chr1.g00930.t01#4    mNeoVis.3.1.1.chr1.g00934.t01#6    mNeoVis.evergladensis.2.1.chr8.g00735.t01#2    mNeoVis.evergladensis.1.1.chr1.g00923.t01#3      2988   125259512:   mNeoVis.evergladensis.2.1.chr1.g00916.t01#2    mNeoVis.vulgivaga.2.1.chr1.g00928.t01#1    mNeoVis.vulgivaga.1.1.chr1.g00931.t01#4    mNeoVis.evergladensis.1.1.chr1.g00924.t01#3    mNeoVis.3.1.1.chr1.g00935.t01#6   mNeoVis.2.1.1.chr1_hap1.g00919.t01#5       2989   125259513:   mNeoVis.vulgivaga.2.1.chr1.g00929.t01#1    mNeoVis.evergladensis.2.1.chr1.g00917.t01#2    mNeoVis.2.1.1.chr1_hap1.g00920.t01#5    mNeoVis.evergladensis.1.1.chr1.g00925.t01#3    mNeoVis.vulgivaga.1.1.chr1.g00932.t01#4   mNeoVis.3.1.1.chr1.g00936.t01#6       2990   125259514:   mNeoVis.evergladensis.2.1.chr7.g01535.t01#2    mNeoVis.vulgivaga.1.1.chr7.g01547.t01#4    mNeoVis.3.1.1.chr7.g01467.t01#6    mNeoVis.vulgivaga.2.1.chr7.g01552.t01#1    mNeoVis.2.1.1.chr7_hap1.g01555.t01#5   mNeoVis.evergladensis.1.1.chr7.g01539.t01#3       2991   125259515:   mNeoVis.evergladensis.1.1.chr1.g00927.t01#3    mNeoVis.evergladensis.2.1.chr1.g00919.t01#2    mNeoVis.vulgivaga.1.1.chr1.g00935.t01#4    mNeoVis.3.1.1.chr1.g00938.t01#6    mNeoVis.2.1.1.chr1_hap1.g00923.t01#5    mNeoVis.vulgivaga.2.1.chr1.g00932.t01#1    mNeoVis.3.1.1.chr1.g00939.t01#6    mNeoVis.vulgivaga.1.1.chr1.g00936.t01#4    mNeoVis.vulgivaga.2.1.chr1.g00931.t01#1      2992   125259516:   mNeoVis.evergladensis.2.1.chr1.g00918.t01#2    mNeoVis.3.1.1.chr1.g00937.t01#6    mNeoVis.vulgivaga.2.1.chr1.g00930.t01#1    mNeoVis.evergladensis.1.1.chr1.g00926.t01#3    mNeoVis.vulgivaga.1.1.chr1.g00933.t01#4   mNeoVis.2.1.1.chr1_hap1.g00921.t01#5       2993   125259517:   mNeoVis.evergladensis.2.1.chr1.g00921.t01#2    mNeoVis.vulgivaga.2.1.chr1.g00933.t01#1    mNeoVis.evergladensis.1.1.chr1.g00928.t01#3    mNeoVis.3.1.1.chr1.g00940.t01#6    mNeoVis.vulgivaga.1.1.chr1.g00937.t01#4   mNeoVis.2.1.1.chr1_hap1.g00925.t01#5       2994   125259518:   mNeoVis.vulgivaga.1.1.chr7.g00093.t01#4    mNeoVis.vulgivaga.2.1.chr7.g00089.t01#1    mNeoVis.evergladensis.1.1.chr7.g00092.t01#3    mNeoVis.2.1.1.chr7_hap1.g00090.t01#5    mNeoVis.3.1.1.chr7.g00075.t01#6   mNeoVis.evergladensis.2.1.chr7.g00090.t01#2       2995   125259519:   mNeoVis.3.1.1.chr1.g00941.t01#6    mNeoVis.2.1.1.chr1_hap1.g00926.t01#5   mNeoVis.vulgivaga.2.1.chr1.g00934.t01#1           2996   125259520:   mNeoVis.vulgivaga.2.1.chr1.g00935.t01#1    mNeoVis.vulgivaga.1.1.chr1.g00938.t01#4    mNeoVis.evergladensis.2.1.chr1.g00922.t01#2    mNeoVis.2.1.1.chr1_hap1.g00927.t01#5   mNeoVis.evergladensis.1.1.chr1.g00929.t01#3       2997   125259521:   mNeoVis.2.1.1.chr1_hap1.g00928.t01#5    mNeoVis.evergladensis.1.1.chr1.g00930.t01#3    mNeoVis.evergladensis.2.1.chr1.g00923.t01#2    mNeoVis.vulgivaga.1.1.chr1.g00939.t01#4    mNeoVis.vulgivaga.2.1.chr1.g00936.t01#1   mNeoVis.3.1.1.chr1.g00942.t01#6       2998   125259522:   mNeoVis.3.1.1.chr1.g00943.t01#6    mNeoVis.vulgivaga.2.1.chr1.g00937.t01#1    mNeoVis.2.1.1.chr1_hap1.g00929.t01#5    mNeoVis.evergladensis.2.1.chr1.g00924.t01#2    mNeoVis.vulgivaga.1.1.chr1.g00940.t01#4   mNeoVis.evergladensis.1.1.chr1.g00931.t01#3       2999   125259523:   mNeoVis.vulgivaga.2.1.chr1.g00938.t01#1    mNeoVis.evergladensis.2.1.chr1.g00925.t01#2    mNeoVis.2.1.1.chr1_hap1.g00930.t01#5    mNeoVis.evergladensis.1.1.chr1.g00932.t01#3   mNeoVis.3.1.1.chr1.g00944.t01#6       3000   125259524:   mNeoVis.vulgivaga.1.1.chr1.g00941.t01#4    mNeoVis.3.1.1.chr1.g00945.t01#6    mNeoVis.evergladensis.2.1.chr1.g00927.t01#2    mNeoVis.vulgivaga.2.1.chr1.g00939.t01#1    mNeoVis.evergladensis.1.1.chr1.g00934.t01#3   mNeoVis.2.1.1.chr1_hap1.g00931.t01#5       3001   125259525:   mNeoVis.vulgivaga.1.1.chr7.g01548.t01#4    mNeoVis.evergladensis.1.1.chr7.g01540.t01#3    mNeoVis.vulgivaga.2.1.chr7.g01553.t01#1    mNeoVis.2.1.1.chr7_hap1.g01556.t01#5    mNeoVis.3.1.1.chr7.g01468.t01#6   mNeoVis.evergladensis.2.1.chr7.g01536.t01#2       3002   125259526:   mNeoVis.vulgivaga.1.1.chr5.g01795.t01#4    mNeoVis.2.1.1.chr9_hap1.g00977.t01#5    mNeoVis.2.1.1.chr2_hap1.g01874.t01#5    mNeoVis.vulgivaga.2.1.chr5.g01815.t01#1    mNeoVis.evergladensis.1.1.chr7.g01541.t01#3    mNeoVis.evergladensis.2.1.chr5.g01812.t01#2    mNeoVis.evergladensis.1.1.chr5.g01822.t01#3    mNeoVis.vulgivaga.2.1.chr7.g01554.t01#1    mNeoVis.vulgivaga.1.1.chr10.g00945.t01#4      3003   125259527:   mNeoVis.evergladensis.2.1.chr4.g00631.t01#2    mNeoVis.2.1.1.chr6_hap1.g00624.t01#5    mNeoVis.vulgivaga.2.1.chr4.g00639.t01#1    mNeoVis.evergladensis.1.1.chr4.g00637.t01#3    mNeoVis.vulgivaga.1.1.chr7.g01549.t01#4   mNeoVis.vulgivaga.1.1.chr4.g00633.t01#4       3004   125259528:   mNeoVis.evergladensis.2.1.chr1.g00928.t01#2    mNeoVis.evergladensis.1.1.chr1.g00935.t01#3    mNeoVis.2.1.1.chr1_hap1.g00932.t01#5    mNeoVis.vulgivaga.1.1.chr1.g00942.t01#4    mNeoVis.vulgivaga.2.1.chr1.g00940.t01#1   mNeoVis.3.1.1.chr1.g00946.t01#6       3005   125259529:   mNeoVis.vulgivaga.2.1.chr1.g00941.t01#1    mNeoVis.vulgivaga.1.1.chr1.g00943.t01#4    mNeoVis.2.1.1.chr1_hap1.g00933.t01#5    mNeoVis.evergladensis.2.1.chr1.g00929.t01#2    mNeoVis.3.1.1.chr1.g00947.t01#6   mNeoVis.evergladensis.1.1.chr1.g00936.t01#3       3006   125259530:   mNeoVis.2.1.1.chr7_hap1.g01557.t01#5    mNeoVis.evergladensis.1.1.chr7.g01542.t01#3    mNeoVis.vulgivaga.1.1.chr7.g01550.t01#4    mNeoVis.evergladensis.2.1.chr7.g01538.t01#2   mNeoVis.vulgivaga.2.1.chr7.g01555.t01#1       3007   125259531:   mNeoVis.3.1.1.chr7.g01469.t01#6    mNeoVis.evergladensis.2.1.chr7.g01539.t01#2    mNeoVis.vulgivaga.2.1.chr7.g01556.t01#1    mNeoVis.2.1.1.chr7_hap1.g01558.t01#5    mNeoVis.evergladensis.1.1.chr7.g01543.t01#3   mNeoVis.vulgivaga.1.1.chr7.g01551.t01#4       3008   125259532:   mNeoVis.evergladensis.2.1.chr1.g00930.t01#2    mNeoVis.2.1.1.chr1_hap1.g00934.t01#5    mNeoVis.vulgivaga.2.1.chr1.g00942.t01#1    mNeoVis.3.1.1.chr1.g00948.t01#6    mNeoVis.vulgivaga.1.1.chr1.g00944.t01#4   mNeoVis.evergladensis.1.1.chr1.g00937.t01#3       3009   125259533:   mNeoVis.vulgivaga.2.1.chr1.g00943.t01#1    mNeoVis.evergladensis.1.1.chr1.g00938.t01#3    mNeoVis.3.1.1.chr1.g00949.t01#6    mNeoVis.evergladensis.2.1.chr1.g00932.t01#2    mNeoVis.vulgivaga.1.1.chr1.g00946.t01#4   mNeoVis.2.1.1.chr1_hap1.g00937.t01#5       3010   125259534:   mNeoVis.evergladensis.2.1.chr1.g00933.t01#2    mNeoVis.vulgivaga.1.1.chr1.g00947.t01#4    mNeoVis.vulgivaga.2.1.chr1.g00944.t01#1    mNeoVis.evergladensis.1.1.chr1.g00939.t01#3   mNeoVis.3.1.1.chr1.g00950.t01#6       3011   125259535:   mNeoVis.vulgivaga.2.1.chr1.g00947.t01#1    mNeoVis.2.1.1.chr1_hap1.g00944.t01#5    mNeoVis.evergladensis.2.1.chr1.g00936.t01#2    mNeoVis.evergladensis.1.1.chr1.g00942.t01#3    mNeoVis.vulgivaga.1.1.chr1.g00950.t01#4    mNeoVis.3.1.1.chr1.g00953.t01#6   mNeoVis.2.1.1.chr1_hap1.g00940.t01#5     3012   125259536:   mNeoVis.evergladensis.1.1.chr1.g00943.t01#3    mNeoVis.2.1.1.chr1_hap1.g00941.t01#5    mNeoVis.vulgivaga.1.1.chr1.g00951.t01#4    mNeoVis.evergladensis.2.1.chr1.g00937.t01#2    mNeoVis.vulgivaga.2.1.chr1.g00948.t01#1   mNeoVis.3.1.1.chr1.g00954.t01#6       3013   125259537:   mNeoVis.2.1.1.chr7_hap1.g01561.t01#5    mNeoVis.evergladensis.2.1.chr7.g01541.t01#2    mNeoVis.evergladensis.1.1.chr7.g01546.t01#3    mNeoVis.vulgivaga.2.1.chr7.g01559.t01#1    mNeoVis.3.1.1.chr7.g01470.t01#6    mNeoVis.vulgivaga.1.1.chr7.g01554.t01#4   mNeoVis.3.1.1.chr7.g01471.t01#6       3014   125259538:   mNeoVis.evergladensis.1.1.chr7.g01544.t01#3    mNeoVis.vulgivaga.2.1.chr7.g01557.t01#1    mNeoVis.vulgivaga.1.1.chr7.g01552.t01#4    mNeoVis.evergladensis.2.1.chr7.g01540.t01#2   mNeoVis.2.1.1.chr7_hap1.g01559.t01#5       3015   125259539:   mNeoVis.2.1.1.chr7_hap1.g01560.t01#5    mNeoVis.vulgivaga.2.1.chr7.g01558.t01#1    mNeoVis.evergladensis.1.1.chr7.g01545.t01#3   mNeoVis.vulgivaga.1.1.chr7.g01553.t01#4         3016   125259540:   mNeoVis.evergladensis.2.1.chr1.g00938.t01#2    mNeoVis.3.1.1.chr1.g00955.t01#6    mNeoVis.2.1.1.chr1_hap1.g00942.t01#5    mNeoVis.vulgivaga.2.1.chr1.g00949.t01#1    mNeoVis.vulgivaga.1.1.chr1.g00952.t01#4   mNeoVis.evergladensis.1.1.chr1.g00944.t01#3       3017   125259541:   mNeoVis.vulgivaga.2.1.chr1.g00950.t01#1    mNeoVis.2.1.1.chr1_hap1.g00943.t01#5    mNeoVis.evergladensis.1.1.chr1.g00945.t01#3    mNeoVis.vulgivaga.1.1.chr1.g00953.t01#4    mNeoVis.evergladensis.2.1.chr1.g00939.t01#2   mNeoVis.3.1.1.chr1.g00956.t01#6       3018   125259542:   mNeoVis.evergladensis.2.1.chr1.g00940.t01#2    mNeoVis.vulgivaga.1.1.chr1.g00954.t01#4    mNeoVis.vulgivaga.2.1.chr1.g00951.t01#1    mNeoVis.2.1.1.chr1_hap1.g00945.t01#5    mNeoVis.evergladensis.1.1.chr1.g00946.t01#3   mNeoVis.3.1.1.chr1.g00957.t01#6       3019   125259543:   mNeoVis.evergladensis.1.1.chr1.g00949.t01#3    mNeoVis.vulgivaga.2.1.chr1.g00954.t01#1    mNeoVis.2.1.1.chr1_hap1.g00948.t01#5    mNeoVis.evergladensis.2.1.chr1.g00943.t01#2   mNeoVis.3.1.1.chr1.g00960.t01#6       3020   125259544:   mNeoVis.evergladensis.2.1.chr7.g00091.t01#2    mNeoVis.2.1.1.chr7_hap1.g00091.t01#5    mNeoVis.3.1.1.chr7.g00076.t01#6    mNeoVis.vulgivaga.1.1.chr7.g00094.t01#4    mNeoVis.vulgivaga.2.1.chr7.g00090.t01#1   mNeoVis.evergladensis.1.1.chr7.g00093.t01#3       3021   125259545:   mNeoVis.vulgivaga.2.1.chr1.g00955.t01#1    mNeoVis.3.1.1.chr1.g00961.t01#6    mNeoVis.evergladensis.2.1.chr1.g00944.t01#2    mNeoVis.vulgivaga.1.1.chr1.g00958.t01#4    mNeoVis.evergladensis.1.1.chr1.g00950.t01#3   mNeoVis.2.1.1.chr1_hap1.g00949.t01#5       3022   125259546:   mNeoVis.evergladensis.2.1.chr7.g01542.t01#2    mNeoVis.vulgivaga.1.1.chr7.g01555.t01#4    mNeoVis.3.1.1.chr7.g01472.t01#6    mNeoVis.vulgivaga.2.1.chr7.g01560.t01#1    mNeoVis.2.1.1.chr7_hap1.g01562.t01#5   mNeoVis.evergladensis.1.1.chr7.g01547.t01#3       3023   125259547:   mNeoVis.vulgivaga.2.1.chr1.g00956.t01#1    mNeoVis.3.1.1.chr1.g00962.t01#6    mNeoVis.vulgivaga.1.1.chr1.g00959.t01#4    mNeoVis.2.1.1.chr1_hap1.g00950.t01#5    mNeoVis.evergladensis.1.1.chr1.g00951.t01#3   mNeoVis.evergladensis.2.1.chr1.g00945.t01#2       3024   125259548:   mNeoVis.evergladensis.1.1.chr1.g00952.t01#3    mNeoVis.vulgivaga.2.1.chr1.g00957.t01#1    mNeoVis.evergladensis.2.1.chr1.g00946.t01#2    mNeoVis.vulgivaga.1.1.chr1.g00960.t01#4    mNeoVis.2.1.1.chr1_hap1.g00951.t01#5   mNeoVis.3.1.1.chr1.g00963.t01#6       3025   125259549:   mNeoVis.2.1.1.chr7_hap1.g01563.t01#5    mNeoVis.vulgivaga.2.1.chr7.g01561.t01#1    mNeoVis.evergladensis.1.1.chr7.g01548.t01#3    mNeoVis.3.1.1.chr7.g01473.t01#6    mNeoVis.vulgivaga.1.1.chr7.g01556.t01#4   mNeoVis.evergladensis.2.1.chr7.g01543.t01#2       3026   125259550:   mNeoVis.vulgivaga.1.1.chr7.g01557.t01#4    mNeoVis.2.1.1.chr7_hap1.g01564.t01#5    mNeoVis.vulgivaga.2.1.chr7.g01562.t01#1    mNeoVis.evergladensis.1.1.chr7.g01549.t01#3    mNeoVis.3.1.1.chr7.g01474.t01#6   mNeoVis.evergladensis.2.1.chr7.g01544.t01#2       3027   125259551:   mNeoVis.evergladensis.1.1.chr7.g01550.t01#3    mNeoVis.3.1.1.chr7.g01475.t01#6    mNeoVis.evergladensis.2.1.chr7.g01545.t01#2    mNeoVis.vulgivaga.1.1.chr7.g01558.t01#4    mNeoVis.2.1.1.chr7_hap1.g01565.t01#5   mNeoVis.vulgivaga.2.1.chr7.g01563.t01#1       3028   125259552:   mNeoVis.evergladensis.1.1.chr1.g00829.t01#3    mNeoVis.2.1.1.chr1_hap1.g00810.t01#5    mNeoVis.evergladensis.2.1.chr1.g00817.t01#2    mNeoVis.evergladensis.2.1.chr1.g00816.t01#2    mNeoVis.3.1.1.chr1.g00836.t01#6    mNeoVis.vulgivaga.2.1.chr1.g00832.t01#1   mNeoVis.vulgivaga.1.1.chr1.g00828.t01#4     3029   125259553:   mNeoVis.2.1.1.chr7_hap1.g01566.t01#5    mNeoVis.vulgivaga.2.1.chr7.g01564.t01#1    mNeoVis.evergladensis.1.1.chr7.g01551.t01#3    mNeoVis.3.1.1.chr7.g01476.t01#6    mNeoVis.vulgivaga.1.1.chr7.g01559.t01#4   mNeoVis.evergladensis.2.1.chr7.g01546.t01#2       3030   125259554:   mNeoVis.2.1.1.chr5_hap1.g01720.t01#5    mNeoVis.2.1.1.chr9_hap1.g00175.t01#5    mNeoVis.evergladensis.2.1.chr9.g00764.t01#2    mNeoVis.evergladensis.2.1.chr10.g00199.t01#2    mNeoVis.vulgivaga.1.1.chr2.g01711.t01#4    mNeoVis.evergladensis.2.1.chr11.g00899.t01#2    mNeoVis.vulgivaga.2.1.chr13.g00605.t01#1    mNeoVis.evergladensis.1.1.chr12.g00096.t01#3    mNeoVis.evergladensis.2.1.chr2.g00319.t01#2      3031   125259555:   mNeoVis.3.1.1.chr6.g00246.t01#6    mNeoVis.vulgivaga.1.1.chr9.g00041.t01#4    mNeoVis.3.1.1.chr5.g01620.t01#6    mNeoVis.vulgivaga.1.1.chr10.g00537.t01#4    mNeoVis.evergladensis.1.1.chr9.g00047.t01#3    mNeoVis.3.1.1.chr1.g01916.t01#6    mNeoVis.vulgivaga.1.1.chr13.g00588.t01#4    mNeoVis.3.1.1.chr5.g00684.t01#6    mNeoVis.2.1.1.chr2_hap1.g01699.t01#5      3032   125259556:   mNeoVis.2.1.1.chr11_hap1.g00268.t01#5   mNeoVis.3.1.1.chr7.g01378.t01#6             3033   125259557:   mNeoVis.3.1.1.chr7.g00251.t01#6   mNeoVis.vulgivaga.2.1.chr7.g00270.t01#1             3034   125259558:   mNeoVis.evergladensis.2.1.chr12.g00096.t01#2   mNeoVis.vulgivaga.2.1.chr7.g00390.t01#1             3035   125259559:   mNeoVis.2.1.1.chr1_hap1.g00812.t01#5    mNeoVis.evergladensis.2.1.chr1.g00819.t01#2    mNeoVis.evergladensis.1.1.chr8.g00156.t01#3    mNeoVis.3.1.1.chr1.g00757.t01#6    mNeoVis.vulgivaga.1.1.chr8.g00155.t01#4    mNeoVis.vulgivaga.2.1.chr1.g00834.t01#1    mNeoVis.vulgivaga.2.1.chr8.g00161.t01#1    mNeoVis.3.1.1.chr9.g00257.t01#6    mNeoVis.vulgivaga.1.1.chr1.g00830.t01#4      3036   125259560:   mNeoVis.evergladensis.2.1.chr10.g01100.t01#2    mNeoVis.evergladensis.1.1.chr10.g01095.t01#3    mNeoVis.vulgivaga.1.1.chr10.g01075.t01#4    mNeoVis.vulgivaga.2.1.chr10.g01089.t01#1   mNeoVis.2.1.1.chr9_hap1.g01110.t01#5       3037   125259561:  mNeoVis.3.1.1.chr8.g00130.t01#6               3038   125259562:   mNeoVis.vulgivaga.1.1.chr1.g00831.t01#4    mNeoVis.3.1.1.chr8.g01148.t01#6    mNeoVis.vulgivaga.2.1.chr1.g00835.t01#1    mNeoVis.2.1.1.chr2_hap1.g01184.t01#5    mNeoVis.3.1.1.chr1.g00839.t01#6    mNeoVis.2.1.1.chr14_hap1.g00064.t01#5    mNeoVis.vulgivaga.1.1.chr8.g01193.t01#4    mNeoVis.vulgivaga.1.1.chr2.g00982.t01#4    mNeoVis.evergladensis.1.1.chr8.g01199.t01#3      3039   125259563:   mNeoVis.evergladensis.2.1.chr7.g01447.t01#2    mNeoVis.3.1.1.chr12.g00077.t01#6    mNeoVis.evergladensis.2.1.chr14.g00342.t01#2    mNeoVis.evergladensis.2.1.chr6.g00691.t01#2    mNeoVis.2.1.1.chr14_hap1.g00342.t01#5    mNeoVis.3.1.1.chr7.g01372.t01#6    mNeoVis.2.1.1.chr12_hap1.g00092.t01#5    mNeoVis.vulgivaga.2.1.chr12.g00082.t01#1    mNeoVis.evergladensis.2.1.chr12.g00078.t01#2      3040   125259564:   mNeoVis.evergladensis.2.1.chr7.g00001.t01#2    mNeoVis.vulgivaga.1.1.chr4.g00817.t01#4    mNeoVis.3.1.1.chr7.g00001.t01#6    mNeoVis.vulgivaga.2.1.chr7.g00001.t01#1    mNeoVis.vulgivaga.1.1.chr7.g00001.t01#4    mNeoVis.vulgivaga.2.1.chr4.g00822.t01#1    mNeoVis.2.1.1.chr7_hap1.g00001.t01#5    mNeoVis.2.1.1.chr6_hap1.g00800.t01#5    mNeoVis.evergladensis.2.1.chr4.g00802.t01#2      3041   125259565:   mNeoVis.vulgivaga.1.1.chr7.g00095.t01#4    mNeoVis.vulgivaga.2.1.chr7.g00091.t01#1    mNeoVis.evergladensis.2.1.chr7.g00092.t01#2    mNeoVis.evergladensis.1.1.chr7.g00094.t01#3    mNeoVis.2.1.1.chr7_hap1.g00092.t01#5   mNeoVis.3.1.1.chr7.g00077.t01#6       3042   125259566:   mNeoVis.3.1.1.chr1.g00840.t01#6    mNeoVis.2.1.1.chr1_hap1.g00813.t01#5    mNeoVis.vulgivaga.1.1.chr1.g00833.t01#4    mNeoVis.evergladensis.2.1.chr1.g00820.t01#2    mNeoVis.evergladensis.1.1.chr1.g00832.t01#3   mNeoVis.vulgivaga.2.1.chr1.g00836.t01#1       3043   125259567:   mNeoVis.evergladensis.2.1.chr7.g01547.t01#2    mNeoVis.3.1.1.chr7.g01477.t01#6    mNeoVis.vulgivaga.2.1.chr7.g01565.t01#1    mNeoVis.vulgivaga.1.1.chr7.g01560.t01#4    mNeoVis.evergladensis.1.1.chr7.g01552.t01#3   mNeoVis.2.1.1.chr7_hap1.g01567.t01#5       3044   125259568:   mNeoVis.vulgivaga.1.1.chr13.g00690.t01#4    mNeoVis.evergladensis.1.1.chr1.g00833.t01#3    mNeoVis.2.1.1.chr13_hap1.g00733.t01#5    mNeoVis.vulgivaga.1.1.chr1.g00834.t01#4    mNeoVis.3.1.1.chr13.g00650.t01#6    mNeoVis.evergladensis.1.1.chr13.g00688.t01#3    mNeoVis.vulgivaga.2.1.chr13.g00696.t01#1    mNeoVis.3.1.1.chr1.g00841.t01#6    mNeoVis.2.1.1.chr1_hap1.g00814.t01#5      3045   125259569:   mNeoVis.evergladensis.1.1.chr7.g01553.t01#3    mNeoVis.2.1.1.chr7_hap1.g01568.t01#5    mNeoVis.evergladensis.2.1.chr7.g01548.t01#2    mNeoVis.3.1.1.chr7.g01478.t01#6    mNeoVis.vulgivaga.2.1.chr7.g01566.t01#1   mNeoVis.vulgivaga.1.1.chr7.g01561.t01#4       3046   125259570:   mNeoVis.3.1.1.chr1.g00843.t01#6    mNeoVis.evergladensis.2.1.chr1.g00823.t01#2    mNeoVis.evergladensis.1.1.chr1.g00835.t01#3    mNeoVis.vulgivaga.1.1.chr1.g00836.t01#4    mNeoVis.vulgivaga.2.1.chr1.g00839.t01#1   mNeoVis.2.1.1.chr1_hap1.g00816.t01#5       3047   125259571:   mNeoVis.vulgivaga.2.1.chr7.g01567.t01#1    mNeoVis.3.1.1.chr7.g01479.t01#6    mNeoVis.evergladensis.2.1.chr7.g01549.t01#2    mNeoVis.vulgivaga.1.1.chr7.g01562.t01#4    mNeoVis.evergladensis.1.1.chr7.g01554.t01#3   mNeoVis.2.1.1.chr7_hap1.g01569.t01#5       3048   125259572:   mNeoVis.vulgivaga.2.1.chr7.g01568.t01#1    mNeoVis.2.1.1.chr7_hap1.g01570.t01#5    mNeoVis.vulgivaga.1.1.chr7.g01563.t01#4    mNeoVis.3.1.1.chr7.g01480.t01#6    mNeoVis.evergladensis.1.1.chr7.g01555.t01#3   mNeoVis.evergladensis.2.1.chr7.g01550.t01#2       3049   125259573:   mNeoVis.vulgivaga.1.1.chr1.g00838.t01#4    mNeoVis.vulgivaga.1.1.chr1.g00837.t01#4    mNeoVis.evergladensis.2.1.chr1.g00824.t01#2    mNeoVis.2.1.1.chr1_hap1.g00817.t01#5    mNeoVis.evergladensis.1.1.chr1.g00836.t01#3    mNeoVis.3.1.1.chr1.g00844.t01#6   mNeoVis.vulgivaga.2.1.chr1.g00840.t01#1     3050   125259574:   mNeoVis.3.1.1.chr1.g00845.t01#6    mNeoVis.vulgivaga.1.1.chr1.g00839.t01#4    mNeoVis.vulgivaga.2.1.chr1.g00841.t01#1    mNeoVis.evergladensis.1.1.chr1.g00837.t01#3    mNeoVis.2.1.1.chr1_hap1.g00818.t01#5   mNeoVis.evergladensis.2.1.chr1.g00825.t01#2       3051   125259575:   mNeoVis.3.1.1.chr7.g01481.t01#6    mNeoVis.evergladensis.2.1.chr7.g01551.t01#2    mNeoVis.vulgivaga.1.1.chr7.g01564.t01#4    mNeoVis.vulgivaga.2.1.chr7.g01569.t01#1    mNeoVis.evergladensis.1.1.chr7.g01556.t01#3   mNeoVis.2.1.1.chr7_hap1.g01571.t01#5       3052   125259576:   mNeoVis.evergladensis.1.1.chr15.g00136.t01#3    mNeoVis.3.1.1.chr15.g00438.t01#6    mNeoVis.evergladensis.2.1.chr15.g00476.t01#2    mNeoVis.evergladensis.1.1.chr1.g00838.t01#3    mNeoVis.vulgivaga.1.1.chr15.g00479.t01#4    mNeoVis.3.1.1.chr1.g00846.t01#6   mNeoVis.vulgivaga.2.1.chr15.g00486.t01#1     3053   125259577:   mNeoVis.vulgivaga.1.1.chr1.g00840.t01#4    mNeoVis.2.1.1.chr1_hap1.g00819.t01#5    mNeoVis.evergladensis.2.1.chr1.g00826.t01#2   mNeoVis.vulgivaga.2.1.chr1.g00842.t01#1         3054   125259578:   mNeoVis.vulgivaga.2.1.chr1.g00843.t01#1    mNeoVis.2.1.1.chr1_hap1.g00820.t01#5    mNeoVis.vulgivaga.1.1.chr1.g00841.t01#4    mNeoVis.evergladensis.2.1.chr1.g00827.t01#2    mNeoVis.evergladensis.1.1.chr1.g00839.t01#3   mNeoVis.3.1.1.chr1.g00847.t01#6       3055   125259579:   mNeoVis.2.1.1.chr7_hap1.g01572.t01#5    mNeoVis.vulgivaga.1.1.chr7.g01565.t01#4    mNeoVis.evergladensis.2.1.chr7.g01552.t01#2    mNeoVis.evergladensis.1.1.chr7.g01557.t01#3    mNeoVis.3.1.1.chr7.g01482.t01#6   mNeoVis.vulgivaga.2.1.chr7.g01570.t01#1       3056   125259580:   mNeoVis.vulgivaga.1.1.chr7.g01566.t01#4    mNeoVis.evergladensis.1.1.chr7.g01558.t01#3    mNeoVis.evergladensis.2.1.chr7.g01553.t01#2    mNeoVis.3.1.1.chr7.g01483.t01#6    mNeoVis.2.1.1.chr7_hap1.g01573.t01#5   mNeoVis.vulgivaga.2.1.chr7.g01571.t01#1       3057   125259581:   mNeoVis.3.1.1.chr7.g01484.t01#6    mNeoVis.evergladensis.2.1.chr7.g01554.t01#2    mNeoVis.vulgivaga.1.1.chr7.g01567.t01#4    mNeoVis.2.1.1.chr7_hap1.g01574.t01#5    mNeoVis.evergladensis.1.1.chr7.g01559.t01#3   mNeoVis.vulgivaga.2.1.chr7.g01572.t01#1       3058   125259582:   mNeoVis.vulgivaga.1.1.chr1.g00842.t01#4    mNeoVis.2.1.1.chr1_hap1.g00821.t01#5    mNeoVis.evergladensis.1.1.chr1.g00840.t01#3    mNeoVis.vulgivaga.2.1.chr1.g00844.t01#1    mNeoVis.3.1.1.chr1.g00848.t01#6   mNeoVis.evergladensis.2.1.chr1.g00828.t01#2       3059   125259583:   mNeoVis.2.1.1.chr1_hap1.g00822.t01#5    mNeoVis.evergladensis.2.1.chr1.g00829.t01#2    mNeoVis.evergladensis.1.1.chr1.g00841.t01#3    mNeoVis.vulgivaga.2.1.chr1.g00845.t01#1    mNeoVis.vulgivaga.1.1.chr1.g00843.t01#4   mNeoVis.3.1.1.chr1.g00849.t01#6       3060   125259584:   mNeoVis.vulgivaga.1.1.chr1.g00844.t01#4    mNeoVis.3.1.1.chr1.g00850.t01#6    mNeoVis.evergladensis.1.1.chr1.g00842.t01#3    mNeoVis.2.1.1.chr1_hap1.g00823.t01#5    mNeoVis.vulgivaga.2.1.chr1.g00846.t01#1   mNeoVis.evergladensis.2.1.chr1.g00830.t01#2       3061   125259585:   mNeoVis.2.1.1.chr1_hap1.g00824.t01#5    mNeoVis.vulgivaga.2.1.chr1.g00847.t01#1    mNeoVis.vulgivaga.1.1.chr1.g00845.t01#4    mNeoVis.3.1.1.chr1.g00851.t01#6    mNeoVis.evergladensis.2.1.chr1.g00831.t01#2   mNeoVis.evergladensis.1.1.chr1.g00843.t01#3       3062   125259586:   mNeoVis.2.1.1.chr1_hap1.g00825.t01#5    mNeoVis.vulgivaga.2.1.chr1.g00848.t01#1    mNeoVis.evergladensis.2.1.chr1.g00832.t01#2    mNeoVis.vulgivaga.1.1.chr1.g00846.t01#4   mNeoVis.evergladensis.1.1.chr1.g00844.t01#3       3063   125259587:   mNeoVis.evergladensis.2.1.chr7.g01490.t01#2    mNeoVis.vulgivaga.2.1.chr7.g01506.t01#1    mNeoVis.evergladensis.1.1.chr7.g01492.t01#3    mNeoVis.3.1.1.chr7.g01417.t01#6    mNeoVis.2.1.1.chr7_hap1.g01508.t01#5   mNeoVis.vulgivaga.1.1.chr7.g01503.t01#4       3064   125259588:  mNeoVis.vulgivaga.2.1.chr1.g00849.t01#1               3065   125259589:   mNeoVis.vulgivaga.2.1.chr7.g01507.t01#1    mNeoVis.evergladensis.1.1.chr7.g01493.t01#3    mNeoVis.2.1.1.chr7_hap1.g01509.t01#5    mNeoVis.vulgivaga.1.1.chr7.g01504.t01#4    mNeoVis.evergladensis.2.1.chr7.g01491.t01#2   mNeoVis.3.1.1.chr7.g01421.t01#6       3066   125259590:   mNeoVis.vulgivaga.1.1.chr1.g00848.t01#4   mNeoVis.vulgivaga.2.1.chr1.g00850.t01#1             3067   125259591:   mNeoVis.3.1.1.chr1.g00856.t01#6    mNeoVis.vulgivaga.1.1.chr1.g00853.t01#4    mNeoVis.vulgivaga.2.1.chr1.g00852.t01#1    mNeoVis.evergladensis.1.1.chr1.g00848.t01#3    mNeoVis.2.1.1.chr1_hap1.g00833.t01#5   mNeoVis.evergladensis.2.1.chr1.g00836.t01#2       3068   125259592:   mNeoVis.vulgivaga.2.1.chr1.g00853.t01#1    mNeoVis.evergladensis.1.1.chr1.g00849.t01#3    mNeoVis.2.1.1.chr1_hap1.g00834.t01#5   mNeoVis.evergladensis.2.1.chr1.g00837.t01#2       3069   125259593:   mNeoVis.vulgivaga.2.1.chr7.g01508.t01#1    mNeoVis.2.1.1.chr7_hap1.g01512.t01#5    mNeoVis.evergladensis.2.1.chr7.g01493.t01#2    mNeoVis.3.1.1.chr7.g01423.t01#6    mNeoVis.vulgivaga.1.1.chr7.g01506.t01#4   mNeoVis.evergladensis.1.1.chr7.g01495.t01#3       3070   125259594:   mNeoVis.vulgivaga.1.1.chr1.g00854.t01#4    mNeoVis.2.1.1.chr8_hap1.g01302.t01#5    mNeoVis.2.1.1.chr1_hap1.g00835.t01#5    mNeoVis.3.1.1.chr8.g01198.t01#6    mNeoVis.evergladensis.1.1.chr8.g01246.t01#3    mNeoVis.evergladensis.2.1.chr9.g01252.t01#2    mNeoVis.evergladensis.1.1.chr1.g00850.t01#3    mNeoVis.vulgivaga.2.1.chr8.g01252.t01#1    mNeoVis.vulgivaga.1.1.chr8.g01238.t01#4      3071   125259595:   mNeoVis.3.1.1.chr1.g00858.t01#6    mNeoVis.vulgivaga.1.1.chr1.g00855.t01#4    mNeoVis.vulgivaga.2.1.chr1.g00855.t01#1    mNeoVis.2.1.1.chr1_hap1.g00836.t01#5    mNeoVis.evergladensis.2.1.chr1.g00839.t01#2   mNeoVis.evergladensis.1.1.chr1.g00851.t01#3       3072   125259596:   mNeoVis.evergladensis.1.1.chr7.g01496.t01#3    mNeoVis.evergladensis.2.1.chr7.g01494.t01#2    mNeoVis.2.1.1.chr7_hap1.g01513.t01#5    mNeoVis.3.1.1.chr7.g01424.t01#6    mNeoVis.vulgivaga.1.1.chr7.g01507.t01#4   mNeoVis.vulgivaga.2.1.chr7.g01509.t01#1       3073   125257549:   mNeoVis.2.1.1.chr1_hap1.g00837.t01#5    mNeoVis.evergladensis.1.1.chr1.g00852.t01#3    mNeoVis.vulgivaga.2.1.chr1.g00856.t01#1    mNeoVis.vulgivaga.1.1.chr1.g00856.t01#4    mNeoVis.evergladensis.2.1.chr1.g00840.t01#2   mNeoVis.3.1.1.chr1.g00859.t01#6       3074   125257550:   mNeoVis.evergladensis.2.1.chr7.g01495.t01#2    mNeoVis.3.1.1.chr7.g01425.t01#6    mNeoVis.2.1.1.chr7_hap1.g01514.t01#5    mNeoVis.vulgivaga.2.1.chr7.g01510.t01#1    mNeoVis.vulgivaga.1.1.chr7.g01508.t01#4   mNeoVis.evergladensis.1.1.chr7.g01497.t01#3       3075   125257551:   mNeoVis.2.1.1.chr7_hap1.g01515.t01#5    mNeoVis.evergladensis.2.1.chr7.g01496.t01#2    mNeoVis.3.1.1.chr7.g01426.t01#6    mNeoVis.vulgivaga.2.1.chr7.g01511.t01#1    mNeoVis.vulgivaga.1.1.chr7.g01509.t01#4   mNeoVis.evergladensis.1.1.chr7.g01498.t01#3       3076   125257552:   mNeoVis.evergladensis.2.1.chr7.g01497.t01#2    mNeoVis.vulgivaga.2.1.chr7.g01513.t01#1    mNeoVis.vulgivaga.2.1.chr7.g01512.t01#1    mNeoVis.2.1.1.chr7_hap1.g01516.t01#5    mNeoVis.3.1.1.chr7.g01427.t01#6    mNeoVis.vulgivaga.1.1.chr7.g01510.t01#4   mNeoVis.evergladensis.1.1.chr7.g01499.t01#3     3077   125257553:   mNeoVis.3.1.1.chr1.g00860.t01#6    mNeoVis.evergladensis.1.1.chr1.g00853.t01#3    mNeoVis.vulgivaga.2.1.chr1.g00857.t01#1    mNeoVis.vulgivaga.1.1.chr1.g00857.t01#4    mNeoVis.2.1.1.chr1_hap1.g00838.t01#5   mNeoVis.evergladensis.2.1.chr1.g00841.t01#2       3078   125257554:   mNeoVis.evergladensis.2.1.chr1.g00842.t01#2    mNeoVis.vulgivaga.1.1.chr1.g00858.t01#4    mNeoVis.3.1.1.chr1.g00862.t01#6    mNeoVis.2.1.1.chr1_hap1.g00839.t01#5    mNeoVis.vulgivaga.2.1.chr1.g00858.t01#1   mNeoVis.evergladensis.1.1.chr1.g00854.t01#3       3079   125257555:   mNeoVis.vulgivaga.1.1.chr7.g01511.t01#4    mNeoVis.vulgivaga.2.1.chr7.g01514.t01#1    mNeoVis.evergladensis.1.1.chr7.g01500.t01#3    mNeoVis.2.1.1.chr7_hap1.g01517.t01#5    mNeoVis.3.1.1.chr7.g01428.t01#6   mNeoVis.evergladensis.2.1.chr7.g01498.t01#2       3080   125257556:   mNeoVis.3.1.1.chr7.g01429.t01#6    mNeoVis.evergladensis.2.1.chr7.g01499.t01#2    mNeoVis.vulgivaga.1.1.chr7.g01512.t01#4    mNeoVis.evergladensis.1.1.chr7.g01501.t01#3   mNeoVis.2.1.1.chr7_hap1.g01518.t01#5       3081   125257557:   mNeoVis.evergladensis.1.1.chr7.g01502.t01#3   mNeoVis.vulgivaga.2.1.chr7.g01515.t01#1             3082   125257558:   mNeoVis.vulgivaga.1.1.chr7.g01513.t01#4    mNeoVis.evergladensis.2.1.chr7.g01500.t01#2    mNeoVis.evergladensis.1.1.chr7.g01503.t01#3    mNeoVis.3.1.1.chr7.g01430.t01#6    mNeoVis.2.1.1.chr7_hap1.g01519.t01#5   mNeoVis.vulgivaga.2.1.chr7.g01516.t01#1       3083   125257559:   mNeoVis.vulgivaga.1.1.chr7.g01514.t01#4    mNeoVis.3.1.1.chr7.g01431.t01#6    mNeoVis.vulgivaga.2.1.chr7.g01517.t01#1    mNeoVis.evergladensis.2.1.chr7.g01501.t01#2    mNeoVis.evergladensis.1.1.chr7.g01504.t01#3   mNeoVis.2.1.1.chr7_hap1.g01520.t01#5       3084   125257560:   mNeoVis.vulgivaga.1.1.chr1.g00859.t01#4   mNeoVis.vulgivaga.2.1.chr1.g00859.t01#1             3085   125257561:   mNeoVis.evergladensis.2.1.chr7.g01502.t01#2    mNeoVis.vulgivaga.1.1.chr7.g01515.t01#4    mNeoVis.vulgivaga.2.1.chr7.g01518.t01#1    mNeoVis.evergladensis.1.1.chr7.g01505.t01#3    mNeoVis.2.1.1.chr7_hap1.g01521.t01#5   mNeoVis.3.1.1.chr7.g01432.t01#6       3086   125257562:   mNeoVis.evergladensis.2.1.chr2.g01042.t01#2    mNeoVis.evergladensis.1.1.chr12.g00102.t01#3    mNeoVis.2.1.1.chr12_hap1.g00128.t01#5    mNeoVis.evergladensis.1.1.chr2.g01040.t01#3    mNeoVis.vulgivaga.2.1.chr12.g00104.t01#1    mNeoVis.3.1.1.chr12.g00099.t01#6    mNeoVis.2.1.1.chr5_hap1.g01049.t01#5    mNeoVis.vulgivaga.1.1.chr1.g00850.t01#4    mNeoVis.vulgivaga.2.1.chr2.g01046.t01#1      3087   125257563:   mNeoVis.evergladensis.2.1.chr1.g00845.t01#2    mNeoVis.vulgivaga.1.1.chr1.g00860.t01#4    mNeoVis.2.1.1.chr1_hap1.g00841.t01#5    mNeoVis.vulgivaga.2.1.chr1.g00860.t01#1    mNeoVis.3.1.1.chr1.g00864.t01#6   mNeoVis.evergladensis.1.1.chr1.g00856.t01#3       3088   125257564:   mNeoVis.2.1.1.chr1_hap1.g00842.t01#5    mNeoVis.evergladensis.1.1.chr1.g00857.t01#3    mNeoVis.vulgivaga.2.1.chr1.g00861.t01#1   mNeoVis.evergladensis.2.1.chr1.g00846.t01#2       3089   125257565:   mNeoVis.vulgivaga.1.1.chr7.g01516.t01#4    mNeoVis.evergladensis.2.1.chr7.g01503.t01#2    mNeoVis.evergladensis.1.1.chr7.g01506.t01#3    mNeoVis.3.1.1.chr7.g01433.t01#6    mNeoVis.vulgivaga.2.1.chr7.g01519.t01#1   mNeoVis.2.1.1.chr7_hap1.g01522.t01#5       3090   125257566:   mNeoVis.evergladensis.1.1.chr1.g00858.t01#3    mNeoVis.evergladensis.2.1.chr1.g00847.t01#2    mNeoVis.vulgivaga.2.1.chr1.g00862.t01#1    mNeoVis.vulgivaga.1.1.chr1.g00861.t01#4    mNeoVis.3.1.1.chr1.g00866.t01#6   mNeoVis.2.1.1.chr1_hap1.g00843.t01#5       3091   125257567:   mNeoVis.evergladensis.1.1.chr7.g01507.t01#3    mNeoVis.3.1.1.chr7.g01434.t01#6    mNeoVis.evergladensis.2.1.chr7.g01504.t01#2    mNeoVis.vulgivaga.2.1.chr7.g01520.t01#1    mNeoVis.2.1.1.chr7_hap1.g01523.t01#5   mNeoVis.vulgivaga.1.1.chr7.g01517.t01#4       3092   125257568:   mNeoVis.evergladensis.1.1.chr7.g01508.t01#3    mNeoVis.vulgivaga.2.1.chr7.g01521.t01#1    mNeoVis.2.1.1.chr7_hap1.g01524.t01#5    mNeoVis.3.1.1.chr7.g01435.t01#6    mNeoVis.evergladensis.2.1.chr7.g01505.t01#2   mNeoVis.vulgivaga.1.1.chr7.g01518.t01#4       3093   125257569:   mNeoVis.3.1.1.chr1.g00867.t01#6    mNeoVis.evergladensis.2.1.chr1.g00848.t01#2    mNeoVis.2.1.1.chr1_hap1.g00844.t01#5    mNeoVis.evergladensis.1.1.chr1.g00859.t01#3    mNeoVis.vulgivaga.2.1.chr1.g00863.t01#1   mNeoVis.vulgivaga.1.1.chr1.g00862.t01#4       3094   125257570:   mNeoVis.evergladensis.2.1.chr1.g00849.t01#2    mNeoVis.3.1.1.chr1.g00868.t01#6    mNeoVis.2.1.1.chr1_hap1.g00845.t01#5    mNeoVis.evergladensis.1.1.chr1.g00860.t01#3    mNeoVis.vulgivaga.2.1.chr1.g00864.t01#1   mNeoVis.vulgivaga.1.1.chr1.g00863.t01#4       3095   125257571:   mNeoVis.3.1.1.chr7.g01436.t01#6    mNeoVis.2.1.1.chr7_hap1.g01525.t01#5    mNeoVis.vulgivaga.1.1.chr7.g01519.t01#4    mNeoVis.vulgivaga.2.1.chr7.g01522.t01#1    mNeoVis.evergladensis.2.1.chr7.g01506.t01#2   mNeoVis.evergladensis.1.1.chr7.g01509.t01#3       3096   125257572:   mNeoVis.vulgivaga.1.1.chr7.g01520.t01#4    mNeoVis.evergladensis.2.1.chr7.g01507.t01#2    mNeoVis.evergladensis.1.1.chr7.g01510.t01#3    mNeoVis.3.1.1.chr7.g01437.t01#6    mNeoVis.vulgivaga.2.1.chr7.g01523.t01#1   mNeoVis.2.1.1.chr7_hap1.g01526.t01#5       3097   125257573:   mNeoVis.evergladensis.2.1.chr7.g01508.t01#2    mNeoVis.2.1.1.chr7_hap1.g01527.t01#5    mNeoVis.vulgivaga.1.1.chr7.g01521.t01#4    mNeoVis.vulgivaga.2.1.chr7.g01524.t01#1    mNeoVis.evergladensis.1.1.chr7.g01511.t01#3   mNeoVis.3.1.1.chr7.g01438.t01#6       3098   125257574:   mNeoVis.2.1.1.chr1_hap1.g00846.t01#5    mNeoVis.evergladensis.1.1.chr1.g00861.t01#3    mNeoVis.vulgivaga.1.1.chr1.g00864.t01#4    mNeoVis.vulgivaga.2.1.chr1.g00865.t01#1    mNeoVis.3.1.1.chr1.g00869.t01#6   mNeoVis.evergladensis.2.1.chr1.g00850.t01#2       3099   125257575:   mNeoVis.vulgivaga.1.1.chr7.g01568.t01#4    mNeoVis.evergladensis.1.1.chr7.g01560.t01#3    mNeoVis.vulgivaga.2.1.chr7.g01573.t01#1    mNeoVis.2.1.1.chr7_hap1.g01575.t01#5    mNeoVis.3.1.1.chr7.g01485.t01#6   mNeoVis.evergladensis.2.1.chr7.g01555.t01#2       3100   125257576:   mNeoVis.evergladensis.2.1.chr7.g01557.t01#2    mNeoVis.2.1.1.chr7_hap1.g01577.t01#5    mNeoVis.vulgivaga.1.1.chr7.g01570.t01#4    mNeoVis.3.1.1.chr7.g01488.t01#6    mNeoVis.vulgivaga.2.1.chr7.g01575.t01#1   mNeoVis.evergladensis.1.1.chr7.g01562.t01#3       3101   125257577:   mNeoVis.2.1.1.chr7_hap1.g00093.t01#5    mNeoVis.vulgivaga.1.1.chr7.g00096.t01#4    mNeoVis.evergladensis.1.1.chr7.g00095.t01#3    mNeoVis.vulgivaga.2.1.chr7.g00092.t01#1   mNeoVis.evergladensis.2.1.chr7.g00093.t01#2       3102   125257578:   mNeoVis.evergladensis.2.1.chr1.g00851.t01#2    mNeoVis.vulgivaga.1.1.chr1.g00865.t01#4    mNeoVis.vulgivaga.2.1.chr1.g00866.t01#1    mNeoVis.2.1.1.chr1_hap1.g00847.t01#5    mNeoVis.3.1.1.chr1.g00870.t01#6    mNeoVis.evergladensis.1.1.chr8.g01196.t01#3    mNeoVis.evergladensis.1.1.chr1.g00862.t01#3   mNeoVis.2.1.1.chr6_hap1.g00249.t01#5     3103   125257579:   mNeoVis.3.1.1.chr9.g01140.t01#6    mNeoVis.2.1.1.chr10_hap1.g01211.t01#5    mNeoVis.vulgivaga.2.1.chr9.g01171.t01#1    mNeoVis.evergladensis.2.1.chr8.g01188.t01#2    mNeoVis.evergladensis.1.1.chr9.g01181.t01#3   mNeoVis.vulgivaga.1.1.chr9.g01187.t01#4       3104   125257580:   mNeoVis.2.1.1.chr10_hap1.g01210.t01#5    mNeoVis.evergladensis.2.1.chr8.g01187.t01#2    mNeoVis.evergladensis.1.1.chr9.g01180.t01#3    mNeoVis.vulgivaga.1.1.chr9.g01186.t01#4   mNeoVis.vulgivaga.2.1.chr9.g01170.t01#1       3105   125257581:   mNeoVis.evergladensis.2.1.chr1.g00852.t01#2   mNeoVis.vulgivaga.2.1.chr1.g00867.t01#1             3106   125257582:   mNeoVis.vulgivaga.1.1.chr7.g01571.t01#4    mNeoVis.3.1.1.chr7.g01489.t01#6    mNeoVis.evergladensis.1.1.chr7.g01563.t01#3    mNeoVis.2.1.1.chr7_hap1.g01578.t01#5    mNeoVis.vulgivaga.2.1.chr7.g01576.t01#1   mNeoVis.evergladensis.2.1.chr7.g01558.t01#2       3107   125257583:   mNeoVis.3.1.1.chr1.g00872.t01#6    mNeoVis.2.1.1.chr1_hap1.g00849.t01#5    mNeoVis.vulgivaga.1.1.chr1.g00867.t01#4    mNeoVis.evergladensis.2.1.chr1.g00853.t01#2    mNeoVis.evergladensis.1.1.chr1.g00864.t01#3   mNeoVis.vulgivaga.2.1.chr1.g00868.t01#1       3108   125257584:   mNeoVis.evergladensis.1.1.chr1.g00865.t01#3    mNeoVis.evergladensis.2.1.chr1.g00855.t01#2    mNeoVis.vulgivaga.1.1.chr1.g00868.t01#4    mNeoVis.3.1.1.chr1.g00874.t01#6    mNeoVis.vulgivaga.2.1.chr1.g00869.t01#1    mNeoVis.2.1.1.chr1_hap1.g00851.t01#5    mNeoVis.vulgivaga.1.1.chr1.g00869.t01#4    mNeoVis.3.1.1.chr1.g00873.t01#6    mNeoVis.evergladensis.2.1.chr1.g00854.t01#2      3109   125257585:   mNeoVis.3.1.1.chr1.g00876.t01#6    mNeoVis.evergladensis.1.1.chr1.g00866.t01#3    mNeoVis.vulgivaga.2.1.chr1.g00871.t01#1    mNeoVis.evergladensis.2.1.chr1.g00856.t01#2    mNeoVis.vulgivaga.1.1.chr1.g00870.t01#4   mNeoVis.2.1.1.chr1_hap1.g00852.t01#5       3110   125257586:   mNeoVis.vulgivaga.2.1.chr1.g00872.t01#1    mNeoVis.vulgivaga.1.1.chr1.g00871.t01#4    mNeoVis.3.1.1.chr1.g00877.t01#6    mNeoVis.evergladensis.2.1.chr1.g00857.t01#2   mNeoVis.evergladensis.1.1.chr1.g00867.t01#3       3111   125257587:   mNeoVis.vulgivaga.2.1.chr7.g01577.t01#1    mNeoVis.vulgivaga.1.1.chr7.g01572.t01#4    mNeoVis.3.1.1.chr7.g01490.t01#6    mNeoVis.evergladensis.1.1.chr7.g01564.t01#3    mNeoVis.evergladensis.2.1.chr7.g01559.t01#2   mNeoVis.2.1.1.chr7_hap1.g01579.t01#5       3112   125257588:   mNeoVis.evergladensis.2.1.chr1.g00859.t01#2    mNeoVis.evergladensis.1.1.chr1.g00869.t01#3    mNeoVis.vulgivaga.1.1.chr1.g00873.t01#4    mNeoVis.vulgivaga.2.1.chr1.g00873.t01#1    mNeoVis.2.1.1.chr1_hap1.g00854.t01#5   mNeoVis.3.1.1.chr1.g00878.t01#6       3113   125257589:   mNeoVis.vulgivaga.1.1.chr7.g01573.t01#4    mNeoVis.3.1.1.chr7.g01491.t01#6    mNeoVis.vulgivaga.2.1.chr7.g01578.t01#1    mNeoVis.evergladensis.2.1.chr7.g01560.t01#2    mNeoVis.evergladensis.1.1.chr7.g01565.t01#3   mNeoVis.2.1.1.chr7_hap1.g01580.t01#5       3114   125257590:   mNeoVis.vulgivaga.1.1.chr7.g01485.t01#4    mNeoVis.3.1.1.chr7.g01399.t01#6    mNeoVis.vulgivaga.2.1.chr7.g01488.t01#1    mNeoVis.2.1.1.chr7_hap1.g01490.t01#5    mNeoVis.evergladensis.1.1.chr7.g01473.t01#3   mNeoVis.evergladensis.2.1.chr7.g01472.t01#2       3115   125257591:   mNeoVis.2.1.1.chr1_hap1.g00855.t01#5    mNeoVis.vulgivaga.2.1.chr1.g00874.t01#1    mNeoVis.evergladensis.2.1.chr1.g00860.t01#2    mNeoVis.3.1.1.chr1.g00879.t01#6    mNeoVis.evergladensis.1.1.chr1.g00870.t01#3   mNeoVis.vulgivaga.1.1.chr1.g00874.t01#4       3116   125257592:   mNeoVis.2.1.1.chr1_hap1.g00856.t01#5    mNeoVis.vulgivaga.2.1.chr1.g00875.t01#1    mNeoVis.3.1.1.chr1.g00880.t01#6   mNeoVis.vulgivaga.1.1.chr1.g00875.t01#4         3117   125257593:   mNeoVis.3.1.1.chr1.g00881.t01#6    mNeoVis.vulgivaga.1.1.chr1.g00876.t01#4    mNeoVis.evergladensis.1.1.chr1.g00871.t01#3    mNeoVis.evergladensis.2.1.chr1.g00861.t01#2    mNeoVis.2.1.1.chr1_hap1.g00857.t01#5   mNeoVis.vulgivaga.2.1.chr1.g00876.t01#1       3118   125257594:   mNeoVis.3.1.1.chr7.g01400.t01#6    mNeoVis.evergladensis.1.1.chr7.g01474.t01#3    mNeoVis.vulgivaga.1.1.chr7.g01486.t01#4    mNeoVis.vulgivaga.2.1.chr7.g01489.t01#1    mNeoVis.evergladensis.2.1.chr7.g01473.t01#2   mNeoVis.2.1.1.chr7_hap1.g01491.t01#5       3119   125257595:   mNeoVis.vulgivaga.2.1.chr7.g00093.t01#1    mNeoVis.2.1.1.chr7_hap1.g00094.t01#5    mNeoVis.evergladensis.1.1.chr7.g00096.t01#3    mNeoVis.vulgivaga.1.1.chr7.g00097.t01#4    mNeoVis.3.1.1.chr7.g00078.t01#6   mNeoVis.evergladensis.2.1.chr7.g00094.t01#2       3120   125257596:   mNeoVis.2.1.1.chr1_hap1.g00858.t01#5    mNeoVis.evergladensis.1.1.chr1.g00873.t01#3    mNeoVis.vulgivaga.2.1.chr1.g00877.t01#1    mNeoVis.3.1.1.chr1.g00882.t01#6    mNeoVis.evergladensis.2.1.chr1.g00862.t01#2   mNeoVis.vulgivaga.1.1.chr1.g00877.t01#4       3121   125257597:   mNeoVis.evergladensis.1.1.chr14.g00690.t01#3    mNeoVis.vulgivaga.1.1.chr14.g00684.t01#4    mNeoVis.3.1.1.chr14.g00663.t01#6    mNeoVis.vulgivaga.2.1.chr14.g00692.t01#1   mNeoVis.evergladensis.2.1.chr14.g00693.t01#2       3122   125257598:   mNeoVis.evergladensis.2.1.chr2.g00498.t01#2    mNeoVis.vulgivaga.1.1.chr2.g00494.t01#4    mNeoVis.vulgivaga.2.1.chr2.g00500.t01#1    mNeoVis.2.1.1.chr5_hap1.g00503.t01#5   mNeoVis.evergladensis.1.1.chr2.g00494.t01#3       3123   125257599:   mNeoVis.3.1.1.chr14.g00662.t01#6   mNeoVis.2.1.1.chr14_hap1.g00693.t01#5             3124   125257600:   mNeoVis.3.1.1.chr1.g00884.t01#6    mNeoVis.evergladensis.2.1.chr1.g00863.t01#2    mNeoVis.vulgivaga.1.1.chr1.g00879.t01#4    mNeoVis.2.1.1.chr1_hap1.g00860.t01#5   mNeoVis.vulgivaga.2.1.chr1.g00878.t01#1       3125   125257601:   mNeoVis.vulgivaga.2.1.chr1.g00879.t01#1    mNeoVis.vulgivaga.1.1.chr1.g00880.t01#4    mNeoVis.evergladensis.2.1.chr1.g00864.t01#2    mNeoVis.2.1.1.chr1_hap1.g00861.t01#5    mNeoVis.evergladensis.1.1.chr1.g00875.t01#3   mNeoVis.3.1.1.chr1.g00885.t01#6       3126   125257602:   mNeoVis.evergladensis.2.1.chr7.g01474.t01#2    mNeoVis.vulgivaga.2.1.chr7.g01490.t01#1    mNeoVis.2.1.1.chr7_hap1.g01492.t01#5    mNeoVis.3.1.1.chr7.g01401.t01#6    mNeoVis.evergladensis.1.1.chr7.g01475.t01#3   mNeoVis.vulgivaga.1.1.chr7.g01487.t01#4       3127   125257603:   mNeoVis.2.1.1.chr7_hap1.g01493.t01#5    mNeoVis.3.1.1.chr7.g01402.t01#6    mNeoVis.evergladensis.1.1.chr7.g01476.t01#3    mNeoVis.vulgivaga.1.1.chr7.g01488.t01#4    mNeoVis.evergladensis.2.1.chr7.g01475.t01#2   mNeoVis.vulgivaga.2.1.chr7.g01491.t01#1       3128   125257604:   mNeoVis.evergladensis.2.1.chr7.g01476.t01#2    mNeoVis.vulgivaga.2.1.chr7.g01492.t01#1    mNeoVis.2.1.1.chr7_hap1.g01494.t01#5    mNeoVis.evergladensis.1.1.chr7.g01477.t01#3    mNeoVis.3.1.1.chr7.g01403.t01#6   mNeoVis.vulgivaga.1.1.chr7.g01489.t01#4       3129   125257605:   mNeoVis.vulgivaga.2.1.chr7.g01493.t01#1    mNeoVis.evergladensis.2.1.chr7.g01477.t01#2    mNeoVis.2.1.1.chr7_hap1.g01495.t01#5    mNeoVis.3.1.1.chr7.g01404.t01#6    mNeoVis.evergladensis.1.1.chr7.g01478.t01#3   mNeoVis.vulgivaga.1.1.chr7.g01490.t01#4       3130   125257606:   mNeoVis.3.1.1.chr7.g01405.t01#6    mNeoVis.evergladensis.2.1.chr7.g01478.t01#2    mNeoVis.2.1.1.chr7_hap1.g01496.t01#5    mNeoVis.evergladensis.1.1.chr7.g01479.t01#3    mNeoVis.vulgivaga.1.1.chr7.g01491.t01#4   mNeoVis.vulgivaga.2.1.chr7.g01494.t01#1       3131   125257607:   mNeoVis.vulgivaga.1.1.chr1.g00882.t01#4    mNeoVis.evergladensis.1.1.chr1.g00877.t01#3    mNeoVis.evergladensis.2.1.chr1.g00867.t01#2    mNeoVis.3.1.1.chr1.g00886.t01#6    mNeoVis.2.1.1.chr1_hap1.g00864.t01#5   mNeoVis.vulgivaga.2.1.chr1.g00880.t01#1       3132   125257608:   mNeoVis.3.1.1.chr6.g00632.t01#6    mNeoVis.vulgivaga.2.1.chr6.g00623.t01#1    mNeoVis.vulgivaga.1.1.chr6.g00625.t01#4    mNeoVis.evergladensis.2.1.chr6.g00626.t01#2    mNeoVis.2.1.1.chr4_hap1.g00617.t01#5   mNeoVis.evergladensis.1.1.chr6.g00628.t01#3       3133   125257609:   mNeoVis.evergladensis.2.1.chr1.g00866.t01#2    mNeoVis.3.1.1.chr1.g00887.t01#6    mNeoVis.vulgivaga.1.1.chr1.g00881.t01#4    mNeoVis.vulgivaga.2.1.chr1.g00881.t01#1    mNeoVis.2.1.1.chr1_hap1.g00863.t01#5   mNeoVis.evergladensis.1.1.chr1.g00876.t01#3       3134   125257610:   mNeoVis.vulgivaga.2.1.chr7.g01495.t01#1    mNeoVis.3.1.1.chr7.g01406.t01#6    mNeoVis.2.1.1.chr7_hap1.g01497.t01#5    mNeoVis.evergladensis.2.1.chr7.g01479.t01#2   mNeoVis.evergladensis.1.1.chr7.g01480.t01#3       3135   125257611:   mNeoVis.vulgivaga.2.1.chr1.g00882.t01#1    mNeoVis.3.1.1.chr1.g00888.t01#6    mNeoVis.evergladensis.2.1.chr1.g00868.t01#2    mNeoVis.2.1.1.chr1_hap1.g00865.t01#5   mNeoVis.evergladensis.1.1.chr1.g00878.t01#3       3136   125257612:   mNeoVis.vulgivaga.1.1.chr7.g01493.t01#4    mNeoVis.vulgivaga.2.1.chr7.g01496.t01#1    mNeoVis.evergladensis.2.1.chr7.g01480.t01#2    mNeoVis.2.1.1.chr7_hap1.g01498.t01#5    mNeoVis.evergladensis.1.1.chr7.g01481.t01#3   mNeoVis.3.1.1.chr7.g01407.t01#6       3137   125257613:   mNeoVis.evergladensis.2.1.chr7.g01481.t01#2    mNeoVis.2.1.1.chr7_hap1.g01499.t01#5    mNeoVis.3.1.1.chr7.g01408.t01#6    mNeoVis.vulgivaga.2.1.chr7.g01498.t01#1    mNeoVis.vulgivaga.1.1.chr7.g01494.t01#4   mNeoVis.evergladensis.1.1.chr7.g01483.t01#3       3138   125257614:   mNeoVis.evergladensis.1.1.chr2.g01239.t01#3    mNeoVis.vulgivaga.2.1.chr2.g01242.t01#1    mNeoVis.evergladensis.2.1.chr2.g01237.t01#2    mNeoVis.2.1.1.chr5_hap1.g01249.t01#5    mNeoVis.vulgivaga.1.1.chr2.g01243.t01#4   mNeoVis.3.1.1.chr2.g01167.t01#6       3139   125257615:   mNeoVis.evergladensis.1.1.chr12.g00255.t01#3    mNeoVis.2.1.1.chr12_hap1.g00283.t01#5    mNeoVis.3.1.1.chr12.g00251.t01#6    mNeoVis.vulgivaga.2.1.chr12.g00258.t01#1    mNeoVis.evergladensis.2.1.chr12.g00251.t01#2   mNeoVis.vulgivaga.1.1.chr12.g00259.t01#4       3140   125257616:   mNeoVis.3.1.1.chr1.g00889.t01#6    mNeoVis.vulgivaga.1.1.chr1.g00884.t01#4    mNeoVis.2.1.1.chr1_hap1.g00866.t01#5    mNeoVis.vulgivaga.2.1.chr1.g00883.t01#1    mNeoVis.evergladensis.2.1.chr1.g00869.t01#2   mNeoVis.evergladensis.1.1.chr1.g00879.t01#3       3141   125257617:   mNeoVis.vulgivaga.1.1.chr1.g00885.t01#4    mNeoVis.vulgivaga.2.1.chr1.g00884.t01#1    mNeoVis.3.1.1.chr1.g00890.t01#6    mNeoVis.evergladensis.2.1.chr1.g00870.t01#2    mNeoVis.2.1.1.chr1_hap1.g00867.t01#5   mNeoVis.evergladensis.1.1.chr1.g00880.t01#3       3142   125257618:   mNeoVis.evergladensis.1.1.chr7.g01484.t01#3    mNeoVis.vulgivaga.2.1.chr7.g01499.t01#1    mNeoVis.3.1.1.chr7.g01409.t01#6    mNeoVis.evergladensis.2.1.chr7.g01482.t01#2   mNeoVis.2.1.1.chr7_hap1.g01500.t01#5       3143   125257619:   mNeoVis.evergladensis.2.1.chr1.g00871.t01#2    mNeoVis.evergladensis.1.1.chr1.g00881.t01#3    mNeoVis.2.1.1.chr1_hap1.g00868.t01#5    mNeoVis.vulgivaga.2.1.chr1.g00885.t01#1    mNeoVis.vulgivaga.1.1.chr1.g00886.t01#4   mNeoVis.3.1.1.chr1.g00891.t01#6       3144   125257620:   mNeoVis.vulgivaga.1.1.chr1.g00887.t01#4    mNeoVis.evergladensis.2.1.chr1.g00872.t01#2    mNeoVis.3.1.1.chr1.g00892.t01#6    mNeoVis.evergladensis.1.1.chr1.g00882.t01#3    mNeoVis.vulgivaga.2.1.chr1.g00886.t01#1   mNeoVis.2.1.1.chr1_hap1.g00869.t01#5       3145   125257621:   mNeoVis.3.1.1.chr15.g00149.t01#6    mNeoVis.vulgivaga.2.1.chr15.g00159.t01#1    mNeoVis.2.1.1.chr15_hap1.g00173.t01#5    mNeoVis.evergladensis.2.1.chr15.g00156.t01#2    mNeoVis.evergladensis.1.1.chr15.g00458.t01#3   mNeoVis.vulgivaga.1.1.chr15.g00155.t01#4       3146   125257622:   mNeoVis.evergladensis.2.1.chr1.g00873.t01#2    mNeoVis.vulgivaga.1.1.chr1.g00888.t01#4    mNeoVis.3.1.1.chr1.g00893.t01#6    mNeoVis.2.1.1.chr1_hap1.g00870.t01#5    mNeoVis.vulgivaga.2.1.chr1.g00887.t01#1   mNeoVis.evergladensis.1.1.chr1.g00883.t01#3       3147   125257623:   mNeoVis.3.1.1.chr7.g01410.t01#6    mNeoVis.vulgivaga.1.1.chr7.g01496.t01#4    mNeoVis.evergladensis.2.1.chr7.g01483.t01#2    mNeoVis.2.1.1.chr7_hap1.g01501.t01#5    mNeoVis.evergladensis.1.1.chr7.g01485.t01#3   mNeoVis.vulgivaga.2.1.chr7.g01500.t01#1       3148   125257624:   mNeoVis.vulgivaga.2.1.chr1.g00789.t01#1    mNeoVis.vulgivaga.1.1.chr1.g00782.t01#4    mNeoVis.3.1.1.chr1.g00791.t01#6    mNeoVis.evergladensis.1.1.chr1.g00783.t01#3    mNeoVis.evergladensis.2.1.chr1.g00773.t01#2   mNeoVis.2.1.1.chr1_hap1.g00765.t01#5       3149   125257625:   mNeoVis.2.1.1.chr1_hap1.g00766.t01#5    mNeoVis.vulgivaga.2.1.chr1.g00790.t01#1    mNeoVis.evergladensis.2.1.chr1.g00774.t01#2    mNeoVis.3.1.1.chr1.g00792.t01#6    mNeoVis.vulgivaga.1.1.chr1.g00783.t01#4   mNeoVis.evergladensis.1.1.chr1.g00784.t01#3       3150   125257626:   mNeoVis.vulgivaga.2.1.chr1.g00791.t01#1    mNeoVis.3.1.1.chr1.g00793.t01#6    mNeoVis.evergladensis.2.1.chr1.g00775.t01#2    mNeoVis.evergladensis.1.1.chr1.g00785.t01#3    mNeoVis.2.1.1.chr1_hap1.g00767.t01#5   mNeoVis.vulgivaga.1.1.chr1.g00784.t01#4       3151   125257627:   mNeoVis.2.1.1.chr1_hap1.g00768.t01#5    mNeoVis.vulgivaga.2.1.chr1.g00792.t01#1    mNeoVis.vulgivaga.1.1.chr1.g00785.t01#4    mNeoVis.evergladensis.2.1.chr1.g00777.t01#2    mNeoVis.evergladensis.1.1.chr1.g00786.t01#3   mNeoVis.3.1.1.chr1.g00794.t01#6       3152   125257628:   mNeoVis.2.1.1.chr7_hap1.g01502.t01#5    mNeoVis.evergladensis.2.1.chr7.g01484.t01#2    mNeoVis.3.1.1.chr7.g01411.t01#6    mNeoVis.vulgivaga.2.1.chr7.g01501.t01#1    mNeoVis.evergladensis.1.1.chr7.g01486.t01#3   mNeoVis.vulgivaga.1.1.chr7.g01497.t01#4       3153   125257629:   mNeoVis.vulgivaga.1.1.chr15.g00158.t01#4    mNeoVis.vulgivaga.2.1.chr15.g00160.t01#1    mNeoVis.3.1.1.chr15.g00154.t01#6    mNeoVis.3.1.1.chr15.g00152.t01#6    mNeoVis.evergladensis.2.1.chr15.g00159.t01#2    mNeoVis.evergladensis.1.1.chr15.g00455.t01#3    mNeoVis.3.1.1.chr15.g00153.t01#6   mNeoVis.2.1.1.chr15_hap1.g00176.t01#5     3154   125257630:   mNeoVis.3.1.1.chr15.g00157.t01#6    mNeoVis.vulgivaga.2.1.chr15.g00164.t01#1   mNeoVis.evergladensis.1.1.chr15.g00451.t01#3         3155   125257631:   mNeoVis.2.1.1.chr7_hap1.g01504.t01#5    mNeoVis.evergladensis.2.1.chr7.g01486.t01#2    mNeoVis.evergladensis.1.1.chr7.g01488.t01#3    mNeoVis.vulgivaga.1.1.chr7.g01499.t01#4   mNeoVis.vulgivaga.2.1.chr7.g01502.t01#1       3156   125257632:   mNeoVis.vulgivaga.2.1.chr7.g01503.t01#1    mNeoVis.3.1.1.chr7.g01414.t01#6    mNeoVis.evergladensis.1.1.chr7.g01489.t01#3    mNeoVis.vulgivaga.1.1.chr7.g01500.t01#4    mNeoVis.evergladensis.2.1.chr7.g01487.t01#2   mNeoVis.2.1.1.chr7_hap1.g01505.t01#5       3157   125257633:   mNeoVis.2.1.1.chr1_hap1.g00770.t01#5    mNeoVis.vulgivaga.1.1.chr1.g00787.t01#4    mNeoVis.3.1.1.chr1.g00796.t01#6    mNeoVis.evergladensis.2.1.chr1.g00779.t01#2    mNeoVis.vulgivaga.2.1.chr1.g00793.t01#1   mNeoVis.evergladensis.1.1.chr1.g00788.t01#3       3158   125257634:   mNeoVis.evergladensis.2.1.chr1.g00780.t01#2    mNeoVis.evergladensis.1.1.chr1.g00789.t01#3    mNeoVis.2.1.1.chr1_hap1.g00771.t01#5    mNeoVis.vulgivaga.2.1.chr1.g00794.t01#1    mNeoVis.vulgivaga.1.1.chr1.g00788.t01#4   mNeoVis.3.1.1.chr1.g00797.t01#6       3159   125257635:   mNeoVis.evergladensis.1.1.chr1.g00792.t01#3    mNeoVis.evergladensis.2.1.chr1.g00781.t01#2    mNeoVis.vulgivaga.2.1.chr1.g00795.t01#1    mNeoVis.2.1.1.chr1_hap1.g00773.t01#5   mNeoVis.vulgivaga.1.1.chr1.g00789.t01#4       3160   125257636:   mNeoVis.evergladensis.2.1.chr1.g00782.t01#2    mNeoVis.2.1.1.chr1_hap1.g00774.t01#5    mNeoVis.vulgivaga.2.1.chr1.g00796.t01#1    mNeoVis.evergladensis.1.1.chr1.g00793.t01#3   mNeoVis.3.1.1.chr1.g00800.t01#6       3161   125257637:   mNeoVis.2.1.1.chr1_hap1.g00775.t01#5    mNeoVis.3.1.1.chr1.g00801.t01#6    mNeoVis.evergladensis.2.1.chr1.g00783.t01#2    mNeoVis.evergladensis.2.1.chr12.g00480.t01#2    mNeoVis.vulgivaga.2.1.chr1.g00797.t01#1   mNeoVis.evergladensis.1.1.chr1.g00794.t01#3       3162   125257638:   mNeoVis.vulgivaga.2.1.chr12.g00492.t01#1    mNeoVis.evergladensis.2.1.chr8.g01096.t01#2    mNeoVis.vulgivaga.1.1.chr12.g00489.t01#4    mNeoVis.3.1.1.chr12.g00487.t01#6    mNeoVis.2.1.1.chr12_hap1.g00536.t01#5   mNeoVis.evergladensis.1.1.chr12.g00487.t01#3       3163   125257639:   mNeoVis.evergladensis.2.1.chr1.g00784.t01#2    mNeoVis.2.1.1.chr1_hap1.g00776.t01#5    mNeoVis.vulgivaga.2.1.chr1.g00798.t01#1    mNeoVis.evergladensis.1.1.chr1.g00795.t01#3    mNeoVis.vulgivaga.1.1.chr1.g00791.t01#4   mNeoVis.3.1.1.chr1.g00802.t01#6       3164   125257640:   mNeoVis.evergladensis.2.1.chr7.g01488.t01#2    mNeoVis.evergladensis.1.1.chr7.g01490.t01#3    mNeoVis.3.1.1.chr7.g01415.t01#6    mNeoVis.vulgivaga.1.1.chr7.g01501.t01#4    mNeoVis.2.1.1.chr7_hap1.g01506.t01#5   mNeoVis.vulgivaga.2.1.chr7.g01504.t01#1       3165   125257641:   mNeoVis.vulgivaga.2.1.chr7.g01505.t01#1    mNeoVis.3.1.1.chr7.g01416.t01#6    mNeoVis.vulgivaga.1.1.chr7.g01502.t01#4    mNeoVis.evergladensis.2.1.chr7.g01489.t01#2    mNeoVis.2.1.1.chr7_hap1.g01507.t01#5   mNeoVis.evergladensis.1.1.chr7.g01491.t01#3       3166   125257642:   mNeoVis.vulgivaga.1.1.chr7.g01522.t01#4    mNeoVis.3.1.1.chr7.g01439.t01#6    mNeoVis.evergladensis.2.1.chr7.g01509.t01#2    mNeoVis.2.1.1.chr7_hap1.g01528.t01#5    mNeoVis.vulgivaga.2.1.chr7.g01525.t01#1   mNeoVis.evergladensis.1.1.chr7.g01512.t01#3       3167   125257643:   mNeoVis.evergladensis.1.1.chr7.g01513.t01#3    mNeoVis.vulgivaga.1.1.chr7.g01523.t01#4    mNeoVis.3.1.1.chr7.g01440.t01#6    mNeoVis.2.1.1.chr7_hap1.g01529.t01#5    mNeoVis.evergladensis.2.1.chr7.g01510.t01#2   mNeoVis.vulgivaga.2.1.chr7.g01526.t01#1       3168   125257644:   mNeoVis.2.1.1.chr7_hap1.g01530.t01#5    mNeoVis.vulgivaga.2.1.chr7.g01527.t01#1    mNeoVis.evergladensis.1.1.chr7.g01514.t01#3    mNeoVis.vulgivaga.1.1.chr7.g01524.t01#4    mNeoVis.evergladensis.2.1.chr7.g01511.t01#2   mNeoVis.3.1.1.chr7.g01441.t01#6       3169   125257645:   mNeoVis.vulgivaga.1.1.chr7.g01525.t01#4    mNeoVis.3.1.1.chr7.g01442.t01#6    mNeoVis.evergladensis.2.1.chr7.g01512.t01#2    mNeoVis.vulgivaga.2.1.chr7.g01529.t01#1    mNeoVis.2.1.1.chr7_hap1.g01531.t01#5    mNeoVis.vulgivaga.2.1.chr7.g01528.t01#1   mNeoVis.evergladensis.1.1.chr7.g01515.t01#3     3170   125257646:   mNeoVis.vulgivaga.2.1.chr7.g01530.t01#1    mNeoVis.2.1.1.chr7_hap1.g01532.t01#5    mNeoVis.evergladensis.2.1.chr7.g01513.t01#2    mNeoVis.evergladensis.1.1.chr7.g01516.t01#3    mNeoVis.vulgivaga.1.1.chr7.g01526.t01#4   mNeoVis.3.1.1.chr7.g01443.t01#6       3171   125257647:   mNeoVis.vulgivaga.2.1.chr1.g00754.t01#1    mNeoVis.3.1.1.chr1.g00755.t01#6    mNeoVis.evergladensis.2.1.chr1.g00743.t01#2    mNeoVis.vulgivaga.1.1.chr1.g00748.t01#4   mNeoVis.2.1.1.chr1_hap1.g00733.t01#5       3172   125257648:   mNeoVis.vulgivaga.1.1.chr1.g00749.t01#4   mNeoVis.vulgivaga.2.1.chr1.g00755.t01#1             3173   125257649:   mNeoVis.evergladensis.1.1.chr7.g01517.t01#3    mNeoVis.3.1.1.chr7.g01444.t01#6    mNeoVis.evergladensis.2.1.chr7.g01514.t01#2    mNeoVis.vulgivaga.1.1.chr7.g01527.t01#4    mNeoVis.vulgivaga.2.1.chr7.g01531.t01#1   mNeoVis.2.1.1.chr7_hap1.g01533.t01#5       3174   125257650:   mNeoVis.evergladensis.1.1.chr1.g00753.t01#3    mNeoVis.vulgivaga.2.1.chr1.g00756.t01#1    mNeoVis.evergladensis.2.1.chr1.g00745.t01#2    mNeoVis.2.1.1.chr1_hap1.g00735.t01#5    mNeoVis.3.1.1.chr1.g00758.t01#6   mNeoVis.vulgivaga.1.1.chr1.g00750.t01#4       3175   125257651:   mNeoVis.evergladensis.2.1.chr1.g00747.t01#2    mNeoVis.evergladensis.1.1.chr1.g00754.t01#3    mNeoVis.vulgivaga.2.1.chr1.g00757.t01#1    mNeoVis.2.1.1.chr1_hap1.g00737.t01#5    mNeoVis.3.1.1.chr1.g00759.t01#6   mNeoVis.vulgivaga.1.1.chr1.g00752.t01#4       3176   125257652:   mNeoVis.vulgivaga.2.1.chr1.g00758.t01#1    mNeoVis.evergladensis.1.1.chr1.g00755.t01#3    mNeoVis.2.1.1.chr1_hap1.g00736.t01#5    mNeoVis.3.1.1.chr10.g00171.t01#6    mNeoVis.evergladensis.2.1.chr10.g00174.t01#2    mNeoVis.vulgivaga.1.1.chr10.g00165.t01#4    mNeoVis.evergladensis.2.1.chr1.g00746.t01#2    mNeoVis.2.1.1.chr9_hap1.g00179.t01#5    mNeoVis.vulgivaga.1.1.chr1.g00751.t01#4      3177   125257653:   mNeoVis.vulgivaga.2.1.chr7.g01532.t01#1    mNeoVis.evergladensis.1.1.chr7.g01518.t01#3    mNeoVis.vulgivaga.1.1.chr7.g01528.t01#4    mNeoVis.3.1.1.chr7.g01445.t01#6    mNeoVis.evergladensis.2.1.chr7.g01515.t01#2   mNeoVis.2.1.1.chr7_hap1.g01534.t01#5       3178   125257654:   mNeoVis.evergladensis.2.1.chr15.g00160.t01#2    mNeoVis.evergladensis.1.1.chr15.g00454.t01#3   mNeoVis.vulgivaga.2.1.chr15.g00161.t01#1         3179   125257655:  mNeoVis.vulgivaga.2.1.chr1.g00759.t01#1               3180   125257656:   mNeoVis.2.1.1.chr7_hap1.g01535.t01#5    mNeoVis.evergladensis.2.1.chr7.g01516.t01#2    mNeoVis.evergladensis.1.1.chr7.g01519.t01#3    mNeoVis.vulgivaga.2.1.chr7.g01533.t01#1    mNeoVis.vulgivaga.1.1.chr7.g01529.t01#4   mNeoVis.3.1.1.chr7.g01446.t01#6       3181   125257657:   mNeoVis.2.1.1.chr1_hap1.g00777.t01#5    mNeoVis.evergladensis.2.1.chr1.g00785.t01#2    mNeoVis.vulgivaga.2.1.chr1.g00799.t01#1    mNeoVis.vulgivaga.1.1.chr1.g00792.t01#4    mNeoVis.3.1.1.chr1.g00803.t01#6   mNeoVis.evergladensis.1.1.chr1.g00796.t01#3       3182   125257658:   mNeoVis.evergladensis.2.1.chr7.g01517.t01#2    mNeoVis.2.1.1.chr7_hap1.g01536.t01#5    mNeoVis.evergladensis.1.1.chr7.g01520.t01#3    mNeoVis.vulgivaga.2.1.chr7.g01534.t01#1    mNeoVis.3.1.1.chr7.g01447.t01#6   mNeoVis.vulgivaga.1.1.chr7.g01530.t01#4       3183   125257659:   mNeoVis.vulgivaga.2.1.chr1.g00800.t01#1    mNeoVis.evergladensis.2.1.chr1.g00786.t01#2    mNeoVis.vulgivaga.1.1.chr1.g00793.t01#4    mNeoVis.2.1.1.chr1_hap1.g00778.t01#5    mNeoVis.3.1.1.chr1.g00804.t01#6   mNeoVis.evergladensis.1.1.chr1.g00797.t01#3       3184   125257660:   mNeoVis.vulgivaga.2.1.chr1.g00760.t01#1    mNeoVis.2.1.1.chr1_hap1.g00739.t01#5    mNeoVis.3.1.1.chr1.g00762.t01#6    mNeoVis.evergladensis.2.1.chr1.g00749.t01#2    mNeoVis.vulgivaga.1.1.chr1.g00754.t01#4   mNeoVis.evergladensis.1.1.chr1.g00757.t01#3       3185   125257661:   mNeoVis.vulgivaga.2.1.chr1.g00801.t01#1    mNeoVis.evergladensis.2.1.chr1.g00787.t01#2    mNeoVis.evergladensis.1.1.chr1.g00798.t01#3    mNeoVis.2.1.1.chr1_hap1.g00779.t01#5    mNeoVis.vulgivaga.1.1.chr1.g00794.t01#4   mNeoVis.3.1.1.chr1.g00805.t01#6       3186   125257662:   mNeoVis.vulgivaga.2.1.chr9.g00415.t01#1    mNeoVis.2.1.1.chr1_hap1.g00281.t01#5    mNeoVis.vulgivaga.1.1.chr1.g00279.t01#4    mNeoVis.vulgivaga.2.1.chr1.g00284.t01#1    mNeoVis.3.1.1.chr6.g00404.t01#6    mNeoVis.vulgivaga.1.1.chr1.g00280.t01#4    mNeoVis.3.1.1.chr1.g00284.t01#6    mNeoVis.evergladensis.2.1.chr6.g00407.t01#2    mNeoVis.evergladensis.2.1.chr1.g00280.t01#2      3187   125257663:   mNeoVis.2.1.1.chr1_hap1.g00235.t01#5    mNeoVis.vulgivaga.1.1.chr1.g00229.t01#4    mNeoVis.evergladensis.2.1.chr1.g00232.t01#2    mNeoVis.evergladensis.1.1.chr1.g00242.t01#3    mNeoVis.vulgivaga.2.1.chr1.g00236.t01#1   mNeoVis.3.1.1.chr1.g00235.t01#6       3188   125257664:   mNeoVis.evergladensis.1.1.chr1.g00799.t01#3    mNeoVis.3.1.1.chr1.g00807.t01#6    mNeoVis.2.1.1.chr1_hap1.g00781.t01#5    mNeoVis.vulgivaga.2.1.chr1.g00803.t01#1    mNeoVis.vulgivaga.1.1.chr1.g00797.t01#4   mNeoVis.evergladensis.2.1.chr1.g00788.t01#2       3189   125257665:   mNeoVis.evergladensis.1.1.chr7.g01424.t01#3    mNeoVis.vulgivaga.2.1.chr7.g01439.t01#1    mNeoVis.vulgivaga.1.1.chr7.g01434.t01#4    mNeoVis.2.1.1.chr7_hap1.g01444.t01#5    mNeoVis.evergladensis.2.1.chr7.g01427.t01#2   mNeoVis.3.1.1.chr7.g01352.t01#6       3190   125257666:   mNeoVis.2.1.1.chr7_hap1.g01445.t01#5    mNeoVis.3.1.1.chr7.g01353.t01#6    mNeoVis.evergladensis.2.1.chr7.g01428.t01#2    mNeoVis.vulgivaga.2.1.chr7.g01440.t01#1    mNeoVis.evergladensis.1.1.chr7.g01425.t01#3   mNeoVis.vulgivaga.1.1.chr7.g01435.t01#4       3191   125257667:   mNeoVis.vulgivaga.2.1.chr7.g01441.t01#1    mNeoVis.vulgivaga.1.1.chr7.g01436.t01#4    mNeoVis.evergladensis.1.1.chr7.g01426.t01#3    mNeoVis.2.1.1.chr7_hap1.g01446.t01#5    mNeoVis.3.1.1.chr7.g01354.t01#6   mNeoVis.evergladensis.2.1.chr7.g01429.t01#2       3192   125257668:   mNeoVis.vulgivaga.1.1.chr1.g00798.t01#4    mNeoVis.vulgivaga.2.1.chr1.g00804.t01#1    mNeoVis.evergladensis.1.1.chr1.g00800.t01#3    mNeoVis.3.1.1.chr1.g00808.t01#6    mNeoVis.2.1.1.chr1_hap1.g00782.t01#5   mNeoVis.evergladensis.2.1.chr1.g00789.t01#2       3193   125257669:  mNeoVis.vulgivaga.2.1.chr1.g00805.t01#1               3194   125257670:  mNeoVis.vulgivaga.2.1.chr1.g00806.t01#1               3195   125257671:   mNeoVis.vulgivaga.2.1.chr7.g01442.t01#1    mNeoVis.2.1.1.chr7_hap1.g01447.t01#5    mNeoVis.evergladensis.2.1.chr7.g01430.t01#2    mNeoVis.evergladensis.1.1.chr7.g01427.t01#3    mNeoVis.vulgivaga.1.1.chr7.g01437.t01#4   mNeoVis.3.1.1.chr7.g01355.t01#6       3196   125257672:   mNeoVis.vulgivaga.1.1.chr7.g01438.t01#4    mNeoVis.evergladensis.1.1.chr7.g01428.t01#3    mNeoVis.2.1.1.chr7_hap1.g01448.t01#5    mNeoVis.3.1.1.chr7.g01356.t01#6    mNeoVis.evergladensis.2.1.chr7.g01431.t01#2   mNeoVis.vulgivaga.2.1.chr7.g01443.t01#1       3197   125257673:   mNeoVis.vulgivaga.2.1.chr1.g00807.t01#1   mNeoVis.evergladensis.1.1.chr1.g00801.t01#3           3198   125257674:   mNeoVis.2.1.1.chr7_hap1.g00095.t01#5    mNeoVis.3.1.1.chr7.g00079.t01#6    mNeoVis.evergladensis.1.1.chr7.g00097.t01#3    mNeoVis.vulgivaga.2.1.chr7.g00094.t01#1    mNeoVis.evergladensis.2.1.chr7.g00095.t01#2   mNeoVis.vulgivaga.1.1.chr7.g00098.t01#4       3199   125257675:   mNeoVis.evergladensis.2.1.chr1.g00791.t01#2    mNeoVis.evergladensis.1.1.chr1.g00802.t01#3    mNeoVis.3.1.1.chr1.g00810.t01#6    mNeoVis.vulgivaga.1.1.chr1.g00800.t01#4    mNeoVis.2.1.1.chr1_hap1.g00785.t01#5   mNeoVis.vulgivaga.2.1.chr1.g00808.t01#1       3200   125257676:   mNeoVis.3.1.1.chr1.g00812.t01#6    mNeoVis.3.1.1.chr1.g00811.t01#6    mNeoVis.evergladensis.1.1.chr1.g00803.t01#3    mNeoVis.evergladensis.2.1.chr1.g00792.t01#2    mNeoVis.2.1.1.chr1_hap1.g00786.t01#5    mNeoVis.vulgivaga.1.1.chr1.g00801.t01#4    mNeoVis.vulgivaga.2.1.chr1.g00809.t01#1   mNeoVis.evergladensis.1.1.chr1.g00804.t01#3     3201   125257677:   mNeoVis.2.1.1.chr1_hap1.g00787.t01#5    mNeoVis.vulgivaga.2.1.chr1.g00810.t01#1    mNeoVis.evergladensis.2.1.chr1.g00793.t01#2    mNeoVis.3.1.1.chr1.g00813.t01#6    mNeoVis.vulgivaga.1.1.chr1.g00802.t01#4   mNeoVis.evergladensis.1.1.chr1.g00805.t01#3       3202   125257678:   mNeoVis.evergladensis.1.1.chr7.g01429.t01#3    mNeoVis.evergladensis.2.1.chr7.g01432.t01#2    mNeoVis.2.1.1.chr7_hap1.g01449.t01#5    mNeoVis.vulgivaga.2.1.chr7.g01444.t01#1    mNeoVis.vulgivaga.1.1.chr7.g01439.t01#4   mNeoVis.3.1.1.chr7.g01357.t01#6       3203   125257679:   mNeoVis.vulgivaga.1.1.chr1.g00803.t01#4    mNeoVis.evergladensis.1.1.chr1.g00806.t01#3    mNeoVis.evergladensis.2.1.chr1.g00794.t01#2    mNeoVis.3.1.1.chr1.g00814.t01#6    mNeoVis.2.1.1.chr1_hap1.g00788.t01#5   mNeoVis.vulgivaga.2.1.chr1.g00811.t01#1       3204   125257680:   mNeoVis.3.1.1.chr7.g01358.t01#6    mNeoVis.vulgivaga.2.1.chr7.g01445.t01#1    mNeoVis.vulgivaga.1.1.chr7.g01440.t01#4    mNeoVis.evergladensis.1.1.chr7.g01430.t01#3    mNeoVis.2.1.1.chr7_hap1.g01450.t01#5   mNeoVis.evergladensis.2.1.chr7.g01433.t01#2       3205   125257681:   mNeoVis.evergladensis.1.1.chr15.g00453.t01#3    mNeoVis.vulgivaga.2.1.chr15.g00162.t01#1    mNeoVis.3.1.1.chr15.g00155.t01#6    mNeoVis.2.1.1.chr15_hap1.g00178.t01#5    mNeoVis.vulgivaga.1.1.chr15.g00160.t01#4   mNeoVis.evergladensis.2.1.chr15.g00161.t01#2       3206   125257682:   mNeoVis.vulgivaga.2.1.chr1.g00812.t01#1    mNeoVis.3.1.1.chr1.g00815.t01#6    mNeoVis.evergladensis.2.1.chr1.g00795.t01#2    mNeoVis.vulgivaga.1.1.chr1.g00804.t01#4    mNeoVis.evergladensis.1.1.chr1.g00807.t01#3   mNeoVis.2.1.1.chr1_hap1.g00789.t01#5       3207   125257683:   mNeoVis.2.1.1.chr7_hap1.g01451.t01#5    mNeoVis.vulgivaga.2.1.chr7.g01446.t01#1    mNeoVis.vulgivaga.1.1.chr7.g01441.t01#4    mNeoVis.3.1.1.chr7.g01359.t01#6    mNeoVis.evergladensis.2.1.chr7.g01434.t01#2   mNeoVis.evergladensis.1.1.chr7.g01431.t01#3       3208   125257684:  mNeoVis.vulgivaga.2.1.chr7.g01447.t01#1               3209   125257685:   mNeoVis.evergladensis.2.1.chr1.g00796.t01#2    mNeoVis.3.1.1.chr1.g00816.t01#6    mNeoVis.vulgivaga.2.1.chr1.g00813.t01#1    mNeoVis.2.1.1.chr1_hap1.g00790.t01#5    mNeoVis.vulgivaga.1.1.chr1.g00805.t01#4   mNeoVis.evergladensis.1.1.chr1.g00808.t01#3       3210   125257686:   mNeoVis.2.1.1.chr1_hap1.g00791.t01#5    mNeoVis.vulgivaga.2.1.chr1.g00814.t01#1    mNeoVis.vulgivaga.1.1.chr1.g00806.t01#4    mNeoVis.evergladensis.1.1.chr1.g00809.t01#3    mNeoVis.3.1.1.chr1.g00817.t01#6   mNeoVis.evergladensis.2.1.chr1.g00797.t01#2       3211   125257687:   mNeoVis.evergladensis.1.1.chr7.g01432.t01#3    mNeoVis.2.1.1.chr7_hap1.g01452.t01#5    mNeoVis.evergladensis.2.1.chr7.g01435.t01#2    mNeoVis.vulgivaga.2.1.chr7.g01448.t01#1    mNeoVis.3.1.1.chr7.g01360.t01#6   mNeoVis.vulgivaga.1.1.chr7.g01442.t01#4       3212   125257688:   mNeoVis.evergladensis.2.1.chr7.g01436.t01#2    mNeoVis.vulgivaga.2.1.chr7.g01449.t01#1    mNeoVis.3.1.1.chr7.g01362.t01#6    mNeoVis.vulgivaga.1.1.chr7.g01443.t01#4    mNeoVis.evergladensis.1.1.chr7.g01433.t01#3   mNeoVis.2.1.1.chr7_hap1.g01454.t01#5       3213   125257689:   mNeoVis.2.1.1.chr1_hap1.g00792.t01#5    mNeoVis.vulgivaga.1.1.chr1.g00807.t01#4    mNeoVis.evergladensis.2.1.chr1.g00798.t01#2    mNeoVis.evergladensis.1.1.chr1.g00810.t01#3    mNeoVis.3.1.1.chr1.g00818.t01#6   mNeoVis.vulgivaga.2.1.chr1.g00815.t01#1       3214   125257690:   mNeoVis.2.1.1.chr1_hap1.g00793.t01#5    mNeoVis.vulgivaga.2.1.chr1.g00816.t01#1    mNeoVis.evergladensis.2.1.chr1.g00799.t01#2    mNeoVis.3.1.1.chr1.g00819.t01#6    mNeoVis.evergladensis.1.1.chr1.g00811.t01#3   mNeoVis.vulgivaga.1.1.chr1.g00808.t01#4       3215   125257691:   mNeoVis.vulgivaga.1.1.chr1.g00809.t01#4    mNeoVis.evergladensis.2.1.chr1.g00800.t01#2    mNeoVis.3.1.1.chr1.g00820.t01#6    mNeoVis.vulgivaga.2.1.chr1.g00817.t01#1    mNeoVis.evergladensis.1.1.chr1.g00812.t01#3   mNeoVis.2.1.1.chr1_hap1.g00794.t01#5       3216   125257692:   mNeoVis.2.1.1.chr7_hap1.g01455.t01#5    mNeoVis.vulgivaga.1.1.chr7.g01444.t01#4    mNeoVis.vulgivaga.2.1.chr7.g01450.t01#1    mNeoVis.evergladensis.1.1.chr7.g01434.t01#3    mNeoVis.evergladensis.2.1.chr7.g01437.t01#2    mNeoVis.vulgivaga.1.1.chr7.g01445.t01#4   mNeoVis.3.1.1.chr7.g01363.t01#6       3217   125257693:   mNeoVis.vulgivaga.1.1.chr7.g01446.t01#4    mNeoVis.3.1.1.chr7.g01364.t01#6    mNeoVis.evergladensis.2.1.chr7.g01438.t01#2    mNeoVis.2.1.1.chr7_hap1.g01456.t01#5    mNeoVis.vulgivaga.2.1.chr7.g01451.t01#1   mNeoVis.evergladensis.1.1.chr7.g01435.t01#3       3218   125257694:   mNeoVis.2.1.1.chr7_hap1.g01457.t01#5   mNeoVis.vulgivaga.2.1.chr7.g01452.t01#1             3219   125257695:   mNeoVis.vulgivaga.2.1.chr15.g00163.t01#1   mNeoVis.3.1.1.chr15.g00156.t01#6             3220   125257696:   mNeoVis.2.1.1.chr7_hap1.g00096.t01#5    mNeoVis.evergladensis.1.1.chr7.g00098.t01#3    mNeoVis.vulgivaga.1.1.chr7.g00099.t01#4    mNeoVis.evergladensis.2.1.chr7.g00096.t01#2    mNeoVis.vulgivaga.2.1.chr7.g00095.t01#1   mNeoVis.3.1.1.chr7.g00080.t01#6       3221   125257697:   mNeoVis.evergladensis.2.1.chr1.g00801.t01#2    mNeoVis.vulgivaga.2.1.chr1.g00818.t01#1    mNeoVis.3.1.1.chr1.g00821.t01#6    mNeoVis.vulgivaga.1.1.chr1.g00811.t01#4    mNeoVis.vulgivaga.1.1.chr1.g00812.t01#4    mNeoVis.evergladensis.1.1.chr1.g00813.t01#3   mNeoVis.2.1.1.chr1_hap1.g00795.t01#5     3222   125257698:   mNeoVis.evergladensis.1.1.chr7.g01436.t01#3    mNeoVis.3.1.1.chr1.g00500.t01#6    mNeoVis.evergladensis.1.1.chr1.g00512.t01#3    mNeoVis.vulgivaga.1.1.chr7.g01447.t01#4    mNeoVis.3.1.1.chr7.g01365.t01#6    mNeoVis.2.1.1.chr1_hap1.g00503.t01#5    mNeoVis.vulgivaga.2.1.chr1.g00511.t01#1    mNeoVis.evergladensis.2.1.chr1.g00499.t01#2    mNeoVis.evergladensis.2.1.chr7.g01439.t01#2      3223   125257699:   mNeoVis.vulgivaga.2.1.chr7.g01454.t01#1    mNeoVis.2.1.1.chr7_hap1.g01459.t01#5    mNeoVis.vulgivaga.1.1.chr7.g01448.t01#4    mNeoVis.3.1.1.chr7.g01366.t01#6    mNeoVis.evergladensis.2.1.chr7.g01440.t01#2   mNeoVis.evergladensis.1.1.chr7.g01438.t01#3       3224   125257700:   mNeoVis.evergladensis.1.1.chr1.g00814.t01#3    mNeoVis.evergladensis.2.1.chr1.g00802.t01#2    mNeoVis.3.1.1.chr1.g00823.t01#6    mNeoVis.vulgivaga.1.1.chr1.g00813.t01#4    mNeoVis.2.1.1.chr1_hap1.g00796.t01#5   mNeoVis.vulgivaga.2.1.chr1.g00819.t01#1       3225   125257701:   mNeoVis.evergladensis.1.1.chr1.g00815.t01#3    mNeoVis.vulgivaga.1.1.chr1.g00814.t01#4    mNeoVis.evergladensis.2.1.chr1.g00803.t01#2    mNeoVis.vulgivaga.2.1.chr1.g00820.t01#1   mNeoVis.2.1.1.chr1_hap1.g00797.t01#5       3226   125257702:   mNeoVis.evergladensis.2.1.chr7.g01441.t01#2    mNeoVis.2.1.1.chr7_hap1.g01460.t01#5    mNeoVis.vulgivaga.2.1.chr7.g01455.t01#1    mNeoVis.vulgivaga.1.1.chr7.g01449.t01#4    mNeoVis.3.1.1.chr7.g01367.t01#6   mNeoVis.evergladensis.1.1.chr7.g01439.t01#3       3227   125257703:   mNeoVis.3.1.1.chr1.g00825.t01#6    mNeoVis.2.1.1.chr1_hap1.g00799.t01#5    mNeoVis.vulgivaga.1.1.chr1.g00815.t01#4    mNeoVis.evergladensis.2.1.chr1.g00805.t01#2    mNeoVis.vulgivaga.2.1.chr1.g00821.t01#1   mNeoVis.evergladensis.1.1.chr1.g00817.t01#3       3228   125257704:   mNeoVis.3.1.1.chr7.g01368.t01#6    mNeoVis.vulgivaga.2.1.chr7.g01457.t01#1    mNeoVis.vulgivaga.1.1.chr7.g01451.t01#4    mNeoVis.2.1.1.chr7_hap1.g01461.t01#5    mNeoVis.evergladensis.2.1.chr7.g01443.t01#2   mNeoVis.evergladensis.1.1.chr7.g01441.t01#3       3229   125257705:   mNeoVis.vulgivaga.1.1.chr7.g01452.t01#4    mNeoVis.3.1.1.chr7.g01369.t01#6    mNeoVis.vulgivaga.2.1.chr7.g01458.t01#1    mNeoVis.2.1.1.chr7_hap1.g01462.t01#5    mNeoVis.evergladensis.1.1.chr7.g01443.t01#3   mNeoVis.evergladensis.2.1.chr7.g01444.t01#2       3230   125257706:   mNeoVis.evergladensis.2.1.chr7.g01445.t01#2    mNeoVis.evergladensis.1.1.chr7.g01444.t01#3    mNeoVis.vulgivaga.2.1.chr7.g01459.t01#1    mNeoVis.3.1.1.chr7.g01370.t01#6    mNeoVis.2.1.1.chr7_hap1.g01464.t01#5   mNeoVis.vulgivaga.1.1.chr7.g01453.t01#4       3231   125257707:   mNeoVis.3.1.1.chr1.g00826.t01#6    mNeoVis.2.1.1.chr1_hap1.g00800.t01#5    mNeoVis.vulgivaga.1.1.chr1.g00817.t01#4    mNeoVis.evergladensis.2.1.chr1.g00806.t01#2    mNeoVis.vulgivaga.2.1.chr1.g00822.t01#1   mNeoVis.evergladensis.1.1.chr1.g00818.t01#3       3232   125257708:   mNeoVis.3.1.1.chr7.g01371.t01#6    mNeoVis.2.1.1.chr7_hap1.g01465.t01#5    mNeoVis.vulgivaga.2.1.chr7.g01460.t01#1    mNeoVis.evergladensis.1.1.chr7.g01445.t01#3    mNeoVis.vulgivaga.1.1.chr7.g01454.t01#4   mNeoVis.evergladensis.2.1.chr7.g01446.t01#2       3233   125257709:   mNeoVis.vulgivaga.2.1.chr7.g01461.t01#1    mNeoVis.3.1.1.chr7.g01373.t01#6    mNeoVis.evergladensis.1.1.chr7.g01446.t01#3    mNeoVis.vulgivaga.1.1.chr7.g01456.t01#4    mNeoVis.2.1.1.chr7_hap1.g01467.t01#5   mNeoVis.evergladensis.2.1.chr7.g01448.t01#2       3234   125257710:   mNeoVis.vulgivaga.1.1.chr1.g00878.t01#4    mNeoVis.2.1.1.chr1_hap1.g00801.t01#5    mNeoVis.evergladensis.1.1.chr1.g00819.t01#3    mNeoVis.vulgivaga.2.1.chr1.g00823.t01#1    mNeoVis.2.1.1.chr1_hap1.g00859.t01#5    mNeoVis.vulgivaga.1.1.chr1.g00819.t01#4    mNeoVis.evergladensis.2.1.chr1.g00807.t01#2    mNeoVis.3.1.1.chr1.g00827.t01#6    mNeoVis.3.1.1.chr1.g00883.t01#6      3235   125257711:   mNeoVis.3.1.1.chr7.g01375.t01#6    mNeoVis.evergladensis.1.1.chr7.g01448.t01#3    mNeoVis.vulgivaga.1.1.chr7.g01458.t01#4    mNeoVis.2.1.1.chr7_hap1.g01469.t01#5    mNeoVis.evergladensis.2.1.chr7.g01450.t01#2   mNeoVis.vulgivaga.2.1.chr7.g01463.t01#1       3236   125257712:   mNeoVis.2.1.1.chr1_hap1.g00802.t01#5    mNeoVis.3.1.1.chr4.g00888.t01#6    mNeoVis.evergladensis.2.1.chr4.g00879.t01#2    mNeoVis.vulgivaga.1.1.chr1.g00820.t01#4    mNeoVis.evergladensis.1.1.chr4.g00897.t01#3   mNeoVis.3.1.1.chr1.g00828.t01#6       3237   125257713:   mNeoVis.evergladensis.1.1.chr1.g00820.t01#3   mNeoVis.evergladensis.2.1.chr1.g00808.t01#2           3238   125257714:  mNeoVis.vulgivaga.2.1.chr1.g00824.t01#1               3239   125257715:   mNeoVis.vulgivaga.1.1.chr1.g00821.t01#4    mNeoVis.vulgivaga.2.1.chr1.g00825.t01#1    mNeoVis.3.1.1.chr1.g00829.t01#6    mNeoVis.evergladensis.2.1.chr1.g00809.t01#2    mNeoVis.2.1.1.chr1_hap1.g00803.t01#5   mNeoVis.evergladensis.1.1.chr1.g00821.t01#3       3240   125257716:   mNeoVis.3.1.1.chr7.g01376.t01#6    mNeoVis.evergladensis.1.1.chr7.g01449.t01#3    mNeoVis.vulgivaga.1.1.chr7.g01459.t01#4    mNeoVis.2.1.1.chr7_hap1.g01470.t01#5    mNeoVis.evergladensis.2.1.chr7.g01451.t01#2   mNeoVis.vulgivaga.2.1.chr7.g01464.t01#1       3241   125257717:   mNeoVis.evergladensis.2.1.chr1.g00810.t01#2    mNeoVis.vulgivaga.2.1.chr1.g00826.t01#1    mNeoVis.evergladensis.1.1.chr1.g00822.t01#3    mNeoVis.2.1.1.chr1_hap1.g00804.t01#5    mNeoVis.vulgivaga.1.1.chr1.g00822.t01#4   mNeoVis.3.1.1.chr1.g00830.t01#6       3242   125257718:   mNeoVis.vulgivaga.1.1.chr1.g00823.t01#4    mNeoVis.vulgivaga.2.1.chr1.g00827.t01#1    mNeoVis.3.1.1.chr1.g00831.t01#6    mNeoVis.evergladensis.1.1.chr1.g00824.t01#3    mNeoVis.evergladensis.2.1.chr1.g00811.t01#2   mNeoVis.2.1.1.chr1_hap1.g00805.t01#5       3243   125257719:   mNeoVis.evergladensis.2.1.chr7.g01452.t01#2    mNeoVis.3.1.1.chr7.g01377.t01#6    mNeoVis.2.1.1.chr7_hap1.g01471.t01#5    mNeoVis.evergladensis.1.1.chr7.g01450.t01#3    mNeoVis.vulgivaga.1.1.chr7.g01460.t01#4   mNeoVis.vulgivaga.2.1.chr7.g01465.t01#1       3244   125257720:   mNeoVis.vulgivaga.2.1.chr7.g01466.t01#1    mNeoVis.3.1.1.chr7.g01379.t01#6    mNeoVis.vulgivaga.1.1.chr7.g01461.t01#4    mNeoVis.2.1.1.chr7_hap1.g01472.t01#5    mNeoVis.evergladensis.2.1.chr7.g01453.t01#2   mNeoVis.evergladensis.1.1.chr7.g01451.t01#3       3245   125257721:   mNeoVis.evergladensis.1.1.chr1.g00825.t01#3    mNeoVis.2.1.1.chr1_hap1.g00806.t01#5    mNeoVis.vulgivaga.2.1.chr1.g00828.t01#1    mNeoVis.3.1.1.chr1.g00832.t01#6    mNeoVis.evergladensis.2.1.chr1.g00812.t01#2   mNeoVis.vulgivaga.1.1.chr1.g00824.t01#4       3246   125257722:   mNeoVis.evergladensis.2.1.chr1.g00813.t01#2    mNeoVis.vulgivaga.1.1.chr1.g00825.t01#4    mNeoVis.2.1.1.chr1_hap1.g00807.t01#5    mNeoVis.vulgivaga.2.1.chr1.g00829.t01#1    mNeoVis.evergladensis.1.1.chr1.g00826.t01#3   mNeoVis.3.1.1.chr1.g00833.t01#6       3247   125257723:   mNeoVis.2.1.1.chr1_hap1.g00808.t01#5    mNeoVis.evergladensis.2.1.chr1.g00814.t01#2    mNeoVis.vulgivaga.2.1.chr1.g00830.t01#1    mNeoVis.evergladensis.1.1.chr1.g00827.t01#3    mNeoVis.3.1.1.chr1.g00834.t01#6   mNeoVis.vulgivaga.1.1.chr1.g00826.t01#4       3248   125257724:   mNeoVis.3.1.1.chr1.g00835.t01#6    mNeoVis.vulgivaga.2.1.chr1.g00831.t01#1    mNeoVis.evergladensis.1.1.chr1.g00828.t01#3    mNeoVis.evergladensis.2.1.chr1.g00815.t01#2    mNeoVis.vulgivaga.1.1.chr1.g00827.t01#4   mNeoVis.2.1.1.chr1_hap1.g00809.t01#5       3249   125257725:   mNeoVis.vulgivaga.1.1.chr7.g01462.t01#4    mNeoVis.vulgivaga.2.1.chr7.g01467.t01#1    mNeoVis.3.1.1.chr7.g01380.t01#6    mNeoVis.evergladensis.2.1.chr7.g01454.t01#2    mNeoVis.2.1.1.chr7_hap1.g01473.t01#5   mNeoVis.evergladensis.1.1.chr7.g01452.t01#3       3250   125257726:   mNeoVis.evergladensis.2.1.chr7.g01455.t01#2    mNeoVis.evergladensis.1.1.chr7.g01453.t01#3    mNeoVis.2.1.1.chr7_hap1.g01474.t01#5    mNeoVis.vulgivaga.2.1.chr7.g01468.t01#1   mNeoVis.vulgivaga.2.1.chr7.g01469.t01#1       3251   125257727:   mNeoVis.vulgivaga.1.1.chr7.g01464.t01#4   mNeoVis.3.1.1.chr7.g01381.t01#6             3252   125257728:   mNeoVis.evergladensis.2.1.chr7.g00097.t01#2    mNeoVis.2.1.1.chr7_hap1.g00097.t01#5    mNeoVis.evergladensis.1.1.chr7.g00099.t01#3    mNeoVis.vulgivaga.2.1.chr7.g00096.t01#1    mNeoVis.vulgivaga.1.1.chr7.g00100.t01#4   mNeoVis.3.1.1.chr7.g00081.t01#6       3253   125257729:   mNeoVis.vulgivaga.2.1.chr1.g00752.t01#1    mNeoVis.2.1.1.chr1_hap1.g00730.t01#5    mNeoVis.evergladensis.2.1.chr1.g00740.t01#2    mNeoVis.evergladensis.1.1.chr1.g00749.t01#3   mNeoVis.vulgivaga.1.1.chr1.g00745.t01#4       3254   125257730:   mNeoVis.evergladensis.2.1.chr7.g01456.t01#2    mNeoVis.3.1.1.chr7.g01382.t01#6    mNeoVis.vulgivaga.1.1.chr7.g01465.t01#4    mNeoVis.vulgivaga.2.1.chr7.g01470.t01#1    mNeoVis.2.1.1.chr7_hap1.g01475.t01#5   mNeoVis.evergladensis.1.1.chr7.g01454.t01#3       3255   125257731:   mNeoVis.evergladensis.2.1.chr1.g00742.t01#2    mNeoVis.3.1.1.chr1.g00754.t01#6    mNeoVis.vulgivaga.2.1.chr1.g00753.t01#1    mNeoVis.vulgivaga.1.1.chr1.g00747.t01#4    mNeoVis.2.1.1.chr1_hap1.g00732.t01#5   mNeoVis.evergladensis.1.1.chr1.g00751.t01#3       3256   125257732:   mNeoVis.vulgivaga.1.1.chr7.g01466.t01#4    mNeoVis.evergladensis.2.1.chr7.g01457.t01#2    mNeoVis.evergladensis.1.1.chr7.g01455.t01#3    mNeoVis.2.1.1.chr7_hap1.g01476.t01#5    mNeoVis.vulgivaga.2.1.chr7.g01471.t01#1   mNeoVis.3.1.1.chr7.g01383.t01#6       3257   125257733:   mNeoVis.vulgivaga.2.1.chr7.g01472.t01#1    mNeoVis.3.1.1.chr7.g01384.t01#6    mNeoVis.2.1.1.chr7_hap1.g01477.t01#5    mNeoVis.evergladensis.2.1.chr7.g01458.t01#2    mNeoVis.vulgivaga.1.1.chr7.g01467.t01#4   mNeoVis.evergladensis.1.1.chr7.g01456.t01#3       3258   125257734:   mNeoVis.evergladensis.2.1.chr7.g01459.t01#2    mNeoVis.evergladensis.1.1.chr7.g01457.t01#3    mNeoVis.vulgivaga.1.1.chr7.g01468.t01#4    mNeoVis.vulgivaga.2.1.chr7.g01473.t01#1    mNeoVis.2.1.1.chr7_hap1.g01478.t01#5   mNeoVis.3.1.1.chr7.g01385.t01#6       3259   125257735:   mNeoVis.3.1.1.chr7.g01386.t01#6    mNeoVis.vulgivaga.2.1.chr7.g01474.t01#1    mNeoVis.evergladensis.1.1.chr7.g01458.t01#3   mNeoVis.vulgivaga.1.1.chr7.g01469.t01#4         3260   125257736:   mNeoVis.2.1.1.chr5_hap1.g01255.t01#5    mNeoVis.vulgivaga.2.1.chr2.g01248.t01#1    mNeoVis.evergladensis.2.1.chr2.g01243.t01#2    mNeoVis.evergladensis.1.1.chr2.g01245.t01#3    mNeoVis.3.1.1.chr2.g01173.t01#6    mNeoVis.vulgivaga.1.1.chr2.g01249.t01#4   mNeoVis.3.1.1.chr2.g01174.t01#6       3261   125257737:   mNeoVis.2.1.1.chr12_hap1.g00495.t01#5    mNeoVis.evergladensis.1.1.chr12.g00450.t01#3    mNeoVis.vulgivaga.2.1.chr12.g00456.t01#1    mNeoVis.vulgivaga.1.1.chr12.g00453.t01#4    mNeoVis.3.1.1.chr12.g00450.t01#6   mNeoVis.evergladensis.2.1.chr12.g00443.t01#2       3262   125257738:   mNeoVis.3.1.1.chr7.g01387.t01#6    mNeoVis.vulgivaga.1.1.chr7.g01470.t01#4    mNeoVis.2.1.1.chr7_hap1.g01479.t01#5   mNeoVis.evergladensis.2.1.chr7.g01460.t01#2       3263   125257739:   mNeoVis.evergladensis.1.1.chr7.g01459.t01#3    mNeoVis.vulgivaga.2.1.chr7.g01475.t01#1   mNeoVis.vulgivaga.1.1.chr7.g01471.t01#4           3264   125257740:   mNeoVis.evergladensis.2.1.chr15.g00091.t01#2    mNeoVis.3.1.1.chr15.g00084.t01#6    mNeoVis.vulgivaga.2.1.chr15.g00092.t01#1    mNeoVis.vulgivaga.1.1.chr15.g00091.t01#4    mNeoVis.evergladensis.1.1.chr15.g00522.t01#3   mNeoVis.2.1.1.chr15_hap1.g00090.t01#5       3265   125257741:   mNeoVis.vulgivaga.2.1.chr7.g01476.t01#1   mNeoVis.evergladensis.2.1.chr7.g01461.t01#2           3266   125257742:   mNeoVis.2.1.1.chr7_hap1.g01481.t01#5    mNeoVis.evergladensis.1.1.chr7.g01461.t01#3    mNeoVis.3.1.1.chr7.g01389.t01#6    mNeoVis.evergladensis.2.1.chr7.g01462.t01#2    mNeoVis.vulgivaga.1.1.chr7.g01473.t01#4   mNeoVis.vulgivaga.2.1.chr7.g01477.t01#1       3267   125257743:   mNeoVis.2.1.1.chr7_hap1.g01482.t01#5    mNeoVis.evergladensis.2.1.chr7.g01463.t01#2    mNeoVis.vulgivaga.2.1.chr7.g01478.t01#1    mNeoVis.3.1.1.chr7.g01390.t01#6    mNeoVis.vulgivaga.1.1.chr7.g01474.t01#4   mNeoVis.evergladensis.1.1.chr7.g01462.t01#3       3268   125257744:   mNeoVis.vulgivaga.2.1.chr7.g01479.t01#1    mNeoVis.evergladensis.2.1.chr7.g01464.t01#2    mNeoVis.vulgivaga.1.1.chr7.g01475.t01#4    mNeoVis.2.1.1.chr7_hap1.g01483.t01#5   mNeoVis.evergladensis.1.1.chr7.g01463.t01#3       3269   125257745:   mNeoVis.3.1.1.chr7.g01391.t01#6    mNeoVis.vulgivaga.2.1.chr7.g01480.t01#1    mNeoVis.vulgivaga.1.1.chr7.g01476.t01#4   mNeoVis.evergladensis.1.1.chr7.g01464.t01#3       3270   125257746:   mNeoVis.vulgivaga.2.1.chr7.g01481.t01#1    mNeoVis.2.1.1.chr7_hap1.g01484.t01#5    mNeoVis.evergladensis.2.1.chr7.g01465.t01#2   mNeoVis.vulgivaga.1.1.chr7.g01477.t01#4         3271   125257747:  mNeoVis.vulgivaga.2.1.chr7.g01482.t01#1               3272   125257748:  mNeoVis.vulgivaga.2.1.chr1.g00761.t01#1               3273   125257749:   mNeoVis.3.1.1.chr1.g00763.t01#6    mNeoVis.vulgivaga.2.1.chr1.g00762.t01#1   mNeoVis.evergladensis.1.1.chr1.g00758.t01#3         3274   125257750:   mNeoVis.evergladensis.1.1.chr1.g00759.t01#3    mNeoVis.evergladensis.2.1.chr1.g00750.t01#2    mNeoVis.vulgivaga.1.1.chr1.g00755.t01#4    mNeoVis.vulgivaga.2.1.chr1.g00763.t01#1    mNeoVis.2.1.1.chr1_hap1.g00741.t01#5   mNeoVis.3.1.1.chr1.g00764.t01#6       3275   125257751:   mNeoVis.vulgivaga.1.1.chr1.g00757.t01#4    mNeoVis.3.1.1.chr1.g00765.t01#6    mNeoVis.vulgivaga.2.1.chr1.g00764.t01#1    mNeoVis.evergladensis.1.1.chr1.g00761.t01#3   mNeoVis.2.1.1.chr1_hap1.g00742.t01#5       3276   125257752:   mNeoVis.vulgivaga.1.1.chr1.g00758.t01#4    mNeoVis.3.1.1.chr1.g00766.t01#6    mNeoVis.evergladensis.2.1.chr1.g00751.t01#2    mNeoVis.evergladensis.1.1.chr1.g00762.t01#3    mNeoVis.vulgivaga.2.1.chr1.g00765.t01#1   mNeoVis.2.1.1.chr1_hap1.g00743.t01#5       3277   125257753:   mNeoVis.evergladensis.1.1.chr9.g01137.t01#3    mNeoVis.evergladensis.2.1.chr8.g01143.t01#2    mNeoVis.3.1.1.chr9.g01097.t01#6    mNeoVis.vulgivaga.1.1.chr9.g01140.t01#4    mNeoVis.vulgivaga.2.1.chr9.g01123.t01#1   mNeoVis.2.1.1.chr10_hap1.g01165.t01#5       3278   125257754:  mNeoVis.3.1.1.chr1.g00767.t01#6               3279   125257755:   mNeoVis.3.1.1.chr1.g00768.t01#6    mNeoVis.evergladensis.2.1.chr1.g00752.t01#2    mNeoVis.evergladensis.1.1.chr1.g00763.t01#3    mNeoVis.vulgivaga.2.1.chr1.g00766.t01#1    mNeoVis.2.1.1.chr1_hap1.g00744.t01#5   mNeoVis.vulgivaga.1.1.chr1.g00759.t01#4       3280   125257756:   mNeoVis.evergladensis.2.1.chr7.g01466.t01#2    mNeoVis.vulgivaga.1.1.chr7.g01479.t01#4    mNeoVis.evergladensis.1.1.chr7.g01466.t01#3    mNeoVis.vulgivaga.2.1.chr7.g01483.t01#1    mNeoVis.2.1.1.chr7_hap1.g01485.t01#5   mNeoVis.3.1.1.chr7.g01393.t01#6       3281   125257757:   mNeoVis.evergladensis.1.1.chr7.g01467.t01#3    mNeoVis.vulgivaga.1.1.chr7.g01480.t01#4    mNeoVis.vulgivaga.2.1.chr7.g01484.t01#1    mNeoVis.3.1.1.chr7.g01394.t01#6    mNeoVis.2.1.1.chr7_hap1.g01486.t01#5   mNeoVis.evergladensis.2.1.chr7.g01467.t01#2       3282   125257758:   mNeoVis.vulgivaga.2.1.chr1.g00767.t01#1    mNeoVis.vulgivaga.1.1.chr1.g00760.t01#4    mNeoVis.evergladensis.2.1.chr1.g00753.t01#2    mNeoVis.3.1.1.chr1.g00769.t01#6    mNeoVis.evergladensis.1.1.chr1.g00764.t01#3   mNeoVis.2.1.1.chr1_hap1.g00745.t01#5       3283   125257759:   mNeoVis.2.1.1.chr1_hap1.g00746.t01#5    mNeoVis.evergladensis.2.1.chr1.g00754.t01#2    mNeoVis.vulgivaga.2.1.chr1.g00768.t01#1    mNeoVis.evergladensis.1.1.chr1.g00765.t01#3    mNeoVis.vulgivaga.1.1.chr1.g00761.t01#4   mNeoVis.3.1.1.chr1.g00770.t01#6       3284   125257760:   mNeoVis.evergladensis.2.1.chr7.g01468.t01#2    mNeoVis.vulgivaga.1.1.chr7.g01481.t01#4    mNeoVis.vulgivaga.2.1.chr7.g01485.t01#1    mNeoVis.evergladensis.1.1.chr7.g01468.t01#3    mNeoVis.3.1.1.chr7.g01395.t01#6   mNeoVis.2.1.1.chr7_hap1.g01487.t01#5       3285   125257761:   mNeoVis.vulgivaga.2.1.chr7.g01486.t01#1    mNeoVis.evergladensis.2.1.chr7.g01470.t01#2    mNeoVis.evergladensis.1.1.chr7.g01470.t01#3   mNeoVis.2.1.1.chr7_hap1.g01488.t01#5         3286   125257762:   mNeoVis.3.1.1.chr7.g01397.t01#6    mNeoVis.evergladensis.1.1.chr7.g01471.t01#3    mNeoVis.vulgivaga.1.1.chr7.g01483.t01#4   mNeoVis.vulgivaga.1.1.chr7.g01482.t01#4         3287   125257763:   mNeoVis.vulgivaga.1.1.chr1.g00763.t01#4    mNeoVis.3.1.1.chr1.g00772.t01#6    mNeoVis.vulgivaga.2.1.chr1.g00769.t01#1    mNeoVis.2.1.1.chr1_hap1.g00747.t01#5    mNeoVis.evergladensis.1.1.chr1.g00766.t01#3   mNeoVis.evergladensis.2.1.chr1.g00755.t01#2       3288   125257764:   mNeoVis.vulgivaga.1.1.chr1.g00764.t01#4    mNeoVis.evergladensis.2.1.chr1.g00756.t01#2    mNeoVis.2.1.1.chr1_hap1.g00748.t01#5    mNeoVis.3.1.1.chr1.g00773.t01#6    mNeoVis.vulgivaga.2.1.chr1.g00770.t01#1   mNeoVis.evergladensis.1.1.chr1.g00767.t01#3       3289   125257765:   mNeoVis.vulgivaga.2.1.chr7.g01487.t01#1    mNeoVis.2.1.1.chr7_hap1.g01489.t01#5    mNeoVis.evergladensis.2.1.chr7.g01471.t01#2    mNeoVis.evergladensis.1.1.chr7.g01472.t01#3    mNeoVis.3.1.1.chr7.g01398.t01#6   mNeoVis.vulgivaga.1.1.chr7.g01484.t01#4       3290   125257766:   mNeoVis.2.1.1.chr7_hap1.g01400.t01#5    mNeoVis.evergladensis.2.1.chr7.g01384.t01#2    mNeoVis.vulgivaga.2.1.chr7.g01394.t01#1    mNeoVis.3.1.1.chr7.g01311.t01#6   mNeoVis.evergladensis.1.1.chr7.g01381.t01#3       3291   125257767:   mNeoVis.vulgivaga.1.1.chr7.g01391.t01#4    mNeoVis.evergladensis.2.1.chr7.g01385.t01#2    mNeoVis.vulgivaga.2.1.chr7.g01395.t01#1    mNeoVis.3.1.1.chr7.g01312.t01#6    mNeoVis.2.1.1.chr7_hap1.g01402.t01#5   mNeoVis.evergladensis.1.1.chr7.g01382.t01#3       3292   125257768:   mNeoVis.evergladensis.2.1.chr7.g01386.t01#2    mNeoVis.3.1.1.chr7.g01313.t01#6    mNeoVis.vulgivaga.1.1.chr7.g01392.t01#4    mNeoVis.vulgivaga.2.1.chr7.g01396.t01#1    mNeoVis.2.1.1.chr7_hap1.g01403.t01#5   mNeoVis.evergladensis.1.1.chr7.g01383.t01#3       3293   125257769:   mNeoVis.vulgivaga.1.1.chr1.g00765.t01#4    mNeoVis.evergladensis.2.1.chr1.g00757.t01#2    mNeoVis.vulgivaga.2.1.chr1.g00771.t01#1    mNeoVis.2.1.1.chr1_hap1.g00749.t01#5    mNeoVis.evergladensis.1.1.chr1.g00768.t01#3   mNeoVis.3.1.1.chr1.g00774.t01#6       3294   125257770:   mNeoVis.3.1.1.chr1.g00775.t01#6    mNeoVis.evergladensis.1.1.chr1.g00769.t01#3    mNeoVis.evergladensis.2.1.chr1.g00758.t01#2    mNeoVis.vulgivaga.1.1.chr1.g00766.t01#4    mNeoVis.2.1.1.chr1_hap1.g00750.t01#5   mNeoVis.vulgivaga.2.1.chr1.g00772.t01#1       3295   125257771:   mNeoVis.evergladensis.1.1.chr7.g01384.t01#3    mNeoVis.3.1.1.chr7.g01314.t01#6    mNeoVis.vulgivaga.1.1.chr7.g01393.t01#4    mNeoVis.2.1.1.chr7_hap1.g01404.t01#5    mNeoVis.vulgivaga.2.1.chr7.g01397.t01#1   mNeoVis.evergladensis.2.1.chr7.g01387.t01#2       3296   125257772:   mNeoVis.evergladensis.1.1.chr7.g01385.t01#3    mNeoVis.vulgivaga.1.1.chr7.g01394.t01#4    mNeoVis.2.1.1.chr7_hap1.g01405.t01#5    mNeoVis.3.1.1.chr7.g01315.t01#6    mNeoVis.vulgivaga.2.1.chr7.g01398.t01#1   mNeoVis.evergladensis.2.1.chr7.g01388.t01#2       3297   125257773:   mNeoVis.3.1.1.chr7.g01316.t01#6    mNeoVis.evergladensis.2.1.chr7.g01389.t01#2    mNeoVis.evergladensis.1.1.chr7.g01386.t01#3    mNeoVis.2.1.1.chr7_hap1.g01406.t01#5    mNeoVis.vulgivaga.1.1.chr7.g01395.t01#4   mNeoVis.vulgivaga.2.1.chr7.g01399.t01#1       3298   125257774:   mNeoVis.evergladensis.1.1.chr1.g00770.t01#3    mNeoVis.2.1.1.chr1_hap1.g00751.t01#5    mNeoVis.evergladensis.2.1.chr1.g00759.t01#2    mNeoVis.vulgivaga.1.1.chr1.g00767.t01#4    mNeoVis.3.1.1.chr1.g00776.t01#6   mNeoVis.vulgivaga.2.1.chr1.g00773.t01#1       3299   125257775:   mNeoVis.3.1.1.chr7.g01317.t01#6    mNeoVis.vulgivaga.1.1.chr7.g01397.t01#4    mNeoVis.vulgivaga.2.1.chr7.g01400.t01#1    mNeoVis.evergladensis.2.1.chr7.g01390.t01#2    mNeoVis.2.1.1.chr7_hap1.g01407.t01#5    mNeoVis.evergladensis.1.1.chr7.g01387.t01#3   mNeoVis.vulgivaga.1.1.chr7.g01396.t01#4     3300   125257776:   mNeoVis.vulgivaga.2.1.chr1.g00774.t01#1    mNeoVis.evergladensis.1.1.chr1.g00771.t01#3    mNeoVis.2.1.1.chr1_hap1.g00752.t01#5    mNeoVis.vulgivaga.1.1.chr1.g00768.t01#4   mNeoVis.evergladensis.2.1.chr1.g00760.t01#2       3301   125257777:   mNeoVis.3.1.1.chr1.g00778.t01#6    mNeoVis.evergladensis.2.1.chr1.g00761.t01#2    mNeoVis.2.1.1.chr1_hap1.g00753.t01#5    mNeoVis.vulgivaga.2.1.chr1.g00775.t01#1    mNeoVis.vulgivaga.1.1.chr1.g00769.t01#4   mNeoVis.evergladensis.1.1.chr1.g00772.t01#3       3302   125257778:   mNeoVis.3.1.1.chr7.g01318.t01#6    mNeoVis.vulgivaga.1.1.chr7.g01398.t01#4    mNeoVis.2.1.1.chr7_hap1.g01408.t01#5    mNeoVis.evergladensis.1.1.chr7.g01388.t01#3    mNeoVis.vulgivaga.2.1.chr7.g01401.t01#1   mNeoVis.evergladensis.2.1.chr7.g01391.t01#2       3303   125257779:   mNeoVis.evergladensis.1.1.chr1.g00773.t01#3    mNeoVis.evergladensis.2.1.chr1.g00762.t01#2    mNeoVis.2.1.1.chr1_hap1.g00754.t01#5    mNeoVis.vulgivaga.2.1.chr1.g00776.t01#1    mNeoVis.vulgivaga.1.1.chr1.g00770.t01#4   mNeoVis.3.1.1.chr1.g00779.t01#6       3304   125257780:   mNeoVis.2.1.1.chr7_hap1.g01409.t01#5    mNeoVis.vulgivaga.2.1.chr7.g01402.t01#1    mNeoVis.evergladensis.1.1.chr7.g01389.t01#3    mNeoVis.evergladensis.2.1.chr7.g01392.t01#2    mNeoVis.3.1.1.chr7.g01319.t01#6   mNeoVis.vulgivaga.1.1.chr7.g01399.t01#4       3305   125257781:   mNeoVis.2.1.1.chr1_hap1.g00755.t01#5    mNeoVis.evergladensis.2.1.chr1.g00763.t01#2    mNeoVis.vulgivaga.1.1.chr1.g00771.t01#4    mNeoVis.evergladensis.1.1.chr1.g00774.t01#3    mNeoVis.3.1.1.chr1.g00780.t01#6   mNeoVis.vulgivaga.2.1.chr1.g00777.t01#1       3306   125257782:   mNeoVis.evergladensis.2.1.chr1.g00764.t01#2    mNeoVis.vulgivaga.1.1.chr1.g00772.t01#4    mNeoVis.vulgivaga.2.1.chr1.g00778.t01#1    mNeoVis.3.1.1.chr1.g00781.t01#6    mNeoVis.2.1.1.chr1_hap1.g00756.t01#5   mNeoVis.evergladensis.1.1.chr1.g00775.t01#3       3307   125257783:   mNeoVis.2.1.1.chr1_hap1.g00757.t01#5    mNeoVis.vulgivaga.2.1.chr1.g00779.t01#1    mNeoVis.evergladensis.1.1.chr1.g00776.t01#3    mNeoVis.vulgivaga.1.1.chr1.g00773.t01#4    mNeoVis.evergladensis.2.1.chr1.g00765.t01#2   mNeoVis.3.1.1.chr1.g00782.t01#6       3308   125257784:   mNeoVis.vulgivaga.1.1.chr7.g01401.t01#4    mNeoVis.2.1.1.chr7_hap1.g01410.t01#5    mNeoVis.evergladensis.1.1.chr7.g01390.t01#3    mNeoVis.evergladensis.2.1.chr7.g01393.t01#2    mNeoVis.3.1.1.chr7.g01320.t01#6   mNeoVis.vulgivaga.2.1.chr7.g01404.t01#1       3309   125257785:   mNeoVis.vulgivaga.2.1.chr1.g00780.t01#1    mNeoVis.vulgivaga.1.1.chr1.g00774.t01#4    mNeoVis.evergladensis.1.1.chr1.g00777.t01#3    mNeoVis.evergladensis.2.1.chr1.g00766.t01#2    mNeoVis.2.1.1.chr1_hap1.g00758.t01#5   mNeoVis.3.1.1.chr1.g00784.t01#6       3310   125257786:   mNeoVis.3.1.1.chr1.g00785.t01#6    mNeoVis.vulgivaga.2.1.chr1.g00781.t01#1    mNeoVis.vulgivaga.1.1.chr1.g00775.t01#4    mNeoVis.2.1.1.chr1_hap1.g00759.t01#5    mNeoVis.evergladensis.2.1.chr1.g00767.t01#2   mNeoVis.evergladensis.1.1.chr1.g00778.t01#3       3311   125257787:   mNeoVis.vulgivaga.1.1.chr7.g01402.t01#4    mNeoVis.3.1.1.chr7.g01321.t01#6    mNeoVis.evergladensis.2.1.chr7.g01394.t01#2    mNeoVis.2.1.1.chr7_hap1.g01411.t01#5    mNeoVis.vulgivaga.2.1.chr7.g01405.t01#1   mNeoVis.evergladensis.1.1.chr7.g01391.t01#3       3312   125257788:   mNeoVis.2.1.1.chr7_hap1.g01412.t01#5    mNeoVis.vulgivaga.2.1.chr7.g01406.t01#1    mNeoVis.evergladensis.2.1.chr7.g01395.t01#2    mNeoVis.evergladensis.1.1.chr7.g01392.t01#3    mNeoVis.vulgivaga.1.1.chr7.g01403.t01#4   mNeoVis.3.1.1.chr7.g01322.t01#6       3313   125257789:   mNeoVis.vulgivaga.2.1.chr1.g00782.t01#1    mNeoVis.vulgivaga.1.1.chr1.g00776.t01#4    mNeoVis.evergladensis.2.1.chr1.g00768.t01#2   mNeoVis.2.1.1.chr1_hap1.g00760.t01#5         3314   125257790:   mNeoVis.vulgivaga.1.1.chr4.g00050.t01#4    mNeoVis.evergladensis.1.1.chr13.g00598.t01#3    mNeoVis.vulgivaga.2.1.chr13.g00602.t01#1    mNeoVis.2.1.1.chr6_hap1.g00748.t01#5    mNeoVis.2.1.1.chr13_hap1.g00630.t01#5    mNeoVis.2.1.1.chr12_hap1.g00516.t01#5    mNeoVis.2.1.1.chr10_hap1.g00987.t01#5    mNeoVis.2.1.1.chr5_hap1.g01180.t01#5    mNeoVis.2.1.1.chr6_hap1.g01074.t01#5      3315   125257791:   mNeoVis.2.1.1.chr12_hap1.g00806.t01#5    mNeoVis.vulgivaga.1.1.chr3.g00846.t01#4    mNeoVis.2.1.1.chr6_hap1.g00626.t01#5    mNeoVis.3.1.1.chr8.g01193.t01#6    mNeoVis.evergladensis.2.1.chr3.g00847.t01#2    mNeoVis.vulgivaga.2.1.chr6.g01038.t01#1    mNeoVis.3.1.1.chr3.g00833.t01#6    mNeoVis.evergladensis.1.1.chr8.g01241.t01#3    mNeoVis.2.1.1.chr12_hap1.g00337.t01#5      3316   125257792:   mNeoVis.2.1.1.chr9_hap1.g00465.t01#5    mNeoVis.evergladensis.1.1.chr10.g00464.t01#3    mNeoVis.evergladensis.2.1.chr2.g01702.t01#2    mNeoVis.vulgivaga.1.1.chr10.g00439.t01#4    mNeoVis.evergladensis.2.1.chr2.g01683.t01#2    mNeoVis.3.1.1.chr10.g00441.t01#6    mNeoVis.vulgivaga.1.1.chr2.g01693.t01#4    mNeoVis.vulgivaga.2.1.chr2.g01699.t01#1    mNeoVis.vulgivaga.2.1.chr10.g00448.t01#1      3317   125257793:   mNeoVis.vulgivaga.2.1.chr6.g00390.t01#1    mNeoVis.3.1.1.chr6.g00391.t01#6    mNeoVis.vulgivaga.1.1.chr6.g00396.t01#4    mNeoVis.evergladensis.2.1.chr6.g00394.t01#2    mNeoVis.evergladensis.1.1.chr6.g00390.t01#3   mNeoVis.2.1.1.chr4_hap1.g00391.t01#5       3318   125257794:   mNeoVis.evergladensis.2.1.chr1.g01466.t01#2    mNeoVis.2.1.1.chr1_hap1.g01502.t01#5    mNeoVis.evergladensis.1.1.chr7.g01393.t01#3   mNeoVis.vulgivaga.2.1.chr7.g01407.t01#1         3319   125257795:   mNeoVis.vulgivaga.2.1.chr5.g01238.t01#1    mNeoVis.2.1.1.chr2_hap1.g01301.t01#5   mNeoVis.3.1.1.chr5.g01226.t01#6           3320   125257796:   mNeoVis.evergladensis.2.1.chr6.g01046.t01#2   mNeoVis.2.1.1.chr4_hap1.g01040.t01#5             3321   125257797:  mNeoVis.evergladensis.1.1.chr1.g00584.t01#3             3322   125257798:  mNeoVis.vulgivaga.2.1.chr11.g00736.t01#1               3323   125257799:   mNeoVis.evergladensis.2.1.chr7.g01396.t01#2    mNeoVis.2.1.1.chr7_hap1.g01413.t01#5    mNeoVis.evergladensis.1.1.chr7.g01394.t01#3    mNeoVis.vulgivaga.2.1.chr7.g01408.t01#1    mNeoVis.vulgivaga.1.1.chr7.g01404.t01#4   mNeoVis.3.1.1.chr7.g01323.t01#6       3324   125257800:   mNeoVis.3.1.1.chr1.g00786.t01#6    mNeoVis.2.1.1.chr1_hap1.g00761.t01#5    mNeoVis.evergladensis.2.1.chr1.g00769.t01#2    mNeoVis.evergladensis.1.1.chr1.g00779.t01#3    mNeoVis.vulgivaga.1.1.chr1.g00777.t01#4   mNeoVis.vulgivaga.2.1.chr1.g00783.t01#1       3325   125257801:   mNeoVis.2.1.1.chr7_hap1.g01414.t01#5    mNeoVis.vulgivaga.2.1.chr7.g01409.t01#1    mNeoVis.evergladensis.1.1.chr7.g01395.t01#3    mNeoVis.3.1.1.chr7.g01324.t01#6    mNeoVis.vulgivaga.1.1.chr7.g01405.t01#4   mNeoVis.evergladensis.2.1.chr7.g01397.t01#2       3326   125257802:   mNeoVis.vulgivaga.1.1.chr1.g00778.t01#4    mNeoVis.3.1.1.chr1.g00787.t01#6   mNeoVis.vulgivaga.2.1.chr1.g00784.t01#1           3327   125257803:   mNeoVis.evergladensis.2.1.chr7.g01398.t01#2    mNeoVis.vulgivaga.1.1.chr7.g01406.t01#4    mNeoVis.evergladensis.1.1.chr7.g01396.t01#3    mNeoVis.3.1.1.chr7.g01325.t01#6    mNeoVis.vulgivaga.2.1.chr7.g01410.t01#1   mNeoVis.2.1.1.chr7_hap1.g01415.t01#5       3328   125257804:  mNeoVis.vulgivaga.2.1.chr15.g00093.t01#1               3329   125257805:   mNeoVis.evergladensis.1.1.chr1.g00780.t01#3    mNeoVis.vulgivaga.1.1.chr1.g00779.t01#4    mNeoVis.vulgivaga.2.1.chr1.g00785.t01#1    mNeoVis.2.1.1.chr1_hap1.g00762.t01#5    mNeoVis.3.1.1.chr1.g00788.t01#6   mNeoVis.evergladensis.2.1.chr1.g00770.t01#2       3330   125257806:   mNeoVis.2.1.1.chr1_hap1.g00763.t01#5    mNeoVis.evergladensis.2.1.chr1.g00771.t01#2    mNeoVis.3.1.1.chr1.g00789.t01#6    mNeoVis.evergladensis.1.1.chr1.g00781.t01#3    mNeoVis.vulgivaga.1.1.chr1.g00780.t01#4   mNeoVis.vulgivaga.2.1.chr1.g00786.t01#1       3331   125257807:   mNeoVis.evergladensis.2.1.chr7.g01399.t01#2    mNeoVis.vulgivaga.1.1.chr7.g01407.t01#4    mNeoVis.vulgivaga.2.1.chr7.g01411.t01#1    mNeoVis.2.1.1.chr7_hap1.g01416.t01#5   mNeoVis.3.1.1.chr7.g01326.t01#6       3332   125257808:   mNeoVis.2.1.1.chr7_hap1.g01417.t01#5    mNeoVis.vulgivaga.1.1.chr7.g01408.t01#4   mNeoVis.vulgivaga.2.1.chr7.g01412.t01#1           3333   125257809:   mNeoVis.vulgivaga.1.1.chr7.g01409.t01#4    mNeoVis.vulgivaga.2.1.chr7.g01413.t01#1    mNeoVis.3.1.1.chr7.g01327.t01#6    mNeoVis.2.1.1.chr7_hap1.g01418.t01#5    mNeoVis.evergladensis.1.1.chr7.g01398.t01#3   mNeoVis.evergladensis.2.1.chr7.g01400.t01#2       3334   125257810:   mNeoVis.3.1.1.chr1.g00790.t01#6    mNeoVis.2.1.1.chr1_hap1.g00764.t01#5    mNeoVis.vulgivaga.1.1.chr1.g00781.t01#4   mNeoVis.vulgivaga.2.1.chr1.g00787.t01#1         3335   125257811:   mNeoVis.evergladensis.2.1.chr1.g00772.t01#2    mNeoVis.evergladensis.1.1.chr1.g00782.t01#3   mNeoVis.vulgivaga.2.1.chr1.g00788.t01#1           3336   125257812:   mNeoVis.3.1.1.chr7.g01328.t01#6    mNeoVis.2.1.1.chr7_hap1.g01419.t01#5    mNeoVis.evergladensis.1.1.chr7.g01399.t01#3    mNeoVis.vulgivaga.2.1.chr7.g01414.t01#1    mNeoVis.vulgivaga.1.1.chr7.g01410.t01#4   mNeoVis.evergladensis.2.1.chr7.g01401.t01#2       3337   125257813:   mNeoVis.vulgivaga.1.1.chr7.g01412.t01#4    mNeoVis.3.1.1.chr7.g01329.t01#6    mNeoVis.vulgivaga.2.1.chr7.g01415.t01#1    mNeoVis.evergladensis.1.1.chr7.g01400.t01#3    mNeoVis.2.1.1.chr7_hap1.g01420.t01#5   mNeoVis.evergladensis.2.1.chr7.g01402.t01#2       3338   125257814:   mNeoVis.evergladensis.2.1.chr12.g00080.t01#2    mNeoVis.3.1.1.chr12.g00079.t01#6    mNeoVis.vulgivaga.1.1.chr12.g00089.t01#4    mNeoVis.vulgivaga.2.1.chr12.g00084.t01#1    mNeoVis.2.1.1.chr12_hap1.g00094.t01#5   mNeoVis.evergladensis.1.1.chr12.g00081.t01#3       3339   125257815:   mNeoVis.vulgivaga.2.1.chr1.g00705.t01#1    mNeoVis.evergladensis.1.1.chr1.g00697.t01#3    mNeoVis.2.1.1.chr1_hap1.g00685.t01#5    mNeoVis.evergladensis.2.1.chr1.g00693.t01#2    mNeoVis.vulgivaga.1.1.chr1.g00697.t01#4   mNeoVis.3.1.1.chr1.g00705.t01#6       3340   125257816:   mNeoVis.vulgivaga.2.1.chr15.g00094.t01#1    mNeoVis.evergladensis.1.1.chr15.g00521.t01#3    mNeoVis.3.1.1.chr15.g00085.t01#6    mNeoVis.evergladensis.2.1.chr15.g00092.t01#2    mNeoVis.vulgivaga.1.1.chr15.g00092.t01#4   mNeoVis.2.1.1.chr15_hap1.g00091.t01#5       3341   125257817:   mNeoVis.vulgivaga.1.1.chr1.g00698.t01#4    mNeoVis.3.1.1.chr1.g00706.t01#6    mNeoVis.vulgivaga.2.1.chr1.g00706.t01#1    mNeoVis.2.1.1.chr1_hap1.g00686.t01#5    mNeoVis.evergladensis.1.1.chr1.g00698.t01#3   mNeoVis.evergladensis.2.1.chr1.g00694.t01#2       3342   125257818:   mNeoVis.vulgivaga.1.1.chr1.g00700.t01#4    mNeoVis.3.1.1.chr1.g00708.t01#6    mNeoVis.2.1.1.chr1_hap1.g00688.t01#5    mNeoVis.evergladensis.1.1.chr1.g00700.t01#3    mNeoVis.vulgivaga.2.1.chr1.g00708.t01#1   mNeoVis.evergladensis.2.1.chr1.g00696.t01#2       3343   125257819:   mNeoVis.vulgivaga.1.1.chr7.g01411.t01#4    mNeoVis.evergladensis.1.1.chr7.g01401.t01#3    mNeoVis.2.1.1.chr7_hap1.g01421.t01#5    mNeoVis.evergladensis.2.1.chr7.g01403.t01#2   mNeoVis.vulgivaga.2.1.chr7.g01416.t01#1       3344   125257820:   mNeoVis.evergladensis.1.1.chr7.g01402.t01#3    mNeoVis.2.1.1.chr7_hap1.g01422.t01#5    mNeoVis.evergladensis.2.1.chr7.g01404.t01#2    mNeoVis.vulgivaga.2.1.chr7.g01417.t01#1   mNeoVis.vulgivaga.1.1.chr7.g01413.t01#4       3345   125257821:  mNeoVis.vulgivaga.2.1.chr7.g01418.t01#1               3346   125257822:   mNeoVis.vulgivaga.2.1.chr7.g01419.t01#1    mNeoVis.3.1.1.chr7.g01333.t01#6    mNeoVis.vulgivaga.1.1.chr7.g01415.t01#4    mNeoVis.evergladensis.2.1.chr7.g01406.t01#2    mNeoVis.2.1.1.chr7_hap1.g01424.t01#5   mNeoVis.evergladensis.1.1.chr7.g01404.t01#3       3347   125257823:   mNeoVis.vulgivaga.1.1.chr6.g00763.t01#4    mNeoVis.evergladensis.1.1.chr6.g00770.t01#3    mNeoVis.evergladensis.2.1.chr6.g00765.t01#2    mNeoVis.3.1.1.chr6.g00764.t01#6    mNeoVis.2.1.1.chr4_hap1.g00758.t01#5   mNeoVis.vulgivaga.2.1.chr6.g00762.t01#1       3348   125257824:   mNeoVis.2.1.1.chr1_hap1.g00689.t01#5    mNeoVis.vulgivaga.2.1.chr1.g00709.t01#1    mNeoVis.evergladensis.1.1.chr1.g00701.t01#3    mNeoVis.vulgivaga.1.1.chr1.g00701.t01#4    mNeoVis.evergladensis.2.1.chr1.g00697.t01#2   mNeoVis.3.1.1.chr1.g00709.t01#6       3349   125257825:   mNeoVis.2.1.1.chr1_hap1.g00690.t01#5    mNeoVis.vulgivaga.2.1.chr1.g00710.t01#1    mNeoVis.evergladensis.1.1.chr1.g00702.t01#3    mNeoVis.evergladensis.2.1.chr1.g00698.t01#2    mNeoVis.vulgivaga.1.1.chr1.g00702.t01#4   mNeoVis.3.1.1.chr1.g00710.t01#6       3350   125257826:   mNeoVis.vulgivaga.1.1.chr7.g01416.t01#4    mNeoVis.2.1.1.chr7_hap1.g01425.t01#5    mNeoVis.evergladensis.2.1.chr7.g01407.t01#2    mNeoVis.evergladensis.1.1.chr7.g01405.t01#3    mNeoVis.vulgivaga.2.1.chr7.g01420.t01#1   mNeoVis.3.1.1.chr7.g01334.t01#6       3351   125257827:   mNeoVis.evergladensis.2.1.chr7.g01408.t01#2    mNeoVis.3.1.1.chr7.g01335.t01#6    mNeoVis.vulgivaga.1.1.chr7.g01417.t01#4   mNeoVis.2.1.1.chr7_hap1.g01427.t01#5         3352   125257828:   mNeoVis.2.1.1.chr7_hap1.g01426.t01#5   mNeoVis.vulgivaga.2.1.chr7.g01421.t01#1             3353   125257829:   mNeoVis.evergladensis.2.1.chr1.g00699.t01#2    mNeoVis.3.1.1.chr1.g00711.t01#6    mNeoVis.vulgivaga.2.1.chr1.g00711.t01#1    mNeoVis.vulgivaga.1.1.chr1.g00703.t01#4    mNeoVis.vulgivaga.2.1.chr1.g00712.t01#1    mNeoVis.2.1.1.chr1_hap1.g00691.t01#5   mNeoVis.evergladensis.1.1.chr1.g00703.t01#3     3354   125257830:   mNeoVis.vulgivaga.2.1.chr7.g01422.t01#1    mNeoVis.evergladensis.1.1.chr7.g01406.t01#3    mNeoVis.vulgivaga.1.1.chr7.g01418.t01#4    mNeoVis.2.1.1.chr7_hap1.g01428.t01#5   mNeoVis.evergladensis.2.1.chr7.g01409.t01#2       3355   125257831:   mNeoVis.2.1.1.chr9_hap1.g01095.t01#5    mNeoVis.3.1.1.chr10.g01054.t01#6    mNeoVis.evergladensis.2.1.chr10.g01085.t01#2   mNeoVis.evergladensis.1.1.chr10.g01082.t01#3       3356   125257832:   mNeoVis.vulgivaga.2.1.chr7.g01423.t01#1    mNeoVis.3.1.1.chr7.g01337.t01#6    mNeoVis.vulgivaga.1.1.chr7.g01419.t01#4    mNeoVis.2.1.1.chr7_hap1.g01429.t01#5    mNeoVis.evergladensis.1.1.chr7.g01407.t01#3   mNeoVis.evergladensis.2.1.chr7.g01410.t01#2       3357   125257833:   mNeoVis.evergladensis.1.1.chr1.g00705.t01#3    mNeoVis.2.1.1.chr1_hap1.g00693.t01#5    mNeoVis.evergladensis.2.1.chr1.g00700.t01#2    mNeoVis.vulgivaga.2.1.chr1.g00713.t01#1    mNeoVis.3.1.1.chr1.g00712.t01#6   mNeoVis.vulgivaga.1.1.chr1.g00704.t01#4       3358   125257834:   mNeoVis.evergladensis.2.1.chr1.g00701.t01#2    mNeoVis.vulgivaga.2.1.chr1.g00714.t01#1    mNeoVis.evergladensis.1.1.chr1.g00706.t01#3    mNeoVis.2.1.1.chr1_hap1.g00694.t01#5    mNeoVis.vulgivaga.1.1.chr1.g00705.t01#4   mNeoVis.3.1.1.chr1.g00713.t01#6       3359   125257835:   mNeoVis.vulgivaga.2.1.chr7.g01424.t01#1    mNeoVis.evergladensis.1.1.chr7.g01408.t01#3    mNeoVis.vulgivaga.1.1.chr7.g01420.t01#4    mNeoVis.evergladensis.2.1.chr7.g01412.t01#2    mNeoVis.2.1.1.chr7_hap1.g01430.t01#5   mNeoVis.3.1.1.chr7.g01338.t01#6       3360   125257836:   mNeoVis.vulgivaga.2.1.chr1.g00715.t01#1    mNeoVis.evergladensis.1.1.chr1.g00707.t01#3    mNeoVis.evergladensis.2.1.chr1.g00702.t01#2    mNeoVis.2.1.1.chr1_hap1.g00695.t01#5    mNeoVis.3.1.1.chr1.g00714.t01#6   mNeoVis.vulgivaga.1.1.chr1.g00706.t01#4       3361   125257837:   mNeoVis.evergladensis.2.1.chr7.g01413.t01#2    mNeoVis.2.1.1.chr7_hap1.g01431.t01#5    mNeoVis.evergladensis.1.1.chr7.g01409.t01#3    mNeoVis.vulgivaga.2.1.chr7.g01425.t01#1   mNeoVis.vulgivaga.1.1.chr7.g01421.t01#4       3362   125257838:   mNeoVis.3.1.1.chr7.g01339.t01#6    mNeoVis.evergladensis.1.1.chr7.g01410.t01#3    mNeoVis.vulgivaga.1.1.chr7.g01422.t01#4    mNeoVis.vulgivaga.2.1.chr7.g01426.t01#1    mNeoVis.evergladensis.2.1.chr7.g01414.t01#2   mNeoVis.2.1.1.chr7_hap1.g01432.t01#5       3363   125257839:   mNeoVis.vulgivaga.1.1.chr3.g01099.t01#4    mNeoVis.2.1.1.chr3_hap1.g01098.t01#5    mNeoVis.3.1.1.chr1.g00715.t01#6    mNeoVis.2.1.1.chr8_hap1.g00154.t01#5    mNeoVis.vulgivaga.2.1.chr3.g01113.t01#1    mNeoVis.evergladensis.2.1.chr9.g00151.t01#2    mNeoVis.evergladensis.2.1.chr1.g00703.t01#2    mNeoVis.vulgivaga.2.1.chr8.g00154.t01#1    mNeoVis.vulgivaga.1.1.chr8.g00148.t01#4      3364   125257840:   mNeoVis.evergladensis.2.1.chr4.g00260.t01#2    mNeoVis.3.1.1.chr4.g00253.t01#6    mNeoVis.evergladensis.1.1.chr4.g00259.t01#3    mNeoVis.vulgivaga.1.1.chr4.g00255.t01#4    mNeoVis.vulgivaga.2.1.chr4.g00259.t01#1   mNeoVis.2.1.1.chr6_hap1.g00254.t01#5       3365   125257841:   mNeoVis.vulgivaga.2.1.chr2.g01900.t01#1    mNeoVis.3.1.1.chr2.g01796.t01#6    mNeoVis.2.1.1.chr5_hap1.g01908.t01#5    mNeoVis.evergladensis.1.1.chr2.g01898.t01#3   mNeoVis.evergladensis.2.1.chr2.g01887.t01#2       3366   125257842:   mNeoVis.vulgivaga.2.1.chr1.g00717.t01#1    mNeoVis.evergladensis.2.1.chr1.g00704.t01#2    mNeoVis.evergladensis.1.1.chr1.g00709.t01#3   mNeoVis.3.1.1.chr1.g00716.t01#6         3367   125257843:   mNeoVis.2.1.1.chr7_hap1.g01433.t01#5    mNeoVis.evergladensis.2.1.chr7.g01415.t01#2    mNeoVis.vulgivaga.2.1.chr7.g01427.t01#1    mNeoVis.evergladensis.1.1.chr7.g01411.t01#3    mNeoVis.3.1.1.chr7.g01340.t01#6    mNeoVis.vulgivaga.1.1.chr1.g00934.t01#4   mNeoVis.2.1.1.chr1_hap1.g00922.t01#5     3368   125257844:   mNeoVis.evergladensis.1.1.chr7.g01412.t01#3    mNeoVis.3.1.1.chr7.g01341.t01#6    mNeoVis.2.1.1.chr7_hap1.g01434.t01#5    mNeoVis.vulgivaga.1.1.chr7.g01423.t01#4    mNeoVis.vulgivaga.2.1.chr7.g01428.t01#1   mNeoVis.evergladensis.2.1.chr7.g01416.t01#2       3369   125257845:   mNeoVis.3.1.1.chr1.g00717.t01#6    mNeoVis.2.1.1.chr1_hap1.g00697.t01#5    mNeoVis.vulgivaga.2.1.chr1.g00718.t01#1    mNeoVis.evergladensis.2.1.chr1.g00705.t01#2    mNeoVis.vulgivaga.1.1.chr1.g00708.t01#4   mNeoVis.evergladensis.1.1.chr1.g00710.t01#3       3370   125257846:   mNeoVis.vulgivaga.2.1.chr1.g00719.t01#1    mNeoVis.2.1.1.chr1_hap1.g00698.t01#5    mNeoVis.evergladensis.1.1.chr1.g00711.t01#3    mNeoVis.evergladensis.2.1.chr1.g00706.t01#2    mNeoVis.3.1.1.chr1.g00718.t01#6   mNeoVis.vulgivaga.1.1.chr1.g00709.t01#4       3371   125257847:   mNeoVis.2.1.1.chr7_hap1.g01435.t01#5    mNeoVis.vulgivaga.1.1.chr7.g01424.t01#4    mNeoVis.3.1.1.chr7.g01342.t01#6    mNeoVis.vulgivaga.2.1.chr7.g01429.t01#1    mNeoVis.evergladensis.1.1.chr7.g01413.t01#3   mNeoVis.evergladensis.2.1.chr7.g01417.t01#2       3372   125257848:   mNeoVis.vulgivaga.2.1.chr1.g00720.t01#1    mNeoVis.evergladensis.1.1.chr1.g00712.t01#3    mNeoVis.2.1.1.chr1_hap1.g00699.t01#5    mNeoVis.vulgivaga.1.1.chr1.g00710.t01#4    mNeoVis.evergladensis.2.1.chr1.g00707.t01#2   mNeoVis.3.1.1.chr1.g00719.t01#6       3373   125257849:   mNeoVis.evergladensis.1.1.chr7.g01414.t01#3    mNeoVis.vulgivaga.2.1.chr7.g01430.t01#1    mNeoVis.evergladensis.2.1.chr7.g01418.t01#2    mNeoVis.vulgivaga.1.1.chr7.g01425.t01#4    mNeoVis.3.1.1.chr8.g00668.t01#6   mNeoVis.2.1.1.chr7_hap1.g01436.t01#5       3374   125257850:   mNeoVis.vulgivaga.1.1.chr14.g00398.t01#4    mNeoVis.evergladensis.2.1.chr14.g00402.t01#2    mNeoVis.vulgivaga.2.1.chr14.g00403.t01#1    mNeoVis.2.1.1.chr14_hap1.g00402.t01#5    mNeoVis.evergladensis.1.1.chr14.g00406.t01#3   mNeoVis.3.1.1.chr14.g00401.t01#6       3375   125257851:   mNeoVis.vulgivaga.1.1.chr1.g00711.t01#4    mNeoVis.evergladensis.1.1.chr1.g00713.t01#3    mNeoVis.2.1.1.chr1_hap1.g00700.t01#5    mNeoVis.vulgivaga.2.1.chr1.g00721.t01#1    mNeoVis.3.1.1.chr1.g00720.t01#6   mNeoVis.evergladensis.2.1.chr1.g00708.t01#2       3376   125257852:   mNeoVis.2.1.1.chr1_hap1.g00701.t01#5    mNeoVis.evergladensis.2.1.chr1.g00709.t01#2    mNeoVis.vulgivaga.1.1.chr1.g00712.t01#4    mNeoVis.vulgivaga.2.1.chr1.g00722.t01#1    mNeoVis.3.1.1.chr1.g00721.t01#6   mNeoVis.evergladensis.1.1.chr1.g00714.t01#3       3377   125257853:   mNeoVis.evergladensis.1.1.chr7.g01415.t01#3    mNeoVis.3.1.1.chr7.g01344.t01#6    mNeoVis.vulgivaga.2.1.chr7.g01431.t01#1    mNeoVis.vulgivaga.1.1.chr7.g01426.t01#4    mNeoVis.evergladensis.2.1.chr7.g01419.t01#2   mNeoVis.2.1.1.chr7_hap1.g01437.t01#5       3378   125257854:   mNeoVis.2.1.1.chr7_hap1.g01438.t01#5    mNeoVis.evergladensis.1.1.chr7.g01416.t01#3    mNeoVis.evergladensis.2.1.chr7.g01420.t01#2    mNeoVis.3.1.1.chr7.g01345.t01#6    mNeoVis.vulgivaga.2.1.chr7.g01432.t01#1   mNeoVis.vulgivaga.1.1.chr7.g01427.t01#4       3379   125257855:   mNeoVis.2.1.1.chr1_hap1.g00702.t01#5    mNeoVis.vulgivaga.1.1.chr1.g00713.t01#4    mNeoVis.evergladensis.2.1.chr1.g00710.t01#2    mNeoVis.3.1.1.chr1.g00722.t01#6    mNeoVis.vulgivaga.2.1.chr1.g00723.t01#1   mNeoVis.evergladensis.1.1.chr1.g00715.t01#3       3380   125257856:   mNeoVis.evergladensis.1.1.chr3.g00222.t01#3    mNeoVis.vulgivaga.1.1.chr3.g00222.t01#4    mNeoVis.2.1.1.chr3_hap1.g00217.t01#5    mNeoVis.vulgivaga.1.1.chr7.g01428.t01#4    mNeoVis.vulgivaga.2.1.chr3.g00219.t01#1    mNeoVis.vulgivaga.2.1.chr7.g01433.t01#1    mNeoVis.evergladensis.2.1.chr3.g00227.t01#2    mNeoVis.evergladensis.2.1.chr7.g01421.t01#2    mNeoVis.evergladensis.1.1.chr7.g01417.t01#3      3381   125257857:   mNeoVis.2.1.1.chr8_hap1.g01436.t01#5    mNeoVis.vulgivaga.1.1.chr8.g01374.t01#4    mNeoVis.evergladensis.2.1.chr9.g01381.t01#2    mNeoVis.3.1.1.chr8.g01333.t01#6   mNeoVis.vulgivaga.2.1.chr8.g01390.t01#1       3382   125257858:   mNeoVis.2.1.1.chr7_hap1.g01439.t01#5    mNeoVis.vulgivaga.2.1.chr7.g01434.t01#1    mNeoVis.evergladensis.1.1.chr7.g01418.t01#3    mNeoVis.3.1.1.chr7.g01347.t01#6    mNeoVis.evergladensis.2.1.chr7.g01422.t01#2   mNeoVis.vulgivaga.1.1.chr7.g01429.t01#4       3383   125257859:   mNeoVis.evergladensis.2.1.chr1.g00711.t01#2    mNeoVis.vulgivaga.1.1.chr1.g00714.t01#4    mNeoVis.3.1.1.chr1.g00724.t01#6    mNeoVis.evergladensis.1.1.chr1.g00717.t01#3    mNeoVis.vulgivaga.2.1.chr1.g00724.t01#1   mNeoVis.2.1.1.chr1_hap1.g00703.t01#5       3384   125257860:   mNeoVis.vulgivaga.1.1.chr7.g01430.t01#4    mNeoVis.evergladensis.2.1.chr7.g01423.t01#2    mNeoVis.evergladensis.1.1.chr7.g01420.t01#3    mNeoVis.2.1.1.chr7_hap1.g01440.t01#5    mNeoVis.vulgivaga.2.1.chr7.g01435.t01#1   mNeoVis.3.1.1.chr7.g01348.t01#6       3385   125257861:   mNeoVis.vulgivaga.2.1.chr1.g00725.t01#1    mNeoVis.2.1.1.chr1_hap1.g00704.t01#5    mNeoVis.evergladensis.1.1.chr1.g00718.t01#3    mNeoVis.vulgivaga.1.1.chr1.g00715.t01#4    mNeoVis.3.1.1.chr1.g00725.t01#6   mNeoVis.evergladensis.2.1.chr1.g00712.t01#2       3386   125257862:   mNeoVis.3.1.1.chr7.g01349.t01#6    mNeoVis.vulgivaga.2.1.chr7.g01436.t01#1    mNeoVis.vulgivaga.1.1.chr7.g01431.t01#4    mNeoVis.evergladensis.1.1.chr7.g01421.t01#3    mNeoVis.evergladensis.2.1.chr7.g01424.t01#2   mNeoVis.2.1.1.chr7_hap1.g01441.t01#5       3387   125257863:   mNeoVis.evergladensis.2.1.chr1.g00713.t01#2    mNeoVis.3.1.1.chr1.g00726.t01#6    mNeoVis.vulgivaga.1.1.chr1.g00716.t01#4    mNeoVis.2.1.1.chr1_hap1.g00705.t01#5    mNeoVis.evergladensis.1.1.chr1.g00719.t01#3   mNeoVis.vulgivaga.2.1.chr1.g00726.t01#1       3388   125257864:   mNeoVis.3.1.1.chr7.g01350.t01#6    mNeoVis.vulgivaga.2.1.chr7.g01437.t01#1    mNeoVis.vulgivaga.1.1.chr7.g01432.t01#4    mNeoVis.evergladensis.1.1.chr7.g01422.t01#3    mNeoVis.evergladensis.2.1.chr7.g01425.t01#2   mNeoVis.2.1.1.chr7_hap1.g01442.t01#5       3389   125257865:   mNeoVis.evergladensis.2.1.chr1.g00714.t01#2    mNeoVis.2.1.1.chr1_hap1.g00706.t01#5    mNeoVis.vulgivaga.2.1.chr1.g00727.t01#1    mNeoVis.vulgivaga.1.1.chr1.g00717.t01#4    mNeoVis.evergladensis.1.1.chr1.g00720.t01#3   mNeoVis.3.1.1.chr1.g00727.t01#6       3390   125257866:   mNeoVis.evergladensis.1.1.chr1.g00721.t01#3    mNeoVis.evergladensis.2.1.chr1.g00715.t01#2    mNeoVis.vulgivaga.2.1.chr1.g00728.t01#1    mNeoVis.2.1.1.chr1_hap1.g00707.t01#5   mNeoVis.vulgivaga.1.1.chr1.g00718.t01#4       3391   125257867:   mNeoVis.vulgivaga.2.1.chr1.g00729.t01#1    mNeoVis.evergladensis.2.1.chr1.g00716.t01#2    mNeoVis.evergladensis.1.1.chr1.g00722.t01#3    mNeoVis.3.1.1.chr1.g00728.t01#6    mNeoVis.vulgivaga.1.1.chr1.g00719.t01#4   mNeoVis.2.1.1.chr1_hap1.g00708.t01#5       3392   125257868:   mNeoVis.3.1.1.chr7.g01351.t01#6    mNeoVis.2.1.1.chr7_hap1.g01443.t01#5    mNeoVis.vulgivaga.1.1.chr7.g01433.t01#4    mNeoVis.vulgivaga.2.1.chr7.g01438.t01#1    mNeoVis.evergladensis.1.1.chr7.g01423.t01#3   mNeoVis.evergladensis.2.1.chr7.g01426.t01#2       3393   125257869:   mNeoVis.vulgivaga.1.1.chr15.g00093.t01#4    mNeoVis.2.1.1.chr15_hap1.g00092.t01#5    mNeoVis.3.1.1.chr15.g00086.t01#6    mNeoVis.evergladensis.2.1.chr15.g00093.t01#2    mNeoVis.vulgivaga.2.1.chr15.g00095.t01#1   mNeoVis.evergladensis.1.1.chr15.g00520.t01#3       3394   125257870:   mNeoVis.2.1.1.chr1_hap1.g00709.t01#5    mNeoVis.vulgivaga.2.1.chr1.g00730.t01#1    mNeoVis.evergladensis.2.1.chr1.g00718.t01#2    mNeoVis.evergladensis.1.1.chr1.g00724.t01#3   mNeoVis.3.1.1.chr1.g00729.t01#6       3395   125257871:   mNeoVis.evergladensis.2.1.chr7.g01334.t01#2    mNeoVis.evergladensis.1.1.chr7.g01334.t01#3    mNeoVis.2.1.1.chr7_hap1.g01351.t01#5    mNeoVis.3.1.1.chr7.g01261.t01#6    mNeoVis.vulgivaga.1.1.chr7.g01341.t01#4   mNeoVis.vulgivaga.2.1.chr7.g01347.t01#1       3396   125257872:   mNeoVis.evergladensis.1.1.chr7.g01335.t01#3    mNeoVis.2.1.1.chr7_hap1.g01352.t01#5    mNeoVis.vulgivaga.2.1.chr7.g01348.t01#1    mNeoVis.3.1.1.chr7.g01262.t01#6    mNeoVis.vulgivaga.1.1.chr7.g01342.t01#4   mNeoVis.evergladensis.2.1.chr7.g01335.t01#2       3397   125257873:   mNeoVis.vulgivaga.2.1.chr7.g01349.t01#1    mNeoVis.3.1.1.chr7.g01263.t01#6    mNeoVis.evergladensis.2.1.chr7.g01336.t01#2    mNeoVis.vulgivaga.1.1.chr7.g01343.t01#4    mNeoVis.evergladensis.1.1.chr7.g01336.t01#3   mNeoVis.2.1.1.chr7_hap1.g01353.t01#5       3398   125257874:   mNeoVis.vulgivaga.2.1.chr7.g00097.t01#1    mNeoVis.2.1.1.chr7_hap1.g00099.t01#5    mNeoVis.evergladensis.1.1.chr7.g00100.t01#3    mNeoVis.vulgivaga.1.1.chr7.g00102.t01#4    mNeoVis.evergladensis.2.1.chr7.g00098.t01#2   mNeoVis.3.1.1.chr7.g00082.t01#6       3399   125257875:   mNeoVis.evergladensis.2.1.chr1.g00719.t01#2    mNeoVis.evergladensis.1.1.chr1.g00726.t01#3    mNeoVis.2.1.1.chr1_hap1.g00710.t01#5    mNeoVis.vulgivaga.2.1.chr1.g00731.t01#1    mNeoVis.vulgivaga.1.1.chr1.g00723.t01#4   mNeoVis.3.1.1.chr1.g00731.t01#6       3400   125257876:   mNeoVis.2.1.1.chr10_hap1.g00678.t01#5    mNeoVis.evergladensis.1.1.chr9.g00667.t01#3    mNeoVis.vulgivaga.1.1.chr9.g00667.t01#4    mNeoVis.vulgivaga.2.1.chr9.g00660.t01#1    mNeoVis.evergladensis.2.1.chr8.g00658.t01#2   mNeoVis.3.1.1.chr9.g00637.t01#6       3401   125257877:   mNeoVis.vulgivaga.1.1.chr1.g00724.t01#4    mNeoVis.3.1.1.chr1.g00732.t01#6    mNeoVis.evergladensis.1.1.chr1.g00727.t01#3    mNeoVis.evergladensis.2.1.chr1.g00721.t01#2    mNeoVis.vulgivaga.2.1.chr1.g00732.t01#1   mNeoVis.2.1.1.chr1_hap1.g00711.t01#5       3402   125257878:   mNeoVis.vulgivaga.2.1.chr7.g01350.t01#1    mNeoVis.evergladensis.1.1.chr7.g01337.t01#3    mNeoVis.2.1.1.chr7_hap1.g01354.t01#5    mNeoVis.vulgivaga.1.1.chr7.g01344.t01#4    mNeoVis.3.1.1.chr7.g01264.t01#6   mNeoVis.evergladensis.2.1.chr7.g01337.t01#2       3403   125257879:   mNeoVis.3.1.1.chr7.g01265.t01#6    mNeoVis.vulgivaga.2.1.chr7.g01351.t01#1    mNeoVis.evergladensis.1.1.chr7.g01338.t01#3    mNeoVis.2.1.1.chr7_hap1.g01355.t01#5    mNeoVis.vulgivaga.1.1.chr7.g01345.t01#4   mNeoVis.evergladensis.2.1.chr7.g01338.t01#2       3404   125257880:   mNeoVis.3.1.1.chr1.g00734.t01#6    mNeoVis.evergladensis.2.1.chr1.g00722.t01#2    mNeoVis.vulgivaga.1.1.chr1.g00726.t01#4    mNeoVis.vulgivaga.2.1.chr1.g00733.t01#1    mNeoVis.evergladensis.1.1.chr1.g00729.t01#3   mNeoVis.2.1.1.chr1_hap1.g00713.t01#5       3405   125257881:   mNeoVis.2.1.1.chr2_hap1.g01261.t01#5    mNeoVis.evergladensis.2.1.chr5.g01205.t01#2    mNeoVis.evergladensis.1.1.chr5.g01201.t01#3    mNeoVis.vulgivaga.2.1.chr5.g01198.t01#1    mNeoVis.vulgivaga.1.1.chr5.g01168.t01#4   mNeoVis.3.1.1.chr5.g01183.t01#6       3406   125257882:   mNeoVis.vulgivaga.1.1.chr2.g00509.t01#4    mNeoVis.evergladensis.1.1.chr1.g00730.t01#3    mNeoVis.vulgivaga.1.1.chr5.g01167.t01#4    mNeoVis.3.1.1.chr2.g00442.t01#6    mNeoVis.3.1.1.chr5.g01182.t01#6   mNeoVis.vulgivaga.2.1.chr2.g00515.t01#1       3407   125257883:   mNeoVis.2.1.1.chr5_hap1.g00519.t01#5    mNeoVis.evergladensis.2.1.chr2.g00513.t01#2   mNeoVis.evergladensis.1.1.chr2.g00510.t01#3         3408   125257884:   mNeoVis.3.1.1.chr7.g01267.t01#6    mNeoVis.vulgivaga.1.1.chr7.g01346.t01#4    mNeoVis.evergladensis.1.1.chr7.g01339.t01#3    mNeoVis.2.1.1.chr7_hap1.g01356.t01#5    mNeoVis.vulgivaga.2.1.chr7.g01352.t01#1   mNeoVis.evergladensis.2.1.chr7.g01339.t01#2       3409   125257885:   mNeoVis.vulgivaga.2.1.chr1.g00683.t01#1    mNeoVis.2.1.1.chr1_hap1.g00664.t01#5    mNeoVis.evergladensis.2.1.chr8.g00669.t01#2    mNeoVis.evergladensis.2.1.chr1.g00670.t01#2    mNeoVis.vulgivaga.2.1.chr1.g00734.t01#1    mNeoVis.vulgivaga.1.1.chr1.g00675.t01#4    mNeoVis.evergladensis.1.1.chr1.g00790.t01#3    mNeoVis.2.1.1.chr10_hap1.g00688.t01#5    mNeoVis.evergladensis.1.1.chr1.g00675.t01#3      3410   125257886:   mNeoVis.vulgivaga.2.1.chr7.g01353.t01#1    mNeoVis.evergladensis.2.1.chr7.g01340.t01#2    mNeoVis.2.1.1.chr7_hap1.g01357.t01#5    mNeoVis.evergladensis.1.1.chr7.g01340.t01#3    mNeoVis.3.1.1.chr7.g01268.t01#6   mNeoVis.vulgivaga.1.1.chr7.g01347.t01#4       3411   125257887:   mNeoVis.vulgivaga.2.1.chr15.g00096.t01#1    mNeoVis.evergladensis.2.1.chr15.g00094.t01#2    mNeoVis.2.1.1.chr15_hap1.g00093.t01#5    mNeoVis.vulgivaga.1.1.chr15.g00094.t01#4    mNeoVis.3.1.1.chr15.g00087.t01#6   mNeoVis.evergladensis.1.1.chr15.g00519.t01#3       3412   125257888:   mNeoVis.evergladensis.1.1.chr3.g00605.t01#3    mNeoVis.2.1.1.chr3_hap1.g00597.t01#5    mNeoVis.3.1.1.chr3.g00589.t01#6    mNeoVis.vulgivaga.2.1.chr3.g00608.t01#1    mNeoVis.evergladensis.2.1.chr3.g00605.t01#2   mNeoVis.vulgivaga.1.1.chr3.g00601.t01#4       3413   125257889:   mNeoVis.3.1.1.chr8.g00875.t01#6    mNeoVis.2.1.1.chr8_hap1.g00943.t01#5    mNeoVis.vulgivaga.1.1.chr8.g00901.t01#4    mNeoVis.vulgivaga.2.1.chr8.g00917.t01#1    mNeoVis.evergladensis.1.1.chr8.g00910.t01#3   mNeoVis.evergladensis.2.1.chr9.g00917.t01#2       3414   125257890:   mNeoVis.vulgivaga.1.1.chr1.g00727.t01#4    mNeoVis.3.1.1.chr1.g00735.t01#6    mNeoVis.evergladensis.2.1.chr1.g00723.t01#2    mNeoVis.evergladensis.1.1.chr1.g00731.t01#3    mNeoVis.2.1.1.chr1_hap1.g00714.t01#5   mNeoVis.vulgivaga.2.1.chr1.g00735.t01#1       3415   125257891:   mNeoVis.vulgivaga.1.1.chr1.g00729.t01#4    mNeoVis.evergladensis.2.1.chr1.g00724.t01#2    mNeoVis.2.1.1.chr1_hap1.g00715.t01#5    mNeoVis.3.1.1.chr1.g00736.t01#6    mNeoVis.vulgivaga.2.1.chr1.g00736.t01#1   mNeoVis.evergladensis.1.1.chr1.g00732.t01#3       3416   125257892:   mNeoVis.evergladensis.2.1.chr7.g00099.t01#2    mNeoVis.vulgivaga.2.1.chr7.g00098.t01#1    mNeoVis.3.1.1.chr7.g00084.t01#6    mNeoVis.evergladensis.2.1.chr7.g00100.t01#2    mNeoVis.evergladensis.1.1.chr7.g00101.t01#3    mNeoVis.2.1.1.chr7_hap1.g00100.t01#5    mNeoVis.vulgivaga.1.1.chr7.g00103.t01#4   mNeoVis.3.1.1.chr7.g00083.t01#6     3417   125257893:   mNeoVis.evergladensis.2.1.chr1.g00725.t01#2    mNeoVis.2.1.1.chr1_hap1.g00716.t01#5    mNeoVis.3.1.1.chr1.g00737.t01#6    mNeoVis.vulgivaga.1.1.chr1.g00730.t01#4    mNeoVis.evergladensis.1.1.chr1.g00733.t01#3   mNeoVis.vulgivaga.2.1.chr1.g00737.t01#1       3418   125257894:   mNeoVis.evergladensis.2.1.chr7.g01341.t01#2    mNeoVis.3.1.1.chr7.g01269.t01#6    mNeoVis.evergladensis.1.1.chr7.g01341.t01#3    mNeoVis.vulgivaga.2.1.chr7.g01354.t01#1    mNeoVis.vulgivaga.1.1.chr7.g01348.t01#4   mNeoVis.2.1.1.chr7_hap1.g01358.t01#5       3419   125257895:  mNeoVis.vulgivaga.2.1.chr1.g00739.t01#1               3420   125257896:   mNeoVis.2.1.1.chr1_hap1.g00717.t01#5    mNeoVis.3.1.1.chr1.g00738.t01#6    mNeoVis.vulgivaga.2.1.chr1.g00738.t01#1    mNeoVis.evergladensis.2.1.chr1.g00727.t01#2    mNeoVis.vulgivaga.1.1.chr1.g00731.t01#4   mNeoVis.evergladensis.1.1.chr1.g00734.t01#3       3421   125257897:   mNeoVis.3.1.1.chr1.g00740.t01#6   mNeoVis.vulgivaga.2.1.chr1.g00740.t01#1             3422   125257898:   mNeoVis.evergladensis.1.1.chr7.g01342.t01#3    mNeoVis.2.1.1.chr7_hap1.g01359.t01#5    mNeoVis.3.1.1.chr7.g01270.t01#6    mNeoVis.vulgivaga.2.1.chr7.g01355.t01#1    mNeoVis.vulgivaga.1.1.chr7.g01349.t01#4   mNeoVis.evergladensis.2.1.chr7.g01342.t01#2       3423   125257899:   mNeoVis.evergladensis.2.1.chr7.g00101.t01#2    mNeoVis.2.1.1.chr7_hap1.g00101.t01#5    mNeoVis.vulgivaga.1.1.chr7.g00104.t01#4    mNeoVis.evergladensis.1.1.chr7.g00102.t01#3    mNeoVis.3.1.1.chr7.g00085.t01#6   mNeoVis.vulgivaga.2.1.chr7.g00099.t01#1       3424   125257900:   mNeoVis.vulgivaga.1.1.chr1.g00735.t01#4    mNeoVis.2.1.1.chr1_hap1.g00720.t01#5    mNeoVis.evergladensis.2.1.chr1.g00729.t01#2    mNeoVis.vulgivaga.2.1.chr1.g00742.t01#1    mNeoVis.evergladensis.1.1.chr1.g00737.t01#3   mNeoVis.3.1.1.chr1.g00742.t01#6       3425   125257901:   mNeoVis.evergladensis.2.1.chr1.g00728.t01#2    mNeoVis.vulgivaga.2.1.chr1.g00741.t01#1    mNeoVis.vulgivaga.1.1.chr1.g00734.t01#4    mNeoVis.evergladensis.1.1.chr1.g00736.t01#3    mNeoVis.2.1.1.chr1_hap1.g00719.t01#5   mNeoVis.3.1.1.chr1.g00741.t01#6       3426   125257902:   mNeoVis.vulgivaga.2.1.chr7.g01356.t01#1    mNeoVis.evergladensis.2.1.chr7.g01343.t01#2    mNeoVis.3.1.1.chr7.g01271.t01#6    mNeoVis.evergladensis.1.1.chr7.g01343.t01#3    mNeoVis.2.1.1.chr7_hap1.g01360.t01#5   mNeoVis.vulgivaga.1.1.chr7.g01350.t01#4       3427   125257903:   mNeoVis.vulgivaga.1.1.chr7.g01351.t01#4    mNeoVis.3.1.1.chr7.g01272.t01#6    mNeoVis.2.1.1.chr7_hap1.g01361.t01#5    mNeoVis.vulgivaga.2.1.chr7.g01357.t01#1    mNeoVis.evergladensis.2.1.chr7.g01344.t01#2   mNeoVis.evergladensis.1.1.chr7.g01344.t01#3       3428   125257904:   mNeoVis.vulgivaga.2.1.chr7.g01358.t01#1    mNeoVis.evergladensis.2.1.chr7.g01346.t01#2    mNeoVis.3.1.1.chr7.g01274.t01#6    mNeoVis.evergladensis.1.1.chr7.g01346.t01#3   mNeoVis.2.1.1.chr7_hap1.g01363.t01#5       3429   125257905:   mNeoVis.vulgivaga.1.1.chr1.g00736.t01#4    mNeoVis.evergladensis.2.1.chr1.g00730.t01#2    mNeoVis.2.1.1.chr1_hap1.g00721.t01#5    mNeoVis.3.1.1.chr1.g00743.t01#6    mNeoVis.vulgivaga.2.1.chr1.g00743.t01#1   mNeoVis.evergladensis.1.1.chr1.g00738.t01#3       3430   125257906:   mNeoVis.2.1.1.chr7_hap1.g01364.t01#5    mNeoVis.evergladensis.2.1.chr7.g01347.t01#2    mNeoVis.vulgivaga.1.1.chr7.g01353.t01#4    mNeoVis.vulgivaga.2.1.chr7.g01359.t01#1   mNeoVis.evergladensis.1.1.chr7.g01347.t01#3       3431   125257907:   mNeoVis.evergladensis.2.1.chr7.g01348.t01#2    mNeoVis.2.1.1.chr7_hap1.g01365.t01#5    mNeoVis.vulgivaga.1.1.chr7.g01354.t01#4    mNeoVis.3.1.1.chr7.g01276.t01#6    mNeoVis.vulgivaga.2.1.chr7.g01360.t01#1   mNeoVis.evergladensis.1.1.chr7.g01348.t01#3       3432   125257908:   mNeoVis.vulgivaga.1.1.chr7.g01355.t01#4    mNeoVis.evergladensis.2.1.chr7.g01349.t01#2    mNeoVis.vulgivaga.2.1.chr7.g01361.t01#1    mNeoVis.evergladensis.1.1.chr7.g01349.t01#3    mNeoVis.3.1.1.chr7.g01277.t01#6   mNeoVis.2.1.1.chr7_hap1.g01366.t01#5       3433   125257909:   mNeoVis.vulgivaga.2.1.chr7.g01362.t01#1    mNeoVis.2.1.1.chr7_hap1.g01367.t01#5    mNeoVis.evergladensis.2.1.chr2.g00707.t01#2    mNeoVis.2.1.1.chr5_hap1.g00707.t01#5    mNeoVis.vulgivaga.1.1.chr7.g01356.t01#4    mNeoVis.vulgivaga.2.1.chr2.g00708.t01#1    mNeoVis.3.1.1.chr2.g00621.t01#6    mNeoVis.3.1.1.chr7.g01278.t01#6    mNeoVis.evergladensis.1.1.chr7.g01350.t01#3      3434   125257910:   mNeoVis.vulgivaga.1.1.chr7.g01357.t01#4    mNeoVis.3.1.1.chr7.g01279.t01#6    mNeoVis.evergladensis.1.1.chr7.g01351.t01#3    mNeoVis.2.1.1.chr7_hap1.g01369.t01#5    mNeoVis.vulgivaga.2.1.chr7.g01363.t01#1   mNeoVis.evergladensis.2.1.chr7.g01352.t01#2       3435   125257911:   mNeoVis.evergladensis.1.1.chr1.g00739.t01#3    mNeoVis.3.1.1.chr1.g00744.t01#6    mNeoVis.evergladensis.2.1.chr1.g00731.t01#2    mNeoVis.2.1.1.chr1_hap1.g00722.t01#5    mNeoVis.vulgivaga.2.1.chr1.g00744.t01#1   mNeoVis.vulgivaga.1.1.chr1.g00737.t01#4       3436   125257912:   mNeoVis.vulgivaga.2.1.chr1.g00745.t01#1    mNeoVis.3.1.1.chr1.g00745.t01#6    mNeoVis.2.1.1.chr1_hap1.g00723.t01#5    mNeoVis.vulgivaga.1.1.chr1.g00738.t01#4    mNeoVis.evergladensis.1.1.chr1.g00740.t01#3   mNeoVis.evergladensis.2.1.chr1.g00732.t01#2       3437   125257913:   mNeoVis.vulgivaga.2.1.chr7.g01364.t01#1    mNeoVis.3.1.1.chr7.g01280.t01#6    mNeoVis.evergladensis.2.1.chr7.g01353.t01#2    mNeoVis.2.1.1.chr7_hap1.g01370.t01#5    mNeoVis.evergladensis.1.1.chr7.g01352.t01#3   mNeoVis.vulgivaga.1.1.chr7.g01359.t01#4       3438   125257914:   mNeoVis.3.1.1.chr1.g00746.t01#6    mNeoVis.2.1.1.chr1_hap1.g00724.t01#5    mNeoVis.vulgivaga.2.1.chr1.g00746.t01#1    mNeoVis.vulgivaga.1.1.chr1.g00739.t01#4    mNeoVis.evergladensis.1.1.chr1.g00741.t01#3   mNeoVis.evergladensis.2.1.chr1.g00733.t01#2       3439   125257915:   mNeoVis.vulgivaga.2.1.chr10.g00177.t01#1    mNeoVis.2.1.1.chr9_hap1.g00184.t01#5    mNeoVis.3.1.1.chr10.g00176.t01#6    mNeoVis.evergladensis.1.1.chr10.g00183.t01#3    mNeoVis.vulgivaga.1.1.chr10.g00170.t01#4   mNeoVis.evergladensis.2.1.chr10.g00179.t01#2       3440   125257916:   mNeoVis.vulgivaga.1.1.chr7.g01360.t01#4    mNeoVis.evergladensis.2.1.chr7.g01354.t01#2    mNeoVis.vulgivaga.2.1.chr7.g01365.t01#1    mNeoVis.evergladensis.1.1.chr7.g01353.t01#3    mNeoVis.2.1.1.chr7_hap1.g01371.t01#5   mNeoVis.3.1.1.chr7.g01281.t01#6       3441   125257917:   mNeoVis.2.1.1.chr1_hap1.g00725.t01#5    mNeoVis.vulgivaga.1.1.chr1.g00740.t01#4    mNeoVis.evergladensis.1.1.chr1.g00743.t01#3    mNeoVis.evergladensis.2.1.chr1.g00735.t01#2   mNeoVis.vulgivaga.2.1.chr1.g00747.t01#1       3442   125257918:   mNeoVis.evergladensis.2.1.chr7.g00014.t01#2    mNeoVis.3.1.1.chr7.g00013.t01#6    mNeoVis.2.1.1.chr7_hap1.g00013.t01#5    mNeoVis.vulgivaga.1.1.chr7.g00015.t01#4    mNeoVis.evergladensis.1.1.chr7.g00015.t01#3   mNeoVis.vulgivaga.2.1.chr7.g00014.t01#1       3443   125257919:   mNeoVis.vulgivaga.2.1.chr7.g01366.t01#1    mNeoVis.3.1.1.chr7.g01282.t01#6    mNeoVis.2.1.1.chr7_hap1.g01372.t01#5    mNeoVis.evergladensis.2.1.chr7.g01355.t01#2    mNeoVis.vulgivaga.1.1.chr7.g01361.t01#4   mNeoVis.evergladensis.1.1.chr7.g01354.t01#3       3444   125257920:   mNeoVis.2.1.1.chr1_hap1.g00726.t01#5    mNeoVis.vulgivaga.1.1.chr1.g00741.t01#4    mNeoVis.vulgivaga.2.1.chr1.g00748.t01#1    mNeoVis.evergladensis.1.1.chr1.g00744.t01#3    mNeoVis.evergladensis.1.1.chr3.g01240.t01#3    mNeoVis.evergladensis.2.1.chr1.g00736.t01#2    mNeoVis.evergladensis.2.1.chr3.g01232.t01#2    mNeoVis.vulgivaga.2.1.chr3.g01246.t01#1    mNeoVis.3.1.1.chr1.g00749.t01#6      3445   125257921:   mNeoVis.evergladensis.1.1.chr12.g00153.t01#3    mNeoVis.vulgivaga.2.1.chr12.g00154.t01#1    mNeoVis.evergladensis.2.1.chr12.g00151.t01#2    mNeoVis.2.1.1.chr12_hap1.g00178.t01#5    mNeoVis.3.1.1.chr12.g00148.t01#6    mNeoVis.evergladensis.1.1.chr12.g00152.t01#3   mNeoVis.vulgivaga.1.1.chr12.g00158.t01#4     3446   125257922:   mNeoVis.vulgivaga.1.1.chr1.g00742.t01#4    mNeoVis.3.1.1.chr1.g00750.t01#6    mNeoVis.2.1.1.chr1_hap1.g00727.t01#5    mNeoVis.evergladensis.2.1.chr1.g00737.t01#2    mNeoVis.vulgivaga.2.1.chr1.g00749.t01#1   mNeoVis.evergladensis.1.1.chr1.g00745.t01#3       3447   125257923:   mNeoVis.vulgivaga.2.1.chr1.g00750.t01#1    mNeoVis.evergladensis.2.1.chr1.g00738.t01#2    mNeoVis.evergladensis.1.1.chr1.g00746.t01#3    mNeoVis.3.1.1.chr1.g00751.t01#6    mNeoVis.vulgivaga.1.1.chr1.g00743.t01#4   mNeoVis.2.1.1.chr1_hap1.g00728.t01#5       3448   125257924:   mNeoVis.vulgivaga.1.1.chr7.g01362.t01#4    mNeoVis.evergladensis.1.1.chr7.g01355.t01#3    mNeoVis.vulgivaga.2.1.chr7.g01367.t01#1    mNeoVis.3.1.1.chr7.g01283.t01#6    mNeoVis.2.1.1.chr7_hap1.g01373.t01#5   mNeoVis.evergladensis.2.1.chr7.g01356.t01#2       3449   125257925:   mNeoVis.vulgivaga.1.1.chr7.g01363.t01#4    mNeoVis.evergladensis.1.1.chr7.g01356.t01#3    mNeoVis.3.1.1.chr7.g01284.t01#6    mNeoVis.evergladensis.2.1.chr7.g01357.t01#2    mNeoVis.vulgivaga.2.1.chr7.g01368.t01#1   mNeoVis.2.1.1.chr7_hap1.g01374.t01#5       3450   125257926:   mNeoVis.3.1.1.chr7.g01285.t01#6    mNeoVis.vulgivaga.1.1.chr7.g01364.t01#4    mNeoVis.vulgivaga.2.1.chr7.g01369.t01#1    mNeoVis.evergladensis.1.1.chr7.g01357.t01#3    mNeoVis.evergladensis.2.1.chr7.g01358.t01#2   mNeoVis.2.1.1.chr7_hap1.g01375.t01#5       3451   125257927:   mNeoVis.vulgivaga.1.1.chr1.g00744.t01#4    mNeoVis.2.1.1.chr1_hap1.g00729.t01#5    mNeoVis.evergladensis.2.1.chr1.g00739.t01#2    mNeoVis.evergladensis.1.1.chr1.g00748.t01#3    mNeoVis.vulgivaga.2.1.chr1.g00751.t01#1   mNeoVis.3.1.1.chr1.g00752.t01#6       3452   125257928:   mNeoVis.vulgivaga.1.1.chr7.g01365.t01#4    mNeoVis.evergladensis.1.1.chr7.g01358.t01#3    mNeoVis.3.1.1.chr7.g01286.t01#6    mNeoVis.2.1.1.chr7_hap1.g01376.t01#5    mNeoVis.evergladensis.2.1.chr7.g01359.t01#2   mNeoVis.vulgivaga.2.1.chr7.g01370.t01#1       3453   125257929:   mNeoVis.vulgivaga.2.1.chr1.g00661.t01#1    mNeoVis.vulgivaga.1.1.chr1.g00654.t01#4    mNeoVis.2.1.1.chr1_hap1.g00645.t01#5    mNeoVis.evergladensis.1.1.chr1.g00656.t01#3    mNeoVis.evergladensis.2.1.chr1.g00650.t01#2   mNeoVis.3.1.1.chr1.g00663.t01#6       3454   125257930:   mNeoVis.evergladensis.1.1.chr2.g01589.t01#3    mNeoVis.vulgivaga.1.1.chr2.g01592.t01#4    mNeoVis.2.1.1.chr5_hap1.g01592.t01#5    mNeoVis.3.1.1.chr2.g01507.t01#6    mNeoVis.evergladensis.2.1.chr2.g01579.t01#2   mNeoVis.vulgivaga.2.1.chr2.g01593.t01#1       3455   125257931:   mNeoVis.evergladensis.1.1.chr1.g00657.t01#3    mNeoVis.evergladensis.2.1.chr1.g00651.t01#2    mNeoVis.3.1.1.chr1.g00664.t01#6    mNeoVis.vulgivaga.1.1.chr1.g00655.t01#4    mNeoVis.2.1.1.chr1_hap1.g00646.t01#5   mNeoVis.vulgivaga.2.1.chr1.g00662.t01#1       3456   125257932:   mNeoVis.2.1.1.chr7_hap1.g01377.t01#5    mNeoVis.vulgivaga.2.1.chr7.g01371.t01#1    mNeoVis.evergladensis.2.1.chr7.g01360.t01#2    mNeoVis.vulgivaga.1.1.chr7.g01366.t01#4    mNeoVis.3.1.1.chr7.g01287.t01#6   mNeoVis.evergladensis.1.1.chr7.g01359.t01#3       3457   125257933:   mNeoVis.3.1.1.chr7.g01288.t01#6    mNeoVis.vulgivaga.1.1.chr7.g01367.t01#4    mNeoVis.evergladensis.2.1.chr7.g01361.t01#2    mNeoVis.evergladensis.1.1.chr7.g01360.t01#3    mNeoVis.2.1.1.chr7_hap1.g01378.t01#5   mNeoVis.vulgivaga.2.1.chr7.g01372.t01#1       3458   125257934:   mNeoVis.2.1.1.chr1_hap1.g00647.t01#5    mNeoVis.3.1.1.chr1.g00665.t01#6    mNeoVis.evergladensis.1.1.chr1.g00658.t01#3    mNeoVis.vulgivaga.1.1.chr1.g00656.t01#4    mNeoVis.vulgivaga.2.1.chr1.g00663.t01#1   mNeoVis.evergladensis.2.1.chr1.g00652.t01#2       3459   125257935:  mNeoVis.vulgivaga.2.1.chr1.g00664.t01#1               3460   125257936:   mNeoVis.vulgivaga.1.1.chr7.g01368.t01#4    mNeoVis.evergladensis.1.1.chr7.g01361.t01#3    mNeoVis.3.1.1.chr7.g01289.t01#6    mNeoVis.evergladensis.2.1.chr7.g01362.t01#2    mNeoVis.vulgivaga.2.1.chr7.g01373.t01#1   mNeoVis.2.1.1.chr7_hap1.g01379.t01#5       3461   125257937:   mNeoVis.vulgivaga.2.1.chr1.g00665.t01#1    mNeoVis.3.1.1.chr1.g00666.t01#6    mNeoVis.evergladensis.1.1.chr1.g00659.t01#3    mNeoVis.vulgivaga.1.1.chr1.g00657.t01#4    mNeoVis.evergladensis.2.1.chr1.g00653.t01#2   mNeoVis.2.1.1.chr1_hap1.g00648.t01#5       3462   125257938:   mNeoVis.vulgivaga.2.1.chr7.g01374.t01#1    mNeoVis.evergladensis.2.1.chr7.g01363.t01#2    mNeoVis.vulgivaga.1.1.chr7.g01369.t01#4    mNeoVis.evergladensis.1.1.chr7.g01362.t01#3    mNeoVis.3.1.1.chr7.g01290.t01#6   mNeoVis.2.1.1.chr7_hap1.g01380.t01#5       3463   125257939:   mNeoVis.evergladensis.1.1.chr1.g00660.t01#3    mNeoVis.vulgivaga.2.1.chr1.g00666.t01#1    mNeoVis.evergladensis.2.1.chr1.g00654.t01#2    mNeoVis.vulgivaga.1.1.chr1.g00658.t01#4    mNeoVis.2.1.1.chr1_hap1.g00649.t01#5   mNeoVis.3.1.1.chr1.g00667.t01#6       3464   125257940:   mNeoVis.vulgivaga.1.1.chr1.g00659.t01#4    mNeoVis.2.1.1.chr1_hap1.g00650.t01#5    mNeoVis.evergladensis.1.1.chr1.g00661.t01#3    mNeoVis.evergladensis.2.1.chr1.g00655.t01#2   mNeoVis.vulgivaga.2.1.chr1.g00667.t01#1       3465   125257941:   mNeoVis.3.1.1.chr1.g00668.t01#6   mNeoVis.vulgivaga.1.1.chr1.g00660.t01#4             3466   125257942:   mNeoVis.evergladensis.2.1.chr1.g00656.t01#2    mNeoVis.vulgivaga.2.1.chr1.g00668.t01#1    mNeoVis.evergladensis.1.1.chr1.g00662.t01#3    mNeoVis.2.1.1.chr1_hap1.g00651.t01#5    mNeoVis.3.1.1.chr1.g00669.t01#6   mNeoVis.vulgivaga.1.1.chr1.g00661.t01#4       3467   125257943:   mNeoVis.evergladensis.2.1.chr7.g01364.t01#2    mNeoVis.vulgivaga.2.1.chr7.g01375.t01#1    mNeoVis.vulgivaga.1.1.chr7.g01370.t01#4    mNeoVis.evergladensis.1.1.chr7.g01363.t01#3    mNeoVis.3.1.1.chr7.g01291.t01#6   mNeoVis.2.1.1.chr7_hap1.g01381.t01#5       3468   125257944:   mNeoVis.2.1.1.chr7_hap1.g01382.t01#5    mNeoVis.3.1.1.chr7.g01292.t01#6    mNeoVis.vulgivaga.2.1.chr7.g01376.t01#1    mNeoVis.vulgivaga.1.1.chr7.g01371.t01#4    mNeoVis.evergladensis.1.1.chr7.g01364.t01#3   mNeoVis.evergladensis.2.1.chr7.g01365.t01#2       3469   125257945:  mNeoVis.vulgivaga.2.1.chr7.g01377.t01#1               3470   125257946:   mNeoVis.2.1.1.chr1_hap1.g00652.t01#5    mNeoVis.evergladensis.2.1.chr1.g00657.t01#2    mNeoVis.3.1.1.chr1.g00670.t01#6    mNeoVis.vulgivaga.2.1.chr1.g00669.t01#1    mNeoVis.vulgivaga.1.1.chr1.g00662.t01#4   mNeoVis.evergladensis.1.1.chr1.g00663.t01#3       3471   125257947:   mNeoVis.vulgivaga.1.1.chr2.g00324.t01#4    mNeoVis.evergladensis.1.1.chr2.g00327.t01#3    mNeoVis.evergladensis.2.1.chr2.g00328.t01#2    mNeoVis.3.1.1.chr2.g00267.t01#6    mNeoVis.vulgivaga.2.1.chr2.g00325.t01#1   mNeoVis.2.1.1.chr5_hap1.g00329.t01#5       3472   125257948:   mNeoVis.3.1.1.chr7.g01293.t01#6    mNeoVis.evergladensis.1.1.chr7.g01365.t01#3    mNeoVis.vulgivaga.2.1.chr7.g01378.t01#1    mNeoVis.vulgivaga.1.1.chr7.g01372.t01#4    mNeoVis.evergladensis.2.1.chr7.g01366.t01#2   mNeoVis.2.1.1.chr7_hap1.g01383.t01#5       3473   125257949:  mNeoVis.vulgivaga.2.1.chr7.g01379.t01#1               3474   125257950:   mNeoVis.vulgivaga.1.1.chr7.g00462.t01#4    mNeoVis.3.1.1.chr7.g00441.t01#6    mNeoVis.2.1.1.chr7_hap1.g00452.t01#5    mNeoVis.vulgivaga.2.1.chr7.g00458.t01#1    mNeoVis.evergladensis.1.1.chr7.g00465.t01#3   mNeoVis.evergladensis.2.1.chr7.g00459.t01#2       3475   125257951:   mNeoVis.2.1.1.chr7_hap1.g01385.t01#5    mNeoVis.vulgivaga.2.1.chr7.g01380.t01#1    mNeoVis.3.1.1.chr7.g01295.t01#6    mNeoVis.evergladensis.1.1.chr7.g01367.t01#3    mNeoVis.evergladensis.2.1.chr7.g01368.t01#2   mNeoVis.vulgivaga.1.1.chr7.g01374.t01#4       3476   125257952:   mNeoVis.3.1.1.chr1.g00671.t01#6    mNeoVis.evergladensis.1.1.chr1.g00664.t01#3    mNeoVis.2.1.1.chr1_hap1.g00653.t01#5    mNeoVis.vulgivaga.2.1.chr1.g00670.t01#1    mNeoVis.evergladensis.2.1.chr1.g00658.t01#2   mNeoVis.vulgivaga.1.1.chr1.g00663.t01#4       3477   125257953:   mNeoVis.evergladensis.1.1.chr7.g01368.t01#3    mNeoVis.3.1.1.chr7.g01296.t01#6    mNeoVis.vulgivaga.2.1.chr7.g01381.t01#1    mNeoVis.vulgivaga.1.1.chr7.g01375.t01#4    mNeoVis.evergladensis.2.1.chr7.g01369.t01#2   mNeoVis.2.1.1.chr7_hap1.g01386.t01#5       3478   125257954:   mNeoVis.vulgivaga.1.1.chr1.g00664.t01#4    mNeoVis.evergladensis.1.1.chr1.g00665.t01#3    mNeoVis.evergladensis.2.1.chr1.g00659.t01#2    mNeoVis.vulgivaga.2.1.chr1.g00671.t01#1    mNeoVis.3.1.1.chr1.g00672.t01#6   mNeoVis.2.1.1.chr1_hap1.g00654.t01#5       3479   125257955:   mNeoVis.evergladensis.2.1.chr7.g01370.t01#2    mNeoVis.3.1.1.chr7.g01297.t01#6    mNeoVis.vulgivaga.1.1.chr7.g01376.t01#4    mNeoVis.2.1.1.chr7_hap1.g01387.t01#5    mNeoVis.vulgivaga.2.1.chr7.g01382.t01#1   mNeoVis.evergladensis.1.1.chr7.g01369.t01#3       3480   125257956:   mNeoVis.2.1.1.chr7_hap1.g01388.t01#5    mNeoVis.vulgivaga.2.1.chr7.g01383.t01#1    mNeoVis.evergladensis.1.1.chr7.g01370.t01#3    mNeoVis.evergladensis.2.1.chr7.g01371.t01#2    mNeoVis.vulgivaga.1.1.chr7.g01377.t01#4   mNeoVis.3.1.1.chr7.g01298.t01#6       3481   125257957:   mNeoVis.3.1.1.chr1.g00673.t01#6    mNeoVis.2.1.1.chr1_hap1.g00655.t01#5    mNeoVis.evergladensis.2.1.chr1.g00660.t01#2    mNeoVis.vulgivaga.1.1.chr1.g00665.t01#4    mNeoVis.evergladensis.1.1.chr1.g00666.t01#3   mNeoVis.vulgivaga.2.1.chr1.g00672.t01#1       3482   125257958:   mNeoVis.evergladensis.2.1.chr1.g00661.t01#2    mNeoVis.evergladensis.1.1.chr1.g00667.t01#3    mNeoVis.vulgivaga.1.1.chr1.g00666.t01#4    mNeoVis.vulgivaga.2.1.chr1.g00673.t01#1   mNeoVis.2.1.1.chr1_hap1.g00656.t01#5       3483   125257959:   mNeoVis.vulgivaga.2.1.chr1.g00674.t01#1    mNeoVis.2.1.1.chr1_hap1.g00657.t01#5    mNeoVis.2.1.1.chr9_hap1.g00734.t01#5    mNeoVis.3.1.1.chr2.g00756.t01#6    mNeoVis.3.1.1.chr1.g00674.t01#6    mNeoVis.evergladensis.2.1.chr1.g01475.t01#2    mNeoVis.evergladensis.1.1.chr10.g00730.t01#3    mNeoVis.evergladensis.2.1.chr1.g00662.t01#2    mNeoVis.evergladensis.1.1.chr1.g00668.t01#3      3484   125257960:   mNeoVis.evergladensis.2.1.chr5.g01238.t01#2    mNeoVis.vulgivaga.1.1.chr5.g01203.t01#4    mNeoVis.vulgivaga.2.1.chr5.g01232.t01#1    mNeoVis.2.1.1.chr2_hap1.g01295.t01#5    mNeoVis.vulgivaga.1.1.chr12.g00352.t01#4    mNeoVis.3.1.1.chr12.g00350.t01#6    mNeoVis.3.1.1.chr5.g01220.t01#6    mNeoVis.evergladensis.1.1.chr5.g01237.t01#3    mNeoVis.evergladensis.1.1.chr12.g00350.t01#3      3485   125257961:   mNeoVis.vulgivaga.2.1.chr12.g00369.t01#1    mNeoVis.vulgivaga.1.1.chr1.g00667.t01#4    mNeoVis.vulgivaga.1.1.chr10.g00712.t01#4    mNeoVis.2.1.1.chr12_hap1.g00405.t01#5    mNeoVis.vulgivaga.1.1.chr12.g00371.t01#4   mNeoVis.3.1.1.chr8.g00593.t01#6       3486   125257962:   mNeoVis.evergladensis.1.1.chr5.g01610.t01#3    mNeoVis.evergladensis.2.1.chr5.g01605.t01#2   mNeoVis.3.1.1.chr5.g01587.t01#6           3487   125257963:   mNeoVis.vulgivaga.2.1.chr5.g00976.t01#1    mNeoVis.2.1.1.chr2_hap1.g01040.t01#5   mNeoVis.3.1.1.chr5.g00966.t01#6           3488   125257964:  mNeoVis.vulgivaga.2.1.chr7.g00459.t01#1               3489   125257965:   mNeoVis.evergladensis.2.1.chr7.g01372.t01#2    mNeoVis.vulgivaga.2.1.chr7.g01384.t01#1    mNeoVis.2.1.1.chr7_hap1.g01389.t01#5    mNeoVis.evergladensis.1.1.chr7.g01371.t01#3    mNeoVis.3.1.1.chr7.g01299.t01#6   mNeoVis.vulgivaga.1.1.chr7.g01378.t01#4       3490   125257966:   mNeoVis.vulgivaga.2.1.chr15.g00524.t01#1    mNeoVis.evergladensis.1.1.chr15.g00099.t01#3    mNeoVis.vulgivaga.1.1.chr15.g00515.t01#4    mNeoVis.evergladensis.2.1.chr15.g00513.t01#2    mNeoVis.2.1.1.chr15_hap1.g00691.t01#5   mNeoVis.3.1.1.chr15.g00472.t01#6       3491   125257967:   mNeoVis.vulgivaga.2.1.chr1.g00675.t01#1    mNeoVis.3.1.1.chr1.g00675.t01#6    mNeoVis.vulgivaga.1.1.chr1.g00668.t01#4    mNeoVis.evergladensis.2.1.chr1.g00663.t01#2    mNeoVis.evergladensis.1.1.chr1.g00669.t01#3   mNeoVis.2.1.1.chr1_hap1.g00658.t01#5       3492   125257968:   mNeoVis.2.1.1.chr1_hap1.g00659.t01#5    mNeoVis.3.1.1.chr1.g00676.t01#6    mNeoVis.evergladensis.2.1.chr1.g00664.t01#2    mNeoVis.vulgivaga.1.1.chr1.g00669.t01#4    mNeoVis.vulgivaga.2.1.chr1.g00676.t01#1   mNeoVis.evergladensis.1.1.chr1.g00670.t01#3       3493   125257969:   mNeoVis.3.1.1.chr1.g00677.t01#6    mNeoVis.vulgivaga.2.1.chr1.g00677.t01#1    mNeoVis.evergladensis.1.1.chr1.g00671.t01#3    mNeoVis.vulgivaga.1.1.chr1.g00670.t01#4    mNeoVis.evergladensis.2.1.chr1.g00665.t01#2   mNeoVis.2.1.1.chr1_hap1.g00660.t01#5       3494   125257970:   mNeoVis.evergladensis.1.1.chr7.g01372.t01#3    mNeoVis.evergladensis.2.1.chr7.g01373.t01#2    mNeoVis.vulgivaga.1.1.chr7.g01379.t01#4    mNeoVis.3.1.1.chr7.g01300.t01#6    mNeoVis.2.1.1.chr7_hap1.g01390.t01#5   mNeoVis.vulgivaga.2.1.chr7.g01385.t01#1       3495   125257971:   mNeoVis.vulgivaga.2.1.chr15.g00525.t01#1    mNeoVis.3.1.1.chr15.g00473.t01#6    mNeoVis.evergladensis.2.1.chr15.g00514.t01#2    mNeoVis.vulgivaga.1.1.chr15.g00516.t01#4    mNeoVis.2.1.1.chr15_hap1.g00692.t01#5   mNeoVis.evergladensis.1.1.chr15.g00098.t01#3       3496   125257972:   mNeoVis.vulgivaga.2.1.chr7.g01387.t01#1    mNeoVis.vulgivaga.1.1.chr7.g01383.t01#4    mNeoVis.evergladensis.2.1.chr7.g01376.t01#2    mNeoVis.3.1.1.chr7.g01303.t01#6    mNeoVis.evergladensis.1.1.chr7.g01374.t01#3   mNeoVis.2.1.1.chr7_hap1.g01392.t01#5       3497   125257973:  mNeoVis.vulgivaga.2.1.chr7.g00460.t01#1               3498   125257974:   mNeoVis.vulgivaga.2.1.chr1.g00678.t01#1    mNeoVis.2.1.1.chr5_hap1.g01830.t01#5    mNeoVis.evergladensis.1.1.chr2.g01826.t01#3    mNeoVis.vulgivaga.2.1.chr2.g01828.t01#1    mNeoVis.3.1.1.chr2.g01732.t01#6    mNeoVis.evergladensis.2.1.chr2.g01813.t01#2    mNeoVis.3.1.1.chr1.g00678.t01#6    mNeoVis.vulgivaga.1.1.chr2.g01819.t01#4    mNeoVis.evergladensis.2.1.chr1.g00666.t01#2      3499   125257975:   mNeoVis.vulgivaga.1.1.chr1.g00671.t01#4   mNeoVis.vulgivaga.2.1.chr1.g00679.t01#1             3500   125257976:   mNeoVis.evergladensis.1.1.chr1.g00672.t01#3    mNeoVis.2.1.1.chr1_hap1.g00661.t01#5    mNeoVis.vulgivaga.1.1.chr1.g00672.t01#4    mNeoVis.vulgivaga.2.1.chr1.g00680.t01#1    mNeoVis.evergladensis.2.1.chr1.g00667.t01#2   mNeoVis.3.1.1.chr10.g00870.t01#6       3501   125257977:   mNeoVis.vulgivaga.2.1.chr9.g01138.t01#1    mNeoVis.3.1.1.chr9.g01113.t01#6    mNeoVis.evergladensis.1.1.chr9.g01154.t01#3    mNeoVis.vulgivaga.1.1.chr9.g01157.t01#4    mNeoVis.evergladensis.2.1.chr8.g01159.t01#2   mNeoVis.2.1.1.chr10_hap1.g01181.t01#5       3502   125257978:   mNeoVis.evergladensis.2.1.chr1.g00668.t01#2    mNeoVis.vulgivaga.2.1.chr1.g00681.t01#1    mNeoVis.2.1.1.chr1_hap1.g00662.t01#5    mNeoVis.vulgivaga.1.1.chr1.g00673.t01#4    mNeoVis.3.1.1.chr1.g00679.t01#6   mNeoVis.evergladensis.1.1.chr1.g00673.t01#3       3503   125257979:   mNeoVis.evergladensis.1.1.chr7.g01375.t01#3    mNeoVis.2.1.1.chr7_hap1.g01393.t01#5    mNeoVis.vulgivaga.2.1.chr7.g01388.t01#1    mNeoVis.evergladensis.2.1.chr7.g01377.t01#2   mNeoVis.vulgivaga.1.1.chr7.g01384.t01#4       3504   125257980:   mNeoVis.evergladensis.2.1.chr1.g00669.t01#2    mNeoVis.2.1.1.chr1_hap1.g00663.t01#5    mNeoVis.vulgivaga.2.1.chr1.g00682.t01#1    mNeoVis.evergladensis.1.1.chr1.g00674.t01#3    mNeoVis.3.1.1.chr1.g00680.t01#6   mNeoVis.vulgivaga.1.1.chr1.g00674.t01#4       3505   125257981:   mNeoVis.evergladensis.1.1.chr7.g01376.t01#3    mNeoVis.2.1.1.chr7_hap1.g01394.t01#5    mNeoVis.vulgivaga.1.1.chr7.g01385.t01#4    mNeoVis.vulgivaga.2.1.chr7.g01389.t01#1    mNeoVis.3.1.1.chr7.g01306.t01#6   mNeoVis.evergladensis.2.1.chr7.g01378.t01#2       3506   125257982:   mNeoVis.evergladensis.2.1.chr7.g01379.t01#2    mNeoVis.evergladensis.1.1.chr7.g01377.t01#3    mNeoVis.2.1.1.chr7_hap1.g01395.t01#5    mNeoVis.vulgivaga.2.1.chr7.g01390.t01#1    mNeoVis.vulgivaga.1.1.chr7.g01386.t01#4   mNeoVis.3.1.1.chr7.g01307.t01#6       3507   125257983:  mNeoVis.vulgivaga.2.1.chr7.g00461.t01#1               3508   125257984:   mNeoVis.2.1.1.chr7_hap1.g01396.t01#5    mNeoVis.vulgivaga.1.1.chr7.g01387.t01#4    mNeoVis.3.1.1.chr7.g01308.t01#6    mNeoVis.evergladensis.2.1.chr7.g01380.t01#2    mNeoVis.vulgivaga.2.1.chr7.g01391.t01#1   mNeoVis.evergladensis.1.1.chr7.g01378.t01#3       3509   125257985:   mNeoVis.evergladensis.1.1.chr1.g00676.t01#3    mNeoVis.2.1.1.chr1_hap1.g00665.t01#5    mNeoVis.evergladensis.2.1.chr1.g00671.t01#2    mNeoVis.3.1.1.chr1.g00682.t01#6    mNeoVis.vulgivaga.1.1.chr1.g00676.t01#4   mNeoVis.vulgivaga.2.1.chr1.g00684.t01#1       3510   125257986:   mNeoVis.3.1.1.chr7.g01309.t01#6    mNeoVis.evergladensis.2.1.chr7.g01381.t01#2    mNeoVis.vulgivaga.1.1.chr7.g01388.t01#4    mNeoVis.2.1.1.chr7_hap1.g01397.t01#5    mNeoVis.evergladensis.1.1.chr7.g01379.t01#3   mNeoVis.vulgivaga.2.1.chr7.g01392.t01#1       3511   125257987:   mNeoVis.evergladensis.2.1.chr1.g00673.t01#2    mNeoVis.2.1.1.chr1_hap1.g00667.t01#5    mNeoVis.evergladensis.1.1.chr1.g00678.t01#3    mNeoVis.vulgivaga.1.1.chr1.g00677.t01#4    mNeoVis.vulgivaga.2.1.chr1.g00686.t01#1   mNeoVis.3.1.1.chr1.g00684.t01#6       3512   125257988:   mNeoVis.evergladensis.1.1.chr1.g00679.t01#3    mNeoVis.vulgivaga.2.1.chr1.g00687.t01#1    mNeoVis.2.1.1.chr1_hap1.g00668.t01#5    mNeoVis.evergladensis.2.1.chr1.g00674.t01#2    mNeoVis.vulgivaga.1.1.chr1.g00678.t01#4   mNeoVis.3.1.1.chr1.g00685.t01#6       3513   125257989:   mNeoVis.vulgivaga.2.1.chr7.g01393.t01#1    mNeoVis.evergladensis.1.1.chr7.g01380.t01#3    mNeoVis.2.1.1.chr7_hap1.g01398.t01#5    mNeoVis.3.1.1.chr7.g01310.t01#6    mNeoVis.evergladensis.2.1.chr7.g01382.t01#2   mNeoVis.vulgivaga.1.1.chr7.g01389.t01#4       3514   125257990:   mNeoVis.2.1.1.chr2_hap1.g00627.t01#5    mNeoVis.vulgivaga.1.1.chr15.g00517.t01#4    mNeoVis.evergladensis.1.1.chr15.g00097.t01#3    mNeoVis.3.1.1.chr5.g00625.t01#6    mNeoVis.evergladensis.2.1.chr12.g00705.t01#2    mNeoVis.vulgivaga.2.1.chr5.g00624.t01#1    mNeoVis.evergladensis.1.1.chr12.g00714.t01#3    mNeoVis.vulgivaga.1.1.chr12.g00729.t01#4    mNeoVis.evergladensis.2.1.chr15.g00515.t01#2      3515   125257991:   mNeoVis.vulgivaga.1.1.chr3.g01134.t01#4    mNeoVis.vulgivaga.2.1.chr3.g01150.t01#1    mNeoVis.evergladensis.1.1.chr3.g01145.t01#3    mNeoVis.evergladensis.2.1.chr3.g01135.t01#2    mNeoVis.3.1.1.chr3.g01124.t01#6   mNeoVis.2.1.1.chr3_hap1.g01136.t01#5       3516   125257992:   mNeoVis.evergladensis.2.1.chr7.g01268.t01#2    mNeoVis.2.1.1.chr7_hap1.g01279.t01#5    mNeoVis.3.1.1.chr7.g01203.t01#6    mNeoVis.vulgivaga.1.1.chr7.g01268.t01#4    mNeoVis.evergladensis.1.1.chr7.g01269.t01#3   mNeoVis.vulgivaga.2.1.chr7.g01281.t01#1       3517   125257993:   mNeoVis.2.1.1.chr7_hap1.g00454.t01#5    mNeoVis.vulgivaga.1.1.chr7.g00464.t01#4   mNeoVis.vulgivaga.2.1.chr7.g00462.t01#1           3518   125257994:   mNeoVis.vulgivaga.1.1.chr7.g01269.t01#4    mNeoVis.evergladensis.1.1.chr7.g01270.t01#3    mNeoVis.evergladensis.2.1.chr7.g01269.t01#2    mNeoVis.vulgivaga.2.1.chr7.g01282.t01#1    mNeoVis.2.1.1.chr7_hap1.g01280.t01#5   mNeoVis.3.1.1.chr7.g01204.t01#6       3519   125257995:   mNeoVis.vulgivaga.2.1.chr1.g00688.t01#1    mNeoVis.evergladensis.2.1.chr1.g00675.t01#2    mNeoVis.evergladensis.1.1.chr1.g00680.t01#3    mNeoVis.3.1.1.chr1.g00686.t01#6    mNeoVis.2.1.1.chr1_hap1.g00669.t01#5   mNeoVis.vulgivaga.1.1.chr1.g00679.t01#4       3520   125257996:   mNeoVis.vulgivaga.2.1.chr7.g01283.t01#1    mNeoVis.2.1.1.chr7_hap1.g01281.t01#5    mNeoVis.vulgivaga.1.1.chr7.g01270.t01#4    mNeoVis.evergladensis.1.1.chr7.g01271.t01#3    mNeoVis.3.1.1.chr7.g01206.t01#6   mNeoVis.evergladensis.2.1.chr7.g01270.t01#2       3521   125257997:   mNeoVis.2.1.1.chr7_hap1.g01282.t01#5    mNeoVis.vulgivaga.2.1.chr7.g01284.t01#1    mNeoVis.3.1.1.chr7.g01207.t01#6    mNeoVis.vulgivaga.1.1.chr7.g01271.t01#4    mNeoVis.evergladensis.2.1.chr7.g01271.t01#2   mNeoVis.evergladensis.1.1.chr7.g01272.t01#3       3522   125257998:   mNeoVis.evergladensis.1.1.chr1.g00681.t01#3    mNeoVis.vulgivaga.1.1.chr1.g00680.t01#4    mNeoVis.evergladensis.2.1.chr1.g00676.t01#2   mNeoVis.vulgivaga.2.1.chr1.g00689.t01#1         3523   125257999:   mNeoVis.3.1.1.chr1.g00687.t01#6    mNeoVis.2.1.1.chr1_hap1.g00670.t01#5   mNeoVis.evergladensis.1.1.chr1.g00682.t01#3         3524   125258000:   mNeoVis.evergladensis.2.1.chr7.g01272.t01#2    mNeoVis.vulgivaga.1.1.chr7.g01273.t01#4    mNeoVis.evergladensis.1.1.chr7.g01273.t01#3    mNeoVis.vulgivaga.2.1.chr7.g01285.t01#1    mNeoVis.3.1.1.chr7.g01208.t01#6   mNeoVis.2.1.1.chr7_hap1.g01283.t01#5       3525   125258001:   mNeoVis.vulgivaga.2.1.chr1.g00690.t01#1    mNeoVis.vulgivaga.1.1.chr1.g00681.t01#4    mNeoVis.3.1.1.chr1.g00688.t01#6    mNeoVis.evergladensis.1.1.chr1.g00683.t01#3    mNeoVis.evergladensis.2.1.chr1.g00677.t01#2   mNeoVis.2.1.1.chr1_hap1.g00671.t01#5       3526   125258002:   mNeoVis.vulgivaga.2.1.chr15.g00527.t01#1    mNeoVis.evergladensis.1.1.chr15.g00096.t01#3    mNeoVis.evergladensis.2.1.chr15.g00516.t01#2    mNeoVis.2.1.1.chr15_hap1.g00694.t01#5   mNeoVis.vulgivaga.1.1.chr15.g00518.t01#4       3527   125258003:   mNeoVis.evergladensis.2.1.chr1.g00678.t01#2    mNeoVis.evergladensis.1.1.chr1.g00684.t01#3    mNeoVis.vulgivaga.1.1.chr1.g00682.t01#4    mNeoVis.2.1.1.chr1_hap1.g00672.t01#5    mNeoVis.vulgivaga.2.1.chr1.g00691.t01#1   mNeoVis.3.1.1.chr1.g00689.t01#6       3528   125258004:   mNeoVis.evergladensis.1.1.chr1.g00685.t01#3    mNeoVis.vulgivaga.1.1.chr1.g00683.t01#4    mNeoVis.vulgivaga.2.1.chr1.g00692.t01#1    mNeoVis.2.1.1.chr1_hap1.g00673.t01#5    mNeoVis.3.1.1.chr1.g00690.t01#6   mNeoVis.evergladensis.2.1.chr1.g00679.t01#2       3529   125258005:   mNeoVis.evergladensis.1.1.chr7.g01274.t01#3    mNeoVis.vulgivaga.2.1.chr7.g01286.t01#1    mNeoVis.3.1.1.chr7.g01209.t01#6    mNeoVis.2.1.1.chr7_hap1.g01284.t01#5    mNeoVis.vulgivaga.1.1.chr7.g01274.t01#4   mNeoVis.evergladensis.2.1.chr7.g01273.t01#2       3530   125258006:   mNeoVis.3.1.1.chr1.g00694.t01#6    mNeoVis.3.1.1.chr1.g00691.t01#6    mNeoVis.2.1.1.chr1_hap1.g00674.t01#5    mNeoVis.3.1.1.chr1.g00693.t01#6    mNeoVis.vulgivaga.1.1.chr1.g00685.t01#4    mNeoVis.evergladensis.2.1.chr1.g00681.t01#2    mNeoVis.vulgivaga.2.1.chr1.g00694.t01#1    mNeoVis.evergladensis.1.1.chr1.g00686.t01#3    mNeoVis.3.1.1.chr1.g00692.t01#6      3531   125258007:   mNeoVis.vulgivaga.2.1.chr1.g00693.t01#1   mNeoVis.vulgivaga.1.1.chr1.g00684.t01#4             3532   125258008:   mNeoVis.2.1.1.chr7_hap1.g01285.t01#5    mNeoVis.vulgivaga.1.1.chr7.g01276.t01#4    mNeoVis.evergladensis.2.1.chr7.g01274.t01#2    mNeoVis.vulgivaga.2.1.chr7.g01287.t01#1   mNeoVis.3.1.1.chr7.g01210.t01#6       3533   125258009:   mNeoVis.evergladensis.2.1.chr1.g00683.t01#2    mNeoVis.2.1.1.chr1_hap1.g00675.t01#5    mNeoVis.vulgivaga.1.1.chr1.g00687.t01#4    mNeoVis.3.1.1.chr1.g00695.t01#6    mNeoVis.vulgivaga.2.1.chr1.g00695.t01#1   mNeoVis.evergladensis.1.1.chr1.g00687.t01#3       3534   125258010:   mNeoVis.vulgivaga.2.1.chr7.g01288.t01#1    mNeoVis.evergladensis.1.1.chr7.g01275.t01#3    mNeoVis.3.1.1.chr7.g01211.t01#6    mNeoVis.evergladensis.2.1.chr7.g01275.t01#2    mNeoVis.2.1.1.chr7_hap1.g01286.t01#5   mNeoVis.vulgivaga.1.1.chr7.g01277.t01#4       3535   125258011:  mNeoVis.vulgivaga.2.1.chr7.g01289.t01#1               3536   125258012:   mNeoVis.2.1.1.chr15_hap1.g00695.t01#5    mNeoVis.3.1.1.chr15.g00476.t01#6    mNeoVis.evergladensis.2.1.chr15.g00517.t01#2    mNeoVis.vulgivaga.1.1.chr15.g00519.t01#4   mNeoVis.vulgivaga.2.1.chr15.g00528.t01#1       3537   125258013:   mNeoVis.vulgivaga.1.1.chr7.g01279.t01#4    mNeoVis.2.1.1.chr7_hap1.g01288.t01#5    mNeoVis.evergladensis.1.1.chr7.g01277.t01#3    mNeoVis.vulgivaga.2.1.chr7.g01290.t01#1    mNeoVis.evergladensis.2.1.chr7.g01277.t01#2   mNeoVis.3.1.1.chr7.g01213.t01#6       3538   125258014:   mNeoVis.evergladensis.2.1.chr7.g00463.t01#2    mNeoVis.3.1.1.chr7.g00443.t01#6    mNeoVis.2.1.1.chr7_hap1.g00455.t01#5    mNeoVis.vulgivaga.1.1.chr7.g00466.t01#4    mNeoVis.vulgivaga.2.1.chr7.g00463.t01#1   mNeoVis.evergladensis.1.1.chr7.g00468.t01#3       3539   125258015:   mNeoVis.evergladensis.1.1.chr1.g00688.t01#3    mNeoVis.evergladensis.2.1.chr1.g00684.t01#2    mNeoVis.2.1.1.chr1_hap1.g00676.t01#5    mNeoVis.3.1.1.chr1.g00696.t01#6    mNeoVis.vulgivaga.1.1.chr1.g00688.t01#4   mNeoVis.vulgivaga.2.1.chr1.g00696.t01#1       3540   125258016:   mNeoVis.vulgivaga.2.1.chr1.g00697.t01#1    mNeoVis.2.1.1.chr1_hap1.g00677.t01#5    mNeoVis.3.1.1.chr1.g00697.t01#6    mNeoVis.evergladensis.2.1.chr1.g00685.t01#2   mNeoVis.evergladensis.1.1.chr1.g00689.t01#3       3541   125258017:   mNeoVis.3.1.1.chr1.g00698.t01#6    mNeoVis.vulgivaga.2.1.chr1.g00698.t01#1    mNeoVis.evergladensis.1.1.chr1.g00690.t01#3    mNeoVis.vulgivaga.1.1.chr1.g00690.t01#4    mNeoVis.evergladensis.2.1.chr1.g00686.t01#2   mNeoVis.2.1.1.chr1_hap1.g00678.t01#5       3542   125258018:   mNeoVis.2.1.1.chr1_hap1.g00679.t01#5    mNeoVis.evergladensis.2.1.chr1.g00687.t01#2    mNeoVis.vulgivaga.1.1.chr1.g00691.t01#4    mNeoVis.3.1.1.chr1.g00699.t01#6    mNeoVis.vulgivaga.2.1.chr1.g00699.t01#1   mNeoVis.evergladensis.1.1.chr1.g00691.t01#3       3543   125258019:   mNeoVis.vulgivaga.1.1.chr1.g00692.t01#4    mNeoVis.vulgivaga.2.1.chr1.g00700.t01#1    mNeoVis.evergladensis.1.1.chr1.g00692.t01#3    mNeoVis.evergladensis.2.1.chr1.g00688.t01#2    mNeoVis.2.1.1.chr1_hap1.g00680.t01#5   mNeoVis.3.1.1.chr1.g00701.t01#6       3544   125258020:   mNeoVis.vulgivaga.1.1.chr1.g00693.t01#4    mNeoVis.vulgivaga.2.1.chr1.g00701.t01#1    mNeoVis.evergladensis.2.1.chr1.g00689.t01#2    mNeoVis.evergladensis.1.1.chr1.g00693.t01#3    mNeoVis.2.1.1.chr1_hap1.g00681.t01#5   mNeoVis.3.1.1.chr1.g00702.t01#6       3545   125258021:   mNeoVis.3.1.1.chr7.g00444.t01#6    mNeoVis.vulgivaga.2.1.chr7.g00464.t01#1   mNeoVis.2.1.1.chr7_hap1.g00456.t01#5           3546   125258022:   mNeoVis.evergladensis.1.1.chr1.g00694.t01#3    mNeoVis.vulgivaga.2.1.chr1.g00702.t01#1    mNeoVis.evergladensis.2.1.chr1.g00690.t01#2    mNeoVis.vulgivaga.1.1.chr1.g00694.t01#4    mNeoVis.3.1.1.chr1.g00703.t01#6   mNeoVis.2.1.1.chr1_hap1.g00682.t01#5       3547   125258023:   mNeoVis.3.1.1.chr1.g00704.t01#6    mNeoVis.2.1.1.chr1_hap1.g00683.t01#5    mNeoVis.vulgivaga.1.1.chr1.g00695.t01#4    mNeoVis.evergladensis.2.1.chr1.g00691.t01#2    mNeoVis.evergladensis.1.1.chr1.g00695.t01#3   mNeoVis.vulgivaga.2.1.chr1.g00703.t01#1       3548   125258024:   mNeoVis.vulgivaga.1.1.chr7.g01280.t01#4    mNeoVis.evergladensis.1.1.chr7.g01278.t01#3    mNeoVis.2.1.1.chr7_hap1.g01290.t01#5    mNeoVis.evergladensis.2.1.chr7.g01279.t01#2    mNeoVis.vulgivaga.2.1.chr7.g01291.t01#1   mNeoVis.3.1.1.chr7.g01214.t01#6       3549   125258025:   mNeoVis.vulgivaga.2.1.chr7.g01292.t01#1    mNeoVis.evergladensis.2.1.chr7.g01280.t01#2    mNeoVis.vulgivaga.1.1.chr7.g01281.t01#4    mNeoVis.evergladensis.1.1.chr7.g01279.t01#3    mNeoVis.2.1.1.chr7_hap1.g01291.t01#5   mNeoVis.3.1.1.chr7.g01215.t01#6       3550   125258026:   mNeoVis.3.1.1.chr1.g00700.t01#6    mNeoVis.2.1.1.chr1_hap1.g00684.t01#5    mNeoVis.vulgivaga.1.1.chr1.g00696.t01#4    mNeoVis.vulgivaga.2.1.chr1.g00704.t01#1    mNeoVis.evergladensis.1.1.chr1.g00696.t01#3   mNeoVis.evergladensis.2.1.chr1.g00692.t01#2       3551   125258027:   mNeoVis.3.1.1.chr1.g00620.t01#6    mNeoVis.2.1.1.chr1_hap1.g00603.t01#5    mNeoVis.vulgivaga.2.1.chr1.g00617.t01#1    mNeoVis.evergladensis.2.1.chr1.g00608.t01#2    mNeoVis.evergladensis.1.1.chr1.g00613.t01#3   mNeoVis.vulgivaga.1.1.chr1.g00613.t01#4       3552   125258028:   mNeoVis.evergladensis.1.1.chr7.g01280.t01#3    mNeoVis.vulgivaga.1.1.chr7.g01283.t01#4    mNeoVis.3.1.1.chr7.g01216.t01#6    mNeoVis.2.1.1.chr7_hap1.g01292.t01#5    mNeoVis.evergladensis.2.1.chr7.g01281.t01#2   mNeoVis.vulgivaga.2.1.chr7.g01293.t01#1       3553   125258029:   mNeoVis.2.1.1.chr7_hap1.g01293.t01#5    mNeoVis.evergladensis.2.1.chr7.g01282.t01#2    mNeoVis.evergladensis.1.1.chr7.g01281.t01#3    mNeoVis.vulgivaga.2.1.chr7.g01294.t01#1   mNeoVis.3.1.1.chr7.g01217.t01#6       3554   125258030:   mNeoVis.vulgivaga.1.1.chr15.g00520.t01#4    mNeoVis.vulgivaga.2.1.chr15.g00529.t01#1    mNeoVis.2.1.1.chr15_hap1.g00696.t01#5    mNeoVis.evergladensis.2.1.chr15.g00518.t01#2    mNeoVis.evergladensis.1.1.chr15.g00095.t01#3   mNeoVis.3.1.1.chr15.g00477.t01#6       3555   125258031:   mNeoVis.vulgivaga.2.1.chr1.g00618.t01#1    mNeoVis.2.1.1.chr1_hap1.g00604.t01#5    mNeoVis.evergladensis.2.1.chr1.g00609.t01#2    mNeoVis.vulgivaga.1.1.chr1.g00614.t01#4    mNeoVis.3.1.1.chr1.g00621.t01#6   mNeoVis.evergladensis.1.1.chr1.g00614.t01#3       3556   125258032:   mNeoVis.vulgivaga.2.1.chr7.g00465.t01#1    mNeoVis.vulgivaga.1.1.chr7.g00468.t01#4   mNeoVis.evergladensis.2.1.chr7.g00465.t01#2         3557   125258033:   mNeoVis.vulgivaga.1.1.chr1.g00616.t01#4    mNeoVis.evergladensis.2.1.chr1.g00610.t01#2    mNeoVis.3.1.1.chr1.g00622.t01#6    mNeoVis.vulgivaga.2.1.chr1.g00619.t01#1   mNeoVis.2.1.1.chr1_hap1.g00605.t01#5       3558   125258034:   mNeoVis.2.1.1.chr7_hap1.g01295.t01#5    mNeoVis.vulgivaga.2.1.chr7.g01296.t01#1    mNeoVis.evergladensis.2.1.chr7.g01284.t01#2    mNeoVis.3.1.1.chr7.g01219.t01#6    mNeoVis.evergladensis.1.1.chr7.g01283.t01#3   mNeoVis.vulgivaga.1.1.chr7.g01286.t01#4       3559   125258035:   mNeoVis.2.1.1.chr7_hap1.g01296.t01#5    mNeoVis.vulgivaga.1.1.chr7.g01288.t01#4    mNeoVis.evergladensis.2.1.chr7.g01285.t01#2    mNeoVis.vulgivaga.2.1.chr7.g01297.t01#1    mNeoVis.evergladensis.1.1.chr7.g01284.t01#3    mNeoVis.vulgivaga.1.1.chr7.g01287.t01#4    mNeoVis.3.1.1.chr7.g01220.t01#6   mNeoVis.3.1.1.chr7.g01221.t01#6     3560   125258036:   mNeoVis.vulgivaga.1.1.chr1.g00617.t01#4    mNeoVis.evergladensis.2.1.chr1.g00611.t01#2    mNeoVis.3.1.1.chr1.g00623.t01#6    mNeoVis.2.1.1.chr1_hap1.g00606.t01#5   mNeoVis.vulgivaga.2.1.chr1.g00620.t01#1       3561   125258037:   mNeoVis.vulgivaga.2.1.chr1.g00621.t01#1    mNeoVis.vulgivaga.1.1.chr1.g00618.t01#4    mNeoVis.evergladensis.2.1.chr1.g00612.t01#2    mNeoVis.3.1.1.chr1.g00624.t01#6    mNeoVis.evergladensis.1.1.chr1.g00616.t01#3   mNeoVis.2.1.1.chr1_hap1.g00607.t01#5       3562   125258038:   mNeoVis.2.1.1.chr7_hap1.g01297.t01#5    mNeoVis.3.1.1.chr7.g01222.t01#6    mNeoVis.vulgivaga.1.1.chr7.g01289.t01#4    mNeoVis.evergladensis.2.1.chr7.g01286.t01#2    mNeoVis.vulgivaga.2.1.chr7.g01298.t01#1   mNeoVis.evergladensis.1.1.chr7.g01285.t01#3       3563   125258039:   mNeoVis.vulgivaga.2.1.chr7.g01299.t01#1    mNeoVis.2.1.1.chr7_hap1.g01298.t01#5    mNeoVis.evergladensis.2.1.chr7.g01287.t01#2    mNeoVis.vulgivaga.1.1.chr7.g01290.t01#4    mNeoVis.evergladensis.1.1.chr7.g01286.t01#3   mNeoVis.3.1.1.chr7.g01223.t01#6       3564   125258040:   mNeoVis.evergladensis.2.1.chr1.g00614.t01#2    mNeoVis.3.1.1.chr1.g00625.t01#6    mNeoVis.2.1.1.chr1_hap1.g00608.t01#5    mNeoVis.vulgivaga.1.1.chr1.g00619.t01#4    mNeoVis.evergladensis.1.1.chr1.g00617.t01#3    mNeoVis.vulgivaga.2.1.chr1.g00623.t01#1    mNeoVis.vulgivaga.2.1.chr1.g00622.t01#1   mNeoVis.evergladensis.2.1.chr1.g00613.t01#2     3565   125258041:   mNeoVis.3.1.1.chr7.g01224.t01#6    mNeoVis.2.1.1.chr7_hap1.g01299.t01#5    mNeoVis.evergladensis.2.1.chr7.g01288.t01#2    mNeoVis.evergladensis.1.1.chr7.g01287.t01#3    mNeoVis.vulgivaga.1.1.chr7.g01291.t01#4   mNeoVis.vulgivaga.2.1.chr7.g01300.t01#1       3566   125258042:   mNeoVis.2.1.1.chr1_hap1.g00609.t01#5    mNeoVis.vulgivaga.2.1.chr1.g00624.t01#1    mNeoVis.evergladensis.2.1.chr1.g00615.t01#2    mNeoVis.evergladensis.1.1.chr1.g00618.t01#3    mNeoVis.vulgivaga.1.1.chr1.g00620.t01#4   mNeoVis.3.1.1.chr1.g00627.t01#6       3567   125258043:   mNeoVis.evergladensis.2.1.chr1.g00616.t01#2    mNeoVis.2.1.1.chr1_hap1.g00610.t01#5    mNeoVis.vulgivaga.2.1.chr1.g00625.t01#1    mNeoVis.evergladensis.1.1.chr1.g00619.t01#3   mNeoVis.vulgivaga.1.1.chr1.g00621.t01#4       3568   125258044:   mNeoVis.vulgivaga.2.1.chr1.g00626.t01#1    mNeoVis.3.1.1.chr1.g00629.t01#6    mNeoVis.evergladensis.1.1.chr1.g00620.t01#3    mNeoVis.vulgivaga.1.1.chr1.g00622.t01#4    mNeoVis.2.1.1.chr1_hap1.g00611.t01#5   mNeoVis.evergladensis.2.1.chr1.g00617.t01#2       3569   125258045:   mNeoVis.2.1.1.chr7_hap1.g01304.t01#5    mNeoVis.evergladensis.1.1.chr7.g01291.t01#3    mNeoVis.evergladensis.2.1.chr7.g01292.t01#2    mNeoVis.vulgivaga.2.1.chr7.g01304.t01#1    mNeoVis.vulgivaga.1.1.chr7.g01295.t01#4   mNeoVis.3.1.1.chr7.g01227.t01#6       3570   125258046:   mNeoVis.evergladensis.1.1.chr7.g01289.t01#3    mNeoVis.3.1.1.chr7.g01226.t01#6    mNeoVis.vulgivaga.1.1.chr7.g01293.t01#4    mNeoVis.vulgivaga.2.1.chr7.g01302.t01#1    mNeoVis.2.1.1.chr7_hap1.g01303.t01#5    mNeoVis.evergladensis.2.1.chr7.g01291.t01#2    mNeoVis.2.1.1.chr7_hap1.g01302.t01#5    mNeoVis.evergladensis.1.1.chr7.g01290.t01#3    mNeoVis.vulgivaga.2.1.chr7.g01303.t01#1      3571   125258047:   mNeoVis.2.1.1.chr7_hap1.g01300.t01#5    mNeoVis.evergladensis.2.1.chr7.g01289.t01#2    mNeoVis.evergladensis.1.1.chr7.g01288.t01#3    mNeoVis.3.1.1.chr7.g01225.t01#6    mNeoVis.vulgivaga.2.1.chr7.g01301.t01#1   mNeoVis.vulgivaga.1.1.chr7.g01292.t01#4       3572   125258048:   mNeoVis.evergladensis.1.1.chr7.g01292.t01#3    mNeoVis.vulgivaga.2.1.chr7.g01305.t01#1    mNeoVis.3.1.1.chr7.g01228.t01#6    mNeoVis.vulgivaga.1.1.chr7.g01296.t01#4    mNeoVis.evergladensis.2.1.chr7.g01293.t01#2   mNeoVis.2.1.1.chr7_hap1.g01305.t01#5       3573   125258049:   mNeoVis.evergladensis.2.1.chr7.g01294.t01#2    mNeoVis.2.1.1.chr7_hap1.g01306.t01#5    mNeoVis.vulgivaga.2.1.chr7.g01306.t01#1    mNeoVis.evergladensis.1.1.chr7.g01293.t01#3    mNeoVis.vulgivaga.1.1.chr7.g01297.t01#4   mNeoVis.3.1.1.chr7.g01229.t01#6       3574   125258050:   mNeoVis.evergladensis.1.1.chr7.g01294.t01#3    mNeoVis.vulgivaga.1.1.chr7.g01298.t01#4    mNeoVis.3.1.1.chr7.g01230.t01#6    mNeoVis.evergladensis.2.1.chr7.g01295.t01#2    mNeoVis.2.1.1.chr7_hap1.g01307.t01#5   mNeoVis.vulgivaga.2.1.chr7.g01307.t01#1       3575   125258051:   mNeoVis.evergladensis.1.1.chr15.g00094.t01#3    mNeoVis.vulgivaga.1.1.chr15.g00522.t01#4    mNeoVis.evergladensis.2.1.chr15.g00519.t01#2    mNeoVis.2.1.1.chr15_hap1.g00697.t01#5   mNeoVis.vulgivaga.2.1.chr15.g00530.t01#1       3576   125258052:   mNeoVis.2.1.1.chr1_hap1.g00612.t01#5    mNeoVis.evergladensis.1.1.chr1.g00621.t01#3    mNeoVis.vulgivaga.2.1.chr1.g00627.t01#1    mNeoVis.vulgivaga.1.1.chr1.g00624.t01#4    mNeoVis.3.1.1.chr1.g00630.t01#6   mNeoVis.evergladensis.2.1.chr1.g00618.t01#2       3577   125258053:   mNeoVis.3.1.1.chr7.g01231.t01#6    mNeoVis.evergladensis.1.1.chr7.g01296.t01#3    mNeoVis.evergladensis.2.1.chr7.g01297.t01#2    mNeoVis.vulgivaga.2.1.chr7.g01308.t01#1    mNeoVis.vulgivaga.1.1.chr7.g01300.t01#4   mNeoVis.2.1.1.chr7_hap1.g01309.t01#5       3578   125258054:   mNeoVis.vulgivaga.1.1.chr7.g01303.t01#4    mNeoVis.vulgivaga.1.1.chr7.g01301.t01#4    mNeoVis.2.1.1.chr7_hap1.g01310.t01#5    mNeoVis.2.1.1.chr7_hap1.g01312.t01#5    mNeoVis.vulgivaga.2.1.chr7.g01310.t01#1   mNeoVis.vulgivaga.2.1.chr7.g01312.t01#1       3579   125258055:   mNeoVis.vulgivaga.2.1.chr7.g01309.t01#1    mNeoVis.vulgivaga.1.1.chr7.g01299.t01#4    mNeoVis.evergladensis.1.1.chr7.g01295.t01#3   mNeoVis.3.1.1.chr7.g01232.t01#6         3580   125258056:   mNeoVis.2.1.1.chr7_hap1.g01308.t01#5   mNeoVis.evergladensis.2.1.chr7.g01296.t01#2           3581   125258057:   mNeoVis.3.1.1.chr7.g01234.t01#6   mNeoVis.vulgivaga.1.1.chr7.g01302.t01#4             3582   125258058:   mNeoVis.2.1.1.chr7_hap1.g01311.t01#5    mNeoVis.vulgivaga.2.1.chr7.g01311.t01#1    mNeoVis.evergladensis.1.1.chr7.g01297.t01#3   mNeoVis.evergladensis.2.1.chr7.g01298.t01#2       3583   125258059:  mNeoVis.vulgivaga.2.1.chr1.g00628.t01#1               3584   125258060:   mNeoVis.evergladensis.2.1.chr7.g01299.t01#2    mNeoVis.vulgivaga.2.1.chr7.g01313.t01#1    mNeoVis.2.1.1.chr7_hap1.g01313.t01#5    mNeoVis.evergladensis.1.1.chr7.g01298.t01#3    mNeoVis.3.1.1.chr7.g01235.t01#6   mNeoVis.vulgivaga.1.1.chr7.g01304.t01#4       3585   125258061:   mNeoVis.3.1.1.chr7.g01236.t01#6    mNeoVis.2.1.1.chr7_hap1.g01314.t01#5    mNeoVis.evergladensis.1.1.chr7.g01299.t01#3    mNeoVis.evergladensis.2.1.chr7.g01300.t01#2    mNeoVis.vulgivaga.1.1.chr7.g01305.t01#4   mNeoVis.vulgivaga.2.1.chr7.g01314.t01#1       3586   125258062:   mNeoVis.vulgivaga.1.1.chr7.g00469.t01#4    mNeoVis.2.1.1.chr7_hap1.g00458.t01#5    mNeoVis.evergladensis.1.1.chr7.g00471.t01#3    mNeoVis.3.1.1.chr7.g00446.t01#6    mNeoVis.evergladensis.2.1.chr7.g00466.t01#2   mNeoVis.vulgivaga.2.1.chr7.g00466.t01#1       3587   125258063:   mNeoVis.vulgivaga.1.1.chr1.g00625.t01#4    mNeoVis.evergladensis.2.1.chr1.g00620.t01#2    mNeoVis.2.1.1.chr1_hap1.g00613.t01#5    mNeoVis.vulgivaga.2.1.chr1.g00629.t01#1    mNeoVis.evergladensis.1.1.chr1.g00622.t01#3   mNeoVis.3.1.1.chr1.g00632.t01#6       3588   125258064:  mNeoVis.vulgivaga.2.1.chr7.g01315.t01#1               3589   125258065:  mNeoVis.vulgivaga.2.1.chr7.g01317.t01#1               3590   125258066:   mNeoVis.evergladensis.2.1.chr7.g01303.t01#2    mNeoVis.vulgivaga.2.1.chr7.g01318.t01#1    mNeoVis.3.1.1.chr7.g01237.t01#6    mNeoVis.vulgivaga.1.1.chr7.g01307.t01#4    mNeoVis.vulgivaga.1.1.chr7.g01306.t01#4    mNeoVis.evergladensis.1.1.chr7.g01301.t01#3    mNeoVis.2.1.1.chr7_hap1.g01316.t01#5   mNeoVis.vulgivaga.2.1.chr7.g01316.t01#1     3591   125258067:   mNeoVis.3.1.1.chr1.g00633.t01#6    mNeoVis.vulgivaga.1.1.chr1.g00626.t01#4    mNeoVis.2.1.1.chr1_hap1.g00614.t01#5    mNeoVis.vulgivaga.2.1.chr1.g00630.t01#1    mNeoVis.evergladensis.1.1.chr1.g00623.t01#3   mNeoVis.evergladensis.2.1.chr1.g00621.t01#2       3592   125258068:   mNeoVis.vulgivaga.2.1.chr7.g01319.t01#1    mNeoVis.2.1.1.chr7_hap1.g01322.t01#5    mNeoVis.evergladensis.1.1.chr7.g01305.t01#3    mNeoVis.evergladensis.2.1.chr7.g01304.t01#2   mNeoVis.2.1.1.chr7_hap1.g01318.t01#5       3593   125258069:   mNeoVis.3.1.1.chr1.g00634.t01#6    mNeoVis.2.1.1.chr1_hap1.g00615.t01#5    mNeoVis.vulgivaga.1.1.chr1.g00627.t01#4    mNeoVis.evergladensis.2.1.chr1.g00622.t01#2    mNeoVis.evergladensis.1.1.chr1.g00624.t01#3   mNeoVis.vulgivaga.2.1.chr1.g00631.t01#1       3594   125258070:   mNeoVis.2.1.1.chr1_hap1.g00616.t01#5    mNeoVis.vulgivaga.1.1.chr1.g00628.t01#4    mNeoVis.vulgivaga.2.1.chr1.g00632.t01#1    mNeoVis.evergladensis.1.1.chr1.g00625.t01#3    mNeoVis.evergladensis.2.1.chr1.g00623.t01#2   mNeoVis.3.1.1.chr1.g00635.t01#6       3595   125258071:   mNeoVis.3.1.1.chr7.g01246.t01#6    mNeoVis.evergladensis.2.1.chr7.g01314.t01#2    mNeoVis.evergladensis.2.1.chr7.g01318.t01#2    mNeoVis.evergladensis.1.1.chr7.g01315.t01#3    mNeoVis.evergladensis.2.1.chr7.g01313.t01#2    mNeoVis.evergladensis.2.1.chr7.g01316.t01#2    mNeoVis.evergladensis.1.1.chr7.g01314.t01#3    mNeoVis.evergladensis.1.1.chr7.g01313.t01#3    mNeoVis.evergladensis.2.1.chr7.g01315.t01#2      3596   125258072:   mNeoVis.evergladensis.1.1.chr7.g01302.t01#3    mNeoVis.vulgivaga.2.1.chr7.g01322.t01#1    mNeoVis.evergladensis.2.1.chr7.g01305.t01#2    mNeoVis.vulgivaga.1.1.chr7.g01309.t01#4    mNeoVis.3.1.1.chr7.g01240.t01#6    mNeoVis.2.1.1.chr7_hap1.g01320.t01#5    mNeoVis.vulgivaga.1.1.chr7.g01308.t01#4    mNeoVis.3.1.1.chr7.g01238.t01#6    mNeoVis.evergladensis.2.1.chr7.g01307.t01#2      3597   125258073:   mNeoVis.evergladensis.2.1.chr7.g01322.t01#2    mNeoVis.2.1.1.chr7_hap1.g01340.t01#5    mNeoVis.vulgivaga.2.1.chr7.g01335.t01#1    mNeoVis.2.1.1.chr7_hap1.g01339.t01#5    mNeoVis.vulgivaga.1.1.chr7.g01329.t01#4    mNeoVis.evergladensis.1.1.chr7.g01321.t01#3    mNeoVis.vulgivaga.1.1.chr7.g01328.t01#4   mNeoVis.evergladensis.2.1.chr7.g01323.t01#2     3598   125258074:   mNeoVis.2.1.1.chr7_hap1.g01323.t01#5    mNeoVis.evergladensis.2.1.chr7.g01310.t01#2    mNeoVis.evergladensis.1.1.chr7.g01304.t01#3    mNeoVis.vulgivaga.1.1.chr7.g01313.t01#4    mNeoVis.evergladensis.2.1.chr7.g01309.t01#2    mNeoVis.3.1.1.chr7.g01242.t01#6   mNeoVis.vulgivaga.2.1.chr7.g01323.t01#1     3599   125258075:   mNeoVis.evergladensis.1.1.chr7.g01306.t01#3    mNeoVis.vulgivaga.2.1.chr7.g01324.t01#1    mNeoVis.2.1.1.chr7_hap1.g01324.t01#5    mNeoVis.3.1.1.chr7.g01243.t01#6    mNeoVis.evergladensis.2.1.chr7.g01311.t01#2   mNeoVis.vulgivaga.1.1.chr7.g01314.t01#4       3600   125258076:   mNeoVis.evergladensis.2.1.chr7.g01306.t01#2    mNeoVis.3.1.1.chr7.g01239.t01#6    mNeoVis.evergladensis.1.1.chr7.g01303.t01#3    mNeoVis.vulgivaga.1.1.chr7.g01311.t01#4   mNeoVis.vulgivaga.1.1.chr7.g01310.t01#4       3601   125258077:   mNeoVis.2.1.1.chr7_hap1.g01334.t01#5    mNeoVis.3.1.1.chr7.g01241.t01#6    mNeoVis.vulgivaga.2.1.chr7.g01332.t01#1   mNeoVis.vulgivaga.1.1.chr7.g01325.t01#4         3602   125258078:   mNeoVis.2.1.1.chr7_hap1.g01336.t01#5    mNeoVis.evergladensis.1.1.chr7.g01318.t01#3    mNeoVis.vulgivaga.1.1.chr7.g01326.t01#4   mNeoVis.evergladensis.2.1.chr7.g01320.t01#2       3603   125258079:   mNeoVis.vulgivaga.2.1.chr7.g01321.t01#1    mNeoVis.evergladensis.2.1.chr7.g01308.t01#2   mNeoVis.2.1.1.chr7_hap1.g01321.t01#5           3604   125258080:   mNeoVis.vulgivaga.2.1.chr7.g01333.t01#1    mNeoVis.2.1.1.chr7_hap1.g01335.t01#5   mNeoVis.evergladensis.1.1.chr7.g01317.t01#3         3605   125258081:   mNeoVis.3.1.1.chr7.g01249.t01#6    mNeoVis.evergladensis.1.1.chr7.g01322.t01#3   mNeoVis.vulgivaga.2.1.chr7.g01336.t01#1           3606   125258082:  mNeoVis.vulgivaga.1.1.chr7.g01312.t01#4               3607   125258083:  mNeoVis.evergladensis.1.1.chr7.g01316.t01#3             3608   125258084:   mNeoVis.vulgivaga.2.1.chr1.g00633.t01#1    mNeoVis.evergladensis.2.1.chr1.g00624.t01#2    mNeoVis.2.1.1.chr1_hap1.g00617.t01#5    mNeoVis.3.1.1.chr1.g00636.t01#6    mNeoVis.vulgivaga.1.1.chr1.g00629.t01#4   mNeoVis.evergladensis.1.1.chr1.g00626.t01#3       3609   125258085:   mNeoVis.2.1.1.chr7_hap1.g01325.t01#5    mNeoVis.evergladensis.2.1.chr7.g01312.t01#2    mNeoVis.3.1.1.chr7.g01244.t01#6    mNeoVis.vulgivaga.1.1.chr7.g01315.t01#4    mNeoVis.evergladensis.1.1.chr7.g01307.t01#3   mNeoVis.vulgivaga.2.1.chr7.g01325.t01#1       3610   125258086:   mNeoVis.evergladensis.1.1.chr1.g00627.t01#3    mNeoVis.2.1.1.chr1_hap1.g00618.t01#5    mNeoVis.3.1.1.chr1.g00637.t01#6    mNeoVis.vulgivaga.2.1.chr1.g00634.t01#1   mNeoVis.evergladensis.2.1.chr1.g00625.t01#2       3611   125258087:   mNeoVis.vulgivaga.1.1.chr1.g00630.t01#4    mNeoVis.evergladensis.2.1.chr1.g00626.t01#2    mNeoVis.vulgivaga.2.1.chr1.g00635.t01#1    mNeoVis.2.1.1.chr1_hap1.g00619.t01#5    mNeoVis.3.1.1.chr1.g00638.t01#6   mNeoVis.evergladensis.1.1.chr1.g00628.t01#3       3612   125258088:   mNeoVis.vulgivaga.2.1.chr1.g00636.t01#1    mNeoVis.3.1.1.chr1.g00639.t01#6    mNeoVis.evergladensis.2.1.chr1.g00627.t01#2    mNeoVis.2.1.1.chr1_hap1.g00620.t01#5    mNeoVis.evergladensis.1.1.chr1.g00629.t01#3   mNeoVis.vulgivaga.1.1.chr1.g00631.t01#4       3613   125258089:   mNeoVis.2.1.1.chr7_hap1.g01337.t01#5    mNeoVis.evergladensis.1.1.chr7.g01319.t01#3    mNeoVis.evergladensis.2.1.chr7.g01321.t01#2    mNeoVis.vulgivaga.2.1.chr7.g01334.t01#1   mNeoVis.vulgivaga.1.1.chr7.g01327.t01#4       3614   125258090:   mNeoVis.2.1.1.chr1_hap1.g00622.t01#5    mNeoVis.evergladensis.2.1.chr1.g00628.t01#2    mNeoVis.3.1.1.chr1.g00640.t01#6    mNeoVis.evergladensis.1.1.chr1.g00632.t01#3    mNeoVis.vulgivaga.1.1.chr1.g00632.t01#4   mNeoVis.vulgivaga.2.1.chr1.g00637.t01#1       3615   125258091:   mNeoVis.vulgivaga.1.1.chr7.g01330.t01#4    mNeoVis.evergladensis.1.1.chr7.g01324.t01#3    mNeoVis.vulgivaga.2.1.chr7.g01337.t01#1    mNeoVis.evergladensis.2.1.chr7.g01324.t01#2    mNeoVis.3.1.1.chr7.g01251.t01#6   mNeoVis.2.1.1.chr7_hap1.g01341.t01#5       3616   125258092:   mNeoVis.2.1.1.chr7_hap1.g01342.t01#5    mNeoVis.3.1.1.chr7.g01252.t01#6    mNeoVis.vulgivaga.1.1.chr7.g01331.t01#4    mNeoVis.evergladensis.1.1.chr7.g01325.t01#3    mNeoVis.vulgivaga.2.1.chr7.g01338.t01#1   mNeoVis.evergladensis.2.1.chr7.g01325.t01#2       3617   125258093:   mNeoVis.vulgivaga.1.1.chr7.g01332.t01#4    mNeoVis.3.1.1.chr7.g01253.t01#6    mNeoVis.evergladensis.1.1.chr7.g01326.t01#3    mNeoVis.evergladensis.2.1.chr7.g01326.t01#2    mNeoVis.2.1.1.chr7_hap1.g01343.t01#5   mNeoVis.vulgivaga.2.1.chr7.g01339.t01#1       3618   125258094:   mNeoVis.evergladensis.2.1.chr7.g01327.t01#2    mNeoVis.vulgivaga.2.1.chr7.g01340.t01#1    mNeoVis.vulgivaga.1.1.chr7.g01333.t01#4    mNeoVis.2.1.1.chr7_hap1.g01344.t01#5    mNeoVis.evergladensis.1.1.chr7.g01327.t01#3   mNeoVis.3.1.1.chr7.g01254.t01#6       3619   125258095:   mNeoVis.evergladensis.1.1.chr1.g00633.t01#3    mNeoVis.vulgivaga.1.1.chr1.g00633.t01#4    mNeoVis.3.1.1.chr1.g00641.t01#6    mNeoVis.2.1.1.chr1_hap1.g00623.t01#5    mNeoVis.vulgivaga.2.1.chr1.g00638.t01#1   mNeoVis.evergladensis.2.1.chr1.g00629.t01#2       3620   125258096:   mNeoVis.3.1.1.chr7.g01255.t01#6    mNeoVis.evergladensis.1.1.chr7.g01328.t01#3    mNeoVis.vulgivaga.1.1.chr7.g01334.t01#4    mNeoVis.evergladensis.2.1.chr7.g01328.t01#2    mNeoVis.2.1.1.chr7_hap1.g01345.t01#5   mNeoVis.vulgivaga.2.1.chr7.g01341.t01#1       3621   125258097:   mNeoVis.vulgivaga.2.1.chr7.g01342.t01#1    mNeoVis.evergladensis.2.1.chr7.g01329.t01#2    mNeoVis.2.1.1.chr7_hap1.g01346.t01#5    mNeoVis.3.1.1.chr7.g01256.t01#6    mNeoVis.vulgivaga.1.1.chr7.g01335.t01#4   mNeoVis.evergladensis.1.1.chr7.g01329.t01#3       3622   125258098:   mNeoVis.3.1.1.chr7.g00447.t01#6    mNeoVis.vulgivaga.2.1.chr7.g00467.t01#1    mNeoVis.vulgivaga.1.1.chr7.g00470.t01#4    mNeoVis.2.1.1.chr7_hap1.g00459.t01#5    mNeoVis.evergladensis.2.1.chr7.g00467.t01#2   mNeoVis.evergladensis.1.1.chr7.g00472.t01#3       3623   125258099:   mNeoVis.vulgivaga.2.1.chr1.g00639.t01#1    mNeoVis.3.1.1.chr8.g01355.t01#6    mNeoVis.3.1.1.chr1.g00642.t01#6    mNeoVis.evergladensis.2.1.chr9.g01406.t01#2    mNeoVis.vulgivaga.2.1.chr8.g01410.t01#1    mNeoVis.2.1.1.chr1_hap1.g00624.t01#5    mNeoVis.evergladensis.1.1.chr1.g00634.t01#3    mNeoVis.evergladensis.1.1.chr8.g01404.t01#3    mNeoVis.2.1.1.chr8_hap1.g01459.t01#5      3624   125258100:   mNeoVis.vulgivaga.2.1.chr1.g00640.t01#1    mNeoVis.3.1.1.chr1.g00643.t01#6   mNeoVis.evergladensis.1.1.chr1.g00635.t01#3         3625   125258101:  mNeoVis.vulgivaga.2.1.chr7.g01343.t01#1               3626   125258102:   mNeoVis.evergladensis.2.1.chr7.g01331.t01#2    mNeoVis.2.1.1.chr7_hap1.g01348.t01#5    mNeoVis.evergladensis.1.1.chr7.g01331.t01#3    mNeoVis.vulgivaga.2.1.chr7.g01344.t01#1    mNeoVis.vulgivaga.1.1.chr7.g01337.t01#4   mNeoVis.3.1.1.chr7.g01258.t01#6       3627   125258103:   mNeoVis.vulgivaga.1.1.chr1.g00635.t01#4    mNeoVis.evergladensis.1.1.chr1.g00636.t01#3    mNeoVis.2.1.1.chr1_hap1.g00625.t01#5    mNeoVis.evergladensis.2.1.chr1.g00631.t01#2   mNeoVis.vulgivaga.2.1.chr1.g00641.t01#1       3628   125258104:   mNeoVis.evergladensis.2.1.chr7.g01332.t01#2    mNeoVis.vulgivaga.2.1.chr7.g01345.t01#1    mNeoVis.evergladensis.1.1.chr7.g01332.t01#3    mNeoVis.3.1.1.chr7.g01259.t01#6   mNeoVis.2.1.1.chr7_hap1.g01349.t01#5       3629   125258105:   mNeoVis.3.1.1.chr1.g00645.t01#6    mNeoVis.vulgivaga.1.1.chr1.g00636.t01#4    mNeoVis.evergladensis.1.1.chr1.g00637.t01#3    mNeoVis.vulgivaga.2.1.chr1.g00642.t01#1    mNeoVis.2.1.1.chr1_hap1.g00626.t01#5   mNeoVis.evergladensis.2.1.chr1.g00632.t01#2       3630   125258106:   mNeoVis.3.1.1.chr7.g01260.t01#6    mNeoVis.vulgivaga.1.1.chr7.g01340.t01#4    mNeoVis.2.1.1.chr7_hap1.g01350.t01#5    mNeoVis.evergladensis.2.1.chr7.g01333.t01#2    mNeoVis.evergladensis.1.1.chr7.g01333.t01#3   mNeoVis.vulgivaga.2.1.chr7.g01346.t01#1       3631   125258107:   mNeoVis.3.1.1.chr1.g00646.t01#6    mNeoVis.vulgivaga.2.1.chr1.g00643.t01#1   mNeoVis.evergladensis.1.1.chr1.g00638.t01#3         3632   125258108:   mNeoVis.evergladensis.1.1.chr1.g00639.t01#3    mNeoVis.2.1.1.chr1_hap1.g00628.t01#5    mNeoVis.vulgivaga.1.1.chr1.g00638.t01#4    mNeoVis.evergladensis.2.1.chr1.g00634.t01#2    mNeoVis.vulgivaga.2.1.chr1.g00644.t01#1   mNeoVis.3.1.1.chr1.g00647.t01#6       3633   125258109:   mNeoVis.evergladensis.2.1.chr5.g01289.t01#2    mNeoVis.evergladensis.1.1.chr5.g01290.t01#3    mNeoVis.3.1.1.chr5.g01273.t01#6    mNeoVis.2.1.1.chr2_hap1.g01346.t01#5    mNeoVis.vulgivaga.1.1.chr5.g01257.t01#4   mNeoVis.vulgivaga.2.1.chr5.g01286.t01#1       3634   125258110:   mNeoVis.vulgivaga.1.1.chr7.g01222.t01#4    mNeoVis.3.1.1.chr7.g01157.t01#6    mNeoVis.2.1.1.chr7_hap1.g01232.t01#5    mNeoVis.vulgivaga.2.1.chr7.g01233.t01#1    mNeoVis.evergladensis.2.1.chr7.g01222.t01#2   mNeoVis.evergladensis.1.1.chr7.g01221.t01#3       3635   125258111:   mNeoVis.vulgivaga.1.1.chr7.g01223.t01#4    mNeoVis.2.1.1.chr7_hap1.g01233.t01#5    mNeoVis.vulgivaga.2.1.chr7.g01234.t01#1    mNeoVis.evergladensis.2.1.chr7.g01223.t01#2   mNeoVis.evergladensis.1.1.chr7.g01222.t01#3       3636   125258112:   mNeoVis.evergladensis.1.1.chr7.g00473.t01#3    mNeoVis.vulgivaga.2.1.chr7.g00468.t01#1    mNeoVis.evergladensis.2.1.chr7.g00468.t01#2    mNeoVis.3.1.1.chr7.g00448.t01#6    mNeoVis.vulgivaga.1.1.chr7.g00471.t01#4   mNeoVis.2.1.1.chr7_hap1.g00460.t01#5       3637   125258113:   mNeoVis.evergladensis.1.1.chr1.g00640.t01#3    mNeoVis.3.1.1.chr1.g00648.t01#6    mNeoVis.vulgivaga.2.1.chr1.g00645.t01#1    mNeoVis.2.1.1.chr1_hap1.g00629.t01#5    mNeoVis.vulgivaga.1.1.chr1.g00639.t01#4   mNeoVis.evergladensis.2.1.chr1.g00635.t01#2       3638   125258114:   mNeoVis.vulgivaga.2.1.chr7.g01235.t01#1    mNeoVis.3.1.1.chr7.g01158.t01#6    mNeoVis.evergladensis.2.1.chr7.g01224.t01#2    mNeoVis.vulgivaga.1.1.chr7.g01224.t01#4    mNeoVis.2.1.1.chr7_hap1.g01234.t01#5   mNeoVis.evergladensis.1.1.chr7.g01223.t01#3       3639   125258115:   mNeoVis.evergladensis.2.1.chr7.g01225.t01#2    mNeoVis.evergladensis.1.1.chr7.g01224.t01#3    mNeoVis.vulgivaga.2.1.chr7.g01236.t01#1    mNeoVis.3.1.1.chr7.g01159.t01#6    mNeoVis.vulgivaga.1.1.chr7.g01225.t01#4   mNeoVis.2.1.1.chr7_hap1.g01235.t01#5       3640   125258116:   mNeoVis.vulgivaga.2.1.chr15.g00011.t01#1    mNeoVis.evergladensis.1.1.chr1.g00641.t01#3    mNeoVis.evergladensis.2.1.chr1.g00636.t01#2    mNeoVis.3.1.1.chr1.g00649.t01#6    mNeoVis.vulgivaga.1.1.chr15.g00011.t01#4    mNeoVis.vulgivaga.2.1.chr1.g00646.t01#1    mNeoVis.evergladensis.2.1.chr15.g00011.t01#2    mNeoVis.2.1.1.chr1_hap1.g00630.t01#5    mNeoVis.evergladensis.2.1.chr4.g00629.t01#2      3641   125258117:   mNeoVis.3.1.1.chr12.g00173.t01#6   mNeoVis.evergladensis.2.1.chr12.g00177.t01#2           3642   125258118:   mNeoVis.vulgivaga.1.1.chr12.g00184.t01#4   mNeoVis.vulgivaga.2.1.chr12.g00181.t01#1           3643   125258119:   mNeoVis.2.1.1.chr7_hap1.g01236.t01#5    mNeoVis.vulgivaga.1.1.chr7.g01226.t01#4    mNeoVis.evergladensis.1.1.chr7.g01225.t01#3    mNeoVis.evergladensis.2.1.chr7.g01226.t01#2    mNeoVis.3.1.1.chr7.g01160.t01#6   mNeoVis.vulgivaga.2.1.chr7.g01237.t01#1       3644   125258120:   mNeoVis.evergladensis.1.1.chr15.g00093.t01#3    mNeoVis.vulgivaga.2.1.chr15.g00531.t01#1    mNeoVis.3.1.1.chr15.g00479.t01#6    mNeoVis.vulgivaga.1.1.chr15.g00523.t01#4    mNeoVis.evergladensis.2.1.chr15.g00520.t01#2   mNeoVis.2.1.1.chr15_hap1.g00698.t01#5       3645   125258121:   mNeoVis.vulgivaga.1.1.chr1.g00641.t01#4    mNeoVis.evergladensis.2.1.chr1.g00637.t01#2    mNeoVis.vulgivaga.2.1.chr1.g00647.t01#1    mNeoVis.evergladensis.1.1.chr1.g00642.t01#3    mNeoVis.3.1.1.chr1.g00650.t01#6   mNeoVis.2.1.1.chr1_hap1.g00631.t01#5       3646   125258122:   mNeoVis.evergladensis.1.1.chr7.g01226.t01#3    mNeoVis.vulgivaga.2.1.chr7.g01238.t01#1    mNeoVis.3.1.1.chr7.g01161.t01#6   mNeoVis.vulgivaga.1.1.chr7.g01227.t01#4         3647   125258123:   mNeoVis.evergladensis.1.1.chr7.g01227.t01#3    mNeoVis.vulgivaga.2.1.chr7.g01239.t01#1    mNeoVis.evergladensis.2.1.chr7.g01227.t01#2    mNeoVis.2.1.1.chr7_hap1.g01237.t01#5    mNeoVis.3.1.1.chr7.g01162.t01#6   mNeoVis.vulgivaga.1.1.chr7.g01228.t01#4       3648   125258124:   mNeoVis.2.1.1.chr1_hap1.g00632.t01#5    mNeoVis.evergladensis.1.1.chr1.g00643.t01#3    mNeoVis.vulgivaga.1.1.chr1.g00642.t01#4    mNeoVis.evergladensis.2.1.chr1.g00638.t01#2    mNeoVis.vulgivaga.2.1.chr1.g00648.t01#1   mNeoVis.3.1.1.chr1.g00651.t01#6       3649   125258125:   mNeoVis.vulgivaga.2.1.chr1.g00649.t01#1    mNeoVis.3.1.1.chr1.g00652.t01#6    mNeoVis.vulgivaga.1.1.chr1.g00643.t01#4    mNeoVis.2.1.1.chr1_hap1.g00633.t01#5    mNeoVis.evergladensis.1.1.chr1.g00644.t01#3   mNeoVis.evergladensis.2.1.chr1.g00639.t01#2       3650   125258126:   mNeoVis.evergladensis.2.1.chr7.g01228.t01#2    mNeoVis.vulgivaga.1.1.chr7.g01229.t01#4    mNeoVis.vulgivaga.2.1.chr7.g01240.t01#1    mNeoVis.evergladensis.1.1.chr7.g01228.t01#3    mNeoVis.3.1.1.chr7.g01163.t01#6   mNeoVis.2.1.1.chr7_hap1.g01238.t01#5       3651   125258127:   mNeoVis.3.1.1.chr1.g00653.t01#6    mNeoVis.evergladensis.1.1.chr1.g00645.t01#3    mNeoVis.evergladensis.2.1.chr1.g00640.t01#2    mNeoVis.vulgivaga.2.1.chr1.g00650.t01#1    mNeoVis.2.1.1.chr1_hap1.g00634.t01#5   mNeoVis.vulgivaga.1.1.chr1.g00644.t01#4       3652   125258128:   mNeoVis.vulgivaga.1.1.chr7.g01230.t01#4    mNeoVis.vulgivaga.2.1.chr7.g01241.t01#1    mNeoVis.evergladensis.1.1.chr7.g01229.t01#3    mNeoVis.2.1.1.chr7_hap1.g01239.t01#5    mNeoVis.3.1.1.chr7.g01164.t01#6   mNeoVis.evergladensis.2.1.chr7.g01229.t01#2       3653   125258129:   mNeoVis.vulgivaga.2.1.chr7.g01242.t01#1    mNeoVis.2.1.1.chr7_hap1.g01240.t01#5    mNeoVis.evergladensis.1.1.chr7.g01230.t01#3    mNeoVis.3.1.1.chr7.g01165.t01#6    mNeoVis.vulgivaga.1.1.chr7.g01231.t01#4   mNeoVis.evergladensis.2.1.chr7.g01230.t01#2       3654   125258130:   mNeoVis.evergladensis.1.1.chr7.g01231.t01#3    mNeoVis.2.1.1.chr7_hap1.g01241.t01#5    mNeoVis.vulgivaga.2.1.chr7.g01243.t01#1    mNeoVis.vulgivaga.1.1.chr7.g01232.t01#4    mNeoVis.3.1.1.chr7.g01166.t01#6   mNeoVis.evergladensis.2.1.chr7.g01231.t01#2       3655   125258131:   mNeoVis.2.1.1.chr1_hap1.g00635.t01#5    mNeoVis.vulgivaga.2.1.chr1.g00651.t01#1    mNeoVis.vulgivaga.1.1.chr1.g00645.t01#4    mNeoVis.3.1.1.chr1.g00654.t01#6   mNeoVis.evergladensis.1.1.chr1.g00646.t01#3       3656   125258132:   mNeoVis.vulgivaga.1.1.chr1.g00646.t01#4    mNeoVis.evergladensis.1.1.chr1.g00648.t01#3    mNeoVis.2.1.1.chr1_hap1.g00637.t01#5    mNeoVis.vulgivaga.2.1.chr1.g00652.t01#1    mNeoVis.3.1.1.chr1.g00655.t01#6   mNeoVis.evergladensis.2.1.chr1.g00642.t01#2       3657   125258133:   mNeoVis.vulgivaga.2.1.chr7.g01244.t01#1    mNeoVis.evergladensis.2.1.chr7.g01232.t01#2    mNeoVis.3.1.1.chr7.g01167.t01#6    mNeoVis.evergladensis.1.1.chr7.g01232.t01#3    mNeoVis.2.1.1.chr7_hap1.g01242.t01#5   mNeoVis.vulgivaga.1.1.chr7.g01233.t01#4       3658   125258134:   mNeoVis.3.1.1.chr7.g01168.t01#6    mNeoVis.evergladensis.2.1.chr7.g01233.t01#2    mNeoVis.2.1.1.chr7_hap1.g01243.t01#5    mNeoVis.evergladensis.1.1.chr7.g01233.t01#3    mNeoVis.vulgivaga.1.1.chr7.g01234.t01#4   mNeoVis.vulgivaga.2.1.chr7.g01245.t01#1       3659   125258135:   mNeoVis.evergladensis.1.1.chr1.g00649.t01#3    mNeoVis.evergladensis.2.1.chr1.g00643.t01#2    mNeoVis.vulgivaga.1.1.chr1.g00647.t01#4    mNeoVis.2.1.1.chr1_hap1.g00638.t01#5    mNeoVis.3.1.1.chr1.g00656.t01#6   mNeoVis.vulgivaga.2.1.chr1.g00653.t01#1       3660   125258136:   mNeoVis.evergladensis.1.1.chr1.g00650.t01#3    mNeoVis.vulgivaga.1.1.chr1.g00648.t01#4    mNeoVis.2.1.1.chr1_hap1.g00639.t01#5    mNeoVis.3.1.1.chr1.g00657.t01#6    mNeoVis.vulgivaga.2.1.chr1.g00654.t01#1    mNeoVis.evergladensis.2.1.chr1.g00644.t01#2   mNeoVis.vulgivaga.2.1.chr1.g00655.t01#1     3661   125258137:   mNeoVis.evergladensis.1.1.chr15.g00092.t01#3    mNeoVis.evergladensis.2.1.chr15.g00521.t01#2    mNeoVis.vulgivaga.1.1.chr15.g00524.t01#4    mNeoVis.2.1.1.chr15_hap1.g00699.t01#5   mNeoVis.vulgivaga.2.1.chr15.g00532.t01#1       3662   125258138:   mNeoVis.2.1.1.chr9_hap1.g00679.t01#5    mNeoVis.evergladensis.1.1.chr10.g00675.t01#3    mNeoVis.vulgivaga.2.1.chr10.g00657.t01#1    mNeoVis.3.1.1.chr10.g00644.t01#6    mNeoVis.evergladensis.2.1.chr10.g00674.t01#2   mNeoVis.vulgivaga.1.1.chr10.g00655.t01#4       3663   125258139:   mNeoVis.vulgivaga.1.1.chr1.g00649.t01#4    mNeoVis.evergladensis.2.1.chr1.g00645.t01#2    mNeoVis.vulgivaga.2.1.chr1.g00656.t01#1    mNeoVis.2.1.1.chr1_hap1.g00640.t01#5   mNeoVis.3.1.1.chr1.g00658.t01#6       3664   125258140:  mNeoVis.evergladensis.1.1.chr1.g00651.t01#3             3665   125258141:   mNeoVis.vulgivaga.1.1.chr1.g00650.t01#4    mNeoVis.2.1.1.chr1_hap1.g00641.t01#5    mNeoVis.3.1.1.chr1.g00659.t01#6    mNeoVis.evergladensis.2.1.chr1.g00646.t01#2   mNeoVis.vulgivaga.2.1.chr1.g00657.t01#1       3666   125258142:   mNeoVis.vulgivaga.2.1.chr1.g00658.t01#1    mNeoVis.evergladensis.2.1.chr1.g00647.t01#2    mNeoVis.evergladensis.1.1.chr1.g00653.t01#3    mNeoVis.vulgivaga.1.1.chr1.g00651.t01#4    mNeoVis.2.1.1.chr1_hap1.g00642.t01#5   mNeoVis.3.1.1.chr1.g00660.t01#6       3667   125258143:   mNeoVis.evergladensis.1.1.chr7.g01234.t01#3    mNeoVis.2.1.1.chr7_hap1.g01244.t01#5    mNeoVis.vulgivaga.1.1.chr7.g01235.t01#4    mNeoVis.vulgivaga.2.1.chr7.g01246.t01#1   mNeoVis.evergladensis.2.1.chr7.g01234.t01#2       3668   125258144:   mNeoVis.vulgivaga.2.1.chr7.g01247.t01#1    mNeoVis.evergladensis.2.1.chr7.g01235.t01#2    mNeoVis.3.1.1.chr7.g01170.t01#6    mNeoVis.evergladensis.1.1.chr7.g01235.t01#3    mNeoVis.2.1.1.chr7_hap1.g01245.t01#5   mNeoVis.vulgivaga.1.1.chr7.g01236.t01#4       3669   125258145:   mNeoVis.3.1.1.chr7.g01171.t01#6    mNeoVis.evergladensis.2.1.chr7.g01236.t01#2    mNeoVis.evergladensis.1.1.chr7.g01236.t01#3    mNeoVis.2.1.1.chr7_hap1.g01246.t01#5    mNeoVis.vulgivaga.1.1.chr7.g01237.t01#4   mNeoVis.vulgivaga.2.1.chr7.g01248.t01#1       3670   125258146:   mNeoVis.evergladensis.1.1.chr1.g00654.t01#3    mNeoVis.vulgivaga.2.1.chr1.g00659.t01#1    mNeoVis.3.1.1.chr1.g00661.t01#6    mNeoVis.evergladensis.2.1.chr1.g00648.t01#2    mNeoVis.vulgivaga.1.1.chr1.g00652.t01#4   mNeoVis.2.1.1.chr1_hap1.g00643.t01#5       3671   125258147:   mNeoVis.evergladensis.2.1.chr1.g00649.t01#2    mNeoVis.vulgivaga.1.1.chr1.g00653.t01#4    mNeoVis.2.1.1.chr1_hap1.g00644.t01#5    mNeoVis.evergladensis.1.1.chr1.g00655.t01#3    mNeoVis.vulgivaga.2.1.chr1.g00660.t01#1   mNeoVis.3.1.1.chr1.g00662.t01#6       3672   125258148:   mNeoVis.vulgivaga.1.1.chr7.g01238.t01#4    mNeoVis.vulgivaga.2.1.chr7.g01249.t01#1    mNeoVis.3.1.1.chr7.g01172.t01#6    mNeoVis.evergladensis.2.1.chr7.g01237.t01#2    mNeoVis.2.1.1.chr7_hap1.g01247.t01#5   mNeoVis.evergladensis.1.1.chr7.g01237.t01#3       3673   125258149:   mNeoVis.evergladensis.1.1.chr7.g01238.t01#3    mNeoVis.evergladensis.2.1.chr7.g01238.t01#2    mNeoVis.2.1.1.chr7_hap1.g01248.t01#5    mNeoVis.vulgivaga.2.1.chr7.g01250.t01#1   mNeoVis.vulgivaga.1.1.chr7.g01239.t01#4       3674   125258150:   mNeoVis.2.1.1.chr7_hap1.g01249.t01#5    mNeoVis.evergladensis.2.1.chr7.g01239.t01#2    mNeoVis.3.1.1.chr7.g01174.t01#6    mNeoVis.evergladensis.1.1.chr7.g01239.t01#3    mNeoVis.vulgivaga.2.1.chr7.g01251.t01#1   mNeoVis.vulgivaga.1.1.chr7.g01240.t01#4       3675   125258151:   mNeoVis.evergladensis.1.1.chr7.g01240.t01#3    mNeoVis.2.1.1.chr7_hap1.g01250.t01#5   mNeoVis.vulgivaga.2.1.chr7.g01252.t01#1           3676   125258152:   mNeoVis.vulgivaga.1.1.chr1.g00568.t01#4    mNeoVis.3.1.1.chr1.g00571.t01#6    mNeoVis.evergladensis.1.1.chr1.g00563.t01#3    mNeoVis.evergladensis.2.1.chr1.g00562.t01#2    mNeoVis.2.1.1.chr1_hap1.g00556.t01#5   mNeoVis.vulgivaga.2.1.chr1.g00572.t01#1       3677   125258153:   mNeoVis.evergladensis.1.1.chr1.g00564.t01#3    mNeoVis.vulgivaga.1.1.chr4.g00327.t01#4    mNeoVis.vulgivaga.2.1.chr5.g00033.t01#1    mNeoVis.2.1.1.chr6_hap1.g00328.t01#5    mNeoVis.3.1.1.chr1.g00572.t01#6    mNeoVis.evergladensis.2.1.chr4.g00334.t01#2    mNeoVis.evergladensis.1.1.chr5.g00035.t01#3    mNeoVis.evergladensis.2.1.chr5.g00035.t01#2    mNeoVis.vulgivaga.2.1.chr1.g00573.t01#1      3678   125258154:   mNeoVis.evergladensis.1.1.chr4.g00824.t01#3    mNeoVis.2.1.1.chr6_hap1.g00808.t01#5    mNeoVis.evergladensis.2.1.chr4.g00810.t01#2    mNeoVis.vulgivaga.2.1.chr4.g00830.t01#1   mNeoVis.3.1.1.chr4.g00818.t01#6       3679   125258155:  mNeoVis.vulgivaga.1.1.chr4.g00825.t01#4               3680   125258156:   mNeoVis.3.1.1.chr15.g00481.t01#6    mNeoVis.2.1.1.chr15_hap1.g00700.t01#5    mNeoVis.vulgivaga.2.1.chr15.g00533.t01#1    mNeoVis.vulgivaga.1.1.chr15.g00525.t01#4    mNeoVis.evergladensis.2.1.chr15.g00522.t01#2   mNeoVis.evergladensis.1.1.chr15.g00091.t01#3       3681   125258157:   mNeoVis.2.1.1.chr1_hap1.g00558.t01#5    mNeoVis.evergladensis.2.1.chr1.g00564.t01#2    mNeoVis.3.1.1.chr1.g00573.t01#6    mNeoVis.vulgivaga.2.1.chr1.g00574.t01#1    mNeoVis.evergladensis.1.1.chr1.g00565.t01#3   mNeoVis.vulgivaga.1.1.chr1.g00570.t01#4       3682   125258158:   mNeoVis.vulgivaga.2.1.chr7.g01253.t01#1    mNeoVis.3.1.1.chr7.g01175.t01#6    mNeoVis.evergladensis.1.1.chr7.g01241.t01#3    mNeoVis.2.1.1.chr7_hap1.g01251.t01#5    mNeoVis.evergladensis.2.1.chr7.g01240.t01#2   mNeoVis.vulgivaga.1.1.chr7.g01241.t01#4       3683   125258159:   mNeoVis.2.1.1.chr1_hap1.g00559.t01#5    mNeoVis.evergladensis.1.1.chr1.g00566.t01#3    mNeoVis.vulgivaga.2.1.chr1.g00575.t01#1    mNeoVis.evergladensis.2.1.chr1.g00565.t01#2    mNeoVis.3.1.1.chr1.g00574.t01#6   mNeoVis.vulgivaga.1.1.chr1.g00571.t01#4       3684   125258160:   mNeoVis.3.1.1.chr4.g00752.t01#6    mNeoVis.evergladensis.2.1.chr4.g00747.t01#2    mNeoVis.vulgivaga.1.1.chr4.g00760.t01#4    mNeoVis.evergladensis.1.1.chr4.g00760.t01#3    mNeoVis.vulgivaga.2.1.chr4.g00766.t01#1   mNeoVis.2.1.1.chr6_hap1.g00743.t01#5       3685   125258161:   mNeoVis.evergladensis.1.1.chr5.g00032.t01#3    mNeoVis.vulgivaga.2.1.chr5.g00030.t01#1    mNeoVis.evergladensis.2.1.chr5.g00032.t01#2    mNeoVis.vulgivaga.1.1.chr5.g00031.t01#4   mNeoVis.2.1.1.chr2_hap1.g00032.t01#5       3686   125258162:   mNeoVis.vulgivaga.1.1.chr7.g01242.t01#4    mNeoVis.3.1.1.chr7.g01176.t01#6    mNeoVis.vulgivaga.2.1.chr7.g01254.t01#1    mNeoVis.evergladensis.1.1.chr7.g01242.t01#3    mNeoVis.2.1.1.chr7_hap1.g01252.t01#5   mNeoVis.evergladensis.2.1.chr7.g01241.t01#2       3687   125258163:   mNeoVis.vulgivaga.2.1.chr7.g01255.t01#1    mNeoVis.evergladensis.1.1.chr7.g01243.t01#3    mNeoVis.evergladensis.2.1.chr7.g01242.t01#2    mNeoVis.2.1.1.chr7_hap1.g01253.t01#5   mNeoVis.vulgivaga.1.1.chr7.g01243.t01#4       3688   125258164:   mNeoVis.vulgivaga.1.1.chr1.g00572.t01#4    mNeoVis.3.1.1.chr1.g00575.t01#6    mNeoVis.evergladensis.2.1.chr1.g00566.t01#2    mNeoVis.evergladensis.1.1.chr1.g00567.t01#3    mNeoVis.vulgivaga.2.1.chr1.g00576.t01#1   mNeoVis.2.1.1.chr1_hap1.g00560.t01#5       3689   125258165:   mNeoVis.evergladensis.1.1.chr7.g01244.t01#3    mNeoVis.evergladensis.2.1.chr7.g01243.t01#2    mNeoVis.3.1.1.chr7.g01178.t01#6    mNeoVis.vulgivaga.1.1.chr7.g01244.t01#4    mNeoVis.vulgivaga.2.1.chr7.g01256.t01#1   mNeoVis.2.1.1.chr7_hap1.g01254.t01#5       3690   125258166:   mNeoVis.evergladensis.1.1.chr15.g00090.t01#3    mNeoVis.2.1.1.chr15_hap1.g00701.t01#5    mNeoVis.vulgivaga.1.1.chr15.g00526.t01#4    mNeoVis.3.1.1.chr15.g00482.t01#6    mNeoVis.evergladensis.2.1.chr15.g00523.t01#2   mNeoVis.vulgivaga.2.1.chr15.g00534.t01#1       3691   125258167:   mNeoVis.vulgivaga.1.1.chr1.g00573.t01#4    mNeoVis.2.1.1.chr1_hap1.g00561.t01#5    mNeoVis.3.1.1.chr1.g00576.t01#6    mNeoVis.evergladensis.1.1.chr1.g00569.t01#3    mNeoVis.vulgivaga.2.1.chr1.g00577.t01#1   mNeoVis.evergladensis.2.1.chr1.g00567.t01#2       3692   125258168:   mNeoVis.evergladensis.1.1.chr7.g01245.t01#3    mNeoVis.vulgivaga.1.1.chr7.g01245.t01#4    mNeoVis.3.1.1.chr7.g01179.t01#6    mNeoVis.vulgivaga.2.1.chr7.g01257.t01#1    mNeoVis.2.1.1.chr7_hap1.g01255.t01#5   mNeoVis.evergladensis.2.1.chr7.g01244.t01#2       3693   125258169:   mNeoVis.vulgivaga.1.1.chr1.g00575.t01#4    mNeoVis.vulgivaga.2.1.chr1.g00578.t01#1    mNeoVis.evergladensis.2.1.chr1.g00569.t01#2    mNeoVis.2.1.1.chr1_hap1.g00563.t01#5    mNeoVis.3.1.1.chr1.g00578.t01#6   mNeoVis.evergladensis.1.1.chr1.g00570.t01#3       3694   125258170:   mNeoVis.2.1.1.chr1_hap1.g00562.t01#5    mNeoVis.evergladensis.2.1.chr1.g00568.t01#2    mNeoVis.3.1.1.chr1.g00577.t01#6    mNeoVis.evergladensis.1.1.chr1.g00571.t01#3    mNeoVis.vulgivaga.1.1.chr1.g00574.t01#4    mNeoVis.vulgivaga.2.1.chr1.g00579.t01#1   mNeoVis.evergladensis.1.1.chr1.g00572.t01#3     3695   125258171:   mNeoVis.vulgivaga.2.1.chr1.g00580.t01#1    mNeoVis.3.1.1.chr1.g00579.t01#6    mNeoVis.evergladensis.2.1.chr1.g00570.t01#2    mNeoVis.2.1.1.chr1_hap1.g00564.t01#5    mNeoVis.evergladensis.1.1.chr1.g00573.t01#3   mNeoVis.vulgivaga.1.1.chr1.g00576.t01#4       3696   125258172:   mNeoVis.evergladensis.2.1.chr7.g01245.t01#2    mNeoVis.vulgivaga.2.1.chr7.g01258.t01#1    mNeoVis.2.1.1.chr7_hap1.g01256.t01#5    mNeoVis.evergladensis.1.1.chr7.g01246.t01#3    mNeoVis.vulgivaga.1.1.chr7.g01246.t01#4   mNeoVis.3.1.1.chr7.g01180.t01#6       3697   125258173:   mNeoVis.evergladensis.1.1.chr7.g01248.t01#3    mNeoVis.vulgivaga.2.1.chr7.g01260.t01#1    mNeoVis.vulgivaga.1.1.chr7.g01248.t01#4    mNeoVis.3.1.1.chr7.g01182.t01#6    mNeoVis.2.1.1.chr7_hap1.g01258.t01#5   mNeoVis.evergladensis.2.1.chr7.g01247.t01#2       3698   125258174:   mNeoVis.evergladensis.2.1.chr11.g00773.t01#2    mNeoVis.3.1.1.chr7.g01181.t01#6    mNeoVis.evergladensis.1.1.chr11.g00781.t01#3    mNeoVis.evergladensis.2.1.chr7.g01246.t01#2    mNeoVis.2.1.1.chr7_hap1.g01257.t01#5    mNeoVis.vulgivaga.2.1.chr7.g01259.t01#1    mNeoVis.vulgivaga.1.1.chr11.g00773.t01#4    mNeoVis.3.1.1.chr1.g01796.t01#6    mNeoVis.2.1.1.chr11_hap1.g00791.t01#5      3699   125258175:   mNeoVis.2.1.1.chr7_hap1.g01259.t01#5    mNeoVis.vulgivaga.2.1.chr7.g01261.t01#1    mNeoVis.evergladensis.1.1.chr7.g01249.t01#3    mNeoVis.vulgivaga.1.1.chr7.g01249.t01#4    mNeoVis.3.1.1.chr7.g01183.t01#6   mNeoVis.evergladensis.2.1.chr7.g01248.t01#2       3700   125258176:   mNeoVis.3.1.1.chr7.g01184.t01#6    mNeoVis.vulgivaga.1.1.chr7.g01250.t01#4    mNeoVis.vulgivaga.2.1.chr7.g01262.t01#1    mNeoVis.evergladensis.2.1.chr7.g01249.t01#2    mNeoVis.2.1.1.chr7_hap1.g01260.t01#5   mNeoVis.evergladensis.1.1.chr7.g01250.t01#3       3701   125258177:   mNeoVis.evergladensis.2.1.chr1.g00572.t01#2    mNeoVis.2.1.1.chr1_hap1.g00566.t01#5    mNeoVis.3.1.1.chr1.g00581.t01#6    mNeoVis.vulgivaga.2.1.chr1.g00581.t01#1   mNeoVis.evergladensis.1.1.chr1.g00575.t01#3       3702   125258178:   mNeoVis.vulgivaga.1.1.chr1.g00578.t01#4    mNeoVis.vulgivaga.2.1.chr1.g00582.t01#1    mNeoVis.2.1.1.chr1_hap1.g00567.t01#5    mNeoVis.3.1.1.chr1.g00582.t01#6    mNeoVis.evergladensis.2.1.chr1.g00573.t01#2   mNeoVis.evergladensis.1.1.chr1.g00576.t01#3       3703   125258179:   mNeoVis.vulgivaga.1.1.chr1.g00579.t01#4    mNeoVis.vulgivaga.2.1.chr1.g00583.t01#1    mNeoVis.2.1.1.chr1_hap1.g00568.t01#5    mNeoVis.3.1.1.chr1.g00583.t01#6    mNeoVis.evergladensis.2.1.chr1.g00574.t01#2   mNeoVis.evergladensis.1.1.chr1.g00577.t01#3       3704   125258180:   mNeoVis.evergladensis.2.1.chr7.g01250.t01#2    mNeoVis.evergladensis.1.1.chr7.g01251.t01#3    mNeoVis.2.1.1.chr7_hap1.g01261.t01#5    mNeoVis.vulgivaga.2.1.chr7.g01263.t01#1    mNeoVis.3.1.1.chr7.g01185.t01#6   mNeoVis.vulgivaga.1.1.chr7.g01251.t01#4       3705   125258181:   mNeoVis.vulgivaga.1.1.chr15.g00527.t01#4    mNeoVis.evergladensis.2.1.chr15.g00524.t01#2    mNeoVis.evergladensis.1.1.chr15.g00089.t01#3    mNeoVis.vulgivaga.2.1.chr15.g00535.t01#1   mNeoVis.2.1.1.chr15_hap1.g00702.t01#5       3706   125258182:   mNeoVis.2.1.1.chr1_hap1.g00569.t01#5    mNeoVis.evergladensis.1.1.chr1.g00578.t01#3    mNeoVis.vulgivaga.2.1.chr1.g00584.t01#1    mNeoVis.3.1.1.chr1.g00584.t01#6    mNeoVis.evergladensis.2.1.chr1.g00575.t01#2   mNeoVis.vulgivaga.1.1.chr1.g00580.t01#4       3707   125258183:   mNeoVis.evergladensis.2.1.chr1.g00576.t01#2    mNeoVis.3.1.1.chr1.g00585.t01#6    mNeoVis.evergladensis.1.1.chr1.g00579.t01#3    mNeoVis.vulgivaga.1.1.chr1.g00581.t01#4    mNeoVis.vulgivaga.2.1.chr1.g00585.t01#1   mNeoVis.2.1.1.chr1_hap1.g00570.t01#5       3708   125258184:   mNeoVis.evergladensis.2.1.chr1.g00577.t01#2    mNeoVis.2.1.1.chr1_hap1.g00571.t01#5    mNeoVis.3.1.1.chr1.g00586.t01#6    mNeoVis.vulgivaga.1.1.chr1.g00582.t01#4    mNeoVis.evergladensis.1.1.chr1.g00580.t01#3   mNeoVis.vulgivaga.2.1.chr1.g00586.t01#1       3709   125258185:   mNeoVis.2.1.1.chr7_hap1.g01262.t01#5    mNeoVis.evergladensis.1.1.chr7.g01252.t01#3    mNeoVis.vulgivaga.2.1.chr7.g01264.t01#1    mNeoVis.vulgivaga.1.1.chr7.g01252.t01#4    mNeoVis.evergladensis.2.1.chr7.g01251.t01#2   mNeoVis.3.1.1.chr7.g01186.t01#6       3710   125258186:   mNeoVis.2.1.1.chr1_hap1.g00572.t01#5    mNeoVis.vulgivaga.2.1.chr1.g00587.t01#1    mNeoVis.evergladensis.2.1.chr1.g00578.t01#2    mNeoVis.3.1.1.chr1.g00587.t01#6    mNeoVis.vulgivaga.1.1.chr1.g00583.t01#4   mNeoVis.evergladensis.1.1.chr1.g00581.t01#3       3711   125258187:  mNeoVis.vulgivaga.2.1.chr15.g00536.t01#1               3712   125258188:   mNeoVis.evergladensis.2.1.chr1.g00579.t01#2    mNeoVis.3.1.1.chr1.g00588.t01#6    mNeoVis.vulgivaga.1.1.chr1.g00584.t01#4    mNeoVis.evergladensis.1.1.chr1.g00582.t01#3    mNeoVis.vulgivaga.2.1.chr1.g00588.t01#1   mNeoVis.2.1.1.chr1_hap1.g00573.t01#5       3713   125258189:   mNeoVis.2.1.1.chr1_hap1.g00574.t01#5    mNeoVis.vulgivaga.2.1.chr1.g00589.t01#1    mNeoVis.evergladensis.2.1.chr1.g00580.t01#2    mNeoVis.3.1.1.chr1.g00589.t01#6    mNeoVis.vulgivaga.1.1.chr1.g00585.t01#4   mNeoVis.evergladensis.1.1.chr1.g00583.t01#3       3714   125258190:   mNeoVis.2.1.1.chr7_hap1.g01264.t01#5    mNeoVis.evergladensis.1.1.chr7.g01254.t01#3    mNeoVis.evergladensis.2.1.chr7.g01253.t01#2    mNeoVis.vulgivaga.2.1.chr7.g01266.t01#1    mNeoVis.vulgivaga.1.1.chr7.g01254.t01#4   mNeoVis.3.1.1.chr7.g01188.t01#6       3715   125258191:   mNeoVis.2.1.1.chr7_hap1.g01265.t01#5    mNeoVis.evergladensis.1.1.chr7.g01255.t01#3    mNeoVis.evergladensis.2.1.chr7.g01254.t01#2    mNeoVis.vulgivaga.2.1.chr7.g01267.t01#1    mNeoVis.vulgivaga.1.1.chr7.g01255.t01#4   mNeoVis.3.1.1.chr7.g01189.t01#6       3716   125258192:   mNeoVis.evergladensis.1.1.chr7.g00474.t01#3    mNeoVis.vulgivaga.2.1.chr7.g00469.t01#1    mNeoVis.vulgivaga.1.1.chr7.g00472.t01#4    mNeoVis.2.1.1.chr7_hap1.g00461.t01#5    mNeoVis.3.1.1.chr7.g00449.t01#6   mNeoVis.evergladensis.2.1.chr7.g00469.t01#2       3717   125258193:   mNeoVis.evergladensis.1.1.chr1.g00585.t01#3    mNeoVis.2.1.1.chr1_hap1.g00576.t01#5    mNeoVis.vulgivaga.1.1.chr1.g00586.t01#4    mNeoVis.3.1.1.chr1.g00590.t01#6    mNeoVis.vulgivaga.2.1.chr1.g00590.t01#1   mNeoVis.evergladensis.2.1.chr1.g00581.t01#2       3718   125258194:   mNeoVis.2.1.1.chr7_hap1.g01266.t01#5    mNeoVis.evergladensis.1.1.chr7.g01256.t01#3    mNeoVis.vulgivaga.2.1.chr7.g01268.t01#1    mNeoVis.vulgivaga.1.1.chr7.g01256.t01#4    mNeoVis.3.1.1.chr7.g01190.t01#6   mNeoVis.evergladensis.2.1.chr7.g01255.t01#2       3719   125258195:   mNeoVis.vulgivaga.1.1.chr7.g01257.t01#4    mNeoVis.evergladensis.2.1.chr7.g01256.t01#2    mNeoVis.3.1.1.chr7.g01191.t01#6    mNeoVis.vulgivaga.2.1.chr7.g01269.t01#1    mNeoVis.evergladensis.1.1.chr7.g01257.t01#3   mNeoVis.2.1.1.chr7_hap1.g01267.t01#5       3720   125258196:  mNeoVis.vulgivaga.2.1.chr15.g00537.t01#1               3721   125258197:   mNeoVis.evergladensis.1.1.chr7.g00475.t01#3    mNeoVis.3.1.1.chr7.g00450.t01#6    mNeoVis.evergladensis.2.1.chr7.g00470.t01#2    mNeoVis.vulgivaga.2.1.chr7.g00470.t01#1    mNeoVis.2.1.1.chr7_hap1.g00462.t01#5   mNeoVis.vulgivaga.1.1.chr7.g00473.t01#4       3722   125258198:   mNeoVis.vulgivaga.2.1.chr1.g00591.t01#1    mNeoVis.evergladensis.2.1.chr1.g00582.t01#2    mNeoVis.2.1.1.chr1_hap1.g00577.t01#5    mNeoVis.vulgivaga.1.1.chr1.g00587.t01#4    mNeoVis.3.1.1.chr1.g00591.t01#6   mNeoVis.evergladensis.1.1.chr1.g00586.t01#3       3723   125258199:   mNeoVis.2.1.1.chr1_hap1.g00578.t01#5    mNeoVis.evergladensis.2.1.chr1.g00583.t01#2    mNeoVis.3.1.1.chr1.g00592.t01#6    mNeoVis.evergladensis.1.1.chr1.g00587.t01#3    mNeoVis.vulgivaga.2.1.chr1.g00592.t01#1   mNeoVis.vulgivaga.1.1.chr1.g00588.t01#4       3724   125258200:   mNeoVis.evergladensis.2.1.chr7.g01257.t01#2    mNeoVis.vulgivaga.2.1.chr7.g01270.t01#1    mNeoVis.vulgivaga.1.1.chr7.g01258.t01#4    mNeoVis.2.1.1.chr7_hap1.g01268.t01#5   mNeoVis.evergladensis.1.1.chr7.g01258.t01#3       3725   125258201:   mNeoVis.vulgivaga.1.1.chr7.g01259.t01#4    mNeoVis.evergladensis.1.1.chr7.g01259.t01#3    mNeoVis.3.1.1.chr7.g01193.t01#6    mNeoVis.2.1.1.chr7_hap1.g01269.t01#5    mNeoVis.vulgivaga.2.1.chr7.g01271.t01#1   mNeoVis.evergladensis.2.1.chr7.g01258.t01#2       3726   125258202:   mNeoVis.vulgivaga.1.1.chr1.g00590.t01#4    mNeoVis.vulgivaga.2.1.chr1.g00594.t01#1    mNeoVis.evergladensis.2.1.chr1.g00585.t01#2    mNeoVis.3.1.1.chr1.g00593.t01#6    mNeoVis.2.1.1.chr1_hap1.g00580.t01#5   mNeoVis.evergladensis.1.1.chr1.g00589.t01#3       3727   125258203:   mNeoVis.2.1.1.chr7_hap1.g01270.t01#5    mNeoVis.evergladensis.2.1.chr7.g01259.t01#2    mNeoVis.vulgivaga.1.1.chr7.g01260.t01#4    mNeoVis.vulgivaga.2.1.chr7.g01272.t01#1    mNeoVis.3.1.1.chr7.g01194.t01#6   mNeoVis.evergladensis.1.1.chr7.g01260.t01#3       3728   125258204:   mNeoVis.2.1.1.chr7_hap1.g01271.t01#5    mNeoVis.evergladensis.2.1.chr7.g01260.t01#2    mNeoVis.vulgivaga.2.1.chr7.g01273.t01#1    mNeoVis.vulgivaga.1.1.chr7.g01261.t01#4    mNeoVis.3.1.1.chr7.g01195.t01#6   mNeoVis.evergladensis.1.1.chr7.g01261.t01#3       3729   125258205:   mNeoVis.vulgivaga.1.1.chr1.g00591.t01#4    mNeoVis.vulgivaga.2.1.chr1.g00595.t01#1    mNeoVis.evergladensis.2.1.chr1.g00586.t01#2    mNeoVis.2.1.1.chr1_hap1.g00581.t01#5    mNeoVis.evergladensis.1.1.chr1.g00591.t01#3    mNeoVis.3.1.1.chr1.g00594.t01#6   mNeoVis.evergladensis.1.1.chr1.g00590.t01#3     3730   125258206:   mNeoVis.vulgivaga.2.1.chr7.g01274.t01#1    mNeoVis.3.1.1.chr7.g01196.t01#6    mNeoVis.vulgivaga.1.1.chr7.g01262.t01#4    mNeoVis.evergladensis.1.1.chr7.g01262.t01#3    mNeoVis.evergladensis.2.1.chr7.g01261.t01#2   mNeoVis.2.1.1.chr7_hap1.g01272.t01#5       3731   125258207:   mNeoVis.vulgivaga.1.1.chr1.g00592.t01#4    mNeoVis.evergladensis.1.1.chr1.g00592.t01#3    mNeoVis.2.1.1.chr1_hap1.g00582.t01#5    mNeoVis.vulgivaga.2.1.chr1.g00596.t01#1    mNeoVis.vulgivaga.1.1.chr1.g00594.t01#4    mNeoVis.3.1.1.chr1.g00595.t01#6    mNeoVis.vulgivaga.1.1.chr1.g00593.t01#4   mNeoVis.evergladensis.2.1.chr1.g00587.t01#2     3732   125258208:   mNeoVis.vulgivaga.2.1.chr7.g01275.t01#1    mNeoVis.evergladensis.2.1.chr7.g01262.t01#2    mNeoVis.vulgivaga.1.1.chr7.g01263.t01#4    mNeoVis.evergladensis.1.1.chr7.g01263.t01#3    mNeoVis.3.1.1.chr7.g01197.t01#6   mNeoVis.2.1.1.chr7_hap1.g01273.t01#5       3733   125258209:   mNeoVis.vulgivaga.1.1.chr7.g01264.t01#4    mNeoVis.vulgivaga.2.1.chr7.g01276.t01#1    mNeoVis.evergladensis.1.1.chr7.g01264.t01#3    mNeoVis.evergladensis.2.1.chr7.g01263.t01#2    mNeoVis.3.1.1.chr7.g01198.t01#6   mNeoVis.2.1.1.chr7_hap1.g01274.t01#5       3734   125258210:   mNeoVis.2.1.1.chr7_hap1.g01275.t01#5    mNeoVis.vulgivaga.2.1.chr7.g01277.t01#1    mNeoVis.3.1.1.chr7.g01199.t01#6    mNeoVis.evergladensis.2.1.chr7.g01264.t01#2    mNeoVis.vulgivaga.1.1.chr7.g01265.t01#4   mNeoVis.evergladensis.1.1.chr7.g01265.t01#3       3735   125258211:   mNeoVis.3.1.1.chr7.g00451.t01#6    mNeoVis.2.1.1.chr7_hap1.g00463.t01#5    mNeoVis.evergladensis.2.1.chr7.g00471.t01#2    mNeoVis.evergladensis.1.1.chr7.g00476.t01#3    mNeoVis.vulgivaga.2.1.chr7.g00471.t01#1   mNeoVis.vulgivaga.1.1.chr7.g00474.t01#4       3736   125258212:   mNeoVis.evergladensis.2.1.chr1.g00588.t01#2    mNeoVis.evergladensis.1.1.chr1.g00593.t01#3    mNeoVis.vulgivaga.2.1.chr1.g00597.t01#1    mNeoVis.2.1.1.chr1_hap1.g00583.t01#5    mNeoVis.vulgivaga.1.1.chr1.g00595.t01#4   mNeoVis.3.1.1.chr1.g00596.t01#6       3737   125258213:   mNeoVis.3.1.1.chr15.g00485.t01#6    mNeoVis.vulgivaga.1.1.chr15.g00529.t01#4    mNeoVis.vulgivaga.2.1.chr15.g00538.t01#1    mNeoVis.2.1.1.chr15_hap1.g00704.t01#5    mNeoVis.evergladensis.2.1.chr15.g00526.t01#2   mNeoVis.evergladensis.1.1.chr15.g00087.t01#3       3738   125258214:   mNeoVis.evergladensis.1.1.chr7.g01266.t01#3    mNeoVis.evergladensis.2.1.chr7.g01265.t01#2    mNeoVis.2.1.1.chr7_hap1.g01276.t01#5    mNeoVis.3.1.1.chr7.g01200.t01#6    mNeoVis.vulgivaga.2.1.chr7.g01278.t01#1   mNeoVis.vulgivaga.1.1.chr7.g01266.t01#4       3739   125258215:   mNeoVis.2.1.1.chr1_hap1.g00585.t01#5    mNeoVis.evergladensis.1.1.chr1.g00595.t01#3    mNeoVis.evergladensis.2.1.chr1.g00590.t01#2    mNeoVis.vulgivaga.2.1.chr1.g00599.t01#1    mNeoVis.vulgivaga.1.1.chr1.g00597.t01#4   mNeoVis.3.1.1.chr1.g00599.t01#6       3740   125258216:   mNeoVis.evergladensis.2.1.chr7.g01266.t01#2    mNeoVis.evergladensis.1.1.chr7.g01267.t01#3    mNeoVis.3.1.1.chr7.g01201.t01#6    mNeoVis.2.1.1.chr7_hap1.g01277.t01#5    mNeoVis.vulgivaga.2.1.chr7.g01279.t01#1   mNeoVis.vulgivaga.1.1.chr7.g01267.t01#4       3741   125258217:   mNeoVis.2.1.1.chr7_hap1.g01278.t01#5    mNeoVis.evergladensis.2.1.chr7.g01267.t01#2    mNeoVis.vulgivaga.2.1.chr7.g01280.t01#1    mNeoVis.evergladensis.1.1.chr7.g01268.t01#3   mNeoVis.3.1.1.chr7.g01202.t01#6       3742   125258218:   mNeoVis.2.1.1.chr1_hap1.g00587.t01#5    mNeoVis.evergladensis.2.1.chr1.g00592.t01#2    mNeoVis.vulgivaga.2.1.chr1.g00600.t01#1    mNeoVis.vulgivaga.1.1.chr1.g00598.t01#4    mNeoVis.3.1.1.chr1.g00601.t01#6   mNeoVis.evergladensis.1.1.chr1.g00596.t01#3       3743   125258219:   mNeoVis.vulgivaga.1.1.chr1.g00599.t01#4    mNeoVis.vulgivaga.2.1.chr1.g00601.t01#1    mNeoVis.3.1.1.chr1.g00602.t01#6    mNeoVis.evergladensis.1.1.chr1.g00597.t01#3    mNeoVis.evergladensis.2.1.chr1.g00591.t01#2   mNeoVis.2.1.1.chr1_hap1.g00586.t01#5       3744   125258220:   mNeoVis.vulgivaga.2.1.chr4.g00539.t01#1    mNeoVis.evergladensis.2.1.chr4.g00530.t01#2    mNeoVis.2.1.1.chr6_hap1.g00525.t01#5    mNeoVis.3.1.1.chr4.g00530.t01#6    mNeoVis.2.1.1.chr7_hap1.g01189.t01#5    mNeoVis.3.1.1.chr7.g01113.t01#6    mNeoVis.evergladensis.2.1.chr7.g01178.t01#2    mNeoVis.evergladensis.1.1.chr7.g01178.t01#3    mNeoVis.vulgivaga.1.1.chr7.g01179.t01#4      3745   125258221:   mNeoVis.vulgivaga.1.1.chr4.g00799.t01#4    mNeoVis.vulgivaga.2.1.chr4.g00803.t01#1    mNeoVis.2.1.1.chr6_hap1.g00781.t01#5    mNeoVis.evergladensis.2.1.chr4.g00784.t01#2    mNeoVis.vulgivaga.2.1.chr7.g01190.t01#1    mNeoVis.3.1.1.chr4.g00789.t01#6   mNeoVis.evergladensis.1.1.chr4.g00797.t01#3     3746   125258222:   mNeoVis.vulgivaga.1.1.chr7.g01180.t01#4    mNeoVis.vulgivaga.2.1.chr7.g01191.t01#1    mNeoVis.3.1.1.chr7.g01114.t01#6    mNeoVis.evergladensis.2.1.chr7.g01179.t01#2    mNeoVis.2.1.1.chr7_hap1.g01190.t01#5   mNeoVis.evergladensis.1.1.chr7.g01179.t01#3       3747   125258223:   mNeoVis.evergladensis.1.1.chr7.g01181.t01#3    mNeoVis.vulgivaga.2.1.chr7.g01192.t01#1    mNeoVis.3.1.1.chr7.g01116.t01#6    mNeoVis.vulgivaga.1.1.chr7.g01182.t01#4    mNeoVis.2.1.1.chr7_hap1.g01192.t01#5   mNeoVis.evergladensis.2.1.chr7.g01181.t01#2       3748   125258224:   mNeoVis.2.1.1.chr7_hap1.g00464.t01#5    mNeoVis.vulgivaga.1.1.chr7.g00475.t01#4    mNeoVis.3.1.1.chr7.g00452.t01#6    mNeoVis.vulgivaga.2.1.chr7.g00472.t01#1    mNeoVis.evergladensis.1.1.chr7.g00477.t01#3   mNeoVis.evergladensis.2.1.chr7.g00472.t01#2       3749   125258225:   mNeoVis.evergladensis.1.1.chr1.g00598.t01#3    mNeoVis.evergladensis.2.1.chr1.g00593.t01#2    mNeoVis.3.1.1.chr1.g00603.t01#6    mNeoVis.vulgivaga.2.1.chr1.g00602.t01#1    mNeoVis.vulgivaga.1.1.chr1.g00600.t01#4   mNeoVis.2.1.1.chr1_hap1.g00588.t01#5       3750   125258226:   mNeoVis.vulgivaga.1.1.chr1.g00602.t01#4    mNeoVis.evergladensis.2.1.chr1.g00595.t01#2    mNeoVis.2.1.1.chr1_hap1.g00592.t01#5    mNeoVis.evergladensis.1.1.chr1.g00601.t01#3    mNeoVis.vulgivaga.2.1.chr1.g00604.t01#1   mNeoVis.3.1.1.chr1.g00606.t01#6       3751   125258227:   mNeoVis.vulgivaga.2.1.chr7.g01194.t01#1    mNeoVis.evergladensis.2.1.chr7.g01183.t01#2    mNeoVis.2.1.1.chr7_hap1.g01194.t01#5    mNeoVis.3.1.1.chr7.g01119.t01#6    mNeoVis.vulgivaga.1.1.chr7.g01184.t01#4   mNeoVis.evergladensis.1.1.chr7.g01183.t01#3       3752   125258228:   mNeoVis.2.1.1.chr1_hap1.g00593.t01#5    mNeoVis.3.1.1.chr1.g00605.t01#6    mNeoVis.evergladensis.2.1.chr1.g00596.t01#2   mNeoVis.vulgivaga.2.1.chr1.g00605.t01#1         3753   125258229:   mNeoVis.vulgivaga.1.1.chr1.g00603.t01#4    mNeoVis.vulgivaga.2.1.chr1.g00606.t01#1    mNeoVis.evergladensis.2.1.chr1.g00597.t01#2    mNeoVis.evergladensis.1.1.chr1.g00600.t01#3    mNeoVis.3.1.1.chr1.g00607.t01#6   mNeoVis.2.1.1.chr1_hap1.g00591.t01#5       3754   125258230:   mNeoVis.vulgivaga.2.1.chr2.g00507.t01#1    mNeoVis.vulgivaga.1.1.chr2.g00501.t01#4    mNeoVis.3.1.1.chr2.g00433.t01#6    mNeoVis.evergladensis.2.1.chr2.g00505.t01#2    mNeoVis.evergladensis.1.1.chr2.g00501.t01#3   mNeoVis.2.1.1.chr5_hap1.g00510.t01#5       3755   125258231:   mNeoVis.vulgivaga.1.1.chr5.g01206.t01#4    mNeoVis.2.1.1.chr2_hap1.g01298.t01#5    mNeoVis.vulgivaga.2.1.chr5.g01235.t01#1    mNeoVis.3.1.1.chr5.g01223.t01#6    mNeoVis.evergladensis.2.1.chr5.g01241.t01#2   mNeoVis.evergladensis.1.1.chr5.g01240.t01#3       3756   125258232:   mNeoVis.vulgivaga.1.1.chr2.g00300.t01#4    mNeoVis.evergladensis.1.1.chr2.g00304.t01#3    mNeoVis.2.1.1.chr5_hap1.g00305.t01#5    mNeoVis.evergladensis.2.1.chr2.g00303.t01#2    mNeoVis.3.1.1.chr2.g00241.t01#6   mNeoVis.vulgivaga.2.1.chr2.g00299.t01#1       3757   125258233:   mNeoVis.vulgivaga.1.1.chr1.g00604.t01#4    mNeoVis.2.1.1.chr1_hap1.g00590.t01#5    mNeoVis.evergladensis.2.1.chr1.g00598.t01#2    mNeoVis.evergladensis.1.1.chr1.g00602.t01#3   mNeoVis.vulgivaga.2.1.chr1.g00607.t01#1       3758   125258234:   mNeoVis.3.1.1.chr1.g00610.t01#6    mNeoVis.evergladensis.1.1.chr1.g00603.t01#3    mNeoVis.vulgivaga.2.1.chr1.g00608.t01#1    mNeoVis.vulgivaga.1.1.chr1.g00605.t01#4    mNeoVis.2.1.1.chr1_hap1.g00594.t01#5   mNeoVis.evergladensis.2.1.chr1.g00599.t01#2       3759   125258235:   mNeoVis.vulgivaga.2.1.chr7.g01195.t01#1    mNeoVis.vulgivaga.1.1.chr7.g01185.t01#4    mNeoVis.evergladensis.1.1.chr7.g01184.t01#3    mNeoVis.2.1.1.chr7_hap1.g01195.t01#5    mNeoVis.evergladensis.2.1.chr7.g01184.t01#2   mNeoVis.3.1.1.chr7.g01120.t01#6       3760   125258236:   mNeoVis.3.1.1.chr1.g00611.t01#6    mNeoVis.evergladensis.1.1.chr1.g00604.t01#3    mNeoVis.vulgivaga.2.1.chr1.g00609.t01#1    mNeoVis.vulgivaga.1.1.chr1.g00606.t01#4    mNeoVis.evergladensis.2.1.chr1.g00600.t01#2   mNeoVis.2.1.1.chr1_hap1.g00595.t01#5       3761   125258237:   mNeoVis.3.1.1.chr7.g01121.t01#6    mNeoVis.evergladensis.1.1.chr7.g01185.t01#3    mNeoVis.vulgivaga.1.1.chr7.g01186.t01#4    mNeoVis.2.1.1.chr7_hap1.g01196.t01#5    mNeoVis.evergladensis.2.1.chr7.g01185.t01#2   mNeoVis.vulgivaga.2.1.chr7.g01196.t01#1       3762   125258238:   mNeoVis.evergladensis.2.1.chr1.g00601.t01#2    mNeoVis.vulgivaga.2.1.chr1.g00610.t01#1    mNeoVis.3.1.1.chr1.g00612.t01#6    mNeoVis.2.1.1.chr1_hap1.g00596.t01#5    mNeoVis.vulgivaga.1.1.chr1.g00608.t01#4   mNeoVis.evergladensis.1.1.chr1.g00605.t01#3       3763   125258239:   mNeoVis.2.1.1.chr7_hap1.g01197.t01#5    mNeoVis.3.1.1.chr7.g01122.t01#6    mNeoVis.evergladensis.2.1.chr7.g01186.t01#2    mNeoVis.evergladensis.1.1.chr7.g01186.t01#3    mNeoVis.vulgivaga.1.1.chr7.g01187.t01#4   mNeoVis.vulgivaga.2.1.chr7.g01197.t01#1       3764   125258240:   mNeoVis.vulgivaga.2.1.chr1.g00611.t01#1    mNeoVis.2.1.1.chr1_hap1.g00597.t01#5    mNeoVis.evergladensis.2.1.chr1.g00602.t01#2    mNeoVis.3.1.1.chr1.g00613.t01#6    mNeoVis.evergladensis.1.1.chr1.g00606.t01#3   mNeoVis.vulgivaga.1.1.chr1.g00609.t01#4       3765   125258241:   mNeoVis.3.1.1.chr7.g01123.t01#6    mNeoVis.evergladensis.2.1.chr7.g01187.t01#2    mNeoVis.evergladensis.1.1.chr7.g01187.t01#3    mNeoVis.vulgivaga.1.1.chr7.g01188.t01#4    mNeoVis.2.1.1.chr7_hap1.g01198.t01#5   mNeoVis.vulgivaga.2.1.chr7.g01198.t01#1       3766   125258242:   mNeoVis.2.1.1.chr1_hap1.g00598.t01#5    mNeoVis.3.1.1.chr1.g00614.t01#6    mNeoVis.evergladensis.2.1.chr1.g00603.t01#2    mNeoVis.vulgivaga.2.1.chr1.g00612.t01#1    mNeoVis.vulgivaga.1.1.chr1.g00607.t01#4   mNeoVis.evergladensis.1.1.chr1.g00607.t01#3       3767   125258243:   mNeoVis.vulgivaga.2.1.chr1.g00613.t01#1    mNeoVis.vulgivaga.1.1.chr1.g00610.t01#4    mNeoVis.3.1.1.chr1.g00616.t01#6    mNeoVis.evergladensis.1.1.chr1.g00608.t01#3    mNeoVis.2.1.1.chr1_hap1.g00599.t01#5   mNeoVis.evergladensis.2.1.chr1.g00604.t01#2       3768   125258244:   mNeoVis.2.1.1.chr1_hap1.g00600.t01#5    mNeoVis.evergladensis.2.1.chr1.g00605.t01#2    mNeoVis.evergladensis.1.1.chr1.g00610.t01#3    mNeoVis.vulgivaga.2.1.chr1.g00614.t01#1    mNeoVis.3.1.1.chr1.g00617.t01#6   mNeoVis.vulgivaga.1.1.chr1.g00611.t01#4       3769   125258245:   mNeoVis.evergladensis.1.1.chr7.g01188.t01#3    mNeoVis.vulgivaga.1.1.chr7.g01189.t01#4    mNeoVis.vulgivaga.2.1.chr7.g01199.t01#1    mNeoVis.evergladensis.2.1.chr7.g01188.t01#2    mNeoVis.3.1.1.chr7.g01124.t01#6   mNeoVis.2.1.1.chr7_hap1.g01199.t01#5       3770   125258246:   mNeoVis.vulgivaga.2.1.chr1.g00616.t01#1    mNeoVis.3.1.1.chr1.g00619.t01#6    mNeoVis.evergladensis.2.1.chr1.g00607.t01#2    mNeoVis.2.1.1.chr1_hap1.g00602.t01#5    mNeoVis.vulgivaga.1.1.chr1.g00612.t01#4   mNeoVis.evergladensis.1.1.chr1.g00612.t01#3       3771   125258247:  mNeoVis.vulgivaga.2.1.chr1.g00507.t01#1               3772   125258248:   mNeoVis.vulgivaga.2.1.chr7.g01200.t01#1    mNeoVis.evergladensis.1.1.chr7.g01189.t01#3    mNeoVis.evergladensis.2.1.chr7.g01189.t01#2    mNeoVis.vulgivaga.1.1.chr7.g01190.t01#4    mNeoVis.3.1.1.chr7.g01125.t01#6   mNeoVis.2.1.1.chr7_hap1.g01200.t01#5       3773   125258249:   mNeoVis.evergladensis.1.1.chr1.g00509.t01#3    mNeoVis.2.1.1.chr1_hap1.g00500.t01#5    mNeoVis.vulgivaga.1.1.chr1.g00493.t01#4    mNeoVis.evergladensis.2.1.chr1.g00496.t01#2    mNeoVis.3.1.1.chr1.g00497.t01#6   mNeoVis.vulgivaga.2.1.chr1.g00508.t01#1       3774   125258250:   mNeoVis.3.1.1.chr7.g01126.t01#6    mNeoVis.evergladensis.1.1.chr7.g01190.t01#3    mNeoVis.vulgivaga.2.1.chr7.g01201.t01#1    mNeoVis.2.1.1.chr7_hap1.g01201.t01#5    mNeoVis.vulgivaga.1.1.chr7.g01191.t01#4   mNeoVis.evergladensis.2.1.chr7.g01190.t01#2       3775   125258251:   mNeoVis.vulgivaga.1.1.chr7.g01192.t01#4    mNeoVis.evergladensis.2.1.chr7.g01191.t01#2    mNeoVis.2.1.1.chr7_hap1.g01202.t01#5    mNeoVis.vulgivaga.2.1.chr7.g01202.t01#1    mNeoVis.3.1.1.chr7.g01127.t01#6   mNeoVis.evergladensis.1.1.chr7.g01191.t01#3       3776   125258252:   mNeoVis.evergladensis.2.1.chr1.g00497.t01#2    mNeoVis.vulgivaga.1.1.chr1.g00494.t01#4    mNeoVis.3.1.1.chr1.g00498.t01#6    mNeoVis.evergladensis.1.1.chr1.g00510.t01#3    mNeoVis.vulgivaga.2.1.chr1.g00509.t01#1   mNeoVis.2.1.1.chr1_hap1.g00501.t01#5       3777   125258253:   mNeoVis.evergladensis.1.1.chr7.g01193.t01#3    mNeoVis.evergladensis.2.1.chr7.g01192.t01#2    mNeoVis.evergladensis.1.1.chr7.g01192.t01#3    mNeoVis.vulgivaga.2.1.chr7.g01203.t01#1    mNeoVis.vulgivaga.1.1.chr7.g01193.t01#4    mNeoVis.2.1.1.chr7_hap1.g01203.t01#5   mNeoVis.3.1.1.chr7.g01128.t01#6       3778   125258254:   mNeoVis.evergladensis.1.1.chr7.g01195.t01#3    mNeoVis.2.1.1.chr7_hap1.g01205.t01#5    mNeoVis.vulgivaga.2.1.chr7.g01204.t01#1    mNeoVis.evergladensis.2.1.chr7.g01194.t01#2    mNeoVis.vulgivaga.1.1.chr7.g01194.t01#4   mNeoVis.3.1.1.chr7.g01129.t01#6       3779   125258255:   mNeoVis.vulgivaga.2.1.chr7.g01205.t01#1    mNeoVis.3.1.1.chr7.g01130.t01#6    mNeoVis.vulgivaga.1.1.chr7.g01195.t01#4    mNeoVis.evergladensis.1.1.chr7.g01196.t01#3    mNeoVis.evergladensis.2.1.chr7.g01195.t01#2   mNeoVis.2.1.1.chr7_hap1.g01206.t01#5       3780   125258256:   mNeoVis.vulgivaga.1.1.chr1.g00495.t01#4    mNeoVis.3.1.1.chr1.g00499.t01#6    mNeoVis.evergladensis.1.1.chr1.g00511.t01#3    mNeoVis.vulgivaga.2.1.chr1.g00510.t01#1    mNeoVis.2.1.1.chr1_hap1.g00502.t01#5   mNeoVis.evergladensis.2.1.chr1.g00498.t01#2       3781   125258257:   mNeoVis.vulgivaga.1.1.chr7.g01196.t01#4    mNeoVis.3.1.1.chr5.g01195.t01#6    mNeoVis.evergladensis.1.1.chr7.g01197.t01#3    mNeoVis.evergladensis.1.1.chr5.g01213.t01#3    mNeoVis.2.1.1.chr2_hap1.g01273.t01#5    mNeoVis.vulgivaga.2.1.chr7.g01206.t01#1    mNeoVis.evergladensis.2.1.chr7.g01196.t01#2    mNeoVis.3.1.1.chr7.g01131.t01#6    mNeoVis.2.1.1.chr7_hap1.g01207.t01#5      3782   125258258:   mNeoVis.evergladensis.2.1.chr7.g00473.t01#2    mNeoVis.vulgivaga.2.1.chr7.g00473.t01#1    mNeoVis.evergladensis.1.1.chr7.g00478.t01#3    mNeoVis.vulgivaga.1.1.chr7.g00476.t01#4   mNeoVis.2.1.1.chr7_hap1.g00465.t01#5       3783   125258259:   mNeoVis.3.1.1.chr7.g01132.t01#6    mNeoVis.vulgivaga.1.1.chr7.g01197.t01#4    mNeoVis.2.1.1.chr7_hap1.g01209.t01#5    mNeoVis.evergladensis.1.1.chr7.g01198.t01#3    mNeoVis.evergladensis.2.1.chr7.g01197.t01#2   mNeoVis.vulgivaga.2.1.chr7.g01207.t01#1       3784   125258260:   mNeoVis.3.1.1.chr1.g00501.t01#6    mNeoVis.2.1.1.chr1_hap1.g00504.t01#5    mNeoVis.evergladensis.1.1.chr1.g00513.t01#3    mNeoVis.vulgivaga.2.1.chr1.g00512.t01#1    mNeoVis.evergladensis.2.1.chr1.g00500.t01#2   mNeoVis.vulgivaga.1.1.chr1.g00497.t01#4       3785   125258261:   mNeoVis.vulgivaga.2.1.chr7.g01208.t01#1    mNeoVis.2.1.1.chr7_hap1.g01210.t01#5    mNeoVis.vulgivaga.1.1.chr7.g01198.t01#4    mNeoVis.evergladensis.1.1.chr7.g01199.t01#3    mNeoVis.3.1.1.chr7.g01134.t01#6   mNeoVis.evergladensis.2.1.chr7.g01199.t01#2       3786   125258262:   mNeoVis.2.1.1.chr7_hap1.g01211.t01#5    mNeoVis.evergladensis.2.1.chr7.g01200.t01#2    mNeoVis.evergladensis.1.1.chr7.g01200.t01#3    mNeoVis.vulgivaga.1.1.chr7.g01199.t01#4   mNeoVis.vulgivaga.2.1.chr7.g01209.t01#1       3787   125258263:   mNeoVis.vulgivaga.2.1.chr1.g00513.t01#1    mNeoVis.vulgivaga.1.1.chr1.g00498.t01#4    mNeoVis.evergladensis.1.1.chr1.g00514.t01#3    mNeoVis.3.1.1.chr1.g00502.t01#6    mNeoVis.evergladensis.2.1.chr1.g00501.t01#2   mNeoVis.2.1.1.chr1_hap1.g00505.t01#5       3788   125258264:   mNeoVis.evergladensis.2.1.chr1.g00504.t01#2    mNeoVis.vulgivaga.2.1.chr1.g00516.t01#1    mNeoVis.vulgivaga.1.1.chr1.g00501.t01#4   mNeoVis.evergladensis.1.1.chr1.g00517.t01#3       3789   125258265:   mNeoVis.vulgivaga.2.1.chr1.g00517.t01#1    mNeoVis.3.1.1.chr1.g00507.t01#6    mNeoVis.evergladensis.2.1.chr1.g00505.t01#2   mNeoVis.evergladensis.2.1.chr1.g00507.t01#2       3790   125258266:   mNeoVis.evergladensis.2.1.chr1.g00502.t01#2    mNeoVis.vulgivaga.2.1.chr1.g00514.t01#1   mNeoVis.3.1.1.chr1.g00503.t01#6           3791   125258267:   mNeoVis.vulgivaga.2.1.chr1.g00518.t01#1    mNeoVis.vulgivaga.1.1.chr1.g00502.t01#4   mNeoVis.evergladensis.1.1.chr1.g00518.t01#3         3792   125258268:   mNeoVis.vulgivaga.1.1.chr1.g00499.t01#4   mNeoVis.3.1.1.chr1.g00504.t01#6             3793   125258269:  mNeoVis.3.1.1.chr1.g00506.t01#6               3794   125258270:  mNeoVis.evergladensis.1.1.chr1.g00515.t01#3             3795   125258271:  mNeoVis.evergladensis.1.1.chr1.g00516.t01#3             3796   125258272:  mNeoVis.vulgivaga.1.1.chr1.g00500.t01#4               3797   125258273:  mNeoVis.evergladensis.2.1.chr1.g00503.t01#2             3798   125258274:  mNeoVis.vulgivaga.2.1.chr1.g00515.t01#1               3799   125258275:   mNeoVis.evergladensis.2.1.chr7.g01201.t01#2    mNeoVis.2.1.1.chr7_hap1.g01212.t01#5   mNeoVis.vulgivaga.2.1.chr7.g01210.t01#1           3800   125258276:   mNeoVis.3.1.1.chr7.g01136.t01#6    mNeoVis.vulgivaga.2.1.chr7.g01211.t01#1    mNeoVis.evergladensis.1.1.chr7.g01201.t01#3    mNeoVis.vulgivaga.1.1.chr7.g01200.t01#4    mNeoVis.evergladensis.2.1.chr7.g01202.t01#2   mNeoVis.2.1.1.chr7_hap1.g01213.t01#5       3801   125258277:   mNeoVis.evergladensis.2.1.chr7.g01203.t01#2    mNeoVis.vulgivaga.2.1.chr7.g01212.t01#1    mNeoVis.evergladensis.1.1.chr7.g01202.t01#3    mNeoVis.3.1.1.chr7.g01137.t01#6    mNeoVis.2.1.1.chr7_hap1.g01214.t01#5   mNeoVis.vulgivaga.1.1.chr7.g01201.t01#4       3802   125258278:   mNeoVis.vulgivaga.2.1.chr1.g00519.t01#1    mNeoVis.evergladensis.1.1.chr1.g00520.t01#3    mNeoVis.evergladensis.2.1.chr1.g00508.t01#2   mNeoVis.vulgivaga.1.1.chr1.g00503.t01#4         3803   125258279:   mNeoVis.vulgivaga.1.1.chr1.g00504.t01#4    mNeoVis.vulgivaga.2.1.chr1.g00520.t01#1    mNeoVis.evergladensis.2.1.chr1.g00509.t01#2   mNeoVis.3.1.1.chr1.g00508.t01#6         3804   125258280:   mNeoVis.3.1.1.chr1.g00510.t01#6    mNeoVis.evergladensis.2.1.chr1.g00511.t01#2    mNeoVis.vulgivaga.1.1.chr1.g00506.t01#4    mNeoVis.2.1.1.chr1_hap1.g00506.t01#5    mNeoVis.evergladensis.1.1.chr1.g00521.t01#3   mNeoVis.vulgivaga.2.1.chr1.g00521.t01#1       3805   125258281:   mNeoVis.2.1.1.chr7_hap1.g01215.t01#5    mNeoVis.evergladensis.1.1.chr7.g01203.t01#3    mNeoVis.3.1.1.chr7.g01138.t01#6    mNeoVis.vulgivaga.1.1.chr7.g01202.t01#4    mNeoVis.vulgivaga.2.1.chr7.g01213.t01#1   mNeoVis.evergladensis.2.1.chr7.g01204.t01#2       3806   125258282:   mNeoVis.vulgivaga.2.1.chr7.g00474.t01#1    mNeoVis.3.1.1.chr7.g00454.t01#6    mNeoVis.2.1.1.chr7_hap1.g00467.t01#5    mNeoVis.2.1.1.chr7_hap1.g00473.t01#5   mNeoVis.evergladensis.2.1.chr7.g00475.t01#2       3807   125258283:  mNeoVis.vulgivaga.2.1.chr1.g00522.t01#1               3808   125258284:   mNeoVis.evergladensis.2.1.chr1.g00512.t01#2    mNeoVis.vulgivaga.2.1.chr1.g00523.t01#1    mNeoVis.vulgivaga.1.1.chr1.g00509.t01#4    mNeoVis.evergladensis.1.1.chr1.g00522.t01#3   mNeoVis.3.1.1.chr1.g00514.t01#6       3809   125258285:   mNeoVis.evergladensis.1.1.chr1.g00523.t01#3    mNeoVis.3.1.1.chr1.g00516.t01#6    mNeoVis.vulgivaga.2.1.chr1.g00525.t01#1   mNeoVis.evergladensis.2.1.chr1.g00514.t01#2       3810   125258286:   mNeoVis.vulgivaga.1.1.chr1.g00512.t01#4    mNeoVis.2.1.1.chr1_hap1.g00510.t01#5   mNeoVis.vulgivaga.1.1.chr1.g00513.t01#4           3811   125258287:   mNeoVis.2.1.1.chr1_hap1.g00511.t01#5   mNeoVis.3.1.1.chr1.g00519.t01#6             3812   125258288:   mNeoVis.2.1.1.chr1_hap1.g00514.t01#5   mNeoVis.evergladensis.1.1.chr1.g00526.t01#3           3813   125258289:   mNeoVis.3.1.1.chr1.g00515.t01#6   mNeoVis.vulgivaga.1.1.chr1.g00510.t01#4             3814   125258290:   mNeoVis.evergladensis.2.1.chr1.g00513.t01#2   mNeoVis.vulgivaga.2.1.chr1.g00524.t01#1             3815   125258291:   mNeoVis.vulgivaga.2.1.chr1.g00526.t01#1   mNeoVis.3.1.1.chr1.g00520.t01#6             3816   125258292:   mNeoVis.evergladensis.2.1.chr1.g00516.t01#2   mNeoVis.vulgivaga.2.1.chr1.g00527.t01#1             3817   125258293:  mNeoVis.2.1.1.chr1_hap1.g00508.t01#5               3818   125258294:  mNeoVis.2.1.1.chr1_hap1.g00509.t01#5               3819   125258295:  mNeoVis.2.1.1.chr1_hap1.g00512.t01#5               3820   125258296:  mNeoVis.2.1.1.chr1_hap1.g00513.t01#5               3821   125258297:  mNeoVis.evergladensis.2.1.chr1.g00515.t01#2             3822   125258298:  mNeoVis.evergladensis.1.1.chr1.g00524.t01#3             3823   125258299:   mNeoVis.3.1.1.chr7.g01139.t01#6    mNeoVis.evergladensis.1.1.chr7.g01204.t01#3    mNeoVis.evergladensis.2.1.chr7.g01205.t01#2    mNeoVis.2.1.1.chr7_hap1.g01216.t01#5    mNeoVis.vulgivaga.1.1.chr7.g01203.t01#4   mNeoVis.vulgivaga.2.1.chr7.g01214.t01#1       3824   125258300:   mNeoVis.2.1.1.chr7_hap1.g01217.t01#5    mNeoVis.3.1.1.chr7.g01140.t01#6    mNeoVis.evergladensis.1.1.chr7.g01205.t01#3    mNeoVis.vulgivaga.1.1.chr7.g01204.t01#4    mNeoVis.vulgivaga.2.1.chr7.g01215.t01#1   mNeoVis.evergladensis.2.1.chr7.g01206.t01#2       3825   125258301:   mNeoVis.2.1.1.chr7_hap1.g01218.t01#5    mNeoVis.vulgivaga.1.1.chr7.g01205.t01#4    mNeoVis.evergladensis.2.1.chr7.g01207.t01#2    mNeoVis.evergladensis.1.1.chr7.g01206.t01#3    mNeoVis.3.1.1.chr7.g01141.t01#6   mNeoVis.vulgivaga.2.1.chr7.g01216.t01#1       3826   125258302:  mNeoVis.vulgivaga.2.1.chr1.g00528.t01#1               3827   125258303:  mNeoVis.vulgivaga.2.1.chr1.g00529.t01#1               3828   125258304:   mNeoVis.evergladensis.2.1.chr7.g01208.t01#2    mNeoVis.evergladensis.1.1.chr7.g01207.t01#3    mNeoVis.3.1.1.chr7.g01142.t01#6    mNeoVis.2.1.1.chr7_hap1.g01219.t01#5    mNeoVis.vulgivaga.1.1.chr7.g01206.t01#4   mNeoVis.vulgivaga.2.1.chr7.g01217.t01#1       3829   125258305:   mNeoVis.vulgivaga.2.1.chr1.g00530.t01#1    mNeoVis.vulgivaga.1.1.chr1.g00515.t01#4    mNeoVis.3.1.1.chr1.g00521.t01#6    mNeoVis.2.1.1.chr1_hap1.g00515.t01#5    mNeoVis.evergladensis.2.1.chr1.g00517.t01#2   mNeoVis.evergladensis.1.1.chr1.g00527.t01#3       3830   125258306:   mNeoVis.2.1.1.chr8_hap1.g01303.t01#5   mNeoVis.evergladensis.1.1.chr8.g01247.t01#3           3831   125258307:   mNeoVis.vulgivaga.1.1.chr7.g01207.t01#4    mNeoVis.2.1.1.chr7_hap1.g01220.t01#5    mNeoVis.3.1.1.chr7.g01143.t01#6    mNeoVis.vulgivaga.2.1.chr7.g01218.t01#1    mNeoVis.evergladensis.1.1.chr7.g01208.t01#3   mNeoVis.evergladensis.2.1.chr7.g01209.t01#2       3832   125258308:   mNeoVis.evergladensis.2.1.chr7.g00476.t01#2    mNeoVis.2.1.1.chr7_hap1.g00468.t01#5    mNeoVis.vulgivaga.1.1.chr7.g00478.t01#4    mNeoVis.vulgivaga.2.1.chr7.g00475.t01#1   mNeoVis.3.1.1.chr7.g00455.t01#6       3833   125258309:   mNeoVis.vulgivaga.2.1.chr1.g00531.t01#1    mNeoVis.evergladensis.2.1.chr1.g00518.t01#2    mNeoVis.evergladensis.1.1.chr1.g00528.t01#3    mNeoVis.3.1.1.chr1.g00522.t01#6    mNeoVis.vulgivaga.1.1.chr1.g00516.t01#4   mNeoVis.2.1.1.chr1_hap1.g00516.t01#5       3834   125258310:   mNeoVis.3.1.1.chr1.g00524.t01#6    mNeoVis.vulgivaga.1.1.chr1.g00517.t01#4    mNeoVis.evergladensis.2.1.chr1.g00519.t01#2   mNeoVis.vulgivaga.2.1.chr1.g00532.t01#1         3835   125258311:   mNeoVis.2.1.1.chr1_hap1.g00517.t01#5    mNeoVis.evergladensis.2.1.chr1.g00520.t01#2    mNeoVis.vulgivaga.1.1.chr1.g00518.t01#4    mNeoVis.3.1.1.chr1.g00525.t01#6   mNeoVis.vulgivaga.2.1.chr1.g00533.t01#1       3836   125258312:   mNeoVis.evergladensis.1.1.chr1.g00532.t01#3    mNeoVis.2.1.1.chr1_hap1.g00521.t01#5    mNeoVis.3.1.1.chr1.g00533.t01#6   mNeoVis.vulgivaga.1.1.chr1.g00527.t01#4         3837   125258313:   mNeoVis.vulgivaga.2.1.chr1.g00540.t01#1   mNeoVis.3.1.1.chr1.g00537.t01#6             3838   125258314:   mNeoVis.evergladensis.1.1.chr1.g00534.t01#3   mNeoVis.vulgivaga.1.1.chr1.g00531.t01#4             3839   125258315:   mNeoVis.evergladensis.2.1.chr1.g00522.t01#2   mNeoVis.vulgivaga.2.1.chr1.g00534.t01#1             3840   125258316:   mNeoVis.evergladensis.2.1.chr1.g00523.t01#2    mNeoVis.vulgivaga.1.1.chr1.g00522.t01#4    mNeoVis.2.1.1.chr1_hap1.g00518.t01#5   mNeoVis.vulgivaga.2.1.chr1.g00535.t01#1         3841   125258317:   mNeoVis.evergladensis.1.1.chr7.g01209.t01#3    mNeoVis.3.1.1.chr7.g01144.t01#6    mNeoVis.vulgivaga.2.1.chr7.g01219.t01#1    mNeoVis.2.1.1.chr7_hap1.g01221.t01#5    mNeoVis.evergladensis.2.1.chr7.g01211.t01#2   mNeoVis.vulgivaga.1.1.chr7.g01208.t01#4       3842   125258318:   mNeoVis.vulgivaga.1.1.chr15.g00530.t01#4    mNeoVis.evergladensis.2.1.chr15.g00527.t01#2    mNeoVis.evergladensis.1.1.chr15.g00086.t01#3    mNeoVis.vulgivaga.2.1.chr15.g00539.t01#1    mNeoVis.2.1.1.chr15_hap1.g00705.t01#5   mNeoVis.3.1.1.chr15.g00486.t01#6       3843   125258319:   mNeoVis.evergladensis.2.1.chr1.g00527.t01#2    mNeoVis.vulgivaga.2.1.chr1.g00536.t01#1    mNeoVis.evergladensis.1.1.chr1.g00533.t01#3    mNeoVis.vulgivaga.1.1.chr1.g00528.t01#4    mNeoVis.2.1.1.chr1_hap1.g00522.t01#5   mNeoVis.3.1.1.chr1.g00534.t01#6       3844   125258320:   mNeoVis.vulgivaga.1.1.chr1.g00526.t01#4    mNeoVis.3.1.1.chr1.g00532.t01#6    mNeoVis.evergladensis.1.1.chr1.g00531.t01#3   mNeoVis.2.1.1.chr1_hap1.g00520.t01#5         3845   125258321:   mNeoVis.vulgivaga.2.1.chr1.g00537.t01#1    mNeoVis.vulgivaga.1.1.chr1.g00529.t01#4   mNeoVis.evergladensis.2.1.chr1.g00528.t01#2         3846   125258322:   mNeoVis.vulgivaga.2.1.chr1.g00538.t01#1    mNeoVis.2.1.1.chr1_hap1.g00523.t01#5    mNeoVis.evergladensis.2.1.chr1.g00529.t01#2    mNeoVis.3.1.1.chr1.g00536.t01#6   mNeoVis.3.1.1.chr1.g00538.t01#6       3847   125258323:   mNeoVis.3.1.1.chr1.g00539.t01#6    mNeoVis.2.1.1.chr1_hap1.g00524.t01#5    mNeoVis.vulgivaga.1.1.chr1.g00532.t01#4    mNeoVis.evergladensis.2.1.chr1.g00530.t01#2   mNeoVis.vulgivaga.2.1.chr1.g00539.t01#1       3848   125258324:   mNeoVis.2.1.1.chr7_hap1.g01222.t01#5    mNeoVis.evergladensis.2.1.chr7.g01212.t01#2    mNeoVis.vulgivaga.1.1.chr7.g01209.t01#4    mNeoVis.vulgivaga.2.1.chr7.g01220.t01#1    mNeoVis.evergladensis.1.1.chr7.g01210.t01#3   mNeoVis.3.1.1.chr7.g01145.t01#6       3849   125258325:   mNeoVis.evergladensis.2.1.chr7.g00477.t01#2    mNeoVis.vulgivaga.2.1.chr7.g00476.t01#1    mNeoVis.2.1.1.chr7_hap1.g00469.t01#5   mNeoVis.vulgivaga.1.1.chr7.g00479.t01#4         3850   125258326:   mNeoVis.vulgivaga.1.1.chr1.g00535.t01#4   mNeoVis.vulgivaga.2.1.chr1.g00541.t01#1             3851   125258327:   mNeoVis.vulgivaga.2.1.chr1.g00542.t01#1    mNeoVis.vulgivaga.1.1.chr1.g00538.t01#4    mNeoVis.evergladensis.2.1.chr1.g00533.t01#2    mNeoVis.evergladensis.1.1.chr1.g00535.t01#3   mNeoVis.2.1.1.chr1_hap1.g00526.t01#5       3852   125258328:   mNeoVis.vulgivaga.2.1.chr1.g00543.t01#1    mNeoVis.2.1.1.chr1_hap1.g00527.t01#5    mNeoVis.vulgivaga.1.1.chr1.g00539.t01#4   mNeoVis.evergladensis.2.1.chr1.g00534.t01#2       3853   125258329:   mNeoVis.3.1.1.chr1.g00543.t01#6    mNeoVis.evergladensis.1.1.chr1.g00536.t01#3    mNeoVis.vulgivaga.2.1.chr1.g00544.t01#1    mNeoVis.evergladensis.2.1.chr1.g00535.t01#2    mNeoVis.2.1.1.chr1_hap1.g00529.t01#5   mNeoVis.vulgivaga.1.1.chr1.g00540.t01#4       3854   125258330:   mNeoVis.evergladensis.2.1.chr7.g01213.t01#2    mNeoVis.evergladensis.1.1.chr7.g01211.t01#3    mNeoVis.3.1.1.chr7.g01146.t01#6    mNeoVis.2.1.1.chr7_hap1.g01223.t01#5    mNeoVis.vulgivaga.2.1.chr7.g01221.t01#1   mNeoVis.vulgivaga.1.1.chr7.g01210.t01#4       3855   125258331:   mNeoVis.vulgivaga.1.1.chr7.g01211.t01#4    mNeoVis.vulgivaga.2.1.chr7.g01222.t01#1    mNeoVis.evergladensis.2.1.chr7.g01214.t01#2    mNeoVis.evergladensis.1.1.chr7.g01212.t01#3    mNeoVis.2.1.1.chr7_hap1.g01224.t01#5   mNeoVis.3.1.1.chr7.g01147.t01#6       3856   125258332:   mNeoVis.3.1.1.chr1.g00544.t01#6    mNeoVis.evergladensis.1.1.chr1.g00537.t01#3    mNeoVis.vulgivaga.2.1.chr1.g00545.t01#1    mNeoVis.evergladensis.2.1.chr1.g00536.t01#2    mNeoVis.vulgivaga.1.1.chr1.g00541.t01#4   mNeoVis.2.1.1.chr1_hap1.g00530.t01#5       3857   125258333:   mNeoVis.vulgivaga.2.1.chr7.g00477.t01#1    mNeoVis.evergladensis.1.1.chr7.g00479.t01#3   mNeoVis.vulgivaga.1.1.chr7.g00480.t01#4           3858   125258334:   mNeoVis.vulgivaga.1.1.chr1.g00542.t01#4    mNeoVis.evergladensis.1.1.chr1.g00538.t01#3    mNeoVis.2.1.1.chr1_hap1.g00531.t01#5    mNeoVis.vulgivaga.2.1.chr1.g00546.t01#1    mNeoVis.3.1.1.chr1.g00545.t01#6   mNeoVis.evergladensis.2.1.chr1.g00537.t01#2       3859   125258335:   mNeoVis.vulgivaga.1.1.chr7.g01212.t01#4    mNeoVis.evergladensis.2.1.chr7.g01215.t01#2    mNeoVis.vulgivaga.2.1.chr7.g01224.t01#1    mNeoVis.evergladensis.1.1.chr7.g01213.t01#3    mNeoVis.vulgivaga.2.1.chr7.g01223.t01#1    mNeoVis.3.1.1.chr7.g01148.t01#6   mNeoVis.2.1.1.chr7_hap1.g01225.t01#5     3860   125258336:   mNeoVis.3.1.1.chr1.g00546.t01#6    mNeoVis.evergladensis.1.1.chr1.g00539.t01#3    mNeoVis.vulgivaga.2.1.chr1.g00547.t01#1    mNeoVis.2.1.1.chr1_hap1.g00532.t01#5    mNeoVis.vulgivaga.1.1.chr1.g00543.t01#4   mNeoVis.evergladensis.2.1.chr1.g00538.t01#2       3861   125258337:   mNeoVis.vulgivaga.1.1.chr7.g01213.t01#4    mNeoVis.vulgivaga.2.1.chr7.g01225.t01#1    mNeoVis.evergladensis.1.1.chr7.g01214.t01#3    mNeoVis.3.1.1.chr7.g01149.t01#6    mNeoVis.evergladensis.2.1.chr7.g01216.t01#2   mNeoVis.2.1.1.chr7_hap1.g01226.t01#5       3862   125258338:   mNeoVis.evergladensis.1.1.chr7.g01215.t01#3   mNeoVis.vulgivaga.2.1.chr7.g01226.t01#1             3863   125258339:   mNeoVis.vulgivaga.2.1.chr7.g01227.t01#1    mNeoVis.2.1.1.chr7_hap1.g01227.t01#5    mNeoVis.3.1.1.chr7.g01150.t01#6    mNeoVis.evergladensis.1.1.chr7.g01216.t01#3    mNeoVis.vulgivaga.1.1.chr7.g01214.t01#4   mNeoVis.evergladensis.2.1.chr7.g01217.t01#2       3864   125258340:   mNeoVis.vulgivaga.2.1.chr15.g00540.t01#1    mNeoVis.2.1.1.chr15_hap1.g00706.t01#5    mNeoVis.3.1.1.chr15.g00487.t01#6    mNeoVis.vulgivaga.1.1.chr15.g00531.t01#4    mNeoVis.evergladensis.1.1.chr15.g00085.t01#3   mNeoVis.evergladensis.2.1.chr15.g00528.t01#2       3865   125258341:   mNeoVis.vulgivaga.1.1.chr1.g00544.t01#4    mNeoVis.vulgivaga.2.1.chr1.g00548.t01#1    mNeoVis.evergladensis.1.1.chr1.g00540.t01#3    mNeoVis.3.1.1.chr1.g00547.t01#6    mNeoVis.2.1.1.chr1_hap1.g00533.t01#5   mNeoVis.evergladensis.2.1.chr1.g00539.t01#2       3866   125258342:   mNeoVis.3.1.1.chr1.g00548.t01#6    mNeoVis.evergladensis.1.1.chr1.g00541.t01#3    mNeoVis.2.1.1.chr1_hap1.g00534.t01#5    mNeoVis.vulgivaga.1.1.chr1.g00545.t01#4    mNeoVis.evergladensis.2.1.chr1.g00541.t01#2   mNeoVis.vulgivaga.2.1.chr1.g00549.t01#1       3867   125258343:   mNeoVis.vulgivaga.2.1.chr7.g00478.t01#1    mNeoVis.2.1.1.chr7_hap1.g00471.t01#5    mNeoVis.evergladensis.1.1.chr7.g00481.t01#3    mNeoVis.evergladensis.2.1.chr7.g00479.t01#2    mNeoVis.vulgivaga.1.1.chr7.g00481.t01#4   mNeoVis.3.1.1.chr7.g00457.t01#6       3868   125258344:   mNeoVis.evergladensis.2.1.chr1.g00542.t01#2    mNeoVis.vulgivaga.2.1.chr1.g00550.t01#1    mNeoVis.2.1.1.chr1_hap1.g00535.t01#5    mNeoVis.evergladensis.1.1.chr1.g00542.t01#3    mNeoVis.vulgivaga.1.1.chr1.g00546.t01#4   mNeoVis.3.1.1.chr1.g00549.t01#6       3869   125258345:   mNeoVis.evergladensis.2.1.chr7.g01218.t01#2    mNeoVis.2.1.1.chr7_hap1.g01228.t01#5    mNeoVis.vulgivaga.2.1.chr7.g01228.t01#1   mNeoVis.evergladensis.1.1.chr7.g01217.t01#3       3870   125258346:   mNeoVis.evergladensis.2.1.chr1.g00543.t01#2    mNeoVis.2.1.1.chr1_hap1.g00536.t01#5    mNeoVis.vulgivaga.1.1.chr1.g00547.t01#4    mNeoVis.vulgivaga.2.1.chr1.g00551.t01#1    mNeoVis.3.1.1.chr1.g00550.t01#6   mNeoVis.evergladensis.1.1.chr1.g00543.t01#3       3871   125258347:  mNeoVis.vulgivaga.2.1.chr1.g00553.t01#1               3872   125258348:   mNeoVis.vulgivaga.2.1.chr1.g00552.t01#1    mNeoVis.evergladensis.2.1.chr1.g00544.t01#2    mNeoVis.3.1.1.chr1.g00551.t01#6    mNeoVis.2.1.1.chr1_hap1.g00537.t01#5    mNeoVis.evergladensis.1.1.chr1.g00544.t01#3   mNeoVis.vulgivaga.1.1.chr1.g00548.t01#4       3873   125258349:   mNeoVis.vulgivaga.1.1.chr4.g00202.t01#4    mNeoVis.2.1.1.chr6_hap1.g00202.t01#5    mNeoVis.3.1.1.chr4.g00198.t01#6    mNeoVis.evergladensis.2.1.chr4.g00207.t01#2   mNeoVis.evergladensis.1.1.chr4.g00203.t01#3       3874   125258350:   mNeoVis.2.1.1.chr1_hap1.g00538.t01#5    mNeoVis.evergladensis.2.1.chr1.g00545.t01#2    mNeoVis.vulgivaga.1.1.chr1.g00549.t01#4    mNeoVis.vulgivaga.2.1.chr1.g00554.t01#1    mNeoVis.3.1.1.chr1.g00552.t01#6   mNeoVis.evergladensis.1.1.chr1.g00545.t01#3       3875   125258351:   mNeoVis.vulgivaga.2.1.chr7.g01229.t01#1   mNeoVis.vulgivaga.1.1.chr7.g01218.t01#4             3876   125258352:   mNeoVis.3.1.1.chr1.g00553.t01#6    mNeoVis.2.1.1.chr1_hap1.g00539.t01#5    mNeoVis.evergladensis.1.1.chr1.g00546.t01#3    mNeoVis.vulgivaga.2.1.chr1.g00555.t01#1    mNeoVis.vulgivaga.1.1.chr1.g00550.t01#4   mNeoVis.evergladensis.2.1.chr1.g00546.t01#2       3877   125258353:   mNeoVis.3.1.1.chr1.g00554.t01#6    mNeoVis.evergladensis.1.1.chr1.g00547.t01#3    mNeoVis.vulgivaga.2.1.chr1.g00556.t01#1    mNeoVis.2.1.1.chr1_hap1.g00540.t01#5    mNeoVis.evergladensis.2.1.chr1.g00547.t01#2   mNeoVis.vulgivaga.1.1.chr1.g00551.t01#4       3878   125258354:   mNeoVis.evergladensis.2.1.chr1.g00548.t01#2    mNeoVis.2.1.1.chr1_hap1.g00541.t01#5    mNeoVis.vulgivaga.1.1.chr1.g00552.t01#4    mNeoVis.vulgivaga.2.1.chr1.g00557.t01#1    mNeoVis.evergladensis.1.1.chr1.g00548.t01#3   mNeoVis.3.1.1.chr1.g00555.t01#6       3879   125258355:   mNeoVis.3.1.1.chr7.g01154.t01#6    mNeoVis.2.1.1.chr7_hap1.g01229.t01#5    mNeoVis.vulgivaga.2.1.chr7.g01230.t01#1    mNeoVis.evergladensis.1.1.chr7.g01218.t01#3    mNeoVis.evergladensis.2.1.chr7.g01219.t01#2   mNeoVis.vulgivaga.1.1.chr7.g01219.t01#4       3880   125258356:   mNeoVis.3.1.1.chr7.g01155.t01#6    mNeoVis.2.1.1.chr7_hap1.g01230.t01#5    mNeoVis.vulgivaga.2.1.chr7.g01231.t01#1    mNeoVis.evergladensis.2.1.chr7.g01220.t01#2    mNeoVis.evergladensis.1.1.chr7.g01219.t01#3   mNeoVis.vulgivaga.1.1.chr7.g01220.t01#4       3881   125258357:   mNeoVis.evergladensis.1.1.chr15.g00084.t01#3    mNeoVis.vulgivaga.2.1.chr15.g00541.t01#1    mNeoVis.2.1.1.chr15_hap1.g00707.t01#5    mNeoVis.3.1.1.chr15.g00488.t01#6    mNeoVis.vulgivaga.1.1.chr15.g00532.t01#4   mNeoVis.vulgivaga.2.1.chr15.g00542.t01#1       3882   125258358:   mNeoVis.vulgivaga.2.1.chr1.g00558.t01#1    mNeoVis.evergladensis.1.1.chr1.g00549.t01#3    mNeoVis.3.1.1.chr1.g00556.t01#6    mNeoVis.evergladensis.2.1.chr1.g00549.t01#2    mNeoVis.2.1.1.chr1_hap1.g00542.t01#5   mNeoVis.vulgivaga.1.1.chr1.g00553.t01#4       3883   125258359:   mNeoVis.2.1.1.chr7_hap1.g00472.t01#5    mNeoVis.vulgivaga.2.1.chr7.g00479.t01#1    mNeoVis.vulgivaga.1.1.chr7.g00483.t01#4    mNeoVis.3.1.1.chr7.g00458.t01#6   mNeoVis.evergladensis.1.1.chr7.g00482.t01#3       3884   125258360:   mNeoVis.vulgivaga.1.1.chr7.g01221.t01#4    mNeoVis.vulgivaga.2.1.chr7.g01232.t01#1    mNeoVis.2.1.1.chr7_hap1.g01231.t01#5    mNeoVis.evergladensis.2.1.chr7.g01221.t01#2    mNeoVis.3.1.1.chr7.g01156.t01#6   mNeoVis.evergladensis.1.1.chr7.g01220.t01#3       3885   125258361:   mNeoVis.vulgivaga.2.1.chr7.g01146.t01#1    mNeoVis.evergladensis.2.1.chr7.g01134.t01#2    mNeoVis.2.1.1.chr7_hap1.g01147.t01#5    mNeoVis.3.1.1.chr7.g01070.t01#6   mNeoVis.evergladensis.1.1.chr7.g01135.t01#3       3886   125258362:   mNeoVis.2.1.1.chr1_hap1.g00544.t01#5    mNeoVis.3.1.1.chr1.g00558.t01#6    mNeoVis.evergladensis.1.1.chr1.g00551.t01#3    mNeoVis.vulgivaga.1.1.chr1.g00555.t01#4    mNeoVis.vulgivaga.2.1.chr1.g00560.t01#1   mNeoVis.evergladensis.2.1.chr1.g00551.t01#2       3887   125258363:   mNeoVis.vulgivaga.2.1.chr1.g00561.t01#1    mNeoVis.3.1.1.chr1.g00559.t01#6    mNeoVis.2.1.1.chr1_hap1.g00545.t01#5    mNeoVis.evergladensis.1.1.chr1.g00552.t01#3    mNeoVis.evergladensis.2.1.chr1.g00552.t01#2   mNeoVis.vulgivaga.1.1.chr1.g00556.t01#4       3888   125258364:   mNeoVis.vulgivaga.1.1.chr7.g01137.t01#4    mNeoVis.evergladensis.1.1.chr7.g01136.t01#3    mNeoVis.vulgivaga.2.1.chr7.g01147.t01#1    mNeoVis.evergladensis.2.1.chr7.g01135.t01#2   mNeoVis.2.1.1.chr7_hap1.g01148.t01#5       3889   125258365:   mNeoVis.evergladensis.1.1.chr1.g00553.t01#3    mNeoVis.2.1.1.chr1_hap1.g00546.t01#5    mNeoVis.vulgivaga.2.1.chr1.g00562.t01#1    mNeoVis.3.1.1.chr1.g00561.t01#6    mNeoVis.vulgivaga.1.1.chr1.g00557.t01#4   mNeoVis.evergladensis.2.1.chr1.g00553.t01#2       3890   125258366:   mNeoVis.2.1.1.chr1_hap1.g00547.t01#5    mNeoVis.vulgivaga.1.1.chr1.g00558.t01#4    mNeoVis.vulgivaga.2.1.chr1.g00563.t01#1   mNeoVis.3.1.1.chr1.g00562.t01#6         3891   125258367:   mNeoVis.vulgivaga.1.1.chr1.g00560.t01#4    mNeoVis.evergladensis.2.1.chr1.g00554.t01#2    mNeoVis.evergladensis.1.1.chr1.g00554.t01#3    mNeoVis.3.1.1.chr1.g00563.t01#6    mNeoVis.vulgivaga.2.1.chr1.g00564.t01#1   mNeoVis.2.1.1.chr1_hap1.g00548.t01#5       3892   125258368:   mNeoVis.3.1.1.chr7.g01072.t01#6    mNeoVis.vulgivaga.2.1.chr7.g01148.t01#1    mNeoVis.2.1.1.chr7_hap1.g01149.t01#5    mNeoVis.vulgivaga.1.1.chr7.g01138.t01#4   mNeoVis.evergladensis.2.1.chr7.g01136.t01#2       3893   125258369:   mNeoVis.vulgivaga.1.1.chr15.g00533.t01#4    mNeoVis.evergladensis.2.1.chr15.g00530.t01#2    mNeoVis.evergladensis.1.1.chr15.g00083.t01#3    mNeoVis.2.1.1.chr15_hap1.g00708.t01#5   mNeoVis.vulgivaga.2.1.chr15.g00543.t01#1       3894   125258370:   mNeoVis.evergladensis.1.1.chr1.g00555.t01#3    mNeoVis.vulgivaga.1.1.chr1.g00561.t01#4    mNeoVis.evergladensis.2.1.chr1.g00555.t01#2    mNeoVis.vulgivaga.2.1.chr1.g00565.t01#1    mNeoVis.3.1.1.chr1.g00564.t01#6   mNeoVis.2.1.1.chr1_hap1.g00549.t01#5       3895   125258371:   mNeoVis.vulgivaga.2.1.chr1.g00566.t01#1    mNeoVis.2.1.1.chr1_hap1.g00550.t01#5    mNeoVis.evergladensis.2.1.chr1.g00556.t01#2    mNeoVis.3.1.1.chr1.g00565.t01#6    mNeoVis.evergladensis.1.1.chr1.g00556.t01#3   mNeoVis.vulgivaga.1.1.chr1.g00562.t01#4       3896   125258372:   mNeoVis.3.1.1.chr7.g01073.t01#6    mNeoVis.vulgivaga.2.1.chr7.g01149.t01#1    mNeoVis.evergladensis.2.1.chr7.g01137.t01#2    mNeoVis.2.1.1.chr7_hap1.g01150.t01#5    mNeoVis.vulgivaga.1.1.chr7.g01139.t01#4   mNeoVis.evergladensis.1.1.chr7.g01137.t01#3       3897   125258373:   mNeoVis.vulgivaga.2.1.chr7.g01150.t01#1    mNeoVis.vulgivaga.1.1.chr7.g01141.t01#4    mNeoVis.evergladensis.1.1.chr7.g01140.t01#3    mNeoVis.2.1.1.chr7_hap1.g01152.t01#5    mNeoVis.3.1.1.chr7.g01074.t01#6   mNeoVis.evergladensis.2.1.chr7.g01139.t01#2       3898   125258374:   mNeoVis.vulgivaga.1.1.chr1.g00563.t01#4    mNeoVis.3.1.1.chr1.g00566.t01#6    mNeoVis.evergladensis.2.1.chr1.g00557.t01#2    mNeoVis.vulgivaga.2.1.chr1.g00567.t01#1    mNeoVis.2.1.1.chr1_hap1.g00551.t01#5   mNeoVis.evergladensis.1.1.chr1.g00558.t01#3       3899   125258375:   mNeoVis.evergladensis.1.1.chr1.g00559.t01#3    mNeoVis.2.1.1.chr1_hap1.g00552.t01#5    mNeoVis.evergladensis.2.1.chr1.g00558.t01#2    mNeoVis.3.1.1.chr1.g00567.t01#6    mNeoVis.vulgivaga.2.1.chr1.g00568.t01#1   mNeoVis.vulgivaga.1.1.chr1.g00564.t01#4       3900   125258376:   mNeoVis.evergladensis.1.1.chr7.g00483.t01#3    mNeoVis.vulgivaga.2.1.chr7.g00481.t01#1    mNeoVis.3.1.1.chr7.g00459.t01#6    mNeoVis.evergladensis.2.1.chr7.g00480.t01#2   mNeoVis.vulgivaga.1.1.chr7.g00485.t01#4       3901   125258377:   mNeoVis.2.1.1.chr15_hap1.g00709.t01#5    mNeoVis.evergladensis.1.1.chr15.g00082.t01#3    mNeoVis.vulgivaga.2.1.chr15.g00544.t01#1    mNeoVis.vulgivaga.1.1.chr15.g00534.t01#4   mNeoVis.evergladensis.2.1.chr15.g00531.t01#2       3902   125258378:   mNeoVis.evergladensis.1.1.chr1.g00560.t01#3    mNeoVis.evergladensis.2.1.chr1.g00559.t01#2    mNeoVis.vulgivaga.2.1.chr1.g00569.t01#1    mNeoVis.2.1.1.chr1_hap1.g00553.t01#5    mNeoVis.vulgivaga.1.1.chr1.g00565.t01#4   mNeoVis.3.1.1.chr1.g00568.t01#6       3903   125258379:   mNeoVis.evergladensis.1.1.chr1.g00561.t01#3    mNeoVis.2.1.1.chr1_hap1.g00554.t01#5    mNeoVis.evergladensis.2.1.chr1.g00560.t01#2    mNeoVis.vulgivaga.1.1.chr1.g00566.t01#4    mNeoVis.3.1.1.chr1.g00569.t01#6   mNeoVis.vulgivaga.2.1.chr1.g00570.t01#1       3904   125258380:   mNeoVis.vulgivaga.2.1.chr1.g00571.t01#1    mNeoVis.vulgivaga.1.1.chr1.g00567.t01#4    mNeoVis.3.1.1.chr1.g00570.t01#6    mNeoVis.evergladensis.1.1.chr1.g00562.t01#3    mNeoVis.2.1.1.chr1_hap1.g00555.t01#5   mNeoVis.evergladensis.2.1.chr1.g00561.t01#2       3905   125258381:   mNeoVis.vulgivaga.2.1.chr1.g00454.t01#1    mNeoVis.evergladensis.1.1.chr1.g00456.t01#3    mNeoVis.evergladensis.2.1.chr1.g00449.t01#2    mNeoVis.vulgivaga.1.1.chr1.g00443.t01#4    mNeoVis.2.1.1.chr1_hap1.g00451.t01#5   mNeoVis.3.1.1.chr1.g00449.t01#6       3906   125258382:   mNeoVis.2.1.1.chr7_hap1.g01153.t01#5    mNeoVis.evergladensis.1.1.chr7.g01141.t01#3    mNeoVis.3.1.1.chr7.g01075.t01#6    mNeoVis.evergladensis.2.1.chr7.g01140.t01#2    mNeoVis.vulgivaga.1.1.chr7.g01142.t01#4   mNeoVis.vulgivaga.2.1.chr7.g01151.t01#1       3907   125258383:   mNeoVis.vulgivaga.2.1.chr7.g01152.t01#1    mNeoVis.evergladensis.1.1.chr7.g01142.t01#3    mNeoVis.3.1.1.chr7.g01076.t01#6    mNeoVis.vulgivaga.1.1.chr7.g01143.t01#4    mNeoVis.2.1.1.chr7_hap1.g01154.t01#5   mNeoVis.evergladensis.2.1.chr7.g01141.t01#2       3908   125258384:   mNeoVis.vulgivaga.1.1.chr7.g00487.t01#4    mNeoVis.3.1.1.chr7.g00461.t01#6    mNeoVis.evergladensis.2.1.chr7.g00481.t01#2    mNeoVis.evergladensis.1.1.chr7.g00485.t01#3    mNeoVis.vulgivaga.2.1.chr7.g00482.t01#1   mNeoVis.2.1.1.chr7_hap1.g00475.t01#5       3909   125258385:   mNeoVis.vulgivaga.1.1.chr1.g00445.t01#4    mNeoVis.evergladensis.1.1.chr1.g00457.t01#3    mNeoVis.vulgivaga.2.1.chr1.g00455.t01#1    mNeoVis.3.1.1.chr1.g00450.t01#6    mNeoVis.evergladensis.2.1.chr1.g00450.t01#2   mNeoVis.2.1.1.chr1_hap1.g00452.t01#5       3910   125258386:   mNeoVis.evergladensis.2.1.chr1.g00451.t01#2    mNeoVis.vulgivaga.1.1.chr1.g00446.t01#4    mNeoVis.3.1.1.chr1.g00451.t01#6    mNeoVis.vulgivaga.2.1.chr1.g00456.t01#1    mNeoVis.evergladensis.1.1.chr1.g00458.t01#3   mNeoVis.2.1.1.chr1_hap1.g00453.t01#5       3911   125258387:   mNeoVis.3.1.1.chr1.g00452.t01#6    mNeoVis.evergladensis.2.1.chr1.g00452.t01#2    mNeoVis.vulgivaga.2.1.chr1.g00457.t01#1    mNeoVis.3.1.1.chr1.g00453.t01#6   mNeoVis.vulgivaga.1.1.chr1.g00447.t01#4       3912   125258388:   mNeoVis.evergladensis.1.1.chr1.g00461.t01#3    mNeoVis.evergladensis.2.1.chr1.g00453.t01#2    mNeoVis.vulgivaga.2.1.chr1.g00460.t01#1   mNeoVis.evergladensis.1.1.chr1.g00460.t01#3       3913   125258389:   mNeoVis.2.1.1.chr1_hap1.g00454.t01#5    mNeoVis.evergladensis.1.1.chr1.g00459.t01#3   mNeoVis.3.1.1.chr1.g00454.t01#6           3914   125258390:   mNeoVis.vulgivaga.2.1.chr1.g00459.t01#1   mNeoVis.vulgivaga.1.1.chr1.g00448.t01#4             3915   125258391:  mNeoVis.vulgivaga.2.1.chr1.g00458.t01#1               3916   125258392:   mNeoVis.evergladensis.2.1.chr7.g01142.t01#2    mNeoVis.vulgivaga.1.1.chr7.g01144.t01#4    mNeoVis.vulgivaga.2.1.chr7.g01153.t01#1    mNeoVis.2.1.1.chr7_hap1.g01155.t01#5    mNeoVis.3.1.1.chr7.g01077.t01#6   mNeoVis.evergladensis.1.1.chr7.g01143.t01#3       3917   125258393:   mNeoVis.vulgivaga.2.1.chr7.g01154.t01#1    mNeoVis.evergladensis.2.1.chr7.g01143.t01#2    mNeoVis.2.1.1.chr7_hap1.g01156.t01#5    mNeoVis.evergladensis.1.1.chr7.g01144.t01#3    mNeoVis.vulgivaga.1.1.chr7.g01145.t01#4   mNeoVis.3.1.1.chr7.g01078.t01#6       3918   125258394:   mNeoVis.2.1.1.chr1_hap1.g00455.t01#5    mNeoVis.vulgivaga.2.1.chr1.g00461.t01#1    mNeoVis.evergladensis.1.1.chr1.g00462.t01#3    mNeoVis.vulgivaga.1.1.chr1.g00449.t01#4    mNeoVis.3.1.1.chr1.g00455.t01#6   mNeoVis.evergladensis.2.1.chr1.g00454.t01#2       3919   125258395:   mNeoVis.evergladensis.1.1.chr7.g01147.t01#3    mNeoVis.vulgivaga.2.1.chr7.g01156.t01#1    mNeoVis.2.1.1.chr7_hap1.g01158.t01#5    mNeoVis.vulgivaga.2.1.chr7.g01157.t01#1    mNeoVis.vulgivaga.1.1.chr7.g01148.t01#4    mNeoVis.3.1.1.chr7.g01081.t01#6   mNeoVis.evergladensis.2.1.chr7.g01146.t01#2     3920   125258396:   mNeoVis.evergladensis.2.1.chr7.g01144.t01#2    mNeoVis.evergladensis.1.1.chr7.g01145.t01#3    mNeoVis.vulgivaga.1.1.chr7.g01146.t01#4    mNeoVis.2.1.1.chr7_hap1.g01157.t01#5    mNeoVis.vulgivaga.2.1.chr7.g01155.t01#1   mNeoVis.3.1.1.chr7.g01079.t01#6       3921   125258397:   mNeoVis.vulgivaga.1.1.chr1.g00450.t01#4    mNeoVis.3.1.1.chr1.g00456.t01#6    mNeoVis.vulgivaga.2.1.chr1.g00462.t01#1    mNeoVis.evergladensis.2.1.chr1.g00455.t01#2    mNeoVis.2.1.1.chr1_hap1.g00456.t01#5   mNeoVis.evergladensis.1.1.chr1.g00463.t01#3       3922   125258398:   mNeoVis.vulgivaga.2.1.chr7.g01158.t01#1    mNeoVis.vulgivaga.1.1.chr7.g01149.t01#4    mNeoVis.3.1.1.chr7.g01082.t01#6    mNeoVis.evergladensis.2.1.chr7.g01147.t01#2    mNeoVis.2.1.1.chr7_hap1.g01159.t01#5   mNeoVis.evergladensis.1.1.chr7.g01148.t01#3       3923   125258399:   mNeoVis.evergladensis.2.1.chr7.g01148.t01#2    mNeoVis.3.1.1.chr7.g01083.t01#6    mNeoVis.vulgivaga.2.1.chr7.g01159.t01#1    mNeoVis.2.1.1.chr7_hap1.g01160.t01#5    mNeoVis.evergladensis.1.1.chr7.g01149.t01#3   mNeoVis.vulgivaga.1.1.chr7.g01150.t01#4       3924   125258400:   mNeoVis.vulgivaga.2.1.chr1.g00463.t01#1    mNeoVis.vulgivaga.1.1.chr1.g00451.t01#4    mNeoVis.evergladensis.2.1.chr1.g00456.t01#2    mNeoVis.3.1.1.chr1.g00457.t01#6    mNeoVis.2.1.1.chr1_hap1.g00457.t01#5   mNeoVis.evergladensis.1.1.chr1.g00464.t01#3       3925   125258401:   mNeoVis.vulgivaga.2.1.chr7.g01160.t01#1    mNeoVis.evergladensis.1.1.chr7.g01150.t01#3    mNeoVis.vulgivaga.1.1.chr7.g01151.t01#4    mNeoVis.evergladensis.2.1.chr7.g01149.t01#2    mNeoVis.3.1.1.chr7.g01084.t01#6   mNeoVis.2.1.1.chr7_hap1.g01161.t01#5       3926   125258402:   mNeoVis.vulgivaga.1.1.chr1.g00452.t01#4    mNeoVis.evergladensis.2.1.chr1.g00457.t01#2    mNeoVis.vulgivaga.2.1.chr1.g00464.t01#1    mNeoVis.evergladensis.1.1.chr1.g00465.t01#3    mNeoVis.3.1.1.chr1.g00458.t01#6   mNeoVis.2.1.1.chr1_hap1.g00458.t01#5       3927   125258403:   mNeoVis.vulgivaga.1.1.chr7.g01152.t01#4    mNeoVis.evergladensis.1.1.chr7.g01151.t01#3    mNeoVis.vulgivaga.2.1.chr7.g01161.t01#1    mNeoVis.evergladensis.2.1.chr7.g01150.t01#2   mNeoVis.3.1.1.chr7.g01085.t01#6       3928   125258404:  mNeoVis.vulgivaga.2.1.chr7.g01162.t01#1               3929   125258405:   mNeoVis.evergladensis.2.1.chr1.g00458.t01#2    mNeoVis.vulgivaga.1.1.chr1.g00453.t01#4    mNeoVis.evergladensis.1.1.chr1.g00466.t01#3    mNeoVis.vulgivaga.2.1.chr1.g00465.t01#1   mNeoVis.2.1.1.chr1_hap1.g00459.t01#5       3930   125258406:   mNeoVis.3.1.1.chr1.g00459.t01#6    mNeoVis.evergladensis.1.1.chr1.g00467.t01#3    mNeoVis.vulgivaga.2.1.chr1.g00466.t01#1   mNeoVis.2.1.1.chr1_hap1.g00460.t01#5         3931   125258407:   mNeoVis.vulgivaga.2.1.chr7.g01163.t01#1    mNeoVis.vulgivaga.1.1.chr7.g01153.t01#4    mNeoVis.2.1.1.chr7_hap1.g01162.t01#5    mNeoVis.evergladensis.2.1.chr7.g01151.t01#2    mNeoVis.evergladensis.1.1.chr7.g01152.t01#3   mNeoVis.3.1.1.chr7.g01086.t01#6       3932   125258408:   mNeoVis.2.1.1.chr1_hap1.g00461.t01#5    mNeoVis.vulgivaga.2.1.chr1.g00467.t01#1    mNeoVis.evergladensis.2.1.chr1.g00459.t01#2    mNeoVis.evergladensis.1.1.chr1.g00468.t01#3    mNeoVis.vulgivaga.1.1.chr1.g00454.t01#4   mNeoVis.3.1.1.chr1.g00460.t01#6       3933   125258409:   mNeoVis.vulgivaga.2.1.chr7.g01164.t01#1    mNeoVis.vulgivaga.1.1.chr7.g01154.t01#4    mNeoVis.3.1.1.chr7.g01087.t01#6    mNeoVis.evergladensis.1.1.chr7.g01153.t01#3    mNeoVis.evergladensis.2.1.chr7.g01152.t01#2   mNeoVis.2.1.1.chr7_hap1.g01163.t01#5       3934   125258410:   mNeoVis.evergladensis.1.1.chr7.g00486.t01#3    mNeoVis.2.1.1.chr7_hap1.g00476.t01#5    mNeoVis.evergladensis.2.1.chr7.g00482.t01#2    mNeoVis.vulgivaga.1.1.chr7.g00488.t01#4   mNeoVis.vulgivaga.2.1.chr7.g00483.t01#1       3935   125258411:   mNeoVis.vulgivaga.1.1.chr7.g01155.t01#4    mNeoVis.2.1.1.chr7_hap1.g01164.t01#5    mNeoVis.evergladensis.1.1.chr7.g01154.t01#3    mNeoVis.vulgivaga.2.1.chr7.g01165.t01#1    mNeoVis.evergladensis.2.1.chr7.g01153.t01#2   mNeoVis.3.1.1.chr7.g01088.t01#6       3936   125258412:   mNeoVis.vulgivaga.2.1.chr15.g00545.t01#1    mNeoVis.evergladensis.1.1.chr15.g00081.t01#3    mNeoVis.2.1.1.chr15_hap1.g00710.t01#5    mNeoVis.vulgivaga.1.1.chr15.g00535.t01#4   mNeoVis.evergladensis.2.1.chr15.g00532.t01#2       3937   125258413:   mNeoVis.vulgivaga.2.1.chr1.g00468.t01#1    mNeoVis.3.1.1.chr1.g00461.t01#6    mNeoVis.evergladensis.1.1.chr1.g00469.t01#3    mNeoVis.2.1.1.chr1_hap1.g00462.t01#5    mNeoVis.evergladensis.2.1.chr1.g00460.t01#2   mNeoVis.vulgivaga.1.1.chr1.g00455.t01#4       3938   125258414:   mNeoVis.evergladensis.2.1.chr7.g00743.t01#2    mNeoVis.3.1.1.chr7.g00706.t01#6    mNeoVis.evergladensis.1.1.chr7.g00753.t01#3    mNeoVis.vulgivaga.1.1.chr7.g00753.t01#4    mNeoVis.vulgivaga.2.1.chr7.g00746.t01#1   mNeoVis.2.1.1.chr7_hap1.g00755.t01#5       3939   125258415:   mNeoVis.evergladensis.2.1.chr7.g01154.t01#2    mNeoVis.2.1.1.chr7_hap1.g01165.t01#5    mNeoVis.vulgivaga.2.1.chr7.g01166.t01#1   mNeoVis.3.1.1.chr7.g01089.t01#6         3940   125258416:   mNeoVis.3.1.1.chr7.g01090.t01#6    mNeoVis.2.1.1.chr7_hap1.g01166.t01#5    mNeoVis.vulgivaga.2.1.chr7.g01167.t01#1    mNeoVis.evergladensis.1.1.chr7.g01155.t01#3    mNeoVis.evergladensis.2.1.chr7.g01155.t01#2   mNeoVis.vulgivaga.1.1.chr7.g01156.t01#4       3941   125258417:   mNeoVis.vulgivaga.2.1.chr1.g00469.t01#1    mNeoVis.vulgivaga.1.1.chr1.g00456.t01#4    mNeoVis.2.1.1.chr1_hap1.g00463.t01#5    mNeoVis.evergladensis.2.1.chr1.g00461.t01#2   mNeoVis.evergladensis.1.1.chr1.g00470.t01#3       3942   125258418:   mNeoVis.3.1.1.chr1.g00462.t01#6    mNeoVis.evergladensis.2.1.chr1.g00462.t01#2    mNeoVis.vulgivaga.1.1.chr1.g00457.t01#4    mNeoVis.vulgivaga.2.1.chr1.g00470.t01#1    mNeoVis.evergladensis.1.1.chr1.g00471.t01#3   mNeoVis.2.1.1.chr1_hap1.g00464.t01#5       3943   125258419:   mNeoVis.evergladensis.2.1.chr7.g01156.t01#2    mNeoVis.evergladensis.1.1.chr7.g01156.t01#3    mNeoVis.3.1.1.chr7.g01091.t01#6    mNeoVis.vulgivaga.1.1.chr7.g01157.t01#4    mNeoVis.2.1.1.chr7_hap1.g01167.t01#5   mNeoVis.vulgivaga.2.1.chr7.g01168.t01#1       3944   125258420:   mNeoVis.evergladensis.1.1.chr7.g01157.t01#3    mNeoVis.3.1.1.chr7.g01092.t01#6    mNeoVis.vulgivaga.1.1.chr7.g01158.t01#4    mNeoVis.vulgivaga.2.1.chr7.g01169.t01#1   mNeoVis.2.1.1.chr7_hap1.g01168.t01#5       3945   125258421:   mNeoVis.evergladensis.1.1.chr1.g00472.t01#3    mNeoVis.vulgivaga.2.1.chr1.g00471.t01#1    mNeoVis.2.1.1.chr1_hap1.g00465.t01#5    mNeoVis.evergladensis.2.1.chr1.g00463.t01#2    mNeoVis.3.1.1.chr1.g00463.t01#6   mNeoVis.vulgivaga.1.1.chr1.g00458.t01#4       3946   125258422:   mNeoVis.3.1.1.chr7.g01093.t01#6    mNeoVis.2.1.1.chr7_hap1.g01169.t01#5    mNeoVis.evergladensis.1.1.chr7.g01158.t01#3    mNeoVis.evergladensis.2.1.chr7.g01158.t01#2    mNeoVis.vulgivaga.2.1.chr7.g01170.t01#1   mNeoVis.vulgivaga.1.1.chr7.g01159.t01#4       3947   125258423:   mNeoVis.2.1.1.chr7_hap1.g01170.t01#5    mNeoVis.evergladensis.2.1.chr7.g01159.t01#2    mNeoVis.vulgivaga.2.1.chr7.g01171.t01#1    mNeoVis.3.1.1.chr7.g01094.t01#6    mNeoVis.evergladensis.1.1.chr7.g01159.t01#3   mNeoVis.vulgivaga.1.1.chr7.g01160.t01#4       3948   125258424:  mNeoVis.vulgivaga.2.1.chr1.g00472.t01#1               3949   125258425:   mNeoVis.evergladensis.2.1.chr15.g00533.t01#2    mNeoVis.evergladensis.1.1.chr15.g00080.t01#3    mNeoVis.vulgivaga.1.1.chr15.g00536.t01#4    mNeoVis.vulgivaga.2.1.chr15.g00546.t01#1   mNeoVis.2.1.1.chr15_hap1.g00711.t01#5       3950   125258426:   mNeoVis.evergladensis.2.1.chr1.g00464.t01#2    mNeoVis.vulgivaga.1.1.chr1.g00460.t01#4    mNeoVis.2.1.1.chr1_hap1.g00466.t01#5    mNeoVis.vulgivaga.2.1.chr1.g00473.t01#1    mNeoVis.vulgivaga.2.1.chr1.g00474.t01#1    mNeoVis.evergladensis.1.1.chr1.g00473.t01#3   mNeoVis.3.1.1.chr1.g00464.t01#6       3951   125258427:   mNeoVis.vulgivaga.2.1.chr11.g00856.t01#1    mNeoVis.2.1.1.chr11_hap1.g00851.t01#5    mNeoVis.evergladensis.1.1.chr11.g00842.t01#3    mNeoVis.vulgivaga.1.1.chr11.g00833.t01#4   mNeoVis.evergladensis.2.1.chr11.g00834.t01#2       3952   125258428:   mNeoVis.vulgivaga.2.1.chr7.g00484.t01#1    mNeoVis.vulgivaga.1.1.chr7.g00491.t01#4    mNeoVis.2.1.1.chr7_hap1.g00477.t01#5   mNeoVis.3.1.1.chr7.g00462.t01#6         3953   125258429:   mNeoVis.evergladensis.2.1.chr7.g00483.t01#2   mNeoVis.vulgivaga.1.1.chr7.g00489.t01#4             3954   125258430:   mNeoVis.vulgivaga.2.1.chr7.g01172.t01#1    mNeoVis.2.1.1.chr7_hap1.g01171.t01#5    mNeoVis.evergladensis.1.1.chr7.g01160.t01#3    mNeoVis.3.1.1.chr7.g01095.t01#6    mNeoVis.vulgivaga.1.1.chr7.g01161.t01#4   mNeoVis.evergladensis.2.1.chr7.g01160.t01#2       3955   125258431:   mNeoVis.3.1.1.chr7.g01096.t01#6    mNeoVis.2.1.1.chr7_hap1.g01172.t01#5    mNeoVis.vulgivaga.2.1.chr7.g01173.t01#1    mNeoVis.evergladensis.1.1.chr7.g01161.t01#3    mNeoVis.vulgivaga.1.1.chr7.g01162.t01#4   mNeoVis.evergladensis.2.1.chr7.g01161.t01#2       3956   125258432:   mNeoVis.vulgivaga.1.1.chr1.g00459.t01#4    mNeoVis.3.1.1.chr1.g00465.t01#6    mNeoVis.evergladensis.2.1.chr1.g00465.t01#2    mNeoVis.evergladensis.1.1.chr1.g00474.t01#3    mNeoVis.vulgivaga.2.1.chr1.g00475.t01#1   mNeoVis.2.1.1.chr1_hap1.g00467.t01#5       3957   125258433:   mNeoVis.vulgivaga.2.1.chr1.g00476.t01#1    mNeoVis.evergladensis.2.1.chr1.g00466.t01#2    mNeoVis.evergladensis.1.1.chr1.g00475.t01#3    mNeoVis.2.1.1.chr1_hap1.g00468.t01#5    mNeoVis.vulgivaga.1.1.chr1.g00461.t01#4   mNeoVis.3.1.1.chr1.g00466.t01#6       3958   125258434:   mNeoVis.3.1.1.chr1.g00467.t01#6    mNeoVis.vulgivaga.1.1.chr1.g00462.t01#4    mNeoVis.evergladensis.2.1.chr1.g00467.t01#2    mNeoVis.vulgivaga.2.1.chr1.g00477.t01#1    mNeoVis.evergladensis.1.1.chr1.g00477.t01#3    mNeoVis.evergladensis.1.1.chr1.g00476.t01#3   mNeoVis.2.1.1.chr1_hap1.g00469.t01#5     3959   125258435:   mNeoVis.evergladensis.2.1.chr7.g01162.t01#2    mNeoVis.2.1.1.chr7_hap1.g02449.t01#5    mNeoVis.2.1.1.chr7_hap1.g01173.t01#5    mNeoVis.evergladensis.1.1.chr7.g01162.t01#3    mNeoVis.vulgivaga.1.1.chr7.g01163.t01#4    mNeoVis.2.1.1.chr7_hap1.g02420.t01#5   mNeoVis.vulgivaga.2.1.chr7.g01174.t01#1     3960   125258436:   mNeoVis.vulgivaga.2.1.chr9.g00465.t01#1    mNeoVis.evergladensis.2.1.chr8.g00461.t01#2    mNeoVis.evergladensis.1.1.chr9.g00474.t01#3    mNeoVis.3.1.1.chr9.g00448.t01#6    mNeoVis.evergladensis.2.1.chr8.g00462.t01#2    mNeoVis.vulgivaga.1.1.chr9.g00467.t01#4   mNeoVis.2.1.1.chr10_hap1.g00483.t01#5     3961   125258437:   mNeoVis.2.1.1.chr7_hap1.g01174.t01#5    mNeoVis.vulgivaga.1.1.chr7.g01164.t01#4    mNeoVis.vulgivaga.2.1.chr7.g01175.t01#1    mNeoVis.evergladensis.1.1.chr7.g01163.t01#3    mNeoVis.3.1.1.chr7.g01097.t01#6   mNeoVis.evergladensis.2.1.chr7.g01163.t01#2       3962   125258438:   mNeoVis.evergladensis.2.1.chr7.g00782.t01#2    mNeoVis.3.1.1.chr7.g00737.t01#6    mNeoVis.vulgivaga.1.1.chr7.g00794.t01#4    mNeoVis.vulgivaga.2.1.chr7.g00787.t01#1    mNeoVis.evergladensis.1.1.chr7.g00793.t01#3   mNeoVis.2.1.1.chr7_hap1.g00794.t01#5       3963   125258439:   mNeoVis.3.1.1.chr1.g00468.t01#6    mNeoVis.evergladensis.1.1.chr1.g00478.t01#3    mNeoVis.vulgivaga.2.1.chr1.g00478.t01#1    mNeoVis.evergladensis.2.1.chr1.g00468.t01#2    mNeoVis.vulgivaga.1.1.chr1.g00463.t01#4   mNeoVis.2.1.1.chr1_hap1.g00470.t01#5       3964   125258440:   mNeoVis.vulgivaga.1.1.chr1.g00464.t01#4    mNeoVis.2.1.1.chr1_hap1.g00471.t01#5    mNeoVis.vulgivaga.2.1.chr1.g00479.t01#1    mNeoVis.3.1.1.chr1.g00469.t01#6    mNeoVis.evergladensis.1.1.chr1.g00479.t01#3   mNeoVis.evergladensis.2.1.chr1.g00469.t01#2       3965   125258441:   mNeoVis.vulgivaga.2.1.chr7.g01176.t01#1    mNeoVis.2.1.1.chr7_hap1.g01175.t01#5    mNeoVis.vulgivaga.1.1.chr7.g01165.t01#4    mNeoVis.evergladensis.1.1.chr7.g01164.t01#3    mNeoVis.evergladensis.2.1.chr7.g01164.t01#2   mNeoVis.3.1.1.chr7.g01098.t01#6       3966   125258442:   mNeoVis.vulgivaga.1.1.chr1.g00465.t01#4    mNeoVis.2.1.1.chr1_hap1.g00472.t01#5    mNeoVis.3.1.1.chr1.g00470.t01#6    mNeoVis.evergladensis.2.1.chr1.g00470.t01#2    mNeoVis.evergladensis.1.1.chr1.g00480.t01#3   mNeoVis.vulgivaga.2.1.chr1.g00480.t01#1       3967   125258443:   mNeoVis.evergladensis.2.1.chr1.g00471.t01#2    mNeoVis.3.1.1.chr1.g00471.t01#6    mNeoVis.vulgivaga.2.1.chr1.g00481.t01#1    mNeoVis.vulgivaga.1.1.chr1.g00466.t01#4    mNeoVis.evergladensis.1.1.chr1.g00481.t01#3   mNeoVis.2.1.1.chr1_hap1.g00473.t01#5       3968   125258444:   mNeoVis.vulgivaga.2.1.chr7.g01177.t01#1    mNeoVis.evergladensis.2.1.chr7.g01165.t01#2    mNeoVis.vulgivaga.1.1.chr7.g01166.t01#4    mNeoVis.2.1.1.chr7_hap1.g01176.t01#5    mNeoVis.3.1.1.chr7.g01099.t01#6   mNeoVis.evergladensis.1.1.chr7.g01165.t01#3       3969   125258445:   mNeoVis.2.1.1.chr1_hap1.g00474.t01#5    mNeoVis.vulgivaga.2.1.chr1.g00482.t01#1    mNeoVis.evergladensis.1.1.chr1.g00482.t01#3    mNeoVis.3.1.1.chr1.g00472.t01#6   mNeoVis.evergladensis.2.1.chr1.g00472.t01#2       3970   125258446:   mNeoVis.2.1.1.chr1_hap1.g00475.t01#5    mNeoVis.3.1.1.chr1.g00473.t01#6    mNeoVis.vulgivaga.2.1.chr1.g00483.t01#1    mNeoVis.vulgivaga.1.1.chr1.g00468.t01#4    mNeoVis.evergladensis.2.1.chr1.g00473.t01#2   mNeoVis.evergladensis.1.1.chr1.g00483.t01#3       3971   125258447:   mNeoVis.3.1.1.chr7.g01100.t01#6    mNeoVis.evergladensis.2.1.chr7.g01166.t01#2    mNeoVis.vulgivaga.2.1.chr7.g01178.t01#1    mNeoVis.vulgivaga.1.1.chr7.g01167.t01#4    mNeoVis.2.1.1.chr7_hap1.g01177.t01#5   mNeoVis.evergladensis.1.1.chr7.g01166.t01#3       3972   125258448:   mNeoVis.2.1.1.chr7_hap1.g01178.t01#5    mNeoVis.vulgivaga.1.1.chr7.g01168.t01#4    mNeoVis.evergladensis.2.1.chr7.g01167.t01#2    mNeoVis.3.1.1.chr7.g01101.t01#6    mNeoVis.evergladensis.1.1.chr7.g01167.t01#3   mNeoVis.vulgivaga.2.1.chr7.g01179.t01#1       3973   125258449:   mNeoVis.2.1.1.chr7_hap1.g01179.t01#5    mNeoVis.vulgivaga.1.1.chr7.g01169.t01#4    mNeoVis.3.1.1.chr7.g01102.t01#6    mNeoVis.evergladensis.1.1.chr7.g01168.t01#3    mNeoVis.vulgivaga.2.1.chr7.g01180.t01#1   mNeoVis.evergladensis.2.1.chr7.g01168.t01#2       3974   125258450:   mNeoVis.3.1.1.chr1.g00475.t01#6    mNeoVis.vulgivaga.2.1.chr1.g00484.t01#1    mNeoVis.evergladensis.2.1.chr1.g00474.t01#2    mNeoVis.2.1.1.chr1_hap1.g00476.t01#5    mNeoVis.evergladensis.1.1.chr1.g00484.t01#3   mNeoVis.vulgivaga.1.1.chr1.g00469.t01#4       3975   125258451:   mNeoVis.vulgivaga.1.1.chr7.g01170.t01#4    mNeoVis.vulgivaga.2.1.chr7.g01181.t01#1    mNeoVis.evergladensis.1.1.chr7.g01169.t01#3    mNeoVis.2.1.1.chr7_hap1.g01180.t01#5    mNeoVis.3.1.1.chr7.g01103.t01#6   mNeoVis.evergladensis.2.1.chr7.g01169.t01#2       3976   125258452:   mNeoVis.evergladensis.2.1.chr7.g01170.t01#2    mNeoVis.2.1.1.chr7_hap1.g01181.t01#5    mNeoVis.evergladensis.1.1.chr7.g01170.t01#3    mNeoVis.3.1.1.chr7.g01104.t01#6    mNeoVis.vulgivaga.1.1.chr7.g01171.t01#4   mNeoVis.vulgivaga.2.1.chr7.g01182.t01#1       3977   125258453:   mNeoVis.vulgivaga.1.1.chr1.g00470.t01#4    mNeoVis.evergladensis.1.1.chr1.g00485.t01#3    mNeoVis.vulgivaga.2.1.chr1.g00485.t01#1    mNeoVis.3.1.1.chr1.g00474.t01#6    mNeoVis.2.1.1.chr1_hap1.g00477.t01#5   mNeoVis.evergladensis.2.1.chr1.g00475.t01#2       3978   125258454:   mNeoVis.2.1.1.chr7_hap1.g00478.t01#5    mNeoVis.3.1.1.chr7.g00463.t01#6    mNeoVis.vulgivaga.2.1.chr7.g00485.t01#1   mNeoVis.evergladensis.1.1.chr7.g00488.t01#3       3979   125258455:   mNeoVis.2.1.1.chr1_hap1.g00478.t01#5    mNeoVis.3.1.1.chr1.g00476.t01#6    mNeoVis.vulgivaga.1.1.chr1.g00471.t01#4    mNeoVis.vulgivaga.2.1.chr1.g00486.t01#1    mNeoVis.evergladensis.2.1.chr1.g00476.t01#2   mNeoVis.evergladensis.1.1.chr1.g00486.t01#3       3980   125258456:   mNeoVis.vulgivaga.2.1.chr4.g00560.t01#1    mNeoVis.2.1.1.chr6_hap1.g00546.t01#5    mNeoVis.vulgivaga.2.1.chr4.g00631.t01#1    mNeoVis.vulgivaga.1.1.chr4.g00624.t01#4    mNeoVis.vulgivaga.1.1.chr4.g00549.t01#4    mNeoVis.3.1.1.chr4.g00551.t01#6    mNeoVis.evergladensis.2.1.chr4.g00551.t01#2   mNeoVis.evergladensis.1.1.chr4.g00556.t01#3     3981   125258457:   mNeoVis.3.1.1.chr5.g00566.t01#6    mNeoVis.vulgivaga.2.1.chr5.g00571.t01#1    mNeoVis.evergladensis.1.1.chr5.g00567.t01#3    mNeoVis.2.1.1.chr2_hap1.g00572.t01#5    mNeoVis.vulgivaga.1.1.chr1.g00472.t01#4    mNeoVis.vulgivaga.1.1.chr5.g00556.t01#4   mNeoVis.evergladensis.2.1.chr5.g00574.t01#2     3982   125258458:   mNeoVis.2.1.1.chr1_hap1.g00479.t01#5    mNeoVis.3.1.1.chr1.g00477.t01#6    mNeoVis.evergladensis.2.1.chr1.g00477.t01#2    mNeoVis.vulgivaga.2.1.chr1.g00487.t01#1   mNeoVis.evergladensis.1.1.chr1.g00487.t01#3       3983   125258459:   mNeoVis.3.1.1.chr7.g01105.t01#6    mNeoVis.vulgivaga.1.1.chr7.g01172.t01#4    mNeoVis.vulgivaga.2.1.chr7.g01183.t01#1    mNeoVis.2.1.1.chr7_hap1.g01182.t01#5    mNeoVis.evergladensis.2.1.chr7.g01171.t01#2   mNeoVis.evergladensis.1.1.chr7.g01171.t01#3       3984   125258460:   mNeoVis.2.1.1.chr1_hap1.g00480.t01#5    mNeoVis.vulgivaga.1.1.chr1.g00473.t01#4    mNeoVis.evergladensis.2.1.chr1.g00478.t01#2    mNeoVis.3.1.1.chr1.g00478.t01#6    mNeoVis.evergladensis.1.1.chr1.g00488.t01#3   mNeoVis.vulgivaga.2.1.chr1.g00488.t01#1       3985   125258461:  mNeoVis.vulgivaga.2.1.chr7.g01184.t01#1               3986   125258462:   mNeoVis.evergladensis.2.1.chr1.g00479.t01#2    mNeoVis.evergladensis.1.1.chr1.g00489.t01#3    mNeoVis.2.1.1.chr1_hap1.g00481.t01#5    mNeoVis.3.1.1.chr1.g00479.t01#6    mNeoVis.vulgivaga.2.1.chr1.g00489.t01#1   mNeoVis.vulgivaga.1.1.chr1.g00474.t01#4       3987   125258463:   mNeoVis.2.1.1.chr1_hap1.g00482.t01#5    mNeoVis.evergladensis.2.1.chr1.g00480.t01#2    mNeoVis.evergladensis.1.1.chr1.g00490.t01#3    mNeoVis.vulgivaga.1.1.chr1.g00475.t01#4    mNeoVis.vulgivaga.2.1.chr1.g00490.t01#1   mNeoVis.3.1.1.chr1.g00480.t01#6       3988   125258464:   mNeoVis.evergladensis.1.1.chr7.g01173.t01#3    mNeoVis.vulgivaga.1.1.chr7.g01174.t01#4    mNeoVis.2.1.1.chr7_hap1.g01184.t01#5    mNeoVis.vulgivaga.2.1.chr7.g01185.t01#1    mNeoVis.evergladensis.2.1.chr7.g01173.t01#2   mNeoVis.3.1.1.chr7.g01107.t01#6       3989   125258465:   mNeoVis.evergladensis.1.1.chr7.g01174.t01#3    mNeoVis.vulgivaga.2.1.chr7.g01186.t01#1    mNeoVis.2.1.1.chr7_hap1.g01185.t01#5    mNeoVis.vulgivaga.1.1.chr7.g01175.t01#4    mNeoVis.evergladensis.2.1.chr7.g01174.t01#2   mNeoVis.3.1.1.chr7.g01108.t01#6       3990   125258466:   mNeoVis.2.1.1.chr7_hap1.g01186.t01#5    mNeoVis.evergladensis.1.1.chr7.g01175.t01#3    mNeoVis.3.1.1.chr7.g01109.t01#6    mNeoVis.vulgivaga.2.1.chr7.g01187.t01#1    mNeoVis.vulgivaga.1.1.chr7.g01176.t01#4   mNeoVis.evergladensis.2.1.chr7.g01175.t01#2       3991   125258467:   mNeoVis.vulgivaga.2.1.chr1.g00491.t01#1    mNeoVis.vulgivaga.1.1.chr5.g01400.t01#4    mNeoVis.vulgivaga.2.1.chr5.g01425.t01#1    mNeoVis.evergladensis.2.1.chr5.g01424.t01#2    mNeoVis.2.1.1.chr2_hap1.g01487.t01#5    mNeoVis.evergladensis.2.1.chr1.g00481.t01#2    mNeoVis.3.1.1.chr1.g00481.t01#6    mNeoVis.3.1.1.chr5.g01408.t01#6    mNeoVis.vulgivaga.1.1.chr1.g00476.t01#4      3992   125258468:   mNeoVis.evergladensis.1.1.chr1.g00492.t01#3    mNeoVis.2.1.1.chr1_hap1.g00484.t01#5    mNeoVis.vulgivaga.2.1.chr1.g00492.t01#1    mNeoVis.3.1.1.chr1.g00482.t01#6    mNeoVis.vulgivaga.1.1.chr1.g00477.t01#4   mNeoVis.evergladensis.2.1.chr1.g00482.t01#2       3993   125258469:   mNeoVis.3.1.1.chr1.g00483.t01#6    mNeoVis.vulgivaga.1.1.chr1.g00478.t01#4    mNeoVis.vulgivaga.2.1.chr1.g00493.t01#1    mNeoVis.2.1.1.chr1_hap1.g00485.t01#5    mNeoVis.evergladensis.2.1.chr1.g00483.t01#2   mNeoVis.evergladensis.1.1.chr1.g00493.t01#3       3994   125258470:   mNeoVis.vulgivaga.2.1.chr1.g00494.t01#1    mNeoVis.3.1.1.chr1.g00484.t01#6    mNeoVis.evergladensis.2.1.chr1.g00484.t01#2    mNeoVis.vulgivaga.1.1.chr1.g00479.t01#4    mNeoVis.evergladensis.1.1.chr1.g00494.t01#3   mNeoVis.2.1.1.chr1_hap1.g00486.t01#5       3995   125258471:   mNeoVis.2.1.1.chr6_hap1.g00802.t01#5    mNeoVis.evergladensis.1.1.chr1.g00495.t01#3    mNeoVis.vulgivaga.2.1.chr4.g00824.t01#1    mNeoVis.vulgivaga.1.1.chr1.g00480.t01#4    mNeoVis.evergladensis.2.1.chr1.g00485.t01#2    mNeoVis.2.1.1.chr1_hap1.g00487.t01#5    mNeoVis.2.1.1.chr9_hap1.g00057.t01#5    mNeoVis.evergladensis.1.1.chr4.g00818.t01#3    mNeoVis.evergladensis.2.1.chr4.g00804.t01#2      3996   125258472:  mNeoVis.vulgivaga.2.1.chr7.g01188.t01#1               3997   125258473:   mNeoVis.3.1.1.chr7.g01112.t01#6    mNeoVis.2.1.1.chr7_hap1.g01188.t01#5    mNeoVis.vulgivaga.2.1.chr7.g01189.t01#1    mNeoVis.evergladensis.2.1.chr7.g01177.t01#2    mNeoVis.vulgivaga.1.1.chr7.g01178.t01#4   mNeoVis.evergladensis.1.1.chr7.g01177.t01#3       3998   125258474:   mNeoVis.vulgivaga.2.1.chr1.g00496.t01#1    mNeoVis.2.1.1.chr1_hap1.g00488.t01#5    mNeoVis.evergladensis.2.1.chr1.g00486.t01#2    mNeoVis.evergladensis.1.1.chr1.g00496.t01#3    mNeoVis.3.1.1.chr1.g00486.t01#6   mNeoVis.vulgivaga.1.1.chr1.g00481.t01#4       3999   125258475:   mNeoVis.vulgivaga.2.1.chr1.g00497.t01#1    mNeoVis.3.1.1.chr1.g00487.t01#6    mNeoVis.vulgivaga.1.1.chr1.g00482.t01#4    mNeoVis.2.1.1.chr1_hap1.g00489.t01#5   mNeoVis.evergladensis.2.1.chr1.g00487.t01#2       4000   125258476:   mNeoVis.vulgivaga.2.1.chr7.g00486.t01#1    mNeoVis.evergladensis.1.1.chr7.g00489.t01#3    mNeoVis.2.1.1.chr7_hap1.g00479.t01#5    mNeoVis.evergladensis.2.1.chr7.g00484.t01#2    mNeoVis.3.1.1.chr7.g00464.t01#6   mNeoVis.evergladensis.1.1.chr7.g00490.t01#3       4001   125258477:   mNeoVis.2.1.1.chr7_hap1.g01098.t01#5    mNeoVis.evergladensis.1.1.chr7.g01094.t01#3    mNeoVis.evergladensis.2.1.chr7.g01090.t01#2    mNeoVis.vulgivaga.2.1.chr7.g01100.t01#1   mNeoVis.vulgivaga.1.1.chr7.g01089.t01#4       4002   125258478:   mNeoVis.vulgivaga.1.1.chr7.g01090.t01#4    mNeoVis.2.1.1.chr7_hap1.g01099.t01#5    mNeoVis.vulgivaga.2.1.chr7.g01101.t01#1    mNeoVis.evergladensis.1.1.chr7.g01095.t01#3    mNeoVis.evergladensis.2.1.chr7.g01091.t01#2   mNeoVis.3.1.1.chr7.g01027.t01#6       4003   125258479:   mNeoVis.3.1.1.chr7.g01028.t01#6    mNeoVis.2.1.1.chr7_hap1.g01100.t01#5    mNeoVis.evergladensis.1.1.chr7.g01096.t01#3    mNeoVis.evergladensis.2.1.chr7.g01092.t01#2    mNeoVis.vulgivaga.1.1.chr7.g01091.t01#4   mNeoVis.vulgivaga.2.1.chr7.g01102.t01#1       4004   125258480:   mNeoVis.evergladensis.1.1.chr15.g00079.t01#3    mNeoVis.2.1.1.chr15_hap1.g00712.t01#5    mNeoVis.vulgivaga.2.1.chr15.g00547.t01#1    mNeoVis.evergladensis.2.1.chr15.g00534.t01#2   mNeoVis.vulgivaga.1.1.chr15.g00537.t01#4       4005   125258481:  mNeoVis.vulgivaga.2.1.chr7.g01103.t01#1               4006   125258482:   mNeoVis.3.1.1.chr1.g00488.t01#6    mNeoVis.vulgivaga.2.1.chr1.g00498.t01#1    mNeoVis.vulgivaga.1.1.chr1.g00483.t01#4    mNeoVis.evergladensis.2.1.chr1.g00488.t01#2   mNeoVis.2.1.1.chr1_hap1.g00490.t01#5       4007   125258483:   mNeoVis.evergladensis.2.1.chr7.g01093.t01#2    mNeoVis.vulgivaga.1.1.chr7.g01092.t01#4    mNeoVis.3.1.1.chr7.g01029.t01#6    mNeoVis.evergladensis.1.1.chr7.g01097.t01#3    mNeoVis.2.1.1.chr7_hap1.g01101.t01#5   mNeoVis.vulgivaga.2.1.chr7.g01104.t01#1       4008   125258484:   mNeoVis.vulgivaga.2.1.chr7.g01105.t01#1    mNeoVis.vulgivaga.1.1.chr7.g01093.t01#4    mNeoVis.evergladensis.2.1.chr7.g01094.t01#2    mNeoVis.2.1.1.chr7_hap1.g01102.t01#5   mNeoVis.evergladensis.1.1.chr7.g01098.t01#3       4009   125258485:   mNeoVis.vulgivaga.1.1.chr7.g00494.t01#4    mNeoVis.evergladensis.1.1.chr7.g00491.t01#3   mNeoVis.vulgivaga.2.1.chr7.g00487.t01#1           4010   125258486:   mNeoVis.vulgivaga.1.1.chr1.g00484.t01#4   mNeoVis.vulgivaga.2.1.chr1.g00499.t01#1             4011   125258487:   mNeoVis.vulgivaga.2.1.chr1.g00500.t01#1    mNeoVis.evergladensis.2.1.chr1.g00489.t01#2    mNeoVis.3.1.1.chr1.g00489.t01#6    mNeoVis.evergladensis.1.1.chr1.g00501.t01#3    mNeoVis.2.1.1.chr1_hap1.g00492.t01#5   mNeoVis.vulgivaga.1.1.chr1.g00485.t01#4       4012   125258488:   mNeoVis.evergladensis.1.1.chr7.g01099.t01#3    mNeoVis.vulgivaga.1.1.chr7.g01094.t01#4    mNeoVis.vulgivaga.2.1.chr7.g01106.t01#1    mNeoVis.2.1.1.chr7_hap1.g01103.t01#5    mNeoVis.evergladensis.2.1.chr7.g01095.t01#2   mNeoVis.3.1.1.chr7.g01031.t01#6       4013   125258489:   mNeoVis.evergladensis.2.1.chr7.g01096.t01#2    mNeoVis.vulgivaga.1.1.chr7.g01095.t01#4    mNeoVis.evergladensis.1.1.chr7.g01100.t01#3    mNeoVis.3.1.1.chr7.g01032.t01#6    mNeoVis.vulgivaga.2.1.chr7.g01107.t01#1   mNeoVis.2.1.1.chr7_hap1.g01104.t01#5       4014   125258490:   mNeoVis.evergladensis.1.1.chr15.g00078.t01#3    mNeoVis.3.1.1.chr15.g00491.t01#6    mNeoVis.2.1.1.chr15_hap1.g00713.t01#5    mNeoVis.vulgivaga.2.1.chr15.g00548.t01#1    mNeoVis.vulgivaga.1.1.chr15.g00538.t01#4   mNeoVis.evergladensis.2.1.chr15.g00535.t01#2       4015   125258491:   mNeoVis.evergladensis.2.1.chr1.g00490.t01#2    mNeoVis.3.1.1.chr1.g00490.t01#6    mNeoVis.2.1.1.chr1_hap1.g00493.t01#5    mNeoVis.vulgivaga.1.1.chr1.g00486.t01#4    mNeoVis.vulgivaga.2.1.chr1.g00501.t01#1   mNeoVis.evergladensis.1.1.chr1.g00502.t01#3       4016   125258492:   mNeoVis.vulgivaga.2.1.chr7.g01108.t01#1    mNeoVis.2.1.1.chr7_hap1.g01105.t01#5    mNeoVis.3.1.1.chr7.g01033.t01#6    mNeoVis.evergladensis.2.1.chr7.g01097.t01#2    mNeoVis.evergladensis.1.1.chr7.g01101.t01#3   mNeoVis.vulgivaga.1.1.chr7.g01096.t01#4       4017   125258493:   mNeoVis.2.1.1.chr1_hap1.g00494.t01#5    mNeoVis.vulgivaga.1.1.chr1.g00487.t01#4    mNeoVis.evergladensis.1.1.chr1.g00503.t01#3    mNeoVis.vulgivaga.2.1.chr1.g00502.t01#1   mNeoVis.3.1.1.chr1.g00491.t01#6       4018   125258494:   mNeoVis.evergladensis.2.1.chr7.g00485.t01#2    mNeoVis.vulgivaga.2.1.chr7.g00488.t01#1   mNeoVis.2.1.1.chr7_hap1.g00481.t01#5           4019   125258495:   mNeoVis.vulgivaga.2.1.chr1.g00503.t01#1    mNeoVis.3.1.1.chr1.g00492.t01#6    mNeoVis.2.1.1.chr1_hap1.g00495.t01#5    mNeoVis.evergladensis.1.1.chr1.g00504.t01#3    mNeoVis.vulgivaga.1.1.chr1.g00488.t01#4   mNeoVis.evergladensis.2.1.chr1.g00492.t01#2       4020   125258496:   mNeoVis.vulgivaga.1.1.chr1.g00489.t01#4    mNeoVis.evergladensis.2.1.chr1.g00493.t01#2    mNeoVis.3.1.1.chr1.g00493.t01#6    mNeoVis.evergladensis.1.1.chr1.g00505.t01#3    mNeoVis.2.1.1.chr1_hap1.g00496.t01#5   mNeoVis.vulgivaga.2.1.chr1.g00504.t01#1       4021   125258497:   mNeoVis.evergladensis.1.1.chr7.g01102.t01#3    mNeoVis.2.1.1.chr7_hap1.g01106.t01#5    mNeoVis.vulgivaga.2.1.chr7.g01109.t01#1    mNeoVis.3.1.1.chr7.g01034.t01#6    mNeoVis.vulgivaga.1.1.chr7.g01097.t01#4   mNeoVis.evergladensis.2.1.chr7.g01098.t01#2       4022   125258498:   mNeoVis.3.1.1.chr1.g00494.t01#6    mNeoVis.2.1.1.chr1_hap1.g00497.t01#5    mNeoVis.evergladensis.1.1.chr1.g00506.t01#3    mNeoVis.evergladensis.2.1.chr1.g00494.t01#2    mNeoVis.vulgivaga.2.1.chr1.g00505.t01#1   mNeoVis.vulgivaga.1.1.chr1.g00490.t01#4       4023   125258499:   mNeoVis.2.1.1.chr1_hap1.g00498.t01#5    mNeoVis.evergladensis.1.1.chr1.g00507.t01#3    mNeoVis.evergladensis.2.1.chr1.g00491.t01#2    mNeoVis.vulgivaga.1.1.chr1.g00491.t01#4    mNeoVis.vulgivaga.2.1.chr1.g00506.t01#1   mNeoVis.3.1.1.chr1.g00495.t01#6       4024   125258500:  mNeoVis.vulgivaga.2.1.chr1.g00396.t01#1               4025   125258501:   mNeoVis.2.1.1.chr1_hap1.g00393.t01#5    mNeoVis.evergladensis.1.1.chr1.g00399.t01#3    mNeoVis.3.1.1.chr1.g00392.t01#6    mNeoVis.vulgivaga.2.1.chr1.g00397.t01#1    mNeoVis.vulgivaga.1.1.chr1.g00390.t01#4   mNeoVis.evergladensis.2.1.chr1.g00391.t01#2       4026   125258502:   mNeoVis.evergladensis.1.1.chr15.g00077.t01#3    mNeoVis.2.1.1.chr15_hap1.g00714.t01#5    mNeoVis.3.1.1.chr15.g00492.t01#6    mNeoVis.evergladensis.2.1.chr15.g00536.t01#2    mNeoVis.vulgivaga.1.1.chr15.g00539.t01#4   mNeoVis.vulgivaga.2.1.chr15.g00549.t01#1       4027   125258503:   mNeoVis.vulgivaga.2.1.chr1.g00398.t01#1    mNeoVis.2.1.1.chr1_hap1.g00394.t01#5    mNeoVis.evergladensis.2.1.chr1.g00392.t01#2   mNeoVis.evergladensis.1.1.chr1.g00401.t01#3       4028   125258504:   mNeoVis.evergladensis.2.1.chr1.g00393.t01#2    mNeoVis.evergladensis.1.1.chr1.g00400.t01#3    mNeoVis.2.1.1.chr1_hap1.g00395.t01#5    mNeoVis.3.1.1.chr1.g00393.t01#6    mNeoVis.vulgivaga.1.1.chr1.g00391.t01#4   mNeoVis.vulgivaga.2.1.chr1.g00399.t01#1       4029   125258505:   mNeoVis.evergladensis.2.1.chr2.g00805.t01#2    mNeoVis.vulgivaga.2.1.chr7.g01110.t01#1    mNeoVis.2.1.1.chr7_hap1.g01109.t01#5    mNeoVis.vulgivaga.2.1.chr5.g00685.t01#1    mNeoVis.evergladensis.1.1.chr2.g00802.t01#3    mNeoVis.evergladensis.2.1.chr5.g00691.t01#2    mNeoVis.vulgivaga.1.1.chr2.g00804.t01#4    mNeoVis.3.1.1.chr2.g00718.t01#6    mNeoVis.vulgivaga.2.1.chr2.g00810.t01#1      4030   125258506:  mNeoVis.2.1.1.chr2_hap1.g00736.t01#5               4031   125258507:   mNeoVis.3.1.1.chr7.g01037.t01#6    mNeoVis.evergladensis.1.1.chr7.g01104.t01#3    mNeoVis.evergladensis.2.1.chr7.g01099.t01#2    mNeoVis.vulgivaga.2.1.chr7.g01112.t01#1    mNeoVis.vulgivaga.1.1.chr7.g01098.t01#4   mNeoVis.2.1.1.chr7_hap1.g01113.t01#5       4032   125258508:   mNeoVis.evergladensis.1.1.chr7.g01105.t01#3    mNeoVis.vulgivaga.1.1.chr7.g01099.t01#4    mNeoVis.3.1.1.chr7.g01036.t01#6    mNeoVis.vulgivaga.2.1.chr7.g01113.t01#1   mNeoVis.2.1.1.chr7_hap1.g01114.t01#5       4033   125258509:   mNeoVis.vulgivaga.2.1.chr7.g00489.t01#1   mNeoVis.evergladensis.1.1.chr7.g00492.t01#3           4034   125258510:  mNeoVis.vulgivaga.2.1.chr7.g01114.t01#1               4035   125258511:   mNeoVis.vulgivaga.1.1.chr1.g00392.t01#4    mNeoVis.2.1.1.chr1_hap1.g00396.t01#5    mNeoVis.3.1.1.chr1.g00395.t01#6    mNeoVis.evergladensis.2.1.chr1.g00394.t01#2    mNeoVis.evergladensis.1.1.chr1.g00402.t01#3   mNeoVis.vulgivaga.2.1.chr1.g00400.t01#1       4036   125258512:   mNeoVis.3.1.1.chr1.g00396.t01#6    mNeoVis.evergladensis.1.1.chr1.g00403.t01#3    mNeoVis.evergladensis.2.1.chr1.g00396.t01#2   mNeoVis.vulgivaga.2.1.chr1.g00401.t01#1         4037   125258513:   mNeoVis.vulgivaga.2.1.chr1.g00402.t01#1    mNeoVis.evergladensis.2.1.chr1.g00397.t01#2    mNeoVis.evergladensis.1.1.chr1.g00404.t01#3    mNeoVis.vulgivaga.1.1.chr1.g00394.t01#4    mNeoVis.2.1.1.chr1_hap1.g00398.t01#5   mNeoVis.3.1.1.chr1.g00397.t01#6       4038   125258514:   mNeoVis.2.1.1.chr1_hap1.g00399.t01#5    mNeoVis.3.1.1.chr1.g00398.t01#6    mNeoVis.evergladensis.1.1.chr1.g00405.t01#3    mNeoVis.evergladensis.2.1.chr1.g00398.t01#2   mNeoVis.vulgivaga.2.1.chr1.g00403.t01#1       4039   125258515:   mNeoVis.vulgivaga.2.1.chr11.g00549.t01#1    mNeoVis.evergladensis.2.1.chr11.g00542.t01#2    mNeoVis.vulgivaga.2.1.chr13.g00393.t01#1    mNeoVis.2.1.1.chr11_hap1.g00555.t01#5    mNeoVis.3.1.1.chr13.g00344.t01#6    mNeoVis.2.1.1.chr13_hap1.g00407.t01#5    mNeoVis.evergladensis.1.1.chr13.g00389.t01#3    mNeoVis.3.1.1.chr11.g00515.t01#6    mNeoVis.vulgivaga.1.1.chr13.g00383.t01#4      4040   125258516:   mNeoVis.vulgivaga.1.1.chr1.g00395.t01#4    mNeoVis.vulgivaga.2.1.chr1.g00404.t01#1    mNeoVis.evergladensis.2.1.chr1.g00399.t01#2    mNeoVis.evergladensis.1.1.chr1.g00406.t01#3    mNeoVis.2.1.1.chr1_hap1.g00400.t01#5   mNeoVis.3.1.1.chr1.g00399.t01#6       4041   125258517:   mNeoVis.3.1.1.chr4.g00043.t01#6    mNeoVis.vulgivaga.1.1.chr4.g00046.t01#4    mNeoVis.evergladensis.1.1.chr4.g00046.t01#3    mNeoVis.vulgivaga.2.1.chr4.g00044.t01#1    mNeoVis.2.1.1.chr6_hap1.g00047.t01#5   mNeoVis.evergladensis.2.1.chr4.g00049.t01#2       4042   125258518:   mNeoVis.2.1.1.chr12_hap1.g00245.t01#5    mNeoVis.evergladensis.2.1.chr12.g00214.t01#2    mNeoVis.3.1.1.chr12.g00212.t01#6    mNeoVis.evergladensis.1.1.chr12.g00216.t01#3    mNeoVis.vulgivaga.1.1.chr12.g00221.t01#4   mNeoVis.vulgivaga.2.1.chr12.g00219.t01#1       4043   125258519:   mNeoVis.evergladensis.1.1.chr7.g01107.t01#3    mNeoVis.evergladensis.2.1.chr7.g01103.t01#2    mNeoVis.vulgivaga.1.1.chr7.g01103.t01#4    mNeoVis.2.1.1.chr7_hap1.g01117.t01#5    mNeoVis.3.1.1.chr7.g01040.t01#6   mNeoVis.vulgivaga.2.1.chr7.g01115.t01#1       4044   125258520:   mNeoVis.vulgivaga.2.1.chr7.g01116.t01#1    mNeoVis.evergladensis.1.1.chr7.g01108.t01#3    mNeoVis.vulgivaga.1.1.chr7.g01104.t01#4    mNeoVis.3.1.1.chr7.g01041.t01#6    mNeoVis.2.1.1.chr7_hap1.g01118.t01#5   mNeoVis.evergladensis.2.1.chr7.g01104.t01#2       4045   125258521:   mNeoVis.3.1.1.chr7.g01042.t01#6    mNeoVis.vulgivaga.1.1.chr7.g01105.t01#4    mNeoVis.2.1.1.chr7_hap1.g01119.t01#5    mNeoVis.vulgivaga.2.1.chr7.g01117.t01#1    mNeoVis.evergladensis.2.1.chr7.g01105.t01#2   mNeoVis.evergladensis.1.1.chr7.g01109.t01#3       4046   125258522:   mNeoVis.vulgivaga.1.1.chr1.g00396.t01#4    mNeoVis.evergladensis.2.1.chr1.g00400.t01#2    mNeoVis.evergladensis.1.1.chr1.g00407.t01#3    mNeoVis.vulgivaga.2.1.chr1.g00405.t01#1   mNeoVis.2.1.1.chr1_hap1.g00401.t01#5       4047   125258523:   mNeoVis.evergladensis.1.1.chr7.g01110.t01#3    mNeoVis.evergladensis.2.1.chr7.g01106.t01#2    mNeoVis.vulgivaga.2.1.chr7.g01118.t01#1    mNeoVis.vulgivaga.1.1.chr7.g01106.t01#4    mNeoVis.2.1.1.chr7_hap1.g01120.t01#5   mNeoVis.3.1.1.chr7.g01043.t01#6       4048   125258524:   mNeoVis.evergladensis.1.1.chr7.g00493.t01#3    mNeoVis.vulgivaga.1.1.chr7.g00497.t01#4    mNeoVis.2.1.1.chr7_hap1.g00482.t01#5    mNeoVis.vulgivaga.2.1.chr7.g00490.t01#1    mNeoVis.3.1.1.chr7.g00465.t01#6   mNeoVis.vulgivaga.1.1.chr7.g00496.t01#4       4049   125258525:   mNeoVis.3.1.1.chr1.g00400.t01#6    mNeoVis.2.1.1.chr1_hap1.g00402.t01#5    mNeoVis.evergladensis.2.1.chr1.g00401.t01#2    mNeoVis.vulgivaga.2.1.chr1.g00406.t01#1    mNeoVis.vulgivaga.1.1.chr1.g00397.t01#4   mNeoVis.evergladensis.1.1.chr1.g00408.t01#3       4050   125258526:   mNeoVis.vulgivaga.2.1.chr7.g01119.t01#1    mNeoVis.vulgivaga.1.1.chr7.g01107.t01#4    mNeoVis.evergladensis.2.1.chr7.g01107.t01#2    mNeoVis.evergladensis.1.1.chr7.g01111.t01#3   mNeoVis.3.1.1.chr7.g01044.t01#6       4051   125258527:   mNeoVis.evergladensis.1.1.chr1.g00409.t01#3    mNeoVis.2.1.1.chr1_hap1.g00403.t01#5    mNeoVis.3.1.1.chr1.g00401.t01#6    mNeoVis.evergladensis.2.1.chr1.g00402.t01#2    mNeoVis.vulgivaga.1.1.chr1.g00398.t01#4   mNeoVis.vulgivaga.2.1.chr1.g00407.t01#1       4052   125258528:   mNeoVis.3.1.1.chr1.g00402.t01#6    mNeoVis.evergladensis.2.1.chr1.g00403.t01#2    mNeoVis.2.1.1.chr6_hap1.g01027.t01#5    mNeoVis.vulgivaga.1.1.chr1.g00399.t01#4    mNeoVis.evergladensis.2.1.chr4.g00970.t01#2    mNeoVis.evergladensis.1.1.chr4.g00993.t01#3    mNeoVis.vulgivaga.1.1.chr4.g00993.t01#4    mNeoVis.vulgivaga.2.1.chr1.g00408.t01#1    mNeoVis.2.1.1.chr1_hap1.g00404.t01#5      4053   125258529:   mNeoVis.vulgivaga.2.1.chr1.g00409.t01#1    mNeoVis.2.1.1.chr1_hap1.g00405.t01#5    mNeoVis.evergladensis.1.1.chr1.g00411.t01#3    mNeoVis.3.1.1.chr1.g00403.t01#6    mNeoVis.vulgivaga.1.1.chr1.g00400.t01#4   mNeoVis.evergladensis.2.1.chr1.g00404.t01#2       4054   125258530:   mNeoVis.vulgivaga.1.1.chr1.g00401.t01#4    mNeoVis.vulgivaga.2.1.chr1.g00410.t01#1    mNeoVis.2.1.1.chr1_hap1.g00406.t01#5    mNeoVis.evergladensis.2.1.chr1.g00405.t01#2    mNeoVis.3.1.1.chr1.g00404.t01#6   mNeoVis.evergladensis.1.1.chr1.g00412.t01#3       4055   125258531:   mNeoVis.vulgivaga.2.1.chr7.g01121.t01#1    mNeoVis.evergladensis.1.1.chr7.g01113.t01#3    mNeoVis.evergladensis.2.1.chr7.g01109.t01#2    mNeoVis.3.1.1.chr7.g01046.t01#6    mNeoVis.2.1.1.chr7_hap1.g01123.t01#5   mNeoVis.vulgivaga.1.1.chr7.g01109.t01#4       4056   125258532:   mNeoVis.3.1.1.chr1.g00405.t01#6    mNeoVis.evergladensis.2.1.chr1.g00406.t01#2    mNeoVis.2.1.1.chr1_hap1.g00408.t01#5    mNeoVis.vulgivaga.2.1.chr1.g00411.t01#1   mNeoVis.evergladensis.1.1.chr1.g00413.t01#3       4057   125258533:   mNeoVis.vulgivaga.2.1.chr1.g00412.t01#1    mNeoVis.evergladensis.2.1.chr1.g00408.t01#2   mNeoVis.evergladensis.1.1.chr1.g00415.t01#3         4058   125258534:   mNeoVis.evergladensis.1.1.chr7.g01114.t01#3    mNeoVis.vulgivaga.2.1.chr7.g01122.t01#1    mNeoVis.vulgivaga.1.1.chr7.g01110.t01#4    mNeoVis.2.1.1.chr7_hap1.g01124.t01#5    mNeoVis.evergladensis.2.1.chr7.g01110.t01#2   mNeoVis.3.1.1.chr7.g01047.t01#6       4059   125258535:   mNeoVis.vulgivaga.2.1.chr7.g01123.t01#1    mNeoVis.vulgivaga.1.1.chr7.g01113.t01#4    mNeoVis.evergladensis.2.1.chr7.g01111.t01#2    mNeoVis.2.1.1.chr7_hap1.g01125.t01#5    mNeoVis.3.1.1.chr7.g01049.t01#6    mNeoVis.vulgivaga.1.1.chr7.g01111.t01#4    mNeoVis.evergladensis.1.1.chr7.g01115.t01#3    mNeoVis.evergladensis.1.1.chr7.g01116.t01#3    mNeoVis.vulgivaga.1.1.chr7.g01112.t01#4      4060   125258536:   mNeoVis.2.1.1.chr1_hap1.g00409.t01#5    mNeoVis.vulgivaga.2.1.chr1.g00413.t01#1    mNeoVis.evergladensis.1.1.chr1.g00414.t01#3    mNeoVis.vulgivaga.1.1.chr1.g00403.t01#4    mNeoVis.evergladensis.2.1.chr1.g00407.t01#2   mNeoVis.3.1.1.chr1.g00406.t01#6       4061   125258537:   mNeoVis.vulgivaga.2.1.chr7.g01124.t01#1    mNeoVis.vulgivaga.1.1.chr7.g01114.t01#4    mNeoVis.3.1.1.chr7.g01050.t01#6    mNeoVis.2.1.1.chr7_hap1.g01126.t01#5    mNeoVis.evergladensis.2.1.chr7.g01112.t01#2   mNeoVis.evergladensis.1.1.chr7.g01117.t01#3       4062   125258538:   mNeoVis.vulgivaga.1.1.chr7.g01115.t01#4    mNeoVis.3.1.1.chr7.g01051.t01#6    mNeoVis.evergladensis.1.1.chr7.g01118.t01#3    mNeoVis.vulgivaga.2.1.chr7.g01125.t01#1    mNeoVis.2.1.1.chr7_hap1.g01127.t01#5   mNeoVis.evergladensis.2.1.chr7.g01113.t01#2       4063   125258539:   mNeoVis.2.1.1.chr6_hap1.g00659.t01#5    mNeoVis.evergladensis.1.1.chr4.g00676.t01#3    mNeoVis.evergladensis.2.1.chr4.g00668.t01#2    mNeoVis.vulgivaga.1.1.chr4.g00674.t01#4    mNeoVis.vulgivaga.2.1.chr4.g00679.t01#1    mNeoVis.3.1.1.chr4.g00666.t01#6   mNeoVis.3.1.1.chr4.g00667.t01#6       4064   125258540:   mNeoVis.evergladensis.1.1.chr1.g00416.t01#3    mNeoVis.vulgivaga.2.1.chr1.g00414.t01#1    mNeoVis.3.1.1.chr1.g00407.t01#6    mNeoVis.2.1.1.chr1_hap1.g00410.t01#5    mNeoVis.evergladensis.2.1.chr1.g00409.t01#2   mNeoVis.vulgivaga.1.1.chr1.g00404.t01#4       4065   125258541:   mNeoVis.evergladensis.2.1.chr7.g01114.t01#2    mNeoVis.evergladensis.1.1.chr7.g01119.t01#3    mNeoVis.vulgivaga.2.1.chr7.g01126.t01#1    mNeoVis.vulgivaga.1.1.chr7.g01116.t01#4   mNeoVis.2.1.1.chr7_hap1.g01128.t01#5       4066   125258542:   mNeoVis.evergladensis.1.1.chr7.g01120.t01#3    mNeoVis.vulgivaga.2.1.chr7.g01127.t01#1    mNeoVis.evergladensis.2.1.chr7.g01115.t01#2    mNeoVis.2.1.1.chr7_hap1.g01129.t01#5    mNeoVis.3.1.1.chr7.g01053.t01#6   mNeoVis.vulgivaga.1.1.chr7.g01117.t01#4       4067   125258543:   mNeoVis.3.1.1.chr7.g00466.t01#6    mNeoVis.evergladensis.2.1.chr7.g00488.t01#2    mNeoVis.vulgivaga.1.1.chr7.g00498.t01#4    mNeoVis.2.1.1.chr7_hap1.g00484.t01#5   mNeoVis.vulgivaga.2.1.chr7.g00491.t01#1       4068   125258544:   mNeoVis.3.1.1.chr1.g00408.t01#6    mNeoVis.2.1.1.chr1_hap1.g00411.t01#5    mNeoVis.vulgivaga.2.1.chr1.g00415.t01#1    mNeoVis.evergladensis.2.1.chr1.g00410.t01#2    mNeoVis.vulgivaga.1.1.chr1.g00405.t01#4   mNeoVis.evergladensis.1.1.chr1.g00417.t01#3       4069   125258545:   mNeoVis.evergladensis.2.1.chr1.g00411.t01#2    mNeoVis.vulgivaga.1.1.chr6.g00449.t01#4    mNeoVis.3.1.1.chr5.g00892.t01#6    mNeoVis.vulgivaga.1.1.chr1.g00406.t01#4    mNeoVis.evergladensis.1.1.chr1.g00418.t01#3    mNeoVis.2.1.1.chr2_hap1.g00959.t01#5    mNeoVis.2.1.1.chr1_hap1.g00412.t01#5    mNeoVis.evergladensis.2.1.chr6.g00449.t01#2    mNeoVis.vulgivaga.2.1.chr1.g00416.t01#1      4070   125258546:   mNeoVis.evergladensis.2.1.chr5.g00454.t01#2    mNeoVis.3.1.1.chr5.g00443.t01#6    mNeoVis.evergladensis.1.1.chr5.g00446.t01#3    mNeoVis.2.1.1.chr2_hap1.g00452.t01#5    mNeoVis.vulgivaga.1.1.chr11.g00702.t01#4    mNeoVis.vulgivaga.2.1.chr5.g00452.t01#1   mNeoVis.vulgivaga.1.1.chr5.g00439.t01#4     4071   125258547:   mNeoVis.2.1.1.chr6_hap1.g00930.t01#5    mNeoVis.vulgivaga.2.1.chr4.g00902.t01#1    mNeoVis.3.1.1.chr4.g00891.t01#6    mNeoVis.evergladensis.2.1.chr4.g00882.t01#2    mNeoVis.vulgivaga.1.1.chr4.g00900.t01#4   mNeoVis.evergladensis.1.1.chr4.g00900.t01#3       4072   125258548:   mNeoVis.evergladensis.2.1.chr15.g00537.t01#2    mNeoVis.vulgivaga.2.1.chr15.g00550.t01#1    mNeoVis.evergladensis.1.1.chr15.g00076.t01#3    mNeoVis.vulgivaga.1.1.chr15.g00540.t01#4    mNeoVis.3.1.1.chr15.g00493.t01#6   mNeoVis.2.1.1.chr15_hap1.g00715.t01#5       4073   125258549:   mNeoVis.evergladensis.1.1.chr1.g00419.t01#3    mNeoVis.vulgivaga.2.1.chr1.g00417.t01#1    mNeoVis.3.1.1.chr1.g00410.t01#6    mNeoVis.evergladensis.2.1.chr1.g00412.t01#2    mNeoVis.2.1.1.chr1_hap1.g00413.t01#5   mNeoVis.vulgivaga.1.1.chr1.g00407.t01#4       4074   125258550:   mNeoVis.evergladensis.1.1.chr4.g00964.t01#3    mNeoVis.vulgivaga.1.1.chr4.g00966.t01#4    mNeoVis.3.1.1.chr1.g00412.t01#6    mNeoVis.3.1.1.chr4.g00954.t01#6    mNeoVis.evergladensis.2.1.chr4.g00942.t01#2    mNeoVis.evergladensis.1.1.chr1.g00421.t01#3    mNeoVis.2.1.1.chr1_hap1.g00415.t01#5    mNeoVis.vulgivaga.2.1.chr4.g00966.t01#1    mNeoVis.2.1.1.chr6_hap1.g00997.t01#5      4075   125258551:   mNeoVis.vulgivaga.1.1.chr1.g00409.t01#4    mNeoVis.3.1.1.chr1.g00413.t01#6    mNeoVis.evergladensis.1.1.chr1.g00422.t01#3    mNeoVis.vulgivaga.2.1.chr1.g00420.t01#1    mNeoVis.2.1.1.chr1_hap1.g00416.t01#5   mNeoVis.evergladensis.2.1.chr1.g00414.t01#2       4076   125258552:   mNeoVis.evergladensis.1.1.chr7.g00496.t01#3    mNeoVis.vulgivaga.1.1.chr7.g00499.t01#4    mNeoVis.vulgivaga.2.1.chr7.g00492.t01#1   mNeoVis.2.1.1.chr7_hap1.g00485.t01#5         4077   125258553:   mNeoVis.evergladensis.2.1.chr12.g00216.t01#2    mNeoVis.2.1.1.chr12_hap1.g00247.t01#5    mNeoVis.3.1.1.chr12.g00214.t01#6    mNeoVis.2.1.1.chr1_hap1.g00417.t01#5    mNeoVis.3.1.1.chr1.g00414.t01#6    mNeoVis.vulgivaga.1.1.chr12.g00223.t01#4    mNeoVis.evergladensis.2.1.chr1.g00415.t01#2    mNeoVis.vulgivaga.2.1.chr1.g00421.t01#1    mNeoVis.evergladensis.1.1.chr1.g00423.t01#3      4078   125258554:   mNeoVis.2.1.1.chr1_hap1.g00418.t01#5    mNeoVis.evergladensis.1.1.chr1.g00424.t01#3    mNeoVis.3.1.1.chr1.g00415.t01#6    mNeoVis.evergladensis.2.1.chr1.g00416.t01#2    mNeoVis.vulgivaga.2.1.chr1.g00422.t01#1   mNeoVis.vulgivaga.1.1.chr1.g00411.t01#4       4079   125258555:   mNeoVis.vulgivaga.1.1.chr1.g00412.t01#4    mNeoVis.vulgivaga.2.1.chr1.g00423.t01#1    mNeoVis.evergladensis.1.1.chr1.g00425.t01#3    mNeoVis.evergladensis.2.1.chr1.g00417.t01#2    mNeoVis.3.1.1.chr1.g00416.t01#6   mNeoVis.2.1.1.chr1_hap1.g00419.t01#5       4080   125258556:   mNeoVis.vulgivaga.1.1.chr7.g01118.t01#4    mNeoVis.evergladensis.2.1.chr7.g01116.t01#2    mNeoVis.3.1.1.chr7.g01054.t01#6    mNeoVis.2.1.1.chr7_hap1.g01130.t01#5    mNeoVis.vulgivaga.2.1.chr7.g01128.t01#1   mNeoVis.evergladensis.1.1.chr7.g01121.t01#3       4081   125258557:   mNeoVis.evergladensis.2.1.chr5.g00513.t01#2    mNeoVis.vulgivaga.2.1.chr5.g00513.t01#1    mNeoVis.vulgivaga.1.1.chr5.g00497.t01#4    mNeoVis.2.1.1.chr2_hap1.g00512.t01#5    mNeoVis.evergladensis.1.1.chr5.g00507.t01#3   mNeoVis.3.1.1.chr5.g00506.t01#6       4082   125258558:   mNeoVis.vulgivaga.1.1.chr1.g00413.t01#4    mNeoVis.evergladensis.2.1.chr1.g00418.t01#2    mNeoVis.vulgivaga.2.1.chr1.g00424.t01#1    mNeoVis.2.1.1.chr1_hap1.g00420.t01#5   mNeoVis.evergladensis.1.1.chr1.g00426.t01#3       4083   125258559:   mNeoVis.evergladensis.2.1.chr1.g00419.t01#2    mNeoVis.vulgivaga.1.1.chr1.g00414.t01#4    mNeoVis.2.1.1.chr1_hap1.g00421.t01#5    mNeoVis.3.1.1.chr1.g00418.t01#6    mNeoVis.vulgivaga.2.1.chr1.g00425.t01#1   mNeoVis.evergladensis.1.1.chr1.g00427.t01#3       4084   125258560:   mNeoVis.vulgivaga.1.1.chr7.g01119.t01#4    mNeoVis.2.1.1.chr7_hap1.g01131.t01#5    mNeoVis.evergladensis.2.1.chr7.g01117.t01#2   mNeoVis.vulgivaga.2.1.chr7.g01129.t01#1         4085   125258561:   mNeoVis.3.1.1.chr7.g01055.t01#6    mNeoVis.2.1.1.chr7_hap1.g01132.t01#5    mNeoVis.vulgivaga.1.1.chr7.g01120.t01#4    mNeoVis.vulgivaga.2.1.chr7.g01130.t01#1    mNeoVis.evergladensis.1.1.chr7.g01122.t01#3   mNeoVis.evergladensis.2.1.chr7.g01118.t01#2       4086   125258562:   mNeoVis.3.1.1.chr7.g01056.t01#6    mNeoVis.evergladensis.2.1.chr7.g01119.t01#2    mNeoVis.vulgivaga.1.1.chr7.g01121.t01#4    mNeoVis.evergladensis.1.1.chr7.g01123.t01#3    mNeoVis.vulgivaga.2.1.chr7.g01131.t01#1   mNeoVis.2.1.1.chr7_hap1.g01133.t01#5       4087   125258563:   mNeoVis.vulgivaga.2.1.chr1.g00426.t01#1    mNeoVis.evergladensis.2.1.chr1.g00420.t01#2    mNeoVis.vulgivaga.1.1.chr1.g00415.t01#4    mNeoVis.2.1.1.chr1_hap1.g00422.t01#5    mNeoVis.3.1.1.chr1.g00419.t01#6   mNeoVis.evergladensis.1.1.chr1.g00428.t01#3       4088   125258564:   mNeoVis.2.1.1.chr10_hap1.g00187.t01#5   mNeoVis.evergladensis.1.1.chr9.g00184.t01#3           4089   125258565:  mNeoVis.vulgivaga.2.1.chr7.g00493.t01#1               4090   125258566:   mNeoVis.vulgivaga.1.1.chr1.g00416.t01#4    mNeoVis.evergladensis.1.1.chr1.g00429.t01#3    mNeoVis.2.1.1.chr1_hap1.g00423.t01#5    mNeoVis.vulgivaga.2.1.chr1.g00427.t01#1    mNeoVis.3.1.1.chr1.g00420.t01#6   mNeoVis.evergladensis.2.1.chr1.g00421.t01#2       4091   125258567:   mNeoVis.vulgivaga.1.1.chr1.g00417.t01#4    mNeoVis.evergladensis.2.1.chr1.g00422.t01#2    mNeoVis.2.1.1.chr1_hap1.g00424.t01#5    mNeoVis.3.1.1.chr1.g00421.t01#6    mNeoVis.vulgivaga.2.1.chr1.g00428.t01#1   mNeoVis.evergladensis.1.1.chr1.g00430.t01#3       4092   125258568:   mNeoVis.vulgivaga.2.1.chr7.g01132.t01#1    mNeoVis.vulgivaga.1.1.chr7.g01122.t01#4    mNeoVis.evergladensis.1.1.chr7.g01124.t01#3    mNeoVis.2.1.1.chr7_hap1.g01134.t01#5    mNeoVis.evergladensis.2.1.chr7.g01120.t01#2   mNeoVis.3.1.1.chr7.g01057.t01#6       4093   125258569:   mNeoVis.3.1.1.chr12.g00004.t01#6    mNeoVis.evergladensis.1.1.chr12.g00004.t01#3    mNeoVis.evergladensis.2.1.chr12.g00005.t01#2    mNeoVis.vulgivaga.2.1.chr12.g00005.t01#1    mNeoVis.2.1.1.chr12_hap1.g00007.t01#5   mNeoVis.vulgivaga.1.1.chr12.g00005.t01#4       4094   125258570:   mNeoVis.vulgivaga.2.1.chr15.g00552.t01#1    mNeoVis.2.1.1.chr15_hap1.g00716.t01#5    mNeoVis.3.1.1.chr15.g00494.t01#6    mNeoVis.evergladensis.1.1.chr15.g00074.t01#3    mNeoVis.vulgivaga.1.1.chr15.g00542.t01#4   mNeoVis.evergladensis.2.1.chr15.g00538.t01#2       4095   125258571:   mNeoVis.vulgivaga.2.1.chr7.g01133.t01#1    mNeoVis.evergladensis.2.1.chr7.g01121.t01#2    mNeoVis.3.1.1.chr7.g01058.t01#6    mNeoVis.2.1.1.chr7_hap1.g01135.t01#5    mNeoVis.vulgivaga.1.1.chr7.g01123.t01#4   mNeoVis.evergladensis.1.1.chr7.g01125.t01#3       4096   125258572:   mNeoVis.evergladensis.2.1.chr7.g01122.t01#2    mNeoVis.vulgivaga.2.1.chr7.g01134.t01#1    mNeoVis.vulgivaga.2.1.chr7.g01135.t01#1    mNeoVis.3.1.1.chr7.g01059.t01#6    mNeoVis.3.1.1.chr7.g01060.t01#6    mNeoVis.vulgivaga.1.1.chr7.g01124.t01#4    mNeoVis.evergladensis.1.1.chr7.g01126.t01#3   mNeoVis.2.1.1.chr7_hap1.g01136.t01#5     4097   125256525:   mNeoVis.vulgivaga.1.1.chr1.g00418.t01#4    mNeoVis.evergladensis.2.1.chr1.g00423.t01#2    mNeoVis.2.1.1.chr1_hap1.g00425.t01#5    mNeoVis.3.1.1.chr1.g00422.t01#6    mNeoVis.vulgivaga.2.1.chr1.g00429.t01#1   mNeoVis.evergladensis.1.1.chr1.g00431.t01#3       4098   125256526:   mNeoVis.3.1.1.chr1.g00423.t01#6    mNeoVis.evergladensis.1.1.chr1.g00432.t01#3    mNeoVis.vulgivaga.1.1.chr1.g00419.t01#4    mNeoVis.2.1.1.chr1_hap1.g00426.t01#5    mNeoVis.vulgivaga.2.1.chr1.g00430.t01#1   mNeoVis.evergladensis.2.1.chr1.g00424.t01#2       4099   125256527:   mNeoVis.evergladensis.2.1.chr7.g01123.t01#2    mNeoVis.vulgivaga.2.1.chr7.g01136.t01#1    mNeoVis.vulgivaga.1.1.chr7.g01125.t01#4    mNeoVis.3.1.1.chr7.g01061.t01#6   mNeoVis.2.1.1.chr7_hap1.g01137.t01#5       4100   125256528:   mNeoVis.vulgivaga.1.1.chr7.g01126.t01#4    mNeoVis.2.1.1.chr7_hap1.g01138.t01#5    mNeoVis.evergladensis.1.1.chr7.g01127.t01#3    mNeoVis.3.1.1.chr7.g01062.t01#6    mNeoVis.vulgivaga.2.1.chr7.g01137.t01#1   mNeoVis.evergladensis.2.1.chr7.g01124.t01#2       4101   125256529:   mNeoVis.evergladensis.1.1.chr1.g00433.t01#3    mNeoVis.vulgivaga.1.1.chr1.g00420.t01#4    mNeoVis.2.1.1.chr1_hap1.g00427.t01#5    mNeoVis.evergladensis.2.1.chr1.g00425.t01#2   mNeoVis.vulgivaga.2.1.chr1.g00431.t01#1       4102   125256530:   mNeoVis.3.1.1.chr1.g00425.t01#6    mNeoVis.vulgivaga.1.1.chr1.g00422.t01#4    mNeoVis.2.1.1.chr1_hap1.g00429.t01#5    mNeoVis.vulgivaga.2.1.chr1.g00432.t01#1    mNeoVis.evergladensis.2.1.chr1.g00426.t01#2   mNeoVis.evergladensis.1.1.chr1.g00434.t01#3       4103   125256531:   mNeoVis.vulgivaga.2.1.chr7.g01138.t01#1    mNeoVis.evergladensis.1.1.chr7.g01128.t01#3    mNeoVis.evergladensis.2.1.chr7.g01126.t01#2    mNeoVis.3.1.1.chr7.g01063.t01#6    mNeoVis.2.1.1.chr7_hap1.g01139.t01#5   mNeoVis.vulgivaga.1.1.chr7.g01127.t01#4       4104   125256532:   mNeoVis.evergladensis.2.1.chr1.g00427.t01#2    mNeoVis.3.1.1.chr1.g00426.t01#6    mNeoVis.evergladensis.1.1.chr1.g00435.t01#3    mNeoVis.2.1.1.chr1_hap1.g00430.t01#5    mNeoVis.vulgivaga.2.1.chr1.g00433.t01#1   mNeoVis.vulgivaga.1.1.chr1.g00423.t01#4       4105   125256533:   mNeoVis.evergladensis.1.1.chr1.g00436.t01#3    mNeoVis.2.1.1.chr1_hap1.g00431.t01#5    mNeoVis.vulgivaga.1.1.chr1.g00424.t01#4    mNeoVis.vulgivaga.2.1.chr1.g00434.t01#1    mNeoVis.evergladensis.2.1.chr1.g00428.t01#2   mNeoVis.3.1.1.chr1.g00427.t01#6       4106   125256534:   mNeoVis.3.1.1.chr1.g00428.t01#6    mNeoVis.vulgivaga.2.1.chr1.g00435.t01#1    mNeoVis.evergladensis.1.1.chr1.g00437.t01#3    mNeoVis.evergladensis.2.1.chr1.g00429.t01#2    mNeoVis.2.1.1.chr1_hap1.g00432.t01#5   mNeoVis.vulgivaga.1.1.chr1.g00425.t01#4       4107   125256535:   mNeoVis.3.1.1.chr7.g01064.t01#6    mNeoVis.2.1.1.chr7_hap1.g01140.t01#5    mNeoVis.evergladensis.1.1.chr7.g01129.t01#3    mNeoVis.vulgivaga.1.1.chr7.g01128.t01#4    mNeoVis.vulgivaga.2.1.chr7.g01139.t01#1   mNeoVis.evergladensis.2.1.chr7.g01127.t01#2       4108   125256536:   mNeoVis.2.1.1.chr7_hap1.g00486.t01#5    mNeoVis.evergladensis.2.1.chr7.g00489.t01#2    mNeoVis.vulgivaga.1.1.chr7.g00501.t01#4    mNeoVis.3.1.1.chr7.g00469.t01#6    mNeoVis.evergladensis.1.1.chr7.g00497.t01#3    mNeoVis.vulgivaga.2.1.chr7.g00494.t01#1   mNeoVis.3.1.1.chr7.g00468.t01#6       4109   125256537:   mNeoVis.evergladensis.1.1.chr7.g00487.t01#3   mNeoVis.vulgivaga.1.1.chr7.g00490.t01#4             4110   125256538:   mNeoVis.vulgivaga.1.1.chr1.g00426.t01#4    mNeoVis.evergladensis.2.1.chr1.g00430.t01#2    mNeoVis.2.1.1.chr1_hap1.g00433.t01#5    mNeoVis.evergladensis.1.1.chr1.g00438.t01#3   mNeoVis.vulgivaga.2.1.chr1.g00436.t01#1       4111   125256539:   mNeoVis.evergladensis.1.1.chr1.g00439.t01#3    mNeoVis.vulgivaga.1.1.chr1.g00427.t01#4    mNeoVis.2.1.1.chr1_hap1.g00434.t01#5    mNeoVis.vulgivaga.2.1.chr1.g00437.t01#1   mNeoVis.evergladensis.2.1.chr1.g00431.t01#2       4112   125256540:   mNeoVis.2.1.1.chr1_hap1.g00435.t01#5    mNeoVis.vulgivaga.2.1.chr1.g00438.t01#1    mNeoVis.vulgivaga.1.1.chr1.g00428.t01#4    mNeoVis.3.1.1.chr1.g00430.t01#6    mNeoVis.evergladensis.2.1.chr1.g00432.t01#2   mNeoVis.evergladensis.1.1.chr1.g00440.t01#3       4113   125256541:   mNeoVis.vulgivaga.2.1.chr1.g00439.t01#1    mNeoVis.vulgivaga.1.1.chr1.g00429.t01#4    mNeoVis.2.1.1.chr1_hap1.g00436.t01#5    mNeoVis.evergladensis.2.1.chr1.g00433.t01#2    mNeoVis.3.1.1.chr1.g00431.t01#6   mNeoVis.evergladensis.1.1.chr1.g00441.t01#3       4114   125256542:   mNeoVis.evergladensis.1.1.chr7.g01130.t01#3    mNeoVis.vulgivaga.1.1.chr7.g01130.t01#4    mNeoVis.evergladensis.2.1.chr7.g01129.t01#2    mNeoVis.2.1.1.chr7_hap1.g01142.t01#5    mNeoVis.vulgivaga.2.1.chr7.g01141.t01#1   mNeoVis.3.1.1.chr7.g01065.t01#6       4115   125256543:   mNeoVis.evergladensis.1.1.chr7.g00498.t01#3    mNeoVis.evergladensis.2.1.chr7.g00491.t01#2    mNeoVis.2.1.1.chr7_hap1.g00488.t01#5    mNeoVis.vulgivaga.2.1.chr7.g00495.t01#1   mNeoVis.3.1.1.chr7.g00470.t01#6       4116   125256544:   mNeoVis.2.1.1.chr1_hap1.g00438.t01#5    mNeoVis.evergladensis.2.1.chr1.g00435.t01#2    mNeoVis.vulgivaga.2.1.chr1.g00440.t01#1    mNeoVis.3.1.1.chr1.g00432.t01#6    mNeoVis.vulgivaga.1.1.chr1.g00431.t01#4   mNeoVis.evergladensis.1.1.chr1.g00443.t01#3       4117   125256545:   mNeoVis.3.1.1.chr7.g01066.t01#6    mNeoVis.vulgivaga.1.1.chr7.g01131.t01#4    mNeoVis.2.1.1.chr7_hap1.g01143.t01#5    mNeoVis.evergladensis.2.1.chr7.g01130.t01#2    mNeoVis.vulgivaga.2.1.chr7.g01142.t01#1   mNeoVis.evergladensis.1.1.chr7.g01131.t01#3       4118   125256546:   mNeoVis.3.1.1.chr8.g01325.t01#6    mNeoVis.vulgivaga.1.1.chr8.g01365.t01#4    mNeoVis.evergladensis.1.1.chr8.g01372.t01#3    mNeoVis.vulgivaga.2.1.chr8.g01382.t01#1    mNeoVis.2.1.1.chr8_hap1.g01428.t01#5    mNeoVis.vulgivaga.1.1.chr8.g01366.t01#4   mNeoVis.evergladensis.2.1.chr9.g01373.t01#2     4119   125256547:   mNeoVis.3.1.1.chr1.g00433.t01#6    mNeoVis.vulgivaga.2.1.chr1.g00441.t01#1    mNeoVis.evergladensis.2.1.chr1.g00434.t01#2    mNeoVis.evergladensis.1.1.chr1.g00442.t01#3   mNeoVis.vulgivaga.1.1.chr1.g00430.t01#4       4120   125256548:   mNeoVis.evergladensis.2.1.chr1.g00436.t01#2    mNeoVis.vulgivaga.2.1.chr1.g00442.t01#1    mNeoVis.3.1.1.chr1.g00434.t01#6    mNeoVis.vulgivaga.1.1.chr1.g00432.t01#4    mNeoVis.evergladensis.1.1.chr1.g00444.t01#3   mNeoVis.2.1.1.chr1_hap1.g00439.t01#5       4121   125256549:   mNeoVis.evergladensis.1.1.chr7.g01132.t01#3    mNeoVis.vulgivaga.1.1.chr7.g01132.t01#4    mNeoVis.vulgivaga.2.1.chr7.g01143.t01#1    mNeoVis.2.1.1.chr7_hap1.g01144.t01#5   mNeoVis.3.1.1.chr7.g01067.t01#6       4122   125256550:   mNeoVis.evergladensis.1.1.chr7.g01133.t01#3    mNeoVis.evergladensis.2.1.chr7.g01132.t01#2    mNeoVis.vulgivaga.1.1.chr7.g01133.t01#4    mNeoVis.3.1.1.chr7.g01068.t01#6    mNeoVis.vulgivaga.2.1.chr7.g01144.t01#1   mNeoVis.2.1.1.chr7_hap1.g01145.t01#5       4123   125256551:   mNeoVis.vulgivaga.1.1.chr7.g01134.t01#4    mNeoVis.evergladensis.1.1.chr7.g01134.t01#3    mNeoVis.evergladensis.2.1.chr7.g01133.t01#2    mNeoVis.2.1.1.chr7_hap1.g01146.t01#5    mNeoVis.3.1.1.chr7.g01069.t01#6    mNeoVis.vulgivaga.2.1.chr7.g01145.t01#1    mNeoVis.vulgivaga.1.1.chr7.g01136.t01#4   mNeoVis.vulgivaga.1.1.chr7.g01135.t01#4     4124   125256552:   mNeoVis.2.1.1.chr12_hap1.g00220.t01#5    mNeoVis.3.1.1.chr12.g00189.t01#6    mNeoVis.evergladensis.2.1.chr12.g00192.t01#2    mNeoVis.vulgivaga.2.1.chr12.g00196.t01#1    mNeoVis.vulgivaga.1.1.chr12.g00198.t01#4    mNeoVis.evergladensis.1.1.chr12.g00194.t01#3   mNeoVis.vulgivaga.1.1.chr12.g00199.t01#4     4125   125256553:   mNeoVis.3.1.1.chr1.g00229.t01#6    mNeoVis.evergladensis.2.1.chr1.g00226.t01#2    mNeoVis.vulgivaga.1.1.chr1.g00222.t01#4    mNeoVis.2.1.1.chr1_hap1.g00229.t01#5    mNeoVis.evergladensis.1.1.chr1.g00236.t01#3   mNeoVis.vulgivaga.2.1.chr1.g00227.t01#1       4126   125256554:   mNeoVis.2.1.1.chr6_hap1.g00649.t01#5    mNeoVis.vulgivaga.1.1.chr4.g00662.t01#4    mNeoVis.evergladensis.2.1.chr4.g00657.t01#2    mNeoVis.vulgivaga.2.1.chr4.g00667.t01#1    mNeoVis.evergladensis.1.1.chr4.g00663.t01#3   mNeoVis.3.1.1.chr4.g00656.t01#6       4127   125256555:   mNeoVis.vulgivaga.1.1.chr12.g00200.t01#4   mNeoVis.3.1.1.chr12.g00190.t01#6             4128   125256556:   mNeoVis.3.1.1.chr1.g00435.t01#6    mNeoVis.vulgivaga.2.1.chr1.g00443.t01#1    mNeoVis.2.1.1.chr1_hap1.g00440.t01#5    mNeoVis.evergladensis.2.1.chr1.g00437.t01#2    mNeoVis.evergladensis.1.1.chr1.g00445.t01#3   mNeoVis.vulgivaga.1.1.chr1.g00433.t01#4       4129   125256557:   mNeoVis.2.1.1.chr7_hap1.g01038.t01#5    mNeoVis.3.1.1.chr7.g00974.t01#6    mNeoVis.vulgivaga.2.1.chr7.g01039.t01#1    mNeoVis.evergladensis.2.1.chr7.g01031.t01#2    mNeoVis.evergladensis.1.1.chr7.g01037.t01#3   mNeoVis.vulgivaga.1.1.chr7.g01033.t01#4       4130   125256558:   mNeoVis.vulgivaga.1.1.chr4.g00749.t01#4    mNeoVis.vulgivaga.1.1.chr1.g00434.t01#4    mNeoVis.evergladensis.1.1.chr4.g00746.t01#3    mNeoVis.evergladensis.2.1.chr4.g00736.t01#2    mNeoVis.3.1.1.chr1.g00436.t01#6    mNeoVis.vulgivaga.2.1.chr1.g00444.t01#1    mNeoVis.2.1.1.chr6_hap1.g00729.t01#5    mNeoVis.vulgivaga.2.1.chr4.g00754.t01#1    mNeoVis.evergladensis.1.1.chr1.g00446.t01#3      4131   125256559:   mNeoVis.evergladensis.2.1.chr9.g01233.t01#2    mNeoVis.2.1.1.chr8_hap1.g01285.t01#5    mNeoVis.3.1.1.chr8.g01181.t01#6    mNeoVis.evergladensis.1.1.chr8.g01230.t01#3    mNeoVis.vulgivaga.1.1.chr8.g01221.t01#4   mNeoVis.vulgivaga.2.1.chr8.g01235.t01#1       4132   125256560:   mNeoVis.vulgivaga.1.1.chr1.g00435.t01#4    mNeoVis.evergladensis.2.1.chr1.g00439.t01#2    mNeoVis.evergladensis.1.1.chr1.g00447.t01#3    mNeoVis.3.1.1.chr1.g00437.t01#6    mNeoVis.vulgivaga.2.1.chr1.g00445.t01#1   mNeoVis.2.1.1.chr1_hap1.g00442.t01#5       4133   125256561:   mNeoVis.evergladensis.1.1.chr7.g01038.t01#3    mNeoVis.2.1.1.chr7_hap1.g01039.t01#5    mNeoVis.3.1.1.chr7.g00975.t01#6    mNeoVis.vulgivaga.2.1.chr7.g01040.t01#1    mNeoVis.evergladensis.2.1.chr7.g01032.t01#2   mNeoVis.vulgivaga.1.1.chr7.g01034.t01#4       4134   125256562:   mNeoVis.vulgivaga.2.1.chr7.g00496.t01#1    mNeoVis.evergladensis.2.1.chr7.g00492.t01#2    mNeoVis.evergladensis.1.1.chr7.g00499.t01#3    mNeoVis.vulgivaga.1.1.chr7.g00502.t01#4    mNeoVis.3.1.1.chr7.g00471.t01#6   mNeoVis.2.1.1.chr7_hap1.g00489.t01#5       4135   125256563:   mNeoVis.evergladensis.1.1.chr1.g00448.t01#3    mNeoVis.3.1.1.chr1.g00439.t01#6    mNeoVis.3.1.1.chr1.g00440.t01#6    mNeoVis.vulgivaga.1.1.chr1.g00436.t01#4    mNeoVis.evergladensis.2.1.chr1.g00440.t01#2    mNeoVis.2.1.1.chr1_hap1.g00443.t01#5   mNeoVis.vulgivaga.2.1.chr1.g00446.t01#1     4136   125256564:   mNeoVis.3.1.1.chr4.g00457.t01#6    mNeoVis.evergladensis.1.1.chr4.g00465.t01#3    mNeoVis.evergladensis.2.1.chr4.g00461.t01#2    mNeoVis.vulgivaga.2.1.chr4.g00467.t01#1    mNeoVis.vulgivaga.1.1.chr4.g00458.t01#4   mNeoVis.2.1.1.chr6_hap1.g00455.t01#5       4137   125256565:   mNeoVis.vulgivaga.1.1.chr7.g01035.t01#4    mNeoVis.vulgivaga.2.1.chr7.g01041.t01#1    mNeoVis.2.1.1.chr7_hap1.g01040.t01#5    mNeoVis.evergladensis.1.1.chr7.g01039.t01#3    mNeoVis.evergladensis.2.1.chr7.g01033.t01#2   mNeoVis.3.1.1.chr7.g00976.t01#6       4138   125256566:   mNeoVis.2.1.1.chr6_hap1.g00913.t01#5    mNeoVis.3.1.1.chr11.g00172.t01#6    mNeoVis.vulgivaga.1.1.chr4.g00882.t01#4    mNeoVis.2.1.1.chr11_hap1.g00203.t01#5    mNeoVis.evergladensis.2.1.chr11.g00200.t01#2    mNeoVis.vulgivaga.2.1.chr11.g00203.t01#1    mNeoVis.evergladensis.1.1.chr11.g00202.t01#3    mNeoVis.2.1.1.chr4_hap1.g00914.t01#5    mNeoVis.evergladensis.2.1.chr6.g00919.t01#2      4139   125256567:   mNeoVis.evergladensis.2.1.chr3.g00399.t01#2    mNeoVis.evergladensis.2.1.chr15.g00540.t01#2    mNeoVis.2.1.1.chr3_hap1.g00392.t01#5    mNeoVis.vulgivaga.1.1.chr3.g00393.t01#4    mNeoVis.vulgivaga.2.1.chr15.g00553.t01#1    mNeoVis.2.1.1.chr15_hap1.g00718.t01#5    mNeoVis.vulgivaga.2.1.chr3.g00393.t01#1    mNeoVis.3.1.1.chr15.g00495.t01#6    mNeoVis.3.1.1.chr3.g00384.t01#6      4140   125256568:   mNeoVis.3.1.1.chr11.g01116.t01#6    mNeoVis.evergladensis.1.1.chr11.g01160.t01#3    mNeoVis.vulgivaga.1.1.chr11.g01140.t01#4    mNeoVis.vulgivaga.2.1.chr11.g01169.t01#1    mNeoVis.vulgivaga.2.1.chr13.g00331.t01#1    mNeoVis.2.1.1.chr11_hap1.g01173.t01#5    mNeoVis.evergladensis.1.1.chr11.g01159.t01#3    mNeoVis.2.1.1.chr13_hap1.g00343.t01#5    mNeoVis.3.1.1.chr13.g00285.t01#6      4141   125256569:   mNeoVis.evergladensis.2.1.chr11.g01142.t01#2    mNeoVis.vulgivaga.1.1.chr6.g00918.t01#4   mNeoVis.vulgivaga.2.1.chr6.g00913.t01#1           4142   125256570:   mNeoVis.vulgivaga.2.1.chr7.g01042.t01#1    mNeoVis.evergladensis.1.1.chr7.g01040.t01#3    mNeoVis.2.1.1.chr7_hap1.g01041.t01#5    mNeoVis.evergladensis.2.1.chr7.g01034.t01#2    mNeoVis.3.1.1.chr7.g00977.t01#6   mNeoVis.vulgivaga.1.1.chr7.g01036.t01#4       4143   125256571:   mNeoVis.evergladensis.2.1.chr1.g00441.t01#2    mNeoVis.2.1.1.chr1_hap1.g00444.t01#5    mNeoVis.vulgivaga.1.1.chr1.g00437.t01#4    mNeoVis.3.1.1.chr1.g00442.t01#6    mNeoVis.evergladensis.1.1.chr1.g00449.t01#3    mNeoVis.vulgivaga.2.1.chr1.g00447.t01#1   mNeoVis.3.1.1.chr1.g00441.t01#6       4144   125256572:   mNeoVis.vulgivaga.1.1.chr7.g01037.t01#4    mNeoVis.evergladensis.2.1.chr7.g01035.t01#2    mNeoVis.2.1.1.chr7_hap1.g01042.t01#5    mNeoVis.evergladensis.1.1.chr7.g01041.t01#3    mNeoVis.vulgivaga.2.1.chr7.g01043.t01#1   mNeoVis.3.1.1.chr7.g00978.t01#6       4145   125256573:   mNeoVis.vulgivaga.2.1.chr7.g01044.t01#1    mNeoVis.2.1.1.chr7_hap1.g01043.t01#5    mNeoVis.vulgivaga.1.1.chr7.g01038.t01#4    mNeoVis.evergladensis.2.1.chr7.g01036.t01#2   mNeoVis.evergladensis.1.1.chr7.g01042.t01#3       4146   125256574:   mNeoVis.evergladensis.2.1.chr7.g01037.t01#2    mNeoVis.vulgivaga.1.1.chr7.g01039.t01#4    mNeoVis.vulgivaga.2.1.chr7.g01045.t01#1    mNeoVis.evergladensis.1.1.chr7.g01043.t01#3    mNeoVis.3.1.1.chr7.g00979.t01#6   mNeoVis.2.1.1.chr7_hap1.g01044.t01#5       4147   125256575:   mNeoVis.3.1.1.chr4.g00685.t01#6    mNeoVis.2.1.1.chr6_hap1.g00677.t01#5    mNeoVis.evergladensis.2.1.chr4.g00683.t01#2    mNeoVis.vulgivaga.1.1.chr4.g00695.t01#4    mNeoVis.evergladensis.1.1.chr4.g00693.t01#3    mNeoVis.vulgivaga.1.1.chr4.g00696.t01#4   mNeoVis.vulgivaga.2.1.chr4.g00698.t01#1     4148   125256576:   mNeoVis.3.1.1.chr1.g00444.t01#6    mNeoVis.vulgivaga.2.1.chr1.g00448.t01#1    mNeoVis.evergladensis.1.1.chr1.g00451.t01#3    mNeoVis.vulgivaga.1.1.chr1.g00438.t01#4    mNeoVis.evergladensis.2.1.chr1.g00443.t01#2   mNeoVis.2.1.1.chr1_hap1.g00446.t01#5       4149   125256577:   mNeoVis.evergladensis.2.1.chr1.g00444.t01#2   mNeoVis.vulgivaga.2.1.chr1.g00449.t01#1             4150   125256578:   mNeoVis.evergladensis.1.1.chr1.g00452.t01#3    mNeoVis.vulgivaga.2.1.chr1.g00450.t01#1    mNeoVis.3.1.1.chr1.g00445.t01#6    mNeoVis.2.1.1.chr1_hap1.g00447.t01#5    mNeoVis.evergladensis.2.1.chr1.g00445.t01#2   mNeoVis.vulgivaga.1.1.chr1.g00439.t01#4       4151   125256579:   mNeoVis.evergladensis.1.1.chr7.g01044.t01#3    mNeoVis.vulgivaga.2.1.chr7.g01046.t01#1    mNeoVis.2.1.1.chr7_hap1.g01045.t01#5    mNeoVis.vulgivaga.1.1.chr7.g01040.t01#4    mNeoVis.evergladensis.2.1.chr7.g01038.t01#2   mNeoVis.3.1.1.chr7.g00980.t01#6       4152   125256580:   mNeoVis.evergladensis.1.1.chr15.g00072.t01#3    mNeoVis.evergladensis.2.1.chr15.g00542.t01#2    mNeoVis.vulgivaga.2.1.chr15.g00554.t01#1    mNeoVis.vulgivaga.1.1.chr15.g00544.t01#4    mNeoVis.2.1.1.chr15_hap1.g00719.t01#5   mNeoVis.3.1.1.chr15.g00496.t01#6       4153   125256581:   mNeoVis.vulgivaga.2.1.chr1.g00451.t01#1    mNeoVis.2.1.1.chr1_hap1.g00448.t01#5    mNeoVis.vulgivaga.1.1.chr1.g00440.t01#4    mNeoVis.3.1.1.chr1.g00446.t01#6    mNeoVis.evergladensis.2.1.chr1.g00446.t01#2   mNeoVis.evergladensis.1.1.chr1.g00453.t01#3       4154   125256582:   mNeoVis.vulgivaga.2.1.chr1.g00452.t01#1    mNeoVis.evergladensis.2.1.chr1.g00447.t01#2    mNeoVis.vulgivaga.1.1.chr1.g00441.t01#4    mNeoVis.2.1.1.chr1_hap1.g00449.t01#5    mNeoVis.evergladensis.1.1.chr1.g00454.t01#3   mNeoVis.3.1.1.chr1.g00447.t01#6       4155   125256583:   mNeoVis.evergladensis.2.1.chr7.g01039.t01#2    mNeoVis.evergladensis.1.1.chr7.g01045.t01#3    mNeoVis.vulgivaga.1.1.chr7.g01041.t01#4    mNeoVis.2.1.1.chr7_hap1.g01046.t01#5    mNeoVis.vulgivaga.2.1.chr7.g01047.t01#1   mNeoVis.3.1.1.chr7.g00981.t01#6       4156   125256584:   mNeoVis.3.1.1.chr1.g00448.t01#6    mNeoVis.evergladensis.1.1.chr1.g00455.t01#3    mNeoVis.2.1.1.chr1_hap1.g00450.t01#5    mNeoVis.vulgivaga.2.1.chr1.g00453.t01#1    mNeoVis.vulgivaga.1.1.chr1.g00442.t01#4   mNeoVis.evergladensis.2.1.chr1.g00448.t01#2       4157   125256585:   mNeoVis.2.1.1.chr7_hap1.g00490.t01#5    mNeoVis.vulgivaga.2.1.chr7.g00497.t01#1    mNeoVis.3.1.1.chr7.g00472.t01#6    mNeoVis.evergladensis.1.1.chr7.g00501.t01#3   mNeoVis.evergladensis.2.1.chr7.g00493.t01#2       4158   125256586:   mNeoVis.evergladensis.2.1.chr1.g00347.t01#2    mNeoVis.2.1.1.chr1_hap1.g00349.t01#5    mNeoVis.vulgivaga.1.1.chr1.g00347.t01#4    mNeoVis.evergladensis.1.1.chr1.g00355.t01#3    mNeoVis.vulgivaga.2.1.chr1.g00355.t01#1   mNeoVis.3.1.1.chr1.g00352.t01#6       4159   125256587:   mNeoVis.2.1.1.chr7_hap1.g01047.t01#5    mNeoVis.evergladensis.2.1.chr7.g01040.t01#2    mNeoVis.vulgivaga.1.1.chr7.g01042.t01#4    mNeoVis.3.1.1.chr7.g00982.t01#6    mNeoVis.vulgivaga.2.1.chr7.g01048.t01#1   mNeoVis.evergladensis.1.1.chr7.g01046.t01#3       4160   125256588:   mNeoVis.evergladensis.2.1.chr7.g01041.t01#2    mNeoVis.vulgivaga.2.1.chr7.g01049.t01#1    mNeoVis.evergladensis.1.1.chr7.g01047.t01#3    mNeoVis.3.1.1.chr7.g00983.t01#6    mNeoVis.2.1.1.chr7_hap1.g01048.t01#5   mNeoVis.vulgivaga.1.1.chr7.g01043.t01#4       4161   125256589:   mNeoVis.vulgivaga.2.1.chr7.g01050.t01#1    mNeoVis.2.1.1.chr7_hap1.g01049.t01#5    mNeoVis.vulgivaga.1.1.chr7.g01044.t01#4    mNeoVis.3.1.1.chr7.g00984.t01#6   mNeoVis.evergladensis.2.1.chr7.g01042.t01#2       4162   125256590:   mNeoVis.vulgivaga.2.1.chr7.g01051.t01#1    mNeoVis.evergladensis.2.1.chr7.g01043.t01#2    mNeoVis.3.1.1.chr7.g00985.t01#6   mNeoVis.2.1.1.chr7_hap1.g01050.t01#5         4163   125256591:   mNeoVis.evergladensis.1.1.chr1.g00356.t01#3    mNeoVis.2.1.1.chr1_hap1.g00350.t01#5    mNeoVis.vulgivaga.2.1.chr1.g00356.t01#1    mNeoVis.evergladensis.2.1.chr1.g00348.t01#2   mNeoVis.vulgivaga.1.1.chr1.g00348.t01#4       4164   125256592:  mNeoVis.vulgivaga.2.1.chr1.g00357.t01#1               4165   125256593:   mNeoVis.evergladensis.1.1.chr15.g00071.t01#3    mNeoVis.2.1.1.chr15_hap1.g00720.t01#5    mNeoVis.vulgivaga.1.1.chr15.g00545.t01#4    mNeoVis.3.1.1.chr15.g00497.t01#6    mNeoVis.vulgivaga.2.1.chr15.g00555.t01#1   mNeoVis.evergladensis.2.1.chr15.g00543.t01#2       4166   125256594:   mNeoVis.evergladensis.2.1.chr1.g00350.t01#2    mNeoVis.3.1.1.chr1.g00355.t01#6    mNeoVis.vulgivaga.1.1.chr1.g00350.t01#4    mNeoVis.evergladensis.1.1.chr1.g00358.t01#3    mNeoVis.vulgivaga.2.1.chr1.g00358.t01#1   mNeoVis.2.1.1.chr1_hap1.g00352.t01#5       4167   125256595:   mNeoVis.vulgivaga.2.1.chr1.g00359.t01#1    mNeoVis.vulgivaga.1.1.chr1.g00351.t01#4    mNeoVis.3.1.1.chr1.g00356.t01#6    mNeoVis.evergladensis.1.1.chr1.g00359.t01#3    mNeoVis.evergladensis.2.1.chr1.g00351.t01#2   mNeoVis.2.1.1.chr1_hap1.g00353.t01#5       4168   125256596:   mNeoVis.vulgivaga.2.1.chr7.g01052.t01#1    mNeoVis.2.1.1.chr7_hap1.g01051.t01#5   mNeoVis.evergladensis.2.1.chr7.g01044.t01#2         4169   125256597:   mNeoVis.2.1.1.chr7_hap1.g01052.t01#5    mNeoVis.evergladensis.2.1.chr7.g01045.t01#2    mNeoVis.vulgivaga.2.1.chr7.g01053.t01#1    mNeoVis.evergladensis.1.1.chr7.g01050.t01#3    mNeoVis.vulgivaga.2.1.chr7.g01054.t01#1   mNeoVis.vulgivaga.1.1.chr7.g01046.t01#4       4170   125256598:   mNeoVis.2.1.1.chr1_hap1.g00354.t01#5    mNeoVis.vulgivaga.2.1.chr1.g00360.t01#1    mNeoVis.vulgivaga.1.1.chr1.g00352.t01#4    mNeoVis.3.1.1.chr1.g00357.t01#6    mNeoVis.evergladensis.1.1.chr1.g00360.t01#3   mNeoVis.evergladensis.2.1.chr1.g00352.t01#2       4171   125256599:   mNeoVis.vulgivaga.2.1.chr7.g01055.t01#1    mNeoVis.vulgivaga.1.1.chr7.g01047.t01#4    mNeoVis.evergladensis.2.1.chr7.g01046.t01#2    mNeoVis.2.1.1.chr7_hap1.g01053.t01#5   mNeoVis.evergladensis.1.1.chr7.g01051.t01#3       4172   125256600:   mNeoVis.3.1.1.chr7.g00987.t01#6    mNeoVis.evergladensis.2.1.chr7.g01047.t01#2    mNeoVis.vulgivaga.1.1.chr7.g01048.t01#4    mNeoVis.2.1.1.chr7_hap1.g01054.t01#5   mNeoVis.vulgivaga.2.1.chr7.g01056.t01#1       4173   125256601:   mNeoVis.2.1.1.chr12_hap1.g00630.t01#5    mNeoVis.evergladensis.2.1.chr12.g00568.t01#2    mNeoVis.vulgivaga.2.1.chr12.g00584.t01#1    mNeoVis.3.1.1.chr12.g00574.t01#6    mNeoVis.evergladensis.1.1.chr12.g00569.t01#3    mNeoVis.vulgivaga.1.1.chr12.g00582.t01#4    mNeoVis.3.1.1.chr12.g00575.t01#6   mNeoVis.vulgivaga.1.1.chr12.g00583.t01#4     4174   125256602:   mNeoVis.evergladensis.1.1.chr7.g00502.t01#3    mNeoVis.evergladensis.2.1.chr7.g00494.t01#2   mNeoVis.2.1.1.chr7_hap1.g00491.t01#5           4175   125256603:   mNeoVis.3.1.1.chr7.g00473.t01#6   mNeoVis.vulgivaga.2.1.chr7.g00498.t01#1             4176   125256604:   mNeoVis.2.1.1.chr7_hap1.g01055.t01#5    mNeoVis.evergladensis.1.1.chr7.g01052.t01#3    mNeoVis.vulgivaga.2.1.chr7.g01057.t01#1   mNeoVis.vulgivaga.1.1.chr7.g01049.t01#4         4177   125256605:   mNeoVis.vulgivaga.1.1.chr1.g00353.t01#4    mNeoVis.evergladensis.2.1.chr1.g00353.t01#2    mNeoVis.vulgivaga.2.1.chr1.g00361.t01#1    mNeoVis.2.1.1.chr1_hap1.g00355.t01#5   mNeoVis.evergladensis.1.1.chr1.g00361.t01#3       4178   125256606:   mNeoVis.evergladensis.2.1.chr1.g00354.t01#2    mNeoVis.vulgivaga.1.1.chr1.g00354.t01#4    mNeoVis.evergladensis.1.1.chr1.g00362.t01#3    mNeoVis.vulgivaga.2.1.chr1.g00362.t01#1    mNeoVis.3.1.1.chr1.g00358.t01#6   mNeoVis.2.1.1.chr1_hap1.g00356.t01#5       4179   125256607:   mNeoVis.2.1.1.chr7_hap1.g01056.t01#5    mNeoVis.vulgivaga.1.1.chr7.g01050.t01#4    mNeoVis.evergladensis.1.1.chr7.g01053.t01#3    mNeoVis.evergladensis.2.1.chr7.g01048.t01#2    mNeoVis.vulgivaga.2.1.chr7.g01058.t01#1   mNeoVis.3.1.1.chr7.g00988.t01#6       4180   125256608:   mNeoVis.vulgivaga.1.1.chr1.g00355.t01#4    mNeoVis.vulgivaga.2.1.chr1.g00363.t01#1    mNeoVis.3.1.1.chr1.g00359.t01#6    mNeoVis.2.1.1.chr1_hap1.g00357.t01#5    mNeoVis.evergladensis.1.1.chr1.g00363.t01#3   mNeoVis.evergladensis.2.1.chr1.g00355.t01#2       4181   125256609:  mNeoVis.vulgivaga.2.1.chr7.g01059.t01#1               4182   125256610:   mNeoVis.2.1.1.chr7_hap1.g01058.t01#5    mNeoVis.vulgivaga.1.1.chr7.g01052.t01#4    mNeoVis.evergladensis.2.1.chr7.g01050.t01#2    mNeoVis.vulgivaga.2.1.chr7.g01060.t01#1   mNeoVis.3.1.1.chr7.g00989.t01#6       4183   125256611:   mNeoVis.evergladensis.1.1.chr7.g01055.t01#3    mNeoVis.evergladensis.2.1.chr7.g01052.t01#2    mNeoVis.vulgivaga.2.1.chr7.g01061.t01#1    mNeoVis.vulgivaga.1.1.chr7.g01054.t01#4   mNeoVis.2.1.1.chr7_hap1.g01060.t01#5       4184   125256612:   mNeoVis.2.1.1.chr7_hap1.g01061.t01#5    mNeoVis.vulgivaga.1.1.chr7.g01055.t01#4    mNeoVis.evergladensis.1.1.chr7.g01056.t01#3    mNeoVis.vulgivaga.2.1.chr7.g01062.t01#1    mNeoVis.evergladensis.2.1.chr7.g01053.t01#2   mNeoVis.3.1.1.chr7.g00991.t01#6       4185   125256613:   mNeoVis.vulgivaga.1.1.chr1.g00356.t01#4    mNeoVis.vulgivaga.2.1.chr1.g00364.t01#1    mNeoVis.evergladensis.1.1.chr1.g00364.t01#3    mNeoVis.3.1.1.chr1.g00360.t01#6    mNeoVis.2.1.1.chr1_hap1.g00358.t01#5   mNeoVis.evergladensis.2.1.chr1.g00356.t01#2       4186   125256614:   mNeoVis.2.1.1.chr1_hap1.g00359.t01#5    mNeoVis.vulgivaga.1.1.chr1.g00357.t01#4    mNeoVis.evergladensis.2.1.chr1.g00357.t01#2    mNeoVis.vulgivaga.2.1.chr1.g00365.t01#1    mNeoVis.3.1.1.chr1.g00361.t01#6   mNeoVis.evergladensis.1.1.chr1.g00365.t01#3       4187   125256615:   mNeoVis.vulgivaga.2.1.chr7.g01063.t01#1   mNeoVis.evergladensis.1.1.chr7.g01057.t01#3           4188   125256616:   mNeoVis.2.1.1.chr1_hap1.g00360.t01#5    mNeoVis.vulgivaga.1.1.chr1.g00358.t01#4    mNeoVis.3.1.1.chr1.g00362.t01#6    mNeoVis.vulgivaga.2.1.chr1.g00366.t01#1    mNeoVis.evergladensis.2.1.chr1.g00358.t01#2   mNeoVis.evergladensis.1.1.chr1.g00366.t01#3       4189   125256617:   mNeoVis.vulgivaga.2.1.chr7.g01064.t01#1    mNeoVis.2.1.1.chr7_hap1.g01063.t01#5    mNeoVis.evergladensis.2.1.chr7.g01055.t01#2    mNeoVis.evergladensis.1.1.chr7.g01058.t01#3   mNeoVis.3.1.1.chr7.g00994.t01#6       4190   125256618:  mNeoVis.vulgivaga.2.1.chr7.g01065.t01#1               4191   125256619:   mNeoVis.vulgivaga.2.1.chr7.g00499.t01#1   mNeoVis.2.1.1.chr7_hap1.g00494.t01#5             4192   125256620:   mNeoVis.vulgivaga.2.1.chr1.g00367.t01#1    mNeoVis.evergladensis.2.1.chr1.g00359.t01#2    mNeoVis.evergladensis.1.1.chr1.g00367.t01#3    mNeoVis.3.1.1.chr1.g00363.t01#6    mNeoVis.vulgivaga.1.1.chr1.g00359.t01#4   mNeoVis.2.1.1.chr1_hap1.g00361.t01#5       4193   125256621:   mNeoVis.evergladensis.2.1.chr7.g01056.t01#2    mNeoVis.evergladensis.1.1.chr7.g01060.t01#3    mNeoVis.vulgivaga.2.1.chr7.g01066.t01#1   mNeoVis.2.1.1.chr7_hap1.g01064.t01#5         4194   125256622:   mNeoVis.2.1.1.chr7_hap1.g01065.t01#5    mNeoVis.vulgivaga.2.1.chr7.g01067.t01#1    mNeoVis.vulgivaga.1.1.chr7.g01057.t01#4    mNeoVis.evergladensis.2.1.chr7.g01057.t01#2   mNeoVis.evergladensis.1.1.chr7.g01061.t01#3       4195   125256623:   mNeoVis.evergladensis.2.1.chr7.g01058.t01#2    mNeoVis.vulgivaga.2.1.chr7.g01068.t01#1    mNeoVis.evergladensis.1.1.chr7.g01062.t01#3    mNeoVis.2.1.1.chr7_hap1.g01066.t01#5    mNeoVis.3.1.1.chr7.g00996.t01#6   mNeoVis.vulgivaga.1.1.chr7.g01058.t01#4       4196   125256624:   mNeoVis.vulgivaga.1.1.chr1.g00360.t01#4    mNeoVis.evergladensis.1.1.chr1.g00368.t01#3    mNeoVis.3.1.1.chr1.g00364.t01#6    mNeoVis.vulgivaga.2.1.chr1.g00368.t01#1    mNeoVis.2.1.1.chr1_hap1.g00362.t01#5   mNeoVis.evergladensis.2.1.chr1.g00360.t01#2       4197   125256625:   mNeoVis.3.1.1.chr1.g00365.t01#6    mNeoVis.2.1.1.chr1_hap1.g00363.t01#5    mNeoVis.evergladensis.2.1.chr1.g00361.t01#2    mNeoVis.vulgivaga.2.1.chr1.g00369.t01#1    mNeoVis.vulgivaga.1.1.chr1.g00361.t01#4   mNeoVis.evergladensis.1.1.chr1.g00369.t01#3       4198   125256626:   mNeoVis.evergladensis.2.1.chr1.g00363.t01#2    mNeoVis.vulgivaga.1.1.chr1.g00362.t01#4    mNeoVis.2.1.1.chr1_hap1.g00366.t01#5    mNeoVis.3.1.1.chr1.g00366.t01#6    mNeoVis.evergladensis.1.1.chr1.g00370.t01#3   mNeoVis.vulgivaga.2.1.chr1.g00370.t01#1       4199   125256627:   mNeoVis.evergladensis.1.1.chr1.g00371.t01#3    mNeoVis.evergladensis.2.1.chr1.g00362.t01#2    mNeoVis.vulgivaga.2.1.chr1.g00371.t01#1    mNeoVis.vulgivaga.1.1.chr1.g00363.t01#4    mNeoVis.3.1.1.chr1.g00367.t01#6   mNeoVis.2.1.1.chr1_hap1.g00365.t01#5       4200   125256628:   mNeoVis.evergladensis.1.1.chr7.g01063.t01#3    mNeoVis.2.1.1.chr7_hap1.g01067.t01#5    mNeoVis.vulgivaga.1.1.chr7.g01059.t01#4    mNeoVis.vulgivaga.2.1.chr7.g01069.t01#1    mNeoVis.3.1.1.chr7.g00997.t01#6   mNeoVis.evergladensis.2.1.chr7.g01059.t01#2       4201   125256629:   mNeoVis.2.1.1.chr7_hap1.g01068.t01#5    mNeoVis.evergladensis.2.1.chr7.g01060.t01#2    mNeoVis.vulgivaga.2.1.chr7.g01070.t01#1   mNeoVis.vulgivaga.1.1.chr7.g01060.t01#4         4202   125256630:   mNeoVis.evergladensis.1.1.chr7.g01064.t01#3    mNeoVis.vulgivaga.1.1.chr7.g01061.t01#4    mNeoVis.evergladensis.2.1.chr7.g01061.t01#2    mNeoVis.2.1.1.chr7_hap1.g01069.t01#5   mNeoVis.vulgivaga.2.1.chr7.g01071.t01#1       4203   125256631:   mNeoVis.vulgivaga.2.1.chr7.g01072.t01#1    mNeoVis.2.1.1.chr7_hap1.g01070.t01#5    mNeoVis.evergladensis.2.1.chr7.g01062.t01#2    mNeoVis.3.1.1.chr7.g00998.t01#6    mNeoVis.evergladensis.1.1.chr7.g01065.t01#3   mNeoVis.vulgivaga.1.1.chr7.g01062.t01#4       4204   125256632:   mNeoVis.vulgivaga.1.1.chr7.g00504.t01#4    mNeoVis.vulgivaga.2.1.chr7.g00500.t01#1    mNeoVis.evergladensis.1.1.chr7.g00504.t01#3    mNeoVis.2.1.1.chr7_hap1.g00495.t01#5   mNeoVis.evergladensis.2.1.chr7.g00496.t01#2       4205   125256633:   mNeoVis.2.1.1.chr15_hap1.g00721.t01#5    mNeoVis.vulgivaga.1.1.chr15.g00546.t01#4    mNeoVis.vulgivaga.2.1.chr15.g00556.t01#1    mNeoVis.3.1.1.chr15.g00498.t01#6   mNeoVis.evergladensis.1.1.chr15.g00070.t01#3       4206   125256634:   mNeoVis.evergladensis.1.1.chr7.g01066.t01#3    mNeoVis.vulgivaga.1.1.chr7.g01063.t01#4    mNeoVis.evergladensis.2.1.chr7.g01063.t01#2    mNeoVis.2.1.1.chr7_hap1.g01071.t01#5    mNeoVis.3.1.1.chr7.g00999.t01#6   mNeoVis.vulgivaga.2.1.chr7.g01073.t01#1       4207   125256635:   mNeoVis.vulgivaga.2.1.chr1.g00372.t01#1    mNeoVis.evergladensis.1.1.chr1.g00372.t01#3    mNeoVis.2.1.1.chr1_hap1.g00367.t01#5    mNeoVis.evergladensis.2.1.chr1.g00364.t01#2    mNeoVis.3.1.1.chr1.g00368.t01#6   mNeoVis.vulgivaga.1.1.chr1.g00365.t01#4       4208   125256636:   mNeoVis.vulgivaga.2.1.chr7.g01074.t01#1    mNeoVis.3.1.1.chr7.g01000.t01#6    mNeoVis.evergladensis.1.1.chr7.g01067.t01#3    mNeoVis.vulgivaga.1.1.chr7.g01064.t01#4    mNeoVis.evergladensis.2.1.chr7.g01064.t01#2   mNeoVis.2.1.1.chr7_hap1.g01072.t01#5       4209   125256637:   mNeoVis.2.1.1.chr1_hap1.g00368.t01#5    mNeoVis.vulgivaga.2.1.chr1.g00373.t01#1    mNeoVis.3.1.1.chr1.g00369.t01#6    mNeoVis.evergladensis.1.1.chr1.g00373.t01#3    mNeoVis.vulgivaga.1.1.chr1.g00366.t01#4   mNeoVis.evergladensis.2.1.chr1.g00365.t01#2       4210   125256638:   mNeoVis.vulgivaga.1.1.chr1.g00367.t01#4    mNeoVis.2.1.1.chr1_hap1.g00369.t01#5    mNeoVis.evergladensis.2.1.chr1.g00366.t01#2    mNeoVis.evergladensis.1.1.chr1.g00374.t01#3    mNeoVis.vulgivaga.2.1.chr1.g00374.t01#1   mNeoVis.3.1.1.chr1.g00370.t01#6       4211   125256639:   mNeoVis.vulgivaga.1.1.chr1.g00368.t01#4    mNeoVis.evergladensis.1.1.chr1.g00375.t01#3    mNeoVis.evergladensis.2.1.chr1.g00367.t01#2    mNeoVis.2.1.1.chr1_hap1.g00370.t01#5    mNeoVis.vulgivaga.2.1.chr1.g00375.t01#1   mNeoVis.3.1.1.chr1.g00371.t01#6       4212   125256640:   mNeoVis.vulgivaga.1.1.chr7.g01065.t01#4    mNeoVis.3.1.1.chr7.g01002.t01#6    mNeoVis.2.1.1.chr7_hap1.g01073.t01#5    mNeoVis.vulgivaga.2.1.chr7.g01075.t01#1   mNeoVis.evergladensis.1.1.chr7.g01068.t01#3       4213   125256641:   mNeoVis.evergladensis.2.1.chr7.g01065.t01#2    mNeoVis.2.1.1.chr7_hap1.g01074.t01#5    mNeoVis.vulgivaga.2.1.chr7.g01076.t01#1    mNeoVis.3.1.1.chr7.g01001.t01#6    mNeoVis.evergladensis.1.1.chr7.g01069.t01#3   mNeoVis.vulgivaga.1.1.chr7.g01066.t01#4       4214   125256642:   mNeoVis.3.1.1.chr7.g01003.t01#6    mNeoVis.2.1.1.chr7_hap1.g01075.t01#5    mNeoVis.evergladensis.2.1.chr7.g01066.t01#2    mNeoVis.evergladensis.1.1.chr7.g01071.t01#3    mNeoVis.vulgivaga.1.1.chr7.g01068.t01#4   mNeoVis.vulgivaga.2.1.chr7.g01077.t01#1       4215   125256643:   mNeoVis.evergladensis.1.1.chr7.g01070.t01#3    mNeoVis.evergladensis.2.1.chr7.g01067.t01#2    mNeoVis.vulgivaga.1.1.chr7.g01067.t01#4    mNeoVis.2.1.1.chr7_hap1.g01076.t01#5    mNeoVis.vulgivaga.2.1.chr7.g01078.t01#1   mNeoVis.3.1.1.chr7.g01004.t01#6       4216   125256644:   mNeoVis.3.1.1.chr7.g01005.t01#6    mNeoVis.evergladensis.2.1.chr7.g01068.t01#2    mNeoVis.vulgivaga.1.1.chr7.g01069.t01#4    mNeoVis.vulgivaga.2.1.chr7.g01079.t01#1   mNeoVis.2.1.1.chr7_hap1.g01077.t01#5       4217   125256645:   mNeoVis.3.1.1.chr7.g01006.t01#6    mNeoVis.vulgivaga.2.1.chr7.g01080.t01#1    mNeoVis.evergladensis.1.1.chr7.g01073.t01#3    mNeoVis.vulgivaga.1.1.chr7.g01070.t01#4    mNeoVis.2.1.1.chr7_hap1.g01078.t01#5   mNeoVis.evergladensis.2.1.chr7.g01069.t01#2       4218   125256646:   mNeoVis.vulgivaga.2.1.chr7.g01082.t01#1    mNeoVis.evergladensis.1.1.chr7.g01075.t01#3    mNeoVis.3.1.1.chr7.g01008.t01#6   mNeoVis.vulgivaga.1.1.chr7.g01072.t01#4         4219   125256647:   mNeoVis.vulgivaga.2.1.chr1.g00376.t01#1    mNeoVis.2.1.1.chr1_hap1.g00371.t01#5    mNeoVis.evergladensis.2.1.chr1.g00368.t01#2   mNeoVis.evergladensis.1.1.chr1.g00376.t01#3       4220   125256648:   mNeoVis.vulgivaga.2.1.chr1.g00377.t01#1    mNeoVis.evergladensis.2.1.chr1.g00369.t01#2    mNeoVis.vulgivaga.1.1.chr1.g00369.t01#4   mNeoVis.2.1.1.chr1_hap1.g00372.t01#5         4221   125256649:   mNeoVis.evergladensis.2.1.chr1.g00370.t01#2    mNeoVis.vulgivaga.1.1.chr1.g00370.t01#4    mNeoVis.3.1.1.chr1.g00372.t01#6    mNeoVis.vulgivaga.2.1.chr1.g00378.t01#1    mNeoVis.2.1.1.chr1_hap1.g00373.t01#5   mNeoVis.evergladensis.1.1.chr1.g00378.t01#3       4222   125256650:   mNeoVis.vulgivaga.2.1.chr7.g01081.t01#1    mNeoVis.evergladensis.1.1.chr7.g01074.t01#3    mNeoVis.vulgivaga.1.1.chr7.g01071.t01#4    mNeoVis.3.1.1.chr7.g01007.t01#6    mNeoVis.evergladensis.2.1.chr7.g01070.t01#2   mNeoVis.2.1.1.chr7_hap1.g01079.t01#5       4223   125256651:   mNeoVis.3.1.1.chr15.g00499.t01#6    mNeoVis.vulgivaga.2.1.chr15.g00557.t01#1    mNeoVis.2.1.1.chr15_hap1.g00722.t01#5    mNeoVis.vulgivaga.1.1.chr15.g00547.t01#4    mNeoVis.evergladensis.1.1.chr15.g00069.t01#3   mNeoVis.evergladensis.2.1.chr15.g00544.t01#2       4224   125256652:   mNeoVis.3.1.1.chr7.g00482.t01#6    mNeoVis.vulgivaga.2.1.chr7.g00501.t01#1    mNeoVis.evergladensis.2.1.chr7.g00498.t01#2    mNeoVis.evergladensis.1.1.chr7.g00506.t01#3    mNeoVis.vulgivaga.1.1.chr7.g00505.t01#4   mNeoVis.2.1.1.chr7_hap1.g00497.t01#5       4225   125256653:   mNeoVis.3.1.1.chr5.g01678.t01#6    mNeoVis.evergladensis.2.1.chr7.g01071.t01#2    mNeoVis.2.1.1.chr7_hap1.g01080.t01#5    mNeoVis.vulgivaga.2.1.chr7.g01083.t01#1    mNeoVis.vulgivaga.1.1.chr7.g01073.t01#4    mNeoVis.evergladensis.1.1.chr7.g01076.t01#3   mNeoVis.3.1.1.chr7.g01009.t01#6       4226   125256654:   mNeoVis.2.1.1.chr7_hap1.g01081.t01#5    mNeoVis.evergladensis.2.1.chr7.g01072.t01#2    mNeoVis.evergladensis.1.1.chr7.g01077.t01#3    mNeoVis.vulgivaga.1.1.chr7.g01074.t01#4    mNeoVis.3.1.1.chr7.g01010.t01#6   mNeoVis.vulgivaga.2.1.chr7.g01084.t01#1       4227   125256655:   mNeoVis.evergladensis.2.1.chr7.g01073.t01#2    mNeoVis.2.1.1.chr7_hap1.g01082.t01#5    mNeoVis.3.1.1.chr7.g01011.t01#6    mNeoVis.vulgivaga.2.1.chr7.g01085.t01#1    mNeoVis.evergladensis.1.1.chr7.g01078.t01#3   mNeoVis.vulgivaga.1.1.chr7.g01075.t01#4       4228   125256656:   mNeoVis.evergladensis.2.1.chr7.g01074.t01#2    mNeoVis.vulgivaga.2.1.chr7.g01086.t01#1    mNeoVis.vulgivaga.1.1.chr7.g01076.t01#4    mNeoVis.evergladensis.1.1.chr7.g01079.t01#3    mNeoVis.3.1.1.chr7.g01012.t01#6   mNeoVis.2.1.1.chr7_hap1.g01083.t01#5       4229   125256657:   mNeoVis.2.1.1.chr7_hap1.g01086.t01#5    mNeoVis.evergladensis.1.1.chr7.g01082.t01#3    mNeoVis.3.1.1.chr7.g01015.t01#6    mNeoVis.vulgivaga.1.1.chr7.g01078.t01#4   mNeoVis.evergladensis.2.1.chr7.g01077.t01#2       4230   125256658:   mNeoVis.2.1.1.chr7_hap1.g01085.t01#5    mNeoVis.evergladensis.2.1.chr7.g01076.t01#2    mNeoVis.evergladensis.1.1.chr7.g01081.t01#3   mNeoVis.vulgivaga.2.1.chr7.g01088.t01#1         4231   125256659:   mNeoVis.3.1.1.chr7.g01014.t01#6   mNeoVis.vulgivaga.2.1.chr7.g01089.t01#1             4232   125256660:   mNeoVis.vulgivaga.2.1.chr1.g00379.t01#1    mNeoVis.evergladensis.2.1.chr1.g00371.t01#2    mNeoVis.3.1.1.chr1.g00373.t01#6    mNeoVis.vulgivaga.1.1.chr1.g00371.t01#4    mNeoVis.evergladensis.1.1.chr1.g00379.t01#3   mNeoVis.2.1.1.chr1_hap1.g00374.t01#5       4233   125256661:   mNeoVis.2.1.1.chr1_hap1.g00375.t01#5    mNeoVis.evergladensis.1.1.chr1.g00380.t01#3    mNeoVis.vulgivaga.2.1.chr1.g00380.t01#1    mNeoVis.vulgivaga.1.1.chr1.g00372.t01#4    mNeoVis.evergladensis.2.1.chr1.g00372.t01#2   mNeoVis.3.1.1.chr1.g00374.t01#6       4234   125256662:   mNeoVis.vulgivaga.1.1.chr1.g00373.t01#4    mNeoVis.evergladensis.1.1.chr1.g00381.t01#3    mNeoVis.vulgivaga.2.1.chr1.g00381.t01#1    mNeoVis.2.1.1.chr1_hap1.g00376.t01#5   mNeoVis.evergladensis.2.1.chr1.g00373.t01#2       4235   125256663:   mNeoVis.evergladensis.1.1.chr7.g00995.t01#3    mNeoVis.3.1.1.chr7.g00935.t01#6    mNeoVis.2.1.1.chr7_hap1.g00996.t01#5    mNeoVis.vulgivaga.1.1.chr7.g00990.t01#4    mNeoVis.vulgivaga.2.1.chr7.g00995.t01#1   mNeoVis.evergladensis.2.1.chr7.g00989.t01#2       4236   125256664:   mNeoVis.vulgivaga.2.1.chr7.g00996.t01#1    mNeoVis.3.1.1.chr7.g00936.t01#6    mNeoVis.evergladensis.1.1.chr7.g00996.t01#3    mNeoVis.evergladensis.2.1.chr7.g00990.t01#2    mNeoVis.vulgivaga.1.1.chr7.g00991.t01#4   mNeoVis.2.1.1.chr7_hap1.g00997.t01#5       4237   125256665:   mNeoVis.vulgivaga.1.1.chr7.g00992.t01#4    mNeoVis.2.1.1.chr7_hap1.g00999.t01#5    mNeoVis.vulgivaga.2.1.chr7.g00997.t01#1    mNeoVis.evergladensis.2.1.chr7.g00991.t01#2    mNeoVis.3.1.1.chr7.g00937.t01#6   mNeoVis.evergladensis.1.1.chr7.g00998.t01#3       4238   125256666:   mNeoVis.evergladensis.2.1.chr7.g00992.t01#2    mNeoVis.3.1.1.chr7.g00938.t01#6    mNeoVis.2.1.1.chr7_hap1.g00998.t01#5    mNeoVis.vulgivaga.2.1.chr7.g00998.t01#1    mNeoVis.vulgivaga.1.1.chr7.g00993.t01#4   mNeoVis.evergladensis.1.1.chr7.g00997.t01#3       4239   125256667:   mNeoVis.vulgivaga.1.1.chr1.g00374.t01#4    mNeoVis.evergladensis.1.1.chr1.g00382.t01#3    mNeoVis.evergladensis.2.1.chr1.g00374.t01#2    mNeoVis.vulgivaga.2.1.chr1.g00382.t01#1    mNeoVis.2.1.1.chr1_hap1.g00377.t01#5   mNeoVis.3.1.1.chr1.g00376.t01#6       4240   125256668:   mNeoVis.evergladensis.2.1.chr1.g00375.t01#2    mNeoVis.vulgivaga.2.1.chr1.g00383.t01#1    mNeoVis.2.1.1.chr1_hap1.g00378.t01#5    mNeoVis.3.1.1.chr1.g00377.t01#6    mNeoVis.evergladensis.1.1.chr1.g00383.t01#3   mNeoVis.vulgivaga.1.1.chr1.g00375.t01#4       4241   125256669:   mNeoVis.vulgivaga.2.1.chr1.g01890.t01#1    mNeoVis.3.1.1.chr1.g00378.t01#6    mNeoVis.evergladensis.1.1.chr1.g00385.t01#3    mNeoVis.vulgivaga.2.1.chr12.g00099.t01#1    mNeoVis.2.1.1.chr12_hap1.g00123.t01#5    mNeoVis.3.1.1.chr15.g00380.t01#6    mNeoVis.vulgivaga.1.1.chr1.g00376.t01#4    mNeoVis.vulgivaga.2.1.chr1.g00384.t01#1    mNeoVis.evergladensis.2.1.chr1.g00377.t01#2      4242   125256670:   mNeoVis.3.1.1.chr7.g00939.t01#6    mNeoVis.vulgivaga.1.1.chr7.g00995.t01#4    mNeoVis.evergladensis.1.1.chr7.g00999.t01#3    mNeoVis.vulgivaga.2.1.chr7.g00999.t01#1    mNeoVis.2.1.1.chr7_hap1.g01000.t01#5   mNeoVis.evergladensis.2.1.chr7.g00993.t01#2       4243   125256671:   mNeoVis.vulgivaga.1.1.chr1.g00377.t01#4    mNeoVis.2.1.1.chr1_hap1.g00381.t01#5    mNeoVis.vulgivaga.2.1.chr1.g00385.t01#1    mNeoVis.evergladensis.2.1.chr1.g00378.t01#2    mNeoVis.3.1.1.chr1.g00379.t01#6   mNeoVis.evergladensis.1.1.chr1.g00386.t01#3       4244   125256672:   mNeoVis.vulgivaga.2.1.chr1.g00386.t01#1    mNeoVis.2.1.1.chr1_hap1.g00382.t01#5    mNeoVis.evergladensis.1.1.chr1.g00387.t01#3    mNeoVis.3.1.1.chr1.g00380.t01#6    mNeoVis.evergladensis.2.1.chr1.g00379.t01#2   mNeoVis.vulgivaga.1.1.chr1.g00378.t01#4       4245   125256673:   mNeoVis.2.1.1.chr7_hap1.g01001.t01#5    mNeoVis.evergladensis.2.1.chr7.g00994.t01#2    mNeoVis.vulgivaga.2.1.chr7.g01000.t01#1    mNeoVis.evergladensis.1.1.chr7.g01000.t01#3    mNeoVis.vulgivaga.1.1.chr7.g00996.t01#4   mNeoVis.3.1.1.chr7.g00940.t01#6       4246   125256674:   mNeoVis.3.1.1.chr7.g00941.t01#6    mNeoVis.evergladensis.2.1.chr7.g00995.t01#2    mNeoVis.evergladensis.1.1.chr7.g01001.t01#3    mNeoVis.vulgivaga.1.1.chr7.g00997.t01#4    mNeoVis.vulgivaga.2.1.chr7.g01001.t01#1   mNeoVis.2.1.1.chr7_hap1.g01002.t01#5       4247   125256675:   mNeoVis.vulgivaga.1.1.chr7.g00998.t01#4    mNeoVis.evergladensis.2.1.chr7.g00996.t01#2    mNeoVis.vulgivaga.2.1.chr7.g01002.t01#1    mNeoVis.3.1.1.chr7.g00942.t01#6    mNeoVis.2.1.1.chr7_hap1.g01003.t01#5   mNeoVis.evergladensis.1.1.chr7.g01002.t01#3       4248   125256676:   mNeoVis.evergladensis.2.1.chr1.g00380.t01#2    mNeoVis.3.1.1.chr1.g00381.t01#6    mNeoVis.evergladensis.1.1.chr1.g00388.t01#3    mNeoVis.2.1.1.chr1_hap1.g00383.t01#5    mNeoVis.vulgivaga.1.1.chr1.g00379.t01#4   mNeoVis.vulgivaga.2.1.chr1.g00387.t01#1       4249   125256677:   mNeoVis.2.1.1.chr7_hap1.g01004.t01#5    mNeoVis.evergladensis.1.1.chr7.g01003.t01#3    mNeoVis.3.1.1.chr7.g00943.t01#6    mNeoVis.vulgivaga.1.1.chr7.g00999.t01#4    mNeoVis.vulgivaga.2.1.chr7.g01003.t01#1   mNeoVis.evergladensis.2.1.chr7.g00997.t01#2       4250   125256678:   mNeoVis.evergladensis.1.1.chr7.g01004.t01#3    mNeoVis.3.1.1.chr7.g00944.t01#6    mNeoVis.vulgivaga.2.1.chr7.g01004.t01#1    mNeoVis.vulgivaga.1.1.chr7.g01000.t01#4    mNeoVis.2.1.1.chr7_hap1.g01005.t01#5   mNeoVis.evergladensis.2.1.chr7.g00998.t01#2       4251   125256679:   mNeoVis.evergladensis.1.1.chr15.g00518.t01#3    mNeoVis.2.1.1.chr15_hap1.g00094.t01#5    mNeoVis.vulgivaga.1.1.chr15.g00095.t01#4    mNeoVis.vulgivaga.2.1.chr15.g00097.t01#1    mNeoVis.3.1.1.chr15.g00088.t01#6   mNeoVis.evergladensis.2.1.chr15.g00095.t01#2       4252   125256680:   mNeoVis.3.1.1.chr1.g00382.t01#6    mNeoVis.vulgivaga.1.1.chr1.g00380.t01#4    mNeoVis.2.1.1.chr1_hap1.g00384.t01#5    mNeoVis.evergladensis.1.1.chr1.g00389.t01#3    mNeoVis.vulgivaga.2.1.chr1.g00388.t01#1   mNeoVis.evergladensis.2.1.chr1.g00381.t01#2       4253   125256681:   mNeoVis.vulgivaga.1.1.chr7.g01001.t01#4    mNeoVis.2.1.1.chr7_hap1.g01006.t01#5    mNeoVis.evergladensis.1.1.chr7.g01005.t01#3    mNeoVis.vulgivaga.2.1.chr7.g01005.t01#1   mNeoVis.evergladensis.2.1.chr7.g00999.t01#2       4254   125256682:  mNeoVis.vulgivaga.2.1.chr7.g01007.t01#1               4255   125256683:   mNeoVis.vulgivaga.1.1.chr7.g01002.t01#4    mNeoVis.evergladensis.2.1.chr7.g01000.t01#2    mNeoVis.evergladensis.1.1.chr7.g01006.t01#3    mNeoVis.3.1.1.chr7.g00946.t01#6    mNeoVis.2.1.1.chr7_hap1.g01007.t01#5   mNeoVis.vulgivaga.2.1.chr7.g01006.t01#1       4256   125256684:   mNeoVis.evergladensis.1.1.chr1.g00390.t01#3    mNeoVis.vulgivaga.1.1.chr1.g00381.t01#4    mNeoVis.vulgivaga.2.1.chr1.g00389.t01#1    mNeoVis.evergladensis.2.1.chr1.g00382.t01#2    mNeoVis.2.1.1.chr1_hap1.g00385.t01#5   mNeoVis.3.1.1.chr1.g00383.t01#6       4257   125256685:   mNeoVis.evergladensis.2.1.chr7.g01001.t01#2    mNeoVis.3.1.1.chr7.g00947.t01#6    mNeoVis.vulgivaga.1.1.chr7.g01003.t01#4    mNeoVis.vulgivaga.2.1.chr7.g01008.t01#1    mNeoVis.vulgivaga.2.1.chr7.g01009.t01#1    mNeoVis.evergladensis.1.1.chr7.g01007.t01#3   mNeoVis.2.1.1.chr7_hap1.g01008.t01#5     4258   125256686:   mNeoVis.2.1.1.chr15_hap1.g00723.t01#5    mNeoVis.vulgivaga.1.1.chr15.g00548.t01#4    mNeoVis.evergladensis.1.1.chr15.g00068.t01#3    mNeoVis.vulgivaga.2.1.chr15.g00558.t01#1   mNeoVis.evergladensis.2.1.chr15.g00545.t01#2       4259   125256687:   mNeoVis.evergladensis.2.1.chr7.g00499.t01#2    mNeoVis.evergladensis.1.1.chr7.g00507.t01#3    mNeoVis.vulgivaga.1.1.chr7.g00506.t01#4    mNeoVis.vulgivaga.2.1.chr7.g00502.t01#1   mNeoVis.2.1.1.chr7_hap1.g00498.t01#5       4260   125256688:   mNeoVis.evergladensis.1.1.chr1.g00391.t01#3    mNeoVis.vulgivaga.2.1.chr1.g00390.t01#1    mNeoVis.2.1.1.chr1_hap1.g00386.t01#5    mNeoVis.3.1.1.chr1.g00384.t01#6    mNeoVis.vulgivaga.1.1.chr1.g00382.t01#4   mNeoVis.evergladensis.2.1.chr1.g00383.t01#2       4261   125256689:   mNeoVis.vulgivaga.2.1.chr7.g01010.t01#1    mNeoVis.evergladensis.2.1.chr7.g01002.t01#2    mNeoVis.vulgivaga.1.1.chr7.g01004.t01#4    mNeoVis.2.1.1.chr7_hap1.g01009.t01#5   mNeoVis.evergladensis.1.1.chr7.g01008.t01#3       4262   125256690:   mNeoVis.3.1.1.chr7.g00948.t01#6    mNeoVis.evergladensis.1.1.chr7.g01009.t01#3    mNeoVis.2.1.1.chr7_hap1.g01010.t01#5    mNeoVis.evergladensis.2.1.chr7.g01003.t01#2    mNeoVis.vulgivaga.1.1.chr7.g01005.t01#4   mNeoVis.vulgivaga.2.1.chr7.g01011.t01#1       4263   125256691:   mNeoVis.3.1.1.chr1.g00386.t01#6    mNeoVis.evergladensis.1.1.chr1.g00393.t01#3    mNeoVis.evergladensis.2.1.chr1.g00385.t01#2    mNeoVis.2.1.1.chr1_hap1.g00388.t01#5   mNeoVis.vulgivaga.2.1.chr1.g00391.t01#1       4264   125256692:   mNeoVis.evergladensis.1.1.chr7.g01010.t01#3    mNeoVis.3.1.1.chr7.g00949.t01#6    mNeoVis.2.1.1.chr7_hap1.g01011.t01#5    mNeoVis.evergladensis.2.1.chr7.g01004.t01#2    mNeoVis.vulgivaga.1.1.chr7.g01006.t01#4   mNeoVis.vulgivaga.2.1.chr7.g01012.t01#1       4265   125256693:   mNeoVis.3.1.1.chr1.g00387.t01#6    mNeoVis.vulgivaga.1.1.chr1.g00385.t01#4    mNeoVis.vulgivaga.2.1.chr1.g00392.t01#1    mNeoVis.2.1.1.chr1_hap1.g00389.t01#5    mNeoVis.evergladensis.1.1.chr1.g00394.t01#3   mNeoVis.evergladensis.2.1.chr1.g00386.t01#2       4266   125256694:   mNeoVis.vulgivaga.1.1.chr7.g01007.t01#4    mNeoVis.evergladensis.2.1.chr7.g01005.t01#2    mNeoVis.vulgivaga.2.1.chr7.g01013.t01#1    mNeoVis.3.1.1.chr7.g00950.t01#6    mNeoVis.evergladensis.1.1.chr7.g01011.t01#3   mNeoVis.2.1.1.chr7_hap1.g01012.t01#5       4267   125256695:   mNeoVis.vulgivaga.2.1.chr7.g01014.t01#1    mNeoVis.evergladensis.2.1.chr7.g01006.t01#2    mNeoVis.evergladensis.1.1.chr7.g01012.t01#3    mNeoVis.3.1.1.chr7.g00951.t01#6    mNeoVis.2.1.1.chr7_hap1.g01013.t01#5   mNeoVis.vulgivaga.1.1.chr7.g01008.t01#4       4268   125256696:   mNeoVis.3.1.1.chr1.g00388.t01#6    mNeoVis.vulgivaga.1.1.chr1.g00386.t01#4    mNeoVis.2.1.1.chr1_hap1.g00390.t01#5    mNeoVis.evergladensis.2.1.chr1.g00387.t01#2    mNeoVis.vulgivaga.2.1.chr1.g00393.t01#1   mNeoVis.evergladensis.1.1.chr1.g00395.t01#3       4269   125256697:   mNeoVis.3.1.1.chr7.g00952.t01#6    mNeoVis.evergladensis.1.1.chr7.g01013.t01#3    mNeoVis.evergladensis.2.1.chr7.g01007.t01#2    mNeoVis.2.1.1.chr7_hap1.g01014.t01#5    mNeoVis.vulgivaga.1.1.chr7.g01009.t01#4   mNeoVis.vulgivaga.2.1.chr7.g01015.t01#1       4270   125256698:   mNeoVis.3.1.1.chr1.g00389.t01#6    mNeoVis.vulgivaga.1.1.chr1.g00387.t01#4    mNeoVis.2.1.1.chr1_hap1.g00391.t01#5    mNeoVis.vulgivaga.2.1.chr1.g00394.t01#1    mNeoVis.evergladensis.2.1.chr1.g00388.t01#2   mNeoVis.evergladensis.1.1.chr1.g00396.t01#3       4271   125256699:   mNeoVis.vulgivaga.1.1.chr1.g00388.t01#4    mNeoVis.evergladensis.1.1.chr1.g00397.t01#3    mNeoVis.3.1.1.chr1.g00390.t01#6    mNeoVis.evergladensis.2.1.chr1.g00389.t01#2    mNeoVis.2.1.1.chr1_hap1.g00392.t01#5   mNeoVis.vulgivaga.2.1.chr1.g00395.t01#1       4272   125256700:   mNeoVis.vulgivaga.2.1.chr7.g01016.t01#1    mNeoVis.evergladensis.2.1.chr7.g01008.t01#2    mNeoVis.3.1.1.chr7.g00953.t01#6    mNeoVis.evergladensis.1.1.chr7.g01014.t01#3    mNeoVis.vulgivaga.1.1.chr7.g01010.t01#4   mNeoVis.2.1.1.chr7_hap1.g01015.t01#5       4273   125256701:   mNeoVis.vulgivaga.1.1.chr1.g00295.t01#4    mNeoVis.vulgivaga.2.1.chr1.g00299.t01#1    mNeoVis.evergladensis.2.1.chr1.g00295.t01#2    mNeoVis.3.1.1.chr1.g00301.t01#6    mNeoVis.evergladensis.1.1.chr1.g00304.t01#3   mNeoVis.2.1.1.chr1_hap1.g00296.t01#5       4274   125256702:   mNeoVis.vulgivaga.2.1.chr7.g01017.t01#1    mNeoVis.evergladensis.2.1.chr7.g01009.t01#2    mNeoVis.3.1.1.chr7.g00954.t01#6    mNeoVis.evergladensis.1.1.chr7.g01015.t01#3    mNeoVis.vulgivaga.1.1.chr7.g01011.t01#4   mNeoVis.2.1.1.chr7_hap1.g01016.t01#5       4275   125256703:   mNeoVis.evergladensis.2.1.chr7.g01010.t01#2    mNeoVis.evergladensis.1.1.chr7.g01016.t01#3    mNeoVis.2.1.1.chr7_hap1.g01017.t01#5    mNeoVis.vulgivaga.1.1.chr7.g01012.t01#4    mNeoVis.3.1.1.chr7.g00955.t01#6   mNeoVis.vulgivaga.2.1.chr7.g01018.t01#1       4276   125256704:   mNeoVis.evergladensis.1.1.chr7.g01017.t01#3    mNeoVis.vulgivaga.1.1.chr7.g01013.t01#4    mNeoVis.2.1.1.chr7_hap1.g01018.t01#5    mNeoVis.evergladensis.2.1.chr7.g01011.t01#2    mNeoVis.3.1.1.chr7.g00956.t01#6   mNeoVis.vulgivaga.2.1.chr7.g01019.t01#1       4277   125256705:   mNeoVis.vulgivaga.1.1.chr1.g00297.t01#4    mNeoVis.evergladensis.1.1.chr1.g00306.t01#3    mNeoVis.evergladensis.2.1.chr1.g00296.t01#2    mNeoVis.3.1.1.chr1.g00303.t01#6    mNeoVis.2.1.1.chr1_hap1.g00298.t01#5   mNeoVis.vulgivaga.2.1.chr1.g00300.t01#1       4278   125256706:   mNeoVis.evergladensis.1.1.chr1.g00305.t01#3    mNeoVis.evergladensis.2.1.chr1.g00297.t01#2    mNeoVis.evergladensis.1.1.chr5.g00222.t01#3    mNeoVis.3.1.1.chr1.g00302.t01#6    mNeoVis.vulgivaga.2.1.chr1.g00301.t01#1    mNeoVis.2.1.1.chr1_hap1.g00297.t01#5   mNeoVis.vulgivaga.1.1.chr1.g00296.t01#4     4279   125256707:   mNeoVis.2.1.1.chr11_hap1.g00938.t01#5    mNeoVis.evergladensis.1.1.chr11.g00930.t01#3    mNeoVis.evergladensis.2.1.chr11.g00916.t01#2    mNeoVis.vulgivaga.1.1.chr11.g00911.t01#4    mNeoVis.vulgivaga.2.1.chr11.g00940.t01#1   mNeoVis.3.1.1.chr11.g00892.t01#6       4280   125256708:   mNeoVis.evergladensis.2.1.chr4.g00281.t01#2    mNeoVis.vulgivaga.1.1.chr4.g00275.t01#4    mNeoVis.vulgivaga.2.1.chr4.g00280.t01#1    mNeoVis.2.1.1.chr6_hap1.g00274.t01#5    mNeoVis.evergladensis.1.1.chr4.g00279.t01#3   mNeoVis.3.1.1.chr4.g00271.t01#6       4281   125256709:   mNeoVis.evergladensis.2.1.chr7.g01012.t01#2    mNeoVis.vulgivaga.1.1.chr7.g01014.t01#4    mNeoVis.evergladensis.1.1.chr7.g01018.t01#3    mNeoVis.3.1.1.chr7.g00957.t01#6   mNeoVis.vulgivaga.2.1.chr7.g01020.t01#1       4282   125256710:   mNeoVis.evergladensis.2.1.chr15.g00546.t01#2    mNeoVis.2.1.1.chr15_hap1.g00724.t01#5    mNeoVis.vulgivaga.1.1.chr15.g00549.t01#4    mNeoVis.vulgivaga.2.1.chr15.g00559.t01#1   mNeoVis.evergladensis.1.1.chr15.g00067.t01#3       4283   125256711:   mNeoVis.evergladensis.2.1.chr1.g00298.t01#2    mNeoVis.2.1.1.chr1_hap1.g00299.t01#5    mNeoVis.3.1.1.chr1.g00304.t01#6    mNeoVis.evergladensis.1.1.chr1.g00307.t01#3    mNeoVis.vulgivaga.1.1.chr1.g00298.t01#4   mNeoVis.vulgivaga.2.1.chr1.g00302.t01#1       4284   125256712:   mNeoVis.3.1.1.chr7.g01016.t01#6    mNeoVis.2.1.1.chr7_hap1.g01087.t01#5    mNeoVis.evergladensis.2.1.chr7.g01078.t01#2    mNeoVis.vulgivaga.1.1.chr7.g01079.t01#4    mNeoVis.vulgivaga.2.1.chr7.g01090.t01#1   mNeoVis.evergladensis.1.1.chr7.g01083.t01#3       4285   125256713:   mNeoVis.vulgivaga.2.1.chr1.g00303.t01#1    mNeoVis.evergladensis.1.1.chr1.g00308.t01#3    mNeoVis.vulgivaga.1.1.chr1.g00299.t01#4    mNeoVis.3.1.1.chr1.g00305.t01#6    mNeoVis.evergladensis.2.1.chr1.g00299.t01#2   mNeoVis.2.1.1.chr1_hap1.g00300.t01#5       4286   125256714:   mNeoVis.2.1.1.chr7_hap1.g01088.t01#5    mNeoVis.vulgivaga.2.1.chr7.g01091.t01#1    mNeoVis.evergladensis.2.1.chr7.g01079.t01#2    mNeoVis.vulgivaga.1.1.chr7.g01080.t01#4    mNeoVis.evergladensis.1.1.chr7.g01084.t01#3   mNeoVis.3.1.1.chr7.g01017.t01#6       4287   125256715:   mNeoVis.3.1.1.chr7.g01018.t01#6    mNeoVis.2.1.1.chr7_hap1.g01089.t01#5    mNeoVis.vulgivaga.1.1.chr7.g01081.t01#4    mNeoVis.evergladensis.1.1.chr7.g01085.t01#3    mNeoVis.vulgivaga.2.1.chr7.g01092.t01#1   mNeoVis.evergladensis.2.1.chr7.g01080.t01#2       4288   125256716:   mNeoVis.3.1.1.chr4.g01071.t01#6    mNeoVis.2.1.1.chr7_hap1.g01090.t01#5    mNeoVis.vulgivaga.1.1.chr7.g01082.t01#4    mNeoVis.vulgivaga.2.1.chr7.g01093.t01#1    mNeoVis.evergladensis.2.1.chr7.g01081.t01#2   mNeoVis.evergladensis.1.1.chr7.g01086.t01#3       4289   125256717:   mNeoVis.vulgivaga.2.1.chr1.g00304.t01#1    mNeoVis.3.1.1.chr1.g00306.t01#6    mNeoVis.evergladensis.1.1.chr1.g00309.t01#3    mNeoVis.2.1.1.chr1_hap1.g00301.t01#5    mNeoVis.evergladensis.2.1.chr1.g00300.t01#2   mNeoVis.vulgivaga.1.1.chr1.g00300.t01#4       4290   125256718:   mNeoVis.3.1.1.chr1.g00855.t01#6    mNeoVis.vulgivaga.1.1.chr1.g00852.t01#4    mNeoVis.vulgivaga.1.1.chr1.g00301.t01#4    mNeoVis.vulgivaga.2.1.chr2.g00691.t01#1    mNeoVis.evergladensis.2.1.chr1.g00301.t01#2    mNeoVis.vulgivaga.2.1.chr1.g00305.t01#1    mNeoVis.2.1.1.chr1_hap1.g00831.t01#5    mNeoVis.3.1.1.chr1.g00307.t01#6    mNeoVis.evergladensis.1.1.chr1.g00847.t01#3      4291   125256719:   mNeoVis.evergladensis.1.1.chr4.g01082.t01#3   mNeoVis.3.1.1.chr4.g01064.t01#6             4292   125256720:   mNeoVis.3.1.1.chr7.g00958.t01#6    mNeoVis.vulgivaga.2.1.chr7.g01021.t01#1    mNeoVis.evergladensis.2.1.chr7.g01013.t01#2    mNeoVis.evergladensis.1.1.chr7.g01019.t01#3    mNeoVis.vulgivaga.1.1.chr7.g01015.t01#4   mNeoVis.2.1.1.chr7_hap1.g01020.t01#5       4293   125256721:   mNeoVis.3.1.1.chr7.g01019.t01#6    mNeoVis.evergladensis.2.1.chr7.g01082.t01#2    mNeoVis.2.1.1.chr7_hap1.g01091.t01#5    mNeoVis.vulgivaga.1.1.chr7.g01083.t01#4    mNeoVis.vulgivaga.2.1.chr7.g01094.t01#1   mNeoVis.evergladensis.1.1.chr7.g01087.t01#3       4294   125256722:   mNeoVis.vulgivaga.2.1.chr1.g00306.t01#1    mNeoVis.3.1.1.chr3.g00362.t01#6    mNeoVis.3.1.1.chr1.g00308.t01#6    mNeoVis.2.1.1.chr1_hap1.g00303.t01#5    mNeoVis.evergladensis.1.1.chr1.g00311.t01#3    mNeoVis.evergladensis.2.1.chr1.g00302.t01#2   mNeoVis.vulgivaga.1.1.chr1.g00302.t01#4     4295   125256723:   mNeoVis.evergladensis.2.1.chr1.g00303.t01#2    mNeoVis.2.1.1.chr1_hap1.g00304.t01#5    mNeoVis.vulgivaga.2.1.chr1.g00307.t01#1    mNeoVis.evergladensis.1.1.chr1.g00312.t01#3    mNeoVis.vulgivaga.1.1.chr1.g00303.t01#4   mNeoVis.3.1.1.chr1.g00309.t01#6       4296   125256724:   mNeoVis.vulgivaga.1.1.chr5.g00222.t01#4    mNeoVis.vulgivaga.1.1.chr4.g00131.t01#4    mNeoVis.evergladensis.2.1.chr4.g00137.t01#2    mNeoVis.evergladensis.1.1.chr4.g00130.t01#3    mNeoVis.evergladensis.1.1.chr5.g00229.t01#3    mNeoVis.vulgivaga.2.1.chr5.g00226.t01#1    mNeoVis.vulgivaga.2.1.chr4.g00130.t01#1    mNeoVis.2.1.1.chr2_hap1.g00225.t01#5    mNeoVis.3.1.1.chr5.g00217.t01#6      4297   125256725:   mNeoVis.vulgivaga.1.1.chr1.g00304.t01#4    mNeoVis.3.1.1.chr1.g00310.t01#6    mNeoVis.evergladensis.1.1.chr1.g00313.t01#3    mNeoVis.evergladensis.2.1.chr1.g00304.t01#2    mNeoVis.2.1.1.chr1_hap1.g00305.t01#5   mNeoVis.vulgivaga.2.1.chr1.g00308.t01#1       4298   125256726:   mNeoVis.evergladensis.1.1.chr1.g00314.t01#3    mNeoVis.3.1.1.chr1.g00311.t01#6    mNeoVis.vulgivaga.1.1.chr1.g00305.t01#4    mNeoVis.2.1.1.chr1_hap1.g00306.t01#5    mNeoVis.vulgivaga.2.1.chr1.g00309.t01#1   mNeoVis.evergladensis.2.1.chr1.g00305.t01#2       4299   125256727:   mNeoVis.evergladensis.1.1.chr7.g01088.t01#3    mNeoVis.evergladensis.2.1.chr7.g01083.t01#2    mNeoVis.vulgivaga.1.1.chr7.g01084.t01#4    mNeoVis.2.1.1.chr7_hap1.g01092.t01#5    mNeoVis.vulgivaga.2.1.chr7.g01095.t01#1   mNeoVis.3.1.1.chr7.g01020.t01#6       4300   125256728:   mNeoVis.evergladensis.2.1.chr7.g01084.t01#2    mNeoVis.vulgivaga.2.1.chr7.g01096.t01#1    mNeoVis.evergladensis.1.1.chr7.g01089.t01#3    mNeoVis.3.1.1.chr7.g01021.t01#6    mNeoVis.vulgivaga.1.1.chr7.g01085.t01#4   mNeoVis.2.1.1.chr7_hap1.g01093.t01#5       4301   125256729:   mNeoVis.2.1.1.chr7_hap1.g00499.t01#5    mNeoVis.evergladensis.1.1.chr7.g00509.t01#3    mNeoVis.vulgivaga.2.1.chr7.g00503.t01#1    mNeoVis.3.1.1.chr7.g00484.t01#6    mNeoVis.evergladensis.2.1.chr7.g00500.t01#2   mNeoVis.vulgivaga.1.1.chr7.g00507.t01#4       4302   125256730:   mNeoVis.vulgivaga.1.1.chr7.g01086.t01#4    mNeoVis.2.1.1.chr7_hap1.g01094.t01#5    mNeoVis.evergladensis.1.1.chr5.g00847.t01#3    mNeoVis.vulgivaga.2.1.chr5.g00845.t01#1    mNeoVis.3.1.1.chr5.g00834.t01#6    mNeoVis.vulgivaga.2.1.chr7.g01097.t01#1    mNeoVis.2.1.1.chr2_hap1.g00901.t01#5    mNeoVis.evergladensis.2.1.chr5.g00856.t01#2    mNeoVis.vulgivaga.1.1.chr5.g00825.t01#4      4303   125256731:   mNeoVis.evergladensis.1.1.chr14.g00527.t01#3    mNeoVis.vulgivaga.2.1.chr14.g00524.t01#1    mNeoVis.2.1.1.chr14_hap1.g00527.t01#5    mNeoVis.evergladensis.2.1.chr14.g00526.t01#2   mNeoVis.vulgivaga.1.1.chr14.g00519.t01#4       4304   125256732:   mNeoVis.evergladensis.1.1.chr14.g00526.t01#3    mNeoVis.2.1.1.chr14_hap1.g00526.t01#5    mNeoVis.vulgivaga.2.1.chr14.g00523.t01#1   mNeoVis.3.1.1.chr14.g00521.t01#6         4305   125256733:   mNeoVis.3.1.1.chr7.g01022.t01#6    mNeoVis.evergladensis.1.1.chr7.g01090.t01#3   mNeoVis.evergladensis.2.1.chr7.g01085.t01#2         4306   125256734:   mNeoVis.vulgivaga.1.1.chr14.g00517.t01#4   mNeoVis.evergladensis.2.1.chr14.g00525.t01#2           4307   125256735:  mNeoVis.3.1.1.chr14.g00522.t01#6               4308   125256736:   mNeoVis.vulgivaga.2.1.chr1.g00310.t01#1   mNeoVis.2.1.1.chr1_hap1.g00307.t01#5             4309   125256737:   mNeoVis.evergladensis.2.1.chr1.g00306.t01#2    mNeoVis.vulgivaga.2.1.chr1.g00311.t01#1    mNeoVis.evergladensis.1.1.chr1.g00315.t01#3    mNeoVis.vulgivaga.1.1.chr1.g00306.t01#4    mNeoVis.2.1.1.chr1_hap1.g00308.t01#5   mNeoVis.3.1.1.chr1.g00312.t01#6       4310   125256738:   mNeoVis.evergladensis.2.1.chr1.g00307.t01#2    mNeoVis.2.1.1.chr1_hap1.g00309.t01#5    mNeoVis.vulgivaga.2.1.chr1.g00312.t01#1   mNeoVis.evergladensis.1.1.chr1.g00316.t01#3       4311   125256739:   mNeoVis.3.1.1.chr7.g01023.t01#6    mNeoVis.2.1.1.chr7_hap1.g01095.t01#5    mNeoVis.evergladensis.2.1.chr7.g01086.t01#2    mNeoVis.vulgivaga.2.1.chr7.g01098.t01#1    mNeoVis.evergladensis.1.1.chr7.g01091.t01#3   mNeoVis.vulgivaga.1.1.chr7.g01087.t01#4       4312   125256740:   mNeoVis.2.1.1.chr1_hap1.g00310.t01#5    mNeoVis.3.1.1.chr1.g00314.t01#6    mNeoVis.evergladensis.1.1.chr1.g00317.t01#3    mNeoVis.vulgivaga.2.1.chr1.g00313.t01#1    mNeoVis.3.1.1.chr5.g00171.t01#6    mNeoVis.vulgivaga.1.1.chr1.g00307.t01#4   mNeoVis.evergladensis.2.1.chr1.g00308.t01#2     4313   125256741:   mNeoVis.vulgivaga.1.1.chr1.g00308.t01#4    mNeoVis.evergladensis.1.1.chr1.g00318.t01#3    mNeoVis.vulgivaga.2.1.chr1.g00314.t01#1    mNeoVis.2.1.1.chr1_hap1.g00311.t01#5    mNeoVis.evergladensis.2.1.chr1.g00309.t01#2   mNeoVis.3.1.1.chr1.g00315.t01#6       4314   125256742:   mNeoVis.evergladensis.2.1.chr7.g01088.t01#2    mNeoVis.vulgivaga.1.1.chr7.g01088.t01#4    mNeoVis.vulgivaga.2.1.chr7.g01099.t01#1    mNeoVis.2.1.1.chr7_hap1.g01097.t01#5    mNeoVis.3.1.1.chr7.g01025.t01#6    mNeoVis.evergladensis.2.1.chr7.g01089.t01#2   mNeoVis.evergladensis.1.1.chr7.g01093.t01#3     4315   125256743:   mNeoVis.vulgivaga.1.1.chr1.g00309.t01#4    mNeoVis.evergladensis.2.1.chr1.g00310.t01#2    mNeoVis.vulgivaga.2.1.chr1.g00315.t01#1    mNeoVis.2.1.1.chr1_hap1.g00312.t01#5    mNeoVis.3.1.1.chr1.g00316.t01#6   mNeoVis.evergladensis.1.1.chr1.g00319.t01#3       4316   125256744:   mNeoVis.evergladensis.1.1.chr1.g00320.t01#3    mNeoVis.vulgivaga.2.1.chr1.g00316.t01#1    mNeoVis.2.1.1.chr1_hap1.g00313.t01#5    mNeoVis.evergladensis.2.1.chr1.g00311.t01#2    mNeoVis.3.1.1.chr1.g00317.t01#6   mNeoVis.vulgivaga.1.1.chr1.g00310.t01#4       4317   125256745:   mNeoVis.evergladensis.1.1.chr7.g00981.t01#3    mNeoVis.2.1.1.chr7_hap1.g00982.t01#5    mNeoVis.vulgivaga.1.1.chr7.g00976.t01#4    mNeoVis.3.1.1.chr7.g00921.t01#6    mNeoVis.evergladensis.2.1.chr7.g00975.t01#2   mNeoVis.vulgivaga.2.1.chr7.g00981.t01#1       4318   125256746:   mNeoVis.vulgivaga.1.1.chr1.g00311.t01#4    mNeoVis.evergladensis.2.1.chr1.g00312.t01#2    mNeoVis.vulgivaga.2.1.chr1.g00317.t01#1    mNeoVis.2.1.1.chr1_hap1.g00314.t01#5    mNeoVis.evergladensis.1.1.chr1.g00321.t01#3   mNeoVis.3.1.1.chr1.g00318.t01#6       4319   125256747:   mNeoVis.3.1.1.chr8.g00939.t01#6    mNeoVis.2.1.1.chr8_hap1.g01007.t01#5    mNeoVis.vulgivaga.2.1.chr8.g00982.t01#1    mNeoVis.evergladensis.2.1.chr9.g00981.t01#2    mNeoVis.vulgivaga.1.1.chr8.g00967.t01#4   mNeoVis.evergladensis.1.1.chr8.g00973.t01#3       4320   125256748:   mNeoVis.vulgivaga.2.1.chr7.g00982.t01#1    mNeoVis.evergladensis.2.1.chr7.g00976.t01#2    mNeoVis.3.1.1.chr7.g00922.t01#6    mNeoVis.evergladensis.1.1.chr7.g00982.t01#3    mNeoVis.2.1.1.chr7_hap1.g00983.t01#5   mNeoVis.vulgivaga.1.1.chr7.g00977.t01#4       4321   125256749:   mNeoVis.3.1.1.chr1.g00320.t01#6    mNeoVis.evergladensis.1.1.chr1.g00323.t01#3    mNeoVis.vulgivaga.1.1.chr1.g00313.t01#4    mNeoVis.evergladensis.2.1.chr1.g00314.t01#2    mNeoVis.2.1.1.chr1_hap1.g00316.t01#5   mNeoVis.vulgivaga.2.1.chr1.g00319.t01#1       4322   125256750:   mNeoVis.2.1.1.chr4_hap1.g00762.t01#5    mNeoVis.evergladensis.2.1.chr6.g00769.t01#2    mNeoVis.evergladensis.1.1.chr1.g00322.t01#3    mNeoVis.3.1.1.chr6.g00768.t01#6    mNeoVis.3.1.1.chr5.g00482.t01#6    mNeoVis.evergladensis.1.1.chr6.g00774.t01#3    mNeoVis.3.1.1.chr1.g00319.t01#6    mNeoVis.evergladensis.1.1.chr5.g00486.t01#3    mNeoVis.vulgivaga.1.1.chr1.g00312.t01#4      4323   125256751:   mNeoVis.3.1.1.chr7.g00923.t01#6    mNeoVis.2.1.1.chr7_hap1.g00984.t01#5    mNeoVis.evergladensis.1.1.chr7.g00983.t01#3    mNeoVis.vulgivaga.1.1.chr7.g00978.t01#4    mNeoVis.evergladensis.2.1.chr7.g00977.t01#2   mNeoVis.vulgivaga.2.1.chr7.g00983.t01#1       4324   125256752:   mNeoVis.vulgivaga.2.1.chr7.g00984.t01#1    mNeoVis.3.1.1.chr7.g00924.t01#6    mNeoVis.evergladensis.2.1.chr7.g00978.t01#2    mNeoVis.evergladensis.1.1.chr7.g00984.t01#3    mNeoVis.2.1.1.chr7_hap1.g00985.t01#5   mNeoVis.vulgivaga.1.1.chr7.g00979.t01#4       4325   125256753:  mNeoVis.vulgivaga.2.1.chr1.g00320.t01#1               4326   125256754:   mNeoVis.evergladensis.2.1.chr1.g00316.t01#2    mNeoVis.evergladensis.1.1.chr1.g00325.t01#3    mNeoVis.3.1.1.chr1.g00322.t01#6    mNeoVis.vulgivaga.2.1.chr1.g00321.t01#1    mNeoVis.vulgivaga.1.1.chr1.g00315.t01#4   mNeoVis.2.1.1.chr1_hap1.g00318.t01#5       4327   125256755:   mNeoVis.vulgivaga.1.1.chr1.g00316.t01#4    mNeoVis.evergladensis.1.1.chr1.g00326.t01#3    mNeoVis.vulgivaga.2.1.chr1.g00322.t01#1    mNeoVis.2.1.1.chr1_hap1.g00319.t01#5    mNeoVis.3.1.1.chr1.g00323.t01#6   mNeoVis.evergladensis.2.1.chr1.g00317.t01#2       4328   125256756:   mNeoVis.evergladensis.2.1.chr1.g00318.t01#2    mNeoVis.evergladensis.1.1.chr1.g00327.t01#3    mNeoVis.vulgivaga.2.1.chr1.g00323.t01#1    mNeoVis.3.1.1.chr1.g00324.t01#6    mNeoVis.vulgivaga.1.1.chr1.g00317.t01#4   mNeoVis.2.1.1.chr1_hap1.g00320.t01#5       4329   125256757:   mNeoVis.vulgivaga.2.1.chr7.g00985.t01#1    mNeoVis.evergladensis.2.1.chr7.g00979.t01#2    mNeoVis.3.1.1.chr7.g00925.t01#6    mNeoVis.2.1.1.chr7_hap1.g00986.t01#5    mNeoVis.evergladensis.1.1.chr7.g00985.t01#3   mNeoVis.vulgivaga.1.1.chr7.g00980.t01#4       4330   125256758:   mNeoVis.evergladensis.1.1.chr15.g00066.t01#3    mNeoVis.3.1.1.chr15.g00502.t01#6    mNeoVis.2.1.1.chr15_hap1.g00725.t01#5    mNeoVis.vulgivaga.1.1.chr15.g00550.t01#4    mNeoVis.vulgivaga.2.1.chr15.g00560.t01#1   mNeoVis.evergladensis.2.1.chr15.g00547.t01#2       4331   125256759:   mNeoVis.3.1.1.chr1.g00325.t01#6    mNeoVis.vulgivaga.2.1.chr1.g00324.t01#1    mNeoVis.2.1.1.chr1_hap1.g00321.t01#5    mNeoVis.vulgivaga.1.1.chr1.g00318.t01#4    mNeoVis.evergladensis.2.1.chr1.g00319.t01#2   mNeoVis.evergladensis.1.1.chr1.g00328.t01#3       4332   125256760:   mNeoVis.evergladensis.1.1.chr1.g00329.t01#3    mNeoVis.3.1.1.chr1.g00326.t01#6    mNeoVis.vulgivaga.2.1.chr1.g00325.t01#1    mNeoVis.vulgivaga.1.1.chr1.g00320.t01#4    mNeoVis.2.1.1.chr1_hap1.g00322.t01#5   mNeoVis.evergladensis.2.1.chr1.g00320.t01#2       4333   125256761:   mNeoVis.evergladensis.2.1.chr7.g00980.t01#2    mNeoVis.vulgivaga.2.1.chr7.g00986.t01#1    mNeoVis.vulgivaga.1.1.chr7.g00981.t01#4    mNeoVis.2.1.1.chr7_hap1.g00987.t01#5   mNeoVis.evergladensis.1.1.chr7.g00986.t01#3       4334   125256762:   mNeoVis.vulgivaga.1.1.chr1.g00321.t01#4    mNeoVis.3.1.1.chr1.g00327.t01#6    mNeoVis.evergladensis.2.1.chr1.g00321.t01#2    mNeoVis.vulgivaga.2.1.chr1.g00326.t01#1    mNeoVis.evergladensis.1.1.chr1.g00330.t01#3   mNeoVis.2.1.1.chr1_hap1.g00323.t01#5       4335   125256763:   mNeoVis.evergladensis.1.1.chr1.g00331.t01#3    mNeoVis.vulgivaga.2.1.chr1.g00327.t01#1    mNeoVis.3.1.1.chr1.g00328.t01#6    mNeoVis.vulgivaga.1.1.chr1.g00322.t01#4    mNeoVis.evergladensis.2.1.chr1.g00322.t01#2   mNeoVis.2.1.1.chr1_hap1.g00324.t01#5       4336   125256764:   mNeoVis.2.1.1.chr7_hap1.g00988.t01#5    mNeoVis.evergladensis.2.1.chr7.g00981.t01#2    mNeoVis.evergladensis.1.1.chr7.g00987.t01#3    mNeoVis.vulgivaga.1.1.chr7.g00982.t01#4    mNeoVis.vulgivaga.2.1.chr7.g00987.t01#1   mNeoVis.3.1.1.chr7.g00927.t01#6       4337   125256765:   mNeoVis.2.1.1.chr1_hap1.g00325.t01#5    mNeoVis.evergladensis.2.1.chr1.g00323.t01#2    mNeoVis.3.1.1.chr1.g00329.t01#6    mNeoVis.evergladensis.1.1.chr1.g00332.t01#3    mNeoVis.vulgivaga.2.1.chr1.g00328.t01#1   mNeoVis.vulgivaga.1.1.chr1.g00323.t01#4       4338   125256766:   mNeoVis.evergladensis.2.1.chr5.g00235.t01#2    mNeoVis.3.1.1.chr5.g00224.t01#6    mNeoVis.vulgivaga.2.1.chr5.g00233.t01#1    mNeoVis.2.1.1.chr2_hap1.g00232.t01#5    mNeoVis.evergladensis.1.1.chr5.g00235.t01#3   mNeoVis.vulgivaga.1.1.chr5.g00227.t01#4       4339   125256767:   mNeoVis.vulgivaga.1.1.chr7.g00983.t01#4    mNeoVis.3.1.1.chr7.g00928.t01#6    mNeoVis.vulgivaga.2.1.chr7.g00988.t01#1    mNeoVis.evergladensis.2.1.chr7.g00982.t01#2    mNeoVis.evergladensis.1.1.chr7.g00988.t01#3   mNeoVis.2.1.1.chr7_hap1.g00989.t01#5       4340   125256768:   mNeoVis.evergladensis.1.1.chr11.g00598.t01#3    mNeoVis.evergladensis.2.1.chr8.g00238.t01#2    mNeoVis.vulgivaga.1.1.chr11.g00589.t01#4    mNeoVis.evergladensis.2.1.chr11.g00595.t01#2    mNeoVis.3.1.1.chr7.g00929.t01#6    mNeoVis.2.1.1.chr11_hap1.g00607.t01#5    mNeoVis.3.1.1.chr9.g00223.t01#6    mNeoVis.evergladensis.2.1.chr7.g00983.t01#2    mNeoVis.vulgivaga.2.1.chr11.g00603.t01#1      4341   125256769:   mNeoVis.2.1.1.chr9_hap1.g00022.t01#5    mNeoVis.vulgivaga.1.1.chr10.g00022.t01#4    mNeoVis.evergladensis.2.1.chr10.g00024.t01#2    mNeoVis.evergladensis.1.1.chr10.g00022.t01#3    mNeoVis.3.1.1.chr10.g00021.t01#6   mNeoVis.vulgivaga.2.1.chr10.g00022.t01#1       4342   125256770:   mNeoVis.evergladensis.1.1.chr1.g00333.t01#3    mNeoVis.evergladensis.2.1.chr1.g00324.t01#2    mNeoVis.vulgivaga.1.1.chr1.g00324.t01#4    mNeoVis.3.1.1.chr1.g00330.t01#6    mNeoVis.vulgivaga.2.1.chr1.g00329.t01#1   mNeoVis.2.1.1.chr1_hap1.g00326.t01#5       4343   125256771:   mNeoVis.evergladensis.2.1.chr1.g00325.t01#2    mNeoVis.2.1.1.chr1_hap1.g00327.t01#5    mNeoVis.vulgivaga.2.1.chr1.g00330.t01#1    mNeoVis.evergladensis.1.1.chr1.g00334.t01#3    mNeoVis.3.1.1.chr1.g00331.t01#6   mNeoVis.vulgivaga.1.1.chr1.g00325.t01#4       4344   125256772:   mNeoVis.vulgivaga.1.1.chr4.g00395.t01#4    mNeoVis.2.1.1.chr6_hap1.g00393.t01#5    mNeoVis.3.1.1.chr4.g00396.t01#6    mNeoVis.vulgivaga.2.1.chr4.g00401.t01#1    mNeoVis.evergladensis.1.1.chr4.g00400.t01#3   mNeoVis.evergladensis.2.1.chr4.g00401.t01#2       4345   125256773:  mNeoVis.vulgivaga.2.1.chr7.g00990.t01#1               4346   125256774:   mNeoVis.evergladensis.1.1.chr7.g00991.t01#3    mNeoVis.3.1.1.chr7.g00931.t01#6    mNeoVis.vulgivaga.1.1.chr7.g00986.t01#4    mNeoVis.2.1.1.chr7_hap1.g00992.t01#5    mNeoVis.vulgivaga.2.1.chr7.g00991.t01#1   mNeoVis.evergladensis.2.1.chr7.g00985.t01#2       4347   125256775:   mNeoVis.vulgivaga.2.1.chr7.g00992.t01#1    mNeoVis.2.1.1.chr7_hap1.g00993.t01#5    mNeoVis.vulgivaga.1.1.chr7.g00987.t01#4    mNeoVis.evergladensis.1.1.chr7.g00992.t01#3    mNeoVis.3.1.1.chr7.g00932.t01#6   mNeoVis.evergladensis.2.1.chr7.g00986.t01#2       4348   125256776:   mNeoVis.evergladensis.1.1.chr7.g00993.t01#3    mNeoVis.3.1.1.chr7.g00933.t01#6    mNeoVis.2.1.1.chr7_hap1.g00994.t01#5    mNeoVis.evergladensis.2.1.chr7.g00987.t01#2    mNeoVis.vulgivaga.1.1.chr7.g00988.t01#4   mNeoVis.vulgivaga.2.1.chr7.g00993.t01#1       4349   125256777:   mNeoVis.vulgivaga.2.1.chr1.g00331.t01#1    mNeoVis.evergladensis.1.1.chr1.g00335.t01#3    mNeoVis.vulgivaga.1.1.chr1.g00326.t01#4    mNeoVis.3.1.1.chr1.g00332.t01#6   mNeoVis.2.1.1.chr1_hap1.g00328.t01#5       4350   125256778:   mNeoVis.2.1.1.chr15_hap1.g00726.t01#5    mNeoVis.evergladensis.2.1.chr15.g00548.t01#2    mNeoVis.vulgivaga.1.1.chr15.g00551.t01#4    mNeoVis.vulgivaga.2.1.chr15.g00561.t01#1    mNeoVis.3.1.1.chr15.g00503.t01#6   mNeoVis.evergladensis.1.1.chr15.g00065.t01#3       4351   125256779:   mNeoVis.vulgivaga.1.1.chr7.g00508.t01#4    mNeoVis.3.1.1.chr7.g00485.t01#6    mNeoVis.evergladensis.1.1.chr7.g00510.t01#3    mNeoVis.vulgivaga.2.1.chr7.g00504.t01#1    mNeoVis.2.1.1.chr7_hap1.g00500.t01#5   mNeoVis.evergladensis.2.1.chr7.g00501.t01#2       4352   125256780:   mNeoVis.evergladensis.1.1.chr7.g00994.t01#3    mNeoVis.evergladensis.2.1.chr7.g00988.t01#2    mNeoVis.2.1.1.chr7_hap1.g00995.t01#5    mNeoVis.vulgivaga.1.1.chr7.g00989.t01#4    mNeoVis.3.1.1.chr7.g00934.t01#6   mNeoVis.vulgivaga.2.1.chr7.g00994.t01#1       4353   125256781:   mNeoVis.evergladensis.2.1.chr7.g01014.t01#2    mNeoVis.evergladensis.1.1.chr7.g01020.t01#3    mNeoVis.3.1.1.chr7.g00959.t01#6    mNeoVis.2.1.1.chr7_hap1.g01021.t01#5    mNeoVis.vulgivaga.2.1.chr7.g01022.t01#1   mNeoVis.vulgivaga.1.1.chr7.g01016.t01#4       4354   125256782:   mNeoVis.evergladensis.2.1.chr1.g00341.t01#2    mNeoVis.2.1.1.chr1_hap1.g00343.t01#5    mNeoVis.evergladensis.1.1.chr1.g00350.t01#3    mNeoVis.vulgivaga.1.1.chr1.g00341.t01#4    mNeoVis.3.1.1.chr1.g00347.t01#6   mNeoVis.vulgivaga.2.1.chr1.g00347.t01#1       4355   125256783:   mNeoVis.vulgivaga.1.1.chr7.g01017.t01#4    mNeoVis.vulgivaga.2.1.chr7.g01023.t01#1    mNeoVis.3.1.1.chr7.g00960.t01#6    mNeoVis.2.1.1.chr7_hap1.g01022.t01#5    mNeoVis.evergladensis.1.1.chr7.g01021.t01#3   mNeoVis.evergladensis.2.1.chr7.g01015.t01#2       4356   125256784:   mNeoVis.3.1.1.chr7.g00961.t01#6    mNeoVis.vulgivaga.2.1.chr7.g01024.t01#1    mNeoVis.evergladensis.1.1.chr7.g01022.t01#3    mNeoVis.2.1.1.chr7_hap1.g01023.t01#5    mNeoVis.vulgivaga.1.1.chr7.g01018.t01#4   mNeoVis.evergladensis.2.1.chr7.g01016.t01#2       4357   125256785:   mNeoVis.vulgivaga.1.1.chr7.g01019.t01#4    mNeoVis.2.1.1.chr7_hap1.g01024.t01#5    mNeoVis.vulgivaga.2.1.chr7.g01025.t01#1    mNeoVis.evergladensis.1.1.chr7.g01023.t01#3    mNeoVis.3.1.1.chr7.g00962.t01#6   mNeoVis.evergladensis.2.1.chr7.g01017.t01#2       4358   125256786:   mNeoVis.3.1.1.chr1.g00348.t01#6    mNeoVis.vulgivaga.2.1.chr1.g00348.t01#1    mNeoVis.2.1.1.chr1_hap1.g00344.t01#5    mNeoVis.vulgivaga.1.1.chr1.g00342.t01#4    mNeoVis.evergladensis.1.1.chr1.g00351.t01#3   mNeoVis.evergladensis.2.1.chr1.g00342.t01#2       4359   125256787:   mNeoVis.evergladensis.1.1.chr7.g01024.t01#3    mNeoVis.vulgivaga.1.1.chr7.g01020.t01#4    mNeoVis.vulgivaga.2.1.chr7.g01026.t01#1    mNeoVis.2.1.1.chr7_hap1.g01025.t01#5    mNeoVis.evergladensis.2.1.chr7.g01018.t01#2   mNeoVis.3.1.1.chr7.g00963.t01#6       4360   125256788:  mNeoVis.vulgivaga.2.1.chr1.g00350.t01#1               4361   125256789:   mNeoVis.evergladensis.1.1.chr15.g00517.t01#3    mNeoVis.vulgivaga.2.1.chr15.g00098.t01#1    mNeoVis.evergladensis.2.1.chr15.g00096.t01#2    mNeoVis.2.1.1.chr15_hap1.g00095.t01#5    mNeoVis.vulgivaga.1.1.chr15.g00096.t01#4   mNeoVis.3.1.1.chr15.g00089.t01#6       4362   125256790:   mNeoVis.vulgivaga.2.1.chr1.g00351.t01#1    mNeoVis.evergladensis.2.1.chr1.g00345.t01#2    mNeoVis.3.1.1.chr4.g00388.t01#6    mNeoVis.evergladensis.1.1.chr1.g00353.t01#3    mNeoVis.vulgivaga.1.1.chr1.g00345.t01#4   mNeoVis.2.1.1.chr1_hap1.g00347.t01#5       4363   125256791:  mNeoVis.vulgivaga.2.1.chr1.g00352.t01#1               4364   125256792:  mNeoVis.vulgivaga.2.1.chr1.g00353.t01#1               4365   125256793:  mNeoVis.vulgivaga.2.1.chr1.g00354.t01#1               4366   125256794:   mNeoVis.vulgivaga.2.1.chr7.g01027.t01#1    mNeoVis.vulgivaga.1.1.chr7.g01021.t01#4    mNeoVis.evergladensis.1.1.chr7.g01025.t01#3    mNeoVis.2.1.1.chr7_hap1.g01026.t01#5   mNeoVis.evergladensis.2.1.chr7.g01019.t01#2       4367   125256795:   mNeoVis.3.1.1.chr7.g00966.t01#6    mNeoVis.vulgivaga.1.1.chr7.g01023.t01#4    mNeoVis.vulgivaga.2.1.chr7.g01029.t01#1    mNeoVis.2.1.1.chr7_hap1.g01028.t01#5    mNeoVis.evergladensis.1.1.chr7.g01027.t01#3   mNeoVis.evergladensis.2.1.chr7.g01021.t01#2       4368   125256796:   mNeoVis.3.1.1.chr1.g00251.t01#6    mNeoVis.vulgivaga.1.1.chr1.g00246.t01#4    mNeoVis.evergladensis.2.1.chr1.g00249.t01#2    mNeoVis.evergladensis.1.1.chr1.g00258.t01#3    mNeoVis.2.1.1.chr1_hap1.g00251.t01#5   mNeoVis.vulgivaga.2.1.chr1.g00253.t01#1       4369   125256797:   mNeoVis.evergladensis.2.1.chr1.g00250.t01#2    mNeoVis.vulgivaga.2.1.chr1.g00254.t01#1    mNeoVis.2.1.1.chr1_hap1.g00252.t01#5    mNeoVis.evergladensis.1.1.chr1.g00259.t01#3    mNeoVis.vulgivaga.1.1.chr1.g00247.t01#4   mNeoVis.3.1.1.chr1.g00252.t01#6       4370   125256798:   mNeoVis.vulgivaga.1.1.chr7.g01024.t01#4    mNeoVis.2.1.1.chr7_hap1.g01029.t01#5    mNeoVis.evergladensis.1.1.chr7.g01028.t01#3    mNeoVis.vulgivaga.2.1.chr7.g01030.t01#1    mNeoVis.evergladensis.2.1.chr7.g01022.t01#2   mNeoVis.3.1.1.chr7.g00967.t01#6       4371   125256799:   mNeoVis.3.1.1.chr1.g00253.t01#6    mNeoVis.vulgivaga.2.1.chr1.g00255.t01#1    mNeoVis.evergladensis.2.1.chr1.g00251.t01#2    mNeoVis.evergladensis.1.1.chr1.g00260.t01#3    mNeoVis.2.1.1.chr1_hap1.g00253.t01#5   mNeoVis.vulgivaga.1.1.chr1.g00248.t01#4       4372   125256800:   mNeoVis.vulgivaga.2.1.chr7.g01031.t01#1    mNeoVis.evergladensis.2.1.chr7.g01023.t01#2    mNeoVis.evergladensis.1.1.chr7.g01029.t01#3    mNeoVis.vulgivaga.1.1.chr7.g01025.t01#4   mNeoVis.2.1.1.chr7_hap1.g01030.t01#5       4373   125256801:   mNeoVis.vulgivaga.1.1.chr7.g01026.t01#4    mNeoVis.evergladensis.1.1.chr7.g01030.t01#3    mNeoVis.2.1.1.chr7_hap1.g01031.t01#5    mNeoVis.evergladensis.2.1.chr7.g01024.t01#2    mNeoVis.vulgivaga.2.1.chr7.g01032.t01#1   mNeoVis.3.1.1.chr2.g00690.t01#6       4374   125256802:   mNeoVis.vulgivaga.1.1.chr7.g01027.t01#4    mNeoVis.evergladensis.1.1.chr7.g01031.t01#3    mNeoVis.2.1.1.chr7_hap1.g01032.t01#5    mNeoVis.3.1.1.chr7.g00968.t01#6    mNeoVis.evergladensis.2.1.chr7.g01025.t01#2   mNeoVis.vulgivaga.2.1.chr7.g01033.t01#1       4375   125256803:   mNeoVis.vulgivaga.2.1.chr1.g00256.t01#1    mNeoVis.3.1.1.chr1.g00254.t01#6    mNeoVis.2.1.1.chr1_hap1.g00254.t01#5    mNeoVis.evergladensis.2.1.chr1.g00252.t01#2    mNeoVis.vulgivaga.1.1.chr1.g00249.t01#4   mNeoVis.evergladensis.1.1.chr1.g00261.t01#3       4376   125256804:   mNeoVis.evergladensis.2.1.chr7.g01026.t01#2    mNeoVis.vulgivaga.1.1.chr7.g01028.t01#4    mNeoVis.evergladensis.1.1.chr7.g01032.t01#3    mNeoVis.3.1.1.chr7.g00969.t01#6    mNeoVis.2.1.1.chr7_hap1.g01033.t01#5   mNeoVis.vulgivaga.2.1.chr7.g01034.t01#1       4377   125256805:   mNeoVis.3.1.1.chr1.g00255.t01#6    mNeoVis.2.1.1.chr1_hap1.g00255.t01#5    mNeoVis.vulgivaga.1.1.chr1.g00250.t01#4    mNeoVis.evergladensis.1.1.chr1.g00262.t01#3    mNeoVis.vulgivaga.2.1.chr1.g00257.t01#1   mNeoVis.evergladensis.2.1.chr1.g00253.t01#2       4378   125256806:   mNeoVis.3.1.1.chr1.g00256.t01#6    mNeoVis.evergladensis.1.1.chr1.g00263.t01#3    mNeoVis.2.1.1.chr1_hap1.g00256.t01#5    mNeoVis.evergladensis.2.1.chr1.g00254.t01#2    mNeoVis.vulgivaga.2.1.chr1.g00258.t01#1   mNeoVis.vulgivaga.1.1.chr1.g00251.t01#4       4379   125256807:   mNeoVis.2.1.1.chr7_hap1.g01034.t01#5    mNeoVis.vulgivaga.2.1.chr7.g01035.t01#1    mNeoVis.evergladensis.2.1.chr7.g01027.t01#2    mNeoVis.3.1.1.chr7.g00970.t01#6    mNeoVis.vulgivaga.1.1.chr7.g01029.t01#4   mNeoVis.evergladensis.1.1.chr7.g01033.t01#3       4380   125256808:   mNeoVis.vulgivaga.1.1.chr7.g01030.t01#4    mNeoVis.vulgivaga.2.1.chr7.g01036.t01#1    mNeoVis.evergladensis.1.1.chr7.g01034.t01#3    mNeoVis.evergladensis.2.1.chr7.g01028.t01#2    mNeoVis.3.1.1.chr7.g00971.t01#6   mNeoVis.2.1.1.chr7_hap1.g01035.t01#5       4381   125256809:   mNeoVis.evergladensis.1.1.chr7.g01035.t01#3    mNeoVis.3.1.1.chr7.g00972.t01#6    mNeoVis.vulgivaga.2.1.chr7.g01037.t01#1    mNeoVis.2.1.1.chr7_hap1.g01036.t01#5    mNeoVis.evergladensis.2.1.chr7.g01029.t01#2   mNeoVis.vulgivaga.1.1.chr7.g01031.t01#4       4382   125256810:   mNeoVis.evergladensis.1.1.chr1.g00264.t01#3    mNeoVis.vulgivaga.2.1.chr1.g00259.t01#1   mNeoVis.vulgivaga.1.1.chr1.g00252.t01#4           4383   125256811:   mNeoVis.vulgivaga.1.1.chr1.g00253.t01#4    mNeoVis.3.1.1.chr1.g00258.t01#6   mNeoVis.vulgivaga.2.1.chr1.g00260.t01#1           4384   125256812:   mNeoVis.3.1.1.chr7.g00973.t01#6    mNeoVis.vulgivaga.2.1.chr7.g01038.t01#1    mNeoVis.evergladensis.1.1.chr7.g01036.t01#3    mNeoVis.vulgivaga.1.1.chr7.g01032.t01#4    mNeoVis.2.1.1.chr7_hap1.g01037.t01#5   mNeoVis.evergladensis.2.1.chr7.g01030.t01#2       4385   125256813:   mNeoVis.vulgivaga.2.1.chr1.g00261.t01#1    mNeoVis.3.1.1.chr1.g00259.t01#6    mNeoVis.evergladensis.1.1.chr1.g00265.t01#3    mNeoVis.2.1.1.chr1_hap1.g00258.t01#5    mNeoVis.evergladensis.2.1.chr1.g00256.t01#2   mNeoVis.vulgivaga.1.1.chr1.g00254.t01#4       4386   125256814:   mNeoVis.3.1.1.chr4.g00819.t01#6    mNeoVis.vulgivaga.1.1.chr4.g00826.t01#4    mNeoVis.evergladensis.2.1.chr4.g00811.t01#2    mNeoVis.2.1.1.chr6_hap1.g00809.t01#5    mNeoVis.evergladensis.1.1.chr4.g00825.t01#3   mNeoVis.vulgivaga.2.1.chr4.g00831.t01#1       4387   125256815:   mNeoVis.3.1.1.chr11.g00369.t01#6    mNeoVis.vulgivaga.2.1.chr11.g00404.t01#1    mNeoVis.vulgivaga.1.1.chr11.g00393.t01#4    mNeoVis.2.1.1.chr11_hap1.g00403.t01#5    mNeoVis.evergladensis.1.1.chr11.g00402.t01#3   mNeoVis.evergladensis.2.1.chr11.g00399.t01#2       4388   125256816:   mNeoVis.3.1.1.chr5.g00034.t01#6    mNeoVis.vulgivaga.1.1.chr5.g00038.t01#4    mNeoVis.2.1.1.chr2_hap1.g00040.t01#5   mNeoVis.vulgivaga.2.1.chr5.g00037.t01#1         4389   125256817:   mNeoVis.evergladensis.1.1.chr5.g00040.t01#3    mNeoVis.vulgivaga.1.1.chr5.g00037.t01#4   mNeoVis.evergladensis.2.1.chr5.g00040.t01#2         4390   125256818:   mNeoVis.vulgivaga.2.1.chr1.g00262.t01#1    mNeoVis.evergladensis.1.1.chr1.g00266.t01#3    mNeoVis.3.1.1.chr1.g00260.t01#6    mNeoVis.evergladensis.2.1.chr1.g00257.t01#2    mNeoVis.2.1.1.chr1_hap1.g00259.t01#5   mNeoVis.vulgivaga.1.1.chr1.g00255.t01#4       4391   125256819:   mNeoVis.vulgivaga.1.1.chr1.g00257.t01#4    mNeoVis.3.1.1.chr1.g00262.t01#6    mNeoVis.2.1.1.chr1_hap1.g00261.t01#5    mNeoVis.evergladensis.2.1.chr1.g00258.t01#2    mNeoVis.vulgivaga.2.1.chr1.g00263.t01#1   mNeoVis.evergladensis.1.1.chr1.g00268.t01#3       4392   125256820:   mNeoVis.vulgivaga.2.1.chr7.g00926.t01#1    mNeoVis.2.1.1.chr7_hap1.g00928.t01#5    mNeoVis.3.1.1.chr7.g00869.t01#6    mNeoVis.evergladensis.2.1.chr7.g00921.t01#2    mNeoVis.vulgivaga.1.1.chr7.g00926.t01#4   mNeoVis.evergladensis.1.1.chr7.g00927.t01#3       4393   125256821:   mNeoVis.vulgivaga.1.1.chr7.g00927.t01#4    mNeoVis.evergladensis.1.1.chr7.g00928.t01#3    mNeoVis.evergladensis.2.1.chr7.g00922.t01#2    mNeoVis.vulgivaga.2.1.chr7.g00927.t01#1    mNeoVis.2.1.1.chr7_hap1.g00929.t01#5   mNeoVis.3.1.1.chr7.g00870.t01#6       4394   125256822:   mNeoVis.3.1.1.chr15.g00090.t01#6    mNeoVis.evergladensis.1.1.chr15.g00516.t01#3    mNeoVis.vulgivaga.2.1.chr15.g00099.t01#1    mNeoVis.evergladensis.2.1.chr15.g00097.t01#2    mNeoVis.vulgivaga.1.1.chr15.g00097.t01#4    mNeoVis.vulgivaga.2.1.chr15.g00100.t01#1   mNeoVis.2.1.1.chr15_hap1.g00096.t01#5     4395   125256823:   mNeoVis.vulgivaga.2.1.chr1.g00264.t01#1    mNeoVis.2.1.1.chr1_hap1.g00262.t01#5    mNeoVis.3.1.1.chr1.g00263.t01#6    mNeoVis.vulgivaga.1.1.chr1.g00258.t01#4    mNeoVis.evergladensis.1.1.chr1.g00269.t01#3   mNeoVis.evergladensis.2.1.chr1.g00259.t01#2       4396   125256824:   mNeoVis.evergladensis.1.1.chr7.g00929.t01#3    mNeoVis.3.1.1.chr7.g00871.t01#6    mNeoVis.evergladensis.2.1.chr7.g00923.t01#2    mNeoVis.2.1.1.chr7_hap1.g00930.t01#5    mNeoVis.vulgivaga.2.1.chr7.g00928.t01#1   mNeoVis.vulgivaga.1.1.chr7.g00928.t01#4       4397   125256825:   mNeoVis.evergladensis.1.1.chr1.g00270.t01#3    mNeoVis.2.1.1.chr1_hap1.g00263.t01#5    mNeoVis.3.1.1.chr1.g00264.t01#6    mNeoVis.vulgivaga.1.1.chr1.g00259.t01#4    mNeoVis.vulgivaga.2.1.chr1.g00265.t01#1   mNeoVis.evergladensis.2.1.chr1.g00260.t01#2       4398   125256826:   mNeoVis.evergladensis.1.1.chr1.g00271.t01#3    mNeoVis.vulgivaga.1.1.chr1.g00260.t01#4    mNeoVis.evergladensis.2.1.chr1.g00261.t01#2    mNeoVis.2.1.1.chr1_hap1.g00264.t01#5    mNeoVis.vulgivaga.2.1.chr1.g00266.t01#1   mNeoVis.3.1.1.chr1.g00265.t01#6       4399   125256827:   mNeoVis.evergladensis.1.1.chr1.g00272.t01#3   mNeoVis.vulgivaga.2.1.chr1.g00267.t01#1             4400   125256828:   mNeoVis.vulgivaga.2.1.chr7.g00929.t01#1    mNeoVis.evergladensis.1.1.chr7.g00930.t01#3    mNeoVis.vulgivaga.1.1.chr7.g00929.t01#4    mNeoVis.evergladensis.2.1.chr7.g00924.t01#2    mNeoVis.3.1.1.chr7.g00872.t01#6   mNeoVis.2.1.1.chr7_hap1.g00931.t01#5       4401   125256829:   mNeoVis.vulgivaga.1.1.chr1.g00262.t01#4    mNeoVis.evergladensis.1.1.chr1.g00273.t01#3    mNeoVis.vulgivaga.2.1.chr1.g00268.t01#1    mNeoVis.2.1.1.chr1_hap1.g00266.t01#5    mNeoVis.evergladensis.2.1.chr1.g00263.t01#2   mNeoVis.3.1.1.chr1.g00267.t01#6       4402   125256830:  mNeoVis.vulgivaga.2.1.chr7.g00931.t01#1               4403   125256831:   mNeoVis.evergladensis.1.1.chr7.g00931.t01#3   mNeoVis.2.1.1.chr7_hap1.g00932.t01#5             4404   125256832:   mNeoVis.evergladensis.1.1.chr7.g00932.t01#3   mNeoVis.evergladensis.2.1.chr7.g00926.t01#2           4405   125256833:   mNeoVis.2.1.1.chr7_hap1.g00933.t01#5   mNeoVis.3.1.1.chr7.g00873.t01#6             4406   125256834:   mNeoVis.vulgivaga.1.1.chr7.g00930.t01#4   mNeoVis.3.1.1.chr7.g00874.t01#6             4407   125256835:  mNeoVis.vulgivaga.2.1.chr7.g00930.t01#1               4408   125256836:  mNeoVis.evergladensis.2.1.chr7.g00925.t01#2             4409   125256837:   mNeoVis.2.1.1.chr7_hap1.g00014.t01#5    mNeoVis.evergladensis.1.1.chr7.g00016.t01#3    mNeoVis.3.1.1.chr7.g00014.t01#6    mNeoVis.vulgivaga.1.1.chr7.g00016.t01#4    mNeoVis.vulgivaga.2.1.chr7.g00015.t01#1   mNeoVis.evergladensis.2.1.chr7.g00015.t01#2       4410   125256838:   mNeoVis.vulgivaga.1.1.chr1.g00263.t01#4    mNeoVis.vulgivaga.2.1.chr1.g00269.t01#1    mNeoVis.evergladensis.2.1.chr1.g00264.t01#2    mNeoVis.2.1.1.chr1_hap1.g00267.t01#5    mNeoVis.3.1.1.chr1.g00268.t01#6   mNeoVis.evergladensis.1.1.chr1.g00274.t01#3       4411   125256839:   mNeoVis.evergladensis.2.1.chr1.g00265.t01#2    mNeoVis.evergladensis.1.1.chr1.g00275.t01#3    mNeoVis.vulgivaga.1.1.chr1.g00264.t01#4    mNeoVis.3.1.1.chr1.g00269.t01#6    mNeoVis.vulgivaga.2.1.chr1.g00270.t01#1   mNeoVis.2.1.1.chr1_hap1.g00268.t01#5       4412   125256840:   mNeoVis.vulgivaga.2.1.chr7.g00932.t01#1   mNeoVis.3.1.1.chr7.g00875.t01#6             4413   125256841:   mNeoVis.vulgivaga.2.1.chr1.g00271.t01#1    mNeoVis.2.1.1.chr1_hap1.g00269.t01#5    mNeoVis.3.1.1.chr1.g00271.t01#6    mNeoVis.evergladensis.2.1.chr1.g00266.t01#2    mNeoVis.vulgivaga.1.1.chr1.g00265.t01#4   mNeoVis.evergladensis.1.1.chr1.g00276.t01#3       4414   125256842:   mNeoVis.evergladensis.1.1.chr7.g00933.t01#3    mNeoVis.vulgivaga.1.1.chr7.g00931.t01#4    mNeoVis.2.1.1.chr7_hap1.g00934.t01#5    mNeoVis.evergladensis.2.1.chr7.g00927.t01#2    mNeoVis.vulgivaga.2.1.chr7.g00933.t01#1   mNeoVis.3.1.1.chr7.g00876.t01#6       4415   125256843:   mNeoVis.vulgivaga.2.1.chr7.g00934.t01#1    mNeoVis.3.1.1.chr7.g00877.t01#6    mNeoVis.2.1.1.chr7_hap1.g00935.t01#5    mNeoVis.evergladensis.1.1.chr7.g00934.t01#3    mNeoVis.vulgivaga.1.1.chr7.g00932.t01#4   mNeoVis.evergladensis.2.1.chr7.g00928.t01#2       4416   125256844:   mNeoVis.2.1.1.chr1_hap1.g00270.t01#5    mNeoVis.3.1.1.chr1.g00272.t01#6    mNeoVis.evergladensis.1.1.chr1.g00277.t01#3    mNeoVis.vulgivaga.2.1.chr1.g00272.t01#1    mNeoVis.evergladensis.2.1.chr1.g00267.t01#2   mNeoVis.vulgivaga.1.1.chr1.g00266.t01#4       4417   125256845:   mNeoVis.vulgivaga.1.1.chr4.g00451.t01#4    mNeoVis.2.1.1.chr6_hap1.g00448.t01#5    mNeoVis.vulgivaga.2.1.chr4.g00460.t01#1    mNeoVis.evergladensis.1.1.chr4.g00458.t01#3    mNeoVis.3.1.1.chr4.g00450.t01#6   mNeoVis.evergladensis.2.1.chr4.g00454.t01#2       4418   125256846:   mNeoVis.evergladensis.1.1.chr7.g00935.t01#3    mNeoVis.vulgivaga.1.1.chr7.g00933.t01#4    mNeoVis.vulgivaga.2.1.chr7.g00935.t01#1    mNeoVis.3.1.1.chr7.g00878.t01#6    mNeoVis.evergladensis.2.1.chr7.g00929.t01#2   mNeoVis.2.1.1.chr7_hap1.g00936.t01#5       4419   125256847:   mNeoVis.vulgivaga.1.1.chr1.g00268.t01#4   mNeoVis.vulgivaga.2.1.chr1.g00273.t01#1             4420   125256848:   mNeoVis.vulgivaga.1.1.chr7.g00934.t01#4    mNeoVis.vulgivaga.2.1.chr7.g00936.t01#1    mNeoVis.3.1.1.chr7.g00879.t01#6    mNeoVis.evergladensis.2.1.chr7.g00930.t01#2    mNeoVis.evergladensis.1.1.chr7.g00936.t01#3   mNeoVis.2.1.1.chr7_hap1.g00937.t01#5       4421   125256849:  mNeoVis.vulgivaga.2.1.chr1.g00274.t01#1               4422   125256850:   mNeoVis.vulgivaga.1.1.chr7.g00935.t01#4    mNeoVis.vulgivaga.2.1.chr7.g00937.t01#1    mNeoVis.3.1.1.chr15.g00501.t01#6    mNeoVis.evergladensis.1.1.chr7.g00937.t01#3    mNeoVis.2.1.1.chr7_hap1.g00938.t01#5   mNeoVis.evergladensis.2.1.chr7.g00931.t01#2       4423   125256851:   mNeoVis.evergladensis.1.1.chr4.g00195.t01#3    mNeoVis.vulgivaga.1.1.chr4.g00195.t01#4    mNeoVis.3.1.1.chr4.g00192.t01#6    mNeoVis.vulgivaga.2.1.chr4.g00196.t01#1    mNeoVis.2.1.1.chr6_hap1.g00196.t01#5    mNeoVis.evergladensis.2.1.chr4.g00201.t01#2   mNeoVis.evergladensis.1.1.chr4.g00196.t01#3     4424   125256852:   mNeoVis.2.1.1.chr1_hap1.g00272.t01#5    mNeoVis.vulgivaga.1.1.chr1.g00270.t01#4    mNeoVis.evergladensis.2.1.chr1.g00270.t01#2    mNeoVis.3.1.1.chr1.g00275.t01#6    mNeoVis.evergladensis.1.1.chr1.g00280.t01#3   mNeoVis.vulgivaga.2.1.chr1.g00275.t01#1       4425   125256853:   mNeoVis.vulgivaga.2.1.chr9.g01075.t01#1    mNeoVis.3.1.1.chr9.g01051.t01#6    mNeoVis.evergladensis.1.1.chr9.g01087.t01#3    mNeoVis.vulgivaga.1.1.chr9.g01089.t01#4    mNeoVis.evergladensis.2.1.chr8.g01099.t01#2   mNeoVis.2.1.1.chr10_hap1.g01121.t01#5       4426   125256854:   mNeoVis.vulgivaga.2.1.chr7.g00938.t01#1    mNeoVis.evergladensis.1.1.chr7.g00938.t01#3    mNeoVis.2.1.1.chr7_hap1.g00939.t01#5    mNeoVis.vulgivaga.1.1.chr7.g00936.t01#4   mNeoVis.evergladensis.2.1.chr7.g00932.t01#2       4427   125256855:   mNeoVis.vulgivaga.1.1.chr7.g00937.t01#4    mNeoVis.3.1.1.chr7.g00880.t01#6    mNeoVis.vulgivaga.2.1.chr7.g00939.t01#1    mNeoVis.evergladensis.1.1.chr7.g00939.t01#3    mNeoVis.2.1.1.chr7_hap1.g00940.t01#5   mNeoVis.evergladensis.2.1.chr7.g00933.t01#2       4428   125256856:  mNeoVis.vulgivaga.2.1.chr1.g00332.t01#1               4429   125256857:  mNeoVis.vulgivaga.2.1.chr1.g00276.t01#1               4430   125256858:   mNeoVis.evergladensis.1.1.chr1.g00336.t01#3    mNeoVis.evergladensis.2.1.chr1.g00327.t01#2    mNeoVis.vulgivaga.1.1.chr1.g00327.t01#4    mNeoVis.3.1.1.chr1.g00333.t01#6    mNeoVis.vulgivaga.2.1.chr1.g00333.t01#1   mNeoVis.2.1.1.chr1_hap1.g00329.t01#5       4431   125256859:   mNeoVis.vulgivaga.2.1.chr7.g00940.t01#1    mNeoVis.vulgivaga.1.1.chr7.g00938.t01#4    mNeoVis.3.1.1.chr7.g00881.t01#6    mNeoVis.evergladensis.2.1.chr7.g00934.t01#2    mNeoVis.evergladensis.1.1.chr7.g00940.t01#3   mNeoVis.2.1.1.chr7_hap1.g00941.t01#5       4432   125256860:   mNeoVis.evergladensis.2.1.chr1.g00271.t01#2    mNeoVis.evergladensis.1.1.chr1.g00281.t01#3    mNeoVis.vulgivaga.2.1.chr1.g00277.t01#1    mNeoVis.2.1.1.chr1_hap1.g00273.t01#5    mNeoVis.vulgivaga.1.1.chr1.g00271.t01#4   mNeoVis.3.1.1.chr1.g00276.t01#6       4433   125256861:   mNeoVis.evergladensis.1.1.chr15.g00515.t01#3    mNeoVis.evergladensis.2.1.chr15.g00098.t01#2    mNeoVis.3.1.1.chr15.g00092.t01#6    mNeoVis.vulgivaga.1.1.chr15.g00098.t01#4    mNeoVis.2.1.1.chr15_hap1.g00097.t01#5   mNeoVis.vulgivaga.2.1.chr15.g00101.t01#1       4434   125256862:   mNeoVis.vulgivaga.1.1.chr1.g00272.t01#4    mNeoVis.3.1.1.chr1.g00277.t01#6    mNeoVis.evergladensis.1.1.chr1.g00282.t01#3    mNeoVis.evergladensis.2.1.chr1.g00272.t01#2    mNeoVis.vulgivaga.2.1.chr1.g00278.t01#1   mNeoVis.2.1.1.chr1_hap1.g00274.t01#5       4435   125256863:   mNeoVis.2.1.1.chr7_hap1.g00943.t01#5    mNeoVis.vulgivaga.1.1.chr7.g00940.t01#4    mNeoVis.3.1.1.chr7.g00883.t01#6    mNeoVis.vulgivaga.2.1.chr7.g00942.t01#1    mNeoVis.evergladensis.2.1.chr7.g00936.t01#2   mNeoVis.evergladensis.1.1.chr7.g00942.t01#3       4436   125256864:   mNeoVis.vulgivaga.2.1.chr1.g00279.t01#1    mNeoVis.2.1.1.chr1_hap1.g00275.t01#5    mNeoVis.vulgivaga.1.1.chr1.g00273.t01#4    mNeoVis.3.1.1.chr1.g00278.t01#6    mNeoVis.evergladensis.1.1.chr1.g00283.t01#3   mNeoVis.evergladensis.2.1.chr1.g00273.t01#2       4437   125256865:   mNeoVis.3.1.1.chr4.g00196.t01#6    mNeoVis.2.1.1.chr6_hap1.g00201.t01#5    mNeoVis.vulgivaga.1.1.chr4.g00200.t01#4    mNeoVis.evergladensis.2.1.chr4.g00206.t01#2    mNeoVis.evergladensis.1.1.chr4.g00201.t01#3   mNeoVis.vulgivaga.2.1.chr4.g00201.t01#1       4438   125256866:   mNeoVis.2.1.1.chr10_hap1.g01118.t01#5    mNeoVis.3.1.1.chr9.g01046.t01#6    mNeoVis.vulgivaga.2.1.chr9.g01073.t01#1    mNeoVis.evergladensis.2.1.chr8.g01094.t01#2   mNeoVis.vulgivaga.1.1.chr9.g01087.t01#4       4439   125256867:   mNeoVis.3.1.1.chr9.g01048.t01#6    mNeoVis.evergladensis.1.1.chr9.g01084.t01#3   mNeoVis.3.1.1.chr9.g01047.t01#6           4440   125256868:   mNeoVis.evergladensis.1.1.chr7.g00943.t01#3    mNeoVis.evergladensis.2.1.chr7.g00937.t01#2    mNeoVis.vulgivaga.2.1.chr7.g00943.t01#1    mNeoVis.vulgivaga.1.1.chr7.g00941.t01#4    mNeoVis.3.1.1.chr7.g00884.t01#6   mNeoVis.2.1.1.chr7_hap1.g00944.t01#5       4441   125256869:   mNeoVis.vulgivaga.2.1.chr7.g00944.t01#1    mNeoVis.evergladensis.1.1.chr7.g00944.t01#3    mNeoVis.evergladensis.2.1.chr7.g00938.t01#2    mNeoVis.vulgivaga.1.1.chr7.g00942.t01#4   mNeoVis.2.1.1.chr7_hap1.g00945.t01#5       4442   125256870:   mNeoVis.3.1.1.chr7.g00015.t01#6    mNeoVis.vulgivaga.2.1.chr7.g00016.t01#1    mNeoVis.2.1.1.chr7_hap1.g00015.t01#5    mNeoVis.vulgivaga.1.1.chr7.g00017.t01#4    mNeoVis.evergladensis.1.1.chr7.g00017.t01#3   mNeoVis.evergladensis.2.1.chr7.g00016.t01#2       4443   125256871:   mNeoVis.vulgivaga.1.1.chr1.g00329.t01#4    mNeoVis.vulgivaga.2.1.chr1.g00335.t01#1    mNeoVis.2.1.1.chr1_hap1.g00331.t01#5    mNeoVis.evergladensis.2.1.chr1.g00329.t01#2    mNeoVis.evergladensis.1.1.chr1.g00338.t01#3   mNeoVis.3.1.1.chr1.g00335.t01#6       4444   125256872:   mNeoVis.evergladensis.2.1.chr1.g00330.t01#2    mNeoVis.evergladensis.1.1.chr1.g00339.t01#3    mNeoVis.3.1.1.chr1.g00336.t01#6    mNeoVis.vulgivaga.1.1.chr1.g00330.t01#4    mNeoVis.vulgivaga.2.1.chr1.g00336.t01#1   mNeoVis.2.1.1.chr1_hap1.g00332.t01#5       4445   125256873:   mNeoVis.evergladensis.1.1.chr3.g00269.t01#3    mNeoVis.vulgivaga.2.1.chr3.g00264.t01#1   mNeoVis.vulgivaga.1.1.chr3.g00265.t01#4           4446   125256874:   mNeoVis.2.1.1.chr7_hap1.g00947.t01#5    mNeoVis.vulgivaga.2.1.chr7.g00946.t01#1    mNeoVis.vulgivaga.1.1.chr7.g00944.t01#4    mNeoVis.3.1.1.chr7.g00887.t01#6    mNeoVis.evergladensis.1.1.chr7.g00946.t01#3   mNeoVis.evergladensis.2.1.chr7.g00940.t01#2       4447   125256875:   mNeoVis.2.1.1.chr7_hap1.g00946.t01#5    mNeoVis.vulgivaga.2.1.chr7.g00945.t01#1    mNeoVis.vulgivaga.1.1.chr7.g00943.t01#4    mNeoVis.evergladensis.1.1.chr7.g00945.t01#3    mNeoVis.3.1.1.chr7.g00886.t01#6   mNeoVis.evergladensis.2.1.chr7.g00939.t01#2       4448   125256876:   mNeoVis.evergladensis.1.1.chr15.g00177.t01#3    mNeoVis.3.1.1.chr15.g00398.t01#6    mNeoVis.evergladensis.2.1.chr15.g00437.t01#2    mNeoVis.2.1.1.chr15_hap1.g00615.t01#5    mNeoVis.vulgivaga.1.1.chr15.g00438.t01#4   mNeoVis.vulgivaga.2.1.chr15.g00445.t01#1       4449   125256877:   mNeoVis.2.1.1.chr9_hap1.g01164.t01#5    mNeoVis.vulgivaga.2.1.chr10.g01144.t01#1    mNeoVis.vulgivaga.1.1.chr10.g01128.t01#4    mNeoVis.evergladensis.1.1.chr10.g01148.t01#3    mNeoVis.3.1.1.chr10.g01117.t01#6   mNeoVis.evergladensis.2.1.chr10.g01152.t01#2       4450   125256878:   mNeoVis.vulgivaga.2.1.chr1.g00337.t01#1    mNeoVis.vulgivaga.1.1.chr1.g00331.t01#4    mNeoVis.evergladensis.1.1.chr1.g00340.t01#3    mNeoVis.3.1.1.chr1.g00337.t01#6    mNeoVis.evergladensis.2.1.chr1.g00331.t01#2   mNeoVis.2.1.1.chr1_hap1.g00333.t01#5       4451   125256879:   mNeoVis.2.1.1.chr1_hap1.g00334.t01#5    mNeoVis.evergladensis.1.1.chr1.g00341.t01#3    mNeoVis.vulgivaga.2.1.chr1.g00338.t01#1    mNeoVis.evergladensis.2.1.chr1.g00332.t01#2    mNeoVis.3.1.1.chr1.g00338.t01#6   mNeoVis.vulgivaga.1.1.chr1.g00332.t01#4       4452   125256880:   mNeoVis.vulgivaga.2.1.chr7.g00947.t01#1    mNeoVis.evergladensis.1.1.chr7.g00947.t01#3    mNeoVis.evergladensis.2.1.chr7.g00941.t01#2    mNeoVis.3.1.1.chr7.g00888.t01#6    mNeoVis.2.1.1.chr7_hap1.g00948.t01#5   mNeoVis.vulgivaga.1.1.chr7.g00945.t01#4       4453   125256881:   mNeoVis.vulgivaga.2.1.chr1.g00339.t01#1    mNeoVis.evergladensis.1.1.chr1.g00342.t01#3    mNeoVis.2.1.1.chr1_hap1.g00335.t01#5    mNeoVis.vulgivaga.1.1.chr1.g00333.t01#4    mNeoVis.evergladensis.2.1.chr1.g00333.t01#2   mNeoVis.3.1.1.chr1.g00339.t01#6       4454   125256882:   mNeoVis.evergladensis.1.1.chr7.g00948.t01#3    mNeoVis.vulgivaga.2.1.chr7.g00948.t01#1    mNeoVis.evergladensis.2.1.chr7.g00942.t01#2    mNeoVis.3.1.1.chr7.g00889.t01#6    mNeoVis.2.1.1.chr7_hap1.g00949.t01#5   mNeoVis.vulgivaga.1.1.chr7.g00946.t01#4       4455   125256883:   mNeoVis.evergladensis.2.1.chr7.g00943.t01#2    mNeoVis.vulgivaga.1.1.chr7.g00947.t01#4    mNeoVis.3.1.1.chr7.g00890.t01#6    mNeoVis.2.1.1.chr7_hap1.g00950.t01#5    mNeoVis.evergladensis.1.1.chr7.g00949.t01#3   mNeoVis.vulgivaga.2.1.chr7.g00949.t01#1       4456   125256884:   mNeoVis.vulgivaga.1.1.chr1.g00334.t01#4    mNeoVis.evergladensis.2.1.chr1.g00334.t01#2    mNeoVis.3.1.1.chr1.g00340.t01#6    mNeoVis.vulgivaga.2.1.chr1.g00340.t01#1    mNeoVis.2.1.1.chr1_hap1.g00336.t01#5   mNeoVis.evergladensis.1.1.chr1.g00343.t01#3       4457   125256885:   mNeoVis.2.1.1.chr1_hap1.g00337.t01#5    mNeoVis.vulgivaga.2.1.chr1.g00341.t01#1    mNeoVis.vulgivaga.1.1.chr1.g00335.t01#4    mNeoVis.3.1.1.chr1.g00341.t01#6    mNeoVis.evergladensis.2.1.chr1.g00335.t01#2   mNeoVis.evergladensis.1.1.chr1.g00344.t01#3       4458   125256886:   mNeoVis.vulgivaga.1.1.chr7.g00948.t01#4    mNeoVis.2.1.1.chr7_hap1.g00951.t01#5    mNeoVis.evergladensis.1.1.chr7.g00950.t01#3    mNeoVis.3.1.1.chr7.g00891.t01#6    mNeoVis.vulgivaga.2.1.chr7.g00950.t01#1   mNeoVis.evergladensis.2.1.chr7.g00944.t01#2       4459   125256887:   mNeoVis.evergladensis.2.1.chr7.g00945.t01#2    mNeoVis.2.1.1.chr7_hap1.g00952.t01#5    mNeoVis.3.1.1.chr7.g00892.t01#6    mNeoVis.vulgivaga.1.1.chr7.g00949.t01#4    mNeoVis.evergladensis.1.1.chr7.g00951.t01#3   mNeoVis.vulgivaga.2.1.chr7.g00951.t01#1       4460   125256888:   mNeoVis.3.1.1.chr7.g00893.t01#6    mNeoVis.evergladensis.1.1.chr7.g00952.t01#3    mNeoVis.vulgivaga.2.1.chr7.g00952.t01#1    mNeoVis.2.1.1.chr7_hap1.g00953.t01#5    mNeoVis.evergladensis.2.1.chr7.g00946.t01#2   mNeoVis.vulgivaga.1.1.chr7.g00950.t01#4       4461   125256889:   mNeoVis.2.1.1.chr1_hap1.g00338.t01#5    mNeoVis.3.1.1.chr1.g00342.t01#6    mNeoVis.vulgivaga.2.1.chr1.g00342.t01#1    mNeoVis.evergladensis.1.1.chr1.g00345.t01#3   mNeoVis.vulgivaga.1.1.chr1.g00336.t01#4       4462   125256890:   mNeoVis.evergladensis.1.1.chr1.g00346.t01#3    mNeoVis.vulgivaga.2.1.chr1.g00343.t01#1    mNeoVis.vulgivaga.1.1.chr1.g00337.t01#4    mNeoVis.2.1.1.chr1_hap1.g00339.t01#5    mNeoVis.evergladensis.2.1.chr1.g00337.t01#2   mNeoVis.3.1.1.chr1.g00343.t01#6       4463   125256891:   mNeoVis.evergladensis.1.1.chr7.g00953.t01#3    mNeoVis.2.1.1.chr7_hap1.g00954.t01#5    mNeoVis.3.1.1.chr7.g00894.t01#6    mNeoVis.evergladensis.2.1.chr7.g00947.t01#2    mNeoVis.vulgivaga.2.1.chr7.g00953.t01#1   mNeoVis.vulgivaga.1.1.chr7.g00951.t01#4       4464   125256892:   mNeoVis.evergladensis.1.1.chr7.g00954.t01#3    mNeoVis.3.1.1.chr7.g00895.t01#6    mNeoVis.2.1.1.chr7_hap1.g00955.t01#5    mNeoVis.vulgivaga.1.1.chr7.g00952.t01#4    mNeoVis.vulgivaga.2.1.chr7.g00954.t01#1   mNeoVis.evergladensis.2.1.chr7.g00948.t01#2       4465   125256893:   mNeoVis.evergladensis.2.1.chr1.g00338.t01#2    mNeoVis.2.1.1.chr1_hap1.g00340.t01#5    mNeoVis.evergladensis.1.1.chr1.g00347.t01#3    mNeoVis.3.1.1.chr1.g00344.t01#6    mNeoVis.vulgivaga.1.1.chr1.g00338.t01#4   mNeoVis.vulgivaga.2.1.chr1.g00344.t01#1       4466   125256894:   mNeoVis.3.1.1.chr15.g00093.t01#6    mNeoVis.evergladensis.2.1.chr15.g00099.t01#2    mNeoVis.vulgivaga.1.1.chr15.g00099.t01#4    mNeoVis.vulgivaga.2.1.chr15.g00102.t01#1    mNeoVis.2.1.1.chr15_hap1.g00098.t01#5   mNeoVis.evergladensis.1.1.chr15.g00514.t01#3       4467   125256895:   mNeoVis.evergladensis.2.1.chr1.g00339.t01#2    mNeoVis.2.1.1.chr1_hap1.g00341.t01#5    mNeoVis.evergladensis.1.1.chr1.g00348.t01#3    mNeoVis.vulgivaga.2.1.chr1.g00345.t01#1    mNeoVis.3.1.1.chr1.g00345.t01#6   mNeoVis.vulgivaga.1.1.chr1.g00339.t01#4       4468   125256896:   mNeoVis.3.1.1.chr4.g00575.t01#6    mNeoVis.vulgivaga.2.1.chr4.g00583.t01#1    mNeoVis.2.1.1.chr6_hap1.g00570.t01#5    mNeoVis.evergladensis.2.1.chr4.g00574.t01#2    mNeoVis.vulgivaga.1.1.chr4.g00573.t01#4   mNeoVis.evergladensis.1.1.chr4.g00581.t01#3       4469   125256897:   mNeoVis.vulgivaga.2.1.chr1.g00346.t01#1    mNeoVis.2.1.1.chr1_hap1.g00342.t01#5    mNeoVis.evergladensis.1.1.chr1.g00349.t01#3   mNeoVis.evergladensis.2.1.chr1.g00340.t01#2       4470   125256898:   mNeoVis.3.1.1.chr2.g01385.t01#6    mNeoVis.evergladensis.1.1.chr2.g01464.t01#3    mNeoVis.evergladensis.1.1.chr1.g00284.t01#3    mNeoVis.evergladensis.2.1.chr1.g00274.t01#2    mNeoVis.3.1.1.chr1.g00279.t01#6    mNeoVis.2.1.1.chr1_hap1.g00276.t01#5    mNeoVis.vulgivaga.1.1.chr2.g01462.t01#4    mNeoVis.evergladensis.2.1.chr2.g01455.t01#2    mNeoVis.2.1.1.chr5_hap1.g01465.t01#5      4471   125256899:   mNeoVis.evergladensis.2.1.chr7.g00949.t01#2    mNeoVis.vulgivaga.2.1.chr7.g00955.t01#1    mNeoVis.3.1.1.chr7.g00896.t01#6    mNeoVis.vulgivaga.1.1.chr7.g00953.t01#4    mNeoVis.evergladensis.1.1.chr7.g00955.t01#3   mNeoVis.2.1.1.chr7_hap1.g00956.t01#5       4472   125256900:   mNeoVis.vulgivaga.2.1.chr1.g00281.t01#1    mNeoVis.evergladensis.2.1.chr1.g00275.t01#2    mNeoVis.evergladensis.1.1.chr1.g00285.t01#3    mNeoVis.2.1.1.chr1_hap1.g00277.t01#5    mNeoVis.vulgivaga.1.1.chr1.g00275.t01#4   mNeoVis.3.1.1.chr1.g00280.t01#6       4473   125256901:   mNeoVis.3.1.1.chr7.g00897.t01#6    mNeoVis.vulgivaga.2.1.chr7.g00956.t01#1    mNeoVis.vulgivaga.1.1.chr7.g00954.t01#4    mNeoVis.evergladensis.2.1.chr7.g00950.t01#2    mNeoVis.2.1.1.chr7_hap1.g00957.t01#5   mNeoVis.evergladensis.1.1.chr7.g00956.t01#3       4474   125256902:   mNeoVis.vulgivaga.1.1.chr1.g00276.t01#4    mNeoVis.2.1.1.chr1_hap1.g00278.t01#5    mNeoVis.3.1.1.chr1.g00281.t01#6    mNeoVis.evergladensis.2.1.chr1.g00276.t01#2    mNeoVis.vulgivaga.2.1.chr1.g00282.t01#1   mNeoVis.evergladensis.1.1.chr1.g00286.t01#3       4475   125256903:   mNeoVis.vulgivaga.1.1.chr7.g00955.t01#4    mNeoVis.2.1.1.chr7_hap1.g00958.t01#5    mNeoVis.evergladensis.1.1.chr7.g00957.t01#3    mNeoVis.3.1.1.chr7.g00898.t01#6    mNeoVis.evergladensis.2.1.chr7.g00951.t01#2   mNeoVis.vulgivaga.2.1.chr7.g00957.t01#1       4476   125256904:   mNeoVis.3.1.1.chr7.g00900.t01#6    mNeoVis.evergladensis.2.1.chr7.g00954.t01#2    mNeoVis.evergladensis.1.1.chr7.g00960.t01#3    mNeoVis.2.1.1.chr7_hap1.g00961.t01#5    mNeoVis.vulgivaga.1.1.chr7.g00957.t01#4   mNeoVis.vulgivaga.2.1.chr7.g00959.t01#1       4477   125256905:   mNeoVis.vulgivaga.2.1.chr7.g00958.t01#1    mNeoVis.2.1.1.chr6_hap1.g00660.t01#5    mNeoVis.evergladensis.1.1.chr7.g00958.t01#3    mNeoVis.evergladensis.2.1.chr15.g00443.t01#2    mNeoVis.evergladensis.2.1.chr2.g01213.t01#2    mNeoVis.2.1.1.chr7_hap1.g00959.t01#5    mNeoVis.vulgivaga.2.1.chr2.g01219.t01#1    mNeoVis.vulgivaga.2.1.chr10.g01078.t01#1    mNeoVis.vulgivaga.2.1.chr2.g01071.t01#1      4478   125256906:   mNeoVis.vulgivaga.1.1.chr5.g00750.t01#4    mNeoVis.vulgivaga.2.1.chr5.g00769.t01#1    mNeoVis.2.1.1.chr2_hap1.g00819.t01#5    mNeoVis.evergladensis.2.1.chr5.g00776.t01#2    mNeoVis.3.1.1.chr5.g00766.t01#6   mNeoVis.evergladensis.1.1.chr5.g00769.t01#3       4479   125256907:   mNeoVis.2.1.1.chr10_hap1.g00497.t01#5    mNeoVis.vulgivaga.1.1.chr9.g00483.t01#4    mNeoVis.evergladensis.2.1.chr8.g00477.t01#2    mNeoVis.vulgivaga.2.1.chr9.g00480.t01#1   mNeoVis.3.1.1.chr9.g00459.t01#6       4480   125256908:   mNeoVis.2.1.1.chr1_hap1.g01517.t01#5    mNeoVis.evergladensis.2.1.chr1.g01483.t01#2    mNeoVis.evergladensis.1.1.chr1.g01482.t01#3   mNeoVis.vulgivaga.1.1.chr1.g01484.t01#4         4481   125256909:  mNeoVis.evergladensis.1.1.chr15.g00172.t01#3             4482   125256910:   mNeoVis.2.1.1.chr7_hap1.g00962.t01#5    mNeoVis.evergladensis.2.1.chr7.g00955.t01#2    mNeoVis.vulgivaga.1.1.chr7.g00958.t01#4    mNeoVis.vulgivaga.2.1.chr7.g00960.t01#1    mNeoVis.3.1.1.chr7.g00901.t01#6   mNeoVis.evergladensis.1.1.chr7.g00961.t01#3       4483   125256911:   mNeoVis.evergladensis.2.1.chr1.g00277.t01#2    mNeoVis.evergladensis.1.1.chr1.g00288.t01#3    mNeoVis.2.1.1.chr1_hap1.g00280.t01#5    mNeoVis.vulgivaga.1.1.chr1.g00278.t01#4    mNeoVis.vulgivaga.2.1.chr1.g00283.t01#1   mNeoVis.3.1.1.chr1.g00282.t01#6       4484   125256912:   mNeoVis.evergladensis.2.1.chr1.g00281.t01#2    mNeoVis.2.1.1.chr1_hap1.g00282.t01#5    mNeoVis.vulgivaga.1.1.chr1.g00281.t01#4    mNeoVis.evergladensis.1.1.chr1.g00290.t01#3    mNeoVis.vulgivaga.2.1.chr1.g00285.t01#1   mNeoVis.3.1.1.chr1.g00286.t01#6       4485   125256913:   mNeoVis.2.1.1.chr7_hap1.g00963.t01#5    mNeoVis.vulgivaga.1.1.chr7.g00959.t01#4    mNeoVis.vulgivaga.2.1.chr7.g00961.t01#1    mNeoVis.3.1.1.chr7.g00902.t01#6    mNeoVis.evergladensis.2.1.chr7.g00956.t01#2   mNeoVis.evergladensis.1.1.chr7.g00962.t01#3       4486   125256914:   mNeoVis.3.1.1.chr7.g00903.t01#6    mNeoVis.evergladensis.1.1.chr7.g00963.t01#3    mNeoVis.2.1.1.chr7_hap1.g00964.t01#5    mNeoVis.vulgivaga.2.1.chr7.g00962.t01#1    mNeoVis.vulgivaga.1.1.chr7.g00960.t01#4   mNeoVis.evergladensis.2.1.chr7.g00957.t01#2       4487   125256915:   mNeoVis.evergladensis.2.1.chr7.g00017.t01#2    mNeoVis.3.1.1.chr7.g00016.t01#6    mNeoVis.evergladensis.1.1.chr7.g00018.t01#3    mNeoVis.vulgivaga.1.1.chr7.g00018.t01#4    mNeoVis.2.1.1.chr7_hap1.g00016.t01#5   mNeoVis.vulgivaga.2.1.chr7.g00017.t01#1       4488   125256916:   mNeoVis.vulgivaga.1.1.chr1.g00283.t01#4    mNeoVis.evergladensis.2.1.chr1.g00283.t01#2    mNeoVis.3.1.1.chr1.g00288.t01#6    mNeoVis.evergladensis.1.1.chr1.g00292.t01#3    mNeoVis.vulgivaga.2.1.chr1.g00287.t01#1   mNeoVis.2.1.1.chr1_hap1.g00284.t01#5       4489   125256917:   mNeoVis.vulgivaga.2.1.chr7.g00963.t01#1    mNeoVis.evergladensis.2.1.chr7.g00958.t01#2    mNeoVis.evergladensis.1.1.chr7.g00964.t01#3    mNeoVis.2.1.1.chr7_hap1.g00965.t01#5    mNeoVis.3.1.1.chr7.g00904.t01#6   mNeoVis.vulgivaga.1.1.chr7.g00961.t01#4       4490   125256918:   mNeoVis.2.1.1.chr7_hap1.g00017.t01#5    mNeoVis.evergladensis.2.1.chr7.g00018.t01#2    mNeoVis.vulgivaga.2.1.chr7.g00018.t01#1    mNeoVis.vulgivaga.1.1.chr7.g00019.t01#4    mNeoVis.evergladensis.1.1.chr7.g00019.t01#3   mNeoVis.3.1.1.chr3.g01152.t01#6       4491   125256919:   mNeoVis.vulgivaga.1.1.chr15.g00100.t01#4    mNeoVis.2.1.1.chr15_hap1.g00099.t01#5    mNeoVis.evergladensis.2.1.chr15.g00100.t01#2    mNeoVis.evergladensis.1.1.chr15.g00513.t01#3   mNeoVis.vulgivaga.2.1.chr15.g00103.t01#1       4492   125256920:   mNeoVis.evergladensis.1.1.chr7.g00965.t01#3    mNeoVis.3.1.1.chr7.g00905.t01#6    mNeoVis.vulgivaga.1.1.chr7.g00962.t01#4    mNeoVis.2.1.1.chr7_hap1.g00966.t01#5    mNeoVis.vulgivaga.2.1.chr7.g00964.t01#1   mNeoVis.evergladensis.2.1.chr7.g00959.t01#2       4493   125256921:   mNeoVis.evergladensis.1.1.chr15.g00512.t01#3    mNeoVis.vulgivaga.1.1.chr15.g00101.t01#4    mNeoVis.2.1.1.chr15_hap1.g00100.t01#5    mNeoVis.evergladensis.2.1.chr15.g00101.t01#2   mNeoVis.vulgivaga.2.1.chr15.g00104.t01#1       4494   125256922:   mNeoVis.vulgivaga.1.1.chr1.g00284.t01#4    mNeoVis.evergladensis.2.1.chr1.g00284.t01#2    mNeoVis.3.1.1.chr1.g00290.t01#6    mNeoVis.evergladensis.1.1.chr1.g00293.t01#3    mNeoVis.vulgivaga.2.1.chr1.g00288.t01#1   mNeoVis.2.1.1.chr1_hap1.g00285.t01#5       4495   125256923:   mNeoVis.evergladensis.1.1.chr11.g00521.t01#3    mNeoVis.vulgivaga.1.1.chr11.g00514.t01#4    mNeoVis.3.1.1.chr11.g00490.t01#6    mNeoVis.2.1.1.chr11_hap1.g00529.t01#5    mNeoVis.evergladensis.2.1.chr11.g00517.t01#2   mNeoVis.vulgivaga.2.1.chr11.g00526.t01#1       4496   125256924:   mNeoVis.2.1.1.chr7_hap1.g00967.t01#5    mNeoVis.evergladensis.1.1.chr7.g00966.t01#3    mNeoVis.evergladensis.2.1.chr7.g00960.t01#2    mNeoVis.vulgivaga.2.1.chr7.g00965.t01#1   mNeoVis.vulgivaga.1.1.chr7.g00963.t01#4       4497   125256925:   mNeoVis.vulgivaga.1.1.chr1.g00285.t01#4    mNeoVis.evergladensis.2.1.chr1.g00285.t01#2    mNeoVis.3.1.1.chr1.g00291.t01#6    mNeoVis.evergladensis.1.1.chr1.g00294.t01#3    mNeoVis.vulgivaga.2.1.chr1.g00289.t01#1   mNeoVis.2.1.1.chr1_hap1.g00286.t01#5       4498   125256926:   mNeoVis.evergladensis.1.1.chr7.g00967.t01#3    mNeoVis.3.1.1.chr7.g00906.t01#6    mNeoVis.evergladensis.2.1.chr7.g00961.t01#2    mNeoVis.vulgivaga.1.1.chr7.g00964.t01#4    mNeoVis.2.1.1.chr7_hap1.g00968.t01#5   mNeoVis.vulgivaga.2.1.chr7.g00966.t01#1       4499   125256927:   mNeoVis.3.1.1.chr7.g00907.t01#6    mNeoVis.2.1.1.chr7_hap1.g00969.t01#5    mNeoVis.evergladensis.2.1.chr7.g00962.t01#2    mNeoVis.evergladensis.1.1.chr7.g00968.t01#3    mNeoVis.vulgivaga.2.1.chr7.g00967.t01#1   mNeoVis.vulgivaga.1.1.chr7.g00965.t01#4       4500   125256928:   mNeoVis.evergladensis.2.1.chr1.g00287.t01#2    mNeoVis.2.1.1.chr1_hap1.g00288.t01#5    mNeoVis.evergladensis.1.1.chr1.g00296.t01#3    mNeoVis.vulgivaga.1.1.chr1.g00287.t01#4    mNeoVis.vulgivaga.2.1.chr1.g00291.t01#1   mNeoVis.3.1.1.chr1.g00293.t01#6       4501   125256929:   mNeoVis.vulgivaga.2.1.chr5.g00344.t01#1    mNeoVis.3.1.1.chr5.g00331.t01#6    mNeoVis.2.1.1.chr2_hap1.g00341.t01#5    mNeoVis.evergladensis.2.1.chr5.g00342.t01#2    mNeoVis.evergladensis.1.1.chr5.g00340.t01#3   mNeoVis.vulgivaga.1.1.chr5.g00335.t01#4       4502   125256930:   mNeoVis.3.1.1.chr11.g00394.t01#6    mNeoVis.evergladensis.2.1.chr11.g00425.t01#2    mNeoVis.evergladensis.1.1.chr11.g00429.t01#3    mNeoVis.2.1.1.chr11_hap1.g00434.t01#5    mNeoVis.vulgivaga.2.1.chr11.g00433.t01#1   mNeoVis.vulgivaga.1.1.chr11.g00422.t01#4       4503   125256931:  mNeoVis.evergladensis.2.1.chr11.g00426.t01#2             4504   125256932:   mNeoVis.evergladensis.1.1.chr7.g00969.t01#3    mNeoVis.3.1.1.chr7.g00909.t01#6    mNeoVis.vulgivaga.2.1.chr7.g00968.t01#1    mNeoVis.2.1.1.chr7_hap1.g00970.t01#5   mNeoVis.evergladensis.2.1.chr7.g00963.t01#2       4505   125256933:   mNeoVis.evergladensis.1.1.chr1.g00297.t01#3    mNeoVis.2.1.1.chr1_hap1.g00289.t01#5    mNeoVis.evergladensis.2.1.chr1.g00288.t01#2    mNeoVis.vulgivaga.1.1.chr1.g00288.t01#4    mNeoVis.vulgivaga.2.1.chr1.g00292.t01#1   mNeoVis.3.1.1.chr1.g00294.t01#6       4506   125256934:   mNeoVis.2.1.1.chr7_hap1.g00971.t01#5    mNeoVis.evergladensis.1.1.chr7.g00970.t01#3    mNeoVis.vulgivaga.2.1.chr7.g00969.t01#1   mNeoVis.evergladensis.2.1.chr7.g00964.t01#2       4507   125256935:   mNeoVis.evergladensis.1.1.chr7.g00020.t01#3    mNeoVis.vulgivaga.2.1.chr7.g00019.t01#1    mNeoVis.vulgivaga.1.1.chr7.g00021.t01#4    mNeoVis.2.1.1.chr7_hap1.g00019.t01#5   mNeoVis.evergladensis.2.1.chr7.g00020.t01#2       4508   125256936:   mNeoVis.evergladensis.2.1.chr7.g00965.t01#2    mNeoVis.3.1.1.chr7.g00911.t01#6    mNeoVis.evergladensis.1.1.chr7.g00971.t01#3    mNeoVis.vulgivaga.1.1.chr7.g00966.t01#4    mNeoVis.vulgivaga.2.1.chr7.g00970.t01#1   mNeoVis.2.1.1.chr7_hap1.g00972.t01#5       4509   125256937:   mNeoVis.vulgivaga.2.1.chr7.g00971.t01#1   mNeoVis.3.1.1.chr7.g00912.t01#6             4510   125256938:   mNeoVis.evergladensis.2.1.chr1.g00289.t01#2    mNeoVis.vulgivaga.1.1.chr1.g00289.t01#4    mNeoVis.vulgivaga.2.1.chr1.g00293.t01#1    mNeoVis.3.1.1.chr1.g00295.t01#6   mNeoVis.2.1.1.chr1_hap1.g00290.t01#5       4511   125256939:   mNeoVis.3.1.1.chr1.g00296.t01#6    mNeoVis.2.1.1.chr1_hap1.g00291.t01#5    mNeoVis.vulgivaga.2.1.chr1.g00294.t01#1    mNeoVis.evergladensis.2.1.chr1.g00290.t01#2    mNeoVis.vulgivaga.1.1.chr1.g00290.t01#4   mNeoVis.evergladensis.1.1.chr1.g00299.t01#3       4512   125256940:   mNeoVis.evergladensis.1.1.chr7.g00972.t01#3    mNeoVis.vulgivaga.2.1.chr7.g00972.t01#1    mNeoVis.vulgivaga.1.1.chr7.g00967.t01#4    mNeoVis.evergladensis.2.1.chr7.g00966.t01#2    mNeoVis.2.1.1.chr7_hap1.g00973.t01#5   mNeoVis.3.1.1.chr7.g00913.t01#6       4513   125256941:   mNeoVis.3.1.1.chr7.g00914.t01#6    mNeoVis.2.1.1.chr7_hap1.g00974.t01#5    mNeoVis.vulgivaga.2.1.chr7.g00973.t01#1    mNeoVis.vulgivaga.1.1.chr7.g00968.t01#4    mNeoVis.evergladensis.1.1.chr7.g00973.t01#3   mNeoVis.evergladensis.2.1.chr7.g00967.t01#2       4514   125256942:   mNeoVis.evergladensis.1.1.chr1.g00300.t01#3    mNeoVis.evergladensis.2.1.chr1.g00291.t01#2    mNeoVis.3.1.1.chr1.g00297.t01#6    mNeoVis.2.1.1.chr1_hap1.g00292.t01#5    mNeoVis.vulgivaga.2.1.chr1.g00295.t01#1   mNeoVis.vulgivaga.1.1.chr1.g00291.t01#4       4515   125256943:   mNeoVis.evergladensis.2.1.chr7.g00968.t01#2    mNeoVis.vulgivaga.1.1.chr7.g00969.t01#4    mNeoVis.vulgivaga.2.1.chr7.g00974.t01#1    mNeoVis.evergladensis.1.1.chr7.g00974.t01#3    mNeoVis.3.1.1.chr7.g00915.t01#6   mNeoVis.2.1.1.chr7_hap1.g00975.t01#5       4516   125256944:   mNeoVis.evergladensis.2.1.chr7.g00969.t01#2    mNeoVis.2.1.1.chr7_hap1.g00976.t01#5    mNeoVis.evergladensis.1.1.chr7.g00975.t01#3    mNeoVis.vulgivaga.2.1.chr7.g00975.t01#1   mNeoVis.vulgivaga.1.1.chr7.g00970.t01#4       4517   125256945:   mNeoVis.vulgivaga.2.1.chr7.g00976.t01#1    mNeoVis.evergladensis.2.1.chr7.g00970.t01#2    mNeoVis.evergladensis.1.1.chr7.g00976.t01#3    mNeoVis.vulgivaga.1.1.chr7.g00971.t01#4    mNeoVis.2.1.1.chr7_hap1.g00977.t01#5   mNeoVis.3.1.1.chr7.g00916.t01#6       4518   125256946:   mNeoVis.3.1.1.chr7.g00917.t01#6    mNeoVis.2.1.1.chr7_hap1.g00978.t01#5    mNeoVis.evergladensis.2.1.chr7.g00971.t01#2    mNeoVis.evergladensis.1.1.chr7.g00977.t01#3    mNeoVis.vulgivaga.2.1.chr7.g00977.t01#1   mNeoVis.vulgivaga.1.1.chr7.g00972.t01#4       4519   125256947:   mNeoVis.3.1.1.chr7.g00918.t01#6    mNeoVis.vulgivaga.2.1.chr7.g00978.t01#1   mNeoVis.evergladensis.1.1.chr7.g00978.t01#3         4520   125256948:   mNeoVis.2.1.1.chr7_hap1.g00980.t01#5    mNeoVis.vulgivaga.2.1.chr7.g00979.t01#1    mNeoVis.evergladensis.2.1.chr7.g00973.t01#2    mNeoVis.3.1.1.chr7.g00919.t01#6    mNeoVis.evergladensis.1.1.chr7.g00979.t01#3   mNeoVis.vulgivaga.1.1.chr7.g00974.t01#4       4521   125256949:   mNeoVis.vulgivaga.2.1.chr1.g00296.t01#1    mNeoVis.3.1.1.chr1.g00298.t01#6    mNeoVis.2.1.1.chr1_hap1.g00293.t01#5    mNeoVis.evergladensis.2.1.chr1.g00292.t01#2    mNeoVis.evergladensis.1.1.chr1.g00301.t01#3   mNeoVis.vulgivaga.1.1.chr1.g00292.t01#4       4522   125256950:   mNeoVis.vulgivaga.1.1.chr1.g00293.t01#4    mNeoVis.evergladensis.2.1.chr1.g00293.t01#2    mNeoVis.vulgivaga.2.1.chr1.g00297.t01#1    mNeoVis.3.1.1.chr1.g00299.t01#6    mNeoVis.2.1.1.chr1_hap1.g00294.t01#5   mNeoVis.evergladensis.1.1.chr1.g00302.t01#3       4523   125256951:   mNeoVis.evergladensis.2.1.chr1.g00294.t01#2    mNeoVis.vulgivaga.2.1.chr1.g00298.t01#1    mNeoVis.3.1.1.chr1.g00300.t01#6    mNeoVis.evergladensis.1.1.chr1.g00303.t01#3    mNeoVis.2.1.1.chr1_hap1.g00295.t01#5   mNeoVis.vulgivaga.1.1.chr1.g00294.t01#4       4524   125256952:   mNeoVis.3.1.1.chr7.g00920.t01#6    mNeoVis.evergladensis.1.1.chr7.g00980.t01#3    mNeoVis.evergladensis.2.1.chr7.g00974.t01#2    mNeoVis.vulgivaga.1.1.chr7.g00975.t01#4    mNeoVis.vulgivaga.2.1.chr7.g00980.t01#1   mNeoVis.2.1.1.chr7_hap1.g00981.t01#5       4525   125256953:   mNeoVis.evergladensis.1.1.chr10.g00056.t01#3    mNeoVis.2.1.1.chr9_hap1.g00053.t01#5    mNeoVis.2.1.1.chr1_hap1.g00204.t01#5    mNeoVis.vulgivaga.2.1.chr10.g00056.t01#1    mNeoVis.3.1.1.chr10.g00057.t01#6    mNeoVis.evergladensis.2.1.chr1.g00199.t01#2    mNeoVis.evergladensis.2.1.chr10.g00059.t01#2    mNeoVis.vulgivaga.1.1.chr1.g00198.t01#4    mNeoVis.vulgivaga.2.1.chr1.g00202.t01#1      4526   125256954:   mNeoVis.vulgivaga.1.1.chr7.g00022.t01#4    mNeoVis.evergladensis.1.1.chr7.g00021.t01#3    mNeoVis.2.1.1.chr7_hap1.g00020.t01#5    mNeoVis.vulgivaga.2.1.chr7.g00020.t01#1   mNeoVis.evergladensis.2.1.chr7.g00021.t01#2       4527   125256955:   mNeoVis.evergladensis.2.1.chr1.g00200.t01#2    mNeoVis.vulgivaga.1.1.chr1.g00199.t01#4    mNeoVis.vulgivaga.2.1.chr10.g00011.t01#1    mNeoVis.3.1.1.chr1.g00206.t01#6    mNeoVis.2.1.1.chr9_hap1.g00009.t01#5    mNeoVis.vulgivaga.1.1.chr10.g00009.t01#4    mNeoVis.vulgivaga.2.1.chr1.g00203.t01#1    mNeoVis.2.1.1.chr1_hap1.g00205.t01#5    mNeoVis.evergladensis.1.1.chr10.g00009.t01#3      4528   125256956:   mNeoVis.2.1.1.chr1_hap1.g00206.t01#5    mNeoVis.vulgivaga.2.1.chr1.g00204.t01#1    mNeoVis.vulgivaga.1.1.chr1.g00200.t01#4    mNeoVis.evergladensis.2.1.chr1.g00201.t01#2    mNeoVis.3.1.1.chr1.g00207.t01#6   mNeoVis.evergladensis.1.1.chr1.g00215.t01#3       4529   125256957:   mNeoVis.vulgivaga.1.1.chr4.g00487.t01#4    mNeoVis.2.1.1.chr6_hap1.g00484.t01#5    mNeoVis.evergladensis.2.1.chr4.g00489.t01#2    mNeoVis.evergladensis.1.1.chr4.g00494.t01#3    mNeoVis.vulgivaga.2.1.chr4.g00497.t01#1   mNeoVis.3.1.1.chr4.g00488.t01#6       4530   125256958:   mNeoVis.vulgivaga.1.1.chr4.g00485.t01#4    mNeoVis.2.1.1.chr6_hap1.g00481.t01#5    mNeoVis.3.1.1.chr4.g00484.t01#6    mNeoVis.evergladensis.1.1.chr4.g00492.t01#3    mNeoVis.evergladensis.2.1.chr4.g00487.t01#2   mNeoVis.vulgivaga.2.1.chr4.g00495.t01#1       4531   125256959:   mNeoVis.3.1.1.chr7.g00826.t01#6    mNeoVis.evergladensis.1.1.chr7.g00884.t01#3    mNeoVis.vulgivaga.2.1.chr7.g00879.t01#1    mNeoVis.vulgivaga.1.1.chr7.g00884.t01#4   mNeoVis.2.1.1.chr7_hap1.g00885.t01#5       4532   125256960:   mNeoVis.evergladensis.2.1.chr7.g00879.t01#2   mNeoVis.vulgivaga.2.1.chr7.g00883.t01#1             4533   125256961:   mNeoVis.2.1.1.chr1_hap1.g00207.t01#5    mNeoVis.evergladensis.2.1.chr1.g00202.t01#2    mNeoVis.vulgivaga.1.1.chr1.g00201.t01#4    mNeoVis.evergladensis.1.1.chr1.g00216.t01#3    mNeoVis.3.1.1.chr1.g00208.t01#6   mNeoVis.vulgivaga.2.1.chr1.g00205.t01#1       4534   125256962:  mNeoVis.vulgivaga.2.1.chr1.g00206.t01#1               4535   125256963:   mNeoVis.3.1.1.chr7.g00823.t01#6    mNeoVis.vulgivaga.1.1.chr7.g00881.t01#4    mNeoVis.2.1.1.chr7_hap1.g00882.t01#5    mNeoVis.evergladensis.2.1.chr7.g00876.t01#2    mNeoVis.vulgivaga.2.1.chr7.g00880.t01#1   mNeoVis.evergladensis.1.1.chr7.g00881.t01#3       4536   125256964:   mNeoVis.3.1.1.chr10.g00217.t01#6    mNeoVis.evergladensis.1.1.chr10.g00223.t01#3    mNeoVis.2.1.1.chr9_hap1.g00224.t01#5    mNeoVis.vulgivaga.1.1.chr10.g00211.t01#4    mNeoVis.vulgivaga.2.1.chr10.g00218.t01#1   mNeoVis.evergladensis.2.1.chr10.g00221.t01#2       4537   125256965:   mNeoVis.vulgivaga.2.1.chr7.g00881.t01#1    mNeoVis.evergladensis.1.1.chr7.g00882.t01#3    mNeoVis.evergladensis.2.1.chr7.g00877.t01#2    mNeoVis.3.1.1.chr7.g00824.t01#6    mNeoVis.vulgivaga.1.1.chr7.g00882.t01#4   mNeoVis.2.1.1.chr7_hap1.g00883.t01#5       4538   125256966:   mNeoVis.3.1.1.chr1.g00210.t01#6    mNeoVis.evergladensis.1.1.chr1.g00218.t01#3    mNeoVis.evergladensis.2.1.chr1.g00204.t01#2    mNeoVis.vulgivaga.1.1.chr1.g00203.t01#4    mNeoVis.2.1.1.chr1_hap1.g00209.t01#5   mNeoVis.vulgivaga.2.1.chr1.g00207.t01#1       4539   125256967:   mNeoVis.vulgivaga.2.1.chr7.g00882.t01#1    mNeoVis.3.1.1.chr7.g00825.t01#6    mNeoVis.evergladensis.1.1.chr7.g00883.t01#3    mNeoVis.evergladensis.2.1.chr7.g00878.t01#2    mNeoVis.vulgivaga.1.1.chr7.g00883.t01#4   mNeoVis.2.1.1.chr7_hap1.g00884.t01#5       4540   125256968:   mNeoVis.evergladensis.2.1.chr1.g00205.t01#2    mNeoVis.vulgivaga.1.1.chr1.g00204.t01#4    mNeoVis.evergladensis.1.1.chr1.g00219.t01#3    mNeoVis.2.1.1.chr1_hap1.g00210.t01#5    mNeoVis.vulgivaga.2.1.chr1.g00208.t01#1   mNeoVis.3.1.1.chr1.g00211.t01#6       4541   125256969:   mNeoVis.vulgivaga.2.1.chr7.g00884.t01#1    mNeoVis.evergladensis.1.1.chr7.g00885.t01#3    mNeoVis.3.1.1.chr7.g00827.t01#6    mNeoVis.2.1.1.chr7_hap1.g00886.t01#5    mNeoVis.evergladensis.2.1.chr7.g00880.t01#2   mNeoVis.vulgivaga.1.1.chr7.g00885.t01#4       4542   125256970:   mNeoVis.vulgivaga.2.1.chr1.g00209.t01#1    mNeoVis.evergladensis.1.1.chr1.g00220.t01#3    mNeoVis.3.1.1.chr1.g00212.t01#6    mNeoVis.evergladensis.2.1.chr1.g00206.t01#2    mNeoVis.vulgivaga.1.1.chr1.g00205.t01#4   mNeoVis.2.1.1.chr1_hap1.g00211.t01#5       4543   125256971:   mNeoVis.2.1.1.chr1_hap1.g00213.t01#5    mNeoVis.3.1.1.chr1.g00214.t01#6    mNeoVis.vulgivaga.2.1.chr1.g00212.t01#1    mNeoVis.evergladensis.2.1.chr1.g00209.t01#2   mNeoVis.vulgivaga.1.1.chr1.g00207.t01#4       4544   125256972:   mNeoVis.vulgivaga.1.1.chr1.g00206.t01#4    mNeoVis.vulgivaga.2.1.chr1.g00211.t01#1    mNeoVis.evergladensis.2.1.chr1.g00207.t01#2    mNeoVis.evergladensis.1.1.chr1.g00221.t01#3   mNeoVis.3.1.1.chr1.g00213.t01#6       4545   125256973:   mNeoVis.2.1.1.chr1_hap1.g00212.t01#5    mNeoVis.evergladensis.2.1.chr1.g00208.t01#2   mNeoVis.vulgivaga.2.1.chr1.g00210.t01#1           4546   125256974:   mNeoVis.evergladensis.2.1.chr7.g00881.t01#2    mNeoVis.vulgivaga.2.1.chr7.g00885.t01#1    mNeoVis.evergladensis.1.1.chr7.g00886.t01#3    mNeoVis.2.1.1.chr7_hap1.g00887.t01#5    mNeoVis.3.1.1.chr7.g00828.t01#6   mNeoVis.vulgivaga.1.1.chr7.g00886.t01#4       4547   125256975:   mNeoVis.evergladensis.1.1.chr15.g00511.t01#3    mNeoVis.3.1.1.chr15.g00095.t01#6    mNeoVis.vulgivaga.2.1.chr15.g00105.t01#1    mNeoVis.evergladensis.2.1.chr15.g00102.t01#2    mNeoVis.2.1.1.chr15_hap1.g00101.t01#5   mNeoVis.vulgivaga.1.1.chr15.g00102.t01#4       4548   125256976:   mNeoVis.vulgivaga.2.1.chr1.g00213.t01#1    mNeoVis.evergladensis.2.1.chr1.g00210.t01#2    mNeoVis.vulgivaga.1.1.chr1.g00208.t01#4    mNeoVis.3.1.1.chr1.g00215.t01#6    mNeoVis.2.1.1.chr1_hap1.g00214.t01#5   mNeoVis.evergladensis.1.1.chr1.g00222.t01#3       4549   125256977:   mNeoVis.vulgivaga.1.1.chr1.g00209.t01#4    mNeoVis.evergladensis.1.1.chr1.g00223.t01#3    mNeoVis.vulgivaga.2.1.chr1.g00214.t01#1    mNeoVis.evergladensis.2.1.chr1.g00211.t01#2    mNeoVis.3.1.1.chr1.g00216.t01#6   mNeoVis.2.1.1.chr1_hap1.g00215.t01#5       4550   125256978:   mNeoVis.evergladensis.1.1.chr7.g00887.t01#3    mNeoVis.vulgivaga.2.1.chr7.g00886.t01#1    mNeoVis.3.1.1.chr7.g00830.t01#6    mNeoVis.vulgivaga.1.1.chr7.g00887.t01#4    mNeoVis.evergladensis.2.1.chr7.g00882.t01#2   mNeoVis.2.1.1.chr7_hap1.g00889.t01#5       4551   125256979:   mNeoVis.vulgivaga.2.1.chr7.g00021.t01#1    mNeoVis.evergladensis.2.1.chr7.g00022.t01#2    mNeoVis.2.1.1.chr7_hap1.g00021.t01#5    mNeoVis.evergladensis.1.1.chr7.g00022.t01#3   mNeoVis.vulgivaga.1.1.chr7.g00023.t01#4       4552   125256980:   mNeoVis.vulgivaga.2.1.chr1.g00215.t01#1    mNeoVis.evergladensis.2.1.chr1.g00212.t01#2    mNeoVis.vulgivaga.1.1.chr1.g00210.t01#4    mNeoVis.3.1.1.chr1.g00217.t01#6    mNeoVis.evergladensis.1.1.chr1.g00224.t01#3   mNeoVis.2.1.1.chr1_hap1.g00216.t01#5       4553   125256981:   mNeoVis.evergladensis.2.1.chr7.g00883.t01#2    mNeoVis.vulgivaga.1.1.chr7.g00888.t01#4    mNeoVis.2.1.1.chr7_hap1.g00890.t01#5    mNeoVis.3.1.1.chr7.g00831.t01#6    mNeoVis.evergladensis.1.1.chr7.g00888.t01#3   mNeoVis.vulgivaga.2.1.chr7.g00887.t01#1       4554   125256982:   mNeoVis.evergladensis.2.1.chr7.g00884.t01#2    mNeoVis.2.1.1.chr7_hap1.g00891.t01#5    mNeoVis.vulgivaga.1.1.chr7.g00889.t01#4    mNeoVis.vulgivaga.2.1.chr7.g00888.t01#1    mNeoVis.evergladensis.1.1.chr7.g00889.t01#3   mNeoVis.3.1.1.chr7.g00832.t01#6       4555   125256983:   mNeoVis.evergladensis.1.1.chr7.g00890.t01#3    mNeoVis.vulgivaga.2.1.chr7.g00889.t01#1    mNeoVis.evergladensis.2.1.chr11.g00590.t01#2    mNeoVis.vulgivaga.1.1.chr7.g00890.t01#4    mNeoVis.vulgivaga.1.1.chr11.g00584.t01#4    mNeoVis.2.1.1.chr7_hap1.g00892.t01#5    mNeoVis.evergladensis.2.1.chr7.g00885.t01#2   mNeoVis.3.1.1.chr7.g00833.t01#6     4556   125256984:   mNeoVis.vulgivaga.1.1.chr7.g00891.t01#4    mNeoVis.evergladensis.1.1.chr7.g00891.t01#3    mNeoVis.vulgivaga.2.1.chr7.g00890.t01#1    mNeoVis.evergladensis.2.1.chr7.g00886.t01#2    mNeoVis.3.1.1.chr7.g00834.t01#6   mNeoVis.2.1.1.chr7_hap1.g00893.t01#5       4557   125256985:   mNeoVis.3.1.1.chr1.g00218.t01#6    mNeoVis.2.1.1.chr1_hap1.g00217.t01#5    mNeoVis.evergladensis.1.1.chr1.g00225.t01#3    mNeoVis.vulgivaga.2.1.chr1.g00216.t01#1    mNeoVis.vulgivaga.1.1.chr1.g00211.t01#4   mNeoVis.evergladensis.2.1.chr1.g00213.t01#2       4558   125256986:   mNeoVis.evergladensis.1.1.chr7.g00892.t01#3    mNeoVis.3.1.1.chr7.g00835.t01#6    mNeoVis.vulgivaga.2.1.chr7.g00891.t01#1    mNeoVis.evergladensis.2.1.chr7.g00887.t01#2    mNeoVis.vulgivaga.1.1.chr7.g00892.t01#4   mNeoVis.2.1.1.chr7_hap1.g00894.t01#5       4559   125256987:   mNeoVis.evergladensis.1.1.chr1.g00226.t01#3    mNeoVis.vulgivaga.1.1.chr1.g00212.t01#4    mNeoVis.3.1.1.chr1.g00219.t01#6    mNeoVis.evergladensis.2.1.chr1.g00214.t01#2    mNeoVis.2.1.1.chr1_hap1.g00218.t01#5   mNeoVis.vulgivaga.2.1.chr1.g00217.t01#1       4560   125256988:   mNeoVis.vulgivaga.2.1.chr1.g00218.t01#1    mNeoVis.2.1.1.chr1_hap1.g00219.t01#5    mNeoVis.evergladensis.2.1.chr1.g00215.t01#2    mNeoVis.evergladensis.1.1.chr1.g00227.t01#3   mNeoVis.vulgivaga.1.1.chr1.g00213.t01#4       4561   125256989:   mNeoVis.evergladensis.2.1.chr7.g00888.t01#2    mNeoVis.evergladensis.1.1.chr7.g00893.t01#3    mNeoVis.vulgivaga.2.1.chr7.g00892.t01#1    mNeoVis.2.1.1.chr7_hap1.g00895.t01#5   mNeoVis.vulgivaga.1.1.chr7.g00893.t01#4       4562   125256990:   mNeoVis.vulgivaga.1.1.chr15.g00103.t01#4    mNeoVis.3.1.1.chr15.g00096.t01#6    mNeoVis.evergladensis.1.1.chr15.g00510.t01#3    mNeoVis.vulgivaga.2.1.chr15.g00106.t01#1    mNeoVis.evergladensis.2.1.chr15.g00103.t01#2   mNeoVis.2.1.1.chr15_hap1.g00102.t01#5       4563   125256991:   mNeoVis.vulgivaga.1.1.chr1.g00214.t01#4    mNeoVis.evergladensis.2.1.chr1.g00216.t01#2    mNeoVis.evergladensis.1.1.chr1.g00228.t01#3    mNeoVis.2.1.1.chr1_hap1.g00220.t01#5    mNeoVis.vulgivaga.2.1.chr1.g00219.t01#1   mNeoVis.3.1.1.chr1.g00220.t01#6       4564   125256992:   mNeoVis.evergladensis.1.1.chr7.g00894.t01#3    mNeoVis.vulgivaga.2.1.chr7.g00893.t01#1    mNeoVis.2.1.1.chr7_hap1.g00896.t01#5    mNeoVis.3.1.1.chr7.g00837.t01#6    mNeoVis.vulgivaga.1.1.chr7.g00894.t01#4   mNeoVis.evergladensis.2.1.chr7.g00889.t01#2       4565   125256993:   mNeoVis.3.1.1.chr7.g00019.t01#6    mNeoVis.2.1.1.chr7_hap1.g00022.t01#5    mNeoVis.vulgivaga.2.1.chr7.g00022.t01#1    mNeoVis.evergladensis.2.1.chr7.g00023.t01#2   mNeoVis.evergladensis.1.1.chr7.g00023.t01#3       4566   125256994:   mNeoVis.vulgivaga.2.1.chr9.g00491.t01#1    mNeoVis.evergladensis.2.1.chr8.g00489.t01#2    mNeoVis.3.1.1.chr9.g00471.t01#6    mNeoVis.vulgivaga.1.1.chr9.g00494.t01#4   mNeoVis.evergladensis.1.1.chr9.g00500.t01#3       4567   125256995:   mNeoVis.2.1.1.chr1_hap1.g00221.t01#5    mNeoVis.3.1.1.chr1.g00221.t01#6    mNeoVis.evergladensis.1.1.chr1.g00229.t01#3    mNeoVis.vulgivaga.2.1.chr1.g00220.t01#1    mNeoVis.vulgivaga.1.1.chr1.g00215.t01#4   mNeoVis.evergladensis.2.1.chr1.g00218.t01#2       4568   125256996:   mNeoVis.evergladensis.2.1.chr1.g00219.t01#2    mNeoVis.vulgivaga.1.1.chr1.g00216.t01#4    mNeoVis.evergladensis.1.1.chr1.g00230.t01#3    mNeoVis.vulgivaga.2.1.chr1.g00221.t01#1    mNeoVis.2.1.1.chr1_hap1.g00222.t01#5   mNeoVis.3.1.1.chr1.g00222.t01#6       4569   125256997:   mNeoVis.evergladensis.1.1.chr7.g00896.t01#3    mNeoVis.2.1.1.chr7_hap1.g00899.t01#5    mNeoVis.vulgivaga.2.1.chr7.g00895.t01#1    mNeoVis.3.1.1.chr7.g00839.t01#6   mNeoVis.evergladensis.2.1.chr7.g00892.t01#2       4570   125256998:   mNeoVis.vulgivaga.2.1.chr7.g00896.t01#1    mNeoVis.3.1.1.chr7.g00840.t01#6    mNeoVis.evergladensis.2.1.chr7.g00891.t01#2    mNeoVis.evergladensis.1.1.chr7.g00897.t01#3    mNeoVis.2.1.1.chr7_hap1.g00898.t01#5   mNeoVis.vulgivaga.1.1.chr7.g00896.t01#4       4571   125256999:   mNeoVis.vulgivaga.1.1.chr7.g00897.t01#4    mNeoVis.vulgivaga.2.1.chr7.g00897.t01#1    mNeoVis.3.1.1.chr7.g00841.t01#6    mNeoVis.2.1.1.chr7_hap1.g00900.t01#5    mNeoVis.evergladensis.2.1.chr7.g00893.t01#2   mNeoVis.evergladensis.1.1.chr7.g00898.t01#3       4572   125257000:   mNeoVis.evergladensis.2.1.chr7.g00024.t01#2    mNeoVis.2.1.1.chr7_hap1.g00023.t01#5    mNeoVis.vulgivaga.2.1.chr7.g00023.t01#1    mNeoVis.vulgivaga.1.1.chr7.g00025.t01#4   mNeoVis.evergladensis.1.1.chr7.g00024.t01#3       4573   125257001:   mNeoVis.vulgivaga.1.1.chr1.g00217.t01#4    mNeoVis.evergladensis.2.1.chr1.g00220.t01#2    mNeoVis.2.1.1.chr1_hap1.g00223.t01#5    mNeoVis.vulgivaga.2.1.chr1.g00222.t01#1    mNeoVis.3.1.1.chr1.g00223.t01#6   mNeoVis.evergladensis.1.1.chr1.g00231.t01#3       4574   125257002:   mNeoVis.3.1.1.chr1.g00224.t01#6    mNeoVis.2.1.1.chr1_hap1.g00224.t01#5    mNeoVis.vulgivaga.1.1.chr1.g00218.t01#4    mNeoVis.evergladensis.2.1.chr1.g00221.t01#2    mNeoVis.evergladensis.1.1.chr1.g00232.t01#3   mNeoVis.vulgivaga.2.1.chr1.g00223.t01#1       4575   125257003:   mNeoVis.evergladensis.2.1.chr1.g00222.t01#2    mNeoVis.2.1.1.chr1_hap1.g00225.t01#5    mNeoVis.vulgivaga.2.1.chr1.g00224.t01#1    mNeoVis.evergladensis.1.1.chr1.g00233.t01#3    mNeoVis.3.1.1.chr1.g00225.t01#6   mNeoVis.vulgivaga.1.1.chr1.g00219.t01#4       4576   125257004:   mNeoVis.evergladensis.1.1.chr7.g00899.t01#3    mNeoVis.3.1.1.chr7.g00842.t01#6    mNeoVis.evergladensis.2.1.chr7.g00894.t01#2    mNeoVis.vulgivaga.1.1.chr7.g00898.t01#4    mNeoVis.2.1.1.chr7_hap1.g00901.t01#5   mNeoVis.vulgivaga.2.1.chr7.g00898.t01#1       4577   125257005:   mNeoVis.3.1.1.chr7.g00843.t01#6    mNeoVis.vulgivaga.2.1.chr7.g00899.t01#1    mNeoVis.2.1.1.chr7_hap1.g00902.t01#5    mNeoVis.evergladensis.1.1.chr7.g00900.t01#3    mNeoVis.evergladensis.2.1.chr7.g00895.t01#2   mNeoVis.vulgivaga.1.1.chr7.g00899.t01#4       4578   125257006:   mNeoVis.vulgivaga.1.1.chr7.g00900.t01#4    mNeoVis.evergladensis.2.1.chr7.g00896.t01#2    mNeoVis.3.1.1.chr7.g00844.t01#6    mNeoVis.2.1.1.chr7_hap1.g00903.t01#5    mNeoVis.vulgivaga.2.1.chr7.g00900.t01#1   mNeoVis.evergladensis.1.1.chr7.g00901.t01#3       4579   125257007:   mNeoVis.evergladensis.1.1.chr1.g00235.t01#3    mNeoVis.evergladensis.2.1.chr1.g00224.t01#2    mNeoVis.vulgivaga.1.1.chr1.g00221.t01#4    mNeoVis.vulgivaga.2.1.chr1.g00226.t01#1    mNeoVis.3.1.1.chr1.g00227.t01#6   mNeoVis.2.1.1.chr1_hap1.g00227.t01#5       4580   125257008:   mNeoVis.vulgivaga.2.1.chr1.g00225.t01#1    mNeoVis.vulgivaga.1.1.chr1.g00220.t01#4    mNeoVis.evergladensis.1.1.chr1.g00234.t01#3    mNeoVis.3.1.1.chr1.g00226.t01#6    mNeoVis.2.1.1.chr1_hap1.g00226.t01#5   mNeoVis.evergladensis.2.1.chr1.g00223.t01#2       4581   125257009:   mNeoVis.evergladensis.1.1.chr12.g00728.t01#3    mNeoVis.evergladensis.2.1.chr12.g00719.t01#2    mNeoVis.2.1.1.chr12_hap1.g00792.t01#5    mNeoVis.3.1.1.chr12.g00723.t01#6    mNeoVis.vulgivaga.2.1.chr12.g00748.t01#1   mNeoVis.vulgivaga.1.1.chr12.g00743.t01#4       4582   125257010:   mNeoVis.evergladensis.2.1.chr12.g00297.t01#2    mNeoVis.vulgivaga.2.1.chr12.g00305.t01#1    mNeoVis.2.1.1.chr12_hap1.g00339.t01#5    mNeoVis.3.1.1.chr12.g00301.t01#6    mNeoVis.vulgivaga.1.1.chr12.g00307.t01#4   mNeoVis.evergladensis.1.1.chr12.g00302.t01#3       4583   125257011:   mNeoVis.evergladensis.2.1.chr4.g00795.t01#2    mNeoVis.vulgivaga.2.1.chr4.g00814.t01#1    mNeoVis.vulgivaga.1.1.chr4.g00810.t01#4    mNeoVis.evergladensis.1.1.chr4.g00808.t01#3   mNeoVis.2.1.1.chr6_hap1.g00792.t01#5       4584   125257012:   mNeoVis.2.1.1.chr7_hap1.g00904.t01#5    mNeoVis.evergladensis.1.1.chr7.g00902.t01#3    mNeoVis.evergladensis.2.1.chr7.g00897.t01#2    mNeoVis.3.1.1.chr7.g00845.t01#6    mNeoVis.vulgivaga.1.1.chr7.g00901.t01#4   mNeoVis.vulgivaga.2.1.chr7.g00901.t01#1       4585   125257013:   mNeoVis.vulgivaga.1.1.chr7.g00902.t01#4    mNeoVis.2.1.1.chr7_hap1.g00905.t01#5    mNeoVis.evergladensis.2.1.chr7.g00898.t01#2    mNeoVis.vulgivaga.2.1.chr7.g00902.t01#1    mNeoVis.evergladensis.1.1.chr7.g00903.t01#3   mNeoVis.3.1.1.chr7.g00846.t01#6       4586   125257014:   mNeoVis.evergladensis.1.1.chr1.g00237.t01#3    mNeoVis.3.1.1.chr1.g00230.t01#6    mNeoVis.vulgivaga.1.1.chr1.g00223.t01#4    mNeoVis.2.1.1.chr1_hap1.g00230.t01#5    mNeoVis.evergladensis.2.1.chr1.g00227.t01#2   mNeoVis.vulgivaga.2.1.chr1.g00228.t01#1       4587   125257015:   mNeoVis.2.1.1.chr7_hap1.g00906.t01#5    mNeoVis.evergladensis.2.1.chr7.g00899.t01#2    mNeoVis.evergladensis.1.1.chr7.g00904.t01#3    mNeoVis.vulgivaga.1.1.chr7.g00903.t01#4    mNeoVis.3.1.1.chr7.g00847.t01#6   mNeoVis.vulgivaga.2.1.chr7.g00903.t01#1       4588   125257016:   mNeoVis.3.1.1.chr1.g00231.t01#6    mNeoVis.evergladensis.1.1.chr1.g00238.t01#3    mNeoVis.vulgivaga.2.1.chr1.g00229.t01#1    mNeoVis.2.1.1.chr1_hap1.g00231.t01#5    mNeoVis.vulgivaga.1.1.chr1.g00225.t01#4   mNeoVis.evergladensis.2.1.chr1.g00228.t01#2       4589   125257017:   mNeoVis.vulgivaga.2.1.chr1.g00230.t01#1    mNeoVis.2.1.1.chr1_hap1.g00232.t01#5    mNeoVis.3.1.1.chr1.g00232.t01#6    mNeoVis.evergladensis.2.1.chr1.g00229.t01#2    mNeoVis.vulgivaga.1.1.chr1.g00226.t01#4   mNeoVis.evergladensis.1.1.chr1.g00239.t01#3       4590   125257018:   mNeoVis.3.1.1.chr7.g00848.t01#6    mNeoVis.2.1.1.chr7_hap1.g00907.t01#5    mNeoVis.evergladensis.1.1.chr7.g00905.t01#3    mNeoVis.vulgivaga.2.1.chr7.g00904.t01#1    mNeoVis.evergladensis.2.1.chr7.g00900.t01#2   mNeoVis.vulgivaga.1.1.chr7.g00904.t01#4       4591   125257019:   mNeoVis.vulgivaga.2.1.chr1.g00232.t01#1    mNeoVis.evergladensis.2.1.chr1.g00230.t01#2    mNeoVis.vulgivaga.1.1.chr1.g00227.t01#4    mNeoVis.vulgivaga.2.1.chr1.g00231.t01#1    mNeoVis.vulgivaga.2.1.chr1.g00233.t01#1    mNeoVis.vulgivaga.2.1.chr1.g00234.t01#1    mNeoVis.3.1.1.chr1.g00233.t01#6    mNeoVis.evergladensis.1.1.chr1.g00240.t01#3    mNeoVis.2.1.1.chr1_hap1.g00233.t01#5      4592   125257020:   mNeoVis.evergladensis.1.1.chr4.g00819.t01#3    mNeoVis.3.1.1.chr4.g00811.t01#6    mNeoVis.vulgivaga.1.1.chr4.g00820.t01#4    mNeoVis.evergladensis.2.1.chr4.g00805.t01#2    mNeoVis.vulgivaga.2.1.chr4.g00825.t01#1    mNeoVis.2.1.1.chr6_hap1.g00803.t01#5   mNeoVis.3.1.1.chr4.g00812.t01#6       4593   125257021:   mNeoVis.evergladensis.1.1.chr1.g00241.t01#3    mNeoVis.vulgivaga.1.1.chr1.g00228.t01#4    mNeoVis.vulgivaga.2.1.chr1.g00235.t01#1    mNeoVis.3.1.1.chr1.g00234.t01#6    mNeoVis.2.1.1.chr1_hap1.g00234.t01#5   mNeoVis.evergladensis.2.1.chr1.g00231.t01#2       4594   125257022:   mNeoVis.3.1.1.chr1.g00236.t01#6    mNeoVis.2.1.1.chr1_hap1.g00236.t01#5    mNeoVis.evergladensis.2.1.chr1.g00233.t01#2    mNeoVis.evergladensis.1.1.chr1.g00243.t01#3    mNeoVis.vulgivaga.2.1.chr1.g00237.t01#1   mNeoVis.vulgivaga.1.1.chr1.g00230.t01#4       4595   125257023:   mNeoVis.2.1.1.chr7_hap1.g00908.t01#5    mNeoVis.vulgivaga.2.1.chr7.g00905.t01#1    mNeoVis.evergladensis.1.1.chr7.g00906.t01#3    mNeoVis.evergladensis.2.1.chr7.g00901.t01#2    mNeoVis.vulgivaga.1.1.chr7.g00905.t01#4   mNeoVis.3.1.1.chr7.g00849.t01#6       4596   125257024:   mNeoVis.vulgivaga.2.1.chr1.g00238.t01#1    mNeoVis.vulgivaga.1.1.chr1.g00231.t01#4    mNeoVis.3.1.1.chr1.g00237.t01#6    mNeoVis.2.1.1.chr1_hap1.g00237.t01#5    mNeoVis.evergladensis.1.1.chr1.g00244.t01#3   mNeoVis.evergladensis.2.1.chr1.g00234.t01#2       4597   125257025:   mNeoVis.evergladensis.1.1.chr7.g00907.t01#3    mNeoVis.2.1.1.chr7_hap1.g00909.t01#5    mNeoVis.evergladensis.2.1.chr7.g00902.t01#2    mNeoVis.vulgivaga.2.1.chr7.g00906.t01#1   mNeoVis.vulgivaga.1.1.chr7.g00906.t01#4       4598   125257026:   mNeoVis.evergladensis.2.1.chr1.g00235.t01#2    mNeoVis.3.1.1.chr1.g00238.t01#6    mNeoVis.2.1.1.chr1_hap1.g00238.t01#5    mNeoVis.vulgivaga.2.1.chr1.g00239.t01#1    mNeoVis.evergladensis.1.1.chr1.g00245.t01#3   mNeoVis.vulgivaga.1.1.chr1.g00232.t01#4       4599   125257027:   mNeoVis.evergladensis.1.1.chr15.g00509.t01#3    mNeoVis.vulgivaga.1.1.chr15.g00104.t01#4    mNeoVis.vulgivaga.2.1.chr15.g00107.t01#1    mNeoVis.evergladensis.2.1.chr15.g00104.t01#2   mNeoVis.2.1.1.chr15_hap1.g00103.t01#5       4600   125257028:   mNeoVis.vulgivaga.1.1.chr1.g00233.t01#4    mNeoVis.evergladensis.1.1.chr1.g00246.t01#3    mNeoVis.3.1.1.chr1.g00239.t01#6    mNeoVis.2.1.1.chr1_hap1.g00239.t01#5    mNeoVis.vulgivaga.2.1.chr1.g00240.t01#1   mNeoVis.evergladensis.2.1.chr1.g00236.t01#2       4601   125257029:   mNeoVis.vulgivaga.2.1.chr4.g00988.t01#1    mNeoVis.evergladensis.1.1.chr4.g00987.t01#3    mNeoVis.2.1.1.chr6_hap1.g01020.t01#5    mNeoVis.vulgivaga.1.1.chr4.g00986.t01#4    mNeoVis.evergladensis.2.1.chr4.g00964.t01#2   mNeoVis.3.1.1.chr4.g00980.t01#6       4602   125257030:   mNeoVis.evergladensis.1.1.chr1.g00247.t01#3    mNeoVis.2.1.1.chr1_hap1.g00240.t01#5    mNeoVis.3.1.1.chr1.g00240.t01#6    mNeoVis.vulgivaga.2.1.chr1.g00241.t01#1    mNeoVis.vulgivaga.1.1.chr1.g00234.t01#4   mNeoVis.evergladensis.2.1.chr1.g00237.t01#2       4603   125257031:   mNeoVis.2.1.1.chr7_hap1.g00911.t01#5    mNeoVis.3.1.1.chr7.g00852.t01#6    mNeoVis.evergladensis.2.1.chr7.g00904.t01#2    mNeoVis.evergladensis.1.1.chr7.g00909.t01#3    mNeoVis.vulgivaga.2.1.chr7.g00908.t01#1   mNeoVis.vulgivaga.1.1.chr7.g00908.t01#4       4604   125257032:   mNeoVis.vulgivaga.2.1.chr7.g00909.t01#1    mNeoVis.3.1.1.chr7.g00853.t01#6    mNeoVis.evergladensis.2.1.chr7.g00905.t01#2    mNeoVis.evergladensis.1.1.chr7.g00910.t01#3   mNeoVis.2.1.1.chr7_hap1.g00912.t01#5       4605   125257033:   mNeoVis.vulgivaga.1.1.chr7.g00910.t01#4    mNeoVis.evergladensis.2.1.chr7.g00906.t01#2    mNeoVis.evergladensis.1.1.chr7.g00911.t01#3    mNeoVis.vulgivaga.2.1.chr7.g00910.t01#1   mNeoVis.2.1.1.chr7_hap1.g00913.t01#5       4606   125257034:   mNeoVis.evergladensis.2.1.chr7.g00907.t01#2    mNeoVis.3.1.1.chr3.g00646.t01#6    mNeoVis.vulgivaga.1.1.chr7.g00911.t01#4    mNeoVis.evergladensis.1.1.chr7.g00912.t01#3    mNeoVis.vulgivaga.2.1.chr7.g00911.t01#1   mNeoVis.2.1.1.chr7_hap1.g00914.t01#5       4607   125257035:   mNeoVis.vulgivaga.2.1.chr7.g00912.t01#1    mNeoVis.2.1.1.chr7_hap1.g00915.t01#5    mNeoVis.evergladensis.2.1.chr7.g00908.t01#2    mNeoVis.vulgivaga.1.1.chr7.g00912.t01#4   mNeoVis.evergladensis.1.1.chr7.g00913.t01#3       4608   125257036:   mNeoVis.evergladensis.2.1.chr7.g00909.t01#2    mNeoVis.vulgivaga.1.1.chr7.g00913.t01#4    mNeoVis.evergladensis.1.1.chr7.g00914.t01#3    mNeoVis.3.1.1.chr7.g00855.t01#6    mNeoVis.evergladensis.1.1.chr7.g00915.t01#3    mNeoVis.vulgivaga.2.1.chr7.g00914.t01#1    mNeoVis.vulgivaga.2.1.chr7.g00913.t01#1    mNeoVis.2.1.1.chr7_hap1.g00916.t01#5    mNeoVis.3.1.1.chr7.g00854.t01#6      4609   125257037:   mNeoVis.2.1.1.chr7_hap1.g00024.t01#5    mNeoVis.vulgivaga.1.1.chr7.g00026.t01#4    mNeoVis.vulgivaga.2.1.chr7.g00024.t01#1    mNeoVis.evergladensis.2.1.chr7.g00025.t01#2    mNeoVis.evergladensis.1.1.chr7.g00025.t01#3   mNeoVis.3.1.1.chr7.g00021.t01#6       4610   125257038:  mNeoVis.vulgivaga.2.1.chr1.g00243.t01#1               4611   125257039:   mNeoVis.vulgivaga.2.1.chr1.g00242.t01#1    mNeoVis.evergladensis.1.1.chr1.g00248.t01#3    mNeoVis.vulgivaga.1.1.chr1.g00235.t01#4    mNeoVis.3.1.1.chr1.g00241.t01#6    mNeoVis.2.1.1.chr1_hap1.g00241.t01#5   mNeoVis.evergladensis.2.1.chr1.g00238.t01#2       4612   125257040:   mNeoVis.3.1.1.chr1.g00242.t01#6    mNeoVis.vulgivaga.2.1.chr1.g00244.t01#1    mNeoVis.vulgivaga.1.1.chr1.g00236.t01#4   mNeoVis.2.1.1.chr1_hap1.g00242.t01#5         4613   125257041:   mNeoVis.vulgivaga.1.1.chr7.g00914.t01#4    mNeoVis.evergladensis.2.1.chr7.g00910.t01#2    mNeoVis.2.1.1.chr7_hap1.g00917.t01#5    mNeoVis.vulgivaga.2.1.chr7.g00915.t01#1    mNeoVis.3.1.1.chr7.g00857.t01#6   mNeoVis.evergladensis.1.1.chr7.g00916.t01#3       4614   125257042:   mNeoVis.evergladensis.1.1.chr1.g00250.t01#3    mNeoVis.vulgivaga.1.1.chr1.g00238.t01#4    mNeoVis.2.1.1.chr1_hap1.g00243.t01#5    mNeoVis.evergladensis.2.1.chr1.g00240.t01#2    mNeoVis.vulgivaga.2.1.chr1.g00245.t01#1   mNeoVis.3.1.1.chr1.g00243.t01#6       4615   125257043:   mNeoVis.3.1.1.chr1.g00244.t01#6    mNeoVis.vulgivaga.1.1.chr1.g00239.t01#4    mNeoVis.evergladensis.2.1.chr1.g00241.t01#2    mNeoVis.2.1.1.chr1_hap1.g00244.t01#5    mNeoVis.vulgivaga.2.1.chr1.g00246.t01#1   mNeoVis.evergladensis.1.1.chr1.g00251.t01#3       4616   125257044:   mNeoVis.3.1.1.chr1.g00245.t01#6    mNeoVis.2.1.1.chr1_hap1.g00245.t01#5    mNeoVis.vulgivaga.1.1.chr1.g00240.t01#4    mNeoVis.evergladensis.1.1.chr1.g00252.t01#3    mNeoVis.evergladensis.2.1.chr1.g00242.t01#2   mNeoVis.vulgivaga.2.1.chr1.g00247.t01#1       4617   125257045:   mNeoVis.evergladensis.2.1.chr1.g00243.t01#2    mNeoVis.3.1.1.chr1.g00246.t01#6    mNeoVis.2.1.1.chr1_hap1.g00246.t01#5    mNeoVis.vulgivaga.1.1.chr1.g00241.t01#4    mNeoVis.evergladensis.1.1.chr1.g00253.t01#3   mNeoVis.vulgivaga.2.1.chr1.g00248.t01#1       4618   125257046:   mNeoVis.evergladensis.2.1.chr3.g00162.t01#2    mNeoVis.2.1.1.chr3_hap1.g00156.t01#5    mNeoVis.3.1.1.chr3.g00159.t01#6    mNeoVis.vulgivaga.1.1.chr3.g00162.t01#4    mNeoVis.evergladensis.1.1.chr3.g00161.t01#3   mNeoVis.vulgivaga.2.1.chr3.g00159.t01#1       4619   125257047:   mNeoVis.evergladensis.2.1.chr1.g00244.t01#2    mNeoVis.vulgivaga.2.1.chr1.g00249.t01#1    mNeoVis.evergladensis.1.1.chr1.g00254.t01#3    mNeoVis.vulgivaga.1.1.chr1.g00242.t01#4   mNeoVis.2.1.1.chr1_hap1.g00247.t01#5       4620   125257048:   mNeoVis.evergladensis.2.1.chr1.g00245.t01#2   mNeoVis.3.1.1.chr1.g00247.t01#6             4621   125257049:   mNeoVis.3.1.1.chr1.g00248.t01#6    mNeoVis.evergladensis.1.1.chr1.g00255.t01#3    mNeoVis.2.1.1.chr1_hap1.g00248.t01#5    mNeoVis.vulgivaga.1.1.chr1.g00243.t01#4    mNeoVis.vulgivaga.2.1.chr1.g00250.t01#1   mNeoVis.evergladensis.2.1.chr1.g00246.t01#2       4622   125257050:   mNeoVis.vulgivaga.2.1.chr7.g00917.t01#1    mNeoVis.evergladensis.1.1.chr7.g00918.t01#3    mNeoVis.evergladensis.2.1.chr7.g00912.t01#2    mNeoVis.vulgivaga.1.1.chr7.g00916.t01#4    mNeoVis.3.1.1.chr7.g00859.t01#6   mNeoVis.2.1.1.chr7_hap1.g00919.t01#5       4623   125257051:   mNeoVis.vulgivaga.2.1.chr7.g00918.t01#1    mNeoVis.evergladensis.1.1.chr7.g00919.t01#3    mNeoVis.vulgivaga.1.1.chr7.g00917.t01#4    mNeoVis.3.1.1.chr7.g00860.t01#6    mNeoVis.evergladensis.2.1.chr7.g00913.t01#2   mNeoVis.2.1.1.chr7_hap1.g00920.t01#5       4624   125257052:   mNeoVis.vulgivaga.2.1.chr7.g00919.t01#1    mNeoVis.evergladensis.1.1.chr7.g00920.t01#3    mNeoVis.vulgivaga.1.1.chr7.g00918.t01#4    mNeoVis.3.1.1.chr7.g00861.t01#6    mNeoVis.evergladensis.2.1.chr7.g00914.t01#2   mNeoVis.2.1.1.chr7_hap1.g00921.t01#5       4625   125257053:   mNeoVis.2.1.1.chr7_hap1.g00922.t01#5    mNeoVis.3.1.1.chr7.g00862.t01#6    mNeoVis.evergladensis.1.1.chr7.g00921.t01#3    mNeoVis.vulgivaga.2.1.chr7.g00920.t01#1    mNeoVis.evergladensis.2.1.chr7.g00915.t01#2   mNeoVis.vulgivaga.1.1.chr7.g00919.t01#4       4626   125257054:   mNeoVis.vulgivaga.2.1.chr1.g00251.t01#1    mNeoVis.2.1.1.chr1_hap1.g00249.t01#5    mNeoVis.vulgivaga.1.1.chr1.g00244.t01#4    mNeoVis.3.1.1.chr1.g00249.t01#6    mNeoVis.evergladensis.2.1.chr1.g00247.t01#2   mNeoVis.evergladensis.1.1.chr1.g00256.t01#3       4627   125257055:   mNeoVis.evergladensis.1.1.chr9.g01076.t01#3    mNeoVis.evergladensis.2.1.chr8.g01087.t01#2    mNeoVis.vulgivaga.2.1.chr9.g01067.t01#1    mNeoVis.3.1.1.chr9.g01039.t01#6    mNeoVis.2.1.1.chr10_hap1.g01111.t01#5   mNeoVis.vulgivaga.1.1.chr9.g01080.t01#4       4628   125257056:   mNeoVis.vulgivaga.1.1.chr7.g00920.t01#4    mNeoVis.3.1.1.chr7.g00863.t01#6    mNeoVis.2.1.1.chr7_hap1.g00923.t01#5    mNeoVis.vulgivaga.2.1.chr7.g00921.t01#1    mNeoVis.evergladensis.2.1.chr7.g00916.t01#2    mNeoVis.vulgivaga.1.1.chr7.g00921.t01#4   mNeoVis.evergladensis.1.1.chr7.g00922.t01#3     4629   125257057:   mNeoVis.evergladensis.2.1.chr1.g00248.t01#2    mNeoVis.2.1.1.chr1_hap1.g00250.t01#5    mNeoVis.vulgivaga.1.1.chr1.g00245.t01#4    mNeoVis.3.1.1.chr1.g00250.t01#6    mNeoVis.evergladensis.1.1.chr1.g00257.t01#3   mNeoVis.vulgivaga.2.1.chr1.g00252.t01#1       4630   125257058:   mNeoVis.2.1.1.chr7_hap1.g00924.t01#5    mNeoVis.evergladensis.1.1.chr7.g00923.t01#3    mNeoVis.vulgivaga.2.1.chr7.g00922.t01#1    mNeoVis.vulgivaga.1.1.chr7.g00922.t01#4    mNeoVis.evergladensis.2.1.chr7.g00917.t01#2   mNeoVis.3.1.1.chr7.g00864.t01#6       4631   125257059:   mNeoVis.2.1.1.chr7_hap1.g00925.t01#5    mNeoVis.3.1.1.chr7.g00865.t01#6    mNeoVis.evergladensis.1.1.chr7.g00924.t01#3    mNeoVis.evergladensis.2.1.chr7.g00918.t01#2    mNeoVis.vulgivaga.1.1.chr7.g00923.t01#4   mNeoVis.vulgivaga.2.1.chr7.g00923.t01#1       4632   125257060:   mNeoVis.3.1.1.chr1.g00163.t01#6    mNeoVis.vulgivaga.1.1.chr1.g00156.t01#4    mNeoVis.evergladensis.2.1.chr1.g00156.t01#2    mNeoVis.evergladensis.1.1.chr1.g00168.t01#3    mNeoVis.vulgivaga.2.1.chr1.g00157.t01#1   mNeoVis.2.1.1.chr1_hap1.g00160.t01#5       4633   125257061:   mNeoVis.vulgivaga.1.1.chr9.g01074.t01#4    mNeoVis.2.1.1.chr10_hap1.g01103.t01#5    mNeoVis.evergladensis.1.1.chr9.g01070.t01#3    mNeoVis.3.1.1.chr9.g01033.t01#6    mNeoVis.vulgivaga.2.1.chr9.g01061.t01#1   mNeoVis.evergladensis.2.1.chr8.g01079.t01#2       4634   125257062:   mNeoVis.2.1.1.chr7_hap1.g00926.t01#5    mNeoVis.evergladensis.2.1.chr7.g00919.t01#2    mNeoVis.evergladensis.1.1.chr7.g00925.t01#3    mNeoVis.vulgivaga.1.1.chr7.g00924.t01#4    mNeoVis.3.1.1.chr7.g00866.t01#6   mNeoVis.vulgivaga.2.1.chr7.g00924.t01#1       4635   125257063:   mNeoVis.2.1.1.chr1_hap1.g00162.t01#5    mNeoVis.vulgivaga.2.1.chr1.g00160.t01#1    mNeoVis.vulgivaga.1.1.chr1.g00159.t01#4    mNeoVis.evergladensis.2.1.chr1.g00158.t01#2    mNeoVis.3.1.1.chr1.g00167.t01#6   mNeoVis.evergladensis.1.1.chr1.g00170.t01#3       4636   125257064:   mNeoVis.vulgivaga.2.1.chr1.g00159.t01#1   mNeoVis.vulgivaga.1.1.chr1.g00158.t01#4             4637   125257065:   mNeoVis.vulgivaga.1.1.chr7.g00925.t01#4    mNeoVis.vulgivaga.2.1.chr7.g00925.t01#1    mNeoVis.2.1.1.chr7_hap1.g00927.t01#5    mNeoVis.3.1.1.chr7.g00868.t01#6    mNeoVis.evergladensis.2.1.chr7.g00920.t01#2   mNeoVis.evergladensis.1.1.chr7.g00926.t01#3       4638   125257066:   mNeoVis.vulgivaga.2.1.chr7.g00828.t01#1    mNeoVis.evergladensis.2.1.chr7.g00822.t01#2    mNeoVis.vulgivaga.1.1.chr7.g00830.t01#4    mNeoVis.2.1.1.chr7_hap1.g00833.t01#5    mNeoVis.3.1.1.chr7.g00777.t01#6   mNeoVis.evergladensis.1.1.chr7.g00830.t01#3       4639   125257067:   mNeoVis.evergladensis.2.1.chr7.g00824.t01#2    mNeoVis.vulgivaga.2.1.chr7.g00829.t01#1    mNeoVis.3.1.1.chr7.g00778.t01#6    mNeoVis.evergladensis.2.1.chr7.g00823.t01#2    mNeoVis.evergladensis.1.1.chr7.g00831.t01#3    mNeoVis.vulgivaga.1.1.chr7.g00831.t01#4   mNeoVis.2.1.1.chr7_hap1.g00834.t01#5     4640   125257068:   mNeoVis.3.1.1.chr7.g00779.t01#6    mNeoVis.evergladensis.1.1.chr7.g00832.t01#3    mNeoVis.vulgivaga.1.1.chr7.g00832.t01#4    mNeoVis.2.1.1.chr7_hap1.g00835.t01#5    mNeoVis.evergladensis.2.1.chr7.g00825.t01#2   mNeoVis.vulgivaga.2.1.chr7.g00830.t01#1       4641   125257069:   mNeoVis.vulgivaga.1.1.chr7.g00833.t01#4    mNeoVis.3.1.1.chr7.g00780.t01#6    mNeoVis.evergladensis.2.1.chr7.g00826.t01#2    mNeoVis.evergladensis.1.1.chr7.g00833.t01#3    mNeoVis.vulgivaga.2.1.chr7.g00831.t01#1   mNeoVis.2.1.1.chr7_hap1.g00836.t01#5       4642   125257070:   mNeoVis.vulgivaga.2.1.chr1.g00161.t01#1    mNeoVis.vulgivaga.1.1.chr1.g00160.t01#4    mNeoVis.3.1.1.chr1.g00166.t01#6    mNeoVis.evergladensis.1.1.chr1.g00171.t01#3    mNeoVis.evergladensis.2.1.chr1.g00159.t01#2   mNeoVis.2.1.1.chr1_hap1.g00163.t01#5       4643   125257071:  mNeoVis.vulgivaga.2.1.chr1.g00162.t01#1               4644   125257072:   mNeoVis.2.1.1.chr7_hap1.g00837.t01#5    mNeoVis.vulgivaga.1.1.chr7.g00834.t01#4    mNeoVis.evergladensis.1.1.chr7.g00834.t01#3    mNeoVis.evergladensis.2.1.chr7.g00827.t01#2    mNeoVis.vulgivaga.2.1.chr7.g00832.t01#1   mNeoVis.3.1.1.chr7.g00781.t01#6       4645   125257073:   mNeoVis.3.1.1.chr1.g00168.t01#6    mNeoVis.2.1.1.chr1_hap1.g00164.t01#5    mNeoVis.evergladensis.2.1.chr1.g00160.t01#2    mNeoVis.evergladensis.1.1.chr1.g00172.t01#3    mNeoVis.vulgivaga.1.1.chr1.g00161.t01#4   mNeoVis.vulgivaga.2.1.chr1.g00163.t01#1       4646   125257074:   mNeoVis.vulgivaga.1.1.chr7.g00835.t01#4    mNeoVis.2.1.1.chr7_hap1.g00838.t01#5    mNeoVis.3.1.1.chr7.g00782.t01#6    mNeoVis.evergladensis.1.1.chr7.g00835.t01#3    mNeoVis.evergladensis.2.1.chr7.g00828.t01#2   mNeoVis.vulgivaga.2.1.chr7.g00833.t01#1       4647   125257075:   mNeoVis.evergladensis.2.1.chr7.g00829.t01#2    mNeoVis.vulgivaga.2.1.chr7.g00834.t01#1    mNeoVis.vulgivaga.1.1.chr7.g00836.t01#4    mNeoVis.3.1.1.chr7.g00783.t01#6    mNeoVis.evergladensis.1.1.chr7.g00836.t01#3   mNeoVis.2.1.1.chr7_hap1.g00839.t01#5       4648   125257076:   mNeoVis.evergladensis.1.1.chr7.g00837.t01#3    mNeoVis.evergladensis.2.1.chr7.g00830.t01#2    mNeoVis.vulgivaga.2.1.chr7.g00835.t01#1    mNeoVis.3.1.1.chr7.g00784.t01#6    mNeoVis.vulgivaga.1.1.chr7.g00837.t01#4   mNeoVis.2.1.1.chr7_hap1.g00840.t01#5       4649   125257077:   mNeoVis.vulgivaga.2.1.chr7.g00836.t01#1    mNeoVis.evergladensis.2.1.chr7.g00831.t01#2    mNeoVis.evergladensis.1.1.chr7.g00838.t01#3    mNeoVis.3.1.1.chr7.g00785.t01#6    mNeoVis.2.1.1.chr7_hap1.g00841.t01#5   mNeoVis.vulgivaga.1.1.chr7.g00838.t01#4       4650   125257078:   mNeoVis.3.1.1.chr7.g00786.t01#6    mNeoVis.vulgivaga.1.1.chr7.g00839.t01#4    mNeoVis.vulgivaga.2.1.chr7.g00837.t01#1    mNeoVis.2.1.1.chr7_hap1.g00842.t01#5    mNeoVis.evergladensis.1.1.chr7.g00839.t01#3   mNeoVis.evergladensis.2.1.chr7.g00832.t01#2       4651   125257079:   mNeoVis.2.1.1.chr7_hap1.g00025.t01#5    mNeoVis.vulgivaga.1.1.chr7.g00027.t01#4    mNeoVis.evergladensis.1.1.chr7.g00026.t01#3   mNeoVis.vulgivaga.2.1.chr7.g00025.t01#1         4652   125257080:  mNeoVis.vulgivaga.2.1.chr1.g00164.t01#1               4653   125257081:  mNeoVis.vulgivaga.2.1.chr1.g00165.t01#1               4654   125257082:   mNeoVis.evergladensis.2.1.chr7.g00833.t01#2    mNeoVis.vulgivaga.1.1.chr7.g00840.t01#4    mNeoVis.2.1.1.chr7_hap1.g00843.t01#5    mNeoVis.vulgivaga.2.1.chr7.g00838.t01#1    mNeoVis.evergladensis.1.1.chr7.g00840.t01#3   mNeoVis.evergladensis.2.1.chr7.g00834.t01#2       4655   125257083:   mNeoVis.evergladensis.1.1.chr7.g00841.t01#3    mNeoVis.evergladensis.2.1.chr7.g00835.t01#2    mNeoVis.2.1.1.chr7_hap1.g00844.t01#5    mNeoVis.3.1.1.chr7.g00787.t01#6    mNeoVis.vulgivaga.2.1.chr7.g00839.t01#1   mNeoVis.vulgivaga.1.1.chr7.g00841.t01#4       4656   125257084:   mNeoVis.evergladensis.1.1.chr1.g00175.t01#3    mNeoVis.3.1.1.chr1.g00170.t01#6    mNeoVis.vulgivaga.2.1.chr1.g00166.t01#1    mNeoVis.vulgivaga.1.1.chr1.g00163.t01#4    mNeoVis.2.1.1.chr1_hap1.g00167.t01#5   mNeoVis.evergladensis.2.1.chr1.g00163.t01#2       4657   125257085:   mNeoVis.vulgivaga.1.1.chr7.g00842.t01#4    mNeoVis.evergladensis.1.1.chr7.g00842.t01#3    mNeoVis.2.1.1.chr7_hap1.g00845.t01#5    mNeoVis.evergladensis.2.1.chr7.g00836.t01#2    mNeoVis.3.1.1.chr7.g00788.t01#6   mNeoVis.vulgivaga.2.1.chr7.g00840.t01#1       4658   125257086:   mNeoVis.vulgivaga.2.1.chr1.g00167.t01#1    mNeoVis.vulgivaga.1.1.chr1.g00164.t01#4    mNeoVis.evergladensis.1.1.chr1.g00176.t01#3    mNeoVis.evergladensis.2.1.chr1.g00164.t01#2    mNeoVis.2.1.1.chr1_hap1.g00168.t01#5   mNeoVis.3.1.1.chr1.g00171.t01#6       4659   125257087:   mNeoVis.2.1.1.chr7_hap1.g00846.t01#5    mNeoVis.evergladensis.2.1.chr7.g00837.t01#2    mNeoVis.vulgivaga.2.1.chr7.g00841.t01#1    mNeoVis.3.1.1.chr7.g00789.t01#6    mNeoVis.evergladensis.1.1.chr7.g00843.t01#3   mNeoVis.vulgivaga.1.1.chr7.g00843.t01#4       4660   125257088:  mNeoVis.vulgivaga.2.1.chr1.g00168.t01#1               4661   125257089:   mNeoVis.vulgivaga.2.1.chr7.g00842.t01#1    mNeoVis.vulgivaga.1.1.chr7.g00844.t01#4    mNeoVis.evergladensis.1.1.chr7.g00844.t01#3    mNeoVis.3.1.1.chr7.g00790.t01#6    mNeoVis.evergladensis.2.1.chr7.g00838.t01#2   mNeoVis.2.1.1.chr7_hap1.g00847.t01#5       4662   125257090:   mNeoVis.evergladensis.1.1.chr1.g00178.t01#3    mNeoVis.2.1.1.chr1_hap1.g00170.t01#5    mNeoVis.vulgivaga.2.1.chr1.g00169.t01#1    mNeoVis.evergladensis.2.1.chr1.g00166.t01#2    mNeoVis.3.1.1.chr1.g00172.t01#6   mNeoVis.vulgivaga.1.1.chr1.g00166.t01#4       4663   125257091:   mNeoVis.2.1.1.chr7_hap1.g00848.t01#5    mNeoVis.evergladensis.2.1.chr7.g00839.t01#2    mNeoVis.vulgivaga.1.1.chr7.g00845.t01#4    mNeoVis.evergladensis.1.1.chr7.g00845.t01#3   mNeoVis.vulgivaga.2.1.chr7.g00843.t01#1       4664   125257092:   mNeoVis.vulgivaga.1.1.chr1.g00168.t01#4    mNeoVis.vulgivaga.2.1.chr1.g00170.t01#1    mNeoVis.evergladensis.1.1.chr1.g00179.t01#3    mNeoVis.3.1.1.chr1.g00173.t01#6    mNeoVis.2.1.1.chr1_hap1.g00172.t01#5   mNeoVis.evergladensis.2.1.chr1.g00167.t01#2       4665   125257093:   mNeoVis.2.1.1.chr1_hap1.g00173.t01#5    mNeoVis.evergladensis.2.1.chr1.g00168.t01#2    mNeoVis.evergladensis.1.1.chr1.g00181.t01#3    mNeoVis.vulgivaga.2.1.chr1.g00171.t01#1   mNeoVis.vulgivaga.1.1.chr1.g00169.t01#4       4666   125257094:   mNeoVis.evergladensis.2.1.chr1.g00169.t01#2    mNeoVis.2.1.1.chr1_hap1.g00174.t01#5    mNeoVis.evergladensis.1.1.chr1.g00182.t01#3    mNeoVis.vulgivaga.1.1.chr1.g00170.t01#4    mNeoVis.3.1.1.chr1.g00175.t01#6   mNeoVis.vulgivaga.2.1.chr1.g00172.t01#1       4667   125257095:   mNeoVis.vulgivaga.2.1.chr1.g00173.t01#1    mNeoVis.evergladensis.2.1.chr1.g00170.t01#2    mNeoVis.3.1.1.chr1.g00176.t01#6    mNeoVis.vulgivaga.1.1.chr1.g00171.t01#4    mNeoVis.2.1.1.chr1_hap1.g00175.t01#5   mNeoVis.evergladensis.1.1.chr1.g00183.t01#3       4668   125257096:   mNeoVis.3.1.1.chr1.g00177.t01#6    mNeoVis.vulgivaga.1.1.chr1.g00172.t01#4    mNeoVis.2.1.1.chr1_hap1.g00176.t01#5    mNeoVis.evergladensis.1.1.chr1.g00184.t01#3    mNeoVis.vulgivaga.2.1.chr1.g00174.t01#1   mNeoVis.evergladensis.2.1.chr1.g00171.t01#2       4669   125257097:   mNeoVis.evergladensis.1.1.chr7.g00846.t01#3    mNeoVis.2.1.1.chr7_hap1.g00849.t01#5    mNeoVis.vulgivaga.1.1.chr7.g00846.t01#4    mNeoVis.evergladensis.2.1.chr7.g00840.t01#2   mNeoVis.vulgivaga.2.1.chr7.g00844.t01#1       4670   125257098:   mNeoVis.evergladensis.2.1.chr1.g00172.t01#2    mNeoVis.2.1.1.chr1_hap1.g00177.t01#5    mNeoVis.vulgivaga.2.1.chr1.g00175.t01#1    mNeoVis.evergladensis.1.1.chr1.g00185.t01#3    mNeoVis.3.1.1.chr1.g00178.t01#6   mNeoVis.vulgivaga.1.1.chr1.g00173.t01#4       4671   125257099:   mNeoVis.evergladensis.1.1.chr7.g00847.t01#3    mNeoVis.vulgivaga.1.1.chr7.g00847.t01#4    mNeoVis.2.1.1.chr7_hap1.g00850.t01#5    mNeoVis.vulgivaga.2.1.chr7.g00845.t01#1    mNeoVis.3.1.1.chr7.g00792.t01#6   mNeoVis.evergladensis.2.1.chr7.g00841.t01#2       4672   125257100:   mNeoVis.2.1.1.chr1_hap1.g00178.t01#5    mNeoVis.vulgivaga.2.1.chr1.g00176.t01#1    mNeoVis.evergladensis.1.1.chr1.g00186.t01#3    mNeoVis.vulgivaga.1.1.chr1.g00174.t01#4    mNeoVis.3.1.1.chr1.g00179.t01#6   mNeoVis.evergladensis.2.1.chr1.g00173.t01#2       4673   125257101:   mNeoVis.evergladensis.2.1.chr7.g00842.t01#2    mNeoVis.vulgivaga.1.1.chr7.g00848.t01#4    mNeoVis.evergladensis.1.1.chr7.g00848.t01#3    mNeoVis.2.1.1.chr7_hap1.g00851.t01#5    mNeoVis.vulgivaga.2.1.chr7.g00846.t01#1   mNeoVis.3.1.1.chr7.g00793.t01#6       4674   125257102:   mNeoVis.evergladensis.2.1.chr1.g00174.t01#2    mNeoVis.3.1.1.chr1.g00180.t01#6    mNeoVis.vulgivaga.2.1.chr1.g00177.t01#1    mNeoVis.vulgivaga.1.1.chr1.g00175.t01#4    mNeoVis.evergladensis.1.1.chr1.g00187.t01#3   mNeoVis.2.1.1.chr1_hap1.g00179.t01#5       4675   125257103:   mNeoVis.evergladensis.1.1.chr7.g00850.t01#3    mNeoVis.evergladensis.2.1.chr7.g00844.t01#2    mNeoVis.2.1.1.chr7_hap1.g00853.t01#5    mNeoVis.3.1.1.chr7.g00795.t01#6    mNeoVis.vulgivaga.1.1.chr7.g00850.t01#4   mNeoVis.vulgivaga.2.1.chr7.g00847.t01#1       4676   125257104:   mNeoVis.vulgivaga.2.1.chr1.g00178.t01#1    mNeoVis.evergladensis.2.1.chr1.g00175.t01#2    mNeoVis.3.1.1.chr1.g00181.t01#6    mNeoVis.evergladensis.1.1.chr1.g00188.t01#3   mNeoVis.2.1.1.chr1_hap1.g00180.t01#5       4677   125257105:   mNeoVis.2.1.1.chr1_hap1.g00181.t01#5    mNeoVis.evergladensis.2.1.chr1.g00176.t01#2    mNeoVis.vulgivaga.2.1.chr1.g00179.t01#1    mNeoVis.evergladensis.1.1.chr1.g00189.t01#3   mNeoVis.3.1.1.chr1.g00182.t01#6       4678   125257106:   mNeoVis.evergladensis.1.1.chr7.g00851.t01#3    mNeoVis.evergladensis.2.1.chr7.g00845.t01#2    mNeoVis.2.1.1.chr7_hap1.g00854.t01#5    mNeoVis.3.1.1.chr7.g00796.t01#6    mNeoVis.vulgivaga.2.1.chr7.g00848.t01#1    mNeoVis.vulgivaga.1.1.chr7.g00851.t01#4   mNeoVis.vulgivaga.2.1.chr7.g00849.t01#1     4679   125257107:   mNeoVis.vulgivaga.1.1.chr1.g00176.t01#4    mNeoVis.evergladensis.1.1.chr1.g00190.t01#3    mNeoVis.3.1.1.chr1.g00183.t01#6    mNeoVis.evergladensis.2.1.chr1.g00177.t01#2    mNeoVis.2.1.1.chr1_hap1.g00182.t01#5   mNeoVis.vulgivaga.2.1.chr1.g00180.t01#1       4680   125257108:   mNeoVis.vulgivaga.2.1.chr7.g00850.t01#1   mNeoVis.vulgivaga.1.1.chr7.g00852.t01#4             4681   125257109:   mNeoVis.evergladensis.1.1.chr7.g00853.t01#3    mNeoVis.vulgivaga.2.1.chr7.g00851.t01#1    mNeoVis.vulgivaga.1.1.chr7.g00853.t01#4    mNeoVis.evergladensis.2.1.chr7.g00847.t01#2   mNeoVis.2.1.1.chr7_hap1.g00855.t01#5       4682   125257110:   mNeoVis.vulgivaga.2.1.chr7.g00026.t01#1    mNeoVis.2.1.1.chr7_hap1.g00026.t01#5    mNeoVis.vulgivaga.1.1.chr7.g00028.t01#4    mNeoVis.evergladensis.1.1.chr7.g00027.t01#3   mNeoVis.evergladensis.2.1.chr7.g00027.t01#2       4683   125257111:   mNeoVis.vulgivaga.2.1.chr1.g00181.t01#1    mNeoVis.evergladensis.2.1.chr1.g00178.t01#2    mNeoVis.3.1.1.chr1.g00184.t01#6    mNeoVis.2.1.1.chr1_hap1.g00183.t01#5    mNeoVis.vulgivaga.1.1.chr1.g00177.t01#4   mNeoVis.evergladensis.1.1.chr1.g00191.t01#3       4684   125257112:   mNeoVis.vulgivaga.2.1.chr7.g00852.t01#1    mNeoVis.vulgivaga.1.1.chr7.g00854.t01#4    mNeoVis.evergladensis.2.1.chr7.g00848.t01#2    mNeoVis.3.1.1.chr7.g00798.t01#6    mNeoVis.2.1.1.chr7_hap1.g00856.t01#5   mNeoVis.evergladensis.1.1.chr7.g00854.t01#3       4685   125257113:   mNeoVis.2.1.1.chr7_hap1.g00857.t01#5    mNeoVis.evergladensis.1.1.chr7.g00855.t01#3    mNeoVis.3.1.1.chr7.g00799.t01#6    mNeoVis.vulgivaga.2.1.chr7.g00853.t01#1    mNeoVis.vulgivaga.1.1.chr7.g00855.t01#4   mNeoVis.evergladensis.2.1.chr7.g00849.t01#2       4686   125257114:   mNeoVis.evergladensis.1.1.chr1.g00192.t01#3    mNeoVis.vulgivaga.1.1.chr1.g00178.t01#4    mNeoVis.3.1.1.chr1.g00185.t01#6    mNeoVis.vulgivaga.2.1.chr1.g00182.t01#1    mNeoVis.2.1.1.chr1_hap1.g00184.t01#5   mNeoVis.evergladensis.2.1.chr1.g00179.t01#2       4687   125257115:   mNeoVis.2.1.1.chr1_hap1.g00185.t01#5    mNeoVis.vulgivaga.2.1.chr1.g00183.t01#1    mNeoVis.evergladensis.2.1.chr1.g00180.t01#2    mNeoVis.vulgivaga.1.1.chr1.g00179.t01#4    mNeoVis.evergladensis.1.1.chr1.g00193.t01#3   mNeoVis.3.1.1.chr1.g00186.t01#6       4688   125257116:   mNeoVis.evergladensis.2.1.chr1.g00181.t01#2    mNeoVis.2.1.1.chr1_hap1.g00186.t01#5    mNeoVis.evergladensis.1.1.chr1.g00194.t01#3    mNeoVis.vulgivaga.1.1.chr1.g00180.t01#4    mNeoVis.3.1.1.chr1.g00187.t01#6   mNeoVis.vulgivaga.2.1.chr1.g00184.t01#1       4689   125257117:   mNeoVis.evergladensis.2.1.chr7.g00850.t01#2    mNeoVis.3.1.1.chr7.g00800.t01#6    mNeoVis.2.1.1.chr7_hap1.g00858.t01#5    mNeoVis.vulgivaga.2.1.chr7.g00854.t01#1    mNeoVis.vulgivaga.2.1.chr7.g00855.t01#1    mNeoVis.evergladensis.1.1.chr7.g00856.t01#3   mNeoVis.vulgivaga.1.1.chr7.g00856.t01#4     4690   125257118:   mNeoVis.vulgivaga.1.1.chr7.g00857.t01#4    mNeoVis.evergladensis.2.1.chr7.g00851.t01#2    mNeoVis.2.1.1.chr7_hap1.g00859.t01#5    mNeoVis.evergladensis.1.1.chr7.g00857.t01#3   mNeoVis.vulgivaga.2.1.chr7.g00856.t01#1       4691   125257119:   mNeoVis.evergladensis.1.1.chr1.g00195.t01#3    mNeoVis.2.1.1.chr1_hap1.g00187.t01#5    mNeoVis.vulgivaga.2.1.chr1.g00185.t01#1    mNeoVis.evergladensis.2.1.chr1.g00182.t01#2    mNeoVis.3.1.1.chr1.g00188.t01#6   mNeoVis.vulgivaga.1.1.chr1.g00181.t01#4       4692   125257120:   mNeoVis.vulgivaga.2.1.chr15.g00108.t01#1    mNeoVis.3.1.1.chr15.g00098.t01#6    mNeoVis.2.1.1.chr15_hap1.g00104.t01#5    mNeoVis.evergladensis.2.1.chr15.g00105.t01#2   mNeoVis.evergladensis.1.1.chr15.g00508.t01#3       4693   125257121:   mNeoVis.evergladensis.2.1.chr1.g00183.t01#2    mNeoVis.3.1.1.chr1.g00189.t01#6    mNeoVis.vulgivaga.2.1.chr1.g00186.t01#1    mNeoVis.vulgivaga.1.1.chr1.g00182.t01#4    mNeoVis.evergladensis.1.1.chr1.g00196.t01#3   mNeoVis.2.1.1.chr1_hap1.g00188.t01#5       4694   125257122:   mNeoVis.2.1.1.chr1_hap1.g00189.t01#5    mNeoVis.evergladensis.2.1.chr1.g00184.t01#2    mNeoVis.vulgivaga.2.1.chr1.g00187.t01#1    mNeoVis.evergladensis.1.1.chr1.g00197.t01#3    mNeoVis.vulgivaga.1.1.chr1.g00183.t01#4   mNeoVis.3.1.1.chr1.g00190.t01#6       4695   125257123:   mNeoVis.vulgivaga.2.1.chr7.g00857.t01#1    mNeoVis.3.1.1.chr7.g00802.t01#6    mNeoVis.2.1.1.chr7_hap1.g00860.t01#5    mNeoVis.evergladensis.1.1.chr7.g00858.t01#3    mNeoVis.evergladensis.2.1.chr7.g00852.t01#2   mNeoVis.vulgivaga.1.1.chr7.g00858.t01#4       4696   125257124:   mNeoVis.vulgivaga.2.1.chr10.g00968.t01#1    mNeoVis.vulgivaga.1.1.chr10.g00242.t01#4    mNeoVis.evergladensis.2.1.chr4.g00644.t01#2    mNeoVis.2.1.1.chr8_hap1.g01193.t01#5    mNeoVis.vulgivaga.2.1.chr4.g00652.t01#1    mNeoVis.evergladensis.2.1.chr12.g00426.t01#2    mNeoVis.vulgivaga.1.1.chr6.g00714.t01#4    mNeoVis.2.1.1.chr4_hap1.g00765.t01#5    mNeoVis.vulgivaga.1.1.chr9.g00963.t01#4      4697   125257125:   mNeoVis.vulgivaga.2.1.chr9.g00744.t01#1    mNeoVis.vulgivaga.2.1.chr5.g01399.t01#1    mNeoVis.evergladensis.2.1.chr6.g00519.t01#2    mNeoVis.vulgivaga.1.1.chr5.g01372.t01#4    mNeoVis.evergladensis.2.1.chr5.g01399.t01#2    mNeoVis.evergladensis.1.1.chr5.g00454.t01#3   mNeoVis.2.1.1.chr4_hap1.g00745.t01#5     4698   125257126:   mNeoVis.3.1.1.chr10.g00246.t01#6    mNeoVis.evergladensis.2.1.chr5.g00462.t01#2    mNeoVis.vulgivaga.2.1.chr5.g00461.t01#1    mNeoVis.3.1.1.chr12.g00433.t01#6   mNeoVis.evergladensis.1.1.chr9.g00753.t01#3       4699   125257127:   mNeoVis.evergladensis.1.1.chr7.g00859.t01#3    mNeoVis.vulgivaga.1.1.chr7.g00859.t01#4   mNeoVis.vulgivaga.2.1.chr7.g00858.t01#1           4700   125257128:   mNeoVis.2.1.1.chr10_hap1.g00478.t01#5    mNeoVis.evergladensis.1.1.chr9.g00470.t01#3   mNeoVis.3.1.1.chr9.g00444.t01#6           4701   125257129:   mNeoVis.2.1.1.chr7_hap1.g00861.t01#5    mNeoVis.vulgivaga.1.1.chr7.g00860.t01#4    mNeoVis.3.1.1.chr7.g00803.t01#6    mNeoVis.vulgivaga.2.1.chr7.g00859.t01#1    mNeoVis.evergladensis.1.1.chr7.g00860.t01#3   mNeoVis.evergladensis.2.1.chr7.g00853.t01#2       4702   125257130:   mNeoVis.vulgivaga.2.1.chr1.g00188.t01#1    mNeoVis.2.1.1.chr1_hap1.g00190.t01#5    mNeoVis.vulgivaga.1.1.chr1.g00184.t01#4    mNeoVis.evergladensis.1.1.chr1.g00198.t01#3    mNeoVis.3.1.1.chr1.g00191.t01#6   mNeoVis.evergladensis.2.1.chr1.g00185.t01#2       4703   125257131:   mNeoVis.evergladensis.1.1.chr7.g00861.t01#3    mNeoVis.vulgivaga.2.1.chr7.g00860.t01#1    mNeoVis.3.1.1.chr7.g00804.t01#6    mNeoVis.evergladensis.2.1.chr7.g00854.t01#2    mNeoVis.vulgivaga.1.1.chr7.g00861.t01#4   mNeoVis.2.1.1.chr7_hap1.g00862.t01#5       4704   125257132:   mNeoVis.vulgivaga.2.1.chr1.g00189.t01#1    mNeoVis.vulgivaga.1.1.chr1.g00185.t01#4    mNeoVis.evergladensis.1.1.chr1.g00199.t01#3    mNeoVis.evergladensis.2.1.chr1.g00186.t01#2    mNeoVis.3.1.1.chr1.g00192.t01#6   mNeoVis.2.1.1.chr1_hap1.g00191.t01#5       4705   125257133:   mNeoVis.2.1.1.chr7_hap1.g00863.t01#5    mNeoVis.evergladensis.2.1.chr7.g00856.t01#2    mNeoVis.3.1.1.chr7.g00805.t01#6    mNeoVis.evergladensis.1.1.chr7.g00862.t01#3    mNeoVis.vulgivaga.2.1.chr7.g00861.t01#1   mNeoVis.vulgivaga.1.1.chr7.g00862.t01#4       4706   125257134:   mNeoVis.evergladensis.2.1.chr1.g00187.t01#2    mNeoVis.2.1.1.chr1_hap1.g00192.t01#5    mNeoVis.vulgivaga.2.1.chr1.g00190.t01#1    mNeoVis.vulgivaga.1.1.chr1.g00186.t01#4    mNeoVis.evergladensis.1.1.chr1.g00200.t01#3   mNeoVis.3.1.1.chr1.g00193.t01#6       4707   125257135:   mNeoVis.evergladensis.1.1.chr1.g00201.t01#3    mNeoVis.2.1.1.chr1_hap1.g00193.t01#5    mNeoVis.evergladensis.2.1.chr1.g00188.t01#2    mNeoVis.3.1.1.chr1.g00194.t01#6    mNeoVis.vulgivaga.1.1.chr1.g00187.t01#4   mNeoVis.vulgivaga.2.1.chr1.g00191.t01#1       4708   125257136:   mNeoVis.vulgivaga.1.1.chr7.g00029.t01#4    mNeoVis.2.1.1.chr7_hap1.g00027.t01#5    mNeoVis.evergladensis.2.1.chr7.g00028.t01#2    mNeoVis.vulgivaga.2.1.chr7.g00027.t01#1   mNeoVis.evergladensis.1.1.chr7.g00028.t01#3       4709   125257137:   mNeoVis.vulgivaga.1.1.chr1.g00188.t01#4    mNeoVis.3.1.1.chr1.g00195.t01#6    mNeoVis.vulgivaga.2.1.chr1.g00192.t01#1    mNeoVis.evergladensis.1.1.chr1.g00202.t01#3    mNeoVis.2.1.1.chr1_hap1.g00194.t01#5   mNeoVis.evergladensis.2.1.chr1.g00189.t01#2       4710   125257138:   mNeoVis.vulgivaga.2.1.chr15.g00109.t01#1    mNeoVis.evergladensis.1.1.chr15.g00507.t01#3    mNeoVis.3.1.1.chr15.g00099.t01#6    mNeoVis.evergladensis.2.1.chr15.g00106.t01#2    mNeoVis.2.1.1.chr15_hap1.g00105.t01#5   mNeoVis.vulgivaga.1.1.chr15.g00107.t01#4       4711   125257139:   mNeoVis.3.1.1.chr1.g00196.t01#6    mNeoVis.vulgivaga.1.1.chr1.g00189.t01#4    mNeoVis.evergladensis.2.1.chr1.g00190.t01#2    mNeoVis.evergladensis.1.1.chr1.g00203.t01#3    mNeoVis.2.1.1.chr1_hap1.g00195.t01#5   mNeoVis.vulgivaga.2.1.chr1.g00193.t01#1       4712   125257140:   mNeoVis.evergladensis.1.1.chr7.g00863.t01#3    mNeoVis.vulgivaga.2.1.chr7.g00862.t01#1    mNeoVis.3.1.1.chr7.g00806.t01#6    mNeoVis.evergladensis.2.1.chr7.g00857.t01#2    mNeoVis.vulgivaga.1.1.chr7.g00863.t01#4   mNeoVis.2.1.1.chr7_hap1.g00864.t01#5       4713   125257141:   mNeoVis.vulgivaga.2.1.chr7.g00863.t01#1    mNeoVis.evergladensis.2.1.chr7.g00858.t01#2    mNeoVis.evergladensis.1.1.chr7.g00864.t01#3    mNeoVis.vulgivaga.1.1.chr7.g00864.t01#4   mNeoVis.2.1.1.chr7_hap1.g00865.t01#5       4714   125257142:   mNeoVis.evergladensis.2.1.chr1.g00191.t01#2    mNeoVis.2.1.1.chr1_hap1.g00196.t01#5    mNeoVis.evergladensis.1.1.chr1.g00204.t01#3    mNeoVis.vulgivaga.1.1.chr1.g00190.t01#4    mNeoVis.3.1.1.chr1.g00197.t01#6   mNeoVis.vulgivaga.2.1.chr1.g00194.t01#1       4715   125257143:   mNeoVis.vulgivaga.2.1.chr7.g00864.t01#1    mNeoVis.2.1.1.chr7_hap1.g00866.t01#5    mNeoVis.evergladensis.2.1.chr7.g00859.t01#2   mNeoVis.vulgivaga.1.1.chr7.g00865.t01#4         4716   125257144:   mNeoVis.vulgivaga.2.1.chr7.g00865.t01#1    mNeoVis.vulgivaga.1.1.chr7.g00866.t01#4    mNeoVis.2.1.1.chr7_hap1.g00867.t01#5    mNeoVis.evergladensis.1.1.chr7.g00865.t01#3    mNeoVis.3.1.1.chr7.g00807.t01#6   mNeoVis.evergladensis.2.1.chr7.g00860.t01#2       4717   125257145:   mNeoVis.evergladensis.2.1.chr1.g00192.t01#2    mNeoVis.vulgivaga.1.1.chr1.g00191.t01#4    mNeoVis.evergladensis.1.1.chr1.g00205.t01#3    mNeoVis.2.1.1.chr1_hap1.g00197.t01#5    mNeoVis.3.1.1.chr1.g00198.t01#6   mNeoVis.vulgivaga.2.1.chr1.g00195.t01#1       4718   125257146:   mNeoVis.vulgivaga.2.1.chr7.g00866.t01#1    mNeoVis.evergladensis.2.1.chr7.g00861.t01#2    mNeoVis.vulgivaga.1.1.chr7.g00867.t01#4    mNeoVis.evergladensis.1.1.chr7.g00866.t01#3   mNeoVis.2.1.1.chr7_hap1.g00868.t01#5       4719   125257147:   mNeoVis.evergladensis.1.1.chr7.g00867.t01#3    mNeoVis.vulgivaga.2.1.chr7.g00867.t01#1   mNeoVis.evergladensis.2.1.chr7.g00862.t01#2         4720   125257148:   mNeoVis.2.1.1.chr7_hap1.g00870.t01#5    mNeoVis.vulgivaga.2.1.chr7.g00868.t01#1    mNeoVis.3.1.1.chr7.g00810.t01#6    mNeoVis.vulgivaga.1.1.chr7.g00869.t01#4    mNeoVis.evergladensis.1.1.chr7.g00869.t01#3   mNeoVis.evergladensis.2.1.chr7.g00863.t01#2       4721   125257149:   mNeoVis.vulgivaga.2.1.chr7.g00869.t01#1    mNeoVis.evergladensis.2.1.chr7.g00864.t01#2    mNeoVis.vulgivaga.1.1.chr7.g00870.t01#4    mNeoVis.2.1.1.chr7_hap1.g00871.t01#5    mNeoVis.3.1.1.chr7.g00811.t01#6   mNeoVis.evergladensis.1.1.chr7.g00868.t01#3       4722   125257150:   mNeoVis.2.1.1.chr1_hap1.g00198.t01#5    mNeoVis.evergladensis.2.1.chr1.g00193.t01#2    mNeoVis.3.1.1.chr1.g00199.t01#6    mNeoVis.vulgivaga.1.1.chr1.g00192.t01#4    mNeoVis.evergladensis.1.1.chr1.g00206.t01#3   mNeoVis.vulgivaga.2.1.chr1.g00196.t01#1       4723   125257151:   mNeoVis.3.1.1.chr7.g00812.t01#6    mNeoVis.evergladensis.1.1.chr7.g00870.t01#3    mNeoVis.vulgivaga.1.1.chr7.g00871.t01#4    mNeoVis.vulgivaga.2.1.chr7.g00870.t01#1    mNeoVis.evergladensis.2.1.chr7.g00865.t01#2   mNeoVis.2.1.1.chr7_hap1.g00872.t01#5       4724   125257152:   mNeoVis.vulgivaga.1.1.chr1.g00193.t01#4    mNeoVis.2.1.1.chr1_hap1.g00199.t01#5    mNeoVis.evergladensis.1.1.chr1.g00208.t01#3    mNeoVis.vulgivaga.2.1.chr1.g00197.t01#1   mNeoVis.evergladensis.2.1.chr1.g00194.t01#2       4725   125257153:   mNeoVis.vulgivaga.2.1.chr1.g00198.t01#1    mNeoVis.evergladensis.2.1.chr1.g00195.t01#2    mNeoVis.evergladensis.1.1.chr1.g00209.t01#3    mNeoVis.vulgivaga.1.1.chr1.g00194.t01#4    mNeoVis.2.1.1.chr1_hap1.g00200.t01#5   mNeoVis.3.1.1.chr1.g00201.t01#6       4726   125257154:   mNeoVis.vulgivaga.1.1.chr7.g00872.t01#4    mNeoVis.evergladensis.1.1.chr7.g00871.t01#3    mNeoVis.3.1.1.chr7.g00813.t01#6    mNeoVis.2.1.1.chr7_hap1.g00873.t01#5    mNeoVis.evergladensis.2.1.chr7.g00866.t01#2   mNeoVis.vulgivaga.2.1.chr7.g00871.t01#1       4727   125257155:   mNeoVis.3.1.1.chr7.g00814.t01#6    mNeoVis.evergladensis.2.1.chr7.g00867.t01#2    mNeoVis.evergladensis.1.1.chr7.g00872.t01#3    mNeoVis.vulgivaga.1.1.chr7.g00873.t01#4    mNeoVis.2.1.1.chr7_hap1.g00874.t01#5   mNeoVis.vulgivaga.2.1.chr7.g00872.t01#1       4728   125257156:   mNeoVis.vulgivaga.2.1.chr1.g00199.t01#1    mNeoVis.evergladensis.2.1.chr1.g00196.t01#2    mNeoVis.3.1.1.chr1.g00202.t01#6    mNeoVis.vulgivaga.1.1.chr1.g00195.t01#4    mNeoVis.2.1.1.chr1_hap1.g00201.t01#5   mNeoVis.evergladensis.1.1.chr1.g00210.t01#3       4729   125257157:   mNeoVis.evergladensis.2.1.chr4.g00494.t01#2    mNeoVis.vulgivaga.1.1.chr4.g00492.t01#4    mNeoVis.2.1.1.chr6_hap1.g00488.t01#5    mNeoVis.3.1.1.chr4.g00493.t01#6    mNeoVis.evergladensis.1.1.chr4.g00499.t01#3   mNeoVis.vulgivaga.2.1.chr4.g00502.t01#1       4730   125257158:   mNeoVis.vulgivaga.1.1.chr6.g00621.t01#4    mNeoVis.2.1.1.chr4_hap1.g00613.t01#5    mNeoVis.vulgivaga.2.1.chr6.g00619.t01#1    mNeoVis.evergladensis.2.1.chr6.g00622.t01#2    mNeoVis.3.1.1.chr6.g00628.t01#6   mNeoVis.evergladensis.1.1.chr6.g00624.t01#3       4731   125257159:   mNeoVis.3.1.1.chr5.g00454.t01#6    mNeoVis.vulgivaga.2.1.chr5.g00463.t01#1    mNeoVis.2.1.1.chr2_hap1.g00461.t01#5    mNeoVis.evergladensis.2.1.chr5.g00464.t01#2    mNeoVis.vulgivaga.1.1.chr5.g00448.t01#4   mNeoVis.evergladensis.1.1.chr5.g00456.t01#3       4732   125257160:   mNeoVis.3.1.1.chr1.g00203.t01#6    mNeoVis.vulgivaga.1.1.chr1.g00196.t01#4    mNeoVis.2.1.1.chr1_hap1.g00202.t01#5    mNeoVis.evergladensis.1.1.chr1.g00211.t01#3    mNeoVis.evergladensis.2.1.chr1.g00197.t01#2   mNeoVis.vulgivaga.2.1.chr1.g00200.t01#1       4733   125257161:   mNeoVis.evergladensis.2.1.chr7.g00868.t01#2    mNeoVis.evergladensis.1.1.chr7.g00873.t01#3    mNeoVis.vulgivaga.2.1.chr7.g00873.t01#1    mNeoVis.evergladensis.1.1.chr3.g01116.t01#3    mNeoVis.evergladensis.2.1.chr3.g01105.t01#2    mNeoVis.2.1.1.chr7_hap1.g00875.t01#5    mNeoVis.vulgivaga.1.1.chr7.g00874.t01#4    mNeoVis.vulgivaga.1.1.chr3.g01107.t01#4    mNeoVis.3.1.1.chr7.g00815.t01#6      4734   125257162:   mNeoVis.evergladensis.2.1.chr1.g00198.t01#2    mNeoVis.vulgivaga.1.1.chr1.g00197.t01#4    mNeoVis.3.1.1.chr1.g00204.t01#6    mNeoVis.2.1.1.chr1_hap1.g00203.t01#5    mNeoVis.vulgivaga.2.1.chr1.g00201.t01#1   mNeoVis.evergladensis.1.1.chr1.g00212.t01#3       4735   125257163:   mNeoVis.vulgivaga.1.1.chr7.g00875.t01#4    mNeoVis.vulgivaga.2.1.chr7.g00874.t01#1    mNeoVis.evergladensis.1.1.chr7.g00874.t01#3    mNeoVis.2.1.1.chr7_hap1.g00876.t01#5   mNeoVis.evergladensis.2.1.chr7.g00869.t01#2       4736   125257164:   mNeoVis.evergladensis.1.1.chr7.g00875.t01#3    mNeoVis.3.1.1.chr7.g00817.t01#6    mNeoVis.vulgivaga.2.1.chr7.g00875.t01#1    mNeoVis.2.1.1.chr7_hap1.g00877.t01#5    mNeoVis.evergladensis.2.1.chr7.g00870.t01#2   mNeoVis.vulgivaga.1.1.chr7.g00876.t01#4       4737   125257165:   mNeoVis.evergladensis.2.1.chr1.g00114.t01#2    mNeoVis.evergladensis.1.1.chr1.g00123.t01#3    mNeoVis.3.1.1.chr1.g00120.t01#6    mNeoVis.vulgivaga.2.1.chr1.g00113.t01#1    mNeoVis.2.1.1.chr1_hap1.g00116.t01#5   mNeoVis.vulgivaga.1.1.chr1.g00116.t01#4       4738   125257166:   mNeoVis.evergladensis.1.1.chr7.g00029.t01#3    mNeoVis.evergladensis.2.1.chr7.g00029.t01#2    mNeoVis.vulgivaga.1.1.chr7.g00030.t01#4    mNeoVis.vulgivaga.2.1.chr7.g00028.t01#1   mNeoVis.2.1.1.chr7_hap1.g00028.t01#5       4739   125257167:   mNeoVis.vulgivaga.2.1.chr15.g00110.t01#1    mNeoVis.evergladensis.2.1.chr15.g00107.t01#2    mNeoVis.vulgivaga.1.1.chr15.g00108.t01#4    mNeoVis.2.1.1.chr15_hap1.g00106.t01#5   mNeoVis.evergladensis.1.1.chr15.g00506.t01#3       4740   125257168:   mNeoVis.2.1.1.chr1_hap1.g00117.t01#5    mNeoVis.evergladensis.2.1.chr1.g00115.t01#2    mNeoVis.vulgivaga.1.1.chr1.g00117.t01#4    mNeoVis.vulgivaga.2.1.chr1.g00114.t01#1    mNeoVis.3.1.1.chr1.g00121.t01#6   mNeoVis.evergladensis.1.1.chr1.g00124.t01#3       4741   125257169:   mNeoVis.evergladensis.2.1.chr1.g01656.t01#2    mNeoVis.2.1.1.chr5_hap1.g01024.t01#5    mNeoVis.evergladensis.1.1.chr1.g01656.t01#3    mNeoVis.vulgivaga.1.1.chr7.g00755.t01#4    mNeoVis.evergladensis.1.1.chr2.g01097.t01#3    mNeoVis.2.1.1.chr7_hap1.g00757.t01#5    mNeoVis.evergladensis.2.1.chr7.g00745.t01#2    mNeoVis.2.1.1.chr1_hap1.g01689.t01#5    mNeoVis.evergladensis.2.1.chr2.g01095.t01#2      4742   125257170:   mNeoVis.vulgivaga.2.1.chr1.g00116.t01#1    mNeoVis.evergladensis.1.1.chr1.g00125.t01#3    mNeoVis.evergladensis.2.1.chr1.g00117.t01#2    mNeoVis.vulgivaga.1.1.chr1.g00118.t01#4    mNeoVis.2.1.1.chr1_hap1.g00118.t01#5   mNeoVis.3.1.1.chr1.g00123.t01#6       4743   125257171:   mNeoVis.evergladensis.2.1.chr7.g00872.t01#2    mNeoVis.evergladensis.1.1.chr7.g00877.t01#3    mNeoVis.vulgivaga.1.1.chr7.g00878.t01#4    mNeoVis.2.1.1.chr7_hap1.g00879.t01#5   mNeoVis.vulgivaga.2.1.chr7.g00877.t01#1       4744   125257172:   mNeoVis.2.1.1.chr7_hap1.g00880.t01#5    mNeoVis.3.1.1.chr7.g00821.t01#6    mNeoVis.vulgivaga.2.1.chr7.g00878.t01#1    mNeoVis.evergladensis.2.1.chr7.g00874.t01#2    mNeoVis.vulgivaga.1.1.chr7.g00880.t01#4   mNeoVis.evergladensis.1.1.chr7.g00879.t01#3       4745   125257173:   mNeoVis.evergladensis.1.1.chr1.g00126.t01#3    mNeoVis.3.1.1.chr1.g00124.t01#6    mNeoVis.evergladensis.2.1.chr1.g00118.t01#2    mNeoVis.vulgivaga.1.1.chr1.g00119.t01#4    mNeoVis.2.1.1.chr1_hap1.g00119.t01#5   mNeoVis.vulgivaga.2.1.chr1.g00117.t01#1       4746   125257174:   mNeoVis.evergladensis.2.1.chr7.g00747.t01#2    mNeoVis.3.1.1.chr7.g00709.t01#6    mNeoVis.2.1.1.chr7_hap1.g00759.t01#5    mNeoVis.evergladensis.1.1.chr7.g00757.t01#3    mNeoVis.vulgivaga.2.1.chr7.g00749.t01#1   mNeoVis.vulgivaga.1.1.chr7.g00758.t01#4       4747   125257175:   mNeoVis.vulgivaga.2.1.chr1.g00118.t01#1    mNeoVis.3.1.1.chr1.g00125.t01#6   mNeoVis.evergladensis.1.1.chr1.g00128.t01#3         4748   125257176:  mNeoVis.vulgivaga.2.1.chr7.g00751.t01#1               4749   125257177:   mNeoVis.vulgivaga.1.1.chr7.g00759.t01#4    mNeoVis.3.1.1.chr7.g00710.t01#6    mNeoVis.2.1.1.chr7_hap1.g00760.t01#5    mNeoVis.evergladensis.2.1.chr7.g00748.t01#2    mNeoVis.vulgivaga.2.1.chr7.g00750.t01#1   mNeoVis.evergladensis.1.1.chr7.g00758.t01#3       4750   125257178:   mNeoVis.vulgivaga.2.1.chr7.g00752.t01#1    mNeoVis.evergladensis.1.1.chr7.g00760.t01#3    mNeoVis.vulgivaga.1.1.chr7.g00761.t01#4   mNeoVis.2.1.1.chr7_hap1.g00762.t01#5         4751   125257179:   mNeoVis.evergladensis.1.1.chr1.g00129.t01#3    mNeoVis.2.1.1.chr1_hap1.g00121.t01#5    mNeoVis.vulgivaga.1.1.chr1.g00120.t01#4    mNeoVis.vulgivaga.2.1.chr1.g00119.t01#1    mNeoVis.3.1.1.chr1.g00126.t01#6   mNeoVis.evergladensis.2.1.chr1.g00119.t01#2       4752   125257180:   mNeoVis.2.1.1.chr7_hap1.g00763.t01#5    mNeoVis.3.1.1.chr7.g00713.t01#6    mNeoVis.vulgivaga.2.1.chr7.g00753.t01#1    mNeoVis.evergladensis.1.1.chr7.g00761.t01#3    mNeoVis.vulgivaga.1.1.chr7.g00762.t01#4   mNeoVis.evergladensis.2.1.chr7.g00751.t01#2       4753   125257181:   mNeoVis.vulgivaga.2.1.chr15.g00111.t01#1    mNeoVis.vulgivaga.1.1.chr15.g00109.t01#4    mNeoVis.evergladensis.1.1.chr15.g00505.t01#3    mNeoVis.2.1.1.chr15_hap1.g00107.t01#5   mNeoVis.evergladensis.2.1.chr15.g00109.t01#2       4754   125257182:   mNeoVis.evergladensis.2.1.chr1.g00120.t01#2    mNeoVis.3.1.1.chr1.g00127.t01#6    mNeoVis.evergladensis.1.1.chr1.g00130.t01#3    mNeoVis.vulgivaga.1.1.chr1.g00121.t01#4    mNeoVis.vulgivaga.2.1.chr1.g00120.t01#1   mNeoVis.2.1.1.chr1_hap1.g00122.t01#5       4755   125257183:   mNeoVis.vulgivaga.2.1.chr7.g00755.t01#1    mNeoVis.evergladensis.1.1.chr7.g00763.t01#3    mNeoVis.evergladensis.2.1.chr7.g00753.t01#2    mNeoVis.3.1.1.chr7.g00715.t01#6    mNeoVis.vulgivaga.1.1.chr7.g00764.t01#4   mNeoVis.2.1.1.chr7_hap1.g00765.t01#5       4756   125257184:   mNeoVis.evergladensis.2.1.chr7.g00754.t01#2    mNeoVis.vulgivaga.1.1.chr7.g00765.t01#4    mNeoVis.2.1.1.chr7_hap1.g00766.t01#5    mNeoVis.3.1.1.chr7.g00716.t01#6    mNeoVis.vulgivaga.2.1.chr7.g00756.t01#1   mNeoVis.evergladensis.1.1.chr7.g00764.t01#3       4757   125257185:   mNeoVis.evergladensis.2.1.chr7.g00031.t01#2    mNeoVis.vulgivaga.1.1.chr7.g00032.t01#4    mNeoVis.vulgivaga.2.1.chr7.g00030.t01#1    mNeoVis.evergladensis.1.1.chr7.g00031.t01#3    mNeoVis.2.1.1.chr7_hap1.g00030.t01#5   mNeoVis.3.1.1.chr7.g00024.t01#6       4758   125257186:   mNeoVis.2.1.1.chr7_hap1.g00767.t01#5    mNeoVis.evergladensis.1.1.chr7.g00765.t01#3    mNeoVis.3.1.1.chr7.g00717.t01#6    mNeoVis.vulgivaga.2.1.chr7.g00757.t01#1   mNeoVis.vulgivaga.1.1.chr7.g00766.t01#4       4759   125257187:  mNeoVis.vulgivaga.2.1.chr1.g00121.t01#1               4760   125257188:   mNeoVis.vulgivaga.2.1.chr1.g00122.t01#1    mNeoVis.vulgivaga.1.1.chr1.g00122.t01#4    mNeoVis.2.1.1.chr1_hap1.g00124.t01#5    mNeoVis.3.1.1.chr1.g00129.t01#6    mNeoVis.evergladensis.2.1.chr1.g00122.t01#2   mNeoVis.evergladensis.1.1.chr1.g00132.t01#3       4761   125257189:   mNeoVis.evergladensis.2.1.chr7.g00755.t01#2    mNeoVis.vulgivaga.2.1.chr7.g00758.t01#1    mNeoVis.evergladensis.1.1.chr7.g00766.t01#3    mNeoVis.2.1.1.chr7_hap1.g00768.t01#5    mNeoVis.vulgivaga.1.1.chr7.g00767.t01#4   mNeoVis.3.1.1.chr7.g00718.t01#6       4762   125257190:   mNeoVis.evergladensis.1.1.chr7.g00767.t01#3    mNeoVis.vulgivaga.1.1.chr7.g00768.t01#4    mNeoVis.evergladensis.2.1.chr7.g00756.t01#2    mNeoVis.vulgivaga.2.1.chr7.g00759.t01#1   mNeoVis.2.1.1.chr7_hap1.g00769.t01#5       4763   125257191:   mNeoVis.3.1.1.chr10.g01138.t01#6    mNeoVis.evergladensis.1.1.chr10.g01169.t01#3    mNeoVis.2.1.1.chr9_hap1.g01188.t01#5    mNeoVis.evergladensis.2.1.chr10.g01176.t01#2   mNeoVis.vulgivaga.1.1.chr10.g01149.t01#4       4764   125257192:   mNeoVis.vulgivaga.2.1.chr7.g00760.t01#1   mNeoVis.evergladensis.1.1.chr7.g00768.t01#3           4765   125257193:   mNeoVis.evergladensis.1.1.chr7.g00775.t01#3    mNeoVis.vulgivaga.1.1.chr7.g00774.t01#4    mNeoVis.2.1.1.chr7_hap1.g00780.t01#5    mNeoVis.evergladensis.2.1.chr7.g00764.t01#2   mNeoVis.vulgivaga.2.1.chr7.g00767.t01#1       4766   125257194:   mNeoVis.evergladensis.1.1.chr7.g00771.t01#3    mNeoVis.2.1.1.chr7_hap1.g00770.t01#5    mNeoVis.evergladensis.2.1.chr7.g00758.t01#2   mNeoVis.vulgivaga.2.1.chr7.g00761.t01#1         4767   125257195:   mNeoVis.2.1.1.chr7_hap1.g00776.t01#5    mNeoVis.vulgivaga.1.1.chr7.g00772.t01#4    mNeoVis.3.1.1.chr7.g00722.t01#6   mNeoVis.vulgivaga.1.1.chr7.g00777.t01#4         4768   125257196:   mNeoVis.evergladensis.1.1.chr7.g00770.t01#3    mNeoVis.vulgivaga.1.1.chr7.g00771.t01#4    mNeoVis.evergladensis.2.1.chr7.g00760.t01#2   mNeoVis.2.1.1.chr7_hap1.g00772.t01#5         4769   125257197:   mNeoVis.2.1.1.chr7_hap1.g00778.t01#5    mNeoVis.evergladensis.1.1.chr7.g00772.t01#3   mNeoVis.evergladensis.2.1.chr7.g00762.t01#2         4770   125257198:   mNeoVis.vulgivaga.1.1.chr7.g00775.t01#4    mNeoVis.evergladensis.2.1.chr7.g00765.t01#2   mNeoVis.evergladensis.1.1.chr7.g00776.t01#3         4771   125257199:   mNeoVis.vulgivaga.2.1.chr7.g00771.t01#1    mNeoVis.vulgivaga.1.1.chr7.g00779.t01#4   mNeoVis.evergladensis.2.1.chr7.g00766.t01#2         4772   125257200:   mNeoVis.vulgivaga.2.1.chr7.g00769.t01#1    mNeoVis.vulgivaga.1.1.chr7.g00778.t01#4   mNeoVis.evergladensis.2.1.chr7.g00767.t01#2         4773   125257201:   mNeoVis.vulgivaga.2.1.chr7.g00772.t01#1   mNeoVis.3.1.1.chr7.g00724.t01#6             4774   125257202:   mNeoVis.2.1.1.chr7_hap1.g00781.t01#5   mNeoVis.vulgivaga.2.1.chr7.g00770.t01#1             4775   125257203:   mNeoVis.2.1.1.chr7_hap1.g00779.t01#5   mNeoVis.evergladensis.2.1.chr7.g00763.t01#2           4776   125257204:   mNeoVis.vulgivaga.2.1.chr1.g00123.t01#1    mNeoVis.evergladensis.1.1.chr1.g00133.t01#3   mNeoVis.evergladensis.2.1.chr1.g00123.t01#2         4777   125257205:  mNeoVis.vulgivaga.2.1.chr7.g00762.t01#1               4778   125257206:   mNeoVis.vulgivaga.2.1.chr1.g00124.t01#1    mNeoVis.3.1.1.chr1.g00130.t01#6    mNeoVis.evergladensis.2.1.chr1.g00124.t01#2    mNeoVis.vulgivaga.1.1.chr1.g00123.t01#4    mNeoVis.evergladensis.1.1.chr1.g00134.t01#3   mNeoVis.2.1.1.chr1_hap1.g00125.t01#5       4779   125257207:   mNeoVis.3.1.1.chr7.g00720.t01#6    mNeoVis.2.1.1.chr7_hap1.g00771.t01#5    mNeoVis.evergladensis.2.1.chr7.g00759.t01#2    mNeoVis.evergladensis.1.1.chr7.g00769.t01#3    mNeoVis.vulgivaga.2.1.chr7.g00763.t01#1   mNeoVis.vulgivaga.1.1.chr7.g00769.t01#4       4780   125257208:  mNeoVis.vulgivaga.2.1.chr7.g00764.t01#1               4781   125257209:  mNeoVis.vulgivaga.2.1.chr7.g00766.t01#1               4782   125257210:   mNeoVis.vulgivaga.1.1.chr1.g00124.t01#4    mNeoVis.2.1.1.chr1_hap1.g00126.t01#5    mNeoVis.evergladensis.2.1.chr1.g00125.t01#2    mNeoVis.evergladensis.1.1.chr1.g00135.t01#3    mNeoVis.3.1.1.chr1.g00131.t01#6   mNeoVis.vulgivaga.2.1.chr1.g00125.t01#1       4783   125257211:   mNeoVis.vulgivaga.2.1.chr15.g00112.t01#1    mNeoVis.3.1.1.chr15.g00101.t01#6   mNeoVis.2.1.1.chr15_hap1.g00108.t01#5           4784   125257212:   mNeoVis.vulgivaga.1.1.chr1.g00125.t01#4    mNeoVis.evergladensis.1.1.chr1.g00136.t01#3    mNeoVis.2.1.1.chr1_hap1.g00127.t01#5    mNeoVis.evergladensis.2.1.chr1.g00126.t01#2    mNeoVis.3.1.1.chr1.g00132.t01#6   mNeoVis.vulgivaga.2.1.chr1.g00126.t01#1       4785   125257213:   mNeoVis.vulgivaga.2.1.chr7.g00768.t01#1   mNeoVis.evergladensis.1.1.chr7.g00777.t01#3           4786   125257214:   mNeoVis.evergladensis.2.1.chr1.g00127.t01#2    mNeoVis.3.1.1.chr1.g00133.t01#6    mNeoVis.vulgivaga.2.1.chr1.g00127.t01#1    mNeoVis.vulgivaga.1.1.chr1.g00126.t01#4    mNeoVis.evergladensis.1.1.chr1.g00137.t01#3   mNeoVis.2.1.1.chr1_hap1.g00128.t01#5       4787   125257215:   mNeoVis.evergladensis.1.1.chr1.g00138.t01#3    mNeoVis.vulgivaga.1.1.chr1.g00127.t01#4    mNeoVis.3.1.1.chr1.g00134.t01#6    mNeoVis.evergladensis.2.1.chr1.g00128.t01#2    mNeoVis.vulgivaga.2.1.chr1.g00129.t01#1    mNeoVis.2.1.1.chr1_hap1.g00129.t01#5   mNeoVis.vulgivaga.2.1.chr1.g00128.t01#1     4788   125257216:   mNeoVis.vulgivaga.2.1.chr5.g01185.t01#1    mNeoVis.vulgivaga.1.1.chr5.g01154.t01#4    mNeoVis.evergladensis.1.1.chr5.g01188.t01#3    mNeoVis.3.1.1.chr5.g01167.t01#6    mNeoVis.evergladensis.2.1.chr5.g01191.t01#2   mNeoVis.2.1.1.chr2_hap1.g01248.t01#5       4789   125257217:   mNeoVis.vulgivaga.1.1.chr1.g00129.t01#4    mNeoVis.3.1.1.chr1.g00136.t01#6    mNeoVis.vulgivaga.2.1.chr1.g00131.t01#1    mNeoVis.2.1.1.chr1_hap1.g00131.t01#5    mNeoVis.evergladensis.1.1.chr1.g00139.t01#3   mNeoVis.evergladensis.2.1.chr1.g00130.t01#2       4790   125257218:   mNeoVis.2.1.1.chr1_hap1.g00132.t01#5    mNeoVis.vulgivaga.2.1.chr1.g00132.t01#1    mNeoVis.evergladensis.2.1.chr1.g00131.t01#2    mNeoVis.evergladensis.1.1.chr1.g00140.t01#3    mNeoVis.3.1.1.chr1.g00137.t01#6   mNeoVis.vulgivaga.1.1.chr1.g00130.t01#4       4791   125257219:   mNeoVis.3.1.1.chr7.g00723.t01#6    mNeoVis.evergladensis.1.1.chr7.g00778.t01#3    mNeoVis.vulgivaga.2.1.chr7.g00773.t01#1    mNeoVis.2.1.1.chr7_hap1.g00782.t01#5   mNeoVis.evergladensis.2.1.chr7.g00769.t01#2       4792   125257220:   mNeoVis.vulgivaga.1.1.chr7.g00781.t01#4    mNeoVis.3.1.1.chr7.g00725.t01#6    mNeoVis.vulgivaga.2.1.chr7.g00774.t01#1   mNeoVis.evergladensis.2.1.chr7.g00768.t01#2       4793   125257221:  mNeoVis.vulgivaga.2.1.chr7.g00775.t01#1               4794   125257222:   mNeoVis.vulgivaga.2.1.chr7.g00776.t01#1    mNeoVis.evergladensis.2.1.chr7.g00771.t01#2    mNeoVis.vulgivaga.1.1.chr7.g00782.t01#4    mNeoVis.3.1.1.chr7.g00726.t01#6    mNeoVis.2.1.1.chr7_hap1.g00784.t01#5   mNeoVis.evergladensis.1.1.chr7.g00780.t01#3       4795   125257223:   mNeoVis.vulgivaga.2.1.chr15.g00129.t01#1    mNeoVis.evergladensis.2.1.chr15.g00126.t01#2    mNeoVis.2.1.1.chr15_hap1.g00139.t01#5    mNeoVis.vulgivaga.1.1.chr15.g00125.t01#4    mNeoVis.3.1.1.chr15.g00117.t01#6    mNeoVis.evergladensis.2.1.chr15.g00125.t01#2   mNeoVis.evergladensis.1.1.chr15.g00490.t01#3     4796   125257224:   mNeoVis.2.1.1.chr15_hap1.g00109.t01#5    mNeoVis.evergladensis.1.1.chr15.g00504.t01#3    mNeoVis.3.1.1.chr15.g00102.t01#6    mNeoVis.vulgivaga.1.1.chr15.g00111.t01#4    mNeoVis.vulgivaga.2.1.chr15.g00113.t01#1   mNeoVis.evergladensis.2.1.chr15.g00110.t01#2       4797   125257225:   mNeoVis.evergladensis.1.1.chr8.g00933.t01#3    mNeoVis.vulgivaga.2.1.chr8.g00940.t01#1    mNeoVis.evergladensis.2.1.chr9.g00942.t01#2    mNeoVis.3.1.1.chr8.g00901.t01#6   mNeoVis.2.1.1.chr8_hap1.g00969.t01#5       4798   125257226:   mNeoVis.2.1.1.chr8_hap1.g00971.t01#5    mNeoVis.vulgivaga.2.1.chr8.g00942.t01#1    mNeoVis.vulgivaga.1.1.chr8.g00929.t01#4    mNeoVis.evergladensis.1.1.chr8.g00935.t01#3   mNeoVis.evergladensis.2.1.chr9.g00944.t01#2       4799   125257227:   mNeoVis.2.1.1.chr15_hap1.g00372.t01#5    mNeoVis.2.1.1.chr15_hap1.g00446.t01#5   mNeoVis.2.1.1.chr15_hap1.g00409.t01#5           4800   125257228:   mNeoVis.3.1.1.chr8.g00903.t01#6   mNeoVis.vulgivaga.1.1.chr8.g00927.t01#4             4801   125257229:   mNeoVis.3.1.1.chr1.g00138.t01#6    mNeoVis.vulgivaga.1.1.chr1.g00131.t01#4    mNeoVis.2.1.1.chr1_hap1.g00133.t01#5    mNeoVis.vulgivaga.2.1.chr1.g00133.t01#1    mNeoVis.evergladensis.1.1.chr1.g00141.t01#3   mNeoVis.evergladensis.2.1.chr1.g00132.t01#2       4802   125257230:   mNeoVis.evergladensis.1.1.chr7.g00781.t01#3    mNeoVis.2.1.1.chr7_hap1.g00785.t01#5    mNeoVis.vulgivaga.1.1.chr7.g00783.t01#4    mNeoVis.evergladensis.2.1.chr7.g00772.t01#2   mNeoVis.vulgivaga.2.1.chr7.g00777.t01#1       4803   125257231:   mNeoVis.3.1.1.chr7.g00728.t01#6   mNeoVis.vulgivaga.2.1.chr7.g00778.t01#1             4804   125257232:   mNeoVis.2.1.1.chr7_hap1.g00787.t01#5    mNeoVis.evergladensis.2.1.chr7.g00774.t01#2    mNeoVis.vulgivaga.2.1.chr7.g00779.t01#1    mNeoVis.evergladensis.1.1.chr7.g00783.t01#3    mNeoVis.vulgivaga.1.1.chr7.g00785.t01#4   mNeoVis.3.1.1.chr7.g00727.t01#6       4805   125257233:   mNeoVis.evergladensis.2.1.chr7.g00775.t01#2    mNeoVis.evergladensis.1.1.chr7.g00784.t01#3    mNeoVis.2.1.1.chr7_hap1.g00788.t01#5    mNeoVis.vulgivaga.2.1.chr7.g00780.t01#1   mNeoVis.vulgivaga.1.1.chr7.g00786.t01#4       4806   125257234:   mNeoVis.vulgivaga.1.1.chr7.g00793.t01#4    mNeoVis.evergladensis.1.1.chr7.g00792.t01#3   mNeoVis.evergladensis.2.1.chr7.g00781.t01#2         4807   125257235:  mNeoVis.3.1.1.chr7.g00729.t01#6               4808   125257236:   mNeoVis.vulgivaga.2.1.chr7.g00781.t01#1    mNeoVis.evergladensis.1.1.chr7.g00785.t01#3    mNeoVis.2.1.1.chr7_hap1.g00789.t01#5    mNeoVis.vulgivaga.1.1.chr7.g00787.t01#4    mNeoVis.3.1.1.chr7.g00730.t01#6   mNeoVis.evergladensis.2.1.chr7.g00776.t01#2       4809   125257237:   mNeoVis.evergladensis.2.1.chr7.g00777.t01#2    mNeoVis.2.1.1.chr7_hap1.g00790.t01#5    mNeoVis.3.1.1.chr7.g00731.t01#6    mNeoVis.evergladensis.1.1.chr7.g00786.t01#3    mNeoVis.vulgivaga.2.1.chr7.g00782.t01#1   mNeoVis.vulgivaga.1.1.chr7.g00788.t01#4       4810   125257238:   mNeoVis.vulgivaga.2.1.chr7.g00783.t01#1   mNeoVis.evergladensis.1.1.chr7.g00788.t01#3           4811   125257239:   mNeoVis.evergladensis.1.1.chr7.g00787.t01#3    mNeoVis.3.1.1.chr7.g00733.t01#6    mNeoVis.vulgivaga.1.1.chr7.g00790.t01#4    mNeoVis.evergladensis.2.1.chr7.g00778.t01#2    mNeoVis.vulgivaga.2.1.chr7.g00784.t01#1   mNeoVis.2.1.1.chr7_hap1.g00791.t01#5       4812   125257240:   mNeoVis.vulgivaga.2.1.chr1.g00134.t01#1    mNeoVis.evergladensis.1.1.chr11.g01005.t01#3    mNeoVis.3.1.1.chr11.g00963.t01#6    mNeoVis.vulgivaga.2.1.chr11.g01011.t01#1    mNeoVis.2.1.1.chr11_hap1.g01013.t01#5    mNeoVis.vulgivaga.1.1.chr9.g01204.t01#4    mNeoVis.evergladensis.1.1.chr1.g00142.t01#3    mNeoVis.2.1.1.chr1_hap1.g00134.t01#5    mNeoVis.vulgivaga.1.1.chr11.g00985.t01#4      4813   125257241:   mNeoVis.2.1.1.chr1_hap1.g00135.t01#5    mNeoVis.evergladensis.2.1.chr1.g00133.t01#2    mNeoVis.vulgivaga.1.1.chr1.g00133.t01#4    mNeoVis.evergladensis.1.1.chr1.g00143.t01#3    mNeoVis.3.1.1.chr1.g00140.t01#6   mNeoVis.vulgivaga.2.1.chr1.g00135.t01#1       4814   125257242:   mNeoVis.2.1.1.chr7_hap1.g00792.t01#5    mNeoVis.evergladensis.2.1.chr7.g00779.t01#2    mNeoVis.vulgivaga.2.1.chr7.g00785.t01#1    mNeoVis.evergladensis.1.1.chr7.g00789.t01#3   mNeoVis.vulgivaga.1.1.chr7.g00791.t01#4       4815   125257243:  mNeoVis.vulgivaga.2.1.chr7.g00786.t01#1               4816   125257244:   mNeoVis.evergladensis.2.1.chr15.g00111.t01#2    mNeoVis.evergladensis.1.1.chr15.g00503.t01#3    mNeoVis.vulgivaga.2.1.chr15.g00114.t01#1    mNeoVis.2.1.1.chr15_hap1.g00110.t01#5    mNeoVis.vulgivaga.1.1.chr15.g00112.t01#4   mNeoVis.3.1.1.chr15.g00103.t01#6       4817   125257245:   mNeoVis.vulgivaga.1.1.chr7.g00031.t01#4    mNeoVis.vulgivaga.2.1.chr7.g00029.t01#1    mNeoVis.2.1.1.chr7_hap1.g00029.t01#5    mNeoVis.evergladensis.2.1.chr7.g00030.t01#2   mNeoVis.evergladensis.1.1.chr7.g00030.t01#3       4818   125257246:   mNeoVis.vulgivaga.2.1.chr1.g00136.t01#1    mNeoVis.evergladensis.2.1.chr1.g00134.t01#2    mNeoVis.evergladensis.1.1.chr1.g00144.t01#3    mNeoVis.2.1.1.chr1_hap1.g00136.t01#5    mNeoVis.vulgivaga.1.1.chr1.g00134.t01#4   mNeoVis.3.1.1.chr1.g00141.t01#6       4819   125257247:   mNeoVis.vulgivaga.2.1.chr1.g00137.t01#1    mNeoVis.evergladensis.2.1.chr1.g00135.t01#2    mNeoVis.evergladensis.1.1.chr1.g00145.t01#3    mNeoVis.3.1.1.chr1.g00142.t01#6    mNeoVis.vulgivaga.1.1.chr1.g00135.t01#4   mNeoVis.2.1.1.chr1_hap1.g00137.t01#5       4820   125257248:   mNeoVis.evergladensis.2.1.chr7.g00784.t01#2    mNeoVis.2.1.1.chr7_hap1.g00796.t01#5    mNeoVis.3.1.1.chr7.g00739.t01#6    mNeoVis.vulgivaga.2.1.chr7.g00788.t01#1   mNeoVis.evergladensis.1.1.chr7.g00795.t01#3       4821   125257249:   mNeoVis.evergladensis.1.1.chr7.g00796.t01#3   mNeoVis.vulgivaga.1.1.chr7.g00796.t01#4             4822   125257250:   mNeoVis.3.1.1.chr7.g00740.t01#6   mNeoVis.evergladensis.2.1.chr7.g00783.t01#2           4823   125257251:   mNeoVis.vulgivaga.2.1.chr7.g00795.t01#1    mNeoVis.3.1.1.chr7.g00746.t01#6    mNeoVis.evergladensis.2.1.chr7.g00790.t01#2    mNeoVis.vulgivaga.1.1.chr7.g00800.t01#4    mNeoVis.2.1.1.chr7_hap1.g00802.t01#5   mNeoVis.evergladensis.1.1.chr7.g00800.t01#3       4824   125257252:   mNeoVis.vulgivaga.2.1.chr7.g00790.t01#1    mNeoVis.3.1.1.chr7.g00742.t01#6    mNeoVis.vulgivaga.1.1.chr7.g00798.t01#4    mNeoVis.evergladensis.2.1.chr7.g00787.t01#2   mNeoVis.2.1.1.chr7_hap1.g00799.t01#5       4825   125257253:   mNeoVis.vulgivaga.2.1.chr7.g00789.t01#1    mNeoVis.evergladensis.2.1.chr7.g00786.t01#2   mNeoVis.2.1.1.chr7_hap1.g00798.t01#5           4826   125257254:   mNeoVis.vulgivaga.1.1.chr7.g00797.t01#4    mNeoVis.vulgivaga.2.1.chr7.g00791.t01#1   mNeoVis.evergladensis.1.1.chr7.g00798.t01#3         4827   125257255:   mNeoVis.vulgivaga.1.1.chr1.g00136.t01#4    mNeoVis.evergladensis.1.1.chr1.g00146.t01#3    mNeoVis.evergladensis.2.1.chr1.g00136.t01#2    mNeoVis.vulgivaga.2.1.chr1.g00138.t01#1   mNeoVis.2.1.1.chr1_hap1.g00138.t01#5       4828   125257256:   mNeoVis.vulgivaga.2.1.chr7.g00792.t01#1    mNeoVis.vulgivaga.1.1.chr7.g00799.t01#4    mNeoVis.evergladensis.1.1.chr7.g00799.t01#3    mNeoVis.3.1.1.chr7.g00743.t01#6    mNeoVis.2.1.1.chr7_hap1.g00800.t01#5   mNeoVis.evergladensis.2.1.chr7.g00788.t01#2       4829   125257257:   mNeoVis.vulgivaga.2.1.chr7.g00031.t01#1    mNeoVis.evergladensis.2.1.chr7.g00032.t01#2   mNeoVis.evergladensis.1.1.chr7.g00032.t01#3         4830   125257258:   mNeoVis.3.1.1.chr1.g00144.t01#6    mNeoVis.vulgivaga.1.1.chr1.g00137.t01#4    mNeoVis.evergladensis.2.1.chr1.g00137.t01#2    mNeoVis.evergladensis.1.1.chr1.g00147.t01#3    mNeoVis.vulgivaga.2.1.chr1.g00139.t01#1   mNeoVis.2.1.1.chr1_hap1.g00139.t01#5       4831   125257259:  mNeoVis.vulgivaga.2.1.chr7.g00794.t01#1               4832   125257260:   mNeoVis.2.1.1.chr7_hap1.g00801.t01#5    mNeoVis.evergladensis.2.1.chr7.g00789.t01#2    mNeoVis.3.1.1.chr7.g00744.t01#6   mNeoVis.vulgivaga.2.1.chr7.g00793.t01#1         4833   125257261:   mNeoVis.evergladensis.1.1.chr1.g00149.t01#3    mNeoVis.3.1.1.chr1.g00145.t01#6    mNeoVis.2.1.1.chr1_hap1.g00141.t01#5    mNeoVis.evergladensis.2.1.chr1.g00139.t01#2    mNeoVis.vulgivaga.1.1.chr1.g00139.t01#4   mNeoVis.vulgivaga.2.1.chr1.g00140.t01#1       4834   125257262:   mNeoVis.evergladensis.2.1.chr1.g00140.t01#2    mNeoVis.3.1.1.chr1.g00146.t01#6    mNeoVis.2.1.1.chr1_hap1.g00142.t01#5    mNeoVis.vulgivaga.2.1.chr1.g00141.t01#1    mNeoVis.vulgivaga.1.1.chr1.g00140.t01#4   mNeoVis.evergladensis.1.1.chr1.g00150.t01#3       4835   125257263:   mNeoVis.evergladensis.2.1.chr6.g00741.t01#2    mNeoVis.vulgivaga.1.1.chr6.g00737.t01#4    mNeoVis.3.1.1.chr6.g00741.t01#6    mNeoVis.vulgivaga.2.1.chr6.g00737.t01#1   mNeoVis.evergladensis.1.1.chr6.g00742.t01#3       4836   125257264:   mNeoVis.vulgivaga.2.1.chr7.g00796.t01#1    mNeoVis.3.1.1.chr7.g00747.t01#6   mNeoVis.2.1.1.chr7_hap1.g00803.t01#5           4837   125257265:   mNeoVis.evergladensis.1.1.chr7.g00801.t01#3    mNeoVis.vulgivaga.2.1.chr7.g00797.t01#1   mNeoVis.evergladensis.2.1.chr7.g00791.t01#2         4838   125257266:   mNeoVis.evergladensis.1.1.chr7.g00802.t01#3    mNeoVis.vulgivaga.2.1.chr7.g00798.t01#1    mNeoVis.3.1.1.chr7.g00748.t01#6   mNeoVis.evergladensis.2.1.chr7.g00792.t01#2       4839   125257267:   mNeoVis.vulgivaga.1.1.chr7.g00802.t01#4    mNeoVis.2.1.1.chr7_hap1.g00804.t01#5   mNeoVis.evergladensis.2.1.chr7.g00793.t01#2         4840   125257268:   mNeoVis.evergladensis.2.1.chr15.g00112.t01#2   mNeoVis.vulgivaga.2.1.chr15.g00115.t01#1           4841   125257269:   mNeoVis.vulgivaga.1.1.chr1.g00141.t01#4    mNeoVis.vulgivaga.2.1.chr1.g00142.t01#1    mNeoVis.evergladensis.2.1.chr1.g00141.t01#2    mNeoVis.evergladensis.1.1.chr1.g00151.t01#3    mNeoVis.3.1.1.chr1.g00147.t01#6   mNeoVis.2.1.1.chr1_hap1.g00143.t01#5       4842   125257270:   mNeoVis.vulgivaga.2.1.chr1.g00143.t01#1    mNeoVis.vulgivaga.1.1.chr12.g00248.t01#4    mNeoVis.evergladensis.1.1.chr1.g00152.t01#3    mNeoVis.evergladensis.2.1.chr1.g00142.t01#2    mNeoVis.evergladensis.2.1.chr12.g00241.t01#2    mNeoVis.2.1.1.chr1_hap1.g00144.t01#5    mNeoVis.3.1.1.chr12.g00239.t01#6    mNeoVis.vulgivaga.1.1.chr1.g00142.t01#4    mNeoVis.2.1.1.chr12_hap1.g00272.t01#5      4843   125257271:   mNeoVis.evergladensis.2.1.chr7.g00794.t01#2    mNeoVis.3.1.1.chr7.g00749.t01#6    mNeoVis.2.1.1.chr7_hap1.g00805.t01#5    mNeoVis.vulgivaga.1.1.chr7.g00803.t01#4    mNeoVis.evergladensis.1.1.chr7.g00803.t01#3    mNeoVis.3.1.1.chr7.g00750.t01#6    mNeoVis.vulgivaga.2.1.chr7.g00799.t01#1   mNeoVis.evergladensis.1.1.chr7.g00804.t01#3     4844   125257272:   mNeoVis.3.1.1.chr7.g00751.t01#6    mNeoVis.vulgivaga.2.1.chr7.g00800.t01#1    mNeoVis.evergladensis.2.1.chr7.g00795.t01#2    mNeoVis.2.1.1.chr7_hap1.g00806.t01#5    mNeoVis.vulgivaga.1.1.chr7.g00804.t01#4   mNeoVis.evergladensis.1.1.chr7.g00805.t01#3       4845   125257273:   mNeoVis.evergladensis.1.1.chr7.g00806.t01#3    mNeoVis.evergladensis.2.1.chr7.g00796.t01#2    mNeoVis.vulgivaga.2.1.chr7.g00801.t01#1    mNeoVis.3.1.1.chr7.g00752.t01#6    mNeoVis.2.1.1.chr7_hap1.g00807.t01#5   mNeoVis.vulgivaga.1.1.chr7.g00805.t01#4       4846   125257274:   mNeoVis.vulgivaga.1.1.chr1.g00143.t01#4    mNeoVis.3.1.1.chr1.g00149.t01#6    mNeoVis.vulgivaga.2.1.chr1.g00144.t01#1    mNeoVis.evergladensis.1.1.chr1.g00153.t01#3    mNeoVis.2.1.1.chr1_hap1.g00145.t01#5   mNeoVis.evergladensis.2.1.chr1.g00143.t01#2       4847   125257275:   mNeoVis.2.1.1.chr7_hap1.g00808.t01#5    mNeoVis.evergladensis.1.1.chr7.g00807.t01#3    mNeoVis.evergladensis.2.1.chr7.g00797.t01#2    mNeoVis.vulgivaga.2.1.chr7.g00802.t01#1    mNeoVis.3.1.1.chr7.g00753.t01#6   mNeoVis.vulgivaga.1.1.chr7.g00806.t01#4       4848   125257276:   mNeoVis.3.1.1.chr1.g00150.t01#6    mNeoVis.evergladensis.1.1.chr1.g00154.t01#3    mNeoVis.evergladensis.2.1.chr1.g00144.t01#2    mNeoVis.vulgivaga.2.1.chr1.g00145.t01#1    mNeoVis.2.1.1.chr1_hap1.g00146.t01#5   mNeoVis.vulgivaga.1.1.chr1.g00144.t01#4       4849   125257277:   mNeoVis.vulgivaga.2.1.chr7.g00803.t01#1    mNeoVis.vulgivaga.1.1.chr7.g00807.t01#4   mNeoVis.3.1.1.chr7.g00755.t01#6           4850   125257278:   mNeoVis.3.1.1.chr7.g00754.t01#6   mNeoVis.vulgivaga.2.1.chr7.g00804.t01#1             4851   125257279:  mNeoVis.evergladensis.2.1.chr7.g00798.t01#2             4852   125257280:  mNeoVis.evergladensis.1.1.chr7.g00808.t01#3             4853   125257281:  mNeoVis.2.1.1.chr7_hap1.g00809.t01#5               4854   125257282:   mNeoVis.vulgivaga.2.1.chr1.g00146.t01#1    mNeoVis.3.1.1.chr1.g00151.t01#6    mNeoVis.evergladensis.1.1.chr1.g00156.t01#3   mNeoVis.evergladensis.1.1.chr1.g00155.t01#3       4855   125257283:   mNeoVis.vulgivaga.2.1.chr1.g00147.t01#1    mNeoVis.evergladensis.2.1.chr1.g00146.t01#2    mNeoVis.2.1.1.chr1_hap1.g00148.t01#5    mNeoVis.3.1.1.chr1.g00152.t01#6    mNeoVis.evergladensis.1.1.chr1.g00157.t01#3   mNeoVis.vulgivaga.1.1.chr1.g00146.t01#4       4856   125257284:   mNeoVis.3.1.1.chr1.g00153.t01#6    mNeoVis.evergladensis.1.1.chr1.g00158.t01#3    mNeoVis.vulgivaga.1.1.chr1.g00147.t01#4    mNeoVis.vulgivaga.2.1.chr1.g00148.t01#1    mNeoVis.evergladensis.2.1.chr1.g00147.t01#2   mNeoVis.2.1.1.chr1_hap1.g00149.t01#5       4857   125257285:   mNeoVis.evergladensis.2.1.chr7.g00817.t01#2    mNeoVis.evergladensis.1.1.chr7.g00809.t01#3    mNeoVis.vulgivaga.1.1.chr7.g00825.t01#4    mNeoVis.2.1.1.chr7_hap1.g00810.t01#5    mNeoVis.2.1.1.chr7_hap1.g00828.t01#5    mNeoVis.3.1.1.chr7.g00756.t01#6    mNeoVis.vulgivaga.2.1.chr7.g00823.t01#1    mNeoVis.evergladensis.1.1.chr7.g00825.t01#3    mNeoVis.3.1.1.chr7.g00772.t01#6      4858   125257286:   mNeoVis.vulgivaga.1.1.chr7.g00809.t01#4    mNeoVis.2.1.1.chr7_hap1.g00811.t01#5    mNeoVis.vulgivaga.2.1.chr7.g00806.t01#1    mNeoVis.evergladensis.2.1.chr7.g00800.t01#2    mNeoVis.evergladensis.1.1.chr7.g00810.t01#3   mNeoVis.3.1.1.chr7.g00757.t01#6       4859   125257287:   mNeoVis.vulgivaga.2.1.chr1.g00149.t01#1    mNeoVis.2.1.1.chr1_hap1.g00150.t01#5    mNeoVis.evergladensis.2.1.chr1.g00148.t01#2    mNeoVis.3.1.1.chr1.g00154.t01#6    mNeoVis.vulgivaga.1.1.chr1.g00148.t01#4   mNeoVis.evergladensis.1.1.chr1.g00159.t01#3       4860   125257288:   mNeoVis.2.1.1.chr7_hap1.g00813.t01#5    mNeoVis.evergladensis.1.1.chr7.g00811.t01#3    mNeoVis.vulgivaga.1.1.chr7.g00811.t01#4   mNeoVis.vulgivaga.2.1.chr7.g00807.t01#1         4861   125257289:   mNeoVis.vulgivaga.2.1.chr7.g00808.t01#1   mNeoVis.3.1.1.chr3.g01196.t01#6             4862   125257290:   mNeoVis.2.1.1.chr7_hap1.g00032.t01#5    mNeoVis.evergladensis.1.1.chr7.g00033.t01#3    mNeoVis.vulgivaga.1.1.chr7.g00034.t01#4    mNeoVis.evergladensis.2.1.chr7.g00033.t01#2    mNeoVis.3.1.1.chr7.g00026.t01#6   mNeoVis.vulgivaga.2.1.chr7.g00032.t01#1       4863   125257291:   mNeoVis.2.1.1.chr1_hap1.g00151.t01#5    mNeoVis.evergladensis.2.1.chr1.g00149.t01#2    mNeoVis.vulgivaga.2.1.chr1.g00150.t01#1    mNeoVis.3.1.1.chr1.g00155.t01#6    mNeoVis.vulgivaga.1.1.chr1.g00149.t01#4   mNeoVis.evergladensis.1.1.chr1.g00160.t01#3       4864   125257292:   mNeoVis.3.1.1.chr6.g01057.t01#6    mNeoVis.vulgivaga.1.1.chr7.g00812.t01#4    mNeoVis.evergladensis.1.1.chr7.g00812.t01#3    mNeoVis.2.1.1.chr7_hap1.g00814.t01#5    mNeoVis.evergladensis.2.1.chr7.g00803.t01#2   mNeoVis.vulgivaga.2.1.chr7.g00809.t01#1       4865   125257293:   mNeoVis.3.1.1.chr15.g00104.t01#6    mNeoVis.2.1.1.chr15_hap1.g00112.t01#5    mNeoVis.evergladensis.1.1.chr15.g00502.t01#3    mNeoVis.vulgivaga.2.1.chr15.g00116.t01#1    mNeoVis.vulgivaga.1.1.chr15.g00113.t01#4   mNeoVis.evergladensis.2.1.chr15.g00113.t01#2       4866   125257294:   mNeoVis.evergladensis.2.1.chr7.g00804.t01#2    mNeoVis.2.1.1.chr7_hap1.g00815.t01#5    mNeoVis.evergladensis.1.1.chr7.g00813.t01#3    mNeoVis.vulgivaga.1.1.chr7.g00813.t01#4    mNeoVis.3.1.1.chr7.g00759.t01#6   mNeoVis.vulgivaga.2.1.chr7.g00810.t01#1       4867   125257295:   mNeoVis.vulgivaga.2.1.chr1.g00151.t01#1   mNeoVis.2.1.1.chr1_hap1.g00152.t01#5             4868   125257296:   mNeoVis.vulgivaga.1.1.chr7.g00814.t01#4    mNeoVis.3.1.1.chr7.g00760.t01#6    mNeoVis.2.1.1.chr7_hap1.g00816.t01#5    mNeoVis.vulgivaga.2.1.chr7.g00811.t01#1    mNeoVis.evergladensis.2.1.chr7.g00805.t01#2   mNeoVis.evergladensis.1.1.chr7.g00814.t01#3       4869   125257297:   mNeoVis.vulgivaga.2.1.chr7.g00812.t01#1    mNeoVis.2.1.1.chr7_hap1.g00817.t01#5    mNeoVis.evergladensis.2.1.chr7.g00806.t01#2    mNeoVis.evergladensis.1.1.chr7.g00815.t01#3    mNeoVis.vulgivaga.1.1.chr7.g00815.t01#4   mNeoVis.3.1.1.chr7.g00761.t01#6       4870   125257298:   mNeoVis.2.1.1.chr1_hap1.g00154.t01#5    mNeoVis.vulgivaga.1.1.chr1.g00150.t01#4    mNeoVis.vulgivaga.2.1.chr1.g00152.t01#1    mNeoVis.evergladensis.1.1.chr1.g00161.t01#3    mNeoVis.3.1.1.chr1.g00157.t01#6   mNeoVis.evergladensis.2.1.chr1.g00151.t01#2       4871   125257299:   mNeoVis.2.1.1.chr7_hap1.g00818.t01#5    mNeoVis.evergladensis.1.1.chr7.g00816.t01#3    mNeoVis.3.1.1.chr7.g00762.t01#6    mNeoVis.evergladensis.2.1.chr7.g00807.t01#2   mNeoVis.vulgivaga.2.1.chr7.g00813.t01#1       4872   125257300:   mNeoVis.3.1.1.chr1.g00158.t01#6    mNeoVis.vulgivaga.2.1.chr1.g00153.t01#1    mNeoVis.2.1.1.chr1_hap1.g00155.t01#5    mNeoVis.vulgivaga.1.1.chr1.g00151.t01#4    mNeoVis.evergladensis.2.1.chr1.g00152.t01#2   mNeoVis.evergladensis.1.1.chr1.g00162.t01#3       4873   125257301:   mNeoVis.evergladensis.2.1.chr7.g00808.t01#2    mNeoVis.vulgivaga.1.1.chr7.g00817.t01#4    mNeoVis.vulgivaga.2.1.chr7.g00814.t01#1    mNeoVis.3.1.1.chr7.g00763.t01#6    mNeoVis.2.1.1.chr7_hap1.g00819.t01#5   mNeoVis.evergladensis.1.1.chr7.g00817.t01#3       4874   125257302:   mNeoVis.evergladensis.1.1.chr1.g00163.t01#3    mNeoVis.2.1.1.chr1_hap1.g00156.t01#5    mNeoVis.vulgivaga.1.1.chr1.g00152.t01#4    mNeoVis.evergladensis.1.1.chr1.g00164.t01#3    mNeoVis.3.1.1.chr1.g00159.t01#6    mNeoVis.vulgivaga.2.1.chr1.g00154.t01#1   mNeoVis.evergladensis.2.1.chr1.g00153.t01#2     4875   125257303:   mNeoVis.evergladensis.2.1.chr7.g00809.t01#2    mNeoVis.vulgivaga.2.1.chr7.g00815.t01#1    mNeoVis.3.1.1.chr7.g00764.t01#6    mNeoVis.evergladensis.1.1.chr7.g00818.t01#3    mNeoVis.vulgivaga.1.1.chr7.g00818.t01#4   mNeoVis.2.1.1.chr7_hap1.g00820.t01#5       4876   125257304:   mNeoVis.evergladensis.1.1.chr1.g00165.t01#3    mNeoVis.2.1.1.chr1_hap1.g00157.t01#5    mNeoVis.evergladensis.2.1.chr1.g00154.t01#2    mNeoVis.vulgivaga.1.1.chr1.g00153.t01#4    mNeoVis.vulgivaga.2.1.chr1.g00155.t01#1   mNeoVis.3.1.1.chr1.g00160.t01#6       4877   125257305:   mNeoVis.evergladensis.2.1.chr7.g00810.t01#2    mNeoVis.2.1.1.chr7_hap1.g00821.t01#5    mNeoVis.vulgivaga.2.1.chr7.g00816.t01#1    mNeoVis.evergladensis.1.1.chr7.g00819.t01#3    mNeoVis.vulgivaga.1.1.chr7.g00819.t01#4   mNeoVis.3.1.1.chr7.g00765.t01#6       4878   125257306:   mNeoVis.2.1.1.chr1_hap1.g00159.t01#5    mNeoVis.vulgivaga.1.1.chr1.g00155.t01#4    mNeoVis.evergladensis.1.1.chr1.g00167.t01#3    mNeoVis.vulgivaga.2.1.chr1.g00156.t01#1    mNeoVis.evergladensis.2.1.chr1.g00155.t01#2   mNeoVis.3.1.1.chr1.g00162.t01#6       4879   125257307:   mNeoVis.2.1.1.chr7_hap1.g00822.t01#5    mNeoVis.evergladensis.1.1.chr7.g00820.t01#3    mNeoVis.vulgivaga.1.1.chr7.g00820.t01#4    mNeoVis.evergladensis.2.1.chr7.g00811.t01#2    mNeoVis.vulgivaga.2.1.chr7.g00817.t01#1   mNeoVis.3.1.1.chr7.g00766.t01#6       4880   125257308:   mNeoVis.3.1.1.chr7.g00767.t01#6    mNeoVis.vulgivaga.1.1.chr7.g00821.t01#4    mNeoVis.2.1.1.chr7_hap1.g00823.t01#5    mNeoVis.evergladensis.1.1.chr7.g00821.t01#3    mNeoVis.evergladensis.2.1.chr7.g00812.t01#2   mNeoVis.vulgivaga.2.1.chr7.g00818.t01#1       4881   125257309:   mNeoVis.vulgivaga.1.1.chr7.g00035.t01#4    mNeoVis.2.1.1.chr7_hap1.g00033.t01#5    mNeoVis.vulgivaga.2.1.chr7.g00033.t01#1    mNeoVis.evergladensis.1.1.chr7.g00034.t01#3   mNeoVis.evergladensis.2.1.chr7.g00034.t01#2       4882   125257310:   mNeoVis.2.1.1.chr1_hap1.g00085.t01#5    mNeoVis.3.1.1.chr1.g00087.t01#6    mNeoVis.evergladensis.1.1.chr1.g00088.t01#3    mNeoVis.evergladensis.2.1.chr1.g00084.t01#2    mNeoVis.vulgivaga.1.1.chr1.g00084.t01#4   mNeoVis.vulgivaga.2.1.chr1.g00080.t01#1       4883   125257311:   mNeoVis.vulgivaga.2.1.chr7.g00820.t01#1    mNeoVis.2.1.1.chr7_hap1.g00825.t01#5   mNeoVis.evergladensis.2.1.chr7.g00814.t01#2         4884   125257312:   mNeoVis.2.1.1.chr7_hap1.g00824.t01#5   mNeoVis.evergladensis.1.1.chr7.g00822.t01#3           4885   125257313:   mNeoVis.3.1.1.chr7.g00768.t01#6   mNeoVis.vulgivaga.1.1.chr7.g00822.t01#4             4886   125257314:  mNeoVis.vulgivaga.2.1.chr7.g00819.t01#1               4887   125257315:  mNeoVis.evergladensis.2.1.chr7.g00813.t01#2             4888   125257316:  mNeoVis.vulgivaga.1.1.chr7.g00823.t01#4               4889   125257317:   mNeoVis.vulgivaga.2.1.chr15.g00117.t01#1   mNeoVis.2.1.1.chr15_hap1.g00128.t01#5             4890   125257318:   mNeoVis.3.1.1.chr1.g00088.t01#6    mNeoVis.evergladensis.2.1.chr1.g00085.t01#2    mNeoVis.vulgivaga.2.1.chr1.g00081.t01#1    mNeoVis.2.1.1.chr1_hap1.g00087.t01#5    mNeoVis.vulgivaga.1.1.chr1.g00085.t01#4   mNeoVis.evergladensis.1.1.chr1.g00090.t01#3       4891   125257319:   mNeoVis.2.1.1.chr1_hap1.g00088.t01#5    mNeoVis.evergladensis.1.1.chr1.g00091.t01#3    mNeoVis.vulgivaga.1.1.chr1.g00086.t01#4    mNeoVis.vulgivaga.2.1.chr1.g00082.t01#1    mNeoVis.evergladensis.2.1.chr1.g00086.t01#2   mNeoVis.3.1.1.chr1.g00089.t01#6       4892   125257320:   mNeoVis.2.1.1.chr7_hap1.g00826.t01#5    mNeoVis.evergladensis.2.1.chr7.g00815.t01#2    mNeoVis.evergladensis.1.1.chr7.g00823.t01#3    mNeoVis.vulgivaga.2.1.chr7.g00821.t01#1    mNeoVis.3.1.1.chr7.g00770.t01#6   mNeoVis.vulgivaga.1.1.chr7.g00824.t01#4       4893   125257321:   mNeoVis.vulgivaga.2.1.chr1.g00083.t01#1   mNeoVis.evergladensis.1.1.chr1.g00092.t01#3           4894   125257322:   mNeoVis.vulgivaga.1.1.chr4.g00110.t01#4    mNeoVis.2.1.1.chr3_hap1.g01199.t01#5    mNeoVis.evergladensis.1.1.chr6.g00332.t01#3    mNeoVis.vulgivaga.2.1.chr5.g01302.t01#1    mNeoVis.evergladensis.1.1.chr10.g00416.t01#3    mNeoVis.evergladensis.2.1.chr4.g00113.t01#2    mNeoVis.evergladensis.2.1.chr10.g00105.t01#2    mNeoVis.evergladensis.1.1.chr3.g01211.t01#3    mNeoVis.3.1.1.chr4.g00107.t01#6      4895   125257323:   mNeoVis.evergladensis.2.1.chr4.g00748.t01#2    mNeoVis.evergladensis.1.1.chr4.g00761.t01#3    mNeoVis.2.1.1.chr6_hap1.g00744.t01#5    mNeoVis.3.1.1.chr4.g00753.t01#6    mNeoVis.vulgivaga.2.1.chr4.g00767.t01#1   mNeoVis.vulgivaga.1.1.chr4.g00761.t01#4       4896   125257324:   mNeoVis.vulgivaga.1.1.chr6.g00804.t01#4    mNeoVis.2.1.1.chr4_hap1.g00802.t01#5    mNeoVis.3.1.1.chr6.g00813.t01#6    mNeoVis.evergladensis.2.1.chr6.g00807.t01#2    mNeoVis.vulgivaga.2.1.chr6.g00801.t01#1   mNeoVis.evergladensis.1.1.chr8.g01117.t01#3       4897   125257325:   mNeoVis.evergladensis.1.1.chr1.g00094.t01#3    mNeoVis.3.1.1.chr1.g00091.t01#6    mNeoVis.2.1.1.chr1_hap1.g00089.t01#5    mNeoVis.vulgivaga.2.1.chr1.g00085.t01#1    mNeoVis.evergladensis.2.1.chr1.g00087.t01#2   mNeoVis.vulgivaga.1.1.chr1.g00088.t01#4       4898   125257326:   mNeoVis.evergladensis.2.1.chr7.g00816.t01#2    mNeoVis.2.1.1.chr7_hap1.g00827.t01#5    mNeoVis.vulgivaga.2.1.chr7.g00822.t01#1    mNeoVis.evergladensis.1.1.chr7.g00824.t01#3   mNeoVis.3.1.1.chr7.g00771.t01#6       4899   125257327:   mNeoVis.evergladensis.2.1.chr7.g00818.t01#2    mNeoVis.evergladensis.1.1.chr7.g00826.t01#3    mNeoVis.vulgivaga.2.1.chr7.g00824.t01#1    mNeoVis.3.1.1.chr7.g00773.t01#6    mNeoVis.vulgivaga.1.1.chr7.g00826.t01#4   mNeoVis.2.1.1.chr7_hap1.g00829.t01#5       4900   125257328:   mNeoVis.3.1.1.chr7.g00774.t01#6    mNeoVis.evergladensis.1.1.chr7.g00827.t01#3    mNeoVis.2.1.1.chr7_hap1.g00830.t01#5    mNeoVis.vulgivaga.2.1.chr7.g00825.t01#1    mNeoVis.vulgivaga.1.1.chr7.g00827.t01#4   mNeoVis.evergladensis.2.1.chr7.g00819.t01#2       4901   125257329:   mNeoVis.vulgivaga.2.1.chr1.g00086.t01#1    mNeoVis.vulgivaga.1.1.chr1.g00089.t01#4    mNeoVis.evergladensis.2.1.chr1.g00088.t01#2    mNeoVis.evergladensis.1.1.chr1.g00095.t01#3    mNeoVis.2.1.1.chr1_hap1.g00090.t01#5   mNeoVis.3.1.1.chr1.g00092.t01#6       4902   125257330:   mNeoVis.vulgivaga.1.1.chr7.g00828.t01#4    mNeoVis.vulgivaga.2.1.chr7.g00826.t01#1    mNeoVis.2.1.1.chr7_hap1.g00831.t01#5    mNeoVis.evergladensis.2.1.chr7.g00820.t01#2    mNeoVis.evergladensis.1.1.chr7.g00828.t01#3   mNeoVis.3.1.1.chr7.g00775.t01#6       4903   125257331:   mNeoVis.vulgivaga.1.1.chr7.g00829.t01#4    mNeoVis.3.1.1.chr7.g00776.t01#6    mNeoVis.evergladensis.1.1.chr7.g00829.t01#3    mNeoVis.2.1.1.chr7_hap1.g00832.t01#5    mNeoVis.vulgivaga.2.1.chr7.g00827.t01#1   mNeoVis.evergladensis.2.1.chr7.g00821.t01#2       4904   125257332:   mNeoVis.vulgivaga.2.1.chr7.g00701.t01#1    mNeoVis.evergladensis.2.1.chr7.g00702.t01#2   mNeoVis.evergladensis.1.1.chr7.g00710.t01#3         4905   125257333:   mNeoVis.vulgivaga.1.1.chr7.g00708.t01#4   mNeoVis.2.1.1.chr7_hap1.g00704.t01#5             4906   125257334:   mNeoVis.vulgivaga.2.1.chr1.g00087.t01#1    mNeoVis.3.1.1.chr1.g00093.t01#6    mNeoVis.evergladensis.2.1.chr1.g00089.t01#2    mNeoVis.vulgivaga.1.1.chr1.g00090.t01#4    mNeoVis.evergladensis.1.1.chr1.g00096.t01#3   mNeoVis.2.1.1.chr1_hap1.g00091.t01#5       4907   125257335:   mNeoVis.2.1.1.chr1_hap1.g00092.t01#5    mNeoVis.evergladensis.2.1.chr1.g00090.t01#2    mNeoVis.evergladensis.1.1.chr1.g00097.t01#3    mNeoVis.vulgivaga.2.1.chr1.g00088.t01#1    mNeoVis.3.1.1.chr1.g00094.t01#6   mNeoVis.vulgivaga.1.1.chr1.g00091.t01#4       4908   125257336:   mNeoVis.evergladensis.1.1.chr7.g00711.t01#3   mNeoVis.vulgivaga.2.1.chr7.g00702.t01#1             4909   125257337:   mNeoVis.vulgivaga.2.1.chr7.g00703.t01#1    mNeoVis.2.1.1.chr7_hap1.g00705.t01#5    mNeoVis.vulgivaga.1.1.chr7.g00709.t01#4    mNeoVis.evergladensis.1.1.chr7.g00712.t01#3   mNeoVis.evergladensis.2.1.chr7.g00703.t01#2       4910   125257338:   mNeoVis.vulgivaga.2.1.chr7.g00704.t01#1    mNeoVis.evergladensis.2.1.chr7.g00704.t01#2    mNeoVis.evergladensis.1.1.chr7.g00713.t01#3    mNeoVis.vulgivaga.1.1.chr7.g00710.t01#4   mNeoVis.2.1.1.chr7_hap1.g00706.t01#5       4911   125257339:   mNeoVis.2.1.1.chr1_hap1.g00093.t01#5    mNeoVis.evergladensis.2.1.chr1.g00091.t01#2    mNeoVis.evergladensis.1.1.chr1.g00098.t01#3    mNeoVis.vulgivaga.2.1.chr1.g00089.t01#1    mNeoVis.3.1.1.chr1.g00095.t01#6   mNeoVis.vulgivaga.1.1.chr1.g00092.t01#4       4912   125257340:   mNeoVis.vulgivaga.2.1.chr1.g00090.t01#1    mNeoVis.evergladensis.2.1.chr1.g00092.t01#2    mNeoVis.vulgivaga.1.1.chr1.g00093.t01#4    mNeoVis.evergladensis.1.1.chr1.g00099.t01#3   mNeoVis.2.1.1.chr1_hap1.g00094.t01#5       4913   125257341:  mNeoVis.vulgivaga.2.1.chr1.g00091.t01#1               4914   125257342:   mNeoVis.3.1.1.chr7.g00667.t01#6    mNeoVis.vulgivaga.2.1.chr7.g00705.t01#1    mNeoVis.evergladensis.1.1.chr7.g00714.t01#3    mNeoVis.2.1.1.chr7_hap1.g00707.t01#5    mNeoVis.evergladensis.2.1.chr7.g00705.t01#2   mNeoVis.vulgivaga.1.1.chr7.g00711.t01#4       4915   125257343:   mNeoVis.evergladensis.1.1.chr1.g00100.t01#3    mNeoVis.evergladensis.2.1.chr1.g00093.t01#2    mNeoVis.3.1.1.chr1.g00097.t01#6    mNeoVis.vulgivaga.1.1.chr1.g00094.t01#4    mNeoVis.vulgivaga.2.1.chr1.g00092.t01#1   mNeoVis.2.1.1.chr1_hap1.g00095.t01#5       4916   125257344:  mNeoVis.vulgivaga.2.1.chr1.g00093.t01#1               4917   125257345:   mNeoVis.3.1.1.chr7.g00668.t01#6    mNeoVis.evergladensis.2.1.chr7.g00706.t01#2    mNeoVis.vulgivaga.2.1.chr7.g00706.t01#1    mNeoVis.2.1.1.chr7_hap1.g00708.t01#5    mNeoVis.evergladensis.1.1.chr7.g00715.t01#3   mNeoVis.vulgivaga.1.1.chr7.g00712.t01#4       4918   125257346:   mNeoVis.evergladensis.1.1.chr7.g00716.t01#3    mNeoVis.vulgivaga.2.1.chr7.g00707.t01#1    mNeoVis.vulgivaga.1.1.chr7.g00713.t01#4    mNeoVis.2.1.1.chr7_hap1.g00709.t01#5    mNeoVis.evergladensis.2.1.chr7.g00707.t01#2   mNeoVis.3.1.1.chr7.g00669.t01#6       4919   125257347:   mNeoVis.vulgivaga.2.1.chr7.g00708.t01#1    mNeoVis.evergladensis.2.1.chr7.g00708.t01#2    mNeoVis.evergladensis.1.1.chr7.g00717.t01#3    mNeoVis.vulgivaga.1.1.chr7.g00714.t01#4   mNeoVis.2.1.1.chr7_hap1.g00710.t01#5       4920   125257348:   mNeoVis.evergladensis.2.1.chr1.g00095.t01#2    mNeoVis.vulgivaga.1.1.chr1.g00096.t01#4    mNeoVis.vulgivaga.2.1.chr1.g00094.t01#1    mNeoVis.2.1.1.chr1_hap1.g00097.t01#5   mNeoVis.3.1.1.chr1.g00099.t01#6       4921   125257349:   mNeoVis.vulgivaga.1.1.chr1.g00097.t01#4    mNeoVis.evergladensis.2.1.chr1.g00096.t01#2    mNeoVis.3.1.1.chr1.g00100.t01#6    mNeoVis.2.1.1.chr1_hap1.g00098.t01#5    mNeoVis.evergladensis.1.1.chr1.g00103.t01#3   mNeoVis.vulgivaga.2.1.chr1.g00095.t01#1       4922   125257350:   mNeoVis.vulgivaga.1.1.chr1.g00098.t01#4    mNeoVis.evergladensis.2.1.chr1.g00097.t01#2    mNeoVis.vulgivaga.2.1.chr1.g00096.t01#1    mNeoVis.evergladensis.1.1.chr1.g00104.t01#3    mNeoVis.3.1.1.chr1.g00101.t01#6   mNeoVis.2.1.1.chr1_hap1.g00099.t01#5       4923   125257351:   mNeoVis.vulgivaga.2.1.chr1.g00097.t01#1    mNeoVis.vulgivaga.1.1.chr1.g00099.t01#4    mNeoVis.3.1.1.chr1.g00102.t01#6    mNeoVis.2.1.1.chr1_hap1.g00100.t01#5    mNeoVis.evergladensis.2.1.chr1.g00098.t01#2   mNeoVis.evergladensis.1.1.chr1.g00105.t01#3       4924   125257352:   mNeoVis.2.1.1.chr7_hap1.g00711.t01#5    mNeoVis.evergladensis.2.1.chr7.g00709.t01#2    mNeoVis.vulgivaga.1.1.chr7.g00715.t01#4    mNeoVis.vulgivaga.2.1.chr7.g00709.t01#1   mNeoVis.evergladensis.1.1.chr7.g00718.t01#3       4925   125257353:   mNeoVis.evergladensis.2.1.chr7.g00710.t01#2    mNeoVis.2.1.1.chr7_hap1.g00712.t01#5    mNeoVis.vulgivaga.1.1.chr7.g00716.t01#4    mNeoVis.3.1.1.chr7.g00671.t01#6    mNeoVis.evergladensis.1.1.chr7.g00719.t01#3   mNeoVis.vulgivaga.2.1.chr7.g00710.t01#1       4926   125257354:   mNeoVis.evergladensis.1.1.chr7.g00720.t01#3    mNeoVis.vulgivaga.1.1.chr7.g00717.t01#4    mNeoVis.vulgivaga.2.1.chr7.g00711.t01#1    mNeoVis.3.1.1.chr7.g00672.t01#6    mNeoVis.evergladensis.2.1.chr7.g00711.t01#2   mNeoVis.2.1.1.chr7_hap1.g00713.t01#5       4927   125257355:   mNeoVis.vulgivaga.1.1.chr1.g00100.t01#4    mNeoVis.evergladensis.1.1.chr1.g00106.t01#3    mNeoVis.evergladensis.2.1.chr1.g00099.t01#2    mNeoVis.3.1.1.chr1.g00103.t01#6    mNeoVis.2.1.1.chr1_hap1.g00101.t01#5   mNeoVis.vulgivaga.2.1.chr1.g00098.t01#1       4928   125257356:   mNeoVis.vulgivaga.1.1.chr15.g00115.t01#4    mNeoVis.evergladensis.1.1.chr15.g00500.t01#3    mNeoVis.vulgivaga.2.1.chr15.g00118.t01#1    mNeoVis.2.1.1.chr15_hap1.g00129.t01#5    mNeoVis.3.1.1.chr15.g00106.t01#6   mNeoVis.evergladensis.2.1.chr15.g00115.t01#2       4929   125257357:   mNeoVis.evergladensis.1.1.chr7.g00721.t01#3    mNeoVis.vulgivaga.1.1.chr7.g00718.t01#4    mNeoVis.evergladensis.2.1.chr7.g00712.t01#2    mNeoVis.2.1.1.chr7_hap1.g00714.t01#5    mNeoVis.vulgivaga.2.1.chr7.g00712.t01#1   mNeoVis.3.1.1.chr7.g00673.t01#6       4930   125257358:   mNeoVis.vulgivaga.2.1.chr1.g00099.t01#1    mNeoVis.evergladensis.2.1.chr1.g00100.t01#2   mNeoVis.evergladensis.1.1.chr1.g00107.t01#3         4931   125257359:   mNeoVis.2.1.1.chr7_hap1.g00715.t01#5    mNeoVis.vulgivaga.2.1.chr7.g00713.t01#1    mNeoVis.vulgivaga.1.1.chr7.g00719.t01#4    mNeoVis.evergladensis.1.1.chr7.g00722.t01#3    mNeoVis.3.1.1.chr7.g00674.t01#6   mNeoVis.evergladensis.2.1.chr7.g00713.t01#2       4932   125257360:   mNeoVis.evergladensis.1.1.chr7.g00723.t01#3    mNeoVis.vulgivaga.1.1.chr7.g00720.t01#4    mNeoVis.3.1.1.chr7.g00675.t01#6    mNeoVis.evergladensis.2.1.chr7.g00714.t01#2    mNeoVis.vulgivaga.2.1.chr7.g00714.t01#1   mNeoVis.2.1.1.chr7_hap1.g00716.t01#5       4933   125257361:   mNeoVis.2.1.1.chr7_hap1.g00717.t01#5    mNeoVis.evergladensis.2.1.chr7.g00715.t01#2    mNeoVis.3.1.1.chr7.g00676.t01#6    mNeoVis.vulgivaga.2.1.chr7.g00715.t01#1    mNeoVis.evergladensis.1.1.chr7.g00724.t01#3   mNeoVis.vulgivaga.1.1.chr7.g00721.t01#4       4934   125257362:   mNeoVis.3.1.1.chr15.g00107.t01#6    mNeoVis.2.1.1.chr15_hap1.g00130.t01#5    mNeoVis.vulgivaga.2.1.chr15.g00119.t01#1    mNeoVis.evergladensis.2.1.chr15.g00116.t01#2    mNeoVis.evergladensis.1.1.chr15.g00499.t01#3   mNeoVis.vulgivaga.1.1.chr15.g00116.t01#4       4935   125257363:   mNeoVis.2.1.1.chr1_hap1.g00103.t01#5    mNeoVis.evergladensis.1.1.chr1.g00108.t01#3   mNeoVis.vulgivaga.2.1.chr1.g00100.t01#1           4936   125257364:   mNeoVis.evergladensis.2.1.chr1.g00101.t01#2    mNeoVis.vulgivaga.2.1.chr1.g00101.t01#1    mNeoVis.evergladensis.1.1.chr1.g00110.t01#3    mNeoVis.vulgivaga.1.1.chr1.g00103.t01#4    mNeoVis.3.1.1.chr1.g00106.t01#6   mNeoVis.2.1.1.chr1_hap1.g00104.t01#5       4937   125257365:   mNeoVis.evergladensis.2.1.chr1.g00102.t01#2    mNeoVis.2.1.1.chr1_hap1.g00105.t01#5    mNeoVis.3.1.1.chr1.g00107.t01#6    mNeoVis.evergladensis.1.1.chr1.g00111.t01#3    mNeoVis.vulgivaga.1.1.chr1.g00104.t01#4   mNeoVis.vulgivaga.2.1.chr1.g00102.t01#1       4938   125257366:   mNeoVis.evergladensis.1.1.chr1.g00112.t01#3    mNeoVis.evergladensis.2.1.chr1.g00103.t01#2    mNeoVis.3.1.1.chr1.g00109.t01#6    mNeoVis.evergladensis.1.1.chr1.g00113.t01#3    mNeoVis.vulgivaga.1.1.chr1.g00106.t01#4    mNeoVis.2.1.1.chr1_hap1.g00106.t01#5    mNeoVis.vulgivaga.1.1.chr1.g00105.t01#4   mNeoVis.3.1.1.chr1.g00108.t01#6     4939   125257367:   mNeoVis.evergladensis.1.1.chr1.g00114.t01#3    mNeoVis.vulgivaga.1.1.chr1.g00107.t01#4    mNeoVis.3.1.1.chr1.g00110.t01#6    mNeoVis.2.1.1.chr1_hap1.g00107.t01#5   mNeoVis.evergladensis.2.1.chr1.g00105.t01#2       4940   125257368:  mNeoVis.vulgivaga.2.1.chr1.g00103.t01#1               4941   125257369:   mNeoVis.vulgivaga.2.1.chr7.g00716.t01#1    mNeoVis.3.1.1.chr7.g00677.t01#6    mNeoVis.2.1.1.chr7_hap1.g00718.t01#5    mNeoVis.evergladensis.1.1.chr7.g00725.t01#3    mNeoVis.evergladensis.2.1.chr7.g00717.t01#2   mNeoVis.vulgivaga.1.1.chr7.g00723.t01#4       4942   125257370:   mNeoVis.vulgivaga.2.1.chr1.g00104.t01#1    mNeoVis.evergladensis.2.1.chr1.g00106.t01#2    mNeoVis.3.1.1.chr1.g00111.t01#6    mNeoVis.evergladensis.1.1.chr1.g00115.t01#3    mNeoVis.2.1.1.chr1_hap1.g00108.t01#5   mNeoVis.vulgivaga.1.1.chr1.g00108.t01#4       4943   125257371:   mNeoVis.3.1.1.chr7.g00678.t01#6    mNeoVis.vulgivaga.1.1.chr7.g00724.t01#4    mNeoVis.evergladensis.2.1.chr7.g00718.t01#2    mNeoVis.evergladensis.1.1.chr7.g00726.t01#3    mNeoVis.2.1.1.chr7_hap1.g00719.t01#5   mNeoVis.vulgivaga.2.1.chr7.g00717.t01#1       4944   125257372:   mNeoVis.3.1.1.chr7.g00679.t01#6    mNeoVis.vulgivaga.2.1.chr7.g00718.t01#1    mNeoVis.vulgivaga.1.1.chr7.g00725.t01#4    mNeoVis.evergladensis.2.1.chr7.g00723.t01#2    mNeoVis.evergladensis.1.1.chr7.g00727.t01#3   mNeoVis.2.1.1.chr7_hap1.g00720.t01#5       4945   125257373:   mNeoVis.2.1.1.chr7_hap1.g00733.t01#5    mNeoVis.vulgivaga.2.1.chr7.g00725.t01#1    mNeoVis.evergladensis.1.1.chr7.g00732.t01#3    mNeoVis.vulgivaga.1.1.chr7.g00732.t01#4   mNeoVis.3.1.1.chr7.g00686.t01#6       4946   125257374:   mNeoVis.evergladensis.1.1.chr15.g00498.t01#3    mNeoVis.vulgivaga.1.1.chr15.g00117.t01#4    mNeoVis.vulgivaga.2.1.chr15.g00120.t01#1    mNeoVis.3.1.1.chr15.g00108.t01#6    mNeoVis.2.1.1.chr15_hap1.g00131.t01#5   mNeoVis.evergladensis.2.1.chr15.g00117.t01#2       4947   125257375:   mNeoVis.evergladensis.1.1.chr1.g00116.t01#3    mNeoVis.vulgivaga.2.1.chr1.g00105.t01#1    mNeoVis.3.1.1.chr1.g00112.t01#6    mNeoVis.vulgivaga.1.1.chr1.g00109.t01#4    mNeoVis.evergladensis.2.1.chr1.g00107.t01#2   mNeoVis.2.1.1.chr1_hap1.g00109.t01#5       4948   125257376:   mNeoVis.vulgivaga.1.1.chr7.g00727.t01#4    mNeoVis.2.1.1.chr7_hap1.g00729.t01#5    mNeoVis.evergladensis.1.1.chr7.g00729.t01#3    mNeoVis.vulgivaga.2.1.chr7.g00721.t01#1    mNeoVis.evergladensis.2.1.chr7.g00720.t01#2   mNeoVis.3.1.1.chr7.g00681.t01#6       4949   125257377:   mNeoVis.3.1.1.chr1.g00113.t01#6    mNeoVis.2.1.1.chr1_hap1.g00110.t01#5    mNeoVis.vulgivaga.1.1.chr1.g00110.t01#4    mNeoVis.evergladensis.1.1.chr1.g00117.t01#3    mNeoVis.vulgivaga.2.1.chr1.g00106.t01#1   mNeoVis.evergladensis.2.1.chr1.g00108.t01#2       4950   125257378:   mNeoVis.2.1.1.chr1_hap1.g00111.t01#5    mNeoVis.vulgivaga.1.1.chr1.g00111.t01#4    mNeoVis.vulgivaga.2.1.chr1.g00107.t01#1    mNeoVis.evergladensis.2.1.chr1.g00109.t01#2    mNeoVis.3.1.1.chr1.g00114.t01#6   mNeoVis.evergladensis.1.1.chr1.g00118.t01#3       4951   125257379:   mNeoVis.2.1.1.chr7_hap1.g00730.t01#5    mNeoVis.evergladensis.1.1.chr7.g00730.t01#3    mNeoVis.evergladensis.2.1.chr7.g00721.t01#2    mNeoVis.3.1.1.chr7.g00682.t01#6    mNeoVis.vulgivaga.2.1.chr7.g00722.t01#1   mNeoVis.vulgivaga.1.1.chr7.g00728.t01#4       4952   125257380:   mNeoVis.2.1.1.chr1_hap1.g00112.t01#5    mNeoVis.vulgivaga.1.1.chr1.g00112.t01#4    mNeoVis.3.1.1.chr1.g00116.t01#6    mNeoVis.evergladensis.2.1.chr1.g00110.t01#2    mNeoVis.vulgivaga.2.1.chr1.g00109.t01#1   mNeoVis.evergladensis.1.1.chr1.g00119.t01#3       4953   125257381:   mNeoVis.3.1.1.chr7.g00683.t01#6    mNeoVis.vulgivaga.2.1.chr7.g00723.t01#1    mNeoVis.vulgivaga.1.1.chr7.g00729.t01#4    mNeoVis.2.1.1.chr7_hap1.g00731.t01#5    mNeoVis.evergladensis.2.1.chr7.g00722.t01#2   mNeoVis.evergladensis.1.1.chr7.g00731.t01#3       4954   125257382:   mNeoVis.evergladensis.2.1.chr1.g00111.t01#2    mNeoVis.evergladensis.1.1.chr1.g00120.t01#3    mNeoVis.vulgivaga.1.1.chr1.g00113.t01#4    mNeoVis.2.1.1.chr1_hap1.g00113.t01#5    mNeoVis.3.1.1.chr1.g00117.t01#6   mNeoVis.vulgivaga.2.1.chr1.g00110.t01#1       4955   125257383:   mNeoVis.3.1.1.chr7.g00687.t01#6    mNeoVis.evergladensis.1.1.chr7.g00733.t01#3    mNeoVis.evergladensis.2.1.chr7.g00724.t01#2    mNeoVis.vulgivaga.1.1.chr7.g00733.t01#4    mNeoVis.vulgivaga.2.1.chr7.g00726.t01#1   mNeoVis.2.1.1.chr7_hap1.g00734.t01#5       4956   125257384:   mNeoVis.2.1.1.chr7_hap1.g00735.t01#5    mNeoVis.vulgivaga.1.1.chr7.g00734.t01#4    mNeoVis.vulgivaga.2.1.chr7.g00727.t01#1    mNeoVis.evergladensis.2.1.chr7.g00725.t01#2    mNeoVis.3.1.1.chr7.g00688.t01#6   mNeoVis.evergladensis.1.1.chr7.g00734.t01#3       4957   125257385:   mNeoVis.2.1.1.chr1_hap1.g00114.t01#5    mNeoVis.evergladensis.1.1.chr1.g00121.t01#3    mNeoVis.3.1.1.chr1.g00118.t01#6    mNeoVis.vulgivaga.1.1.chr1.g00114.t01#4    mNeoVis.evergladensis.2.1.chr1.g00112.t01#2   mNeoVis.vulgivaga.2.1.chr1.g00111.t01#1       4958   125257386:   mNeoVis.vulgivaga.2.1.chr7.g00728.t01#1    mNeoVis.evergladensis.1.1.chr7.g00735.t01#3    mNeoVis.evergladensis.2.1.chr7.g00726.t01#2    mNeoVis.vulgivaga.1.1.chr7.g00735.t01#4   mNeoVis.2.1.1.chr7_hap1.g00736.t01#5       4959   125257387:   mNeoVis.evergladensis.2.1.chr7.g00727.t01#2    mNeoVis.2.1.1.chr7_hap1.g00737.t01#5    mNeoVis.vulgivaga.1.1.chr7.g00736.t01#4    mNeoVis.3.1.1.chr7.g00690.t01#6    mNeoVis.vulgivaga.2.1.chr7.g00729.t01#1   mNeoVis.evergladensis.1.1.chr7.g00736.t01#3       4960   125257388:   mNeoVis.2.1.1.chr7_hap1.g00738.t01#5    mNeoVis.3.1.1.chr7.g02153.t01#6    mNeoVis.vulgivaga.1.1.chr7.g00737.t01#4    mNeoVis.vulgivaga.2.1.chr7.g00730.t01#1    mNeoVis.evergladensis.1.1.chr7.g00737.t01#3   mNeoVis.evergladensis.2.1.chr7.g00728.t01#2       4961   125257389:   mNeoVis.vulgivaga.1.1.chr7.g00738.t01#4    mNeoVis.evergladensis.2.1.chr7.g00729.t01#2    mNeoVis.2.1.1.chr7_hap1.g00739.t01#5    mNeoVis.evergladensis.1.1.chr7.g00738.t01#3   mNeoVis.vulgivaga.2.1.chr7.g00731.t01#1       4962   125257390:   mNeoVis.evergladensis.1.1.chr7.g00739.t01#3    mNeoVis.evergladensis.2.1.chr7.g00730.t01#2    mNeoVis.3.1.1.chr7.g00692.t01#6    mNeoVis.2.1.1.chr7_hap1.g00740.t01#5   mNeoVis.vulgivaga.2.1.chr7.g00732.t01#1       4963   125257391:   mNeoVis.vulgivaga.2.1.chr7.g00733.t01#1    mNeoVis.evergladensis.2.1.chr7.g00731.t01#2   mNeoVis.evergladensis.1.1.chr7.g00740.t01#3         4964   125257392:   mNeoVis.3.1.1.chr1.g00119.t01#6    mNeoVis.vulgivaga.1.1.chr1.g00115.t01#4    mNeoVis.evergladensis.2.1.chr1.g00113.t01#2    mNeoVis.vulgivaga.2.1.chr1.g00112.t01#1    mNeoVis.2.1.1.chr1_hap1.g00115.t01#5   mNeoVis.evergladensis.1.1.chr1.g00122.t01#3       4965   125257393:   mNeoVis.vulgivaga.2.1.chr7.g00734.t01#1    mNeoVis.evergladensis.1.1.chr7.g00741.t01#3    mNeoVis.2.1.1.chr7_hap1.g00743.t01#5    mNeoVis.vulgivaga.1.1.chr7.g00741.t01#4   mNeoVis.3.1.1.chr7.g00694.t01#6       4966   125257394:  mNeoVis.vulgivaga.2.1.chr7.g00736.t01#1               4967   125257395:   mNeoVis.2.1.1.chr7_hap1.g00747.t01#5    mNeoVis.vulgivaga.1.1.chr7.g00745.t01#4    mNeoVis.2.1.1.chr7_hap1.g00742.t01#5    mNeoVis.vulgivaga.2.1.chr7.g00738.t01#1    mNeoVis.evergladensis.2.1.chr7.g00735.t01#2    mNeoVis.3.1.1.chr7.g00698.t01#6   mNeoVis.evergladensis.1.1.chr7.g00745.t01#3     4968   125257396:   mNeoVis.evergladensis.2.1.chr7.g00732.t01#2    mNeoVis.2.1.1.chr7_hap1.g00744.t01#5    mNeoVis.vulgivaga.2.1.chr7.g00735.t01#1    mNeoVis.3.1.1.chr7.g00695.t01#6    mNeoVis.vulgivaga.1.1.chr7.g00743.t01#4   mNeoVis.evergladensis.1.1.chr7.g00742.t01#3       4969   125257397:   mNeoVis.vulgivaga.2.1.chr7.g00737.t01#1    mNeoVis.3.1.1.chr7.g00697.t01#6    mNeoVis.evergladensis.2.1.chr7.g00734.t01#2    mNeoVis.evergladensis.1.1.chr7.g00744.t01#3    mNeoVis.vulgivaga.1.1.chr7.g00744.t01#4   mNeoVis.2.1.1.chr7_hap1.g00746.t01#5       4970   125257398:   mNeoVis.evergladensis.1.1.chr1.g00046.t01#3    mNeoVis.vulgivaga.1.1.chr1.g00043.t01#4    mNeoVis.3.1.1.chr1.g00049.t01#6    mNeoVis.2.1.1.chr1_hap1.g00046.t01#5   mNeoVis.vulgivaga.2.1.chr1.g00045.t01#1       4971   125257399:   mNeoVis.vulgivaga.2.1.chr1.g00046.t01#1    mNeoVis.3.1.1.chr1.g00050.t01#6    mNeoVis.evergladensis.1.1.chr1.g00047.t01#3    mNeoVis.vulgivaga.1.1.chr1.g00044.t01#4    mNeoVis.evergladensis.2.1.chr1.g00044.t01#2   mNeoVis.2.1.1.chr1_hap1.g00047.t01#5       4972   125257400:   mNeoVis.vulgivaga.2.1.chr7.g00739.t01#1    mNeoVis.evergladensis.1.1.chr7.g00746.t01#3    mNeoVis.2.1.1.chr7_hap1.g00748.t01#5    mNeoVis.vulgivaga.1.1.chr7.g00746.t01#4    mNeoVis.3.1.1.chr7.g00699.t01#6   mNeoVis.evergladensis.2.1.chr7.g00736.t01#2       4973   125257401:   mNeoVis.3.1.1.chr7.g00700.t01#6    mNeoVis.vulgivaga.2.1.chr7.g00740.t01#1    mNeoVis.evergladensis.1.1.chr7.g00747.t01#3    mNeoVis.evergladensis.2.1.chr7.g00737.t01#2    mNeoVis.vulgivaga.1.1.chr7.g00747.t01#4   mNeoVis.2.1.1.chr7_hap1.g00749.t01#5       4974   125257402:   mNeoVis.vulgivaga.2.1.chr7.g00741.t01#1    mNeoVis.3.1.1.chr7.g00701.t01#6    mNeoVis.evergladensis.1.1.chr7.g00748.t01#3    mNeoVis.2.1.1.chr7_hap1.g00750.t01#5    mNeoVis.evergladensis.2.1.chr7.g00738.t01#2   mNeoVis.vulgivaga.1.1.chr7.g00748.t01#4       4975   125257403:   mNeoVis.2.1.1.chr1_hap1.g00048.t01#5    mNeoVis.evergladensis.2.1.chr1.g00045.t01#2    mNeoVis.3.1.1.chr1.g00051.t01#6    mNeoVis.vulgivaga.1.1.chr1.g00045.t01#4    mNeoVis.evergladensis.1.1.chr1.g00048.t01#3   mNeoVis.vulgivaga.2.1.chr1.g00047.t01#1       4976   125257404:   mNeoVis.vulgivaga.2.1.chr1.g00048.t01#1    mNeoVis.3.1.1.chr1.g00052.t01#6    mNeoVis.vulgivaga.1.1.chr1.g00046.t01#4    mNeoVis.evergladensis.2.1.chr1.g00046.t01#2    mNeoVis.2.1.1.chr1_hap1.g00049.t01#5   mNeoVis.evergladensis.1.1.chr1.g00049.t01#3       4977   125257405:   mNeoVis.evergladensis.2.1.chr1.g00047.t01#2    mNeoVis.evergladensis.1.1.chr1.g00050.t01#3    mNeoVis.vulgivaga.2.1.chr1.g00049.t01#1    mNeoVis.vulgivaga.1.1.chr1.g00047.t01#4    mNeoVis.2.1.1.chr1_hap1.g00050.t01#5   mNeoVis.3.1.1.chr1.g00053.t01#6       4978   125257406:   mNeoVis.2.1.1.chr1_hap1.g00051.t01#5    mNeoVis.evergladensis.2.1.chr1.g00048.t01#2    mNeoVis.3.1.1.chr1.g00054.t01#6   mNeoVis.vulgivaga.2.1.chr1.g00050.t01#1         4979   125257407:   mNeoVis.evergladensis.1.1.chr1.g00052.t01#3    mNeoVis.evergladensis.2.1.chr1.g00049.t01#2    mNeoVis.vulgivaga.2.1.chr1.g00051.t01#1    mNeoVis.2.1.1.chr1_hap1.g00052.t01#5   mNeoVis.vulgivaga.1.1.chr1.g00048.t01#4       4980   125257408:   mNeoVis.evergladensis.2.1.chr7.g00739.t01#2    mNeoVis.vulgivaga.2.1.chr7.g00742.t01#1    mNeoVis.3.1.1.chr7.g00702.t01#6    mNeoVis.evergladensis.1.1.chr7.g00749.t01#3    mNeoVis.vulgivaga.1.1.chr7.g00749.t01#4   mNeoVis.2.1.1.chr7_hap1.g00751.t01#5       4981   125257409:   mNeoVis.evergladensis.2.1.chr7.g00740.t01#2    mNeoVis.vulgivaga.2.1.chr7.g00743.t01#1    mNeoVis.evergladensis.1.1.chr7.g00750.t01#3    mNeoVis.vulgivaga.1.1.chr7.g00750.t01#4   mNeoVis.2.1.1.chr7_hap1.g00752.t01#5       4982   125257410:   mNeoVis.vulgivaga.2.1.chr7.g00744.t01#1    mNeoVis.evergladensis.1.1.chr7.g00751.t01#3    mNeoVis.2.1.1.chr7_hap1.g00753.t01#5    mNeoVis.vulgivaga.1.1.chr7.g00751.t01#4   mNeoVis.evergladensis.2.1.chr7.g00741.t01#2       4983   125257411:   mNeoVis.vulgivaga.2.1.chr1.g00052.t01#1   mNeoVis.2.1.1.chr1_hap1.g00053.t01#5             4984   125257412:   mNeoVis.evergladensis.2.1.chr7.g00742.t01#2    mNeoVis.vulgivaga.2.1.chr7.g00745.t01#1    mNeoVis.evergladensis.1.1.chr7.g00752.t01#3    mNeoVis.3.1.1.chr7.g00705.t01#6    mNeoVis.vulgivaga.1.1.chr7.g00752.t01#4   mNeoVis.2.1.1.chr7_hap1.g00754.t01#5       4985   125257413:   mNeoVis.vulgivaga.2.1.chr1.g00053.t01#1    mNeoVis.2.1.1.chr1_hap1.g00054.t01#5    mNeoVis.vulgivaga.1.1.chr1.g00050.t01#4    mNeoVis.3.1.1.chr1.g00056.t01#6    mNeoVis.evergladensis.1.1.chr1.g00053.t01#3   mNeoVis.evergladensis.2.1.chr1.g00050.t01#2       4986   125257414:   mNeoVis.evergladensis.1.1.chr7.g00754.t01#3    mNeoVis.vulgivaga.2.1.chr7.g00747.t01#1    mNeoVis.vulgivaga.1.1.chr7.g00754.t01#4    mNeoVis.evergladensis.2.1.chr7.g00744.t01#2    mNeoVis.2.1.1.chr7_hap1.g00756.t01#5   mNeoVis.3.1.1.chr7.g00707.t01#6       4987   125257415:   mNeoVis.evergladensis.2.1.chr7.g00746.t01#2    mNeoVis.evergladensis.1.1.chr7.g00756.t01#3    mNeoVis.2.1.1.chr7_hap1.g00758.t01#5    mNeoVis.vulgivaga.1.1.chr7.g00757.t01#4   mNeoVis.vulgivaga.2.1.chr7.g00748.t01#1       4988   125257416:   mNeoVis.2.1.1.chr7_hap1.g00606.t01#5    mNeoVis.vulgivaga.2.1.chr7.g00612.t01#1    mNeoVis.evergladensis.2.1.chr7.g00608.t01#2    mNeoVis.vulgivaga.1.1.chr7.g00615.t01#4    mNeoVis.evergladensis.1.1.chr7.g00616.t01#3   mNeoVis.3.1.1.chr7.g00589.t01#6       4989   125257417:   mNeoVis.vulgivaga.1.1.chr7.g00616.t01#4    mNeoVis.evergladensis.1.1.chr7.g00617.t01#3    mNeoVis.3.1.1.chr7.g00590.t01#6    mNeoVis.2.1.1.chr7_hap1.g00607.t01#5    mNeoVis.vulgivaga.2.1.chr7.g00613.t01#1   mNeoVis.evergladensis.2.1.chr7.g00609.t01#2       4990   125257418:   mNeoVis.vulgivaga.2.1.chr7.g00614.t01#1    mNeoVis.3.1.1.chr7.g00591.t01#6   mNeoVis.evergladensis.1.1.chr7.g00618.t01#3         4991   125257419:   mNeoVis.vulgivaga.1.1.chr7.g00618.t01#4    mNeoVis.vulgivaga.2.1.chr7.g00615.t01#1    mNeoVis.evergladensis.1.1.chr7.g00619.t01#3    mNeoVis.2.1.1.chr7_hap1.g00609.t01#5   mNeoVis.evergladensis.2.1.chr7.g00611.t01#2       4992   125257420:   mNeoVis.evergladensis.2.1.chr7.g00613.t01#2    mNeoVis.2.1.1.chr7_hap1.g00611.t01#5    mNeoVis.vulgivaga.1.1.chr7.g00619.t01#4    mNeoVis.vulgivaga.2.1.chr7.g00616.t01#1    mNeoVis.evergladensis.1.1.chr7.g00620.t01#3   mNeoVis.3.1.1.chr7.g00592.t01#6       4993   125257421:   mNeoVis.evergladensis.2.1.chr7.g00614.t01#2    mNeoVis.vulgivaga.1.1.chr7.g00620.t01#4    mNeoVis.vulgivaga.2.1.chr7.g00617.t01#1   mNeoVis.2.1.1.chr7_hap1.g00612.t01#5         4994   125257422:   mNeoVis.evergladensis.1.1.chr7.g00621.t01#3    mNeoVis.vulgivaga.1.1.chr7.g00621.t01#4    mNeoVis.3.1.1.chr4.g00622.t01#6    mNeoVis.evergladensis.2.1.chr7.g00615.t01#2    mNeoVis.vulgivaga.2.1.chr7.g00618.t01#1   mNeoVis.2.1.1.chr7_hap1.g00613.t01#5       4995   125257423:   mNeoVis.vulgivaga.1.1.chr7.g00622.t01#4    mNeoVis.evergladensis.2.1.chr7.g00616.t01#2    mNeoVis.2.1.1.chr7_hap1.g00614.t01#5    mNeoVis.vulgivaga.2.1.chr7.g00619.t01#1   mNeoVis.3.1.1.chr7.g00593.t01#6       4996   125257424:   mNeoVis.2.1.1.chr7_hap1.g00034.t01#5    mNeoVis.vulgivaga.2.1.chr7.g00034.t01#1    mNeoVis.evergladensis.2.1.chr7.g00035.t01#2    mNeoVis.evergladensis.1.1.chr7.g00035.t01#3   mNeoVis.vulgivaga.1.1.chr7.g00036.t01#4       4997   125257425:   mNeoVis.vulgivaga.2.1.chr1.g00054.t01#1    mNeoVis.3.1.1.chr1.g00057.t01#6    mNeoVis.2.1.1.chr1_hap1.g00055.t01#5    mNeoVis.vulgivaga.1.1.chr1.g00051.t01#4   mNeoVis.evergladensis.1.1.chr1.g00054.t01#3       4998   125257426:   mNeoVis.vulgivaga.1.1.chr7.g00623.t01#4    mNeoVis.vulgivaga.2.1.chr7.g00620.t01#1    mNeoVis.2.1.1.chr7_hap1.g00615.t01#5    mNeoVis.3.1.1.chr7.g00594.t01#6   mNeoVis.evergladensis.2.1.chr7.g00617.t01#2       4999   125257427:   mNeoVis.evergladensis.2.1.chr7.g00618.t01#2    mNeoVis.evergladensis.1.1.chr7.g00624.t01#3    mNeoVis.vulgivaga.1.1.chr7.g00624.t01#4    mNeoVis.vulgivaga.2.1.chr7.g00621.t01#1   mNeoVis.2.1.1.chr7_hap1.g00616.t01#5       5000   125257428:   mNeoVis.vulgivaga.1.1.chr7.g00625.t01#4    mNeoVis.vulgivaga.2.1.chr7.g00622.t01#1    mNeoVis.3.1.1.chr7.g00595.t01#6    mNeoVis.2.1.1.chr7_hap1.g00617.t01#5   mNeoVis.evergladensis.1.1.chr7.g00625.t01#3       5001   125257429:   mNeoVis.evergladensis.1.1.chr7.g00626.t01#3    mNeoVis.vulgivaga.1.1.chr7.g00626.t01#4    mNeoVis.evergladensis.2.1.chr7.g00619.t01#2    mNeoVis.3.1.1.chr7.g00596.t01#6    mNeoVis.2.1.1.chr7_hap1.g00618.t01#5   mNeoVis.vulgivaga.2.1.chr7.g00623.t01#1       5002   125257430:   mNeoVis.evergladensis.1.1.chr7.g00627.t01#3    mNeoVis.evergladensis.2.1.chr7.g00620.t01#2    mNeoVis.vulgivaga.1.1.chr7.g00627.t01#4    mNeoVis.vulgivaga.2.1.chr7.g00624.t01#1   mNeoVis.2.1.1.chr7_hap1.g00619.t01#5       5003   125257431:   mNeoVis.3.1.1.chr7.g00597.t01#6    mNeoVis.vulgivaga.2.1.chr7.g00625.t01#1    mNeoVis.evergladensis.1.1.chr7.g00628.t01#3   mNeoVis.evergladensis.2.1.chr7.g00621.t01#2       5004   125257432:   mNeoVis.vulgivaga.1.1.chr7.g00628.t01#4    mNeoVis.vulgivaga.2.1.chr7.g00626.t01#1    mNeoVis.3.1.1.chr7.g00598.t01#6    mNeoVis.2.1.1.chr7_hap1.g00620.t01#5    mNeoVis.evergladensis.1.1.chr7.g00629.t01#3   mNeoVis.evergladensis.2.1.chr7.g00622.t01#2       5005   125257433:   mNeoVis.evergladensis.1.1.chr7.g00630.t01#3    mNeoVis.vulgivaga.2.1.chr7.g00627.t01#1    mNeoVis.3.1.1.chr7.g00599.t01#6    mNeoVis.vulgivaga.1.1.chr7.g00629.t01#4    mNeoVis.2.1.1.chr7_hap1.g00621.t01#5   mNeoVis.evergladensis.2.1.chr7.g00623.t01#2       5006   125257434:   mNeoVis.evergladensis.2.1.chr7.g00624.t01#2    mNeoVis.vulgivaga.1.1.chr7.g00630.t01#4    mNeoVis.evergladensis.1.1.chr7.g00631.t01#3    mNeoVis.vulgivaga.2.1.chr7.g00628.t01#1   mNeoVis.2.1.1.chr7_hap1.g00622.t01#5       5007   125257435:   mNeoVis.2.1.1.chr15_hap1.g00132.t01#5    mNeoVis.vulgivaga.2.1.chr15.g00121.t01#1    mNeoVis.vulgivaga.1.1.chr15.g00118.t01#4    mNeoVis.evergladensis.2.1.chr15.g00118.t01#2   mNeoVis.evergladensis.1.1.chr15.g00497.t01#3       5008   125257436:   mNeoVis.evergladensis.2.1.chr7.g00625.t01#2    mNeoVis.vulgivaga.1.1.chr7.g00631.t01#4    mNeoVis.3.1.1.chr7.g00601.t01#6    mNeoVis.evergladensis.1.1.chr7.g00632.t01#3    mNeoVis.vulgivaga.2.1.chr7.g00629.t01#1   mNeoVis.2.1.1.chr7_hap1.g00623.t01#5       5009   125257437:   mNeoVis.vulgivaga.1.1.chr1.g00052.t01#4    mNeoVis.vulgivaga.2.1.chr1.g00055.t01#1    mNeoVis.2.1.1.chr1_hap1.g00056.t01#5    mNeoVis.3.1.1.chr1.g00058.t01#6    mNeoVis.evergladensis.2.1.chr1.g00054.t01#2   mNeoVis.evergladensis.1.1.chr1.g00055.t01#3       5010   125257438:   mNeoVis.2.1.1.chr7_hap1.g00625.t01#5    mNeoVis.vulgivaga.2.1.chr7.g00630.t01#1   mNeoVis.vulgivaga.1.1.chr7.g00633.t01#4           5011   125257439:   mNeoVis.vulgivaga.2.1.chr1.g00056.t01#1    mNeoVis.vulgivaga.1.1.chr1.g00053.t01#4    mNeoVis.evergladensis.2.1.chr1.g00055.t01#2    mNeoVis.evergladensis.1.1.chr1.g00056.t01#3    mNeoVis.2.1.1.chr1_hap1.g00057.t01#5   mNeoVis.3.1.1.chr1.g00059.t01#6       5012   125257440:  mNeoVis.vulgivaga.2.1.chr7.g00632.t01#1               5013   125257441:   mNeoVis.vulgivaga.1.1.chr7.g00635.t01#4    mNeoVis.2.1.1.chr7_hap1.g00630.t01#5    mNeoVis.evergladensis.1.1.chr7.g00636.t01#3    mNeoVis.evergladensis.2.1.chr7.g00629.t01#2    mNeoVis.3.1.1.chr7.g00602.t01#6   mNeoVis.vulgivaga.2.1.chr7.g00631.t01#1       5014   125257442:   mNeoVis.3.1.1.chr7.g00606.t01#6    mNeoVis.2.1.1.chr7_hap1.g00632.t01#5    mNeoVis.vulgivaga.2.1.chr7.g00633.t01#1    mNeoVis.vulgivaga.1.1.chr7.g00637.t01#4    mNeoVis.evergladensis.1.1.chr7.g00638.t01#3   mNeoVis.evergladensis.2.1.chr7.g00631.t01#2       5015   125257443:   mNeoVis.vulgivaga.2.1.chr1.g00057.t01#1    mNeoVis.vulgivaga.1.1.chr1.g00054.t01#4    mNeoVis.evergladensis.1.1.chr1.g00057.t01#3    mNeoVis.3.1.1.chr1.g00060.t01#6    mNeoVis.2.1.1.chr1_hap1.g00058.t01#5   mNeoVis.evergladensis.2.1.chr1.g00056.t01#2       5016   125257444:   mNeoVis.evergladensis.2.1.chr7.g00633.t01#2    mNeoVis.evergladensis.1.1.chr7.g00640.t01#3    mNeoVis.2.1.1.chr7_hap1.g00634.t01#5    mNeoVis.vulgivaga.1.1.chr7.g00639.t01#4    mNeoVis.vulgivaga.2.1.chr7.g00634.t01#1   mNeoVis.3.1.1.chr7.g00607.t01#6       5017   125257445:   mNeoVis.vulgivaga.2.1.chr7.g00635.t01#1    mNeoVis.evergladensis.1.1.chr7.g00641.t01#3   mNeoVis.vulgivaga.1.1.chr7.g00640.t01#4           5018   125257446:   mNeoVis.3.1.1.chr1.g00061.t01#6    mNeoVis.2.1.1.chr1_hap1.g00059.t01#5    mNeoVis.vulgivaga.2.1.chr1.g00058.t01#1    mNeoVis.evergladensis.2.1.chr1.g00057.t01#2    mNeoVis.vulgivaga.1.1.chr1.g00055.t01#4   mNeoVis.evergladensis.1.1.chr1.g00058.t01#3       5019   125257447:   mNeoVis.3.1.1.chr1.g00062.t01#6    mNeoVis.2.1.1.chr1_hap1.g00060.t01#5    mNeoVis.evergladensis.2.1.chr1.g00058.t01#2    mNeoVis.vulgivaga.2.1.chr1.g00059.t01#1    mNeoVis.evergladensis.1.1.chr1.g00060.t01#3    mNeoVis.evergladensis.1.1.chr1.g00059.t01#3   mNeoVis.vulgivaga.1.1.chr1.g00056.t01#4     5020   125257448:   mNeoVis.vulgivaga.1.1.chr1.g00057.t01#4    mNeoVis.vulgivaga.2.1.chr1.g00060.t01#1    mNeoVis.evergladensis.1.1.chr1.g00061.t01#3    mNeoVis.2.1.1.chr1_hap1.g00061.t01#5   mNeoVis.evergladensis.2.1.chr1.g00059.t01#2       5021   125257449:   mNeoVis.vulgivaga.2.1.chr1.g00061.t01#1    mNeoVis.2.1.1.chr1_hap1.g00063.t01#5    mNeoVis.3.1.1.chr1.g00064.t01#6    mNeoVis.evergladensis.2.1.chr1.g00061.t01#2    mNeoVis.evergladensis.1.1.chr1.g00063.t01#3   mNeoVis.vulgivaga.1.1.chr1.g00059.t01#4       5022   125257450:   mNeoVis.2.1.1.chr7_hap1.g00636.t01#5    mNeoVis.vulgivaga.2.1.chr7.g00636.t01#1    mNeoVis.evergladensis.1.1.chr7.g00643.t01#3    mNeoVis.vulgivaga.1.1.chr7.g00642.t01#4   mNeoVis.evergladensis.2.1.chr7.g00635.t01#2       5023   125257451:  mNeoVis.vulgivaga.2.1.chr7.g00637.t01#1               5024   125257452:  mNeoVis.vulgivaga.2.1.chr7.g00638.t01#1               5025   125257453:   mNeoVis.2.1.1.chr7_hap1.g00639.t01#5    mNeoVis.vulgivaga.2.1.chr7.g00639.t01#1   mNeoVis.evergladensis.2.1.chr7.g00637.t01#2         5026   125257454:   mNeoVis.vulgivaga.1.1.chr7.g00646.t01#4    mNeoVis.vulgivaga.2.1.chr7.g00640.t01#1    mNeoVis.evergladensis.1.1.chr7.g00647.t01#3    mNeoVis.3.1.1.chr7.g00610.t01#6    mNeoVis.evergladensis.2.1.chr7.g00638.t01#2   mNeoVis.2.1.1.chr7_hap1.g00640.t01#5       5027   125257455:   mNeoVis.vulgivaga.2.1.chr7.g00641.t01#1    mNeoVis.2.1.1.chr7_hap1.g00641.t01#5    mNeoVis.vulgivaga.1.1.chr7.g00647.t01#4   mNeoVis.evergladensis.1.1.chr7.g00648.t01#3       5028   125257456:   mNeoVis.3.1.1.chr1.g00065.t01#6    mNeoVis.evergladensis.2.1.chr1.g00062.t01#2    mNeoVis.vulgivaga.1.1.chr1.g00060.t01#4    mNeoVis.vulgivaga.2.1.chr1.g00062.t01#1    mNeoVis.evergladensis.1.1.chr1.g00064.t01#3   mNeoVis.2.1.1.chr1_hap1.g00064.t01#5       5029   125257457:   mNeoVis.evergladensis.1.1.chr1.g00065.t01#3    mNeoVis.vulgivaga.1.1.chr1.g00061.t01#4    mNeoVis.vulgivaga.2.1.chr1.g00063.t01#1    mNeoVis.2.1.1.chr1_hap1.g00065.t01#5    mNeoVis.3.1.1.chr1.g00066.t01#6   mNeoVis.evergladensis.2.1.chr1.g00063.t01#2       5030   125257458:   mNeoVis.vulgivaga.1.1.chr7.g00648.t01#4    mNeoVis.vulgivaga.2.1.chr7.g00642.t01#1    mNeoVis.evergladensis.1.1.chr7.g00649.t01#3    mNeoVis.evergladensis.2.1.chr7.g00639.t01#2   mNeoVis.2.1.1.chr7_hap1.g00642.t01#5       5031   125257459:   mNeoVis.vulgivaga.1.1.chr7.g00649.t01#4    mNeoVis.vulgivaga.2.1.chr7.g00643.t01#1    mNeoVis.3.1.1.chr7.g00612.t01#6    mNeoVis.evergladensis.2.1.chr7.g00640.t01#2    mNeoVis.2.1.1.chr7_hap1.g00643.t01#5   mNeoVis.evergladensis.1.1.chr7.g00650.t01#3       5032   125257460:   mNeoVis.vulgivaga.1.1.chr1.g00062.t01#4    mNeoVis.2.1.1.chr1_hap1.g00066.t01#5    mNeoVis.vulgivaga.2.1.chr1.g00064.t01#1    mNeoVis.3.1.1.chr1.g00067.t01#6    mNeoVis.evergladensis.2.1.chr1.g00064.t01#2   mNeoVis.evergladensis.1.1.chr1.g00066.t01#3       5033   125257461:   mNeoVis.vulgivaga.2.1.chr7.g00644.t01#1    mNeoVis.evergladensis.2.1.chr7.g00642.t01#2   mNeoVis.2.1.1.chr7_hap1.g00645.t01#5           5034   125257462:   mNeoVis.evergladensis.1.1.chr7.g00652.t01#3    mNeoVis.vulgivaga.2.1.chr7.g00645.t01#1    mNeoVis.evergladensis.2.1.chr7.g00643.t01#2   mNeoVis.2.1.1.chr7_hap1.g00646.t01#5         5035   125257463:   mNeoVis.3.1.1.chr7.g00614.t01#6    mNeoVis.2.1.1.chr7_hap1.g00647.t01#5    mNeoVis.evergladensis.1.1.chr7.g00653.t01#3    mNeoVis.vulgivaga.1.1.chr7.g00651.t01#4    mNeoVis.vulgivaga.2.1.chr7.g00646.t01#1   mNeoVis.evergladensis.2.1.chr7.g00644.t01#2       5036   125257464:   mNeoVis.vulgivaga.1.1.chr7.g00652.t01#4    mNeoVis.vulgivaga.2.1.chr7.g00647.t01#1    mNeoVis.2.1.1.chr7_hap1.g00648.t01#5    mNeoVis.evergladensis.1.1.chr7.g00654.t01#3   mNeoVis.evergladensis.2.1.chr7.g00645.t01#2       5037   125257465:   mNeoVis.2.1.1.chr7_hap1.g00650.t01#5   mNeoVis.evergladensis.2.1.chr7.g00647.t01#2           5038   125257466:   mNeoVis.2.1.1.chr7_hap1.g00649.t01#5    mNeoVis.evergladensis.1.1.chr7.g00655.t01#3    mNeoVis.vulgivaga.1.1.chr7.g00653.t01#4    mNeoVis.vulgivaga.2.1.chr7.g00648.t01#1    mNeoVis.3.1.1.chr7.g00615.t01#6   mNeoVis.evergladensis.2.1.chr7.g00646.t01#2       5039   125257467:   mNeoVis.evergladensis.1.1.chr7.g00656.t01#3    mNeoVis.evergladensis.2.1.chr7.g00648.t01#2    mNeoVis.vulgivaga.2.1.chr7.g00649.t01#1    mNeoVis.2.1.1.chr7_hap1.g00651.t01#5    mNeoVis.3.1.1.chr7.g00616.t01#6   mNeoVis.vulgivaga.1.1.chr7.g00654.t01#4       5040   125257468:   mNeoVis.3.1.1.chr7.g00617.t01#6    mNeoVis.vulgivaga.2.1.chr7.g00650.t01#1    mNeoVis.evergladensis.1.1.chr7.g00657.t01#3    mNeoVis.evergladensis.2.1.chr7.g00649.t01#2    mNeoVis.2.1.1.chr7_hap1.g00652.t01#5   mNeoVis.vulgivaga.1.1.chr7.g00655.t01#4       5041   125257469:   mNeoVis.2.1.1.chr1_hap1.g00067.t01#5    mNeoVis.evergladensis.1.1.chr1.g00069.t01#3    mNeoVis.vulgivaga.2.1.chr1.g00065.t01#1    mNeoVis.evergladensis.2.1.chr1.g00065.t01#2    mNeoVis.vulgivaga.1.1.chr1.g00064.t01#4   mNeoVis.3.1.1.chr1.g00070.t01#6       5042   125257470:   mNeoVis.2.1.1.chr1_hap1.g00069.t01#5    mNeoVis.vulgivaga.2.1.chr1.g00066.t01#1    mNeoVis.evergladensis.2.1.chr1.g00066.t01#2    mNeoVis.evergladensis.1.1.chr1.g00070.t01#3    mNeoVis.3.1.1.chr1.g00072.t01#6   mNeoVis.vulgivaga.1.1.chr1.g00065.t01#4       5043   125257471:   mNeoVis.2.1.1.chr8_hap1.g01107.t01#5    mNeoVis.3.1.1.chr8.g01028.t01#6    mNeoVis.evergladensis.2.1.chr9.g01085.t01#2    mNeoVis.vulgivaga.2.1.chr8.g01084.t01#1    mNeoVis.evergladensis.1.1.chr8.g01077.t01#3   mNeoVis.vulgivaga.1.1.chr8.g01072.t01#4       5044   125257472:   mNeoVis.evergladensis.2.1.chr1.g00068.t01#2    mNeoVis.vulgivaga.1.1.chr1.g00066.t01#4    mNeoVis.evergladensis.2.1.chr1.g00067.t01#2    mNeoVis.2.1.1.chr1_hap1.g00070.t01#5    mNeoVis.3.1.1.chr1.g00074.t01#6    mNeoVis.evergladensis.1.1.chr1.g00071.t01#3   mNeoVis.vulgivaga.2.1.chr1.g00067.t01#1     5045   125257473:   mNeoVis.2.1.1.chr7_hap1.g00653.t01#5    mNeoVis.vulgivaga.1.1.chr7.g00656.t01#4    mNeoVis.evergladensis.2.1.chr7.g00650.t01#2   mNeoVis.vulgivaga.2.1.chr7.g00652.t01#1         5046   125257474:   mNeoVis.vulgivaga.2.1.chr7.g00651.t01#1   mNeoVis.evergladensis.1.1.chr7.g00658.t01#3           5047   125257475:   mNeoVis.2.1.1.chr7_hap1.g00035.t01#5    mNeoVis.vulgivaga.2.1.chr7.g00035.t01#1    mNeoVis.evergladensis.1.1.chr7.g00036.t01#3    mNeoVis.evergladensis.2.1.chr7.g00036.t01#2   mNeoVis.vulgivaga.1.1.chr7.g00037.t01#4       5048   125257476:   mNeoVis.evergladensis.1.1.chr1.g00072.t01#3    mNeoVis.3.1.1.chr1.g00075.t01#6    mNeoVis.vulgivaga.1.1.chr1.g00068.t01#4    mNeoVis.vulgivaga.2.1.chr1.g00068.t01#1    mNeoVis.vulgivaga.1.1.chr1.g00067.t01#4    mNeoVis.evergladensis.2.1.chr1.g00069.t01#2    mNeoVis.vulgivaga.1.1.chr1.g00069.t01#4   mNeoVis.2.1.1.chr1_hap1.g00071.t01#5     5049   125257477:   mNeoVis.2.1.1.chr1_hap1.g00079.t01#5    mNeoVis.3.1.1.chr1.g00080.t01#6    mNeoVis.evergladensis.1.1.chr1.g00081.t01#3    mNeoVis.vulgivaga.2.1.chr1.g00073.t01#1    mNeoVis.vulgivaga.1.1.chr1.g00078.t01#4   mNeoVis.evergladensis.2.1.chr1.g00078.t01#2       5050   125257478:   mNeoVis.vulgivaga.1.1.chr7.g00657.t01#4    mNeoVis.vulgivaga.2.1.chr7.g00653.t01#1    mNeoVis.3.1.1.chr7.g00619.t01#6    mNeoVis.2.1.1.chr7_hap1.g00654.t01#5    mNeoVis.evergladensis.2.1.chr7.g00651.t01#2   mNeoVis.evergladensis.1.1.chr7.g00659.t01#3       5051   125257479:   mNeoVis.vulgivaga.1.1.chr15.g00119.t01#4    mNeoVis.vulgivaga.2.1.chr15.g00122.t01#1    mNeoVis.evergladensis.1.1.chr15.g00496.t01#3    mNeoVis.3.1.1.chr15.g00110.t01#6    mNeoVis.2.1.1.chr15_hap1.g00133.t01#5   mNeoVis.evergladensis.2.1.chr15.g00119.t01#2       5052   125257480:   mNeoVis.evergladensis.1.1.chr7.g00660.t01#3    mNeoVis.vulgivaga.2.1.chr7.g00654.t01#1    mNeoVis.vulgivaga.1.1.chr7.g00658.t01#4    mNeoVis.2.1.1.chr7_hap1.g00655.t01#5   mNeoVis.evergladensis.2.1.chr7.g00652.t01#2       5053   125257481:   mNeoVis.vulgivaga.2.1.chr7.g00655.t01#1    mNeoVis.2.1.1.chr7_hap1.g00656.t01#5    mNeoVis.3.1.1.chr7.g00620.t01#6    mNeoVis.vulgivaga.1.1.chr7.g00659.t01#4    mNeoVis.evergladensis.2.1.chr7.g00653.t01#2   mNeoVis.evergladensis.1.1.chr7.g00661.t01#3       5054   125257482:   mNeoVis.2.1.1.chr7_hap1.g00661.t01#5    mNeoVis.evergladensis.2.1.chr7.g00657.t01#2    mNeoVis.3.1.1.chr7.g00624.t01#6    mNeoVis.vulgivaga.2.1.chr7.g00660.t01#1    mNeoVis.vulgivaga.1.1.chr7.g00664.t01#4   mNeoVis.evergladensis.1.1.chr7.g00666.t01#3       5055   125257483:   mNeoVis.evergladensis.2.1.chr7.g00656.t01#2    mNeoVis.vulgivaga.1.1.chr7.g00663.t01#4    mNeoVis.evergladensis.1.1.chr7.g00665.t01#3    mNeoVis.2.1.1.chr7_hap1.g00660.t01#5    mNeoVis.3.1.1.chr7.g00622.t01#6    mNeoVis.2.1.1.chr7_hap1.g00659.t01#5    mNeoVis.3.1.1.chr7.g00623.t01#6    mNeoVis.evergladensis.1.1.chr7.g00664.t01#3    mNeoVis.vulgivaga.2.1.chr7.g00656.t01#1      5056   125257484:   mNeoVis.evergladensis.1.1.chr7.g00663.t01#3    mNeoVis.vulgivaga.2.1.chr7.g00657.t01#1    mNeoVis.evergladensis.2.1.chr7.g00654.t01#2    mNeoVis.3.1.1.chr7.g00621.t01#6    mNeoVis.vulgivaga.1.1.chr7.g00661.t01#4   mNeoVis.2.1.1.chr7_hap1.g00658.t01#5       5057   125257485:   mNeoVis.evergladensis.2.1.chr1.g00070.t01#2    mNeoVis.vulgivaga.2.1.chr1.g00069.t01#1    mNeoVis.2.1.1.chr1_hap1.g00072.t01#5    mNeoVis.evergladensis.1.1.chr1.g00073.t01#3    mNeoVis.vulgivaga.1.1.chr1.g00070.t01#4   mNeoVis.3.1.1.chr1.g00076.t01#6       5058   125257486:   mNeoVis.vulgivaga.2.1.chr5.g01604.t01#1    mNeoVis.evergladensis.2.1.chr5.g01601.t01#2    mNeoVis.2.1.1.chr2_hap1.g01661.t01#5    mNeoVis.3.1.1.chr5.g01583.t01#6    mNeoVis.vulgivaga.1.1.chr5.g01581.t01#4   mNeoVis.evergladensis.1.1.chr5.g01606.t01#3       5059   125257487:   mNeoVis.3.1.1.chr1.g00077.t01#6    mNeoVis.vulgivaga.2.1.chr1.g00070.t01#1    mNeoVis.evergladensis.1.1.chr1.g00074.t01#3    mNeoVis.vulgivaga.1.1.chr1.g00072.t01#4    mNeoVis.2.1.1.chr1_hap1.g00073.t01#5   mNeoVis.evergladensis.2.1.chr1.g00071.t01#2       5060   125257488:   mNeoVis.vulgivaga.1.1.chr1.g00073.t01#4    mNeoVis.2.1.1.chr1_hap1.g00074.t01#5    mNeoVis.vulgivaga.2.1.chr1.g00071.t01#1    mNeoVis.evergladensis.1.1.chr1.g00075.t01#3   mNeoVis.evergladensis.2.1.chr1.g00072.t01#2       5061   125257489:   mNeoVis.3.1.1.chr12.g00015.t01#6    mNeoVis.2.1.1.chr7_hap1.g02461.t01#5   mNeoVis.2.1.1.chr7_hap1.g02432.t01#5           5062   125257490:   mNeoVis.evergladensis.2.1.chr1.g00076.t01#2    mNeoVis.vulgivaga.2.1.chr1.g00072.t01#1    mNeoVis.2.1.1.chr1_hap1.g00077.t01#5    mNeoVis.3.1.1.chr1.g00078.t01#6    mNeoVis.evergladensis.1.1.chr1.g00079.t01#3   mNeoVis.vulgivaga.1.1.chr1.g00076.t01#4       5063   125257491:   mNeoVis.3.1.1.chr7.g00626.t01#6    mNeoVis.vulgivaga.1.1.chr7.g00666.t01#4    mNeoVis.evergladensis.2.1.chr7.g00658.t01#2    mNeoVis.vulgivaga.2.1.chr7.g00661.t01#1    mNeoVis.2.1.1.chr7_hap1.g00663.t01#5   mNeoVis.evergladensis.1.1.chr7.g00667.t01#3       5064   125257492:   mNeoVis.vulgivaga.2.1.chr7.g00663.t01#1    mNeoVis.vulgivaga.1.1.chr7.g00668.t01#4    mNeoVis.evergladensis.1.1.chr7.g00669.t01#3    mNeoVis.3.1.1.chr7.g00628.t01#6    mNeoVis.2.1.1.chr7_hap1.g00665.t01#5   mNeoVis.evergladensis.2.1.chr7.g00660.t01#2       5065   125257493:   mNeoVis.2.1.1.chr1_hap1.g00878.t01#5    mNeoVis.vulgivaga.1.1.chr1.g00894.t01#4   mNeoVis.evergladensis.2.1.chr1.g00881.t01#2         5066   125257494:   mNeoVis.3.1.1.chr7.g00630.t01#6    mNeoVis.vulgivaga.2.1.chr7.g00664.t01#1    mNeoVis.vulgivaga.1.1.chr7.g00671.t01#4    mNeoVis.evergladensis.2.1.chr7.g00663.t01#2   mNeoVis.evergladensis.1.1.chr7.g00671.t01#3       5067   125257495:   mNeoVis.evergladensis.1.1.chr1.g00082.t01#3    mNeoVis.3.1.1.chr1.g00081.t01#6    mNeoVis.vulgivaga.2.1.chr1.g00074.t01#1    mNeoVis.vulgivaga.1.1.chr1.g00079.t01#4    mNeoVis.evergladensis.2.1.chr1.g00079.t01#2   mNeoVis.2.1.1.chr1_hap1.g00080.t01#5       5068   125257496:   mNeoVis.2.1.1.chr7_hap1.g00667.t01#5    mNeoVis.vulgivaga.1.1.chr7.g00670.t01#4    mNeoVis.3.1.1.chr7.g00629.t01#6    mNeoVis.evergladensis.2.1.chr7.g00662.t01#2    mNeoVis.evergladensis.1.1.chr7.g00672.t01#3   mNeoVis.evergladensis.1.1.chr7.g00670.t01#3       5069   125257497:   mNeoVis.vulgivaga.1.1.chr7.g00672.t01#4    mNeoVis.evergladensis.2.1.chr7.g00665.t01#2    mNeoVis.evergladensis.1.1.chr7.g00673.t01#3   mNeoVis.vulgivaga.2.1.chr7.g00665.t01#1         5070   125257498:   mNeoVis.2.1.1.chr7_hap1.g00668.t01#5   mNeoVis.3.1.1.chr7.g00631.t01#6             5071   125257499:  mNeoVis.evergladensis.2.1.chr7.g00664.t01#2             5072   125257500:   mNeoVis.3.1.1.chr7.g00633.t01#6    mNeoVis.vulgivaga.2.1.chr7.g00666.t01#1    mNeoVis.evergladensis.1.1.chr7.g00677.t01#3   mNeoVis.2.1.1.chr7_hap1.g00671.t01#5         5073   125257501:   mNeoVis.3.1.1.chr7.g00634.t01#6    mNeoVis.vulgivaga.2.1.chr7.g00667.t01#1    mNeoVis.vulgivaga.1.1.chr7.g00676.t01#4    mNeoVis.evergladensis.1.1.chr7.g00678.t01#3    mNeoVis.2.1.1.chr7_hap1.g00672.t01#5   mNeoVis.evergladensis.2.1.chr7.g00670.t01#2       5074   125257502:   mNeoVis.vulgivaga.2.1.chr7.g00668.t01#1    mNeoVis.evergladensis.2.1.chr7.g00671.t01#2    mNeoVis.vulgivaga.1.1.chr7.g00677.t01#4    mNeoVis.evergladensis.1.1.chr7.g00679.t01#3    mNeoVis.2.1.1.chr7_hap1.g00673.t01#5   mNeoVis.3.1.1.chr7.g00635.t01#6       5075   125257503:   mNeoVis.2.1.1.chr7_hap1.g00674.t01#5    mNeoVis.3.1.1.chr7.g00638.t01#6    mNeoVis.vulgivaga.2.1.chr7.g00669.t01#1    mNeoVis.vulgivaga.1.1.chr7.g00678.t01#4    mNeoVis.evergladensis.1.1.chr7.g00680.t01#3   mNeoVis.evergladensis.2.1.chr7.g00672.t01#2       5076   125257504:   mNeoVis.2.1.1.chr1_hap1.g00081.t01#5    mNeoVis.evergladensis.2.1.chr1.g00080.t01#2    mNeoVis.vulgivaga.2.1.chr1.g00075.t01#1    mNeoVis.evergladensis.1.1.chr1.g00083.t01#3    mNeoVis.3.1.1.chr1.g00082.t01#6   mNeoVis.vulgivaga.1.1.chr1.g00080.t01#4       5077   125257505:   mNeoVis.vulgivaga.1.1.chr7.g00679.t01#4    mNeoVis.evergladensis.1.1.chr7.g00681.t01#3    mNeoVis.evergladensis.2.1.chr7.g00673.t01#2    mNeoVis.2.1.1.chr7_hap1.g00675.t01#5   mNeoVis.vulgivaga.2.1.chr7.g00670.t01#1       5078   125257506:   mNeoVis.evergladensis.2.1.chr7.g00674.t01#2    mNeoVis.2.1.1.chr7_hap1.g00676.t01#5    mNeoVis.vulgivaga.2.1.chr7.g00671.t01#1    mNeoVis.vulgivaga.1.1.chr7.g00680.t01#4    mNeoVis.evergladensis.1.1.chr7.g00682.t01#3   mNeoVis.3.1.1.chr7.g00640.t01#6       5079   125257507:   mNeoVis.vulgivaga.2.1.chr1.g00076.t01#1    mNeoVis.evergladensis.1.1.chr1.g00084.t01#3    mNeoVis.2.1.1.chr1_hap1.g00082.t01#5    mNeoVis.3.1.1.chr1.g00083.t01#6   mNeoVis.evergladensis.2.1.chr1.g00081.t01#2       5080   125257508:   mNeoVis.vulgivaga.1.1.chr7.g00681.t01#4    mNeoVis.evergladensis.1.1.chr7.g00683.t01#3    mNeoVis.3.1.1.chr7.g00641.t01#6    mNeoVis.evergladensis.2.1.chr7.g00675.t01#2    mNeoVis.2.1.1.chr7_hap1.g00677.t01#5   mNeoVis.vulgivaga.2.1.chr7.g00672.t01#1       5081   125257509:   mNeoVis.vulgivaga.1.1.chr7.g00038.t01#4    mNeoVis.evergladensis.2.1.chr7.g00037.t01#2    mNeoVis.2.1.1.chr7_hap1.g00036.t01#5    mNeoVis.vulgivaga.2.1.chr7.g00036.t01#1   mNeoVis.evergladensis.1.1.chr7.g00037.t01#3       5082   125257510:   mNeoVis.evergladensis.1.1.chr7.g00684.t01#3    mNeoVis.3.1.1.chr7.g00642.t01#6    mNeoVis.evergladensis.2.1.chr7.g00676.t01#2    mNeoVis.2.1.1.chr7_hap1.g00678.t01#5    mNeoVis.vulgivaga.2.1.chr7.g00673.t01#1   mNeoVis.vulgivaga.1.1.chr7.g00682.t01#4       5083   125257511:   mNeoVis.3.1.1.chr1.g00084.t01#6    mNeoVis.evergladensis.1.1.chr1.g00085.t01#3    mNeoVis.vulgivaga.2.1.chr1.g00077.t01#1    mNeoVis.2.1.1.chr1_hap1.g00083.t01#5    mNeoVis.vulgivaga.1.1.chr1.g00082.t01#4   mNeoVis.evergladensis.2.1.chr1.g00082.t01#2       5084   125257512:   mNeoVis.evergladensis.1.1.chr1.g00086.t01#3    mNeoVis.3.1.1.chr1.g00085.t01#6   mNeoVis.vulgivaga.2.1.chr1.g00078.t01#1           5085   125257513:   mNeoVis.vulgivaga.2.1.chr1.g00079.t01#1    mNeoVis.2.1.1.chr1_hap1.g00084.t01#5    mNeoVis.3.1.1.chr1.g00086.t01#6    mNeoVis.evergladensis.2.1.chr1.g00083.t01#2    mNeoVis.vulgivaga.1.1.chr1.g00083.t01#4   mNeoVis.evergladensis.1.1.chr1.g00087.t01#3       5086   125257514:   mNeoVis.3.1.1.chr1.g00001.t01#6    mNeoVis.vulgivaga.2.1.chr1.g00001.t01#1    mNeoVis.vulgivaga.1.1.chr1.g00001.t01#4   mNeoVis.evergladensis.1.1.chr1.g00001.t01#3       5087   125257515:  mNeoVis.vulgivaga.2.1.chr7.g00675.t01#1               5088   125257516:   mNeoVis.vulgivaga.2.1.chr7.g00674.t01#1    mNeoVis.evergladensis.2.1.chr7.g00677.t01#2    mNeoVis.vulgivaga.1.1.chr7.g00683.t01#4   mNeoVis.evergladensis.1.1.chr7.g00685.t01#3       5089   125257517:   mNeoVis.vulgivaga.2.1.chr7.g00676.t01#1   mNeoVis.2.1.1.chr7_hap1.g00680.t01#5             5090   125257518:   mNeoVis.vulgivaga.1.1.chr7.g00685.t01#4    mNeoVis.2.1.1.chr7_hap1.g00681.t01#5    mNeoVis.evergladensis.1.1.chr7.g00686.t01#3    mNeoVis.evergladensis.2.1.chr7.g00679.t01#2    mNeoVis.vulgivaga.2.1.chr7.g00677.t01#1   mNeoVis.3.1.1.chr7.g00646.t01#6       5091   125257519:   mNeoVis.vulgivaga.2.1.chr7.g00678.t01#1    mNeoVis.3.1.1.chr7.g00647.t01#6    mNeoVis.vulgivaga.1.1.chr7.g00686.t01#4    mNeoVis.evergladensis.1.1.chr7.g00687.t01#3    mNeoVis.2.1.1.chr7_hap1.g00682.t01#5   mNeoVis.evergladensis.2.1.chr7.g00680.t01#2       5092   125257520:   mNeoVis.vulgivaga.1.1.chr7.g00687.t01#4    mNeoVis.evergladensis.2.1.chr7.g00681.t01#2    mNeoVis.vulgivaga.2.1.chr7.g00679.t01#1    mNeoVis.2.1.1.chr7_hap1.g00683.t01#5    mNeoVis.evergladensis.1.1.chr7.g00688.t01#3   mNeoVis.3.1.1.chr7.g00648.t01#6       5093   125257521:   mNeoVis.vulgivaga.2.1.chr7.g00037.t01#1   mNeoVis.evergladensis.2.1.chr7.g00038.t01#2           5094   125257522:   mNeoVis.2.1.1.chr1_hap1.g00003.t01#5    mNeoVis.3.1.1.chr1.g00002.t01#6    mNeoVis.evergladensis.2.1.chr1.g00001.t01#2    mNeoVis.vulgivaga.2.1.chr1.g00002.t01#1    mNeoVis.vulgivaga.1.1.chr1.g00002.t01#4   mNeoVis.evergladensis.1.1.chr1.g00002.t01#3       5095   125257523:   mNeoVis.evergladensis.1.1.chr1.g00003.t01#3    mNeoVis.vulgivaga.1.1.chr1.g00003.t01#4    mNeoVis.2.1.1.chr1_hap1.g00004.t01#5    mNeoVis.3.1.1.chr1.g00003.t01#6    mNeoVis.evergladensis.2.1.chr1.g00002.t01#2   mNeoVis.vulgivaga.2.1.chr1.g00003.t01#1       5096   125257524:   mNeoVis.vulgivaga.2.1.chr7.g00680.t01#1    mNeoVis.vulgivaga.1.1.chr7.g00688.t01#4    mNeoVis.3.1.1.chr7.g00649.t01#6    mNeoVis.2.1.1.chr7_hap1.g00684.t01#5    mNeoVis.evergladensis.1.1.chr7.g00689.t01#3   mNeoVis.evergladensis.2.1.chr7.g00682.t01#2       5097   125257525:   mNeoVis.2.1.1.chr1_hap1.g00005.t01#5    mNeoVis.vulgivaga.2.1.chr1.g00004.t01#1    mNeoVis.vulgivaga.1.1.chr1.g00004.t01#4    mNeoVis.3.1.1.chr1.g00004.t01#6    mNeoVis.evergladensis.1.1.chr1.g00004.t01#3   mNeoVis.evergladensis.2.1.chr1.g00003.t01#2       5098   125257526:   mNeoVis.3.1.1.chr7.g00650.t01#6    mNeoVis.vulgivaga.1.1.chr7.g00689.t01#4    mNeoVis.vulgivaga.2.1.chr7.g00681.t01#1    mNeoVis.evergladensis.2.1.chr7.g00683.t01#2    mNeoVis.2.1.1.chr7_hap1.g00685.t01#5   mNeoVis.evergladensis.1.1.chr7.g00690.t01#3       5099   125257527:   mNeoVis.vulgivaga.1.1.chr7.g00690.t01#4    mNeoVis.3.1.1.chr7.g00651.t01#6    mNeoVis.vulgivaga.2.1.chr7.g00682.t01#1    mNeoVis.2.1.1.chr7_hap1.g00686.t01#5    mNeoVis.evergladensis.1.1.chr7.g00691.t01#3   mNeoVis.evergladensis.2.1.chr7.g00684.t01#2       5100   125257528:   mNeoVis.2.1.1.chr7_hap1.g00687.t01#5    mNeoVis.evergladensis.1.1.chr7.g00692.t01#3    mNeoVis.evergladensis.2.1.chr7.g00685.t01#2    mNeoVis.3.1.1.chr7.g00652.t01#6    mNeoVis.vulgivaga.2.1.chr7.g00683.t01#1   mNeoVis.vulgivaga.1.1.chr7.g00691.t01#4       5101   125257529:   mNeoVis.evergladensis.2.1.chr1.g00004.t01#2    mNeoVis.2.1.1.chr1_hap1.g00006.t01#5    mNeoVis.evergladensis.2.1.chr1.g00005.t01#2    mNeoVis.vulgivaga.1.1.chr1.g00005.t01#4    mNeoVis.vulgivaga.2.1.chr1.g00005.t01#1    mNeoVis.3.1.1.chr1.g00005.t01#6   mNeoVis.evergladensis.1.1.chr1.g00005.t01#3     5102   125257530:   mNeoVis.vulgivaga.2.1.chr7.g00684.t01#1    mNeoVis.3.1.1.chr7.g00654.t01#6    mNeoVis.evergladensis.2.1.chr7.g00686.t01#2    mNeoVis.vulgivaga.1.1.chr7.g00692.t01#4    mNeoVis.evergladensis.1.1.chr7.g00693.t01#3   mNeoVis.2.1.1.chr7_hap1.g00688.t01#5       5103   125257531:   mNeoVis.vulgivaga.1.1.chr7.g00693.t01#4    mNeoVis.evergladensis.1.1.chr7.g00694.t01#3    mNeoVis.evergladensis.2.1.chr7.g00687.t01#2    mNeoVis.vulgivaga.2.1.chr7.g00685.t01#1    mNeoVis.2.1.1.chr7_hap1.g00689.t01#5   mNeoVis.3.1.1.chr7.g00655.t01#6       5104   125257532:   mNeoVis.evergladensis.1.1.chr7.g00695.t01#3    mNeoVis.2.1.1.chr7_hap1.g00690.t01#5    mNeoVis.vulgivaga.2.1.chr7.g00686.t01#1    mNeoVis.evergladensis.2.1.chr7.g00688.t01#2   mNeoVis.vulgivaga.1.1.chr7.g00694.t01#4       5105   125257533:   mNeoVis.evergladensis.1.1.chr1.g00006.t01#3    mNeoVis.3.1.1.chr1.g00006.t01#6    mNeoVis.evergladensis.2.1.chr1.g00006.t01#2    mNeoVis.vulgivaga.2.1.chr1.g00006.t01#1    mNeoVis.vulgivaga.1.1.chr1.g00006.t01#4   mNeoVis.2.1.1.chr1_hap1.g00007.t01#5       5106   125257534:   mNeoVis.evergladensis.2.1.chr7.g00689.t01#2    mNeoVis.vulgivaga.2.1.chr7.g00687.t01#1    mNeoVis.vulgivaga.1.1.chr7.g00695.t01#4    mNeoVis.evergladensis.1.1.chr7.g00696.t01#3   mNeoVis.2.1.1.chr7_hap1.g00691.t01#5       5107   125257535:   mNeoVis.vulgivaga.1.1.chr7.g00696.t01#4    mNeoVis.evergladensis.1.1.chr7.g00697.t01#3    mNeoVis.3.1.1.chr7.g00656.t01#6    mNeoVis.2.1.1.chr7_hap1.g00692.t01#5    mNeoVis.evergladensis.2.1.chr7.g00690.t01#2   mNeoVis.vulgivaga.2.1.chr7.g00688.t01#1       5108   125257536:   mNeoVis.evergladensis.1.1.chr7.g00698.t01#3    mNeoVis.vulgivaga.1.1.chr7.g00697.t01#4    mNeoVis.vulgivaga.2.1.chr7.g00689.t01#1    mNeoVis.3.1.1.chr7.g00657.t01#6    mNeoVis.2.1.1.chr7_hap1.g00693.t01#5   mNeoVis.evergladensis.2.1.chr7.g00691.t01#2       5109   125257537:   mNeoVis.3.1.1.chr1.g00008.t01#6    mNeoVis.2.1.1.chr1_hap1.g00008.t01#5    mNeoVis.evergladensis.2.1.chr1.g00007.t01#2    mNeoVis.evergladensis.1.1.chr1.g00007.t01#3    mNeoVis.vulgivaga.1.1.chr1.g00007.t01#4   mNeoVis.vulgivaga.2.1.chr1.g00007.t01#1       5110   125257538:   mNeoVis.evergladensis.2.1.chr7.g00692.t01#2    mNeoVis.2.1.1.chr7_hap1.g00694.t01#5    mNeoVis.vulgivaga.2.1.chr7.g00690.t01#1    mNeoVis.evergladensis.1.1.chr7.g00699.t01#3   mNeoVis.vulgivaga.1.1.chr7.g00698.t01#4       5111   125257539:   mNeoVis.2.1.1.chr1_hap1.g00009.t01#5    mNeoVis.3.1.1.chr1.g00010.t01#6    mNeoVis.evergladensis.2.1.chr1.g00008.t01#2    mNeoVis.vulgivaga.1.1.chr1.g00008.t01#4    mNeoVis.vulgivaga.2.1.chr1.g00008.t01#1   mNeoVis.evergladensis.1.1.chr1.g00008.t01#3       5112   125257540:   mNeoVis.vulgivaga.2.1.chr7.g00691.t01#1    mNeoVis.evergladensis.2.1.chr7.g00693.t01#2    mNeoVis.evergladensis.1.1.chr7.g00700.t01#3   mNeoVis.vulgivaga.1.1.chr7.g00699.t01#4         5113   125257541:   mNeoVis.3.1.1.chr7.g00660.t01#6    mNeoVis.2.1.1.chr7_hap1.g00695.t01#5    mNeoVis.evergladensis.1.1.chr7.g00701.t01#3   mNeoVis.vulgivaga.2.1.chr7.g00692.t01#1         5114   125257542:   mNeoVis.vulgivaga.2.1.chr7.g00693.t01#1    mNeoVis.2.1.1.chr7_hap1.g00696.t01#5    mNeoVis.vulgivaga.1.1.chr7.g00700.t01#4    mNeoVis.evergladensis.2.1.chr7.g00694.t01#2    mNeoVis.3.1.1.chr7.g00661.t01#6   mNeoVis.evergladensis.1.1.chr7.g00702.t01#3       5115   125257543:   mNeoVis.evergladensis.1.1.chr7.g00703.t01#3    mNeoVis.3.1.1.chr7.g00662.t01#6    mNeoVis.2.1.1.chr7_hap1.g00697.t01#5    mNeoVis.vulgivaga.1.1.chr7.g00701.t01#4    mNeoVis.vulgivaga.2.1.chr7.g00694.t01#1   mNeoVis.evergladensis.2.1.chr7.g00695.t01#2       5116   125257544:   mNeoVis.evergladensis.2.1.chr1.g00009.t01#2    mNeoVis.2.1.1.chr1_hap1.g00010.t01#5    mNeoVis.vulgivaga.1.1.chr1.g00009.t01#4    mNeoVis.3.1.1.chr1.g00012.t01#6    mNeoVis.vulgivaga.2.1.chr1.g00009.t01#1   mNeoVis.evergladensis.1.1.chr1.g00009.t01#3       5117   125257545:   mNeoVis.evergladensis.1.1.chr7.g00704.t01#3    mNeoVis.vulgivaga.2.1.chr7.g00695.t01#1    mNeoVis.vulgivaga.1.1.chr7.g00702.t01#4    mNeoVis.2.1.1.chr7_hap1.g00698.t01#5   mNeoVis.evergladensis.2.1.chr7.g00696.t01#2       5118   125257546:   mNeoVis.vulgivaga.2.1.chr1.g00010.t01#1    mNeoVis.vulgivaga.1.1.chr1.g00010.t01#4    mNeoVis.evergladensis.1.1.chr1.g00010.t01#3    mNeoVis.evergladensis.2.1.chr1.g00010.t01#2   mNeoVis.2.1.1.chr1_hap1.g00011.t01#5       5119   125257547:   mNeoVis.evergladensis.1.1.chr1.g00011.t01#3    mNeoVis.2.1.1.chr1_hap1.g00012.t01#5    mNeoVis.vulgivaga.2.1.chr1.g00011.t01#1   mNeoVis.vulgivaga.1.1.chr1.g00011.t01#4         5120   125257548:   mNeoVis.vulgivaga.2.1.chr7.g00696.t01#1    mNeoVis.vulgivaga.1.1.chr7.g00703.t01#4    mNeoVis.evergladensis.1.1.chr7.g00705.t01#3    mNeoVis.evergladensis.2.1.chr7.g00697.t01#2    mNeoVis.3.1.1.chr7.g00664.t01#6   mNeoVis.2.1.1.chr7_hap1.g00699.t01#5       5121   125255501:   mNeoVis.evergladensis.2.1.chr7.g00698.t01#2    mNeoVis.2.1.1.chr7_hap1.g00700.t01#5    mNeoVis.vulgivaga.2.1.chr7.g00697.t01#1    mNeoVis.vulgivaga.1.1.chr7.g00704.t01#4    mNeoVis.evergladensis.1.1.chr7.g00706.t01#3   mNeoVis.3.1.1.chr2.g01616.t01#6       5122   125255502:   mNeoVis.evergladensis.1.1.chr7.g00707.t01#3    mNeoVis.evergladensis.2.1.chr7.g00699.t01#2    mNeoVis.vulgivaga.1.1.chr7.g00705.t01#4    mNeoVis.2.1.1.chr7_hap1.g00701.t01#5   mNeoVis.vulgivaga.2.1.chr7.g00698.t01#1       5123   125255503:   mNeoVis.evergladensis.2.1.chr7.g00700.t01#2    mNeoVis.vulgivaga.1.1.chr7.g00706.t01#4    mNeoVis.2.1.1.chr7_hap1.g00702.t01#5    mNeoVis.vulgivaga.2.1.chr7.g00699.t01#1   mNeoVis.evergladensis.1.1.chr7.g00708.t01#3       5124   125255504:   mNeoVis.2.1.1.chr1_hap1.g00013.t01#5    mNeoVis.3.1.1.chr1.g00015.t01#6    mNeoVis.evergladensis.2.1.chr1.g00012.t01#2    mNeoVis.vulgivaga.1.1.chr1.g00012.t01#4    mNeoVis.evergladensis.1.1.chr1.g00012.t01#3   mNeoVis.vulgivaga.2.1.chr1.g00012.t01#1       5125   125255505:   mNeoVis.vulgivaga.2.1.chr7.g00700.t01#1    mNeoVis.vulgivaga.1.1.chr7.g00707.t01#4    mNeoVis.2.1.1.chr7_hap1.g00703.t01#5    mNeoVis.evergladensis.2.1.chr7.g00701.t01#2   mNeoVis.evergladensis.1.1.chr7.g00709.t01#3       5126   125255506:   mNeoVis.vulgivaga.1.1.chr14.g00399.t01#4    mNeoVis.evergladensis.2.1.chr14.g00403.t01#2    mNeoVis.evergladensis.1.1.chr14.g00407.t01#3    mNeoVis.2.1.1.chr14_hap1.g00403.t01#5    mNeoVis.vulgivaga.2.1.chr14.g00404.t01#1   mNeoVis.3.1.1.chr14.g00402.t01#6       5127   125255507:   mNeoVis.evergladensis.1.1.chr7.g00564.t01#3    mNeoVis.vulgivaga.1.1.chr7.g00563.t01#4    mNeoVis.vulgivaga.2.1.chr7.g00558.t01#1    mNeoVis.evergladensis.2.1.chr7.g00556.t01#2    mNeoVis.2.1.1.chr7_hap1.g00555.t01#5   mNeoVis.3.1.1.chr7.g00538.t01#6       5128   125255508:   mNeoVis.vulgivaga.1.1.chr1.g00013.t01#4    mNeoVis.vulgivaga.2.1.chr1.g00013.t01#1    mNeoVis.evergladensis.2.1.chr1.g00013.t01#2    mNeoVis.2.1.1.chr1_hap1.g00014.t01#5    mNeoVis.3.1.1.chr1.g00016.t01#6   mNeoVis.evergladensis.1.1.chr1.g00013.t01#3       5129   125255509:   mNeoVis.evergladensis.2.1.chr7.g00557.t01#2    mNeoVis.2.1.1.chr7_hap1.g00556.t01#5    mNeoVis.vulgivaga.1.1.chr7.g00565.t01#4    mNeoVis.3.1.1.chr7.g00539.t01#6    mNeoVis.evergladensis.1.1.chr7.g00565.t01#3   mNeoVis.vulgivaga.2.1.chr7.g00559.t01#1       5130   125255510:   mNeoVis.2.1.1.chr1_hap1.g00015.t01#5    mNeoVis.evergladensis.1.1.chr1.g00014.t01#3    mNeoVis.evergladensis.2.1.chr9.g01187.t01#2    mNeoVis.evergladensis.2.1.chr1.g00014.t01#2    mNeoVis.vulgivaga.2.1.chr1.g00014.t01#1    mNeoVis.2.1.1.chr8_hap1.g01239.t01#5    mNeoVis.vulgivaga.1.1.chr8.g01176.t01#4   mNeoVis.vulgivaga.1.1.chr1.g00014.t01#4     5131   125255511:   mNeoVis.evergladensis.2.1.chr1.g00015.t01#2    mNeoVis.vulgivaga.2.1.chr1.g00015.t01#1    mNeoVis.2.1.1.chr1_hap1.g00016.t01#5   mNeoVis.evergladensis.1.1.chr1.g00015.t01#3       5132   125255512:   mNeoVis.vulgivaga.2.1.chr7.g00560.t01#1    mNeoVis.2.1.1.chr7_hap1.g00557.t01#5    mNeoVis.vulgivaga.1.1.chr7.g00566.t01#4    mNeoVis.evergladensis.1.1.chr7.g00567.t01#3    mNeoVis.evergladensis.2.1.chr7.g00558.t01#2   mNeoVis.3.1.1.chr7.g00540.t01#6       5133   125255513:   mNeoVis.3.1.1.chr15.g00111.t01#6    mNeoVis.evergladensis.2.1.chr15.g00120.t01#2    mNeoVis.vulgivaga.2.1.chr15.g00123.t01#1    mNeoVis.evergladensis.1.1.chr15.g00495.t01#3    mNeoVis.vulgivaga.1.1.chr15.g00120.t01#4   mNeoVis.2.1.1.chr15_hap1.g00134.t01#5       5134   125255514:   mNeoVis.vulgivaga.2.1.chr7.g00561.t01#1    mNeoVis.evergladensis.2.1.chr7.g00559.t01#2    mNeoVis.3.1.1.chr7.g00541.t01#6    mNeoVis.2.1.1.chr7_hap1.g00558.t01#5    mNeoVis.evergladensis.1.1.chr7.g00568.t01#3   mNeoVis.vulgivaga.1.1.chr7.g00567.t01#4       5135   125255515:   mNeoVis.2.1.1.chr7_hap1.g00559.t01#5    mNeoVis.evergladensis.1.1.chr7.g00569.t01#3    mNeoVis.vulgivaga.1.1.chr7.g00568.t01#4    mNeoVis.evergladensis.2.1.chr7.g00560.t01#2    mNeoVis.3.1.1.chr7.g00542.t01#6   mNeoVis.vulgivaga.2.1.chr7.g00562.t01#1       5136   125255516:   mNeoVis.evergladensis.2.1.chr7.g00039.t01#2    mNeoVis.vulgivaga.2.1.chr7.g00038.t01#1    mNeoVis.vulgivaga.1.1.chr7.g00040.t01#4    mNeoVis.evergladensis.1.1.chr7.g00039.t01#3   mNeoVis.2.1.1.chr7_hap1.g00037.t01#5       5137   125255517:   mNeoVis.3.1.1.chr1.g00017.t01#6    mNeoVis.evergladensis.2.1.chr1.g00016.t01#2    mNeoVis.evergladensis.1.1.chr1.g00016.t01#3    mNeoVis.2.1.1.chr1_hap1.g00017.t01#5    mNeoVis.vulgivaga.1.1.chr1.g00017.t01#4   mNeoVis.vulgivaga.2.1.chr1.g00016.t01#1       5138   125255518:   mNeoVis.evergladensis.1.1.chr1.g00017.t01#3    mNeoVis.2.1.1.chr1_hap1.g00018.t01#5    mNeoVis.vulgivaga.2.1.chr1.g00017.t01#1   mNeoVis.vulgivaga.1.1.chr1.g00018.t01#4         5139   125255519:   mNeoVis.3.1.1.chr14.g00072.t01#6    mNeoVis.evergladensis.1.1.chr14.g00081.t01#3    mNeoVis.2.1.1.chr14_hap1.g00076.t01#5    mNeoVis.vulgivaga.1.1.chr14.g00078.t01#4    mNeoVis.vulgivaga.2.1.chr14.g00076.t01#1   mNeoVis.evergladensis.2.1.chr14.g00077.t01#2       5140   125255520:   mNeoVis.vulgivaga.1.1.chr9.g00021.t01#4    mNeoVis.2.1.1.chr10_hap1.g00028.t01#5    mNeoVis.vulgivaga.2.1.chr9.g00027.t01#1    mNeoVis.evergladensis.2.1.chr8.g00026.t01#2   mNeoVis.evergladensis.1.1.chr9.g00024.t01#3       5141   125255521:   mNeoVis.vulgivaga.2.1.chr7.g00563.t01#1    mNeoVis.evergladensis.1.1.chr7.g00570.t01#3    mNeoVis.3.1.1.chr7.g00543.t01#6   mNeoVis.evergladensis.2.1.chr7.g00561.t01#2       5142   125255522:   mNeoVis.2.1.1.chr7_hap1.g00560.t01#5   mNeoVis.vulgivaga.1.1.chr7.g00569.t01#4             5143   125255523:   mNeoVis.vulgivaga.2.1.chr7.g00564.t01#1    mNeoVis.vulgivaga.1.1.chr7.g00570.t01#4    mNeoVis.3.1.1.chr7.g00544.t01#6    mNeoVis.evergladensis.2.1.chr7.g00562.t01#2    mNeoVis.2.1.1.chr7_hap1.g00561.t01#5   mNeoVis.evergladensis.1.1.chr7.g00571.t01#3       5144   125255524:   mNeoVis.vulgivaga.2.1.chr1.g00018.t01#1    mNeoVis.evergladensis.1.1.chr1.g00018.t01#3    mNeoVis.2.1.1.chr1_hap1.g00019.t01#5   mNeoVis.evergladensis.2.1.chr1.g00017.t01#2       5145   125255525:   mNeoVis.vulgivaga.2.1.chr7.g00565.t01#1    mNeoVis.2.1.1.chr7_hap1.g00562.t01#5    mNeoVis.evergladensis.1.1.chr7.g00572.t01#3    mNeoVis.3.1.1.chr7.g00545.t01#6    mNeoVis.vulgivaga.1.1.chr7.g00571.t01#4   mNeoVis.evergladensis.2.1.chr7.g00563.t01#2       5146   125255526:   mNeoVis.vulgivaga.2.1.chr1.g00019.t01#1    mNeoVis.evergladensis.1.1.chr1.g00019.t01#3    mNeoVis.vulgivaga.1.1.chr1.g00020.t01#4    mNeoVis.evergladensis.2.1.chr1.g00018.t01#2   mNeoVis.2.1.1.chr1_hap1.g00020.t01#5       5147   125255527:   mNeoVis.3.1.1.chr7.g00546.t01#6    mNeoVis.vulgivaga.1.1.chr7.g00572.t01#4    mNeoVis.evergladensis.2.1.chr7.g00564.t01#2    mNeoVis.2.1.1.chr7_hap1.g00563.t01#5    mNeoVis.evergladensis.1.1.chr7.g00573.t01#3   mNeoVis.vulgivaga.2.1.chr7.g00566.t01#1       5148   125255528:   mNeoVis.evergladensis.1.1.chr7.g00574.t01#3    mNeoVis.vulgivaga.2.1.chr7.g00567.t01#1    mNeoVis.evergladensis.2.1.chr7.g00565.t01#2    mNeoVis.vulgivaga.1.1.chr7.g00573.t01#4   mNeoVis.2.1.1.chr7_hap1.g00564.t01#5       5149   125255529:   mNeoVis.vulgivaga.2.1.chr7.g00568.t01#1   mNeoVis.3.1.1.chr7.g00547.t01#6             5150   125255530:   mNeoVis.vulgivaga.2.1.chr7.g00569.t01#1    mNeoVis.2.1.1.chr7_hap1.g00565.t01#5    mNeoVis.evergladensis.1.1.chr7.g00575.t01#3    mNeoVis.evergladensis.2.1.chr7.g00567.t01#2    mNeoVis.vulgivaga.1.1.chr7.g00574.t01#4    mNeoVis.evergladensis.2.1.chr7.g00566.t01#2   mNeoVis.3.1.1.chr7.g00548.t01#6       5151   125255531:  mNeoVis.vulgivaga.2.1.chr7.g00570.t01#1               5152   125255532:   mNeoVis.2.1.1.chr1_hap1.g00021.t01#5    mNeoVis.vulgivaga.2.1.chr1.g00020.t01#1    mNeoVis.vulgivaga.1.1.chr1.g00021.t01#4    mNeoVis.evergladensis.2.1.chr1.g00019.t01#2   mNeoVis.evergladensis.1.1.chr1.g00020.t01#3       5153   125255533:   mNeoVis.vulgivaga.2.1.chr7.g00571.t01#1    mNeoVis.vulgivaga.1.1.chr7.g00576.t01#4    mNeoVis.2.1.1.chr7_hap1.g00567.t01#5    mNeoVis.3.1.1.chr7.g00550.t01#6    mNeoVis.evergladensis.1.1.chr7.g00577.t01#3   mNeoVis.evergladensis.2.1.chr7.g00569.t01#2       5154   125255534:   mNeoVis.vulgivaga.2.1.chr7.g00572.t01#1    mNeoVis.vulgivaga.1.1.chr7.g00577.t01#4    mNeoVis.3.1.1.chr7.g00551.t01#6    mNeoVis.2.1.1.chr7_hap1.g00568.t01#5    mNeoVis.evergladensis.2.1.chr7.g00570.t01#2   mNeoVis.evergladensis.1.1.chr7.g00578.t01#3       5155   125255535:   mNeoVis.2.1.1.chr1_hap1.g00022.t01#5    mNeoVis.evergladensis.2.1.chr1.g00020.t01#2    mNeoVis.vulgivaga.2.1.chr1.g00021.t01#1    mNeoVis.evergladensis.1.1.chr1.g00021.t01#3   mNeoVis.3.1.1.chr1.g00020.t01#6       5156   125255536:   mNeoVis.2.1.1.chr1_hap1.g00024.t01#5    mNeoVis.vulgivaga.2.1.chr1.g00023.t01#1    mNeoVis.evergladensis.2.1.chr1.g00022.t01#2    mNeoVis.vulgivaga.1.1.chr1.g00024.t01#4    mNeoVis.evergladensis.1.1.chr1.g00023.t01#3   mNeoVis.3.1.1.chr1.g00022.t01#6       5157   125255537:   mNeoVis.vulgivaga.1.1.chr1.g00023.t01#4    mNeoVis.evergladensis.2.1.chr1.g00021.t01#2    mNeoVis.3.1.1.chr1.g00021.t01#6    mNeoVis.evergladensis.1.1.chr1.g00022.t01#3    mNeoVis.2.1.1.chr1_hap1.g00023.t01#5   mNeoVis.vulgivaga.2.1.chr1.g00022.t01#1       5158   125255538:   mNeoVis.vulgivaga.1.1.chr7.g00041.t01#4    mNeoVis.evergladensis.1.1.chr7.g00040.t01#3    mNeoVis.2.1.1.chr7_hap1.g00038.t01#5    mNeoVis.vulgivaga.2.1.chr7.g00039.t01#1   mNeoVis.evergladensis.2.1.chr7.g00040.t01#2       5159   125255539:   mNeoVis.vulgivaga.2.1.chr1.g00024.t01#1    mNeoVis.evergladensis.1.1.chr1.g00024.t01#3    mNeoVis.evergladensis.2.1.chr1.g00023.t01#2    mNeoVis.vulgivaga.1.1.chr1.g00025.t01#4   mNeoVis.2.1.1.chr1_hap1.g00025.t01#5       5160   125255540:   mNeoVis.vulgivaga.1.1.chr1.g00026.t01#4    mNeoVis.vulgivaga.2.1.chr1.g00025.t01#1    mNeoVis.evergladensis.2.1.chr1.g00024.t01#2    mNeoVis.2.1.1.chr1_hap1.g00026.t01#5   mNeoVis.evergladensis.1.1.chr1.g00026.t01#3       5161   125255541:   mNeoVis.3.1.1.chr7.g00552.t01#6    mNeoVis.2.1.1.chr7_hap1.g00569.t01#5    mNeoVis.evergladensis.2.1.chr7.g00571.t01#2    mNeoVis.vulgivaga.2.1.chr7.g00573.t01#1    mNeoVis.vulgivaga.1.1.chr7.g00578.t01#4   mNeoVis.evergladensis.1.1.chr7.g00579.t01#3       5162   125255542:  mNeoVis.vulgivaga.2.1.chr1.g00027.t01#1               5163   125255543:  mNeoVis.vulgivaga.2.1.chr7.g00574.t01#1               5164   125255544:   mNeoVis.evergladensis.1.1.chr1.g00027.t01#3    mNeoVis.evergladensis.2.1.chr1.g00025.t01#2    mNeoVis.3.1.1.chr1.g00023.t01#6    mNeoVis.2.1.1.chr1_hap1.g00027.t01#5    mNeoVis.vulgivaga.2.1.chr1.g00026.t01#1   mNeoVis.vulgivaga.1.1.chr1.g00027.t01#4       5165   125255545:   mNeoVis.3.1.1.chr7.g00553.t01#6    mNeoVis.vulgivaga.1.1.chr7.g00579.t01#4    mNeoVis.evergladensis.2.1.chr7.g00572.t01#2    mNeoVis.2.1.1.chr7_hap1.g00570.t01#5    mNeoVis.evergladensis.1.1.chr7.g00580.t01#3   mNeoVis.vulgivaga.2.1.chr7.g00575.t01#1       5166   125255546:   mNeoVis.evergladensis.1.1.chr1.g00028.t01#3    mNeoVis.2.1.1.chr1_hap1.g00028.t01#5    mNeoVis.vulgivaga.1.1.chr1.g00028.t01#4   mNeoVis.evergladensis.2.1.chr1.g00026.t01#2       5167   125255547:   mNeoVis.vulgivaga.2.1.chr1.g00028.t01#1   mNeoVis.3.1.1.chr1.g00026.t01#6             5168   125255548:   mNeoVis.evergladensis.1.1.chr1.g00029.t01#3    mNeoVis.3.1.1.chr1.g00027.t01#6    mNeoVis.vulgivaga.2.1.chr1.g00029.t01#1    mNeoVis.vulgivaga.1.1.chr1.g00029.t01#4    mNeoVis.evergladensis.2.1.chr1.g00027.t01#2   mNeoVis.2.1.1.chr1_hap1.g00029.t01#5       5169   125255549:   mNeoVis.evergladensis.1.1.chr1.g00030.t01#3    mNeoVis.2.1.1.chr1_hap1.g00030.t01#5    mNeoVis.vulgivaga.2.1.chr1.g00030.t01#1   mNeoVis.3.1.1.chr1.g00028.t01#6         5170   125255550:   mNeoVis.evergladensis.2.1.chr1.g00029.t01#2    mNeoVis.2.1.1.chr1_hap1.g00031.t01#5    mNeoVis.evergladensis.1.1.chr1.g00031.t01#3    mNeoVis.3.1.1.chr1.g00029.t01#6    mNeoVis.vulgivaga.1.1.chr1.g00031.t01#4    mNeoVis.evergladensis.2.1.chr1.g00030.t01#2   mNeoVis.vulgivaga.2.1.chr1.g00031.t01#1     5171   125255551:   mNeoVis.vulgivaga.1.1.chr1.g00032.t01#4    mNeoVis.2.1.1.chr1_hap1.g00032.t01#5    mNeoVis.vulgivaga.2.1.chr1.g00032.t01#1    mNeoVis.3.1.1.chr1.g00030.t01#6    mNeoVis.evergladensis.1.1.chr1.g00032.t01#3   mNeoVis.evergladensis.2.1.chr1.g00031.t01#2       5172   125255552:   mNeoVis.3.1.1.chr7.g00554.t01#6    mNeoVis.2.1.1.chr7_hap1.g00571.t01#5    mNeoVis.vulgivaga.2.1.chr7.g00576.t01#1    mNeoVis.evergladensis.2.1.chr7.g00573.t01#2    mNeoVis.evergladensis.1.1.chr7.g00581.t01#3   mNeoVis.vulgivaga.1.1.chr7.g00580.t01#4       5173   125255553:   mNeoVis.3.1.1.chr7.g00555.t01#6    mNeoVis.2.1.1.chr7_hap1.g00572.t01#5    mNeoVis.evergladensis.2.1.chr7.g00574.t01#2    mNeoVis.evergladensis.1.1.chr7.g00582.t01#3    mNeoVis.vulgivaga.1.1.chr7.g00581.t01#4   mNeoVis.vulgivaga.2.1.chr7.g00577.t01#1       5174   125255554:   mNeoVis.2.1.1.chr7_hap1.g00573.t01#5    mNeoVis.3.1.1.chr7.g00556.t01#6    mNeoVis.vulgivaga.2.1.chr7.g00578.t01#1    mNeoVis.evergladensis.2.1.chr7.g00575.t01#2    mNeoVis.evergladensis.1.1.chr7.g00583.t01#3   mNeoVis.vulgivaga.1.1.chr7.g00582.t01#4       5175   125255555:   mNeoVis.vulgivaga.1.1.chr7.g00042.t01#4    mNeoVis.vulgivaga.2.1.chr7.g00040.t01#1    mNeoVis.evergladensis.1.1.chr7.g00041.t01#3    mNeoVis.2.1.1.chr7_hap1.g00039.t01#5   mNeoVis.evergladensis.2.1.chr7.g00041.t01#2       5176   125255556:   mNeoVis.vulgivaga.2.1.chr7.g00579.t01#1    mNeoVis.3.1.1.chr7.g00557.t01#6    mNeoVis.2.1.1.chr7_hap1.g00574.t01#5    mNeoVis.evergladensis.2.1.chr7.g00576.t01#2    mNeoVis.evergladensis.1.1.chr7.g00584.t01#3   mNeoVis.vulgivaga.1.1.chr7.g00583.t01#4       5177   125255557:   mNeoVis.vulgivaga.2.1.chr15.g00124.t01#1    mNeoVis.3.1.1.chr15.g00112.t01#6    mNeoVis.2.1.1.chr15_hap1.g00135.t01#5    mNeoVis.evergladensis.2.1.chr15.g00121.t01#2    mNeoVis.evergladensis.1.1.chr15.g00494.t01#3   mNeoVis.vulgivaga.1.1.chr15.g00121.t01#4       5178   125255558:   mNeoVis.evergladensis.1.1.chr1.g00033.t01#3    mNeoVis.vulgivaga.1.1.chr1.g00033.t01#4    mNeoVis.vulgivaga.2.1.chr1.g00033.t01#1    mNeoVis.evergladensis.2.1.chr1.g00032.t01#2    mNeoVis.3.1.1.chr1.g00032.t01#6   mNeoVis.2.1.1.chr1_hap1.g00033.t01#5       5179   125255559:   mNeoVis.evergladensis.2.1.chr1.g00033.t01#2    mNeoVis.vulgivaga.2.1.chr1.g00034.t01#1    mNeoVis.vulgivaga.1.1.chr1.g00034.t01#4    mNeoVis.3.1.1.chr1.g00033.t01#6    mNeoVis.2.1.1.chr1_hap1.g00034.t01#5   mNeoVis.evergladensis.1.1.chr1.g00034.t01#3       5180   125255560:   mNeoVis.vulgivaga.2.1.chr7.g00580.t01#1    mNeoVis.2.1.1.chr7_hap1.g00575.t01#5    mNeoVis.evergladensis.2.1.chr12.g00356.t01#2    mNeoVis.vulgivaga.1.1.chr7.g00584.t01#4    mNeoVis.evergladensis.2.1.chr7.g00577.t01#2    mNeoVis.vulgivaga.1.1.chr12.g00363.t01#4    mNeoVis.evergladensis.1.1.chr12.g00363.t01#3    mNeoVis.evergladensis.1.1.chr7.g00585.t01#3    mNeoVis.3.1.1.chr7.g00558.t01#6      5181   125255561:   mNeoVis.vulgivaga.2.1.chr1.g00035.t01#1    mNeoVis.evergladensis.1.1.chr1.g00035.t01#3    mNeoVis.3.1.1.chr1.g00034.t01#6    mNeoVis.2.1.1.chr1_hap1.g00035.t01#5   mNeoVis.evergladensis.2.1.chr1.g00034.t01#2       5182   125255562:   mNeoVis.evergladensis.2.1.chr7.g00578.t01#2    mNeoVis.vulgivaga.1.1.chr7.g00585.t01#4    mNeoVis.2.1.1.chr7_hap1.g00576.t01#5    mNeoVis.vulgivaga.2.1.chr7.g00581.t01#1    mNeoVis.evergladensis.1.1.chr7.g00586.t01#3   mNeoVis.3.1.1.chr7.g00559.t01#6       5183   125255563:   mNeoVis.vulgivaga.2.1.chr7.g00582.t01#1    mNeoVis.evergladensis.1.1.chr7.g00587.t01#3    mNeoVis.evergladensis.2.1.chr7.g00579.t01#2    mNeoVis.vulgivaga.1.1.chr7.g00586.t01#4    mNeoVis.3.1.1.chr7.g00560.t01#6   mNeoVis.2.1.1.chr7_hap1.g00577.t01#5       5184   125255564:   mNeoVis.3.1.1.chr5.g00678.t01#6    mNeoVis.evergladensis.1.1.chr5.g00681.t01#3    mNeoVis.evergladensis.2.1.chr5.g00687.t01#2    mNeoVis.vulgivaga.2.1.chr5.g00678.t01#1    mNeoVis.2.1.1.chr2_hap1.g00728.t01#5   mNeoVis.vulgivaga.1.1.chr5.g00664.t01#4       5185   125255565:   mNeoVis.2.1.1.chr1_hap1.g00036.t01#5    mNeoVis.evergladensis.2.1.chr1.g00035.t01#2    mNeoVis.evergladensis.1.1.chr1.g00036.t01#3   mNeoVis.vulgivaga.2.1.chr1.g00036.t01#1         5186   125255566:   mNeoVis.evergladensis.1.1.chr7.g00588.t01#3    mNeoVis.evergladensis.2.1.chr7.g00580.t01#2    mNeoVis.vulgivaga.1.1.chr7.g00587.t01#4    mNeoVis.2.1.1.chr7_hap1.g00578.t01#5    mNeoVis.3.1.1.chr7.g00561.t01#6   mNeoVis.vulgivaga.2.1.chr7.g00583.t01#1       5187   125255567:   mNeoVis.vulgivaga.1.1.chr1.g00035.t01#4    mNeoVis.vulgivaga.2.1.chr1.g00037.t01#1    mNeoVis.evergladensis.1.1.chr1.g00037.t01#3    mNeoVis.2.1.1.chr1_hap1.g00038.t01#5   mNeoVis.evergladensis.2.1.chr1.g00036.t01#2       5188   125255568:   mNeoVis.3.1.1.chr7.g00031.t01#6    mNeoVis.vulgivaga.1.1.chr7.g00043.t01#4    mNeoVis.vulgivaga.2.1.chr7.g00041.t01#1    mNeoVis.evergladensis.2.1.chr7.g00042.t01#2    mNeoVis.evergladensis.1.1.chr7.g00042.t01#3   mNeoVis.2.1.1.chr7_hap1.g00040.t01#5       5189   125255569:   mNeoVis.vulgivaga.2.1.chr7.g00584.t01#1    mNeoVis.evergladensis.1.1.chr7.g00589.t01#3    mNeoVis.3.1.1.chr7.g00562.t01#6    mNeoVis.vulgivaga.1.1.chr7.g00588.t01#4    mNeoVis.2.1.1.chr7_hap1.g00579.t01#5   mNeoVis.evergladensis.2.1.chr7.g00581.t01#2       5190   125255570:   mNeoVis.evergladensis.1.1.chr7.g00590.t01#3    mNeoVis.vulgivaga.1.1.chr7.g00589.t01#4    mNeoVis.vulgivaga.2.1.chr7.g00585.t01#1    mNeoVis.evergladensis.2.1.chr7.g00582.t01#2    mNeoVis.2.1.1.chr7_hap1.g00580.t01#5   mNeoVis.3.1.1.chr7.g00563.t01#6       5191   125255571:   mNeoVis.3.1.1.chr15.g00113.t01#6   mNeoVis.vulgivaga.2.1.chr15.g00125.t01#1           5192   125255572:   mNeoVis.2.1.1.chr7_hap1.g00581.t01#5    mNeoVis.evergladensis.2.1.chr7.g00583.t01#2    mNeoVis.vulgivaga.2.1.chr7.g00587.t01#1    mNeoVis.evergladensis.1.1.chr7.g00592.t01#3    mNeoVis.2.1.1.chr7_hap1.g00582.t01#5    mNeoVis.evergladensis.1.1.chr7.g00591.t01#3   mNeoVis.vulgivaga.1.1.chr7.g00590.t01#4     5193   125255573:   mNeoVis.vulgivaga.2.1.chr7.g00586.t01#1   mNeoVis.3.1.1.chr7.g00564.t01#6             5194   125255574:   mNeoVis.3.1.1.chr1.g00039.t01#6    mNeoVis.evergladensis.1.1.chr1.g00038.t01#3    mNeoVis.vulgivaga.1.1.chr1.g00036.t01#4    mNeoVis.vulgivaga.2.1.chr1.g00038.t01#1   mNeoVis.2.1.1.chr1_hap1.g00039.t01#5       5195   125255575:   mNeoVis.evergladensis.2.1.chr1.g00037.t01#2    mNeoVis.2.1.1.chr1_hap1.g00040.t01#5    mNeoVis.vulgivaga.2.1.chr1.g00039.t01#1    mNeoVis.3.1.1.chr1.g00040.t01#6    mNeoVis.evergladensis.1.1.chr1.g00039.t01#3   mNeoVis.vulgivaga.1.1.chr1.g00037.t01#4       5196   125255576:   mNeoVis.evergladensis.2.1.chr1.g00038.t01#2    mNeoVis.vulgivaga.2.1.chr1.g00040.t01#1    mNeoVis.2.1.1.chr1_hap1.g00041.t01#5    mNeoVis.vulgivaga.1.1.chr1.g00038.t01#4    mNeoVis.3.1.1.chr1.g00041.t01#6   mNeoVis.evergladensis.1.1.chr1.g00040.t01#3       5197   125255577:   mNeoVis.vulgivaga.1.1.chr7.g00591.t01#4    mNeoVis.vulgivaga.2.1.chr7.g00588.t01#1    mNeoVis.evergladensis.2.1.chr7.g00584.t01#2    mNeoVis.2.1.1.chr7_hap1.g00583.t01#5    mNeoVis.3.1.1.chr7.g00565.t01#6   mNeoVis.evergladensis.1.1.chr7.g00593.t01#3       5198   125255578:   mNeoVis.vulgivaga.1.1.chr7.g00592.t01#4    mNeoVis.3.1.1.chr7.g00566.t01#6   mNeoVis.evergladensis.2.1.chr7.g00585.t01#2         5199   125255579:   mNeoVis.2.1.1.chr15_hap1.g00136.t01#5    mNeoVis.evergladensis.2.1.chr15.g00122.t01#2    mNeoVis.3.1.1.chr15.g00114.t01#6    mNeoVis.vulgivaga.2.1.chr15.g00126.t01#1    mNeoVis.vulgivaga.1.1.chr15.g00122.t01#4   mNeoVis.evergladensis.1.1.chr15.g00492.t01#3       5200   125255580:   mNeoVis.2.1.1.chr1_hap1.g00042.t01#5    mNeoVis.vulgivaga.1.1.chr1.g00039.t01#4    mNeoVis.evergladensis.2.1.chr1.g00039.t01#2    mNeoVis.vulgivaga.2.1.chr1.g00041.t01#1    mNeoVis.evergladensis.1.1.chr1.g00041.t01#3   mNeoVis.3.1.1.chr1.g00045.t01#6       5201   125255581:   mNeoVis.evergladensis.2.1.chr7.g00586.t01#2    mNeoVis.2.1.1.chr7_hap1.g00584.t01#5    mNeoVis.evergladensis.1.1.chr7.g00594.t01#3    mNeoVis.vulgivaga.2.1.chr7.g00589.t01#1    mNeoVis.3.1.1.chr7.g00567.t01#6   mNeoVis.vulgivaga.1.1.chr7.g00593.t01#4       5202   125255582:   mNeoVis.2.1.1.chr7_hap1.g00585.t01#5    mNeoVis.vulgivaga.2.1.chr7.g00590.t01#1    mNeoVis.evergladensis.1.1.chr7.g00595.t01#3    mNeoVis.vulgivaga.1.1.chr7.g00594.t01#4    mNeoVis.3.1.1.chr7.g00568.t01#6   mNeoVis.evergladensis.2.1.chr7.g00587.t01#2       5203   125255583:   mNeoVis.vulgivaga.2.1.chr1.g00042.t01#1    mNeoVis.2.1.1.chr1_hap1.g00043.t01#5    mNeoVis.evergladensis.2.1.chr1.g00040.t01#2    mNeoVis.vulgivaga.1.1.chr1.g00040.t01#4    mNeoVis.evergladensis.1.1.chr1.g00043.t01#3   mNeoVis.3.1.1.chr1.g00046.t01#6       5204   125255584:   mNeoVis.vulgivaga.1.1.chr1.g00041.t01#4    mNeoVis.3.1.1.chr1.g00047.t01#6    mNeoVis.evergladensis.2.1.chr1.g00041.t01#2    mNeoVis.2.1.1.chr1_hap1.g00044.t01#5    mNeoVis.evergladensis.1.1.chr1.g00044.t01#3   mNeoVis.vulgivaga.2.1.chr1.g00043.t01#1       5205   125255585:   mNeoVis.3.1.1.chr1.g00048.t01#6    mNeoVis.vulgivaga.1.1.chr1.g00042.t01#4    mNeoVis.2.1.1.chr1_hap1.g00045.t01#5    mNeoVis.evergladensis.1.1.chr1.g00045.t01#3    mNeoVis.evergladensis.2.1.chr1.g00042.t01#2   mNeoVis.vulgivaga.2.1.chr1.g00044.t01#1       5206   125255586:   mNeoVis.vulgivaga.1.1.chr7.g00595.t01#4    mNeoVis.3.1.1.chr7.g00569.t01#6    mNeoVis.2.1.1.chr7_hap1.g00586.t01#5    mNeoVis.vulgivaga.2.1.chr7.g00591.t01#1    mNeoVis.evergladensis.2.1.chr7.g00588.t01#2   mNeoVis.evergladensis.1.1.chr7.g00596.t01#3       5207   125255587:   mNeoVis.vulgivaga.2.1.chr6.g00769.t01#1    mNeoVis.3.1.1.chr6.g00772.t01#6    mNeoVis.2.1.1.chr4_hap1.g00767.t01#5    mNeoVis.evergladensis.2.1.chr6.g00773.t01#2    mNeoVis.evergladensis.1.1.chr6.g00778.t01#3   mNeoVis.vulgivaga.1.1.chr6.g00771.t01#4       5208   125255588:   mNeoVis.vulgivaga.2.1.chr14.g00412.t01#1    mNeoVis.3.1.1.chr14.g00411.t01#6    mNeoVis.vulgivaga.1.1.chr14.g00406.t01#4    mNeoVis.2.1.1.chr14_hap1.g00411.t01#5    mNeoVis.evergladensis.1.1.chr14.g00414.t01#3   mNeoVis.evergladensis.2.1.chr14.g00410.t01#2       5209   125255589:   mNeoVis.evergladensis.1.1.chr14.g00415.t01#3    mNeoVis.evergladensis.2.1.chr14.g00411.t01#2    mNeoVis.3.1.1.chr14.g00412.t01#6    mNeoVis.vulgivaga.2.1.chr14.g00413.t01#1    mNeoVis.vulgivaga.1.1.chr14.g00407.t01#4   mNeoVis.2.1.1.chr14_hap1.g00412.t01#5       5210   125255590:   mNeoVis.vulgivaga.1.1.chr6.g00773.t01#4    mNeoVis.2.1.1.chr4_hap1.g00772.t01#5    mNeoVis.vulgivaga.2.1.chr6.g00770.t01#1   mNeoVis.evergladensis.1.1.chr6.g00780.t01#3       5211   125255591:   mNeoVis.2.1.1.chr14_hap1.g00414.t01#5    mNeoVis.vulgivaga.1.1.chr14.g00408.t01#4    mNeoVis.3.1.1.chr14.g00413.t01#6    mNeoVis.vulgivaga.2.1.chr14.g00414.t01#1    mNeoVis.evergladensis.2.1.chr14.g00412.t01#2   mNeoVis.evergladensis.1.1.chr14.g00416.t01#3       5212   125255592:   mNeoVis.evergladensis.2.1.chr14.g00413.t01#2    mNeoVis.2.1.1.chr14_hap1.g00415.t01#5    mNeoVis.evergladensis.1.1.chr14.g00417.t01#3    mNeoVis.vulgivaga.2.1.chr14.g00415.t01#1    mNeoVis.vulgivaga.1.1.chr14.g00409.t01#4   mNeoVis.3.1.1.chr14.g00414.t01#6       5213   125255593:   mNeoVis.3.1.1.chr14.g00416.t01#6    mNeoVis.vulgivaga.2.1.chr14.g00416.t01#1    mNeoVis.vulgivaga.1.1.chr14.g00410.t01#4    mNeoVis.evergladensis.2.1.chr14.g00415.t01#2    mNeoVis.2.1.1.chr14_hap1.g00417.t01#5   mNeoVis.evergladensis.1.1.chr14.g00418.t01#3       5214   125255594:   mNeoVis.vulgivaga.2.1.chr14.g00417.t01#1    mNeoVis.3.1.1.chr14.g00415.t01#6    mNeoVis.evergladensis.2.1.chr14.g00414.t01#2   mNeoVis.2.1.1.chr14_hap1.g00416.t01#5         5215   125255595:   mNeoVis.vulgivaga.1.1.chr15.g00553.t01#4    mNeoVis.2.1.1.chr15_hap1.g00728.t01#5    mNeoVis.evergladensis.2.1.chr15.g00550.t01#2    mNeoVis.vulgivaga.2.1.chr15.g00562.t01#1    mNeoVis.evergladensis.1.1.chr15.g00063.t01#3   mNeoVis.3.1.1.chr15.g00505.t01#6       5216   125255596:   mNeoVis.vulgivaga.1.1.chr7.g00509.t01#4    mNeoVis.vulgivaga.2.1.chr7.g00505.t01#1    mNeoVis.3.1.1.chr7.g00486.t01#6    mNeoVis.2.1.1.chr7_hap1.g00501.t01#5    mNeoVis.evergladensis.1.1.chr7.g00511.t01#3   mNeoVis.evergladensis.2.1.chr7.g00502.t01#2       5217   125255597:   mNeoVis.3.1.1.chr6.g00779.t01#6    mNeoVis.evergladensis.2.1.chr6.g00777.t01#2    mNeoVis.vulgivaga.1.1.chr6.g00774.t01#4    mNeoVis.vulgivaga.2.1.chr6.g00771.t01#1    mNeoVis.3.1.1.chr6.g00780.t01#6    mNeoVis.evergladensis.1.1.chr6.g00782.t01#3    mNeoVis.evergladensis.1.1.chr6.g00781.t01#3   mNeoVis.2.1.1.chr4_hap1.g00773.t01#5     5218   125255598:   mNeoVis.vulgivaga.1.1.chr14.g00411.t01#4    mNeoVis.evergladensis.2.1.chr14.g00416.t01#2    mNeoVis.3.1.1.chr14.g00417.t01#6    mNeoVis.2.1.1.chr14_hap1.g00418.t01#5    mNeoVis.evergladensis.1.1.chr14.g00419.t01#3   mNeoVis.vulgivaga.2.1.chr14.g00418.t01#1       5219   125255599:   mNeoVis.evergladensis.2.1.chr14.g00417.t01#2    mNeoVis.vulgivaga.1.1.chr14.g00412.t01#4    mNeoVis.2.1.1.chr14_hap1.g00419.t01#5    mNeoVis.evergladensis.1.1.chr14.g00420.t01#3   mNeoVis.vulgivaga.2.1.chr14.g00419.t01#1       5220   125255600:   mNeoVis.vulgivaga.2.1.chr14.g00420.t01#1    mNeoVis.evergladensis.2.1.chr14.g00418.t01#2    mNeoVis.vulgivaga.1.1.chr14.g00413.t01#4    mNeoVis.3.1.1.chr14.g00418.t01#6    mNeoVis.2.1.1.chr14_hap1.g00420.t01#5   mNeoVis.evergladensis.1.1.chr14.g00421.t01#3       5221   125255601:   mNeoVis.vulgivaga.2.1.chr6.g00772.t01#1    mNeoVis.vulgivaga.1.1.chr6.g00775.t01#4    mNeoVis.evergladensis.2.1.chr6.g00778.t01#2    mNeoVis.3.1.1.chr6.g00781.t01#6   mNeoVis.evergladensis.1.1.chr6.g00783.t01#3       5222   125255602:   mNeoVis.3.1.1.chr11.g00031.t01#6    mNeoVis.2.1.1.chr11_hap1.g00055.t01#5   mNeoVis.vulgivaga.2.1.chr11.g00054.t01#1         5223   125255603:   mNeoVis.2.1.1.chr4_hap1.g00774.t01#5    mNeoVis.vulgivaga.1.1.chr6.g00776.t01#4    mNeoVis.vulgivaga.2.1.chr6.g00773.t01#1    mNeoVis.evergladensis.1.1.chr6.g00784.t01#3    mNeoVis.evergladensis.2.1.chr6.g00779.t01#2   mNeoVis.3.1.1.chr6.g00782.t01#6       5224   125255604:   mNeoVis.vulgivaga.2.1.chr14.g00422.t01#1    mNeoVis.3.1.1.chr14.g00420.t01#6    mNeoVis.evergladensis.2.1.chr14.g00420.t01#2    mNeoVis.vulgivaga.1.1.chr14.g00415.t01#4    mNeoVis.evergladensis.1.1.chr14.g00424.t01#3   mNeoVis.2.1.1.chr14_hap1.g00422.t01#5       5225   125255605:   mNeoVis.3.1.1.chr14.g00421.t01#6    mNeoVis.evergladensis.1.1.chr14.g00425.t01#3    mNeoVis.2.1.1.chr14_hap1.g00423.t01#5    mNeoVis.vulgivaga.2.1.chr14.g00423.t01#1   mNeoVis.evergladensis.2.1.chr14.g00421.t01#2       5226   125255606:   mNeoVis.evergladensis.2.1.chr14.g00422.t01#2    mNeoVis.evergladensis.1.1.chr14.g00426.t01#3    mNeoVis.vulgivaga.1.1.chr14.g00416.t01#4    mNeoVis.2.1.1.chr14_hap1.g00424.t01#5    mNeoVis.vulgivaga.2.1.chr14.g00424.t01#1   mNeoVis.3.1.1.chr14.g00423.t01#6       5227   125255607:   mNeoVis.evergladensis.1.1.chr6.g00785.t01#3    mNeoVis.evergladensis.2.1.chr6.g00780.t01#2    mNeoVis.vulgivaga.1.1.chr6.g00777.t01#4    mNeoVis.vulgivaga.2.1.chr6.g00774.t01#1    mNeoVis.3.1.1.chr6.g00783.t01#6   mNeoVis.2.1.1.chr4_hap1.g00775.t01#5       5228   125255608:   mNeoVis.2.1.1.chr4_hap1.g00776.t01#5    mNeoVis.3.1.1.chr6.g00784.t01#6    mNeoVis.vulgivaga.1.1.chr6.g00778.t01#4    mNeoVis.evergladensis.2.1.chr6.g00781.t01#2    mNeoVis.evergladensis.1.1.chr6.g00786.t01#3   mNeoVis.vulgivaga.2.1.chr6.g00775.t01#1       5229   125255609:   mNeoVis.3.1.1.chr14.g00424.t01#6    mNeoVis.vulgivaga.1.1.chr14.g00417.t01#4    mNeoVis.vulgivaga.2.1.chr14.g00425.t01#1    mNeoVis.2.1.1.chr14_hap1.g00425.t01#5    mNeoVis.evergladensis.2.1.chr14.g00423.t01#2   mNeoVis.evergladensis.1.1.chr14.g00427.t01#3       5230   125255610:   mNeoVis.2.1.1.chr4_hap1.g00777.t01#5    mNeoVis.evergladensis.2.1.chr6.g00782.t01#2    mNeoVis.vulgivaga.1.1.chr6.g00779.t01#4    mNeoVis.evergladensis.1.1.chr6.g00787.t01#3    mNeoVis.vulgivaga.2.1.chr6.g00776.t01#1   mNeoVis.3.1.1.chr6.g00786.t01#6       5231   125255611:   mNeoVis.evergladensis.1.1.chr6.g00788.t01#3    mNeoVis.vulgivaga.2.1.chr6.g00777.t01#1    mNeoVis.3.1.1.chr6.g00788.t01#6    mNeoVis.evergladensis.2.1.chr6.g00783.t01#2    mNeoVis.2.1.1.chr4_hap1.g00778.t01#5    mNeoVis.3.1.1.chr6.g00789.t01#6   mNeoVis.vulgivaga.1.1.chr6.g00780.t01#4     5232   125255612:   mNeoVis.evergladensis.2.1.chr7.g00503.t01#2    mNeoVis.vulgivaga.2.1.chr7.g00506.t01#1    mNeoVis.3.1.1.chr7.g00487.t01#6    mNeoVis.vulgivaga.1.1.chr7.g00510.t01#4    mNeoVis.2.1.1.chr7_hap1.g00502.t01#5   mNeoVis.evergladensis.1.1.chr7.g00512.t01#3       5233   125255613:   mNeoVis.vulgivaga.1.1.chr6.g00781.t01#4    mNeoVis.2.1.1.chr4_hap1.g00779.t01#5    mNeoVis.evergladensis.1.1.chr6.g00789.t01#3    mNeoVis.vulgivaga.2.1.chr6.g00778.t01#1   mNeoVis.evergladensis.2.1.chr6.g00784.t01#2       5234   125255614:   mNeoVis.evergladensis.2.1.chr6.g00785.t01#2    mNeoVis.2.1.1.chr4_hap1.g00780.t01#5    mNeoVis.3.1.1.chr6.g00790.t01#6    mNeoVis.vulgivaga.1.1.chr6.g00783.t01#4    mNeoVis.evergladensis.1.1.chr6.g00790.t01#3   mNeoVis.vulgivaga.2.1.chr6.g00779.t01#1       5235   125255615:   mNeoVis.evergladensis.1.1.chr14.g00428.t01#3    mNeoVis.3.1.1.chr14.g00425.t01#6    mNeoVis.vulgivaga.2.1.chr14.g00426.t01#1    mNeoVis.vulgivaga.1.1.chr14.g00419.t01#4    mNeoVis.vulgivaga.1.1.chr14.g00418.t01#4    mNeoVis.evergladensis.2.1.chr14.g00424.t01#2   mNeoVis.2.1.1.chr14_hap1.g00426.t01#5     5236   125255616:   mNeoVis.evergladensis.1.1.chr9.g00197.t01#3    mNeoVis.vulgivaga.1.1.chr9.g00194.t01#4    mNeoVis.2.1.1.chr10_hap1.g00200.t01#5    mNeoVis.vulgivaga.1.1.chr9.g00193.t01#4    mNeoVis.evergladensis.2.1.chr8.g00192.t01#2    mNeoVis.3.1.1.chr9.g00175.t01#6   mNeoVis.vulgivaga.2.1.chr9.g00194.t01#1     5237   125255617:   mNeoVis.evergladensis.1.1.chr6.g00791.t01#3    mNeoVis.vulgivaga.1.1.chr6.g00784.t01#4    mNeoVis.evergladensis.2.1.chr6.g00786.t01#2    mNeoVis.3.1.1.chr6.g00791.t01#6    mNeoVis.2.1.1.chr4_hap1.g00781.t01#5   mNeoVis.vulgivaga.2.1.chr6.g00780.t01#1       5238   125255618:  mNeoVis.vulgivaga.2.1.chr14.g00427.t01#1               5239   125255619:   mNeoVis.vulgivaga.2.1.chr6.g00781.t01#1    mNeoVis.2.1.1.chr4_hap1.g00782.t01#5    mNeoVis.vulgivaga.1.1.chr6.g00785.t01#4    mNeoVis.evergladensis.2.1.chr6.g00787.t01#2    mNeoVis.evergladensis.1.1.chr6.g00792.t01#3   mNeoVis.3.1.1.chr6.g00792.t01#6       5240   125255620:   mNeoVis.3.1.1.chr7.g00488.t01#6    mNeoVis.evergladensis.1.1.chr7.g00513.t01#3    mNeoVis.vulgivaga.2.1.chr7.g00507.t01#1    mNeoVis.2.1.1.chr7_hap1.g00503.t01#5    mNeoVis.vulgivaga.1.1.chr7.g00511.t01#4   mNeoVis.evergladensis.2.1.chr7.g00504.t01#2       5241   125255621:   mNeoVis.evergladensis.2.1.chr14.g00337.t01#2    mNeoVis.3.1.1.chr14.g00336.t01#6    mNeoVis.vulgivaga.1.1.chr14.g00333.t01#4    mNeoVis.2.1.1.chr14_hap1.g00337.t01#5    mNeoVis.evergladensis.1.1.chr14.g00342.t01#3   mNeoVis.vulgivaga.2.1.chr14.g00339.t01#1       5242   125255622:   mNeoVis.evergladensis.2.1.chr6.g00788.t01#2    mNeoVis.vulgivaga.1.1.chr6.g00786.t01#4    mNeoVis.evergladensis.1.1.chr6.g00793.t01#3    mNeoVis.vulgivaga.2.1.chr6.g00782.t01#1    mNeoVis.3.1.1.chr6.g00793.t01#6   mNeoVis.2.1.1.chr4_hap1.g00783.t01#5       5243   125255623:   mNeoVis.2.1.1.chr11_hap1.g00140.t01#5    mNeoVis.evergladensis.2.1.chr11.g00137.t01#2    mNeoVis.vulgivaga.1.1.chr11.g00134.t01#4    mNeoVis.evergladensis.1.1.chr11.g00138.t01#3    mNeoVis.3.1.1.chr11.g00111.t01#6   mNeoVis.vulgivaga.2.1.chr11.g00140.t01#1       5244   125255624:   mNeoVis.vulgivaga.2.1.chr14.g00340.t01#1    mNeoVis.evergladensis.1.1.chr14.g00343.t01#3    mNeoVis.evergladensis.2.1.chr14.g00339.t01#2    mNeoVis.2.1.1.chr14_hap1.g00338.t01#5    mNeoVis.3.1.1.chr14.g00337.t01#6   mNeoVis.vulgivaga.1.1.chr14.g00335.t01#4       5245   125255625:   mNeoVis.evergladensis.2.1.chr6.g00790.t01#2    mNeoVis.evergladensis.1.1.chr6.g00794.t01#3    mNeoVis.2.1.1.chr4_hap1.g00785.t01#5    mNeoVis.vulgivaga.1.1.chr6.g00787.t01#4    mNeoVis.vulgivaga.2.1.chr6.g00783.t01#1   mNeoVis.3.1.1.chr6.g00794.t01#6       5246   125255626:   mNeoVis.3.1.1.chr6.g00795.t01#6    mNeoVis.vulgivaga.1.1.chr6.g00788.t01#4    mNeoVis.vulgivaga.2.1.chr6.g00784.t01#1    mNeoVis.evergladensis.2.1.chr6.g00791.t01#2    mNeoVis.evergladensis.1.1.chr6.g00795.t01#3   mNeoVis.2.1.1.chr4_hap1.g00786.t01#5       5247   125255627:   mNeoVis.3.1.1.chr6.g00796.t01#6    mNeoVis.vulgivaga.1.1.chr6.g00789.t01#4    mNeoVis.2.1.1.chr4_hap1.g00787.t01#5    mNeoVis.vulgivaga.2.1.chr6.g00785.t01#1    mNeoVis.evergladensis.1.1.chr6.g00796.t01#3   mNeoVis.evergladensis.2.1.chr6.g00792.t01#2       5248   125255628:   mNeoVis.vulgivaga.1.1.chr4.g00592.t01#4    mNeoVis.2.1.1.chr6_hap1.g00586.t01#5    mNeoVis.3.1.1.chr6.g00797.t01#6    mNeoVis.vulgivaga.1.1.chr6.g00790.t01#4    mNeoVis.vulgivaga.2.1.chr6.g00786.t01#1    mNeoVis.2.1.1.chr4_hap1.g00788.t01#5    mNeoVis.evergladensis.2.1.chr4.g00590.t01#2    mNeoVis.evergladensis.1.1.chr4.g00599.t01#3    mNeoVis.3.1.1.chr4.g00589.t01#6      5249   125255629:   mNeoVis.vulgivaga.2.1.chr14.g00341.t01#1    mNeoVis.3.1.1.chr14.g00338.t01#6    mNeoVis.evergladensis.2.1.chr14.g00340.t01#2    mNeoVis.evergladensis.1.1.chr14.g00344.t01#3    mNeoVis.2.1.1.chr14_hap1.g00339.t01#5   mNeoVis.vulgivaga.1.1.chr14.g00336.t01#4       5250   125255630:   mNeoVis.evergladensis.1.1.chr10.g00940.t01#3    mNeoVis.evergladensis.2.1.chr10.g00939.t01#2    mNeoVis.vulgivaga.1.1.chr10.g00921.t01#4    mNeoVis.2.1.1.chr9_hap1.g00954.t01#5    mNeoVis.3.1.1.chr10.g00909.t01#6   mNeoVis.vulgivaga.2.1.chr10.g00930.t01#1       5251   125255631:   mNeoVis.vulgivaga.2.1.chr6.g00787.t01#1    mNeoVis.2.1.1.chr4_hap1.g00789.t01#5    mNeoVis.vulgivaga.1.1.chr6.g00791.t01#4    mNeoVis.evergladensis.2.1.chr6.g00794.t01#2    mNeoVis.evergladensis.1.1.chr6.g00798.t01#3   mNeoVis.3.1.1.chr6.g00798.t01#6       5252   125255632:   mNeoVis.3.1.1.chr14.g00339.t01#6    mNeoVis.vulgivaga.1.1.chr14.g00337.t01#4    mNeoVis.evergladensis.2.1.chr14.g00341.t01#2    mNeoVis.evergladensis.1.1.chr14.g00345.t01#3    mNeoVis.vulgivaga.2.1.chr14.g00342.t01#1   mNeoVis.2.1.1.chr14_hap1.g00340.t01#5       5253   125255633:   mNeoVis.evergladensis.1.1.chr6.g00799.t01#3    mNeoVis.vulgivaga.2.1.chr6.g00788.t01#1    mNeoVis.vulgivaga.1.1.chr6.g00792.t01#4    mNeoVis.2.1.1.chr4_hap1.g00790.t01#5    mNeoVis.3.1.1.chr6.g00799.t01#6   mNeoVis.evergladensis.2.1.chr6.g00795.t01#2       5254   125255634:   mNeoVis.vulgivaga.1.1.chr6.g00698.t01#4    mNeoVis.vulgivaga.2.1.chr6.g00697.t01#1    mNeoVis.evergladensis.2.1.chr6.g00702.t01#2    mNeoVis.evergladensis.1.1.chr6.g00701.t01#3    mNeoVis.3.1.1.chr6.g00702.t01#6   mNeoVis.2.1.1.chr4_hap1.g00698.t01#5       5255   125255635:   mNeoVis.evergladensis.1.1.chr14.g00346.t01#3    mNeoVis.vulgivaga.2.1.chr14.g00343.t01#1    mNeoVis.2.1.1.chr14_hap1.g00341.t01#5    mNeoVis.vulgivaga.1.1.chr14.g00338.t01#4    mNeoVis.3.1.1.chr14.g00340.t01#6   mNeoVis.evergladensis.2.1.chr6.g00690.t01#2       5256   125255636:   mNeoVis.2.1.1.chr7_hap1.g00504.t01#5    mNeoVis.evergladensis.2.1.chr7.g00505.t01#2    mNeoVis.3.1.1.chr7.g00489.t01#6    mNeoVis.vulgivaga.2.1.chr7.g00508.t01#1    mNeoVis.evergladensis.1.1.chr7.g00514.t01#3   mNeoVis.vulgivaga.1.1.chr7.g00512.t01#4       5257   125255637:   mNeoVis.evergladensis.1.1.chr14.g00348.t01#3    mNeoVis.evergladensis.2.1.chr14.g00344.t01#2    mNeoVis.3.1.1.chr14.g00342.t01#6    mNeoVis.vulgivaga.1.1.chr14.g00340.t01#4    mNeoVis.2.1.1.chr14_hap1.g00344.t01#5   mNeoVis.vulgivaga.2.1.chr14.g00345.t01#1       5258   125255638:   mNeoVis.evergladensis.2.1.chr3.g01165.t01#2    mNeoVis.vulgivaga.2.1.chr10.g00173.t01#1    mNeoVis.vulgivaga.2.1.chr10.g00568.t01#1    mNeoVis.evergladensis.2.1.chr14.g00343.t01#2    mNeoVis.evergladensis.2.1.chr10.g00176.t01#2    mNeoVis.evergladensis.2.1.chr5.g01032.t01#2    mNeoVis.vulgivaga.2.1.chr14.g00344.t01#1    mNeoVis.evergladensis.1.1.chr3.g00336.t01#3    mNeoVis.vulgivaga.1.1.chr10.g00167.t01#4      5259   125255639:   mNeoVis.2.1.1.chr9_hap1.g00181.t01#5    mNeoVis.2.1.1.chr9_hap1.g00589.t01#5    mNeoVis.2.1.1.chr3_hap1.g00333.t01#5    mNeoVis.2.1.1.chr9_hap1.g00587.t01#5    mNeoVis.2.1.1.chr14_hap1.g00343.t01#5    mNeoVis.2.1.1.chr3_hap1.g00329.t01#5    mNeoVis.vulgivaga.1.1.chr5.g00995.t01#4    mNeoVis.2.1.1.chr9_hap1.g00588.t01#5    mNeoVis.evergladensis.1.1.chr5.g01023.t01#3      5260   125255640:   mNeoVis.evergladensis.1.1.chr10.g00585.t01#3    mNeoVis.3.1.1.chr8.g01303.t01#6    mNeoVis.2.1.1.chr9_hap1.g00775.t01#5    mNeoVis.vulgivaga.2.1.chr4.g00743.t01#1    mNeoVis.evergladensis.2.1.chr3.g00341.t01#2    mNeoVis.vulgivaga.1.1.chr10.g00750.t01#4   mNeoVis.vulgivaga.1.1.chr3.g00331.t01#4     5261   125255641:   mNeoVis.evergladensis.1.1.chr10.g00018.t01#3    mNeoVis.vulgivaga.2.1.chr10.g00020.t01#1    mNeoVis.3.1.1.chr10.g00018.t01#6    mNeoVis.2.1.1.chr9_hap1.g00019.t01#5    mNeoVis.evergladensis.2.1.chr10.g00021.t01#2   mNeoVis.vulgivaga.1.1.chr10.g00019.t01#4       5262   125255642:   mNeoVis.2.1.1.chr6_hap1.g00271.t01#5    mNeoVis.evergladensis.2.1.chr4.g00277.t01#2    mNeoVis.evergladensis.1.1.chr4.g00276.t01#3   mNeoVis.vulgivaga.1.1.chr4.g00270.t01#4         5263   125255643:   mNeoVis.vulgivaga.2.1.chr4.g00277.t01#1   mNeoVis.evergladensis.1.1.chr10.g00765.t01#3           5264   125255644:   mNeoVis.evergladensis.1.1.chr3.g01176.t01#3   mNeoVis.3.1.1.chr3.g01150.t01#6             5265   125255645:   mNeoVis.evergladensis.2.1.chr10.g00580.t01#2   mNeoVis.3.1.1.chr5.g01011.t01#6             5266   125255646:   mNeoVis.3.1.1.chr10.g00557.t01#6   mNeoVis.vulgivaga.1.1.chr10.g00564.t01#4           5267   125255647:   mNeoVis.evergladensis.1.1.chr3.g00340.t01#3   mNeoVis.vulgivaga.1.1.chr3.g00335.t01#4             5268   125255648:  mNeoVis.vulgivaga.2.1.chr3.g01180.t01#1               5269   125255649:  mNeoVis.vulgivaga.2.1.chr10.g00569.t01#1               5270   125255650:  mNeoVis.vulgivaga.2.1.chr3.g00332.t01#1               5271   125255651:   mNeoVis.vulgivaga.1.1.chr14.g00341.t01#4    mNeoVis.evergladensis.1.1.chr14.g00349.t01#3    mNeoVis.evergladensis.2.1.chr14.g00345.t01#2    mNeoVis.2.1.1.chr14_hap1.g00345.t01#5    mNeoVis.vulgivaga.2.1.chr14.g00346.t01#1   mNeoVis.3.1.1.chr14.g00343.t01#6       5272   125255652:   mNeoVis.evergladensis.1.1.chr6.g00702.t01#3    mNeoVis.vulgivaga.2.1.chr6.g00698.t01#1    mNeoVis.evergladensis.2.1.chr6.g00703.t01#2    mNeoVis.vulgivaga.1.1.chr6.g00699.t01#4    mNeoVis.3.1.1.chr6.g00703.t01#6   mNeoVis.2.1.1.chr4_hap1.g00699.t01#5       5273   125255653:   mNeoVis.vulgivaga.2.1.chr6.g00699.t01#1   mNeoVis.3.1.1.chr6.g00704.t01#6             5274   125255654:   mNeoVis.evergladensis.1.1.chr14.g00350.t01#3    mNeoVis.3.1.1.chr14.g00344.t01#6    mNeoVis.vulgivaga.1.1.chr14.g00342.t01#4    mNeoVis.2.1.1.chr14_hap1.g00346.t01#5    mNeoVis.vulgivaga.2.1.chr14.g00347.t01#1   mNeoVis.evergladensis.2.1.chr14.g00346.t01#2       5275   125255655:   mNeoVis.evergladensis.2.1.chr6.g00705.t01#2    mNeoVis.vulgivaga.2.1.chr6.g00700.t01#1    mNeoVis.3.1.1.chr6.g00705.t01#6    mNeoVis.evergladensis.1.1.chr6.g00704.t01#3    mNeoVis.2.1.1.chr4_hap1.g00701.t01#5   mNeoVis.vulgivaga.1.1.chr6.g00701.t01#4       5276   125255656:   mNeoVis.evergladensis.1.1.chr14.g00351.t01#3    mNeoVis.3.1.1.chr14.g00345.t01#6    mNeoVis.2.1.1.chr14_hap1.g00347.t01#5    mNeoVis.vulgivaga.1.1.chr14.g00343.t01#4    mNeoVis.vulgivaga.2.1.chr14.g00348.t01#1   mNeoVis.evergladensis.2.1.chr14.g00347.t01#2       5277   125255657:   mNeoVis.vulgivaga.1.1.chr7.g00513.t01#4    mNeoVis.evergladensis.2.1.chr7.g00506.t01#2    mNeoVis.2.1.1.chr7_hap1.g00505.t01#5    mNeoVis.evergladensis.1.1.chr7.g00515.t01#3    mNeoVis.vulgivaga.2.1.chr7.g00509.t01#1   mNeoVis.3.1.1.chr7.g00490.t01#6       5278   125255658:   mNeoVis.3.1.1.chr14.g00346.t01#6    mNeoVis.evergladensis.1.1.chr14.g00352.t01#3    mNeoVis.2.1.1.chr14_hap1.g00348.t01#5    mNeoVis.vulgivaga.1.1.chr14.g00344.t01#4    mNeoVis.vulgivaga.2.1.chr14.g00349.t01#1   mNeoVis.evergladensis.2.1.chr14.g00348.t01#2       5279   125255659:   mNeoVis.evergladensis.1.1.chr14.g00353.t01#3    mNeoVis.vulgivaga.1.1.chr14.g00345.t01#4    mNeoVis.2.1.1.chr14_hap1.g00349.t01#5    mNeoVis.3.1.1.chr14.g00347.t01#6    mNeoVis.vulgivaga.2.1.chr14.g00350.t01#1   mNeoVis.evergladensis.2.1.chr14.g00349.t01#2       5280   125255660:   mNeoVis.vulgivaga.1.1.chr6.g00702.t01#4    mNeoVis.2.1.1.chr4_hap1.g00702.t01#5    mNeoVis.evergladensis.2.1.chr6.g00706.t01#2    mNeoVis.vulgivaga.2.1.chr6.g00701.t01#1   mNeoVis.evergladensis.1.1.chr6.g00705.t01#3       5281   125255661:   mNeoVis.evergladensis.1.1.chr6.g00706.t01#3    mNeoVis.2.1.1.chr4_hap1.g00703.t01#5    mNeoVis.3.1.1.chr6.g00706.t01#6    mNeoVis.evergladensis.2.1.chr6.g00707.t01#2    mNeoVis.vulgivaga.1.1.chr6.g00703.t01#4   mNeoVis.vulgivaga.2.1.chr6.g00702.t01#1       5282   125255662:   mNeoVis.evergladensis.1.1.chr14.g00354.t01#3    mNeoVis.vulgivaga.1.1.chr14.g00346.t01#4    mNeoVis.2.1.1.chr14_hap1.g00350.t01#5    mNeoVis.3.1.1.chr14.g00348.t01#6    mNeoVis.evergladensis.2.1.chr14.g00350.t01#2   mNeoVis.vulgivaga.2.1.chr14.g00351.t01#1       5283   125255663:   mNeoVis.vulgivaga.2.1.chr6.g00703.t01#1    mNeoVis.evergladensis.1.1.chr6.g00707.t01#3    mNeoVis.3.1.1.chr6.g00707.t01#6    mNeoVis.vulgivaga.1.1.chr6.g00704.t01#4    mNeoVis.2.1.1.chr4_hap1.g00704.t01#5   mNeoVis.evergladensis.2.1.chr6.g00708.t01#2       5284   125255664:   mNeoVis.2.1.1.chr15_hap1.g00729.t01#5    mNeoVis.vulgivaga.1.1.chr15.g00554.t01#4    mNeoVis.evergladensis.2.1.chr15.g00551.t01#2    mNeoVis.3.1.1.chr15.g00506.t01#6    mNeoVis.evergladensis.1.1.chr15.g00062.t01#3   mNeoVis.vulgivaga.2.1.chr15.g00563.t01#1       5285   125255665:   mNeoVis.3.1.1.chr6.g00708.t01#6    mNeoVis.evergladensis.2.1.chr6.g00709.t01#2    mNeoVis.evergladensis.1.1.chr6.g00708.t01#3    mNeoVis.2.1.1.chr4_hap1.g00705.t01#5    mNeoVis.vulgivaga.2.1.chr6.g00704.t01#1   mNeoVis.vulgivaga.1.1.chr6.g00705.t01#4       5286   125255666:   mNeoVis.evergladensis.1.1.chr6.g00709.t01#3    mNeoVis.vulgivaga.1.1.chr6.g00706.t01#4    mNeoVis.vulgivaga.2.1.chr6.g00705.t01#1    mNeoVis.evergladensis.2.1.chr6.g00710.t01#2    mNeoVis.2.1.1.chr4_hap1.g00706.t01#5   mNeoVis.3.1.1.chr6.g00709.t01#6       5287   125255667:   mNeoVis.vulgivaga.2.1.chr14.g00352.t01#1    mNeoVis.evergladensis.2.1.chr14.g00351.t01#2    mNeoVis.vulgivaga.1.1.chr14.g00347.t01#4    mNeoVis.evergladensis.1.1.chr14.g00355.t01#3    mNeoVis.2.1.1.chr14_hap1.g00351.t01#5   mNeoVis.3.1.1.chr14.g00349.t01#6       5288   125255668:   mNeoVis.evergladensis.2.1.chr14.g00352.t01#2    mNeoVis.3.1.1.chr14.g00350.t01#6    mNeoVis.vulgivaga.2.1.chr14.g00353.t01#1    mNeoVis.vulgivaga.1.1.chr14.g00348.t01#4    mNeoVis.evergladensis.1.1.chr14.g00356.t01#3   mNeoVis.2.1.1.chr14_hap1.g00352.t01#5       5289   125255669:   mNeoVis.evergladensis.1.1.chr14.g00357.t01#3    mNeoVis.evergladensis.2.1.chr14.g00353.t01#2    mNeoVis.2.1.1.chr14_hap1.g00353.t01#5    mNeoVis.vulgivaga.1.1.chr14.g00349.t01#4    mNeoVis.vulgivaga.2.1.chr14.g00354.t01#1   mNeoVis.3.1.1.chr14.g00351.t01#6       5290   125255670:   mNeoVis.vulgivaga.2.1.chr6.g00706.t01#1    mNeoVis.vulgivaga.1.1.chr6.g00707.t01#4    mNeoVis.2.1.1.chr4_hap1.g00707.t01#5    mNeoVis.evergladensis.1.1.chr6.g00710.t01#3   mNeoVis.evergladensis.2.1.chr6.g00711.t01#2       5291   125255671:   mNeoVis.evergladensis.1.1.chr14.g00358.t01#3    mNeoVis.vulgivaga.2.1.chr14.g00355.t01#1    mNeoVis.2.1.1.chr14_hap1.g00354.t01#5    mNeoVis.vulgivaga.1.1.chr14.g00350.t01#4    mNeoVis.evergladensis.2.1.chr14.g00354.t01#2   mNeoVis.3.1.1.chr5.g01632.t01#6       5292   125255672:   mNeoVis.2.1.1.chr4_hap1.g00708.t01#5    mNeoVis.vulgivaga.2.1.chr6.g00707.t01#1    mNeoVis.3.1.1.chr6.g00711.t01#6    mNeoVis.evergladensis.1.1.chr6.g00711.t01#3    mNeoVis.vulgivaga.1.1.chr6.g00708.t01#4   mNeoVis.evergladensis.2.1.chr6.g00712.t01#2       5293   125255673:   mNeoVis.3.1.1.chr6.g00713.t01#6   mNeoVis.evergladensis.1.1.chr6.g00713.t01#3           5294   125255674:   mNeoVis.vulgivaga.2.1.chr14.g00356.t01#1    mNeoVis.3.1.1.chr14.g00354.t01#6    mNeoVis.evergladensis.2.1.chr14.g00355.t01#2    mNeoVis.evergladensis.1.1.chr14.g00359.t01#3    mNeoVis.vulgivaga.1.1.chr14.g00351.t01#4   mNeoVis.2.1.1.chr14_hap1.g00355.t01#5       5295   125255675:   mNeoVis.vulgivaga.1.1.chr6.g00709.t01#4    mNeoVis.2.1.1.chr4_hap1.g00709.t01#5    mNeoVis.vulgivaga.2.1.chr6.g00708.t01#1    mNeoVis.evergladensis.2.1.chr6.g00714.t01#2    mNeoVis.3.1.1.chr6.g00712.t01#6   mNeoVis.evergladensis.1.1.chr6.g00712.t01#3       5296   125255676:   mNeoVis.3.1.1.chr5.g01668.t01#6    mNeoVis.evergladensis.2.1.chr5.g01684.t01#2    mNeoVis.vulgivaga.2.1.chr5.g01685.t01#1    mNeoVis.2.1.1.chr2_hap1.g01744.t01#5    mNeoVis.evergladensis.1.1.chr5.g01690.t01#3   mNeoVis.vulgivaga.1.1.chr5.g01664.t01#4       5297   125255677:   mNeoVis.evergladensis.2.1.chr6.g00715.t01#2    mNeoVis.2.1.1.chr4_hap1.g00710.t01#5    mNeoVis.vulgivaga.2.1.chr6.g00709.t01#1    mNeoVis.vulgivaga.1.1.chr6.g00710.t01#4   mNeoVis.evergladensis.1.1.chr6.g00714.t01#3       5298   125255678:   mNeoVis.3.1.1.chr14.g00355.t01#6    mNeoVis.evergladensis.1.1.chr14.g00360.t01#3    mNeoVis.vulgivaga.1.1.chr14.g00352.t01#4    mNeoVis.evergladensis.2.1.chr14.g00356.t01#2    mNeoVis.2.1.1.chr14_hap1.g00356.t01#5   mNeoVis.vulgivaga.2.1.chr14.g00357.t01#1       5299   125255679:   mNeoVis.vulgivaga.2.1.chr14.g00358.t01#1    mNeoVis.evergladensis.2.1.chr14.g00357.t01#2    mNeoVis.vulgivaga.1.1.chr14.g00353.t01#4    mNeoVis.3.1.1.chr14.g00356.t01#6    mNeoVis.evergladensis.1.1.chr14.g00361.t01#3   mNeoVis.2.1.1.chr14_hap1.g00357.t01#5       5300   125255680:   mNeoVis.vulgivaga.2.1.chr6.g00710.t01#1   mNeoVis.3.1.1.chr6.g00715.t01#6             5301   125255681:   mNeoVis.evergladensis.1.1.chr14.g00362.t01#3    mNeoVis.vulgivaga.2.1.chr14.g00359.t01#1    mNeoVis.2.1.1.chr14_hap1.g00358.t01#5    mNeoVis.vulgivaga.1.1.chr14.g00354.t01#4    mNeoVis.3.1.1.chr14.g00357.t01#6   mNeoVis.evergladensis.2.1.chr14.g00358.t01#2       5302   125255682:   mNeoVis.vulgivaga.1.1.chr6.g00711.t01#4    mNeoVis.evergladensis.1.1.chr6.g00715.t01#3    mNeoVis.2.1.1.chr4_hap1.g00711.t01#5    mNeoVis.3.1.1.chr6.g00716.t01#6    mNeoVis.evergladensis.2.1.chr6.g00716.t01#2   mNeoVis.vulgivaga.2.1.chr6.g00711.t01#1       5303   125255683:   mNeoVis.vulgivaga.2.1.chr14.g00360.t01#1    mNeoVis.evergladensis.2.1.chr14.g00359.t01#2    mNeoVis.2.1.1.chr14_hap1.g00359.t01#5    mNeoVis.evergladensis.1.1.chr14.g00363.t01#3    mNeoVis.vulgivaga.1.1.chr14.g00355.t01#4   mNeoVis.3.1.1.chr14.g00358.t01#6       5304   125255684:   mNeoVis.vulgivaga.2.1.chr6.g00712.t01#1    mNeoVis.vulgivaga.1.1.chr6.g00712.t01#4   mNeoVis.3.1.1.chr6.g00717.t01#6           5305   125255685:   mNeoVis.vulgivaga.2.1.chr14.g00361.t01#1    mNeoVis.evergladensis.1.1.chr14.g00364.t01#3    mNeoVis.evergladensis.2.1.chr14.g00360.t01#2    mNeoVis.2.1.1.chr14_hap1.g00360.t01#5   mNeoVis.vulgivaga.1.1.chr14.g00356.t01#4       5306   125255686:   mNeoVis.vulgivaga.2.1.chr14.g00362.t01#1    mNeoVis.2.1.1.chr14_hap1.g00361.t01#5    mNeoVis.evergladensis.1.1.chr14.g00365.t01#3    mNeoVis.vulgivaga.1.1.chr14.g00357.t01#4    mNeoVis.3.1.1.chr14.g00360.t01#6   mNeoVis.evergladensis.2.1.chr14.g00361.t01#2       5307   125255687:   mNeoVis.2.1.1.chr4_hap1.g00713.t01#5    mNeoVis.evergladensis.1.1.chr6.g00721.t01#3    mNeoVis.vulgivaga.1.1.chr6.g00715.t01#4    mNeoVis.evergladensis.2.1.chr6.g00719.t01#2    mNeoVis.3.1.1.chr6.g00719.t01#6   mNeoVis.vulgivaga.2.1.chr6.g00713.t01#1       5308   125255688:   mNeoVis.evergladensis.2.1.chr6.g00720.t01#2    mNeoVis.2.1.1.chr4_hap1.g00714.t01#5    mNeoVis.vulgivaga.1.1.chr6.g00716.t01#4    mNeoVis.vulgivaga.2.1.chr6.g00715.t01#1    mNeoVis.vulgivaga.2.1.chr6.g00714.t01#1   mNeoVis.evergladensis.1.1.chr6.g00722.t01#3       5309   125255689:   mNeoVis.evergladensis.2.1.chr14.g00363.t01#2    mNeoVis.evergladensis.1.1.chr14.g00367.t01#3    mNeoVis.vulgivaga.2.1.chr14.g00363.t01#1    mNeoVis.2.1.1.chr14_hap1.g00363.t01#5    mNeoVis.vulgivaga.1.1.chr14.g00359.t01#4   mNeoVis.3.1.1.chr14.g00362.t01#6       5310   125255690:   mNeoVis.2.1.1.chr4_hap1.g00715.t01#5    mNeoVis.evergladensis.2.1.chr6.g00721.t01#2    mNeoVis.vulgivaga.2.1.chr6.g00716.t01#1    mNeoVis.3.1.1.chr6.g00721.t01#6    mNeoVis.vulgivaga.1.1.chr6.g00717.t01#4   mNeoVis.evergladensis.1.1.chr6.g00723.t01#3       5311   125255691:   mNeoVis.vulgivaga.2.1.chr14.g00364.t01#1    mNeoVis.3.1.1.chr14.g00361.t01#6    mNeoVis.evergladensis.1.1.chr14.g00366.t01#3    mNeoVis.2.1.1.chr14_hap1.g00362.t01#5    mNeoVis.evergladensis.2.1.chr14.g00362.t01#2   mNeoVis.vulgivaga.1.1.chr14.g00358.t01#4       5312   125255692:   mNeoVis.vulgivaga.1.1.chr14.g00360.t01#4    mNeoVis.3.1.1.chr11.g01040.t01#6    mNeoVis.evergladensis.2.1.chr14.g00364.t01#2    mNeoVis.vulgivaga.2.1.chr11.g01094.t01#1    mNeoVis.vulgivaga.1.1.chr11.g01065.t01#4    mNeoVis.3.1.1.chr14.g00363.t01#6    mNeoVis.evergladensis.2.1.chr11.g01066.t01#2    mNeoVis.2.1.1.chr14_hap1.g00364.t01#5    mNeoVis.evergladensis.1.1.chr14.g00368.t01#3      5313   125255693:   mNeoVis.vulgivaga.1.1.chr6.g00745.t01#4    mNeoVis.evergladensis.2.1.chr6.g00750.t01#2    mNeoVis.evergladensis.1.1.chr6.g00754.t01#3    mNeoVis.2.1.1.chr4_hap1.g00741.t01#5   mNeoVis.vulgivaga.2.1.chr6.g00746.t01#1       5314   125255694:   mNeoVis.evergladensis.1.1.chr6.g00724.t01#3    mNeoVis.evergladensis.2.1.chr6.g00722.t01#2    mNeoVis.vulgivaga.1.1.chr6.g00719.t01#4    mNeoVis.2.1.1.chr4_hap1.g00716.t01#5    mNeoVis.vulgivaga.2.1.chr6.g00717.t01#1   mNeoVis.3.1.1.chr6.g00722.t01#6       5315   125255695:   mNeoVis.2.1.1.chr14_hap1.g00365.t01#5    mNeoVis.evergladensis.2.1.chr14.g00365.t01#2    mNeoVis.evergladensis.1.1.chr14.g00369.t01#3    mNeoVis.vulgivaga.1.1.chr14.g00361.t01#4   mNeoVis.vulgivaga.2.1.chr14.g00366.t01#1       5316   125255696:   mNeoVis.evergladensis.2.1.chr6.g00723.t01#2    mNeoVis.evergladensis.1.1.chr6.g00725.t01#3    mNeoVis.vulgivaga.1.1.chr6.g00720.t01#4    mNeoVis.vulgivaga.2.1.chr6.g00718.t01#1   mNeoVis.2.1.1.chr4_hap1.g00717.t01#5       5317   125255697:   mNeoVis.evergladensis.1.1.chr14.g00370.t01#3    mNeoVis.2.1.1.chr14_hap1.g00366.t01#5    mNeoVis.3.1.1.chr14.g00365.t01#6    mNeoVis.vulgivaga.2.1.chr14.g00367.t01#1    mNeoVis.evergladensis.2.1.chr14.g00366.t01#2   mNeoVis.vulgivaga.1.1.chr14.g00362.t01#4       5318   125255698:   mNeoVis.2.1.1.chr4_hap1.g00718.t01#5    mNeoVis.evergladensis.2.1.chr6.g00724.t01#2    mNeoVis.evergladensis.1.1.chr6.g00726.t01#3    mNeoVis.vulgivaga.2.1.chr6.g00719.t01#1    mNeoVis.3.1.1.chr6.g00724.t01#6   mNeoVis.vulgivaga.1.1.chr6.g00721.t01#4       5319   125255699:   mNeoVis.vulgivaga.2.1.chr6.g00720.t01#1    mNeoVis.3.1.1.chr6.g00725.t01#6    mNeoVis.evergladensis.1.1.chr6.g00727.t01#3    mNeoVis.2.1.1.chr4_hap1.g00719.t01#5    mNeoVis.evergladensis.2.1.chr6.g00725.t01#2   mNeoVis.vulgivaga.1.1.chr6.g00722.t01#4       5320   125255700:   mNeoVis.3.1.1.chr14.g00366.t01#6    mNeoVis.vulgivaga.1.1.chr14.g00363.t01#4    mNeoVis.evergladensis.1.1.chr14.g00371.t01#3    mNeoVis.evergladensis.2.1.chr14.g00367.t01#2    mNeoVis.vulgivaga.2.1.chr14.g00368.t01#1   mNeoVis.2.1.1.chr14_hap1.g00367.t01#5       5321   125255701:   mNeoVis.evergladensis.2.1.chr6.g00726.t01#2    mNeoVis.vulgivaga.2.1.chr6.g00721.t01#1    mNeoVis.2.1.1.chr4_hap1.g00720.t01#5    mNeoVis.evergladensis.1.1.chr6.g00728.t01#3    mNeoVis.vulgivaga.1.1.chr6.g00723.t01#4   mNeoVis.3.1.1.chr6.g00726.t01#6       5322   125255702:   mNeoVis.evergladensis.2.1.chr6.g00727.t01#2    mNeoVis.evergladensis.1.1.chr6.g00729.t01#3    mNeoVis.vulgivaga.2.1.chr6.g00722.t01#1    mNeoVis.vulgivaga.1.1.chr6.g00724.t01#4    mNeoVis.3.1.1.chr6.g00727.t01#6   mNeoVis.2.1.1.chr4_hap1.g00721.t01#5       5323   125255703:   mNeoVis.evergladensis.2.1.chr14.g00368.t01#2    mNeoVis.vulgivaga.2.1.chr14.g00369.t01#1    mNeoVis.vulgivaga.1.1.chr14.g00364.t01#4    mNeoVis.evergladensis.1.1.chr14.g00372.t01#3    mNeoVis.2.1.1.chr14_hap1.g00368.t01#5   mNeoVis.3.1.1.chr14.g00367.t01#6       5324   125255704:   mNeoVis.evergladensis.1.1.chr6.g00730.t01#3    mNeoVis.vulgivaga.1.1.chr6.g00725.t01#4    mNeoVis.3.1.1.chr6.g00728.t01#6   mNeoVis.vulgivaga.2.1.chr6.g00723.t01#1         5325   125255705:   mNeoVis.vulgivaga.1.1.chr6.g00726.t01#4    mNeoVis.2.1.1.chr4_hap1.g00722.t01#5    mNeoVis.3.1.1.chr6.g00729.t01#6    mNeoVis.evergladensis.2.1.chr6.g00728.t01#2    mNeoVis.evergladensis.1.1.chr6.g00731.t01#3   mNeoVis.vulgivaga.2.1.chr6.g00724.t01#1       5326   125255706:   mNeoVis.evergladensis.1.1.chr14.g00373.t01#3    mNeoVis.2.1.1.chr14_hap1.g00369.t01#5    mNeoVis.3.1.1.chr14.g00368.t01#6    mNeoVis.vulgivaga.2.1.chr14.g00370.t01#1    mNeoVis.vulgivaga.1.1.chr14.g00365.t01#4   mNeoVis.evergladensis.2.1.chr14.g00369.t01#2       5327   125255707:   mNeoVis.3.1.1.chr6.g00730.t01#6    mNeoVis.2.1.1.chr4_hap1.g00723.t01#5    mNeoVis.evergladensis.2.1.chr6.g00729.t01#2    mNeoVis.evergladensis.1.1.chr6.g00732.t01#3    mNeoVis.vulgivaga.1.1.chr6.g00727.t01#4   mNeoVis.vulgivaga.2.1.chr6.g00725.t01#1       5328   125255708:   mNeoVis.vulgivaga.2.1.chr6.g00726.t01#1   mNeoVis.3.1.1.chr6.g00731.t01#6             5329   125255709:   mNeoVis.vulgivaga.1.1.chr14.g00366.t01#4    mNeoVis.3.1.1.chr14.g00369.t01#6    mNeoVis.evergladensis.2.1.chr14.g00370.t01#2    mNeoVis.vulgivaga.2.1.chr14.g00371.t01#1    mNeoVis.evergladensis.1.1.chr14.g00374.t01#3   mNeoVis.2.1.1.chr14_hap1.g00370.t01#5       5330   125255710:   mNeoVis.evergladensis.2.1.chr15.g00552.t01#2    mNeoVis.3.1.1.chr15.g00507.t01#6    mNeoVis.vulgivaga.1.1.chr15.g00555.t01#4    mNeoVis.evergladensis.1.1.chr15.g00061.t01#3    mNeoVis.vulgivaga.2.1.chr15.g00564.t01#1   mNeoVis.2.1.1.chr15_hap1.g00730.t01#5       5331   125255711:   mNeoVis.vulgivaga.2.1.chr6.g00727.t01#1   mNeoVis.evergladensis.1.1.chr6.g00733.t01#3           5332   125255712:   mNeoVis.vulgivaga.2.1.chr6.g00728.t01#1    mNeoVis.evergladensis.2.1.chr6.g00730.t01#2    mNeoVis.3.1.1.chr6.g00732.t01#6    mNeoVis.evergladensis.1.1.chr6.g00734.t01#3    mNeoVis.vulgivaga.1.1.chr6.g00728.t01#4   mNeoVis.2.1.1.chr4_hap1.g00724.t01#5       5333   125255713:   mNeoVis.vulgivaga.1.1.chr2.g01674.t01#4    mNeoVis.evergladensis.2.1.chr2.g01663.t01#2    mNeoVis.3.1.1.chr6.g00733.t01#6    mNeoVis.evergladensis.1.1.chr2.g01677.t01#3    mNeoVis.vulgivaga.1.1.chr6.g00729.t01#4    mNeoVis.evergladensis.2.1.chr6.g00731.t01#2    mNeoVis.2.1.1.chr4_hap1.g00725.t01#5    mNeoVis.2.1.1.chr5_hap1.g01680.t01#5    mNeoVis.vulgivaga.2.1.chr2.g01680.t01#1      5334   125255714:   mNeoVis.3.1.1.chr6.g00734.t01#6    mNeoVis.vulgivaga.2.1.chr6.g00730.t01#1    mNeoVis.evergladensis.2.1.chr6.g00732.t01#2    mNeoVis.2.1.1.chr4_hap1.g00726.t01#5    mNeoVis.evergladensis.1.1.chr6.g00735.t01#3   mNeoVis.vulgivaga.1.1.chr6.g00730.t01#4       5335   125255715:   mNeoVis.vulgivaga.2.1.chr14.g00372.t01#1    mNeoVis.evergladensis.2.1.chr14.g00371.t01#2    mNeoVis.2.1.1.chr14_hap1.g00371.t01#5    mNeoVis.3.1.1.chr14.g00370.t01#6    mNeoVis.evergladensis.1.1.chr14.g00375.t01#3   mNeoVis.vulgivaga.1.1.chr14.g00367.t01#4       5336   125255716:   mNeoVis.2.1.1.chr14_hap1.g00372.t01#5    mNeoVis.evergladensis.1.1.chr14.g00376.t01#3    mNeoVis.vulgivaga.2.1.chr14.g00373.t01#1    mNeoVis.vulgivaga.1.1.chr14.g00368.t01#4    mNeoVis.3.1.1.chr14.g00371.t01#6   mNeoVis.evergladensis.2.1.chr14.g00372.t01#2       5337   125255717:   mNeoVis.3.1.1.chr6.g00735.t01#6    mNeoVis.evergladensis.1.1.chr6.g00736.t01#3    mNeoVis.evergladensis.2.1.chr6.g00733.t01#2    mNeoVis.vulgivaga.2.1.chr6.g00731.t01#1    mNeoVis.vulgivaga.1.1.chr6.g00731.t01#4   mNeoVis.2.1.1.chr4_hap1.g00727.t01#5       5338   125255718:   mNeoVis.3.1.1.chr6.g00736.t01#6    mNeoVis.2.1.1.chr11_hap1.g00486.t01#5    mNeoVis.evergladensis.2.1.chr6.g00736.t01#2    mNeoVis.evergladensis.2.1.chr6.g00735.t01#2    mNeoVis.2.1.1.chr8_hap1.g00655.t01#5    mNeoVis.evergladensis.1.1.chr6.g00737.t01#3    mNeoVis.vulgivaga.2.1.chr6.g00732.t01#1    mNeoVis.2.1.1.chr4_hap1.g00728.t01#5    mNeoVis.vulgivaga.1.1.chr6.g00732.t01#4      5339   125255719:   mNeoVis.vulgivaga.1.1.chr2.g01355.t01#4    mNeoVis.3.1.1.chr2.g01280.t01#6    mNeoVis.evergladensis.2.1.chr2.g01343.t01#2    mNeoVis.vulgivaga.2.1.chr2.g01353.t01#1   mNeoVis.evergladensis.1.1.chr2.g01352.t01#3       5340   125255720:   mNeoVis.evergladensis.1.1.chr6.g00738.t01#3    mNeoVis.3.1.1.chr6.g00737.t01#6    mNeoVis.vulgivaga.1.1.chr6.g00733.t01#4    mNeoVis.evergladensis.2.1.chr6.g00737.t01#2    mNeoVis.vulgivaga.2.1.chr6.g00733.t01#1   mNeoVis.2.1.1.chr4_hap1.g00729.t01#5       5341   125255721:   mNeoVis.vulgivaga.2.1.chr14.g00374.t01#1    mNeoVis.3.1.1.chr14.g00372.t01#6    mNeoVis.2.1.1.chr14_hap1.g00373.t01#5    mNeoVis.vulgivaga.1.1.chr14.g00369.t01#4    mNeoVis.evergladensis.1.1.chr14.g00377.t01#3   mNeoVis.evergladensis.2.1.chr14.g00373.t01#2       5342   125255722:   mNeoVis.evergladensis.1.1.chr14.g00378.t01#3    mNeoVis.evergladensis.2.1.chr14.g00374.t01#2    mNeoVis.vulgivaga.1.1.chr14.g00370.t01#4    mNeoVis.3.1.1.chr14.g00373.t01#6    mNeoVis.vulgivaga.2.1.chr14.g00375.t01#1   mNeoVis.2.1.1.chr14_hap1.g00374.t01#5       5343   125255723:   mNeoVis.evergladensis.2.1.chr7.g00507.t01#2    mNeoVis.3.1.1.chr7.g00491.t01#6    mNeoVis.vulgivaga.1.1.chr7.g00514.t01#4    mNeoVis.2.1.1.chr7_hap1.g00506.t01#5    mNeoVis.vulgivaga.2.1.chr7.g00510.t01#1   mNeoVis.evergladensis.1.1.chr7.g00516.t01#3       5344   125255724:   mNeoVis.vulgivaga.2.1.chr6.g00734.t01#1    mNeoVis.evergladensis.1.1.chr6.g00739.t01#3    mNeoVis.3.1.1.chr6.g00738.t01#6    mNeoVis.vulgivaga.1.1.chr6.g00734.t01#4    mNeoVis.2.1.1.chr4_hap1.g00730.t01#5   mNeoVis.evergladensis.2.1.chr6.g00738.t01#2       5345   125255725:   mNeoVis.2.1.1.chr14_hap1.g00375.t01#5    mNeoVis.3.1.1.chr14.g00374.t01#6    mNeoVis.vulgivaga.2.1.chr14.g00376.t01#1    mNeoVis.vulgivaga.1.1.chr14.g00371.t01#4    mNeoVis.evergladensis.1.1.chr14.g00379.t01#3   mNeoVis.evergladensis.2.1.chr14.g00375.t01#2       5346   125255726:   mNeoVis.2.1.1.chr4_hap1.g00731.t01#5    mNeoVis.evergladensis.2.1.chr6.g00739.t01#2    mNeoVis.vulgivaga.2.1.chr6.g00735.t01#1    mNeoVis.vulgivaga.1.1.chr6.g00735.t01#4    mNeoVis.evergladensis.1.1.chr6.g00740.t01#3   mNeoVis.3.1.1.chr6.g00739.t01#6       5347   125255727:   mNeoVis.vulgivaga.1.1.chr6.g00736.t01#4    mNeoVis.3.1.1.chr6.g00740.t01#6    mNeoVis.2.1.1.chr4_hap1.g00732.t01#5    mNeoVis.vulgivaga.2.1.chr6.g00736.t01#1    mNeoVis.evergladensis.2.1.chr6.g00740.t01#2   mNeoVis.evergladensis.1.1.chr6.g00741.t01#3       5348   125255728:   mNeoVis.vulgivaga.2.1.chr9.g00856.t01#1    mNeoVis.evergladensis.2.1.chr8.g00869.t01#2    mNeoVis.evergladensis.1.1.chr9.g00863.t01#3    mNeoVis.vulgivaga.1.1.chr9.g00869.t01#4    mNeoVis.3.1.1.chr9.g00827.t01#6   mNeoVis.2.1.1.chr10_hap1.g00899.t01#5       5349   125255729:   mNeoVis.2.1.1.chr14_hap1.g00376.t01#5    mNeoVis.vulgivaga.1.1.chr14.g00372.t01#4    mNeoVis.evergladensis.2.1.chr14.g00376.t01#2    mNeoVis.evergladensis.1.1.chr14.g00380.t01#3   mNeoVis.vulgivaga.2.1.chr14.g00377.t01#1       5350   125255730:   mNeoVis.evergladensis.1.1.chr6.g00743.t01#3    mNeoVis.evergladensis.2.1.chr6.g00742.t01#2    mNeoVis.vulgivaga.2.1.chr6.g00738.t01#1    mNeoVis.3.1.1.chr6.g00742.t01#6    mNeoVis.vulgivaga.1.1.chr6.g00738.t01#4   mNeoVis.2.1.1.chr4_hap1.g00733.t01#5       5351   125255731:   mNeoVis.3.1.1.chr15.g00508.t01#6    mNeoVis.vulgivaga.2.1.chr15.g00565.t01#1    mNeoVis.2.1.1.chr15_hap1.g00731.t01#5    mNeoVis.evergladensis.1.1.chr15.g00060.t01#3    mNeoVis.vulgivaga.1.1.chr15.g00556.t01#4   mNeoVis.evergladensis.2.1.chr15.g00553.t01#2       5352   125255732:   mNeoVis.3.1.1.chr14.g00376.t01#6    mNeoVis.vulgivaga.1.1.chr14.g00373.t01#4    mNeoVis.vulgivaga.2.1.chr14.g00378.t01#1    mNeoVis.evergladensis.2.1.chr14.g00377.t01#2    mNeoVis.evergladensis.1.1.chr14.g00381.t01#3   mNeoVis.2.1.1.chr14_hap1.g00377.t01#5       5353   125255733:   mNeoVis.2.1.1.chr14_hap1.g00378.t01#5    mNeoVis.vulgivaga.1.1.chr14.g00374.t01#4    mNeoVis.evergladensis.2.1.chr14.g00378.t01#2    mNeoVis.vulgivaga.2.1.chr14.g00379.t01#1    mNeoVis.3.1.1.chr14.g00377.t01#6   mNeoVis.evergladensis.1.1.chr14.g00382.t01#3       5354   125255734:   mNeoVis.evergladensis.2.1.chr7.g00508.t01#2    mNeoVis.vulgivaga.1.1.chr7.g00515.t01#4    mNeoVis.3.1.1.chr7.g00492.t01#6    mNeoVis.2.1.1.chr7_hap1.g00507.t01#5    mNeoVis.evergladensis.1.1.chr7.g00517.t01#3   mNeoVis.vulgivaga.2.1.chr7.g00511.t01#1       5355   125255735:   mNeoVis.evergladensis.1.1.chr6.g00744.t01#3   mNeoVis.vulgivaga.2.1.chr6.g00739.t01#1             5356   125255736:   mNeoVis.vulgivaga.2.1.chr14.g00380.t01#1    mNeoVis.2.1.1.chr14_hap1.g00379.t01#5    mNeoVis.3.1.1.chr14.g00378.t01#6    mNeoVis.vulgivaga.1.1.chr14.g00375.t01#4    mNeoVis.evergladensis.1.1.chr14.g00383.t01#3   mNeoVis.evergladensis.2.1.chr14.g00379.t01#2       5357   125255737:   mNeoVis.vulgivaga.2.1.chr6.g00741.t01#1    mNeoVis.3.1.1.chr6.g00744.t01#6    mNeoVis.vulgivaga.1.1.chr6.g00740.t01#4    mNeoVis.evergladensis.2.1.chr6.g00744.t01#2    mNeoVis.2.1.1.chr4_hap1.g00735.t01#5    mNeoVis.evergladensis.1.1.chr6.g00746.t01#3   mNeoVis.evergladensis.1.1.chr6.g00747.t01#3     5358   125255738:   mNeoVis.3.1.1.chr14.g00379.t01#6    mNeoVis.vulgivaga.1.1.chr14.g00376.t01#4    mNeoVis.vulgivaga.2.1.chr14.g00381.t01#1    mNeoVis.evergladensis.1.1.chr14.g00384.t01#3    mNeoVis.2.1.1.chr14_hap1.g00380.t01#5   mNeoVis.evergladensis.2.1.chr14.g00380.t01#2       5359   125255739:   mNeoVis.2.1.1.chr14_hap1.g00301.t01#5    mNeoVis.evergladensis.2.1.chr14.g00302.t01#2    mNeoVis.vulgivaga.2.1.chr14.g00302.t01#1   mNeoVis.evergladensis.1.1.chr14.g00307.t01#3       5360   125255740:   mNeoVis.3.1.1.chr14.g00302.t01#6    mNeoVis.evergladensis.1.1.chr14.g00308.t01#3    mNeoVis.evergladensis.2.1.chr14.g00303.t01#2    mNeoVis.vulgivaga.1.1.chr14.g00298.t01#4    mNeoVis.vulgivaga.2.1.chr14.g00303.t01#1   mNeoVis.2.1.1.chr14_hap1.g00302.t01#5       5361   125255741:   mNeoVis.evergladensis.2.1.chr13.g00242.t01#2    mNeoVis.vulgivaga.2.1.chr13.g00239.t01#1    mNeoVis.evergladensis.1.1.chr13.g00241.t01#3    mNeoVis.vulgivaga.1.1.chr13.g00236.t01#4    mNeoVis.3.1.1.chr13.g00194.t01#6   mNeoVis.2.1.1.chr13_hap1.g00244.t01#5       5362   125255742:   mNeoVis.3.1.1.chr7.g00493.t01#6    mNeoVis.evergladensis.1.1.chr7.g00518.t01#3    mNeoVis.2.1.1.chr7_hap1.g00508.t01#5    mNeoVis.evergladensis.2.1.chr7.g00509.t01#2    mNeoVis.vulgivaga.1.1.chr7.g00516.t01#4   mNeoVis.vulgivaga.2.1.chr7.g00512.t01#1       5363   125255743:   mNeoVis.2.1.1.chr14_hap1.g00303.t01#5    mNeoVis.3.1.1.chr14.g00303.t01#6    mNeoVis.vulgivaga.2.1.chr14.g00304.t01#1    mNeoVis.evergladensis.2.1.chr14.g00304.t01#2   mNeoVis.evergladensis.1.1.chr14.g00309.t01#3       5364   125255744:   mNeoVis.evergladensis.2.1.chr6.g00745.t01#2    mNeoVis.vulgivaga.1.1.chr6.g00741.t01#4    mNeoVis.evergladensis.1.1.chr6.g00749.t01#3    mNeoVis.vulgivaga.2.1.chr6.g00742.t01#1    mNeoVis.2.1.1.chr4_hap1.g00736.t01#5   mNeoVis.3.1.1.chr6.g00745.t01#6       5365   125255745:   mNeoVis.vulgivaga.1.1.chr14.g00300.t01#4    mNeoVis.3.1.1.chr14.g00304.t01#6    mNeoVis.evergladensis.1.1.chr14.g00310.t01#3    mNeoVis.evergladensis.2.1.chr14.g00305.t01#2    mNeoVis.2.1.1.chr14_hap1.g00304.t01#5   mNeoVis.vulgivaga.2.1.chr14.g00305.t01#1       5366   125255746:   mNeoVis.evergladensis.1.1.chr15.g00059.t01#3    mNeoVis.3.1.1.chr15.g00509.t01#6    mNeoVis.2.1.1.chr15_hap1.g00732.t01#5    mNeoVis.vulgivaga.2.1.chr15.g00566.t01#1    mNeoVis.evergladensis.2.1.chr15.g00554.t01#2   mNeoVis.vulgivaga.1.1.chr15.g00557.t01#4       5367   125255747:   mNeoVis.vulgivaga.1.1.chr6.g00742.t01#4    mNeoVis.evergladensis.1.1.chr6.g00750.t01#3    mNeoVis.vulgivaga.2.1.chr6.g00743.t01#1    mNeoVis.2.1.1.chr4_hap1.g00737.t01#5    mNeoVis.evergladensis.2.1.chr6.g00746.t01#2   mNeoVis.3.1.1.chr6.g00746.t01#6       5368   125255748:   mNeoVis.evergladensis.2.1.chr6.g00748.t01#2    mNeoVis.2.1.1.chr4_hap1.g00739.t01#5    mNeoVis.vulgivaga.2.1.chr6.g00744.t01#1    mNeoVis.3.1.1.chr6.g00747.t01#6    mNeoVis.evergladensis.1.1.chr6.g00751.t01#3   mNeoVis.vulgivaga.1.1.chr6.g00743.t01#4       5369   125255749:   mNeoVis.vulgivaga.2.1.chr6.g00745.t01#1    mNeoVis.2.1.1.chr4_hap1.g00740.t01#5    mNeoVis.vulgivaga.1.1.chr6.g00744.t01#4    mNeoVis.3.1.1.chr6.g00749.t01#6    mNeoVis.evergladensis.2.1.chr6.g00749.t01#2   mNeoVis.evergladensis.1.1.chr6.g00753.t01#3       5370   125255750:   mNeoVis.2.1.1.chr4_hap1.g00647.t01#5    mNeoVis.vulgivaga.2.1.chr6.g00651.t01#1    mNeoVis.vulgivaga.1.1.chr6.g00652.t01#4    mNeoVis.evergladensis.2.1.chr6.g00654.t01#2    mNeoVis.evergladensis.1.1.chr6.g00657.t01#3   mNeoVis.3.1.1.chr6.g00659.t01#6       5371   125255751:   mNeoVis.3.1.1.chr7.g00494.t01#6    mNeoVis.evergladensis.2.1.chr7.g00510.t01#2    mNeoVis.2.1.1.chr7_hap1.g00509.t01#5    mNeoVis.evergladensis.1.1.chr7.g00519.t01#3    mNeoVis.vulgivaga.1.1.chr7.g00517.t01#4   mNeoVis.vulgivaga.2.1.chr7.g00513.t01#1       5372   125255752:   mNeoVis.vulgivaga.2.1.chr6.g00652.t01#1    mNeoVis.vulgivaga.1.1.chr6.g00653.t01#4    mNeoVis.2.1.1.chr4_hap1.g00648.t01#5    mNeoVis.evergladensis.2.1.chr6.g00655.t01#2    mNeoVis.evergladensis.1.1.chr6.g00658.t01#3   mNeoVis.3.1.1.chr6.g00660.t01#6       5373   125255753:   mNeoVis.3.1.1.chr6.g00661.t01#6    mNeoVis.vulgivaga.1.1.chr6.g00654.t01#4    mNeoVis.evergladensis.2.1.chr6.g00656.t01#2    mNeoVis.vulgivaga.2.1.chr6.g00653.t01#1    mNeoVis.evergladensis.1.1.chr6.g00659.t01#3   mNeoVis.2.1.1.chr4_hap1.g00649.t01#5       5374   125255754:   mNeoVis.vulgivaga.2.1.chr6.g00654.t01#1    mNeoVis.vulgivaga.1.1.chr6.g00655.t01#4    mNeoVis.evergladensis.1.1.chr6.g00660.t01#3    mNeoVis.3.1.1.chr6.g00662.t01#6    mNeoVis.2.1.1.chr4_hap1.g00650.t01#5   mNeoVis.evergladensis.2.1.chr6.g00657.t01#2       5375   125255755:   mNeoVis.2.1.1.chr4_hap1.g00651.t01#5    mNeoVis.evergladensis.1.1.chr6.g00661.t01#3    mNeoVis.vulgivaga.2.1.chr6.g00655.t01#1    mNeoVis.vulgivaga.1.1.chr6.g00656.t01#4    mNeoVis.evergladensis.2.1.chr6.g00658.t01#2   mNeoVis.3.1.1.chr6.g00663.t01#6       5376   125255756:   mNeoVis.vulgivaga.2.1.chr14.g00306.t01#1    mNeoVis.vulgivaga.1.1.chr14.g00301.t01#4    mNeoVis.evergladensis.2.1.chr14.g00306.t01#2    mNeoVis.3.1.1.chr14.g00305.t01#6    mNeoVis.evergladensis.1.1.chr14.g00311.t01#3   mNeoVis.2.1.1.chr14_hap1.g00305.t01#5       5377   125255757:   mNeoVis.evergladensis.1.1.chr14.g00312.t01#3    mNeoVis.3.1.1.chr14.g00306.t01#6    mNeoVis.2.1.1.chr14_hap1.g00306.t01#5    mNeoVis.vulgivaga.2.1.chr14.g00307.t01#1    mNeoVis.vulgivaga.1.1.chr14.g00302.t01#4   mNeoVis.evergladensis.2.1.chr14.g00307.t01#2       5378   125255758:   mNeoVis.evergladensis.1.1.chr14.g00313.t01#3    mNeoVis.3.1.1.chr14.g00307.t01#6    mNeoVis.evergladensis.2.1.chr14.g00308.t01#2    mNeoVis.2.1.1.chr14_hap1.g00307.t01#5    mNeoVis.vulgivaga.2.1.chr14.g00308.t01#1   mNeoVis.vulgivaga.1.1.chr14.g00303.t01#4       5379   125255759:   mNeoVis.vulgivaga.2.1.chr14.g00309.t01#1    mNeoVis.vulgivaga.1.1.chr14.g00304.t01#4    mNeoVis.3.1.1.chr14.g00308.t01#6    mNeoVis.evergladensis.1.1.chr14.g00314.t01#3    mNeoVis.evergladensis.2.1.chr14.g00309.t01#2   mNeoVis.2.1.1.chr14_hap1.g00308.t01#5       5380   125255760:   mNeoVis.2.1.1.chr4_hap1.g00652.t01#5    mNeoVis.evergladensis.2.1.chr6.g00659.t01#2    mNeoVis.evergladensis.1.1.chr6.g00662.t01#3    mNeoVis.3.1.1.chr6.g00664.t01#6    mNeoVis.vulgivaga.2.1.chr6.g00656.t01#1   mNeoVis.vulgivaga.1.1.chr6.g00657.t01#4       5381   125255761:   mNeoVis.evergladensis.1.1.chr6.g00663.t01#3    mNeoVis.2.1.1.chr4_hap1.g00653.t01#5    mNeoVis.evergladensis.2.1.chr6.g00660.t01#2    mNeoVis.3.1.1.chr6.g00665.t01#6    mNeoVis.vulgivaga.1.1.chr6.g00658.t01#4   mNeoVis.vulgivaga.2.1.chr6.g00657.t01#1       5382   125255762:   mNeoVis.evergladensis.2.1.chr14.g00310.t01#2    mNeoVis.vulgivaga.2.1.chr14.g00310.t01#1    mNeoVis.3.1.1.chr14.g00309.t01#6    mNeoVis.2.1.1.chr14_hap1.g00309.t01#5   mNeoVis.evergladensis.1.1.chr14.g00315.t01#3       5383   125255763:   mNeoVis.3.1.1.chr14.g00310.t01#6    mNeoVis.vulgivaga.1.1.chr14.g00306.t01#4    mNeoVis.evergladensis.2.1.chr14.g00311.t01#2    mNeoVis.vulgivaga.2.1.chr14.g00311.t01#1    mNeoVis.2.1.1.chr14_hap1.g00310.t01#5   mNeoVis.evergladensis.1.1.chr14.g00316.t01#3       5384   125255764:   mNeoVis.vulgivaga.2.1.chr6.g00659.t01#1    mNeoVis.evergladensis.1.1.chr6.g00665.t01#3    mNeoVis.2.1.1.chr4_hap1.g00655.t01#5    mNeoVis.3.1.1.chr6.g00666.t01#6    mNeoVis.evergladensis.2.1.chr6.g00662.t01#2   mNeoVis.vulgivaga.1.1.chr6.g00660.t01#4       5385   125255765:   mNeoVis.2.1.1.chr4_hap1.g00657.t01#5    mNeoVis.evergladensis.1.1.chr6.g00667.t01#3    mNeoVis.vulgivaga.2.1.chr6.g00661.t01#1    mNeoVis.vulgivaga.1.1.chr6.g00662.t01#4   mNeoVis.evergladensis.2.1.chr6.g00664.t01#2       5386   125255766:   mNeoVis.vulgivaga.2.1.chr14.g00312.t01#1    mNeoVis.2.1.1.chr14_hap1.g00311.t01#5    mNeoVis.vulgivaga.1.1.chr14.g00307.t01#4    mNeoVis.evergladensis.2.1.chr14.g00312.t01#2   mNeoVis.evergladensis.1.1.chr14.g00317.t01#3       5387   125255767:   mNeoVis.vulgivaga.1.1.chr14.g00308.t01#4    mNeoVis.2.1.1.chr14_hap1.g00312.t01#5    mNeoVis.3.1.1.chr14.g00311.t01#6    mNeoVis.vulgivaga.2.1.chr14.g00313.t01#1    mNeoVis.evergladensis.1.1.chr14.g00318.t01#3   mNeoVis.evergladensis.2.1.chr14.g00313.t01#2       5388   125255768:   mNeoVis.vulgivaga.2.1.chr6.g00662.t01#1    mNeoVis.evergladensis.1.1.chr6.g00668.t01#3    mNeoVis.3.1.1.chr6.g00669.t01#6    mNeoVis.evergladensis.2.1.chr6.g00665.t01#2    mNeoVis.vulgivaga.1.1.chr6.g00663.t01#4   mNeoVis.2.1.1.chr4_hap1.g00658.t01#5       5389   125255769:   mNeoVis.vulgivaga.1.1.chr6.g00664.t01#4    mNeoVis.2.1.1.chr4_hap1.g00659.t01#5    mNeoVis.vulgivaga.2.1.chr6.g00663.t01#1    mNeoVis.evergladensis.2.1.chr6.g00666.t01#2    mNeoVis.3.1.1.chr6.g00670.t01#6   mNeoVis.evergladensis.1.1.chr6.g00669.t01#3       5390   125255770:   mNeoVis.evergladensis.2.1.chr14.g00314.t01#2    mNeoVis.vulgivaga.2.1.chr14.g00314.t01#1    mNeoVis.vulgivaga.1.1.chr14.g00309.t01#4    mNeoVis.evergladensis.1.1.chr14.g00319.t01#3    mNeoVis.2.1.1.chr14_hap1.g00313.t01#5   mNeoVis.3.1.1.chr14.g00312.t01#6       5391   125255771:   mNeoVis.2.1.1.chr15_hap1.g00637.t01#5    mNeoVis.evergladensis.2.1.chr15.g00459.t01#2    mNeoVis.evergladensis.1.1.chr15.g00155.t01#3    mNeoVis.3.1.1.chr15.g00418.t01#6    mNeoVis.vulgivaga.2.1.chr15.g00466.t01#1   mNeoVis.vulgivaga.1.1.chr15.g00460.t01#4       5392   125255772:   mNeoVis.2.1.1.chr4_hap1.g00660.t01#5    mNeoVis.evergladensis.1.1.chr6.g00670.t01#3    mNeoVis.vulgivaga.1.1.chr6.g00665.t01#4    mNeoVis.evergladensis.2.1.chr6.g00667.t01#2    mNeoVis.vulgivaga.2.1.chr6.g00664.t01#1   mNeoVis.3.1.1.chr6.g00671.t01#6       5393   125255773:   mNeoVis.vulgivaga.1.1.chr14.g00310.t01#4    mNeoVis.2.1.1.chr14_hap1.g00314.t01#5    mNeoVis.vulgivaga.2.1.chr14.g00315.t01#1    mNeoVis.evergladensis.1.1.chr14.g00320.t01#3    mNeoVis.evergladensis.2.1.chr14.g00315.t01#2   mNeoVis.3.1.1.chr14.g00313.t01#6       5394   125255774:   mNeoVis.3.1.1.chr7.g00495.t01#6    mNeoVis.evergladensis.1.1.chr7.g00520.t01#3    mNeoVis.2.1.1.chr7_hap1.g00510.t01#5    mNeoVis.vulgivaga.1.1.chr7.g00518.t01#4    mNeoVis.vulgivaga.2.1.chr7.g00514.t01#1   mNeoVis.evergladensis.2.1.chr7.g00511.t01#2       5395   125255775:   mNeoVis.vulgivaga.2.1.chr6.g00665.t01#1    mNeoVis.evergladensis.1.1.chr6.g00671.t01#3    mNeoVis.vulgivaga.1.1.chr6.g00666.t01#4    mNeoVis.2.1.1.chr4_hap1.g00661.t01#5    mNeoVis.evergladensis.2.1.chr6.g00668.t01#2   mNeoVis.3.1.1.chr6.g00672.t01#6       5396   125255776:   mNeoVis.2.1.1.chr4_hap1.g00662.t01#5    mNeoVis.evergladensis.1.1.chr6.g00672.t01#3    mNeoVis.vulgivaga.1.1.chr6.g00667.t01#4    mNeoVis.3.1.1.chr6.g00673.t01#6    mNeoVis.vulgivaga.2.1.chr6.g00666.t01#1   mNeoVis.evergladensis.2.1.chr6.g00669.t01#2       5397   125255777:   mNeoVis.2.1.1.chr4_hap1.g00663.t01#5    mNeoVis.evergladensis.1.1.chr6.g00673.t01#3    mNeoVis.vulgivaga.1.1.chr6.g00668.t01#4    mNeoVis.vulgivaga.2.1.chr6.g00667.t01#1   mNeoVis.evergladensis.2.1.chr6.g00670.t01#2       5398   125255778:   mNeoVis.evergladensis.2.1.chr6.g00671.t01#2    mNeoVis.3.1.1.chr6.g00675.t01#6    mNeoVis.evergladensis.1.1.chr6.g00674.t01#3    mNeoVis.vulgivaga.2.1.chr6.g00668.t01#1    mNeoVis.vulgivaga.1.1.chr6.g00669.t01#4   mNeoVis.2.1.1.chr4_hap1.g00664.t01#5       5399   125255779:   mNeoVis.3.1.1.chr6.g00676.t01#6    mNeoVis.vulgivaga.1.1.chr6.g00670.t01#4    mNeoVis.evergladensis.1.1.chr6.g00676.t01#3    mNeoVis.2.1.1.chr4_hap1.g00665.t01#5    mNeoVis.vulgivaga.2.1.chr6.g00669.t01#1   mNeoVis.evergladensis.2.1.chr6.g00672.t01#2       5400   125255780:   mNeoVis.vulgivaga.1.1.chr14.g00311.t01#4    mNeoVis.2.1.1.chr14_hap1.g00315.t01#5    mNeoVis.evergladensis.2.1.chr14.g00316.t01#2    mNeoVis.evergladensis.1.1.chr14.g00321.t01#3    mNeoVis.3.1.1.chr14.g00314.t01#6   mNeoVis.vulgivaga.2.1.chr14.g00316.t01#1       5401   125255781:   mNeoVis.vulgivaga.2.1.chr6.g00670.t01#1    mNeoVis.evergladensis.2.1.chr6.g00673.t01#2    mNeoVis.3.1.1.chr6.g00677.t01#6    mNeoVis.2.1.1.chr4_hap1.g00666.t01#5    mNeoVis.evergladensis.1.1.chr6.g00677.t01#3   mNeoVis.vulgivaga.1.1.chr6.g00671.t01#4       5402   125255782:   mNeoVis.3.1.1.chr6.g00678.t01#6    mNeoVis.2.1.1.chr4_hap1.g00667.t01#5    mNeoVis.evergladensis.1.1.chr6.g00678.t01#3    mNeoVis.vulgivaga.2.1.chr6.g00671.t01#1    mNeoVis.vulgivaga.1.1.chr6.g00672.t01#4   mNeoVis.evergladensis.2.1.chr6.g00674.t01#2       5403   125255783:   mNeoVis.2.1.1.chr4_hap1.g00668.t01#5    mNeoVis.2.1.1.chr13_hap1.g00419.t01#5    mNeoVis.vulgivaga.1.1.chr6.g00673.t01#4   mNeoVis.evergladensis.2.1.chr13.g00402.t01#2       5404   125255784:   mNeoVis.vulgivaga.2.1.chr6.g00672.t01#1   mNeoVis.evergladensis.2.1.chr6.g00675.t01#2           5405   125255785:   mNeoVis.evergladensis.1.1.chr6.g00679.t01#3    mNeoVis.3.1.1.chr6.g00679.t01#6    mNeoVis.vulgivaga.1.1.chr6.g00675.t01#4    mNeoVis.evergladensis.2.1.chr6.g00676.t01#2    mNeoVis.vulgivaga.2.1.chr6.g00673.t01#1   mNeoVis.2.1.1.chr4_hap1.g00669.t01#5       5406   125255786:   mNeoVis.evergladensis.1.1.chr14.g00322.t01#3    mNeoVis.evergladensis.2.1.chr14.g00317.t01#2    mNeoVis.vulgivaga.2.1.chr14.g00317.t01#1    mNeoVis.vulgivaga.1.1.chr14.g00312.t01#4   mNeoVis.2.1.1.chr14_hap1.g00316.t01#5       5407   125255787:   mNeoVis.vulgivaga.1.1.chr14.g00313.t01#4    mNeoVis.2.1.1.chr14_hap1.g00317.t01#5    mNeoVis.vulgivaga.2.1.chr14.g00318.t01#1    mNeoVis.3.1.1.chr14.g00315.t01#6    mNeoVis.evergladensis.1.1.chr14.g00323.t01#3   mNeoVis.evergladensis.2.1.chr14.g00318.t01#2       5408   125255788:   mNeoVis.evergladensis.1.1.chr14.g00324.t01#3    mNeoVis.vulgivaga.2.1.chr14.g00319.t01#1   mNeoVis.vulgivaga.1.1.chr14.g00314.t01#4         5409   125255789:   mNeoVis.3.1.1.chr14.g00316.t01#6    mNeoVis.evergladensis.2.1.chr14.g00319.t01#2   mNeoVis.2.1.1.chr14_hap1.g00318.t01#5           5410   125255790:   mNeoVis.vulgivaga.2.1.chr6.g00674.t01#1    mNeoVis.evergladensis.1.1.chr6.g00680.t01#3    mNeoVis.evergladensis.2.1.chr6.g00677.t01#2    mNeoVis.vulgivaga.1.1.chr6.g00676.t01#4   mNeoVis.2.1.1.chr4_hap1.g00670.t01#5       5411   125255791:   mNeoVis.evergladensis.2.1.chr6.g00678.t01#2   mNeoVis.vulgivaga.2.1.chr6.g00675.t01#1             5412   125255792:   mNeoVis.vulgivaga.1.1.chr6.g00677.t01#4    mNeoVis.vulgivaga.2.1.chr6.g00676.t01#1    mNeoVis.evergladensis.2.1.chr6.g00679.t01#2    mNeoVis.2.1.1.chr4_hap1.g00671.t01#5    mNeoVis.3.1.1.chr6.g00680.t01#6   mNeoVis.evergladensis.1.1.chr6.g00681.t01#3       5413   125255793:   mNeoVis.evergladensis.2.1.chr6.g00680.t01#2    mNeoVis.vulgivaga.2.1.chr6.g00677.t01#1    mNeoVis.vulgivaga.1.1.chr6.g00678.t01#4    mNeoVis.3.1.1.chr6.g00681.t01#6    mNeoVis.evergladensis.1.1.chr6.g00682.t01#3   mNeoVis.2.1.1.chr4_hap1.g00672.t01#5       5414   125255794:   mNeoVis.evergladensis.1.1.chr14.g00325.t01#3    mNeoVis.2.1.1.chr14_hap1.g00319.t01#5    mNeoVis.evergladensis.2.1.chr14.g00320.t01#2    mNeoVis.vulgivaga.2.1.chr14.g00320.t01#1    mNeoVis.3.1.1.chr14.g00317.t01#6   mNeoVis.vulgivaga.1.1.chr14.g00315.t01#4       5415   125255795:   mNeoVis.3.1.1.chr6.g00682.t01#6    mNeoVis.vulgivaga.2.1.chr6.g00678.t01#1    mNeoVis.evergladensis.1.1.chr6.g00683.t01#3    mNeoVis.2.1.1.chr4_hap1.g00673.t01#5    mNeoVis.evergladensis.2.1.chr6.g00681.t01#2   mNeoVis.vulgivaga.1.1.chr6.g00679.t01#4       5416   125255796:   mNeoVis.vulgivaga.2.1.chr14.g00321.t01#1    mNeoVis.3.1.1.chr14.g00318.t01#6    mNeoVis.vulgivaga.1.1.chr14.g00316.t01#4   mNeoVis.2.1.1.chr14_hap1.g00320.t01#5         5417   125255797:   mNeoVis.vulgivaga.1.1.chr6.g00681.t01#4    mNeoVis.3.1.1.chr6.g00684.t01#6    mNeoVis.evergladensis.2.1.chr6.g00683.t01#2    mNeoVis.2.1.1.chr4_hap1.g00676.t01#5    mNeoVis.vulgivaga.2.1.chr6.g00680.t01#1   mNeoVis.evergladensis.1.1.chr6.g00685.t01#3       5418   125255798:   mNeoVis.3.1.1.chr6.g00683.t01#6    mNeoVis.vulgivaga.2.1.chr6.g00679.t01#1    mNeoVis.evergladensis.1.1.chr6.g00684.t01#3    mNeoVis.2.1.1.chr4_hap1.g00675.t01#5    mNeoVis.evergladensis.2.1.chr6.g00682.t01#2   mNeoVis.vulgivaga.1.1.chr6.g00680.t01#4       5419   125255799:   mNeoVis.2.1.1.chr3_hap1.g00744.t01#5    mNeoVis.vulgivaga.2.1.chr3.g00751.t01#1    mNeoVis.evergladensis.1.1.chr3.g00745.t01#3    mNeoVis.3.1.1.chr3.g00728.t01#6    mNeoVis.evergladensis.2.1.chr3.g00744.t01#2   mNeoVis.vulgivaga.1.1.chr3.g00740.t01#4       5420   125255800:   mNeoVis.evergladensis.1.1.chr6.g00686.t01#3    mNeoVis.vulgivaga.2.1.chr6.g00681.t01#1    mNeoVis.vulgivaga.1.1.chr6.g00682.t01#4    mNeoVis.evergladensis.2.1.chr6.g00684.t01#2    mNeoVis.3.1.1.chr6.g00685.t01#6   mNeoVis.2.1.1.chr4_hap1.g00677.t01#5       5421   125255801:   mNeoVis.vulgivaga.1.1.chr14.g00317.t01#4    mNeoVis.evergladensis.1.1.chr14.g00326.t01#3    mNeoVis.3.1.1.chr14.g00319.t01#6    mNeoVis.evergladensis.2.1.chr14.g00321.t01#2    mNeoVis.2.1.1.chr14_hap1.g00321.t01#5   mNeoVis.vulgivaga.2.1.chr14.g00322.t01#1       5422   125255802:   mNeoVis.2.1.1.chr4_hap1.g00678.t01#5    mNeoVis.evergladensis.1.1.chr6.g00688.t01#3    mNeoVis.3.1.1.chr6.g00686.t01#6    mNeoVis.vulgivaga.2.1.chr6.g00682.t01#1    mNeoVis.vulgivaga.1.1.chr6.g00684.t01#4   mNeoVis.evergladensis.2.1.chr6.g00685.t01#2       5423   125255803:   mNeoVis.2.1.1.chr14_hap1.g00322.t01#5    mNeoVis.3.1.1.chr14.g00320.t01#6    mNeoVis.evergladensis.1.1.chr14.g00327.t01#3    mNeoVis.vulgivaga.1.1.chr14.g00318.t01#4    mNeoVis.evergladensis.2.1.chr14.g00322.t01#2   mNeoVis.vulgivaga.2.1.chr14.g00323.t01#1       5424   125255804:  mNeoVis.vulgivaga.2.1.chr14.g00324.t01#1               5425   125255805:   mNeoVis.2.1.1.chr14_hap1.g00323.t01#5    mNeoVis.vulgivaga.1.1.chr14.g00319.t01#4    mNeoVis.evergladensis.2.1.chr14.g00323.t01#2    mNeoVis.vulgivaga.2.1.chr14.g00325.t01#1    mNeoVis.3.1.1.chr14.g00321.t01#6   mNeoVis.evergladensis.1.1.chr14.g00328.t01#3       5426   125255806:   mNeoVis.evergladensis.2.1.chr14.g00324.t01#2    mNeoVis.2.1.1.chr14_hap1.g00324.t01#5    mNeoVis.vulgivaga.2.1.chr14.g00326.t01#1    mNeoVis.evergladensis.1.1.chr14.g00329.t01#3    mNeoVis.vulgivaga.1.1.chr14.g00320.t01#4   mNeoVis.3.1.1.chr14.g00322.t01#6       5427   125255807:   mNeoVis.evergladensis.1.1.chr10.g00936.t01#3    mNeoVis.vulgivaga.2.1.chr10.g00925.t01#1    mNeoVis.vulgivaga.1.1.chr10.g00917.t01#4    mNeoVis.evergladensis.2.1.chr10.g00935.t01#2   mNeoVis.2.1.1.chr9_hap1.g00950.t01#5       5428   125255808:   mNeoVis.2.1.1.chr4_hap1.g00680.t01#5    mNeoVis.vulgivaga.2.1.chr6.g00683.t01#1    mNeoVis.evergladensis.1.1.chr6.g00690.t01#3    mNeoVis.vulgivaga.1.1.chr6.g00685.t01#4    mNeoVis.evergladensis.2.1.chr6.g00687.t01#2   mNeoVis.3.1.1.chr6.g00687.t01#6       5429   125255809:   mNeoVis.2.1.1.chr14_hap1.g00325.t01#5    mNeoVis.3.1.1.chr14.g00323.t01#6    mNeoVis.vulgivaga.1.1.chr14.g00321.t01#4    mNeoVis.evergladensis.1.1.chr14.g00330.t01#3    mNeoVis.vulgivaga.2.1.chr14.g00327.t01#1   mNeoVis.evergladensis.2.1.chr14.g00325.t01#2       5430   125255810:   mNeoVis.vulgivaga.1.1.chr5.g00220.t01#4    mNeoVis.2.1.1.chr2_hap1.g00223.t01#5    mNeoVis.evergladensis.1.1.chr5.g00227.t01#3    mNeoVis.2.1.1.chr4_hap1.g00684.t01#5    mNeoVis.3.1.1.chr6.g00690.t01#6    mNeoVis.vulgivaga.1.1.chr6.g00686.t01#4    mNeoVis.vulgivaga.2.1.chr5.g00224.t01#1    mNeoVis.3.1.1.chr5.g00215.t01#6    mNeoVis.evergladensis.2.1.chr5.g00227.t01#2      5431   125255811:   mNeoVis.2.1.1.chr4_hap1.g00686.t01#5    mNeoVis.vulgivaga.2.1.chr6.g00686.t01#1    mNeoVis.vulgivaga.1.1.chr6.g00687.t01#4    mNeoVis.evergladensis.1.1.chr6.g00691.t01#3    mNeoVis.evergladensis.2.1.chr6.g00689.t01#2   mNeoVis.3.1.1.chr6.g00692.t01#6       5432   125255812:   mNeoVis.vulgivaga.2.1.chr14.g00328.t01#1    mNeoVis.evergladensis.1.1.chr14.g00331.t01#3    mNeoVis.2.1.1.chr14_hap1.g00326.t01#5    mNeoVis.vulgivaga.1.1.chr14.g00322.t01#4    mNeoVis.3.1.1.chr14.g00324.t01#6   mNeoVis.evergladensis.2.1.chr14.g00326.t01#2       5433   125255813:   mNeoVis.vulgivaga.2.1.chr6.g00687.t01#1    mNeoVis.vulgivaga.1.1.chr6.g00688.t01#4    mNeoVis.3.1.1.chr6.g00693.t01#6    mNeoVis.2.1.1.chr4_hap1.g00688.t01#5    mNeoVis.evergladensis.1.1.chr6.g00692.t01#3   mNeoVis.evergladensis.2.1.chr6.g00692.t01#2       5434   125255814:   mNeoVis.2.1.1.chr4_hap1.g00689.t01#5    mNeoVis.vulgivaga.2.1.chr6.g00688.t01#1    mNeoVis.vulgivaga.1.1.chr6.g00689.t01#4    mNeoVis.evergladensis.1.1.chr6.g00693.t01#3    mNeoVis.evergladensis.2.1.chr6.g00693.t01#2   mNeoVis.3.1.1.chr6.g00694.t01#6       5435   125255815:   mNeoVis.vulgivaga.2.1.chr14.g00329.t01#1    mNeoVis.2.1.1.chr14_hap1.g00327.t01#5    mNeoVis.vulgivaga.1.1.chr14.g00323.t01#4    mNeoVis.3.1.1.chr14.g00325.t01#6    mNeoVis.evergladensis.1.1.chr14.g00332.t01#3   mNeoVis.evergladensis.2.1.chr14.g00327.t01#2       5436   125255816:   mNeoVis.vulgivaga.2.1.chr14.g00330.t01#1    mNeoVis.2.1.1.chr14_hap1.g00328.t01#5    mNeoVis.vulgivaga.1.1.chr14.g00324.t01#4    mNeoVis.evergladensis.1.1.chr14.g00333.t01#3    mNeoVis.evergladensis.2.1.chr14.g00328.t01#2   mNeoVis.3.1.1.chr14.g00326.t01#6       5437   125255817:   mNeoVis.vulgivaga.2.1.chr14.g00331.t01#1    mNeoVis.3.1.1.chr14.g00327.t01#6    mNeoVis.evergladensis.2.1.chr14.g00329.t01#2    mNeoVis.2.1.1.chr14_hap1.g00329.t01#5    mNeoVis.vulgivaga.1.1.chr14.g00325.t01#4   mNeoVis.evergladensis.1.1.chr14.g00334.t01#3       5438   125255818:   mNeoVis.vulgivaga.1.1.chr12.g00562.t01#4    mNeoVis.evergladensis.1.1.chr12.g00552.t01#3    mNeoVis.vulgivaga.2.1.chr12.g00564.t01#1    mNeoVis.2.1.1.chr12_hap1.g00611.t01#5    mNeoVis.3.1.1.chr12.g00557.t01#6   mNeoVis.evergladensis.2.1.chr12.g00548.t01#2       5439   125255819:   mNeoVis.evergladensis.1.1.chr7.g00521.t01#3    mNeoVis.evergladensis.2.1.chr7.g00512.t01#2    mNeoVis.vulgivaga.2.1.chr7.g00515.t01#1    mNeoVis.2.1.1.chr7_hap1.g00511.t01#5   mNeoVis.vulgivaga.1.1.chr7.g00519.t01#4       5440   125255820:   mNeoVis.3.1.1.chr15.g00419.t01#6    mNeoVis.vulgivaga.1.1.chr15.g00461.t01#4    mNeoVis.2.1.1.chr15_hap1.g00638.t01#5    mNeoVis.evergladensis.2.1.chr15.g00460.t01#2    mNeoVis.vulgivaga.2.1.chr15.g00467.t01#1   mNeoVis.evergladensis.1.1.chr15.g00154.t01#3       5441   125255821:   mNeoVis.vulgivaga.2.1.chr14.g00332.t01#1    mNeoVis.evergladensis.2.1.chr14.g00330.t01#2    mNeoVis.3.1.1.chr14.g00328.t01#6    mNeoVis.2.1.1.chr14_hap1.g00330.t01#5    mNeoVis.evergladensis.1.1.chr14.g00335.t01#3   mNeoVis.vulgivaga.1.1.chr14.g00326.t01#4       5442   125255822:   mNeoVis.3.1.1.chr14.g00329.t01#6    mNeoVis.evergladensis.1.1.chr14.g00336.t01#3    mNeoVis.2.1.1.chr14_hap1.g00331.t01#5    mNeoVis.vulgivaga.1.1.chr14.g00327.t01#4    mNeoVis.evergladensis.2.1.chr14.g00331.t01#2   mNeoVis.vulgivaga.2.1.chr14.g00333.t01#1       5443   125255823:   mNeoVis.evergladensis.2.1.chr14.g00332.t01#2    mNeoVis.3.1.1.chr14.g00331.t01#6    mNeoVis.2.1.1.chr14_hap1.g00332.t01#5    mNeoVis.vulgivaga.1.1.chr14.g00328.t01#4    mNeoVis.evergladensis.1.1.chr14.g00337.t01#3   mNeoVis.vulgivaga.2.1.chr14.g00334.t01#1       5444   125255824:   mNeoVis.vulgivaga.2.1.chr14.g00335.t01#1    mNeoVis.evergladensis.1.1.chr14.g00338.t01#3    mNeoVis.2.1.1.chr14_hap1.g00333.t01#5    mNeoVis.evergladensis.2.1.chr14.g00333.t01#2    mNeoVis.vulgivaga.1.1.chr14.g00329.t01#4   mNeoVis.3.1.1.chr14.g00332.t01#6       5445   125255825:   mNeoVis.3.1.1.chr2.g00749.t01#6    mNeoVis.evergladensis.2.1.chr2.g00840.t01#2    mNeoVis.vulgivaga.1.1.chr2.g00841.t01#4    mNeoVis.2.1.1.chr5_hap1.g00839.t01#5    mNeoVis.vulgivaga.2.1.chr2.g00842.t01#1   mNeoVis.evergladensis.1.1.chr2.g00835.t01#3       5446   125255826:   mNeoVis.evergladensis.1.1.chr14.g00339.t01#3    mNeoVis.2.1.1.chr14_hap1.g00334.t01#5    mNeoVis.evergladensis.2.1.chr14.g00334.t01#2    mNeoVis.vulgivaga.1.1.chr14.g00330.t01#4    mNeoVis.3.1.1.chr14.g00333.t01#6   mNeoVis.vulgivaga.2.1.chr14.g00336.t01#1       5447   125255827:   mNeoVis.3.1.1.chr14.g00334.t01#6    mNeoVis.vulgivaga.2.1.chr14.g00337.t01#1    mNeoVis.evergladensis.1.1.chr14.g00340.t01#3    mNeoVis.vulgivaga.1.1.chr14.g00331.t01#4    mNeoVis.evergladensis.2.1.chr14.g00335.t01#2   mNeoVis.2.1.1.chr14_hap1.g00335.t01#5       5448   125255828:   mNeoVis.vulgivaga.1.1.chr6.g00690.t01#4    mNeoVis.evergladensis.2.1.chr6.g00694.t01#2    mNeoVis.3.1.1.chr6.g00695.t01#6    mNeoVis.2.1.1.chr4_hap1.g00690.t01#5    mNeoVis.evergladensis.1.1.chr6.g00694.t01#3   mNeoVis.vulgivaga.2.1.chr6.g00689.t01#1       5449   125255829:   mNeoVis.2.1.1.chr14_hap1.g00336.t01#5    mNeoVis.vulgivaga.1.1.chr14.g00332.t01#4    mNeoVis.evergladensis.1.1.chr14.g00341.t01#3    mNeoVis.evergladensis.2.1.chr14.g00336.t01#2    mNeoVis.3.1.1.chr14.g00335.t01#6   mNeoVis.vulgivaga.2.1.chr14.g00338.t01#1       5450   125255830:   mNeoVis.vulgivaga.2.1.chr6.g00691.t01#1    mNeoVis.vulgivaga.1.1.chr6.g00692.t01#4    mNeoVis.evergladensis.2.1.chr6.g00696.t01#2    mNeoVis.2.1.1.chr4_hap1.g00692.t01#5    mNeoVis.3.1.1.chr6.g00697.t01#6   mNeoVis.evergladensis.1.1.chr6.g00696.t01#3       5451   125255831:   mNeoVis.evergladensis.2.1.chr14.g00254.t01#2    mNeoVis.vulgivaga.1.1.chr10.g00972.t01#4    mNeoVis.2.1.1.chr9_hap1.g01004.t01#5    mNeoVis.vulgivaga.2.1.chr14.g00255.t01#1    mNeoVis.2.1.1.chr14_hap1.g00254.t01#5    mNeoVis.3.1.1.chr14.g00256.t01#6    mNeoVis.evergladensis.1.1.chr14.g00260.t01#3    mNeoVis.vulgivaga.2.1.chr10.g00983.t01#1    mNeoVis.evergladensis.2.1.chr10.g00991.t01#2      5452   125255832:   mNeoVis.vulgivaga.1.1.chr14.g00251.t01#4    mNeoVis.3.1.1.chr14.g00257.t01#6    mNeoVis.evergladensis.2.1.chr14.g00255.t01#2    mNeoVis.2.1.1.chr14_hap1.g00255.t01#5    mNeoVis.evergladensis.1.1.chr14.g00261.t01#3   mNeoVis.vulgivaga.2.1.chr14.g00256.t01#1       5453   125255833:   mNeoVis.vulgivaga.1.1.chr6.g00693.t01#4    mNeoVis.evergladensis.2.1.chr6.g00697.t01#2    mNeoVis.vulgivaga.2.1.chr6.g00692.t01#1    mNeoVis.2.1.1.chr4_hap1.g00693.t01#5    mNeoVis.3.1.1.chr6.g00698.t01#6   mNeoVis.evergladensis.1.1.chr6.g00697.t01#3       5454   125255834:   mNeoVis.evergladensis.1.1.chr6.g00698.t01#3    mNeoVis.2.1.1.chr4_hap1.g00694.t01#5    mNeoVis.3.1.1.chr6.g00699.t01#6    mNeoVis.vulgivaga.2.1.chr6.g00693.t01#1    mNeoVis.evergladensis.2.1.chr6.g00698.t01#2   mNeoVis.vulgivaga.1.1.chr6.g00694.t01#4       5455   125255835:   mNeoVis.vulgivaga.1.1.chr6.g00695.t01#4    mNeoVis.vulgivaga.2.1.chr6.g00694.t01#1    mNeoVis.evergladensis.2.1.chr6.g00699.t01#2    mNeoVis.evergladensis.1.1.chr6.g00699.t01#3    mNeoVis.2.1.1.chr4_hap1.g00695.t01#5   mNeoVis.3.1.1.chr6.g00700.t01#6       5456   125255836:   mNeoVis.vulgivaga.2.1.chr14.g00257.t01#1    mNeoVis.2.1.1.chr14_hap1.g00256.t01#5    mNeoVis.evergladensis.2.1.chr14.g00256.t01#2    mNeoVis.3.1.1.chr14.g00258.t01#6    mNeoVis.vulgivaga.1.1.chr14.g00252.t01#4   mNeoVis.evergladensis.1.1.chr14.g00262.t01#3       5457   125255837:   mNeoVis.2.1.1.chr11_hap1.g00726.t01#5    mNeoVis.evergladensis.2.1.chr11.g00709.t01#2    mNeoVis.vulgivaga.2.1.chr6.g00695.t01#1    mNeoVis.2.1.1.chr4_hap1.g00696.t01#5   mNeoVis.vulgivaga.1.1.chr11.g00708.t01#4       5458   125255838:   mNeoVis.3.1.1.chr11.g00688.t01#6   mNeoVis.evergladensis.2.1.chr6.g00700.t01#2           5459   125255839:   mNeoVis.vulgivaga.1.1.chr6.g00696.t01#4    mNeoVis.vulgivaga.2.1.chr6.g00696.t01#1    mNeoVis.evergladensis.2.1.chr6.g00701.t01#2    mNeoVis.3.1.1.chr6.g00701.t01#6    mNeoVis.2.1.1.chr4_hap1.g00697.t01#5   mNeoVis.evergladensis.1.1.chr6.g00700.t01#3       5460   125255840:   mNeoVis.3.1.1.chr6.g00625.t01#6    mNeoVis.evergladensis.1.1.chr6.g00619.t01#3    mNeoVis.evergladensis.2.1.chr6.g00617.t01#2    mNeoVis.vulgivaga.1.1.chr6.g00616.t01#4    mNeoVis.vulgivaga.2.1.chr6.g00614.t01#1   mNeoVis.2.1.1.chr4_hap1.g00608.t01#5       5461   125255841:   mNeoVis.vulgivaga.2.1.chr14.g00258.t01#1    mNeoVis.vulgivaga.1.1.chr14.g00253.t01#4    mNeoVis.evergladensis.1.1.chr14.g00263.t01#3    mNeoVis.evergladensis.2.1.chr14.g00257.t01#2    mNeoVis.3.1.1.chr14.g00259.t01#6   mNeoVis.2.1.1.chr14_hap1.g00257.t01#5       5462   125255842:   mNeoVis.3.1.1.chr14.g00260.t01#6    mNeoVis.evergladensis.2.1.chr14.g00258.t01#2    mNeoVis.vulgivaga.2.1.chr14.g00259.t01#1    mNeoVis.2.1.1.chr14_hap1.g00258.t01#5    mNeoVis.vulgivaga.1.1.chr14.g00255.t01#4   mNeoVis.evergladensis.1.1.chr14.g00264.t01#3       5463   125255843:   mNeoVis.2.1.1.chr14_hap1.g00259.t01#5    mNeoVis.3.1.1.chr14.g00261.t01#6    mNeoVis.evergladensis.1.1.chr14.g00265.t01#3    mNeoVis.vulgivaga.2.1.chr14.g00260.t01#1   mNeoVis.evergladensis.2.1.chr14.g00259.t01#2       5464   125255844:   mNeoVis.3.1.1.chr6.g00626.t01#6    mNeoVis.2.1.1.chr4_hap1.g00609.t01#5    mNeoVis.evergladensis.1.1.chr6.g00620.t01#3    mNeoVis.evergladensis.2.1.chr6.g00618.t01#2    mNeoVis.vulgivaga.1.1.chr6.g00617.t01#4   mNeoVis.vulgivaga.2.1.chr6.g00615.t01#1       5465   125255845:   mNeoVis.2.1.1.chr14_hap1.g00260.t01#5    mNeoVis.vulgivaga.1.1.chr14.g00257.t01#4    mNeoVis.vulgivaga.2.1.chr14.g00261.t01#1    mNeoVis.3.1.1.chr14.g00262.t01#6    mNeoVis.evergladensis.2.1.chr14.g00260.t01#2   mNeoVis.evergladensis.1.1.chr14.g00266.t01#3       5466   125255846:   mNeoVis.evergladensis.2.1.chr15.g00461.t01#2    mNeoVis.vulgivaga.1.1.chr15.g00462.t01#4    mNeoVis.evergladensis.1.1.chr15.g00153.t01#3    mNeoVis.3.1.1.chr15.g00420.t01#6    mNeoVis.vulgivaga.2.1.chr15.g00468.t01#1   mNeoVis.2.1.1.chr15_hap1.g00639.t01#5       5467   125255847:   mNeoVis.2.1.1.chr4_hap1.g00610.t01#5    mNeoVis.evergladensis.1.1.chr6.g00621.t01#3    mNeoVis.vulgivaga.1.1.chr6.g00618.t01#4    mNeoVis.vulgivaga.2.1.chr6.g00616.t01#1   mNeoVis.evergladensis.2.1.chr6.g00619.t01#2       5468   125255848:   mNeoVis.vulgivaga.1.1.chr14.g00258.t01#4    mNeoVis.3.1.1.chr14.g00263.t01#6    mNeoVis.evergladensis.1.1.chr14.g00267.t01#3    mNeoVis.vulgivaga.2.1.chr14.g00262.t01#1    mNeoVis.2.1.1.chr14_hap1.g00261.t01#5   mNeoVis.evergladensis.2.1.chr14.g00261.t01#2       5469   125255849:   mNeoVis.2.1.1.chr4_hap1.g00611.t01#5    mNeoVis.evergladensis.1.1.chr6.g00622.t01#3    mNeoVis.evergladensis.2.1.chr6.g00620.t01#2    mNeoVis.vulgivaga.2.1.chr6.g00617.t01#1   mNeoVis.vulgivaga.1.1.chr6.g00619.t01#4       5470   125255850:   mNeoVis.vulgivaga.1.1.chr6.g00620.t01#4    mNeoVis.3.1.1.chr6.g00627.t01#6    mNeoVis.2.1.1.chr4_hap1.g00612.t01#5    mNeoVis.vulgivaga.2.1.chr6.g00618.t01#1    mNeoVis.evergladensis.2.1.chr6.g00621.t01#2   mNeoVis.evergladensis.1.1.chr6.g00623.t01#3       5471   125255851:   mNeoVis.3.1.1.chr14.g00264.t01#6    mNeoVis.2.1.1.chr14_hap1.g00262.t01#5    mNeoVis.vulgivaga.2.1.chr14.g00263.t01#1    mNeoVis.evergladensis.2.1.chr14.g00262.t01#2    mNeoVis.vulgivaga.1.1.chr14.g00259.t01#4   mNeoVis.evergladensis.1.1.chr14.g00268.t01#3       5472   125255852:   mNeoVis.2.1.1.chr14_hap1.g00263.t01#5    mNeoVis.vulgivaga.2.1.chr14.g00264.t01#1    mNeoVis.vulgivaga.1.1.chr14.g00260.t01#4    mNeoVis.evergladensis.2.1.chr14.g00263.t01#2    mNeoVis.3.1.1.chr14.g00265.t01#6   mNeoVis.evergladensis.1.1.chr14.g00269.t01#3       5473   125255853:   mNeoVis.2.1.1.chr14_hap1.g00264.t01#5    mNeoVis.vulgivaga.2.1.chr14.g00265.t01#1    mNeoVis.evergladensis.2.1.chr14.g00264.t01#2    mNeoVis.evergladensis.1.1.chr14.g00270.t01#3   mNeoVis.vulgivaga.1.1.chr14.g00261.t01#4       5474   125255854:   mNeoVis.vulgivaga.2.1.chr14.g00266.t01#1    mNeoVis.2.1.1.chr14_hap1.g00266.t01#5    mNeoVis.evergladensis.2.1.chr14.g00265.t01#2    mNeoVis.3.1.1.chr14.g00266.t01#6    mNeoVis.vulgivaga.1.1.chr14.g00262.t01#4   mNeoVis.evergladensis.1.1.chr14.g00271.t01#3       5475   125255855:   mNeoVis.vulgivaga.1.1.chr14.g00263.t01#4    mNeoVis.2.1.1.chr14_hap1.g00265.t01#5    mNeoVis.evergladensis.1.1.chr14.g00272.t01#3    mNeoVis.vulgivaga.2.1.chr14.g00267.t01#1   mNeoVis.evergladensis.2.1.chr14.g00266.t01#2       5476   125255856:   mNeoVis.vulgivaga.2.1.chr6.g00620.t01#1    mNeoVis.evergladensis.1.1.chr6.g00625.t01#3    mNeoVis.3.1.1.chr6.g00630.t01#6    mNeoVis.vulgivaga.1.1.chr6.g00622.t01#4    mNeoVis.evergladensis.2.1.chr6.g00623.t01#2   mNeoVis.2.1.1.chr4_hap1.g00614.t01#5       5477   125255857:   mNeoVis.2.1.1.chr4_hap1.g00615.t01#5    mNeoVis.evergladensis.1.1.chr6.g00626.t01#3    mNeoVis.vulgivaga.1.1.chr6.g00623.t01#4    mNeoVis.3.1.1.chr6.g00631.t01#6    mNeoVis.vulgivaga.2.1.chr6.g00621.t01#1   mNeoVis.evergladensis.2.1.chr6.g00624.t01#2       5478   125255858:   mNeoVis.vulgivaga.1.1.chr14.g00264.t01#4    mNeoVis.evergladensis.2.1.chr14.g00267.t01#2    mNeoVis.vulgivaga.2.1.chr14.g00268.t01#1    mNeoVis.2.1.1.chr14_hap1.g00267.t01#5    mNeoVis.evergladensis.1.1.chr14.g00273.t01#3   mNeoVis.3.1.1.chr14.g00268.t01#6       5479   125255859:   mNeoVis.vulgivaga.2.1.chr6.g00622.t01#1    mNeoVis.2.1.1.chr4_hap1.g00616.t01#5    mNeoVis.vulgivaga.1.1.chr6.g00624.t01#4    mNeoVis.evergladensis.2.1.chr6.g00625.t01#2   mNeoVis.evergladensis.1.1.chr6.g00627.t01#3       5480   125255860:   mNeoVis.3.1.1.chr14.g00269.t01#6    mNeoVis.2.1.1.chr14_hap1.g00268.t01#5    mNeoVis.evergladensis.2.1.chr14.g00268.t01#2    mNeoVis.evergladensis.1.1.chr14.g00274.t01#3    mNeoVis.vulgivaga.1.1.chr14.g00265.t01#4   mNeoVis.vulgivaga.2.1.chr14.g00269.t01#1       5481   125255861:   mNeoVis.3.1.1.chr5.g00368.t01#6    mNeoVis.evergladensis.2.1.chr5.g00381.t01#2    mNeoVis.vulgivaga.2.1.chr5.g00380.t01#1    mNeoVis.2.1.1.chr2_hap1.g00382.t01#5    mNeoVis.evergladensis.1.1.chr5.g00377.t01#3   mNeoVis.vulgivaga.1.1.chr5.g00368.t01#4       5482   125255862:   mNeoVis.2.1.1.chr4_hap1.g00618.t01#5    mNeoVis.evergladensis.2.1.chr6.g00627.t01#2    mNeoVis.vulgivaga.2.1.chr6.g00624.t01#1    mNeoVis.vulgivaga.1.1.chr6.g00626.t01#4   mNeoVis.evergladensis.1.1.chr6.g00629.t01#3       5483   125255863:  mNeoVis.3.1.1.chr6.g00633.t01#6               5484   125255864:   mNeoVis.2.1.1.chr4_hap1.g00619.t01#5    mNeoVis.evergladensis.1.1.chr6.g00630.t01#3    mNeoVis.vulgivaga.2.1.chr6.g00625.t01#1    mNeoVis.evergladensis.2.1.chr6.g00628.t01#2   mNeoVis.vulgivaga.1.1.chr6.g00627.t01#4       5485   125255865:  mNeoVis.vulgivaga.2.1.chr14.g00270.t01#1               5486   125255866:   mNeoVis.vulgivaga.2.1.chr14.g00271.t01#1    mNeoVis.evergladensis.2.1.chr14.g00270.t01#2    mNeoVis.2.1.1.chr14_hap1.g00270.t01#5    mNeoVis.evergladensis.1.1.chr14.g00276.t01#3   mNeoVis.3.1.1.chr14.g00271.t01#6       5487   125255867:   mNeoVis.3.1.1.chr8.g00749.t01#6    mNeoVis.2.1.1.chr8_hap1.g00799.t01#5    mNeoVis.vulgivaga.1.1.chr8.g00773.t01#4    mNeoVis.evergladensis.2.1.chr9.g00786.t01#2    mNeoVis.evergladensis.1.1.chr8.g00780.t01#3    mNeoVis.vulgivaga.2.1.chr8.g00792.t01#1   mNeoVis.evergladensis.1.1.chr8.g00781.t01#3     5488   125255868:   mNeoVis.evergladensis.2.1.chr6.g00629.t01#2    mNeoVis.evergladensis.1.1.chr6.g00631.t01#3    mNeoVis.vulgivaga.1.1.chr6.g00628.t01#4    mNeoVis.vulgivaga.2.1.chr6.g00626.t01#1    mNeoVis.3.1.1.chr6.g00634.t01#6   mNeoVis.2.1.1.chr4_hap1.g00620.t01#5       5489   125255869:   mNeoVis.evergladensis.1.1.chr5.g00437.t01#3    mNeoVis.evergladensis.2.1.chr5.g00442.t01#2    mNeoVis.vulgivaga.1.1.chr5.g00428.t01#4    mNeoVis.3.1.1.chr5.g00433.t01#6    mNeoVis.vulgivaga.2.1.chr5.g00442.t01#1   mNeoVis.2.1.1.chr2_hap1.g00443.t01#5       5490   125255870:   mNeoVis.evergladensis.1.1.chr5.g01399.t01#3    mNeoVis.evergladensis.2.1.chr5.g01395.t01#2    mNeoVis.2.1.1.chr2_hap1.g01457.t01#5    mNeoVis.vulgivaga.1.1.chr5.g01368.t01#4    mNeoVis.3.1.1.chr5.g01378.t01#6   mNeoVis.vulgivaga.2.1.chr5.g01395.t01#1       5491   125255871:   mNeoVis.evergladensis.1.1.chr4.g00566.t01#3    mNeoVis.vulgivaga.2.1.chr4.g00569.t01#1    mNeoVis.vulgivaga.2.1.chr6.g00627.t01#1    mNeoVis.3.1.1.chr4.g00561.t01#6    mNeoVis.evergladensis.2.1.chr6.g00630.t01#2    mNeoVis.3.1.1.chr6.g00635.t01#6    mNeoVis.evergladensis.1.1.chr6.g00632.t01#3    mNeoVis.evergladensis.2.1.chr4.g00559.t01#2    mNeoVis.2.1.1.chr4_hap1.g00621.t01#5      5492   125255872:   mNeoVis.vulgivaga.2.1.chr6.g00628.t01#1    mNeoVis.evergladensis.2.1.chr6.g00631.t01#2    mNeoVis.3.1.1.chr6.g00636.t01#6    mNeoVis.2.1.1.chr4_hap1.g00622.t01#5    mNeoVis.evergladensis.1.1.chr6.g00633.t01#3   mNeoVis.vulgivaga.1.1.chr6.g00630.t01#4       5493   125255873:   mNeoVis.vulgivaga.2.1.chr6.g00629.t01#1    mNeoVis.3.1.1.chr6.g00637.t01#6    mNeoVis.evergladensis.2.1.chr6.g00632.t01#2    mNeoVis.2.1.1.chr4_hap1.g00623.t01#5   mNeoVis.vulgivaga.1.1.chr6.g00631.t01#4       5494   125255874:   mNeoVis.3.1.1.chr14.g00274.t01#6    mNeoVis.vulgivaga.1.1.chr14.g00268.t01#4    mNeoVis.vulgivaga.2.1.chr14.g00272.t01#1   mNeoVis.evergladensis.1.1.chr14.g00277.t01#3       5495   125255875:   mNeoVis.evergladensis.2.1.chr14.g00271.t01#2    mNeoVis.2.1.1.chr14_hap1.g00272.t01#5   mNeoVis.3.1.1.chr14.g00275.t01#6           5496   125255876:  mNeoVis.vulgivaga.2.1.chr14.g00274.t01#1               5497   125255877:   mNeoVis.vulgivaga.1.1.chr14.g00269.t01#4    mNeoVis.3.1.1.chr14.g00272.t01#6    mNeoVis.3.1.1.chr14.g00273.t01#6    mNeoVis.2.1.1.chr14_hap1.g00271.t01#5    mNeoVis.evergladensis.2.1.chr14.g00272.t01#2    mNeoVis.evergladensis.1.1.chr14.g00278.t01#3   mNeoVis.vulgivaga.2.1.chr14.g00273.t01#1     5498   125255878:   mNeoVis.evergladensis.2.1.chr6.g00633.t01#2    mNeoVis.vulgivaga.2.1.chr6.g00630.t01#1    mNeoVis.2.1.1.chr4_hap1.g00624.t01#5    mNeoVis.3.1.1.chr6.g00638.t01#6    mNeoVis.evergladensis.1.1.chr6.g00634.t01#3   mNeoVis.vulgivaga.1.1.chr6.g00632.t01#4       5499   125255879:   mNeoVis.2.1.1.chr4_hap1.g00625.t01#5    mNeoVis.evergladensis.2.1.chr6.g00634.t01#2    mNeoVis.vulgivaga.2.1.chr6.g00631.t01#1    mNeoVis.vulgivaga.1.1.chr6.g00633.t01#4   mNeoVis.evergladensis.1.1.chr6.g00635.t01#3       5500   125255880:   mNeoVis.3.1.1.chr2.g00374.t01#6    mNeoVis.vulgivaga.1.1.chr14.g00672.t01#4    mNeoVis.2.1.1.chr5_hap1.g00451.t01#5    mNeoVis.evergladensis.1.1.chr2.g00442.t01#3    mNeoVis.evergladensis.2.1.chr2.g00445.t01#2    mNeoVis.vulgivaga.2.1.chr14.g00680.t01#1    mNeoVis.evergladensis.2.1.chr14.g00680.t01#2    mNeoVis.vulgivaga.1.1.chr2.g00443.t01#4    mNeoVis.2.1.1.chr14_hap1.g00682.t01#5      5501   125255881:   mNeoVis.2.1.1.chr10_hap1.g01115.t01#5    mNeoVis.vulgivaga.2.1.chr9.g01070.t01#1    mNeoVis.evergladensis.2.1.chr8.g01090.t01#2    mNeoVis.evergladensis.1.1.chr9.g01080.t01#3    mNeoVis.3.1.1.chr9.g01042.t01#6    mNeoVis.3.1.1.chr9.g00907.t01#6   mNeoVis.vulgivaga.1.1.chr9.g01083.t01#4     5502   125255882:   mNeoVis.vulgivaga.1.1.chr14.g00271.t01#4    mNeoVis.3.1.1.chr14.g00276.t01#6    mNeoVis.evergladensis.2.1.chr14.g00273.t01#2    mNeoVis.2.1.1.chr14_hap1.g00273.t01#5    mNeoVis.evergladensis.1.1.chr14.g00279.t01#3   mNeoVis.vulgivaga.2.1.chr14.g00275.t01#1       5503   125255883:   mNeoVis.vulgivaga.2.1.chr9.g00933.t01#1    mNeoVis.evergladensis.1.1.chr9.g00943.t01#3    mNeoVis.evergladensis.2.1.chr8.g00947.t01#2    mNeoVis.vulgivaga.1.1.chr9.g00947.t01#4   mNeoVis.2.1.1.chr10_hap1.g00977.t01#5       5504   125255884:   mNeoVis.vulgivaga.2.1.chr13.g00482.t01#1    mNeoVis.evergladensis.1.1.chr13.g00480.t01#3    mNeoVis.2.1.1.chr13_hap1.g00500.t01#5    mNeoVis.3.1.1.chr13.g00435.t01#6   mNeoVis.vulgivaga.1.1.chr13.g00475.t01#4       5505   125255885:  mNeoVis.vulgivaga.2.1.chr2.g00447.t01#1               5506   125255886:   mNeoVis.vulgivaga.2.1.chr14.g00276.t01#1    mNeoVis.evergladensis.2.1.chr14.g00275.t01#2    mNeoVis.vulgivaga.1.1.chr14.g00272.t01#4    mNeoVis.evergladensis.1.1.chr14.g00281.t01#3    mNeoVis.3.1.1.chr14.g00277.t01#6   mNeoVis.2.1.1.chr14_hap1.g00274.t01#5       5507   125255887:   mNeoVis.3.1.1.chr6.g00639.t01#6    mNeoVis.evergladensis.1.1.chr6.g00636.t01#3    mNeoVis.vulgivaga.2.1.chr6.g00632.t01#1    mNeoVis.2.1.1.chr4_hap1.g00626.t01#5    mNeoVis.vulgivaga.1.1.chr6.g00634.t01#4   mNeoVis.evergladensis.2.1.chr6.g00635.t01#2       5508   125255888:   mNeoVis.vulgivaga.1.1.chr6.g00635.t01#4    mNeoVis.vulgivaga.2.1.chr6.g00633.t01#1    mNeoVis.2.1.1.chr4_hap1.g00627.t01#5    mNeoVis.evergladensis.1.1.chr6.g00637.t01#3   mNeoVis.evergladensis.2.1.chr6.g00636.t01#2       5509   125255889:   mNeoVis.evergladensis.2.1.chr6.g00637.t01#2    mNeoVis.evergladensis.1.1.chr6.g00638.t01#3    mNeoVis.3.1.1.chr6.g00640.t01#6    mNeoVis.2.1.1.chr4_hap1.g00628.t01#5    mNeoVis.vulgivaga.2.1.chr6.g00634.t01#1   mNeoVis.vulgivaga.1.1.chr6.g00636.t01#4       5510   125255890:   mNeoVis.vulgivaga.2.1.chr14.g00277.t01#1    mNeoVis.evergladensis.1.1.chr14.g00282.t01#3    mNeoVis.2.1.1.chr14_hap1.g00275.t01#5    mNeoVis.evergladensis.2.1.chr14.g00278.t01#2    mNeoVis.vulgivaga.1.1.chr14.g00273.t01#4   mNeoVis.3.1.1.chr14.g00278.t01#6       5511   125255891:   mNeoVis.vulgivaga.2.1.chr6.g00635.t01#1    mNeoVis.vulgivaga.1.1.chr6.g00637.t01#4    mNeoVis.evergladensis.2.1.chr6.g00638.t01#2    mNeoVis.evergladensis.1.1.chr6.g00639.t01#3   mNeoVis.2.1.1.chr4_hap1.g00629.t01#5       5512   125255892:   mNeoVis.2.1.1.chr14_hap1.g00277.t01#5    mNeoVis.vulgivaga.2.1.chr14.g00278.t01#1    mNeoVis.vulgivaga.1.1.chr14.g00275.t01#4    mNeoVis.evergladensis.2.1.chr14.g00280.t01#2   mNeoVis.evergladensis.1.1.chr14.g00284.t01#3       5513   125255893:   mNeoVis.2.1.1.chr14_hap1.g00278.t01#5    mNeoVis.evergladensis.1.1.chr14.g00285.t01#3    mNeoVis.vulgivaga.2.1.chr14.g00279.t01#1    mNeoVis.vulgivaga.1.1.chr14.g00276.t01#4   mNeoVis.evergladensis.2.1.chr14.g00281.t01#2       5514   125255894:  mNeoVis.3.1.1.chr14.g00280.t01#6               5515   125255895:   mNeoVis.2.1.1.chr4_hap1.g00630.t01#5    mNeoVis.evergladensis.2.1.chr6.g00639.t01#2    mNeoVis.evergladensis.1.1.chr6.g00640.t01#3    mNeoVis.vulgivaga.2.1.chr6.g00636.t01#1    mNeoVis.vulgivaga.1.1.chr6.g00638.t01#4   mNeoVis.3.1.1.chr6.g00641.t01#6       5516   125255896:   mNeoVis.evergladensis.1.1.chr6.g00641.t01#3    mNeoVis.evergladensis.2.1.chr6.g00640.t01#2    mNeoVis.vulgivaga.2.1.chr6.g00637.t01#1    mNeoVis.vulgivaga.1.1.chr6.g00639.t01#4   mNeoVis.2.1.1.chr4_hap1.g00631.t01#5       5517   125255897:   mNeoVis.evergladensis.1.1.chr14.g00286.t01#3    mNeoVis.evergladensis.2.1.chr14.g00282.t01#2    mNeoVis.3.1.1.chr14.g00281.t01#6    mNeoVis.2.1.1.chr14_hap1.g00279.t01#5    mNeoVis.vulgivaga.2.1.chr14.g00280.t01#1   mNeoVis.vulgivaga.1.1.chr14.g00277.t01#4       5518   125255898:   mNeoVis.evergladensis.1.1.chr14.g00287.t01#3    mNeoVis.vulgivaga.2.1.chr14.g00281.t01#1    mNeoVis.2.1.1.chr14_hap1.g00280.t01#5    mNeoVis.3.1.1.chr14.g00282.t01#6    mNeoVis.evergladensis.2.1.chr14.g00283.t01#2   mNeoVis.vulgivaga.1.1.chr14.g00278.t01#4       5519   125255899:   mNeoVis.vulgivaga.2.1.chr6.g00638.t01#1    mNeoVis.evergladensis.2.1.chr6.g00641.t01#2   mNeoVis.2.1.1.chr4_hap1.g00632.t01#5           5520   125255900:   mNeoVis.evergladensis.1.1.chr6.g00643.t01#3    mNeoVis.evergladensis.2.1.chr6.g00643.t01#2    mNeoVis.vulgivaga.1.1.chr6.g00641.t01#4    mNeoVis.2.1.1.chr4_hap1.g00634.t01#5    mNeoVis.vulgivaga.2.1.chr6.g00639.t01#1   mNeoVis.3.1.1.chr6.g00644.t01#6       5521   125255901:   mNeoVis.vulgivaga.2.1.chr14.g00282.t01#1    mNeoVis.evergladensis.1.1.chr14.g00288.t01#3    mNeoVis.3.1.1.chr14.g00283.t01#6   mNeoVis.2.1.1.chr14_hap1.g00281.t01#5         5522   125255902:   mNeoVis.2.1.1.chr4_hap1.g00635.t01#5    mNeoVis.evergladensis.2.1.chr6.g00644.t01#2    mNeoVis.vulgivaga.1.1.chr6.g00642.t01#4    mNeoVis.vulgivaga.2.1.chr6.g00640.t01#1   mNeoVis.evergladensis.1.1.chr6.g00644.t01#3       5523   125255903:   mNeoVis.vulgivaga.2.1.chr14.g00283.t01#1    mNeoVis.evergladensis.1.1.chr14.g00289.t01#3    mNeoVis.evergladensis.2.1.chr14.g00284.t01#2    mNeoVis.3.1.1.chr14.g00284.t01#6    mNeoVis.2.1.1.chr14_hap1.g00282.t01#5   mNeoVis.vulgivaga.1.1.chr14.g00279.t01#4       5524   125255904:   mNeoVis.evergladensis.2.1.chr6.g00645.t01#2    mNeoVis.vulgivaga.1.1.chr6.g00643.t01#4    mNeoVis.vulgivaga.2.1.chr6.g00641.t01#1    mNeoVis.evergladensis.1.1.chr6.g00645.t01#3   mNeoVis.2.1.1.chr4_hap1.g00636.t01#5       5525   125255905:   mNeoVis.evergladensis.1.1.chr14.g00290.t01#3    mNeoVis.3.1.1.chr14.g00285.t01#6    mNeoVis.evergladensis.2.1.chr14.g00285.t01#2    mNeoVis.2.1.1.chr14_hap1.g00283.t01#5    mNeoVis.vulgivaga.2.1.chr14.g00284.t01#1   mNeoVis.vulgivaga.1.1.chr14.g00280.t01#4       5526   125255906:   mNeoVis.evergladensis.1.1.chr14.g00291.t01#3    mNeoVis.2.1.1.chr14_hap1.g00284.t01#5    mNeoVis.3.1.1.chr14.g00286.t01#6    mNeoVis.vulgivaga.1.1.chr14.g00281.t01#4    mNeoVis.evergladensis.2.1.chr14.g00286.t01#2   mNeoVis.vulgivaga.2.1.chr14.g00285.t01#1       5527   125255907:   mNeoVis.vulgivaga.1.1.chr14.g00282.t01#4    mNeoVis.evergladensis.2.1.chr14.g00287.t01#2    mNeoVis.vulgivaga.2.1.chr14.g00286.t01#1    mNeoVis.2.1.1.chr14_hap1.g00285.t01#5    mNeoVis.3.1.1.chr14.g00287.t01#6   mNeoVis.evergladensis.1.1.chr14.g00292.t01#3       5528   125255908:   mNeoVis.3.1.1.chr6.g00647.t01#6    mNeoVis.vulgivaga.2.1.chr6.g00642.t01#1    mNeoVis.vulgivaga.1.1.chr6.g00644.t01#4    mNeoVis.evergladensis.2.1.chr6.g00646.t01#2    mNeoVis.2.1.1.chr4_hap1.g00637.t01#5   mNeoVis.evergladensis.1.1.chr6.g00646.t01#3       5529   125255909:   mNeoVis.2.1.1.chr14_hap1.g00286.t01#5    mNeoVis.vulgivaga.2.1.chr14.g00287.t01#1    mNeoVis.vulgivaga.1.1.chr14.g00283.t01#4    mNeoVis.3.1.1.chr14.g00288.t01#6    mNeoVis.evergladensis.1.1.chr14.g00293.t01#3   mNeoVis.evergladensis.2.1.chr14.g00288.t01#2       5530   125255910:   mNeoVis.3.1.1.chr14.g00289.t01#6    mNeoVis.vulgivaga.1.1.chr14.g00284.t01#4    mNeoVis.evergladensis.1.1.chr14.g00294.t01#3    mNeoVis.evergladensis.2.1.chr14.g00289.t01#2    mNeoVis.2.1.1.chr14_hap1.g00287.t01#5   mNeoVis.vulgivaga.2.1.chr14.g00288.t01#1       5531   125255911:   mNeoVis.vulgivaga.1.1.chr14.g00285.t01#4    mNeoVis.evergladensis.2.1.chr14.g00290.t01#2    mNeoVis.evergladensis.1.1.chr14.g00295.t01#3    mNeoVis.3.1.1.chr14.g00290.t01#6    mNeoVis.2.1.1.chr14_hap1.g00288.t01#5   mNeoVis.vulgivaga.2.1.chr14.g00289.t01#1       5532   125255912:   mNeoVis.evergladensis.1.1.chr14.g00296.t01#3    mNeoVis.evergladensis.2.1.chr14.g00291.t01#2    mNeoVis.2.1.1.chr14_hap1.g00289.t01#5    mNeoVis.vulgivaga.2.1.chr14.g00290.t01#1    mNeoVis.3.1.1.chr14.g00291.t01#6   mNeoVis.vulgivaga.1.1.chr14.g00286.t01#4       5533   125255913:   mNeoVis.3.1.1.chr14.g00292.t01#6    mNeoVis.2.1.1.chr14_hap1.g00290.t01#5    mNeoVis.evergladensis.2.1.chr14.g00292.t01#2    mNeoVis.evergladensis.1.1.chr14.g00297.t01#3    mNeoVis.vulgivaga.2.1.chr14.g00291.t01#1   mNeoVis.vulgivaga.1.1.chr14.g00287.t01#4       5534   125255914:   mNeoVis.evergladensis.1.1.chr2.g00685.t01#3    mNeoVis.2.1.1.chr5_hap1.g00689.t01#5    mNeoVis.vulgivaga.2.1.chr2.g00689.t01#1    mNeoVis.vulgivaga.1.1.chr2.g00680.t01#4   mNeoVis.evergladensis.2.1.chr2.g00686.t01#2       5535   125255915:   mNeoVis.evergladensis.2.1.chr14.g00293.t01#2    mNeoVis.2.1.1.chr14_hap1.g00291.t01#5    mNeoVis.vulgivaga.2.1.chr14.g00292.t01#1    mNeoVis.evergladensis.1.1.chr14.g00298.t01#3    mNeoVis.3.1.1.chr14.g00293.t01#6   mNeoVis.vulgivaga.1.1.chr14.g00288.t01#4       5536   125255916:   mNeoVis.2.1.1.chr4_hap1.g00638.t01#5   mNeoVis.vulgivaga.2.1.chr6.g00643.t01#1             5537   125255917:   mNeoVis.2.1.1.chr4_hap1.g00639.t01#5    mNeoVis.evergladensis.2.1.chr6.g00647.t01#2    mNeoVis.evergladensis.1.1.chr6.g00647.t01#3    mNeoVis.vulgivaga.2.1.chr6.g00644.t01#1    mNeoVis.3.1.1.chr6.g00648.t01#6   mNeoVis.vulgivaga.1.1.chr6.g00645.t01#4       5538   125255918:   mNeoVis.evergladensis.1.1.chr14.g00299.t01#3    mNeoVis.vulgivaga.2.1.chr14.g00293.t01#1    mNeoVis.vulgivaga.1.1.chr14.g00289.t01#4    mNeoVis.evergladensis.2.1.chr14.g00294.t01#2    mNeoVis.3.1.1.chr14.g00294.t01#6   mNeoVis.2.1.1.chr14_hap1.g00292.t01#5       5539   125255919:   mNeoVis.evergladensis.1.1.chr14.g00300.t01#3    mNeoVis.3.1.1.chr14.g00295.t01#6    mNeoVis.2.1.1.chr14_hap1.g00293.t01#5    mNeoVis.vulgivaga.2.1.chr14.g00294.t01#1    mNeoVis.evergladensis.2.1.chr14.g00295.t01#2   mNeoVis.vulgivaga.1.1.chr14.g00290.t01#4       5540   125255920:   mNeoVis.evergladensis.2.1.chr14.g00296.t01#2    mNeoVis.vulgivaga.2.1.chr14.g00295.t01#1    mNeoVis.vulgivaga.1.1.chr14.g00291.t01#4    mNeoVis.evergladensis.1.1.chr14.g00301.t01#3    mNeoVis.3.1.1.chr14.g00296.t01#6   mNeoVis.2.1.1.chr14_hap1.g00294.t01#5       5541   125255921:   mNeoVis.vulgivaga.2.1.chr6.g00645.t01#1    mNeoVis.evergladensis.1.1.chr6.g00648.t01#3    mNeoVis.2.1.1.chr4_hap1.g00640.t01#5    mNeoVis.vulgivaga.1.1.chr6.g00646.t01#4    mNeoVis.evergladensis.2.1.chr6.g00648.t01#2   mNeoVis.3.1.1.chr6.g00649.t01#6       5542   125255922:   mNeoVis.vulgivaga.1.1.chr6.g00647.t01#4    mNeoVis.evergladensis.1.1.chr6.g00649.t01#3    mNeoVis.2.1.1.chr4_hap1.g00641.t01#5    mNeoVis.vulgivaga.2.1.chr6.g00646.t01#1    mNeoVis.3.1.1.chr6.g00651.t01#6   mNeoVis.evergladensis.2.1.chr6.g00649.t01#2       5543   125255923:  mNeoVis.vulgivaga.2.1.chr6.g00647.t01#1               5544   125255924:   mNeoVis.vulgivaga.1.1.chr6.g00649.t01#4    mNeoVis.evergladensis.2.1.chr6.g00651.t01#2   mNeoVis.vulgivaga.2.1.chr6.g00648.t01#1           5545   125255925:   mNeoVis.evergladensis.1.1.chr14.g00302.t01#3    mNeoVis.evergladensis.2.1.chr14.g00297.t01#2    mNeoVis.vulgivaga.1.1.chr14.g00292.t01#4    mNeoVis.vulgivaga.2.1.chr14.g00296.t01#1    mNeoVis.2.1.1.chr14_hap1.g00295.t01#5   mNeoVis.3.1.1.chr14.g00297.t01#6       5546   125255926:   mNeoVis.vulgivaga.1.1.chr14.g00293.t01#4    mNeoVis.2.1.1.chr14_hap1.g00296.t01#5    mNeoVis.3.1.1.chr14.g00298.t01#6    mNeoVis.vulgivaga.2.1.chr14.g00297.t01#1    mNeoVis.evergladensis.2.1.chr14.g00298.t01#2   mNeoVis.evergladensis.1.1.chr14.g00303.t01#3       5547   125255927:   mNeoVis.2.1.1.chr14_hap1.g00297.t01#5    mNeoVis.evergladensis.1.1.chr14.g00304.t01#3    mNeoVis.evergladensis.2.1.chr14.g00299.t01#2    mNeoVis.vulgivaga.1.1.chr14.g00294.t01#4    mNeoVis.vulgivaga.2.1.chr14.g00298.t01#1   mNeoVis.3.1.1.chr14.g00299.t01#6       5548   125255928:   mNeoVis.evergladensis.2.1.chr14.g00300.t01#2    mNeoVis.evergladensis.1.1.chr14.g00305.t01#3    mNeoVis.vulgivaga.1.1.chr14.g00295.t01#4    mNeoVis.2.1.1.chr14_hap1.g00298.t01#5    mNeoVis.3.1.1.chr14.g00300.t01#6   mNeoVis.vulgivaga.2.1.chr14.g00299.t01#1       5549   125255929:   mNeoVis.evergladensis.1.1.chr14.g00306.t01#3    mNeoVis.evergladensis.2.1.chr14.g00301.t01#2    mNeoVis.vulgivaga.1.1.chr14.g00296.t01#4    mNeoVis.2.1.1.chr14_hap1.g00299.t01#5   mNeoVis.vulgivaga.2.1.chr14.g00300.t01#1       5550   125255930:   mNeoVis.2.1.1.chr4_hap1.g00645.t01#5    mNeoVis.vulgivaga.2.1.chr6.g00649.t01#1    mNeoVis.vulgivaga.1.1.chr6.g00650.t01#4    mNeoVis.evergladensis.2.1.chr6.g00652.t01#2    mNeoVis.evergladensis.1.1.chr6.g00655.t01#3   mNeoVis.3.1.1.chr6.g00657.t01#6       5551   125255931:   mNeoVis.vulgivaga.2.1.chr6.g00650.t01#1    mNeoVis.vulgivaga.1.1.chr6.g00651.t01#4    mNeoVis.evergladensis.1.1.chr6.g00656.t01#3    mNeoVis.3.1.1.chr6.g00658.t01#6    mNeoVis.evergladensis.2.1.chr6.g00653.t01#2   mNeoVis.2.1.1.chr4_hap1.g00646.t01#5       5552   125255932:   mNeoVis.vulgivaga.1.1.chr14.g00297.t01#4    mNeoVis.2.1.1.chr14_hap1.g00300.t01#5   mNeoVis.vulgivaga.2.1.chr14.g00301.t01#1         5553   125255933:   mNeoVis.3.1.1.chr7.g00388.t01#6    mNeoVis.2.1.1.chr7_hap1.g00402.t01#5    mNeoVis.vulgivaga.1.1.chr7.g00413.t01#4    mNeoVis.evergladensis.1.1.chr7.g00413.t01#3    mNeoVis.vulgivaga.2.1.chr7.g00406.t01#1   mNeoVis.evergladensis.2.1.chr7.g00406.t01#2       5554   125255934:   mNeoVis.vulgivaga.2.1.chr14.g00216.t01#1    mNeoVis.evergladensis.1.1.chr14.g00221.t01#3    mNeoVis.2.1.1.chr14_hap1.g00213.t01#5    mNeoVis.vulgivaga.1.1.chr14.g00211.t01#4    mNeoVis.evergladensis.2.1.chr14.g00214.t01#2   mNeoVis.3.1.1.chr14.g00212.t01#6       5555   125255935:   mNeoVis.evergladensis.2.1.chr6.g00573.t01#2    mNeoVis.3.1.1.chr6.g00577.t01#6    mNeoVis.2.1.1.chr4_hap1.g00566.t01#5    mNeoVis.vulgivaga.2.1.chr6.g00573.t01#1    mNeoVis.vulgivaga.1.1.chr6.g00573.t01#4    mNeoVis.vulgivaga.2.1.chr6.g00572.t01#1   mNeoVis.evergladensis.1.1.chr6.g00572.t01#3     5556   125255936:   mNeoVis.vulgivaga.2.1.chr14.g00217.t01#1    mNeoVis.evergladensis.1.1.chr14.g00222.t01#3    mNeoVis.2.1.1.chr14_hap1.g00214.t01#5    mNeoVis.3.1.1.chr14.g00213.t01#6    mNeoVis.evergladensis.2.1.chr14.g00215.t01#2   mNeoVis.vulgivaga.1.1.chr14.g00212.t01#4       5557   125255937:   mNeoVis.3.1.1.chr14.g00214.t01#6    mNeoVis.evergladensis.2.1.chr14.g00216.t01#2    mNeoVis.vulgivaga.1.1.chr14.g00213.t01#4    mNeoVis.evergladensis.1.1.chr14.g00223.t01#3    mNeoVis.vulgivaga.2.1.chr14.g00218.t01#1   mNeoVis.2.1.1.chr14_hap1.g00215.t01#5       5558   125255938:   mNeoVis.evergladensis.2.1.chr6.g00574.t01#2    mNeoVis.vulgivaga.1.1.chr6.g00574.t01#4    mNeoVis.3.1.1.chr6.g00578.t01#6    mNeoVis.evergladensis.1.1.chr6.g00573.t01#3    mNeoVis.2.1.1.chr4_hap1.g00567.t01#5   mNeoVis.vulgivaga.2.1.chr6.g00574.t01#1       5559   125255939:   mNeoVis.evergladensis.2.1.chr14.g00217.t01#2    mNeoVis.evergladensis.1.1.chr14.g00224.t01#3    mNeoVis.2.1.1.chr14_hap1.g00216.t01#5    mNeoVis.vulgivaga.1.1.chr14.g00214.t01#4    mNeoVis.vulgivaga.2.1.chr14.g00219.t01#1   mNeoVis.3.1.1.chr14.g00215.t01#6       5560   125255940:   mNeoVis.3.1.1.chr6.g00579.t01#6    mNeoVis.vulgivaga.1.1.chr6.g00575.t01#4    mNeoVis.vulgivaga.2.1.chr6.g00575.t01#1    mNeoVis.evergladensis.1.1.chr6.g00574.t01#3    mNeoVis.2.1.1.chr4_hap1.g00568.t01#5   mNeoVis.evergladensis.2.1.chr6.g00575.t01#2       5561   125255941:   mNeoVis.3.1.1.chr7.g00820.t01#6    mNeoVis.evergladensis.2.1.chr7.g00873.t01#2    mNeoVis.evergladensis.1.1.chr7.g00878.t01#3   mNeoVis.vulgivaga.1.1.chr7.g00879.t01#4         5562   125255942:   mNeoVis.vulgivaga.2.1.chr6.g00576.t01#1    mNeoVis.3.1.1.chr6.g00581.t01#6    mNeoVis.2.1.1.chr4_hap1.g00569.t01#5    mNeoVis.evergladensis.2.1.chr6.g00576.t01#2    mNeoVis.vulgivaga.1.1.chr6.g00576.t01#4    mNeoVis.evergladensis.1.1.chr6.g00575.t01#3   mNeoVis.3.1.1.chr6.g00580.t01#6       5563   125255943:   mNeoVis.3.1.1.chr14.g00216.t01#6    mNeoVis.2.1.1.chr14_hap1.g00217.t01#5    mNeoVis.vulgivaga.2.1.chr14.g00220.t01#1    mNeoVis.evergladensis.2.1.chr14.g00218.t01#2    mNeoVis.vulgivaga.1.1.chr14.g00215.t01#4   mNeoVis.evergladensis.1.1.chr14.g00225.t01#3       5564   125255944:   mNeoVis.vulgivaga.1.1.chr14.g00216.t01#4    mNeoVis.evergladensis.1.1.chr14.g00226.t01#3    mNeoVis.3.1.1.chr14.g00217.t01#6    mNeoVis.evergladensis.2.1.chr14.g00219.t01#2    mNeoVis.2.1.1.chr14_hap1.g00218.t01#5   mNeoVis.vulgivaga.2.1.chr14.g00221.t01#1       5565   125255945:   mNeoVis.evergladensis.1.1.chr14.g00227.t01#3    mNeoVis.2.1.1.chr14_hap1.g00219.t01#5    mNeoVis.3.1.1.chr14.g00218.t01#6    mNeoVis.evergladensis.2.1.chr14.g00220.t01#2    mNeoVis.vulgivaga.2.1.chr14.g00222.t01#1   mNeoVis.vulgivaga.1.1.chr14.g00217.t01#4       5566   125255946:   mNeoVis.vulgivaga.1.1.chr14.g00218.t01#4    mNeoVis.evergladensis.2.1.chr14.g00221.t01#2    mNeoVis.evergladensis.1.1.chr14.g00228.t01#3    mNeoVis.2.1.1.chr14_hap1.g00220.t01#5   mNeoVis.vulgivaga.2.1.chr14.g00223.t01#1       5567   125255947:   mNeoVis.evergladensis.2.1.chr6.g00577.t01#2    mNeoVis.evergladensis.1.1.chr6.g00576.t01#3    mNeoVis.vulgivaga.1.1.chr6.g00577.t01#4    mNeoVis.2.1.1.chr4_hap1.g00570.t01#5    mNeoVis.vulgivaga.2.1.chr6.g00577.t01#1   mNeoVis.3.1.1.chr6.g00583.t01#6       5568   125255948:   mNeoVis.3.1.1.chr6.g00584.t01#6    mNeoVis.vulgivaga.2.1.chr6.g00578.t01#1    mNeoVis.vulgivaga.1.1.chr6.g00578.t01#4    mNeoVis.evergladensis.2.1.chr6.g00578.t01#2    mNeoVis.evergladensis.1.1.chr6.g00577.t01#3    mNeoVis.3.1.1.chr6.g00585.t01#6   mNeoVis.2.1.1.chr4_hap1.g00571.t01#5     5569   125255949:   mNeoVis.3.1.1.chr6.g00586.t01#6    mNeoVis.evergladensis.2.1.chr6.g00579.t01#2    mNeoVis.evergladensis.1.1.chr6.g00579.t01#3    mNeoVis.vulgivaga.2.1.chr6.g00579.t01#1    mNeoVis.2.1.1.chr4_hap1.g00574.t01#5   mNeoVis.vulgivaga.1.1.chr6.g00581.t01#4       5570   125255950:   mNeoVis.vulgivaga.2.1.chr14.g00224.t01#1    mNeoVis.evergladensis.2.1.chr14.g00223.t01#2    mNeoVis.evergladensis.1.1.chr14.g00229.t01#3    mNeoVis.vulgivaga.1.1.chr14.g00220.t01#4    mNeoVis.3.1.1.chr14.g00223.t01#6   mNeoVis.2.1.1.chr14_hap1.g00221.t01#5       5571   125255951:   mNeoVis.vulgivaga.2.1.chr14.g00225.t01#1    mNeoVis.2.1.1.chr14_hap1.g00222.t01#5    mNeoVis.evergladensis.2.1.chr14.g00224.t01#2    mNeoVis.evergladensis.1.1.chr14.g00230.t01#3   mNeoVis.vulgivaga.1.1.chr14.g00221.t01#4       5572   125255952:   mNeoVis.3.1.1.chr6.g00587.t01#6   mNeoVis.vulgivaga.2.1.chr6.g00580.t01#1             5573   125255953:   mNeoVis.evergladensis.1.1.chr6.g00580.t01#3    mNeoVis.evergladensis.2.1.chr6.g00581.t01#2    mNeoVis.vulgivaga.1.1.chr6.g00582.t01#4    mNeoVis.evergladensis.2.1.chr6.g00580.t01#2    mNeoVis.vulgivaga.2.1.chr6.g00581.t01#1   mNeoVis.2.1.1.chr4_hap1.g00576.t01#5       5574   125255954:   mNeoVis.vulgivaga.2.1.chr6.g00582.t01#1    mNeoVis.2.1.1.chr4_hap1.g00577.t01#5    mNeoVis.evergladensis.1.1.chr6.g00582.t01#3    mNeoVis.vulgivaga.1.1.chr6.g00583.t01#4    mNeoVis.3.1.1.chr6.g00588.t01#6   mNeoVis.evergladensis.2.1.chr6.g00582.t01#2       5575   125255955:   mNeoVis.3.1.1.chr6.g00589.t01#6    mNeoVis.vulgivaga.1.1.chr6.g00586.t01#4    mNeoVis.2.1.1.chr4_hap1.g00578.t01#5    mNeoVis.vulgivaga.2.1.chr6.g00583.t01#1    mNeoVis.evergladensis.2.1.chr6.g00583.t01#2   mNeoVis.evergladensis.1.1.chr6.g00584.t01#3       5576   125255956:   mNeoVis.vulgivaga.1.1.chr14.g00222.t01#4    mNeoVis.2.1.1.chr14_hap1.g00223.t01#5    mNeoVis.evergladensis.1.1.chr14.g00231.t01#3    mNeoVis.3.1.1.chr14.g00225.t01#6    mNeoVis.vulgivaga.2.1.chr14.g00226.t01#1   mNeoVis.evergladensis.2.1.chr14.g00225.t01#2       5577   125255957:   mNeoVis.evergladensis.2.1.chr14.g00226.t01#2    mNeoVis.vulgivaga.2.1.chr14.g00227.t01#1    mNeoVis.vulgivaga.1.1.chr14.g00223.t01#4    mNeoVis.2.1.1.chr14_hap1.g00224.t01#5    mNeoVis.evergladensis.1.1.chr14.g00232.t01#3   mNeoVis.3.1.1.chr14.g00226.t01#6       5578   125255958:   mNeoVis.2.1.1.chr4_hap1.g00579.t01#5    mNeoVis.vulgivaga.2.1.chr6.g00584.t01#1    mNeoVis.evergladensis.1.1.chr6.g00585.t01#3    mNeoVis.3.1.1.chr6.g00590.t01#6    mNeoVis.vulgivaga.1.1.chr6.g00587.t01#4   mNeoVis.evergladensis.2.1.chr6.g00584.t01#2       5579   125255959:   mNeoVis.vulgivaga.2.1.chr14.g00228.t01#1    mNeoVis.evergladensis.2.1.chr14.g00227.t01#2    mNeoVis.evergladensis.1.1.chr14.g00233.t01#3    mNeoVis.vulgivaga.1.1.chr14.g00224.t01#4    mNeoVis.2.1.1.chr14_hap1.g00225.t01#5   mNeoVis.3.1.1.chr14.g00227.t01#6       5580   125255960:   mNeoVis.evergladensis.1.1.chr6.g00586.t01#3    mNeoVis.vulgivaga.1.1.chr6.g00588.t01#4    mNeoVis.3.1.1.chr6.g00591.t01#6    mNeoVis.evergladensis.2.1.chr6.g00585.t01#2    mNeoVis.2.1.1.chr4_hap1.g00580.t01#5   mNeoVis.vulgivaga.2.1.chr6.g00585.t01#1       5581   125255961:   mNeoVis.evergladensis.1.1.chr14.g00234.t01#3    mNeoVis.vulgivaga.2.1.chr14.g00229.t01#1    mNeoVis.evergladensis.2.1.chr14.g00228.t01#2    mNeoVis.3.1.1.chr14.g00228.t01#6    mNeoVis.2.1.1.chr14_hap1.g00226.t01#5   mNeoVis.vulgivaga.1.1.chr14.g00225.t01#4       5582   125255962:   mNeoVis.2.1.1.chr14_hap1.g00227.t01#5    mNeoVis.evergladensis.2.1.chr14.g00229.t01#2    mNeoVis.vulgivaga.1.1.chr14.g00226.t01#4    mNeoVis.evergladensis.1.1.chr14.g00235.t01#3    mNeoVis.vulgivaga.2.1.chr14.g00230.t01#1   mNeoVis.3.1.1.chr14.g00229.t01#6       5583   125255963:   mNeoVis.vulgivaga.2.1.chr6.g00586.t01#1    mNeoVis.vulgivaga.1.1.chr6.g00589.t01#4    mNeoVis.evergladensis.1.1.chr6.g00587.t01#3    mNeoVis.3.1.1.chr6.g00592.t01#6    mNeoVis.2.1.1.chr4_hap1.g00581.t01#5   mNeoVis.evergladensis.2.1.chr6.g00586.t01#2       5584   125255964:   mNeoVis.3.1.1.chr6.g00593.t01#6    mNeoVis.vulgivaga.1.1.chr6.g00590.t01#4    mNeoVis.evergladensis.1.1.chr6.g00588.t01#3   mNeoVis.2.1.1.chr4_hap1.g00582.t01#5         5585   125255965:   mNeoVis.3.1.1.chr6.g00594.t01#6    mNeoVis.evergladensis.2.1.chr6.g00587.t01#2   mNeoVis.vulgivaga.2.1.chr6.g00587.t01#1           5586   125255966:   mNeoVis.vulgivaga.1.1.chr6.g00591.t01#4    mNeoVis.evergladensis.1.1.chr6.g00589.t01#3   mNeoVis.2.1.1.chr4_hap1.g00583.t01#5           5587   125255967:   mNeoVis.3.1.1.chr14.g00231.t01#6    mNeoVis.evergladensis.2.1.chr14.g00231.t01#2    mNeoVis.vulgivaga.2.1.chr14.g00232.t01#1    mNeoVis.evergladensis.1.1.chr14.g00237.t01#3    mNeoVis.2.1.1.chr14_hap1.g00229.t01#5   mNeoVis.vulgivaga.1.1.chr14.g00228.t01#4       5588   125255968:   mNeoVis.evergladensis.1.1.chr14.g00238.t01#3    mNeoVis.vulgivaga.2.1.chr14.g00233.t01#1    mNeoVis.2.1.1.chr14_hap1.g00230.t01#5    mNeoVis.evergladensis.2.1.chr14.g00232.t01#2    mNeoVis.3.1.1.chr14.g00232.t01#6   mNeoVis.vulgivaga.1.1.chr14.g00229.t01#4       5589   125255969:   mNeoVis.3.1.1.chr6.g00595.t01#6    mNeoVis.vulgivaga.2.1.chr6.g00588.t01#1    mNeoVis.evergladensis.2.1.chr6.g00588.t01#2    mNeoVis.vulgivaga.1.1.chr6.g00592.t01#4    mNeoVis.evergladensis.1.1.chr6.g00590.t01#3   mNeoVis.2.1.1.chr4_hap1.g00584.t01#5       5590   125255970:   mNeoVis.2.1.1.chr14_hap1.g00232.t01#5    mNeoVis.evergladensis.2.1.chr14.g00234.t01#2    mNeoVis.vulgivaga.2.1.chr14.g00235.t01#1   mNeoVis.evergladensis.1.1.chr14.g00240.t01#3       5591   125255971:   mNeoVis.evergladensis.2.1.chr14.g00235.t01#2    mNeoVis.3.1.1.chr14.g00235.t01#6    mNeoVis.vulgivaga.2.1.chr14.g00236.t01#1    mNeoVis.vulgivaga.1.1.chr14.g00231.t01#4    mNeoVis.evergladensis.1.1.chr14.g00241.t01#3   mNeoVis.2.1.1.chr14_hap1.g00233.t01#5       5592   125255972:   mNeoVis.evergladensis.1.1.chr7.g00414.t01#3    mNeoVis.vulgivaga.1.1.chr7.g00414.t01#4    mNeoVis.3.1.1.chr7.g00389.t01#6    mNeoVis.evergladensis.2.1.chr7.g00407.t01#2    mNeoVis.vulgivaga.2.1.chr7.g00407.t01#1   mNeoVis.2.1.1.chr7_hap1.g00403.t01#5       5593   125255973:   mNeoVis.vulgivaga.2.1.chr6.g00589.t01#1    mNeoVis.evergladensis.2.1.chr6.g00589.t01#2    mNeoVis.2.1.1.chr4_hap1.g00585.t01#5    mNeoVis.3.1.1.chr6.g00596.t01#6    mNeoVis.vulgivaga.1.1.chr6.g00593.t01#4   mNeoVis.evergladensis.1.1.chr6.g00591.t01#3       5594   125255974:   mNeoVis.vulgivaga.2.1.chr6.g00590.t01#1    mNeoVis.evergladensis.1.1.chr6.g00592.t01#3    mNeoVis.vulgivaga.1.1.chr6.g00594.t01#4    mNeoVis.evergladensis.2.1.chr6.g00590.t01#2    mNeoVis.3.1.1.chr6.g00597.t01#6   mNeoVis.2.1.1.chr4_hap1.g00586.t01#5       5595   125255975:   mNeoVis.3.1.1.chr14.g00236.t01#6    mNeoVis.vulgivaga.1.1.chr14.g00232.t01#4    mNeoVis.vulgivaga.2.1.chr14.g00237.t01#1    mNeoVis.evergladensis.2.1.chr14.g00236.t01#2    mNeoVis.evergladensis.1.1.chr14.g00242.t01#3   mNeoVis.2.1.1.chr14_hap1.g00234.t01#5       5596   125255976:   mNeoVis.2.1.1.chr14_hap1.g00235.t01#5    mNeoVis.vulgivaga.2.1.chr14.g00238.t01#1    mNeoVis.3.1.1.chr14.g00237.t01#6    mNeoVis.evergladensis.2.1.chr14.g00237.t01#2    mNeoVis.vulgivaga.1.1.chr14.g00233.t01#4   mNeoVis.evergladensis.1.1.chr14.g00243.t01#3       5597   125255977:   mNeoVis.2.1.1.chr4_hap1.g00587.t01#5    mNeoVis.vulgivaga.1.1.chr6.g00595.t01#4    mNeoVis.3.1.1.chr6.g00598.t01#6    mNeoVis.evergladensis.1.1.chr6.g00593.t01#3    mNeoVis.evergladensis.2.1.chr6.g00591.t01#2   mNeoVis.vulgivaga.2.1.chr6.g00591.t01#1       5598   125255978:   mNeoVis.evergladensis.2.1.chr14.g00238.t01#2    mNeoVis.vulgivaga.1.1.chr14.g00234.t01#4    mNeoVis.vulgivaga.2.1.chr14.g00239.t01#1    mNeoVis.3.1.1.chr14.g00238.t01#6   mNeoVis.2.1.1.chr14_hap1.g00236.t01#5       5599   125255979:   mNeoVis.3.1.1.chr15.g00422.t01#6    mNeoVis.vulgivaga.2.1.chr15.g00470.t01#1    mNeoVis.vulgivaga.1.1.chr15.g00464.t01#4    mNeoVis.evergladensis.1.1.chr15.g00151.t01#3    mNeoVis.evergladensis.2.1.chr15.g00463.t01#2   mNeoVis.2.1.1.chr15_hap1.g00641.t01#5       5600   125255980:   mNeoVis.3.1.1.chr6.g00599.t01#6    mNeoVis.2.1.1.chr4_hap1.g00588.t01#5    mNeoVis.vulgivaga.2.1.chr6.g00592.t01#1    mNeoVis.evergladensis.1.1.chr6.g00594.t01#3    mNeoVis.vulgivaga.1.1.chr6.g00596.t01#4   mNeoVis.evergladensis.2.1.chr6.g00592.t01#2       5601   125255981:   mNeoVis.vulgivaga.2.1.chr14.g00240.t01#1    mNeoVis.3.1.1.chr14.g00239.t01#6    mNeoVis.evergladensis.1.1.chr14.g00245.t01#3    mNeoVis.2.1.1.chr14_hap1.g00237.t01#5    mNeoVis.vulgivaga.1.1.chr14.g00235.t01#4   mNeoVis.evergladensis.2.1.chr14.g00239.t01#2       5602   125255982:   mNeoVis.evergladensis.1.1.chr14.g00246.t01#3    mNeoVis.2.1.1.chr14_hap1.g00238.t01#5    mNeoVis.vulgivaga.1.1.chr14.g00236.t01#4    mNeoVis.3.1.1.chr14.g00240.t01#6    mNeoVis.evergladensis.2.1.chr14.g00240.t01#2   mNeoVis.vulgivaga.2.1.chr14.g00241.t01#1       5603   125255983:   mNeoVis.evergladensis.1.1.chr6.g00595.t01#3    mNeoVis.2.1.1.chr4_hap1.g00589.t01#5    mNeoVis.vulgivaga.1.1.chr6.g00597.t01#4    mNeoVis.3.1.1.chr6.g00600.t01#6    mNeoVis.vulgivaga.2.1.chr6.g00593.t01#1   mNeoVis.evergladensis.2.1.chr6.g00593.t01#2       5604   125255984:   mNeoVis.3.1.1.chr6.g00601.t01#6    mNeoVis.vulgivaga.2.1.chr6.g00594.t01#1    mNeoVis.evergladensis.2.1.chr6.g00594.t01#2    mNeoVis.evergladensis.1.1.chr6.g00596.t01#3    mNeoVis.vulgivaga.1.1.chr6.g00598.t01#4   mNeoVis.2.1.1.chr4_hap1.g00590.t01#5       5605   125255985:   mNeoVis.evergladensis.1.1.chr7.g00415.t01#3    mNeoVis.vulgivaga.2.1.chr7.g00408.t01#1    mNeoVis.evergladensis.2.1.chr7.g00408.t01#2    mNeoVis.2.1.1.chr7_hap1.g00405.t01#5   mNeoVis.3.1.1.chr7.g00391.t01#6       5606   125255986:   mNeoVis.evergladensis.2.1.chr6.g00595.t01#2    mNeoVis.vulgivaga.1.1.chr6.g00599.t01#4    mNeoVis.3.1.1.chr6.g00603.t01#6    mNeoVis.2.1.1.chr4_hap1.g00591.t01#5    mNeoVis.evergladensis.1.1.chr6.g00597.t01#3   mNeoVis.vulgivaga.2.1.chr6.g00595.t01#1       5607   125255987:   mNeoVis.vulgivaga.2.1.chr6.g00596.t01#1    mNeoVis.evergladensis.1.1.chr6.g00598.t01#3    mNeoVis.evergladensis.2.1.chr6.g00596.t01#2    mNeoVis.3.1.1.chr6.g00604.t01#6    mNeoVis.vulgivaga.1.1.chr6.g00600.t01#4   mNeoVis.2.1.1.chr4_hap1.g00592.t01#5       5608   125255988:   mNeoVis.vulgivaga.1.1.chr6.g00601.t01#4    mNeoVis.3.1.1.chr6.g00605.t01#6    mNeoVis.2.1.1.chr4_hap1.g00593.t01#5    mNeoVis.vulgivaga.2.1.chr6.g00597.t01#1    mNeoVis.evergladensis.2.1.chr6.g00597.t01#2   mNeoVis.evergladensis.1.1.chr6.g00599.t01#3       5609   125255989:   mNeoVis.evergladensis.2.1.chr14.g00242.t01#2    mNeoVis.evergladensis.1.1.chr14.g00248.t01#3    mNeoVis.3.1.1.chr14.g00243.t01#6    mNeoVis.2.1.1.chr14_hap1.g00241.t01#5    mNeoVis.vulgivaga.2.1.chr14.g00243.t01#1   mNeoVis.vulgivaga.1.1.chr14.g00238.t01#4       5610   125255990:   mNeoVis.evergladensis.2.1.chr15.g00464.t01#2    mNeoVis.2.1.1.chr15_hap1.g00642.t01#5    mNeoVis.vulgivaga.1.1.chr15.g00465.t01#4    mNeoVis.vulgivaga.2.1.chr15.g00471.t01#1    mNeoVis.3.1.1.chr15.g00423.t01#6   mNeoVis.evergladensis.1.1.chr15.g00150.t01#3       5611   125255991:   mNeoVis.vulgivaga.1.1.chr7.g00415.t01#4    mNeoVis.2.1.1.chr7_hap1.g00406.t01#5    mNeoVis.3.1.1.chr7.g00392.t01#6    mNeoVis.evergladensis.1.1.chr7.g00416.t01#3    mNeoVis.vulgivaga.2.1.chr7.g00409.t01#1   mNeoVis.evergladensis.2.1.chr7.g00409.t01#2       5612   125255992:   mNeoVis.evergladensis.2.1.chr6.g00599.t01#2    mNeoVis.vulgivaga.2.1.chr6.g00599.t01#1    mNeoVis.3.1.1.chr6.g00607.t01#6    mNeoVis.2.1.1.chr4_hap1.g00595.t01#5    mNeoVis.vulgivaga.1.1.chr6.g00603.t01#4   mNeoVis.evergladensis.1.1.chr6.g00601.t01#3       5613   125255993:   mNeoVis.vulgivaga.2.1.chr6.g00598.t01#1    mNeoVis.3.1.1.chr6.g00606.t01#6    mNeoVis.2.1.1.chr4_hap1.g00594.t01#5    mNeoVis.evergladensis.1.1.chr6.g00600.t01#3    mNeoVis.vulgivaga.1.1.chr6.g00602.t01#4   mNeoVis.evergladensis.2.1.chr6.g00598.t01#2       5614   125255994:   mNeoVis.evergladensis.1.1.chr6.g00602.t01#3    mNeoVis.evergladensis.2.1.chr6.g00600.t01#2    mNeoVis.vulgivaga.1.1.chr6.g00604.t01#4    mNeoVis.vulgivaga.2.1.chr6.g00600.t01#1    mNeoVis.2.1.1.chr4_hap1.g00596.t01#5   mNeoVis.3.1.1.chr6.g00608.t01#6       5615   125255995:   mNeoVis.evergladensis.1.1.chr14.g00249.t01#3    mNeoVis.evergladensis.2.1.chr14.g00243.t01#2    mNeoVis.2.1.1.chr14_hap1.g00242.t01#5    mNeoVis.vulgivaga.2.1.chr14.g00244.t01#1    mNeoVis.3.1.1.chr14.g00244.t01#6   mNeoVis.vulgivaga.1.1.chr14.g00239.t01#4       5616   125255996:   mNeoVis.2.1.1.chr4_hap1.g00597.t01#5    mNeoVis.vulgivaga.2.1.chr6.g00601.t01#1    mNeoVis.3.1.1.chr6.g00609.t01#6   mNeoVis.evergladensis.2.1.chr6.g00601.t01#2       5617   125255997:   mNeoVis.2.1.1.chr2_hap1.g00633.t01#5    mNeoVis.2.1.1.chr9_hap1.g00309.t01#5    mNeoVis.2.1.1.chr2_hap1.g00628.t01#5    mNeoVis.evergladensis.2.1.chr10.g00306.t01#2    mNeoVis.vulgivaga.2.1.chr10.g00301.t01#1   mNeoVis.evergladensis.1.1.chr10.g00309.t01#3       5618   125255998:   mNeoVis.3.1.1.chr10.g00301.t01#6    mNeoVis.2.1.1.chr2_hap1.g00635.t01#5    mNeoVis.vulgivaga.1.1.chr10.g00297.t01#4   mNeoVis.3.1.1.chr10.g00300.t01#6         5619   125255999:   mNeoVis.vulgivaga.1.1.chr6.g00607.t01#4    mNeoVis.2.1.1.chr4_hap1.g00598.t01#5   mNeoVis.evergladensis.1.1.chr6.g00605.t01#3         5620   125256000:  mNeoVis.vulgivaga.2.1.chr6.g00602.t01#1               5621   125256001:  mNeoVis.3.1.1.chr6.g00614.t01#6               5622   125256002:  mNeoVis.evergladensis.2.1.chr6.g00603.t01#2             5623   125256003:   mNeoVis.vulgivaga.2.1.chr14.g00245.t01#1    mNeoVis.evergladensis.2.1.chr14.g00244.t01#2    mNeoVis.2.1.1.chr14_hap1.g00243.t01#5    mNeoVis.vulgivaga.1.1.chr14.g00240.t01#4    mNeoVis.evergladensis.1.1.chr14.g00250.t01#3   mNeoVis.3.1.1.chr14.g00245.t01#6       5624   125256004:   mNeoVis.evergladensis.2.1.chr6.g00604.t01#2    mNeoVis.2.1.1.chr4_hap1.g00599.t01#5    mNeoVis.evergladensis.1.1.chr6.g00606.t01#3    mNeoVis.vulgivaga.1.1.chr6.g00608.t01#4    mNeoVis.3.1.1.chr6.g00615.t01#6   mNeoVis.vulgivaga.2.1.chr6.g00603.t01#1       5625   125256005:   mNeoVis.evergladensis.2.1.chr5.g00631.t01#2    mNeoVis.3.1.1.chr5.g00626.t01#6    mNeoVis.evergladensis.1.1.chr5.g00624.t01#3    mNeoVis.vulgivaga.1.1.chr5.g00608.t01#4   mNeoVis.vulgivaga.2.1.chr5.g00625.t01#1       5626   125256006:  mNeoVis.2.1.1.chr2_hap1.g00630.t01#5               5627   125256007:   mNeoVis.evergladensis.1.1.chr7.g00417.t01#3    mNeoVis.vulgivaga.2.1.chr7.g00410.t01#1    mNeoVis.2.1.1.chr7_hap1.g00404.t01#5   mNeoVis.evergladensis.2.1.chr7.g00410.t01#2       5628   125256008:   mNeoVis.2.1.1.chr2_hap1.g00652.t01#5    mNeoVis.2.1.1.chr2_hap1.g00649.t01#5    mNeoVis.2.1.1.chr2_hap1.g00642.t01#5    mNeoVis.2.1.1.chr2_hap1.g00638.t01#5   mNeoVis.2.1.1.chr2_hap1.g00631.t01#5       5629   125256009:   mNeoVis.evergladensis.2.1.chr6.g00605.t01#2    mNeoVis.vulgivaga.1.1.chr6.g00609.t01#4    mNeoVis.2.1.1.chr4_hap1.g00600.t01#5   mNeoVis.vulgivaga.2.1.chr6.g00604.t01#1         5630   125256010:   mNeoVis.2.1.1.chr2_hap1.g00717.t01#5    mNeoVis.evergladensis.2.1.chr6.g00609.t01#2    mNeoVis.evergladensis.1.1.chr6.g00608.t01#3   mNeoVis.vulgivaga.2.1.chr6.g00607.t01#1         5631   125256011:   mNeoVis.evergladensis.1.1.chr5.g00626.t01#3    mNeoVis.2.1.1.chr2_hap1.g00718.t01#5    mNeoVis.2.1.1.chr2_hap1.g00648.t01#5   mNeoVis.2.1.1.chr2_hap1.g00641.t01#5         5632   125256012:   mNeoVis.2.1.1.chr2_hap1.g00636.t01#5    mNeoVis.2.1.1.chr2_hap1.g00647.t01#5   mNeoVis.2.1.1.chr2_hap1.g00640.t01#5           5633   125256013:   mNeoVis.2.1.1.chr2_hap1.g00637.t01#5    mNeoVis.vulgivaga.2.1.chr5.g00626.t01#1   mNeoVis.2.1.1.chr2_hap1.g00595.t01#5           5634   125256014:   mNeoVis.2.1.1.chr2_hap1.g00711.t01#5    mNeoVis.2.1.1.chr2_hap1.g00713.t01#5   mNeoVis.2.1.1.chr2_hap1.g00716.t01#5           5635   125256015:   mNeoVis.3.1.1.chr6.g00617.t01#6   mNeoVis.evergladensis.1.1.chr6.g00612.t01#3           5636   125256016:   mNeoVis.3.1.1.chr6.g00616.t01#6   mNeoVis.evergladensis.1.1.chr6.g00607.t01#3           5637   125256017:   mNeoVis.vulgivaga.2.1.chr6.g00606.t01#1   mNeoVis.evergladensis.2.1.chr6.g00608.t01#2           5638   125256018:   mNeoVis.2.1.1.chr2_hap1.g00712.t01#5   mNeoVis.2.1.1.chr2_hap1.g00639.t01#5             5639   125256019:   mNeoVis.vulgivaga.2.1.chr14.g00246.t01#1    mNeoVis.evergladensis.2.1.chr14.g00245.t01#2    mNeoVis.2.1.1.chr14_hap1.g00244.t01#5    mNeoVis.vulgivaga.1.1.chr14.g00241.t01#4    mNeoVis.evergladensis.1.1.chr14.g00251.t01#3   mNeoVis.3.1.1.chr14.g00246.t01#6       5640   125256020:   mNeoVis.vulgivaga.1.1.chr14.g00242.t01#4    mNeoVis.3.1.1.chr14.g00248.t01#6    mNeoVis.2.1.1.chr14_hap1.g00246.t01#5    mNeoVis.evergladensis.1.1.chr14.g00252.t01#3    mNeoVis.evergladensis.2.1.chr14.g00246.t01#2   mNeoVis.vulgivaga.2.1.chr14.g00247.t01#1       5641   125256021:   mNeoVis.evergladensis.1.1.chr14.g00253.t01#3    mNeoVis.vulgivaga.2.1.chr14.g00248.t01#1    mNeoVis.evergladensis.2.1.chr14.g00247.t01#2    mNeoVis.3.1.1.chr14.g00250.t01#6    mNeoVis.2.1.1.chr14_hap1.g00247.t01#5   mNeoVis.vulgivaga.1.1.chr14.g00243.t01#4       5642   125256022:   mNeoVis.evergladensis.1.1.chr13.g00790.t01#3    mNeoVis.2.1.1.chr13_hap1.g00835.t01#5    mNeoVis.vulgivaga.2.1.chr13.g00793.t01#1    mNeoVis.vulgivaga.1.1.chr13.g00786.t01#4    mNeoVis.evergladensis.2.1.chr13.g00797.t01#2   mNeoVis.3.1.1.chr13.g00754.t01#6       5643   125256023:   mNeoVis.vulgivaga.2.1.chr6.g00605.t01#1    mNeoVis.evergladensis.2.1.chr6.g00607.t01#2    mNeoVis.vulgivaga.2.1.chr10.g00303.t01#1   mNeoVis.evergladensis.1.1.chr6.g00611.t01#3       5644   125256024:   mNeoVis.2.1.1.chr15_hap1.g00643.t01#5    mNeoVis.evergladensis.1.1.chr15.g00149.t01#3    mNeoVis.vulgivaga.2.1.chr15.g00472.t01#1    mNeoVis.vulgivaga.1.1.chr15.g00466.t01#4    mNeoVis.evergladensis.2.1.chr15.g00465.t01#2   mNeoVis.3.1.1.chr15.g00424.t01#6       5645   125256025:   mNeoVis.vulgivaga.2.1.chr14.g00249.t01#1    mNeoVis.evergladensis.2.1.chr14.g00248.t01#2    mNeoVis.2.1.1.chr14_hap1.g00248.t01#5    mNeoVis.vulgivaga.1.1.chr14.g00244.t01#4   mNeoVis.evergladensis.1.1.chr14.g00254.t01#3       5646   125256026:   mNeoVis.3.1.1.chr6.g00620.t01#6    mNeoVis.vulgivaga.1.1.chr6.g00611.t01#4    mNeoVis.evergladensis.1.1.chr6.g00614.t01#3    mNeoVis.2.1.1.chr4_hap1.g00603.t01#5    mNeoVis.vulgivaga.2.1.chr6.g00609.t01#1   mNeoVis.evergladensis.2.1.chr6.g00611.t01#2       5647   125256027:   mNeoVis.evergladensis.1.1.chr7.g00419.t01#3    mNeoVis.2.1.1.chr7_hap1.g00408.t01#5    mNeoVis.vulgivaga.1.1.chr7.g00416.t01#4    mNeoVis.vulgivaga.2.1.chr7.g00411.t01#1    mNeoVis.evergladensis.2.1.chr7.g00412.t01#2    mNeoVis.evergladensis.2.1.chr7.g00411.t01#2    mNeoVis.evergladensis.1.1.chr7.g00418.t01#3   mNeoVis.3.1.1.chr7.g00393.t01#6     5648   125256028:   mNeoVis.evergladensis.1.1.chr6.g00615.t01#3    mNeoVis.evergladensis.2.1.chr6.g00612.t01#2    mNeoVis.vulgivaga.1.1.chr6.g00612.t01#4    mNeoVis.3.1.1.chr6.g00621.t01#6    mNeoVis.2.1.1.chr4_hap1.g00604.t01#5   mNeoVis.vulgivaga.2.1.chr6.g00610.t01#1       5649   125256029:   mNeoVis.2.1.1.chr4_hap1.g00605.t01#5    mNeoVis.evergladensis.1.1.chr6.g00616.t01#3    mNeoVis.3.1.1.chr6.g00622.t01#6    mNeoVis.vulgivaga.2.1.chr6.g00611.t01#1    mNeoVis.vulgivaga.1.1.chr6.g00613.t01#4   mNeoVis.evergladensis.2.1.chr6.g00613.t01#2       5650   125256030:   mNeoVis.2.1.1.chr14_hap1.g00249.t01#5    mNeoVis.evergladensis.2.1.chr14.g00249.t01#2    mNeoVis.vulgivaga.1.1.chr14.g00245.t01#4    mNeoVis.evergladensis.1.1.chr14.g00255.t01#3    mNeoVis.vulgivaga.2.1.chr14.g00250.t01#1   mNeoVis.3.1.1.chr14.g00252.t01#6       5651   125256031:   mNeoVis.evergladensis.2.1.chr14.g00250.t01#2    mNeoVis.vulgivaga.2.1.chr14.g00251.t01#1    mNeoVis.2.1.1.chr14_hap1.g00250.t01#5    mNeoVis.vulgivaga.1.1.chr14.g00246.t01#4   mNeoVis.evergladensis.1.1.chr14.g00256.t01#3       5652   125256032:   mNeoVis.3.1.1.chr15.g00425.t01#6    mNeoVis.vulgivaga.1.1.chr15.g00467.t01#4    mNeoVis.vulgivaga.2.1.chr15.g00473.t01#1    mNeoVis.evergladensis.2.1.chr15.g00466.t01#2    mNeoVis.evergladensis.1.1.chr15.g00148.t01#3   mNeoVis.2.1.1.chr15_hap1.g00644.t01#5       5653   125256033:   mNeoVis.evergladensis.2.1.chr6.g00614.t01#2    mNeoVis.vulgivaga.1.1.chr6.g00614.t01#4    mNeoVis.3.1.1.chr6.g00623.t01#6    mNeoVis.evergladensis.1.1.chr6.g00617.t01#3    mNeoVis.evergladensis.2.1.chr6.g00615.t01#2    mNeoVis.vulgivaga.2.1.chr6.g00612.t01#1   mNeoVis.2.1.1.chr4_hap1.g00606.t01#5     5654   125256034:   mNeoVis.2.1.1.chr2_hap1.g00281.t01#5    mNeoVis.evergladensis.1.1.chr5.g00285.t01#3    mNeoVis.vulgivaga.2.1.chr5.g00284.t01#1    mNeoVis.vulgivaga.1.1.chr5.g00275.t01#4    mNeoVis.evergladensis.2.1.chr5.g00285.t01#2   mNeoVis.3.1.1.chr5.g00273.t01#6       5655   125256035:   mNeoVis.vulgivaga.2.1.chr14.g00252.t01#1    mNeoVis.evergladensis.2.1.chr14.g00251.t01#2    mNeoVis.2.1.1.chr14_hap1.g00251.t01#5   mNeoVis.vulgivaga.1.1.chr14.g00247.t01#4       5656   125256036:   mNeoVis.2.1.1.chr4_hap1.g00607.t01#5    mNeoVis.evergladensis.2.1.chr6.g00616.t01#2    mNeoVis.vulgivaga.2.1.chr6.g00613.t01#1    mNeoVis.vulgivaga.1.1.chr6.g00615.t01#4   mNeoVis.evergladensis.1.1.chr6.g00618.t01#3       5657   125256037:   mNeoVis.evergladensis.1.1.chr14.g00258.t01#3    mNeoVis.vulgivaga.2.1.chr14.g00253.t01#1    mNeoVis.3.1.1.chr14.g00254.t01#6    mNeoVis.evergladensis.2.1.chr14.g00252.t01#2    mNeoVis.2.1.1.chr14_hap1.g00252.t01#5   mNeoVis.vulgivaga.1.1.chr14.g00248.t01#4       5658   125256038:   mNeoVis.2.1.1.chr14_hap1.g00253.t01#5    mNeoVis.vulgivaga.1.1.chr14.g00249.t01#4    mNeoVis.3.1.1.chr14.g00255.t01#6    mNeoVis.evergladensis.1.1.chr14.g00259.t01#3    mNeoVis.vulgivaga.2.1.chr14.g00254.t01#1   mNeoVis.evergladensis.2.1.chr14.g00253.t01#2       5659   125256039:   mNeoVis.vulgivaga.2.1.chr6.g00543.t01#1    mNeoVis.vulgivaga.1.1.chr6.g00544.t01#4    mNeoVis.2.1.1.chr4_hap1.g00534.t01#5    mNeoVis.evergladensis.1.1.chr6.g00540.t01#3    mNeoVis.3.1.1.chr6.g00543.t01#6   mNeoVis.evergladensis.2.1.chr6.g00540.t01#2       5660   125256040:   mNeoVis.evergladensis.2.1.chr7.g00413.t01#2    mNeoVis.vulgivaga.2.1.chr7.g00412.t01#1   mNeoVis.evergladensis.1.1.chr7.g00420.t01#3         5661   125256041:   mNeoVis.vulgivaga.1.1.chr7.g00418.t01#4    mNeoVis.2.1.1.chr7_hap1.g00409.t01#5   mNeoVis.2.1.1.chr7_hap1.g00410.t01#5           5662   125256042:   mNeoVis.evergladensis.1.1.chr7.g00421.t01#3   mNeoVis.3.1.1.chr7.g00394.t01#6             5663   125256043:   mNeoVis.3.1.1.chr14.g00175.t01#6    mNeoVis.vulgivaga.1.1.chr14.g00175.t01#4    mNeoVis.evergladensis.1.1.chr14.g00179.t01#3    mNeoVis.2.1.1.chr14_hap1.g00177.t01#5    mNeoVis.evergladensis.2.1.chr14.g00174.t01#2   mNeoVis.vulgivaga.2.1.chr14.g00178.t01#1       5664   125256044:   mNeoVis.vulgivaga.2.1.chr6.g00544.t01#1    mNeoVis.3.1.1.chr6.g00544.t01#6    mNeoVis.evergladensis.1.1.chr6.g00541.t01#3    mNeoVis.2.1.1.chr4_hap1.g00535.t01#5    mNeoVis.evergladensis.2.1.chr6.g00541.t01#2   mNeoVis.vulgivaga.1.1.chr6.g00545.t01#4       5665   125256045:   mNeoVis.evergladensis.1.1.chr14.g00183.t01#3    mNeoVis.vulgivaga.1.1.chr14.g00178.t01#4    mNeoVis.2.1.1.chr14_hap1.g00178.t01#5    mNeoVis.3.1.1.chr14.g00178.t01#6    mNeoVis.evergladensis.2.1.chr14.g00178.t01#2   mNeoVis.vulgivaga.2.1.chr14.g00179.t01#1       5666   125256046:   mNeoVis.evergladensis.1.1.chr6.g00542.t01#3    mNeoVis.vulgivaga.1.1.chr10.g00726.t01#4    mNeoVis.vulgivaga.1.1.chr6.g00546.t01#4    mNeoVis.vulgivaga.2.1.chr6.g00545.t01#1    mNeoVis.3.1.1.chr10.g00716.t01#6    mNeoVis.vulgivaga.2.1.chr10.g00730.t01#1    mNeoVis.evergladensis.2.1.chr10.g00739.t01#2    mNeoVis.3.1.1.chr6.g00545.t01#6    mNeoVis.evergladensis.1.1.chr10.g00743.t01#3      5667   125256047:   mNeoVis.2.1.1.chr4_hap1.g00537.t01#5    mNeoVis.vulgivaga.1.1.chr6.g00547.t01#4    mNeoVis.vulgivaga.2.1.chr6.g00546.t01#1    mNeoVis.evergladensis.1.1.chr6.g00543.t01#3    mNeoVis.3.1.1.chr6.g00546.t01#6   mNeoVis.evergladensis.2.1.chr6.g00543.t01#2       5668   125256048:   mNeoVis.vulgivaga.2.1.chr14.g00180.t01#1    mNeoVis.evergladensis.1.1.chr14.g00181.t01#3    mNeoVis.vulgivaga.1.1.chr14.g00176.t01#4    mNeoVis.3.1.1.chr14.g00176.t01#6    mNeoVis.evergladensis.2.1.chr14.g00176.t01#2   mNeoVis.2.1.1.chr14_hap1.g00179.t01#5       5669   125256049:   mNeoVis.vulgivaga.2.1.chr14.g00181.t01#1    mNeoVis.2.1.1.chr14_hap1.g00180.t01#5    mNeoVis.evergladensis.1.1.chr14.g00182.t01#3    mNeoVis.evergladensis.2.1.chr14.g00177.t01#2    mNeoVis.3.1.1.chr14.g00177.t01#6   mNeoVis.vulgivaga.1.1.chr14.g00177.t01#4       5670   125256050:   mNeoVis.3.1.1.chr6.g00547.t01#6    mNeoVis.evergladensis.1.1.chr6.g00544.t01#3    mNeoVis.evergladensis.2.1.chr6.g00544.t01#2    mNeoVis.vulgivaga.2.1.chr6.g00547.t01#1    mNeoVis.vulgivaga.1.1.chr6.g00548.t01#4   mNeoVis.2.1.1.chr4_hap1.g00538.t01#5       5671   125256051:   mNeoVis.3.1.1.chr14.g00179.t01#6    mNeoVis.vulgivaga.2.1.chr14.g00182.t01#1    mNeoVis.evergladensis.1.1.chr14.g00184.t01#3    mNeoVis.evergladensis.2.1.chr14.g00179.t01#2    mNeoVis.vulgivaga.1.1.chr14.g00179.t01#4   mNeoVis.2.1.1.chr14_hap1.g00181.t01#5       5672   125256052:   mNeoVis.evergladensis.2.1.chr14.g00180.t01#2    mNeoVis.evergladensis.1.1.chr14.g00185.t01#3    mNeoVis.vulgivaga.2.1.chr14.g00183.t01#1    mNeoVis.3.1.1.chr14.g00180.t01#6    mNeoVis.vulgivaga.1.1.chr14.g00180.t01#4   mNeoVis.2.1.1.chr14_hap1.g00182.t01#5       5673   125256053:   mNeoVis.evergladensis.2.1.chr14.g00181.t01#2    mNeoVis.vulgivaga.2.1.chr14.g00184.t01#1    mNeoVis.evergladensis.1.1.chr14.g00186.t01#3    mNeoVis.vulgivaga.1.1.chr14.g00181.t01#4    mNeoVis.3.1.1.chr14.g00181.t01#6   mNeoVis.2.1.1.chr14_hap1.g00183.t01#5       5674   125256054:   mNeoVis.evergladensis.2.1.chr6.g00545.t01#2    mNeoVis.3.1.1.chr6.g00548.t01#6    mNeoVis.evergladensis.1.1.chr6.g00545.t01#3    mNeoVis.2.1.1.chr4_hap1.g00539.t01#5    mNeoVis.vulgivaga.1.1.chr6.g00549.t01#4   mNeoVis.vulgivaga.2.1.chr6.g00548.t01#1       5675   125256055:   mNeoVis.2.1.1.chr2_hap1.g00982.t01#5    mNeoVis.evergladensis.2.1.chr5.g00935.t01#2    mNeoVis.evergladensis.1.1.chr5.g00926.t01#3    mNeoVis.vulgivaga.2.1.chr5.g00922.t01#1    mNeoVis.3.1.1.chr5.g00915.t01#6   mNeoVis.vulgivaga.1.1.chr5.g00903.t01#4       5676   125256056:   mNeoVis.evergladensis.1.1.chr2.g00665.t01#3    mNeoVis.evergladensis.2.1.chr2.g00666.t01#2    mNeoVis.vulgivaga.2.1.chr2.g00669.t01#1    mNeoVis.vulgivaga.1.1.chr2.g00660.t01#4    mNeoVis.3.1.1.chr2.g00586.t01#6   mNeoVis.2.1.1.chr5_hap1.g00669.t01#5       5677   125256057:   mNeoVis.evergladensis.2.1.chr7.g00414.t01#2    mNeoVis.2.1.1.chr7_hap1.g00411.t01#5    mNeoVis.evergladensis.1.1.chr7.g00422.t01#3    mNeoVis.vulgivaga.1.1.chr7.g00419.t01#4    mNeoVis.vulgivaga.2.1.chr7.g00413.t01#1   mNeoVis.3.1.1.chr7.g00395.t01#6       5678   125256058:   mNeoVis.2.1.1.chr15_hap1.g00645.t01#5    mNeoVis.vulgivaga.2.1.chr15.g00474.t01#1    mNeoVis.vulgivaga.1.1.chr15.g00468.t01#4    mNeoVis.evergladensis.1.1.chr15.g00147.t01#3    mNeoVis.evergladensis.2.1.chr15.g00467.t01#2   mNeoVis.3.1.1.chr15.g00426.t01#6       5679   125256059:   mNeoVis.3.1.1.chr6.g00549.t01#6    mNeoVis.2.1.1.chr4_hap1.g00540.t01#5    mNeoVis.vulgivaga.2.1.chr6.g00549.t01#1   mNeoVis.evergladensis.2.1.chr6.g00546.t01#2       5680   125256060:   mNeoVis.vulgivaga.1.1.chr5.g00904.t01#4    mNeoVis.vulgivaga.2.1.chr6.g00550.t01#1    mNeoVis.evergladensis.2.1.chr5.g00936.t01#2    mNeoVis.2.1.1.chr4_hap1.g00541.t01#5    mNeoVis.evergladensis.1.1.chr6.g00546.t01#3    mNeoVis.3.1.1.chr5.g00916.t01#6    mNeoVis.2.1.1.chr2_hap1.g00983.t01#5    mNeoVis.evergladensis.1.1.chr5.g00927.t01#3    mNeoVis.evergladensis.2.1.chr6.g00547.t01#2      5681   125256061:   mNeoVis.2.1.1.chr5_hap1.g00671.t01#5    mNeoVis.vulgivaga.1.1.chr2.g00662.t01#4    mNeoVis.evergladensis.2.1.chr2.g00668.t01#2    mNeoVis.vulgivaga.2.1.chr2.g00671.t01#1    mNeoVis.evergladensis.1.1.chr2.g00667.t01#3   mNeoVis.3.1.1.chr2.g00588.t01#6       5682   125256062:   mNeoVis.vulgivaga.2.1.chr14.g00185.t01#1    mNeoVis.2.1.1.chr14_hap1.g00184.t01#5    mNeoVis.evergladensis.1.1.chr14.g00188.t01#3    mNeoVis.evergladensis.2.1.chr14.g00182.t01#2    mNeoVis.3.1.1.chr14.g00183.t01#6   mNeoVis.vulgivaga.1.1.chr14.g00182.t01#4       5683   125256063:   mNeoVis.vulgivaga.2.1.chr6.g00551.t01#1    mNeoVis.evergladensis.1.1.chr6.g00549.t01#3    mNeoVis.3.1.1.chr6.g00552.t01#6    mNeoVis.evergladensis.2.1.chr6.g00551.t01#2    mNeoVis.vulgivaga.1.1.chr6.g00552.t01#4   mNeoVis.2.1.1.chr4_hap1.g00544.t01#5       5684   125256064:   mNeoVis.vulgivaga.1.1.chr14.g00183.t01#4    mNeoVis.evergladensis.2.1.chr14.g00183.t01#2    mNeoVis.3.1.1.chr14.g00182.t01#6    mNeoVis.vulgivaga.2.1.chr14.g00186.t01#1   mNeoVis.evergladensis.1.1.chr14.g00187.t01#3       5685   125256065:   mNeoVis.vulgivaga.2.1.chr14.g00187.t01#1    mNeoVis.evergladensis.1.1.chr14.g00189.t01#3    mNeoVis.vulgivaga.1.1.chr14.g00184.t01#4    mNeoVis.evergladensis.2.1.chr14.g00184.t01#2    mNeoVis.2.1.1.chr14_hap1.g00185.t01#5   mNeoVis.3.1.1.chr14.g00184.t01#6       5686   125256066:   mNeoVis.vulgivaga.2.1.chr6.g00552.t01#1    mNeoVis.evergladensis.2.1.chr6.g00552.t01#2    mNeoVis.evergladensis.1.1.chr6.g00550.t01#3    mNeoVis.2.1.1.chr4_hap1.g00545.t01#5    mNeoVis.3.1.1.chr6.g00553.t01#6   mNeoVis.vulgivaga.1.1.chr6.g00553.t01#4       5687   125256067:   mNeoVis.evergladensis.2.1.chr6.g00554.t01#2    mNeoVis.3.1.1.chr6.g00554.t01#6    mNeoVis.vulgivaga.2.1.chr6.g00553.t01#1    mNeoVis.2.1.1.chr4_hap1.g00547.t01#5    mNeoVis.vulgivaga.1.1.chr6.g00554.t01#4   mNeoVis.evergladensis.1.1.chr6.g00552.t01#3       5688   125256068:   mNeoVis.evergladensis.2.1.chr6.g00555.t01#2    mNeoVis.2.1.1.chr4_hap1.g00548.t01#5    mNeoVis.vulgivaga.2.1.chr6.g00554.t01#1    mNeoVis.vulgivaga.1.1.chr6.g00555.t01#4    mNeoVis.evergladensis.1.1.chr6.g00553.t01#3   mNeoVis.3.1.1.chr6.g00555.t01#6       5689   125256069:   mNeoVis.vulgivaga.1.1.chr14.g00185.t01#4    mNeoVis.evergladensis.2.1.chr14.g00185.t01#2    mNeoVis.3.1.1.chr14.g00185.t01#6    mNeoVis.evergladensis.1.1.chr14.g00190.t01#3    mNeoVis.vulgivaga.2.1.chr14.g00188.t01#1   mNeoVis.2.1.1.chr14_hap1.g00186.t01#5       5690   125256070:   mNeoVis.vulgivaga.1.1.chr14.g00186.t01#4    mNeoVis.3.1.1.chr14.g00186.t01#6    mNeoVis.vulgivaga.2.1.chr14.g00189.t01#1    mNeoVis.evergladensis.1.1.chr14.g00191.t01#3    mNeoVis.evergladensis.2.1.chr14.g00186.t01#2   mNeoVis.2.1.1.chr14_hap1.g00187.t01#5       5691   125256071:   mNeoVis.3.1.1.chr6.g00556.t01#6    mNeoVis.evergladensis.1.1.chr6.g00554.t01#3    mNeoVis.evergladensis.2.1.chr6.g00556.t01#2    mNeoVis.vulgivaga.2.1.chr6.g00555.t01#1    mNeoVis.vulgivaga.1.1.chr6.g00556.t01#4   mNeoVis.2.1.1.chr4_hap1.g00549.t01#5       5692   125256072:  mNeoVis.vulgivaga.2.1.chr14.g00191.t01#1               5693   125256073:   mNeoVis.2.1.1.chr9_hap1.g00302.t01#5    mNeoVis.evergladensis.1.1.chr10.g00302.t01#3    mNeoVis.evergladensis.1.1.chr14.g00192.t01#3    mNeoVis.3.1.1.chr14.g00187.t01#6    mNeoVis.2.1.1.chr14_hap1.g00188.t01#5    mNeoVis.evergladensis.2.1.chr14.g00187.t01#2    mNeoVis.vulgivaga.2.1.chr14.g00190.t01#1    mNeoVis.3.1.1.chr10.g00290.t01#6    mNeoVis.vulgivaga.1.1.chr14.g00187.t01#4      5694   125256074:   mNeoVis.evergladensis.2.1.chr14.g00188.t01#2    mNeoVis.evergladensis.1.1.chr14.g00193.t01#3    mNeoVis.2.1.1.chr14_hap1.g00189.t01#5    mNeoVis.vulgivaga.2.1.chr14.g00192.t01#1    mNeoVis.vulgivaga.1.1.chr14.g00188.t01#4   mNeoVis.3.1.1.chr14.g00188.t01#6       5695   125256075:   mNeoVis.evergladensis.1.1.chr14.g00194.t01#3    mNeoVis.2.1.1.chr14_hap1.g00190.t01#5    mNeoVis.vulgivaga.2.1.chr14.g00193.t01#1    mNeoVis.3.1.1.chr14.g00189.t01#6    mNeoVis.evergladensis.2.1.chr14.g00189.t01#2   mNeoVis.vulgivaga.1.1.chr14.g00189.t01#4       5696   125256076:   mNeoVis.evergladensis.1.1.chr6.g00555.t01#3    mNeoVis.vulgivaga.2.1.chr6.g00556.t01#1    mNeoVis.3.1.1.chr6.g00557.t01#6    mNeoVis.vulgivaga.1.1.chr6.g00557.t01#4    mNeoVis.evergladensis.2.1.chr6.g00557.t01#2   mNeoVis.2.1.1.chr4_hap1.g00550.t01#5       5697   125256077:   mNeoVis.evergladensis.1.1.chr14.g00195.t01#3    mNeoVis.2.1.1.chr14_hap1.g00191.t01#5    mNeoVis.3.1.1.chr14.g00190.t01#6    mNeoVis.vulgivaga.1.1.chr14.g00190.t01#4    mNeoVis.vulgivaga.2.1.chr14.g00194.t01#1   mNeoVis.evergladensis.2.1.chr14.g00190.t01#2       5698   125256078:   mNeoVis.vulgivaga.2.1.chr15.g00475.t01#1    mNeoVis.3.1.1.chr15.g00427.t01#6    mNeoVis.evergladensis.1.1.chr15.g00146.t01#3    mNeoVis.2.1.1.chr15_hap1.g00646.t01#5    mNeoVis.vulgivaga.1.1.chr15.g00469.t01#4    mNeoVis.evergladensis.2.1.chr15.g00468.t01#2   mNeoVis.vulgivaga.2.1.chr15.g00476.t01#1     5699   125256079:   mNeoVis.3.1.1.chr11.g01062.t01#6    mNeoVis.vulgivaga.2.1.chr11.g01115.t01#1    mNeoVis.evergladensis.1.1.chr11.g01105.t01#3    mNeoVis.2.1.1.chr11_hap1.g01120.t01#5    mNeoVis.vulgivaga.1.1.chr11.g01088.t01#4    mNeoVis.evergladensis.2.1.chr11.g01089.t01#2   mNeoVis.vulgivaga.2.1.chr11.g01116.t01#1     5700   125256080:   mNeoVis.vulgivaga.1.1.chr12.g00102.t01#4    mNeoVis.evergladensis.2.1.chr12.g00095.t01#2    mNeoVis.evergladensis.1.1.chr12.g00095.t01#3   mNeoVis.2.1.1.chr12_hap1.g00120.t01#5         5701   125256081:   mNeoVis.vulgivaga.2.1.chr6.g00557.t01#1    mNeoVis.3.1.1.chr6.g00558.t01#6    mNeoVis.2.1.1.chr4_hap1.g00551.t01#5    mNeoVis.vulgivaga.1.1.chr6.g00558.t01#4    mNeoVis.evergladensis.1.1.chr6.g00556.t01#3   mNeoVis.evergladensis.2.1.chr6.g00558.t01#2       5702   125256082:   mNeoVis.vulgivaga.1.1.chr6.g00559.t01#4    mNeoVis.evergladensis.1.1.chr6.g00557.t01#3    mNeoVis.evergladensis.2.1.chr6.g00559.t01#2    mNeoVis.2.1.1.chr4_hap1.g00552.t01#5    mNeoVis.vulgivaga.2.1.chr6.g00558.t01#1   mNeoVis.3.1.1.chr6.g00559.t01#6       5703   125256083:   mNeoVis.vulgivaga.1.1.chr14.g00191.t01#4    mNeoVis.evergladensis.1.1.chr14.g00196.t01#3    mNeoVis.3.1.1.chr14.g00191.t01#6    mNeoVis.vulgivaga.2.1.chr14.g00195.t01#1    mNeoVis.2.1.1.chr14_hap1.g00192.t01#5   mNeoVis.evergladensis.2.1.chr14.g00191.t01#2       5704   125256084:   mNeoVis.evergladensis.1.1.chr6.g00558.t01#3    mNeoVis.evergladensis.2.1.chr6.g00560.t01#2    mNeoVis.vulgivaga.1.1.chr6.g00561.t01#4    mNeoVis.2.1.1.chr4_hap1.g00553.t01#5    mNeoVis.3.1.1.chr6.g00560.t01#6   mNeoVis.vulgivaga.2.1.chr6.g00559.t01#1       5705   125256085:   mNeoVis.evergladensis.1.1.chr14.g00197.t01#3    mNeoVis.3.1.1.chr14.g00192.t01#6    mNeoVis.vulgivaga.1.1.chr14.g00192.t01#4    mNeoVis.evergladensis.2.1.chr14.g00192.t01#2    mNeoVis.vulgivaga.2.1.chr14.g00196.t01#1   mNeoVis.2.1.1.chr14_hap1.g00193.t01#5       5706   125256086:   mNeoVis.2.1.1.chr14_hap1.g00194.t01#5    mNeoVis.evergladensis.2.1.chr14.g00193.t01#2    mNeoVis.vulgivaga.1.1.chr14.g00193.t01#4    mNeoVis.evergladensis.1.1.chr14.g00198.t01#3    mNeoVis.vulgivaga.2.1.chr14.g00197.t01#1   mNeoVis.3.1.1.chr14.g00193.t01#6       5707   125256087:   mNeoVis.3.1.1.chr6.g00561.t01#6    mNeoVis.evergladensis.1.1.chr6.g00559.t01#3    mNeoVis.vulgivaga.2.1.chr6.g00560.t01#1    mNeoVis.vulgivaga.1.1.chr6.g00560.t01#4    mNeoVis.evergladensis.2.1.chr6.g00561.t01#2   mNeoVis.2.1.1.chr4_hap1.g00554.t01#5       5708   125256088:   mNeoVis.vulgivaga.2.1.chr6.g00561.t01#1    mNeoVis.2.1.1.chr4_hap1.g00555.t01#5    mNeoVis.evergladensis.2.1.chr6.g00562.t01#2    mNeoVis.vulgivaga.1.1.chr6.g00562.t01#4    mNeoVis.evergladensis.1.1.chr6.g00560.t01#3   mNeoVis.3.1.1.chr6.g00562.t01#6       5709   125256089:   mNeoVis.evergladensis.2.1.chr14.g00194.t01#2    mNeoVis.evergladensis.1.1.chr14.g00199.t01#3   mNeoVis.vulgivaga.2.1.chr14.g00198.t01#1         5710   125256090:   mNeoVis.evergladensis.1.1.chr6.g00561.t01#3    mNeoVis.2.1.1.chr4_hap1.g00556.t01#5    mNeoVis.3.1.1.chr6.g00563.t01#6    mNeoVis.evergladensis.2.1.chr6.g00563.t01#2    mNeoVis.vulgivaga.2.1.chr6.g00562.t01#1   mNeoVis.vulgivaga.1.1.chr6.g00563.t01#4       5711   125256091:   mNeoVis.vulgivaga.1.1.chr14.g00195.t01#4    mNeoVis.3.1.1.chr14.g00194.t01#6    mNeoVis.evergladensis.2.1.chr14.g00196.t01#2    mNeoVis.vulgivaga.2.1.chr14.g00200.t01#1    mNeoVis.evergladensis.1.1.chr14.g00201.t01#3   mNeoVis.2.1.1.chr14_hap1.g00196.t01#5       5712   125256092:   mNeoVis.evergladensis.1.1.chr6.g00562.t01#3    mNeoVis.evergladensis.2.1.chr6.g00564.t01#2    mNeoVis.3.1.1.chr6.g00564.t01#6    mNeoVis.vulgivaga.2.1.chr6.g00563.t01#1    mNeoVis.vulgivaga.1.1.chr6.g00564.t01#4   mNeoVis.2.1.1.chr4_hap1.g00557.t01#5       5713   125256093:   mNeoVis.3.1.1.chr6.g00565.t01#6    mNeoVis.vulgivaga.1.1.chr6.g00565.t01#4    mNeoVis.evergladensis.1.1.chr6.g00563.t01#3    mNeoVis.vulgivaga.2.1.chr6.g00564.t01#1    mNeoVis.2.1.1.chr4_hap1.g00558.t01#5   mNeoVis.evergladensis.2.1.chr6.g00565.t01#2       5714   125256094:   mNeoVis.2.1.1.chr14_hap1.g00197.t01#5    mNeoVis.evergladensis.2.1.chr14.g00197.t01#2    mNeoVis.vulgivaga.1.1.chr14.g00196.t01#4    mNeoVis.vulgivaga.2.1.chr14.g00201.t01#1    mNeoVis.3.1.1.chr14.g00195.t01#6   mNeoVis.evergladensis.1.1.chr14.g00202.t01#3       5715   125256095:   mNeoVis.vulgivaga.1.1.chr15.g00470.t01#4    mNeoVis.evergladensis.1.1.chr15.g00144.t01#3    mNeoVis.3.1.1.chr15.g00428.t01#6    mNeoVis.evergladensis.2.1.chr15.g00469.t01#2   mNeoVis.2.1.1.chr15_hap1.g00647.t01#5       5716   125256096:   mNeoVis.vulgivaga.2.1.chr11.g00938.t01#1    mNeoVis.evergladensis.1.1.chr11.g00928.t01#3    mNeoVis.2.1.1.chr11_hap1.g00936.t01#5   mNeoVis.evergladensis.2.1.chr11.g00914.t01#2       5717   125256097:   mNeoVis.evergladensis.1.1.chr15.g00145.t01#3   mNeoVis.vulgivaga.2.1.chr15.g00477.t01#1           5718   125256098:   mNeoVis.evergladensis.1.1.chr6.g00564.t01#3    mNeoVis.vulgivaga.1.1.chr6.g00566.t01#4    mNeoVis.3.1.1.chr6.g00566.t01#6    mNeoVis.vulgivaga.2.1.chr6.g00565.t01#1    mNeoVis.evergladensis.2.1.chr6.g00566.t01#2   mNeoVis.2.1.1.chr4_hap1.g00559.t01#5       5719   125256099:   mNeoVis.vulgivaga.2.1.chr3.g00934.t01#1    mNeoVis.3.1.1.chr1.g00905.t01#6    mNeoVis.2.1.1.chr1_hap1.g00884.t01#5    mNeoVis.evergladensis.2.1.chr3.g00917.t01#2   mNeoVis.vulgivaga.1.1.chr1.g00901.t01#4       5720   125256100:   mNeoVis.3.1.1.chr14.g00197.t01#6    mNeoVis.2.1.1.chr14_hap1.g00199.t01#5    mNeoVis.evergladensis.1.1.chr14.g00205.t01#3    mNeoVis.evergladensis.2.1.chr14.g00199.t01#2    mNeoVis.vulgivaga.1.1.chr14.g00198.t01#4   mNeoVis.vulgivaga.2.1.chr14.g00203.t01#1       5721   125256101:   mNeoVis.vulgivaga.2.1.chr14.g00202.t01#1    mNeoVis.vulgivaga.1.1.chr12.g00756.t01#4    mNeoVis.evergladensis.1.1.chr12.g00739.t01#3    mNeoVis.evergladensis.2.1.chr6.g00434.t01#2    mNeoVis.evergladensis.2.1.chr12.g00731.t01#2    mNeoVis.vulgivaga.1.1.chr12.g00279.t01#4    mNeoVis.evergladensis.1.1.chr12.g00275.t01#3    mNeoVis.evergladensis.1.1.chr14.g00204.t01#3    mNeoVis.vulgivaga.2.1.chr12.g00761.t01#1      5722   125256102:   mNeoVis.evergladensis.1.1.chr6.g00565.t01#3    mNeoVis.vulgivaga.1.1.chr6.g00567.t01#4    mNeoVis.evergladensis.2.1.chr6.g00567.t01#2    mNeoVis.2.1.1.chr4_hap1.g00560.t01#5    mNeoVis.vulgivaga.2.1.chr6.g00566.t01#1   mNeoVis.3.1.1.chr6.g00567.t01#6       5723   125256103:   mNeoVis.2.1.1.chr14_hap1.g00200.t01#5    mNeoVis.3.1.1.chr14.g00198.t01#6    mNeoVis.vulgivaga.1.1.chr14.g00199.t01#4    mNeoVis.evergladensis.1.1.chr14.g00206.t01#3    mNeoVis.vulgivaga.2.1.chr14.g00204.t01#1   mNeoVis.evergladensis.2.1.chr14.g00200.t01#2       5724   125256104:   mNeoVis.evergladensis.2.1.chr6.g00568.t01#2    mNeoVis.2.1.1.chr4_hap1.g00561.t01#5    mNeoVis.vulgivaga.2.1.chr6.g00567.t01#1    mNeoVis.evergladensis.1.1.chr6.g00566.t01#3   mNeoVis.vulgivaga.1.1.chr6.g00568.t01#4       5725   125256105:   mNeoVis.2.1.1.chr14_hap1.g00201.t01#5    mNeoVis.evergladensis.1.1.chr14.g00207.t01#3    mNeoVis.evergladensis.2.1.chr14.g00201.t01#2    mNeoVis.vulgivaga.2.1.chr14.g00205.t01#1   mNeoVis.3.1.1.chr14.g00199.t01#6       5726   125256106:   mNeoVis.evergladensis.1.1.chr7.g00423.t01#3    mNeoVis.3.1.1.chr7.g00396.t01#6    mNeoVis.2.1.1.chr7_hap1.g00412.t01#5    mNeoVis.vulgivaga.2.1.chr7.g00414.t01#1    mNeoVis.vulgivaga.1.1.chr7.g00420.t01#4   mNeoVis.evergladensis.2.1.chr7.g00415.t01#2       5727   125256107:   mNeoVis.evergladensis.1.1.chr14.g00209.t01#3    mNeoVis.vulgivaga.1.1.chr14.g00202.t01#4    mNeoVis.2.1.1.chr14_hap1.g00203.t01#5    mNeoVis.vulgivaga.2.1.chr14.g00206.t01#1    mNeoVis.evergladensis.2.1.chr14.g00202.t01#2   mNeoVis.3.1.1.chr14.g00200.t01#6       5728   125256108:   mNeoVis.evergladensis.2.1.chr15.g00470.t01#2    mNeoVis.vulgivaga.2.1.chr15.g00478.t01#1    mNeoVis.vulgivaga.1.1.chr15.g00471.t01#4    mNeoVis.evergladensis.1.1.chr15.g00143.t01#3   mNeoVis.2.1.1.chr15_hap1.g00648.t01#5       5729   125256109:   mNeoVis.2.1.1.chr14_hap1.g00204.t01#5    mNeoVis.3.1.1.chr14.g00201.t01#6    mNeoVis.vulgivaga.2.1.chr14.g00207.t01#1    mNeoVis.evergladensis.1.1.chr14.g00210.t01#3    mNeoVis.evergladensis.2.1.chr14.g00204.t01#2   mNeoVis.vulgivaga.1.1.chr14.g00203.t01#4       5730   125256110:   mNeoVis.2.1.1.chr4_hap1.g00562.t01#5    mNeoVis.evergladensis.1.1.chr6.g00567.t01#3    mNeoVis.vulgivaga.1.1.chr6.g00569.t01#4    mNeoVis.vulgivaga.2.1.chr6.g00568.t01#1    mNeoVis.evergladensis.2.1.chr6.g00569.t01#2   mNeoVis.3.1.1.chr6.g00573.t01#6       5731   125256111:   mNeoVis.evergladensis.2.1.chr6.g00570.t01#2    mNeoVis.vulgivaga.2.1.chr6.g00569.t01#1    mNeoVis.evergladensis.1.1.chr6.g00568.t01#3    mNeoVis.2.1.1.chr4_hap1.g00563.t01#5    mNeoVis.vulgivaga.1.1.chr6.g00570.t01#4   mNeoVis.3.1.1.chr6.g00574.t01#6       5732   125256112:   mNeoVis.3.1.1.chr6.g00575.t01#6    mNeoVis.vulgivaga.1.1.chr6.g00571.t01#4    mNeoVis.evergladensis.2.1.chr6.g00571.t01#2    mNeoVis.vulgivaga.2.1.chr6.g00570.t01#1    mNeoVis.evergladensis.1.1.chr6.g00569.t01#3   mNeoVis.2.1.1.chr4_hap1.g00564.t01#5       5733   125256113:   mNeoVis.2.1.1.chr4_hap1.g00565.t01#5    mNeoVis.vulgivaga.2.1.chr6.g00571.t01#1    mNeoVis.evergladensis.1.1.chr6.g00570.t01#3    mNeoVis.evergladensis.2.1.chr6.g00572.t01#2    mNeoVis.vulgivaga.1.1.chr6.g00572.t01#4   mNeoVis.3.1.1.chr6.g00576.t01#6       5734   125256114:   mNeoVis.vulgivaga.2.1.chr6.g00503.t01#1   mNeoVis.evergladensis.1.1.chr6.g00502.t01#3           5735   125256115:   mNeoVis.vulgivaga.1.1.chr6.g00506.t01#4    mNeoVis.evergladensis.1.1.chr6.g00503.t01#3    mNeoVis.3.1.1.chr6.g00500.t01#6    mNeoVis.evergladensis.2.1.chr6.g00504.t01#2    mNeoVis.2.1.1.chr4_hap1.g00498.t01#5   mNeoVis.vulgivaga.2.1.chr6.g00504.t01#1       5736   125256116:   mNeoVis.vulgivaga.2.1.chr6.g00505.t01#1    mNeoVis.evergladensis.1.1.chr6.g00504.t01#3    mNeoVis.evergladensis.2.1.chr6.g00505.t01#2    mNeoVis.vulgivaga.1.1.chr6.g00507.t01#4    mNeoVis.3.1.1.chr6.g00501.t01#6   mNeoVis.2.1.1.chr4_hap1.g00499.t01#5       5737   125256117:   mNeoVis.vulgivaga.1.1.chr6.g00508.t01#4    mNeoVis.vulgivaga.2.1.chr6.g00506.t01#1    mNeoVis.evergladensis.2.1.chr6.g00506.t01#2    mNeoVis.2.1.1.chr4_hap1.g00500.t01#5   mNeoVis.evergladensis.1.1.chr6.g00505.t01#3       5738   125256118:   mNeoVis.3.1.1.chr6.g00509.t01#6    mNeoVis.vulgivaga.2.1.chr6.g00507.t01#1    mNeoVis.evergladensis.1.1.chr6.g00506.t01#3    mNeoVis.2.1.1.chr4_hap1.g00501.t01#5    mNeoVis.evergladensis.2.1.chr6.g00507.t01#2   mNeoVis.vulgivaga.1.1.chr6.g00509.t01#4       5739   125256119:   mNeoVis.2.1.1.chr4_hap1.g00503.t01#5    mNeoVis.vulgivaga.1.1.chr6.g00511.t01#4    mNeoVis.vulgivaga.2.1.chr6.g00508.t01#1   mNeoVis.3.1.1.chr6.g00510.t01#6         5740   125256120:   mNeoVis.vulgivaga.2.1.chr6.g00509.t01#1    mNeoVis.2.1.1.chr4_hap1.g00504.t01#5    mNeoVis.evergladensis.1.1.chr6.g00508.t01#3    mNeoVis.evergladensis.2.1.chr6.g00508.t01#2   mNeoVis.vulgivaga.1.1.chr6.g00512.t01#4       5741   125256121:   mNeoVis.3.1.1.chr6.g00512.t01#6    mNeoVis.evergladensis.1.1.chr6.g00509.t01#3    mNeoVis.vulgivaga.2.1.chr6.g00510.t01#1    mNeoVis.evergladensis.2.1.chr6.g00509.t01#2    mNeoVis.vulgivaga.1.1.chr6.g00513.t01#4   mNeoVis.2.1.1.chr4_hap1.g00505.t01#5       5742   125256122:   mNeoVis.evergladensis.1.1.chr6.g00510.t01#3    mNeoVis.vulgivaga.2.1.chr6.g00511.t01#1    mNeoVis.evergladensis.2.1.chr6.g00510.t01#2    mNeoVis.2.1.1.chr4_hap1.g00506.t01#5   mNeoVis.vulgivaga.1.1.chr6.g00514.t01#4       5743   125256123:   mNeoVis.3.1.1.chr6.g00513.t01#6    mNeoVis.vulgivaga.2.1.chr6.g00512.t01#1    mNeoVis.evergladensis.1.1.chr6.g00511.t01#3    mNeoVis.evergladensis.2.1.chr6.g00511.t01#2    mNeoVis.2.1.1.chr4_hap1.g00507.t01#5   mNeoVis.vulgivaga.1.1.chr6.g00515.t01#4       5744   125256124:   mNeoVis.evergladensis.1.1.chr6.g00512.t01#3    mNeoVis.2.1.1.chr4_hap1.g00508.t01#5    mNeoVis.vulgivaga.1.1.chr6.g00516.t01#4    mNeoVis.evergladensis.2.1.chr6.g00512.t01#2    mNeoVis.3.1.1.chr6.g00514.t01#6   mNeoVis.vulgivaga.2.1.chr6.g00513.t01#1       5745   125256125:   mNeoVis.2.1.1.chr14_hap1.g00205.t01#5    mNeoVis.evergladensis.2.1.chr14.g00205.t01#2    mNeoVis.vulgivaga.2.1.chr14.g00208.t01#1   mNeoVis.evergladensis.1.1.chr14.g00211.t01#3       5746   125256126:   mNeoVis.evergladensis.2.1.chr14.g00206.t01#2    mNeoVis.evergladensis.1.1.chr14.g00212.t01#3    mNeoVis.3.1.1.chr14.g00202.t01#6    mNeoVis.2.1.1.chr14_hap1.g00206.t01#5    mNeoVis.vulgivaga.1.1.chr14.g00204.t01#4   mNeoVis.vulgivaga.2.1.chr14.g00209.t01#1       5747   125256127:   mNeoVis.2.1.1.chr14_hap1.g00207.t01#5    mNeoVis.3.1.1.chr14.g00203.t01#6    mNeoVis.vulgivaga.2.1.chr14.g00210.t01#1    mNeoVis.vulgivaga.1.1.chr14.g00205.t01#4    mNeoVis.evergladensis.1.1.chr14.g00213.t01#3   mNeoVis.evergladensis.2.1.chr14.g00207.t01#2       5748   125256128:   mNeoVis.evergladensis.1.1.chr14.g00214.t01#3    mNeoVis.3.1.1.chr14.g00204.t01#6    mNeoVis.vulgivaga.1.1.chr14.g00206.t01#4    mNeoVis.2.1.1.chr14_hap1.g00208.t01#5    mNeoVis.evergladensis.2.1.chr14.g00208.t01#2   mNeoVis.vulgivaga.2.1.chr14.g00211.t01#1       5749   125256129:   mNeoVis.vulgivaga.1.1.chr14.g00207.t01#4    mNeoVis.2.1.1.chr14_hap1.g00209.t01#5    mNeoVis.vulgivaga.2.1.chr14.g00212.t01#1    mNeoVis.evergladensis.1.1.chr14.g00215.t01#3    mNeoVis.evergladensis.2.1.chr14.g00209.t01#2   mNeoVis.3.1.1.chr14.g00205.t01#6       5750   125256130:   mNeoVis.vulgivaga.1.1.chr14.g00208.t01#4    mNeoVis.2.1.1.chr14_hap1.g00210.t01#5   mNeoVis.vulgivaga.2.1.chr14.g00213.t01#1         5751   125256131:   mNeoVis.evergladensis.1.1.chr14.g00217.t01#3    mNeoVis.3.1.1.chr14.g00208.t01#6    mNeoVis.2.1.1.chr14_hap1.g00211.t01#5    mNeoVis.evergladensis.2.1.chr14.g00211.t01#2    mNeoVis.vulgivaga.1.1.chr14.g00209.t01#4   mNeoVis.vulgivaga.2.1.chr14.g00214.t01#1       5752   125256132:   mNeoVis.vulgivaga.1.1.chr14.g00210.t01#4    mNeoVis.vulgivaga.2.1.chr14.g00215.t01#1    mNeoVis.3.1.1.chr14.g00211.t01#6    mNeoVis.evergladensis.1.1.chr14.g00220.t01#3    mNeoVis.2.1.1.chr14_hap1.g00212.t01#5   mNeoVis.evergladensis.2.1.chr14.g00213.t01#2       5753   125256133:   mNeoVis.3.1.1.chr14.g00144.t01#6    mNeoVis.evergladensis.1.1.chr14.g00144.t01#3    mNeoVis.vulgivaga.1.1.chr14.g00142.t01#4    mNeoVis.vulgivaga.2.1.chr14.g00144.t01#1    mNeoVis.evergladensis.2.1.chr14.g00142.t01#2   mNeoVis.2.1.1.chr14_hap1.g00146.t01#5       5754   125256134:  mNeoVis.vulgivaga.2.1.chr14.g00146.t01#1               5755   125256135:   mNeoVis.evergladensis.2.1.chr14.g00143.t01#2    mNeoVis.vulgivaga.2.1.chr14.g00145.t01#1    mNeoVis.2.1.1.chr14_hap1.g00147.t01#5    mNeoVis.vulgivaga.1.1.chr14.g00145.t01#4    mNeoVis.3.1.1.chr14.g00145.t01#6   mNeoVis.evergladensis.1.1.chr14.g00148.t01#3       5756   125256136:   mNeoVis.2.1.1.chr14_hap1.g00148.t01#5    mNeoVis.vulgivaga.1.1.chr14.g00146.t01#4    mNeoVis.vulgivaga.2.1.chr14.g00147.t01#1    mNeoVis.evergladensis.2.1.chr14.g00144.t01#2    mNeoVis.3.1.1.chr14.g00146.t01#6   mNeoVis.evergladensis.1.1.chr14.g00149.t01#3       5757   125256137:   mNeoVis.3.1.1.chr14.g00147.t01#6    mNeoVis.evergladensis.2.1.chr14.g00145.t01#2    mNeoVis.2.1.1.chr14_hap1.g00149.t01#5    mNeoVis.vulgivaga.1.1.chr14.g00147.t01#4    mNeoVis.vulgivaga.2.1.chr14.g00148.t01#1   mNeoVis.evergladensis.1.1.chr14.g00150.t01#3       5758   125256138:   mNeoVis.3.1.1.chr14.g00148.t01#6    mNeoVis.2.1.1.chr14_hap1.g00150.t01#5    mNeoVis.vulgivaga.1.1.chr14.g00148.t01#4    mNeoVis.vulgivaga.2.1.chr14.g00149.t01#1    mNeoVis.evergladensis.2.1.chr14.g00146.t01#2   mNeoVis.evergladensis.1.1.chr14.g00151.t01#3       5759   125256139:   mNeoVis.evergladensis.2.1.chr14.g00147.t01#2    mNeoVis.evergladensis.1.1.chr14.g00152.t01#3    mNeoVis.3.1.1.chr14.g00149.t01#6    mNeoVis.vulgivaga.1.1.chr14.g00149.t01#4    mNeoVis.2.1.1.chr14_hap1.g00151.t01#5   mNeoVis.vulgivaga.2.1.chr14.g00150.t01#1       5760   125256140:   mNeoVis.evergladensis.1.1.chr14.g00153.t01#3    mNeoVis.vulgivaga.2.1.chr14.g00151.t01#1    mNeoVis.2.1.1.chr14_hap1.g00152.t01#5    mNeoVis.vulgivaga.1.1.chr14.g00150.t01#4    mNeoVis.evergladensis.2.1.chr14.g00148.t01#2   mNeoVis.3.1.1.chr14.g00150.t01#6       5761   125256141:   mNeoVis.vulgivaga.1.1.chr7.g00421.t01#4    mNeoVis.evergladensis.1.1.chr7.g00424.t01#3    mNeoVis.3.1.1.chr7.g00397.t01#6    mNeoVis.evergladensis.1.1.chr7.g00425.t01#3    mNeoVis.evergladensis.2.1.chr7.g00416.t01#2    mNeoVis.2.1.1.chr7_hap1.g00413.t01#5   mNeoVis.vulgivaga.2.1.chr7.g00415.t01#1     5762   125256142:   mNeoVis.vulgivaga.2.1.chr14.g00152.t01#1    mNeoVis.3.1.1.chr14.g00151.t01#6    mNeoVis.2.1.1.chr14_hap1.g00153.t01#5    mNeoVis.vulgivaga.1.1.chr14.g00151.t01#4    mNeoVis.evergladensis.1.1.chr14.g00154.t01#3   mNeoVis.evergladensis.2.1.chr14.g00149.t01#2       5763   125256143:  mNeoVis.vulgivaga.2.1.chr6.g00514.t01#1               5764   125256144:   mNeoVis.vulgivaga.1.1.chr6.g00517.t01#4    mNeoVis.evergladensis.2.1.chr6.g00513.t01#2    mNeoVis.vulgivaga.2.1.chr6.g00515.t01#1    mNeoVis.evergladensis.1.1.chr6.g00513.t01#3    mNeoVis.3.1.1.chr6.g00515.t01#6   mNeoVis.2.1.1.chr4_hap1.g00509.t01#5       5765   125256145:   mNeoVis.vulgivaga.2.1.chr15.g00479.t01#1    mNeoVis.vulgivaga.1.1.chr15.g00472.t01#4    mNeoVis.evergladensis.1.1.chr15.g00142.t01#3    mNeoVis.3.1.1.chr15.g00430.t01#6    mNeoVis.2.1.1.chr15_hap1.g00649.t01#5   mNeoVis.evergladensis.2.1.chr15.g00471.t01#2       5766   125256146:  mNeoVis.vulgivaga.2.1.chr6.g00516.t01#1               5767   125256147:   mNeoVis.evergladensis.1.1.chr6.g00514.t01#3    mNeoVis.2.1.1.chr4_hap1.g00510.t01#5    mNeoVis.3.1.1.chr6.g00516.t01#6    mNeoVis.vulgivaga.2.1.chr6.g00517.t01#1    mNeoVis.evergladensis.2.1.chr6.g00514.t01#2   mNeoVis.vulgivaga.1.1.chr6.g00518.t01#4       5768   125256148:   mNeoVis.evergladensis.2.1.chr14.g00150.t01#2    mNeoVis.vulgivaga.2.1.chr14.g00153.t01#1    mNeoVis.2.1.1.chr14_hap1.g00154.t01#5    mNeoVis.vulgivaga.1.1.chr14.g00152.t01#4    mNeoVis.3.1.1.chr14.g00152.t01#6   mNeoVis.evergladensis.1.1.chr14.g00155.t01#3       5769   125256149:  mNeoVis.vulgivaga.2.1.chr6.g00518.t01#1               5770   125256150:   mNeoVis.vulgivaga.2.1.chr14.g00154.t01#1    mNeoVis.2.1.1.chr14_hap1.g00155.t01#5    mNeoVis.evergladensis.1.1.chr14.g00156.t01#3    mNeoVis.3.1.1.chr14.g00153.t01#6    mNeoVis.vulgivaga.1.1.chr14.g00153.t01#4   mNeoVis.evergladensis.2.1.chr14.g00151.t01#2       5771   125256151:   mNeoVis.evergladensis.1.1.chr6.g00515.t01#3    mNeoVis.2.1.1.chr4_hap1.g00511.t01#5    mNeoVis.3.1.1.chr6.g00518.t01#6    mNeoVis.evergladensis.2.1.chr6.g00515.t01#2    mNeoVis.vulgivaga.1.1.chr6.g00519.t01#4   mNeoVis.vulgivaga.2.1.chr6.g00519.t01#1       5772   125256152:   mNeoVis.evergladensis.1.1.chr6.g00516.t01#3    mNeoVis.evergladensis.2.1.chr6.g00516.t01#2    mNeoVis.vulgivaga.2.1.chr6.g00520.t01#1    mNeoVis.vulgivaga.1.1.chr6.g00520.t01#4    mNeoVis.3.1.1.chr6.g00519.t01#6   mNeoVis.2.1.1.chr4_hap1.g00512.t01#5       5773   125256153:   mNeoVis.vulgivaga.1.1.chr14.g00154.t01#4    mNeoVis.vulgivaga.2.1.chr14.g00155.t01#1    mNeoVis.2.1.1.chr14_hap1.g00156.t01#5    mNeoVis.evergladensis.2.1.chr14.g00152.t01#2    mNeoVis.evergladensis.1.1.chr14.g00158.t01#3   mNeoVis.3.1.1.chr14.g00154.t01#6       5774   125256154:   mNeoVis.evergladensis.2.1.chr6.g00517.t01#2    mNeoVis.vulgivaga.1.1.chr6.g00521.t01#4    mNeoVis.2.1.1.chr4_hap1.g00513.t01#5    mNeoVis.vulgivaga.2.1.chr6.g00521.t01#1   mNeoVis.evergladensis.1.1.chr6.g00517.t01#3       5775   125256155:   mNeoVis.2.1.1.chr4_hap1.g00514.t01#5    mNeoVis.vulgivaga.2.1.chr6.g00522.t01#1    mNeoVis.vulgivaga.1.1.chr6.g00522.t01#4    mNeoVis.3.1.1.chr6.g00520.t01#6    mNeoVis.evergladensis.1.1.chr6.g00518.t01#3   mNeoVis.evergladensis.2.1.chr6.g00518.t01#2       5776   125256156:   mNeoVis.vulgivaga.2.1.chr15.g00480.t01#1   mNeoVis.2.1.1.chr15_hap1.g00650.t01#5             5777   125256157:   mNeoVis.vulgivaga.1.1.chr6.g00523.t01#4    mNeoVis.evergladensis.2.1.chr6.g00520.t01#2    mNeoVis.2.1.1.chr4_hap1.g00515.t01#5    mNeoVis.evergladensis.1.1.chr6.g00519.t01#3    mNeoVis.vulgivaga.2.1.chr6.g00523.t01#1   mNeoVis.3.1.1.chr6.g00521.t01#6       5778   125256158:   mNeoVis.vulgivaga.2.1.chr6.g00524.t01#1    mNeoVis.2.1.1.chr4_hap1.g00516.t01#5    mNeoVis.evergladensis.2.1.chr6.g00521.t01#2    mNeoVis.vulgivaga.1.1.chr6.g00524.t01#4    mNeoVis.3.1.1.chr6.g00523.t01#6   mNeoVis.evergladensis.1.1.chr6.g00520.t01#3       5779   125256159:   mNeoVis.evergladensis.2.1.chr7.g00417.t01#2    mNeoVis.vulgivaga.1.1.chr7.g00422.t01#4    mNeoVis.vulgivaga.2.1.chr7.g00416.t01#1    mNeoVis.3.1.1.chr7.g00398.t01#6    mNeoVis.evergladensis.1.1.chr7.g00426.t01#3   mNeoVis.2.1.1.chr7_hap1.g00414.t01#5       5780   125256160:   mNeoVis.2.1.1.chr4_hap1.g00517.t01#5    mNeoVis.vulgivaga.1.1.chr6.g00525.t01#4    mNeoVis.vulgivaga.2.1.chr6.g00525.t01#1    mNeoVis.evergladensis.2.1.chr6.g00522.t01#2    mNeoVis.3.1.1.chr6.g00524.t01#6   mNeoVis.evergladensis.1.1.chr6.g00521.t01#3       5781   125256161:   mNeoVis.vulgivaga.1.1.chr14.g00155.t01#4    mNeoVis.2.1.1.chr14_hap1.g00157.t01#5    mNeoVis.evergladensis.2.1.chr14.g00153.t01#2    mNeoVis.vulgivaga.2.1.chr14.g00156.t01#1    mNeoVis.3.1.1.chr14.g00155.t01#6   mNeoVis.evergladensis.1.1.chr14.g00159.t01#3       5782   125256162:   mNeoVis.vulgivaga.2.1.chr15.g00481.t01#1    mNeoVis.evergladensis.1.1.chr15.g00140.t01#3    mNeoVis.3.1.1.chr15.g00432.t01#6    mNeoVis.vulgivaga.1.1.chr15.g00474.t01#4    mNeoVis.evergladensis.2.1.chr15.g00473.t01#2   mNeoVis.2.1.1.chr15_hap1.g00651.t01#5       5783   125256163:   mNeoVis.vulgivaga.1.1.chr14.g00156.t01#4    mNeoVis.evergladensis.2.1.chr14.g00154.t01#2    mNeoVis.2.1.1.chr14_hap1.g00158.t01#5    mNeoVis.vulgivaga.2.1.chr14.g00157.t01#1    mNeoVis.3.1.1.chr14.g00156.t01#6   mNeoVis.evergladensis.1.1.chr14.g00160.t01#3       5784   125256164:   mNeoVis.vulgivaga.1.1.chr13.g00228.t01#4    mNeoVis.2.1.1.chr13_hap1.g00236.t01#5    mNeoVis.3.1.1.chr13.g00187.t01#6    mNeoVis.vulgivaga.2.1.chr13.g00231.t01#1    mNeoVis.evergladensis.1.1.chr13.g00233.t01#3   mNeoVis.evergladensis.2.1.chr13.g00234.t01#2       5785   125256165:   mNeoVis.2.1.1.chr14_hap1.g00159.t01#5    mNeoVis.3.1.1.chr14.g00157.t01#6    mNeoVis.evergladensis.1.1.chr14.g00161.t01#3    mNeoVis.evergladensis.2.1.chr14.g00155.t01#2    mNeoVis.vulgivaga.1.1.chr14.g00157.t01#4   mNeoVis.vulgivaga.2.1.chr14.g00158.t01#1       5786   125256166:   mNeoVis.vulgivaga.2.1.chr6.g00526.t01#1    mNeoVis.2.1.1.chr4_hap1.g00518.t01#5    mNeoVis.evergladensis.1.1.chr6.g00522.t01#3    mNeoVis.evergladensis.2.1.chr6.g00523.t01#2    mNeoVis.3.1.1.chr6.g00525.t01#6   mNeoVis.vulgivaga.1.1.chr6.g00526.t01#4       5787   125256167:   mNeoVis.2.1.1.chr4_hap1.g00519.t01#5    mNeoVis.3.1.1.chr6.g00526.t01#6    mNeoVis.evergladensis.2.1.chr6.g00524.t01#2    mNeoVis.vulgivaga.2.1.chr6.g00527.t01#1    mNeoVis.vulgivaga.1.1.chr6.g00527.t01#4   mNeoVis.evergladensis.1.1.chr6.g00523.t01#3       5788   125256168:   mNeoVis.vulgivaga.1.1.chr14.g00158.t01#4    mNeoVis.2.1.1.chr14_hap1.g00160.t01#5    mNeoVis.3.1.1.chr14.g00158.t01#6    mNeoVis.evergladensis.2.1.chr14.g00156.t01#2    mNeoVis.evergladensis.1.1.chr14.g00162.t01#3   mNeoVis.vulgivaga.2.1.chr14.g00159.t01#1       5789   125256169:   mNeoVis.3.1.1.chr14.g00159.t01#6    mNeoVis.vulgivaga.2.1.chr14.g00160.t01#1    mNeoVis.evergladensis.1.1.chr14.g00163.t01#3    mNeoVis.vulgivaga.1.1.chr14.g00159.t01#4    mNeoVis.evergladensis.2.1.chr14.g00157.t01#2   mNeoVis.2.1.1.chr14_hap1.g00161.t01#5       5790   125256170:   mNeoVis.evergladensis.2.1.chr7.g00418.t01#2    mNeoVis.3.1.1.chr7.g00399.t01#6    mNeoVis.vulgivaga.2.1.chr7.g00417.t01#1    mNeoVis.vulgivaga.1.1.chr7.g00423.t01#4    mNeoVis.evergladensis.1.1.chr7.g00427.t01#3   mNeoVis.2.1.1.chr7_hap1.g00415.t01#5       5791   125256171:   mNeoVis.evergladensis.1.1.chr6.g00524.t01#3    mNeoVis.2.1.1.chr4_hap1.g00520.t01#5    mNeoVis.vulgivaga.2.1.chr6.g00528.t01#1    mNeoVis.3.1.1.chr6.g00527.t01#6    mNeoVis.evergladensis.2.1.chr6.g00525.t01#2   mNeoVis.vulgivaga.1.1.chr6.g00528.t01#4       5792   125256172:   mNeoVis.vulgivaga.1.1.chr8.g01357.t01#4    mNeoVis.vulgivaga.1.1.chr14.g00160.t01#4    mNeoVis.2.1.1.chr8_hap1.g01420.t01#5    mNeoVis.2.1.1.chr14_hap1.g00162.t01#5    mNeoVis.3.1.1.chr14.g00160.t01#6    mNeoVis.3.1.1.chr8.g01318.t01#6    mNeoVis.vulgivaga.2.1.chr14.g00161.t01#1    mNeoVis.vulgivaga.2.1.chr8.g01374.t01#1    mNeoVis.evergladensis.1.1.chr14.g00164.t01#3      5793   125256173:   mNeoVis.3.1.1.chr14.g00161.t01#6    mNeoVis.vulgivaga.2.1.chr14.g00162.t01#1   mNeoVis.vulgivaga.1.1.chr14.g00161.t01#4         5794   125256174:   mNeoVis.2.1.1.chr4_hap1.g00521.t01#5    mNeoVis.vulgivaga.1.1.chr6.g00529.t01#4    mNeoVis.vulgivaga.2.1.chr6.g00529.t01#1    mNeoVis.evergladensis.2.1.chr6.g00526.t01#2    mNeoVis.evergladensis.1.1.chr6.g00525.t01#3   mNeoVis.3.1.1.chr6.g00528.t01#6       5795   125256175:   mNeoVis.vulgivaga.2.1.chr6.g00530.t01#1    mNeoVis.evergladensis.1.1.chr6.g00526.t01#3    mNeoVis.3.1.1.chr6.g00529.t01#6    mNeoVis.evergladensis.2.1.chr6.g00527.t01#2    mNeoVis.vulgivaga.1.1.chr6.g00530.t01#4   mNeoVis.2.1.1.chr4_hap1.g00522.t01#5       5796   125256176:   mNeoVis.vulgivaga.2.1.chr15.g00482.t01#1    mNeoVis.vulgivaga.1.1.chr15.g00475.t01#4    mNeoVis.evergladensis.1.1.chr15.g00139.t01#3    mNeoVis.2.1.1.chr15_hap1.g00652.t01#5    mNeoVis.evergladensis.2.1.chr15.g00474.t01#2   mNeoVis.3.1.1.chr15.g00433.t01#6       5797   125256177:   mNeoVis.3.1.1.chr14.g00162.t01#6    mNeoVis.2.1.1.chr14_hap1.g00164.t01#5   mNeoVis.vulgivaga.2.1.chr14.g00163.t01#1         5798   125256178:   mNeoVis.evergladensis.2.1.chr14.g00161.t01#2    mNeoVis.2.1.1.chr14_hap1.g00165.t01#5   mNeoVis.vulgivaga.2.1.chr14.g00164.t01#1         5799   125256179:   mNeoVis.vulgivaga.2.1.chr14.g00165.t01#1    mNeoVis.evergladensis.1.1.chr14.g00167.t01#3    mNeoVis.evergladensis.2.1.chr14.g00162.t01#2    mNeoVis.vulgivaga.1.1.chr14.g00163.t01#4    mNeoVis.3.1.1.chr14.g00163.t01#6   mNeoVis.2.1.1.chr14_hap1.g00166.t01#5       5800   125256180:   mNeoVis.evergladensis.2.1.chr14.g00163.t01#2    mNeoVis.vulgivaga.1.1.chr14.g00164.t01#4    mNeoVis.3.1.1.chr14.g00164.t01#6    mNeoVis.2.1.1.chr14_hap1.g00167.t01#5    mNeoVis.vulgivaga.2.1.chr14.g00166.t01#1   mNeoVis.evergladensis.1.1.chr14.g00168.t01#3       5801   125256181:   mNeoVis.3.1.1.chr6.g00530.t01#6    mNeoVis.evergladensis.2.1.chr6.g00528.t01#2    mNeoVis.2.1.1.chr4_hap1.g00523.t01#5    mNeoVis.evergladensis.1.1.chr6.g00527.t01#3    mNeoVis.vulgivaga.2.1.chr6.g00531.t01#1   mNeoVis.vulgivaga.1.1.chr6.g00531.t01#4       5802   125256182:  mNeoVis.vulgivaga.2.1.chr14.g00167.t01#1               5803   125256183:   mNeoVis.evergladensis.1.1.chr14.g00170.t01#3    mNeoVis.3.1.1.chr14.g00166.t01#6    mNeoVis.evergladensis.2.1.chr14.g00165.t01#2    mNeoVis.vulgivaga.2.1.chr14.g00168.t01#1    mNeoVis.vulgivaga.1.1.chr14.g00166.t01#4   mNeoVis.2.1.1.chr14_hap1.g00169.t01#5       5804   125256184:   mNeoVis.2.1.1.chr14_hap1.g00170.t01#5    mNeoVis.evergladensis.1.1.chr14.g00171.t01#3    mNeoVis.vulgivaga.2.1.chr14.g00169.t01#1    mNeoVis.3.1.1.chr14.g00167.t01#6    mNeoVis.evergladensis.2.1.chr14.g00166.t01#2   mNeoVis.vulgivaga.1.1.chr14.g00167.t01#4       5805   125256185:   mNeoVis.evergladensis.1.1.chr14.g00172.t01#3    mNeoVis.2.1.1.chr14_hap1.g00171.t01#5    mNeoVis.vulgivaga.1.1.chr14.g00168.t01#4    mNeoVis.evergladensis.2.1.chr14.g00167.t01#2    mNeoVis.vulgivaga.2.1.chr14.g00170.t01#1   mNeoVis.3.1.1.chr14.g00168.t01#6       5806   125256186:   mNeoVis.vulgivaga.1.1.chr6.g00533.t01#4    mNeoVis.evergladensis.1.1.chr6.g00528.t01#3    mNeoVis.vulgivaga.2.1.chr6.g00532.t01#1    mNeoVis.2.1.1.chr4_hap1.g00524.t01#5    mNeoVis.evergladensis.2.1.chr6.g00529.t01#2   mNeoVis.3.1.1.chr6.g00532.t01#6       5807   125256187:   mNeoVis.vulgivaga.2.1.chr6.g00533.t01#1    mNeoVis.evergladensis.2.1.chr6.g00530.t01#2    mNeoVis.2.1.1.chr4_hap1.g00525.t01#5    mNeoVis.3.1.1.chr6.g00533.t01#6    mNeoVis.vulgivaga.1.1.chr6.g00534.t01#4   mNeoVis.evergladensis.1.1.chr6.g00529.t01#3       5808   125256188:   mNeoVis.2.1.1.chr14_hap1.g00172.t01#5    mNeoVis.vulgivaga.1.1.chr14.g00169.t01#4    mNeoVis.evergladensis.1.1.chr14.g00173.t01#3    mNeoVis.vulgivaga.2.1.chr14.g00171.t01#1    mNeoVis.evergladensis.2.1.chr14.g00168.t01#2   mNeoVis.3.1.1.chr14.g00169.t01#6       5809   125256189:   mNeoVis.evergladensis.1.1.chr15.g00138.t01#3    mNeoVis.3.1.1.chr15.g00434.t01#6    mNeoVis.vulgivaga.1.1.chr15.g00476.t01#4   mNeoVis.vulgivaga.2.1.chr15.g00483.t01#1       5810   125256190:   mNeoVis.vulgivaga.1.1.chr6.g00535.t01#4    mNeoVis.3.1.1.chr6.g00534.t01#6    mNeoVis.evergladensis.2.1.chr6.g00531.t01#2    mNeoVis.evergladensis.1.1.chr6.g00530.t01#3    mNeoVis.vulgivaga.2.1.chr6.g00534.t01#1   mNeoVis.2.1.1.chr4_hap1.g00526.t01#5       5811   125256191:   mNeoVis.vulgivaga.2.1.chr14.g00172.t01#1    mNeoVis.3.1.1.chr14.g00170.t01#6    mNeoVis.evergladensis.1.1.chr14.g00174.t01#3    mNeoVis.vulgivaga.1.1.chr14.g00170.t01#4    mNeoVis.2.1.1.chr14_hap1.g00173.t01#5   mNeoVis.evergladensis.2.1.chr14.g00169.t01#2       5812   125256192:   mNeoVis.vulgivaga.1.1.chr7.g00424.t01#4    mNeoVis.3.1.1.chr7.g00400.t01#6    mNeoVis.evergladensis.2.1.chr7.g00419.t01#2    mNeoVis.evergladensis.1.1.chr7.g00428.t01#3    mNeoVis.2.1.1.chr7_hap1.g00416.t01#5   mNeoVis.vulgivaga.2.1.chr7.g00418.t01#1       5813   125256193:  mNeoVis.vulgivaga.2.1.chr14.g00173.t01#1               5814   125256194:   mNeoVis.evergladensis.2.1.chr6.g00532.t01#2    mNeoVis.vulgivaga.2.1.chr6.g00535.t01#1    mNeoVis.2.1.1.chr4_hap1.g00527.t01#5    mNeoVis.3.1.1.chr6.g00535.t01#6    mNeoVis.vulgivaga.1.1.chr6.g00536.t01#4   mNeoVis.evergladensis.1.1.chr6.g00531.t01#3       5815   125256195:   mNeoVis.evergladensis.2.1.chr14.g00170.t01#2    mNeoVis.3.1.1.chr14.g00171.t01#6    mNeoVis.vulgivaga.2.1.chr14.g00174.t01#1    mNeoVis.vulgivaga.1.1.chr14.g00171.t01#4   mNeoVis.evergladensis.1.1.chr14.g00175.t01#3       5816   125256196:   mNeoVis.vulgivaga.2.1.chr14.g00175.t01#1    mNeoVis.3.1.1.chr14.g00172.t01#6    mNeoVis.2.1.1.chr14_hap1.g00174.t01#5    mNeoVis.evergladensis.1.1.chr14.g00176.t01#3    mNeoVis.vulgivaga.1.1.chr14.g00172.t01#4   mNeoVis.evergladensis.2.1.chr14.g00171.t01#2       5817   125256197:   mNeoVis.2.1.1.chr4_hap1.g00528.t01#5    mNeoVis.evergladensis.2.1.chr6.g00533.t01#2    mNeoVis.vulgivaga.2.1.chr6.g00536.t01#1    mNeoVis.3.1.1.chr6.g00537.t01#6    mNeoVis.evergladensis.1.1.chr6.g00533.t01#3   mNeoVis.vulgivaga.1.1.chr6.g00537.t01#4       5818   125256198:   mNeoVis.vulgivaga.1.1.chr6.g00538.t01#4    mNeoVis.2.1.1.chr4_hap1.g00529.t01#5    mNeoVis.vulgivaga.2.1.chr6.g00537.t01#1   mNeoVis.evergladensis.2.1.chr6.g00534.t01#2       5819   125256199:   mNeoVis.vulgivaga.1.1.chr15.g00482.t01#4    mNeoVis.3.1.1.chr15.g00436.t01#6    mNeoVis.3.1.1.chr15.g00435.t01#6    mNeoVis.vulgivaga.2.1.chr15.g00484.t01#1    mNeoVis.evergladensis.2.1.chr15.g00475.t01#2    mNeoVis.2.1.1.chr15_hap1.g00653.t01#5    mNeoVis.evergladensis.1.1.chr15.g00137.t01#3    mNeoVis.2.1.1.chr15_hap1.g00654.t01#5    mNeoVis.evergladensis.1.1.chr15.g00135.t01#3      5820   125256200:   mNeoVis.2.1.1.chr15_hap1.g00656.t01#5    mNeoVis.evergladensis.2.1.chr15.g00478.t01#2    mNeoVis.3.1.1.chr15.g00440.t01#6    mNeoVis.vulgivaga.2.1.chr15.g00488.t01#1    mNeoVis.evergladensis.1.1.chr15.g00134.t01#3   mNeoVis.vulgivaga.1.1.chr15.g00481.t01#4       5821   125256201:   mNeoVis.evergladensis.2.1.chr15.g00477.t01#2    mNeoVis.vulgivaga.1.1.chr15.g00480.t01#4    mNeoVis.vulgivaga.2.1.chr15.g00487.t01#1    mNeoVis.2.1.1.chr15_hap1.g00655.t01#5   mNeoVis.3.1.1.chr15.g00439.t01#6       5822   125256202:   mNeoVis.evergladensis.2.1.chr6.g00535.t01#2    mNeoVis.evergladensis.1.1.chr6.g00534.t01#3    mNeoVis.vulgivaga.1.1.chr6.g00539.t01#4    mNeoVis.vulgivaga.2.1.chr6.g00538.t01#1   mNeoVis.3.1.1.chr6.g00538.t01#6       5823   125256203:   mNeoVis.2.1.1.chr4_hap1.g00531.t01#5    mNeoVis.vulgivaga.2.1.chr6.g00539.t01#1    mNeoVis.3.1.1.chr6.g00539.t01#6    mNeoVis.vulgivaga.1.1.chr6.g00540.t01#4    mNeoVis.evergladensis.1.1.chr6.g00535.t01#3   mNeoVis.evergladensis.2.1.chr6.g00536.t01#2       5824   125256204:   mNeoVis.2.1.1.chr4_hap1.g00530.t01#5    mNeoVis.3.1.1.chr6.g00540.t01#6    mNeoVis.evergladensis.2.1.chr6.g00537.t01#2    mNeoVis.vulgivaga.1.1.chr6.g00541.t01#4    mNeoVis.vulgivaga.2.1.chr6.g00540.t01#1   mNeoVis.evergladensis.1.1.chr6.g00536.t01#3       5825   125256205:   mNeoVis.evergladensis.2.1.chr6.g00538.t01#2    mNeoVis.3.1.1.chr6.g00541.t01#6    mNeoVis.vulgivaga.1.1.chr6.g00542.t01#4    mNeoVis.vulgivaga.2.1.chr6.g00541.t01#1    mNeoVis.evergladensis.1.1.chr6.g00537.t01#3   mNeoVis.2.1.1.chr4_hap1.g00532.t01#5       5826   125256206:   mNeoVis.vulgivaga.1.1.chr6.g00543.t01#4    mNeoVis.2.1.1.chr4_hap1.g00533.t01#5    mNeoVis.evergladensis.1.1.chr6.g00538.t01#3    mNeoVis.evergladensis.2.1.chr6.g00539.t01#2    mNeoVis.vulgivaga.2.1.chr6.g00542.t01#1   mNeoVis.3.1.1.chr6.g00542.t01#6       5827   125256207:   mNeoVis.3.1.1.chr6.g00457.t01#6    mNeoVis.vulgivaga.2.1.chr6.g00457.t01#1    mNeoVis.vulgivaga.1.1.chr6.g00464.t01#4    mNeoVis.evergladensis.1.1.chr6.g00462.t01#3    mNeoVis.2.1.1.chr4_hap1.g00457.t01#5   mNeoVis.evergladensis.2.1.chr6.g00464.t01#2       5828   125256208:   mNeoVis.vulgivaga.1.1.chr6.g00465.t01#4    mNeoVis.vulgivaga.2.1.chr6.g00458.t01#1    mNeoVis.3.1.1.chr6.g00458.t01#6    mNeoVis.2.1.1.chr4_hap1.g00458.t01#5    mNeoVis.vulgivaga.2.1.chr6.g00459.t01#1    mNeoVis.evergladensis.1.1.chr6.g00463.t01#3   mNeoVis.evergladensis.2.1.chr6.g00465.t01#2     5829   125256209:   mNeoVis.2.1.1.chr14_hap1.g00175.t01#5    mNeoVis.vulgivaga.2.1.chr14.g00176.t01#1    mNeoVis.3.1.1.chr14.g00173.t01#6    mNeoVis.vulgivaga.1.1.chr14.g00173.t01#4    mNeoVis.evergladensis.1.1.chr14.g00177.t01#3   mNeoVis.evergladensis.2.1.chr14.g00172.t01#2       5830   125256210:   mNeoVis.2.1.1.chr4_hap1.g00459.t01#5    mNeoVis.3.1.1.chr6.g00459.t01#6    mNeoVis.evergladensis.2.1.chr6.g00466.t01#2    mNeoVis.vulgivaga.1.1.chr6.g00466.t01#4    mNeoVis.vulgivaga.2.1.chr6.g00460.t01#1   mNeoVis.evergladensis.1.1.chr6.g00464.t01#3       5831   125256211:   mNeoVis.vulgivaga.1.1.chr7.g00425.t01#4    mNeoVis.vulgivaga.2.1.chr7.g00419.t01#1    mNeoVis.3.1.1.chr7.g00401.t01#6    mNeoVis.evergladensis.2.1.chr7.g00420.t01#2    mNeoVis.evergladensis.1.1.chr7.g00429.t01#3   mNeoVis.2.1.1.chr7_hap1.g00417.t01#5       5832   125256212:   mNeoVis.vulgivaga.2.1.chr6.g00461.t01#1    mNeoVis.evergladensis.1.1.chr6.g00465.t01#3    mNeoVis.2.1.1.chr4_hap1.g00460.t01#5    mNeoVis.3.1.1.chr6.g00460.t01#6    mNeoVis.vulgivaga.1.1.chr6.g00467.t01#4   mNeoVis.evergladensis.2.1.chr6.g00467.t01#2       5833   125256213:   mNeoVis.vulgivaga.2.1.chr14.g00177.t01#1    mNeoVis.3.1.1.chr14.g00174.t01#6    mNeoVis.vulgivaga.1.1.chr14.g00174.t01#4    mNeoVis.evergladensis.1.1.chr14.g00178.t01#3    mNeoVis.2.1.1.chr14_hap1.g00176.t01#5   mNeoVis.evergladensis.2.1.chr14.g00173.t01#2       5834   125256214:   mNeoVis.3.1.1.chr6.g00461.t01#6    mNeoVis.vulgivaga.1.1.chr6.g00468.t01#4    mNeoVis.evergladensis.2.1.chr6.g00468.t01#2    mNeoVis.evergladensis.1.1.chr6.g00466.t01#3    mNeoVis.vulgivaga.2.1.chr6.g00462.t01#1   mNeoVis.2.1.1.chr4_hap1.g00461.t01#5       5835   125256215:   mNeoVis.2.1.1.chr15_hap1.g00137.t01#5    mNeoVis.evergladensis.2.1.chr15.g00123.t01#2    mNeoVis.vulgivaga.1.1.chr15.g00123.t01#4    mNeoVis.vulgivaga.2.1.chr15.g00127.t01#1    mNeoVis.evergladensis.1.1.chr15.g00493.t01#3   mNeoVis.3.1.1.chr15.g00115.t01#6       5836   125256216:   mNeoVis.3.1.1.chr14.g00091.t01#6    mNeoVis.vulgivaga.2.1.chr14.g00097.t01#1    mNeoVis.3.1.1.chr7.g01266.t01#6    mNeoVis.vulgivaga.1.1.chr14.g00097.t01#4    mNeoVis.2.1.1.chr14_hap1.g00098.t01#5    mNeoVis.evergladensis.2.1.chr14.g00097.t01#2   mNeoVis.evergladensis.1.1.chr14.g00102.t01#3     5837   125256217:   mNeoVis.2.1.1.chr4_hap1.g00784.t01#5    mNeoVis.2.1.1.chr12_hap1.g00702.t01#5    mNeoVis.2.1.1.chr1_hap1.g01956.t01#5    mNeoVis.vulgivaga.1.1.chr12.g00153.t01#4    mNeoVis.evergladensis.1.1.chr13.g00533.t01#3    mNeoVis.evergladensis.1.1.chr12.g00641.t01#3    mNeoVis.2.1.1.chr5_hap1.g01589.t01#5    mNeoVis.vulgivaga.2.1.chr13.g00540.t01#1    mNeoVis.vulgivaga.2.1.chr12.g00149.t01#1      5838   125256218:   mNeoVis.vulgivaga.2.1.chr6.g00463.t01#1   mNeoVis.vulgivaga.1.1.chr6.g00470.t01#4             5839   125256219:   mNeoVis.evergladensis.2.1.chr14.g00098.t01#2    mNeoVis.vulgivaga.1.1.chr14.g00098.t01#4    mNeoVis.2.1.1.chr14_hap1.g00099.t01#5    mNeoVis.vulgivaga.2.1.chr14.g00098.t01#1    mNeoVis.evergladensis.1.1.chr14.g00103.t01#3   mNeoVis.3.1.1.chr14.g00092.t01#6       5840   125256220:   mNeoVis.3.1.1.chr14.g00093.t01#6    mNeoVis.evergladensis.2.1.chr14.g00099.t01#2    mNeoVis.vulgivaga.1.1.chr14.g00099.t01#4    mNeoVis.evergladensis.1.1.chr14.g00104.t01#3    mNeoVis.2.1.1.chr14_hap1.g00100.t01#5   mNeoVis.vulgivaga.2.1.chr14.g00099.t01#1       5841   125256221:  mNeoVis.vulgivaga.2.1.chr6.g00465.t01#1               5842   125256222:   mNeoVis.evergladensis.1.1.chr6.g00467.t01#3    mNeoVis.3.1.1.chr6.g00462.t01#6    mNeoVis.2.1.1.chr4_hap1.g00462.t01#5    mNeoVis.vulgivaga.2.1.chr6.g00464.t01#1    mNeoVis.evergladensis.2.1.chr6.g00469.t01#2   mNeoVis.vulgivaga.1.1.chr6.g00469.t01#4       5843   125256223:   mNeoVis.evergladensis.1.1.chr6.g00468.t01#3    mNeoVis.evergladensis.2.1.chr6.g00470.t01#2    mNeoVis.vulgivaga.2.1.chr6.g00466.t01#1    mNeoVis.2.1.1.chr4_hap1.g00463.t01#5    mNeoVis.vulgivaga.1.1.chr6.g00471.t01#4   mNeoVis.3.1.1.chr6.g00463.t01#6       5844   125256224:   mNeoVis.evergladensis.2.1.chr14.g00100.t01#2    mNeoVis.vulgivaga.1.1.chr14.g00100.t01#4    mNeoVis.3.1.1.chr14.g00095.t01#6    mNeoVis.vulgivaga.2.1.chr14.g00100.t01#1    mNeoVis.2.1.1.chr14_hap1.g00101.t01#5   mNeoVis.evergladensis.1.1.chr14.g00105.t01#3       5845   125256225:   mNeoVis.evergladensis.2.1.chr14.g00101.t01#2    mNeoVis.vulgivaga.1.1.chr14.g00101.t01#4    mNeoVis.evergladensis.1.1.chr14.g00106.t01#3    mNeoVis.vulgivaga.2.1.chr14.g00101.t01#1    mNeoVis.3.1.1.chr14.g00096.t01#6   mNeoVis.2.1.1.chr14_hap1.g00102.t01#5       5846   125256226:   mNeoVis.evergladensis.1.1.chr6.g00469.t01#3    mNeoVis.vulgivaga.1.1.chr6.g00472.t01#4    mNeoVis.vulgivaga.2.1.chr6.g00467.t01#1    mNeoVis.2.1.1.chr4_hap1.g00464.t01#5    mNeoVis.evergladensis.2.1.chr6.g00471.t01#2   mNeoVis.3.1.1.chr15.g00376.t01#6       5847   125256227:   mNeoVis.evergladensis.1.1.chr14.g00107.t01#3    mNeoVis.2.1.1.chr14_hap1.g00103.t01#5    mNeoVis.evergladensis.2.1.chr14.g00102.t01#2    mNeoVis.vulgivaga.1.1.chr14.g00102.t01#4    mNeoVis.3.1.1.chr14.g00097.t01#6   mNeoVis.vulgivaga.2.1.chr14.g00102.t01#1       5848   125256228:   mNeoVis.vulgivaga.2.1.chr14.g00103.t01#1    mNeoVis.2.1.1.chr14_hap1.g00104.t01#5    mNeoVis.evergladensis.2.1.chr14.g00103.t01#2    mNeoVis.evergladensis.1.1.chr14.g00108.t01#3    mNeoVis.3.1.1.chr14.g00098.t01#6   mNeoVis.vulgivaga.1.1.chr14.g00103.t01#4       5849   125256229:   mNeoVis.evergladensis.2.1.chr6.g00472.t01#2   mNeoVis.vulgivaga.2.1.chr6.g00468.t01#1             5850   125256230:   mNeoVis.vulgivaga.1.1.chr6.g00474.t01#4    mNeoVis.evergladensis.1.1.chr6.g00470.t01#3    mNeoVis.2.1.1.chr4_hap1.g00466.t01#5    mNeoVis.3.1.1.chr6.g00465.t01#6    mNeoVis.vulgivaga.2.1.chr6.g00469.t01#1   mNeoVis.evergladensis.2.1.chr6.g00473.t01#2       5851   125256231:   mNeoVis.vulgivaga.2.1.chr6.g00470.t01#1    mNeoVis.3.1.1.chr6.g00466.t01#6    mNeoVis.evergladensis.2.1.chr6.g00474.t01#2    mNeoVis.vulgivaga.1.1.chr6.g00475.t01#4    mNeoVis.evergladensis.1.1.chr6.g00471.t01#3   mNeoVis.2.1.1.chr4_hap1.g00467.t01#5       5852   125256232:   mNeoVis.evergladensis.1.1.chr6.g00472.t01#3    mNeoVis.vulgivaga.1.1.chr6.g00476.t01#4    mNeoVis.vulgivaga.2.1.chr6.g00471.t01#1    mNeoVis.2.1.1.chr4_hap1.g00468.t01#5    mNeoVis.3.1.1.chr6.g00467.t01#6   mNeoVis.evergladensis.2.1.chr6.g00475.t01#2       5853   125256233:   mNeoVis.evergladensis.2.1.chr6.g00476.t01#2    mNeoVis.3.1.1.chr6.g00468.t01#6    mNeoVis.vulgivaga.1.1.chr6.g00477.t01#4    mNeoVis.vulgivaga.2.1.chr6.g00472.t01#1    mNeoVis.evergladensis.1.1.chr6.g00473.t01#3   mNeoVis.2.1.1.chr4_hap1.g00469.t01#5       5854   125256234:   mNeoVis.vulgivaga.2.1.chr14.g00104.t01#1    mNeoVis.3.1.1.chr14.g00099.t01#6    mNeoVis.evergladensis.2.1.chr14.g00104.t01#2    mNeoVis.vulgivaga.1.1.chr14.g00104.t01#4    mNeoVis.evergladensis.1.1.chr14.g00109.t01#3   mNeoVis.2.1.1.chr14_hap1.g00105.t01#5       5855   125256235:   mNeoVis.evergladensis.2.1.chr3.g00885.t01#2    mNeoVis.evergladensis.1.1.chr3.g00892.t01#3    mNeoVis.vulgivaga.2.1.chr3.g00901.t01#1    mNeoVis.3.1.1.chr3.g00874.t01#6    mNeoVis.vulgivaga.1.1.chr3.g00889.t01#4    mNeoVis.2.1.1.chr3_hap1.g00891.t01#5   mNeoVis.3.1.1.chr3.g00875.t01#6       5856   125256236:   mNeoVis.vulgivaga.1.1.chr6.g00478.t01#4    mNeoVis.evergladensis.2.1.chr6.g00477.t01#2    mNeoVis.evergladensis.1.1.chr6.g00474.t01#3    mNeoVis.2.1.1.chr4_hap1.g00470.t01#5    mNeoVis.3.1.1.chr6.g00469.t01#6   mNeoVis.vulgivaga.2.1.chr6.g00473.t01#1       5857   125256237:   mNeoVis.vulgivaga.2.1.chr14.g00105.t01#1   mNeoVis.vulgivaga.1.1.chr14.g00105.t01#4           5858   125256238:   mNeoVis.3.1.1.chr6.g00472.t01#6    mNeoVis.3.1.1.chr6.g00471.t01#6    mNeoVis.evergladensis.2.1.chr6.g00478.t01#2    mNeoVis.3.1.1.chr6.g00470.t01#6    mNeoVis.vulgivaga.1.1.chr6.g00479.t01#4    mNeoVis.evergladensis.1.1.chr6.g00475.t01#3    mNeoVis.2.1.1.chr4_hap1.g00471.t01#5   mNeoVis.vulgivaga.2.1.chr6.g00474.t01#1     5859   125256239:   mNeoVis.vulgivaga.2.1.chr3.g00902.t01#1    mNeoVis.2.1.1.chr3_hap1.g00892.t01#5    mNeoVis.evergladensis.1.1.chr3.g00893.t01#3    mNeoVis.evergladensis.2.1.chr3.g00886.t01#2    mNeoVis.3.1.1.chr3.g00876.t01#6   mNeoVis.vulgivaga.1.1.chr3.g00890.t01#4       5860   125256240:   mNeoVis.evergladensis.2.1.chr14.g00106.t01#2    mNeoVis.3.1.1.chr14.g00101.t01#6    mNeoVis.2.1.1.chr14_hap1.g00107.t01#5    mNeoVis.vulgivaga.2.1.chr14.g00107.t01#1    mNeoVis.vulgivaga.1.1.chr14.g00107.t01#4   mNeoVis.evergladensis.1.1.chr14.g00112.t01#3       5861   125256241:   mNeoVis.vulgivaga.2.1.chr6.g00475.t01#1    mNeoVis.3.1.1.chr6.g00473.t01#6    mNeoVis.2.1.1.chr4_hap1.g00472.t01#5    mNeoVis.evergladensis.2.1.chr6.g00479.t01#2    mNeoVis.vulgivaga.1.1.chr6.g00480.t01#4   mNeoVis.evergladensis.1.1.chr6.g00476.t01#3       5862   125256242:   mNeoVis.3.1.1.chr14.g00102.t01#6    mNeoVis.2.1.1.chr14_hap1.g00108.t01#5    mNeoVis.evergladensis.1.1.chr14.g00113.t01#3    mNeoVis.evergladensis.2.1.chr14.g00107.t01#2    mNeoVis.vulgivaga.2.1.chr14.g00108.t01#1   mNeoVis.vulgivaga.1.1.chr14.g00108.t01#4       5863   125256243:   mNeoVis.evergladensis.2.1.chr14.g00108.t01#2    mNeoVis.2.1.1.chr14_hap1.g00109.t01#5    mNeoVis.vulgivaga.2.1.chr14.g00109.t01#1    mNeoVis.vulgivaga.1.1.chr14.g00109.t01#4    mNeoVis.3.1.1.chr14.g00103.t01#6   mNeoVis.evergladensis.1.1.chr14.g00114.t01#3       5864   125256244:   mNeoVis.vulgivaga.1.1.chr14.g00110.t01#4    mNeoVis.evergladensis.1.1.chr14.g00115.t01#3    mNeoVis.vulgivaga.2.1.chr14.g00110.t01#1    mNeoVis.evergladensis.2.1.chr14.g00109.t01#2    mNeoVis.2.1.1.chr14_hap1.g00112.t01#5   mNeoVis.3.1.1.chr14.g00105.t01#6       5865   125256245:   mNeoVis.evergladensis.1.1.chr6.g00477.t01#3    mNeoVis.evergladensis.2.1.chr6.g00480.t01#2    mNeoVis.2.1.1.chr4_hap1.g00473.t01#5    mNeoVis.3.1.1.chr6.g00474.t01#6    mNeoVis.vulgivaga.2.1.chr6.g00476.t01#1    mNeoVis.evergladensis.2.1.chr6.g00481.t01#2   mNeoVis.vulgivaga.1.1.chr6.g00481.t01#4     5866   125256246:   mNeoVis.evergladensis.2.1.chr14.g00110.t01#2    mNeoVis.3.1.1.chr14.g00106.t01#6    mNeoVis.vulgivaga.1.1.chr14.g00111.t01#4    mNeoVis.vulgivaga.2.1.chr14.g00111.t01#1   mNeoVis.2.1.1.chr14_hap1.g00113.t01#5       5867   125256247:   mNeoVis.vulgivaga.2.1.chr6.g00477.t01#1    mNeoVis.evergladensis.1.1.chr6.g00478.t01#3   mNeoVis.3.1.1.chr6.g00475.t01#6           5868   125256248:   mNeoVis.vulgivaga.2.1.chr14.g00112.t01#1    mNeoVis.vulgivaga.1.1.chr14.g00112.t01#4    mNeoVis.evergladensis.1.1.chr14.g00116.t01#3    mNeoVis.3.1.1.chr14.g00107.t01#6    mNeoVis.2.1.1.chr14_hap1.g00114.t01#5   mNeoVis.evergladensis.2.1.chr14.g00111.t01#2       5869   125256249:   mNeoVis.evergladensis.1.1.chr6.g00479.t01#3    mNeoVis.3.1.1.chr6.g00477.t01#6    mNeoVis.2.1.1.chr4_hap1.g00475.t01#5    mNeoVis.evergladensis.2.1.chr6.g00483.t01#2    mNeoVis.vulgivaga.1.1.chr6.g00483.t01#4   mNeoVis.vulgivaga.2.1.chr6.g00478.t01#1       5870   125256250:   mNeoVis.vulgivaga.2.1.chr14.g00113.t01#1   mNeoVis.2.1.1.chr14_hap1.g00115.t01#5             5871   125256251:   mNeoVis.evergladensis.2.1.chr14.g00113.t01#2    mNeoVis.vulgivaga.2.1.chr14.g00114.t01#1    mNeoVis.evergladensis.1.1.chr14.g00118.t01#3    mNeoVis.vulgivaga.1.1.chr14.g00114.t01#4    mNeoVis.2.1.1.chr14_hap1.g00116.t01#5   mNeoVis.3.1.1.chr14.g00110.t01#6       5872   125256252:   mNeoVis.vulgivaga.1.1.chr6.g00484.t01#4    mNeoVis.vulgivaga.2.1.chr6.g00479.t01#1    mNeoVis.3.1.1.chr6.g00478.t01#6    mNeoVis.2.1.1.chr4_hap1.g00476.t01#5    mNeoVis.evergladensis.1.1.chr6.g00480.t01#3   mNeoVis.evergladensis.2.1.chr6.g00484.t01#2       5873   125256253:   mNeoVis.vulgivaga.1.1.chr14.g00115.t01#4    mNeoVis.evergladensis.1.1.chr14.g00119.t01#3    mNeoVis.3.1.1.chr14.g00111.t01#6    mNeoVis.2.1.1.chr14_hap1.g00117.t01#5    mNeoVis.evergladensis.2.1.chr14.g00114.t01#2   mNeoVis.vulgivaga.2.1.chr14.g00115.t01#1       5874   125256254:  mNeoVis.vulgivaga.2.1.chr6.g00480.t01#1               5875   125256255:   mNeoVis.vulgivaga.2.1.chr6.g00481.t01#1   mNeoVis.3.1.1.chr6.g00479.t01#6             5876   125256256:   mNeoVis.vulgivaga.2.1.chr6.g00482.t01#1    mNeoVis.evergladensis.1.1.chr6.g00481.t01#3    mNeoVis.3.1.1.chr6.g00480.t01#6    mNeoVis.2.1.1.chr4_hap1.g00477.t01#5   mNeoVis.vulgivaga.1.1.chr6.g00485.t01#4       5877   125256257:   mNeoVis.2.1.1.chr14_hap1.g00118.t01#5    mNeoVis.vulgivaga.2.1.chr14.g00116.t01#1    mNeoVis.evergladensis.1.1.chr14.g00120.t01#3    mNeoVis.3.1.1.chr14.g00112.t01#6    mNeoVis.vulgivaga.1.1.chr14.g00116.t01#4   mNeoVis.evergladensis.2.1.chr14.g00115.t01#2       5878   125256258:   mNeoVis.2.1.1.chr4_hap1.g00478.t01#5    mNeoVis.vulgivaga.2.1.chr6.g00483.t01#1    mNeoVis.vulgivaga.1.1.chr6.g00486.t01#4    mNeoVis.3.1.1.chr6.g00481.t01#6    mNeoVis.evergladensis.2.1.chr6.g00485.t01#2   mNeoVis.evergladensis.1.1.chr6.g00482.t01#3       5879   125256259:   mNeoVis.2.1.1.chr14_hap1.g00119.t01#5    mNeoVis.vulgivaga.2.1.chr14.g00117.t01#1   mNeoVis.evergladensis.1.1.chr14.g00121.t01#3         5880   125256260:   mNeoVis.3.1.1.chr14.g00114.t01#6    mNeoVis.vulgivaga.2.1.chr14.g00118.t01#1    mNeoVis.vulgivaga.1.1.chr14.g00117.t01#4    mNeoVis.evergladensis.1.1.chr14.g00122.t01#3    mNeoVis.2.1.1.chr14_hap1.g00120.t01#5   mNeoVis.evergladensis.2.1.chr14.g00116.t01#2       5881   125256261:  mNeoVis.vulgivaga.2.1.chr7.g00420.t01#1               5882   125256262:   mNeoVis.evergladensis.1.1.chr6.g00483.t01#3    mNeoVis.evergladensis.2.1.chr6.g00486.t01#2    mNeoVis.vulgivaga.2.1.chr6.g00484.t01#1    mNeoVis.2.1.1.chr4_hap1.g00479.t01#5    mNeoVis.3.1.1.chr6.g00482.t01#6   mNeoVis.vulgivaga.1.1.chr6.g00487.t01#4       5883   125256263:   mNeoVis.evergladensis.1.1.chr14.g00124.t01#3    mNeoVis.evergladensis.2.1.chr14.g00117.t01#2    mNeoVis.vulgivaga.1.1.chr14.g00120.t01#4    mNeoVis.vulgivaga.2.1.chr14.g00119.t01#1   mNeoVis.3.1.1.chr14.g00116.t01#6       5884   125256264:  mNeoVis.vulgivaga.2.1.chr14.g00120.t01#1               5885   125256265:   mNeoVis.vulgivaga.2.1.chr6.g00485.t01#1    mNeoVis.2.1.1.chr4_hap1.g00480.t01#5    mNeoVis.evergladensis.2.1.chr6.g00487.t01#2    mNeoVis.vulgivaga.1.1.chr6.g00488.t01#4    mNeoVis.evergladensis.1.1.chr6.g00484.t01#3   mNeoVis.3.1.1.chr6.g00483.t01#6       5886   125256266:   mNeoVis.3.1.1.chr14.g00118.t01#6    mNeoVis.vulgivaga.1.1.chr14.g00121.t01#4    mNeoVis.vulgivaga.2.1.chr14.g00121.t01#1    mNeoVis.evergladensis.2.1.chr14.g00118.t01#2    mNeoVis.2.1.1.chr14_hap1.g00122.t01#5   mNeoVis.evergladensis.1.1.chr14.g00126.t01#3       5887   125256267:   mNeoVis.evergladensis.1.1.chr14.g00127.t01#3    mNeoVis.2.1.1.chr14_hap1.g00123.t01#5    mNeoVis.evergladensis.2.1.chr14.g00119.t01#2    mNeoVis.3.1.1.chr14.g00119.t01#6    mNeoVis.vulgivaga.2.1.chr14.g00122.t01#1   mNeoVis.vulgivaga.1.1.chr14.g00122.t01#4       5888   125256268:   mNeoVis.vulgivaga.1.1.chr14.g00123.t01#4    mNeoVis.3.1.1.chr14.g00121.t01#6    mNeoVis.evergladensis.2.1.chr14.g00120.t01#2    mNeoVis.2.1.1.chr14_hap1.g00124.t01#5    mNeoVis.vulgivaga.2.1.chr14.g00123.t01#1   mNeoVis.evergladensis.1.1.chr14.g00128.t01#3       5889   125256269:   mNeoVis.evergladensis.1.1.chr6.g00485.t01#3    mNeoVis.3.1.1.chr6.g00485.t01#6    mNeoVis.2.1.1.chr4_hap1.g00481.t01#5    mNeoVis.vulgivaga.2.1.chr6.g00486.t01#1    mNeoVis.evergladensis.2.1.chr6.g00488.t01#2   mNeoVis.vulgivaga.1.1.chr6.g00489.t01#4       5890   125256270:   mNeoVis.2.1.1.chr4_hap1.g00483.t01#5    mNeoVis.vulgivaga.1.1.chr6.g00490.t01#4    mNeoVis.vulgivaga.2.1.chr6.g00487.t01#1    mNeoVis.evergladensis.2.1.chr6.g00490.t01#2   mNeoVis.evergladensis.1.1.chr6.g00487.t01#3       5891   125256271:   mNeoVis.2.1.1.chr4_hap1.g00482.t01#5    mNeoVis.evergladensis.1.1.chr6.g00486.t01#3    mNeoVis.3.1.1.chr6.g00484.t01#6    mNeoVis.vulgivaga.2.1.chr6.g00488.t01#1    mNeoVis.evergladensis.2.1.chr6.g00489.t01#2   mNeoVis.vulgivaga.1.1.chr6.g00491.t01#4       5892   125256272:   mNeoVis.vulgivaga.2.1.chr6.g00489.t01#1    mNeoVis.evergladensis.1.1.chr6.g00488.t01#3    mNeoVis.vulgivaga.1.1.chr6.g00492.t01#4    mNeoVis.evergladensis.2.1.chr6.g00491.t01#2   mNeoVis.2.1.1.chr4_hap1.g00484.t01#5       5893   125256273:   mNeoVis.evergladensis.2.1.chr6.g00492.t01#2    mNeoVis.3.1.1.chr6.g00487.t01#6    mNeoVis.evergladensis.1.1.chr6.g00489.t01#3    mNeoVis.2.1.1.chr4_hap1.g00485.t01#5    mNeoVis.vulgivaga.1.1.chr6.g00493.t01#4   mNeoVis.vulgivaga.2.1.chr6.g00490.t01#1       5894   125256274:   mNeoVis.3.1.1.chr14.g00122.t01#6    mNeoVis.vulgivaga.1.1.chr14.g00124.t01#4    mNeoVis.2.1.1.chr14_hap1.g00125.t01#5    mNeoVis.evergladensis.1.1.chr14.g00129.t01#3    mNeoVis.vulgivaga.2.1.chr14.g00124.t01#1   mNeoVis.evergladensis.2.1.chr14.g00121.t01#2       5895   125256275:   mNeoVis.3.1.1.chr14.g00123.t01#6    mNeoVis.2.1.1.chr14_hap1.g00126.t01#5   mNeoVis.vulgivaga.2.1.chr14.g00125.t01#1         5896   125256276:   mNeoVis.vulgivaga.2.1.chr6.g00491.t01#1    mNeoVis.evergladensis.2.1.chr6.g00493.t01#2    mNeoVis.3.1.1.chr6.g00488.t01#6    mNeoVis.evergladensis.1.1.chr6.g00490.t01#3    mNeoVis.vulgivaga.1.1.chr6.g00494.t01#4   mNeoVis.2.1.1.chr4_hap1.g00486.t01#5       5897   125256277:   mNeoVis.vulgivaga.2.1.chr14.g00126.t01#1    mNeoVis.vulgivaga.1.1.chr14.g00125.t01#4    mNeoVis.evergladensis.2.1.chr14.g00122.t01#2    mNeoVis.3.1.1.chr14.g00124.t01#6    mNeoVis.evergladensis.1.1.chr14.g00130.t01#3   mNeoVis.2.1.1.chr14_hap1.g00127.t01#5       5898   125256278:   mNeoVis.evergladensis.2.1.chr14.g00123.t01#2    mNeoVis.vulgivaga.1.1.chr14.g00126.t01#4    mNeoVis.evergladensis.2.1.chr14.g00124.t01#2    mNeoVis.evergladensis.1.1.chr14.g00131.t01#3    mNeoVis.3.1.1.chr14.g00125.t01#6    mNeoVis.2.1.1.chr14_hap1.g00128.t01#5   mNeoVis.vulgivaga.2.1.chr14.g00127.t01#1     5899   125256279:   mNeoVis.evergladensis.2.1.chr14.g00125.t01#2    mNeoVis.vulgivaga.2.1.chr14.g00128.t01#1   mNeoVis.3.1.1.chr14.g00126.t01#6           5900   125256280:   mNeoVis.evergladensis.2.1.chr14.g00126.t01#2    mNeoVis.3.1.1.chr14.g00127.t01#6    mNeoVis.evergladensis.1.1.chr14.g00132.t01#3    mNeoVis.2.1.1.chr14_hap1.g00129.t01#5    mNeoVis.vulgivaga.2.1.chr14.g00129.t01#1   mNeoVis.vulgivaga.1.1.chr14.g00127.t01#4       5901   125256281:   mNeoVis.evergladensis.1.1.chr6.g00491.t01#3    mNeoVis.2.1.1.chr4_hap1.g00487.t01#5    mNeoVis.vulgivaga.1.1.chr6.g00495.t01#4    mNeoVis.vulgivaga.2.1.chr6.g00492.t01#1    mNeoVis.3.1.1.chr6.g00489.t01#6   mNeoVis.evergladensis.2.1.chr6.g00494.t01#2       5902   125256282:   mNeoVis.2.1.1.chr14_hap1.g00130.t01#5    mNeoVis.vulgivaga.2.1.chr14.g00130.t01#1    mNeoVis.evergladensis.2.1.chr14.g00127.t01#2    mNeoVis.vulgivaga.1.1.chr14.g00128.t01#4    mNeoVis.3.1.1.chr14.g00128.t01#6   mNeoVis.evergladensis.1.1.chr14.g00133.t01#3       5903   125256283:   mNeoVis.vulgivaga.2.1.chr6.g00493.t01#1    mNeoVis.evergladensis.2.1.chr6.g00495.t01#2    mNeoVis.vulgivaga.1.1.chr6.g00496.t01#4    mNeoVis.evergladensis.1.1.chr6.g00492.t01#3    mNeoVis.2.1.1.chr4_hap1.g00488.t01#5   mNeoVis.3.1.1.chr6.g00490.t01#6       5904   125256284:   mNeoVis.2.1.1.chr4_hap1.g00489.t01#5    mNeoVis.vulgivaga.2.1.chr6.g00494.t01#1    mNeoVis.evergladensis.2.1.chr6.g00496.t01#2    mNeoVis.vulgivaga.1.1.chr6.g00497.t01#4    mNeoVis.evergladensis.1.1.chr6.g00493.t01#3   mNeoVis.3.1.1.chr6.g00491.t01#6       5905   125256285:   mNeoVis.vulgivaga.2.1.chr6.g00495.t01#1    mNeoVis.2.1.1.chr4_hap1.g00490.t01#5    mNeoVis.evergladensis.2.1.chr6.g00497.t01#2    mNeoVis.evergladensis.1.1.chr6.g00494.t01#3    mNeoVis.vulgivaga.1.1.chr6.g00498.t01#4   mNeoVis.3.1.1.chr6.g00492.t01#6       5906   125256286:   mNeoVis.vulgivaga.1.1.chr6.g00499.t01#4    mNeoVis.vulgivaga.2.1.chr6.g00496.t01#1   mNeoVis.evergladensis.1.1.chr6.g00495.t01#3         5907   125256287:  mNeoVis.vulgivaga.2.1.chr14.g00131.t01#1               5908   125256288:   mNeoVis.evergladensis.1.1.chr7.g00431.t01#3    mNeoVis.2.1.1.chr7_hap1.g00419.t01#5    mNeoVis.3.1.1.chr7.g00405.t01#6    mNeoVis.evergladensis.2.1.chr7.g00423.t01#2    mNeoVis.vulgivaga.1.1.chr7.g00427.t01#4   mNeoVis.vulgivaga.2.1.chr7.g00421.t01#1       5909   125256289:   mNeoVis.vulgivaga.1.1.chr6.g00500.t01#4    mNeoVis.vulgivaga.2.1.chr6.g00497.t01#1    mNeoVis.3.1.1.chr6.g00494.t01#6    mNeoVis.2.1.1.chr4_hap1.g00492.t01#5    mNeoVis.evergladensis.1.1.chr6.g00496.t01#3   mNeoVis.evergladensis.2.1.chr6.g00499.t01#2       5910   125256290:   mNeoVis.vulgivaga.2.1.chr6.g00498.t01#1    mNeoVis.vulgivaga.1.1.chr6.g00501.t01#4    mNeoVis.3.1.1.chr6.g00495.t01#6    mNeoVis.evergladensis.1.1.chr6.g00497.t01#3    mNeoVis.2.1.1.chr4_hap1.g00491.t01#5   mNeoVis.evergladensis.2.1.chr6.g00498.t01#2       5911   125256291:   mNeoVis.evergladensis.2.1.chr14.g00128.t01#2    mNeoVis.vulgivaga.1.1.chr14.g00130.t01#4    mNeoVis.evergladensis.1.1.chr14.g00134.t01#3    mNeoVis.vulgivaga.2.1.chr14.g00132.t01#1   mNeoVis.2.1.1.chr14_hap1.g00132.t01#5       5912   125256292:   mNeoVis.evergladensis.2.1.chr14.g00129.t01#2    mNeoVis.3.1.1.chr14.g00130.t01#6    mNeoVis.evergladensis.1.1.chr14.g00135.t01#3    mNeoVis.2.1.1.chr14_hap1.g00133.t01#5    mNeoVis.vulgivaga.2.1.chr14.g00133.t01#1   mNeoVis.vulgivaga.1.1.chr14.g00131.t01#4       5913   125256293:   mNeoVis.evergladensis.2.1.chr7.g00424.t01#2    mNeoVis.vulgivaga.1.1.chr7.g00428.t01#4    mNeoVis.3.1.1.chr7.g00406.t01#6    mNeoVis.2.1.1.chr7_hap1.g00420.t01#5    mNeoVis.vulgivaga.2.1.chr7.g00422.t01#1   mNeoVis.evergladensis.1.1.chr7.g00432.t01#3       5914   125256294:   mNeoVis.vulgivaga.1.1.chr6.g00502.t01#4    mNeoVis.vulgivaga.2.1.chr6.g00499.t01#1    mNeoVis.evergladensis.1.1.chr6.g00498.t01#3    mNeoVis.2.1.1.chr4_hap1.g00494.t01#5    mNeoVis.evergladensis.2.1.chr6.g00500.t01#2   mNeoVis.3.1.1.chr6.g00496.t01#6       5915   125256295:   mNeoVis.vulgivaga.1.1.chr6.g00503.t01#4    mNeoVis.vulgivaga.2.1.chr6.g00500.t01#1    mNeoVis.2.1.1.chr4_hap1.g00495.t01#5    mNeoVis.evergladensis.2.1.chr6.g00501.t01#2    mNeoVis.3.1.1.chr6.g00497.t01#6   mNeoVis.evergladensis.1.1.chr6.g00499.t01#3       5916   125256296:   mNeoVis.evergladensis.1.1.chr14.g00136.t01#3    mNeoVis.vulgivaga.2.1.chr14.g00134.t01#1    mNeoVis.2.1.1.chr14_hap1.g00134.t01#5    mNeoVis.3.1.1.chr14.g00131.t01#6    mNeoVis.evergladensis.2.1.chr14.g00130.t01#2   mNeoVis.vulgivaga.1.1.chr14.g00132.t01#4       5917   125256297:   mNeoVis.vulgivaga.1.1.chr6.g00504.t01#4    mNeoVis.3.1.1.chr6.g00498.t01#6    mNeoVis.evergladensis.1.1.chr6.g00500.t01#3    mNeoVis.evergladensis.2.1.chr6.g00502.t01#2    mNeoVis.2.1.1.chr4_hap1.g00496.t01#5   mNeoVis.vulgivaga.2.1.chr6.g00501.t01#1       5918   125256298:   mNeoVis.evergladensis.2.1.chr14.g00132.t01#2    mNeoVis.3.1.1.chr14.g00132.t01#6    mNeoVis.evergladensis.1.1.chr14.g00137.t01#3    mNeoVis.vulgivaga.2.1.chr14.g00135.t01#1    mNeoVis.2.1.1.chr14_hap1.g00136.t01#5   mNeoVis.vulgivaga.1.1.chr14.g00134.t01#4       5919   125256299:   mNeoVis.vulgivaga.1.1.chr14.g00135.t01#4    mNeoVis.3.1.1.chr14.g00133.t01#6    mNeoVis.evergladensis.1.1.chr14.g00138.t01#3    mNeoVis.2.1.1.chr14_hap1.g00138.t01#5    mNeoVis.vulgivaga.2.1.chr14.g00136.t01#1   mNeoVis.evergladensis.2.1.chr14.g00133.t01#2       5920   125256300:   mNeoVis.3.1.1.chr6.g00499.t01#6    mNeoVis.vulgivaga.1.1.chr6.g00505.t01#4    mNeoVis.2.1.1.chr4_hap1.g00497.t01#5    mNeoVis.vulgivaga.2.1.chr6.g00502.t01#1    mNeoVis.evergladensis.1.1.chr6.g00501.t01#3   mNeoVis.evergladensis.2.1.chr6.g00503.t01#2       5921   125256301:   mNeoVis.evergladensis.1.1.chr6.g00416.t01#3    mNeoVis.vulgivaga.1.1.chr6.g00423.t01#4    mNeoVis.3.1.1.chr6.g00417.t01#6    mNeoVis.evergladensis.2.1.chr6.g00421.t01#2    mNeoVis.2.1.1.chr4_hap1.g00418.t01#5   mNeoVis.vulgivaga.2.1.chr6.g00415.t01#1       5922   125256302:   mNeoVis.evergladensis.2.1.chr14.g00134.t01#2    mNeoVis.2.1.1.chr14_hap1.g00139.t01#5    mNeoVis.vulgivaga.1.1.chr14.g00136.t01#4    mNeoVis.3.1.1.chr14.g00134.t01#6    mNeoVis.vulgivaga.2.1.chr14.g00137.t01#1   mNeoVis.evergladensis.1.1.chr14.g00139.t01#3       5923   125256303:   mNeoVis.evergladensis.2.1.chr14.g00135.t01#2    mNeoVis.2.1.1.chr14_hap1.g00140.t01#5    mNeoVis.vulgivaga.1.1.chr14.g00137.t01#4    mNeoVis.3.1.1.chr14.g00135.t01#6    mNeoVis.evergladensis.1.1.chr14.g00140.t01#3   mNeoVis.vulgivaga.2.1.chr14.g00138.t01#1       5924   125256304:   mNeoVis.vulgivaga.1.1.chr14.g00138.t01#4    mNeoVis.evergladensis.2.1.chr14.g00136.t01#2    mNeoVis.evergladensis.1.1.chr14.g00141.t01#3    mNeoVis.vulgivaga.2.1.chr14.g00139.t01#1    mNeoVis.3.1.1.chr14.g00136.t01#6   mNeoVis.2.1.1.chr14_hap1.g00141.t01#5       5925   125256305:   mNeoVis.evergladensis.1.1.chr14.g00142.t01#3    mNeoVis.vulgivaga.1.1.chr14.g00139.t01#4    mNeoVis.evergladensis.2.1.chr14.g00137.t01#2    mNeoVis.2.1.1.chr14_hap1.g00142.t01#5   mNeoVis.vulgivaga.2.1.chr14.g00140.t01#1       5926   125256306:   mNeoVis.evergladensis.1.1.chr14.g00143.t01#3    mNeoVis.vulgivaga.2.1.chr14.g00141.t01#1    mNeoVis.evergladensis.2.1.chr14.g00139.t01#2    mNeoVis.vulgivaga.1.1.chr14.g00140.t01#4    mNeoVis.2.1.1.chr14_hap1.g00143.t01#5    mNeoVis.3.1.1.chr12.g00389.t01#6    mNeoVis.evergladensis.2.1.chr12.g00386.t01#2    mNeoVis.3.1.1.chr14.g00141.t01#6    mNeoVis.vulgivaga.2.1.chr12.g00394.t01#1      5927   125256307:   mNeoVis.evergladensis.2.1.chr14.g00141.t01#2    mNeoVis.vulgivaga.2.1.chr14.g00142.t01#1    mNeoVis.3.1.1.chr14.g00143.t01#6    mNeoVis.evergladensis.1.1.chr14.g00145.t01#3    mNeoVis.vulgivaga.1.1.chr14.g00141.t01#4   mNeoVis.2.1.1.chr14_hap1.g00145.t01#5       5928   125256308:   mNeoVis.evergladensis.1.1.chr7.g00043.t01#3   mNeoVis.vulgivaga.2.1.chr7.g00042.t01#1             5929   125256309:   mNeoVis.vulgivaga.1.1.chr6.g00425.t01#4    mNeoVis.2.1.1.chr4_hap1.g00419.t01#5    mNeoVis.3.1.1.chr6.g00418.t01#6    mNeoVis.evergladensis.2.1.chr6.g00422.t01#2    mNeoVis.evergladensis.1.1.chr6.g00419.t01#3   mNeoVis.vulgivaga.2.1.chr6.g00416.t01#1       5930   125256310:   mNeoVis.vulgivaga.2.1.chr14.g00143.t01#1    mNeoVis.vulgivaga.1.1.chr14.g00143.t01#4    mNeoVis.2.1.1.chr14_hap1.g00144.t01#5    mNeoVis.evergladensis.2.1.chr14.g00140.t01#2    mNeoVis.evergladensis.1.1.chr14.g00146.t01#3   mNeoVis.3.1.1.chr14.g00142.t01#6       5931   125256311:   mNeoVis.vulgivaga.2.1.chr14.g00057.t01#1    mNeoVis.2.1.1.chr14_hap1.g00058.t01#5    mNeoVis.vulgivaga.1.1.chr14.g00056.t01#4    mNeoVis.evergladensis.1.1.chr14.g00062.t01#3    mNeoVis.evergladensis.2.1.chr14.g00058.t01#2   mNeoVis.3.1.1.chr14.g00053.t01#6       5932   125256312:  mNeoVis.vulgivaga.2.1.chr6.g00417.t01#1               5933   125256313:   mNeoVis.evergladensis.2.1.chr15.g00124.t01#2    mNeoVis.vulgivaga.2.1.chr15.g00128.t01#1    mNeoVis.evergladensis.1.1.chr15.g00491.t01#3    mNeoVis.vulgivaga.1.1.chr15.g00124.t01#4    mNeoVis.3.1.1.chr15.g00116.t01#6   mNeoVis.2.1.1.chr15_hap1.g00138.t01#5       5934   125256314:   mNeoVis.vulgivaga.2.1.chr6.g00419.t01#1    mNeoVis.3.1.1.chr6.g00421.t01#6    mNeoVis.evergladensis.2.1.chr6.g00425.t01#2    mNeoVis.2.1.1.chr4_hap1.g00421.t01#5    mNeoVis.evergladensis.1.1.chr6.g00422.t01#3   mNeoVis.vulgivaga.1.1.chr6.g00427.t01#4       5935   125256315:   mNeoVis.vulgivaga.1.1.chr14.g00057.t01#4    mNeoVis.evergladensis.1.1.chr14.g00064.t01#3    mNeoVis.vulgivaga.2.1.chr14.g00058.t01#1    mNeoVis.evergladensis.2.1.chr14.g00059.t01#2    mNeoVis.3.1.1.chr14.g00054.t01#6   mNeoVis.2.1.1.chr14_hap1.g00059.t01#5       5936   125256316:   mNeoVis.vulgivaga.1.1.chr6.g00428.t01#4    mNeoVis.evergladensis.2.1.chr6.g00426.t01#2    mNeoVis.evergladensis.1.1.chr6.g00423.t01#3    mNeoVis.3.1.1.chr6.g00422.t01#6    mNeoVis.2.1.1.chr4_hap1.g00422.t01#5   mNeoVis.vulgivaga.2.1.chr6.g00420.t01#1       5937   125256317:   mNeoVis.evergladensis.2.1.chr14.g00060.t01#2    mNeoVis.vulgivaga.2.1.chr14.g00059.t01#1    mNeoVis.evergladensis.1.1.chr14.g00065.t01#3   mNeoVis.2.1.1.chr14_hap1.g00060.t01#5         5938   125256318:   mNeoVis.evergladensis.2.1.chr14.g00061.t01#2    mNeoVis.vulgivaga.1.1.chr14.g00059.t01#4    mNeoVis.evergladensis.1.1.chr14.g00066.t01#3    mNeoVis.2.1.1.chr14_hap1.g00061.t01#5   mNeoVis.vulgivaga.2.1.chr14.g00060.t01#1       5939   125256319:   mNeoVis.3.1.1.chr6.g00423.t01#6    mNeoVis.2.1.1.chr4_hap1.g00423.t01#5    mNeoVis.evergladensis.1.1.chr6.g00424.t01#3    mNeoVis.vulgivaga.2.1.chr6.g00421.t01#1    mNeoVis.vulgivaga.1.1.chr6.g00429.t01#4   mNeoVis.evergladensis.2.1.chr6.g00427.t01#2       5940   125256320:   mNeoVis.vulgivaga.2.1.chr14.g00061.t01#1    mNeoVis.evergladensis.1.1.chr14.g00067.t01#3    mNeoVis.evergladensis.2.1.chr14.g00062.t01#2   mNeoVis.vulgivaga.1.1.chr14.g00060.t01#4       5941   125256321:   mNeoVis.3.1.1.chr14.g00058.t01#6    mNeoVis.vulgivaga.1.1.chr14.g00061.t01#4    mNeoVis.vulgivaga.2.1.chr14.g00062.t01#1    mNeoVis.evergladensis.2.1.chr14.g00063.t01#2    mNeoVis.evergladensis.1.1.chr14.g00068.t01#3   mNeoVis.2.1.1.chr14_hap1.g00063.t01#5       5942   125256322:   mNeoVis.2.1.1.chr4_hap1.g00424.t01#5    mNeoVis.vulgivaga.2.1.chr6.g00422.t01#1    mNeoVis.evergladensis.2.1.chr6.g00428.t01#2    mNeoVis.3.1.1.chr6.g00424.t01#6    mNeoVis.evergladensis.1.1.chr6.g00425.t01#3   mNeoVis.vulgivaga.1.1.chr6.g00430.t01#4       5943   125256323:   mNeoVis.2.1.1.chr14_hap1.g00065.t01#5    mNeoVis.evergladensis.1.1.chr14.g00069.t01#3    mNeoVis.vulgivaga.1.1.chr14.g00063.t01#4    mNeoVis.evergladensis.2.1.chr14.g00064.t01#2    mNeoVis.vulgivaga.2.1.chr14.g00063.t01#1   mNeoVis.3.1.1.chr14.g00059.t01#6       5944   125256324:   mNeoVis.2.1.1.chr5_hap1.g00798.t01#5    mNeoVis.3.1.1.chr2.g00711.t01#6    mNeoVis.evergladensis.2.1.chr2.g00799.t01#2    mNeoVis.vulgivaga.1.1.chr2.g00797.t01#4    mNeoVis.vulgivaga.2.1.chr2.g00803.t01#1   mNeoVis.evergladensis.1.1.chr2.g00796.t01#3       5945   125256325:   mNeoVis.evergladensis.1.1.chr6.g00426.t01#3    mNeoVis.3.1.1.chr6.g00425.t01#6    mNeoVis.vulgivaga.2.1.chr6.g00423.t01#1   mNeoVis.2.1.1.chr4_hap1.g00425.t01#5         5946   125256326:   mNeoVis.evergladensis.1.1.chr6.g00427.t01#3    mNeoVis.3.1.1.chr6.g00426.t01#6    mNeoVis.vulgivaga.2.1.chr6.g00424.t01#1    mNeoVis.2.1.1.chr4_hap1.g00426.t01#5   mNeoVis.evergladensis.2.1.chr6.g00430.t01#2       5947   125256327:   mNeoVis.vulgivaga.2.1.chr6.g00425.t01#1    mNeoVis.3.1.1.chr6.g00427.t01#6    mNeoVis.evergladensis.1.1.chr6.g00428.t01#3    mNeoVis.vulgivaga.1.1.chr6.g00433.t01#4    mNeoVis.2.1.1.chr4_hap1.g00427.t01#5   mNeoVis.evergladensis.2.1.chr6.g00431.t01#2       5948   125256328:   mNeoVis.evergladensis.1.1.chr6.g00429.t01#3    mNeoVis.vulgivaga.2.1.chr6.g00426.t01#1    mNeoVis.vulgivaga.1.1.chr6.g00434.t01#4    mNeoVis.evergladensis.2.1.chr6.g00432.t01#2   mNeoVis.2.1.1.chr4_hap1.g00428.t01#5       5949   125256329:   mNeoVis.2.1.1.chr4_hap1.g00429.t01#5    mNeoVis.vulgivaga.2.1.chr6.g00427.t01#1    mNeoVis.3.1.1.chr6.g00428.t01#6    mNeoVis.evergladensis.2.1.chr6.g00433.t01#2    mNeoVis.vulgivaga.1.1.chr6.g00435.t01#4   mNeoVis.evergladensis.1.1.chr6.g00430.t01#3       5950   125256330:   mNeoVis.evergladensis.2.1.chr14.g00065.t01#2    mNeoVis.evergladensis.1.1.chr14.g00070.t01#3    mNeoVis.2.1.1.chr14_hap1.g00066.t01#5    mNeoVis.vulgivaga.2.1.chr14.g00064.t01#1    mNeoVis.vulgivaga.1.1.chr14.g00064.t01#4   mNeoVis.3.1.1.chr14.g00060.t01#6       5951   125256331:   mNeoVis.vulgivaga.1.1.chr14.g00065.t01#4    mNeoVis.3.1.1.chr14.g00061.t01#6    mNeoVis.evergladensis.2.1.chr14.g00066.t01#2    mNeoVis.evergladensis.1.1.chr14.g00071.t01#3   mNeoVis.vulgivaga.2.1.chr14.g00065.t01#1       5952   125256332:   mNeoVis.vulgivaga.2.1.chr14.g00067.t01#1    mNeoVis.evergladensis.2.1.chr14.g00068.t01#2    mNeoVis.2.1.1.chr14_hap1.g00067.t01#5   mNeoVis.vulgivaga.1.1.chr14.g00067.t01#4       5953   125256333:  mNeoVis.vulgivaga.2.1.chr7.g00043.t01#1               5954   125256334:  mNeoVis.vulgivaga.2.1.chr6.g00428.t01#1               5955   125256335:   mNeoVis.vulgivaga.2.1.chr14.g00066.t01#1    mNeoVis.vulgivaga.1.1.chr14.g00066.t01#4    mNeoVis.evergladensis.2.1.chr14.g00067.t01#2    mNeoVis.3.1.1.chr14.g00063.t01#6    mNeoVis.evergladensis.1.1.chr14.g00072.t01#3   mNeoVis.2.1.1.chr14_hap1.g00068.t01#5       5956   125256336:   mNeoVis.vulgivaga.1.1.chr6.g00437.t01#4    mNeoVis.3.1.1.chr6.g00430.t01#6    mNeoVis.evergladensis.2.1.chr6.g00435.t01#2    mNeoVis.2.1.1.chr4_hap1.g00431.t01#5    mNeoVis.vulgivaga.2.1.chr6.g00430.t01#1   mNeoVis.evergladensis.1.1.chr6.g00433.t01#3       5957   125256337:   mNeoVis.vulgivaga.1.1.chr14.g00068.t01#4    mNeoVis.evergladensis.1.1.chr14.g00073.t01#3    mNeoVis.vulgivaga.2.1.chr14.g00068.t01#1    mNeoVis.evergladensis.2.1.chr14.g00069.t01#2   mNeoVis.3.1.1.chr14.g00064.t01#6       5958   125256338:   mNeoVis.evergladensis.1.1.chr14.g00074.t01#3    mNeoVis.3.1.1.chr14.g00065.t01#6    mNeoVis.vulgivaga.1.1.chr14.g00069.t01#4    mNeoVis.2.1.1.chr14_hap1.g00069.t01#5    mNeoVis.vulgivaga.2.1.chr14.g00069.t01#1   mNeoVis.evergladensis.2.1.chr14.g00070.t01#2       5959   125256339:   mNeoVis.2.1.1.chr4_hap1.g00432.t01#5    mNeoVis.evergladensis.1.1.chr6.g00434.t01#3    mNeoVis.3.1.1.chr6.g00431.t01#6    mNeoVis.vulgivaga.2.1.chr6.g00431.t01#1    mNeoVis.vulgivaga.1.1.chr6.g00438.t01#4   mNeoVis.evergladensis.2.1.chr6.g00436.t01#2       5960   125256340:   mNeoVis.3.1.1.chr6.g00432.t01#6    mNeoVis.evergladensis.2.1.chr6.g00437.t01#2    mNeoVis.vulgivaga.2.1.chr6.g00432.t01#1    mNeoVis.vulgivaga.1.1.chr6.g00439.t01#4    mNeoVis.evergladensis.1.1.chr6.g00435.t01#3   mNeoVis.2.1.1.chr4_hap1.g00433.t01#5       5961   125256341:   mNeoVis.evergladensis.1.1.chr7.g00044.t01#3   mNeoVis.vulgivaga.2.1.chr7.g00044.t01#1             5962   125256342:   mNeoVis.vulgivaga.2.1.chr6.g00433.t01#1    mNeoVis.evergladensis.1.1.chr6.g00436.t01#3    mNeoVis.2.1.1.chr4_hap1.g00434.t01#5    mNeoVis.evergladensis.2.1.chr6.g00438.t01#2    mNeoVis.3.1.1.chr6.g00433.t01#6   mNeoVis.vulgivaga.1.1.chr6.g00440.t01#4       5963   125256343:   mNeoVis.vulgivaga.1.1.chr14.g00070.t01#4    mNeoVis.2.1.1.chr14_hap1.g00070.t01#5    mNeoVis.evergladensis.1.1.chr14.g00075.t01#3    mNeoVis.evergladensis.2.1.chr14.g00071.t01#2    mNeoVis.3.1.1.chr14.g00066.t01#6   mNeoVis.vulgivaga.2.1.chr14.g00070.t01#1       5964   125256344:   mNeoVis.vulgivaga.1.1.chr6.g00441.t01#4    mNeoVis.3.1.1.chr6.g00434.t01#6    mNeoVis.2.1.1.chr4_hap1.g00435.t01#5    mNeoVis.evergladensis.1.1.chr6.g00437.t01#3    mNeoVis.vulgivaga.2.1.chr6.g00434.t01#1   mNeoVis.evergladensis.2.1.chr6.g00439.t01#2       5965   125256345:   mNeoVis.2.1.1.chr14_hap1.g00071.t01#5    mNeoVis.vulgivaga.2.1.chr14.g00071.t01#1    mNeoVis.evergladensis.2.1.chr14.g00072.t01#2    mNeoVis.evergladensis.1.1.chr14.g00076.t01#3    mNeoVis.3.1.1.chr14.g00067.t01#6   mNeoVis.vulgivaga.1.1.chr14.g00071.t01#4       5966   125256346:   mNeoVis.evergladensis.1.1.chr14.g00077.t01#3    mNeoVis.vulgivaga.1.1.chr14.g00072.t01#4    mNeoVis.2.1.1.chr14_hap1.g00072.t01#5    mNeoVis.vulgivaga.2.1.chr14.g00072.t01#1    mNeoVis.3.1.1.chr14.g00068.t01#6   mNeoVis.evergladensis.2.1.chr14.g00073.t01#2       5967   125256347:   mNeoVis.2.1.1.chr4_hap1.g00436.t01#5    mNeoVis.evergladensis.2.1.chr6.g00440.t01#2    mNeoVis.vulgivaga.1.1.chr6.g00442.t01#4    mNeoVis.3.1.1.chr6.g00435.t01#6    mNeoVis.vulgivaga.2.1.chr6.g00435.t01#1   mNeoVis.evergladensis.1.1.chr6.g00438.t01#3       5968   125256348:   mNeoVis.evergladensis.2.1.chr6.g00441.t01#2    mNeoVis.evergladensis.1.1.chr6.g00439.t01#3    mNeoVis.3.1.1.chr6.g00436.t01#6    mNeoVis.vulgivaga.1.1.chr6.g00443.t01#4    mNeoVis.2.1.1.chr4_hap1.g00437.t01#5   mNeoVis.vulgivaga.2.1.chr6.g00436.t01#1       5969   125256349:   mNeoVis.vulgivaga.2.1.chr14.g00073.t01#1    mNeoVis.evergladensis.2.1.chr14.g00074.t01#2    mNeoVis.evergladensis.1.1.chr14.g00078.t01#3    mNeoVis.3.1.1.chr14.g00069.t01#6   mNeoVis.2.1.1.chr14_hap1.g00073.t01#5       5970   125256350:   mNeoVis.evergladensis.1.1.chr14.g00079.t01#3    mNeoVis.2.1.1.chr14_hap1.g00074.t01#5    mNeoVis.evergladensis.2.1.chr14.g00075.t01#2    mNeoVis.3.1.1.chr14.g00070.t01#6    mNeoVis.vulgivaga.2.1.chr14.g00074.t01#1   mNeoVis.vulgivaga.1.1.chr14.g00076.t01#4       5971   125256351:   mNeoVis.vulgivaga.2.1.chr7.g00045.t01#1   mNeoVis.evergladensis.1.1.chr7.g00045.t01#3           5972   125256352:   mNeoVis.vulgivaga.1.1.chr6.g00444.t01#4    mNeoVis.2.1.1.chr4_hap1.g00438.t01#5    mNeoVis.evergladensis.2.1.chr6.g00442.t01#2    mNeoVis.3.1.1.chr6.g00437.t01#6    mNeoVis.evergladensis.1.1.chr6.g00440.t01#3   mNeoVis.vulgivaga.2.1.chr6.g00437.t01#1       5973   125256353:   mNeoVis.vulgivaga.1.1.chr14.g00077.t01#4    mNeoVis.vulgivaga.2.1.chr14.g00075.t01#1    mNeoVis.evergladensis.1.1.chr14.g00080.t01#3    mNeoVis.2.1.1.chr14_hap1.g00075.t01#5    mNeoVis.3.1.1.chr14.g00071.t01#6   mNeoVis.evergladensis.2.1.chr14.g00076.t01#2       5974   125256354:   mNeoVis.vulgivaga.2.1.chr6.g00438.t01#1    mNeoVis.evergladensis.2.1.chr6.g00444.t01#2    mNeoVis.2.1.1.chr4_hap1.g00439.t01#5    mNeoVis.vulgivaga.1.1.chr6.g00445.t01#4    mNeoVis.evergladensis.1.1.chr6.g00442.t01#3   mNeoVis.3.1.1.chr6.g00439.t01#6       5975   125256355:   mNeoVis.3.1.1.chr6.g00440.t01#6    mNeoVis.evergladensis.1.1.chr6.g00443.t01#3    mNeoVis.vulgivaga.2.1.chr6.g00439.t01#1    mNeoVis.evergladensis.2.1.chr6.g00445.t01#2    mNeoVis.vulgivaga.1.1.chr6.g00446.t01#4   mNeoVis.2.1.1.chr4_hap1.g00440.t01#5       5976   125256356:   mNeoVis.vulgivaga.2.1.chr6.g00440.t01#1    mNeoVis.2.1.1.chr4_hap1.g00441.t01#5    mNeoVis.evergladensis.1.1.chr6.g00444.t01#3    mNeoVis.evergladensis.2.1.chr6.g00446.t01#2    mNeoVis.3.1.1.chr6.g00441.t01#6   mNeoVis.vulgivaga.1.1.chr6.g00447.t01#4       5977   125256357:   mNeoVis.vulgivaga.1.1.chr6.g00448.t01#4    mNeoVis.2.1.1.chr4_hap1.g00442.t01#5    mNeoVis.evergladensis.2.1.chr6.g00447.t01#2    mNeoVis.evergladensis.1.1.chr6.g00445.t01#3    mNeoVis.vulgivaga.2.1.chr6.g00441.t01#1   mNeoVis.3.1.1.chr6.g00442.t01#6       5978   125256358:   mNeoVis.evergladensis.2.1.chr5.g00623.t01#2    mNeoVis.vulgivaga.1.1.chr2.g00873.t01#4    mNeoVis.evergladensis.2.1.chr2.g00871.t01#2    mNeoVis.vulgivaga.2.1.chr6.g00442.t01#1    mNeoVis.evergladensis.1.1.chr2.g01594.t01#3    mNeoVis.3.1.1.chr2.g00783.t01#6    mNeoVis.3.1.1.chr2.g01512.t01#6    mNeoVis.2.1.1.chr4_hap1.g00443.t01#5    mNeoVis.vulgivaga.1.1.chr2.g01597.t01#4      5979   125256359:   mNeoVis.3.1.1.chr6.g00443.t01#6    mNeoVis.vulgivaga.2.1.chr6.g00443.t01#1    mNeoVis.vulgivaga.1.1.chr6.g00450.t01#4    mNeoVis.2.1.1.chr4_hap1.g00444.t01#5    mNeoVis.evergladensis.1.1.chr6.g00448.t01#3   mNeoVis.evergladensis.2.1.chr6.g00450.t01#2       5980   125256360:   mNeoVis.vulgivaga.2.1.chr14.g00077.t01#1    mNeoVis.2.1.1.chr14_hap1.g00077.t01#5    mNeoVis.vulgivaga.1.1.chr14.g00079.t01#4    mNeoVis.3.1.1.chr14.g00073.t01#6    mNeoVis.evergladensis.1.1.chr14.g00082.t01#3   mNeoVis.evergladensis.2.1.chr14.g00078.t01#2       5981   125256361:   mNeoVis.evergladensis.2.1.chr14.g00079.t01#2    mNeoVis.3.1.1.chr14.g00074.t01#6    mNeoVis.vulgivaga.1.1.chr14.g00080.t01#4    mNeoVis.evergladensis.1.1.chr14.g00083.t01#3    mNeoVis.vulgivaga.2.1.chr14.g00078.t01#1   mNeoVis.2.1.1.chr14_hap1.g00078.t01#5       5982   125256362:   mNeoVis.evergladensis.2.1.chr14.g00080.t01#2    mNeoVis.vulgivaga.2.1.chr14.g00079.t01#1    mNeoVis.evergladensis.1.1.chr14.g00084.t01#3    mNeoVis.2.1.1.chr14_hap1.g00079.t01#5    mNeoVis.3.1.1.chr14.g00075.t01#6   mNeoVis.vulgivaga.1.1.chr14.g00081.t01#4       5983   125256363:   mNeoVis.evergladensis.1.1.chr7.g00047.t01#3    mNeoVis.vulgivaga.1.1.chr7.g00046.t01#4    mNeoVis.2.1.1.chr7_hap1.g00043.t01#5    mNeoVis.vulgivaga.2.1.chr7.g00046.t01#1   mNeoVis.evergladensis.2.1.chr7.g00045.t01#2       5984   125256364:   mNeoVis.vulgivaga.1.1.chr14.g00082.t01#4    mNeoVis.vulgivaga.2.1.chr14.g00080.t01#1    mNeoVis.evergladensis.2.1.chr14.g00081.t01#2    mNeoVis.3.1.1.chr14.g00076.t01#6    mNeoVis.evergladensis.1.1.chr14.g00085.t01#3   mNeoVis.2.1.1.chr14_hap1.g00080.t01#5       5985   125256365:   mNeoVis.evergladensis.1.1.chr9.g00026.t01#3    mNeoVis.vulgivaga.1.1.chr9.g00023.t01#4    mNeoVis.2.1.1.chr10_hap1.g00030.t01#5    mNeoVis.evergladensis.2.1.chr8.g00028.t01#2   mNeoVis.vulgivaga.2.1.chr9.g00029.t01#1       5986   125256366:   mNeoVis.3.1.1.chr14.g00077.t01#6    mNeoVis.vulgivaga.1.1.chr14.g00083.t01#4    mNeoVis.evergladensis.2.1.chr14.g00082.t01#2    mNeoVis.2.1.1.chr14_hap1.g00081.t01#5    mNeoVis.evergladensis.1.1.chr14.g00086.t01#3   mNeoVis.vulgivaga.2.1.chr14.g00081.t01#1       5987   125256367:   mNeoVis.2.1.1.chr4_hap1.g00445.t01#5    mNeoVis.vulgivaga.2.1.chr6.g00444.t01#1    mNeoVis.3.1.1.chr6.g00444.t01#6    mNeoVis.vulgivaga.1.1.chr6.g00451.t01#4    mNeoVis.evergladensis.2.1.chr6.g00451.t01#2   mNeoVis.evergladensis.1.1.chr6.g00449.t01#3       5988   125256368:   mNeoVis.evergladensis.1.1.chr6.g00450.t01#3    mNeoVis.vulgivaga.2.1.chr6.g00445.t01#1    mNeoVis.3.1.1.chr6.g00445.t01#6    mNeoVis.evergladensis.2.1.chr6.g00453.t01#2    mNeoVis.2.1.1.chr4_hap1.g00446.t01#5   mNeoVis.vulgivaga.1.1.chr6.g00452.t01#4       5989   125256369:   mNeoVis.2.1.1.chr4_hap1.g00447.t01#5    mNeoVis.evergladensis.2.1.chr6.g00454.t01#2    mNeoVis.vulgivaga.2.1.chr6.g00446.t01#1    mNeoVis.vulgivaga.1.1.chr6.g00453.t01#4    mNeoVis.evergladensis.1.1.chr6.g00451.t01#3   mNeoVis.3.1.1.chr6.g00446.t01#6       5990   125256370:  mNeoVis.vulgivaga.2.1.chr14.g00082.t01#1               5991   125256371:   mNeoVis.2.1.1.chr14_hap1.g00083.t01#5    mNeoVis.vulgivaga.1.1.chr14.g00084.t01#4    mNeoVis.evergladensis.1.1.chr14.g00087.t01#3    mNeoVis.evergladensis.2.1.chr14.g00083.t01#2   mNeoVis.vulgivaga.2.1.chr14.g00083.t01#1       5992   125256372:   mNeoVis.2.1.1.chr14_hap1.g00085.t01#5    mNeoVis.vulgivaga.2.1.chr14.g00084.t01#1    mNeoVis.evergladensis.1.1.chr14.g00089.t01#3   mNeoVis.evergladensis.2.1.chr14.g00084.t01#2       5993   125256373:   mNeoVis.3.1.1.chr14.g00080.t01#6    mNeoVis.evergladensis.2.1.chr14.g00085.t01#2    mNeoVis.vulgivaga.2.1.chr14.g00085.t01#1    mNeoVis.2.1.1.chr14_hap1.g00086.t01#5   mNeoVis.evergladensis.1.1.chr14.g00090.t01#3       5994   125256374:   mNeoVis.3.1.1.chr6.g00447.t01#6    mNeoVis.evergladensis.2.1.chr6.g00455.t01#2    mNeoVis.2.1.1.chr4_hap1.g00448.t01#5    mNeoVis.vulgivaga.1.1.chr6.g00454.t01#4    mNeoVis.evergladensis.1.1.chr6.g00452.t01#3   mNeoVis.vulgivaga.2.1.chr6.g00447.t01#1       5995   125256375:   mNeoVis.vulgivaga.2.1.chr6.g00448.t01#1    mNeoVis.evergladensis.2.1.chr6.g00456.t01#2    mNeoVis.2.1.1.chr4_hap1.g00449.t01#5    mNeoVis.3.1.1.chr6.g00448.t01#6    mNeoVis.vulgivaga.1.1.chr6.g00455.t01#4   mNeoVis.evergladensis.1.1.chr6.g00453.t01#3       5996   125256376:   mNeoVis.2.1.1.chr14_hap1.g00087.t01#5    mNeoVis.vulgivaga.2.1.chr14.g00086.t01#1    mNeoVis.evergladensis.1.1.chr14.g00091.t01#3    mNeoVis.3.1.1.chr14.g00081.t01#6    mNeoVis.vulgivaga.1.1.chr14.g00086.t01#4   mNeoVis.evergladensis.2.1.chr14.g00086.t01#2       5997   125256377:  mNeoVis.vulgivaga.2.1.chr14.g00087.t01#1               5998   125256378:   mNeoVis.evergladensis.2.1.chr6.g00457.t01#2    mNeoVis.vulgivaga.2.1.chr6.g00449.t01#1    mNeoVis.3.1.1.chr6.g00449.t01#6    mNeoVis.2.1.1.chr4_hap1.g00450.t01#5    mNeoVis.vulgivaga.1.1.chr6.g00456.t01#4   mNeoVis.evergladensis.1.1.chr6.g00454.t01#3       5999   125256379:   mNeoVis.vulgivaga.2.1.chr15.g00130.t01#1    mNeoVis.3.1.1.chr15.g00118.t01#6    mNeoVis.evergladensis.2.1.chr15.g00127.t01#2    mNeoVis.2.1.1.chr15_hap1.g00141.t01#5    mNeoVis.vulgivaga.1.1.chr15.g00127.t01#4   mNeoVis.evergladensis.1.1.chr15.g00489.t01#3       6000   125256380:   mNeoVis.2.1.1.chr14_hap1.g00089.t01#5    mNeoVis.evergladensis.1.1.chr14.g00093.t01#3    mNeoVis.vulgivaga.2.1.chr14.g00088.t01#1    mNeoVis.vulgivaga.1.1.chr14.g00088.t01#4   mNeoVis.evergladensis.2.1.chr14.g00088.t01#2       6001   125256381:   mNeoVis.vulgivaga.1.1.chr6.g00457.t01#4    mNeoVis.2.1.1.chr4_hap1.g00451.t01#5    mNeoVis.evergladensis.1.1.chr6.g00455.t01#3    mNeoVis.evergladensis.2.1.chr6.g00458.t01#2    mNeoVis.vulgivaga.2.1.chr6.g00450.t01#1   mNeoVis.3.1.1.chr6.g00450.t01#6       6002   125256382:   mNeoVis.2.1.1.chr4_hap1.g00452.t01#5    mNeoVis.vulgivaga.1.1.chr6.g00458.t01#4    mNeoVis.evergladensis.1.1.chr6.g00456.t01#3    mNeoVis.evergladensis.2.1.chr6.g00459.t01#2    mNeoVis.vulgivaga.2.1.chr6.g00451.t01#1   mNeoVis.3.1.1.chr6.g00451.t01#6       6003   125256383:   mNeoVis.2.1.1.chr14_hap1.g00090.t01#5    mNeoVis.evergladensis.1.1.chr14.g00094.t01#3    mNeoVis.vulgivaga.2.1.chr14.g00089.t01#1    mNeoVis.vulgivaga.1.1.chr14.g00089.t01#4    mNeoVis.3.1.1.chr14.g00084.t01#6   mNeoVis.evergladensis.2.1.chr14.g00089.t01#2       6004   125256384:   mNeoVis.3.1.1.chr14.g00085.t01#6    mNeoVis.evergladensis.1.1.chr14.g00095.t01#3    mNeoVis.vulgivaga.1.1.chr14.g00090.t01#4    mNeoVis.evergladensis.2.1.chr14.g00090.t01#2    mNeoVis.2.1.1.chr14_hap1.g00091.t01#5   mNeoVis.vulgivaga.2.1.chr14.g00090.t01#1       6005   125256385:   mNeoVis.3.1.1.chr14.g00086.t01#6    mNeoVis.vulgivaga.1.1.chr14.g00091.t01#4    mNeoVis.evergladensis.1.1.chr14.g00096.t01#3    mNeoVis.evergladensis.2.1.chr14.g00091.t01#2    mNeoVis.vulgivaga.2.1.chr14.g00091.t01#1   mNeoVis.2.1.1.chr14_hap1.g00092.t01#5       6006   125256386:   mNeoVis.evergladensis.1.1.chr6.g00457.t01#3    mNeoVis.evergladensis.2.1.chr6.g00460.t01#2    mNeoVis.vulgivaga.2.1.chr6.g00452.t01#1    mNeoVis.3.1.1.chr6.g00452.t01#6    mNeoVis.2.1.1.chr4_hap1.g00453.t01#5   mNeoVis.vulgivaga.1.1.chr6.g00459.t01#4       6007   125256387:   mNeoVis.vulgivaga.2.1.chr14.g00092.t01#1    mNeoVis.2.1.1.chr14_hap1.g00093.t01#5    mNeoVis.3.1.1.chr14.g00087.t01#6    mNeoVis.evergladensis.2.1.chr14.g00092.t01#2    mNeoVis.vulgivaga.1.1.chr14.g00092.t01#4   mNeoVis.evergladensis.1.1.chr14.g00097.t01#3       6008   125256388:   mNeoVis.3.1.1.chr6.g00453.t01#6   mNeoVis.vulgivaga.2.1.chr6.g00453.t01#1             6009   125256389:   mNeoVis.3.1.1.chr6.g00454.t01#6    mNeoVis.vulgivaga.1.1.chr6.g00460.t01#4    mNeoVis.vulgivaga.2.1.chr6.g00454.t01#1    mNeoVis.2.1.1.chr4_hap1.g00454.t01#5    mNeoVis.evergladensis.1.1.chr6.g00458.t01#3   mNeoVis.evergladensis.2.1.chr6.g00461.t01#2       6010   125256390:   mNeoVis.evergladensis.2.1.chr12.g00310.t01#2    mNeoVis.2.1.1.chr12_hap1.g00351.t01#5    mNeoVis.vulgivaga.1.1.chr12.g00320.t01#4    mNeoVis.3.1.1.chr12.g00316.t01#6    mNeoVis.evergladensis.2.1.chr12.g00311.t01#2   mNeoVis.evergladensis.1.1.chr12.g00317.t01#3       6011   125256391:   mNeoVis.vulgivaga.2.1.chr14.g00093.t01#1    mNeoVis.2.1.1.chr14_hap1.g00094.t01#5    mNeoVis.evergladensis.1.1.chr14.g00098.t01#3    mNeoVis.vulgivaga.1.1.chr14.g00093.t01#4    mNeoVis.evergladensis.2.1.chr14.g00093.t01#2   mNeoVis.3.1.1.chr14.g00088.t01#6       6012   125256392:   mNeoVis.vulgivaga.2.1.chr7.g00047.t01#1    mNeoVis.evergladensis.1.1.chr7.g00049.t01#3    mNeoVis.2.1.1.chr7_hap1.g00044.t01#5    mNeoVis.vulgivaga.1.1.chr7.g00047.t01#4   mNeoVis.evergladensis.2.1.chr7.g00046.t01#2       6013   125256393:   mNeoVis.evergladensis.1.1.chr6.g00459.t01#3    mNeoVis.3.1.1.chr6.g00455.t01#6    mNeoVis.vulgivaga.2.1.chr6.g00455.t01#1    mNeoVis.2.1.1.chr4_hap1.g00455.t01#5    mNeoVis.evergladensis.2.1.chr6.g00462.t01#2   mNeoVis.vulgivaga.1.1.chr6.g00461.t01#4       6014   125256394:   mNeoVis.vulgivaga.1.1.chr6.g00463.t01#4    mNeoVis.2.1.1.chr4_hap1.g00456.t01#5    mNeoVis.3.1.1.chr6.g00456.t01#6    mNeoVis.vulgivaga.2.1.chr6.g00456.t01#1    mNeoVis.evergladensis.2.1.chr6.g00463.t01#2   mNeoVis.evergladensis.1.1.chr6.g00461.t01#3       6015   125256395:   mNeoVis.evergladensis.1.1.chr6.g00355.t01#3    mNeoVis.vulgivaga.1.1.chr6.g00362.t01#4    mNeoVis.evergladensis.2.1.chr6.g00361.t01#2    mNeoVis.2.1.1.chr4_hap1.g00357.t01#5   mNeoVis.vulgivaga.2.1.chr6.g00354.t01#1       6016   125256396:   mNeoVis.2.1.1.chr14_hap1.g00095.t01#5    mNeoVis.evergladensis.2.1.chr14.g00094.t01#2    mNeoVis.3.1.1.chr14.g00089.t01#6    mNeoVis.evergladensis.1.1.chr14.g00099.t01#3    mNeoVis.vulgivaga.1.1.chr14.g00094.t01#4   mNeoVis.vulgivaga.2.1.chr14.g00094.t01#1       6017   125256397:  mNeoVis.vulgivaga.2.1.chr6.g00355.t01#1               6018   125256398:   mNeoVis.3.1.1.chr6.g00360.t01#6    mNeoVis.2.1.1.chr4_hap1.g00358.t01#5    mNeoVis.vulgivaga.2.1.chr6.g00356.t01#1    mNeoVis.evergladensis.1.1.chr6.g00356.t01#3    mNeoVis.evergladensis.2.1.chr6.g00362.t01#2   mNeoVis.vulgivaga.1.1.chr6.g00363.t01#4       6019   125256399:   mNeoVis.vulgivaga.2.1.chr6.g00357.t01#1    mNeoVis.vulgivaga.1.1.chr6.g00364.t01#4    mNeoVis.evergladensis.2.1.chr6.g00363.t01#2    mNeoVis.evergladensis.1.1.chr6.g00357.t01#3   mNeoVis.2.1.1.chr4_hap1.g00359.t01#5       6020   125256400:   mNeoVis.evergladensis.2.1.chr6.g00364.t01#2    mNeoVis.evergladensis.1.1.chr6.g00358.t01#3    mNeoVis.vulgivaga.1.1.chr6.g00365.t01#4    mNeoVis.2.1.1.chr4_hap1.g00360.t01#5   mNeoVis.vulgivaga.2.1.chr6.g00358.t01#1       6021   125256401:   mNeoVis.vulgivaga.1.1.chr6.g00366.t01#4    mNeoVis.vulgivaga.2.1.chr6.g00359.t01#1    mNeoVis.3.1.1.chr6.g00362.t01#6    mNeoVis.evergladensis.2.1.chr6.g00365.t01#2    mNeoVis.evergladensis.1.1.chr6.g00359.t01#3   mNeoVis.2.1.1.chr4_hap1.g00361.t01#5       6022   125256402:   mNeoVis.evergladensis.1.1.chr6.g00360.t01#3    mNeoVis.vulgivaga.2.1.chr6.g00360.t01#1    mNeoVis.evergladensis.2.1.chr6.g00366.t01#2    mNeoVis.vulgivaga.1.1.chr6.g00367.t01#4    mNeoVis.3.1.1.chr6.g00363.t01#6   mNeoVis.2.1.1.chr4_hap1.g00362.t01#5       6023   125256403:   mNeoVis.evergladensis.2.1.chr14.g00095.t01#2    mNeoVis.vulgivaga.2.1.chr14.g00095.t01#1    mNeoVis.evergladensis.1.1.chr14.g00100.t01#3    mNeoVis.vulgivaga.1.1.chr14.g00095.t01#4   mNeoVis.2.1.1.chr14_hap1.g00096.t01#5       6024   125256404:   mNeoVis.3.1.1.chr6.g00364.t01#6    mNeoVis.evergladensis.2.1.chr6.g00367.t01#2    mNeoVis.evergladensis.1.1.chr6.g00361.t01#3    mNeoVis.2.1.1.chr4_hap1.g00363.t01#5    mNeoVis.vulgivaga.1.1.chr6.g00368.t01#4   mNeoVis.vulgivaga.2.1.chr6.g00361.t01#1       6025   125256405:   mNeoVis.2.1.1.chr4_hap1.g00364.t01#5    mNeoVis.evergladensis.1.1.chr6.g00362.t01#3    mNeoVis.vulgivaga.1.1.chr6.g00369.t01#4    mNeoVis.evergladensis.2.1.chr6.g00368.t01#2    mNeoVis.3.1.1.chr6.g00365.t01#6   mNeoVis.vulgivaga.2.1.chr6.g00362.t01#1       6026   125256406:   mNeoVis.evergladensis.2.1.chr6.g00369.t01#2   mNeoVis.vulgivaga.2.1.chr6.g00363.t01#1             6027   125256407:   mNeoVis.2.1.1.chr4_hap1.g00366.t01#5    mNeoVis.3.1.1.chr6.g00367.t01#6    mNeoVis.vulgivaga.2.1.chr6.g00364.t01#1    mNeoVis.evergladensis.2.1.chr6.g00370.t01#2    mNeoVis.vulgivaga.1.1.chr6.g00371.t01#4   mNeoVis.evergladensis.1.1.chr6.g00364.t01#3       6028   125256408:   mNeoVis.evergladensis.2.1.chr14.g00096.t01#2    mNeoVis.evergladensis.1.1.chr14.g00101.t01#3    mNeoVis.2.1.1.chr14_hap1.g00097.t01#5    mNeoVis.vulgivaga.1.1.chr14.g00096.t01#4   mNeoVis.vulgivaga.2.1.chr14.g00096.t01#1       6029   125256409:   mNeoVis.vulgivaga.2.1.chr7.g00048.t01#1    mNeoVis.vulgivaga.1.1.chr7.g00048.t01#4    mNeoVis.evergladensis.1.1.chr7.g00050.t01#3    mNeoVis.evergladensis.2.1.chr7.g00047.t01#2    mNeoVis.2.1.1.chr7_hap1.g00045.t01#5   mNeoVis.3.1.1.chr7.g00036.t01#6       6030   125256410:   mNeoVis.evergladensis.2.1.chr3.g00863.t01#2    mNeoVis.vulgivaga.2.1.chr3.g00877.t01#1    mNeoVis.vulgivaga.1.1.chr3.g00865.t01#4    mNeoVis.2.1.1.chr3_hap1.g00867.t01#5    mNeoVis.vulgivaga.1.1.chr3.g00864.t01#4    mNeoVis.evergladensis.1.1.chr3.g00870.t01#3   mNeoVis.3.1.1.chr3.g00850.t01#6       6031   125256411:   mNeoVis.evergladensis.2.1.chr6.g00371.t01#2    mNeoVis.vulgivaga.2.1.chr6.g00365.t01#1    mNeoVis.2.1.1.chr4_hap1.g00367.t01#5    mNeoVis.3.1.1.chr6.g00368.t01#6    mNeoVis.vulgivaga.1.1.chr6.g00372.t01#4   mNeoVis.evergladensis.1.1.chr6.g00365.t01#3       6032   125256412:   mNeoVis.vulgivaga.2.1.chr14.g00010.t01#1    mNeoVis.vulgivaga.1.1.chr14.g00010.t01#4    mNeoVis.2.1.1.chr14_hap1.g00010.t01#5    mNeoVis.evergladensis.2.1.chr14.g00010.t01#2    mNeoVis.3.1.1.chr14.g00009.t01#6   mNeoVis.evergladensis.1.1.chr14.g00010.t01#3       6033   125256413:   mNeoVis.2.1.1.chr4_hap1.g00368.t01#5    mNeoVis.3.1.1.chr6.g00369.t01#6    mNeoVis.evergladensis.2.1.chr6.g00372.t01#2    mNeoVis.vulgivaga.2.1.chr6.g00366.t01#1    mNeoVis.vulgivaga.1.1.chr6.g00373.t01#4   mNeoVis.evergladensis.1.1.chr6.g00366.t01#3       6034   125256414:   mNeoVis.2.1.1.chr14_hap1.g00011.t01#5    mNeoVis.vulgivaga.1.1.chr14.g00011.t01#4    mNeoVis.3.1.1.chr14.g00010.t01#6    mNeoVis.evergladensis.2.1.chr14.g00011.t01#2    mNeoVis.vulgivaga.2.1.chr14.g00011.t01#1   mNeoVis.evergladensis.1.1.chr14.g00011.t01#3       6035   125256415:   mNeoVis.evergladensis.2.1.chr6.g00373.t01#2    mNeoVis.3.1.1.chr6.g00370.t01#6    mNeoVis.vulgivaga.1.1.chr6.g00374.t01#4    mNeoVis.evergladensis.1.1.chr6.g00367.t01#3    mNeoVis.2.1.1.chr4_hap1.g00369.t01#5   mNeoVis.vulgivaga.2.1.chr6.g00367.t01#1       6036   125256416:   mNeoVis.vulgivaga.2.1.chr6.g00368.t01#1    mNeoVis.3.1.1.chr6.g00371.t01#6    mNeoVis.vulgivaga.1.1.chr6.g00375.t01#4    mNeoVis.2.1.1.chr4_hap1.g00370.t01#5    mNeoVis.evergladensis.1.1.chr6.g00368.t01#3    mNeoVis.evergladensis.1.1.chr6.g00369.t01#3   mNeoVis.evergladensis.2.1.chr6.g00374.t01#2     6037   125256417:   mNeoVis.vulgivaga.1.1.chr14.g00012.t01#4    mNeoVis.3.1.1.chr14.g00011.t01#6    mNeoVis.2.1.1.chr14_hap1.g00012.t01#5    mNeoVis.vulgivaga.2.1.chr14.g00012.t01#1    mNeoVis.evergladensis.1.1.chr14.g00012.t01#3   mNeoVis.evergladensis.2.1.chr14.g00012.t01#2       6038   125256418:   mNeoVis.evergladensis.1.1.chr14.g00013.t01#3    mNeoVis.vulgivaga.2.1.chr14.g00013.t01#1    mNeoVis.2.1.1.chr14_hap1.g00013.t01#5    mNeoVis.evergladensis.1.1.chr14.g00015.t01#3    mNeoVis.3.1.1.chr14.g00012.t01#6    mNeoVis.vulgivaga.1.1.chr14.g00013.t01#4   mNeoVis.evergladensis.2.1.chr14.g00013.t01#2     6039   125256419:   mNeoVis.evergladensis.2.1.chr6.g00375.t01#2    mNeoVis.evergladensis.1.1.chr6.g00370.t01#3    mNeoVis.vulgivaga.1.1.chr6.g00376.t01#4    mNeoVis.2.1.1.chr4_hap1.g00371.t01#5    mNeoVis.vulgivaga.2.1.chr6.g00369.t01#1   mNeoVis.3.1.1.chr6.g00372.t01#6       6040   125256420:   mNeoVis.vulgivaga.1.1.chr6.g00377.t01#4    mNeoVis.vulgivaga.2.1.chr6.g00370.t01#1    mNeoVis.evergladensis.1.1.chr6.g00371.t01#3    mNeoVis.evergladensis.2.1.chr6.g00376.t01#2    mNeoVis.3.1.1.chr6.g00373.t01#6   mNeoVis.2.1.1.chr4_hap1.g00372.t01#5       6041   125256421:   mNeoVis.evergladensis.2.1.chr14.g00014.t01#2    mNeoVis.vulgivaga.2.1.chr14.g00014.t01#1    mNeoVis.vulgivaga.1.1.chr14.g00014.t01#4    mNeoVis.2.1.1.chr14_hap1.g00014.t01#5    mNeoVis.evergladensis.1.1.chr14.g00014.t01#3   mNeoVis.3.1.1.chr14.g00013.t01#6       6042   125256422:   mNeoVis.vulgivaga.1.1.chr14.g00015.t01#4    mNeoVis.evergladensis.2.1.chr14.g00015.t01#2    mNeoVis.2.1.1.chr14_hap1.g00015.t01#5    mNeoVis.vulgivaga.2.1.chr14.g00015.t01#1    mNeoVis.evergladensis.1.1.chr14.g00016.t01#3   mNeoVis.3.1.1.chr14.g00014.t01#6       6043   125256423:  mNeoVis.vulgivaga.2.1.chr6.g00371.t01#1               6044   125256424:   mNeoVis.evergladensis.2.1.chr14.g00016.t01#2    mNeoVis.vulgivaga.1.1.chr14.g00016.t01#4    mNeoVis.vulgivaga.2.1.chr14.g00016.t01#1    mNeoVis.evergladensis.1.1.chr14.g00017.t01#3    mNeoVis.3.1.1.chr14.g00015.t01#6   mNeoVis.2.1.1.chr14_hap1.g00016.t01#5       6045   125256425:   mNeoVis.2.1.1.chr14_hap1.g00017.t01#5    mNeoVis.evergladensis.1.1.chr14.g00018.t01#3    mNeoVis.evergladensis.2.1.chr14.g00017.t01#2    mNeoVis.vulgivaga.2.1.chr14.g00017.t01#1    mNeoVis.3.1.1.chr14.g00016.t01#6   mNeoVis.vulgivaga.1.1.chr14.g00017.t01#4       6046   125256426:   mNeoVis.2.1.1.chr4_hap1.g00373.t01#5    mNeoVis.evergladensis.2.1.chr6.g00377.t01#2    mNeoVis.vulgivaga.2.1.chr6.g00372.t01#1    mNeoVis.vulgivaga.1.1.chr6.g00378.t01#4    mNeoVis.3.1.1.chr6.g00374.t01#6   mNeoVis.evergladensis.1.1.chr6.g00372.t01#3       6047   125256427:   mNeoVis.3.1.1.chr6.g00375.t01#6    mNeoVis.vulgivaga.1.1.chr6.g00379.t01#4    mNeoVis.2.1.1.chr4_hap1.g00374.t01#5    mNeoVis.evergladensis.2.1.chr6.g00378.t01#2    mNeoVis.evergladensis.1.1.chr6.g00373.t01#3   mNeoVis.vulgivaga.2.1.chr6.g00373.t01#1       6048   125256428:   mNeoVis.evergladensis.2.1.chr6.g00379.t01#2    mNeoVis.evergladensis.1.1.chr6.g00374.t01#3    mNeoVis.vulgivaga.2.1.chr6.g00374.t01#1    mNeoVis.vulgivaga.1.1.chr6.g00380.t01#4   mNeoVis.2.1.1.chr4_hap1.g00375.t01#5       6049   125256429:   mNeoVis.2.1.1.chr14_hap1.g00018.t01#5    mNeoVis.evergladensis.2.1.chr14.g00018.t01#2    mNeoVis.3.1.1.chr14.g00017.t01#6    mNeoVis.vulgivaga.2.1.chr14.g00018.t01#1    mNeoVis.evergladensis.1.1.chr14.g00020.t01#3   mNeoVis.vulgivaga.1.1.chr14.g00018.t01#4       6050   125256430:   mNeoVis.evergladensis.1.1.chr15.g00581.t01#3    mNeoVis.evergladensis.2.1.chr15.g00034.t01#2    mNeoVis.3.1.1.chr15.g00030.t01#6    mNeoVis.vulgivaga.2.1.chr15.g00036.t01#1    mNeoVis.2.1.1.chr15_hap1.g00033.t01#5   mNeoVis.vulgivaga.1.1.chr15.g00035.t01#4       6051   125256431:   mNeoVis.2.1.1.chr4_hap1.g00376.t01#5    mNeoVis.3.1.1.chr6.g00377.t01#6    mNeoVis.vulgivaga.1.1.chr6.g00381.t01#4    mNeoVis.evergladensis.2.1.chr6.g00380.t01#2    mNeoVis.evergladensis.1.1.chr6.g00375.t01#3   mNeoVis.vulgivaga.2.1.chr6.g00375.t01#1       6052   125256432:   mNeoVis.vulgivaga.2.1.chr6.g00377.t01#1    mNeoVis.2.1.1.chr4_hap1.g00378.t01#5    mNeoVis.evergladensis.2.1.chr6.g00382.t01#2    mNeoVis.vulgivaga.1.1.chr6.g00383.t01#4    mNeoVis.3.1.1.chr6.g00378.t01#6   mNeoVis.evergladensis.1.1.chr6.g00377.t01#3       6053   125256433:   mNeoVis.evergladensis.2.1.chr6.g00381.t01#2    mNeoVis.vulgivaga.1.1.chr6.g00382.t01#4    mNeoVis.evergladensis.1.1.chr6.g00376.t01#3    mNeoVis.vulgivaga.2.1.chr6.g00376.t01#1    mNeoVis.2.1.1.chr4_hap1.g00377.t01#5   mNeoVis.3.1.1.chr6.g00379.t01#6       6054   125256434:   mNeoVis.vulgivaga.1.1.chr6.g00384.t01#4    mNeoVis.evergladensis.2.1.chr6.g00383.t01#2    mNeoVis.2.1.1.chr4_hap1.g00379.t01#5    mNeoVis.3.1.1.chr6.g00380.t01#6    mNeoVis.evergladensis.1.1.chr6.g00378.t01#3   mNeoVis.vulgivaga.2.1.chr6.g00378.t01#1       6055   125256435:   mNeoVis.evergladensis.1.1.chr6.g00379.t01#3    mNeoVis.vulgivaga.2.1.chr6.g00379.t01#1    mNeoVis.3.1.1.chr6.g00381.t01#6    mNeoVis.evergladensis.2.1.chr6.g00384.t01#2    mNeoVis.vulgivaga.1.1.chr6.g00388.t01#4   mNeoVis.2.1.1.chr4_hap1.g00380.t01#5       6056   125256436:   mNeoVis.3.1.1.chr7.g00037.t01#6    mNeoVis.evergladensis.1.1.chr7.g00051.t01#3    mNeoVis.vulgivaga.2.1.chr7.g00049.t01#1    mNeoVis.2.1.1.chr7_hap1.g00046.t01#5   mNeoVis.evergladensis.2.1.chr7.g00048.t01#2       6057   125256437:   mNeoVis.vulgivaga.2.1.chr6.g00380.t01#1    mNeoVis.2.1.1.chr4_hap1.g00381.t01#5    mNeoVis.3.1.1.chr6.g00382.t01#6    mNeoVis.evergladensis.1.1.chr6.g00380.t01#3   mNeoVis.evergladensis.2.1.chr6.g00385.t01#2       6058   125256438:   mNeoVis.evergladensis.2.1.chr6.g00386.t01#2    mNeoVis.evergladensis.1.1.chr6.g00381.t01#3    mNeoVis.vulgivaga.1.1.chr6.g00386.t01#4    mNeoVis.2.1.1.chr4_hap1.g00382.t01#5    mNeoVis.vulgivaga.2.1.chr6.g00381.t01#1   mNeoVis.3.1.1.chr6.g00383.t01#6       6059   125256439:   mNeoVis.evergladensis.1.1.chr14.g00021.t01#3    mNeoVis.vulgivaga.2.1.chr14.g00019.t01#1    mNeoVis.vulgivaga.1.1.chr14.g00019.t01#4    mNeoVis.evergladensis.2.1.chr14.g00019.t01#2    mNeoVis.3.1.1.chr14.g00018.t01#6   mNeoVis.2.1.1.chr14_hap1.g00019.t01#5       6060   125256440:   mNeoVis.evergladensis.2.1.chr12.g00406.t01#2    mNeoVis.vulgivaga.1.1.chr12.g00416.t01#4    mNeoVis.evergladensis.1.1.chr12.g00413.t01#3    mNeoVis.2.1.1.chr12_hap1.g00449.t01#5    mNeoVis.3.1.1.chr12.g00411.t01#6   mNeoVis.vulgivaga.2.1.chr12.g00414.t01#1       6061   125256441:   mNeoVis.2.1.1.chr4_hap1.g00383.t01#5    mNeoVis.evergladensis.2.1.chr6.g00387.t01#2    mNeoVis.evergladensis.1.1.chr6.g00382.t01#3    mNeoVis.vulgivaga.2.1.chr6.g00382.t01#1   mNeoVis.vulgivaga.1.1.chr6.g00387.t01#4       6062   125256442:   mNeoVis.3.1.1.chr6.g00384.t01#6    mNeoVis.vulgivaga.1.1.chr6.g00389.t01#4    mNeoVis.2.1.1.chr4_hap1.g00384.t01#5    mNeoVis.evergladensis.1.1.chr6.g00383.t01#3    mNeoVis.vulgivaga.2.1.chr6.g00383.t01#1   mNeoVis.evergladensis.2.1.chr6.g00388.t01#2       6063   125256443:   mNeoVis.evergladensis.1.1.chr6.g00384.t01#3    mNeoVis.evergladensis.2.1.chr6.g00389.t01#2    mNeoVis.vulgivaga.1.1.chr6.g00390.t01#4    mNeoVis.vulgivaga.2.1.chr6.g00384.t01#1    mNeoVis.2.1.1.chr4_hap1.g00385.t01#5   mNeoVis.3.1.1.chr6.g00385.t01#6       6064   125256444:   mNeoVis.vulgivaga.2.1.chr6.g00385.t01#1   mNeoVis.evergladensis.1.1.chr6.g00385.t01#3           6065   125256445:   mNeoVis.evergladensis.2.1.chr6.g00391.t01#2    mNeoVis.vulgivaga.1.1.chr6.g00393.t01#4    mNeoVis.2.1.1.chr4_hap1.g00387.t01#5    mNeoVis.3.1.1.chr6.g00387.t01#6    mNeoVis.evergladensis.1.1.chr6.g00386.t01#3   mNeoVis.vulgivaga.2.1.chr6.g00386.t01#1       6066   125256446:   mNeoVis.vulgivaga.1.1.chr14.g00020.t01#4    mNeoVis.3.1.1.chr14.g00019.t01#6    mNeoVis.vulgivaga.2.1.chr14.g00020.t01#1    mNeoVis.evergladensis.2.1.chr14.g00020.t01#2   mNeoVis.2.1.1.chr14_hap1.g00020.t01#5       6067   125256447:   mNeoVis.vulgivaga.1.1.chr6.g00394.t01#4    mNeoVis.evergladensis.2.1.chr6.g00392.t01#2    mNeoVis.2.1.1.chr4_hap1.g00389.t01#5    mNeoVis.3.1.1.chr6.g00389.t01#6    mNeoVis.evergladensis.1.1.chr6.g00388.t01#3   mNeoVis.vulgivaga.2.1.chr6.g00387.t01#1       6068   125256448:  mNeoVis.vulgivaga.2.1.chr6.g00388.t01#1               6069   125256449:   mNeoVis.vulgivaga.1.1.chr14.g00022.t01#4    mNeoVis.2.1.1.chr14_hap1.g00022.t01#5    mNeoVis.3.1.1.chr14.g00020.t01#6    mNeoVis.evergladensis.1.1.chr14.g00024.t01#3    mNeoVis.evergladensis.2.1.chr14.g00022.t01#2   mNeoVis.vulgivaga.2.1.chr14.g00021.t01#1       6070   125256450:   mNeoVis.2.1.1.chr4_hap1.g00390.t01#5    mNeoVis.evergladensis.2.1.chr6.g00393.t01#2    mNeoVis.evergladensis.1.1.chr5.g00460.t01#3    mNeoVis.evergladensis.1.1.chr6.g00389.t01#3    mNeoVis.vulgivaga.2.1.chr5.g00467.t01#1    mNeoVis.3.1.1.chr6.g00390.t01#6    mNeoVis.evergladensis.2.1.chr5.g00468.t01#2    mNeoVis.2.1.1.chr12_hap1.g00271.t01#5    mNeoVis.vulgivaga.1.1.chr6.g00395.t01#4      6071   125256451:  mNeoVis.vulgivaga.2.1.chr14.g00023.t01#1               6072   125256452:   mNeoVis.vulgivaga.2.1.chr14.g00022.t01#1    mNeoVis.vulgivaga.1.1.chr14.g00023.t01#4    mNeoVis.evergladensis.1.1.chr14.g00025.t01#3    mNeoVis.2.1.1.chr14_hap1.g00023.t01#5    mNeoVis.3.1.1.chr14.g00021.t01#6   mNeoVis.evergladensis.2.1.chr14.g00023.t01#2       6073   125256453:   mNeoVis.vulgivaga.2.1.chr6.g00391.t01#1    mNeoVis.2.1.1.chr4_hap1.g00392.t01#5    mNeoVis.3.1.1.chr6.g00392.t01#6    mNeoVis.vulgivaga.1.1.chr6.g00397.t01#4    mNeoVis.evergladensis.2.1.chr6.g00395.t01#2   mNeoVis.evergladensis.1.1.chr6.g00391.t01#3       6074   125256454:   mNeoVis.evergladensis.1.1.chr6.g00392.t01#3    mNeoVis.vulgivaga.2.1.chr6.g00392.t01#1    mNeoVis.evergladensis.2.1.chr6.g00396.t01#2    mNeoVis.3.1.1.chr6.g00393.t01#6    mNeoVis.2.1.1.chr4_hap1.g00393.t01#5   mNeoVis.vulgivaga.1.1.chr6.g00398.t01#4       6075   125256455:   mNeoVis.3.1.1.chr6.g00394.t01#6    mNeoVis.vulgivaga.1.1.chr6.g00399.t01#4    mNeoVis.evergladensis.2.1.chr6.g00397.t01#2    mNeoVis.2.1.1.chr4_hap1.g00394.t01#5    mNeoVis.vulgivaga.2.1.chr6.g00393.t01#1   mNeoVis.evergladensis.1.1.chr6.g00393.t01#3       6076   125256456:   mNeoVis.evergladensis.2.1.chr14.g00024.t01#2    mNeoVis.vulgivaga.2.1.chr14.g00024.t01#1    mNeoVis.2.1.1.chr14_hap1.g00024.t01#5    mNeoVis.vulgivaga.1.1.chr14.g00024.t01#4   mNeoVis.evergladensis.1.1.chr14.g00026.t01#3       6077   125256457:   mNeoVis.2.1.1.chr14_hap1.g00025.t01#5    mNeoVis.3.1.1.chr14.g00023.t01#6    mNeoVis.vulgivaga.2.1.chr14.g00025.t01#1    mNeoVis.evergladensis.2.1.chr14.g00025.t01#2    mNeoVis.evergladensis.1.1.chr14.g00027.t01#3   mNeoVis.vulgivaga.1.1.chr14.g00025.t01#4       6078   125256458:   mNeoVis.3.1.1.chr15.g00031.t01#6    mNeoVis.vulgivaga.1.1.chr15.g00036.t01#4    mNeoVis.evergladensis.2.1.chr15.g00035.t01#2    mNeoVis.vulgivaga.2.1.chr15.g00037.t01#1    mNeoVis.evergladensis.1.1.chr15.g00580.t01#3   mNeoVis.2.1.1.chr15_hap1.g00034.t01#5       6079   125256459:   mNeoVis.vulgivaga.2.1.chr6.g00394.t01#1    mNeoVis.evergladensis.2.1.chr6.g00398.t01#2    mNeoVis.vulgivaga.1.1.chr6.g00400.t01#4    mNeoVis.2.1.1.chr4_hap1.g00395.t01#5    mNeoVis.3.1.1.chr6.g00395.t01#6    mNeoVis.3.1.1.chr1.g01736.t01#6   mNeoVis.evergladensis.1.1.chr1.g01744.t01#3     6080   125256460:   mNeoVis.evergladensis.2.1.chr3.g00840.t01#2    mNeoVis.vulgivaga.1.1.chr3.g00839.t01#4    mNeoVis.vulgivaga.2.1.chr3.g00851.t01#1    mNeoVis.3.1.1.chr3.g00826.t01#6    mNeoVis.evergladensis.1.1.chr3.g00844.t01#3   mNeoVis.2.1.1.chr3_hap1.g00843.t01#5       6081   125256461:   mNeoVis.evergladensis.2.1.chr6.g00399.t01#2    mNeoVis.vulgivaga.2.1.chr6.g00395.t01#1    mNeoVis.2.1.1.chr4_hap1.g00396.t01#5    mNeoVis.3.1.1.chr6.g00396.t01#6    mNeoVis.evergladensis.1.1.chr6.g00394.t01#3   mNeoVis.vulgivaga.1.1.chr6.g00401.t01#4       6082   125256462:   mNeoVis.vulgivaga.2.1.chr14.g00026.t01#1    mNeoVis.3.1.1.chr14.g00024.t01#6    mNeoVis.2.1.1.chr14_hap1.g00026.t01#5    mNeoVis.vulgivaga.1.1.chr14.g00026.t01#4    mNeoVis.evergladensis.2.1.chr14.g00026.t01#2   mNeoVis.evergladensis.1.1.chr14.g00028.t01#3       6083   125256463:   mNeoVis.evergladensis.2.1.chr6.g00400.t01#2    mNeoVis.2.1.1.chr4_hap1.g00397.t01#5    mNeoVis.evergladensis.1.1.chr6.g00395.t01#3    mNeoVis.vulgivaga.2.1.chr6.g00396.t01#1    mNeoVis.3.1.1.chr6.g00397.t01#6   mNeoVis.vulgivaga.1.1.chr6.g00402.t01#4       6084   125256464:   mNeoVis.evergladensis.1.1.chr7.g00052.t01#3    mNeoVis.2.1.1.chr7_hap1.g00047.t01#5   mNeoVis.vulgivaga.2.1.chr7.g00050.t01#1           6085   125256465:  mNeoVis.vulgivaga.2.1.chr14.g00027.t01#1               6086   125256466:   mNeoVis.3.1.1.chr6.g00398.t01#6    mNeoVis.evergladensis.1.1.chr6.g00396.t01#3    mNeoVis.vulgivaga.2.1.chr6.g00397.t01#1    mNeoVis.vulgivaga.1.1.chr6.g00403.t01#4    mNeoVis.2.1.1.chr4_hap1.g00398.t01#5   mNeoVis.evergladensis.2.1.chr6.g00401.t01#2       6087   125256467:   mNeoVis.vulgivaga.1.1.chr14.g00027.t01#4    mNeoVis.2.1.1.chr14_hap1.g00027.t01#5    mNeoVis.evergladensis.2.1.chr14.g00027.t01#2   mNeoVis.vulgivaga.2.1.chr14.g00028.t01#1       6088   125256468:   mNeoVis.vulgivaga.2.1.chr14.g00029.t01#1    mNeoVis.evergladensis.2.1.chr14.g00029.t01#2   mNeoVis.2.1.1.chr14_hap1.g00028.t01#5           6089   125256469:   mNeoVis.3.1.1.chr6.g00399.t01#6    mNeoVis.vulgivaga.2.1.chr6.g00398.t01#1    mNeoVis.evergladensis.2.1.chr6.g00402.t01#2    mNeoVis.2.1.1.chr4_hap1.g00399.t01#5    mNeoVis.evergladensis.1.1.chr6.g00397.t01#3   mNeoVis.vulgivaga.1.1.chr6.g00404.t01#4       6090   125256470:   mNeoVis.vulgivaga.1.1.chr6.g00405.t01#4    mNeoVis.2.1.1.chr4_hap1.g00400.t01#5    mNeoVis.3.1.1.chr6.g00400.t01#6    mNeoVis.evergladensis.2.1.chr6.g00403.t01#2    mNeoVis.vulgivaga.2.1.chr6.g00399.t01#1   mNeoVis.evergladensis.1.1.chr6.g00398.t01#3       6091   125256471:   mNeoVis.vulgivaga.1.1.chr14.g00029.t01#4   mNeoVis.vulgivaga.2.1.chr14.g00030.t01#1           6092   125256472:   mNeoVis.vulgivaga.2.1.chr14.g00031.t01#1    mNeoVis.2.1.1.chr14_hap1.g00030.t01#5    mNeoVis.evergladensis.2.1.chr14.g00030.t01#2    mNeoVis.evergladensis.1.1.chr14.g00031.t01#3    mNeoVis.vulgivaga.1.1.chr14.g00030.t01#4   mNeoVis.3.1.1.chr14.g00027.t01#6       6093   125256473:   mNeoVis.vulgivaga.2.1.chr14.g00032.t01#1    mNeoVis.evergladensis.1.1.chr14.g00032.t01#3    mNeoVis.evergladensis.2.1.chr14.g00031.t01#2    mNeoVis.vulgivaga.1.1.chr14.g00031.t01#4    mNeoVis.2.1.1.chr14_hap1.g00031.t01#5   mNeoVis.3.1.1.chr14.g00028.t01#6       6094   125256474:  mNeoVis.vulgivaga.2.1.chr14.g00033.t01#1               6095   125256475:   mNeoVis.evergladensis.1.1.chr14.g00034.t01#3    mNeoVis.evergladensis.2.1.chr14.g00032.t01#2    mNeoVis.3.1.1.chr14.g00029.t01#6    mNeoVis.2.1.1.chr14_hap1.g00033.t01#5    mNeoVis.vulgivaga.1.1.chr14.g00033.t01#4   mNeoVis.vulgivaga.2.1.chr14.g00034.t01#1       6096   125256476:   mNeoVis.2.1.1.chr4_hap1.g00401.t01#5    mNeoVis.evergladensis.2.1.chr6.g00404.t01#2    mNeoVis.vulgivaga.2.1.chr6.g00400.t01#1    mNeoVis.evergladensis.1.1.chr6.g00399.t01#3    mNeoVis.vulgivaga.1.1.chr6.g00406.t01#4   mNeoVis.3.1.1.chr6.g00401.t01#6       6097   125256477:   mNeoVis.2.1.1.chr4_hap1.g00402.t01#5    mNeoVis.vulgivaga.2.1.chr6.g00401.t01#1    mNeoVis.evergladensis.1.1.chr6.g00400.t01#3    mNeoVis.vulgivaga.1.1.chr6.g00407.t01#4    mNeoVis.3.1.1.chr6.g00402.t01#6   mNeoVis.evergladensis.2.1.chr6.g00405.t01#2       6098   125256478:   mNeoVis.evergladensis.1.1.chr6.g00401.t01#3    mNeoVis.vulgivaga.1.1.chr6.g00408.t01#4    mNeoVis.3.1.1.chr6.g00403.t01#6    mNeoVis.2.1.1.chr4_hap1.g00403.t01#5    mNeoVis.vulgivaga.2.1.chr6.g00402.t01#1   mNeoVis.evergladensis.2.1.chr6.g00406.t01#2       6099   125256479:   mNeoVis.2.1.1.chr14_hap1.g00034.t01#5    mNeoVis.3.1.1.chr14.g00030.t01#6    mNeoVis.evergladensis.2.1.chr14.g00033.t01#2    mNeoVis.vulgivaga.2.1.chr14.g00035.t01#1   mNeoVis.evergladensis.1.1.chr14.g00035.t01#3       6100   125256480:   mNeoVis.evergladensis.1.1.chr14.g00036.t01#3    mNeoVis.2.1.1.chr14_hap1.g00035.t01#5    mNeoVis.evergladensis.2.1.chr14.g00034.t01#2    mNeoVis.3.1.1.chr14.g00031.t01#6   mNeoVis.vulgivaga.2.1.chr14.g00036.t01#1       6101   125256481:  mNeoVis.vulgivaga.1.1.chr14.g00034.t01#4               6102   125256482:   mNeoVis.evergladensis.2.1.chr6.g00408.t01#2    mNeoVis.vulgivaga.1.1.chr6.g00410.t01#4    mNeoVis.3.1.1.chr6.g00405.t01#6    mNeoVis.vulgivaga.2.1.chr6.g00404.t01#1    mNeoVis.evergladensis.1.1.chr6.g00403.t01#3   mNeoVis.2.1.1.chr4_hap1.g00405.t01#5       6103   125256483:   mNeoVis.vulgivaga.1.1.chr6.g00411.t01#4    mNeoVis.vulgivaga.2.1.chr6.g00405.t01#1    mNeoVis.evergladensis.2.1.chr6.g00409.t01#2    mNeoVis.evergladensis.1.1.chr6.g00404.t01#3    mNeoVis.2.1.1.chr4_hap1.g00406.t01#5   mNeoVis.3.1.1.chr6.g00406.t01#6       6104   125256484:   mNeoVis.evergladensis.2.1.chr6.g00410.t01#2    mNeoVis.vulgivaga.2.1.chr6.g00406.t01#1    mNeoVis.3.1.1.chr6.g00407.t01#6    mNeoVis.evergladensis.1.1.chr6.g00406.t01#3    mNeoVis.vulgivaga.1.1.chr6.g00412.t01#4   mNeoVis.2.1.1.chr4_hap1.g00407.t01#5       6105   125256485:   mNeoVis.vulgivaga.1.1.chr6.g00413.t01#4    mNeoVis.2.1.1.chr4_hap1.g00408.t01#5    mNeoVis.vulgivaga.2.1.chr6.g00407.t01#1    mNeoVis.evergladensis.2.1.chr6.g00411.t01#2    mNeoVis.3.1.1.chr6.g00408.t01#6   mNeoVis.evergladensis.1.1.chr6.g00407.t01#3       6106   125256486:   mNeoVis.vulgivaga.1.1.chr14.g00035.t01#4    mNeoVis.evergladensis.2.1.chr14.g00035.t01#2    mNeoVis.3.1.1.chr14.g00032.t01#6    mNeoVis.vulgivaga.2.1.chr14.g00037.t01#1    mNeoVis.evergladensis.1.1.chr14.g00037.t01#3   mNeoVis.2.1.1.chr14_hap1.g00036.t01#5       6107   125256487:   mNeoVis.3.1.1.chr14.g00033.t01#6    mNeoVis.evergladensis.2.1.chr14.g00036.t01#2    mNeoVis.vulgivaga.2.1.chr14.g00038.t01#1   mNeoVis.vulgivaga.1.1.chr14.g00036.t01#4       6108   125256488:   mNeoVis.evergladensis.1.1.chr15.g00579.t01#3    mNeoVis.2.1.1.chr15_hap1.g00035.t01#5    mNeoVis.vulgivaga.2.1.chr15.g00038.t01#1    mNeoVis.vulgivaga.1.1.chr15.g00037.t01#4    mNeoVis.evergladensis.2.1.chr15.g00036.t01#2   mNeoVis.3.1.1.chr15.g00032.t01#6       6109   125256489:   mNeoVis.vulgivaga.1.1.chr6.g00414.t01#4    mNeoVis.2.1.1.chr4_hap1.g00409.t01#5    mNeoVis.3.1.1.chr6.g00409.t01#6    mNeoVis.vulgivaga.2.1.chr6.g00408.t01#1    mNeoVis.evergladensis.2.1.chr6.g00412.t01#2   mNeoVis.evergladensis.1.1.chr6.g00408.t01#3       6110   125256490:   mNeoVis.evergladensis.2.1.chr14.g00037.t01#2    mNeoVis.vulgivaga.2.1.chr14.g00039.t01#1    mNeoVis.vulgivaga.1.1.chr14.g00037.t01#4   mNeoVis.3.1.1.chr14.g00034.t01#6         6111   125256491:   mNeoVis.vulgivaga.2.1.chr7.g00051.t01#1    mNeoVis.evergladensis.2.1.chr7.g00050.t01#2    mNeoVis.vulgivaga.1.1.chr7.g00052.t01#4   mNeoVis.2.1.1.chr7_hap1.g00049.t01#5         6112   125256492:   mNeoVis.2.1.1.chr14_hap1.g00038.t01#5    mNeoVis.3.1.1.chr14.g00035.t01#6    mNeoVis.evergladensis.1.1.chr14.g00040.t01#3    mNeoVis.evergladensis.2.1.chr14.g00039.t01#2    mNeoVis.vulgivaga.1.1.chr14.g00038.t01#4   mNeoVis.vulgivaga.2.1.chr14.g00040.t01#1       6113   125256493:   mNeoVis.vulgivaga.1.1.chr6.g00416.t01#4    mNeoVis.evergladensis.1.1.chr6.g00409.t01#3    mNeoVis.3.1.1.chr6.g00410.t01#6    mNeoVis.vulgivaga.1.1.chr6.g00415.t01#4    mNeoVis.2.1.1.chr4_hap1.g00410.t01#5    mNeoVis.vulgivaga.2.1.chr6.g00409.t01#1   mNeoVis.evergladensis.2.1.chr6.g00413.t01#2     6114   125256494:   mNeoVis.3.1.1.chr6.g00411.t01#6    mNeoVis.evergladensis.2.1.chr6.g00414.t01#2    mNeoVis.evergladensis.1.1.chr6.g00410.t01#3    mNeoVis.2.1.1.chr4_hap1.g00411.t01#5    mNeoVis.vulgivaga.2.1.chr6.g00410.t01#1   mNeoVis.vulgivaga.1.1.chr6.g00417.t01#4       6115   125256495:   mNeoVis.vulgivaga.1.1.chr6.g00419.t01#4    mNeoVis.evergladensis.1.1.chr6.g00411.t01#3    mNeoVis.vulgivaga.2.1.chr6.g00411.t01#1    mNeoVis.2.1.1.chr4_hap1.g00413.t01#5    mNeoVis.3.1.1.chr6.g00412.t01#6   mNeoVis.evergladensis.2.1.chr6.g00416.t01#2       6116   125256496:   mNeoVis.vulgivaga.2.1.chr6.g00412.t01#1    mNeoVis.evergladensis.1.1.chr6.g00413.t01#3    mNeoVis.evergladensis.2.1.chr6.g00417.t01#2    mNeoVis.vulgivaga.1.1.chr6.g00420.t01#4    mNeoVis.3.1.1.chr6.g00413.t01#6    mNeoVis.2.1.1.chr4_hap1.g00414.t01#5    mNeoVis.evergladensis.1.1.chr6.g00412.t01#3   mNeoVis.3.1.1.chr6.g00414.t01#6     6117   125256497:   mNeoVis.2.1.1.chr4_hap1.g00321.t01#5    mNeoVis.evergladensis.1.1.chr6.g00320.t01#3    mNeoVis.evergladensis.2.1.chr6.g00325.t01#2    mNeoVis.vulgivaga.1.1.chr6.g00328.t01#4    mNeoVis.3.1.1.chr6.g00326.t01#6   mNeoVis.vulgivaga.2.1.chr6.g00318.t01#1       6118   125256498:   mNeoVis.3.1.1.chr6.g00327.t01#6    mNeoVis.evergladensis.2.1.chr6.g00326.t01#2    mNeoVis.vulgivaga.1.1.chr6.g00329.t01#4    mNeoVis.vulgivaga.2.1.chr6.g00319.t01#1    mNeoVis.2.1.1.chr4_hap1.g00323.t01#5   mNeoVis.evergladensis.1.1.chr6.g00321.t01#3       6119   125256499:   mNeoVis.3.1.1.chr6.g00328.t01#6    mNeoVis.2.1.1.chr4_hap1.g00324.t01#5    mNeoVis.vulgivaga.2.1.chr6.g00320.t01#1    mNeoVis.vulgivaga.1.1.chr6.g00330.t01#4    mNeoVis.evergladensis.2.1.chr6.g00327.t01#2   mNeoVis.evergladensis.1.1.chr6.g00322.t01#3       6120   125256500:   mNeoVis.2.1.1.chr13_hap1.g00811.t01#5    mNeoVis.evergladensis.1.1.chr13.g00769.t01#3    mNeoVis.vulgivaga.1.1.chr13.g00766.t01#4    mNeoVis.evergladensis.2.1.chr13.g00776.t01#2   mNeoVis.vulgivaga.2.1.chr13.g00774.t01#1       6121   125256501:   mNeoVis.vulgivaga.2.1.chr13.g00775.t01#1    mNeoVis.2.1.1.chr13_hap1.g00812.t01#5    mNeoVis.3.1.1.chr13.g00735.t01#6    mNeoVis.vulgivaga.1.1.chr13.g00767.t01#4    mNeoVis.evergladensis.1.1.chr13.g00770.t01#3   mNeoVis.evergladensis.2.1.chr13.g00777.t01#2       6122   125256502:   mNeoVis.vulgivaga.2.1.chr6.g00321.t01#1    mNeoVis.3.1.1.chr6.g00329.t01#6    mNeoVis.2.1.1.chr4_hap1.g00325.t01#5    mNeoVis.vulgivaga.1.1.chr6.g00331.t01#4    mNeoVis.evergladensis.1.1.chr6.g00323.t01#3   mNeoVis.evergladensis.2.1.chr6.g00328.t01#2       6123   125256503:   mNeoVis.evergladensis.1.1.chr6.g00324.t01#3    mNeoVis.3.1.1.chr6.g00330.t01#6    mNeoVis.evergladensis.2.1.chr6.g00329.t01#2    mNeoVis.2.1.1.chr4_hap1.g00326.t01#5    mNeoVis.vulgivaga.1.1.chr6.g00332.t01#4   mNeoVis.vulgivaga.2.1.chr6.g00322.t01#1       6124   125256504:   mNeoVis.2.1.1.chr13_hap1.g00813.t01#5    mNeoVis.vulgivaga.1.1.chr13.g00768.t01#4    mNeoVis.evergladensis.2.1.chr13.g00778.t01#2   mNeoVis.3.1.1.chr13.g00737.t01#6         6125   125256505:   mNeoVis.3.1.1.chr13.g00736.t01#6    mNeoVis.evergladensis.1.1.chr13.g00771.t01#3   mNeoVis.vulgivaga.2.1.chr13.g00776.t01#1         6126   125256506:   mNeoVis.vulgivaga.2.1.chr6.g00323.t01#1    mNeoVis.vulgivaga.1.1.chr6.g00333.t01#4    mNeoVis.2.1.1.chr4_hap1.g00327.t01#5    mNeoVis.evergladensis.2.1.chr6.g00330.t01#2    mNeoVis.evergladensis.1.1.chr6.g00325.t01#3   mNeoVis.3.1.1.chr6.g00331.t01#6       6127   125256507:   mNeoVis.evergladensis.2.1.chr13.g00779.t01#2    mNeoVis.evergladensis.1.1.chr13.g00772.t01#3    mNeoVis.2.1.1.chr13_hap1.g00814.t01#5    mNeoVis.3.1.1.chr13.g00738.t01#6    mNeoVis.vulgivaga.2.1.chr13.g00777.t01#1   mNeoVis.vulgivaga.1.1.chr13.g00769.t01#4       6128   125256508:   mNeoVis.vulgivaga.2.1.chr11.g01036.t01#1    mNeoVis.3.1.1.chr11.g00988.t01#6    mNeoVis.2.1.1.chr11_hap1.g01037.t01#5    mNeoVis.vulgivaga.1.1.chr11.g01010.t01#4    mNeoVis.evergladensis.1.1.chr11.g01030.t01#3   mNeoVis.evergladensis.2.1.chr11.g01014.t01#2       6129   125256509:   mNeoVis.vulgivaga.2.1.chr13.g00778.t01#1    mNeoVis.vulgivaga.1.1.chr13.g00770.t01#4    mNeoVis.evergladensis.1.1.chr13.g00773.t01#3    mNeoVis.evergladensis.2.1.chr13.g00780.t01#2   mNeoVis.2.1.1.chr13_hap1.g00816.t01#5       6130   125256510:   mNeoVis.2.1.1.chr4_hap1.g00328.t01#5    mNeoVis.evergladensis.2.1.chr6.g00331.t01#2    mNeoVis.vulgivaga.1.1.chr6.g00334.t01#4    mNeoVis.evergladensis.1.1.chr6.g00326.t01#3    mNeoVis.3.1.1.chr6.g00333.t01#6   mNeoVis.vulgivaga.2.1.chr6.g00324.t01#1       6131   125256511:   mNeoVis.2.1.1.chr4_hap1.g00329.t01#5    mNeoVis.evergladensis.2.1.chr6.g00332.t01#2    mNeoVis.vulgivaga.2.1.chr6.g00325.t01#1    mNeoVis.vulgivaga.1.1.chr6.g00335.t01#4    mNeoVis.evergladensis.1.1.chr6.g00327.t01#3   mNeoVis.3.1.1.chr6.g00334.t01#6       6132   125256512:   mNeoVis.evergladensis.1.1.chr14.g00041.t01#3    mNeoVis.vulgivaga.2.1.chr14.g00041.t01#1    mNeoVis.evergladensis.2.1.chr14.g00040.t01#2    mNeoVis.2.1.1.chr14_hap1.g00039.t01#5    mNeoVis.3.1.1.chr14.g00036.t01#6   mNeoVis.vulgivaga.1.1.chr14.g00039.t01#4       6133   125256513:   mNeoVis.2.1.1.chr14_hap1.g00040.t01#5    mNeoVis.vulgivaga.1.1.chr14.g00040.t01#4    mNeoVis.evergladensis.1.1.chr14.g00042.t01#3    mNeoVis.3.1.1.chr14.g00037.t01#6    mNeoVis.vulgivaga.2.1.chr14.g00042.t01#1   mNeoVis.evergladensis.2.1.chr14.g00041.t01#2       6134   125256514:   mNeoVis.2.1.1.chr4_hap1.g00330.t01#5    mNeoVis.3.1.1.chr6.g00335.t01#6    mNeoVis.vulgivaga.1.1.chr6.g00336.t01#4    mNeoVis.vulgivaga.2.1.chr6.g00326.t01#1    mNeoVis.evergladensis.2.1.chr6.g00333.t01#2   mNeoVis.evergladensis.1.1.chr6.g00328.t01#3       6135   125256515:   mNeoVis.vulgivaga.2.1.chr13.g00779.t01#1    mNeoVis.vulgivaga.1.1.chr13.g00771.t01#4    mNeoVis.2.1.1.chr13_hap1.g00819.t01#5    mNeoVis.evergladensis.1.1.chr13.g00774.t01#3    mNeoVis.evergladensis.2.1.chr13.g00781.t01#2    mNeoVis.evergladensis.1.1.chr13.g00775.t01#3   mNeoVis.3.1.1.chr13.g00739.t01#6       6136   125256516:   mNeoVis.3.1.1.chr5.g01253.t01#6    mNeoVis.vulgivaga.1.1.chr13.g00772.t01#4    mNeoVis.vulgivaga.2.1.chr13.g00780.t01#1    mNeoVis.3.1.1.chr13.g00740.t01#6    mNeoVis.evergladensis.2.1.chr13.g00782.t01#2    mNeoVis.evergladensis.1.1.chr13.g00776.t01#3   mNeoVis.2.1.1.chr13_hap1.g00820.t01#5     6137   125256517:   mNeoVis.2.1.1.chr12_hap1.g00561.t01#5    mNeoVis.evergladensis.2.1.chr12.g00504.t01#2    mNeoVis.2.1.1.chr12_hap1.g00560.t01#5   mNeoVis.vulgivaga.2.1.chr12.g00518.t01#1       6138   125256518:   mNeoVis.3.1.1.chr13.g00742.t01#6    mNeoVis.2.1.1.chr13_hap1.g00822.t01#5    mNeoVis.vulgivaga.1.1.chr13.g00774.t01#4    mNeoVis.vulgivaga.2.1.chr13.g00781.t01#1    mNeoVis.evergladensis.2.1.chr13.g00783.t01#2   mNeoVis.evergladensis.1.1.chr13.g00777.t01#3       6139   125256519:   mNeoVis.2.1.1.chr13_hap1.g00823.t01#5    mNeoVis.evergladensis.1.1.chr13.g00778.t01#3    mNeoVis.vulgivaga.2.1.chr13.g00782.t01#1    mNeoVis.3.1.1.chr13.g00743.t01#6    mNeoVis.evergladensis.2.1.chr13.g00784.t01#2   mNeoVis.vulgivaga.1.1.chr13.g00775.t01#4       6140   125256520:   mNeoVis.vulgivaga.1.1.chr13.g00776.t01#4    mNeoVis.3.1.1.chr13.g00744.t01#6    mNeoVis.evergladensis.2.1.chr13.g00785.t01#2    mNeoVis.evergladensis.1.1.chr13.g00779.t01#3    mNeoVis.2.1.1.chr13_hap1.g00824.t01#5   mNeoVis.vulgivaga.2.1.chr13.g00783.t01#1       6141   125256521:   mNeoVis.evergladensis.1.1.chr6.g00329.t01#3    mNeoVis.3.1.1.chr6.g00336.t01#6    mNeoVis.evergladensis.2.1.chr6.g00334.t01#2    mNeoVis.vulgivaga.2.1.chr6.g00327.t01#1    mNeoVis.vulgivaga.1.1.chr6.g00337.t01#4   mNeoVis.2.1.1.chr4_hap1.g00331.t01#5       6142   125256522:   mNeoVis.vulgivaga.2.1.chr14.g00043.t01#1    mNeoVis.evergladensis.2.1.chr14.g00042.t01#2    mNeoVis.vulgivaga.1.1.chr14.g00041.t01#4    mNeoVis.3.1.1.chr14.g00038.t01#6    mNeoVis.2.1.1.chr14_hap1.g00041.t01#5   mNeoVis.evergladensis.1.1.chr14.g00043.t01#3       6143   125256523:   mNeoVis.2.1.1.chr4_hap1.g00332.t01#5    mNeoVis.vulgivaga.2.1.chr6.g00328.t01#1    mNeoVis.3.1.1.chr6.g00337.t01#6    mNeoVis.vulgivaga.1.1.chr6.g00338.t01#4   mNeoVis.evergladensis.1.1.chr6.g00330.t01#3       6144   125256524:   mNeoVis.evergladensis.1.1.chr14.g00047.t01#3    mNeoVis.vulgivaga.1.1.chr14.g00042.t01#4    mNeoVis.2.1.1.chr14_hap1.g00042.t01#5    mNeoVis.evergladensis.1.1.chr14.g00046.t01#3    mNeoVis.vulgivaga.2.1.chr14.g00044.t01#1    mNeoVis.evergladensis.2.1.chr14.g00043.t01#2    mNeoVis.3.1.1.chr14.g00039.t01#6   mNeoVis.evergladensis.1.1.chr14.g00044.t01#3     6145   125254477:   mNeoVis.3.1.1.chr13.g00145.t01#6    mNeoVis.vulgivaga.2.1.chr13.g00188.t01#1    mNeoVis.2.1.1.chr13_hap1.g00193.t01#5    mNeoVis.evergladensis.1.1.chr13.g00189.t01#3    mNeoVis.vulgivaga.1.1.chr13.g00185.t01#4   mNeoVis.evergladensis.2.1.chr13.g00191.t01#2       6146   125254478:   mNeoVis.3.1.1.chr6.g00338.t01#6    mNeoVis.2.1.1.chr4_hap1.g00333.t01#5    mNeoVis.vulgivaga.1.1.chr6.g00339.t01#4    mNeoVis.vulgivaga.2.1.chr6.g00329.t01#1    mNeoVis.evergladensis.2.1.chr6.g00335.t01#2   mNeoVis.evergladensis.1.1.chr6.g00331.t01#3       6147   125254479:   mNeoVis.3.1.1.chr6.g00339.t01#6    mNeoVis.vulgivaga.1.1.chr6.g00340.t01#4    mNeoVis.evergladensis.1.1.chr6.g00333.t01#3    mNeoVis.evergladensis.2.1.chr6.g00337.t01#2   mNeoVis.vulgivaga.2.1.chr6.g00331.t01#1       6148   125254480:   mNeoVis.evergladensis.1.1.chr6.g00334.t01#3    mNeoVis.3.1.1.chr6.g00340.t01#6    mNeoVis.2.1.1.chr4_hap1.g00335.t01#5    mNeoVis.vulgivaga.2.1.chr6.g00332.t01#1    mNeoVis.vulgivaga.1.1.chr6.g00341.t01#4   mNeoVis.evergladensis.2.1.chr6.g00338.t01#2       6149   125254481:  mNeoVis.vulgivaga.2.1.chr14.g00046.t01#1               6150   125254482:   mNeoVis.vulgivaga.2.1.chr14.g00045.t01#1    mNeoVis.evergladensis.2.1.chr14.g00045.t01#2    mNeoVis.vulgivaga.1.1.chr14.g00044.t01#4    mNeoVis.3.1.1.chr14.g00040.t01#6    mNeoVis.evergladensis.1.1.chr14.g00049.t01#3   mNeoVis.2.1.1.chr14_hap1.g00046.t01#5       6151   125254483:   mNeoVis.2.1.1.chr14_hap1.g00047.t01#5    mNeoVis.evergladensis.2.1.chr14.g00047.t01#2    mNeoVis.vulgivaga.1.1.chr14.g00046.t01#4    mNeoVis.vulgivaga.2.1.chr14.g00047.t01#1   mNeoVis.3.1.1.chr14.g00042.t01#6       6152   125254484:   mNeoVis.vulgivaga.1.1.chr11.g00294.t01#4    mNeoVis.3.1.1.chr11.g00271.t01#6    mNeoVis.vulgivaga.2.1.chr11.g00299.t01#1    mNeoVis.2.1.1.chr11_hap1.g00301.t01#5   mNeoVis.evergladensis.2.1.chr11.g00298.t01#2       6153   125254485:   mNeoVis.vulgivaga.1.1.chr7.g00053.t01#4    mNeoVis.vulgivaga.2.1.chr7.g00052.t01#1   mNeoVis.2.1.1.chr7_hap1.g00050.t01#5           6154   125254486:   mNeoVis.evergladensis.1.1.chr7.g00054.t01#3   mNeoVis.evergladensis.2.1.chr7.g00051.t01#2           6155   125254487:   mNeoVis.2.1.1.chr4_hap1.g00336.t01#5   mNeoVis.vulgivaga.2.1.chr6.g00333.t01#1             6156   125254488:  mNeoVis.vulgivaga.2.1.chr14.g00048.t01#1               6157   125254489:   mNeoVis.evergladensis.2.1.chr6.g00345.t01#2    mNeoVis.evergladensis.1.1.chr6.g00336.t01#3    mNeoVis.2.1.1.chr4_hap1.g00341.t01#5    mNeoVis.vulgivaga.2.1.chr6.g00338.t01#1   mNeoVis.vulgivaga.1.1.chr6.g00347.t01#4       6158   125254490:   mNeoVis.evergladensis.2.1.chr6.g00340.t01#2   mNeoVis.vulgivaga.2.1.chr6.g00334.t01#1             6159   125254491:   mNeoVis.vulgivaga.2.1.chr6.g00336.t01#1    mNeoVis.evergladensis.2.1.chr6.g00342.t01#2    mNeoVis.2.1.1.chr4_hap1.g00338.t01#5    mNeoVis.evergladensis.1.1.chr6.g00338.t01#3    mNeoVis.vulgivaga.1.1.chr6.g00345.t01#4   mNeoVis.3.1.1.chr6.g00342.t01#6       6160   125254492:   mNeoVis.3.1.1.chr14.g00044.t01#6    mNeoVis.vulgivaga.1.1.chr14.g00048.t01#4    mNeoVis.evergladensis.2.1.chr14.g00049.t01#2    mNeoVis.2.1.1.chr14_hap1.g00049.t01#5    mNeoVis.vulgivaga.2.1.chr14.g00049.t01#1   mNeoVis.evergladensis.1.1.chr14.g00053.t01#3       6161   125254493:   mNeoVis.2.1.1.chr14_hap1.g00050.t01#5    mNeoVis.evergladensis.2.1.chr14.g00050.t01#2    mNeoVis.vulgivaga.2.1.chr14.g00050.t01#1    mNeoVis.3.1.1.chr14.g00045.t01#6    mNeoVis.evergladensis.1.1.chr14.g00054.t01#3   mNeoVis.vulgivaga.1.1.chr14.g00049.t01#4       6162   125254494:   mNeoVis.vulgivaga.2.1.chr6.g00337.t01#1    mNeoVis.3.1.1.chr6.g00343.t01#6    mNeoVis.vulgivaga.1.1.chr6.g00346.t01#4    mNeoVis.evergladensis.1.1.chr6.g00339.t01#3    mNeoVis.evergladensis.2.1.chr6.g00343.t01#2   mNeoVis.2.1.1.chr4_hap1.g00339.t01#5       6163   125254495:   mNeoVis.evergladensis.1.1.chr14.g00055.t01#3    mNeoVis.vulgivaga.1.1.chr14.g00050.t01#4    mNeoVis.vulgivaga.2.1.chr14.g00051.t01#1    mNeoVis.2.1.1.chr14_hap1.g00051.t01#5    mNeoVis.evergladensis.2.1.chr14.g00051.t01#2   mNeoVis.3.1.1.chr14.g00046.t01#6       6164   125254496:   mNeoVis.evergladensis.1.1.chr14.g00057.t01#3    mNeoVis.vulgivaga.2.1.chr14.g00052.t01#1    mNeoVis.2.1.1.chr14_hap1.g00052.t01#5    mNeoVis.evergladensis.2.1.chr14.g00053.t01#2    mNeoVis.vulgivaga.1.1.chr14.g00051.t01#4   mNeoVis.3.1.1.chr14.g00048.t01#6       6165   125254497:   mNeoVis.evergladensis.2.1.chr6.g00346.t01#2    mNeoVis.vulgivaga.2.1.chr6.g00339.t01#1    mNeoVis.3.1.1.chr6.g00345.t01#6    mNeoVis.2.1.1.chr4_hap1.g00342.t01#5    mNeoVis.vulgivaga.1.1.chr6.g00348.t01#4   mNeoVis.evergladensis.1.1.chr6.g00340.t01#3       6166   125254498:   mNeoVis.evergladensis.1.1.chr14.g00058.t01#3    mNeoVis.evergladensis.2.1.chr14.g00054.t01#2    mNeoVis.2.1.1.chr14_hap1.g00053.t01#5    mNeoVis.3.1.1.chr14.g00049.t01#6    mNeoVis.vulgivaga.2.1.chr14.g00053.t01#1   mNeoVis.vulgivaga.1.1.chr14.g00052.t01#4       6167   125254499:   mNeoVis.vulgivaga.2.1.chr7.g00053.t01#1    mNeoVis.2.1.1.chr7_hap1.g00051.t01#5    mNeoVis.vulgivaga.1.1.chr7.g00054.t01#4    mNeoVis.evergladensis.1.1.chr7.g00055.t01#3    mNeoVis.evergladensis.2.1.chr7.g00052.t01#2   mNeoVis.3.1.1.chr7.g00041.t01#6       6168   125254500:  mNeoVis.vulgivaga.2.1.chr6.g00341.t01#1               6169   125254501:   mNeoVis.vulgivaga.1.1.chr6.g00349.t01#4    mNeoVis.evergladensis.2.1.chr6.g00347.t01#2    mNeoVis.vulgivaga.2.1.chr6.g00340.t01#1    mNeoVis.3.1.1.chr6.g00346.t01#6   mNeoVis.2.1.1.chr4_hap1.g00343.t01#5       6170   125254502:   mNeoVis.2.1.1.chr4_hap1.g00345.t01#5    mNeoVis.evergladensis.2.1.chr6.g00349.t01#2    mNeoVis.vulgivaga.1.1.chr6.g00350.t01#4    mNeoVis.3.1.1.chr6.g00347.t01#6    mNeoVis.evergladensis.1.1.chr6.g00342.t01#3   mNeoVis.vulgivaga.2.1.chr6.g00342.t01#1       6171   125254503:   mNeoVis.vulgivaga.1.1.chr14.g00053.t01#4    mNeoVis.vulgivaga.2.1.chr14.g00054.t01#1    mNeoVis.evergladensis.1.1.chr14.g00059.t01#3    mNeoVis.evergladensis.2.1.chr14.g00055.t01#2    mNeoVis.3.1.1.chr14.g00050.t01#6   mNeoVis.2.1.1.chr14_hap1.g00055.t01#5       6172   125254504:   mNeoVis.evergladensis.2.1.chr14.g00056.t01#2    mNeoVis.evergladensis.1.1.chr14.g00060.t01#3    mNeoVis.vulgivaga.2.1.chr14.g00055.t01#1    mNeoVis.2.1.1.chr14_hap1.g00056.t01#5    mNeoVis.3.1.1.chr14.g00051.t01#6   mNeoVis.vulgivaga.1.1.chr14.g00054.t01#4       6173   125254505:   mNeoVis.evergladensis.2.1.chr6.g00350.t01#2    mNeoVis.evergladensis.1.1.chr6.g00343.t01#3    mNeoVis.vulgivaga.1.1.chr6.g00351.t01#4    mNeoVis.2.1.1.chr4_hap1.g00346.t01#5    mNeoVis.vulgivaga.2.1.chr6.g00343.t01#1   mNeoVis.3.1.1.chr6.g00348.t01#6       6174   125254506:   mNeoVis.vulgivaga.2.1.chr14.g00056.t01#1    mNeoVis.evergladensis.1.1.chr14.g00061.t01#3    mNeoVis.3.1.1.chr14.g00052.t01#6    mNeoVis.2.1.1.chr14_hap1.g00057.t01#5    mNeoVis.evergladensis.2.1.chr14.g00057.t01#2   mNeoVis.vulgivaga.1.1.chr14.g00055.t01#4       6175   125254507:   mNeoVis.vulgivaga.2.1.chr6.g00344.t01#1    mNeoVis.2.1.1.chr4_hap1.g00347.t01#5    mNeoVis.3.1.1.chr6.g00349.t01#6    mNeoVis.evergladensis.1.1.chr6.g00344.t01#3    mNeoVis.vulgivaga.1.1.chr6.g00352.t01#4   mNeoVis.evergladensis.2.1.chr6.g00351.t01#2       6176   125254508:   mNeoVis.vulgivaga.1.1.chr13.g00751.t01#4    mNeoVis.2.1.1.chr13_hap1.g00795.t01#5    mNeoVis.evergladensis.1.1.chr13.g00752.t01#3    mNeoVis.3.1.1.chr13.g00712.t01#6    mNeoVis.evergladensis.2.1.chr13.g00759.t01#2   mNeoVis.vulgivaga.2.1.chr13.g00758.t01#1       6177   125254509:   mNeoVis.vulgivaga.2.1.chr6.g00345.t01#1    mNeoVis.3.1.1.chr6.g00350.t01#6    mNeoVis.evergladensis.2.1.chr6.g00352.t01#2    mNeoVis.evergladensis.1.1.chr6.g00345.t01#3    mNeoVis.vulgivaga.1.1.chr6.g00353.t01#4   mNeoVis.2.1.1.chr4_hap1.g00348.t01#5       6178   125254510:   mNeoVis.evergladensis.2.1.chr6.g00353.t01#2    mNeoVis.2.1.1.chr4_hap1.g00349.t01#5    mNeoVis.3.1.1.chr6.g00351.t01#6    mNeoVis.vulgivaga.1.1.chr6.g00354.t01#4    mNeoVis.evergladensis.1.1.chr6.g00346.t01#3   mNeoVis.vulgivaga.2.1.chr6.g00346.t01#1       6179   125254511:   mNeoVis.evergladensis.2.1.chr6.g00354.t01#2    mNeoVis.2.1.1.chr4_hap1.g00350.t01#5    mNeoVis.vulgivaga.1.1.chr6.g00355.t01#4    mNeoVis.3.1.1.chr6.g00352.t01#6    mNeoVis.vulgivaga.2.1.chr6.g00347.t01#1   mNeoVis.evergladensis.1.1.chr6.g00347.t01#3       6180   125254512:   mNeoVis.vulgivaga.1.1.chr13.g00752.t01#4    mNeoVis.3.1.1.chr13.g00713.t01#6    mNeoVis.vulgivaga.2.1.chr13.g00759.t01#1    mNeoVis.2.1.1.chr13_hap1.g00796.t01#5    mNeoVis.evergladensis.2.1.chr13.g00760.t01#2   mNeoVis.evergladensis.1.1.chr13.g00753.t01#3       6181   125254513:   mNeoVis.vulgivaga.1.1.chr13.g00753.t01#4    mNeoVis.evergladensis.2.1.chr13.g00761.t01#2    mNeoVis.vulgivaga.2.1.chr13.g00760.t01#1    mNeoVis.2.1.1.chr13_hap1.g00797.t01#5    mNeoVis.3.1.1.chr13.g00714.t01#6   mNeoVis.evergladensis.1.1.chr13.g00754.t01#3       6182   125254514:   mNeoVis.3.1.1.chr15.g00033.t01#6    mNeoVis.2.1.1.chr15_hap1.g00036.t01#5    mNeoVis.vulgivaga.1.1.chr15.g00038.t01#4    mNeoVis.evergladensis.1.1.chr15.g00578.t01#3    mNeoVis.vulgivaga.2.1.chr15.g00039.t01#1   mNeoVis.evergladensis.2.1.chr15.g00037.t01#2       6183   125254515:   mNeoVis.evergladensis.1.1.chr6.g00414.t01#3    mNeoVis.2.1.1.chr4_hap1.g00415.t01#5    mNeoVis.vulgivaga.2.1.chr6.g00413.t01#1    mNeoVis.evergladensis.2.1.chr6.g00418.t01#2   mNeoVis.3.1.1.chr6.g00415.t01#6       6184   125254516:   mNeoVis.vulgivaga.2.1.chr6.g00414.t01#1    mNeoVis.evergladensis.1.1.chr6.g00415.t01#3    mNeoVis.2.1.1.chr4_hap1.g00417.t01#5    mNeoVis.evergladensis.2.1.chr6.g00420.t01#2    mNeoVis.vulgivaga.1.1.chr6.g00422.t01#4   mNeoVis.3.1.1.chr6.g00416.t01#6       6185   125254517:   mNeoVis.3.1.1.chr6.g00353.t01#6    mNeoVis.vulgivaga.1.1.chr6.g00356.t01#4    mNeoVis.2.1.1.chr4_hap1.g00351.t01#5    mNeoVis.evergladensis.1.1.chr6.g00348.t01#3    mNeoVis.evergladensis.2.1.chr6.g00355.t01#2   mNeoVis.vulgivaga.2.1.chr6.g00348.t01#1       6186   125254518:   mNeoVis.evergladensis.1.1.chr13.g00755.t01#3    mNeoVis.vulgivaga.2.1.chr13.g00761.t01#1    mNeoVis.2.1.1.chr13_hap1.g00798.t01#5    mNeoVis.3.1.1.chr13.g00715.t01#6   mNeoVis.evergladensis.2.1.chr13.g00762.t01#2       6187   125254519:   mNeoVis.evergladensis.2.1.chr13.g00763.t01#2    mNeoVis.evergladensis.1.1.chr13.g00756.t01#3    mNeoVis.2.1.1.chr13_hap1.g00799.t01#5    mNeoVis.vulgivaga.1.1.chr13.g00754.t01#4    mNeoVis.3.1.1.chr13.g00716.t01#6   mNeoVis.vulgivaga.2.1.chr13.g00762.t01#1       6188   125254520:   mNeoVis.vulgivaga.1.1.chr13.g00755.t01#4    mNeoVis.2.1.1.chr13_hap1.g00800.t01#5    mNeoVis.3.1.1.chr13.g00717.t01#6    mNeoVis.evergladensis.2.1.chr13.g00764.t01#2    mNeoVis.evergladensis.1.1.chr13.g00757.t01#3   mNeoVis.vulgivaga.2.1.chr13.g00763.t01#1       6189   125254521:   mNeoVis.vulgivaga.1.1.chr7.g00055.t01#4    mNeoVis.evergladensis.1.1.chr7.g00056.t01#3    mNeoVis.2.1.1.chr7_hap1.g00052.t01#5    mNeoVis.3.1.1.chr7.g00042.t01#6    mNeoVis.evergladensis.2.1.chr7.g00053.t01#2   mNeoVis.vulgivaga.2.1.chr7.g00054.t01#1       6190   125254522:   mNeoVis.vulgivaga.2.1.chr13.g00764.t01#1    mNeoVis.3.1.1.chr13.g00718.t01#6    mNeoVis.evergladensis.2.1.chr13.g00765.t01#2    mNeoVis.vulgivaga.1.1.chr13.g00756.t01#4    mNeoVis.evergladensis.1.1.chr13.g00758.t01#3   mNeoVis.2.1.1.chr13_hap1.g00801.t01#5       6191   125254523:   mNeoVis.vulgivaga.2.1.chr6.g00307.t01#1    mNeoVis.3.1.1.chr6.g00314.t01#6   mNeoVis.2.1.1.chr4_hap1.g00311.t01#5           6192   125254524:   mNeoVis.evergladensis.1.1.chr13.g00759.t01#3    mNeoVis.vulgivaga.1.1.chr13.g00757.t01#4    mNeoVis.2.1.1.chr13_hap1.g00802.t01#5    mNeoVis.evergladensis.2.1.chr13.g00766.t01#2   mNeoVis.vulgivaga.2.1.chr13.g00765.t01#1       6193   125254525:   mNeoVis.vulgivaga.1.1.chr13.g00758.t01#4    mNeoVis.vulgivaga.2.1.chr13.g00766.t01#1    mNeoVis.evergladensis.2.1.chr13.g00767.t01#2    mNeoVis.evergladensis.1.1.chr13.g00760.t01#3   mNeoVis.2.1.1.chr13_hap1.g00803.t01#5       6194   125254526:   mNeoVis.evergladensis.1.1.chr6.g00311.t01#3    mNeoVis.vulgivaga.2.1.chr6.g00308.t01#1    mNeoVis.evergladensis.2.1.chr6.g00316.t01#2   mNeoVis.2.1.1.chr4_hap1.g00312.t01#5         6195   125254527:   mNeoVis.evergladensis.1.1.chr6.g00312.t01#3    mNeoVis.vulgivaga.2.1.chr6.g00309.t01#1    mNeoVis.3.1.1.chr6.g00316.t01#6    mNeoVis.vulgivaga.1.1.chr6.g00319.t01#4    mNeoVis.evergladensis.2.1.chr6.g00317.t01#2   mNeoVis.2.1.1.chr4_hap1.g00313.t01#5       6196   125254528:  mNeoVis.vulgivaga.2.1.chr6.g00310.t01#1               6197   125254529:   mNeoVis.3.1.1.chr6.g00318.t01#6    mNeoVis.2.1.1.chr4_hap1.g00314.t01#5    mNeoVis.vulgivaga.1.1.chr6.g00321.t01#4    mNeoVis.evergladensis.1.1.chr6.g00313.t01#3    mNeoVis.vulgivaga.2.1.chr6.g00311.t01#1   mNeoVis.evergladensis.2.1.chr6.g00318.t01#2       6198   125254530:   mNeoVis.vulgivaga.2.1.chr6.g00312.t01#1    mNeoVis.evergladensis.1.1.chr6.g00314.t01#3    mNeoVis.vulgivaga.1.1.chr6.g00322.t01#4    mNeoVis.evergladensis.2.1.chr6.g00319.t01#2   mNeoVis.2.1.1.chr4_hap1.g00315.t01#5       6199   125254531:   mNeoVis.2.1.1.chr4_hap1.g00317.t01#5    mNeoVis.3.1.1.chr6.g00320.t01#6    mNeoVis.vulgivaga.1.1.chr6.g00324.t01#4    mNeoVis.evergladensis.2.1.chr6.g00321.t01#2    mNeoVis.vulgivaga.2.1.chr6.g00313.t01#1   mNeoVis.evergladensis.1.1.chr6.g00316.t01#3       6200   125254532:   mNeoVis.vulgivaga.2.1.chr6.g00314.t01#1    mNeoVis.vulgivaga.1.1.chr6.g00323.t01#4    mNeoVis.2.1.1.chr4_hap1.g00316.t01#5    mNeoVis.evergladensis.1.1.chr6.g00315.t01#3   mNeoVis.evergladensis.2.1.chr6.g00320.t01#2       6201   125254533:   mNeoVis.vulgivaga.2.1.chr13.g00767.t01#1    mNeoVis.vulgivaga.1.1.chr13.g00759.t01#4    mNeoVis.evergladensis.2.1.chr13.g00768.t01#2    mNeoVis.evergladensis.1.1.chr13.g00762.t01#3   mNeoVis.2.1.1.chr13_hap1.g00804.t01#5       6202   125254534:   mNeoVis.3.1.1.chr13.g00721.t01#6   mNeoVis.evergladensis.1.1.chr13.g00761.t01#3           6203   125254535:   mNeoVis.vulgivaga.1.1.chr13.g00760.t01#4    mNeoVis.2.1.1.chr13_hap1.g00805.t01#5    mNeoVis.evergladensis.1.1.chr13.g00763.t01#3    mNeoVis.vulgivaga.2.1.chr13.g00768.t01#1   mNeoVis.evergladensis.2.1.chr13.g00769.t01#2       6204   125254536:   mNeoVis.vulgivaga.2.1.chr13.g00769.t01#1    mNeoVis.evergladensis.1.1.chr13.g00764.t01#3    mNeoVis.3.1.1.chr13.g00723.t01#6    mNeoVis.evergladensis.2.1.chr13.g00770.t01#2    mNeoVis.2.1.1.chr13_hap1.g00806.t01#5   mNeoVis.vulgivaga.1.1.chr13.g00761.t01#4       6205   125254537:   mNeoVis.vulgivaga.2.1.chr6.g00315.t01#1    mNeoVis.vulgivaga.1.1.chr6.g00325.t01#4    mNeoVis.2.1.1.chr4_hap1.g00318.t01#5    mNeoVis.3.1.1.chr6.g00322.t01#6    mNeoVis.evergladensis.1.1.chr6.g00317.t01#3    mNeoVis.3.1.1.chr6.g00321.t01#6   mNeoVis.evergladensis.2.1.chr6.g00322.t01#2     6206   125254538:   mNeoVis.2.1.1.chr11_hap1.g00160.t01#5    mNeoVis.vulgivaga.2.1.chr11.g00159.t01#1    mNeoVis.evergladensis.1.1.chr11.g00157.t01#3    mNeoVis.evergladensis.2.1.chr11.g00157.t01#2    mNeoVis.vulgivaga.1.1.chr11.g00154.t01#4   mNeoVis.3.1.1.chr11.g00132.t01#6       6207   125254539:   mNeoVis.evergladensis.2.1.chr13.g00771.t01#2    mNeoVis.evergladensis.1.1.chr13.g00765.t01#3    mNeoVis.3.1.1.chr13.g00724.t01#6    mNeoVis.2.1.1.chr13_hap1.g00807.t01#5    mNeoVis.vulgivaga.2.1.chr13.g00770.t01#1   mNeoVis.vulgivaga.1.1.chr13.g00762.t01#4       6208   125254540:   mNeoVis.2.1.1.chr13_hap1.g00808.t01#5    mNeoVis.vulgivaga.2.1.chr13.g00771.t01#1    mNeoVis.vulgivaga.1.1.chr13.g00763.t01#4   mNeoVis.evergladensis.1.1.chr13.g00766.t01#3       6209   125254541:   mNeoVis.2.1.1.chr4_hap1.g00319.t01#5    mNeoVis.3.1.1.chr6.g00323.t01#6    mNeoVis.evergladensis.1.1.chr6.g00318.t01#3    mNeoVis.vulgivaga.2.1.chr6.g00316.t01#1    mNeoVis.evergladensis.2.1.chr6.g00323.t01#2   mNeoVis.vulgivaga.1.1.chr6.g00326.t01#4       6210   125254542:   mNeoVis.evergladensis.2.1.chr6.g00324.t01#2    mNeoVis.vulgivaga.2.1.chr6.g00317.t01#1    mNeoVis.3.1.1.chr6.g00324.t01#6    mNeoVis.evergladensis.1.1.chr6.g00319.t01#3    mNeoVis.2.1.1.chr4_hap1.g00320.t01#5   mNeoVis.vulgivaga.1.1.chr6.g00327.t01#4       6211   125254543:   mNeoVis.evergladensis.1.1.chr13.g00767.t01#3    mNeoVis.vulgivaga.1.1.chr13.g00764.t01#4    mNeoVis.vulgivaga.2.1.chr13.g00772.t01#1    mNeoVis.evergladensis.2.1.chr13.g00774.t01#2    mNeoVis.2.1.1.chr13_hap1.g00809.t01#5   mNeoVis.3.1.1.chr13.g00728.t01#6       6212   125254544:   mNeoVis.3.1.1.chr13.g00729.t01#6    mNeoVis.evergladensis.1.1.chr13.g00768.t01#3    mNeoVis.evergladensis.2.1.chr13.g00775.t01#2    mNeoVis.vulgivaga.1.1.chr13.g00765.t01#4    mNeoVis.vulgivaga.2.1.chr13.g00773.t01#1   mNeoVis.2.1.1.chr13_hap1.g00810.t01#5       6213   125254545:   mNeoVis.vulgivaga.2.1.chr13.g00785.t01#1    mNeoVis.evergladensis.1.1.chr13.g00781.t01#3    mNeoVis.2.1.1.chr13_hap1.g00826.t01#5   mNeoVis.evergladensis.2.1.chr13.g00787.t01#2       6214   125254546:   mNeoVis.evergladensis.1.1.chr13.g00780.t01#3    mNeoVis.vulgivaga.1.1.chr8.g00649.t01#4    mNeoVis.2.1.1.chr13_hap1.g00825.t01#5    mNeoVis.vulgivaga.2.1.chr13.g00784.t01#1    mNeoVis.2.1.1.chr8_hap1.g00671.t01#5    mNeoVis.evergladensis.1.1.chr8.g00653.t01#3    mNeoVis.vulgivaga.1.1.chr13.g00777.t01#4    mNeoVis.evergladensis.2.1.chr5.g00290.t01#2    mNeoVis.vulgivaga.2.1.chr8.g00666.t01#1      6215   125254547:   mNeoVis.vulgivaga.1.1.chr14.g00689.t01#4    mNeoVis.evergladensis.1.1.chr14.g00695.t01#3    mNeoVis.evergladensis.2.1.chr14.g00698.t01#2    mNeoVis.2.1.1.chr14_hap1.g00698.t01#5    mNeoVis.3.1.1.chr14.g00667.t01#6   mNeoVis.vulgivaga.2.1.chr14.g00697.t01#1       6216   125254548:   mNeoVis.vulgivaga.2.1.chr6.g00349.t01#1    mNeoVis.vulgivaga.1.1.chr6.g00357.t01#4    mNeoVis.2.1.1.chr4_hap1.g00352.t01#5    mNeoVis.3.1.1.chr6.g00354.t01#6    mNeoVis.evergladensis.1.1.chr6.g00349.t01#3   mNeoVis.evergladensis.2.1.chr6.g00356.t01#2       6217   125254549:   mNeoVis.3.1.1.chr6.g00355.t01#6    mNeoVis.evergladensis.2.1.chr6.g00357.t01#2    mNeoVis.vulgivaga.1.1.chr6.g00358.t01#4    mNeoVis.evergladensis.1.1.chr6.g00350.t01#3    mNeoVis.vulgivaga.2.1.chr6.g00350.t01#1   mNeoVis.2.1.1.chr4_hap1.g00353.t01#5       6218   125254550:   mNeoVis.evergladensis.1.1.chr6.g00351.t01#3    mNeoVis.2.1.1.chr4_hap1.g00354.t01#5    mNeoVis.evergladensis.2.1.chr6.g00358.t01#2    mNeoVis.vulgivaga.2.1.chr6.g00351.t01#1    mNeoVis.3.1.1.chr6.g00356.t01#6   mNeoVis.vulgivaga.1.1.chr6.g00359.t01#4       6219   125254551:   mNeoVis.vulgivaga.2.1.chr13.g00786.t01#1    mNeoVis.evergladensis.2.1.chr13.g00789.t01#2    mNeoVis.2.1.1.chr13_hap1.g00827.t01#5    mNeoVis.3.1.1.chr13.g00747.t01#6    mNeoVis.vulgivaga.1.1.chr13.g00779.t01#4   mNeoVis.evergladensis.1.1.chr13.g00782.t01#3       6220   125254552:   mNeoVis.vulgivaga.2.1.chr13.g00787.t01#1    mNeoVis.2.1.1.chr13_hap1.g00828.t01#5    mNeoVis.evergladensis.1.1.chr13.g00784.t01#3    mNeoVis.vulgivaga.1.1.chr13.g00780.t01#4   mNeoVis.evergladensis.2.1.chr13.g00790.t01#2       6221   125254553:   mNeoVis.vulgivaga.2.1.chr6.g00352.t01#1    mNeoVis.vulgivaga.1.1.chr6.g00360.t01#4    mNeoVis.evergladensis.2.1.chr6.g00359.t01#2    mNeoVis.evergladensis.1.1.chr6.g00352.t01#3    mNeoVis.3.1.1.chr6.g00357.t01#6   mNeoVis.2.1.1.chr4_hap1.g00355.t01#5       6222   125254554:   mNeoVis.3.1.1.chr13.g00750.t01#6    mNeoVis.vulgivaga.2.1.chr13.g00789.t01#1    mNeoVis.2.1.1.chr13_hap1.g00830.t01#5    mNeoVis.evergladensis.1.1.chr13.g00786.t01#3    mNeoVis.evergladensis.2.1.chr13.g00792.t01#2   mNeoVis.vulgivaga.1.1.chr13.g00782.t01#4       6223   125254555:   mNeoVis.vulgivaga.2.1.chr6.g00353.t01#1    mNeoVis.2.1.1.chr4_hap1.g00356.t01#5    mNeoVis.vulgivaga.1.1.chr6.g00361.t01#4    mNeoVis.3.1.1.chr6.g00358.t01#6    mNeoVis.evergladensis.2.1.chr6.g00360.t01#2   mNeoVis.evergladensis.1.1.chr6.g00353.t01#3       6224   125254556:   mNeoVis.evergladensis.1.1.chr13.g00787.t01#3    mNeoVis.vulgivaga.1.1.chr13.g00783.t01#4    mNeoVis.2.1.1.chr13_hap1.g00831.t01#5    mNeoVis.evergladensis.2.1.chr13.g00793.t01#2    mNeoVis.3.1.1.chr13.g00751.t01#6   mNeoVis.vulgivaga.2.1.chr13.g00790.t01#1       6225   125254557:   mNeoVis.vulgivaga.2.1.chr13.g00791.t01#1    mNeoVis.evergladensis.1.1.chr13.g00788.t01#3    mNeoVis.3.1.1.chr13.g00752.t01#6    mNeoVis.vulgivaga.1.1.chr13.g00784.t01#4    mNeoVis.2.1.1.chr13_hap1.g00833.t01#5   mNeoVis.evergladensis.2.1.chr13.g00795.t01#2       6226   125254558:  mNeoVis.vulgivaga.2.1.chr6.g00260.t01#1               6227   125254559:   mNeoVis.vulgivaga.1.1.chr13.g00785.t01#4    mNeoVis.3.1.1.chr13.g00753.t01#6    mNeoVis.evergladensis.2.1.chr13.g00796.t01#2    mNeoVis.evergladensis.1.1.chr13.g00789.t01#3    mNeoVis.vulgivaga.2.1.chr13.g00792.t01#1   mNeoVis.2.1.1.chr13_hap1.g00834.t01#5       6228   125254560:   mNeoVis.vulgivaga.2.1.chr6.g01065.t01#1    mNeoVis.evergladensis.2.1.chr6.g01072.t01#2    mNeoVis.evergladensis.1.1.chr6.g01078.t01#3   mNeoVis.2.1.1.chr4_hap1.g01071.t01#5         6229   125254561:   mNeoVis.evergladensis.1.1.chr6.g00264.t01#3    mNeoVis.vulgivaga.2.1.chr6.g00261.t01#1    mNeoVis.evergladensis.2.1.chr6.g00263.t01#2    mNeoVis.2.1.1.chr4_hap1.g00263.t01#5    mNeoVis.3.1.1.chr6.g00265.t01#6   mNeoVis.vulgivaga.1.1.chr6.g00269.t01#4       6230   125254562:   mNeoVis.vulgivaga.2.1.chr15.g00040.t01#1    mNeoVis.vulgivaga.1.1.chr15.g00039.t01#4    mNeoVis.evergladensis.1.1.chr15.g00577.t01#3    mNeoVis.evergladensis.2.1.chr15.g00038.t01#2   mNeoVis.2.1.1.chr15_hap1.g00037.t01#5       6231   125254563:   mNeoVis.evergladensis.1.1.chr14.g00001.t01#3    mNeoVis.vulgivaga.1.1.chr14.g00001.t01#4    mNeoVis.evergladensis.2.1.chr14.g00001.t01#2    mNeoVis.3.1.1.chr14.g00001.t01#6    mNeoVis.2.1.1.chr14_hap1.g00001.t01#5   mNeoVis.vulgivaga.2.1.chr14.g00001.t01#1       6232   125254564:   mNeoVis.vulgivaga.1.1.chr6.g01067.t01#4    mNeoVis.evergladensis.1.1.chr6.g01079.t01#3    mNeoVis.vulgivaga.2.1.chr6.g01066.t01#1    mNeoVis.evergladensis.2.1.chr6.g01073.t01#2   mNeoVis.2.1.1.chr4_hap1.g01072.t01#5       6233   125254565:   mNeoVis.vulgivaga.2.1.chr14.g00002.t01#1    mNeoVis.3.1.1.chr14.g00002.t01#6    mNeoVis.vulgivaga.1.1.chr14.g00002.t01#4    mNeoVis.evergladensis.1.1.chr14.g00002.t01#3    mNeoVis.evergladensis.2.1.chr14.g00002.t01#2   mNeoVis.2.1.1.chr14_hap1.g00002.t01#5       6234   125254566:   mNeoVis.2.1.1.chr14_hap1.g00003.t01#5    mNeoVis.vulgivaga.2.1.chr14.g00003.t01#1    mNeoVis.vulgivaga.1.1.chr14.g00003.t01#4    mNeoVis.evergladensis.2.1.chr14.g00003.t01#2   mNeoVis.evergladensis.1.1.chr14.g00003.t01#3       6235   125254567:   mNeoVis.vulgivaga.1.1.chr6.g00271.t01#4    mNeoVis.3.1.1.chr6.g00263.t01#6    mNeoVis.evergladensis.2.1.chr6.g00266.t01#2    mNeoVis.evergladensis.1.1.chr6.g00266.t01#3    mNeoVis.evergladensis.2.1.chr6.g00265.t01#2    mNeoVis.2.1.1.chr4_hap1.g00265.t01#5   mNeoVis.vulgivaga.2.1.chr6.g00262.t01#1     6236   125254568:   mNeoVis.vulgivaga.2.1.chr6.g00263.t01#1    mNeoVis.evergladensis.1.1.chr6.g00267.t01#3    mNeoVis.evergladensis.2.1.chr6.g00267.t01#2    mNeoVis.3.1.1.chr6.g00264.t01#6    mNeoVis.2.1.1.chr4_hap1.g00266.t01#5   mNeoVis.vulgivaga.1.1.chr6.g00272.t01#4       6237   125254569:   mNeoVis.3.1.1.chr6.g00266.t01#6    mNeoVis.vulgivaga.2.1.chr6.g00264.t01#1    mNeoVis.vulgivaga.1.1.chr6.g00273.t01#4    mNeoVis.evergladensis.1.1.chr6.g00268.t01#3    mNeoVis.2.1.1.chr4_hap1.g00267.t01#5   mNeoVis.evergladensis.2.1.chr6.g00268.t01#2       6238   125254570:   mNeoVis.vulgivaga.1.1.chr14.g00004.t01#4    mNeoVis.2.1.1.chr14_hap1.g00004.t01#5    mNeoVis.evergladensis.1.1.chr14.g00004.t01#3    mNeoVis.evergladensis.2.1.chr14.g00004.t01#2    mNeoVis.vulgivaga.2.1.chr14.g00004.t01#1   mNeoVis.3.1.1.chr14.g00004.t01#6       6239   125254571:   mNeoVis.evergladensis.1.1.chr6.g00269.t01#3    mNeoVis.2.1.1.chr4_hap1.g00268.t01#5    mNeoVis.evergladensis.2.1.chr6.g00269.t01#2    mNeoVis.3.1.1.chr6.g00268.t01#6    mNeoVis.vulgivaga.1.1.chr6.g00274.t01#4   mNeoVis.vulgivaga.2.1.chr6.g00265.t01#1       6240   125254572:   mNeoVis.2.1.1.chr4_hap1.g01073.t01#5    mNeoVis.evergladensis.2.1.chr6.g01074.t01#2    mNeoVis.evergladensis.1.1.chr6.g01080.t01#3    mNeoVis.vulgivaga.2.1.chr6.g01067.t01#1   mNeoVis.vulgivaga.1.1.chr6.g01068.t01#4       6241   125254573:   mNeoVis.vulgivaga.2.1.chr6.g00266.t01#1    mNeoVis.evergladensis.1.1.chr6.g00270.t01#3    mNeoVis.evergladensis.2.1.chr6.g00270.t01#2    mNeoVis.2.1.1.chr4_hap1.g00269.t01#5    mNeoVis.3.1.1.chr6.g00269.t01#6   mNeoVis.vulgivaga.1.1.chr6.g00275.t01#4       6242   125254574:   mNeoVis.vulgivaga.1.1.chr14.g00006.t01#4    mNeoVis.2.1.1.chr14_hap1.g00006.t01#5    mNeoVis.evergladensis.1.1.chr14.g00006.t01#3    mNeoVis.evergladensis.2.1.chr14.g00006.t01#2    mNeoVis.vulgivaga.2.1.chr14.g00006.t01#1   mNeoVis.3.1.1.chr14.g00006.t01#6       6243   125254575:   mNeoVis.2.1.1.chr4_hap1.g00270.t01#5    mNeoVis.evergladensis.2.1.chr6.g00271.t01#2    mNeoVis.vulgivaga.2.1.chr6.g00267.t01#1   mNeoVis.evergladensis.1.1.chr6.g00271.t01#3       6244   125254576:   mNeoVis.2.1.1.chr4_hap1.g00271.t01#5    mNeoVis.3.1.1.chr6.g00270.t01#6    mNeoVis.evergladensis.2.1.chr6.g00272.t01#2    mNeoVis.evergladensis.1.1.chr6.g00272.t01#3    mNeoVis.vulgivaga.2.1.chr6.g00268.t01#1   mNeoVis.vulgivaga.1.1.chr6.g00276.t01#4       6245   125254577:   mNeoVis.3.1.1.chr6.g00271.t01#6    mNeoVis.vulgivaga.1.1.chr6.g00277.t01#4    mNeoVis.vulgivaga.2.1.chr6.g00269.t01#1    mNeoVis.2.1.1.chr4_hap1.g00272.t01#5    mNeoVis.evergladensis.1.1.chr6.g00273.t01#3   mNeoVis.evergladensis.2.1.chr6.g00273.t01#2       6246   125254578:   mNeoVis.vulgivaga.2.1.chr6.g00270.t01#1    mNeoVis.evergladensis.2.1.chr6.g00274.t01#2    mNeoVis.evergladensis.1.1.chr6.g00274.t01#3    mNeoVis.2.1.1.chr4_hap1.g00273.t01#5   mNeoVis.vulgivaga.1.1.chr6.g00278.t01#4       6247   125254579:   mNeoVis.3.1.1.chr14.g00007.t01#6    mNeoVis.2.1.1.chr14_hap1.g00008.t01#5    mNeoVis.evergladensis.2.1.chr14.g00007.t01#2    mNeoVis.evergladensis.1.1.chr14.g00007.t01#3    mNeoVis.vulgivaga.2.1.chr14.g00007.t01#1   mNeoVis.vulgivaga.1.1.chr14.g00007.t01#4       6248   125254580:   mNeoVis.vulgivaga.2.1.chr14.g00008.t01#1    mNeoVis.2.1.1.chr14_hap1.g00009.t01#5    mNeoVis.evergladensis.1.1.chr14.g00008.t01#3    mNeoVis.3.1.1.chr14.g00008.t01#6    mNeoVis.vulgivaga.1.1.chr14.g00008.t01#4   mNeoVis.evergladensis.2.1.chr14.g00008.t01#2       6249   125254581:   mNeoVis.vulgivaga.2.1.chr6.g01068.t01#1    mNeoVis.3.1.1.chr3.g01014.t01#6    mNeoVis.evergladensis.1.1.chr6.g01081.t01#3    mNeoVis.vulgivaga.1.1.chr6.g01069.t01#4    mNeoVis.evergladensis.2.1.chr6.g01075.t01#2   mNeoVis.2.1.1.chr4_hap1.g01074.t01#5       6250   125254582:   mNeoVis.evergladensis.2.1.chr15.g00039.t01#2    mNeoVis.2.1.1.chr15_hap1.g00038.t01#5    mNeoVis.vulgivaga.2.1.chr15.g00041.t01#1    mNeoVis.evergladensis.1.1.chr15.g00575.t01#3    mNeoVis.vulgivaga.1.1.chr15.g00040.t01#4   mNeoVis.3.1.1.chr15.g00035.t01#6       6251   125254583:   mNeoVis.vulgivaga.2.1.chr14.g00009.t01#1    mNeoVis.evergladensis.2.1.chr14.g00009.t01#2    mNeoVis.2.1.1.chr14_hap1.g00007.t01#5    mNeoVis.evergladensis.1.1.chr14.g00009.t01#3   mNeoVis.vulgivaga.1.1.chr14.g00009.t01#4       6252   125254584:   mNeoVis.vulgivaga.2.1.chr13.g00715.t01#1    mNeoVis.2.1.1.chr13_hap1.g00752.t01#5    mNeoVis.3.1.1.chr13.g00670.t01#6    mNeoVis.vulgivaga.1.1.chr13.g00708.t01#4    mNeoVis.evergladensis.1.1.chr13.g00710.t01#3   mNeoVis.evergladensis.2.1.chr13.g00717.t01#2       6253   125254585:   mNeoVis.vulgivaga.2.1.chr6.g00271.t01#1    mNeoVis.evergladensis.2.1.chr6.g00275.t01#2    mNeoVis.3.1.1.chr6.g00274.t01#6    mNeoVis.evergladensis.1.1.chr6.g00275.t01#3    mNeoVis.vulgivaga.1.1.chr6.g00279.t01#4   mNeoVis.2.1.1.chr4_hap1.g00274.t01#5       6254   125254586:   mNeoVis.vulgivaga.1.1.chr13.g00710.t01#4    mNeoVis.2.1.1.chr13_hap1.g00754.t01#5    mNeoVis.evergladensis.2.1.chr13.g00718.t01#2    mNeoVis.vulgivaga.2.1.chr13.g00716.t01#1    mNeoVis.3.1.1.chr13.g00671.t01#6   mNeoVis.evergladensis.1.1.chr13.g00711.t01#3       6255   125254587:   mNeoVis.evergladensis.1.1.chr6.g00276.t01#3    mNeoVis.3.1.1.chr6.g00275.t01#6    mNeoVis.2.1.1.chr4_hap1.g00275.t01#5    mNeoVis.vulgivaga.1.1.chr6.g00280.t01#4    mNeoVis.evergladensis.2.1.chr6.g00276.t01#2   mNeoVis.vulgivaga.2.1.chr6.g00272.t01#1       6256   125254588:   mNeoVis.vulgivaga.1.1.chr6.g00281.t01#4    mNeoVis.evergladensis.2.1.chr6.g00277.t01#2    mNeoVis.3.1.1.chr6.g00276.t01#6    mNeoVis.2.1.1.chr4_hap1.g00276.t01#5    mNeoVis.evergladensis.1.1.chr6.g00277.t01#3   mNeoVis.vulgivaga.2.1.chr6.g00273.t01#1       6257   125254589:   mNeoVis.evergladensis.1.1.chr6.g00278.t01#3    mNeoVis.2.1.1.chr4_hap1.g00277.t01#5    mNeoVis.evergladensis.2.1.chr6.g00278.t01#2   mNeoVis.vulgivaga.2.1.chr6.g00274.t01#1         6258   125254590:   mNeoVis.2.1.1.chr4_hap1.g00278.t01#5    mNeoVis.vulgivaga.2.1.chr6.g00275.t01#1    mNeoVis.vulgivaga.1.1.chr6.g00283.t01#4    mNeoVis.evergladensis.1.1.chr6.g00279.t01#3    mNeoVis.evergladensis.2.1.chr6.g00279.t01#2   mNeoVis.3.1.1.chr6.g00277.t01#6       6259   125254591:   mNeoVis.evergladensis.1.1.chr13.g00712.t01#3    mNeoVis.2.1.1.chr13_hap1.g00755.t01#5    mNeoVis.evergladensis.2.1.chr13.g00719.t01#2    mNeoVis.vulgivaga.2.1.chr13.g00717.t01#1    mNeoVis.3.1.1.chr13.g00672.t01#6   mNeoVis.vulgivaga.1.1.chr13.g00711.t01#4       6260   125254592:   mNeoVis.vulgivaga.2.1.chr13.g00718.t01#1    mNeoVis.3.1.1.chr13.g00673.t01#6    mNeoVis.vulgivaga.1.1.chr13.g00712.t01#4    mNeoVis.2.1.1.chr13_hap1.g00756.t01#5    mNeoVis.evergladensis.1.1.chr13.g00713.t01#3   mNeoVis.evergladensis.2.1.chr13.g00720.t01#2       6261   125254593:   mNeoVis.2.1.1.chr4_hap1.g00279.t01#5    mNeoVis.vulgivaga.1.1.chr6.g00284.t01#4    mNeoVis.vulgivaga.2.1.chr6.g00276.t01#1    mNeoVis.evergladensis.1.1.chr6.g00280.t01#3    mNeoVis.evergladensis.2.1.chr6.g00280.t01#2   mNeoVis.3.1.1.chr6.g00278.t01#6       6262   125254594:   mNeoVis.evergladensis.1.1.chr6.g01082.t01#3    mNeoVis.evergladensis.2.1.chr6.g01076.t01#2    mNeoVis.2.1.1.chr4_hap1.g01075.t01#5    mNeoVis.vulgivaga.1.1.chr6.g01070.t01#4   mNeoVis.vulgivaga.2.1.chr6.g01069.t01#1       6263   125254595:   mNeoVis.3.1.1.chr6.g00279.t01#6   mNeoVis.vulgivaga.2.1.chr6.g00277.t01#1             6264   125254596:   mNeoVis.vulgivaga.2.1.chr15.g00042.t01#1    mNeoVis.3.1.1.chr15.g00036.t01#6    mNeoVis.evergladensis.2.1.chr15.g00040.t01#2    mNeoVis.evergladensis.1.1.chr15.g00574.t01#3   mNeoVis.2.1.1.chr15_hap1.g00039.t01#5       6265   125254597:  mNeoVis.vulgivaga.2.1.chr13.g00719.t01#1               6266   125254598:   mNeoVis.3.1.1.chr6.g00280.t01#6    mNeoVis.evergladensis.2.1.chr6.g00284.t01#2    mNeoVis.vulgivaga.1.1.chr6.g00286.t01#4    mNeoVis.2.1.1.chr4_hap1.g00282.t01#5   mNeoVis.vulgivaga.2.1.chr6.g00278.t01#1       6267   125254599:   mNeoVis.3.1.1.chr13.g00674.t01#6    mNeoVis.vulgivaga.2.1.chr13.g00720.t01#1    mNeoVis.2.1.1.chr13_hap1.g00758.t01#5    mNeoVis.evergladensis.2.1.chr13.g00722.t01#2    mNeoVis.vulgivaga.1.1.chr13.g00715.t01#4   mNeoVis.evergladensis.1.1.chr13.g00714.t01#3       6268   125254600:   mNeoVis.vulgivaga.2.1.chr13.g00721.t01#1    mNeoVis.evergladensis.2.1.chr13.g00723.t01#2    mNeoVis.3.1.1.chr13.g00675.t01#6    mNeoVis.evergladensis.1.1.chr13.g00715.t01#3    mNeoVis.vulgivaga.1.1.chr13.g00716.t01#4   mNeoVis.2.1.1.chr13_hap1.g00759.t01#5       6269   125254601:   mNeoVis.evergladensis.1.1.chr6.g00284.t01#3    mNeoVis.vulgivaga.1.1.chr6.g00287.t01#4    mNeoVis.vulgivaga.2.1.chr6.g00279.t01#1    mNeoVis.evergladensis.2.1.chr6.g00286.t01#2    mNeoVis.3.1.1.chr6.g00282.t01#6   mNeoVis.2.1.1.chr4_hap1.g00283.t01#5       6270   125254602:   mNeoVis.3.1.1.chr13.g00677.t01#6    mNeoVis.2.1.1.chr13_hap1.g00760.t01#5    mNeoVis.evergladensis.1.1.chr13.g00716.t01#3    mNeoVis.vulgivaga.2.1.chr13.g00722.t01#1    mNeoVis.evergladensis.2.1.chr13.g00724.t01#2   mNeoVis.vulgivaga.1.1.chr13.g00717.t01#4       6271   125254603:   mNeoVis.evergladensis.2.1.chr13.g00725.t01#2    mNeoVis.vulgivaga.2.1.chr13.g00723.t01#1    mNeoVis.vulgivaga.1.1.chr13.g00718.t01#4    mNeoVis.2.1.1.chr13_hap1.g00761.t01#5    mNeoVis.3.1.1.chr13.g00678.t01#6   mNeoVis.evergladensis.1.1.chr13.g00717.t01#3       6272   125254604:   mNeoVis.vulgivaga.2.1.chr13.g00724.t01#1    mNeoVis.vulgivaga.1.1.chr13.g00719.t01#4    mNeoVis.evergladensis.1.1.chr13.g00718.t01#3    mNeoVis.evergladensis.2.1.chr13.g00726.t01#2    mNeoVis.2.1.1.chr13_hap1.g00762.t01#5   mNeoVis.3.1.1.chr13.g00679.t01#6       6273   125254605:   mNeoVis.vulgivaga.2.1.chr15.g00043.t01#1    mNeoVis.evergladensis.2.1.chr15.g00041.t01#2    mNeoVis.vulgivaga.1.1.chr15.g00041.t01#4    mNeoVis.evergladensis.1.1.chr15.g00573.t01#3    mNeoVis.2.1.1.chr15_hap1.g00040.t01#5   mNeoVis.3.1.1.chr15.g00037.t01#6       6274   125254606:   mNeoVis.2.1.1.chr13_hap1.g00763.t01#5    mNeoVis.evergladensis.2.1.chr13.g00727.t01#2    mNeoVis.vulgivaga.1.1.chr13.g00720.t01#4    mNeoVis.vulgivaga.2.1.chr13.g00725.t01#1   mNeoVis.evergladensis.1.1.chr13.g00719.t01#3       6275   125254607:   mNeoVis.3.1.1.chr6.g00283.t01#6    mNeoVis.2.1.1.chr4_hap1.g00284.t01#5    mNeoVis.evergladensis.2.1.chr6.g00287.t01#2    mNeoVis.vulgivaga.2.1.chr6.g00280.t01#1    mNeoVis.evergladensis.1.1.chr6.g00285.t01#3   mNeoVis.vulgivaga.1.1.chr6.g00288.t01#4       6276   125254608:   mNeoVis.3.1.1.chr13.g00680.t01#6    mNeoVis.evergladensis.1.1.chr13.g00720.t01#3    mNeoVis.evergladensis.2.1.chr13.g00728.t01#2    mNeoVis.vulgivaga.2.1.chr13.g00726.t01#1    mNeoVis.2.1.1.chr13_hap1.g00764.t01#5   mNeoVis.vulgivaga.1.1.chr13.g00721.t01#4       6277   125254609:   mNeoVis.2.1.1.chr4_hap1.g00285.t01#5    mNeoVis.vulgivaga.2.1.chr6.g00281.t01#1    mNeoVis.vulgivaga.1.1.chr6.g00289.t01#4    mNeoVis.3.1.1.chr6.g00284.t01#6    mNeoVis.evergladensis.1.1.chr6.g00286.t01#3   mNeoVis.evergladensis.2.1.chr6.g00288.t01#2       6278   125254610:   mNeoVis.3.1.1.chr13.g00681.t01#6    mNeoVis.evergladensis.1.1.chr13.g00721.t01#3    mNeoVis.vulgivaga.2.1.chr13.g00727.t01#1    mNeoVis.vulgivaga.1.1.chr13.g00722.t01#4    mNeoVis.2.1.1.chr13_hap1.g00765.t01#5   mNeoVis.evergladensis.2.1.chr13.g00730.t01#2       6279   125254611:   mNeoVis.3.1.1.chr6.g00286.t01#6    mNeoVis.2.1.1.chr4_hap1.g00287.t01#5    mNeoVis.evergladensis.1.1.chr6.g00288.t01#3    mNeoVis.vulgivaga.1.1.chr6.g00291.t01#4    mNeoVis.vulgivaga.2.1.chr6.g00282.t01#1   mNeoVis.evergladensis.2.1.chr6.g00290.t01#2       6280   125254612:   mNeoVis.vulgivaga.1.1.chr6.g00293.t01#4    mNeoVis.evergladensis.1.1.chr6.g00290.t01#3    mNeoVis.vulgivaga.2.1.chr6.g00284.t01#1    mNeoVis.evergladensis.2.1.chr6.g00292.t01#2    mNeoVis.3.1.1.chr6.g00289.t01#6   mNeoVis.2.1.1.chr4_hap1.g00289.t01#5       6281   125254613:   mNeoVis.vulgivaga.2.1.chr5.g00881.t01#1    mNeoVis.evergladensis.1.1.chr5.g00883.t01#3    mNeoVis.3.1.1.chr5.g00870.t01#6    mNeoVis.2.1.1.chr4_hap1.g00288.t01#5    mNeoVis.3.1.1.chr6.g00287.t01#6    mNeoVis.vulgivaga.1.1.chr5.g00863.t01#4    mNeoVis.evergladensis.1.1.chr6.g00289.t01#3    mNeoVis.2.1.1.chr2_hap1.g00938.t01#5    mNeoVis.evergladensis.2.1.chr6.g00291.t01#2      6282   125254614:   mNeoVis.vulgivaga.1.1.chr6.g00292.t01#4   mNeoVis.3.1.1.chr6.g00288.t01#6             6283   125254615:  mNeoVis.vulgivaga.2.1.chr6.g00283.t01#1               6284   125254616:   mNeoVis.evergladensis.1.1.chr15.g00571.t01#3    mNeoVis.2.1.1.chr15_hap1.g00042.t01#5    mNeoVis.vulgivaga.1.1.chr15.g00043.t01#4    mNeoVis.3.1.1.chr15.g00039.t01#6    mNeoVis.vulgivaga.2.1.chr15.g00044.t01#1   mNeoVis.evergladensis.2.1.chr15.g00043.t01#2       6285   125254617:   mNeoVis.evergladensis.1.1.chr6.g01083.t01#3    mNeoVis.2.1.1.chr4_hap1.g01076.t01#5    mNeoVis.evergladensis.2.1.chr6.g01077.t01#2    mNeoVis.vulgivaga.1.1.chr6.g01071.t01#4   mNeoVis.vulgivaga.2.1.chr6.g01070.t01#1       6286   125254618:   mNeoVis.evergladensis.1.1.chr13.g00722.t01#3    mNeoVis.evergladensis.2.1.chr13.g00731.t01#2    mNeoVis.2.1.1.chr13_hap1.g00766.t01#5    mNeoVis.vulgivaga.1.1.chr13.g00723.t01#4    mNeoVis.3.1.1.chr13.g00682.t01#6   mNeoVis.vulgivaga.2.1.chr13.g00728.t01#1       6287   125254619:   mNeoVis.3.1.1.chr6.g00292.t01#6    mNeoVis.evergladensis.2.1.chr6.g00293.t01#2    mNeoVis.vulgivaga.2.1.chr6.g00285.t01#1    mNeoVis.evergladensis.1.1.chr6.g00291.t01#3    mNeoVis.2.1.1.chr4_hap1.g00290.t01#5   mNeoVis.vulgivaga.1.1.chr6.g00295.t01#4       6288   125254620:   mNeoVis.evergladensis.1.1.chr6.g00292.t01#3    mNeoVis.vulgivaga.1.1.chr6.g00296.t01#4    mNeoVis.2.1.1.chr4_hap1.g00291.t01#5    mNeoVis.vulgivaga.2.1.chr6.g00286.t01#1    mNeoVis.3.1.1.chr6.g00291.t01#6   mNeoVis.evergladensis.2.1.chr6.g00295.t01#2       6289   125254621:   mNeoVis.vulgivaga.2.1.chr6.g00287.t01#1    mNeoVis.evergladensis.2.1.chr6.g00296.t01#2    mNeoVis.evergladensis.1.1.chr6.g00293.t01#3    mNeoVis.3.1.1.chr6.g00293.t01#6    mNeoVis.2.1.1.chr4_hap1.g00292.t01#5   mNeoVis.vulgivaga.1.1.chr6.g00297.t01#4       6290   125254622:   mNeoVis.2.1.1.chr13_hap1.g00767.t01#5    mNeoVis.evergladensis.2.1.chr13.g00732.t01#2    mNeoVis.vulgivaga.2.1.chr13.g00729.t01#1    mNeoVis.3.1.1.chr13.g00683.t01#6    mNeoVis.vulgivaga.1.1.chr13.g00724.t01#4   mNeoVis.evergladensis.1.1.chr13.g00723.t01#3       6291   125254623:   mNeoVis.vulgivaga.2.1.chr6.g00288.t01#1    mNeoVis.evergladensis.2.1.chr6.g00297.t01#2    mNeoVis.evergladensis.1.1.chr6.g00294.t01#3    mNeoVis.2.1.1.chr4_hap1.g00293.t01#5    mNeoVis.vulgivaga.1.1.chr6.g00298.t01#4   mNeoVis.3.1.1.chr6.g00294.t01#6       6292   125254624:   mNeoVis.2.1.1.chr13_hap1.g00768.t01#5    mNeoVis.vulgivaga.1.1.chr13.g00725.t01#4    mNeoVis.evergladensis.2.1.chr13.g00733.t01#2    mNeoVis.vulgivaga.2.1.chr13.g00730.t01#1    mNeoVis.3.1.1.chr13.g00684.t01#6   mNeoVis.evergladensis.1.1.chr13.g00724.t01#3       6293   125254625:   mNeoVis.vulgivaga.2.1.chr6.g00289.t01#1    mNeoVis.evergladensis.1.1.chr6.g00296.t01#3    mNeoVis.2.1.1.chr4_hap1.g00295.t01#5    mNeoVis.3.1.1.chr6.g00295.t01#6    mNeoVis.vulgivaga.1.1.chr6.g00300.t01#4   mNeoVis.evergladensis.2.1.chr6.g00298.t01#2       6294   125254626:   mNeoVis.evergladensis.2.1.chr13.g00734.t01#2    mNeoVis.evergladensis.1.1.chr13.g00725.t01#3    mNeoVis.2.1.1.chr13_hap1.g00769.t01#5    mNeoVis.vulgivaga.1.1.chr13.g00726.t01#4    mNeoVis.vulgivaga.2.1.chr13.g00731.t01#1   mNeoVis.3.1.1.chr13.g00685.t01#6       6295   125254627:   mNeoVis.3.1.1.chr13.g00686.t01#6    mNeoVis.vulgivaga.1.1.chr13.g00727.t01#4    mNeoVis.vulgivaga.2.1.chr13.g00732.t01#1    mNeoVis.evergladensis.2.1.chr13.g00736.t01#2    mNeoVis.evergladensis.1.1.chr13.g00726.t01#3   mNeoVis.2.1.1.chr13_hap1.g00770.t01#5       6296   125254628:   mNeoVis.2.1.1.chr4_hap1.g00296.t01#5    mNeoVis.vulgivaga.2.1.chr6.g00290.t01#1    mNeoVis.vulgivaga.1.1.chr6.g00301.t01#4    mNeoVis.evergladensis.2.1.chr6.g00299.t01#2    mNeoVis.3.1.1.chr6.g00296.t01#6   mNeoVis.evergladensis.1.1.chr6.g00297.t01#3       6297   125254629:   mNeoVis.evergladensis.1.1.chr6.g00295.t01#3    mNeoVis.evergladensis.2.1.chr6.g00300.t01#2    mNeoVis.3.1.1.chr6.g00297.t01#6    mNeoVis.vulgivaga.1.1.chr6.g00299.t01#4    mNeoVis.2.1.1.chr4_hap1.g00294.t01#5   mNeoVis.vulgivaga.2.1.chr6.g00291.t01#1       6298   125254630:   mNeoVis.vulgivaga.2.1.chr6.g00292.t01#1    mNeoVis.evergladensis.1.1.chr6.g00298.t01#3    mNeoVis.vulgivaga.1.1.chr6.g00302.t01#4    mNeoVis.evergladensis.2.1.chr6.g00301.t01#2    mNeoVis.3.1.1.chr6.g00299.t01#6   mNeoVis.2.1.1.chr4_hap1.g00297.t01#5       6299   125254631:   mNeoVis.vulgivaga.1.1.chr6.g00304.t01#4    mNeoVis.evergladensis.1.1.chr6.g00299.t01#3    mNeoVis.vulgivaga.2.1.chr6.g00294.t01#1    mNeoVis.3.1.1.chr6.g00300.t01#6    mNeoVis.evergladensis.2.1.chr6.g00302.t01#2    mNeoVis.2.1.1.chr4_hap1.g00298.t01#5    mNeoVis.vulgivaga.2.1.chr6.g00293.t01#1   mNeoVis.vulgivaga.1.1.chr6.g00303.t01#4     6300   125254632:   mNeoVis.evergladensis.2.1.chr13.g00737.t01#2    mNeoVis.3.1.1.chr13.g00688.t01#6    mNeoVis.evergladensis.1.1.chr13.g00727.t01#3    mNeoVis.2.1.1.chr13_hap1.g00771.t01#5    mNeoVis.vulgivaga.2.1.chr13.g00733.t01#1   mNeoVis.vulgivaga.1.1.chr13.g00728.t01#4       6301   125254633:   mNeoVis.vulgivaga.1.1.chr6.g01072.t01#4    mNeoVis.evergladensis.1.1.chr6.g01084.t01#3    mNeoVis.3.1.1.chr6.g01076.t01#6    mNeoVis.vulgivaga.2.1.chr6.g01071.t01#1    mNeoVis.evergladensis.2.1.chr6.g01078.t01#2   mNeoVis.2.1.1.chr4_hap1.g01077.t01#5       6302   125254634:   mNeoVis.vulgivaga.2.1.chr6.g00295.t01#1   mNeoVis.vulgivaga.1.1.chr6.g00305.t01#4             6303   125254635:   mNeoVis.evergladensis.1.1.chr6.g00300.t01#3    mNeoVis.vulgivaga.2.1.chr6.g00296.t01#1    mNeoVis.evergladensis.2.1.chr6.g00303.t01#2   mNeoVis.2.1.1.chr4_hap1.g00299.t01#5         6304   125254636:   mNeoVis.3.1.1.chr6.g00301.t01#6   mNeoVis.vulgivaga.1.1.chr6.g00306.t01#4             6305   125254637:   mNeoVis.vulgivaga.1.1.chr6.g00308.t01#4    mNeoVis.vulgivaga.2.1.chr6.g00298.t01#1    mNeoVis.evergladensis.2.1.chr6.g00305.t01#2    mNeoVis.3.1.1.chr6.g00303.t01#6    mNeoVis.evergladensis.1.1.chr6.g00302.t01#3   mNeoVis.2.1.1.chr4_hap1.g00301.t01#5       6306   125254638:   mNeoVis.2.1.1.chr4_hap1.g00302.t01#5    mNeoVis.evergladensis.2.1.chr6.g00306.t01#2    mNeoVis.evergladensis.1.1.chr6.g00303.t01#3    mNeoVis.3.1.1.chr6.g00304.t01#6    mNeoVis.vulgivaga.2.1.chr6.g00299.t01#1   mNeoVis.vulgivaga.1.1.chr6.g00309.t01#4       6307   125254639:   mNeoVis.3.1.1.chr6.g00305.t01#6    mNeoVis.evergladensis.2.1.chr6.g00307.t01#2    mNeoVis.3.1.1.chr6.g00307.t01#6    mNeoVis.vulgivaga.1.1.chr6.g00310.t01#4    mNeoVis.2.1.1.chr4_hap1.g00303.t01#5    mNeoVis.vulgivaga.2.1.chr6.g00300.t01#1   mNeoVis.evergladensis.1.1.chr6.g00304.t01#3     6308   125254640:   mNeoVis.vulgivaga.1.1.chr13.g00729.t01#4    mNeoVis.2.1.1.chr13_hap1.g00772.t01#5    mNeoVis.evergladensis.1.1.chr13.g00728.t01#3    mNeoVis.vulgivaga.2.1.chr13.g00734.t01#1    mNeoVis.evergladensis.2.1.chr13.g00738.t01#2   mNeoVis.3.1.1.chr13.g00689.t01#6       6309   125254641:   mNeoVis.evergladensis.2.1.chr13.g00739.t01#2    mNeoVis.evergladensis.1.1.chr13.g00729.t01#3    mNeoVis.3.1.1.chr13.g00690.t01#6    mNeoVis.vulgivaga.1.1.chr13.g00730.t01#4    mNeoVis.2.1.1.chr13_hap1.g00773.t01#5   mNeoVis.vulgivaga.2.1.chr13.g00735.t01#1       6310   125254642:   mNeoVis.vulgivaga.2.1.chr6.g00301.t01#1    mNeoVis.vulgivaga.1.1.chr6.g00313.t01#4    mNeoVis.2.1.1.chr4_hap1.g00306.t01#5    mNeoVis.evergladensis.1.1.chr6.g00305.t01#3   mNeoVis.evergladensis.2.1.chr6.g00311.t01#2       6311   125254643:   mNeoVis.vulgivaga.2.1.chr6.g00302.t01#1    mNeoVis.evergladensis.1.1.chr6.g00306.t01#3    mNeoVis.2.1.1.chr4_hap1.g00305.t01#5    mNeoVis.vulgivaga.1.1.chr6.g00311.t01#4    mNeoVis.3.1.1.chr6.g00308.t01#6   mNeoVis.evergladensis.2.1.chr6.g00308.t01#2       6312   125254644:   mNeoVis.evergladensis.2.1.chr13.g00740.t01#2    mNeoVis.2.1.1.chr13_hap1.g00774.t01#5    mNeoVis.vulgivaga.1.1.chr13.g00731.t01#4    mNeoVis.vulgivaga.2.1.chr13.g00736.t01#1    mNeoVis.evergladensis.1.1.chr13.g00730.t01#3   mNeoVis.3.1.1.chr13.g00691.t01#6       6313   125254645:   mNeoVis.evergladensis.1.1.chr13.g00732.t01#3   mNeoVis.3.1.1.chr13.g00692.t01#6             6314   125254646:   mNeoVis.vulgivaga.2.1.chr6.g00303.t01#1    mNeoVis.evergladensis.2.1.chr6.g00312.t01#2    mNeoVis.evergladensis.1.1.chr6.g00307.t01#3    mNeoVis.2.1.1.chr4_hap1.g00307.t01#5    mNeoVis.vulgivaga.1.1.chr6.g00314.t01#4   mNeoVis.3.1.1.chr6.g00309.t01#6       6315   125254647:   mNeoVis.vulgivaga.2.1.chr6.g00304.t01#1    mNeoVis.2.1.1.chr4_hap1.g00308.t01#5    mNeoVis.3.1.1.chr6.g00310.t01#6    mNeoVis.evergladensis.2.1.chr6.g00313.t01#2    mNeoVis.evergladensis.1.1.chr6.g00308.t01#3   mNeoVis.vulgivaga.1.1.chr6.g00315.t01#4       6316   125254648:   mNeoVis.vulgivaga.2.1.chr6.g00305.t01#1    mNeoVis.evergladensis.2.1.chr6.g00314.t01#2    mNeoVis.2.1.1.chr4_hap1.g00309.t01#5    mNeoVis.3.1.1.chr6.g00311.t01#6    mNeoVis.evergladensis.1.1.chr6.g00309.t01#3   mNeoVis.vulgivaga.1.1.chr6.g00316.t01#4       6317   125254649:   mNeoVis.3.1.1.chr13.g00693.t01#6    mNeoVis.2.1.1.chr13_hap1.g00775.t01#5    mNeoVis.evergladensis.1.1.chr13.g00733.t01#3    mNeoVis.vulgivaga.2.1.chr13.g00737.t01#1    mNeoVis.evergladensis.2.1.chr13.g00741.t01#2   mNeoVis.vulgivaga.1.1.chr13.g00732.t01#4       6318   125254650:   mNeoVis.3.1.1.chr6.g00312.t01#6    mNeoVis.2.1.1.chr4_hap1.g00310.t01#5    mNeoVis.evergladensis.2.1.chr6.g00315.t01#2    mNeoVis.vulgivaga.2.1.chr6.g00306.t01#1    mNeoVis.evergladensis.1.1.chr6.g00310.t01#3   mNeoVis.vulgivaga.1.1.chr6.g00317.t01#4       6319   125254651:   mNeoVis.3.1.1.chr13.g00694.t01#6    mNeoVis.vulgivaga.1.1.chr13.g00733.t01#4    mNeoVis.evergladensis.2.1.chr13.g00742.t01#2    mNeoVis.vulgivaga.2.1.chr13.g00738.t01#1    mNeoVis.2.1.1.chr13_hap1.g00776.t01#5   mNeoVis.evergladensis.1.1.chr13.g00734.t01#3       6320   125254652:   mNeoVis.3.1.1.chr13.g00695.t01#6    mNeoVis.evergladensis.1.1.chr13.g00735.t01#3    mNeoVis.2.1.1.chr13_hap1.g00777.t01#5    mNeoVis.vulgivaga.2.1.chr13.g00739.t01#1    mNeoVis.evergladensis.2.1.chr13.g00743.t01#2   mNeoVis.vulgivaga.1.1.chr13.g00735.t01#4       6321   125254653:   mNeoVis.evergladensis.2.1.chr6.g00213.t01#2    mNeoVis.vulgivaga.1.1.chr6.g00221.t01#4    mNeoVis.evergladensis.1.1.chr6.g00215.t01#3    mNeoVis.3.1.1.chr6.g00214.t01#6    mNeoVis.2.1.1.chr4_hap1.g00214.t01#5   mNeoVis.vulgivaga.2.1.chr6.g00214.t01#1       6322   125254654:   mNeoVis.3.1.1.chr6.g00215.t01#6    mNeoVis.vulgivaga.1.1.chr6.g00222.t01#4    mNeoVis.evergladensis.1.1.chr6.g00216.t01#3    mNeoVis.evergladensis.2.1.chr6.g00214.t01#2    mNeoVis.vulgivaga.2.1.chr6.g00215.t01#1   mNeoVis.2.1.1.chr4_hap1.g00215.t01#5       6323   125254655:   mNeoVis.vulgivaga.2.1.chr13.g00740.t01#1    mNeoVis.evergladensis.2.1.chr13.g00744.t01#2    mNeoVis.3.1.1.chr13.g00696.t01#6    mNeoVis.vulgivaga.1.1.chr13.g00734.t01#4    mNeoVis.2.1.1.chr13_hap1.g00778.t01#5   mNeoVis.evergladensis.1.1.chr13.g00736.t01#3       6324   125254656:   mNeoVis.3.1.1.chr6.g00217.t01#6    mNeoVis.vulgivaga.1.1.chr6.g00224.t01#4    mNeoVis.evergladensis.2.1.chr6.g00216.t01#2    mNeoVis.evergladensis.1.1.chr6.g00218.t01#3    mNeoVis.vulgivaga.2.1.chr6.g00217.t01#1   mNeoVis.2.1.1.chr4_hap1.g00217.t01#5       6325   125254657:   mNeoVis.evergladensis.2.1.chr3.g00831.t01#2    mNeoVis.evergladensis.1.1.chr3.g00836.t01#3    mNeoVis.vulgivaga.1.1.chr3.g00832.t01#4    mNeoVis.2.1.1.chr12_hap1.g00580.t01#5    mNeoVis.3.1.1.chr7.g00703.t01#6    mNeoVis.vulgivaga.2.1.chr12.g00533.t01#1    mNeoVis.evergladensis.2.1.chr15.g00385.t01#2    mNeoVis.evergladensis.2.1.chr12.g00518.t01#2    mNeoVis.2.1.1.chr3_hap1.g00836.t01#5      6326   125254658:   mNeoVis.vulgivaga.2.1.chr6.g00216.t01#1    mNeoVis.evergladensis.2.1.chr6.g00215.t01#2    mNeoVis.3.1.1.chr3.g00816.t01#6    mNeoVis.2.1.1.chr4_hap1.g00216.t01#5    mNeoVis.vulgivaga.2.1.chr3.g00843.t01#1    mNeoVis.vulgivaga.1.1.chr6.g00223.t01#4    mNeoVis.evergladensis.1.1.chr6.g00217.t01#3   mNeoVis.3.1.1.chr6.g00216.t01#6     6327   125254659:   mNeoVis.evergladensis.2.1.chr13.g00745.t01#2    mNeoVis.evergladensis.1.1.chr13.g00737.t01#3    mNeoVis.2.1.1.chr13_hap1.g00779.t01#5    mNeoVis.3.1.1.chr13.g00697.t01#6    mNeoVis.vulgivaga.2.1.chr13.g00741.t01#1   mNeoVis.vulgivaga.1.1.chr13.g00736.t01#4       6328   125254660:   mNeoVis.evergladensis.2.1.chr13.g00746.t01#2    mNeoVis.vulgivaga.1.1.chr13.g00737.t01#4    mNeoVis.vulgivaga.2.1.chr13.g00742.t01#1    mNeoVis.2.1.1.chr13_hap1.g00780.t01#5    mNeoVis.3.1.1.chr13.g00698.t01#6   mNeoVis.evergladensis.1.1.chr13.g00738.t01#3       6329   125254661:   mNeoVis.vulgivaga.1.1.chr13.g00738.t01#4    mNeoVis.evergladensis.2.1.chr13.g00747.t01#2    mNeoVis.2.1.1.chr13_hap1.g00781.t01#5    mNeoVis.3.1.1.chr13.g00699.t01#6    mNeoVis.vulgivaga.2.1.chr13.g00743.t01#1   mNeoVis.evergladensis.1.1.chr13.g00739.t01#3       6330   125254662:   mNeoVis.evergladensis.2.1.chr13.g00748.t01#2    mNeoVis.evergladensis.1.1.chr13.g00740.t01#3    mNeoVis.vulgivaga.2.1.chr13.g00744.t01#1    mNeoVis.2.1.1.chr13_hap1.g00782.t01#5    mNeoVis.3.1.1.chr13.g00700.t01#6   mNeoVis.vulgivaga.1.1.chr13.g00739.t01#4       6331   125254663:   mNeoVis.vulgivaga.2.1.chr5.g00916.t01#1    mNeoVis.3.1.1.chr5.g00908.t01#6    mNeoVis.2.1.1.chr2_hap1.g00975.t01#5    mNeoVis.vulgivaga.1.1.chr5.g00896.t01#4    mNeoVis.vulgivaga.1.1.chr5.g00897.t01#4    mNeoVis.evergladensis.2.1.chr5.g00929.t01#2   mNeoVis.evergladensis.1.1.chr5.g00920.t01#3     6332   125254664:   mNeoVis.evergladensis.2.1.chr6.g00217.t01#2    mNeoVis.vulgivaga.1.1.chr6.g00225.t01#4    mNeoVis.evergladensis.1.1.chr6.g00219.t01#3    mNeoVis.2.1.1.chr4_hap1.g00218.t01#5    mNeoVis.3.1.1.chr6.g00218.t01#6   mNeoVis.vulgivaga.2.1.chr6.g00218.t01#1       6333   125254665:   mNeoVis.evergladensis.2.1.chr13.g00749.t01#2    mNeoVis.vulgivaga.1.1.chr13.g00740.t01#4    mNeoVis.3.1.1.chr13.g00701.t01#6    mNeoVis.2.1.1.chr13_hap1.g00783.t01#5    mNeoVis.evergladensis.1.1.chr13.g00741.t01#3   mNeoVis.vulgivaga.2.1.chr13.g00745.t01#1       6334   125254666:   mNeoVis.vulgivaga.2.1.chr6.g00219.t01#1    mNeoVis.3.1.1.chr6.g00219.t01#6    mNeoVis.vulgivaga.1.1.chr6.g00226.t01#4    mNeoVis.evergladensis.1.1.chr6.g00220.t01#3    mNeoVis.evergladensis.2.1.chr6.g00218.t01#2   mNeoVis.2.1.1.chr4_hap1.g00219.t01#5       6335   125254667:   mNeoVis.vulgivaga.1.1.chr13.g00741.t01#4    mNeoVis.3.1.1.chr13.g00702.t01#6    mNeoVis.2.1.1.chr13_hap1.g00784.t01#5    mNeoVis.evergladensis.2.1.chr13.g00750.t01#2    mNeoVis.evergladensis.1.1.chr13.g00742.t01#3   mNeoVis.vulgivaga.2.1.chr13.g00746.t01#1       6336   125254668:   mNeoVis.vulgivaga.2.1.chr6.g00220.t01#1    mNeoVis.2.1.1.chr4_hap1.g00220.t01#5    mNeoVis.3.1.1.chr6.g00220.t01#6    mNeoVis.evergladensis.1.1.chr6.g00221.t01#3    mNeoVis.evergladensis.2.1.chr6.g00219.t01#2   mNeoVis.vulgivaga.1.1.chr6.g00227.t01#4       6337   125254669:   mNeoVis.vulgivaga.2.1.chr6.g00221.t01#1    mNeoVis.evergladensis.2.1.chr6.g00220.t01#2    mNeoVis.vulgivaga.1.1.chr6.g00228.t01#4    mNeoVis.evergladensis.1.1.chr6.g00222.t01#3    mNeoVis.2.1.1.chr4_hap1.g00221.t01#5   mNeoVis.3.1.1.chr6.g00221.t01#6       6338   125254670:   mNeoVis.evergladensis.1.1.chr13.g00745.t01#3    mNeoVis.evergladensis.2.1.chr13.g00752.t01#2    mNeoVis.vulgivaga.2.1.chr13.g00747.t01#1    mNeoVis.vulgivaga.1.1.chr13.g00744.t01#4    mNeoVis.3.1.1.chr13.g00705.t01#6   mNeoVis.2.1.1.chr13_hap1.g00785.t01#5       6339   125254671:   mNeoVis.vulgivaga.2.1.chr13.g00748.t01#1    mNeoVis.vulgivaga.1.1.chr13.g00742.t01#4    mNeoVis.3.1.1.chr13.g00703.t01#6    mNeoVis.evergladensis.1.1.chr13.g00743.t01#3    mNeoVis.2.1.1.chr13_hap1.g00786.t01#5   mNeoVis.evergladensis.2.1.chr13.g00751.t01#2       6340   125254672:   mNeoVis.3.1.1.chr13.g00704.t01#6    mNeoVis.evergladensis.1.1.chr13.g00744.t01#3    mNeoVis.2.1.1.chr13_hap1.g00787.t01#5    mNeoVis.vulgivaga.2.1.chr13.g00749.t01#1    mNeoVis.evergladensis.2.1.chr13.g00753.t01#2   mNeoVis.vulgivaga.1.1.chr13.g00743.t01#4       6341   125254673:   mNeoVis.evergladensis.1.1.chr13.g00746.t01#3    mNeoVis.vulgivaga.2.1.chr13.g00750.t01#1    mNeoVis.vulgivaga.1.1.chr13.g00745.t01#4    mNeoVis.3.1.1.chr13.g00706.t01#6    mNeoVis.evergladensis.2.1.chr13.g00754.t01#2   mNeoVis.2.1.1.chr13_hap1.g00788.t01#5       6342   125254674:   mNeoVis.evergladensis.2.1.chr13.g00755.t01#2    mNeoVis.evergladensis.1.1.chr13.g00747.t01#3    mNeoVis.vulgivaga.2.1.chr13.g00751.t01#1    mNeoVis.vulgivaga.1.1.chr13.g00746.t01#4    mNeoVis.3.1.1.chr13.g00707.t01#6   mNeoVis.2.1.1.chr13_hap1.g00789.t01#5       6343   125254675:   mNeoVis.2.1.1.chr4_hap1.g00222.t01#5    mNeoVis.vulgivaga.2.1.chr6.g00222.t01#1    mNeoVis.3.1.1.chr6.g00222.t01#6    mNeoVis.evergladensis.1.1.chr6.g00223.t01#3    mNeoVis.evergladensis.2.1.chr6.g00221.t01#2   mNeoVis.vulgivaga.1.1.chr6.g00229.t01#4       6344   125254676:   mNeoVis.evergladensis.1.1.chr6.g01086.t01#3    mNeoVis.vulgivaga.1.1.chr6.g01073.t01#4    mNeoVis.vulgivaga.2.1.chr6.g01072.t01#1    mNeoVis.3.1.1.chr6.g01077.t01#6    mNeoVis.evergladensis.2.1.chr6.g01079.t01#2   mNeoVis.2.1.1.chr4_hap1.g01078.t01#5       6345   125254677:   mNeoVis.vulgivaga.2.1.chr6.g00223.t01#1    mNeoVis.evergladensis.1.1.chr6.g00224.t01#3    mNeoVis.2.1.1.chr4_hap1.g00223.t01#5    mNeoVis.evergladensis.2.1.chr6.g00222.t01#2   mNeoVis.vulgivaga.1.1.chr6.g00230.t01#4       6346   125254678:   mNeoVis.evergladensis.2.1.chr6.g01080.t01#2    mNeoVis.2.1.1.chr4_hap1.g01079.t01#5    mNeoVis.evergladensis.1.1.chr6.g01085.t01#3    mNeoVis.3.1.1.chr6.g01078.t01#6    mNeoVis.vulgivaga.1.1.chr6.g01074.t01#4   mNeoVis.vulgivaga.2.1.chr6.g01073.t01#1       6347   125254679:   mNeoVis.vulgivaga.2.1.chr13.g00753.t01#1    mNeoVis.vulgivaga.1.1.chr13.g00747.t01#4    mNeoVis.evergladensis.2.1.chr13.g00756.t01#2    mNeoVis.2.1.1.chr13_hap1.g00791.t01#5    mNeoVis.3.1.1.chr13.g00708.t01#6   mNeoVis.evergladensis.1.1.chr13.g00748.t01#3       6348   125254680:   mNeoVis.2.1.1.chr13_hap1.g00792.t01#5    mNeoVis.vulgivaga.1.1.chr13.g00748.t01#4    mNeoVis.3.1.1.chr13.g00709.t01#6    mNeoVis.vulgivaga.2.1.chr13.g00754.t01#1    mNeoVis.evergladensis.2.1.chr13.g00757.t01#2   mNeoVis.evergladensis.1.1.chr13.g00749.t01#3       6349   125254681:   mNeoVis.2.1.1.chr5_hap1.g01020.t01#5   mNeoVis.2.1.1.chr5_hap1.g01019.t01#5             6350   125254682:   mNeoVis.2.1.1.chr7_hap1.g01111.t01#5   mNeoVis.evergladensis.2.1.chr2.g01015.t01#2           6351   125254683:   mNeoVis.vulgivaga.2.1.chr2.g01020.t01#1   mNeoVis.vulgivaga.1.1.chr2.g01019.t01#4             6352   125254684:   mNeoVis.evergladensis.1.1.chr6.g00225.t01#3   mNeoVis.vulgivaga.2.1.chr6.g00224.t01#1             6353   125254685:  mNeoVis.vulgivaga.2.1.chr13.g00755.t01#1               6354   125254686:   mNeoVis.vulgivaga.1.1.chr6.g00231.t01#4    mNeoVis.3.1.1.chr6.g00224.t01#6    mNeoVis.vulgivaga.2.1.chr6.g00225.t01#1    mNeoVis.2.1.1.chr4_hap1.g00225.t01#5    mNeoVis.evergladensis.2.1.chr6.g00224.t01#2   mNeoVis.evergladensis.1.1.chr6.g00226.t01#3       6355   125254687:   mNeoVis.2.1.1.chr4_hap1.g00226.t01#5    mNeoVis.3.1.1.chr6.g00225.t01#6    mNeoVis.evergladensis.1.1.chr6.g00227.t01#3    mNeoVis.evergladensis.2.1.chr6.g00225.t01#2    mNeoVis.vulgivaga.1.1.chr6.g00232.t01#4   mNeoVis.vulgivaga.2.1.chr6.g00226.t01#1       6356   125254688:   mNeoVis.2.1.1.chr13_hap1.g00793.t01#5    mNeoVis.vulgivaga.2.1.chr13.g00756.t01#1    mNeoVis.vulgivaga.1.1.chr13.g00749.t01#4   mNeoVis.evergladensis.1.1.chr13.g00750.t01#3       6357   125254689:   mNeoVis.vulgivaga.2.1.chr13.g00757.t01#1    mNeoVis.3.1.1.chr13.g00711.t01#6    mNeoVis.vulgivaga.1.1.chr13.g00750.t01#4    mNeoVis.evergladensis.2.1.chr13.g00758.t01#2    mNeoVis.2.1.1.chr13_hap1.g00794.t01#5   mNeoVis.evergladensis.1.1.chr13.g00751.t01#3       6358   125254690:   mNeoVis.evergladensis.1.1.chr6.g00228.t01#3    mNeoVis.vulgivaga.1.1.chr6.g00233.t01#4    mNeoVis.2.1.1.chr4_hap1.g00227.t01#5    mNeoVis.evergladensis.2.1.chr6.g00226.t01#2   mNeoVis.vulgivaga.2.1.chr6.g00227.t01#1       6359   125254691:   mNeoVis.3.1.1.chr13.g00631.t01#6    mNeoVis.evergladensis.1.1.chr13.g00671.t01#3    mNeoVis.vulgivaga.1.1.chr13.g00672.t01#4    mNeoVis.evergladensis.2.1.chr13.g00679.t01#2    mNeoVis.2.1.1.chr13_hap1.g00716.t01#5   mNeoVis.vulgivaga.2.1.chr13.g00679.t01#1       6360   125254692:   mNeoVis.evergladensis.2.1.chr13.g00680.t01#2    mNeoVis.evergladensis.1.1.chr13.g00672.t01#3    mNeoVis.2.1.1.chr13_hap1.g00717.t01#5    mNeoVis.vulgivaga.1.1.chr13.g00673.t01#4    mNeoVis.3.1.1.chr13.g00632.t01#6   mNeoVis.vulgivaga.2.1.chr13.g00680.t01#1       6361   125254693:   mNeoVis.vulgivaga.1.1.chr15.g00044.t01#4    mNeoVis.2.1.1.chr15_hap1.g00043.t01#5    mNeoVis.3.1.1.chr15.g00040.t01#6    mNeoVis.vulgivaga.2.1.chr15.g00045.t01#1    mNeoVis.evergladensis.2.1.chr15.g00044.t01#2   mNeoVis.evergladensis.1.1.chr15.g00570.t01#3       6362   125254694:   mNeoVis.evergladensis.2.1.chr6.g00227.t01#2    mNeoVis.evergladensis.1.1.chr6.g00229.t01#3    mNeoVis.vulgivaga.1.1.chr6.g00234.t01#4    mNeoVis.2.1.1.chr4_hap1.g00228.t01#5   mNeoVis.vulgivaga.2.1.chr6.g00228.t01#1       6363   125254695:   mNeoVis.3.1.1.chr13.g00634.t01#6    mNeoVis.evergladensis.2.1.chr13.g00683.t01#2    mNeoVis.vulgivaga.2.1.chr13.g00682.t01#1    mNeoVis.3.1.1.chr13.g00633.t01#6    mNeoVis.2.1.1.chr13_hap1.g00720.t01#5    mNeoVis.vulgivaga.1.1.chr13.g00676.t01#4   mNeoVis.evergladensis.1.1.chr13.g00674.t01#3     6364   125254696:   mNeoVis.2.1.1.chr13_hap1.g00718.t01#5    mNeoVis.vulgivaga.2.1.chr13.g00681.t01#1    mNeoVis.evergladensis.1.1.chr13.g00673.t01#3    mNeoVis.vulgivaga.1.1.chr13.g00674.t01#4   mNeoVis.evergladensis.2.1.chr13.g00681.t01#2       6365   125254697:   mNeoVis.evergladensis.2.1.chr6.g00228.t01#2    mNeoVis.3.1.1.chr6.g00228.t01#6    mNeoVis.2.1.1.chr4_hap1.g00229.t01#5    mNeoVis.evergladensis.1.1.chr6.g00230.t01#3    mNeoVis.vulgivaga.1.1.chr6.g00235.t01#4   mNeoVis.vulgivaga.2.1.chr6.g00229.t01#1       6366   125254698:   mNeoVis.3.1.1.chr13.g00635.t01#6    mNeoVis.vulgivaga.1.1.chr13.g00677.t01#4    mNeoVis.2.1.1.chr13_hap1.g00721.t01#5    mNeoVis.evergladensis.2.1.chr13.g00684.t01#2    mNeoVis.evergladensis.1.1.chr13.g00675.t01#3   mNeoVis.vulgivaga.2.1.chr13.g00683.t01#1       6367   125254699:   mNeoVis.vulgivaga.1.1.chr6.g00236.t01#4    mNeoVis.evergladensis.1.1.chr6.g00231.t01#3    mNeoVis.vulgivaga.2.1.chr6.g00230.t01#1    mNeoVis.evergladensis.2.1.chr6.g00229.t01#2    mNeoVis.3.1.1.chr6.g00229.t01#6   mNeoVis.2.1.1.chr4_hap1.g00230.t01#5       6368   125254700:   mNeoVis.evergladensis.2.1.chr15.g00045.t01#2    mNeoVis.3.1.1.chr15.g00041.t01#6    mNeoVis.evergladensis.1.1.chr15.g00569.t01#3    mNeoVis.2.1.1.chr15_hap1.g00044.t01#5    mNeoVis.vulgivaga.2.1.chr15.g00046.t01#1   mNeoVis.vulgivaga.1.1.chr15.g00045.t01#4       6369   125254701:   mNeoVis.vulgivaga.1.1.chr6.g00238.t01#4    mNeoVis.3.1.1.chr6.g00231.t01#6    mNeoVis.evergladensis.1.1.chr6.g00232.t01#3    mNeoVis.vulgivaga.2.1.chr6.g00231.t01#1    mNeoVis.2.1.1.chr4_hap1.g00232.t01#5    mNeoVis.vulgivaga.2.1.chr6.g00232.t01#1   mNeoVis.evergladensis.2.1.chr6.g00231.t01#2     6370   125254702:   mNeoVis.vulgivaga.1.1.chr6.g00237.t01#4    mNeoVis.evergladensis.2.1.chr6.g00230.t01#2    mNeoVis.2.1.1.chr4_hap1.g00231.t01#5   mNeoVis.3.1.1.chr6.g00230.t01#6         6371   125254703:   mNeoVis.vulgivaga.2.1.chr6.g00233.t01#1    mNeoVis.evergladensis.1.1.chr6.g00233.t01#3    mNeoVis.vulgivaga.1.1.chr6.g00239.t01#4    mNeoVis.2.1.1.chr4_hap1.g00233.t01#5    mNeoVis.evergladensis.2.1.chr6.g00232.t01#2   mNeoVis.3.1.1.chr6.g00232.t01#6       6372   125254704:   mNeoVis.evergladensis.2.1.chr6.g00233.t01#2    mNeoVis.3.1.1.chr6.g00233.t01#6    mNeoVis.2.1.1.chr4_hap1.g00234.t01#5    mNeoVis.vulgivaga.2.1.chr6.g00234.t01#1    mNeoVis.evergladensis.1.1.chr6.g00234.t01#3   mNeoVis.vulgivaga.1.1.chr6.g00240.t01#4       6373   125254705:   mNeoVis.evergladensis.2.1.chr2.g00633.t01#2    mNeoVis.evergladensis.1.1.chr2.g00632.t01#3    mNeoVis.2.1.1.chr5_hap1.g00636.t01#5    mNeoVis.vulgivaga.1.1.chr2.g00627.t01#4    mNeoVis.3.1.1.chr2.g00555.t01#6   mNeoVis.vulgivaga.2.1.chr2.g00634.t01#1       6374   125254706:   mNeoVis.vulgivaga.2.1.chr6.g00235.t01#1    mNeoVis.3.1.1.chr6.g00234.t01#6    mNeoVis.evergladensis.1.1.chr6.g00235.t01#3    mNeoVis.2.1.1.chr4_hap1.g00235.t01#5   mNeoVis.evergladensis.2.1.chr6.g00234.t01#2       6375   125254707:   mNeoVis.3.1.1.chr6.g00235.t01#6    mNeoVis.vulgivaga.1.1.chr6.g00242.t01#4    mNeoVis.evergladensis.2.1.chr6.g00235.t01#2    mNeoVis.2.1.1.chr4_hap1.g00236.t01#5    mNeoVis.vulgivaga.2.1.chr6.g00236.t01#1   mNeoVis.evergladensis.1.1.chr6.g00236.t01#3       6376   125254708:   mNeoVis.evergladensis.2.1.chr6.g00236.t01#2    mNeoVis.evergladensis.1.1.chr6.g00237.t01#3    mNeoVis.3.1.1.chr6.g00236.t01#6    mNeoVis.vulgivaga.1.1.chr6.g00243.t01#4    mNeoVis.2.1.1.chr4_hap1.g00237.t01#5   mNeoVis.vulgivaga.2.1.chr6.g00237.t01#1       6377   125254709:   mNeoVis.2.1.1.chr13_hap1.g00722.t01#5    mNeoVis.vulgivaga.2.1.chr13.g00684.t01#1    mNeoVis.3.1.1.chr13.g00636.t01#6    mNeoVis.evergladensis.1.1.chr13.g00676.t01#3    mNeoVis.vulgivaga.1.1.chr13.g00678.t01#4   mNeoVis.evergladensis.2.1.chr13.g00685.t01#2       6378   125254710:   mNeoVis.vulgivaga.1.1.chr13.g00679.t01#4    mNeoVis.evergladensis.2.1.chr13.g00687.t01#2    mNeoVis.vulgivaga.2.1.chr13.g00685.t01#1    mNeoVis.evergladensis.1.1.chr13.g00677.t01#3   mNeoVis.2.1.1.chr13_hap1.g00723.t01#5       6379   125254711:  mNeoVis.vulgivaga.2.1.chr6.g01074.t01#1               6380   125254712:   mNeoVis.evergladensis.1.1.chr6.g00240.t01#3    mNeoVis.vulgivaga.1.1.chr6.g00244.t01#4    mNeoVis.3.1.1.chr6.g00237.t01#6    mNeoVis.evergladensis.2.1.chr6.g00237.t01#2    mNeoVis.vulgivaga.1.1.chr6.g00246.t01#4    mNeoVis.vulgivaga.2.1.chr6.g00238.t01#1    mNeoVis.2.1.1.chr4_hap1.g00238.t01#5   mNeoVis.evergladensis.1.1.chr6.g00238.t01#3     6381   125254713:   mNeoVis.vulgivaga.2.1.chr15.g00047.t01#1    mNeoVis.2.1.1.chr15_hap1.g00045.t01#5    mNeoVis.vulgivaga.1.1.chr15.g00046.t01#4    mNeoVis.evergladensis.2.1.chr15.g00046.t01#2    mNeoVis.3.1.1.chr15.g00042.t01#6   mNeoVis.evergladensis.1.1.chr15.g00568.t01#3       6382   125254714:   mNeoVis.3.1.1.chr6.g00238.t01#6    mNeoVis.2.1.1.chr4_hap1.g00239.t01#5    mNeoVis.vulgivaga.2.1.chr6.g00239.t01#1    mNeoVis.evergladensis.1.1.chr6.g00239.t01#3    mNeoVis.evergladensis.2.1.chr6.g00238.t01#2   mNeoVis.vulgivaga.1.1.chr6.g00245.t01#4       6383   125254715:   mNeoVis.vulgivaga.2.1.chr15.g00048.t01#1   mNeoVis.2.1.1.chr15_hap1.g00046.t01#5             6384   125254716:   mNeoVis.vulgivaga.2.1.chr6.g00240.t01#1    mNeoVis.vulgivaga.1.1.chr6.g00247.t01#4    mNeoVis.2.1.1.chr4_hap1.g00240.t01#5    mNeoVis.evergladensis.1.1.chr6.g00241.t01#3    mNeoVis.3.1.1.chr6.g00239.t01#6   mNeoVis.evergladensis.2.1.chr6.g00239.t01#2       6385   125254717:   mNeoVis.2.1.1.chr13_hap1.g00724.t01#5    mNeoVis.evergladensis.2.1.chr13.g00688.t01#2    mNeoVis.vulgivaga.2.1.chr13.g00686.t01#1    mNeoVis.evergladensis.1.1.chr13.g00678.t01#3    mNeoVis.3.1.1.chr13.g00640.t01#6   mNeoVis.vulgivaga.1.1.chr13.g00680.t01#4       6386   125254718:   mNeoVis.3.1.1.chr13.g00641.t01#6    mNeoVis.evergladensis.1.1.chr13.g00679.t01#3    mNeoVis.vulgivaga.1.1.chr13.g00681.t01#4    mNeoVis.vulgivaga.2.1.chr13.g00687.t01#1    mNeoVis.2.1.1.chr13_hap1.g00725.t01#5   mNeoVis.evergladensis.2.1.chr13.g00689.t01#2       6387   125254719:   mNeoVis.vulgivaga.1.1.chr13.g00682.t01#4    mNeoVis.evergladensis.1.1.chr13.g00680.t01#3    mNeoVis.2.1.1.chr13_hap1.g00726.t01#5    mNeoVis.vulgivaga.2.1.chr13.g00688.t01#1    mNeoVis.3.1.1.chr13.g00642.t01#6   mNeoVis.evergladensis.2.1.chr13.g00690.t01#2       6388   125254720:   mNeoVis.vulgivaga.2.1.chr6.g00241.t01#1    mNeoVis.2.1.1.chr4_hap1.g00241.t01#5    mNeoVis.evergladensis.1.1.chr6.g00242.t01#3    mNeoVis.3.1.1.chr6.g00240.t01#6    mNeoVis.vulgivaga.1.1.chr6.g00248.t01#4   mNeoVis.evergladensis.2.1.chr6.g00240.t01#2       6389   125254721:   mNeoVis.vulgivaga.2.1.chr6.g00243.t01#1    mNeoVis.evergladensis.1.1.chr6.g00244.t01#3    mNeoVis.2.1.1.chr4_hap1.g00243.t01#5    mNeoVis.vulgivaga.1.1.chr6.g00250.t01#4    mNeoVis.3.1.1.chr6.g00242.t01#6   mNeoVis.evergladensis.2.1.chr6.g00242.t01#2       6390   125254722:   mNeoVis.2.1.1.chr4_hap1.g00244.t01#5    mNeoVis.evergladensis.2.1.chr6.g00243.t01#2    mNeoVis.vulgivaga.1.1.chr6.g00251.t01#4    mNeoVis.vulgivaga.2.1.chr6.g00244.t01#1    mNeoVis.3.1.1.chr6.g00243.t01#6   mNeoVis.evergladensis.1.1.chr6.g00245.t01#3       6391   125254723:   mNeoVis.evergladensis.2.1.chr15.g00047.t01#2    mNeoVis.2.1.1.chr15_hap1.g00047.t01#5    mNeoVis.evergladensis.1.1.chr15.g00567.t01#3    mNeoVis.vulgivaga.2.1.chr15.g00049.t01#1    mNeoVis.vulgivaga.1.1.chr15.g00047.t01#4   mNeoVis.3.1.1.chr15.g00044.t01#6       6392   125254724:   mNeoVis.vulgivaga.1.1.chr6.g01076.t01#4    mNeoVis.vulgivaga.2.1.chr6.g01075.t01#1    mNeoVis.evergladensis.1.1.chr6.g01087.t01#3   mNeoVis.evergladensis.2.1.chr6.g01081.t01#2       6393   125254725:   mNeoVis.2.1.1.chr4_hap1.g01084.t01#5   mNeoVis.2.1.1.chr4_hap1.g01082.t01#5             6394   125254726:   mNeoVis.2.1.1.chr15_hap1.g00369.t01#5    mNeoVis.2.1.1.chr15_hap1.g00443.t01#5    mNeoVis.evergladensis.2.1.chr6.g00245.t01#2    mNeoVis.vulgivaga.2.1.chr6.g00246.t01#1    mNeoVis.vulgivaga.1.1.chr6.g00253.t01#4    mNeoVis.2.1.1.chr4_hap1.g00246.t01#5    mNeoVis.evergladensis.1.1.chr6.g00247.t01#3    mNeoVis.2.1.1.chr15_hap1.g00406.t01#5    mNeoVis.3.1.1.chr6.g00245.t01#6      6395   125254727:   mNeoVis.2.1.1.chr13_hap1.g00727.t01#5    mNeoVis.vulgivaga.2.1.chr13.g00689.t01#1    mNeoVis.3.1.1.chr13.g00643.t01#6    mNeoVis.evergladensis.2.1.chr13.g00691.t01#2    mNeoVis.evergladensis.1.1.chr13.g00681.t01#3   mNeoVis.vulgivaga.1.1.chr13.g00683.t01#4       6396   125254728:   mNeoVis.evergladensis.1.1.chr6.g00249.t01#3    mNeoVis.evergladensis.2.1.chr6.g00247.t01#2    mNeoVis.vulgivaga.1.1.chr6.g00255.t01#4    mNeoVis.2.1.1.chr4_hap1.g00248.t01#5    mNeoVis.vulgivaga.2.1.chr6.g00248.t01#1   mNeoVis.3.1.1.chr6.g00247.t01#6       6397   125254729:   mNeoVis.evergladensis.1.1.chr13.g00682.t01#3    mNeoVis.vulgivaga.2.1.chr13.g00690.t01#1    mNeoVis.vulgivaga.1.1.chr13.g00684.t01#4    mNeoVis.3.1.1.chr13.g00644.t01#6    mNeoVis.evergladensis.2.1.chr13.g00692.t01#2   mNeoVis.2.1.1.chr13_hap1.g00728.t01#5       6398   125254730:   mNeoVis.evergladensis.2.1.chr13.g00693.t01#2    mNeoVis.3.1.1.chr13.g00645.t01#6    mNeoVis.vulgivaga.2.1.chr13.g00691.t01#1    mNeoVis.vulgivaga.1.1.chr13.g00685.t01#4    mNeoVis.evergladensis.1.1.chr13.g00683.t01#3   mNeoVis.2.1.1.chr13_hap1.g00729.t01#5       6399   125254731:   mNeoVis.vulgivaga.2.1.chr13.g00692.t01#1    mNeoVis.3.1.1.chr13.g00646.t01#6    mNeoVis.evergladensis.2.1.chr13.g00694.t01#2    mNeoVis.vulgivaga.1.1.chr13.g00686.t01#4    mNeoVis.2.1.1.chr13_hap1.g00730.t01#5   mNeoVis.evergladensis.1.1.chr13.g00684.t01#3       6400   125254732:   mNeoVis.evergladensis.2.1.chr6.g00248.t01#2    mNeoVis.3.1.1.chr6.g00248.t01#6    mNeoVis.vulgivaga.2.1.chr6.g00249.t01#1    mNeoVis.evergladensis.1.1.chr6.g00250.t01#3    mNeoVis.vulgivaga.1.1.chr6.g00256.t01#4   mNeoVis.2.1.1.chr4_hap1.g00249.t01#5       6401   125254733:   mNeoVis.2.1.1.chr4_hap1.g00250.t01#5    mNeoVis.3.1.1.chr6.g00249.t01#6    mNeoVis.vulgivaga.2.1.chr6.g00250.t01#1   mNeoVis.evergladensis.1.1.chr6.g00251.t01#3       6402   125254734:   mNeoVis.evergladensis.2.1.chr13.g00695.t01#2    mNeoVis.vulgivaga.2.1.chr13.g00693.t01#1    mNeoVis.2.1.1.chr13_hap1.g00731.t01#5    mNeoVis.3.1.1.chr13.g00647.t01#6    mNeoVis.evergladensis.1.1.chr13.g00685.t01#3   mNeoVis.vulgivaga.1.1.chr13.g00687.t01#4       6403   125254735:   mNeoVis.2.1.1.chr4_hap1.g01085.t01#5    mNeoVis.vulgivaga.1.1.chr6.g01077.t01#4    mNeoVis.vulgivaga.2.1.chr6.g01076.t01#1    mNeoVis.evergladensis.1.1.chr6.g01088.t01#3   mNeoVis.evergladensis.2.1.chr6.g01082.t01#2       6404   125254736:   mNeoVis.vulgivaga.1.1.chr6.g00257.t01#4    mNeoVis.evergladensis.1.1.chr6.g00253.t01#3    mNeoVis.2.1.1.chr4_hap1.g00252.t01#5    mNeoVis.evergladensis.2.1.chr6.g00251.t01#2    mNeoVis.3.1.1.chr6.g00250.t01#6   mNeoVis.vulgivaga.2.1.chr6.g00251.t01#1       6405   125254737:   mNeoVis.evergladensis.2.1.chr6.g00252.t01#2    mNeoVis.2.1.1.chr4_hap1.g00253.t01#5    mNeoVis.vulgivaga.2.1.chr6.g00252.t01#1    mNeoVis.3.1.1.chr6.g00251.t01#6    mNeoVis.vulgivaga.1.1.chr6.g00258.t01#4    mNeoVis.evergladensis.2.1.chr6.g00253.t01#2   mNeoVis.evergladensis.1.1.chr6.g00254.t01#3     6406   125254738:   mNeoVis.3.1.1.chr6.g00253.t01#6    mNeoVis.vulgivaga.2.1.chr6.g00253.t01#1    mNeoVis.vulgivaga.1.1.chr6.g00260.t01#4    mNeoVis.2.1.1.chr4_hap1.g00254.t01#5   mNeoVis.evergladensis.1.1.chr6.g00256.t01#3       6407   125254739:   mNeoVis.3.1.1.chr6.g00252.t01#6    mNeoVis.evergladensis.2.1.chr6.g00254.t01#2    mNeoVis.vulgivaga.1.1.chr6.g00259.t01#4   mNeoVis.evergladensis.1.1.chr6.g00255.t01#3       6408   125254740:   mNeoVis.vulgivaga.2.1.chr6.g00254.t01#1    mNeoVis.evergladensis.1.1.chr6.g00257.t01#3    mNeoVis.evergladensis.2.1.chr6.g00255.t01#2    mNeoVis.vulgivaga.1.1.chr6.g00261.t01#4    mNeoVis.2.1.1.chr4_hap1.g00255.t01#5   mNeoVis.3.1.1.chr6.g00254.t01#6       6409   125254741:   mNeoVis.vulgivaga.2.1.chr6.g00255.t01#1    mNeoVis.evergladensis.2.1.chr6.g00256.t01#2    mNeoVis.3.1.1.chr6.g00255.t01#6    mNeoVis.2.1.1.chr4_hap1.g00256.t01#5    mNeoVis.vulgivaga.1.1.chr6.g00262.t01#4   mNeoVis.evergladensis.1.1.chr6.g00258.t01#3       6410   125254742:   mNeoVis.3.1.1.chr6.g00256.t01#6    mNeoVis.evergladensis.1.1.chr6.g00259.t01#3    mNeoVis.evergladensis.2.1.chr6.g00257.t01#2    mNeoVis.vulgivaga.2.1.chr6.g00256.t01#1    mNeoVis.vulgivaga.1.1.chr6.g00263.t01#4   mNeoVis.2.1.1.chr4_hap1.g00257.t01#5       6411   125254743:   mNeoVis.evergladensis.1.1.chr13.g00687.t01#3    mNeoVis.vulgivaga.1.1.chr13.g00689.t01#4    mNeoVis.2.1.1.chr13_hap1.g00732.t01#5    mNeoVis.evergladensis.2.1.chr13.g00698.t01#2    mNeoVis.3.1.1.chr13.g00648.t01#6   mNeoVis.vulgivaga.2.1.chr13.g00694.t01#1       6412   125254744:   mNeoVis.2.1.1.chr4_hap1.g00259.t01#5    mNeoVis.3.1.1.chr6.g00257.t01#6    mNeoVis.vulgivaga.1.1.chr6.g00265.t01#4    mNeoVis.vulgivaga.2.1.chr6.g00257.t01#1    mNeoVis.evergladensis.1.1.chr6.g00260.t01#3   mNeoVis.evergladensis.2.1.chr6.g00259.t01#2       6413   125254745:   mNeoVis.vulgivaga.1.1.chr15.g00048.t01#4    mNeoVis.3.1.1.chr15.g00045.t01#6    mNeoVis.evergladensis.1.1.chr15.g00566.t01#3    mNeoVis.evergladensis.2.1.chr15.g00048.t01#2    mNeoVis.2.1.1.chr15_hap1.g00048.t01#5   mNeoVis.vulgivaga.2.1.chr15.g00050.t01#1       6414   125254746:   mNeoVis.evergladensis.1.1.chr3.g00593.t01#3    mNeoVis.evergladensis.2.1.chr3.g00593.t01#2    mNeoVis.3.1.1.chr3.g00577.t01#6    mNeoVis.vulgivaga.1.1.chr3.g00589.t01#4    mNeoVis.2.1.1.chr3_hap1.g00585.t01#5   mNeoVis.vulgivaga.2.1.chr3.g00596.t01#1       6415   125254747:   mNeoVis.3.1.1.chr15.g00484.t01#6    mNeoVis.evergladensis.1.1.chr15.g00088.t01#3    mNeoVis.2.1.1.chr15_hap1.g00703.t01#5    mNeoVis.vulgivaga.1.1.chr15.g00528.t01#4   mNeoVis.evergladensis.2.1.chr15.g00525.t01#2       6416   125254748:   mNeoVis.vulgivaga.2.1.chr6.g00258.t01#1    mNeoVis.3.1.1.chr6.g00258.t01#6    mNeoVis.2.1.1.chr4_hap1.g00260.t01#5    mNeoVis.evergladensis.2.1.chr6.g00260.t01#2    mNeoVis.vulgivaga.1.1.chr6.g00266.t01#4   mNeoVis.evergladensis.1.1.chr6.g00261.t01#3       6417   125254749:   mNeoVis.3.1.1.chr13.g00649.t01#6    mNeoVis.evergladensis.1.1.chr13.g00686.t01#3    mNeoVis.vulgivaga.2.1.chr13.g00695.t01#1   mNeoVis.evergladensis.2.1.chr13.g00697.t01#2       6418   125254750:   mNeoVis.evergladensis.1.1.chr6.g00262.t01#3    mNeoVis.2.1.1.chr4_hap1.g00261.t01#5    mNeoVis.vulgivaga.2.1.chr6.g00259.t01#1    mNeoVis.vulgivaga.1.1.chr6.g00267.t01#4    mNeoVis.3.1.1.chr6.g00259.t01#6   mNeoVis.evergladensis.2.1.chr6.g00261.t01#2       6419   125254751:   mNeoVis.vulgivaga.2.1.chr6.g00173.t01#1    mNeoVis.2.1.1.chr4_hap1.g00173.t01#5    mNeoVis.evergladensis.2.1.chr6.g00174.t01#2    mNeoVis.evergladensis.1.1.chr6.g00175.t01#3    mNeoVis.vulgivaga.1.1.chr6.g00175.t01#4   mNeoVis.3.1.1.chr6.g00173.t01#6       6420   125254752:   mNeoVis.vulgivaga.1.1.chr6.g00176.t01#4    mNeoVis.3.1.1.chr6.g00175.t01#6   mNeoVis.vulgivaga.2.1.chr6.g00174.t01#1           6421   125254753:   mNeoVis.evergladensis.1.1.chr6.g00177.t01#3    mNeoVis.vulgivaga.2.1.chr6.g00175.t01#1    mNeoVis.vulgivaga.1.1.chr6.g00177.t01#4    mNeoVis.evergladensis.2.1.chr6.g00175.t01#2    mNeoVis.3.1.1.chr6.g00176.t01#6   mNeoVis.2.1.1.chr4_hap1.g00174.t01#5       6422   125254754:   mNeoVis.evergladensis.2.1.chr6.g00176.t01#2    mNeoVis.3.1.1.chr6.g00177.t01#6    mNeoVis.2.1.1.chr4_hap1.g00175.t01#5    mNeoVis.evergladensis.1.1.chr6.g00178.t01#3    mNeoVis.vulgivaga.2.1.chr6.g00176.t01#1   mNeoVis.vulgivaga.1.1.chr6.g00178.t01#4       6423   125254755:   mNeoVis.evergladensis.1.1.chr13.g00690.t01#3    mNeoVis.vulgivaga.2.1.chr13.g00697.t01#1    mNeoVis.evergladensis.2.1.chr13.g00701.t01#2    mNeoVis.vulgivaga.1.1.chr13.g00692.t01#4    mNeoVis.2.1.1.chr13_hap1.g00736.t01#5   mNeoVis.3.1.1.chr13.g00651.t01#6       6424   125254756:   mNeoVis.3.1.1.chr13.g00652.t01#6    mNeoVis.vulgivaga.2.1.chr13.g00698.t01#1    mNeoVis.evergladensis.1.1.chr13.g00691.t01#3    mNeoVis.2.1.1.chr13_hap1.g00735.t01#5    mNeoVis.evergladensis.2.1.chr13.g00700.t01#2   mNeoVis.vulgivaga.1.1.chr13.g00691.t01#4       6425   125254757:   mNeoVis.vulgivaga.2.1.chr13.g00699.t01#1    mNeoVis.evergladensis.1.1.chr13.g00692.t01#3    mNeoVis.3.1.1.chr13.g00653.t01#6    mNeoVis.evergladensis.2.1.chr13.g00702.t01#2    mNeoVis.vulgivaga.1.1.chr13.g00693.t01#4   mNeoVis.2.1.1.chr13_hap1.g00737.t01#5       6426   125254758:   mNeoVis.vulgivaga.1.1.chr6.g01078.t01#4    mNeoVis.evergladensis.2.1.chr6.g01083.t01#2    mNeoVis.vulgivaga.2.1.chr6.g01077.t01#1    mNeoVis.2.1.1.chr4_hap1.g01086.t01#5   mNeoVis.evergladensis.1.1.chr6.g01089.t01#3       6427   125254759:   mNeoVis.vulgivaga.1.1.chr6.g00179.t01#4    mNeoVis.evergladensis.2.1.chr6.g00177.t01#2    mNeoVis.vulgivaga.2.1.chr6.g00177.t01#1    mNeoVis.3.1.1.chr6.g00178.t01#6    mNeoVis.2.1.1.chr4_hap1.g00176.t01#5   mNeoVis.evergladensis.1.1.chr6.g00179.t01#3       6428   125254760:   mNeoVis.3.1.1.chr6.g00179.t01#6    mNeoVis.2.1.1.chr4_hap1.g00177.t01#5    mNeoVis.evergladensis.1.1.chr6.g00180.t01#3    mNeoVis.vulgivaga.1.1.chr6.g00180.t01#4    mNeoVis.vulgivaga.2.1.chr6.g00178.t01#1   mNeoVis.evergladensis.2.1.chr6.g00178.t01#2       6429   125254761:   mNeoVis.evergladensis.1.1.chr13.g00693.t01#3    mNeoVis.vulgivaga.2.1.chr13.g00700.t01#1    mNeoVis.evergladensis.2.1.chr13.g00703.t01#2    mNeoVis.3.1.1.chr13.g00654.t01#6    mNeoVis.vulgivaga.1.1.chr13.g00694.t01#4   mNeoVis.2.1.1.chr13_hap1.g00738.t01#5       6430   125254762:   mNeoVis.2.1.1.chr2_hap1.g00483.t01#5    mNeoVis.evergladensis.2.1.chr5.g00486.t01#2    mNeoVis.vulgivaga.1.1.chr6.g00181.t01#4    mNeoVis.vulgivaga.1.1.chr5.g00470.t01#4    mNeoVis.evergladensis.2.1.chr9.g01067.t01#2    mNeoVis.2.1.1.chr8_hap1.g01091.t01#5    mNeoVis.3.1.1.chr8.g01013.t01#6    mNeoVis.vulgivaga.2.1.chr6.g00179.t01#1    mNeoVis.3.1.1.chr5.g00476.t01#6      6431   125254763:   mNeoVis.evergladensis.2.1.chr6.g00179.t01#2   mNeoVis.2.1.1.chr4_hap1.g00178.t01#5             6432   125254764:   mNeoVis.vulgivaga.1.1.chr3.g01116.t01#4    mNeoVis.vulgivaga.1.1.chr13.g00695.t01#4    mNeoVis.evergladensis.2.1.chr3.g01115.t01#2    mNeoVis.evergladensis.2.1.chr13.g00704.t01#2    mNeoVis.vulgivaga.2.1.chr13.g00701.t01#1    mNeoVis.2.1.1.chr3_hap1.g01114.t01#5    mNeoVis.vulgivaga.2.1.chr3.g01130.t01#1    mNeoVis.3.1.1.chr3.g01105.t01#6    mNeoVis.3.1.1.chr13.g00655.t01#6      6433   125254765:   mNeoVis.2.1.1.chr3_hap1.g01116.t01#5    mNeoVis.evergladensis.1.1.chr3.g01126.t01#3    mNeoVis.vulgivaga.1.1.chr3.g01117.t01#4    mNeoVis.3.1.1.chr3.g01104.t01#6    mNeoVis.evergladensis.2.1.chr3.g01116.t01#2   mNeoVis.vulgivaga.2.1.chr3.g01132.t01#1       6434   125254766:   mNeoVis.2.1.1.chr4_hap1.g00179.t01#5    mNeoVis.vulgivaga.2.1.chr6.g00180.t01#1    mNeoVis.evergladensis.2.1.chr6.g00180.t01#2    mNeoVis.evergladensis.1.1.chr6.g00182.t01#3    mNeoVis.3.1.1.chr6.g00181.t01#6   mNeoVis.vulgivaga.1.1.chr6.g00182.t01#4       6435   125254767:   mNeoVis.3.1.1.chr13.g00656.t01#6    mNeoVis.vulgivaga.2.1.chr13.g00702.t01#1    mNeoVis.evergladensis.1.1.chr13.g00695.t01#3    mNeoVis.2.1.1.chr13_hap1.g00740.t01#5    mNeoVis.evergladensis.2.1.chr13.g00705.t01#2   mNeoVis.vulgivaga.1.1.chr13.g00696.t01#4       6436   125254768:   mNeoVis.vulgivaga.2.1.chr13.g00704.t01#1    mNeoVis.2.1.1.chr13_hap1.g00742.t01#5    mNeoVis.evergladensis.1.1.chr13.g00696.t01#3    mNeoVis.vulgivaga.1.1.chr13.g00697.t01#4   mNeoVis.3.1.1.chr13.g00658.t01#6       6437   125254769:   mNeoVis.evergladensis.1.1.chr13.g00366.t01#3    mNeoVis.3.1.1.chr6.g00801.t01#6    mNeoVis.2.1.1.chr13_hap1.g00377.t01#5    mNeoVis.evergladensis.1.1.chr6.g00801.t01#3    mNeoVis.2.1.1.chr4_hap1.g00792.t01#5    mNeoVis.evergladensis.2.1.chr6.g00797.t01#2    mNeoVis.vulgivaga.1.1.chr6.g00794.t01#4    mNeoVis.2.1.1.chr8_hap1.g01488.t01#5    mNeoVis.vulgivaga.2.1.chr6.g00790.t01#1      6438   125254770:   mNeoVis.evergladensis.2.1.chr13.g00706.t01#2    mNeoVis.3.1.1.chr13.g00657.t01#6    mNeoVis.2.1.1.chr13_hap1.g00741.t01#5    mNeoVis.vulgivaga.2.1.chr13.g00703.t01#1   mNeoVis.vulgivaga.1.1.chr13.g00698.t01#4       6439   125254771:   mNeoVis.vulgivaga.1.1.chr15.g00049.t01#4    mNeoVis.evergladensis.1.1.chr15.g00565.t01#3    mNeoVis.3.1.1.chr15.g00046.t01#6    mNeoVis.vulgivaga.2.1.chr15.g00051.t01#1    mNeoVis.2.1.1.chr15_hap1.g00049.t01#5   mNeoVis.evergladensis.2.1.chr15.g00049.t01#2       6440   125254772:   mNeoVis.vulgivaga.2.1.chr13.g00706.t01#1    mNeoVis.vulgivaga.1.1.chr13.g00700.t01#4    mNeoVis.3.1.1.chr13.g00660.t01#6    mNeoVis.evergladensis.1.1.chr13.g00698.t01#3    mNeoVis.2.1.1.chr13_hap1.g00744.t01#5   mNeoVis.evergladensis.2.1.chr13.g00709.t01#2       6441   125254773:   mNeoVis.3.1.1.chr6.g01081.t01#6    mNeoVis.vulgivaga.1.1.chr6.g01079.t01#4    mNeoVis.2.1.1.chr4_hap1.g01087.t01#5    mNeoVis.evergladensis.1.1.chr6.g01090.t01#3    mNeoVis.vulgivaga.2.1.chr6.g01078.t01#1   mNeoVis.evergladensis.2.1.chr6.g01084.t01#2       6442   125254774:   mNeoVis.vulgivaga.1.1.chr6.g00183.t01#4    mNeoVis.3.1.1.chr6.g00182.t01#6    mNeoVis.vulgivaga.2.1.chr6.g00181.t01#1    mNeoVis.evergladensis.2.1.chr6.g00181.t01#2    mNeoVis.2.1.1.chr4_hap1.g00180.t01#5   mNeoVis.evergladensis.1.1.chr6.g00183.t01#3       6443   125254775:   mNeoVis.vulgivaga.2.1.chr6.g00182.t01#1    mNeoVis.2.1.1.chr4_hap1.g00181.t01#5    mNeoVis.evergladensis.2.1.chr6.g00182.t01#2   mNeoVis.vulgivaga.1.1.chr6.g00185.t01#4         6444   125254776:   mNeoVis.evergladensis.1.1.chr6.g00184.t01#3    mNeoVis.vulgivaga.1.1.chr6.g00184.t01#4   mNeoVis.3.1.1.chr6.g00183.t01#6           6445   125254777:   mNeoVis.2.1.1.chr2_hap1.g00173.t01#5    mNeoVis.2.1.1.chr2_hap1.g00174.t01#5    mNeoVis.evergladensis.1.1.chr5.g00180.t01#3    mNeoVis.3.1.1.chr5.g00164.t01#6    mNeoVis.evergladensis.1.1.chr5.g00179.t01#3    mNeoVis.evergladensis.2.1.chr5.g00177.t01#2    mNeoVis.vulgivaga.2.1.chr5.g00174.t01#1    mNeoVis.vulgivaga.2.1.chr5.g00173.t01#1    mNeoVis.evergladensis.2.1.chr5.g00178.t01#2      6446   125254778:   mNeoVis.vulgivaga.2.1.chr13.g00707.t01#1    mNeoVis.3.1.1.chr13.g00661.t01#6    mNeoVis.evergladensis.2.1.chr13.g00710.t01#2    mNeoVis.vulgivaga.1.1.chr13.g00701.t01#4   mNeoVis.evergladensis.1.1.chr13.g00699.t01#3       6447   125254779:   mNeoVis.3.1.1.chr13.g00662.t01#6    mNeoVis.vulgivaga.2.1.chr13.g00708.t01#1    mNeoVis.2.1.1.chr13_hap1.g00746.t01#5    mNeoVis.evergladensis.2.1.chr13.g00711.t01#2    mNeoVis.vulgivaga.1.1.chr13.g00702.t01#4   mNeoVis.evergladensis.1.1.chr13.g00700.t01#3       6448   125254780:   mNeoVis.evergladensis.2.1.chr13.g00712.t01#2    mNeoVis.vulgivaga.1.1.chr13.g00703.t01#4    mNeoVis.3.1.1.chr13.g00663.t01#6    mNeoVis.2.1.1.chr13_hap1.g00747.t01#5   mNeoVis.vulgivaga.2.1.chr13.g00709.t01#1       6449   125254781:   mNeoVis.vulgivaga.2.1.chr6.g00183.t01#1    mNeoVis.evergladensis.1.1.chr6.g00185.t01#3    mNeoVis.3.1.1.chr6.g00184.t01#6    mNeoVis.2.1.1.chr4_hap1.g00182.t01#5    mNeoVis.evergladensis.2.1.chr6.g00183.t01#2   mNeoVis.vulgivaga.1.1.chr6.g00186.t01#4       6450   125254782:   mNeoVis.3.1.1.chr6.g00185.t01#6    mNeoVis.2.1.1.chr4_hap1.g00183.t01#5    mNeoVis.vulgivaga.2.1.chr6.g00184.t01#1    mNeoVis.evergladensis.1.1.chr6.g00186.t01#3    mNeoVis.evergladensis.2.1.chr6.g00184.t01#2   mNeoVis.vulgivaga.1.1.chr6.g00187.t01#4       6451   125254783:   mNeoVis.vulgivaga.2.1.chr13.g00710.t01#1   mNeoVis.3.1.1.chr13.g00664.t01#6             6452   125254784:   mNeoVis.2.1.1.chr4_hap1.g00184.t01#5    mNeoVis.3.1.1.chr6.g00186.t01#6    mNeoVis.evergladensis.1.1.chr6.g00187.t01#3    mNeoVis.vulgivaga.1.1.chr6.g00188.t01#4    mNeoVis.evergladensis.2.1.chr6.g00185.t01#2   mNeoVis.vulgivaga.2.1.chr6.g00185.t01#1       6453   125254785:   mNeoVis.vulgivaga.2.1.chr13.g00711.t01#1   mNeoVis.evergladensis.2.1.chr13.g00713.t01#2           6454   125254786:   mNeoVis.vulgivaga.2.1.chr6.g00186.t01#1    mNeoVis.evergladensis.2.1.chr6.g00186.t01#2    mNeoVis.vulgivaga.1.1.chr6.g00189.t01#4    mNeoVis.evergladensis.1.1.chr6.g00188.t01#3    mNeoVis.3.1.1.chr6.g00187.t01#6   mNeoVis.2.1.1.chr4_hap1.g00185.t01#5       6455   125254787:   mNeoVis.3.1.1.chr13.g00667.t01#6    mNeoVis.vulgivaga.1.1.chr13.g00705.t01#4    mNeoVis.evergladensis.1.1.chr13.g00707.t01#3    mNeoVis.2.1.1.chr13_hap1.g00749.t01#5    mNeoVis.evergladensis.2.1.chr13.g00714.t01#2   mNeoVis.vulgivaga.2.1.chr13.g00712.t01#1       6456   125254788:   mNeoVis.3.1.1.chr6.g00188.t01#6    mNeoVis.vulgivaga.1.1.chr6.g00190.t01#4    mNeoVis.2.1.1.chr4_hap1.g00186.t01#5    mNeoVis.evergladensis.1.1.chr6.g00189.t01#3    mNeoVis.vulgivaga.2.1.chr6.g00187.t01#1   mNeoVis.evergladensis.2.1.chr6.g00187.t01#2       6457   125254789:   mNeoVis.evergladensis.2.1.chr15.g00050.t01#2    mNeoVis.3.1.1.chr15.g00047.t01#6    mNeoVis.vulgivaga.2.1.chr15.g00052.t01#1    mNeoVis.vulgivaga.1.1.chr15.g00050.t01#4    mNeoVis.2.1.1.chr15_hap1.g00050.t01#5   mNeoVis.evergladensis.1.1.chr15.g00564.t01#3       6458   125254790:   mNeoVis.evergladensis.1.1.chr6.g00190.t01#3    mNeoVis.vulgivaga.1.1.chr6.g00191.t01#4    mNeoVis.3.1.1.chr6.g00189.t01#6    mNeoVis.evergladensis.2.1.chr6.g00188.t01#2    mNeoVis.vulgivaga.2.1.chr6.g00188.t01#1   mNeoVis.2.1.1.chr4_hap1.g00187.t01#5       6459   125254791:   mNeoVis.vulgivaga.2.1.chr15.g00053.t01#1    mNeoVis.2.1.1.chr15_hap1.g00051.t01#5    mNeoVis.evergladensis.2.1.chr15.g00051.t01#2    mNeoVis.3.1.1.chr15.g00048.t01#6    mNeoVis.vulgivaga.1.1.chr15.g00051.t01#4   mNeoVis.evergladensis.1.1.chr15.g00563.t01#3       6460   125254792:   mNeoVis.evergladensis.1.1.chr13.g00708.t01#3    mNeoVis.vulgivaga.1.1.chr13.g00706.t01#4    mNeoVis.2.1.1.chr13_hap1.g00750.t01#5    mNeoVis.evergladensis.2.1.chr13.g00715.t01#2    mNeoVis.vulgivaga.2.1.chr13.g00713.t01#1   mNeoVis.3.1.1.chr13.g00668.t01#6       6461   125254793:   mNeoVis.2.1.1.chr4_hap1.g01089.t01#5    mNeoVis.evergladensis.2.1.chr6.g01086.t01#2    mNeoVis.vulgivaga.2.1.chr6.g01079.t01#1    mNeoVis.vulgivaga.1.1.chr6.g01080.t01#4   mNeoVis.evergladensis.1.1.chr6.g01091.t01#3       6462   125254794:   mNeoVis.evergladensis.1.1.chr6.g00191.t01#3    mNeoVis.2.1.1.chr4_hap1.g00188.t01#5    mNeoVis.3.1.1.chr6.g00190.t01#6    mNeoVis.vulgivaga.2.1.chr6.g00189.t01#1    mNeoVis.evergladensis.2.1.chr6.g00189.t01#2   mNeoVis.vulgivaga.1.1.chr6.g00192.t01#4       6463   125254795:   mNeoVis.vulgivaga.1.1.chr6.g00193.t01#4    mNeoVis.vulgivaga.2.1.chr6.g00190.t01#1    mNeoVis.evergladensis.1.1.chr6.g00192.t01#3   mNeoVis.2.1.1.chr4_hap1.g00189.t01#5         6464   125254796:   mNeoVis.vulgivaga.1.1.chr13.g00707.t01#4    mNeoVis.vulgivaga.2.1.chr13.g00714.t01#1    mNeoVis.2.1.1.chr13_hap1.g00751.t01#5    mNeoVis.3.1.1.chr13.g00669.t01#6    mNeoVis.evergladensis.1.1.chr13.g00709.t01#3   mNeoVis.evergladensis.2.1.chr13.g00716.t01#2       6465   125254797:   mNeoVis.vulgivaga.2.1.chr6.g00191.t01#1    mNeoVis.3.1.1.chr6.g00191.t01#6    mNeoVis.vulgivaga.1.1.chr6.g00195.t01#4    mNeoVis.2.1.1.chr4_hap1.g00191.t01#5    mNeoVis.evergladensis.1.1.chr6.g00193.t01#3   mNeoVis.evergladensis.2.1.chr6.g00190.t01#2       6466   125254798:   mNeoVis.evergladensis.2.1.chr13.g00632.t01#2    mNeoVis.3.1.1.chr13.g00584.t01#6    mNeoVis.2.1.1.chr13_hap1.g00659.t01#5    mNeoVis.evergladensis.1.1.chr13.g00631.t01#3    mNeoVis.vulgivaga.2.1.chr13.g00635.t01#1   mNeoVis.vulgivaga.1.1.chr13.g00620.t01#4       6467   125254799:   mNeoVis.vulgivaga.1.1.chr6.g01081.t01#4    mNeoVis.2.1.1.chr4_hap1.g01090.t01#5    mNeoVis.evergladensis.1.1.chr6.g01092.t01#3    mNeoVis.vulgivaga.2.1.chr6.g01080.t01#1   mNeoVis.evergladensis.2.1.chr6.g01087.t01#2       6468   125254800:   mNeoVis.evergladensis.1.1.chr6.g00194.t01#3    mNeoVis.evergladensis.2.1.chr6.g00191.t01#2    mNeoVis.vulgivaga.1.1.chr6.g00196.t01#4    mNeoVis.3.1.1.chr6.g00192.t01#6    mNeoVis.vulgivaga.2.1.chr6.g00192.t01#1   mNeoVis.2.1.1.chr4_hap1.g00192.t01#5       6469   125254801:   mNeoVis.vulgivaga.1.1.chr3.g01048.t01#4    mNeoVis.3.1.1.chr3.g01033.t01#6    mNeoVis.evergladensis.2.1.chr3.g01044.t01#2    mNeoVis.evergladensis.1.1.chr3.g01055.t01#3    mNeoVis.2.1.1.chr3_hap1.g01045.t01#5   mNeoVis.vulgivaga.2.1.chr3.g01062.t01#1       6470   125254802:   mNeoVis.2.1.1.chr4_hap1.g00193.t01#5    mNeoVis.evergladensis.1.1.chr6.g00195.t01#3    mNeoVis.vulgivaga.1.1.chr6.g00197.t01#4    mNeoVis.evergladensis.2.1.chr6.g00192.t01#2    mNeoVis.3.1.1.chr6.g00193.t01#6   mNeoVis.vulgivaga.2.1.chr6.g00193.t01#1       6471   125254803:   mNeoVis.2.1.1.chr13_hap1.g00660.t01#5    mNeoVis.evergladensis.2.1.chr13.g00633.t01#2    mNeoVis.3.1.1.chr13.g00585.t01#6    mNeoVis.vulgivaga.2.1.chr13.g00636.t01#1    mNeoVis.evergladensis.1.1.chr13.g00632.t01#3   mNeoVis.vulgivaga.1.1.chr13.g00621.t01#4       6472   125254804:   mNeoVis.evergladensis.2.1.chr13.g00634.t01#2    mNeoVis.vulgivaga.2.1.chr13.g00637.t01#1    mNeoVis.2.1.1.chr13_hap1.g00661.t01#5    mNeoVis.3.1.1.chr13.g00586.t01#6    mNeoVis.evergladensis.1.1.chr13.g00633.t01#3   mNeoVis.vulgivaga.1.1.chr13.g00622.t01#4       6473   125254805:   mNeoVis.vulgivaga.2.1.chr6.g00194.t01#1   mNeoVis.3.1.1.chr6.g00194.t01#6             6474   125254806:   mNeoVis.evergladensis.1.1.chr13.g00634.t01#3    mNeoVis.vulgivaga.2.1.chr13.g00638.t01#1    mNeoVis.vulgivaga.1.1.chr13.g00623.t01#4    mNeoVis.evergladensis.2.1.chr13.g00635.t01#2    mNeoVis.2.1.1.chr13_hap1.g00662.t01#5   mNeoVis.3.1.1.chr13.g00587.t01#6       6475   125254807:   mNeoVis.vulgivaga.2.1.chr6.g00195.t01#1    mNeoVis.3.1.1.chr6.g00195.t01#6    mNeoVis.vulgivaga.1.1.chr6.g00201.t01#4    mNeoVis.vulgivaga.1.1.chr6.g00200.t01#4    mNeoVis.evergladensis.2.1.chr6.g00194.t01#2    mNeoVis.2.1.1.chr4_hap1.g00195.t01#5   mNeoVis.evergladensis.1.1.chr6.g00197.t01#3     6476   125254808:   mNeoVis.2.1.1.chr15_hap1.g00052.t01#5    mNeoVis.evergladensis.1.1.chr15.g00562.t01#3    mNeoVis.vulgivaga.2.1.chr15.g00054.t01#1    mNeoVis.evergladensis.2.1.chr15.g00052.t01#2    mNeoVis.vulgivaga.1.1.chr15.g00052.t01#4   mNeoVis.3.1.1.chr15.g00049.t01#6       6477   125254809:   mNeoVis.evergladensis.2.1.chr13.g00636.t01#2    mNeoVis.2.1.1.chr13_hap1.g00663.t01#5    mNeoVis.3.1.1.chr13.g00588.t01#6    mNeoVis.evergladensis.1.1.chr13.g00635.t01#3    mNeoVis.vulgivaga.2.1.chr13.g00639.t01#1   mNeoVis.vulgivaga.1.1.chr13.g00624.t01#4       6478   125254810:   mNeoVis.3.1.1.chr6.g00197.t01#6    mNeoVis.2.1.1.chr4_hap1.g00196.t01#5    mNeoVis.evergladensis.1.1.chr6.g00198.t01#3    mNeoVis.evergladensis.2.1.chr6.g00195.t01#2    mNeoVis.vulgivaga.2.1.chr6.g00196.t01#1   mNeoVis.vulgivaga.1.1.chr6.g00203.t01#4       6479   125254811:   mNeoVis.evergladensis.2.1.chr13.g00637.t01#2    mNeoVis.evergladensis.1.1.chr13.g00636.t01#3    mNeoVis.3.1.1.chr13.g00589.t01#6    mNeoVis.vulgivaga.2.1.chr13.g00640.t01#1    mNeoVis.vulgivaga.1.1.chr13.g00625.t01#4   mNeoVis.2.1.1.chr13_hap1.g00664.t01#5       6480   125254812:   mNeoVis.evergladensis.1.1.chr6.g00199.t01#3    mNeoVis.2.1.1.chr4_hap1.g00197.t01#5    mNeoVis.3.1.1.chr6.g00198.t01#6    mNeoVis.vulgivaga.2.1.chr6.g00197.t01#1    mNeoVis.evergladensis.2.1.chr6.g00196.t01#2   mNeoVis.vulgivaga.1.1.chr6.g00204.t01#4       6481   125254813:   mNeoVis.evergladensis.1.1.chr13.g00637.t01#3    mNeoVis.vulgivaga.2.1.chr13.g00641.t01#1    mNeoVis.3.1.1.chr13.g00590.t01#6    mNeoVis.vulgivaga.1.1.chr13.g00626.t01#4    mNeoVis.evergladensis.2.1.chr13.g00638.t01#2   mNeoVis.2.1.1.chr13_hap1.g00665.t01#5       6482   125254814:   mNeoVis.vulgivaga.1.1.chr15.g00053.t01#4    mNeoVis.vulgivaga.2.1.chr15.g00055.t01#1    mNeoVis.2.1.1.chr15_hap1.g00053.t01#5    mNeoVis.evergladensis.1.1.chr15.g00561.t01#3    mNeoVis.3.1.1.chr15.g00050.t01#6   mNeoVis.evergladensis.2.1.chr15.g00053.t01#2       6483   125254815:   mNeoVis.vulgivaga.1.1.chr6.g01082.t01#4    mNeoVis.2.1.1.chr4_hap1.g01091.t01#5    mNeoVis.evergladensis.1.1.chr6.g01093.t01#3    mNeoVis.vulgivaga.2.1.chr6.g01081.t01#1   mNeoVis.evergladensis.2.1.chr6.g01088.t01#2       6484   125254816:  mNeoVis.vulgivaga.2.1.chr6.g00198.t01#1               6485   125254817:  mNeoVis.vulgivaga.2.1.chr13.g00643.t01#1               6486   125254818:   mNeoVis.evergladensis.1.1.chr13.g00638.t01#3    mNeoVis.vulgivaga.1.1.chr13.g00628.t01#4    mNeoVis.3.1.1.chr13.g00591.t01#6    mNeoVis.vulgivaga.2.1.chr13.g00642.t01#1    mNeoVis.evergladensis.2.1.chr13.g00639.t01#2   mNeoVis.2.1.1.chr13_hap1.g00666.t01#5       6487   125254819:   mNeoVis.vulgivaga.1.1.chr6.g00206.t01#4    mNeoVis.3.1.1.chr6.g00200.t01#6    mNeoVis.evergladensis.2.1.chr6.g00198.t01#2    mNeoVis.evergladensis.1.1.chr6.g00201.t01#3    mNeoVis.2.1.1.chr4_hap1.g00199.t01#5   mNeoVis.vulgivaga.2.1.chr6.g00199.t01#1       6488   125254820:  mNeoVis.vulgivaga.2.1.chr13.g00645.t01#1               6489   125254821:   mNeoVis.3.1.1.chr13.g00594.t01#6    mNeoVis.evergladensis.1.1.chr13.g00640.t01#3    mNeoVis.vulgivaga.1.1.chr13.g00630.t01#4    mNeoVis.evergladensis.2.1.chr13.g00641.t01#2    mNeoVis.vulgivaga.2.1.chr13.g00644.t01#1   mNeoVis.2.1.1.chr13_hap1.g00668.t01#5       6490   125254822:   mNeoVis.3.1.1.chr13.g00593.t01#6    mNeoVis.evergladensis.2.1.chr13.g00640.t01#2    mNeoVis.evergladensis.1.1.chr13.g00639.t01#3    mNeoVis.vulgivaga.1.1.chr13.g00629.t01#4   mNeoVis.2.1.1.chr13_hap1.g00667.t01#5       6491   125254823:   mNeoVis.vulgivaga.1.1.chr15.g00054.t01#4    mNeoVis.evergladensis.1.1.chr15.g00560.t01#3    mNeoVis.2.1.1.chr15_hap1.g00054.t01#5    mNeoVis.evergladensis.2.1.chr15.g00054.t01#2   mNeoVis.vulgivaga.2.1.chr15.g00056.t01#1       6492   125254824:   mNeoVis.2.1.1.chr4_hap1.g00200.t01#5    mNeoVis.3.1.1.chr6.g00201.t01#6   mNeoVis.vulgivaga.2.1.chr6.g00200.t01#1           6493   125254825:   mNeoVis.vulgivaga.2.1.chr6.g00201.t01#1    mNeoVis.evergladensis.1.1.chr6.g00202.t01#3    mNeoVis.2.1.1.chr4_hap1.g00201.t01#5    mNeoVis.vulgivaga.1.1.chr6.g00208.t01#4   mNeoVis.evergladensis.2.1.chr6.g00200.t01#2       6494   125254826:  mNeoVis.vulgivaga.2.1.chr13.g00646.t01#1               6495   125254827:   mNeoVis.2.1.1.chr4_hap1.g00202.t01#5    mNeoVis.vulgivaga.2.1.chr6.g00202.t01#1    mNeoVis.vulgivaga.1.1.chr6.g00209.t01#4    mNeoVis.evergladensis.1.1.chr6.g00203.t01#3    mNeoVis.evergladensis.2.1.chr6.g00201.t01#2   mNeoVis.3.1.1.chr6.g00202.t01#6       6496   125254828:   mNeoVis.3.1.1.chr13.g00595.t01#6    mNeoVis.evergladensis.2.1.chr13.g00644.t01#2    mNeoVis.evergladensis.1.1.chr13.g00642.t01#3    mNeoVis.2.1.1.chr13_hap1.g00672.t01#5   mNeoVis.vulgivaga.2.1.chr13.g00647.t01#1       6497   125254829:   mNeoVis.vulgivaga.2.1.chr15.g00057.t01#1    mNeoVis.vulgivaga.1.1.chr15.g00055.t01#4    mNeoVis.2.1.1.chr15_hap1.g00055.t01#5    mNeoVis.evergladensis.1.1.chr15.g00559.t01#3    mNeoVis.3.1.1.chr15.g00051.t01#6   mNeoVis.evergladensis.2.1.chr15.g00055.t01#2       6498   125254830:   mNeoVis.2.1.1.chr4_hap1.g00203.t01#5    mNeoVis.vulgivaga.1.1.chr6.g00210.t01#4    mNeoVis.vulgivaga.2.1.chr6.g00203.t01#1    mNeoVis.evergladensis.1.1.chr6.g00204.t01#3    mNeoVis.evergladensis.2.1.chr6.g00202.t01#2   mNeoVis.3.1.1.chr6.g00203.t01#6       6499   125254831:   mNeoVis.3.1.1.chr2.g00843.t01#6    mNeoVis.evergladensis.1.1.chr2.g00926.t01#3    mNeoVis.vulgivaga.2.1.chr2.g00932.t01#1    mNeoVis.evergladensis.2.1.chr2.g00928.t01#2    mNeoVis.vulgivaga.1.1.chr2.g00932.t01#4   mNeoVis.2.1.1.chr5_hap1.g00931.t01#5       6500   125254832:   mNeoVis.2.1.1.chr13_hap1.g00673.t01#5    mNeoVis.vulgivaga.2.1.chr13.g00648.t01#1    mNeoVis.evergladensis.2.1.chr13.g00645.t01#2    mNeoVis.evergladensis.1.1.chr13.g00643.t01#3   mNeoVis.3.1.1.chr13.g00596.t01#6       6501   125254833:   mNeoVis.3.1.1.chr6.g00204.t01#6    mNeoVis.evergladensis.2.1.chr6.g00203.t01#2    mNeoVis.2.1.1.chr4_hap1.g00204.t01#5    mNeoVis.evergladensis.1.1.chr6.g00205.t01#3    mNeoVis.vulgivaga.1.1.chr6.g00211.t01#4   mNeoVis.vulgivaga.2.1.chr6.g00204.t01#1       6502   125254834:   mNeoVis.evergladensis.2.1.chr13.g00646.t01#2    mNeoVis.vulgivaga.1.1.chr13.g00637.t01#4    mNeoVis.3.1.1.chr13.g00597.t01#6    mNeoVis.evergladensis.1.1.chr13.g00644.t01#3    mNeoVis.2.1.1.chr13_hap1.g00674.t01#5   mNeoVis.vulgivaga.2.1.chr13.g00649.t01#1       6503   125254835:   mNeoVis.2.1.1.chr13_hap1.g00675.t01#5    mNeoVis.vulgivaga.1.1.chr13.g00638.t01#4    mNeoVis.vulgivaga.2.1.chr13.g00650.t01#1   mNeoVis.3.1.1.chr13.g00598.t01#6         6504   125254836:   mNeoVis.vulgivaga.2.1.chr6.g00205.t01#1    mNeoVis.2.1.1.chr4_hap1.g00205.t01#5    mNeoVis.evergladensis.1.1.chr6.g00206.t01#3    mNeoVis.evergladensis.2.1.chr6.g00204.t01#2    mNeoVis.vulgivaga.1.1.chr6.g00212.t01#4   mNeoVis.3.1.1.chr6.g00205.t01#6       6505   125254837:   mNeoVis.vulgivaga.1.1.chr13.g00639.t01#4    mNeoVis.3.1.1.chr13.g00599.t01#6    mNeoVis.2.1.1.chr13_hap1.g00676.t01#5    mNeoVis.vulgivaga.2.1.chr13.g00651.t01#1    mNeoVis.evergladensis.1.1.chr13.g00645.t01#3   mNeoVis.evergladensis.2.1.chr13.g00647.t01#2       6506   125254838:   mNeoVis.evergladensis.1.1.chr15.g00557.t01#3    mNeoVis.2.1.1.chr15_hap1.g00056.t01#5    mNeoVis.vulgivaga.1.1.chr15.g00057.t01#4    mNeoVis.evergladensis.2.1.chr15.g00056.t01#2    mNeoVis.3.1.1.chr15.g00052.t01#6   mNeoVis.vulgivaga.2.1.chr15.g00058.t01#1       6507   125254839:   mNeoVis.3.1.1.chr6.g00206.t01#6    mNeoVis.vulgivaga.2.1.chr6.g00206.t01#1    mNeoVis.evergladensis.1.1.chr6.g00207.t01#3    mNeoVis.2.1.1.chr4_hap1.g00206.t01#5    mNeoVis.evergladensis.2.1.chr6.g00205.t01#2   mNeoVis.vulgivaga.1.1.chr6.g00213.t01#4       6508   125254840:   mNeoVis.evergladensis.2.1.chr6.g00206.t01#2    mNeoVis.vulgivaga.1.1.chr6.g00214.t01#4    mNeoVis.evergladensis.1.1.chr6.g00208.t01#3    mNeoVis.3.1.1.chr6.g00207.t01#6    mNeoVis.vulgivaga.2.1.chr6.g00207.t01#1   mNeoVis.2.1.1.chr4_hap1.g00207.t01#5       6509   125254841:   mNeoVis.evergladensis.1.1.chr6.g00209.t01#3    mNeoVis.vulgivaga.2.1.chr6.g00208.t01#1    mNeoVis.3.1.1.chr6.g00208.t01#6    mNeoVis.2.1.1.chr4_hap1.g00208.t01#5    mNeoVis.vulgivaga.1.1.chr6.g00215.t01#4   mNeoVis.evergladensis.2.1.chr6.g00207.t01#2       6510   125254842:   mNeoVis.evergladensis.1.1.chr13.g00646.t01#3    mNeoVis.evergladensis.2.1.chr13.g00648.t01#2    mNeoVis.vulgivaga.2.1.chr13.g00652.t01#1    mNeoVis.vulgivaga.1.1.chr13.g00640.t01#4    mNeoVis.2.1.1.chr13_hap1.g00677.t01#5   mNeoVis.3.1.1.chr13.g00600.t01#6       6511   125254843:   mNeoVis.vulgivaga.2.1.chr5.g00166.t01#1    mNeoVis.2.1.1.chr2_hap1.g00166.t01#5    mNeoVis.evergladensis.1.1.chr5.g00171.t01#3   mNeoVis.evergladensis.2.1.chr5.g00168.t01#2       6512   125254844:   mNeoVis.evergladensis.1.1.chr13.g00648.t01#3    mNeoVis.vulgivaga.2.1.chr13.g00653.t01#1    mNeoVis.2.1.1.chr13_hap1.g00679.t01#5    mNeoVis.evergladensis.2.1.chr13.g00650.t01#2    mNeoVis.3.1.1.chr13.g00602.t01#6   mNeoVis.vulgivaga.1.1.chr13.g00642.t01#4       6513   125254845:   mNeoVis.3.1.1.chr13.g00603.t01#6    mNeoVis.vulgivaga.2.1.chr13.g00654.t01#1    mNeoVis.vulgivaga.1.1.chr13.g00643.t01#4    mNeoVis.evergladensis.1.1.chr13.g00649.t01#3    mNeoVis.evergladensis.2.1.chr13.g00651.t01#2   mNeoVis.2.1.1.chr13_hap1.g00680.t01#5       6514   125254846:   mNeoVis.vulgivaga.1.1.chr6.g00216.t01#4    mNeoVis.evergladensis.1.1.chr6.g00210.t01#3    mNeoVis.vulgivaga.2.1.chr6.g00209.t01#1    mNeoVis.3.1.1.chr6.g00209.t01#6    mNeoVis.2.1.1.chr4_hap1.g00209.t01#5   mNeoVis.evergladensis.2.1.chr6.g00208.t01#2       6515   125254847:   mNeoVis.vulgivaga.1.1.chr6.g01083.t01#4    mNeoVis.2.1.1.chr4_hap1.g01092.t01#5    mNeoVis.evergladensis.1.1.chr6.g01094.t01#3    mNeoVis.vulgivaga.2.1.chr6.g01082.t01#1   mNeoVis.evergladensis.2.1.chr6.g01089.t01#2       6516   125254848:   mNeoVis.3.1.1.chr6.g00210.t01#6    mNeoVis.vulgivaga.1.1.chr6.g00217.t01#4    mNeoVis.2.1.1.chr4_hap1.g00210.t01#5    mNeoVis.vulgivaga.2.1.chr6.g00210.t01#1    mNeoVis.evergladensis.2.1.chr6.g00209.t01#2   mNeoVis.evergladensis.1.1.chr6.g00211.t01#3       6517   125254849:   mNeoVis.vulgivaga.2.1.chr13.g00655.t01#1    mNeoVis.vulgivaga.1.1.chr13.g00644.t01#4    mNeoVis.evergladensis.2.1.chr13.g00652.t01#2    mNeoVis.evergladensis.1.1.chr13.g00650.t01#3    mNeoVis.3.1.1.chr13.g00604.t01#6   mNeoVis.2.1.1.chr13_hap1.g00681.t01#5       6518   125254850:   mNeoVis.2.1.1.chr4_hap1.g00211.t01#5    mNeoVis.vulgivaga.1.1.chr6.g00218.t01#4    mNeoVis.evergladensis.2.1.chr6.g00210.t01#2    mNeoVis.evergladensis.1.1.chr6.g00212.t01#3    mNeoVis.vulgivaga.2.1.chr6.g00211.t01#1   mNeoVis.3.1.1.chr6.g00211.t01#6       6519   125254851:   mNeoVis.2.1.1.chr4_hap1.g00212.t01#5    mNeoVis.vulgivaga.1.1.chr6.g00219.t01#4    mNeoVis.vulgivaga.2.1.chr6.g00212.t01#1    mNeoVis.evergladensis.1.1.chr6.g00213.t01#3    mNeoVis.evergladensis.2.1.chr6.g00211.t01#2   mNeoVis.3.1.1.chr6.g00212.t01#6       6520   125254852:   mNeoVis.3.1.1.chr6.g00213.t01#6    mNeoVis.vulgivaga.1.1.chr6.g00220.t01#4    mNeoVis.vulgivaga.2.1.chr6.g00213.t01#1    mNeoVis.2.1.1.chr4_hap1.g00213.t01#5    mNeoVis.evergladensis.1.1.chr6.g00214.t01#3   mNeoVis.evergladensis.2.1.chr6.g00212.t01#2       6521   125254853:   mNeoVis.evergladensis.2.1.chr13.g00654.t01#2    mNeoVis.vulgivaga.2.1.chr13.g00656.t01#1    mNeoVis.vulgivaga.1.1.chr13.g00647.t01#4    mNeoVis.3.1.1.chr13.g00605.t01#6    mNeoVis.evergladensis.1.1.chr13.g00652.t01#3   mNeoVis.2.1.1.chr13_hap1.g00683.t01#5       6522   125254854:   mNeoVis.vulgivaga.2.1.chr6.g00127.t01#1    mNeoVis.evergladensis.2.1.chr6.g00129.t01#2    mNeoVis.2.1.1.chr4_hap1.g00127.t01#5    mNeoVis.vulgivaga.1.1.chr6.g00130.t01#4    mNeoVis.evergladensis.1.1.chr6.g00131.t01#3   mNeoVis.3.1.1.chr6.g00127.t01#6       6523   125254855:  mNeoVis.vulgivaga.2.1.chr6.g01083.t01#1               6524   125254856:   mNeoVis.evergladensis.1.1.chr13.g00653.t01#3    mNeoVis.vulgivaga.2.1.chr13.g00657.t01#1   mNeoVis.2.1.1.chr13_hap1.g00684.t01#5           6525   125254857:   mNeoVis.vulgivaga.2.1.chr13.g00658.t01#1    mNeoVis.3.1.1.chr13.g00606.t01#6    mNeoVis.2.1.1.chr13_hap1.g00685.t01#5    mNeoVis.evergladensis.2.1.chr13.g00655.t01#2    mNeoVis.evergladensis.1.1.chr13.g00654.t01#3   mNeoVis.vulgivaga.1.1.chr13.g00648.t01#4       6526   125254858:   mNeoVis.2.1.1.chr13_hap1.g00686.t01#5    mNeoVis.vulgivaga.2.1.chr13.g00659.t01#1    mNeoVis.vulgivaga.1.1.chr13.g00649.t01#4    mNeoVis.evergladensis.1.1.chr13.g00655.t01#3    mNeoVis.evergladensis.2.1.chr13.g00656.t01#2   mNeoVis.3.1.1.chr13.g00607.t01#6       6527   125254859:   mNeoVis.3.1.1.chr13.g00608.t01#6    mNeoVis.vulgivaga.1.1.chr13.g00651.t01#4    mNeoVis.vulgivaga.2.1.chr13.g00660.t01#1    mNeoVis.2.1.1.chr13_hap1.g00687.t01#5    mNeoVis.evergladensis.1.1.chr13.g00656.t01#3   mNeoVis.evergladensis.2.1.chr13.g00657.t01#2       6528   125254860:   mNeoVis.vulgivaga.2.1.chr6.g00128.t01#1    mNeoVis.evergladensis.2.1.chr6.g00130.t01#2    mNeoVis.2.1.1.chr4_hap1.g00128.t01#5    mNeoVis.evergladensis.1.1.chr6.g00132.t01#3   mNeoVis.vulgivaga.1.1.chr6.g00131.t01#4       6529   125254861:   mNeoVis.evergladensis.2.1.chr6.g00131.t01#2    mNeoVis.evergladensis.1.1.chr6.g00133.t01#3    mNeoVis.3.1.1.chr6.g00129.t01#6    mNeoVis.vulgivaga.2.1.chr6.g00129.t01#1    mNeoVis.vulgivaga.1.1.chr6.g00132.t01#4   mNeoVis.2.1.1.chr4_hap1.g00129.t01#5       6530   125254862:   mNeoVis.3.1.1.chr13.g00609.t01#6    mNeoVis.evergladensis.2.1.chr13.g00658.t01#2    mNeoVis.vulgivaga.2.1.chr13.g00661.t01#1    mNeoVis.evergladensis.1.1.chr13.g00658.t01#3    mNeoVis.2.1.1.chr13_hap1.g00688.t01#5   mNeoVis.vulgivaga.1.1.chr13.g00652.t01#4       6531   125254863:   mNeoVis.evergladensis.1.1.chr6.g00135.t01#3    mNeoVis.vulgivaga.2.1.chr6.g00130.t01#1    mNeoVis.3.1.1.chr6.g00130.t01#6    mNeoVis.vulgivaga.1.1.chr6.g00133.t01#4    mNeoVis.evergladensis.1.1.chr6.g00134.t01#3    mNeoVis.vulgivaga.2.1.chr6.g00131.t01#1    mNeoVis.2.1.1.chr4_hap1.g00130.t01#5    mNeoVis.evergladensis.2.1.chr6.g00133.t01#2    mNeoVis.evergladensis.2.1.chr6.g00132.t01#2      6532   125254864:   mNeoVis.3.1.1.chr6.g00132.t01#6    mNeoVis.vulgivaga.2.1.chr6.g00133.t01#1    mNeoVis.2.1.1.chr4_hap1.g00132.t01#5    mNeoVis.evergladensis.2.1.chr6.g00135.t01#2    mNeoVis.evergladensis.2.1.chr6.g00134.t01#2    mNeoVis.2.1.1.chr4_hap1.g00133.t01#5    mNeoVis.vulgivaga.1.1.chr6.g00135.t01#4    mNeoVis.3.1.1.chr6.g00131.t01#6    mNeoVis.vulgivaga.2.1.chr6.g00132.t01#1      6533   125254865:   mNeoVis.evergladensis.1.1.chr6.g00116.t01#3    mNeoVis.2.1.1.chr4_hap1.g00112.t01#5    mNeoVis.vulgivaga.2.1.chr6.g00111.t01#1    mNeoVis.evergladensis.2.1.chr6.g00114.t01#2    mNeoVis.2.1.1.chr4_hap1.g00113.t01#5    mNeoVis.3.1.1.chr6.g00112.t01#6    mNeoVis.vulgivaga.2.1.chr6.g00112.t01#1    mNeoVis.vulgivaga.1.1.chr6.g00117.t01#4    mNeoVis.evergladensis.2.1.chr6.g00115.t01#2      6534   125254866:  mNeoVis.vulgivaga.1.1.chr6.g00134.t01#4               6535   125254867:   mNeoVis.2.1.1.chr15_hap1.g00057.t01#5    mNeoVis.vulgivaga.1.1.chr15.g00058.t01#4    mNeoVis.evergladensis.2.1.chr15.g00057.t01#2    mNeoVis.evergladensis.1.1.chr15.g00556.t01#3    mNeoVis.3.1.1.chr15.g00053.t01#6   mNeoVis.vulgivaga.2.1.chr15.g00059.t01#1       6536   125254868:   mNeoVis.evergladensis.2.1.chr6.g01090.t01#2    mNeoVis.2.1.1.chr4_hap1.g01093.t01#5    mNeoVis.vulgivaga.1.1.chr6.g01084.t01#4   mNeoVis.vulgivaga.2.1.chr6.g01084.t01#1         6537   125254869:   mNeoVis.vulgivaga.1.1.chr6.g00136.t01#4    mNeoVis.3.1.1.chr6.g00133.t01#6    mNeoVis.evergladensis.1.1.chr6.g00137.t01#3    mNeoVis.evergladensis.2.1.chr6.g00136.t01#2    mNeoVis.vulgivaga.2.1.chr6.g00134.t01#1   mNeoVis.2.1.1.chr4_hap1.g00134.t01#5       6538   125254870:   mNeoVis.vulgivaga.1.1.chr6.g00137.t01#4    mNeoVis.3.1.1.chr6.g00134.t01#6    mNeoVis.evergladensis.2.1.chr6.g00137.t01#2    mNeoVis.vulgivaga.2.1.chr6.g00135.t01#1    mNeoVis.2.1.1.chr4_hap1.g00135.t01#5   mNeoVis.evergladensis.1.1.chr6.g00138.t01#3       6539   125254871:   mNeoVis.evergladensis.2.1.chr13.g00659.t01#2    mNeoVis.evergladensis.1.1.chr13.g00659.t01#3    mNeoVis.vulgivaga.1.1.chr13.g00653.t01#4    mNeoVis.3.1.1.chr13.g00610.t01#6    mNeoVis.vulgivaga.2.1.chr13.g00662.t01#1   mNeoVis.2.1.1.chr13_hap1.g00689.t01#5       6540   125254872:   mNeoVis.evergladensis.1.1.chr6.g00139.t01#3    mNeoVis.evergladensis.2.1.chr6.g00138.t01#2    mNeoVis.3.1.1.chr6.g00135.t01#6    mNeoVis.vulgivaga.2.1.chr6.g00136.t01#1    mNeoVis.2.1.1.chr4_hap1.g00136.t01#5   mNeoVis.vulgivaga.1.1.chr6.g00138.t01#4       6541   125254873:   mNeoVis.evergladensis.2.1.chr6.g00140.t01#2    mNeoVis.vulgivaga.1.1.chr6.g00140.t01#4    mNeoVis.vulgivaga.2.1.chr6.g00137.t01#1    mNeoVis.evergladensis.1.1.chr6.g00140.t01#3    mNeoVis.3.1.1.chr6.g00138.t01#6   mNeoVis.2.1.1.chr4_hap1.g00138.t01#5       6542   125254874:   mNeoVis.2.1.1.chr4_hap1.g00139.t01#5    mNeoVis.evergladensis.1.1.chr6.g00141.t01#3    mNeoVis.evergladensis.2.1.chr6.g00141.t01#2    mNeoVis.vulgivaga.1.1.chr6.g00141.t01#4    mNeoVis.vulgivaga.2.1.chr6.g00138.t01#1   mNeoVis.3.1.1.chr6.g00137.t01#6       6543   125254875:   mNeoVis.evergladensis.1.1.chr13.g00660.t01#3    mNeoVis.vulgivaga.2.1.chr13.g00663.t01#1    mNeoVis.2.1.1.chr13_hap1.g00690.t01#5    mNeoVis.vulgivaga.1.1.chr13.g00654.t01#4   mNeoVis.evergladensis.2.1.chr13.g00660.t01#2       6544   125254876:   mNeoVis.3.1.1.chr6.g00139.t01#6    mNeoVis.2.1.1.chr4_hap1.g00140.t01#5    mNeoVis.vulgivaga.2.1.chr6.g00139.t01#1    mNeoVis.evergladensis.1.1.chr6.g00142.t01#3    mNeoVis.evergladensis.2.1.chr6.g00142.t01#2   mNeoVis.vulgivaga.1.1.chr6.g00142.t01#4       6545   125254877:   mNeoVis.3.1.1.chr13.g00612.t01#6    mNeoVis.evergladensis.1.1.chr13.g00661.t01#3    mNeoVis.2.1.1.chr13_hap1.g00691.t01#5    mNeoVis.vulgivaga.1.1.chr13.g00655.t01#4    mNeoVis.vulgivaga.2.1.chr13.g00664.t01#1   mNeoVis.evergladensis.2.1.chr13.g00661.t01#2       6546   125254878:   mNeoVis.vulgivaga.1.1.chr6.g00143.t01#4    mNeoVis.evergladensis.2.1.chr6.g00143.t01#2    mNeoVis.3.1.1.chr6.g00140.t01#6    mNeoVis.vulgivaga.2.1.chr6.g00140.t01#1    mNeoVis.2.1.1.chr4_hap1.g00141.t01#5    mNeoVis.3.1.1.chr6.g00141.t01#6   mNeoVis.evergladensis.1.1.chr6.g00143.t01#3     6547   125254879:   mNeoVis.vulgivaga.1.1.chr13.g00656.t01#4    mNeoVis.vulgivaga.2.1.chr13.g00665.t01#1    mNeoVis.evergladensis.2.1.chr13.g00662.t01#2    mNeoVis.3.1.1.chr13.g00613.t01#6    mNeoVis.2.1.1.chr13_hap1.g00692.t01#5   mNeoVis.evergladensis.1.1.chr13.g00662.t01#3       6548   125254880:   mNeoVis.vulgivaga.1.1.chr6.g00144.t01#4    mNeoVis.vulgivaga.2.1.chr6.g00141.t01#1    mNeoVis.3.1.1.chr6.g00142.t01#6    mNeoVis.evergladensis.1.1.chr6.g00144.t01#3    mNeoVis.2.1.1.chr4_hap1.g00142.t01#5   mNeoVis.evergladensis.2.1.chr6.g00145.t01#2       6549   125254881:   mNeoVis.evergladensis.2.1.chr6.g00146.t01#2    mNeoVis.vulgivaga.1.1.chr6.g00145.t01#4    mNeoVis.3.1.1.chr6.g00143.t01#6    mNeoVis.evergladensis.1.1.chr6.g00145.t01#3    mNeoVis.vulgivaga.2.1.chr6.g00142.t01#1   mNeoVis.2.1.1.chr4_hap1.g00143.t01#5       6550   125254882:   mNeoVis.vulgivaga.2.1.chr13.g00666.t01#1    mNeoVis.3.1.1.chr13.g00614.t01#6    mNeoVis.2.1.1.chr13_hap1.g00693.t01#5    mNeoVis.evergladensis.2.1.chr13.g00663.t01#2   mNeoVis.vulgivaga.1.1.chr13.g00657.t01#4       6551   125254883:   mNeoVis.vulgivaga.2.1.chr13.g00667.t01#1    mNeoVis.vulgivaga.1.1.chr13.g00658.t01#4    mNeoVis.evergladensis.2.1.chr13.g00664.t01#2    mNeoVis.2.1.1.chr13_hap1.g00694.t01#5   mNeoVis.evergladensis.1.1.chr13.g00664.t01#3       6552   125254884:   mNeoVis.3.1.1.chr6.g00144.t01#6    mNeoVis.vulgivaga.2.1.chr6.g00143.t01#1    mNeoVis.evergladensis.2.1.chr6.g00147.t01#2    mNeoVis.2.1.1.chr4_hap1.g00144.t01#5    mNeoVis.vulgivaga.1.1.chr6.g00146.t01#4   mNeoVis.evergladensis.1.1.chr6.g00146.t01#3       6553   125254885:   mNeoVis.vulgivaga.1.1.chr6.g00147.t01#4    mNeoVis.2.1.1.chr4_hap1.g00145.t01#5    mNeoVis.3.1.1.chr6.g00145.t01#6    mNeoVis.evergladensis.1.1.chr6.g00147.t01#3    mNeoVis.evergladensis.2.1.chr6.g00144.t01#2   mNeoVis.vulgivaga.2.1.chr6.g00144.t01#1       6554   125254886:   mNeoVis.3.1.1.chr13.g00618.t01#6    mNeoVis.vulgivaga.1.1.chr13.g00661.t01#4    mNeoVis.evergladensis.1.1.chr13.g00666.t01#3   mNeoVis.2.1.1.chr13_hap1.g00705.t01#5         6555   125254887:   mNeoVis.evergladensis.2.1.chr13.g00668.t01#2    mNeoVis.2.1.1.chr13_hap1.g00699.t01#5   mNeoVis.vulgivaga.2.1.chr13.g00668.t01#1         6556   125254888:   mNeoVis.3.1.1.chr6.g00146.t01#6   mNeoVis.vulgivaga.2.1.chr6.g00145.t01#1             6557   125254889:   mNeoVis.vulgivaga.1.1.chr15.g00059.t01#4    mNeoVis.vulgivaga.2.1.chr15.g00060.t01#1    mNeoVis.evergladensis.2.1.chr15.g00058.t01#2    mNeoVis.3.1.1.chr15.g00054.t01#6    mNeoVis.2.1.1.chr15_hap1.g00058.t01#5   mNeoVis.evergladensis.1.1.chr15.g00555.t01#3       6558   125254890:   mNeoVis.evergladensis.1.1.chr6.g00149.t01#3    mNeoVis.3.1.1.chr6.g00147.t01#6    mNeoVis.2.1.1.chr4_hap1.g00147.t01#5    mNeoVis.vulgivaga.2.1.chr6.g00146.t01#1    mNeoVis.evergladensis.2.1.chr6.g00149.t01#2   mNeoVis.vulgivaga.1.1.chr6.g00149.t01#4       6559   125254891:   mNeoVis.evergladensis.2.1.chr13.g00669.t01#2   mNeoVis.vulgivaga.2.1.chr13.g00670.t01#1           6560   125254892:  mNeoVis.vulgivaga.2.1.chr13.g00671.t01#1               6561   125254893:   mNeoVis.evergladensis.1.1.chr6.g00150.t01#3    mNeoVis.3.1.1.chr6.g00148.t01#6    mNeoVis.2.1.1.chr4_hap1.g00148.t01#5    mNeoVis.evergladensis.2.1.chr6.g00150.t01#2    mNeoVis.vulgivaga.2.1.chr6.g00147.t01#1   mNeoVis.vulgivaga.1.1.chr6.g00150.t01#4       6562   125254894:   mNeoVis.vulgivaga.1.1.chr13.g00663.t01#4   mNeoVis.vulgivaga.2.1.chr13.g00672.t01#1           6563   125254895:   mNeoVis.evergladensis.2.1.chr13.g00671.t01#2    mNeoVis.2.1.1.chr13_hap1.g00708.t01#5   mNeoVis.vulgivaga.2.1.chr13.g00673.t01#1         6564   125254896:   mNeoVis.vulgivaga.2.1.chr13.g00674.t01#1    mNeoVis.3.1.1.chr13.g00623.t01#6    mNeoVis.evergladensis.2.1.chr13.g00672.t01#2   mNeoVis.vulgivaga.1.1.chr13.g00664.t01#4       6565   125254897:   mNeoVis.vulgivaga.1.1.chr13.g00665.t01#4   mNeoVis.vulgivaga.2.1.chr13.g00675.t01#1           6566   125254898:   mNeoVis.3.1.1.chr15.g00055.t01#6    mNeoVis.2.1.1.chr15_hap1.g00059.t01#5    mNeoVis.evergladensis.1.1.chr15.g00554.t01#3    mNeoVis.vulgivaga.2.1.chr15.g00061.t01#1    mNeoVis.vulgivaga.1.1.chr15.g00060.t01#4   mNeoVis.evergladensis.2.1.chr15.g00059.t01#2       6567   125254899:   mNeoVis.2.1.1.chr4_hap1.g00149.t01#5    mNeoVis.vulgivaga.1.1.chr6.g00151.t01#4    mNeoVis.evergladensis.1.1.chr6.g00151.t01#3    mNeoVis.3.1.1.chr6.g00149.t01#6    mNeoVis.vulgivaga.2.1.chr6.g00148.t01#1   mNeoVis.evergladensis.2.1.chr6.g00151.t01#2       6568   125254900:   mNeoVis.2.1.1.chr13_hap1.g00709.t01#5    mNeoVis.evergladensis.2.1.chr13.g00673.t01#2    mNeoVis.vulgivaga.1.1.chr13.g00666.t01#4    mNeoVis.evergladensis.1.1.chr13.g00667.t01#3    mNeoVis.3.1.1.chr13.g00624.t01#6   mNeoVis.vulgivaga.2.1.chr13.g00676.t01#1       6569   125254901:   mNeoVis.2.1.1.chr13_hap1.g00714.t01#5    mNeoVis.vulgivaga.1.1.chr13.g00670.t01#4    mNeoVis.evergladensis.2.1.chr13.g00677.t01#2    mNeoVis.vulgivaga.2.1.chr13.g00678.t01#1    mNeoVis.3.1.1.chr13.g00630.t01#6   mNeoVis.evergladensis.1.1.chr13.g00670.t01#3       6570   125254902:   mNeoVis.2.1.1.chr10_hap1.g01030.t01#5    mNeoVis.3.1.1.chr9.g00963.t01#6    mNeoVis.evergladensis.2.1.chr8.g01003.t01#2    mNeoVis.vulgivaga.2.1.chr9.g00987.t01#1    mNeoVis.evergladensis.1.1.chr9.g00997.t01#3   mNeoVis.vulgivaga.1.1.chr9.g01002.t01#4       6571   125254903:   mNeoVis.evergladensis.2.1.chr6.g00152.t01#2   mNeoVis.vulgivaga.2.1.chr6.g00149.t01#1             6572   125254904:   mNeoVis.vulgivaga.2.1.chr6.g00150.t01#1    mNeoVis.vulgivaga.1.1.chr6.g00153.t01#4    mNeoVis.evergladensis.1.1.chr6.g00153.t01#3    mNeoVis.evergladensis.2.1.chr6.g00153.t01#2    mNeoVis.2.1.1.chr4_hap1.g00151.t01#5   mNeoVis.3.1.1.chr6.g00151.t01#6       6573   125254905:   mNeoVis.2.1.1.chr4_hap1.g01094.t01#5    mNeoVis.evergladensis.2.1.chr6.g01091.t01#2    mNeoVis.evergladensis.1.1.chr6.g01096.t01#3    mNeoVis.vulgivaga.2.1.chr6.g01085.t01#1    mNeoVis.3.1.1.chr6.g01083.t01#6   mNeoVis.vulgivaga.1.1.chr6.g01085.t01#4       6574   125254906:   mNeoVis.vulgivaga.1.1.chr13.g00669.t01#4    mNeoVis.evergladensis.2.1.chr13.g00676.t01#2    mNeoVis.vulgivaga.2.1.chr13.g00677.t01#1   mNeoVis.evergladensis.1.1.chr13.g00668.t01#3       6575   125254907:   mNeoVis.evergladensis.2.1.chr6.g00154.t01#2    mNeoVis.3.1.1.chr6.g00152.t01#6    mNeoVis.evergladensis.1.1.chr6.g00154.t01#3    mNeoVis.2.1.1.chr4_hap1.g00152.t01#5    mNeoVis.vulgivaga.1.1.chr6.g00154.t01#4   mNeoVis.vulgivaga.2.1.chr6.g00151.t01#1       6576   125254908:   mNeoVis.vulgivaga.1.1.chr15.g00061.t01#4    mNeoVis.vulgivaga.2.1.chr15.g00062.t01#1    mNeoVis.evergladensis.1.1.chr15.g00553.t01#3    mNeoVis.evergladensis.2.1.chr15.g00060.t01#2    mNeoVis.3.1.1.chr15.g00056.t01#6   mNeoVis.2.1.1.chr15_hap1.g00060.t01#5       6577   125254909:   mNeoVis.3.1.1.chr6.g00153.t01#6    mNeoVis.evergladensis.1.1.chr6.g00155.t01#3    mNeoVis.evergladensis.2.1.chr6.g00155.t01#2    mNeoVis.2.1.1.chr4_hap1.g00153.t01#5    mNeoVis.vulgivaga.2.1.chr6.g00152.t01#1   mNeoVis.vulgivaga.1.1.chr6.g00155.t01#4       6578   125254910:   mNeoVis.vulgivaga.2.1.chr6.g00153.t01#1    mNeoVis.vulgivaga.1.1.chr6.g00156.t01#4    mNeoVis.3.1.1.chr6.g00154.t01#6    mNeoVis.evergladensis.2.1.chr6.g00156.t01#2    mNeoVis.evergladensis.1.1.chr6.g00156.t01#3   mNeoVis.2.1.1.chr4_hap1.g00154.t01#5       6579   125254911:   mNeoVis.evergladensis.1.1.chr13.g00592.t01#3    mNeoVis.vulgivaga.1.1.chr13.g00580.t01#4    mNeoVis.evergladensis.2.1.chr13.g00596.t01#2    mNeoVis.vulgivaga.2.1.chr13.g00596.t01#1   mNeoVis.2.1.1.chr13_hap1.g00620.t01#5       6580   125254912:   mNeoVis.vulgivaga.1.1.chr13.g00579.t01#4    mNeoVis.evergladensis.2.1.chr8.g01095.t01#2    mNeoVis.vulgivaga.2.1.chr12.g00493.t01#1    mNeoVis.evergladensis.1.1.chr9.g01085.t01#3    mNeoVis.vulgivaga.1.1.chr12.g00490.t01#4    mNeoVis.2.1.1.chr13_hap1.g00619.t01#5    mNeoVis.evergladensis.2.1.chr8.g01097.t01#2    mNeoVis.2.1.1.chr12_hap1.g00537.t01#5    mNeoVis.3.1.1.chr12.g00488.t01#6      6581   125254913:   mNeoVis.evergladensis.2.1.chr13.g00595.t01#2   mNeoVis.vulgivaga.2.1.chr13.g00595.t01#1           6582   125254914:   mNeoVis.evergladensis.1.1.chr7.g02491.t01#3   mNeoVis.evergladensis.2.1.chr7.g02494.t01#2           6583   125254915:  mNeoVis.vulgivaga.2.1.chr6.g00155.t01#1               6584   125254916:   mNeoVis.3.1.1.chr6.g00155.t01#6    mNeoVis.evergladensis.2.1.chr6.g00157.t01#2    mNeoVis.vulgivaga.1.1.chr6.g00157.t01#4    mNeoVis.2.1.1.chr4_hap1.g00155.t01#5    mNeoVis.evergladensis.1.1.chr6.g00157.t01#3   mNeoVis.vulgivaga.2.1.chr6.g00154.t01#1       6585   125254917:   mNeoVis.3.1.1.chr13.g00546.t01#6    mNeoVis.vulgivaga.1.1.chr13.g00581.t01#4    mNeoVis.evergladensis.1.1.chr13.g00593.t01#3    mNeoVis.vulgivaga.2.1.chr13.g00597.t01#1   mNeoVis.2.1.1.chr13_hap1.g00621.t01#5       6586   125254918:   mNeoVis.evergladensis.2.1.chr6.g00158.t01#2    mNeoVis.3.1.1.chr6.g00156.t01#6    mNeoVis.evergladensis.1.1.chr6.g00158.t01#3    mNeoVis.2.1.1.chr4_hap1.g00156.t01#5    mNeoVis.vulgivaga.1.1.chr6.g00160.t01#4   mNeoVis.vulgivaga.2.1.chr6.g00156.t01#1       6587   125254919:   mNeoVis.2.1.1.chr13_hap1.g00622.t01#5    mNeoVis.3.1.1.chr13.g00547.t01#6    mNeoVis.evergladensis.1.1.chr13.g00594.t01#3    mNeoVis.vulgivaga.1.1.chr13.g00582.t01#4    mNeoVis.vulgivaga.2.1.chr13.g00598.t01#1   mNeoVis.evergladensis.2.1.chr13.g00597.t01#2       6588   125254920:  mNeoVis.vulgivaga.2.1.chr6.g00157.t01#1               6589   125254921:   mNeoVis.vulgivaga.1.1.chr13.g00583.t01#4    mNeoVis.evergladensis.1.1.chr13.g00595.t01#3    mNeoVis.evergladensis.2.1.chr13.g00598.t01#2    mNeoVis.2.1.1.chr13_hap1.g00623.t01#5   mNeoVis.vulgivaga.2.1.chr13.g00599.t01#1       6590   125254922:   mNeoVis.evergladensis.1.1.chr6.g00159.t01#3    mNeoVis.vulgivaga.1.1.chr6.g00161.t01#4    mNeoVis.evergladensis.2.1.chr6.g00159.t01#2    mNeoVis.2.1.1.chr4_hap1.g00157.t01#5    mNeoVis.3.1.1.chr6.g00157.t01#6   mNeoVis.vulgivaga.2.1.chr6.g00158.t01#1       6591   125254923:   mNeoVis.vulgivaga.2.1.chr15.g00391.t01#1    mNeoVis.evergladensis.2.1.chr15.g00382.t01#2    mNeoVis.evergladensis.1.1.chr15.g00231.t01#3    mNeoVis.2.1.1.chr15_hap1.g00516.t01#5    mNeoVis.3.1.1.chr15.g00345.t01#6   mNeoVis.vulgivaga.1.1.chr15.g00383.t01#4       6592   125254924:   mNeoVis.2.1.1.chr4_hap1.g00159.t01#5    mNeoVis.vulgivaga.2.1.chr6.g00160.t01#1    mNeoVis.3.1.1.chr6.g00159.t01#6    mNeoVis.evergladensis.1.1.chr6.g00161.t01#3   mNeoVis.evergladensis.2.1.chr6.g00161.t01#2       6593   125254925:   mNeoVis.vulgivaga.1.1.chr6.g00162.t01#4    mNeoVis.vulgivaga.2.1.chr6.g00159.t01#1   mNeoVis.evergladensis.2.1.chr6.g00160.t01#2         6594   125254926:   mNeoVis.evergladensis.1.1.chr6.g00160.t01#3    mNeoVis.2.1.1.chr4_hap1.g00158.t01#5   mNeoVis.3.1.1.chr6.g00158.t01#6           6595   125254927:   mNeoVis.evergladensis.2.1.chr6.g00162.t01#2    mNeoVis.vulgivaga.1.1.chr6.g00163.t01#4    mNeoVis.evergladensis.1.1.chr6.g00162.t01#3    mNeoVis.2.1.1.chr4_hap1.g00160.t01#5    mNeoVis.3.1.1.chr6.g00160.t01#6   mNeoVis.vulgivaga.2.1.chr6.g00161.t01#1       6596   125254928:   mNeoVis.evergladensis.2.1.chr13.g00599.t01#2    mNeoVis.evergladensis.1.1.chr13.g00596.t01#3    mNeoVis.vulgivaga.1.1.chr13.g00584.t01#4    mNeoVis.vulgivaga.2.1.chr13.g00600.t01#1    mNeoVis.2.1.1.chr13_hap1.g00624.t01#5   mNeoVis.3.1.1.chr13.g00550.t01#6       6597   125254929:   mNeoVis.evergladensis.1.1.chr6.g00771.t01#3    mNeoVis.evergladensis.2.1.chr6.g00766.t01#2    mNeoVis.vulgivaga.1.1.chr6.g00764.t01#4    mNeoVis.vulgivaga.2.1.chr6.g00763.t01#1    mNeoVis.3.1.1.chr6.g00765.t01#6   mNeoVis.2.1.1.chr4_hap1.g00759.t01#5       6598   125254930:   mNeoVis.2.1.1.chr13_hap1.g00625.t01#5    mNeoVis.evergladensis.1.1.chr13.g00597.t01#3    mNeoVis.evergladensis.2.1.chr13.g00600.t01#2    mNeoVis.3.1.1.chr13.g00551.t01#6    mNeoVis.vulgivaga.1.1.chr13.g00585.t01#4   mNeoVis.vulgivaga.2.1.chr13.g00601.t01#1       6599   125254931:   mNeoVis.vulgivaga.1.1.chr6.g00164.t01#4    mNeoVis.3.1.1.chr6.g00161.t01#6    mNeoVis.vulgivaga.2.1.chr6.g00162.t01#1    mNeoVis.evergladensis.1.1.chr6.g00164.t01#3    mNeoVis.2.1.1.chr4_hap1.g00162.t01#5   mNeoVis.evergladensis.2.1.chr6.g00163.t01#2       6600   125254932:   mNeoVis.evergladensis.1.1.chr6.g00165.t01#3    mNeoVis.vulgivaga.2.1.chr6.g00163.t01#1    mNeoVis.evergladensis.2.1.chr6.g00164.t01#2    mNeoVis.2.1.1.chr4_hap1.g00163.t01#5    mNeoVis.3.1.1.chr6.g00162.t01#6   mNeoVis.vulgivaga.1.1.chr6.g00165.t01#4       6601   125254933:   mNeoVis.vulgivaga.2.1.chr13.g00603.t01#1    mNeoVis.evergladensis.1.1.chr13.g00599.t01#3    mNeoVis.vulgivaga.1.1.chr13.g00586.t01#4    mNeoVis.3.1.1.chr13.g00552.t01#6    mNeoVis.2.1.1.chr13_hap1.g00626.t01#5   mNeoVis.evergladensis.2.1.chr13.g00601.t01#2       6602   125254934:   mNeoVis.vulgivaga.1.1.chr6.g00166.t01#4    mNeoVis.2.1.1.chr4_hap1.g00164.t01#5    mNeoVis.evergladensis.1.1.chr6.g00166.t01#3    mNeoVis.evergladensis.2.1.chr6.g00165.t01#2   mNeoVis.vulgivaga.2.1.chr6.g00164.t01#1       6603   125254935:   mNeoVis.evergladensis.2.1.chr6.g00166.t01#2    mNeoVis.vulgivaga.1.1.chr6.g00167.t01#4    mNeoVis.vulgivaga.2.1.chr6.g00165.t01#1    mNeoVis.evergladensis.1.1.chr6.g00167.t01#3    mNeoVis.3.1.1.chr6.g00164.t01#6   mNeoVis.2.1.1.chr4_hap1.g00165.t01#5       6604   125254936:   mNeoVis.vulgivaga.1.1.chr13.g00587.t01#4    mNeoVis.3.1.1.chr13.g00553.t01#6    mNeoVis.vulgivaga.2.1.chr13.g00604.t01#1    mNeoVis.evergladensis.2.1.chr13.g00602.t01#2    mNeoVis.evergladensis.1.1.chr13.g00600.t01#3   mNeoVis.2.1.1.chr13_hap1.g00627.t01#5       6605   125254937:   mNeoVis.2.1.1.chr4_hap1.g00166.t01#5    mNeoVis.vulgivaga.1.1.chr6.g00168.t01#4    mNeoVis.evergladensis.1.1.chr6.g00168.t01#3    mNeoVis.vulgivaga.2.1.chr6.g00166.t01#1    mNeoVis.3.1.1.chr6.g00165.t01#6   mNeoVis.evergladensis.2.1.chr6.g00167.t01#2       6606   125254938:   mNeoVis.evergladensis.1.1.chr13.g00602.t01#3    mNeoVis.2.1.1.chr13_hap1.g00629.t01#5    mNeoVis.evergladensis.2.1.chr13.g00604.t01#2    mNeoVis.3.1.1.chr13.g00555.t01#6    mNeoVis.vulgivaga.2.1.chr13.g00606.t01#1   mNeoVis.vulgivaga.1.1.chr13.g00589.t01#4       6607   125254939:   mNeoVis.3.1.1.chr6.g00166.t01#6    mNeoVis.vulgivaga.2.1.chr6.g00167.t01#1    mNeoVis.vulgivaga.1.1.chr6.g00169.t01#4    mNeoVis.2.1.1.chr4_hap1.g00167.t01#5    mNeoVis.evergladensis.1.1.chr6.g00169.t01#3   mNeoVis.evergladensis.2.1.chr6.g00168.t01#2       6608   125254940:   mNeoVis.vulgivaga.2.1.chr6.g00168.t01#1    mNeoVis.evergladensis.1.1.chr6.g00170.t01#3    mNeoVis.3.1.1.chr6.g00167.t01#6    mNeoVis.evergladensis.2.1.chr6.g00169.t01#2    mNeoVis.vulgivaga.1.1.chr6.g00170.t01#4   mNeoVis.2.1.1.chr4_hap1.g00168.t01#5       6609   125254941:   mNeoVis.2.1.1.chr13_hap1.g00631.t01#5    mNeoVis.evergladensis.2.1.chr13.g00606.t01#2    mNeoVis.3.1.1.chr13.g00556.t01#6    mNeoVis.evergladensis.1.1.chr13.g00603.t01#3   mNeoVis.vulgivaga.2.1.chr13.g00607.t01#1       6610   125254942:   mNeoVis.evergladensis.2.1.chr6.g00170.t01#2    mNeoVis.3.1.1.chr6.g00168.t01#6    mNeoVis.2.1.1.chr4_hap1.g00169.t01#5    mNeoVis.evergladensis.1.1.chr6.g00171.t01#3    mNeoVis.vulgivaga.2.1.chr6.g00169.t01#1   mNeoVis.vulgivaga.1.1.chr6.g00171.t01#4       6611   125254943:   mNeoVis.evergladensis.1.1.chr6.g00172.t01#3    mNeoVis.vulgivaga.1.1.chr6.g00172.t01#4    mNeoVis.evergladensis.2.1.chr6.g00171.t01#2    mNeoVis.3.1.1.chr6.g00169.t01#6    mNeoVis.vulgivaga.2.1.chr6.g00170.t01#1   mNeoVis.2.1.1.chr4_hap1.g00170.t01#5       6612   125254944:   mNeoVis.evergladensis.2.1.chr13.g00607.t01#2    mNeoVis.evergladensis.2.1.chr13.g00729.t01#2    mNeoVis.vulgivaga.2.1.chr13.g00608.t01#1    mNeoVis.2.1.1.chr13_hap1.g00632.t01#5    mNeoVis.evergladensis.1.1.chr13.g00604.t01#3    mNeoVis.vulgivaga.1.1.chr13.g00592.t01#4   mNeoVis.3.1.1.chr13.g00557.t01#6       6613   125254945:   mNeoVis.2.1.1.chr13_hap1.g00633.t01#5    mNeoVis.evergladensis.1.1.chr13.g00605.t01#3    mNeoVis.3.1.1.chr13.g00558.t01#6    mNeoVis.evergladensis.2.1.chr13.g00608.t01#2    mNeoVis.vulgivaga.1.1.chr13.g00593.t01#4   mNeoVis.vulgivaga.2.1.chr13.g00609.t01#1       6614   125254946:   mNeoVis.3.1.1.chr6.g00170.t01#6    mNeoVis.2.1.1.chr4_hap1.g00171.t01#5    mNeoVis.evergladensis.2.1.chr6.g00172.t01#2    mNeoVis.vulgivaga.2.1.chr6.g00171.t01#1    mNeoVis.evergladensis.1.1.chr6.g00173.t01#3   mNeoVis.vulgivaga.1.1.chr6.g00173.t01#4       6615   125254947:   mNeoVis.evergladensis.1.1.chr13.g00606.t01#3    mNeoVis.vulgivaga.2.1.chr13.g00610.t01#1   mNeoVis.3.1.1.chr13.g00559.t01#6           6616   125254948:   mNeoVis.vulgivaga.1.1.chr6.g00174.t01#4    mNeoVis.3.1.1.chr6.g00172.t01#6    mNeoVis.evergladensis.1.1.chr6.g00174.t01#3    mNeoVis.evergladensis.2.1.chr6.g00173.t01#2    mNeoVis.vulgivaga.2.1.chr6.g00172.t01#1   mNeoVis.2.1.1.chr4_hap1.g00172.t01#5       6617   125254949:   mNeoVis.2.1.1.chr3_hap1.g00618.t01#5    mNeoVis.3.1.1.chr3.g00611.t01#6    mNeoVis.evergladensis.1.1.chr3.g00625.t01#3    mNeoVis.evergladensis.2.1.chr3.g00626.t01#2    mNeoVis.vulgivaga.1.1.chr3.g00621.t01#4   mNeoVis.vulgivaga.2.1.chr3.g00629.t01#1       6618   125254950:   mNeoVis.3.1.1.chr13.g00562.t01#6    mNeoVis.vulgivaga.2.1.chr13.g00612.t01#1    mNeoVis.vulgivaga.1.1.chr13.g00595.t01#4    mNeoVis.2.1.1.chr13_hap1.g00635.t01#5   mNeoVis.evergladensis.1.1.chr13.g00608.t01#3       6619   125254951:   mNeoVis.3.1.1.chr13.g00563.t01#6    mNeoVis.vulgivaga.1.1.chr13.g00596.t01#4    mNeoVis.vulgivaga.2.1.chr13.g00613.t01#1    mNeoVis.evergladensis.2.1.chr13.g00610.t01#2    mNeoVis.evergladensis.1.1.chr13.g00609.t01#3   mNeoVis.2.1.1.chr13_hap1.g00636.t01#5       6620   125254952:   mNeoVis.vulgivaga.1.1.chr6.g00071.t01#4    mNeoVis.3.1.1.chr6.g00068.t01#6    mNeoVis.2.1.1.chr4_hap1.g00065.t01#5    mNeoVis.evergladensis.1.1.chr6.g00067.t01#3    mNeoVis.vulgivaga.2.1.chr6.g00070.t01#1   mNeoVis.evergladensis.2.1.chr6.g00070.t01#2       6621   125254953:   mNeoVis.evergladensis.2.1.chr6.g00071.t01#2    mNeoVis.evergladensis.1.1.chr6.g00068.t01#3    mNeoVis.2.1.1.chr4_hap1.g00066.t01#5   mNeoVis.vulgivaga.2.1.chr6.g00071.t01#1         6622   125254954:   mNeoVis.evergladensis.2.1.chr6.g00072.t01#2    mNeoVis.evergladensis.1.1.chr6.g00069.t01#3    mNeoVis.2.1.1.chr4_hap1.g00067.t01#5    mNeoVis.3.1.1.chr6.g00070.t01#6    mNeoVis.vulgivaga.2.1.chr6.g00072.t01#1   mNeoVis.vulgivaga.1.1.chr6.g00073.t01#4       6623   125254955:   mNeoVis.evergladensis.2.1.chr6.g00073.t01#2    mNeoVis.evergladensis.1.1.chr6.g00070.t01#3    mNeoVis.3.1.1.chr6.g00071.t01#6    mNeoVis.2.1.1.chr4_hap1.g00068.t01#5    mNeoVis.vulgivaga.2.1.chr6.g00073.t01#1   mNeoVis.vulgivaga.1.1.chr6.g00074.t01#4       6624   125254956:   mNeoVis.vulgivaga.2.1.chr6.g00074.t01#1    mNeoVis.vulgivaga.1.1.chr6.g00075.t01#4    mNeoVis.2.1.1.chr4_hap1.g00069.t01#5    mNeoVis.3.1.1.chr6.g00072.t01#6    mNeoVis.evergladensis.1.1.chr6.g00071.t01#3   mNeoVis.evergladensis.2.1.chr6.g00074.t01#2       6625   125254957:   mNeoVis.3.1.1.chr13.g00564.t01#6    mNeoVis.vulgivaga.1.1.chr13.g00597.t01#4    mNeoVis.evergladensis.1.1.chr13.g00610.t01#3    mNeoVis.evergladensis.2.1.chr13.g00611.t01#2    mNeoVis.vulgivaga.2.1.chr13.g00614.t01#1   mNeoVis.2.1.1.chr13_hap1.g00637.t01#5       6626   125254958:   mNeoVis.evergladensis.2.1.chr15.g00061.t01#2    mNeoVis.vulgivaga.1.1.chr15.g00062.t01#4    mNeoVis.3.1.1.chr15.g00057.t01#6    mNeoVis.evergladensis.1.1.chr15.g00552.t01#3    mNeoVis.vulgivaga.2.1.chr15.g00063.t01#1   mNeoVis.2.1.1.chr15_hap1.g00061.t01#5       6627   125254959:   mNeoVis.evergladensis.2.1.chr13.g00612.t01#2    mNeoVis.vulgivaga.1.1.chr13.g00598.t01#4    mNeoVis.2.1.1.chr13_hap1.g00638.t01#5    mNeoVis.evergladensis.1.1.chr13.g00611.t01#3    mNeoVis.3.1.1.chr13.g00565.t01#6   mNeoVis.vulgivaga.2.1.chr13.g00615.t01#1       6628   125254960:   mNeoVis.evergladensis.1.1.chr6.g00072.t01#3    mNeoVis.evergladensis.2.1.chr6.g00075.t01#2    mNeoVis.3.1.1.chr6.g00074.t01#6    mNeoVis.vulgivaga.2.1.chr6.g00075.t01#1    mNeoVis.3.1.1.chr6.g00073.t01#6    mNeoVis.2.1.1.chr4_hap1.g00070.t01#5    mNeoVis.evergladensis.1.1.chr6.g00073.t01#3   mNeoVis.vulgivaga.1.1.chr6.g00076.t01#4     6629   125254961:   mNeoVis.vulgivaga.1.1.chr6.g00078.t01#4    mNeoVis.2.1.1.chr4_hap1.g00072.t01#5    mNeoVis.vulgivaga.2.1.chr12.g00453.t01#1    mNeoVis.vulgivaga.2.1.chr6.g00076.t01#1    mNeoVis.evergladensis.1.1.chr6.g00075.t01#3    mNeoVis.3.1.1.chr6.g00076.t01#6    mNeoVis.2.1.1.chr12_hap1.g00491.t01#5    mNeoVis.evergladensis.1.1.chr12.g00447.t01#3    mNeoVis.vulgivaga.1.1.chr12.g00450.t01#4      6630   125254962:   mNeoVis.vulgivaga.1.1.chr13.g00599.t01#4    mNeoVis.evergladensis.1.1.chr13.g00612.t01#3    mNeoVis.vulgivaga.2.1.chr13.g00616.t01#1    mNeoVis.2.1.1.chr13_hap1.g00639.t01#5    mNeoVis.3.1.1.chr13.g00566.t01#6   mNeoVis.evergladensis.2.1.chr13.g00613.t01#2       6631   125254963:   mNeoVis.vulgivaga.2.1.chr6.g00077.t01#1    mNeoVis.evergladensis.1.1.chr6.g00076.t01#3    mNeoVis.2.1.1.chr4_hap1.g00073.t01#5    mNeoVis.evergladensis.2.1.chr9.g01086.t01#2    mNeoVis.3.1.1.chr8.g01029.t01#6    mNeoVis.2.1.1.chr8_hap1.g01108.t01#5    mNeoVis.vulgivaga.1.1.chr8.g01073.t01#4    mNeoVis.evergladensis.1.1.chr8.g01078.t01#3    mNeoVis.vulgivaga.1.1.chr6.g00079.t01#4      6632   125254964:   mNeoVis.3.1.1.chr6.g00077.t01#6    mNeoVis.evergladensis.2.1.chr6.g00078.t01#2    mNeoVis.vulgivaga.1.1.chr6.g00080.t01#4    mNeoVis.2.1.1.chr4_hap1.g00074.t01#5    mNeoVis.vulgivaga.2.1.chr6.g00078.t01#1   mNeoVis.evergladensis.1.1.chr6.g00077.t01#3       6633   125254965:   mNeoVis.vulgivaga.1.1.chr6.g00081.t01#4    mNeoVis.3.1.1.chr6.g00078.t01#6    mNeoVis.evergladensis.1.1.chr6.g00078.t01#3    mNeoVis.evergladensis.2.1.chr6.g00080.t01#2    mNeoVis.vulgivaga.2.1.chr6.g00079.t01#1   mNeoVis.2.1.1.chr4_hap1.g00076.t01#5       6634   125254966:   mNeoVis.vulgivaga.1.1.chr6.g00082.t01#4    mNeoVis.3.1.1.chr6.g00079.t01#6    mNeoVis.evergladensis.2.1.chr6.g00081.t01#2    mNeoVis.2.1.1.chr4_hap1.g00077.t01#5    mNeoVis.vulgivaga.2.1.chr6.g00080.t01#1   mNeoVis.evergladensis.1.1.chr6.g00079.t01#3       6635   125254967:   mNeoVis.evergladensis.2.1.chr13.g00614.t01#2    mNeoVis.vulgivaga.2.1.chr13.g00617.t01#1    mNeoVis.vulgivaga.1.1.chr13.g00600.t01#4    mNeoVis.3.1.1.chr13.g00567.t01#6    mNeoVis.2.1.1.chr13_hap1.g00640.t01#5   mNeoVis.evergladensis.1.1.chr13.g00613.t01#3       6636   125254968:   mNeoVis.evergladensis.2.1.chr13.g00615.t01#2    mNeoVis.3.1.1.chr13.g00568.t01#6    mNeoVis.2.1.1.chr13_hap1.g00641.t01#5    mNeoVis.vulgivaga.1.1.chr13.g00601.t01#4    mNeoVis.evergladensis.1.1.chr13.g00614.t01#3   mNeoVis.vulgivaga.2.1.chr13.g00618.t01#1       6637   125254969:   mNeoVis.vulgivaga.1.1.chr15.g00063.t01#4    mNeoVis.vulgivaga.2.1.chr15.g00064.t01#1    mNeoVis.evergladensis.2.1.chr15.g00062.t01#2    mNeoVis.3.1.1.chr15.g00058.t01#6    mNeoVis.evergladensis.1.1.chr15.g00551.t01#3   mNeoVis.2.1.1.chr15_hap1.g00062.t01#5       6638   125254970:   mNeoVis.2.1.1.chr4_hap1.g00078.t01#5    mNeoVis.evergladensis.2.1.chr6.g00082.t01#2    mNeoVis.vulgivaga.2.1.chr6.g00081.t01#1    mNeoVis.vulgivaga.1.1.chr6.g00083.t01#4   mNeoVis.evergladensis.1.1.chr6.g00080.t01#3       6639   125254971:   mNeoVis.vulgivaga.2.1.chr6.g00082.t01#1    mNeoVis.evergladensis.1.1.chr6.g00081.t01#3    mNeoVis.3.1.1.chr6.g00080.t01#6    mNeoVis.vulgivaga.1.1.chr6.g00084.t01#4    mNeoVis.2.1.1.chr4_hap1.g00079.t01#5   mNeoVis.evergladensis.2.1.chr6.g00083.t01#2       6640   125254972:   mNeoVis.evergladensis.1.1.chr13.g00616.t01#3    mNeoVis.vulgivaga.1.1.chr13.g00603.t01#4    mNeoVis.evergladensis.1.1.chr13.g00615.t01#3    mNeoVis.vulgivaga.2.1.chr13.g00619.t01#1   mNeoVis.evergladensis.2.1.chr13.g00617.t01#2       6641   125254973:   mNeoVis.vulgivaga.2.1.chr13.g00620.t01#1    mNeoVis.evergladensis.1.1.chr13.g00617.t01#3   mNeoVis.vulgivaga.1.1.chr13.g00604.t01#4         6642   125254974:  mNeoVis.3.1.1.chr13.g00569.t01#6               6643   125254975:  mNeoVis.2.1.1.chr13_hap1.g00643.t01#5               6644   125254976:   mNeoVis.evergladensis.1.1.chr6.g00082.t01#3    mNeoVis.evergladensis.2.1.chr6.g00084.t01#2    mNeoVis.vulgivaga.2.1.chr6.g00083.t01#1    mNeoVis.vulgivaga.1.1.chr6.g00085.t01#4    mNeoVis.2.1.1.chr4_hap1.g00080.t01#5   mNeoVis.3.1.1.chr6.g00082.t01#6       6645   125254977:   mNeoVis.3.1.1.chr12.g00432.t01#6    mNeoVis.evergladensis.1.1.chr12.g00431.t01#3    mNeoVis.vulgivaga.1.1.chr12.g00436.t01#4    mNeoVis.2.1.1.chr12_hap1.g00474.t01#5    mNeoVis.vulgivaga.2.1.chr12.g00435.t01#1   mNeoVis.evergladensis.2.1.chr12.g00425.t01#2       6646   125254978:   mNeoVis.evergladensis.1.1.chr13.g00618.t01#3    mNeoVis.3.1.1.chr13.g00570.t01#6    mNeoVis.2.1.1.chr13_hap1.g00644.t01#5    mNeoVis.evergladensis.2.1.chr13.g00618.t01#2    mNeoVis.vulgivaga.2.1.chr13.g00621.t01#1   mNeoVis.vulgivaga.1.1.chr13.g00605.t01#4       6647   125254979:   mNeoVis.evergladensis.2.1.chr6.g00086.t01#2    mNeoVis.vulgivaga.2.1.chr6.g00084.t01#1    mNeoVis.vulgivaga.1.1.chr6.g00087.t01#4    mNeoVis.evergladensis.1.1.chr6.g00085.t01#3   mNeoVis.2.1.1.chr4_hap1.g00082.t01#5       6648   125254980:   mNeoVis.evergladensis.2.1.chr6.g00087.t01#2    mNeoVis.2.1.1.chr4_hap1.g00083.t01#5    mNeoVis.vulgivaga.1.1.chr6.g00088.t01#4    mNeoVis.evergladensis.1.1.chr6.g00086.t01#3    mNeoVis.vulgivaga.2.1.chr6.g00085.t01#1   mNeoVis.3.1.1.chr6.g00084.t01#6       6649   125254981:   mNeoVis.evergladensis.2.1.chr15.g00063.t01#2    mNeoVis.vulgivaga.2.1.chr15.g00065.t01#1    mNeoVis.evergladensis.1.1.chr15.g00550.t01#3    mNeoVis.3.1.1.chr15.g00059.t01#6    mNeoVis.2.1.1.chr15_hap1.g00063.t01#5   mNeoVis.vulgivaga.1.1.chr15.g00064.t01#4       6650   125254982:   mNeoVis.3.1.1.chr13.g00571.t01#6    mNeoVis.2.1.1.chr13_hap1.g00645.t01#5    mNeoVis.evergladensis.1.1.chr13.g00619.t01#3    mNeoVis.vulgivaga.2.1.chr13.g00622.t01#1    mNeoVis.evergladensis.2.1.chr13.g00619.t01#2   mNeoVis.vulgivaga.1.1.chr13.g00606.t01#4       6651   125254983:   mNeoVis.evergladensis.1.1.chr13.g00620.t01#3    mNeoVis.2.1.1.chr13_hap1.g00646.t01#5    mNeoVis.evergladensis.2.1.chr13.g00620.t01#2    mNeoVis.vulgivaga.2.1.chr13.g00623.t01#1    mNeoVis.3.1.1.chr13.g00572.t01#6   mNeoVis.vulgivaga.1.1.chr13.g00607.t01#4       6652   125254984:   mNeoVis.vulgivaga.1.1.chr8.g01407.t01#4    mNeoVis.vulgivaga.2.1.chr8.g01421.t01#1    mNeoVis.evergladensis.1.1.chr8.g01414.t01#3    mNeoVis.evergladensis.2.1.chr9.g01417.t01#2    mNeoVis.2.1.1.chr8_hap1.g01469.t01#5   mNeoVis.3.1.1.chr8.g01364.t01#6       6653   125254985:   mNeoVis.2.1.1.chr4_hap1.g00087.t01#5    mNeoVis.vulgivaga.1.1.chr6.g00092.t01#4    mNeoVis.evergladensis.1.1.chr6.g00090.t01#3    mNeoVis.3.1.1.chr6.g00088.t01#6    mNeoVis.evergladensis.2.1.chr6.g00089.t01#2   mNeoVis.vulgivaga.2.1.chr6.g00087.t01#1       6654   125254986:   mNeoVis.vulgivaga.1.1.chr6.g00091.t01#4    mNeoVis.3.1.1.chr6.g00090.t01#6    mNeoVis.evergladensis.1.1.chr6.g00092.t01#3    mNeoVis.vulgivaga.2.1.chr6.g00089.t01#1    mNeoVis.evergladensis.2.1.chr6.g00092.t01#2   mNeoVis.2.1.1.chr4_hap1.g00089.t01#5       6655   125254987:   mNeoVis.2.1.1.chr4_hap1.g00084.t01#5    mNeoVis.3.1.1.chr6.g00087.t01#6    mNeoVis.3.1.1.chr6.g00085.t01#6    mNeoVis.evergladensis.1.1.chr6.g00087.t01#3    mNeoVis.evergladensis.1.1.chr6.g00089.t01#3   mNeoVis.2.1.1.chr4_hap1.g00086.t01#5       6656   125254988:   mNeoVis.vulgivaga.1.1.chr6.g00095.t01#4    mNeoVis.2.1.1.chr4_hap1.g00090.t01#5    mNeoVis.evergladensis.1.1.chr6.g00093.t01#3    mNeoVis.evergladensis.2.1.chr6.g00093.t01#2    mNeoVis.3.1.1.chr6.g00091.t01#6   mNeoVis.vulgivaga.2.1.chr6.g00090.t01#1       6657   125254989:   mNeoVis.evergladensis.1.1.chr6.g00088.t01#3    mNeoVis.vulgivaga.1.1.chr6.g00090.t01#4    mNeoVis.2.1.1.chr4_hap1.g00085.t01#5    mNeoVis.3.1.1.chr6.g00086.t01#6   mNeoVis.evergladensis.2.1.chr6.g00091.t01#2       6658   125254990:   mNeoVis.vulgivaga.2.1.chr6.g00088.t01#1    mNeoVis.3.1.1.chr6.g00089.t01#6    mNeoVis.2.1.1.chr4_hap1.g00088.t01#5   mNeoVis.vulgivaga.1.1.chr6.g00094.t01#4         6659   125254991:   mNeoVis.vulgivaga.2.1.chr6.g00086.t01#1    mNeoVis.vulgivaga.1.1.chr6.g00089.t01#4   mNeoVis.evergladensis.2.1.chr6.g00088.t01#2         6660   125254992:   mNeoVis.evergladensis.2.1.chr6.g00090.t01#2    mNeoVis.vulgivaga.1.1.chr6.g00093.t01#4   mNeoVis.evergladensis.1.1.chr6.g00091.t01#3         6661   125254993:   mNeoVis.vulgivaga.2.1.chr13.g00624.t01#1    mNeoVis.2.1.1.chr13_hap1.g00647.t01#5    mNeoVis.vulgivaga.1.1.chr13.g00608.t01#4    mNeoVis.3.1.1.chr13.g00573.t01#6    mNeoVis.evergladensis.2.1.chr13.g00621.t01#2   mNeoVis.evergladensis.1.1.chr13.g00621.t01#3       6662   125254994:   mNeoVis.vulgivaga.1.1.chr13.g00609.t01#4    mNeoVis.evergladensis.1.1.chr13.g00622.t01#3    mNeoVis.vulgivaga.2.1.chr13.g00625.t01#1    mNeoVis.evergladensis.2.1.chr13.g00622.t01#2    mNeoVis.3.1.1.chr13.g00574.t01#6   mNeoVis.2.1.1.chr13_hap1.g00648.t01#5       6663   125254995:   mNeoVis.3.1.1.chr6.g00092.t01#6    mNeoVis.evergladensis.2.1.chr6.g00094.t01#2    mNeoVis.vulgivaga.2.1.chr6.g00091.t01#1    mNeoVis.2.1.1.chr4_hap1.g00091.t01#5    mNeoVis.vulgivaga.1.1.chr6.g00096.t01#4   mNeoVis.evergladensis.1.1.chr6.g00094.t01#3       6664   125254996:   mNeoVis.evergladensis.2.1.chr6.g00095.t01#2    mNeoVis.3.1.1.chr6.g00093.t01#6    mNeoVis.vulgivaga.1.1.chr6.g00097.t01#4    mNeoVis.2.1.1.chr4_hap1.g00092.t01#5    mNeoVis.evergladensis.1.1.chr6.g00095.t01#3   mNeoVis.vulgivaga.2.1.chr6.g00092.t01#1       6665   125254997:   mNeoVis.vulgivaga.2.1.chr13.g00626.t01#1   mNeoVis.vulgivaga.1.1.chr13.g00610.t01#4           6666   125254998:   mNeoVis.3.1.1.chr6.g00095.t01#6    mNeoVis.vulgivaga.2.1.chr6.g00094.t01#1    mNeoVis.evergladensis.1.1.chr6.g00097.t01#3    mNeoVis.vulgivaga.1.1.chr6.g00098.t01#4    mNeoVis.evergladensis.1.1.chr6.g00096.t01#3    mNeoVis.2.1.1.chr4_hap1.g00096.t01#5    mNeoVis.evergladensis.2.1.chr6.g00097.t01#2    mNeoVis.vulgivaga.1.1.chr6.g00099.t01#4    mNeoVis.vulgivaga.2.1.chr6.g00095.t01#1      6667   125254999:   mNeoVis.evergladensis.1.1.chr13.g00623.t01#3    mNeoVis.evergladensis.2.1.chr13.g00624.t01#2    mNeoVis.vulgivaga.2.1.chr13.g00627.t01#1    mNeoVis.3.1.1.chr13.g00576.t01#6    mNeoVis.2.1.1.chr13_hap1.g00651.t01#5   mNeoVis.vulgivaga.1.1.chr13.g00611.t01#4       6668   125255000:   mNeoVis.2.1.1.chr13_hap1.g00652.t01#5    mNeoVis.evergladensis.1.1.chr13.g00624.t01#3    mNeoVis.vulgivaga.2.1.chr13.g00628.t01#1    mNeoVis.evergladensis.2.1.chr13.g00625.t01#2    mNeoVis.3.1.1.chr13.g00577.t01#6   mNeoVis.vulgivaga.1.1.chr13.g00612.t01#4       6669   125255001:   mNeoVis.vulgivaga.2.1.chr6.g00097.t01#1    mNeoVis.3.1.1.chr6.g00096.t01#6   mNeoVis.vulgivaga.1.1.chr6.g00102.t01#4           6670   125255002:   mNeoVis.evergladensis.2.1.chr13.g00626.t01#2    mNeoVis.2.1.1.chr13_hap1.g00653.t01#5    mNeoVis.vulgivaga.1.1.chr13.g00614.t01#4    mNeoVis.evergladensis.1.1.chr13.g00625.t01#3    mNeoVis.vulgivaga.2.1.chr13.g00629.t01#1   mNeoVis.3.1.1.chr13.g00578.t01#6       6671   125255003:   mNeoVis.evergladensis.1.1.chr13.g00641.t01#3    mNeoVis.3.1.1.chr13.g00592.t01#6    mNeoVis.vulgivaga.1.1.chr13.g00631.t01#4    mNeoVis.evergladensis.2.1.chr13.g00643.t01#2   mNeoVis.2.1.1.chr13_hap1.g00670.t01#5       6672   125255004:   mNeoVis.evergladensis.2.1.chr13.g00642.t01#2    mNeoVis.2.1.1.chr13_hap1.g00671.t01#5   mNeoVis.2.1.1.chr13_hap1.g00669.t01#5           6673   125255005:   mNeoVis.3.1.1.chr6.g01084.t01#6    mNeoVis.vulgivaga.1.1.chr6.g01086.t01#4    mNeoVis.evergladensis.2.1.chr6.g01092.t01#2    mNeoVis.evergladensis.1.1.chr6.g01097.t01#3    mNeoVis.vulgivaga.2.1.chr6.g01086.t01#1   mNeoVis.2.1.1.chr4_hap1.g01095.t01#5       6674   125255006:   mNeoVis.2.1.1.chr4_hap1.g00098.t01#5    mNeoVis.evergladensis.2.1.chr6.g00099.t01#2    mNeoVis.evergladensis.1.1.chr6.g00101.t01#3    mNeoVis.3.1.1.chr6.g00098.t01#6    mNeoVis.vulgivaga.1.1.chr6.g00103.t01#4   mNeoVis.vulgivaga.2.1.chr6.g00098.t01#1       6675   125255007:   mNeoVis.evergladensis.2.1.chr6.g00102.t01#2    mNeoVis.vulgivaga.2.1.chr6.g00099.t01#1    mNeoVis.vulgivaga.1.1.chr6.g00105.t01#4    mNeoVis.2.1.1.chr4_hap1.g00100.t01#5    mNeoVis.3.1.1.chr6.g00100.t01#6   mNeoVis.evergladensis.1.1.chr6.g00104.t01#3       6676   125255008:   mNeoVis.evergladensis.1.1.chr6.g00105.t01#3    mNeoVis.vulgivaga.2.1.chr6.g00100.t01#1    mNeoVis.evergladensis.2.1.chr6.g00103.t01#2    mNeoVis.2.1.1.chr4_hap1.g00101.t01#5    mNeoVis.vulgivaga.1.1.chr6.g00106.t01#4   mNeoVis.3.1.1.chr6.g00101.t01#6       6677   125255009:  mNeoVis.vulgivaga.2.1.chr13.g00631.t01#1               6678   125255010:   mNeoVis.vulgivaga.2.1.chr13.g00632.t01#1    mNeoVis.2.1.1.chr13_hap1.g00656.t01#5    mNeoVis.vulgivaga.1.1.chr13.g00617.t01#4    mNeoVis.3.1.1.chr13.g00581.t01#6    mNeoVis.evergladensis.2.1.chr13.g00629.t01#2   mNeoVis.evergladensis.1.1.chr13.g00628.t01#3       6679   125255011:   mNeoVis.evergladensis.1.1.chr6.g00106.t01#3    mNeoVis.2.1.1.chr4_hap1.g00102.t01#5    mNeoVis.vulgivaga.2.1.chr6.g00101.t01#1    mNeoVis.3.1.1.chr6.g00102.t01#6    mNeoVis.vulgivaga.1.1.chr6.g00107.t01#4   mNeoVis.evergladensis.2.1.chr6.g00104.t01#2       6680   125255012:   mNeoVis.2.1.1.chr13_hap1.g00657.t01#5    mNeoVis.3.1.1.chr13.g00582.t01#6    mNeoVis.evergladensis.2.1.chr13.g00630.t01#2    mNeoVis.vulgivaga.2.1.chr13.g00633.t01#1    mNeoVis.vulgivaga.1.1.chr13.g00618.t01#4   mNeoVis.evergladensis.1.1.chr13.g00629.t01#3       6681   125255013:   mNeoVis.vulgivaga.1.1.chr6.g00108.t01#4    mNeoVis.2.1.1.chr4_hap1.g00103.t01#5    mNeoVis.3.1.1.chr6.g00103.t01#6    mNeoVis.evergladensis.1.1.chr6.g00107.t01#3    mNeoVis.evergladensis.2.1.chr6.g00105.t01#2   mNeoVis.vulgivaga.2.1.chr6.g00102.t01#1       6682   125255014:   mNeoVis.vulgivaga.1.1.chr6.g00109.t01#4    mNeoVis.evergladensis.2.1.chr6.g00106.t01#2    mNeoVis.2.1.1.chr4_hap1.g00104.t01#5    mNeoVis.3.1.1.chr6.g00104.t01#6    mNeoVis.evergladensis.1.1.chr6.g00108.t01#3   mNeoVis.vulgivaga.2.1.chr6.g00103.t01#1       6683   125255015:   mNeoVis.2.1.1.chr13_hap1.g00658.t01#5    mNeoVis.3.1.1.chr13.g00583.t01#6    mNeoVis.evergladensis.2.1.chr13.g00631.t01#2    mNeoVis.vulgivaga.2.1.chr13.g00634.t01#1    mNeoVis.evergladensis.1.1.chr13.g00630.t01#3   mNeoVis.vulgivaga.1.1.chr13.g00619.t01#4       6684   125255016:   mNeoVis.3.1.1.chr13.g00488.t01#6    mNeoVis.2.1.1.chr13_hap1.g00547.t01#5    mNeoVis.vulgivaga.2.1.chr13.g00533.t01#1    mNeoVis.evergladensis.1.1.chr13.g00527.t01#3    mNeoVis.evergladensis.2.1.chr13.g00530.t01#2   mNeoVis.vulgivaga.1.1.chr13.g00522.t01#4       6685   125255017:   mNeoVis.vulgivaga.1.1.chr6.g00110.t01#4    mNeoVis.vulgivaga.2.1.chr6.g00104.t01#1    mNeoVis.3.1.1.chr6.g00105.t01#6   mNeoVis.2.1.1.chr4_hap1.g00105.t01#5         6686   125255018:   mNeoVis.vulgivaga.1.1.chr6.g00111.t01#4    mNeoVis.vulgivaga.2.1.chr6.g00105.t01#1    mNeoVis.evergladensis.1.1.chr6.g00110.t01#3    mNeoVis.evergladensis.2.1.chr6.g00108.t01#2    mNeoVis.3.1.1.chr6.g00106.t01#6   mNeoVis.2.1.1.chr4_hap1.g00106.t01#5       6687   125255019:   mNeoVis.vulgivaga.1.1.chr13.g00523.t01#4    mNeoVis.2.1.1.chr13_hap1.g00548.t01#5    mNeoVis.evergladensis.1.1.chr13.g00528.t01#3    mNeoVis.evergladensis.2.1.chr13.g00531.t01#2    mNeoVis.vulgivaga.2.1.chr13.g00534.t01#1   mNeoVis.3.1.1.chr13.g00489.t01#6       6688   125255020:  mNeoVis.vulgivaga.2.1.chr13.g00535.t01#1               6689   125255021:   mNeoVis.2.1.1.chr4_hap1.g00107.t01#5    mNeoVis.3.1.1.chr6.g00107.t01#6    mNeoVis.vulgivaga.2.1.chr6.g00106.t01#1    mNeoVis.vulgivaga.1.1.chr6.g00112.t01#4    mNeoVis.evergladensis.2.1.chr6.g00109.t01#2   mNeoVis.evergladensis.1.1.chr6.g00111.t01#3       6690   125255022:   mNeoVis.2.1.1.chr13_hap1.g00549.t01#5    mNeoVis.vulgivaga.1.1.chr13.g00524.t01#4    mNeoVis.vulgivaga.2.1.chr13.g00536.t01#1    mNeoVis.evergladensis.1.1.chr13.g00529.t01#3    mNeoVis.evergladensis.2.1.chr13.g00532.t01#2   mNeoVis.3.1.1.chr13.g00490.t01#6       6691   125255023:   mNeoVis.evergladensis.2.1.chr6.g00110.t01#2    mNeoVis.vulgivaga.1.1.chr6.g00113.t01#4    mNeoVis.2.1.1.chr4_hap1.g00108.t01#5    mNeoVis.3.1.1.chr6.g00108.t01#6    mNeoVis.evergladensis.1.1.chr6.g00112.t01#3   mNeoVis.vulgivaga.2.1.chr6.g00107.t01#1       6692   125255024:   mNeoVis.2.1.1.chr13_hap1.g00550.t01#5    mNeoVis.vulgivaga.2.1.chr13.g00537.t01#1    mNeoVis.evergladensis.1.1.chr13.g00530.t01#3    mNeoVis.evergladensis.2.1.chr13.g00533.t01#2   mNeoVis.3.1.1.chr13.g00491.t01#6       6693   125255025:   mNeoVis.evergladensis.1.1.chr6.g00113.t01#3    mNeoVis.vulgivaga.2.1.chr6.g00108.t01#1    mNeoVis.evergladensis.2.1.chr6.g00111.t01#2    mNeoVis.2.1.1.chr4_hap1.g00109.t01#5    mNeoVis.vulgivaga.1.1.chr6.g00114.t01#4   mNeoVis.3.1.1.chr6.g00109.t01#6       6694   125255026:   mNeoVis.evergladensis.1.1.chr13.g00531.t01#3    mNeoVis.3.1.1.chr13.g00492.t01#6    mNeoVis.evergladensis.2.1.chr13.g00534.t01#2    mNeoVis.2.1.1.chr13_hap1.g00551.t01#5    mNeoVis.vulgivaga.1.1.chr13.g00526.t01#4   mNeoVis.vulgivaga.2.1.chr13.g00538.t01#1       6695   125255027:   mNeoVis.2.1.1.chr4_hap1.g00110.t01#5    mNeoVis.evergladensis.2.1.chr6.g00112.t01#2    mNeoVis.vulgivaga.2.1.chr6.g00109.t01#1    mNeoVis.3.1.1.chr6.g00110.t01#6    mNeoVis.vulgivaga.1.1.chr6.g00115.t01#4   mNeoVis.evergladensis.1.1.chr6.g00114.t01#3       6696   125255028:   mNeoVis.evergladensis.1.1.chr6.g00115.t01#3    mNeoVis.3.1.1.chr6.g00111.t01#6    mNeoVis.vulgivaga.1.1.chr6.g00116.t01#4    mNeoVis.2.1.1.chr4_hap1.g00111.t01#5    mNeoVis.vulgivaga.2.1.chr6.g00110.t01#1   mNeoVis.evergladensis.2.1.chr6.g00113.t01#2       6697   125255029:   mNeoVis.evergladensis.1.1.chr13.g00532.t01#3    mNeoVis.evergladensis.2.1.chr13.g00535.t01#2    mNeoVis.3.1.1.chr13.g00493.t01#6    mNeoVis.2.1.1.chr13_hap1.g00552.t01#5    mNeoVis.vulgivaga.1.1.chr13.g00527.t01#4   mNeoVis.vulgivaga.2.1.chr13.g00539.t01#1       6698   125255030:   mNeoVis.vulgivaga.2.1.chr6.g00113.t01#1    mNeoVis.vulgivaga.1.1.chr6.g00118.t01#4    mNeoVis.evergladensis.2.1.chr6.g00116.t01#2    mNeoVis.2.1.1.chr4_hap1.g00114.t01#5    mNeoVis.3.1.1.chr6.g00113.t01#6   mNeoVis.evergladensis.1.1.chr6.g00117.t01#3       6699   125255031:   mNeoVis.2.1.1.chr4_hap1.g00115.t01#5    mNeoVis.vulgivaga.1.1.chr6.g00119.t01#4    mNeoVis.evergladensis.1.1.chr6.g00118.t01#3    mNeoVis.evergladensis.2.1.chr6.g00117.t01#2    mNeoVis.3.1.1.chr6.g00115.t01#6   mNeoVis.vulgivaga.2.1.chr6.g00114.t01#1       6700   125255032:   mNeoVis.vulgivaga.2.1.chr13.g00541.t01#1    mNeoVis.evergladensis.2.1.chr13.g00537.t01#2    mNeoVis.3.1.1.chr13.g00495.t01#6    mNeoVis.vulgivaga.1.1.chr13.g00529.t01#4    mNeoVis.2.1.1.chr13_hap1.g00554.t01#5   mNeoVis.evergladensis.1.1.chr13.g00534.t01#3       6701   125255033:   mNeoVis.vulgivaga.2.1.chr6.g00116.t01#1    mNeoVis.evergladensis.2.1.chr6.g00119.t01#2    mNeoVis.evergladensis.1.1.chr6.g00121.t01#3    mNeoVis.3.1.1.chr6.g00117.t01#6    mNeoVis.vulgivaga.1.1.chr6.g00121.t01#4    mNeoVis.2.1.1.chr4_hap1.g00117.t01#5   mNeoVis.evergladensis.1.1.chr6.g00120.t01#3     6702   125255034:   mNeoVis.vulgivaga.2.1.chr6.g00115.t01#1    mNeoVis.vulgivaga.1.1.chr6.g00120.t01#4    mNeoVis.2.1.1.chr4_hap1.g00116.t01#5   mNeoVis.evergladensis.1.1.chr6.g00119.t01#3       6703   125255035:   mNeoVis.3.1.1.chr6.g00116.t01#6   mNeoVis.evergladensis.2.1.chr6.g00118.t01#2           6704   125255036:   mNeoVis.evergladensis.1.1.chr13.g00535.t01#3    mNeoVis.3.1.1.chr13.g00496.t01#6    mNeoVis.vulgivaga.2.1.chr13.g00542.t01#1    mNeoVis.2.1.1.chr13_hap1.g00555.t01#5    mNeoVis.vulgivaga.1.1.chr13.g00530.t01#4   mNeoVis.evergladensis.2.1.chr13.g00538.t01#2       6705   125255037:   mNeoVis.vulgivaga.1.1.chr13.g00531.t01#4    mNeoVis.3.1.1.chr13.g00497.t01#6    mNeoVis.2.1.1.chr13_hap1.g00556.t01#5    mNeoVis.evergladensis.1.1.chr13.g00536.t01#3    mNeoVis.vulgivaga.2.1.chr13.g00543.t01#1   mNeoVis.evergladensis.2.1.chr13.g00539.t01#2       6706   125255038:   mNeoVis.evergladensis.1.1.chr13.g00537.t01#3    mNeoVis.3.1.1.chr13.g00498.t01#6    mNeoVis.vulgivaga.2.1.chr13.g00544.t01#1    mNeoVis.2.1.1.chr13_hap1.g00557.t01#5    mNeoVis.vulgivaga.1.1.chr13.g00532.t01#4   mNeoVis.evergladensis.2.1.chr13.g00540.t01#2       6707   125255039:   mNeoVis.vulgivaga.1.1.chr13.g00533.t01#4    mNeoVis.3.1.1.chr13.g00499.t01#6    mNeoVis.2.1.1.chr13_hap1.g00558.t01#5    mNeoVis.evergladensis.1.1.chr13.g00538.t01#3    mNeoVis.evergladensis.2.1.chr13.g00541.t01#2   mNeoVis.vulgivaga.2.1.chr13.g00545.t01#1       6708   125255040:   mNeoVis.3.1.1.chr13.g00500.t01#6    mNeoVis.vulgivaga.2.1.chr13.g00546.t01#1    mNeoVis.vulgivaga.1.1.chr13.g00534.t01#4    mNeoVis.2.1.1.chr13_hap1.g00559.t01#5    mNeoVis.evergladensis.1.1.chr13.g00539.t01#3   mNeoVis.evergladensis.2.1.chr13.g00542.t01#2       6709   125255041:   mNeoVis.evergladensis.1.1.chr6.g00122.t01#3    mNeoVis.3.1.1.chr6.g00118.t01#6    mNeoVis.vulgivaga.2.1.chr6.g00117.t01#1    mNeoVis.2.1.1.chr4_hap1.g00118.t01#5    mNeoVis.evergladensis.2.1.chr6.g00120.t01#2   mNeoVis.vulgivaga.1.1.chr6.g00122.t01#4       6710   125255042:   mNeoVis.vulgivaga.1.1.chr13.g00535.t01#4    mNeoVis.2.1.1.chr13_hap1.g00560.t01#5    mNeoVis.evergladensis.1.1.chr13.g00540.t01#3    mNeoVis.evergladensis.2.1.chr13.g00543.t01#2    mNeoVis.3.1.1.chr13.g00501.t01#6   mNeoVis.vulgivaga.2.1.chr13.g00547.t01#1       6711   125255043:   mNeoVis.evergladensis.2.1.chr7.g00425.t01#2    mNeoVis.vulgivaga.1.1.chr7.g00429.t01#4    mNeoVis.evergladensis.1.1.chr7.g00433.t01#3    mNeoVis.vulgivaga.2.1.chr7.g00423.t01#1    mNeoVis.2.1.1.chr7_hap1.g00421.t01#5   mNeoVis.3.1.1.chr7.g00407.t01#6       6712   125255044:   mNeoVis.evergladensis.2.1.chr6.g00122.t01#2    mNeoVis.evergladensis.1.1.chr6.g00125.t01#3    mNeoVis.2.1.1.chr4_hap1.g00121.t01#5    mNeoVis.vulgivaga.2.1.chr6.g00119.t01#1    mNeoVis.2.1.1.chr4_hap1.g00120.t01#5    mNeoVis.3.1.1.chr6.g00121.t01#6    mNeoVis.vulgivaga.1.1.chr6.g00124.t01#4   mNeoVis.evergladensis.1.1.chr6.g00124.t01#3     6713   125255045:   mNeoVis.vulgivaga.2.1.chr6.g00118.t01#1    mNeoVis.vulgivaga.1.1.chr6.g00123.t01#4    mNeoVis.evergladensis.1.1.chr6.g00123.t01#3    mNeoVis.evergladensis.2.1.chr6.g00121.t01#2    mNeoVis.2.1.1.chr4_hap1.g00119.t01#5   mNeoVis.3.1.1.chr6.g00120.t01#6       6714   125255046:   mNeoVis.vulgivaga.1.1.chr6.g00125.t01#4    mNeoVis.3.1.1.chr6.g00122.t01#6    mNeoVis.2.1.1.chr4_hap1.g00122.t01#5    mNeoVis.evergladensis.1.1.chr6.g00126.t01#3   mNeoVis.evergladensis.2.1.chr6.g00123.t01#2       6715   125255047:  mNeoVis.vulgivaga.2.1.chr6.g00120.t01#1               6716   125255048:  mNeoVis.vulgivaga.2.1.chr13.g00548.t01#1               6717   125255049:  mNeoVis.vulgivaga.2.1.chr6.g00121.t01#1               6718   125255050:   mNeoVis.vulgivaga.1.1.chr6.g00126.t01#4    mNeoVis.evergladensis.2.1.chr6.g00124.t01#2    mNeoVis.evergladensis.1.1.chr6.g00127.t01#3    mNeoVis.2.1.1.chr4_hap1.g00123.t01#5    mNeoVis.3.1.1.chr6.g00123.t01#6   mNeoVis.vulgivaga.2.1.chr6.g00122.t01#1       6719   125255051:   mNeoVis.vulgivaga.2.1.chr13.g00549.t01#1    mNeoVis.evergladensis.1.1.chr13.g00542.t01#3    mNeoVis.evergladensis.2.1.chr13.g00546.t01#2    mNeoVis.2.1.1.chr13_hap1.g00562.t01#5   mNeoVis.3.1.1.chr13.g00502.t01#6       6720   125255052:   mNeoVis.vulgivaga.1.1.chr6.g00127.t01#4    mNeoVis.3.1.1.chr6.g00124.t01#6    mNeoVis.evergladensis.1.1.chr6.g00128.t01#3    mNeoVis.evergladensis.2.1.chr6.g00125.t01#2    mNeoVis.vulgivaga.2.1.chr6.g00123.t01#1   mNeoVis.2.1.1.chr4_hap1.g00124.t01#5       6721   125255053:   mNeoVis.3.1.1.chr13.g00503.t01#6    mNeoVis.vulgivaga.2.1.chr13.g00550.t01#1    mNeoVis.2.1.1.chr13_hap1.g00563.t01#5    mNeoVis.evergladensis.2.1.chr13.g00547.t01#2    mNeoVis.evergladensis.1.1.chr13.g00543.t01#3   mNeoVis.vulgivaga.1.1.chr13.g00538.t01#4       6722   125255054:   mNeoVis.3.1.1.chr6.g00125.t01#6    mNeoVis.2.1.1.chr4_hap1.g00125.t01#5    mNeoVis.evergladensis.1.1.chr6.g00129.t01#3    mNeoVis.evergladensis.2.1.chr6.g00126.t01#2    mNeoVis.vulgivaga.2.1.chr6.g00124.t01#1   mNeoVis.vulgivaga.1.1.chr6.g00128.t01#4       6723   125255055:   mNeoVis.evergladensis.1.1.chr13.g00545.t01#3    mNeoVis.evergladensis.2.1.chr13.g00549.t01#2    mNeoVis.3.1.1.chr13.g00504.t01#6    mNeoVis.vulgivaga.1.1.chr13.g00539.t01#4    mNeoVis.2.1.1.chr13_hap1.g00565.t01#5   mNeoVis.vulgivaga.2.1.chr13.g00551.t01#1       6724   125255056:   mNeoVis.evergladensis.2.1.chr6.g00127.t01#2   mNeoVis.vulgivaga.2.1.chr6.g00125.t01#1             6725   125255057:   mNeoVis.vulgivaga.2.1.chr6.g00126.t01#1   mNeoVis.evergladensis.2.1.chr6.g00128.t01#2           6726   125255058:   mNeoVis.vulgivaga.2.1.chr13.g00552.t01#1    mNeoVis.2.1.1.chr13_hap1.g00566.t01#5   mNeoVis.evergladensis.2.1.chr13.g00550.t01#2         6727   125255059:   mNeoVis.evergladensis.1.1.chr13.g00547.t01#3    mNeoVis.2.1.1.chr13_hap1.g00568.t01#5    mNeoVis.evergladensis.2.1.chr13.g00552.t01#2    mNeoVis.vulgivaga.1.1.chr13.g00541.t01#4    mNeoVis.vulgivaga.2.1.chr13.g00554.t01#1   mNeoVis.3.1.1.chr13.g00507.t01#6       6728   125255060:   mNeoVis.evergladensis.2.1.chr13.g00554.t01#2    mNeoVis.vulgivaga.2.1.chr13.g00556.t01#1    mNeoVis.evergladensis.1.1.chr13.g00549.t01#3    mNeoVis.3.1.1.chr13.g00508.t01#6    mNeoVis.2.1.1.chr13_hap1.g00570.t01#5   mNeoVis.vulgivaga.1.1.chr13.g00542.t01#4       6729   125255061:   mNeoVis.vulgivaga.1.1.chr2.g01892.t01#4    mNeoVis.vulgivaga.1.1.chr13.g00543.t01#4    mNeoVis.3.1.1.chr13.g00509.t01#6    mNeoVis.evergladensis.1.1.chr13.g00550.t01#3    mNeoVis.evergladensis.2.1.chr13.g00555.t01#2    mNeoVis.2.1.1.chr13_hap1.g00571.t01#5   mNeoVis.vulgivaga.2.1.chr13.g00557.t01#1     6730   125255062:   mNeoVis.evergladensis.1.1.chr2.g01892.t01#3    mNeoVis.vulgivaga.2.1.chr2.g01895.t01#1    mNeoVis.2.1.1.chr5_hap1.g01899.t01#5    mNeoVis.vulgivaga.1.1.chr2.g01893.t01#4    mNeoVis.vulgivaga.2.1.chr2.g01894.t01#1   mNeoVis.evergladensis.2.1.chr2.g01879.t01#2       6731   125255063:   mNeoVis.evergladensis.2.1.chr2.g01877.t01#2    mNeoVis.vulgivaga.2.1.chr2.g01892.t01#1    mNeoVis.evergladensis.1.1.chr2.g01891.t01#3   mNeoVis.2.1.1.chr5_hap1.g01897.t01#5         6732   125255064:   mNeoVis.2.1.1.chr5_hap1.g01904.t01#5    mNeoVis.2.1.1.chr5_hap1.g01855.t01#5    mNeoVis.vulgivaga.1.1.chr2.g01848.t01#4   mNeoVis.evergladensis.1.1.chr2.g01848.t01#3       6733   125255065:   mNeoVis.evergladensis.2.1.chr2.g01884.t01#2    mNeoVis.evergladensis.1.1.chr2.g01896.t01#3   mNeoVis.2.1.1.chr5_hap1.g01906.t01#5           6734   125255066:   mNeoVis.3.1.1.chr2.g01791.t01#6    mNeoVis.evergladensis.2.1.chr2.g01885.t01#2   mNeoVis.2.1.1.chr5_hap1.g01898.t01#5           6735   125255067:   mNeoVis.vulgivaga.2.1.chr2.g01893.t01#1   mNeoVis.evergladensis.2.1.chr2.g01878.t01#2           6736   125255068:   mNeoVis.3.1.1.chr2.g01792.t01#6   mNeoVis.2.1.1.chr5_hap1.g01900.t01#5             6737   125255069:   mNeoVis.evergladensis.1.1.chr2.g01894.t01#3   mNeoVis.2.1.1.chr5_hap1.g01902.t01#5             6738   125255070:   mNeoVis.vulgivaga.2.1.chr2.g01851.t01#1   mNeoVis.vulgivaga.1.1.chr2.g01847.t01#4             6739   125255071:  mNeoVis.2.1.1.chr5_hap1.g01853.t01#5               6740   125255072:  mNeoVis.evergladensis.2.1.chr2.g01880.t01#2             6741   125255073:  mNeoVis.evergladensis.2.1.chr2.g01882.t01#2             6742   125255074:  mNeoVis.vulgivaga.1.1.chr2.g01896.t01#4               6743   125255075:   mNeoVis.vulgivaga.1.1.chr13.g00544.t01#4    mNeoVis.vulgivaga.2.1.chr13.g00558.t01#1    mNeoVis.evergladensis.2.1.chr13.g00556.t01#2    mNeoVis.evergladensis.1.1.chr13.g00551.t01#3    mNeoVis.3.1.1.chr13.g00510.t01#6   mNeoVis.2.1.1.chr13_hap1.g00572.t01#5       6744   125255076:   mNeoVis.vulgivaga.2.1.chr6.g00029.t01#1   mNeoVis.3.1.1.chr6.g00027.t01#6             6745   125255077:   mNeoVis.3.1.1.chr6.g00028.t01#6    mNeoVis.evergladensis.1.1.chr6.g00025.t01#3    mNeoVis.evergladensis.2.1.chr6.g00025.t01#2    mNeoVis.vulgivaga.1.1.chr6.g00028.t01#4    mNeoVis.vulgivaga.2.1.chr6.g00030.t01#1   mNeoVis.2.1.1.chr4_hap1.g00025.t01#5       6746   125255078:   mNeoVis.evergladensis.2.1.chr13.g00557.t01#2    mNeoVis.vulgivaga.1.1.chr13.g00545.t01#4    mNeoVis.2.1.1.chr13_hap1.g00573.t01#5    mNeoVis.3.1.1.chr13.g00511.t01#6    mNeoVis.evergladensis.1.1.chr13.g00553.t01#3    mNeoVis.vulgivaga.2.1.chr13.g00559.t01#1   mNeoVis.evergladensis.1.1.chr13.g00552.t01#3     6747   125255079:   mNeoVis.evergladensis.2.1.chr7.g00426.t01#2    mNeoVis.3.1.1.chr7.g00408.t01#6    mNeoVis.2.1.1.chr7_hap1.g00422.t01#5    mNeoVis.vulgivaga.2.1.chr7.g00424.t01#1   mNeoVis.vulgivaga.1.1.chr7.g00430.t01#4       6748   125255080:   mNeoVis.evergladensis.2.1.chr6.g00026.t01#2    mNeoVis.vulgivaga.1.1.chr13.g00573.t01#4    mNeoVis.2.1.1.chr13_hap1.g00613.t01#5    mNeoVis.evergladensis.1.1.chr6.g00026.t01#3    mNeoVis.vulgivaga.2.1.chr13.g00589.t01#1    mNeoVis.evergladensis.2.1.chr4.g00149.t01#2    mNeoVis.3.1.1.chr13.g00537.t01#6    mNeoVis.vulgivaga.1.1.chr3.g00863.t01#4    mNeoVis.evergladensis.1.1.chr13.g00585.t01#3      6749   125255081:   mNeoVis.vulgivaga.2.1.chr6.g00032.t01#1    mNeoVis.vulgivaga.1.1.chr6.g00029.t01#4    mNeoVis.2.1.1.chr4_hap1.g00026.t01#5    mNeoVis.3.1.1.chr6.g00030.t01#6    mNeoVis.evergladensis.2.1.chr6.g00027.t01#2    mNeoVis.evergladensis.1.1.chr6.g00027.t01#3   mNeoVis.3.1.1.chr6.g00029.t01#6       6750   125255082:   mNeoVis.vulgivaga.2.1.chr6.g00033.t01#1    mNeoVis.evergladensis.1.1.chr6.g00028.t01#3    mNeoVis.2.1.1.chr4_hap1.g00027.t01#5    mNeoVis.evergladensis.2.1.chr6.g00030.t01#2    mNeoVis.vulgivaga.1.1.chr6.g00030.t01#4   mNeoVis.3.1.1.chr6.g00032.t01#6       6751   125255083:   mNeoVis.vulgivaga.1.1.chr10.g00569.t01#4    mNeoVis.2.1.1.chr9_hap1.g00594.t01#5    mNeoVis.evergladensis.2.1.chr10.g00588.t01#2    mNeoVis.evergladensis.1.1.chr10.g00590.t01#3    mNeoVis.3.1.1.chr10.g00562.t01#6   mNeoVis.vulgivaga.2.1.chr10.g00574.t01#1       6752   125255084:   mNeoVis.2.1.1.chr13_hap1.g00574.t01#5    mNeoVis.evergladensis.1.1.chr13.g00554.t01#3    mNeoVis.vulgivaga.1.1.chr13.g00546.t01#4    mNeoVis.evergladensis.2.1.chr13.g00558.t01#2    mNeoVis.3.1.1.chr13.g00512.t01#6   mNeoVis.vulgivaga.2.1.chr13.g00560.t01#1       6753   125255085:   mNeoVis.evergladensis.1.1.chr13.g00555.t01#3    mNeoVis.3.1.1.chr13.g00513.t01#6    mNeoVis.evergladensis.2.1.chr13.g00559.t01#2    mNeoVis.vulgivaga.1.1.chr13.g00547.t01#4    mNeoVis.2.1.1.chr13_hap1.g00575.t01#5   mNeoVis.vulgivaga.2.1.chr13.g00561.t01#1       6754   125255086:   mNeoVis.2.1.1.chr13_hap1.g00576.t01#5    mNeoVis.vulgivaga.1.1.chr13.g00548.t01#4    mNeoVis.evergladensis.1.1.chr13.g00556.t01#3    mNeoVis.evergladensis.2.1.chr13.g00560.t01#2    mNeoVis.3.1.1.chr13.g00514.t01#6   mNeoVis.vulgivaga.2.1.chr13.g00562.t01#1       6755   125255087:   mNeoVis.vulgivaga.2.1.chr7.g00425.t01#1    mNeoVis.vulgivaga.1.1.chr7.g00431.t01#4    mNeoVis.evergladensis.2.1.chr7.g00427.t01#2    mNeoVis.2.1.1.chr7_hap1.g00423.t01#5    mNeoVis.3.1.1.chr7.g00409.t01#6   mNeoVis.evergladensis.1.1.chr7.g00434.t01#3       6756   125255088:   mNeoVis.evergladensis.1.1.chr6.g00029.t01#3    mNeoVis.vulgivaga.2.1.chr6.g00034.t01#1   mNeoVis.evergladensis.2.1.chr6.g00029.t01#2         6757   125255089:   mNeoVis.3.1.1.chr6.g00033.t01#6    mNeoVis.2.1.1.chr4_hap1.g00029.t01#5    mNeoVis.evergladensis.1.1.chr6.g00030.t01#3    mNeoVis.vulgivaga.2.1.chr6.g00035.t01#1    mNeoVis.evergladensis.2.1.chr6.g00031.t01#2   mNeoVis.vulgivaga.1.1.chr6.g00032.t01#4       6758   125255090:   mNeoVis.vulgivaga.2.1.chr13.g00563.t01#1    mNeoVis.vulgivaga.1.1.chr13.g00549.t01#4   mNeoVis.3.1.1.chr13.g00515.t01#6           6759   125255091:   mNeoVis.2.1.1.chr13_hap1.g00577.t01#5    mNeoVis.evergladensis.1.1.chr13.g00557.t01#3   mNeoVis.evergladensis.2.1.chr13.g00561.t01#2         6760   125255092:   mNeoVis.vulgivaga.2.1.chr13.g00564.t01#1    mNeoVis.evergladensis.1.1.chr13.g00558.t01#3    mNeoVis.vulgivaga.1.1.chr13.g00550.t01#4    mNeoVis.2.1.1.chr13_hap1.g00578.t01#5   mNeoVis.evergladensis.2.1.chr13.g00562.t01#2       6761   125255093:   mNeoVis.3.1.1.chr13.g00516.t01#6    mNeoVis.evergladensis.2.1.chr13.g00563.t01#2    mNeoVis.2.1.1.chr13_hap1.g00579.t01#5    mNeoVis.evergladensis.1.1.chr13.g00559.t01#3    mNeoVis.vulgivaga.1.1.chr13.g00551.t01#4   mNeoVis.vulgivaga.2.1.chr13.g00565.t01#1       6762   125255094:   mNeoVis.vulgivaga.2.1.chr6.g00036.t01#1    mNeoVis.evergladensis.2.1.chr6.g00032.t01#2    mNeoVis.2.1.1.chr4_hap1.g00030.t01#5    mNeoVis.evergladensis.1.1.chr6.g00031.t01#3    mNeoVis.3.1.1.chr6.g00034.t01#6   mNeoVis.vulgivaga.1.1.chr6.g00033.t01#4       6763   125255095:   mNeoVis.vulgivaga.1.1.chr6.g00034.t01#4    mNeoVis.evergladensis.2.1.chr6.g00033.t01#2    mNeoVis.vulgivaga.2.1.chr6.g00037.t01#1    mNeoVis.2.1.1.chr4_hap1.g00031.t01#5   mNeoVis.evergladensis.1.1.chr6.g00032.t01#3       6764   125255096:   mNeoVis.vulgivaga.2.1.chr6.g00038.t01#1    mNeoVis.3.1.1.chr6.g00036.t01#6    mNeoVis.vulgivaga.1.1.chr6.g00035.t01#4    mNeoVis.evergladensis.2.1.chr6.g00034.t01#2    mNeoVis.2.1.1.chr4_hap1.g00032.t01#5   mNeoVis.evergladensis.1.1.chr6.g00033.t01#3       6765   125255097:   mNeoVis.3.1.1.chr13.g00517.t01#6    mNeoVis.vulgivaga.2.1.chr13.g00566.t01#1    mNeoVis.2.1.1.chr13_hap1.g00580.t01#5    mNeoVis.evergladensis.2.1.chr13.g00564.t01#2    mNeoVis.evergladensis.1.1.chr13.g00560.t01#3   mNeoVis.vulgivaga.1.1.chr13.g00552.t01#4       6766   125255098:   mNeoVis.vulgivaga.2.1.chr5.g01248.t01#1    mNeoVis.vulgivaga.1.1.chr5.g01219.t01#4    mNeoVis.2.1.1.chr2_hap1.g01311.t01#5    mNeoVis.evergladensis.1.1.chr5.g01254.t01#3    mNeoVis.evergladensis.2.1.chr5.g01252.t01#2   mNeoVis.3.1.1.chr5.g01236.t01#6       6767   125255099:   mNeoVis.evergladensis.2.1.chr2.g00312.t01#2    mNeoVis.vulgivaga.1.1.chr2.g00309.t01#4    mNeoVis.2.1.1.chr5_hap1.g00313.t01#5    mNeoVis.3.1.1.chr2.g00251.t01#6    mNeoVis.vulgivaga.2.1.chr2.g00308.t01#1   mNeoVis.evergladensis.1.1.chr2.g00312.t01#3       6768   125255100:   mNeoVis.vulgivaga.2.1.chr13.g00567.t01#1    mNeoVis.2.1.1.chr13_hap1.g00581.t01#5    mNeoVis.evergladensis.1.1.chr13.g00561.t01#3    mNeoVis.vulgivaga.1.1.chr13.g00553.t01#4    mNeoVis.evergladensis.2.1.chr13.g00565.t01#2   mNeoVis.3.1.1.chr13.g00518.t01#6       6769   125255101:   mNeoVis.vulgivaga.1.1.chr13.g00554.t01#4    mNeoVis.2.1.1.chr13_hap1.g00583.t01#5    mNeoVis.evergladensis.2.1.chr13.g00566.t01#2    mNeoVis.evergladensis.1.1.chr13.g00563.t01#3    mNeoVis.vulgivaga.2.1.chr13.g00568.t01#1   mNeoVis.3.1.1.chr13.g00520.t01#6       6770   125255102:   mNeoVis.evergladensis.2.1.chr13.g00567.t01#2    mNeoVis.vulgivaga.2.1.chr13.g00569.t01#1    mNeoVis.vulgivaga.1.1.chr13.g00555.t01#4    mNeoVis.3.1.1.chr13.g00521.t01#6    mNeoVis.evergladensis.1.1.chr13.g00564.t01#3   mNeoVis.2.1.1.chr13_hap1.g00584.t01#5       6771   125255103:   mNeoVis.vulgivaga.2.1.chr6.g00039.t01#1    mNeoVis.3.1.1.chr6.g00037.t01#6    mNeoVis.evergladensis.2.1.chr6.g00035.t01#2    mNeoVis.vulgivaga.1.1.chr6.g00037.t01#4    mNeoVis.2.1.1.chr4_hap1.g00033.t01#5   mNeoVis.evergladensis.1.1.chr6.g00034.t01#3       6772   125255104:   mNeoVis.evergladensis.1.1.chr13.g00565.t01#3    mNeoVis.2.1.1.chr13_hap1.g00585.t01#5    mNeoVis.vulgivaga.2.1.chr13.g00570.t01#1    mNeoVis.3.1.1.chr13.g00523.t01#6    mNeoVis.evergladensis.2.1.chr13.g00568.t01#2   mNeoVis.vulgivaga.1.1.chr13.g00557.t01#4       6773   125255105:   mNeoVis.2.1.1.chr13_hap1.g00587.t01#5    mNeoVis.evergladensis.2.1.chr13.g00569.t01#2    mNeoVis.3.1.1.chr13.g00524.t01#6    mNeoVis.vulgivaga.2.1.chr13.g00571.t01#1    mNeoVis.evergladensis.1.1.chr13.g00566.t01#3   mNeoVis.vulgivaga.1.1.chr13.g00558.t01#4       6774   125255106:   mNeoVis.evergladensis.1.1.chr6.g00035.t01#3    mNeoVis.vulgivaga.2.1.chr6.g00040.t01#1    mNeoVis.2.1.1.chr4_hap1.g00034.t01#5    mNeoVis.vulgivaga.1.1.chr6.g00038.t01#4   mNeoVis.evergladensis.2.1.chr6.g00036.t01#2       6775   125255107:   mNeoVis.vulgivaga.2.1.chr13.g00574.t01#1    mNeoVis.3.1.1.chr12.g00227.t01#6    mNeoVis.2.1.1.chr13_hap1.g00597.t01#5    mNeoVis.vulgivaga.1.1.chr12.g00236.t01#4    mNeoVis.evergladensis.2.1.chr12.g00228.t01#2    mNeoVis.evergladensis.2.1.chr13.g00574.t01#2    mNeoVis.3.1.1.chr13.g00527.t01#6    mNeoVis.vulgivaga.1.1.chr13.g00561.t01#4    mNeoVis.evergladensis.1.1.chr13.g00571.t01#3      6776   125255108:   mNeoVis.2.1.1.chr13_hap1.g00589.t01#5    mNeoVis.2.1.1.chr13_hap1.g00604.t01#5    mNeoVis.2.1.1.chr13_hap1.g00596.t01#5    mNeoVis.2.1.1.chr13_hap1.g00588.t01#5    mNeoVis.2.1.1.chr13_hap1.g00603.t01#5    mNeoVis.2.1.1.chr13_hap1.g00595.t01#5    mNeoVis.2.1.1.chr13_hap1.g00602.t01#5    mNeoVis.2.1.1.chr13_hap1.g00594.t01#5    mNeoVis.2.1.1.chr13_hap1.g00601.t01#5      6777   125255109:   mNeoVis.evergladensis.2.1.chr13.g00571.t01#2    mNeoVis.evergladensis.2.1.chr13.g00579.t01#2    mNeoVis.evergladensis.2.1.chr13.g00576.t01#2    mNeoVis.evergladensis.2.1.chr13.g00572.t01#2    mNeoVis.evergladensis.2.1.chr13.g00570.t01#2    mNeoVis.evergladensis.2.1.chr13.g00581.t01#2    mNeoVis.evergladensis.2.1.chr13.g00578.t01#2    mNeoVis.evergladensis.2.1.chr13.g00573.t01#2    mNeoVis.evergladensis.2.1.chr13.g00577.t01#2      6778   125255110:   mNeoVis.evergladensis.1.1.chr13.g00573.t01#3    mNeoVis.evergladensis.1.1.chr13.g00569.t01#3    mNeoVis.vulgivaga.1.1.chr13.g00559.t01#4    mNeoVis.evergladensis.1.1.chr13.g00570.t01#3    mNeoVis.evergladensis.1.1.chr13.g00575.t01#3    mNeoVis.evergladensis.1.1.chr13.g00568.t01#3    mNeoVis.evergladensis.1.1.chr13.g00576.t01#3    mNeoVis.vulgivaga.1.1.chr13.g00560.t01#4    mNeoVis.evergladensis.1.1.chr13.g00567.t01#3      6779   125255111:   mNeoVis.vulgivaga.2.1.chr13.g00578.t01#1    mNeoVis.vulgivaga.2.1.chr13.g00576.t01#1    mNeoVis.vulgivaga.2.1.chr13.g00579.t01#1    mNeoVis.vulgivaga.2.1.chr13.g00572.t01#1    mNeoVis.vulgivaga.2.1.chr13.g00580.t01#1    mNeoVis.vulgivaga.2.1.chr13.g00573.t01#1   mNeoVis.vulgivaga.2.1.chr13.g00577.t01#1     6780   125255112:   mNeoVis.3.1.1.chr13.g00528.t01#6    mNeoVis.vulgivaga.1.1.chr13.g00562.t01#4    mNeoVis.evergladensis.2.1.chr13.g00575.t01#2    mNeoVis.2.1.1.chr13_hap1.g00598.t01#5    mNeoVis.vulgivaga.2.1.chr13.g00575.t01#1   mNeoVis.evergladensis.1.1.chr13.g00572.t01#3       6781   125255113:   mNeoVis.2.1.1.chr13_hap1.g00605.t01#5    mNeoVis.evergladensis.2.1.chr13.g00582.t01#2    mNeoVis.vulgivaga.1.1.chr13.g00563.t01#4    mNeoVis.evergladensis.1.1.chr13.g00574.t01#3    mNeoVis.vulgivaga.2.1.chr13.g00581.t01#1   mNeoVis.3.1.1.chr13.g00529.t01#6       6782   125255114:   mNeoVis.vulgivaga.1.1.chr6.g00039.t01#4    mNeoVis.2.1.1.chr4_hap1.g00035.t01#5    mNeoVis.evergladensis.2.1.chr6.g00037.t01#2    mNeoVis.evergladensis.1.1.chr6.g00036.t01#3    mNeoVis.vulgivaga.2.1.chr6.g00041.t01#1   mNeoVis.3.1.1.chr6.g00038.t01#6       6783   125255115:   mNeoVis.vulgivaga.2.1.chr13.g00582.t01#1    mNeoVis.evergladensis.2.1.chr13.g00583.t01#2    mNeoVis.evergladensis.1.1.chr13.g00577.t01#3    mNeoVis.3.1.1.chr13.g00530.t01#6    mNeoVis.vulgivaga.1.1.chr13.g00564.t01#4   mNeoVis.2.1.1.chr13_hap1.g00606.t01#5       6784   125255116:   mNeoVis.vulgivaga.2.1.chr6.g00043.t01#1    mNeoVis.2.1.1.chr4_hap1.g00037.t01#5    mNeoVis.vulgivaga.1.1.chr6.g00041.t01#4    mNeoVis.evergladensis.2.1.chr6.g00039.t01#2   mNeoVis.evergladensis.1.1.chr6.g00038.t01#3       6785   125255117:   mNeoVis.vulgivaga.1.1.chr6.g00042.t01#4   mNeoVis.3.1.1.chr6.g00040.t01#6             6786   125255118:   mNeoVis.vulgivaga.2.1.chr13.g00583.t01#1    mNeoVis.vulgivaga.1.1.chr13.g00565.t01#4    mNeoVis.evergladensis.1.1.chr13.g00578.t01#3    mNeoVis.3.1.1.chr13.g00531.t01#6    mNeoVis.2.1.1.chr13_hap1.g00607.t01#5   mNeoVis.evergladensis.2.1.chr13.g00584.t01#2       6787   125255119:   mNeoVis.2.1.1.chr15_hap1.g00657.t01#5    mNeoVis.vulgivaga.2.1.chr15.g00489.t01#1    mNeoVis.evergladensis.2.1.chr15.g00479.t01#2    mNeoVis.evergladensis.1.1.chr15.g00133.t01#3    mNeoVis.vulgivaga.1.1.chr15.g00483.t01#4   mNeoVis.3.1.1.chr15.g00441.t01#6       6788   125255120:   mNeoVis.vulgivaga.2.1.chr13.g00584.t01#1    mNeoVis.3.1.1.chr13.g00532.t01#6    mNeoVis.evergladensis.1.1.chr13.g00579.t01#3    mNeoVis.2.1.1.chr13_hap1.g00608.t01#5    mNeoVis.vulgivaga.1.1.chr13.g00567.t01#4   mNeoVis.evergladensis.2.1.chr13.g00585.t01#2       6789   125255121:   mNeoVis.2.1.1.chr4_hap1.g00038.t01#5    mNeoVis.evergladensis.2.1.chr6.g00040.t01#2    mNeoVis.evergladensis.1.1.chr6.g00039.t01#3    mNeoVis.vulgivaga.2.1.chr6.g00044.t01#1    mNeoVis.vulgivaga.1.1.chr6.g00043.t01#4   mNeoVis.3.1.1.chr6.g00041.t01#6       6790   125255122:   mNeoVis.vulgivaga.1.1.chr15.g00484.t01#4    mNeoVis.2.1.1.chr15_hap1.g00658.t01#5    mNeoVis.vulgivaga.2.1.chr15.g00490.t01#1    mNeoVis.evergladensis.1.1.chr15.g00132.t01#3    mNeoVis.3.1.1.chr15.g00442.t01#6   mNeoVis.evergladensis.2.1.chr15.g00480.t01#2       6791   125255123:   mNeoVis.vulgivaga.1.1.chr6.g00044.t01#4    mNeoVis.2.1.1.chr4_hap1.g00039.t01#5    mNeoVis.vulgivaga.2.1.chr6.g00045.t01#1    mNeoVis.evergladensis.1.1.chr6.g00040.t01#3    mNeoVis.3.1.1.chr6.g00042.t01#6   mNeoVis.evergladensis.2.1.chr6.g00041.t01#2       6792   125255124:   mNeoVis.3.1.1.chr6.g00043.t01#6    mNeoVis.evergladensis.2.1.chr6.g00042.t01#2    mNeoVis.vulgivaga.2.1.chr6.g00046.t01#1    mNeoVis.2.1.1.chr4_hap1.g00040.t01#5   mNeoVis.evergladensis.1.1.chr6.g00041.t01#3       6793   125255125:   mNeoVis.3.1.1.chr7.g00410.t01#6    mNeoVis.vulgivaga.1.1.chr7.g00432.t01#4    mNeoVis.2.1.1.chr7_hap1.g00424.t01#5    mNeoVis.evergladensis.2.1.chr7.g00428.t01#2    mNeoVis.vulgivaga.2.1.chr7.g00426.t01#1   mNeoVis.evergladensis.1.1.chr7.g00435.t01#3       6794   125255126:   mNeoVis.vulgivaga.2.1.chr6.g00047.t01#1    mNeoVis.evergladensis.2.1.chr6.g00043.t01#2    mNeoVis.3.1.1.chr6.g00044.t01#6    mNeoVis.evergladensis.1.1.chr6.g00042.t01#3    mNeoVis.vulgivaga.1.1.chr6.g00047.t01#4   mNeoVis.2.1.1.chr4_hap1.g00041.t01#5       6795   125255127:   mNeoVis.vulgivaga.1.1.chr6.g00048.t01#4    mNeoVis.evergladensis.2.1.chr6.g00044.t01#2    mNeoVis.2.1.1.chr4_hap1.g00042.t01#5    mNeoVis.vulgivaga.2.1.chr6.g00048.t01#1    mNeoVis.3.1.1.chr6.g00045.t01#6   mNeoVis.evergladensis.1.1.chr6.g00043.t01#3       6796   125255128:   mNeoVis.3.1.1.chr13.g00533.t01#6    mNeoVis.vulgivaga.1.1.chr13.g00569.t01#4    mNeoVis.vulgivaga.2.1.chr13.g00585.t01#1    mNeoVis.evergladensis.1.1.chr1.g01541.t01#3    mNeoVis.evergladensis.1.1.chr13.g00581.t01#3    mNeoVis.evergladensis.2.1.chr13.g00586.t01#2   mNeoVis.2.1.1.chr13_hap1.g00609.t01#5     6797   125255129:   mNeoVis.3.1.1.chr6.g00046.t01#6    mNeoVis.vulgivaga.2.1.chr6.g00049.t01#1    mNeoVis.2.1.1.chr4_hap1.g00043.t01#5    mNeoVis.evergladensis.1.1.chr6.g00044.t01#3    mNeoVis.vulgivaga.1.1.chr6.g00049.t01#4   mNeoVis.evergladensis.2.1.chr6.g00045.t01#2       6798   125255130:   mNeoVis.2.1.1.chr6_hap1.g01068.t01#5    mNeoVis.3.1.1.chr12.g00771.t01#6    mNeoVis.vulgivaga.2.1.chr12.g00799.t01#1    mNeoVis.vulgivaga.1.1.chr4.g01032.t01#4    mNeoVis.evergladensis.2.1.chr12.g00770.t01#2    mNeoVis.vulgivaga.2.1.chr13.g00586.t01#1    mNeoVis.2.1.1.chr12_hap1.g00842.t01#5    mNeoVis.3.1.1.chr4.g01023.t01#6    mNeoVis.vulgivaga.1.1.chr13.g00570.t01#4      6799   125255131:   mNeoVis.evergladensis.2.1.chr12.g00218.t01#2    mNeoVis.evergladensis.1.1.chr13.g00582.t01#3    mNeoVis.2.1.1.chr12_hap1.g00249.t01#5    mNeoVis.2.1.1.chr13_hap1.g00610.t01#5    mNeoVis.evergladensis.1.1.chr12.g00220.t01#3   mNeoVis.3.1.1.chr13.g00534.t01#6       6800   125255132:   mNeoVis.vulgivaga.1.1.chr4.g00441.t01#4    mNeoVis.evergladensis.1.1.chr4.g00447.t01#3    mNeoVis.2.1.1.chr6_hap1.g00439.t01#5    mNeoVis.3.1.1.chr4.g00439.t01#6    mNeoVis.vulgivaga.2.1.chr4.g00448.t01#1   mNeoVis.evergladensis.2.1.chr4.g00446.t01#2       6801   125255133:   mNeoVis.3.1.1.chr12.g00216.t01#6   mNeoVis.vulgivaga.1.1.chr12.g00225.t01#4           6802   125255134:   mNeoVis.2.1.1.chr13_hap1.g00611.t01#5    mNeoVis.vulgivaga.1.1.chr13.g00571.t01#4    mNeoVis.evergladensis.1.1.chr13.g00583.t01#3    mNeoVis.vulgivaga.2.1.chr13.g00587.t01#1   mNeoVis.evergladensis.2.1.chr13.g00587.t01#2       6803   125255135:   mNeoVis.evergladensis.2.1.chr13.g00588.t01#2    mNeoVis.3.1.1.chr13.g00536.t01#6    mNeoVis.vulgivaga.2.1.chr13.g00588.t01#1    mNeoVis.2.1.1.chr13_hap1.g00612.t01#5    mNeoVis.vulgivaga.1.1.chr13.g00572.t01#4   mNeoVis.evergladensis.1.1.chr13.g00584.t01#3       6804   125255136:   mNeoVis.3.1.1.chr6.g00047.t01#6    mNeoVis.vulgivaga.1.1.chr6.g00050.t01#4    mNeoVis.evergladensis.1.1.chr6.g00045.t01#3    mNeoVis.vulgivaga.2.1.chr6.g00050.t01#1    mNeoVis.evergladensis.2.1.chr6.g00046.t01#2   mNeoVis.2.1.1.chr4_hap1.g00044.t01#5       6805   125255137:   mNeoVis.3.1.1.chr13.g00538.t01#6    mNeoVis.vulgivaga.1.1.chr13.g00574.t01#4    mNeoVis.evergladensis.2.1.chr13.g00590.t01#2    mNeoVis.evergladensis.1.1.chr13.g00586.t01#3    mNeoVis.vulgivaga.2.1.chr13.g00590.t01#1   mNeoVis.2.1.1.chr13_hap1.g00614.t01#5       6806   125255138:   mNeoVis.2.1.1.chr4_hap1.g00045.t01#5    mNeoVis.vulgivaga.2.1.chr6.g00051.t01#1    mNeoVis.evergladensis.2.1.chr6.g00047.t01#2    mNeoVis.vulgivaga.1.1.chr6.g00051.t01#4    mNeoVis.evergladensis.1.1.chr6.g00046.t01#3   mNeoVis.3.1.1.chr6.g00048.t01#6       6807   125255139:   mNeoVis.evergladensis.1.1.chr6.g00047.t01#3    mNeoVis.3.1.1.chr6.g00049.t01#6    mNeoVis.vulgivaga.2.1.chr6.g00052.t01#1    mNeoVis.2.1.1.chr4_hap1.g00046.t01#5    mNeoVis.evergladensis.2.1.chr6.g00048.t01#2   mNeoVis.vulgivaga.1.1.chr6.g00052.t01#4       6808   125255140:   mNeoVis.evergladensis.1.1.chr6.g00048.t01#3    mNeoVis.3.1.1.chr6.g00050.t01#6    mNeoVis.vulgivaga.2.1.chr6.g00053.t01#1    mNeoVis.vulgivaga.1.1.chr6.g00053.t01#4    mNeoVis.evergladensis.2.1.chr6.g00049.t01#2   mNeoVis.2.1.1.chr4_hap1.g00047.t01#5       6809   125255141:   mNeoVis.vulgivaga.2.1.chr13.g00591.t01#1    mNeoVis.evergladensis.1.1.chr13.g00588.t01#3    mNeoVis.evergladensis.2.1.chr13.g00591.t01#2    mNeoVis.3.1.1.chr13.g00539.t01#6    mNeoVis.2.1.1.chr13_hap1.g00615.t01#5   mNeoVis.vulgivaga.1.1.chr13.g00575.t01#4       6810   125255142:   mNeoVis.vulgivaga.1.1.chr15.g00485.t01#4    mNeoVis.evergladensis.1.1.chr15.g00131.t01#3    mNeoVis.2.1.1.chr15_hap1.g00659.t01#5    mNeoVis.vulgivaga.2.1.chr15.g00491.t01#1    mNeoVis.3.1.1.chr15.g00443.t01#6   mNeoVis.evergladensis.2.1.chr15.g00481.t01#2       6811   125255143:   mNeoVis.vulgivaga.2.1.chr13.g00592.t01#1    mNeoVis.2.1.1.chr13_hap1.g00616.t01#5    mNeoVis.evergladensis.1.1.chr13.g00589.t01#3    mNeoVis.evergladensis.2.1.chr13.g00592.t01#2    mNeoVis.3.1.1.chr13.g00540.t01#6   mNeoVis.vulgivaga.1.1.chr13.g00576.t01#4       6812   125255144:   mNeoVis.evergladensis.2.1.chr6.g00050.t01#2    mNeoVis.evergladensis.1.1.chr6.g00049.t01#3    mNeoVis.2.1.1.chr4_hap1.g00048.t01#5    mNeoVis.vulgivaga.2.1.chr6.g00054.t01#1   mNeoVis.vulgivaga.1.1.chr6.g00054.t01#4       6813   125255145:   mNeoVis.evergladensis.1.1.chr12.g00507.t01#3    mNeoVis.evergladensis.2.1.chr12.g00500.t01#2    mNeoVis.vulgivaga.1.1.chr13.g00577.t01#4    mNeoVis.vulgivaga.2.1.chr12.g00513.t01#1    mNeoVis.2.1.1.chr13_hap1.g00617.t01#5    mNeoVis.3.1.1.chr13.g00541.t01#6    mNeoVis.vulgivaga.1.1.chr12.g00513.t01#4    mNeoVis.2.1.1.chr12_hap1.g00555.t01#5    mNeoVis.3.1.1.chr12.g00513.t01#6      6814   125255146:   mNeoVis.vulgivaga.2.1.chr13.g00594.t01#1    mNeoVis.evergladensis.1.1.chr13.g00591.t01#3    mNeoVis.vulgivaga.1.1.chr13.g00578.t01#4    mNeoVis.evergladensis.2.1.chr13.g00594.t01#2    mNeoVis.3.1.1.chr13.g00542.t01#6   mNeoVis.2.1.1.chr13_hap1.g00618.t01#5       6815   125255147:   mNeoVis.vulgivaga.2.1.chr13.g00483.t01#1    mNeoVis.vulgivaga.1.1.chr13.g00476.t01#4    mNeoVis.2.1.1.chr13_hap1.g00501.t01#5    mNeoVis.3.1.1.chr13.g00436.t01#6    mNeoVis.evergladensis.2.1.chr13.g00482.t01#2   mNeoVis.evergladensis.1.1.chr13.g00481.t01#3       6816   125255148:   mNeoVis.evergladensis.2.1.chr6.g00052.t01#2    mNeoVis.3.1.1.chr6.g00052.t01#6    mNeoVis.vulgivaga.2.1.chr6.g00055.t01#1    mNeoVis.vulgivaga.1.1.chr6.g00055.t01#4    mNeoVis.2.1.1.chr4_hap1.g00049.t01#5   mNeoVis.evergladensis.1.1.chr6.g00050.t01#3       6817   125255149:   mNeoVis.3.1.1.chr13.g00437.t01#6    mNeoVis.evergladensis.1.1.chr13.g00482.t01#3    mNeoVis.vulgivaga.2.1.chr13.g00484.t01#1    mNeoVis.evergladensis.2.1.chr13.g00483.t01#2    mNeoVis.vulgivaga.1.1.chr13.g00477.t01#4   mNeoVis.2.1.1.chr13_hap1.g00502.t01#5       6818   125255150:   mNeoVis.vulgivaga.1.1.chr13.g00478.t01#4    mNeoVis.evergladensis.1.1.chr13.g00483.t01#3    mNeoVis.evergladensis.2.1.chr13.g00484.t01#2    mNeoVis.3.1.1.chr13.g00438.t01#6    mNeoVis.2.1.1.chr13_hap1.g00503.t01#5   mNeoVis.vulgivaga.2.1.chr13.g00485.t01#1       6819   125255151:   mNeoVis.vulgivaga.2.1.chr6.g00056.t01#1    mNeoVis.evergladensis.2.1.chr6.g00054.t01#2    mNeoVis.evergladensis.1.1.chr6.g00052.t01#3    mNeoVis.vulgivaga.1.1.chr6.g00056.t01#4   mNeoVis.2.1.1.chr4_hap1.g00050.t01#5       6820   125255152:   mNeoVis.evergladensis.1.1.chr13.g00484.t01#3    mNeoVis.vulgivaga.2.1.chr13.g00486.t01#1    mNeoVis.2.1.1.chr13_hap1.g00504.t01#5    mNeoVis.3.1.1.chr13.g00439.t01#6    mNeoVis.evergladensis.2.1.chr13.g00485.t01#2   mNeoVis.vulgivaga.1.1.chr13.g00479.t01#4       6821   125255153:   mNeoVis.2.1.1.chr15_hap1.g00661.t01#5    mNeoVis.evergladensis.1.1.chr15.g00130.t01#3    mNeoVis.vulgivaga.2.1.chr15.g00493.t01#1    mNeoVis.vulgivaga.1.1.chr15.g00487.t01#4    mNeoVis.evergladensis.2.1.chr15.g00483.t01#2   mNeoVis.3.1.1.chr15.g00445.t01#6       6822   125255154:   mNeoVis.3.1.1.chr6.g00053.t01#6    mNeoVis.vulgivaga.2.1.chr6.g00057.t01#1    mNeoVis.evergladensis.1.1.chr6.g00053.t01#3    mNeoVis.2.1.1.chr4_hap1.g00051.t01#5    mNeoVis.vulgivaga.1.1.chr6.g00057.t01#4   mNeoVis.evergladensis.2.1.chr6.g00055.t01#2       6823   125255155:   mNeoVis.2.1.1.chr13_hap1.g00505.t01#5    mNeoVis.3.1.1.chr13.g00440.t01#6    mNeoVis.evergladensis.1.1.chr13.g00485.t01#3    mNeoVis.vulgivaga.1.1.chr13.g00480.t01#4    mNeoVis.evergladensis.2.1.chr13.g00486.t01#2   mNeoVis.vulgivaga.2.1.chr13.g00487.t01#1       6824   125255156:   mNeoVis.3.1.1.chr7.g00411.t01#6    mNeoVis.evergladensis.2.1.chr7.g00429.t01#2    mNeoVis.vulgivaga.1.1.chr7.g00433.t01#4    mNeoVis.vulgivaga.2.1.chr7.g00427.t01#1    mNeoVis.2.1.1.chr7_hap1.g00425.t01#5   mNeoVis.evergladensis.1.1.chr7.g00436.t01#3       6825   125255157:   mNeoVis.evergladensis.2.1.chr6.g00056.t01#2    mNeoVis.evergladensis.1.1.chr6.g00054.t01#3    mNeoVis.2.1.1.chr4_hap1.g00052.t01#5    mNeoVis.3.1.1.chr6.g00054.t01#6    mNeoVis.vulgivaga.2.1.chr6.g00058.t01#1   mNeoVis.vulgivaga.1.1.chr6.g00058.t01#4       6826   125255158:   mNeoVis.evergladensis.2.1.chr13.g00487.t01#2    mNeoVis.evergladensis.1.1.chr13.g00486.t01#3    mNeoVis.vulgivaga.1.1.chr13.g00481.t01#4    mNeoVis.vulgivaga.2.1.chr13.g00488.t01#1    mNeoVis.2.1.1.chr13_hap1.g00506.t01#5   mNeoVis.3.1.1.chr13.g00441.t01#6       6827   125255159:   mNeoVis.vulgivaga.2.1.chr6.g00059.t01#1    mNeoVis.3.1.1.chr6.g00055.t01#6    mNeoVis.evergladensis.1.1.chr6.g00055.t01#3    mNeoVis.2.1.1.chr4_hap1.g00053.t01#5    mNeoVis.vulgivaga.1.1.chr6.g00059.t01#4   mNeoVis.evergladensis.2.1.chr6.g00057.t01#2       6828   125255160:   mNeoVis.3.1.1.chr6.g00057.t01#6    mNeoVis.evergladensis.1.1.chr6.g00056.t01#3    mNeoVis.vulgivaga.1.1.chr6.g00060.t01#4    mNeoVis.2.1.1.chr4_hap1.g00054.t01#5    mNeoVis.evergladensis.2.1.chr6.g00059.t01#2   mNeoVis.vulgivaga.2.1.chr6.g00060.t01#1       6829   125255161:   mNeoVis.evergladensis.1.1.chr13.g00487.t01#3    mNeoVis.vulgivaga.2.1.chr13.g00489.t01#1    mNeoVis.3.1.1.chr13.g00442.t01#6    mNeoVis.2.1.1.chr13_hap1.g00507.t01#5    mNeoVis.vulgivaga.1.1.chr13.g00482.t01#4   mNeoVis.evergladensis.2.1.chr13.g00488.t01#2       6830   125255162:   mNeoVis.vulgivaga.1.1.chr7.g00434.t01#4    mNeoVis.3.1.1.chr7.g00412.t01#6    mNeoVis.2.1.1.chr7_hap1.g00426.t01#5    mNeoVis.evergladensis.1.1.chr7.g00437.t01#3    mNeoVis.evergladensis.2.1.chr7.g00430.t01#2   mNeoVis.vulgivaga.2.1.chr7.g00428.t01#1       6831   125255163:   mNeoVis.vulgivaga.2.1.chr6.g00061.t01#1    mNeoVis.evergladensis.1.1.chr6.g00057.t01#3    mNeoVis.evergladensis.2.1.chr6.g00060.t01#2    mNeoVis.vulgivaga.1.1.chr6.g00061.t01#4    mNeoVis.2.1.1.chr4_hap1.g00055.t01#5   mNeoVis.3.1.1.chr6.g00058.t01#6       6832   125255164:   mNeoVis.evergladensis.1.1.chr13.g00488.t01#3    mNeoVis.3.1.1.chr13.g00443.t01#6    mNeoVis.2.1.1.chr13_hap1.g00508.t01#5    mNeoVis.vulgivaga.2.1.chr13.g00490.t01#1    mNeoVis.vulgivaga.1.1.chr13.g00483.t01#4   mNeoVis.evergladensis.2.1.chr13.g00489.t01#2       6833   125255165:   mNeoVis.evergladensis.2.1.chr15.g00484.t01#2    mNeoVis.3.1.1.chr15.g00446.t01#6    mNeoVis.vulgivaga.1.1.chr15.g00488.t01#4    mNeoVis.evergladensis.1.1.chr15.g00129.t01#3    mNeoVis.2.1.1.chr15_hap1.g00662.t01#5   mNeoVis.vulgivaga.2.1.chr15.g00494.t01#1       6834   125255166:   mNeoVis.evergladensis.1.1.chr6.g00058.t01#3    mNeoVis.3.1.1.chr6.g00059.t01#6    mNeoVis.vulgivaga.2.1.chr6.g00062.t01#1    mNeoVis.evergladensis.2.1.chr6.g00061.t01#2    mNeoVis.vulgivaga.1.1.chr6.g00062.t01#4   mNeoVis.2.1.1.chr4_hap1.g00056.t01#5       6835   125255167:   mNeoVis.vulgivaga.1.1.chr6.g00064.t01#4    mNeoVis.3.1.1.chr6.g00061.t01#6    mNeoVis.vulgivaga.2.1.chr6.g00063.t01#1    mNeoVis.evergladensis.2.1.chr6.g00063.t01#2    mNeoVis.2.1.1.chr4_hap1.g00058.t01#5   mNeoVis.evergladensis.1.1.chr6.g00060.t01#3       6836   125255168:   mNeoVis.evergladensis.1.1.chr13.g00489.t01#3    mNeoVis.vulgivaga.2.1.chr13.g00491.t01#1    mNeoVis.vulgivaga.1.1.chr13.g00484.t01#4   mNeoVis.3.1.1.chr13.g00444.t01#6         6837   125255169:   mNeoVis.evergladensis.1.1.chr13.g00490.t01#3    mNeoVis.vulgivaga.2.1.chr13.g00493.t01#1    mNeoVis.3.1.1.chr13.g00445.t01#6   mNeoVis.vulgivaga.1.1.chr13.g00485.t01#4       6838   125255170:   mNeoVis.evergladensis.2.1.chr13.g00490.t01#2   mNeoVis.2.1.1.chr13_hap1.g00509.t01#5             6839   125255171:  mNeoVis.vulgivaga.2.1.chr13.g00492.t01#1               6840   125255172:   mNeoVis.evergladensis.1.1.chr7.g00438.t01#3    mNeoVis.2.1.1.chr7_hap1.g00427.t01#5    mNeoVis.3.1.1.chr7.g00413.t01#6    mNeoVis.vulgivaga.1.1.chr7.g00435.t01#4    mNeoVis.evergladensis.2.1.chr7.g00431.t01#2   mNeoVis.vulgivaga.2.1.chr7.g00429.t01#1       6841   125255173:   mNeoVis.evergladensis.1.1.chr6.g00061.t01#3    mNeoVis.vulgivaga.1.1.chr6.g00065.t01#4   mNeoVis.vulgivaga.2.1.chr6.g00064.t01#1           6842   125255174:   mNeoVis.vulgivaga.1.1.chr6.g00066.t01#4    mNeoVis.3.1.1.chr6.g00063.t01#6    mNeoVis.evergladensis.2.1.chr6.g00065.t01#2    mNeoVis.2.1.1.chr4_hap1.g00060.t01#5    mNeoVis.vulgivaga.2.1.chr6.g00065.t01#1   mNeoVis.evergladensis.1.1.chr6.g00062.t01#3       6843   125255175:   mNeoVis.evergladensis.1.1.chr6.g00063.t01#3    mNeoVis.vulgivaga.1.1.chr6.g00067.t01#4    mNeoVis.evergladensis.2.1.chr6.g00066.t01#2    mNeoVis.vulgivaga.2.1.chr6.g00066.t01#1   mNeoVis.3.1.1.chr6.g00064.t01#6       6844   125255176:   mNeoVis.evergladensis.2.1.chr6.g00067.t01#2    mNeoVis.vulgivaga.1.1.chr6.g00068.t01#4    mNeoVis.vulgivaga.2.1.chr6.g00067.t01#1    mNeoVis.2.1.1.chr4_hap1.g00062.t01#5   mNeoVis.evergladensis.1.1.chr6.g00064.t01#3       6845   125255177:   mNeoVis.2.1.1.chr13_hap1.g00510.t01#5    mNeoVis.vulgivaga.1.1.chr13.g00486.t01#4    mNeoVis.3.1.1.chr13.g00446.t01#6    mNeoVis.evergladensis.1.1.chr13.g00491.t01#3    mNeoVis.vulgivaga.2.1.chr13.g00494.t01#1   mNeoVis.evergladensis.2.1.chr13.g00491.t01#2       6846   125255178:   mNeoVis.vulgivaga.1.1.chr7.g00436.t01#4    mNeoVis.vulgivaga.2.1.chr7.g00430.t01#1    mNeoVis.3.1.1.chr7.g00414.t01#6    mNeoVis.evergladensis.1.1.chr7.g00439.t01#3    mNeoVis.2.1.1.chr7_hap1.g00428.t01#5   mNeoVis.evergladensis.2.1.chr7.g00432.t01#2       6847   125255179:   mNeoVis.evergladensis.2.1.chr15.g00485.t01#2    mNeoVis.evergladensis.1.1.chr15.g00128.t01#3    mNeoVis.2.1.1.chr15_hap1.g00663.t01#5    mNeoVis.3.1.1.chr15.g00447.t01#6   mNeoVis.vulgivaga.2.1.chr15.g00495.t01#1       6848   125255180:   mNeoVis.2.1.1.chr15_hap1.g00664.t01#5    mNeoVis.vulgivaga.1.1.chr15.g00489.t01#4    mNeoVis.evergladensis.1.1.chr15.g00127.t01#3    mNeoVis.evergladensis.2.1.chr15.g00486.t01#2   mNeoVis.vulgivaga.2.1.chr15.g00496.t01#1       6849   125255181:   mNeoVis.evergladensis.2.1.chr7.g00433.t01#2    mNeoVis.2.1.1.chr7_hap1.g00429.t01#5    mNeoVis.3.1.1.chr7.g00415.t01#6    mNeoVis.evergladensis.1.1.chr7.g00440.t01#3    mNeoVis.vulgivaga.2.1.chr7.g00431.t01#1   mNeoVis.vulgivaga.1.1.chr7.g00437.t01#4       6850   125255182:  mNeoVis.vulgivaga.2.1.chr13.g00496.t01#1               6851   125255183:   mNeoVis.vulgivaga.2.1.chr13.g00495.t01#1   mNeoVis.3.1.1.chr13.g00452.t01#6             6852   125255184:   mNeoVis.evergladensis.1.1.chr13.g00494.t01#3    mNeoVis.3.1.1.chr13.g00453.t01#6    mNeoVis.vulgivaga.2.1.chr13.g00498.t01#1    mNeoVis.2.1.1.chr13_hap1.g00512.t01#5    mNeoVis.evergladensis.2.1.chr13.g00493.t01#2   mNeoVis.vulgivaga.1.1.chr13.g00488.t01#4       6853   125255185:   mNeoVis.vulgivaga.1.1.chr13.g00489.t01#4    mNeoVis.3.1.1.chr13.g00454.t01#6    mNeoVis.2.1.1.chr13_hap1.g00513.t01#5    mNeoVis.evergladensis.1.1.chr13.g00495.t01#3    mNeoVis.evergladensis.2.1.chr13.g00494.t01#2   mNeoVis.vulgivaga.2.1.chr13.g00499.t01#1       6854   125255186:   mNeoVis.vulgivaga.1.1.chr6.g00069.t01#4    mNeoVis.3.1.1.chr6.g00066.t01#6    mNeoVis.vulgivaga.2.1.chr6.g00068.t01#1    mNeoVis.evergladensis.1.1.chr6.g00065.t01#3    mNeoVis.2.1.1.chr4_hap1.g00063.t01#5   mNeoVis.evergladensis.2.1.chr6.g00068.t01#2       6855   125255187:   mNeoVis.2.1.1.chr13_hap1.g00514.t01#5    mNeoVis.vulgivaga.2.1.chr13.g00500.t01#1    mNeoVis.evergladensis.2.1.chr13.g00495.t01#2    mNeoVis.vulgivaga.1.1.chr13.g00490.t01#4    mNeoVis.evergladensis.1.1.chr13.g00496.t01#3   mNeoVis.3.1.1.chr13.g00455.t01#6       6856   125255188:   mNeoVis.vulgivaga.2.1.chr6.g00069.t01#1    mNeoVis.2.1.1.chr4_hap1.g00064.t01#5    mNeoVis.evergladensis.2.1.chr6.g00069.t01#2    mNeoVis.vulgivaga.1.1.chr6.g00070.t01#4   mNeoVis.evergladensis.1.1.chr6.g00066.t01#3       6857   125255189:   mNeoVis.vulgivaga.2.1.chr7.g00432.t01#1    mNeoVis.vulgivaga.1.1.chr7.g00438.t01#4    mNeoVis.evergladensis.2.1.chr7.g00434.t01#2    mNeoVis.3.1.1.chr7.g00417.t01#6    mNeoVis.evergladensis.1.1.chr7.g00441.t01#3   mNeoVis.2.1.1.chr7_hap1.g00430.t01#5       6858   125255190:   mNeoVis.evergladensis.1.1.chr13.g00497.t01#3    mNeoVis.3.1.1.chr13.g00456.t01#6    mNeoVis.vulgivaga.2.1.chr13.g00501.t01#1    mNeoVis.2.1.1.chr13_hap1.g00515.t01#5    mNeoVis.vulgivaga.1.1.chr13.g00491.t01#4   mNeoVis.evergladensis.2.1.chr13.g00496.t01#2       6859   125255191:   mNeoVis.vulgivaga.1.1.chr5.g01863.t01#4    mNeoVis.evergladensis.2.1.chr5.g01881.t01#2    mNeoVis.vulgivaga.2.1.chr5.g01883.t01#1    mNeoVis.2.1.1.chr2_hap1.g01942.t01#5   mNeoVis.evergladensis.1.1.chr5.g01890.t01#3       6860   125255192:   mNeoVis.evergladensis.1.1.chr5.g01891.t01#3    mNeoVis.2.1.1.chr2_hap1.g01944.t01#5    mNeoVis.evergladensis.2.1.chr5.g01883.t01#2    mNeoVis.vulgivaga.1.1.chr5.g01864.t01#4   mNeoVis.vulgivaga.2.1.chr5.g01884.t01#1       6861   125255193:   mNeoVis.vulgivaga.2.1.chr13.g00502.t01#1    mNeoVis.3.1.1.chr13.g00457.t01#6    mNeoVis.evergladensis.2.1.chr13.g00497.t01#2    mNeoVis.2.1.1.chr13_hap1.g00516.t01#5    mNeoVis.vulgivaga.1.1.chr13.g00492.t01#4   mNeoVis.evergladensis.1.1.chr13.g00498.t01#3       6862   125255194:   mNeoVis.vulgivaga.2.1.chr15.g00497.t01#1    mNeoVis.vulgivaga.1.1.chr15.g00490.t01#4    mNeoVis.evergladensis.2.1.chr15.g00487.t01#2    mNeoVis.2.1.1.chr15_hap1.g00665.t01#5   mNeoVis.evergladensis.1.1.chr15.g00126.t01#3       6863   125255195:   mNeoVis.3.1.1.chr13.g00458.t01#6    mNeoVis.vulgivaga.2.1.chr13.g00503.t01#1   mNeoVis.evergladensis.2.1.chr13.g00498.t01#2         6864   125255196:   mNeoVis.evergladensis.1.1.chr13.g00499.t01#3    mNeoVis.2.1.1.chr13_hap1.g00517.t01#5    mNeoVis.evergladensis.2.1.chr13.g00499.t01#2    mNeoVis.vulgivaga.1.1.chr13.g00493.t01#4    mNeoVis.vulgivaga.2.1.chr13.g00504.t01#1   mNeoVis.3.1.1.chr13.g00459.t01#6       6865   125255197:   mNeoVis.vulgivaga.1.1.chr5.g01058.t01#4    mNeoVis.2.1.1.chr2_hap1.g01157.t01#5    mNeoVis.evergladensis.1.1.chr5.g01091.t01#3    mNeoVis.evergladensis.2.1.chr5.g01100.t01#2    mNeoVis.3.1.1.chr5.g01074.t01#6    mNeoVis.vulgivaga.2.1.chr5.g01087.t01#1   mNeoVis.evergladensis.2.1.chr5.g01099.t01#2     6866   125255198:   mNeoVis.2.1.1.chr13_hap1.g00518.t01#5    mNeoVis.vulgivaga.2.1.chr13.g00505.t01#1    mNeoVis.vulgivaga.1.1.chr13.g00494.t01#4    mNeoVis.evergladensis.1.1.chr13.g00500.t01#3    mNeoVis.3.1.1.chr13.g00460.t01#6   mNeoVis.evergladensis.2.1.chr13.g00500.t01#2       6867   125255199:   mNeoVis.vulgivaga.1.1.chr13.g00495.t01#4    mNeoVis.2.1.1.chr8_hap1.g01208.t01#5    mNeoVis.evergladensis.2.1.chr13.g00501.t01#2    mNeoVis.2.1.1.chr13_hap1.g00519.t01#5    mNeoVis.evergladensis.1.1.chr13.g00501.t01#3    mNeoVis.2.1.1.chr8_hap1.g01205.t01#5    mNeoVis.vulgivaga.2.1.chr13.g00506.t01#1    mNeoVis.2.1.1.chr8_hap1.g01203.t01#5    mNeoVis.2.1.1.chr8_hap1.g01192.t01#5      6868   125255200:   mNeoVis.2.1.1.chr2_hap1.g01948.t01#5    mNeoVis.vulgivaga.2.1.chr5.g01885.t01#1    mNeoVis.evergladensis.1.1.chr5.g01894.t01#3    mNeoVis.vulgivaga.1.1.chr5.g01867.t01#4   mNeoVis.evergladensis.2.1.chr5.g01885.t01#2       6869   125255201:   mNeoVis.vulgivaga.1.1.chr5.g01868.t01#4    mNeoVis.evergladensis.1.1.chr5.g01895.t01#3    mNeoVis.3.1.1.chr5.g01845.t01#6    mNeoVis.2.1.1.chr2_hap1.g01949.t01#5    mNeoVis.vulgivaga.2.1.chr5.g01886.t01#1   mNeoVis.evergladensis.2.1.chr5.g01886.t01#2       6870   125255202:   mNeoVis.vulgivaga.2.1.chr5.g01887.t01#1    mNeoVis.vulgivaga.1.1.chr5.g01869.t01#4    mNeoVis.evergladensis.1.1.chr5.g01896.t01#3    mNeoVis.3.1.1.chr5.g01846.t01#6    mNeoVis.evergladensis.2.1.chr5.g01887.t01#2   mNeoVis.2.1.1.chr2_hap1.g01950.t01#5       6871   125255203:   mNeoVis.evergladensis.1.1.chr13.g00502.t01#3    mNeoVis.vulgivaga.2.1.chr13.g00507.t01#1   mNeoVis.3.1.1.chr13.g00462.t01#6           6872   125255204:   mNeoVis.3.1.1.chr13.g00463.t01#6    mNeoVis.evergladensis.2.1.chr13.g00505.t01#2    mNeoVis.2.1.1.chr13_hap1.g00523.t01#5    mNeoVis.evergladensis.1.1.chr13.g00503.t01#3    mNeoVis.vulgivaga.2.1.chr13.g00508.t01#1   mNeoVis.vulgivaga.1.1.chr13.g00498.t01#4       6873   125255205:   mNeoVis.vulgivaga.1.1.chr5.g01870.t01#4    mNeoVis.evergladensis.1.1.chr5.g01897.t01#3    mNeoVis.evergladensis.2.1.chr5.g01888.t01#2    mNeoVis.3.1.1.chr5.g01847.t01#6    mNeoVis.vulgivaga.2.1.chr5.g01888.t01#1   mNeoVis.2.1.1.chr2_hap1.g01951.t01#5       6874   125255206:   mNeoVis.evergladensis.1.1.chr13.g00504.t01#3    mNeoVis.vulgivaga.2.1.chr13.g00509.t01#1    mNeoVis.evergladensis.2.1.chr13.g00506.t01#2    mNeoVis.3.1.1.chr13.g00464.t01#6    mNeoVis.vulgivaga.1.1.chr13.g00499.t01#4   mNeoVis.2.1.1.chr13_hap1.g00524.t01#5       6875   125255207:   mNeoVis.2.1.1.chr13_hap1.g00525.t01#5    mNeoVis.vulgivaga.1.1.chr13.g00500.t01#4    mNeoVis.evergladensis.2.1.chr13.g00507.t01#2    mNeoVis.evergladensis.1.1.chr13.g00506.t01#3    mNeoVis.vulgivaga.2.1.chr13.g00510.t01#1   mNeoVis.3.1.1.chr13.g00465.t01#6       6876   125255208:   mNeoVis.2.1.1.chr2_hap1.g01952.t01#5    mNeoVis.3.1.1.chr5.g01848.t01#6    mNeoVis.evergladensis.2.1.chr5.g01889.t01#2    mNeoVis.vulgivaga.2.1.chr5.g01889.t01#1    mNeoVis.evergladensis.1.1.chr5.g01898.t01#3   mNeoVis.vulgivaga.1.1.chr5.g01871.t01#4       6877   125255209:   mNeoVis.vulgivaga.2.1.chr13.g00512.t01#1    mNeoVis.vulgivaga.1.1.chr13.g00502.t01#4    mNeoVis.evergladensis.2.1.chr13.g00509.t01#2    mNeoVis.3.1.1.chr13.g00467.t01#6    mNeoVis.2.1.1.chr13_hap1.g00527.t01#5   mNeoVis.evergladensis.1.1.chr13.g00505.t01#3       6878   125255210:   mNeoVis.3.1.1.chr13.g00466.t01#6    mNeoVis.vulgivaga.2.1.chr13.g00511.t01#1    mNeoVis.2.1.1.chr13_hap1.g00526.t01#5    mNeoVis.evergladensis.2.1.chr13.g00508.t01#2   mNeoVis.vulgivaga.1.1.chr13.g00501.t01#4       6879   125255211:   mNeoVis.vulgivaga.2.1.chr13.g00513.t01#1    mNeoVis.vulgivaga.1.1.chr13.g00503.t01#4    mNeoVis.evergladensis.2.1.chr13.g00510.t01#2    mNeoVis.3.1.1.chr13.g00468.t01#6    mNeoVis.2.1.1.chr13_hap1.g00528.t01#5    mNeoVis.evergladensis.1.1.chr13.g00507.t01#3   mNeoVis.evergladensis.1.1.chr13.g00508.t01#3     6880   125255212:   mNeoVis.2.1.1.chr15_hap1.g00666.t01#5    mNeoVis.vulgivaga.1.1.chr15.g00491.t01#4    mNeoVis.evergladensis.2.1.chr15.g00488.t01#2    mNeoVis.vulgivaga.2.1.chr15.g00498.t01#1   mNeoVis.evergladensis.1.1.chr15.g00125.t01#3       6881   125255213:   mNeoVis.evergladensis.2.1.chr5.g01890.t01#2    mNeoVis.2.1.1.chr2_hap1.g01953.t01#5    mNeoVis.vulgivaga.1.1.chr5.g01872.t01#4    mNeoVis.evergladensis.1.1.chr5.g01899.t01#3    mNeoVis.3.1.1.chr5.g01849.t01#6   mNeoVis.vulgivaga.2.1.chr5.g01890.t01#1       6882   125255214:   mNeoVis.vulgivaga.2.1.chr13.g00514.t01#1    mNeoVis.evergladensis.1.1.chr13.g00509.t01#3    mNeoVis.vulgivaga.1.1.chr13.g00504.t01#4    mNeoVis.3.1.1.chr13.g00469.t01#6    mNeoVis.2.1.1.chr13_hap1.g00529.t01#5    mNeoVis.evergladensis.2.1.chr13.g00511.t01#2   mNeoVis.evergladensis.2.1.chr13.g00512.t01#2     6883   125255215:   mNeoVis.vulgivaga.1.1.chr13.g00505.t01#4    mNeoVis.evergladensis.2.1.chr13.g00513.t01#2    mNeoVis.vulgivaga.2.1.chr13.g00515.t01#1    mNeoVis.evergladensis.1.1.chr13.g00510.t01#3    mNeoVis.2.1.1.chr13_hap1.g00530.t01#5   mNeoVis.3.1.1.chr13.g00470.t01#6       6884   125255216:   mNeoVis.2.1.1.chr4_hap1.g00001.t01#5    mNeoVis.vulgivaga.1.1.chr6.g00001.t01#4    mNeoVis.evergladensis.1.1.chr6.g00001.t01#3    mNeoVis.vulgivaga.2.1.chr6.g00001.t01#1   mNeoVis.evergladensis.2.1.chr6.g00001.t01#2       6885   125255217:   mNeoVis.vulgivaga.1.1.chr13.g00506.t01#4    mNeoVis.evergladensis.2.1.chr13.g00514.t01#2    mNeoVis.3.1.1.chr13.g00471.t01#6    mNeoVis.vulgivaga.2.1.chr13.g00516.t01#1    mNeoVis.evergladensis.1.1.chr13.g00511.t01#3   mNeoVis.2.1.1.chr13_hap1.g00531.t01#5       6886   125255218:   mNeoVis.vulgivaga.1.1.chr6.g00002.t01#4   mNeoVis.vulgivaga.2.1.chr6.g00002.t01#1             6887   125255219:   mNeoVis.evergladensis.2.1.chr13.g00515.t01#2    mNeoVis.vulgivaga.2.1.chr13.g00517.t01#1    mNeoVis.vulgivaga.1.1.chr13.g00507.t01#4    mNeoVis.3.1.1.chr13.g00472.t01#6    mNeoVis.evergladensis.1.1.chr13.g00512.t01#3   mNeoVis.2.1.1.chr13_hap1.g00532.t01#5       6888   125255220:   mNeoVis.evergladensis.1.1.chr13.g00513.t01#3    mNeoVis.3.1.1.chr13.g00473.t01#6    mNeoVis.2.1.1.chr13_hap1.g00533.t01#5    mNeoVis.vulgivaga.2.1.chr13.g00518.t01#1    mNeoVis.vulgivaga.1.1.chr13.g00508.t01#4   mNeoVis.evergladensis.2.1.chr13.g00516.t01#2       6889   125255221:   mNeoVis.evergladensis.2.1.chr6.g00002.t01#2    mNeoVis.vulgivaga.2.1.chr6.g00003.t01#1    mNeoVis.vulgivaga.1.1.chr6.g00003.t01#4    mNeoVis.3.1.1.chr6.g00002.t01#6    mNeoVis.evergladensis.1.1.chr6.g00002.t01#3   mNeoVis.2.1.1.chr4_hap1.g00002.t01#5       6890   125255222:   mNeoVis.vulgivaga.2.1.chr13.g00519.t01#1    mNeoVis.3.1.1.chr13.g00474.t01#6    mNeoVis.2.1.1.chr13_hap1.g00534.t01#5    mNeoVis.evergladensis.1.1.chr13.g00514.t01#3    mNeoVis.evergladensis.2.1.chr13.g00517.t01#2   mNeoVis.vulgivaga.1.1.chr13.g00509.t01#4       6891   125255223:   mNeoVis.vulgivaga.1.1.chr13.g00510.t01#4    mNeoVis.evergladensis.2.1.chr13.g00518.t01#2    mNeoVis.vulgivaga.2.1.chr13.g00520.t01#1    mNeoVis.3.1.1.chr13.g00475.t01#6    mNeoVis.2.1.1.chr13_hap1.g00535.t01#5   mNeoVis.evergladensis.1.1.chr13.g00515.t01#3       6892   125255224:   mNeoVis.3.1.1.chr7.g00418.t01#6    mNeoVis.vulgivaga.1.1.chr7.g00439.t01#4    mNeoVis.2.1.1.chr7_hap1.g00431.t01#5    mNeoVis.vulgivaga.2.1.chr7.g00433.t01#1   mNeoVis.evergladensis.1.1.chr7.g00442.t01#3       6893   125255225:   mNeoVis.2.1.1.chr4_hap1.g00003.t01#5    mNeoVis.vulgivaga.2.1.chr6.g00004.t01#1    mNeoVis.evergladensis.2.1.chr6.g00003.t01#2    mNeoVis.vulgivaga.1.1.chr6.g00004.t01#4    mNeoVis.evergladensis.1.1.chr6.g00003.t01#3   mNeoVis.3.1.1.chr6.g00003.t01#6       6894   125255226:   mNeoVis.evergladensis.2.1.chr6.g00004.t01#2    mNeoVis.vulgivaga.1.1.chr6.g00005.t01#4    mNeoVis.vulgivaga.2.1.chr6.g00005.t01#1    mNeoVis.2.1.1.chr4_hap1.g00004.t01#5    mNeoVis.evergladensis.1.1.chr6.g00004.t01#3   mNeoVis.3.1.1.chr6.g00004.t01#6       6895   125255227:   mNeoVis.vulgivaga.2.1.chr13.g00521.t01#1    mNeoVis.2.1.1.chr13_hap1.g00536.t01#5    mNeoVis.3.1.1.chr13.g00476.t01#6    mNeoVis.evergladensis.1.1.chr13.g00516.t01#3    mNeoVis.evergladensis.2.1.chr13.g00519.t01#2   mNeoVis.vulgivaga.1.1.chr13.g00511.t01#4       6896   125255228:   mNeoVis.evergladensis.1.1.chr13.g00517.t01#3    mNeoVis.2.1.1.chr13_hap1.g00537.t01#5    mNeoVis.3.1.1.chr13.g00477.t01#6    mNeoVis.evergladensis.2.1.chr13.g00520.t01#2    mNeoVis.vulgivaga.1.1.chr13.g00512.t01#4   mNeoVis.vulgivaga.2.1.chr13.g00522.t01#1       6897   125255229:   mNeoVis.evergladensis.1.1.chr13.g00518.t01#3    mNeoVis.2.1.1.chr13_hap1.g00538.t01#5    mNeoVis.3.1.1.chr13.g00478.t01#6    mNeoVis.vulgivaga.2.1.chr13.g00523.t01#1   mNeoVis.evergladensis.2.1.chr13.g00521.t01#2       6898   125255230:   mNeoVis.2.1.1.chr13_hap1.g00539.t01#5    mNeoVis.vulgivaga.1.1.chr13.g00514.t01#4    mNeoVis.vulgivaga.2.1.chr13.g00524.t01#1    mNeoVis.evergladensis.2.1.chr13.g00522.t01#2    mNeoVis.evergladensis.1.1.chr13.g00519.t01#3   mNeoVis.3.1.1.chr13.g00479.t01#6       6899   125255231:   mNeoVis.vulgivaga.2.1.chr13.g00525.t01#1   mNeoVis.vulgivaga.1.1.chr13.g00515.t01#4           6900   125255232:   mNeoVis.vulgivaga.2.1.chr13.g00526.t01#1    mNeoVis.evergladensis.1.1.chr13.g00521.t01#3    mNeoVis.2.1.1.chr13_hap1.g00541.t01#5    mNeoVis.evergladensis.2.1.chr13.g00524.t01#2    mNeoVis.3.1.1.chr13.g00481.t01#6   mNeoVis.vulgivaga.1.1.chr13.g00516.t01#4       6901   125255233:  mNeoVis.vulgivaga.2.1.chr13.g00527.t01#1               6902   125255234:   mNeoVis.vulgivaga.2.1.chr6.g00006.t01#1    mNeoVis.evergladensis.2.1.chr6.g00005.t01#2   mNeoVis.2.1.1.chr4_hap1.g00005.t01#5           6903   125255235:   mNeoVis.vulgivaga.1.1.chr13.g00517.t01#4    mNeoVis.3.1.1.chr13.g00483.t01#6    mNeoVis.vulgivaga.2.1.chr13.g00528.t01#1    mNeoVis.evergladensis.1.1.chr13.g00522.t01#3    mNeoVis.2.1.1.chr13_hap1.g00542.t01#5   mNeoVis.evergladensis.2.1.chr13.g00525.t01#2       6904   125255236:   mNeoVis.evergladensis.1.1.chr6.g00006.t01#3    mNeoVis.evergladensis.2.1.chr6.g00006.t01#2    mNeoVis.vulgivaga.1.1.chr6.g00008.t01#4    mNeoVis.vulgivaga.2.1.chr6.g00007.t01#1   mNeoVis.2.1.1.chr4_hap1.g00006.t01#5       6905   125255237:   mNeoVis.evergladensis.1.1.chr6.g00007.t01#3    mNeoVis.vulgivaga.2.1.chr6.g00008.t01#1    mNeoVis.vulgivaga.1.1.chr6.g00009.t01#4    mNeoVis.2.1.1.chr4_hap1.g00007.t01#5    mNeoVis.3.1.1.chr6.g00009.t01#6   mNeoVis.evergladensis.2.1.chr6.g00007.t01#2       6906   125255238:   mNeoVis.vulgivaga.1.1.chr13.g00518.t01#4    mNeoVis.2.1.1.chr13_hap1.g00543.t01#5    mNeoVis.vulgivaga.2.1.chr13.g00529.t01#1    mNeoVis.evergladensis.1.1.chr13.g00523.t01#3    mNeoVis.3.1.1.chr13.g00484.t01#6   mNeoVis.evergladensis.2.1.chr13.g00526.t01#2       6907   125255239:   mNeoVis.vulgivaga.2.1.chr7.g00434.t01#1    mNeoVis.evergladensis.1.1.chr7.g00443.t01#3    mNeoVis.2.1.1.chr7_hap1.g00432.t01#5    mNeoVis.vulgivaga.1.1.chr7.g00440.t01#4    mNeoVis.evergladensis.2.1.chr7.g00436.t01#2   mNeoVis.3.1.1.chr7.g00419.t01#6       6908   125255240:   mNeoVis.evergladensis.1.1.chr13.g00524.t01#3    mNeoVis.3.1.1.chr13.g00485.t01#6    mNeoVis.vulgivaga.2.1.chr13.g00530.t01#1    mNeoVis.2.1.1.chr13_hap1.g00544.t01#5    mNeoVis.evergladensis.2.1.chr13.g00527.t01#2   mNeoVis.vulgivaga.1.1.chr13.g00519.t01#4       6909   125255241:   mNeoVis.vulgivaga.1.1.chr6.g00010.t01#4    mNeoVis.2.1.1.chr4_hap1.g00008.t01#5    mNeoVis.vulgivaga.2.1.chr6.g00009.t01#1    mNeoVis.evergladensis.2.1.chr6.g00008.t01#2   mNeoVis.evergladensis.1.1.chr6.g00008.t01#3       6910   125255242:   mNeoVis.vulgivaga.2.1.chr15.g00499.t01#1    mNeoVis.2.1.1.chr15_hap1.g00667.t01#5    mNeoVis.vulgivaga.1.1.chr15.g00492.t01#4    mNeoVis.evergladensis.2.1.chr15.g00489.t01#2   mNeoVis.evergladensis.1.1.chr15.g00124.t01#3       6911   125255243:   mNeoVis.evergladensis.1.1.chr6.g00009.t01#3    mNeoVis.evergladensis.2.1.chr6.g00009.t01#2    mNeoVis.vulgivaga.2.1.chr6.g00010.t01#1    mNeoVis.vulgivaga.1.1.chr6.g00011.t01#4   mNeoVis.2.1.1.chr4_hap1.g00009.t01#5       6912   125255244:  mNeoVis.vulgivaga.2.1.chr6.g00011.t01#1               6913   125255245:   mNeoVis.2.1.1.chr13_hap1.g00545.t01#5    mNeoVis.vulgivaga.1.1.chr13.g00520.t01#4    mNeoVis.vulgivaga.2.1.chr13.g00531.t01#1    mNeoVis.evergladensis.1.1.chr13.g00525.t01#3    mNeoVis.evergladensis.2.1.chr13.g00528.t01#2   mNeoVis.3.1.1.chr13.g00486.t01#6       6914   125255246:   mNeoVis.2.1.1.chr4_hap1.g00010.t01#5    mNeoVis.vulgivaga.1.1.chr6.g00012.t01#4    mNeoVis.evergladensis.1.1.chr6.g00010.t01#3    mNeoVis.evergladensis.2.1.chr6.g00010.t01#2    mNeoVis.vulgivaga.2.1.chr6.g00012.t01#1   mNeoVis.3.1.1.chr6.g00011.t01#6       6915   125255247:   mNeoVis.evergladensis.2.1.chr13.g00529.t01#2    mNeoVis.vulgivaga.1.1.chr13.g00521.t01#4    mNeoVis.vulgivaga.2.1.chr13.g00532.t01#1    mNeoVis.2.1.1.chr13_hap1.g00546.t01#5    mNeoVis.evergladensis.1.1.chr13.g00526.t01#3   mNeoVis.3.1.1.chr13.g00487.t01#6       6916   125255248:   mNeoVis.vulgivaga.1.1.chr13.g00424.t01#4    mNeoVis.3.1.1.chr13.g00385.t01#6    mNeoVis.evergladensis.1.1.chr13.g00428.t01#3    mNeoVis.2.1.1.chr13_hap1.g00447.t01#5    mNeoVis.evergladensis.2.1.chr13.g00429.t01#2   mNeoVis.vulgivaga.2.1.chr13.g00432.t01#1       6917   125255249:   mNeoVis.3.1.1.chr6.g00012.t01#6    mNeoVis.vulgivaga.2.1.chr6.g00013.t01#1   mNeoVis.vulgivaga.1.1.chr6.g00013.t01#4           6918   125255250:   mNeoVis.evergladensis.1.1.chr6.g00012.t01#3    mNeoVis.evergladensis.2.1.chr6.g00012.t01#2    mNeoVis.vulgivaga.1.1.chr6.g00014.t01#4    mNeoVis.vulgivaga.2.1.chr6.g00014.t01#1   mNeoVis.2.1.1.chr4_hap1.g00012.t01#5       6919   125255251:   mNeoVis.3.1.1.chr13.g00386.t01#6    mNeoVis.evergladensis.1.1.chr13.g00429.t01#3    mNeoVis.2.1.1.chr13_hap1.g00448.t01#5    mNeoVis.evergladensis.2.1.chr13.g00430.t01#2    mNeoVis.vulgivaga.2.1.chr13.g00433.t01#1   mNeoVis.vulgivaga.1.1.chr13.g00425.t01#4       6920   125255252:   mNeoVis.evergladensis.2.1.chr6.g00013.t01#2    mNeoVis.vulgivaga.1.1.chr6.g00015.t01#4    mNeoVis.2.1.1.chr4_hap1.g00013.t01#5    mNeoVis.evergladensis.1.1.chr6.g00013.t01#3    mNeoVis.vulgivaga.2.1.chr6.g00015.t01#1   mNeoVis.3.1.1.chr6.g00014.t01#6       6921   125255253:   mNeoVis.vulgivaga.1.1.chr13.g00426.t01#4    mNeoVis.evergladensis.1.1.chr13.g00430.t01#3    mNeoVis.vulgivaga.2.1.chr13.g00434.t01#1    mNeoVis.3.1.1.chr13.g00387.t01#6    mNeoVis.2.1.1.chr13_hap1.g00449.t01#5   mNeoVis.evergladensis.2.1.chr13.g00431.t01#2       6922   125255254:   mNeoVis.evergladensis.1.1.chr6.g00014.t01#3    mNeoVis.2.1.1.chr4_hap1.g00014.t01#5    mNeoVis.vulgivaga.1.1.chr6.g00016.t01#4    mNeoVis.3.1.1.chr6.g00015.t01#6    mNeoVis.vulgivaga.2.1.chr6.g00016.t01#1   mNeoVis.evergladensis.2.1.chr6.g00014.t01#2       6923   125255255:   mNeoVis.2.1.1.chr4_hap1.g00015.t01#5    mNeoVis.vulgivaga.1.1.chr6.g00017.t01#4    mNeoVis.vulgivaga.2.1.chr6.g00017.t01#1    mNeoVis.evergladensis.1.1.chr6.g00015.t01#3   mNeoVis.evergladensis.2.1.chr6.g00015.t01#2       6924   125255256:   mNeoVis.vulgivaga.2.1.chr13.g00435.t01#1    mNeoVis.vulgivaga.1.1.chr13.g00427.t01#4    mNeoVis.evergladensis.1.1.chr13.g00431.t01#3    mNeoVis.3.1.1.chr13.g00388.t01#6    mNeoVis.evergladensis.2.1.chr13.g00432.t01#2   mNeoVis.2.1.1.chr13_hap1.g00450.t01#5       6925   125255257:   mNeoVis.evergladensis.1.1.chr6.g00016.t01#3    mNeoVis.2.1.1.chr4_hap1.g00016.t01#5    mNeoVis.evergladensis.2.1.chr6.g00016.t01#2    mNeoVis.vulgivaga.2.1.chr6.g00018.t01#1   mNeoVis.vulgivaga.1.1.chr6.g00018.t01#4       6926   125255258:   mNeoVis.evergladensis.1.1.chr3.g00759.t01#3    mNeoVis.evergladensis.2.1.chr3.g00758.t01#2    mNeoVis.vulgivaga.1.1.chr3.g00755.t01#4    mNeoVis.2.1.1.chr3_hap1.g00760.t01#5   mNeoVis.vulgivaga.2.1.chr3.g00766.t01#1       6927   125255259:   mNeoVis.evergladensis.1.1.chr6.g00017.t01#3    mNeoVis.vulgivaga.1.1.chr6.g00019.t01#4    mNeoVis.3.1.1.chr6.g00018.t01#6    mNeoVis.2.1.1.chr4_hap1.g00017.t01#5    mNeoVis.evergladensis.2.1.chr6.g00017.t01#2   mNeoVis.vulgivaga.2.1.chr6.g00019.t01#1       6928   125255260:   mNeoVis.vulgivaga.2.1.chr6.g00020.t01#1    mNeoVis.evergladensis.2.1.chr6.g00018.t01#2    mNeoVis.vulgivaga.1.1.chr6.g00020.t01#4    mNeoVis.evergladensis.1.1.chr6.g00018.t01#3    mNeoVis.3.1.1.chr6.g00019.t01#6   mNeoVis.2.1.1.chr4_hap1.g00018.t01#5       6929   125255261:   mNeoVis.vulgivaga.1.1.chr7.g00441.t01#4    mNeoVis.vulgivaga.2.1.chr7.g00435.t01#1    mNeoVis.3.1.1.chr7.g00420.t01#6    mNeoVis.evergladensis.1.1.chr7.g00444.t01#3    mNeoVis.evergladensis.2.1.chr7.g00437.t01#2   mNeoVis.2.1.1.chr7_hap1.g00433.t01#5       6930   125255262:   mNeoVis.3.1.1.chr6.g00020.t01#6    mNeoVis.vulgivaga.2.1.chr6.g00021.t01#1    mNeoVis.2.1.1.chr4_hap1.g00019.t01#5    mNeoVis.evergladensis.1.1.chr6.g00019.t01#3   mNeoVis.evergladensis.2.1.chr6.g00019.t01#2       6931   125255263:   mNeoVis.vulgivaga.1.1.chr13.g00428.t01#4    mNeoVis.vulgivaga.2.1.chr13.g00436.t01#1    mNeoVis.3.1.1.chr13.g00389.t01#6    mNeoVis.evergladensis.2.1.chr13.g00433.t01#2    mNeoVis.evergladensis.1.1.chr13.g00432.t01#3   mNeoVis.2.1.1.chr13_hap1.g00451.t01#5       6932   125255264:   mNeoVis.vulgivaga.1.1.chr6.g00023.t01#4    mNeoVis.3.1.1.chr6.g00022.t01#6   mNeoVis.vulgivaga.2.1.chr6.g00023.t01#1           6933   125255265:   mNeoVis.2.1.1.chr4_hap1.g00021.t01#5    mNeoVis.vulgivaga.2.1.chr6.g00024.t01#1    mNeoVis.evergladensis.1.1.chr6.g00021.t01#3    mNeoVis.3.1.1.chr6.g00023.t01#6    mNeoVis.vulgivaga.1.1.chr6.g00024.t01#4   mNeoVis.evergladensis.2.1.chr6.g00021.t01#2       6934   125255266:   mNeoVis.3.1.1.chr3.g00749.t01#6    mNeoVis.evergladensis.2.1.chr3.g00764.t01#2    mNeoVis.evergladensis.1.1.chr3.g00767.t01#3    mNeoVis.vulgivaga.2.1.chr3.g00772.t01#1    mNeoVis.2.1.1.chr3_hap1.g00767.t01#5   mNeoVis.vulgivaga.1.1.chr3.g00761.t01#4       6935   125255267:   mNeoVis.vulgivaga.1.1.chr6.g00025.t01#4    mNeoVis.vulgivaga.2.1.chr6.g00025.t01#1    mNeoVis.evergladensis.1.1.chr6.g00022.t01#3    mNeoVis.3.1.1.chr6.g00024.t01#6    mNeoVis.evergladensis.2.1.chr6.g00022.t01#2   mNeoVis.2.1.1.chr4_hap1.g00022.t01#5       6936   125255268:   mNeoVis.evergladensis.1.1.chr13.g00433.t01#3    mNeoVis.2.1.1.chr13_hap1.g00452.t01#5    mNeoVis.evergladensis.2.1.chr13.g00434.t01#2    mNeoVis.vulgivaga.1.1.chr13.g00429.t01#4    mNeoVis.3.1.1.chr13.g00390.t01#6   mNeoVis.vulgivaga.2.1.chr13.g00437.t01#1       6937   125255269:   mNeoVis.evergladensis.2.1.chr13.g00435.t01#2    mNeoVis.3.1.1.chr13.g00391.t01#6    mNeoVis.vulgivaga.1.1.chr13.g00430.t01#4    mNeoVis.vulgivaga.2.1.chr13.g00438.t01#1    mNeoVis.2.1.1.chr13_hap1.g00453.t01#5   mNeoVis.evergladensis.1.1.chr13.g00434.t01#3       6938   125255270:   mNeoVis.vulgivaga.2.1.chr13.g00439.t01#1    mNeoVis.3.1.1.chr13.g00392.t01#6    mNeoVis.2.1.1.chr13_hap1.g00454.t01#5    mNeoVis.evergladensis.2.1.chr13.g00436.t01#2   mNeoVis.vulgivaga.1.1.chr13.g00431.t01#4       6939   125255271:   mNeoVis.evergladensis.1.1.chr6.g00023.t01#3    mNeoVis.vulgivaga.2.1.chr6.g00026.t01#1    mNeoVis.2.1.1.chr4_hap1.g00023.t01#5    mNeoVis.evergladensis.2.1.chr6.g00023.t01#2   mNeoVis.vulgivaga.1.1.chr6.g00026.t01#4       6940   125255272:   mNeoVis.3.1.1.chr6.g00025.t01#6   mNeoVis.vulgivaga.2.1.chr6.g00027.t01#1             6941   125255273:   mNeoVis.evergladensis.2.1.chr13.g00437.t01#2    mNeoVis.3.1.1.chr13.g00393.t01#6    mNeoVis.vulgivaga.2.1.chr13.g00440.t01#1    mNeoVis.evergladensis.1.1.chr13.g00436.t01#3    mNeoVis.2.1.1.chr13_hap1.g00455.t01#5   mNeoVis.vulgivaga.1.1.chr13.g00432.t01#4       6942   125255274:   mNeoVis.vulgivaga.1.1.chr13.g00433.t01#4    mNeoVis.evergladensis.2.1.chr13.g00438.t01#2    mNeoVis.3.1.1.chr13.g00394.t01#6    mNeoVis.2.1.1.chr13_hap1.g00456.t01#5    mNeoVis.evergladensis.1.1.chr13.g00437.t01#3   mNeoVis.vulgivaga.2.1.chr13.g00441.t01#1       6943   125255275:   mNeoVis.3.1.1.chr6.g00026.t01#6    mNeoVis.vulgivaga.2.1.chr6.g00028.t01#1    mNeoVis.evergladensis.2.1.chr6.g00024.t01#2    mNeoVis.vulgivaga.1.1.chr6.g00027.t01#4    mNeoVis.2.1.1.chr4_hap1.g00024.t01#5   mNeoVis.evergladensis.1.1.chr6.g00024.t01#3       6944   125255276:   mNeoVis.vulgivaga.1.1.chr13.g00434.t01#4    mNeoVis.2.1.1.chr13_hap1.g00457.t01#5    mNeoVis.vulgivaga.2.1.chr13.g00442.t01#1    mNeoVis.evergladensis.1.1.chr13.g00438.t01#3    mNeoVis.evergladensis.2.1.chr13.g00439.t01#2   mNeoVis.3.1.1.chr13.g00395.t01#6       6945   125255277:   mNeoVis.3.1.1.chr15.g00449.t01#6    mNeoVis.2.1.1.chr15_hap1.g00668.t01#5    mNeoVis.vulgivaga.1.1.chr15.g00493.t01#4    mNeoVis.evergladensis.2.1.chr15.g00490.t01#2    mNeoVis.vulgivaga.2.1.chr15.g00500.t01#1   mNeoVis.evergladensis.1.1.chr15.g00122.t01#3       6946   125255278:   mNeoVis.3.1.1.chr13.g00396.t01#6    mNeoVis.vulgivaga.2.1.chr13.g00443.t01#1    mNeoVis.2.1.1.chr13_hap1.g00458.t01#5    mNeoVis.evergladensis.1.1.chr13.g00439.t01#3    mNeoVis.vulgivaga.1.1.chr13.g00435.t01#4   mNeoVis.evergladensis.2.1.chr13.g00440.t01#2       6947   125255279:   mNeoVis.3.1.1.chr13.g00397.t01#6    mNeoVis.2.1.1.chr13_hap1.g00459.t01#5    mNeoVis.evergladensis.2.1.chr13.g00441.t01#2    mNeoVis.evergladensis.1.1.chr13.g00440.t01#3    mNeoVis.vulgivaga.1.1.chr13.g00436.t01#4   mNeoVis.vulgivaga.2.1.chr13.g00444.t01#1       6948   125255280:   mNeoVis.vulgivaga.1.1.chr7.g00442.t01#4    mNeoVis.3.1.1.chr7.g00421.t01#6    mNeoVis.vulgivaga.2.1.chr7.g00436.t01#1    mNeoVis.evergladensis.2.1.chr7.g00438.t01#2    mNeoVis.evergladensis.1.1.chr7.g00445.t01#3   mNeoVis.2.1.1.chr7_hap1.g00434.t01#5       6949   125255281:   mNeoVis.vulgivaga.1.1.chr13.g00437.t01#4    mNeoVis.2.1.1.chr13_hap1.g00460.t01#5    mNeoVis.evergladensis.1.1.chr13.g00441.t01#3    mNeoVis.vulgivaga.2.1.chr13.g00445.t01#1    mNeoVis.evergladensis.2.1.chr13.g00442.t01#2   mNeoVis.3.1.1.chr13.g00398.t01#6       6950   125255282:   mNeoVis.vulgivaga.2.1.chr13.g00446.t01#1    mNeoVis.evergladensis.2.1.chr13.g00443.t01#2    mNeoVis.vulgivaga.1.1.chr13.g00438.t01#4    mNeoVis.evergladensis.1.1.chr13.g00442.t01#3    mNeoVis.2.1.1.chr13_hap1.g00461.t01#5   mNeoVis.3.1.1.chr13.g00399.t01#6       6951   125255283:   mNeoVis.vulgivaga.2.1.chr5.g01837.t01#1    mNeoVis.evergladensis.2.1.chr5.g01834.t01#2    mNeoVis.vulgivaga.1.1.chr5.g01817.t01#4    mNeoVis.2.1.1.chr2_hap1.g01896.t01#5   mNeoVis.evergladensis.1.1.chr5.g01843.t01#3       6952   125255284:   mNeoVis.vulgivaga.2.1.chr13.g00447.t01#1    mNeoVis.vulgivaga.1.1.chr13.g00439.t01#4    mNeoVis.evergladensis.2.1.chr13.g00444.t01#2    mNeoVis.3.1.1.chr13.g00400.t01#6    mNeoVis.2.1.1.chr13_hap1.g00462.t01#5   mNeoVis.evergladensis.1.1.chr13.g00443.t01#3       6953   125255285:   mNeoVis.evergladensis.2.1.chr5.g01835.t01#2    mNeoVis.vulgivaga.2.1.chr5.g01838.t01#1    mNeoVis.3.1.1.chr5.g01817.t01#6    mNeoVis.evergladensis.1.1.chr5.g01844.t01#3    mNeoVis.vulgivaga.1.1.chr5.g01818.t01#4   mNeoVis.2.1.1.chr2_hap1.g01897.t01#5       6954   125255286:   mNeoVis.evergladensis.2.1.chr5.g01836.t01#2    mNeoVis.evergladensis.1.1.chr5.g01845.t01#3    mNeoVis.vulgivaga.1.1.chr5.g01819.t01#4    mNeoVis.vulgivaga.2.1.chr5.g01839.t01#1   mNeoVis.2.1.1.chr2_hap1.g01898.t01#5       6955   125255287:   mNeoVis.2.1.1.chr13_hap1.g00463.t01#5    mNeoVis.3.1.1.chr13.g00401.t01#6    mNeoVis.evergladensis.2.1.chr13.g00445.t01#2    mNeoVis.vulgivaga.1.1.chr13.g00440.t01#4    mNeoVis.vulgivaga.2.1.chr13.g00448.t01#1   mNeoVis.evergladensis.1.1.chr13.g00444.t01#3       6956   125255288:   mNeoVis.3.1.1.chr13.g00402.t01#6    mNeoVis.vulgivaga.1.1.chr13.g00441.t01#4    mNeoVis.vulgivaga.2.1.chr13.g00449.t01#1    mNeoVis.evergladensis.1.1.chr13.g00445.t01#3    mNeoVis.2.1.1.chr13_hap1.g00464.t01#5   mNeoVis.evergladensis.2.1.chr13.g00446.t01#2       6957   125255289:   mNeoVis.evergladensis.2.1.chr5.g01837.t01#2    mNeoVis.vulgivaga.2.1.chr5.g01840.t01#1    mNeoVis.vulgivaga.1.1.chr5.g01820.t01#4    mNeoVis.evergladensis.1.1.chr5.g01846.t01#3   mNeoVis.2.1.1.chr2_hap1.g01899.t01#5       6958   125255290:   mNeoVis.vulgivaga.1.1.chr13.g00442.t01#4    mNeoVis.2.1.1.chr13_hap1.g00465.t01#5    mNeoVis.evergladensis.1.1.chr13.g00446.t01#3    mNeoVis.3.1.1.chr13.g00403.t01#6    mNeoVis.evergladensis.2.1.chr13.g00447.t01#2   mNeoVis.vulgivaga.2.1.chr13.g00450.t01#1       6959   125255291:   mNeoVis.vulgivaga.2.1.chr5.g00241.t01#1    mNeoVis.vulgivaga.1.1.chr5.g00234.t01#4    mNeoVis.2.1.1.chr2_hap1.g00239.t01#5    mNeoVis.evergladensis.2.1.chr5.g00243.t01#2    mNeoVis.evergladensis.1.1.chr5.g00242.t01#3    mNeoVis.vulgivaga.2.1.chr5.g00240.t01#1   mNeoVis.3.1.1.chr5.g00231.t01#6       6960   125255292:   mNeoVis.vulgivaga.2.1.chr13.g00451.t01#1    mNeoVis.evergladensis.2.1.chr13.g00448.t01#2    mNeoVis.2.1.1.chr13_hap1.g00466.t01#5    mNeoVis.vulgivaga.1.1.chr13.g00443.t01#4    mNeoVis.3.1.1.chr13.g00404.t01#6   mNeoVis.evergladensis.1.1.chr13.g00447.t01#3       6961   125255293:   mNeoVis.evergladensis.1.1.chr11.g00893.t01#3    mNeoVis.vulgivaga.1.1.chr11.g00877.t01#4    mNeoVis.vulgivaga.2.1.chr11.g00903.t01#1    mNeoVis.evergladensis.2.1.chr11.g00878.t01#2    mNeoVis.2.1.1.chr11_hap1.g00900.t01#5   mNeoVis.3.1.1.chr11.g00855.t01#6       6962   125255294:   mNeoVis.evergladensis.2.1.chr13.g00449.t01#2    mNeoVis.evergladensis.1.1.chr13.g00448.t01#3    mNeoVis.3.1.1.chr13.g00405.t01#6    mNeoVis.vulgivaga.2.1.chr13.g00452.t01#1    mNeoVis.vulgivaga.1.1.chr13.g00444.t01#4   mNeoVis.2.1.1.chr13_hap1.g00467.t01#5       6963   125255295:   mNeoVis.3.1.1.chr5.g01814.t01#6    mNeoVis.vulgivaga.1.1.chr5.g01821.t01#4    mNeoVis.2.1.1.chr2_hap1.g01900.t01#5    mNeoVis.vulgivaga.2.1.chr5.g01841.t01#1    mNeoVis.evergladensis.1.1.chr5.g01847.t01#3   mNeoVis.evergladensis.2.1.chr5.g01838.t01#2       6964   125255296:   mNeoVis.vulgivaga.2.1.chr5.g01842.t01#1    mNeoVis.2.1.1.chr2_hap1.g01901.t01#5    mNeoVis.3.1.1.chr5.g01815.t01#6    mNeoVis.evergladensis.1.1.chr5.g01848.t01#3    mNeoVis.evergladensis.2.1.chr5.g01839.t01#2   mNeoVis.vulgivaga.1.1.chr5.g01822.t01#4       6965   125255297:   mNeoVis.vulgivaga.2.1.chr5.g01843.t01#1    mNeoVis.evergladensis.2.1.chr5.g01840.t01#2    mNeoVis.2.1.1.chr2_hap1.g01902.t01#5    mNeoVis.vulgivaga.1.1.chr5.g01823.t01#4    mNeoVis.evergladensis.1.1.chr5.g01849.t01#3   mNeoVis.3.1.1.chr5.g01816.t01#6       6966   125255298:   mNeoVis.evergladensis.2.1.chr13.g00450.t01#2    mNeoVis.vulgivaga.2.1.chr13.g00453.t01#1    mNeoVis.evergladensis.1.1.chr13.g00449.t01#3    mNeoVis.2.1.1.chr13_hap1.g00468.t01#5    mNeoVis.3.1.1.chr13.g00406.t01#6   mNeoVis.vulgivaga.1.1.chr13.g00445.t01#4       6967   125255299:   mNeoVis.2.1.1.chr13_hap1.g00470.t01#5    mNeoVis.evergladensis.1.1.chr13.g00450.t01#3    mNeoVis.3.1.1.chr13.g00407.t01#6    mNeoVis.vulgivaga.2.1.chr13.g00454.t01#1    mNeoVis.vulgivaga.1.1.chr13.g00447.t01#4   mNeoVis.evergladensis.2.1.chr13.g00451.t01#2       6968   125255300:   mNeoVis.2.1.1.chr15_hap1.g00669.t01#5    mNeoVis.vulgivaga.1.1.chr15.g00494.t01#4    mNeoVis.evergladensis.2.1.chr15.g00491.t01#2    mNeoVis.3.1.1.chr15.g00450.t01#6    mNeoVis.evergladensis.1.1.chr15.g00123.t01#3   mNeoVis.vulgivaga.2.1.chr15.g00501.t01#1       6969   125255301:   mNeoVis.vulgivaga.1.1.chr5.g01824.t01#4    mNeoVis.2.1.1.chr2_hap1.g01903.t01#5    mNeoVis.vulgivaga.2.1.chr5.g01844.t01#1    mNeoVis.evergladensis.2.1.chr5.g01841.t01#2   mNeoVis.evergladensis.1.1.chr5.g01850.t01#3       6970   125255302:   mNeoVis.evergladensis.2.1.chr5.g01842.t01#2    mNeoVis.vulgivaga.1.1.chr5.g01825.t01#4    mNeoVis.2.1.1.chr2_hap1.g01904.t01#5    mNeoVis.vulgivaga.2.1.chr5.g01845.t01#1   mNeoVis.evergladensis.1.1.chr5.g01851.t01#3       6971   125255303:   mNeoVis.2.1.1.chr2_hap1.g01905.t01#5    mNeoVis.evergladensis.1.1.chr5.g01852.t01#3    mNeoVis.vulgivaga.1.1.chr5.g01826.t01#4    mNeoVis.vulgivaga.2.1.chr5.g01846.t01#1    mNeoVis.evergladensis.2.1.chr5.g01843.t01#2   mNeoVis.3.1.1.chr5.g01819.t01#6       6972   125255304:   mNeoVis.3.1.1.chr5.g00096.t01#6    mNeoVis.vulgivaga.1.1.chr5.g00101.t01#4    mNeoVis.evergladensis.2.1.chr5.g00105.t01#2    mNeoVis.evergladensis.2.1.chr13.g00452.t01#2    mNeoVis.vulgivaga.1.1.chr13.g00448.t01#4    mNeoVis.2.1.1.chr13_hap1.g00471.t01#5    mNeoVis.evergladensis.1.1.chr13.g00451.t01#3    mNeoVis.evergladensis.1.1.chr5.g00104.t01#3    mNeoVis.3.1.1.chr13.g00408.t01#6      6973   125255305:   mNeoVis.vulgivaga.2.1.chr13.g00456.t01#1    mNeoVis.2.1.1.chr13_hap1.g00472.t01#5    mNeoVis.3.1.1.chr13.g00409.t01#6    mNeoVis.vulgivaga.1.1.chr13.g00449.t01#4    mNeoVis.evergladensis.2.1.chr13.g00453.t01#2   mNeoVis.evergladensis.1.1.chr13.g00452.t01#3       6974   125255306:   mNeoVis.vulgivaga.2.1.chr13.g00457.t01#1    mNeoVis.2.1.1.chr13_hap1.g00473.t01#5    mNeoVis.evergladensis.2.1.chr13.g00454.t01#2    mNeoVis.3.1.1.chr13.g00410.t01#6    mNeoVis.vulgivaga.1.1.chr13.g00450.t01#4   mNeoVis.evergladensis.1.1.chr13.g00453.t01#3       6975   125255307:   mNeoVis.3.1.1.chr13.g00411.t01#6    mNeoVis.evergladensis.1.1.chr13.g00454.t01#3    mNeoVis.evergladensis.2.1.chr13.g00455.t01#2    mNeoVis.vulgivaga.2.1.chr13.g00458.t01#1    mNeoVis.vulgivaga.1.1.chr13.g00451.t01#4   mNeoVis.2.1.1.chr13_hap1.g00474.t01#5       6976   125255308:   mNeoVis.evergladensis.1.1.chr13.g00455.t01#3    mNeoVis.evergladensis.2.1.chr13.g00456.t01#2    mNeoVis.2.1.1.chr13_hap1.g00475.t01#5    mNeoVis.vulgivaga.2.1.chr13.g00459.t01#1    mNeoVis.3.1.1.chr13.g00412.t01#6   mNeoVis.vulgivaga.1.1.chr13.g00452.t01#4       6977   125255309:   mNeoVis.2.1.1.chr2_hap1.g01906.t01#5    mNeoVis.vulgivaga.1.1.chr5.g01827.t01#4    mNeoVis.3.1.1.chr5.g01820.t01#6    mNeoVis.evergladensis.1.1.chr5.g01853.t01#3    mNeoVis.vulgivaga.2.1.chr5.g01847.t01#1   mNeoVis.evergladensis.2.1.chr5.g01844.t01#2       6978   125255310:   mNeoVis.vulgivaga.1.1.chr5.g01828.t01#4    mNeoVis.vulgivaga.2.1.chr5.g01848.t01#1    mNeoVis.evergladensis.1.1.chr5.g01854.t01#3    mNeoVis.2.1.1.chr2_hap1.g01907.t01#5   mNeoVis.evergladensis.2.1.chr5.g01845.t01#2       6979   125255311:   mNeoVis.2.1.1.chr13_hap1.g00487.t01#5    mNeoVis.evergladensis.2.1.chr13.g00467.t01#2    mNeoVis.evergladensis.1.1.chr13.g00465.t01#3    mNeoVis.vulgivaga.2.1.chr13.g00469.t01#1    mNeoVis.3.1.1.chr13.g00421.t01#6   mNeoVis.vulgivaga.1.1.chr13.g00462.t01#4       6980   125255312:   mNeoVis.3.1.1.chr13.g00415.t01#6    mNeoVis.vulgivaga.1.1.chr13.g00456.t01#4    mNeoVis.2.1.1.chr13_hap1.g00476.t01#5    mNeoVis.vulgivaga.1.1.chr13.g00457.t01#4    mNeoVis.vulgivaga.2.1.chr13.g00461.t01#1    mNeoVis.3.1.1.chr13.g00416.t01#6    mNeoVis.evergladensis.2.1.chr13.g00463.t01#2    mNeoVis.vulgivaga.1.1.chr13.g00455.t01#4    mNeoVis.2.1.1.chr13_hap1.g00481.t01#5      6981   125255313:   mNeoVis.evergladensis.2.1.chr13.g00388.t01#2    mNeoVis.3.1.1.chr13.g00342.t01#6    mNeoVis.2.1.1.chr13_hap1.g00404.t01#5    mNeoVis.2.1.1.chr13_hap1.g00403.t01#5    mNeoVis.vulgivaga.2.1.chr13.g00390.t01#1    mNeoVis.vulgivaga.2.1.chr13.g00389.t01#1    mNeoVis.vulgivaga.1.1.chr13.g00379.t01#4    mNeoVis.vulgivaga.1.1.chr13.g00380.t01#4    mNeoVis.evergladensis.2.1.chr13.g00387.t01#2      6982   125255314:   mNeoVis.2.1.1.chr13_hap1.g00396.t01#5    mNeoVis.vulgivaga.1.1.chr13.g00375.t01#4    mNeoVis.evergladensis.2.1.chr13.g00383.t01#2    mNeoVis.vulgivaga.2.1.chr13.g00383.t01#1    mNeoVis.3.1.1.chr13.g00337.t01#6   mNeoVis.evergladensis.1.1.chr13.g00382.t01#3       6983   125255315:   mNeoVis.vulgivaga.1.1.chr13.g00373.t01#4    mNeoVis.3.1.1.chr13.g00341.t01#6    mNeoVis.2.1.1.chr13_hap1.g00401.t01#5    mNeoVis.vulgivaga.2.1.chr13.g00388.t01#1    mNeoVis.evergladensis.1.1.chr13.g00387.t01#3   mNeoVis.evergladensis.2.1.chr13.g00381.t01#2       6984   125255316:   mNeoVis.evergladensis.1.1.chr13.g00457.t01#3    mNeoVis.2.1.1.chr13_hap1.g00477.t01#5    mNeoVis.evergladensis.2.1.chr13.g00458.t01#2    mNeoVis.3.1.1.chr13.g00414.t01#6   mNeoVis.vulgivaga.1.1.chr13.g00454.t01#4       6985   125255317:   mNeoVis.2.1.1.chr13_hap1.g00405.t01#5    mNeoVis.vulgivaga.1.1.chr13.g00381.t01#4    mNeoVis.evergladensis.2.1.chr13.g00389.t01#2    mNeoVis.3.1.1.chr13.g00343.t01#6   mNeoVis.vulgivaga.2.1.chr13.g00391.t01#1       6986   125255318:   mNeoVis.2.1.1.chr13_hap1.g00399.t01#5    mNeoVis.vulgivaga.2.1.chr13.g00386.t01#1    mNeoVis.evergladensis.2.1.chr13.g00384.t01#2    mNeoVis.evergladensis.1.1.chr13.g00385.t01#3   mNeoVis.vulgivaga.1.1.chr13.g00377.t01#4       6987   125255319:   mNeoVis.3.1.1.chr13.g00420.t01#6   mNeoVis.vulgivaga.2.1.chr13.g00468.t01#1           6988   125255320:   mNeoVis.evergladensis.1.1.chr13.g00466.t01#3   mNeoVis.vulgivaga.2.1.chr13.g00470.t01#1           6989   125255321:   mNeoVis.vulgivaga.1.1.chr13.g00463.t01#4    mNeoVis.2.1.1.chr13_hap1.g00488.t01#5    mNeoVis.evergladensis.1.1.chr13.g00467.t01#3    mNeoVis.vulgivaga.2.1.chr13.g00471.t01#1    mNeoVis.3.1.1.chr13.g00423.t01#6   mNeoVis.evergladensis.2.1.chr13.g00468.t01#2       6990   125255322:   mNeoVis.evergladensis.2.1.chr13.g00470.t01#2    mNeoVis.evergladensis.1.1.chr13.g00469.t01#3    mNeoVis.3.1.1.chr13.g00422.t01#6    mNeoVis.2.1.1.chr13_hap1.g00490.t01#5    mNeoVis.vulgivaga.2.1.chr13.g00472.t01#1   mNeoVis.vulgivaga.1.1.chr13.g00465.t01#4       6991   125255323:   mNeoVis.vulgivaga.1.1.chr5.g01829.t01#4    mNeoVis.evergladensis.1.1.chr5.g01855.t01#3    mNeoVis.2.1.1.chr2_hap1.g01908.t01#5   mNeoVis.vulgivaga.2.1.chr5.g01849.t01#1         6992   125255324:   mNeoVis.2.1.1.chr15_hap1.g00670.t01#5    mNeoVis.evergladensis.1.1.chr15.g00120.t01#3    mNeoVis.vulgivaga.1.1.chr15.g00495.t01#4    mNeoVis.vulgivaga.2.1.chr15.g00502.t01#1    mNeoVis.evergladensis.2.1.chr15.g00492.t01#2   mNeoVis.3.1.1.chr15.g00451.t01#6       6993   125255325:   mNeoVis.vulgivaga.2.1.chr5.g01850.t01#1    mNeoVis.evergladensis.2.1.chr5.g01847.t01#2    mNeoVis.vulgivaga.1.1.chr5.g01830.t01#4    mNeoVis.2.1.1.chr2_hap1.g01909.t01#5   mNeoVis.evergladensis.1.1.chr5.g01856.t01#3       6994   125255326:   mNeoVis.3.1.1.chr13.g00425.t01#6    mNeoVis.vulgivaga.1.1.chr13.g00466.t01#4    mNeoVis.2.1.1.chr13_hap1.g00491.t01#5    mNeoVis.evergladensis.2.1.chr13.g00471.t01#2   mNeoVis.vulgivaga.2.1.chr13.g00473.t01#1       6995   125255327:   mNeoVis.3.1.1.chr2.g01290.t01#6    mNeoVis.2.1.1.chr15_hap1.g00678.t01#5    mNeoVis.2.1.1.chr5_hap1.g01367.t01#5    mNeoVis.2.1.1.chr2_hap1.g01910.t01#5    mNeoVis.evergladensis.1.1.chr2.g01364.t01#3    mNeoVis.3.1.1.chr15.g00459.t01#6    mNeoVis.vulgivaga.1.1.chr5.g01831.t01#4    mNeoVis.vulgivaga.1.1.chr2.g01367.t01#4    mNeoVis.evergladensis.2.1.chr5.g01848.t01#2      6996   125255328:   mNeoVis.evergladensis.2.1.chr10.g00268.t01#2    mNeoVis.evergladensis.1.1.chr10.g00269.t01#3   mNeoVis.vulgivaga.2.1.chr10.g00263.t01#1         6997   125255329:   mNeoVis.3.1.1.chr13.g00426.t01#6    mNeoVis.vulgivaga.2.1.chr13.g00474.t01#1    mNeoVis.evergladensis.1.1.chr13.g00472.t01#3    mNeoVis.vulgivaga.1.1.chr13.g00467.t01#4    mNeoVis.evergladensis.2.1.chr13.g00472.t01#2   mNeoVis.2.1.1.chr13_hap1.g00492.t01#5       6998   125255330:   mNeoVis.2.1.1.chr15_hap1.g00672.t01#5    mNeoVis.3.1.1.chr15.g00453.t01#6    mNeoVis.vulgivaga.2.1.chr15.g00503.t01#1    mNeoVis.evergladensis.1.1.chr15.g00119.t01#3    mNeoVis.vulgivaga.1.1.chr15.g00497.t01#4   mNeoVis.evergladensis.2.1.chr15.g00493.t01#2       6999   125255331:   mNeoVis.3.1.1.chr7.g00422.t01#6    mNeoVis.vulgivaga.1.1.chr7.g00443.t01#4    mNeoVis.vulgivaga.2.1.chr7.g00437.t01#1    mNeoVis.evergladensis.2.1.chr7.g00439.t01#2    mNeoVis.evergladensis.1.1.chr7.g00446.t01#3   mNeoVis.2.1.1.chr7_hap1.g00435.t01#5       7000   125255332:   mNeoVis.vulgivaga.2.1.chr5.g01852.t01#1    mNeoVis.2.1.1.chr2_hap1.g01911.t01#5    mNeoVis.evergladensis.1.1.chr5.g01858.t01#3    mNeoVis.vulgivaga.1.1.chr5.g01832.t01#4   mNeoVis.evergladensis.2.1.chr5.g01849.t01#2       7001   125255333:   mNeoVis.2.1.1.chr13_hap1.g00493.t01#5    mNeoVis.evergladensis.1.1.chr13.g00473.t01#3    mNeoVis.3.1.1.chr13.g00427.t01#6    mNeoVis.vulgivaga.1.1.chr13.g00468.t01#4    mNeoVis.evergladensis.2.1.chr13.g00473.t01#2   mNeoVis.vulgivaga.2.1.chr13.g00475.t01#1       7002   125255334:   mNeoVis.vulgivaga.2.1.chr5.g01853.t01#1    mNeoVis.3.1.1.chr5.g01824.t01#6   mNeoVis.evergladensis.1.1.chr5.g01859.t01#3         7003   125255335:   mNeoVis.2.1.1.chr13_hap1.g00494.t01#5    mNeoVis.evergladensis.1.1.chr13.g00474.t01#3    mNeoVis.vulgivaga.1.1.chr13.g00469.t01#4    mNeoVis.3.1.1.chr13.g00428.t01#6    mNeoVis.vulgivaga.2.1.chr13.g00476.t01#1   mNeoVis.evergladensis.2.1.chr13.g00474.t01#2       7004   125255336:   mNeoVis.vulgivaga.2.1.chr5.g01854.t01#1    mNeoVis.evergladensis.1.1.chr5.g01860.t01#3    mNeoVis.2.1.1.chr2_hap1.g01913.t01#5    mNeoVis.evergladensis.2.1.chr5.g01851.t01#2    mNeoVis.vulgivaga.1.1.chr5.g01834.t01#4   mNeoVis.3.1.1.chr5.g01825.t01#6       7005   125255337:   mNeoVis.2.1.1.chr13_hap1.g00495.t01#5    mNeoVis.vulgivaga.1.1.chr13.g00470.t01#4    mNeoVis.vulgivaga.2.1.chr13.g00477.t01#1    mNeoVis.evergladensis.1.1.chr13.g00475.t01#3    mNeoVis.evergladensis.2.1.chr13.g00475.t01#2   mNeoVis.3.1.1.chr13.g00430.t01#6       7006   125255338:   mNeoVis.vulgivaga.2.1.chr13.g00478.t01#1    mNeoVis.evergladensis.2.1.chr13.g00476.t01#2    mNeoVis.evergladensis.1.1.chr13.g00476.t01#3    mNeoVis.3.1.1.chr13.g00431.t01#6    mNeoVis.2.1.1.chr13_hap1.g00496.t01#5   mNeoVis.vulgivaga.1.1.chr13.g00471.t01#4       7007   125255339:   mNeoVis.evergladensis.2.1.chr7.g00440.t01#2    mNeoVis.evergladensis.1.1.chr7.g00447.t01#3    mNeoVis.3.1.1.chr7.g00423.t01#6    mNeoVis.2.1.1.chr7_hap1.g00436.t01#5    mNeoVis.vulgivaga.1.1.chr7.g00444.t01#4   mNeoVis.vulgivaga.2.1.chr7.g00438.t01#1       7008   125255340:   mNeoVis.3.1.1.chr13.g00432.t01#6    mNeoVis.vulgivaga.2.1.chr13.g00479.t01#1    mNeoVis.evergladensis.1.1.chr13.g00477.t01#3    mNeoVis.evergladensis.2.1.chr13.g00477.t01#2    mNeoVis.2.1.1.chr13_hap1.g00497.t01#5    mNeoVis.evergladensis.2.1.chr13.g00478.t01#2   mNeoVis.vulgivaga.1.1.chr13.g00472.t01#4     7009   125255341:   mNeoVis.evergladensis.2.1.chr5.g01852.t01#2    mNeoVis.evergladensis.1.1.chr5.g01861.t01#3    mNeoVis.vulgivaga.2.1.chr5.g01855.t01#1    mNeoVis.vulgivaga.1.1.chr5.g01835.t01#4   mNeoVis.2.1.1.chr2_hap1.g01914.t01#5       7010   125255342:   mNeoVis.vulgivaga.2.1.chr5.g01856.t01#1    mNeoVis.evergladensis.1.1.chr5.g01862.t01#3    mNeoVis.evergladensis.2.1.chr5.g01853.t01#2    mNeoVis.2.1.1.chr2_hap1.g01915.t01#5   mNeoVis.vulgivaga.1.1.chr5.g01836.t01#4       7011   125255343:   mNeoVis.vulgivaga.2.1.chr13.g00480.t01#1    mNeoVis.evergladensis.1.1.chr13.g00478.t01#3    mNeoVis.2.1.1.chr13_hap1.g00498.t01#5    mNeoVis.vulgivaga.1.1.chr13.g00473.t01#4    mNeoVis.evergladensis.2.1.chr13.g00479.t01#2   mNeoVis.3.1.1.chr13.g00433.t01#6       7012   125255344:   mNeoVis.evergladensis.2.1.chr5.g01854.t01#2    mNeoVis.evergladensis.1.1.chr5.g01864.t01#3    mNeoVis.2.1.1.chr2_hap1.g01916.t01#5    mNeoVis.vulgivaga.1.1.chr5.g01837.t01#4   mNeoVis.vulgivaga.2.1.chr5.g01857.t01#1       7013   125255345:   mNeoVis.vulgivaga.2.1.chr5.g01858.t01#1    mNeoVis.vulgivaga.1.1.chr12.g00625.t01#4    mNeoVis.evergladensis.2.1.chr12.g00609.t01#2    mNeoVis.3.1.1.chr5.g01826.t01#6    mNeoVis.evergladensis.1.1.chr12.g00613.t01#3    mNeoVis.2.1.1.chr2_hap1.g01917.t01#5    mNeoVis.evergladensis.2.1.chr5.g01855.t01#2    mNeoVis.2.1.1.chr12_hap1.g00671.t01#5    mNeoVis.vulgivaga.2.1.chr12.g00628.t01#1      7014   125255346:   mNeoVis.vulgivaga.1.1.chr9.g00948.t01#4    mNeoVis.3.1.1.chr9.g00908.t01#6    mNeoVis.2.1.1.chr10_hap1.g00978.t01#5    mNeoVis.vulgivaga.2.1.chr9.g00934.t01#1    mNeoVis.evergladensis.2.1.chr8.g00949.t01#2   mNeoVis.evergladensis.1.1.chr9.g00944.t01#3       7015   125255347:   mNeoVis.vulgivaga.2.1.chr13.g00481.t01#1    mNeoVis.evergladensis.1.1.chr13.g00479.t01#3    mNeoVis.vulgivaga.1.1.chr13.g00474.t01#4    mNeoVis.evergladensis.2.1.chr13.g00480.t01#2   mNeoVis.2.1.1.chr13_hap1.g00499.t01#5       7016   125255348:   mNeoVis.2.1.1.chr15_hap1.g00673.t01#5    mNeoVis.vulgivaga.2.1.chr15.g00504.t01#1    mNeoVis.3.1.1.chr15.g00454.t01#6    mNeoVis.evergladensis.1.1.chr15.g00118.t01#3    mNeoVis.evergladensis.2.1.chr15.g00494.t01#2   mNeoVis.vulgivaga.1.1.chr15.g00498.t01#4       7017   125255349:   mNeoVis.evergladensis.1.1.chr5.g01866.t01#3    mNeoVis.2.1.1.chr2_hap1.g01918.t01#5    mNeoVis.vulgivaga.1.1.chr5.g01839.t01#4    mNeoVis.evergladensis.2.1.chr5.g01856.t01#2   mNeoVis.vulgivaga.2.1.chr5.g01859.t01#1       7018   125255350:   mNeoVis.2.1.1.chr2_hap1.g01919.t01#5    mNeoVis.evergladensis.1.1.chr5.g01867.t01#3    mNeoVis.vulgivaga.1.1.chr5.g01840.t01#4    mNeoVis.vulgivaga.2.1.chr5.g01860.t01#1   mNeoVis.evergladensis.2.1.chr5.g01857.t01#2       7019   125255351:   mNeoVis.2.1.1.chr2_hap1.g01920.t01#5    mNeoVis.evergladensis.2.1.chr5.g01858.t01#2    mNeoVis.evergladensis.1.1.chr5.g01868.t01#3    mNeoVis.vulgivaga.2.1.chr5.g01861.t01#1   mNeoVis.vulgivaga.1.1.chr5.g01841.t01#4       7020   125255352:   mNeoVis.vulgivaga.2.1.chr5.g01862.t01#1    mNeoVis.3.1.1.chr5.g01828.t01#6    mNeoVis.vulgivaga.1.1.chr5.g01842.t01#4    mNeoVis.evergladensis.2.1.chr5.g01859.t01#2    mNeoVis.evergladensis.1.1.chr5.g01869.t01#3   mNeoVis.2.1.1.chr2_hap1.g01921.t01#5       7021   125255353:   mNeoVis.2.1.1.chr2_hap1.g01922.t01#5    mNeoVis.vulgivaga.2.1.chr5.g01863.t01#1    mNeoVis.evergladensis.1.1.chr5.g01870.t01#3    mNeoVis.3.1.1.chr5.g01829.t01#6    mNeoVis.vulgivaga.1.1.chr5.g01843.t01#4   mNeoVis.evergladensis.2.1.chr5.g01860.t01#2       7022   125255354:   mNeoVis.vulgivaga.1.1.chr5.g01844.t01#4    mNeoVis.evergladensis.2.1.chr5.g01861.t01#2    mNeoVis.3.1.1.chr5.g01830.t01#6    mNeoVis.2.1.1.chr2_hap1.g01923.t01#5    mNeoVis.vulgivaga.2.1.chr5.g01864.t01#1   mNeoVis.evergladensis.1.1.chr5.g01871.t01#3       7023   125255355:   mNeoVis.evergladensis.1.1.chr15.g00117.t01#3    mNeoVis.evergladensis.2.1.chr15.g00495.t01#2    mNeoVis.vulgivaga.2.1.chr15.g00505.t01#1    mNeoVis.vulgivaga.1.1.chr15.g00499.t01#4    mNeoVis.2.1.1.chr15_hap1.g00674.t01#5   mNeoVis.3.1.1.chr15.g00455.t01#6       7024   125255356:   mNeoVis.evergladensis.1.1.chr5.g01872.t01#3    mNeoVis.evergladensis.2.1.chr5.g01862.t01#2    mNeoVis.2.1.1.chr2_hap1.g01924.t01#5    mNeoVis.vulgivaga.1.1.chr5.g01845.t01#4   mNeoVis.vulgivaga.2.1.chr5.g01865.t01#1       7025   125255357:   mNeoVis.vulgivaga.1.1.chr5.g01846.t01#4    mNeoVis.vulgivaga.2.1.chr5.g01866.t01#1    mNeoVis.evergladensis.2.1.chr5.g01863.t01#2    mNeoVis.evergladensis.2.1.chr5.g01864.t01#2    mNeoVis.evergladensis.1.1.chr5.g01873.t01#3   mNeoVis.2.1.1.chr2_hap1.g01925.t01#5       7026   125255358:   mNeoVis.3.1.1.chr5.g01832.t01#6    mNeoVis.vulgivaga.1.1.chr5.g01848.t01#4    mNeoVis.evergladensis.1.1.chr5.g01875.t01#3    mNeoVis.vulgivaga.2.1.chr5.g01868.t01#1    mNeoVis.evergladensis.2.1.chr5.g01866.t01#2   mNeoVis.2.1.1.chr2_hap1.g01927.t01#5       7027   125255359:   mNeoVis.evergladensis.2.1.chr5.g01865.t01#2    mNeoVis.vulgivaga.2.1.chr5.g01867.t01#1    mNeoVis.2.1.1.chr2_hap1.g01926.t01#5    mNeoVis.evergladensis.1.1.chr5.g01874.t01#3   mNeoVis.vulgivaga.1.1.chr5.g01847.t01#4       7028   125255360:   mNeoVis.2.1.1.chr9_hap1.g00620.t01#5    mNeoVis.vulgivaga.2.1.chr10.g00596.t01#1    mNeoVis.vulgivaga.1.1.chr10.g00593.t01#4    mNeoVis.evergladensis.2.1.chr13.g00377.t01#2    mNeoVis.evergladensis.1.1.chr13.g00376.t01#3    mNeoVis.3.1.1.chr13.g00332.t01#6    mNeoVis.vulgivaga.1.1.chr13.g00369.t01#4    mNeoVis.vulgivaga.2.1.chr13.g00377.t01#1    mNeoVis.2.1.1.chr13_hap1.g00388.t01#5      7029   125255361:   mNeoVis.vulgivaga.2.1.chr5.g01869.t01#1    mNeoVis.vulgivaga.1.1.chr5.g01849.t01#4    mNeoVis.evergladensis.1.1.chr5.g01876.t01#3    mNeoVis.evergladensis.2.1.chr5.g01867.t01#2   mNeoVis.2.1.1.chr2_hap1.g01928.t01#5       7030   125255362:  mNeoVis.vulgivaga.2.1.chr13.g00378.t01#1               7031   125255363:   mNeoVis.vulgivaga.1.1.chr5.g01850.t01#4    mNeoVis.vulgivaga.2.1.chr5.g01870.t01#1    mNeoVis.evergladensis.2.1.chr5.g01868.t01#2    mNeoVis.2.1.1.chr2_hap1.g01929.t01#5   mNeoVis.evergladensis.1.1.chr5.g01877.t01#3       7032   125255364:   mNeoVis.evergladensis.1.1.chr5.g01878.t01#3    mNeoVis.2.1.1.chr2_hap1.g01930.t01#5    mNeoVis.vulgivaga.1.1.chr5.g01851.t01#4    mNeoVis.vulgivaga.2.1.chr5.g01871.t01#1    mNeoVis.3.1.1.chr5.g01833.t01#6   mNeoVis.evergladensis.2.1.chr5.g01869.t01#2       7033   125255365:   mNeoVis.vulgivaga.1.1.chr5.g01852.t01#4    mNeoVis.evergladensis.2.1.chr5.g01870.t01#2    mNeoVis.2.1.1.chr2_hap1.g01931.t01#5    mNeoVis.3.1.1.chr5.g01834.t01#6    mNeoVis.vulgivaga.2.1.chr5.g01872.t01#1   mNeoVis.evergladensis.1.1.chr5.g01879.t01#3       7034   125255366:   mNeoVis.vulgivaga.1.1.chr5.g01853.t01#4    mNeoVis.evergladensis.2.1.chr5.g01871.t01#2    mNeoVis.evergladensis.1.1.chr5.g01880.t01#3    mNeoVis.2.1.1.chr2_hap1.g01932.t01#5    mNeoVis.3.1.1.chr5.g01835.t01#6   mNeoVis.vulgivaga.2.1.chr5.g01873.t01#1       7035   125255367:   mNeoVis.evergladensis.1.1.chr13.g00379.t01#3    mNeoVis.vulgivaga.1.1.chr13.g00372.t01#4    mNeoVis.3.1.1.chr13.g00335.t01#6    mNeoVis.2.1.1.chr13_hap1.g00392.t01#5    mNeoVis.evergladensis.2.1.chr13.g00380.t01#2   mNeoVis.vulgivaga.2.1.chr13.g00379.t01#1       7036   125255368:   mNeoVis.2.1.1.chr13_hap1.g00390.t01#5    mNeoVis.evergladensis.1.1.chr13.g00378.t01#3    mNeoVis.3.1.1.chr13.g00334.t01#6    mNeoVis.vulgivaga.1.1.chr13.g00371.t01#4   mNeoVis.evergladensis.2.1.chr13.g00379.t01#2       7037   125255369:   mNeoVis.2.1.1.chr13_hap1.g00402.t01#5    mNeoVis.evergladensis.2.1.chr13.g00386.t01#2    mNeoVis.vulgivaga.1.1.chr13.g00382.t01#4   mNeoVis.vulgivaga.2.1.chr13.g00380.t01#1       7038   125255370:   mNeoVis.evergladensis.1.1.chr7.g00448.t01#3    mNeoVis.vulgivaga.1.1.chr7.g00445.t01#4    mNeoVis.2.1.1.chr7_hap1.g00437.t01#5    mNeoVis.evergladensis.2.1.chr7.g00441.t01#2    mNeoVis.3.1.1.chr7.g00424.t01#6   mNeoVis.vulgivaga.2.1.chr7.g00439.t01#1       7039   125255371:   mNeoVis.vulgivaga.1.1.chr5.g01854.t01#4    mNeoVis.vulgivaga.2.1.chr5.g01874.t01#1    mNeoVis.3.1.1.chr5.g01836.t01#6    mNeoVis.evergladensis.2.1.chr5.g01872.t01#2    mNeoVis.2.1.1.chr2_hap1.g01933.t01#5   mNeoVis.evergladensis.1.1.chr5.g01881.t01#3       7040   125255372:   mNeoVis.vulgivaga.2.1.chr13.g00381.t01#1    mNeoVis.evergladensis.1.1.chr13.g00380.t01#3    mNeoVis.2.1.1.chr13_hap1.g00395.t01#5    mNeoVis.vulgivaga.2.1.chr13.g00382.t01#1    mNeoVis.evergladensis.2.1.chr13.g00382.t01#2    mNeoVis.2.1.1.chr13_hap1.g00394.t01#5    mNeoVis.3.1.1.chr13.g00336.t01#6    mNeoVis.evergladensis.1.1.chr13.g00381.t01#3    mNeoVis.vulgivaga.1.1.chr13.g00374.t01#4      7041   125255373:   mNeoVis.evergladensis.1.1.chr13.g00383.t01#3    mNeoVis.vulgivaga.2.1.chr13.g00384.t01#1    mNeoVis.2.1.1.chr13_hap1.g00398.t01#5    mNeoVis.vulgivaga.1.1.chr13.g00376.t01#4    mNeoVis.evergladensis.1.1.chr13.g00384.t01#3   mNeoVis.2.1.1.chr13_hap1.g00397.t01#5       7042   125255374:   mNeoVis.3.1.1.chr13.g00338.t01#6   mNeoVis.vulgivaga.2.1.chr13.g00385.t01#1           7043   125255375:   mNeoVis.vulgivaga.1.1.chr13.g00378.t01#4    mNeoVis.evergladensis.2.1.chr13.g00385.t01#2    mNeoVis.vulgivaga.2.1.chr13.g00387.t01#1    mNeoVis.evergladensis.1.1.chr13.g00386.t01#3    mNeoVis.2.1.1.chr13_hap1.g00400.t01#5   mNeoVis.3.1.1.chr13.g00340.t01#6       7044   125255376:   mNeoVis.vulgivaga.1.1.chr15.g00500.t01#4    mNeoVis.evergladensis.2.1.chr15.g00496.t01#2    mNeoVis.2.1.1.chr15_hap1.g00675.t01#5    mNeoVis.3.1.1.chr15.g00456.t01#6    mNeoVis.vulgivaga.2.1.chr15.g00506.t01#1   mNeoVis.evergladensis.1.1.chr15.g00116.t01#3       7045   125255377:   mNeoVis.vulgivaga.1.1.chr5.g01855.t01#4    mNeoVis.2.1.1.chr2_hap1.g01934.t01#5    mNeoVis.evergladensis.2.1.chr5.g01873.t01#2    mNeoVis.evergladensis.1.1.chr5.g01882.t01#3    mNeoVis.vulgivaga.2.1.chr5.g01875.t01#1   mNeoVis.3.1.1.chr5.g01837.t01#6       7046   125255378:   mNeoVis.evergladensis.1.1.chr5.g01883.t01#3    mNeoVis.vulgivaga.1.1.chr5.g01856.t01#4    mNeoVis.vulgivaga.2.1.chr5.g01876.t01#1    mNeoVis.evergladensis.2.1.chr5.g01874.t01#2    mNeoVis.2.1.1.chr2_hap1.g01935.t01#5   mNeoVis.3.1.1.chr5.g01838.t01#6       7047   125255379:   mNeoVis.vulgivaga.2.1.chr13.g00392.t01#1    mNeoVis.evergladensis.1.1.chr13.g00388.t01#3    mNeoVis.2.1.1.chr13_hap1.g00406.t01#5   mNeoVis.evergladensis.2.1.chr13.g00390.t01#2       7048   125255380:   mNeoVis.vulgivaga.2.1.chr5.g01877.t01#1    mNeoVis.2.1.1.chr2_hap1.g01936.t01#5    mNeoVis.3.1.1.chr5.g01839.t01#6    mNeoVis.evergladensis.2.1.chr5.g01875.t01#2    mNeoVis.vulgivaga.1.1.chr5.g01857.t01#4   mNeoVis.evergladensis.1.1.chr5.g01884.t01#3       7049   125255381:   mNeoVis.2.1.1.chr13_hap1.g00408.t01#5    mNeoVis.vulgivaga.1.1.chr13.g00384.t01#4    mNeoVis.vulgivaga.2.1.chr13.g00394.t01#1    mNeoVis.evergladensis.1.1.chr13.g00390.t01#3    mNeoVis.3.1.1.chr13.g00345.t01#6   mNeoVis.evergladensis.2.1.chr13.g00391.t01#2       7050   125255382:   mNeoVis.evergladensis.2.1.chr5.g01876.t01#2    mNeoVis.vulgivaga.2.1.chr5.g01878.t01#1   mNeoVis.vulgivaga.1.1.chr5.g01858.t01#4           7051   125255383:   mNeoVis.evergladensis.1.1.chr5.g01885.t01#3    mNeoVis.2.1.1.chr2_hap1.g01937.t01#5   mNeoVis.3.1.1.chr5.g01840.t01#6           7052   125255384:   mNeoVis.vulgivaga.2.1.chr5.g01879.t01#1    mNeoVis.evergladensis.2.1.chr5.g01877.t01#2    mNeoVis.evergladensis.1.1.chr5.g01886.t01#3    mNeoVis.3.1.1.chr5.g01841.t01#6    mNeoVis.vulgivaga.1.1.chr5.g01859.t01#4   mNeoVis.2.1.1.chr2_hap1.g01938.t01#5       7053   125255385:   mNeoVis.evergladensis.1.1.chr7.g00449.t01#3    mNeoVis.2.1.1.chr7_hap1.g00438.t01#5    mNeoVis.evergladensis.2.1.chr7.g00442.t01#2    mNeoVis.vulgivaga.2.1.chr7.g00440.t01#1   mNeoVis.vulgivaga.1.1.chr7.g00446.t01#4       7054   125255386:   mNeoVis.vulgivaga.2.1.chr5.g01880.t01#1    mNeoVis.evergladensis.2.1.chr5.g01878.t01#2    mNeoVis.vulgivaga.1.1.chr5.g01860.t01#4    mNeoVis.evergladensis.1.1.chr5.g01887.t01#3    mNeoVis.3.1.1.chr5.g01842.t01#6   mNeoVis.2.1.1.chr2_hap1.g01939.t01#5       7055   125255387:   mNeoVis.vulgivaga.2.1.chr5.g01881.t01#1    mNeoVis.2.1.1.chr2_hap1.g01940.t01#5    mNeoVis.vulgivaga.1.1.chr5.g01861.t01#4    mNeoVis.evergladensis.2.1.chr5.g01879.t01#2   mNeoVis.evergladensis.1.1.chr5.g01888.t01#3       7056   125255388:   mNeoVis.evergladensis.2.1.chr13.g00392.t01#2    mNeoVis.2.1.1.chr13_hap1.g00409.t01#5    mNeoVis.evergladensis.1.1.chr13.g00391.t01#3    mNeoVis.vulgivaga.1.1.chr13.g00385.t01#4    mNeoVis.vulgivaga.2.1.chr13.g00395.t01#1   mNeoVis.3.1.1.chr13.g00347.t01#6       7057   125255389:   mNeoVis.vulgivaga.1.1.chr15.g00501.t01#4    mNeoVis.evergladensis.1.1.chr15.g00115.t01#3    mNeoVis.evergladensis.2.1.chr15.g00497.t01#2    mNeoVis.2.1.1.chr15_hap1.g00676.t01#5    mNeoVis.3.1.1.chr15.g00457.t01#6   mNeoVis.vulgivaga.2.1.chr15.g00507.t01#1       7058   125255390:   mNeoVis.evergladensis.1.1.chr13.g00392.t01#3    mNeoVis.evergladensis.2.1.chr13.g00393.t01#2    mNeoVis.2.1.1.chr13_hap1.g00410.t01#5    mNeoVis.vulgivaga.2.1.chr13.g00397.t01#1    mNeoVis.vulgivaga.1.1.chr13.g00386.t01#4   mNeoVis.3.1.1.chr13.g00348.t01#6       7059   125255391:   mNeoVis.evergladensis.1.1.chr5.g01889.t01#3    mNeoVis.vulgivaga.2.1.chr5.g01882.t01#1    mNeoVis.vulgivaga.1.1.chr5.g01862.t01#4    mNeoVis.2.1.1.chr2_hap1.g01941.t01#5   mNeoVis.evergladensis.2.1.chr5.g01880.t01#2       7060   125255392:   mNeoVis.2.1.1.chr2_hap1.g01856.t01#5    mNeoVis.evergladensis.2.1.chr5.g01794.t01#2    mNeoVis.vulgivaga.2.1.chr5.g01798.t01#1    mNeoVis.evergladensis.1.1.chr5.g01804.t01#3   mNeoVis.vulgivaga.1.1.chr5.g01777.t01#4       7061   125255393:   mNeoVis.vulgivaga.2.1.chr5.g01799.t01#1    mNeoVis.vulgivaga.1.1.chr5.g01778.t01#4    mNeoVis.evergladensis.2.1.chr5.g01795.t01#2    mNeoVis.evergladensis.1.1.chr5.g01805.t01#3   mNeoVis.2.1.1.chr2_hap1.g01857.t01#5       7062   125255394:   mNeoVis.2.1.1.chr13_hap1.g00411.t01#5    mNeoVis.vulgivaga.1.1.chr13.g00387.t01#4    mNeoVis.evergladensis.2.1.chr13.g00394.t01#2    mNeoVis.vulgivaga.2.1.chr13.g00398.t01#1    mNeoVis.3.1.1.chr13.g00349.t01#6   mNeoVis.evergladensis.1.1.chr13.g00393.t01#3       7063   125255395:   mNeoVis.evergladensis.1.1.chr11.g00744.t01#3    mNeoVis.vulgivaga.2.1.chr11.g00756.t01#1    mNeoVis.vulgivaga.1.1.chr11.g00736.t01#4    mNeoVis.2.1.1.chr11_hap1.g00754.t01#5   mNeoVis.3.1.1.chr11.g00717.t01#6       7064   125255396:   mNeoVis.3.1.1.chr13.g00350.t01#6    mNeoVis.vulgivaga.1.1.chr13.g00389.t01#4    mNeoVis.vulgivaga.1.1.chr13.g00388.t01#4    mNeoVis.evergladensis.2.1.chr13.g00395.t01#2    mNeoVis.vulgivaga.2.1.chr13.g00399.t01#1    mNeoVis.2.1.1.chr13_hap1.g00412.t01#5   mNeoVis.evergladensis.1.1.chr13.g00394.t01#3     7065   125255397:   mNeoVis.2.1.1.chr2_hap1.g01858.t01#5    mNeoVis.evergladensis.1.1.chr5.g01806.t01#3    mNeoVis.evergladensis.2.1.chr5.g01796.t01#2    mNeoVis.vulgivaga.2.1.chr5.g01800.t01#1    mNeoVis.vulgivaga.1.1.chr5.g01779.t01#4   mNeoVis.3.1.1.chr5.g01786.t01#6       7066   125255398:   mNeoVis.2.1.1.chr13_hap1.g00413.t01#5    mNeoVis.vulgivaga.1.1.chr13.g00390.t01#4    mNeoVis.evergladensis.1.1.chr13.g00395.t01#3    mNeoVis.vulgivaga.2.1.chr13.g00400.t01#1    mNeoVis.evergladensis.2.1.chr13.g00396.t01#2   mNeoVis.3.1.1.chr13.g00352.t01#6       7067   125255399:   mNeoVis.3.1.1.chr13.g00354.t01#6    mNeoVis.vulgivaga.1.1.chr13.g00391.t01#4    mNeoVis.vulgivaga.2.1.chr13.g00401.t01#1    mNeoVis.evergladensis.2.1.chr13.g00397.t01#2    mNeoVis.evergladensis.1.1.chr13.g00396.t01#3   mNeoVis.2.1.1.chr13_hap1.g00414.t01#5       7068   125255400:   mNeoVis.vulgivaga.1.1.chr13.g00393.t01#4    mNeoVis.2.1.1.chr13_hap1.g00416.t01#5    mNeoVis.evergladensis.1.1.chr13.g00398.t01#3    mNeoVis.evergladensis.2.1.chr13.g00399.t01#2    mNeoVis.3.1.1.chr13.g00355.t01#6   mNeoVis.vulgivaga.2.1.chr13.g00402.t01#1       7069   125255401:   mNeoVis.vulgivaga.2.1.chr13.g00403.t01#1   mNeoVis.3.1.1.chr13.g00356.t01#6             7070   125255402:   mNeoVis.evergladensis.1.1.chr5.g01807.t01#3    mNeoVis.2.1.1.chr2_hap1.g01859.t01#5    mNeoVis.vulgivaga.1.1.chr5.g01780.t01#4    mNeoVis.evergladensis.2.1.chr5.g01797.t01#2   mNeoVis.vulgivaga.2.1.chr5.g01801.t01#1       7071   125255403:   mNeoVis.evergladensis.2.1.chr13.g00401.t01#2    mNeoVis.2.1.1.chr13_hap1.g00418.t01#5    mNeoVis.vulgivaga.1.1.chr13.g00395.t01#4    mNeoVis.vulgivaga.2.1.chr13.g00404.t01#1    mNeoVis.3.1.1.chr13.g00357.t01#6   mNeoVis.evergladensis.1.1.chr13.g00400.t01#3       7072   125255404:   mNeoVis.vulgivaga.2.1.chr5.g01802.t01#1    mNeoVis.2.1.1.chr2_hap1.g01860.t01#5    mNeoVis.vulgivaga.1.1.chr5.g01781.t01#4    mNeoVis.evergladensis.1.1.chr5.g01808.t01#3   mNeoVis.evergladensis.2.1.chr5.g01798.t01#2       7073   125255405:   mNeoVis.evergladensis.2.1.chr5.g01799.t01#2    mNeoVis.vulgivaga.2.1.chr5.g01803.t01#1    mNeoVis.2.1.1.chr2_hap1.g01861.t01#5    mNeoVis.vulgivaga.1.1.chr5.g01782.t01#4    mNeoVis.evergladensis.1.1.chr5.g01809.t01#3   mNeoVis.3.1.1.chr5.g01787.t01#6       7074   125255406:   mNeoVis.vulgivaga.1.1.chr5.g01783.t01#4    mNeoVis.vulgivaga.2.1.chr5.g01804.t01#1    mNeoVis.2.1.1.chr2_hap1.g01862.t01#5    mNeoVis.evergladensis.2.1.chr5.g01800.t01#2   mNeoVis.evergladensis.1.1.chr5.g01810.t01#3       7075   125255407:   mNeoVis.evergladensis.1.1.chr5.g01811.t01#3    mNeoVis.vulgivaga.1.1.chr5.g01784.t01#4    mNeoVis.vulgivaga.2.1.chr5.g01805.t01#1    mNeoVis.2.1.1.chr2_hap1.g01863.t01#5   mNeoVis.evergladensis.2.1.chr5.g01801.t01#2       7076   125255408:   mNeoVis.3.1.1.chr5.g01788.t01#6    mNeoVis.evergladensis.1.1.chr5.g01812.t01#3    mNeoVis.2.1.1.chr2_hap1.g01864.t01#5    mNeoVis.vulgivaga.2.1.chr5.g01806.t01#1    mNeoVis.evergladensis.2.1.chr5.g01802.t01#2   mNeoVis.vulgivaga.1.1.chr5.g01785.t01#4       7077   125255409:   mNeoVis.evergladensis.2.1.chr13.g00403.t01#2    mNeoVis.2.1.1.chr13_hap1.g00420.t01#5    mNeoVis.3.1.1.chr13.g00358.t01#6    mNeoVis.vulgivaga.1.1.chr13.g00396.t01#4    mNeoVis.vulgivaga.2.1.chr13.g00405.t01#1   mNeoVis.evergladensis.1.1.chr13.g00401.t01#3       7078   125255410:   mNeoVis.evergladensis.1.1.chr13.g00402.t01#3    mNeoVis.vulgivaga.2.1.chr13.g00406.t01#1    mNeoVis.vulgivaga.1.1.chr13.g00397.t01#4    mNeoVis.evergladensis.2.1.chr13.g00404.t01#2    mNeoVis.3.1.1.chr13.g00359.t01#6   mNeoVis.2.1.1.chr13_hap1.g00421.t01#5       7079   125255411:   mNeoVis.3.1.1.chr13.g00360.t01#6    mNeoVis.2.1.1.chr13_hap1.g00422.t01#5    mNeoVis.evergladensis.1.1.chr13.g00403.t01#3   mNeoVis.vulgivaga.2.1.chr13.g00407.t01#1       7080   125255412:   mNeoVis.evergladensis.2.1.chr13.g00405.t01#2    mNeoVis.evergladensis.1.1.chr13.g00404.t01#3    mNeoVis.vulgivaga.2.1.chr13.g00408.t01#1    mNeoVis.vulgivaga.1.1.chr13.g00398.t01#4    mNeoVis.3.1.1.chr13.g00361.t01#6   mNeoVis.2.1.1.chr13_hap1.g00423.t01#5       7081   125255413:   mNeoVis.3.1.1.chr5.g01789.t01#6    mNeoVis.2.1.1.chr2_hap1.g01865.t01#5    mNeoVis.evergladensis.2.1.chr5.g01803.t01#2    mNeoVis.vulgivaga.2.1.chr5.g01807.t01#1    mNeoVis.evergladensis.1.1.chr5.g01813.t01#3   mNeoVis.vulgivaga.1.1.chr5.g01786.t01#4       7082   125255414:   mNeoVis.3.1.1.chr13.g00362.t01#6    mNeoVis.vulgivaga.2.1.chr13.g00409.t01#1    mNeoVis.evergladensis.1.1.chr13.g00405.t01#3    mNeoVis.evergladensis.2.1.chr13.g00406.t01#2    mNeoVis.vulgivaga.1.1.chr13.g00399.t01#4   mNeoVis.2.1.1.chr13_hap1.g00424.t01#5       7083   125255415:   mNeoVis.3.1.1.chr5.g01790.t01#6    mNeoVis.vulgivaga.2.1.chr5.g01808.t01#1    mNeoVis.2.1.1.chr2_hap1.g01866.t01#5    mNeoVis.evergladensis.2.1.chr5.g01804.t01#2   mNeoVis.evergladensis.1.1.chr5.g01814.t01#3       7084   125255416:   mNeoVis.evergladensis.1.1.chr13.g00406.t01#3    mNeoVis.evergladensis.2.1.chr13.g00407.t01#2    mNeoVis.3.1.1.chr13.g00363.t01#6    mNeoVis.vulgivaga.1.1.chr13.g00402.t01#4    mNeoVis.2.1.1.chr13_hap1.g00425.t01#5   mNeoVis.vulgivaga.2.1.chr13.g00410.t01#1       7085   125255417:   mNeoVis.2.1.1.chr13_hap1.g00426.t01#5    mNeoVis.vulgivaga.1.1.chr13.g00403.t01#4    mNeoVis.vulgivaga.2.1.chr13.g00411.t01#1    mNeoVis.evergladensis.2.1.chr13.g00408.t01#2   mNeoVis.evergladensis.1.1.chr13.g00407.t01#3       7086   125255418:   mNeoVis.2.1.1.chr15_hap1.g00677.t01#5    mNeoVis.3.1.1.chr15.g00458.t01#6    mNeoVis.evergladensis.1.1.chr15.g00114.t01#3    mNeoVis.vulgivaga.2.1.chr15.g00508.t01#1    mNeoVis.evergladensis.2.1.chr15.g00498.t01#2   mNeoVis.vulgivaga.1.1.chr15.g00502.t01#4       7087   125255419:   mNeoVis.vulgivaga.1.1.chr5.g01788.t01#4    mNeoVis.evergladensis.1.1.chr5.g01815.t01#3   mNeoVis.vulgivaga.2.1.chr5.g01809.t01#1           7088   125255420:   mNeoVis.2.1.1.chr13_hap1.g00427.t01#5    mNeoVis.evergladensis.2.1.chr13.g00409.t01#2    mNeoVis.vulgivaga.1.1.chr13.g00404.t01#4    mNeoVis.vulgivaga.2.1.chr13.g00412.t01#1    mNeoVis.3.1.1.chr13.g00365.t01#6   mNeoVis.evergladensis.1.1.chr13.g00408.t01#3       7089   125255421:  mNeoVis.vulgivaga.2.1.chr13.g00413.t01#1               7090   125255422:   mNeoVis.evergladensis.2.1.chr5.g01806.t01#2    mNeoVis.evergladensis.1.1.chr5.g01816.t01#3    mNeoVis.vulgivaga.2.1.chr5.g01810.t01#1    mNeoVis.2.1.1.chr2_hap1.g01868.t01#5   mNeoVis.vulgivaga.1.1.chr5.g01789.t01#4       7091   125255423:   mNeoVis.2.1.1.chr2_hap1.g01818.t01#5    mNeoVis.vulgivaga.1.1.chr5.g01737.t01#4    mNeoVis.3.1.1.chr5.g01743.t01#6    mNeoVis.evergladensis.1.1.chr5.g01766.t01#3    mNeoVis.vulgivaga.2.1.chr5.g01761.t01#1   mNeoVis.evergladensis.2.1.chr5.g01757.t01#2       7092   125255424:   mNeoVis.vulgivaga.1.1.chr13.g00406.t01#4    mNeoVis.3.1.1.chr13.g00367.t01#6    mNeoVis.evergladensis.2.1.chr13.g00410.t01#2    mNeoVis.2.1.1.chr13_hap1.g00429.t01#5    mNeoVis.vulgivaga.2.1.chr13.g00414.t01#1   mNeoVis.evergladensis.1.1.chr13.g00410.t01#3       7093   125255425:   mNeoVis.evergladensis.1.1.chr11.g00467.t01#3    mNeoVis.evergladensis.2.1.chr11.g00462.t01#2    mNeoVis.3.1.1.chr11.g00436.t01#6    mNeoVis.vulgivaga.1.1.chr11.g00460.t01#4   mNeoVis.2.1.1.chr11_hap1.g00472.t01#5       7094   125255426:   mNeoVis.evergladensis.1.1.chr5.g01767.t01#3    mNeoVis.3.1.1.chr5.g01744.t01#6    mNeoVis.2.1.1.chr2_hap1.g01819.t01#5    mNeoVis.vulgivaga.1.1.chr5.g01738.t01#4    mNeoVis.evergladensis.2.1.chr5.g01758.t01#2   mNeoVis.vulgivaga.2.1.chr5.g01762.t01#1       7095   125255427:   mNeoVis.evergladensis.1.1.chr13.g00411.t01#3    mNeoVis.3.1.1.chr13.g00368.t01#6    mNeoVis.vulgivaga.2.1.chr13.g00415.t01#1    mNeoVis.2.1.1.chr13_hap1.g00430.t01#5    mNeoVis.vulgivaga.1.1.chr13.g00407.t01#4   mNeoVis.evergladensis.2.1.chr13.g00411.t01#2       7096   125255428:   mNeoVis.3.1.1.chr13.g00369.t01#6    mNeoVis.2.1.1.chr13_hap1.g00431.t01#5    mNeoVis.vulgivaga.1.1.chr13.g00408.t01#4    mNeoVis.evergladensis.2.1.chr13.g00412.t01#2    mNeoVis.evergladensis.1.1.chr13.g00412.t01#3   mNeoVis.vulgivaga.2.1.chr13.g00416.t01#1       7097   125255429:   mNeoVis.vulgivaga.2.1.chr13.g00417.t01#1    mNeoVis.evergladensis.2.1.chr13.g00413.t01#2    mNeoVis.vulgivaga.1.1.chr13.g00409.t01#4    mNeoVis.3.1.1.chr13.g00370.t01#6    mNeoVis.2.1.1.chr13_hap1.g00432.t01#5   mNeoVis.evergladensis.1.1.chr13.g00413.t01#3       7098   125255430:   mNeoVis.evergladensis.1.1.chr5.g01817.t01#3    mNeoVis.vulgivaga.1.1.chr5.g01790.t01#4    mNeoVis.evergladensis.2.1.chr5.g01807.t01#2    mNeoVis.vulgivaga.2.1.chr5.g01811.t01#1    mNeoVis.2.1.1.chr2_hap1.g01869.t01#5   mNeoVis.3.1.1.chr5.g01792.t01#6       7099   125255431:   mNeoVis.vulgivaga.1.1.chr5.g01739.t01#4    mNeoVis.evergladensis.2.1.chr5.g01759.t01#2    mNeoVis.vulgivaga.2.1.chr5.g01763.t01#1    mNeoVis.3.1.1.chr5.g01745.t01#6    mNeoVis.2.1.1.chr2_hap1.g01820.t01#5   mNeoVis.evergladensis.1.1.chr5.g01768.t01#3       7100   125255432:   mNeoVis.evergladensis.2.1.chr5.g01760.t01#2    mNeoVis.evergladensis.1.1.chr5.g01769.t01#3    mNeoVis.vulgivaga.1.1.chr5.g01740.t01#4    mNeoVis.vulgivaga.2.1.chr5.g01764.t01#1    mNeoVis.2.1.1.chr2_hap1.g01821.t01#5   mNeoVis.3.1.1.chr14.g00330.t01#6       7101   125255433:   mNeoVis.vulgivaga.2.1.chr13.g00418.t01#1    mNeoVis.3.1.1.chr13.g00371.t01#6    mNeoVis.2.1.1.chr13_hap1.g00433.t01#5    mNeoVis.vulgivaga.1.1.chr13.g00410.t01#4    mNeoVis.evergladensis.2.1.chr13.g00414.t01#2   mNeoVis.evergladensis.1.1.chr13.g00414.t01#3       7102   125255434:   mNeoVis.vulgivaga.2.1.chr5.g01812.t01#1    mNeoVis.vulgivaga.1.1.chr5.g01791.t01#4    mNeoVis.evergladensis.1.1.chr5.g01818.t01#3    mNeoVis.2.1.1.chr2_hap1.g01870.t01#5   mNeoVis.evergladensis.2.1.chr5.g01808.t01#2       7103   125255435:   mNeoVis.vulgivaga.1.1.chr13.g00411.t01#4    mNeoVis.3.1.1.chr13.g00372.t01#6    mNeoVis.2.1.1.chr13_hap1.g00434.t01#5    mNeoVis.evergladensis.1.1.chr13.g00415.t01#3    mNeoVis.evergladensis.2.1.chr13.g00415.t01#2    mNeoVis.3.1.1.chr3.g00011.t01#6   mNeoVis.vulgivaga.2.1.chr13.g00419.t01#1     7104   125255436:   mNeoVis.evergladensis.2.1.chr13.g00416.t01#2    mNeoVis.2.1.1.chr13_hap1.g00435.t01#5    mNeoVis.vulgivaga.1.1.chr13.g00412.t01#4    mNeoVis.evergladensis.1.1.chr13.g00416.t01#3    mNeoVis.3.1.1.chr13.g00373.t01#6   mNeoVis.vulgivaga.2.1.chr13.g00420.t01#1       7105   125255437:   mNeoVis.vulgivaga.1.1.chr15.g00504.t01#4    mNeoVis.evergladensis.2.1.chr15.g00501.t01#2    mNeoVis.2.1.1.chr15_hap1.g00679.t01#5    mNeoVis.evergladensis.1.1.chr15.g00112.t01#3   mNeoVis.vulgivaga.2.1.chr15.g00510.t01#1       7106   125255438:   mNeoVis.evergladensis.1.1.chr5.g01770.t01#3    mNeoVis.vulgivaga.1.1.chr5.g01741.t01#4    mNeoVis.vulgivaga.2.1.chr5.g01765.t01#1    mNeoVis.3.1.1.chr5.g01747.t01#6    mNeoVis.evergladensis.2.1.chr5.g01761.t01#2   mNeoVis.2.1.1.chr2_hap1.g01822.t01#5       7107   125255439:   mNeoVis.evergladensis.1.1.chr13.g00417.t01#3    mNeoVis.evergladensis.2.1.chr13.g00417.t01#2    mNeoVis.3.1.1.chr13.g00374.t01#6    mNeoVis.2.1.1.chr13_hap1.g00436.t01#5    mNeoVis.vulgivaga.2.1.chr13.g00421.t01#1   mNeoVis.vulgivaga.1.1.chr13.g00413.t01#4       7108   125255440:   mNeoVis.3.1.1.chr5.g01793.t01#6    mNeoVis.evergladensis.1.1.chr5.g01820.t01#3    mNeoVis.vulgivaga.1.1.chr5.g01793.t01#4    mNeoVis.vulgivaga.2.1.chr5.g01813.t01#1    mNeoVis.2.1.1.chr2_hap1.g01872.t01#5   mNeoVis.evergladensis.2.1.chr5.g01810.t01#2       7109   125255441:   mNeoVis.vulgivaga.2.1.chr5.g01814.t01#1    mNeoVis.2.1.1.chr2_hap1.g01873.t01#5    mNeoVis.vulgivaga.1.1.chr5.g01794.t01#4    mNeoVis.evergladensis.1.1.chr5.g01821.t01#3   mNeoVis.evergladensis.2.1.chr5.g01811.t01#2       7110   125255442:   mNeoVis.evergladensis.2.1.chr13.g00418.t01#2    mNeoVis.evergladensis.1.1.chr13.g00418.t01#3    mNeoVis.3.1.1.chr13.g00375.t01#6    mNeoVis.vulgivaga.1.1.chr13.g00414.t01#4    mNeoVis.vulgivaga.2.1.chr13.g00422.t01#1   mNeoVis.2.1.1.chr13_hap1.g00437.t01#5       7111   125255443:   mNeoVis.evergladensis.2.1.chr13.g00419.t01#2    mNeoVis.vulgivaga.1.1.chr13.g00415.t01#4    mNeoVis.vulgivaga.2.1.chr13.g00423.t01#1    mNeoVis.2.1.1.chr13_hap1.g00438.t01#5    mNeoVis.evergladensis.1.1.chr13.g00419.t01#3   mNeoVis.3.1.1.chr13.g00376.t01#6       7112   125255444:   mNeoVis.evergladensis.2.1.chr5.g01813.t01#2    mNeoVis.vulgivaga.1.1.chr5.g01796.t01#4    mNeoVis.vulgivaga.2.1.chr5.g01816.t01#1    mNeoVis.3.1.1.chr5.g01796.t01#6    mNeoVis.2.1.1.chr2_hap1.g01875.t01#5   mNeoVis.evergladensis.1.1.chr5.g01823.t01#3       7113   125255445:   mNeoVis.vulgivaga.2.1.chr13.g00424.t01#1    mNeoVis.vulgivaga.1.1.chr13.g00416.t01#4    mNeoVis.evergladensis.2.1.chr13.g00420.t01#2    mNeoVis.2.1.1.chr13_hap1.g00439.t01#5    mNeoVis.evergladensis.1.1.chr13.g00420.t01#3   mNeoVis.3.1.1.chr13.g00377.t01#6       7114   125255446:   mNeoVis.evergladensis.1.1.chr13.g00421.t01#3    mNeoVis.vulgivaga.2.1.chr13.g00425.t01#1    mNeoVis.2.1.1.chr13_hap1.g00440.t01#5   mNeoVis.3.1.1.chr13.g00378.t01#6         7115   125255447:   mNeoVis.vulgivaga.1.1.chr13.g00418.t01#4    mNeoVis.evergladensis.1.1.chr13.g00422.t01#3    mNeoVis.3.1.1.chr13.g00379.t01#6    mNeoVis.evergladensis.2.1.chr13.g00423.t01#2    mNeoVis.2.1.1.chr13_hap1.g00441.t01#5    mNeoVis.evergladensis.2.1.chr13.g00422.t01#2   mNeoVis.vulgivaga.2.1.chr13.g00426.t01#1     7116   125255448:   mNeoVis.vulgivaga.1.1.chr13.g00419.t01#4    mNeoVis.vulgivaga.2.1.chr13.g00427.t01#1    mNeoVis.3.1.1.chr13.g00380.t01#6    mNeoVis.evergladensis.1.1.chr13.g00423.t01#3    mNeoVis.evergladensis.2.1.chr13.g00424.t01#2   mNeoVis.2.1.1.chr13_hap1.g00442.t01#5       7117   125255449:   mNeoVis.2.1.1.chr13_hap1.g00443.t01#5    mNeoVis.evergladensis.1.1.chr13.g00424.t01#3    mNeoVis.evergladensis.2.1.chr13.g00425.t01#2    mNeoVis.vulgivaga.2.1.chr13.g00428.t01#1    mNeoVis.3.1.1.chr13.g00381.t01#6   mNeoVis.vulgivaga.1.1.chr13.g00420.t01#4       7118   125255450:   mNeoVis.vulgivaga.2.1.chr5.g01817.t01#1    mNeoVis.evergladensis.2.1.chr5.g01814.t01#2    mNeoVis.2.1.1.chr2_hap1.g01876.t01#5    mNeoVis.vulgivaga.1.1.chr5.g01797.t01#4   mNeoVis.evergladensis.1.1.chr5.g01824.t01#3       7119   125255451:   mNeoVis.3.1.1.chr13.g00383.t01#6    mNeoVis.vulgivaga.2.1.chr13.g00430.t01#1    mNeoVis.evergladensis.2.1.chr13.g00427.t01#2    mNeoVis.vulgivaga.1.1.chr13.g00422.t01#4    mNeoVis.2.1.1.chr13_hap1.g00445.t01#5   mNeoVis.evergladensis.1.1.chr13.g00426.t01#3       7120   125255452:   mNeoVis.vulgivaga.2.1.chr12.g00361.t01#1    mNeoVis.vulgivaga.1.1.chr12.g00361.t01#4    mNeoVis.2.1.1.chr10_hap1.g00380.t01#5    mNeoVis.vulgivaga.2.1.chr9.g00364.t01#1    mNeoVis.evergladensis.2.1.chr12.g00354.t01#2    mNeoVis.2.1.1.chr12_hap1.g00397.t01#5    mNeoVis.evergladensis.1.1.chr9.g00369.t01#3    mNeoVis.vulgivaga.1.1.chr9.g00363.t01#4    mNeoVis.evergladensis.1.1.chr12.g00361.t01#3      7121   125255453:   mNeoVis.evergladensis.2.1.chr13.g00426.t01#2    mNeoVis.vulgivaga.1.1.chr13.g00421.t01#4    mNeoVis.3.1.1.chr13.g00382.t01#6    mNeoVis.evergladensis.1.1.chr13.g00425.t01#3    mNeoVis.vulgivaga.2.1.chr13.g00429.t01#1   mNeoVis.2.1.1.chr13_hap1.g00444.t01#5       7122   125255454:   mNeoVis.vulgivaga.2.1.chr7.g00441.t01#1    mNeoVis.evergladensis.2.1.chr7.g00443.t01#2    mNeoVis.3.1.1.chr7.g00426.t01#6    mNeoVis.2.1.1.chr7_hap1.g00439.t01#5    mNeoVis.evergladensis.1.1.chr7.g00450.t01#3   mNeoVis.vulgivaga.1.1.chr7.g00447.t01#4       7123   125255455:   mNeoVis.evergladensis.2.1.chr5.g01815.t01#2    mNeoVis.vulgivaga.2.1.chr5.g01818.t01#1    mNeoVis.3.1.1.chr5.g01798.t01#6    mNeoVis.2.1.1.chr2_hap1.g01877.t01#5    mNeoVis.vulgivaga.1.1.chr5.g01798.t01#4   mNeoVis.evergladensis.1.1.chr5.g01825.t01#3       7124   125255456:   mNeoVis.vulgivaga.2.1.chr13.g00431.t01#1    mNeoVis.3.1.1.chr13.g00384.t01#6    mNeoVis.vulgivaga.1.1.chr13.g00423.t01#4    mNeoVis.evergladensis.1.1.chr13.g00427.t01#3    mNeoVis.2.1.1.chr13_hap1.g00446.t01#5   mNeoVis.evergladensis.2.1.chr13.g00428.t01#2       7125   125255457:   mNeoVis.evergladensis.2.1.chr5.g01816.t01#2    mNeoVis.vulgivaga.1.1.chr5.g01799.t01#4    mNeoVis.2.1.1.chr2_hap1.g01878.t01#5    mNeoVis.3.1.1.chr5.g01800.t01#6   mNeoVis.vulgivaga.2.1.chr5.g01819.t01#1       7126   125255458:   mNeoVis.vulgivaga.1.1.chr13.g00328.t01#4    mNeoVis.3.1.1.chr13.g00290.t01#6    mNeoVis.evergladensis.1.1.chr13.g00336.t01#3    mNeoVis.evergladensis.2.1.chr13.g00337.t01#2    mNeoVis.2.1.1.chr13_hap1.g00348.t01#5   mNeoVis.vulgivaga.2.1.chr13.g00335.t01#1       7127   125255459:   mNeoVis.vulgivaga.2.1.chr13.g00336.t01#1   mNeoVis.3.1.1.chr13.g00291.t01#6             7128   125255460:   mNeoVis.evergladensis.1.1.chr13.g00337.t01#3    mNeoVis.2.1.1.chr13_hap1.g00349.t01#5    mNeoVis.3.1.1.chr13.g00292.t01#6    mNeoVis.vulgivaga.2.1.chr13.g00337.t01#1    mNeoVis.vulgivaga.1.1.chr13.g00329.t01#4   mNeoVis.evergladensis.2.1.chr13.g00338.t01#2       7129   125255461:   mNeoVis.evergladensis.2.1.chr5.g01818.t01#2    mNeoVis.2.1.1.chr2_hap1.g01880.t01#5    mNeoVis.vulgivaga.2.1.chr5.g01820.t01#1    mNeoVis.evergladensis.1.1.chr5.g01827.t01#3    mNeoVis.vulgivaga.1.1.chr5.g01800.t01#4   mNeoVis.3.1.1.chr5.g01804.t01#6       7130   125255462:   mNeoVis.evergladensis.2.1.chr5.g01819.t01#2    mNeoVis.vulgivaga.2.1.chr5.g01821.t01#1    mNeoVis.evergladensis.1.1.chr5.g01828.t01#3    mNeoVis.vulgivaga.1.1.chr5.g01801.t01#4   mNeoVis.2.1.1.chr2_hap1.g01881.t01#5       7131   125255463:   mNeoVis.vulgivaga.2.1.chr13.g00338.t01#1    mNeoVis.vulgivaga.1.1.chr13.g00331.t01#4    mNeoVis.3.1.1.chr13.g00293.t01#6    mNeoVis.evergladensis.2.1.chr13.g00339.t01#2    mNeoVis.evergladensis.1.1.chr13.g00338.t01#3   mNeoVis.2.1.1.chr13_hap1.g00350.t01#5       7132   125255464:   mNeoVis.3.1.1.chr5.g01805.t01#6    mNeoVis.vulgivaga.2.1.chr5.g01822.t01#1    mNeoVis.evergladensis.1.1.chr5.g01829.t01#3    mNeoVis.evergladensis.2.1.chr5.g01820.t01#2    mNeoVis.vulgivaga.1.1.chr5.g01802.t01#4   mNeoVis.2.1.1.chr2_hap1.g01882.t01#5       7133   125255465:   mNeoVis.evergladensis.2.1.chr2.g01857.t01#2    mNeoVis.3.1.1.chr2.g01775.t01#6    mNeoVis.evergladensis.1.1.chr2.g01870.t01#3    mNeoVis.vulgivaga.2.1.chr2.g01872.t01#1    mNeoVis.vulgivaga.1.1.chr2.g01871.t01#4   mNeoVis.2.1.1.chr7_hap1.g00440.t01#5       7134   125255466:   mNeoVis.evergladensis.2.1.chr7.g00444.t01#2    mNeoVis.vulgivaga.2.1.chr7.g00442.t01#1    mNeoVis.3.1.1.chr7.g00427.t01#6    mNeoVis.vulgivaga.1.1.chr7.g00448.t01#4   mNeoVis.evergladensis.1.1.chr7.g00451.t01#3       7135   125255467:  mNeoVis.2.1.1.chr5_hap1.g01877.t01#5               7136   125255468:   mNeoVis.3.1.1.chr13.g00294.t01#6    mNeoVis.evergladensis.2.1.chr13.g00340.t01#2    mNeoVis.2.1.1.chr13_hap1.g00351.t01#5    mNeoVis.vulgivaga.2.1.chr13.g00339.t01#1    mNeoVis.vulgivaga.1.1.chr13.g00332.t01#4   mNeoVis.evergladensis.1.1.chr13.g00339.t01#3       7137   125255469:   mNeoVis.vulgivaga.1.1.chr5.g01803.t01#4    mNeoVis.evergladensis.1.1.chr5.g01830.t01#3    mNeoVis.evergladensis.2.1.chr5.g01821.t01#2    mNeoVis.vulgivaga.2.1.chr5.g01823.t01#1   mNeoVis.2.1.1.chr2_hap1.g01883.t01#5       7138   125255470:   mNeoVis.evergladensis.1.1.chr5.g01831.t01#3    mNeoVis.2.1.1.chr2_hap1.g01884.t01#5    mNeoVis.vulgivaga.1.1.chr5.g01804.t01#4    mNeoVis.vulgivaga.2.1.chr5.g01824.t01#1   mNeoVis.evergladensis.2.1.chr5.g01822.t01#2       7139   125255471:   mNeoVis.vulgivaga.1.1.chr13.g00333.t01#4    mNeoVis.2.1.1.chr13_hap1.g00353.t01#5    mNeoVis.evergladensis.1.1.chr13.g00340.t01#3    mNeoVis.vulgivaga.2.1.chr13.g00341.t01#1    mNeoVis.3.1.1.chr13.g00296.t01#6   mNeoVis.evergladensis.2.1.chr13.g00341.t01#2       7140   125255472:   mNeoVis.2.1.1.chr2_hap1.g01885.t01#5    mNeoVis.vulgivaga.2.1.chr5.g01825.t01#1    mNeoVis.3.1.1.chr5.g01806.t01#6    mNeoVis.vulgivaga.1.1.chr5.g01805.t01#4    mNeoVis.evergladensis.1.1.chr5.g01832.t01#3   mNeoVis.evergladensis.2.1.chr5.g01823.t01#2       7141   125255473:   mNeoVis.vulgivaga.1.1.chr13.g00335.t01#4    mNeoVis.evergladensis.1.1.chr13.g00342.t01#3    mNeoVis.2.1.1.chr13_hap1.g00355.t01#5    mNeoVis.evergladensis.2.1.chr13.g00343.t01#2   mNeoVis.vulgivaga.2.1.chr13.g00342.t01#1       7142   125255474:   mNeoVis.vulgivaga.1.1.chr13.g00334.t01#4    mNeoVis.evergladensis.1.1.chr13.g00341.t01#3    mNeoVis.2.1.1.chr13_hap1.g00354.t01#5    mNeoVis.vulgivaga.2.1.chr13.g00343.t01#1   mNeoVis.evergladensis.2.1.chr13.g00342.t01#2       7143   125255475:   mNeoVis.vulgivaga.2.1.chr5.g01826.t01#1    mNeoVis.vulgivaga.1.1.chr5.g01806.t01#4    mNeoVis.evergladensis.2.1.chr5.g01824.t01#2    mNeoVis.evergladensis.1.1.chr5.g01833.t01#3   mNeoVis.2.1.1.chr2_hap1.g01886.t01#5       7144   125255476:   mNeoVis.evergladensis.2.1.chr5.g01825.t01#2    mNeoVis.3.1.1.chr5.g01807.t01#6    mNeoVis.vulgivaga.2.1.chr5.g01827.t01#1    mNeoVis.vulgivaga.1.1.chr5.g01807.t01#4    mNeoVis.2.1.1.chr2_hap1.g01887.t01#5   mNeoVis.evergladensis.1.1.chr5.g01834.t01#3       7145   125255477:   mNeoVis.evergladensis.2.1.chr13.g00344.t01#2    mNeoVis.3.1.1.chr13.g00299.t01#6    mNeoVis.evergladensis.1.1.chr13.g00343.t01#3    mNeoVis.vulgivaga.1.1.chr13.g00336.t01#4    mNeoVis.vulgivaga.2.1.chr13.g00344.t01#1   mNeoVis.2.1.1.chr13_hap1.g00356.t01#5       7146   125255478:   mNeoVis.evergladensis.2.1.chr5.g01826.t01#2    mNeoVis.vulgivaga.1.1.chr5.g01808.t01#4    mNeoVis.2.1.1.chr2_hap1.g01888.t01#5    mNeoVis.vulgivaga.2.1.chr5.g01828.t01#1    mNeoVis.evergladensis.1.1.chr5.g01835.t01#3   mNeoVis.3.1.1.chr5.g01808.t01#6       7147   125255479:   mNeoVis.2.1.1.chr13_hap1.g00357.t01#5    mNeoVis.vulgivaga.1.1.chr13.g00337.t01#4    mNeoVis.vulgivaga.2.1.chr12.g00091.t01#1    mNeoVis.evergladensis.1.1.chr13.g00344.t01#3    mNeoVis.vulgivaga.2.1.chr13.g00345.t01#1    mNeoVis.2.1.1.chr12_hap1.g00112.t01#5    mNeoVis.evergladensis.2.1.chr12.g00087.t01#2    mNeoVis.3.1.1.chr13.g00300.t01#6    mNeoVis.evergladensis.1.1.chr12.g00087.t01#3      7148   125255480:   mNeoVis.evergladensis.2.1.chr5.g01827.t01#2    mNeoVis.vulgivaga.1.1.chr5.g01809.t01#4    mNeoVis.evergladensis.1.1.chr5.g01836.t01#3    mNeoVis.vulgivaga.2.1.chr5.g01829.t01#1   mNeoVis.2.1.1.chr2_hap1.g01889.t01#5       7149   125255481:   mNeoVis.vulgivaga.2.1.chr15.g00511.t01#1    mNeoVis.evergladensis.2.1.chr15.g00502.t01#2    mNeoVis.vulgivaga.1.1.chr15.g00505.t01#4    mNeoVis.3.1.1.chr15.g00461.t01#6    mNeoVis.2.1.1.chr15_hap1.g00680.t01#5   mNeoVis.evergladensis.1.1.chr15.g00111.t01#3       7150   125255482:  mNeoVis.vulgivaga.2.1.chr13.g00347.t01#1               7151   125255483:   mNeoVis.evergladensis.1.1.chr13.g00345.t01#3    mNeoVis.3.1.1.chr13.g00301.t01#6    mNeoVis.vulgivaga.2.1.chr13.g00346.t01#1    mNeoVis.vulgivaga.1.1.chr13.g00338.t01#4    mNeoVis.evergladensis.2.1.chr13.g00346.t01#2    mNeoVis.2.1.1.chr13_hap1.g00358.t01#5   mNeoVis.evergladensis.1.1.chr13.g00346.t01#3     7152   125255484:   mNeoVis.vulgivaga.2.1.chr7.g00443.t01#1    mNeoVis.evergladensis.2.1.chr7.g00445.t01#2    mNeoVis.3.1.1.chr7.g00428.t01#6    mNeoVis.vulgivaga.1.1.chr7.g00449.t01#4    mNeoVis.evergladensis.1.1.chr7.g00452.t01#3   mNeoVis.2.1.1.chr7_hap1.g00441.t01#5       7153   125255485:   mNeoVis.3.1.1.chr13.g00302.t01#6    mNeoVis.vulgivaga.1.1.chr13.g00339.t01#4    mNeoVis.evergladensis.2.1.chr13.g00347.t01#2    mNeoVis.evergladensis.1.1.chr13.g00347.t01#3    mNeoVis.2.1.1.chr13_hap1.g00359.t01#5   mNeoVis.vulgivaga.2.1.chr13.g00348.t01#1       7154   125255486:   mNeoVis.evergladensis.1.1.chr15.g00110.t01#3    mNeoVis.3.1.1.chr15.g00462.t01#6    mNeoVis.vulgivaga.2.1.chr15.g00512.t01#1    mNeoVis.evergladensis.2.1.chr15.g00503.t01#2    mNeoVis.vulgivaga.1.1.chr15.g00506.t01#4   mNeoVis.2.1.1.chr15_hap1.g00681.t01#5       7155   125255487:   mNeoVis.vulgivaga.2.1.chr5.g01830.t01#1    mNeoVis.evergladensis.2.1.chr5.g01828.t01#2    mNeoVis.2.1.1.chr2_hap1.g01890.t01#5   mNeoVis.evergladensis.1.1.chr5.g01837.t01#3       7156   125255488:   mNeoVis.vulgivaga.2.1.chr5.g01831.t01#1    mNeoVis.evergladensis.1.1.chr5.g01838.t01#3    mNeoVis.vulgivaga.1.1.chr5.g01811.t01#4    mNeoVis.2.1.1.chr2_hap1.g01891.t01#5   mNeoVis.evergladensis.2.1.chr5.g01829.t01#2       7157   125255489:  mNeoVis.vulgivaga.2.1.chr5.g01832.t01#1               7158   125255490:   mNeoVis.3.1.1.chr13.g00303.t01#6    mNeoVis.evergladensis.1.1.chr13.g00348.t01#3    mNeoVis.vulgivaga.2.1.chr13.g00349.t01#1    mNeoVis.2.1.1.chr13_hap1.g00360.t01#5    mNeoVis.vulgivaga.1.1.chr13.g00340.t01#4   mNeoVis.evergladensis.2.1.chr13.g00348.t01#2       7159   125255491:   mNeoVis.vulgivaga.1.1.chr7.g00450.t01#4    mNeoVis.evergladensis.2.1.chr7.g00446.t01#2    mNeoVis.evergladensis.1.1.chr7.g00453.t01#3    mNeoVis.2.1.1.chr7_hap1.g00442.t01#5   mNeoVis.vulgivaga.2.1.chr7.g00444.t01#1       7160   125255492:   mNeoVis.vulgivaga.2.1.chr5.g01833.t01#1    mNeoVis.vulgivaga.1.1.chr5.g01813.t01#4    mNeoVis.evergladensis.1.1.chr5.g01839.t01#3    mNeoVis.evergladensis.2.1.chr5.g01830.t01#2   mNeoVis.2.1.1.chr2_hap1.g01892.t01#5       7161   125255493:   mNeoVis.vulgivaga.1.1.chr12.g00188.t01#4    mNeoVis.2.1.1.chr12_hap1.g00208.t01#5    mNeoVis.vulgivaga.2.1.chr12.g00185.t01#1    mNeoVis.evergladensis.2.1.chr12.g00181.t01#2   mNeoVis.evergladensis.1.1.chr12.g00183.t01#3       7162   125255494:   mNeoVis.vulgivaga.1.1.chr5.g01814.t01#4    mNeoVis.3.1.1.chr5.g01812.t01#6    mNeoVis.2.1.1.chr2_hap1.g01893.t01#5    mNeoVis.vulgivaga.2.1.chr5.g01834.t01#1    mNeoVis.evergladensis.1.1.chr5.g01840.t01#3   mNeoVis.evergladensis.2.1.chr5.g01831.t01#2       7163   125255495:   mNeoVis.vulgivaga.2.1.chr13.g00350.t01#1    mNeoVis.evergladensis.1.1.chr13.g00349.t01#3    mNeoVis.2.1.1.chr13_hap1.g00361.t01#5    mNeoVis.3.1.1.chr13.g00304.t01#6    mNeoVis.vulgivaga.1.1.chr13.g00341.t01#4   mNeoVis.evergladensis.2.1.chr13.g00349.t01#2       7164   125255496:   mNeoVis.3.1.1.chr13.g00305.t01#6    mNeoVis.vulgivaga.1.1.chr13.g00342.t01#4    mNeoVis.vulgivaga.2.1.chr13.g00351.t01#1    mNeoVis.2.1.1.chr13_hap1.g00362.t01#5    mNeoVis.evergladensis.2.1.chr13.g00350.t01#2   mNeoVis.evergladensis.1.1.chr13.g00350.t01#3       7165   125255497:   mNeoVis.2.1.1.chr2_hap1.g01894.t01#5    mNeoVis.evergladensis.2.1.chr5.g01832.t01#2    mNeoVis.evergladensis.1.1.chr5.g01841.t01#3    mNeoVis.vulgivaga.2.1.chr5.g01835.t01#1    mNeoVis.3.1.1.chr5.g01813.t01#6   mNeoVis.vulgivaga.1.1.chr5.g01815.t01#4       7166   125255498:  mNeoVis.vulgivaga.2.1.chr15.g00513.t01#1               7167   125255499:  mNeoVis.vulgivaga.2.1.chr5.g01836.t01#1               7168   125255500:   mNeoVis.3.1.1.chr13.g00306.t01#6    mNeoVis.evergladensis.1.1.chr13.g00351.t01#3    mNeoVis.vulgivaga.1.1.chr13.g00343.t01#4    mNeoVis.vulgivaga.2.1.chr13.g00352.t01#1    mNeoVis.2.1.1.chr13_hap1.g00363.t01#5   mNeoVis.evergladensis.2.1.chr13.g00351.t01#2       7169   125253453:   mNeoVis.vulgivaga.2.1.chr5.g01766.t01#1    mNeoVis.vulgivaga.1.1.chr5.g01742.t01#4    mNeoVis.3.1.1.chr5.g01748.t01#6    mNeoVis.evergladensis.2.1.chr5.g01762.t01#2    mNeoVis.2.1.1.chr2_hap1.g01823.t01#5   mNeoVis.evergladensis.1.1.chr5.g01771.t01#3       7170   125253454:   mNeoVis.3.1.1.chr13.g00256.t01#6    mNeoVis.evergladensis.2.1.chr13.g00303.t01#2    mNeoVis.vulgivaga.2.1.chr13.g00300.t01#1    mNeoVis.evergladensis.1.1.chr13.g00300.t01#3    mNeoVis.vulgivaga.1.1.chr13.g00296.t01#4   mNeoVis.2.1.1.chr13_hap1.g00312.t01#5       7171   125253455:   mNeoVis.evergladensis.2.1.chr13.g00304.t01#2    mNeoVis.vulgivaga.1.1.chr13.g00297.t01#4    mNeoVis.vulgivaga.2.1.chr13.g00301.t01#1    mNeoVis.2.1.1.chr13_hap1.g00313.t01#5   mNeoVis.evergladensis.1.1.chr13.g00301.t01#3       7172   125253456:   mNeoVis.vulgivaga.2.1.chr5.g01767.t01#1    mNeoVis.2.1.1.chr2_hap1.g01824.t01#5    mNeoVis.evergladensis.1.1.chr5.g01772.t01#3    mNeoVis.evergladensis.2.1.chr5.g01763.t01#2    mNeoVis.vulgivaga.1.1.chr5.g01743.t01#4   mNeoVis.3.1.1.chr5.g01749.t01#6       7173   125253457:   mNeoVis.2.1.1.chr2_hap1.g01825.t01#5    mNeoVis.evergladensis.1.1.chr5.g01773.t01#3    mNeoVis.3.1.1.chr5.g01750.t01#6    mNeoVis.evergladensis.2.1.chr5.g01764.t01#2    mNeoVis.vulgivaga.2.1.chr5.g01768.t01#1   mNeoVis.vulgivaga.1.1.chr5.g01744.t01#4       7174   125253458:   mNeoVis.evergladensis.1.1.chr13.g00303.t01#3    mNeoVis.evergladensis.2.1.chr13.g00305.t01#2    mNeoVis.3.1.1.chr13.g00258.t01#6    mNeoVis.vulgivaga.1.1.chr13.g00298.t01#4    mNeoVis.2.1.1.chr13_hap1.g00314.t01#5   mNeoVis.vulgivaga.2.1.chr13.g00302.t01#1       7175   125253459:   mNeoVis.vulgivaga.2.1.chr5.g01769.t01#1    mNeoVis.vulgivaga.1.1.chr5.g01745.t01#4    mNeoVis.evergladensis.2.1.chr5.g01765.t01#2    mNeoVis.evergladensis.1.1.chr5.g01774.t01#3   mNeoVis.2.1.1.chr2_hap1.g01826.t01#5       7176   125253460:   mNeoVis.evergladensis.1.1.chr7.g00454.t01#3    mNeoVis.vulgivaga.2.1.chr7.g00445.t01#1    mNeoVis.2.1.1.chr7_hap1.g00443.t01#5    mNeoVis.evergladensis.2.1.chr7.g00447.t01#2    mNeoVis.3.1.1.chr7.g00430.t01#6    mNeoVis.evergladensis.2.1.chr7.g00448.t01#2   mNeoVis.vulgivaga.1.1.chr7.g00451.t01#4     7177   125253461:   mNeoVis.evergladensis.1.1.chr5.g01775.t01#3    mNeoVis.3.1.1.chr5.g01752.t01#6    mNeoVis.vulgivaga.2.1.chr5.g01770.t01#1    mNeoVis.2.1.1.chr2_hap1.g01827.t01#5    mNeoVis.vulgivaga.1.1.chr5.g01746.t01#4   mNeoVis.evergladensis.2.1.chr5.g01766.t01#2       7178   125253462:   mNeoVis.vulgivaga.2.1.chr5.g01771.t01#1    mNeoVis.3.1.1.chr5.g01753.t01#6    mNeoVis.evergladensis.1.1.chr5.g01776.t01#3    mNeoVis.vulgivaga.1.1.chr5.g01747.t01#4    mNeoVis.2.1.1.chr2_hap1.g01828.t01#5   mNeoVis.evergladensis.2.1.chr5.g01767.t01#2       7179   125253463:   mNeoVis.evergladensis.1.1.chr5.g01777.t01#3    mNeoVis.2.1.1.chr2_hap1.g01829.t01#5    mNeoVis.3.1.1.chr5.g01754.t01#6    mNeoVis.vulgivaga.1.1.chr5.g01748.t01#4    mNeoVis.evergladensis.2.1.chr5.g01768.t01#2   mNeoVis.vulgivaga.2.1.chr5.g01772.t01#1       7180   125253464:   mNeoVis.vulgivaga.2.1.chr5.g01773.t01#1    mNeoVis.2.1.1.chr2_hap1.g01830.t01#5    mNeoVis.evergladensis.1.1.chr5.g01778.t01#3    mNeoVis.evergladensis.2.1.chr5.g01769.t01#2    mNeoVis.3.1.1.chr5.g01755.t01#6   mNeoVis.vulgivaga.1.1.chr5.g01749.t01#4       7181   125253465:   mNeoVis.vulgivaga.1.1.chr13.g00345.t01#4    mNeoVis.evergladensis.2.1.chr13.g00352.t01#2    mNeoVis.3.1.1.chr13.g00307.t01#6    mNeoVis.2.1.1.chr13_hap1.g00364.t01#5    mNeoVis.evergladensis.1.1.chr13.g00352.t01#3   mNeoVis.vulgivaga.2.1.chr13.g00353.t01#1       7182   125253466:   mNeoVis.3.1.1.chr5.g01756.t01#6    mNeoVis.evergladensis.1.1.chr5.g01779.t01#3    mNeoVis.vulgivaga.2.1.chr5.g01774.t01#1    mNeoVis.evergladensis.2.1.chr5.g01770.t01#2    mNeoVis.vulgivaga.1.1.chr5.g01750.t01#4   mNeoVis.2.1.1.chr2_hap1.g01831.t01#5       7183   125253467:   mNeoVis.vulgivaga.1.1.chr13.g00346.t01#4    mNeoVis.3.1.1.chr13.g00308.t01#6    mNeoVis.2.1.1.chr13_hap1.g00365.t01#5    mNeoVis.evergladensis.1.1.chr13.g00353.t01#3    mNeoVis.vulgivaga.2.1.chr13.g00354.t01#1   mNeoVis.evergladensis.2.1.chr13.g00353.t01#2       7184   125253468:   mNeoVis.vulgivaga.2.1.chr7.g00446.t01#1    mNeoVis.3.1.1.chr7.g00431.t01#6    mNeoVis.evergladensis.2.1.chr7.g00449.t01#2    mNeoVis.evergladensis.1.1.chr7.g00455.t01#3    mNeoVis.vulgivaga.1.1.chr7.g00452.t01#4   mNeoVis.2.1.1.chr7_hap1.g00444.t01#5       7185   125253469:   mNeoVis.2.1.1.chr13_hap1.g00315.t01#5    mNeoVis.vulgivaga.1.1.chr13.g00299.t01#4    mNeoVis.3.1.1.chr13.g00259.t01#6    mNeoVis.vulgivaga.2.1.chr13.g00303.t01#1    mNeoVis.evergladensis.2.1.chr13.g00306.t01#2   mNeoVis.evergladensis.1.1.chr13.g00304.t01#3       7186   125253470:   mNeoVis.evergladensis.1.1.chr13.g00305.t01#3    mNeoVis.evergladensis.2.1.chr13.g00307.t01#2    mNeoVis.2.1.1.chr13_hap1.g00316.t01#5    mNeoVis.3.1.1.chr13.g00260.t01#6    mNeoVis.vulgivaga.1.1.chr13.g00300.t01#4   mNeoVis.vulgivaga.2.1.chr13.g00304.t01#1       7187   125253471:   mNeoVis.evergladensis.2.1.chr13.g00308.t01#2    mNeoVis.3.1.1.chr13.g00261.t01#6    mNeoVis.evergladensis.1.1.chr13.g00307.t01#3    mNeoVis.vulgivaga.1.1.chr13.g00301.t01#4    mNeoVis.vulgivaga.2.1.chr13.g00305.t01#1   mNeoVis.2.1.1.chr13_hap1.g00317.t01#5       7188   125253472:  mNeoVis.vulgivaga.2.1.chr13.g00355.t01#1               7189   125253473:   mNeoVis.2.1.1.chr13_hap1.g00318.t01#5    mNeoVis.vulgivaga.1.1.chr13.g00302.t01#4    mNeoVis.3.1.1.chr13.g00262.t01#6    mNeoVis.vulgivaga.2.1.chr13.g00306.t01#1    mNeoVis.evergladensis.2.1.chr13.g00309.t01#2   mNeoVis.evergladensis.1.1.chr13.g00308.t01#3       7190   125253474:   mNeoVis.vulgivaga.1.1.chr13.g00348.t01#4    mNeoVis.2.1.1.chr13_hap1.g00367.t01#5    mNeoVis.evergladensis.2.1.chr13.g00355.t01#2    mNeoVis.vulgivaga.2.1.chr13.g00356.t01#1   mNeoVis.evergladensis.1.1.chr13.g00355.t01#3       7191   125253475:   mNeoVis.evergladensis.2.1.chr5.g01771.t01#2    mNeoVis.3.1.1.chr5.g01758.t01#6    mNeoVis.2.1.1.chr2_hap1.g01832.t01#5    mNeoVis.vulgivaga.1.1.chr5.g01752.t01#4    mNeoVis.evergladensis.1.1.chr5.g01781.t01#3   mNeoVis.vulgivaga.2.1.chr5.g01775.t01#1       7192   125253476:   mNeoVis.evergladensis.2.1.chr13.g00356.t01#2    mNeoVis.vulgivaga.2.1.chr13.g00357.t01#1    mNeoVis.evergladensis.1.1.chr13.g00356.t01#3    mNeoVis.3.1.1.chr13.g00312.t01#6    mNeoVis.vulgivaga.1.1.chr13.g00349.t01#4   mNeoVis.2.1.1.chr13_hap1.g00368.t01#5       7193   125253477:   mNeoVis.vulgivaga.1.1.chr5.g01753.t01#4    mNeoVis.evergladensis.1.1.chr5.g01782.t01#3    mNeoVis.evergladensis.2.1.chr5.g01772.t01#2    mNeoVis.3.1.1.chr5.g01759.t01#6    mNeoVis.vulgivaga.2.1.chr5.g01776.t01#1   mNeoVis.2.1.1.chr2_hap1.g01833.t01#5       7194   125253478:   mNeoVis.3.1.1.chr13.g00313.t01#6    mNeoVis.2.1.1.chr13_hap1.g00369.t01#5    mNeoVis.vulgivaga.2.1.chr13.g00358.t01#1    mNeoVis.evergladensis.2.1.chr13.g00357.t01#2    mNeoVis.evergladensis.1.1.chr13.g00358.t01#3   mNeoVis.vulgivaga.1.1.chr13.g00350.t01#4       7195   125253479:   mNeoVis.2.1.1.chr2_hap1.g01834.t01#5    mNeoVis.evergladensis.1.1.chr5.g01783.t01#3    mNeoVis.vulgivaga.1.1.chr5.g01754.t01#4    mNeoVis.vulgivaga.2.1.chr5.g01777.t01#1    mNeoVis.evergladensis.2.1.chr5.g01773.t01#2   mNeoVis.3.1.1.chr5.g01760.t01#6       7196   125253480:   mNeoVis.2.1.1.chr2_hap1.g01835.t01#5    mNeoVis.vulgivaga.2.1.chr5.g01778.t01#1    mNeoVis.3.1.1.chr5.g01761.t01#6    mNeoVis.vulgivaga.1.1.chr5.g01755.t01#4   mNeoVis.evergladensis.1.1.chr5.g01784.t01#3       7197   125253481:   mNeoVis.2.1.1.chr2_hap1.g01836.t01#5    mNeoVis.3.1.1.chr5.g01762.t01#6    mNeoVis.vulgivaga.2.1.chr5.g01779.t01#1    mNeoVis.evergladensis.1.1.chr5.g01785.t01#3    mNeoVis.evergladensis.2.1.chr5.g01774.t01#2   mNeoVis.vulgivaga.1.1.chr5.g01756.t01#4       7198   125253482:   mNeoVis.2.1.1.chr2_hap1.g01837.t01#5    mNeoVis.vulgivaga.1.1.chr5.g01757.t01#4    mNeoVis.3.1.1.chr5.g01763.t01#6    mNeoVis.evergladensis.2.1.chr5.g01775.t01#2    mNeoVis.evergladensis.1.1.chr5.g01786.t01#3   mNeoVis.vulgivaga.2.1.chr5.g01780.t01#1       7199   125253483:   mNeoVis.3.1.1.chr5.g01764.t01#6    mNeoVis.evergladensis.1.1.chr5.g01787.t01#3    mNeoVis.vulgivaga.2.1.chr5.g01781.t01#1    mNeoVis.evergladensis.2.1.chr5.g01776.t01#2    mNeoVis.vulgivaga.1.1.chr5.g01758.t01#4   mNeoVis.2.1.1.chr2_hap1.g01838.t01#5       7200   125253484:   mNeoVis.evergladensis.1.1.chr13.g00359.t01#3    mNeoVis.evergladensis.2.1.chr13.g00359.t01#2    mNeoVis.vulgivaga.1.1.chr13.g00351.t01#4    mNeoVis.vulgivaga.2.1.chr13.g00359.t01#1    mNeoVis.3.1.1.chr13.g00314.t01#6   mNeoVis.2.1.1.chr13_hap1.g00370.t01#5       7201   125253485:   mNeoVis.vulgivaga.2.1.chr5.g01782.t01#1    mNeoVis.2.1.1.chr2_hap1.g01839.t01#5    mNeoVis.vulgivaga.1.1.chr5.g01759.t01#4    mNeoVis.evergladensis.1.1.chr5.g01788.t01#3   mNeoVis.evergladensis.2.1.chr5.g01777.t01#2       7202   125253486:   mNeoVis.vulgivaga.2.1.chr13.g00360.t01#1    mNeoVis.3.1.1.chr13.g00315.t01#6    mNeoVis.evergladensis.2.1.chr13.g00360.t01#2    mNeoVis.2.1.1.chr13_hap1.g00371.t01#5    mNeoVis.vulgivaga.1.1.chr13.g00352.t01#4   mNeoVis.evergladensis.1.1.chr13.g00360.t01#3       7203   125253487:   mNeoVis.evergladensis.2.1.chr13.g00361.t01#2    mNeoVis.evergladensis.1.1.chr13.g00361.t01#3    mNeoVis.vulgivaga.2.1.chr13.g00361.t01#1    mNeoVis.vulgivaga.1.1.chr13.g00353.t01#4    mNeoVis.2.1.1.chr13_hap1.g00372.t01#5   mNeoVis.3.1.1.chr13.g00316.t01#6       7204   125253488:   mNeoVis.3.1.1.chr7.g00432.t01#6    mNeoVis.2.1.1.chr7_hap1.g00445.t01#5    mNeoVis.evergladensis.1.1.chr7.g00456.t01#3    mNeoVis.vulgivaga.1.1.chr7.g00453.t01#4    mNeoVis.evergladensis.2.1.chr7.g00450.t01#2   mNeoVis.vulgivaga.2.1.chr7.g00447.t01#1       7205   125253489:   mNeoVis.2.1.1.chr2_hap1.g01840.t01#5    mNeoVis.3.1.1.chr5.g01766.t01#6    mNeoVis.evergladensis.1.1.chr5.g01789.t01#3    mNeoVis.vulgivaga.2.1.chr5.g01783.t01#1    mNeoVis.vulgivaga.1.1.chr5.g01760.t01#4   mNeoVis.evergladensis.2.1.chr5.g01778.t01#2       7206   125253490:   mNeoVis.vulgivaga.2.1.chr13.g00362.t01#1    mNeoVis.3.1.1.chr13.g00318.t01#6    mNeoVis.2.1.1.chr13_hap1.g00373.t01#5    mNeoVis.evergladensis.2.1.chr13.g00362.t01#2    mNeoVis.vulgivaga.1.1.chr13.g00354.t01#4   mNeoVis.evergladensis.1.1.chr13.g00362.t01#3       7207   125253491:   mNeoVis.vulgivaga.2.1.chr13.g00363.t01#1    mNeoVis.2.1.1.chr13_hap1.g00374.t01#5    mNeoVis.3.1.1.chr13.g00319.t01#6    mNeoVis.evergladensis.2.1.chr13.g00363.t01#2    mNeoVis.vulgivaga.1.1.chr13.g00355.t01#4   mNeoVis.evergladensis.1.1.chr13.g00363.t01#3       7208   125253492:   mNeoVis.2.1.1.chr2_hap1.g01841.t01#5    mNeoVis.evergladensis.1.1.chr5.g01790.t01#3    mNeoVis.3.1.1.chr5.g01767.t01#6    mNeoVis.vulgivaga.2.1.chr5.g01784.t01#1    mNeoVis.evergladensis.2.1.chr5.g01779.t01#2   mNeoVis.vulgivaga.1.1.chr5.g01761.t01#4       7209   125253493:   mNeoVis.vulgivaga.1.1.chr5.g01763.t01#4    mNeoVis.vulgivaga.1.1.chr5.g01764.t01#4    mNeoVis.evergladensis.1.1.chr5.g01792.t01#3    mNeoVis.2.1.1.chr2_hap1.g01843.t01#5    mNeoVis.vulgivaga.2.1.chr5.g01786.t01#1    mNeoVis.evergladensis.2.1.chr5.g01781.t01#2   mNeoVis.3.1.1.chr5.g01769.t01#6       7210   125253494:   mNeoVis.3.1.1.chr5.g01768.t01#6    mNeoVis.evergladensis.2.1.chr5.g01780.t01#2    mNeoVis.2.1.1.chr2_hap1.g01842.t01#5    mNeoVis.vulgivaga.2.1.chr5.g01785.t01#1    mNeoVis.vulgivaga.1.1.chr5.g01762.t01#4   mNeoVis.evergladensis.1.1.chr5.g01791.t01#3       7211   125253495:   mNeoVis.evergladensis.2.1.chr2.g01523.t01#2    mNeoVis.3.1.1.chr2.g01455.t01#6    mNeoVis.2.1.1.chr5_hap1.g01537.t01#5    mNeoVis.evergladensis.1.1.chr2.g01533.t01#3    mNeoVis.vulgivaga.2.1.chr2.g01537.t01#1   mNeoVis.vulgivaga.1.1.chr2.g01534.t01#4       7212   125253496:   mNeoVis.vulgivaga.2.1.chr5.g01787.t01#1   mNeoVis.3.1.1.chr5.g01770.t01#6             7213   125253497:   mNeoVis.vulgivaga.1.1.chr5.g01766.t01#4    mNeoVis.2.1.1.chr2_hap1.g01845.t01#5    mNeoVis.3.1.1.chr5.g01772.t01#6    mNeoVis.evergladensis.2.1.chr5.g01783.t01#2    mNeoVis.evergladensis.1.1.chr5.g01794.t01#3   mNeoVis.vulgivaga.2.1.chr5.g01788.t01#1       7214   125253498:   mNeoVis.evergladensis.2.1.chr5.g01784.t01#2    mNeoVis.vulgivaga.2.1.chr5.g01789.t01#1    mNeoVis.2.1.1.chr2_hap1.g01846.t01#5    mNeoVis.evergladensis.1.1.chr5.g01795.t01#3    mNeoVis.3.1.1.chr5.g01773.t01#6   mNeoVis.vulgivaga.1.1.chr5.g01767.t01#4       7215   125253499:   mNeoVis.evergladensis.1.1.chr13.g00364.t01#3    mNeoVis.evergladensis.2.1.chr13.g00364.t01#2    mNeoVis.vulgivaga.1.1.chr13.g00356.t01#4    mNeoVis.3.1.1.chr13.g00320.t01#6    mNeoVis.2.1.1.chr13_hap1.g00375.t01#5   mNeoVis.vulgivaga.2.1.chr13.g00364.t01#1       7216   125253500:   mNeoVis.3.1.1.chr13.g00321.t01#6    mNeoVis.2.1.1.chr13_hap1.g00376.t01#5    mNeoVis.vulgivaga.1.1.chr13.g00357.t01#4    mNeoVis.evergladensis.2.1.chr13.g00365.t01#2    mNeoVis.vulgivaga.2.1.chr13.g00365.t01#1   mNeoVis.evergladensis.1.1.chr13.g00365.t01#3       7217   125253501:   mNeoVis.vulgivaga.2.1.chr13.g00366.t01#1    mNeoVis.vulgivaga.1.1.chr13.g00358.t01#4    mNeoVis.evergladensis.2.1.chr13.g00366.t01#2   mNeoVis.3.1.1.chr13.g00322.t01#6         7218   125253502:   mNeoVis.vulgivaga.2.1.chr13.g00367.t01#1    mNeoVis.evergladensis.2.1.chr13.g00367.t01#2    mNeoVis.3.1.1.chr13.g00324.t01#6   mNeoVis.2.1.1.chr13_hap1.g00378.t01#5         7219   125253503:   mNeoVis.evergladensis.1.1.chr5.g01796.t01#3    mNeoVis.evergladensis.2.1.chr5.g01785.t01#2    mNeoVis.2.1.1.chr2_hap1.g01847.t01#5    mNeoVis.vulgivaga.1.1.chr5.g01768.t01#4    mNeoVis.vulgivaga.2.1.chr5.g01790.t01#1   mNeoVis.3.1.1.chr5.g01774.t01#6       7220   125253504:   mNeoVis.2.1.1.chr8_hap1.g01109.t01#5    mNeoVis.vulgivaga.1.1.chr8.g01074.t01#4    mNeoVis.evergladensis.1.1.chr8.g01079.t01#3    mNeoVis.vulgivaga.2.1.chr8.g01086.t01#1   mNeoVis.evergladensis.2.1.chr9.g01087.t01#2       7221   125253505:   mNeoVis.evergladensis.2.1.chr10.g00274.t01#2    mNeoVis.vulgivaga.2.1.chr10.g00270.t01#1    mNeoVis.evergladensis.1.1.chr10.g00275.t01#3    mNeoVis.vulgivaga.1.1.chr10.g00264.t01#4   mNeoVis.2.1.1.chr9_hap1.g00275.t01#5       7222   125253506:   mNeoVis.vulgivaga.1.1.chr13.g00360.t01#4    mNeoVis.evergladensis.1.1.chr13.g00367.t01#3    mNeoVis.evergladensis.2.1.chr13.g00368.t01#2    mNeoVis.2.1.1.chr13_hap1.g00379.t01#5   mNeoVis.vulgivaga.2.1.chr13.g00368.t01#1       7223   125253507:   mNeoVis.vulgivaga.1.1.chr5.g01770.t01#4    mNeoVis.evergladensis.2.1.chr5.g01787.t01#2    mNeoVis.evergladensis.1.1.chr5.g01797.t01#3    mNeoVis.2.1.1.chr2_hap1.g01849.t01#5   mNeoVis.vulgivaga.2.1.chr5.g01791.t01#1       7224   125253508:   mNeoVis.vulgivaga.1.1.chr13.g00361.t01#4    mNeoVis.evergladensis.1.1.chr13.g00368.t01#3    mNeoVis.evergladensis.2.1.chr13.g00369.t01#2    mNeoVis.3.1.1.chr13.g00326.t01#6    mNeoVis.vulgivaga.2.1.chr13.g00369.t01#1   mNeoVis.2.1.1.chr13_hap1.g00380.t01#5       7225   125253509:   mNeoVis.vulgivaga.2.1.chr13.g00370.t01#1    mNeoVis.2.1.1.chr13_hap1.g00381.t01#5    mNeoVis.vulgivaga.1.1.chr13.g00362.t01#4    mNeoVis.evergladensis.1.1.chr13.g00369.t01#3   mNeoVis.evergladensis.2.1.chr13.g00370.t01#2       7226   125253510:   mNeoVis.vulgivaga.1.1.chr5.g01771.t01#4    mNeoVis.vulgivaga.2.1.chr5.g01792.t01#1    mNeoVis.evergladensis.2.1.chr5.g01788.t01#2    mNeoVis.evergladensis.1.1.chr5.g01798.t01#3    mNeoVis.2.1.1.chr2_hap1.g01850.t01#5   mNeoVis.3.1.1.chr5.g01778.t01#6       7227   125253511:   mNeoVis.evergladensis.2.1.chr13.g00371.t01#2    mNeoVis.2.1.1.chr13_hap1.g00382.t01#5   mNeoVis.vulgivaga.2.1.chr13.g00371.t01#1         7228   125253512:   mNeoVis.vulgivaga.1.1.chr13.g00363.t01#4   mNeoVis.evergladensis.1.1.chr13.g00370.t01#3           7229   125253513:   mNeoVis.3.1.1.chr5.g01779.t01#6    mNeoVis.vulgivaga.2.1.chr5.g01793.t01#1    mNeoVis.2.1.1.chr2_hap1.g01851.t01#5    mNeoVis.evergladensis.2.1.chr5.g01789.t01#2    mNeoVis.vulgivaga.1.1.chr5.g01772.t01#4   mNeoVis.evergladensis.1.1.chr5.g01799.t01#3       7230   125253514:   mNeoVis.vulgivaga.2.1.chr13.g00372.t01#1    mNeoVis.evergladensis.1.1.chr13.g00371.t01#3    mNeoVis.vulgivaga.1.1.chr13.g00364.t01#4   mNeoVis.evergladensis.2.1.chr13.g00372.t01#2       7231   125253515:   mNeoVis.vulgivaga.2.1.chr5.g01794.t01#1    mNeoVis.2.1.1.chr2_hap1.g01852.t01#5    mNeoVis.evergladensis.1.1.chr5.g01800.t01#3    mNeoVis.vulgivaga.1.1.chr5.g01773.t01#4    mNeoVis.3.1.1.chr5.g01780.t01#6   mNeoVis.evergladensis.2.1.chr5.g01790.t01#2       7232   125253516:   mNeoVis.2.1.1.chr2_hap1.g01853.t01#5    mNeoVis.vulgivaga.1.1.chr5.g01774.t01#4    mNeoVis.evergladensis.1.1.chr5.g01801.t01#3    mNeoVis.3.1.1.chr5.g01781.t01#6    mNeoVis.evergladensis.2.1.chr5.g01791.t01#2   mNeoVis.vulgivaga.2.1.chr5.g01795.t01#1       7233   125253517:   mNeoVis.3.1.1.chr13.g00328.t01#6    mNeoVis.2.1.1.chr13_hap1.g00384.t01#5    mNeoVis.evergladensis.1.1.chr13.g00372.t01#3    mNeoVis.evergladensis.2.1.chr13.g00373.t01#2    mNeoVis.vulgivaga.2.1.chr13.g00373.t01#1   mNeoVis.vulgivaga.1.1.chr13.g00365.t01#4       7234   125253518:   mNeoVis.3.1.1.chr13.g00329.t01#6    mNeoVis.2.1.1.chr13_hap1.g00385.t01#5    mNeoVis.vulgivaga.2.1.chr13.g00374.t01#1    mNeoVis.evergladensis.1.1.chr13.g00373.t01#3    mNeoVis.vulgivaga.1.1.chr13.g00366.t01#4   mNeoVis.evergladensis.2.1.chr13.g00374.t01#2       7235   125253519:   mNeoVis.vulgivaga.1.1.chr13.g00367.t01#4    mNeoVis.vulgivaga.2.1.chr13.g00375.t01#1    mNeoVis.3.1.1.chr13.g00330.t01#6    mNeoVis.evergladensis.2.1.chr13.g00375.t01#2    mNeoVis.evergladensis.1.1.chr13.g00374.t01#3   mNeoVis.2.1.1.chr13_hap1.g00386.t01#5       7236   125253520:   mNeoVis.2.1.1.chr2_hap1.g01854.t01#5    mNeoVis.vulgivaga.2.1.chr5.g01796.t01#1    mNeoVis.vulgivaga.1.1.chr5.g01775.t01#4   mNeoVis.evergladensis.1.1.chr5.g01802.t01#3       7237   125253521:   mNeoVis.2.1.1.chr2_hap1.g01855.t01#5    mNeoVis.vulgivaga.2.1.chr5.g01797.t01#1    mNeoVis.evergladensis.2.1.chr5.g01793.t01#2    mNeoVis.3.1.1.chr5.g01782.t01#6    mNeoVis.evergladensis.1.1.chr5.g01803.t01#3   mNeoVis.vulgivaga.1.1.chr5.g01776.t01#4       7238   125253522:   mNeoVis.evergladensis.1.1.chr13.g00375.t01#3    mNeoVis.3.1.1.chr13.g00331.t01#6    mNeoVis.vulgivaga.1.1.chr13.g00368.t01#4    mNeoVis.2.1.1.chr13_hap1.g00387.t01#5    mNeoVis.vulgivaga.2.1.chr13.g00376.t01#1   mNeoVis.evergladensis.2.1.chr13.g00376.t01#2       7239   125253523:   mNeoVis.3.1.1.chr13.g00248.t01#6    mNeoVis.vulgivaga.2.1.chr13.g00292.t01#1    mNeoVis.evergladensis.1.1.chr13.g00293.t01#3    mNeoVis.evergladensis.2.1.chr13.g00296.t01#2    mNeoVis.2.1.1.chr13_hap1.g00305.t01#5   mNeoVis.vulgivaga.1.1.chr13.g00289.t01#4       7240   125253524:   mNeoVis.vulgivaga.2.1.chr13.g00293.t01#1    mNeoVis.evergladensis.1.1.chr13.g00294.t01#3    mNeoVis.3.1.1.chr13.g00249.t01#6    mNeoVis.evergladensis.2.1.chr13.g00297.t01#2    mNeoVis.vulgivaga.1.1.chr13.g00290.t01#4   mNeoVis.2.1.1.chr13_hap1.g00306.t01#5       7241   125253525:   mNeoVis.vulgivaga.1.1.chr5.g01697.t01#4    mNeoVis.3.1.1.chr5.g01702.t01#6    mNeoVis.2.1.1.chr2_hap1.g01777.t01#5    mNeoVis.vulgivaga.2.1.chr5.g01721.t01#1   mNeoVis.evergladensis.2.1.chr5.g01717.t01#2       7242   125253526:   mNeoVis.vulgivaga.2.1.chr5.g01722.t01#1    mNeoVis.evergladensis.2.1.chr5.g01718.t01#2    mNeoVis.2.1.1.chr2_hap1.g01778.t01#5    mNeoVis.evergladensis.1.1.chr5.g01726.t01#3    mNeoVis.3.1.1.chr5.g01703.t01#6   mNeoVis.vulgivaga.1.1.chr5.g01698.t01#4       7243   125253527:   mNeoVis.vulgivaga.2.1.chr13.g00294.t01#1    mNeoVis.evergladensis.2.1.chr13.g00298.t01#2    mNeoVis.evergladensis.1.1.chr13.g00295.t01#3    mNeoVis.3.1.1.chr13.g00250.t01#6    mNeoVis.vulgivaga.1.1.chr13.g00291.t01#4   mNeoVis.2.1.1.chr13_hap1.g00307.t01#5       7244   125253528:   mNeoVis.vulgivaga.2.1.chr7.g00448.t01#1    mNeoVis.vulgivaga.1.1.chr7.g00454.t01#4   mNeoVis.3.1.1.chr7.g00433.t01#6           7245   125253529:   mNeoVis.evergladensis.1.1.chr7.g00457.t01#3    mNeoVis.2.1.1.chr7_hap1.g00446.t01#5   mNeoVis.evergladensis.2.1.chr7.g00451.t01#2         7246   125253530:   mNeoVis.3.1.1.chr7.g00434.t01#6   mNeoVis.vulgivaga.2.1.chr7.g00449.t01#1             7247   125253531:   mNeoVis.vulgivaga.2.1.chr5.g01723.t01#1    mNeoVis.evergladensis.2.1.chr5.g01719.t01#2    mNeoVis.evergladensis.1.1.chr5.g01727.t01#3    mNeoVis.vulgivaga.1.1.chr5.g01699.t01#4   mNeoVis.2.1.1.chr2_hap1.g01779.t01#5       7248   125253532:   mNeoVis.evergladensis.1.1.chr13.g00296.t01#3    mNeoVis.vulgivaga.2.1.chr13.g00295.t01#1    mNeoVis.3.1.1.chr13.g00251.t01#6    mNeoVis.evergladensis.2.1.chr13.g00299.t01#2    mNeoVis.2.1.1.chr13_hap1.g00308.t01#5   mNeoVis.vulgivaga.1.1.chr13.g00292.t01#4       7249   125253533:   mNeoVis.vulgivaga.2.1.chr5.g01724.t01#1    mNeoVis.evergladensis.1.1.chr5.g01728.t01#3    mNeoVis.evergladensis.2.1.chr5.g01720.t01#2    mNeoVis.vulgivaga.1.1.chr5.g01700.t01#4   mNeoVis.2.1.1.chr2_hap1.g01780.t01#5       7250   125253534:   mNeoVis.vulgivaga.2.1.chr13.g00296.t01#1    mNeoVis.3.1.1.chr13.g00252.t01#6    mNeoVis.evergladensis.2.1.chr13.g00300.t01#2   mNeoVis.evergladensis.1.1.chr13.g00297.t01#3       7251   125253535:   mNeoVis.vulgivaga.1.1.chr5.g01701.t01#4    mNeoVis.evergladensis.2.1.chr5.g01721.t01#2    mNeoVis.vulgivaga.2.1.chr5.g01725.t01#1    mNeoVis.2.1.1.chr2_hap1.g01781.t01#5    mNeoVis.3.1.1.chr5.g01704.t01#6   mNeoVis.evergladensis.1.1.chr5.g01729.t01#3       7252   125253536:   mNeoVis.2.1.1.chr13_hap1.g00309.t01#5    mNeoVis.3.1.1.chr13.g00253.t01#6    mNeoVis.vulgivaga.1.1.chr13.g00293.t01#4   mNeoVis.vulgivaga.2.1.chr13.g00297.t01#1       7253   125253537:   mNeoVis.vulgivaga.2.1.chr13.g00298.t01#1    mNeoVis.vulgivaga.1.1.chr13.g00294.t01#4    mNeoVis.2.1.1.chr13_hap1.g00310.t01#5    mNeoVis.evergladensis.2.1.chr13.g00301.t01#2    mNeoVis.evergladensis.1.1.chr13.g00298.t01#3   mNeoVis.3.1.1.chr13.g00254.t01#6       7254   125253538:   mNeoVis.2.1.1.chr13_hap1.g00311.t01#5    mNeoVis.3.1.1.chr13.g00255.t01#6    mNeoVis.vulgivaga.2.1.chr13.g00299.t01#1    mNeoVis.evergladensis.1.1.chr13.g00299.t01#3    mNeoVis.evergladensis.2.1.chr13.g00302.t01#2   mNeoVis.vulgivaga.1.1.chr13.g00295.t01#4       7255   125253539:   mNeoVis.2.1.1.chr2_hap1.g01782.t01#5    mNeoVis.vulgivaga.2.1.chr5.g01726.t01#1   mNeoVis.evergladensis.2.1.chr5.g01722.t01#2         7256   125253540:   mNeoVis.vulgivaga.2.1.chr5.g01727.t01#1    mNeoVis.evergladensis.1.1.chr5.g01731.t01#3    mNeoVis.evergladensis.2.1.chr5.g01723.t01#2    mNeoVis.2.1.1.chr2_hap1.g01783.t01#5    mNeoVis.vulgivaga.1.1.chr5.g01703.t01#4   mNeoVis.3.1.1.chr5.g01707.t01#6       7257   125253541:   mNeoVis.2.1.1.chr5_hap1.g00825.t01#5    mNeoVis.3.1.1.chr13.g00263.t01#6    mNeoVis.evergladensis.2.1.chr2.g00826.t01#2    mNeoVis.vulgivaga.2.1.chr2.g00829.t01#1    mNeoVis.vulgivaga.2.1.chr13.g00307.t01#1    mNeoVis.vulgivaga.1.1.chr13.g00303.t01#4    mNeoVis.3.1.1.chr2.g00736.t01#6    mNeoVis.evergladensis.2.1.chr13.g00310.t01#2    mNeoVis.2.1.1.chr13_hap1.g00319.t01#5      7258   125253542:   mNeoVis.evergladensis.2.1.chr13.g00311.t01#2    mNeoVis.vulgivaga.2.1.chr13.g00309.t01#1    mNeoVis.2.1.1.chr13_hap1.g00320.t01#5    mNeoVis.evergladensis.1.1.chr13.g00310.t01#3    mNeoVis.3.1.1.chr13.g00264.t01#6    mNeoVis.vulgivaga.2.1.chr13.g00308.t01#1   mNeoVis.vulgivaga.1.1.chr13.g00304.t01#4     7259   125253543:   mNeoVis.evergladensis.1.1.chr5.g01732.t01#3    mNeoVis.2.1.1.chr2_hap1.g01784.t01#5    mNeoVis.3.1.1.chr5.g01708.t01#6    mNeoVis.vulgivaga.1.1.chr5.g01704.t01#4    mNeoVis.vulgivaga.2.1.chr5.g01728.t01#1   mNeoVis.evergladensis.2.1.chr5.g01724.t01#2       7260   125253544:   mNeoVis.evergladensis.1.1.chr5.g01733.t01#3    mNeoVis.2.1.1.chr2_hap1.g01785.t01#5    mNeoVis.vulgivaga.1.1.chr5.g01705.t01#4    mNeoVis.vulgivaga.2.1.chr5.g01729.t01#1    mNeoVis.3.1.1.chr5.g01709.t01#6   mNeoVis.evergladensis.2.1.chr5.g01725.t01#2       7261   125253545:   mNeoVis.evergladensis.2.1.chr13.g00312.t01#2    mNeoVis.vulgivaga.2.1.chr13.g00310.t01#1    mNeoVis.2.1.1.chr13_hap1.g00321.t01#5    mNeoVis.evergladensis.1.1.chr13.g00311.t01#3    mNeoVis.3.1.1.chr13.g00265.t01#6   mNeoVis.vulgivaga.1.1.chr13.g00305.t01#4       7262   125253546:   mNeoVis.3.1.1.chr5.g01710.t01#6    mNeoVis.evergladensis.2.1.chr5.g01726.t01#2    mNeoVis.vulgivaga.1.1.chr5.g01706.t01#4    mNeoVis.vulgivaga.2.1.chr5.g01730.t01#1    mNeoVis.2.1.1.chr2_hap1.g01786.t01#5   mNeoVis.evergladensis.1.1.chr5.g01734.t01#3       7263   125253547:   mNeoVis.vulgivaga.1.1.chr15.g00418.t01#4    mNeoVis.evergladensis.2.1.chr15.g00418.t01#2    mNeoVis.vulgivaga.2.1.chr15.g00425.t01#1    mNeoVis.2.1.1.chr15_hap1.g00596.t01#5    mNeoVis.3.1.1.chr15.g00378.t01#6   mNeoVis.evergladensis.1.1.chr15.g00198.t01#3       7264   125253548:   mNeoVis.3.1.1.chr5.g01711.t01#6    mNeoVis.vulgivaga.2.1.chr5.g01731.t01#1    mNeoVis.evergladensis.2.1.chr5.g01727.t01#2    mNeoVis.vulgivaga.1.1.chr5.g01707.t01#4    mNeoVis.2.1.1.chr2_hap1.g01787.t01#5   mNeoVis.evergladensis.1.1.chr5.g01735.t01#3       7265   125253549:   mNeoVis.evergladensis.1.1.chr3.g01079.t01#3    mNeoVis.vulgivaga.2.1.chr3.g01086.t01#1    mNeoVis.3.1.1.chr3.g01060.t01#6    mNeoVis.vulgivaga.1.1.chr3.g01073.t01#4    mNeoVis.2.1.1.chr3_hap1.g01069.t01#5   mNeoVis.evergladensis.2.1.chr3.g01069.t01#2       7266   125253550:   mNeoVis.2.1.1.chr2_hap1.g01788.t01#5    mNeoVis.evergladensis.1.1.chr5.g01736.t01#3    mNeoVis.vulgivaga.1.1.chr5.g01708.t01#4    mNeoVis.vulgivaga.2.1.chr5.g01732.t01#1    mNeoVis.evergladensis.2.1.chr5.g01728.t01#2   mNeoVis.3.1.1.chr5.g01712.t01#6       7267   125253551:   mNeoVis.vulgivaga.2.1.chr13.g00311.t01#1   mNeoVis.evergladensis.2.1.chr13.g00313.t01#2           7268   125253552:   mNeoVis.vulgivaga.2.1.chr13.g00312.t01#1    mNeoVis.vulgivaga.1.1.chr13.g00306.t01#4    mNeoVis.evergladensis.2.1.chr13.g00314.t01#2    mNeoVis.evergladensis.1.1.chr13.g00313.t01#3    mNeoVis.3.1.1.chr13.g00267.t01#6   mNeoVis.2.1.1.chr13_hap1.g00325.t01#5       7269   125253553:   mNeoVis.vulgivaga.1.1.chr5.g01709.t01#4    mNeoVis.3.1.1.chr5.g01713.t01#6    mNeoVis.vulgivaga.2.1.chr5.g01733.t01#1    mNeoVis.2.1.1.chr2_hap1.g01789.t01#5    mNeoVis.evergladensis.2.1.chr5.g01729.t01#2   mNeoVis.evergladensis.1.1.chr5.g01737.t01#3       7270   125253554:   mNeoVis.vulgivaga.2.1.chr5.g01734.t01#1    mNeoVis.2.1.1.chr2_hap1.g01790.t01#5    mNeoVis.evergladensis.1.1.chr5.g01738.t01#3    mNeoVis.vulgivaga.1.1.chr5.g01710.t01#4   mNeoVis.3.1.1.chr5.g01714.t01#6       7271   125253555:   mNeoVis.2.1.1.chr2_hap1.g01791.t01#5    mNeoVis.evergladensis.1.1.chr5.g01739.t01#3    mNeoVis.3.1.1.chr5.g01715.t01#6   mNeoVis.evergladensis.2.1.chr5.g01730.t01#2       7272   125253556:   mNeoVis.3.1.1.chr7.g00435.t01#6    mNeoVis.evergladensis.2.1.chr7.g00452.t01#2    mNeoVis.vulgivaga.2.1.chr7.g00450.t01#1    mNeoVis.vulgivaga.1.1.chr7.g00455.t01#4   mNeoVis.evergladensis.1.1.chr7.g00458.t01#3       7273   125253557:   mNeoVis.evergladensis.2.1.chr5.g01731.t01#2    mNeoVis.vulgivaga.1.1.chr5.g01711.t01#4    mNeoVis.evergladensis.1.1.chr5.g01740.t01#3    mNeoVis.vulgivaga.2.1.chr5.g01735.t01#1   mNeoVis.2.1.1.chr2_hap1.g01792.t01#5       7274   125253558:   mNeoVis.2.1.1.chr13_hap1.g00326.t01#5    mNeoVis.evergladensis.2.1.chr13.g00316.t01#2    mNeoVis.3.1.1.chr13.g00268.t01#6    mNeoVis.evergladensis.1.1.chr13.g00315.t01#3    mNeoVis.vulgivaga.2.1.chr13.g00314.t01#1    mNeoVis.3.1.1.chr13.g00269.t01#6   mNeoVis.vulgivaga.1.1.chr13.g00308.t01#4     7275   125253559:   mNeoVis.vulgivaga.1.1.chr13.g00307.t01#4    mNeoVis.vulgivaga.2.1.chr13.g00313.t01#1   mNeoVis.evergladensis.1.1.chr13.g00314.t01#3         7276   125253560:   mNeoVis.evergladensis.1.1.chr5.g01741.t01#3    mNeoVis.2.1.1.chr2_hap1.g01793.t01#5    mNeoVis.evergladensis.2.1.chr5.g01732.t01#2    mNeoVis.vulgivaga.2.1.chr5.g01736.t01#1    mNeoVis.vulgivaga.1.1.chr5.g01712.t01#4   mNeoVis.3.1.1.chr5.g01719.t01#6       7277   125253561:   mNeoVis.vulgivaga.1.1.chr5.g01713.t01#4    mNeoVis.evergladensis.1.1.chr5.g01742.t01#3    mNeoVis.vulgivaga.2.1.chr5.g01737.t01#1    mNeoVis.evergladensis.2.1.chr5.g01733.t01#2    mNeoVis.3.1.1.chr5.g01720.t01#6   mNeoVis.2.1.1.chr2_hap1.g01794.t01#5       7278   125253562:   mNeoVis.2.1.1.chr2_hap1.g01795.t01#5    mNeoVis.vulgivaga.1.1.chr5.g01714.t01#4    mNeoVis.3.1.1.chr5.g01721.t01#6    mNeoVis.vulgivaga.2.1.chr5.g01738.t01#1    mNeoVis.evergladensis.1.1.chr5.g01743.t01#3   mNeoVis.evergladensis.2.1.chr5.g01734.t01#2       7279   125253563:   mNeoVis.evergladensis.2.1.chr5.g01735.t01#2    mNeoVis.evergladensis.1.1.chr5.g01744.t01#3    mNeoVis.3.1.1.chr5.g01722.t01#6    mNeoVis.vulgivaga.1.1.chr5.g01715.t01#4    mNeoVis.2.1.1.chr2_hap1.g01796.t01#5   mNeoVis.vulgivaga.2.1.chr5.g01739.t01#1       7280   125253564:   mNeoVis.evergladensis.2.1.chr5.g01736.t01#2    mNeoVis.vulgivaga.1.1.chr5.g01716.t01#4    mNeoVis.vulgivaga.2.1.chr5.g01740.t01#1    mNeoVis.evergladensis.1.1.chr5.g01745.t01#3   mNeoVis.2.1.1.chr2_hap1.g01797.t01#5       7281   125253565:   mNeoVis.2.1.1.chr2_hap1.g01798.t01#5    mNeoVis.evergladensis.2.1.chr5.g01737.t01#2    mNeoVis.vulgivaga.1.1.chr5.g01717.t01#4    mNeoVis.vulgivaga.2.1.chr5.g01741.t01#1    mNeoVis.3.1.1.chr5.g01724.t01#6   mNeoVis.evergladensis.1.1.chr5.g01746.t01#3       7282   125253566:   mNeoVis.2.1.1.chr13_hap1.g00327.t01#5    mNeoVis.3.1.1.chr13.g00270.t01#6    mNeoVis.evergladensis.2.1.chr13.g00317.t01#2    mNeoVis.evergladensis.1.1.chr13.g00316.t01#3    mNeoVis.vulgivaga.2.1.chr13.g00315.t01#1   mNeoVis.vulgivaga.1.1.chr13.g00309.t01#4       7283   125253567:   mNeoVis.evergladensis.1.1.chr5.g01747.t01#3    mNeoVis.3.1.1.chr5.g01725.t01#6    mNeoVis.2.1.1.chr2_hap1.g01799.t01#5    mNeoVis.vulgivaga.1.1.chr5.g01718.t01#4    mNeoVis.vulgivaga.2.1.chr5.g01742.t01#1   mNeoVis.evergladensis.2.1.chr5.g01738.t01#2       7284   125253568:   mNeoVis.3.1.1.chr13.g00271.t01#6    mNeoVis.evergladensis.2.1.chr13.g00318.t01#2    mNeoVis.vulgivaga.2.1.chr13.g00316.t01#1    mNeoVis.vulgivaga.1.1.chr13.g00310.t01#4    mNeoVis.evergladensis.1.1.chr13.g00317.t01#3   mNeoVis.2.1.1.chr13_hap1.g00328.t01#5       7285   125253569:   mNeoVis.3.1.1.chr13.g00272.t01#6    mNeoVis.evergladensis.1.1.chr13.g00318.t01#3    mNeoVis.vulgivaga.1.1.chr13.g00311.t01#4    mNeoVis.evergladensis.2.1.chr13.g00319.t01#2    mNeoVis.vulgivaga.2.1.chr13.g00317.t01#1   mNeoVis.2.1.1.chr13_hap1.g00329.t01#5       7286   125253570:   mNeoVis.vulgivaga.2.1.chr7.g00451.t01#1   mNeoVis.3.1.1.chr7.g00436.t01#6             7287   125253571:   mNeoVis.vulgivaga.2.1.chr13.g00318.t01#1    mNeoVis.2.1.1.chr13_hap1.g00330.t01#5    mNeoVis.vulgivaga.1.1.chr13.g00312.t01#4    mNeoVis.evergladensis.2.1.chr13.g00320.t01#2    mNeoVis.evergladensis.1.1.chr13.g00319.t01#3   mNeoVis.3.1.1.chr13.g00273.t01#6       7288   125253572:   mNeoVis.vulgivaga.1.1.chr13.g00313.t01#4    mNeoVis.vulgivaga.2.1.chr13.g00319.t01#1    mNeoVis.3.1.1.chr13.g00274.t01#6    mNeoVis.2.1.1.chr13_hap1.g00331.t01#5    mNeoVis.evergladensis.1.1.chr13.g00320.t01#3   mNeoVis.evergladensis.2.1.chr13.g00321.t01#2       7289   125253573:   mNeoVis.vulgivaga.1.1.chr15.g00419.t01#4    mNeoVis.evergladensis.1.1.chr15.g00196.t01#3    mNeoVis.evergladensis.2.1.chr15.g00419.t01#2    mNeoVis.vulgivaga.2.1.chr15.g00426.t01#1   mNeoVis.2.1.1.chr15_hap1.g00597.t01#5       7290   125253574:   mNeoVis.evergladensis.1.1.chr13.g00321.t01#3    mNeoVis.vulgivaga.2.1.chr13.g00320.t01#1    mNeoVis.evergladensis.2.1.chr13.g00322.t01#2    mNeoVis.vulgivaga.1.1.chr13.g00314.t01#4    mNeoVis.3.1.1.chr13.g00275.t01#6   mNeoVis.2.1.1.chr13_hap1.g00332.t01#5       7291   125253575:   mNeoVis.evergladensis.2.1.chr13.g00323.t01#2    mNeoVis.3.1.1.chr13.g00276.t01#6    mNeoVis.vulgivaga.2.1.chr13.g00321.t01#1    mNeoVis.2.1.1.chr13_hap1.g00333.t01#5    mNeoVis.evergladensis.1.1.chr13.g00322.t01#3   mNeoVis.vulgivaga.1.1.chr13.g00315.t01#4       7292   125253576:   mNeoVis.vulgivaga.2.1.chr5.g01743.t01#1    mNeoVis.evergladensis.2.1.chr5.g01739.t01#2    mNeoVis.3.1.1.chr5.g01726.t01#6    mNeoVis.vulgivaga.1.1.chr5.g01719.t01#4    mNeoVis.evergladensis.1.1.chr5.g01748.t01#3   mNeoVis.2.1.1.chr2_hap1.g01800.t01#5       7293   125253577:   mNeoVis.vulgivaga.2.1.chr13.g00322.t01#1    mNeoVis.evergladensis.1.1.chr13.g00323.t01#3    mNeoVis.evergladensis.2.1.chr13.g00324.t01#2    mNeoVis.3.1.1.chr13.g00277.t01#6    mNeoVis.2.1.1.chr13_hap1.g00334.t01#5   mNeoVis.vulgivaga.1.1.chr13.g00316.t01#4       7294   125253578:   mNeoVis.vulgivaga.2.1.chr5.g01744.t01#1    mNeoVis.2.1.1.chr2_hap1.g01801.t01#5    mNeoVis.3.1.1.chr5.g01727.t01#6    mNeoVis.evergladensis.2.1.chr5.g01740.t01#2    mNeoVis.vulgivaga.1.1.chr5.g01720.t01#4   mNeoVis.evergladensis.1.1.chr5.g01749.t01#3       7295   125253579:   mNeoVis.2.1.1.chr7_hap1.g00447.t01#5    mNeoVis.evergladensis.2.1.chr7.g00453.t01#2    mNeoVis.vulgivaga.1.1.chr7.g00456.t01#4    mNeoVis.vulgivaga.2.1.chr7.g00452.t01#1   mNeoVis.evergladensis.1.1.chr7.g00459.t01#3       7296   125253580:   mNeoVis.vulgivaga.1.1.chr13.g00317.t01#4    mNeoVis.evergladensis.2.1.chr13.g00325.t01#2    mNeoVis.2.1.1.chr13_hap1.g00335.t01#5    mNeoVis.vulgivaga.2.1.chr13.g00323.t01#1    mNeoVis.3.1.1.chr13.g00278.t01#6   mNeoVis.evergladensis.1.1.chr13.g00324.t01#3       7297   125253581:   mNeoVis.3.1.1.chr13.g00279.t01#6    mNeoVis.2.1.1.chr13_hap1.g00336.t01#5    mNeoVis.evergladensis.1.1.chr13.g00325.t01#3    mNeoVis.vulgivaga.1.1.chr13.g00318.t01#4    mNeoVis.evergladensis.2.1.chr13.g00326.t01#2   mNeoVis.vulgivaga.2.1.chr13.g00324.t01#1       7298   125253582:   mNeoVis.2.1.1.chr2_hap1.g01802.t01#5    mNeoVis.vulgivaga.1.1.chr5.g01721.t01#4    mNeoVis.evergladensis.2.1.chr5.g01741.t01#2    mNeoVis.vulgivaga.2.1.chr5.g01745.t01#1   mNeoVis.evergladensis.1.1.chr5.g01750.t01#3       7299   125253583:   mNeoVis.vulgivaga.2.1.chr13.g00325.t01#1    mNeoVis.3.1.1.chr13.g00280.t01#6    mNeoVis.2.1.1.chr13_hap1.g00337.t01#5    mNeoVis.vulgivaga.1.1.chr13.g00319.t01#4    mNeoVis.evergladensis.2.1.chr13.g00327.t01#2   mNeoVis.evergladensis.1.1.chr13.g00326.t01#3       7300   125253584:   mNeoVis.evergladensis.2.1.chr13.g00328.t01#2    mNeoVis.vulgivaga.2.1.chr13.g00326.t01#1    mNeoVis.2.1.1.chr13_hap1.g00338.t01#5    mNeoVis.evergladensis.1.1.chr13.g00327.t01#3    mNeoVis.3.1.1.chr13.g00281.t01#6   mNeoVis.vulgivaga.1.1.chr13.g00320.t01#4       7301   125253585:   mNeoVis.3.1.1.chr5.g01728.t01#6    mNeoVis.evergladensis.1.1.chr5.g01751.t01#3    mNeoVis.vulgivaga.2.1.chr5.g01746.t01#1    mNeoVis.2.1.1.chr2_hap1.g01803.t01#5    mNeoVis.evergladensis.2.1.chr5.g01742.t01#2   mNeoVis.vulgivaga.1.1.chr5.g01722.t01#4       7302   125253586:   mNeoVis.2.1.1.chr13_hap1.g00339.t01#5    mNeoVis.vulgivaga.1.1.chr13.g00321.t01#4    mNeoVis.evergladensis.1.1.chr13.g00328.t01#3    mNeoVis.vulgivaga.2.1.chr13.g00327.t01#1    mNeoVis.3.1.1.chr13.g00282.t01#6   mNeoVis.evergladensis.2.1.chr13.g00329.t01#2       7303   125253587:   mNeoVis.vulgivaga.1.1.chr5.g01723.t01#4    mNeoVis.3.1.1.chr5.g01729.t01#6    mNeoVis.evergladensis.2.1.chr5.g01743.t01#2    mNeoVis.vulgivaga.2.1.chr5.g01747.t01#1    mNeoVis.evergladensis.1.1.chr5.g01752.t01#3   mNeoVis.2.1.1.chr2_hap1.g01804.t01#5       7304   125253588:   mNeoVis.3.1.1.chr13.g00283.t01#6    mNeoVis.vulgivaga.1.1.chr13.g00322.t01#4    mNeoVis.2.1.1.chr13_hap1.g00340.t01#5    mNeoVis.evergladensis.1.1.chr13.g00329.t01#3    mNeoVis.vulgivaga.2.1.chr13.g00328.t01#1   mNeoVis.evergladensis.2.1.chr13.g00330.t01#2       7305   125253589:   mNeoVis.3.1.1.chr5.g01730.t01#6    mNeoVis.evergladensis.1.1.chr5.g01753.t01#3    mNeoVis.vulgivaga.2.1.chr5.g01748.t01#1    mNeoVis.2.1.1.chr2_hap1.g01805.t01#5    mNeoVis.vulgivaga.1.1.chr5.g01724.t01#4   mNeoVis.evergladensis.2.1.chr5.g01744.t01#2       7306   125253590:   mNeoVis.evergladensis.2.1.chr13.g00331.t01#2    mNeoVis.vulgivaga.2.1.chr13.g00329.t01#1    mNeoVis.vulgivaga.1.1.chr13.g00323.t01#4   mNeoVis.3.1.1.chr13.g00284.t01#6         7307   125253591:   mNeoVis.2.1.1.chr13_hap1.g00341.t01#5   mNeoVis.evergladensis.1.1.chr13.g00330.t01#3           7308   125253592:   mNeoVis.vulgivaga.2.1.chr5.g01749.t01#1    mNeoVis.vulgivaga.1.1.chr5.g01725.t01#4    mNeoVis.evergladensis.1.1.chr5.g01754.t01#3    mNeoVis.evergladensis.2.1.chr5.g01745.t01#2    mNeoVis.2.1.1.chr2_hap1.g01806.t01#5   mNeoVis.3.1.1.chr5.g01731.t01#6       7309   125253593:   mNeoVis.vulgivaga.1.1.chr13.g00324.t01#4    mNeoVis.2.1.1.chr13_hap1.g00344.t01#5    mNeoVis.vulgivaga.2.1.chr13.g00330.t01#1    mNeoVis.evergladensis.1.1.chr13.g00332.t01#3    mNeoVis.3.1.1.chr13.g00286.t01#6   mNeoVis.evergladensis.2.1.chr13.g00333.t01#2       7310   125253594:   mNeoVis.vulgivaga.2.1.chr5.g01750.t01#1    mNeoVis.evergladensis.2.1.chr5.g01746.t01#2    mNeoVis.evergladensis.1.1.chr5.g01755.t01#3    mNeoVis.3.1.1.chr5.g01732.t01#6    mNeoVis.2.1.1.chr2_hap1.g01807.t01#5   mNeoVis.vulgivaga.1.1.chr5.g01726.t01#4       7311   125253595:   mNeoVis.vulgivaga.1.1.chr5.g01727.t01#4    mNeoVis.vulgivaga.2.1.chr5.g01751.t01#1    mNeoVis.evergladensis.2.1.chr5.g01747.t01#2    mNeoVis.evergladensis.1.1.chr5.g01756.t01#3    mNeoVis.2.1.1.chr2_hap1.g01808.t01#5   mNeoVis.3.1.1.chr5.g01733.t01#6       7312   125253596:   mNeoVis.evergladensis.2.1.chr13.g00334.t01#2    mNeoVis.2.1.1.chr13_hap1.g00345.t01#5    mNeoVis.3.1.1.chr13.g00287.t01#6    mNeoVis.vulgivaga.1.1.chr13.g00325.t01#4    mNeoVis.evergladensis.1.1.chr13.g00333.t01#3   mNeoVis.vulgivaga.2.1.chr13.g00332.t01#1       7313   125253597:   mNeoVis.vulgivaga.2.1.chr13.g00333.t01#1    mNeoVis.vulgivaga.1.1.chr13.g00326.t01#4    mNeoVis.evergladensis.1.1.chr13.g00334.t01#3    mNeoVis.3.1.1.chr13.g00288.t01#6    mNeoVis.2.1.1.chr13_hap1.g00346.t01#5   mNeoVis.evergladensis.2.1.chr13.g00335.t01#2       7314   125253598:   mNeoVis.evergladensis.2.1.chr13.g00336.t01#2    mNeoVis.3.1.1.chr13.g00289.t01#6    mNeoVis.2.1.1.chr13_hap1.g00347.t01#5    mNeoVis.evergladensis.1.1.chr13.g00335.t01#3   mNeoVis.vulgivaga.2.1.chr13.g00334.t01#1       7315   125253599:   mNeoVis.vulgivaga.2.1.chr5.g01752.t01#1    mNeoVis.vulgivaga.1.1.chr5.g01728.t01#4    mNeoVis.evergladensis.2.1.chr5.g01748.t01#2    mNeoVis.evergladensis.1.1.chr5.g01757.t01#3    mNeoVis.2.1.1.chr2_hap1.g01809.t01#5   mNeoVis.3.1.1.chr5.g01734.t01#6       7316   125253600:   mNeoVis.vulgivaga.1.1.chr13.g00233.t01#4    mNeoVis.evergladensis.1.1.chr13.g00238.t01#3    mNeoVis.2.1.1.chr13_hap1.g00241.t01#5    mNeoVis.vulgivaga.2.1.chr13.g00236.t01#1   mNeoVis.evergladensis.2.1.chr13.g00239.t01#2       7317   125253601:   mNeoVis.evergladensis.1.1.chr13.g00239.t01#3    mNeoVis.vulgivaga.1.1.chr13.g00234.t01#4    mNeoVis.evergladensis.2.1.chr13.g00240.t01#2    mNeoVis.2.1.1.chr13_hap1.g00242.t01#5   mNeoVis.vulgivaga.2.1.chr13.g00237.t01#1       7318   125253602:   mNeoVis.vulgivaga.2.1.chr5.g01753.t01#1    mNeoVis.3.1.1.chr5.g01736.t01#6    mNeoVis.evergladensis.2.1.chr5.g01749.t01#2    mNeoVis.vulgivaga.1.1.chr5.g01729.t01#4    mNeoVis.2.1.1.chr2_hap1.g01810.t01#5   mNeoVis.evergladensis.1.1.chr5.g01758.t01#3       7319   125253603:   mNeoVis.evergladensis.2.1.chr5.g01750.t01#2    mNeoVis.2.1.1.chr2_hap1.g01811.t01#5    mNeoVis.evergladensis.1.1.chr5.g01759.t01#3    mNeoVis.vulgivaga.2.1.chr5.g01754.t01#1   mNeoVis.vulgivaga.1.1.chr5.g01730.t01#4       7320   125253604:   mNeoVis.2.1.1.chr2_hap1.g01812.t01#5    mNeoVis.evergladensis.1.1.chr5.g01760.t01#3    mNeoVis.evergladensis.2.1.chr5.g01751.t01#2    mNeoVis.vulgivaga.2.1.chr5.g01755.t01#1    mNeoVis.3.1.1.chr5.g01737.t01#6   mNeoVis.vulgivaga.1.1.chr5.g01731.t01#4       7321   125253605:   mNeoVis.3.1.1.chr5.g01739.t01#6    mNeoVis.evergladensis.1.1.chr5.g01762.t01#3    mNeoVis.vulgivaga.1.1.chr5.g01733.t01#4    mNeoVis.2.1.1.chr2_hap1.g01814.t01#5    mNeoVis.evergladensis.2.1.chr5.g01753.t01#2   mNeoVis.vulgivaga.2.1.chr5.g01756.t01#1       7322   125253606:   mNeoVis.2.1.1.chr13_hap1.g00243.t01#5    mNeoVis.3.1.1.chr13.g00193.t01#6    mNeoVis.vulgivaga.1.1.chr13.g00235.t01#4    mNeoVis.evergladensis.1.1.chr13.g00240.t01#3    mNeoVis.vulgivaga.2.1.chr13.g00238.t01#1   mNeoVis.evergladensis.2.1.chr13.g00241.t01#2       7323   125253607:   mNeoVis.2.1.1.chr2_hap1.g01813.t01#5    mNeoVis.3.1.1.chr5.g01738.t01#6    mNeoVis.evergladensis.1.1.chr5.g01761.t01#3    mNeoVis.vulgivaga.2.1.chr5.g01757.t01#1    mNeoVis.evergladensis.2.1.chr5.g01752.t01#2   mNeoVis.vulgivaga.1.1.chr5.g01732.t01#4       7324   125253608:   mNeoVis.evergladensis.1.1.chr13.g00242.t01#3    mNeoVis.evergladensis.2.1.chr13.g00243.t01#2    mNeoVis.2.1.1.chr13_hap1.g00245.t01#5    mNeoVis.vulgivaga.2.1.chr13.g00240.t01#1    mNeoVis.3.1.1.chr13.g00195.t01#6   mNeoVis.vulgivaga.1.1.chr13.g00237.t01#4       7325   125253609:   mNeoVis.evergladensis.2.1.chr5.g01754.t01#2    mNeoVis.evergladensis.1.1.chr5.g01763.t01#3    mNeoVis.vulgivaga.1.1.chr5.g01734.t01#4    mNeoVis.3.1.1.chr5.g01740.t01#6   mNeoVis.vulgivaga.2.1.chr5.g01758.t01#1       7326   125253610:   mNeoVis.evergladensis.2.1.chr5.g01755.t01#2    mNeoVis.evergladensis.1.1.chr5.g01764.t01#3    mNeoVis.vulgivaga.2.1.chr5.g01759.t01#1    mNeoVis.vulgivaga.1.1.chr5.g01735.t01#4    mNeoVis.2.1.1.chr2_hap1.g01816.t01#5   mNeoVis.3.1.1.chr5.g01741.t01#6       7327   125253611:   mNeoVis.evergladensis.2.1.chr13.g00244.t01#2    mNeoVis.vulgivaga.2.1.chr13.g00241.t01#1    mNeoVis.vulgivaga.1.1.chr13.g00238.t01#4    mNeoVis.2.1.1.chr13_hap1.g00246.t01#5    mNeoVis.evergladensis.1.1.chr13.g00243.t01#3   mNeoVis.3.1.1.chr13.g00196.t01#6       7328   125253612:   mNeoVis.evergladensis.2.1.chr8.g00090.t01#2    mNeoVis.3.1.1.chr9.g00069.t01#6    mNeoVis.vulgivaga.2.1.chr9.g00090.t01#1    mNeoVis.vulgivaga.1.1.chr9.g00086.t01#4    mNeoVis.3.1.1.chr9.g00068.t01#6    mNeoVis.evergladensis.1.1.chr9.g00091.t01#3   mNeoVis.2.1.1.chr10_hap1.g00095.t01#5     7329   125253613:   mNeoVis.vulgivaga.1.1.chr15.g00420.t01#4    mNeoVis.3.1.1.chr15.g00381.t01#6    mNeoVis.evergladensis.1.1.chr15.g00197.t01#3    mNeoVis.vulgivaga.2.1.chr15.g00427.t01#1    mNeoVis.evergladensis.2.1.chr15.g00420.t01#2   mNeoVis.2.1.1.chr15_hap1.g00598.t01#5       7330   125253614:   mNeoVis.2.1.1.chr2_hap1.g01817.t01#5    mNeoVis.vulgivaga.2.1.chr5.g01760.t01#1    mNeoVis.evergladensis.1.1.chr5.g01765.t01#3    mNeoVis.3.1.1.chr5.g01742.t01#6    mNeoVis.evergladensis.2.1.chr5.g01756.t01#2   mNeoVis.vulgivaga.1.1.chr5.g01736.t01#4       7331   125253615:   mNeoVis.evergladensis.2.1.chr13.g00245.t01#2    mNeoVis.evergladensis.1.1.chr13.g00244.t01#3    mNeoVis.2.1.1.chr13_hap1.g00247.t01#5    mNeoVis.vulgivaga.2.1.chr13.g00242.t01#1    mNeoVis.vulgivaga.1.1.chr13.g00239.t01#4   mNeoVis.3.1.1.chr13.g00197.t01#6       7332   125253616:   mNeoVis.2.1.1.chr13_hap1.g00248.t01#5    mNeoVis.evergladensis.1.1.chr13.g00245.t01#3    mNeoVis.vulgivaga.1.1.chr13.g00240.t01#4    mNeoVis.3.1.1.chr13.g00198.t01#6    mNeoVis.vulgivaga.2.1.chr13.g00243.t01#1   mNeoVis.evergladensis.2.1.chr13.g00246.t01#2       7333   125253617:   mNeoVis.3.1.1.chr15.g00382.t01#6   mNeoVis.vulgivaga.2.1.chr15.g00428.t01#1           7334   125253618:   mNeoVis.vulgivaga.2.1.chr13.g00244.t01#1    mNeoVis.2.1.1.chr13_hap1.g00250.t01#5    mNeoVis.evergladensis.2.1.chr13.g00248.t01#2    mNeoVis.evergladensis.1.1.chr13.g00247.t01#3    mNeoVis.vulgivaga.1.1.chr13.g00241.t01#4   mNeoVis.3.1.1.chr13.g00200.t01#6       7335   125253619:   mNeoVis.vulgivaga.2.1.chr13.g00245.t01#1    mNeoVis.evergladensis.2.1.chr13.g00249.t01#2    mNeoVis.3.1.1.chr13.g00201.t01#6    mNeoVis.vulgivaga.1.1.chr13.g00242.t01#4    mNeoVis.evergladensis.1.1.chr13.g00248.t01#3   mNeoVis.2.1.1.chr13_hap1.g00251.t01#5       7336   125253620:   mNeoVis.evergladensis.1.1.chr13.g00249.t01#3    mNeoVis.3.1.1.chr13.g00202.t01#6    mNeoVis.evergladensis.2.1.chr13.g00250.t01#2    mNeoVis.2.1.1.chr13_hap1.g00252.t01#5    mNeoVis.vulgivaga.1.1.chr13.g00244.t01#4   mNeoVis.vulgivaga.2.1.chr13.g00246.t01#1       7337   125253621:   mNeoVis.vulgivaga.1.1.chr13.g00245.t01#4    mNeoVis.evergladensis.2.1.chr13.g00251.t01#2    mNeoVis.vulgivaga.2.1.chr13.g00247.t01#1    mNeoVis.2.1.1.chr13_hap1.g00253.t01#5    mNeoVis.evergladensis.1.1.chr13.g00250.t01#3   mNeoVis.3.1.1.chr13.g00203.t01#6       7338   125253622:   mNeoVis.2.1.1.chr13_hap1.g00254.t01#5    mNeoVis.vulgivaga.2.1.chr13.g00248.t01#1    mNeoVis.evergladensis.2.1.chr13.g00252.t01#2    mNeoVis.3.1.1.chr13.g00204.t01#6    mNeoVis.vulgivaga.1.1.chr13.g00246.t01#4   mNeoVis.evergladensis.1.1.chr13.g00251.t01#3       7339   125253623:   mNeoVis.vulgivaga.2.1.chr5.g01688.t01#1    mNeoVis.3.1.1.chr5.g01671.t01#6    mNeoVis.evergladensis.1.1.chr5.g01693.t01#3    mNeoVis.2.1.1.chr2_hap1.g01747.t01#5    mNeoVis.evergladensis.2.1.chr5.g01687.t01#2   mNeoVis.vulgivaga.1.1.chr5.g01667.t01#4       7340   125253624:   mNeoVis.vulgivaga.1.1.chr13.g00247.t01#4    mNeoVis.3.1.1.chr13.g00205.t01#6    mNeoVis.evergladensis.1.1.chr13.g00252.t01#3    mNeoVis.vulgivaga.2.1.chr13.g00249.t01#1    mNeoVis.evergladensis.2.1.chr13.g00253.t01#2   mNeoVis.2.1.1.chr13_hap1.g00255.t01#5       7341   125253625:   mNeoVis.3.1.1.chr13.g00211.t01#6    mNeoVis.2.1.1.chr13_hap1.g00266.t01#5    mNeoVis.2.1.1.chr13_hap1.g00265.t01#5    mNeoVis.evergladensis.2.1.chr13.g00258.t01#2    mNeoVis.2.1.1.chr13_hap1.g00267.t01#5    mNeoVis.vulgivaga.1.1.chr13.g00253.t01#4    mNeoVis.evergladensis.1.1.chr13.g00258.t01#3   mNeoVis.vulgivaga.2.1.chr13.g00255.t01#1     7342   125253626:   mNeoVis.evergladensis.1.1.chr13.g00254.t01#3    mNeoVis.vulgivaga.2.1.chr13.g00250.t01#1    mNeoVis.evergladensis.2.1.chr13.g00254.t01#2    mNeoVis.3.1.1.chr13.g00206.t01#6    mNeoVis.vulgivaga.1.1.chr13.g00248.t01#4   mNeoVis.2.1.1.chr13_hap1.g00256.t01#5       7343   125253627:   mNeoVis.2.1.1.chr13_hap1.g00262.t01#5    mNeoVis.vulgivaga.1.1.chr13.g00251.t01#4    mNeoVis.evergladensis.1.1.chr13.g00257.t01#3    mNeoVis.vulgivaga.2.1.chr13.g00252.t01#1    mNeoVis.evergladensis.1.1.chr13.g00255.t01#3   mNeoVis.3.1.1.chr13.g00208.t01#6       7344   125253628:   mNeoVis.vulgivaga.2.1.chr13.g00254.t01#1    mNeoVis.3.1.1.chr13.g00210.t01#6    mNeoVis.evergladensis.2.1.chr13.g00259.t01#2    mNeoVis.2.1.1.chr13_hap1.g00264.t01#5   mNeoVis.vulgivaga.1.1.chr13.g00252.t01#4       7345   125253629:   mNeoVis.2.1.1.chr13_hap1.g00261.t01#5    mNeoVis.evergladensis.2.1.chr13.g00257.t01#2    mNeoVis.2.1.1.chr13_hap1.g00260.t01#5   mNeoVis.2.1.1.chr13_hap1.g00259.t01#5         7346   125253630:   mNeoVis.evergladensis.1.1.chr13.g00256.t01#3    mNeoVis.3.1.1.chr13.g00209.t01#6    mNeoVis.2.1.1.chr13_hap1.g00263.t01#5   mNeoVis.vulgivaga.2.1.chr13.g00253.t01#1       7347   125253631:   mNeoVis.evergladensis.2.1.chr13.g00256.t01#2    mNeoVis.2.1.1.chr13_hap1.g00258.t01#5   mNeoVis.vulgivaga.1.1.chr13.g00250.t01#4         7348   125253632:   mNeoVis.evergladensis.1.1.chr5.g01694.t01#3    mNeoVis.3.1.1.chr5.g01672.t01#6    mNeoVis.vulgivaga.1.1.chr5.g01668.t01#4    mNeoVis.vulgivaga.2.1.chr5.g01689.t01#1    mNeoVis.2.1.1.chr2_hap1.g01748.t01#5   mNeoVis.evergladensis.2.1.chr5.g01688.t01#2       7349   125253633:   mNeoVis.3.1.1.chr13.g00207.t01#6    mNeoVis.vulgivaga.1.1.chr13.g00249.t01#4    mNeoVis.2.1.1.chr13_hap1.g00257.t01#5    mNeoVis.vulgivaga.2.1.chr13.g00251.t01#1    mNeoVis.evergladensis.2.1.chr13.g00255.t01#2   mNeoVis.evergladensis.1.1.chr13.g00253.t01#3       7350   125253634:   mNeoVis.vulgivaga.1.1.chr13.g00254.t01#4    mNeoVis.evergladensis.1.1.chr13.g00259.t01#3    mNeoVis.vulgivaga.2.1.chr13.g00256.t01#1    mNeoVis.evergladensis.2.1.chr13.g00260.t01#2    mNeoVis.2.1.1.chr13_hap1.g00268.t01#5   mNeoVis.3.1.1.chr13.g00212.t01#6       7351   125253635:   mNeoVis.vulgivaga.2.1.chr13.g00257.t01#1    mNeoVis.2.1.1.chr13_hap1.g00269.t01#5    mNeoVis.3.1.1.chr13.g00213.t01#6    mNeoVis.vulgivaga.1.1.chr13.g00255.t01#4    mNeoVis.evergladensis.1.1.chr13.g00260.t01#3   mNeoVis.evergladensis.2.1.chr13.g00261.t01#2       7352   125253636:   mNeoVis.2.1.1.chr13_hap1.g00270.t01#5    mNeoVis.3.1.1.chr13.g00214.t01#6    mNeoVis.vulgivaga.2.1.chr13.g00258.t01#1    mNeoVis.vulgivaga.1.1.chr13.g00256.t01#4    mNeoVis.evergladensis.1.1.chr13.g00261.t01#3   mNeoVis.evergladensis.2.1.chr13.g00262.t01#2       7353   125253637:   mNeoVis.evergladensis.1.1.chr5.g01695.t01#3    mNeoVis.evergladensis.2.1.chr5.g01689.t01#2    mNeoVis.vulgivaga.1.1.chr5.g01669.t01#4    mNeoVis.3.1.1.chr5.g01673.t01#6    mNeoVis.2.1.1.chr2_hap1.g01749.t01#5   mNeoVis.vulgivaga.2.1.chr5.g01690.t01#1       7354   125253638:   mNeoVis.3.1.1.chr13.g00216.t01#6    mNeoVis.2.1.1.chr13_hap1.g00273.t01#5    mNeoVis.evergladensis.2.1.chr13.g00264.t01#2    mNeoVis.vulgivaga.1.1.chr13.g00258.t01#4   mNeoVis.vulgivaga.2.1.chr13.g00261.t01#1       7355   125253639:   mNeoVis.2.1.1.chr13_hap1.g00272.t01#5    mNeoVis.vulgivaga.2.1.chr13.g00260.t01#1    mNeoVis.3.1.1.chr13.g00215.t01#6   mNeoVis.vulgivaga.1.1.chr13.g00257.t01#4       7356   125253640:   mNeoVis.evergladensis.2.1.chr13.g00263.t01#2    mNeoVis.vulgivaga.2.1.chr13.g00259.t01#1   mNeoVis.2.1.1.chr13_hap1.g00271.t01#5           7357   125253641:  mNeoVis.evergladensis.1.1.chr13.g00262.t01#3             7358   125253642:   mNeoVis.vulgivaga.2.1.chr15.g00066.t01#1    mNeoVis.vulgivaga.1.1.chr15.g00065.t01#4    mNeoVis.evergladensis.2.1.chr15.g00064.t01#2    mNeoVis.evergladensis.1.1.chr15.g00549.t01#3   mNeoVis.2.1.1.chr15_hap1.g00064.t01#5       7359   125253643:   mNeoVis.vulgivaga.2.1.chr6.g01087.t01#1    mNeoVis.evergladensis.2.1.chr6.g01095.t01#2    mNeoVis.evergladensis.2.1.chr6.g01093.t01#2    mNeoVis.2.1.1.chr4_hap1.g01096.t01#5    mNeoVis.3.1.1.chr6.g01085.t01#6    mNeoVis.vulgivaga.1.1.chr6.g01087.t01#4   mNeoVis.evergladensis.1.1.chr6.g01098.t01#3     7360   125253644:   mNeoVis.2.1.1.chr13_hap1.g00274.t01#5    mNeoVis.3.1.1.chr13.g00217.t01#6    mNeoVis.vulgivaga.2.1.chr13.g00262.t01#1    mNeoVis.evergladensis.2.1.chr13.g00265.t01#2    mNeoVis.vulgivaga.2.1.chr13.g00263.t01#1    mNeoVis.evergladensis.1.1.chr13.g00263.t01#3    mNeoVis.vulgivaga.1.1.chr13.g00259.t01#4    mNeoVis.2.1.1.chr13_hap1.g00275.t01#5    mNeoVis.3.1.1.chr13.g00218.t01#6      7361   125253645:   mNeoVis.vulgivaga.2.1.chr13.g00266.t01#1    mNeoVis.2.1.1.chr13_hap1.g00279.t01#5    mNeoVis.vulgivaga.1.1.chr13.g00264.t01#4    mNeoVis.evergladensis.1.1.chr13.g00268.t01#3    mNeoVis.evergladensis.2.1.chr13.g00269.t01#2   mNeoVis.3.1.1.chr13.g00221.t01#6       7362   125253646:   mNeoVis.vulgivaga.1.1.chr13.g00263.t01#4    mNeoVis.3.1.1.chr13.g00220.t01#6    mNeoVis.evergladensis.2.1.chr13.g00268.t01#2    mNeoVis.evergladensis.1.1.chr13.g00267.t01#3    mNeoVis.vulgivaga.2.1.chr13.g00265.t01#1   mNeoVis.2.1.1.chr13_hap1.g00278.t01#5       7363   125253647:   mNeoVis.evergladensis.2.1.chr13.g00270.t01#2    mNeoVis.2.1.1.chr13_hap1.g00280.t01#5    mNeoVis.3.1.1.chr13.g00222.t01#6    mNeoVis.evergladensis.1.1.chr13.g00269.t01#3    mNeoVis.vulgivaga.2.1.chr13.g00267.t01#1   mNeoVis.vulgivaga.1.1.chr13.g00265.t01#4       7364   125253648:   mNeoVis.evergladensis.2.1.chr13.g00266.t01#2    mNeoVis.evergladensis.1.1.chr13.g00265.t01#3    mNeoVis.vulgivaga.1.1.chr13.g00261.t01#4    mNeoVis.2.1.1.chr13_hap1.g00276.t01#5   mNeoVis.3.1.1.chr13.g00219.t01#6       7365   125253649:   mNeoVis.2.1.1.chr13_hap1.g00277.t01#5    mNeoVis.evergladensis.2.1.chr13.g00267.t01#2    mNeoVis.evergladensis.1.1.chr13.g00266.t01#3    mNeoVis.vulgivaga.1.1.chr13.g00262.t01#4   mNeoVis.vulgivaga.2.1.chr13.g00264.t01#1       7366   125253650:   mNeoVis.vulgivaga.2.1.chr5.g01691.t01#1    mNeoVis.evergladensis.1.1.chr5.g01696.t01#3    mNeoVis.3.1.1.chr5.g01674.t01#6    mNeoVis.evergladensis.2.1.chr5.g01690.t01#2    mNeoVis.2.1.1.chr2_hap1.g01750.t01#5   mNeoVis.vulgivaga.1.1.chr5.g01670.t01#4       7367   125253651:   mNeoVis.3.1.1.chr5.g01675.t01#6    mNeoVis.evergladensis.2.1.chr5.g01691.t01#2    mNeoVis.2.1.1.chr2_hap1.g01751.t01#5    mNeoVis.vulgivaga.2.1.chr5.g01692.t01#1    mNeoVis.evergladensis.1.1.chr5.g01698.t01#3   mNeoVis.vulgivaga.1.1.chr5.g01671.t01#4       7368   125253652:   mNeoVis.evergladensis.1.1.chr5.g01699.t01#3    mNeoVis.3.1.1.chr5.g01676.t01#6    mNeoVis.evergladensis.2.1.chr5.g01692.t01#2    mNeoVis.vulgivaga.1.1.chr5.g01672.t01#4    mNeoVis.2.1.1.chr2_hap1.g01752.t01#5   mNeoVis.vulgivaga.2.1.chr5.g01693.t01#1       7369   125253653:   mNeoVis.vulgivaga.2.1.chr13.g00268.t01#1    mNeoVis.vulgivaga.1.1.chr13.g00266.t01#4    mNeoVis.evergladensis.1.1.chr13.g00270.t01#3    mNeoVis.3.1.1.chr13.g00223.t01#6    mNeoVis.evergladensis.2.1.chr13.g00271.t01#2   mNeoVis.2.1.1.chr13_hap1.g00281.t01#5       7370   125253654:   mNeoVis.evergladensis.1.1.chr13.g00271.t01#3    mNeoVis.evergladensis.2.1.chr13.g00273.t01#2    mNeoVis.evergladensis.2.1.chr13.g00272.t01#2    mNeoVis.vulgivaga.2.1.chr13.g00269.t01#1    mNeoVis.3.1.1.chr13.g00224.t01#6    mNeoVis.2.1.1.chr13_hap1.g00282.t01#5   mNeoVis.vulgivaga.1.1.chr13.g00267.t01#4     7371   125253655:   mNeoVis.3.1.1.chr13.g00226.t01#6    mNeoVis.evergladensis.1.1.chr13.g00272.t01#3    mNeoVis.2.1.1.chr13_hap1.g00283.t01#5    mNeoVis.vulgivaga.2.1.chr13.g00270.t01#1   mNeoVis.evergladensis.2.1.chr13.g00274.t01#2       7372   125253656:   mNeoVis.vulgivaga.2.1.chr5.g01694.t01#1    mNeoVis.evergladensis.1.1.chr5.g01700.t01#3    mNeoVis.evergladensis.2.1.chr5.g01693.t01#2    mNeoVis.2.1.1.chr2_hap1.g01753.t01#5    mNeoVis.3.1.1.chr5.g01677.t01#6   mNeoVis.vulgivaga.1.1.chr5.g01673.t01#4       7373   125253657:   mNeoVis.2.1.1.chr2_hap1.g01754.t01#5    mNeoVis.evergladensis.2.1.chr5.g01694.t01#2    mNeoVis.evergladensis.1.1.chr5.g01701.t01#3    mNeoVis.vulgivaga.1.1.chr5.g01674.t01#4   mNeoVis.vulgivaga.2.1.chr5.g01695.t01#1       7374   125253658:   mNeoVis.vulgivaga.1.1.chr13.g00268.t01#4    mNeoVis.evergladensis.2.1.chr13.g00275.t01#2    mNeoVis.vulgivaga.2.1.chr13.g00271.t01#1    mNeoVis.3.1.1.chr13.g00227.t01#6    mNeoVis.evergladensis.1.1.chr13.g00273.t01#3   mNeoVis.2.1.1.chr13_hap1.g00284.t01#5       7375   125253659:   mNeoVis.3.1.1.chr13.g00228.t01#6    mNeoVis.2.1.1.chr13_hap1.g00285.t01#5    mNeoVis.vulgivaga.1.1.chr13.g00269.t01#4    mNeoVis.evergladensis.1.1.chr13.g00274.t01#3    mNeoVis.evergladensis.2.1.chr13.g00276.t01#2   mNeoVis.vulgivaga.2.1.chr13.g00272.t01#1       7376   125253660:   mNeoVis.vulgivaga.2.1.chr13.g00273.t01#1    mNeoVis.evergladensis.1.1.chr13.g00275.t01#3    mNeoVis.3.1.1.chr13.g00229.t01#6    mNeoVis.2.1.1.chr13_hap1.g00286.t01#5    mNeoVis.vulgivaga.1.1.chr13.g00270.t01#4   mNeoVis.evergladensis.2.1.chr13.g00277.t01#2       7377   125253661:   mNeoVis.vulgivaga.2.1.chr13.g00274.t01#1    mNeoVis.evergladensis.1.1.chr13.g00276.t01#3    mNeoVis.vulgivaga.1.1.chr13.g00271.t01#4    mNeoVis.2.1.1.chr13_hap1.g00287.t01#5    mNeoVis.evergladensis.2.1.chr13.g00278.t01#2   mNeoVis.3.1.1.chr13.g00230.t01#6       7378   125253662:   mNeoVis.evergladensis.1.1.chr5.g01702.t01#3    mNeoVis.vulgivaga.1.1.chr5.g01675.t01#4    mNeoVis.3.1.1.chr5.g01681.t01#6    mNeoVis.vulgivaga.2.1.chr5.g01696.t01#1    mNeoVis.evergladensis.2.1.chr5.g01695.t01#2   mNeoVis.2.1.1.chr2_hap1.g01755.t01#5       7379   125253663:   mNeoVis.evergladensis.2.1.chr5.g01696.t01#2    mNeoVis.3.1.1.chr5.g01682.t01#6    mNeoVis.2.1.1.chr2_hap1.g01756.t01#5    mNeoVis.vulgivaga.1.1.chr5.g01676.t01#4    mNeoVis.vulgivaga.2.1.chr5.g01697.t01#1   mNeoVis.evergladensis.1.1.chr5.g01703.t01#3       7380   125253664:   mNeoVis.2.1.1.chr13_hap1.g00288.t01#5    mNeoVis.3.1.1.chr13.g00231.t01#6    mNeoVis.vulgivaga.2.1.chr13.g00275.t01#1    mNeoVis.evergladensis.2.1.chr13.g00279.t01#2    mNeoVis.evergladensis.1.1.chr13.g00277.t01#3   mNeoVis.vulgivaga.1.1.chr13.g00272.t01#4       7381   125253665:   mNeoVis.vulgivaga.2.1.chr13.g00276.t01#1    mNeoVis.evergladensis.1.1.chr13.g00278.t01#3    mNeoVis.2.1.1.chr13_hap1.g00289.t01#5    mNeoVis.vulgivaga.1.1.chr13.g00273.t01#4    mNeoVis.evergladensis.2.1.chr13.g00280.t01#2   mNeoVis.3.1.1.chr13.g00232.t01#6       7382   125253666:   mNeoVis.vulgivaga.1.1.chr13.g00274.t01#4    mNeoVis.evergladensis.2.1.chr13.g00281.t01#2    mNeoVis.2.1.1.chr13_hap1.g00290.t01#5    mNeoVis.evergladensis.1.1.chr13.g00279.t01#3    mNeoVis.3.1.1.chr13.g00233.t01#6   mNeoVis.vulgivaga.2.1.chr13.g00277.t01#1       7383   125253667:   mNeoVis.evergladensis.1.1.chr15.g00195.t01#3    mNeoVis.vulgivaga.2.1.chr15.g00429.t01#1    mNeoVis.vulgivaga.1.1.chr15.g00421.t01#4    mNeoVis.evergladensis.2.1.chr15.g00421.t01#2    mNeoVis.3.1.1.chr15.g00383.t01#6   mNeoVis.2.1.1.chr15_hap1.g00599.t01#5       7384   125253668:   mNeoVis.evergladensis.2.1.chr7.g00454.t01#2    mNeoVis.vulgivaga.2.1.chr7.g00453.t01#1    mNeoVis.vulgivaga.1.1.chr7.g00457.t01#4    mNeoVis.2.1.1.chr7_hap1.g00448.t01#5   mNeoVis.evergladensis.1.1.chr7.g00460.t01#3       7385   125253669:   mNeoVis.evergladensis.1.1.chr13.g00280.t01#3    mNeoVis.2.1.1.chr13_hap1.g00291.t01#5    mNeoVis.evergladensis.2.1.chr13.g00282.t01#2    mNeoVis.vulgivaga.1.1.chr13.g00275.t01#4    mNeoVis.3.1.1.chr13.g00234.t01#6   mNeoVis.vulgivaga.2.1.chr13.g00278.t01#1       7386   125253670:   mNeoVis.3.1.1.chr2.g00824.t01#6    mNeoVis.2.1.1.chr2_hap1.g01757.t01#5    mNeoVis.vulgivaga.2.1.chr5.g01698.t01#1    mNeoVis.evergladensis.1.1.chr5.g01704.t01#3    mNeoVis.evergladensis.2.1.chr5.g01697.t01#2    mNeoVis.vulgivaga.1.1.chr5.g01677.t01#4   mNeoVis.3.1.1.chr5.g01683.t01#6       7387   125253671:   mNeoVis.evergladensis.2.1.chr5.g01698.t01#2    mNeoVis.vulgivaga.1.1.chr5.g01678.t01#4    mNeoVis.3.1.1.chr5.g01684.t01#6    mNeoVis.evergladensis.1.1.chr5.g01705.t01#3    mNeoVis.vulgivaga.2.1.chr5.g01699.t01#1   mNeoVis.2.1.1.chr2_hap1.g01758.t01#5       7388   125253672:   mNeoVis.2.1.1.chr13_hap1.g00292.t01#5    mNeoVis.evergladensis.1.1.chr1.g01756.t01#3    mNeoVis.vulgivaga.2.1.chr13.g00279.t01#1    mNeoVis.evergladensis.2.1.chr1.g01755.t01#2    mNeoVis.vulgivaga.1.1.chr1.g01760.t01#4    mNeoVis.3.1.1.chr13.g00235.t01#6    mNeoVis.2.1.1.chr1_hap1.g01788.t01#5   mNeoVis.3.1.1.chr1.g01746.t01#6     7389   125253673:   mNeoVis.2.1.1.chr13_hap1.g00293.t01#5    mNeoVis.3.1.1.chr13.g00236.t01#6   mNeoVis.vulgivaga.2.1.chr13.g00280.t01#1         7390   125253674:   mNeoVis.vulgivaga.1.1.chr13.g00276.t01#4   mNeoVis.evergladensis.2.1.chr13.g00283.t01#2           7391   125253675:  mNeoVis.evergladensis.1.1.chr13.g00281.t01#3             7392   125253676:   mNeoVis.vulgivaga.2.1.chr13.g00281.t01#1    mNeoVis.2.1.1.chr13_hap1.g00294.t01#5    mNeoVis.evergladensis.2.1.chr13.g00284.t01#2    mNeoVis.vulgivaga.1.1.chr13.g00277.t01#4    mNeoVis.evergladensis.1.1.chr13.g00282.t01#3   mNeoVis.3.1.1.chr13.g00237.t01#6       7393   125253677:   mNeoVis.3.1.1.chr5.g01685.t01#6    mNeoVis.evergladensis.2.1.chr5.g01699.t01#2    mNeoVis.evergladensis.1.1.chr5.g01706.t01#3    mNeoVis.vulgivaga.1.1.chr5.g01679.t01#4    mNeoVis.2.1.1.chr2_hap1.g01759.t01#5   mNeoVis.vulgivaga.2.1.chr5.g01700.t01#1       7394   125253678:   mNeoVis.vulgivaga.2.1.chr13.g00282.t01#1    mNeoVis.2.1.1.chr13_hap1.g00295.t01#5    mNeoVis.evergladensis.2.1.chr13.g00285.t01#2    mNeoVis.vulgivaga.1.1.chr13.g00278.t01#4    mNeoVis.evergladensis.1.1.chr13.g00283.t01#3   mNeoVis.3.1.1.chr13.g00238.t01#6       7395   125253679:   mNeoVis.2.1.1.chr2_hap1.g01760.t01#5    mNeoVis.3.1.1.chr5.g01686.t01#6    mNeoVis.evergladensis.1.1.chr5.g01707.t01#3    mNeoVis.evergladensis.2.1.chr5.g01700.t01#2    mNeoVis.vulgivaga.1.1.chr5.g01680.t01#4   mNeoVis.vulgivaga.2.1.chr5.g01701.t01#1       7396   125253680:   mNeoVis.2.1.1.chr13_hap1.g00296.t01#5    mNeoVis.evergladensis.2.1.chr13.g00287.t01#2    mNeoVis.vulgivaga.2.1.chr13.g00283.t01#1    mNeoVis.3.1.1.chr13.g00239.t01#6    mNeoVis.evergladensis.1.1.chr13.g00284.t01#3   mNeoVis.vulgivaga.1.1.chr13.g00279.t01#4       7397   125253681:   mNeoVis.3.1.1.chr13.g00240.t01#6    mNeoVis.vulgivaga.2.1.chr13.g00284.t01#1    mNeoVis.vulgivaga.1.1.chr13.g00280.t01#4    mNeoVis.evergladensis.1.1.chr13.g00285.t01#3    mNeoVis.evergladensis.2.1.chr13.g00288.t01#2   mNeoVis.2.1.1.chr13_hap1.g00297.t01#5       7398   125253682:   mNeoVis.vulgivaga.1.1.chr5.g01681.t01#4    mNeoVis.2.1.1.chr2_hap1.g01761.t01#5    mNeoVis.evergladensis.2.1.chr5.g01701.t01#2    mNeoVis.evergladensis.1.1.chr5.g01708.t01#3    mNeoVis.vulgivaga.2.1.chr5.g01702.t01#1   mNeoVis.3.1.1.chr5.g01687.t01#6       7399   125253683:   mNeoVis.2.1.1.chr2_hap1.g01762.t01#5    mNeoVis.evergladensis.2.1.chr5.g01702.t01#2    mNeoVis.vulgivaga.2.1.chr5.g01703.t01#1    mNeoVis.evergladensis.1.1.chr5.g01709.t01#3   mNeoVis.vulgivaga.1.1.chr5.g01682.t01#4       7400   125253684:   mNeoVis.evergladensis.2.1.chr15.g00422.t01#2    mNeoVis.2.1.1.chr15_hap1.g00600.t01#5    mNeoVis.3.1.1.chr15.g00385.t01#6    mNeoVis.vulgivaga.2.1.chr15.g00430.t01#1    mNeoVis.evergladensis.1.1.chr15.g00194.t01#3   mNeoVis.vulgivaga.1.1.chr15.g00422.t01#4       7401   125253685:   mNeoVis.evergladensis.1.1.chr5.g01710.t01#3    mNeoVis.evergladensis.2.1.chr5.g01703.t01#2    mNeoVis.2.1.1.chr2_hap1.g01763.t01#5    mNeoVis.vulgivaga.2.1.chr5.g01704.t01#1   mNeoVis.vulgivaga.1.1.chr5.g01683.t01#4       7402   125253686:   mNeoVis.vulgivaga.2.1.chr13.g00285.t01#1    mNeoVis.evergladensis.1.1.chr13.g00286.t01#3    mNeoVis.2.1.1.chr13_hap1.g00298.t01#5    mNeoVis.3.1.1.chr13.g00241.t01#6    mNeoVis.vulgivaga.1.1.chr13.g00281.t01#4   mNeoVis.evergladensis.2.1.chr13.g00289.t01#2       7403   125253687:   mNeoVis.2.1.1.chr2_hap1.g01764.t01#5    mNeoVis.evergladensis.2.1.chr5.g01704.t01#2    mNeoVis.vulgivaga.2.1.chr5.g01705.t01#1    mNeoVis.vulgivaga.1.1.chr5.g01684.t01#4   mNeoVis.evergladensis.1.1.chr5.g01711.t01#3       7404   125253688:   mNeoVis.3.1.1.chr7.g00437.t01#6    mNeoVis.evergladensis.2.1.chr7.g00455.t01#2    mNeoVis.vulgivaga.1.1.chr7.g00458.t01#4    mNeoVis.vulgivaga.2.1.chr7.g00454.t01#1    mNeoVis.2.1.1.chr7_hap1.g00449.t01#5   mNeoVis.evergladensis.1.1.chr7.g00461.t01#3       7405   125253689:  mNeoVis.vulgivaga.2.1.chr5.g01706.t01#1               7406   125253690:  mNeoVis.vulgivaga.2.1.chr5.g01707.t01#1               7407   125253691:   mNeoVis.3.1.1.chr5.g01690.t01#6    mNeoVis.vulgivaga.1.1.chr5.g01685.t01#4    mNeoVis.evergladensis.1.1.chr5.g01712.t01#3    mNeoVis.vulgivaga.2.1.chr5.g01708.t01#1    mNeoVis.2.1.1.chr2_hap1.g01765.t01#5   mNeoVis.evergladensis.2.1.chr5.g01705.t01#2       7408   125253692:   mNeoVis.2.1.1.chr2_hap1.g01766.t01#5    mNeoVis.evergladensis.2.1.chr5.g01706.t01#2    mNeoVis.evergladensis.1.1.chr5.g01713.t01#3    mNeoVis.3.1.1.chr5.g01691.t01#6    mNeoVis.vulgivaga.2.1.chr5.g01709.t01#1   mNeoVis.vulgivaga.1.1.chr5.g01686.t01#4       7409   125253693:   mNeoVis.2.1.1.chr13_hap1.g00299.t01#5    mNeoVis.evergladensis.1.1.chr13.g00287.t01#3    mNeoVis.vulgivaga.2.1.chr13.g00286.t01#1    mNeoVis.vulgivaga.1.1.chr13.g00282.t01#4    mNeoVis.3.1.1.chr13.g00242.t01#6   mNeoVis.evergladensis.2.1.chr13.g00290.t01#2       7410   125253694:   mNeoVis.vulgivaga.1.1.chr13.g00284.t01#4    mNeoVis.evergladensis.2.1.chr13.g00291.t01#2    mNeoVis.3.1.1.chr13.g00243.t01#6    mNeoVis.vulgivaga.2.1.chr13.g00287.t01#1   mNeoVis.2.1.1.chr13_hap1.g00300.t01#5       7411   125253695:   mNeoVis.evergladensis.1.1.chr13.g00288.t01#3   mNeoVis.vulgivaga.1.1.chr13.g00283.t01#4           7412   125253696:   mNeoVis.vulgivaga.2.1.chr5.g01710.t01#1    mNeoVis.vulgivaga.1.1.chr5.g01687.t01#4    mNeoVis.evergladensis.1.1.chr5.g01714.t01#3    mNeoVis.3.1.1.chr5.g01692.t01#6    mNeoVis.2.1.1.chr2_hap1.g01767.t01#5   mNeoVis.evergladensis.2.1.chr5.g01707.t01#2       7413   125253697:   mNeoVis.vulgivaga.2.1.chr5.g01711.t01#1    mNeoVis.vulgivaga.1.1.chr5.g01688.t01#4    mNeoVis.evergladensis.2.1.chr5.g01708.t01#2    mNeoVis.2.1.1.chr2_hap1.g01768.t01#5    mNeoVis.3.1.1.chr5.g01693.t01#6    mNeoVis.vulgivaga.2.1.chr5.g01712.t01#1   mNeoVis.evergladensis.1.1.chr5.g01715.t01#3     7414   125253698:   mNeoVis.vulgivaga.2.1.chr13.g00288.t01#1    mNeoVis.vulgivaga.1.1.chr13.g00285.t01#4   mNeoVis.3.1.1.chr13.g00244.t01#6           7415   125253699:   mNeoVis.evergladensis.1.1.chr13.g00290.t01#3    mNeoVis.2.1.1.chr13_hap1.g00302.t01#5    mNeoVis.vulgivaga.2.1.chr13.g00289.t01#1    mNeoVis.evergladensis.2.1.chr13.g00293.t01#2    mNeoVis.vulgivaga.1.1.chr13.g00286.t01#4   mNeoVis.3.1.1.chr13.g00245.t01#6       7416   125253700:   mNeoVis.vulgivaga.2.1.chr5.g01713.t01#1    mNeoVis.vulgivaga.1.1.chr5.g01689.t01#4    mNeoVis.evergladensis.2.1.chr5.g01709.t01#2    mNeoVis.evergladensis.1.1.chr5.g01716.t01#3    mNeoVis.3.1.1.chr5.g01694.t01#6   mNeoVis.2.1.1.chr2_hap1.g01769.t01#5       7417   125253701:   mNeoVis.evergladensis.2.1.chr5.g01710.t01#2    mNeoVis.2.1.1.chr2_hap1.g01770.t01#5    mNeoVis.vulgivaga.1.1.chr5.g01690.t01#4    mNeoVis.evergladensis.1.1.chr5.g01717.t01#3    mNeoVis.vulgivaga.2.1.chr5.g01714.t01#1   mNeoVis.3.1.1.chr5.g01695.t01#6       7418   125253702:   mNeoVis.2.1.1.chr13_hap1.g00303.t01#5    mNeoVis.vulgivaga.1.1.chr13.g00287.t01#4    mNeoVis.evergladensis.2.1.chr13.g00294.t01#2    mNeoVis.3.1.1.chr13.g00246.t01#6    mNeoVis.evergladensis.1.1.chr13.g00291.t01#3   mNeoVis.vulgivaga.2.1.chr13.g00290.t01#1       7419   125253703:   mNeoVis.3.1.1.chr13.g00247.t01#6    mNeoVis.evergladensis.1.1.chr13.g00292.t01#3    mNeoVis.vulgivaga.2.1.chr13.g00291.t01#1    mNeoVis.2.1.1.chr13_hap1.g00304.t01#5    mNeoVis.vulgivaga.1.1.chr13.g00288.t01#4   mNeoVis.evergladensis.2.1.chr13.g00295.t01#2       7420   125253704:   mNeoVis.3.1.1.chr5.g01696.t01#6    mNeoVis.2.1.1.chr2_hap1.g01771.t01#5    mNeoVis.evergladensis.2.1.chr5.g01711.t01#2    mNeoVis.evergladensis.1.1.chr5.g01718.t01#3    mNeoVis.vulgivaga.1.1.chr5.g01691.t01#4   mNeoVis.vulgivaga.2.1.chr5.g01715.t01#1       7421   125253705:   mNeoVis.evergladensis.2.1.chr13.g00193.t01#2    mNeoVis.2.1.1.chr13_hap1.g00195.t01#5    mNeoVis.vulgivaga.1.1.chr13.g00187.t01#4    mNeoVis.evergladensis.1.1.chr13.g00191.t01#3   mNeoVis.vulgivaga.2.1.chr13.g00190.t01#1       7422   125253706:   mNeoVis.3.1.1.chr13.g00147.t01#6    mNeoVis.vulgivaga.2.1.chr13.g00191.t01#1    mNeoVis.evergladensis.2.1.chr13.g00194.t01#2    mNeoVis.evergladensis.1.1.chr13.g00192.t01#3    mNeoVis.vulgivaga.1.1.chr13.g00188.t01#4   mNeoVis.2.1.1.chr13_hap1.g00196.t01#5       7423   125253707:   mNeoVis.vulgivaga.1.1.chr15.g00423.t01#4    mNeoVis.vulgivaga.2.1.chr15.g00431.t01#1    mNeoVis.evergladensis.2.1.chr15.g00423.t01#2    mNeoVis.evergladensis.1.1.chr15.g00193.t01#3   mNeoVis.2.1.1.chr15_hap1.g00601.t01#5       7424   125253708:   mNeoVis.2.1.1.chr13_hap1.g00197.t01#5    mNeoVis.3.1.1.chr13.g00148.t01#6    mNeoVis.evergladensis.2.1.chr13.g00195.t01#2    mNeoVis.vulgivaga.2.1.chr13.g00192.t01#1    mNeoVis.vulgivaga.1.1.chr13.g00189.t01#4   mNeoVis.evergladensis.1.1.chr13.g00193.t01#3       7425   125253709:   mNeoVis.evergladensis.1.1.chr5.g01719.t01#3    mNeoVis.vulgivaga.1.1.chr5.g01692.t01#4    mNeoVis.2.1.1.chr2_hap1.g01772.t01#5    mNeoVis.evergladensis.2.1.chr5.g01712.t01#2    mNeoVis.vulgivaga.2.1.chr5.g01716.t01#1   mNeoVis.3.1.1.chr5.g01697.t01#6       7426   125253710:   mNeoVis.evergladensis.1.1.chr5.g01720.t01#3    mNeoVis.vulgivaga.2.1.chr5.g01717.t01#1    mNeoVis.vulgivaga.1.1.chr5.g01693.t01#4    mNeoVis.2.1.1.chr2_hap1.g01773.t01#5    mNeoVis.evergladensis.2.1.chr5.g01713.t01#2   mNeoVis.3.1.1.chr5.g01698.t01#6       7427   125253711:   mNeoVis.2.1.1.chr13_hap1.g00198.t01#5    mNeoVis.3.1.1.chr13.g00149.t01#6    mNeoVis.vulgivaga.2.1.chr13.g00193.t01#1    mNeoVis.vulgivaga.1.1.chr13.g00190.t01#4    mNeoVis.evergladensis.1.1.chr13.g00194.t01#3   mNeoVis.evergladensis.2.1.chr13.g00196.t01#2       7428   125253712:   mNeoVis.vulgivaga.2.1.chr13.g00194.t01#1    mNeoVis.evergladensis.2.1.chr13.g00197.t01#2    mNeoVis.vulgivaga.1.1.chr13.g00191.t01#4    mNeoVis.evergladensis.1.1.chr13.g00195.t01#3    mNeoVis.2.1.1.chr13_hap1.g00199.t01#5   mNeoVis.3.1.1.chr13.g00150.t01#6       7429   125253713:   mNeoVis.2.1.1.chr13_hap1.g00200.t01#5    mNeoVis.vulgivaga.2.1.chr13.g00195.t01#1    mNeoVis.vulgivaga.1.1.chr13.g00192.t01#4    mNeoVis.evergladensis.1.1.chr13.g00196.t01#3   mNeoVis.evergladensis.2.1.chr13.g00198.t01#2       7430   125253714:   mNeoVis.vulgivaga.2.1.chr5.g01718.t01#1    mNeoVis.evergladensis.2.1.chr5.g01714.t01#2    mNeoVis.evergladensis.1.1.chr5.g01721.t01#3    mNeoVis.vulgivaga.1.1.chr5.g01694.t01#4    mNeoVis.3.1.1.chr5.g01699.t01#6   mNeoVis.2.1.1.chr2_hap1.g01774.t01#5       7431   125253715:   mNeoVis.evergladensis.1.1.chr5.g01722.t01#3    mNeoVis.vulgivaga.2.1.chr5.g01719.t01#1    mNeoVis.vulgivaga.1.1.chr5.g01695.t01#4    mNeoVis.evergladensis.2.1.chr5.g01715.t01#2    mNeoVis.2.1.1.chr2_hap1.g01775.t01#5   mNeoVis.3.1.1.chr5.g01700.t01#6       7432   125253716:   mNeoVis.vulgivaga.2.1.chr5.g01720.t01#1    mNeoVis.vulgivaga.1.1.chr5.g01696.t01#4    mNeoVis.evergladensis.1.1.chr5.g01723.t01#3    mNeoVis.3.1.1.chr5.g01701.t01#6    mNeoVis.2.1.1.chr2_hap1.g01776.t01#5   mNeoVis.evergladensis.2.1.chr5.g01716.t01#2       7433   125253717:   mNeoVis.vulgivaga.1.1.chr13.g00193.t01#4    mNeoVis.evergladensis.1.1.chr13.g00197.t01#3    mNeoVis.3.1.1.chr13.g00152.t01#6    mNeoVis.2.1.1.chr13_hap1.g00201.t01#5    mNeoVis.evergladensis.2.1.chr13.g00199.t01#2   mNeoVis.vulgivaga.2.1.chr13.g00196.t01#1       7434   125253718:   mNeoVis.2.1.1.chr13_hap1.g00202.t01#5    mNeoVis.evergladensis.2.1.chr13.g00200.t01#2    mNeoVis.vulgivaga.2.1.chr13.g00197.t01#1    mNeoVis.evergladensis.1.1.chr13.g00198.t01#3    mNeoVis.3.1.1.chr13.g00153.t01#6   mNeoVis.vulgivaga.1.1.chr13.g00194.t01#4       7435   125253719:   mNeoVis.vulgivaga.2.1.chr5.g01646.t01#1    mNeoVis.evergladensis.2.1.chr5.g01644.t01#2    mNeoVis.3.1.1.chr5.g01627.t01#6    mNeoVis.vulgivaga.1.1.chr5.g01624.t01#4    mNeoVis.evergladensis.1.1.chr5.g01649.t01#3   mNeoVis.2.1.1.chr2_hap1.g01705.t01#5       7436   125253720:   mNeoVis.3.1.1.chr13.g00154.t01#6    mNeoVis.vulgivaga.1.1.chr13.g00195.t01#4    mNeoVis.evergladensis.1.1.chr13.g00199.t01#3    mNeoVis.2.1.1.chr13_hap1.g00203.t01#5    mNeoVis.evergladensis.2.1.chr13.g00201.t01#2   mNeoVis.vulgivaga.2.1.chr13.g00198.t01#1       7437   125253721:   mNeoVis.evergladensis.2.1.chr5.g01645.t01#2    mNeoVis.vulgivaga.2.1.chr3.g00993.t01#1    mNeoVis.evergladensis.2.1.chr3.g00974.t01#2    mNeoVis.vulgivaga.1.1.chr4.g00885.t01#4    mNeoVis.evergladensis.1.1.chr3.g00987.t01#3    mNeoVis.vulgivaga.1.1.chr5.g01625.t01#4    mNeoVis.vulgivaga.1.1.chr3.g00981.t01#4    mNeoVis.2.1.1.chr3_hap1.g00979.t01#5    mNeoVis.3.1.1.chr3.g00969.t01#6      7438   125253722:   mNeoVis.evergladensis.2.1.chr5.g01646.t01#2    mNeoVis.evergladensis.1.1.chr5.g01651.t01#3    mNeoVis.vulgivaga.1.1.chr5.g01626.t01#4    mNeoVis.2.1.1.chr2_hap1.g01707.t01#5    mNeoVis.3.1.1.chr5.g01629.t01#6   mNeoVis.vulgivaga.2.1.chr5.g01648.t01#1       7439   125253723:   mNeoVis.evergladensis.2.1.chr15.g00065.t01#2    mNeoVis.3.1.1.chr15.g00060.t01#6    mNeoVis.vulgivaga.2.1.chr15.g00067.t01#1    mNeoVis.2.1.1.chr15_hap1.g00065.t01#5    mNeoVis.evergladensis.1.1.chr15.g00548.t01#3   mNeoVis.vulgivaga.1.1.chr15.g00066.t01#4       7440   125253724:   mNeoVis.evergladensis.1.1.chr5.g01652.t01#3    mNeoVis.evergladensis.2.1.chr5.g01647.t01#2    mNeoVis.2.1.1.chr2_hap1.g01708.t01#5    mNeoVis.vulgivaga.1.1.chr5.g01627.t01#4    mNeoVis.vulgivaga.2.1.chr5.g01649.t01#1   mNeoVis.3.1.1.chr5.g01630.t01#6       7441   125253725:   mNeoVis.vulgivaga.2.1.chr13.g00199.t01#1    mNeoVis.vulgivaga.1.1.chr13.g00196.t01#4    mNeoVis.evergladensis.1.1.chr13.g00200.t01#3    mNeoVis.evergladensis.2.1.chr13.g00202.t01#2    mNeoVis.3.1.1.chr13.g00155.t01#6   mNeoVis.2.1.1.chr13_hap1.g00204.t01#5       7442   125253726:   mNeoVis.evergladensis.2.1.chr5.g01648.t01#2    mNeoVis.vulgivaga.1.1.chr5.g01628.t01#4    mNeoVis.evergladensis.1.1.chr5.g01653.t01#3    mNeoVis.2.1.1.chr2_hap1.g01709.t01#5   mNeoVis.vulgivaga.2.1.chr5.g01650.t01#1       7443   125253727:   mNeoVis.evergladensis.2.1.chr13.g00203.t01#2    mNeoVis.vulgivaga.2.1.chr13.g00200.t01#1    mNeoVis.3.1.1.chr13.g00156.t01#6    mNeoVis.evergladensis.1.1.chr13.g00201.t01#3    mNeoVis.2.1.1.chr13_hap1.g00205.t01#5   mNeoVis.vulgivaga.1.1.chr13.g00197.t01#4       7444   125253728:   mNeoVis.vulgivaga.2.1.chr13.g00201.t01#1    mNeoVis.vulgivaga.1.1.chr13.g00198.t01#4    mNeoVis.evergladensis.2.1.chr13.g00204.t01#2    mNeoVis.evergladensis.1.1.chr13.g00202.t01#3   mNeoVis.2.1.1.chr13_hap1.g00206.t01#5       7445   125253729:   mNeoVis.2.1.1.chr4_hap1.g00812.t01#5    mNeoVis.vulgivaga.2.1.chr13.g00202.t01#1    mNeoVis.evergladensis.2.1.chr13.g00205.t01#2    mNeoVis.vulgivaga.2.1.chr6.g00812.t01#1    mNeoVis.evergladensis.2.1.chr6.g00816.t01#2    mNeoVis.vulgivaga.1.1.chr13.g00199.t01#4    mNeoVis.vulgivaga.1.1.chr6.g00814.t01#4    mNeoVis.3.1.1.chr13.g00158.t01#6    mNeoVis.evergladensis.1.1.chr13.g00203.t01#3      7446   125253730:   mNeoVis.3.1.1.chr13.g00159.t01#6    mNeoVis.evergladensis.2.1.chr13.g00206.t01#2    mNeoVis.vulgivaga.1.1.chr13.g00200.t01#4    mNeoVis.evergladensis.1.1.chr13.g00204.t01#3    mNeoVis.2.1.1.chr13_hap1.g00208.t01#5   mNeoVis.vulgivaga.2.1.chr13.g00203.t01#1       7447   125253731:   mNeoVis.3.1.1.chr13.g00160.t01#6    mNeoVis.evergladensis.2.1.chr13.g00207.t01#2    mNeoVis.vulgivaga.1.1.chr13.g00201.t01#4    mNeoVis.evergladensis.1.1.chr13.g00205.t01#3    mNeoVis.2.1.1.chr13_hap1.g00209.t01#5   mNeoVis.vulgivaga.2.1.chr13.g00204.t01#1       7448   125253732:   mNeoVis.2.1.1.chr13_hap1.g00210.t01#5    mNeoVis.vulgivaga.1.1.chr13.g00202.t01#4    mNeoVis.3.1.1.chr13.g00161.t01#6    mNeoVis.evergladensis.1.1.chr13.g00206.t01#3    mNeoVis.evergladensis.2.1.chr13.g00208.t01#2   mNeoVis.vulgivaga.2.1.chr13.g00205.t01#1       7449   125253733:   mNeoVis.vulgivaga.1.1.chr13.g00203.t01#4    mNeoVis.vulgivaga.2.1.chr13.g00206.t01#1    mNeoVis.evergladensis.2.1.chr13.g00209.t01#2    mNeoVis.evergladensis.1.1.chr13.g00207.t01#3    mNeoVis.2.1.1.chr13_hap1.g00211.t01#5   mNeoVis.3.1.1.chr13.g00162.t01#6       7450   125253734:   mNeoVis.evergladensis.1.1.chr15.g00547.t01#3    mNeoVis.evergladensis.2.1.chr15.g00066.t01#2    mNeoVis.2.1.1.chr15_hap1.g00066.t01#5    mNeoVis.vulgivaga.2.1.chr15.g00068.t01#1    mNeoVis.3.1.1.chr15.g00061.t01#6   mNeoVis.vulgivaga.1.1.chr15.g00067.t01#4       7451   125253735:   mNeoVis.vulgivaga.2.1.chr5.g01651.t01#1    mNeoVis.3.1.1.chr5.g01633.t01#6   mNeoVis.evergladensis.1.1.chr5.g01654.t01#3         7452   125253736:   mNeoVis.2.1.1.chr2_hap1.g01710.t01#5    mNeoVis.evergladensis.1.1.chr5.g01655.t01#3    mNeoVis.3.1.1.chr5.g01634.t01#6    mNeoVis.vulgivaga.1.1.chr5.g01629.t01#4    mNeoVis.evergladensis.2.1.chr5.g01649.t01#2   mNeoVis.vulgivaga.2.1.chr5.g01652.t01#1       7453   125253737:   mNeoVis.evergladensis.1.1.chr5.g01656.t01#3    mNeoVis.evergladensis.2.1.chr5.g01650.t01#2    mNeoVis.vulgivaga.2.1.chr5.g01653.t01#1    mNeoVis.2.1.1.chr2_hap1.g01711.t01#5   mNeoVis.vulgivaga.1.1.chr5.g01630.t01#4       7454   125253738:   mNeoVis.vulgivaga.2.1.chr13.g00207.t01#1   mNeoVis.evergladensis.2.1.chr13.g00210.t01#2           7455   125253739:   mNeoVis.3.1.1.chr13.g00164.t01#6    mNeoVis.2.1.1.chr13_hap1.g00213.t01#5    mNeoVis.evergladensis.2.1.chr13.g00211.t01#2    mNeoVis.evergladensis.1.1.chr13.g00210.t01#3    mNeoVis.vulgivaga.1.1.chr13.g00205.t01#4   mNeoVis.vulgivaga.2.1.chr13.g00208.t01#1       7456   125253740:   mNeoVis.vulgivaga.2.1.chr13.g00209.t01#1    mNeoVis.evergladensis.2.1.chr13.g00212.t01#2    mNeoVis.evergladensis.1.1.chr13.g00211.t01#3    mNeoVis.3.1.1.chr13.g00165.t01#6    mNeoVis.2.1.1.chr13_hap1.g00214.t01#5   mNeoVis.vulgivaga.1.1.chr13.g00206.t01#4       7457   125253741:   mNeoVis.evergladensis.2.1.chr6.g01096.t01#2    mNeoVis.evergladensis.1.1.chr6.g01099.t01#3   mNeoVis.vulgivaga.2.1.chr6.g01088.t01#1           7458   125253742:   mNeoVis.2.1.1.chr15_hap1.g00067.t01#5    mNeoVis.evergladensis.1.1.chr15.g00546.t01#3    mNeoVis.3.1.1.chr15.g00062.t01#6    mNeoVis.vulgivaga.1.1.chr15.g00068.t01#4   mNeoVis.vulgivaga.2.1.chr15.g00069.t01#1       7459   125253743:   mNeoVis.3.1.1.chr5.g01636.t01#6    mNeoVis.vulgivaga.1.1.chr5.g01631.t01#4    mNeoVis.evergladensis.1.1.chr5.g01657.t01#3    mNeoVis.2.1.1.chr2_hap1.g01712.t01#5    mNeoVis.evergladensis.2.1.chr5.g01651.t01#2   mNeoVis.vulgivaga.2.1.chr5.g01654.t01#1       7460   125253744:   mNeoVis.evergladensis.1.1.chr13.g00212.t01#3    mNeoVis.2.1.1.chr13_hap1.g00215.t01#5    mNeoVis.vulgivaga.2.1.chr13.g00210.t01#1    mNeoVis.3.1.1.chr13.g00166.t01#6    mNeoVis.vulgivaga.1.1.chr13.g00207.t01#4   mNeoVis.evergladensis.2.1.chr13.g00213.t01#2       7461   125253745:   mNeoVis.evergladensis.2.1.chr6.g01097.t01#2    mNeoVis.vulgivaga.2.1.chr6.g01089.t01#1    mNeoVis.vulgivaga.1.1.chr6.g01090.t01#4    mNeoVis.evergladensis.1.1.chr6.g01100.t01#3   mNeoVis.2.1.1.chr4_hap1.g01099.t01#5       7462   125253746:   mNeoVis.3.1.1.chr13.g00167.t01#6    mNeoVis.evergladensis.1.1.chr13.g00213.t01#3    mNeoVis.vulgivaga.1.1.chr13.g00208.t01#4    mNeoVis.2.1.1.chr13_hap1.g00216.t01#5    mNeoVis.evergladensis.2.1.chr13.g00214.t01#2   mNeoVis.vulgivaga.2.1.chr13.g00211.t01#1       7463   125253747:   mNeoVis.vulgivaga.2.1.chr5.g01655.t01#1    mNeoVis.3.1.1.chr5.g01637.t01#6    mNeoVis.evergladensis.1.1.chr5.g01658.t01#3    mNeoVis.evergladensis.2.1.chr5.g01652.t01#2    mNeoVis.2.1.1.chr2_hap1.g01713.t01#5   mNeoVis.vulgivaga.1.1.chr5.g01632.t01#4       7464   125253748:   mNeoVis.evergladensis.1.1.chr5.g01659.t01#3    mNeoVis.vulgivaga.1.1.chr5.g01633.t01#4    mNeoVis.evergladensis.2.1.chr5.g01653.t01#2    mNeoVis.2.1.1.chr2_hap1.g01714.t01#5    mNeoVis.vulgivaga.2.1.chr5.g01656.t01#1   mNeoVis.3.1.1.chr5.g01638.t01#6       7465   125253749:   mNeoVis.3.1.1.chr9.g00434.t01#6    mNeoVis.evergladensis.2.1.chr8.g00447.t01#2    mNeoVis.evergladensis.1.1.chr9.g00458.t01#3    mNeoVis.2.1.1.chr10_hap1.g00468.t01#5    mNeoVis.vulgivaga.2.1.chr9.g00451.t01#1   mNeoVis.vulgivaga.1.1.chr9.g00452.t01#4       7466   125253750:   mNeoVis.2.1.1.chr13_hap1.g00217.t01#5    mNeoVis.evergladensis.1.1.chr13.g00214.t01#3    mNeoVis.vulgivaga.2.1.chr13.g00212.t01#1    mNeoVis.vulgivaga.1.1.chr13.g00209.t01#4   mNeoVis.evergladensis.2.1.chr13.g00215.t01#2       7467   125253751:   mNeoVis.2.1.1.chr13_hap1.g00218.t01#5    mNeoVis.evergladensis.1.1.chr13.g00215.t01#3    mNeoVis.vulgivaga.2.1.chr13.g00213.t01#1    mNeoVis.vulgivaga.1.1.chr13.g00210.t01#4   mNeoVis.evergladensis.2.1.chr13.g00216.t01#2       7468   125253752:   mNeoVis.vulgivaga.1.1.chr5.g01635.t01#4    mNeoVis.2.1.1.chr2_hap1.g01716.t01#5    mNeoVis.evergladensis.1.1.chr5.g01661.t01#3    mNeoVis.evergladensis.2.1.chr5.g01655.t01#2    mNeoVis.3.1.1.chr5.g01640.t01#6   mNeoVis.vulgivaga.2.1.chr5.g01657.t01#1       7469   125253753:   mNeoVis.evergladensis.1.1.chr13.g00216.t01#3    mNeoVis.vulgivaga.1.1.chr13.g00211.t01#4    mNeoVis.2.1.1.chr13_hap1.g00219.t01#5    mNeoVis.evergladensis.2.1.chr13.g00217.t01#2    mNeoVis.3.1.1.chr13.g00170.t01#6   mNeoVis.vulgivaga.2.1.chr13.g00214.t01#1       7470   125253754:   mNeoVis.evergladensis.2.1.chr13.g00218.t01#2    mNeoVis.3.1.1.chr13.g00171.t01#6    mNeoVis.evergladensis.1.1.chr13.g00217.t01#3    mNeoVis.vulgivaga.1.1.chr13.g00212.t01#4    mNeoVis.vulgivaga.2.1.chr13.g00215.t01#1   mNeoVis.2.1.1.chr13_hap1.g00220.t01#5       7471   125253755:   mNeoVis.evergladensis.2.1.chr5.g01657.t01#2    mNeoVis.3.1.1.chr5.g01642.t01#6    mNeoVis.vulgivaga.1.1.chr5.g01637.t01#4    mNeoVis.evergladensis.1.1.chr5.g01663.t01#3    mNeoVis.vulgivaga.2.1.chr5.g01659.t01#1   mNeoVis.2.1.1.chr2_hap1.g01718.t01#5       7472   125253756:   mNeoVis.2.1.1.chr13_hap1.g00221.t01#5    mNeoVis.evergladensis.1.1.chr13.g00218.t01#3    mNeoVis.vulgivaga.2.1.chr13.g00216.t01#1    mNeoVis.3.1.1.chr13.g00173.t01#6    mNeoVis.vulgivaga.1.1.chr13.g00213.t01#4   mNeoVis.evergladensis.2.1.chr13.g00219.t01#2       7473   125253757:  mNeoVis.vulgivaga.2.1.chr5.g01661.t01#1               7474   125253758:   mNeoVis.vulgivaga.2.1.chr5.g01660.t01#1    mNeoVis.2.1.1.chr2_hap1.g01719.t01#5    mNeoVis.3.1.1.chr5.g01643.t01#6    mNeoVis.evergladensis.1.1.chr5.g01664.t01#3    mNeoVis.vulgivaga.1.1.chr5.g01638.t01#4   mNeoVis.evergladensis.2.1.chr5.g01658.t01#2       7475   125253759:   mNeoVis.vulgivaga.2.1.chr13.g00217.t01#1    mNeoVis.3.1.1.chr13.g00174.t01#6    mNeoVis.evergladensis.2.1.chr13.g00220.t01#2    mNeoVis.2.1.1.chr13_hap1.g00222.t01#5    mNeoVis.vulgivaga.1.1.chr13.g00214.t01#4   mNeoVis.evergladensis.1.1.chr13.g00219.t01#3       7476   125253760:   mNeoVis.evergladensis.1.1.chr15.g00545.t01#3    mNeoVis.2.1.1.chr15_hap1.g00068.t01#5    mNeoVis.vulgivaga.2.1.chr15.g00070.t01#1    mNeoVis.vulgivaga.1.1.chr15.g00069.t01#4   mNeoVis.evergladensis.2.1.chr15.g00069.t01#2       7477   125253761:   mNeoVis.3.1.1.chr13.g00175.t01#6    mNeoVis.2.1.1.chr13_hap1.g00223.t01#5    mNeoVis.evergladensis.2.1.chr13.g00221.t01#2    mNeoVis.evergladensis.1.1.chr13.g00220.t01#3    mNeoVis.vulgivaga.2.1.chr13.g00218.t01#1   mNeoVis.vulgivaga.1.1.chr13.g00215.t01#4       7478   125253762:   mNeoVis.evergladensis.2.1.chr13.g00222.t01#2    mNeoVis.2.1.1.chr13_hap1.g00224.t01#5    mNeoVis.3.1.1.chr13.g00176.t01#6    mNeoVis.evergladensis.1.1.chr13.g00221.t01#3    mNeoVis.vulgivaga.2.1.chr13.g00219.t01#1   mNeoVis.vulgivaga.1.1.chr13.g00216.t01#4       7479   125253763:   mNeoVis.vulgivaga.2.1.chr5.g01662.t01#1    mNeoVis.2.1.1.chr2_hap1.g01721.t01#5    mNeoVis.evergladensis.2.1.chr5.g01661.t01#2    mNeoVis.evergladensis.1.1.chr5.g01666.t01#3    mNeoVis.vulgivaga.1.1.chr5.g01640.t01#4   mNeoVis.3.1.1.chr5.g01645.t01#6       7480   125253764:   mNeoVis.vulgivaga.2.1.chr13.g00220.t01#1    mNeoVis.evergladensis.2.1.chr13.g00223.t01#2    mNeoVis.evergladensis.1.1.chr13.g00222.t01#3    mNeoVis.2.1.1.chr13_hap1.g00225.t01#5   mNeoVis.vulgivaga.1.1.chr13.g00217.t01#4       7481   125253765:   mNeoVis.vulgivaga.1.1.chr13.g00218.t01#4    mNeoVis.vulgivaga.2.1.chr13.g00221.t01#1    mNeoVis.evergladensis.1.1.chr13.g00223.t01#3    mNeoVis.2.1.1.chr13_hap1.g00226.t01#5    mNeoVis.3.1.1.chr13.g00177.t01#6   mNeoVis.evergladensis.2.1.chr13.g00224.t01#2       7482   125253766:   mNeoVis.vulgivaga.2.1.chr5.g01663.t01#1    mNeoVis.2.1.1.chr2_hap1.g01722.t01#5    mNeoVis.evergladensis.2.1.chr5.g01662.t01#2    mNeoVis.evergladensis.1.1.chr5.g01667.t01#3    mNeoVis.3.1.1.chr5.g01646.t01#6   mNeoVis.vulgivaga.1.1.chr5.g01641.t01#4       7483   125253767:   mNeoVis.vulgivaga.1.1.chr5.g01642.t01#4    mNeoVis.2.1.1.chr2_hap1.g01723.t01#5    mNeoVis.evergladensis.1.1.chr5.g01668.t01#3    mNeoVis.3.1.1.chr5.g01647.t01#6    mNeoVis.vulgivaga.2.1.chr5.g01664.t01#1   mNeoVis.evergladensis.2.1.chr5.g01663.t01#2       7484   125253768:   mNeoVis.evergladensis.1.1.chr5.g01669.t01#3    mNeoVis.vulgivaga.1.1.chr5.g01643.t01#4    mNeoVis.vulgivaga.2.1.chr5.g01665.t01#1    mNeoVis.2.1.1.chr2_hap1.g01724.t01#5    mNeoVis.evergladensis.2.1.chr5.g01664.t01#2   mNeoVis.3.1.1.chr5.g01648.t01#6       7485   125253769:   mNeoVis.vulgivaga.2.1.chr5.g01666.t01#1    mNeoVis.2.1.1.chr2_hap1.g01725.t01#5    mNeoVis.evergladensis.1.1.chr5.g01670.t01#3    mNeoVis.evergladensis.2.1.chr5.g01665.t01#2    mNeoVis.3.1.1.chr5.g01649.t01#6   mNeoVis.vulgivaga.1.1.chr5.g01644.t01#4       7486   125253770:   mNeoVis.2.1.1.chr15_hap1.g00069.t01#5    mNeoVis.evergladensis.1.1.chr15.g00544.t01#3    mNeoVis.vulgivaga.1.1.chr15.g00070.t01#4    mNeoVis.3.1.1.chr15.g00063.t01#6   mNeoVis.vulgivaga.2.1.chr15.g00071.t01#1       7487   125253771:   mNeoVis.2.1.1.chr2_hap1.g01726.t01#5    mNeoVis.evergladensis.1.1.chr5.g01671.t01#3    mNeoVis.vulgivaga.1.1.chr5.g01645.t01#4    mNeoVis.3.1.1.chr5.g01650.t01#6    mNeoVis.vulgivaga.2.1.chr5.g01667.t01#1   mNeoVis.evergladensis.2.1.chr5.g01666.t01#2       7488   125253772:   mNeoVis.vulgivaga.2.1.chr5.g01668.t01#1   mNeoVis.evergladensis.1.1.chr5.g01672.t01#3           7489   125253773:   mNeoVis.2.1.1.chr13_hap1.g00228.t01#5    mNeoVis.vulgivaga.2.1.chr13.g00223.t01#1    mNeoVis.3.1.1.chr13.g00179.t01#6    mNeoVis.vulgivaga.1.1.chr13.g00220.t01#4    mNeoVis.evergladensis.1.1.chr13.g00225.t01#3   mNeoVis.evergladensis.2.1.chr13.g00226.t01#2       7490   125253774:   mNeoVis.evergladensis.1.1.chr5.g01673.t01#3    mNeoVis.vulgivaga.1.1.chr5.g01646.t01#4    mNeoVis.2.1.1.chr2_hap1.g01727.t01#5    mNeoVis.vulgivaga.2.1.chr5.g01669.t01#1    mNeoVis.3.1.1.chr5.g01651.t01#6   mNeoVis.evergladensis.2.1.chr5.g01667.t01#2       7491   125253775:   mNeoVis.3.1.1.chr13.g00180.t01#6    mNeoVis.evergladensis.1.1.chr13.g00226.t01#3    mNeoVis.2.1.1.chr13_hap1.g00229.t01#5    mNeoVis.vulgivaga.1.1.chr13.g00221.t01#4    mNeoVis.vulgivaga.2.1.chr13.g00224.t01#1   mNeoVis.evergladensis.2.1.chr13.g00227.t01#2       7492   125253776:   mNeoVis.evergladensis.2.1.chr5.g01668.t01#2    mNeoVis.2.1.1.chr2_hap1.g01728.t01#5    mNeoVis.evergladensis.1.1.chr5.g01674.t01#3    mNeoVis.vulgivaga.2.1.chr5.g01670.t01#1    mNeoVis.3.1.1.chr5.g01652.t01#6   mNeoVis.vulgivaga.1.1.chr5.g01647.t01#4       7493   125253777:   mNeoVis.vulgivaga.2.1.chr5.g01671.t01#1    mNeoVis.evergladensis.1.1.chr5.g01675.t01#3    mNeoVis.3.1.1.chr5.g01653.t01#6    mNeoVis.vulgivaga.1.1.chr5.g01648.t01#4   mNeoVis.2.1.1.chr2_hap1.g01729.t01#5       7494   125253778:   mNeoVis.3.1.1.chr5.g01654.t01#6    mNeoVis.evergladensis.1.1.chr5.g01676.t01#3    mNeoVis.evergladensis.2.1.chr5.g01670.t01#2    mNeoVis.2.1.1.chr2_hap1.g01730.t01#5    mNeoVis.vulgivaga.2.1.chr5.g01672.t01#1   mNeoVis.vulgivaga.1.1.chr5.g01649.t01#4       7495   125253779:   mNeoVis.2.1.1.chr13_hap1.g00230.t01#5    mNeoVis.evergladensis.1.1.chr13.g00227.t01#3    mNeoVis.vulgivaga.2.1.chr13.g00225.t01#1    mNeoVis.vulgivaga.1.1.chr13.g00222.t01#4    mNeoVis.3.1.1.chr13.g00181.t01#6   mNeoVis.evergladensis.2.1.chr13.g00228.t01#2       7496   125253780:   mNeoVis.vulgivaga.1.1.chr12.g00149.t01#4    mNeoVis.3.1.1.chr12.g00137.t01#6    mNeoVis.evergladensis.1.1.chr12.g00143.t01#3    mNeoVis.evergladensis.2.1.chr12.g00144.t01#2    mNeoVis.vulgivaga.2.1.chr12.g00145.t01#1   mNeoVis.2.1.1.chr12_hap1.g00170.t01#5       7497   125253781:   mNeoVis.evergladensis.2.1.chr5.g01671.t01#2    mNeoVis.vulgivaga.2.1.chr5.g01673.t01#1    mNeoVis.vulgivaga.1.1.chr5.g01650.t01#4    mNeoVis.2.1.1.chr2_hap1.g01731.t01#5    mNeoVis.evergladensis.1.1.chr5.g01677.t01#3   mNeoVis.3.1.1.chr5.g01655.t01#6       7498   125253782:   mNeoVis.vulgivaga.1.1.chr5.g01651.t01#4    mNeoVis.3.1.1.chr5.g01656.t01#6    mNeoVis.evergladensis.2.1.chr5.g01672.t01#2    mNeoVis.vulgivaga.2.1.chr5.g01674.t01#1    mNeoVis.2.1.1.chr2_hap1.g01732.t01#5   mNeoVis.evergladensis.1.1.chr5.g01678.t01#3       7499   125253783:   mNeoVis.3.1.1.chr5.g01657.t01#6    mNeoVis.evergladensis.2.1.chr5.g01673.t01#2    mNeoVis.evergladensis.1.1.chr5.g01679.t01#3    mNeoVis.vulgivaga.2.1.chr5.g01675.t01#1    mNeoVis.2.1.1.chr2_hap1.g01733.t01#5   mNeoVis.vulgivaga.1.1.chr5.g01652.t01#4       7500   125253784:   mNeoVis.2.1.1.chr2_hap1.g01734.t01#5    mNeoVis.vulgivaga.2.1.chr5.g01676.t01#1    mNeoVis.evergladensis.1.1.chr5.g01680.t01#3    mNeoVis.evergladensis.2.1.chr5.g01674.t01#2    mNeoVis.vulgivaga.1.1.chr5.g01653.t01#4   mNeoVis.3.1.1.chr5.g01658.t01#6       7501   125253785:   mNeoVis.3.1.1.chr13.g00183.t01#6    mNeoVis.evergladensis.1.1.chr13.g00229.t01#3    mNeoVis.2.1.1.chr13_hap1.g00232.t01#5    mNeoVis.vulgivaga.2.1.chr13.g00227.t01#1    mNeoVis.evergladensis.2.1.chr13.g00230.t01#2   mNeoVis.vulgivaga.1.1.chr13.g00224.t01#4       7502   125253786:   mNeoVis.3.1.1.chr13.g00182.t01#6    mNeoVis.vulgivaga.2.1.chr13.g00226.t01#1    mNeoVis.evergladensis.2.1.chr13.g00229.t01#2    mNeoVis.2.1.1.chr13_hap1.g00231.t01#5    mNeoVis.vulgivaga.1.1.chr13.g00223.t01#4   mNeoVis.evergladensis.1.1.chr13.g00228.t01#3       7503   125253787:   mNeoVis.evergladensis.1.1.chr13.g00230.t01#3    mNeoVis.2.1.1.chr13_hap1.g00233.t01#5    mNeoVis.evergladensis.2.1.chr13.g00231.t01#2    mNeoVis.vulgivaga.2.1.chr13.g00228.t01#1    mNeoVis.3.1.1.chr13.g00184.t01#6   mNeoVis.vulgivaga.1.1.chr13.g00225.t01#4       7504   125253788:   mNeoVis.vulgivaga.2.1.chr13.g00229.t01#1    mNeoVis.2.1.1.chr13_hap1.g00234.t01#5    mNeoVis.evergladensis.1.1.chr13.g00231.t01#3    mNeoVis.evergladensis.2.1.chr13.g00232.t01#2    mNeoVis.3.1.1.chr13.g00185.t01#6   mNeoVis.vulgivaga.1.1.chr13.g00226.t01#4       7505   125253789:   mNeoVis.3.1.1.chr5.g01659.t01#6    mNeoVis.evergladensis.1.1.chr5.g01681.t01#3    mNeoVis.vulgivaga.1.1.chr5.g01655.t01#4    mNeoVis.vulgivaga.2.1.chr5.g01677.t01#1    mNeoVis.evergladensis.2.1.chr5.g01675.t01#2   mNeoVis.2.1.1.chr2_hap1.g01735.t01#5       7506   125253790:   mNeoVis.evergladensis.1.1.chr5.g01682.t01#3    mNeoVis.vulgivaga.1.1.chr5.g01656.t01#4    mNeoVis.vulgivaga.2.1.chr5.g01678.t01#1    mNeoVis.evergladensis.2.1.chr5.g01676.t01#2   mNeoVis.2.1.1.chr2_hap1.g01736.t01#5       7507   125253791:   mNeoVis.evergladensis.1.1.chr5.g01683.t01#3    mNeoVis.vulgivaga.1.1.chr5.g01657.t01#4    mNeoVis.evergladensis.2.1.chr5.g01677.t01#2    mNeoVis.3.1.1.chr5.g01661.t01#6    mNeoVis.2.1.1.chr2_hap1.g01737.t01#5   mNeoVis.vulgivaga.2.1.chr5.g01679.t01#1       7508   125253792:   mNeoVis.evergladensis.2.1.chr13.g00233.t01#2    mNeoVis.2.1.1.chr13_hap1.g00235.t01#5    mNeoVis.3.1.1.chr13.g00186.t01#6    mNeoVis.vulgivaga.1.1.chr13.g00227.t01#4    mNeoVis.evergladensis.1.1.chr13.g00232.t01#3   mNeoVis.vulgivaga.2.1.chr13.g00230.t01#1       7509   125253793:   mNeoVis.evergladensis.2.1.chr5.g01678.t01#2    mNeoVis.vulgivaga.1.1.chr5.g01658.t01#4    mNeoVis.vulgivaga.2.1.chr5.g01680.t01#1    mNeoVis.2.1.1.chr2_hap1.g01738.t01#5    mNeoVis.evergladensis.1.1.chr5.g01684.t01#3   mNeoVis.3.1.1.chr5.g01662.t01#6       7510   125253794:   mNeoVis.vulgivaga.2.1.chr5.g01681.t01#1    mNeoVis.evergladensis.2.1.chr5.g01679.t01#2    mNeoVis.3.1.1.chr5.g01663.t01#6    mNeoVis.vulgivaga.1.1.chr5.g01659.t01#4    mNeoVis.2.1.1.chr2_hap1.g01739.t01#5   mNeoVis.evergladensis.1.1.chr5.g01685.t01#3       7511   125253795:   mNeoVis.3.1.1.chr5.g01664.t01#6    mNeoVis.evergladensis.1.1.chr5.g01686.t01#3    mNeoVis.vulgivaga.1.1.chr5.g01660.t01#4    mNeoVis.evergladensis.2.1.chr5.g01680.t01#2    mNeoVis.vulgivaga.2.1.chr5.g01682.t01#1   mNeoVis.2.1.1.chr2_hap1.g01740.t01#5       7512   125253796:   mNeoVis.2.1.1.chr15_hap1.g00070.t01#5    mNeoVis.evergladensis.2.1.chr15.g00071.t01#2    mNeoVis.vulgivaga.1.1.chr15.g00071.t01#4    mNeoVis.evergladensis.1.1.chr15.g00543.t01#3   mNeoVis.vulgivaga.2.1.chr15.g00072.t01#1       7513   125253797:   mNeoVis.evergladensis.2.1.chr5.g01681.t01#2    mNeoVis.3.1.1.chr5.g01665.t01#6    mNeoVis.vulgivaga.1.1.chr5.g01661.t01#4    mNeoVis.evergladensis.1.1.chr5.g01687.t01#3    mNeoVis.vulgivaga.2.1.chr5.g01683.t01#1   mNeoVis.2.1.1.chr2_hap1.g01741.t01#5       7514   125253798:   mNeoVis.vulgivaga.2.1.chr13.g00232.t01#1    mNeoVis.evergladensis.2.1.chr13.g00235.t01#2    mNeoVis.evergladensis.1.1.chr13.g00234.t01#3    mNeoVis.2.1.1.chr13_hap1.g00237.t01#5    mNeoVis.3.1.1.chr13.g00188.t01#6   mNeoVis.vulgivaga.1.1.chr13.g00229.t01#4       7515   125253799:   mNeoVis.2.1.1.chr2_hap1.g01742.t01#5    mNeoVis.3.1.1.chr5.g01666.t01#6    mNeoVis.evergladensis.1.1.chr5.g01688.t01#3    mNeoVis.evergladensis.1.1.chr5.g01689.t01#3    mNeoVis.vulgivaga.2.1.chr5.g01684.t01#1    mNeoVis.evergladensis.2.1.chr5.g01682.t01#2   mNeoVis.vulgivaga.1.1.chr5.g01662.t01#4     7516   125253800:   mNeoVis.2.1.1.chr2_hap1.g01743.t01#5    mNeoVis.evergladensis.2.1.chr5.g01683.t01#2   mNeoVis.vulgivaga.1.1.chr5.g01663.t01#4           7517   125253801:   mNeoVis.3.1.1.chr13.g00189.t01#6    mNeoVis.evergladensis.2.1.chr13.g00236.t01#2    mNeoVis.vulgivaga.2.1.chr13.g00233.t01#1    mNeoVis.vulgivaga.1.1.chr13.g00230.t01#4    mNeoVis.evergladensis.1.1.chr13.g00235.t01#3   mNeoVis.2.1.1.chr13_hap1.g00238.t01#5       7518   125253802:   mNeoVis.3.1.1.chr13.g00190.t01#6    mNeoVis.evergladensis.2.1.chr13.g00237.t01#2    mNeoVis.vulgivaga.1.1.chr13.g00231.t01#4    mNeoVis.evergladensis.1.1.chr13.g00236.t01#3    mNeoVis.2.1.1.chr13_hap1.g00239.t01#5   mNeoVis.vulgivaga.2.1.chr13.g00234.t01#1       7519   125253803:   mNeoVis.evergladensis.2.1.chr5.g01685.t01#2    mNeoVis.2.1.1.chr2_hap1.g01745.t01#5    mNeoVis.vulgivaga.2.1.chr5.g01686.t01#1    mNeoVis.vulgivaga.1.1.chr5.g01665.t01#4    mNeoVis.3.1.1.chr5.g01669.t01#6   mNeoVis.evergladensis.1.1.chr5.g01691.t01#3       7520   125253804:   mNeoVis.evergladensis.2.1.chr13.g00238.t01#2    mNeoVis.vulgivaga.2.1.chr13.g00235.t01#1    mNeoVis.evergladensis.1.1.chr13.g00237.t01#3    mNeoVis.2.1.1.chr13_hap1.g00240.t01#5    mNeoVis.vulgivaga.1.1.chr13.g00232.t01#4   mNeoVis.3.1.1.chr13.g00191.t01#6       7521   125253805:   mNeoVis.2.1.1.chr13_hap1.g00157.t01#5    mNeoVis.evergladensis.1.1.chr13.g00153.t01#3    mNeoVis.evergladensis.2.1.chr13.g00155.t01#2    mNeoVis.vulgivaga.1.1.chr13.g00149.t01#4    mNeoVis.vulgivaga.2.1.chr13.g00153.t01#1   mNeoVis.3.1.1.chr13.g00109.t01#6       7522   125253806:   mNeoVis.2.1.1.chr2_hap1.g01746.t01#5    mNeoVis.vulgivaga.2.1.chr5.g01687.t01#1    mNeoVis.evergladensis.2.1.chr5.g01686.t01#2    mNeoVis.3.1.1.chr5.g01670.t01#6    mNeoVis.vulgivaga.1.1.chr5.g01666.t01#4   mNeoVis.evergladensis.1.1.chr5.g01692.t01#3       7523   125253807:   mNeoVis.vulgivaga.2.1.chr13.g00154.t01#1    mNeoVis.2.1.1.chr13_hap1.g00158.t01#5    mNeoVis.vulgivaga.1.1.chr13.g00150.t01#4    mNeoVis.evergladensis.2.1.chr13.g00156.t01#2    mNeoVis.evergladensis.1.1.chr13.g00154.t01#3   mNeoVis.3.1.1.chr13.g00110.t01#6       7524   125253808:   mNeoVis.vulgivaga.2.1.chr13.g00155.t01#1    mNeoVis.vulgivaga.1.1.chr13.g00151.t01#4    mNeoVis.evergladensis.1.1.chr13.g00155.t01#3    mNeoVis.evergladensis.2.1.chr13.g00157.t01#2    mNeoVis.3.1.1.chr13.g00111.t01#6   mNeoVis.2.1.1.chr13_hap1.g00159.t01#5       7525   125253809:   mNeoVis.2.1.1.chr2_hap1.g01651.t01#5    mNeoVis.3.1.1.chr5.g01573.t01#6    mNeoVis.vulgivaga.2.1.chr5.g01593.t01#1    mNeoVis.evergladensis.1.1.chr5.g01596.t01#3    mNeoVis.vulgivaga.1.1.chr5.g01569.t01#4   mNeoVis.evergladensis.2.1.chr5.g01591.t01#2       7526   125253810:   mNeoVis.3.1.1.chr5.g01574.t01#6    mNeoVis.2.1.1.chr2_hap1.g01652.t01#5    mNeoVis.vulgivaga.1.1.chr5.g01570.t01#4   mNeoVis.vulgivaga.2.1.chr5.g01594.t01#1         7527   125253811:   mNeoVis.3.1.1.chr5.g01575.t01#6    mNeoVis.2.1.1.chr2_hap1.g01653.t01#5    mNeoVis.evergladensis.1.1.chr5.g01598.t01#3    mNeoVis.vulgivaga.2.1.chr5.g01595.t01#1    mNeoVis.vulgivaga.1.1.chr5.g01571.t01#4   mNeoVis.evergladensis.2.1.chr5.g01593.t01#2       7528   125253812:   mNeoVis.vulgivaga.1.1.chr13.g00152.t01#4    mNeoVis.evergladensis.1.1.chr13.g00156.t01#3    mNeoVis.vulgivaga.2.1.chr13.g00156.t01#1    mNeoVis.3.1.1.chr13.g00112.t01#6    mNeoVis.evergladensis.2.1.chr13.g00158.t01#2   mNeoVis.2.1.1.chr13_hap1.g00160.t01#5       7529   125253813:   mNeoVis.evergladensis.1.1.chr13.g00157.t01#3    mNeoVis.3.1.1.chr13.g00113.t01#6    mNeoVis.vulgivaga.1.1.chr13.g00153.t01#4    mNeoVis.2.1.1.chr13_hap1.g00161.t01#5    mNeoVis.vulgivaga.2.1.chr13.g00157.t01#1   mNeoVis.evergladensis.2.1.chr13.g00159.t01#2       7530   125253814:   mNeoVis.vulgivaga.1.1.chr13.g00154.t01#4    mNeoVis.evergladensis.1.1.chr13.g00158.t01#3    mNeoVis.3.1.1.chr13.g00114.t01#6    mNeoVis.evergladensis.2.1.chr13.g00160.t01#2    mNeoVis.vulgivaga.2.1.chr13.g00158.t01#1   mNeoVis.2.1.1.chr13_hap1.g00162.t01#5       7531   125253815:   mNeoVis.vulgivaga.2.1.chr13.g00159.t01#1    mNeoVis.3.1.1.chr13.g00115.t01#6    mNeoVis.evergladensis.1.1.chr13.g00159.t01#3    mNeoVis.vulgivaga.1.1.chr13.g00155.t01#4    mNeoVis.2.1.1.chr13_hap1.g00163.t01#5   mNeoVis.evergladensis.2.1.chr13.g00161.t01#2       7532   125253816:   mNeoVis.vulgivaga.2.1.chr5.g01596.t01#1    mNeoVis.evergladensis.1.1.chr5.g01599.t01#3    mNeoVis.vulgivaga.1.1.chr5.g01572.t01#4    mNeoVis.2.1.1.chr2_hap1.g01654.t01#5    mNeoVis.3.1.1.chr5.g01576.t01#6   mNeoVis.evergladensis.2.1.chr5.g01594.t01#2       7533   125253817:   mNeoVis.vulgivaga.2.1.chr5.g01597.t01#1    mNeoVis.2.1.1.chr2_hap1.g01655.t01#5    mNeoVis.3.1.1.chr5.g01577.t01#6    mNeoVis.evergladensis.2.1.chr5.g01595.t01#2    mNeoVis.evergladensis.1.1.chr5.g01600.t01#3   mNeoVis.vulgivaga.1.1.chr5.g01573.t01#4       7534   125253818:   mNeoVis.evergladensis.1.1.chr13.g00160.t01#3    mNeoVis.evergladensis.2.1.chr13.g00162.t01#2    mNeoVis.3.1.1.chr13.g00116.t01#6    mNeoVis.vulgivaga.2.1.chr13.g00160.t01#1    mNeoVis.2.1.1.chr13_hap1.g00164.t01#5   mNeoVis.vulgivaga.1.1.chr13.g00156.t01#4       7535   125253819:   mNeoVis.vulgivaga.2.1.chr13.g00161.t01#1    mNeoVis.evergladensis.1.1.chr13.g00161.t01#3   mNeoVis.evergladensis.2.1.chr13.g00163.t01#2         7536   125253820:   mNeoVis.2.1.1.chr2_hap1.g01656.t01#5    mNeoVis.evergladensis.2.1.chr5.g01596.t01#2    mNeoVis.evergladensis.1.1.chr5.g01601.t01#3    mNeoVis.vulgivaga.2.1.chr5.g01598.t01#1   mNeoVis.vulgivaga.1.1.chr5.g01574.t01#4       7537   125253821:   mNeoVis.vulgivaga.2.1.chr13.g00162.t01#1    mNeoVis.evergladensis.2.1.chr13.g00164.t01#2    mNeoVis.2.1.1.chr13_hap1.g00166.t01#5    mNeoVis.3.1.1.chr13.g00118.t01#6    mNeoVis.vulgivaga.1.1.chr13.g00158.t01#4   mNeoVis.evergladensis.1.1.chr13.g00162.t01#3       7538   125253822:   mNeoVis.evergladensis.1.1.chr5.g01602.t01#3    mNeoVis.evergladensis.2.1.chr5.g01597.t01#2    mNeoVis.vulgivaga.1.1.chr5.g01576.t01#4    mNeoVis.3.1.1.chr5.g01579.t01#6    mNeoVis.2.1.1.chr2_hap1.g01657.t01#5    mNeoVis.vulgivaga.1.1.chr5.g01575.t01#4    mNeoVis.vulgivaga.2.1.chr5.g01600.t01#1   mNeoVis.vulgivaga.2.1.chr5.g01599.t01#1     7539   125253823:   mNeoVis.2.1.1.chr13_hap1.g00167.t01#5    mNeoVis.vulgivaga.2.1.chr13.g00163.t01#1    mNeoVis.vulgivaga.1.1.chr13.g00159.t01#4    mNeoVis.3.1.1.chr13.g00119.t01#6    mNeoVis.evergladensis.1.1.chr13.g00163.t01#3   mNeoVis.evergladensis.2.1.chr13.g00165.t01#2       7540   125253824:   mNeoVis.vulgivaga.2.1.chr5.g01601.t01#1    mNeoVis.3.1.1.chr5.g01580.t01#6    mNeoVis.evergladensis.1.1.chr5.g01603.t01#3    mNeoVis.2.1.1.chr2_hap1.g01658.t01#5    mNeoVis.evergladensis.2.1.chr5.g01598.t01#2   mNeoVis.vulgivaga.1.1.chr5.g01577.t01#4       7541   125253825:   mNeoVis.2.1.1.chr13_hap1.g00168.t01#5    mNeoVis.evergladensis.2.1.chr13.g00166.t01#2    mNeoVis.3.1.1.chr13.g00120.t01#6    mNeoVis.vulgivaga.1.1.chr13.g00160.t01#4    mNeoVis.vulgivaga.2.1.chr13.g00164.t01#1   mNeoVis.evergladensis.1.1.chr13.g00164.t01#3       7542   125253826:   mNeoVis.evergladensis.2.1.chr15.g00072.t01#2    mNeoVis.3.1.1.chr15.g00065.t01#6    mNeoVis.2.1.1.chr15_hap1.g00071.t01#5    mNeoVis.vulgivaga.2.1.chr15.g00073.t01#1    mNeoVis.evergladensis.1.1.chr15.g00542.t01#3   mNeoVis.vulgivaga.1.1.chr15.g00072.t01#4       7543   125253827:   mNeoVis.2.1.1.chr4_hap1.g01100.t01#5    mNeoVis.evergladensis.1.1.chr6.g01101.t01#3    mNeoVis.3.1.1.chr6.g01090.t01#6    mNeoVis.vulgivaga.1.1.chr6.g01092.t01#4    mNeoVis.evergladensis.2.1.chr6.g01098.t01#2   mNeoVis.vulgivaga.2.1.chr6.g01090.t01#1       7544   125253828:   mNeoVis.3.1.1.chr5.g01581.t01#6    mNeoVis.evergladensis.1.1.chr5.g01604.t01#3    mNeoVis.evergladensis.2.1.chr5.g01599.t01#2    mNeoVis.vulgivaga.1.1.chr5.g01578.t01#4    mNeoVis.2.1.1.chr2_hap1.g01659.t01#5    mNeoVis.vulgivaga.1.1.chr5.g01579.t01#4   mNeoVis.vulgivaga.2.1.chr5.g01602.t01#1     7545   125253829:   mNeoVis.2.1.1.chr13_hap1.g00169.t01#5    mNeoVis.3.1.1.chr13.g00121.t01#6    mNeoVis.vulgivaga.1.1.chr13.g00161.t01#4    mNeoVis.vulgivaga.2.1.chr13.g00165.t01#1    mNeoVis.evergladensis.2.1.chr13.g00167.t01#2   mNeoVis.evergladensis.1.1.chr13.g00165.t01#3       7546   125253830:   mNeoVis.2.1.1.chr2_hap1.g01660.t01#5    mNeoVis.vulgivaga.2.1.chr5.g01603.t01#1    mNeoVis.evergladensis.2.1.chr5.g01600.t01#2    mNeoVis.3.1.1.chr5.g01582.t01#6    mNeoVis.evergladensis.1.1.chr5.g01605.t01#3   mNeoVis.vulgivaga.1.1.chr5.g01580.t01#4       7547   125253831:   mNeoVis.vulgivaga.1.1.chr13.g00162.t01#4    mNeoVis.evergladensis.1.1.chr13.g00166.t01#3    mNeoVis.2.1.1.chr13_hap1.g00170.t01#5    mNeoVis.vulgivaga.2.1.chr13.g00166.t01#1   mNeoVis.evergladensis.2.1.chr13.g00168.t01#2       7548   125253832:   mNeoVis.vulgivaga.2.1.chr5.g01605.t01#1    mNeoVis.evergladensis.2.1.chr5.g01602.t01#2    mNeoVis.2.1.1.chr2_hap1.g01662.t01#5    mNeoVis.vulgivaga.1.1.chr5.g01582.t01#4   mNeoVis.evergladensis.1.1.chr5.g01607.t01#3       7549   125253833:   mNeoVis.evergladensis.1.1.chr5.g01608.t01#3    mNeoVis.evergladensis.1.1.chr5.g01609.t01#3    mNeoVis.3.1.1.chr5.g01585.t01#6    mNeoVis.2.1.1.chr2_hap1.g01663.t01#5    mNeoVis.evergladensis.2.1.chr5.g01604.t01#2    mNeoVis.evergladensis.2.1.chr5.g01603.t01#2    mNeoVis.vulgivaga.2.1.chr5.g01606.t01#1    mNeoVis.2.1.1.chr2_hap1.g01664.t01#5    mNeoVis.3.1.1.chr5.g01586.t01#6      7550   125253834:   mNeoVis.3.1.1.chr5.g01588.t01#6    mNeoVis.evergladensis.2.1.chr5.g01606.t01#2    mNeoVis.vulgivaga.2.1.chr5.g01608.t01#1    mNeoVis.2.1.1.chr2_hap1.g01665.t01#5    mNeoVis.vulgivaga.1.1.chr5.g01585.t01#4   mNeoVis.evergladensis.1.1.chr5.g01611.t01#3       7551   125253835:   mNeoVis.2.1.1.chr15_hap1.g00072.t01#5    mNeoVis.3.1.1.chr15.g00066.t01#6    mNeoVis.vulgivaga.1.1.chr15.g00073.t01#4    mNeoVis.evergladensis.2.1.chr15.g00073.t01#2    mNeoVis.evergladensis.1.1.chr15.g00541.t01#3   mNeoVis.vulgivaga.2.1.chr15.g00074.t01#1       7552   125253836:   mNeoVis.vulgivaga.2.1.chr6.g01091.t01#1    mNeoVis.2.1.1.chr4_hap1.g01101.t01#5    mNeoVis.evergladensis.1.1.chr6.g01102.t01#3    mNeoVis.vulgivaga.1.1.chr6.g01093.t01#4    mNeoVis.evergladensis.2.1.chr6.g01099.t01#2   mNeoVis.3.1.1.chr6.g01091.t01#6       7553   125253837:   mNeoVis.3.1.1.chr5.g01589.t01#6    mNeoVis.evergladensis.1.1.chr5.g01612.t01#3    mNeoVis.vulgivaga.1.1.chr5.g01586.t01#4    mNeoVis.evergladensis.2.1.chr5.g01607.t01#2    mNeoVis.2.1.1.chr2_hap1.g01666.t01#5   mNeoVis.vulgivaga.2.1.chr5.g01609.t01#1       7554   125253838:   mNeoVis.vulgivaga.1.1.chr2.g01807.t01#4    mNeoVis.2.1.1.chr5_hap1.g01815.t01#5    mNeoVis.3.1.1.chr2.g01721.t01#6    mNeoVis.evergladensis.2.1.chr2.g01801.t01#2    mNeoVis.evergladensis.1.1.chr2.g01813.t01#3   mNeoVis.vulgivaga.2.1.chr2.g01815.t01#1       7555   125253839:   mNeoVis.vulgivaga.2.1.chr5.g01610.t01#1    mNeoVis.evergladensis.1.1.chr5.g01613.t01#3    mNeoVis.vulgivaga.1.1.chr5.g01587.t01#4    mNeoVis.evergladensis.2.1.chr5.g01608.t01#2    mNeoVis.2.1.1.chr2_hap1.g01667.t01#5   mNeoVis.3.1.1.chr5.g01590.t01#6       7556   125253840:   mNeoVis.evergladensis.1.1.chr13.g00167.t01#3    mNeoVis.3.1.1.chr13.g00123.t01#6    mNeoVis.vulgivaga.1.1.chr13.g00163.t01#4    mNeoVis.evergladensis.2.1.chr13.g00169.t01#2    mNeoVis.2.1.1.chr13_hap1.g00171.t01#5   mNeoVis.vulgivaga.2.1.chr13.g00167.t01#1       7557   125253841:   mNeoVis.3.1.1.chr13.g00124.t01#6    mNeoVis.evergladensis.1.1.chr13.g00168.t01#3    mNeoVis.2.1.1.chr13_hap1.g00172.t01#5    mNeoVis.vulgivaga.2.1.chr13.g00168.t01#1    mNeoVis.vulgivaga.1.1.chr13.g00164.t01#4   mNeoVis.evergladensis.2.1.chr13.g00170.t01#2       7558   125253842:   mNeoVis.evergladensis.2.1.chr5.g01609.t01#2    mNeoVis.vulgivaga.2.1.chr5.g01611.t01#1    mNeoVis.evergladensis.1.1.chr5.g01614.t01#3    mNeoVis.3.1.1.chr5.g01591.t01#6    mNeoVis.2.1.1.chr2_hap1.g01668.t01#5   mNeoVis.vulgivaga.1.1.chr5.g01588.t01#4       7559   125253843:   mNeoVis.vulgivaga.2.1.chr5.g01612.t01#1    mNeoVis.3.1.1.chr5.g01592.t01#6    mNeoVis.evergladensis.1.1.chr5.g01615.t01#3    mNeoVis.vulgivaga.1.1.chr5.g01589.t01#4    mNeoVis.2.1.1.chr2_hap1.g01669.t01#5   mNeoVis.evergladensis.2.1.chr5.g01610.t01#2       7560   125253844:   mNeoVis.2.1.1.chr2_hap1.g01670.t01#5    mNeoVis.evergladensis.2.1.chr5.g01611.t01#2    mNeoVis.vulgivaga.1.1.chr5.g01590.t01#4    mNeoVis.evergladensis.1.1.chr5.g01616.t01#3    mNeoVis.vulgivaga.2.1.chr5.g01613.t01#1   mNeoVis.3.1.1.chr5.g01593.t01#6       7561   125253845:   mNeoVis.2.1.1.chr13_hap1.g00173.t01#5    mNeoVis.vulgivaga.2.1.chr13.g00169.t01#1    mNeoVis.vulgivaga.1.1.chr13.g00165.t01#4    mNeoVis.evergladensis.1.1.chr13.g00169.t01#3    mNeoVis.evergladensis.2.1.chr13.g00171.t01#2   mNeoVis.vulgivaga.1.1.chr13.g00166.t01#4       7562   125253846:   mNeoVis.vulgivaga.2.1.chr13.g00170.t01#1    mNeoVis.evergladensis.1.1.chr13.g00170.t01#3    mNeoVis.evergladensis.1.1.chr13.g00171.t01#3    mNeoVis.vulgivaga.1.1.chr13.g00167.t01#4    mNeoVis.evergladensis.2.1.chr13.g00173.t01#2    mNeoVis.2.1.1.chr13_hap1.g00174.t01#5    mNeoVis.3.1.1.chr13.g00125.t01#6    mNeoVis.2.1.1.chr13_hap1.g00175.t01#5    mNeoVis.evergladensis.2.1.chr13.g00172.t01#2      7563   125253847:   mNeoVis.vulgivaga.1.1.chr5.g01591.t01#4    mNeoVis.evergladensis.1.1.chr5.g01617.t01#3    mNeoVis.3.1.1.chr5.g01594.t01#6    mNeoVis.vulgivaga.2.1.chr5.g01614.t01#1    mNeoVis.2.1.1.chr2_hap1.g01671.t01#5   mNeoVis.evergladensis.2.1.chr5.g01612.t01#2       7564   125253848:   mNeoVis.evergladensis.1.1.chr5.g01618.t01#3    mNeoVis.3.1.1.chr5.g01595.t01#6    mNeoVis.evergladensis.2.1.chr5.g01613.t01#2    mNeoVis.2.1.1.chr2_hap1.g01672.t01#5    mNeoVis.vulgivaga.2.1.chr5.g01615.t01#1   mNeoVis.vulgivaga.1.1.chr5.g01592.t01#4       7565   125253849:   mNeoVis.2.1.1.chr13_hap1.g00176.t01#5    mNeoVis.3.1.1.chr13.g00126.t01#6    mNeoVis.evergladensis.2.1.chr13.g00174.t01#2    mNeoVis.vulgivaga.1.1.chr13.g00168.t01#4    mNeoVis.evergladensis.1.1.chr13.g00172.t01#3   mNeoVis.vulgivaga.2.1.chr13.g00171.t01#1       7566   125253850:  mNeoVis.vulgivaga.2.1.chr15.g00075.t01#1               7567   125253851:   mNeoVis.evergladensis.1.1.chr13.g00173.t01#3    mNeoVis.vulgivaga.1.1.chr13.g00169.t01#4    mNeoVis.vulgivaga.2.1.chr13.g00172.t01#1    mNeoVis.evergladensis.2.1.chr13.g00175.t01#2    mNeoVis.2.1.1.chr13_hap1.g00177.t01#5   mNeoVis.3.1.1.chr13.g00127.t01#6       7568   125253852:   mNeoVis.evergladensis.2.1.chr6.g01100.t01#2    mNeoVis.vulgivaga.2.1.chr6.g01092.t01#1    mNeoVis.vulgivaga.1.1.chr6.g01094.t01#4    mNeoVis.2.1.1.chr4_hap1.g01102.t01#5   mNeoVis.evergladensis.1.1.chr6.g01103.t01#3       7569   125253853:  mNeoVis.vulgivaga.2.1.chr5.g01616.t01#1               7570   125253854:   mNeoVis.evergladensis.2.1.chr13.g00176.t01#2    mNeoVis.3.1.1.chr13.g00128.t01#6    mNeoVis.vulgivaga.2.1.chr13.g00173.t01#1    mNeoVis.vulgivaga.1.1.chr13.g00170.t01#4    mNeoVis.evergladensis.1.1.chr13.g00174.t01#3   mNeoVis.2.1.1.chr13_hap1.g00178.t01#5       7571   125253855:   mNeoVis.vulgivaga.2.1.chr5.g01617.t01#1    mNeoVis.2.1.1.chr2_hap1.g01674.t01#5    mNeoVis.evergladensis.1.1.chr5.g01620.t01#3    mNeoVis.evergladensis.2.1.chr5.g01615.t01#2   mNeoVis.vulgivaga.1.1.chr5.g01594.t01#4       7572   125253856:   mNeoVis.3.1.1.chr13.g00130.t01#6    mNeoVis.vulgivaga.1.1.chr13.g00171.t01#4    mNeoVis.vulgivaga.2.1.chr13.g00174.t01#1    mNeoVis.2.1.1.chr13_hap1.g00179.t01#5    mNeoVis.evergladensis.1.1.chr13.g00175.t01#3   mNeoVis.evergladensis.2.1.chr13.g00177.t01#2       7573   125253857:   mNeoVis.2.1.1.chr2_hap1.g01675.t01#5    mNeoVis.3.1.1.chr5.g01598.t01#6    mNeoVis.evergladensis.2.1.chr5.g01616.t01#2    mNeoVis.evergladensis.1.1.chr5.g01621.t01#3    mNeoVis.vulgivaga.2.1.chr5.g01618.t01#1   mNeoVis.vulgivaga.1.1.chr5.g01595.t01#4       7574   125253858:   mNeoVis.2.1.1.chr13_hap1.g00180.t01#5    mNeoVis.3.1.1.chr2.g00799.t01#6    mNeoVis.vulgivaga.2.1.chr13.g00175.t01#1    mNeoVis.vulgivaga.1.1.chr13.g00172.t01#4    mNeoVis.3.1.1.chr13.g00131.t01#6    mNeoVis.evergladensis.2.1.chr13.g00178.t01#2   mNeoVis.evergladensis.1.1.chr13.g00176.t01#3     7575   125253859:   mNeoVis.vulgivaga.2.1.chr15.g00076.t01#1    mNeoVis.vulgivaga.1.1.chr15.g00075.t01#4    mNeoVis.evergladensis.2.1.chr15.g00075.t01#2    mNeoVis.2.1.1.chr15_hap1.g00074.t01#5    mNeoVis.evergladensis.1.1.chr15.g00539.t01#3   mNeoVis.3.1.1.chr15.g00068.t01#6       7576   125253860:   mNeoVis.evergladensis.2.1.chr9.g00756.t01#2    mNeoVis.evergladensis.1.1.chr8.g00750.t01#3    mNeoVis.vulgivaga.1.1.chr8.g00744.t01#4    mNeoVis.vulgivaga.2.1.chr8.g00760.t01#1    mNeoVis.2.1.1.chr8_hap1.g00769.t01#5   mNeoVis.3.1.1.chr8.g00719.t01#6       7577   125253861:   mNeoVis.vulgivaga.2.1.chr6.g01093.t01#1    mNeoVis.2.1.1.chr4_hap1.g01125.t01#5    mNeoVis.evergladensis.1.1.chr6.g01104.t01#3    mNeoVis.evergladensis.2.1.chr6.g01101.t01#2    mNeoVis.3.1.1.chr6.g01092.t01#6   mNeoVis.vulgivaga.1.1.chr6.g01091.t01#4       7578   125253862:   mNeoVis.3.1.1.chr5.g01599.t01#6    mNeoVis.evergladensis.1.1.chr5.g01622.t01#3    mNeoVis.vulgivaga.2.1.chr5.g01619.t01#1    mNeoVis.evergladensis.2.1.chr5.g01617.t01#2    mNeoVis.2.1.1.chr2_hap1.g01676.t01#5   mNeoVis.vulgivaga.1.1.chr5.g01596.t01#4       7579   125253863:   mNeoVis.evergladensis.2.1.chr11.g00692.t01#2    mNeoVis.evergladensis.1.1.chr11.g00700.t01#3    mNeoVis.3.1.1.chr11.g00671.t01#6    mNeoVis.vulgivaga.2.1.chr11.g00709.t01#1   mNeoVis.vulgivaga.1.1.chr11.g00690.t01#4       7580   125253864:   mNeoVis.evergladensis.2.1.chr1.g01910.t01#2    mNeoVis.vulgivaga.1.1.chr1.g01916.t01#4    mNeoVis.evergladensis.1.1.chr1.g01910.t01#3    mNeoVis.2.1.1.chr1_hap1.g01944.t01#5   mNeoVis.vulgivaga.2.1.chr1.g01907.t01#1       7581   125253865:   mNeoVis.evergladensis.1.1.chr1.g01909.t01#3   mNeoVis.3.1.1.chr1.g01893.t01#6             7582   125253866:   mNeoVis.evergladensis.1.1.chr13.g00178.t01#3    mNeoVis.2.1.1.chr13_hap1.g00182.t01#5    mNeoVis.evergladensis.2.1.chr13.g00180.t01#2    mNeoVis.vulgivaga.1.1.chr13.g00174.t01#4   mNeoVis.vulgivaga.2.1.chr13.g00177.t01#1       7583   125253867:   mNeoVis.2.1.1.chr2_hap1.g01678.t01#5    mNeoVis.vulgivaga.2.1.chr5.g01621.t01#1    mNeoVis.vulgivaga.1.1.chr5.g01598.t01#4    mNeoVis.evergladensis.2.1.chr5.g01619.t01#2    mNeoVis.evergladensis.1.1.chr5.g01624.t01#3   mNeoVis.3.1.1.chr5.g01601.t01#6       7584   125253868:   mNeoVis.2.1.1.chr13_hap1.g00183.t01#5    mNeoVis.evergladensis.2.1.chr13.g00181.t01#2    mNeoVis.vulgivaga.2.1.chr13.g00178.t01#1    mNeoVis.evergladensis.1.1.chr13.g00179.t01#3    mNeoVis.3.1.1.chr13.g00134.t01#6   mNeoVis.vulgivaga.1.1.chr13.g00175.t01#4       7585   125253869:   mNeoVis.evergladensis.2.1.chr5.g01620.t01#2    mNeoVis.vulgivaga.1.1.chr5.g01599.t01#4    mNeoVis.evergladensis.1.1.chr5.g01625.t01#3    mNeoVis.3.1.1.chr5.g01602.t01#6    mNeoVis.2.1.1.chr2_hap1.g01679.t01#5   mNeoVis.vulgivaga.2.1.chr5.g01622.t01#1       7586   125253870:   mNeoVis.vulgivaga.2.1.chr5.g01623.t01#1    mNeoVis.evergladensis.1.1.chr5.g01626.t01#3    mNeoVis.2.1.1.chr2_hap1.g01680.t01#5    mNeoVis.evergladensis.2.1.chr5.g01621.t01#2    mNeoVis.vulgivaga.1.1.chr5.g01600.t01#4   mNeoVis.3.1.1.chr3.g00418.t01#6       7587   125253871:   mNeoVis.vulgivaga.2.1.chr5.g01624.t01#1    mNeoVis.evergladensis.1.1.chr5.g01627.t01#3    mNeoVis.vulgivaga.1.1.chr5.g01601.t01#4    mNeoVis.2.1.1.chr2_hap1.g01681.t01#5    mNeoVis.3.1.1.chr5.g01604.t01#6   mNeoVis.evergladensis.2.1.chr5.g01622.t01#2       7588   125253872:   mNeoVis.vulgivaga.1.1.chr5.g01602.t01#4    mNeoVis.2.1.1.chr2_hap1.g01682.t01#5    mNeoVis.3.1.1.chr5.g01605.t01#6    mNeoVis.evergladensis.1.1.chr5.g01628.t01#3    mNeoVis.vulgivaga.2.1.chr5.g01625.t01#1   mNeoVis.evergladensis.2.1.chr5.g01623.t01#2       7589   125253873:   mNeoVis.evergladensis.1.1.chr5.g01629.t01#3    mNeoVis.3.1.1.chr5.g01606.t01#6    mNeoVis.2.1.1.chr2_hap1.g01683.t01#5    mNeoVis.evergladensis.2.1.chr5.g01624.t01#2    mNeoVis.vulgivaga.2.1.chr5.g01626.t01#1   mNeoVis.vulgivaga.1.1.chr5.g01603.t01#4       7590   125253874:   mNeoVis.evergladensis.1.1.chr5.g01630.t01#3    mNeoVis.vulgivaga.2.1.chr5.g01627.t01#1    mNeoVis.evergladensis.2.1.chr5.g01625.t01#2    mNeoVis.3.1.1.chr5.g01607.t01#6    mNeoVis.2.1.1.chr2_hap1.g01684.t01#5   mNeoVis.vulgivaga.1.1.chr5.g01604.t01#4       7591   125253875:   mNeoVis.evergladensis.1.1.chr13.g00180.t01#3    mNeoVis.evergladensis.2.1.chr13.g00182.t01#2    mNeoVis.vulgivaga.2.1.chr13.g00179.t01#1    mNeoVis.2.1.1.chr13_hap1.g00184.t01#5    mNeoVis.3.1.1.chr13.g00136.t01#6   mNeoVis.vulgivaga.1.1.chr13.g00176.t01#4       7592   125253876:   mNeoVis.3.1.1.chr5.g01608.t01#6    mNeoVis.vulgivaga.2.1.chr5.g01628.t01#1    mNeoVis.evergladensis.1.1.chr1.g01854.t01#3    mNeoVis.vulgivaga.2.1.chr3.g01228.t01#1    mNeoVis.vulgivaga.1.1.chr1.g01859.t01#4    mNeoVis.2.1.1.chr1_hap1.g01887.t01#5    mNeoVis.evergladensis.2.1.chr5.g01626.t01#2    mNeoVis.vulgivaga.1.1.chr2.g00814.t01#4    mNeoVis.2.1.1.chr2_hap1.g01685.t01#5      7593   125253877:   mNeoVis.3.1.1.chr5.g01609.t01#6    mNeoVis.evergladensis.2.1.chr5.g01627.t01#2    mNeoVis.vulgivaga.2.1.chr5.g01629.t01#1    mNeoVis.vulgivaga.1.1.chr5.g01606.t01#4    mNeoVis.evergladensis.1.1.chr5.g01632.t01#3   mNeoVis.2.1.1.chr2_hap1.g01686.t01#5       7594   125253878:   mNeoVis.evergladensis.1.1.chr5.g01633.t01#3    mNeoVis.vulgivaga.1.1.chr5.g01607.t01#4    mNeoVis.2.1.1.chr2_hap1.g01687.t01#5    mNeoVis.vulgivaga.2.1.chr5.g01630.t01#1    mNeoVis.3.1.1.chr5.g01610.t01#6   mNeoVis.evergladensis.2.1.chr5.g01628.t01#2       7595   125253879:   mNeoVis.vulgivaga.1.1.chr13.g00177.t01#4    mNeoVis.vulgivaga.2.1.chr13.g00180.t01#1    mNeoVis.evergladensis.2.1.chr13.g00183.t01#2    mNeoVis.2.1.1.chr13_hap1.g00185.t01#5    mNeoVis.3.1.1.chr13.g00137.t01#6   mNeoVis.evergladensis.1.1.chr13.g00181.t01#3       7596   125253880:   mNeoVis.vulgivaga.2.1.chr5.g01631.t01#1    mNeoVis.evergladensis.1.1.chr5.g01634.t01#3    mNeoVis.vulgivaga.1.1.chr5.g01608.t01#4    mNeoVis.2.1.1.chr2_hap1.g01688.t01#5    mNeoVis.evergladensis.2.1.chr5.g01629.t01#2   mNeoVis.3.1.1.chr5.g01611.t01#6       7597   125253881:   mNeoVis.2.1.1.chr15_hap1.g00075.t01#5    mNeoVis.vulgivaga.2.1.chr15.g00077.t01#1    mNeoVis.vulgivaga.1.1.chr15.g00076.t01#4    mNeoVis.evergladensis.1.1.chr15.g00538.t01#3    mNeoVis.3.1.1.chr15.g00069.t01#6   mNeoVis.evergladensis.2.1.chr15.g00076.t01#2       7598   125253882:   mNeoVis.3.1.1.chr13.g00138.t01#6    mNeoVis.evergladensis.2.1.chr13.g00184.t01#2    mNeoVis.evergladensis.1.1.chr13.g00182.t01#3    mNeoVis.vulgivaga.2.1.chr13.g00181.t01#1    mNeoVis.vulgivaga.1.1.chr13.g00178.t01#4   mNeoVis.2.1.1.chr13_hap1.g00186.t01#5       7599   125253883:   mNeoVis.vulgivaga.2.1.chr5.g01632.t01#1    mNeoVis.vulgivaga.1.1.chr5.g01609.t01#4    mNeoVis.3.1.1.chr5.g01612.t01#6    mNeoVis.evergladensis.2.1.chr5.g01630.t01#2    mNeoVis.evergladensis.1.1.chr5.g01635.t01#3   mNeoVis.2.1.1.chr2_hap1.g01689.t01#5       7600   125253884:   mNeoVis.evergladensis.1.1.chr13.g00183.t01#3    mNeoVis.2.1.1.chr13_hap1.g00187.t01#5    mNeoVis.vulgivaga.2.1.chr13.g00182.t01#1    mNeoVis.evergladensis.2.1.chr13.g00185.t01#2    mNeoVis.vulgivaga.1.1.chr13.g00179.t01#4   mNeoVis.3.1.1.chr13.g00139.t01#6       7601   125253885:   mNeoVis.vulgivaga.2.1.chr13.g00183.t01#1    mNeoVis.vulgivaga.1.1.chr13.g00180.t01#4    mNeoVis.evergladensis.2.1.chr13.g00186.t01#2    mNeoVis.2.1.1.chr13_hap1.g00188.t01#5    mNeoVis.3.1.1.chr13.g00140.t01#6   mNeoVis.evergladensis.1.1.chr13.g00184.t01#3       7602   125253886:   mNeoVis.2.1.1.chr2_hap1.g01690.t01#5    mNeoVis.vulgivaga.2.1.chr5.g01633.t01#1   mNeoVis.vulgivaga.1.1.chr5.g01610.t01#4           7603   125253887:   mNeoVis.vulgivaga.2.1.chr13.g00184.t01#1    mNeoVis.evergladensis.2.1.chr13.g00187.t01#2    mNeoVis.evergladensis.1.1.chr13.g00185.t01#3    mNeoVis.2.1.1.chr13_hap1.g00189.t01#5   mNeoVis.vulgivaga.1.1.chr13.g00181.t01#4       7604   125253888:   mNeoVis.vulgivaga.1.1.chr5.g01611.t01#4    mNeoVis.evergladensis.1.1.chr5.g01636.t01#3    mNeoVis.evergladensis.2.1.chr5.g01631.t01#2    mNeoVis.vulgivaga.2.1.chr5.g01634.t01#1    mNeoVis.3.1.1.chr5.g01613.t01#6   mNeoVis.2.1.1.chr2_hap1.g01691.t01#5       7605   125253889:   mNeoVis.evergladensis.2.1.chr1.g01891.t01#2    mNeoVis.2.1.1.chr1_hap1.g01924.t01#5    mNeoVis.3.1.1.chr1.g01874.t01#6    mNeoVis.vulgivaga.2.1.chr1.g01888.t01#1    mNeoVis.evergladensis.1.1.chr1.g01893.t01#3   mNeoVis.vulgivaga.1.1.chr1.g01898.t01#4       7606   125253890:   mNeoVis.evergladensis.2.1.chr1.g00225.t01#2    mNeoVis.3.1.1.chr1.g00228.t01#6   mNeoVis.2.1.1.chr1_hap1.g00228.t01#5           7607   125253891:   mNeoVis.vulgivaga.1.1.chr5.g01612.t01#4    mNeoVis.evergladensis.1.1.chr5.g01637.t01#3    mNeoVis.2.1.1.chr2_hap1.g01692.t01#5    mNeoVis.evergladensis.2.1.chr5.g01632.t01#2   mNeoVis.vulgivaga.2.1.chr5.g01635.t01#1       7608   125253892:   mNeoVis.vulgivaga.2.1.chr5.g01636.t01#1    mNeoVis.vulgivaga.1.1.chr5.g01613.t01#4    mNeoVis.evergladensis.1.1.chr5.g01638.t01#3    mNeoVis.3.1.1.chr5.g01614.t01#6    mNeoVis.evergladensis.2.1.chr5.g01633.t01#2   mNeoVis.2.1.1.chr2_hap1.g01693.t01#5       7609   125253893:   mNeoVis.3.1.1.chr13.g00142.t01#6    mNeoVis.evergladensis.2.1.chr13.g00188.t01#2    mNeoVis.evergladensis.1.1.chr13.g00186.t01#3    mNeoVis.vulgivaga.1.1.chr13.g00182.t01#4    mNeoVis.2.1.1.chr13_hap1.g00190.t01#5   mNeoVis.vulgivaga.2.1.chr13.g00185.t01#1       7610   125253894:   mNeoVis.2.1.1.chr2_hap1.g01694.t01#5    mNeoVis.evergladensis.2.1.chr5.g01634.t01#2    mNeoVis.evergladensis.1.1.chr5.g01639.t01#3    mNeoVis.3.1.1.chr5.g01615.t01#6    mNeoVis.vulgivaga.1.1.chr5.g01614.t01#4   mNeoVis.vulgivaga.2.1.chr5.g01637.t01#1       7611   125253895:   mNeoVis.vulgivaga.2.1.chr5.g01638.t01#1    mNeoVis.evergladensis.2.1.chr5.g01635.t01#2    mNeoVis.2.1.1.chr2_hap1.g01695.t01#5    mNeoVis.evergladensis.1.1.chr5.g01640.t01#3    mNeoVis.3.1.1.chr5.g01616.t01#6   mNeoVis.vulgivaga.1.1.chr5.g01615.t01#4       7612   125253896:   mNeoVis.3.1.1.chr13.g00144.t01#6    mNeoVis.evergladensis.1.1.chr13.g00188.t01#3    mNeoVis.2.1.1.chr13_hap1.g00192.t01#5   mNeoVis.vulgivaga.1.1.chr13.g00184.t01#4       7613   125253897:   mNeoVis.vulgivaga.2.1.chr13.g00186.t01#1   mNeoVis.evergladensis.2.1.chr13.g00189.t01#2           7614   125253898:   mNeoVis.evergladensis.2.1.chr6.g01102.t01#2    mNeoVis.2.1.1.chr4_hap1.g01126.t01#5    mNeoVis.evergladensis.1.1.chr6.g01105.t01#3    mNeoVis.vulgivaga.1.1.chr6.g01095.t01#4    mNeoVis.3.1.1.chr6.g01093.t01#6   mNeoVis.vulgivaga.2.1.chr6.g01094.t01#1       7615   125253899:   mNeoVis.vulgivaga.1.1.chr5.g01616.t01#4    mNeoVis.2.1.1.chr2_hap1.g01696.t01#5    mNeoVis.evergladensis.1.1.chr5.g01641.t01#3    mNeoVis.evergladensis.2.1.chr5.g01636.t01#2    mNeoVis.3.1.1.chr5.g01617.t01#6   mNeoVis.vulgivaga.2.1.chr5.g01639.t01#1       7616   125253900:   mNeoVis.vulgivaga.2.1.chr5.g01640.t01#1    mNeoVis.evergladensis.2.1.chr5.g01637.t01#2    mNeoVis.2.1.1.chr2_hap1.g01697.t01#5    mNeoVis.evergladensis.1.1.chr5.g01642.t01#3    mNeoVis.3.1.1.chr5.g01618.t01#6   mNeoVis.vulgivaga.1.1.chr5.g01617.t01#4       7617   125253901:   mNeoVis.2.1.1.chr15_hap1.g00076.t01#5    mNeoVis.vulgivaga.2.1.chr15.g00078.t01#1    mNeoVis.3.1.1.chr15.g00070.t01#6    mNeoVis.vulgivaga.1.1.chr15.g00077.t01#4    mNeoVis.evergladensis.1.1.chr15.g00537.t01#3   mNeoVis.evergladensis.2.1.chr15.g00077.t01#2       7618   125253902:   mNeoVis.vulgivaga.1.1.chr13.g00183.t01#4    mNeoVis.3.1.1.chr13.g00143.t01#6    mNeoVis.vulgivaga.2.1.chr13.g00187.t01#1    mNeoVis.evergladensis.2.1.chr13.g00190.t01#2    mNeoVis.2.1.1.chr13_hap1.g00191.t01#5   mNeoVis.evergladensis.1.1.chr13.g00187.t01#3       7619   125253903:   mNeoVis.3.1.1.chr5.g01621.t01#6    mNeoVis.evergladensis.2.1.chr5.g01639.t01#2    mNeoVis.evergladensis.1.1.chr5.g01644.t01#3    mNeoVis.vulgivaga.1.1.chr5.g01619.t01#4    mNeoVis.vulgivaga.2.1.chr5.g01641.t01#1   mNeoVis.2.1.1.chr2_hap1.g01700.t01#5       7620   125253904:   mNeoVis.vulgivaga.1.1.chr5.g01620.t01#4    mNeoVis.evergladensis.1.1.chr5.g01645.t01#3    mNeoVis.vulgivaga.2.1.chr5.g01642.t01#1    mNeoVis.2.1.1.chr2_hap1.g01701.t01#5    mNeoVis.3.1.1.chr5.g01623.t01#6   mNeoVis.evergladensis.2.1.chr5.g01640.t01#2       7621   125253905:   mNeoVis.vulgivaga.2.1.chr5.g01643.t01#1    mNeoVis.evergladensis.2.1.chr5.g01641.t01#2    mNeoVis.3.1.1.chr5.g01624.t01#6    mNeoVis.vulgivaga.1.1.chr5.g01621.t01#4    mNeoVis.evergladensis.1.1.chr5.g01646.t01#3   mNeoVis.2.1.1.chr2_hap1.g01702.t01#5       7622   125253906:   mNeoVis.evergladensis.1.1.chr5.g01647.t01#3    mNeoVis.2.1.1.chr2_hap1.g01703.t01#5    mNeoVis.vulgivaga.1.1.chr5.g01622.t01#4    mNeoVis.evergladensis.2.1.chr5.g01642.t01#2    mNeoVis.vulgivaga.2.1.chr5.g01644.t01#1   mNeoVis.3.1.1.chr5.g01625.t01#6       7623   125253907:   mNeoVis.vulgivaga.1.1.chr6.g01096.t01#4    mNeoVis.vulgivaga.2.1.chr6.g01095.t01#1    mNeoVis.evergladensis.2.1.chr6.g01103.t01#2    mNeoVis.evergladensis.1.1.chr6.g01106.t01#3   mNeoVis.2.1.1.chr4_hap1.g01127.t01#5       7624   125253908:   mNeoVis.evergladensis.1.1.chr5.g01648.t01#3    mNeoVis.vulgivaga.1.1.chr5.g01623.t01#4    mNeoVis.vulgivaga.2.1.chr5.g01645.t01#1    mNeoVis.2.1.1.chr2_hap1.g01704.t01#5    mNeoVis.evergladensis.2.1.chr5.g01643.t01#2   mNeoVis.3.1.1.chr5.g01626.t01#6       7625   125253909:   mNeoVis.evergladensis.1.1.chr13.g00190.t01#3    mNeoVis.vulgivaga.1.1.chr13.g00186.t01#4    mNeoVis.vulgivaga.2.1.chr13.g00189.t01#1    mNeoVis.2.1.1.chr13_hap1.g00194.t01#5   mNeoVis.evergladensis.2.1.chr13.g00192.t01#2       7626   125253910:   mNeoVis.evergladensis.1.1.chr13.g00120.t01#3    mNeoVis.vulgivaga.1.1.chr13.g00118.t01#4    mNeoVis.3.1.1.chr13.g00071.t01#6    mNeoVis.vulgivaga.2.1.chr13.g00121.t01#1    mNeoVis.2.1.1.chr13_hap1.g00127.t01#5   mNeoVis.evergladensis.2.1.chr13.g00124.t01#2       7627   125253911:   mNeoVis.3.1.1.chr5.g01517.t01#6    mNeoVis.vulgivaga.1.1.chr5.g01513.t01#4    mNeoVis.2.1.1.chr2_hap1.g01596.t01#5    mNeoVis.evergladensis.1.1.chr11.g00800.t01#3    mNeoVis.vulgivaga.2.1.chr5.g01538.t01#1    mNeoVis.evergladensis.1.1.chr5.g01541.t01#3    mNeoVis.3.1.1.chr11.g00772.t01#6   mNeoVis.evergladensis.2.1.chr5.g01535.t01#2     7628   125253912:   mNeoVis.vulgivaga.1.1.chr11.g00791.t01#4    mNeoVis.evergladensis.2.1.chr11.g00792.t01#2    mNeoVis.2.1.1.chr11_hap1.g00809.t01#5   mNeoVis.vulgivaga.2.1.chr11.g00811.t01#1       7629   125253913:   mNeoVis.evergladensis.2.1.chr5.g01536.t01#2    mNeoVis.vulgivaga.1.1.chr5.g01514.t01#4    mNeoVis.vulgivaga.2.1.chr5.g01539.t01#1    mNeoVis.evergladensis.1.1.chr5.g01542.t01#3    mNeoVis.2.1.1.chr2_hap1.g01597.t01#5   mNeoVis.3.1.1.chr5.g01518.t01#6       7630   125253914:   mNeoVis.vulgivaga.2.1.chr5.g01540.t01#1    mNeoVis.3.1.1.chr5.g01519.t01#6    mNeoVis.vulgivaga.1.1.chr5.g01515.t01#4    mNeoVis.evergladensis.2.1.chr5.g01537.t01#2    mNeoVis.evergladensis.1.1.chr5.g01543.t01#3   mNeoVis.2.1.1.chr2_hap1.g01598.t01#5       7631   125253915:   mNeoVis.2.1.1.chr13_hap1.g00128.t01#5    mNeoVis.vulgivaga.1.1.chr13.g00119.t01#4    mNeoVis.3.1.1.chr13.g00072.t01#6    mNeoVis.evergladensis.2.1.chr13.g00125.t01#2    mNeoVis.evergladensis.1.1.chr13.g00121.t01#3   mNeoVis.vulgivaga.2.1.chr13.g00122.t01#1       7632   125253916:   mNeoVis.evergladensis.2.1.chr13.g00126.t01#2    mNeoVis.vulgivaga.2.1.chr13.g00123.t01#1    mNeoVis.evergladensis.1.1.chr13.g00122.t01#3    mNeoVis.2.1.1.chr13_hap1.g00129.t01#5    mNeoVis.vulgivaga.1.1.chr13.g00120.t01#4   mNeoVis.3.1.1.chr13.g00073.t01#6       7633   125253917:   mNeoVis.vulgivaga.2.1.chr5.g01541.t01#1    mNeoVis.evergladensis.2.1.chr5.g01538.t01#2    mNeoVis.vulgivaga.1.1.chr5.g01516.t01#4    mNeoVis.evergladensis.1.1.chr5.g01544.t01#3    mNeoVis.2.1.1.chr2_hap1.g01599.t01#5   mNeoVis.3.1.1.chr5.g01520.t01#6       7634   125253918:   mNeoVis.evergladensis.2.1.chr13.g00127.t01#2    mNeoVis.3.1.1.chr13.g00075.t01#6    mNeoVis.evergladensis.1.1.chr13.g00123.t01#3    mNeoVis.vulgivaga.1.1.chr13.g00121.t01#4    mNeoVis.2.1.1.chr13_hap1.g00130.t01#5    mNeoVis.vulgivaga.2.1.chr13.g00124.t01#1   mNeoVis.3.1.1.chr13.g00074.t01#6       7635   125253919:   mNeoVis.vulgivaga.2.1.chr5.g01542.t01#1    mNeoVis.3.1.1.chr5.g01521.t01#6    mNeoVis.evergladensis.2.1.chr5.g01539.t01#2    mNeoVis.evergladensis.1.1.chr5.g01545.t01#3    mNeoVis.vulgivaga.1.1.chr5.g01517.t01#4   mNeoVis.2.1.1.chr2_hap1.g01600.t01#5       7636   125253920:   mNeoVis.2.1.1.chr2_hap1.g01601.t01#5    mNeoVis.evergladensis.1.1.chr5.g01546.t01#3    mNeoVis.vulgivaga.1.1.chr5.g01518.t01#4    mNeoVis.3.1.1.chr5.g01522.t01#6    mNeoVis.evergladensis.2.1.chr5.g01540.t01#2   mNeoVis.vulgivaga.2.1.chr5.g01543.t01#1       7637   125253921:   mNeoVis.evergladensis.2.1.chr5.g01541.t01#2    mNeoVis.evergladensis.1.1.chr5.g01547.t01#3    mNeoVis.3.1.1.chr5.g01523.t01#6    mNeoVis.vulgivaga.2.1.chr5.g01544.t01#1    mNeoVis.2.1.1.chr2_hap1.g01602.t01#5   mNeoVis.vulgivaga.1.1.chr5.g01519.t01#4       7638   125253922:   mNeoVis.vulgivaga.2.1.chr13.g00125.t01#1   mNeoVis.3.1.1.chr13.g00082.t01#6             7639   125253923:   mNeoVis.vulgivaga.2.1.chr5.g01545.t01#1    mNeoVis.evergladensis.2.1.chr5.g01542.t01#2    mNeoVis.3.1.1.chr5.g01524.t01#6    mNeoVis.2.1.1.chr2_hap1.g01603.t01#5    mNeoVis.vulgivaga.1.1.chr5.g01520.t01#4   mNeoVis.evergladensis.1.1.chr5.g01548.t01#3       7640   125253924:   mNeoVis.vulgivaga.2.1.chr13.g00126.t01#1    mNeoVis.evergladensis.1.1.chr13.g00125.t01#3    mNeoVis.evergladensis.2.1.chr13.g00129.t01#2    mNeoVis.3.1.1.chr13.g00083.t01#6   mNeoVis.2.1.1.chr13_hap1.g00132.t01#5       7641   125253925:   mNeoVis.2.1.1.chr2_hap1.g01604.t01#5    mNeoVis.evergladensis.2.1.chr5.g01543.t01#2    mNeoVis.evergladensis.1.1.chr5.g01549.t01#3    mNeoVis.vulgivaga.1.1.chr5.g01521.t01#4    mNeoVis.vulgivaga.2.1.chr5.g01546.t01#1   mNeoVis.3.1.1.chr5.g01525.t01#6       7642   125253926:   mNeoVis.evergladensis.1.1.chr13.g00127.t01#3    mNeoVis.2.1.1.chr13_hap1.g00133.t01#5    mNeoVis.evergladensis.2.1.chr13.g00130.t01#2    mNeoVis.3.1.1.chr13.g00084.t01#6    mNeoVis.evergladensis.1.1.chr13.g00126.t01#3    mNeoVis.vulgivaga.1.1.chr13.g00124.t01#4   mNeoVis.vulgivaga.2.1.chr13.g00127.t01#1     7643   125253927:   mNeoVis.2.1.1.chr4_hap1.g01129.t01#5    mNeoVis.vulgivaga.1.1.chr6.g01097.t01#4    mNeoVis.vulgivaga.2.1.chr6.g01096.t01#1    mNeoVis.evergladensis.2.1.chr6.g01105.t01#2   mNeoVis.evergladensis.1.1.chr6.g01107.t01#3       7644   125253928:   mNeoVis.3.1.1.chr5.g01526.t01#6    mNeoVis.vulgivaga.2.1.chr5.g01547.t01#1    mNeoVis.evergladensis.2.1.chr5.g01544.t01#2    mNeoVis.evergladensis.1.1.chr5.g01550.t01#3    mNeoVis.2.1.1.chr2_hap1.g01605.t01#5   mNeoVis.vulgivaga.1.1.chr5.g01522.t01#4       7645   125253929:   mNeoVis.vulgivaga.2.1.chr5.g01548.t01#1    mNeoVis.3.1.1.chr5.g01527.t01#6    mNeoVis.evergladensis.1.1.chr5.g01551.t01#3    mNeoVis.vulgivaga.1.1.chr5.g01523.t01#4    mNeoVis.evergladensis.2.1.chr5.g01545.t01#2   mNeoVis.2.1.1.chr2_hap1.g01606.t01#5       7646   125253930:   mNeoVis.3.1.1.chr13.g00085.t01#6    mNeoVis.vulgivaga.2.1.chr13.g00128.t01#1    mNeoVis.2.1.1.chr13_hap1.g00134.t01#5    mNeoVis.evergladensis.2.1.chr13.g00131.t01#2    mNeoVis.vulgivaga.1.1.chr13.g00125.t01#4   mNeoVis.evergladensis.1.1.chr13.g00128.t01#3       7647   125253931:   mNeoVis.3.1.1.chr5.g01528.t01#6    mNeoVis.evergladensis.1.1.chr5.g01552.t01#3    mNeoVis.vulgivaga.2.1.chr5.g01549.t01#1    mNeoVis.2.1.1.chr2_hap1.g01607.t01#5    mNeoVis.vulgivaga.1.1.chr5.g01524.t01#4   mNeoVis.evergladensis.2.1.chr5.g01546.t01#2       7648   125253932:   mNeoVis.vulgivaga.2.1.chr5.g01550.t01#1    mNeoVis.3.1.1.chr5.g01529.t01#6    mNeoVis.evergladensis.1.1.chr5.g01553.t01#3    mNeoVis.evergladensis.2.1.chr5.g01547.t01#2    mNeoVis.2.1.1.chr2_hap1.g01608.t01#5   mNeoVis.vulgivaga.1.1.chr5.g01525.t01#4       7649   125253933:   mNeoVis.vulgivaga.1.1.chr5.g01526.t01#4    mNeoVis.vulgivaga.2.1.chr5.g01551.t01#1    mNeoVis.3.1.1.chr5.g01530.t01#6    mNeoVis.evergladensis.2.1.chr5.g01548.t01#2    mNeoVis.2.1.1.chr2_hap1.g01609.t01#5   mNeoVis.evergladensis.1.1.chr5.g01554.t01#3       7650   125253934:   mNeoVis.evergladensis.1.1.chr13.g00129.t01#3    mNeoVis.vulgivaga.1.1.chr13.g00126.t01#4    mNeoVis.vulgivaga.2.1.chr13.g00129.t01#1    mNeoVis.2.1.1.chr13_hap1.g00135.t01#5    mNeoVis.3.1.1.chr13.g00087.t01#6   mNeoVis.evergladensis.2.1.chr13.g00132.t01#2       7651   125253935:   mNeoVis.vulgivaga.2.1.chr13.g00130.t01#1    mNeoVis.evergladensis.1.1.chr13.g00130.t01#3    mNeoVis.evergladensis.2.1.chr13.g00133.t01#2    mNeoVis.vulgivaga.1.1.chr13.g00127.t01#4    mNeoVis.3.1.1.chr13.g00088.t01#6   mNeoVis.2.1.1.chr13_hap1.g00136.t01#5       7652   125253936:   mNeoVis.evergladensis.2.1.chr5.g01549.t01#2    mNeoVis.evergladensis.1.1.chr5.g01555.t01#3    mNeoVis.vulgivaga.2.1.chr5.g01552.t01#1   mNeoVis.2.1.1.chr2_hap1.g01610.t01#5         7653   125253937:   mNeoVis.evergladensis.1.1.chr5.g01556.t01#3    mNeoVis.vulgivaga.2.1.chr5.g01553.t01#1    mNeoVis.vulgivaga.1.1.chr5.g01528.t01#4    mNeoVis.evergladensis.2.1.chr5.g01550.t01#2    mNeoVis.2.1.1.chr2_hap1.g01611.t01#5   mNeoVis.3.1.1.chr5.g01532.t01#6       7654   125253938:   mNeoVis.evergladensis.1.1.chr13.g00131.t01#3    mNeoVis.evergladensis.2.1.chr13.g00134.t01#2    mNeoVis.vulgivaga.1.1.chr13.g00128.t01#4    mNeoVis.vulgivaga.2.1.chr13.g00131.t01#1    mNeoVis.2.1.1.chr13_hap1.g00137.t01#5   mNeoVis.3.1.1.chr13.g00089.t01#6       7655   125253939:   mNeoVis.evergladensis.2.1.chr5.g01551.t01#2    mNeoVis.2.1.1.chr2_hap1.g01612.t01#5    mNeoVis.evergladensis.1.1.chr5.g01557.t01#3    mNeoVis.vulgivaga.2.1.chr5.g01554.t01#1   mNeoVis.vulgivaga.1.1.chr5.g01529.t01#4       7656   125253940:   mNeoVis.2.1.1.chr15_hap1.g00078.t01#5    mNeoVis.vulgivaga.2.1.chr15.g00080.t01#1    mNeoVis.vulgivaga.1.1.chr15.g00079.t01#4    mNeoVis.evergladensis.2.1.chr15.g00079.t01#2    mNeoVis.3.1.1.chr15.g00072.t01#6   mNeoVis.evergladensis.1.1.chr15.g00535.t01#3       7657   125253941:   mNeoVis.vulgivaga.1.1.chr5.g01530.t01#4    mNeoVis.evergladensis.1.1.chr5.g01558.t01#3    mNeoVis.3.1.1.chr5.g01534.t01#6    mNeoVis.vulgivaga.2.1.chr5.g01555.t01#1    mNeoVis.evergladensis.2.1.chr5.g01552.t01#2   mNeoVis.2.1.1.chr2_hap1.g01613.t01#5       7658   125253942:   mNeoVis.3.1.1.chr13.g00090.t01#6    mNeoVis.2.1.1.chr13_hap1.g00138.t01#5    mNeoVis.2.1.1.chr9_hap1.g00131.t01#5    mNeoVis.vulgivaga.1.1.chr13.g00129.t01#4    mNeoVis.evergladensis.1.1.chr13.g00132.t01#3    mNeoVis.evergladensis.2.1.chr13.g00135.t01#2   mNeoVis.vulgivaga.2.1.chr13.g00132.t01#1     7659   125253943:   mNeoVis.vulgivaga.2.1.chr5.g01556.t01#1    mNeoVis.evergladensis.1.1.chr5.g01559.t01#3    mNeoVis.3.1.1.chr5.g01535.t01#6    mNeoVis.evergladensis.2.1.chr5.g01553.t01#2    mNeoVis.vulgivaga.1.1.chr5.g01531.t01#4   mNeoVis.2.1.1.chr2_hap1.g01614.t01#5       7660   125253944:   mNeoVis.3.1.1.chr5.g01536.t01#6    mNeoVis.2.1.1.chr2_hap1.g01615.t01#5    mNeoVis.evergladensis.1.1.chr5.g01560.t01#3    mNeoVis.vulgivaga.2.1.chr5.g01557.t01#1    mNeoVis.evergladensis.2.1.chr5.g01554.t01#2   mNeoVis.vulgivaga.1.1.chr5.g01532.t01#4       7661   125253945:   mNeoVis.2.1.1.chr2_hap1.g01616.t01#5    mNeoVis.evergladensis.2.1.chr5.g01555.t01#2    mNeoVis.vulgivaga.2.1.chr5.g01558.t01#1   mNeoVis.evergladensis.1.1.chr5.g01561.t01#3       7662   125253946:   mNeoVis.2.1.1.chr2_hap1.g01617.t01#5    mNeoVis.vulgivaga.2.1.chr5.g01559.t01#1    mNeoVis.evergladensis.1.1.chr5.g01562.t01#3    mNeoVis.vulgivaga.1.1.chr5.g01534.t01#4   mNeoVis.evergladensis.2.1.chr5.g01556.t01#2       7663   125253947:  mNeoVis.vulgivaga.2.1.chr13.g00133.t01#1               7664   125253948:   mNeoVis.evergladensis.2.1.chr13.g00136.t01#2    mNeoVis.2.1.1.chr13_hap1.g00139.t01#5    mNeoVis.vulgivaga.1.1.chr13.g00130.t01#4   mNeoVis.evergladensis.1.1.chr13.g00133.t01#3       7665   125253949:   mNeoVis.vulgivaga.2.1.chr13.g00134.t01#1   mNeoVis.evergladensis.1.1.chr13.g00134.t01#3           7666   125253950:   mNeoVis.vulgivaga.2.1.chr13.g00135.t01#1    mNeoVis.vulgivaga.1.1.chr13.g00131.t01#4    mNeoVis.3.1.1.chr13.g00086.t01#6    mNeoVis.2.1.1.chr13_hap1.g00140.t01#5    mNeoVis.evergladensis.1.1.chr13.g00135.t01#3   mNeoVis.evergladensis.2.1.chr13.g00137.t01#2       7667   125253951:   mNeoVis.vulgivaga.1.1.chr6.g01098.t01#4    mNeoVis.vulgivaga.2.1.chr6.g01097.t01#1    mNeoVis.2.1.1.chr4_hap1.g01128.t01#5    mNeoVis.evergladensis.1.1.chr6.g01108.t01#3   mNeoVis.evergladensis.2.1.chr6.g01104.t01#2       7668   125253952:   mNeoVis.2.1.1.chr2_hap1.g01618.t01#5    mNeoVis.vulgivaga.1.1.chr5.g01535.t01#4    mNeoVis.vulgivaga.2.1.chr5.g01560.t01#1    mNeoVis.evergladensis.1.1.chr5.g01563.t01#3   mNeoVis.evergladensis.2.1.chr5.g01557.t01#2       7669   125253953:   mNeoVis.2.1.1.chr13_hap1.g00141.t01#5    mNeoVis.3.1.1.chr13.g00092.t01#6    mNeoVis.evergladensis.2.1.chr13.g00138.t01#2    mNeoVis.vulgivaga.1.1.chr13.g00132.t01#4    mNeoVis.vulgivaga.2.1.chr13.g00136.t01#1   mNeoVis.evergladensis.1.1.chr13.g00136.t01#3       7670   125253954:   mNeoVis.vulgivaga.1.1.chr5.g01536.t01#4    mNeoVis.evergladensis.2.1.chr5.g01558.t01#2    mNeoVis.3.1.1.chr5.g01540.t01#6    mNeoVis.vulgivaga.2.1.chr5.g01561.t01#1    mNeoVis.evergladensis.1.1.chr5.g01564.t01#3   mNeoVis.2.1.1.chr2_hap1.g01619.t01#5       7671   125253955:   mNeoVis.vulgivaga.1.1.chr5.g01537.t01#4    mNeoVis.3.1.1.chr5.g01541.t01#6    mNeoVis.vulgivaga.2.1.chr5.g01562.t01#1    mNeoVis.evergladensis.1.1.chr5.g01565.t01#3    mNeoVis.evergladensis.2.1.chr5.g01559.t01#2   mNeoVis.2.1.1.chr2_hap1.g01620.t01#5       7672   125253956:   mNeoVis.vulgivaga.1.1.chr13.g00133.t01#4    mNeoVis.2.1.1.chr13_hap1.g00142.t01#5    mNeoVis.3.1.1.chr13.g00093.t01#6    mNeoVis.vulgivaga.2.1.chr13.g00137.t01#1    mNeoVis.evergladensis.2.1.chr13.g00139.t01#2   mNeoVis.evergladensis.1.1.chr13.g00137.t01#3       7673   125253957:   mNeoVis.evergladensis.2.1.chr13.g00140.t01#2    mNeoVis.vulgivaga.1.1.chr13.g00134.t01#4    mNeoVis.vulgivaga.2.1.chr13.g00138.t01#1    mNeoVis.evergladensis.1.1.chr13.g00138.t01#3    mNeoVis.2.1.1.chr13_hap1.g00143.t01#5   mNeoVis.3.1.1.chr13.g00094.t01#6       7674   125253958:   mNeoVis.vulgivaga.2.1.chr5.g01563.t01#1    mNeoVis.evergladensis.2.1.chr5.g01561.t01#2    mNeoVis.vulgivaga.1.1.chr5.g01538.t01#4    mNeoVis.2.1.1.chr2_hap1.g01621.t01#5   mNeoVis.evergladensis.1.1.chr5.g01566.t01#3       7675   125253959:   mNeoVis.evergladensis.2.1.chr5.g01560.t01#2   mNeoVis.3.1.1.chr5.g01542.t01#6             7676   125253960:   mNeoVis.evergladensis.2.1.chr5.g01562.t01#2    mNeoVis.evergladensis.1.1.chr5.g01567.t01#3    mNeoVis.vulgivaga.1.1.chr5.g01540.t01#4    mNeoVis.2.1.1.chr2_hap1.g01622.t01#5    mNeoVis.vulgivaga.2.1.chr5.g01564.t01#1    mNeoVis.3.1.1.chr5.g01543.t01#6   mNeoVis.vulgivaga.1.1.chr5.g01539.t01#4     7677   125253961:   mNeoVis.vulgivaga.1.1.chr5.g01541.t01#4    mNeoVis.vulgivaga.2.1.chr5.g01565.t01#1    mNeoVis.evergladensis.2.1.chr5.g01563.t01#2    mNeoVis.2.1.1.chr2_hap1.g01623.t01#5    mNeoVis.3.1.1.chr5.g01544.t01#6   mNeoVis.evergladensis.1.1.chr5.g01568.t01#3       7678   125253962:   mNeoVis.2.1.1.chr2_hap1.g01624.t01#5    mNeoVis.evergladensis.1.1.chr5.g01569.t01#3    mNeoVis.vulgivaga.2.1.chr5.g01566.t01#1    mNeoVis.evergladensis.2.1.chr5.g01564.t01#2   mNeoVis.vulgivaga.1.1.chr5.g01542.t01#4       7679   125253963:   mNeoVis.vulgivaga.1.1.chr5.g01543.t01#4    mNeoVis.3.1.1.chr5.g01546.t01#6    mNeoVis.evergladensis.2.1.chr5.g01565.t01#2    mNeoVis.vulgivaga.2.1.chr5.g01567.t01#1    mNeoVis.evergladensis.1.1.chr5.g01570.t01#3   mNeoVis.2.1.1.chr2_hap1.g01625.t01#5       7680   125253964:   mNeoVis.2.1.1.chr13_hap1.g00144.t01#5    mNeoVis.vulgivaga.2.1.chr13.g00139.t01#1    mNeoVis.evergladensis.1.1.chr13.g00139.t01#3    mNeoVis.evergladensis.2.1.chr13.g00141.t01#2    mNeoVis.3.1.1.chr13.g00095.t01#6   mNeoVis.vulgivaga.1.1.chr13.g00135.t01#4       7681   125253965:   mNeoVis.evergladensis.2.1.chr5.g01566.t01#2    mNeoVis.2.1.1.chr2_hap1.g01626.t01#5    mNeoVis.vulgivaga.2.1.chr5.g01568.t01#1    mNeoVis.3.1.1.chr5.g01547.t01#6    mNeoVis.evergladensis.1.1.chr5.g01571.t01#3   mNeoVis.vulgivaga.1.1.chr5.g01544.t01#4       7682   125253966:   mNeoVis.evergladensis.1.1.chr5.g01572.t01#3    mNeoVis.evergladensis.2.1.chr5.g01567.t01#2    mNeoVis.vulgivaga.1.1.chr5.g01545.t01#4    mNeoVis.2.1.1.chr2_hap1.g01627.t01#5    mNeoVis.vulgivaga.2.1.chr5.g01569.t01#1   mNeoVis.3.1.1.chr5.g01548.t01#6       7683   125253967:   mNeoVis.evergladensis.1.1.chr13.g00140.t01#3    mNeoVis.vulgivaga.2.1.chr13.g00140.t01#1   mNeoVis.evergladensis.2.1.chr13.g00142.t01#2         7684   125253968:   mNeoVis.evergladensis.2.1.chr13.g00143.t01#2    mNeoVis.2.1.1.chr13_hap1.g00145.t01#5    mNeoVis.evergladensis.1.1.chr13.g00141.t01#3    mNeoVis.3.1.1.chr13.g00096.t01#6    mNeoVis.vulgivaga.2.1.chr13.g00141.t01#1   mNeoVis.vulgivaga.1.1.chr13.g00136.t01#4       7685   125253969:   mNeoVis.evergladensis.1.1.chr5.g01573.t01#3    mNeoVis.vulgivaga.1.1.chr5.g01546.t01#4    mNeoVis.vulgivaga.2.1.chr5.g01570.t01#1    mNeoVis.evergladensis.2.1.chr5.g01568.t01#2    mNeoVis.2.1.1.chr2_hap1.g01628.t01#5   mNeoVis.3.1.1.chr5.g01549.t01#6       7686   125253970:   mNeoVis.evergladensis.1.1.chr5.g01574.t01#3    mNeoVis.2.1.1.chr2_hap1.g01629.t01#5    mNeoVis.vulgivaga.1.1.chr5.g01547.t01#4    mNeoVis.evergladensis.2.1.chr5.g01569.t01#2    mNeoVis.vulgivaga.2.1.chr5.g01571.t01#1   mNeoVis.3.1.1.chr5.g01551.t01#6       7687   125253971:   mNeoVis.3.1.1.chr13.g00097.t01#6    mNeoVis.2.1.1.chr13_hap1.g00146.t01#5    mNeoVis.vulgivaga.2.1.chr13.g00142.t01#1    mNeoVis.vulgivaga.1.1.chr13.g00137.t01#4    mNeoVis.evergladensis.1.1.chr13.g00142.t01#3    mNeoVis.vulgivaga.1.1.chr13.g00138.t01#4   mNeoVis.evergladensis.2.1.chr13.g00144.t01#2     7688   125253972:   mNeoVis.evergladensis.2.1.chr13.g00145.t01#2    mNeoVis.vulgivaga.1.1.chr13.g00139.t01#4    mNeoVis.2.1.1.chr13_hap1.g00147.t01#5    mNeoVis.vulgivaga.2.1.chr13.g00143.t01#1    mNeoVis.3.1.1.chr13.g00098.t01#6   mNeoVis.evergladensis.1.1.chr13.g00143.t01#3       7689   125253973:   mNeoVis.vulgivaga.2.1.chr6.g01098.t01#1    mNeoVis.2.1.1.chr4_hap1.g01130.t01#5    mNeoVis.vulgivaga.1.1.chr6.g01099.t01#4    mNeoVis.evergladensis.2.1.chr6.g01106.t01#2   mNeoVis.evergladensis.1.1.chr6.g01109.t01#3       7690   125253974:   mNeoVis.evergladensis.1.1.chr5.g01575.t01#3    mNeoVis.vulgivaga.2.1.chr5.g01572.t01#1    mNeoVis.vulgivaga.1.1.chr5.g01548.t01#4    mNeoVis.3.1.1.chr5.g01550.t01#6    mNeoVis.2.1.1.chr2_hap1.g01630.t01#5   mNeoVis.evergladensis.2.1.chr5.g01570.t01#2       7691   125253975:   mNeoVis.3.1.1.chr3.g00344.t01#6    mNeoVis.vulgivaga.1.1.chr3.g00353.t01#4    mNeoVis.vulgivaga.2.1.chr3.g00354.t01#1    mNeoVis.3.1.1.chr15.g00073.t01#6    mNeoVis.2.1.1.chr15_hap1.g00079.t01#5    mNeoVis.evergladensis.1.1.chr3.g00358.t01#3    mNeoVis.evergladensis.2.1.chr3.g00359.t01#2    mNeoVis.2.1.1.chr3_hap1.g00351.t01#5    mNeoVis.evergladensis.2.1.chr15.g00080.t01#2      7692   125253976:   mNeoVis.vulgivaga.1.1.chr15.g00080.t01#4    mNeoVis.vulgivaga.2.1.chr15.g00081.t01#1   mNeoVis.evergladensis.1.1.chr15.g00534.t01#3         7693   125253977:   mNeoVis.vulgivaga.1.1.chr5.g01549.t01#4    mNeoVis.3.1.1.chr5.g01552.t01#6    mNeoVis.evergladensis.1.1.chr5.g01576.t01#3    mNeoVis.vulgivaga.2.1.chr5.g01573.t01#1    mNeoVis.evergladensis.2.1.chr5.g01571.t01#2   mNeoVis.2.1.1.chr2_hap1.g01631.t01#5       7694   125253978:   mNeoVis.evergladensis.1.1.chr5.g01577.t01#3    mNeoVis.vulgivaga.1.1.chr5.g01550.t01#4    mNeoVis.2.1.1.chr2_hap1.g01632.t01#5    mNeoVis.evergladensis.2.1.chr5.g01572.t01#2    mNeoVis.vulgivaga.2.1.chr5.g01574.t01#1   mNeoVis.3.1.1.chr5.g01553.t01#6       7695   125253979:   mNeoVis.2.1.1.chr13_hap1.g00148.t01#5    mNeoVis.vulgivaga.1.1.chr13.g00140.t01#4    mNeoVis.3.1.1.chr13.g00099.t01#6    mNeoVis.evergladensis.1.1.chr13.g00144.t01#3    mNeoVis.evergladensis.2.1.chr13.g00146.t01#2   mNeoVis.vulgivaga.2.1.chr13.g00144.t01#1       7696   125253980:   mNeoVis.vulgivaga.1.1.chr13.g00141.t01#4    mNeoVis.2.1.1.chr13_hap1.g00149.t01#5    mNeoVis.evergladensis.2.1.chr13.g00147.t01#2    mNeoVis.evergladensis.1.1.chr13.g00145.t01#3    mNeoVis.vulgivaga.2.1.chr13.g00145.t01#1   mNeoVis.3.1.1.chr13.g00100.t01#6       7697   125253981:   mNeoVis.vulgivaga.2.1.chr13.g00146.t01#1    mNeoVis.evergladensis.1.1.chr13.g00146.t01#3    mNeoVis.3.1.1.chr13.g00101.t01#6    mNeoVis.evergladensis.2.1.chr13.g00148.t01#2    mNeoVis.vulgivaga.1.1.chr13.g00142.t01#4   mNeoVis.2.1.1.chr13_hap1.g00150.t01#5       7698   125253982:   mNeoVis.evergladensis.1.1.chr13.g00147.t01#3    mNeoVis.vulgivaga.2.1.chr13.g00147.t01#1    mNeoVis.evergladensis.2.1.chr13.g00149.t01#2    mNeoVis.2.1.1.chr13_hap1.g00151.t01#5   mNeoVis.vulgivaga.1.1.chr13.g00143.t01#4       7699   125253983:   mNeoVis.vulgivaga.1.1.chr5.g01553.t01#4    mNeoVis.evergladensis.2.1.chr5.g01575.t01#2    mNeoVis.3.1.1.chr5.g01557.t01#6    mNeoVis.vulgivaga.2.1.chr5.g01577.t01#1    mNeoVis.2.1.1.chr2_hap1.g01635.t01#5   mNeoVis.evergladensis.1.1.chr5.g01580.t01#3       7700   125253984:   mNeoVis.evergladensis.2.1.chr5.g01576.t01#2    mNeoVis.vulgivaga.1.1.chr5.g01554.t01#4    mNeoVis.vulgivaga.2.1.chr5.g01578.t01#1    mNeoVis.evergladensis.1.1.chr5.g01581.t01#3    mNeoVis.2.1.1.chr2_hap1.g01636.t01#5   mNeoVis.3.1.1.chr5.g01558.t01#6       7701   125253985:   mNeoVis.evergladensis.1.1.chr5.g01582.t01#3    mNeoVis.vulgivaga.2.1.chr5.g01579.t01#1    mNeoVis.evergladensis.2.1.chr5.g01577.t01#2    mNeoVis.2.1.1.chr2_hap1.g01637.t01#5    mNeoVis.vulgivaga.1.1.chr5.g01555.t01#4   mNeoVis.3.1.1.chr5.g01559.t01#6       7702   125253986:   mNeoVis.vulgivaga.2.1.chr5.g01580.t01#1    mNeoVis.evergladensis.1.1.chr5.g01583.t01#3    mNeoVis.evergladensis.2.1.chr5.g01578.t01#2    mNeoVis.2.1.1.chr2_hap1.g01638.t01#5    mNeoVis.3.1.1.chr5.g01560.t01#6   mNeoVis.vulgivaga.1.1.chr5.g01556.t01#4       7703   125253987:   mNeoVis.vulgivaga.2.1.chr15.g00082.t01#1    mNeoVis.3.1.1.chr15.g00074.t01#6    mNeoVis.vulgivaga.1.1.chr15.g00081.t01#4    mNeoVis.evergladensis.2.1.chr15.g00081.t01#2    mNeoVis.2.1.1.chr15_hap1.g00080.t01#5   mNeoVis.evergladensis.1.1.chr15.g00532.t01#3       7704   125253988:   mNeoVis.2.1.1.chr2_hap1.g01639.t01#5    mNeoVis.vulgivaga.2.1.chr5.g01581.t01#1    mNeoVis.evergladensis.2.1.chr5.g01579.t01#2    mNeoVis.3.1.1.chr5.g01561.t01#6   mNeoVis.evergladensis.1.1.chr5.g01584.t01#3       7705   125253989:   mNeoVis.2.1.1.chr2_hap1.g01640.t01#5    mNeoVis.vulgivaga.2.1.chr5.g01582.t01#1    mNeoVis.evergladensis.2.1.chr5.g01580.t01#2    mNeoVis.evergladensis.1.1.chr5.g01585.t01#3    mNeoVis.3.1.1.chr5.g01562.t01#6   mNeoVis.vulgivaga.1.1.chr5.g01558.t01#4       7706   125253990:   mNeoVis.2.1.1.chr2_hap1.g01641.t01#5    mNeoVis.vulgivaga.2.1.chr5.g01583.t01#1    mNeoVis.evergladensis.2.1.chr5.g01581.t01#2    mNeoVis.evergladensis.1.1.chr5.g01586.t01#3    mNeoVis.3.1.1.chr5.g01563.t01#6   mNeoVis.vulgivaga.1.1.chr5.g01559.t01#4       7707   125253991:   mNeoVis.evergladensis.1.1.chr5.g01587.t01#3    mNeoVis.vulgivaga.2.1.chr5.g01584.t01#1    mNeoVis.vulgivaga.1.1.chr5.g01560.t01#4    mNeoVis.3.1.1.chr5.g01564.t01#6    mNeoVis.evergladensis.2.1.chr5.g01582.t01#2   mNeoVis.2.1.1.chr2_hap1.g01642.t01#5       7708   125253992:   mNeoVis.vulgivaga.2.1.chr6.g01099.t01#1    mNeoVis.evergladensis.1.1.chr6.g01110.t01#3    mNeoVis.vulgivaga.1.1.chr6.g01100.t01#4    mNeoVis.2.1.1.chr4_hap1.g01131.t01#5   mNeoVis.evergladensis.2.1.chr6.g01107.t01#2       7709   125253993:   mNeoVis.3.1.1.chr13.g00103.t01#6    mNeoVis.evergladensis.1.1.chr13.g00148.t01#3    mNeoVis.2.1.1.chr13_hap1.g00152.t01#5    mNeoVis.vulgivaga.2.1.chr13.g00148.t01#1    mNeoVis.evergladensis.2.1.chr13.g00150.t01#2   mNeoVis.vulgivaga.1.1.chr13.g00144.t01#4       7710   125253994:   mNeoVis.3.1.1.chr5.g01565.t01#6    mNeoVis.2.1.1.chr2_hap1.g01643.t01#5    mNeoVis.evergladensis.1.1.chr5.g01588.t01#3    mNeoVis.vulgivaga.1.1.chr5.g01561.t01#4    mNeoVis.evergladensis.2.1.chr5.g01583.t01#2   mNeoVis.vulgivaga.2.1.chr5.g01585.t01#1       7711   125253995:   mNeoVis.3.1.1.chr13.g00104.t01#6    mNeoVis.evergladensis.2.1.chr13.g00151.t01#2    mNeoVis.2.1.1.chr13_hap1.g00153.t01#5    mNeoVis.vulgivaga.2.1.chr13.g00149.t01#1    mNeoVis.evergladensis.1.1.chr13.g00149.t01#3    mNeoVis.3.1.1.chr13.g00105.t01#6   mNeoVis.vulgivaga.1.1.chr13.g00145.t01#4     7712   125253996:   mNeoVis.evergladensis.2.1.chr5.g01584.t01#2    mNeoVis.2.1.1.chr2_hap1.g01644.t01#5    mNeoVis.vulgivaga.1.1.chr5.g01562.t01#4    mNeoVis.evergladensis.1.1.chr5.g01589.t01#3    mNeoVis.vulgivaga.2.1.chr5.g01586.t01#1   mNeoVis.3.1.1.chr5.g01566.t01#6       7713   125253997:   mNeoVis.evergladensis.1.1.chr5.g01590.t01#3    mNeoVis.vulgivaga.1.1.chr5.g01563.t01#4    mNeoVis.2.1.1.chr2_hap1.g01645.t01#5    mNeoVis.evergladensis.2.1.chr5.g01585.t01#2    mNeoVis.3.1.1.chr5.g01567.t01#6   mNeoVis.vulgivaga.2.1.chr5.g01587.t01#1       7714   125253998:   mNeoVis.3.1.1.chr13.g00106.t01#6    mNeoVis.vulgivaga.1.1.chr13.g00146.t01#4    mNeoVis.vulgivaga.2.1.chr13.g00150.t01#1    mNeoVis.evergladensis.2.1.chr13.g00152.t01#2    mNeoVis.evergladensis.1.1.chr13.g00150.t01#3   mNeoVis.2.1.1.chr13_hap1.g00154.t01#5       7715   125253999:   mNeoVis.3.1.1.chr5.g01568.t01#6    mNeoVis.vulgivaga.2.1.chr5.g01588.t01#1    mNeoVis.vulgivaga.1.1.chr5.g01564.t01#4    mNeoVis.2.1.1.chr2_hap1.g01646.t01#5    mNeoVis.evergladensis.1.1.chr5.g01591.t01#3   mNeoVis.evergladensis.2.1.chr5.g01586.t01#2       7716   125254000:   mNeoVis.vulgivaga.1.1.chr5.g01565.t01#4    mNeoVis.evergladensis.2.1.chr5.g01587.t01#2    mNeoVis.vulgivaga.2.1.chr5.g01589.t01#1    mNeoVis.3.1.1.chr5.g01569.t01#6    mNeoVis.2.1.1.chr2_hap1.g01647.t01#5   mNeoVis.evergladensis.1.1.chr5.g01592.t01#3       7717   125254001:   mNeoVis.2.1.1.chr2_hap1.g01648.t01#5    mNeoVis.3.1.1.chr5.g01570.t01#6    mNeoVis.vulgivaga.1.1.chr5.g01566.t01#4    mNeoVis.evergladensis.1.1.chr5.g01593.t01#3    mNeoVis.evergladensis.2.1.chr5.g01588.t01#2   mNeoVis.vulgivaga.2.1.chr5.g01590.t01#1       7718   125254002:   mNeoVis.3.1.1.chr13.g00107.t01#6    mNeoVis.evergladensis.2.1.chr13.g00153.t01#2    mNeoVis.2.1.1.chr13_hap1.g00155.t01#5    mNeoVis.vulgivaga.1.1.chr13.g00147.t01#4    mNeoVis.vulgivaga.2.1.chr13.g00151.t01#1   mNeoVis.evergladensis.1.1.chr13.g00151.t01#3       7719   125254003:   mNeoVis.vulgivaga.2.1.chr5.g01591.t01#1    mNeoVis.vulgivaga.1.1.chr5.g01567.t01#4    mNeoVis.evergladensis.2.1.chr5.g01589.t01#2    mNeoVis.2.1.1.chr2_hap1.g01649.t01#5    mNeoVis.evergladensis.1.1.chr5.g01594.t01#3   mNeoVis.3.1.1.chr5.g01571.t01#6       7720   125254004:   mNeoVis.evergladensis.2.1.chr15.g00082.t01#2    mNeoVis.evergladensis.1.1.chr15.g00531.t01#3    mNeoVis.vulgivaga.2.1.chr15.g00083.t01#1    mNeoVis.2.1.1.chr15_hap1.g00081.t01#5    mNeoVis.3.1.1.chr15.g00075.t01#6   mNeoVis.vulgivaga.1.1.chr15.g00082.t01#4       7721   125254005:   mNeoVis.evergladensis.1.1.chr5.g01595.t01#3    mNeoVis.2.1.1.chr2_hap1.g01650.t01#5    mNeoVis.3.1.1.chr5.g01572.t01#6    mNeoVis.vulgivaga.1.1.chr5.g01568.t01#4    mNeoVis.vulgivaga.2.1.chr5.g01592.t01#1   mNeoVis.evergladensis.2.1.chr5.g01590.t01#2       7722   125254006:   mNeoVis.evergladensis.2.1.chr5.g01491.t01#2    mNeoVis.2.1.1.chr2_hap1.g01553.t01#5    mNeoVis.vulgivaga.2.1.chr5.g01495.t01#1    mNeoVis.vulgivaga.1.1.chr5.g01470.t01#4   mNeoVis.evergladensis.1.1.chr5.g01498.t01#3       7723   125254007:   mNeoVis.evergladensis.1.1.chr15.g00530.t01#3    mNeoVis.evergladensis.2.1.chr15.g00083.t01#2    mNeoVis.vulgivaga.2.1.chr15.g00084.t01#1    mNeoVis.2.1.1.chr15_hap1.g00082.t01#5   mNeoVis.3.1.1.chr15.g00076.t01#6       7724   125254008:   mNeoVis.evergladensis.1.1.chr13.g00152.t01#3    mNeoVis.evergladensis.2.1.chr13.g00154.t01#2    mNeoVis.2.1.1.chr13_hap1.g00156.t01#5    mNeoVis.vulgivaga.2.1.chr13.g00152.t01#1    mNeoVis.vulgivaga.1.1.chr13.g00148.t01#4   mNeoVis.3.1.1.chr13.g00108.t01#6       7725   125254009:   mNeoVis.evergladensis.1.1.chr5.g01499.t01#3    mNeoVis.vulgivaga.2.1.chr5.g01496.t01#1    mNeoVis.evergladensis.2.1.chr5.g01492.t01#2    mNeoVis.3.1.1.chr5.g01475.t01#6    mNeoVis.vulgivaga.1.1.chr5.g01471.t01#4   mNeoVis.2.1.1.chr2_hap1.g01554.t01#5       7726   125254010:   mNeoVis.2.1.1.chr2_hap1.g01555.t01#5    mNeoVis.3.1.1.chr5.g01476.t01#6    mNeoVis.vulgivaga.2.1.chr5.g01497.t01#1    mNeoVis.evergladensis.1.1.chr5.g01500.t01#3    mNeoVis.evergladensis.2.1.chr5.g01493.t01#2   mNeoVis.vulgivaga.1.1.chr5.g01472.t01#4       7727   125254011:  mNeoVis.vulgivaga.2.1.chr13.g00081.t01#1               7728   125254012:   mNeoVis.vulgivaga.2.1.chr5.g01498.t01#1    mNeoVis.3.1.1.chr5.g01477.t01#6    mNeoVis.evergladensis.1.1.chr5.g01501.t01#3    mNeoVis.evergladensis.2.1.chr5.g01494.t01#2    mNeoVis.vulgivaga.1.1.chr5.g01473.t01#4   mNeoVis.2.1.1.chr2_hap1.g01556.t01#5       7729   125254013:   mNeoVis.evergladensis.2.1.chr5.g01495.t01#2    mNeoVis.2.1.1.chr2_hap1.g01557.t01#5    mNeoVis.vulgivaga.2.1.chr5.g01499.t01#1    mNeoVis.evergladensis.1.1.chr5.g01502.t01#3    mNeoVis.vulgivaga.1.1.chr5.g01474.t01#4   mNeoVis.3.1.1.chr5.g01478.t01#6       7730   125254014:   mNeoVis.2.1.1.chr2_hap1.g01558.t01#5    mNeoVis.vulgivaga.1.1.chr5.g01475.t01#4    mNeoVis.vulgivaga.2.1.chr5.g01500.t01#1    mNeoVis.3.1.1.chr5.g01479.t01#6   mNeoVis.evergladensis.1.1.chr5.g01503.t01#3       7731   125254015:   mNeoVis.2.1.1.chr13_hap1.g00083.t01#5    mNeoVis.evergladensis.1.1.chr13.g00080.t01#3    mNeoVis.evergladensis.2.1.chr13.g00084.t01#2    mNeoVis.vulgivaga.2.1.chr13.g00082.t01#1   mNeoVis.2.1.1.chr13_hap1.g00084.t01#5       7732   125254016:   mNeoVis.vulgivaga.2.1.chr13.g00083.t01#1    mNeoVis.evergladensis.1.1.chr13.g00081.t01#3    mNeoVis.vulgivaga.1.1.chr13.g00082.t01#4    mNeoVis.evergladensis.2.1.chr13.g00085.t01#2    mNeoVis.3.1.1.chr13.g00047.t01#6   mNeoVis.2.1.1.chr13_hap1.g00085.t01#5       7733   125254017:   mNeoVis.evergladensis.2.1.chr5.g01497.t01#2    mNeoVis.2.1.1.chr2_hap1.g01559.t01#5    mNeoVis.evergladensis.1.1.chr5.g01504.t01#3    mNeoVis.3.1.1.chr5.g01480.t01#6    mNeoVis.vulgivaga.1.1.chr5.g01476.t01#4   mNeoVis.vulgivaga.2.1.chr5.g01501.t01#1       7734   125254018:   mNeoVis.2.1.1.chr2_hap1.g01560.t01#5    mNeoVis.3.1.1.chr5.g01481.t01#6    mNeoVis.vulgivaga.2.1.chr5.g01502.t01#1    mNeoVis.evergladensis.2.1.chr5.g01498.t01#2    mNeoVis.evergladensis.1.1.chr5.g01505.t01#3   mNeoVis.vulgivaga.1.1.chr5.g01477.t01#4       7735   125254019:   mNeoVis.2.1.1.chr13_hap1.g00086.t01#5    mNeoVis.evergladensis.2.1.chr13.g00086.t01#2    mNeoVis.evergladensis.1.1.chr13.g00082.t01#3    mNeoVis.vulgivaga.1.1.chr13.g00083.t01#4   mNeoVis.vulgivaga.2.1.chr13.g00084.t01#1       7736   125254020:   mNeoVis.3.1.1.chr5.g01482.t01#6    mNeoVis.vulgivaga.1.1.chr5.g01478.t01#4    mNeoVis.evergladensis.2.1.chr5.g01499.t01#2    mNeoVis.vulgivaga.2.1.chr5.g01503.t01#1    mNeoVis.evergladensis.1.1.chr5.g01506.t01#3   mNeoVis.2.1.1.chr2_hap1.g01561.t01#5       7737   125254021:   mNeoVis.vulgivaga.2.1.chr13.g00085.t01#1    mNeoVis.evergladensis.2.1.chr13.g00087.t01#2    mNeoVis.evergladensis.1.1.chr13.g00083.t01#3    mNeoVis.vulgivaga.1.1.chr13.g00084.t01#4   mNeoVis.2.1.1.chr13_hap1.g00087.t01#5       7738   125254022:   mNeoVis.2.1.1.chr13_hap1.g00088.t01#5    mNeoVis.evergladensis.1.1.chr13.g00084.t01#3    mNeoVis.vulgivaga.1.1.chr13.g00085.t01#4    mNeoVis.vulgivaga.2.1.chr13.g00086.t01#1   mNeoVis.evergladensis.2.1.chr13.g00088.t01#2       7739   125254023:   mNeoVis.2.1.1.chr13_hap1.g00089.t01#5    mNeoVis.evergladensis.2.1.chr13.g00089.t01#2    mNeoVis.vulgivaga.2.1.chr13.g00087.t01#1    mNeoVis.vulgivaga.1.1.chr13.g00086.t01#4   mNeoVis.evergladensis.1.1.chr13.g00085.t01#3       7740   125254024:   mNeoVis.2.1.1.chr13_hap1.g00094.t01#5    mNeoVis.vulgivaga.2.1.chr13.g00088.t01#1    mNeoVis.evergladensis.2.1.chr13.g00091.t01#2    mNeoVis.vulgivaga.1.1.chr13.g00088.t01#4   mNeoVis.evergladensis.1.1.chr13.g00087.t01#3       7741   125254025:   mNeoVis.evergladensis.1.1.chr15.g00529.t01#3    mNeoVis.3.1.1.chr15.g00077.t01#6    mNeoVis.vulgivaga.1.1.chr15.g00084.t01#4    mNeoVis.vulgivaga.2.1.chr15.g00085.t01#1    mNeoVis.2.1.1.chr15_hap1.g00083.t01#5   mNeoVis.evergladensis.2.1.chr15.g00084.t01#2       7742   125254026:   mNeoVis.evergladensis.1.1.chr13.g00088.t01#3    mNeoVis.2.1.1.chr13_hap1.g00093.t01#5    mNeoVis.vulgivaga.2.1.chr13.g00089.t01#1   mNeoVis.evergladensis.2.1.chr13.g00092.t01#2       7743   125254027:   mNeoVis.vulgivaga.2.1.chr5.g01505.t01#1    mNeoVis.2.1.1.chr2_hap1.g01563.t01#5    mNeoVis.evergladensis.1.1.chr5.g01508.t01#3    mNeoVis.evergladensis.2.1.chr5.g01501.t01#2    mNeoVis.3.1.1.chr5.g01484.t01#6   mNeoVis.vulgivaga.1.1.chr5.g01480.t01#4       7744   125254028:   mNeoVis.2.1.1.chr2_hap1.g01564.t01#5    mNeoVis.evergladensis.1.1.chr5.g01509.t01#3    mNeoVis.vulgivaga.2.1.chr5.g01506.t01#1    mNeoVis.vulgivaga.1.1.chr5.g01481.t01#4    mNeoVis.3.1.1.chr5.g01485.t01#6   mNeoVis.evergladensis.2.1.chr5.g01502.t01#2       7745   125254029:   mNeoVis.evergladensis.2.1.chr13.g00093.t01#2    mNeoVis.2.1.1.chr13_hap1.g00095.t01#5    mNeoVis.vulgivaga.2.1.chr13.g00090.t01#1    mNeoVis.3.1.1.chr13.g00049.t01#6   mNeoVis.evergladensis.1.1.chr13.g00089.t01#3       7746   125254030:   mNeoVis.vulgivaga.1.1.chr5.g01483.t01#4    mNeoVis.evergladensis.1.1.chr5.g01511.t01#3    mNeoVis.3.1.1.chr5.g01487.t01#6    mNeoVis.2.1.1.chr2_hap1.g01566.t01#5   mNeoVis.vulgivaga.2.1.chr5.g01508.t01#1       7747   125254031:   mNeoVis.evergladensis.1.1.chr5.g01512.t01#3    mNeoVis.vulgivaga.2.1.chr5.g01509.t01#1    mNeoVis.2.1.1.chr2_hap1.g01567.t01#5    mNeoVis.vulgivaga.1.1.chr5.g01484.t01#4    mNeoVis.evergladensis.2.1.chr5.g01506.t01#2   mNeoVis.3.1.1.chr5.g01488.t01#6       7748   125254032:   mNeoVis.evergladensis.2.1.chr13.g00094.t01#2    mNeoVis.2.1.1.chr13_hap1.g00096.t01#5    mNeoVis.vulgivaga.2.1.chr13.g00091.t01#1    mNeoVis.vulgivaga.1.1.chr13.g00090.t01#4   mNeoVis.evergladensis.1.1.chr13.g00090.t01#3       7749   125254033:   mNeoVis.evergladensis.1.1.chr13.g00091.t01#3    mNeoVis.evergladensis.2.1.chr13.g00095.t01#2    mNeoVis.vulgivaga.2.1.chr13.g00092.t01#1    mNeoVis.vulgivaga.1.1.chr13.g00091.t01#4   mNeoVis.2.1.1.chr13_hap1.g00097.t01#5       7750   125254034:   mNeoVis.evergladensis.1.1.chr5.g01513.t01#3    mNeoVis.vulgivaga.2.1.chr5.g01510.t01#1    mNeoVis.vulgivaga.1.1.chr5.g01485.t01#4    mNeoVis.2.1.1.chr2_hap1.g01568.t01#5    mNeoVis.evergladensis.2.1.chr5.g01507.t01#2   mNeoVis.3.1.1.chr5.g01489.t01#6       7751   125254035:   mNeoVis.evergladensis.2.1.chr13.g00096.t01#2    mNeoVis.vulgivaga.2.1.chr13.g00093.t01#1    mNeoVis.vulgivaga.1.1.chr13.g00092.t01#4    mNeoVis.2.1.1.chr13_hap1.g00098.t01#5   mNeoVis.evergladensis.1.1.chr13.g00092.t01#3       7752   125254036:   mNeoVis.3.1.1.chr13.g00053.t01#6    mNeoVis.evergladensis.1.1.chr13.g00093.t01#3    mNeoVis.2.1.1.chr13_hap1.g00099.t01#5    mNeoVis.evergladensis.2.1.chr13.g00097.t01#2    mNeoVis.vulgivaga.2.1.chr13.g00094.t01#1   mNeoVis.vulgivaga.1.1.chr13.g00093.t01#4       7753   125254037:   mNeoVis.vulgivaga.1.1.chr13.g00094.t01#4    mNeoVis.2.1.1.chr13_hap1.g00100.t01#5    mNeoVis.vulgivaga.2.1.chr13.g00095.t01#1    mNeoVis.evergladensis.1.1.chr13.g00094.t01#3    mNeoVis.evergladensis.2.1.chr13.g00098.t01#2   mNeoVis.3.1.1.chr13.g00054.t01#6       7754   125254038:   mNeoVis.evergladensis.2.1.chr13.g00099.t01#2    mNeoVis.2.1.1.chr13_hap1.g00101.t01#5    mNeoVis.evergladensis.1.1.chr13.g00095.t01#3   mNeoVis.vulgivaga.2.1.chr13.g00096.t01#1       7755   125254039:   mNeoVis.vulgivaga.2.1.chr15.g00086.t01#1    mNeoVis.vulgivaga.1.1.chr15.g00085.t01#4    mNeoVis.evergladensis.2.1.chr15.g00085.t01#2    mNeoVis.2.1.1.chr15_hap1.g00084.t01#5    mNeoVis.3.1.1.chr15.g00078.t01#6   mNeoVis.evergladensis.1.1.chr15.g00528.t01#3       7756   125254040:   mNeoVis.evergladensis.1.1.chr13.g00096.t01#3    mNeoVis.vulgivaga.2.1.chr13.g00097.t01#1    mNeoVis.evergladensis.2.1.chr13.g00100.t01#2    mNeoVis.vulgivaga.1.1.chr13.g00096.t01#4   mNeoVis.2.1.1.chr13_hap1.g00102.t01#5       7757   125254041:   mNeoVis.vulgivaga.1.1.chr13.g00097.t01#4    mNeoVis.3.1.1.chr15.g00238.t01#6    mNeoVis.2.1.1.chr13_hap1.g00103.t01#5    mNeoVis.evergladensis.2.1.chr13.g00101.t01#2    mNeoVis.vulgivaga.2.1.chr13.g00098.t01#1   mNeoVis.evergladensis.1.1.chr13.g00097.t01#3       7758   125254042:   mNeoVis.evergladensis.1.1.chr5.g00946.t01#3    mNeoVis.evergladensis.2.1.chr5.g00954.t01#2    mNeoVis.3.1.1.chr5.g00933.t01#6    mNeoVis.2.1.1.chr2_hap1.g01001.t01#5    mNeoVis.vulgivaga.1.1.chr5.g00921.t01#4   mNeoVis.vulgivaga.2.1.chr5.g00940.t01#1       7759   125254043:   mNeoVis.evergladensis.2.1.chr5.g01508.t01#2    mNeoVis.vulgivaga.1.1.chr5.g01486.t01#4    mNeoVis.evergladensis.1.1.chr5.g01514.t01#3    mNeoVis.2.1.1.chr2_hap1.g01569.t01#5    mNeoVis.3.1.1.chr5.g01490.t01#6   mNeoVis.vulgivaga.2.1.chr5.g01511.t01#1       7760   125254044:   mNeoVis.vulgivaga.1.1.chr13.g00098.t01#4    mNeoVis.evergladensis.2.1.chr13.g00102.t01#2    mNeoVis.vulgivaga.2.1.chr13.g00099.t01#1    mNeoVis.evergladensis.1.1.chr13.g00098.t01#3   mNeoVis.2.1.1.chr13_hap1.g00104.t01#5       7761   125254045:   mNeoVis.vulgivaga.2.1.chr13.g00100.t01#1    mNeoVis.evergladensis.2.1.chr13.g00103.t01#2    mNeoVis.2.1.1.chr13_hap1.g00105.t01#5   mNeoVis.vulgivaga.1.1.chr13.g00099.t01#4       7762   125254046:   mNeoVis.vulgivaga.1.1.chr5.g01487.t01#4    mNeoVis.2.1.1.chr2_hap1.g01570.t01#5    mNeoVis.vulgivaga.2.1.chr5.g01512.t01#1    mNeoVis.3.1.1.chr5.g01491.t01#6    mNeoVis.evergladensis.1.1.chr5.g01515.t01#3   mNeoVis.evergladensis.2.1.chr5.g01509.t01#2       7763   125254047:   mNeoVis.vulgivaga.2.1.chr5.g01513.t01#1    mNeoVis.3.1.1.chr5.g01492.t01#6    mNeoVis.2.1.1.chr2_hap1.g01571.t01#5    mNeoVis.vulgivaga.1.1.chr5.g01488.t01#4    mNeoVis.evergladensis.1.1.chr5.g01516.t01#3   mNeoVis.evergladensis.2.1.chr5.g01510.t01#2       7764   125254048:   mNeoVis.2.1.1.chr4_hap1.g01104.t01#5    mNeoVis.vulgivaga.2.1.chr6.g01100.t01#1   mNeoVis.2.1.1.chr4_hap1.g01152.t01#5           7765   125254049:   mNeoVis.evergladensis.1.1.chr5.g01517.t01#3    mNeoVis.evergladensis.2.1.chr5.g01511.t01#2    mNeoVis.vulgivaga.1.1.chr5.g01489.t01#4    mNeoVis.2.1.1.chr2_hap1.g01572.t01#5    mNeoVis.vulgivaga.2.1.chr5.g01514.t01#1   mNeoVis.3.1.1.chr5.g01493.t01#6       7766   125254050:   mNeoVis.evergladensis.2.1.chr13.g00104.t01#2    mNeoVis.vulgivaga.2.1.chr13.g00101.t01#1    mNeoVis.vulgivaga.1.1.chr13.g00100.t01#4   mNeoVis.evergladensis.1.1.chr13.g00099.t01#3       7767   125254051:   mNeoVis.2.1.1.chr2_hap1.g01573.t01#5    mNeoVis.3.1.1.chr5.g01494.t01#6    mNeoVis.vulgivaga.2.1.chr5.g01515.t01#1    mNeoVis.vulgivaga.1.1.chr5.g01490.t01#4    mNeoVis.evergladensis.1.1.chr5.g01518.t01#3   mNeoVis.evergladensis.2.1.chr5.g01512.t01#2       7768   125254052:   mNeoVis.vulgivaga.2.1.chr5.g01516.t01#1    mNeoVis.evergladensis.2.1.chr5.g01513.t01#2    mNeoVis.evergladensis.1.1.chr5.g01519.t01#3    mNeoVis.2.1.1.chr2_hap1.g01574.t01#5    mNeoVis.3.1.1.chr5.g01495.t01#6   mNeoVis.vulgivaga.1.1.chr5.g01491.t01#4       7769   125254053:   mNeoVis.3.1.1.chr5.g01496.t01#6    mNeoVis.2.1.1.chr2_hap1.g01575.t01#5    mNeoVis.evergladensis.1.1.chr5.g01520.t01#3    mNeoVis.vulgivaga.2.1.chr5.g01517.t01#1    mNeoVis.vulgivaga.1.1.chr5.g01492.t01#4   mNeoVis.evergladensis.2.1.chr5.g01514.t01#2       7770   125254054:   mNeoVis.evergladensis.2.1.chr13.g00105.t01#2    mNeoVis.vulgivaga.2.1.chr13.g00102.t01#1    mNeoVis.vulgivaga.1.1.chr13.g00101.t01#4    mNeoVis.3.1.1.chr13.g00057.t01#6    mNeoVis.evergladensis.1.1.chr13.g00100.t01#3   mNeoVis.2.1.1.chr13_hap1.g00107.t01#5       7771   125254055:   mNeoVis.evergladensis.2.1.chr13.g00106.t01#2    mNeoVis.3.1.1.chr13.g00058.t01#6    mNeoVis.evergladensis.1.1.chr13.g00101.t01#3    mNeoVis.2.1.1.chr13_hap1.g00108.t01#5    mNeoVis.vulgivaga.2.1.chr13.g00103.t01#1   mNeoVis.vulgivaga.1.1.chr13.g00102.t01#4       7772   125254056:   mNeoVis.2.1.1.chr15_hap1.g00085.t01#5    mNeoVis.evergladensis.2.1.chr15.g00086.t01#2    mNeoVis.evergladensis.1.1.chr15.g00527.t01#3    mNeoVis.3.1.1.chr15.g00079.t01#6    mNeoVis.vulgivaga.1.1.chr15.g00086.t01#4   mNeoVis.vulgivaga.2.1.chr15.g00087.t01#1       7773   125254057:   mNeoVis.evergladensis.2.1.chr5.g01515.t01#2    mNeoVis.2.1.1.chr2_hap1.g01576.t01#5    mNeoVis.vulgivaga.2.1.chr5.g01518.t01#1    mNeoVis.3.1.1.chr5.g01497.t01#6    mNeoVis.evergladensis.1.1.chr5.g01521.t01#3   mNeoVis.vulgivaga.1.1.chr5.g01493.t01#4       7774   125254058:  mNeoVis.vulgivaga.2.1.chr5.g01519.t01#1               7775   125254059:   mNeoVis.3.1.1.chr5.g01501.t01#6   mNeoVis.vulgivaga.2.1.chr5.g01520.t01#1             7776   125254060:   mNeoVis.vulgivaga.2.1.chr13.g00104.t01#1    mNeoVis.evergladensis.1.1.chr13.g00102.t01#3    mNeoVis.evergladensis.2.1.chr13.g00107.t01#2    mNeoVis.vulgivaga.1.1.chr13.g00103.t01#4    mNeoVis.3.1.1.chr13.g00059.t01#6   mNeoVis.2.1.1.chr13_hap1.g00109.t01#5       7777   125254061:   mNeoVis.3.1.1.chr13.g00060.t01#6    mNeoVis.evergladensis.1.1.chr13.g00103.t01#3    mNeoVis.2.1.1.chr13_hap1.g00110.t01#5    mNeoVis.vulgivaga.2.1.chr13.g00105.t01#1    mNeoVis.evergladensis.2.1.chr13.g00108.t01#2   mNeoVis.vulgivaga.1.1.chr13.g00104.t01#4       7778   125254062:   mNeoVis.vulgivaga.1.1.chr15.g00087.t01#4    mNeoVis.3.1.1.chr15.g00080.t01#6    mNeoVis.evergladensis.1.1.chr15.g00526.t01#3    mNeoVis.2.1.1.chr15_hap1.g00086.t01#5    mNeoVis.evergladensis.2.1.chr15.g00087.t01#2   mNeoVis.vulgivaga.2.1.chr15.g00088.t01#1       7779   125254063:   mNeoVis.evergladensis.1.1.chr5.g01525.t01#3    mNeoVis.vulgivaga.1.1.chr5.g01497.t01#4    mNeoVis.evergladensis.2.1.chr5.g01519.t01#2    mNeoVis.vulgivaga.2.1.chr5.g01521.t01#1    mNeoVis.3.1.1.chr5.g01502.t01#6   mNeoVis.2.1.1.chr2_hap1.g01580.t01#5       7780   125254064:   mNeoVis.evergladensis.1.1.chr5.g01526.t01#3    mNeoVis.evergladensis.2.1.chr5.g01520.t01#2    mNeoVis.vulgivaga.1.1.chr5.g01498.t01#4    mNeoVis.3.1.1.chr5.g01503.t01#6    mNeoVis.2.1.1.chr2_hap1.g01581.t01#5   mNeoVis.vulgivaga.2.1.chr5.g01522.t01#1       7781   125254065:   mNeoVis.vulgivaga.2.1.chr7.g00002.t01#1    mNeoVis.evergladensis.1.1.chr7.g00002.t01#3    mNeoVis.evergladensis.2.1.chr7.g00002.t01#2    mNeoVis.vulgivaga.1.1.chr7.g00002.t01#4   mNeoVis.2.1.1.chr7_hap1.g00002.t01#5       7782   125254066:   mNeoVis.vulgivaga.1.1.chr5.g01499.t01#4    mNeoVis.vulgivaga.2.1.chr5.g01523.t01#1    mNeoVis.evergladensis.1.1.chr5.g01527.t01#3    mNeoVis.evergladensis.2.1.chr5.g01521.t01#2    mNeoVis.2.1.1.chr2_hap1.g01582.t01#5   mNeoVis.3.1.1.chr5.g01504.t01#6       7783   125254067:   mNeoVis.evergladensis.2.1.chr13.g00109.t01#2    mNeoVis.vulgivaga.1.1.chr13.g00105.t01#4    mNeoVis.evergladensis.1.1.chr13.g00104.t01#3    mNeoVis.2.1.1.chr13_hap1.g00111.t01#5   mNeoVis.vulgivaga.2.1.chr13.g00106.t01#1       7784   125254068:   mNeoVis.evergladensis.1.1.chr5.g01528.t01#3    mNeoVis.vulgivaga.1.1.chr5.g01500.t01#4    mNeoVis.2.1.1.chr2_hap1.g01583.t01#5    mNeoVis.3.1.1.chr5.g01505.t01#6    mNeoVis.vulgivaga.2.1.chr5.g01524.t01#1   mNeoVis.evergladensis.2.1.chr5.g01522.t01#2       7785   125254069:   mNeoVis.evergladensis.2.1.chr5.g01523.t01#2    mNeoVis.2.1.1.chr2_hap1.g01584.t01#5    mNeoVis.vulgivaga.2.1.chr5.g01525.t01#1    mNeoVis.evergladensis.1.1.chr5.g01529.t01#3    mNeoVis.vulgivaga.1.1.chr5.g01501.t01#4   mNeoVis.3.1.1.chr5.g01506.t01#6       7786   125254070:   mNeoVis.2.1.1.chr13_hap1.g00112.t01#5    mNeoVis.evergladensis.2.1.chr13.g00110.t01#2    mNeoVis.vulgivaga.2.1.chr13.g00107.t01#1    mNeoVis.vulgivaga.1.1.chr13.g00106.t01#4   mNeoVis.evergladensis.1.1.chr13.g00105.t01#3       7787   125254071:   mNeoVis.evergladensis.2.1.chr5.g01524.t01#2    mNeoVis.3.1.1.chr5.g01507.t01#6    mNeoVis.2.1.1.chr2_hap1.g01585.t01#5    mNeoVis.evergladensis.1.1.chr5.g01530.t01#3    mNeoVis.vulgivaga.1.1.chr5.g01502.t01#4   mNeoVis.vulgivaga.2.1.chr5.g01526.t01#1       7788   125254072:   mNeoVis.vulgivaga.1.1.chr13.g00107.t01#4    mNeoVis.vulgivaga.2.1.chr13.g00108.t01#1    mNeoVis.evergladensis.2.1.chr13.g00111.t01#2    mNeoVis.2.1.1.chr13_hap1.g00113.t01#5   mNeoVis.evergladensis.1.1.chr13.g00106.t01#3       7789   125254073:   mNeoVis.2.1.1.chr15_hap1.g00087.t01#5    mNeoVis.evergladensis.2.1.chr15.g00088.t01#2    mNeoVis.evergladensis.1.1.chr15.g00525.t01#3    mNeoVis.vulgivaga.2.1.chr15.g00089.t01#1   mNeoVis.vulgivaga.1.1.chr15.g00088.t01#4       7790   125254074:   mNeoVis.vulgivaga.1.1.chr13.g00108.t01#4    mNeoVis.evergladensis.1.1.chr13.g00108.t01#3    mNeoVis.evergladensis.2.1.chr13.g00112.t01#2    mNeoVis.vulgivaga.2.1.chr13.g00109.t01#1   mNeoVis.2.1.1.chr13_hap1.g00114.t01#5       7791   125254075:  mNeoVis.vulgivaga.2.1.chr5.g01527.t01#1               7792   125254076:   mNeoVis.vulgivaga.1.1.chr5.g01503.t01#4    mNeoVis.vulgivaga.2.1.chr5.g01528.t01#1    mNeoVis.evergladensis.1.1.chr5.g01531.t01#3    mNeoVis.2.1.1.chr2_hap1.g01586.t01#5   mNeoVis.evergladensis.2.1.chr5.g01525.t01#2       7793   125254077:   mNeoVis.vulgivaga.1.1.chr5.g01504.t01#4    mNeoVis.evergladensis.2.1.chr5.g01526.t01#2    mNeoVis.evergladensis.1.1.chr5.g01532.t01#3    mNeoVis.3.1.1.chr5.g01508.t01#6    mNeoVis.2.1.1.chr2_hap1.g01587.t01#5   mNeoVis.vulgivaga.2.1.chr5.g01529.t01#1       7794   125254078:   mNeoVis.evergladensis.2.1.chr13.g00113.t01#2   mNeoVis.vulgivaga.2.1.chr13.g00110.t01#1           7795   125254079:   mNeoVis.evergladensis.1.1.chr13.g00110.t01#3    mNeoVis.vulgivaga.2.1.chr13.g00111.t01#1    mNeoVis.2.1.1.chr13_hap1.g00117.t01#5    mNeoVis.vulgivaga.1.1.chr13.g00109.t01#4   mNeoVis.3.1.1.chr13.g00061.t01#6       7796   125254080:   mNeoVis.vulgivaga.2.1.chr13.g00112.t01#1    mNeoVis.evergladensis.1.1.chr13.g00111.t01#3    mNeoVis.evergladensis.2.1.chr13.g00115.t01#2    mNeoVis.vulgivaga.1.1.chr13.g00110.t01#4   mNeoVis.2.1.1.chr13_hap1.g00118.t01#5       7797   125254081:   mNeoVis.evergladensis.1.1.chr13.g00112.t01#3    mNeoVis.3.1.1.chr13.g00063.t01#6    mNeoVis.evergladensis.2.1.chr13.g00116.t01#2    mNeoVis.vulgivaga.2.1.chr13.g00113.t01#1    mNeoVis.vulgivaga.1.1.chr13.g00111.t01#4   mNeoVis.2.1.1.chr13_hap1.g00119.t01#5       7798   125254082:   mNeoVis.vulgivaga.1.1.chr5.g01505.t01#4    mNeoVis.3.1.1.chr5.g01509.t01#6    mNeoVis.evergladensis.1.1.chr5.g01533.t01#3    mNeoVis.2.1.1.chr2_hap1.g01588.t01#5    mNeoVis.evergladensis.2.1.chr5.g01527.t01#2   mNeoVis.vulgivaga.2.1.chr5.g01530.t01#1       7799   125254083:   mNeoVis.2.1.1.chr2_hap1.g01589.t01#5    mNeoVis.evergladensis.2.1.chr5.g01528.t01#2    mNeoVis.vulgivaga.2.1.chr5.g01531.t01#1    mNeoVis.vulgivaga.1.1.chr5.g01506.t01#4    mNeoVis.3.1.1.chr5.g01510.t01#6   mNeoVis.evergladensis.1.1.chr5.g01534.t01#3       7800   125254084:  mNeoVis.vulgivaga.2.1.chr7.g00003.t01#1               7801   125254085:   mNeoVis.evergladensis.1.1.chr5.g01535.t01#3    mNeoVis.evergladensis.2.1.chr5.g01529.t01#2    mNeoVis.vulgivaga.2.1.chr5.g01532.t01#1    mNeoVis.2.1.1.chr2_hap1.g01590.t01#5    mNeoVis.vulgivaga.1.1.chr5.g01507.t01#4   mNeoVis.3.1.1.chr5.g01511.t01#6       7802   125254086:   mNeoVis.vulgivaga.1.1.chr13.g00112.t01#4    mNeoVis.evergladensis.1.1.chr13.g00113.t01#3    mNeoVis.2.1.1.chr13_hap1.g00120.t01#5    mNeoVis.evergladensis.2.1.chr13.g00117.t01#2   mNeoVis.vulgivaga.2.1.chr13.g00114.t01#1       7803   125254087:   mNeoVis.evergladensis.2.1.chr5.g01530.t01#2    mNeoVis.3.1.1.chr5.g01512.t01#6    mNeoVis.2.1.1.chr2_hap1.g01591.t01#5    mNeoVis.vulgivaga.2.1.chr5.g01533.t01#1    mNeoVis.evergladensis.1.1.chr5.g01536.t01#3   mNeoVis.vulgivaga.1.1.chr5.g01508.t01#4       7804   125254088:   mNeoVis.evergladensis.2.1.chr5.g01531.t01#2    mNeoVis.2.1.1.chr2_hap1.g01592.t01#5    mNeoVis.3.1.1.chr5.g01513.t01#6    mNeoVis.vulgivaga.2.1.chr5.g01534.t01#1    mNeoVis.evergladensis.1.1.chr5.g01537.t01#3   mNeoVis.vulgivaga.1.1.chr5.g01509.t01#4       7805   125254089:   mNeoVis.vulgivaga.2.1.chr5.g01535.t01#1    mNeoVis.evergladensis.2.1.chr5.g01532.t01#2    mNeoVis.evergladensis.1.1.chr5.g01538.t01#3    mNeoVis.2.1.1.chr11_hap1.g00414.t01#5    mNeoVis.vulgivaga.1.1.chr5.g01510.t01#4    mNeoVis.2.1.1.chr2_hap1.g01593.t01#5   mNeoVis.3.1.1.chr5.g01514.t01#6       7806   125254090:   mNeoVis.2.1.1.chr13_hap1.g00121.t01#5    mNeoVis.evergladensis.1.1.chr13.g00114.t01#3    mNeoVis.3.1.1.chr13.g00065.t01#6    mNeoVis.vulgivaga.1.1.chr13.g00113.t01#4    mNeoVis.evergladensis.2.1.chr13.g00118.t01#2   mNeoVis.vulgivaga.2.1.chr13.g00115.t01#1       7807   125254091:   mNeoVis.evergladensis.2.1.chr5.g01534.t01#2    mNeoVis.vulgivaga.2.1.chr5.g01537.t01#1    mNeoVis.vulgivaga.1.1.chr5.g01512.t01#4    mNeoVis.evergladensis.1.1.chr5.g01540.t01#3    mNeoVis.3.1.1.chr5.g01516.t01#6   mNeoVis.2.1.1.chr2_hap1.g01595.t01#5       7808   125254092:   mNeoVis.3.1.1.chr5.g01515.t01#6    mNeoVis.evergladensis.2.1.chr5.g01533.t01#2    mNeoVis.vulgivaga.2.1.chr5.g01536.t01#1    mNeoVis.evergladensis.1.1.chr5.g01539.t01#3    mNeoVis.vulgivaga.1.1.chr5.g01511.t01#4   mNeoVis.2.1.1.chr2_hap1.g01594.t01#5       7809   125254093:   mNeoVis.evergladensis.1.1.chr13.g00115.t01#3    mNeoVis.3.1.1.chr13.g00066.t01#6    mNeoVis.vulgivaga.2.1.chr13.g00116.t01#1    mNeoVis.evergladensis.2.1.chr13.g00119.t01#2    mNeoVis.2.1.1.chr13_hap1.g00122.t01#5   mNeoVis.vulgivaga.1.1.chr13.g00114.t01#4       7810   125254094:   mNeoVis.evergladensis.1.1.chr13.g00116.t01#3    mNeoVis.3.1.1.chr13.g00067.t01#6    mNeoVis.vulgivaga.2.1.chr13.g00117.t01#1    mNeoVis.evergladensis.2.1.chr13.g00120.t01#2    mNeoVis.2.1.1.chr13_hap1.g00123.t01#5   mNeoVis.vulgivaga.1.1.chr13.g00115.t01#4       7811   125254095:   mNeoVis.vulgivaga.2.1.chr13.g00118.t01#1    mNeoVis.evergladensis.2.1.chr13.g00121.t01#2    mNeoVis.3.1.1.chr13.g00068.t01#6    mNeoVis.evergladensis.1.1.chr13.g00117.t01#3    mNeoVis.vulgivaga.1.1.chr13.g00116.t01#4   mNeoVis.2.1.1.chr13_hap1.g00124.t01#5       7812   125254096:   mNeoVis.2.1.1.chr2_hap1.g01505.t01#5    mNeoVis.evergladensis.1.1.chr5.g01446.t01#3    mNeoVis.evergladensis.2.1.chr5.g01442.t01#2    mNeoVis.vulgivaga.2.1.chr5.g01443.t01#1    mNeoVis.vulgivaga.1.1.chr5.g01419.t01#4   mNeoVis.3.1.1.chr5.g01427.t01#6       7813   125254097:   mNeoVis.vulgivaga.1.1.chr5.g01420.t01#4    mNeoVis.3.1.1.chr5.g01428.t01#6    mNeoVis.evergladensis.1.1.chr5.g01447.t01#3    mNeoVis.2.1.1.chr2_hap1.g01506.t01#5    mNeoVis.evergladensis.2.1.chr5.g01443.t01#2   mNeoVis.vulgivaga.2.1.chr5.g01444.t01#1       7814   125254098:   mNeoVis.3.1.1.chr5.g01426.t01#6    mNeoVis.vulgivaga.1.1.chr5.g01418.t01#4    mNeoVis.2.1.1.chr2_hap1.g01504.t01#5    mNeoVis.evergladensis.2.1.chr5.g01441.t01#2    mNeoVis.evergladensis.1.1.chr5.g01445.t01#3   mNeoVis.vulgivaga.2.1.chr5.g01442.t01#1       7815   125254099:   mNeoVis.2.1.1.chr4_hap1.g01043.t01#5    mNeoVis.vulgivaga.1.1.chr6.g01040.t01#4    mNeoVis.vulgivaga.2.1.chr6.g01040.t01#1    mNeoVis.evergladensis.2.1.chr6.g01048.t01#2   mNeoVis.evergladensis.1.1.chr6.g01050.t01#3       7816   125254100:   mNeoVis.3.1.1.chr13.g00069.t01#6    mNeoVis.evergladensis.2.1.chr13.g00122.t01#2    mNeoVis.2.1.1.chr13_hap1.g00125.t01#5    mNeoVis.vulgivaga.2.1.chr13.g00119.t01#1    mNeoVis.vulgivaga.1.1.chr13.g00117.t01#4   mNeoVis.evergladensis.1.1.chr13.g00118.t01#3       7817   125254101:   mNeoVis.evergladensis.2.1.chr15.g00089.t01#2    mNeoVis.evergladensis.1.1.chr15.g00524.t01#3    mNeoVis.vulgivaga.2.1.chr15.g00090.t01#1    mNeoVis.vulgivaga.1.1.chr15.g00089.t01#4    mNeoVis.3.1.1.chr15.g00082.t01#6   mNeoVis.2.1.1.chr15_hap1.g00088.t01#5       7818   125254102:   mNeoVis.3.1.1.chr5.g01430.t01#6    mNeoVis.vulgivaga.2.1.chr5.g01445.t01#1    mNeoVis.evergladensis.1.1.chr5.g01449.t01#3    mNeoVis.evergladensis.2.1.chr5.g01444.t01#2    mNeoVis.vulgivaga.1.1.chr5.g01421.t01#4   mNeoVis.2.1.1.chr2_hap1.g01507.t01#5       7819   125254103:   mNeoVis.3.1.1.chr13.g00070.t01#6    mNeoVis.evergladensis.2.1.chr13.g00123.t01#2    mNeoVis.evergladensis.1.1.chr13.g00119.t01#3    mNeoVis.2.1.1.chr13_hap1.g00126.t01#5   mNeoVis.vulgivaga.2.1.chr13.g00120.t01#1       7820   125254104:   mNeoVis.vulgivaga.1.1.chr13.g00038.t01#4    mNeoVis.evergladensis.2.1.chr13.g00039.t01#2    mNeoVis.evergladensis.1.1.chr13.g00036.t01#3    mNeoVis.2.1.1.chr13_hap1.g00040.t01#5    mNeoVis.3.1.1.chr13.g00021.t01#6   mNeoVis
